# A draft of human N-glycans of glycoRNA

**DOI:** 10.1101/2023.09.18.558371

**Authors:** Ming Bi, Zirui Zhang, Tao Wang, Hongwei Liang, Zhixin Tian

**Author notes:** These authors made equal contributions (Ming Bi, Zirui Zhang).

## Abstract

In addition to the backbone molecules of proteins and lipids, RNAs have recently been found to be N-glycosylated as well in cell models. Some overlap of N-glycans between RNA and protein exist in terms of monosaccharide composition. Here we report a draft of human tissue N-glycans of glycoRNA covering 12 typical organs as characterized by mass spectrometry-based N-glycomics. RNAs were first prepared, N-glycans were then enzymatically released, hydrophilically enriched, permethylated, analyzed by RPLC-MS/MS, and finally identified by N-glycan search engine GlySeeker. A total of 676 putative sequence structures with 236 monosaccharide compositions were identified across the 12 organs. Organ-specific similarity and heterogeneity of N-glycosylation in glycoRNAs were annotated. This first comprehensive draft of human glycoRNAs serves a foundation for future structural and functional studies.

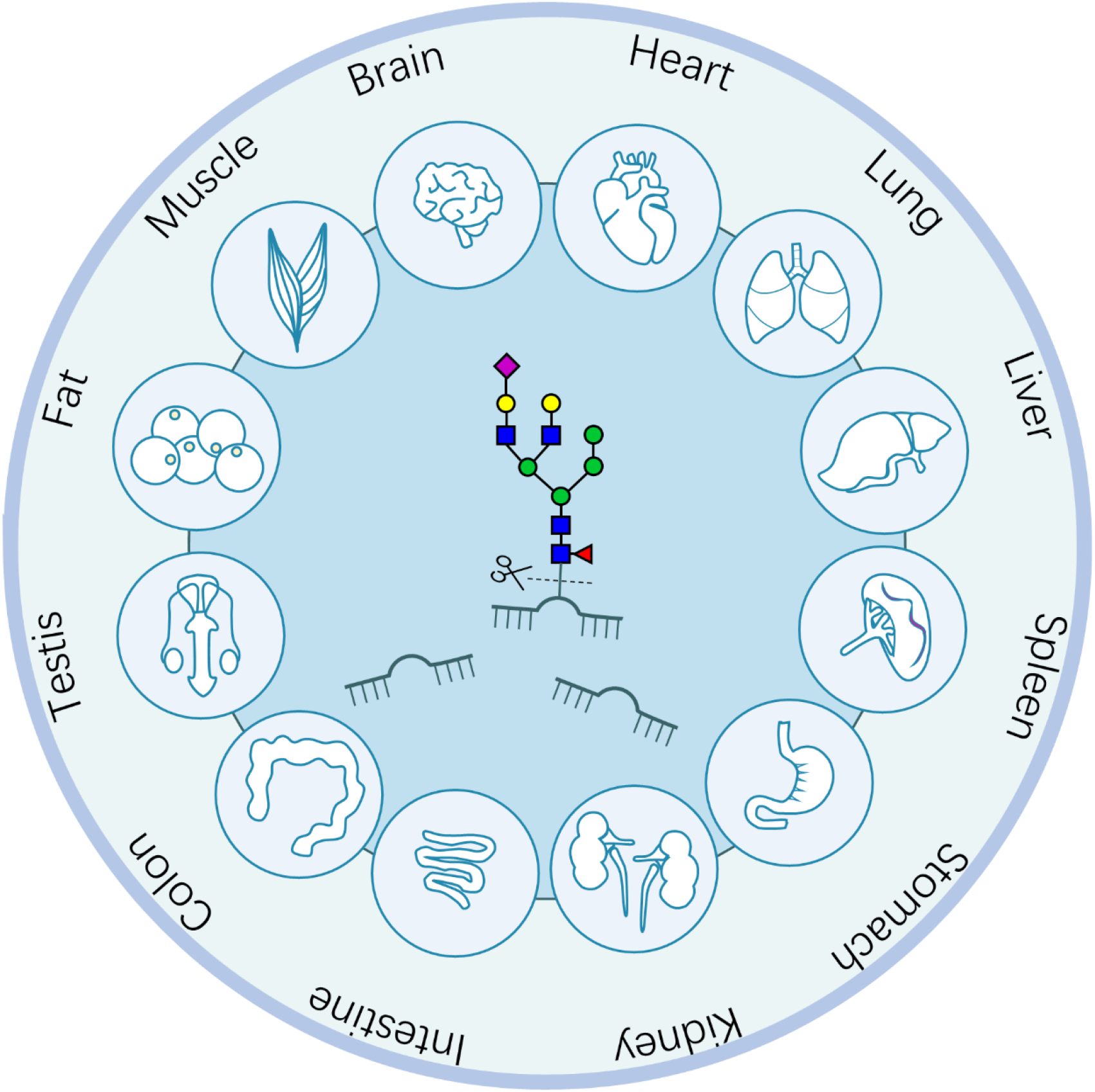
TOC Graphic.

## 1 INTRODUCTION

Glycobiology is defined as the study of the monosaccharide composition, structure and biological function of carbohydrates, primarily those associated with proteins or lipids^1^. However, the concept of glycobiology has undergone a significant expansion with the finding of glycoRNA, a term coined to describe ribonucleic acid modified with N-glycans. In the initial discovery of glycoRNA, Flynn et al ^2^ extracted small RNAs from 293T, H9 and Hela cells, uncovering 260 distinct monosaccharide compositions using PGC-LC-MS/MS analysis and GlycoNote identification based on exact mass^3^. Notably, an increase in the fucosylated N-glycans in glycoRNAs was observed, compared to glycopeptides in 293T and H9 cells, while HeLa cells exhibited a higher degree of sialylated N-glycans in glycoRNAs than in glycopeptides. This discovery, highlighting the predominance of sialic acid residues in glycoRNA, led to the development of techniques such as the sialic acid aptamer and RNA in situ hybridization-mediated proximity ligation assay (ARPLA)^4^; the monocyte–endothelial cell interactions suggested that glycoRNAs may mediate cell–cell interactions during the immune response. A periodate oxidation and aldehyde ligation method named “rPAL” was developed to label native sialoglycoRNAs from isolated RNA samples^5^. A chemoenzymatic method, termed solid-phase capture of oxidized N-linked glycoRNAs (SPCgRNA), was developed to capture glycoRNA by hydrazide chemistry^6^. The captured glycoRNAs were digested with PNGase F, confirming glycosylation in various RNA substrates including miRNA, snoRNA (small nucleolar RNA), snRNA (small nuclear RNA), rRNA (rRNA), and Y_RNA. With LC-MS/MS analysis and GlycoWorkBench identification, 15 and 9 N-glycan compositions in hTERT-HPNE and MIA PaCa-2 cancer cells were found^7^.

Here we report a draft of human tissue N-glycans of glycoRNA covering 12 typical organs as characterized by mass spectrometry-based N-glycomics. RNAs were first prepared, N-glycans were then enzymatically released, hydrophilically enriched, permethylated, analyzed by RPLC-MS/MS, and finally identified by N-glycan search engine GlySeeker^8^. A total of 676 putative sequence structures with 236 monosaccharide compositions were identified across the 12 organs. Organ-specific similarity and heterogeneity of N-glycosylation in glycoRNAs were annotated. This first comprehensive draft of human glycoRNAs serves a foundation for future structural and functional studies.

## 2 MATERIALS AND METHODS

### Chemicals and reagents

TRIzol, ethanol (EtOH), dimethyl sulfoxide (DMSO) and Proteinase K were purchased from Thermo Fisher Scientific (San Jose, CA, USA). PNGase F from Elizabethkingia meningoseptica (NEB), acetonitrile (ACN), trifluoroacetic acid (TFA) and formic acid (FA) were purchased from Sigma-Aldrich (St. Louis, USA). Ammonium bicarbonate (ABC) was bought from Sangon Biotech (Shanghai, China). Iodomethane was brought from TCI (Shanghai, China) and chloroform was from Sinopharm Chemical Reagent (Shanghai, China). Ultrapure water was produced onsite with a Millipore Simplicity System (Billerica, MA, USA).

### Tissues samples

In total, 12 diverse organ tissues were collected from Biobank of Nanjing Drum Tower Hospital (Nanjing, China). The organs/tissues samples were immediately frozen in liquid nitrogen when they were collected and stored at −80 °C. The experiments were performed in accordance with the Code of Ethics of the World Medical Association (Declaration of Helsinki) and the guidelines of Nanjing University and approved by the ethics committee of Nanjing University Medical School (2020-158-01).

### Preparation of N-glycans from glycoRNA

The RNAs were isolated from 12 diverse organs/tissues following a previous report^2^. Initially, TRIzol reagent (Thermo Fisher Scientific) was used to lyse and denature tissues. Homogenization was performed in TRIzol by pipetting, followed by an incubation step at 37 ℃ to further denature non-covalent interactions. Phase separation was initiated by adding 0.2x volumes of 100% chloroform, which was mixed by vortexing and then spun at 12,000x g for 15 minutes at 4 ℃. The resulting aqueous phase was carefully transferred to a fresh tube and mixed with 2x volumes of 100% ethanol (EtOH). This solution was purified using a Zymo RNA cleaning and concentrating column (Zymo Research): the sample solution was added to Zymo columns and spun at 10,000x g for 20 seconds, discarding the flow-through. Three separate washes were performed using 1x 400 mL of RNA Prep Buffer (Zymo Research) and 2x 400uL RNA Wash Buffer (Zymo Research), spinning each time at 10,000x g for 20 seconds. To elute the RNAs, two volumes of pure water were used. Subsequently, the RNA underwent protein digestion by adding 1 mg of Proteinase K (PK, Thermo Fisher Scientific) to 25 mg of purified RNA and incubating it at 37 ℃ for 45 minutes. After PK digestion, the RNAs were purified again using a Zymo RNA clean and concentrator as described above. Next, the RNA samples were sequentially digested with two glycosidases. Typically, for experimental samples, 25 mg of small RNA from the 12 diverse organs/tissues was resuspended in 10 mL of 1x GlycoBuffer 2 (NEB) and 7.5 mL PNGaseF (NEB), and the final reaction volume was adjusted to 100 mL with water. PNGaseF cleavage occurred overnight at 37 ℃. After digestion, the released glycans were subjected to enrichment and permethylation.

### Enrichment and permethylation of N-glycans

The PNGaseF digestion solution was subjected to purification using HyperCarb columns (Thermo Scientific, USA). In brief, Porous Graphitic Carbon (PGC) columns were first activated with ACN and equilibrated with 0.1% TFA solution. The N-glycan digestion solution was repeatedly loaded onto the column for five times; after washing with 0.1% TFA, N-glycans were eluted using 50% ACN and 0.1% TFA; the eluant was dried in a Speed-Vac (Thermo Fisher). The dried N-glycans were resuspended in a solution consisting of 2 μL water, 50 μL dimethyl sulfoxide (DMSO), and 30 μL iodomethane. The mixture was loaded onto a homemade sodium hydroxide (0.1 g) column five times. Subsequently, an additional 30 μL of iodomethane was added to the remaining solution, which was allowed to stand for 30 minutes. After undergoing ACN washing thrice and quenching the reaction with 2 μL acetic acid, the glycan elution was mixed with 200 μL chloroform. The mixture was washed ten times with water, and the chloroform was subsequently evaporated under vacuum. The resulting samples were re-suspended in 30 μL 0.1% TFA for subsequent LC-MS/MS analysis.

### C18-RPLC-ESI-MS/MS analysis of the permethylated N-glycans

C18-RPLC-ESI-MS/MS analysis of the permethylated N-glycans were carried out on a Dionex UltiMate 3000 nano-HPLC system coupled with a nano-electrospray ionization (nano-ESI) Q Exactive mass spectrometer (Thermo Fisher Scientific, San Jose, CA, USA).

Permethylated N-glycans were trapped on a C18 (5 μm, 300 Å, Jupiter, Phenomenex) trap column and separated on a C18 analytical column (75 μid, 60 cm long). Buffer A is 0.1% TFA, and Buffer B is 95% ACN with 0.2% TFA. The flow rate of the mobile phase was 300 nL/min. The elution gradient is as follows: 1-25% B for 20 min, up to 60% B in 190 min, up to 95% B in 10 min, kept at 95% B for 10 min, down to 1% B in 5 min and kept at 1% B for 15 min for equilibrium.

Mass spectra were acquired in the *m/z* range 500-2500 using a mass resolution 70 k. For MS/MS spectra, the mass resolution was set at 17,500. Fragmentation was obtained in a data-dependent mode (Top20) with normalized collisional energy of 10%. The automatic gain control (AGC) target values for MS and MS/MS were placed at 2 × 10^5^ and 2 × 10^5^, respectively; the corresponding maximum injection time were 50 and 250 ms. The isolation window and dynamic exclusion time window are 3.0 *m/z* and 20.0 s. The temperature of the ion transfer capillary was set to 280 °C and the spray voltage was set to 2.8 kV.

With the aforementioned instrumental parameters, three technical replicates of RPLC-MS/MS (HCD) analysis of N-glycans from each organ were carried out.

### Database search and identification of N-glycans by GlySeeker and GlycoNote

For identification by GlySeeker^8-10^, the search parameters of isotopic peak *m/z* deviation (IPMD), isotopic peak abundance cutoff (IPACO) and isotopic peak abundance deviation (IPAD) were set at 15 ppm/40%/ 50% for precursor ions and 15 ppm/20%/ 30% for fragment ions. A/B/C/X/Y/Z ions were comprehensively interpreted. Monosaccharide sequence isomers (i.e., same composition but different sequences) were distinguished and confirmed with structure-diagnostic fragment ions (i.e., glycoform score).

## 3 RESULTS

### Overall identification

The glycoRNA N-glycans from 12 human healthy organs were extracted, analyzed by RPLC-MS/MS(HCD) in triplicates, and identified by N-glycan search engine GlySeeker. Across the 12 different organs studied, a total of 677 N-glycan linkages were identified, corresponding to 236 distinct compositions (**Figure 1a**). Remarkably, the linkages exhibited significant variability among the different organs. Specifically, the number of linkages ranged from 20 in the heart to 253 in the colon, with intermediate values for brain (65), lung (59), liver (68), spleen (183), stomach (199), kidney (185), intestine (185), testis (75), fat (110), and muscle (24). The detailed MS spectral and identification information for the N-glycans identified by GlySeeker from the 12 human organs is provided in **Supplemental Information 1** (12 sheets, one for each organ). The isotopic envelope fingerprinting map, annotated MS/MS spectrum with the matched fragment ions, and graphical fragmentation map of each N-glycan identified from the 12 human organs are provided in **Supplementary information 2-13** (PDF files, 12 in total, one for each organ). Furthermore, the number of distinct compositions for these N-glycan linkages varied as well, ranging from 17 in the heart to 118 in the spleen, and with different numbers for other organs, including brain (32), lung (39), liver (43), stomach (79), kidney (89), intestine (89), colon (75), testis (60), fat (20), and muscle (20). These variations can be attributed to the presence of sequence isomers, wherein the same composition is linked to different structural arrangements. The percentage of isomer varied across the organs, with heart exhibiting the lowest percentage of isomers at 11.85% (comprising 2 isomers), while colon displayed the highest isomerism at 59.6% (comprising 53 isomers).

**Figure 1.**
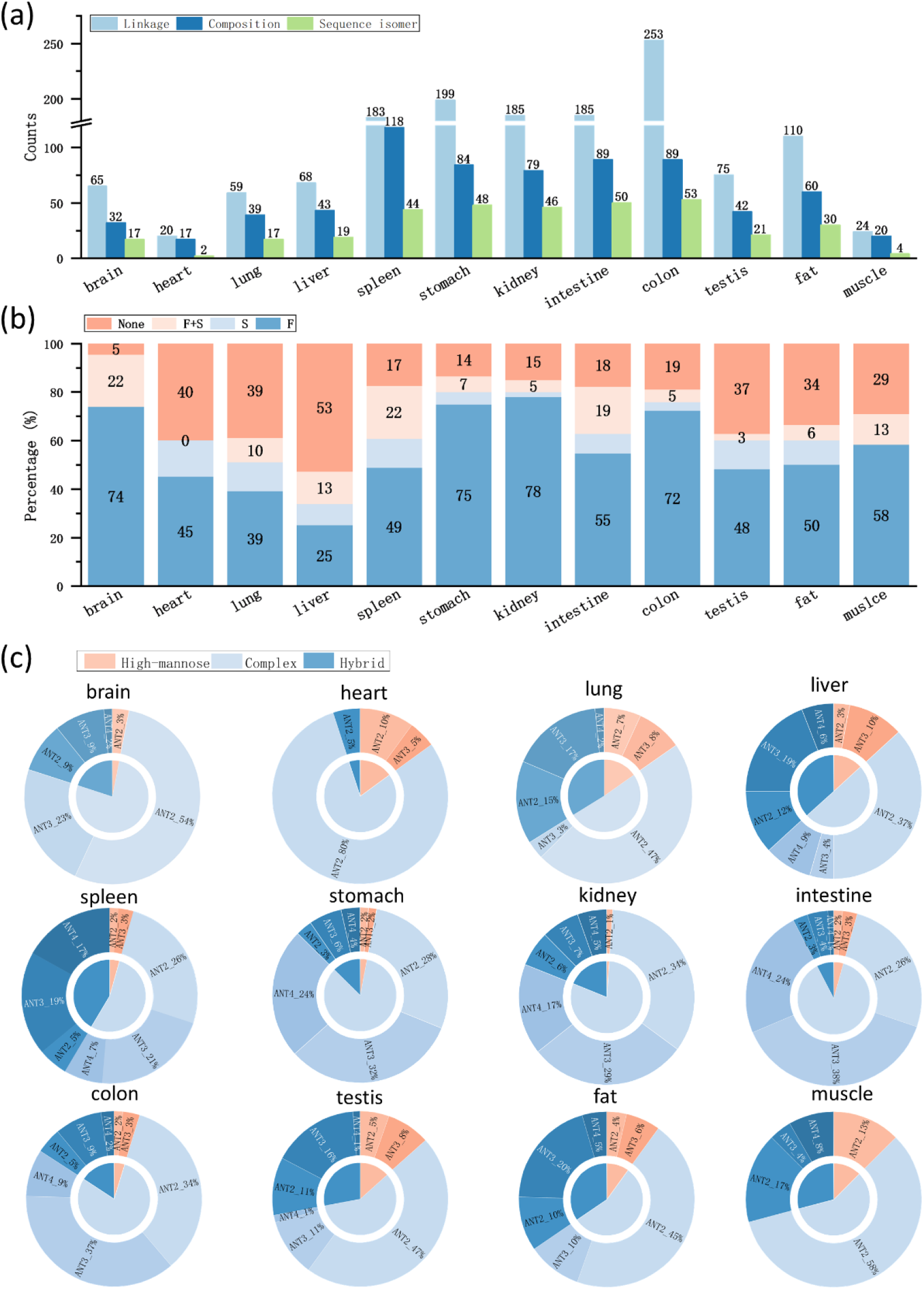
Overview of the glycoRNA N-glycans identified from 12 organs of human. **(a)** Number of linkages, compositions and sequence isomers. **(b)** Percentages of N-glycans without any fucosylation (F) or sialylation (S) (None), with both F and S (F+S), with S only (S) and with F only (F). **(c)** Distribution of N-glycan types (high-mannose, complex, hybrid) and number of antenna.

In terms of glycan composition, our human N-glycan database comprises seven monosaccharides: fucose (F), glucose (G), mannose (M), galactose (G), N-acetylglucosamine (Y), N-acetylgalactosamine (V) and N-acetylneuraminic acid (S). Notably, the database excludes N-glycolylneuraminic acid (Neu5Gc) due to the absence of cytidine-5’-monophosphate-N-acetylneuraminic acid (CMAH) in healthy human body ^11^. In the context of N-glycans attached to RNA, it is notable that these glycans predominantly contain fucose, with a significant presence, while sialic acid occupies a smaller proportion (**Figure 1b**). Among the 12 examined organs, the majority of them exhibited over 50% of N-glycan linkages containing fucose. Liver displayed the lowest percentage, with approximately 38% of N-glycan linkages containing fucose, while the brain stood out with approximately 96% of its N-glycans containing fucose. In terms of sialic acid content, it ranged from 7% to 34% in different organs. While N-glycans modified with both fucose and sialic acid simultaneously were relatively uncommon in RNA.

The presence of fucose and sialic acid in these N-glycans suggests a complexity in the N-glycan structures attached to RNA. The identified N-glycans were further analyzed in terms of types and antennas (**Figure 1c**). High-mannose, complex and hybrid types of N-glycan occupied distinct proportions. Within our database, we cataloged 20 different types of high-mannose N-glycans, although they constituted a relatively small portion of the overall database. Consequently, the number of high-mannose N-glycans identified in our results was proportionately limited. Notably, the kidney exhibited the lowest percentage of high-mannose N-glycans, accounting for only 1% of the N-glycan content. In contrast, complex N-glycan types displayed a clear predominance over hybrid types. The intestine, heart, and stomach stood out with over 80% of their N-glycan content being of the complex type, while all examined organs contained more than 50% complex N-glycans. Additionally, we observed variations in the antenna structures corresponding to different N-glycan types (**Figure 1c**). Tetra-antenna types of N-glycans were present in small percentage across all examined organs.

For the heterogeneity of RNA glycans across different organs (**Figure 2a-2b**), out of the 677 identified N-glycan structures, 431 were unique to a single organ, constituting 63.8% of the total, while 109 linkages were observed in two organs, representing 16.1%. The remaining N-glycans were identified in more than two organs. Only six N-glycans were found consistently across all the 12 organs, indicating limited overlap and highlighting the organ-specific nature of RNA glycans. The 109 N-glycans shared by two organs are detailed in **Figure 2c**. Notably, spleen, stomach, kidney, intestine, and colon exhibited a higher number of unique N-glycans, ranging from 64 to 92 unique structures. In contrast, there was relatively little overlap between pairs of organs. Specifically, the stomach shared 15 and 22 N-glycans with the kidney and colon, respectively, while the spleen shared 10 N-glycans with the intestine. **Figure 2d** displays N-glycans that were shared by more than three organs, highlighting the diversity of glycan structures that are common across multiple organs.

**Figure 2.**
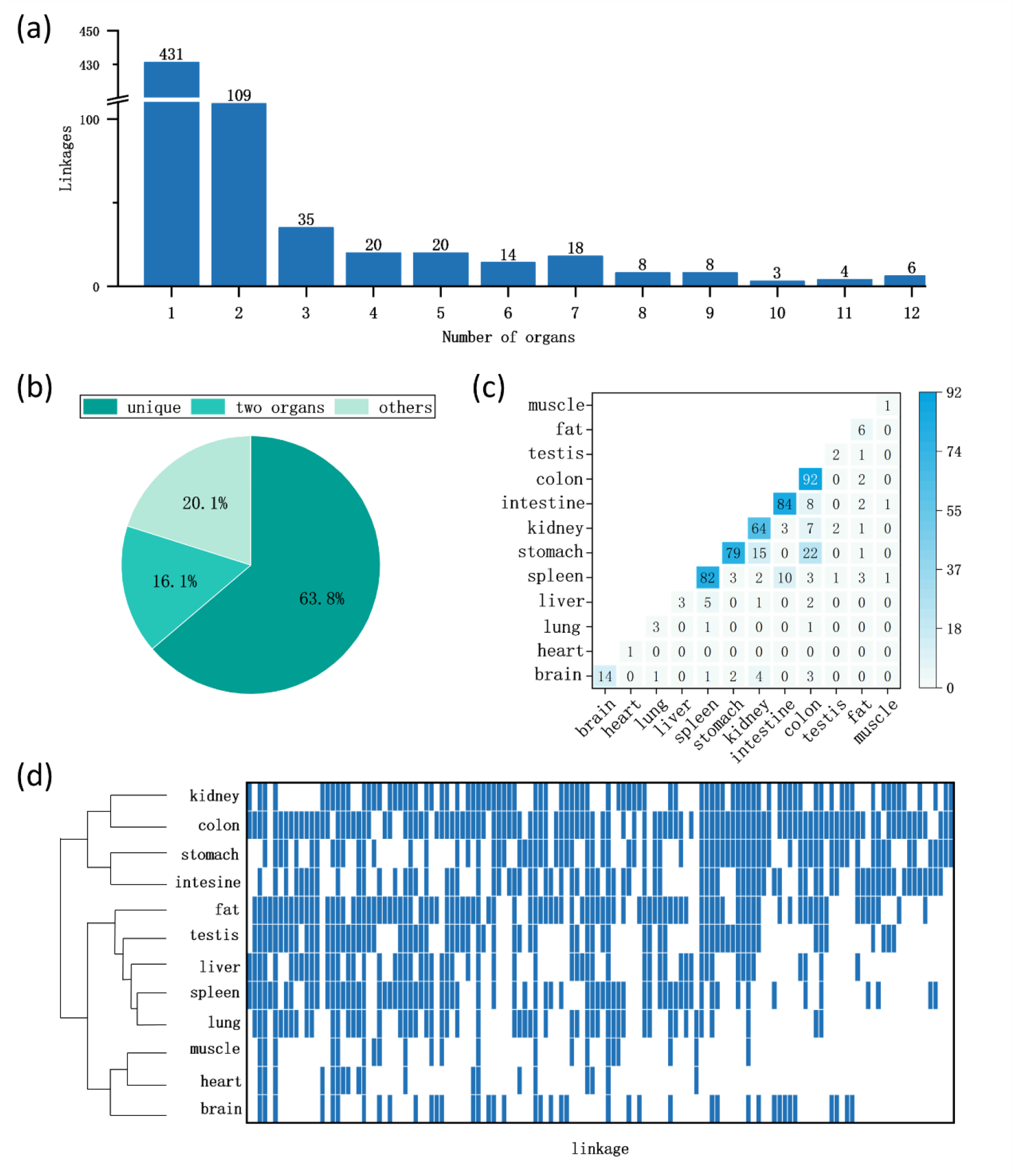
Overlap of the identified glycoRNA N-glycans among the 12 organs of human. **(a)** Distribution of organ-specific occurrence. **(b)** Percentages of unique, two and more than two occurrences. **(c)** Crosstalk between every two organs. **(d)** Heatmap of N-glycans with no less than three occurrences across the 12 organs.

### High-mannose N-glycans

The identification and quantification of 14 high-mannose N-glycans across the 12 organs were illustrated in **Figure 3a**. Despite the identification of only 3 high-mannose N-glycans in the heart, their abundance was relatively high, especially 01Y41Y41M(31M21M21M31G31G21G)61M(31M)61M. N-glycan N2H4F0S0 (Man4) was exclusively identified in muscle, whereas N-glycan N2H5F0S0 (Man5) was quantified in all examined organs, with relative abundances ranging from 0.14% to 3.42%. It’s worth noting that Man5 corresponds to a single structure, namely, 01Y41Y41M(31M)61M(31M)61M. Therapeutic antibodies with Man5 which exhibited enhanced antibody-dependent cell-mediated cytotoxicity (ADCC) compared with antibodies with fucosylated complex or hybrid glycans and increased binding affinity to the Fc gamma RIIIA ^12^. Furthermore, faster clearance of antibody with high-mannose glycoform increased the advantage of therapy ^12^.

**Figure 3.**
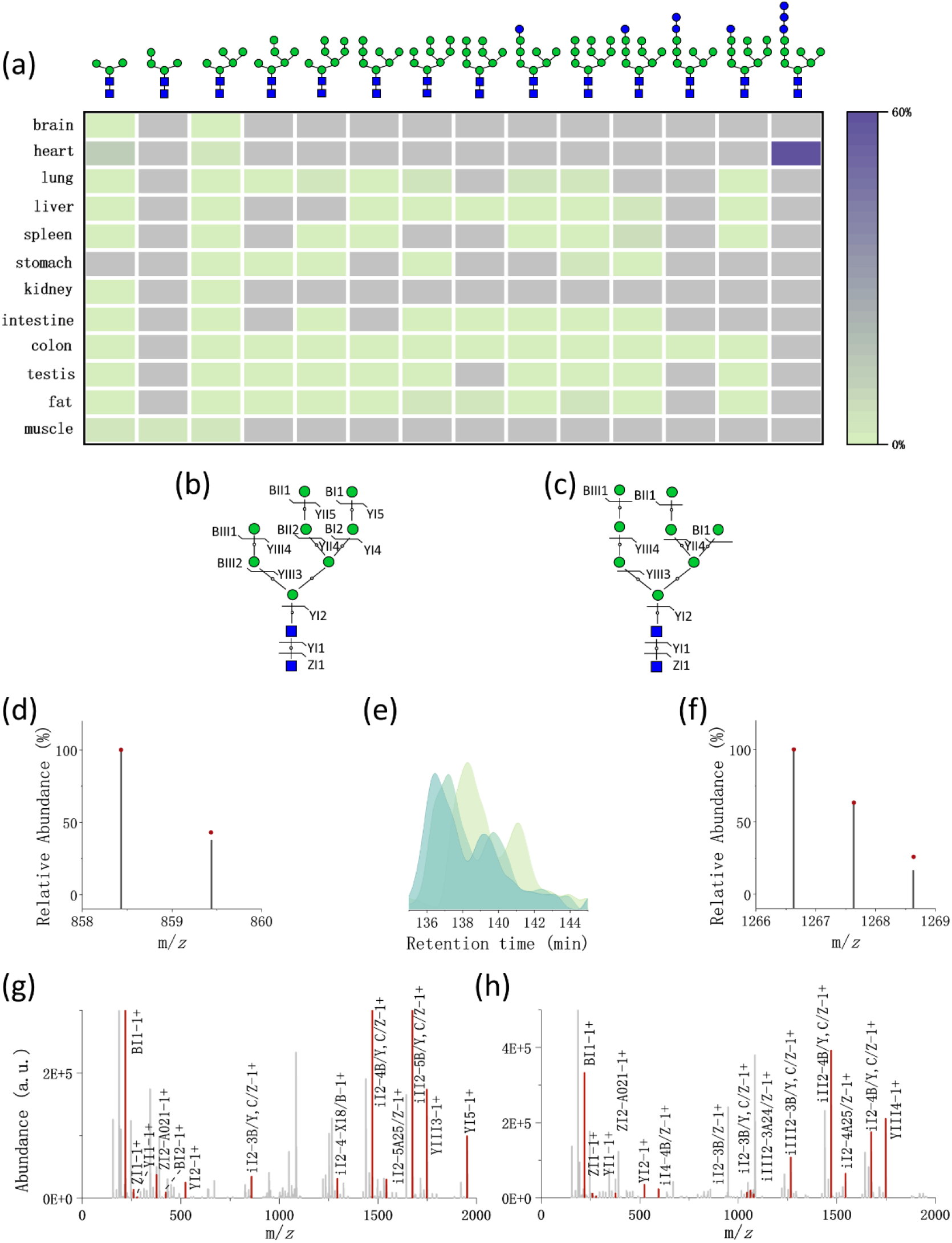
High-mannose N-glycans identified from the glycoRNAs of the 12 organs of human. **(a)** Heatmap of relative abundance. **(b, c)** Graphical fragmentation map of the two sequence isomers 01Y41Y41M(31M21M)61M(31M21M)61M21M and 01Y41Y41M(31M21M21M)61M(31M21M)61M for M8 (N2H8). **(d, f)** Isotopic envelope fingerprinting maps (bar-experimental, dot-theoretical) of the two representative structure-diagnostic fragment ions iI2-3-B/Y,C/Z and iIII2-3-B/Y,C/Z for the two structures in (b) and (c). **(e)** Extract ion chromatograms of monosaccharide composition N2H8 (*m/z* = 1086.06) in the three technical replications. **(g, h)** Annotated MS/MS spectra with the matched fragment ions for the two structures in (b) and (c). Symbols: blue square=N-acetylglucosamine, green circle=mannose, green circle=glucose.

Fourteen distinct high-mannose N-glycan structures, corresponding to eight compositions, were differentiated through the use of structure diagnostic ions. Taking the N-glycan N2H8F0S0 as an illustrative example, the two sequence isomers were separately identified with the retention times (**Figure 3e**) and fragment ions (**Figure 3b**, **Figure 3c**). The annotated MS/MS spectra corresponding to the two sequence structures were visualized in **Figure 3g** and **Figure 3h**. Chromatographic separation of the two sequence structures was also achieved reproductively in the three technical replications (**Figure 3e**). One sequence structure elutes at 136-138 min and the other at 139-141 min, although there is still some overlap. Structure diagnostic-ion iI2-3-B/Y, C/Z (*m/z*= 858.43, *z*=1, **Figure 3d**) determined the earlier eluted N-glycan structure of 01Y41Y41M(31M21M)61M(31M21M)61M21M, and the iIII2-3-B/Y, C/Z (*m/z*= 1266.63, *z*=1, **Figure 3f**) ion determined the second structure of 01Y41Y41M(31M21M21M)61M(31M21M)61M.

### Fucosylated N-glycans

The addition of fucose to glycan structures can vary depending on the specific fucosyltransferases involved, leading to fucose modifications at different positions including the core, branch and or terminal. Fucose has been found to play a significant role in various biological processes, including growth, immune responses and disease such as inflammation and cancer in mammals ^13^.

Figure 4a depicts the fucosylation levels, represented as the sum of abundance percentages for all the N-glycans containing fucose residues, within the RNA. It is noteworthy that the majority of RNA N-glycans were found to be modified with at least one fucose residue, with fucosylation exceeding 13% across all examined organs. Certain organs exhibited particularly elevated levels of fucosylation, exemplified by the stomach, where N-glycans with 6 fucoses accounted for 0.76% of the total abundance. Regarding the position of fucose residues, our analysis emphasizes the quantification of core fucosylation. With the exception of the liver, all other organs displayed core fucosylation levels exceeding 40%. Further classification of the 667 RNA N-glycans revealed that 557 of them were modified with fucose. These fucose-modified N-glycans were categorized based on the position of the fucose residues, as illustrated in Figure 4b. Specifically, there were 253 N-glycans with core fucose, 266 with branch fucose, and 216 with terminal fucose. Intriguingly, 139 N-glycans exhibited modifications with both core and branch fucoses, while 70 N-glycans featured both core and terminal fucose. Additionally, 26 N-glycans bore modifications of both branch and terminal fucose, and 22 N-glycans displayed fucose residues at all three positions.

**Figure 4.**
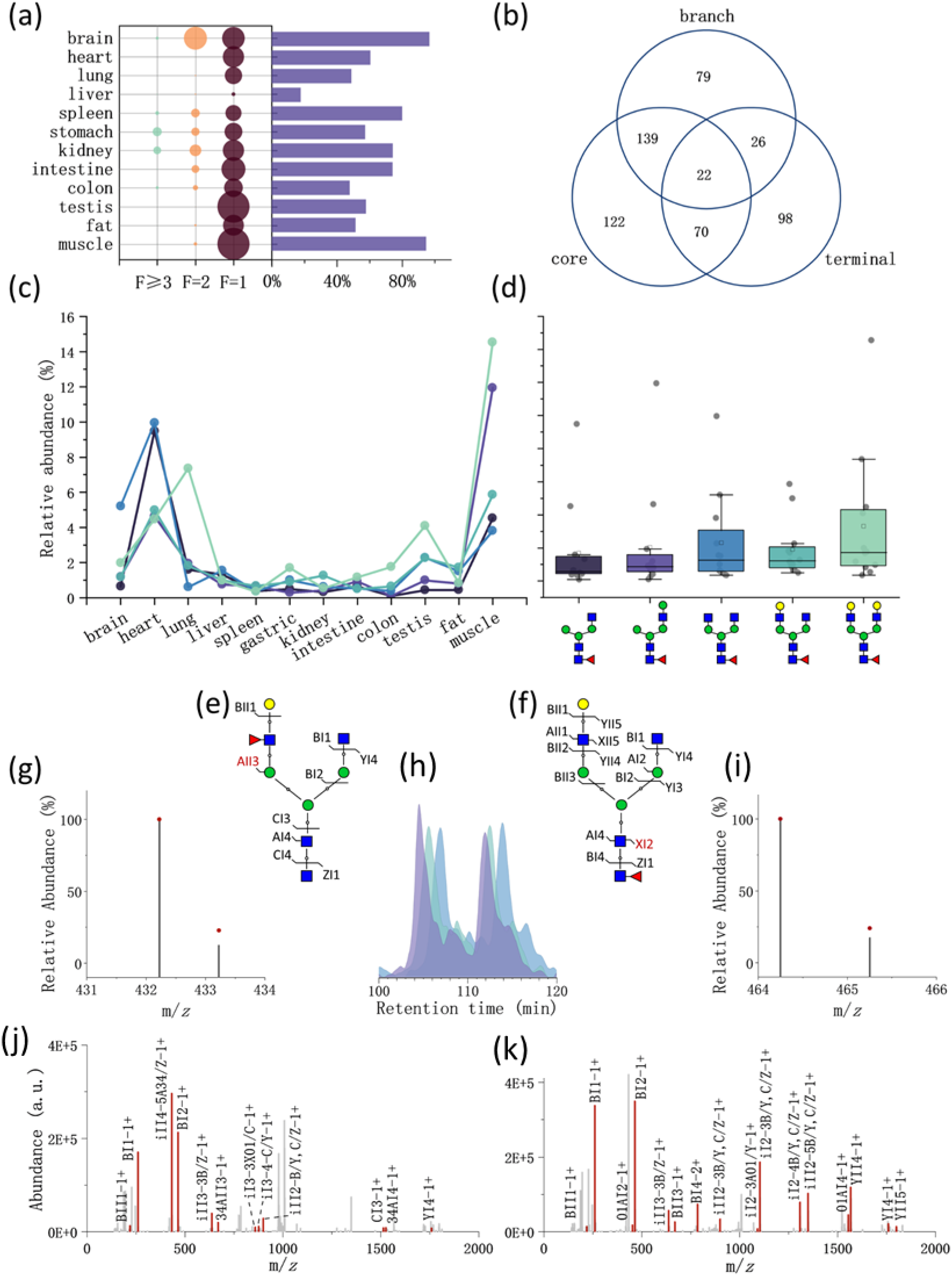
Fucosylated N-glycans identified from the glycoRNAs of the 12 organs of human. **(a)** Bubble chart of the sum of relative abundance of N-glycans with one, two, no less than three fucoses alongside histogram of abundance of N-glycans with the core fucose. **(b)** Venn diagram of N-glycans modified with fucose at different positions of the core, branch and terminal. **(c, d)** Distributions of five common N-glycans among the 12 organs. **(e, f)** Graphical fragmentation map of two representative fucose sequence (position) isomers 01Y41Y41M(31M41Y(31F)41L)61M61Y and 01Y(61F)41Y41M(31M41Y41L)61M61Y with the same monosaccharide composition of N4H4F1. **(g, j)** Isotopic envelope fingerprinting maps (bar-experimental, dot-theoretical) of the two representative structure-diagnostic fragment ions AII3 and XI2 for the two structures in (e) and (f). **(i)** Extract ion chromatograms of the monosaccharide composition N4H4F1S0 (*m/z* = 1010.03) in the three technical replications. **(k, l)** Annotated MS/MS spectra with the matched fragment ions for the two structures in (e) and (f). Symbols: blue square=N-acetylglucosamine, green circle=mannose, yellow circle=galactose, red triangle=fucose.

Among the six N-glycans shared across all 12 organs, one belonged to the high-mannose type, while the remaining five shared a common feature of core fucosylation. The relative abundances of these five N-glycans in each organ are graphically presented in Figure 4c. Notably, these five N-glycans exhibited similar abundance patterns within the same organs. In heart and muscle, these N-glycans showed relative high abundance. The average relative abundances of these five N-glycans across all the 12 organs were summarized in Figure 4d. These values were found to be 1.7%, 2.1%, 2.3%, 1.9%, and 3.3%, corresponding to the N-glycan structures of 01Y(61F)41Y41M(31M)61M61Y, 01Y(61F)41Y41M(31M)61M61Y41L, 01Y(61F)41Y41M(31M41Y)61M61Y, 01Y(61F)41Y41M(31M41Y41L)61M61Y, and 01Y(61F)41Y41M(31M41Y41L)61M61Y41L, respectively.

The position isomers of fucosylated N-glycans are distinguished with diagnostic fragment ions (Figure 3e-3k). The N-glycan composition N4H4F1S0 (*m/z*=1010.02694, *z*=2) was identified with the fucose core and branch isomers. The branch isomer eluted at approximately 107.2 minutes, as evidenced by the extracted ion chromatogram displaying two separate chromatographic peaks. The core isomer was identified at an elution time of 113.6 minutes. The branch and core isomers are confirmed with the structure diagnostic fragment ions iII4-5-A34/Z (*m/z*= 432.22, *z*=1) and XI2 (*m/z*=464.24, *z*=1), respectively.

### Sialylated N-glycans

Sialic acid stands as a common terminal monosaccharide known for its unique negative charge. N-glycoproteins with or without sialic acid have different conformations^14^. Terminal sialic acid significantly influences signal transmission, receptor recognition in both health and disease contexts^15^. In this study of RNA, sialylated N-glycans were not big in number (Figure 1b), but high in abundance (Figure 5a). This quantification was derived from the sum of abundance multiplied by a sialylation factor, considering all glycans containing NeuAc (sialic acid), where the factor accounts for the fraction of antennae that are sialylated. It’s noteworthy that all sialylated N-glycans across various organs featured at least one NeuAc modification, with eight organs showcasing two NeuAc modifications. In specific cases, a select few N-glycans in the brain, spleen, and colon exhibited three NeuAc modifications. Notably, the organs with the highest sialylation levels were the lung (27.6%), spleen (27.2%), and liver (25.2%), whereas the testis (0.8%) and kidney (2.7%) exhibited the lowest levels of sialylation. The shared linkages among the organs with the highest sialylation levels are illustrated in Figure 5b. The spleen featured 161 types of N-glycans with sialic acid and shared 18 linkages with the liver and lung. Among these shared linkages, three specific ones, namely 01Y(61F)41Y41M(31M)61M61Y41L32S, 01Y41Y41M(31M)61M(21Y41L32S)61Y41L, and 01Y41Y41M(31M41Y41L32S)61M61M, were observed in all the three organs.

The linkage types of sialic acid with other monosaccharides are α2,3-linked and α2,6-linked. Mass spectrometry distinguishes the linkage isomer through chemical derivatization methods. Nonetheless, we employed a lengthy RPLC approach, which has proven capable of separating isomeric permethylated N-glycan, including sialylated N-glycan linkage isomer^16^. Manual inspection of the chromatogram identified separated sialic acid linkage isomers (Figure 5c-5i). For instance, the extracted ion chromatogram of N-glycan N3H6F0S1 (*m/z*= 1185.10579, *z*=2) have two separated peaks (Figure 5f). The fragment ions of the N-glycan were showed in Figure 5h and **5i** and among those, structure diagnostic ions YIII4-A02 and ^3,5^AIII3 confirmed the existence of sialic acid. Based on prior research findings, it is known that α2,6-linked N-glycans tend to elute earlier than their α2,3-linked N-glycans. Consequently, we deduced that the left peak represented the glycoform 01Y41Y41M(31M41Y41L62S)61M(31M)61M, while the right peak corresponded to 01Y41Y41M(31M41Y41L32S)61M(31M)61M.

### Antigen T, Lewis X, and sialyl Lewis X

Galactosylated N-glycans comprised a significant portion of RNA glycans, with terminal galactose modifications being prevalent in nearly 30% of N-glycans. Galactosylation, defined similarly to sialylation, exhibited a range of abundance, spanning from 13.4% to 51.1% (Figure 6a).

**Figure5.**
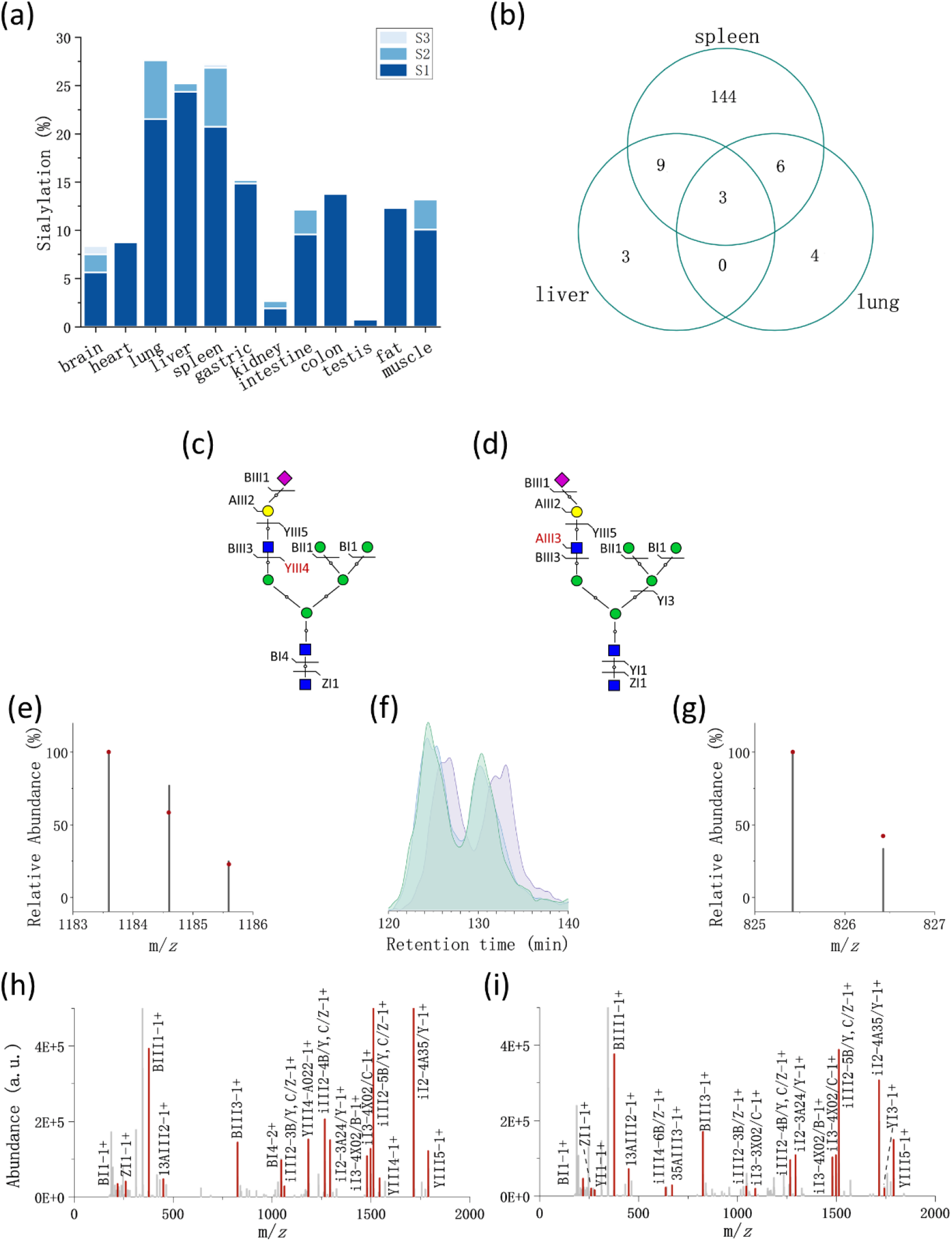
Sialylated N-glycans identified from the glycoRNAs of the 12 organs of human. **(a)** Stacked histogram of relative abundance of sialylated N-glycan with different number of sialic acid residues across the 12 organs. **(b)** Venn diagram of the sialylated N-glycans identified in spleen, liver and lung. **(c, d)** Graphical fragmentation map of two representative sialic acid linkage isomers 01Y41Y41M(31M41Y41L62S)61M(31M)61M (α2,6) and 01Y41Y41M(31M41Y41L32S)61M(31M)61M (α2,3). **(e, g)** Isotopic envelope fingerprinting maps (bar-experimental, dot-theoretical) of the two representative structure-diagnostic fragment ions YIII4 and AIII3 for the two structures in (e) and (g). **(f)** Extract ion chromatograms of the monosaccharide composition N3H6F0S1 (*m/z* = 1185.11) in the three technical replications. (h, i) Annotated MS/MS spectra with the matched fragment ions for the two structures in (e) and (g). Symbols: blue square=N-acetylglucosamine, green circle=mannose, yellow circle=galactose, purple diamond= N-Acetylneuraminic acid.

**Figure 6.**
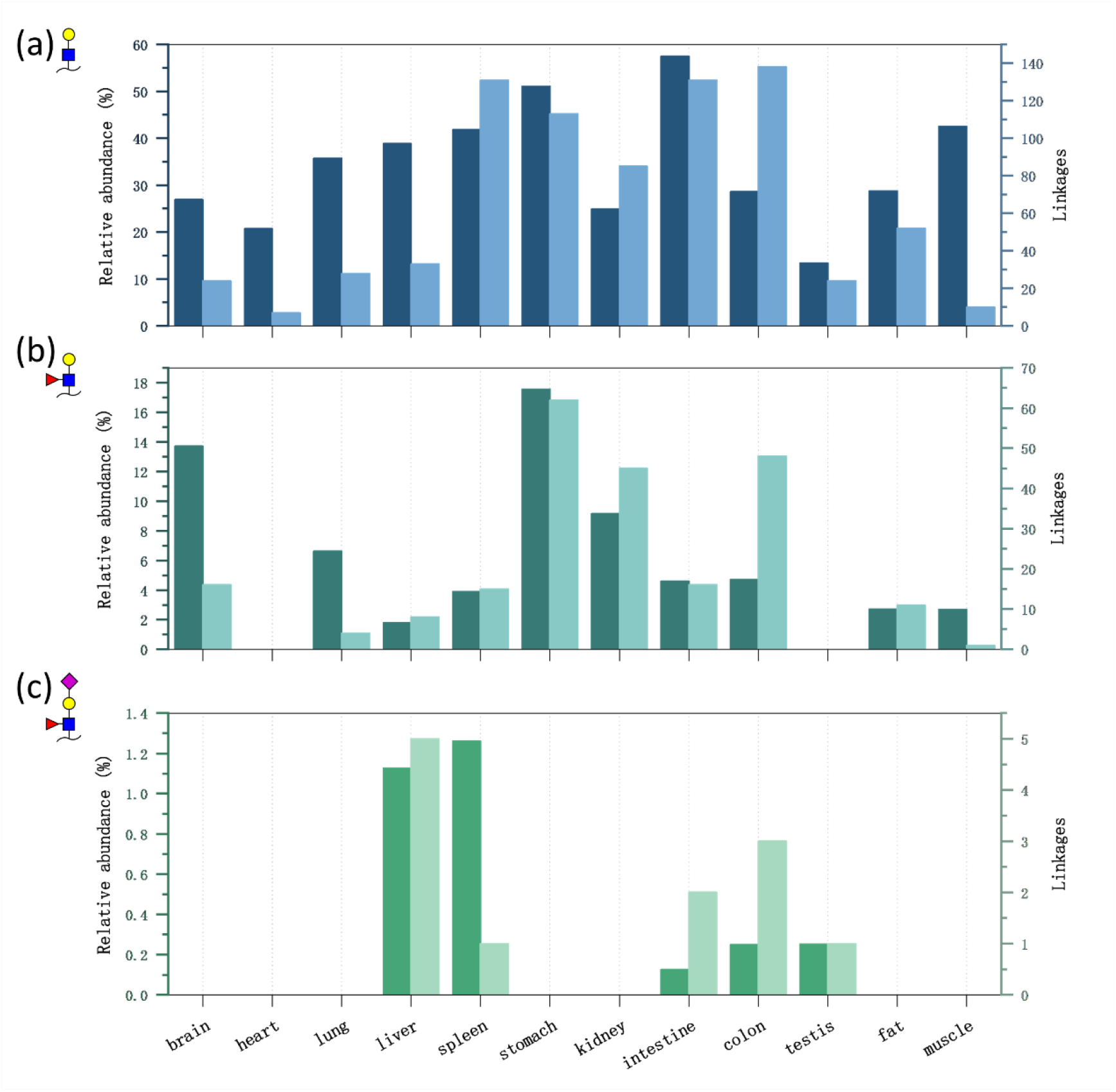
Relative abundance of antigens T (a), Lewis X (b), and sialyl Lewis X (c) identified from the glycoRNAs of the 12 organs of human.

The epitope Lewis X is the structure of fucosylated GlcNAc linked a terminal galactose. The Lewis X epitope, identified in 10 out of the 12 organs (absent in heart and testis), is less abundant than galactosylation (Figure 6b). The Lewis X epitope is rich in stomach, where 62 N-glycans have the structure with 17.6% relative abundance. α1,3-fucosyltransferase 9 (FUT9), unique responsible for the transfer of fucose for the synthesis of the Lewis x (Lex) structure, is expressed in stomach, kidney and brain^17, 18^. Among the 12 organs, stomach, brain and kidney have the Top 3 most IDs with relative abundance ratios of 17.6%, 13.7% and 9.1% respectively.

Sialylated Lewis X was identified in liver, spleen, intestine, colon and testis; and the relative abundance was below 1.3% (Figure 6c).

### Bisecting N-glycans

Bisect GlcNAc was frequently observed in RNA glycan. The percentage of numbers and sum of relative abundance were showed in Figure 7a and **7b**. About half of N-glycans were bisect in intestine and the least percentage of bisect N-glycan was 12.5% in muscle. Some organs did not have so much bisect N-glycan but showed great abundance, such as N-glycan in brain occupied 42.7% relative abundance. The special position of bisect GlcNAc influences the configuration of N-glycan. The bisect GlcNAc N-glycans were mostly fucosylated and but least sialylated (Figure 7c). There were 28.2% sialylated ones in all of the bisecting N-glycans, which is consistent with the fact that bisect GlcNAc would suppress terminal modification ^19^.

**Figure 7.**
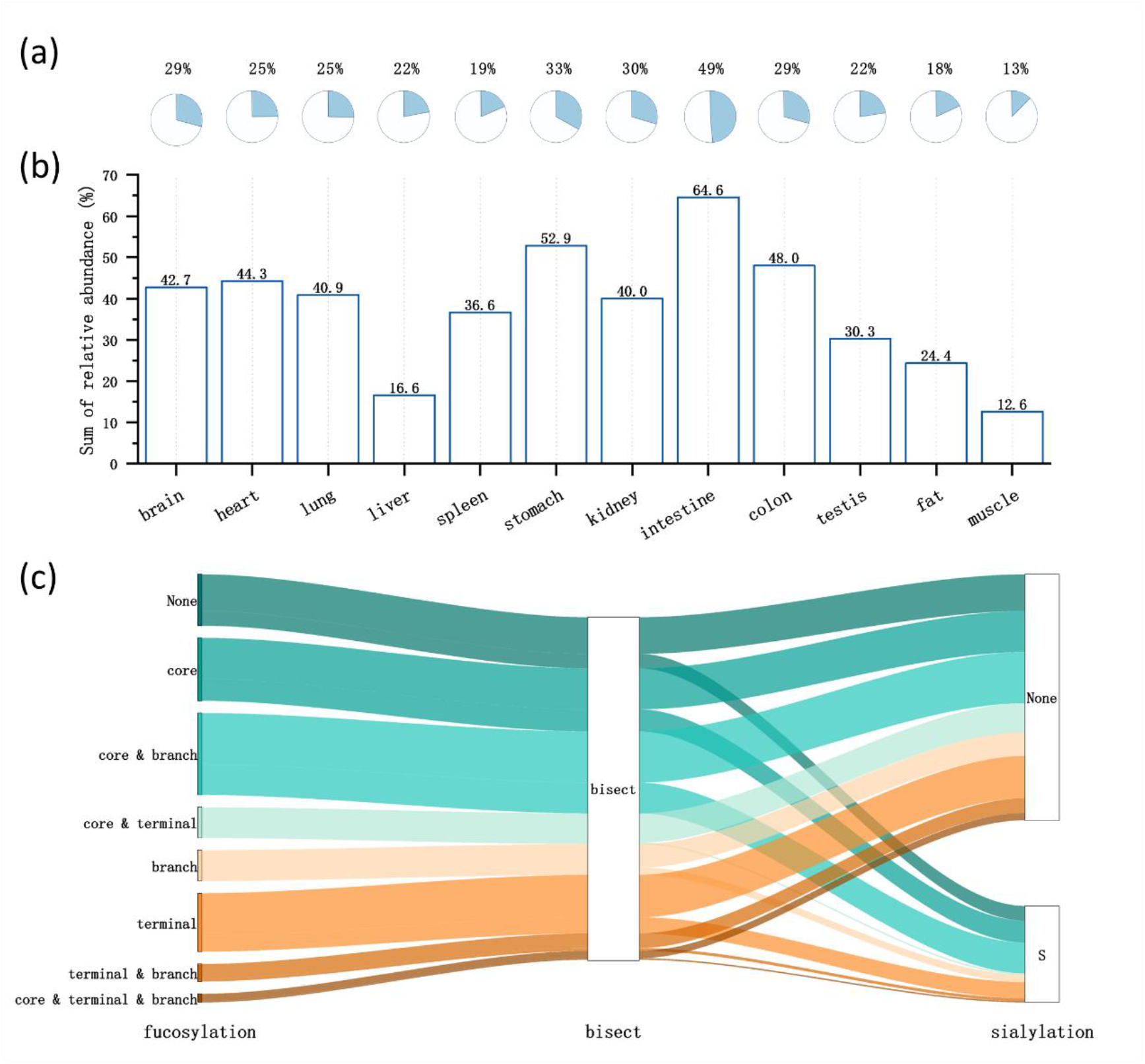
Bisect N-glycans identified from the glycoRNAs of the 12 organs of human. (a) percentages, (b) relative abundance, and (c) alluvial chart of proportional relationship of bisect N-glycans with fucosylated and sialylated ones.

## 4 DISCUSSION

The biofunction of glycoRNA remains a puzzle, and the intricacies of glycosylation in this context have yet to be fully comprehended. While glycosylation of proteins and lipids has been a focus of research, with a predominant emphasis on comparing disease and healthy states, it is crucial to acknowledge that each organ serves unique functions, and glycosylation in different organs may exhibit distinct characteristics. Our study aims to shed light on this aspect by profiling the N-glycans of glycoRNA across 12 human organs.

We have collated and compared our findings with these glycoRNA results to elucidate potential relationships between RNA and protein glycosylation in brain. The human brain is a remarkably complex organ with a dynamic N-glycome. By comparison with the brain N-glycome of protein in rat, macaque, chimpanzee and human, Klaric et al ^20^ found that a progressively more intricate N-glycan landscape and α2,6-linked N-acetylneuraminic acid increased with the evolution. Our glycoRNA analysis in humans mirrors these findings, as complex and hybrid types of N-glycans dominate, comprising 96% of the glycoRNA N-glycome. Sialylation, fucosylation, bisect and LDNF (structures with fucose and N-acetylgalactosamine on the same arm) were major characteristics in N-glycome in protein ^21, 22^. 96% of glycoRNA N-glycans in brain were modified with fucose and 22% N-glycan modified with sialic acid. The fucosylation is the predominated feature. Interestingly, the presence of N-glycans with up to three sialic acids was identified in the brain, constituting 0.87% of the glycoRNA N-glycome. This finding parallels the presence of poly-sialic acid residues as a distinct feature in glycoproteins of the brain, hinting at the possibility that this characteristic may also extend to glycoRNA. Furthermore, our results revealed the presence of 19 bisect N-glycans in the brain, accounting for a relative abundance of 42.7%.

GlycoRNA N-glycans are mostly complex and hybrid with bi/tri-antenna. The level of fucosylation is high, and core fucose is predominant in comparison to the branch and terminal positions. Sialylation is relatively less compared with galactosylation which occupied relative abundance of 10-60%. Bisect N-glycan took a relatively high proportion and had a strong relationship with fucosylation.

## 5. N-GLYCAN IDENTIFICATION BY GLYCONOTE

Besides identification by GlySeeker, the raw RPCL-MS/MS datasets from the 12 organs were also searched by GlycoNote. Raw mass spectra were first converted into the .mgf format, and the mass tolerance for the precursor and fragment ions was 15 ppm. Monosaccharides Hex, HexNAc, dHex and NeuAc were considered with the residue number ranges of 2-15, 2-10, 0-5 and 0-4, respectively, per N-glycan.

A total of 590 N-glycans in terms of unique monosaccharide compositions were identified among 11 out of the 12 organs, where 86 compositions were shared by the GlySeeker IDs; i.e., GlycoNote and GlySeeker have 504 and 150 unique compositions, respectively.

Among the 504 unique compositions of GlycoNote, 55 are present in the theoretical DB adopted by GlySeeker, which has 857 unique compositions constructed specifically for proteins with the known synthetic rules. Shared compositions (Figure 8b) are categorized mainly into two clusters, higher stoichiometry of either N or H with lower stoichiometry of F and S; Unique compositions of GlySeeker (Figure 8a) tend to have quite even stoichiometry of N and H with medium stoichiometry of F and lower stoichiometry of S; Unique compositions of GlycoNote (Figure 8c) have higher stoichiometry of F and S.

**Figure 8.**
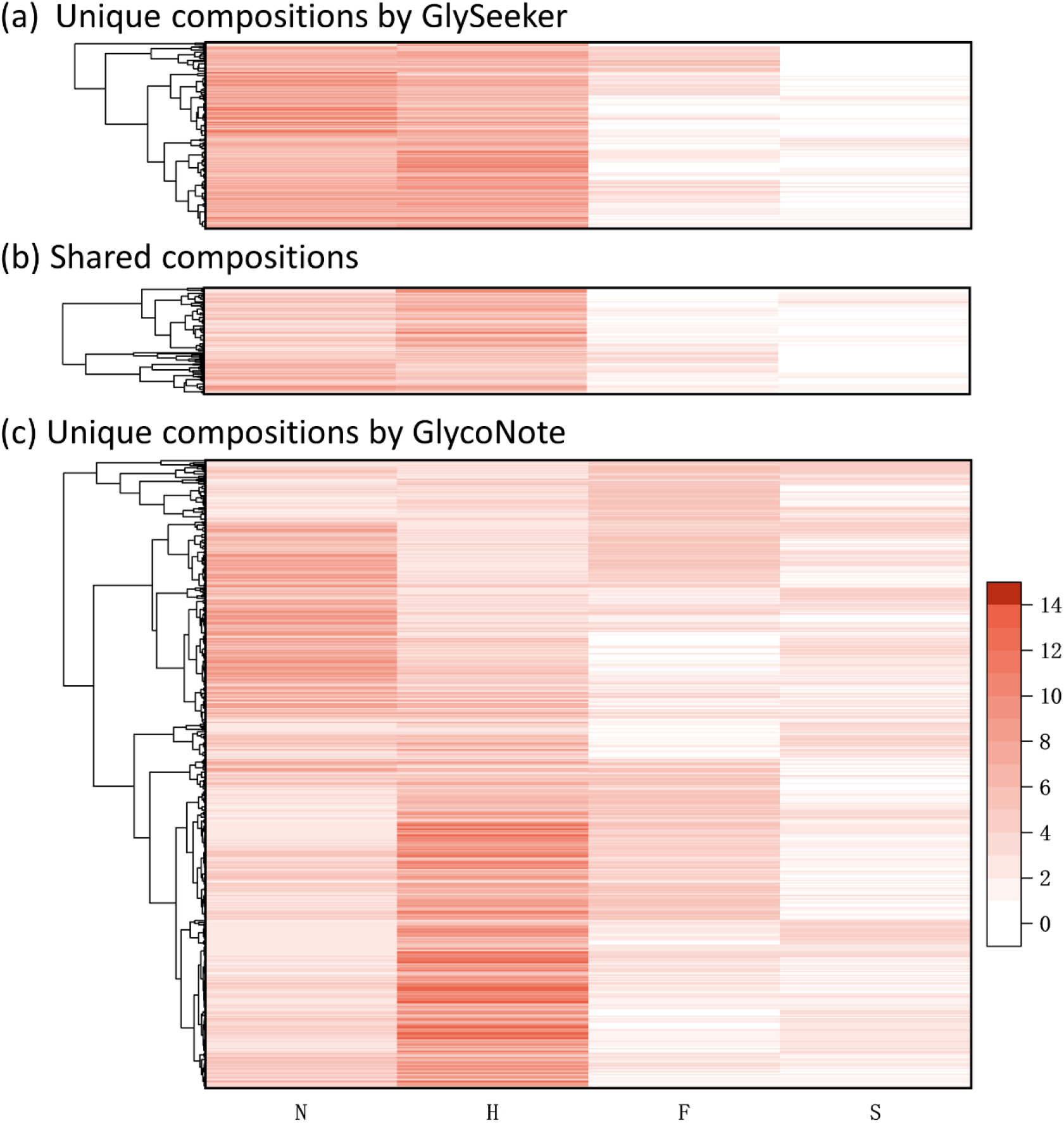
Comparison of the identified monosaccharide compositions by N-glycan search engines GlySeeker and GlycoNote; (a) unique IDs of GlySeeker, (b) shared IDs by the two engines, (c) unique IDs of GlycoNote.

## 6. LIMITATION OF THIS STUDY

Only 12 organs are adopted because of limited access to all the other organs. The theoretical N-glycan DB adopted by GlySeeker, originally built for N-glycoproteins with known biosynthetic rules, is assumed to be fully shared by glycoRNAs. Only partial sequence structures are distinguished from their isomers with structure-diagnostic fragment ions because of intrinsic low concentration, insufficient fragmentation efficiency, too many isomers per composition among others. Automatic interpretation of the linkage isomers of sialic acid separated chromatographically is not available, and only some representative ones are manually explored.

### Author contributions

LHW and ZXT designed the study; ZRZ and TW collected the tissue samples; LHW prepared the glycoRNA; MB did sample cleanup, permethylation, LC-MS/MS and data analysis; MB, LHW and ZXT wrote the manuscript.

## Supplementary information

**Supplementary information 1:** The detailed MS spectral and identification information for the N-glycans identified by GlySeeker from the 12 human organs (MS Excel files, 12 sheets, one for each organ).

**Supplementary information 2-13:** The isotopic envelope fingerprinting map, annotated MS/MS spectrum with the matched fragment ions, and graphical fragmentation map of each N-glycans identified from the 12 human organs of (PDF files, 12 in total, one for each organ)

**Supplementary information 14:** The monosaccharide compositions for the N-glycans identified by GlycoNote from 11 human organs (MS Excel files, 11 sheets, one for each organ; no glycan was identified by GlycoNote in the other one organs).

## Acknowledgements

This research was financially supported by the National Natural Science Foundation of China (22074105 and 21775110) and the Shanghai Science and Technology Commission (14DZ2261100).

## References

1. In Essentials of Glycobiology, Varki, A.; Cummings, R. D.; Esko, J. D.; Stanley, P.; Hart, G. W.; Aebi, M.; Darvill, A. G.; Kinoshita, T.; Packer, N. H.; Prestegard, J. H.; Schnaar, R. L.; Seeberger, P. H., Eds. Cold Spring Harbor Laboratory Press Copyright 2015-2017 by The Consortium of Glycobiology Editors, La Jolla, California. All rights reserved.: Cold Spring Harbor (NY), 2015.

2. Flynn, R. A.; Pedram, K.; Malaker, S. A.; Batista, P. J.; Smith, B. A. H.; Johnson, A. G.; George, B. M.; Majzoub, K.; Villalta, P. W.; Carette, J. E.; Bertozzi, C. R., Small RNAs are modified with N-glycans and displayed on the surface of living cells. Cell 2021, 184 (12), 3109-+.

3. Liu, M. Q.; Treves, G.; Amicucci, M.; Guerrero, A.; Xu, G. G.; Gong, T. Q.; Davis, J.; Park, D.; Galermo, A.; Wu, L. R.; Cao, W. Q.; Lebrilla, C. B., GlycoNote with Iterative Decoy Searching and Open-Search Component Analysis for High-Throughput and Reliable Glycan Spectral Interpretation. Analytical Chemistry 2023, 95 (21), 8223–8231.

4. Ma, Y.; Guo, W. J.; Mou, Q. B.; Shao, X. L.; Lyu, M.; Garcia, V.; Kong, L. G.; Lewis, W.; Ward, C.; Yang, Z. L.; Pan, X. X.; Yi, S. S.; Lu, Y., Spatial imaging of glycoRNA in single cells with ARPLA. Nature Biotechnology 2023.

5. Hemberger, H.; Chai, P.; Lebedenko, C. G.; Caldwell, R. M.; George, B. M.; Flynn, R. A., Rapid and sensitive detection of native glycoRNAs. bioRxiv 2023.

6. Li, J. J.; Yue, S.; Gao, Z. Y.; Hu, W. H.; Liu, Z. L.; Xu, G. Q.; Wu, Z.; Zhang, X. M.; Zhang, G. L.; Qian, F. L.; Jiang, J. H.; Yang, S., Novel Approach to Enriching Glycosylated RNAs: Specific Capture of GlycoRNAs via Solid-Phase Chemistry. Analytical Chemistry 2023, 95 (32), 11969–11977.

7. Damerell, D.; Ceroni, A.; Maass, K.; Ranzinger, R.; Dell, A.; Haslam, S. M., Annotation of glycomics MS and MS/MS spectra using the GlycoWorkbench software tool. *Methods in molecular biology (Clifton*, N.J.) 2015, 1273, 3–15.

8. Xiao, K. J.; Wang, Y.; Shen, Y.; Han, Y. Y.; Tian, Z. X., Large-scale identification and visualization of N-glycans with primary structures using GlySeeker. Rapid Communications in Mass Spectrometry 2018, 32 (2), 142–148.

9. Han, Y. Y.; Xiao, K. J.; Tian, Z. X., Comparative Glycomics Study of Cell-Surface N-Glycomes of HepG2 versus LO2 Cell Lines. Journal of Proteome Research 2019, 18 (1), 372–379.

10. Xiao, K. J.; Han, Y. Y.; Tian, Z. X., Large-scale identification and visualization of human liver N-glycome enriched from LO2 cells. Analytical and Bioanalytical Chemistry 2018, 410 (17), 4195–4202.

11. Wang, J.; Shewell, L. K.; Day, C. J.; Jennings, M. P., N-glycolylneuraminic acid as a carbohydrate cancer biomarker. Translational Oncology 2023, 31.

12. Yu, M.; Brown, D.; Reed, C.; Chung, S.; Lutman, J.; Stefanich, E.; Wong, A.; Stephan, J. P.; Bayer, R., Production, characterization and pharmacokinetic properties of antibodies with N-linked Mannose-5 glycans. Mabs 2012, 4 (4), 475–487.

13. Schneider, M.; Al-Shareffi, E.; Haltiwanger, R. S., Biological functions of fucose in mammals. Glycobiology 2017, 27 (7), 601–618.

14. Guillot, A.; Dauchez, M.; Belloy, N.; Jonquet, J.; Duca, L.; Romier, B.; Maurice, P.; Debelle, L.; Martiny, L.; Durlach, V.; Baud, S.; Blaise, S., Impact of sialic acids on the molecular dynamic of bi-antennary and tri-antennary glycans. Scientific Reports 2016, 6.

15. Bhide, G. P.; Colley, K. J., Sialylation of N-glycans: mechanism, cellular compartmentalization and function. Histochemistry and Cell Biology 2017, 147 (2), 149–174.

16. Wang, J. Y.; Dong, X.; Yu, A. Y.; Huang, Y. F.; Peng, W. J.; Mechref, Y., Isomeric separation of permethylated glycans by extra-long reversed-phase liquid chromatography (RPLC)-MS/MS. Analyst 2022, 147 (10), 2048–2059.

17. Comelli, E. M.; Head, S. R.; Gilmartin, T.; Whisenant, T.; Haslam, S. M.; North, S. J.; Wong, N. K.; Kudo, T.; Narimatsu, H.; Esko, J. D.; Drickamer, K.; Dell, A.; Paulson, J. C., A focused microarray approach to functional glycomics: transcriptional regulation of the glycome. Glycobiology 2006, 16 (2), 117–131.

18. Noro, E.; Togayachi, A.; Sato, T.; Tomioka, A.; Fujita, M.; Sukegawa, M.; Suzuki, N.; Kaji, H.; Narimatsu, H., Large-Scale Identification of N-Glycan Glycoproteins Carrying Lewis x and Site-Specific N-Glycan Alterations in Fut9 Knockout Mice. Journal of Proteome Research 2015, 14 (9), 3823–3834.

19. Nakano, M.; Mishra, S. K.; Tokoro, Y.; Sato, K.; Nakajima, K.; Yamaguchi, Y.; Taniguchi, N.; Kizuka, Y., Bisecting GlcNAc Is a General Suppressor of Terminal Modification of N-glycan. Molecular & cellular proteomics : MCP 2019, 18 (10), 2044–2057.

20. Klaric, T. S.; Gudelj, I.; Santpere, G.; Sousa, A. M. M.; Novokmet, M.; Vuckovic, F.; Ma, S.; Beceheli, I.; Sherwood, C. C.; Ely, J. J.; Hof, P. R.; Josic, D.; Lauc, G.; Sestan, N., Human-specific features and developmental dynamics of the brain N-glycome. bioRxiv : the preprint server for biology 2023.

21. Helm, J.; Hirtler, L.; Altmann, F., Towards Mapping of the Human Brain N-Glycome with Standardized Graphitic Carbon Chromatography. Biomolecules 2022, 12 (1).

22. Klaric, T. S.; Lauc, G., The dynamic brain N-glycome. Glycoconjugate Journal 2022, 39 (3), 443–471.

