## Supplemental Information 11 for "A draft of human N-glycans of glycoRNA"

Spectrum Index: 17084(NoBHs:1)

#### Precursor iEF Map

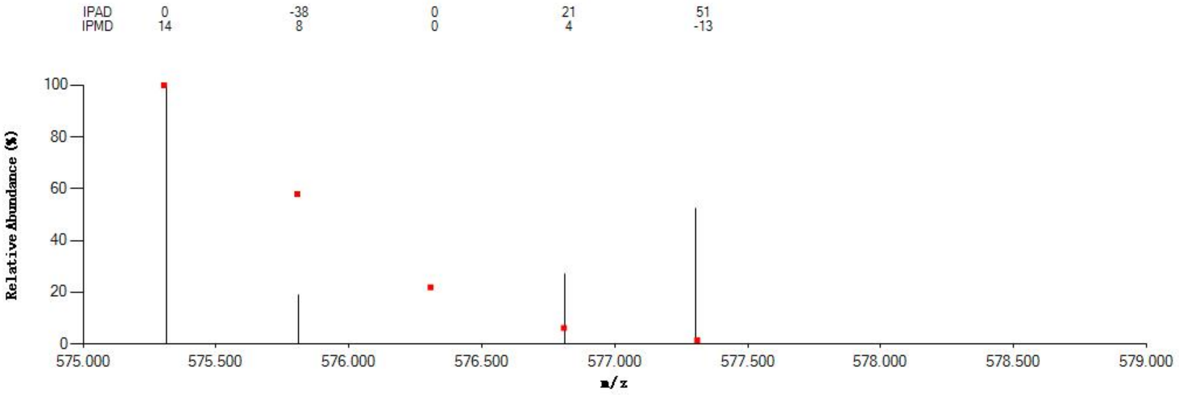


#### Annotated MS/MS Spectrum

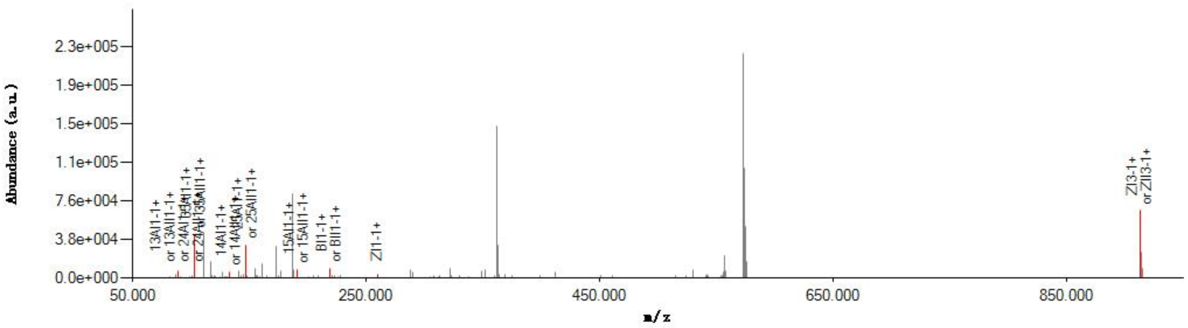


#### Graphical Fragmentation Map

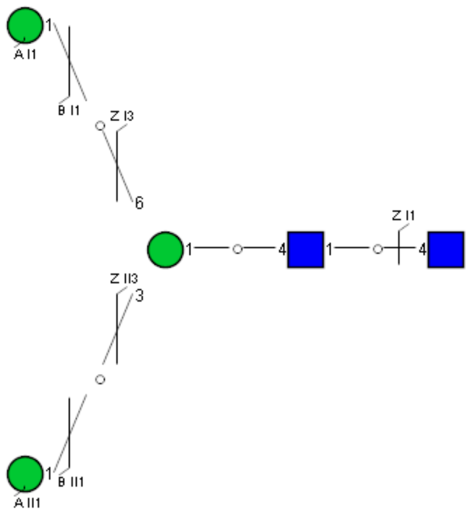

Spectrum Index: 20565(NoBHs:1)

#### Precursor iEF Map

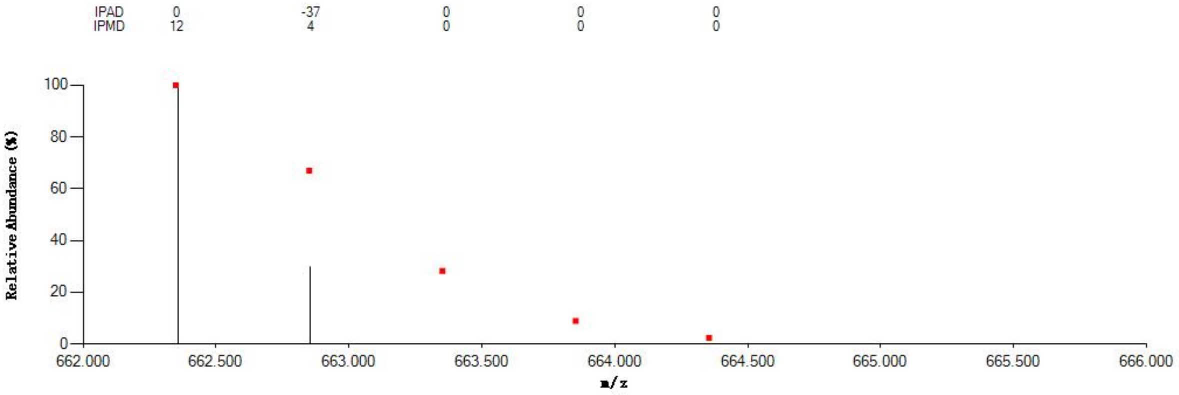


#### Annotated MS/MS Spectrum

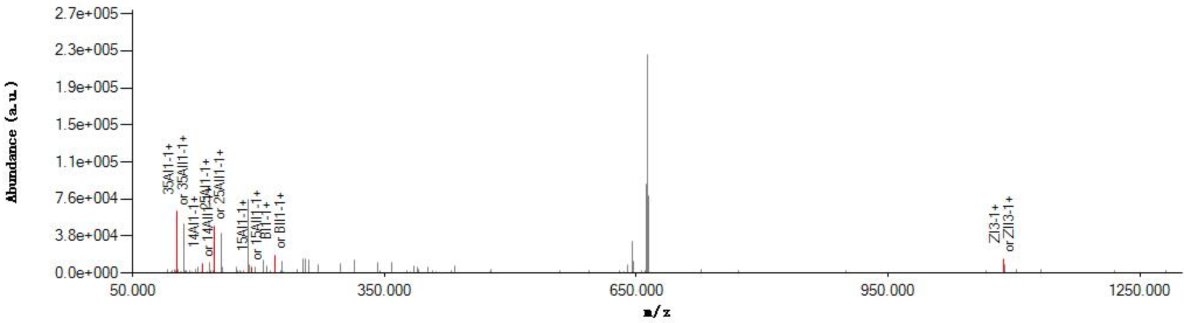


#### Graphical Fragmentation Map

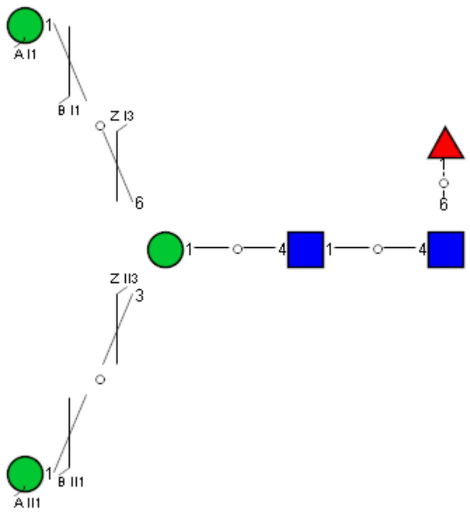

Spectrum Index: 23914(NoBHs:1)

#### Precursor iEF Map

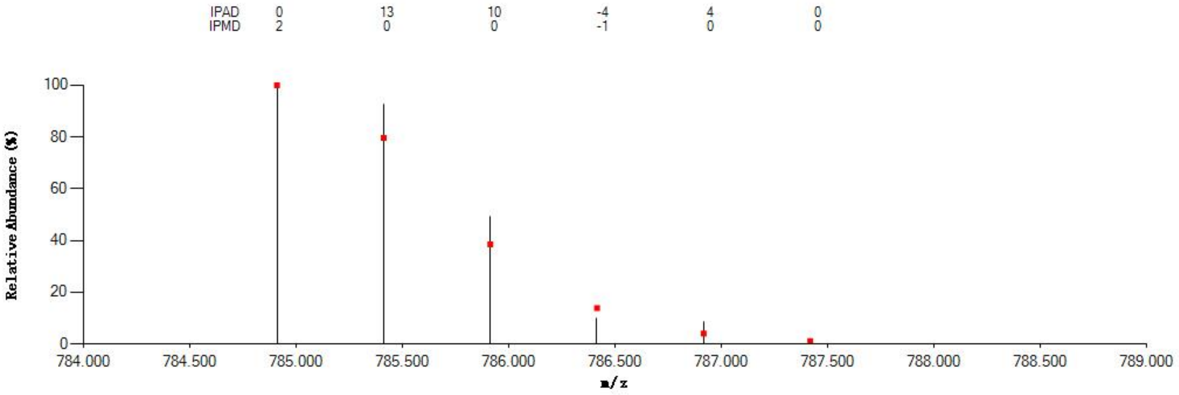


#### Annotated MS/MS Spectrum

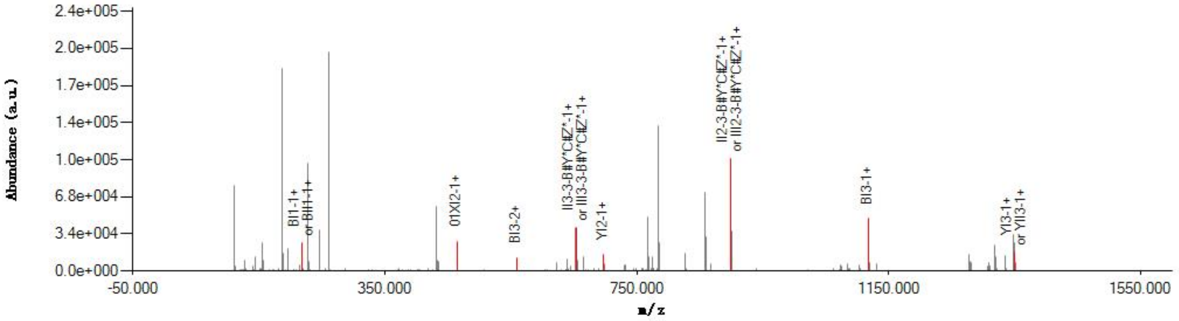


#### Graphical Fragmentation Map

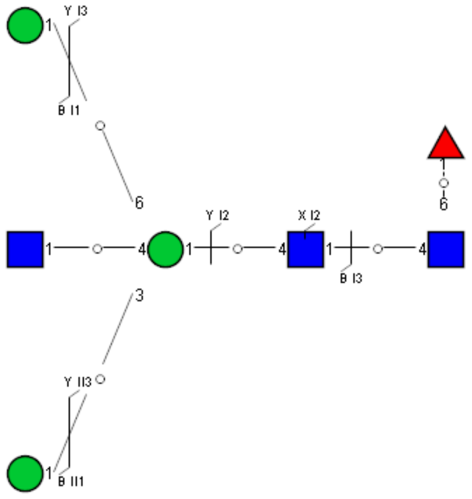

Spectrum Index: 24828(NoBHs:1)

#### Precursor iEF Map

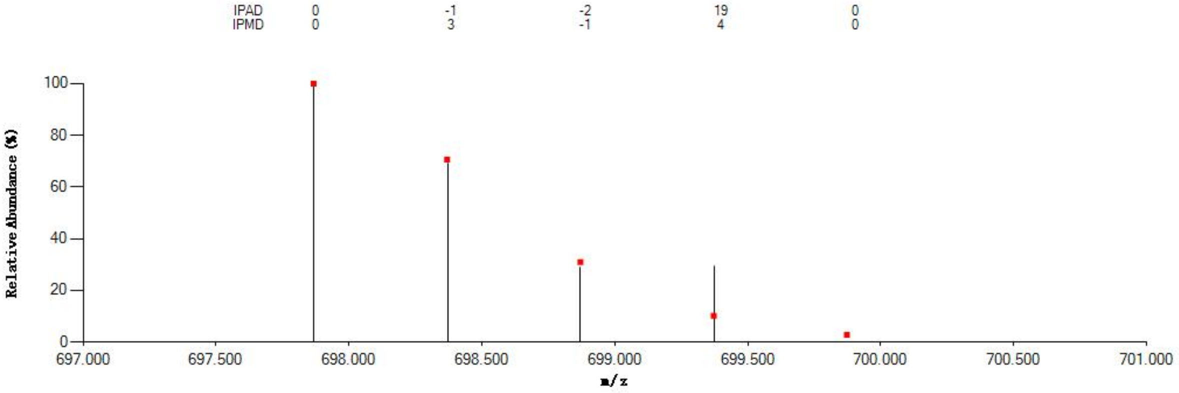


#### Annotated MS/MS Spectrum

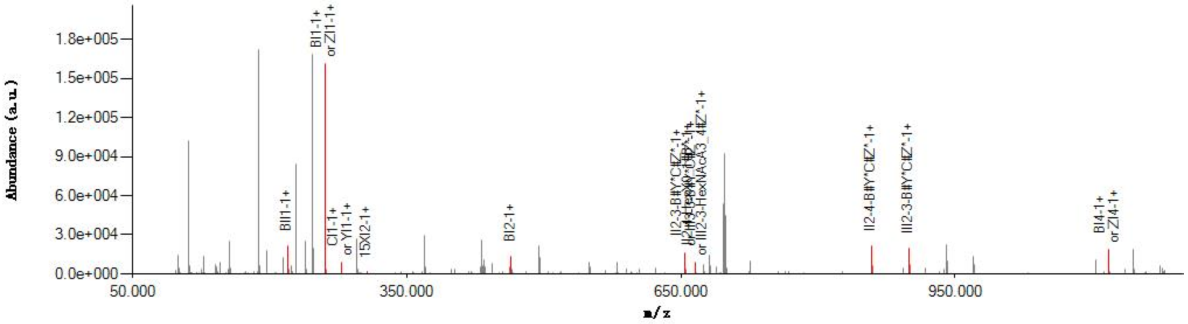


#### Graphical Fragmentation Map

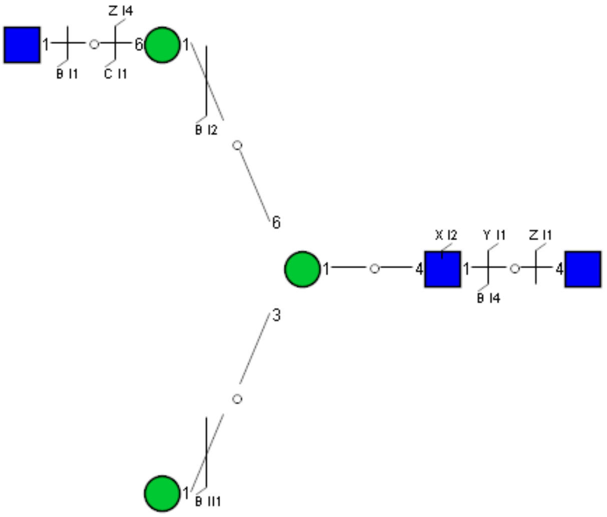

Spectrum Index: 25196(NoBHs:1)

#### Precursor iEF Map

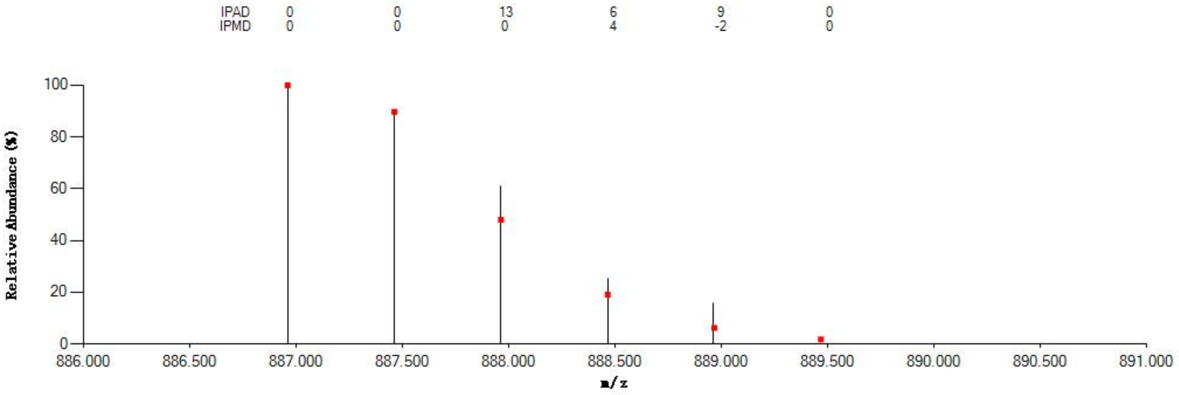


#### Annotated MS/MS Spectrum

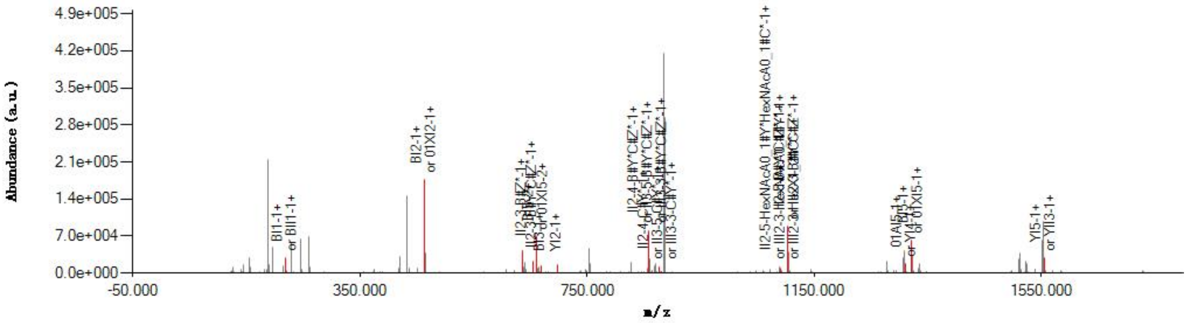


#### Graphical Fragmentation Map

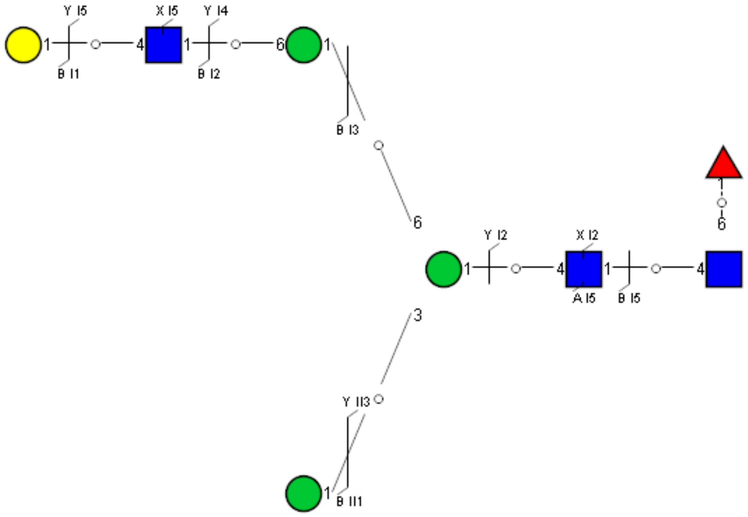

Spectrum Index: 26037(NoBHs:1)

#### Precursor iEF Map

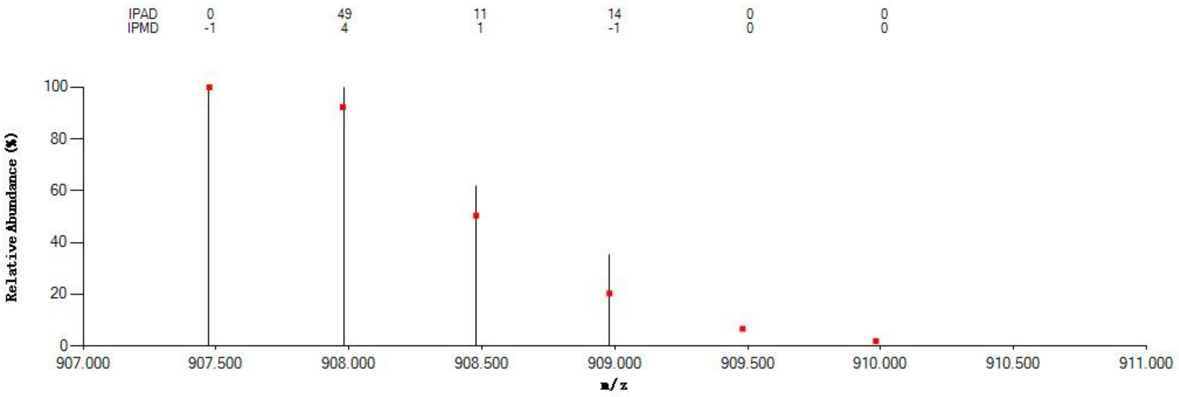


#### Annotated MS/MS Spectrum

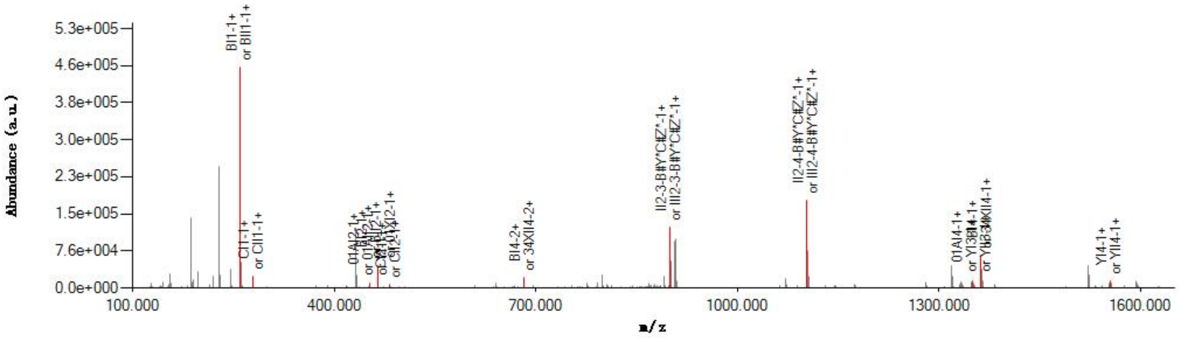


#### Graphical Fragmentation Map

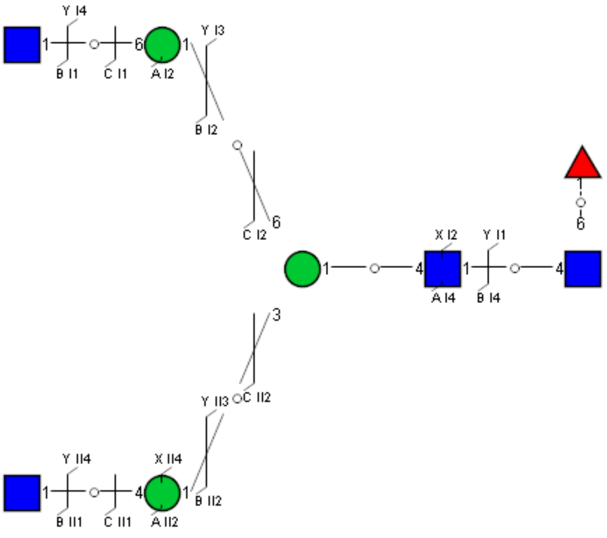

Spectrum Index: 26697(NoBHs:1)

#### Precursor iEF Map

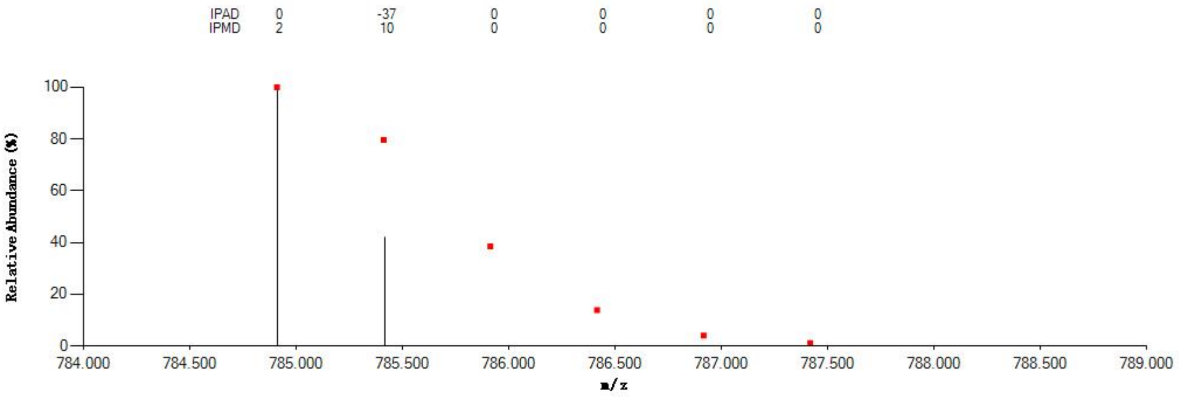


#### Annotated MS/MS Spectrum

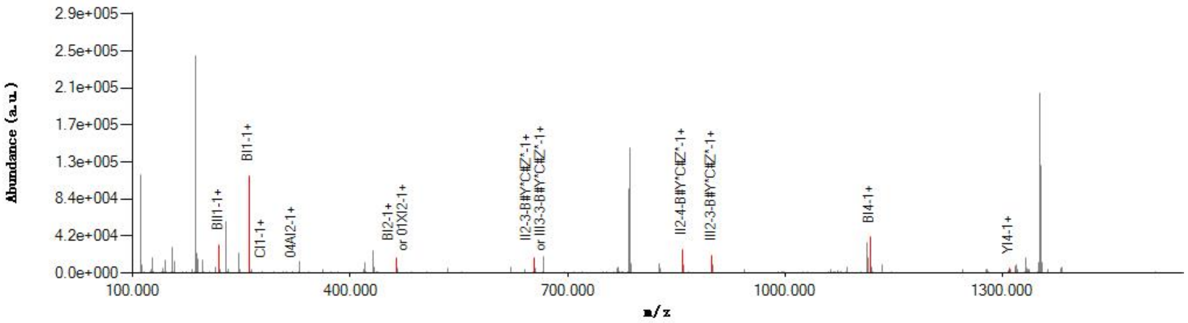


#### Graphical Fragmentation Map

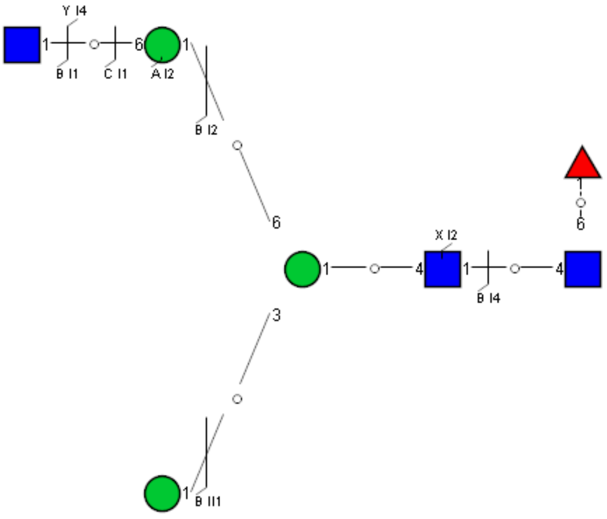

Spectrum Index: 26713(NoBHs:1)

#### Precursor iEF Map

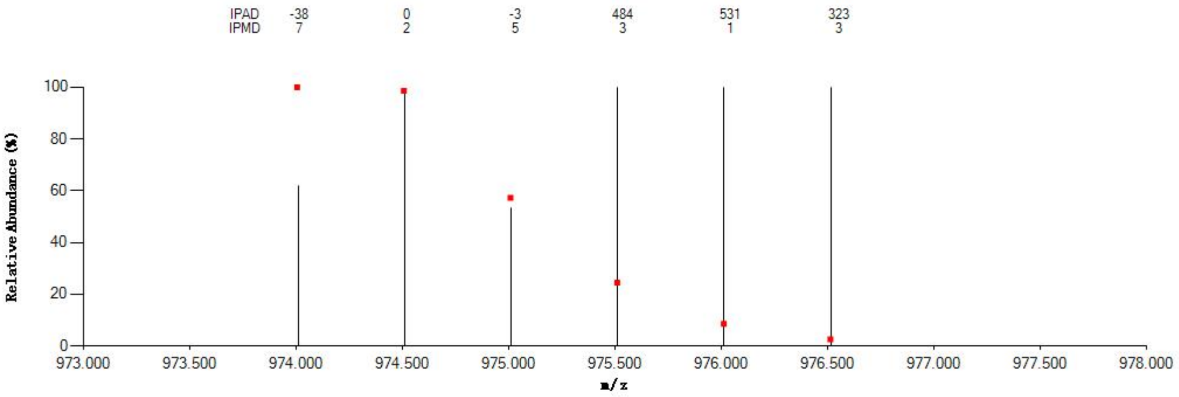


#### Annotated MS/MS Spectrum

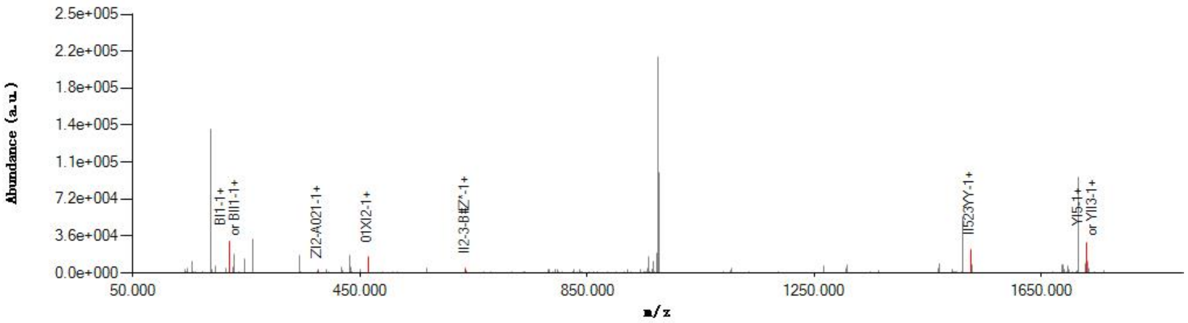


#### Graphical Fragmentation Map

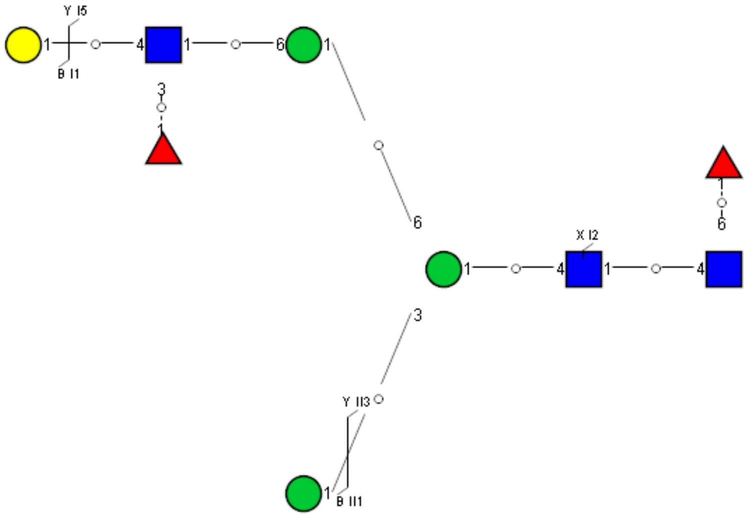

### Spectrum Index: 26797(NoBHs:1)

#### Precursor iEF Map

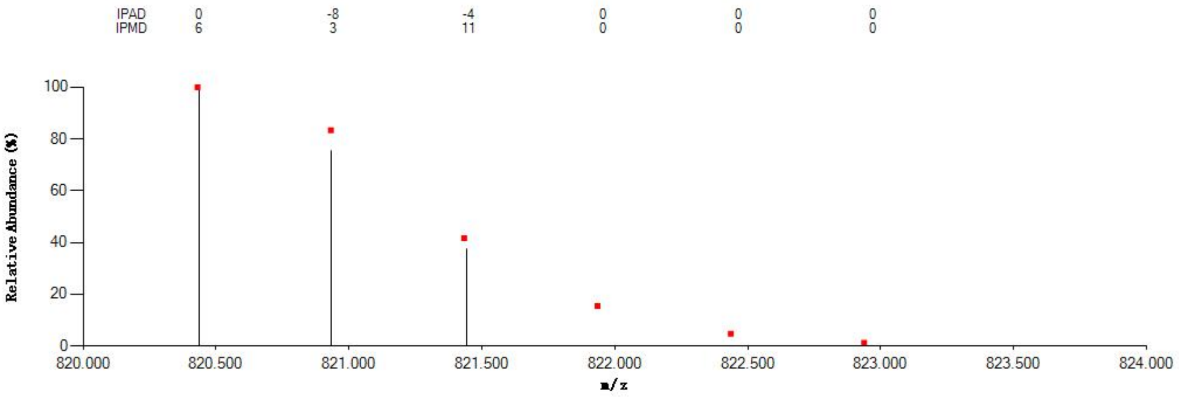


#### Annotated MS/MS Spectrum

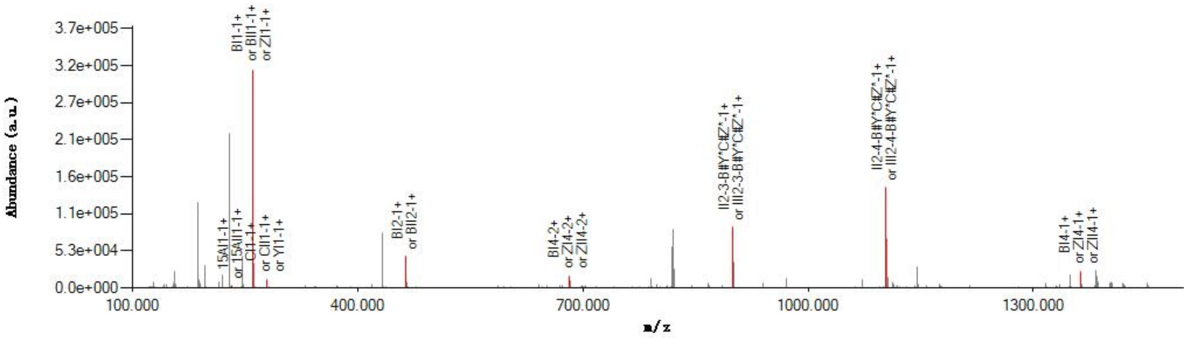


#### Graphical Fragmentation Map

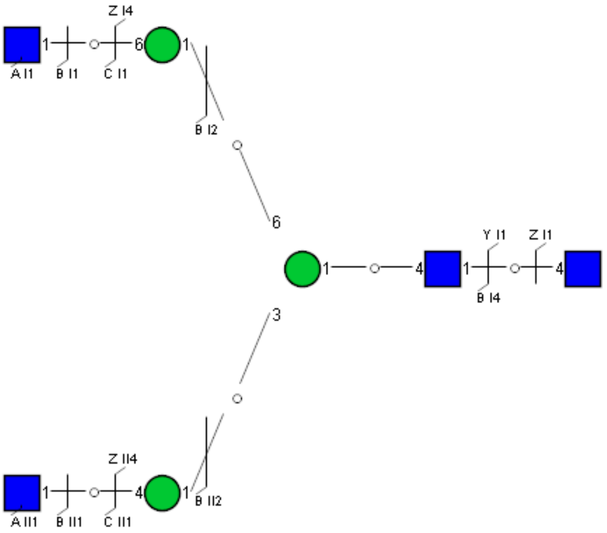

Spectrum Index: 27012(NoBHs:1)

#### Precursor iEF Map

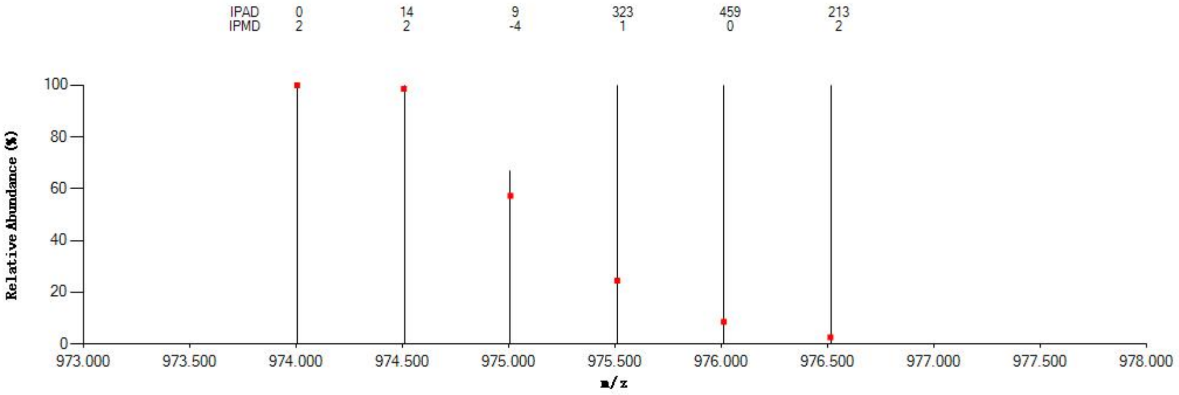


#### Annotated MS/MS Spectrum

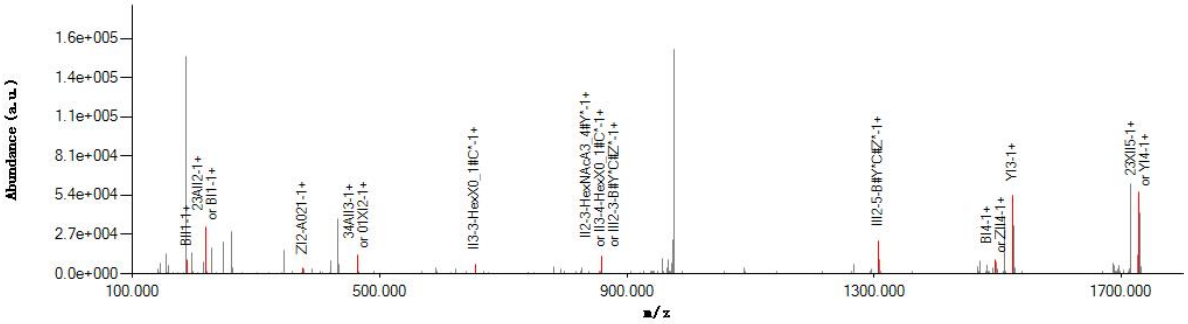


#### Graphical Fragmentation Map

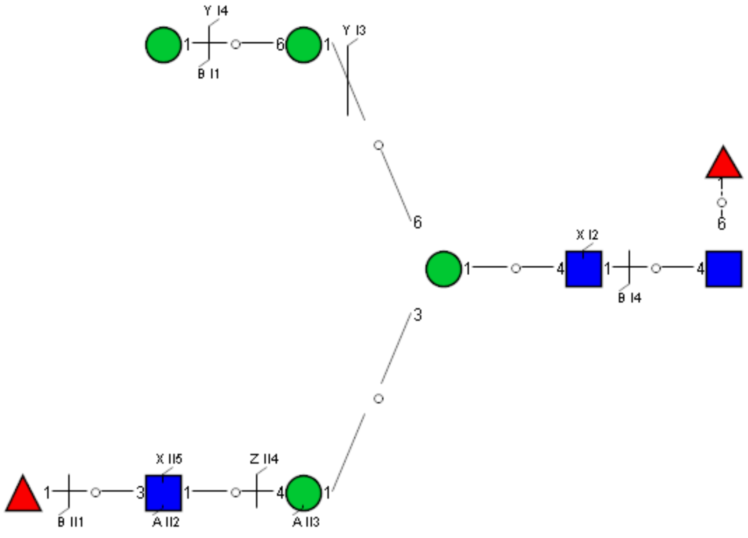

Spectrum Index: 27850(NoBHs:1)

#### Precursor iEF Map

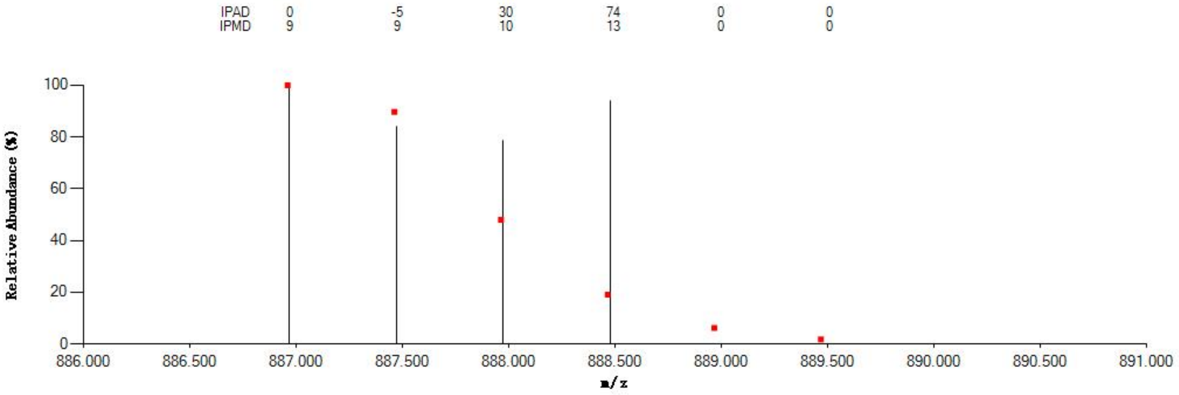


#### Annotated MS/MS Spectrum

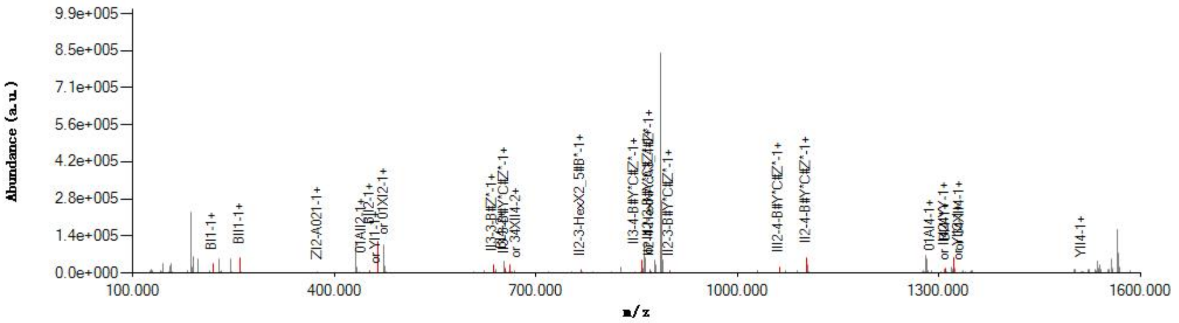


#### Graphical Fragmentation Map

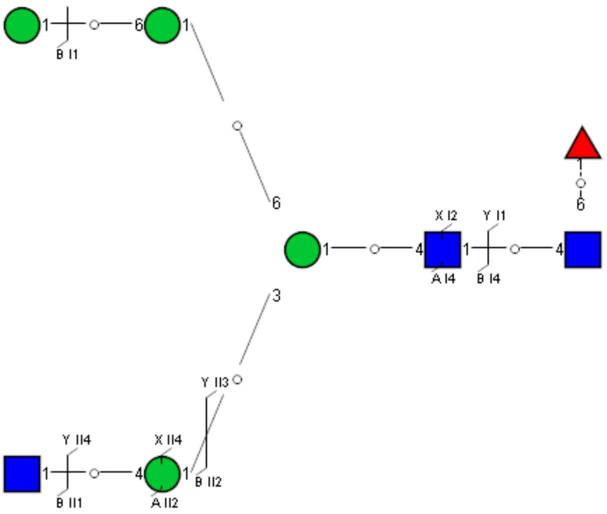

Spectrum Index: 27873(NoBHs:1)

#### Precursor iEF Map

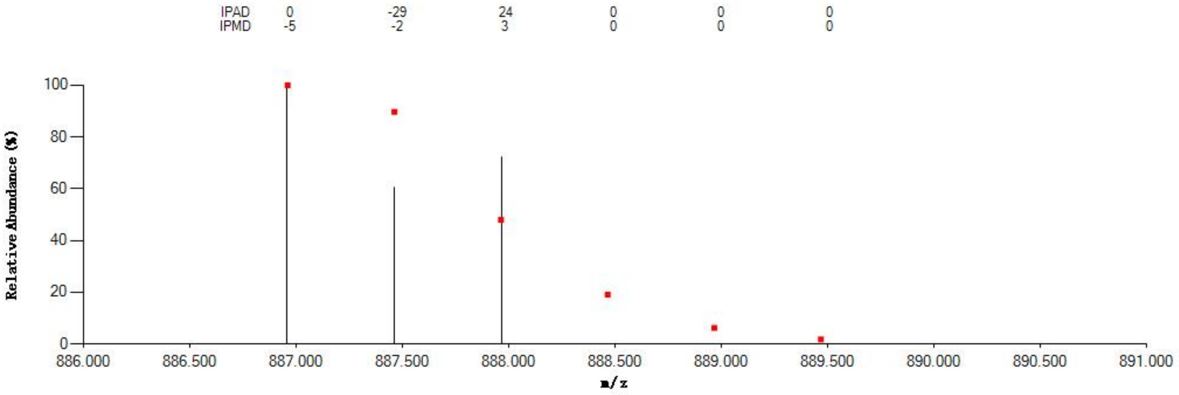


#### Annotated MS/MS Spectrum

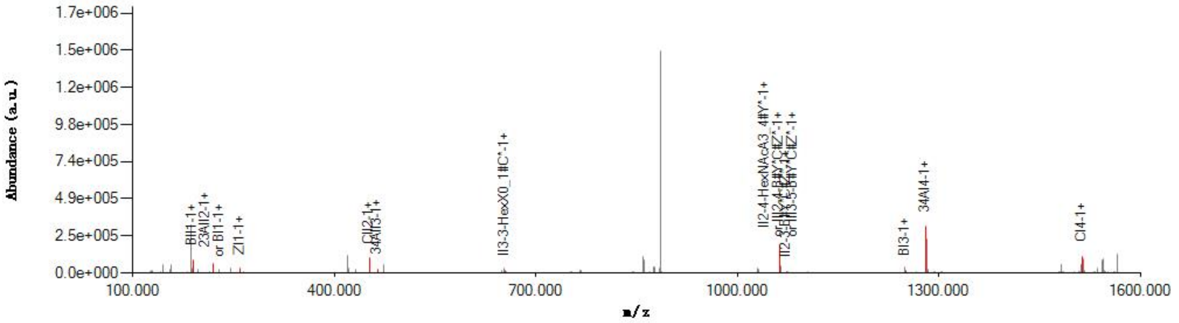


#### Graphical Fragmentation Map

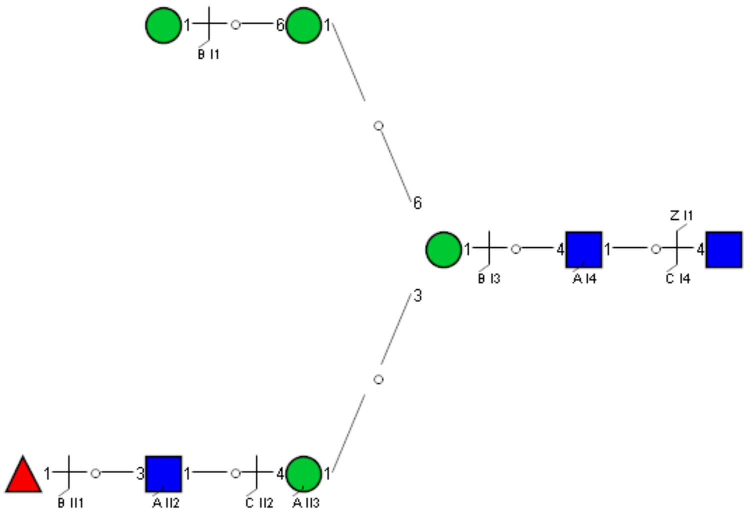

### Spectrum Index: 27904(NoBHs:1)

#### Precursor iEF Map

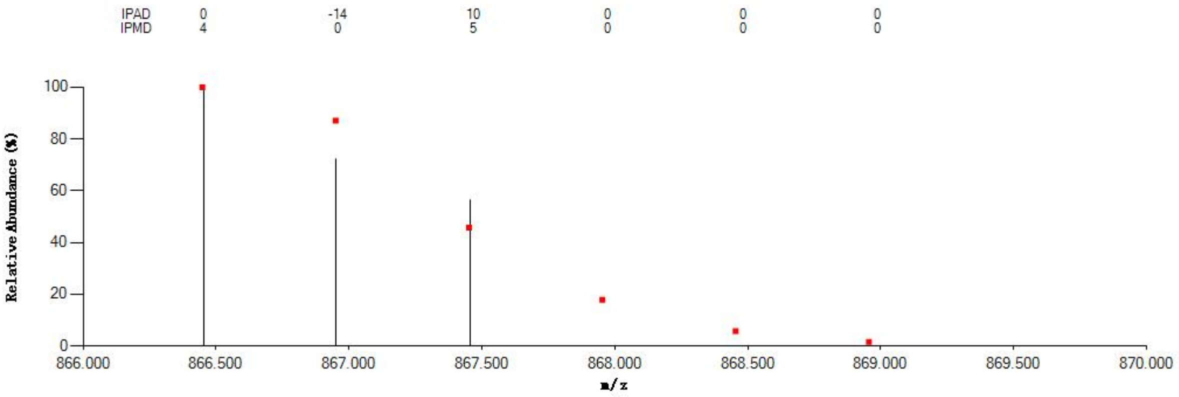


#### Annotated MS/MS Spectrum

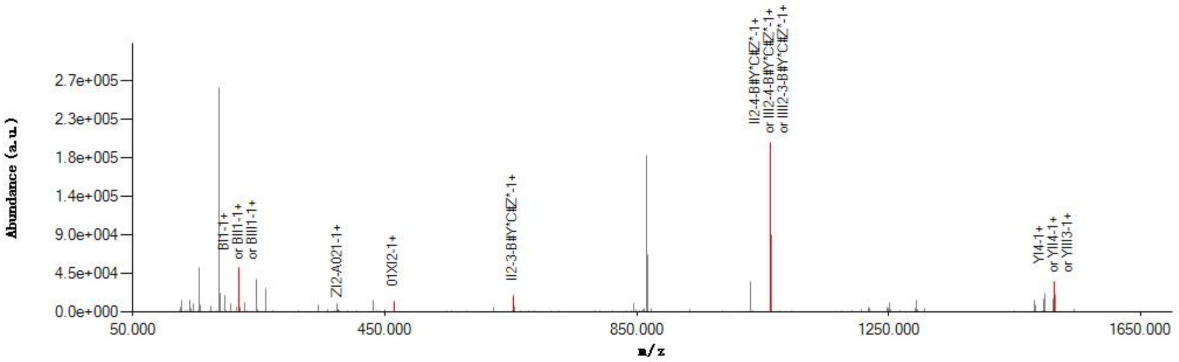


#### Graphical Fragmentation Map

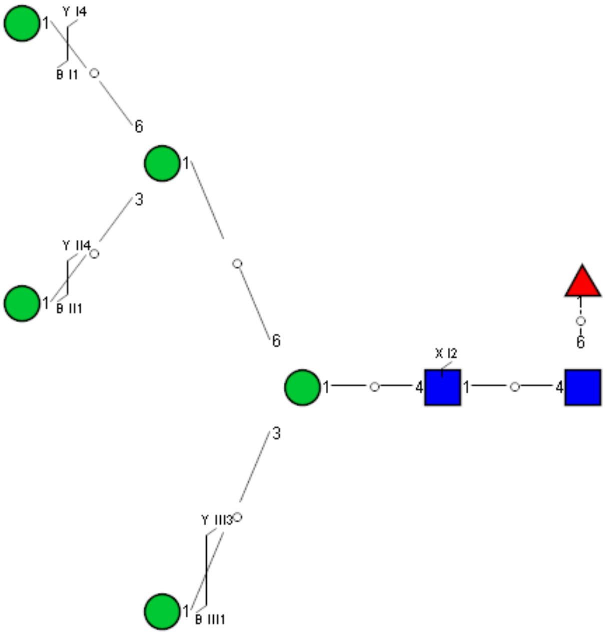

Spectrum Index: 28315(NoBHs:1)

#### Precursor iEF Map

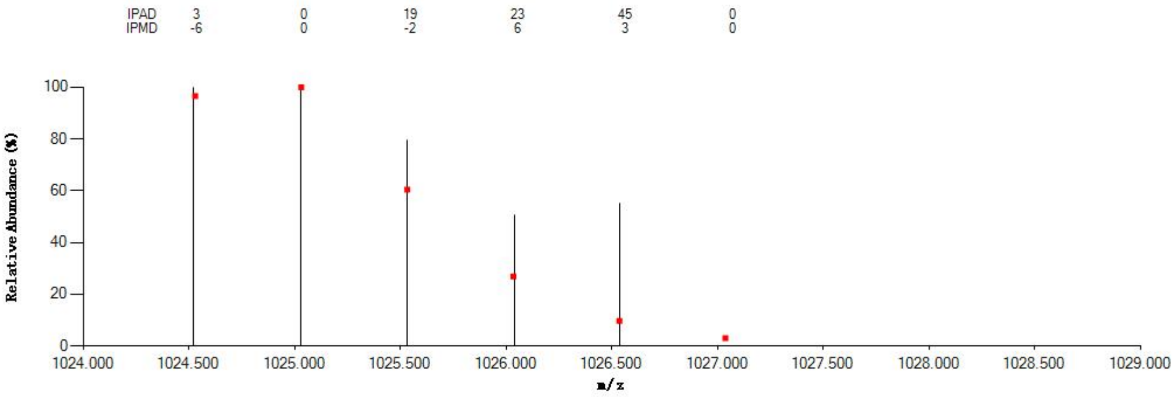


#### Annotated MS/MS Spectrum

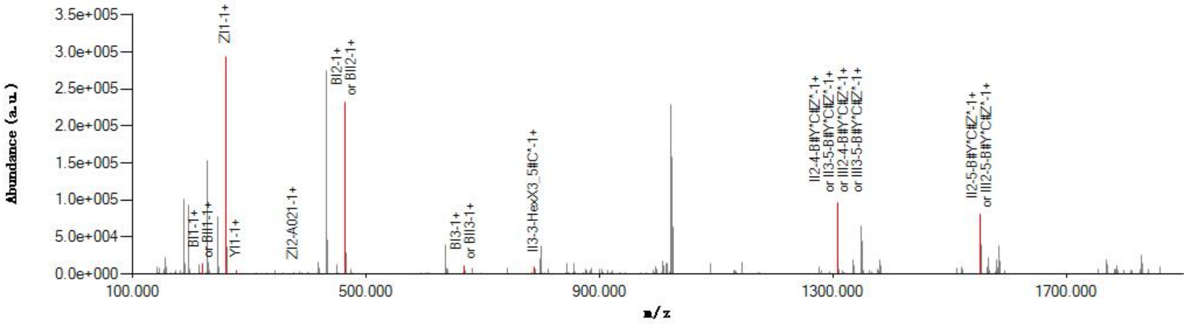


#### Graphical Fragmentation Map

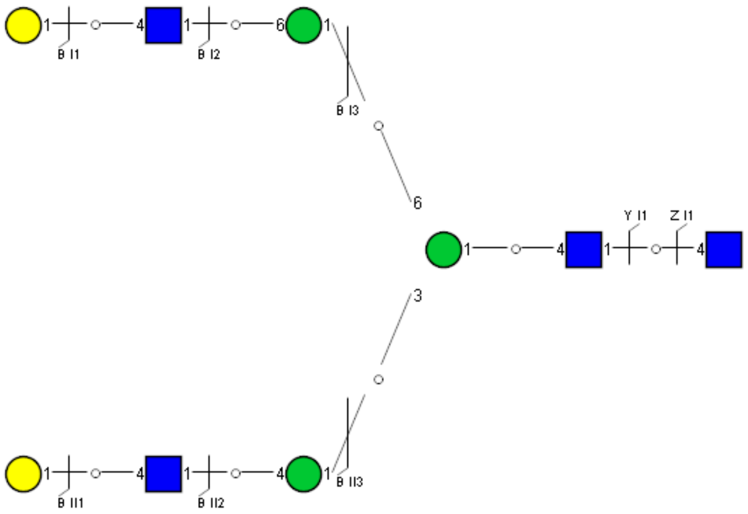

Spectrum Index: 28452(NoBHs:1)

#### Precursor iEF Map

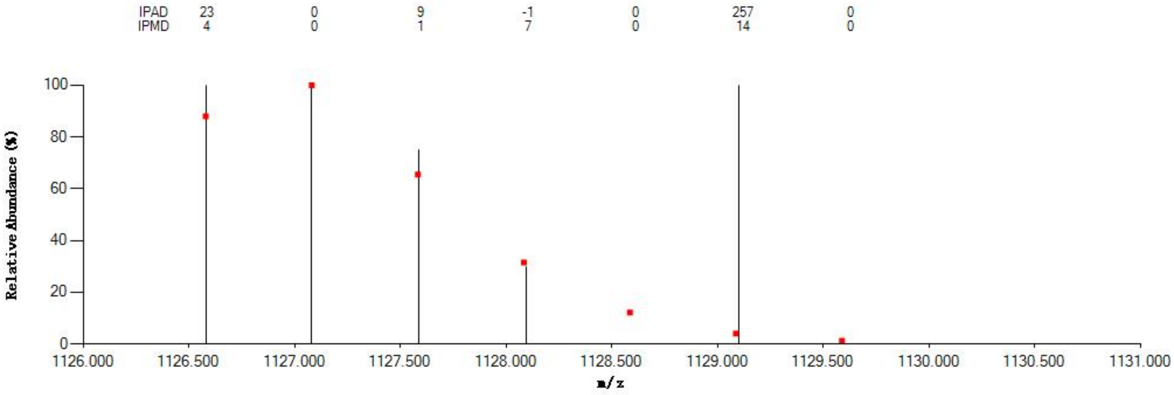


#### Annotated MS/MS Spectrum

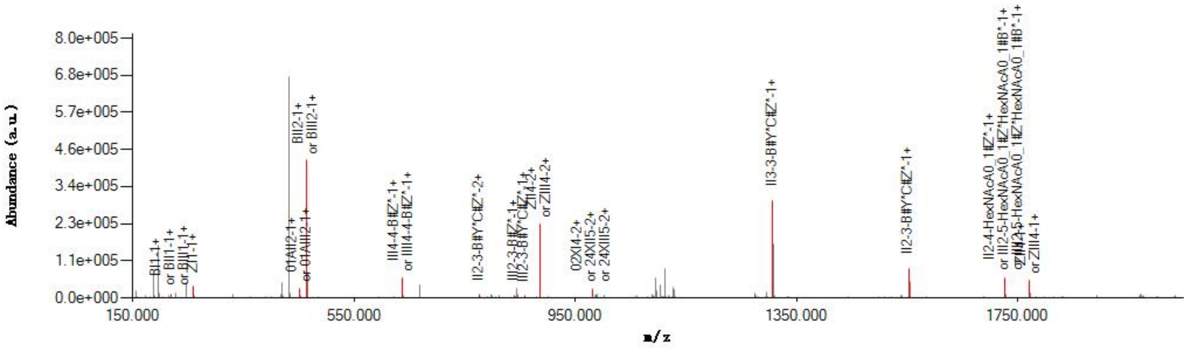


#### Graphical Fragmentation Map

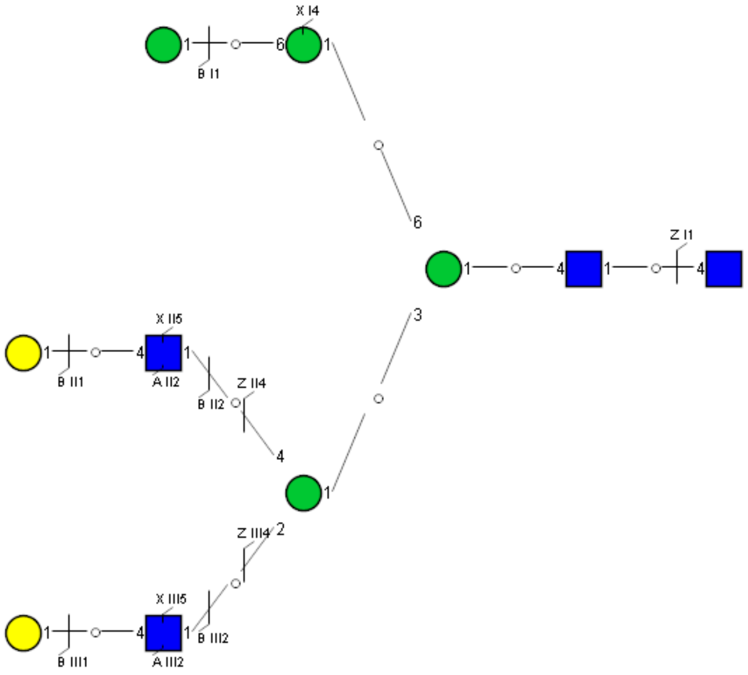

### Spectrum Index: 28455(NoBHs:1)

#### Precursor iEF Map

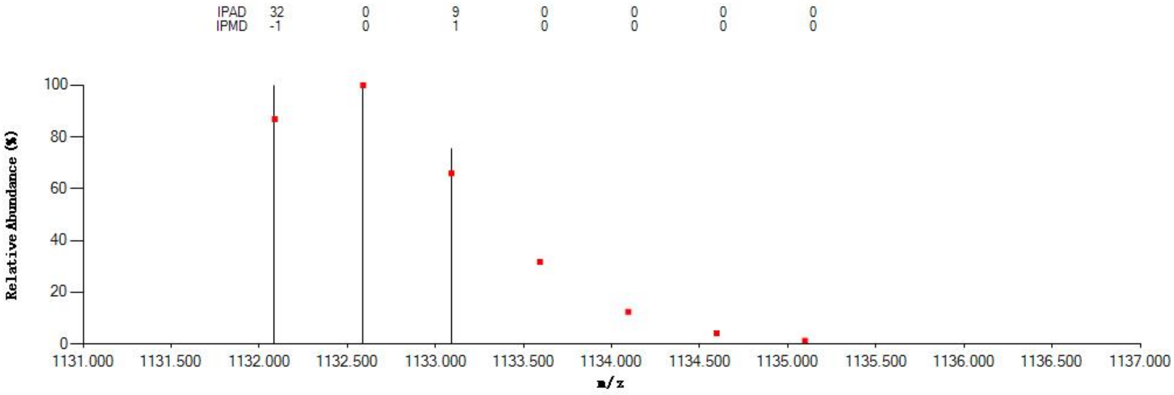


#### Annotated MS/MS Spectrum

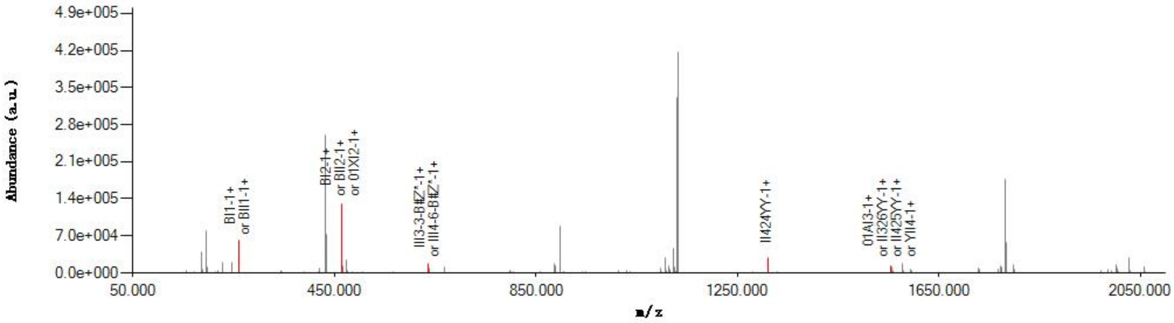


#### Graphical Fragmentation Map

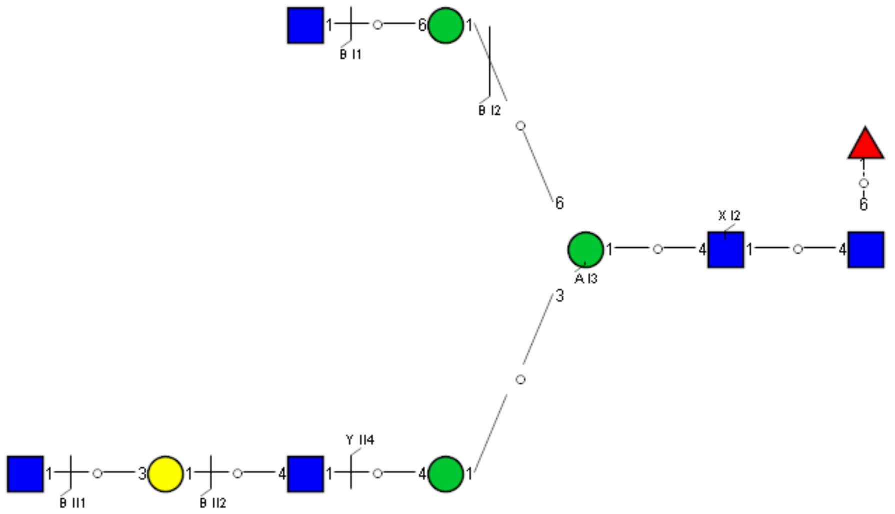

Spectrum Index: 28585(NoBHs:1)

#### Precursor iEF Map

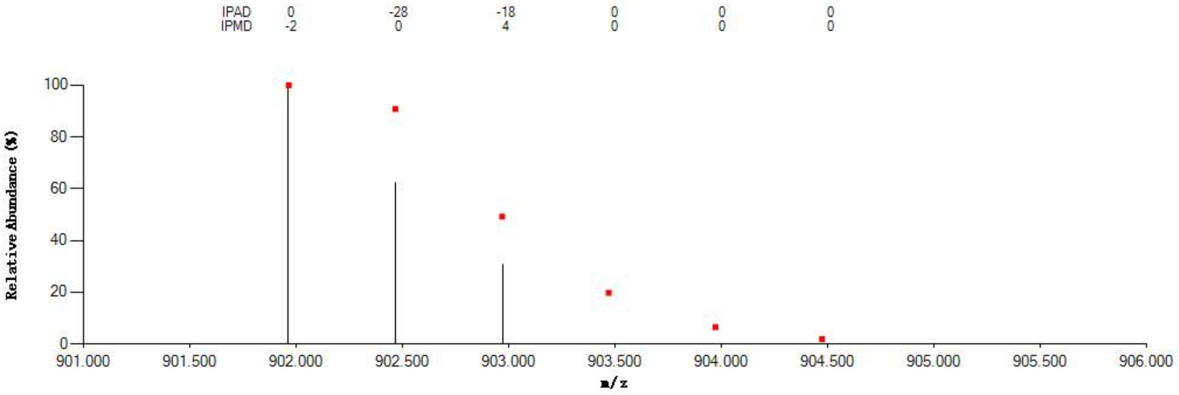


#### Annotated MS/MS Spectrum

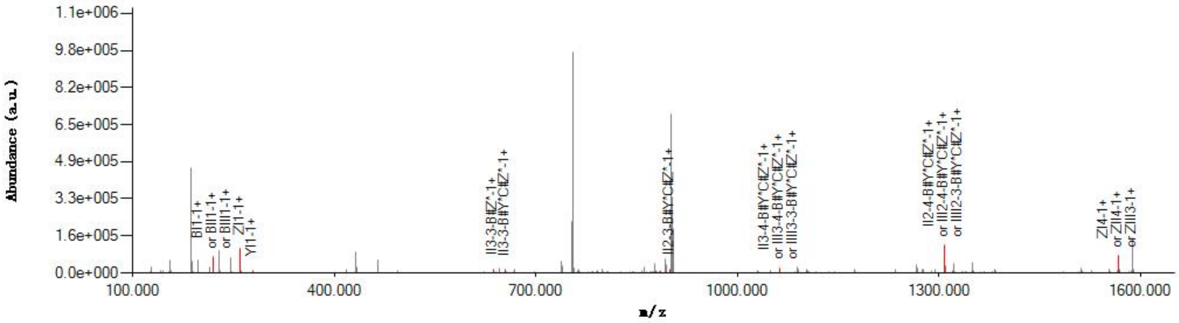


#### Graphical Fragmentation Map

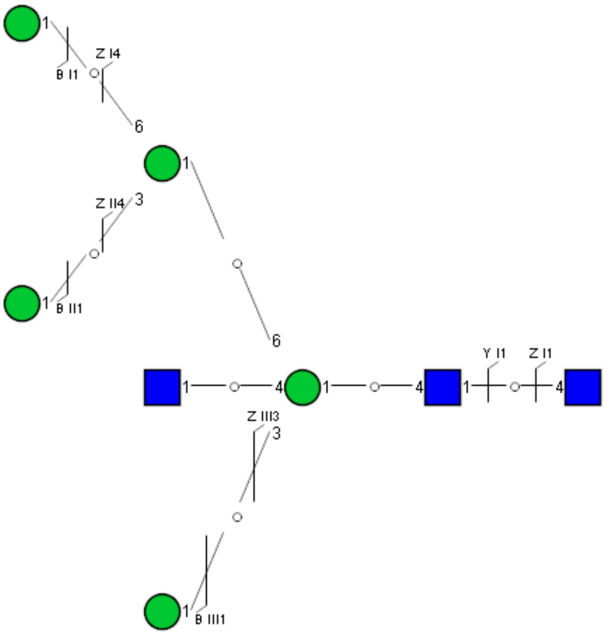

Spectrum Index: 29080(NoBHs:1)

#### Precursor iEF Map

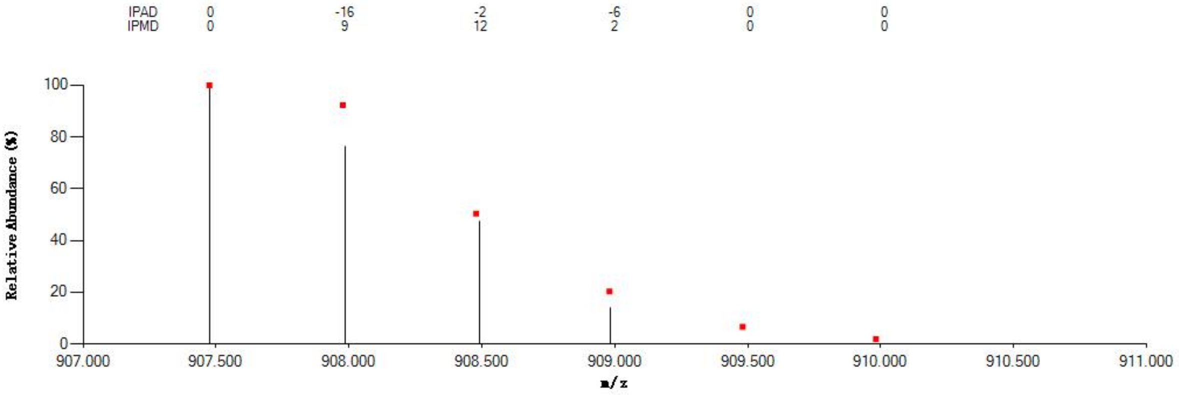


#### Annotated MS/MS Spectrum

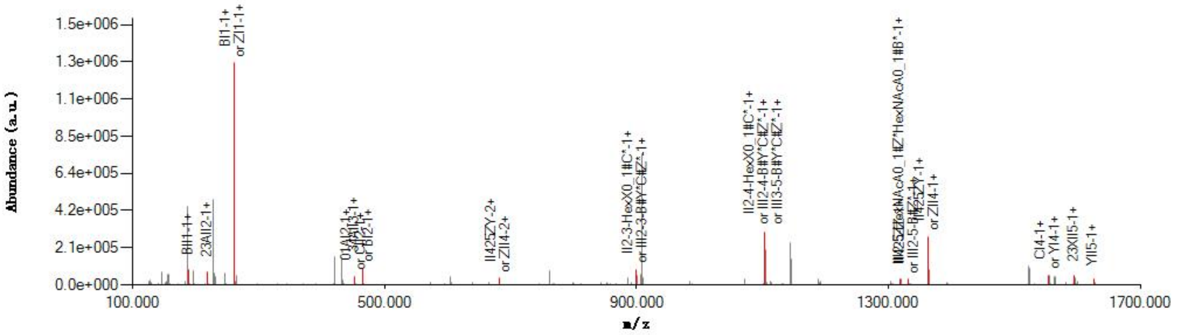


#### Graphical Fragmentation Map

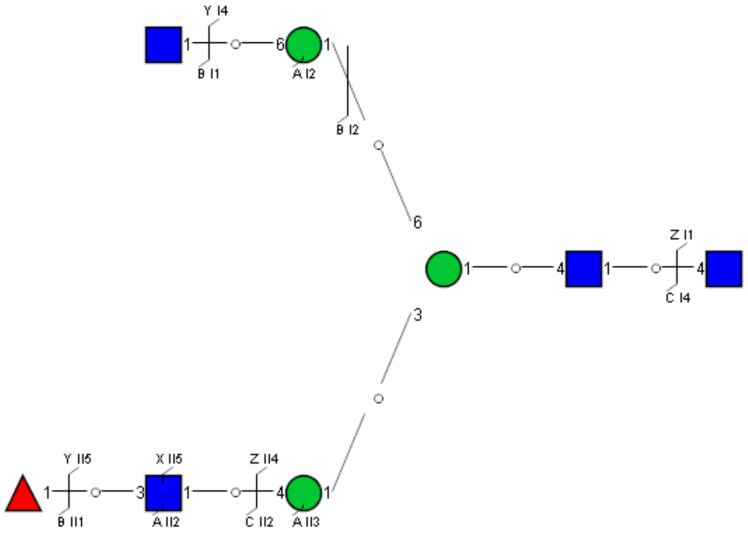

Spectrum Index: 29094(NoBHs:1)

#### Precursor iEF Map

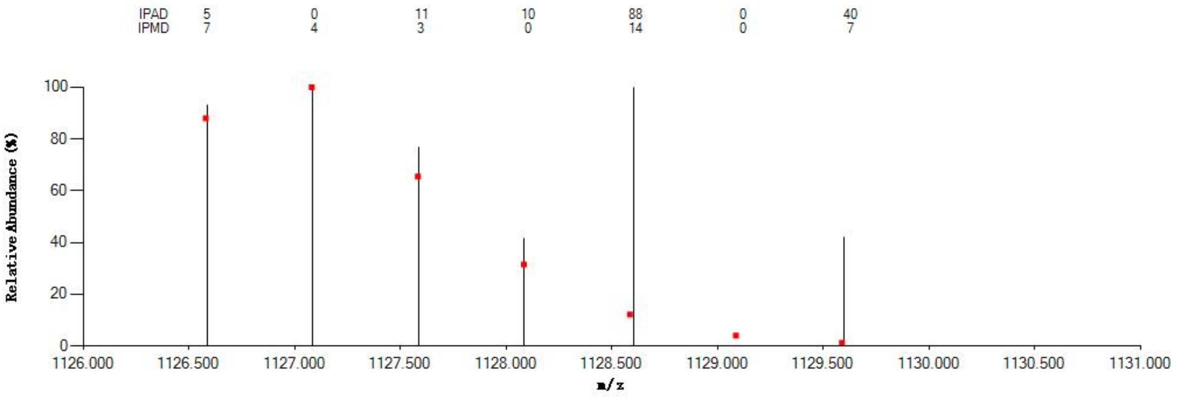

#### Annotated MS/MS Spectrum

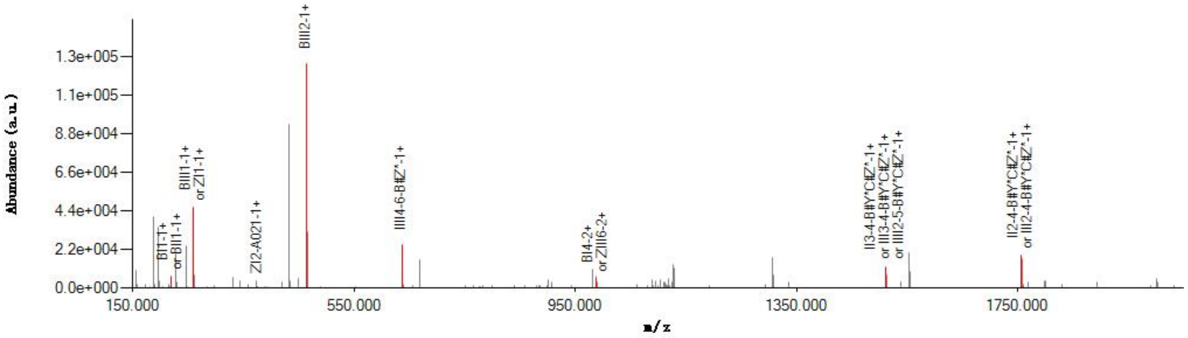

#### Graphical Fragmentation Map

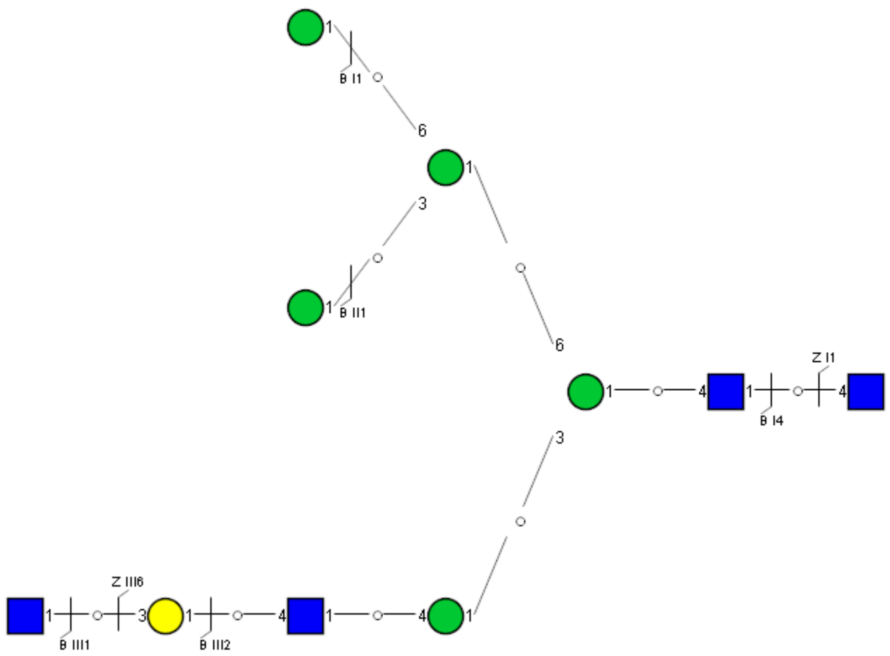

Spectrum Index: 29130(NoBHs:1)

#### Precursor iEF Map

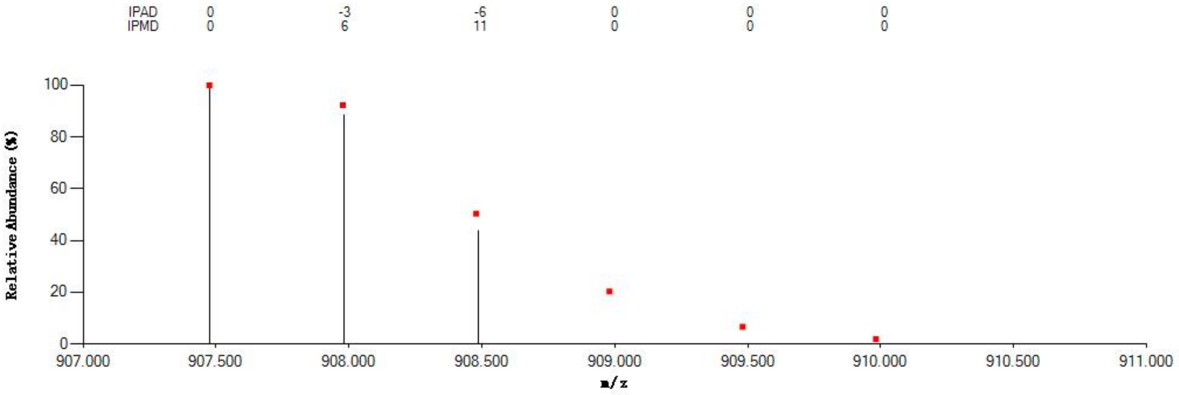

#### Annotated MS/MS Spectrum

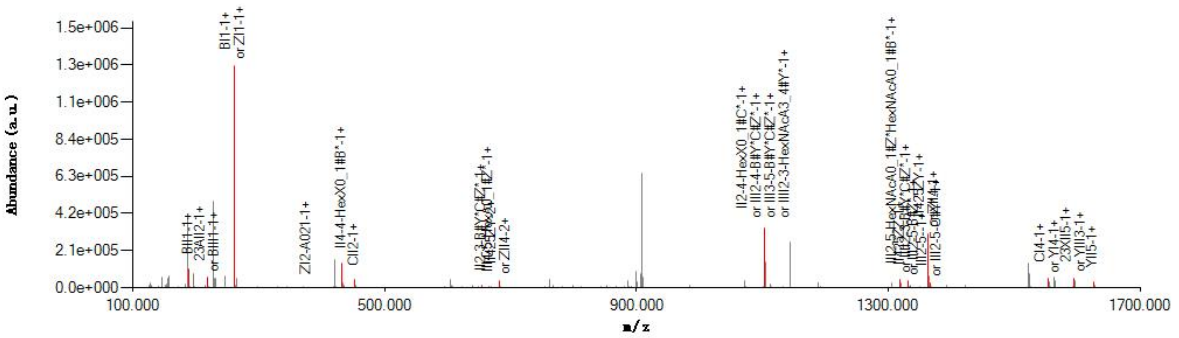

#### Graphical Fragmentation Map

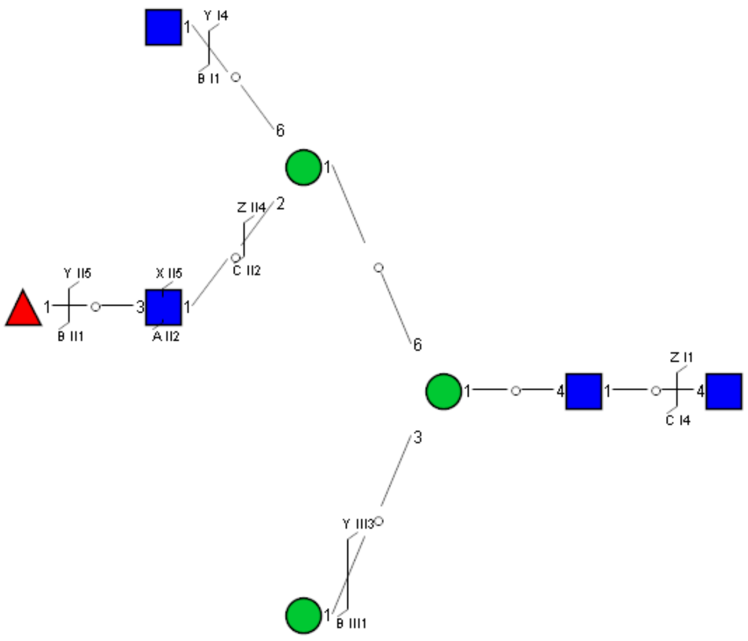

Spectrum Index: 29247(NoBHs:1)

#### Precursor iEF Map

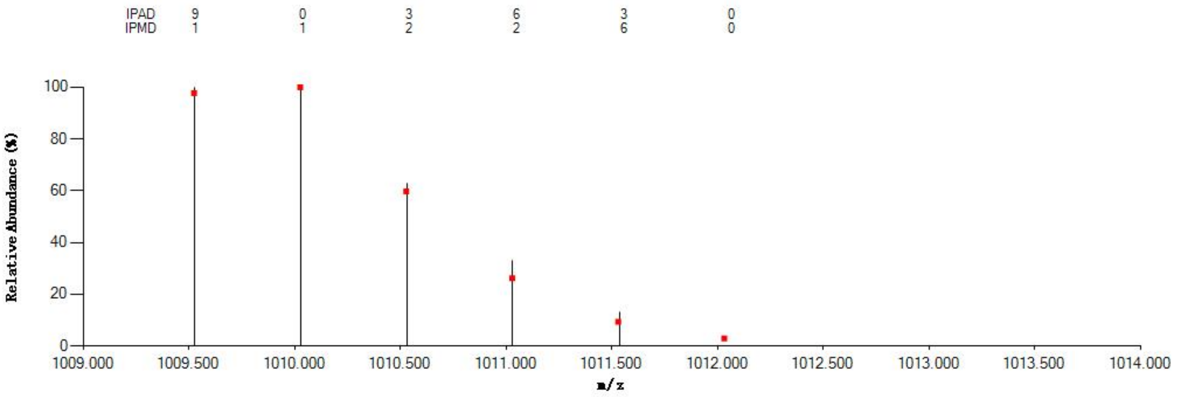

#### Annotated MS/MS Spectrum

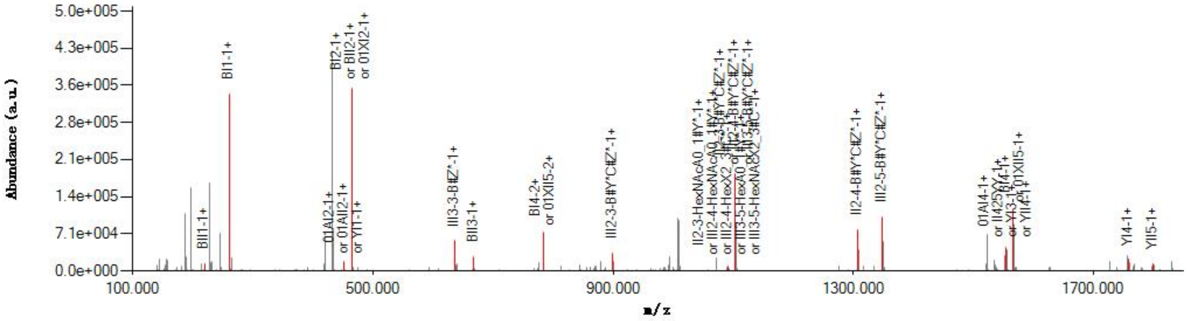

#### Graphical Fragmentation Map

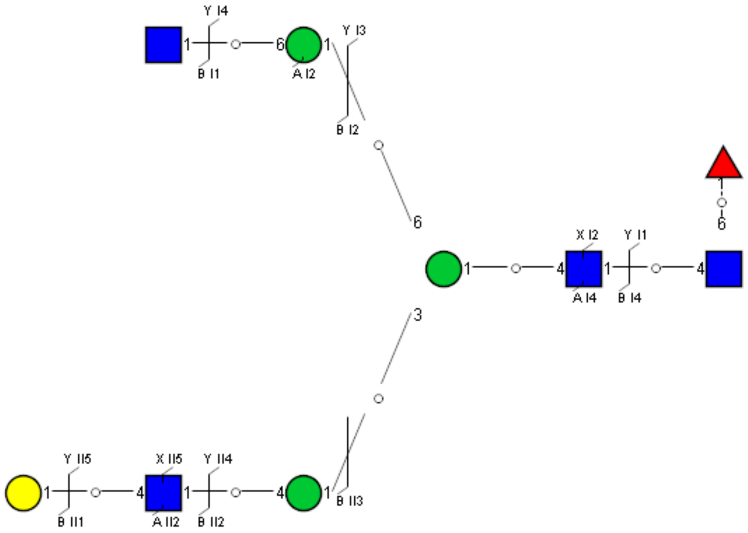

### Spectrum Index: 29252(NoBHs:1)

#### Precursor iEF Map

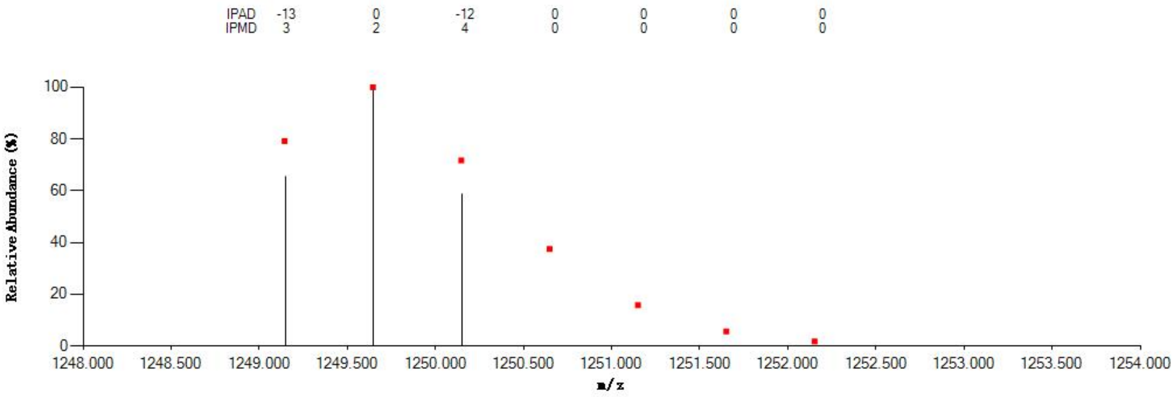


#### Annotated MS/MS Spectrum

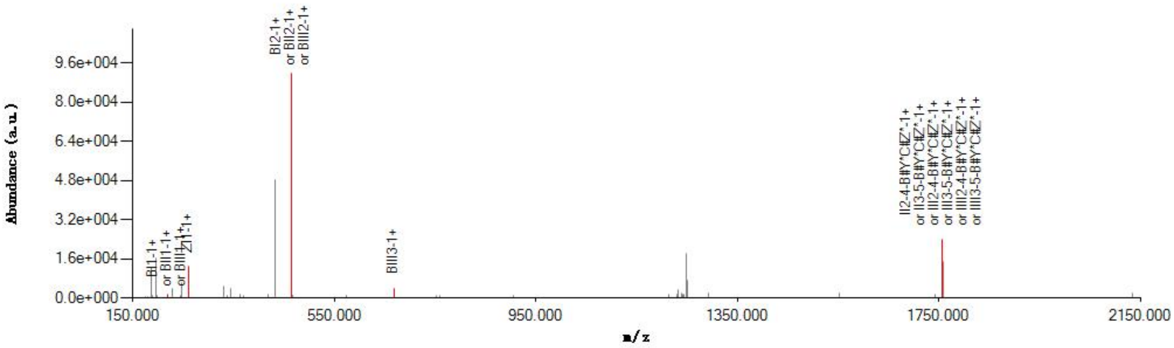


#### Graphical Fragmentation Map

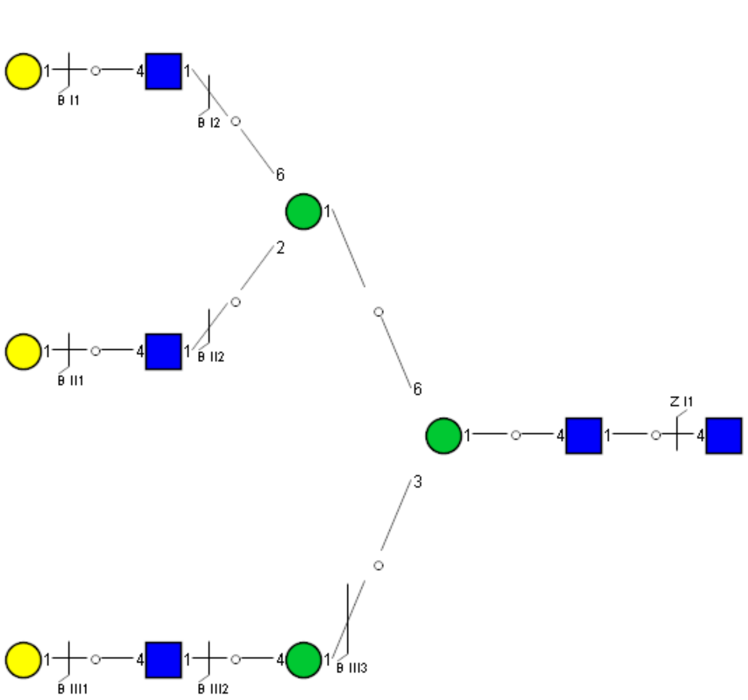

### Spectrum Index: 29319(NoBHs:1)

#### Precursor iEF Map

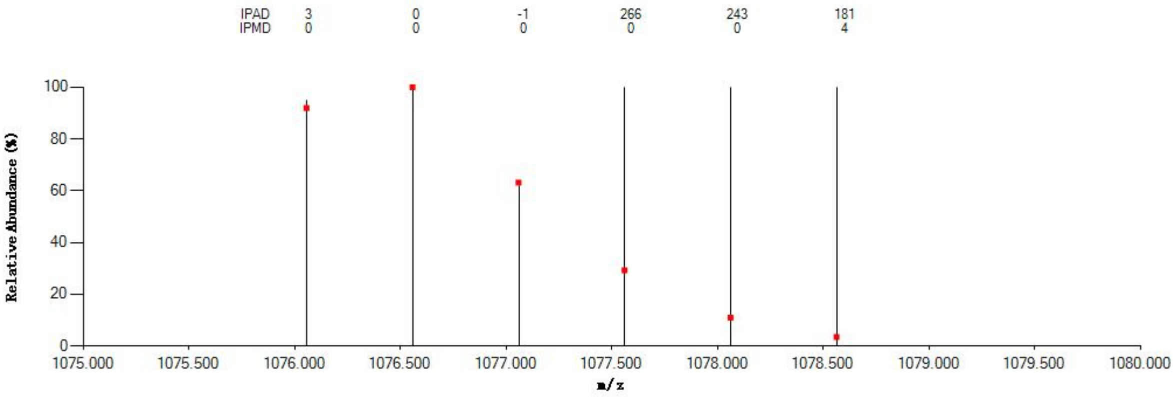


#### Annotated MS/MS Spectrum

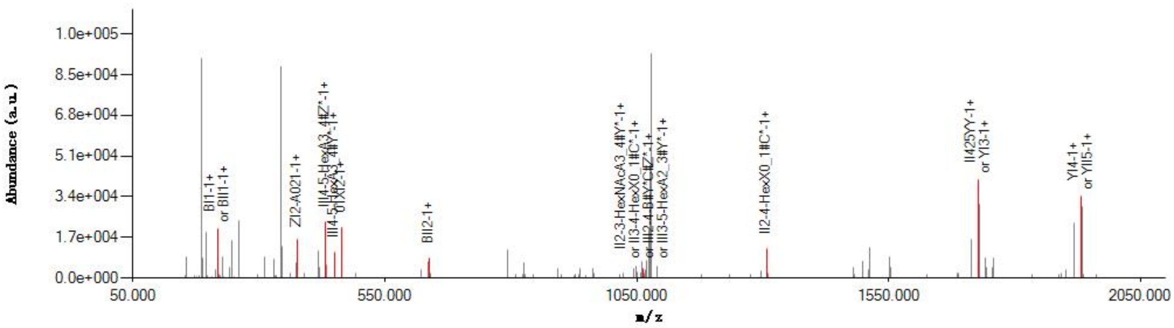


#### Graphical Fragmentation Map

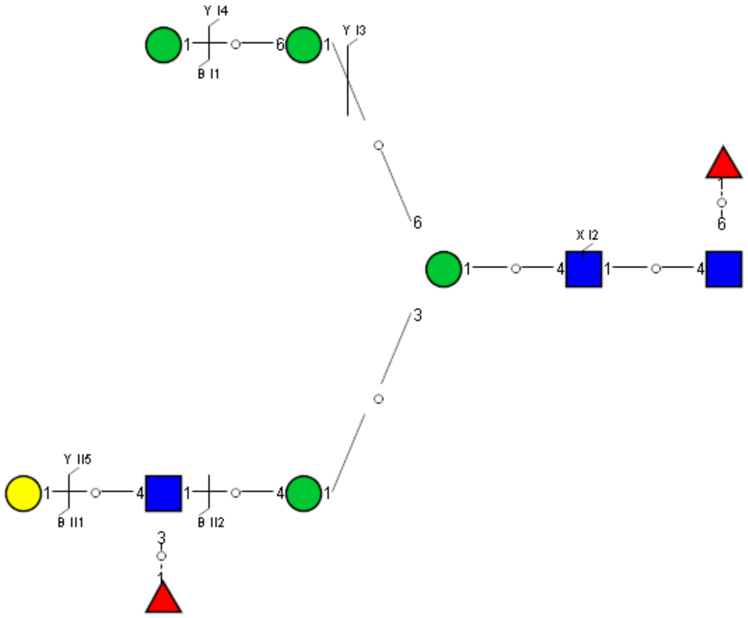

Spectrum Index: 29472(NoBHs:1)

#### Precursor iEF Map

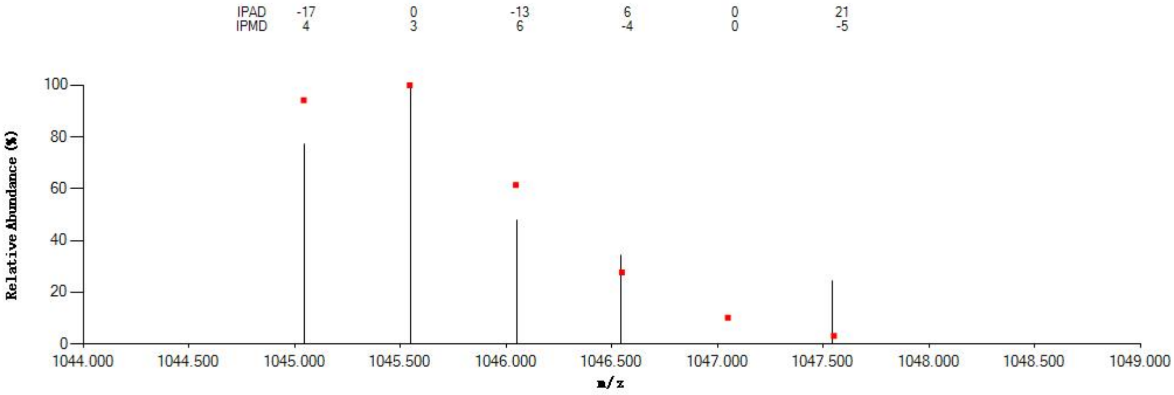


#### Annotated MS/MS Spectrum

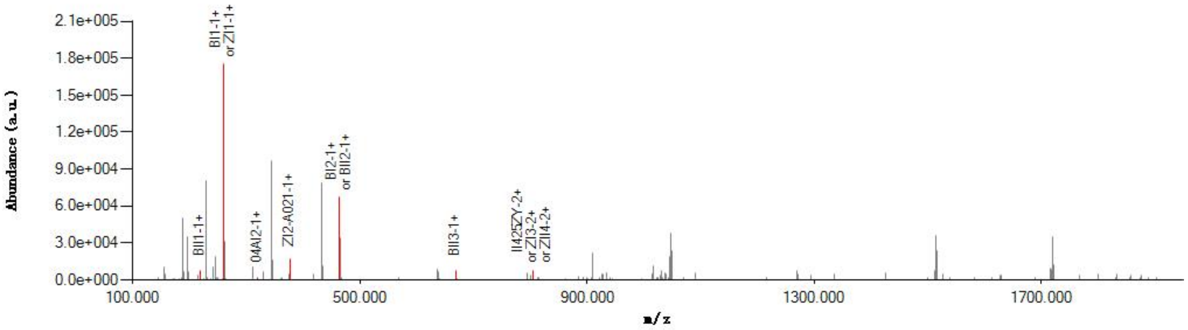


#### Graphical Fragmentation Map

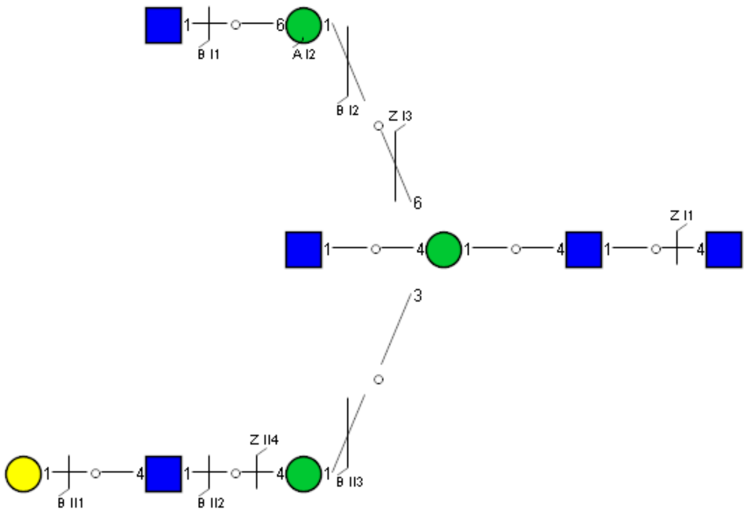

Spectrum Index: 29646(NoBHs:1)

#### Precursor iEF Map

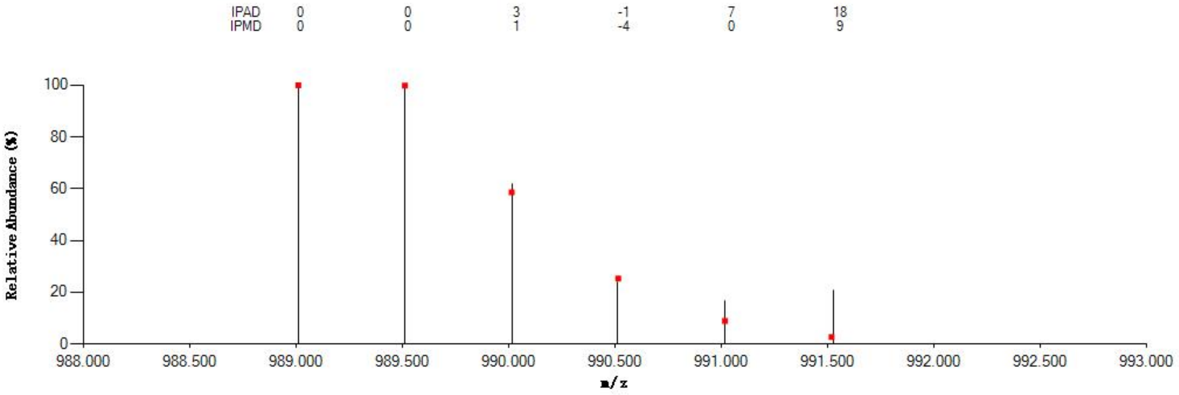


#### Annotated MS/MS Spectrum

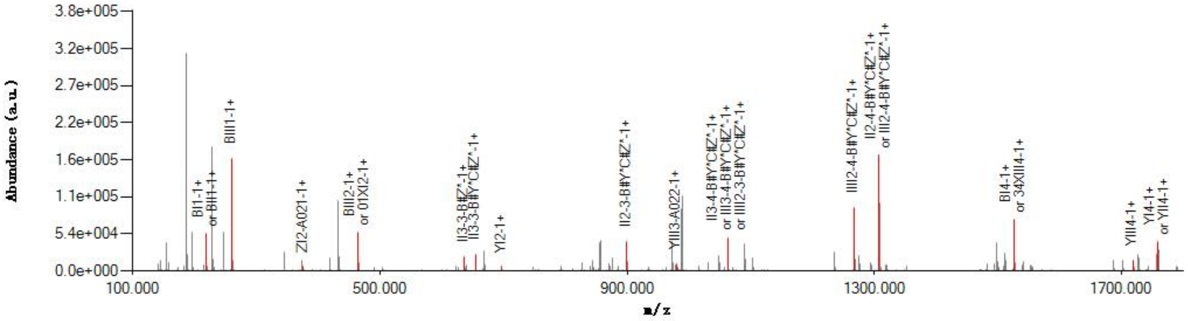


#### Graphical Fragmentation Map

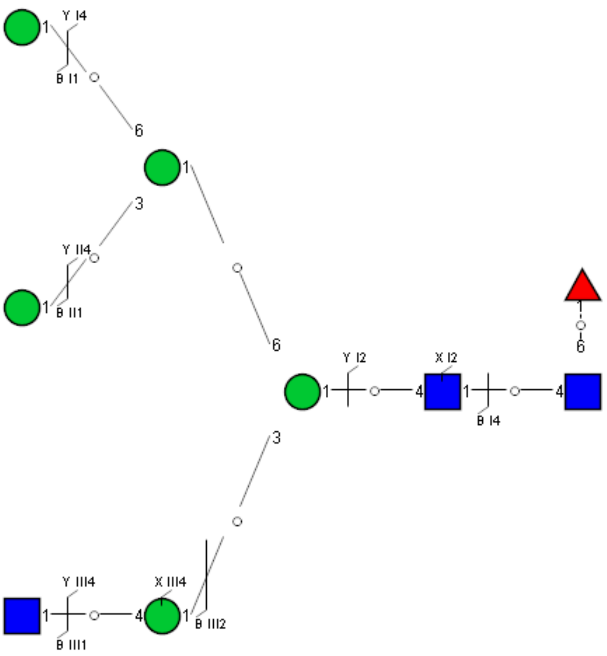

Spectrum Index: 29944(NoBHs:1)

#### Precursor iEF Map

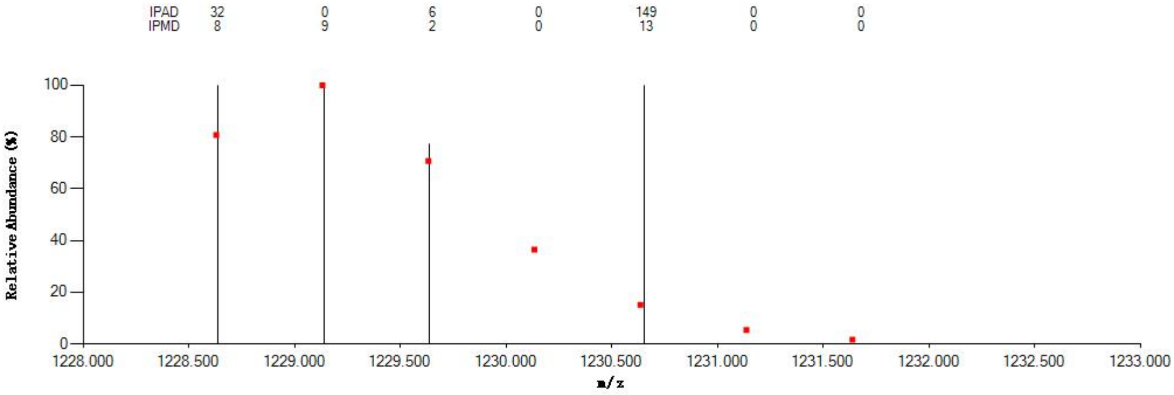


#### Annotated MS/MS Spectrum

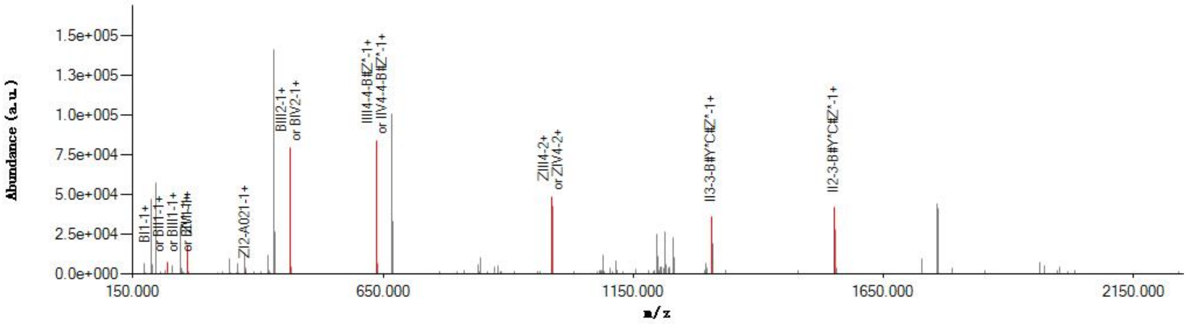


#### Graphical Fragmentation Map

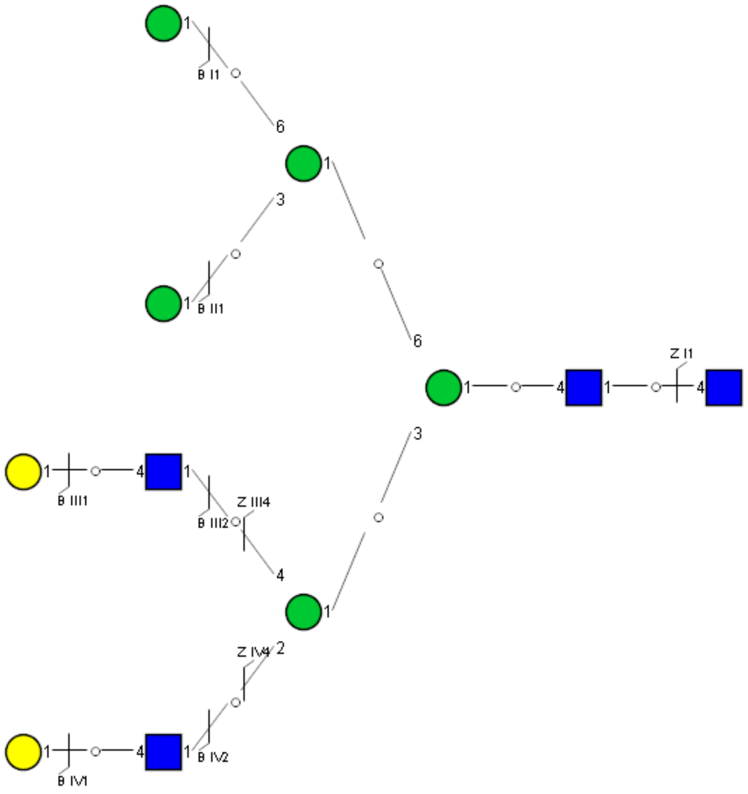

Spectrum Index: 30004(NoBHs:1)

Precursor iEF Map

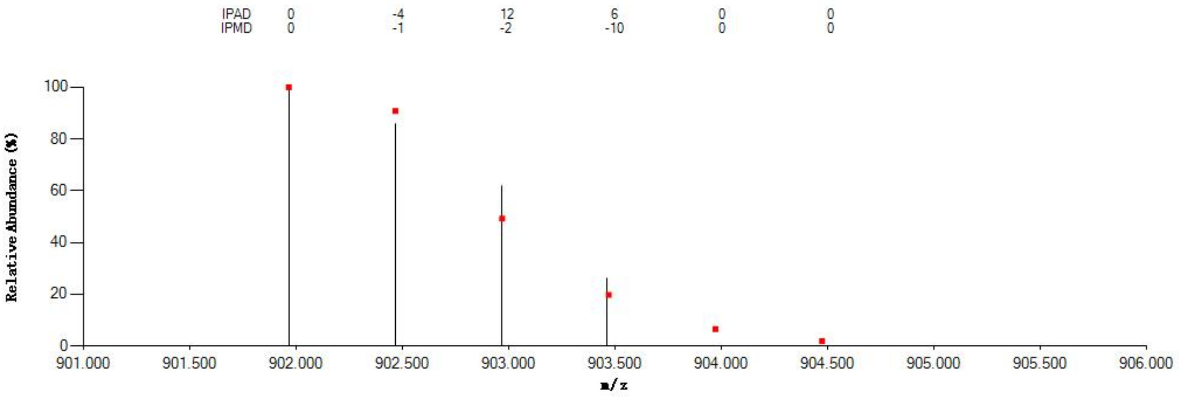


Annotated MS/MS Spectrum

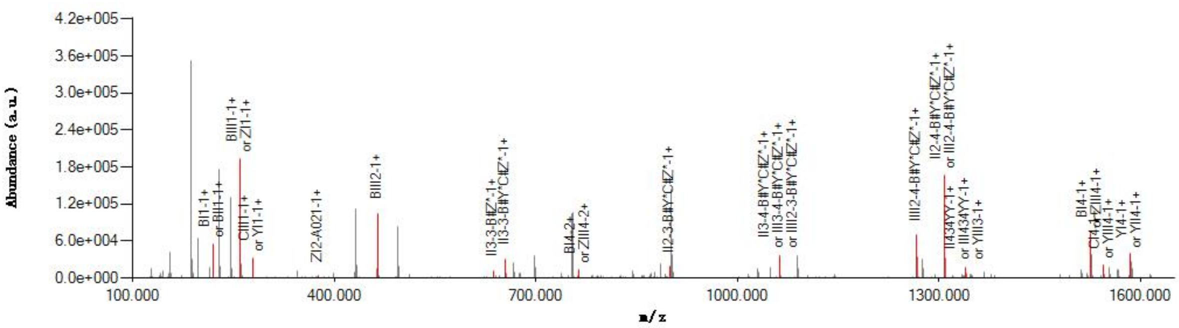


Graphical Fragmentation Map

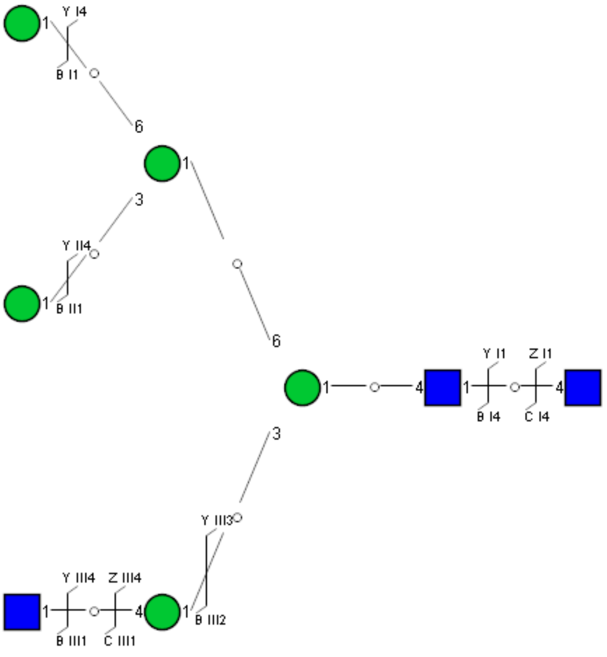

Spectrum Index: 30256(NoBHs:1)

#### Precursor iEF Map

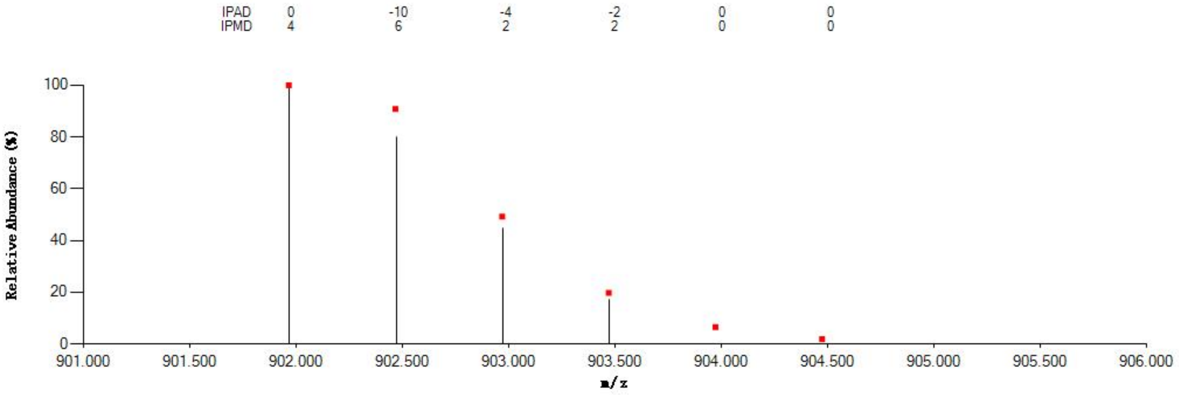


#### Annotated MS/MS Spectrum

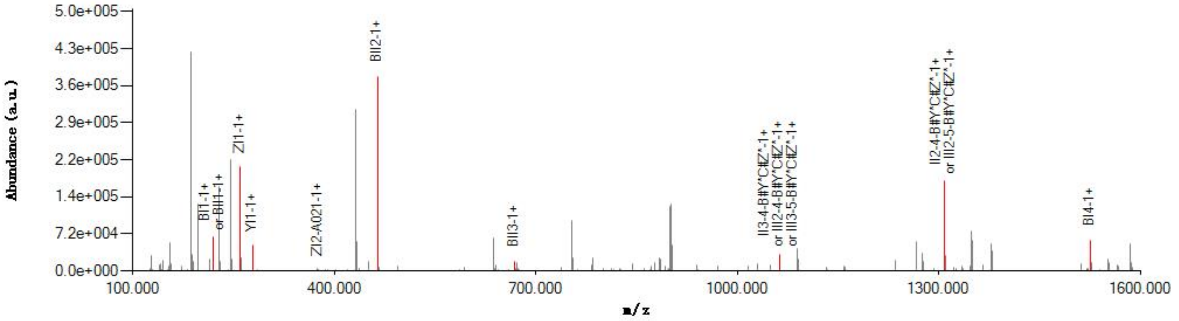


#### Graphical Fragmentation Map

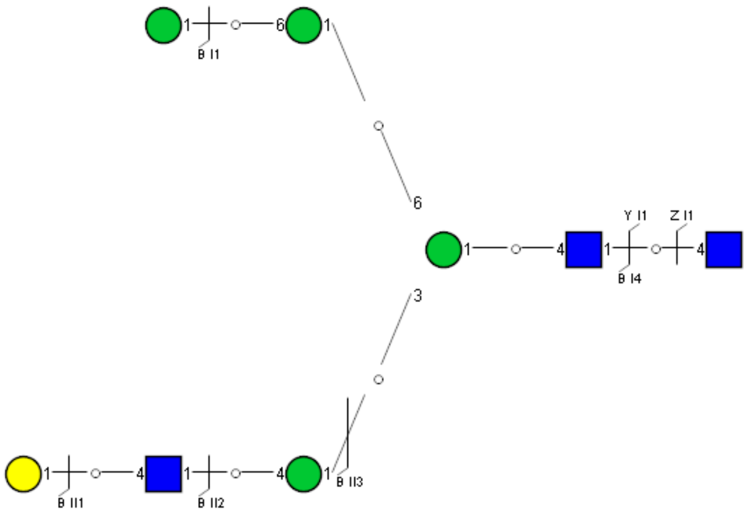

Spectrum Index: 30675(NoBHs:1)

#### Precursor iEF Map

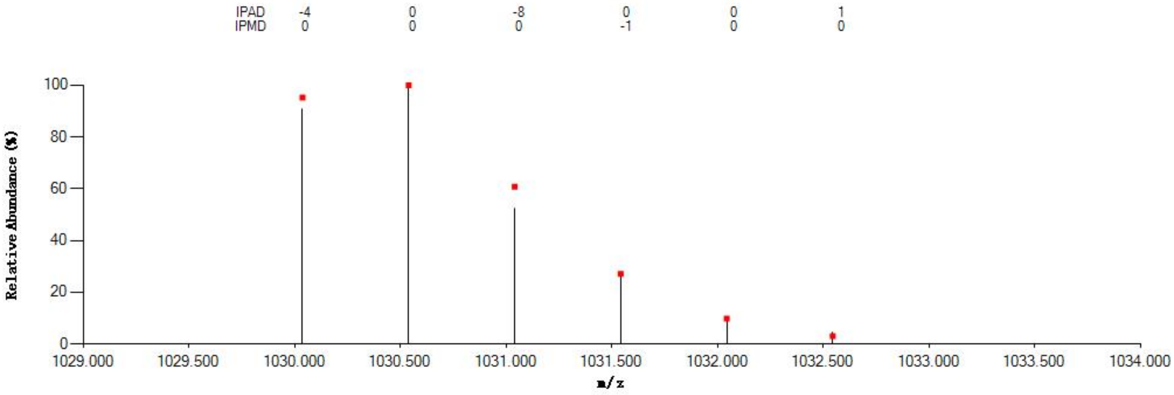

#### Annotated MS/MS Spectrum

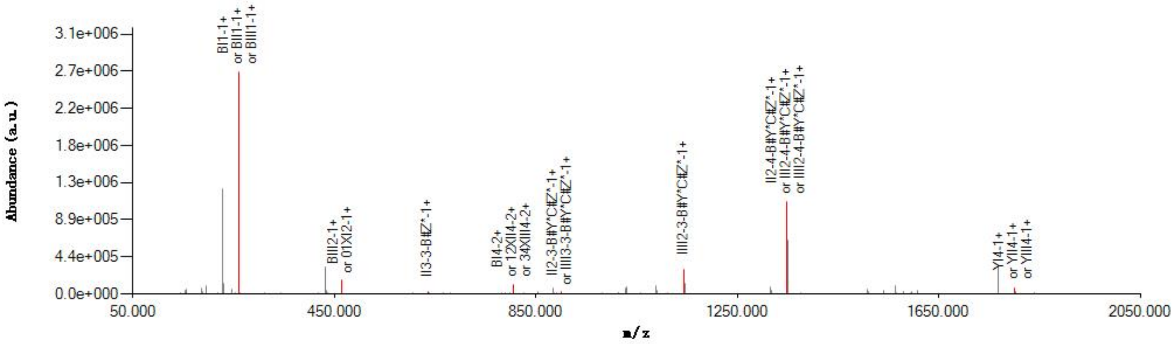

#### Graphical Fragmentation Map

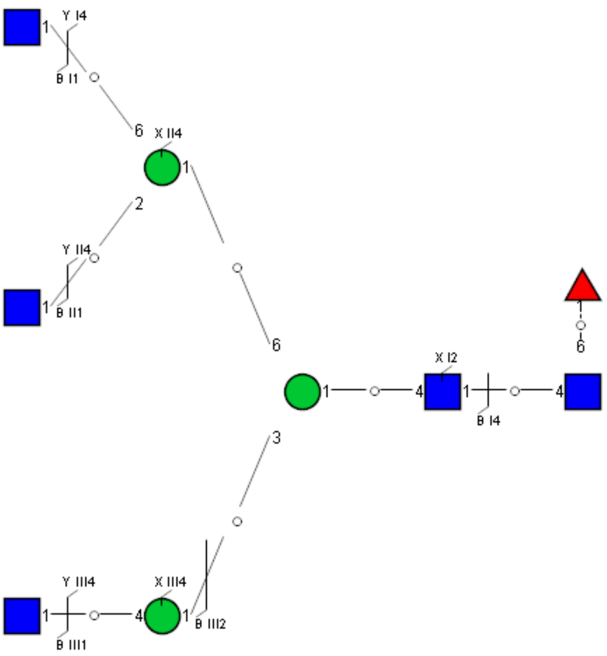

Spectrum Index: 30741(NoBHs:1)

#### Precursor iEF Map

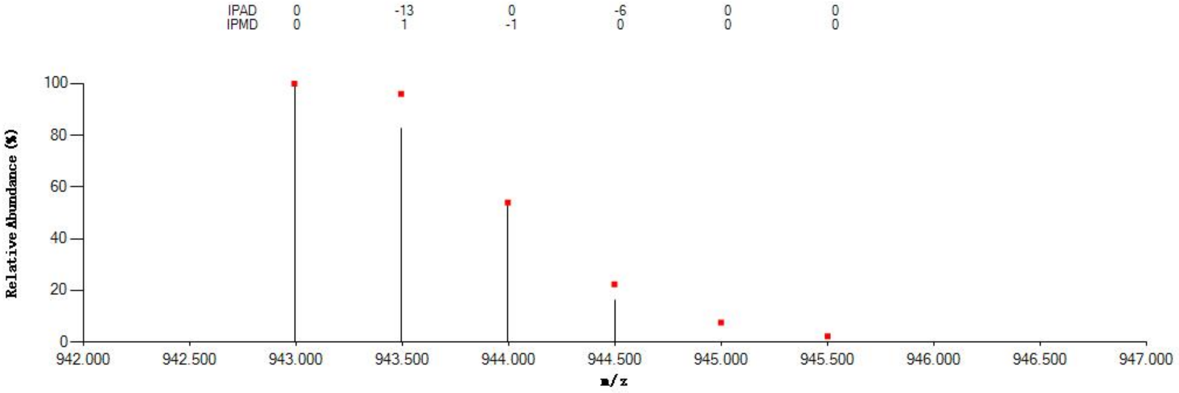


#### Annotated MS/MS Spectrum

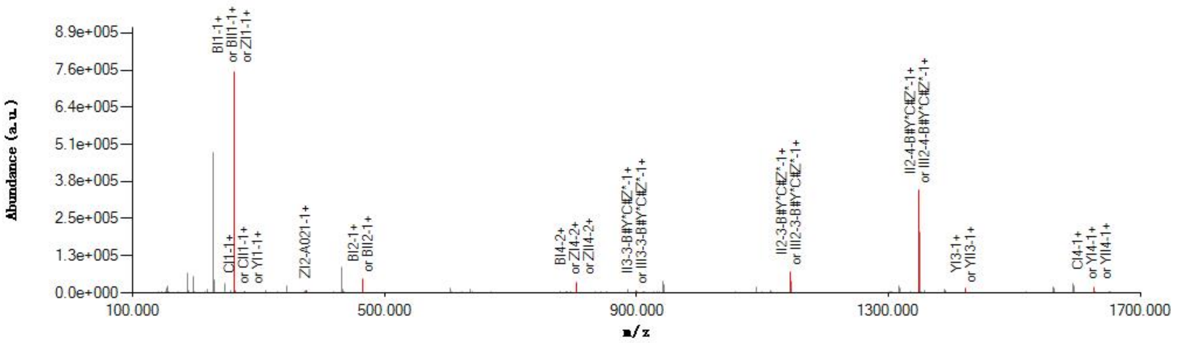


#### Graphical Fragmentation Map

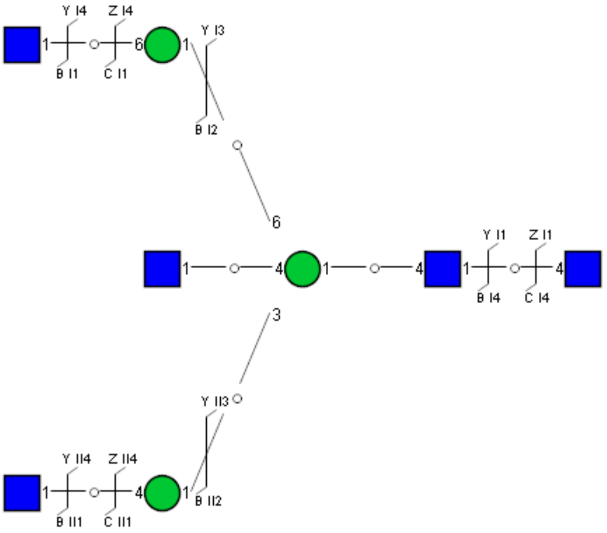

Spectrum Index: 30822(NoBHs:1)

#### Precursor iEF Map

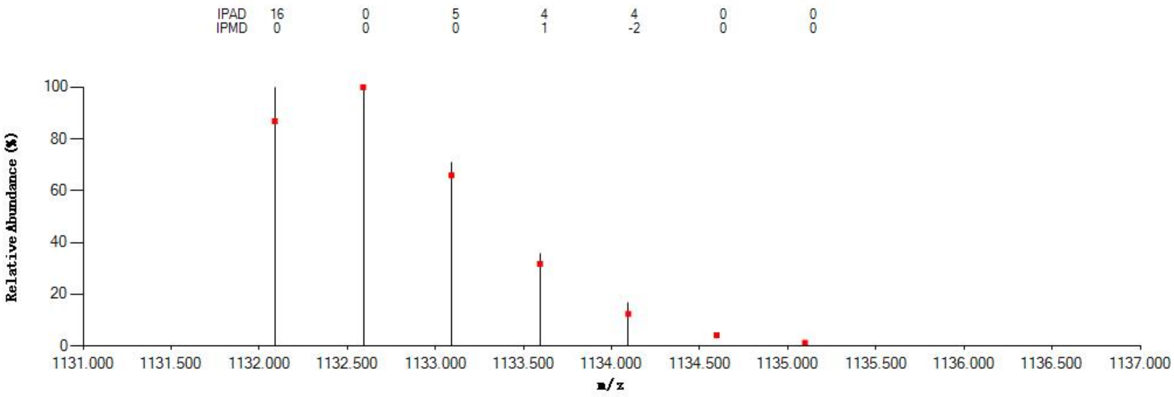


#### Annotated MS/MS Spectrum

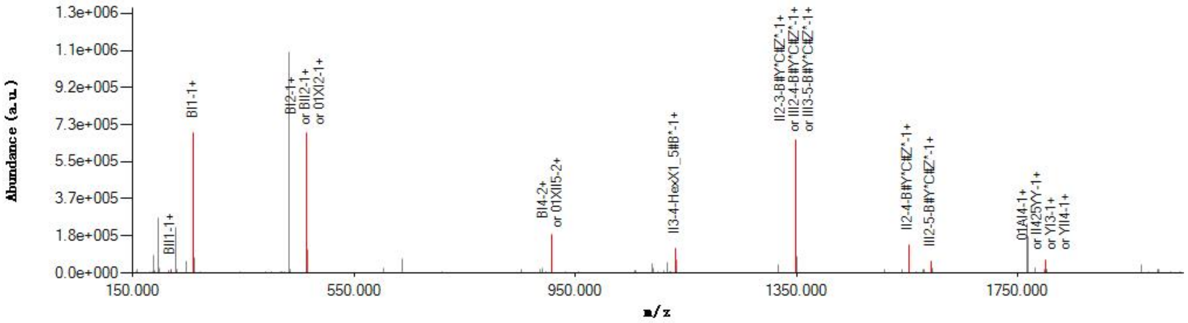


#### Graphical Fragmentation Map

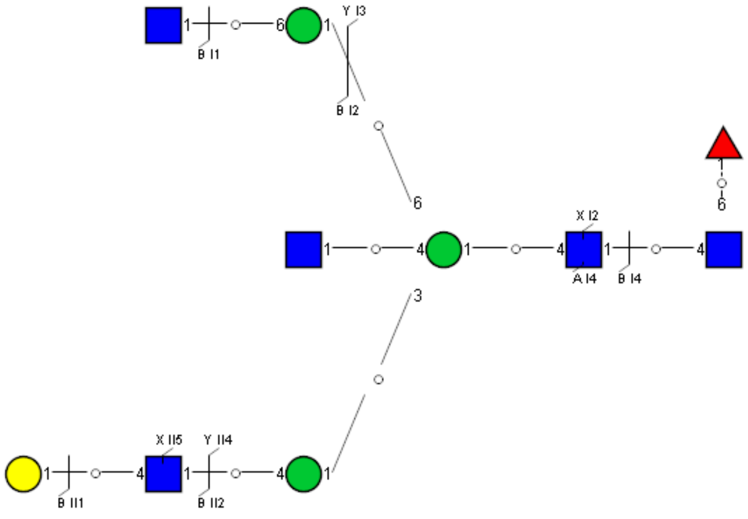

### Spectrum Index: 30904(NoBHs:1)

#### Precursor iEF Map

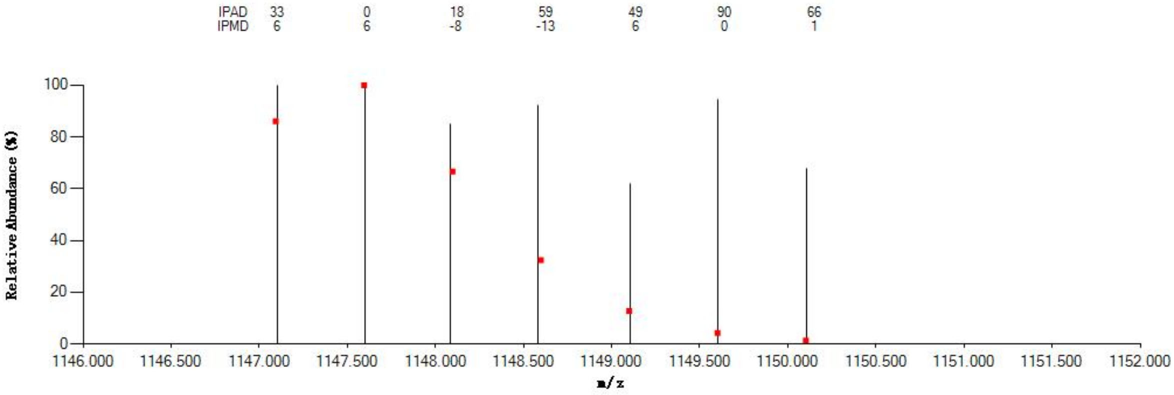

#### Annotated MS/MS Spectrum

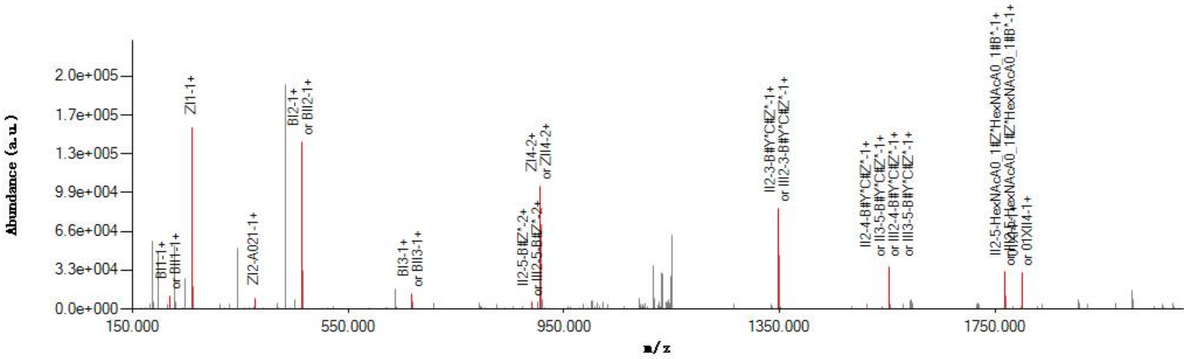

#### Graphical Fragmentation Map

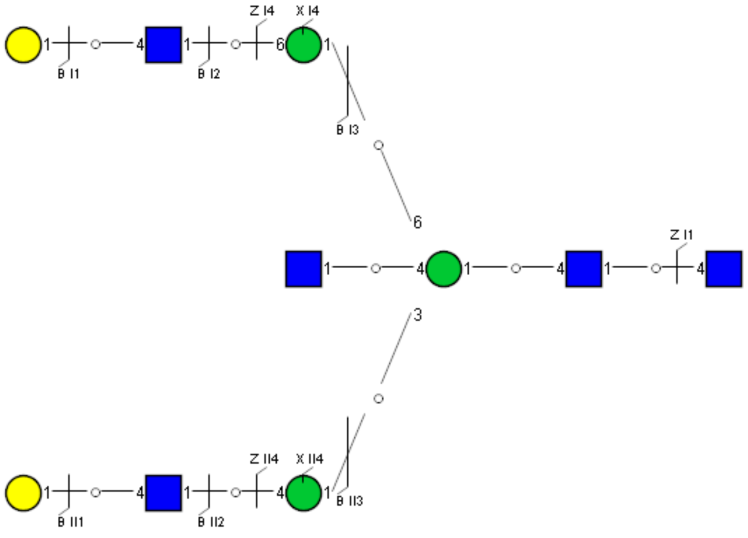

Spectrum Index: 30971(NoBHs:1)

#### Precursor iEF Map

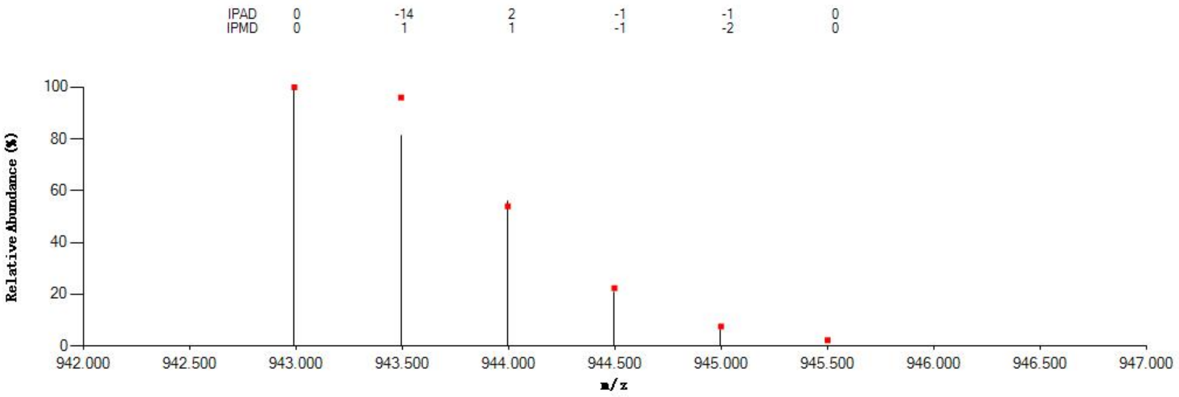


#### Annotated MS/MS Spectrum

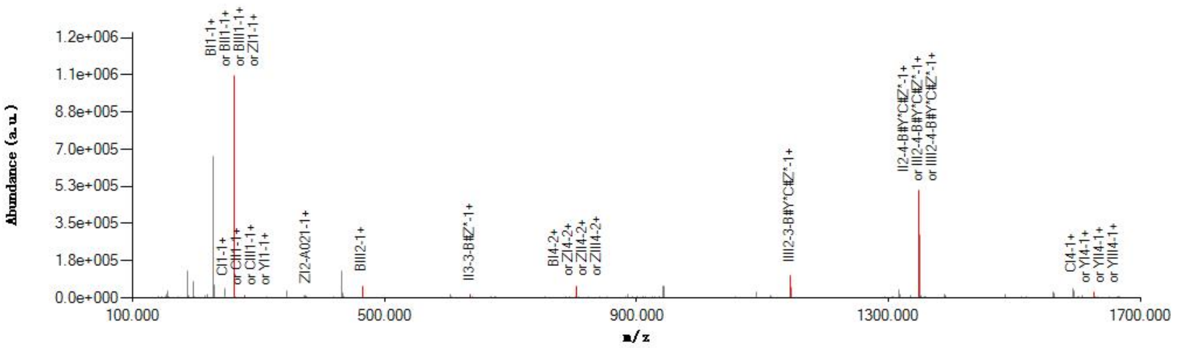


#### Graphical Fragmentation Map

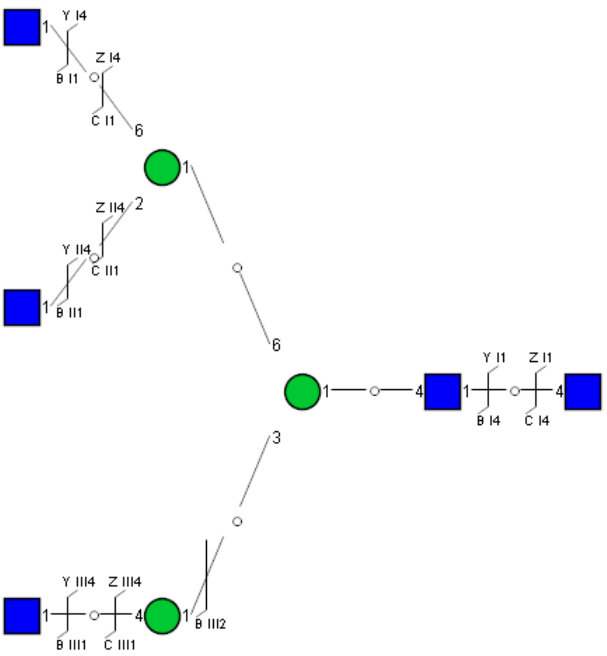

Spectrum Index: 31434(NoBHs:1)

#### Precursor iEF Map

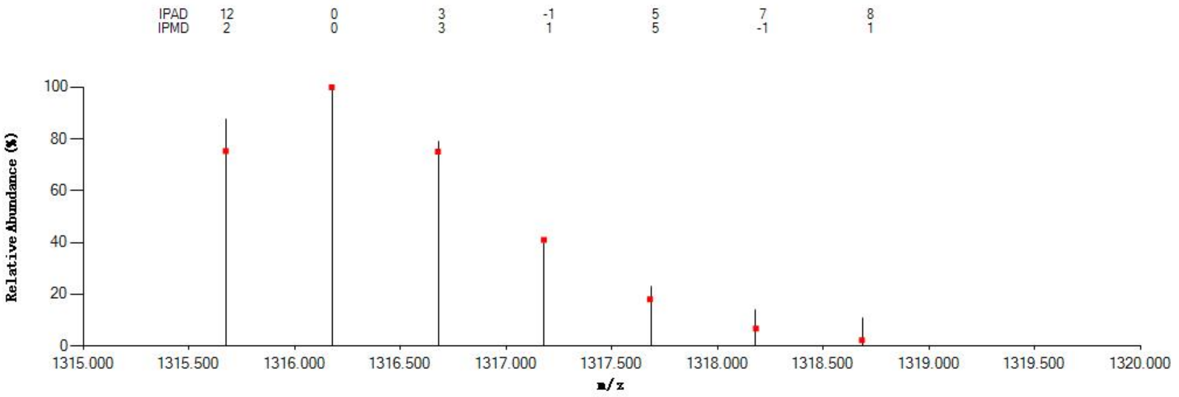

#### Annotated MS/MS Spectrum

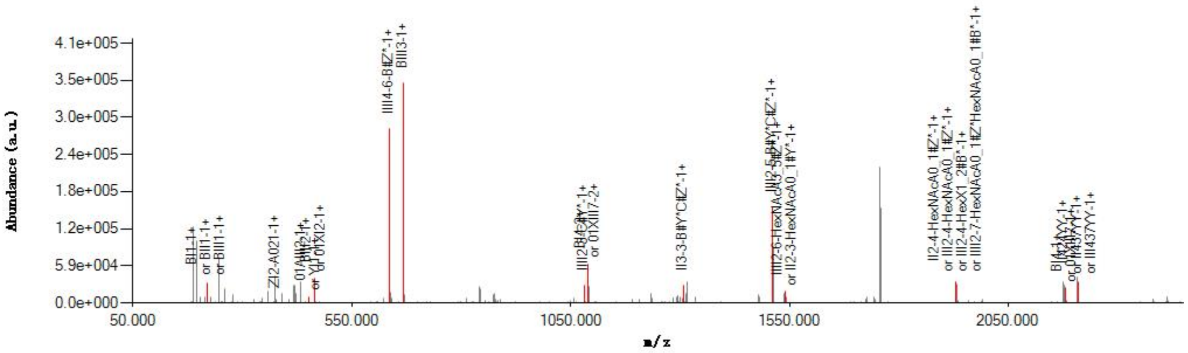

#### Graphical Fragmentation Map

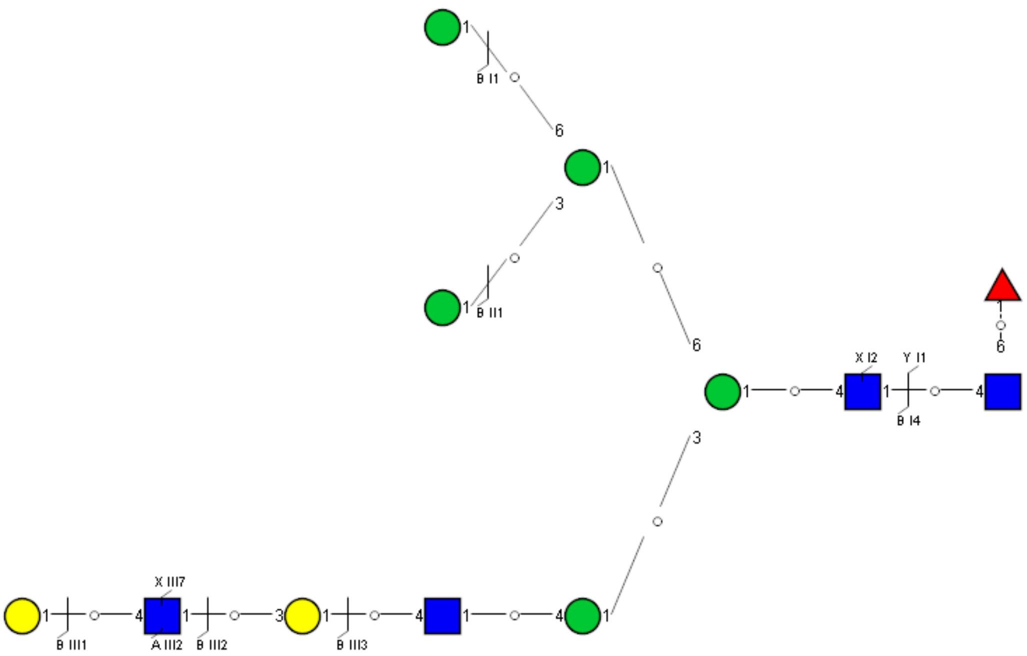

Spectrum Index: 31473(NoBHs:1)

#### Precursor iEF Map

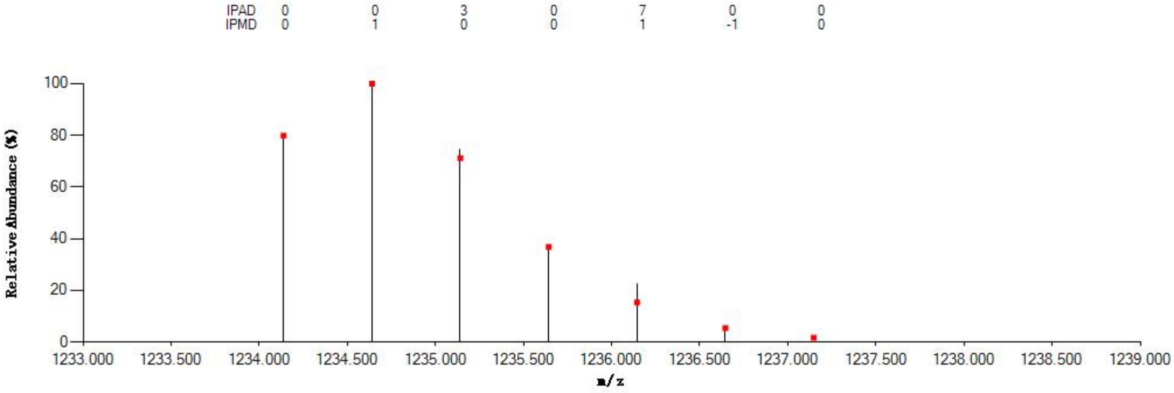


#### Annotated MS/MS Spectrum

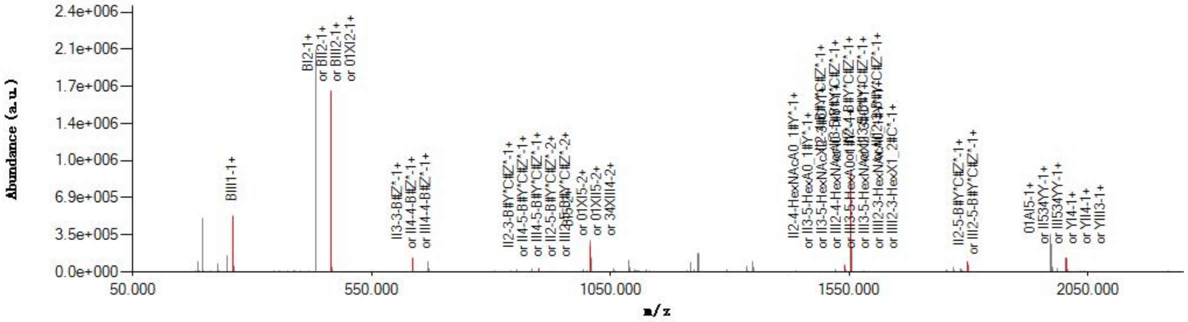


#### Graphical Fragmentation Map

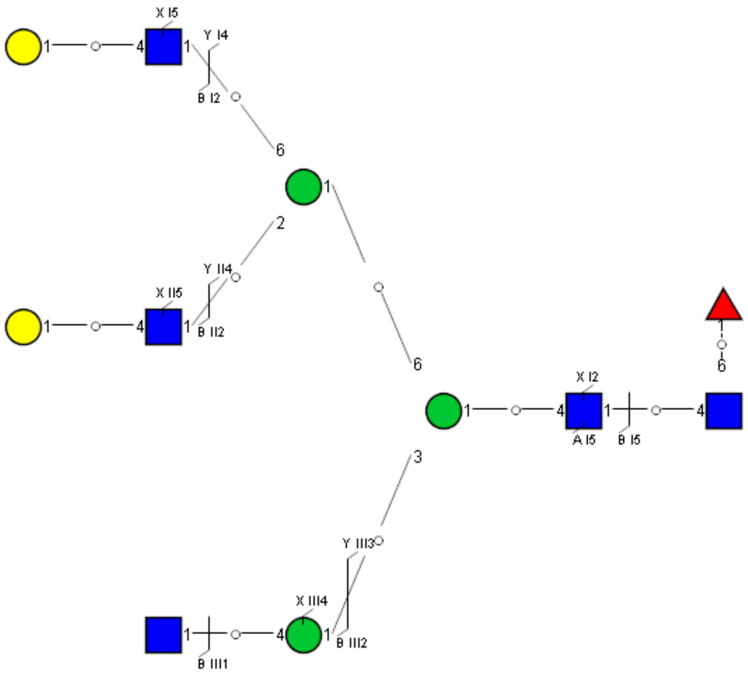

### Spectrum Index: 31519(NoBHs:1)

#### Precursor iEF Map

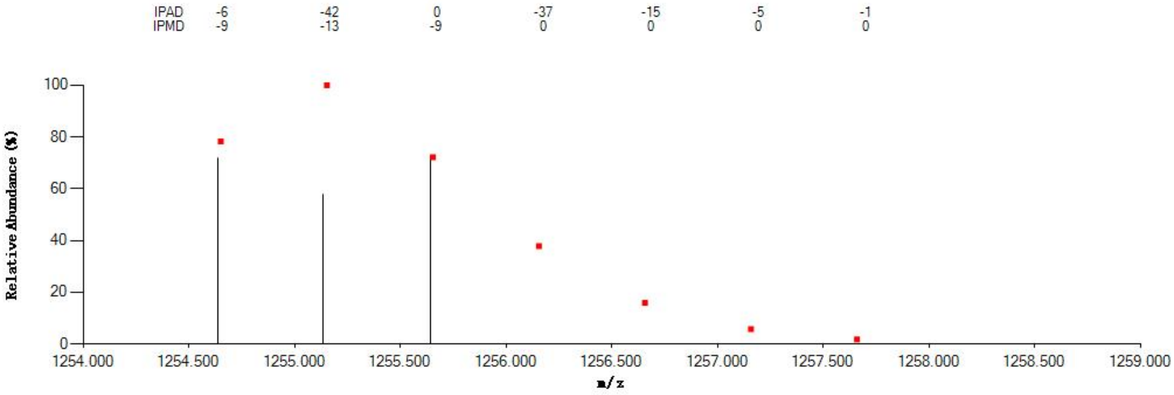

#### Annotated MS/MS Spectrum

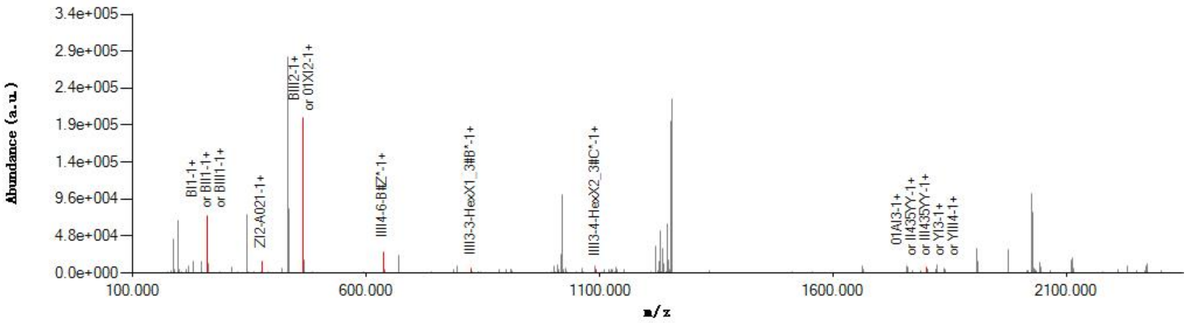

#### Graphical Fragmentation Map

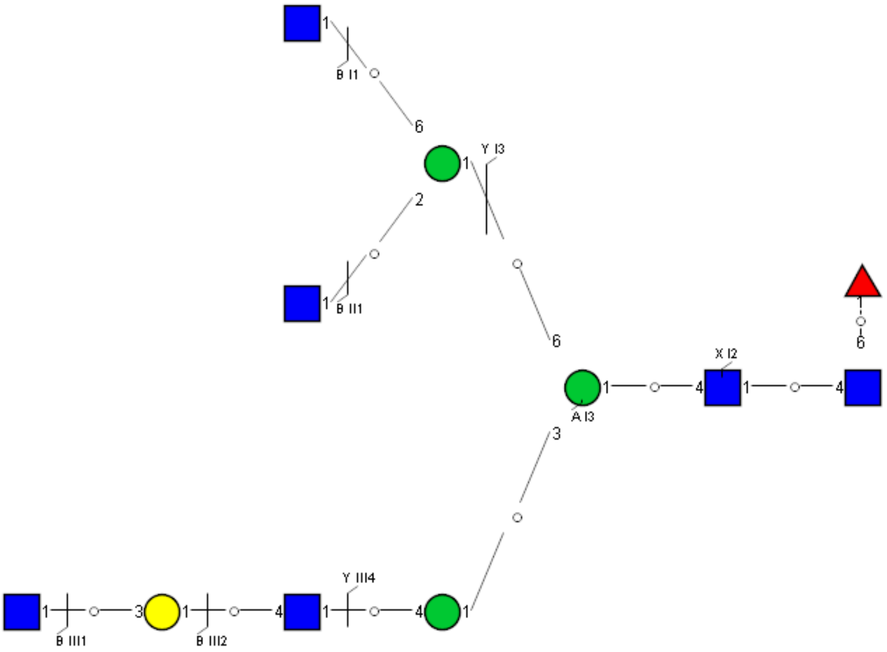

Spectrum Index: 31525(NoBHs:1)

#### Precursor iEF Map

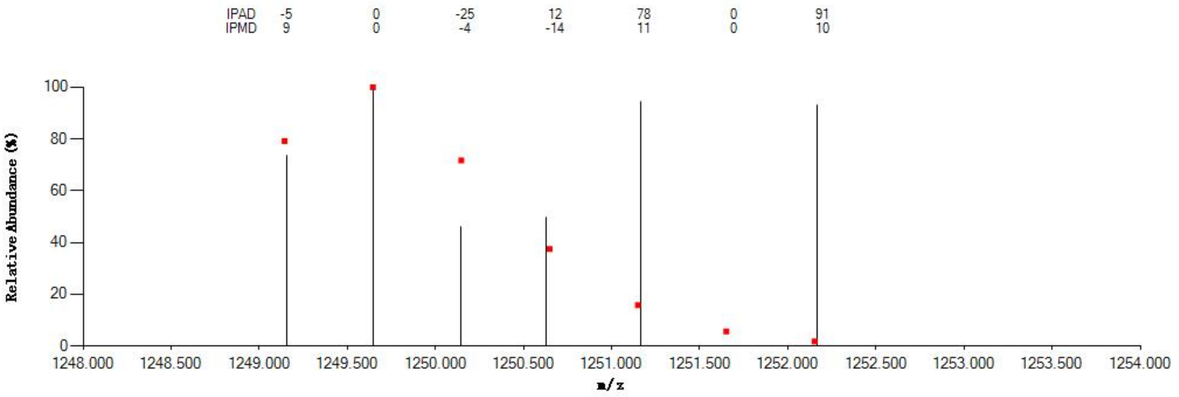


#### Annotated MS/MS Spectrum

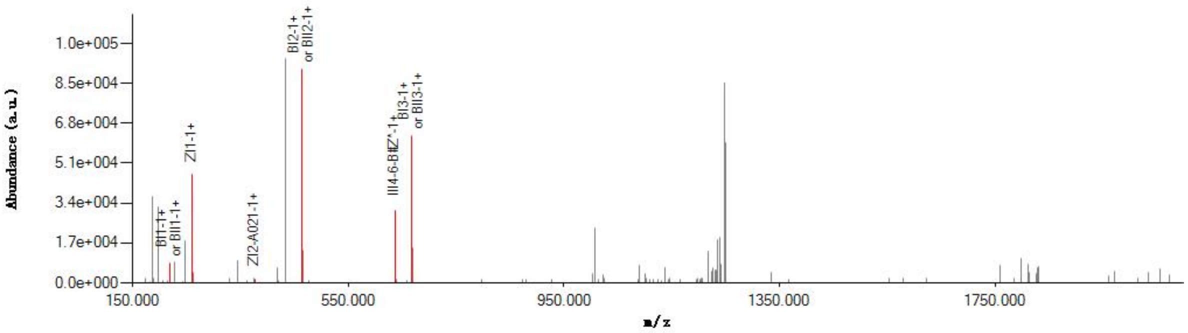


#### Graphical Fragmentation Map

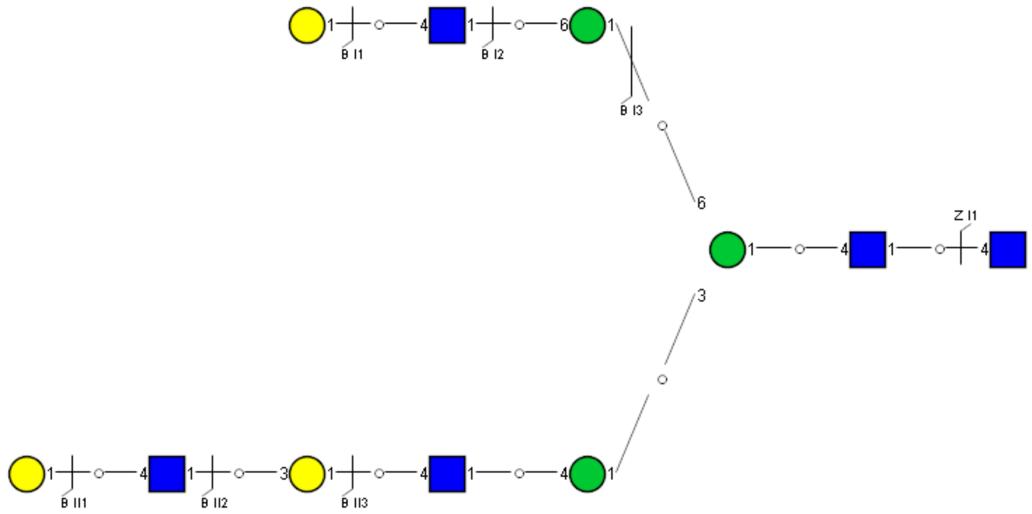

### Spectrum Index: 31620(NoBHs:1)

#### Precursor iEF Map

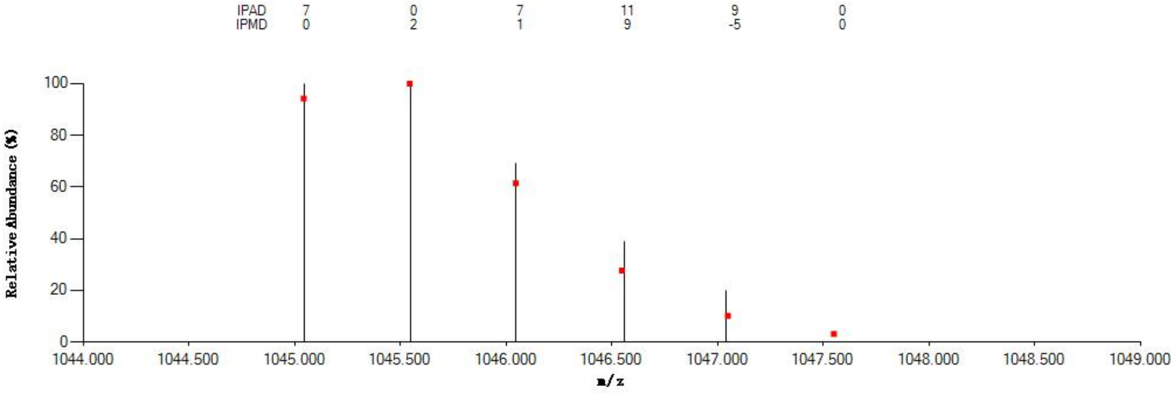


#### Annotated MS/MS Spectrum

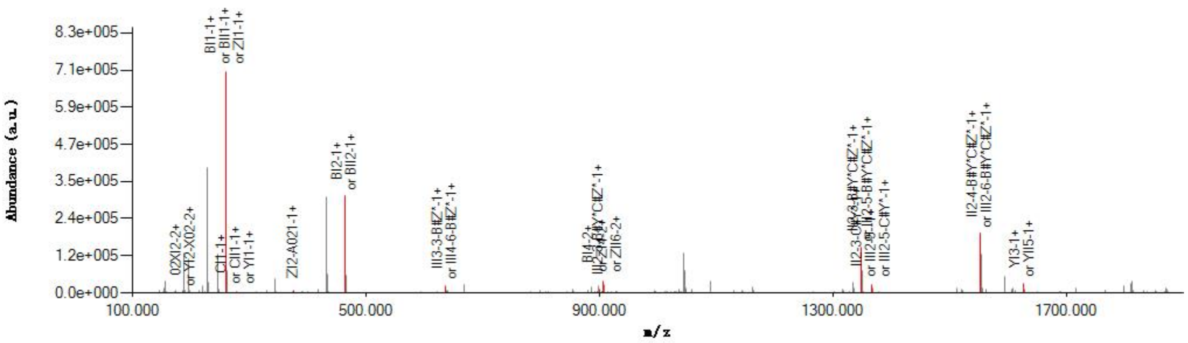


#### Graphical Fragmentation Map

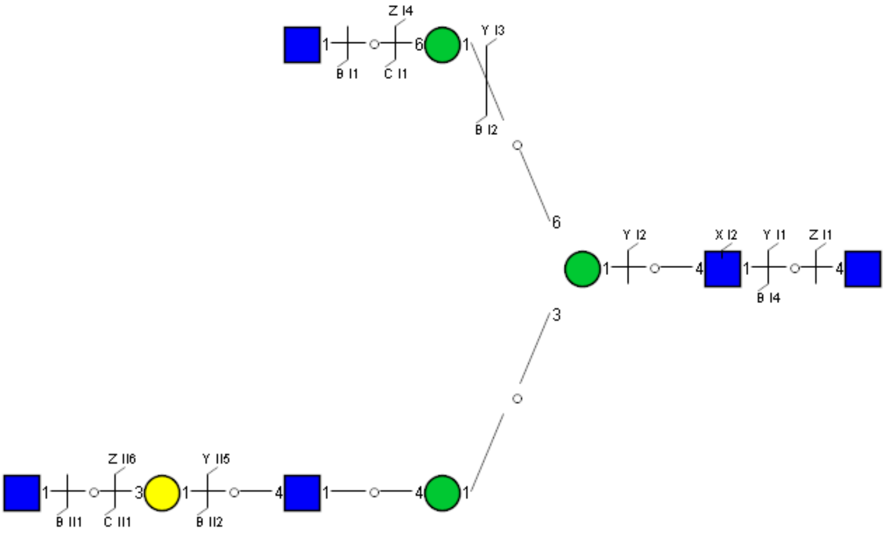

Spectrum Index: 31657(NoBHs:1)

Precursor iEF Map

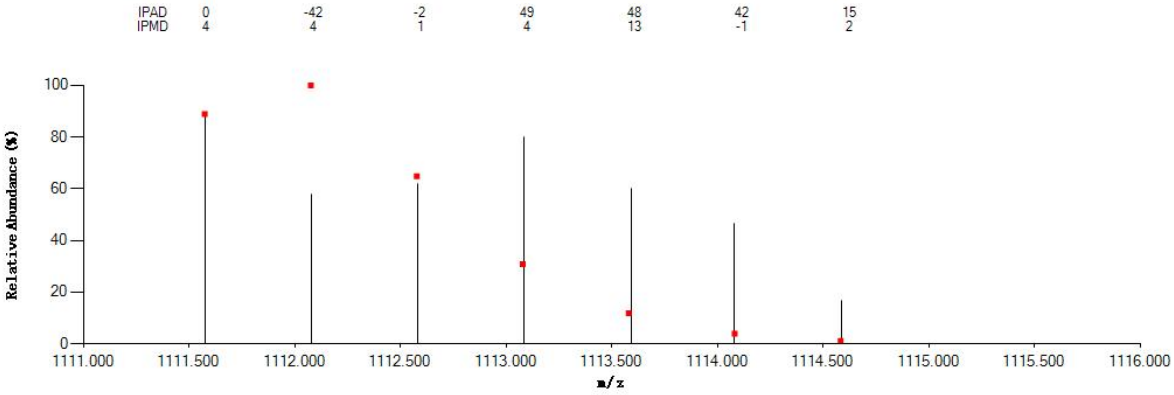


Annotated MS/MS Spectrum

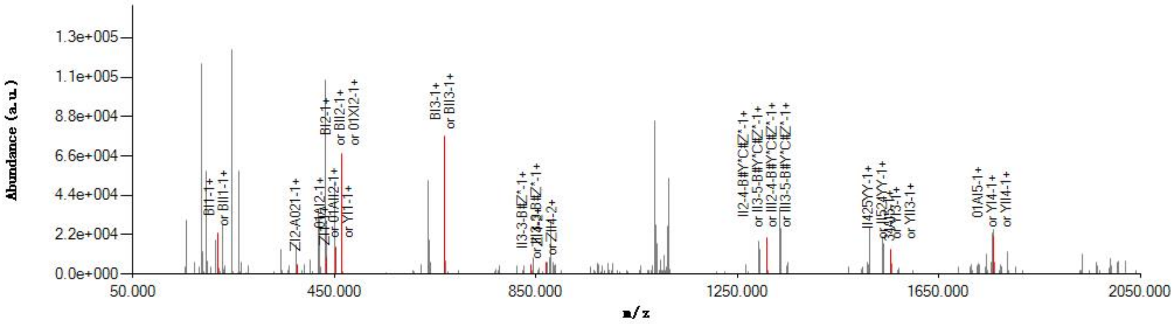


Graphical Fragmentation Map

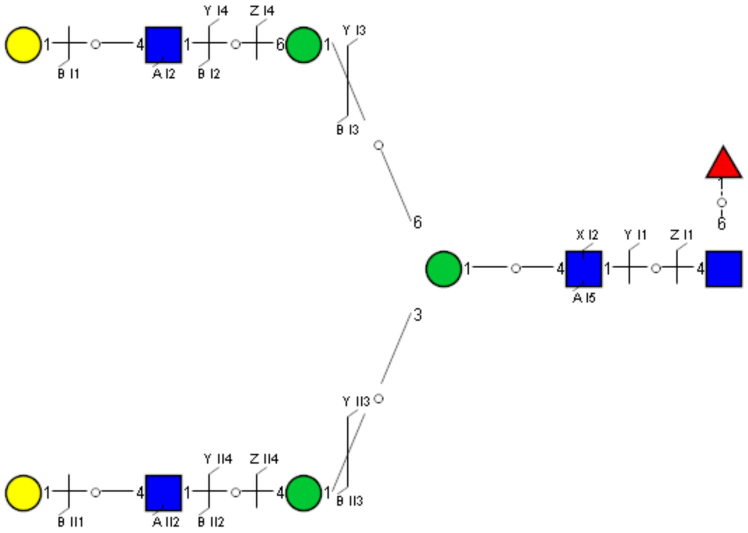

Spectrum Index: 31663(NoBHs:1)

#### Precursor iEF Map

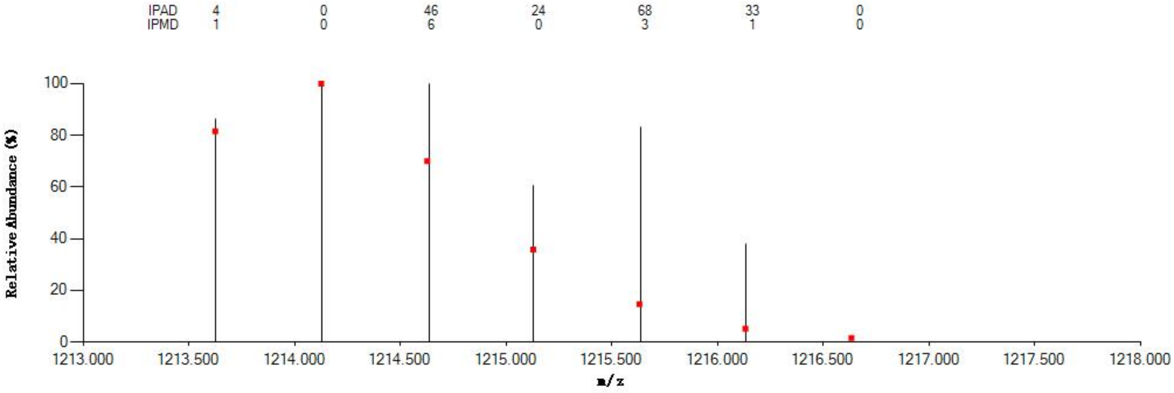


#### Annotated MS/MS Spectrum

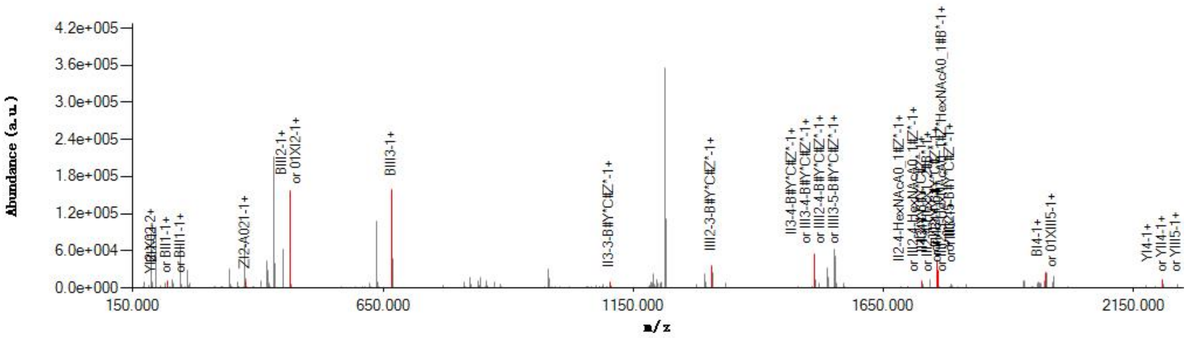


#### Graphical Fragmentation Map

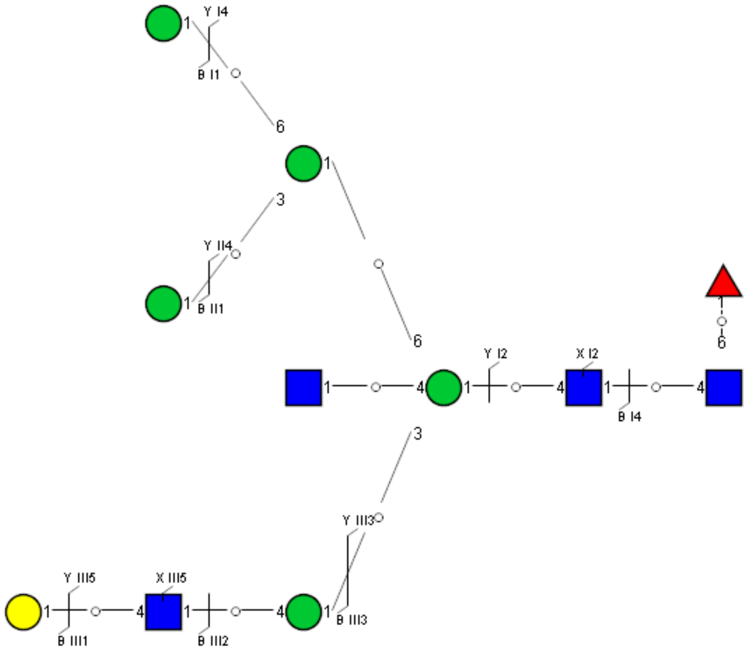

Spectrum Index: 31810(NoBHs:1)

#### Precursor iEF Map

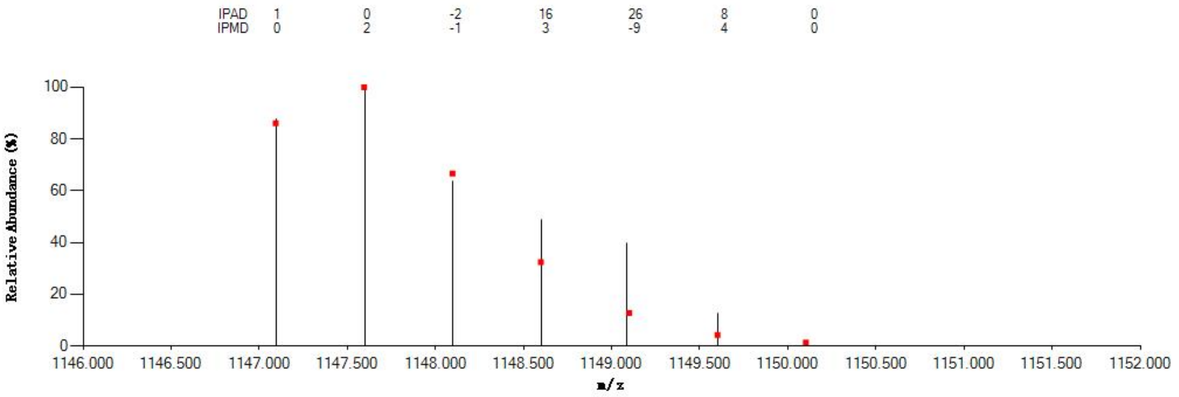


#### Annotated MS/MS Spectrum

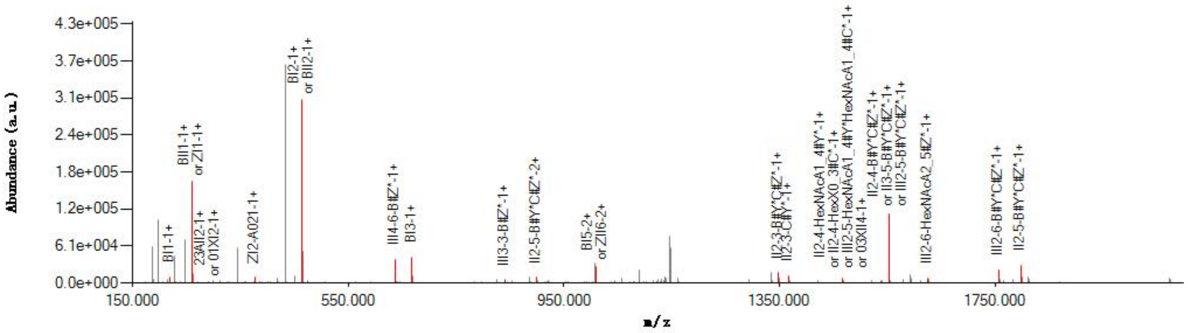


#### Graphical Fragmentation Map

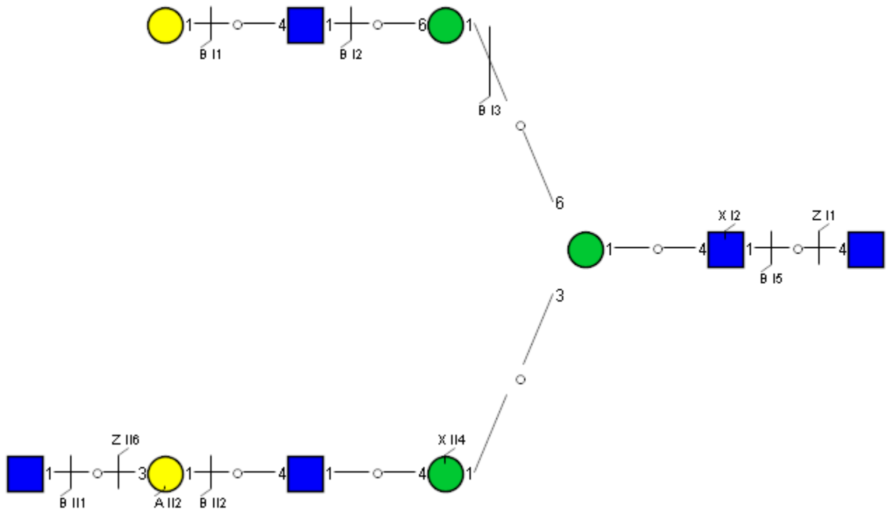

Spectrum Index: 31917(NoBHs:1)

#### Precursor iEF Map

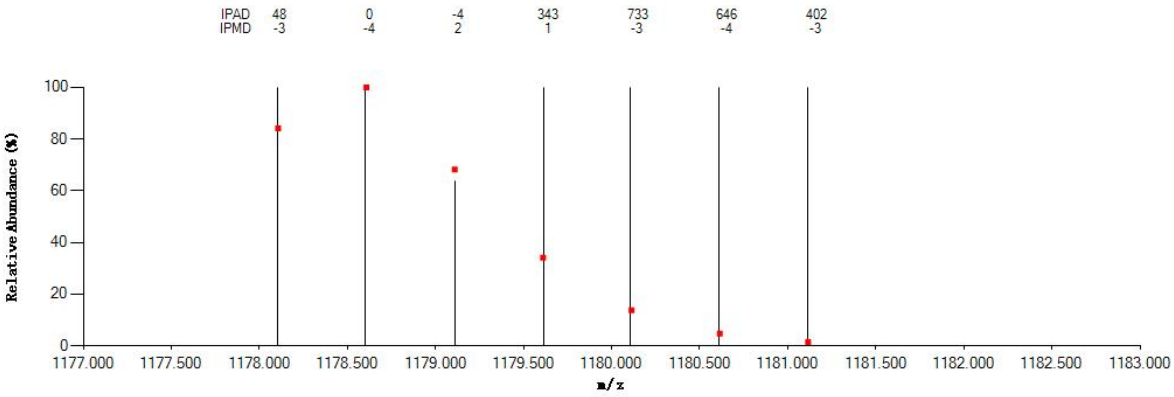


#### Annotated MS/MS Spectrum

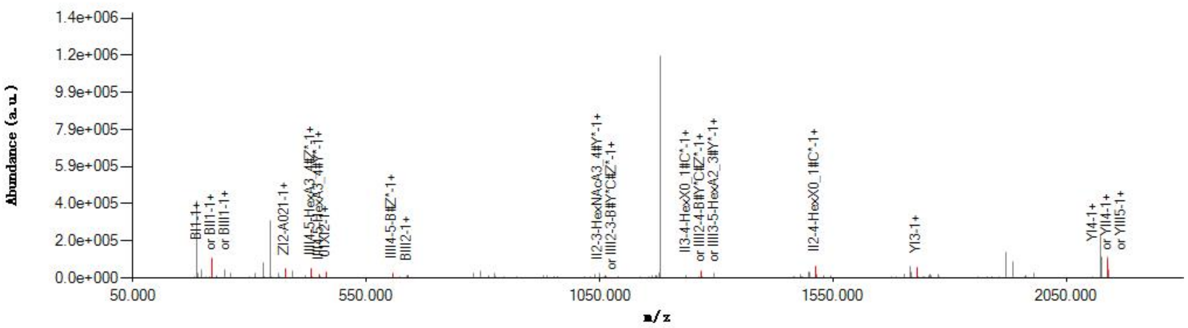


#### Graphical Fragmentation Map

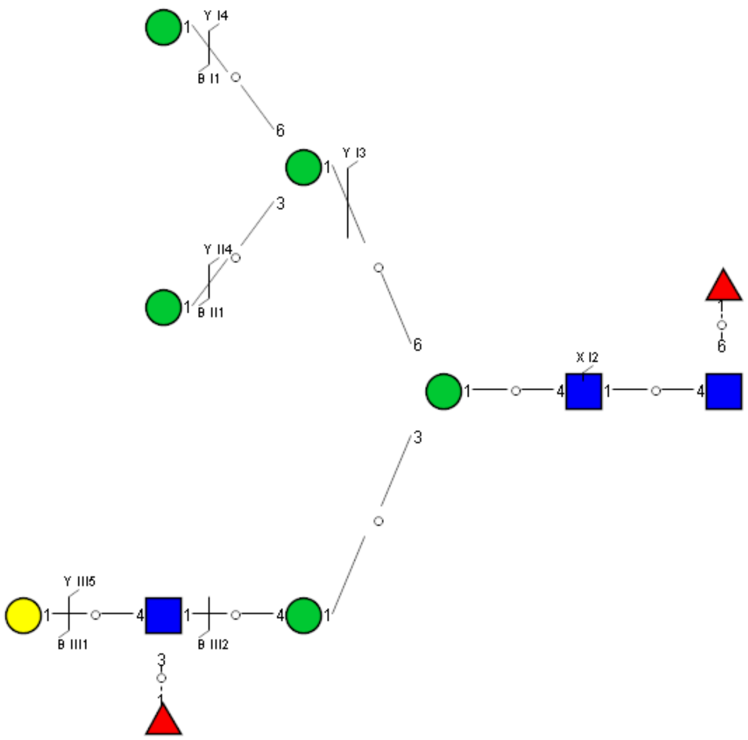

Spectrum Index: 31950(NoBHs:1)

#### Precursor iEF Map

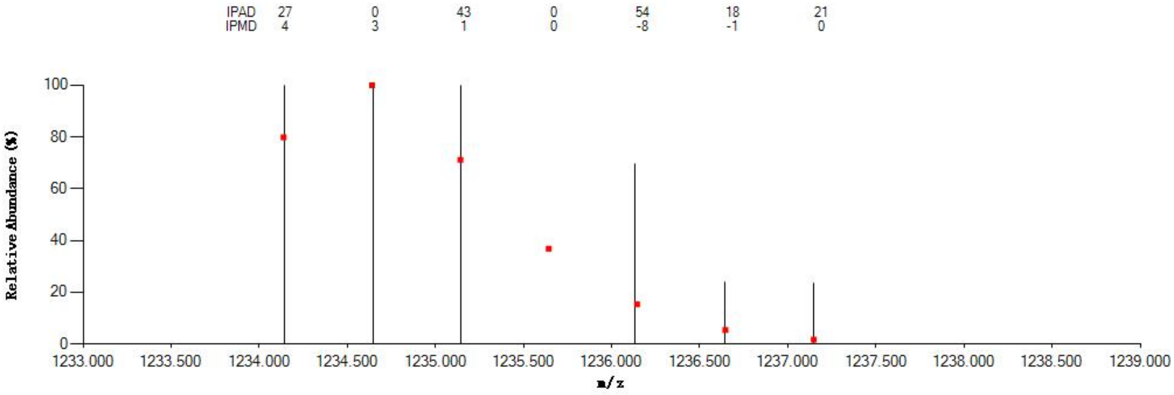

#### Annotated MS/MS Spectrum

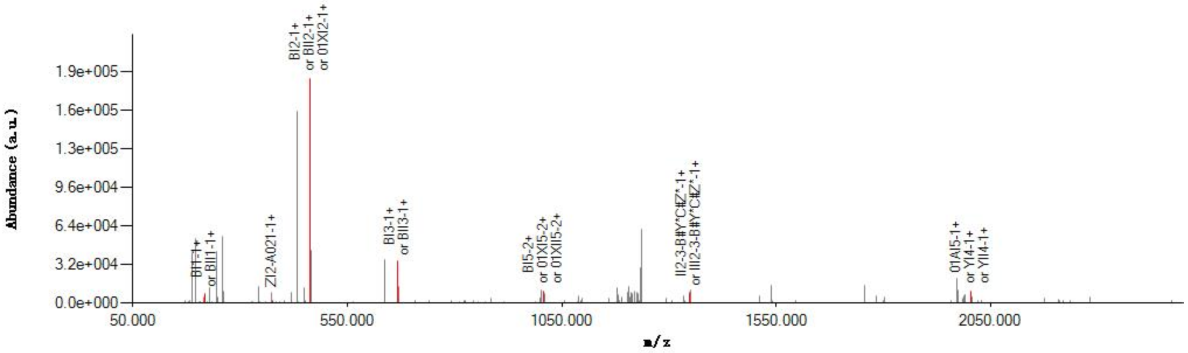

#### Graphical Fragmentation Map

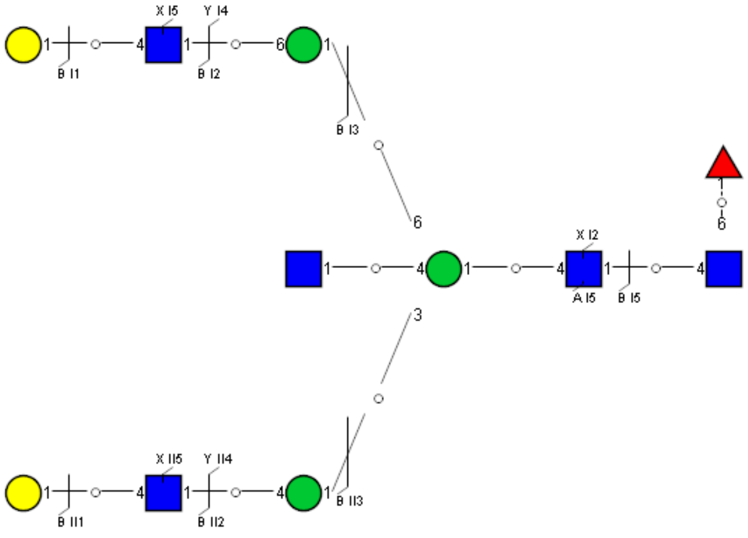

Spectrum Index: 32103(NoBHs:1)

#### Precursor iEF Map

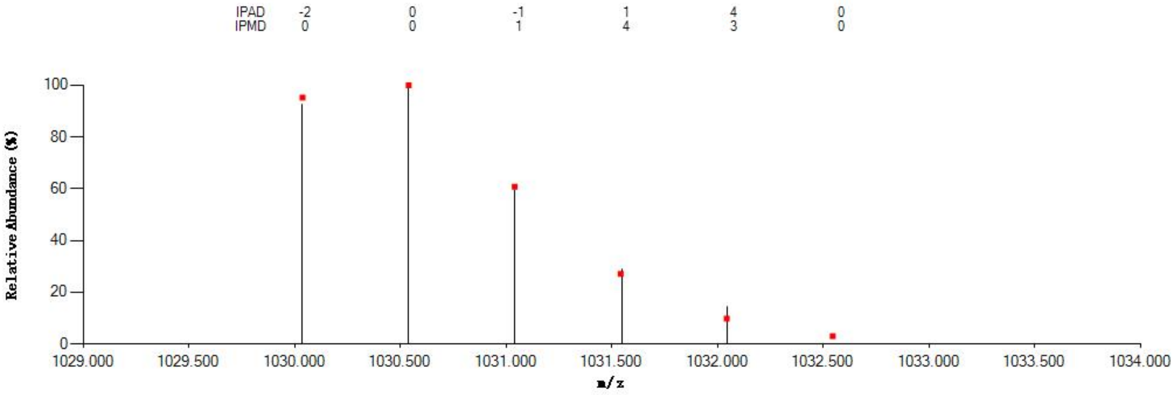

#### Annotated MS/MS Spectrum

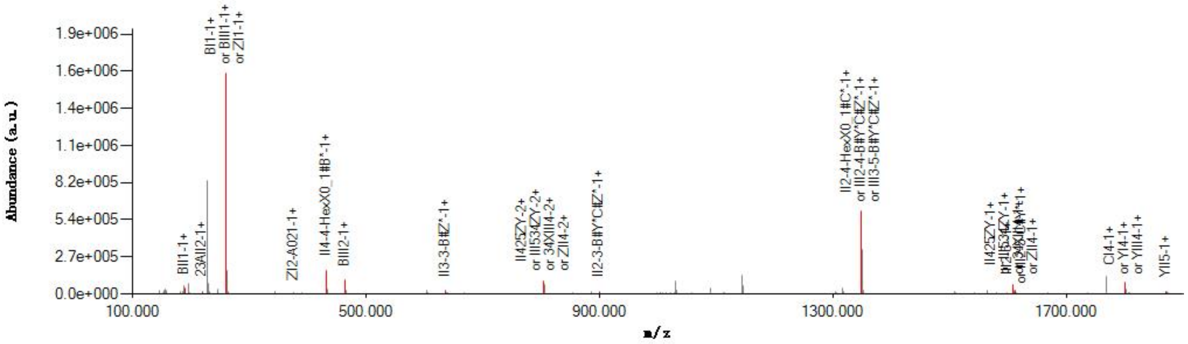

#### Graphical Fragmentation Map

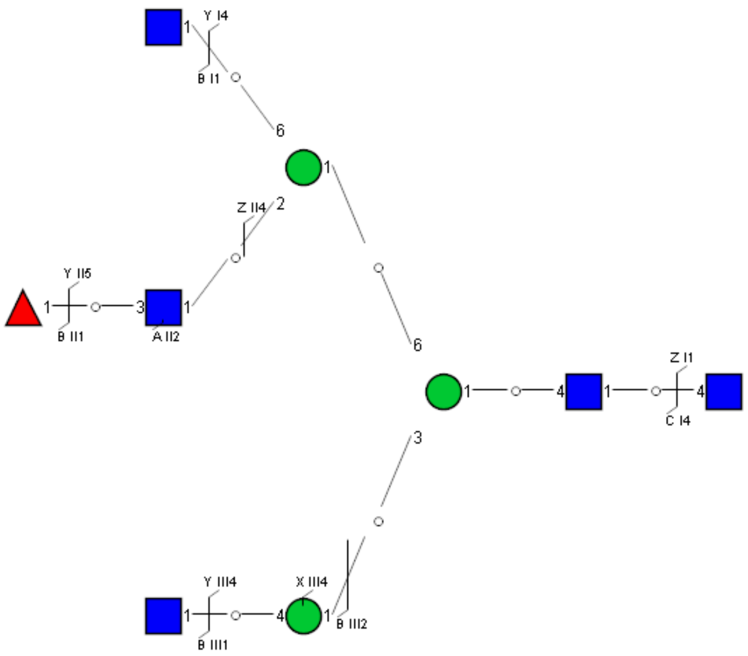

Spectrum Index: 32776(NoBHs:1)

#### Precursor iEF Map

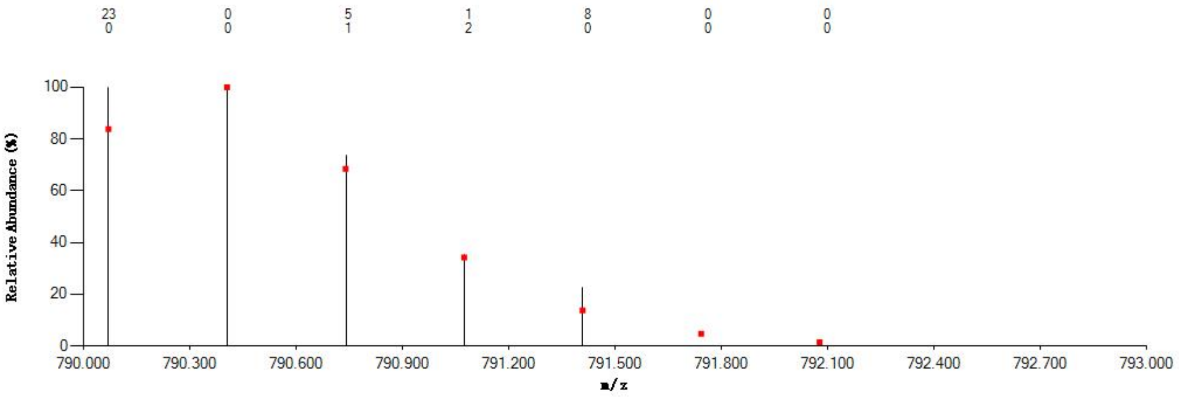

#### Annotated MS/MS Spectrum

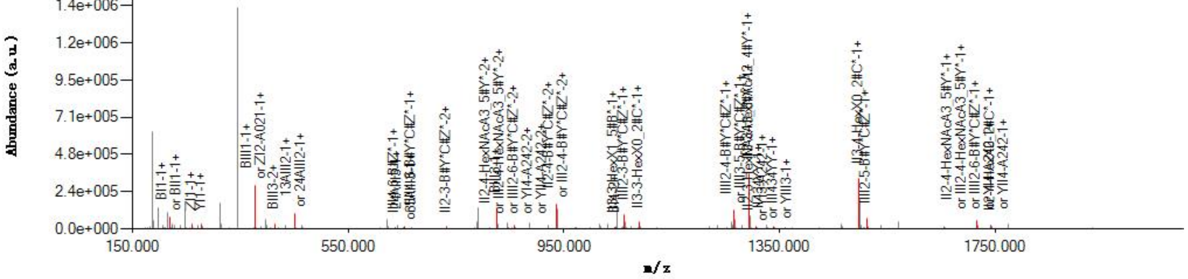

#### Graphical Fragmentation Map

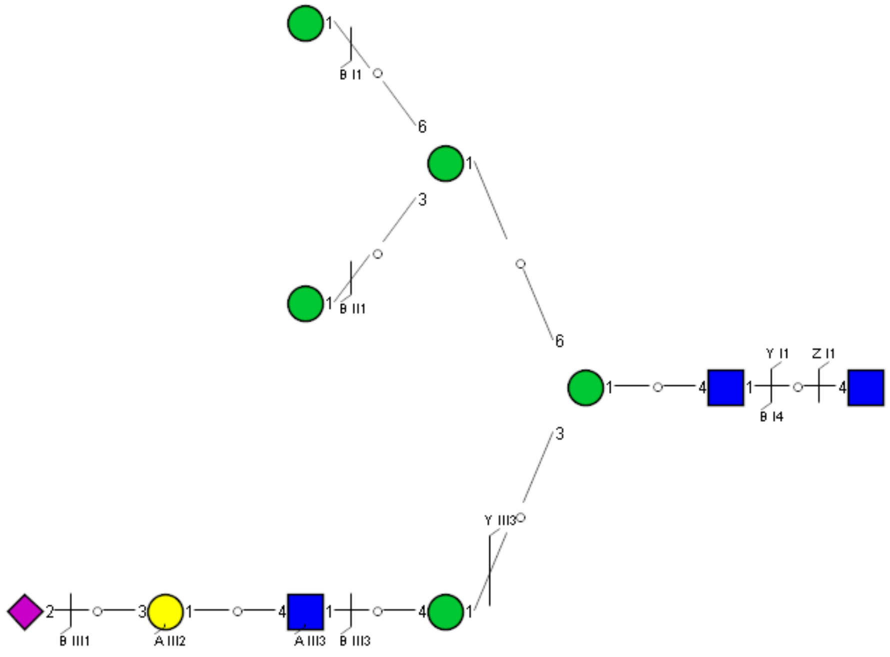

Spectrum Index: 32777(NoBHs:1)

#### Precursor iEF Map

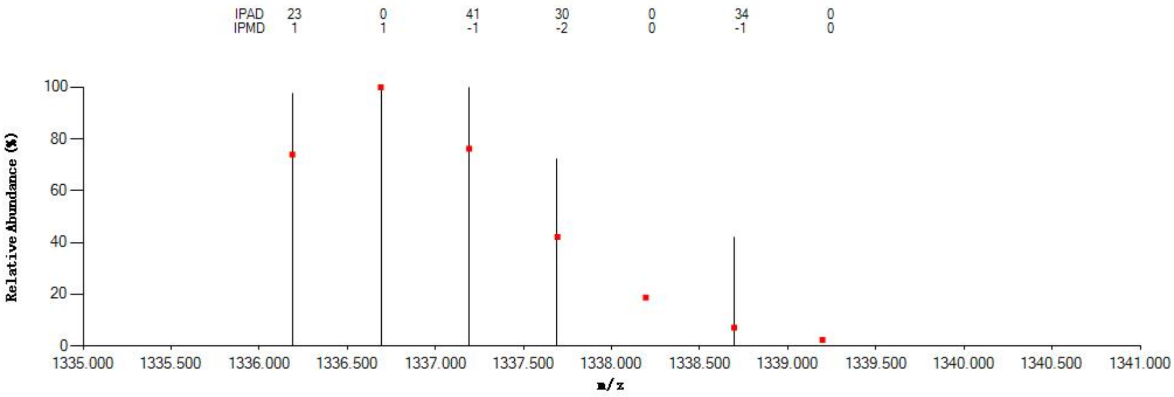


#### Annotated MS/MS Spectrum

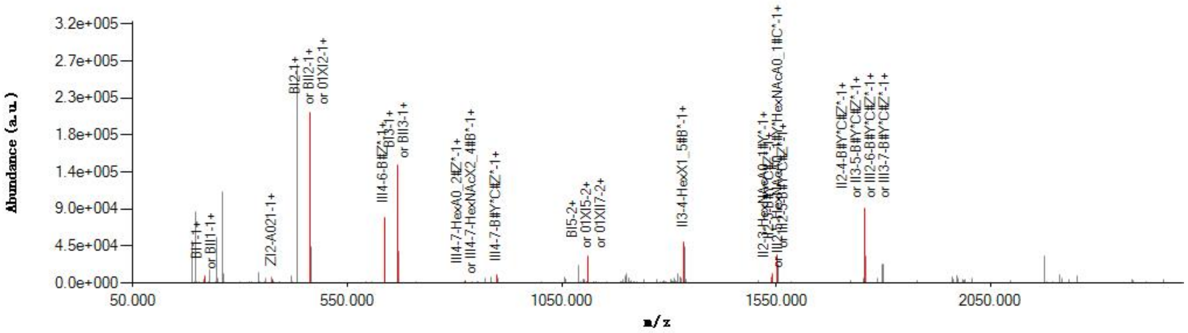


#### Graphical Fragmentation Map

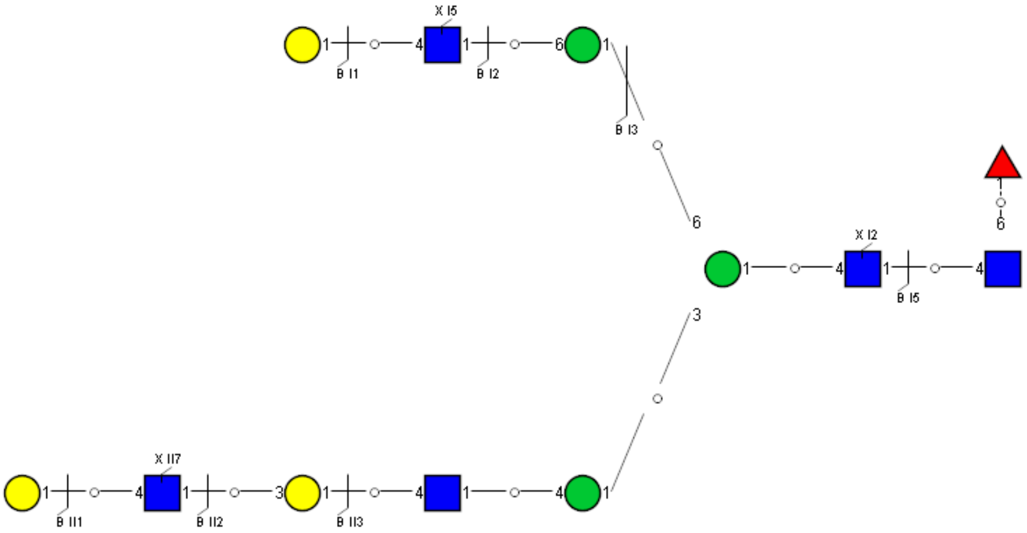

Spectrum Index: 33533(NoBHs:1)

#### Precursor iEF Map

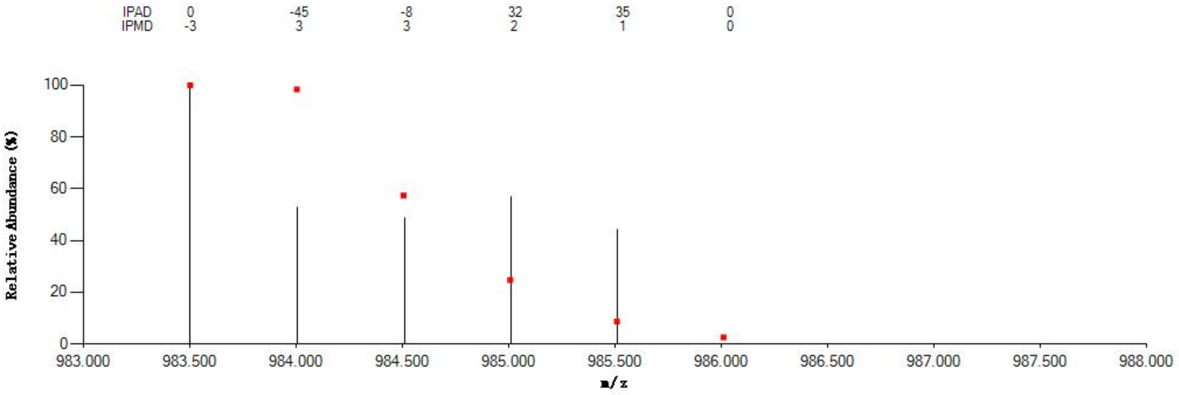


#### Annotated MS/MS Spectrum

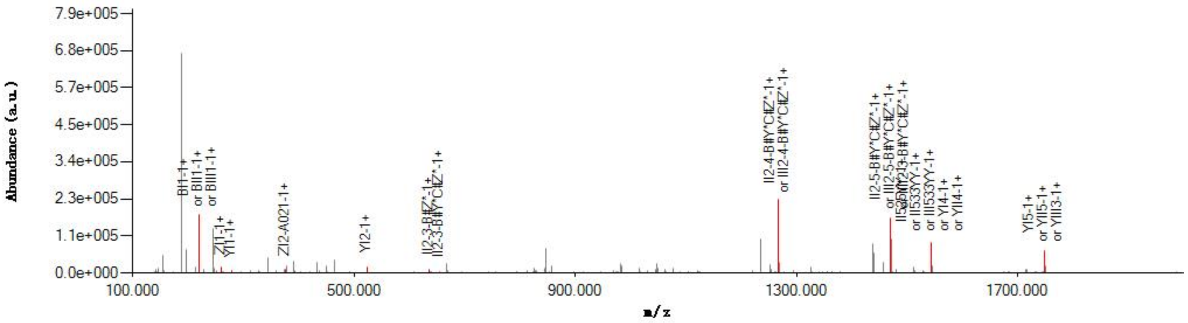


#### Graphical Fragmentation Map

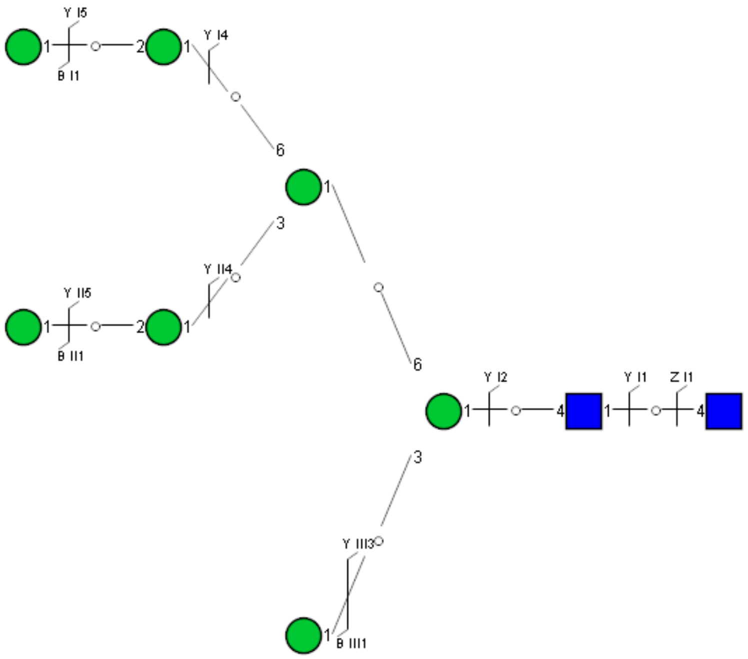

Spectrum Index: 33616(NoBHs:1)

#### Precursor iEF Map

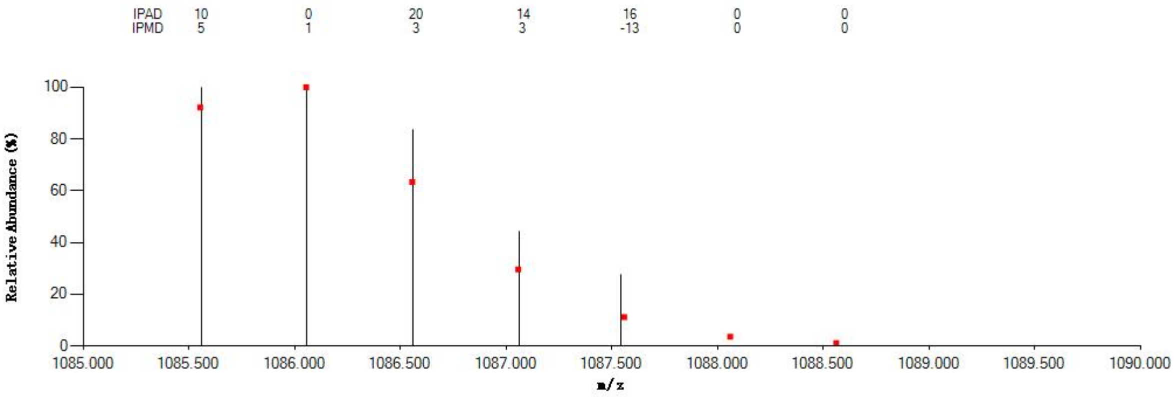


#### Annotated MS/MS Spectrum

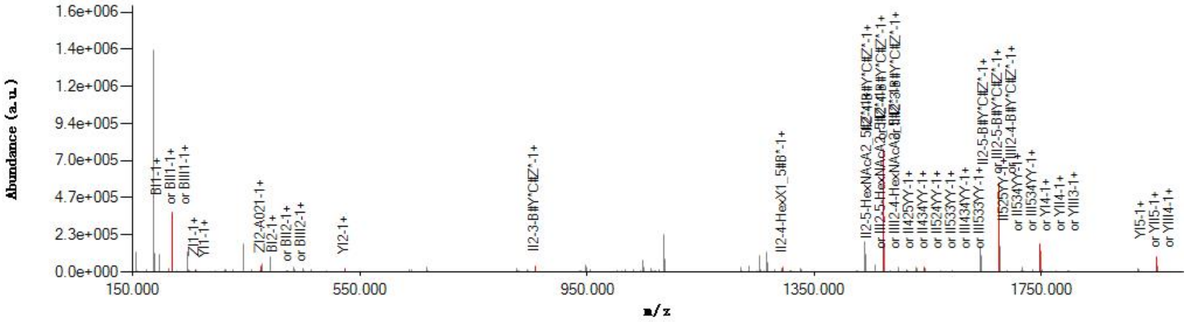


#### Graphical Fragmentation Map

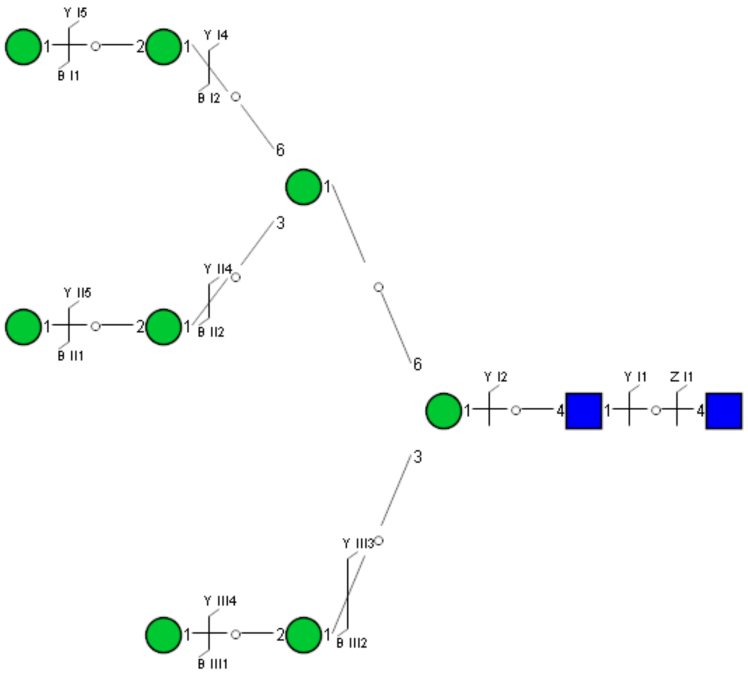

Spectrum Index: 33900(NoBHs:1)

#### Precursor iEF Map

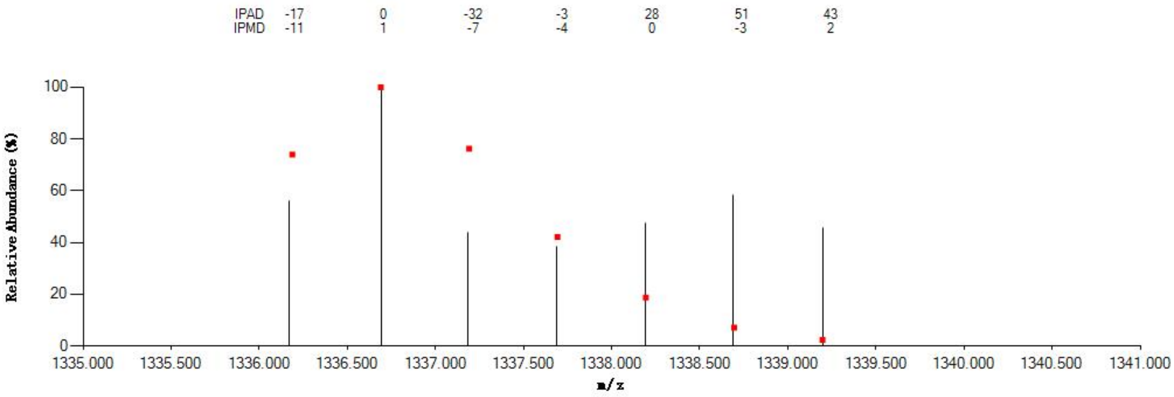


#### Annotated MS/MS Spectrum

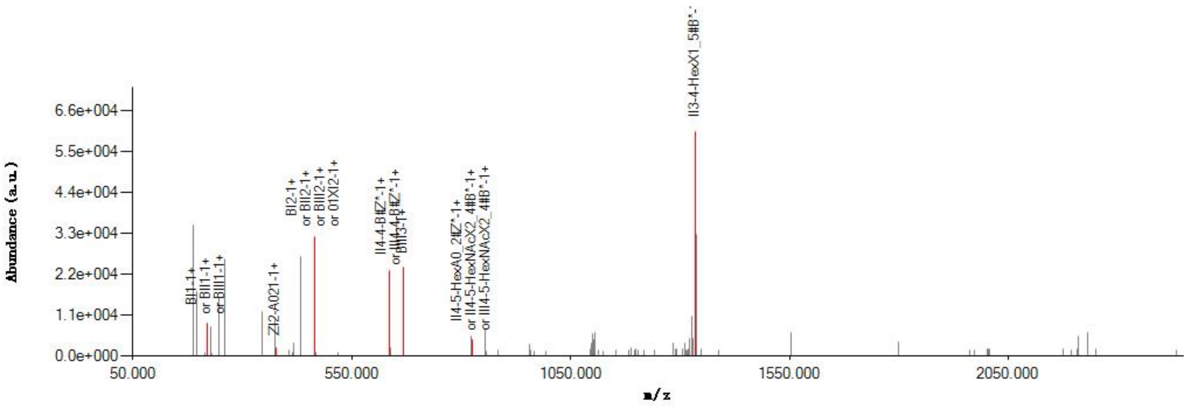


#### Graphical Fragmentation Map

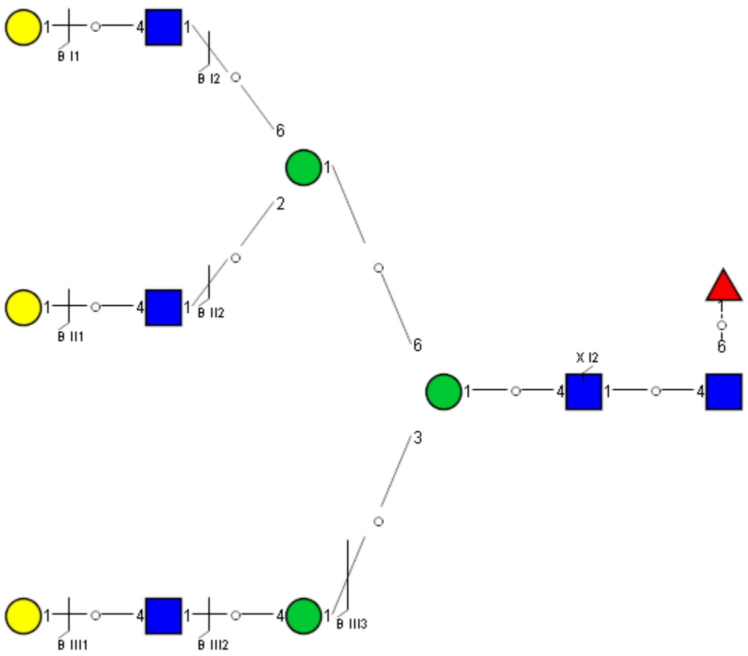

Spectrum Index: 34065(NoBHs:1)

#### Precursor iEF Map

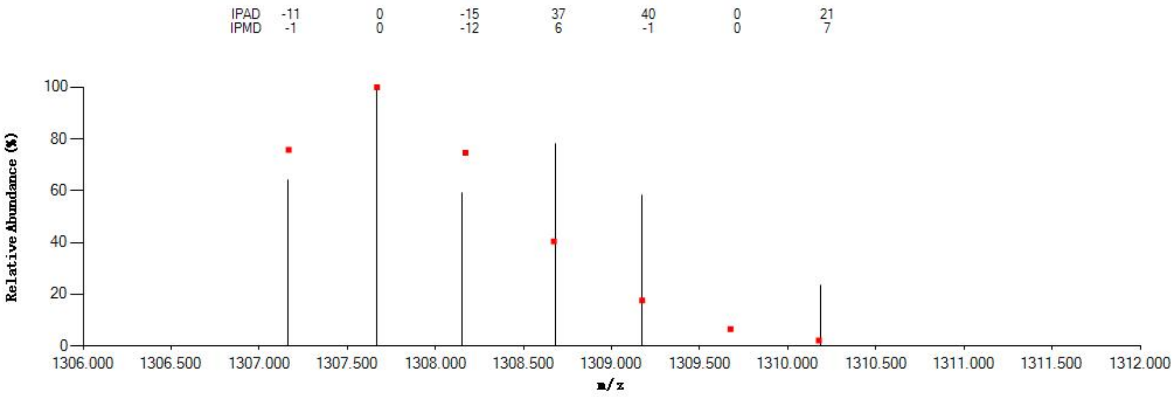


#### Annotated MS/MS Spectrum

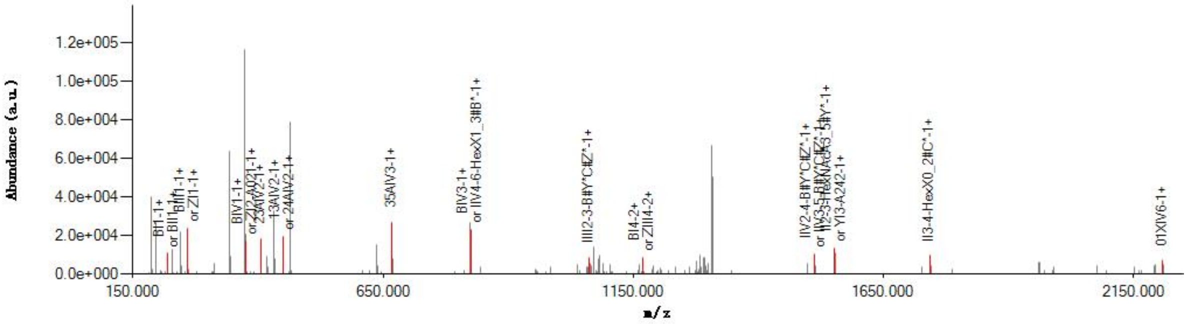


#### Graphical Fragmentation Map

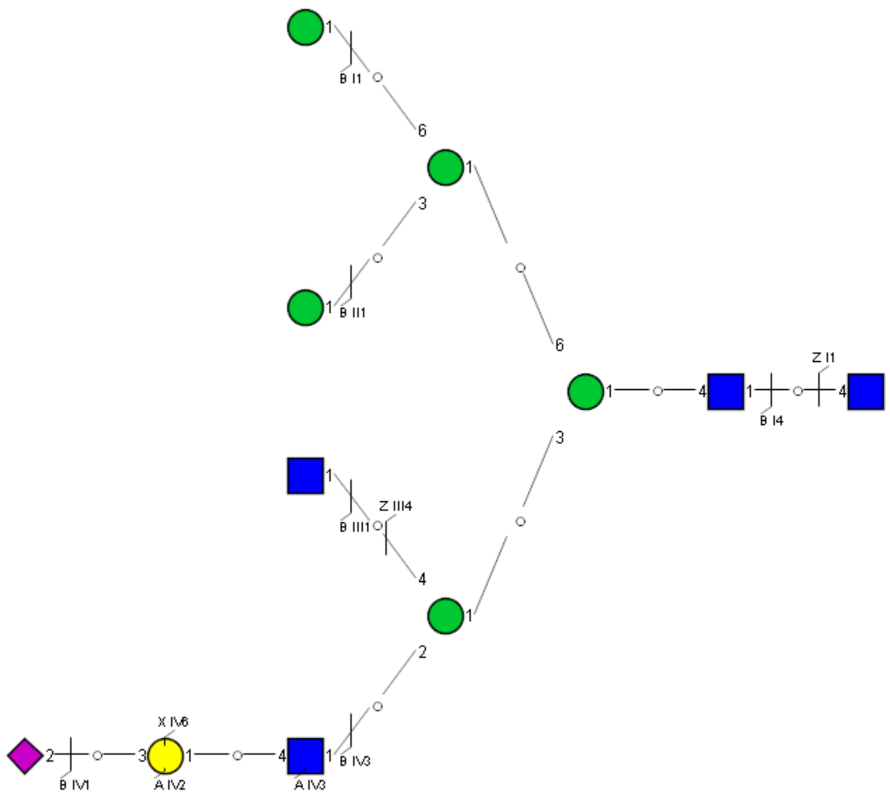

Spectrum Index: 34380(NoBHs:1)

#### Precursor iEF Map

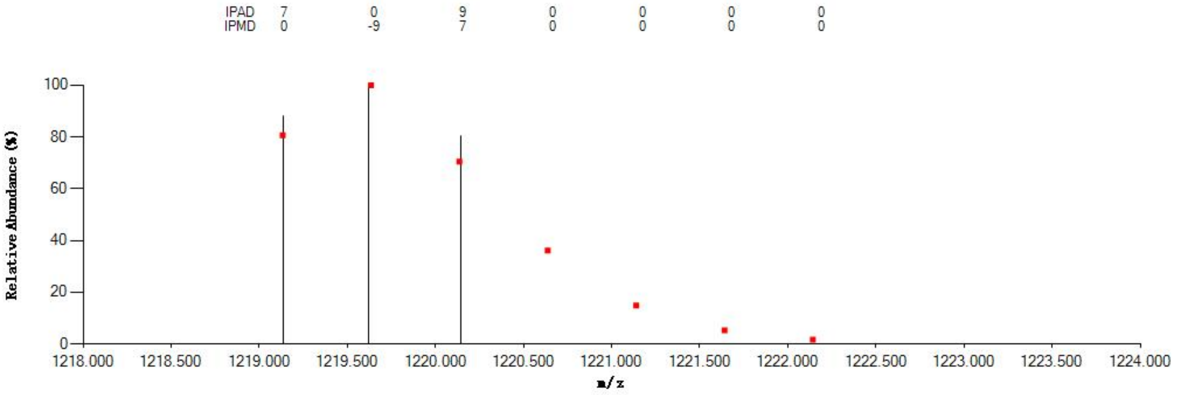


#### Annotated MS/MS Spectrum

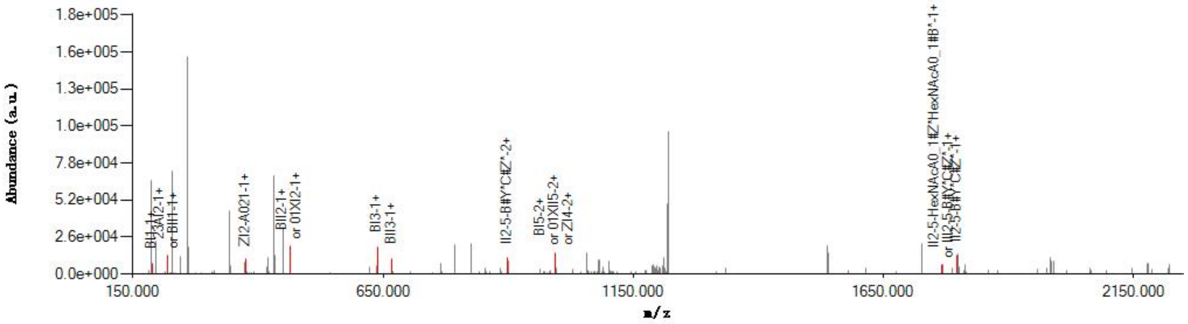


#### Graphical Fragmentation Map

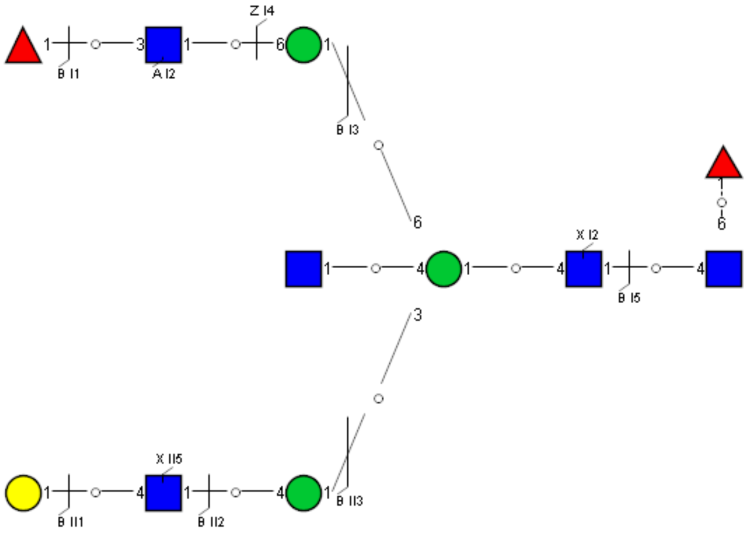

Spectrum Index: 34390(NoBHs:1)

#### Precursor iEF Map

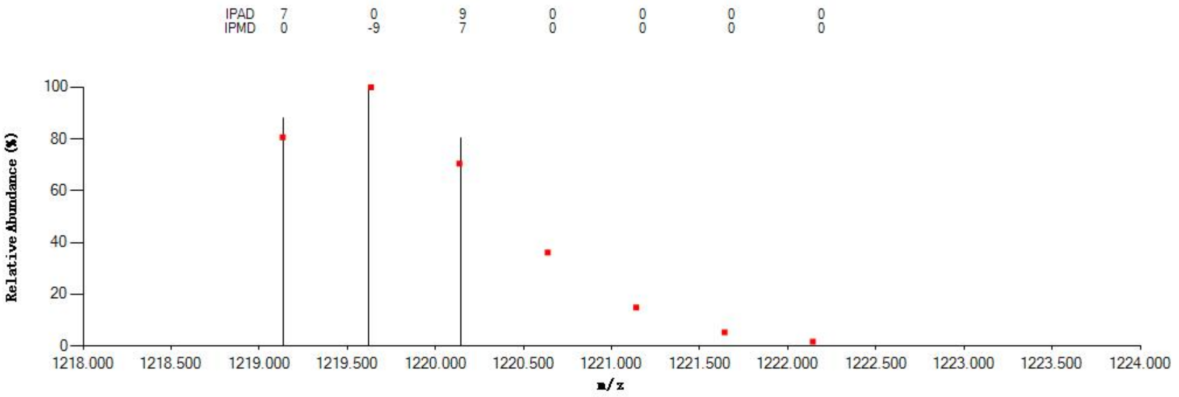


#### Annotated MS/MS Spectrum

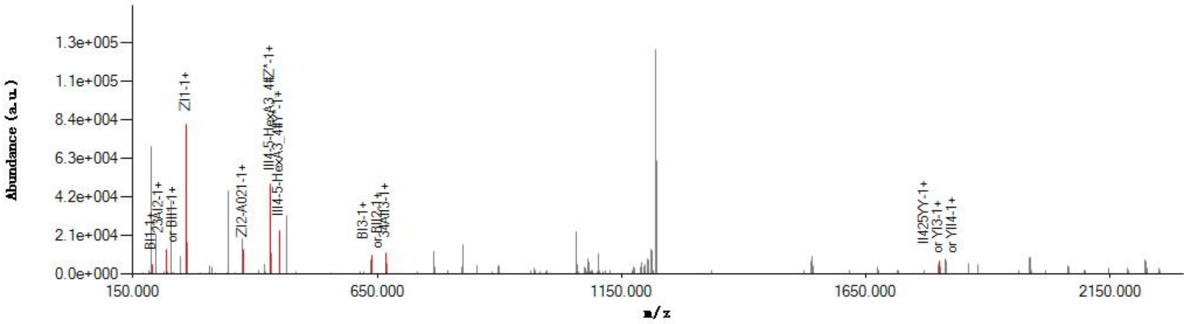


#### Graphical Fragmentation Map

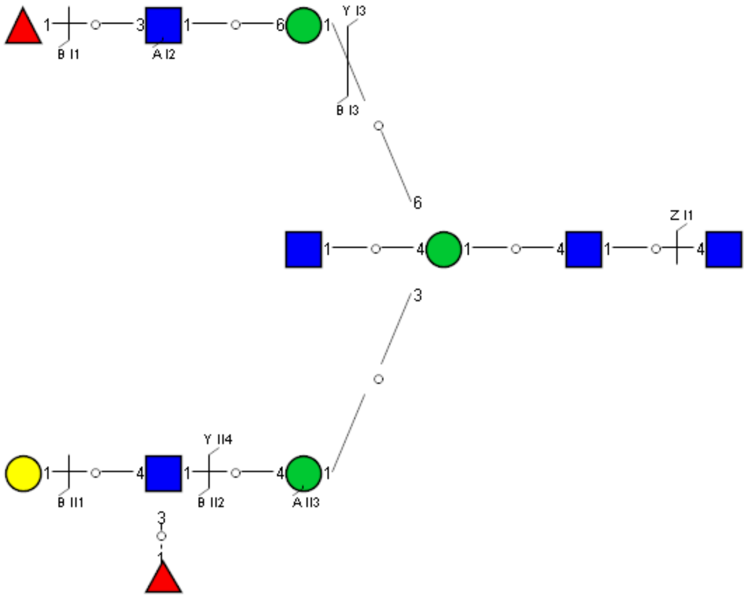

### Spectrum Index: 34709(NoBHs:1)

#### Precursor iEF Map

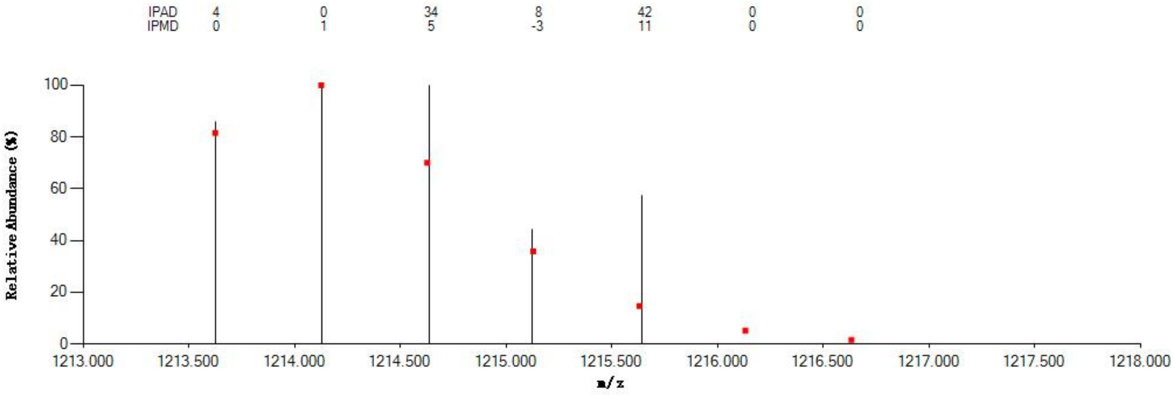


#### Annotated MS/MS Spectrum

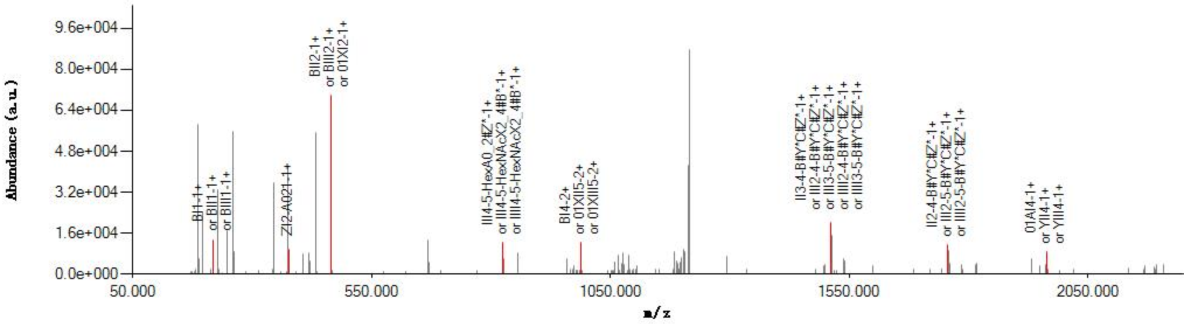


#### Graphical Fragmentation Map

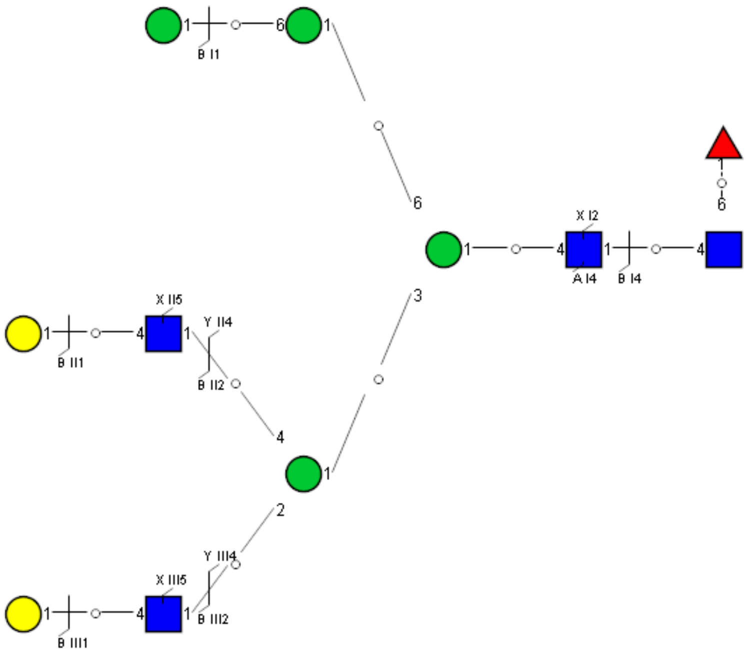

Spectrum Index: 35724(NoBHs:1)

#### Precursor iEF Map

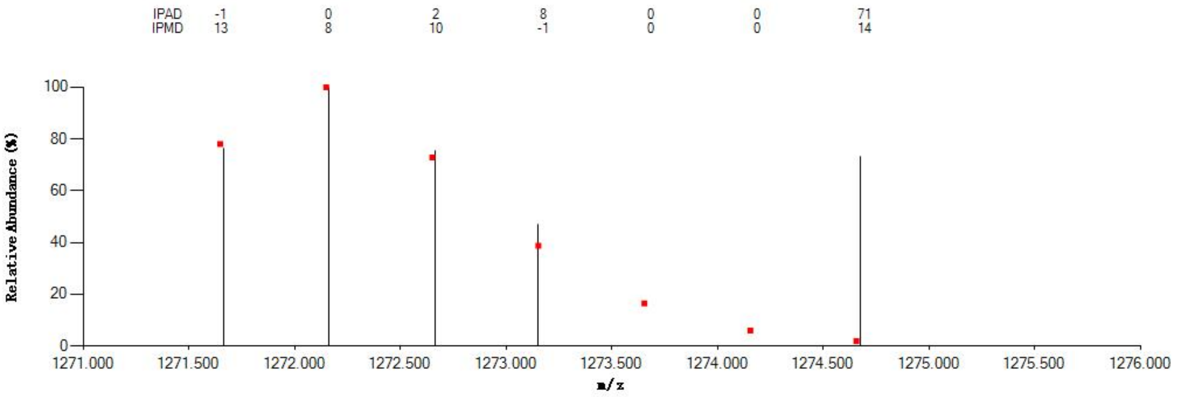


#### Annotated MS/MS Spectrum

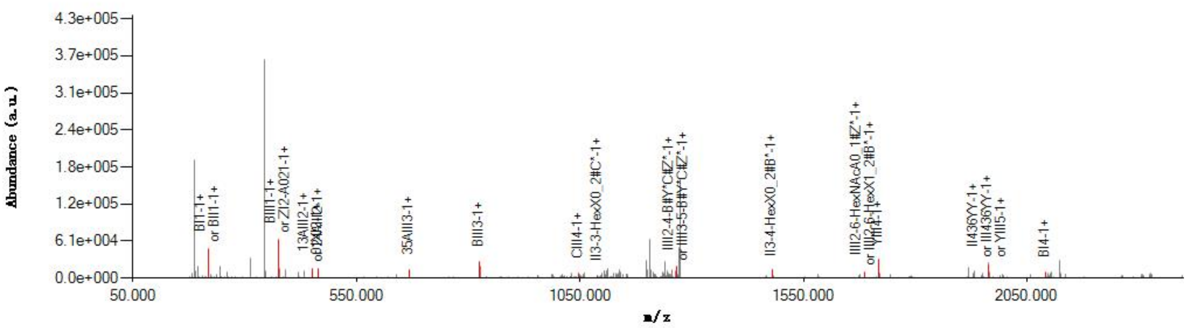


#### Graphical Fragmentation Map

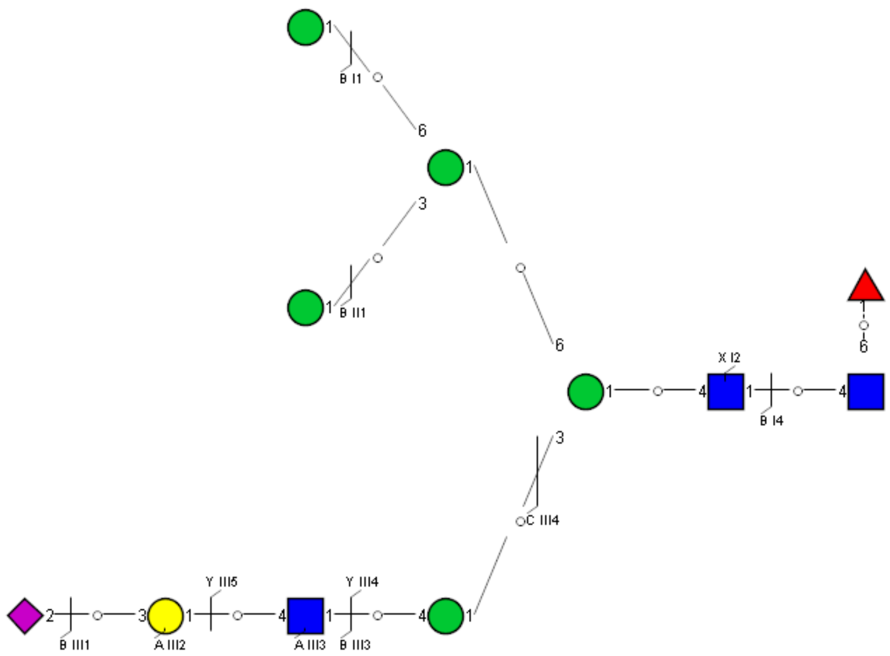

Spectrum Index: 35968(NoBHs:1)

#### Precursor iEF Map

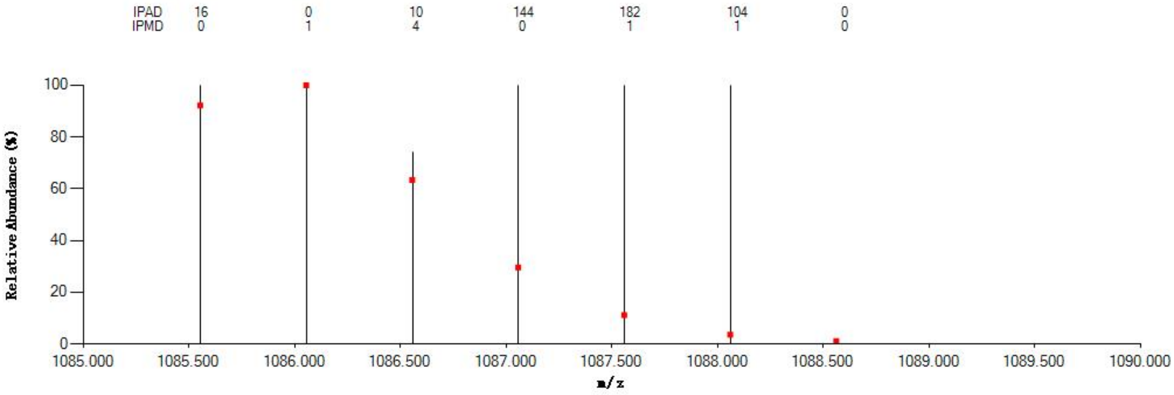


#### Annotated MS/MS Spectrum

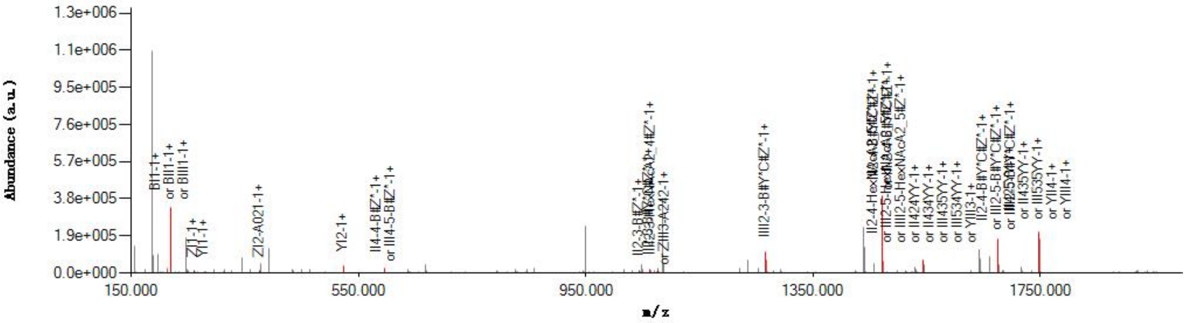


#### Graphical Fragmentation Map

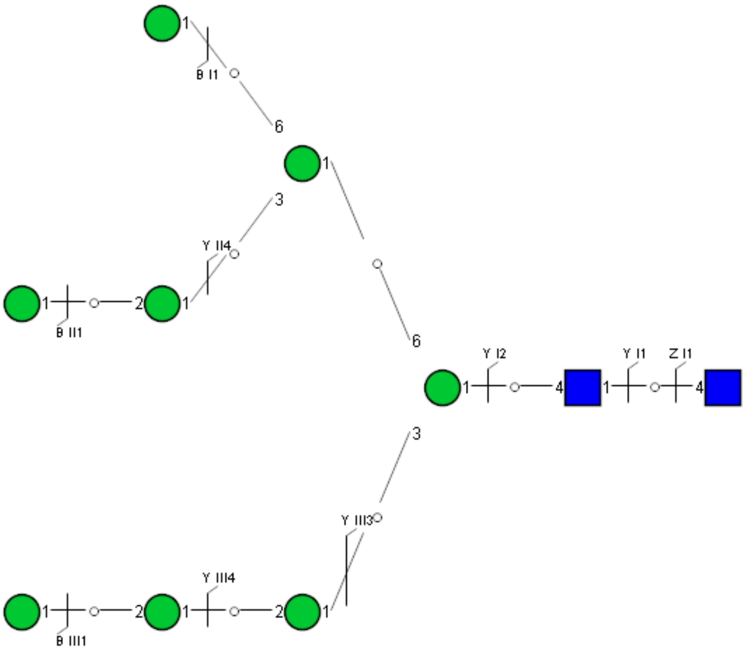

Spectrum Index: 38169(NoBHs:1)

#### Precursor iEF Map

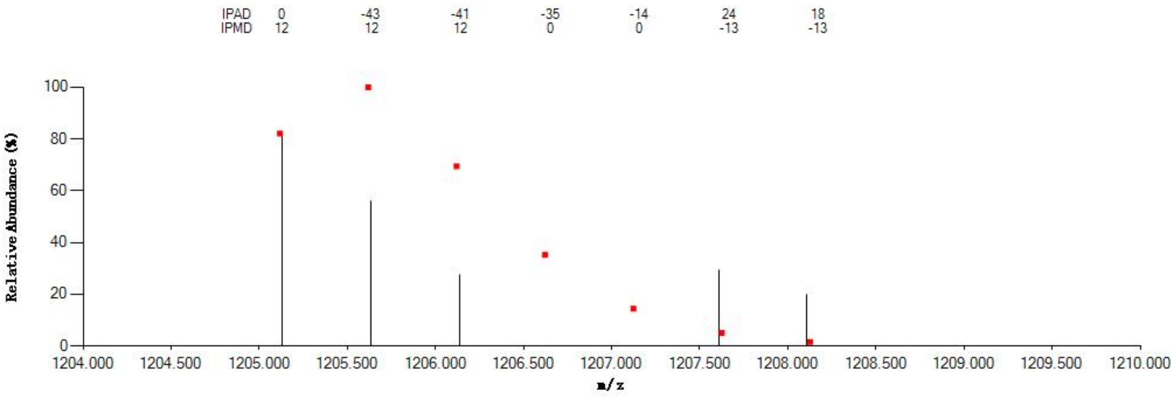


#### Annotated MS/MS Spectrum

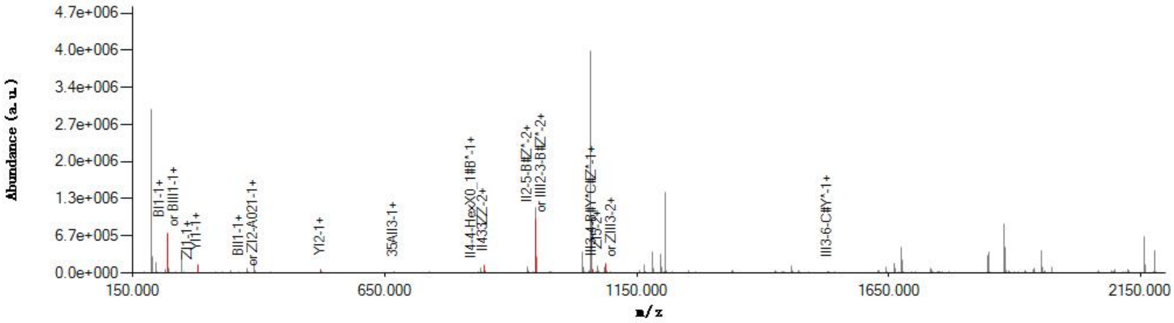


#### Graphical Fragmentation Map

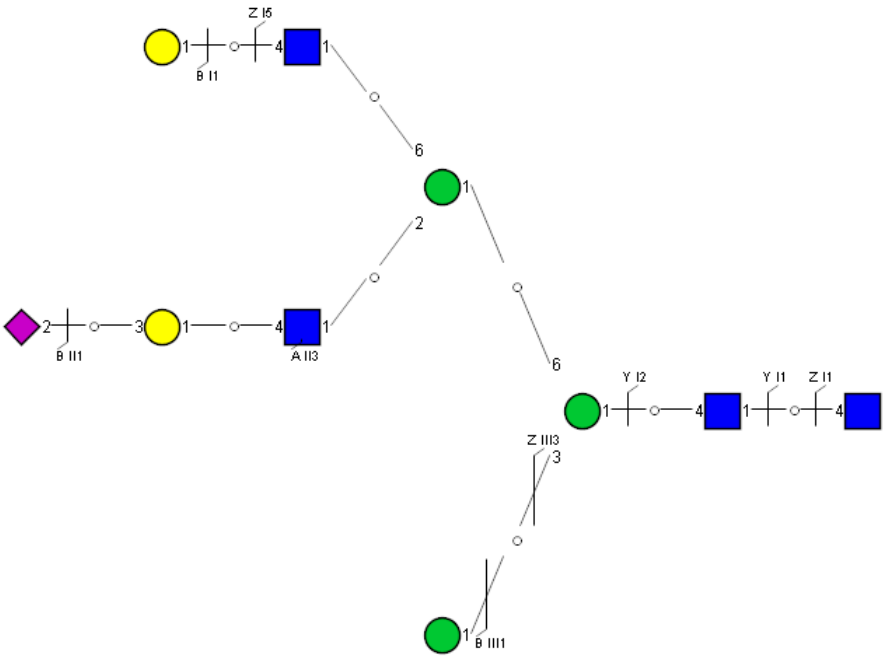

Spectrum Index: 39873(NoBHs:1)

#### Precursor iEF Map

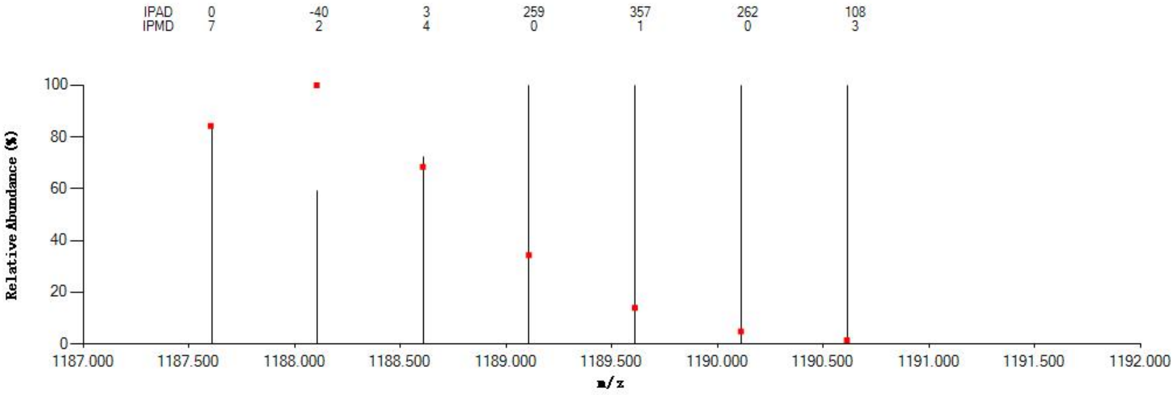

#### Annotated MS/MS Spectrum

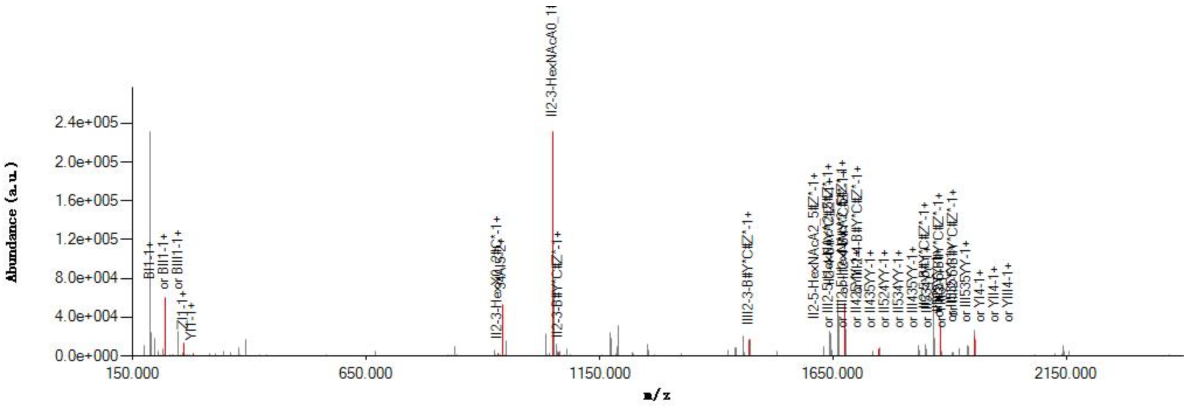

#### Graphical Fragmentation Map

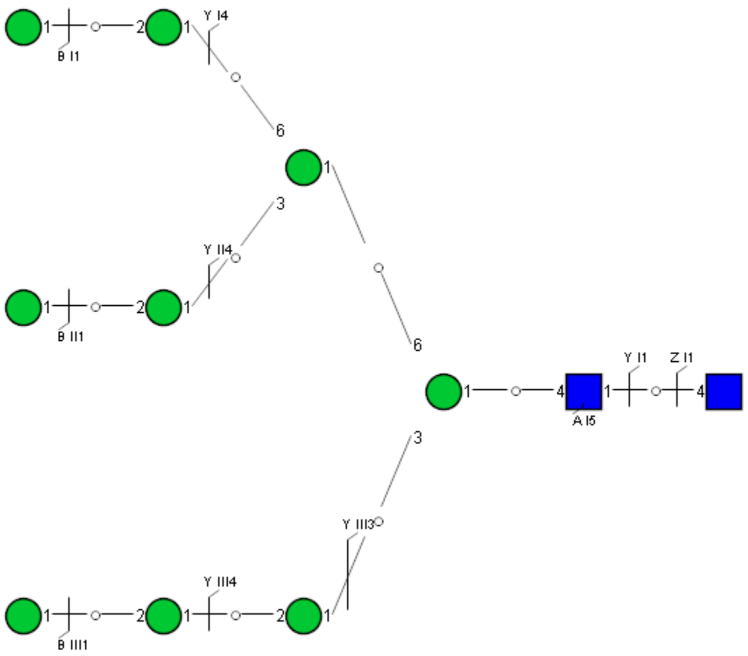

Spectrum Index: 23448(NoBHs:1)

#### Precursor iEF Map

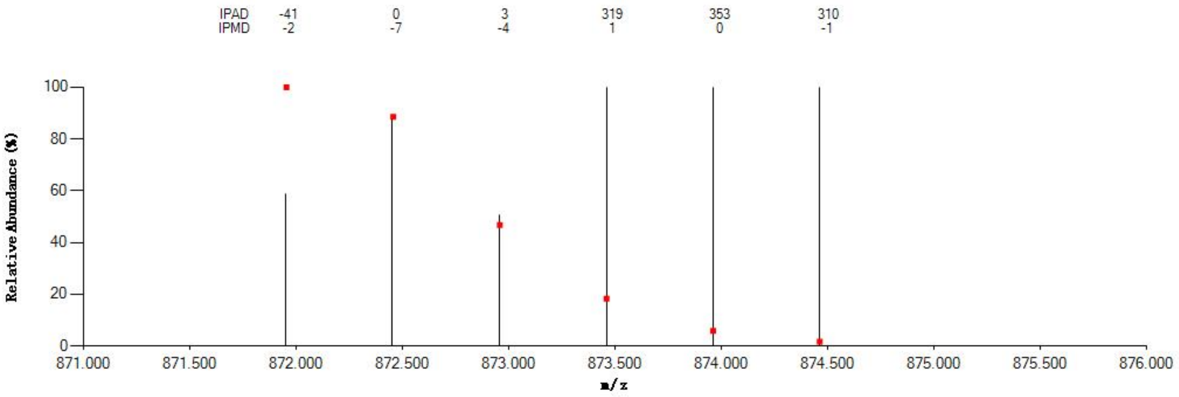


#### Annotated MS/MS Spectrum

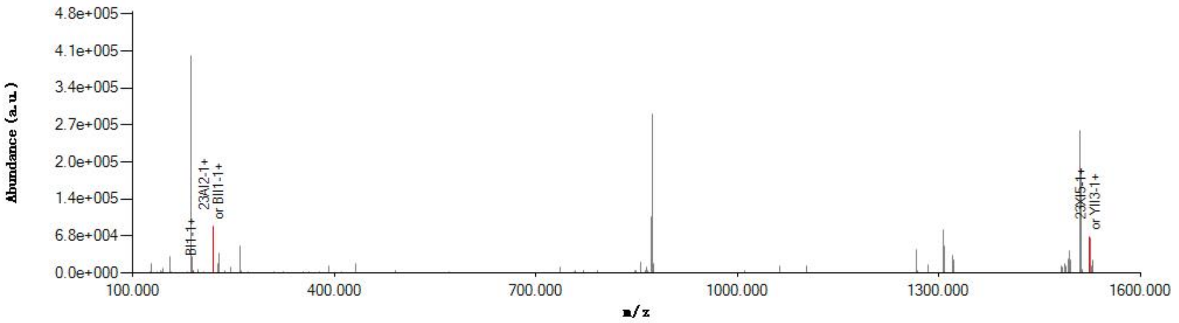


#### Graphical Fragmentation Map

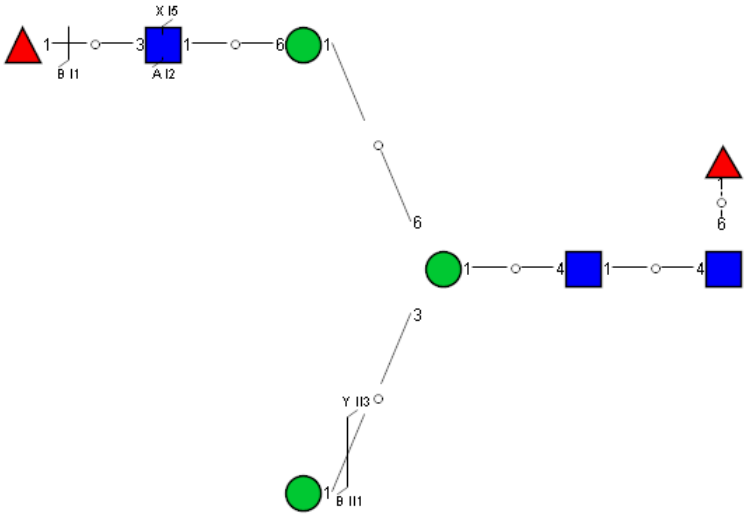

Spectrum Index: 23715(NoBHs:1)

#### Precursor iEF Map

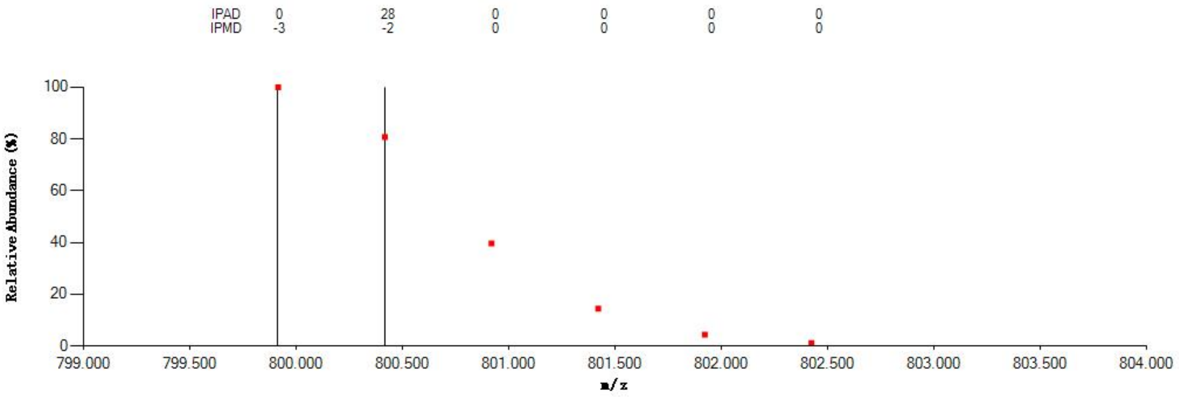


#### Annotated MS/MS Spectrum

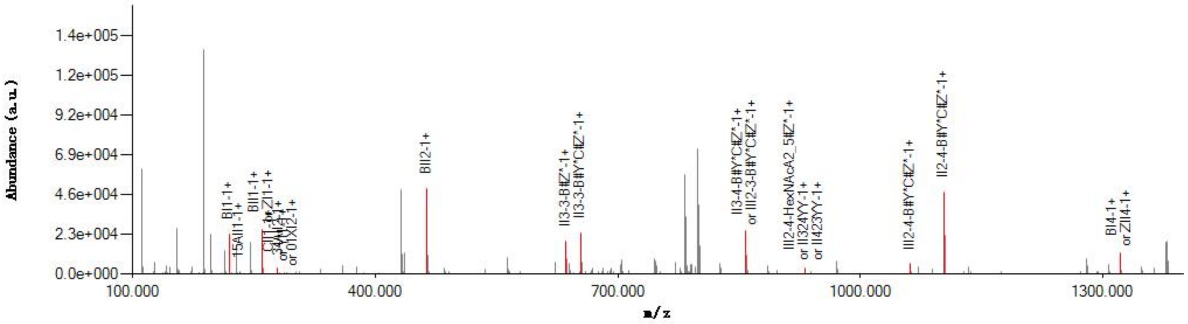


#### Graphical Fragmentation Map

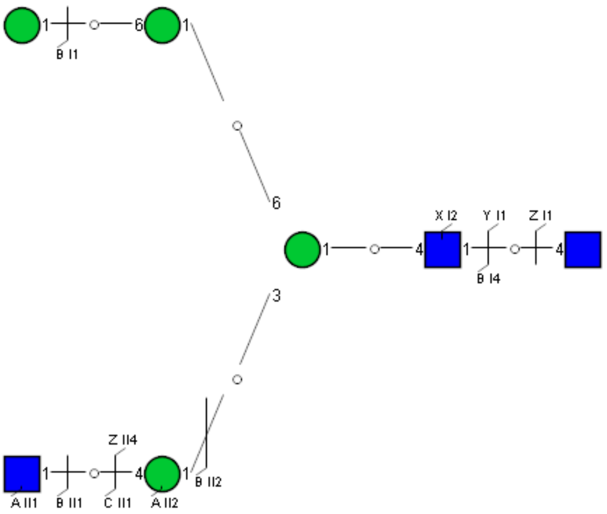

Spectrum Index: 27062(NoBHs:1)

#### Precursor iEF Map

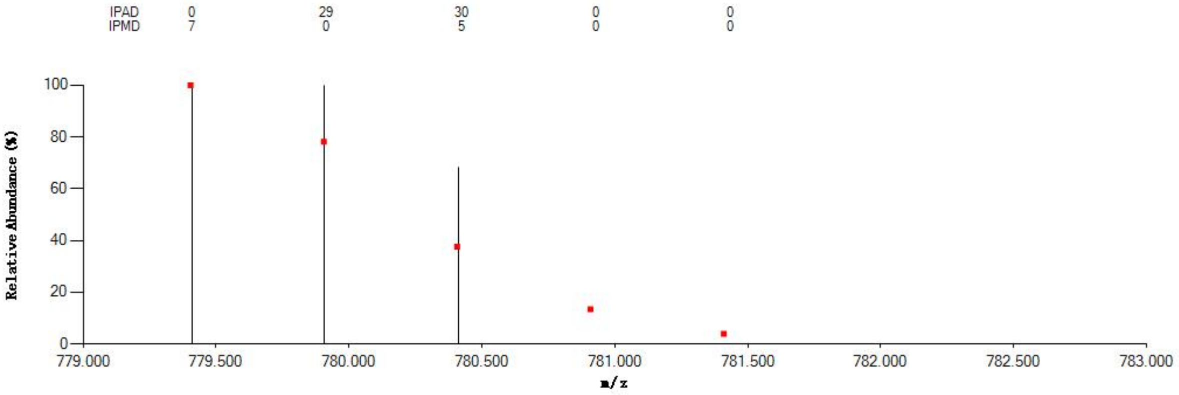

#### Annotated MS/MS Spectrum

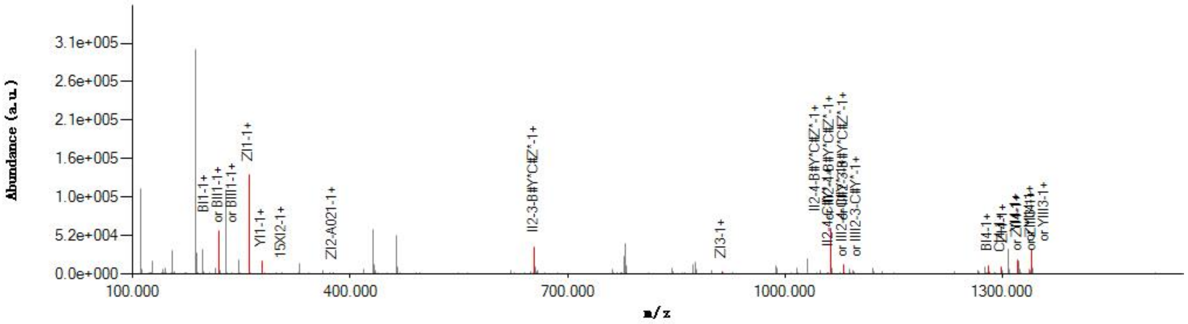

#### Graphical Fragmentation Map

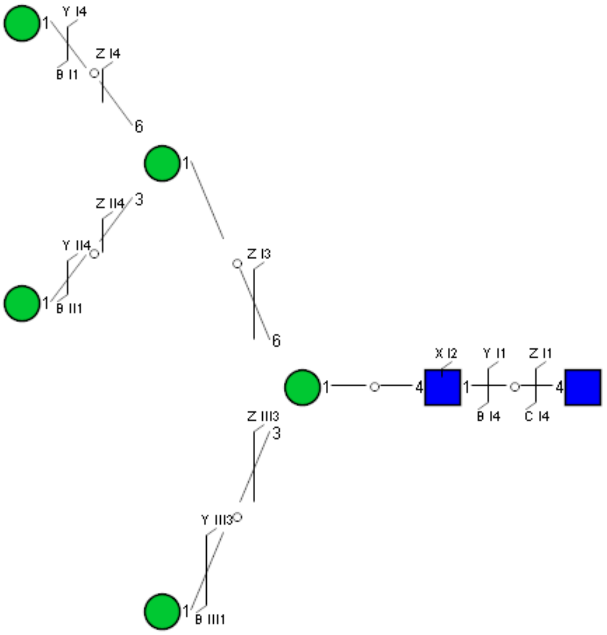

Spectrum Index: 27262(NoBHs:1)

#### Precursor iEF Map

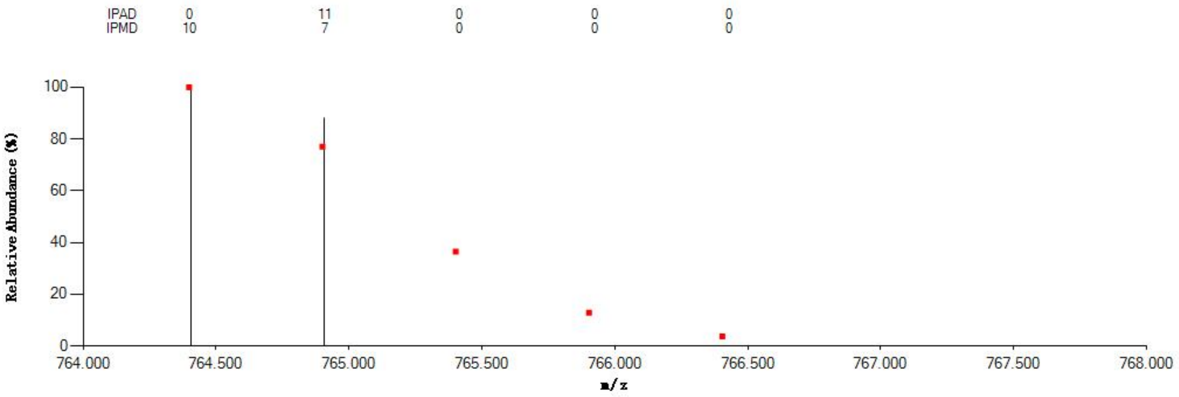


#### Annotated MS/MS Spectrum

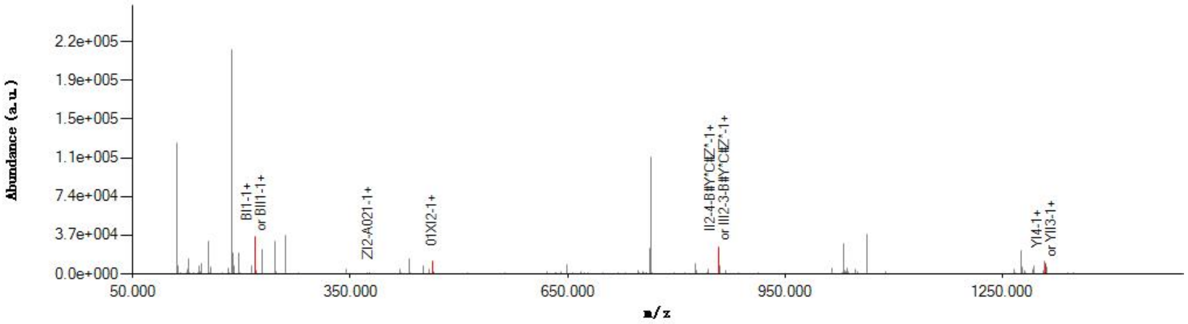


#### Graphical Fragmentation Map

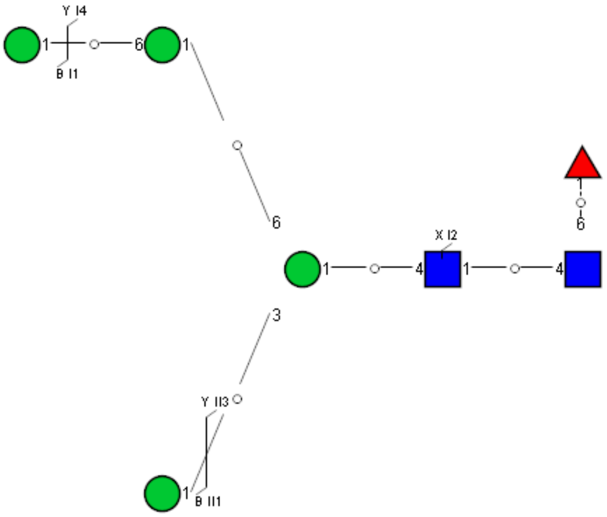

Spectrum Index: 27380(NoBHs:1)

#### Precursor iEF Map

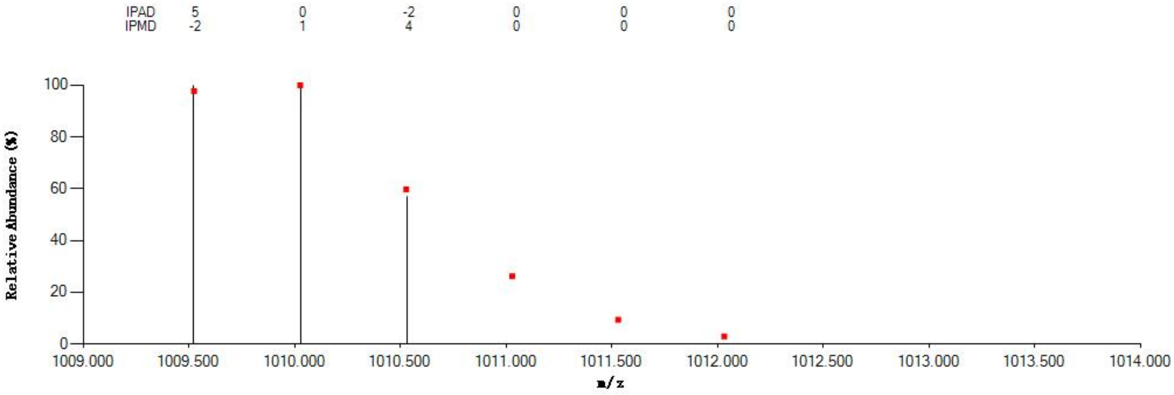


#### Annotated MS/MS Spectrum

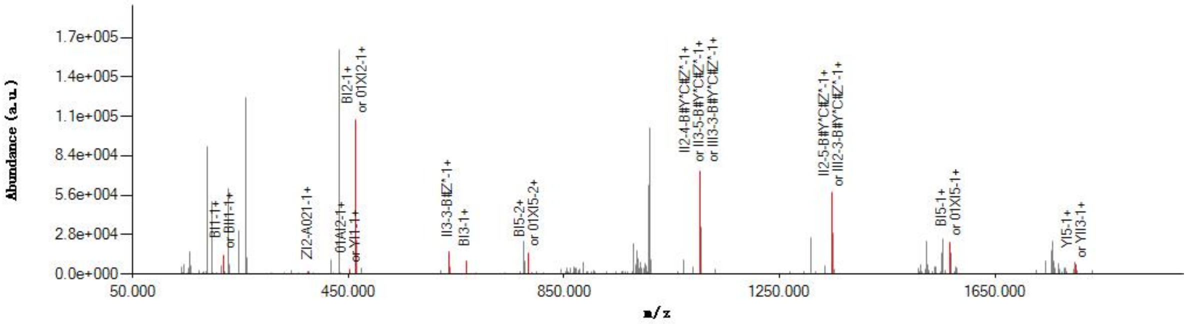


#### Graphical Fragmentation Map

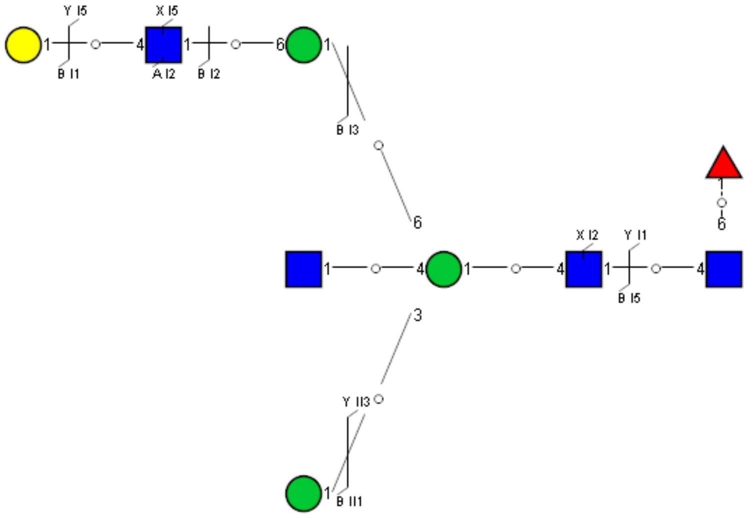

### Spectrum Index: 27826(NoBHs:1)

#### Precursor iEF Map

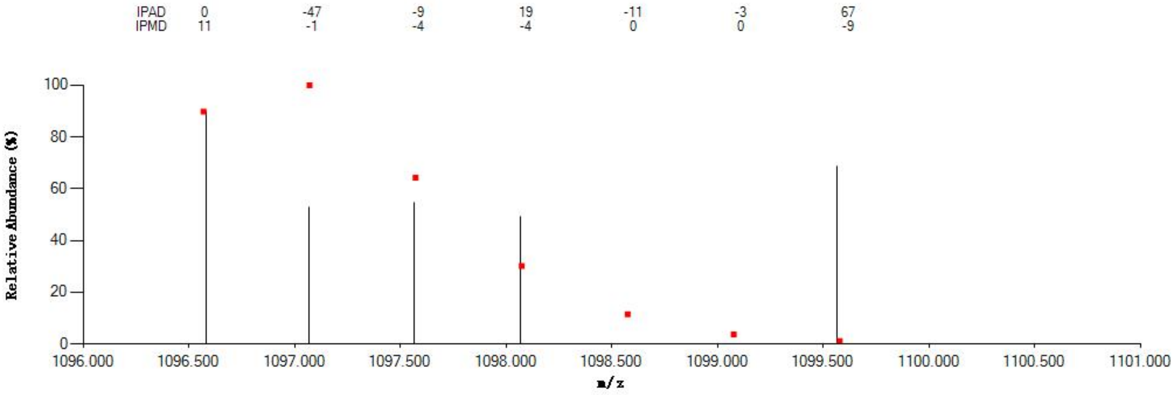


#### Annotated MS/MS Spectrum

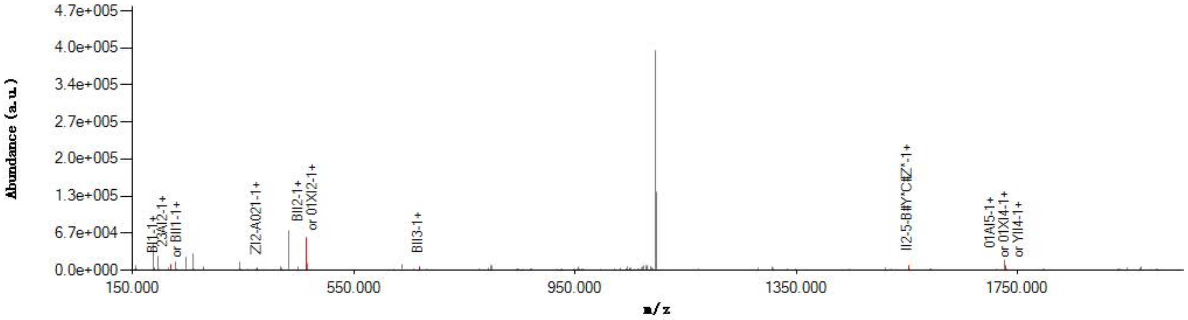


#### Graphical Fragmentation Map

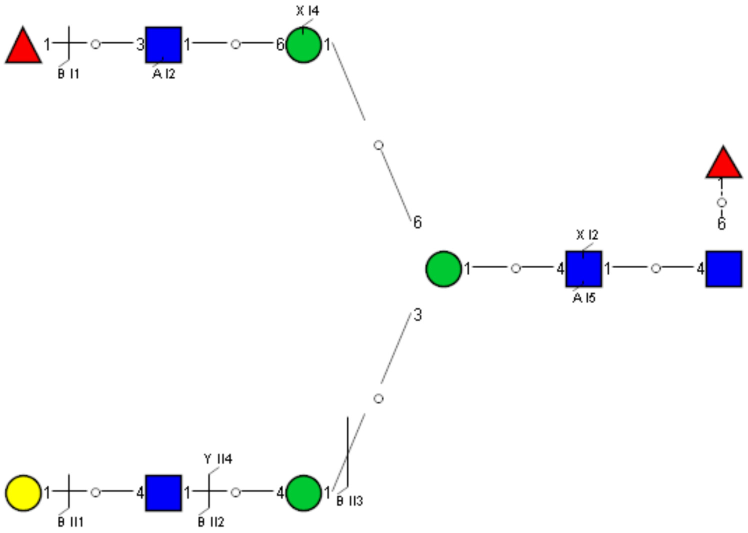

Spectrum Index: 28034(NoBHs:1)

#### Precursor iEF Map

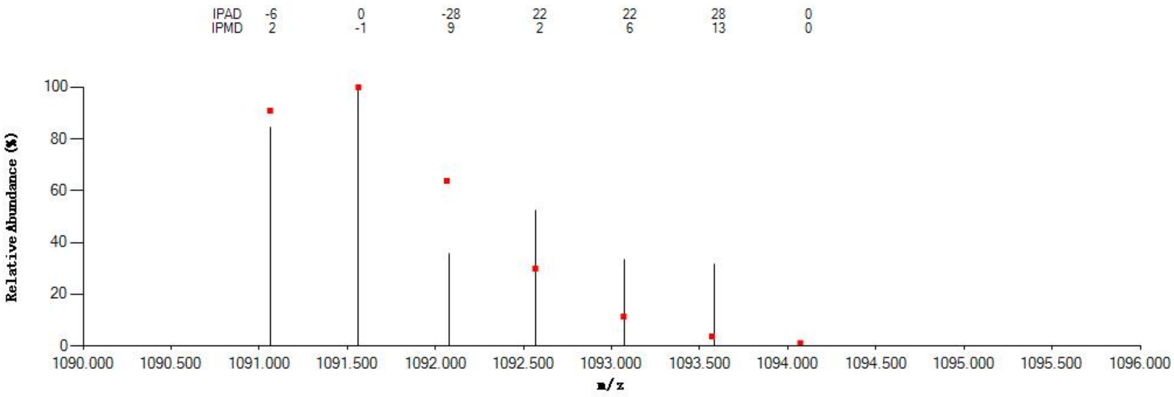


#### Annotated MS/MS Spectrum

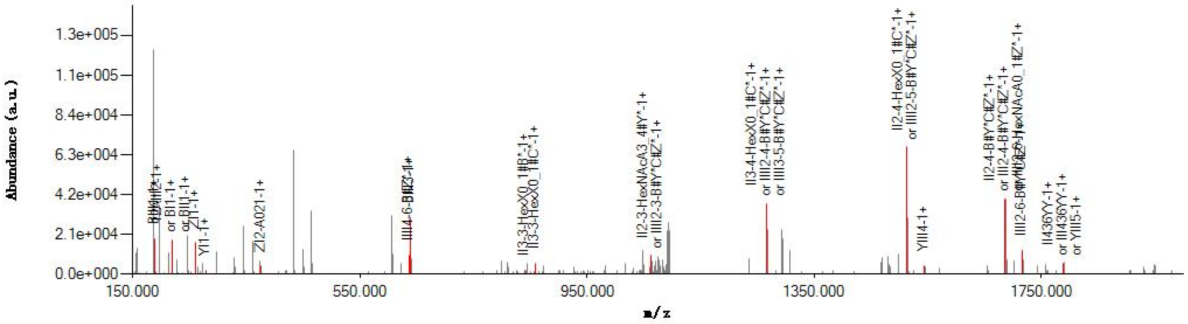


#### Graphical Fragmentation Map

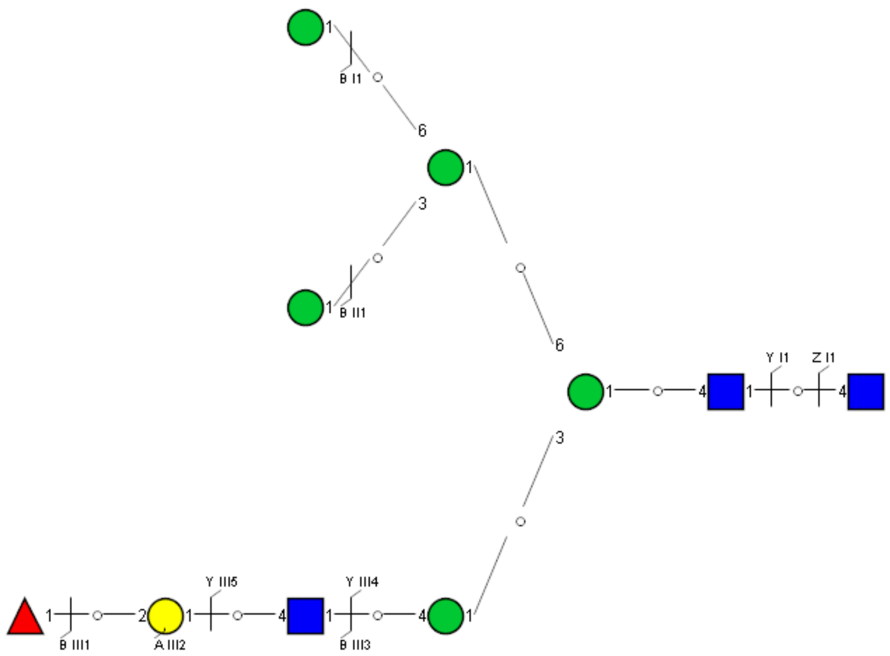

Spectrum Index: 28606(NoBHs:1)

#### Precursor iEF Map

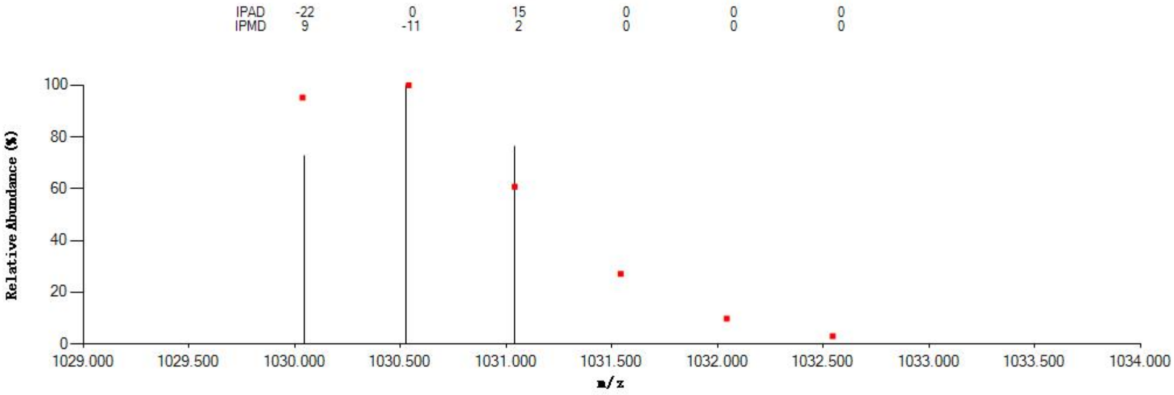


#### Annotated MS/MS Spectrum

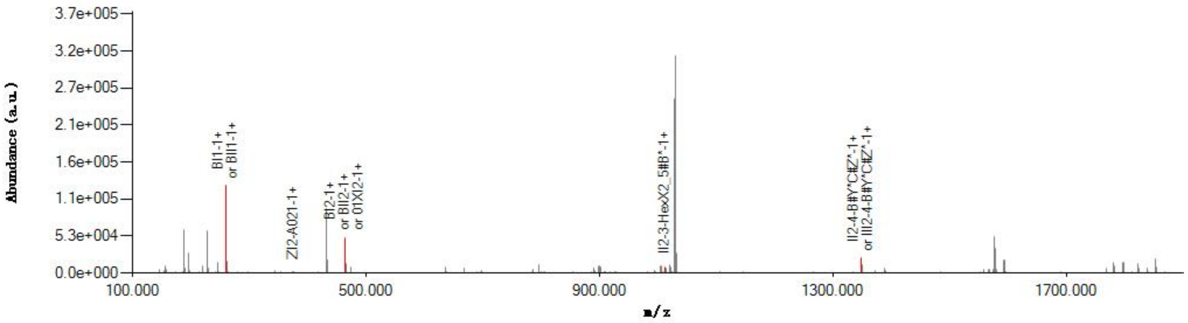


#### Graphical Fragmentation Map

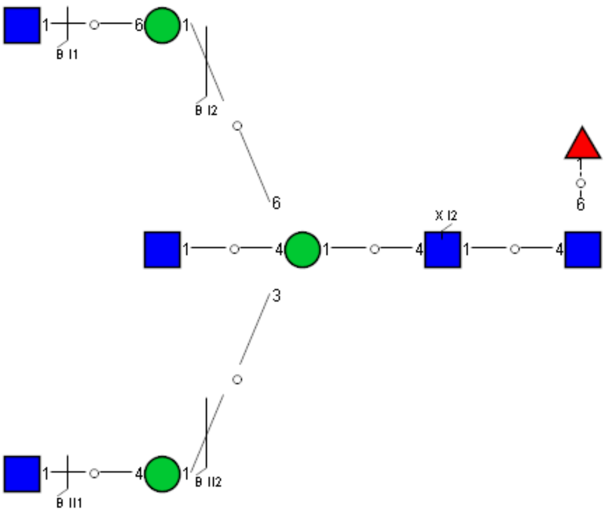

### Spectrum Index: 28765(NoBHs:1)

#### Precursor iEF Map

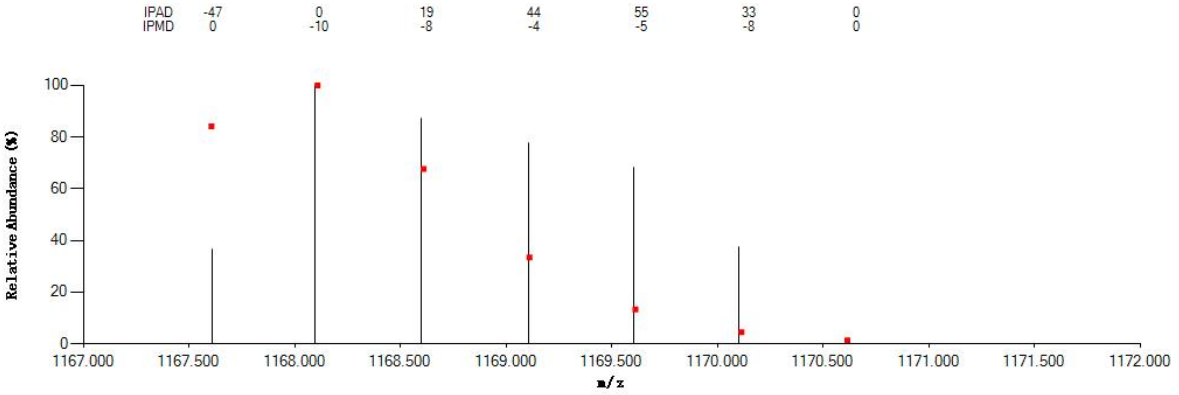


#### Annotated MS/MS Spectrum

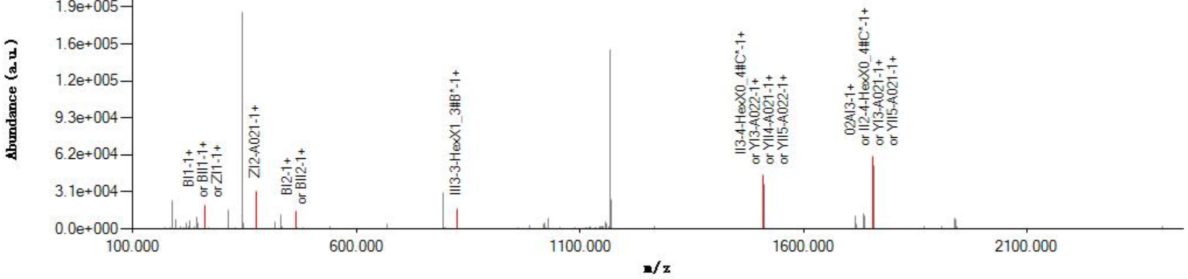


#### Graphical Fragmentation Map

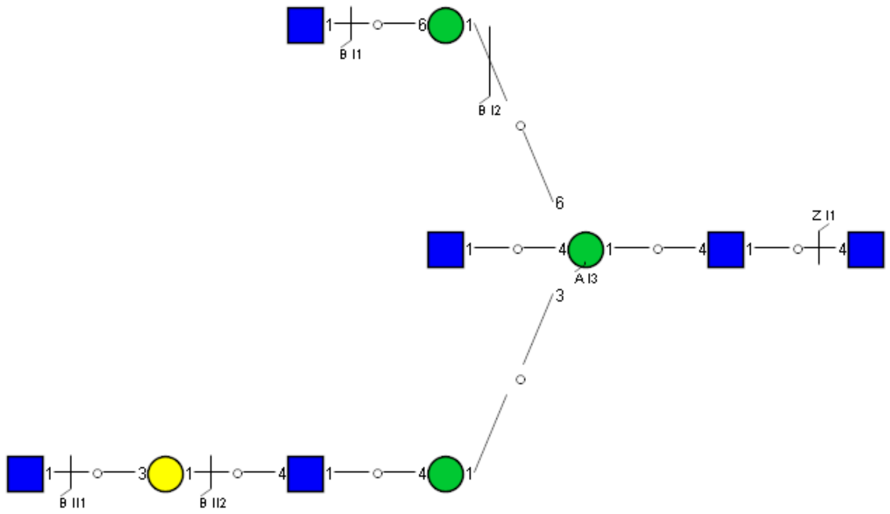

### Spectrum Index: 28841(NoBHs:1)

#### Precursor iEF Map

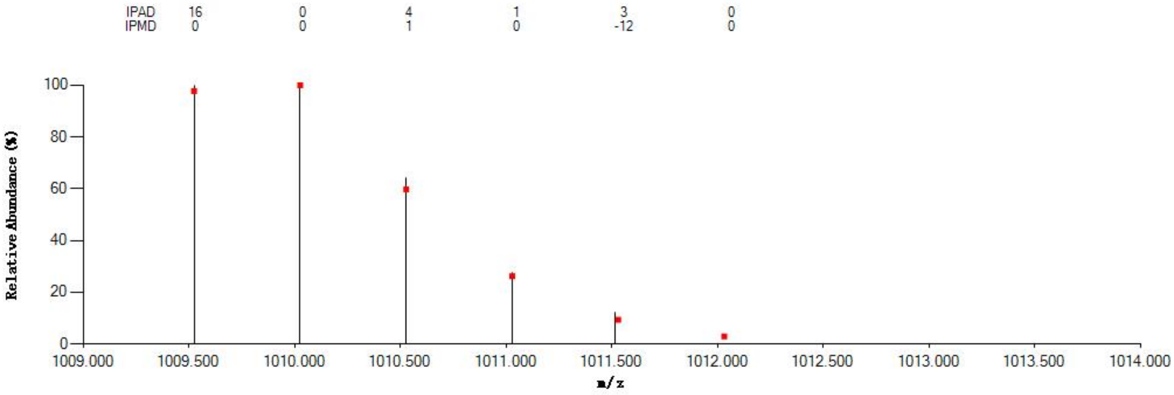

#### Annotated MS/MS Spectrum

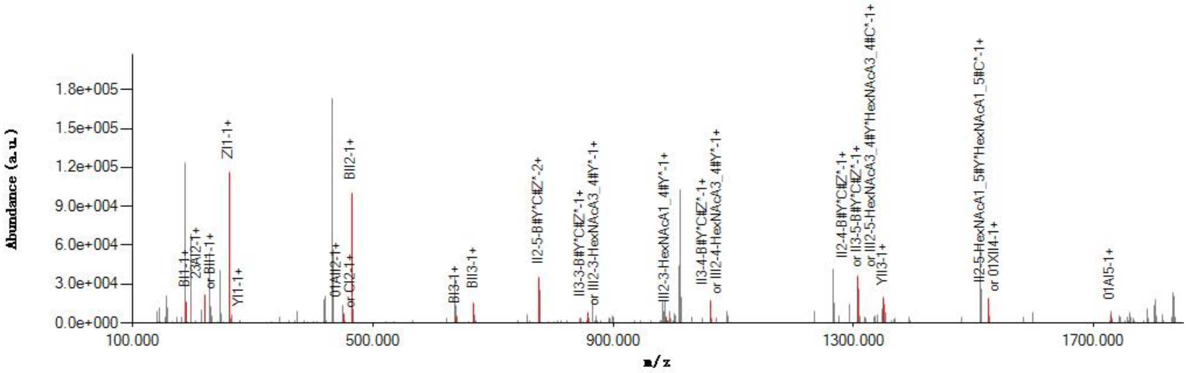

#### Graphical Fragmentation Map

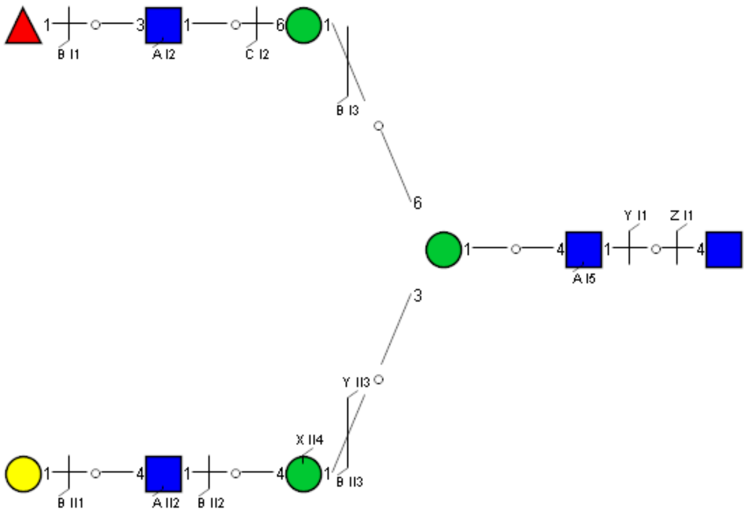

Spectrum Index: 29220(NoBHs:1)

#### Precursor iEF Map

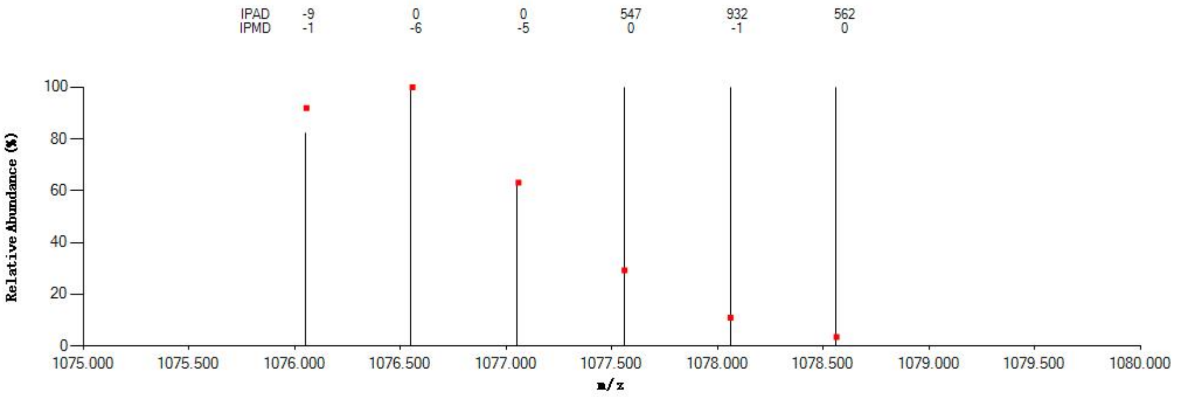


#### Annotated MS/MS Spectrum

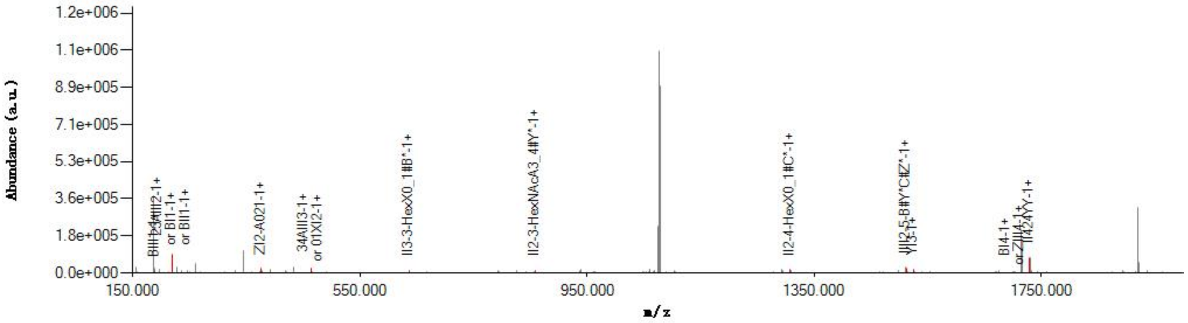


#### Graphical Fragmentation Map

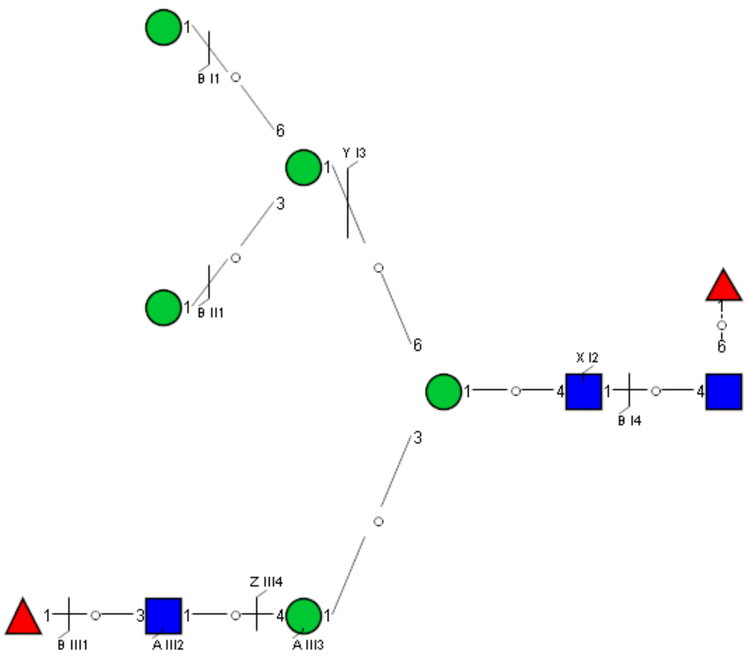

Spectrum Index: 29500(NoBHs:1)

#### Precursor iEF Map

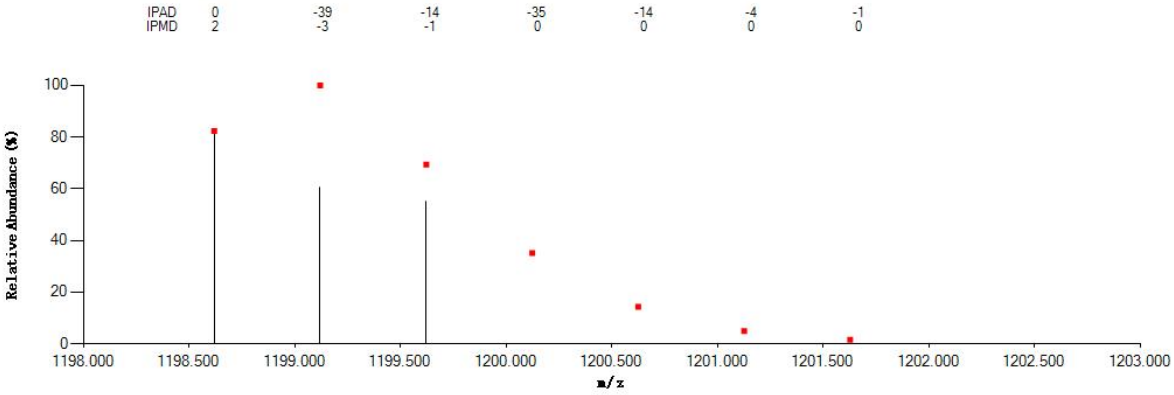


#### Annotated MS/MS Spectrum

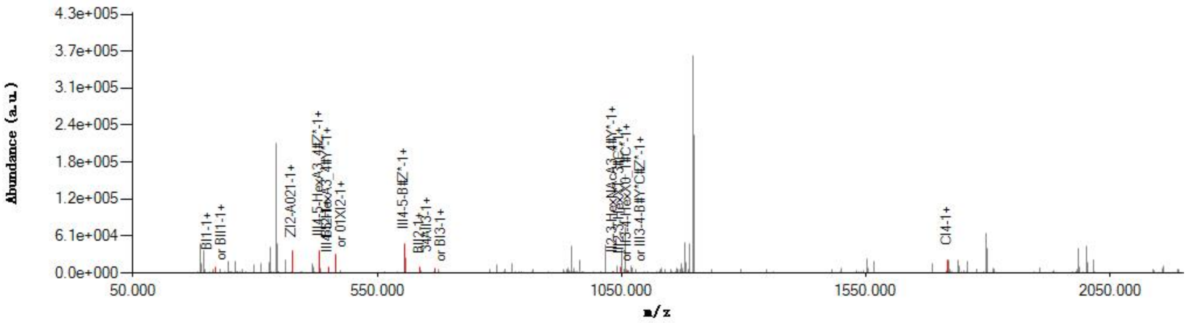


#### Graphical Fragmentation Map

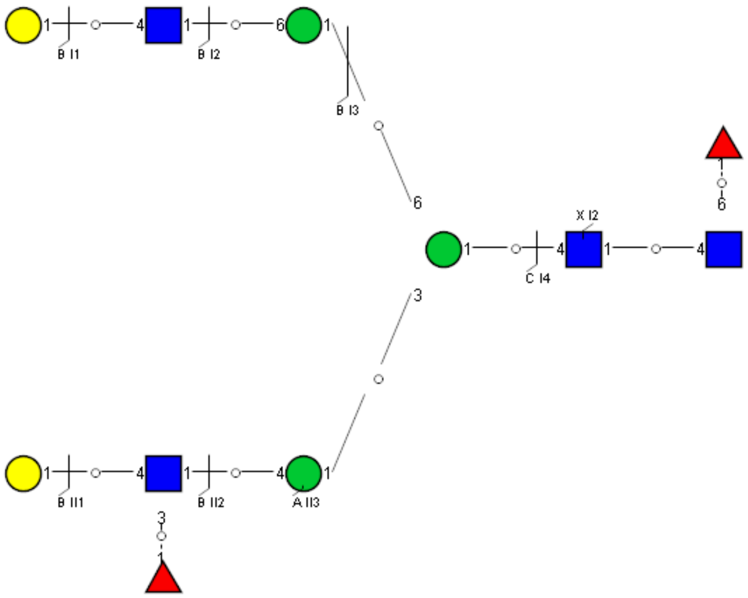

Spectrum Index: 29609(NoBHs:1)

#### Precursor iEF Map

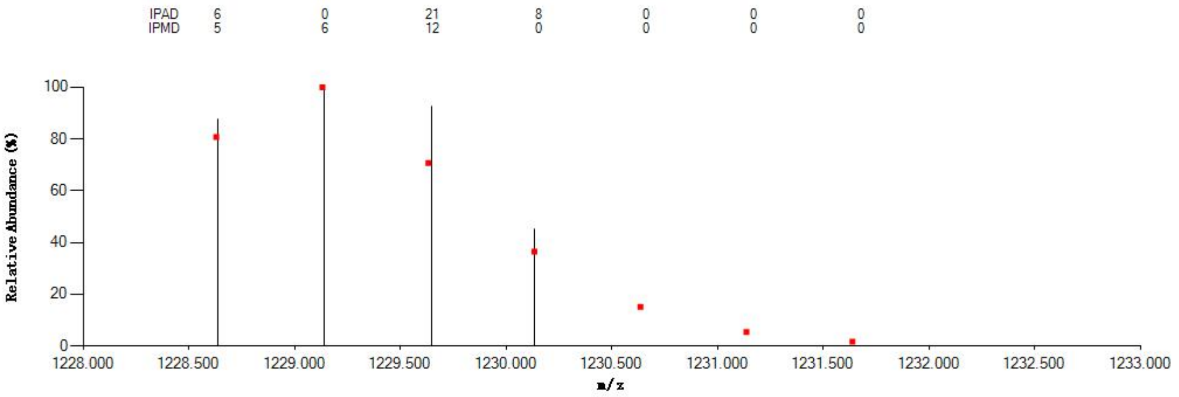


#### Annotated MS/MS Spectrum

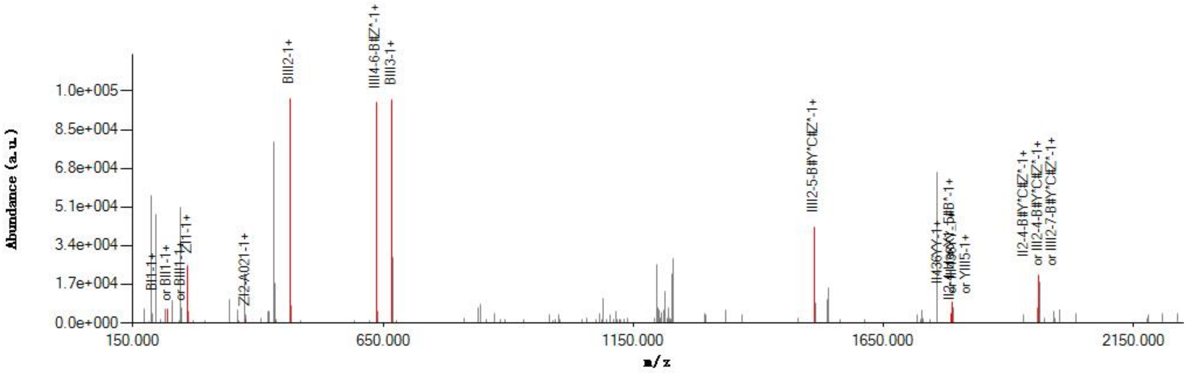


#### Graphical Fragmentation Map

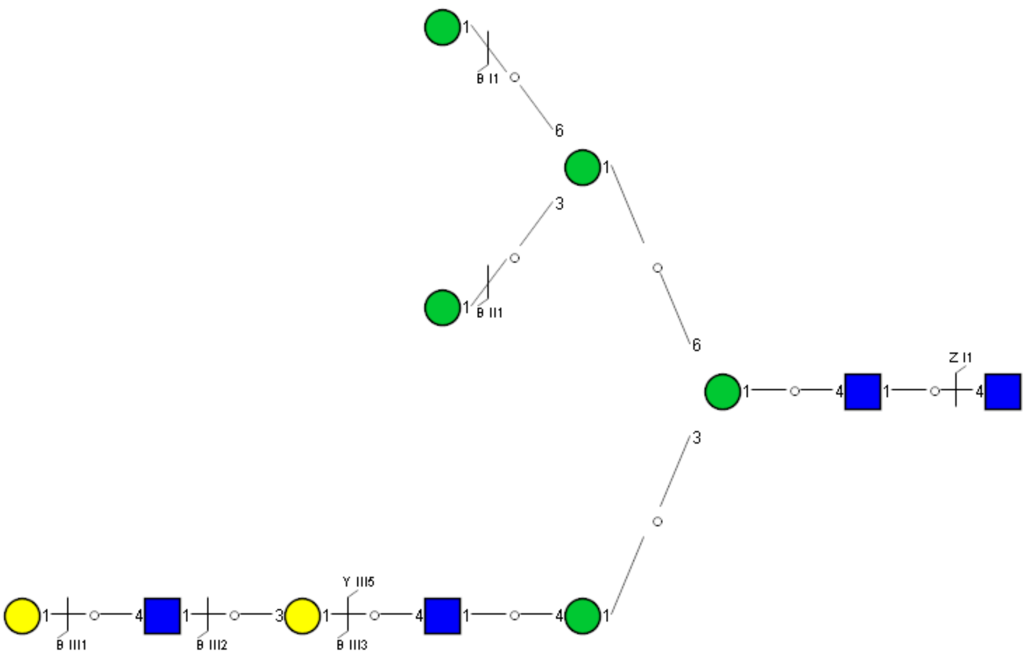

### Spectrum Index: 30642(NoBHs:1)

#### Precursor iEF Map

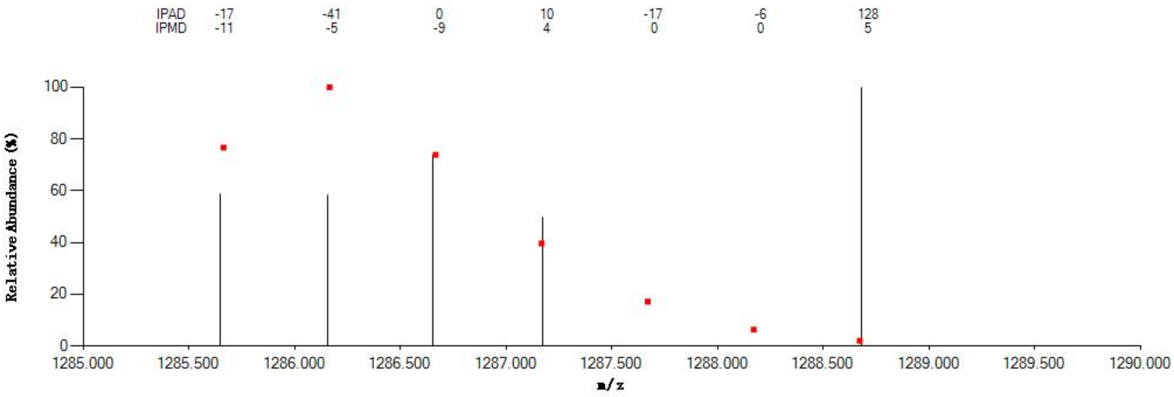


#### Annotated MS/MS Spectrum

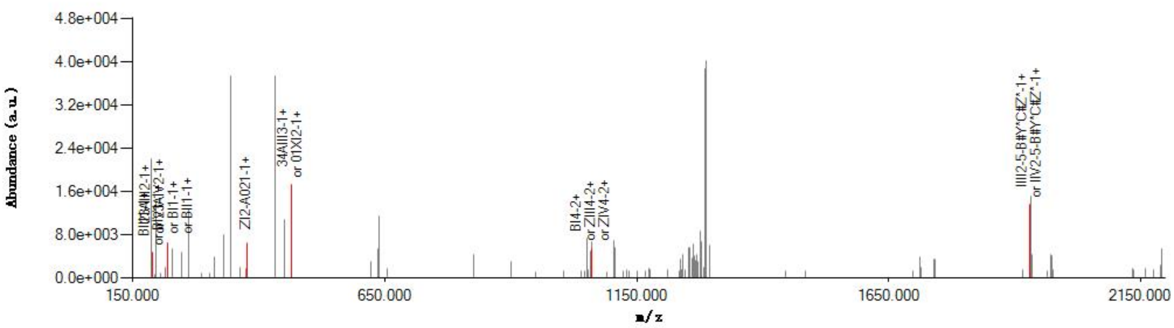


#### Graphical Fragmentation Map

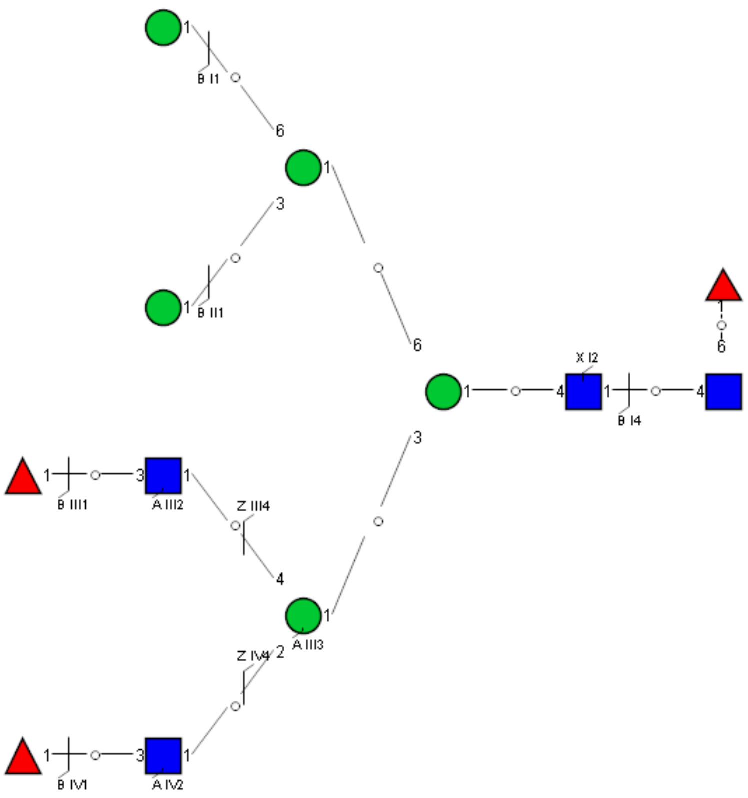

Spectrum Index: 31001(NoBHs:1)

#### Precursor iEF Map

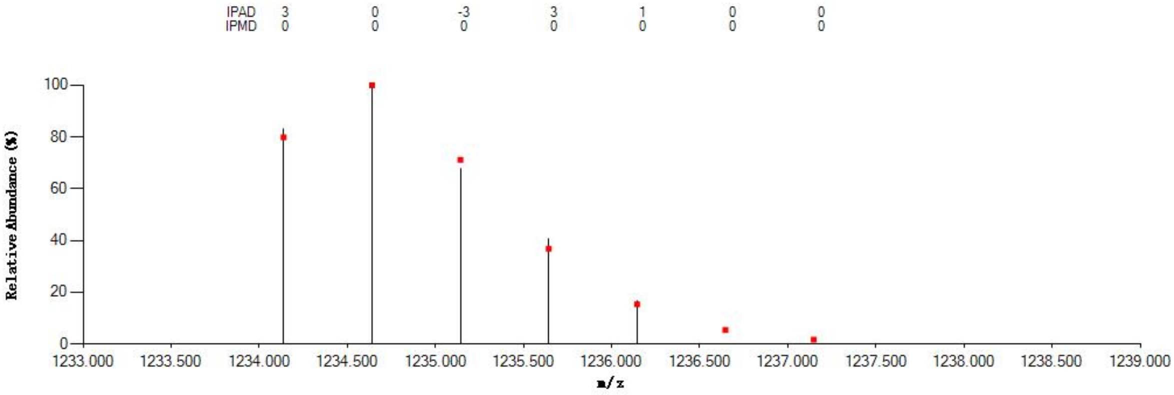


#### Annotated MS/MS Spectrum

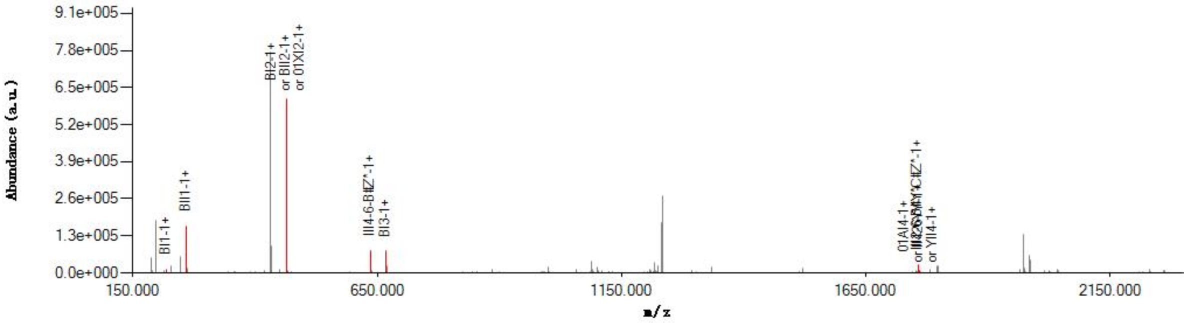


#### Graphical Fragmentation Map

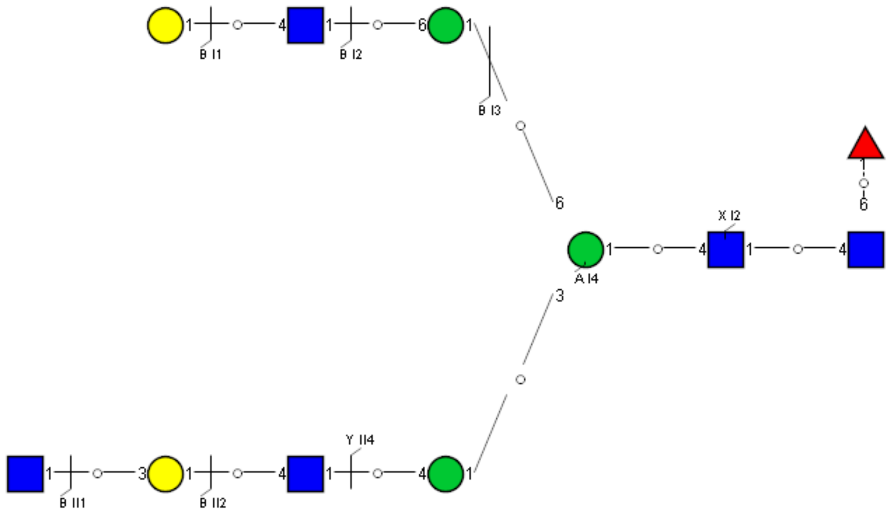

Spectrum Index: 31498(NoBHs:1)

#### Precursor iEF Map

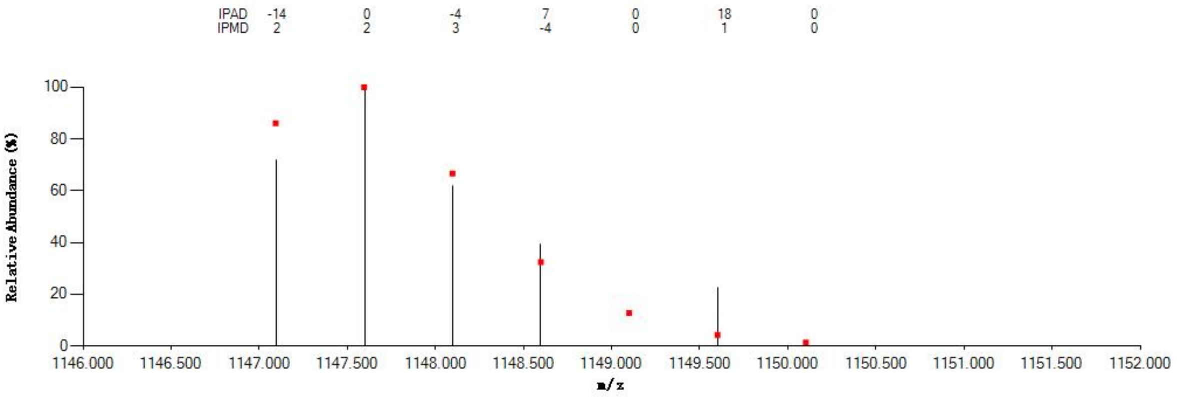

#### Annotated MS/MS Spectrum

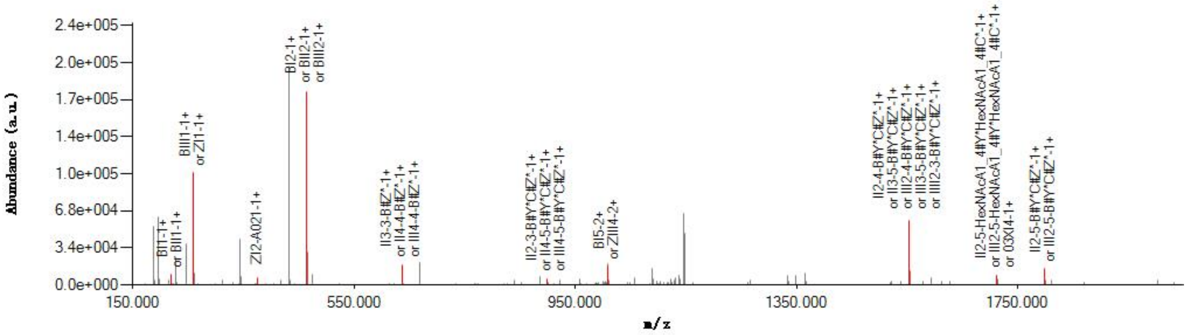

#### Graphical Fragmentation Map

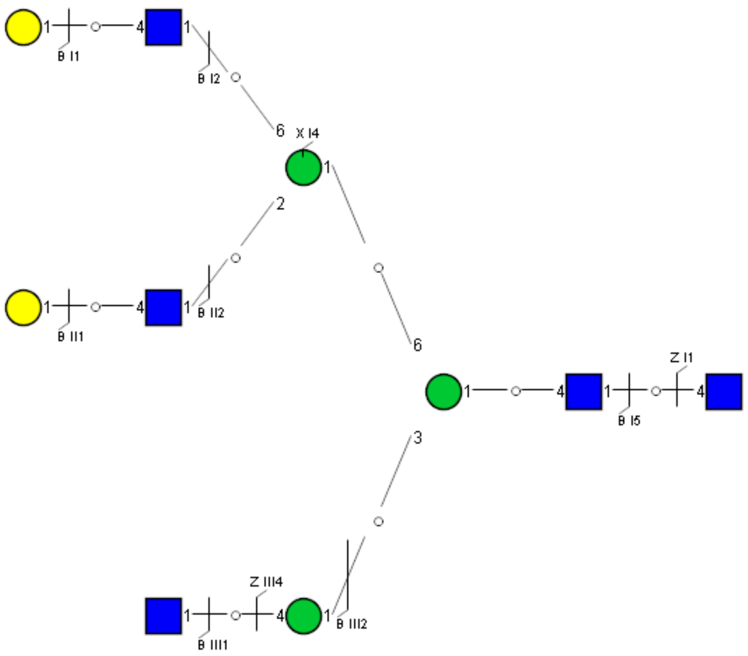

Spectrum Index: 31554(NoBHs:1)

Precursor iEF Map

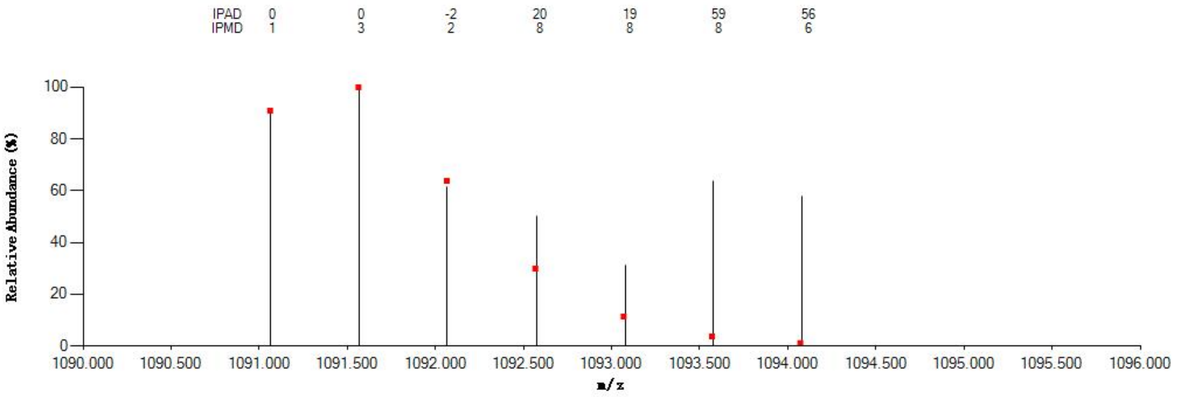


Annotated MS/MS Spectrum

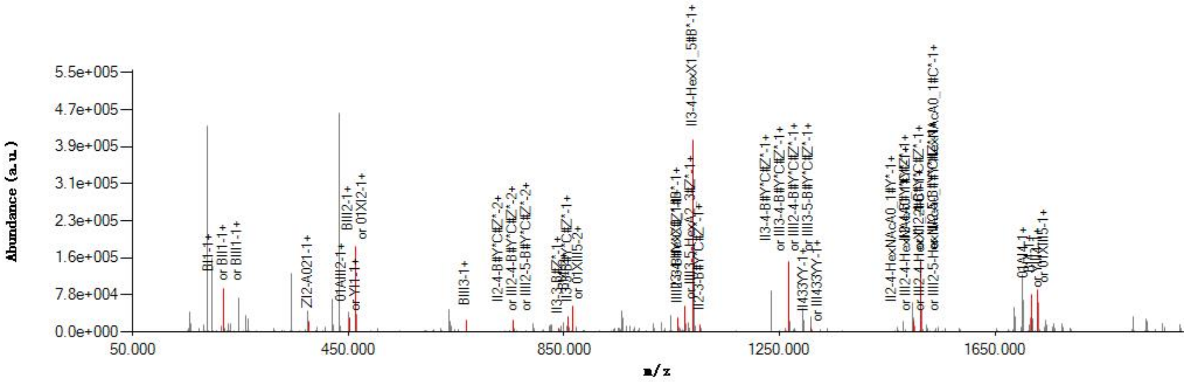


Graphical Fragmentation Map

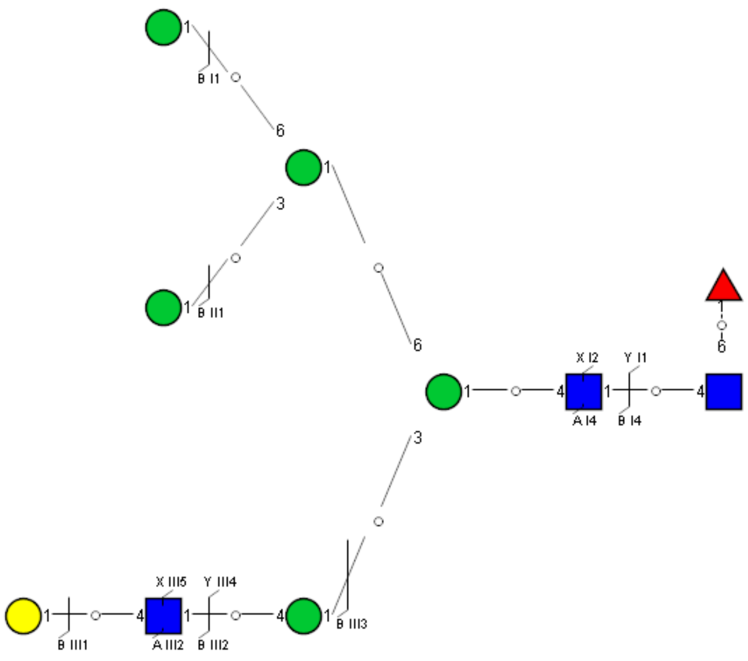

Spectrum Index: 31566(NoBHs:1)

#### Precursor iEF Map

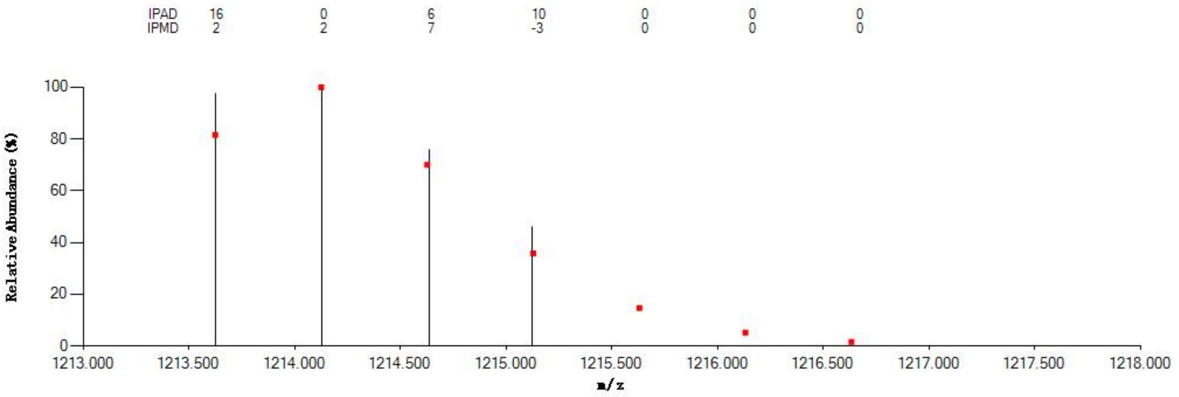

#### Annotated MS/MS Spectrum

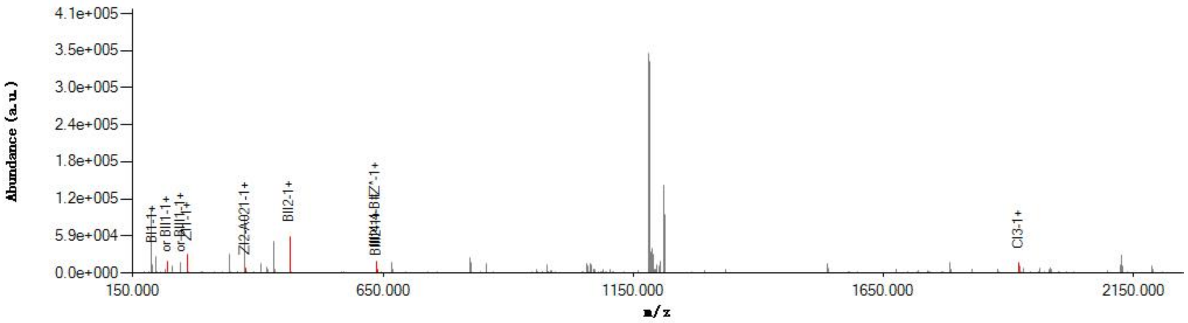

#### Graphical Fragmentation Map

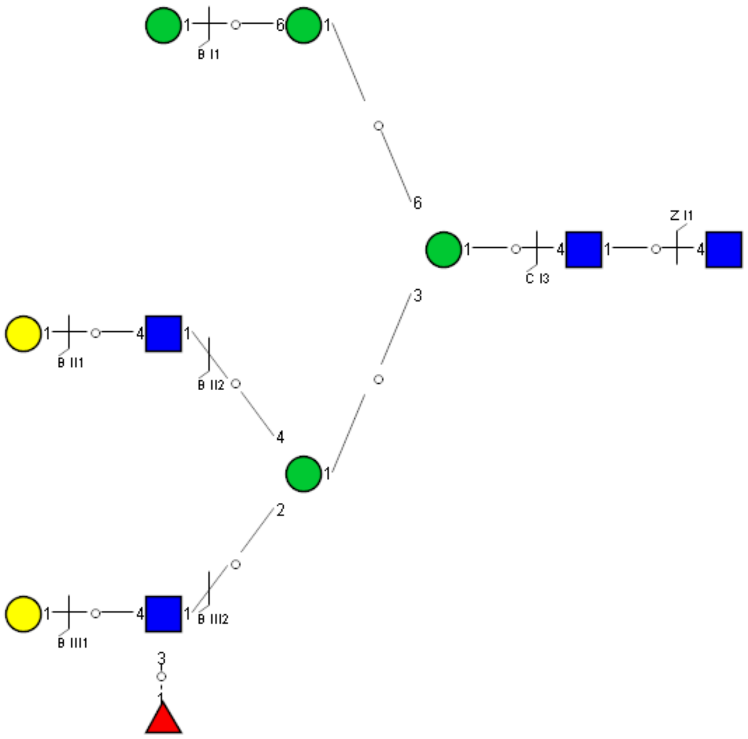

Spectrum Index: 31859(NoBHs:1)

#### Precursor iEF Map

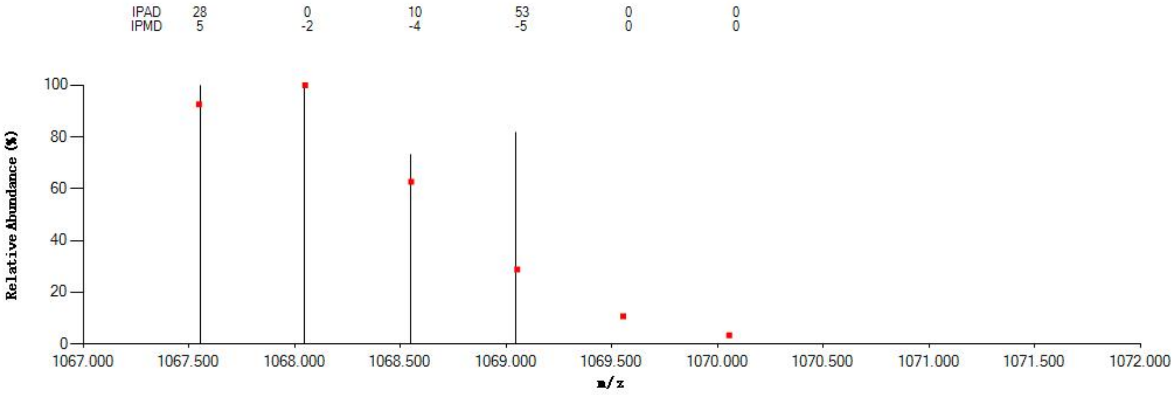

#### Annotated MS/MS Spectrum

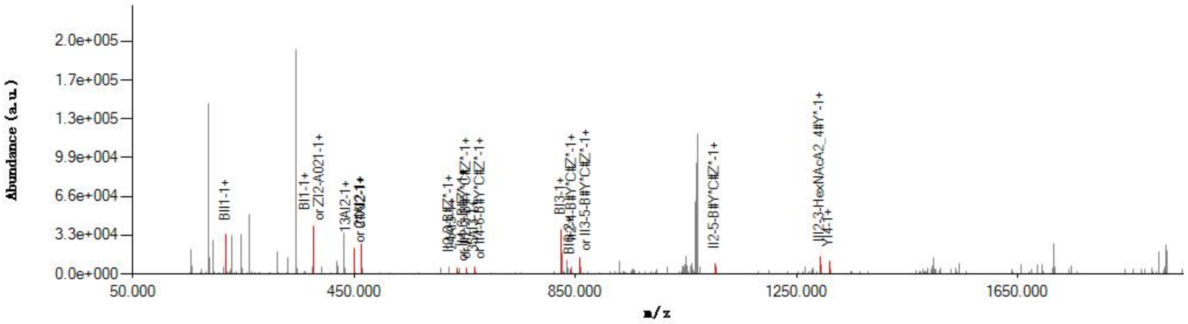

#### Graphical Fragmentation Map

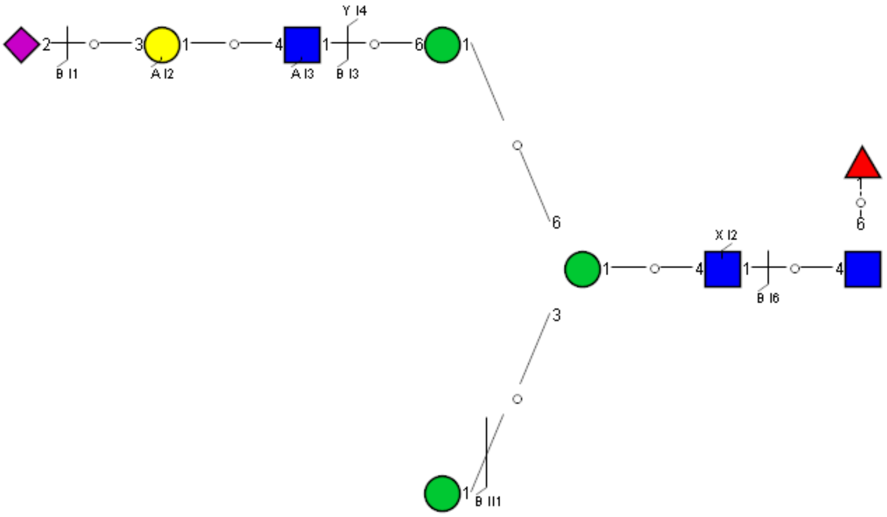

Spectrum Index: 31977(NoBHs:1)

Precursor iEF Map

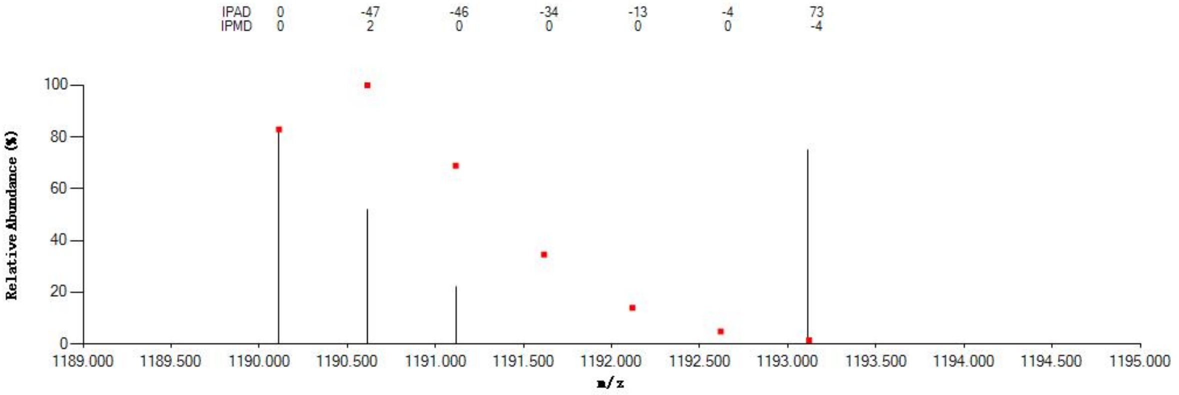

Annotated MS/MS Spectrum

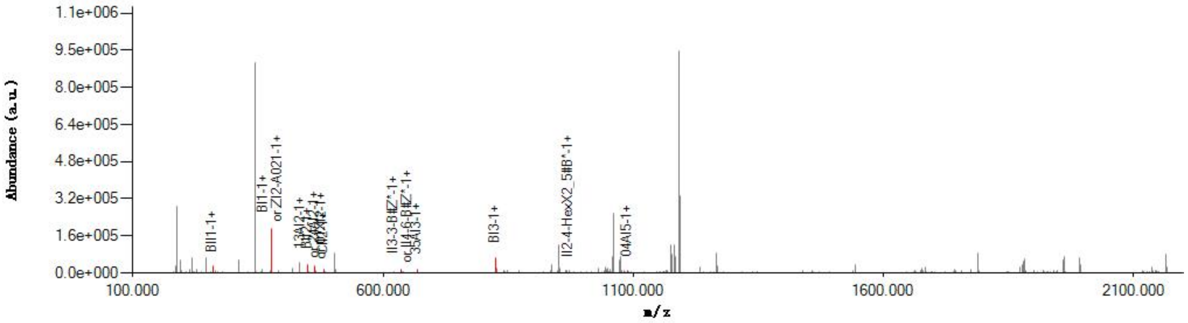

Graphical Fragmentation Map

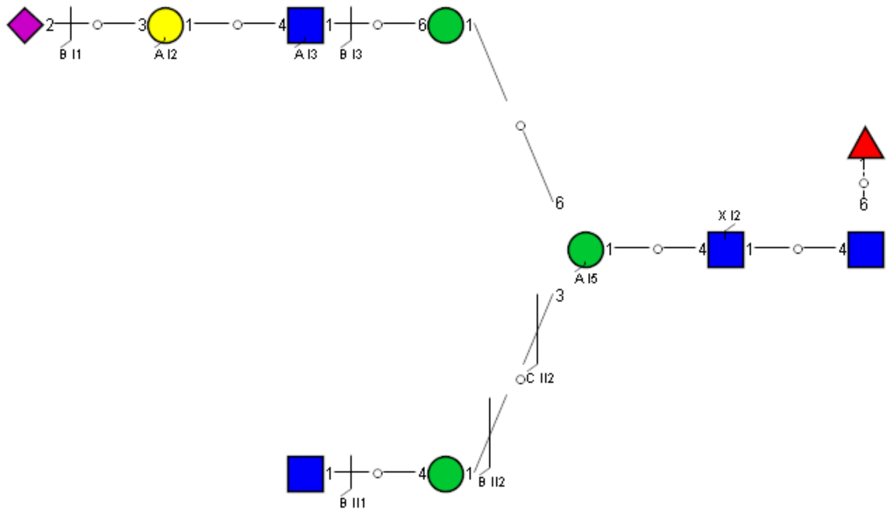

Spectrum Index: 33406(NoBHs:1)

#### Precursor iEF Map

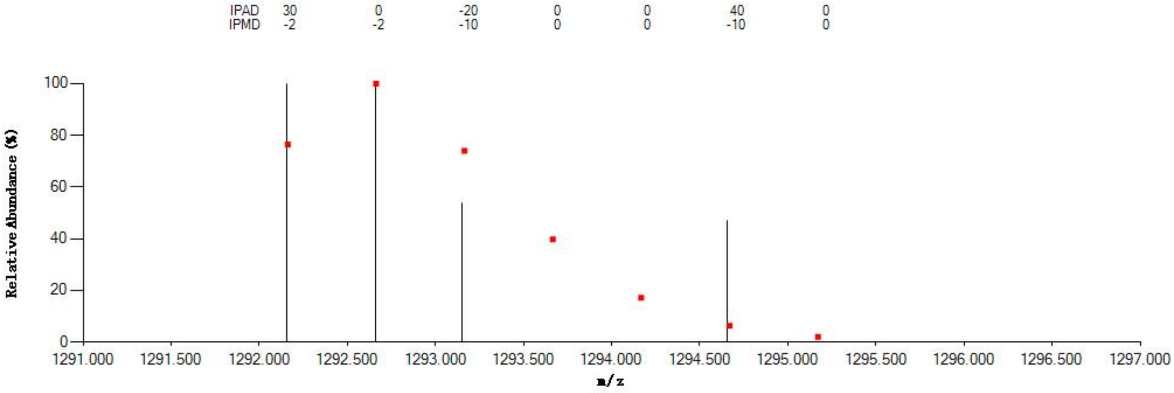


#### Annotated MS/MS Spectrum

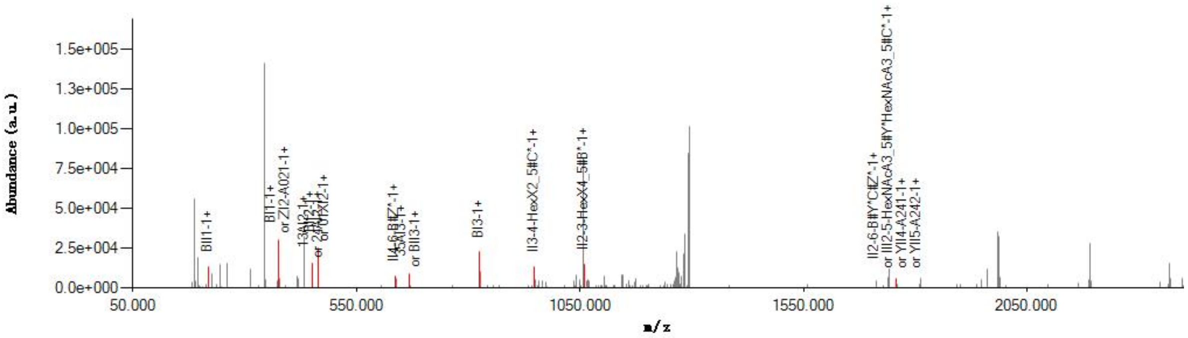


#### Graphical Fragmentation Map

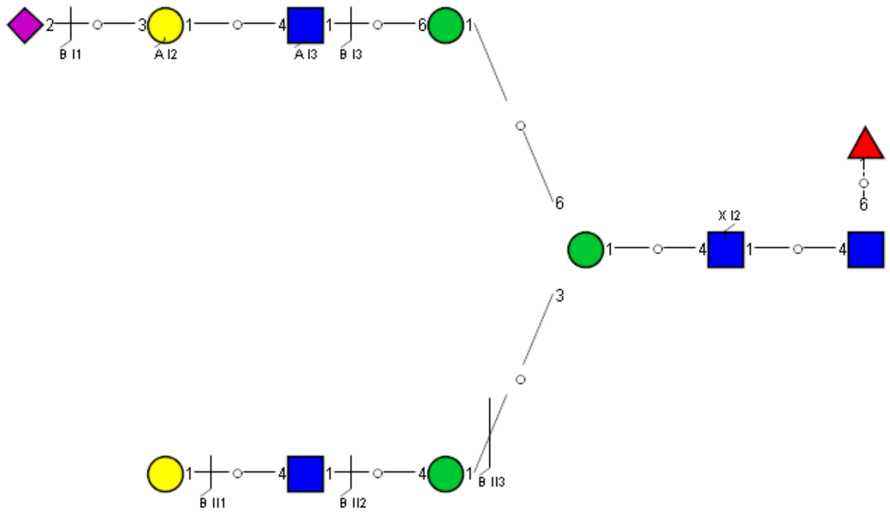

Spectrum Index: 33879(NoBHs:1)

#### Precursor iEF Map

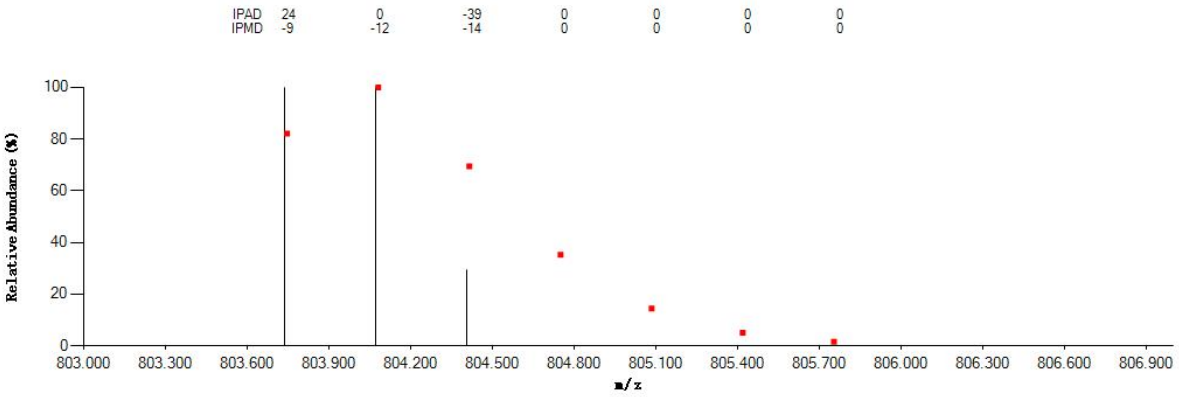

#### Annotated MS/MS Spectrum

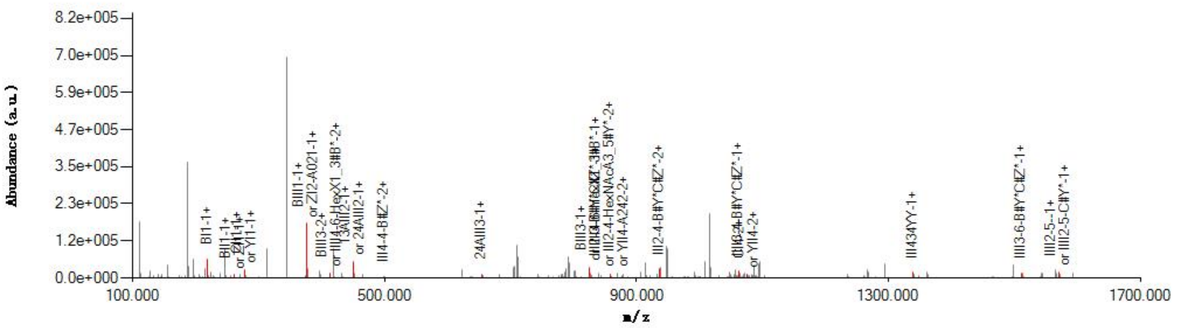

#### Graphical Fragmentation Map

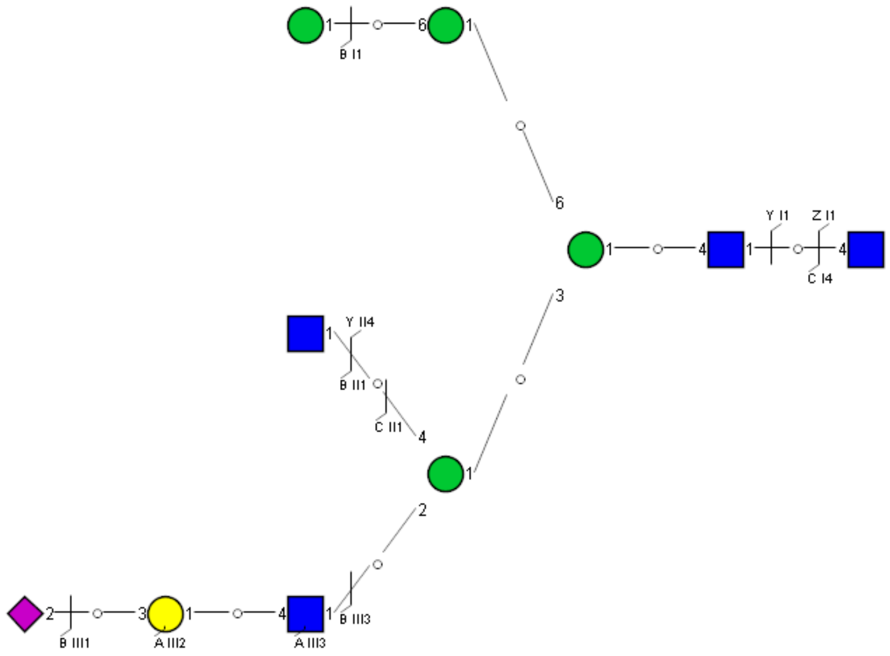

Spectrum Index: 34379(NoBHs:1)

#### Precursor iEF Map

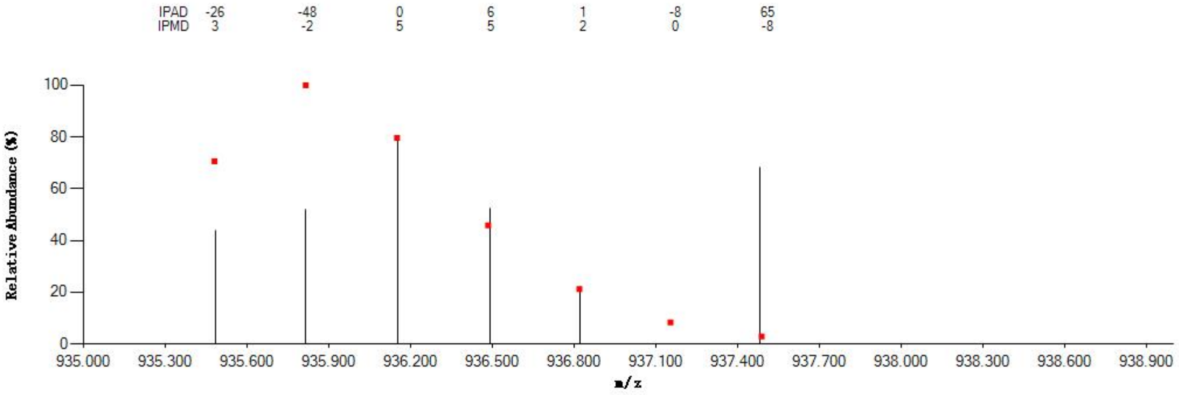


#### Annotated MS/MS Spectrum

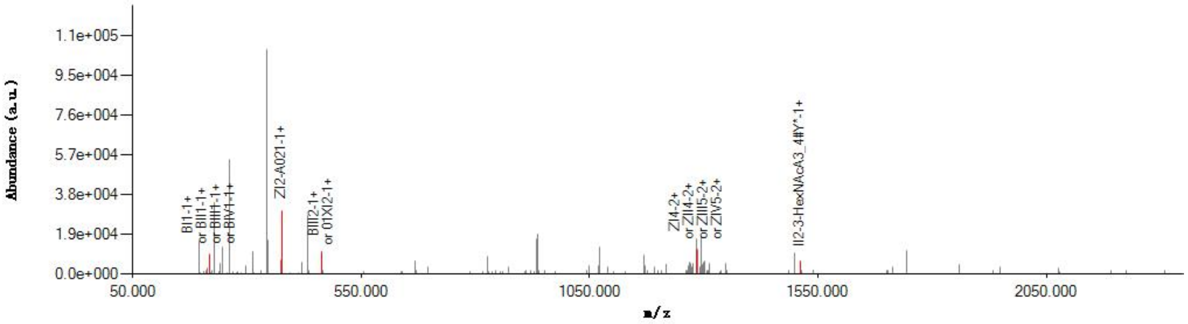


#### Graphical Fragmentation Map

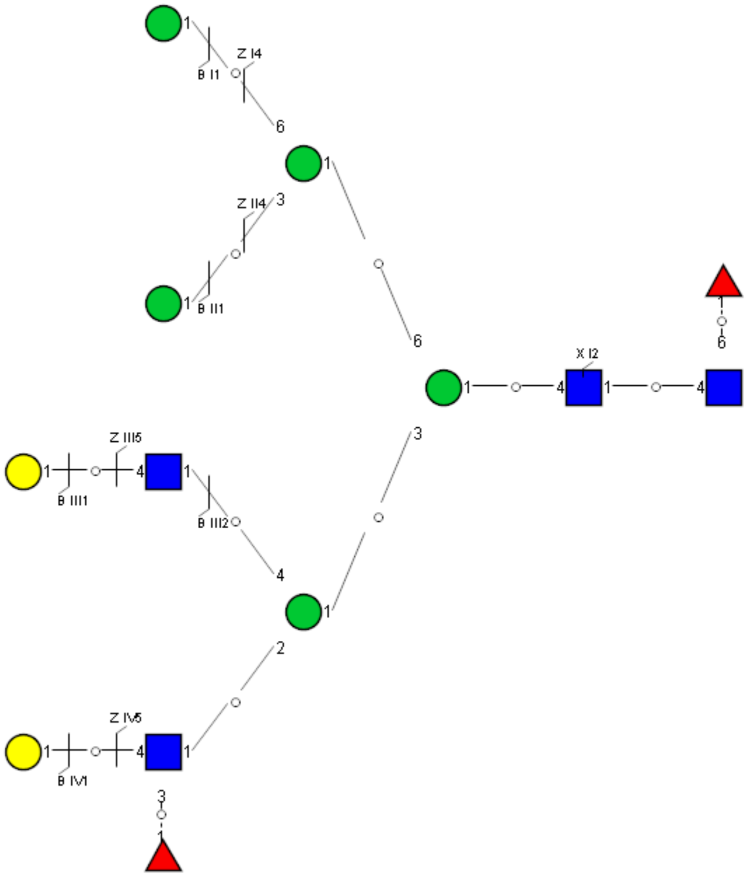

Spectrum Index: 34688(NoBHs:1)

#### Precursor iEF Map

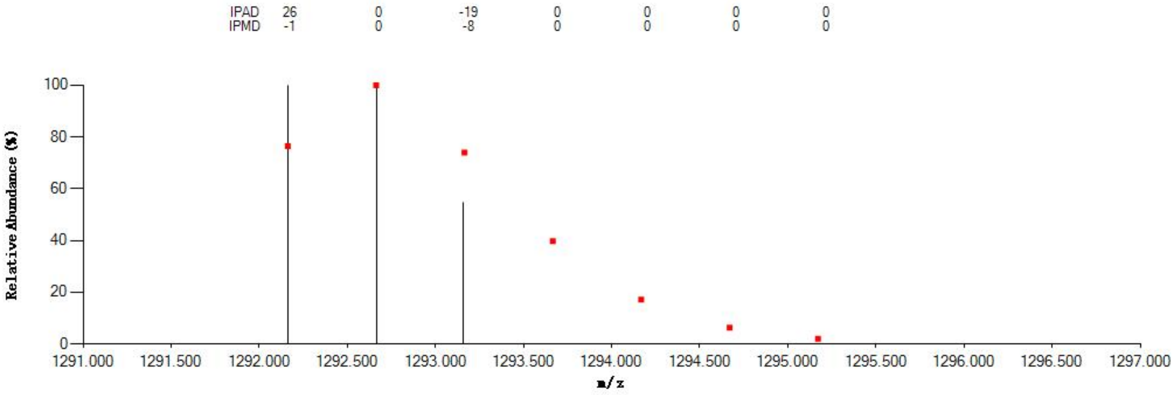


#### Annotated MS/MS Spectrum

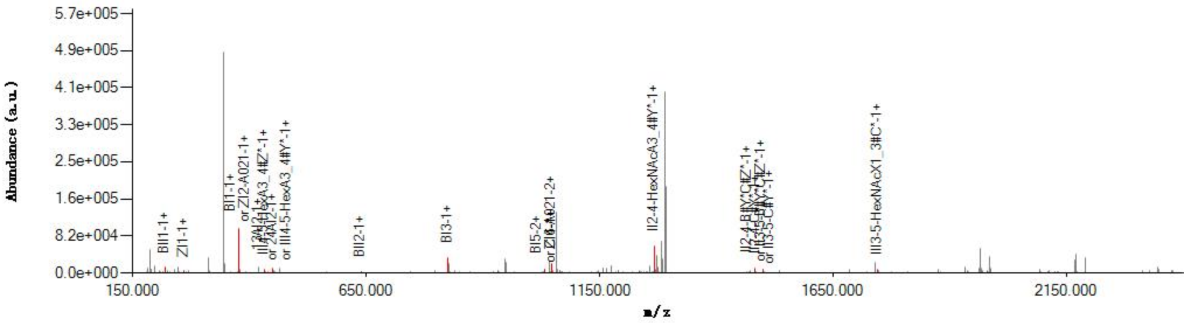


#### Graphical Fragmentation Map

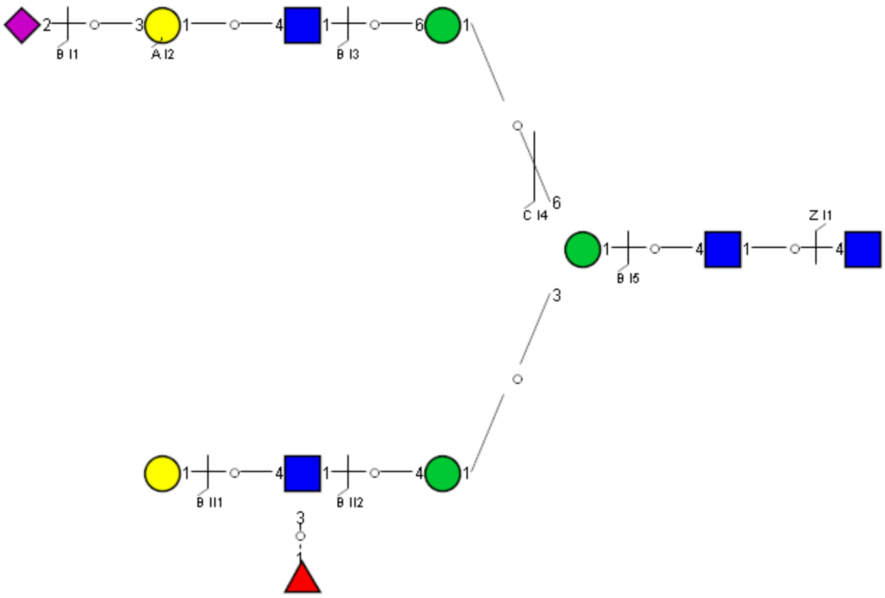

Spectrum Index: 34720(NoBHs:1)

#### Precursor iEF Map

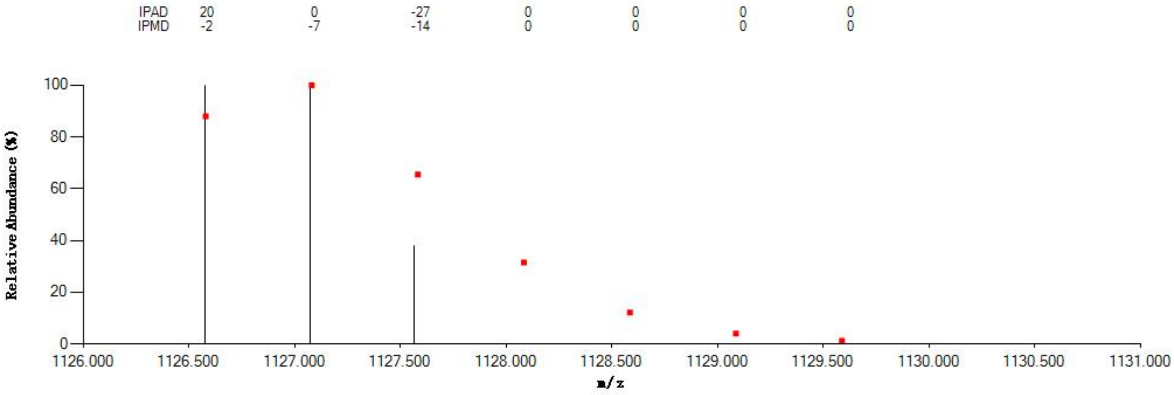


#### Annotated MS/MS Spectrum

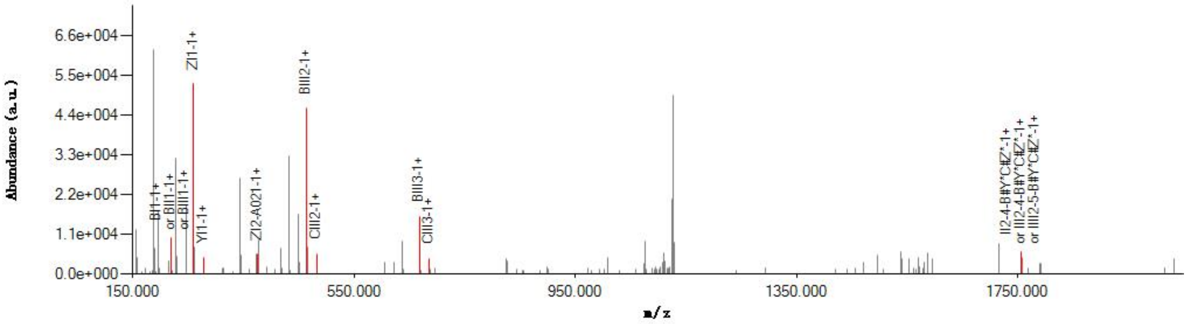


#### Graphical Fragmentation Map

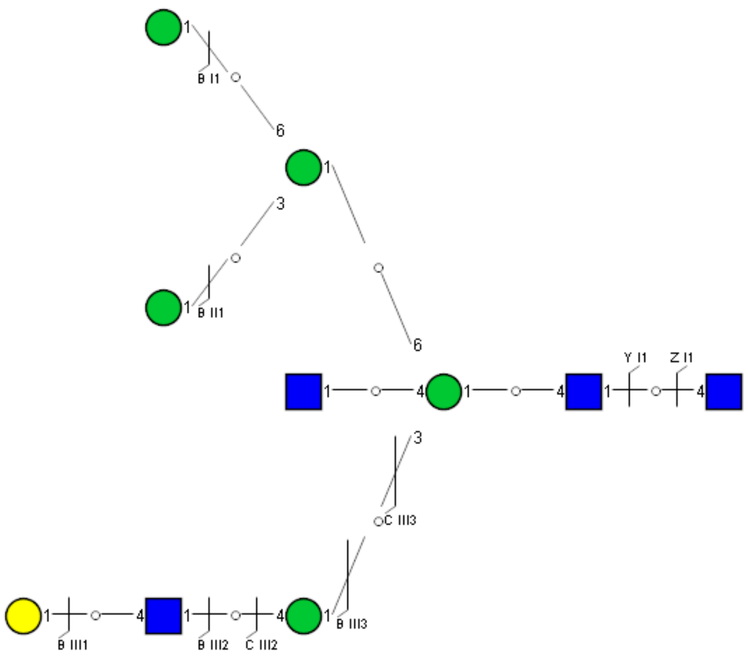

Spectrum Index: 34817(NoBHs:1)

#### Precursor iEF Map

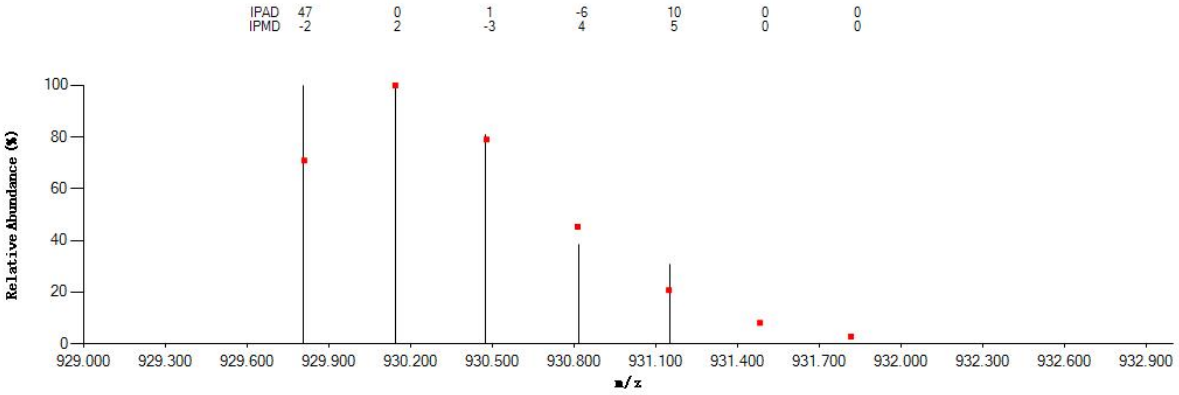


#### Annotated MS/MS Spectrum

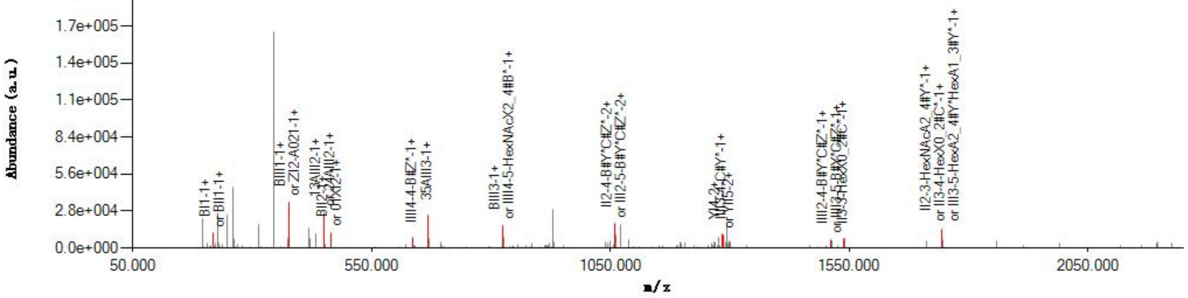


#### Graphical Fragmentation Map

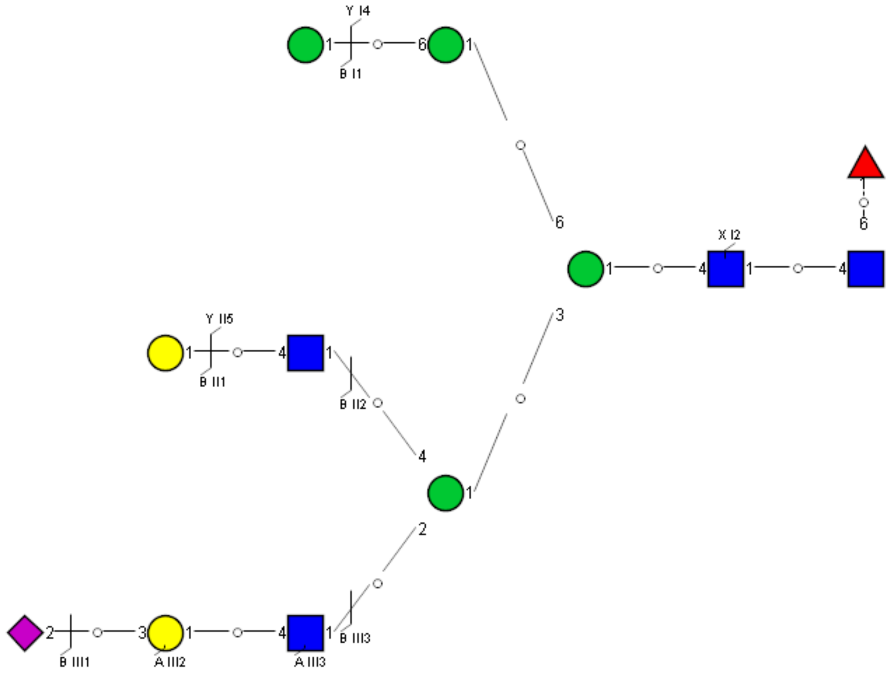

Spectrum Index: 34830(NoBHs:1)

#### Precursor iEF Map

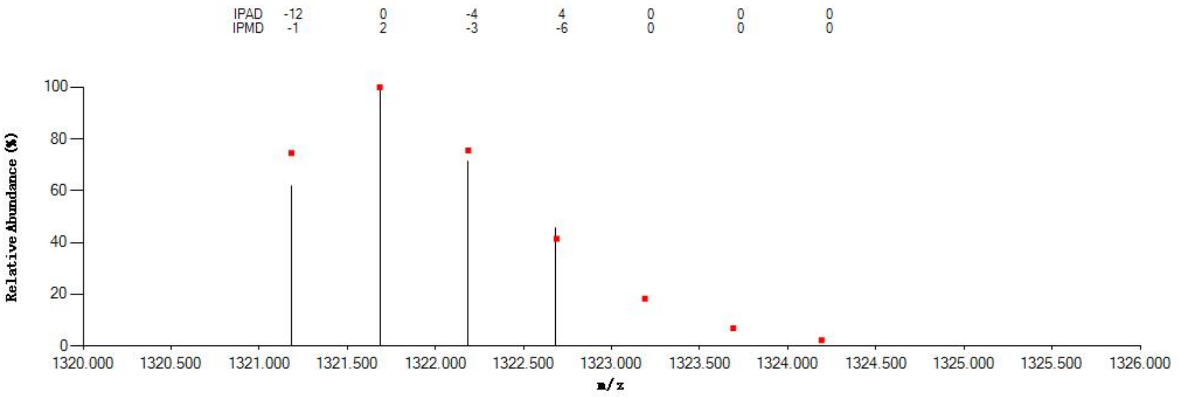


#### Annotated MS/MS Spectrum

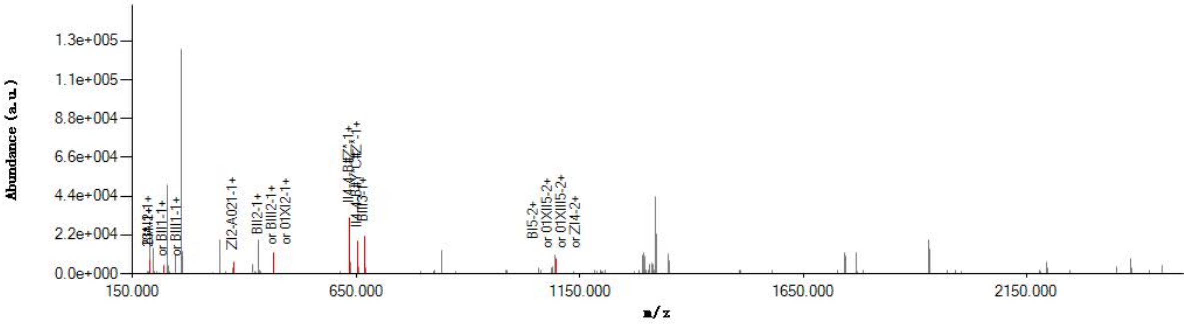


#### Graphical Fragmentation Map

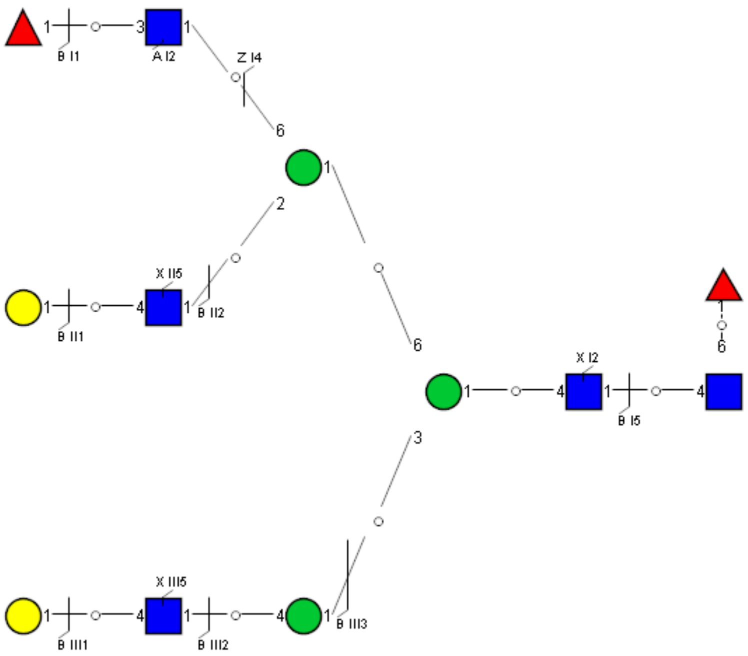

Spectrum Index: 35105(NoBHs:1)

Precursor iEF Map

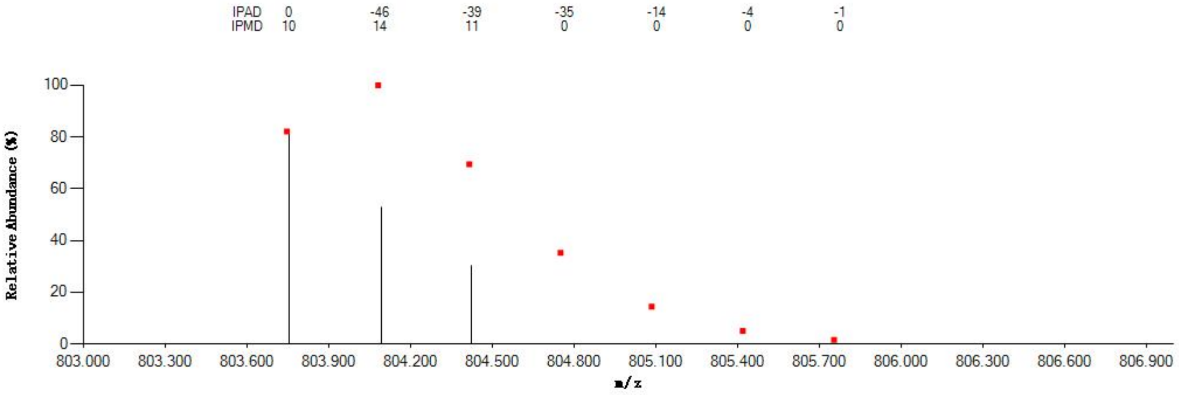


Annotated MS/MS Spectrum

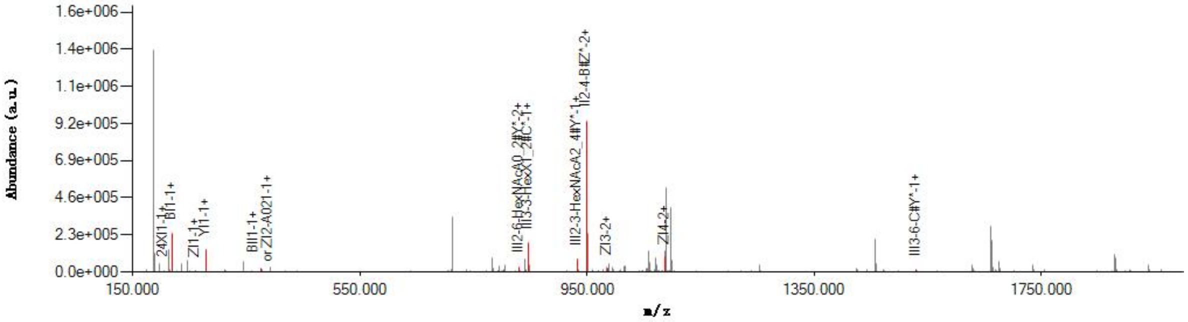


Graphical Fragmentation Map

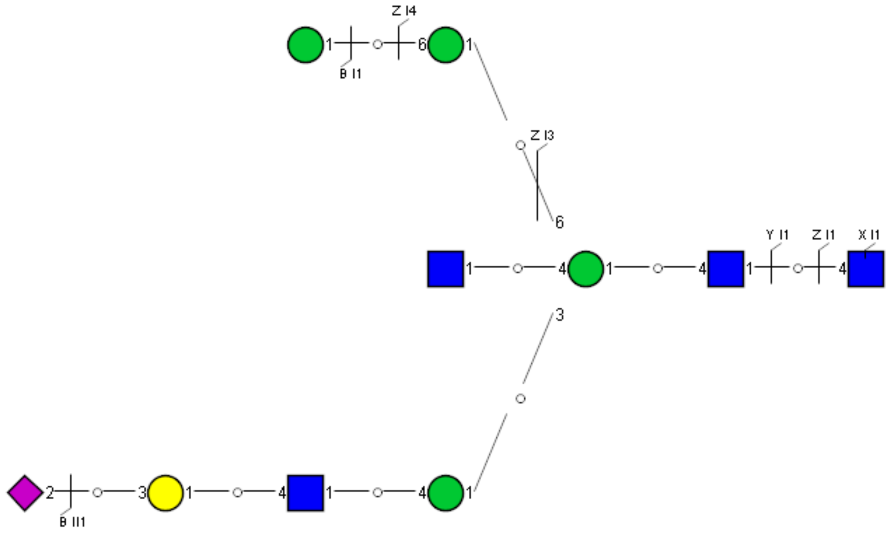

Spectrum Index: 35828(NoBHs:1)

#### Precursor iEF Map

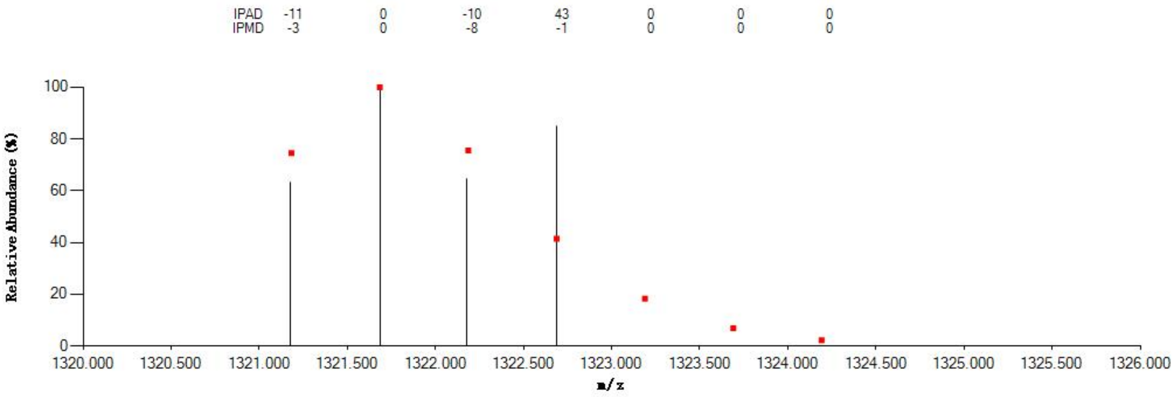

#### Annotated MS/MS Spectrum

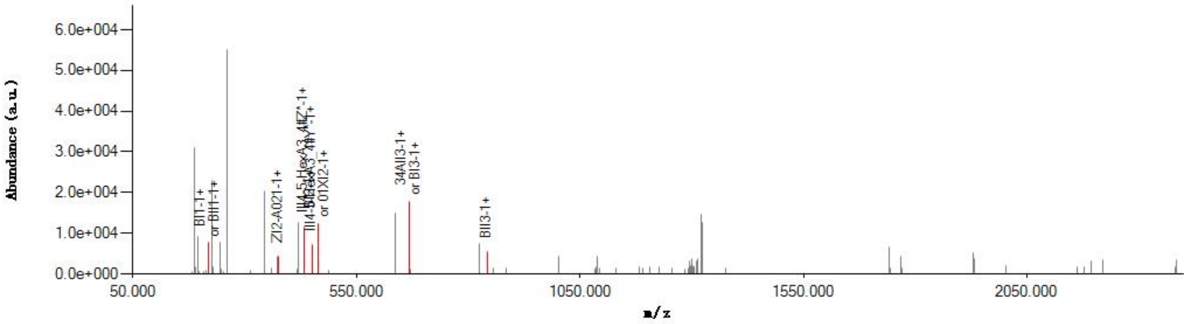

#### Graphical Fragmentation Map

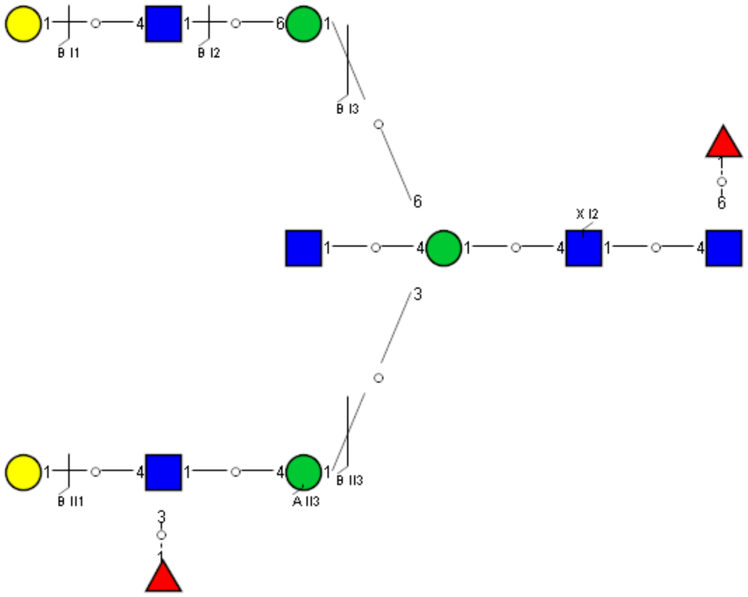

Spectrum Index: 36031(NoBHs:1)

#### Precursor iEF Map

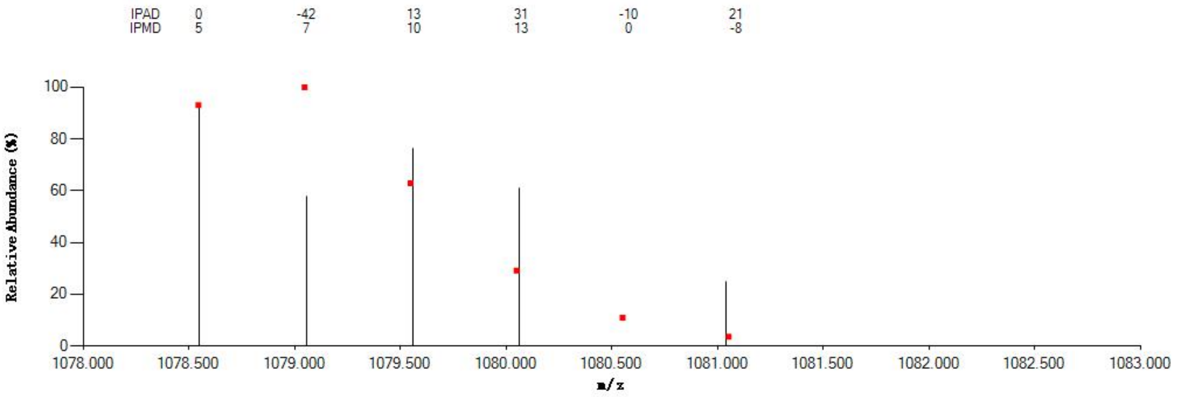


#### Annotated MS/MS Spectrum

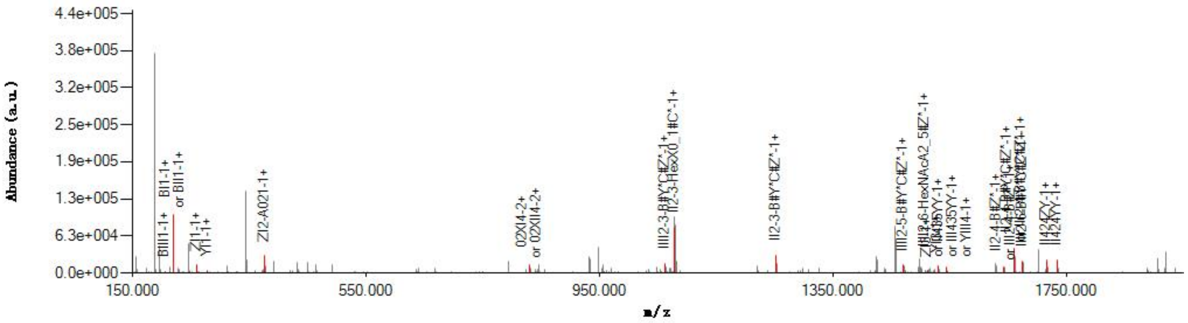


#### Graphical Fragmentation Map

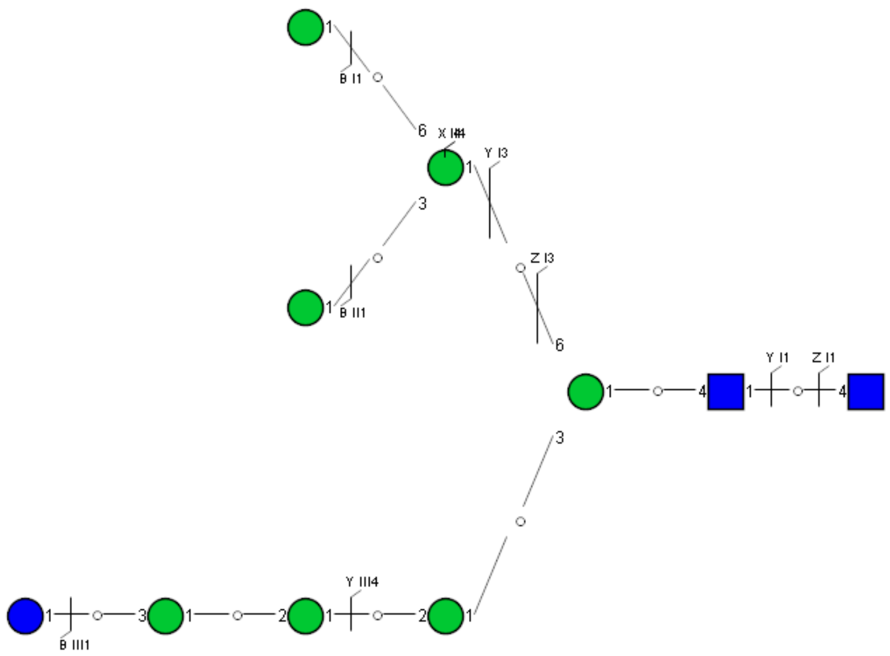

### Spectrum Index: 37640(NoBHs:1)

#### Precursor iEF Map

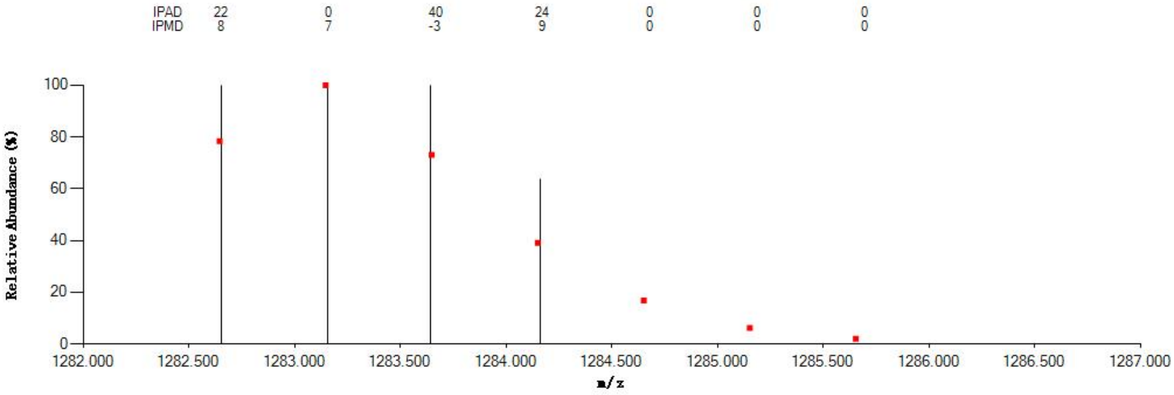


#### Annotated MS/MS Spectrum

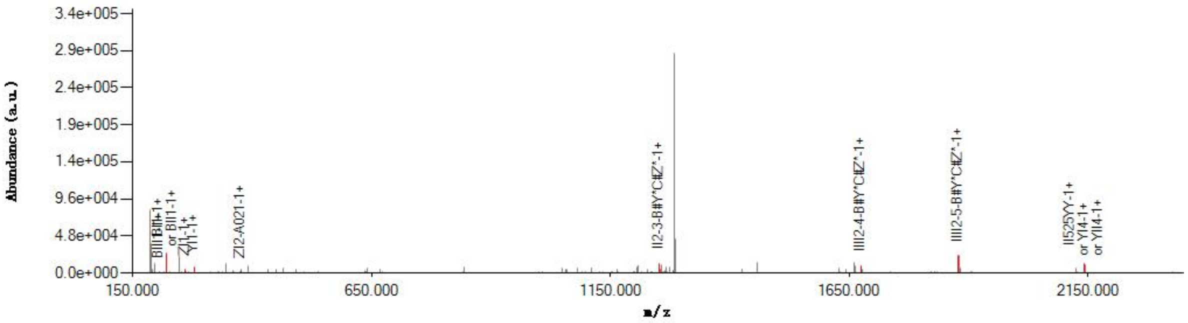


#### Graphical Fragmentation Map

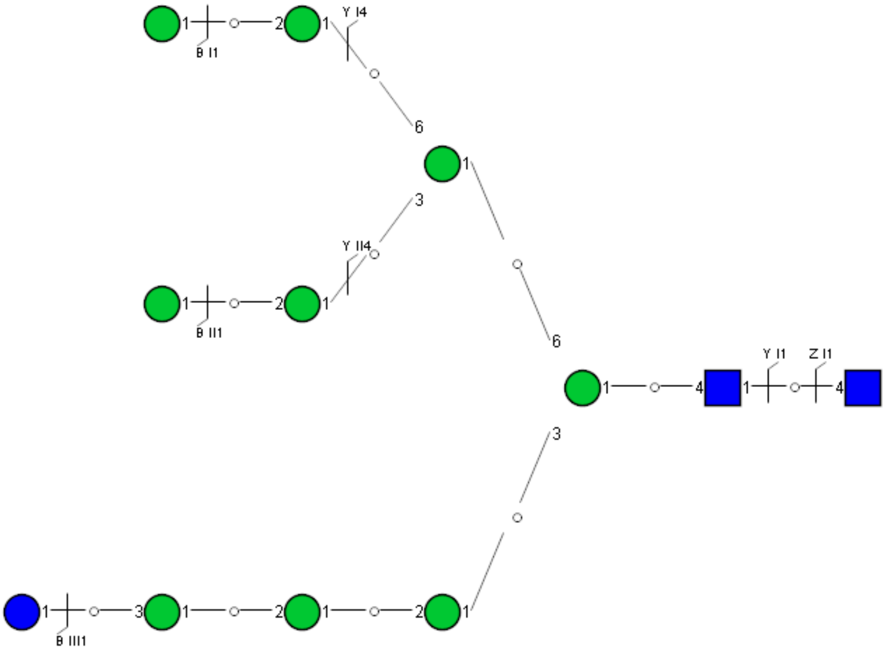

Spectrum Index: 39345(NoBHs:1)

#### Precursor iEF Map

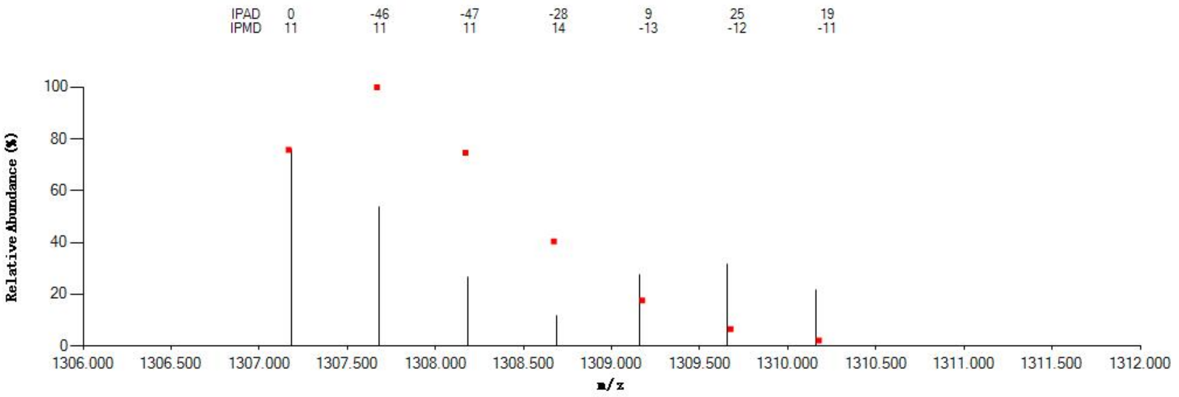


#### Annotated MS/MS Spectrum

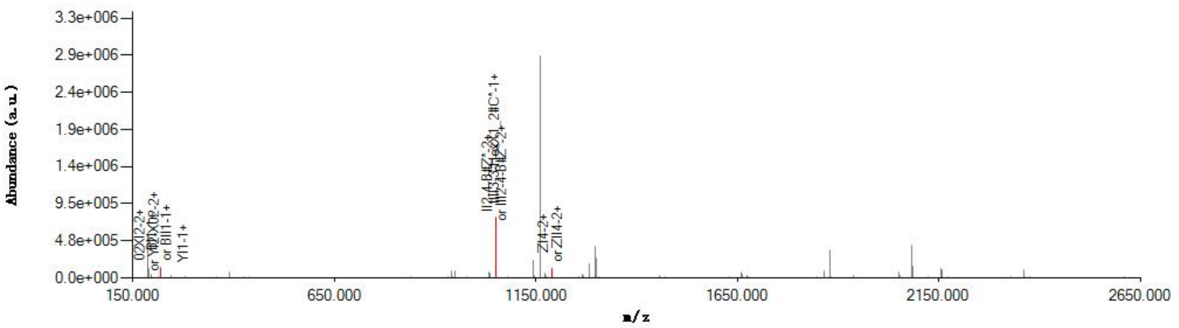


#### Graphical Fragmentation Map

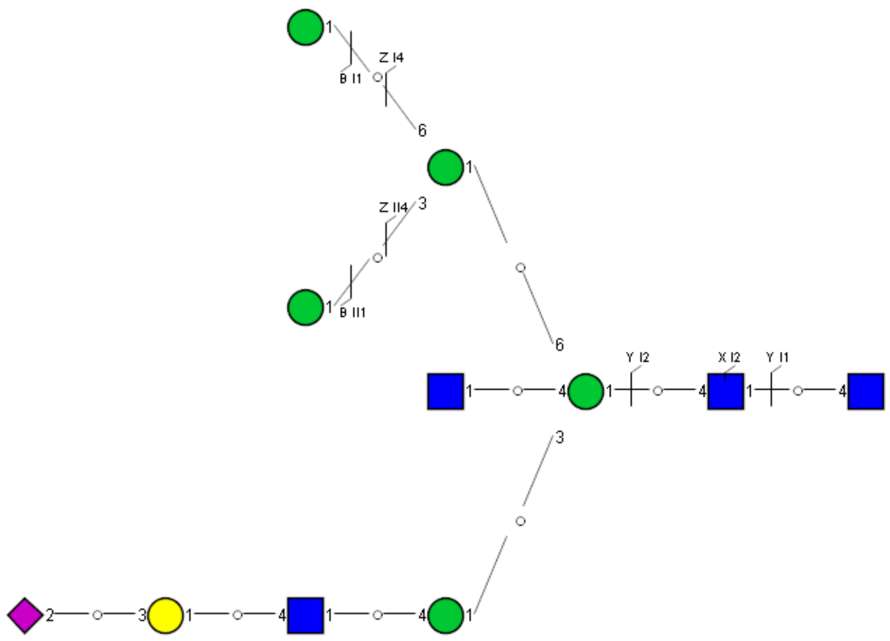

Spectrum Index: 41235(NoBHs:1)

Precursor iEF Map

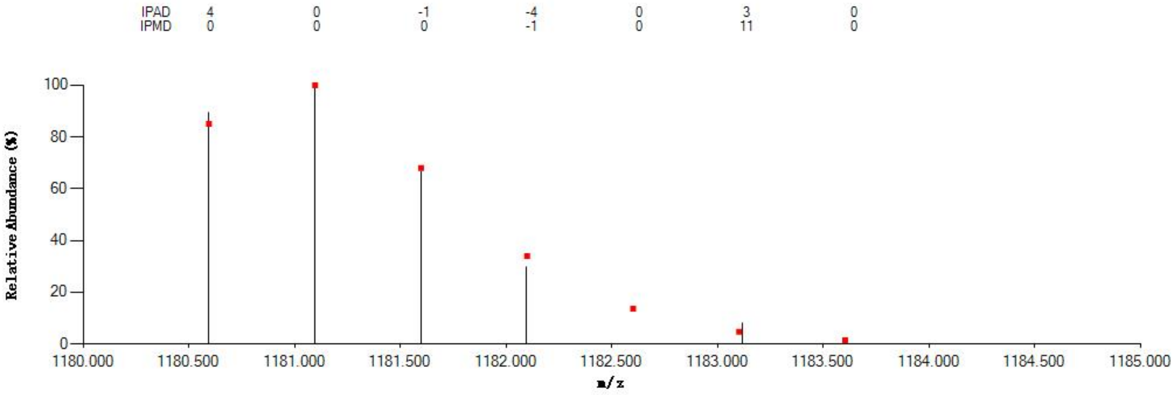


Annotated MS/MS Spectrum

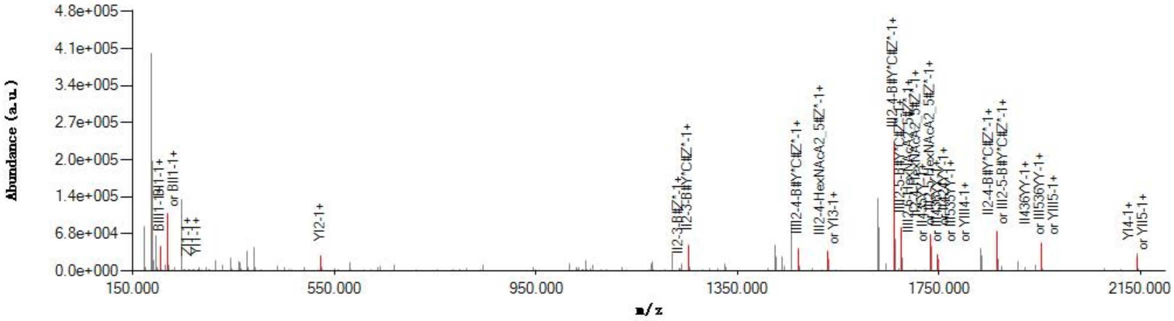


Graphical Fragmentation Map

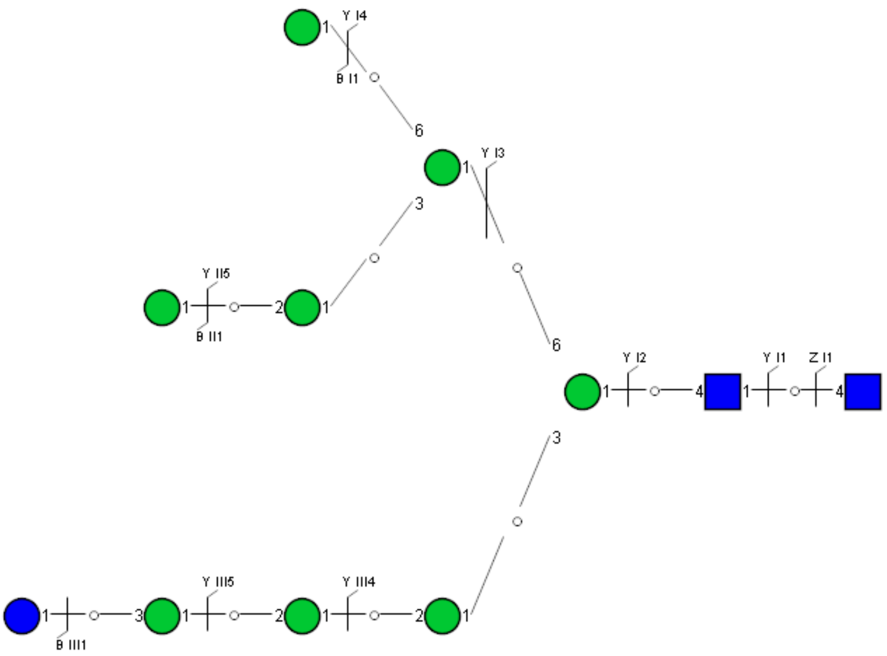

Spectrum Index: 26416(NoBHs:1)

#### Precursor iEF Map

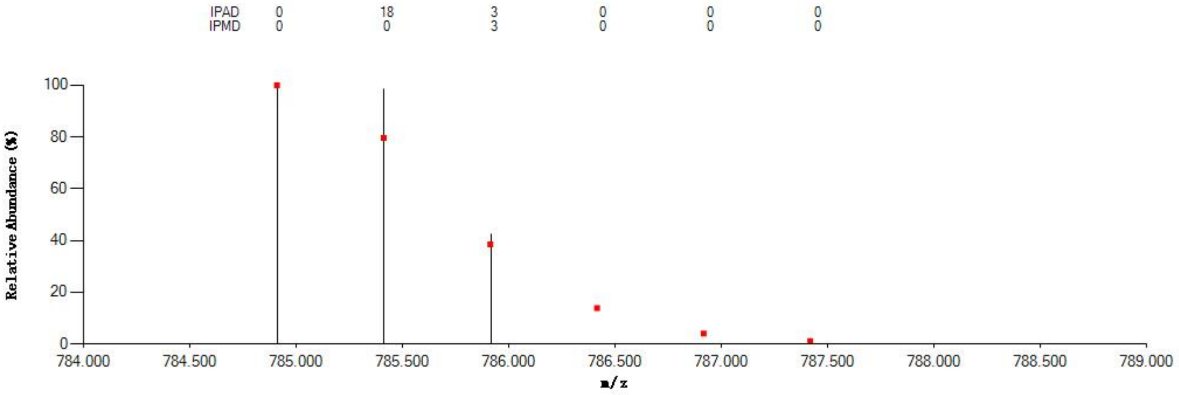


#### Annotated MS/MS Spectrum

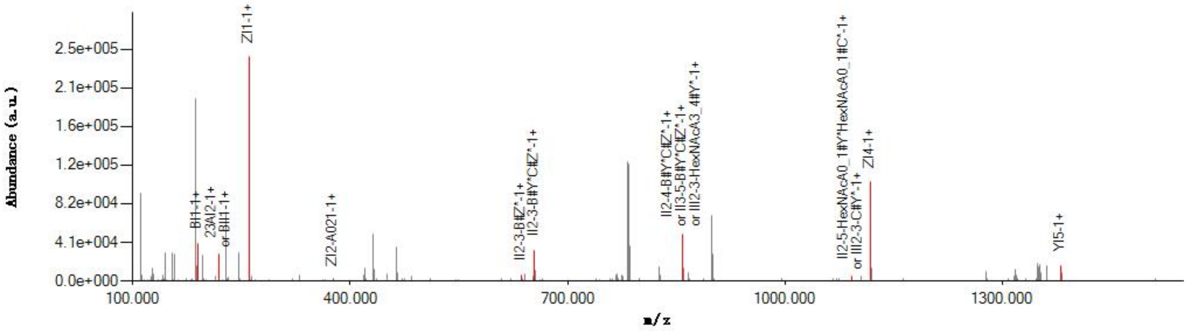


#### Graphical Fragmentation Map

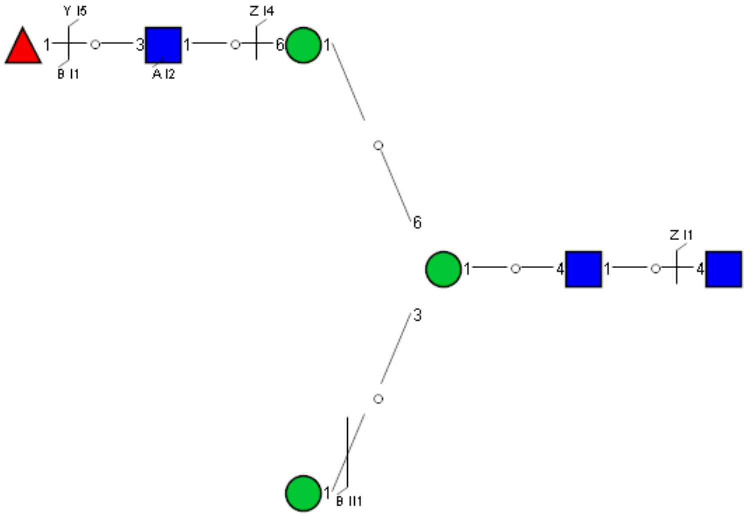

Spectrum Index: 26946(NoBHs:1)

#### Precursor iEF Map

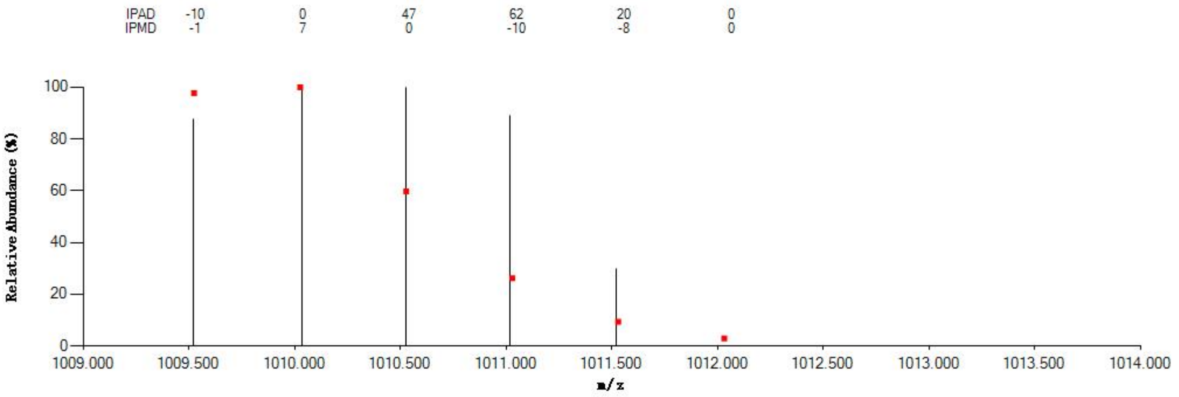


#### Annotated MS/MS Spectrum

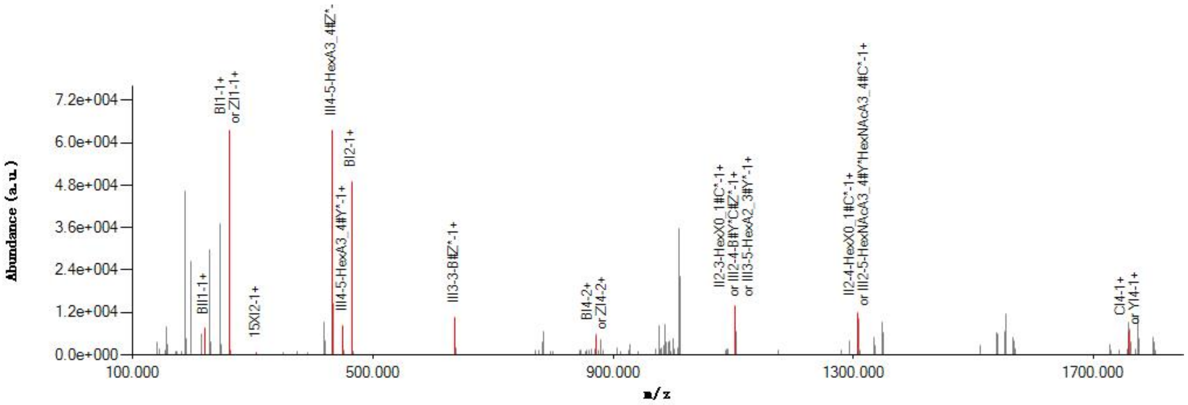


#### Graphical Fragmentation Map

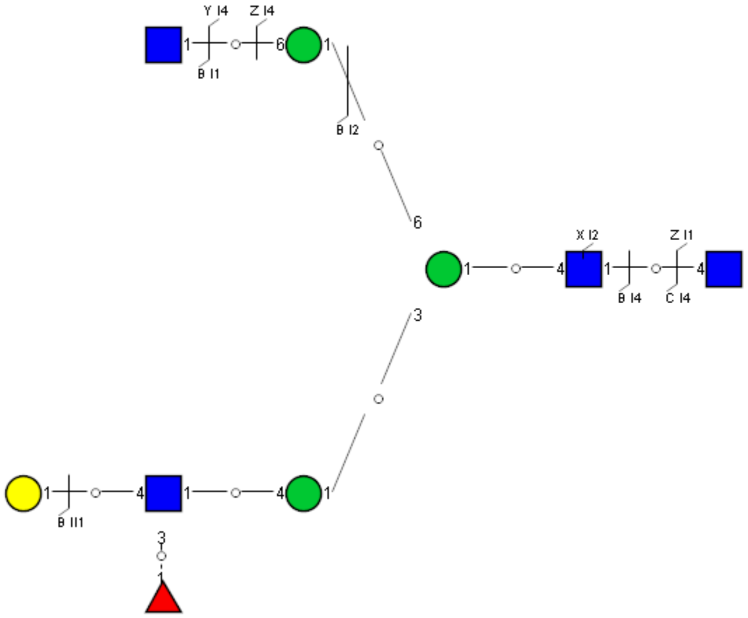

Spectrum Index: 27365(NoBHs:1)

#### Precursor iEF Map

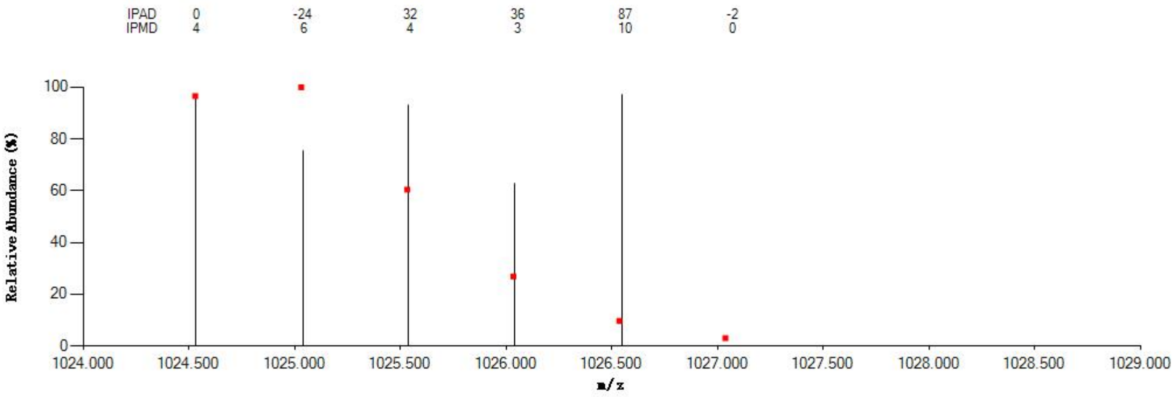

#### Annotated MS/MS Spectrum

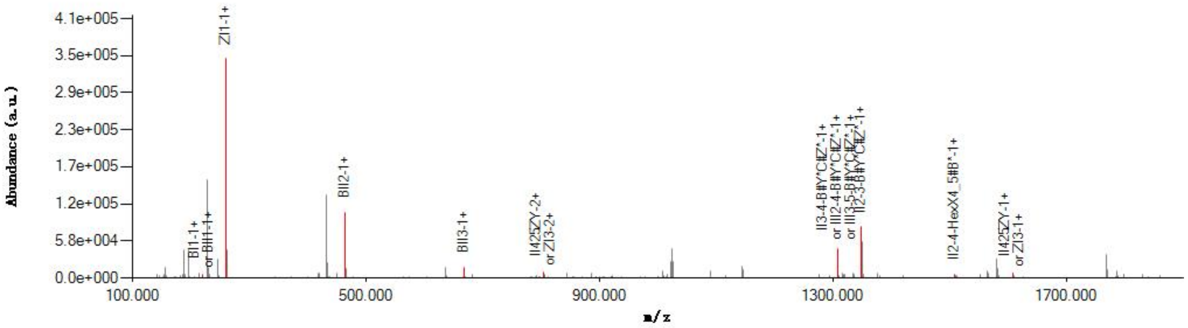

#### Graphical Fragmentation Map

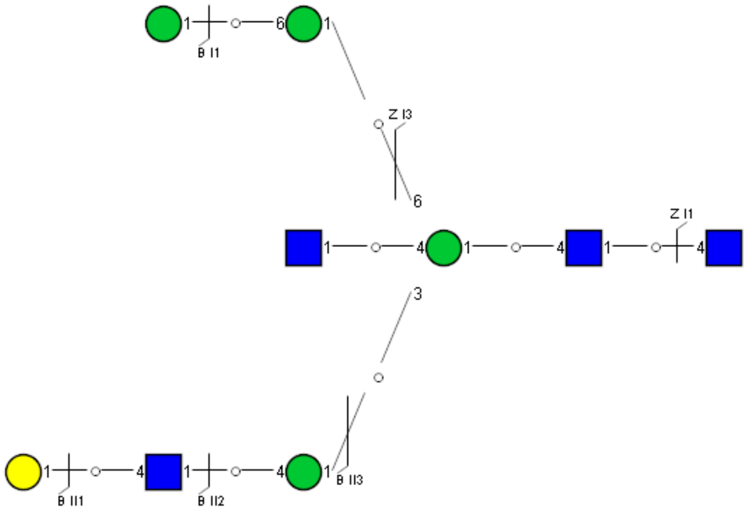

Spectrum Index: 27546(NoBHs:1)

#### Precursor iEF Map

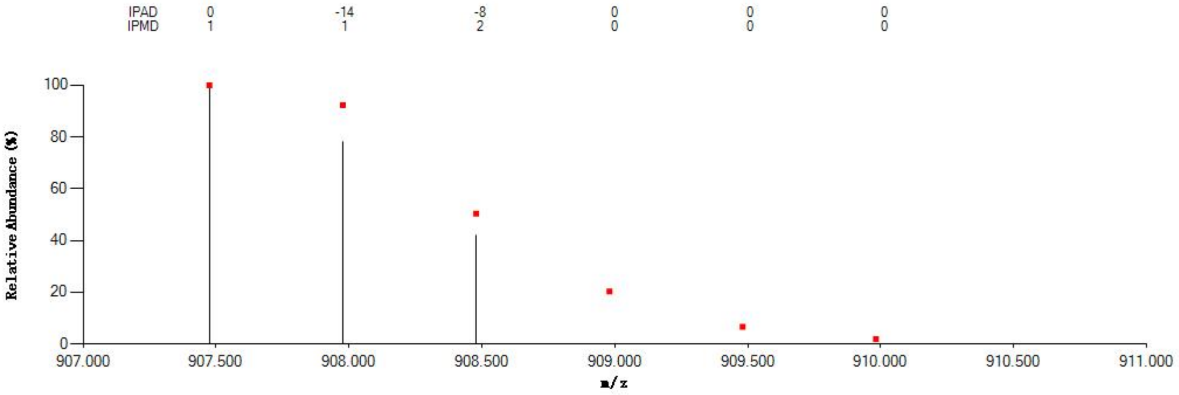


#### Annotated MS/MS Spectrum

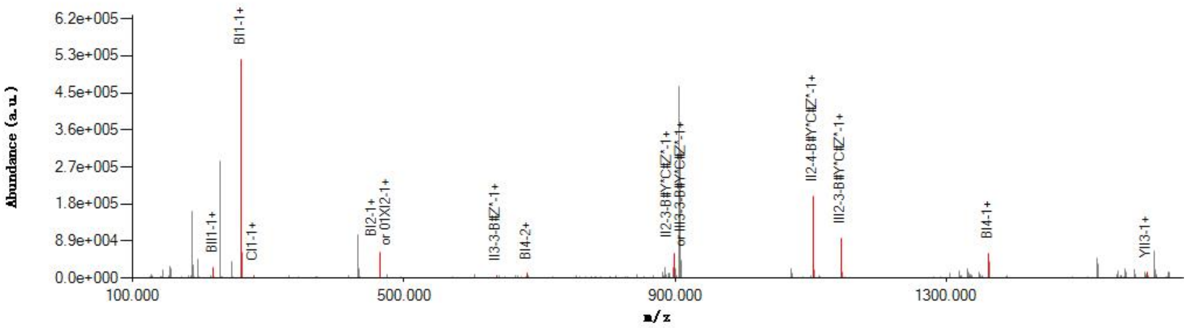


#### Graphical Fragmentation Map

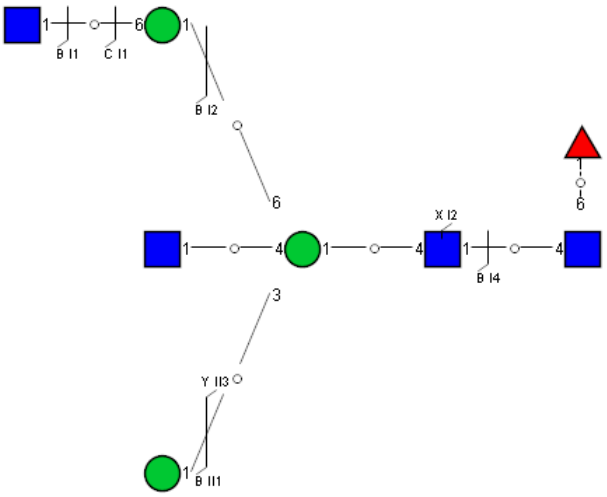

Spectrum Index: 28244(NoBHs:1)

#### Precursor iEF Map

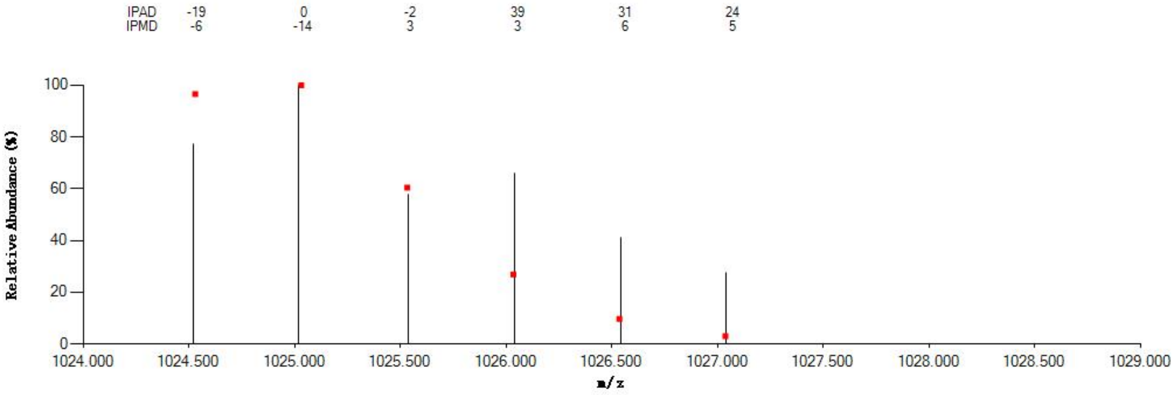


#### Annotated MS/MS Spectrum

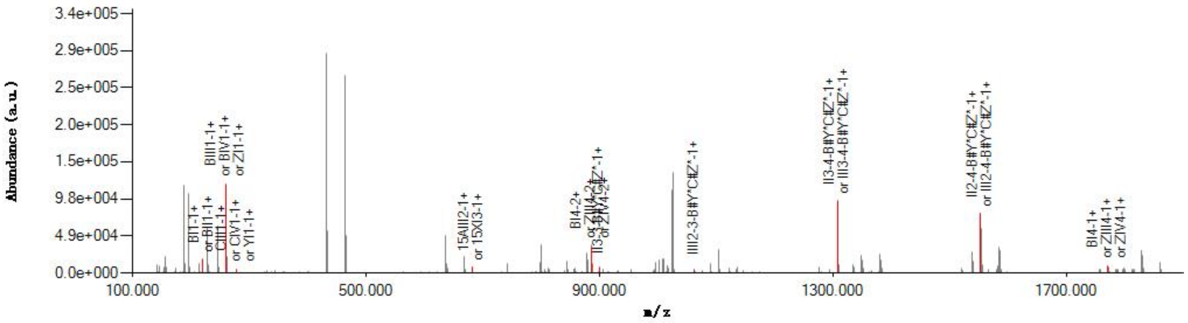


#### Graphical Fragmentation Map

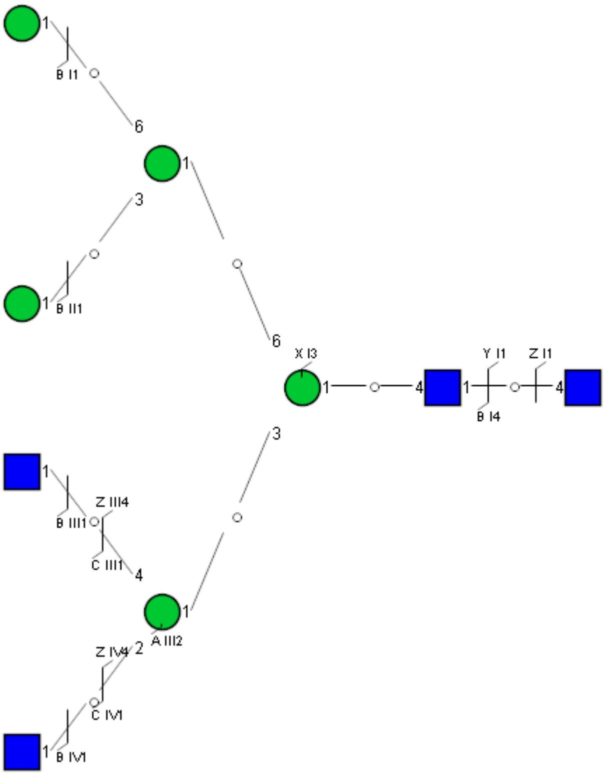

Spectrum Index: 28537(NoBHs:1)

#### Precursor iEF Map

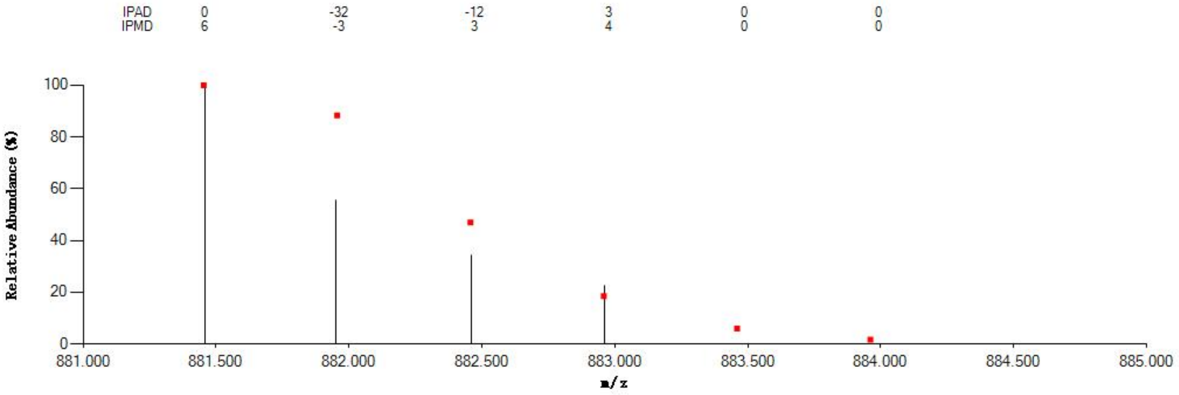


#### Annotated MS/MS Spectrum

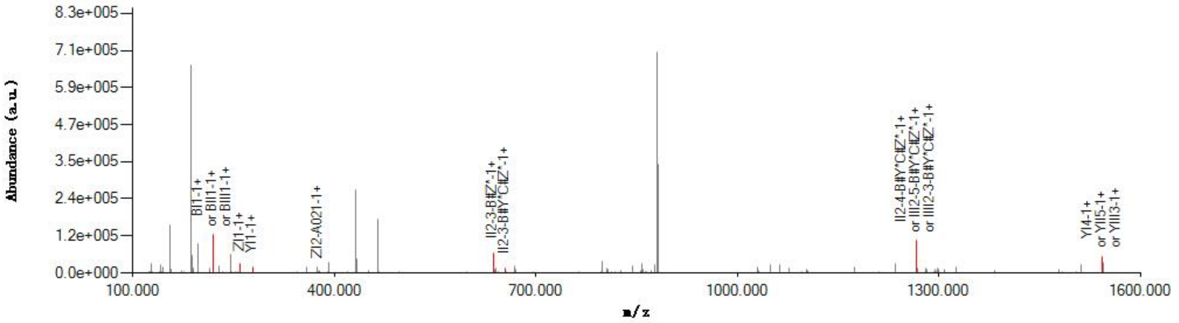


#### Graphical Fragmentation Map

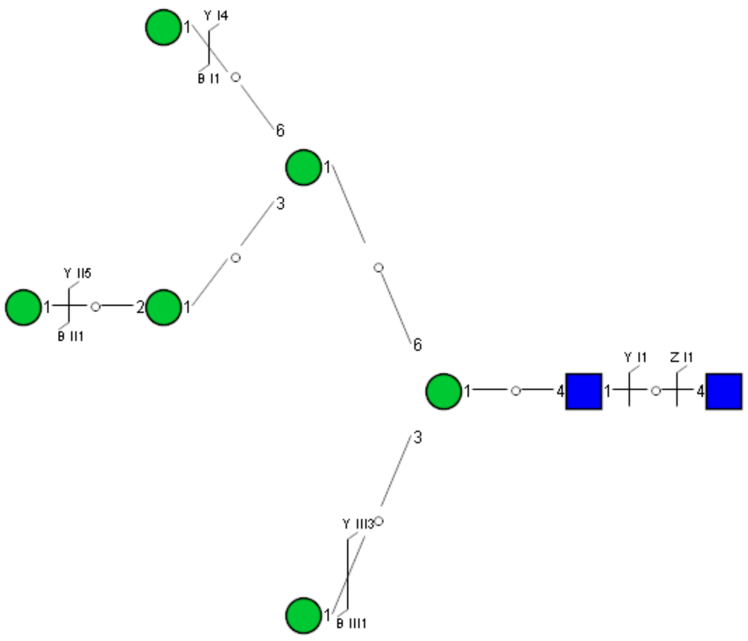

Spectrum Index: 29479(NoBHs:1)

#### Precursor iEF Map

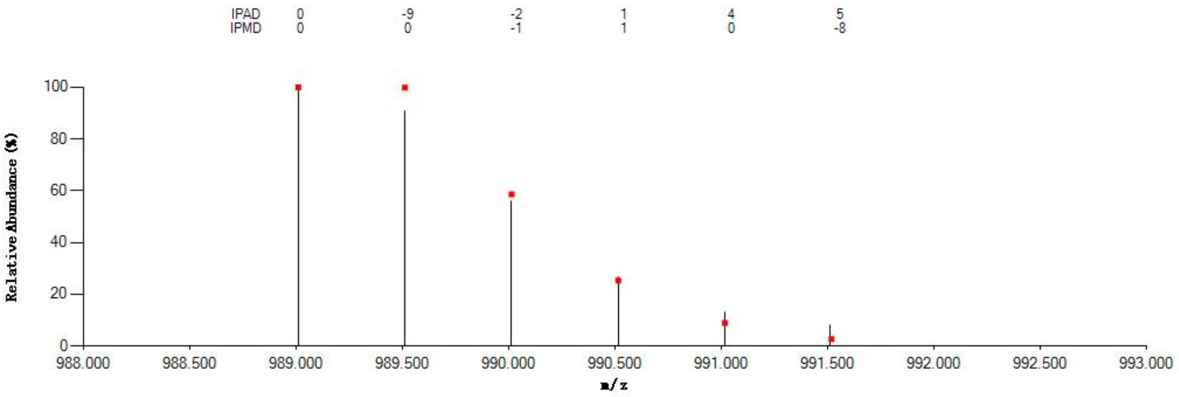


#### Annotated MS/MS Spectrum

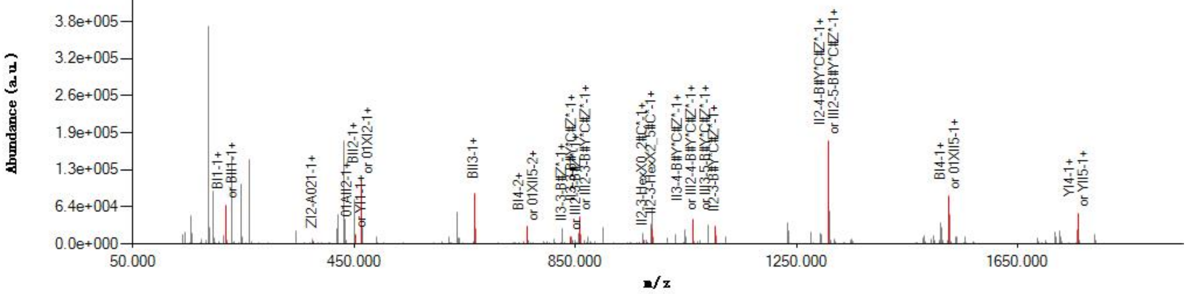


#### Graphical Fragmentation Map

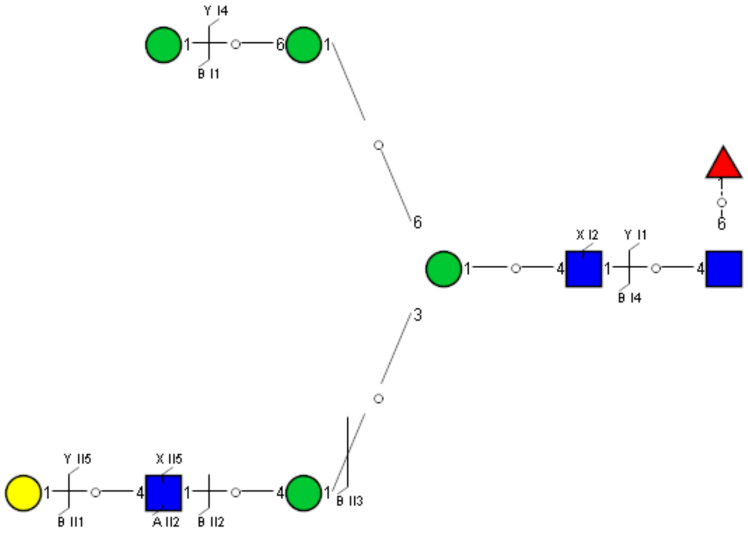

Spectrum Index: 29589(NoBHs:1)

#### Precursor iEF Map

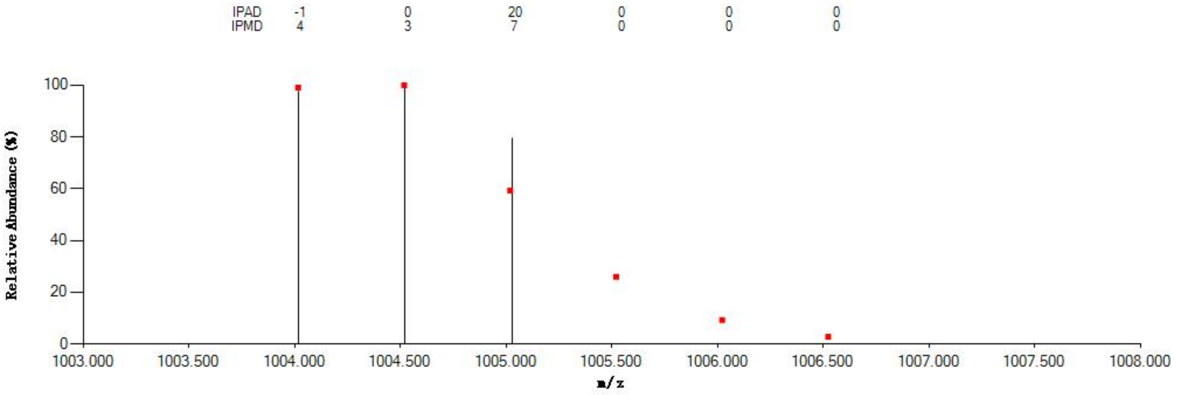


#### Annotated MS/MS Spectrum

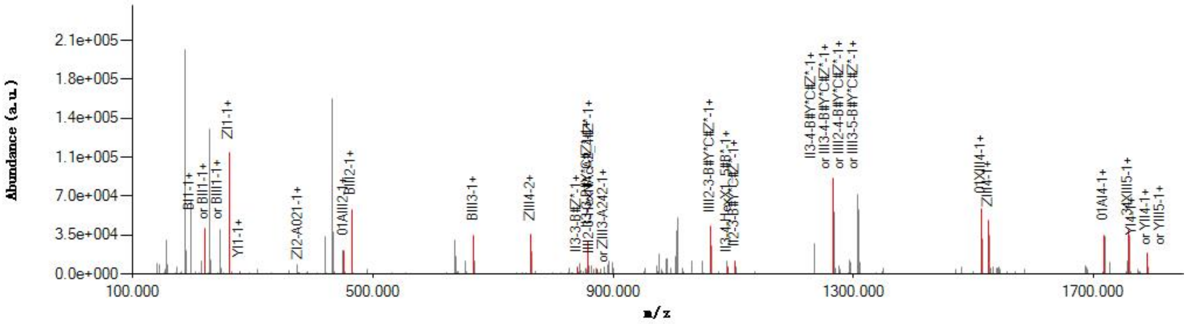


#### Graphical Fragmentation Map

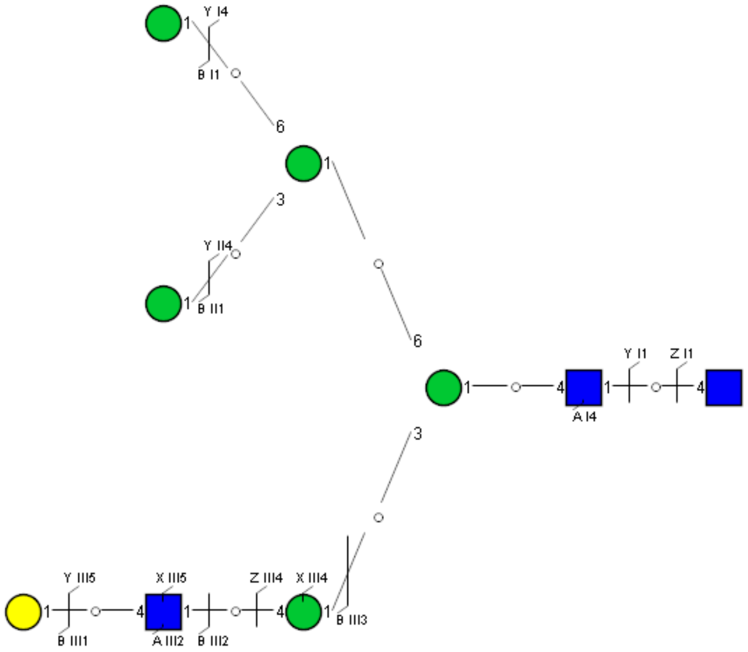

Spectrum Index: 29937(NoBHs:1)

#### Precursor iEF Map

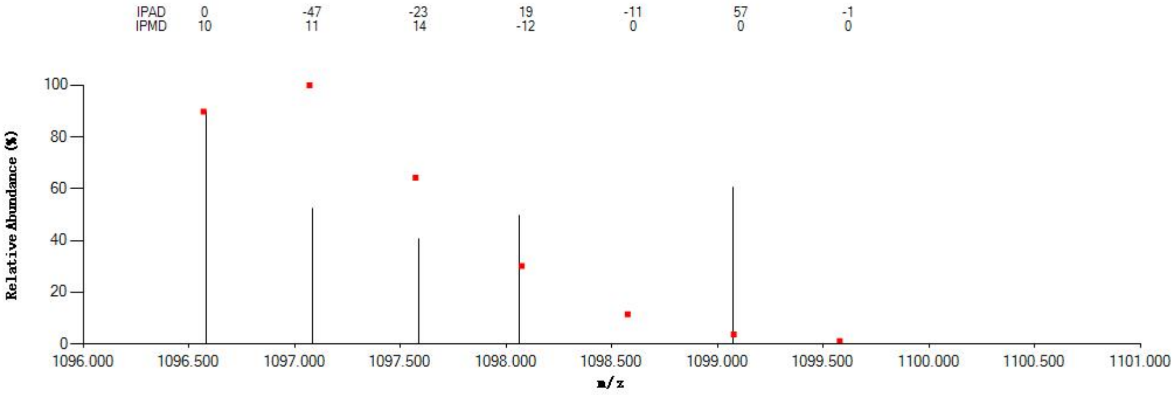

#### Annotated MS/MS Spectrum

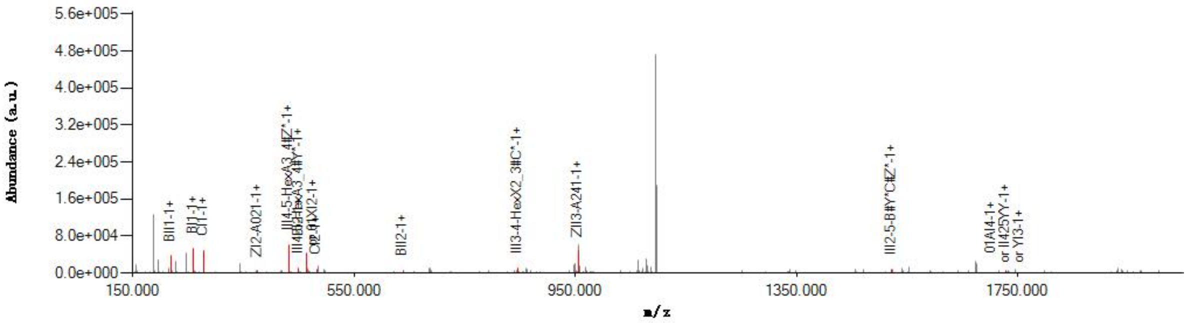

#### Graphical Fragmentation Map

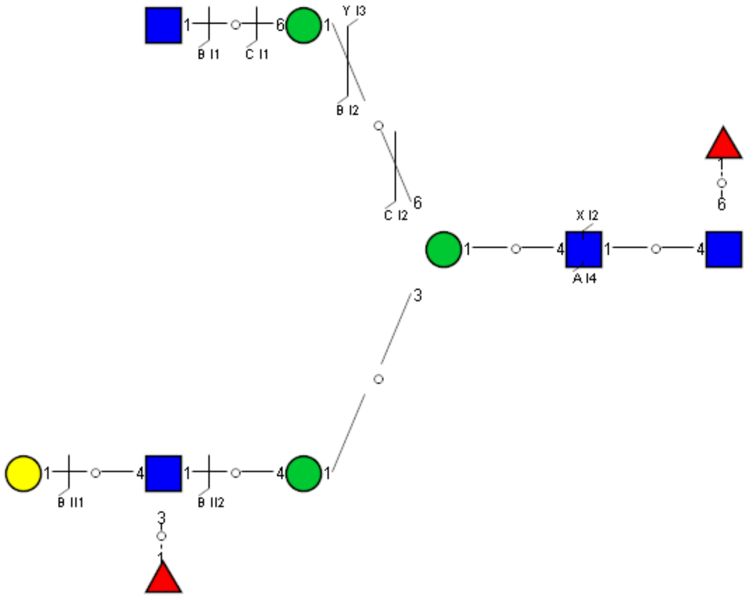

Spectrum Index: 31474(NoBHs:1)

#### Precursor iEF Map

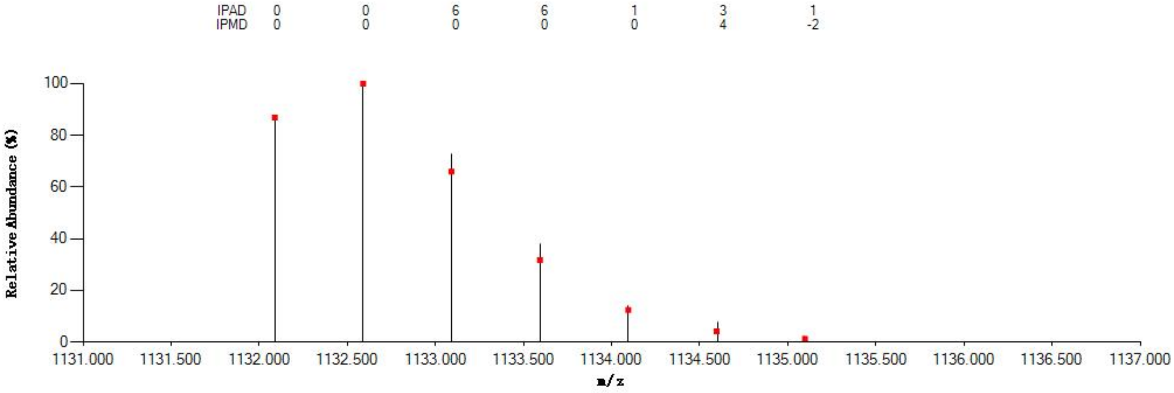


#### Annotated MS/MS Spectrum

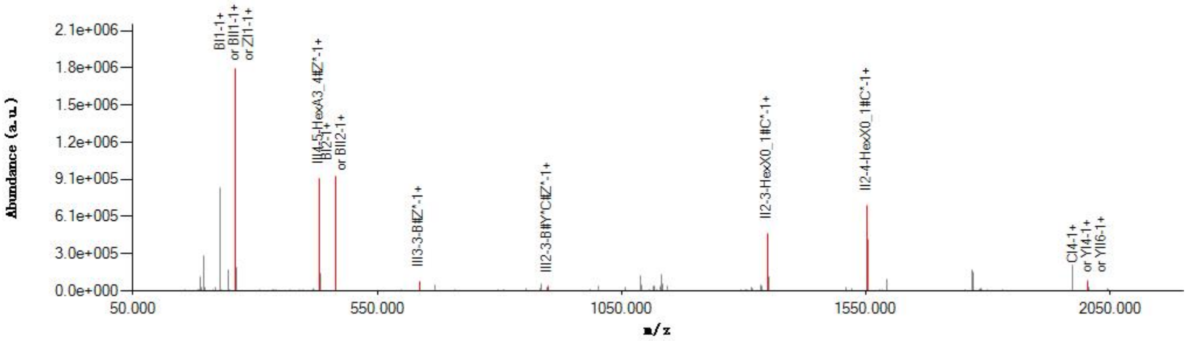


#### Graphical Fragmentation Map

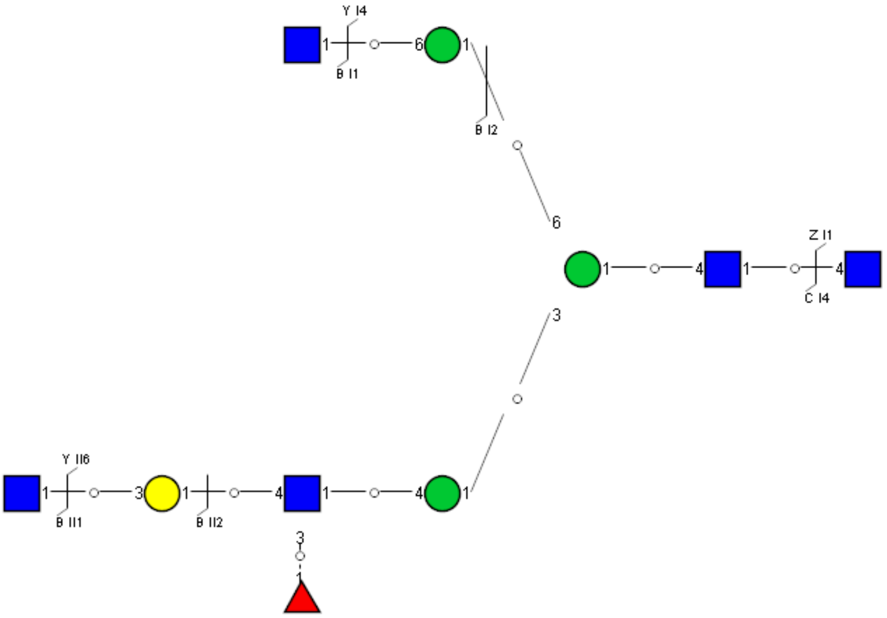

Spectrum Index: 32092(NoBHs:2)

#### Precursor iEF Map

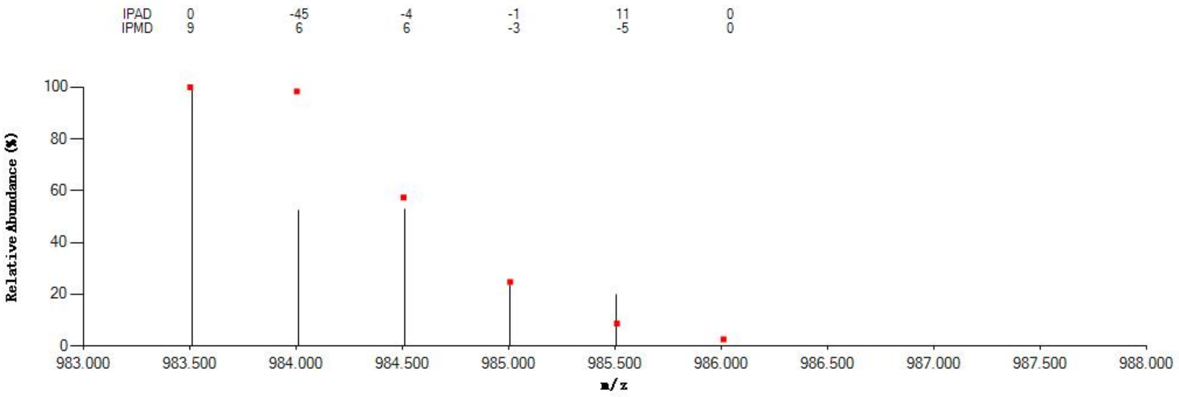


#### Annotated MS/MS Spectrum

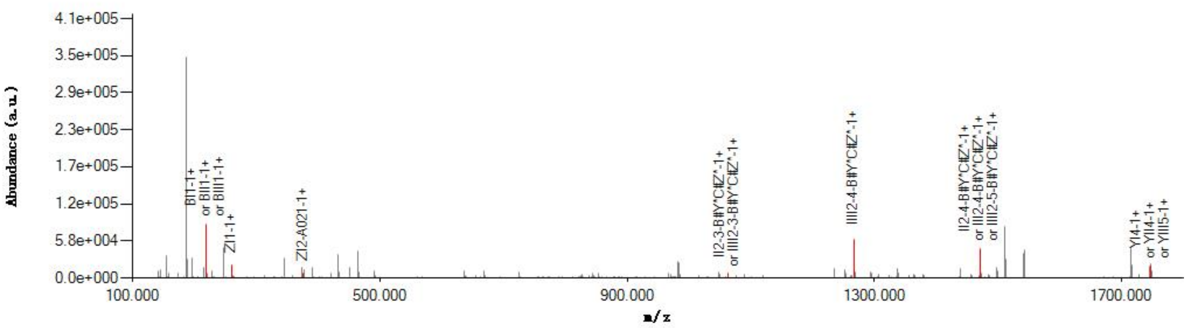


#### Graphical Fragmentation Map

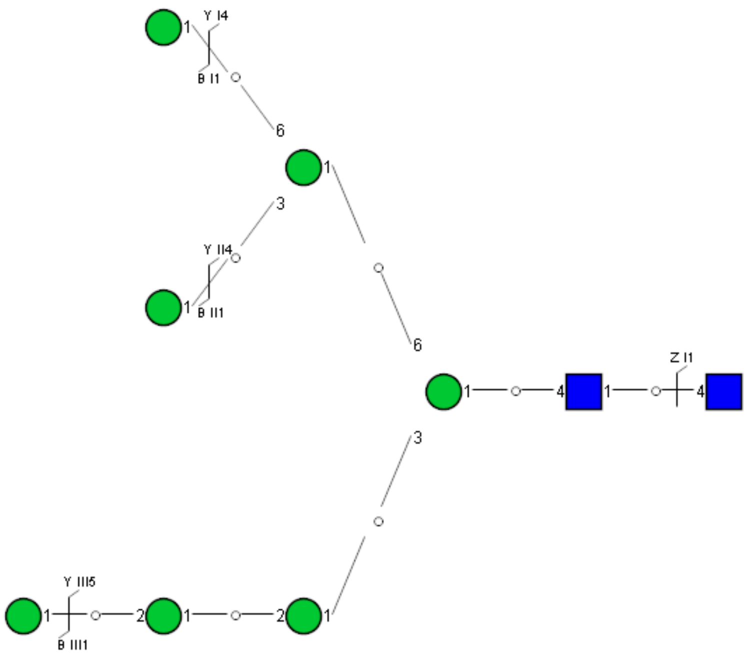

Spectrum Index: 32790(NoBHs:1)

#### Precursor iEF Map

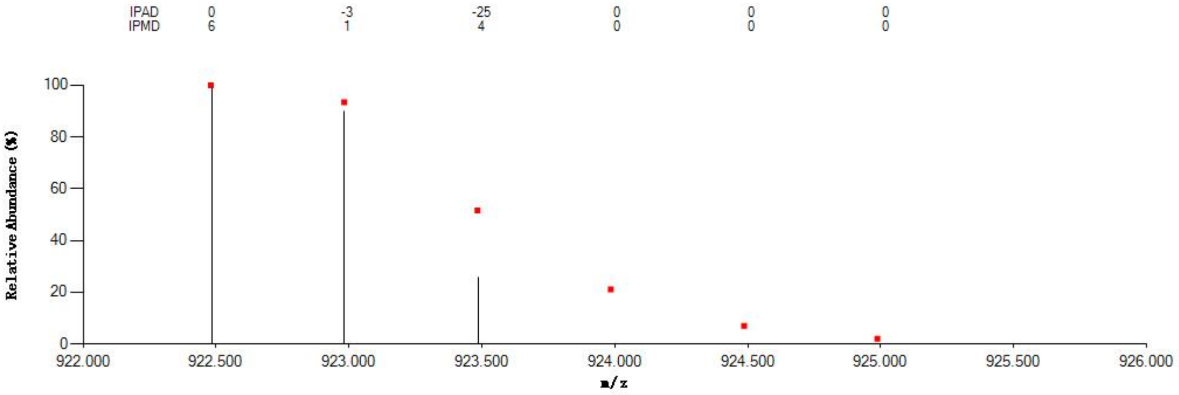


#### Annotated MS/MS Spectrum

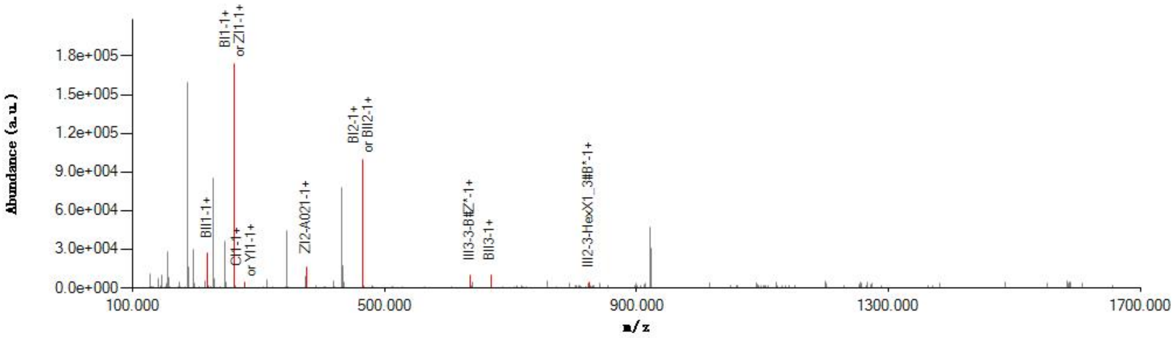


#### Graphical Fragmentation Map

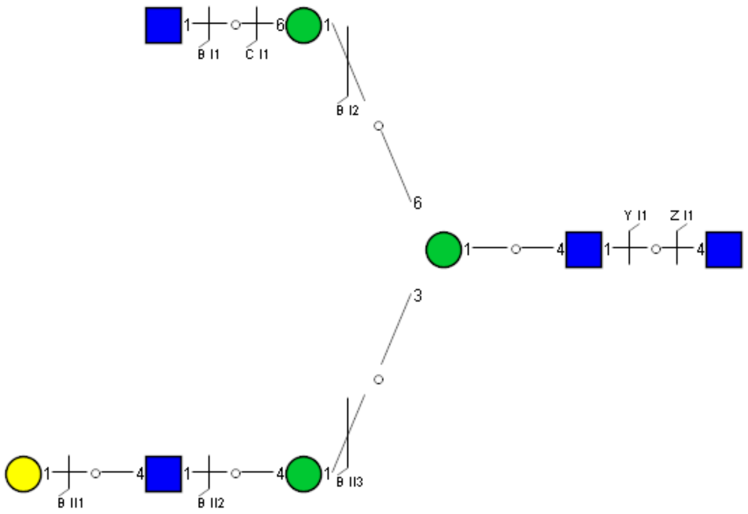

Spectrum Index: 32907(NoBHs:1)

#### Precursor iEF Map

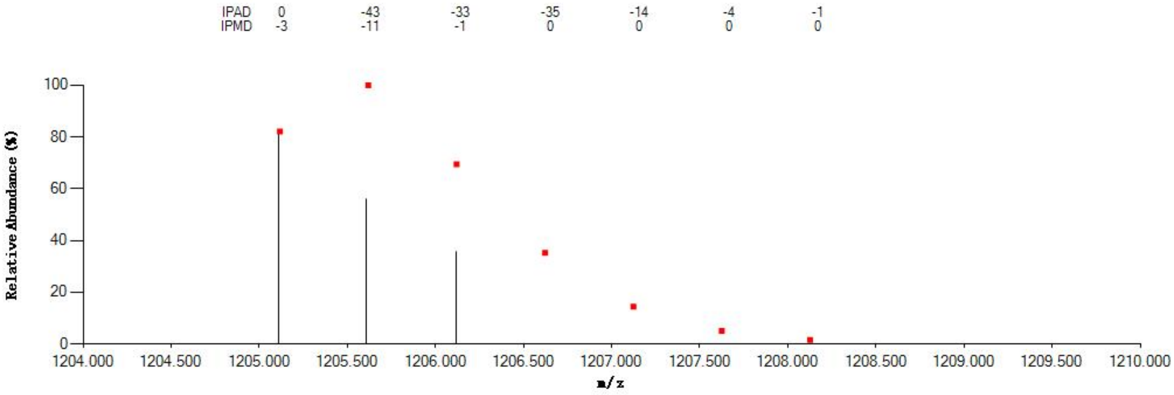

#### Annotated MS/MS Spectrum

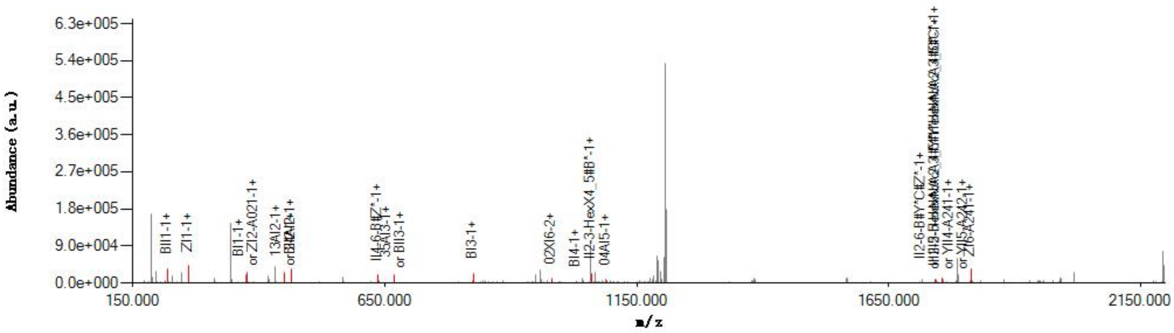

#### Graphical Fragmentation Map

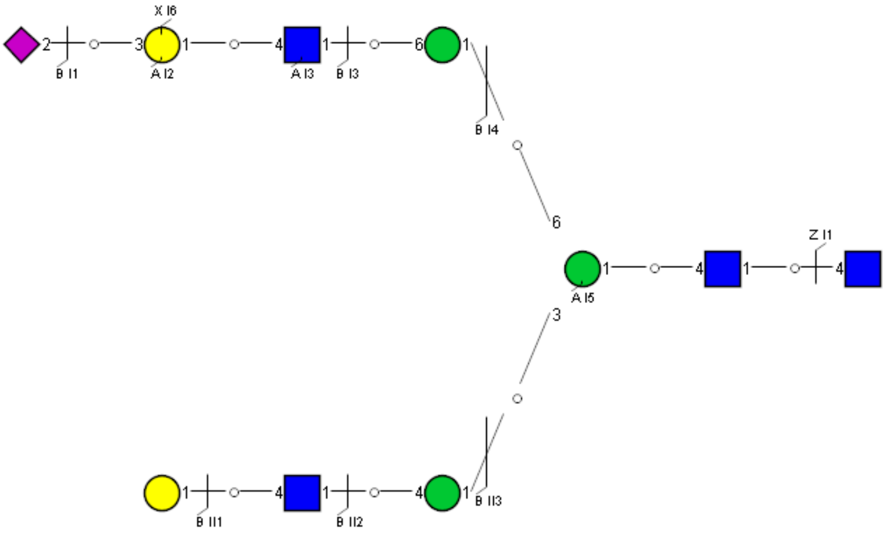

Spectrum Index: 33707(NoBHs:1)

#### Precursor iEF Map

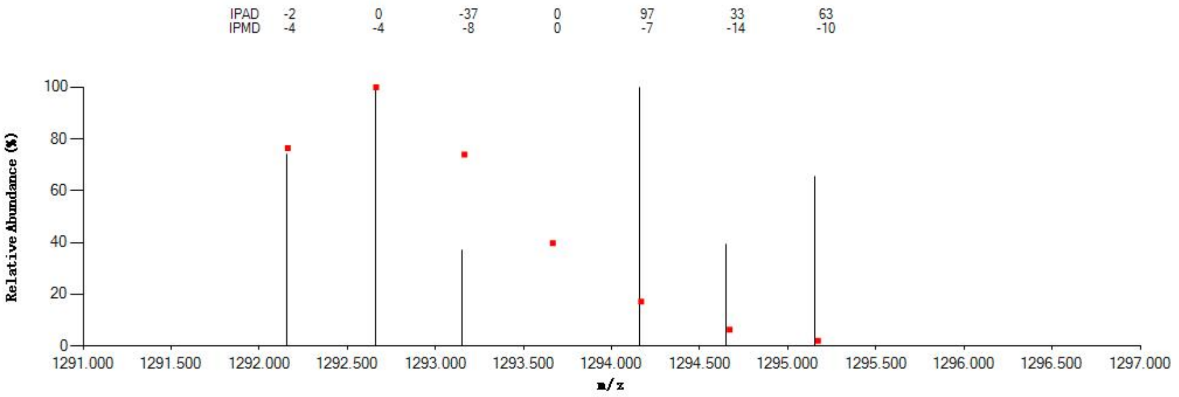


#### Annotated MS/MS Spectrum

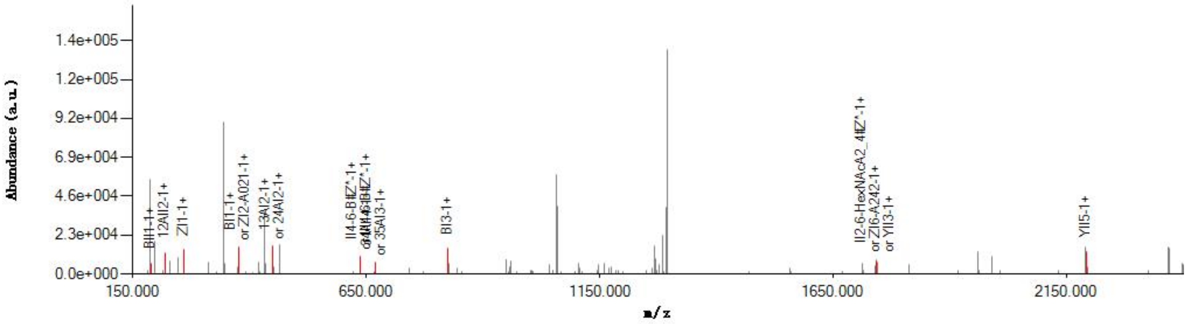


#### Graphical Fragmentation Map

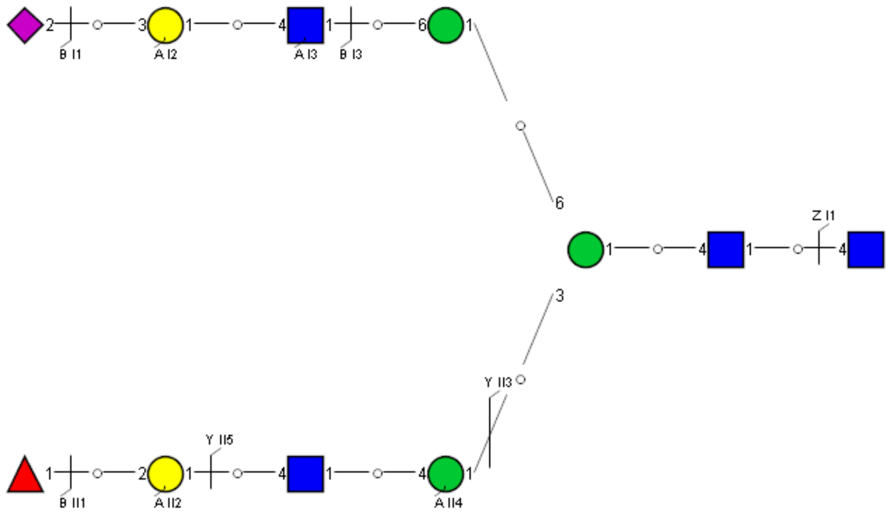

Spectrum Index: 34476(NoBHs:1)

#### Precursor iEF Map

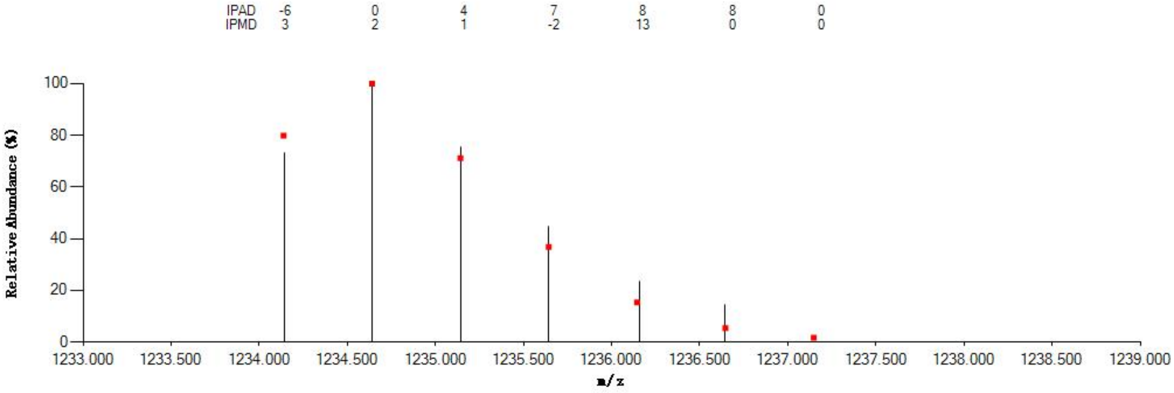

#### Annotated MS/MS Spectrum

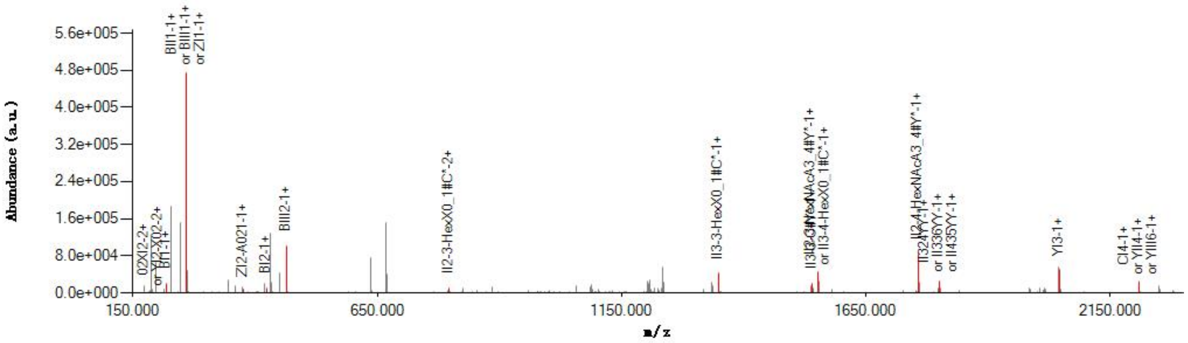

#### Graphical Fragmentation Map

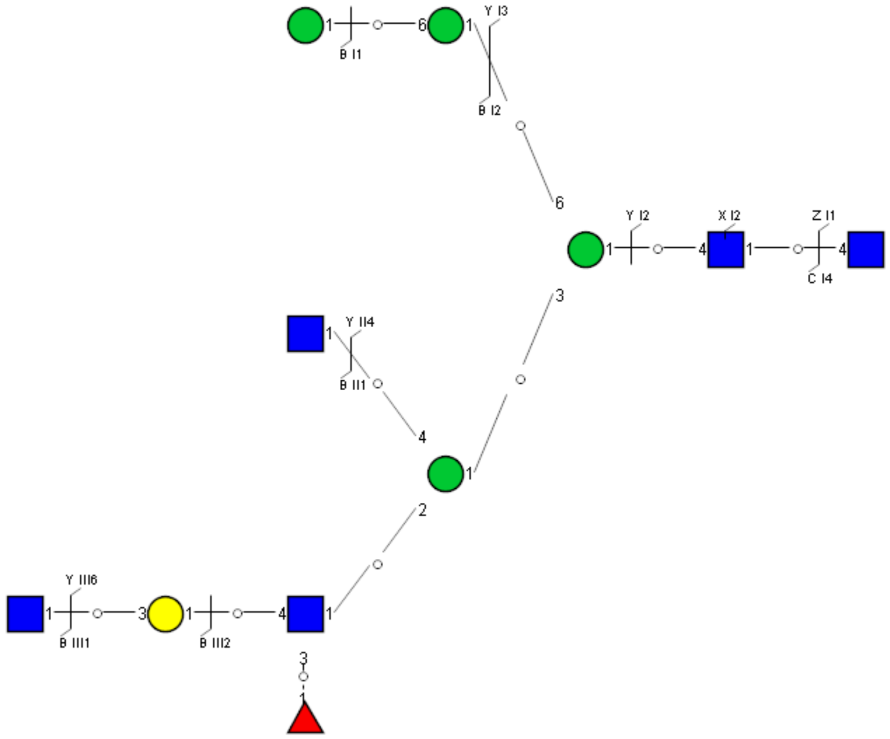

Spectrum Index: 34487(NoBHs:1)

#### Precursor iEF Map

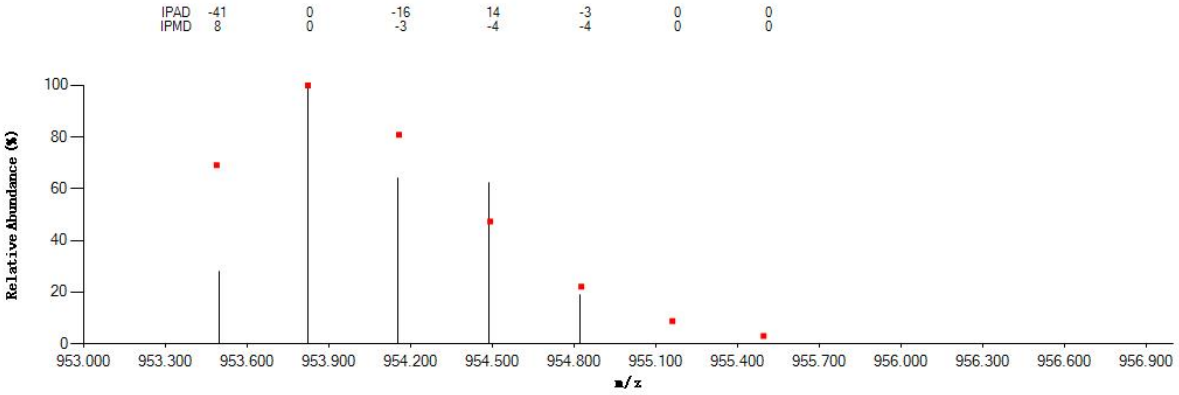

#### Annotated MS/MS Spectrum

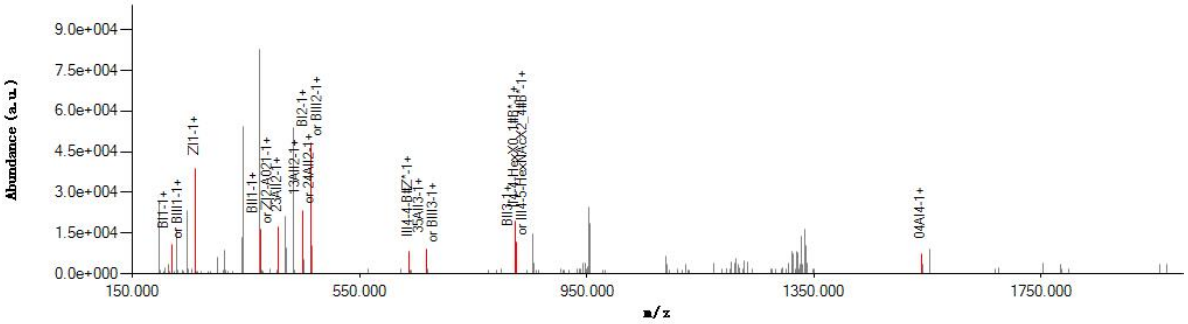

#### Graphical Fragmentation Map

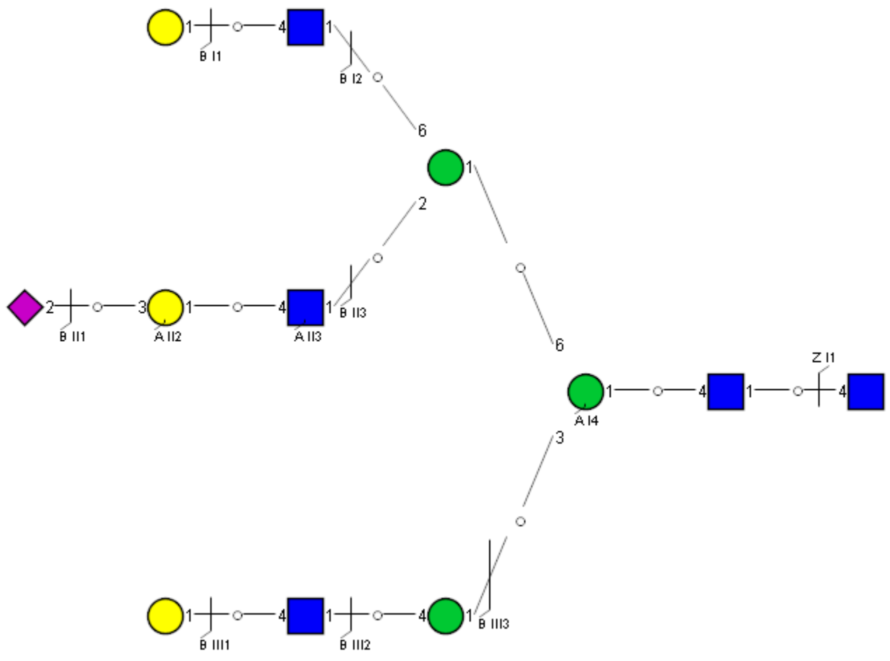

Spectrum Index: 34768(NoBHs:1)

Precursor iEF Map

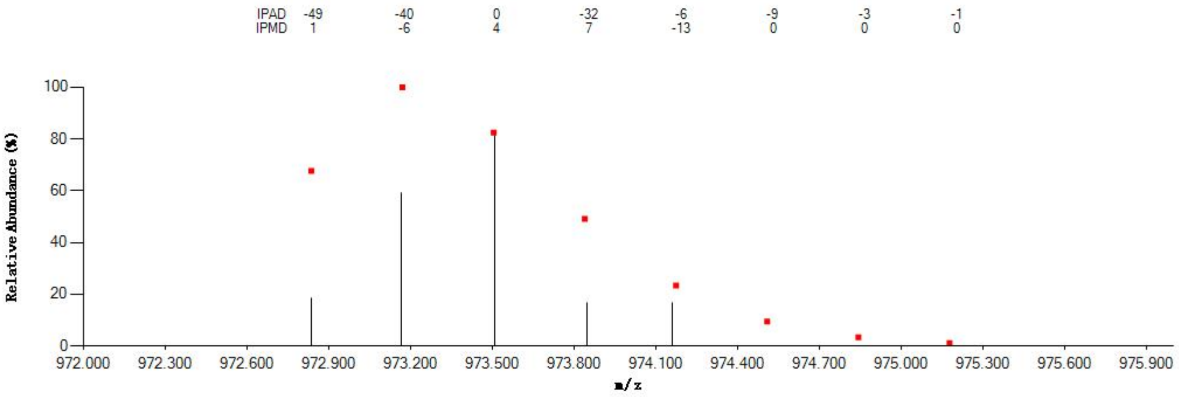


Annotated MS/MS Spectrum

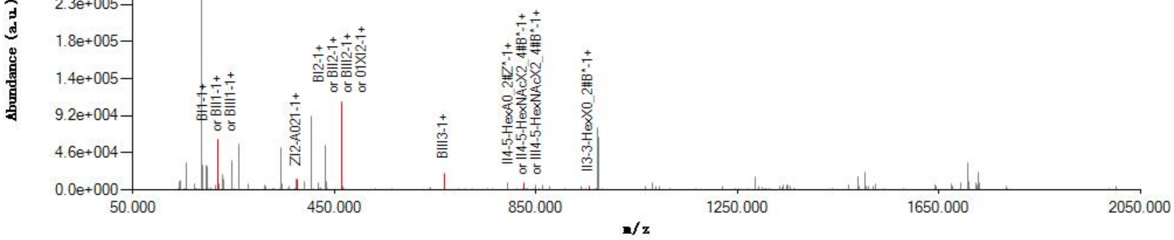


Graphical Fragmentation Map

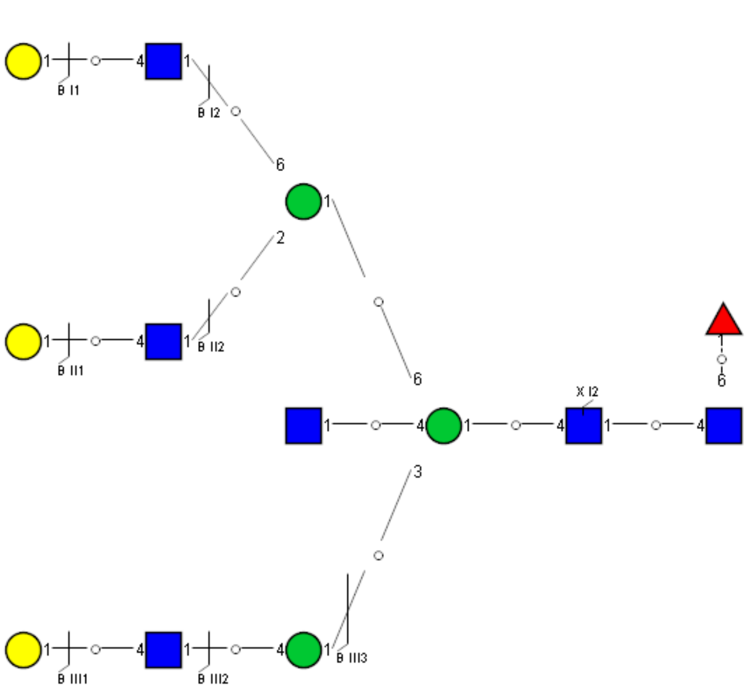

Spectrum Index: 36367(NoBHs:1)

#### Precursor iEF Map

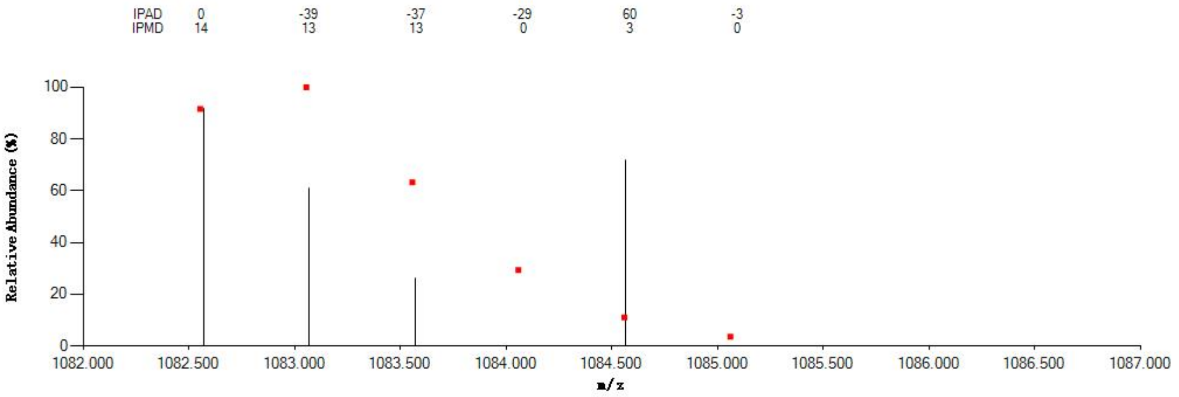


#### Annotated MS/MS Spectrum

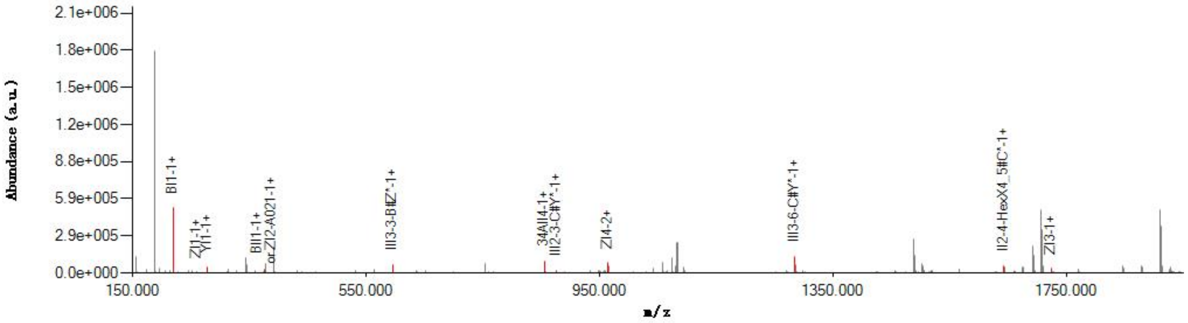


#### Graphical Fragmentation Map

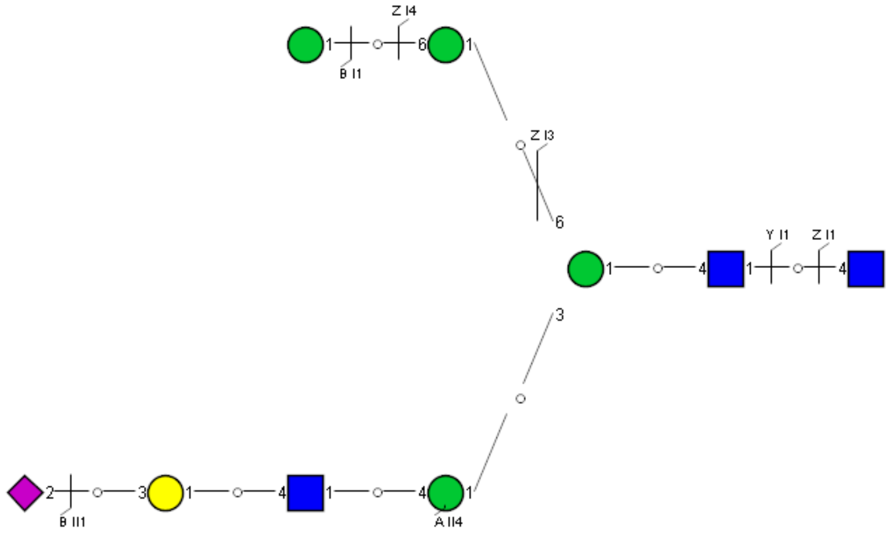

Spectrum Index: 38320(NoBHs:1)

#### Precursor iEF Map

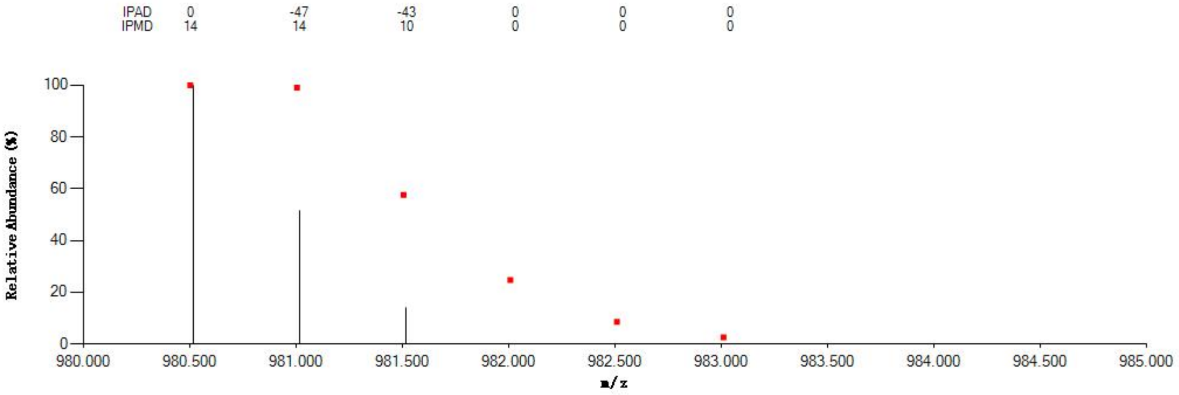


#### Annotated MS/MS Spectrum

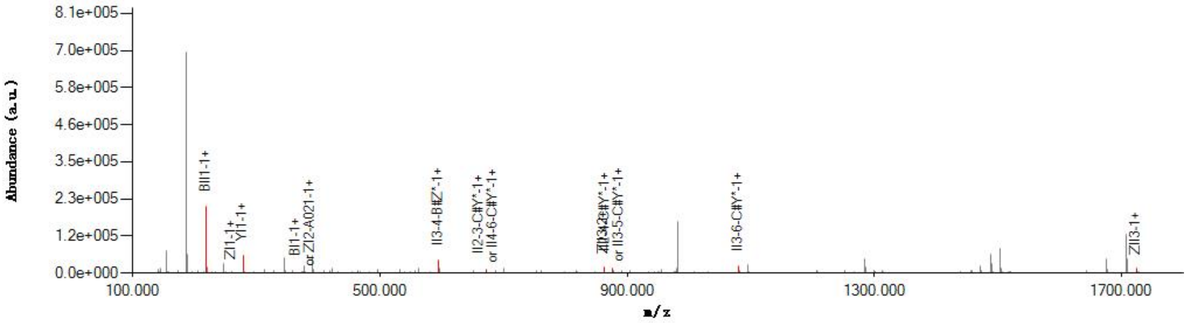


#### Graphical Fragmentation Map

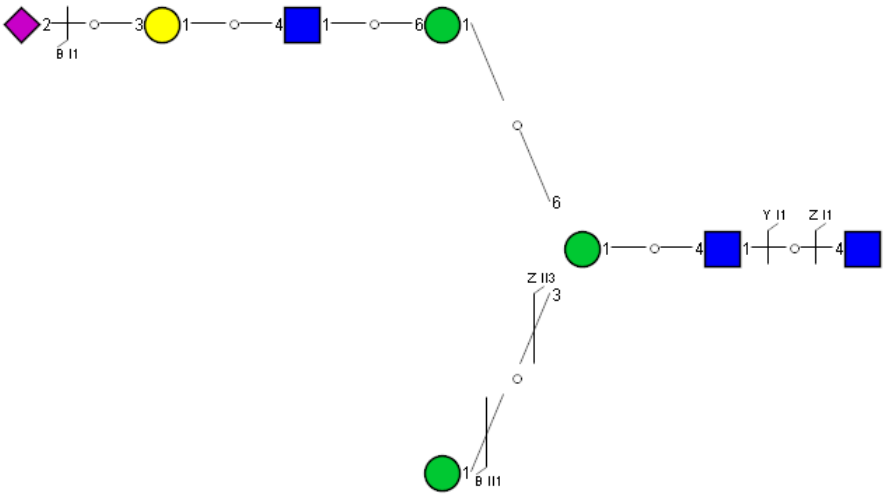

Spectrum Index: 42096(NoBHs:1)

#### Precursor iEF Map

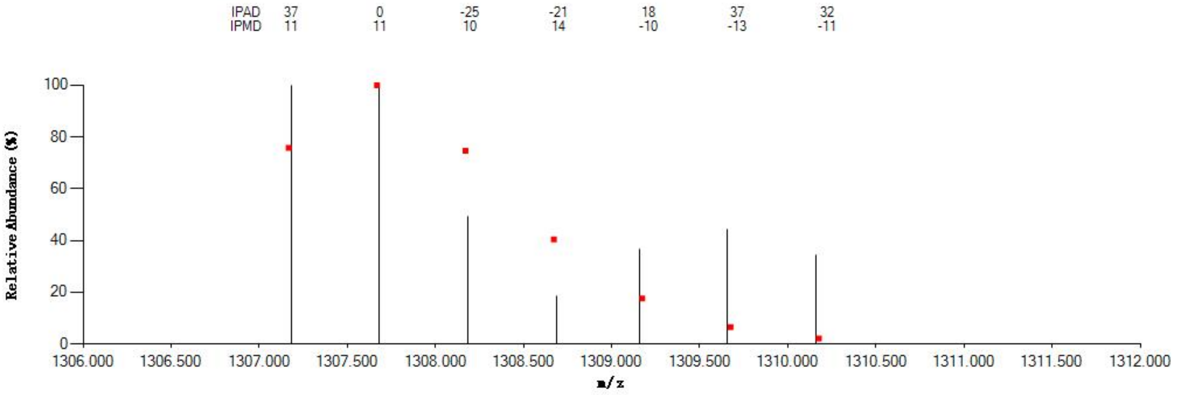


#### Annotated MS/MS Spectrum

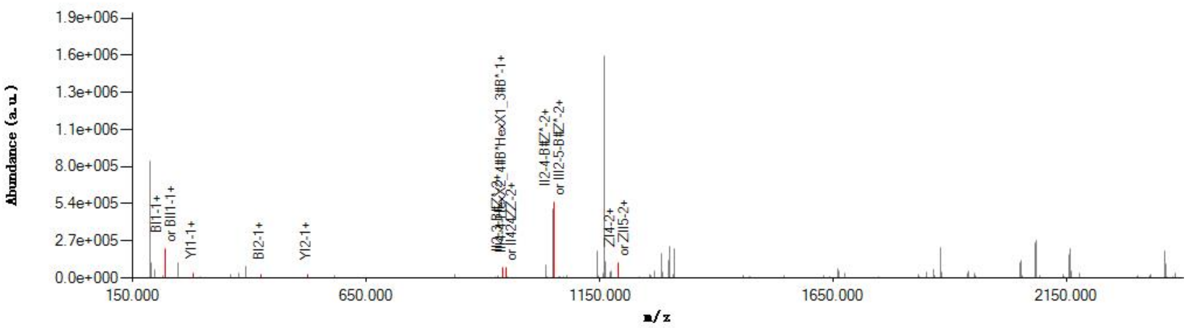


#### Graphical Fragmentation Map

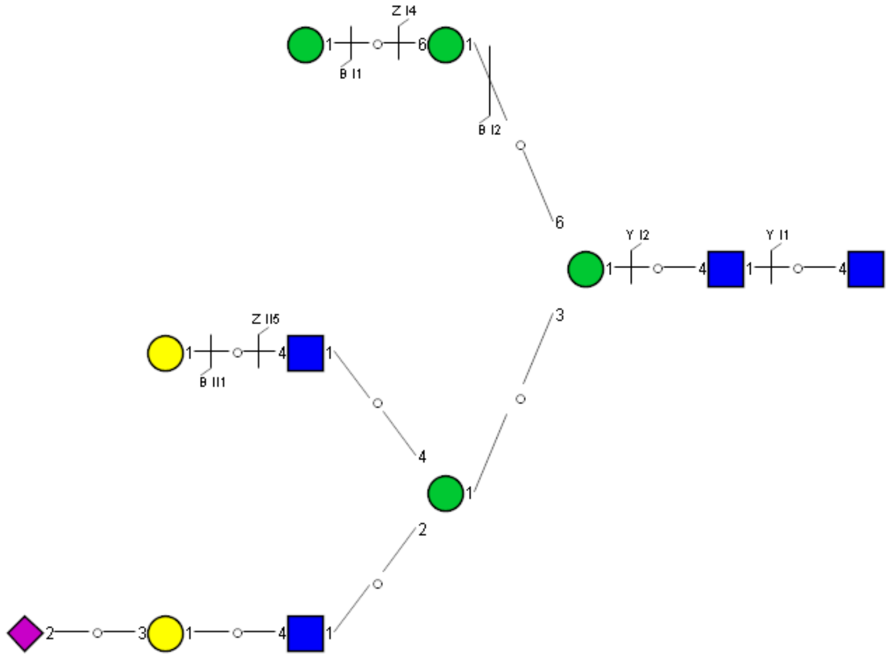
