## Supplemental Information 2 for "A draft of human N-glycans of glycoRNA"

### Spectrum Index: 23610(NoBHs:1)

#### Precursor iEF Map

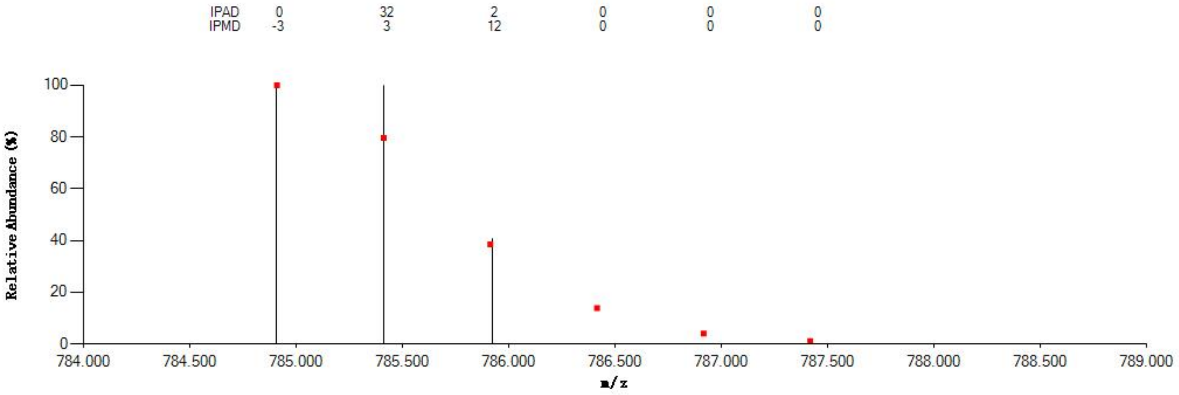


#### Annotated MS/MS Spectrum

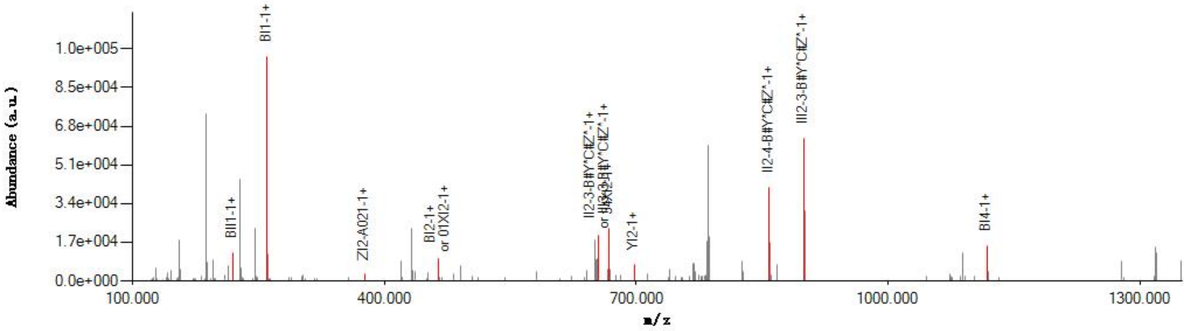


#### Graphical Fragmentation Map

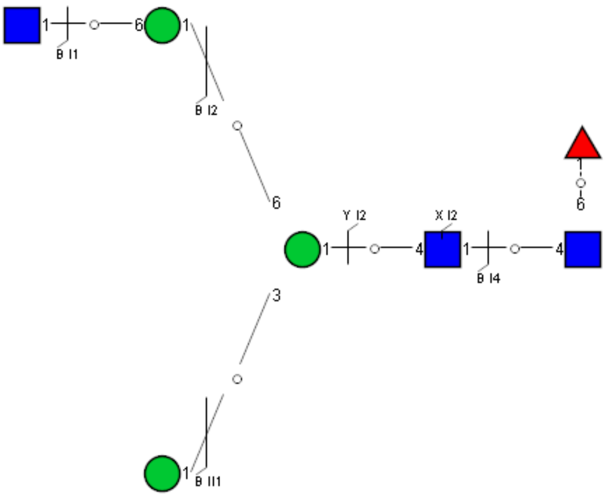

Spectrum Index: 23845(NoBHs:1)

Precursor iEF Map

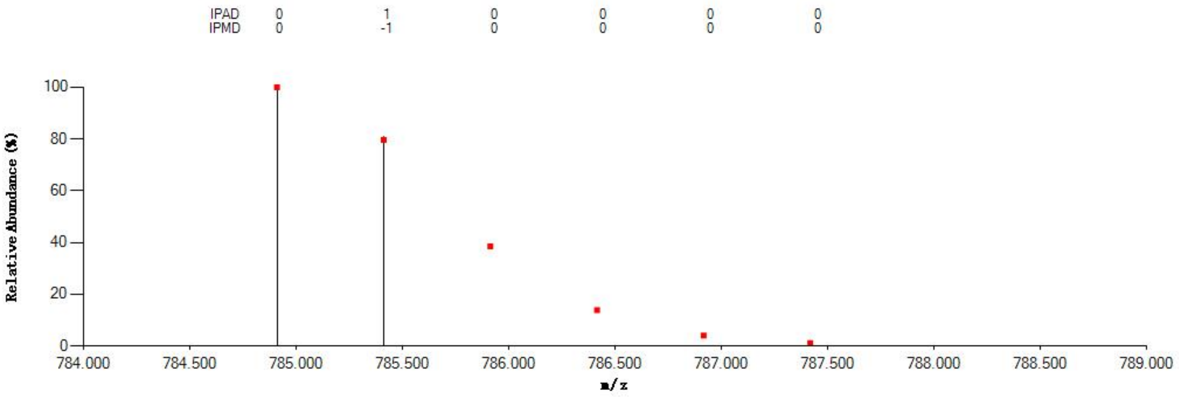


Annotated MS/MS Spectrum

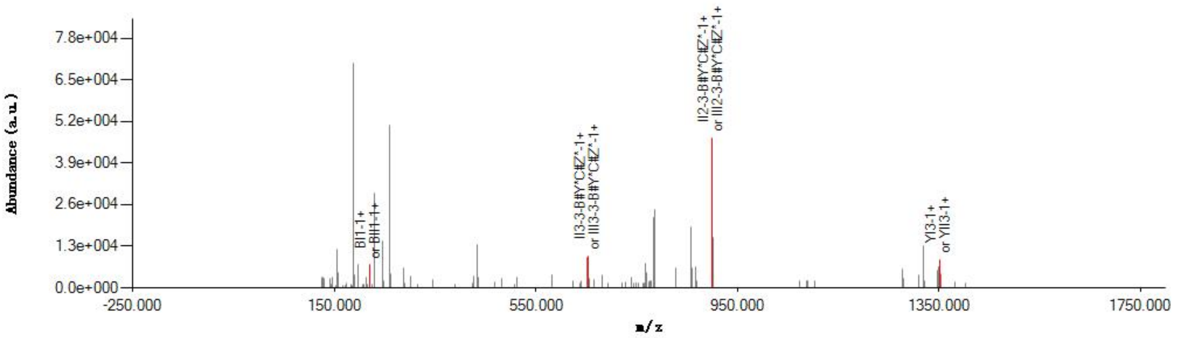


Graphical Fragmentation Map

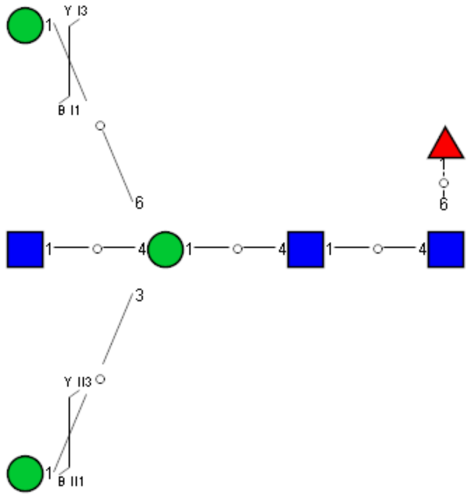

Spectrum Index: 25538(NoBHs:1)

#### Precursor iEF Map

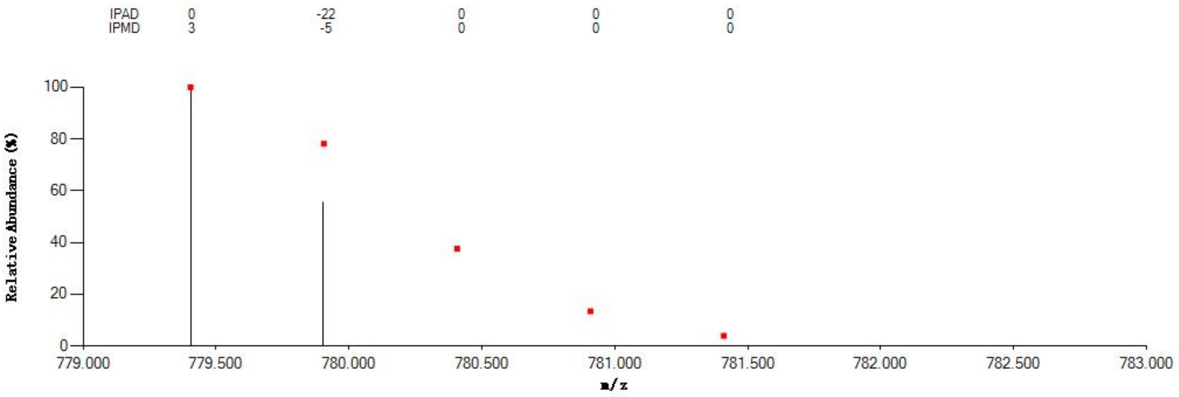


#### Annotated MS/MS Spectrum

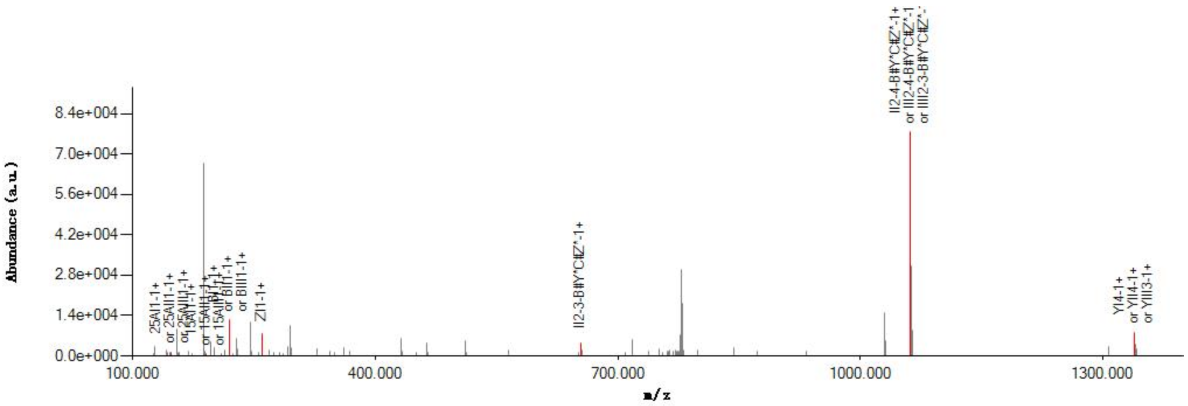


#### Graphical Fragmentation Map

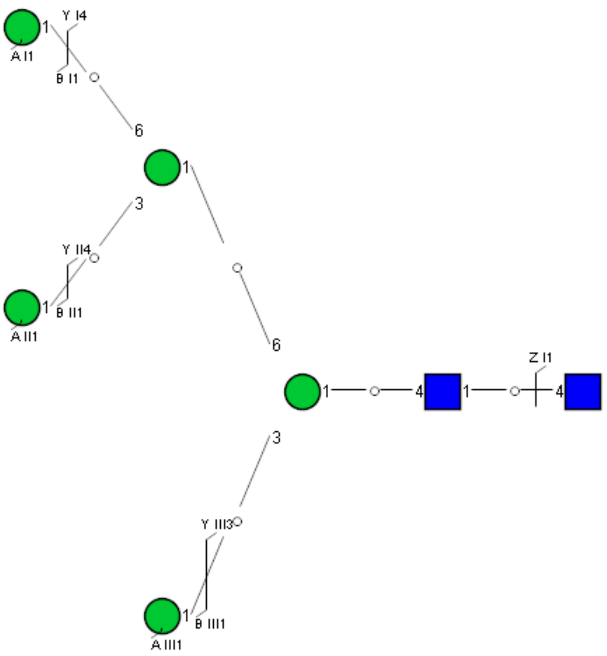

Spectrum Index: 26123(NoBHs:1)

#### Precursor iEF Map

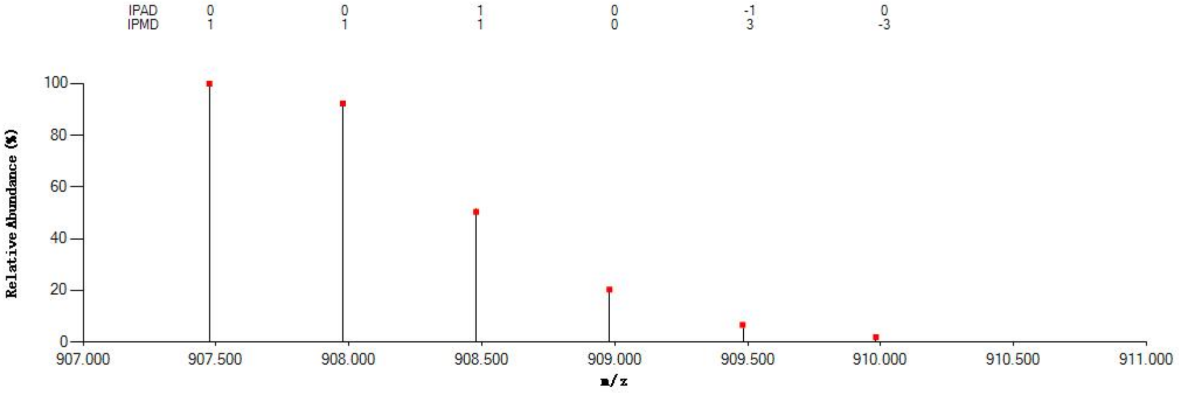


#### Annotated MS/MS Spectrum

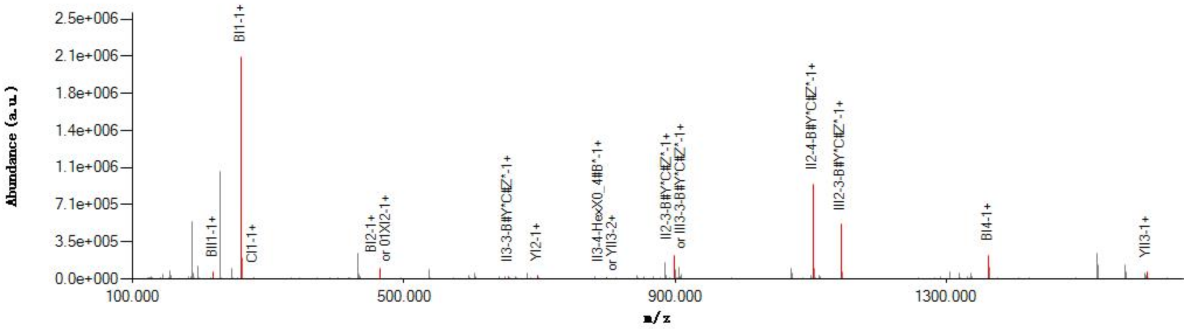


#### Graphical Fragmentation Map

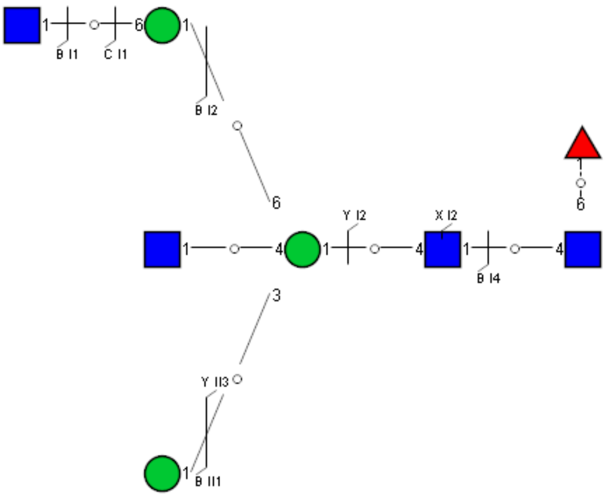

Spectrum Index: 26517(NoBHs:1)

#### Precursor iEF Map

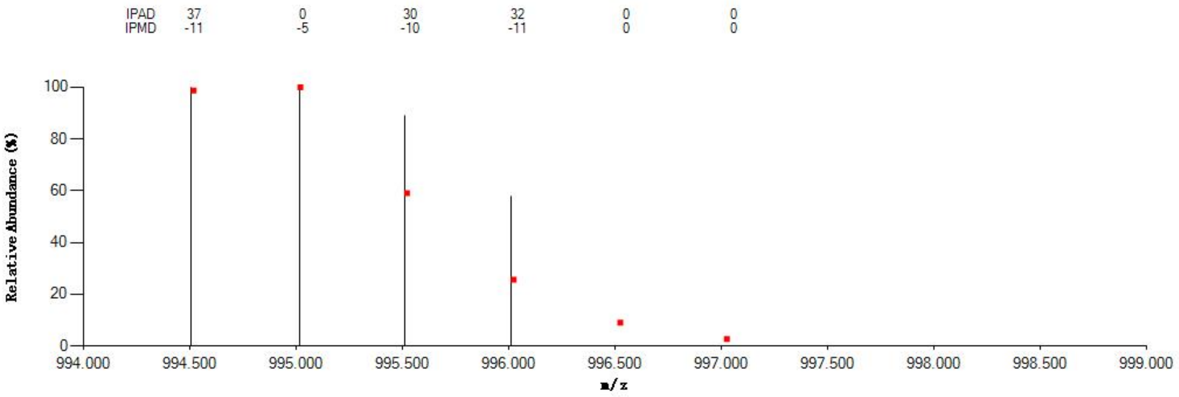


#### Annotated MS/MS Spectrum

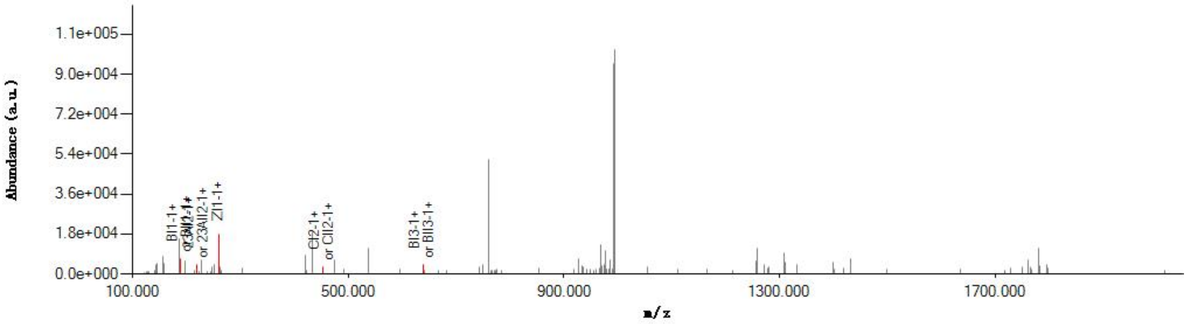


#### Graphical Fragmentation Map

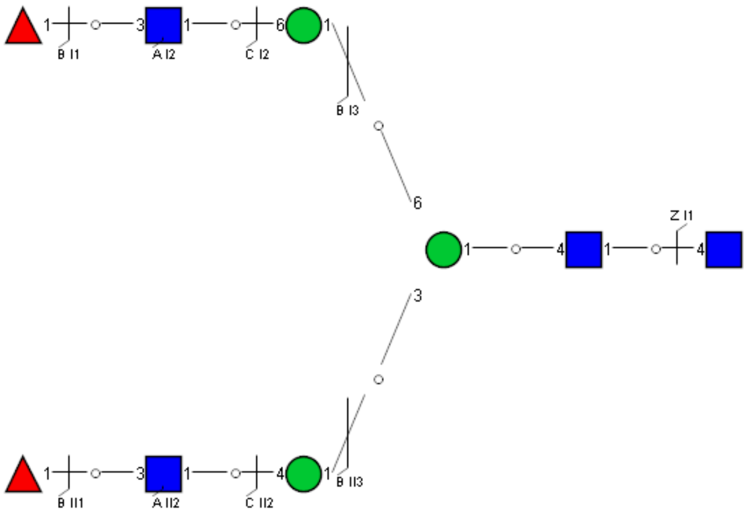

Spectrum Index: 27047(NoBHs:1)

#### Precursor iEF Map

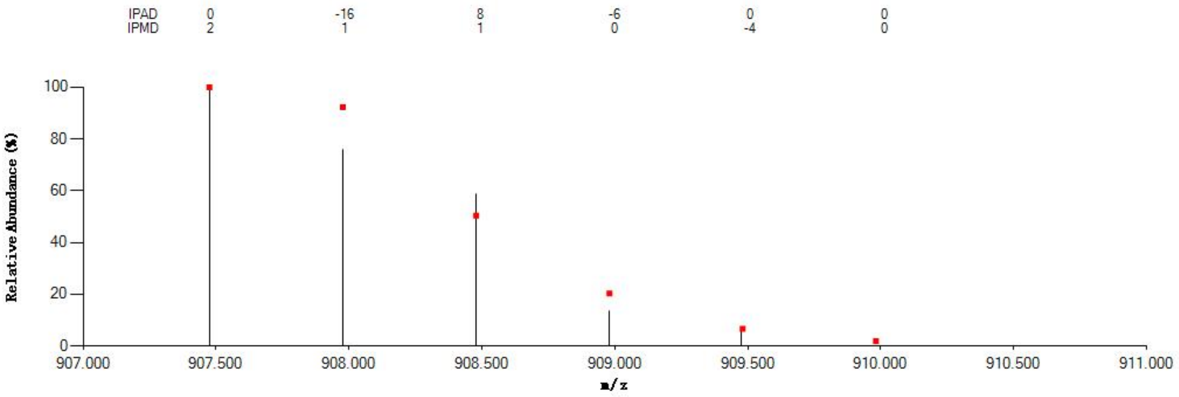

#### Annotated MS/MS Spectrum

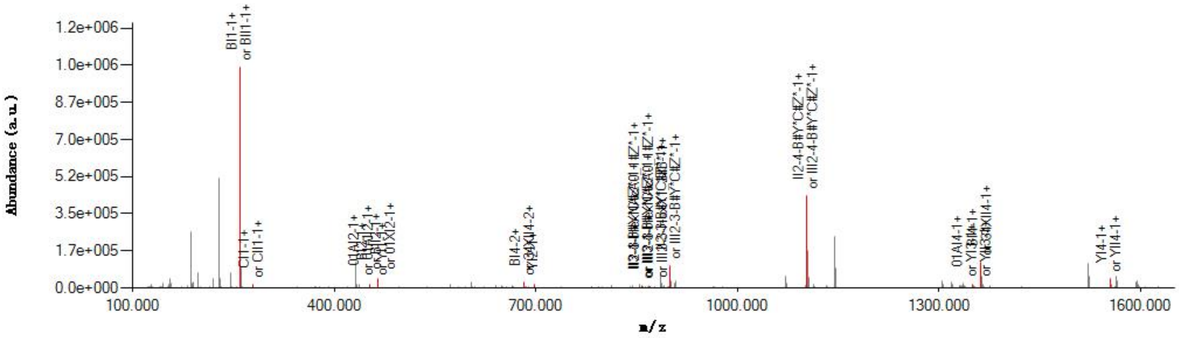

#### Graphical Fragmentation Map

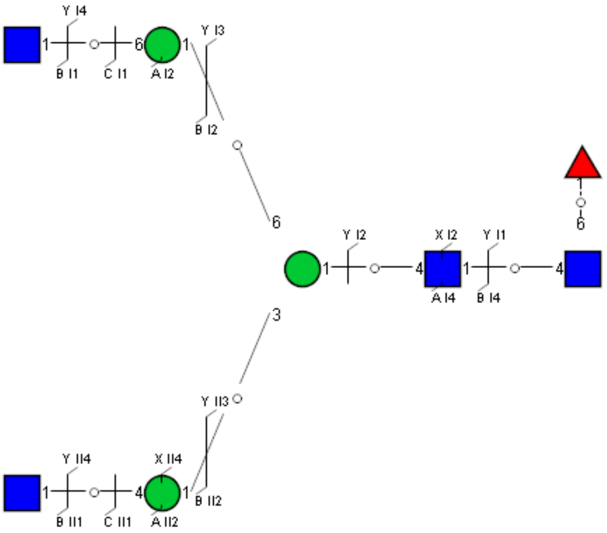

Spectrum Index: 27133(NoBHs:1)

#### Precursor iEF Map

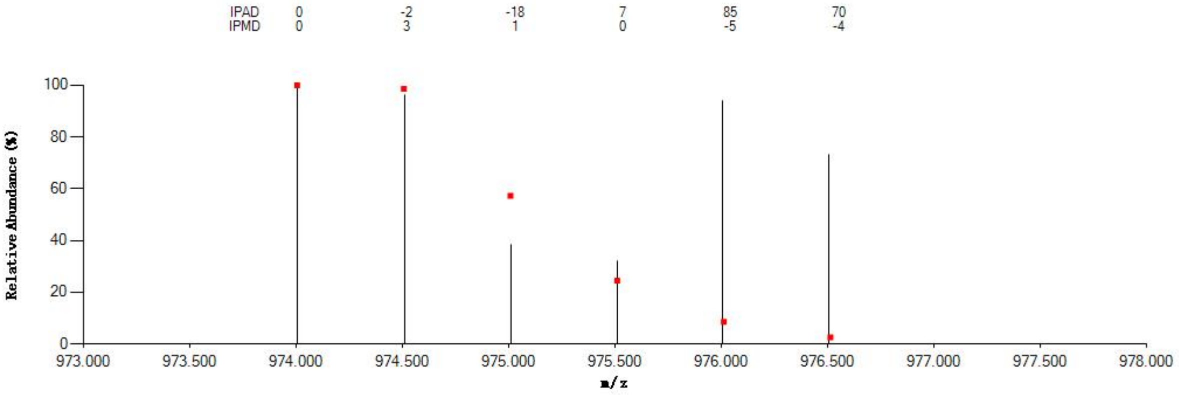


#### Annotated MS/MS Spectrum

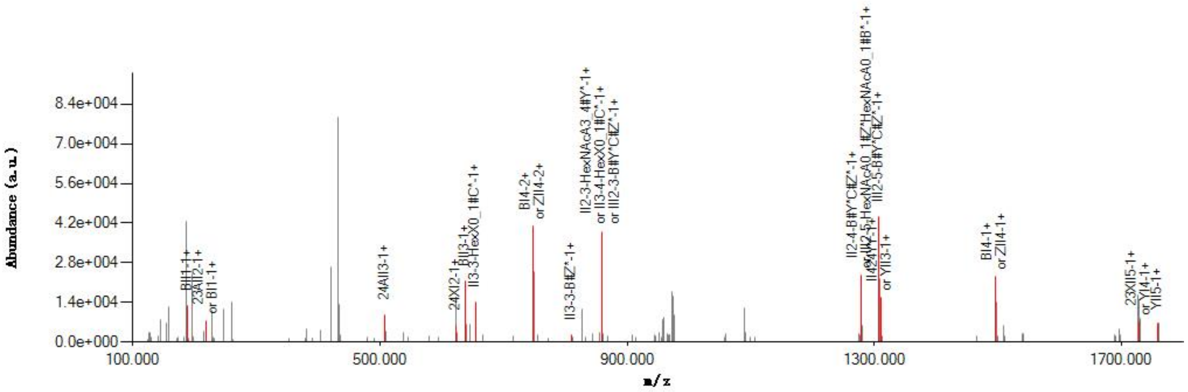


#### Graphical Fragmentation Map

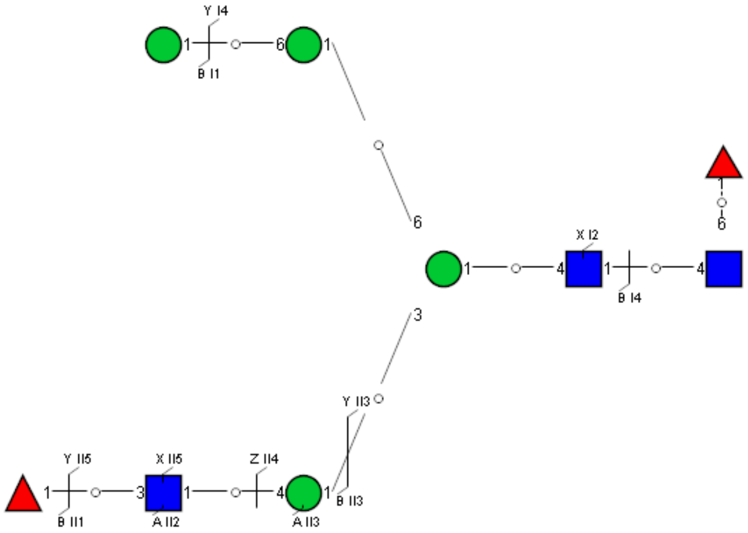

Spectrum Index: 27911(NoBHs:1)

#### Precursor iEF Map

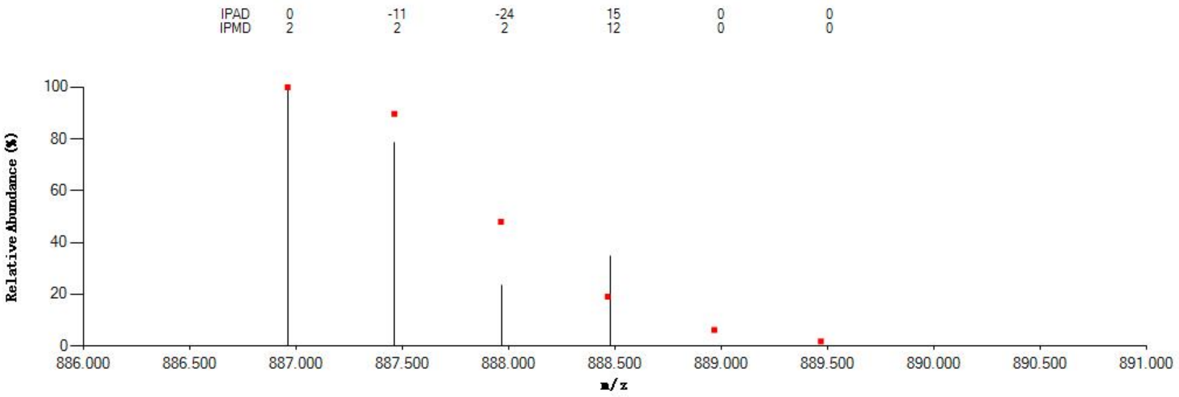


#### Annotated MS/MS Spectrum

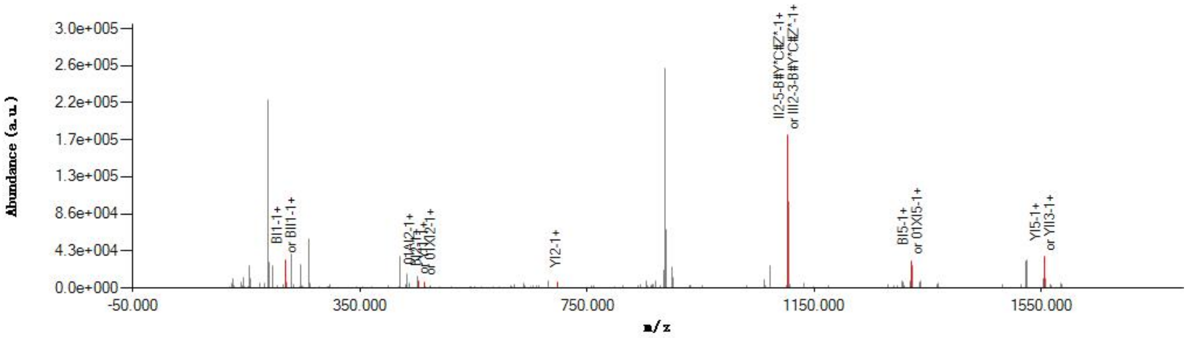


#### Graphical Fragmentation Map

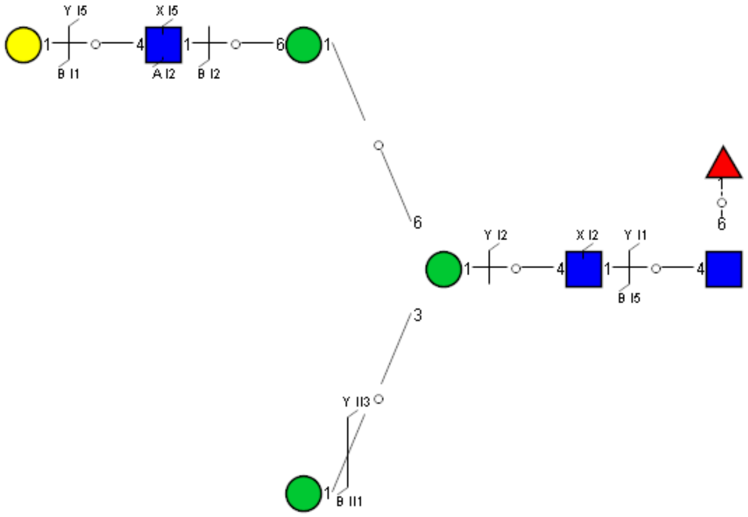

Spectrum Index: 28146(NoBHs:1)

#### Precursor iEF Map

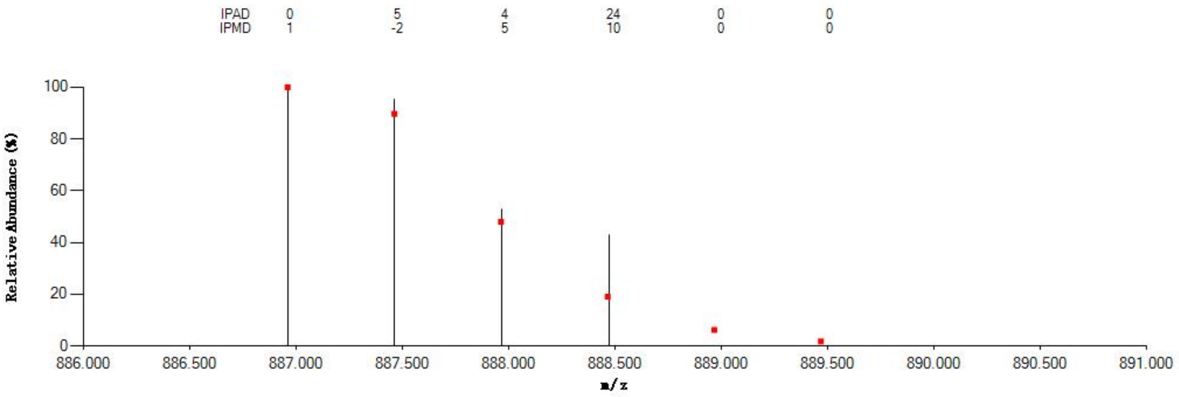

#### Annotated MS/MS Spectrum

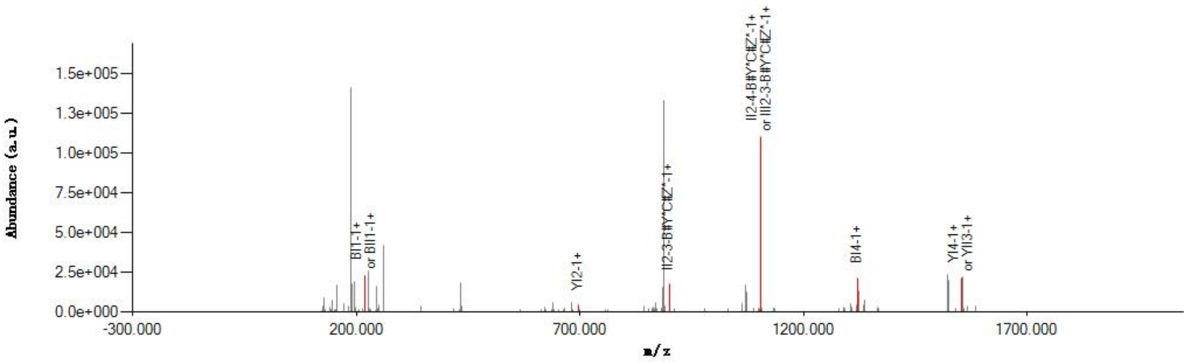

#### Graphical Fragmentation Map

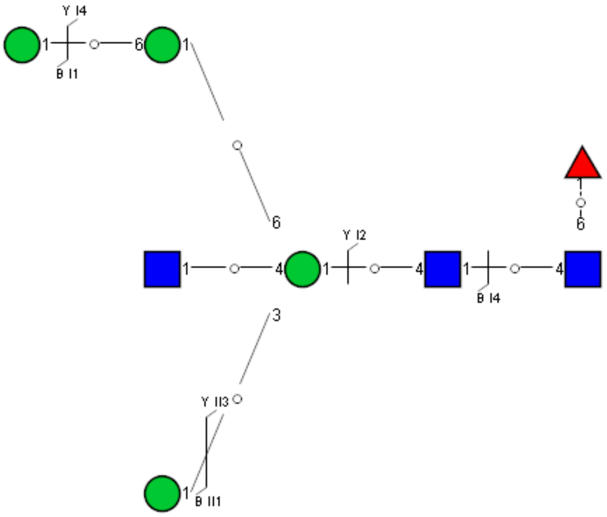

Spectrum Index: 28433(NoBHs:1)

#### Precursor iEF Map

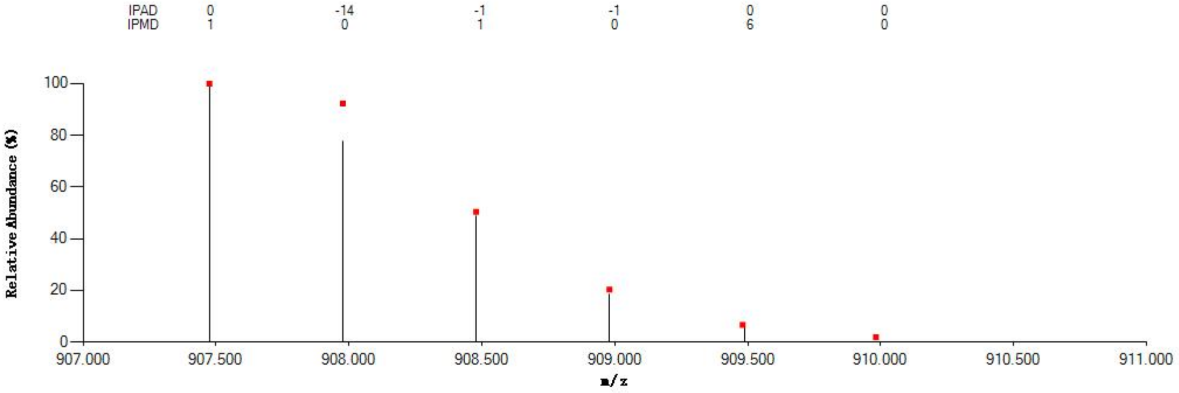

#### Annotated MS/MS Spectrum

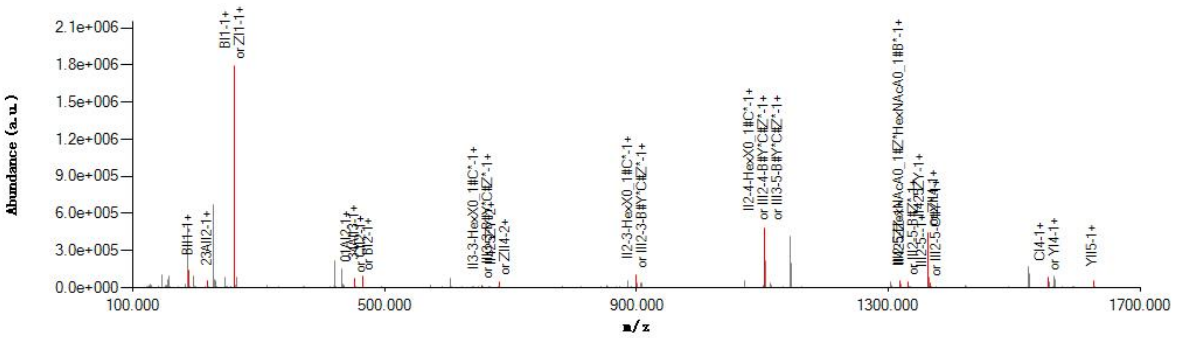

#### Graphical Fragmentation Map

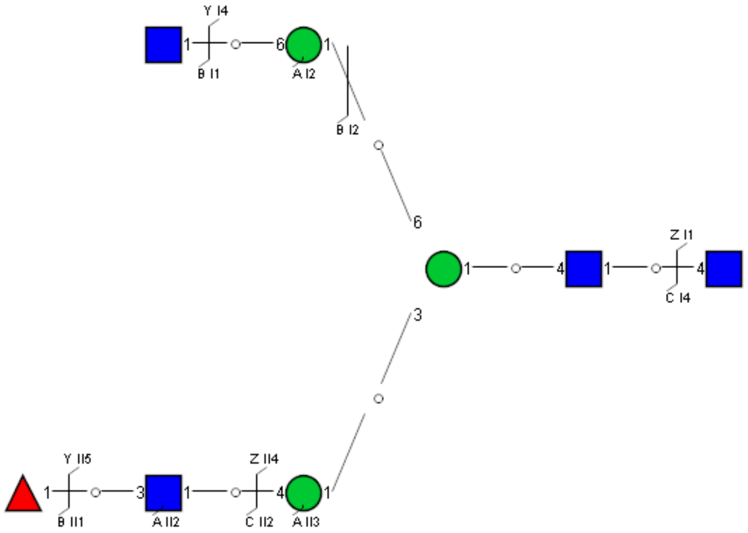

Spectrum Index: 28858(NoBHs:1)

#### Precursor iEF Map

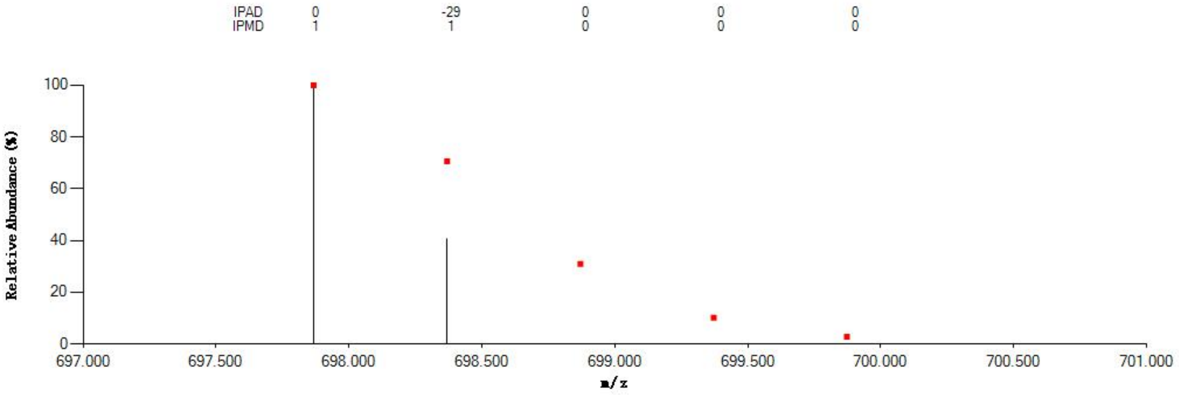


#### Annotated MS/MS Spectrum

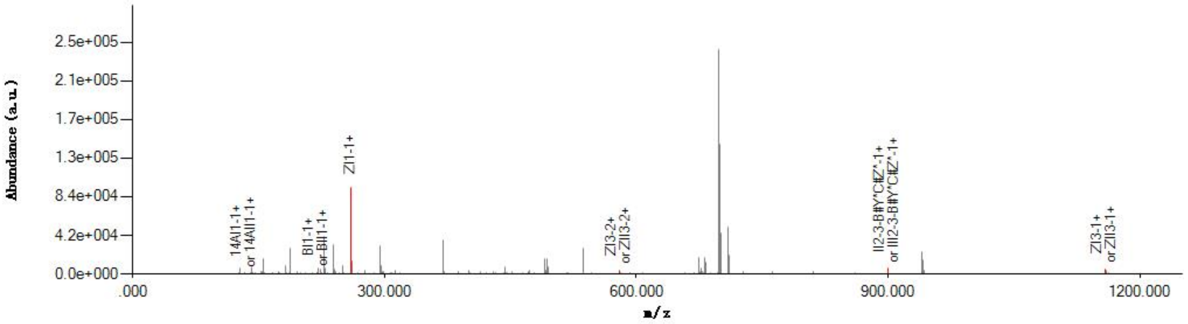


#### Graphical Fragmentation Map

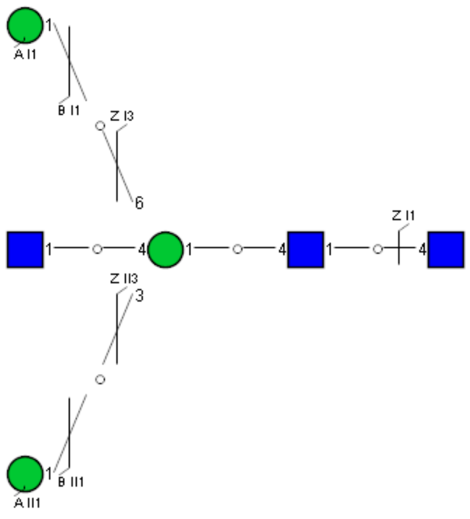

Spectrum Index: 28941(NoBHs:1)

#### Precursor iEF Map

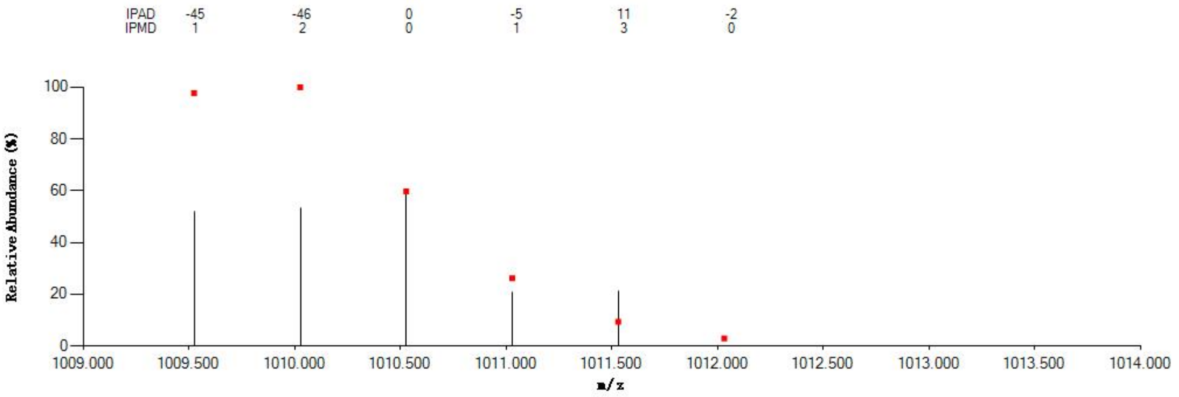

#### Annotated MS/MS Spectrum

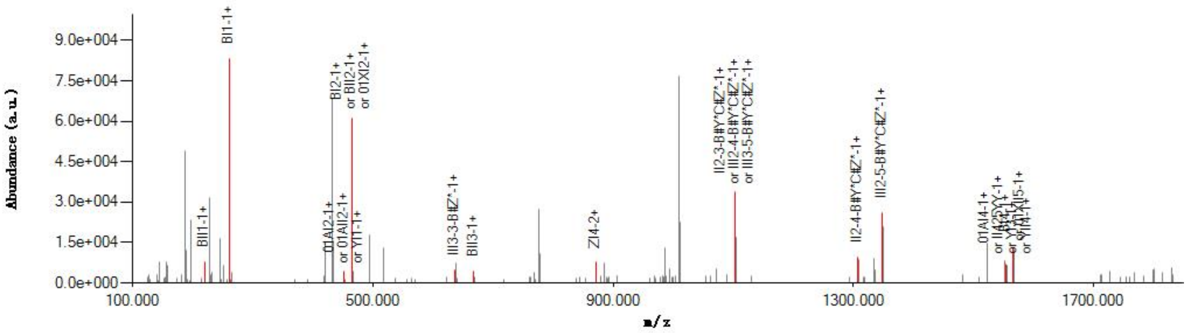

#### Graphical Fragmentation Map

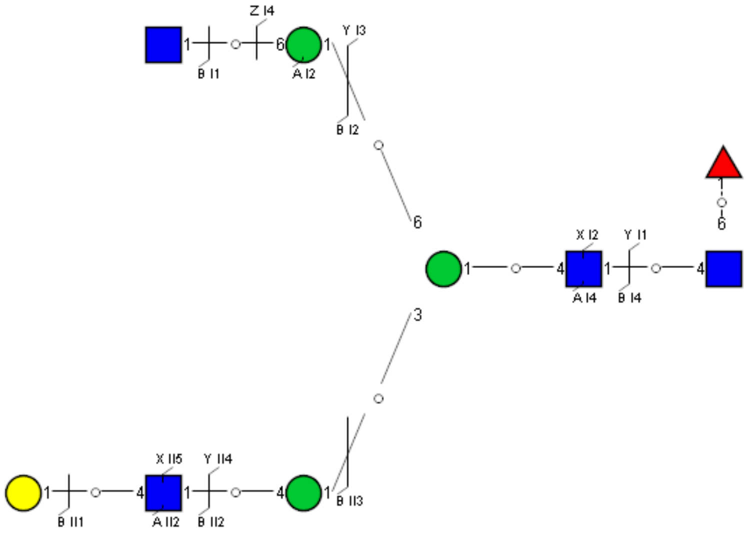

Spectrum Index: 29263(NoBHs:1)

#### Precursor iEF Map

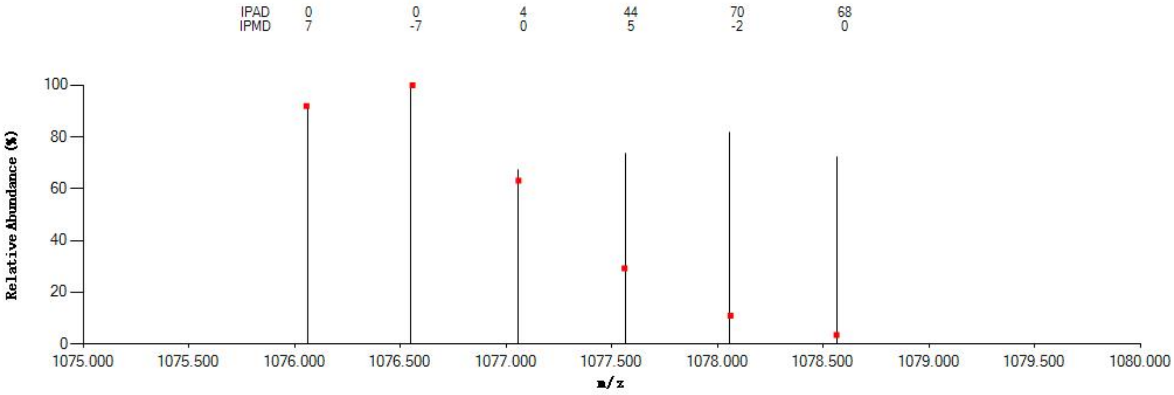


#### Annotated MS/MS Spectrum

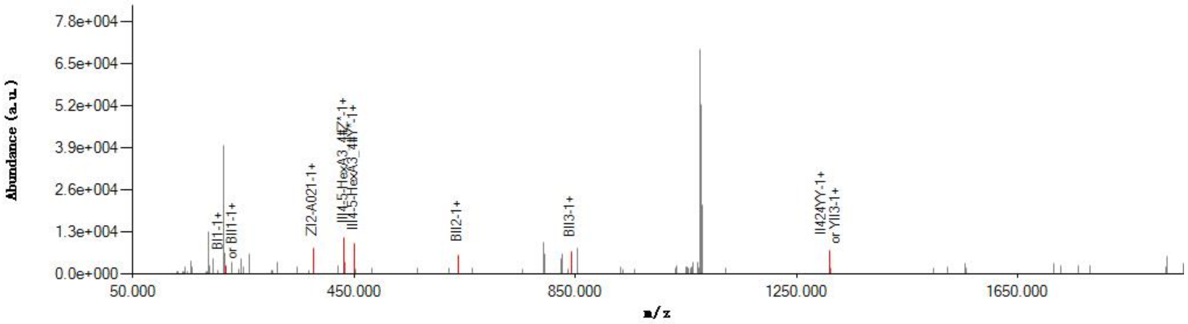


#### Graphical Fragmentation Map

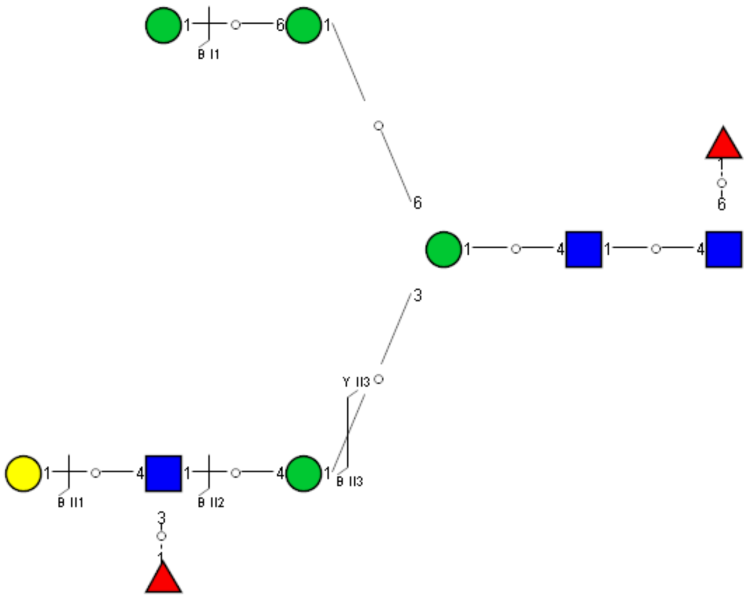

Spectrum Index: 29317(NoBHs:1)

#### Precursor iEF Map

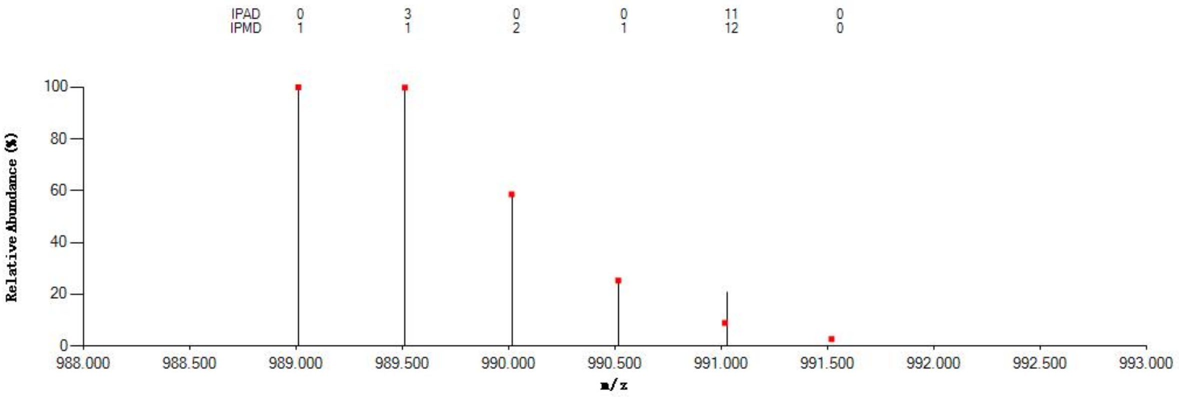


#### Annotated MS/MS Spectrum

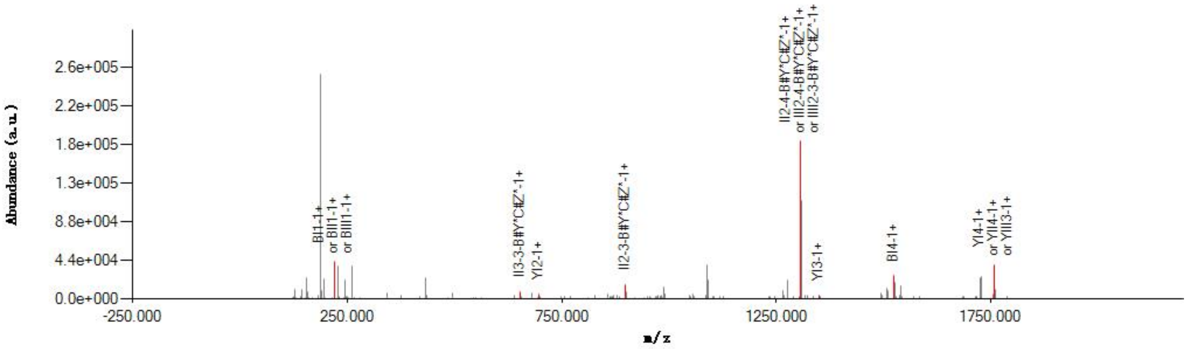


#### Graphical Fragmentation Map

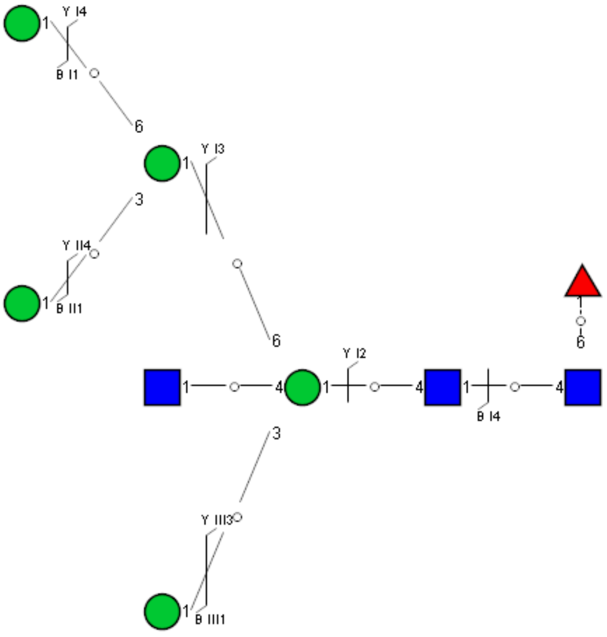

Spectrum Index: 29672(NoBHs:1)

#### Precursor iEF Map

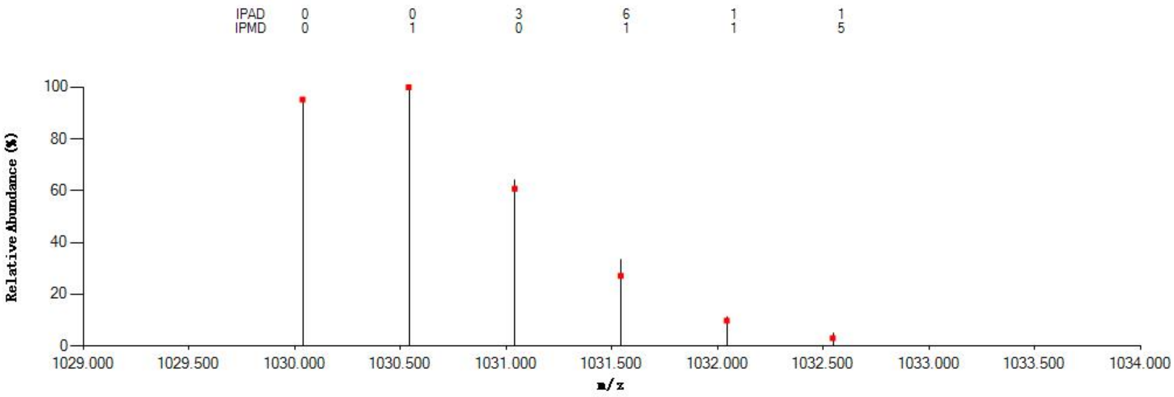


#### Annotated MS/MS Spectrum

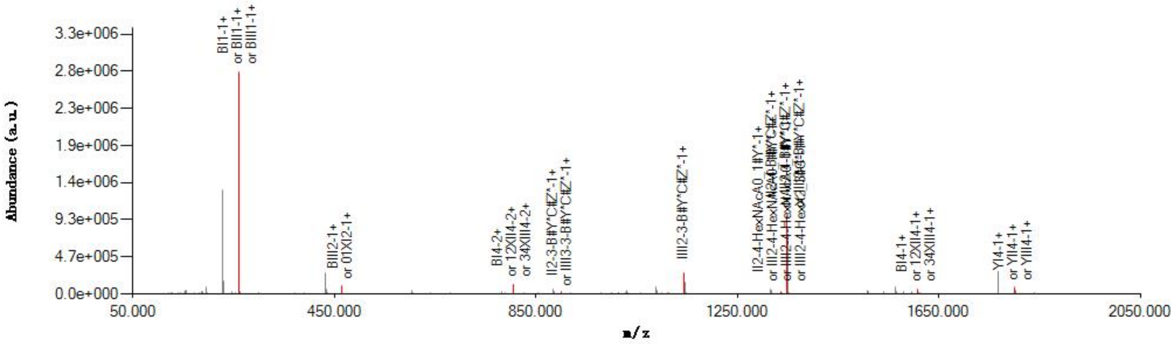


#### Graphical Fragmentation Map

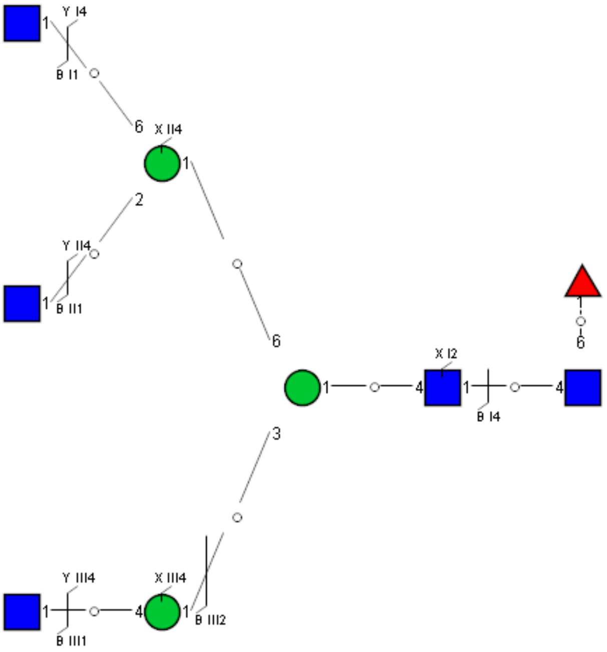

Spectrum Index: 30409(NoBHs:1)

#### Precursor iEF Map

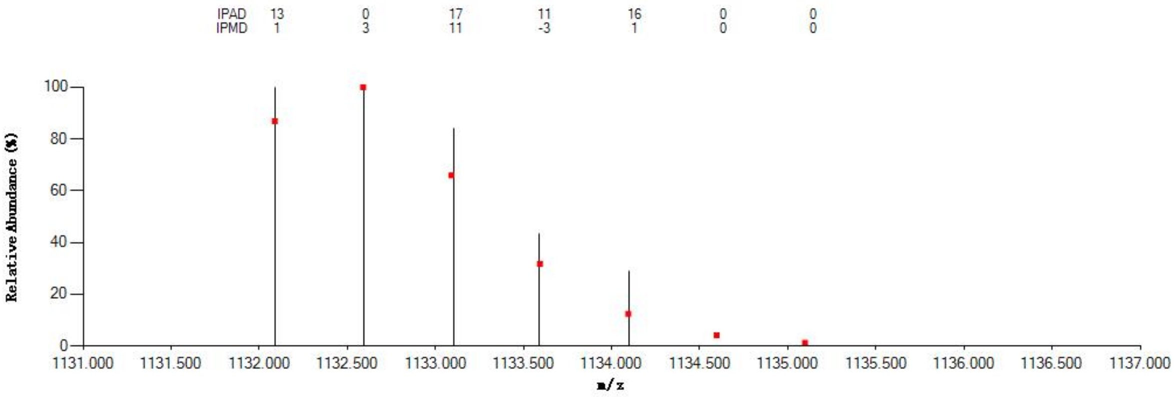


#### Annotated MS/MS Spectrum

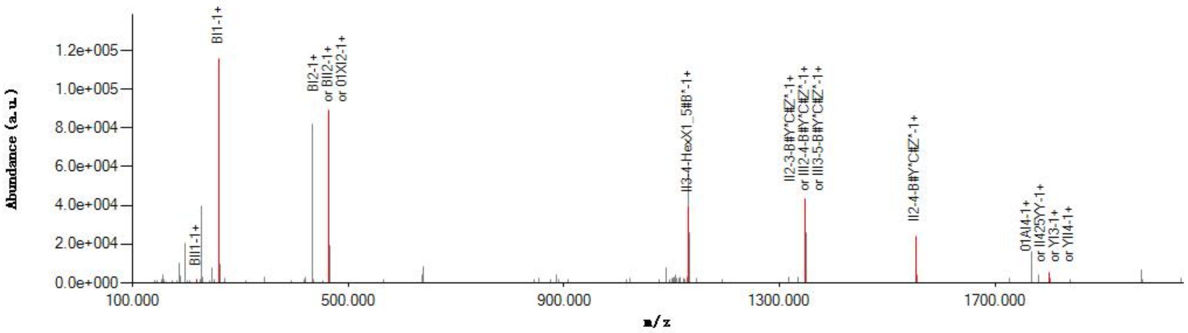


#### Graphical Fragmentation Map

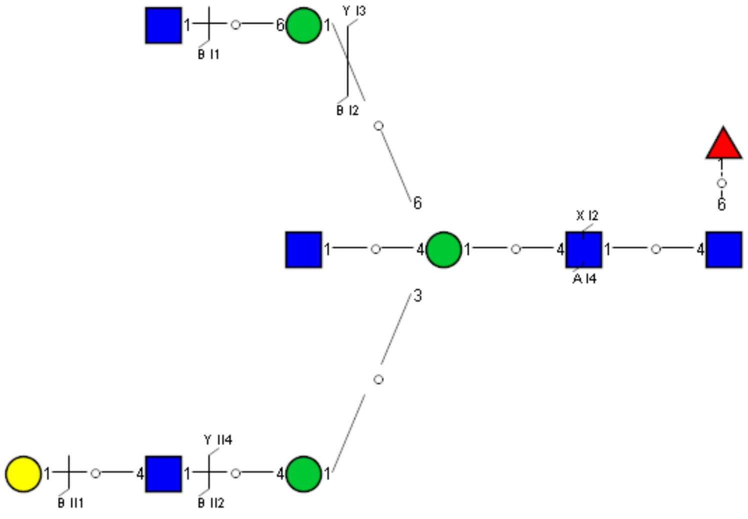

Spectrum Index: 30456(NoBHs:1)

#### Precursor iEF Map

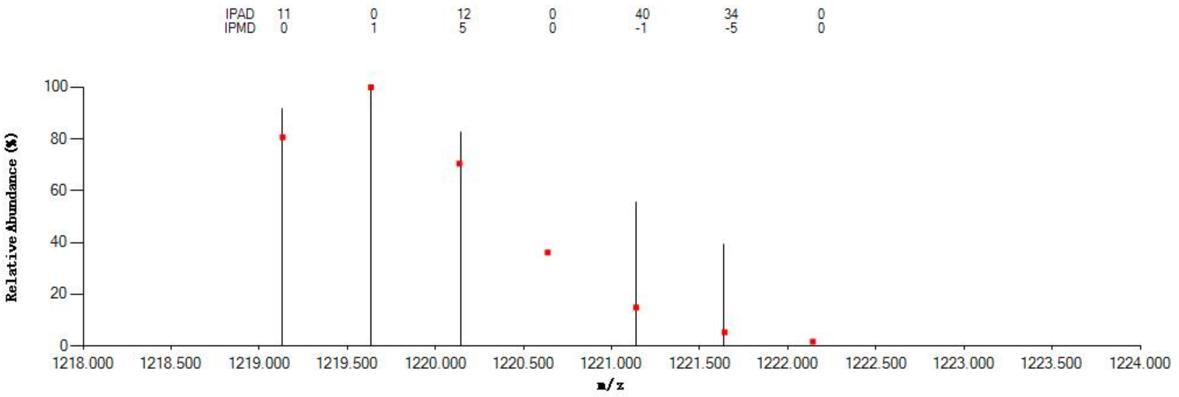


#### Annotated MS/MS Spectrum

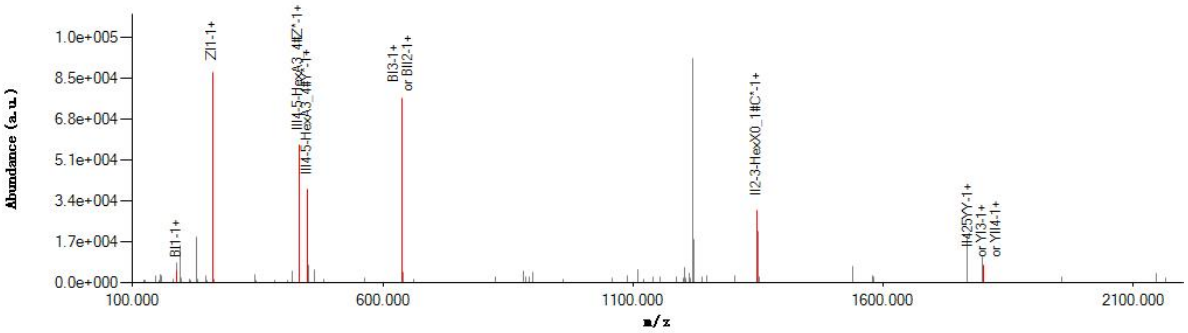


#### Graphical Fragmentation Map

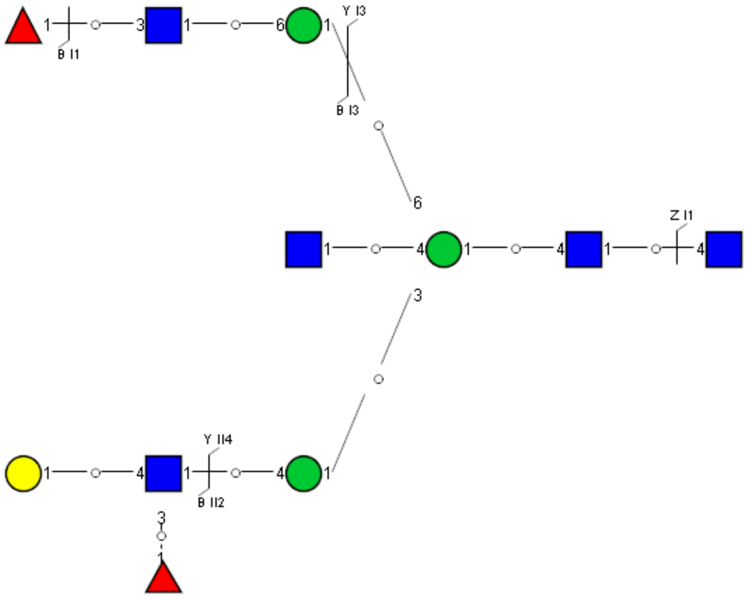

Spectrum Index: 30623(NoBHs:1)

#### Precursor iEF Map

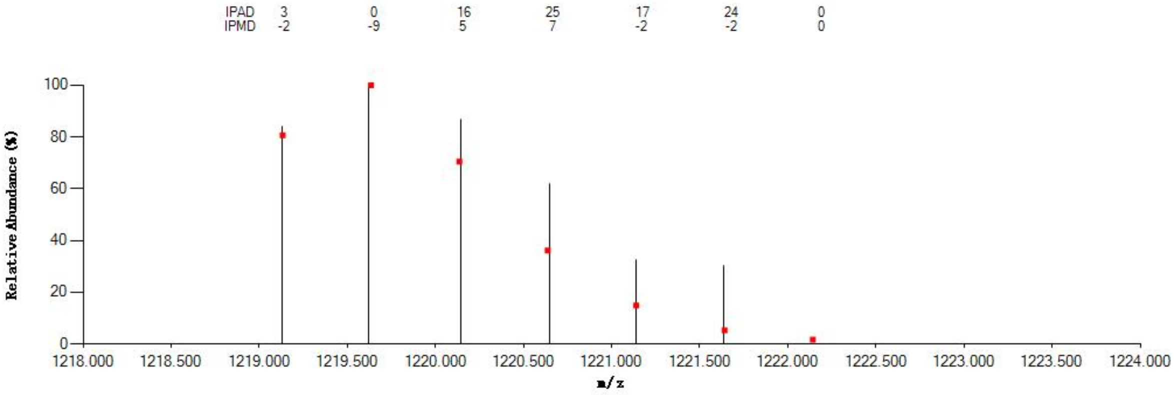


#### Annotated MS/MS Spectrum

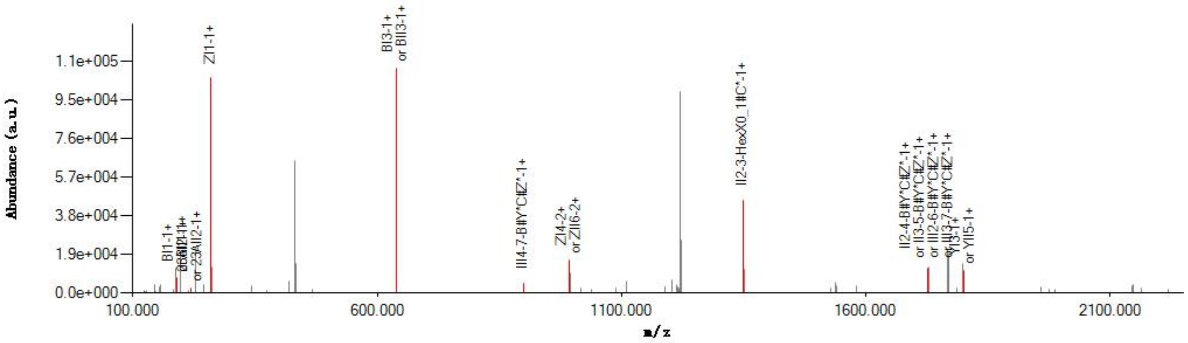


#### Graphical Fragmentation Map

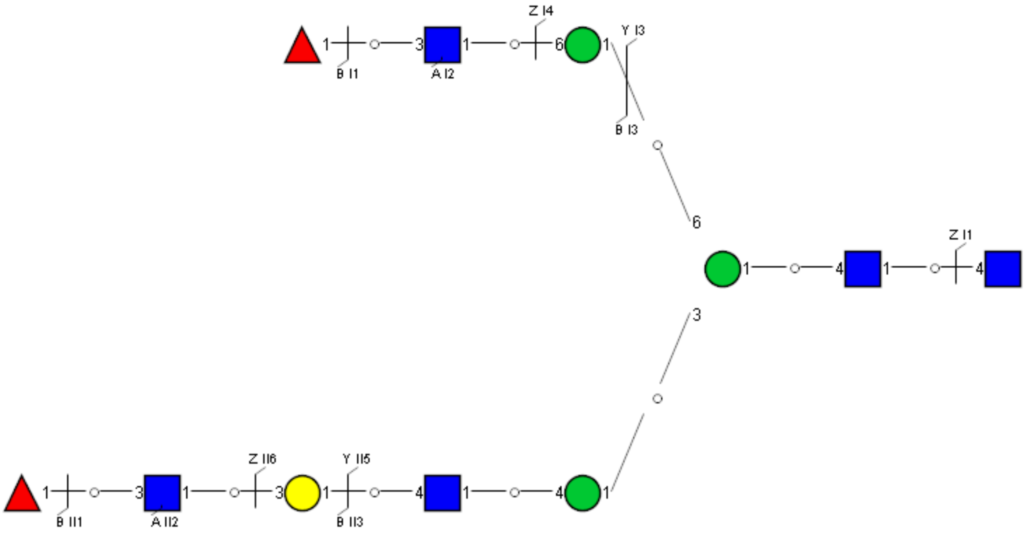

Spectrum Index: 30624(NoBHs:1)

#### Precursor iEF Map

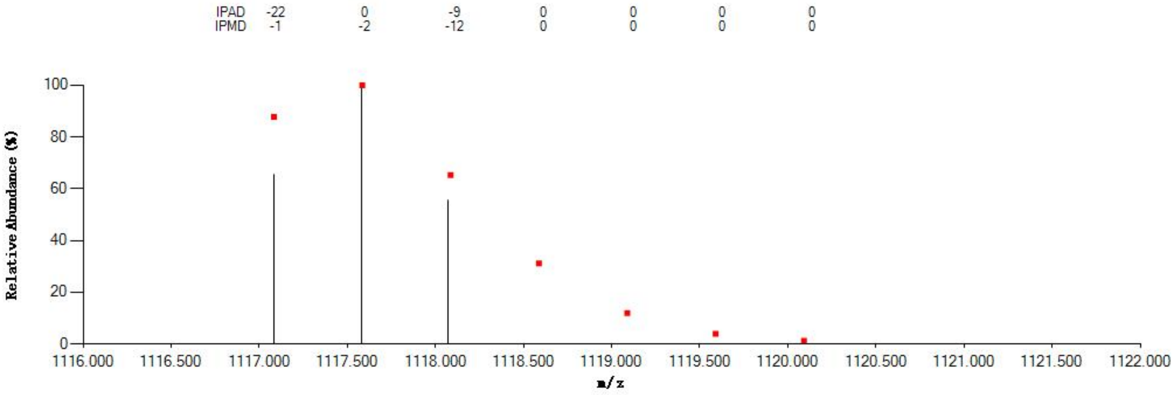


#### Annotated MS/MS Spectrum

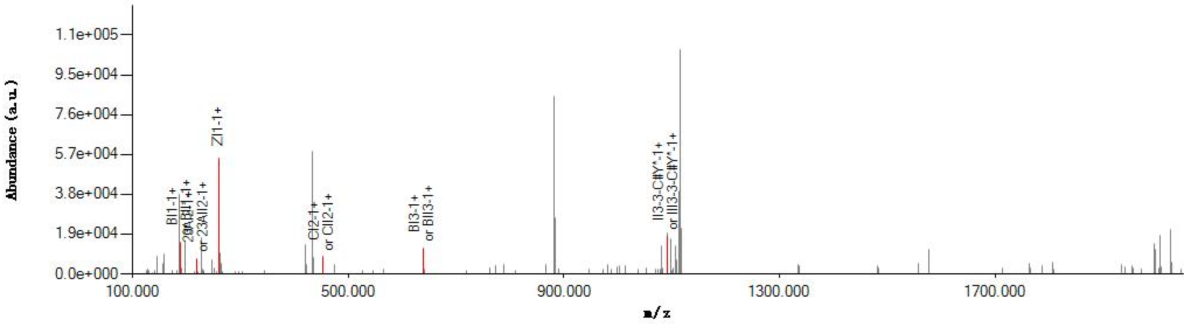


#### Graphical Fragmentation Map

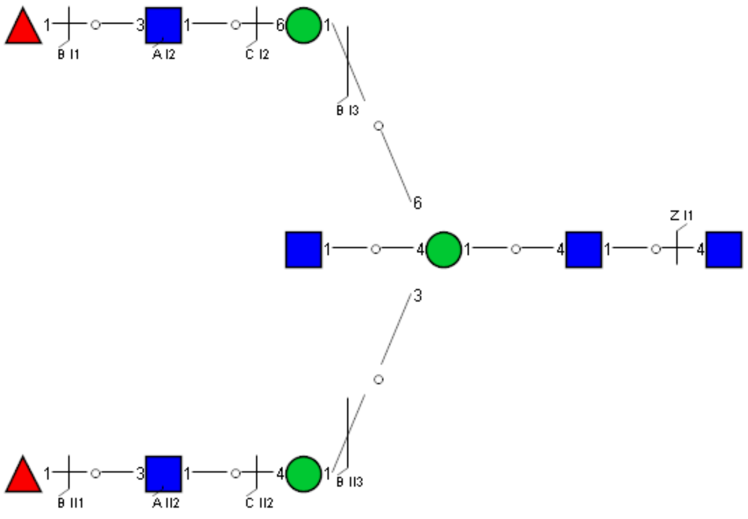

Spectrum Index: 30995(NoBHs:1)

#### Precursor iEF Map

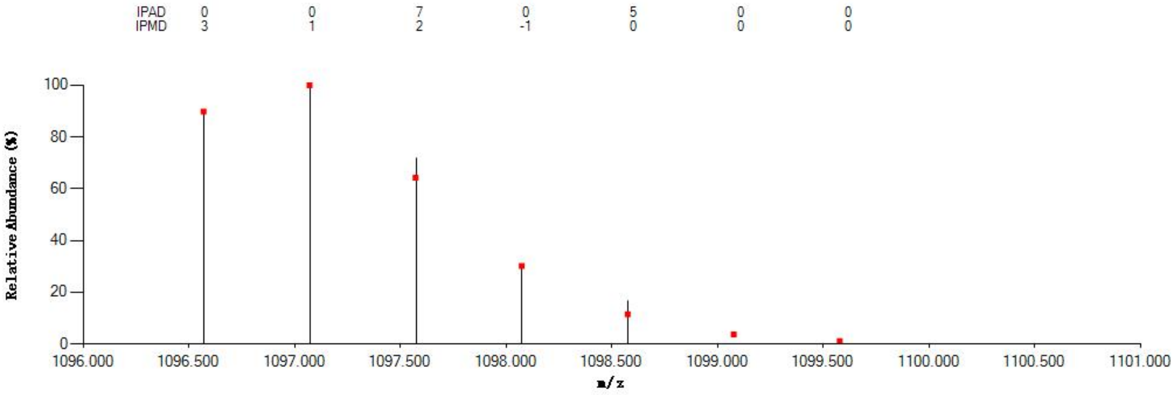

#### Annotated MS/MS Spectrum

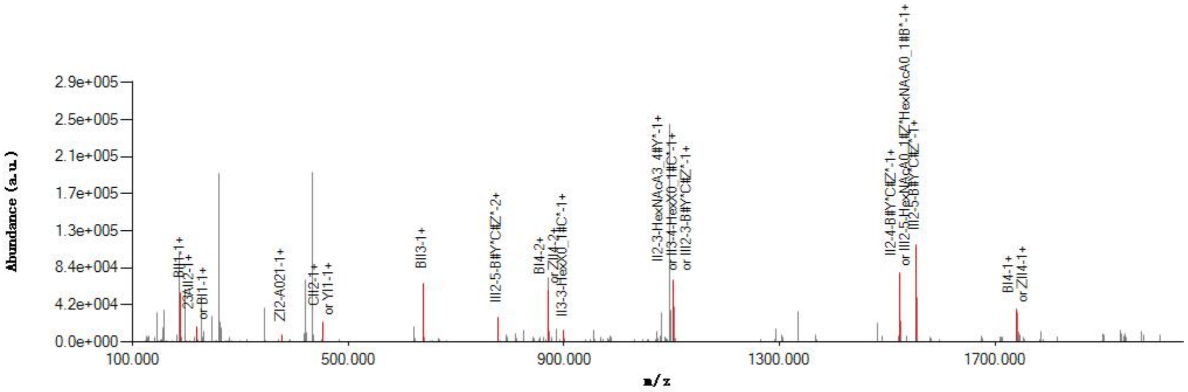

#### Graphical Fragmentation Map

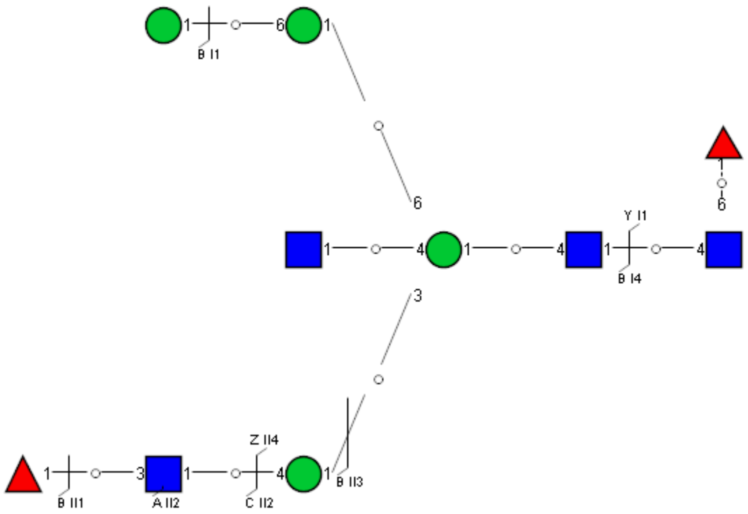

Spectrum Index: 31271(NoBHs:1)

#### Precursor iEF Map

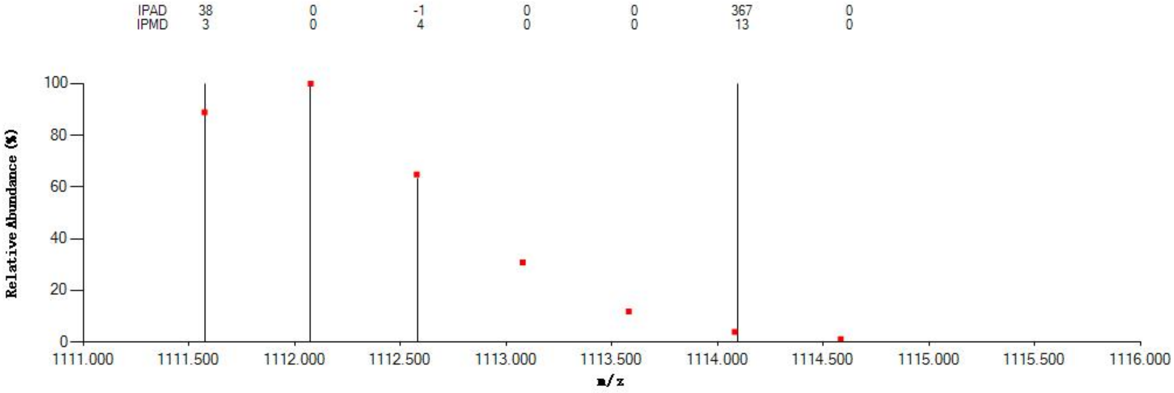


#### Annotated MS/MS Spectrum

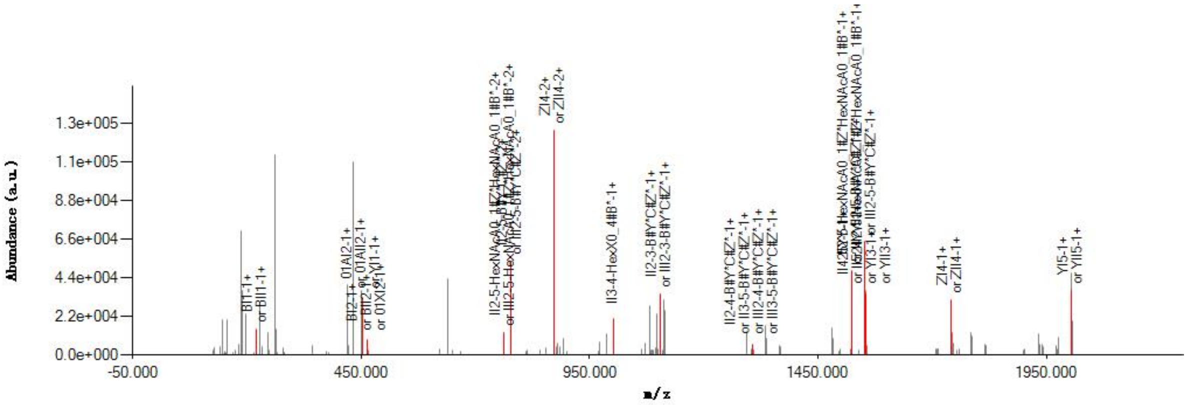


#### Graphical Fragmentation Map

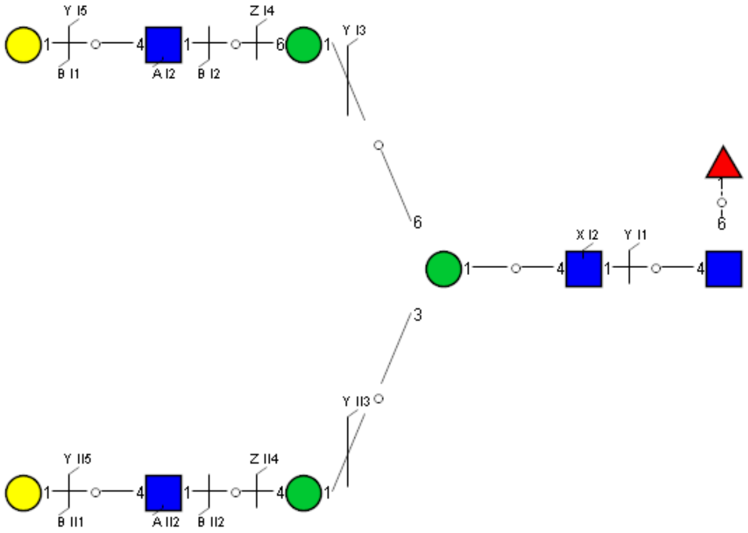

Spectrum Index: 31541(NoBHs:1)

#### Precursor iEF Map

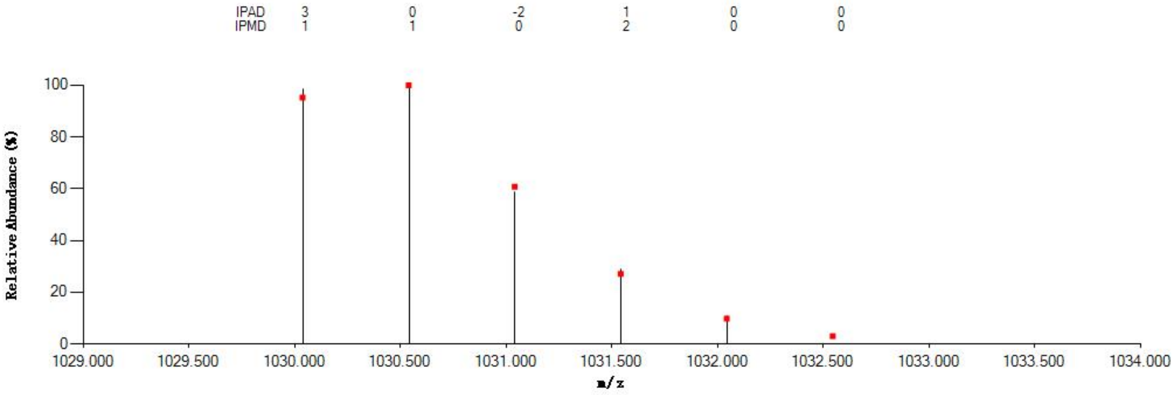


#### Annotated MS/MS Spectrum

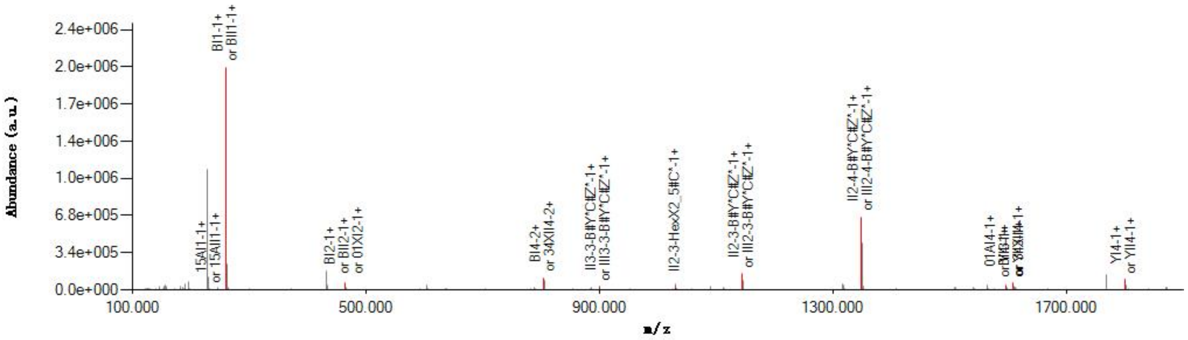


#### Graphical Fragmentation Map

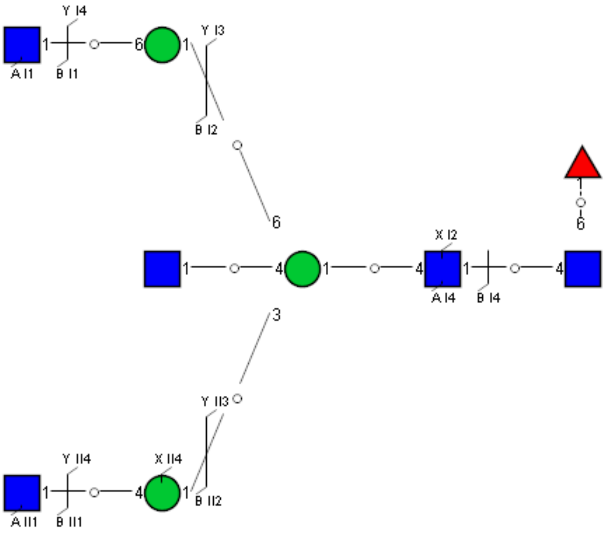

Spectrum Index: 31562(NoBHs:1)

#### Precursor iEF Map

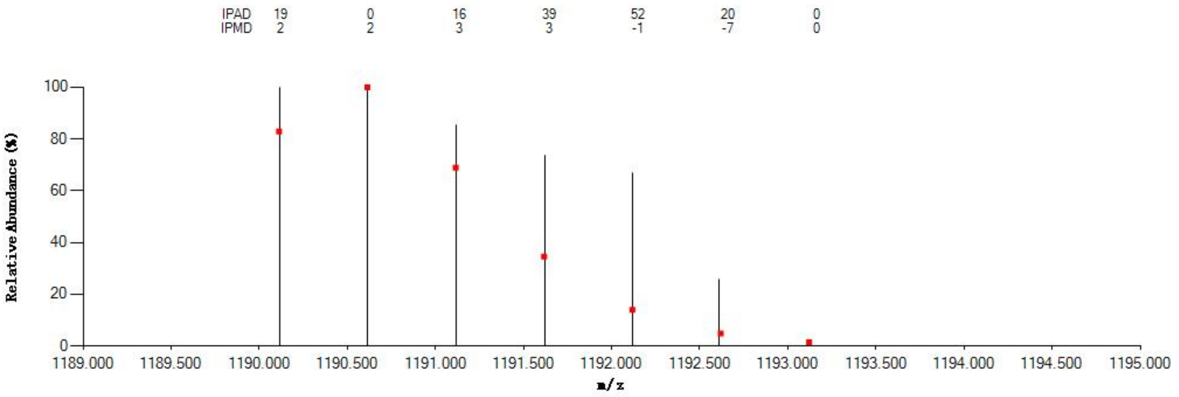


#### Annotated MS/MS Spectrum

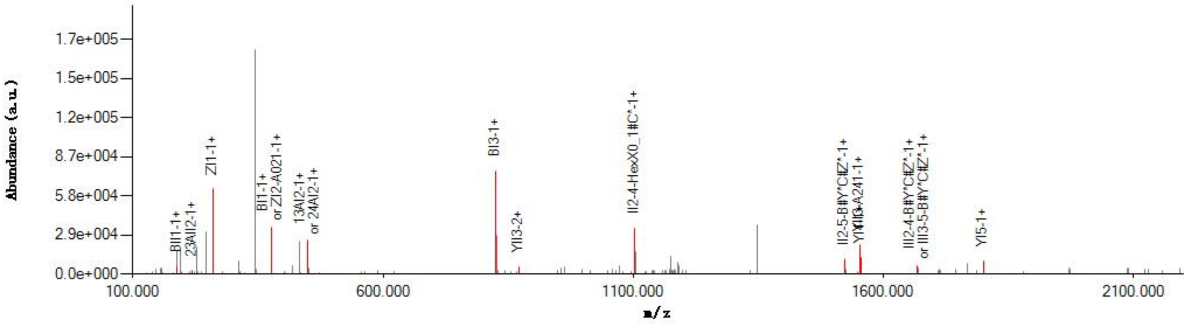


#### Graphical Fragmentation Map

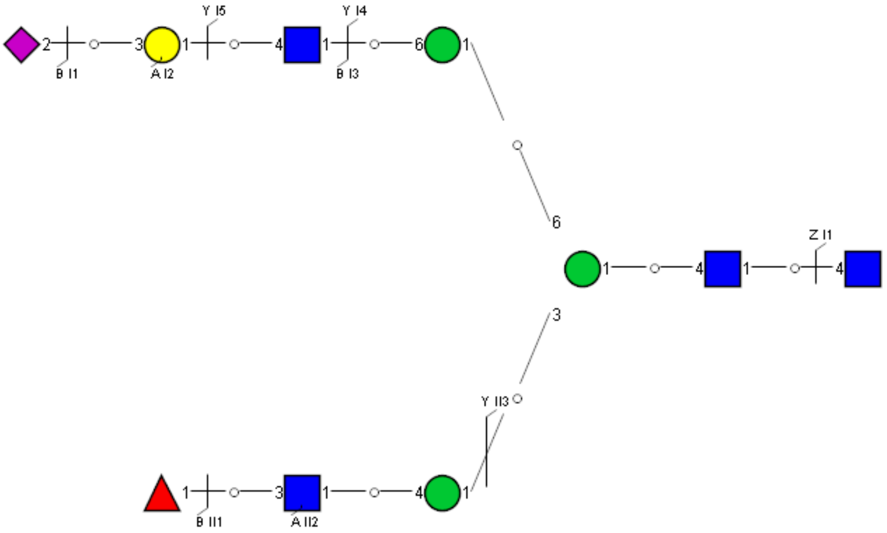

Spectrum Index: 31919(NoBHs:1)

#### Precursor iEF Map

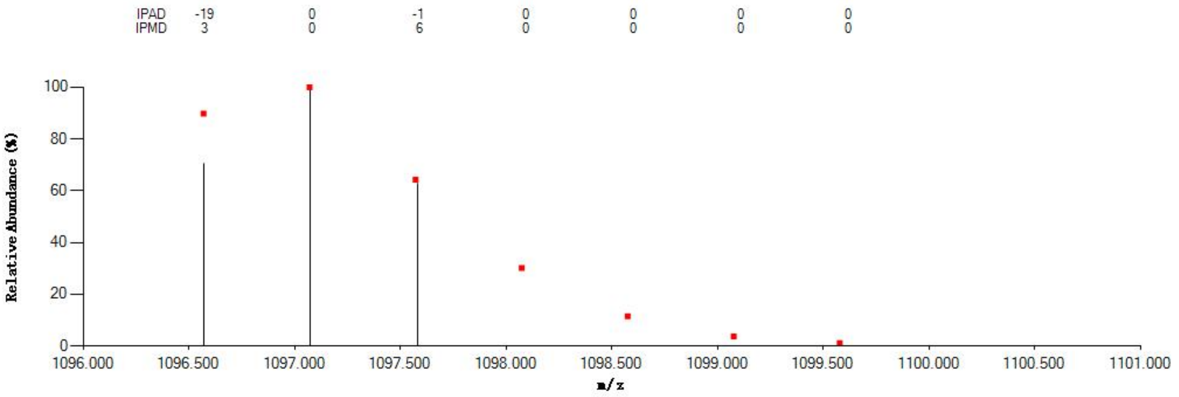

#### Annotated MS/MS Spectrum

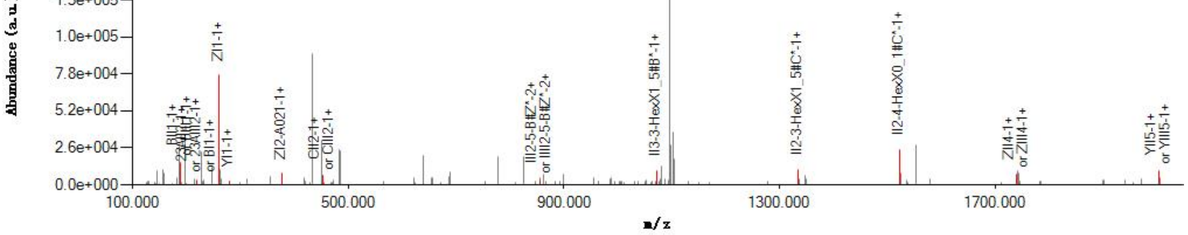

#### Graphical Fragmentation Map

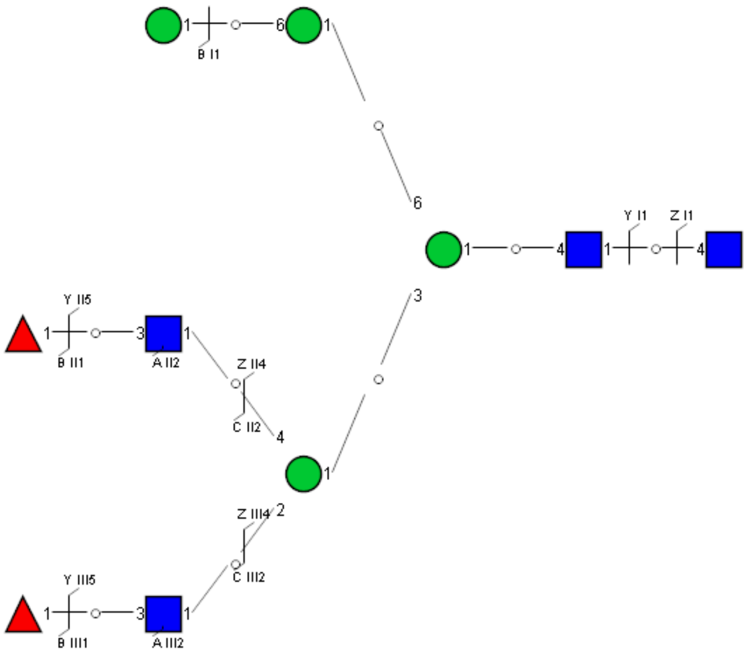

Spectrum Index: 31976(NoBHs:1)

#### Precursor iEF Map

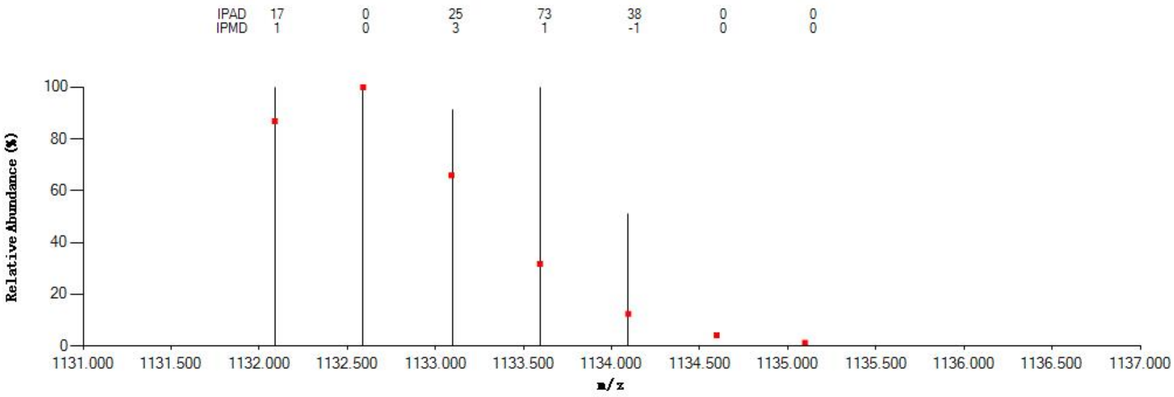


#### Annotated MS/MS Spectrum

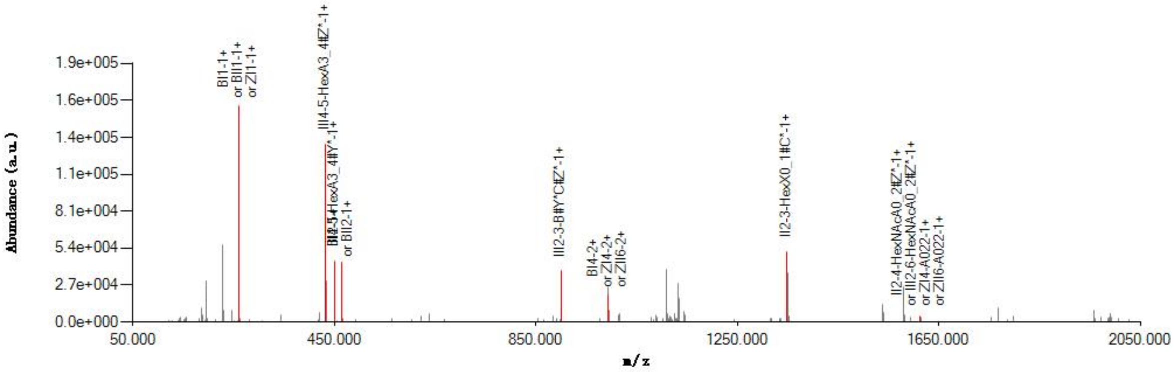


#### Graphical Fragmentation Map

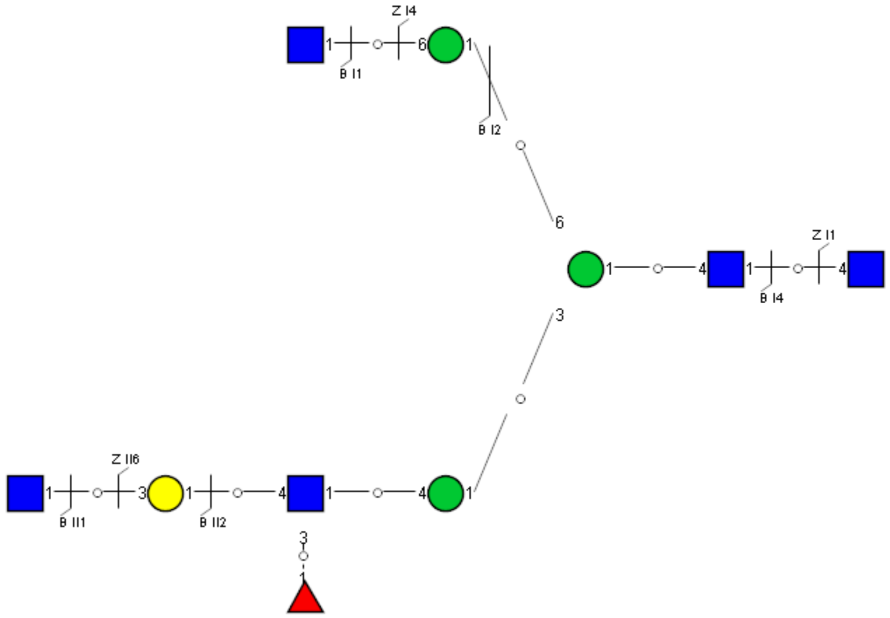

Spectrum Index: 32056(NoBHs:1)

#### Precursor iEF Map

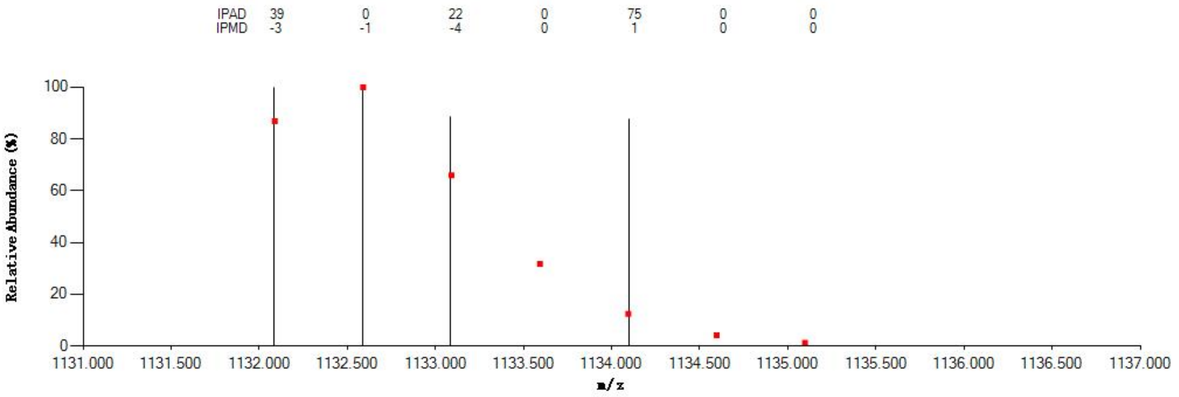


#### Annotated MS/MS Spectrum

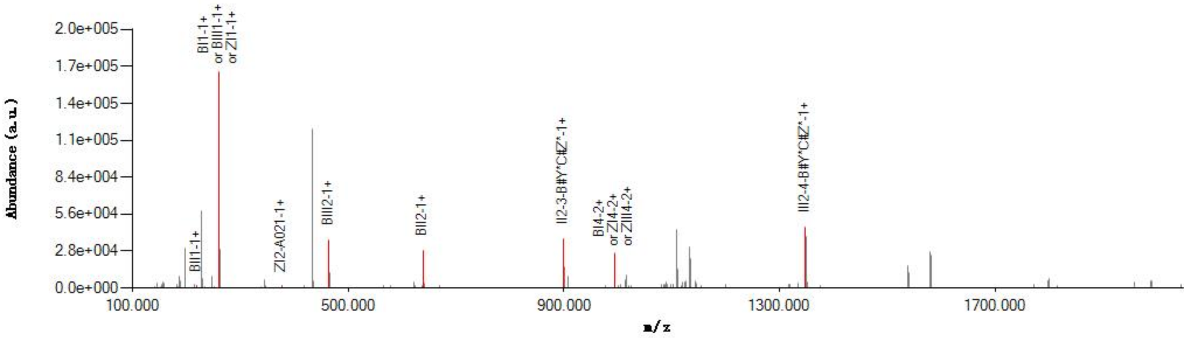


#### Graphical Fragmentation Map

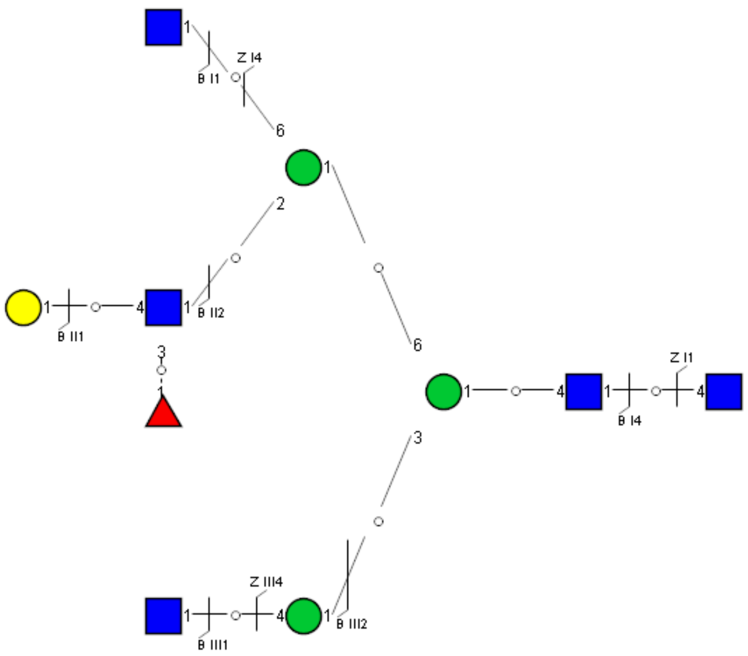

Spectrum Index: 32293(NoBHs:1)

#### Precursor iEF Map

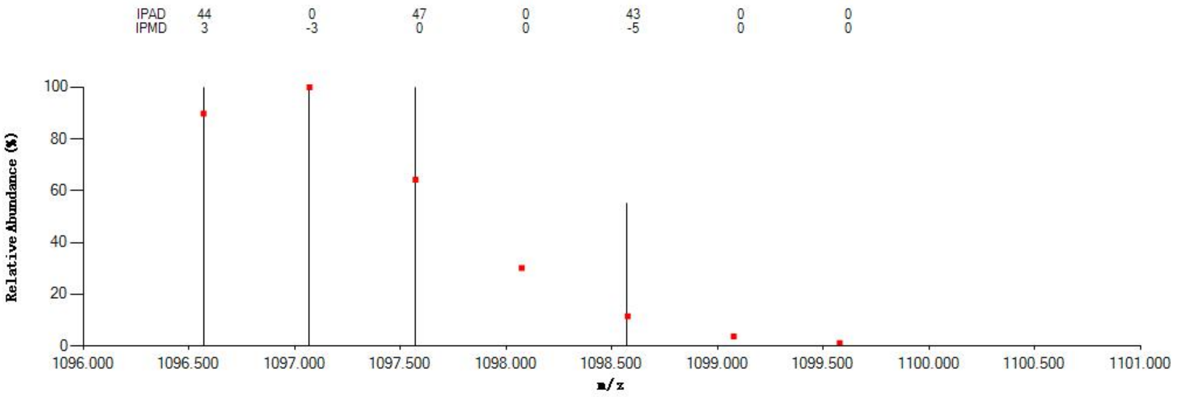


#### Annotated MS/MS Spectrum

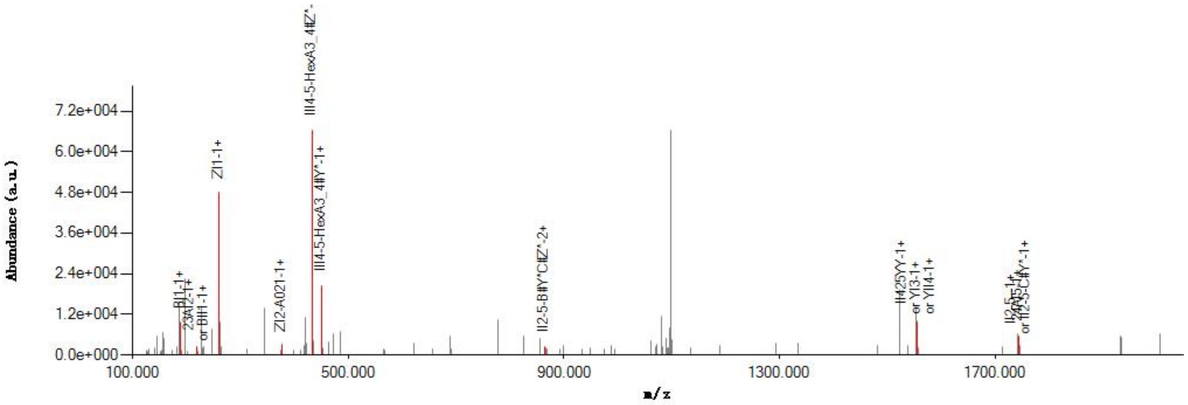


#### Graphical Fragmentation Map

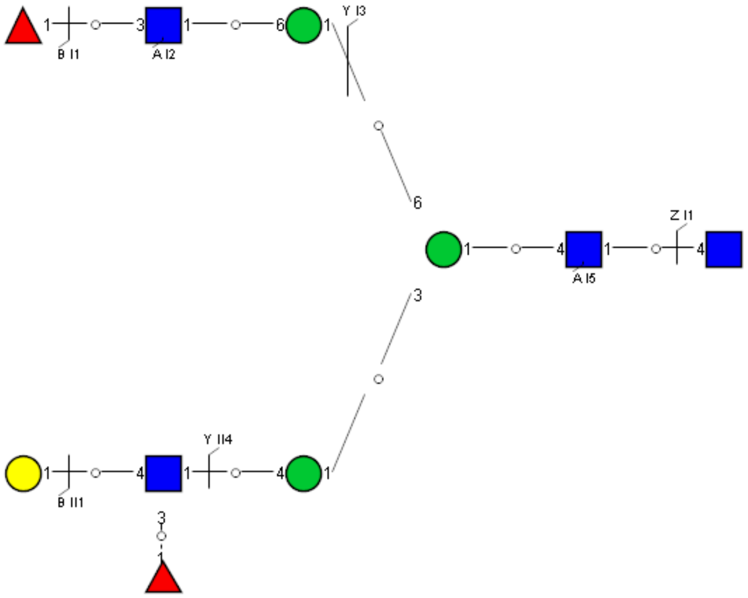

Spectrum Index: 32318(NoBHs:1)

#### Precursor iEF Map

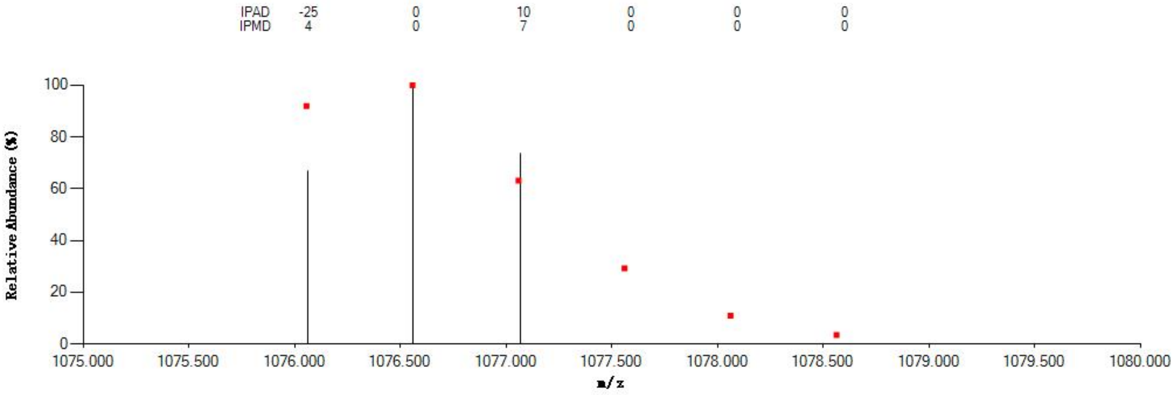

#### Annotated MS/MS Spectrum

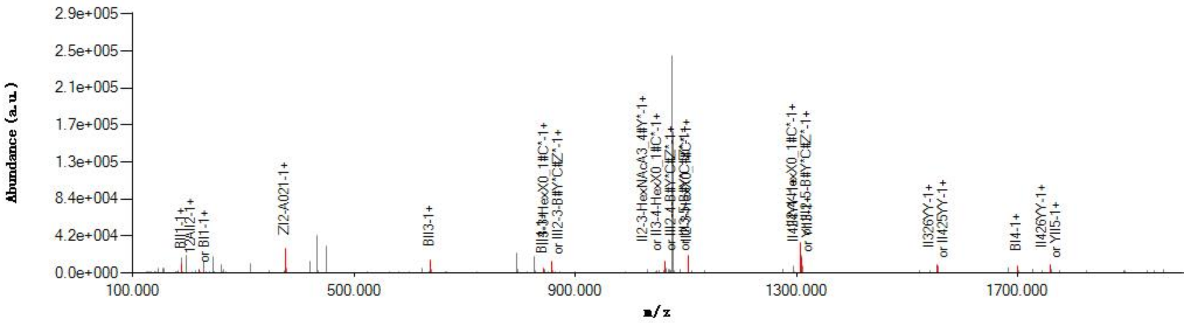

#### Graphical Fragmentation Map

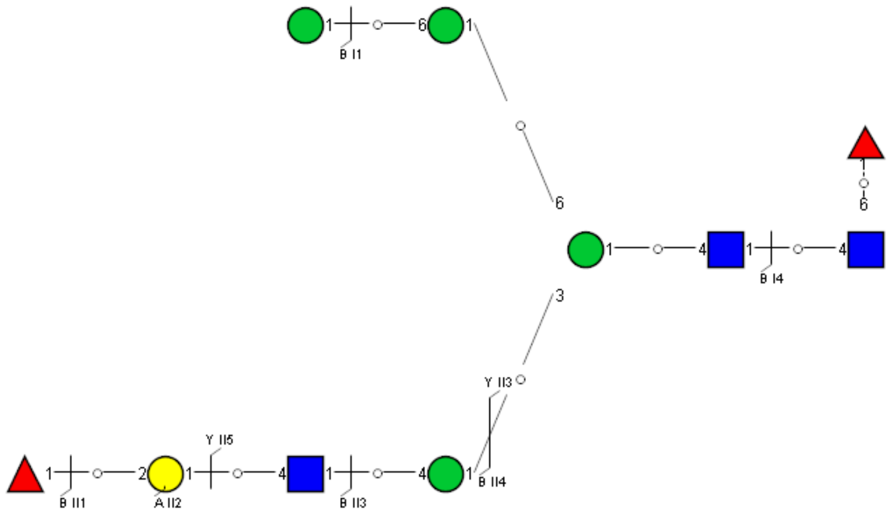

Spectrum Index: 32507(NoBHs:1)

#### Precursor iEF Map

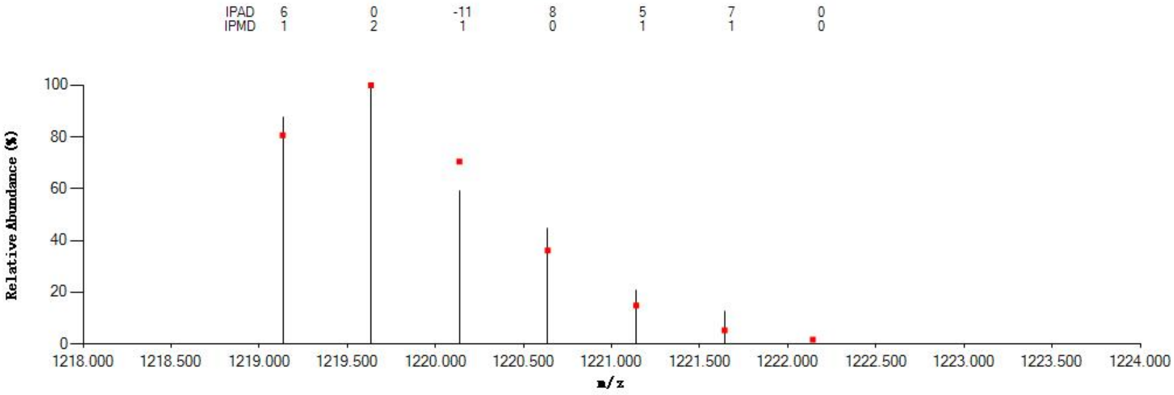


#### Annotated MS/MS Spectrum

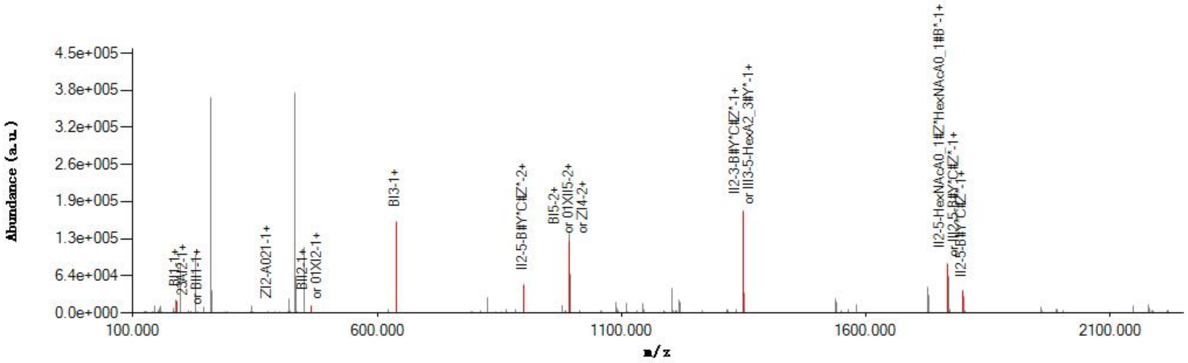


#### Graphical Fragmentation Map

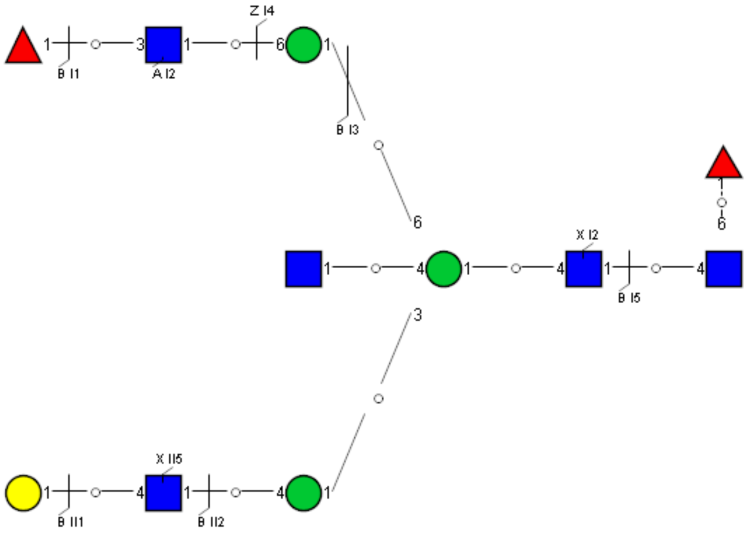

Spectrum Index: 32512(NoBHs:1)

#### Precursor iEF Map

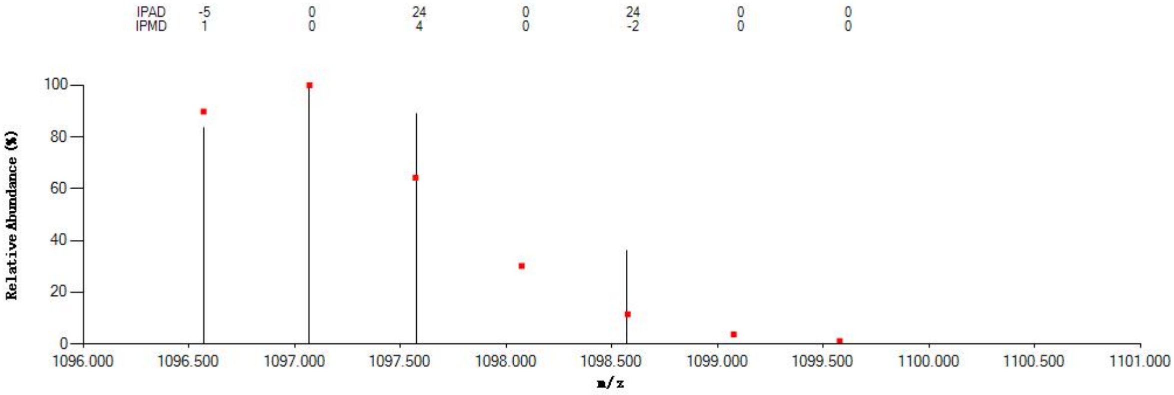


#### Annotated MS/MS Spectrum

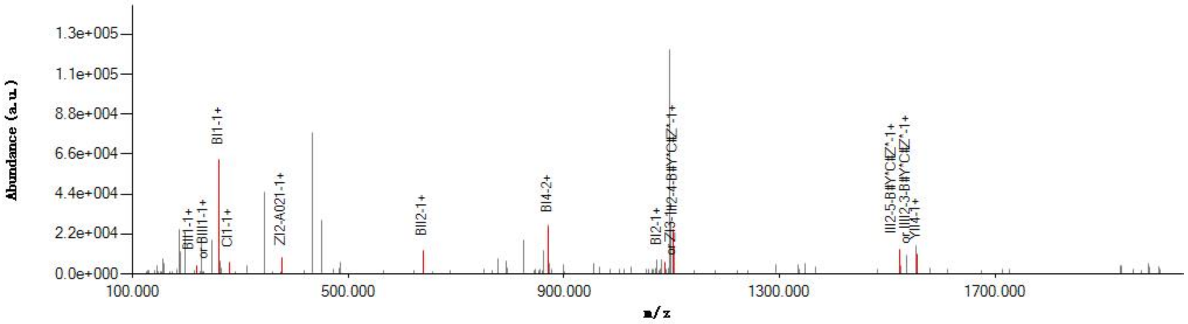


#### Graphical Fragmentation Map

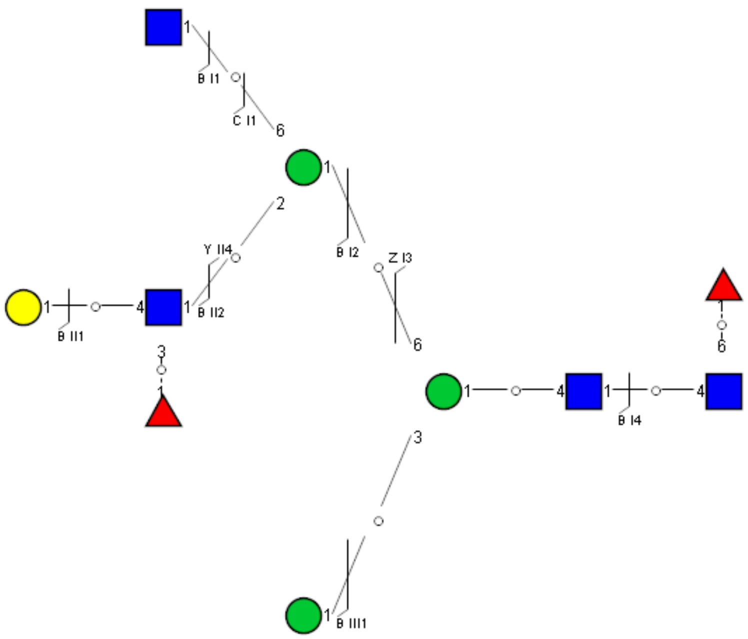

Spectrum Index: 32659(NoBHs:1)

#### Precursor iEF Map

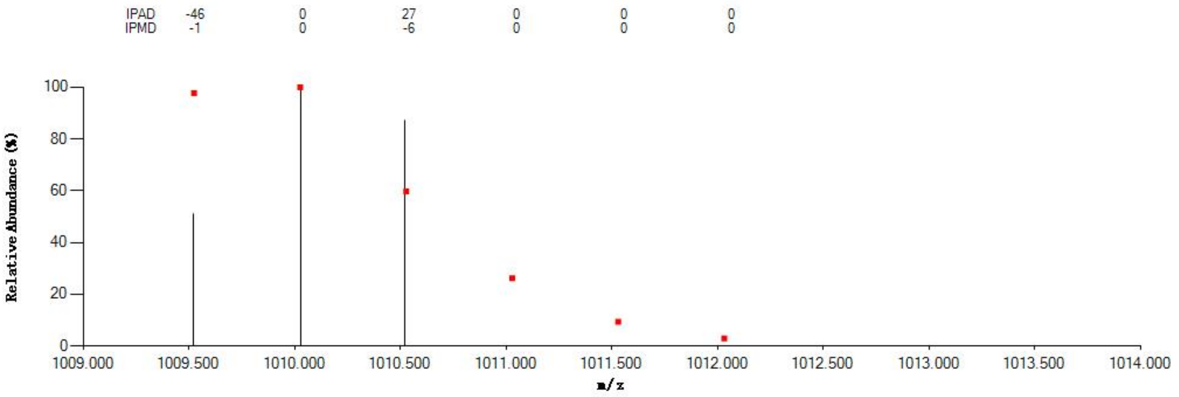


#### Annotated MS/MS Spectrum

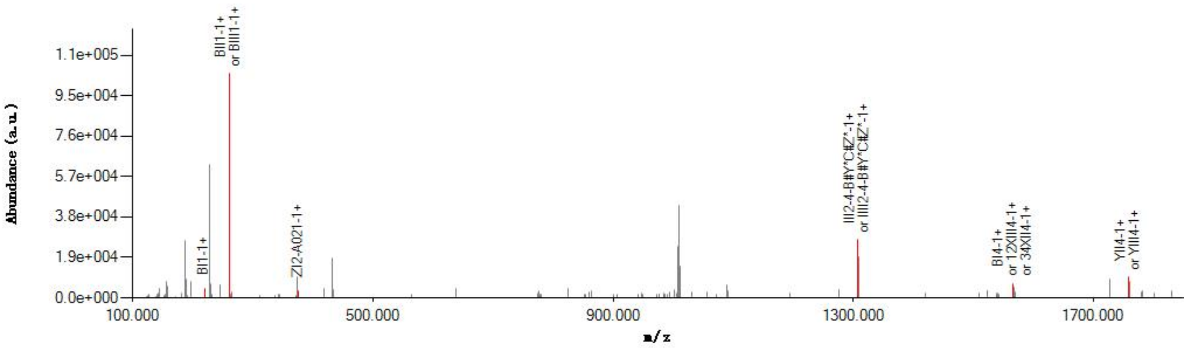


#### Graphical Fragmentation Map

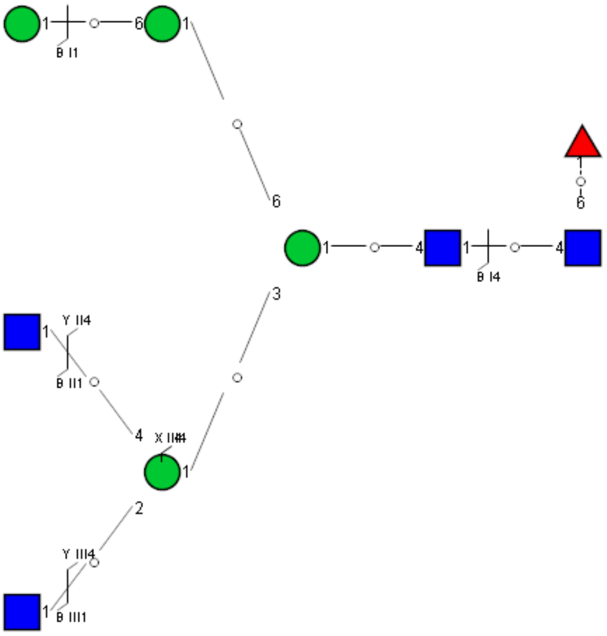

Spectrum Index: 32866(NoBHs:1)

#### Precursor iEF Map

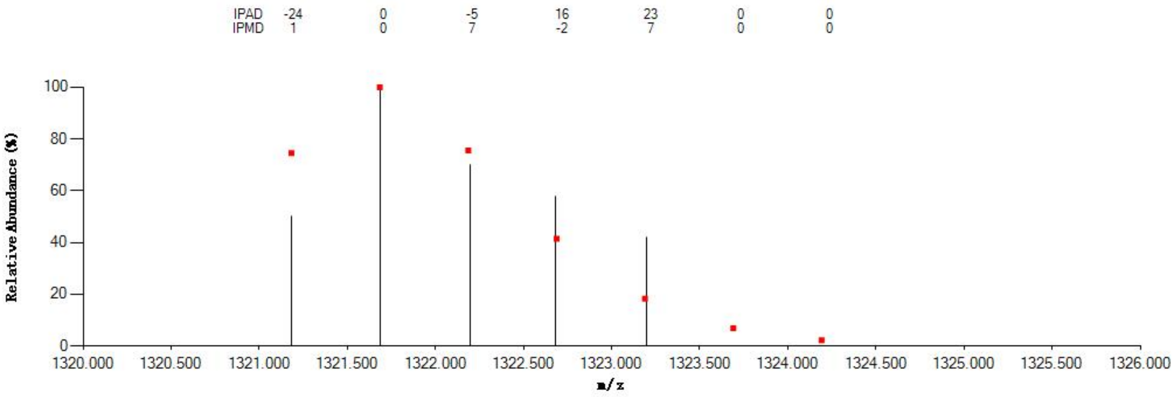


#### Annotated MS/MS Spectrum

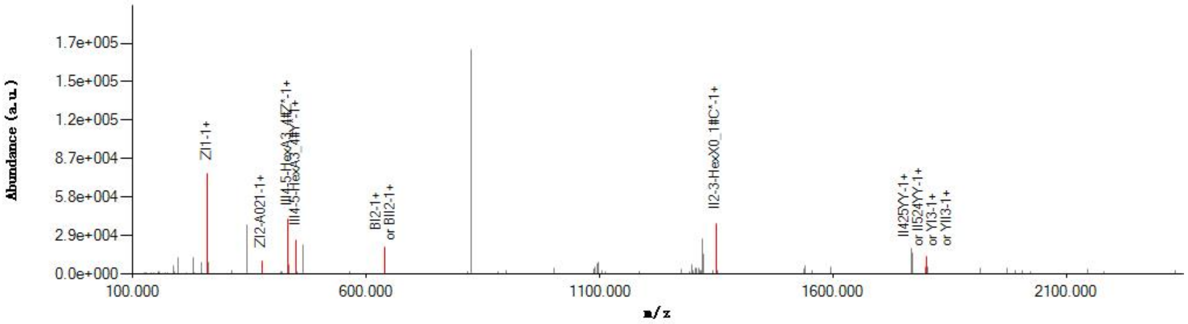


#### Graphical Fragmentation Map

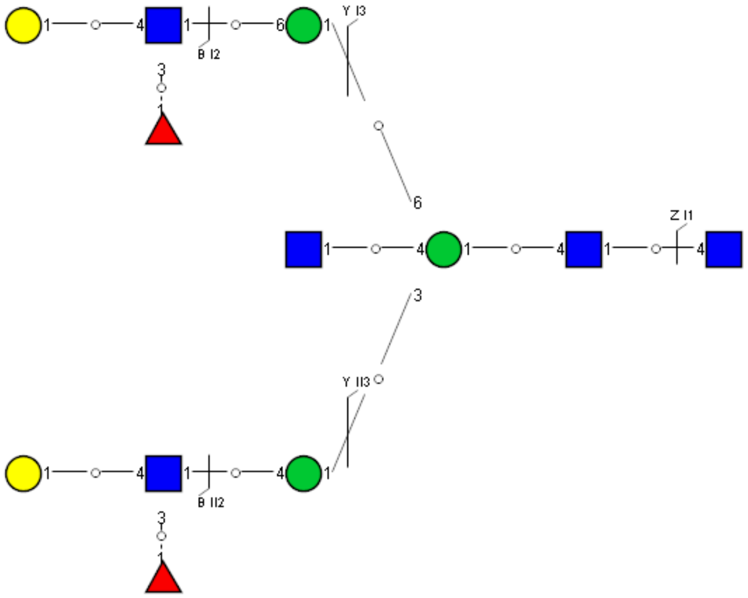

Spectrum Index: 33012(NoBHs:1)

#### Precursor iEF Map

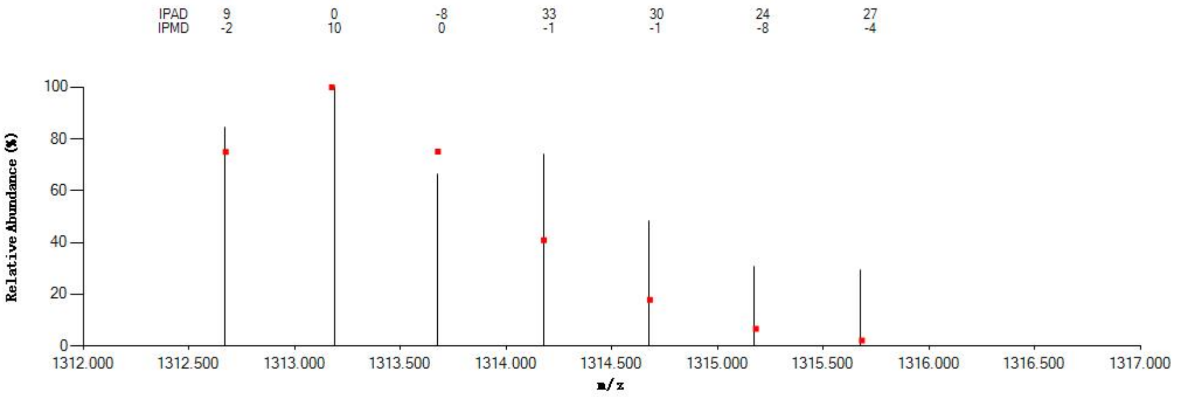


#### Annotated MS/MS Spectrum

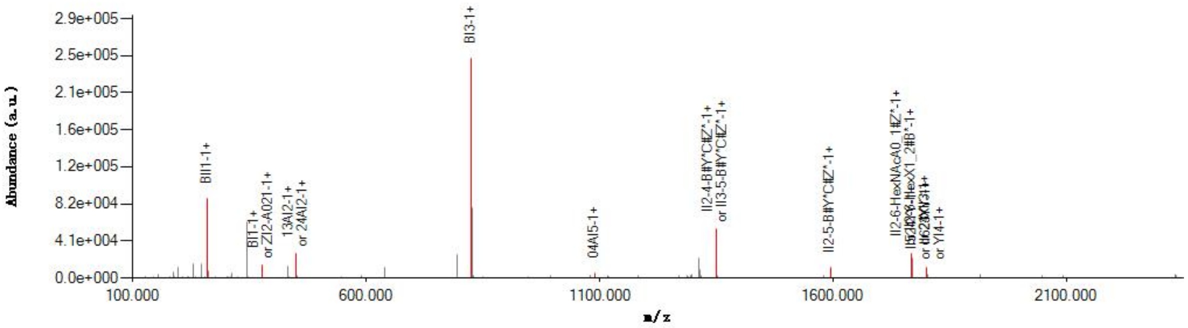


#### Graphical Fragmentation Map

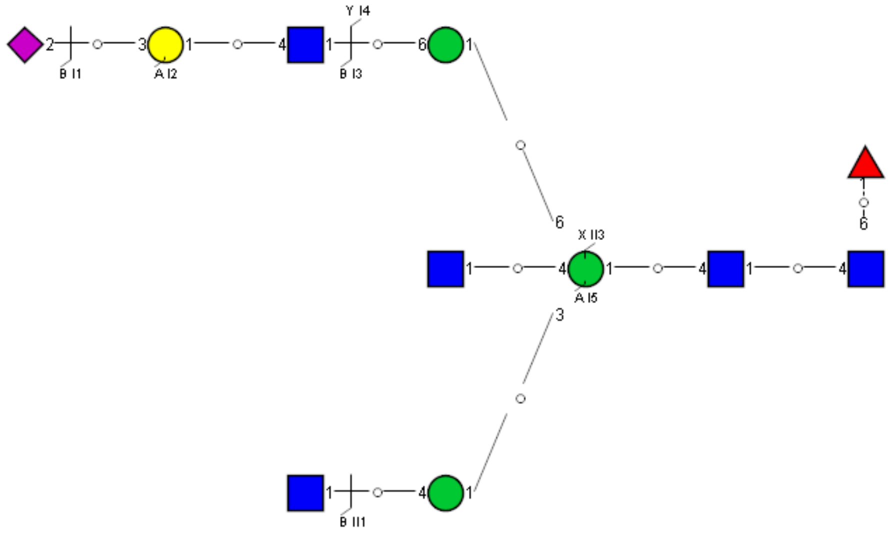

Spectrum Index: 33098(NoBHs:1)

#### Precursor iEF Map

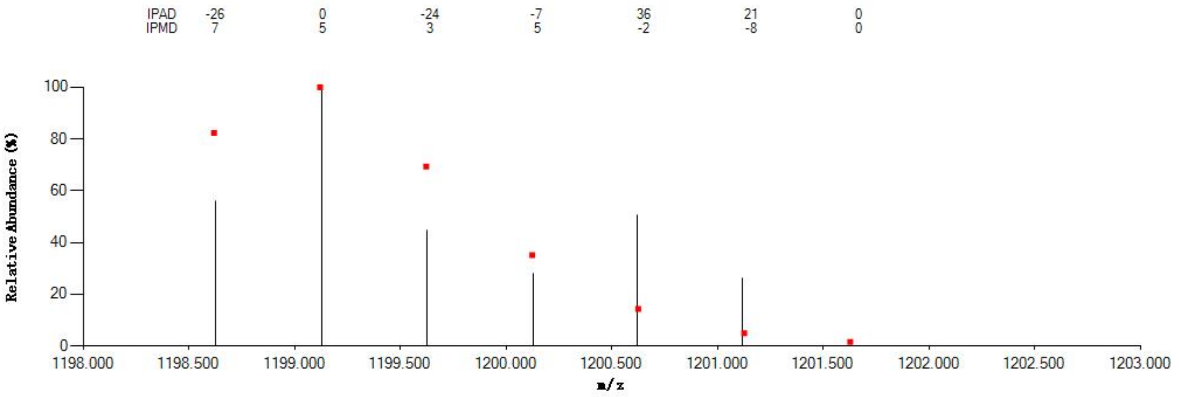

#### Annotated MS/MS Spectrum

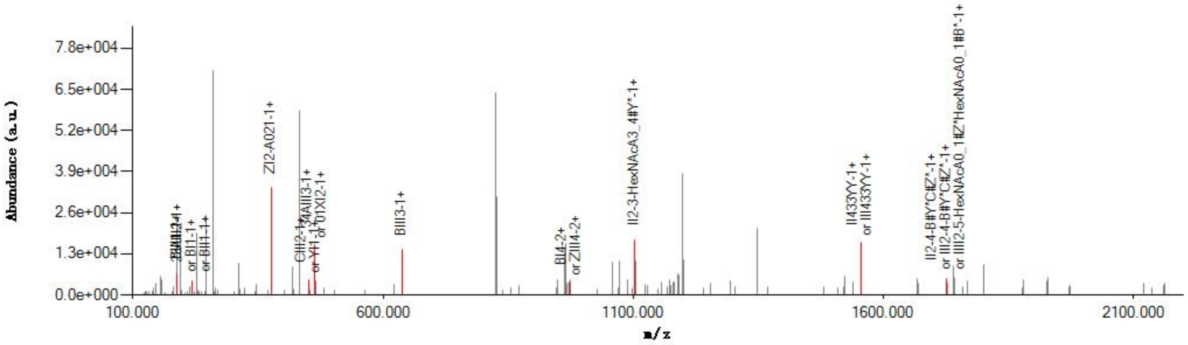

#### Graphical Fragmentation Map

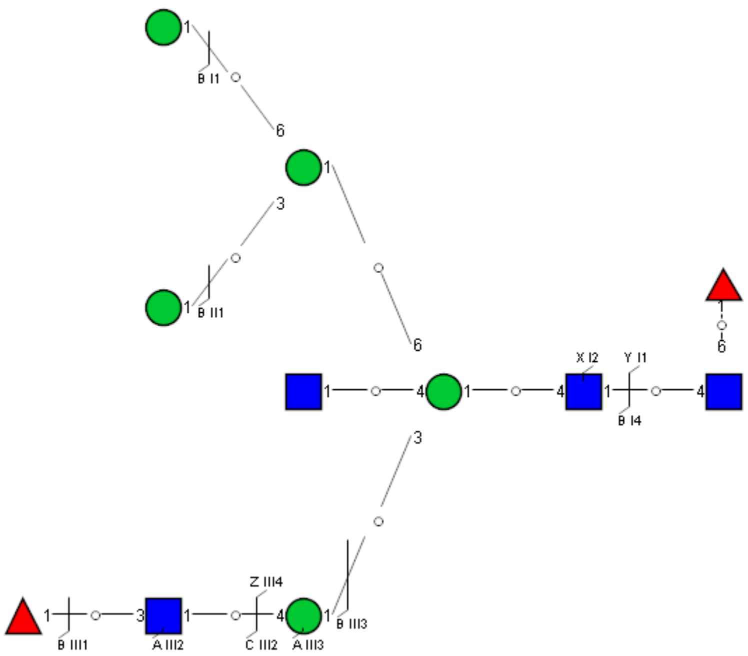

Spectrum Index: 33139(NoBHs:1)

#### Precursor iEF Map

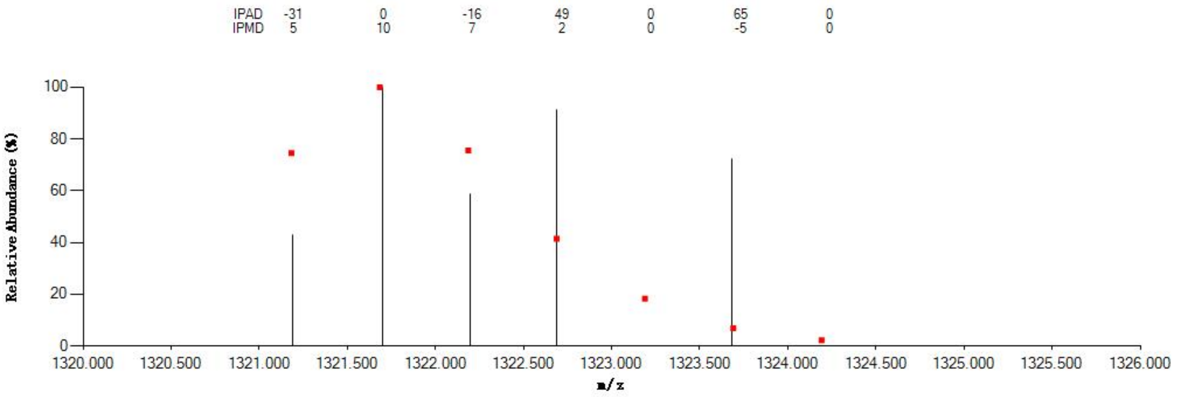


#### Annotated MS/MS Spectrum

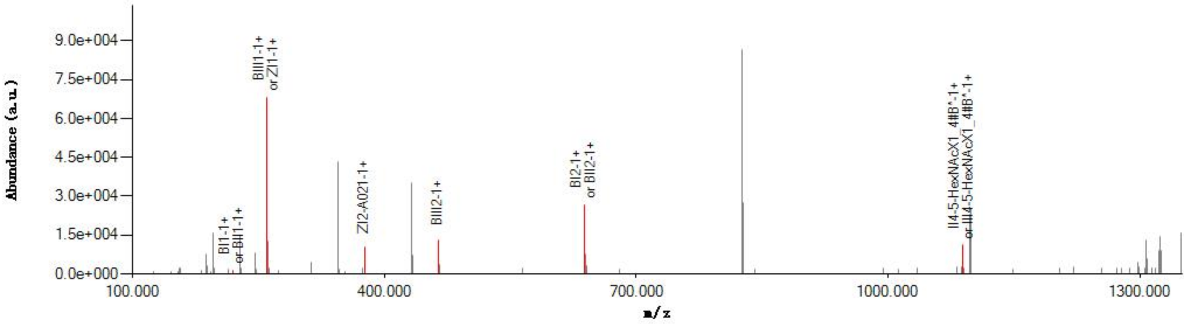


#### Graphical Fragmentation Map

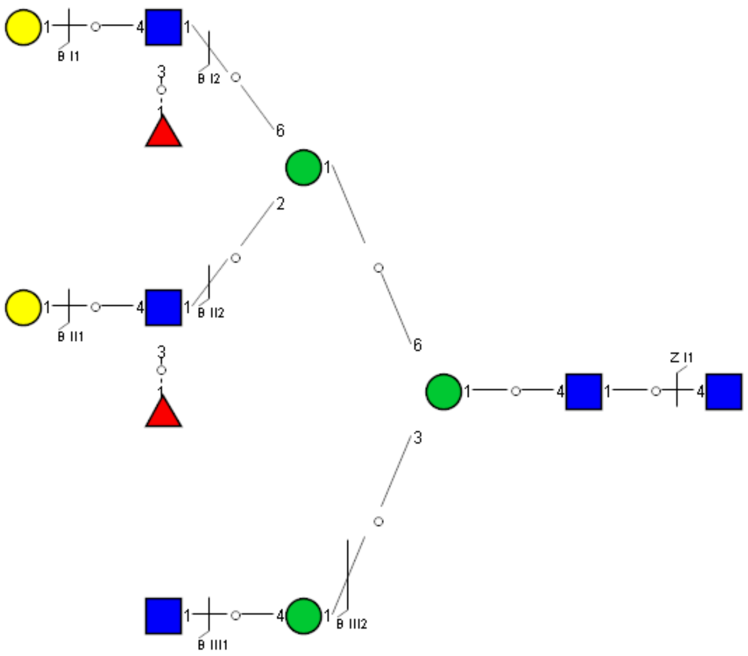

Spectrum Index: 33242(NoBHs:1)

#### Precursor iEF Map

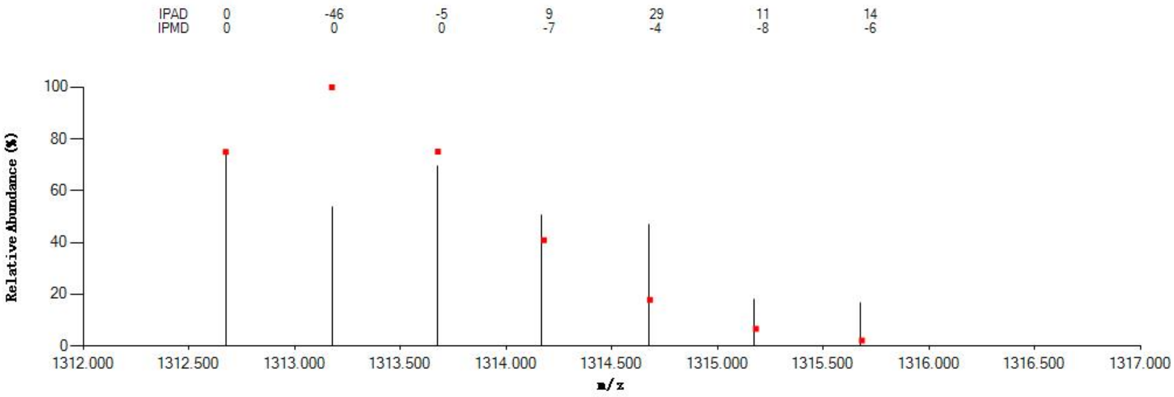


#### Annotated MS/MS Spectrum

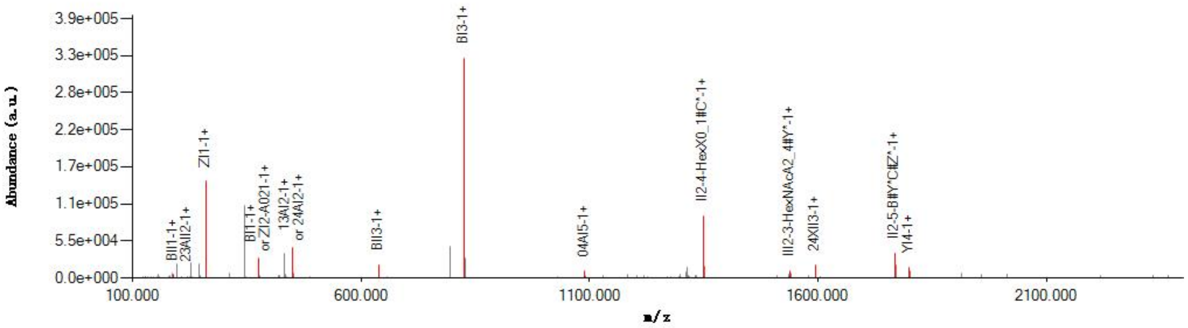


#### Graphical Fragmentation Map

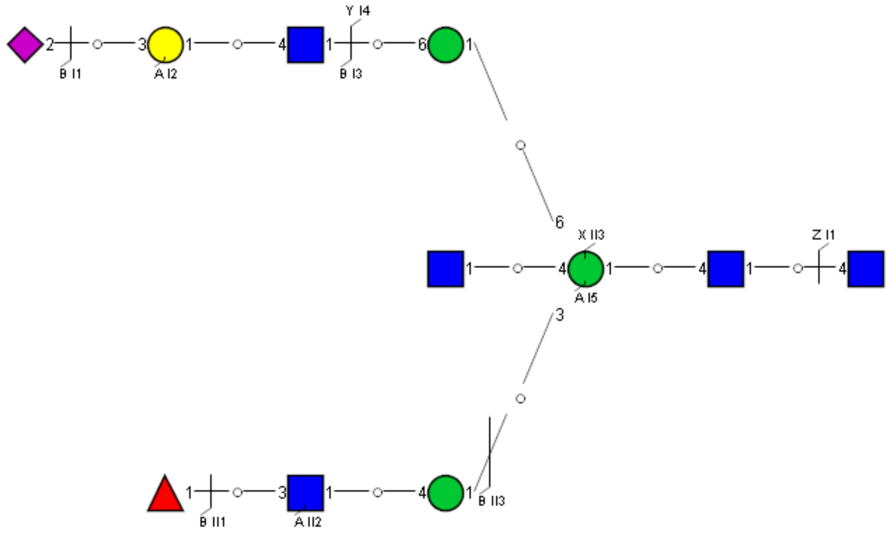

Spectrum Index: 33328(NoBHs:1)

#### Precursor iEF Map

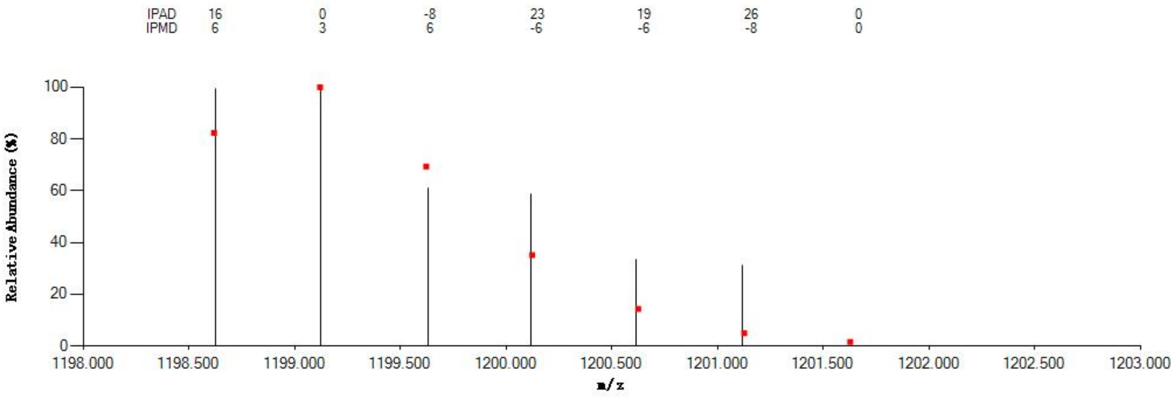


#### Annotated MS/MS Spectrum

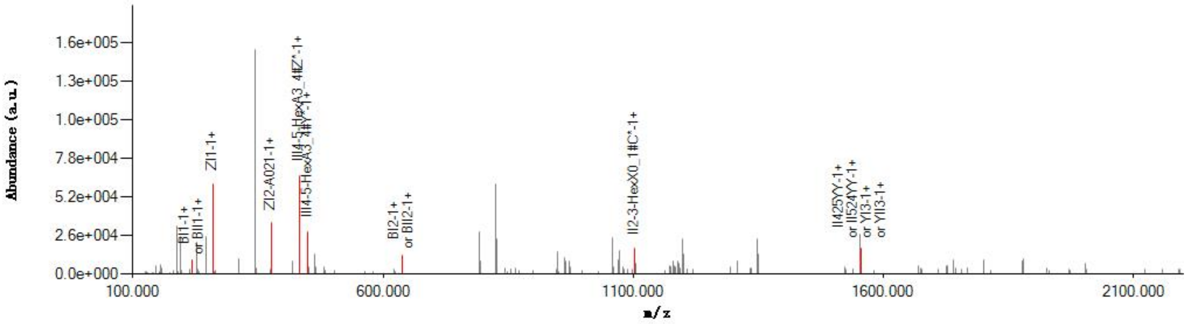


#### Graphical Fragmentation Map

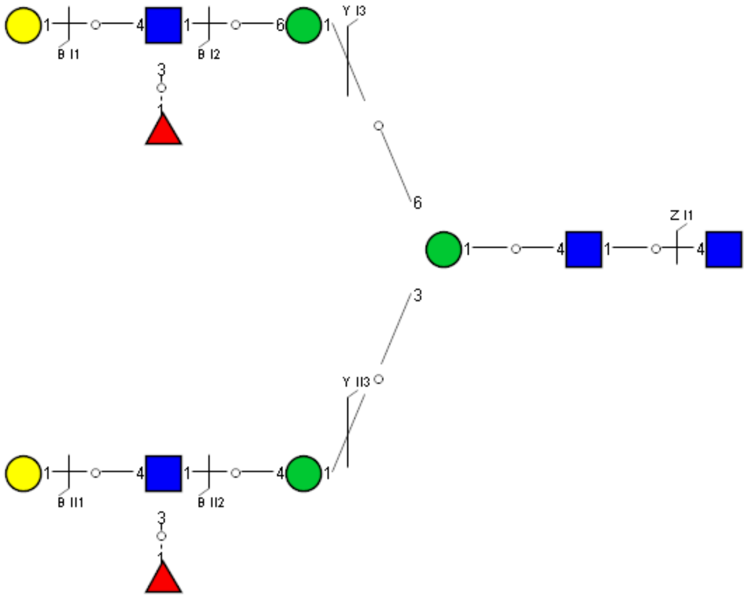

### Spectrum Index: 34084(NoBHs:1)

#### Precursor iEF Map

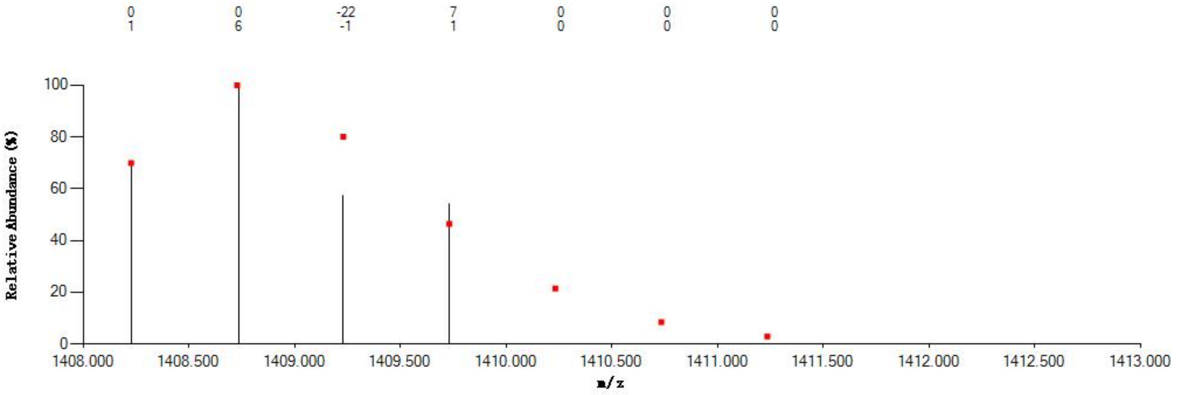

#### Annotated MS/MS Spectrum

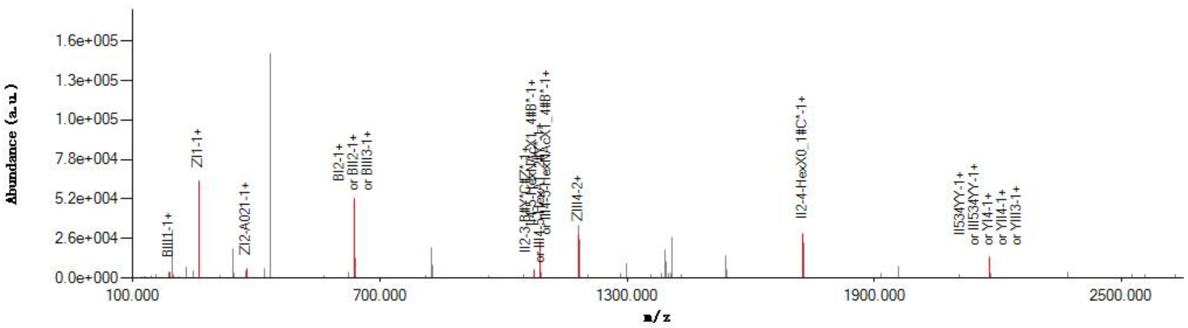

#### Graphical Fragmentation Map

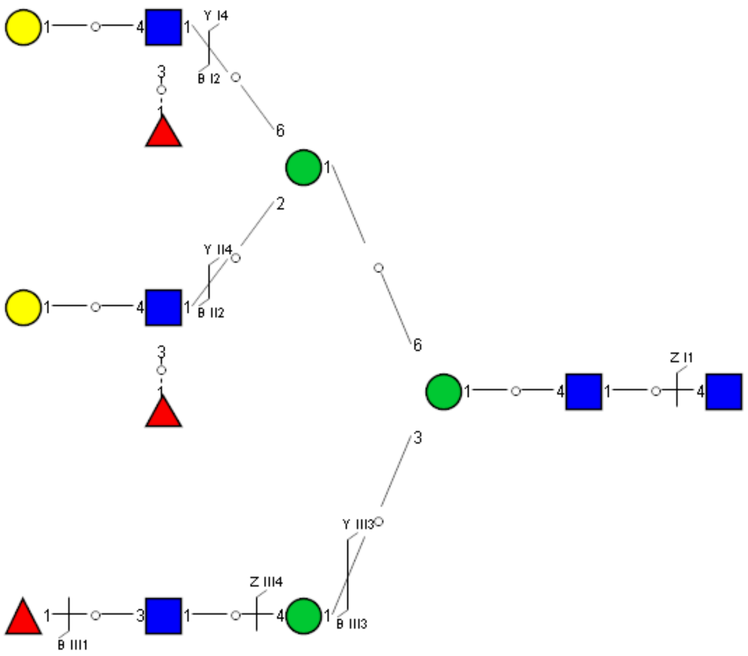

Spectrum Index: 35595(NoBHs:1)

#### Precursor iEF Map

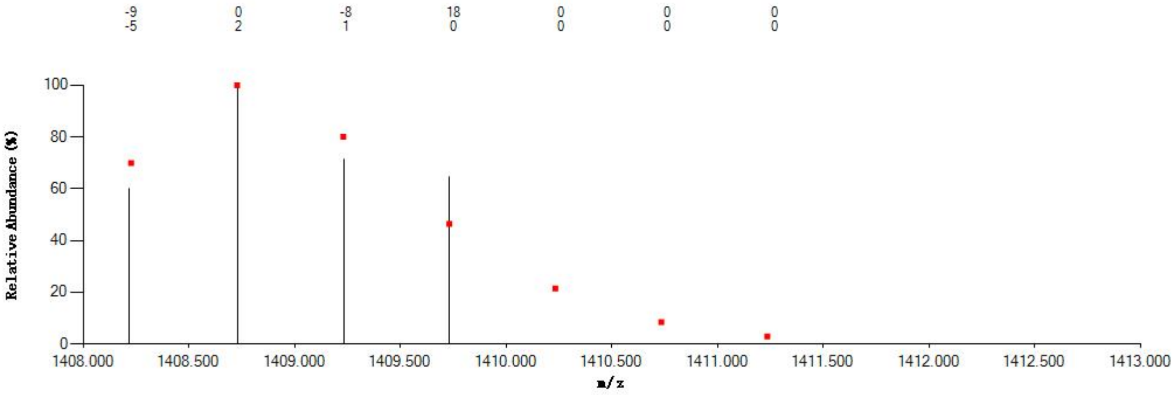

#### Annotated MS/MS Spectrum

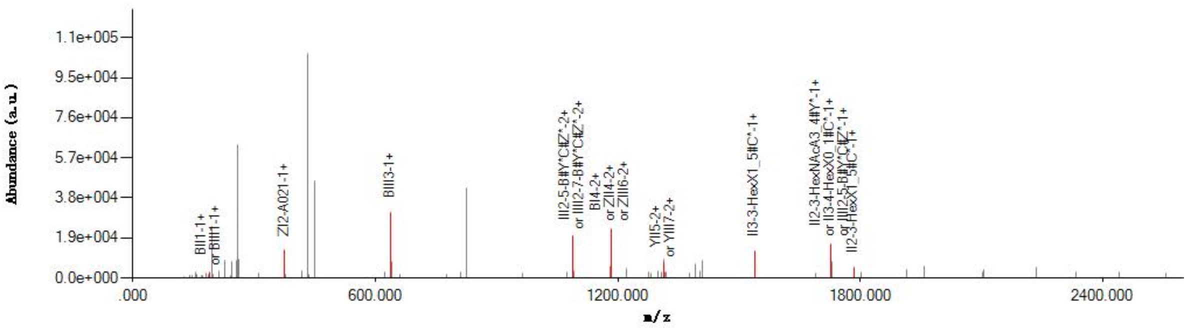

#### Graphical Fragmentation Map

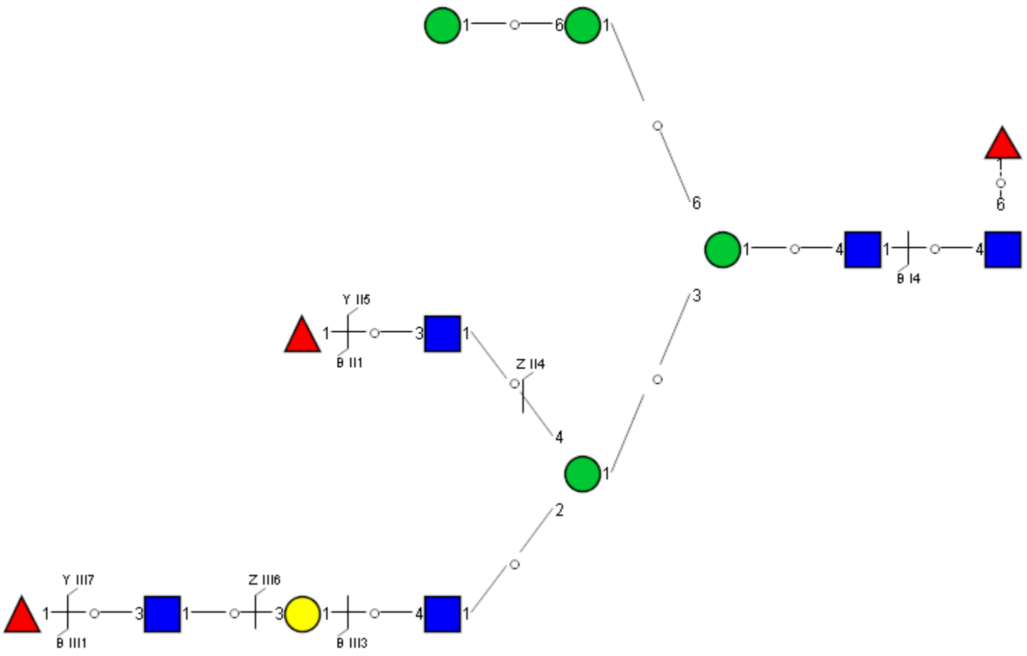

Spectrum Index: 36354(NoBHs:1)

#### Precursor iEF Map

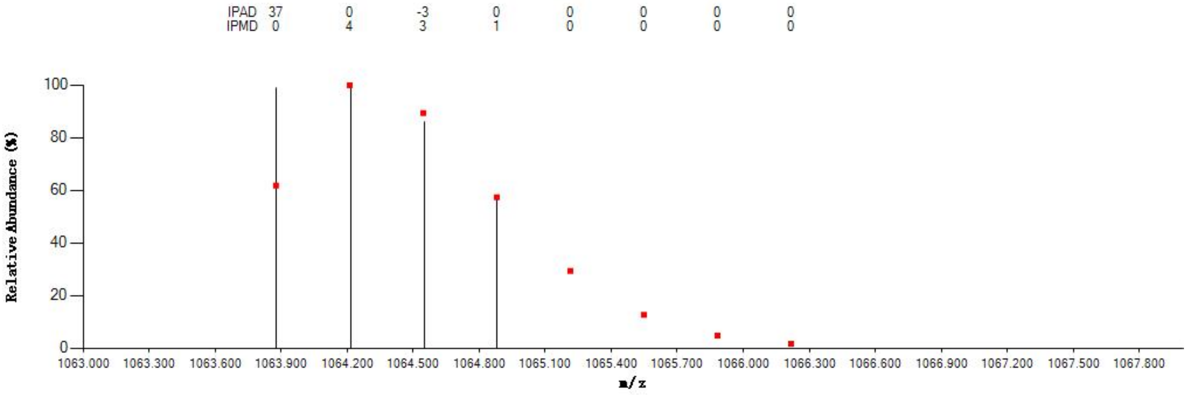


#### Annotated MS/MS Spectrum

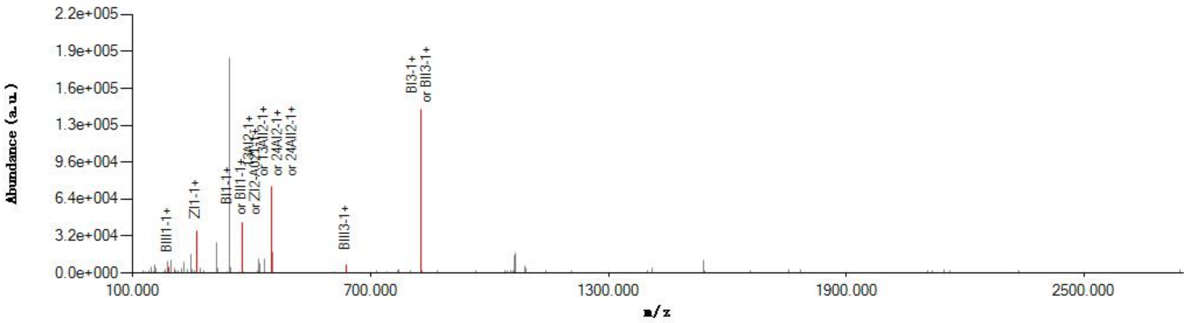


#### Graphical Fragmentation Map

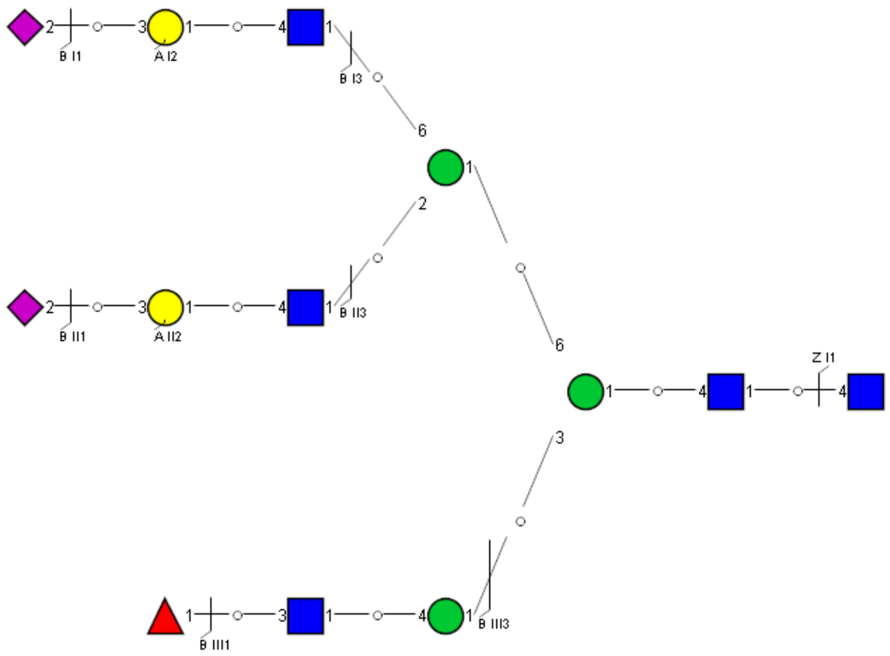

Spectrum Index: 37425(NoBHs:2)

#### Precursor iEF Map

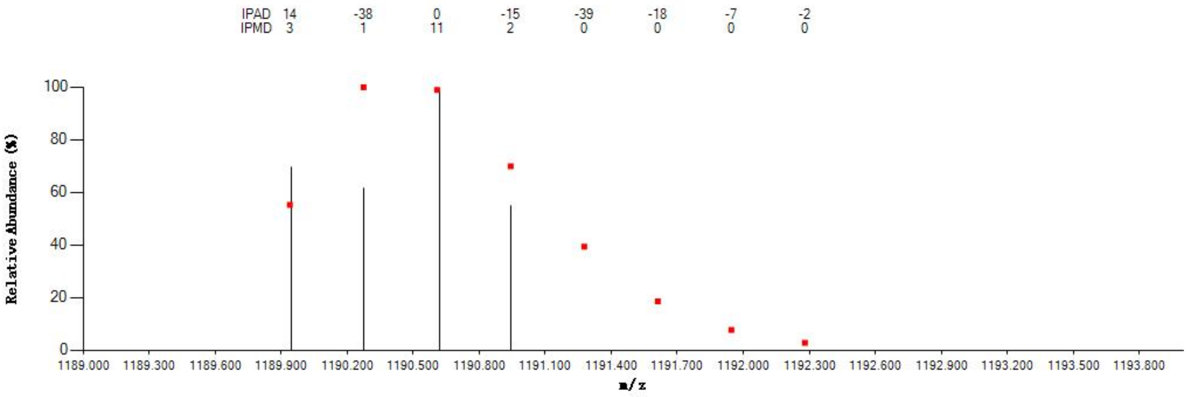


#### Annotated MS/MS Spectrum

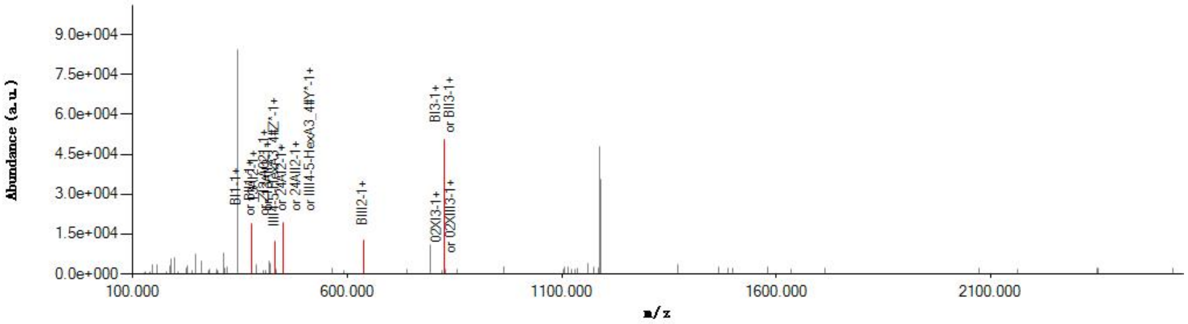


#### Graphical Fragmentation Map

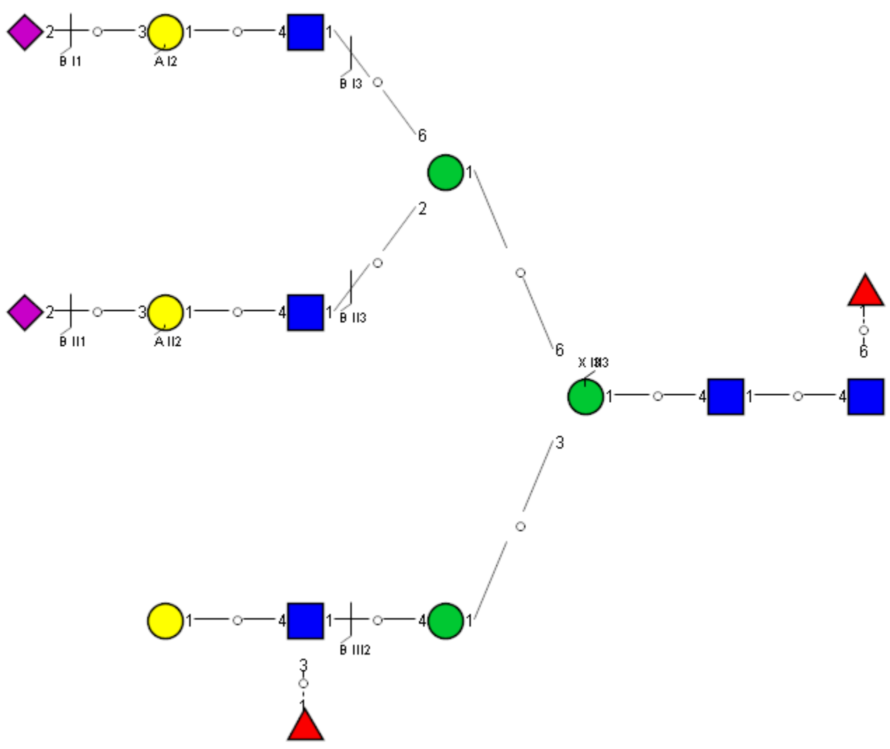

Spectrum Index: 37425(NoBHs:2)

#### Precursor iEF Map

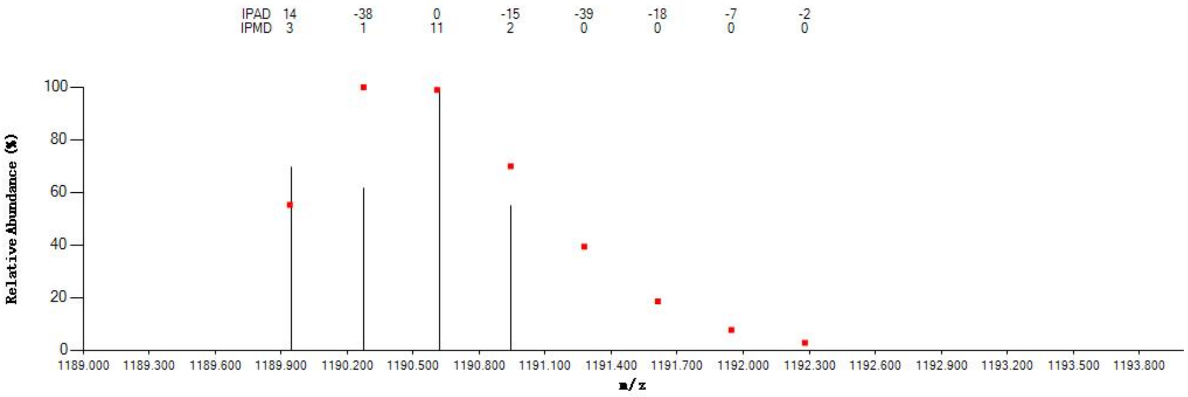


#### Annotated MS/MS Spectrum

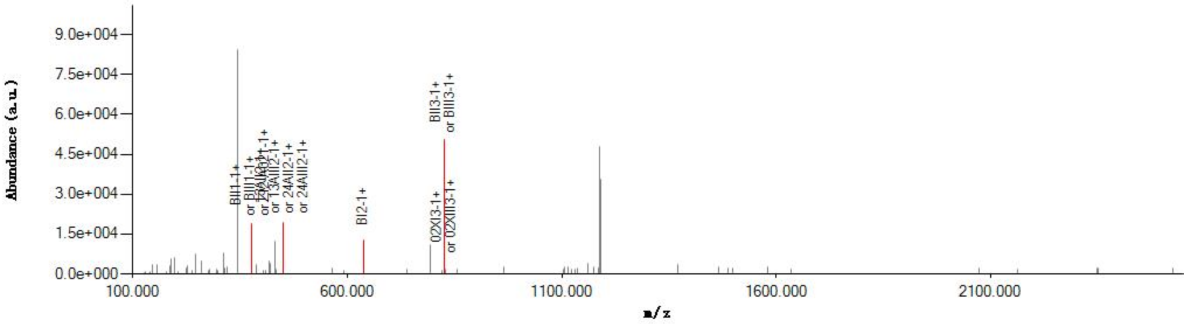


#### Graphical Fragmentation Map

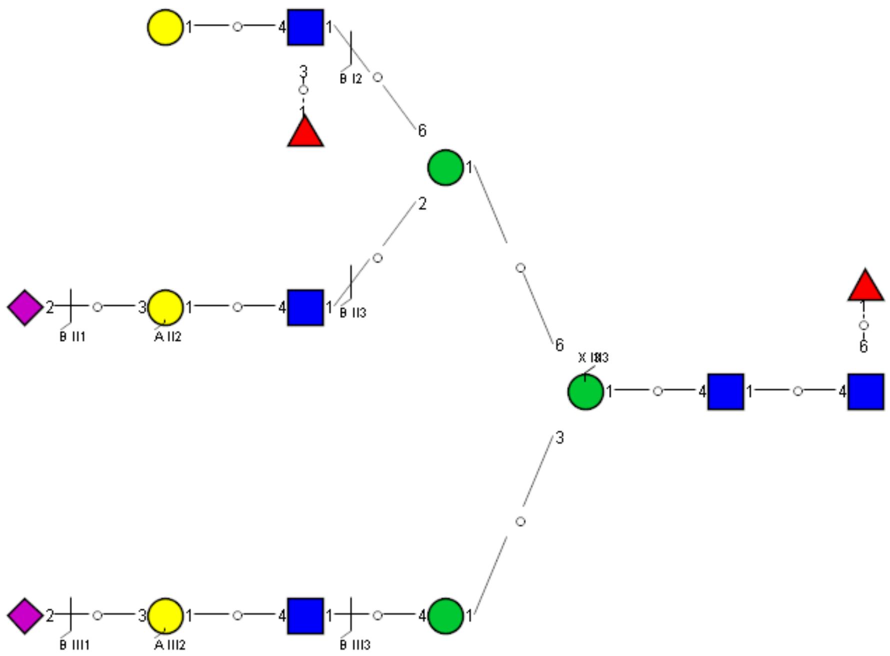

Spectrum Index: 37762(NoBHs:1)

#### Precursor iEF Map

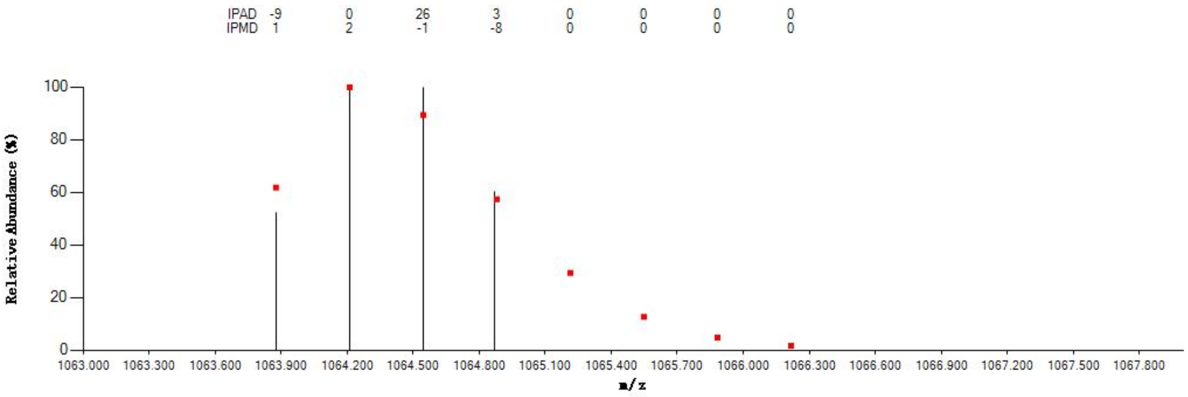


#### Annotated MS/MS Spectrum

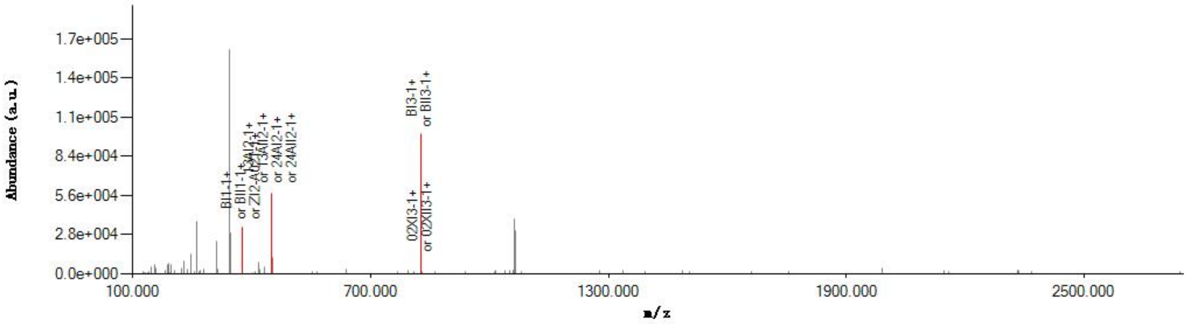


#### Graphical Fragmentation Map

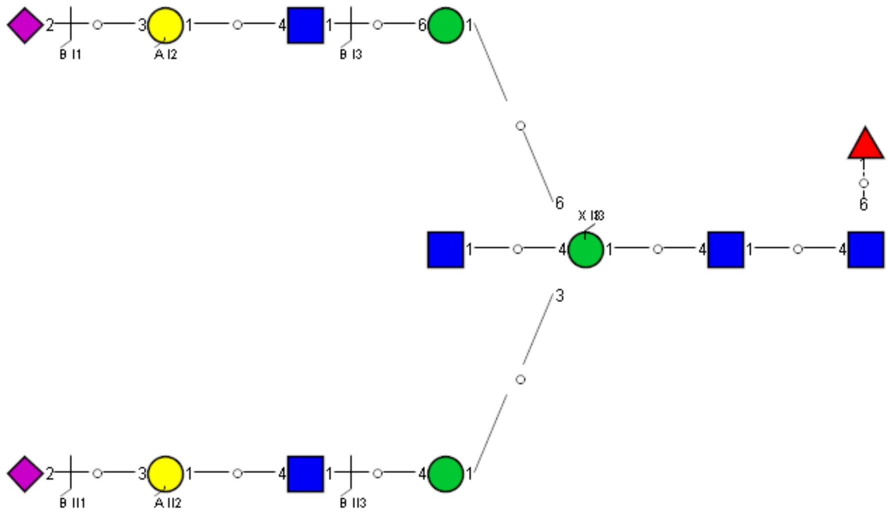

Spectrum Index: 38400(NoBHs:1)

#### Precursor iEF Map

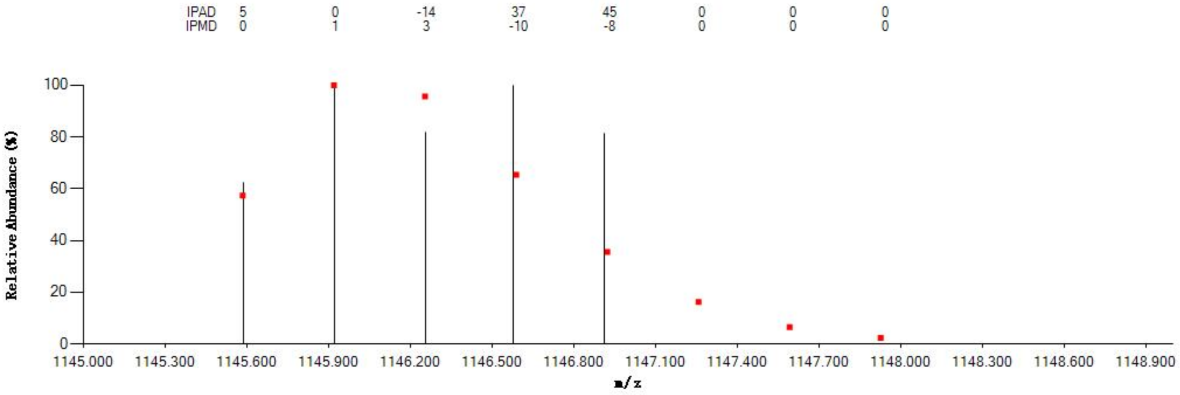


#### Annotated MS/MS Spectrum

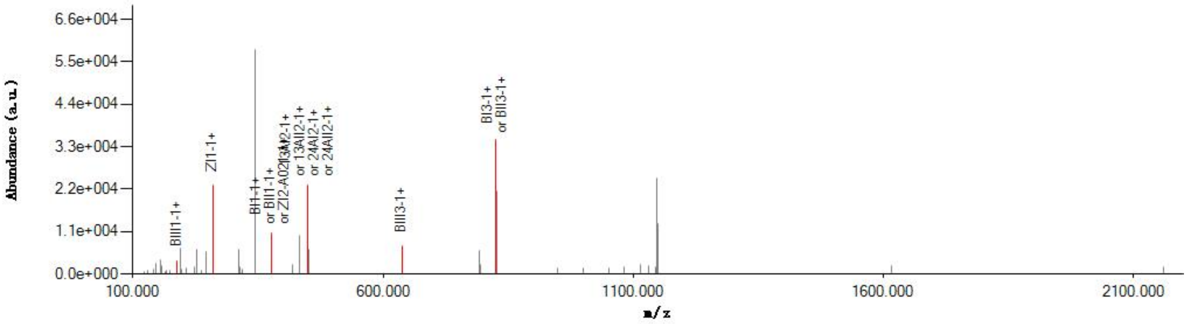


#### Graphical Fragmentation Map

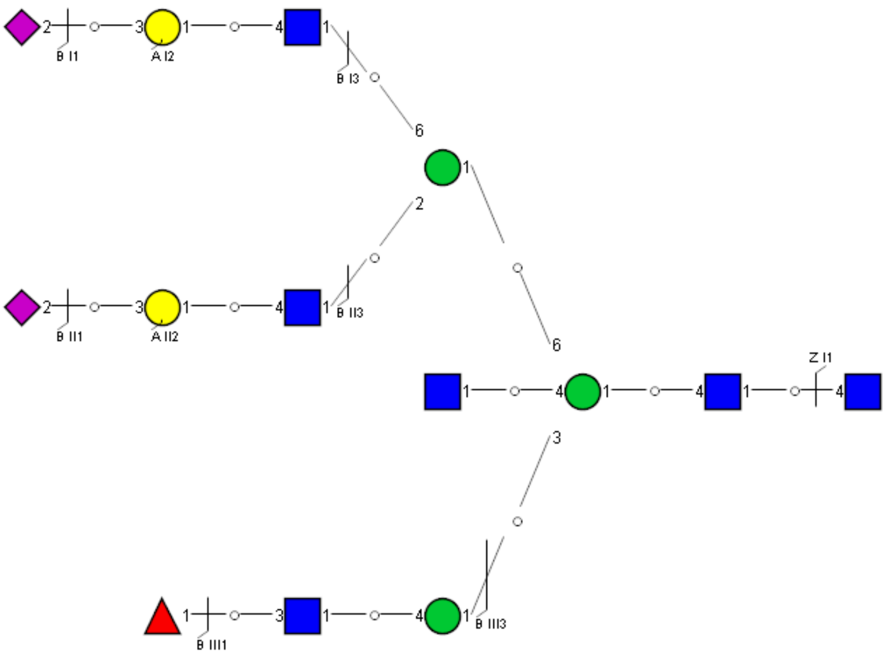

Spectrum Index: 18603(NoBHs:1)

#### Precursor iEF Map

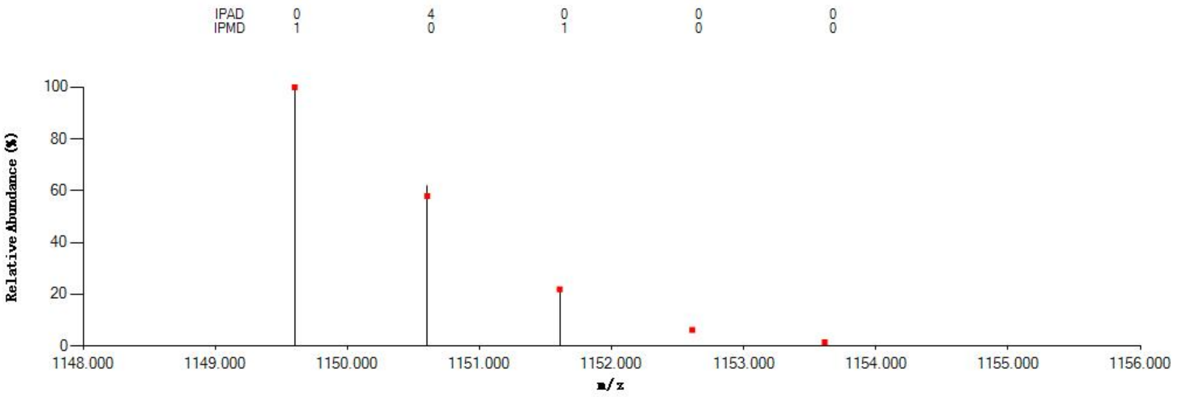


#### Annotated MS/MS Spectrum

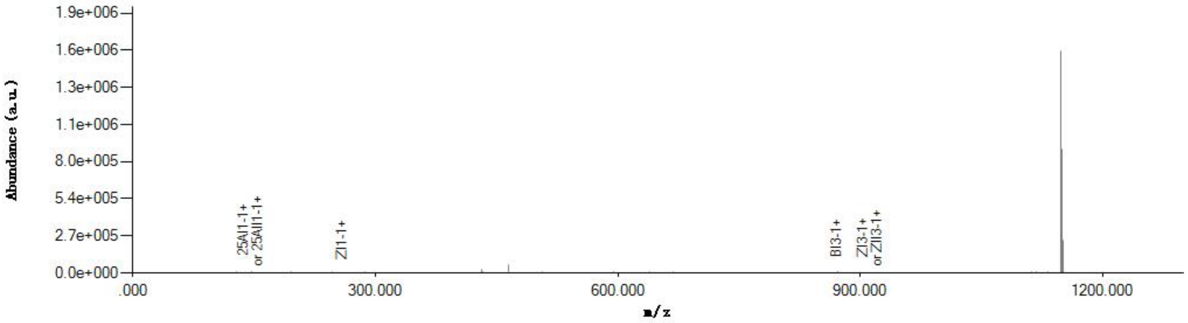


#### Graphical Fragmentation Map

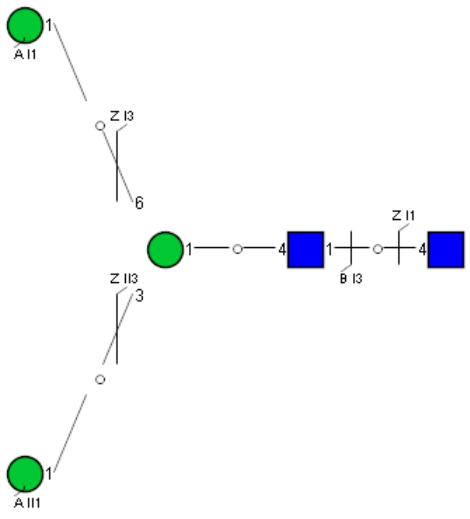

Spectrum Index: 27670(NoBHs:1)

#### Precursor iEF Map

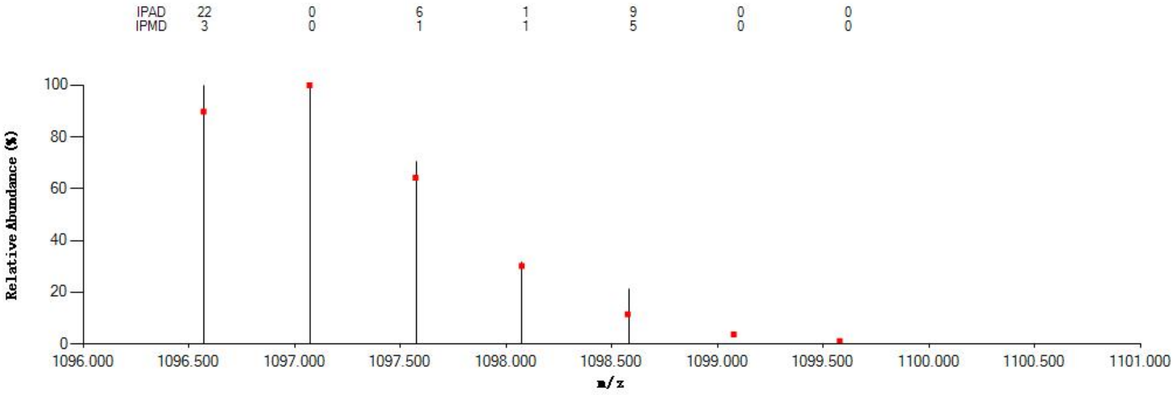


#### Annotated MS/MS Spectrum

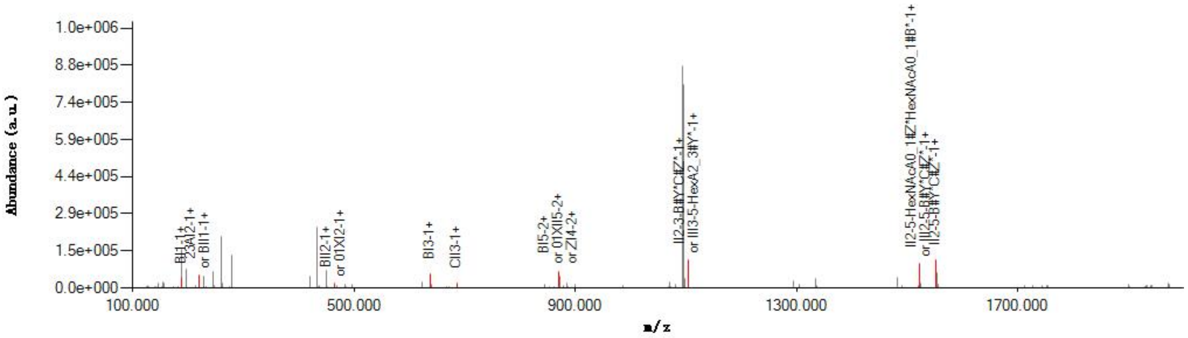


#### Graphical Fragmentation Map

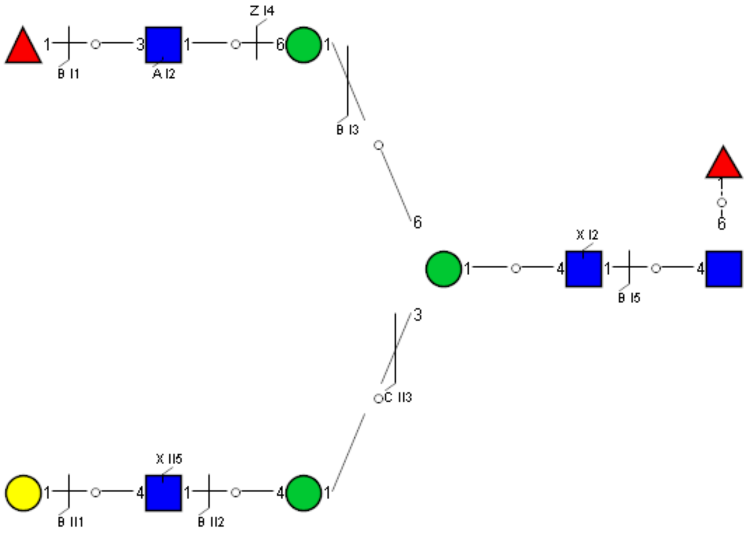

Spectrum Index: 28140(NoBHs:1)

#### Precursor iEF Map

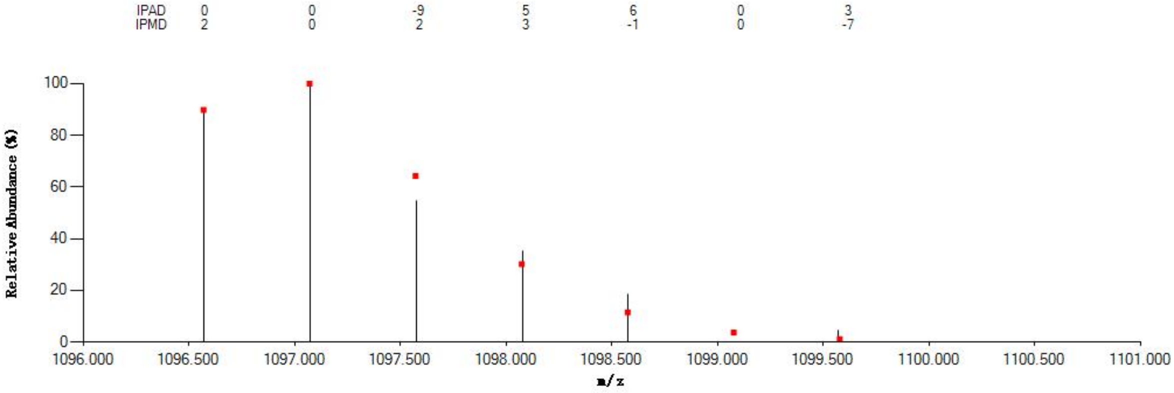


#### Annotated MS/MS Spectrum

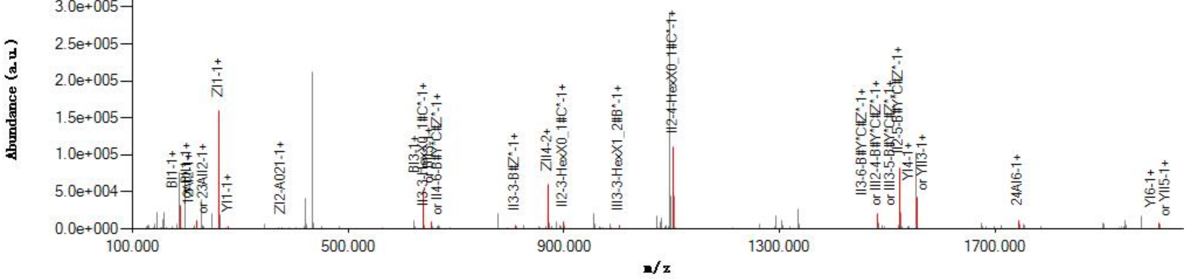


#### Graphical Fragmentation Map

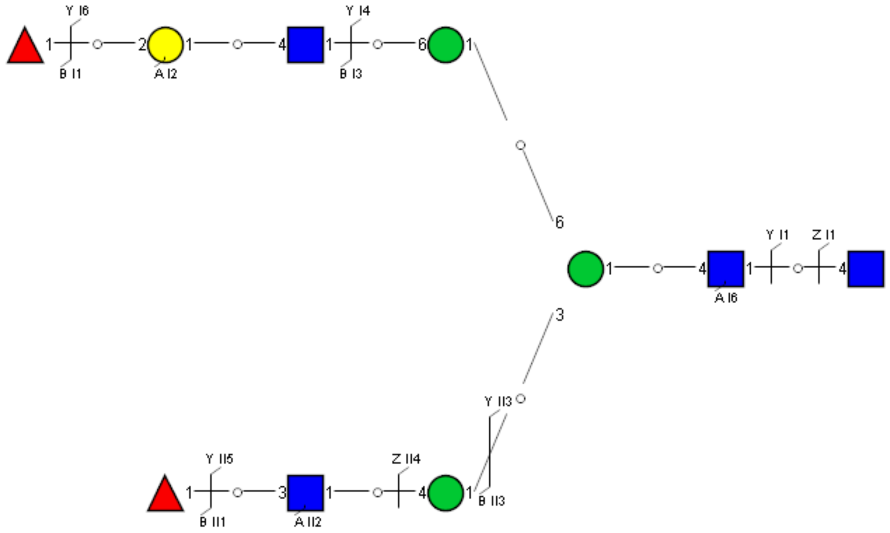

Spectrum Index: 29254(NoBHs:1)

#### Precursor iEF Map

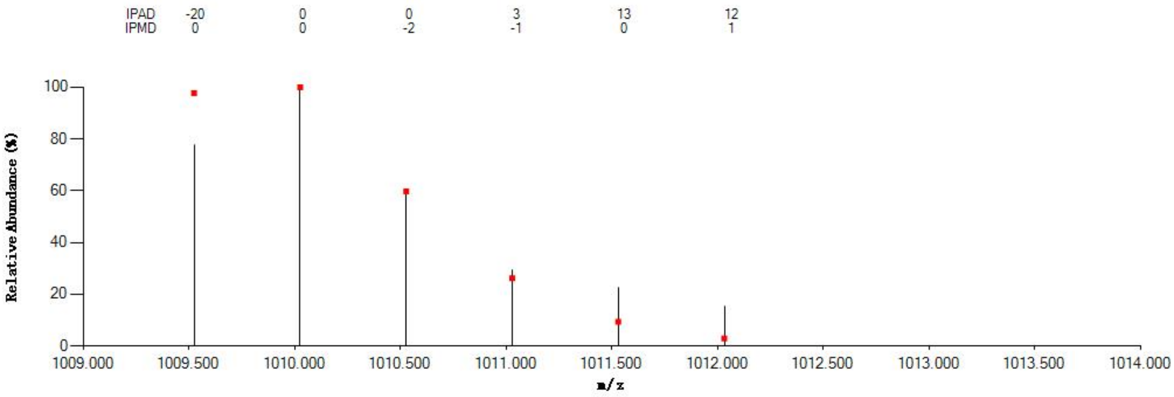


#### Annotated MS/MS Spectrum

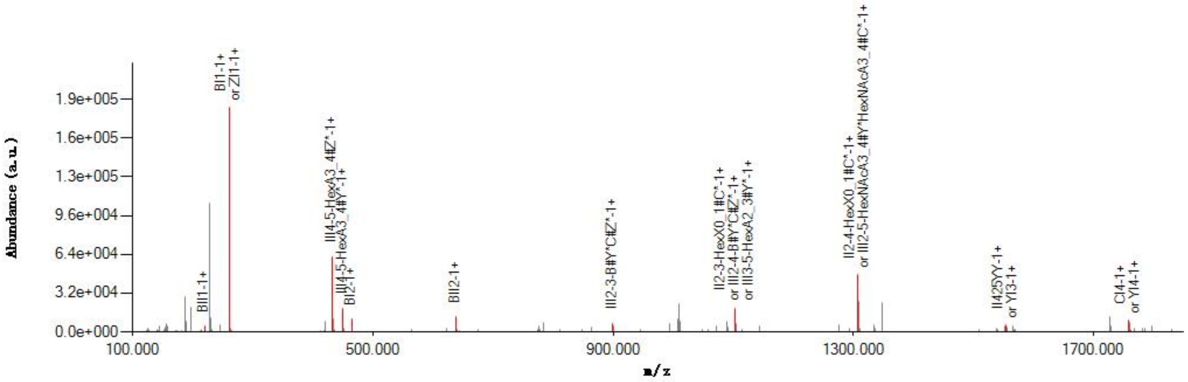


#### Graphical Fragmentation Map

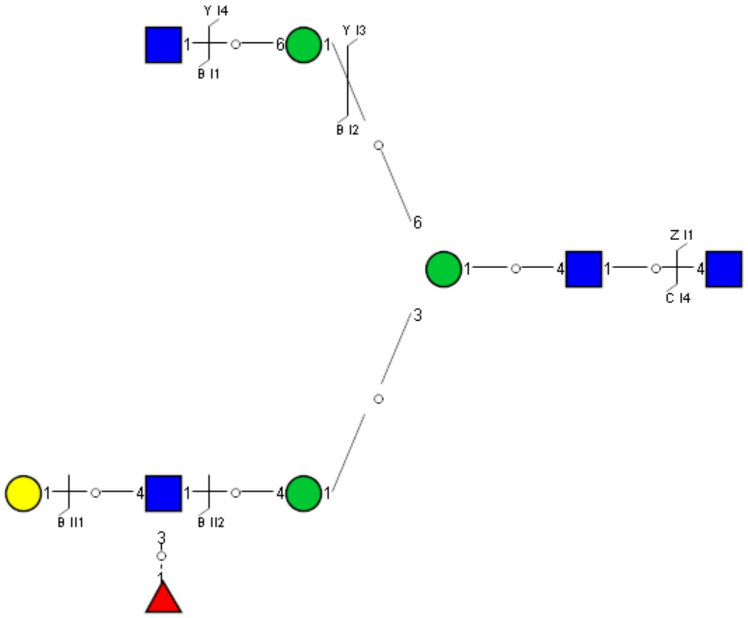

Spectrum Index: 30074(NoBHs:1)

#### Precursor iEF Map

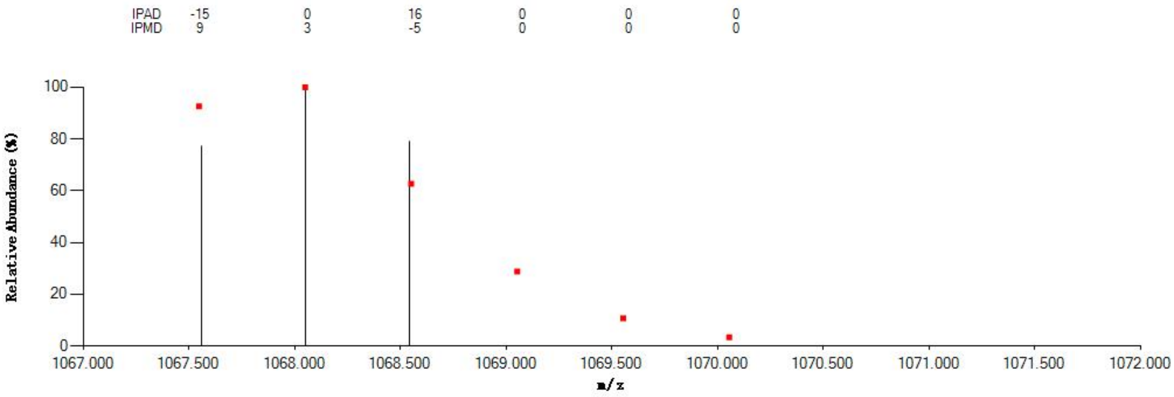


#### Annotated MS/MS Spectrum

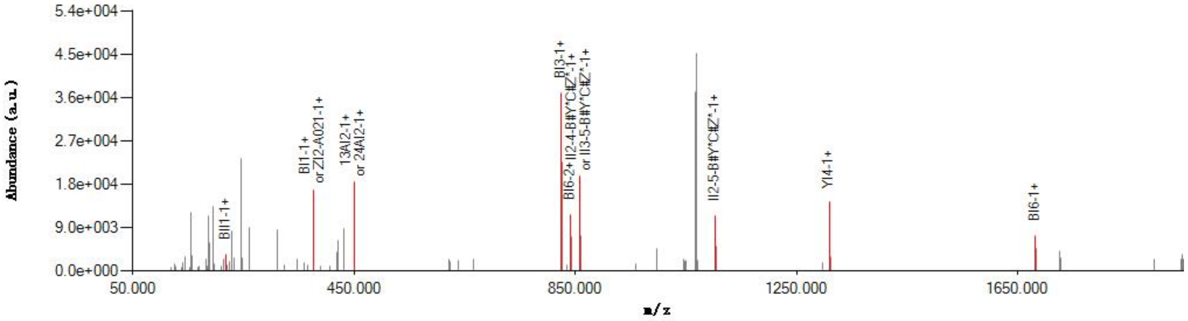


#### Graphical Fragmentation Map

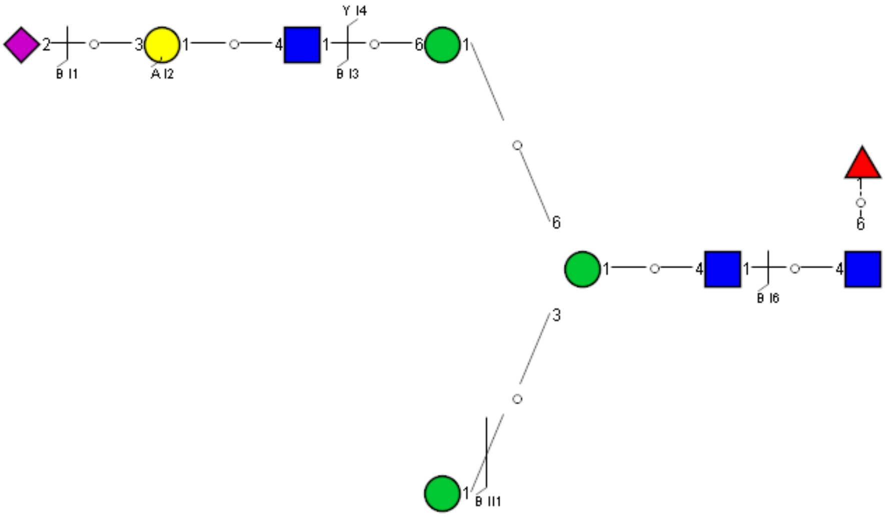

Spectrum Index: 30093(NoBHs:1)

Precursor iEF Map

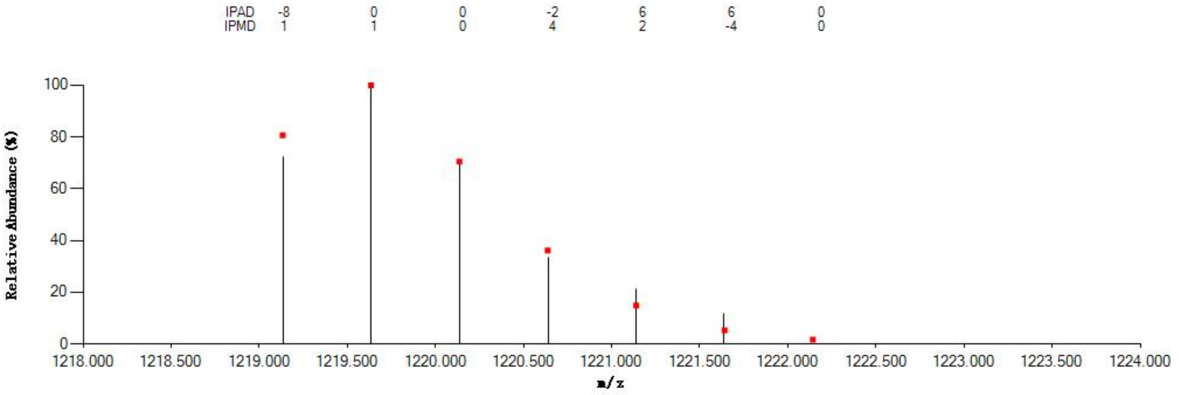


Annotated MS/MS Spectrum

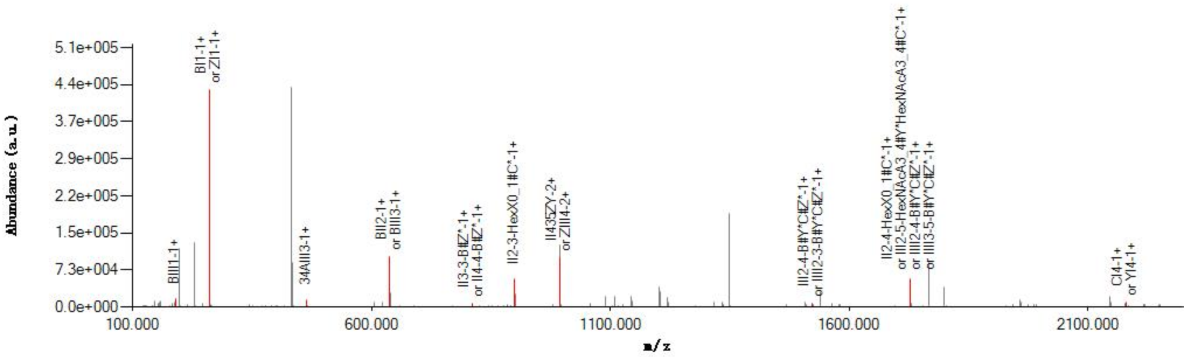


Graphical Fragmentation Map

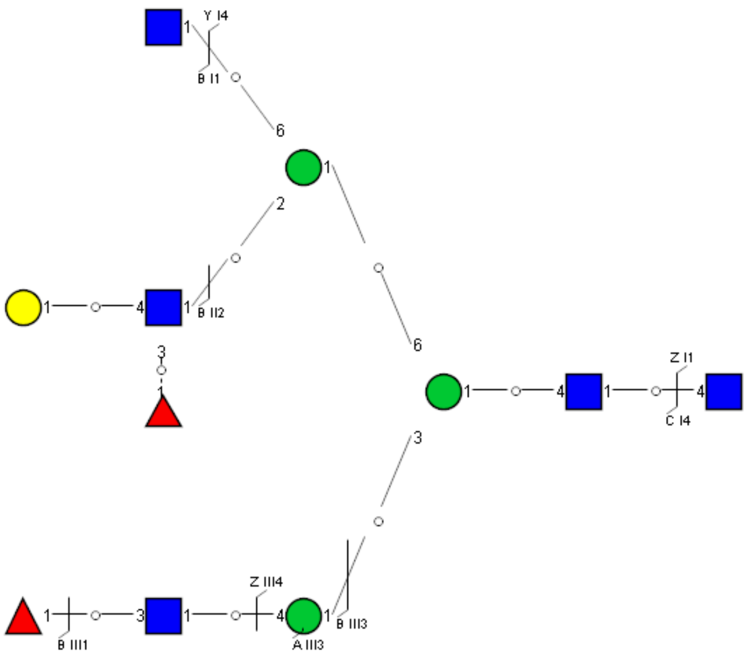

Spectrum Index: 31839(NoBHs:1)

#### Precursor iEF Map

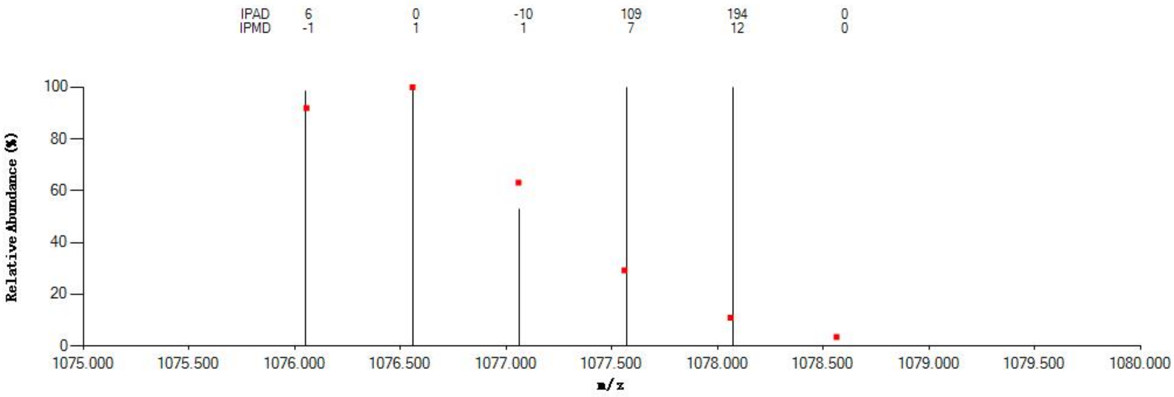


#### Annotated MS/MS Spectrum

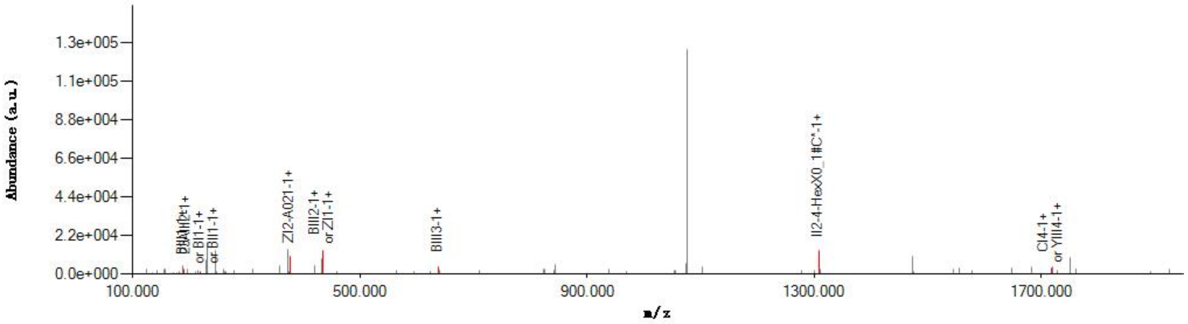


#### Graphical Fragmentation Map

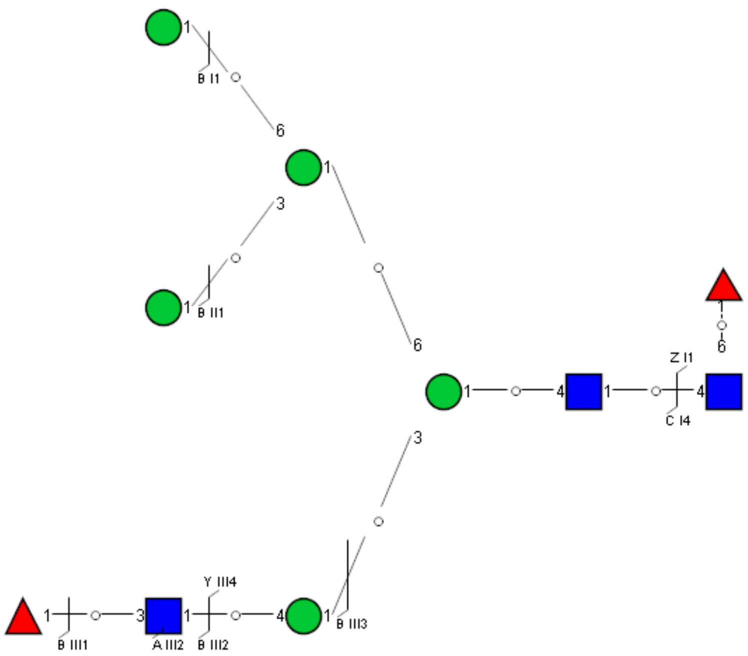

Spectrum Index: 32009(NoBHs:1)

#### Precursor iEF Map

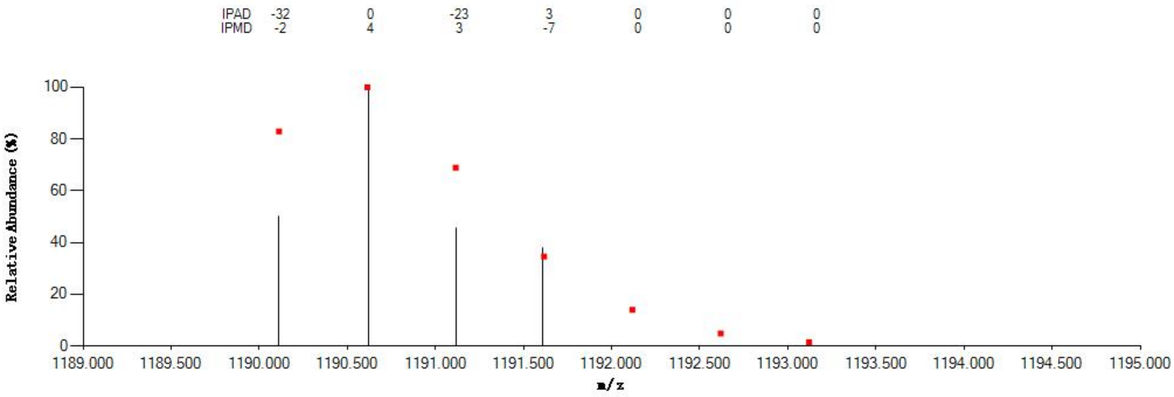


#### Annotated MS/MS Spectrum

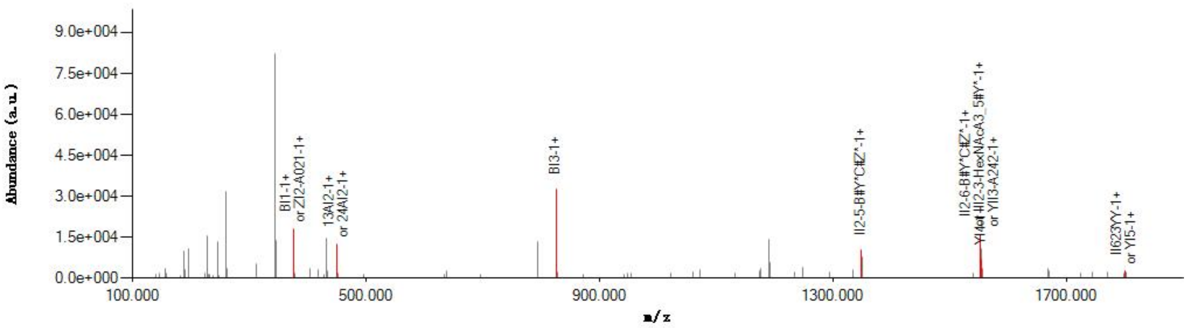


#### Graphical Fragmentation Map

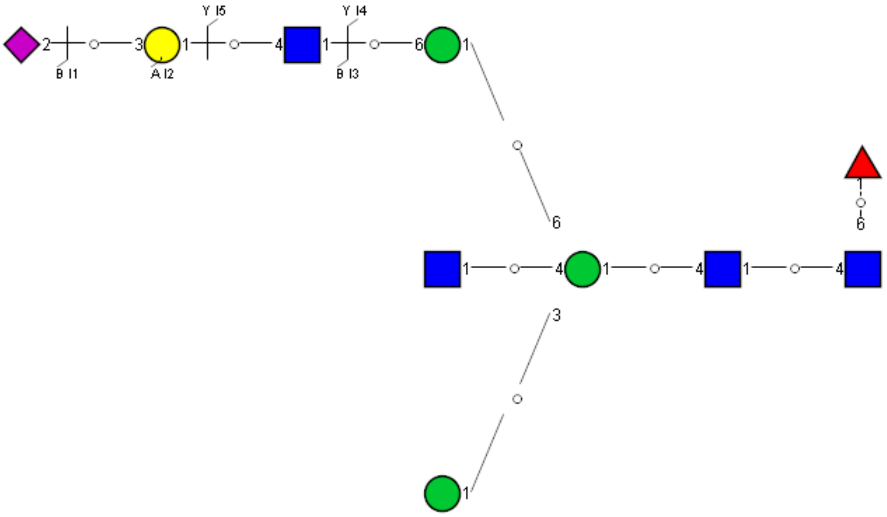

Spectrum Index: 32026(NoBHs:1)

#### Precursor iEF Map

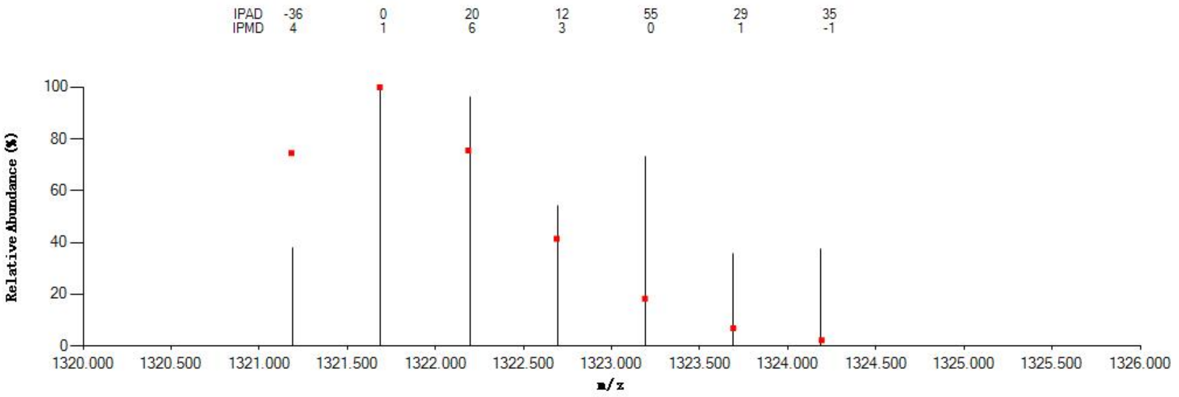


#### Annotated MS/MS Spectrum

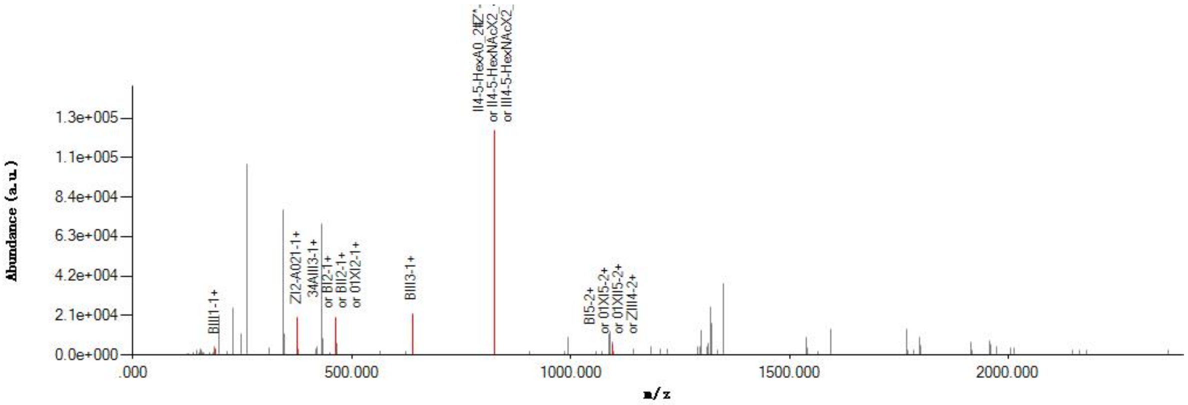


#### Graphical Fragmentation Map

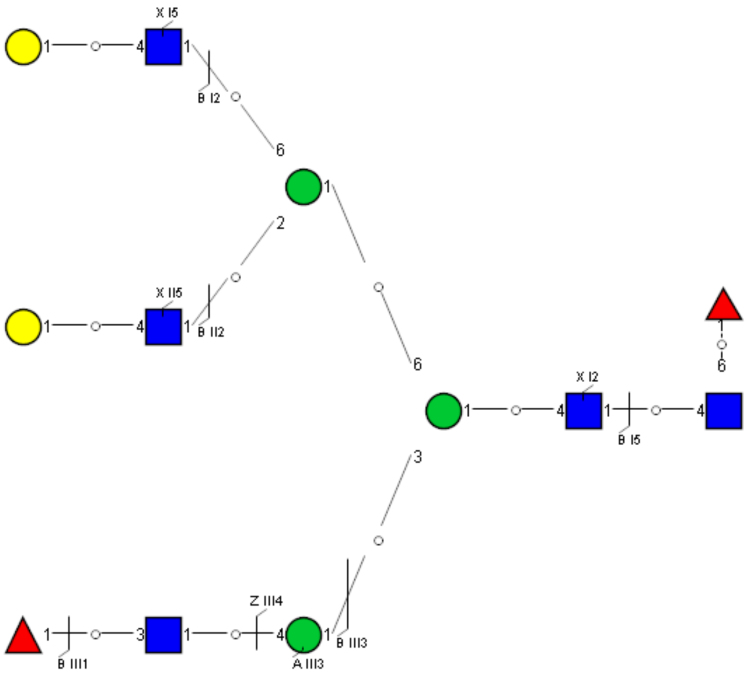

Spectrum Index: 32972(NoBHs:1)

#### Precursor iEF Map

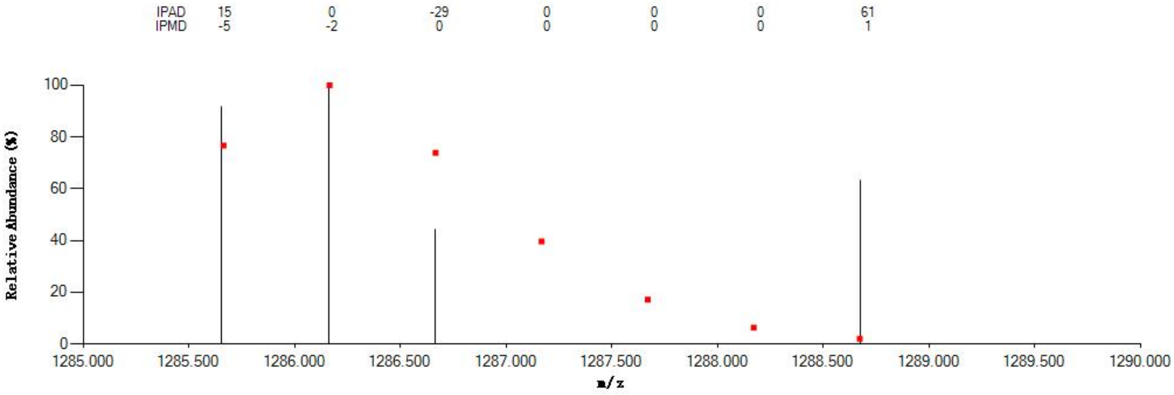


#### Annotated MS/MS Spectrum

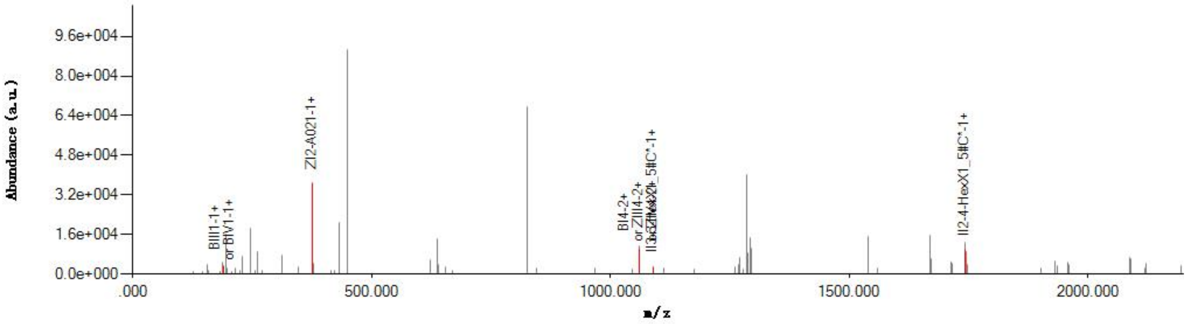


#### Graphical Fragmentation Map

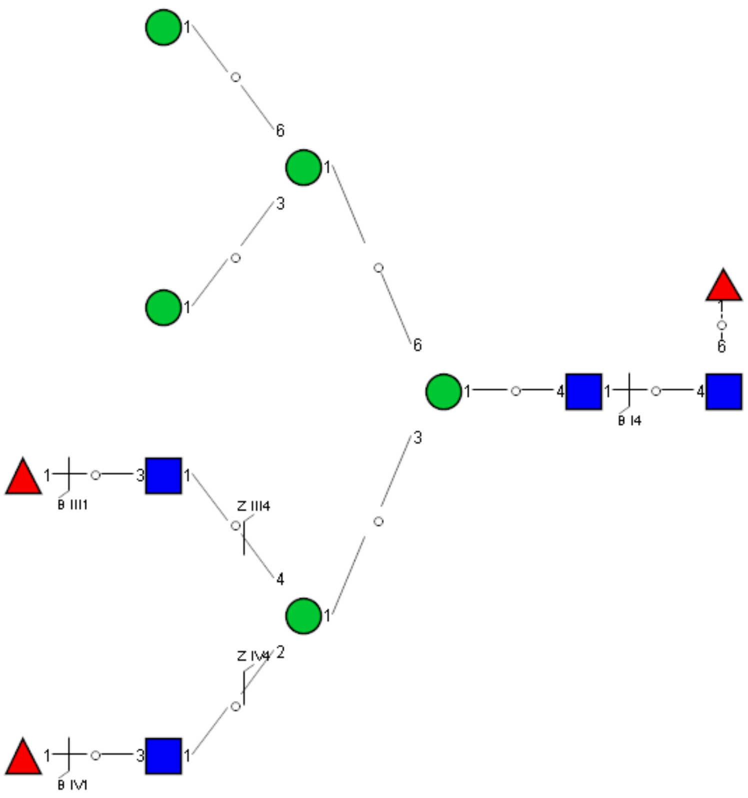

Spectrum Index: 33035(NoBHs:1)

#### Precursor iEF Map

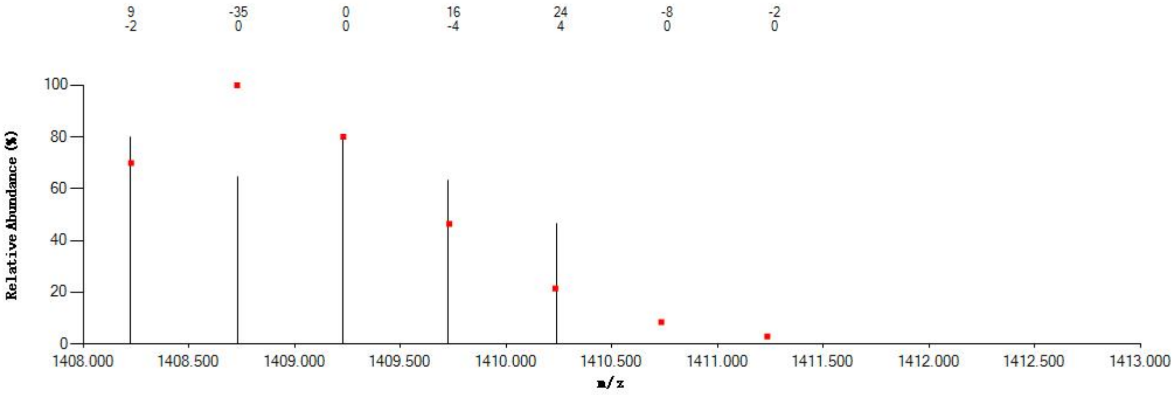

#### Annotated MS/MS Spectrum

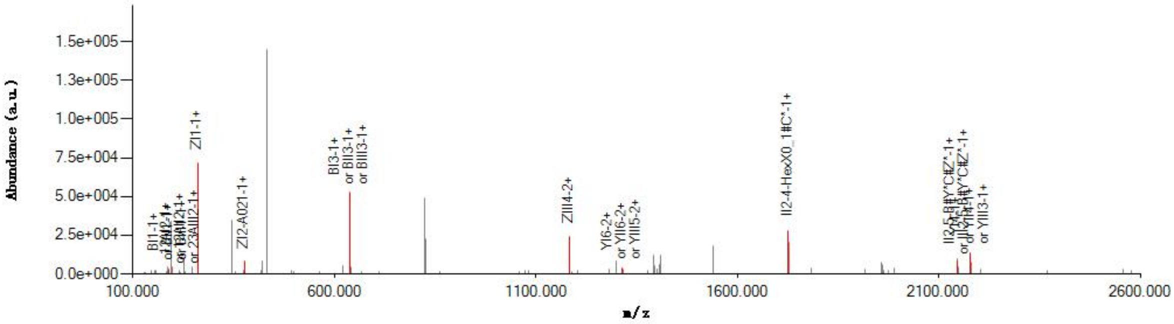

#### Graphical Fragmentation Map

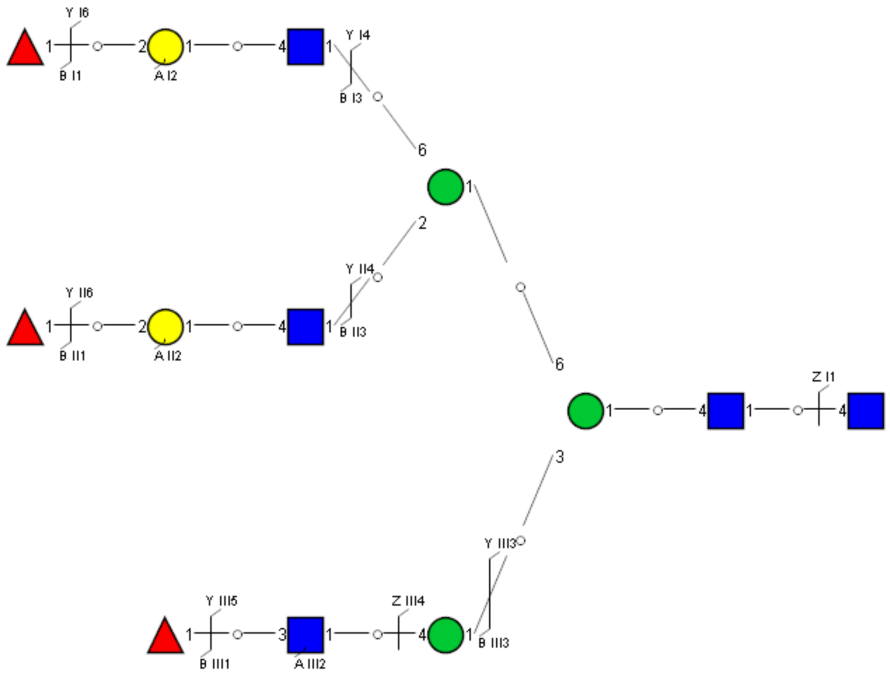

Spectrum Index: 33207(NoBHs:1)

#### Precursor iEF Map

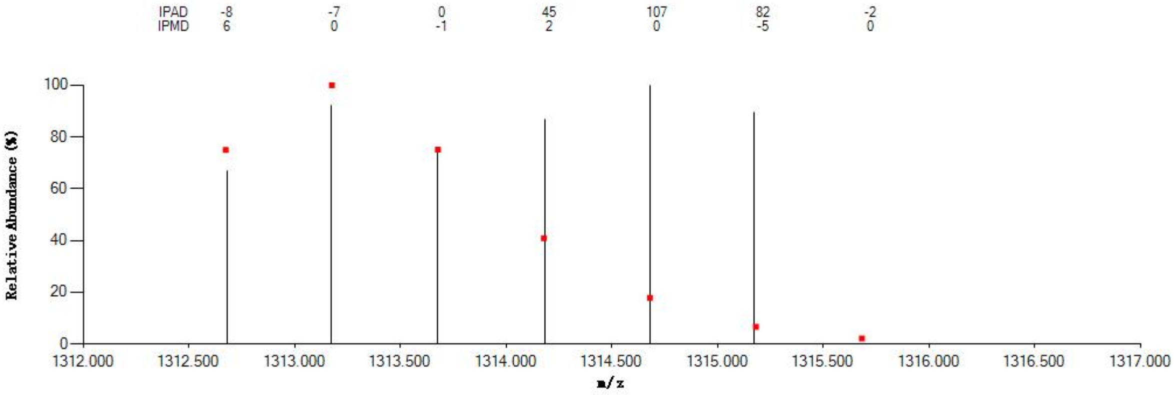


#### Annotated MS/MS Spectrum

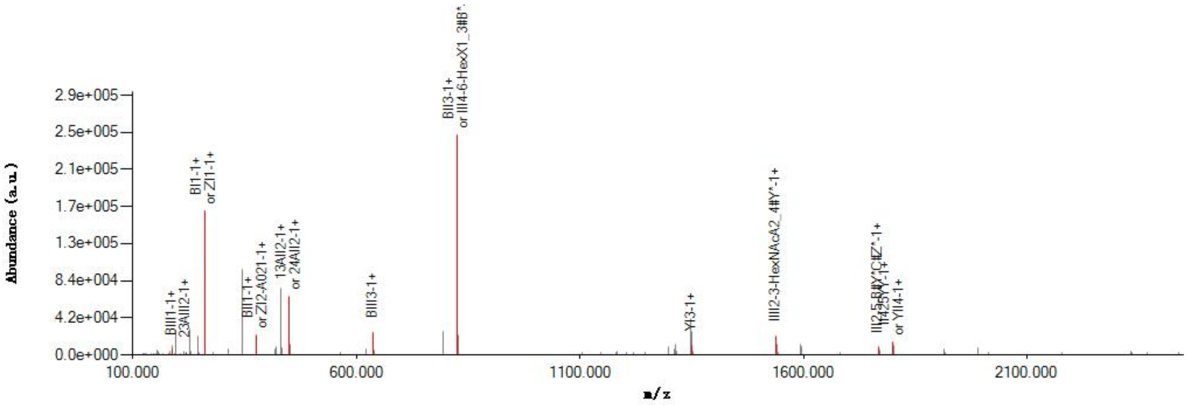


#### Graphical Fragmentation Map

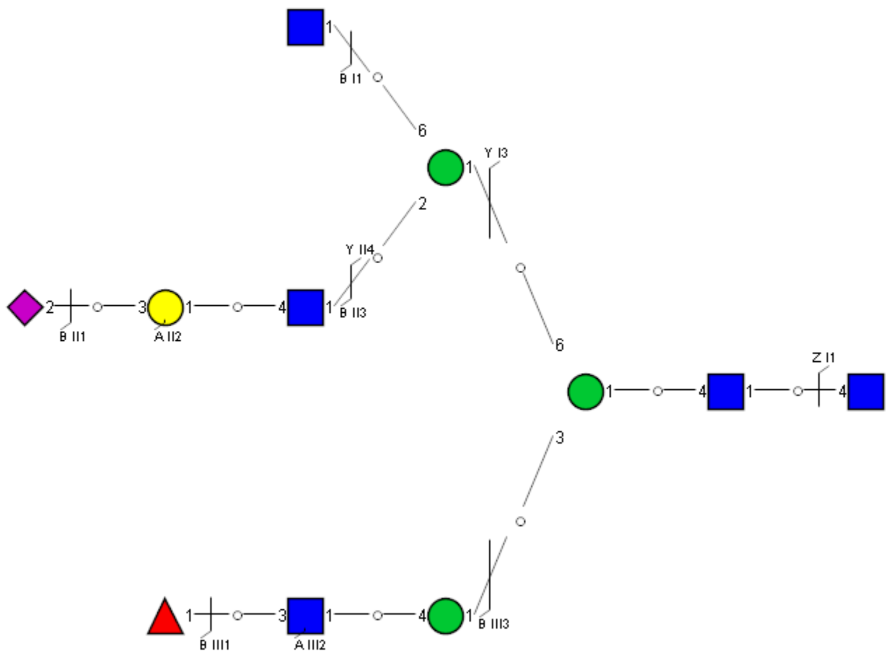

Spectrum Index: 35162(NoBHs:1)

#### Precursor iEF Map

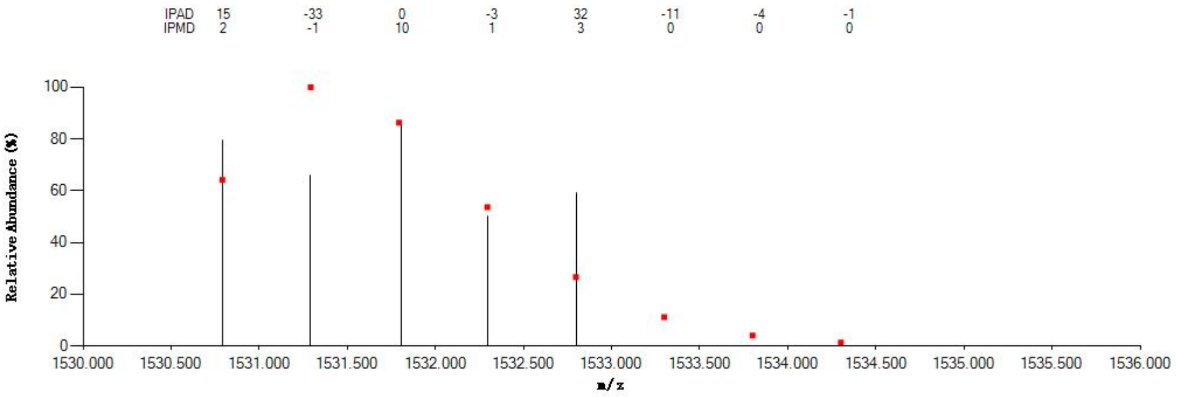


#### Annotated MS/MS Spectrum

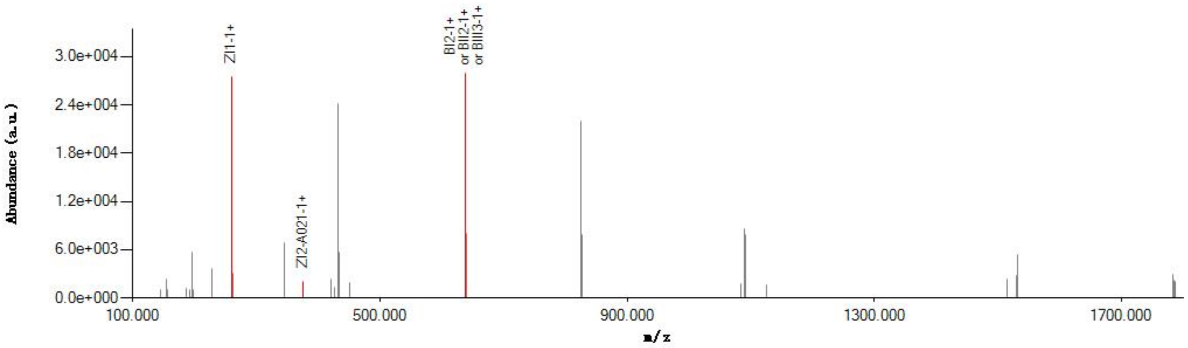


#### Graphical Fragmentation Map

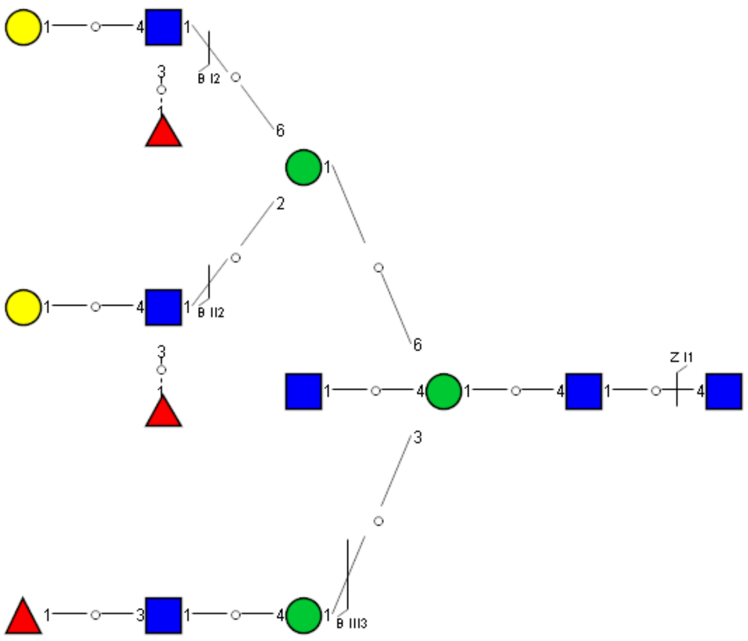

Spectrum Index: 19253(NoBHs:1)

#### Precursor iEF Map

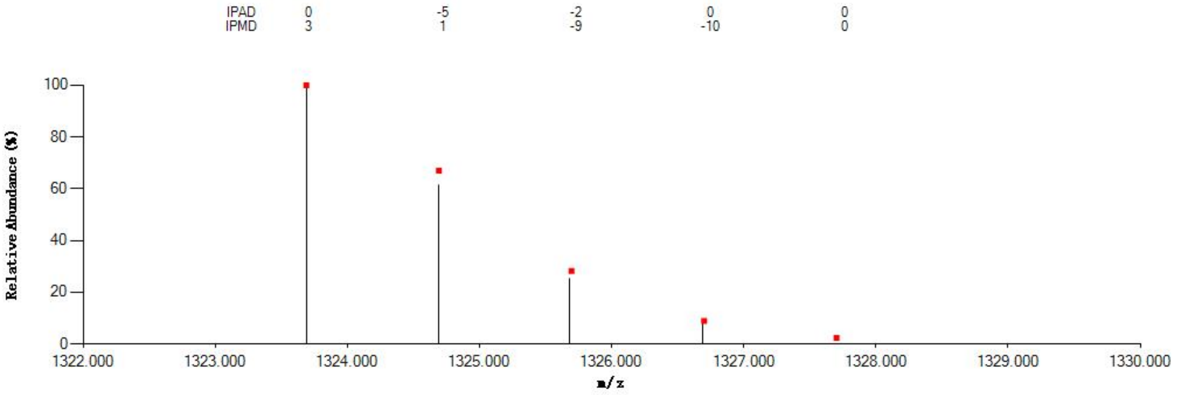


#### Annotated MS/MS Spectrum

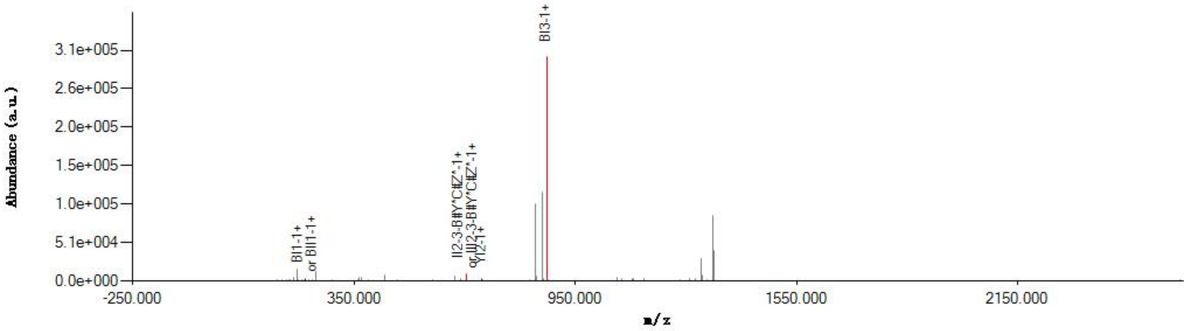


#### Graphical Fragmentation Map

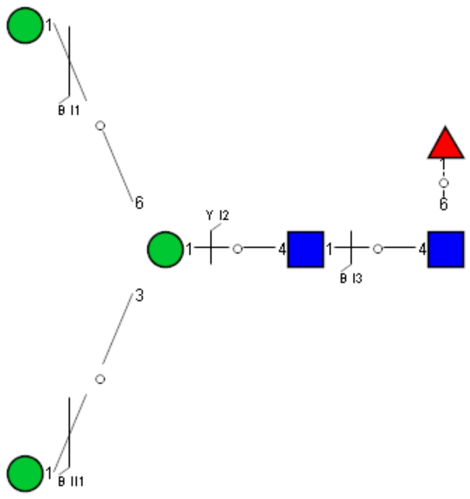

Spectrum Index: 25647(NoBHs:1)

#### Precursor iEF Map

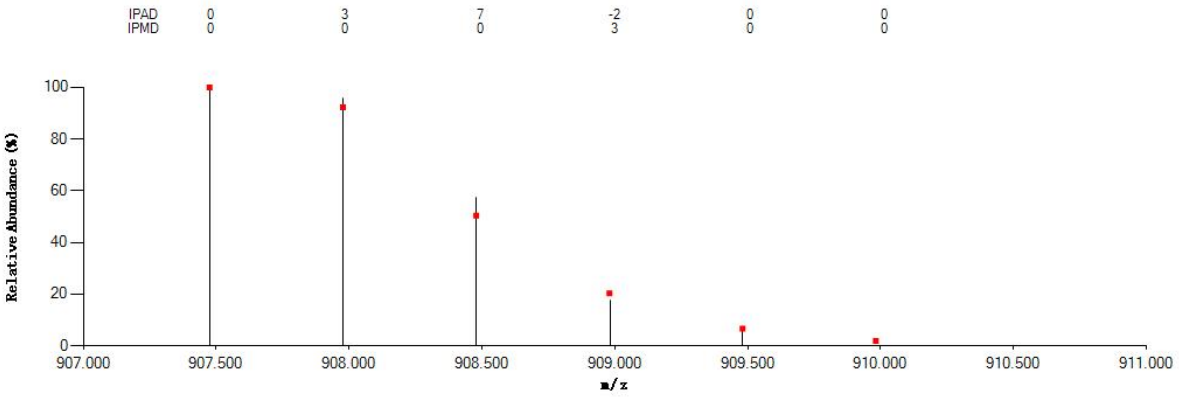


#### Annotated MS/MS Spectrum

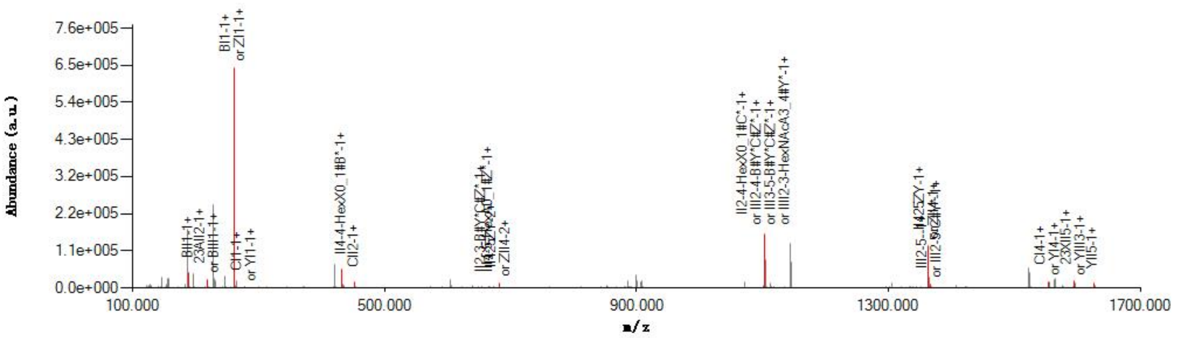


#### Graphical Fragmentation Map

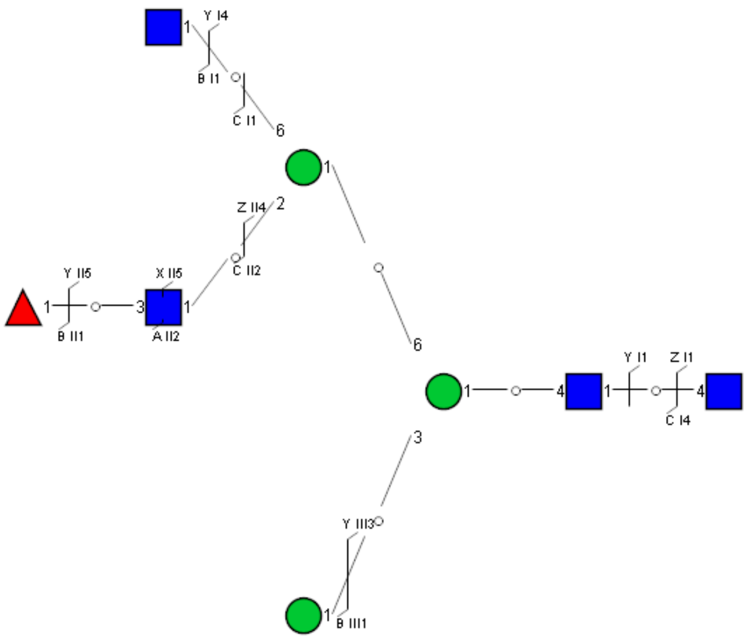

Spectrum Index: 28315(NoBHs:1)

#### Precursor iEF Map

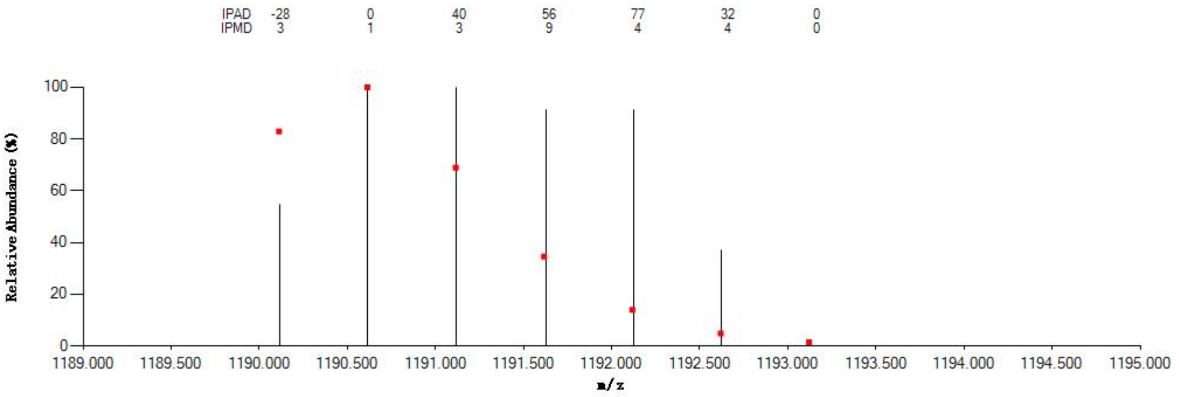


#### Annotated MS/MS Spectrum

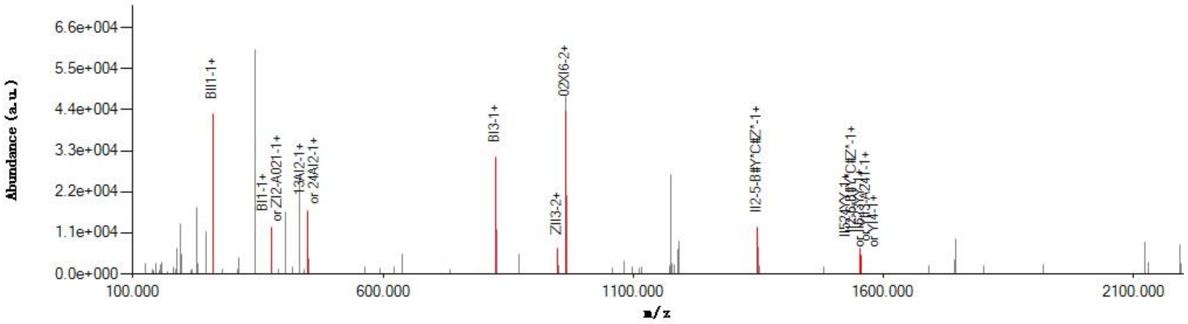


#### Graphical Fragmentation Map

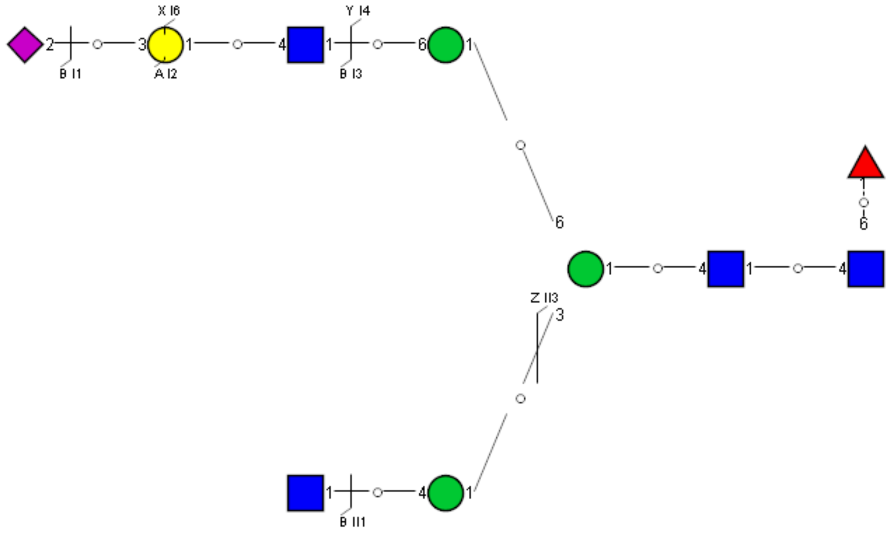

Spectrum Index: 29551(NoBHs:1)

#### Precursor iEF Map

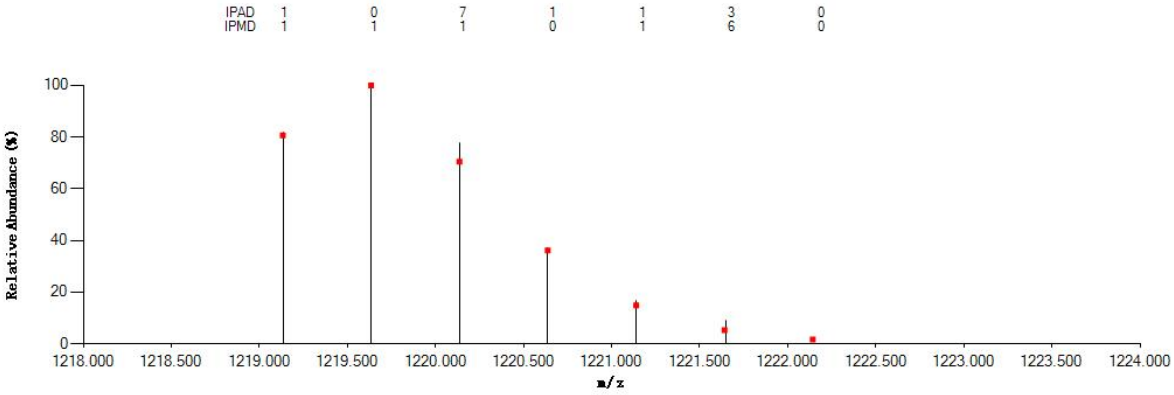


#### Annotated MS/MS Spectrum

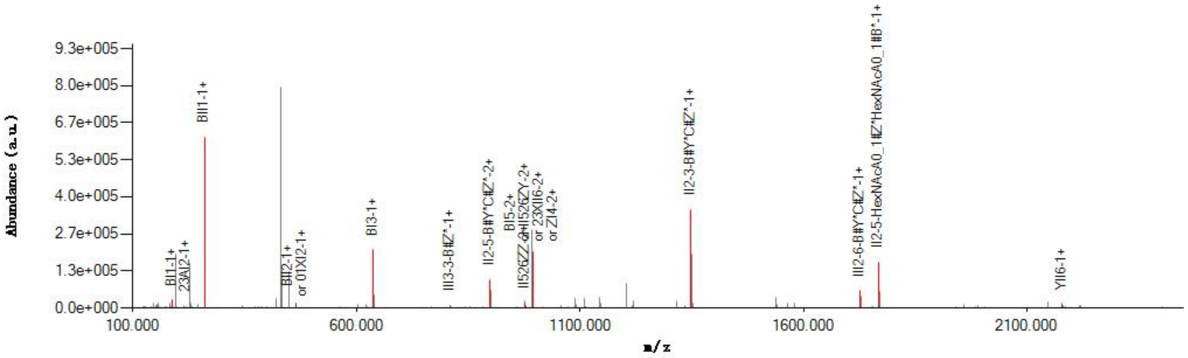


#### Graphical Fragmentation Map

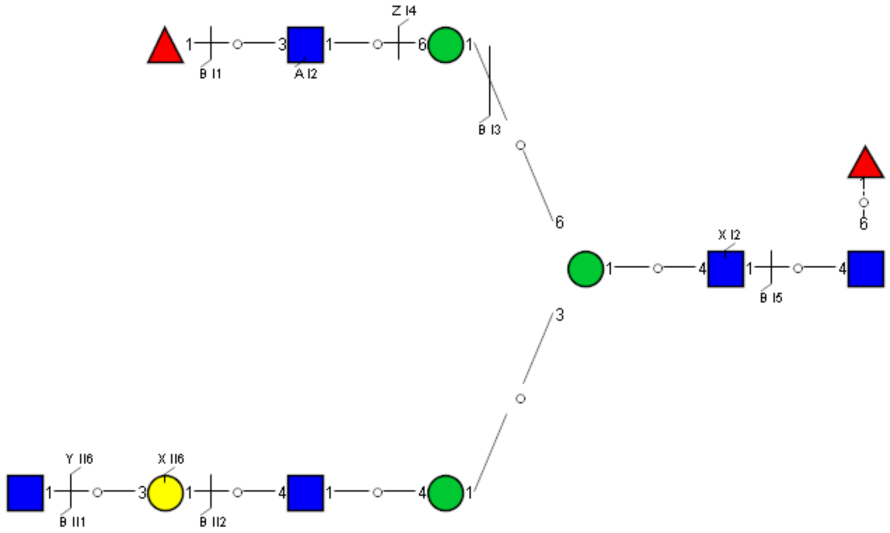

Spectrum Index: 30814(NoBHs:1)

#### Precursor iEF Map

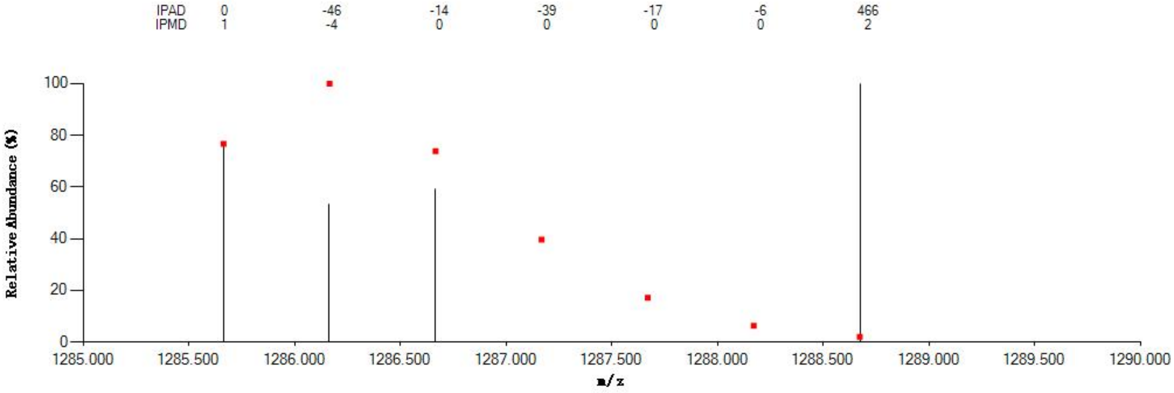


#### Annotated MS/MS Spectrum

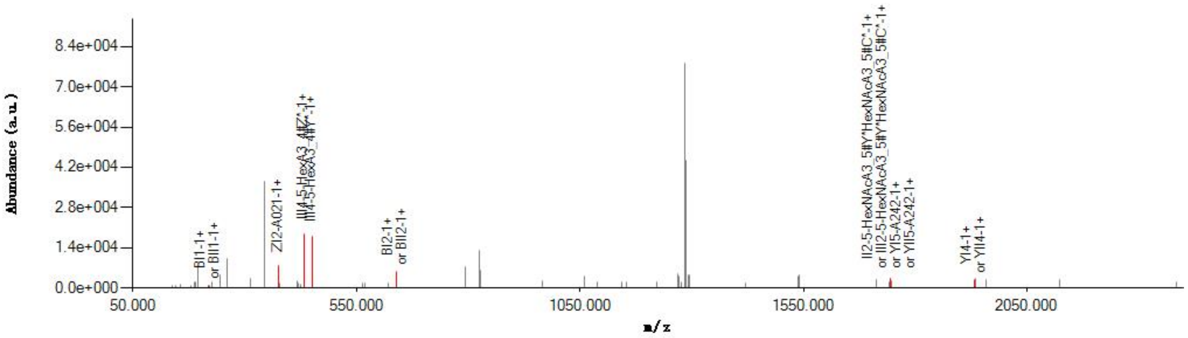


#### Graphical Fragmentation Map

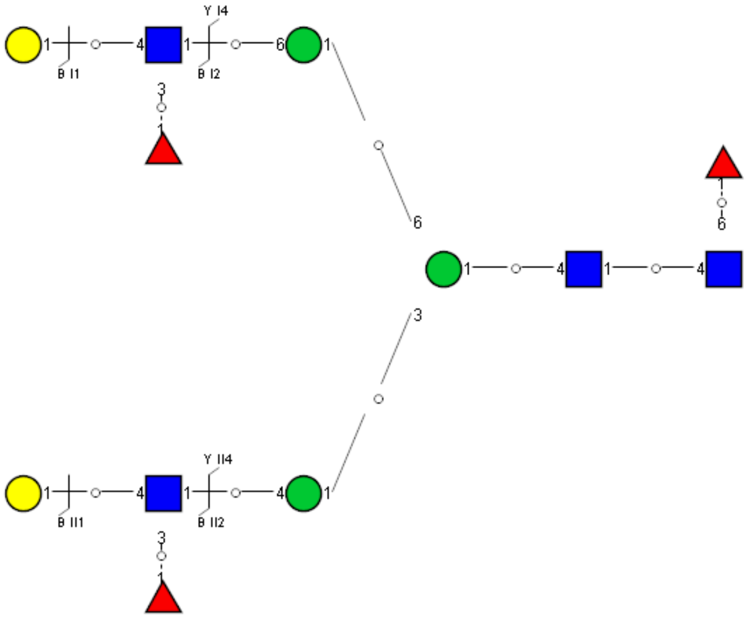

Spectrum Index: 32727(NoBHs:1)

Precursor iEF Map

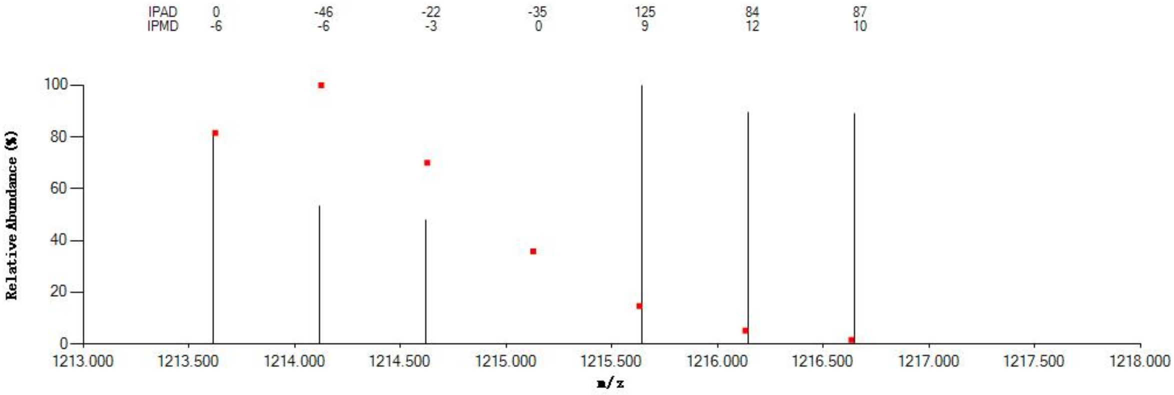

Annotated MS/MS Spectrum

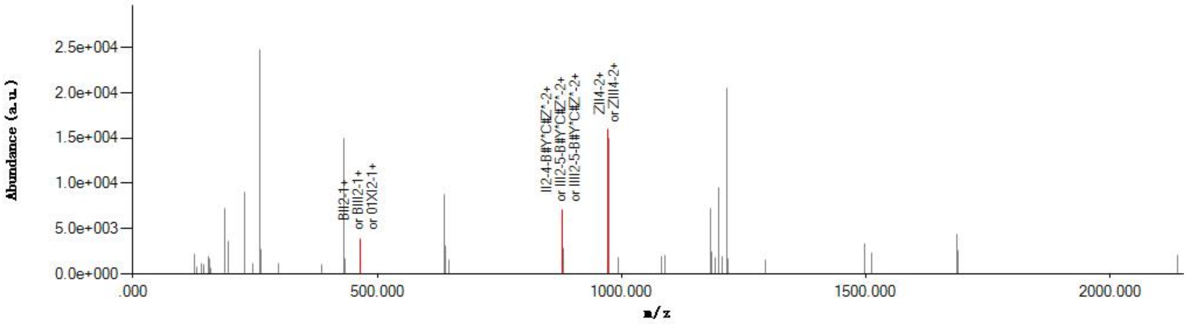

Graphical Fragmentation Map

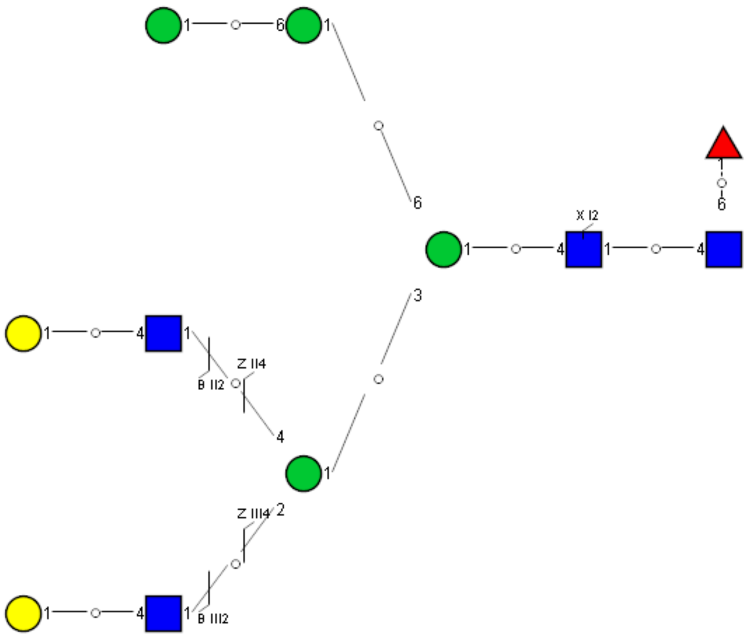

Spectrum Index: 33459(NoBHs:1)

#### Precursor iEF Map

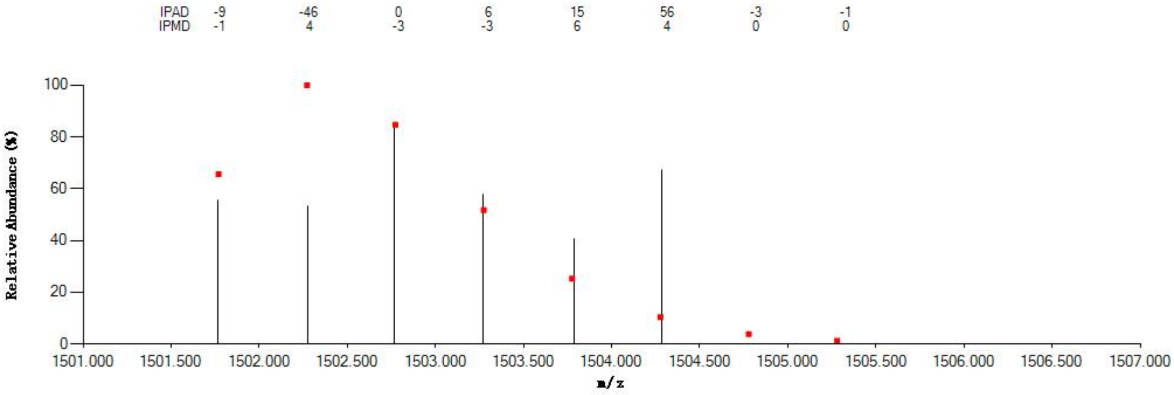


#### Annotated MS/MS Spectrum

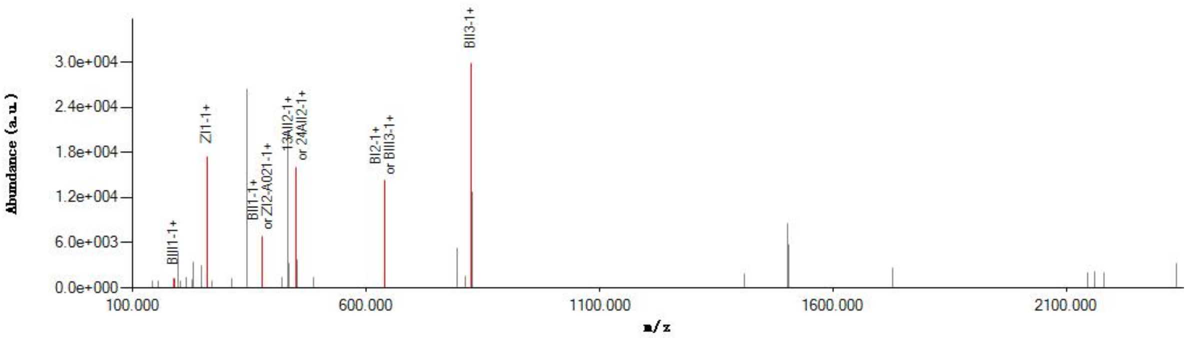


#### Graphical Fragmentation Map

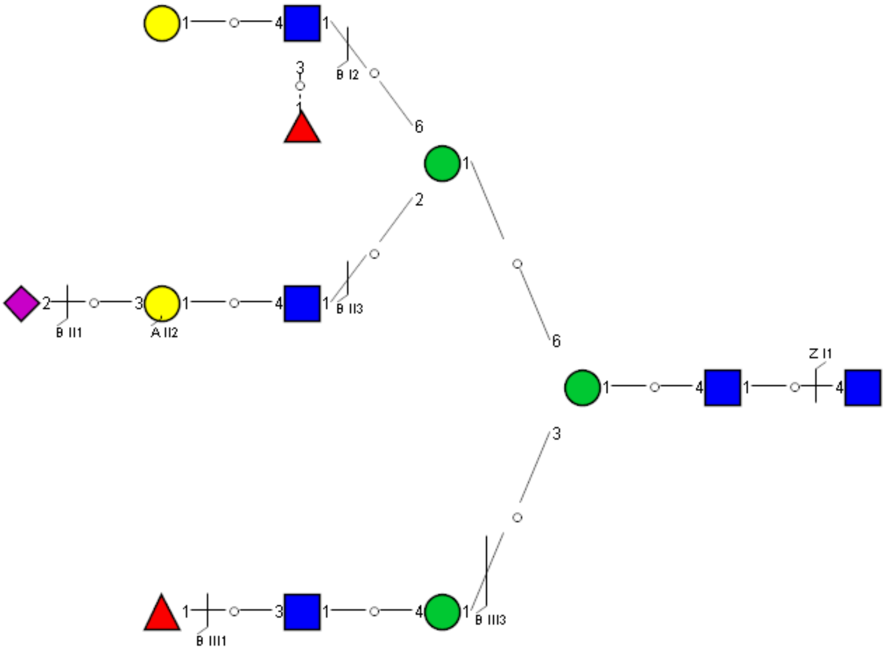

Spectrum Index: 35666(NoBHs:1)

Precursor iEF Map

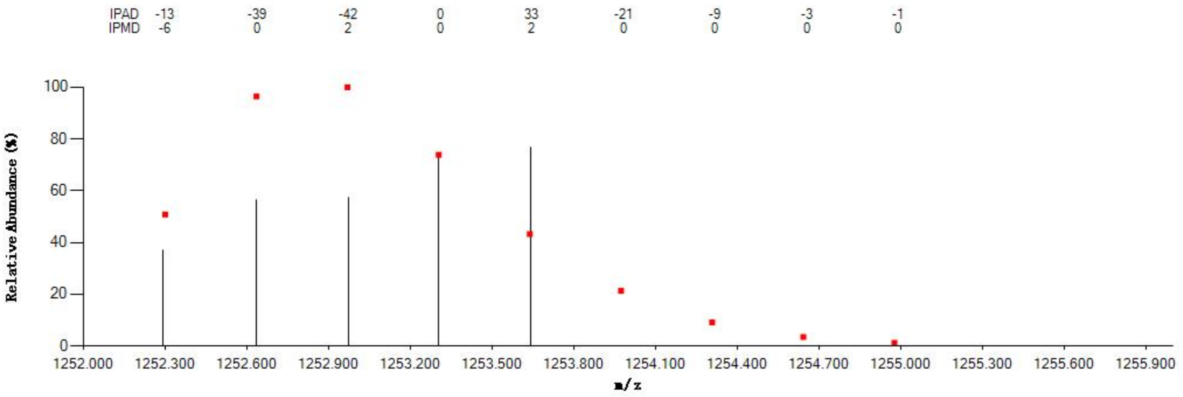


Annotated MS/MS Spectrum

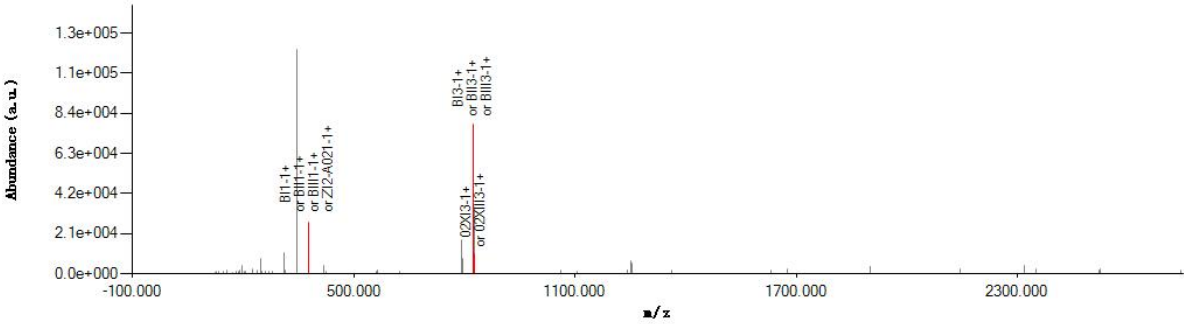


Graphical Fragmentation Map

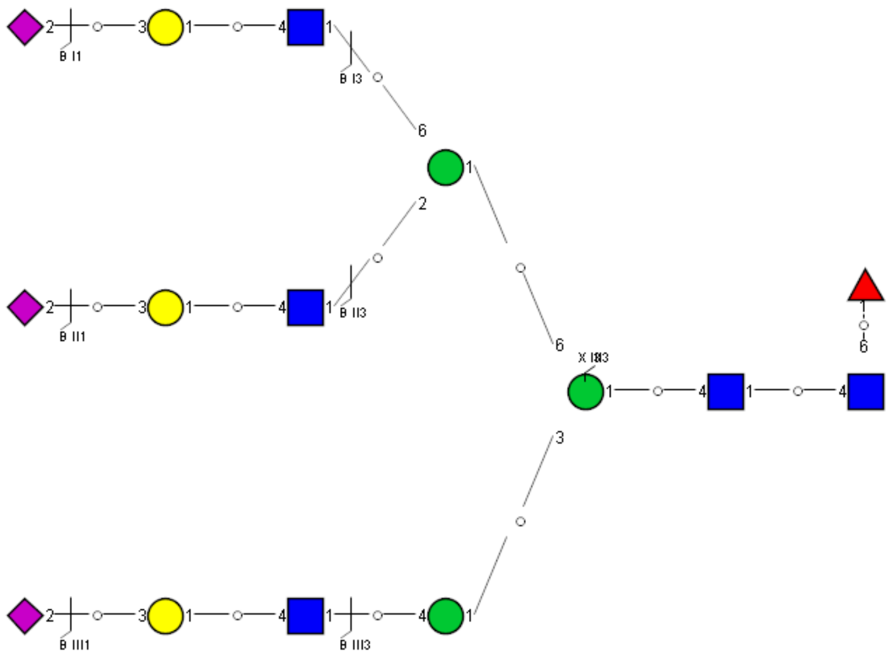
