## Supplemental Information 8 for "A draft of human N-glycans of glycoRNA"

Spectrum Index: 27962(NoBHs:1)

#### Precursor iEF Map

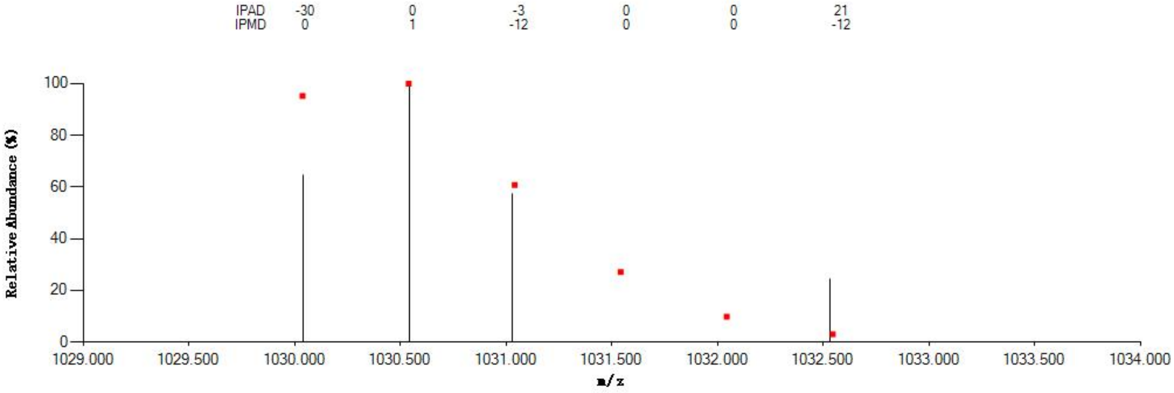


#### Annotated MS/MS Spectrum

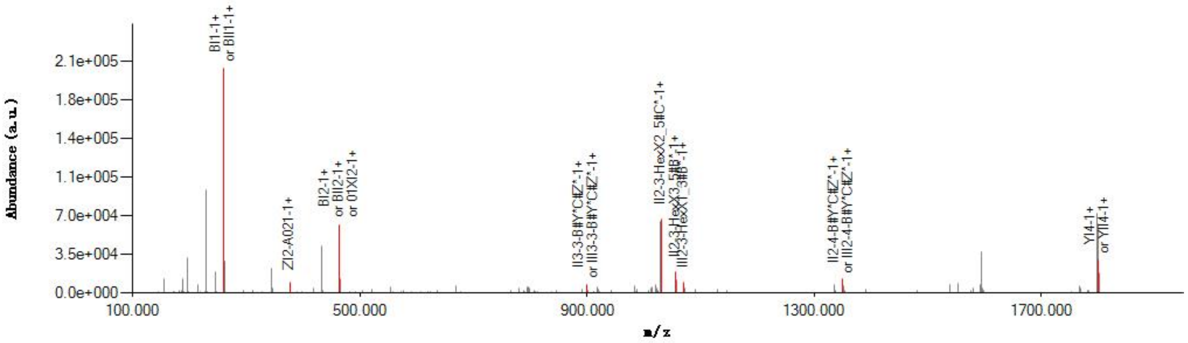


#### Graphical Fragmentation Map

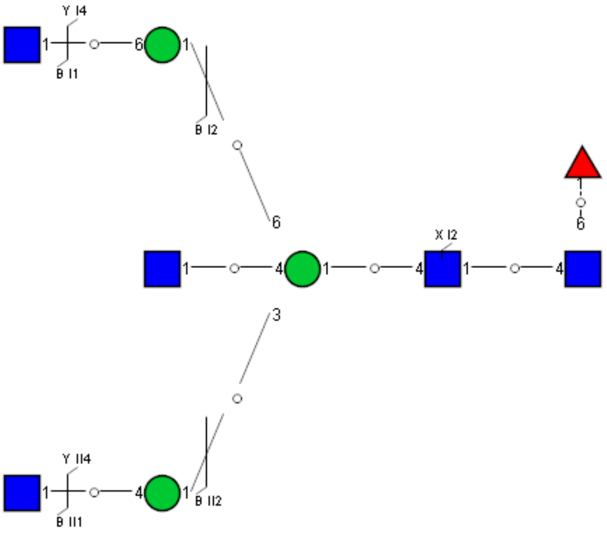

Spectrum Index: 29047(NoBHs:1)

#### Precursor iEF Map

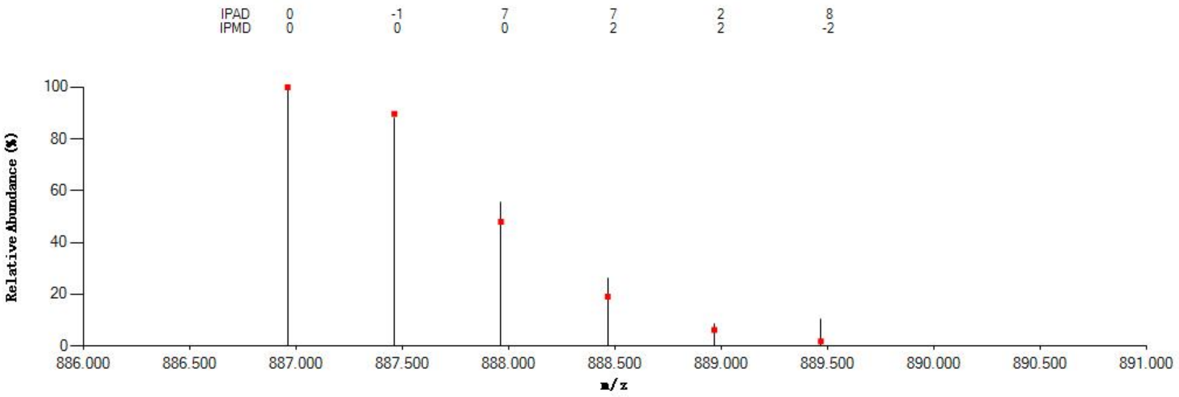


#### Annotated MS/MS Spectrum

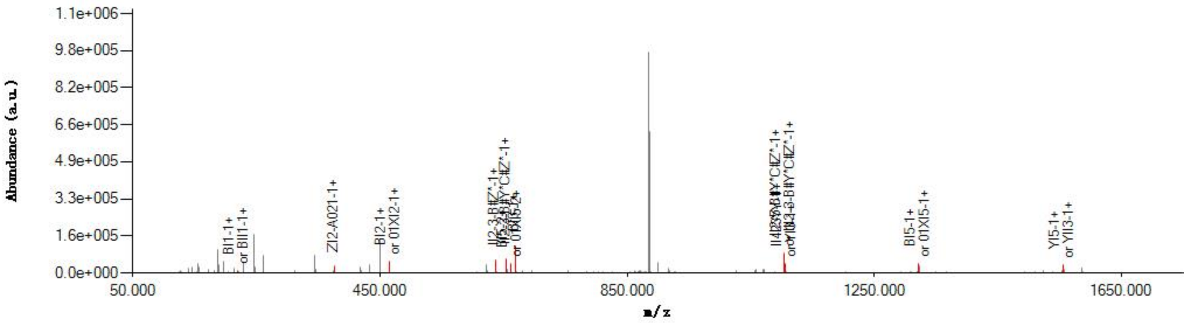


#### Graphical Fragmentation Map

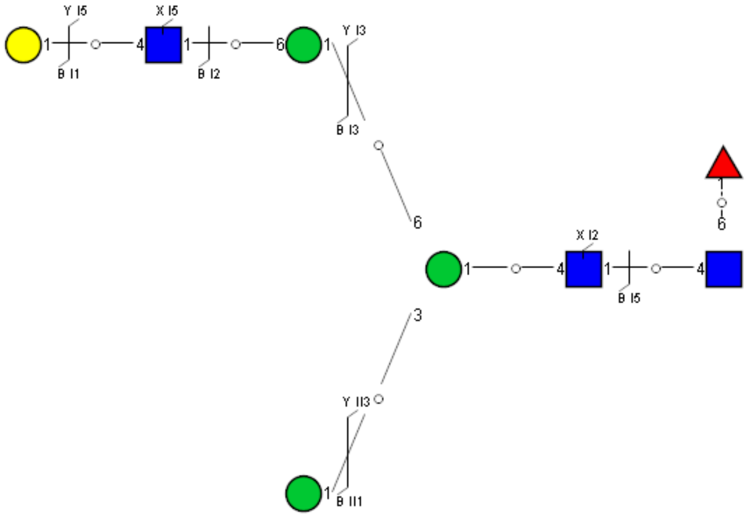

Spectrum Index: 29341(NoBHs:1)

#### Precursor iEF Map

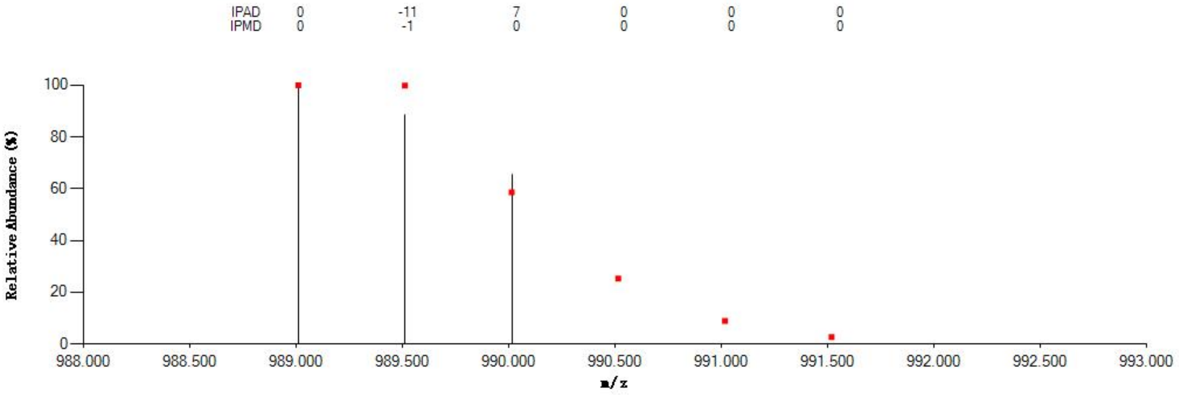


#### Annotated MS/MS Spectrum

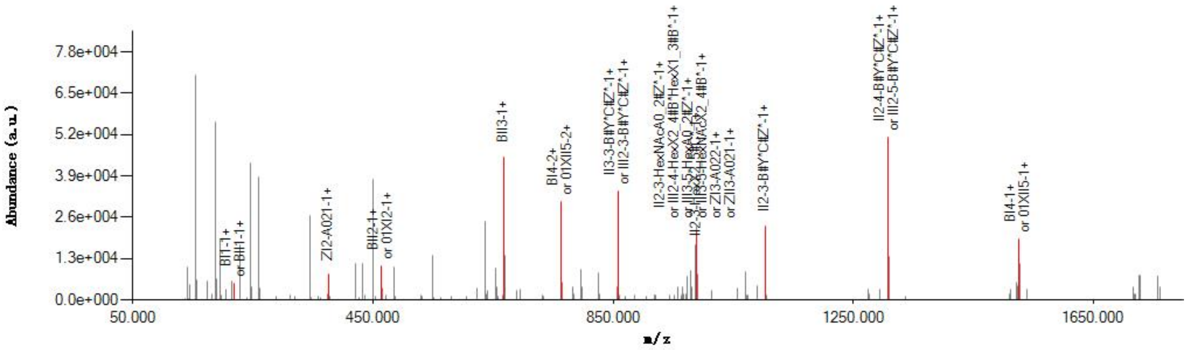


#### Graphical Fragmentation Map

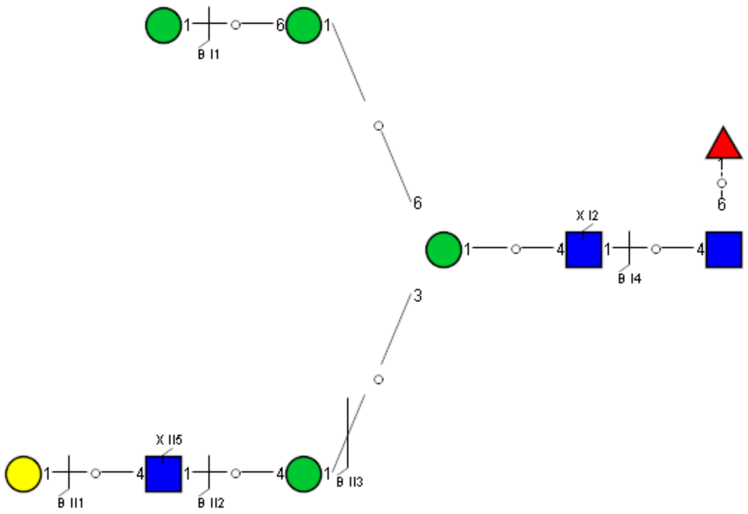

Spectrum Index: 29615(NoBHs:1)

#### Precursor iEF Map

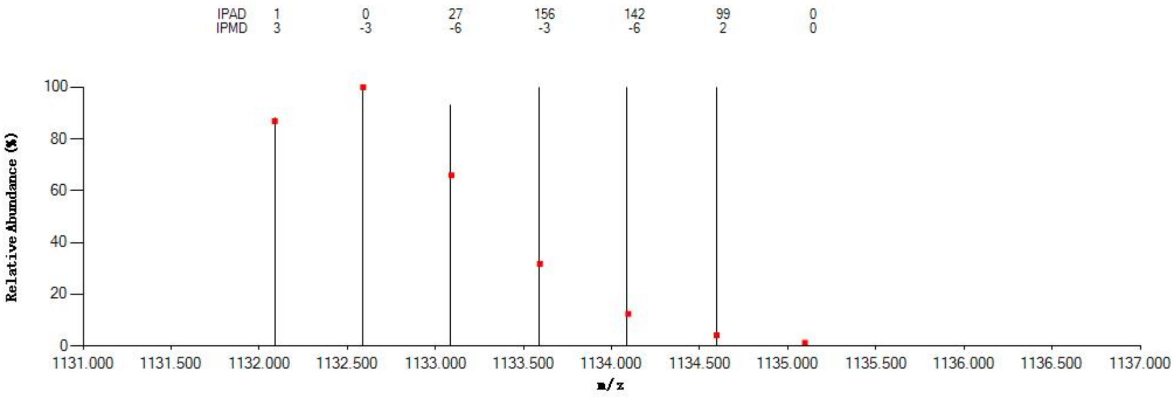

#### Annotated MS/MS Spectrum

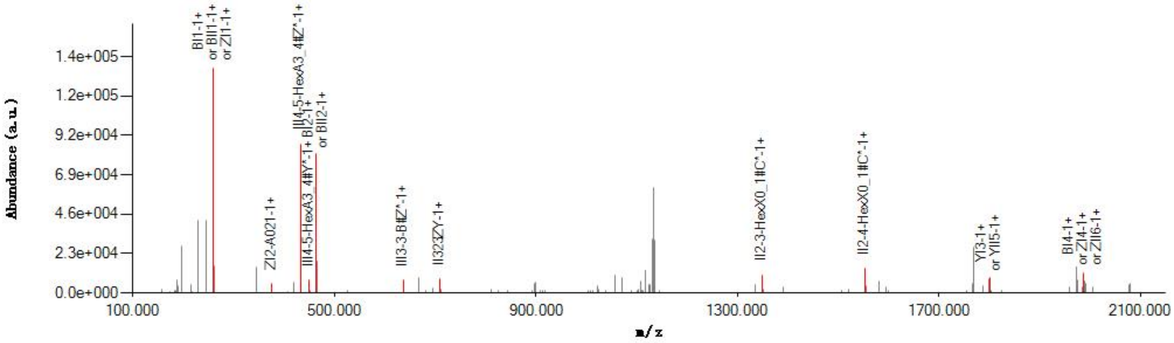

#### Graphical Fragmentation Map

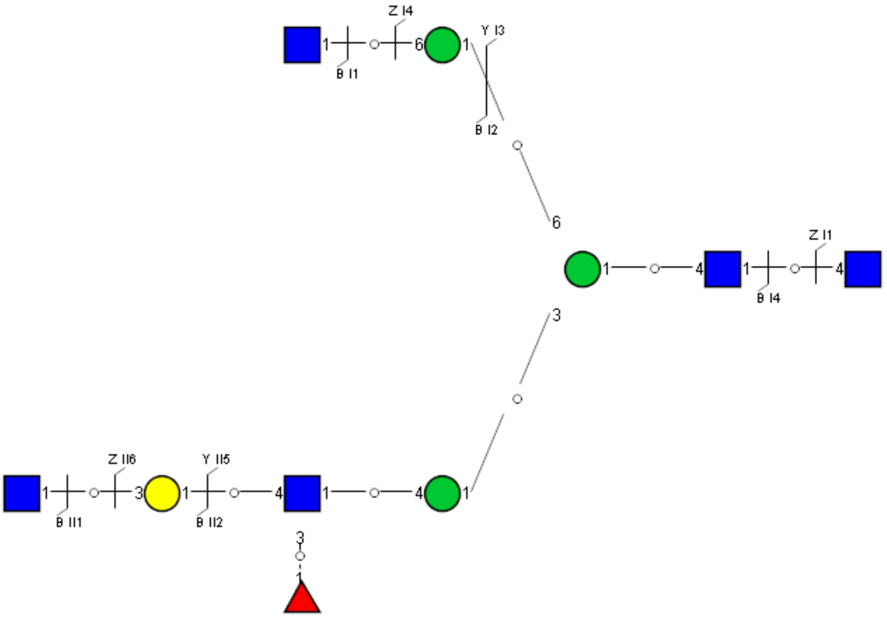

Spectrum Index: 29986(NoBHs:1)

#### Precursor iEF Map

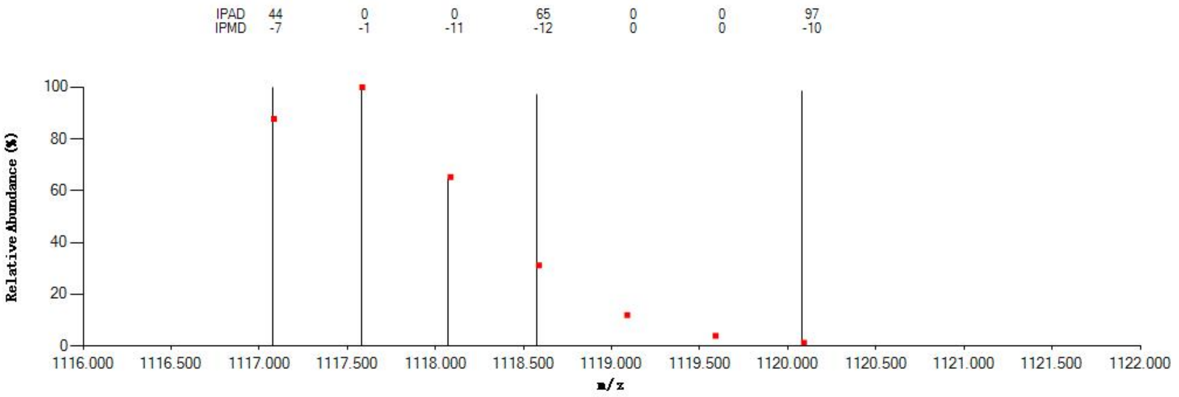


#### Annotated MS/MS Spectrum

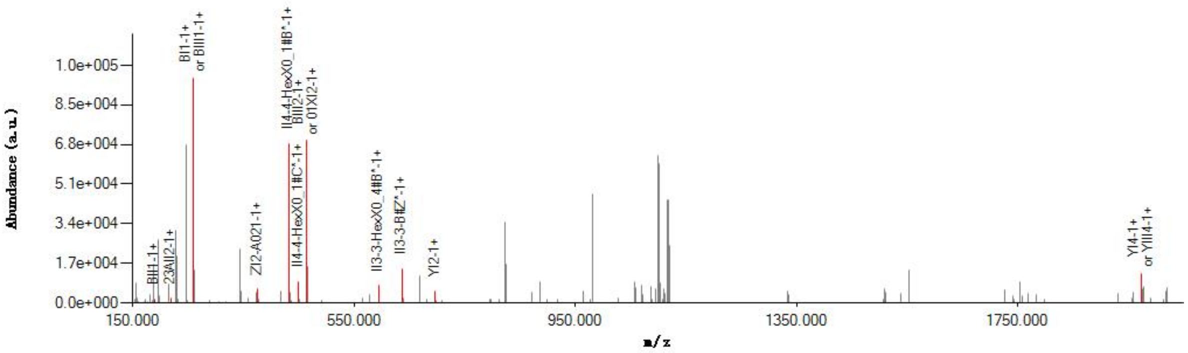


#### Graphical Fragmentation Map

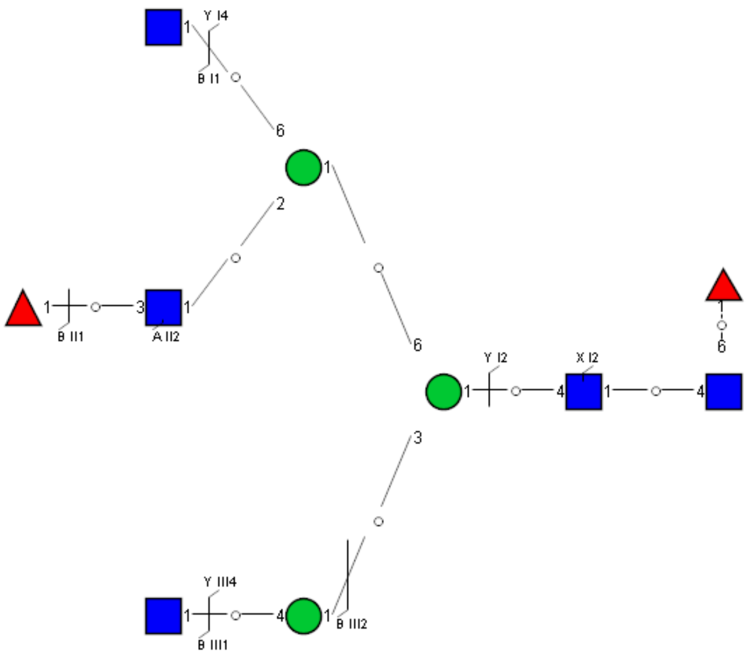

Spectrum Index: 30243(NoBHs:1)

#### Precursor iEF Map

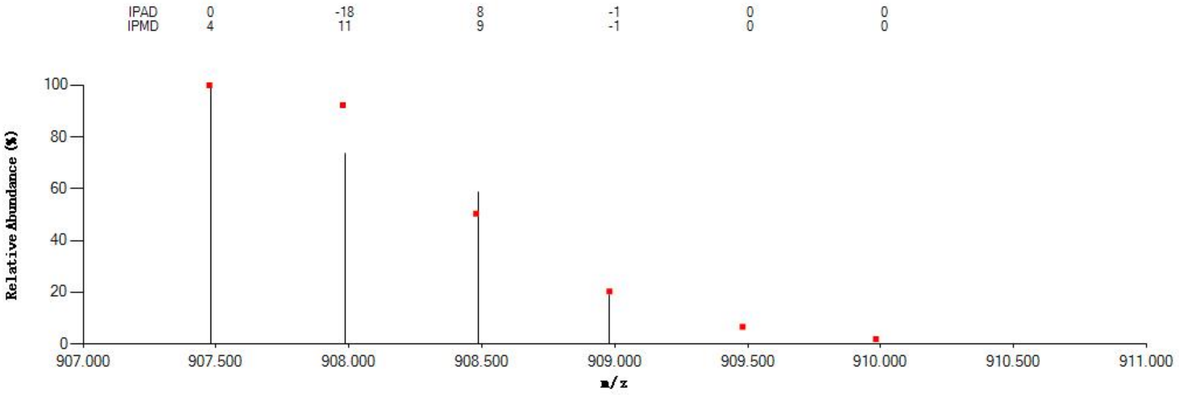


#### Annotated MS/MS Spectrum

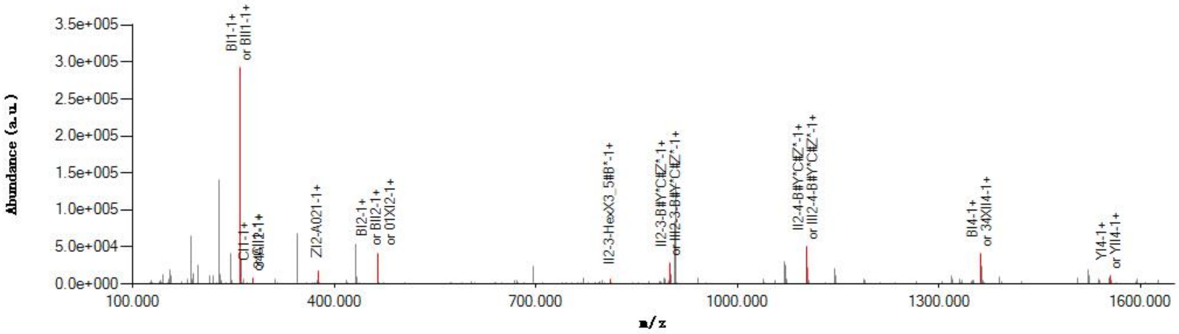


#### Graphical Fragmentation Map

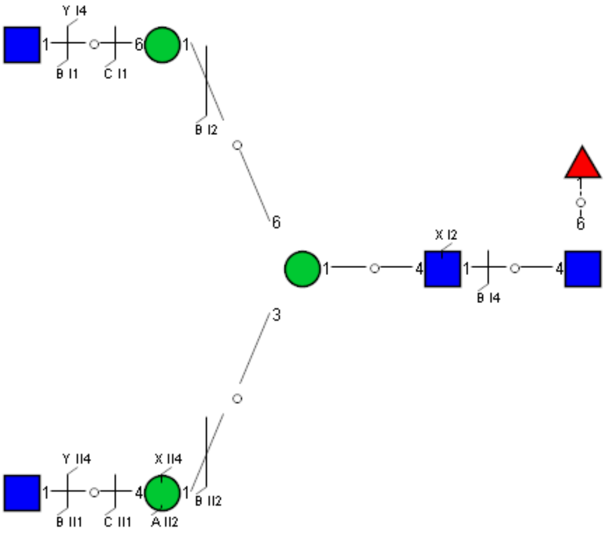

### Spectrum Index: 30622(NoBHs:1)

#### Precursor iEF Map

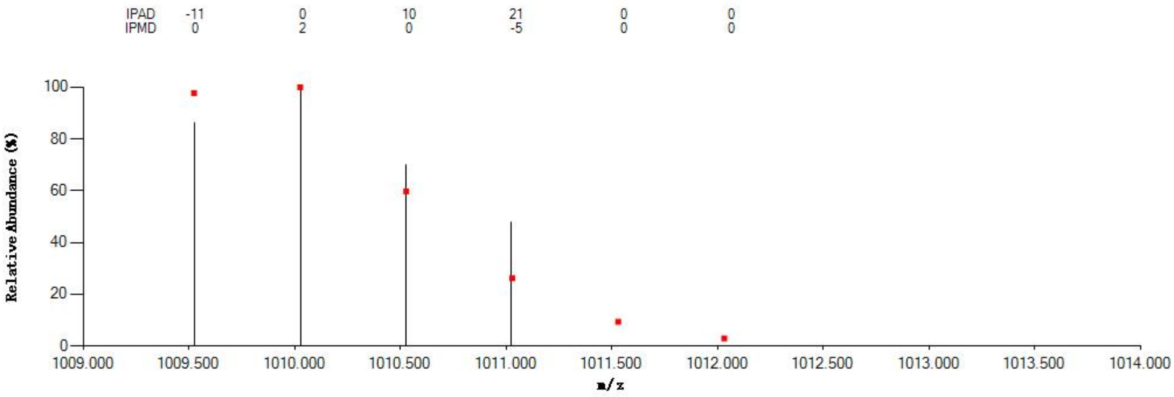


#### Annotated MS/MS Spectrum

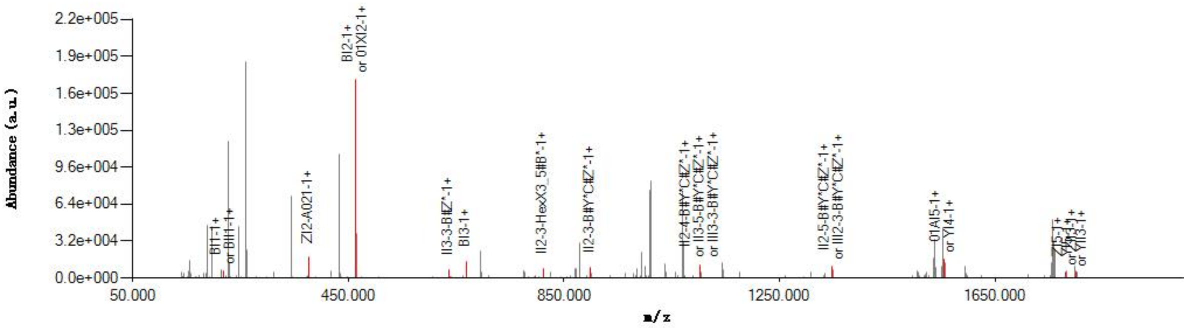


#### Graphical Fragmentation Map

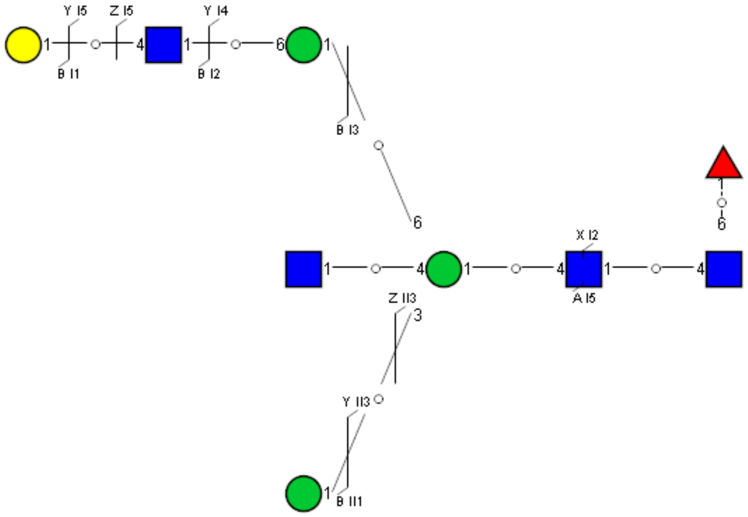

Spectrum Index: 31439(NoBHs:1)

#### Precursor iEF Map

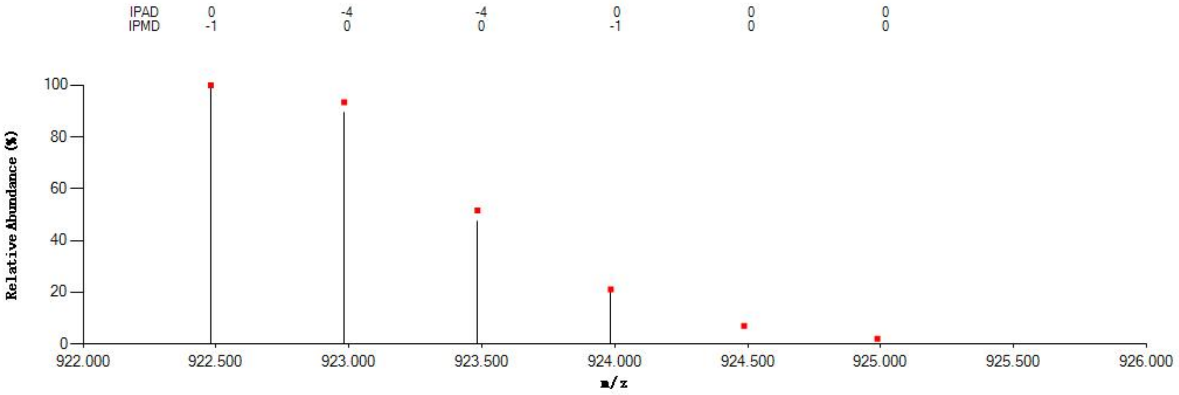

#### Annotated MS/MS Spectrum

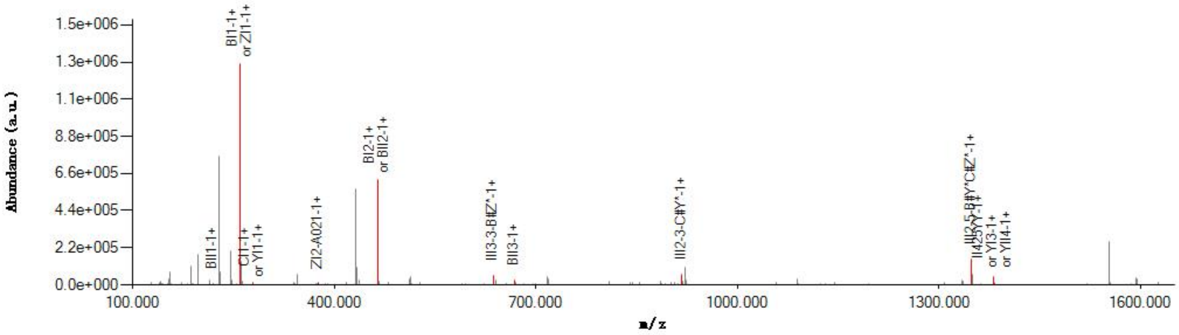

#### Graphical Fragmentation Map

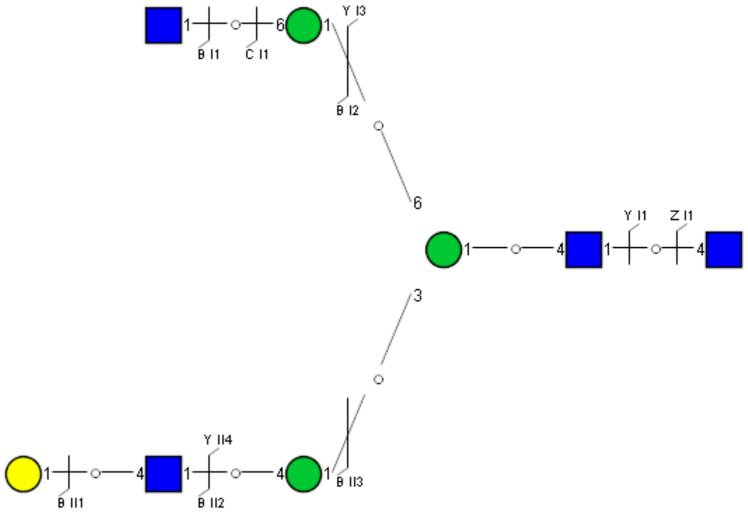

Spectrum Index: 31716(NoBHs:1)

#### Precursor iEF Map

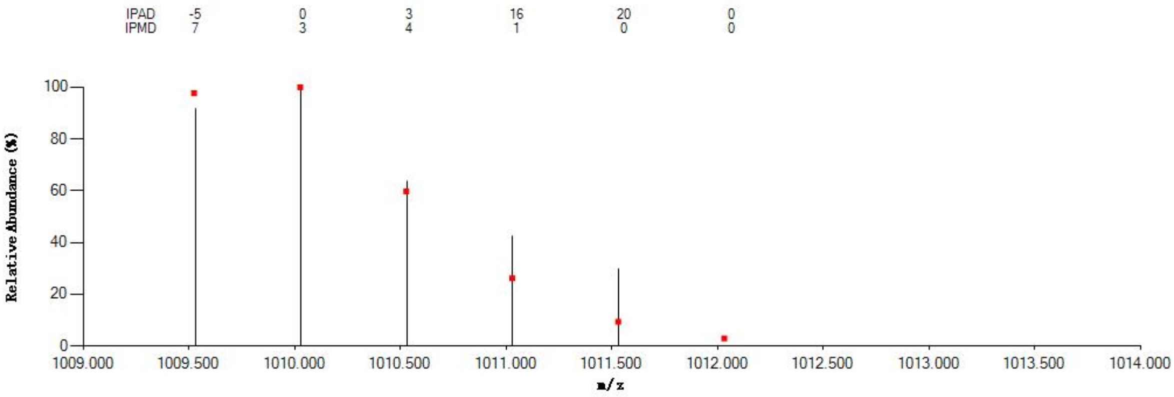

#### Annotated MS/MS Spectrum

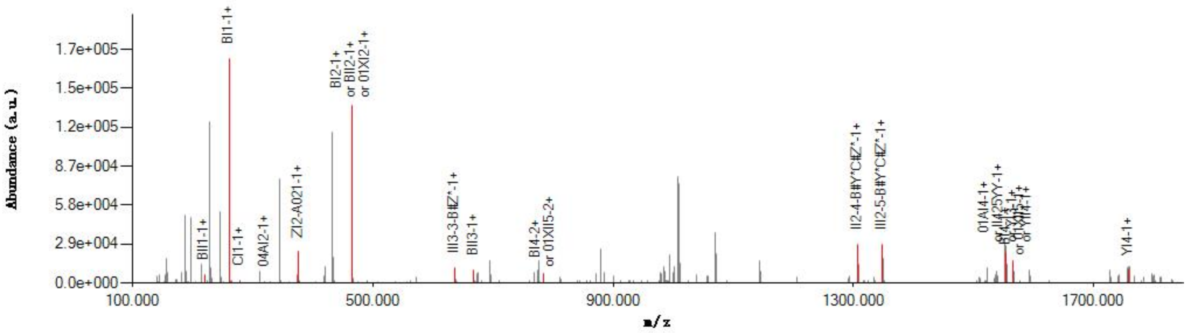

#### Graphical Fragmentation Map

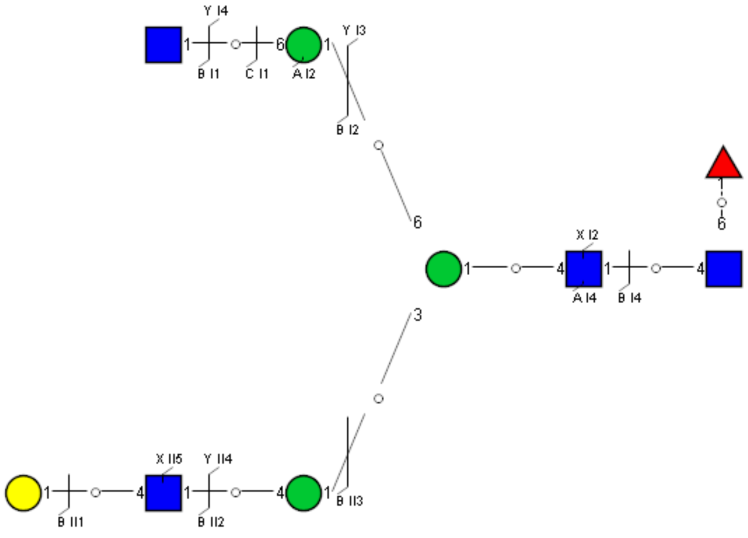

Spectrum Index: 31795(NoBHs:1)

#### Precursor iEF Map

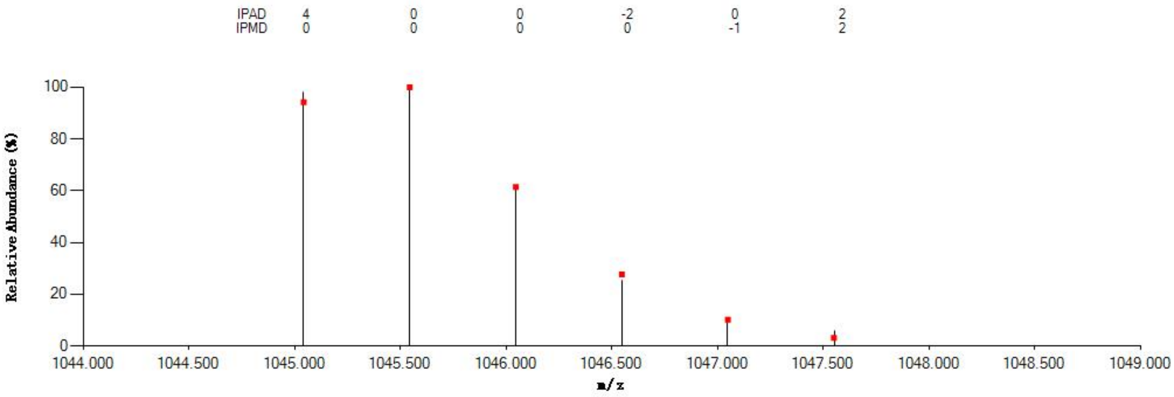


#### Annotated MS/MS Spectrum

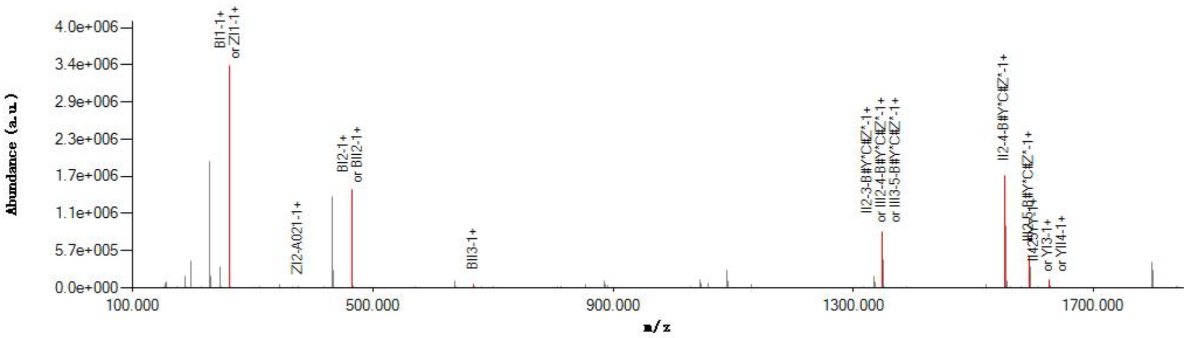


#### Graphical Fragmentation Map

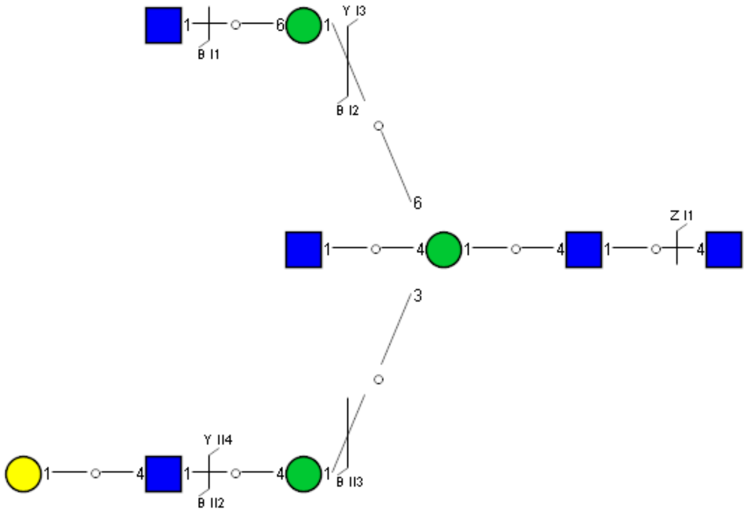

Spectrum Index: 31837(NoBHs:1)

#### Precursor iEF Map

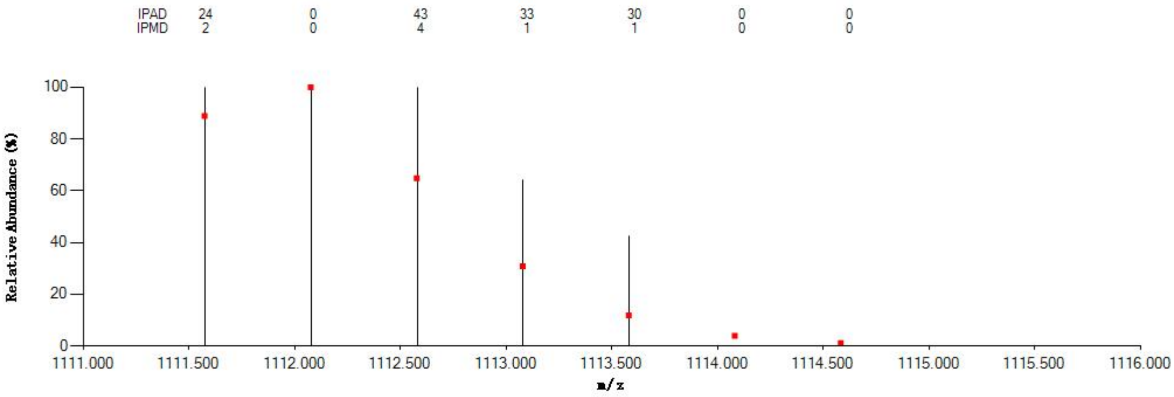


#### Annotated MS/MS Spectrum

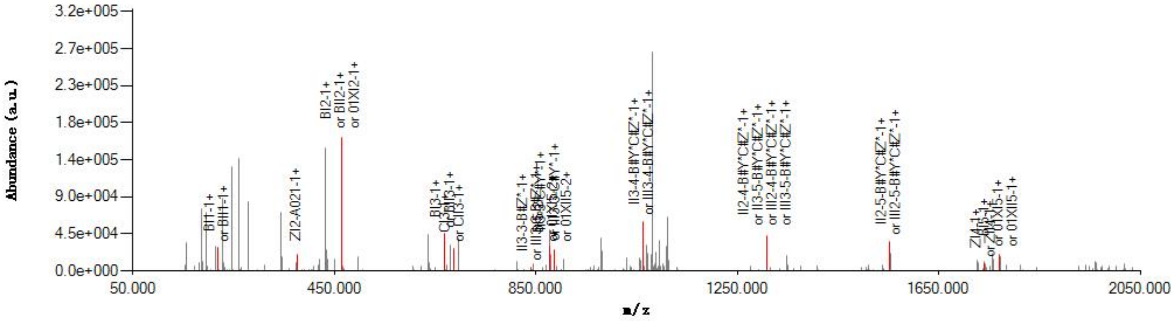


#### Graphical Fragmentation Map

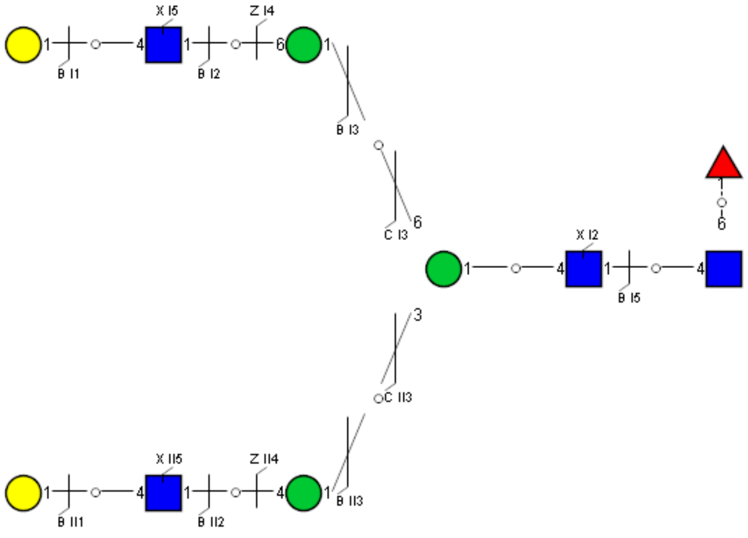

Spectrum Index: 31922(NoBHs:1)

#### Precursor iEF Map

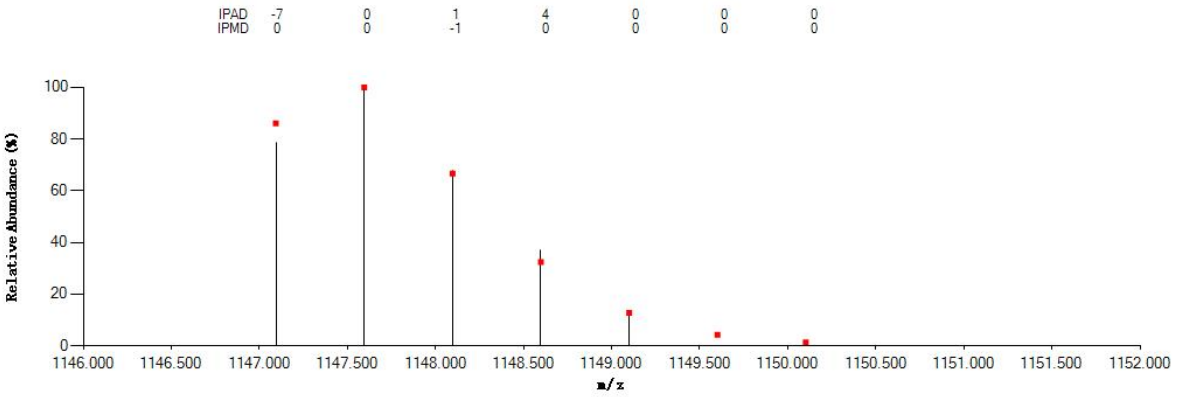

#### Annotated MS/MS Spectrum

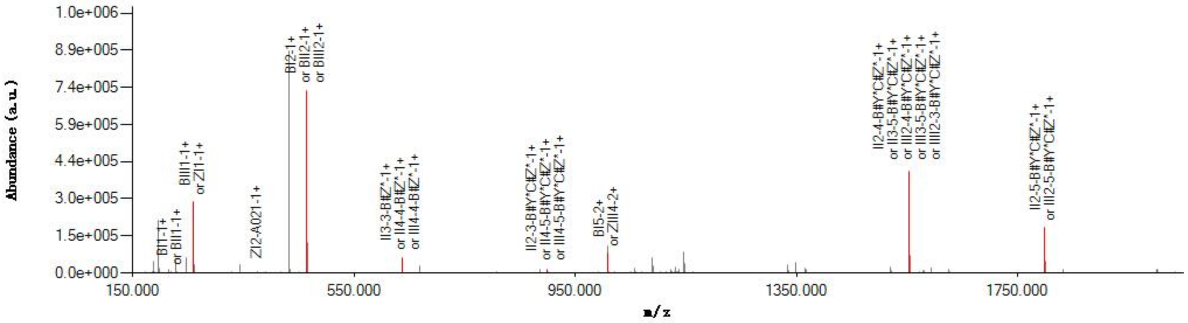

#### Graphical Fragmentation Map

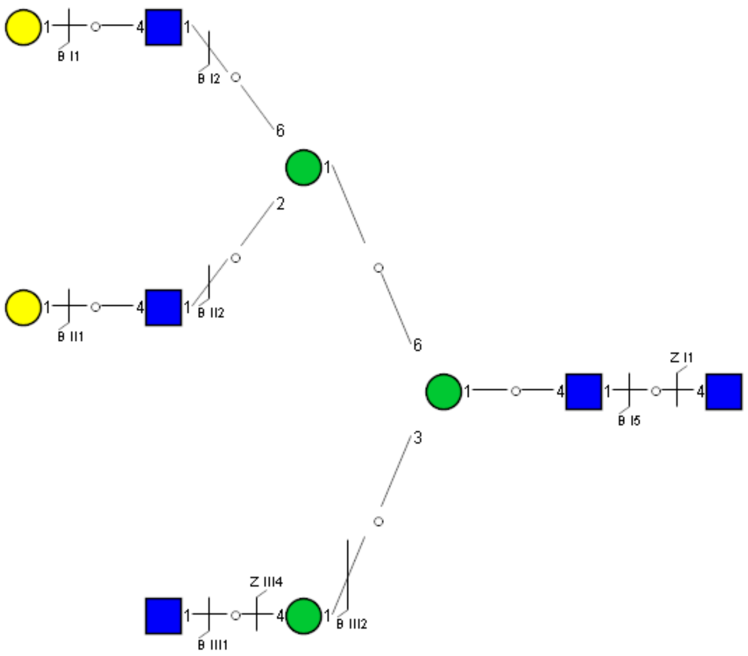

Spectrum Index: 32030(NoBHs:1)

#### Precursor iEF Map

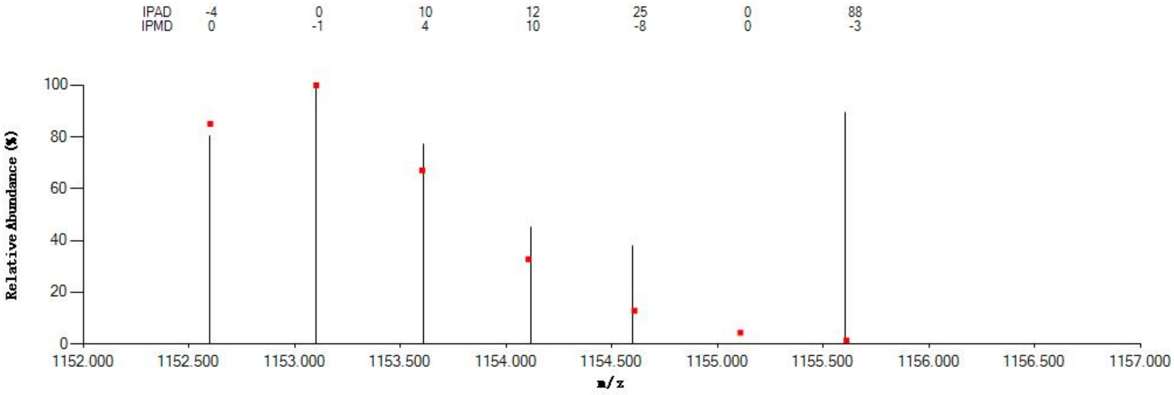


#### Annotated MS/MS Spectrum

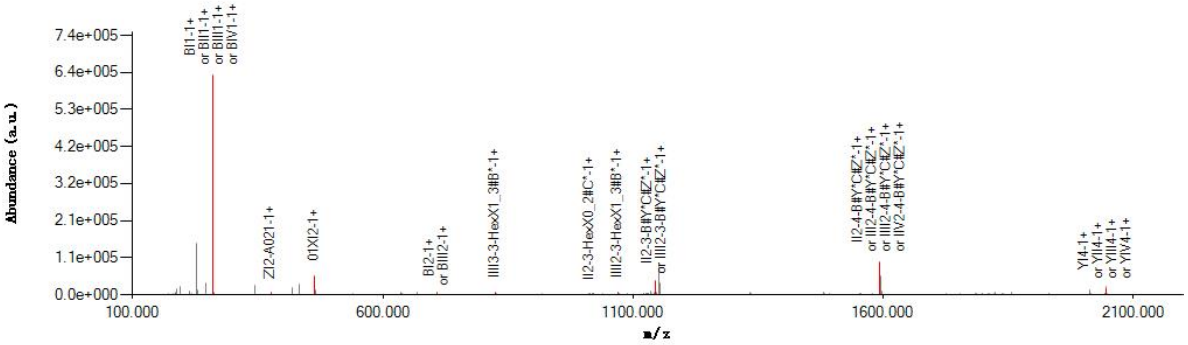


#### Graphical Fragmentation Map

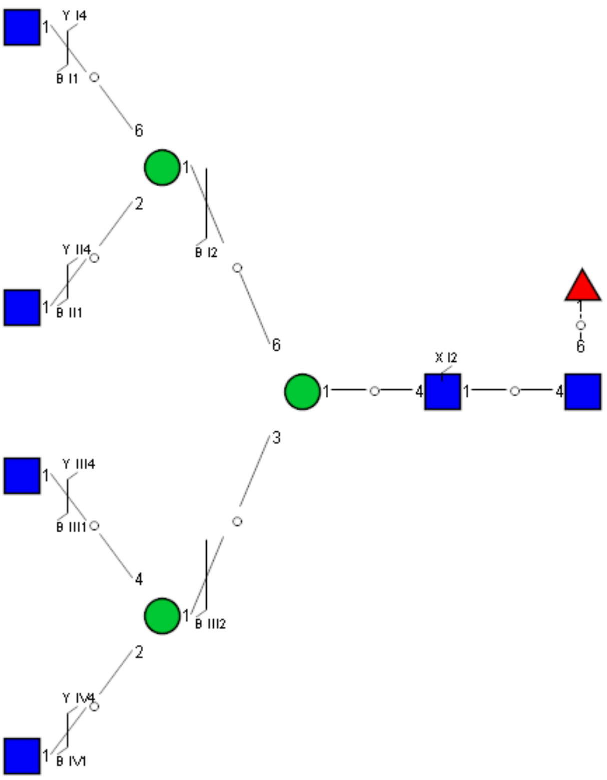

Spectrum Index: 32486(NoBHs:1)

#### Precursor iEF Map

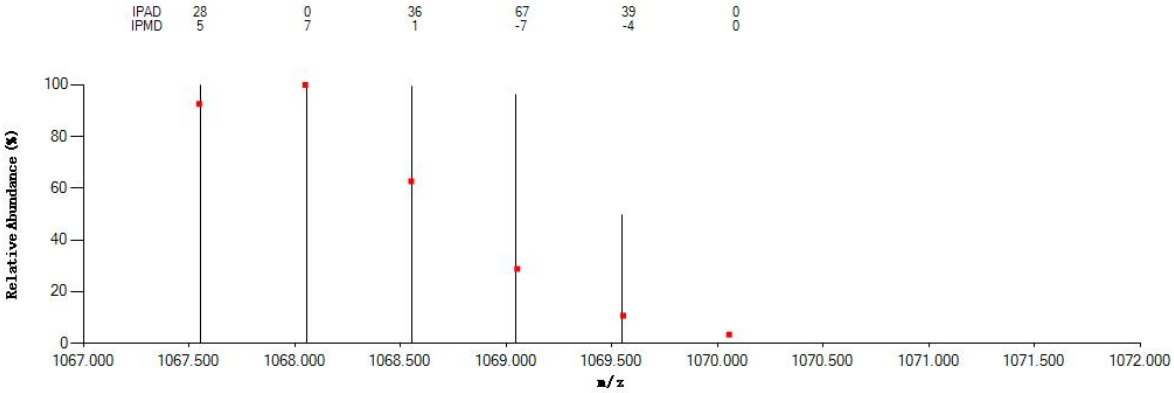


#### Annotated MS/MS Spectrum

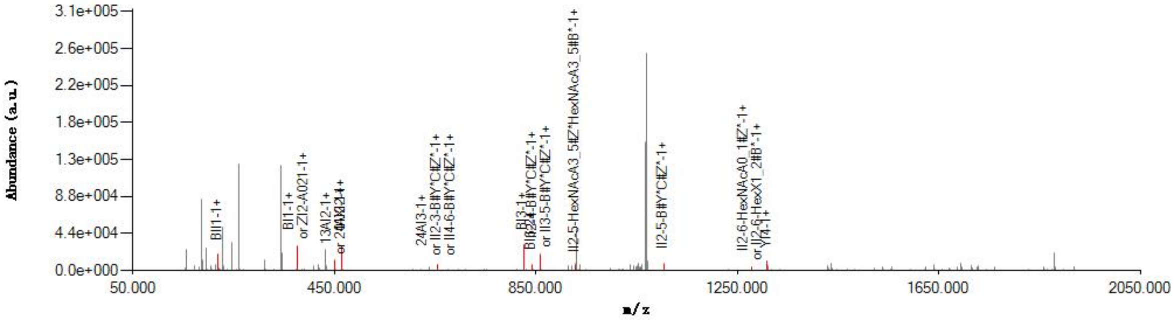


#### Graphical Fragmentation Map

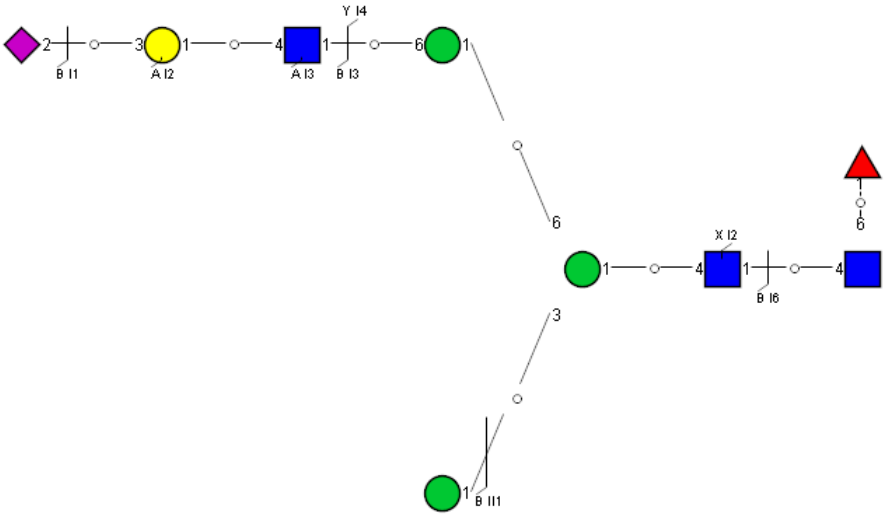

Spectrum Index: 32493(NoBHs:1)

#### Precursor iEF Map

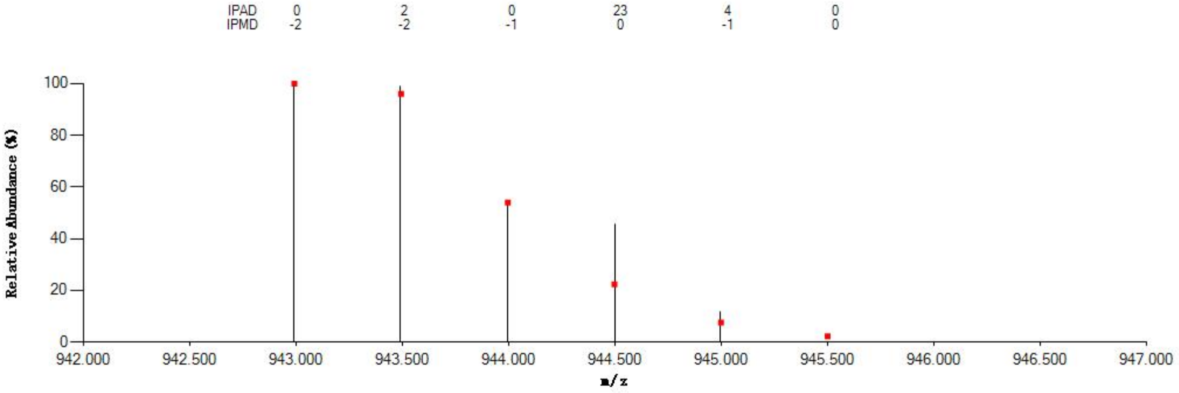


#### Annotated MS/MS Spectrum

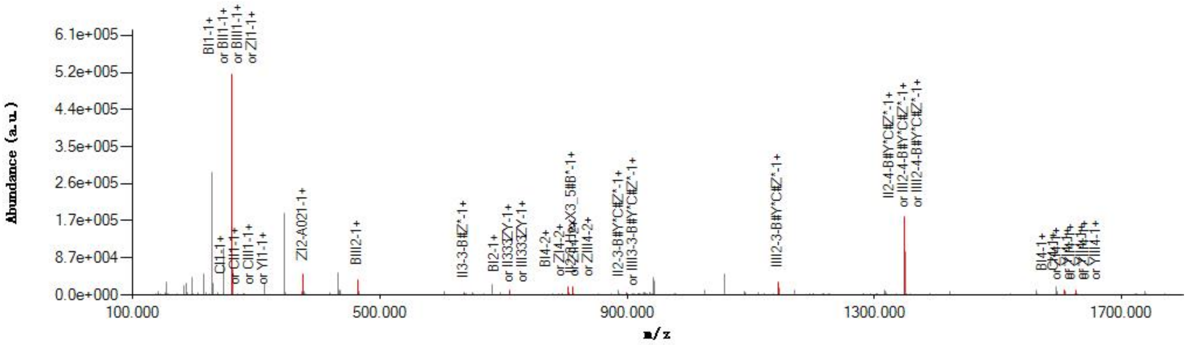


#### Graphical Fragmentation Map

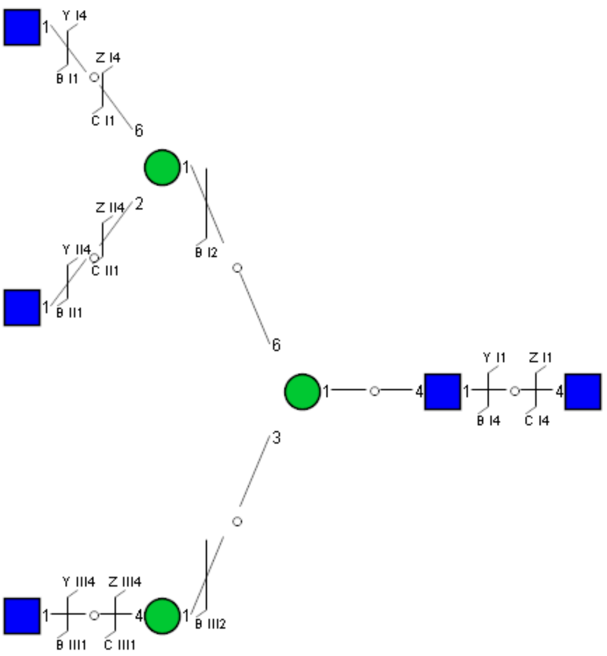

Spectrum Index: 32552(NoBHs:1)

#### Precursor iEF Map

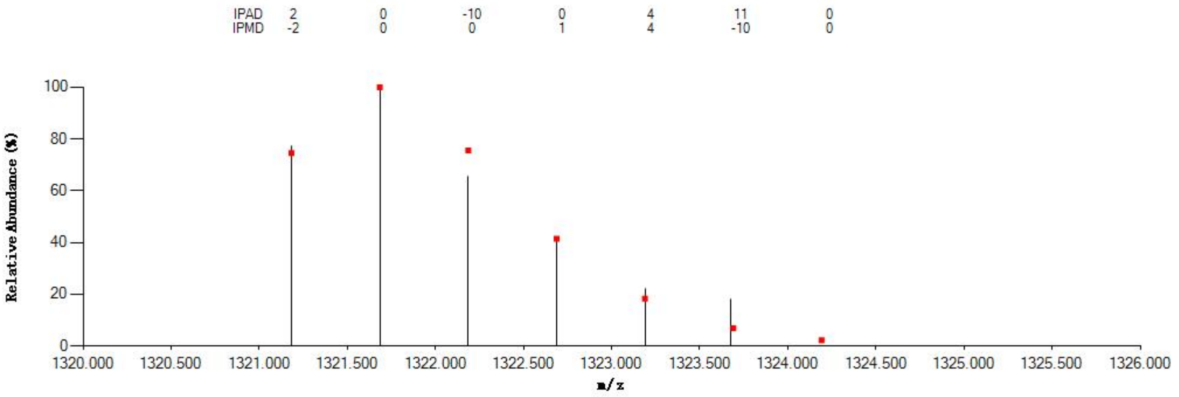

#### Annotated MS/MS Spectrum

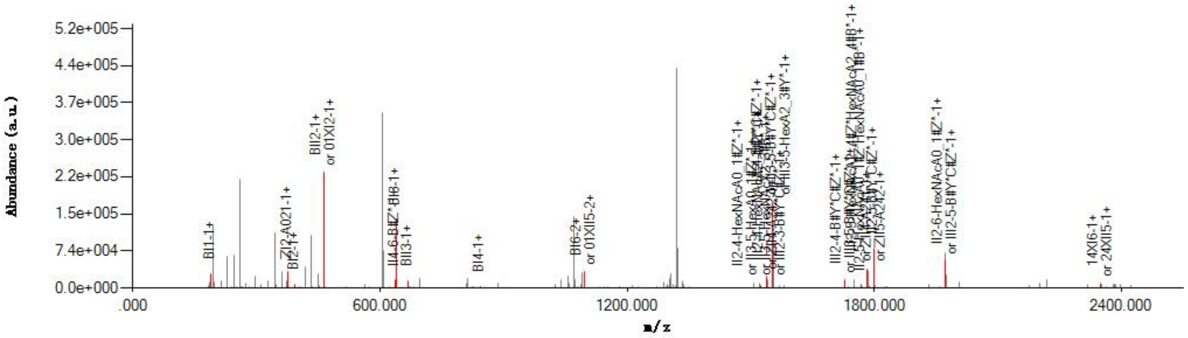

#### Graphical Fragmentation Map

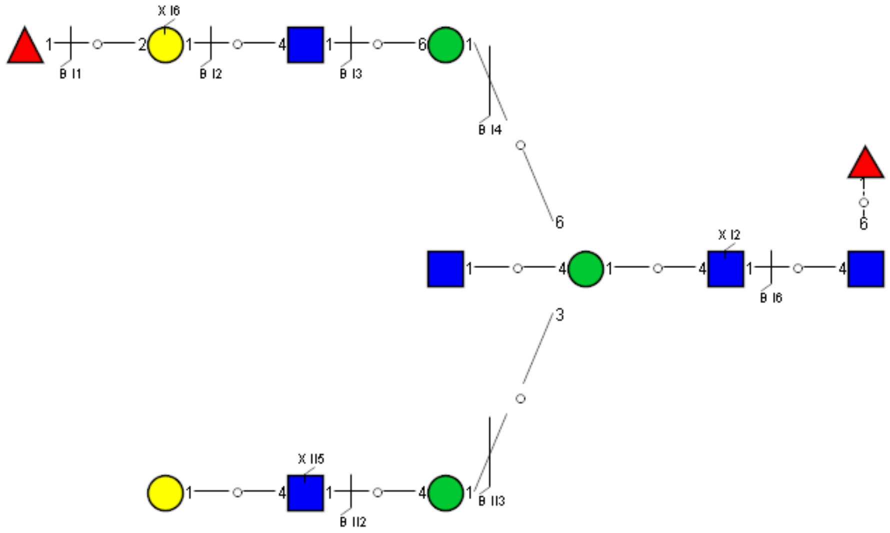

### Spectrum Index: 32614(NoBHs:1)

#### Precursor iEF Map

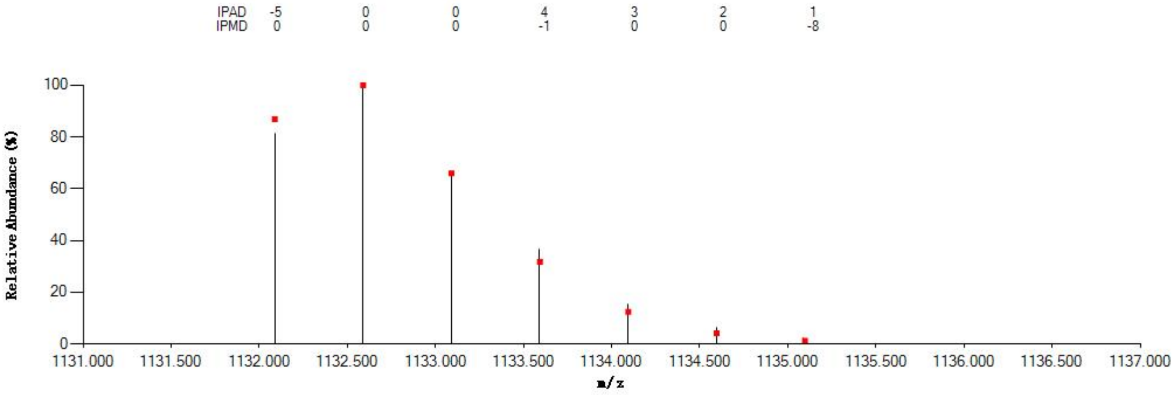


#### Annotated MS/MS Spectrum

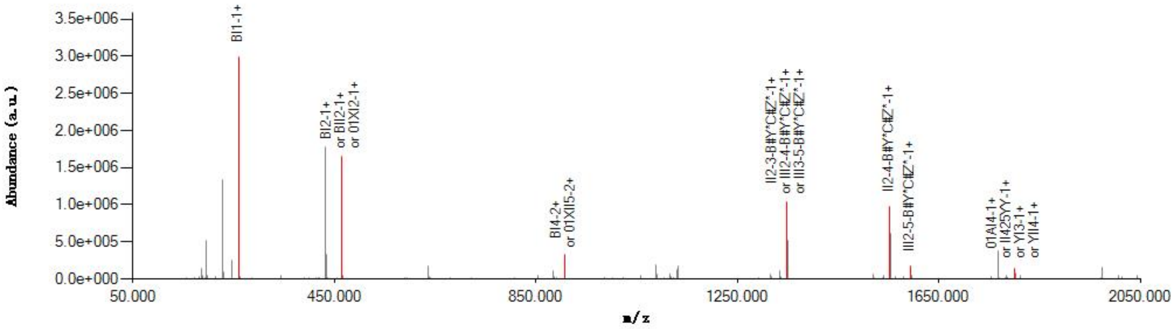


#### Graphical Fragmentation Map

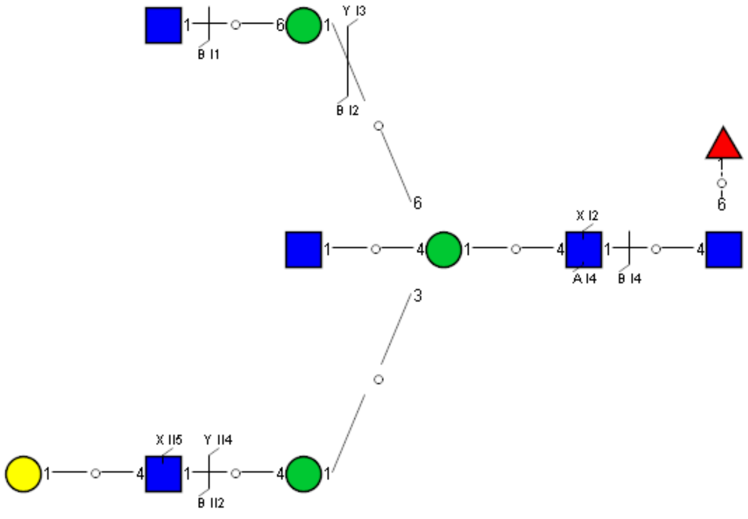

Spectrum Index: 33000(NoBHs:1)

#### Precursor iEF Map

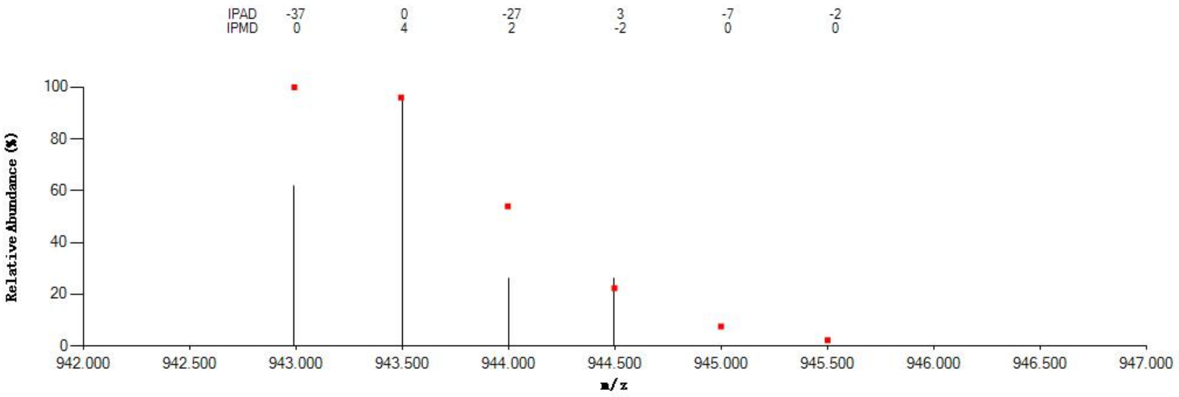

#### Annotated MS/MS Spectrum

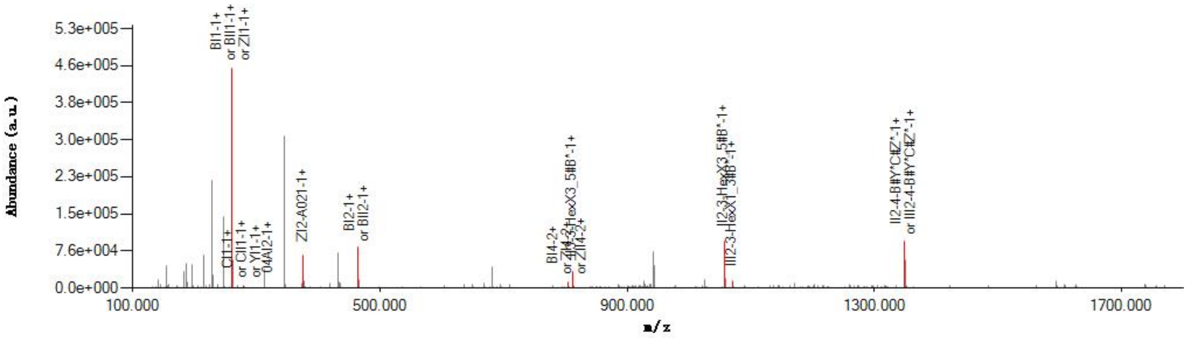

#### Graphical Fragmentation Map

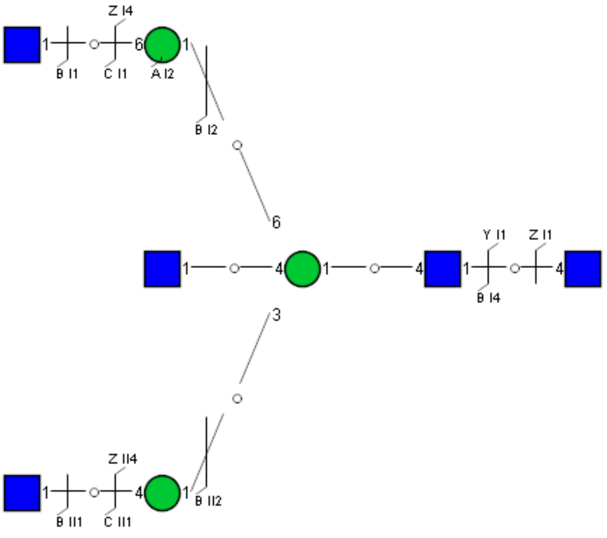

Spectrum Index: 33206(NoBHs:1)

#### Precursor iEF Map

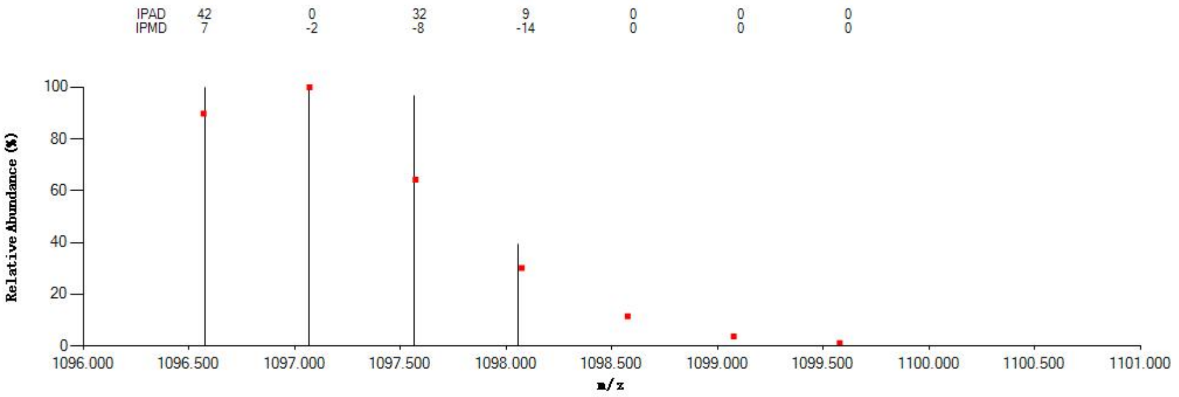


#### Annotated MS/MS Spectrum

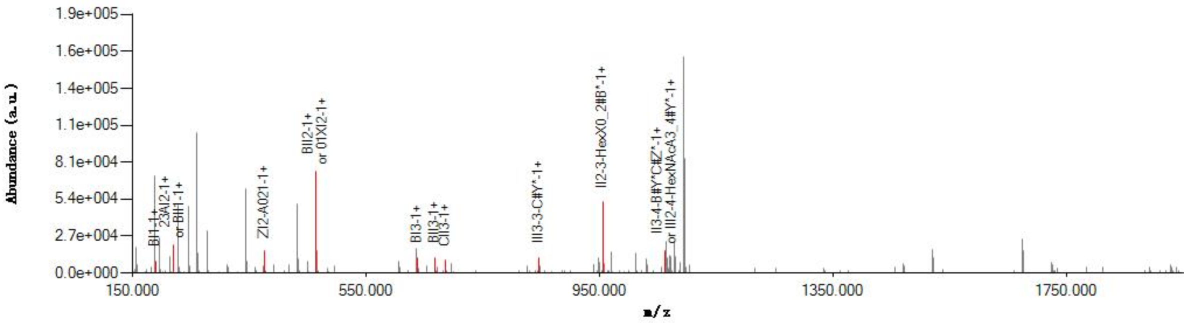


#### Graphical Fragmentation Map

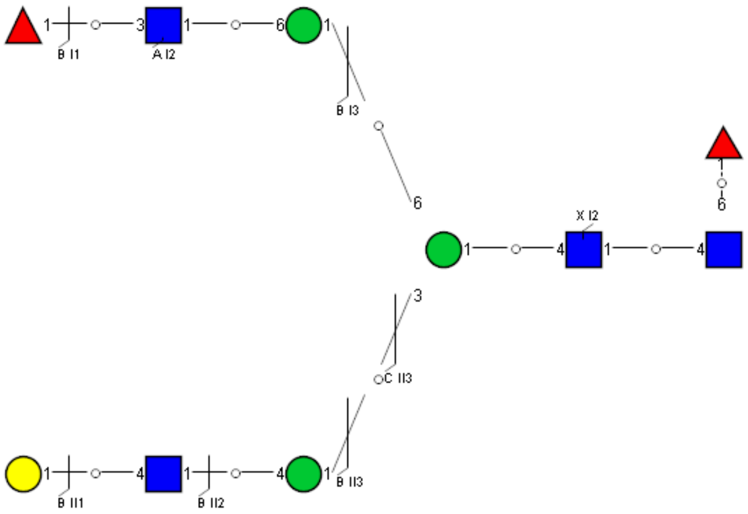

Spectrum Index: 33349(NoBHs:1)

#### Precursor iEF Map

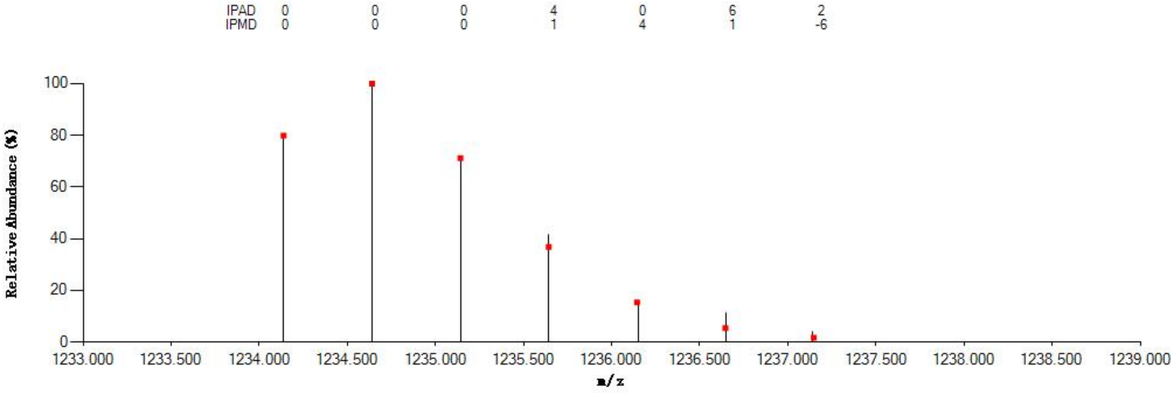


#### Annotated MS/MS Spectrum

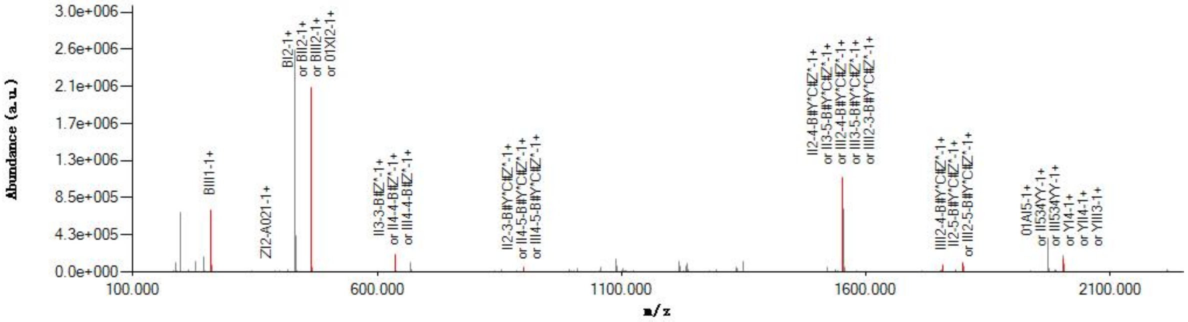


#### Graphical Fragmentation Map

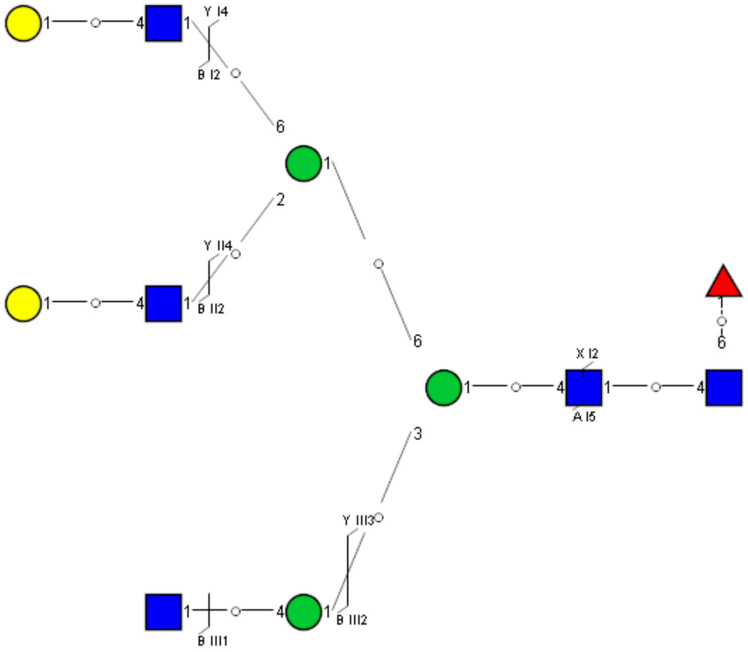

Spectrum Index: 33408(NoBHs:1)

#### Precursor iEF Map

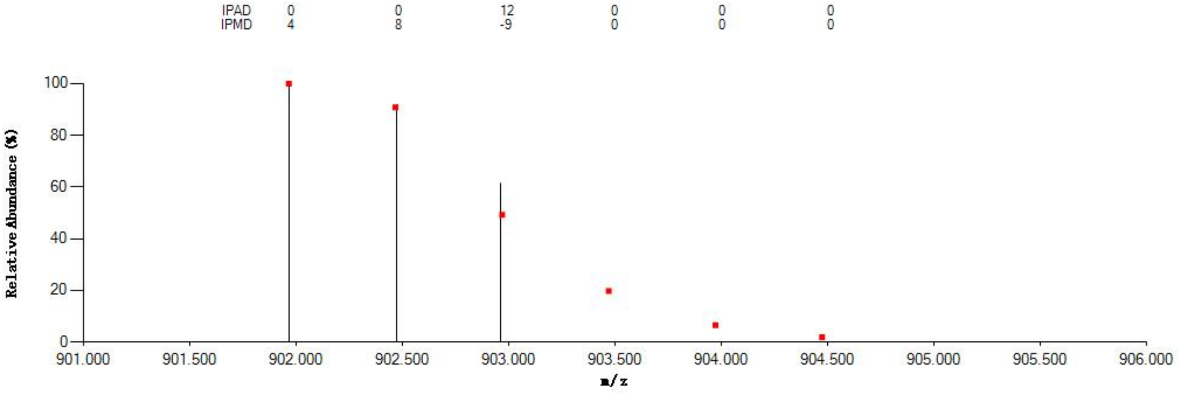


#### Annotated MS/MS Spectrum

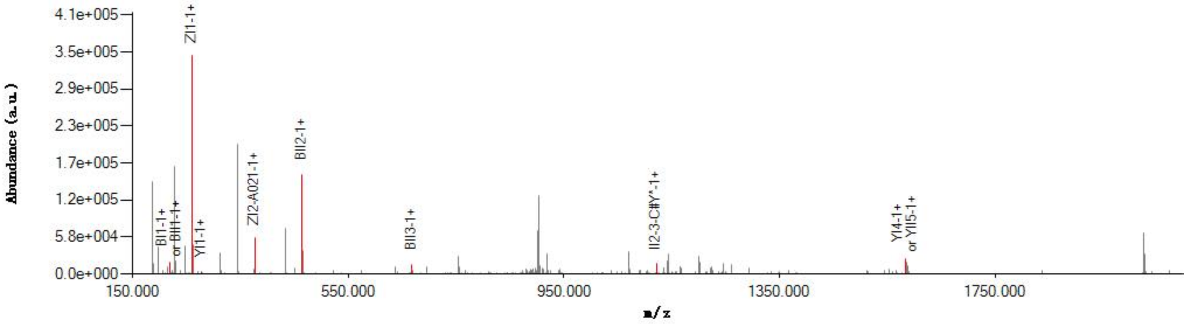


#### Graphical Fragmentation Map

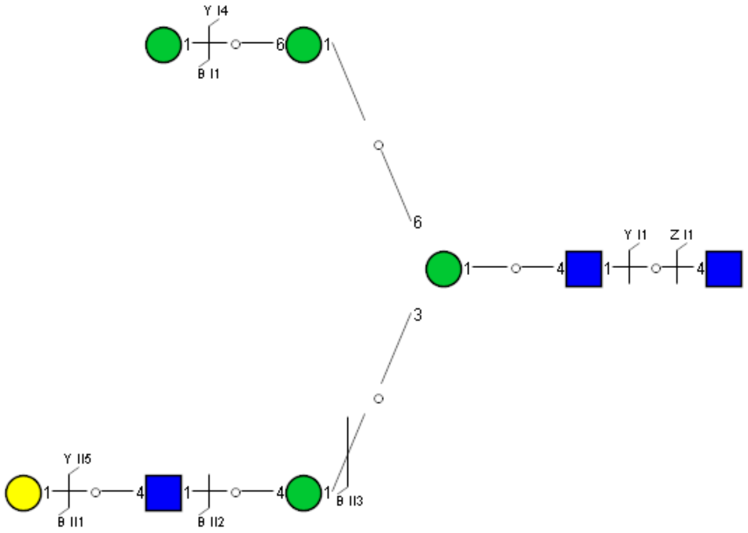

Spectrum Index: 33475(NoBHs:1)

#### Precursor iEF Map

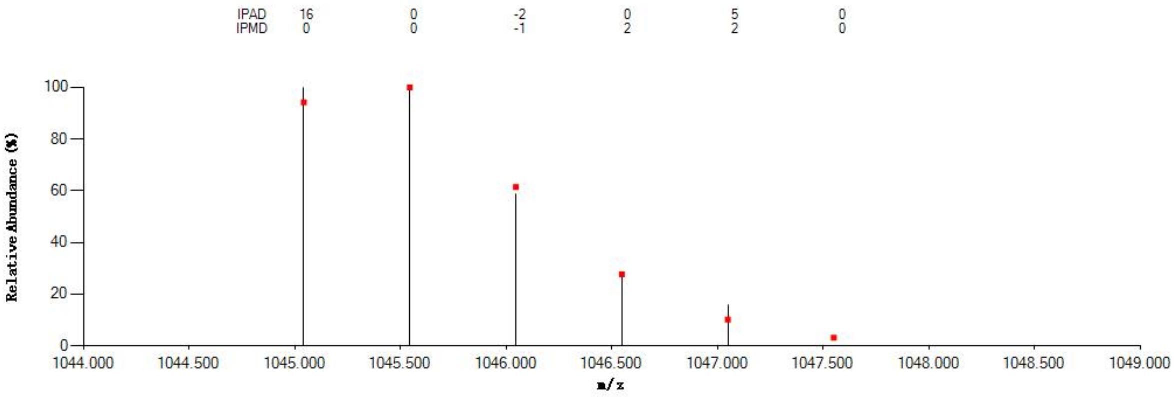


#### Annotated MS/MS Spectrum

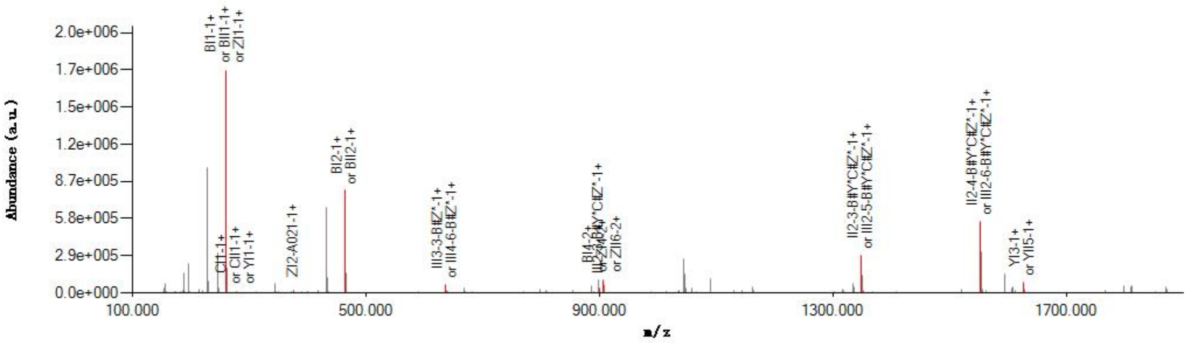


#### Graphical Fragmentation Map

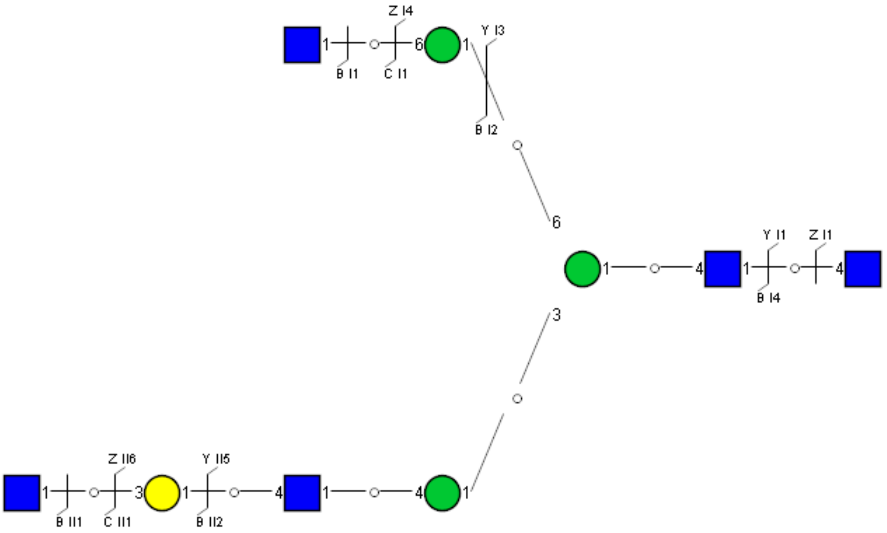

Spectrum Index: 33492(NoBHs:1)

#### Precursor iEF Map

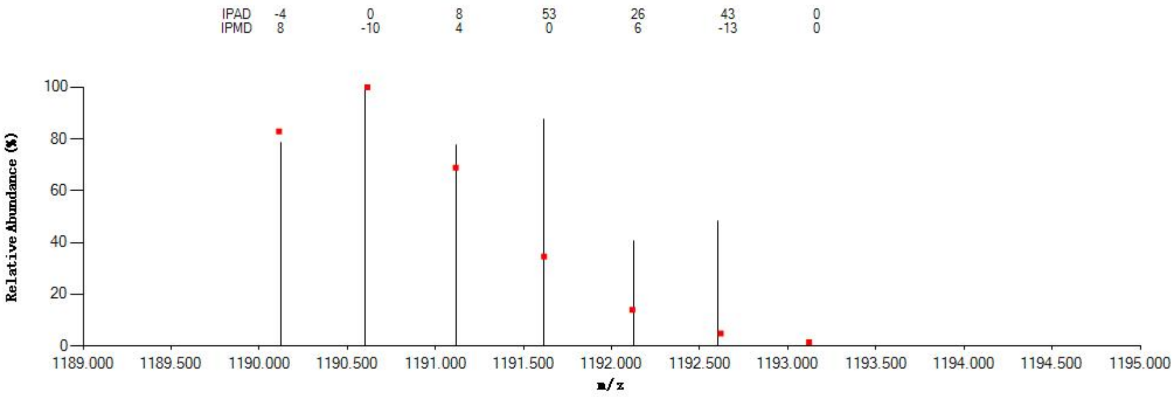


#### Annotated MS/MS Spectrum

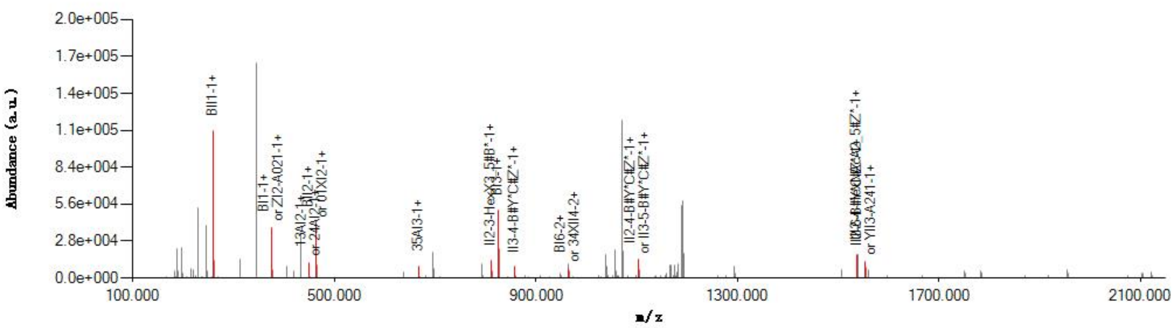


#### Graphical Fragmentation Map

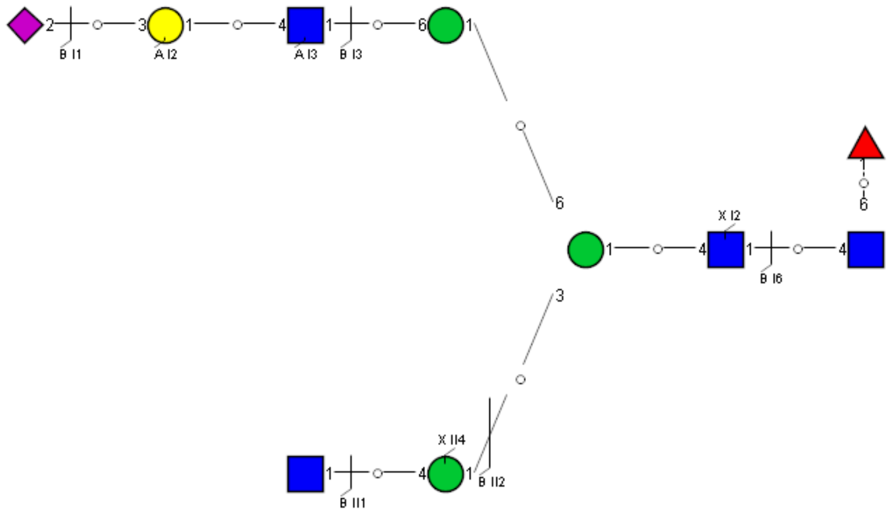

### Spectrum Index: 33518(NoBHs:1)

#### Precursor iEF Map

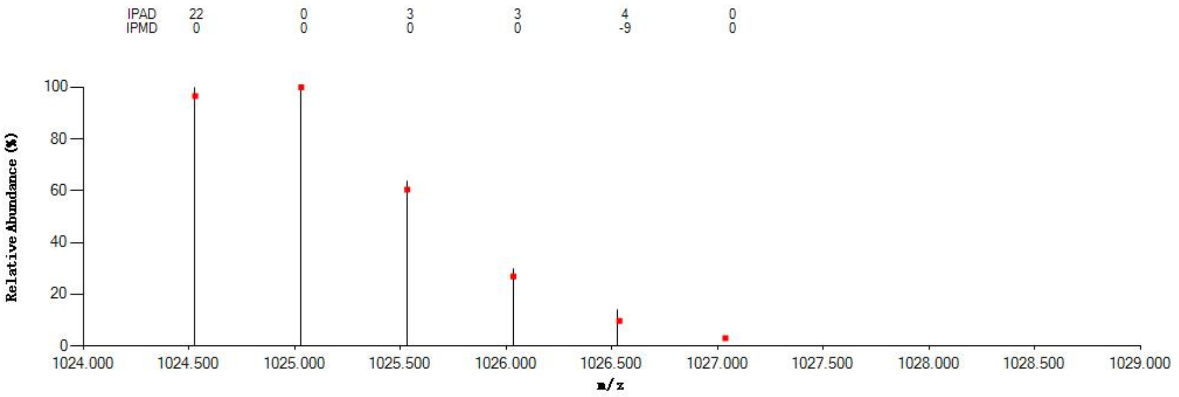


#### Annotated MS/MS Spectrum

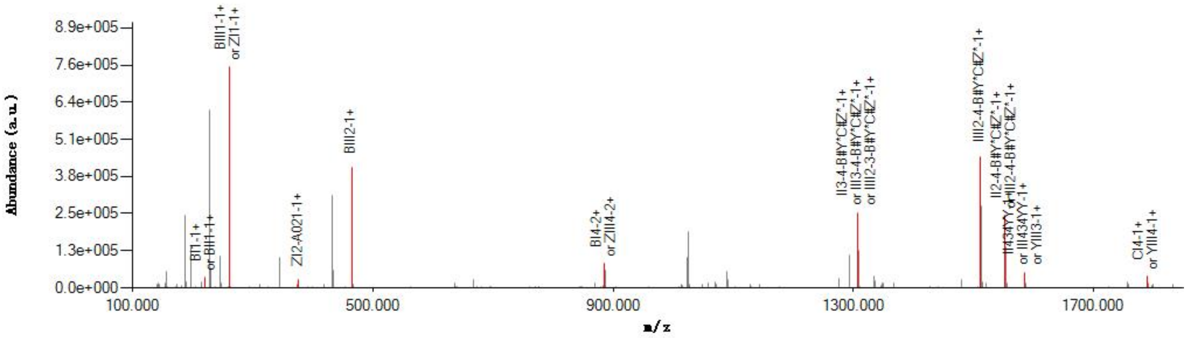


#### Graphical Fragmentation Map

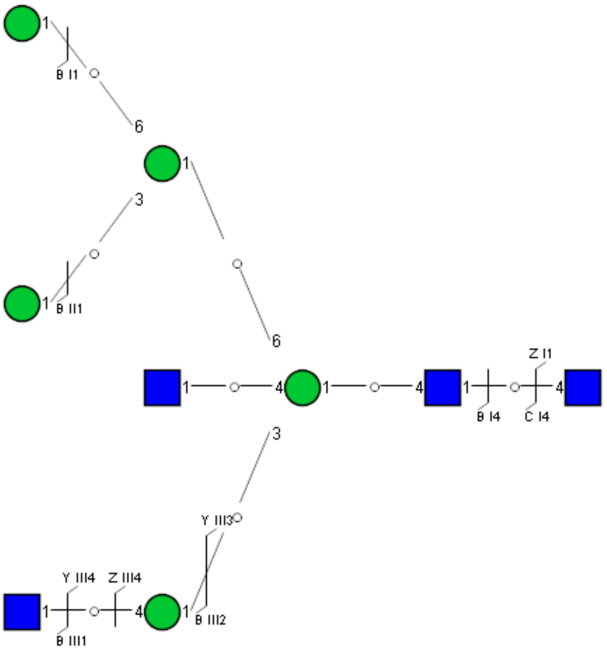

Spectrum Index: 33604(NoBHs:1)

#### Precursor iEF Map

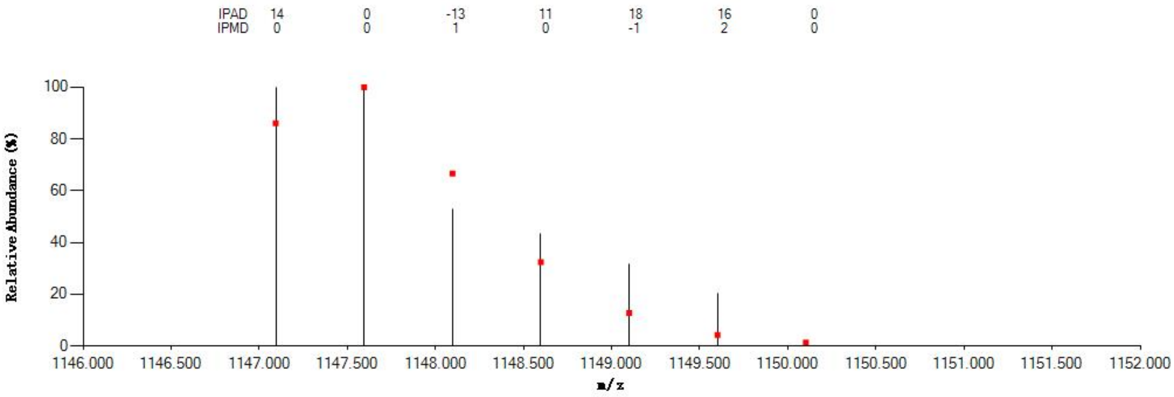

#### Annotated MS/MS Spectrum

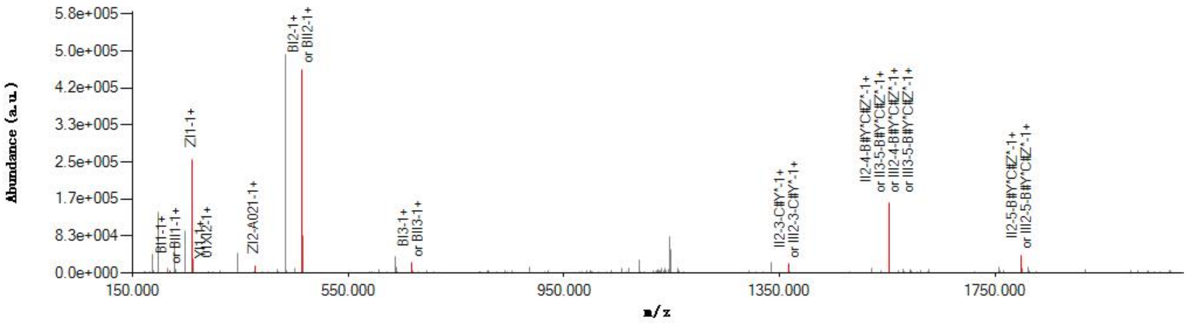

#### Graphical Fragmentation Map

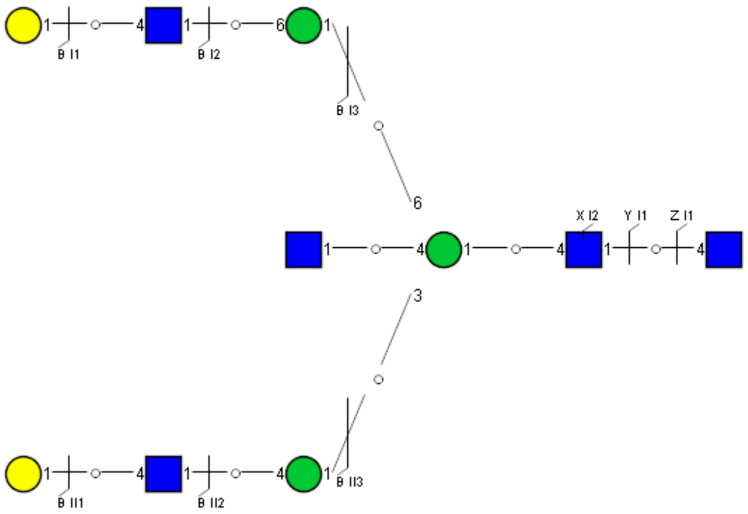

Spectrum Index: 33952(NoBHs:1)

#### Precursor iEF Map

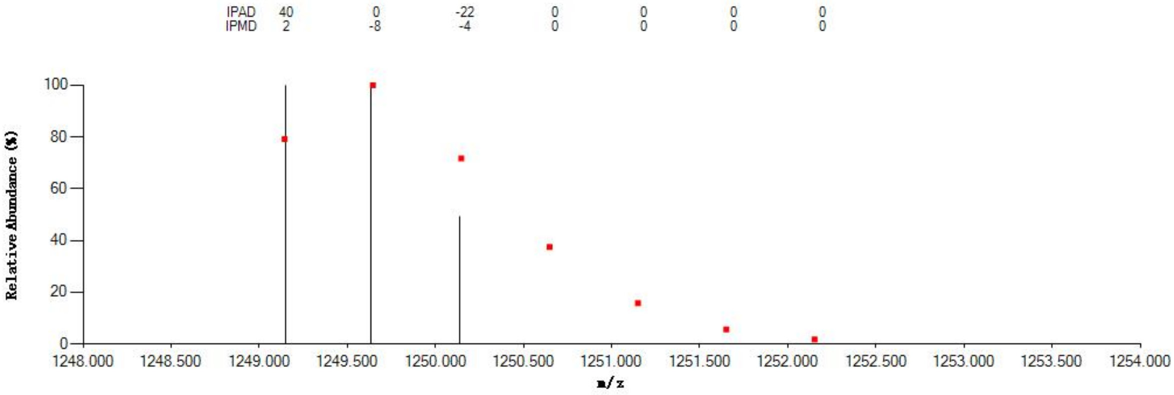


#### Annotated MS/MS Spectrum

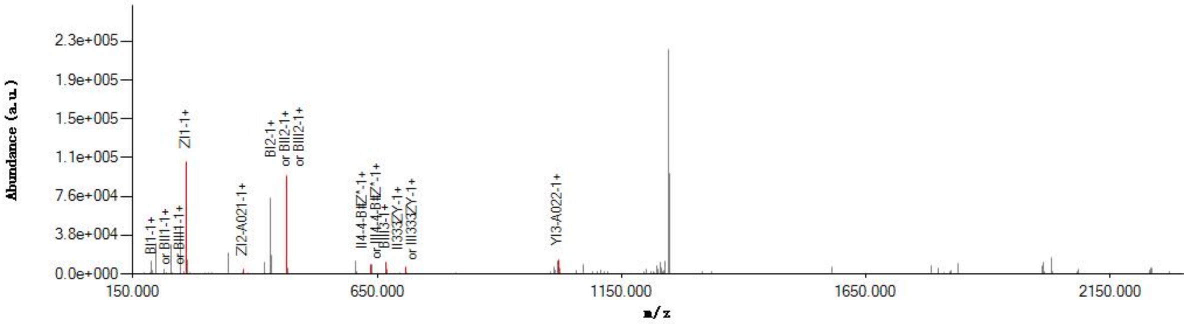


#### Graphical Fragmentation Map

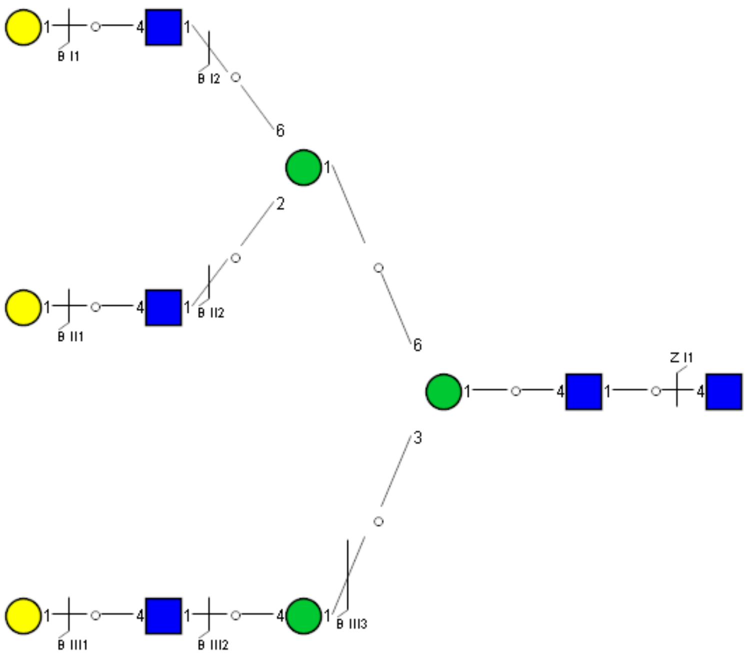

Spectrum Index: 34215(NoBHs:1)

#### Precursor iEF Map

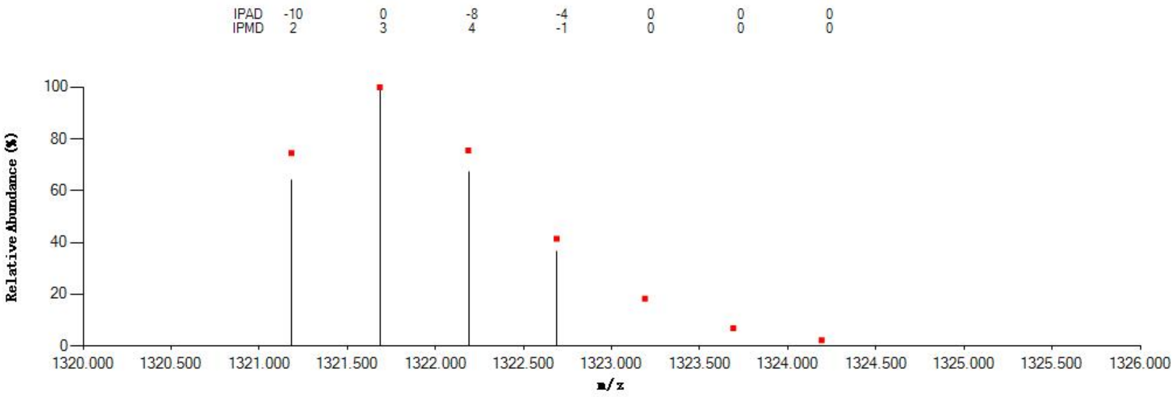


#### Annotated MS/MS Spectrum

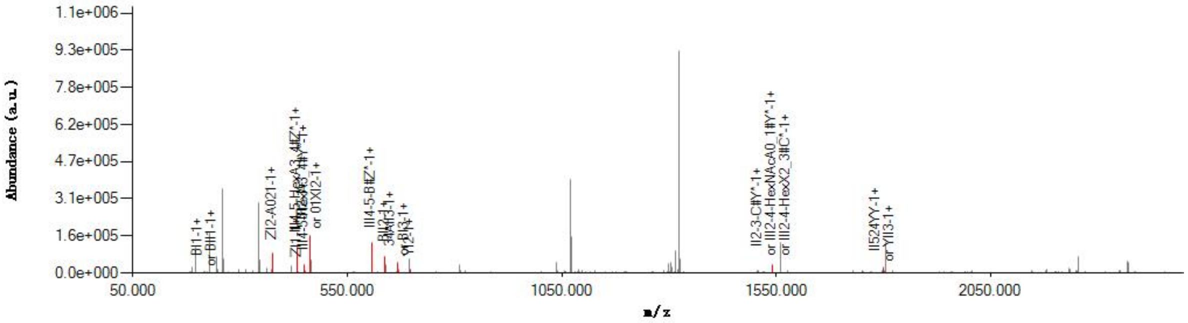


#### Graphical Fragmentation Map

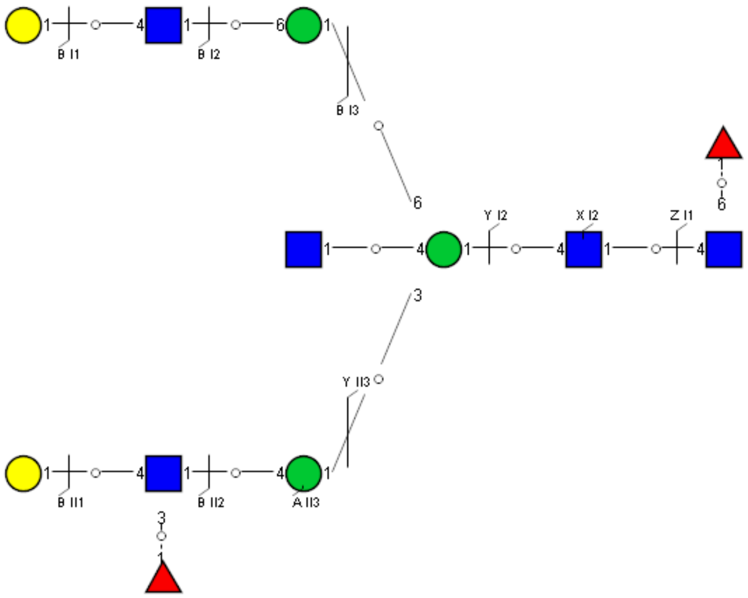

Spectrum Index: 34225(NoBHs:1)

#### Precursor iEF Map

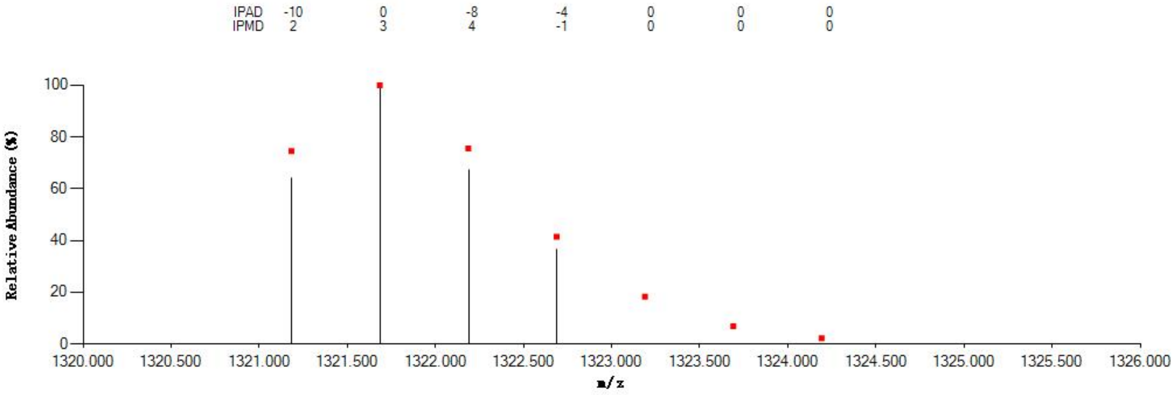


#### Annotated MS/MS Spectrum

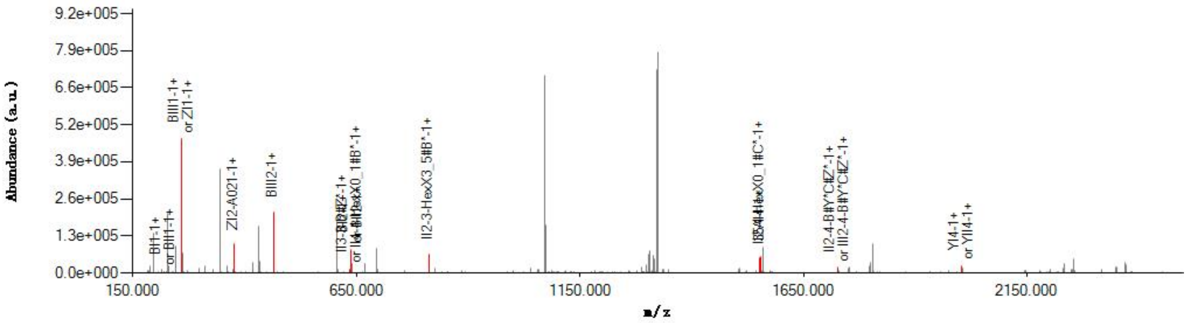


#### Graphical Fragmentation Map

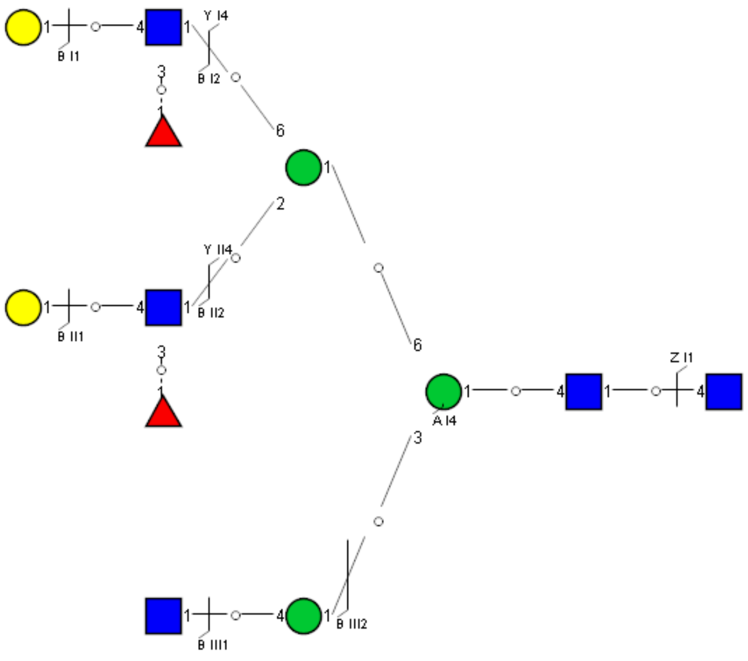

Spectrum Index: 34441(NoBHs:1)

#### Precursor iEF Map

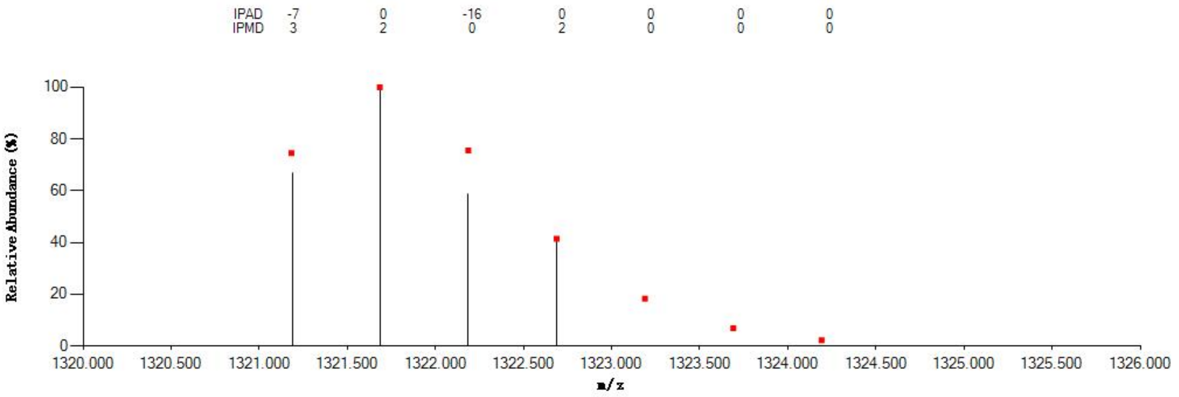


#### Annotated MS/MS Spectrum

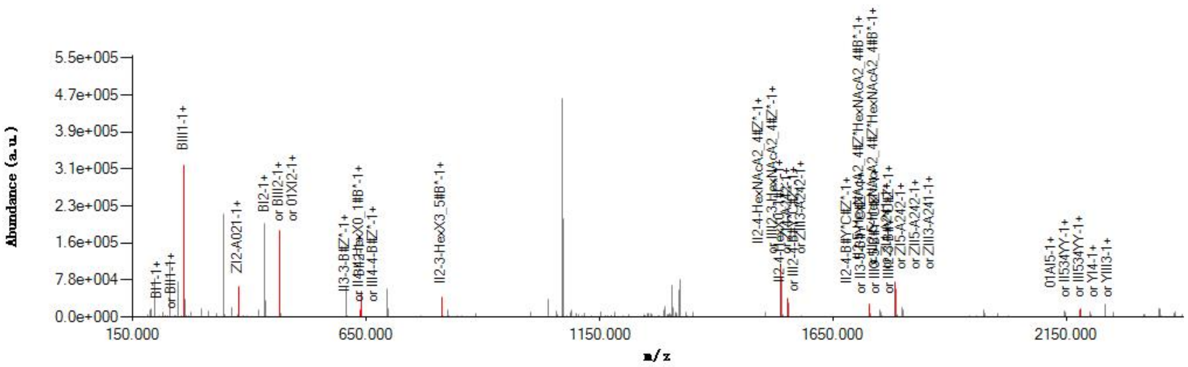


#### Graphical Fragmentation Map

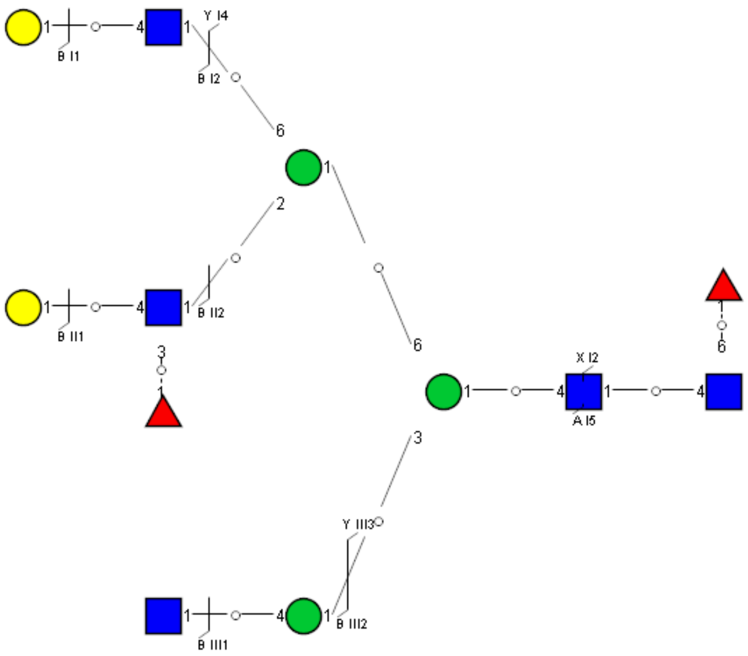

Spectrum Index: 34526(NoBHs:1)

#### Precursor iEF Map

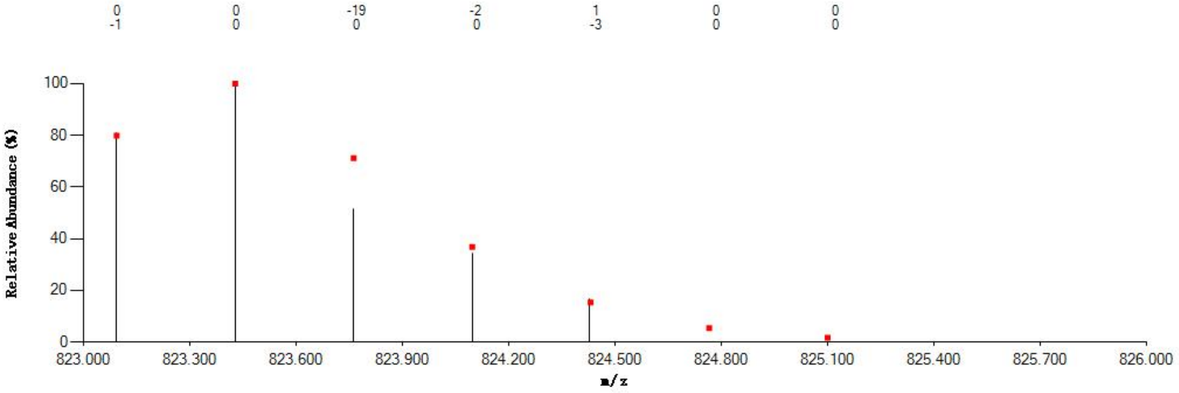


#### Annotated MS/MS Spectrum

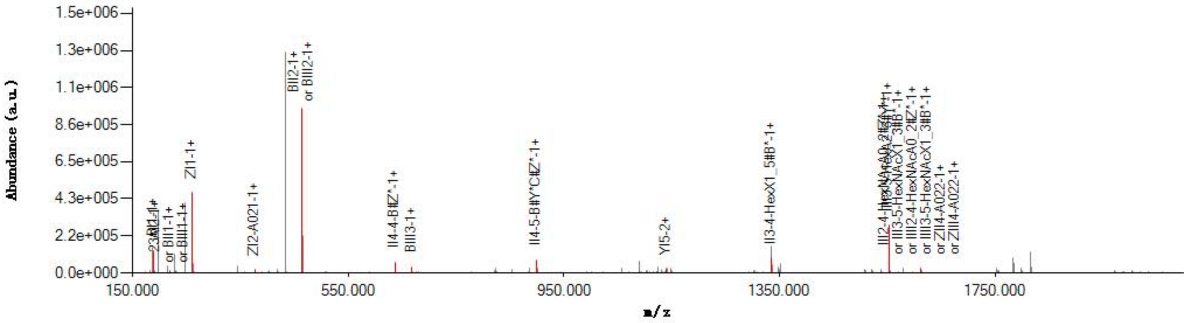


#### Graphical Fragmentation Map

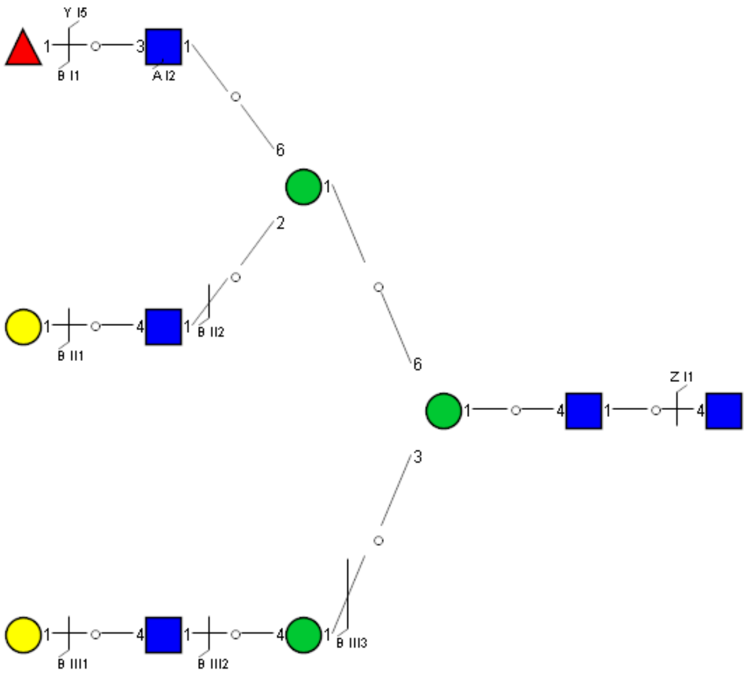

Spectrum Index: 34542(NoBHs:1)

#### Precursor iEF Map

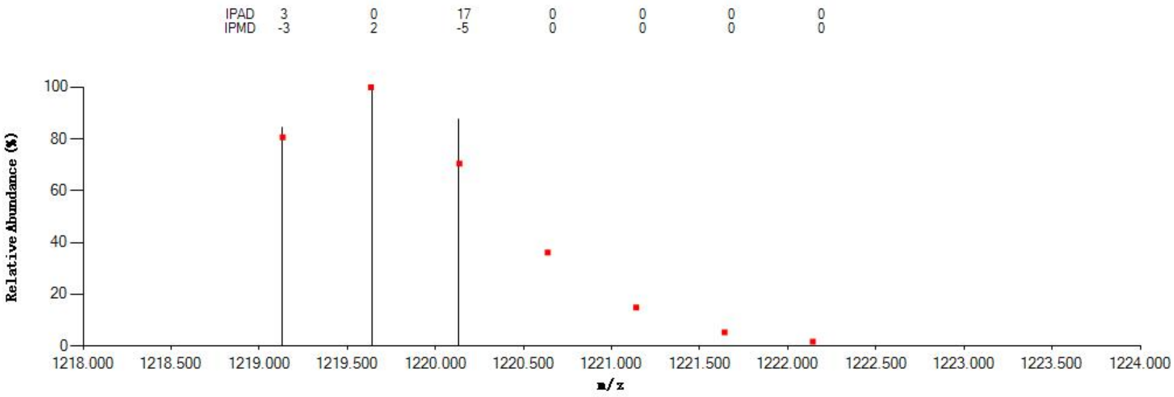


#### Annotated MS/MS Spectrum

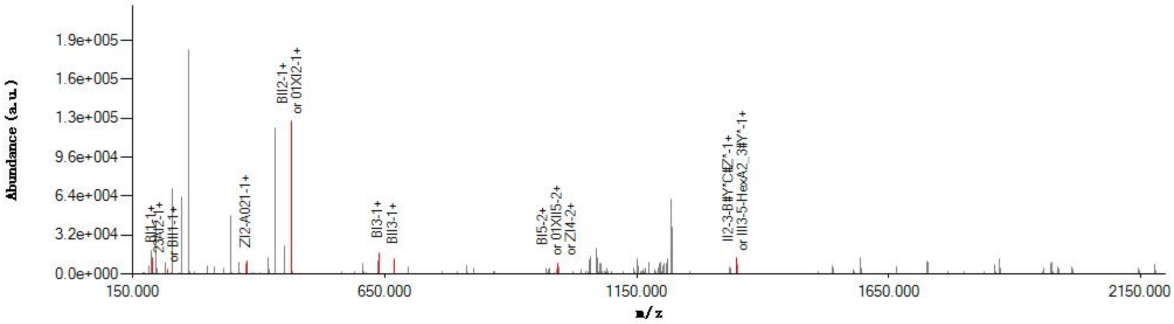


#### Graphical Fragmentation Map

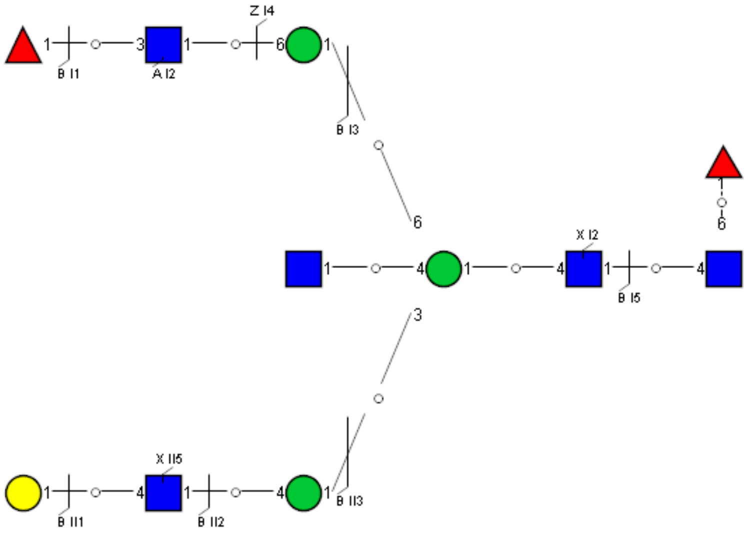

### Spectrum Index: 34737(NoBHs:1)

#### Precursor iEF Map

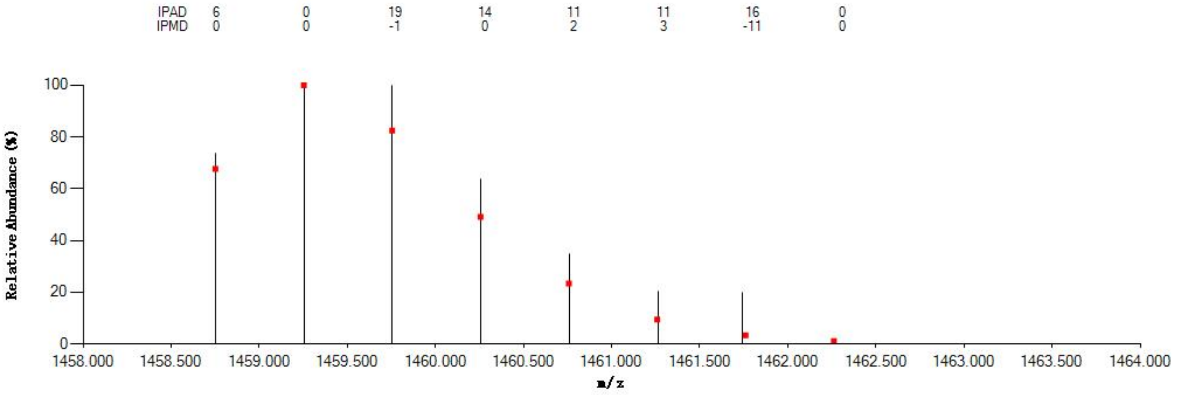


#### Annotated MS/MS Spectrum

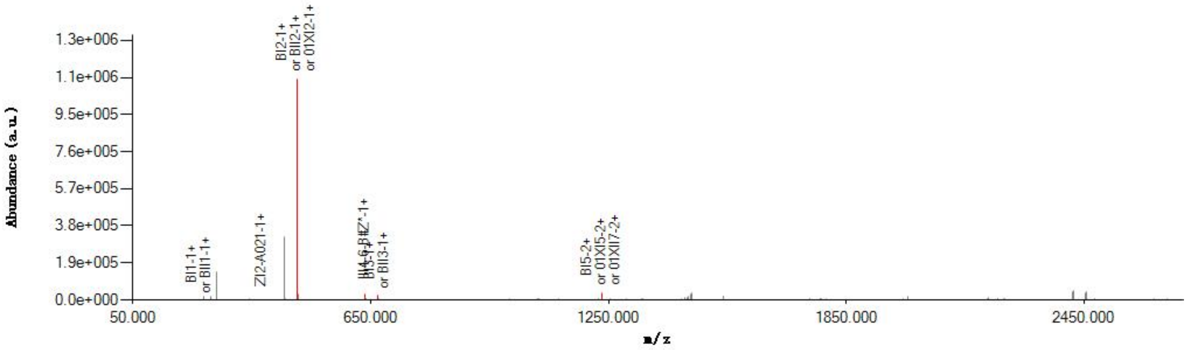


#### Graphical Fragmentation Map

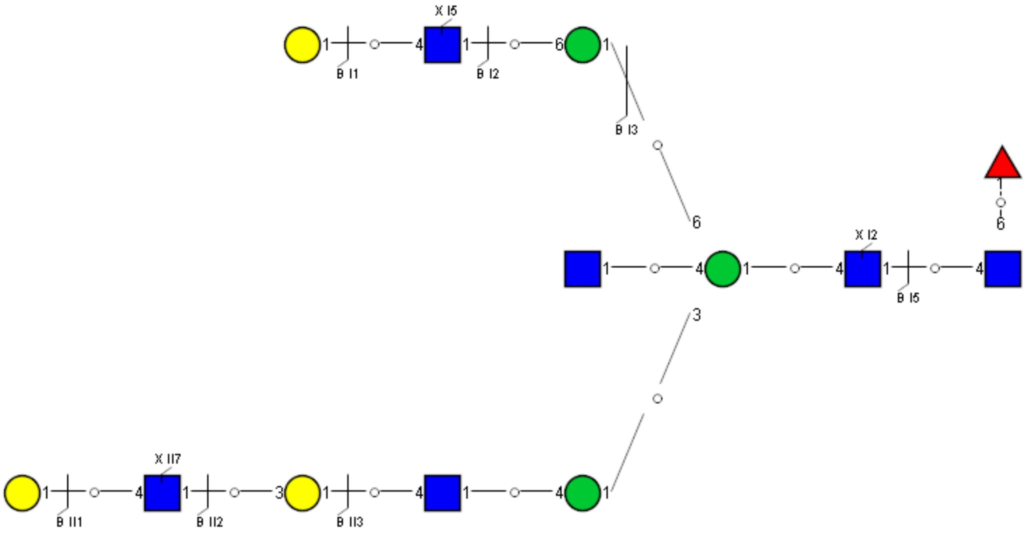

Spectrum Index: 34743(NoBHs:1)

#### Precursor iEF Map

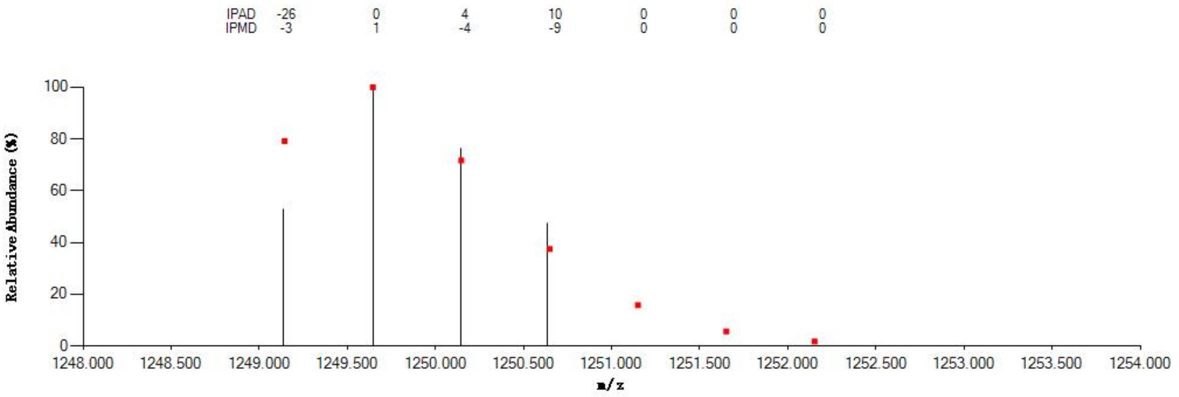

#### Annotated MS/MS Spectrum

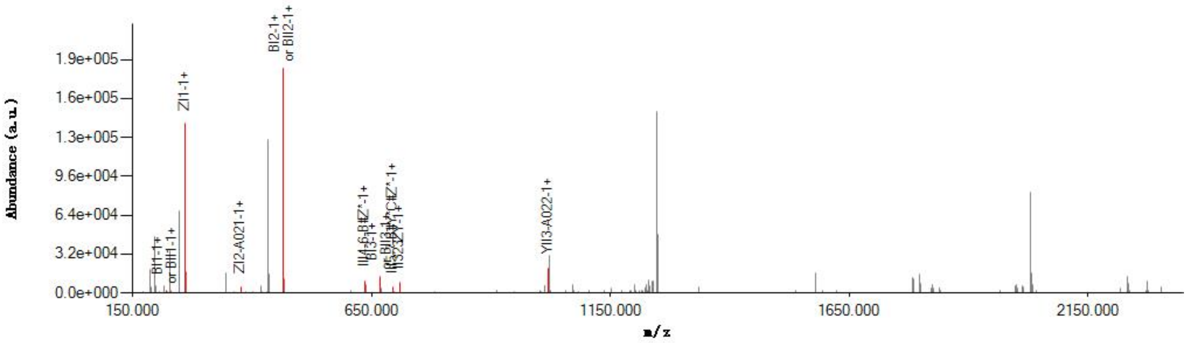

#### Graphical Fragmentation Map

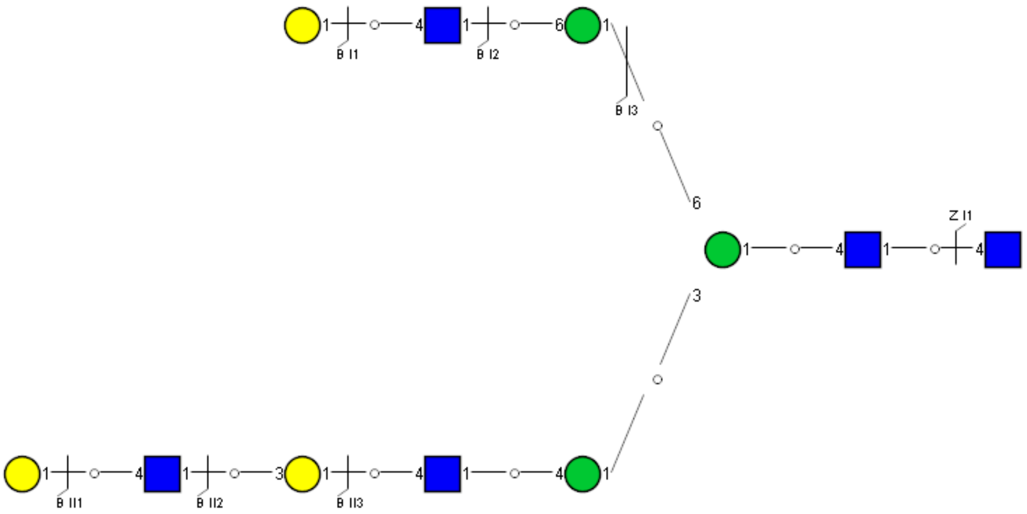

Spectrum Index: 34887(NoBHs:1)

#### Precursor iEF Map

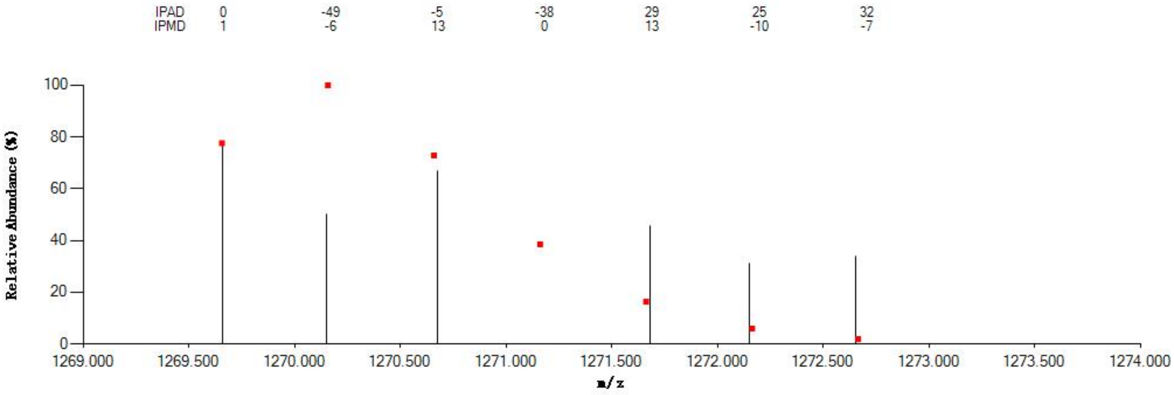

#### Annotated MS/MS Spectrum

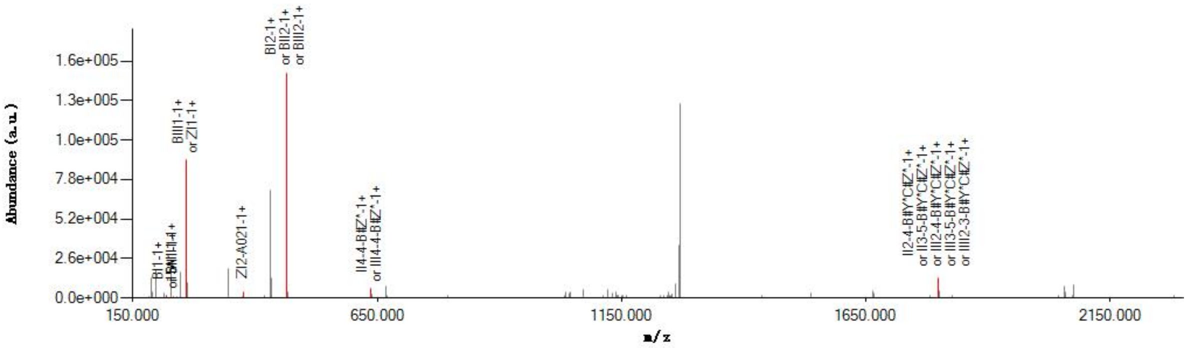

#### Graphical Fragmentation Map

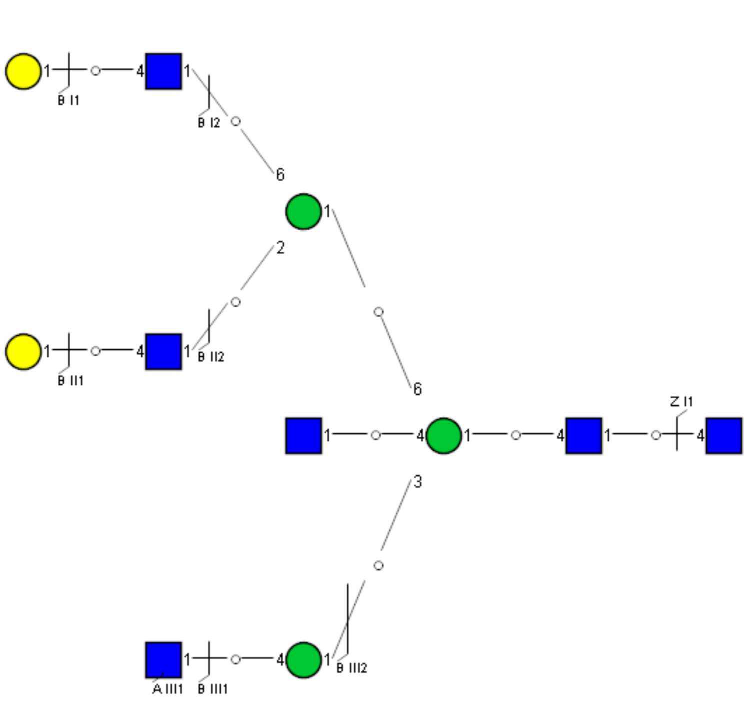

Spectrum Index: 34925(NoBHs:1)

#### Precursor iEF Map

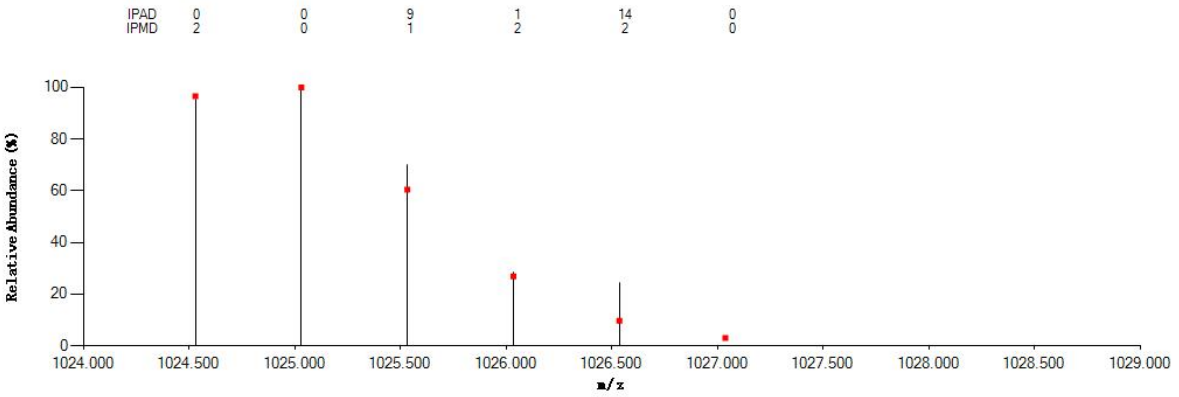


#### Annotated MS/MS Spectrum

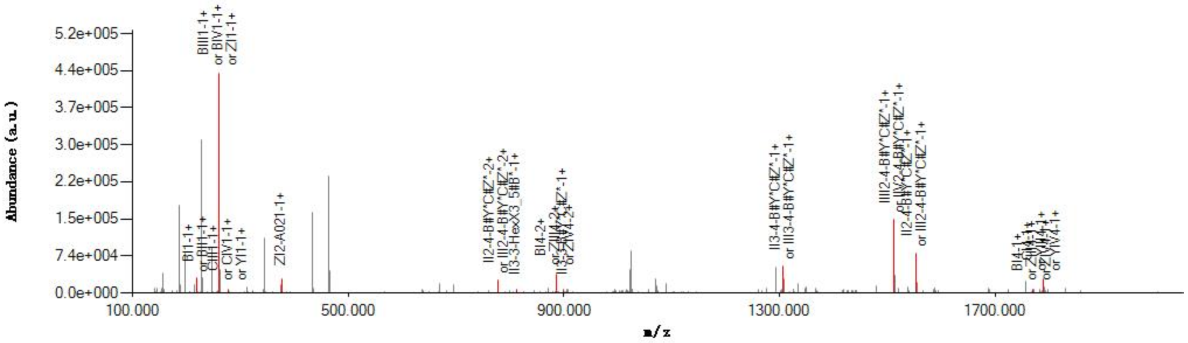


#### Graphical Fragmentation Map

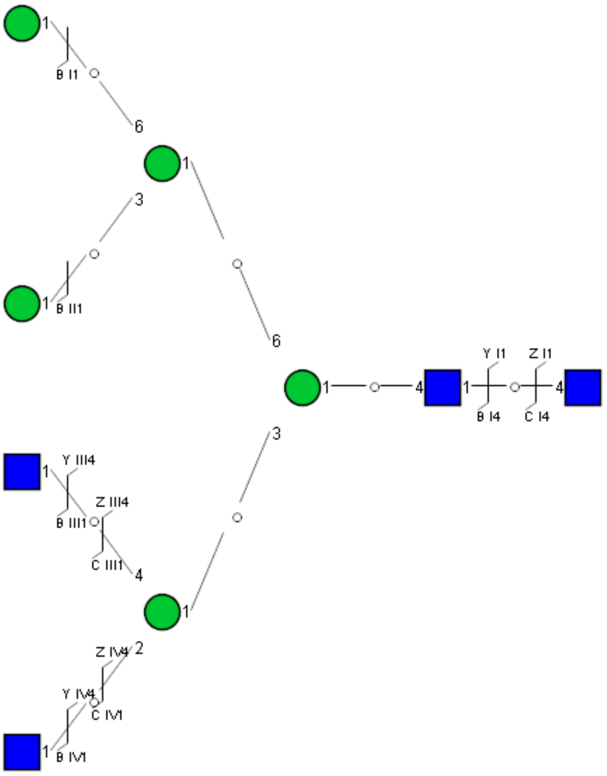

Spectrum Index: 35013(NoBHs:1)

#### Precursor iEF Map

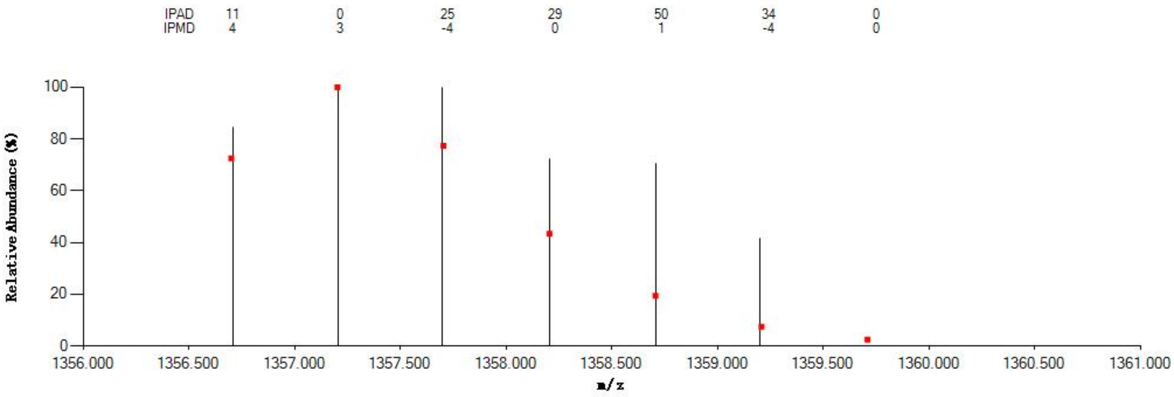

#### Annotated MS/MS Spectrum

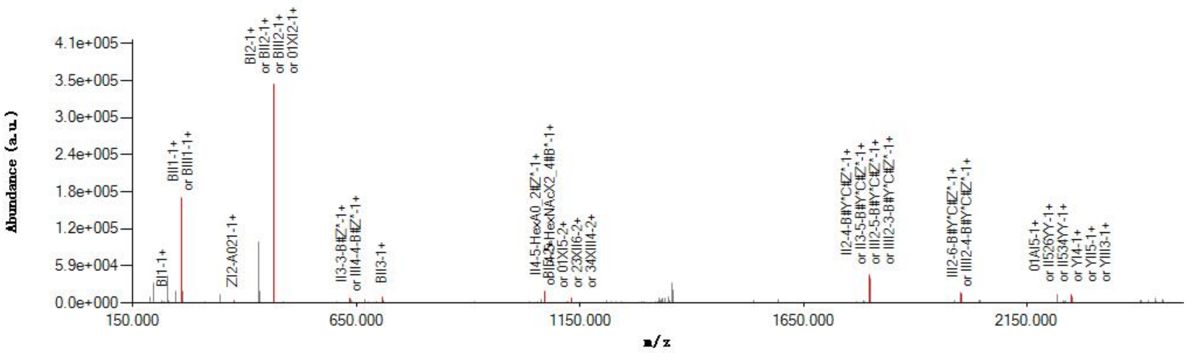

#### Graphical Fragmentation Map

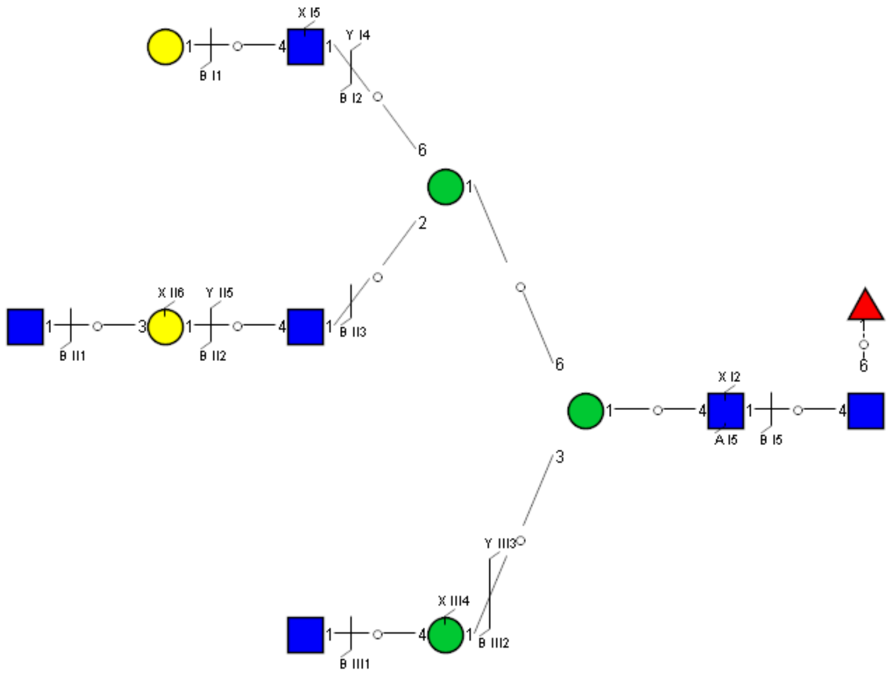

Spectrum Index: 35067(NoBHs:1)

#### Precursor iEF Map

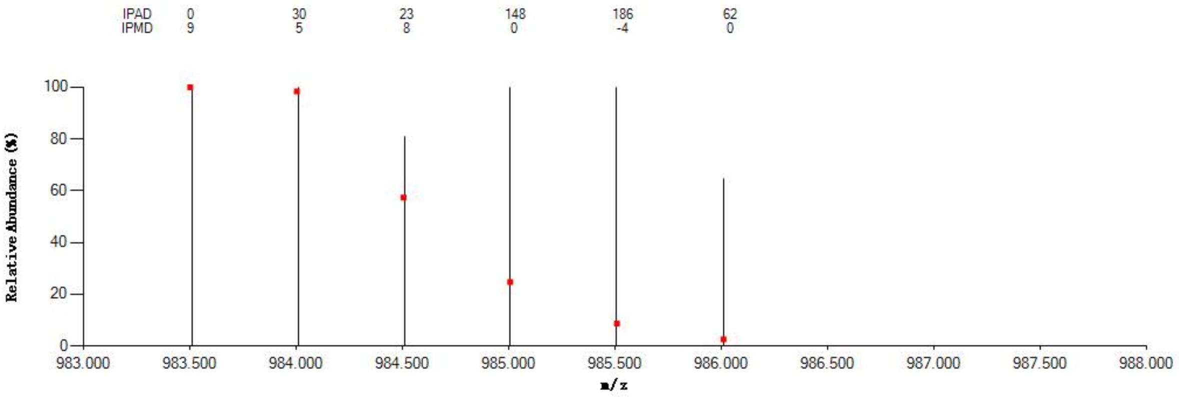


#### Annotated MS/MS Spectrum

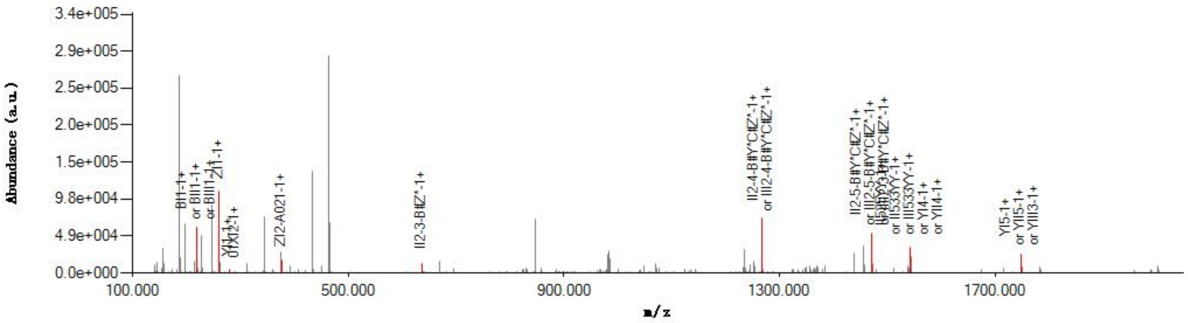


#### Graphical Fragmentation Map

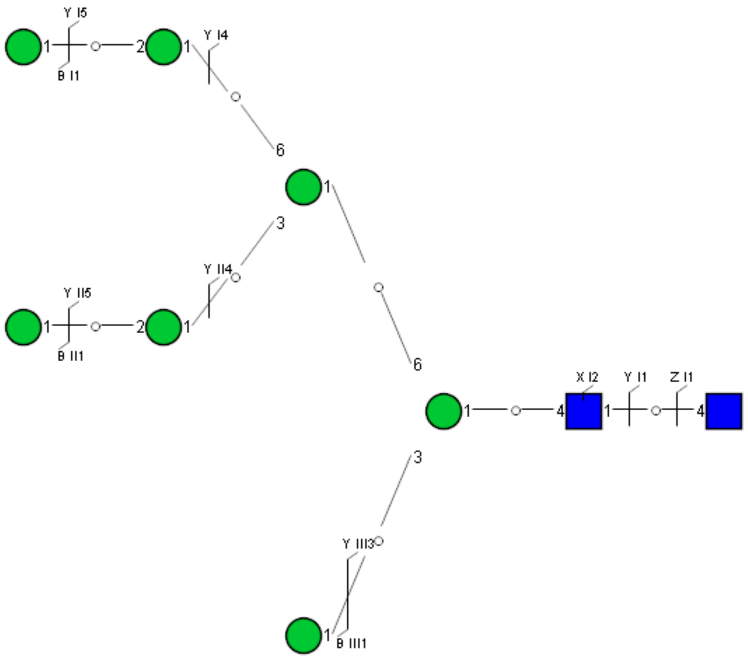

Spectrum Index: 35082(NoBHs:1)

#### Precursor iEF Map

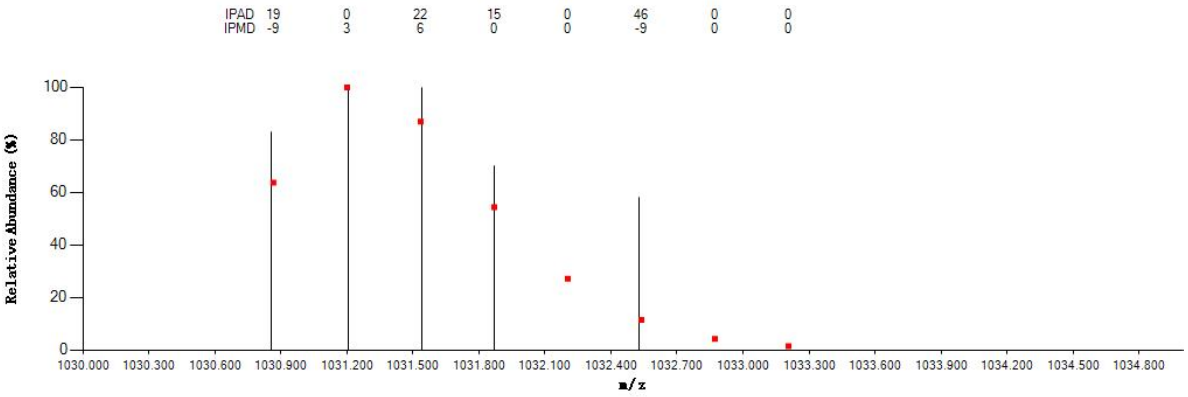


#### Annotated MS/MS Spectrum

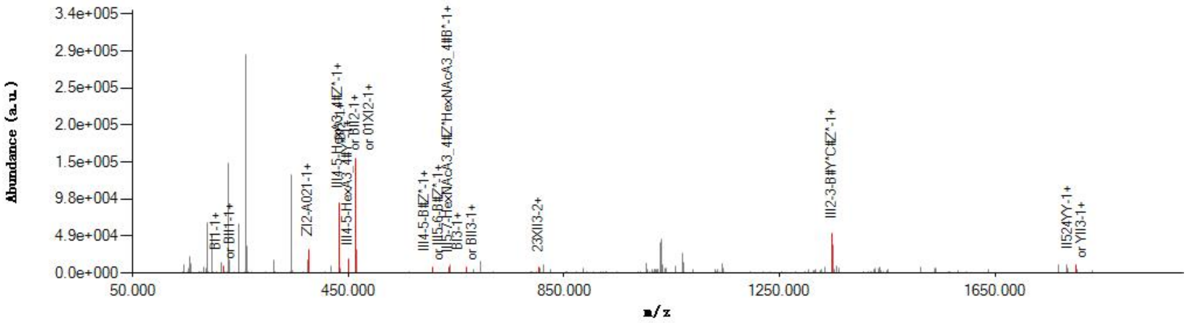


#### Graphical Fragmentation Map

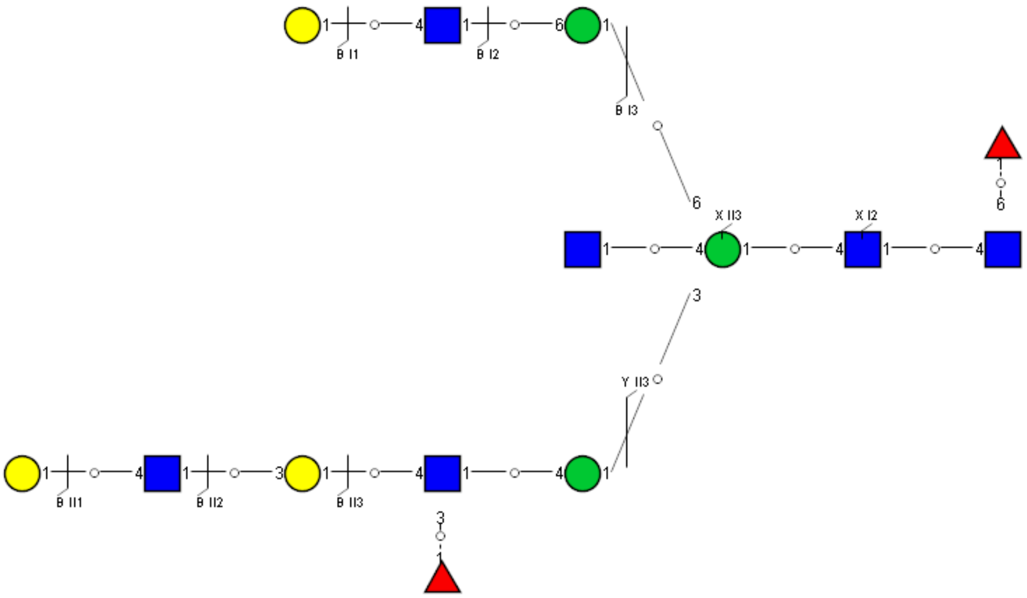

Spectrum Index: 35114(NoBHs:1)

#### Precursor iEF Map

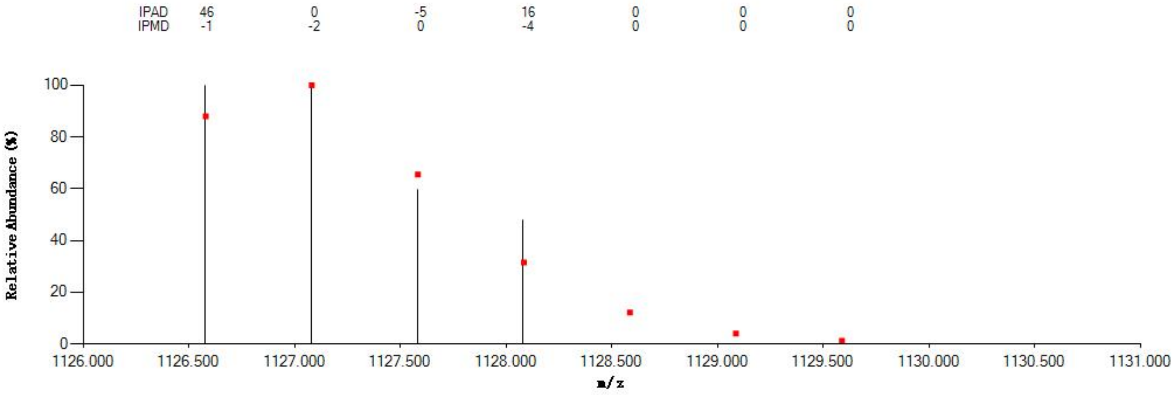

#### Annotated MS/MS Spectrum

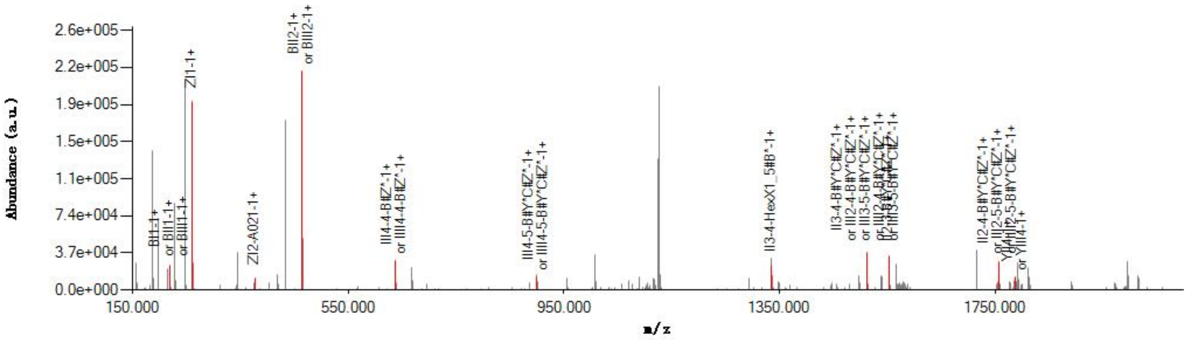

#### Graphical Fragmentation Map

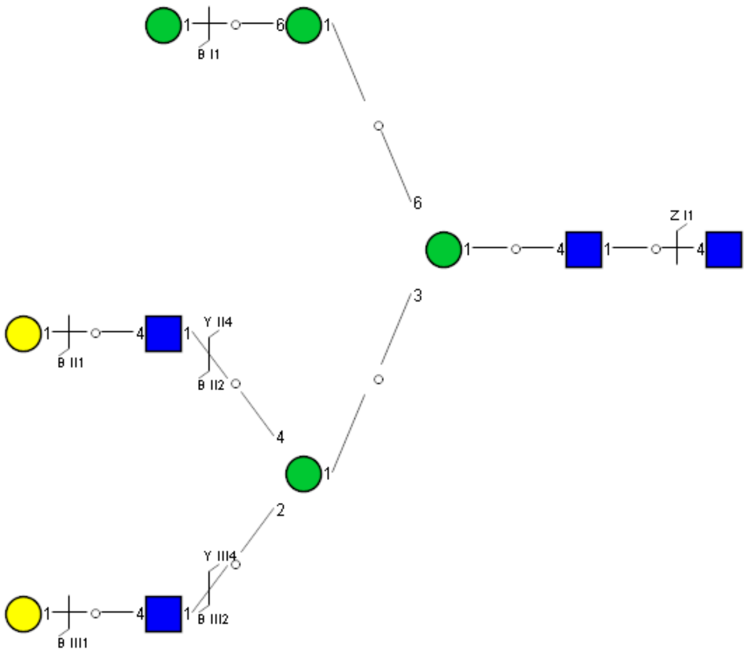

### Spectrum Index: 35371(NoBHs:2)

#### Precursor iEF Map

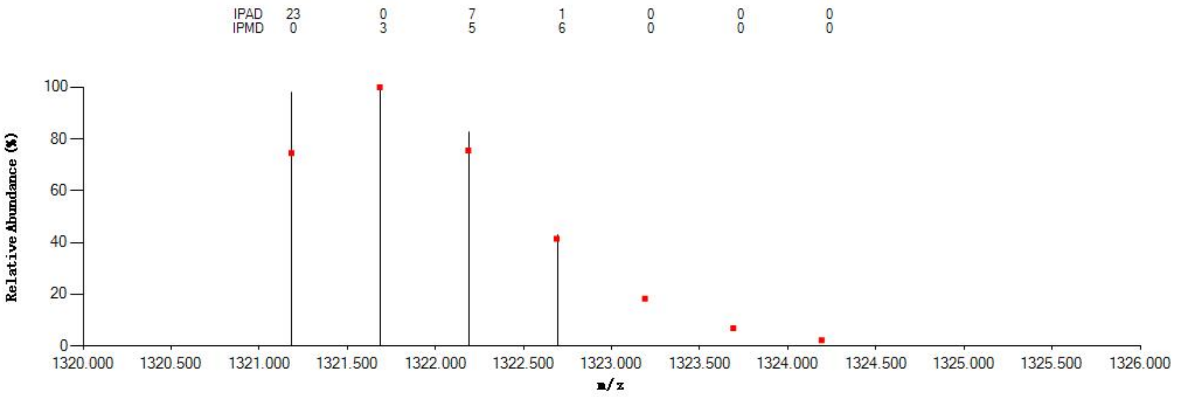

#### Annotated MS/MS Spectrum

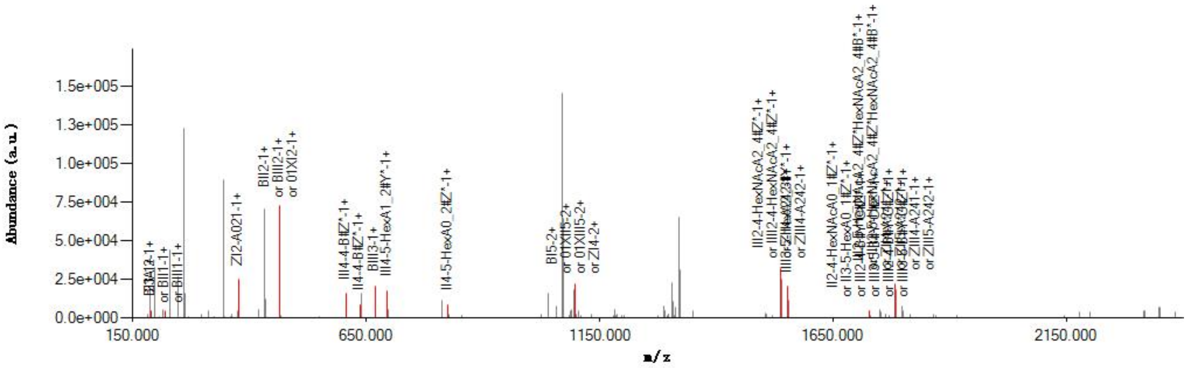

#### Graphical Fragmentation Map

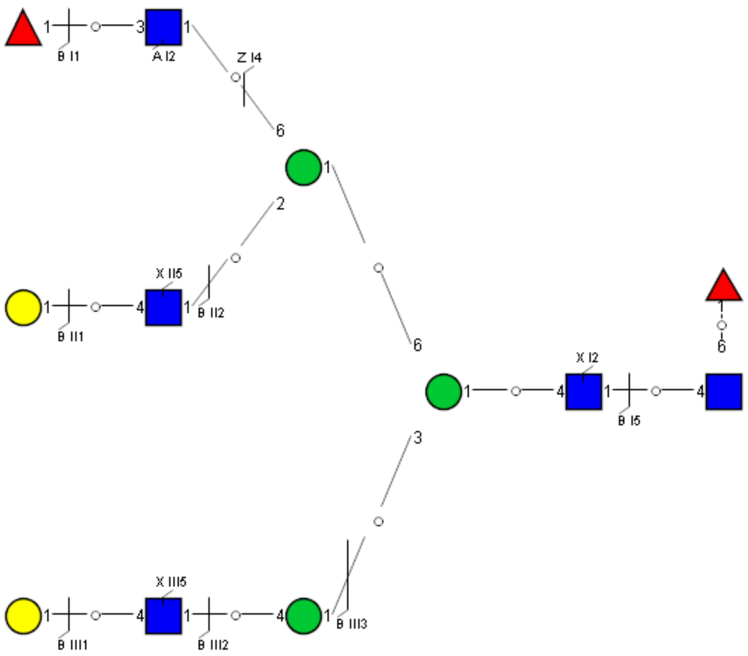

Spectrum Index: 35371(NoBHs:2)

#### Precursor iEF Map

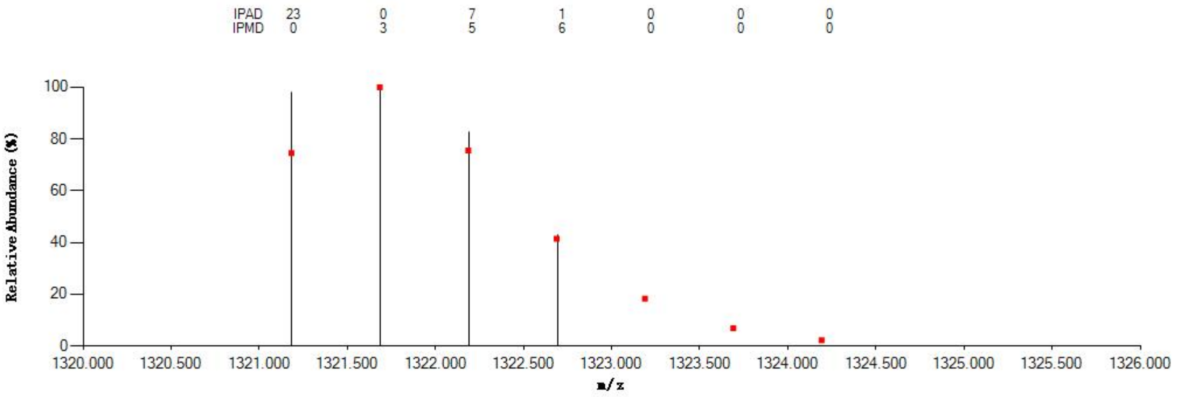

#### Annotated MS/MS Spectrum

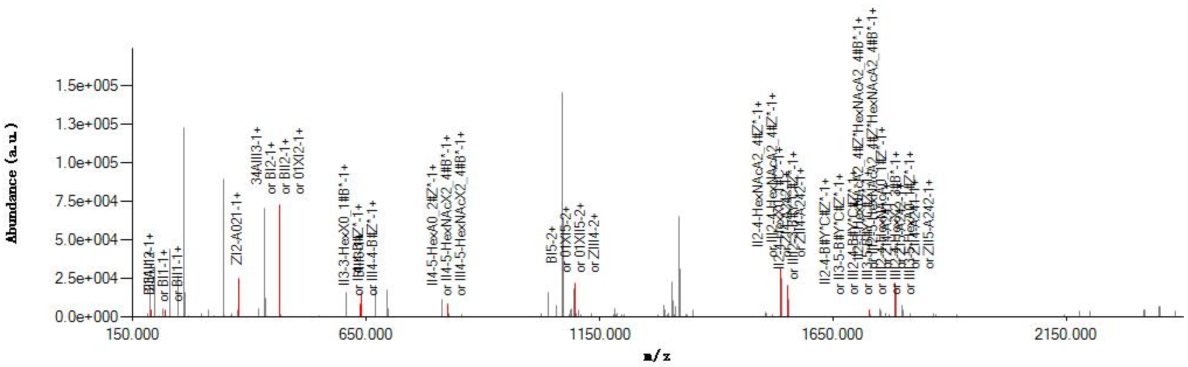

#### Graphical Fragmentation Map

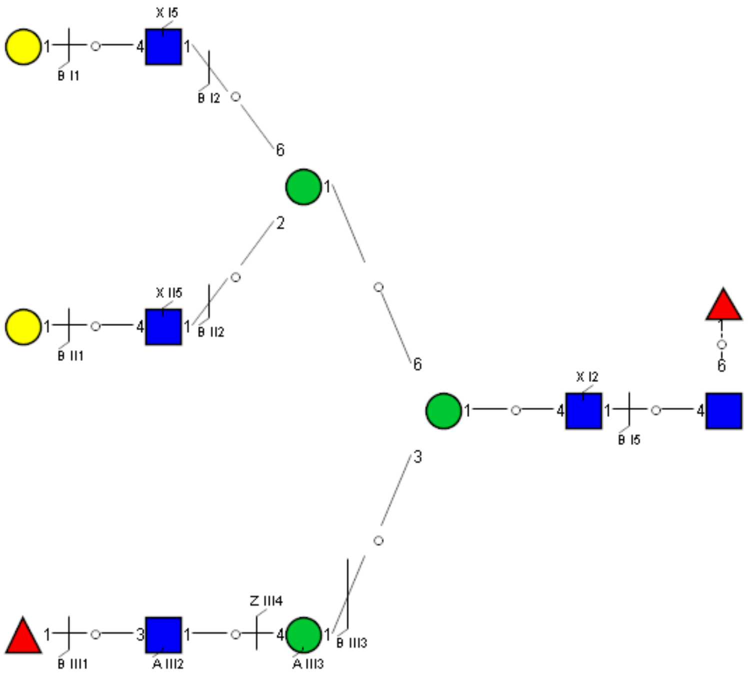

Spectrum Index: 35392(NoBHs:1)

#### Precursor iEF Map

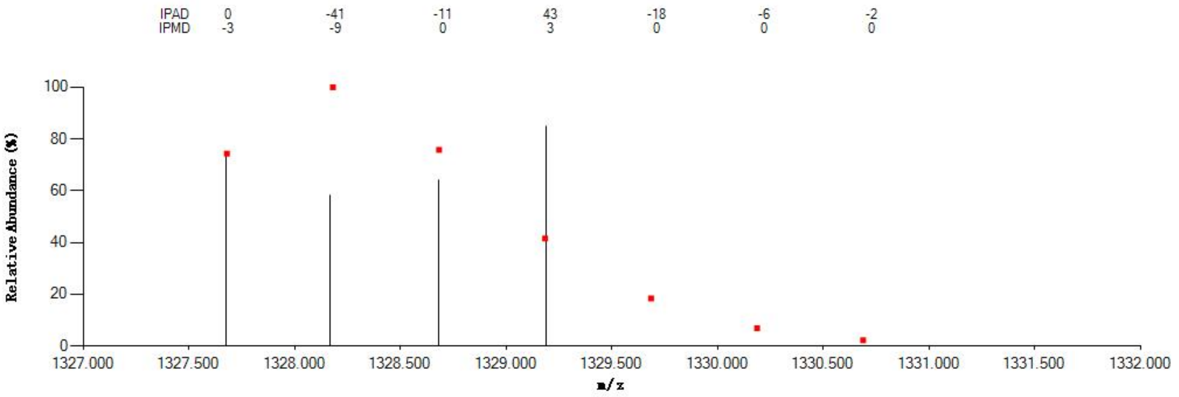


#### Annotated MS/MS Spectrum

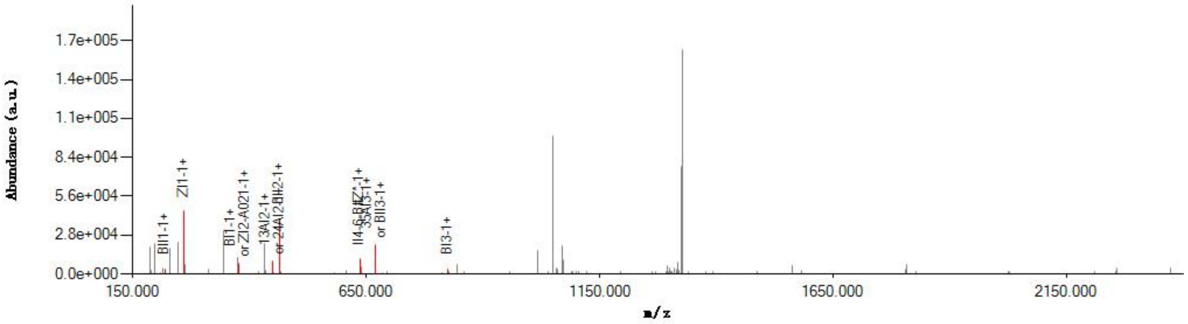


#### Graphical Fragmentation Map

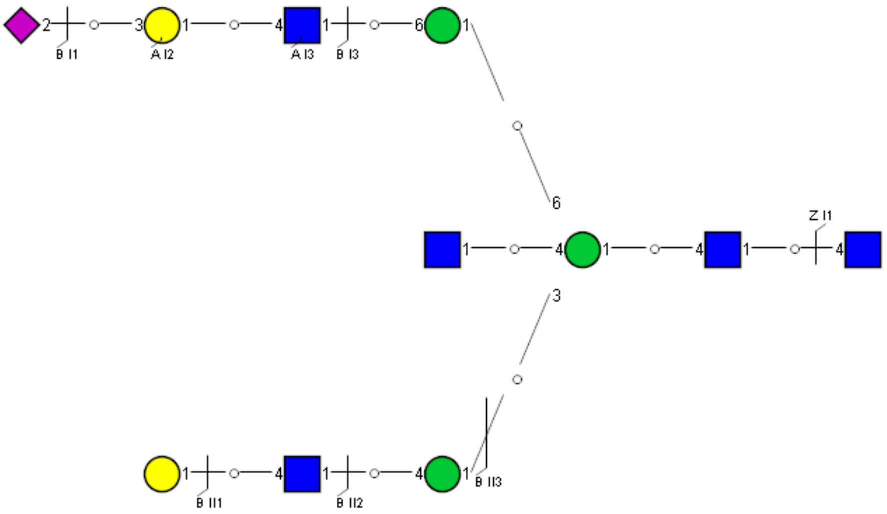

Spectrum Index: 35482(NoBHs:1)

#### Precursor iEF Map

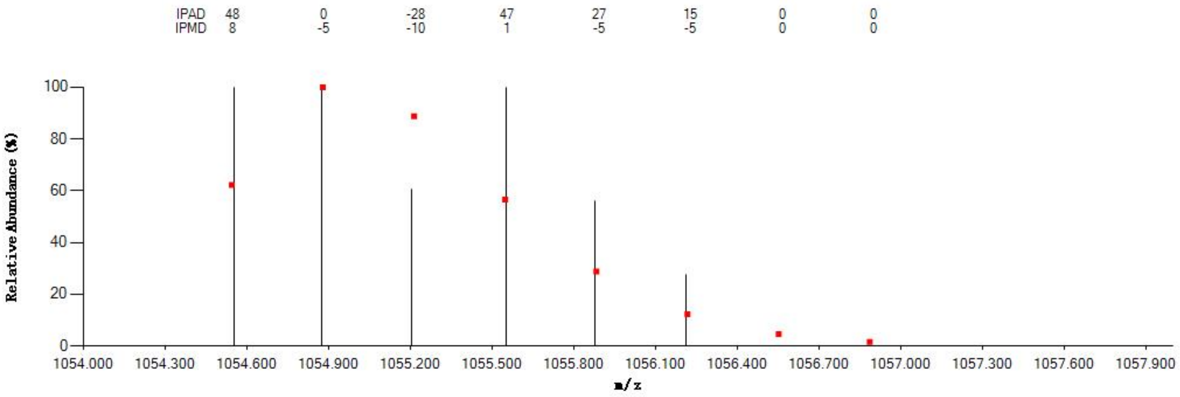


#### Annotated MS/MS Spectrum

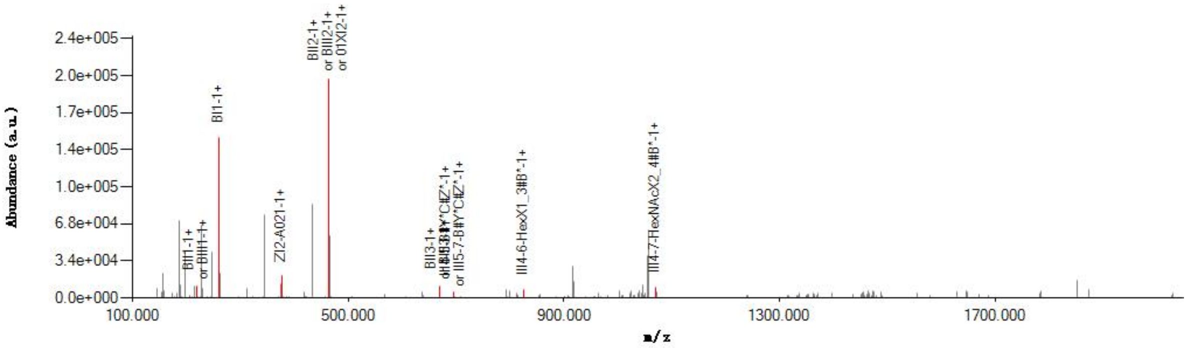


#### Graphical Fragmentation Map

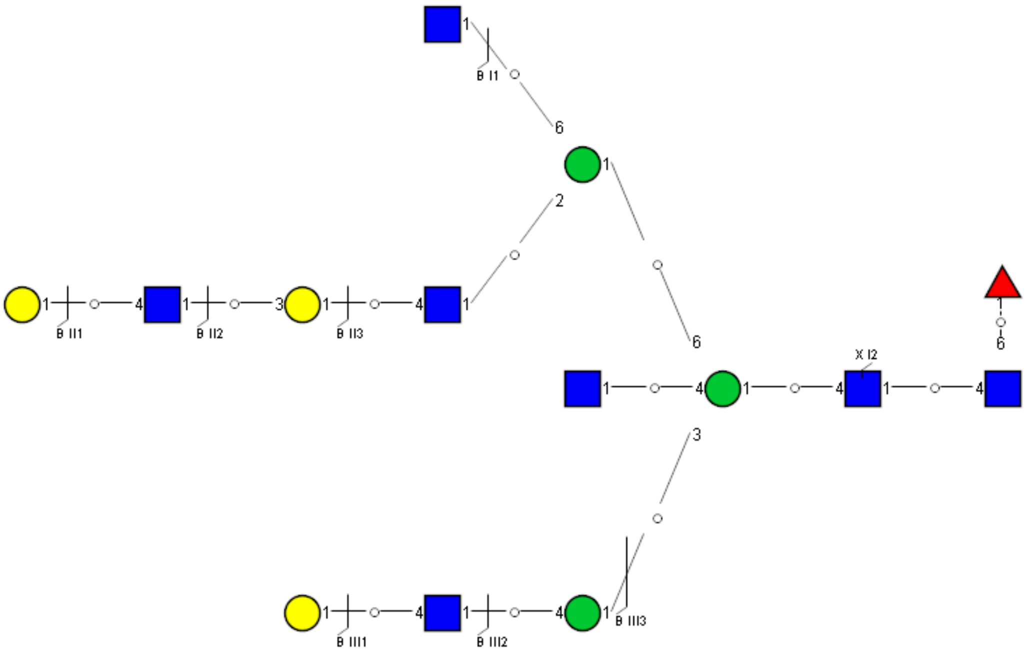

Spectrum Index: 35524(NoBHs:1)

#### Precursor iEF Map

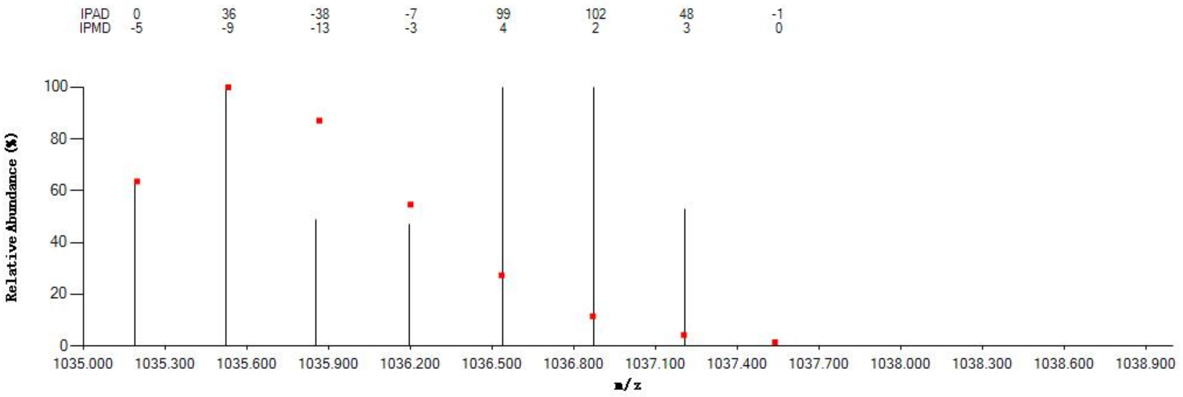

#### Annotated MS/MS Spectrum

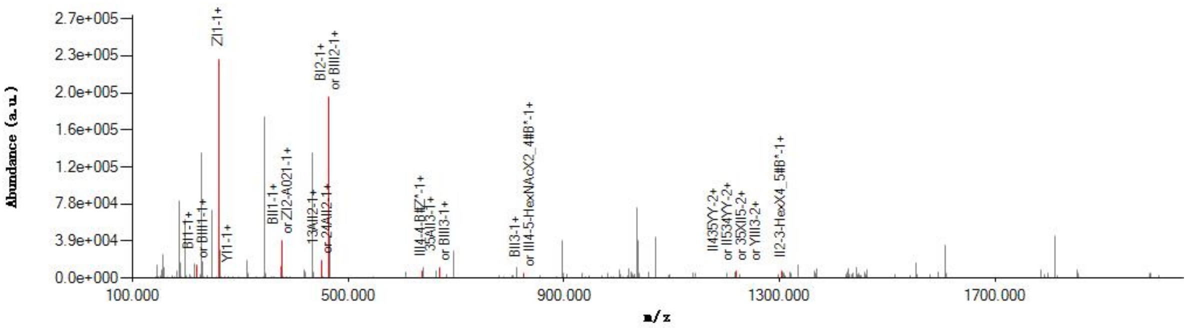

#### Graphical Fragmentation Map

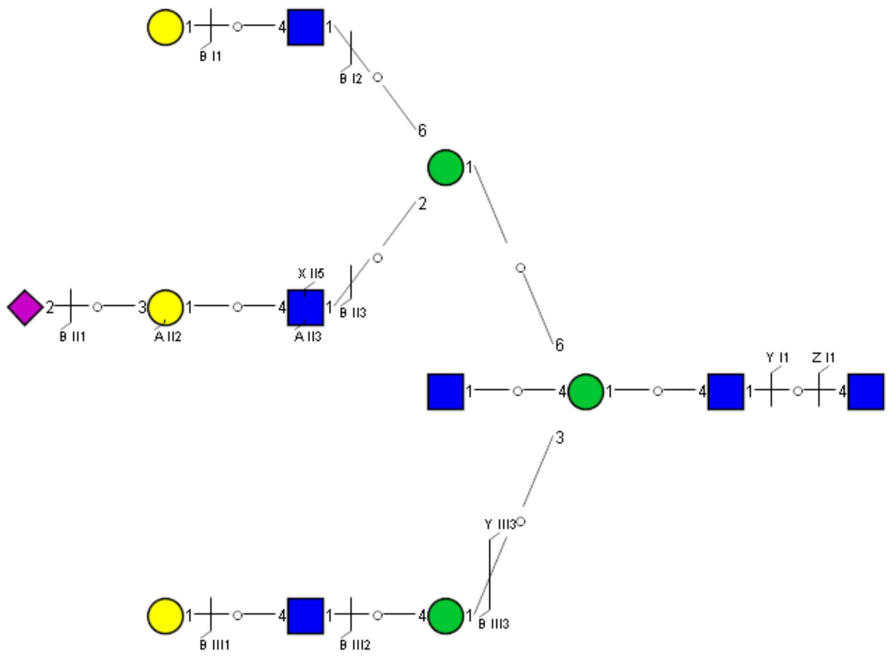

Spectrum Index: 35659(NoBHs:1)

#### Precursor iEF Map

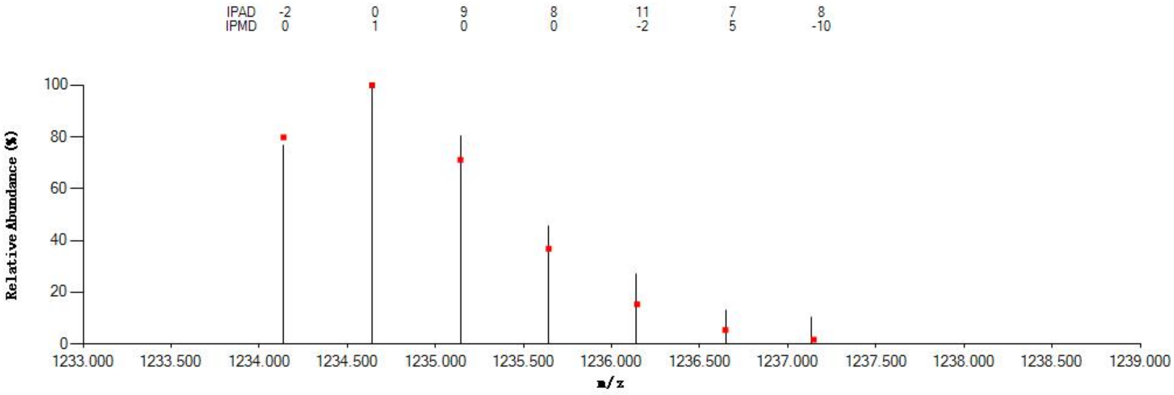

#### Annotated MS/MS Spectrum

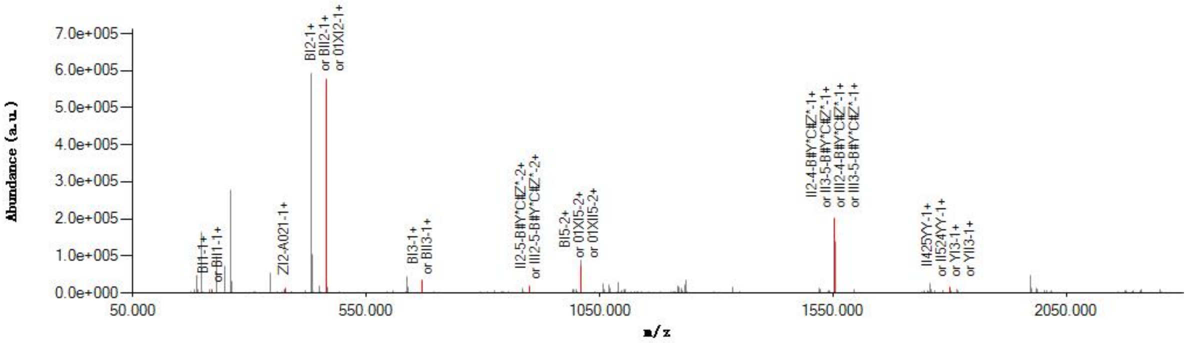

#### Graphical Fragmentation Map

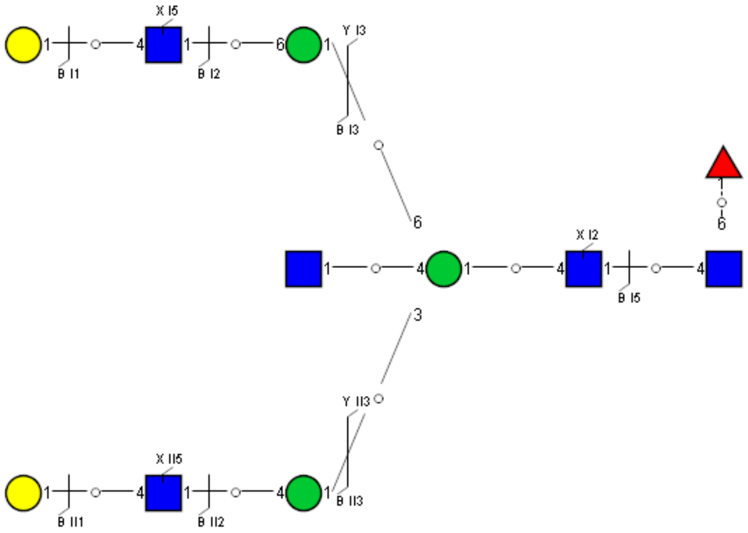

Spectrum Index: 35677(NoBHs:1)

#### Precursor iEF Map

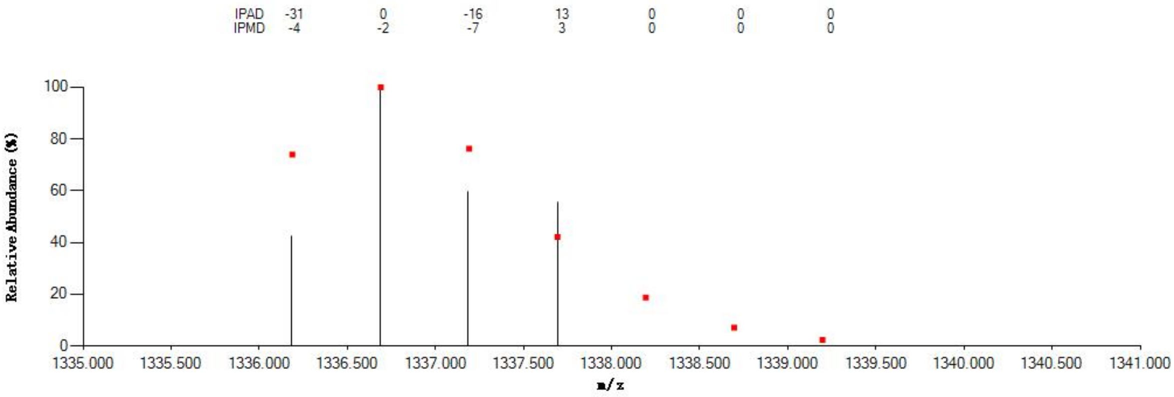


#### Annotated MS/MS Spectrum

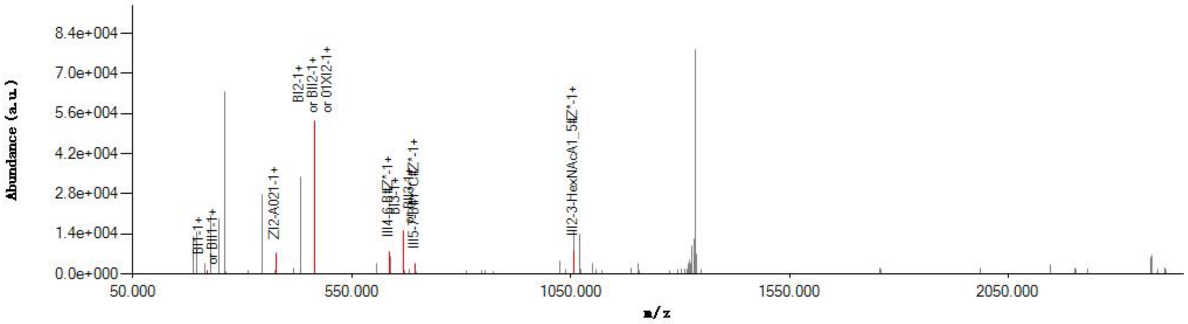


#### Graphical Fragmentation Map

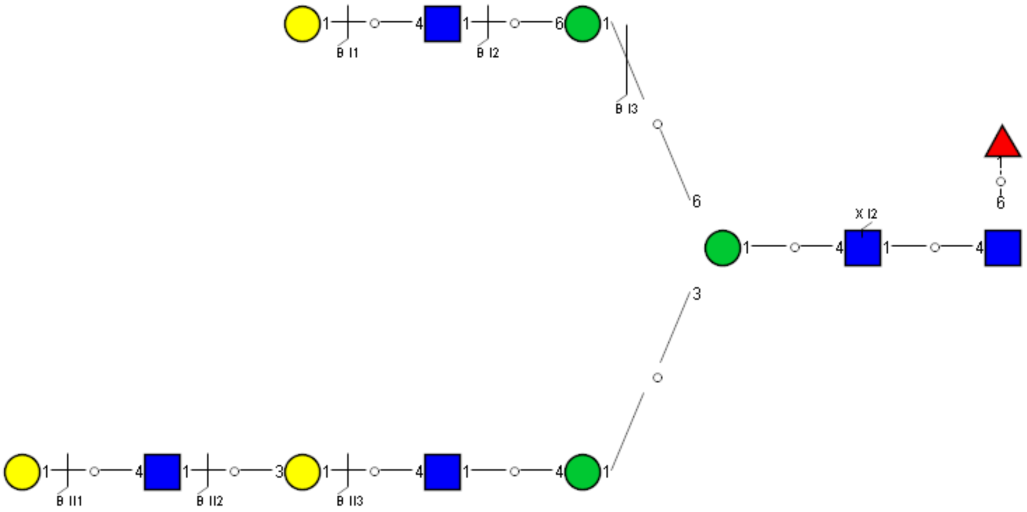

### Spectrum Index: 35964(NoBHs:1)

#### Precursor iEF Map

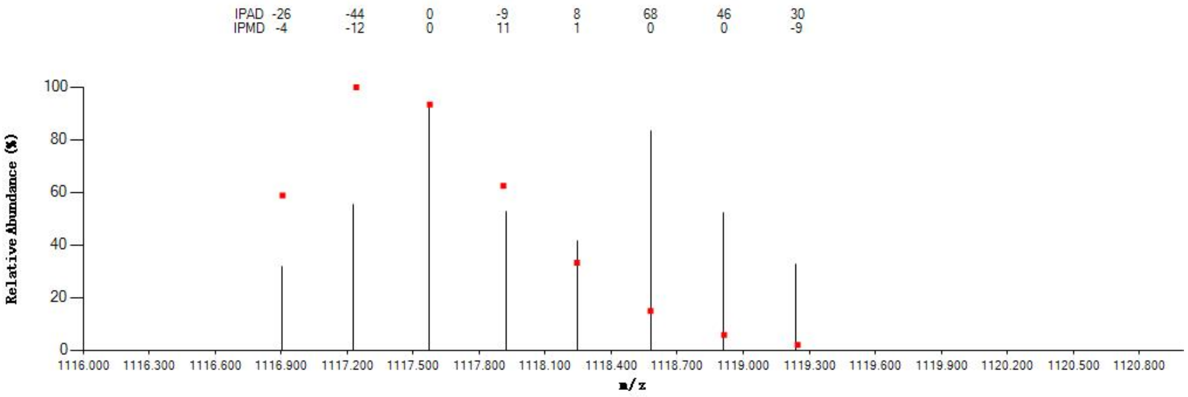

#### Annotated MS/MS Spectrum

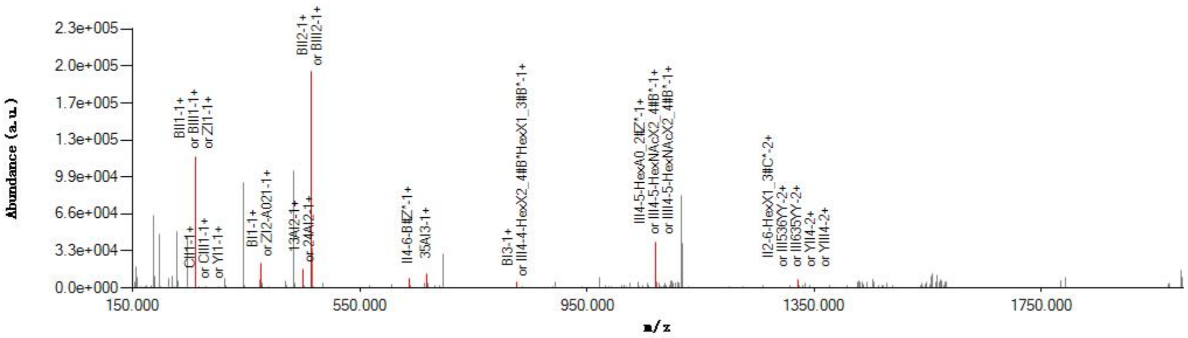

#### Graphical Fragmentation Map

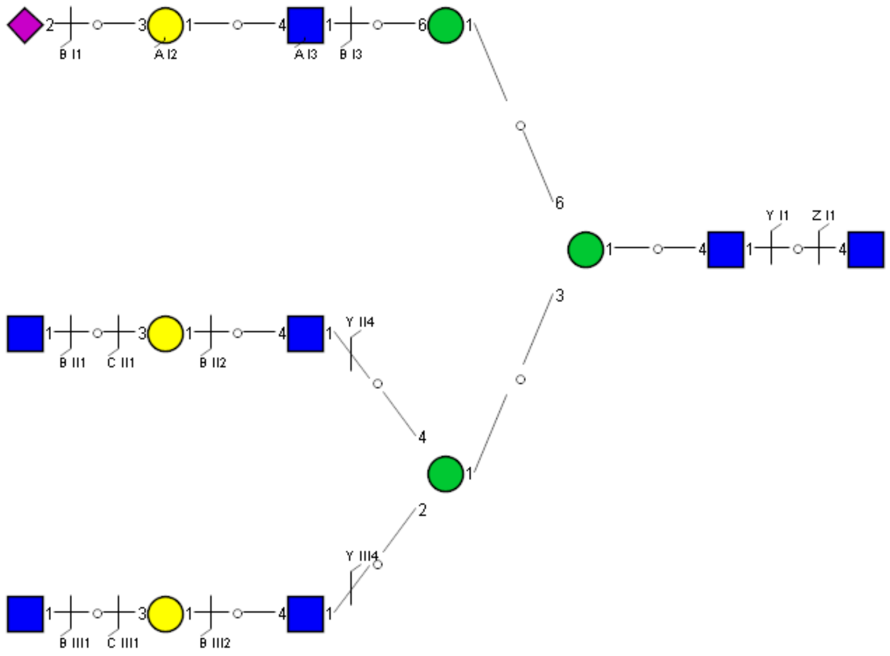

Spectrum Index: 35989(NoBHs:1)

#### Precursor iEF Map

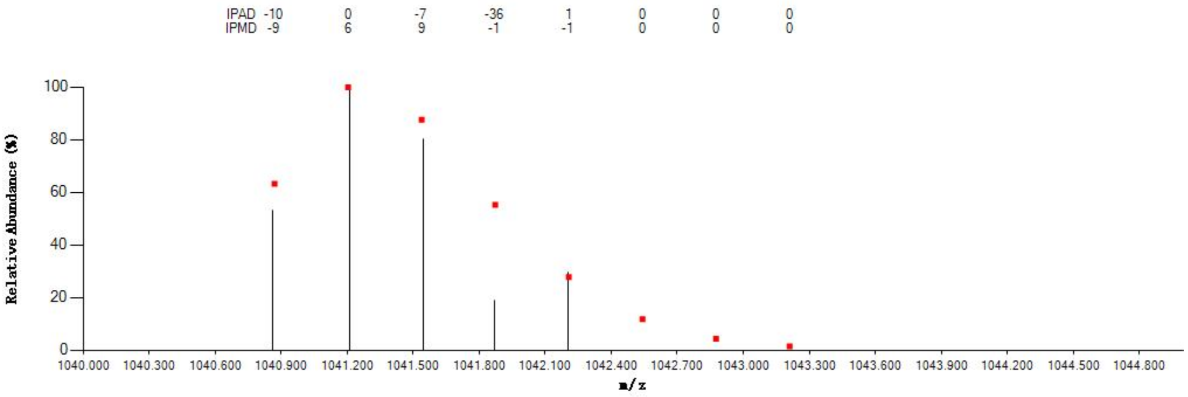


#### Annotated MS/MS Spectrum

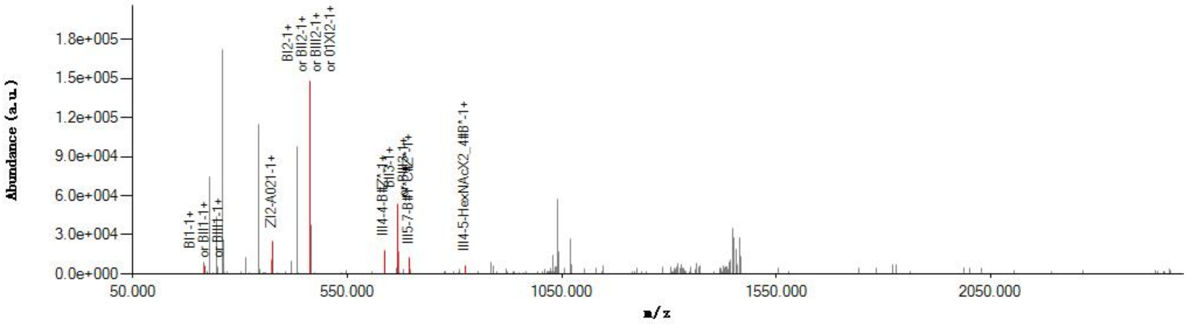


#### Graphical Fragmentation Map

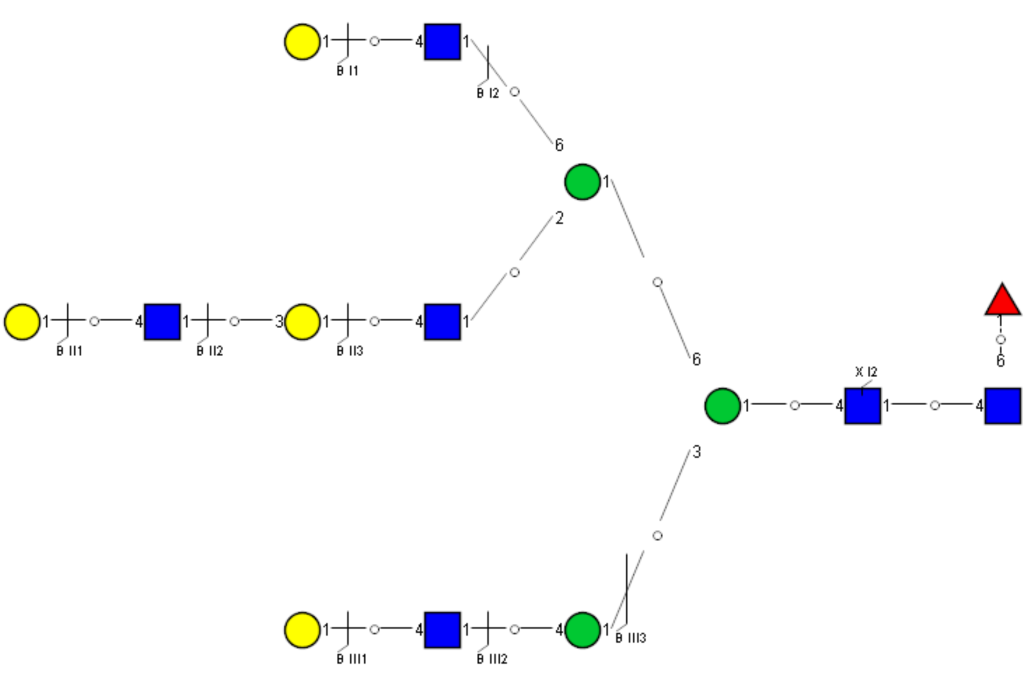

Spectrum Index: 36035(NoBHs:1)

#### Precursor iEF Map

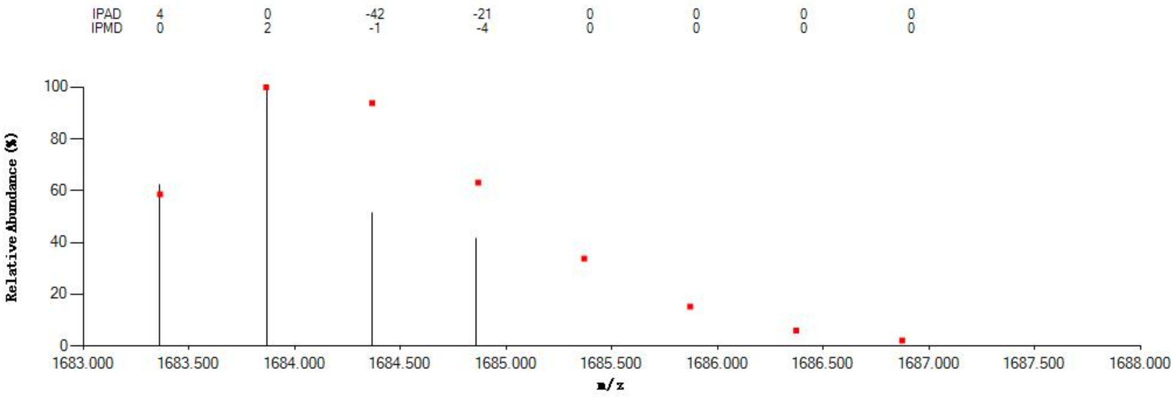


#### Annotated MS/MS Spectrum

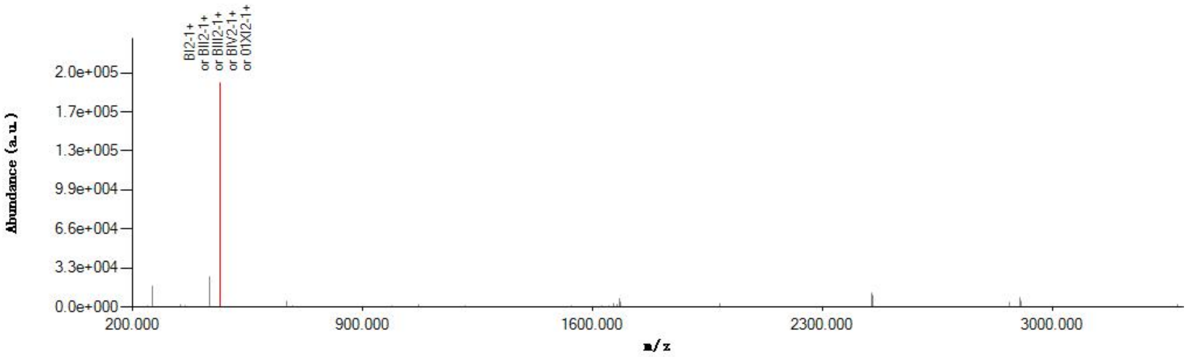


#### Graphical Fragmentation Map

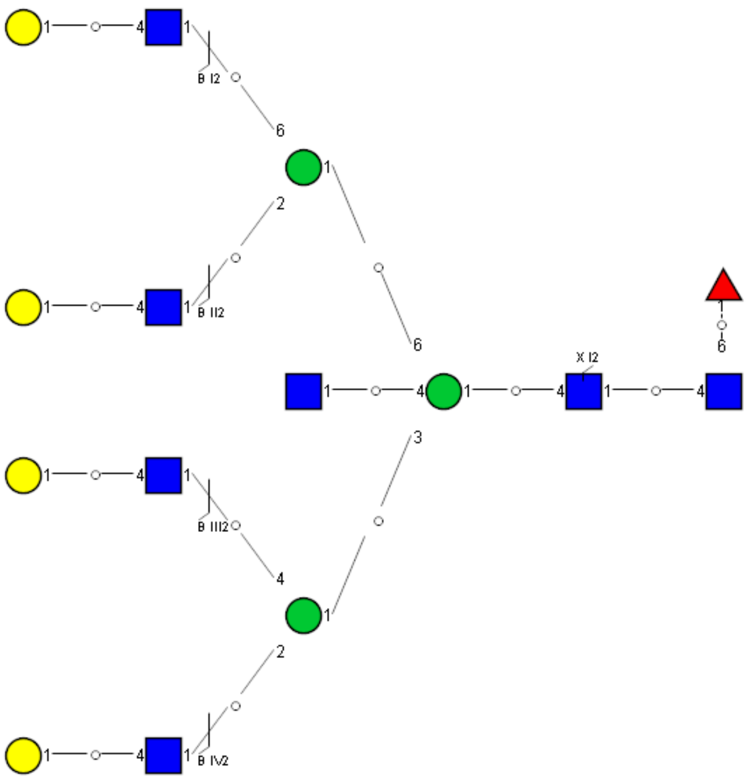

Spectrum Index: 36037(NoBHs:1)

#### Precursor iEF Map

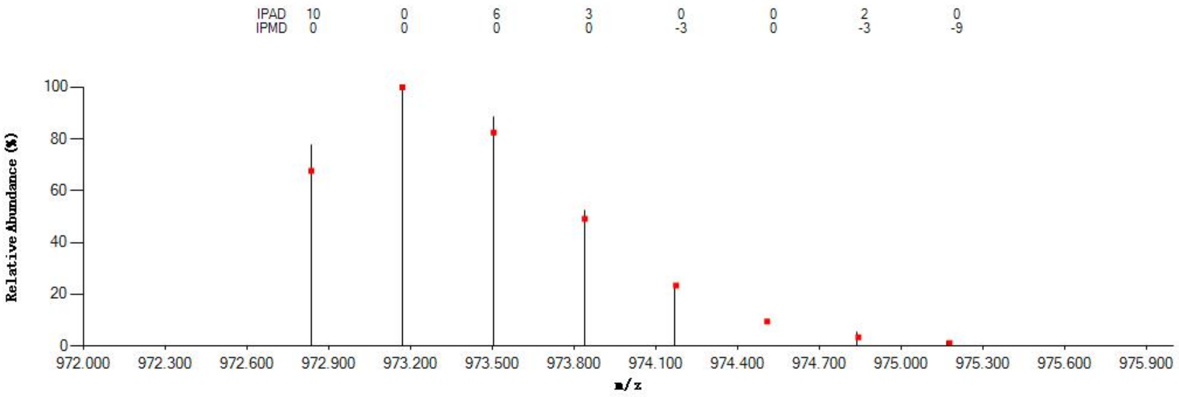


#### Annotated MS/MS Spectrum

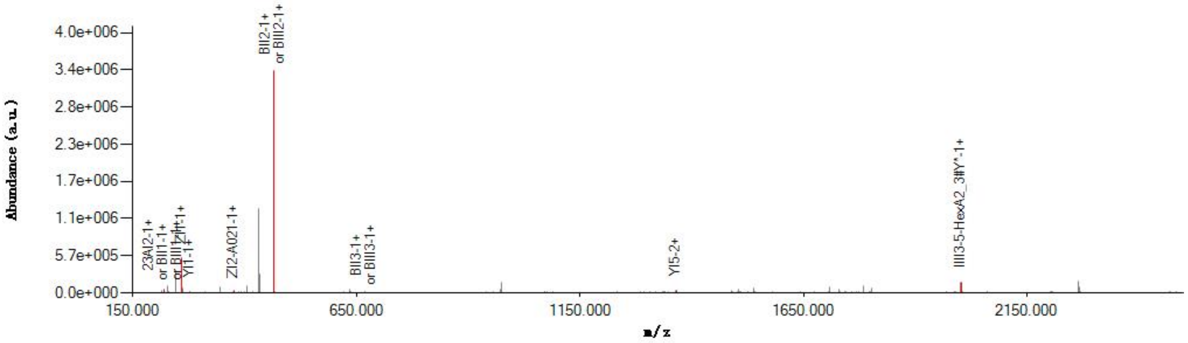


#### Graphical Fragmentation Map

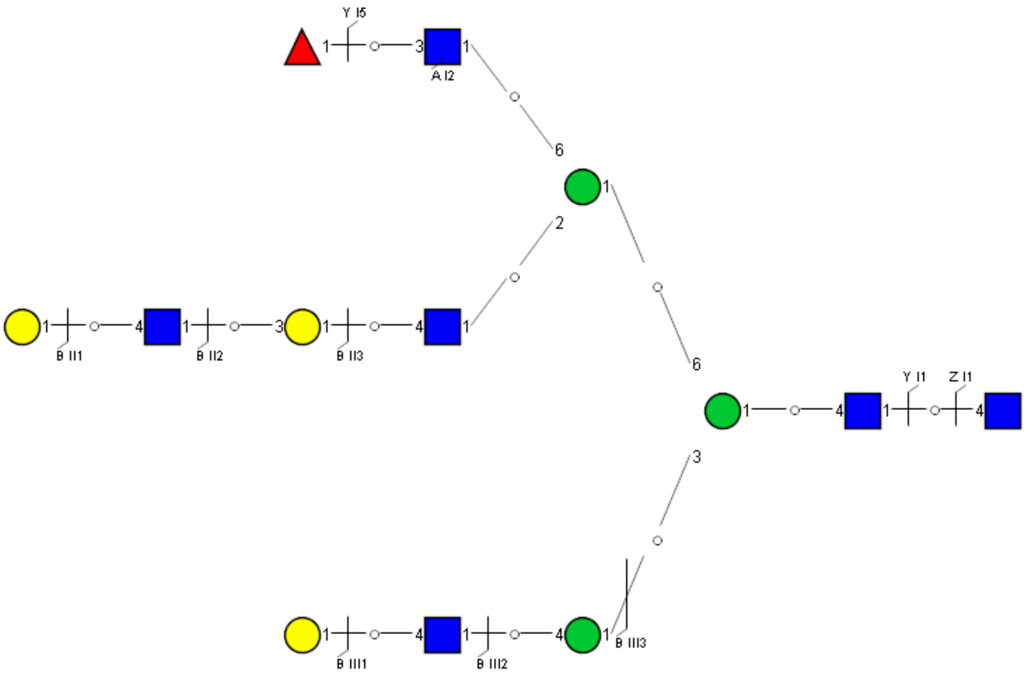

Spectrum Index: 36083(NoBHs:1)

#### Precursor iEF Map

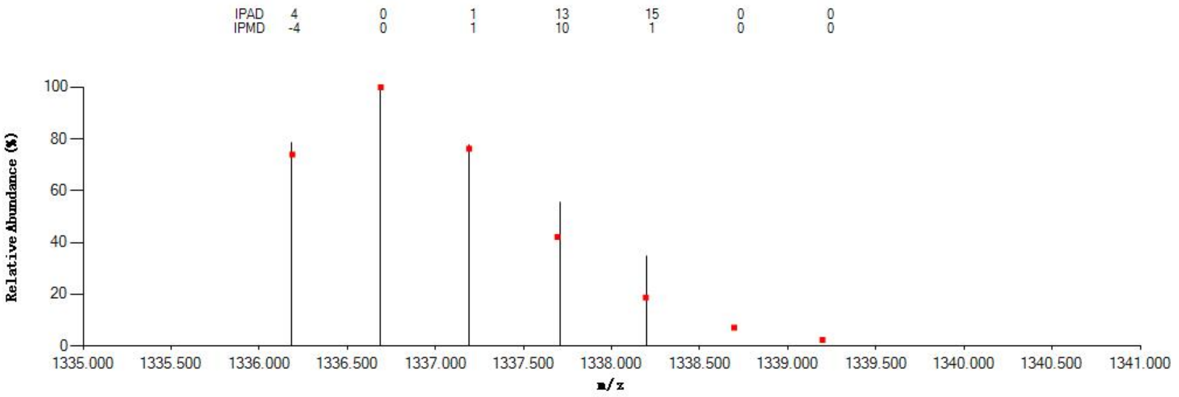

#### Annotated MS/MS Spectrum

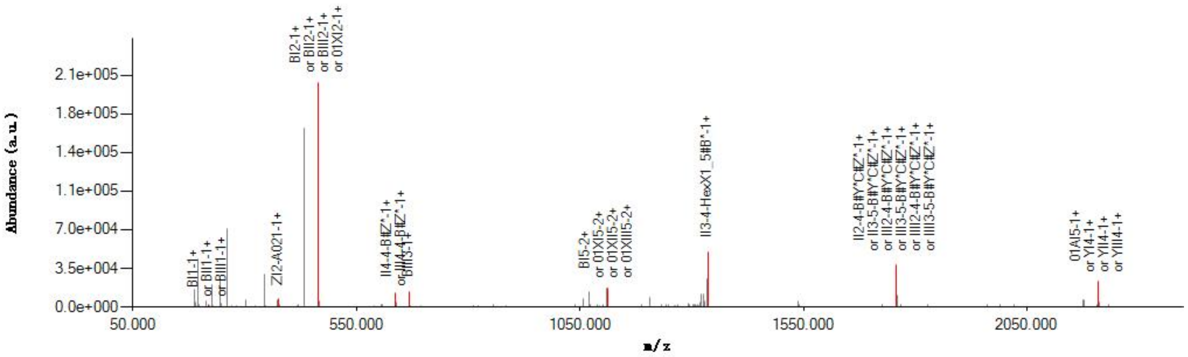

#### Graphical Fragmentation Map

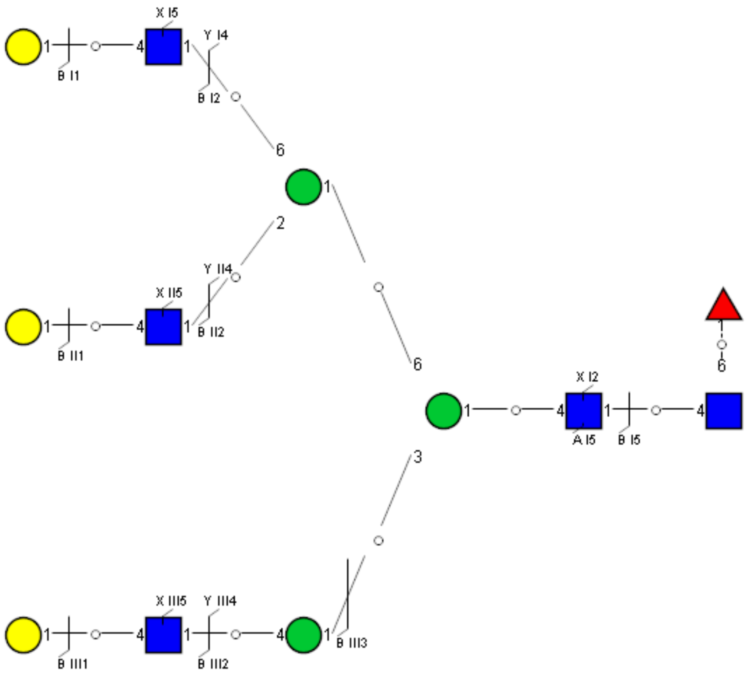

Spectrum Index: 36158(NoBHs:1)

#### Precursor iEF Map

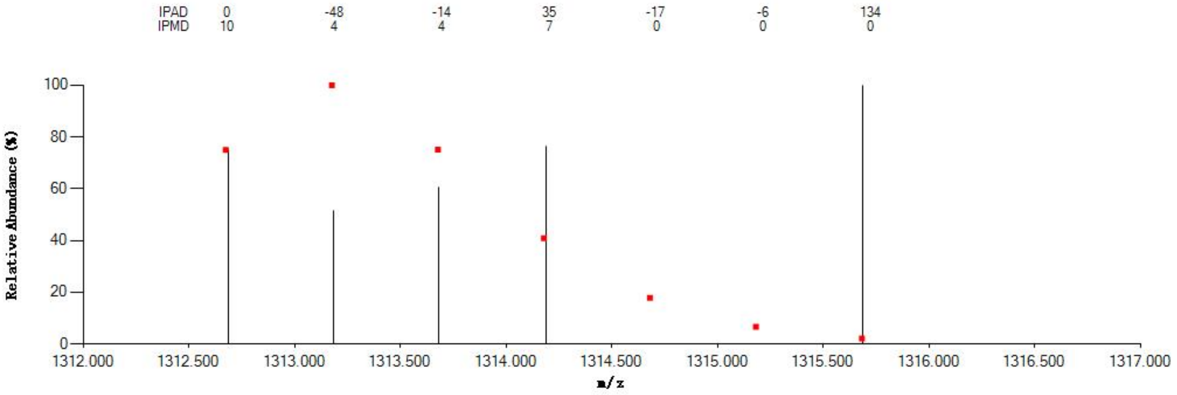


#### Annotated MS/MS Spectrum

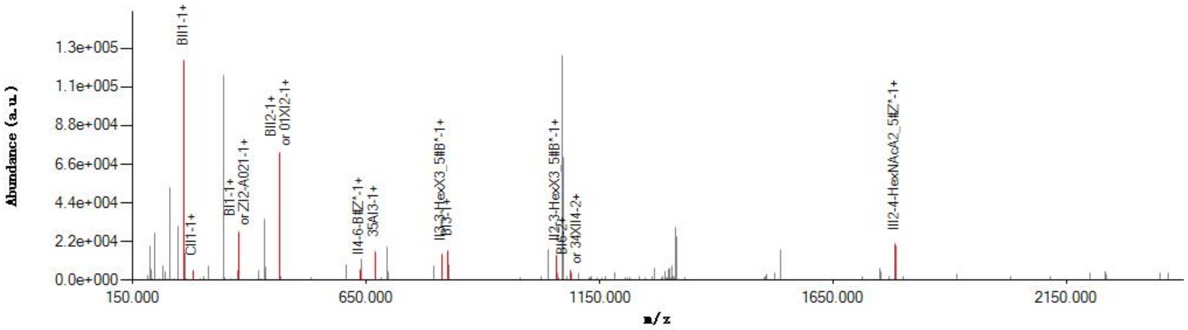


#### Graphical Fragmentation Map

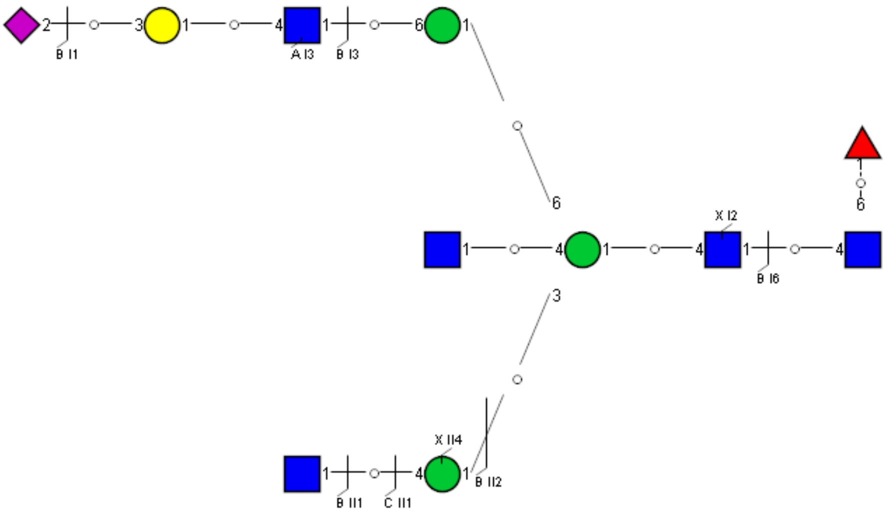

### Spectrum Index: 36165(NoBHs:1)

#### Precursor iEF Map

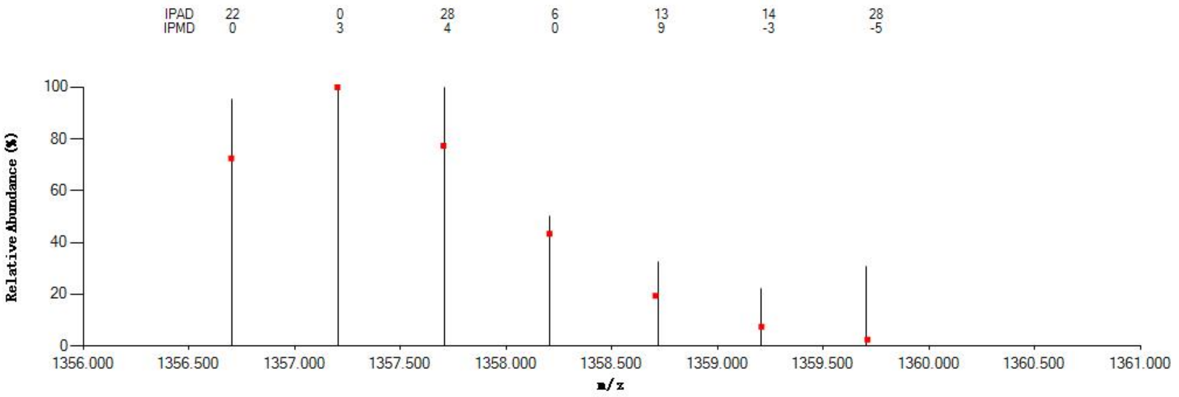


#### Annotated MS/MS Spectrum

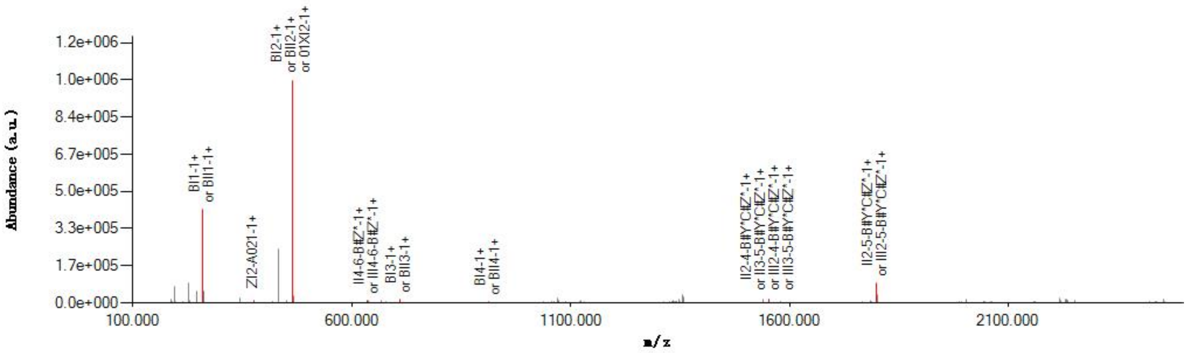


#### Graphical Fragmentation Map

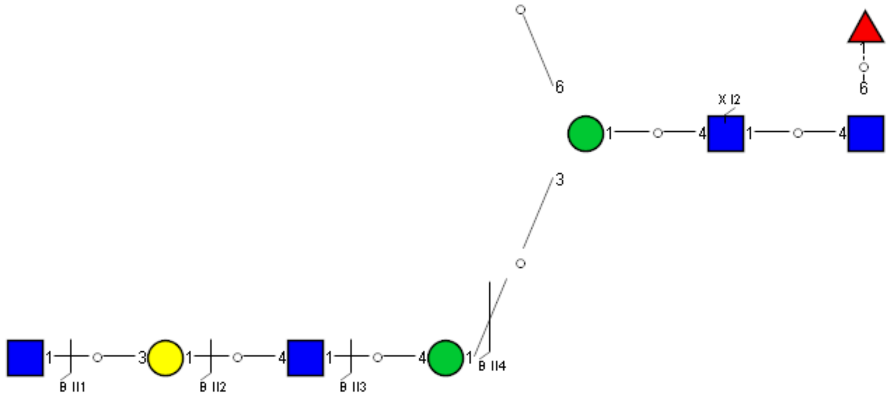

Spectrum Index: 36257(NoBHs:1)

#### Precursor iEF Map

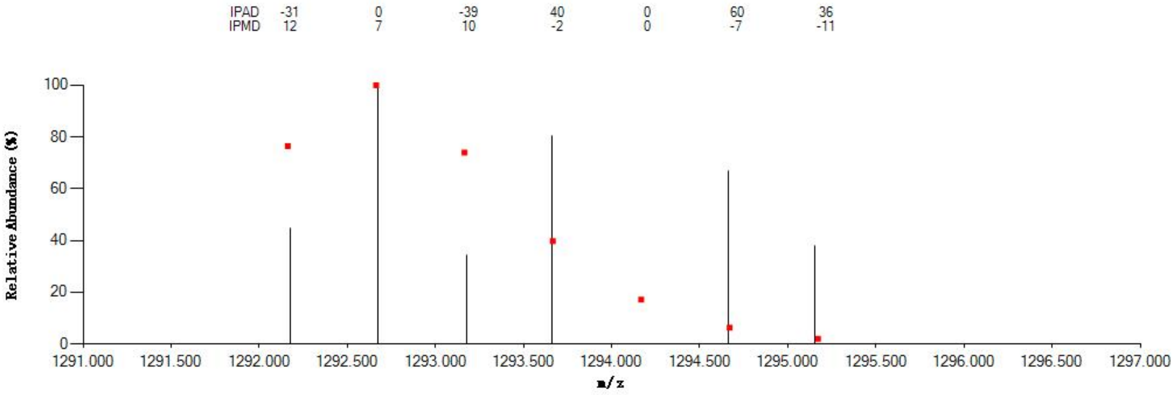

#### Annotated MS/MS Spectrum

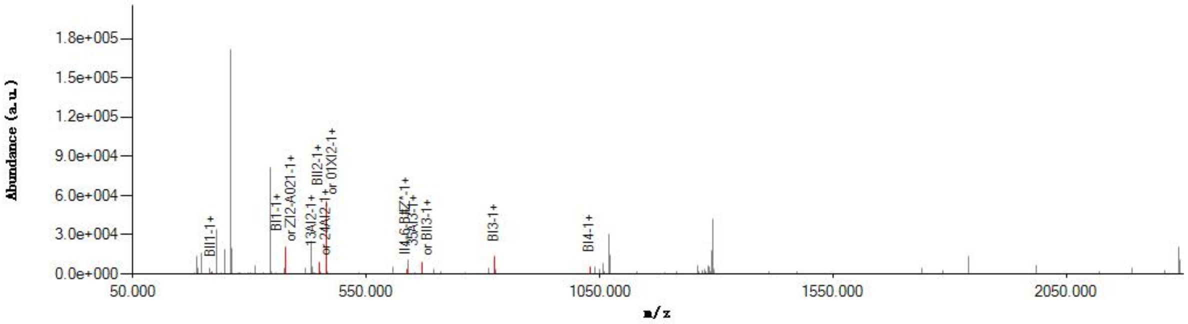

#### Graphical Fragmentation Map

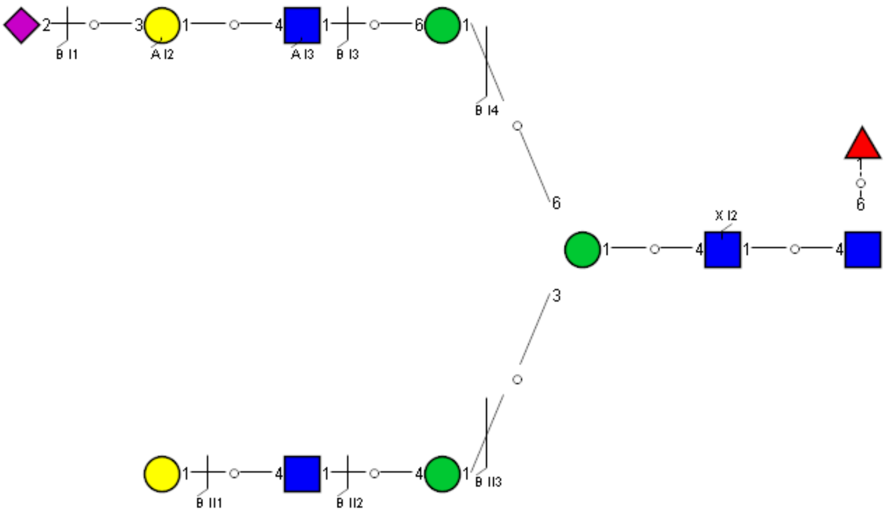

Spectrum Index: 36375(NoBHs:1)

#### Precursor iEF Map

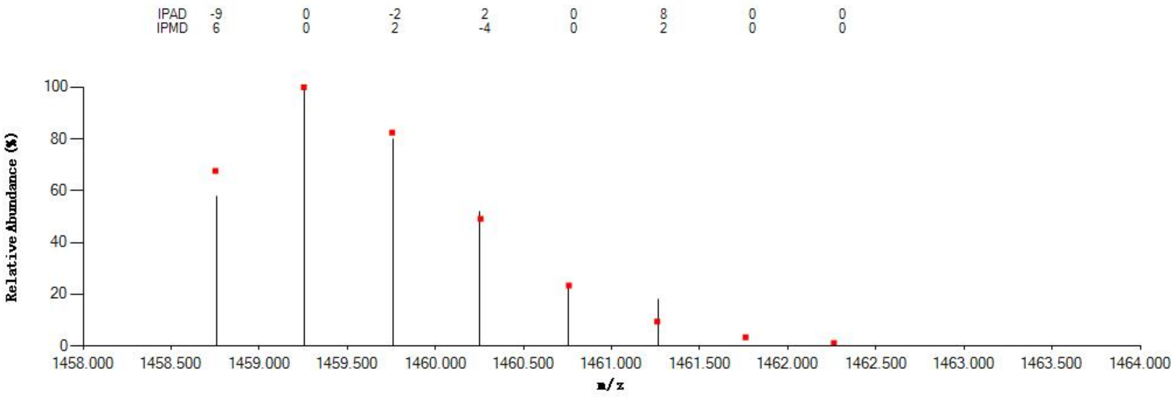


#### Annotated MS/MS Spectrum

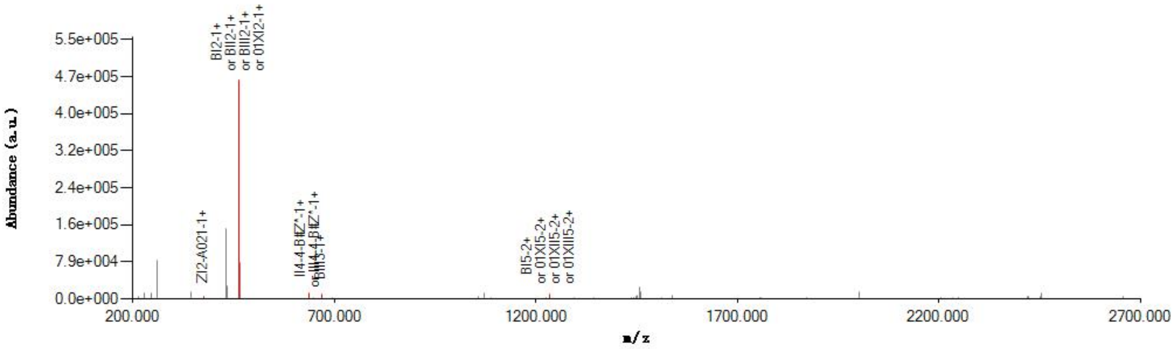


#### Graphical Fragmentation Map

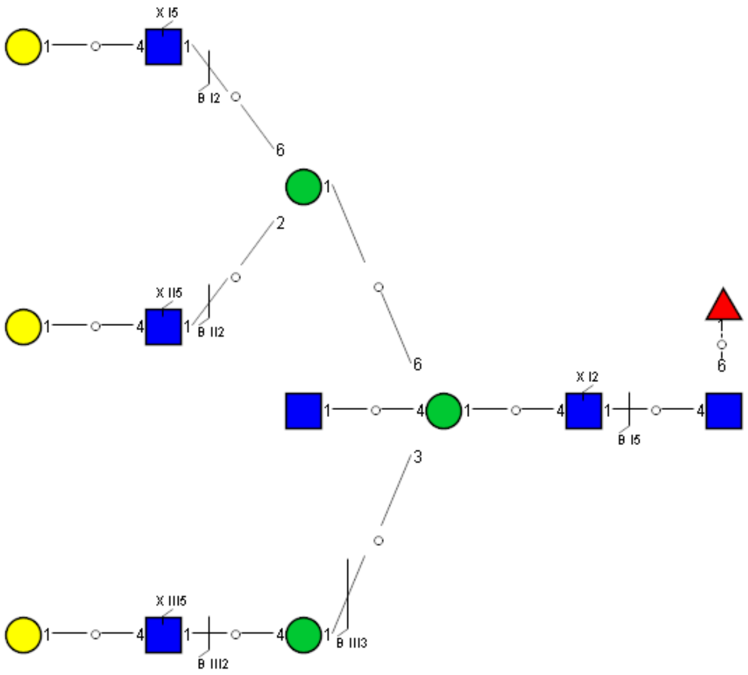

Spectrum Index: 36453(NoBHs:1)

#### Precursor iEF Map

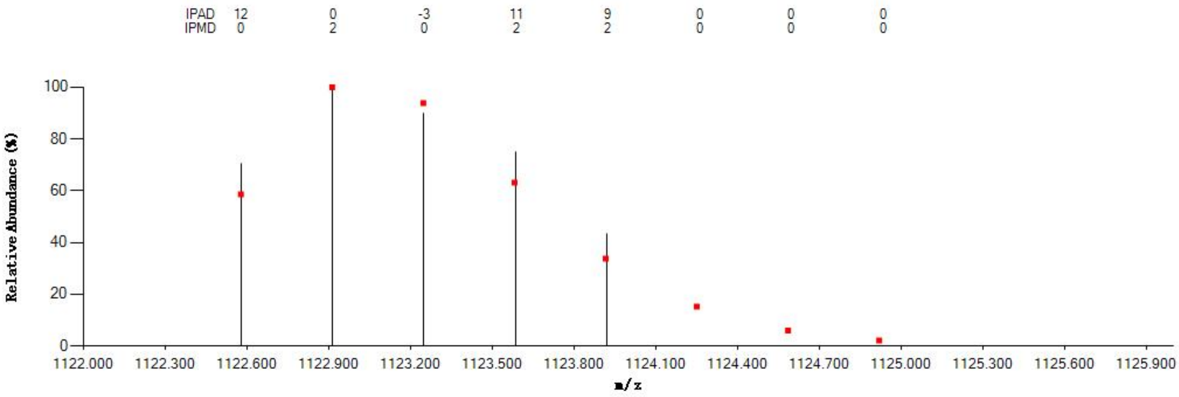


#### Annotated MS/MS Spectrum

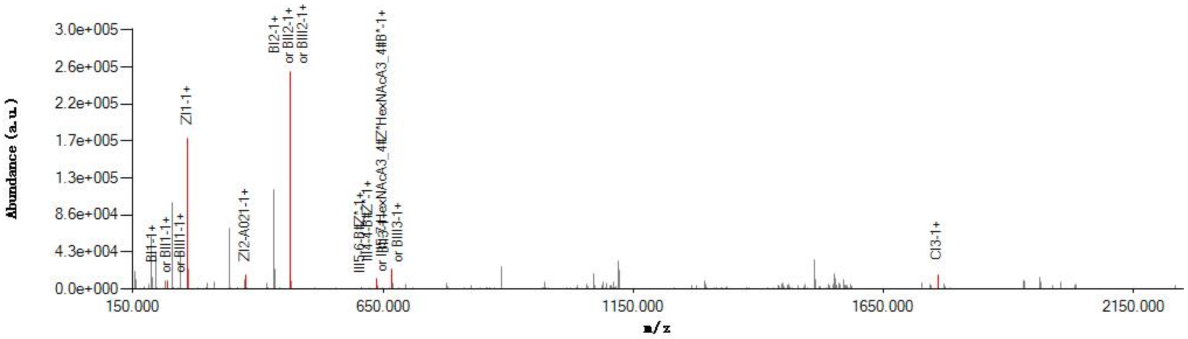


#### Graphical Fragmentation Map

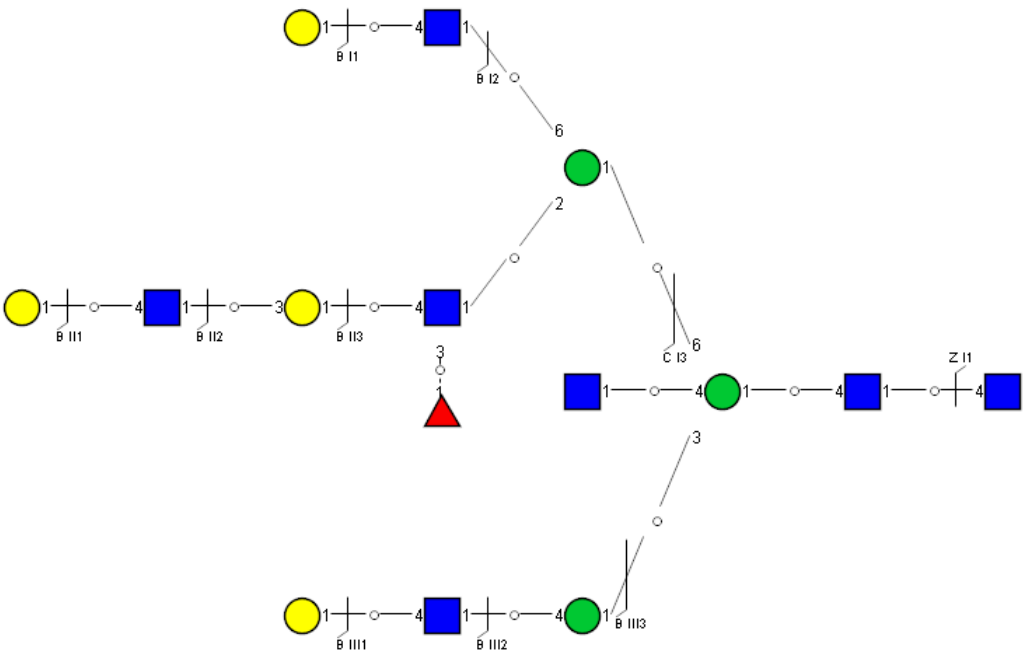

Spectrum Index: 36678(NoBHs:1)

#### Precursor iEF Map

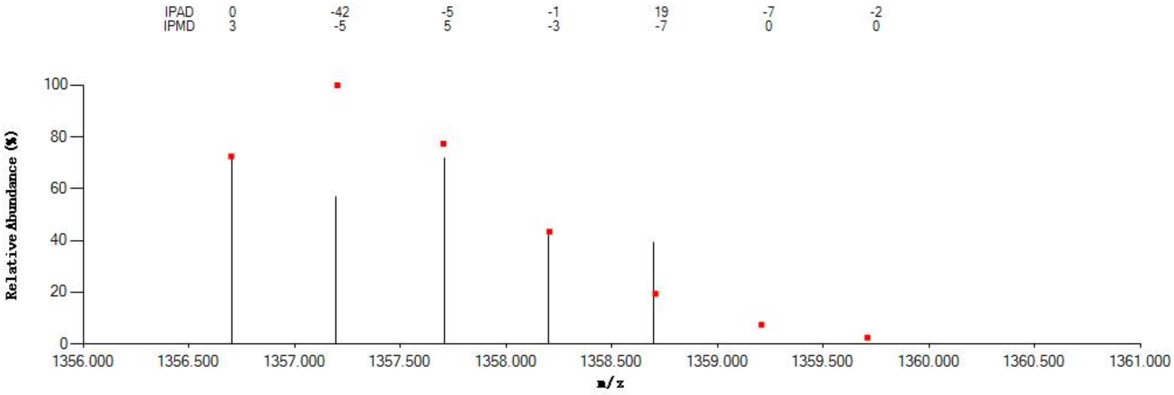


#### Annotated MS/MS Spectrum

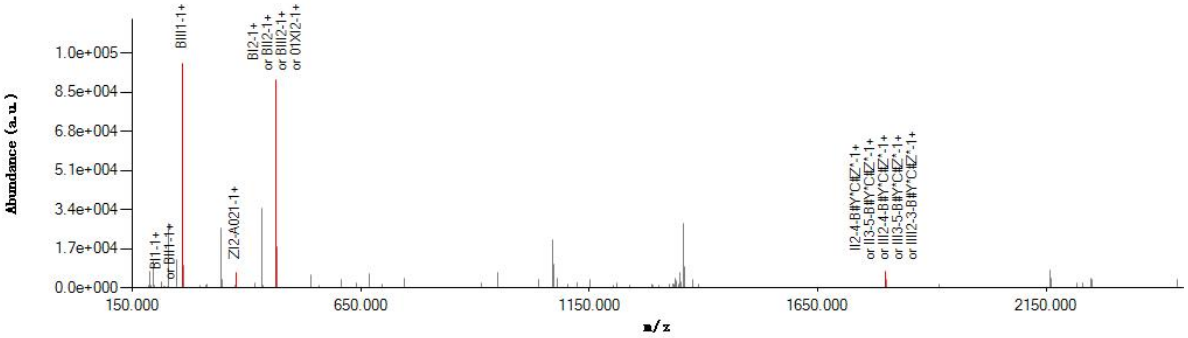


#### Graphical Fragmentation Map

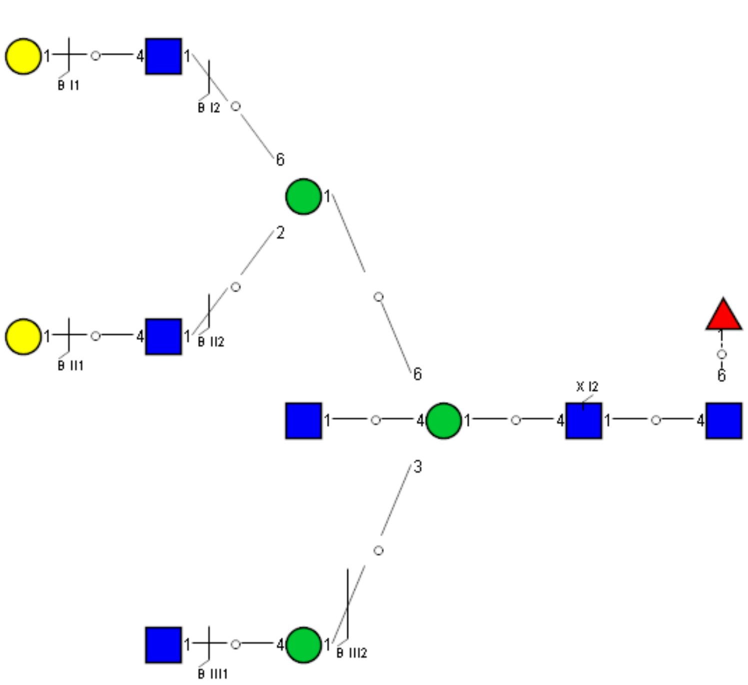

Spectrum Index: 36839(NoBHs:1)

#### Precursor iEF Map

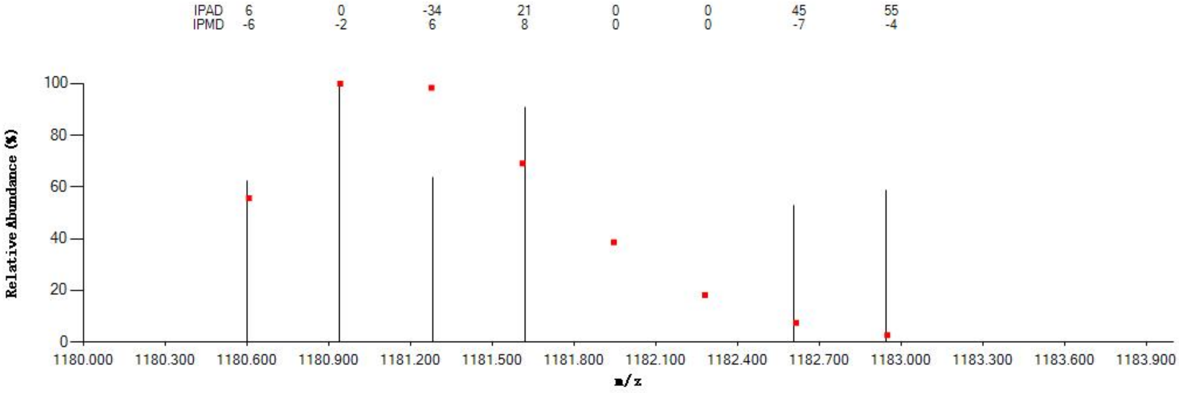


#### Annotated MS/MS Spectrum

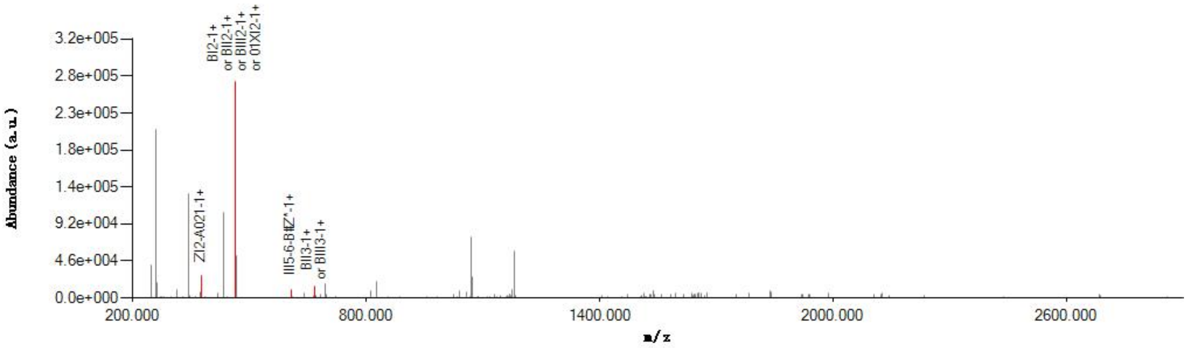


#### Graphical Fragmentation Map

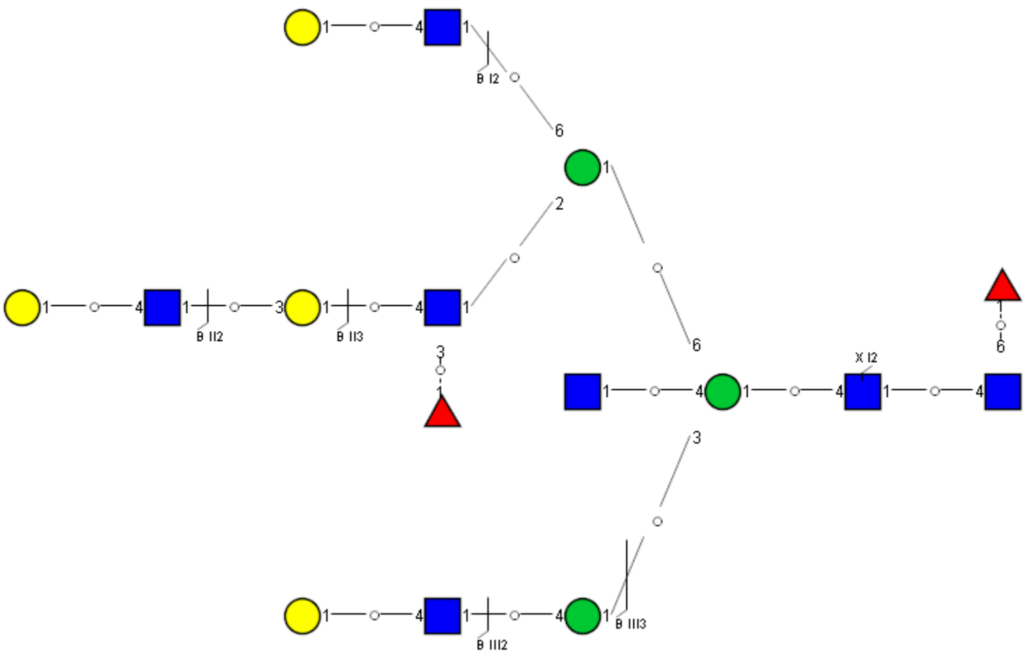

Spectrum Index: 37058(NoBHs:1)

#### Precursor iEF Map

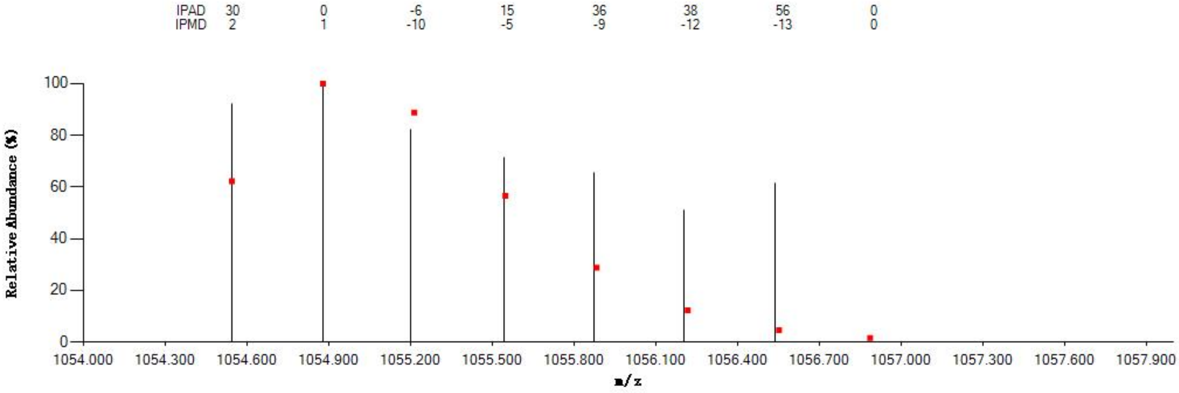


#### Annotated MS/MS Spectrum

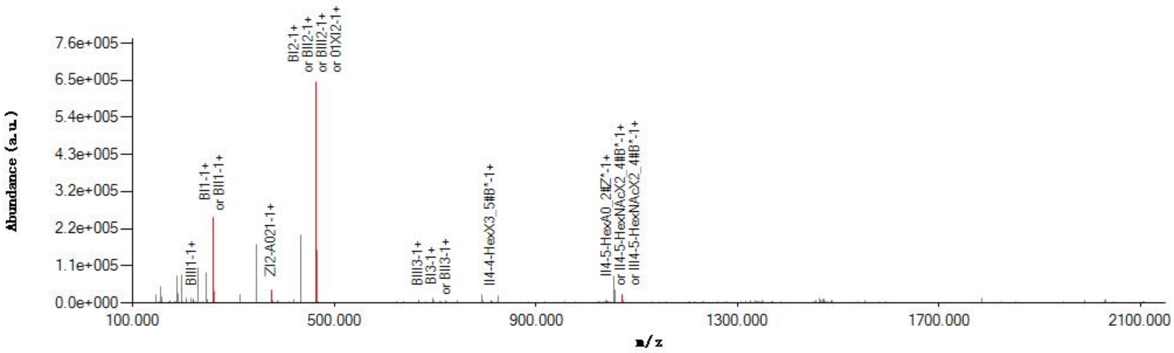


#### Graphical Fragmentation Map

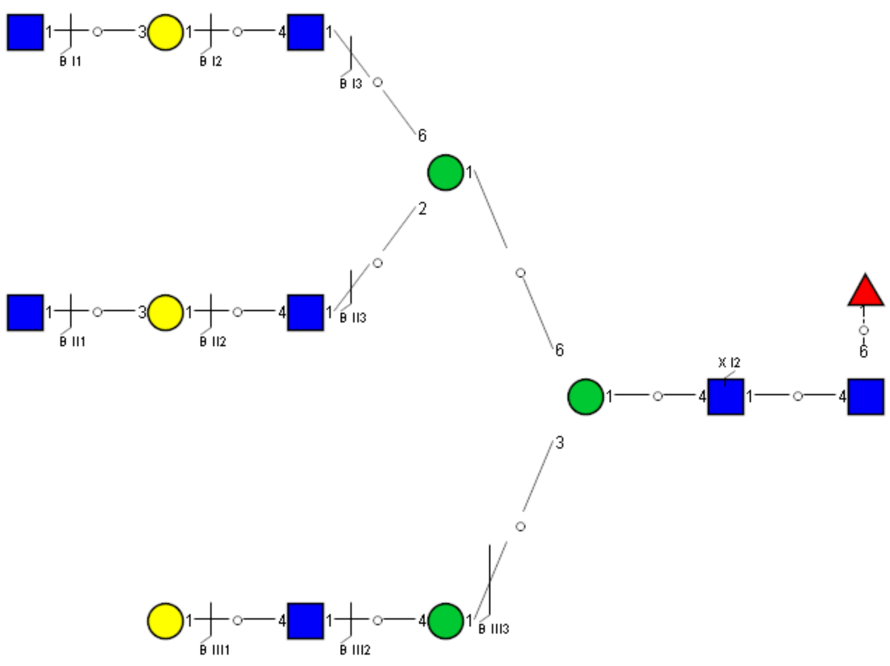

Spectrum Index: 37114(NoBHs:1)

#### Precursor iEF Map

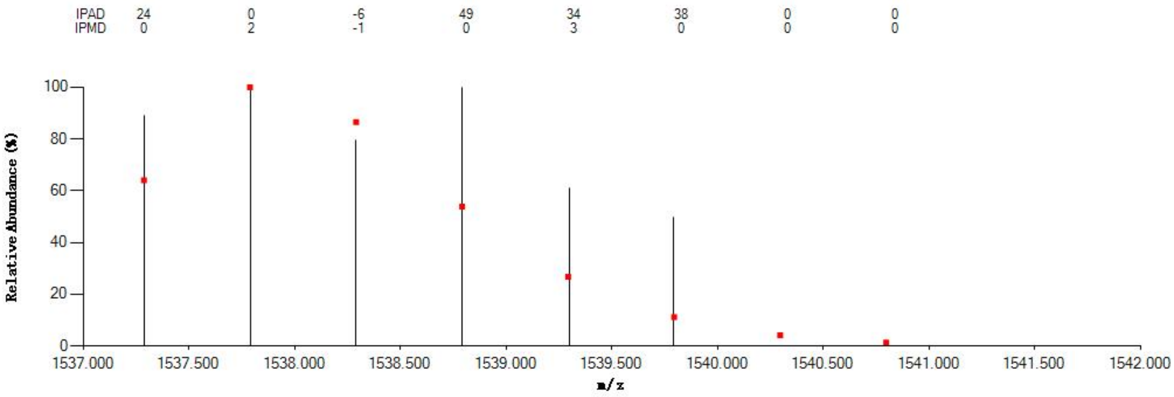


#### Annotated MS/MS Spectrum

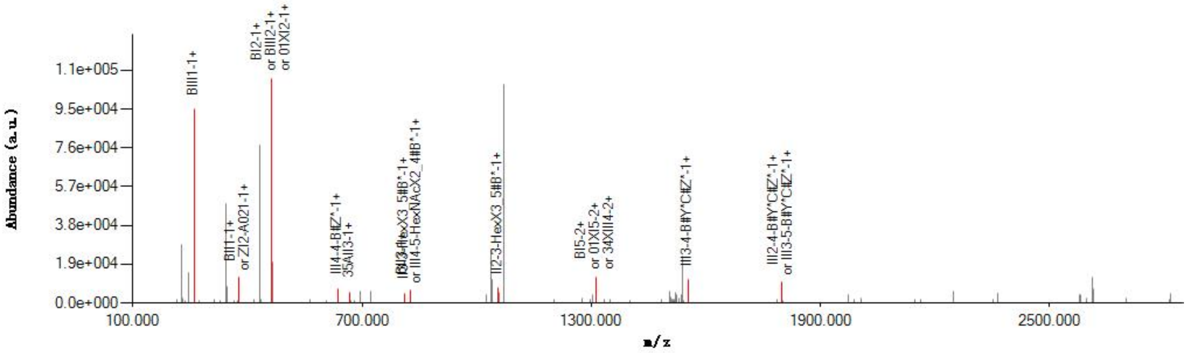


#### Graphical Fragmentation Map

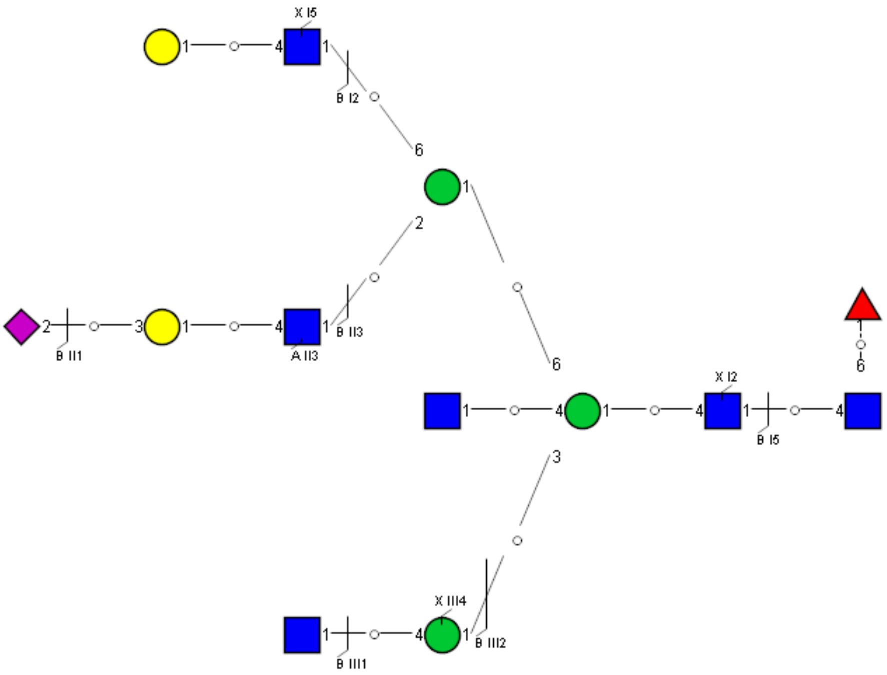

Spectrum Index: 37196(NoBHs:1)

#### Precursor iEF Map

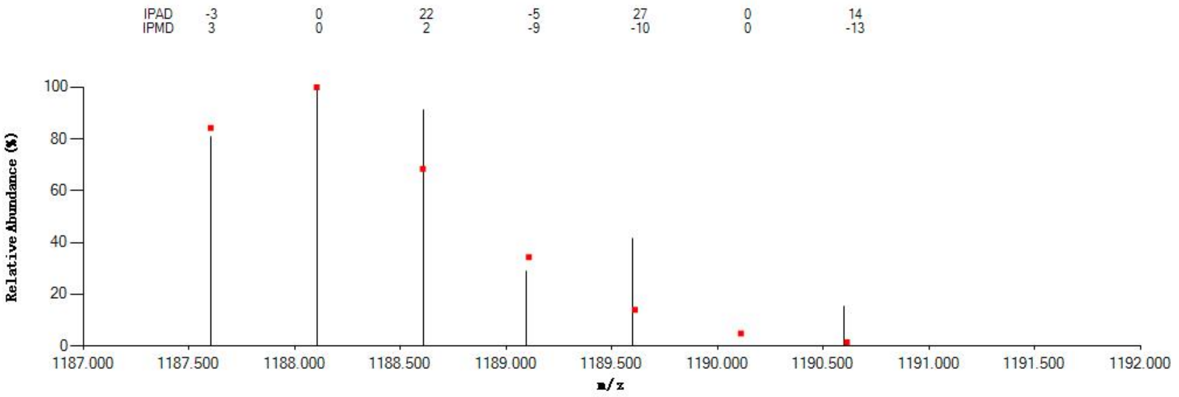


#### Annotated MS/MS Spectrum

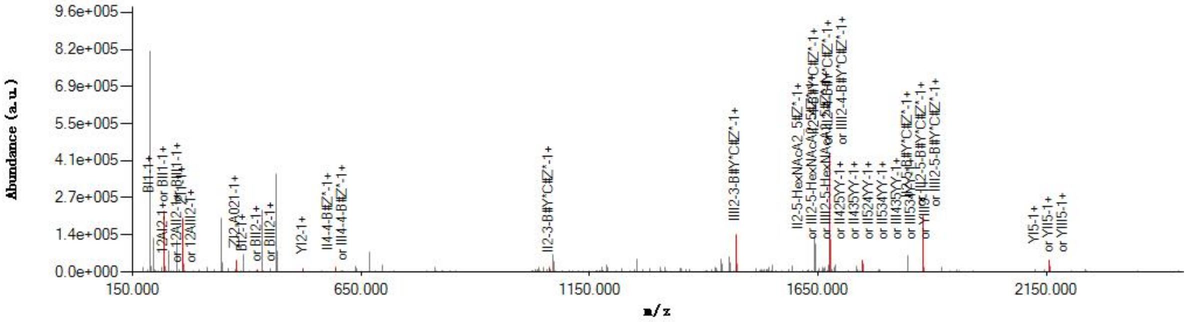


#### Graphical Fragmentation Map

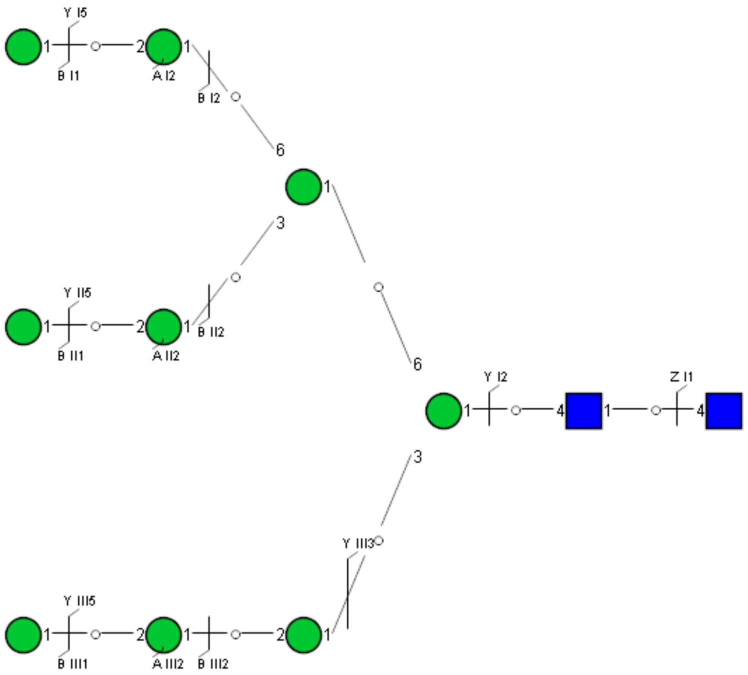

Spectrum Index: 37226(NoBHs:1)

#### Precursor iEF Map

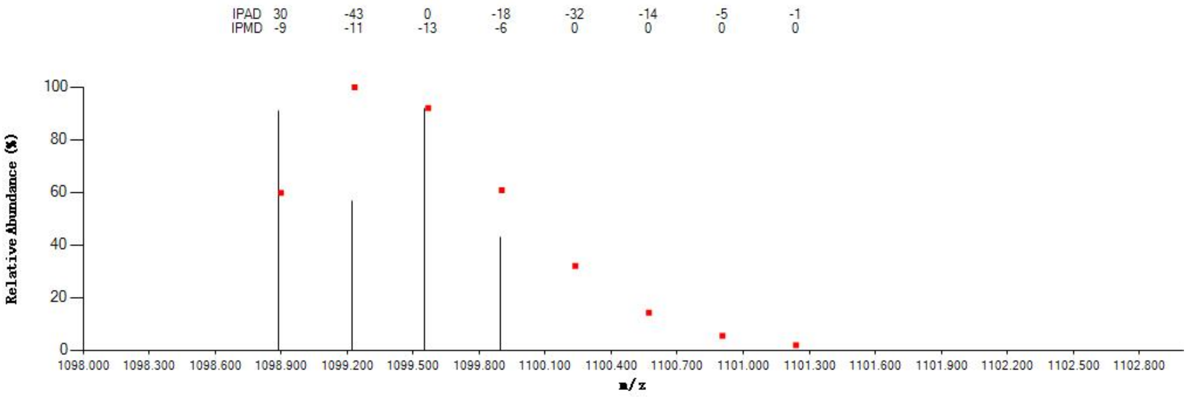


#### Annotated MS/MS Spectrum

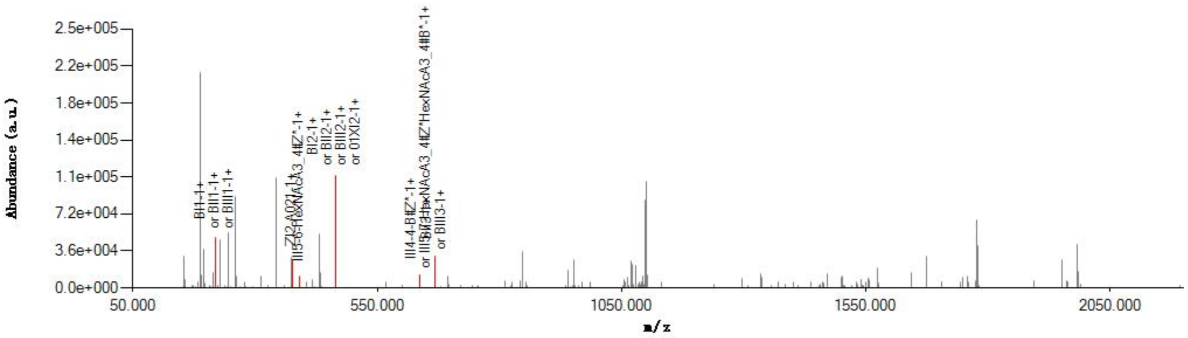


#### Graphical Fragmentation Map

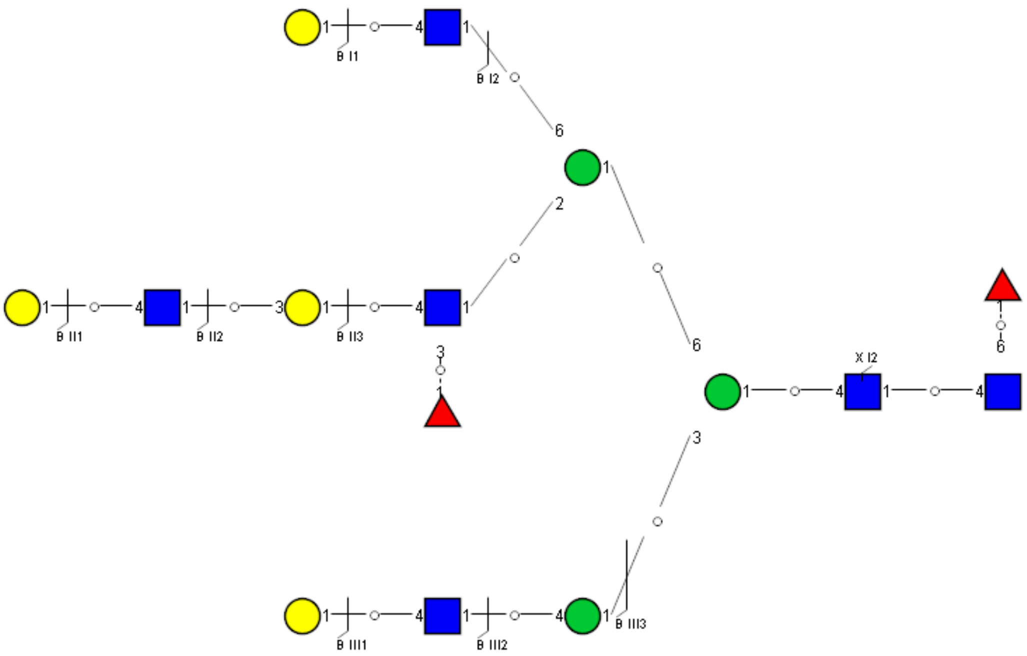

Spectrum Index: 37887(NoBHs:1)

#### Precursor iEF Map

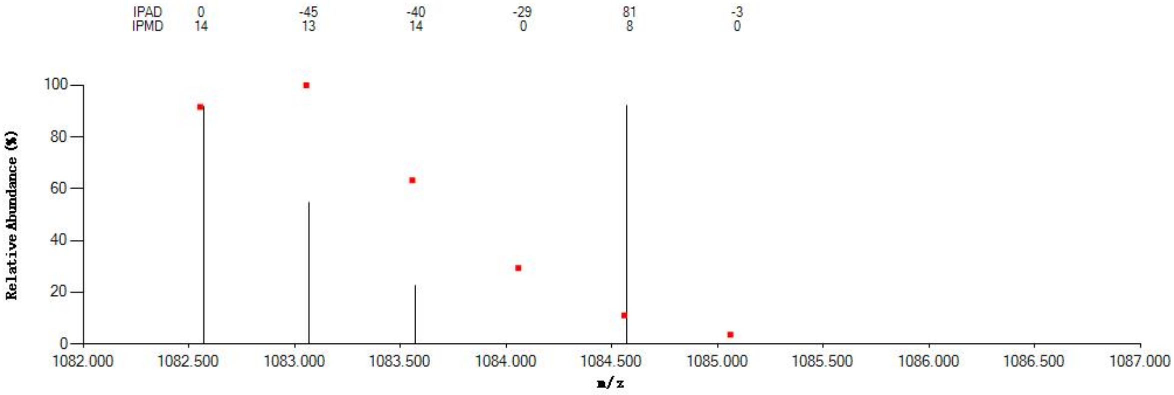


#### Annotated MS/MS Spectrum

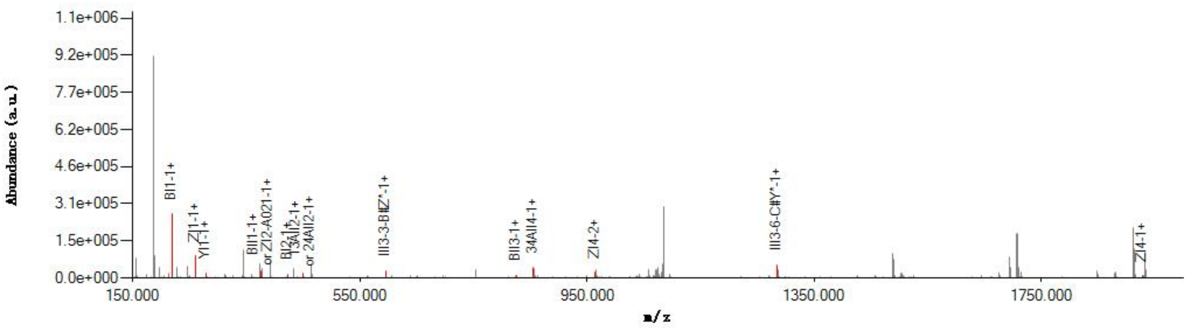


#### Graphical Fragmentation Map

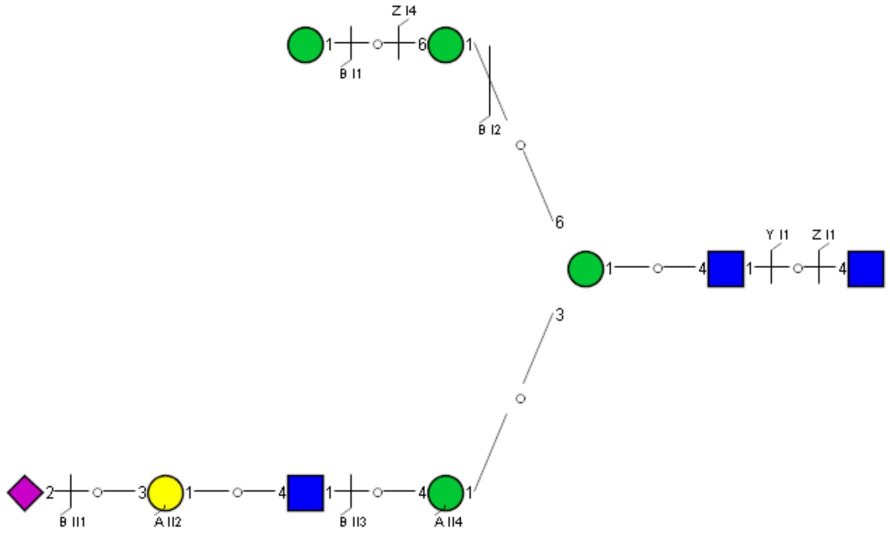

Spectrum Index: 37902(NoBHs:1)

#### Precursor iEF Map

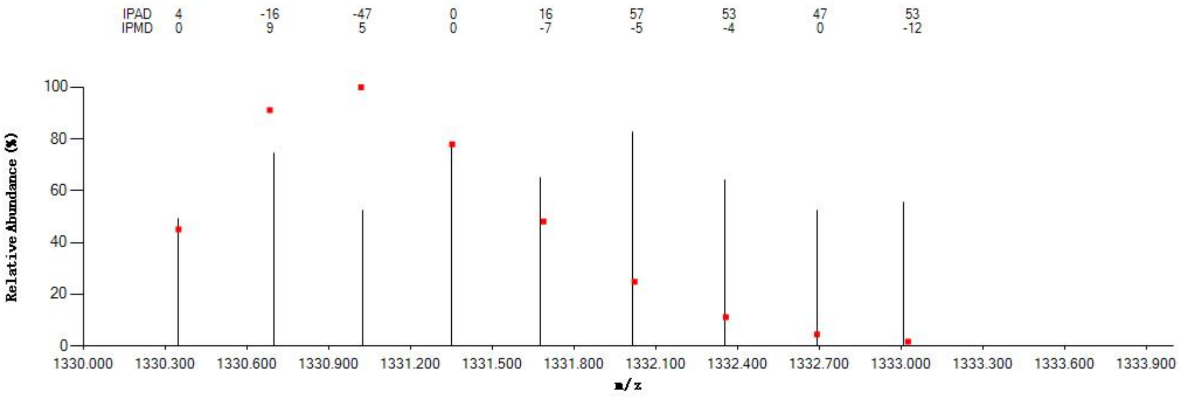

#### Annotated MS/MS Spectrum

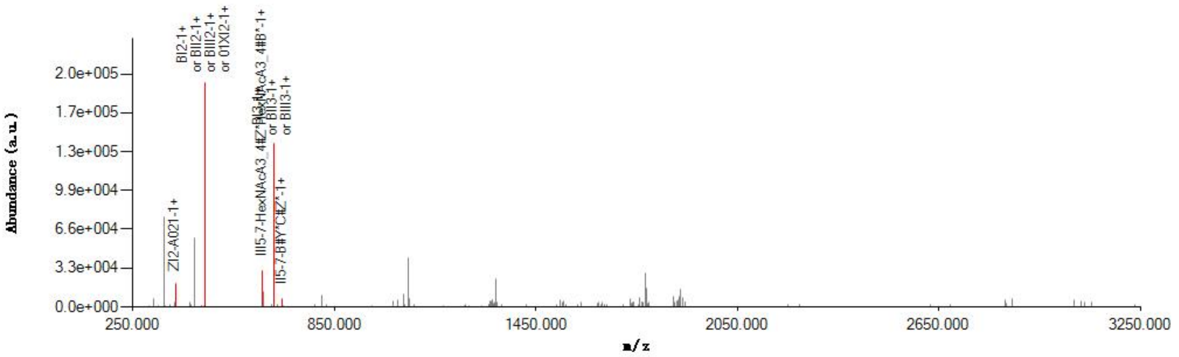

#### Graphical Fragmentation Map

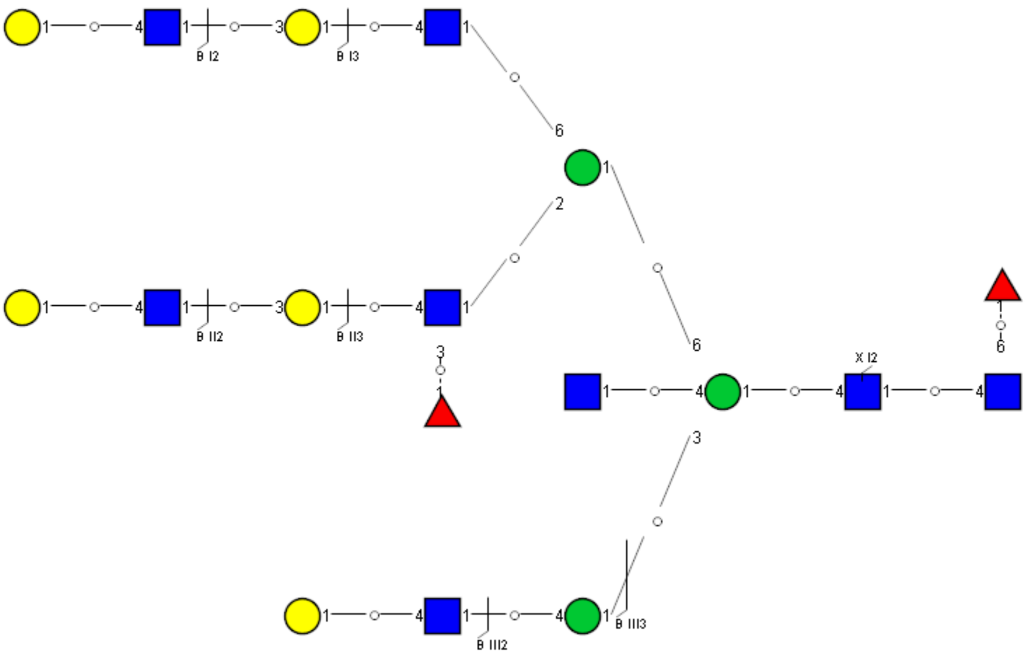

Spectrum Index: 38001(NoBHs:1)

#### Precursor iEF Map

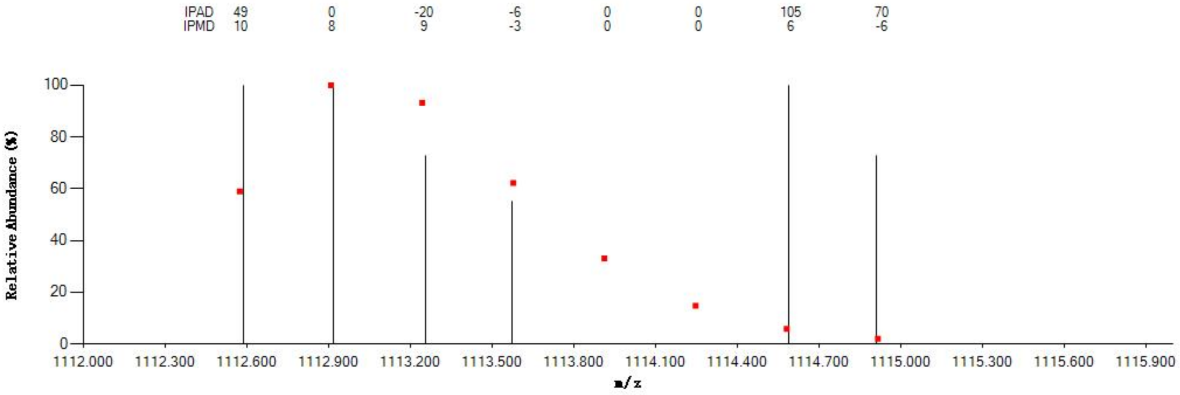


#### Annotated MS/MS Spectrum

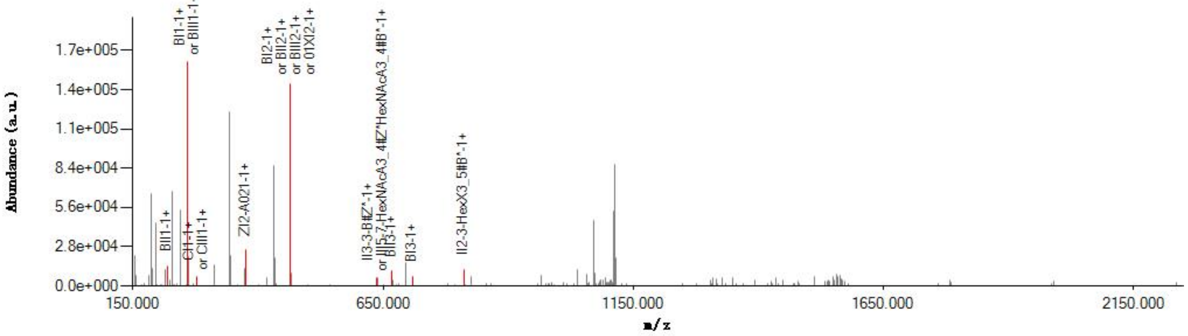


#### Graphical Fragmentation Map

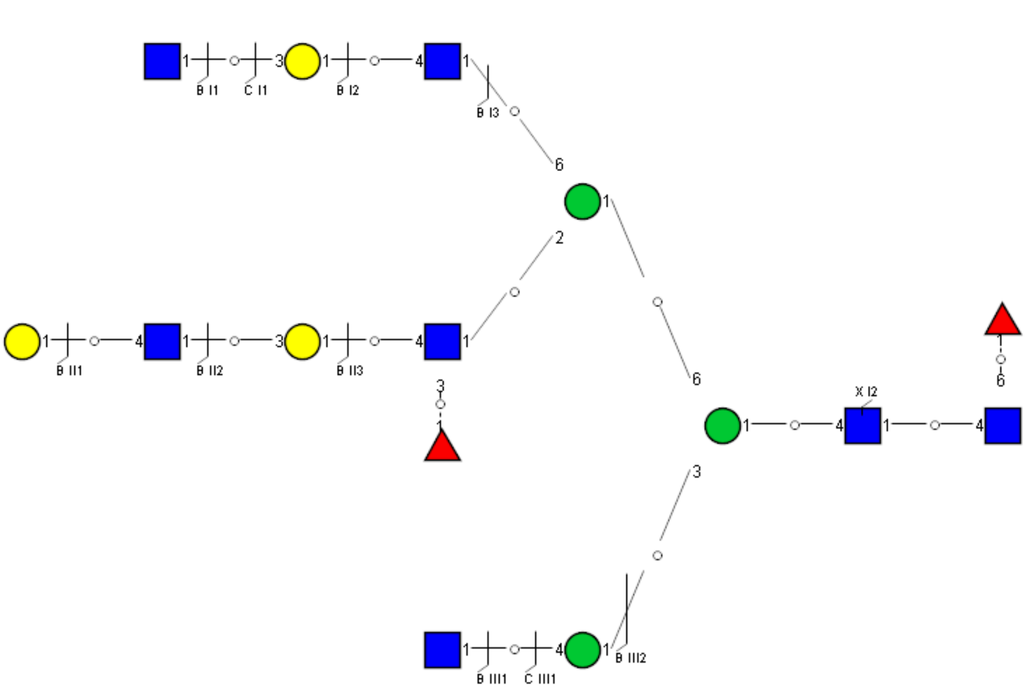

Spectrum Index: 38040(NoBHs:1)

#### Precursor iEF Map

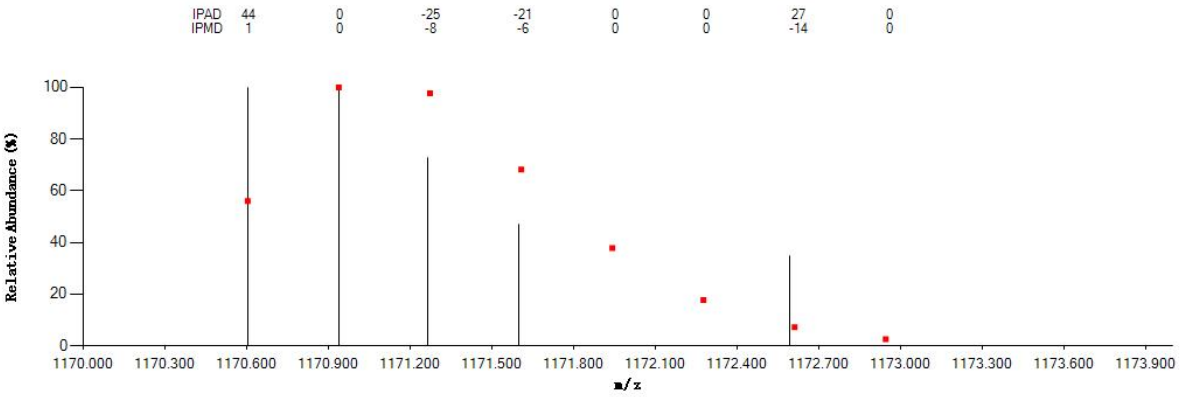


#### Annotated MS/MS Spectrum

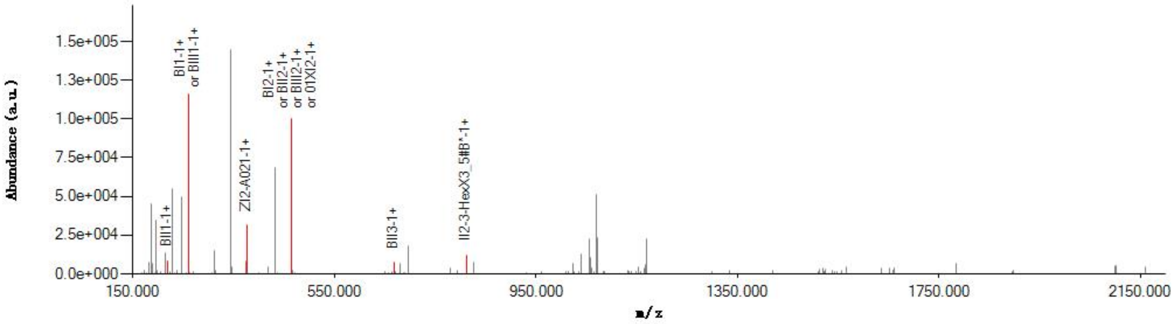


#### Graphical Fragmentation Map

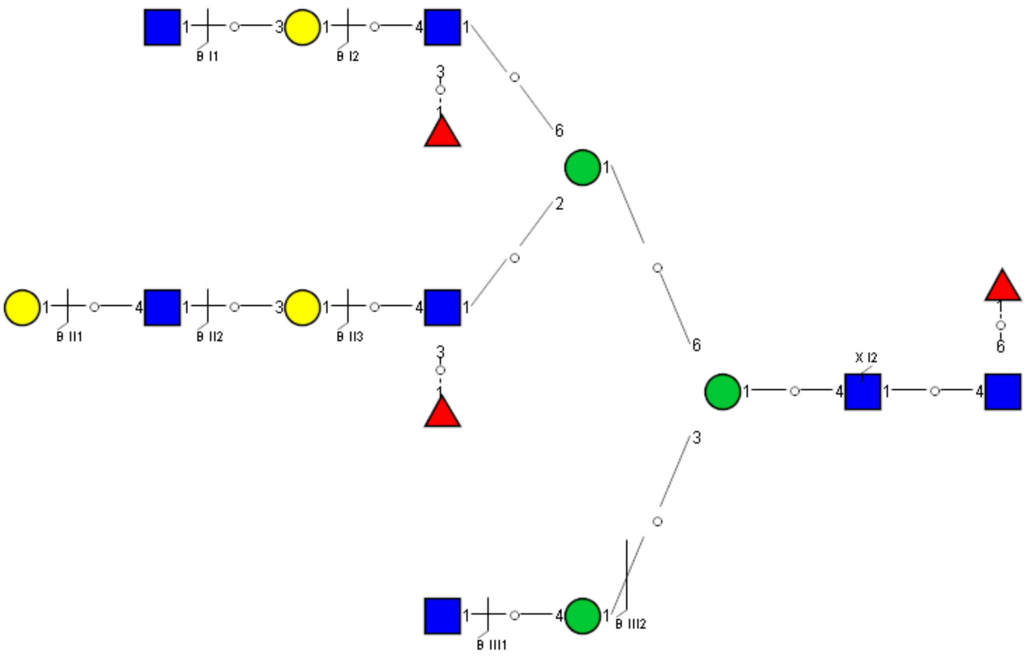

Spectrum Index: 38055(NoBHs:1)

#### Precursor iEF Map

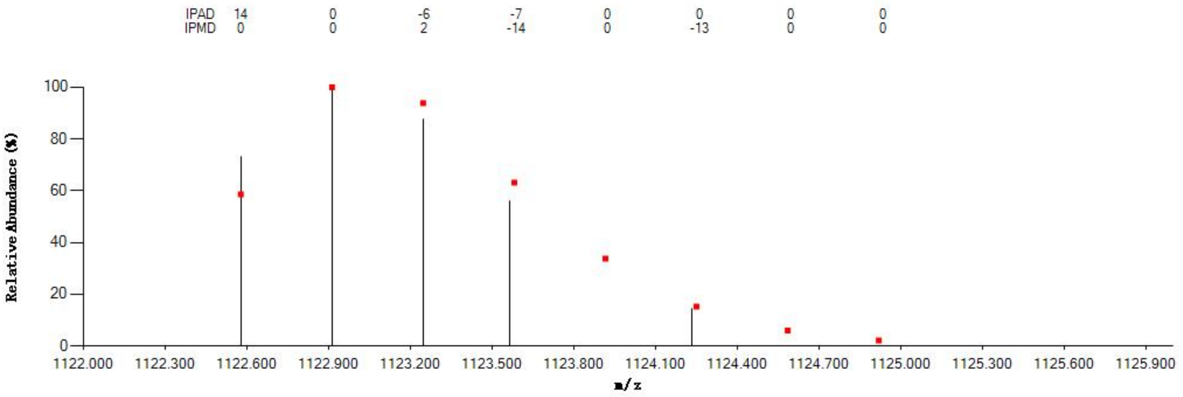


#### Annotated MS/MS Spectrum

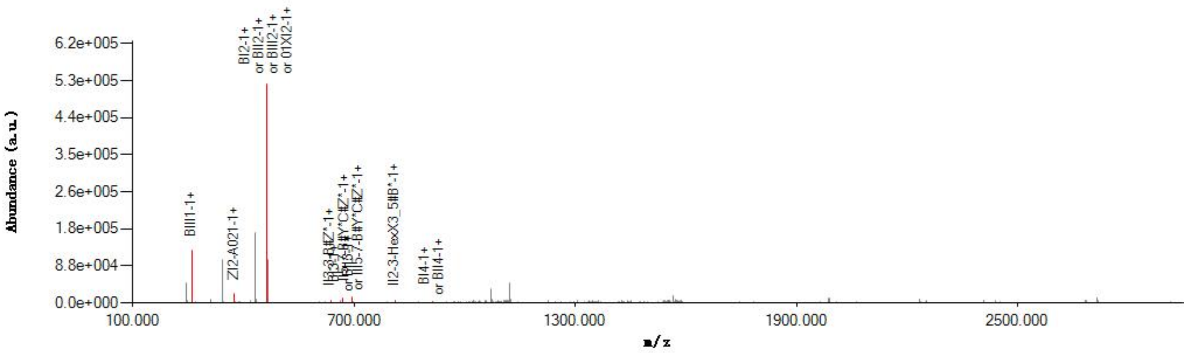


#### Graphical Fragmentation Map

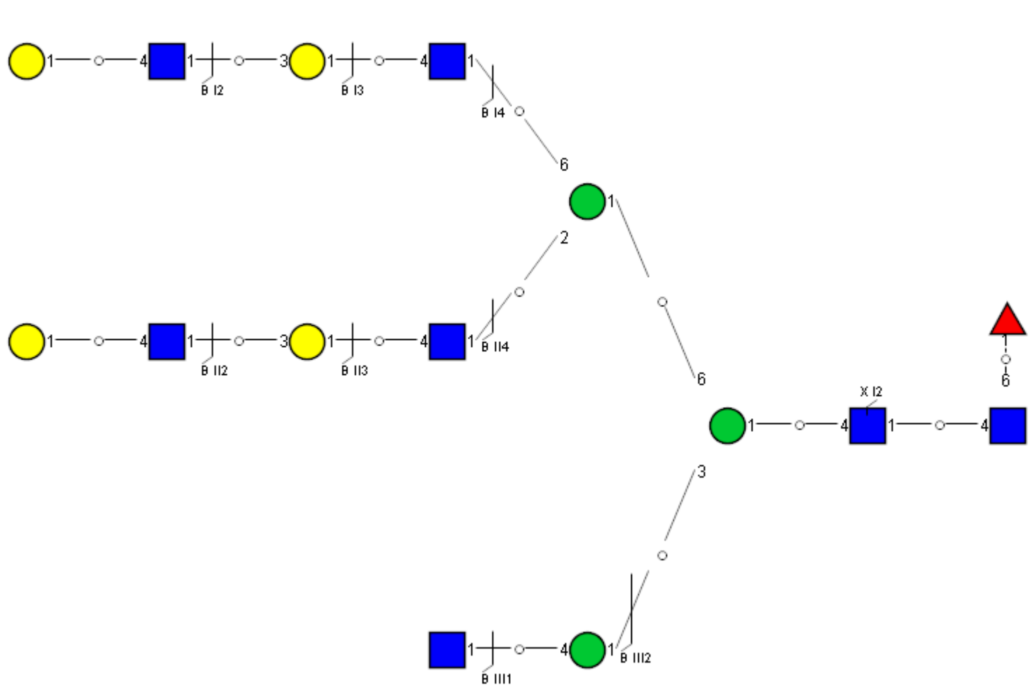

### Spectrum Index: 38099(NoBHs:1)

#### Precursor iEF Map

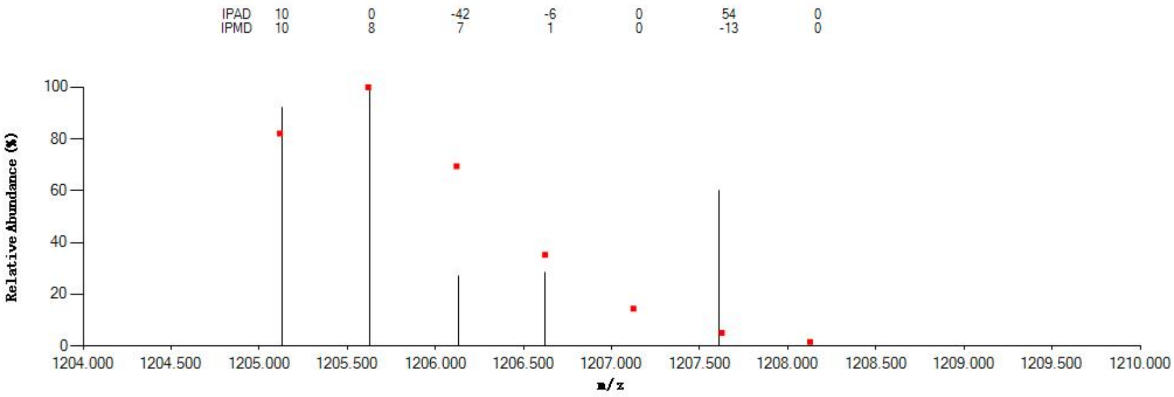


#### Annotated MS/MS Spectrum

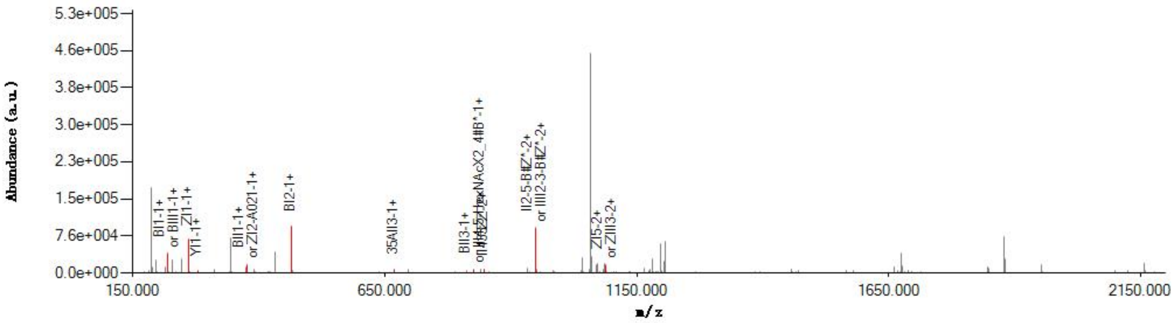


#### Graphical Fragmentation Map

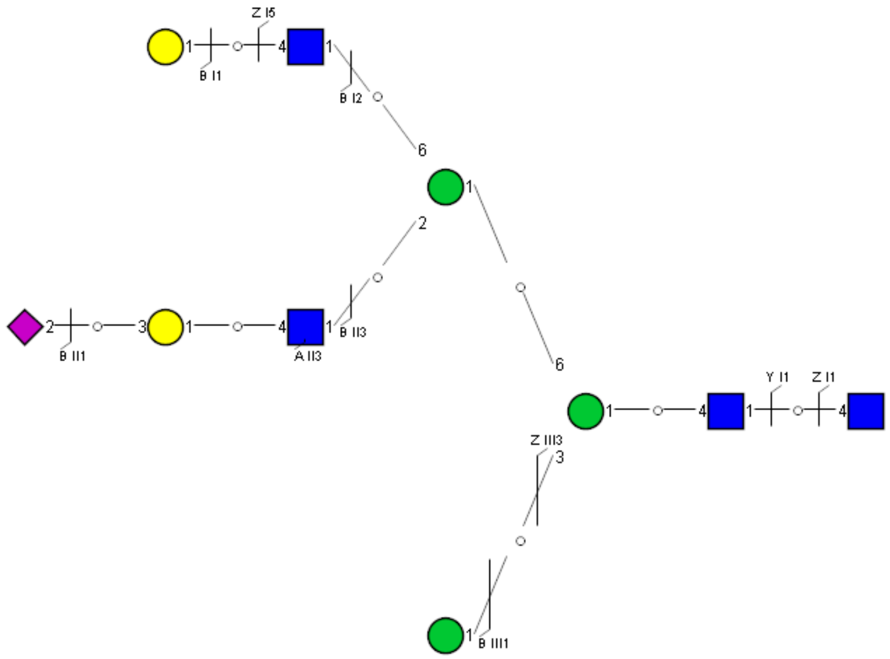

Spectrum Index: 38107(NoBHs:2)

#### Precursor iEF Map

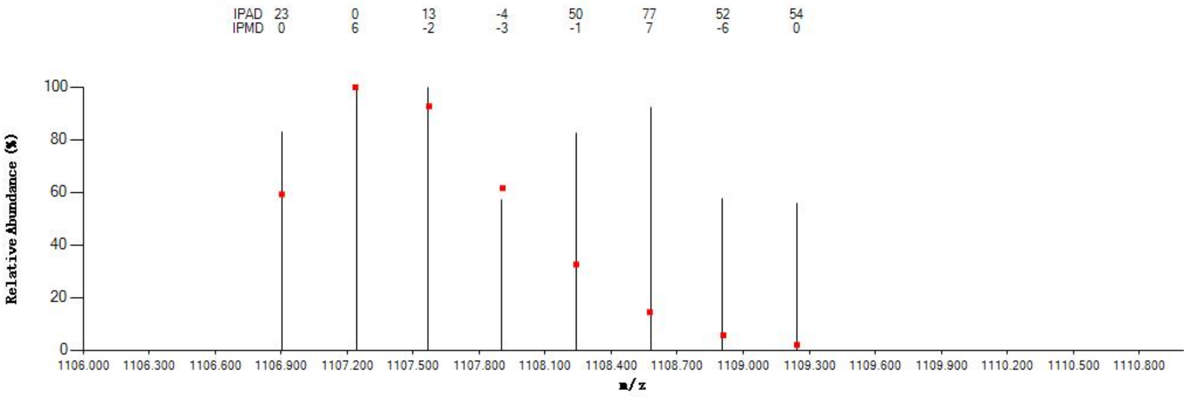


#### Annotated MS/MS Spectrum

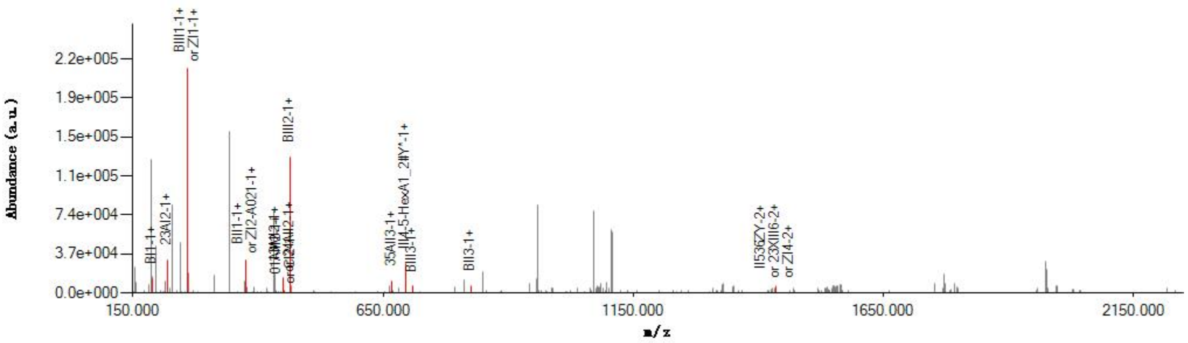


#### Graphical Fragmentation Map

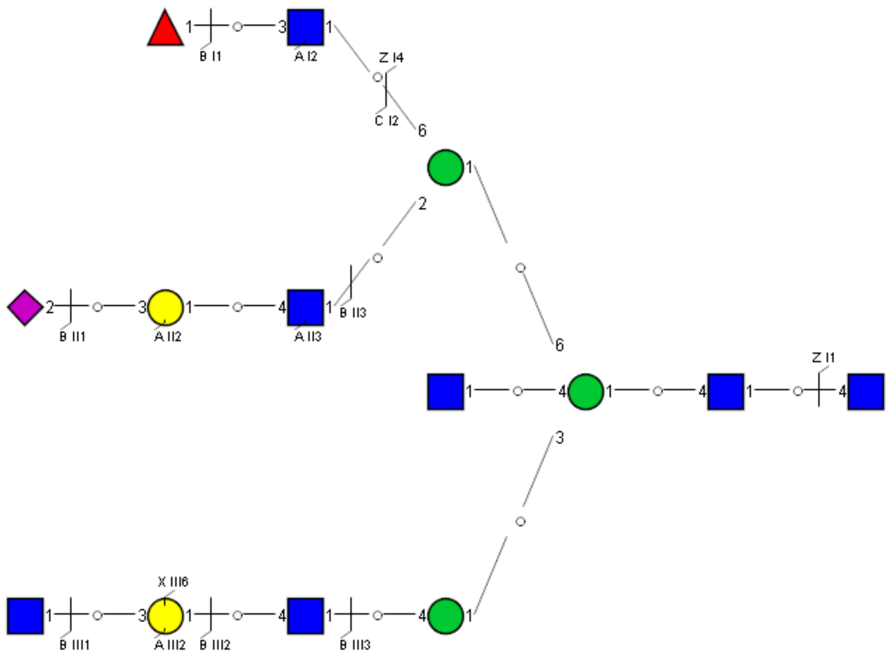

Spectrum Index: 38107(NoBHs:2)

#### Precursor iEF Map

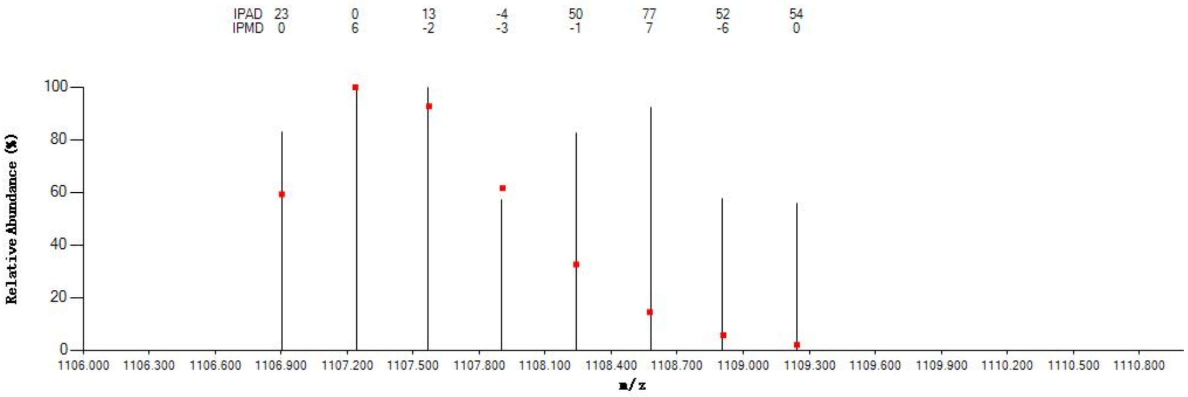

#### Annotated MS/MS Spectrum

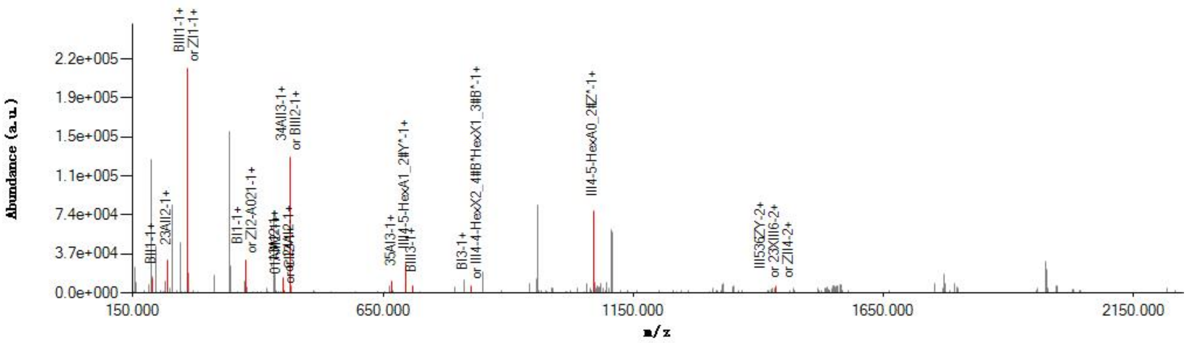

#### Graphical Fragmentation Map

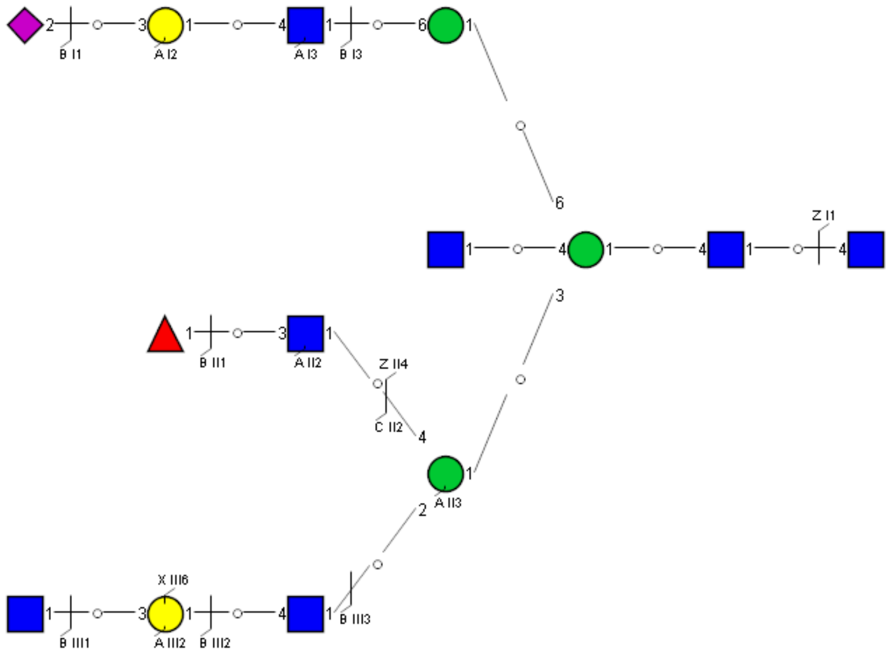

Spectrum Index: 38182(NoBHs:1)

#### Precursor iEF Map

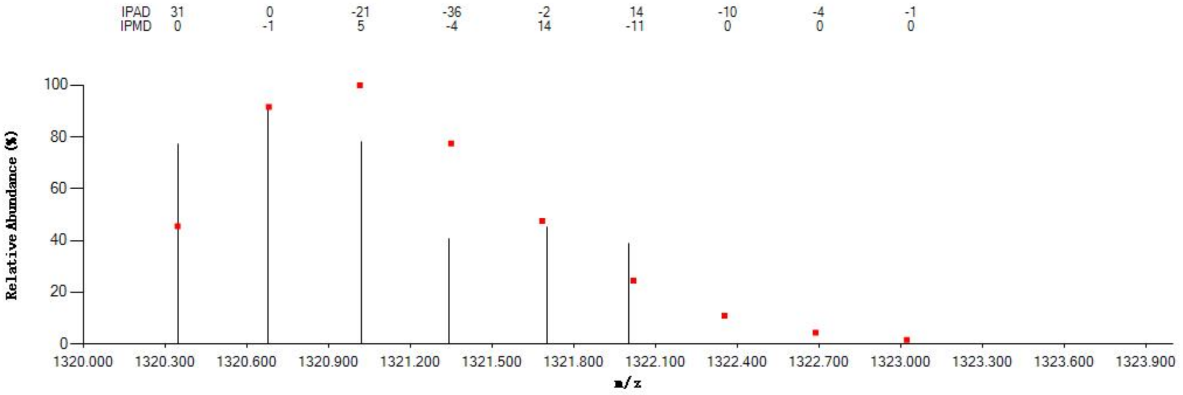


#### Annotated MS/MS Spectrum

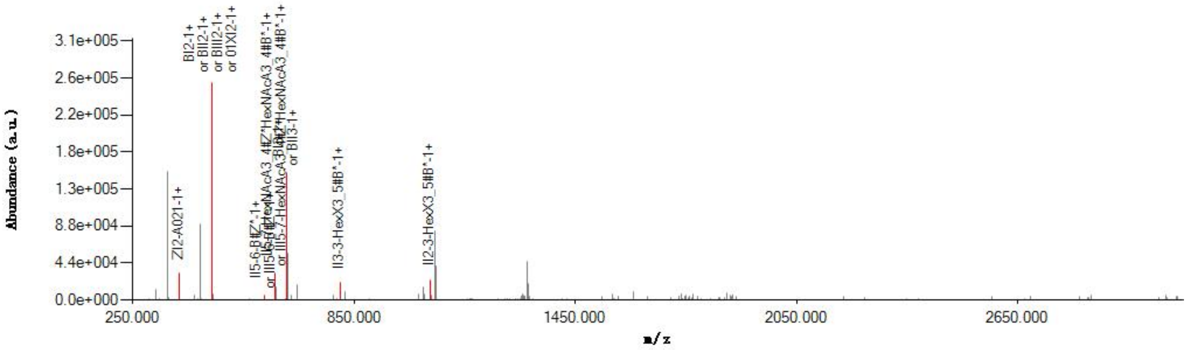


#### Graphical Fragmentation Map

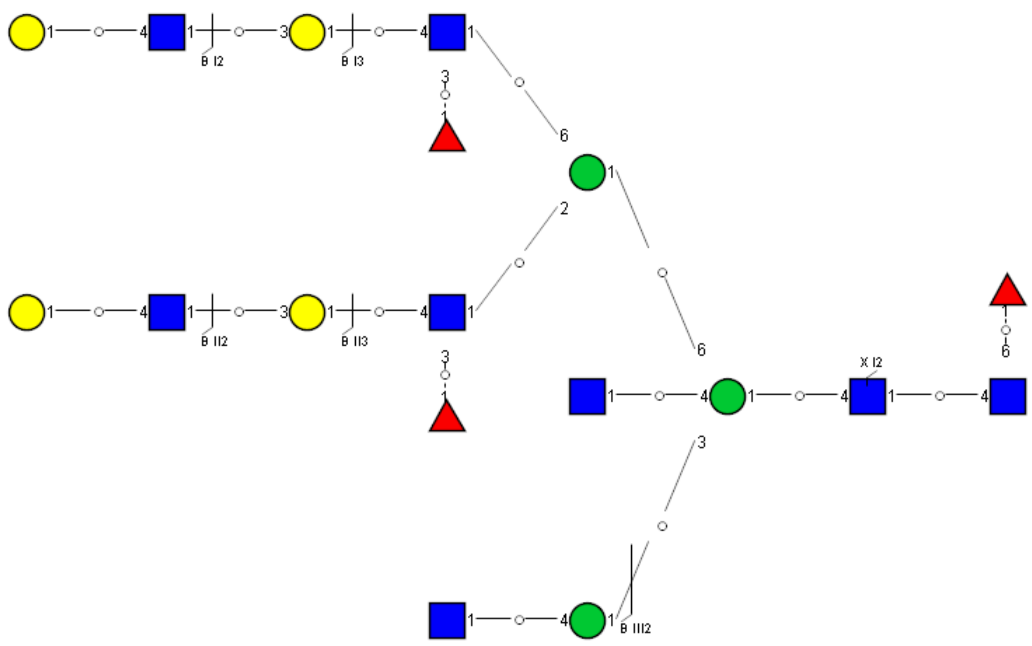

Spectrum Index: 38228(NoBHs:2)

#### Precursor iEF Map

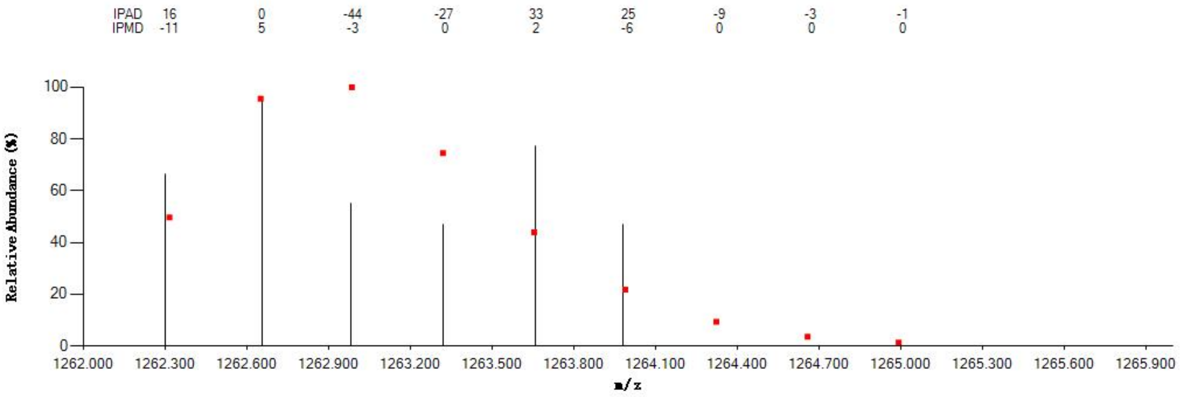


#### Annotated MS/MS Spectrum

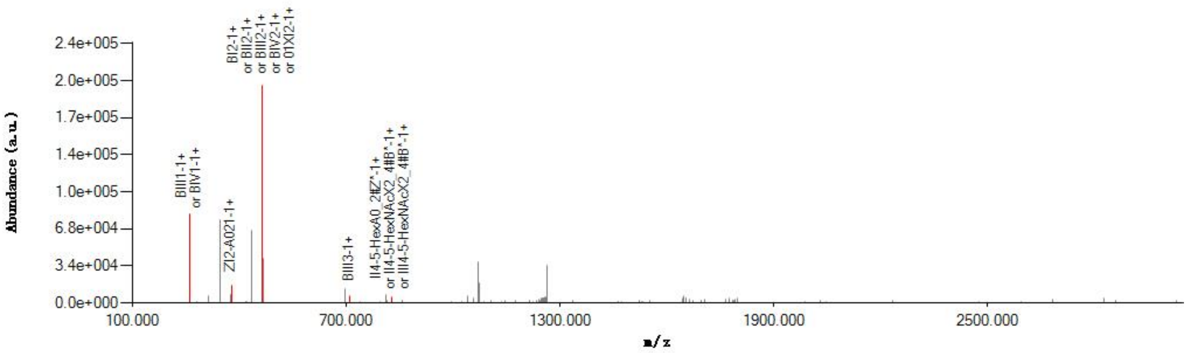


#### Graphical Fragmentation Map

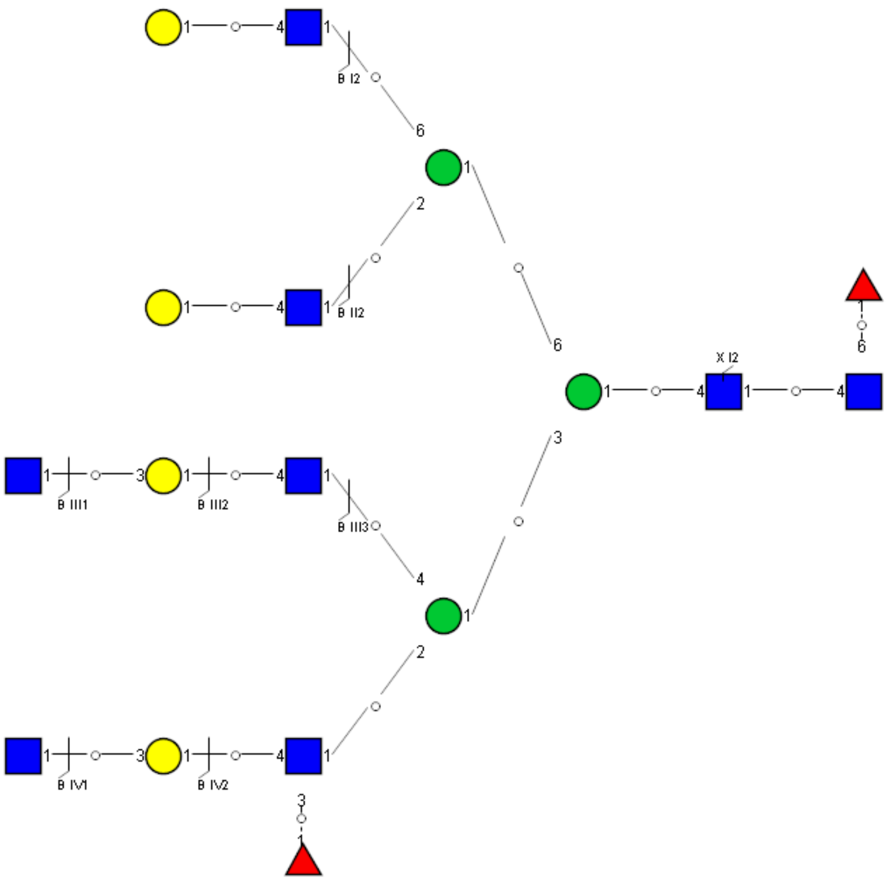

Spectrum Index: 38228(NoBHs:2)

#### Precursor iEF Map

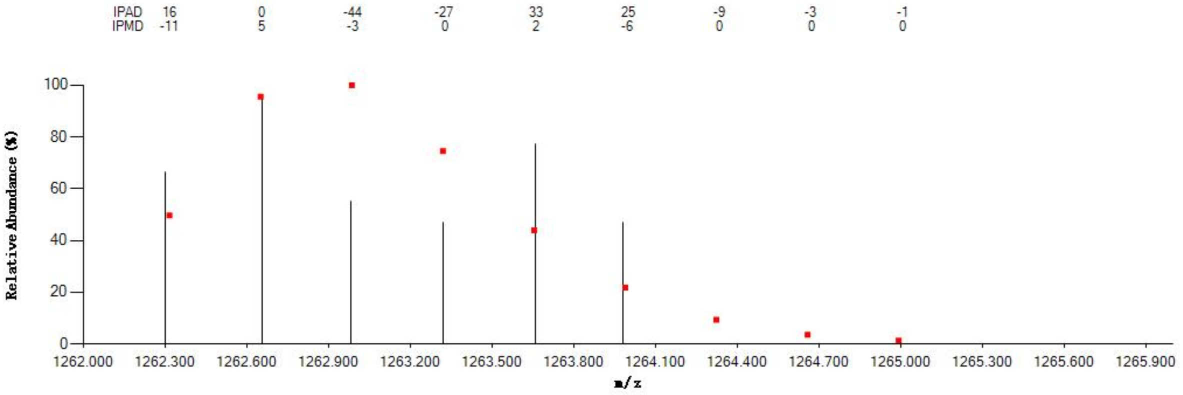

#### Annotated MS/MS Spectrum

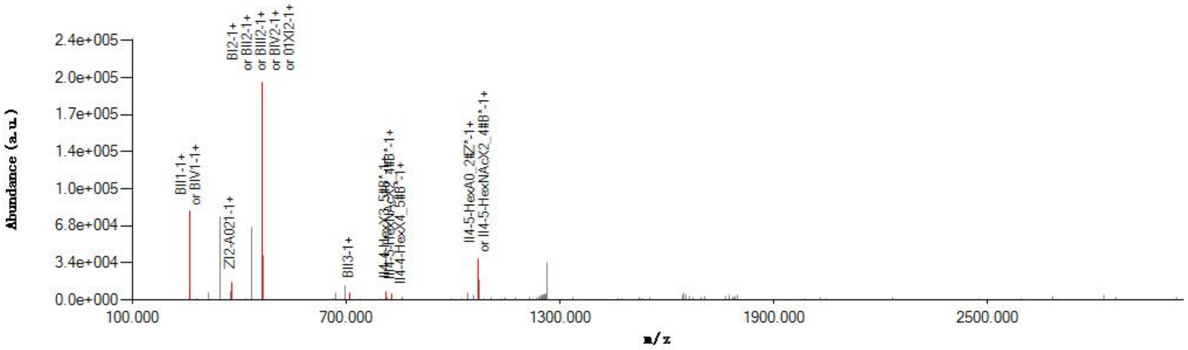

#### Graphical Fragmentation Map

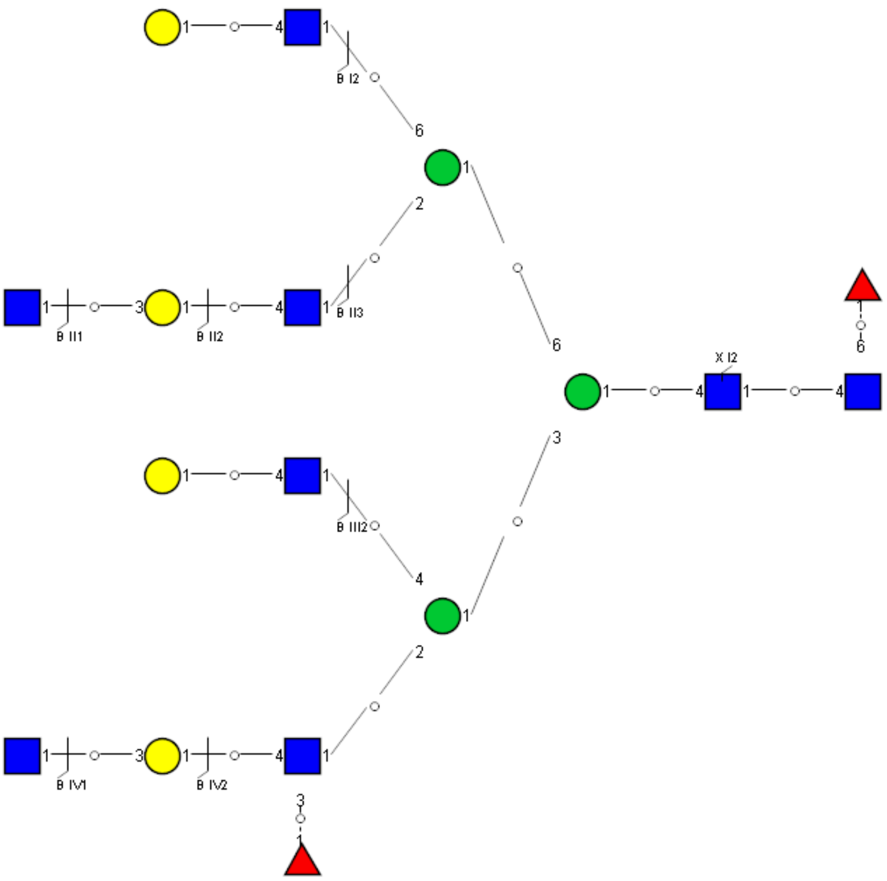

Spectrum Index: 38296(NoBHs:1)

#### Precursor iEF Map

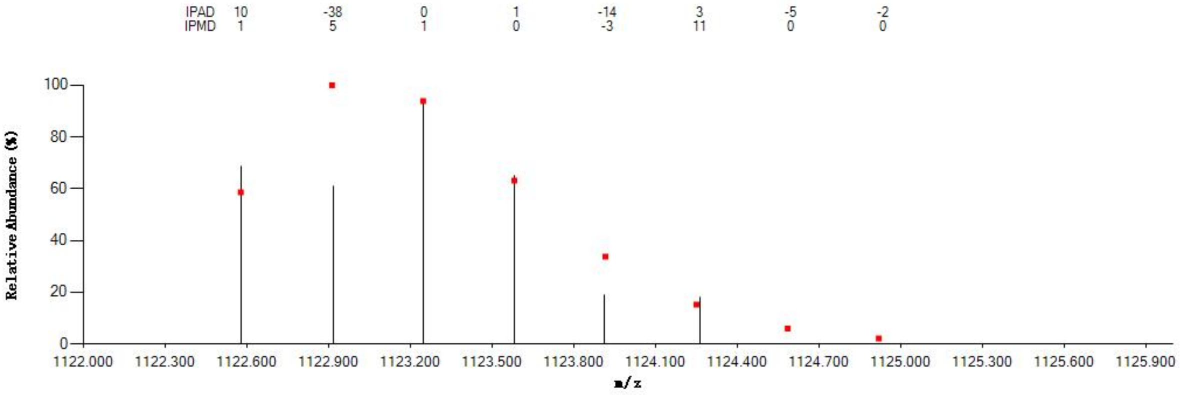


#### Annotated MS/MS Spectrum

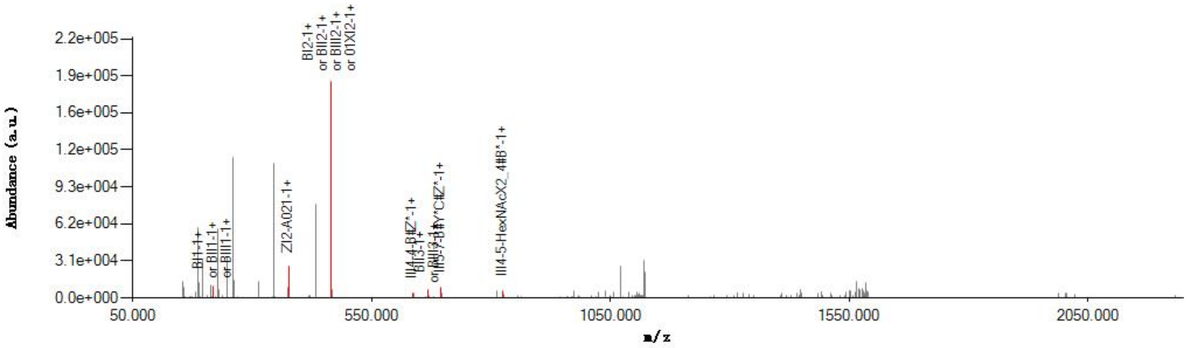


#### Graphical Fragmentation Map

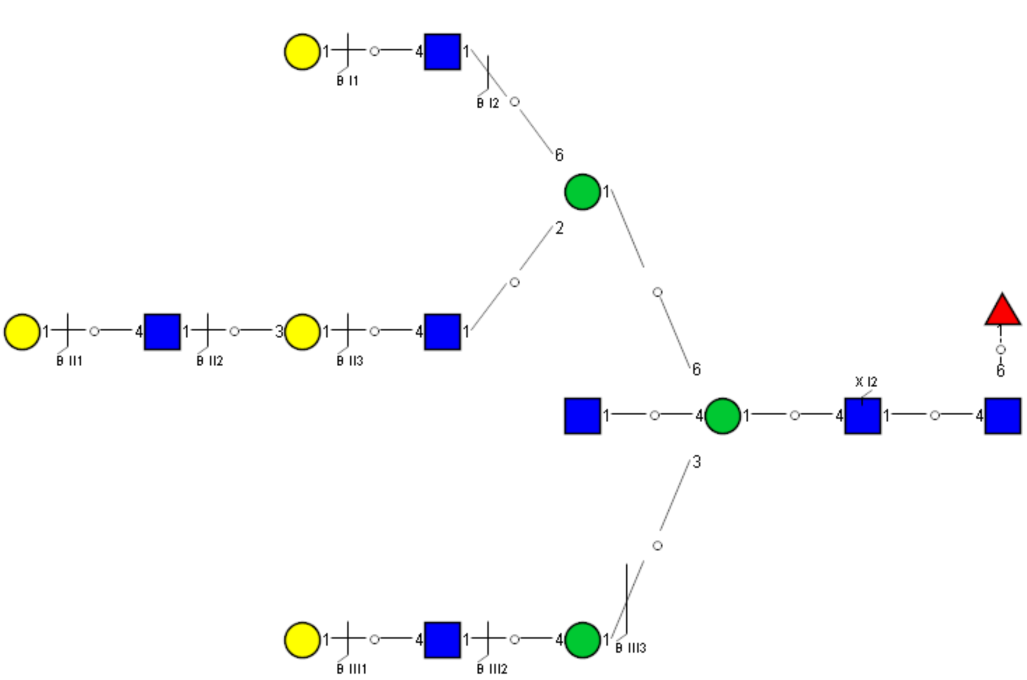

Spectrum Index: 38523(NoBHs:1)

#### Precursor iEF Map

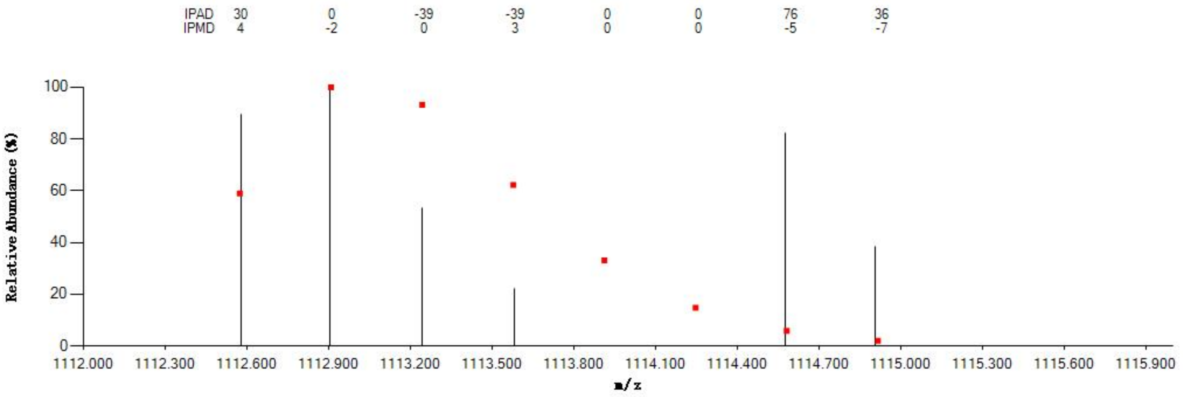


#### Annotated MS/MS Spectrum

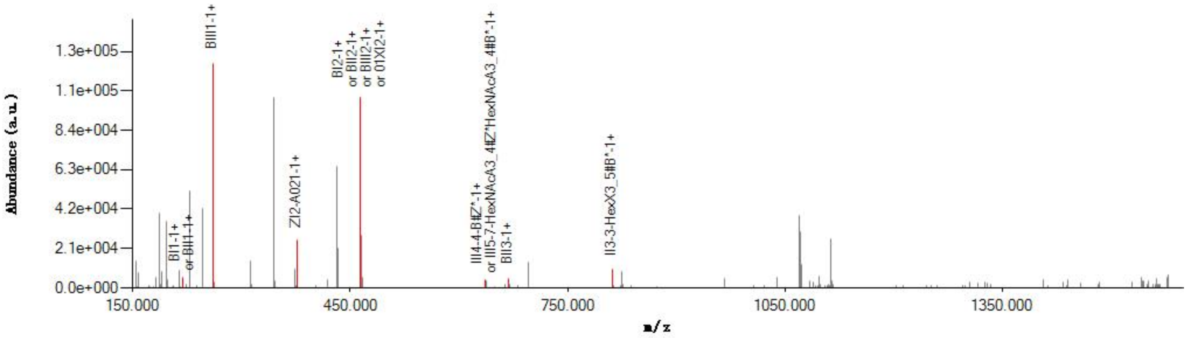


#### Graphical Fragmentation Map

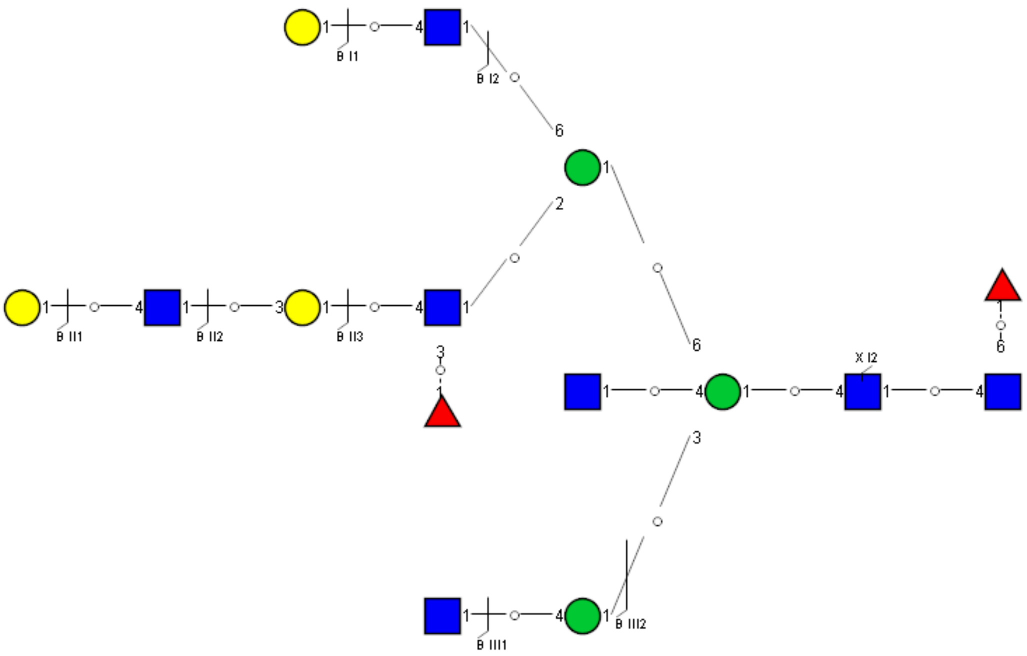

Spectrum Index: 38644(NoBHs:1)

#### Precursor iEF Map

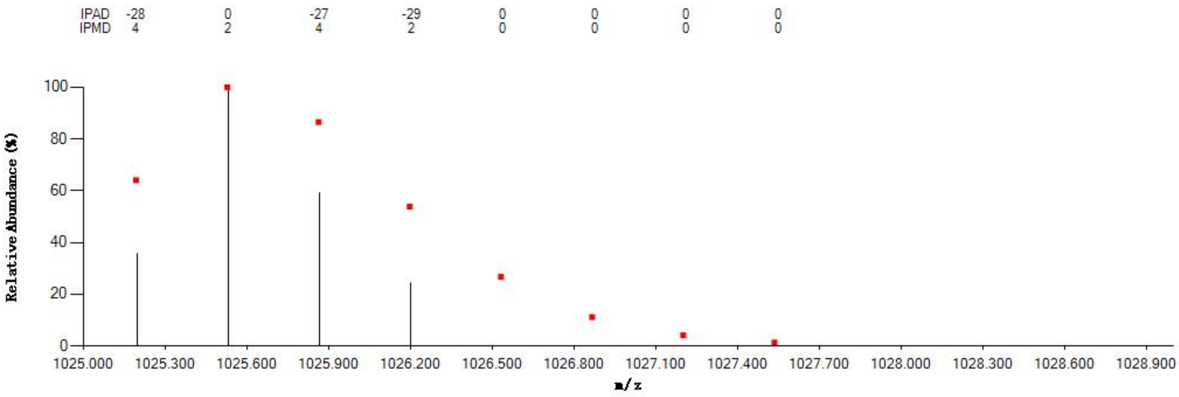


#### Annotated MS/MS Spectrum

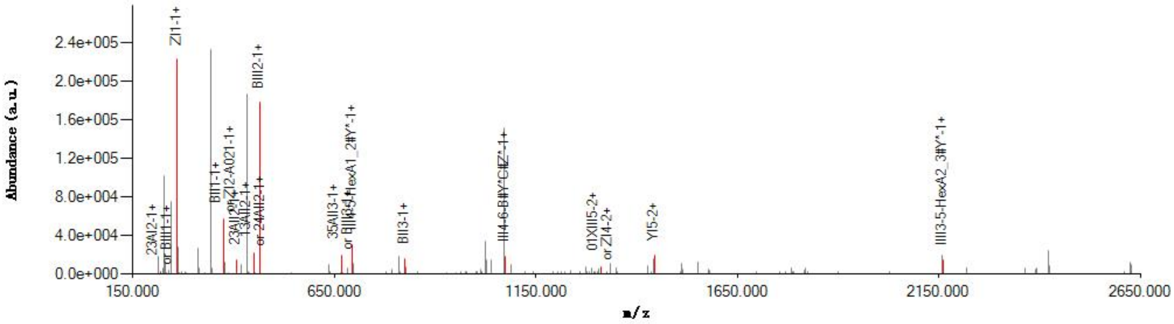


#### Graphical Fragmentation Map

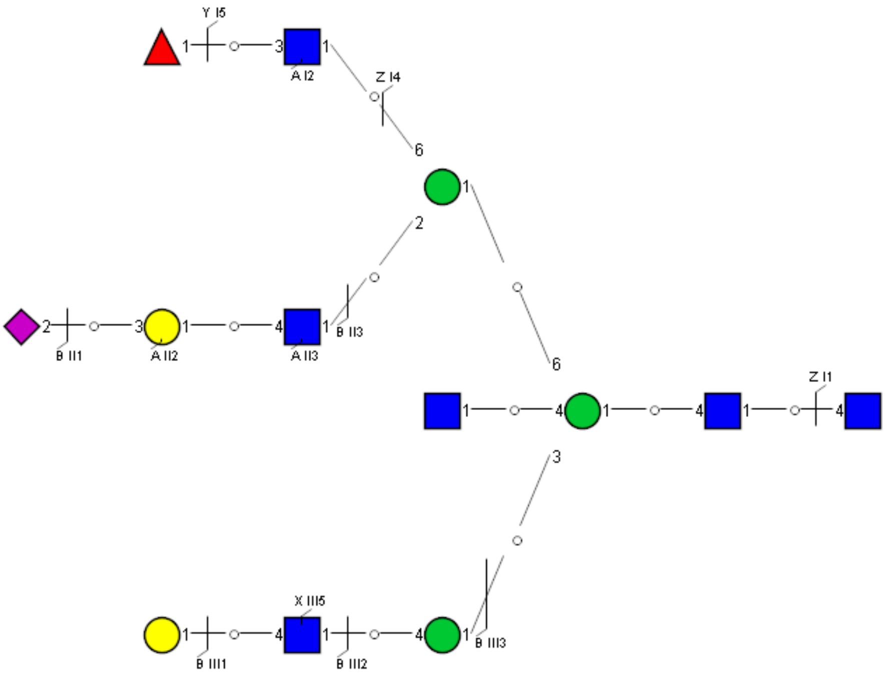

Spectrum Index: 38788(NoBHs:1)

#### Precursor iEF Map

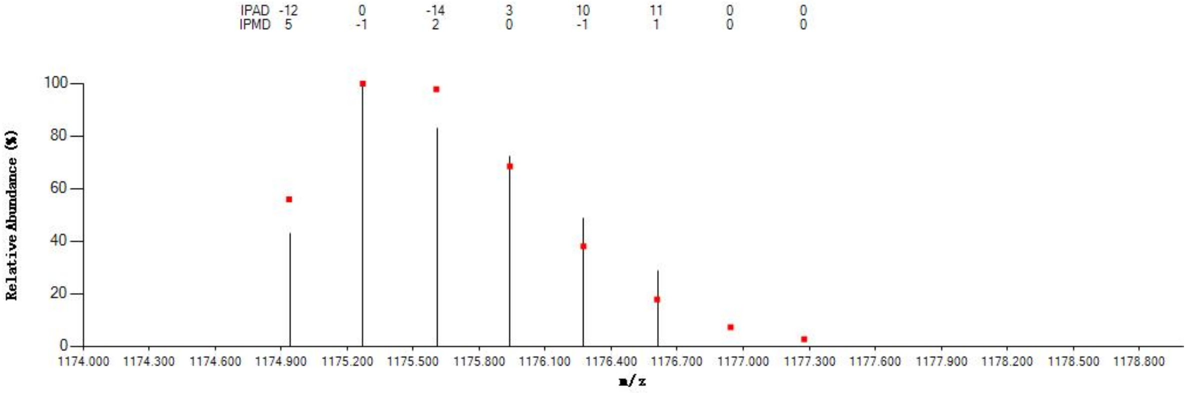


#### Annotated MS/MS Spectrum

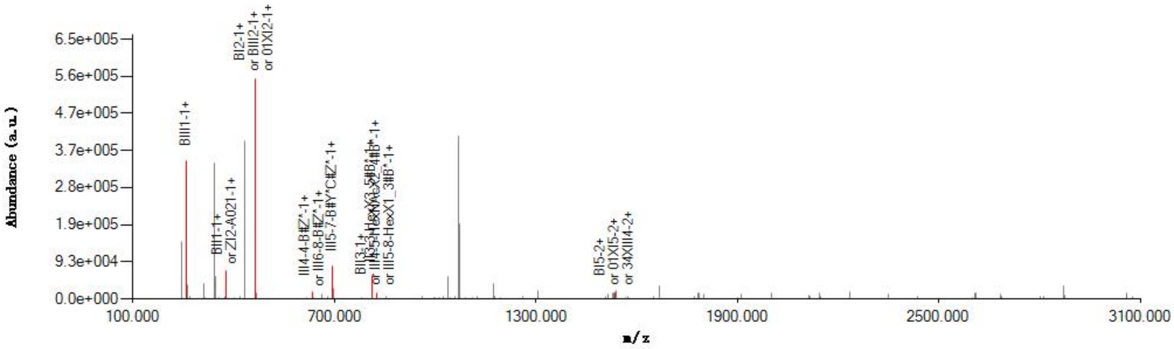


#### Graphical Fragmentation Map

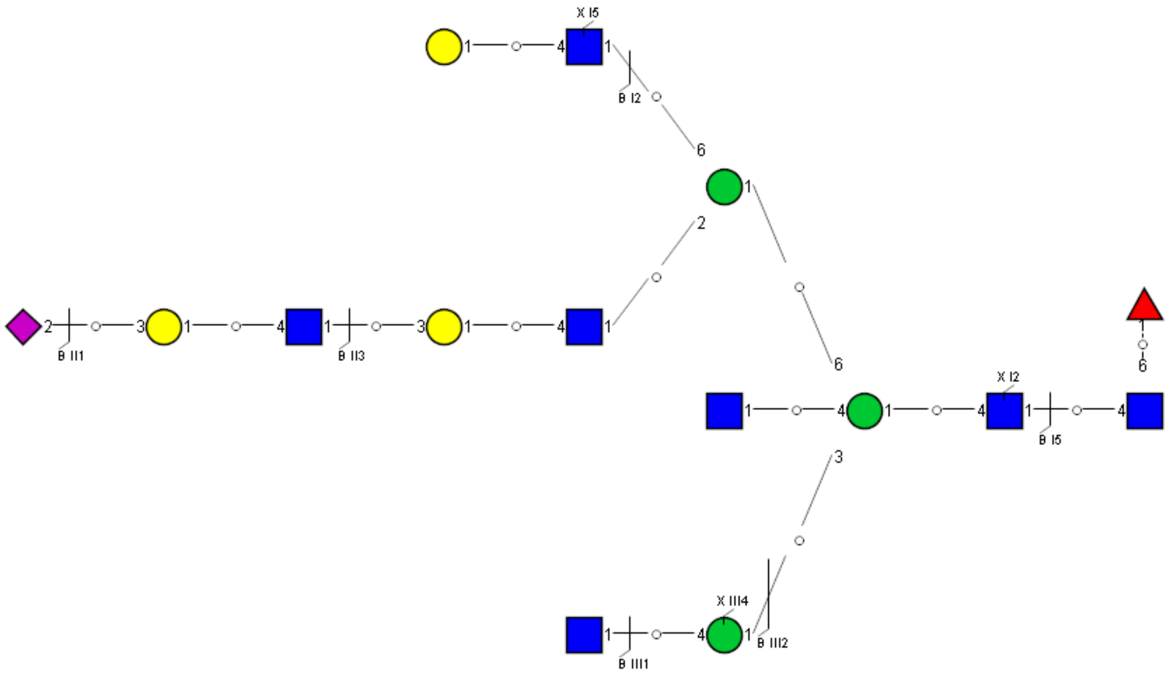

Spectrum Index: 39104(NoBHs:2)

#### Precursor iEF Map

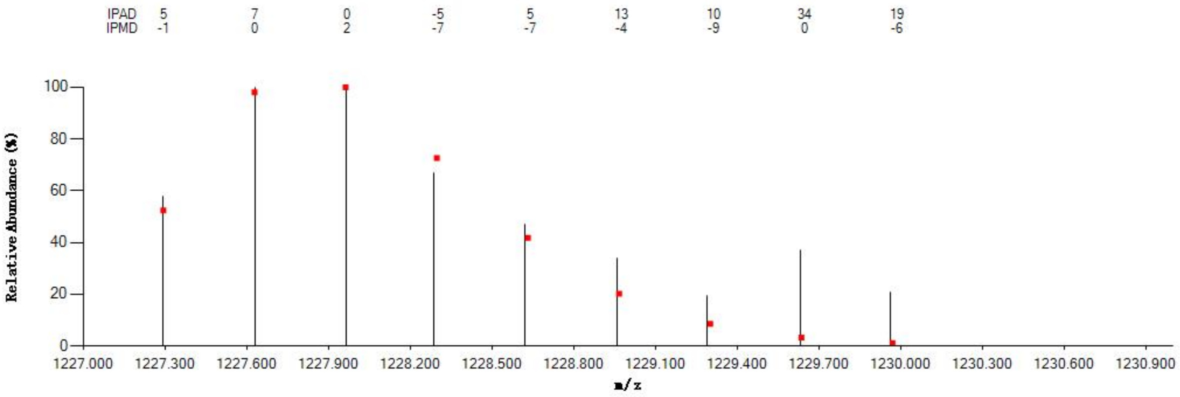


#### Annotated MS/MS Spectrum

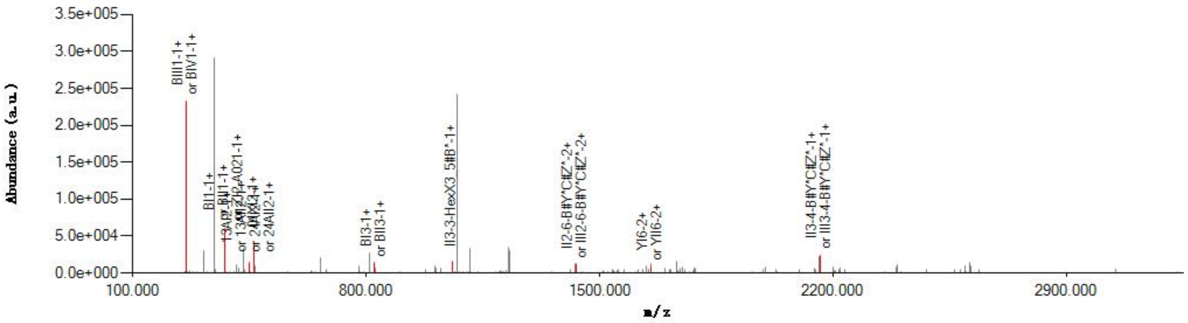


#### Graphical Fragmentation Map

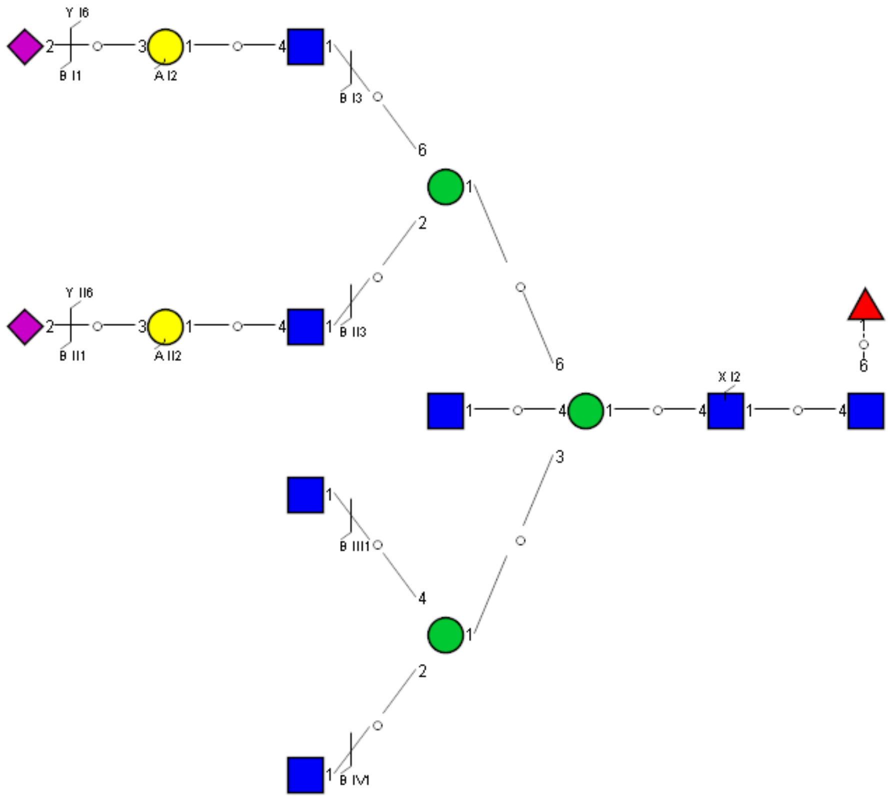

### Spectrum Index: 39104(NoBHs:2)

#### Precursor iEF Map

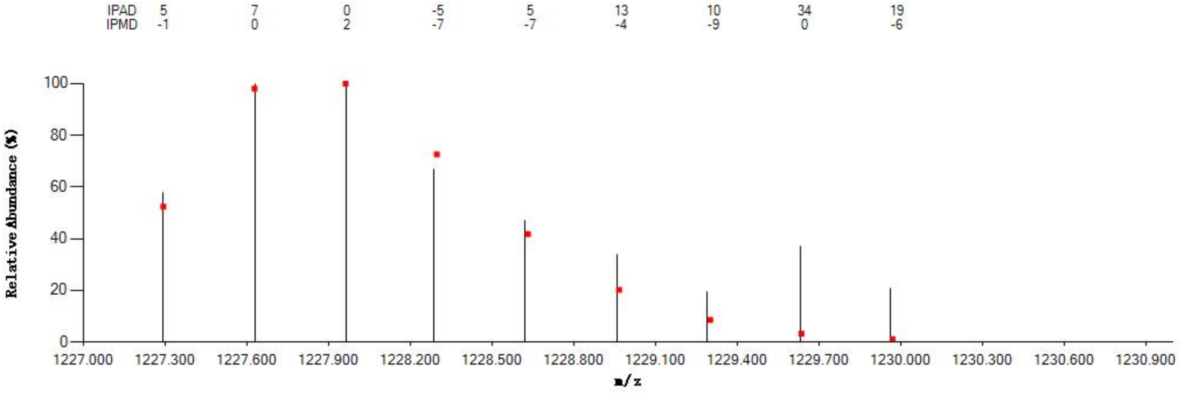


#### Annotated MS/MS Spectrum

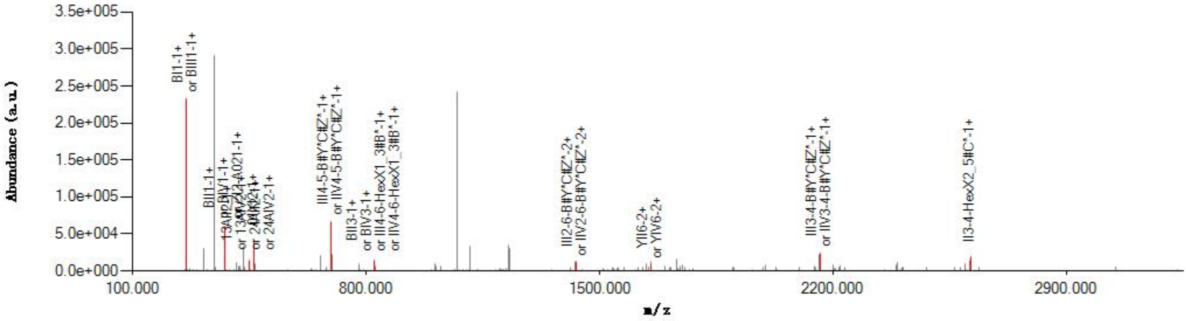


#### Graphical Fragmentation Map

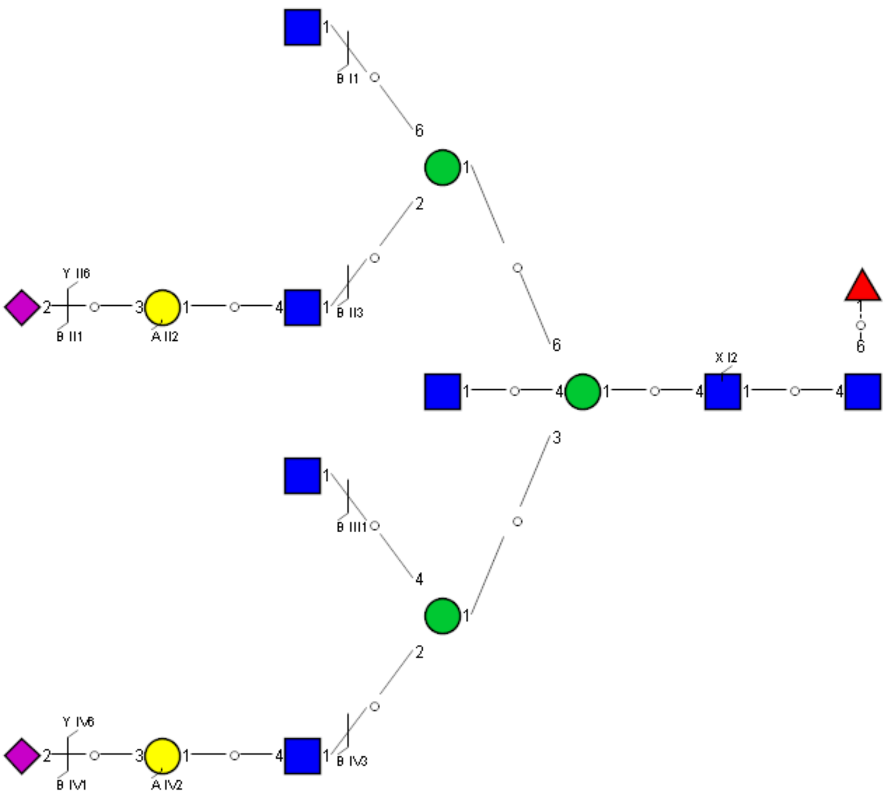

Spectrum Index: 39405(NoBHs:1)

#### Precursor iEF Map

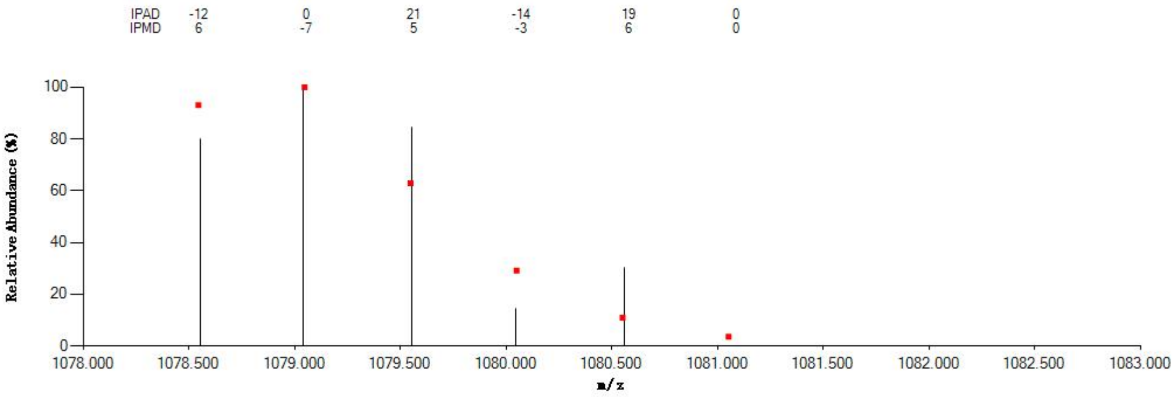


#### Annotated MS/MS Spectrum

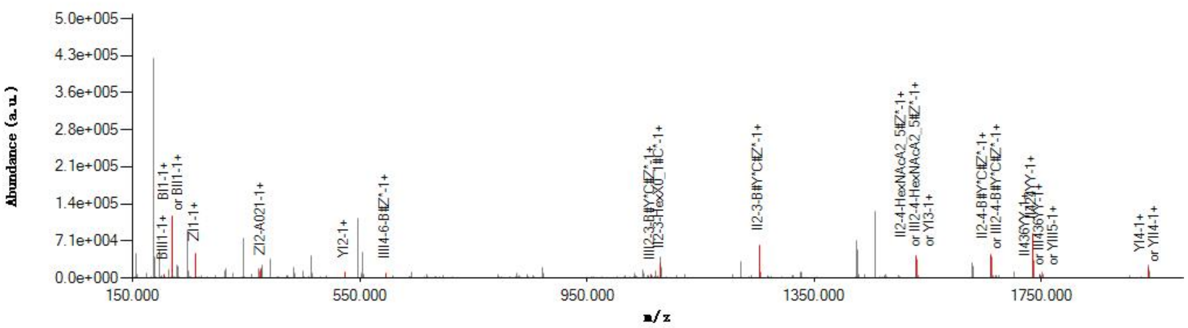


#### Graphical Fragmentation Map

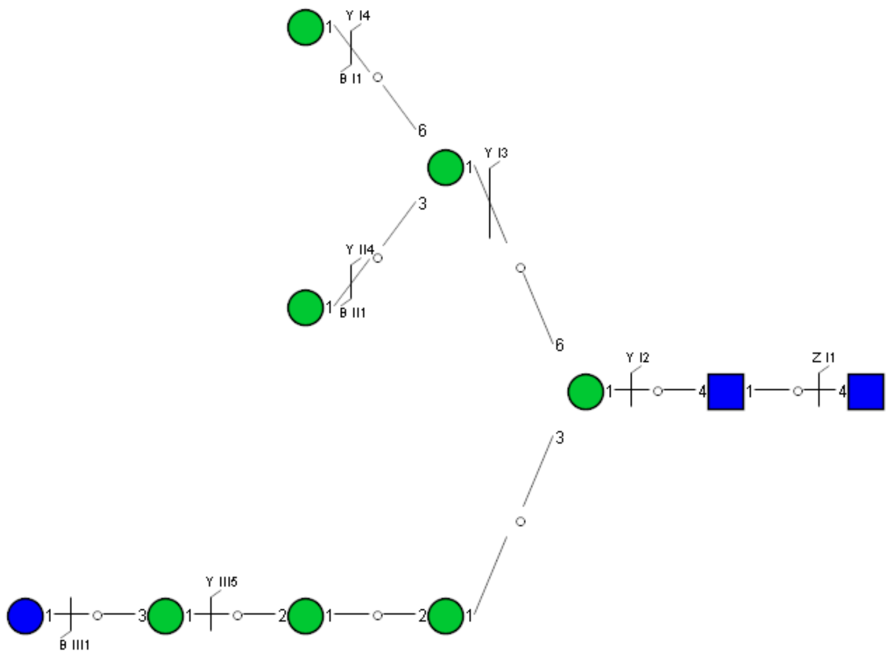

### Spectrum Index: 39674(NoBHs:1)

#### Precursor iEF Map

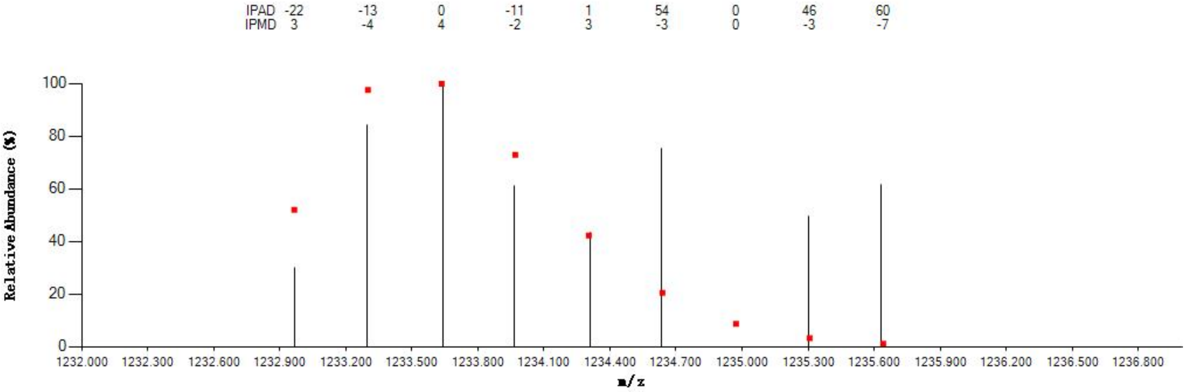


#### Annotated MS/MS Spectrum

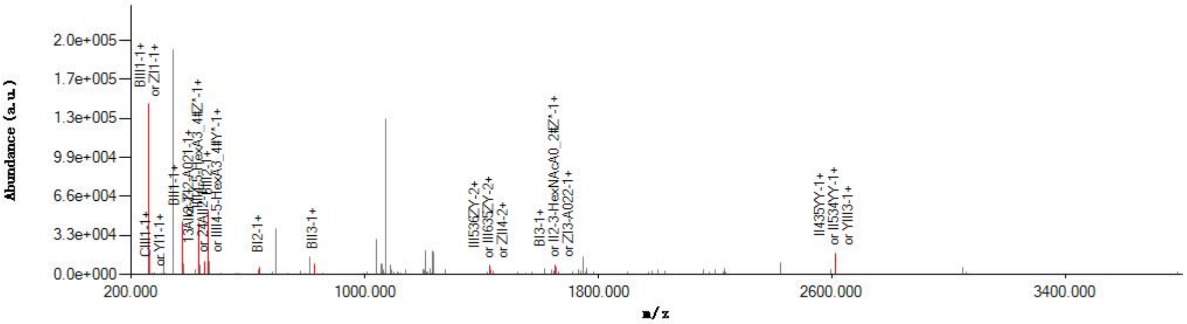


#### Graphical Fragmentation Map

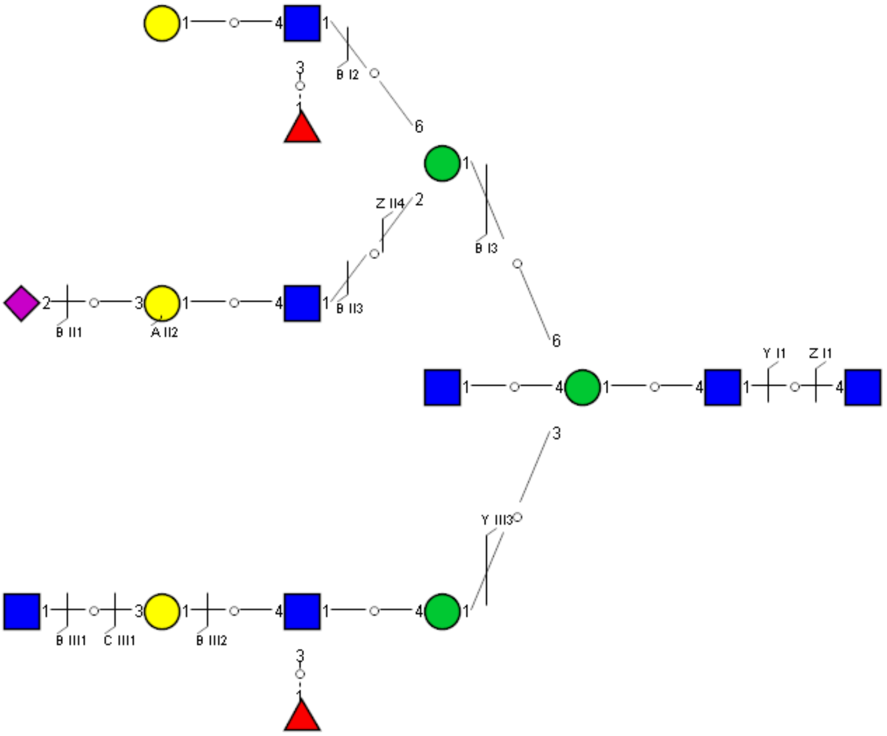

### Spectrum Index: 39733(NoBHs:1)

#### Precursor iEF Map

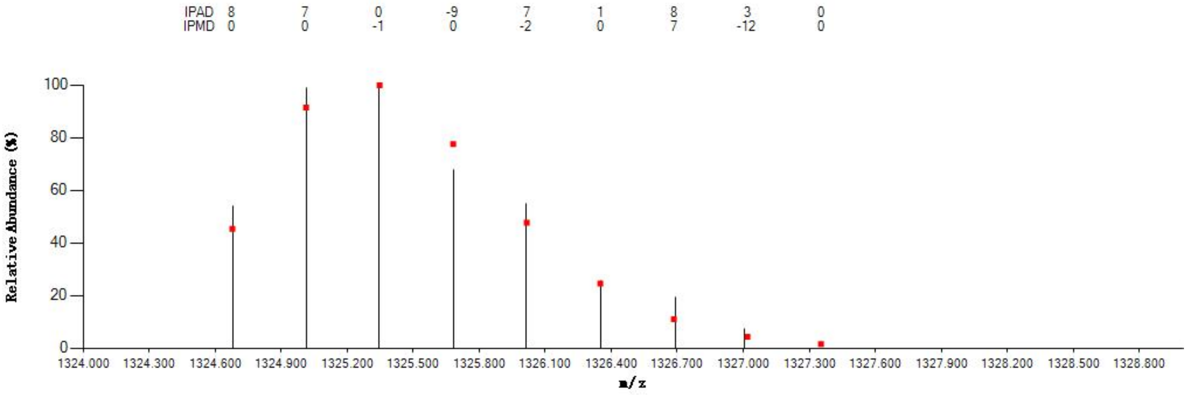


#### Annotated MS/MS Spectrum

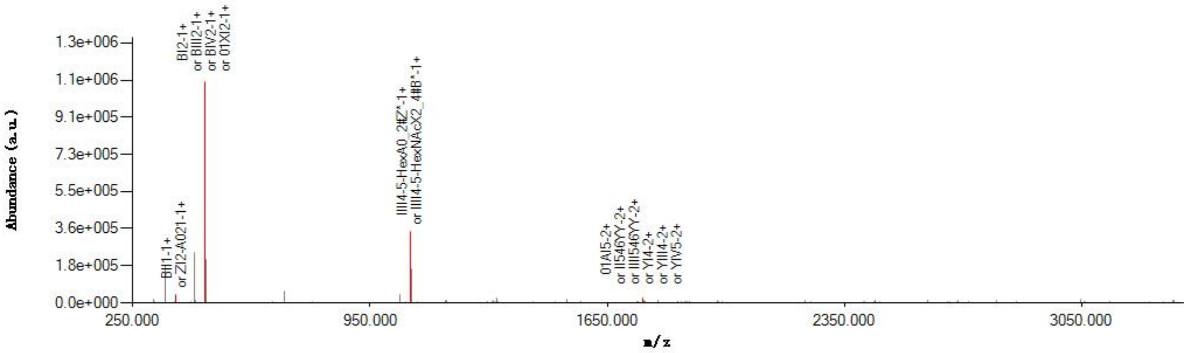


#### Graphical Fragmentation Map

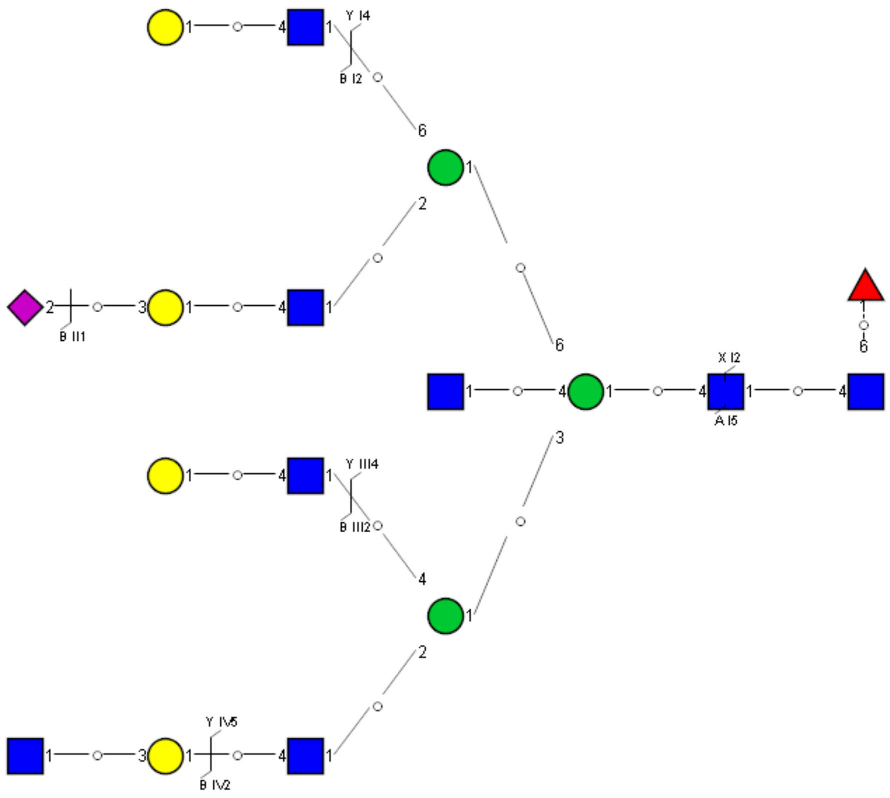

Spectrum Index: 39805(NoBHs:1)

#### Precursor iEF Map

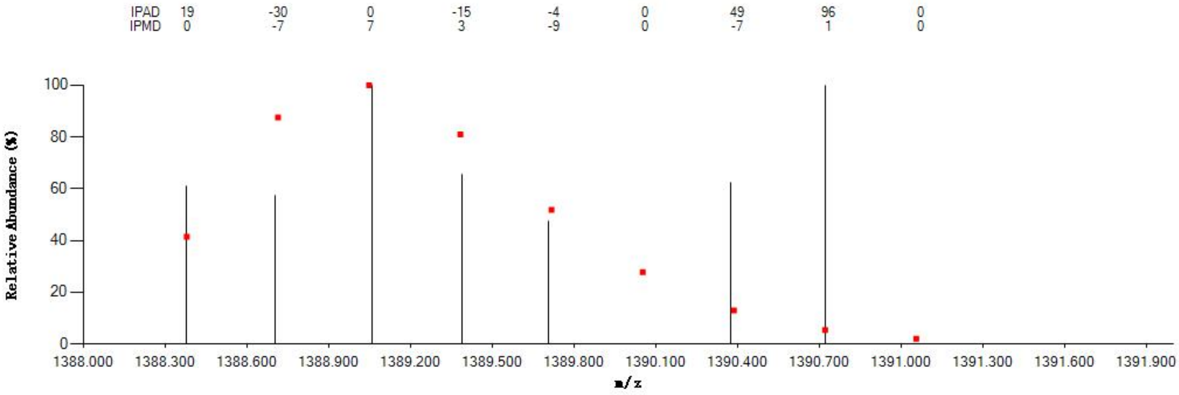


#### Annotated MS/MS Spectrum

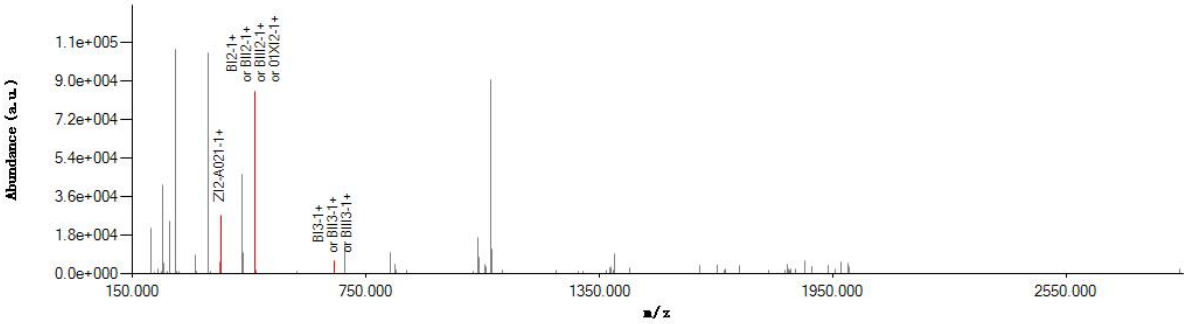


#### Graphical Fragmentation Map

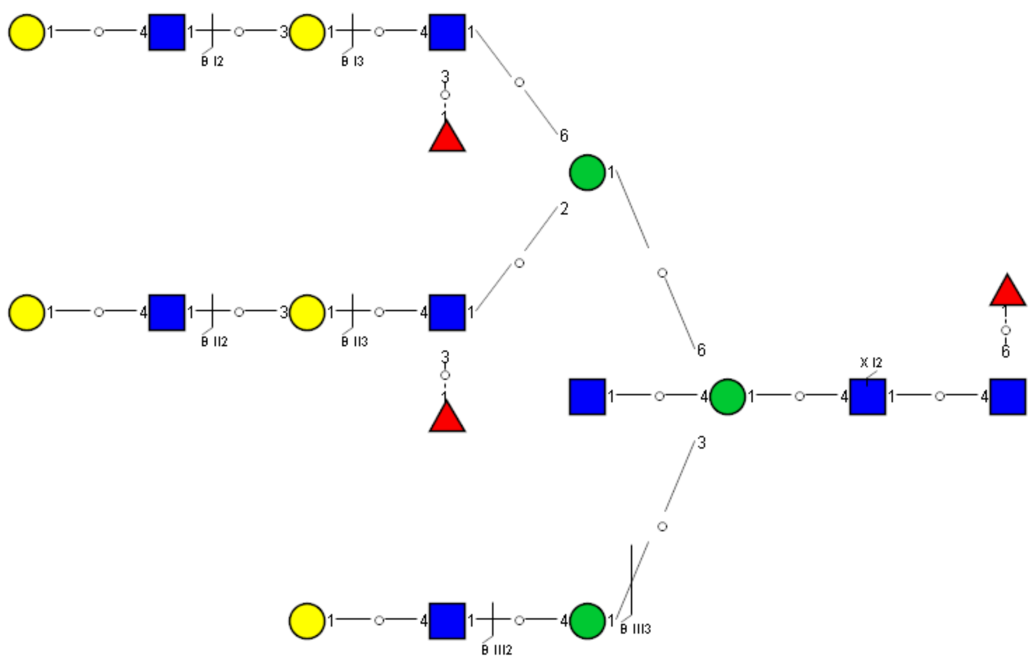

Spectrum Index: 39833(NoBHs:2)

#### Precursor iEF Map

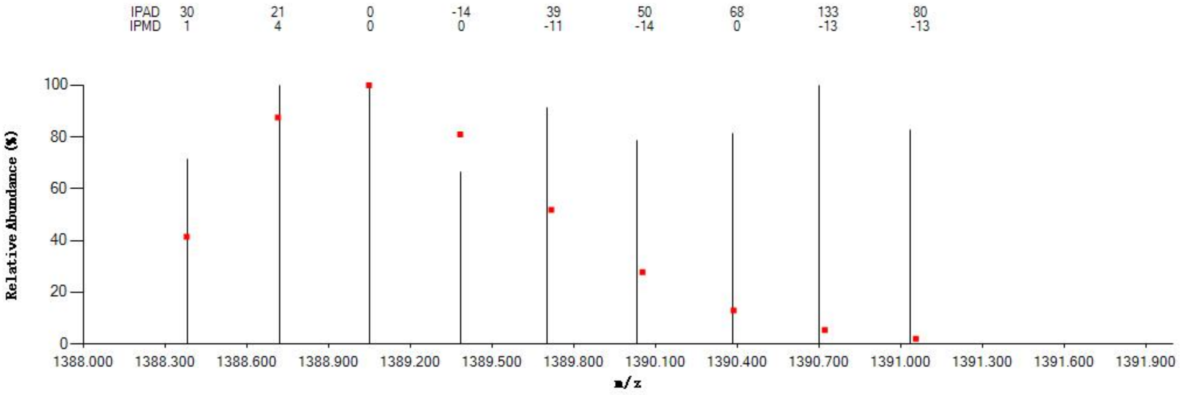


#### Annotated MS/MS Spectrum

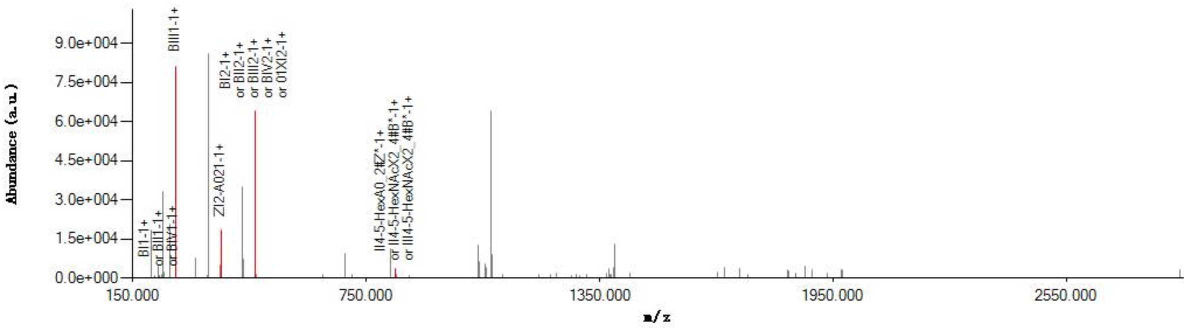


#### Graphical Fragmentation Map

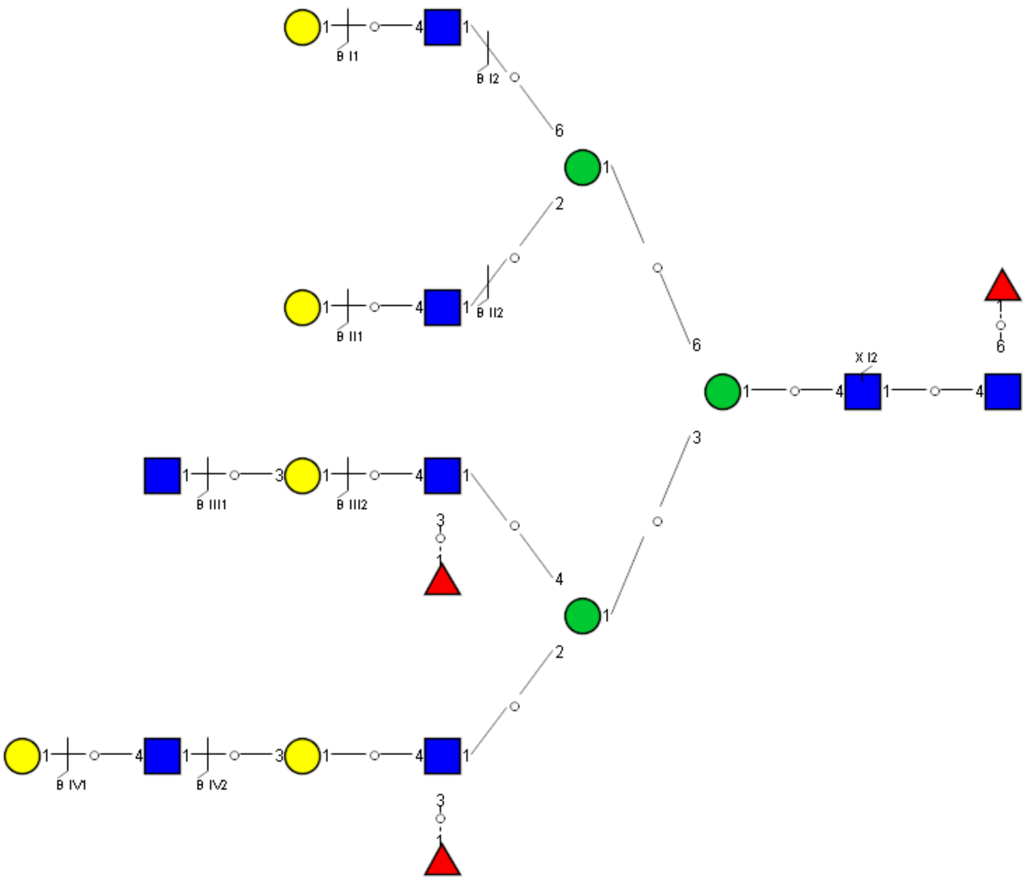

Spectrum Index: 39833(NoBHs:2)

#### Precursor iEF Map

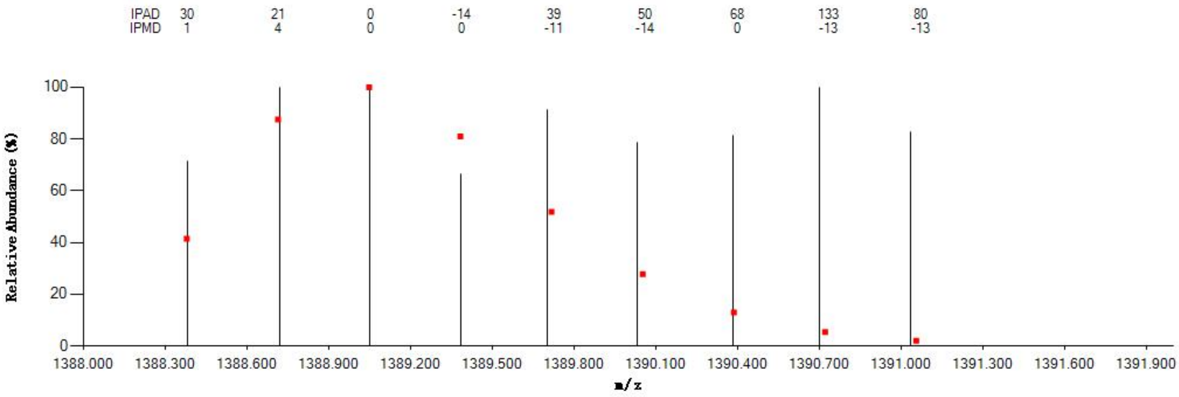


#### Annotated MS/MS Spectrum

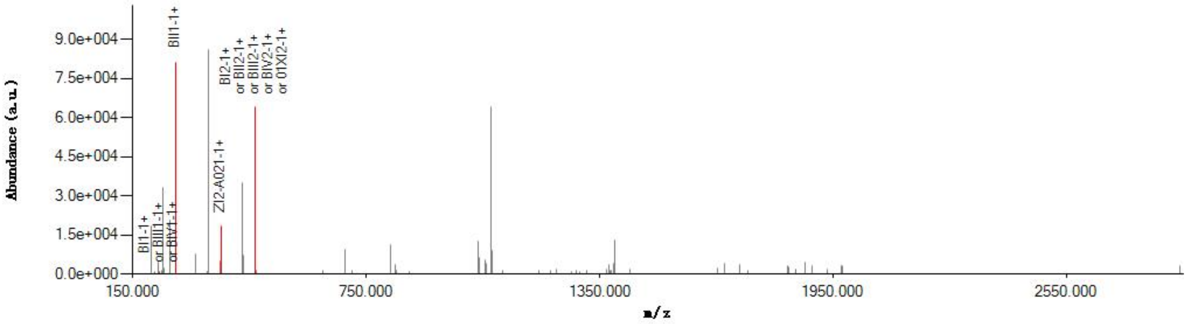


#### Graphical Fragmentation Map

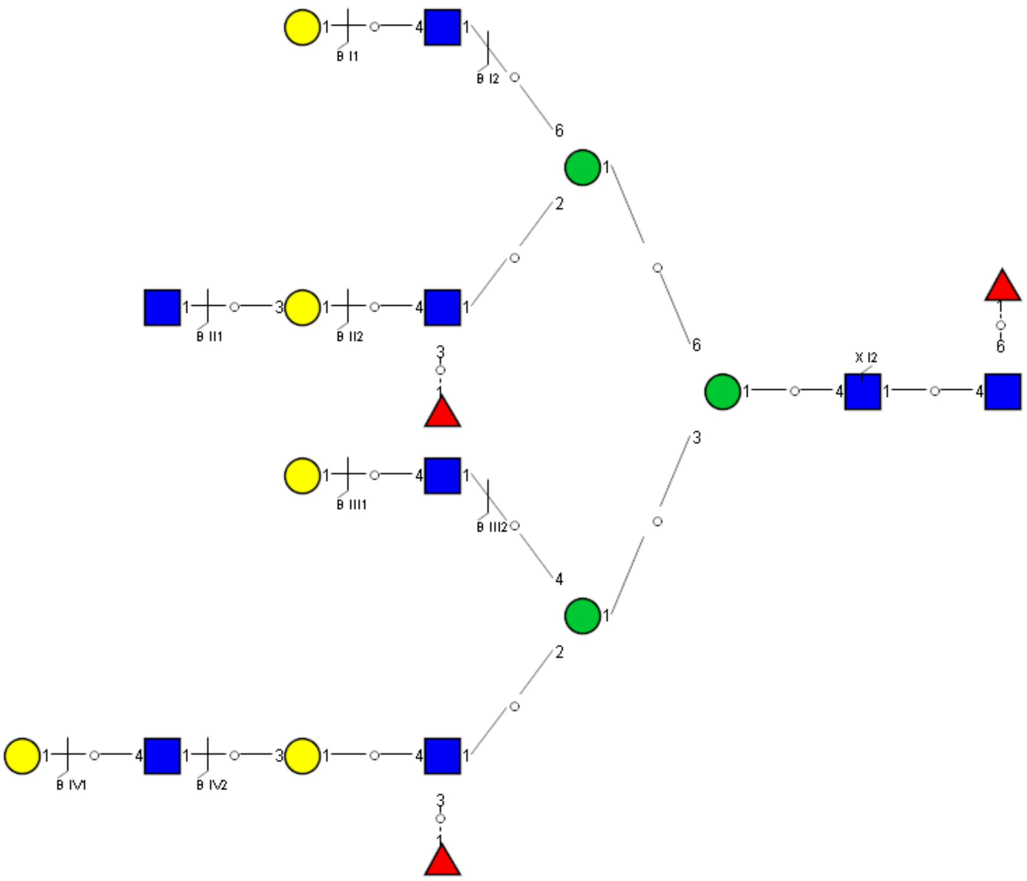

### Spectrum Index: 40219(NoBHs:1)

#### Precursor iEF Map

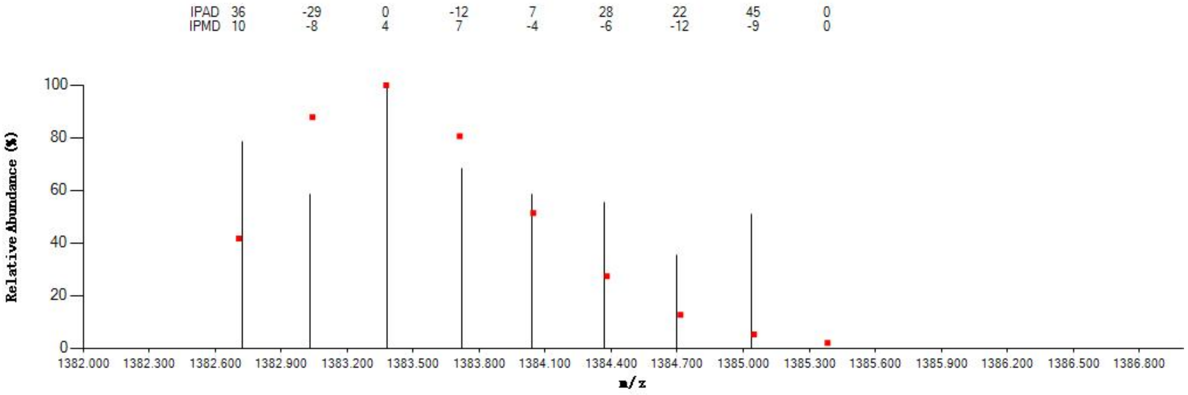


#### Annotated MS/MS Spectrum

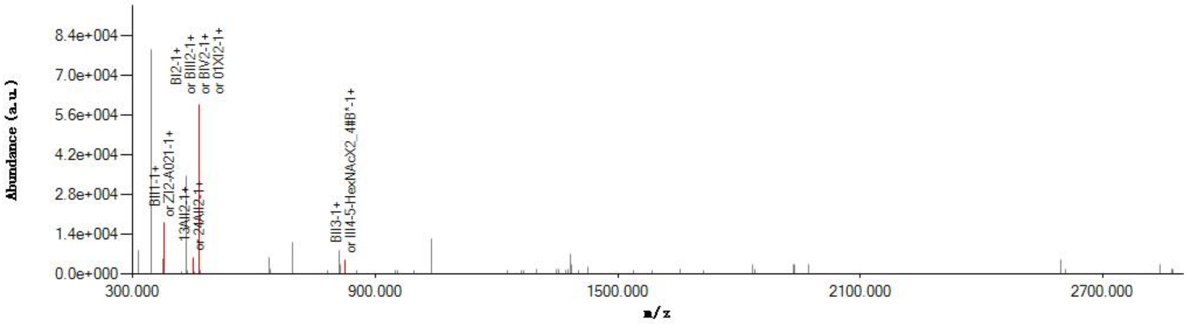


#### Graphical Fragmentation Map

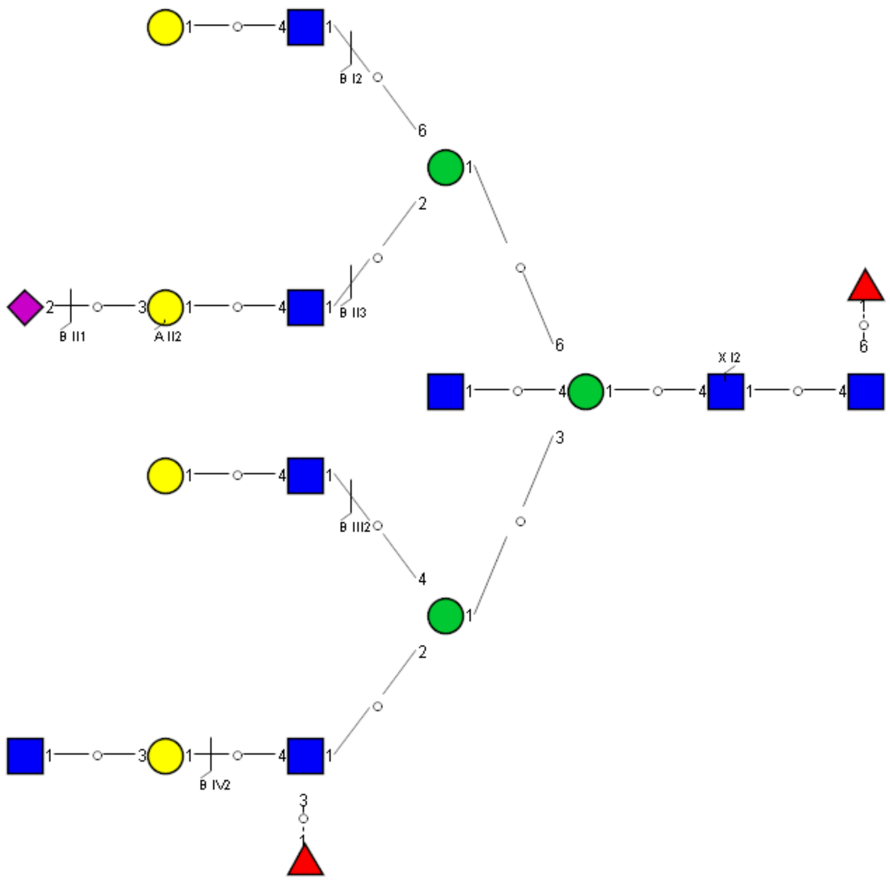

### Spectrum Index: 40279(NoBHs:2)

#### Precursor iEF Map

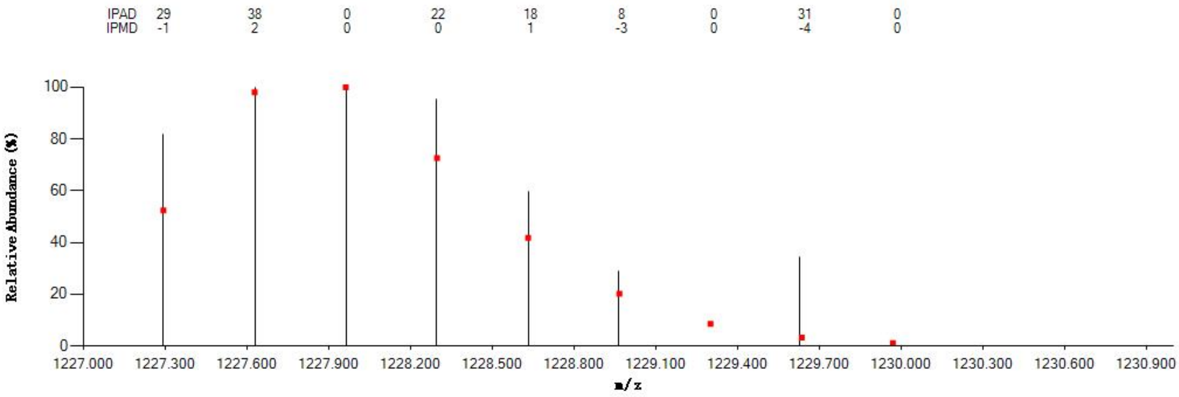

#### Annotated MS/MS Spectrum

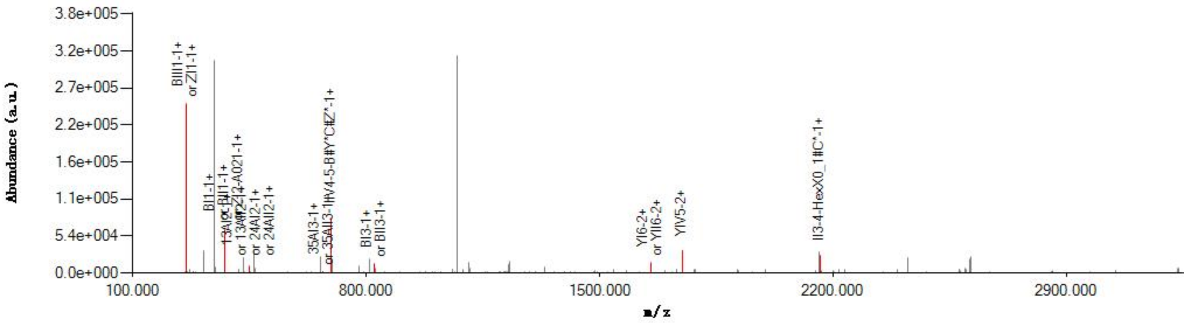

#### Graphical Fragmentation Map

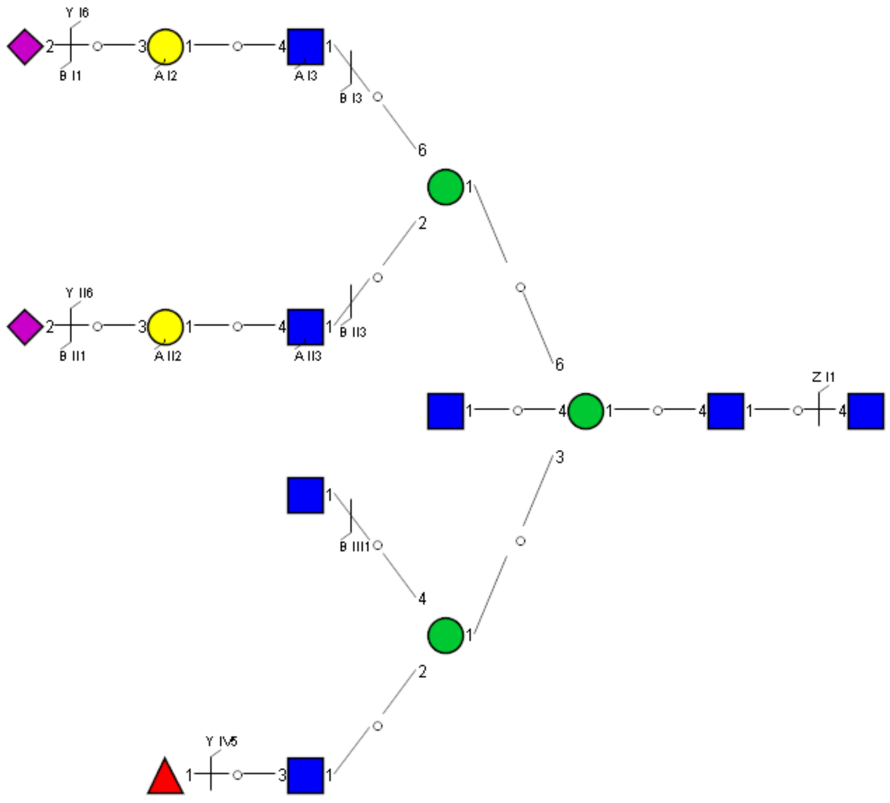

Spectrum Index: 40279(NoBHs:2)

#### Precursor iEF Map

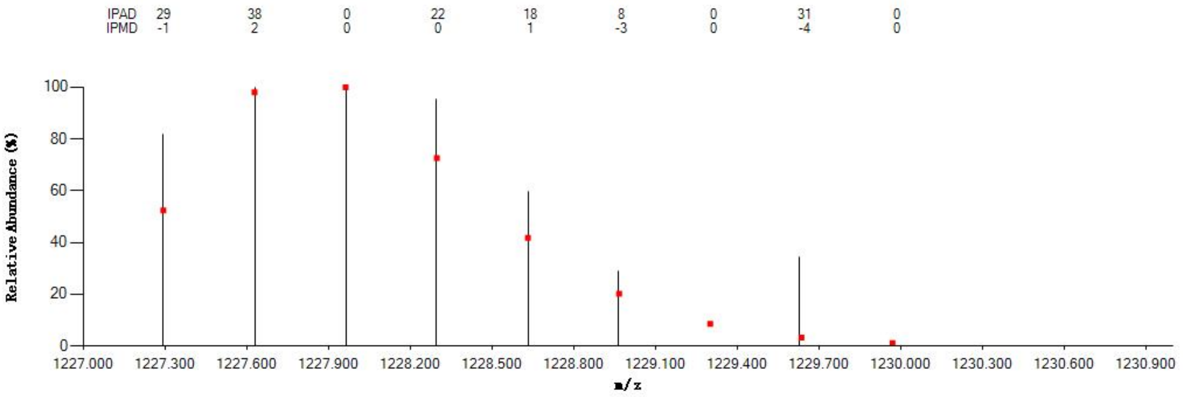


#### Annotated MS/MS Spectrum

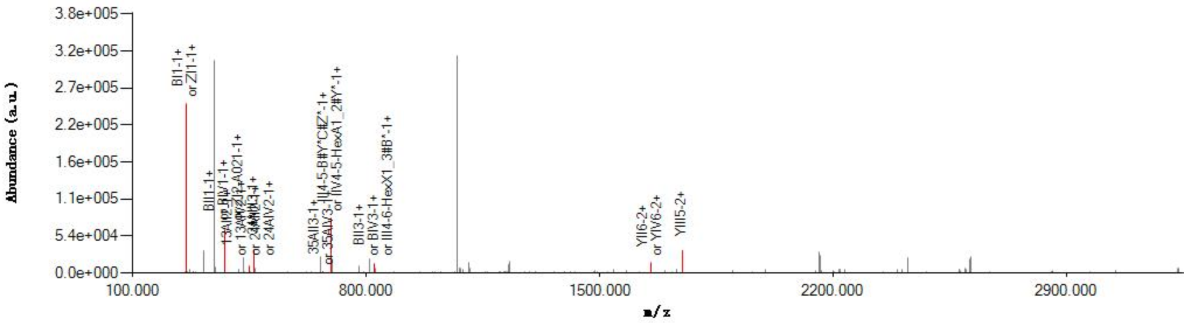


#### Graphical Fragmentation Map

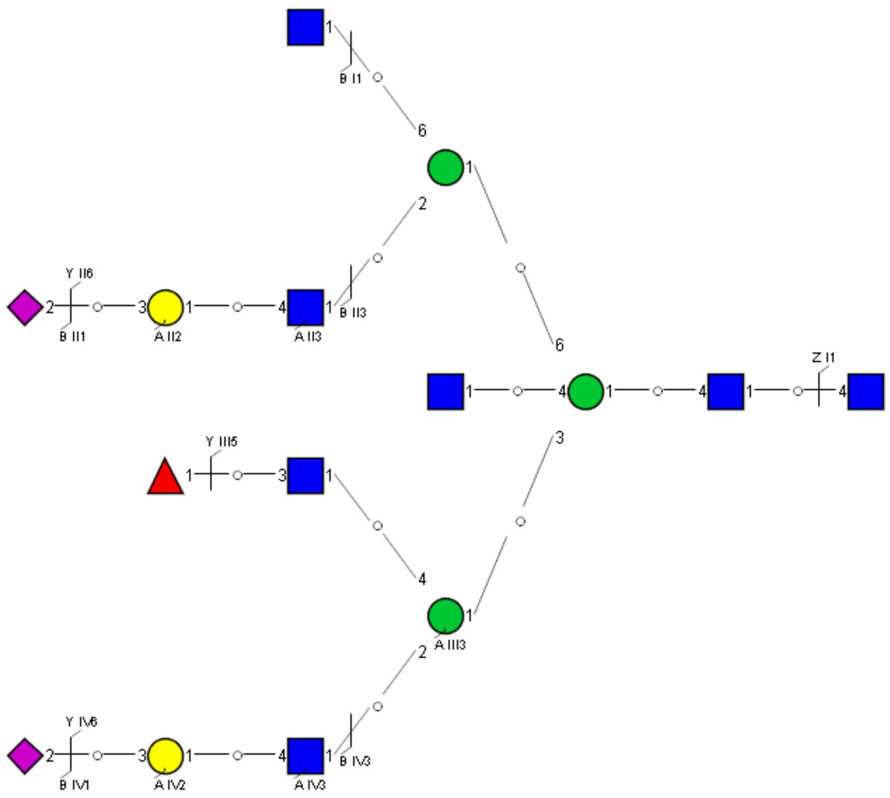

Spectrum Index: 40490(NoBHs:2)

#### Precursor iEF Map

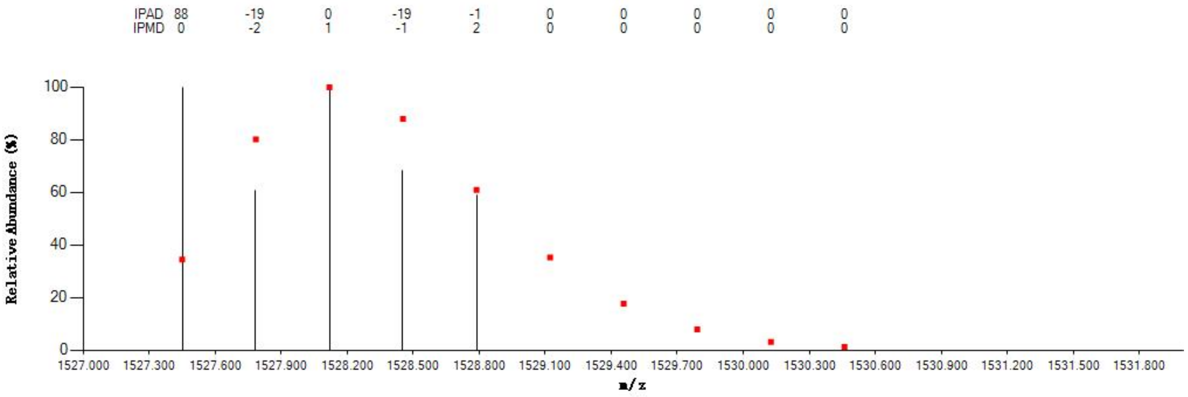


#### Annotated MS/MS Spectrum

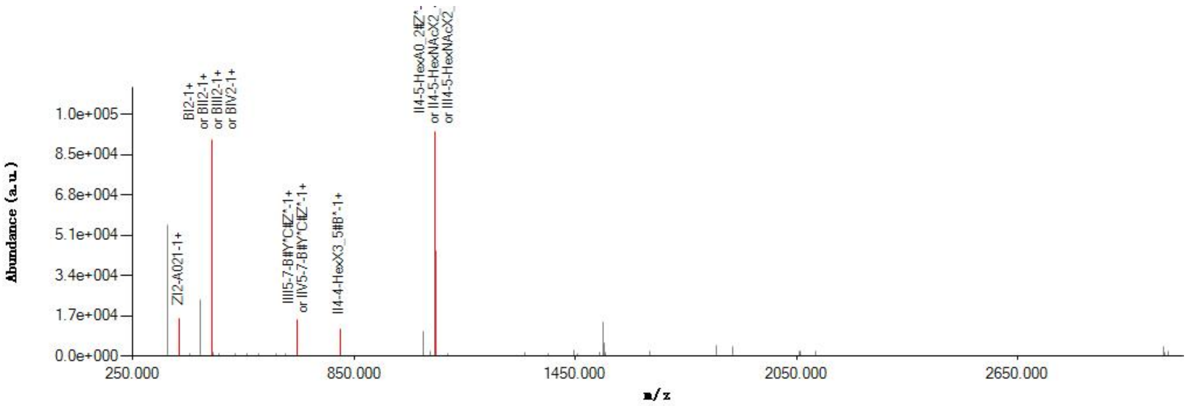


#### Graphical Fragmentation Map

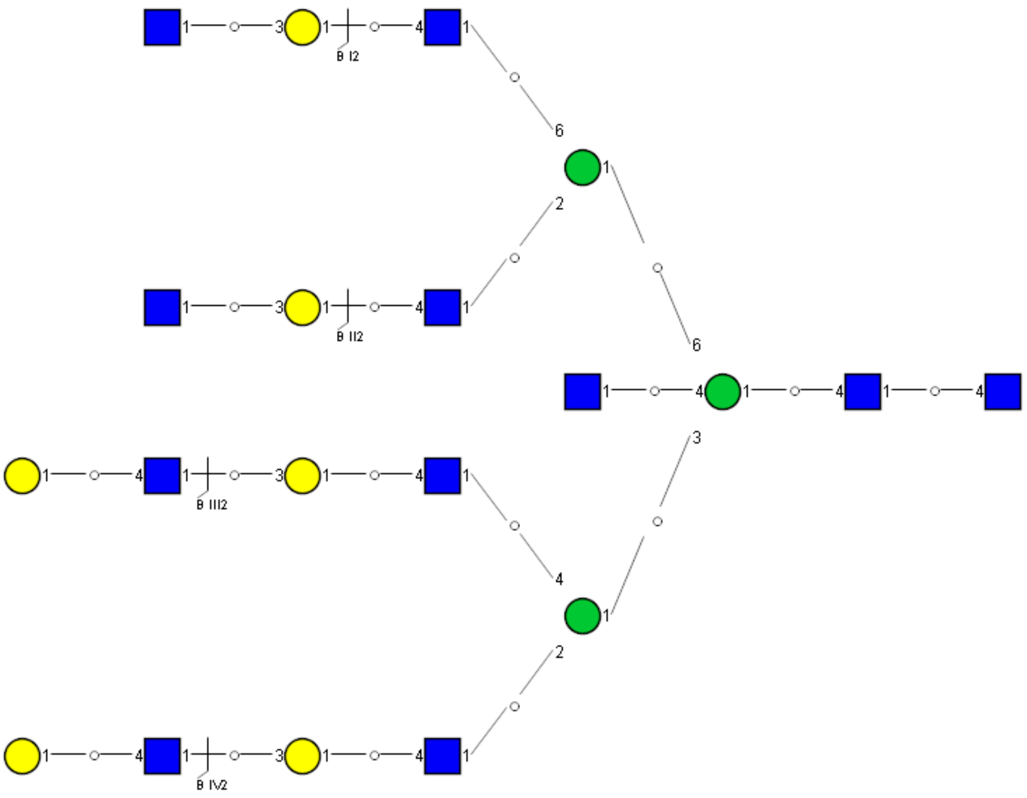

### Spectrum Index: 40490(NoBHs:2)

#### Precursor iEF Map

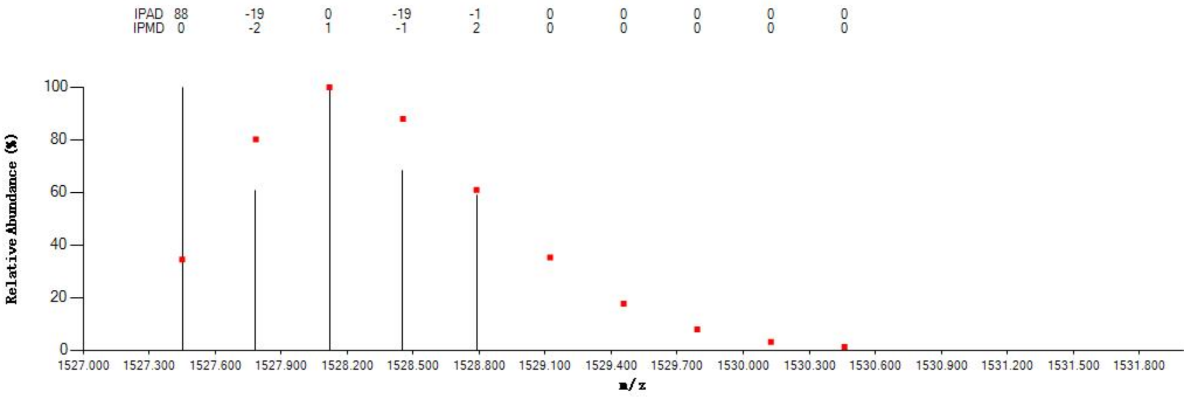


#### Annotated MS/MS Spectrum

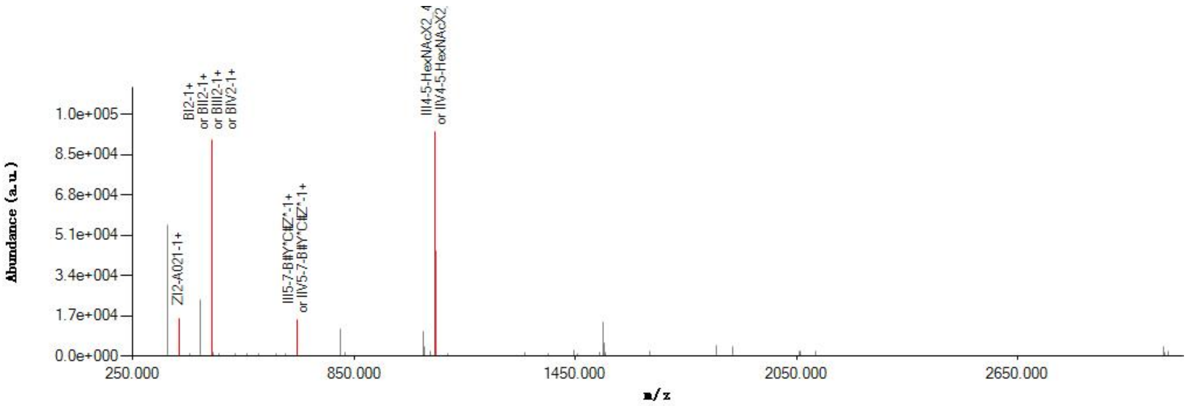


#### Graphical Fragmentation Map

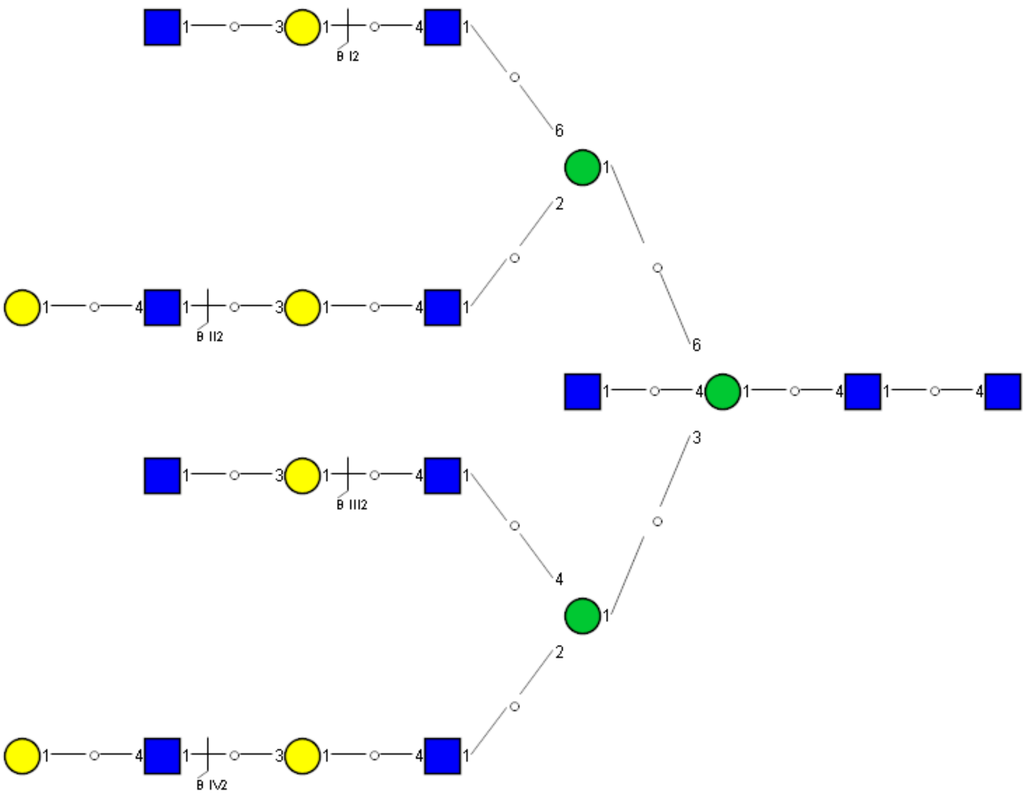

Spectrum Index: 27224(NoBHs:1)

#### Precursor iEF Map

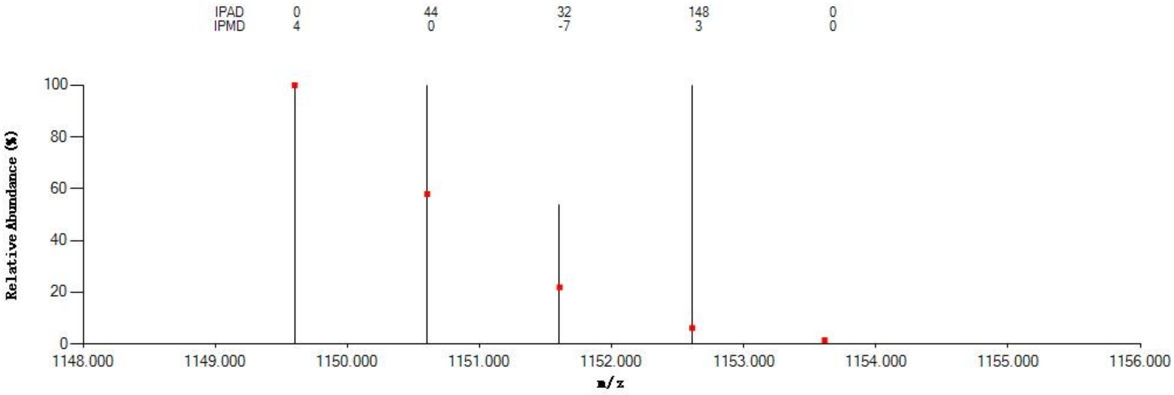


#### Annotated MS/MS Spectrum

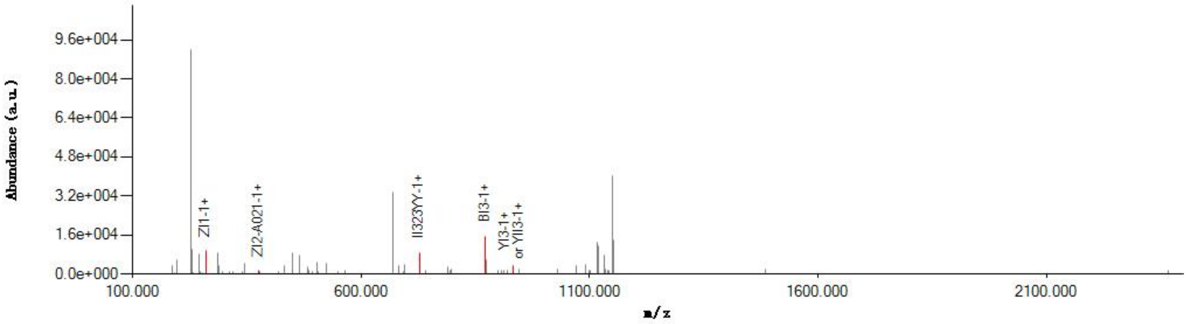


#### Graphical Fragmentation Map

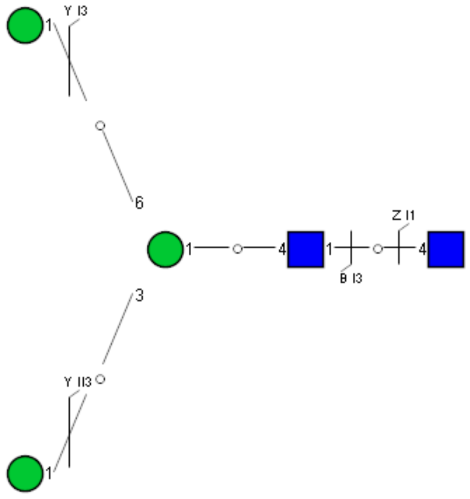

Spectrum Index: 29603(NoBHs:1)

#### Precursor iEF Map

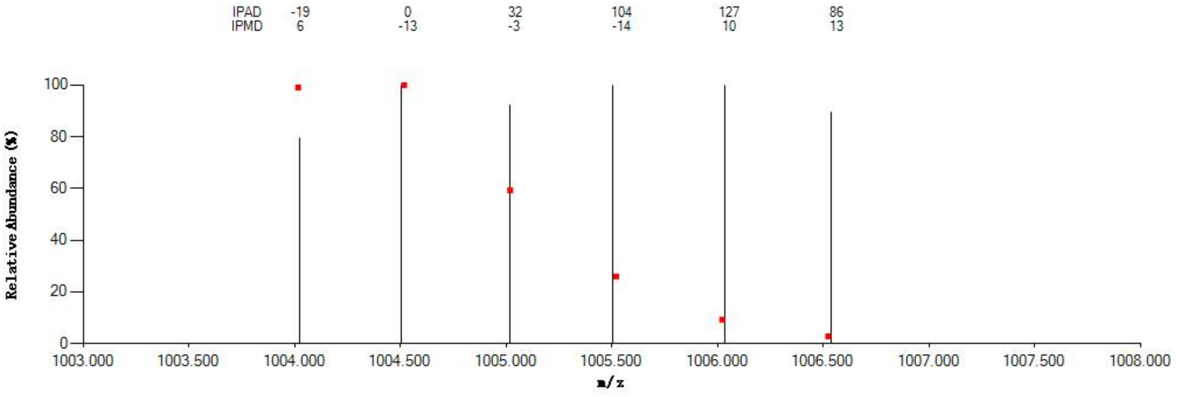


#### Annotated MS/MS Spectrum

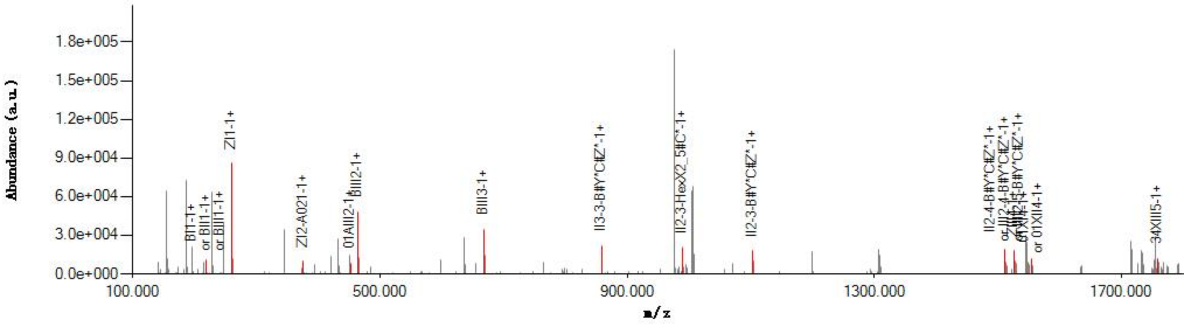


#### Graphical Fragmentation Map

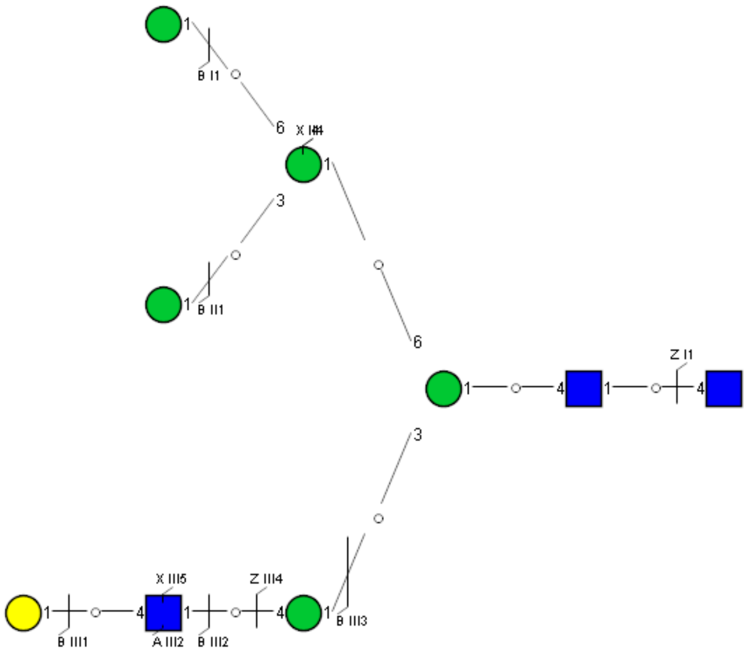

Spectrum Index: 31251(NoBHs:1)

#### Precursor iEF Map

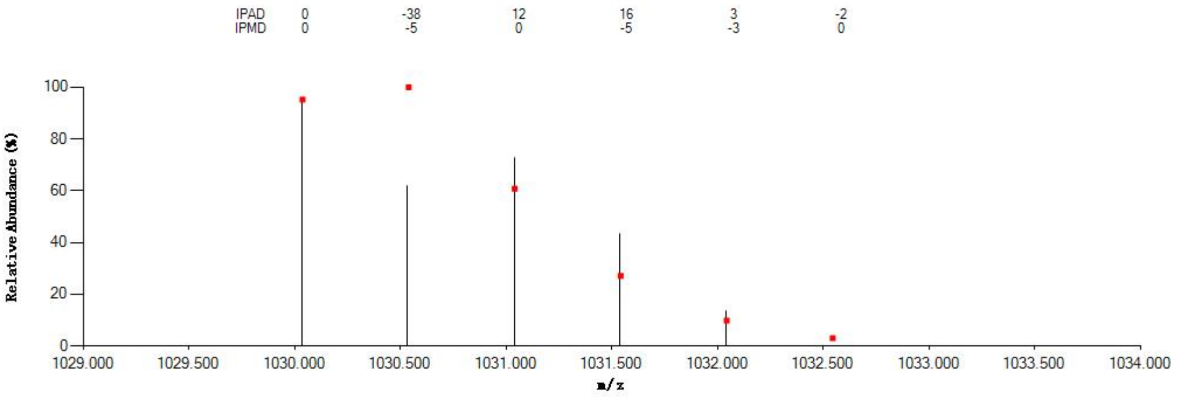


#### Annotated MS/MS Spectrum

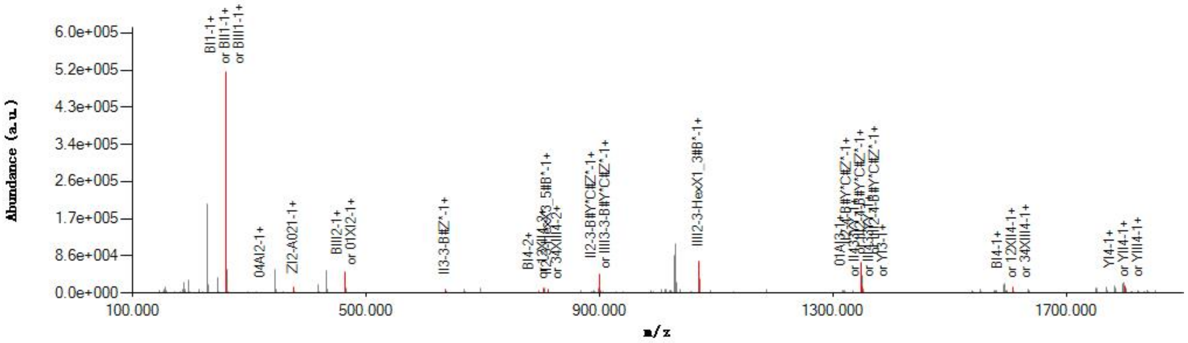


#### Graphical Fragmentation Map

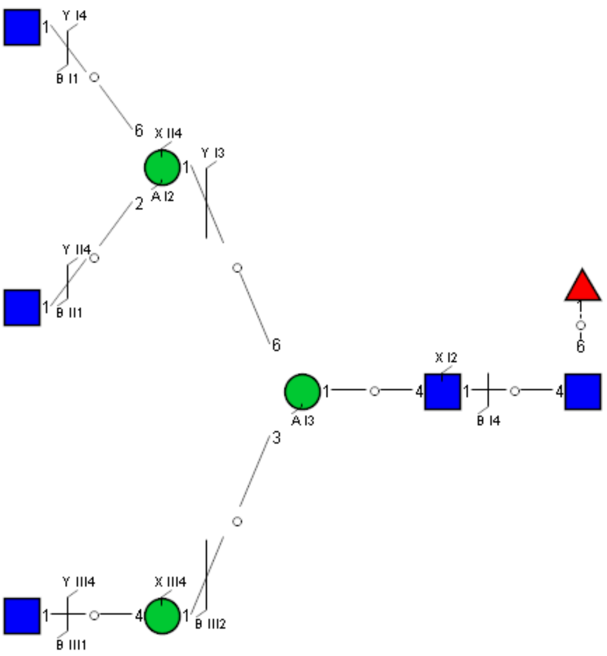

Spectrum Index: 33179(NoBHs:1)

#### Precursor iEF Map

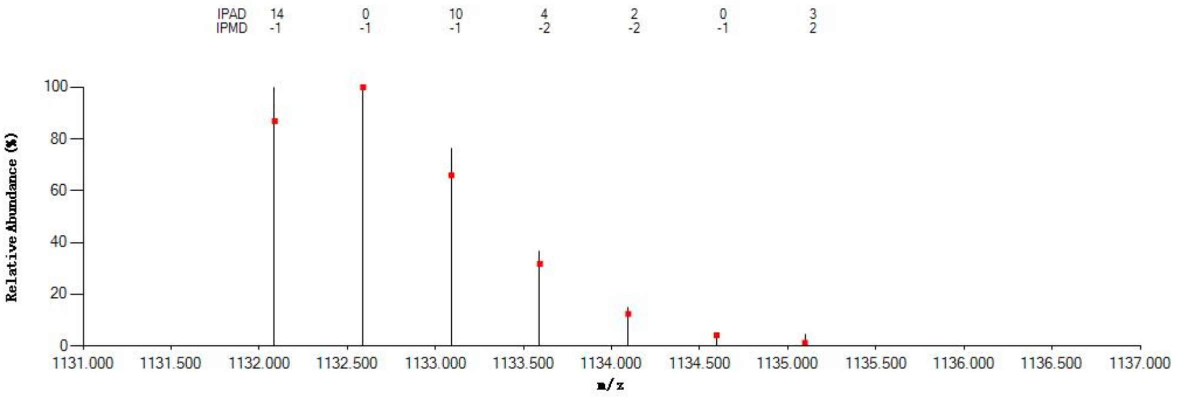


#### Annotated MS/MS Spectrum

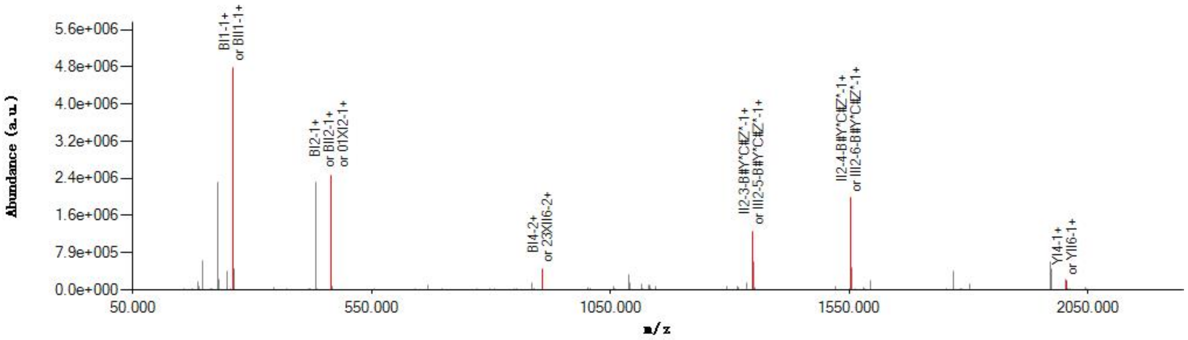


#### Graphical Fragmentation Map

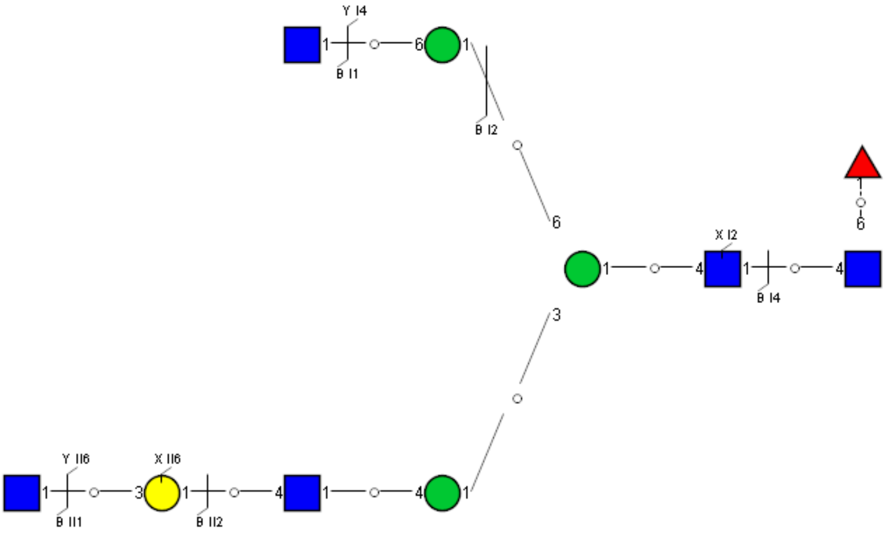

Spectrum Index: 33538(NoBHs:1)

#### Precursor iEF Map

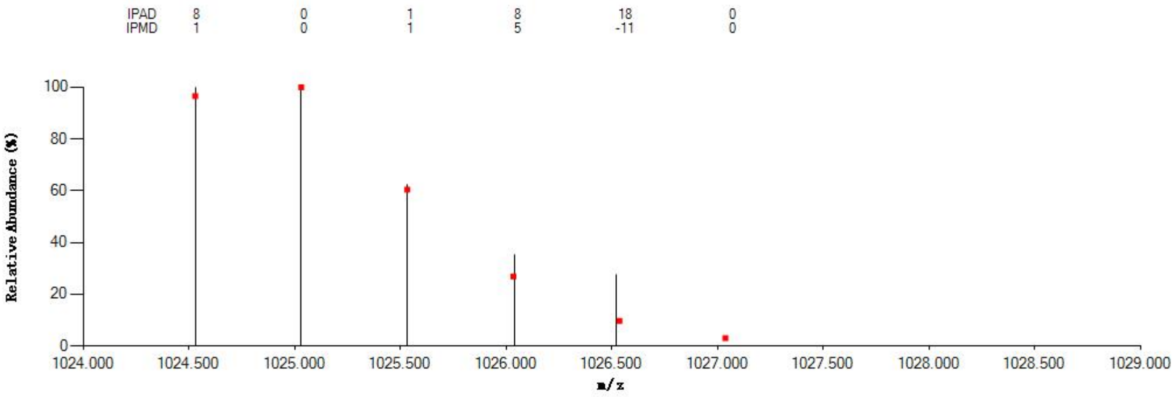


#### Annotated MS/MS Spectrum

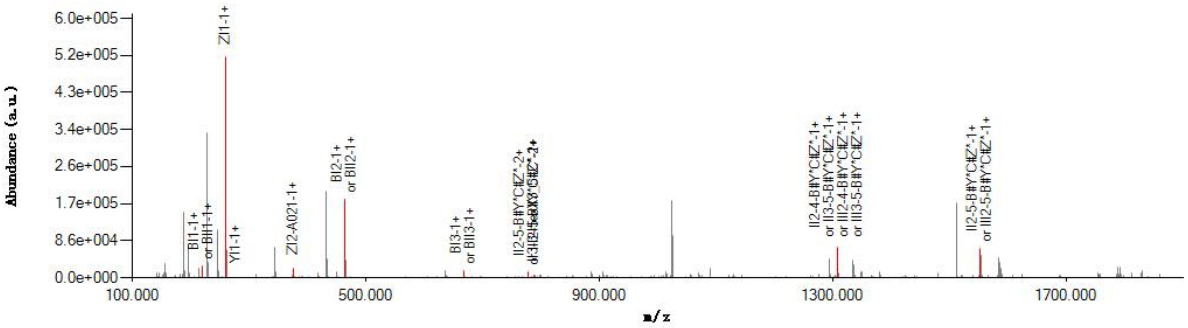


#### Graphical Fragmentation Map

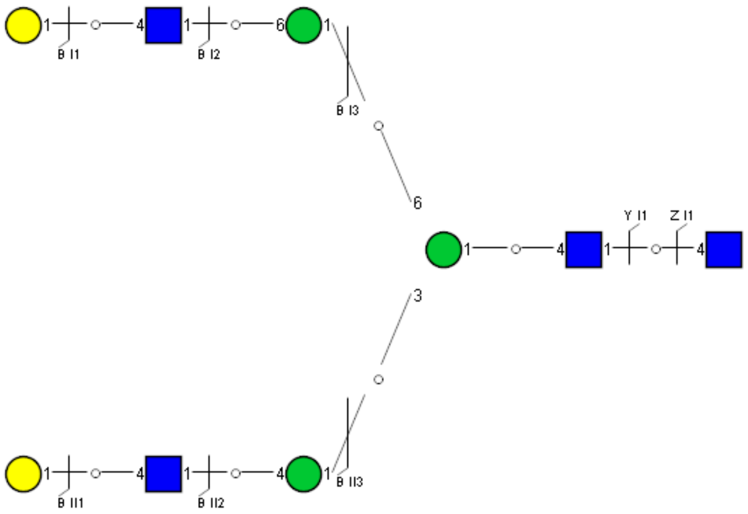

Spectrum Index: 33927(NoBHs:1)

#### Precursor iEF Map

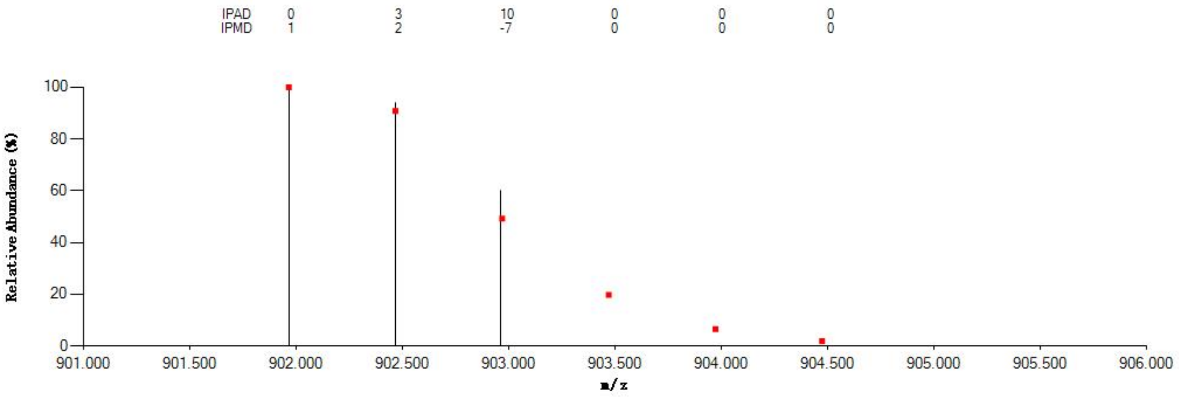


#### Annotated MS/MS Spectrum

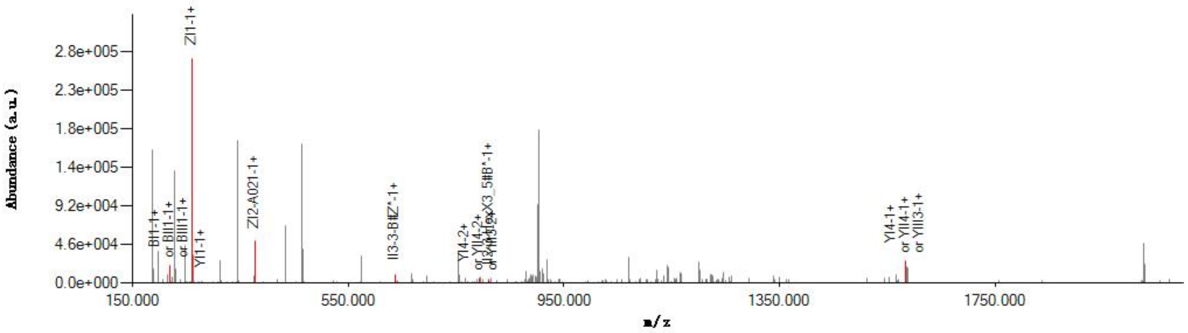


#### Graphical Fragmentation Map

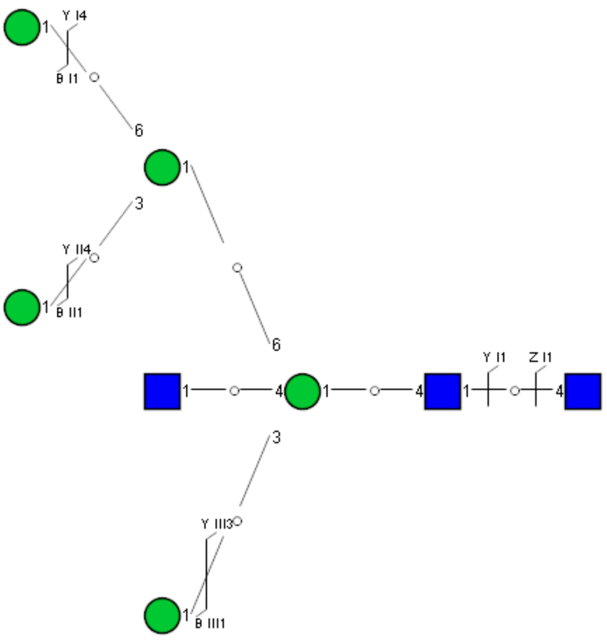

Spectrum Index: 34096(NoBHs:1)

#### Precursor iEF Map

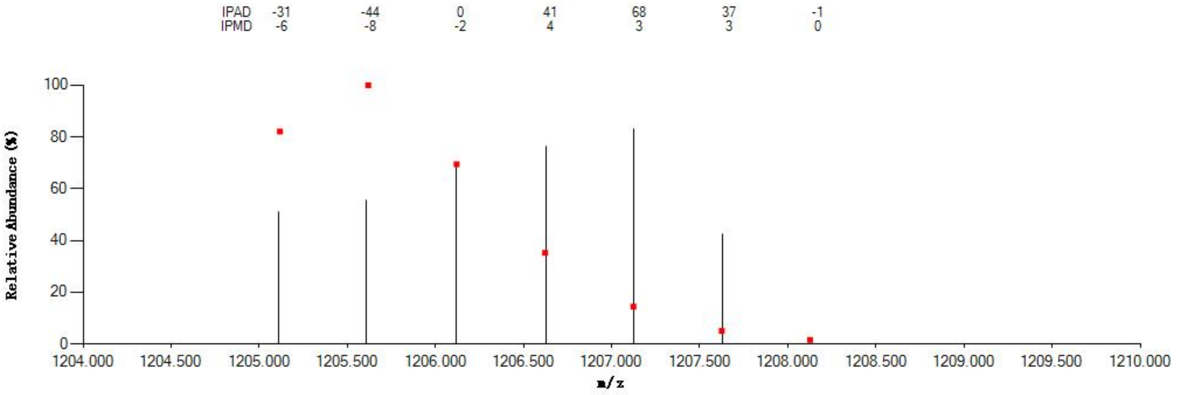


#### Annotated MS/MS Spectrum

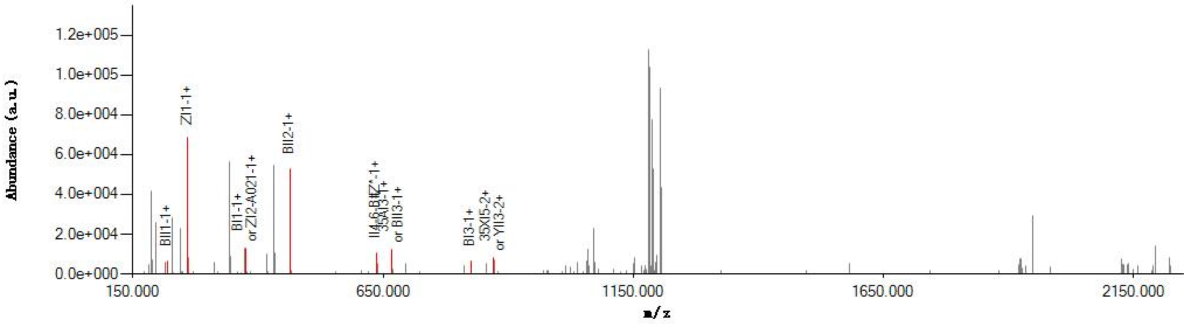


#### Graphical Fragmentation Map

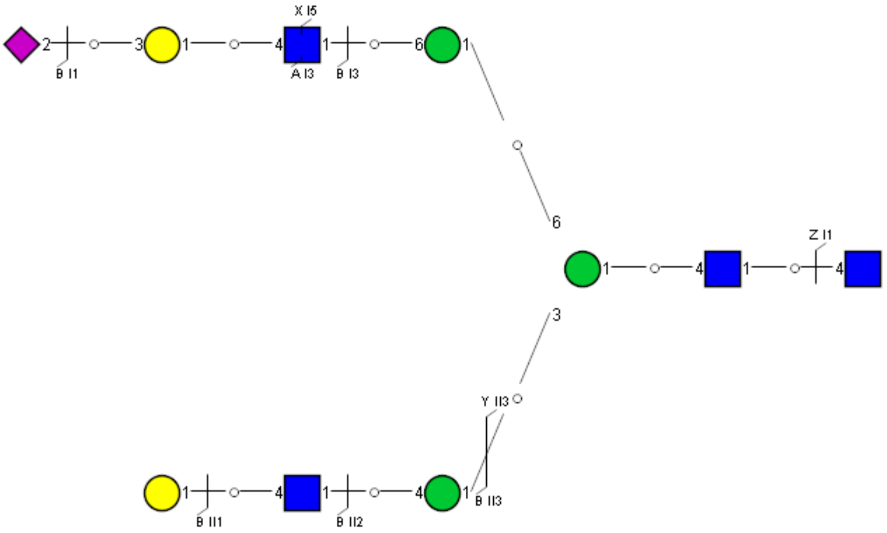

Spectrum Index: 34237(NoBHs:1)

#### Precursor iEF Map

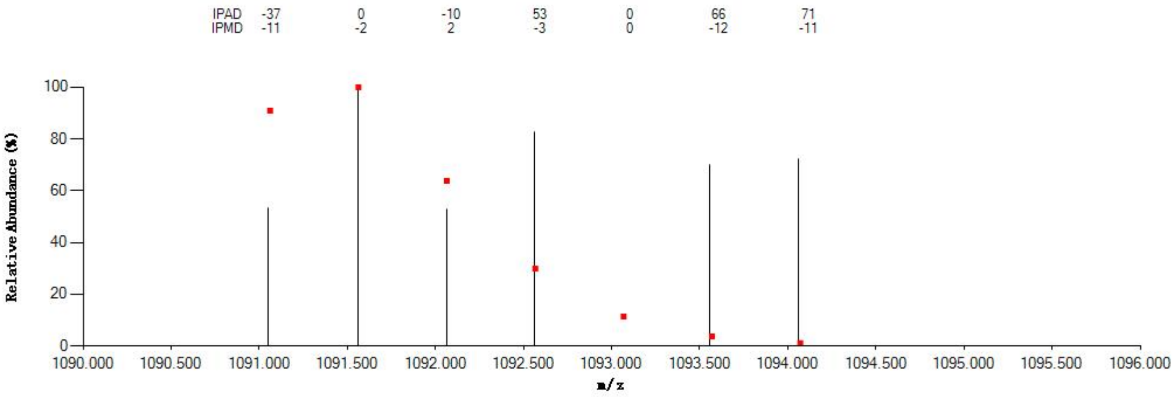


#### Annotated MS/MS Spectrum

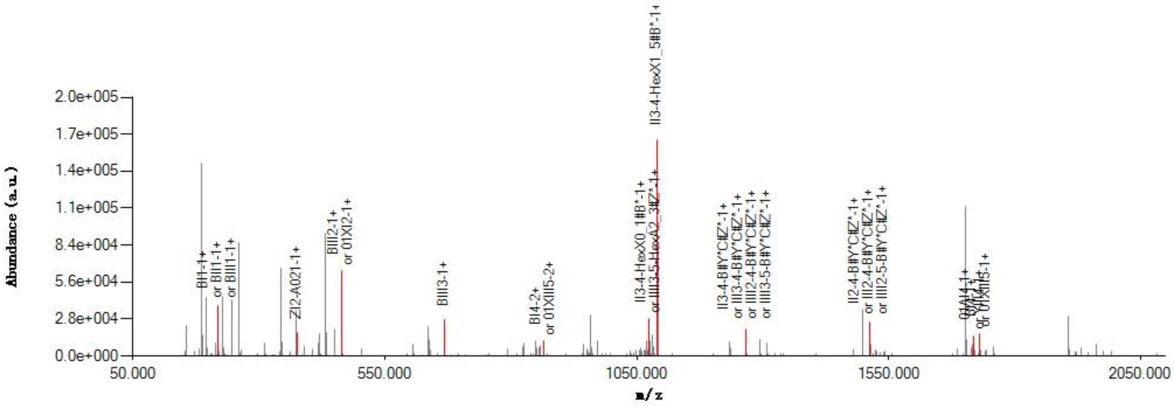


#### Graphical Fragmentation Map

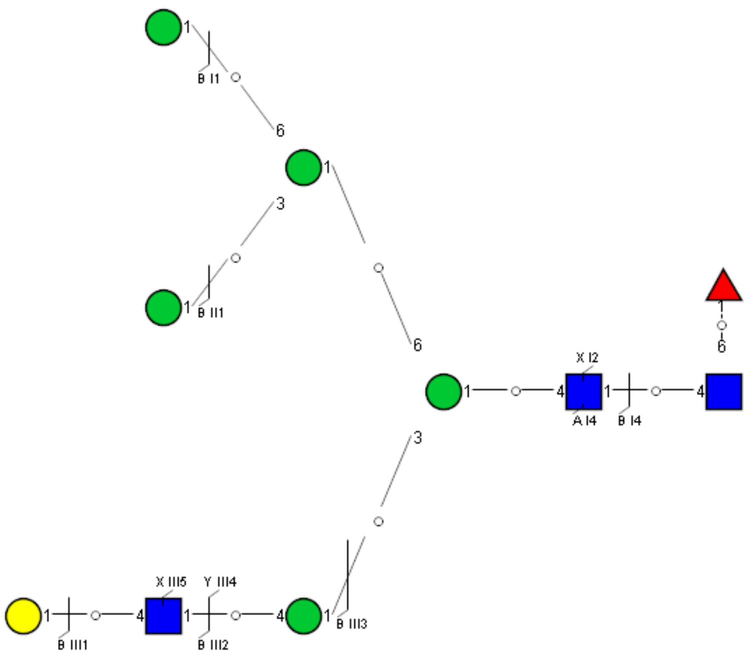

Spectrum Index: 34633(NoBHs:1)

#### Precursor iEF Map

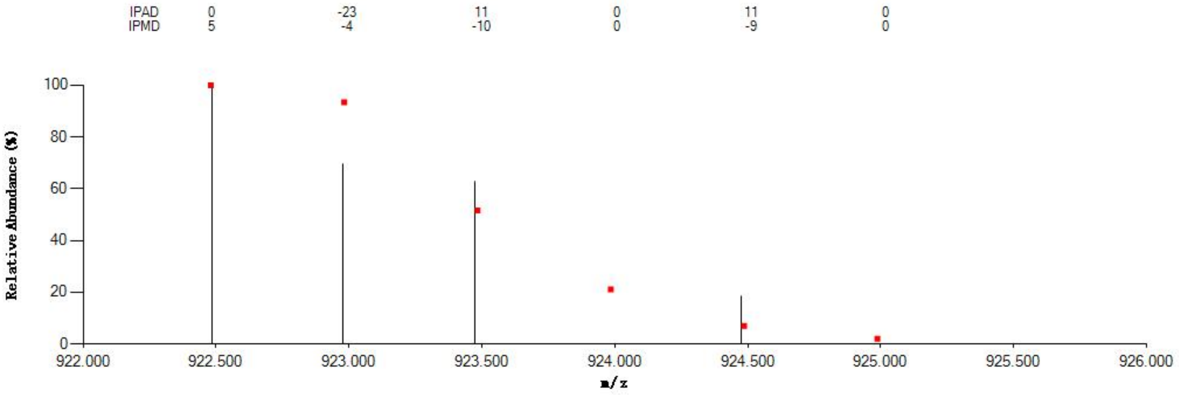


#### Annotated MS/MS Spectrum

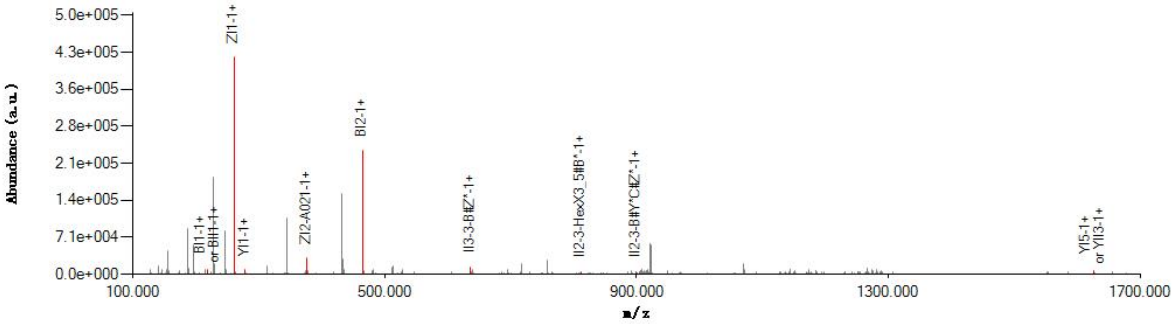


#### Graphical Fragmentation Map

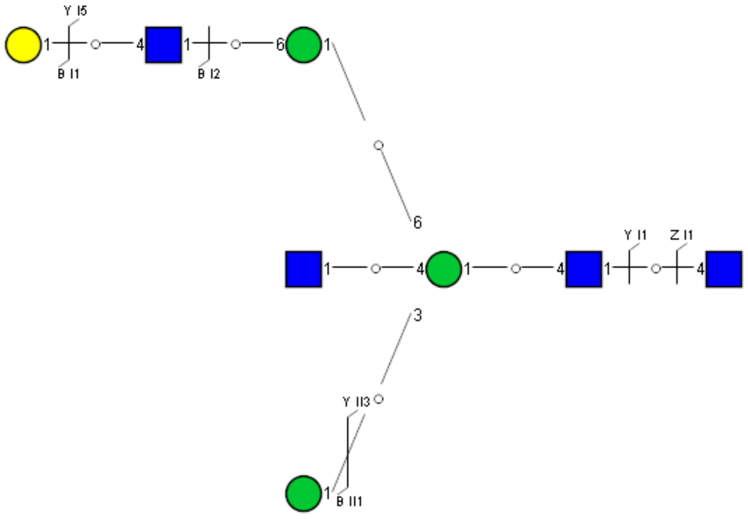

Spectrum Index: 34986(NoBHs:1)

#### Precursor iEF Map

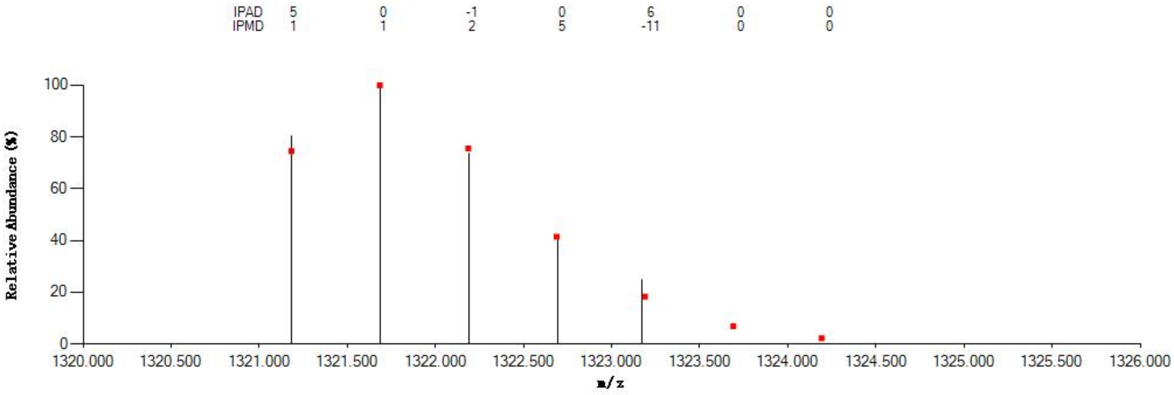


#### Annotated MS/MS Spectrum

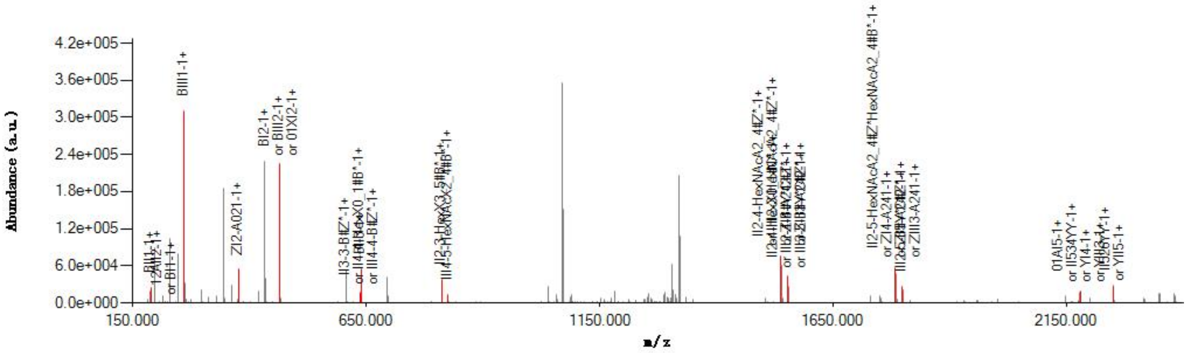


#### Graphical Fragmentation Map

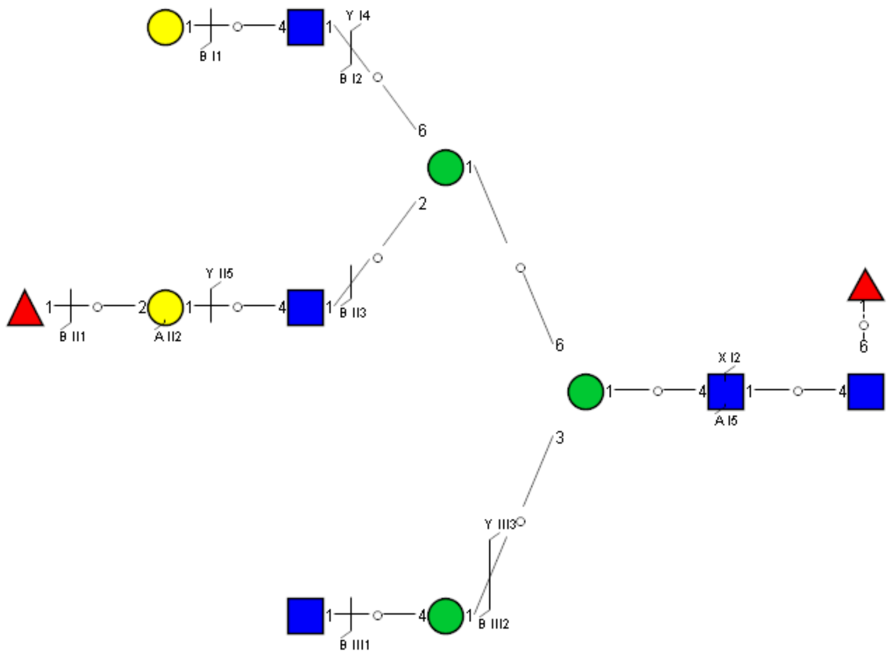

Spectrum Index: 35346(NoBHs:1)

#### Precursor iEF Map

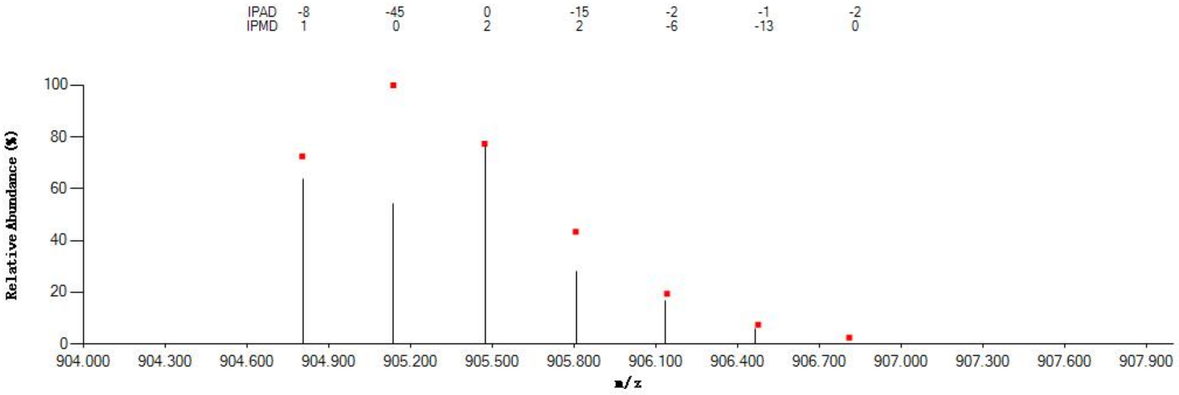


#### Annotated MS/MS Spectrum

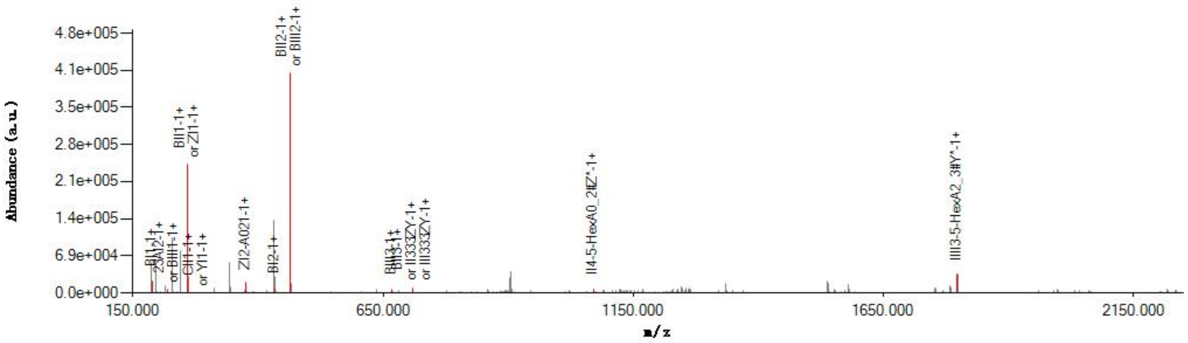


#### Graphical Fragmentation Map

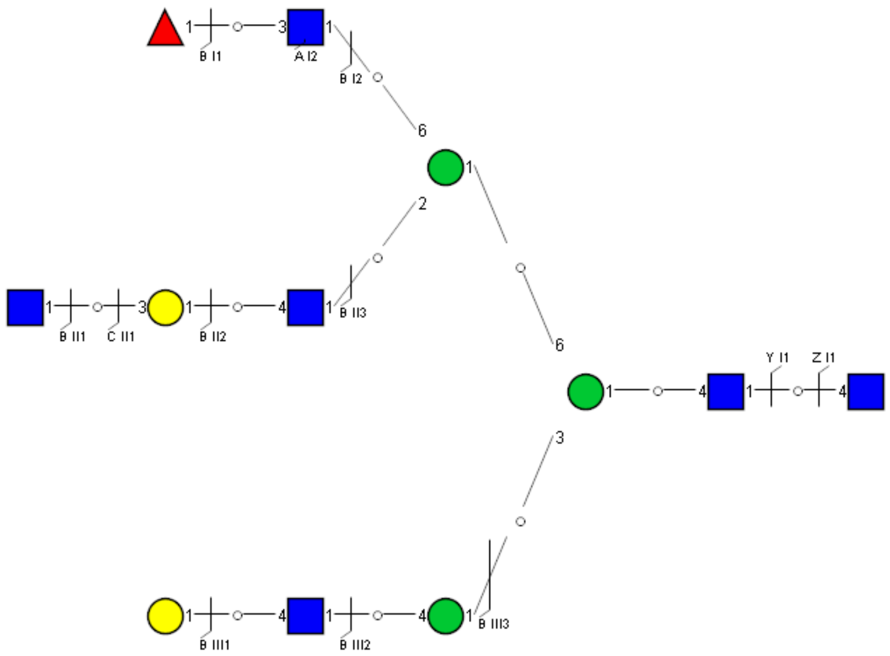

Spectrum Index: 35423(NoBHs:1)

#### Precursor iEF Map

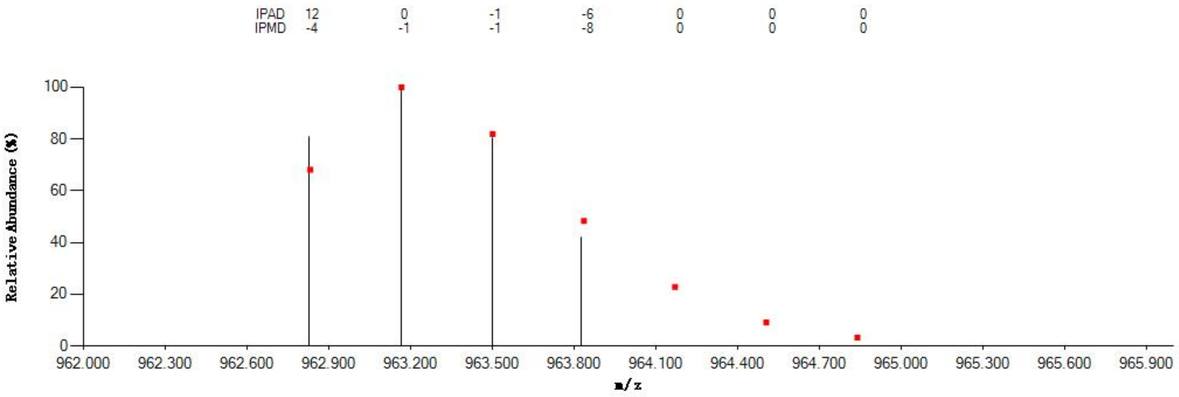


#### Annotated MS/MS Spectrum

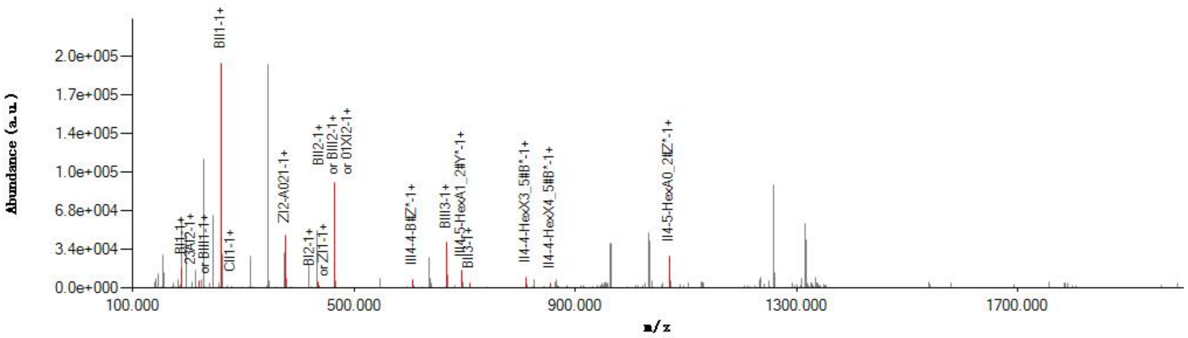


#### Graphical Fragmentation Map

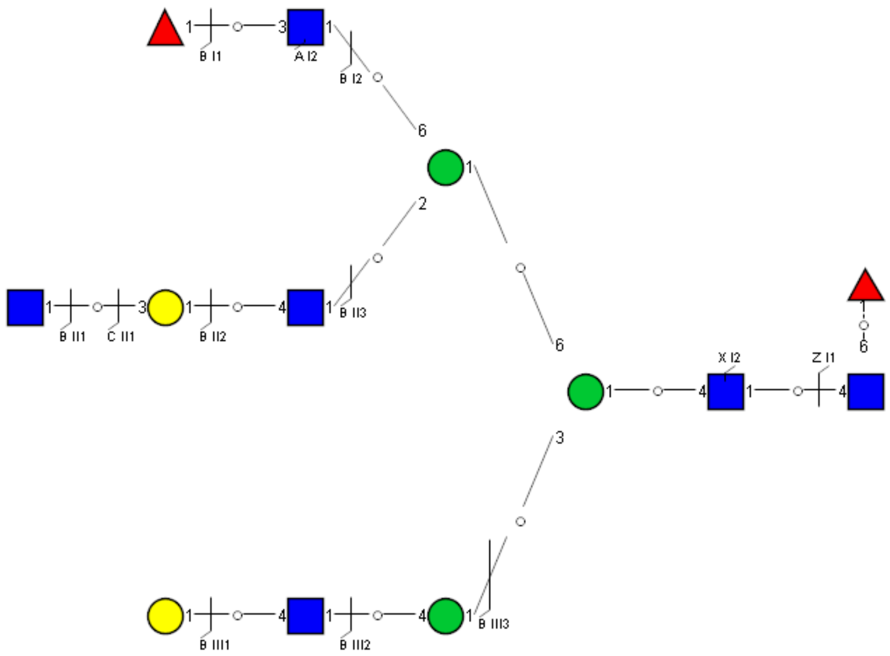

Spectrum Index: 35465(NoBHs:1)

#### Precursor iEF Map

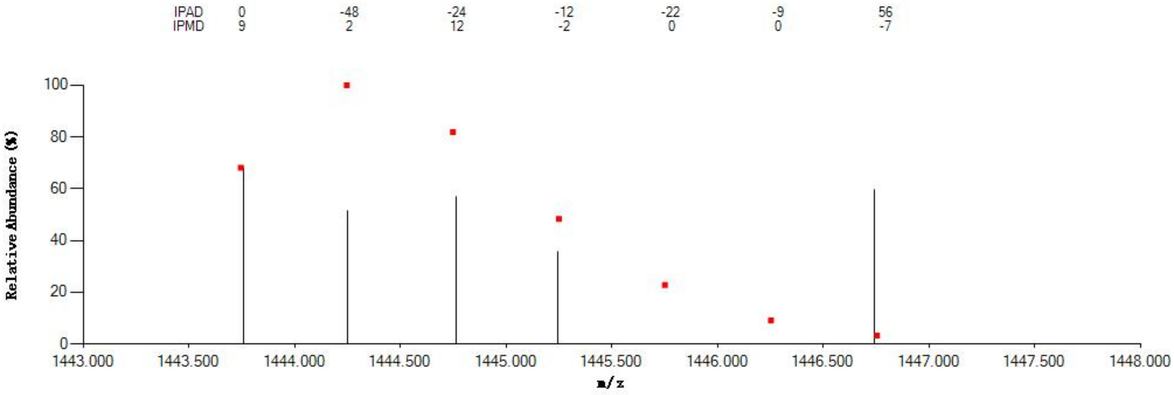

#### Annotated MS/MS Spectrum

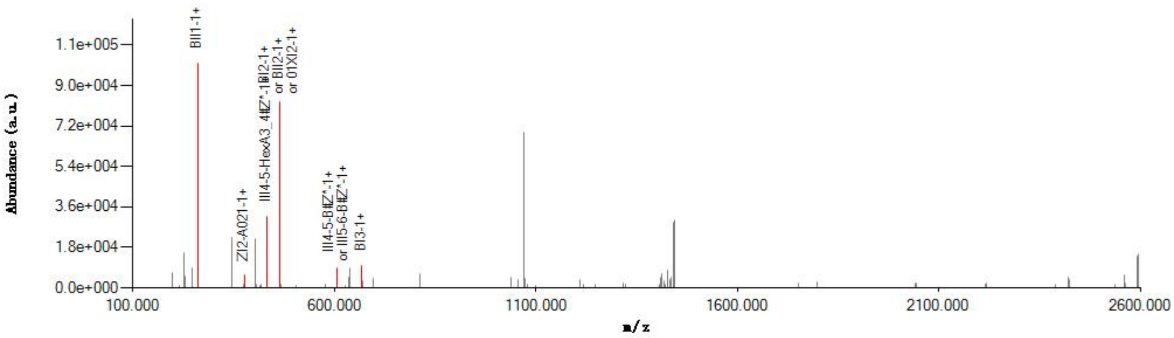

#### Graphical Fragmentation Map

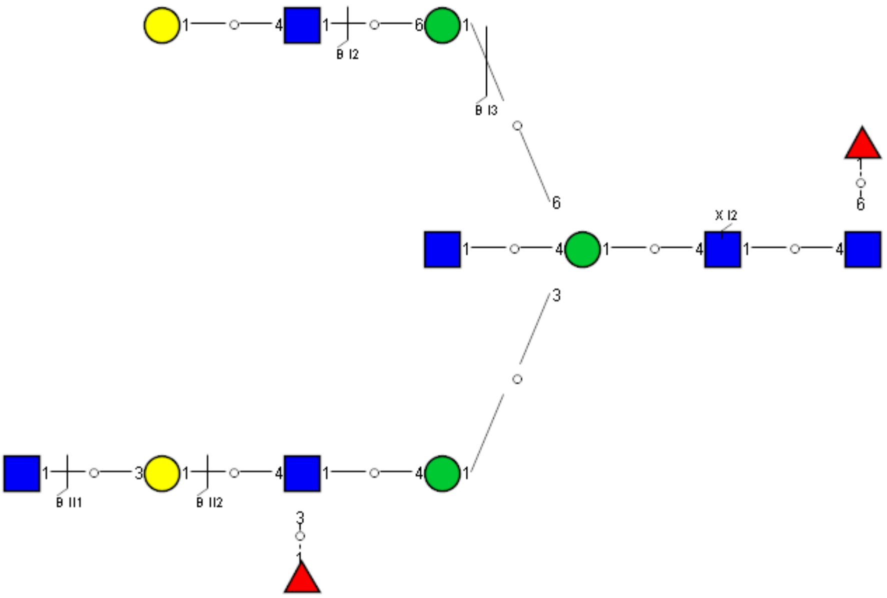

Spectrum Index: 35760(NoBHs:1)

#### Precursor iEF Map

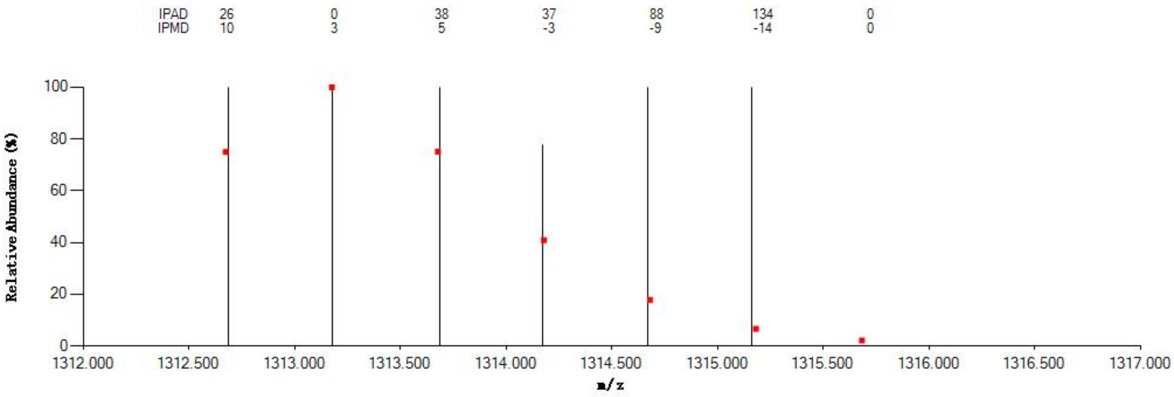

#### Annotated MS/MS Spectrum

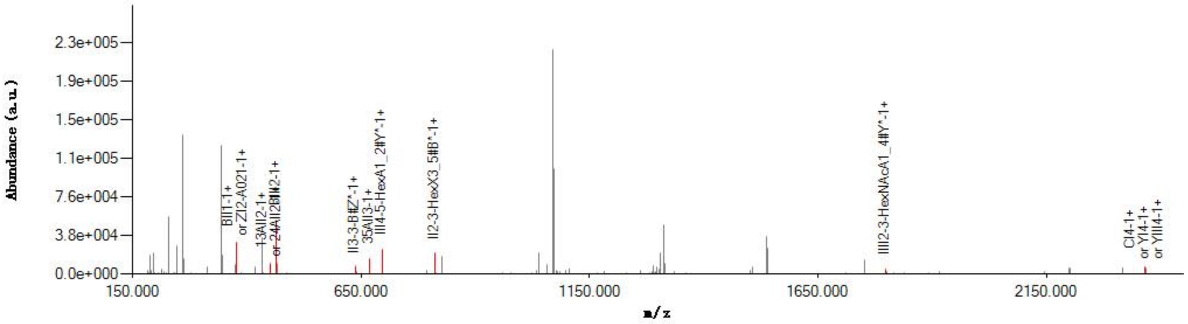

#### Graphical Fragmentation Map

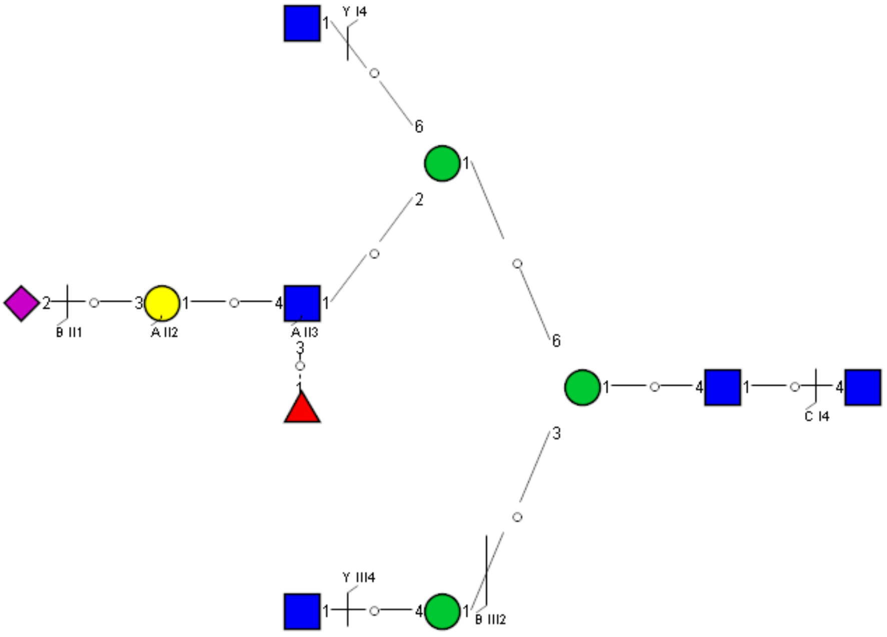

Spectrum Index: 35976(NoBHs:2)

#### Precursor iEF Map

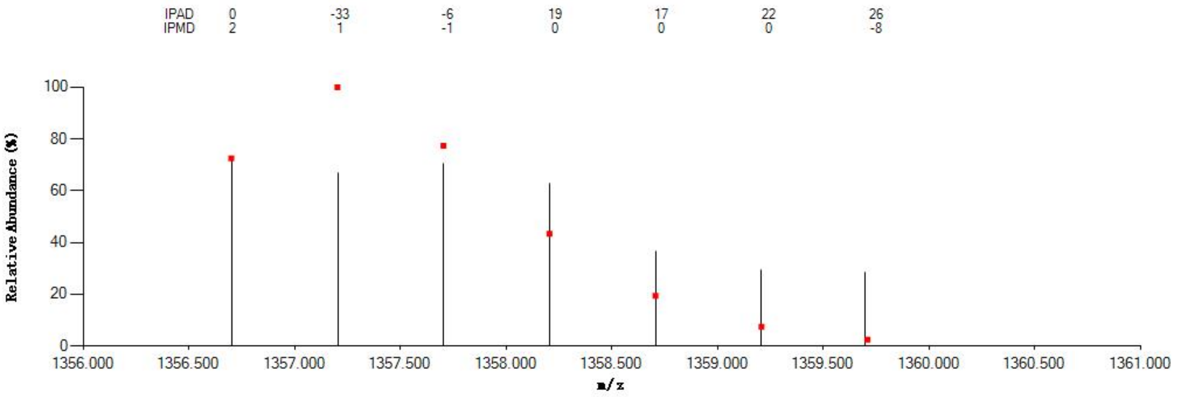

#### Annotated MS/MS Spectrum

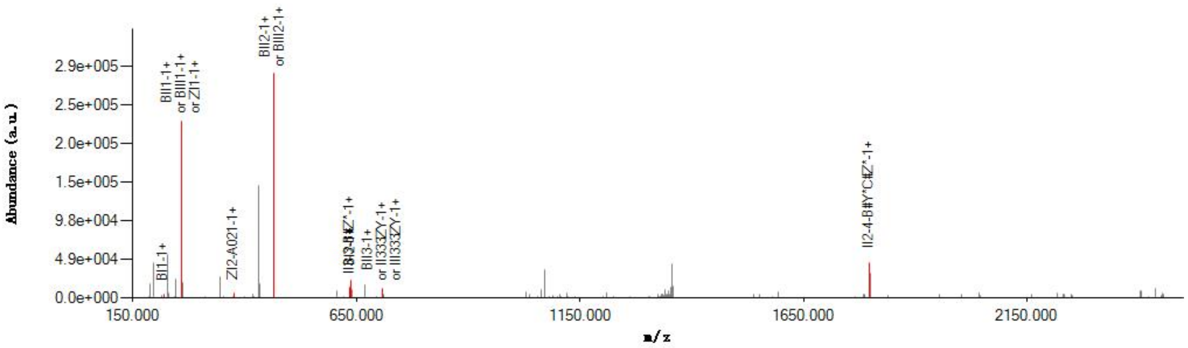

#### Graphical Fragmentation Map

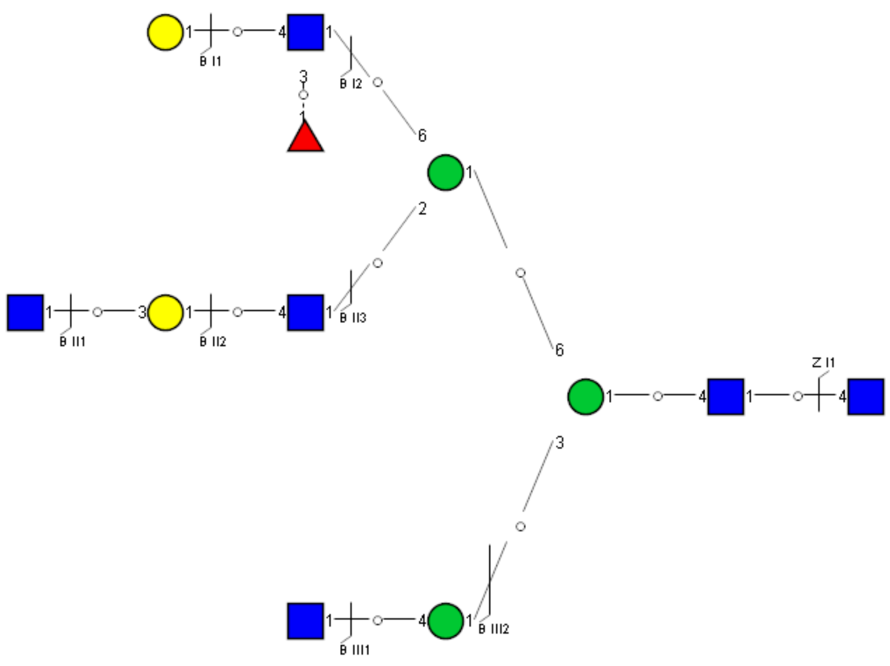

Spectrum Index: 35977(NoBHs:1)

#### Precursor iEF Map

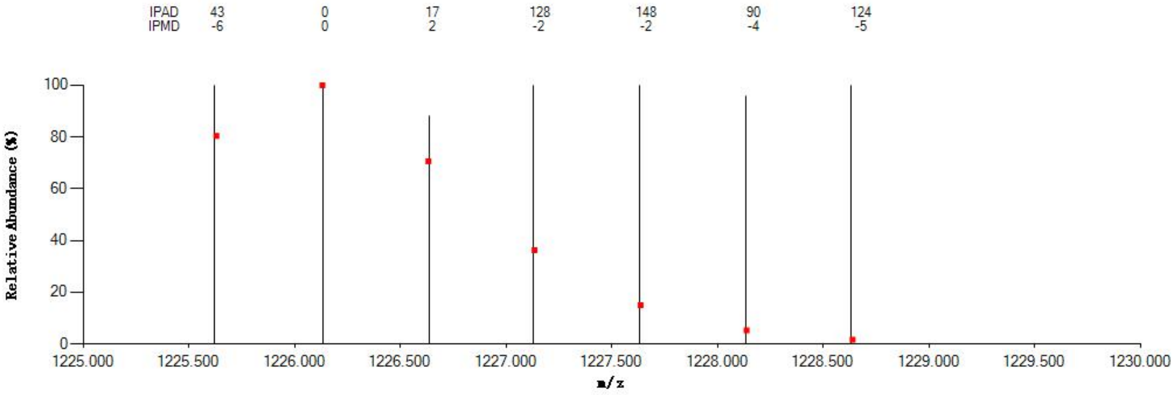


#### Annotated MS/MS Spectrum

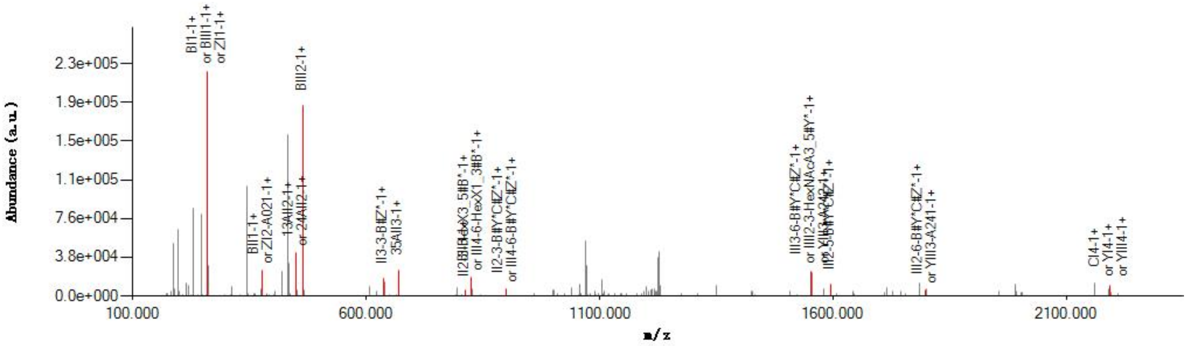


#### Graphical Fragmentation Map

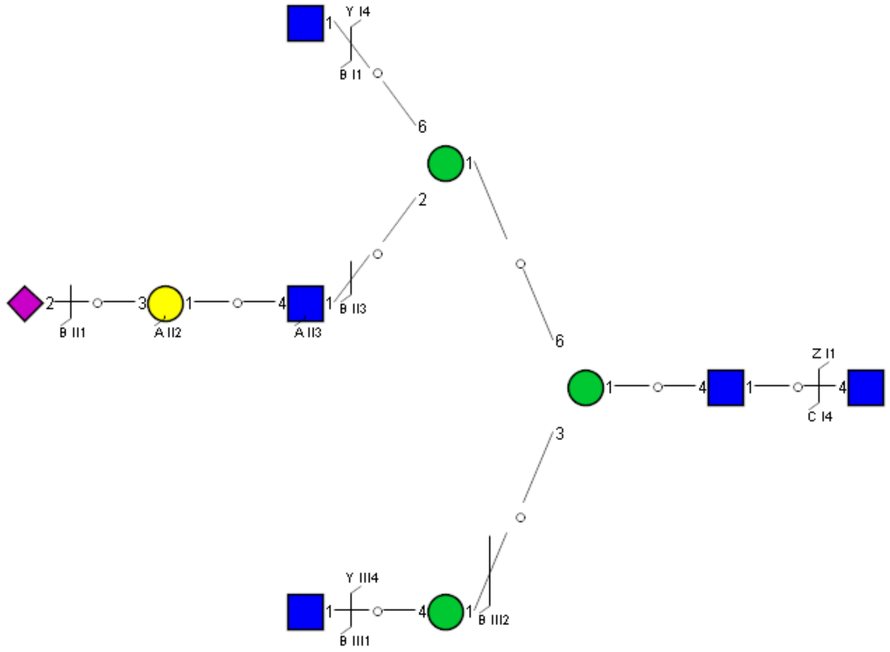

Spectrum Index: 36142(NoBHs:1)

#### Precursor iEF Map

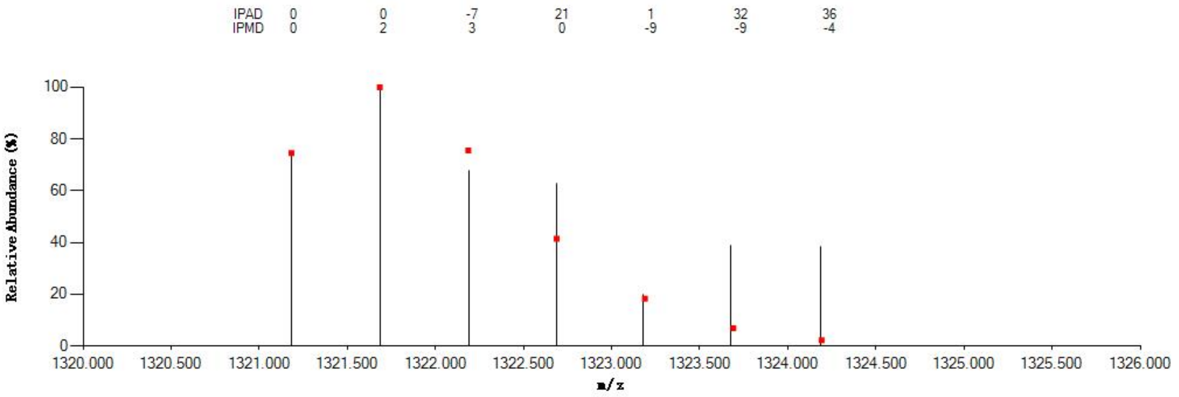


#### Annotated MS/MS Spectrum

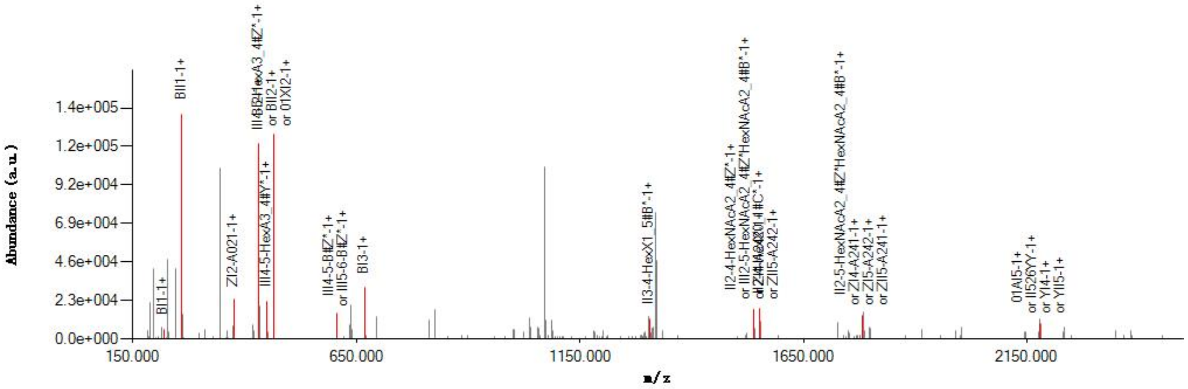


#### Graphical Fragmentation Map

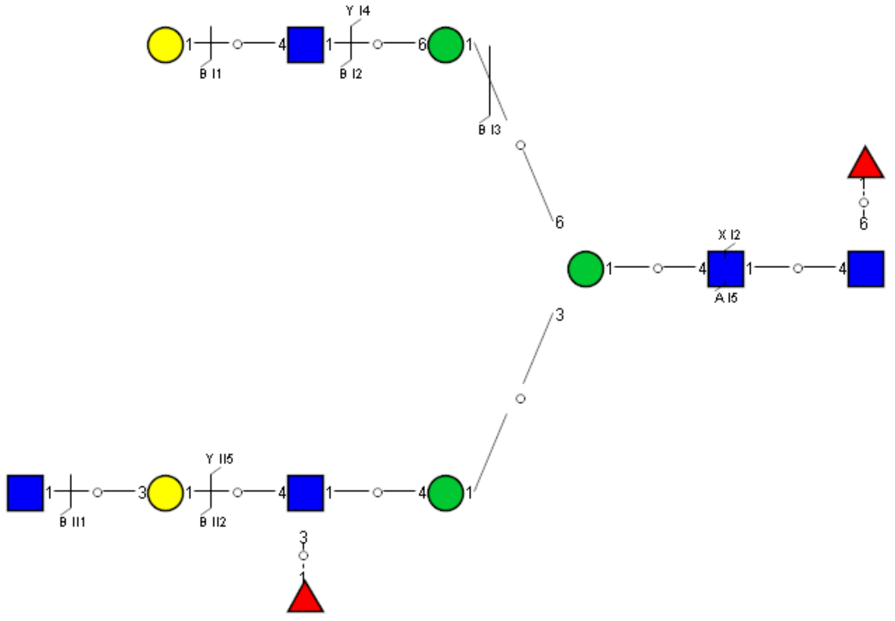

Spectrum Index: 36262(NoBHs:1)

#### Precursor iEF Map

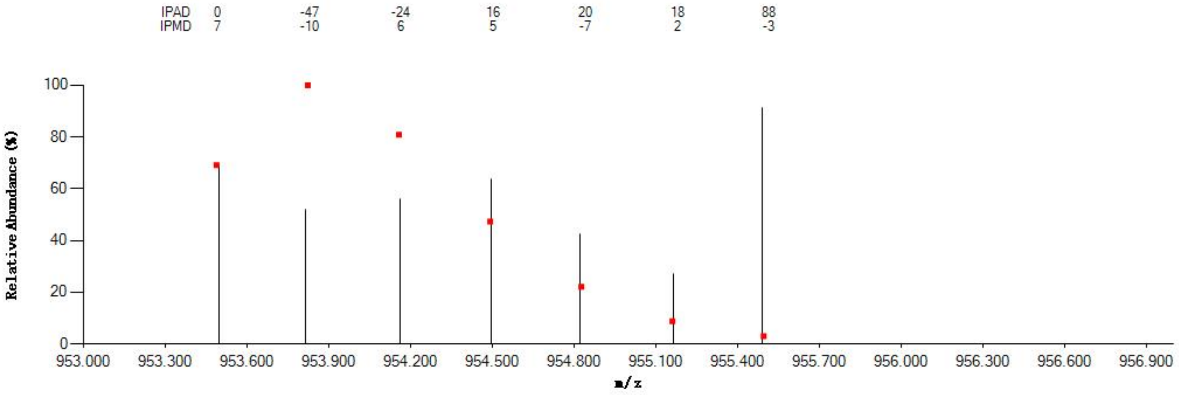


#### Annotated MS/MS Spectrum

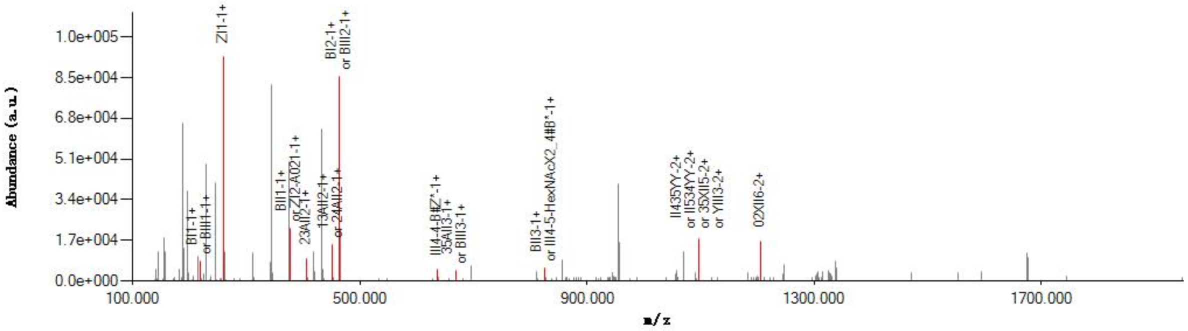


#### Graphical Fragmentation Map

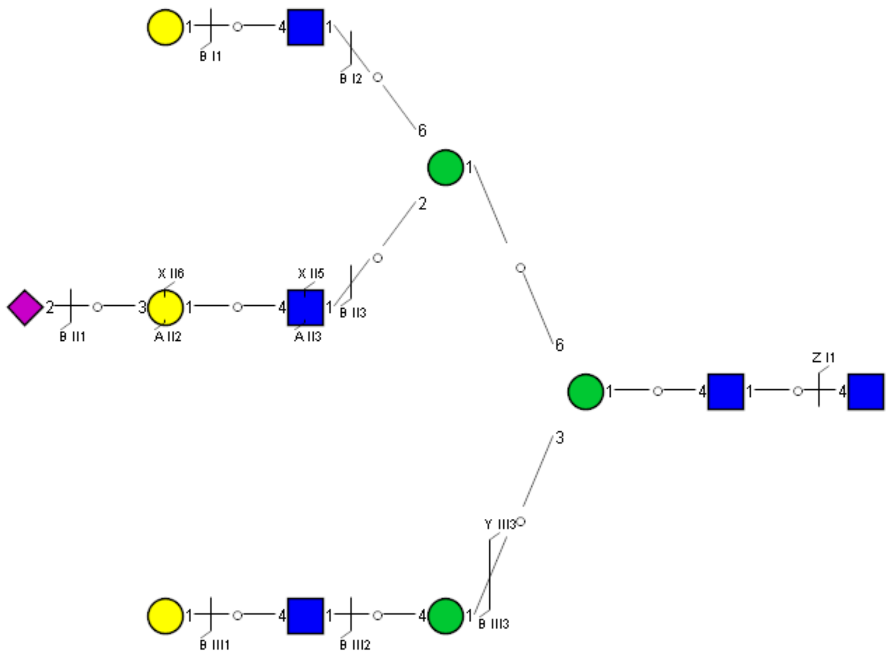

Spectrum Index: 36271(NoBHs:1)

#### Precursor iEF Map

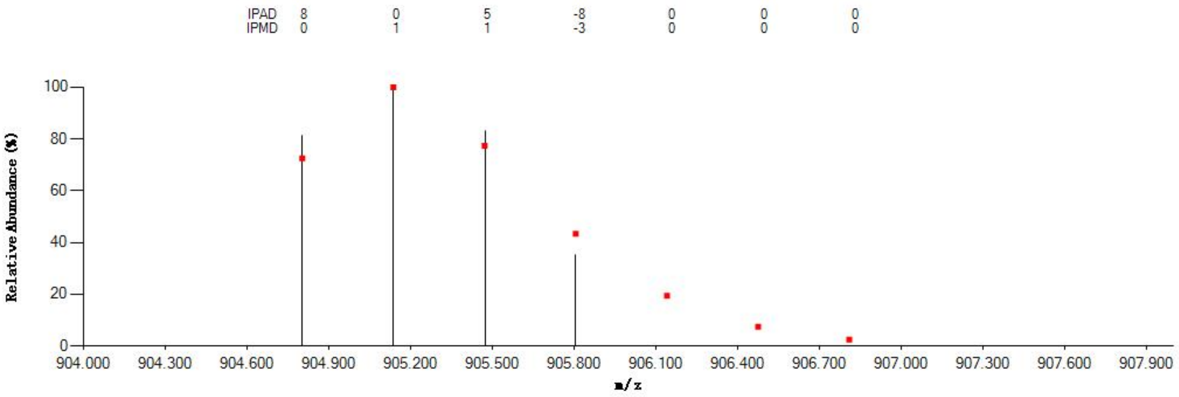


#### Annotated MS/MS Spectrum

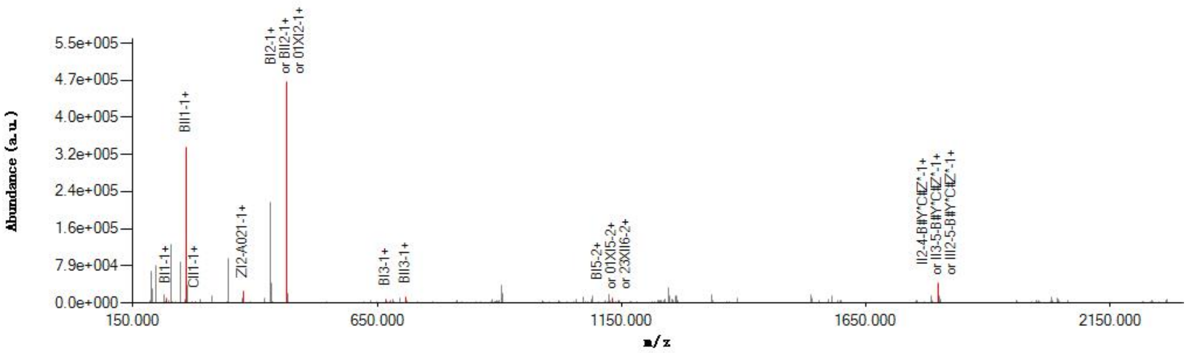


#### Graphical Fragmentation Map

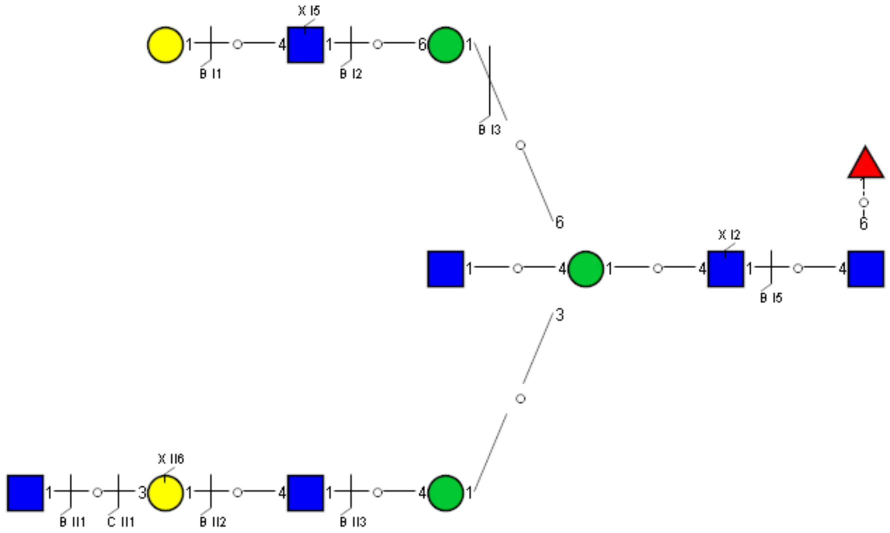

### Spectrum Index: 36306(NoBHs:1)

#### Precursor iEF Map

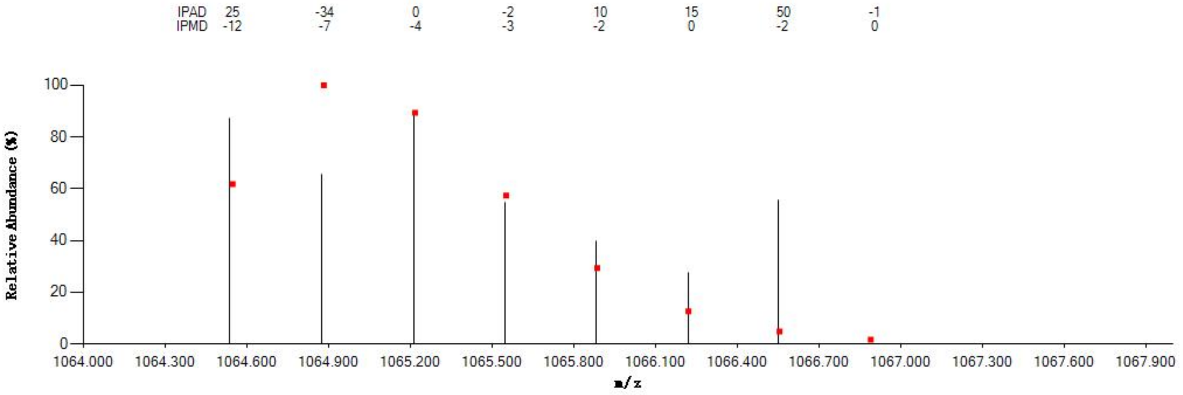


#### Annotated MS/MS Spectrum

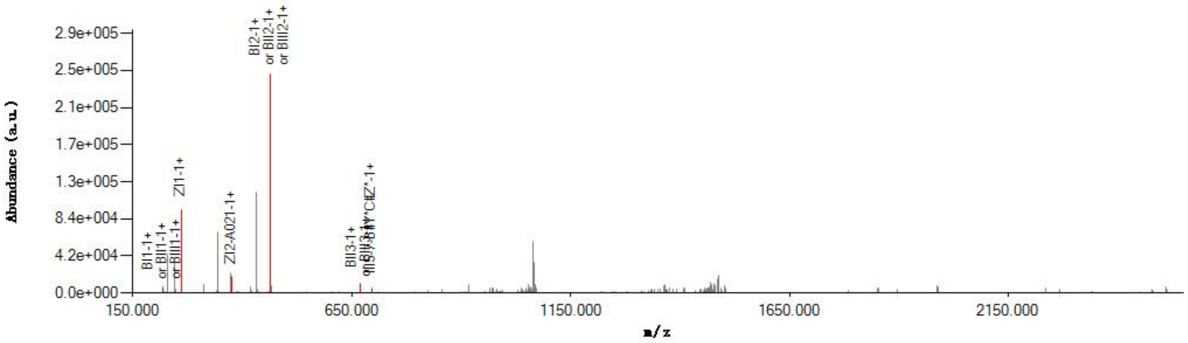


#### Graphical Fragmentation Map

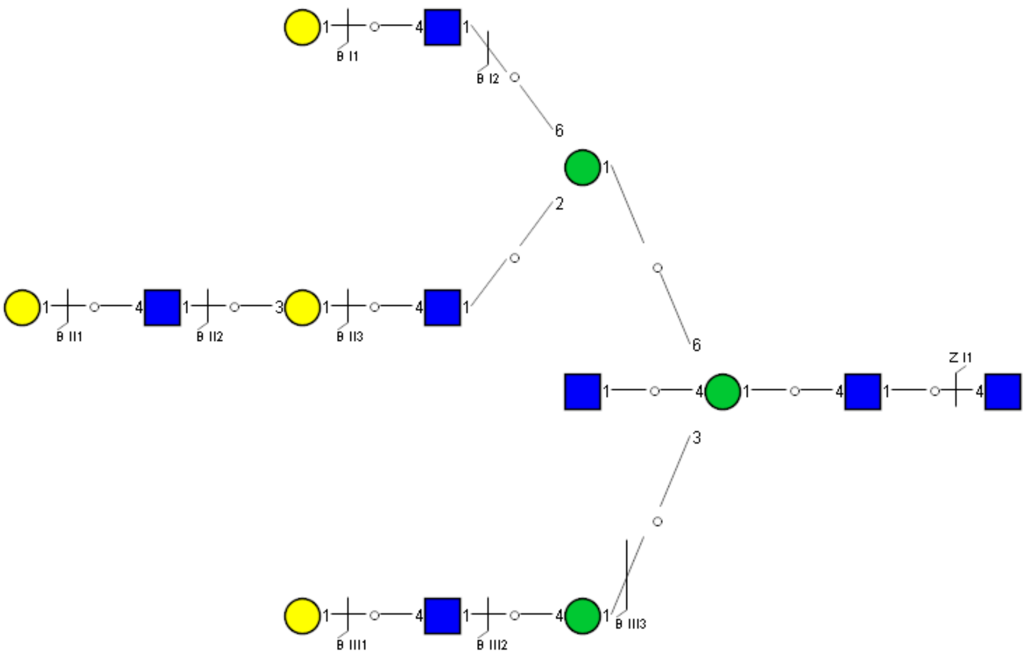

Spectrum Index: 36351(NoBHs:1)

#### Precursor iEF Map

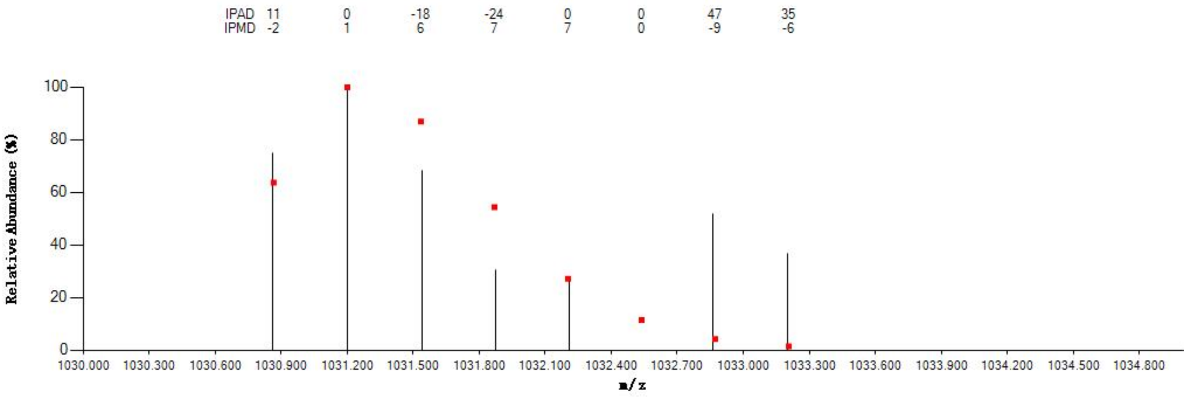

#### Annotated MS/MS Spectrum

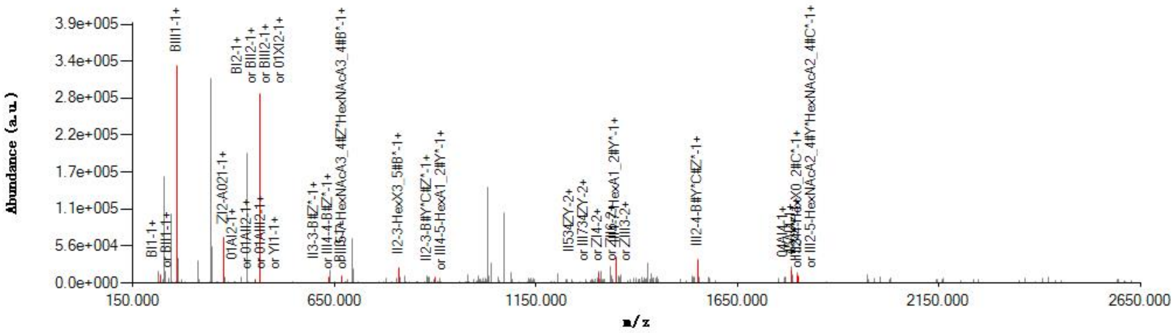

#### Graphical Fragmentation Map

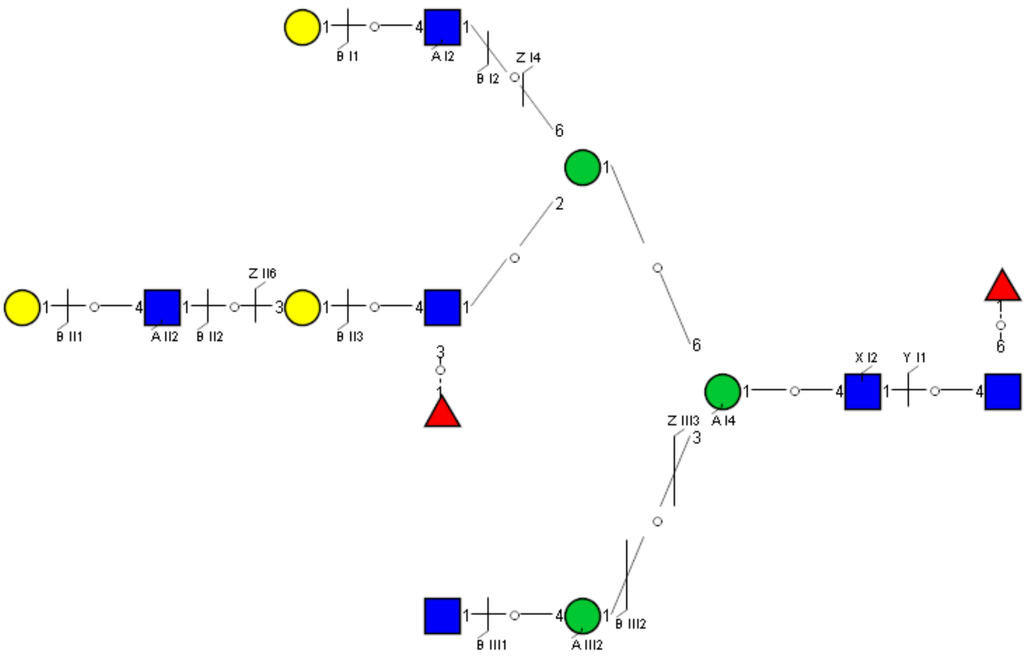

Spectrum Index: 36378(NoBHs:1)

#### Precursor iEF Map

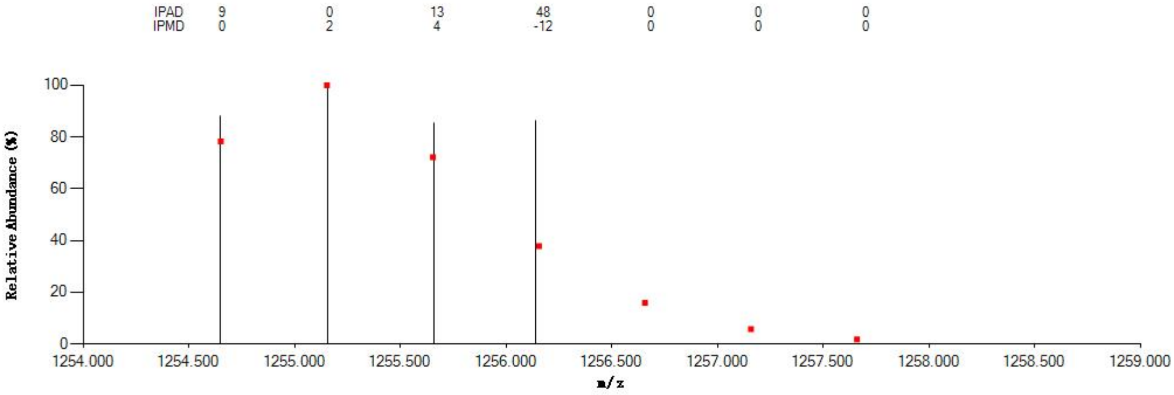


#### Annotated MS/MS Spectrum

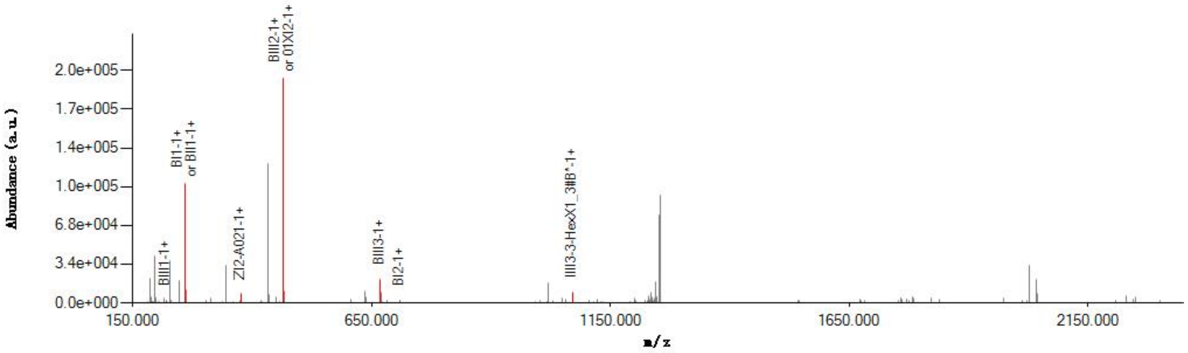


#### Graphical Fragmentation Map

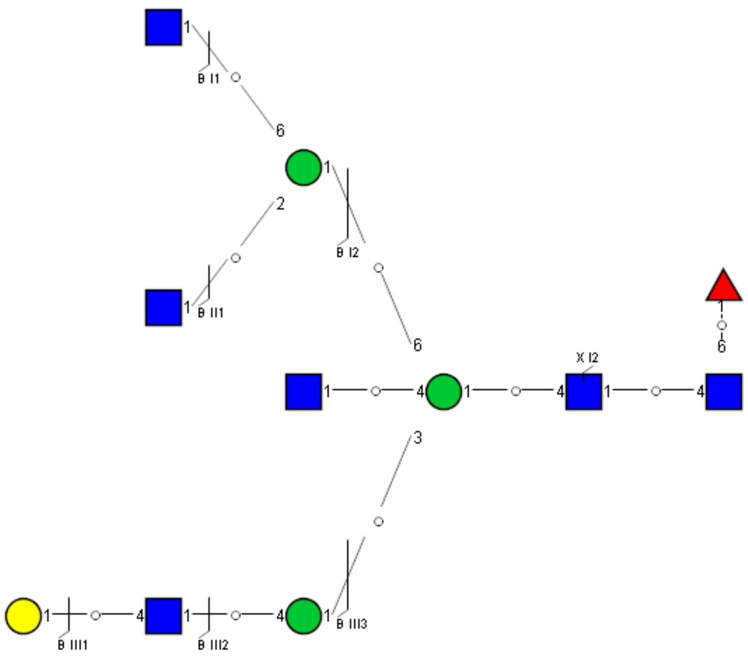

### Spectrum Index: 36485(NoBHs:1)

#### Precursor iEF Map

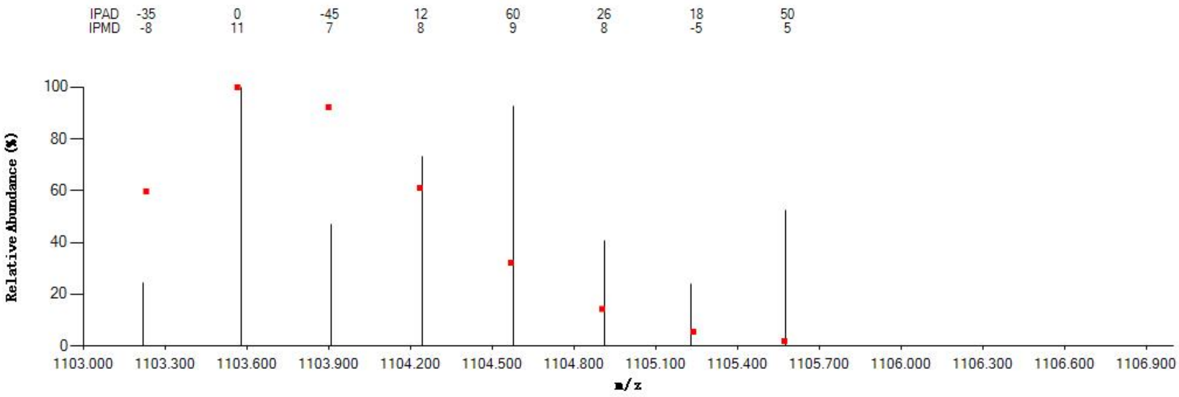


#### Annotated MS/MS Spectrum

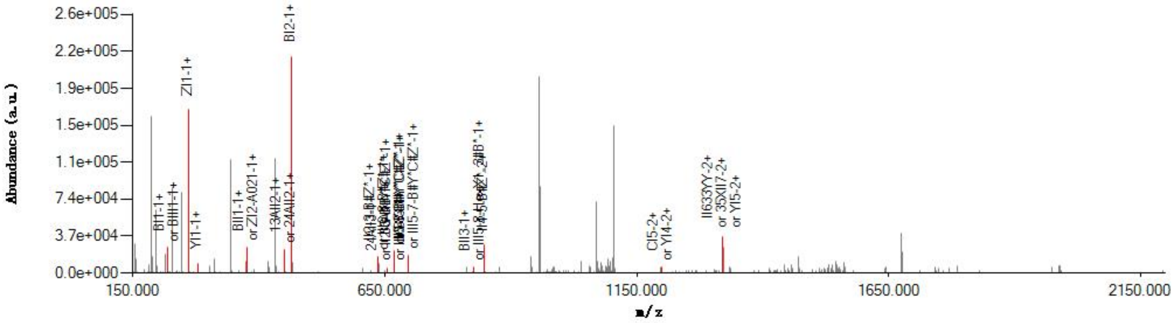


#### Graphical Fragmentation Map

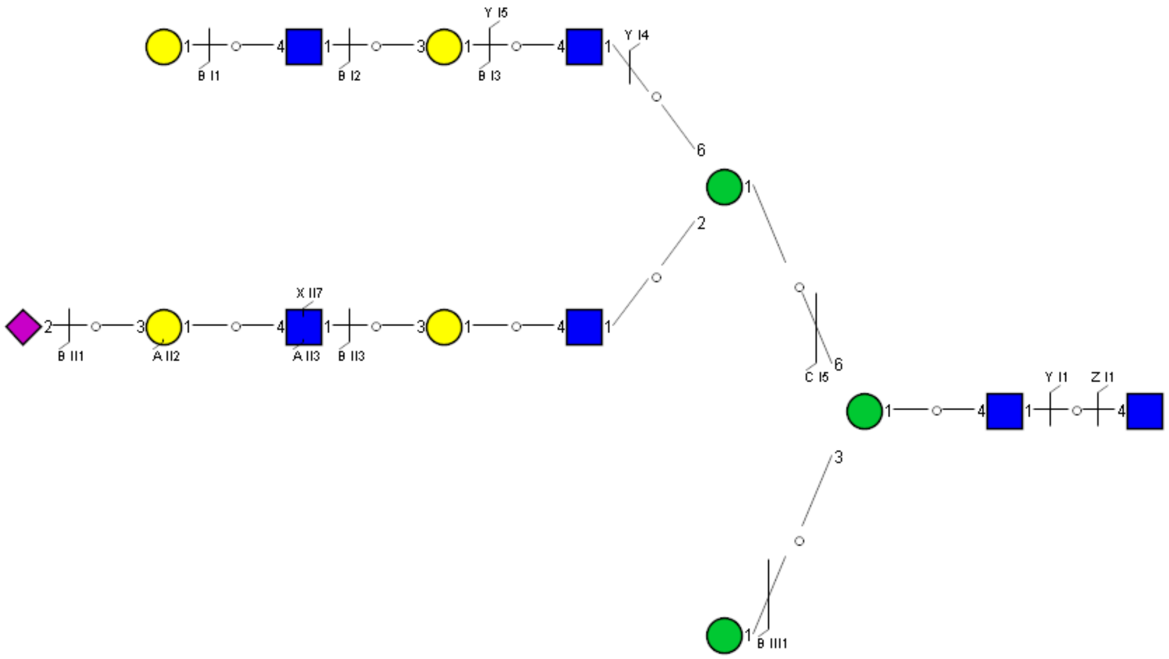

Spectrum Index: 36556(NoBHs:1)

#### Precursor iEF Map

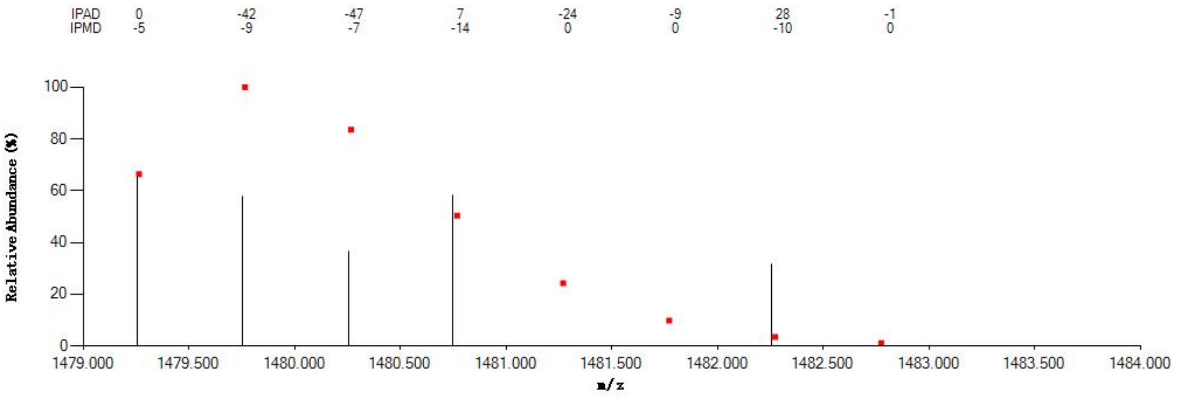


#### Annotated MS/MS Spectrum

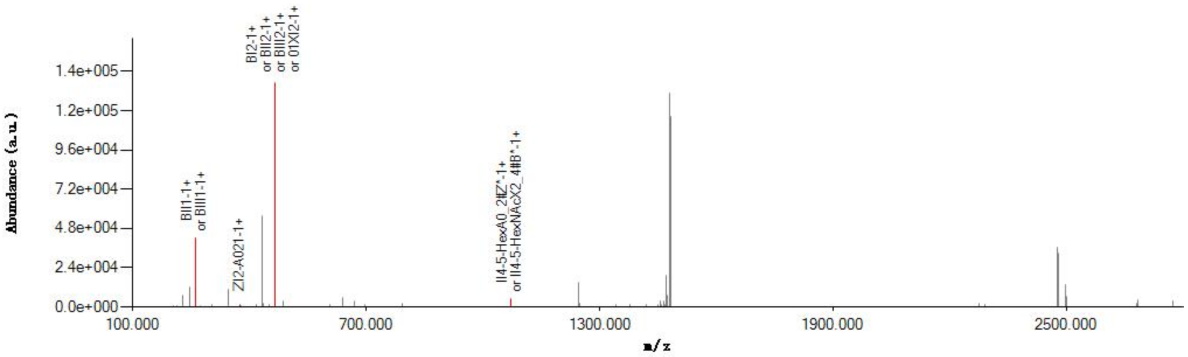


#### Graphical Fragmentation Map

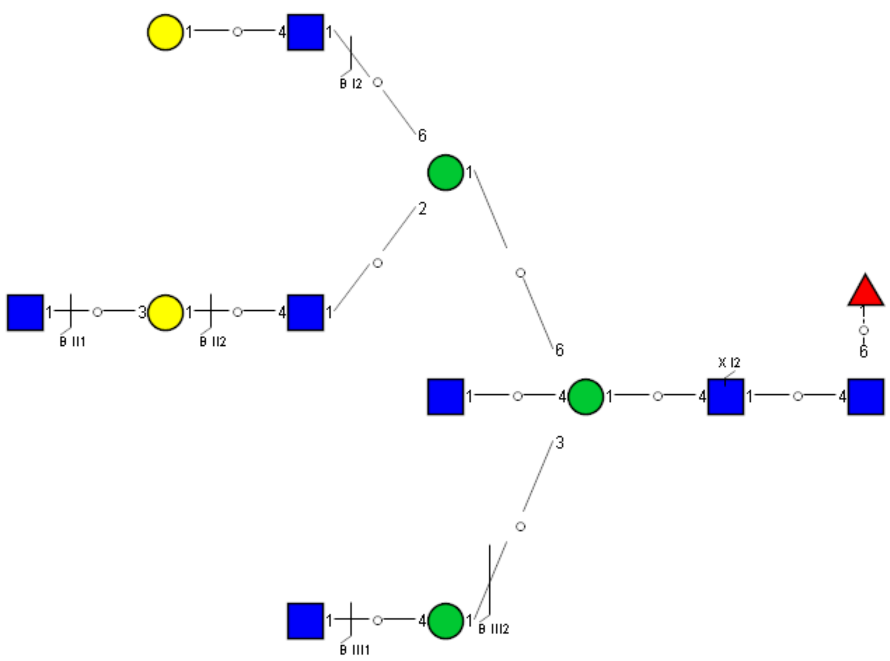

### Spectrum Index: 36724(NoBHs:1)

#### Precursor iEF Map

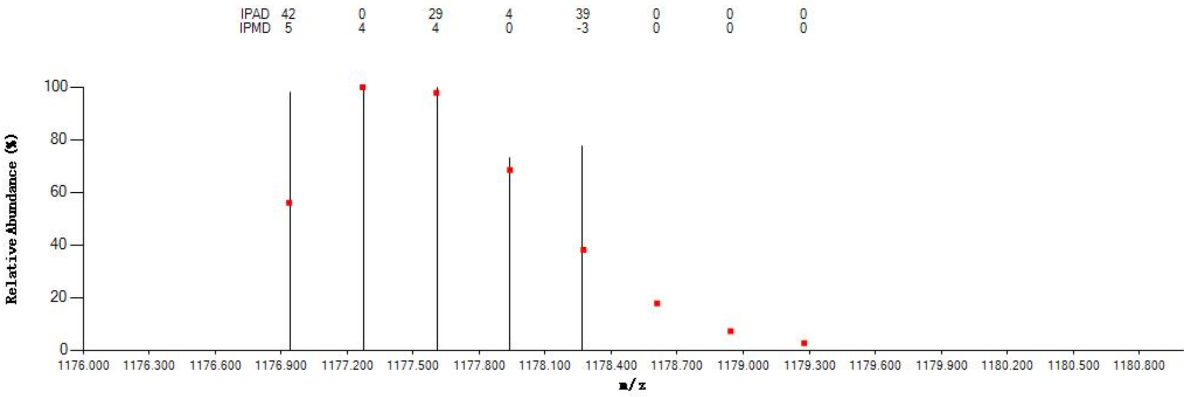

#### Annotated MS/MS Spectrum

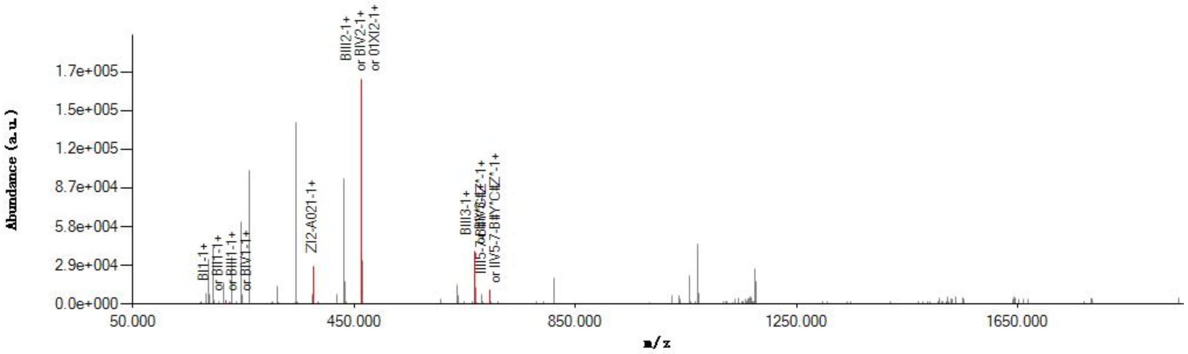

#### Graphical Fragmentation Map

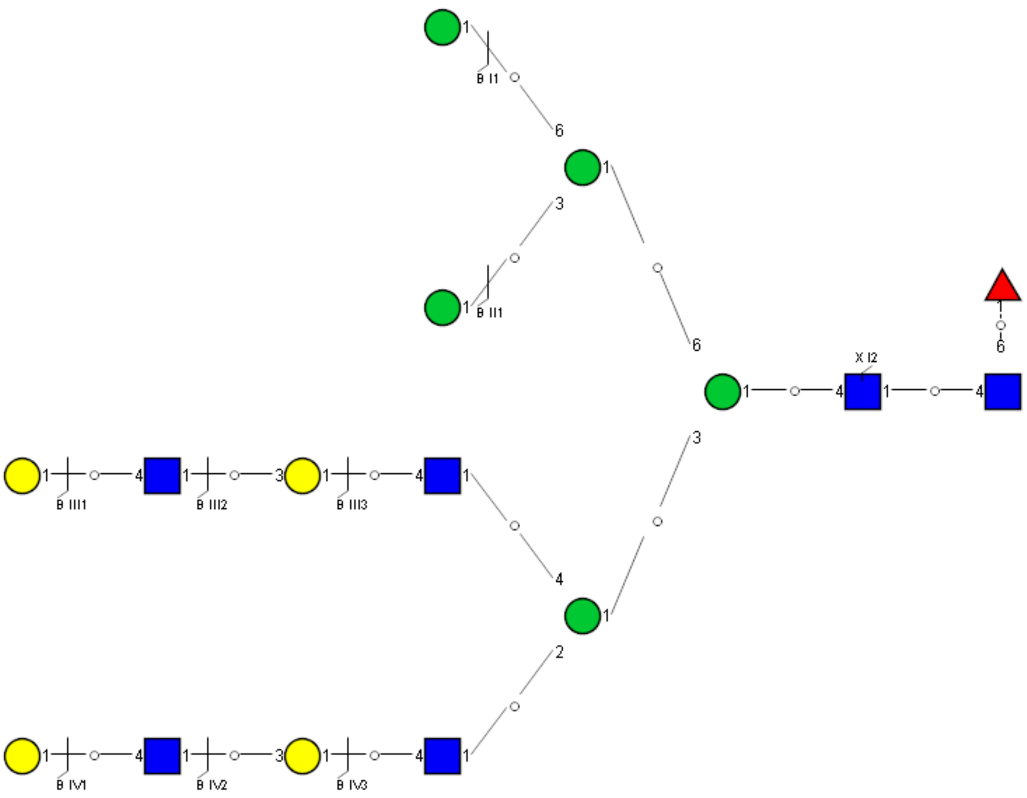

Spectrum Index: 37003(NoBHs:1)

#### Precursor iEF Map

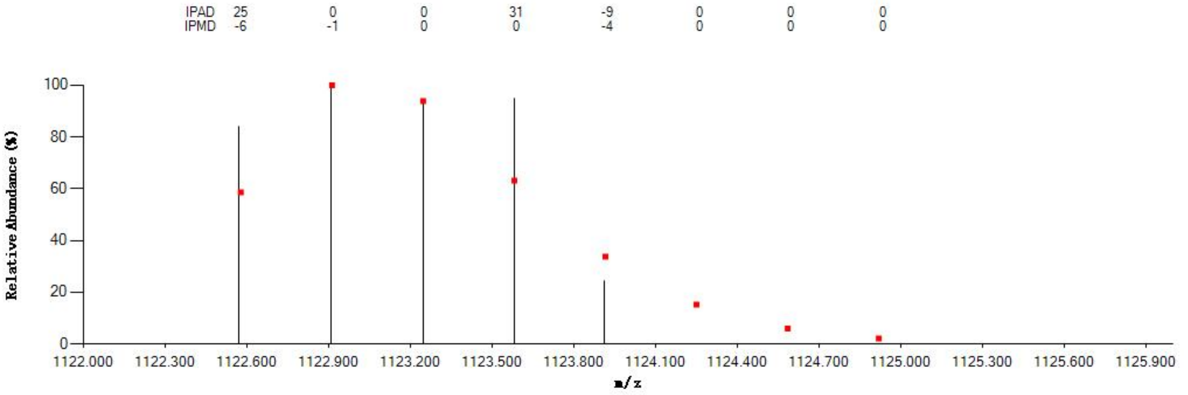

#### Annotated MS/MS Spectrum

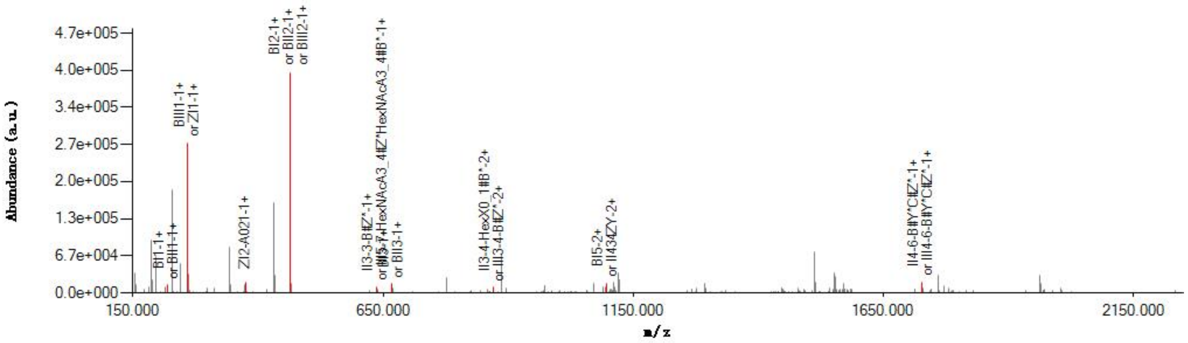

#### Graphical Fragmentation Map

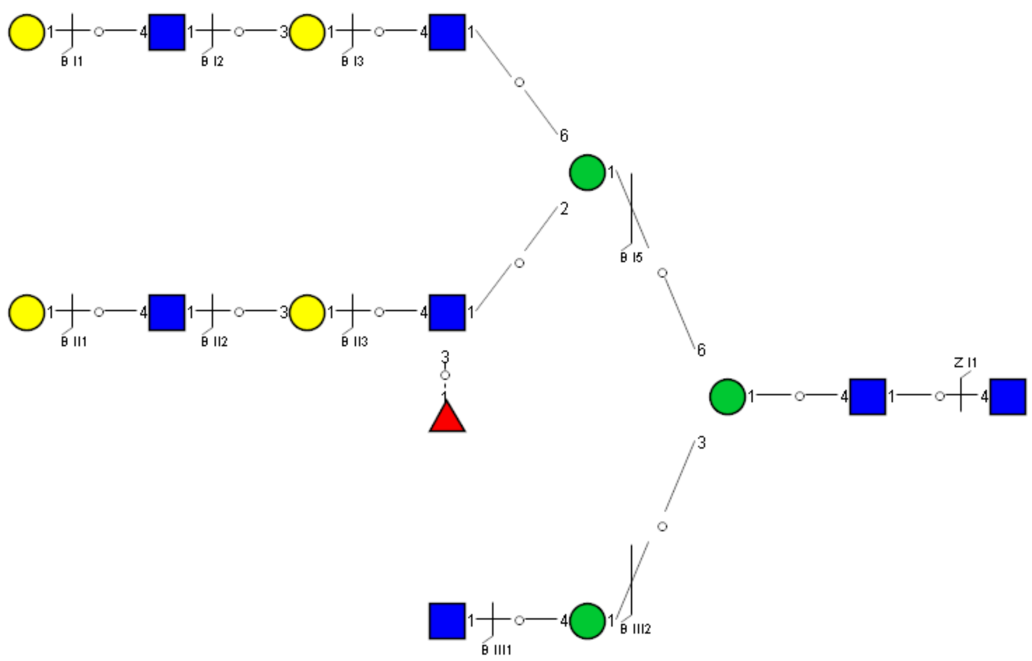

### Spectrum Index: 37094(NoBHs:1)

#### Precursor iEF Map

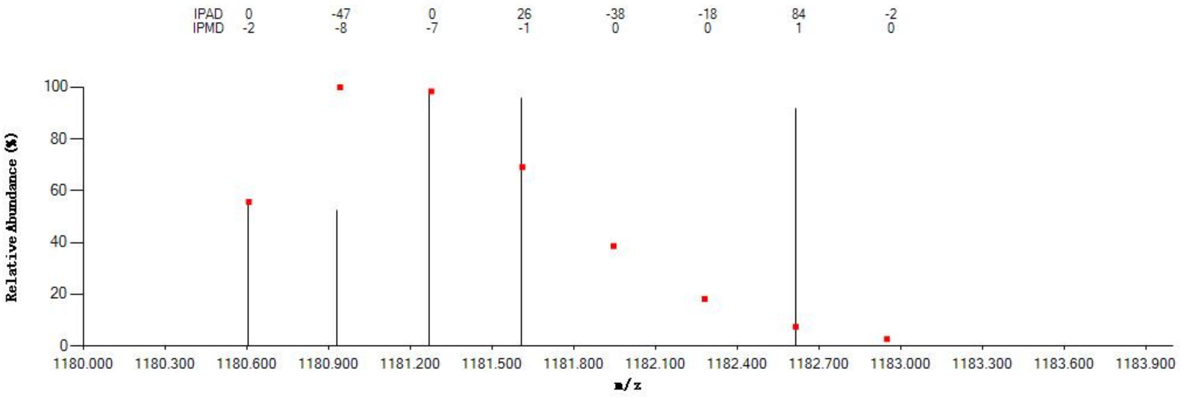


#### Annotated MS/MS Spectrum

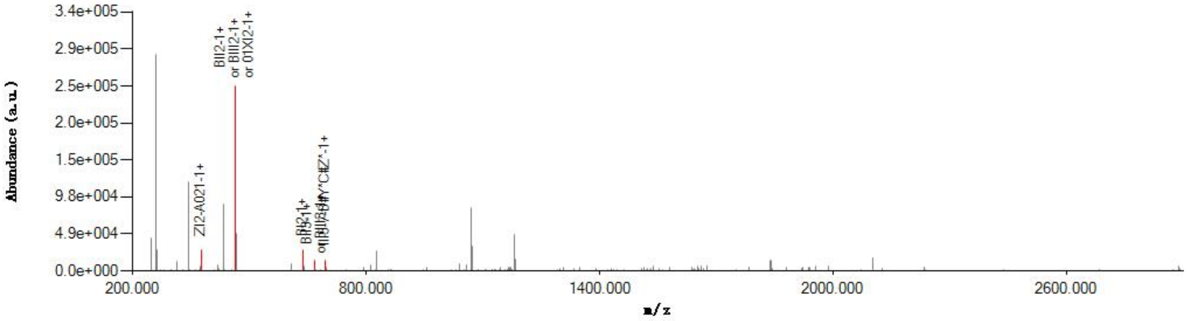


#### Graphical Fragmentation Map

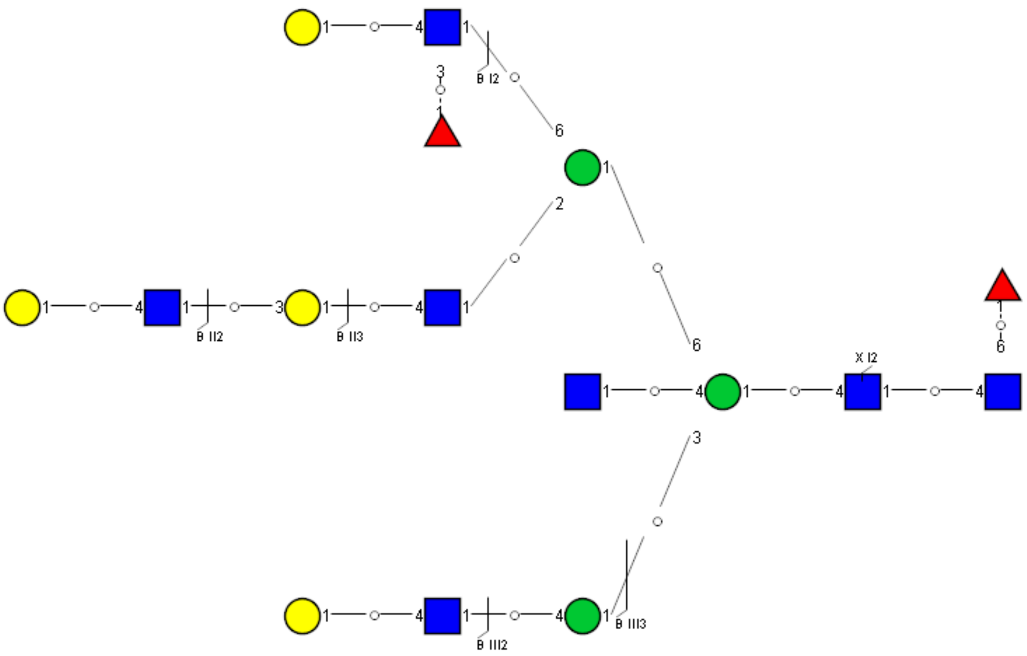

Spectrum Index: 37114(NoBHs:1)

#### Precursor iEF Map

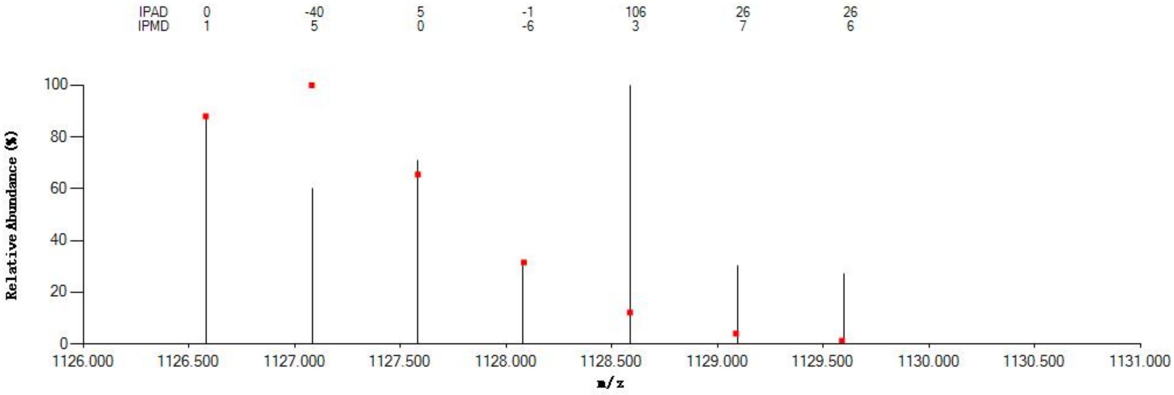

#### Annotated MS/MS Spectrum

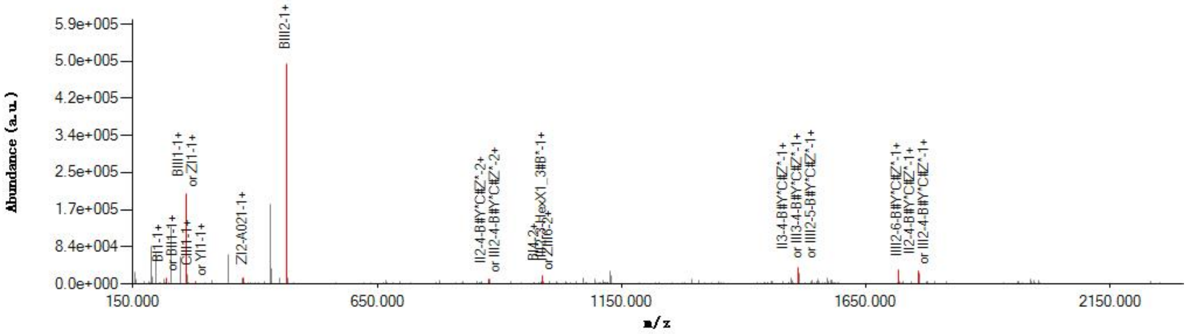

#### Graphical Fragmentation Map

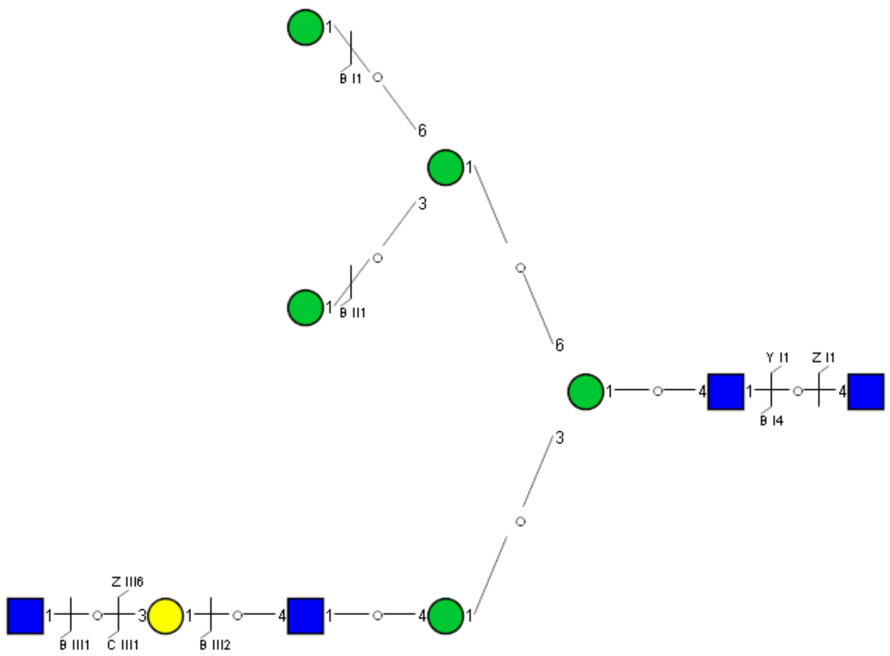

Spectrum Index: 37214(NoBHs:1)

#### Precursor iEF Map

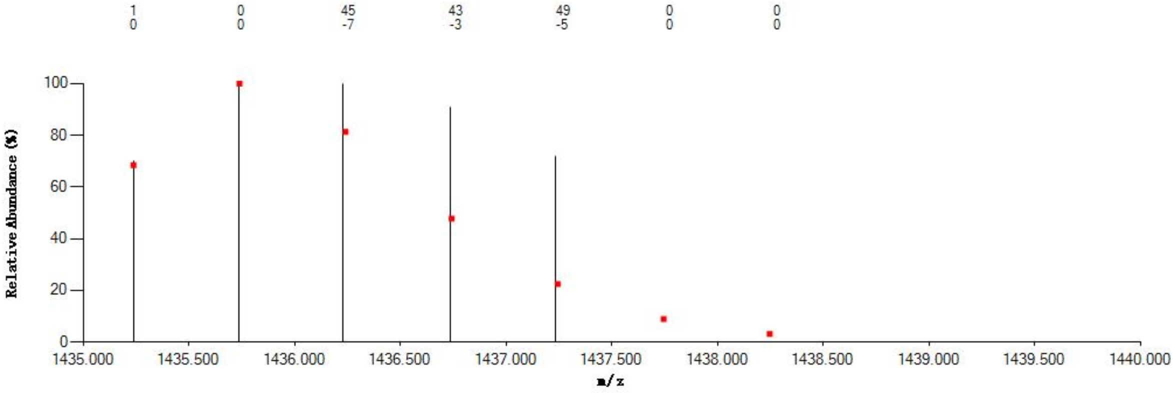


#### Annotated MS/MS Spectrum

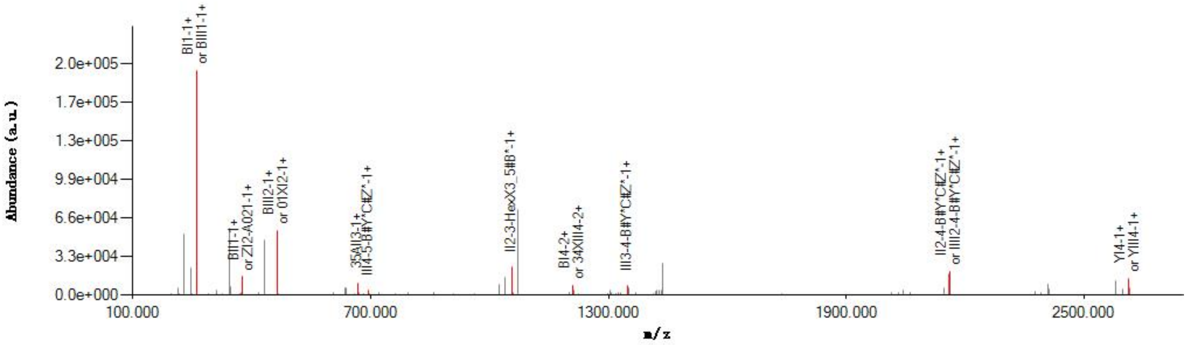


#### Graphical Fragmentation Map

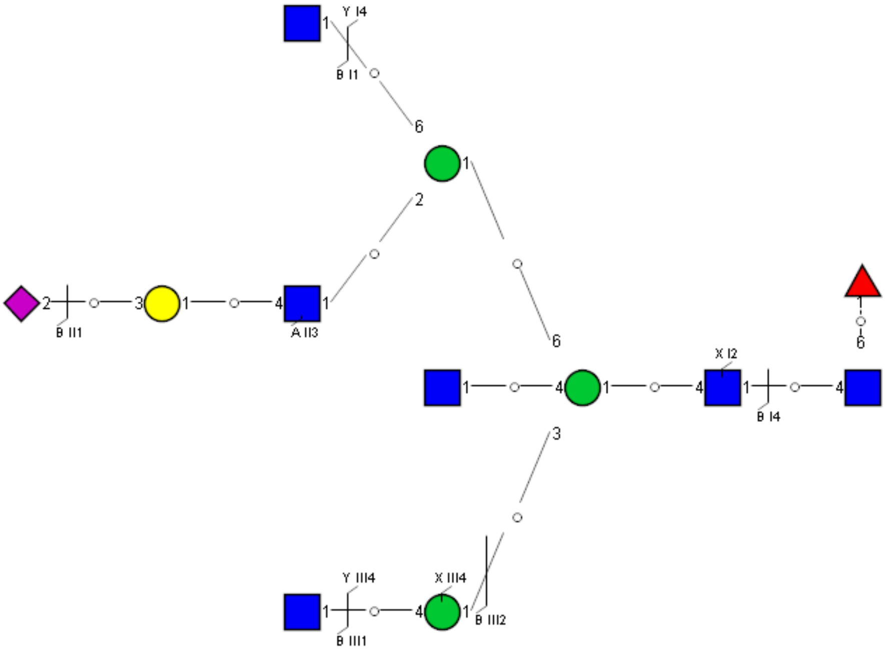

Spectrum Index: 37408(NoBHs:1)

#### Precursor iEF Map

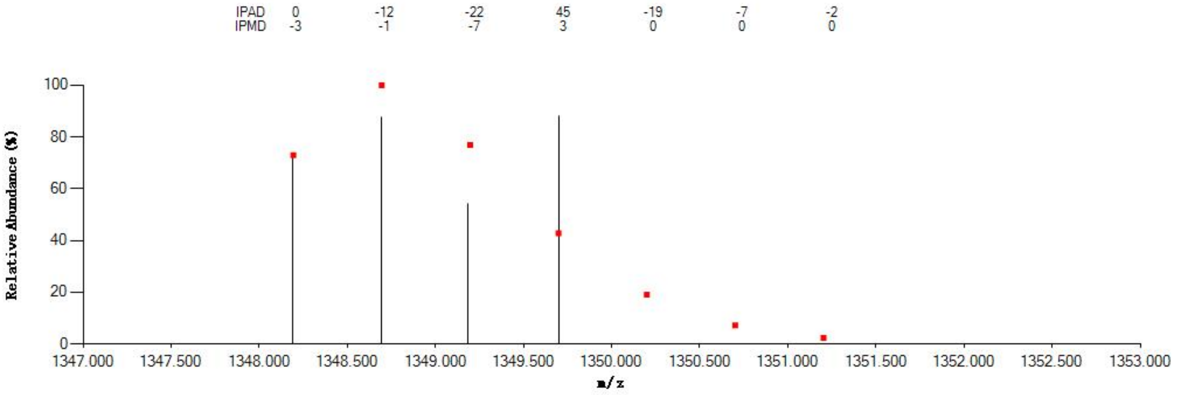


#### Annotated MS/MS Spectrum

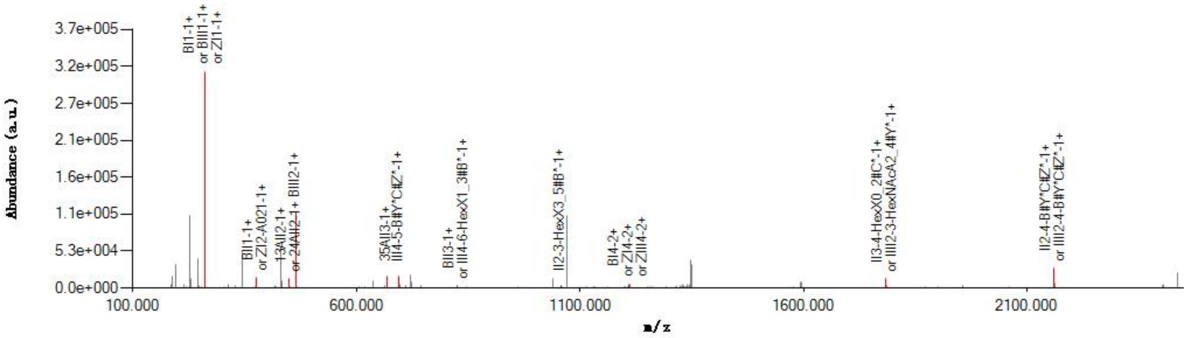


#### Graphical Fragmentation Map

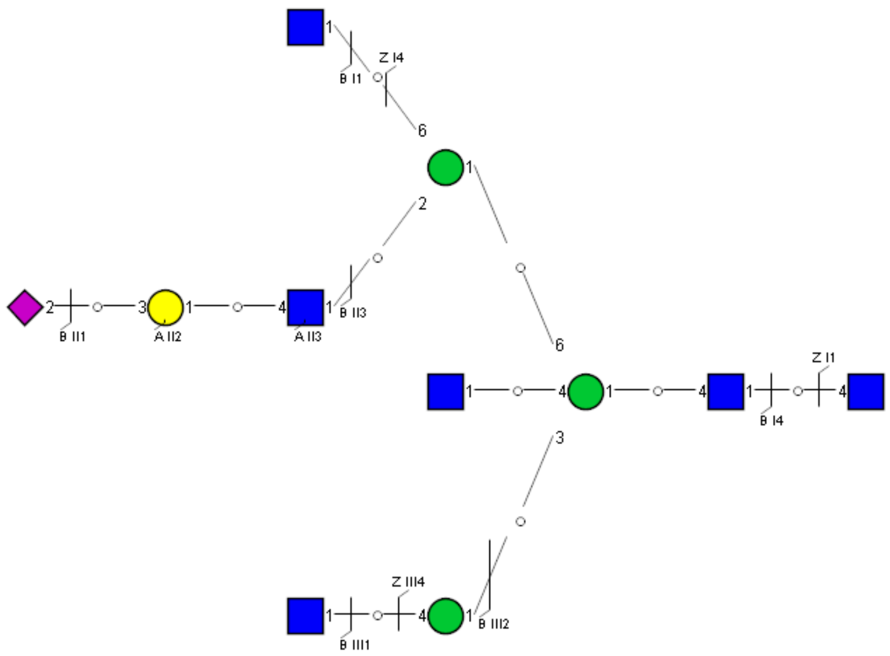

Spectrum Index: 37823(NoBHs:1)

#### Precursor iEF Map

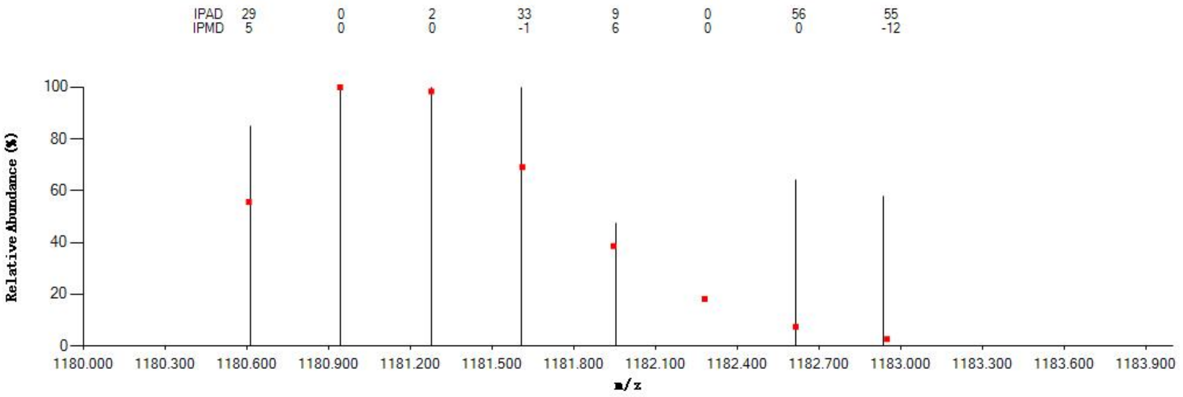


#### Annotated MS/MS Spectrum

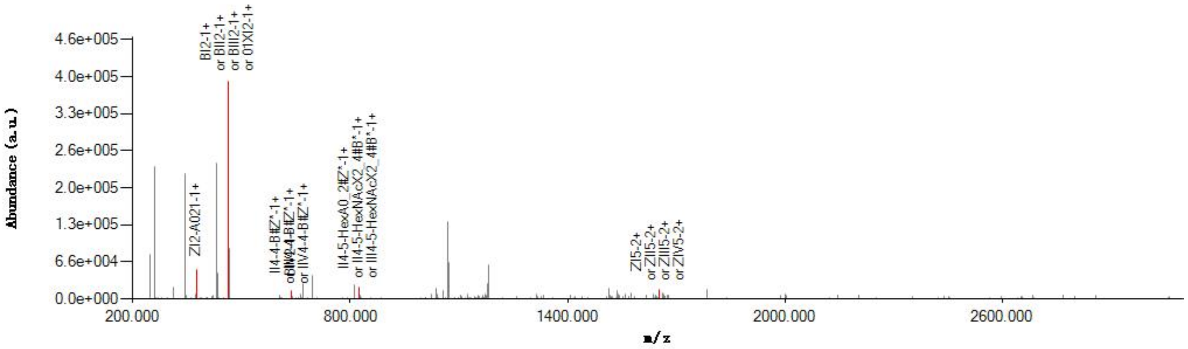


#### Graphical Fragmentation Map

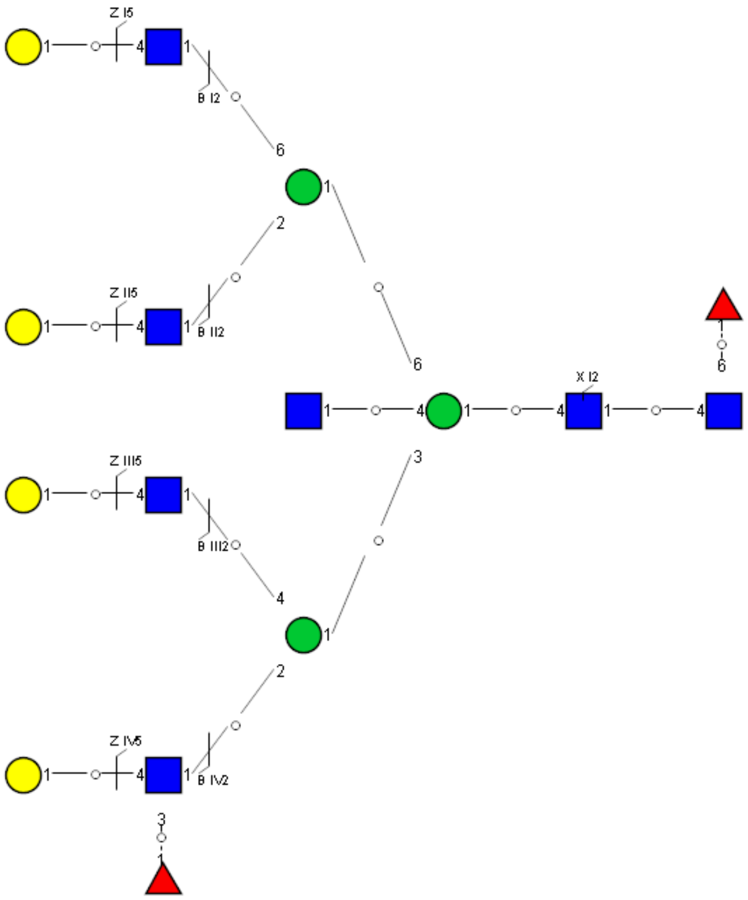

Spectrum Index: 37948(NoBHs:1)

#### Precursor iEF Map

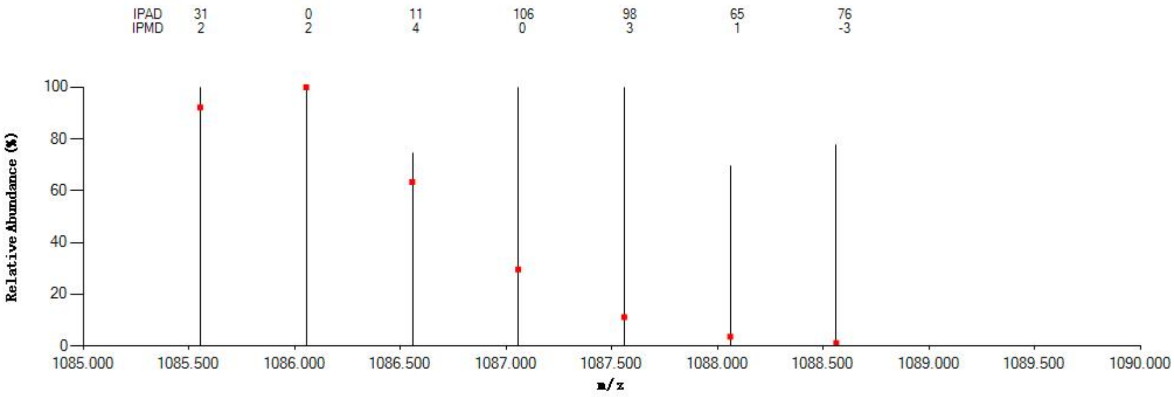


#### Annotated MS/MS Spectrum

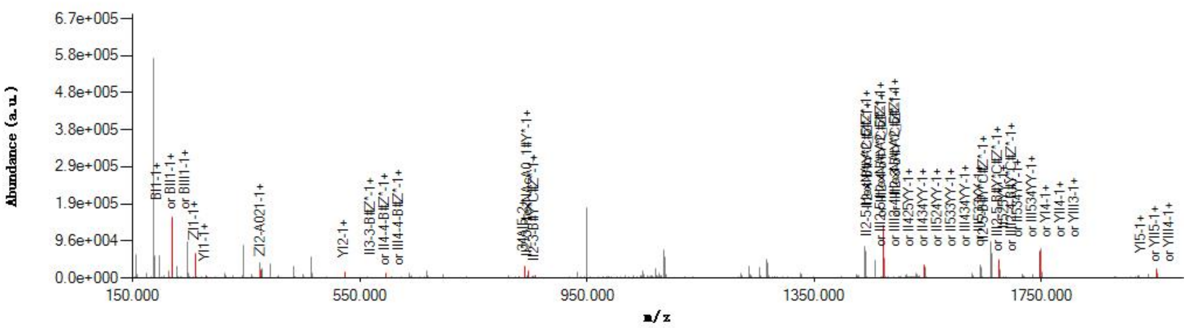


#### Graphical Fragmentation Map

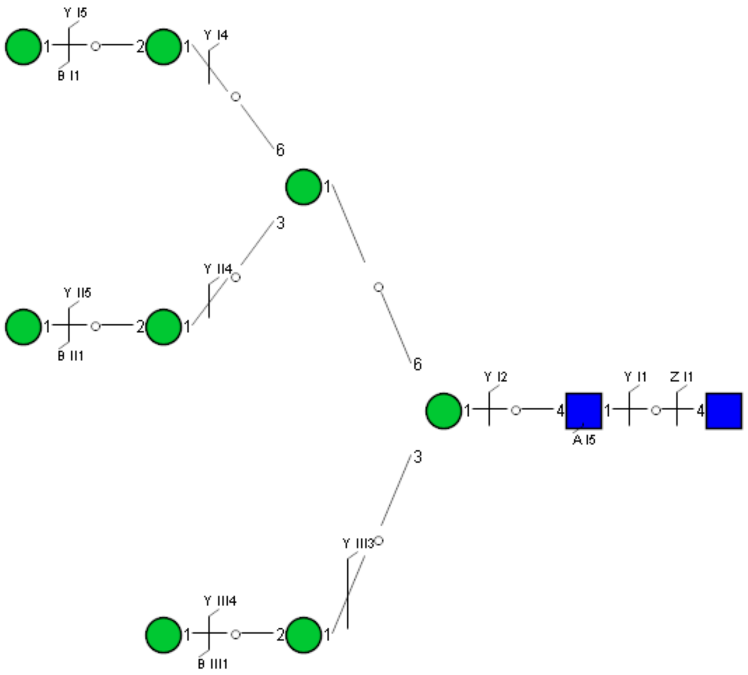

Spectrum Index: 37984(NoBHs:1)

#### Precursor iEF Map

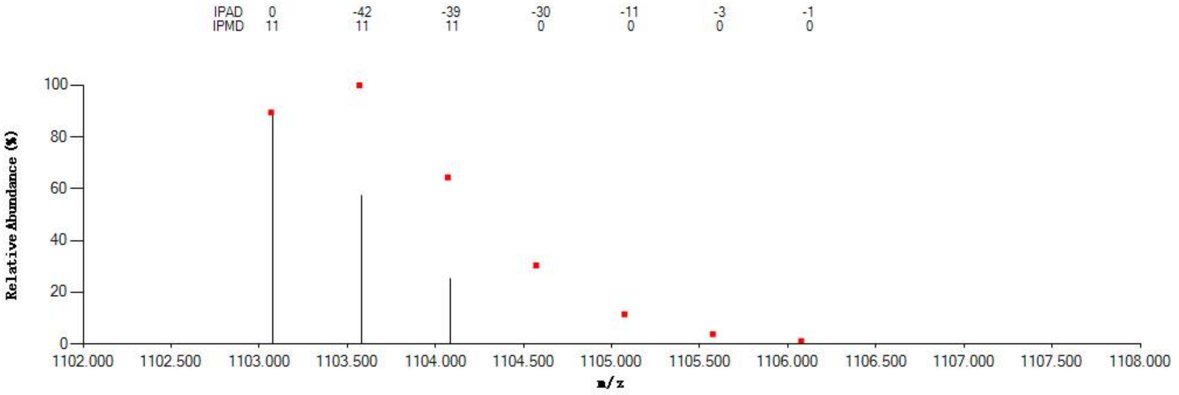


#### Annotated MS/MS Spectrum

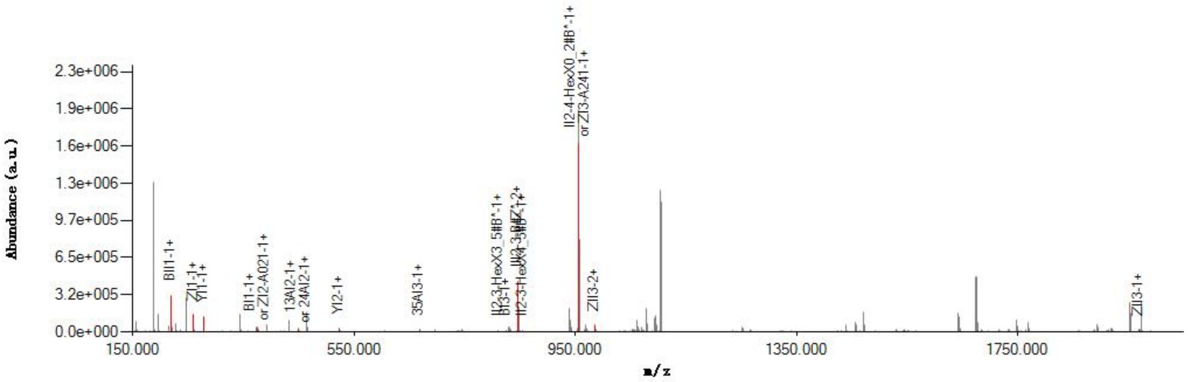


#### Graphical Fragmentation Map

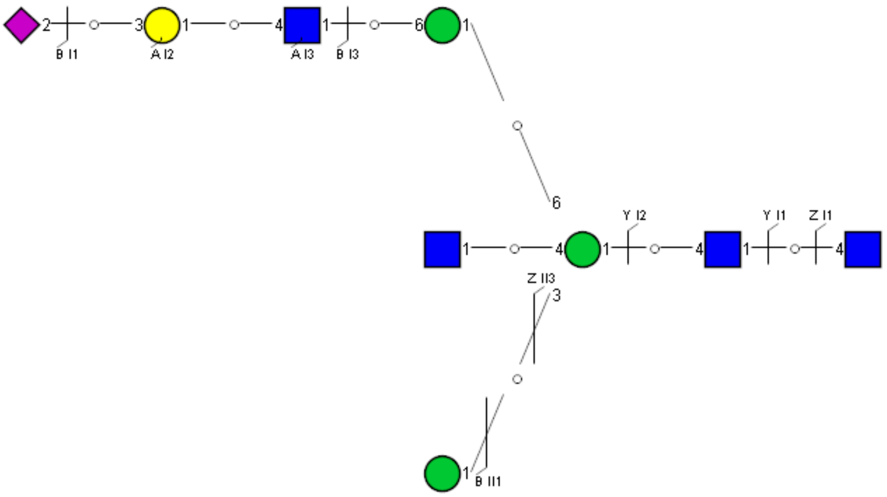

Spectrum Index: 38205(NoBHs:2)

#### Precursor iEF Map

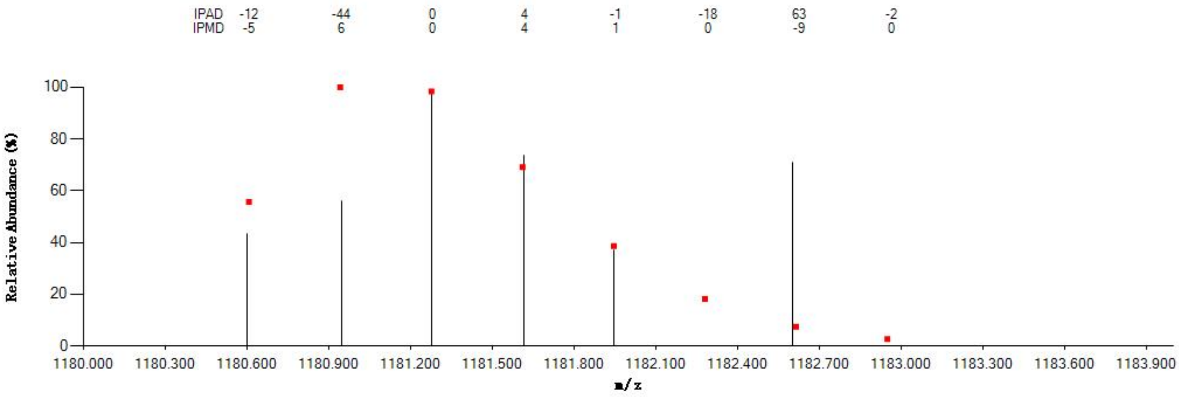


#### Annotated MS/MS Spectrum

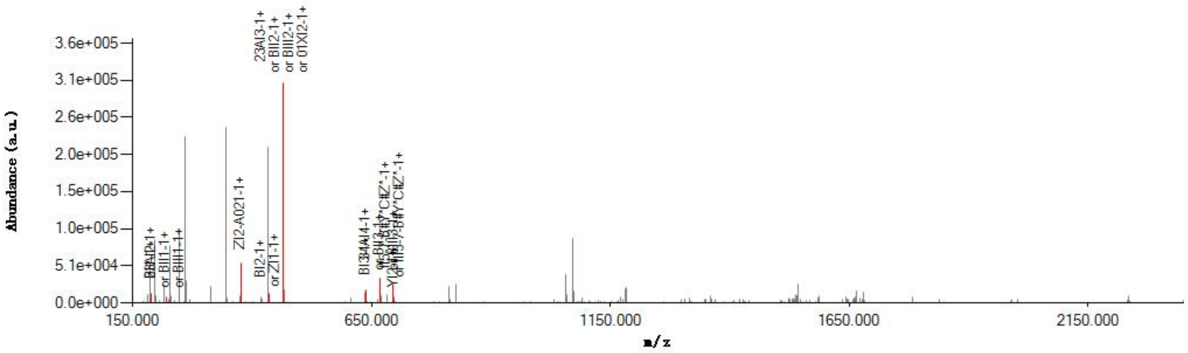


#### Graphical Fragmentation Map

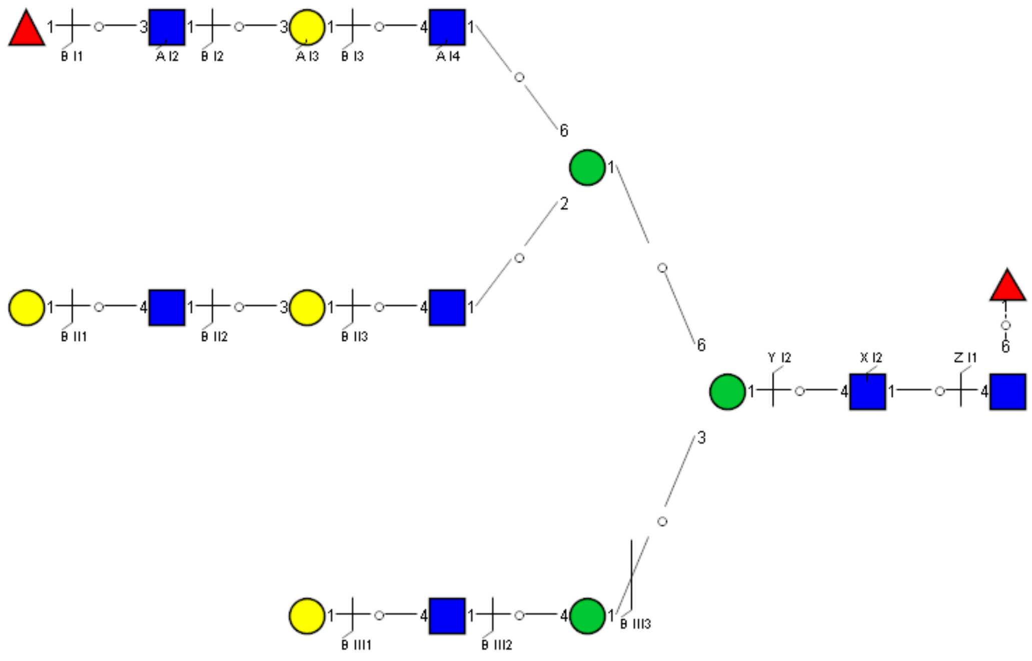

Spectrum Index: 38205(NoBHs:2)

#### Precursor iEF Map

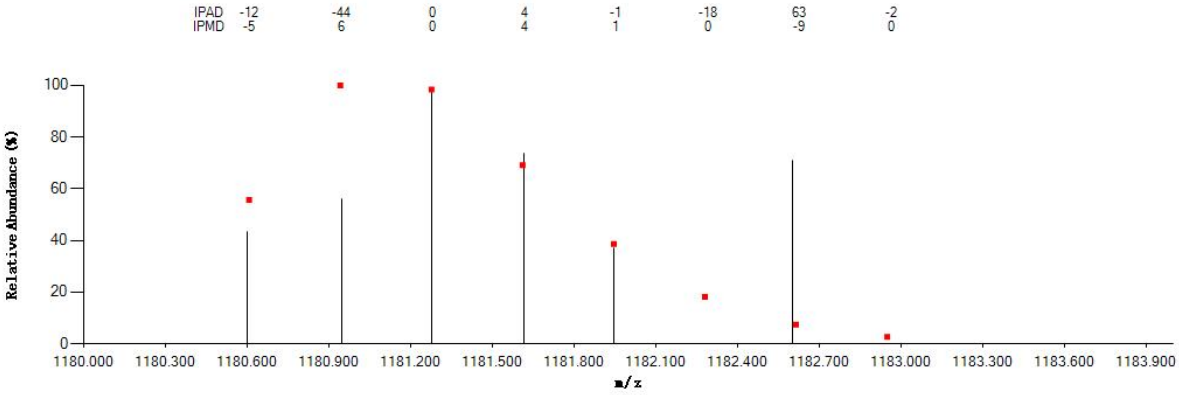


#### Annotated MS/MS Spectrum

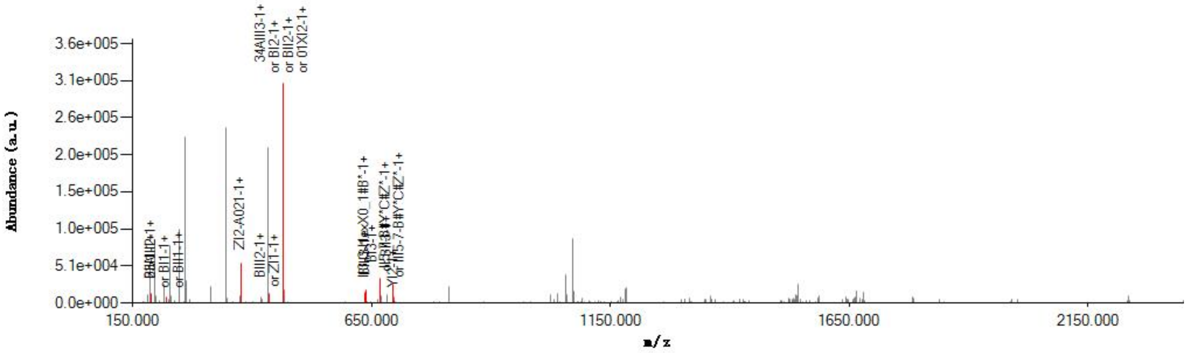


#### Graphical Fragmentation Map

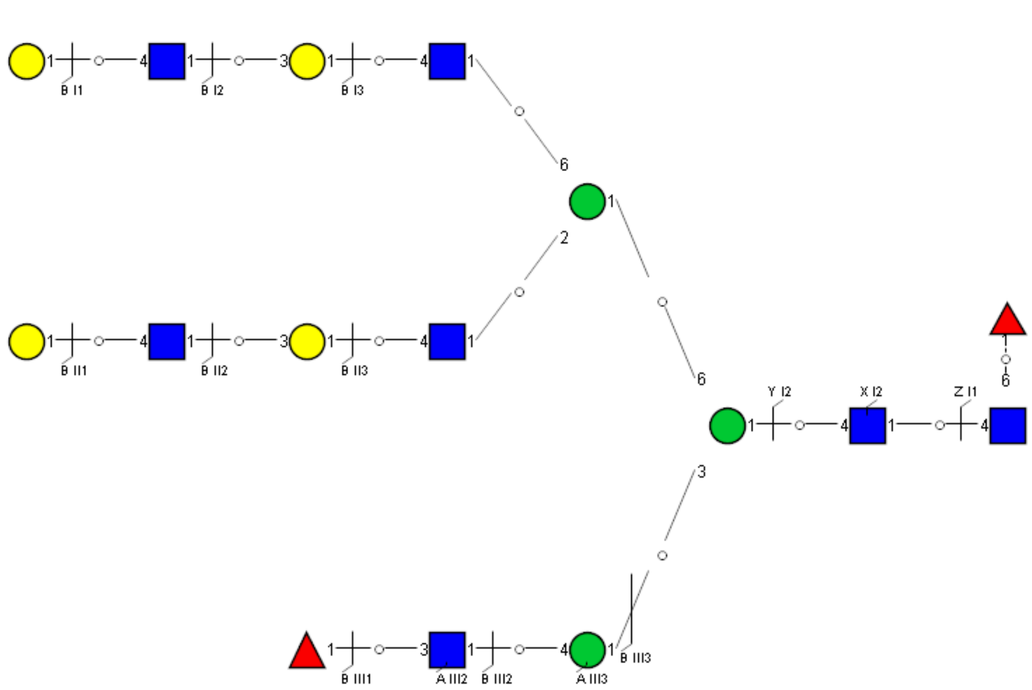

Spectrum Index: 38332(NoBHs:1)

#### Precursor iEF Map

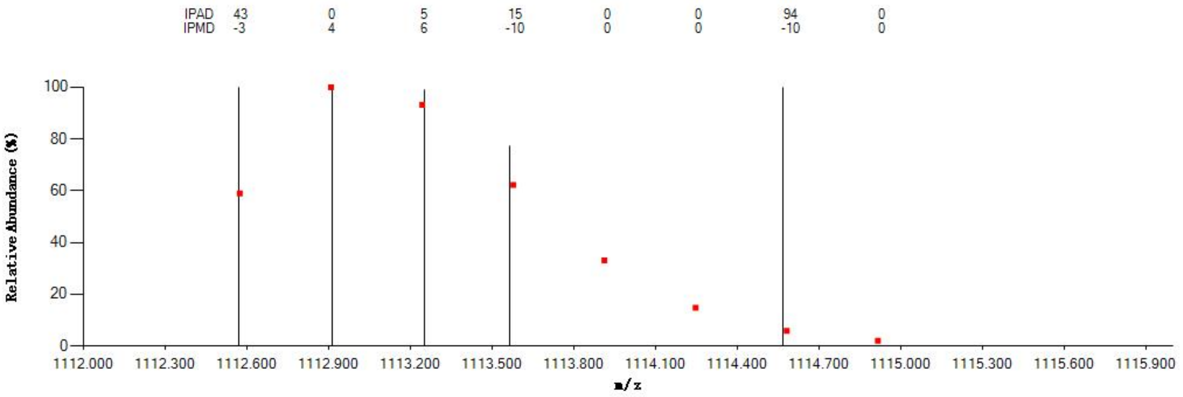


#### Annotated MS/MS Spectrum

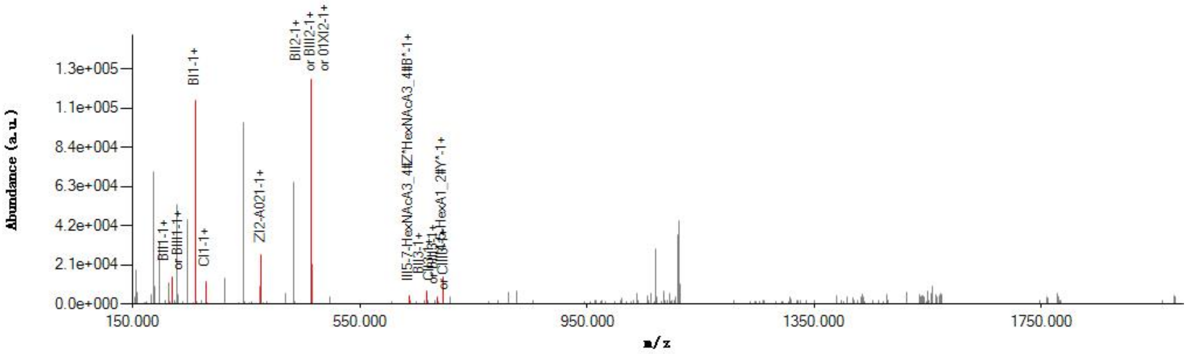


#### Graphical Fragmentation Map

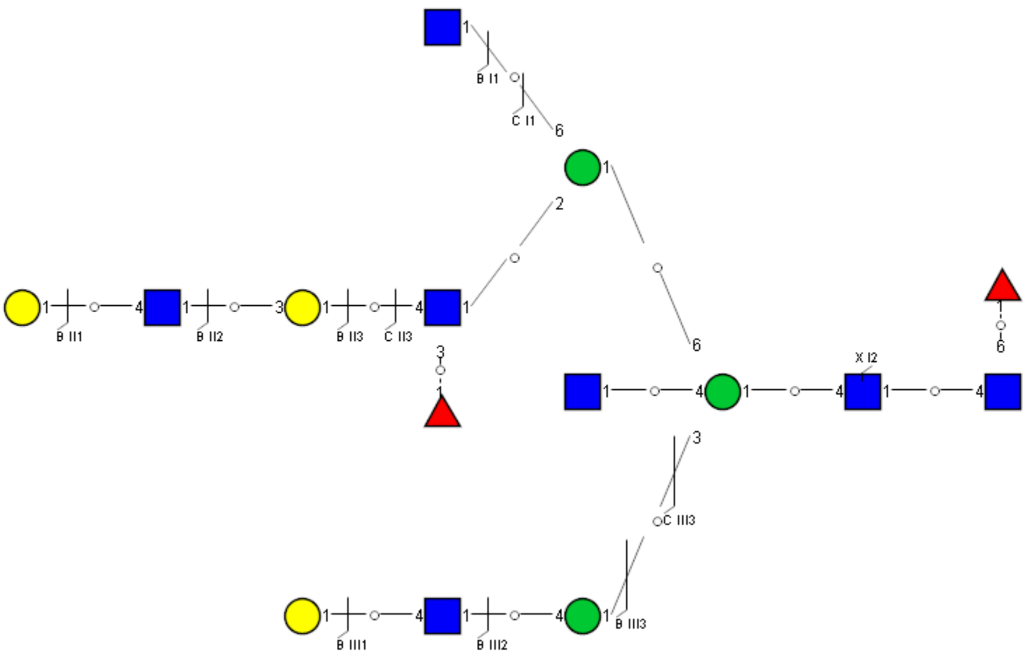

Spectrum Index: 38597(NoBHs:2)

#### Precursor iEF Map

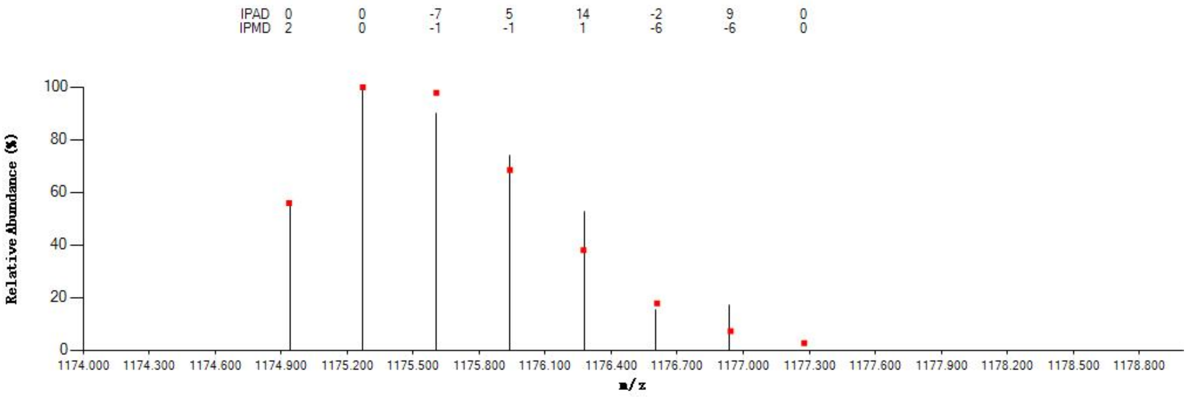


#### Annotated MS/MS Spectrum

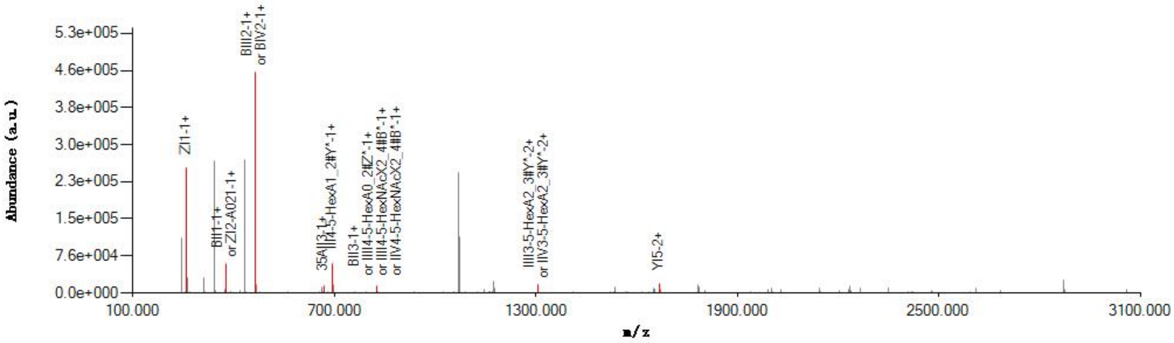


#### Graphical Fragmentation Map

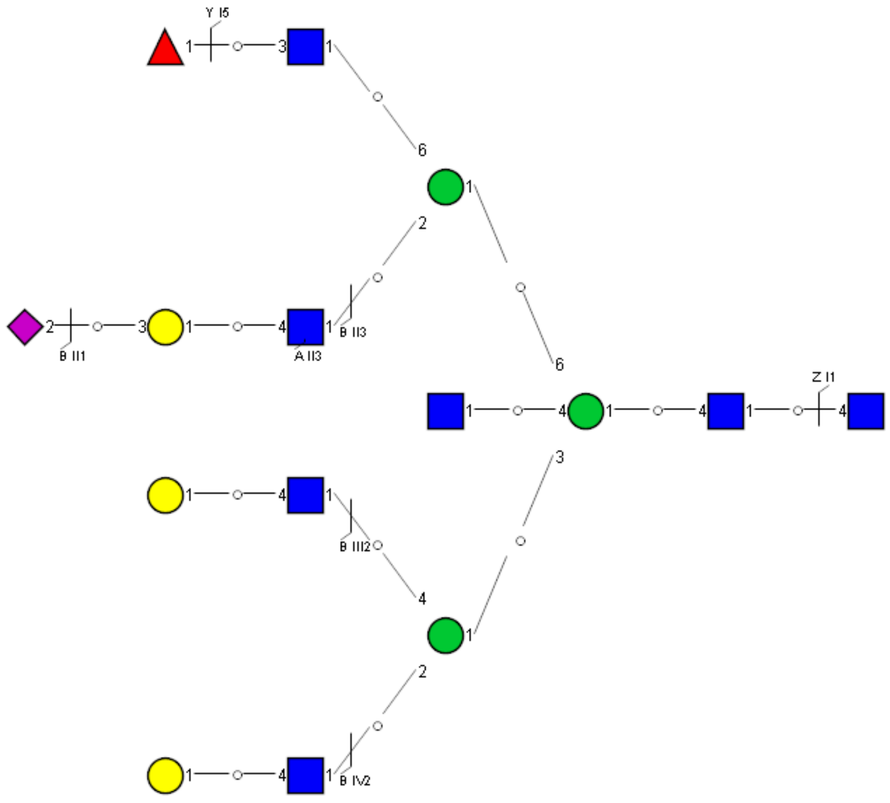

### Spectrum Index: 38597(NoBHs:2)

#### Precursor iEF Map

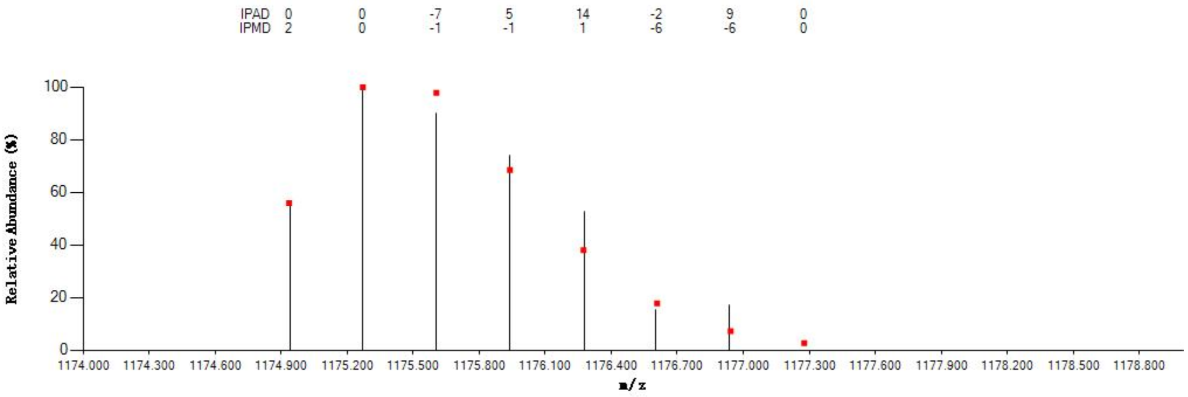


#### Annotated MS/MS Spectrum

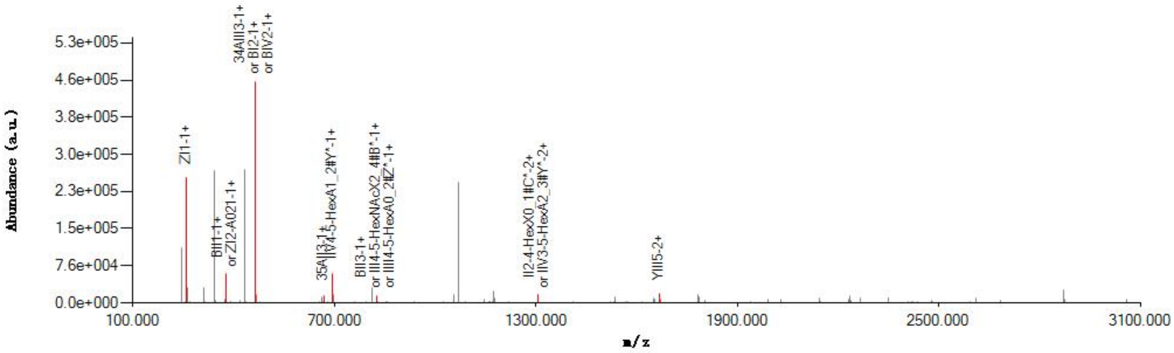


#### Graphical Fragmentation Map

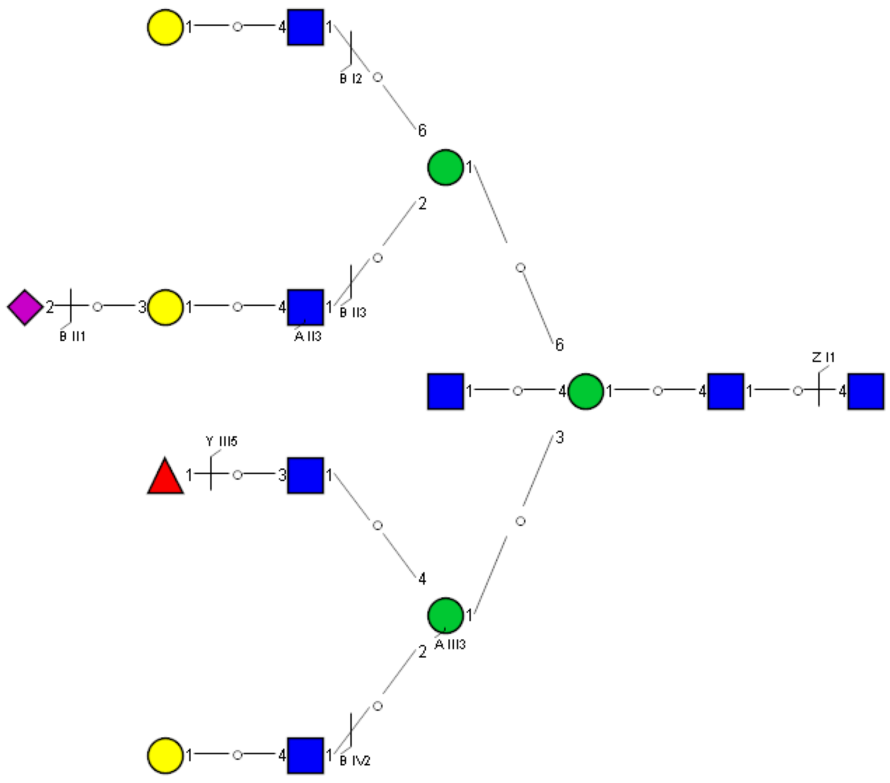

### Spectrum Index: 38859(NoBHs:1)

#### Precursor iEF Map

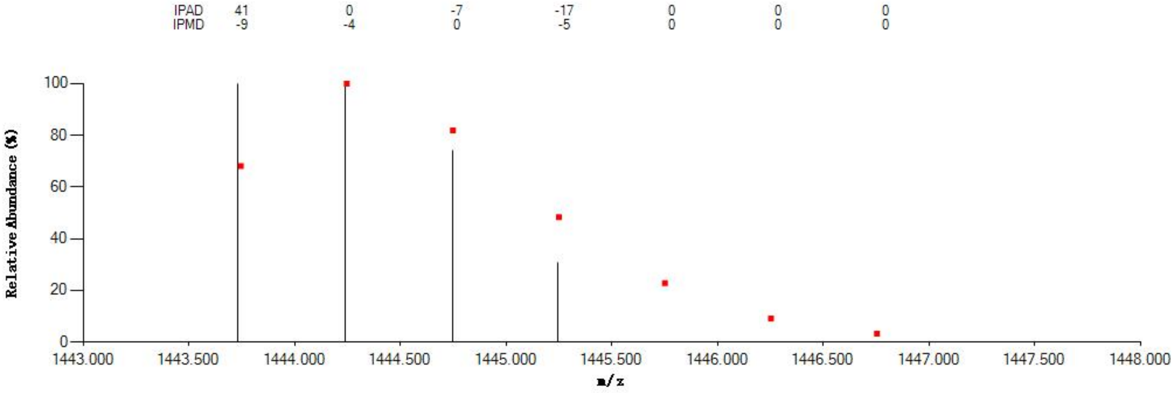


#### Annotated MS/MS Spectrum

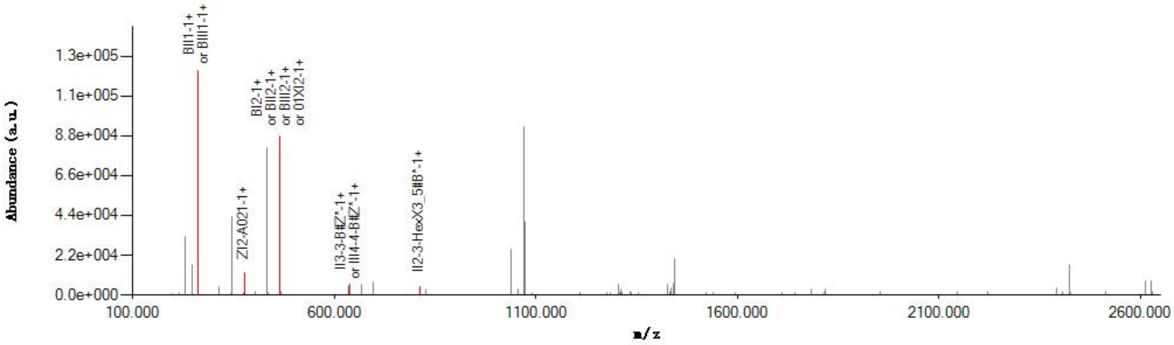


#### Graphical Fragmentation Map

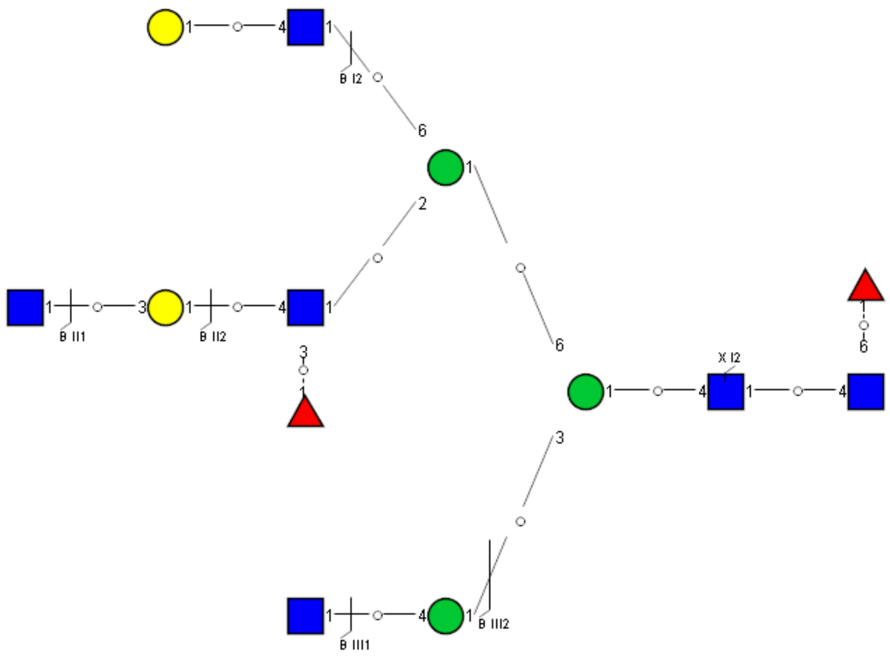

Spectrum Index: 38935(NoBHs:1)

#### Precursor iEF Map

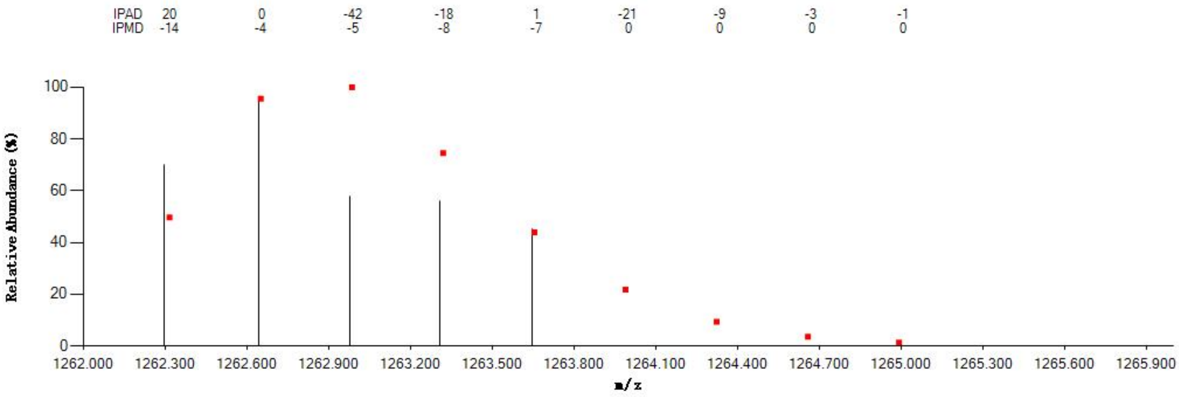


#### Annotated MS/MS Spectrum

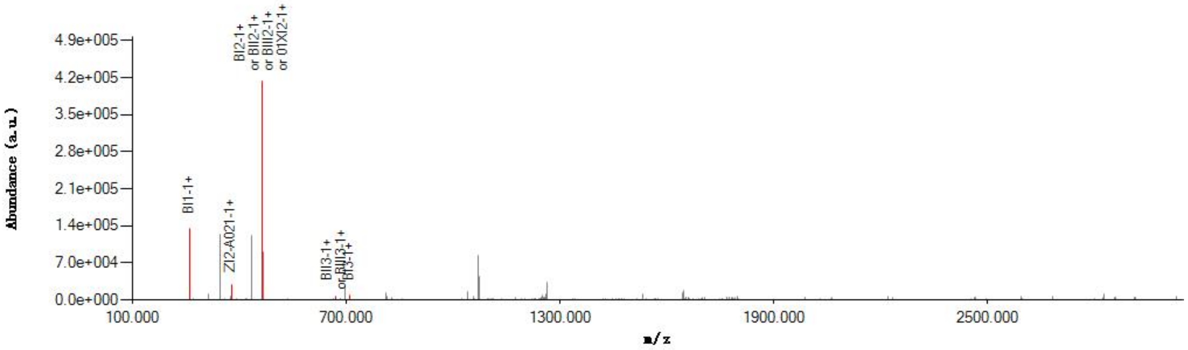


#### Graphical Fragmentation Map

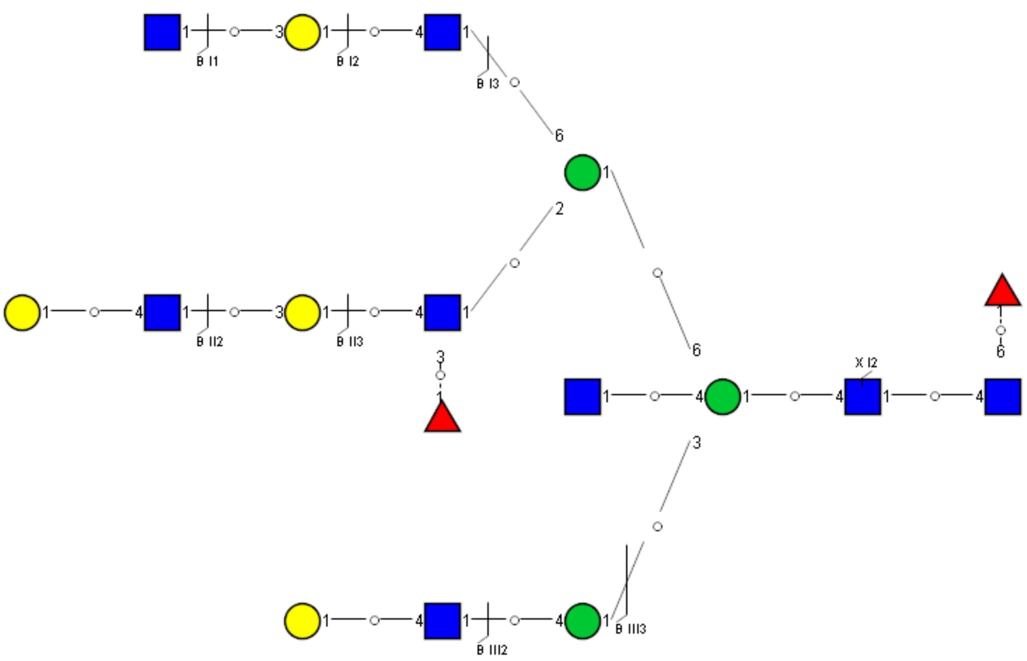

Spectrum Index: 39173(NoBHs:2)

Precursor iEF Map

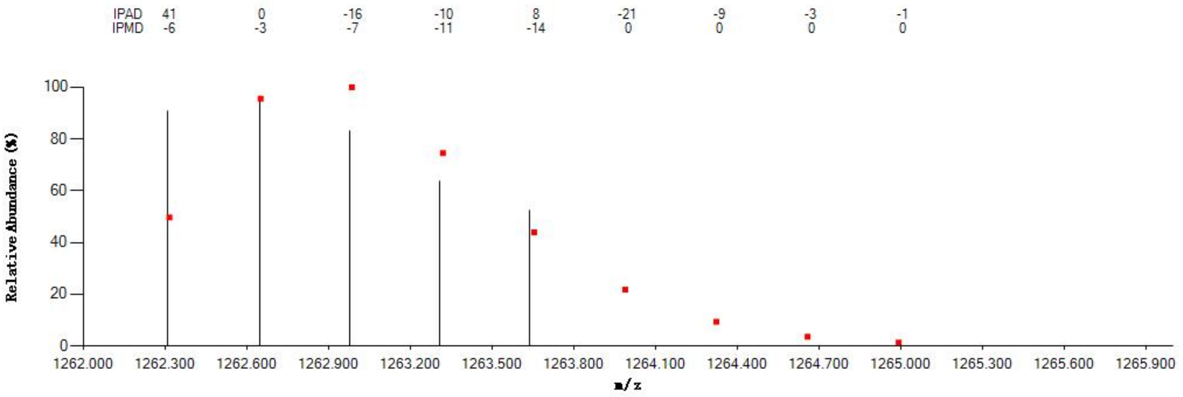

Annotated MS/MS Spectrum

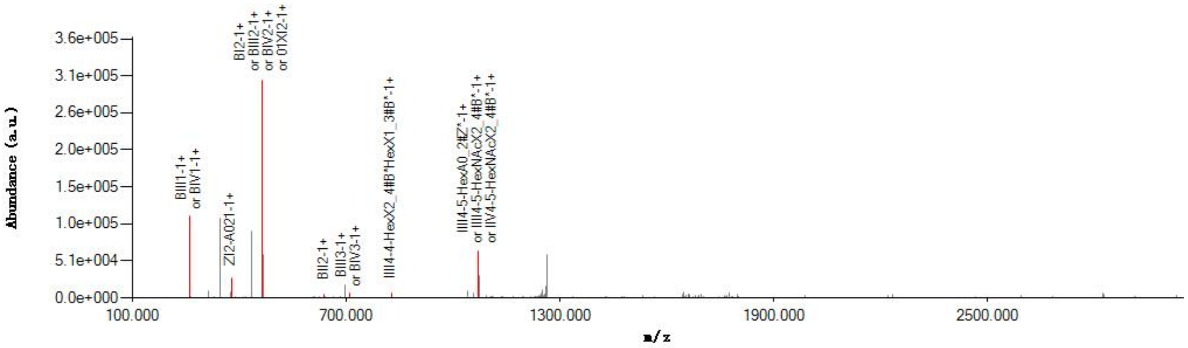

Graphical Fragmentation Map

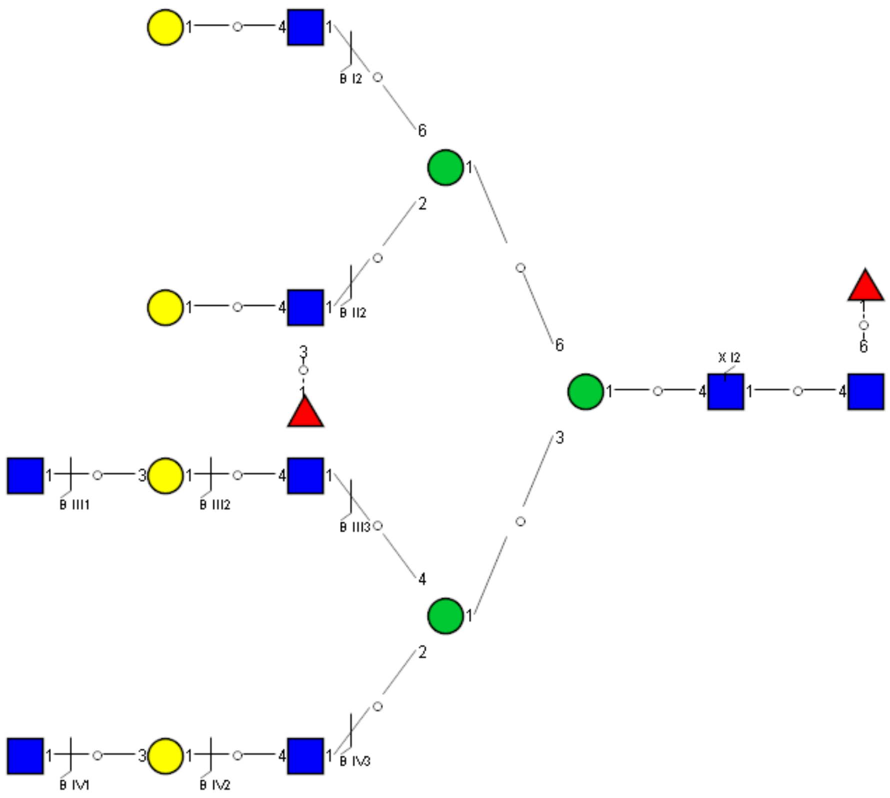

Spectrum Index: 39173(NoBHs:2)

#### Precursor iEF Map

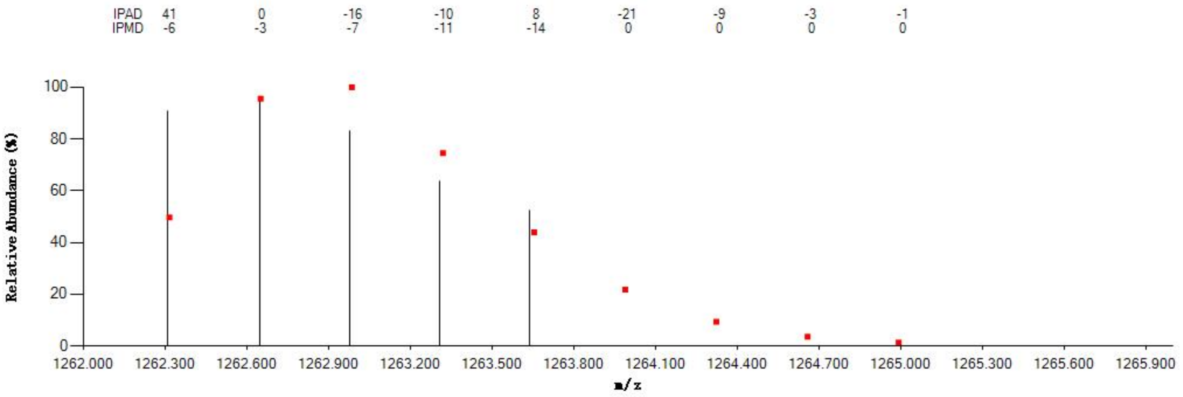

#### Annotated MS/MS Spectrum

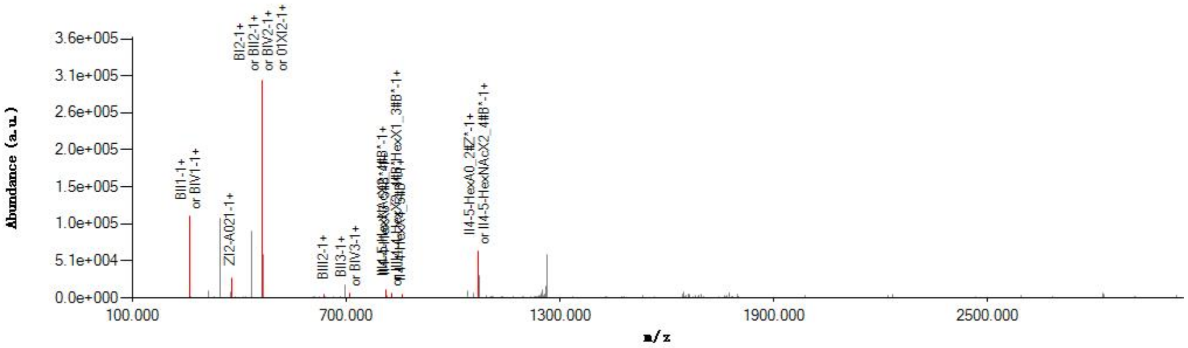

#### Graphical Fragmentation Map

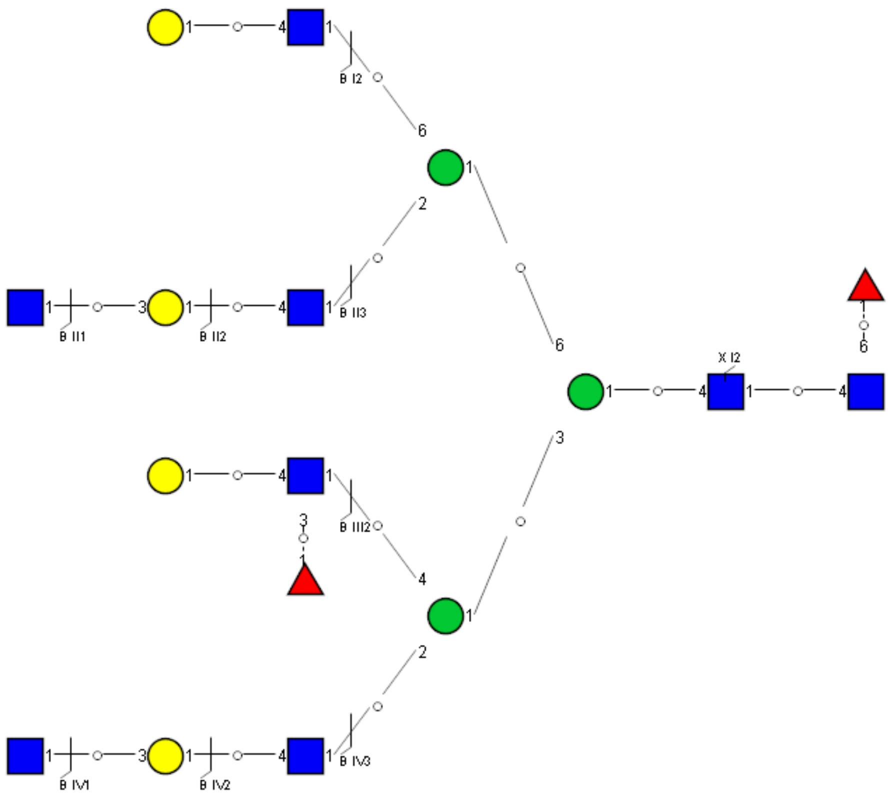

### Spectrum Index: 39249(NoBHs:1)

#### Precursor iEF Map

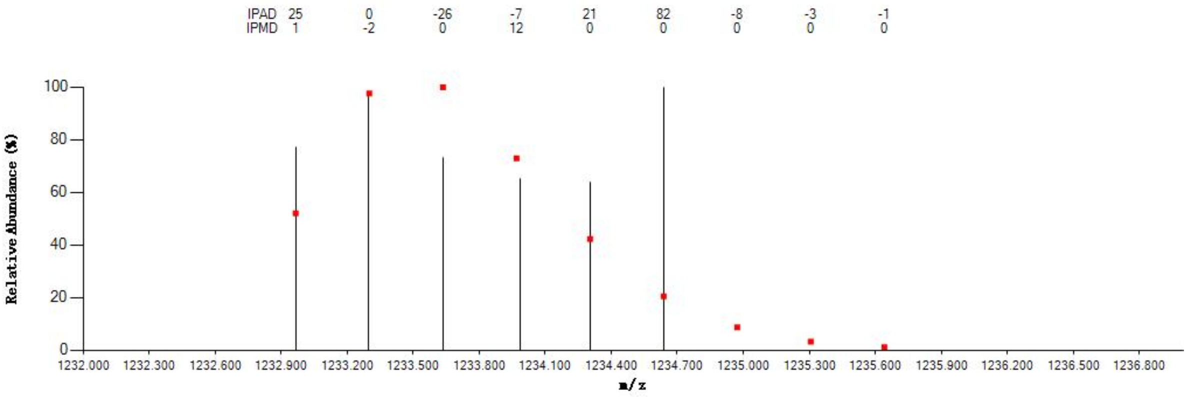

#### Annotated MS/MS Spectrum

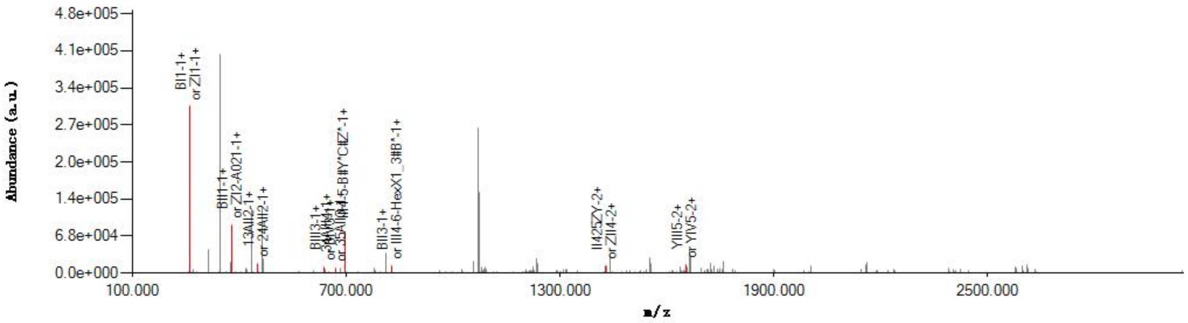

#### Graphical Fragmentation Map

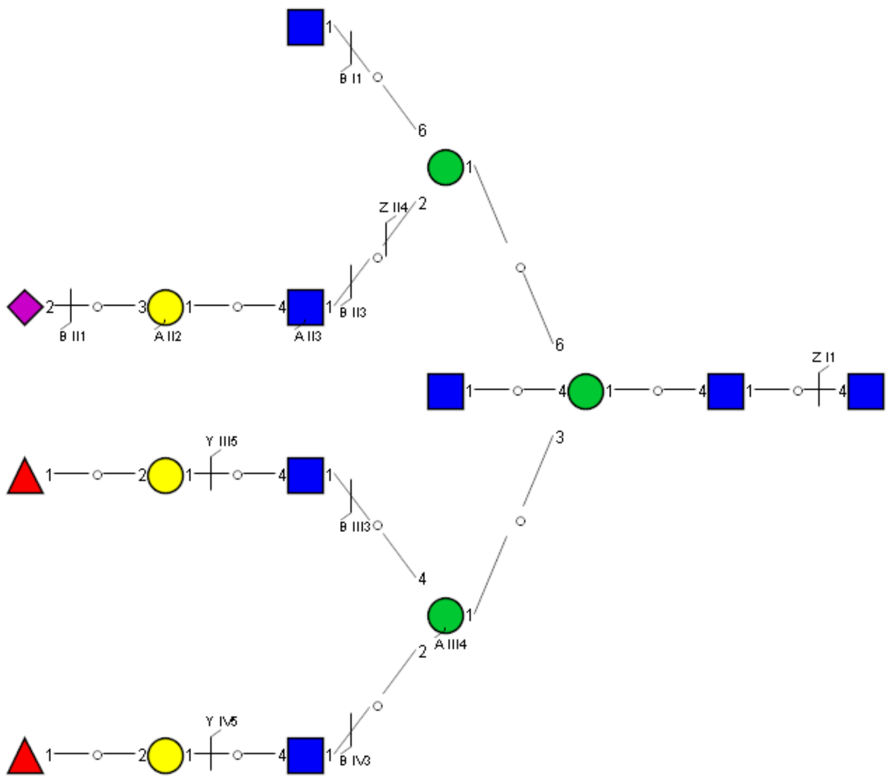

### Spectrum Index: 39455(NoBHs:2)

#### Precursor iEF Map

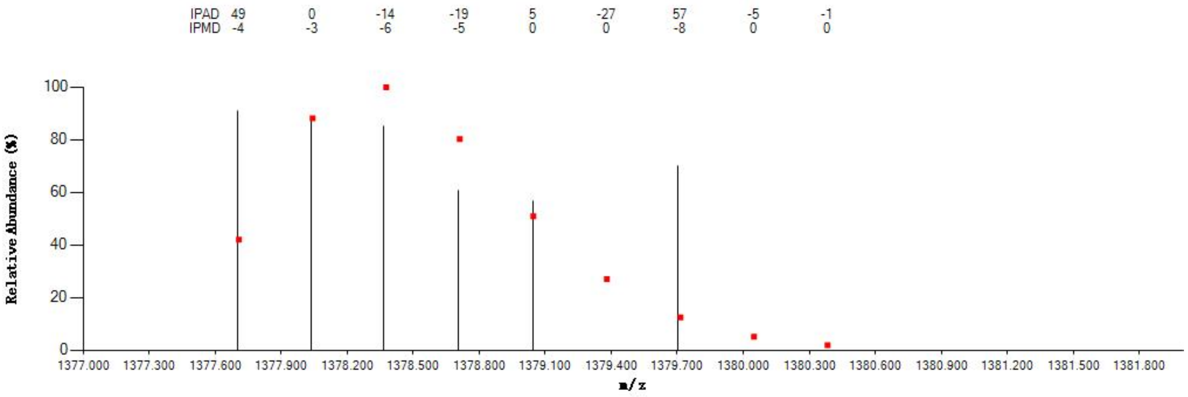

#### Annotated MS/MS Spectrum

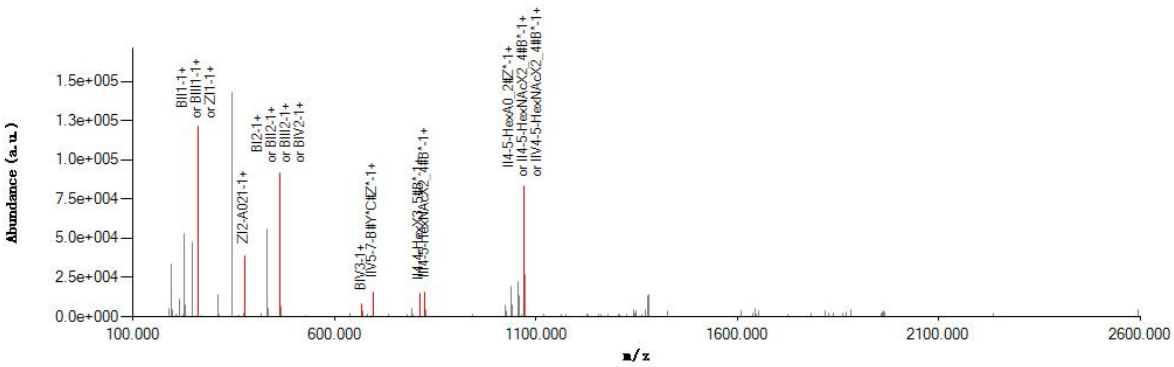

#### Graphical Fragmentation Map

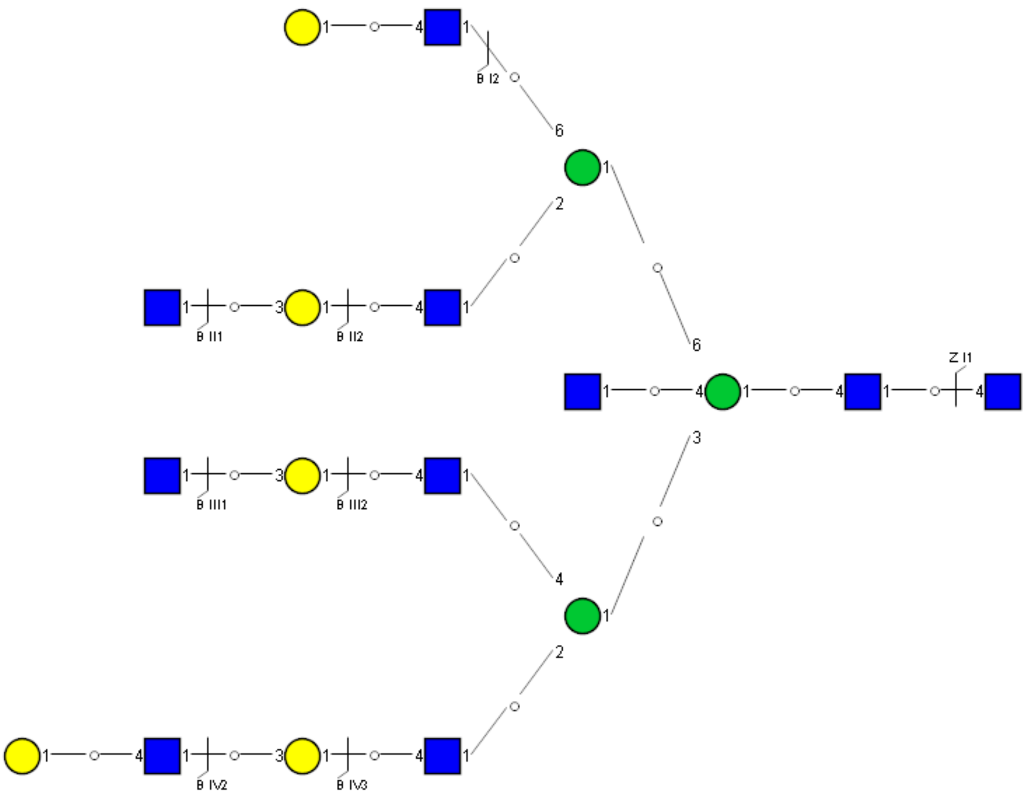

Spectrum Index: 39455(NoBHs:2)

#### Precursor iEF Map

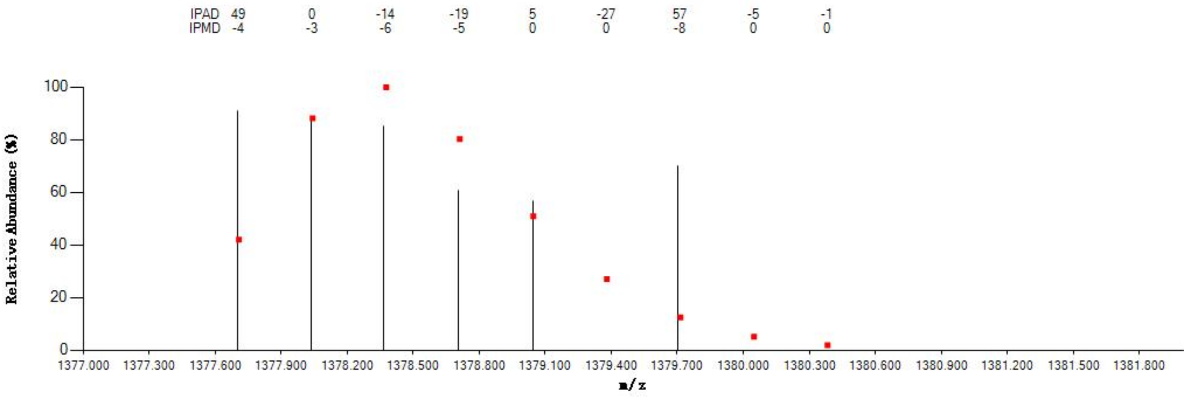


#### Annotated MS/MS Spectrum

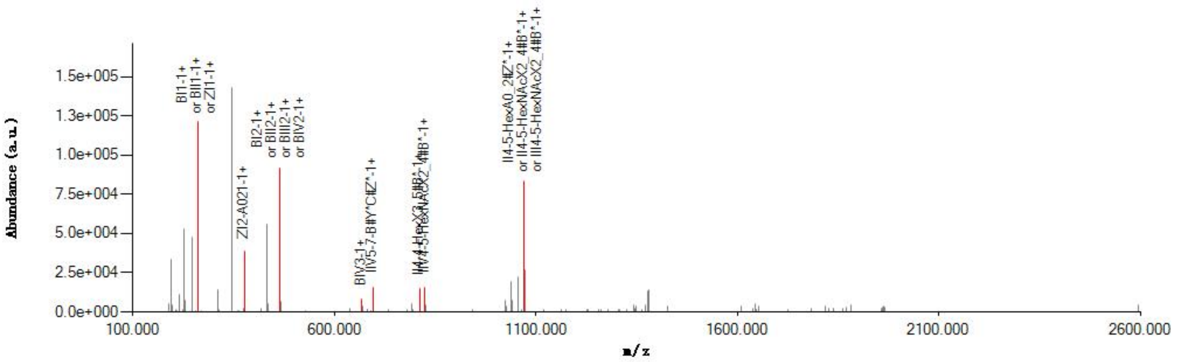


#### Graphical Fragmentation Map

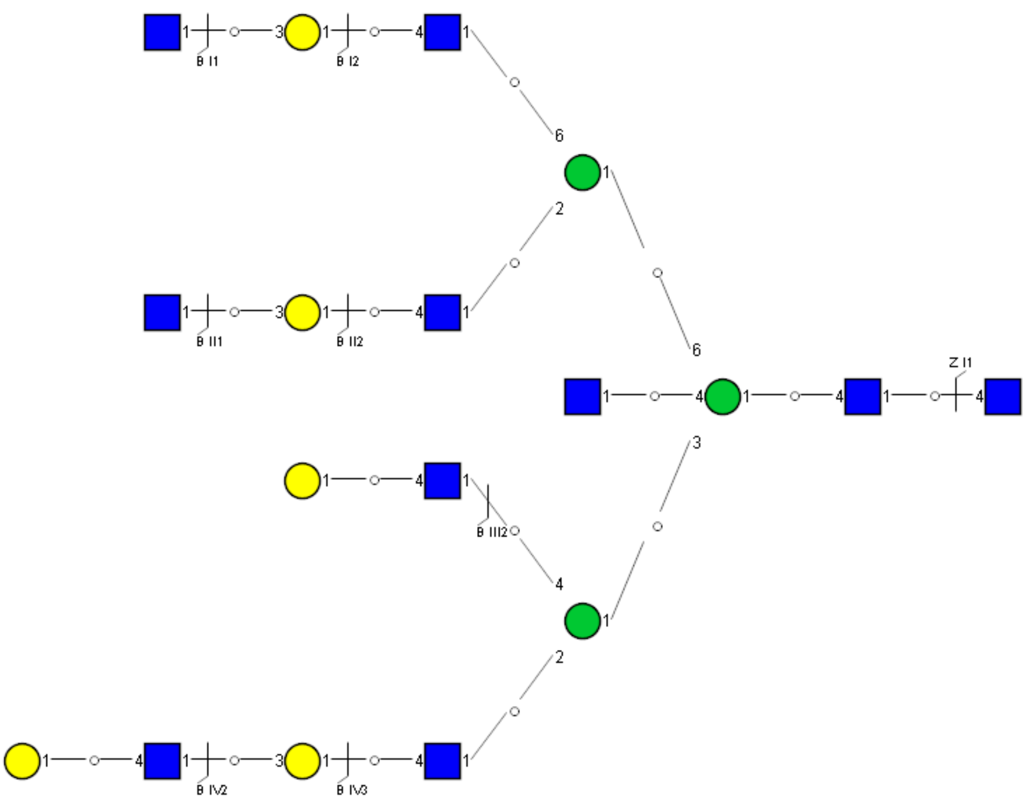

Spectrum Index: 39968(NoBHs:1)

#### Precursor iEF Map

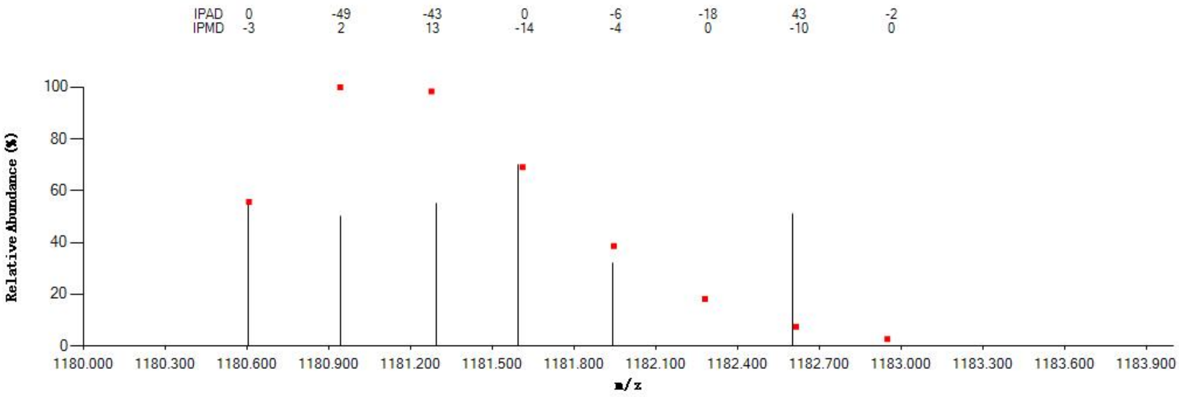


#### Annotated MS/MS Spectrum

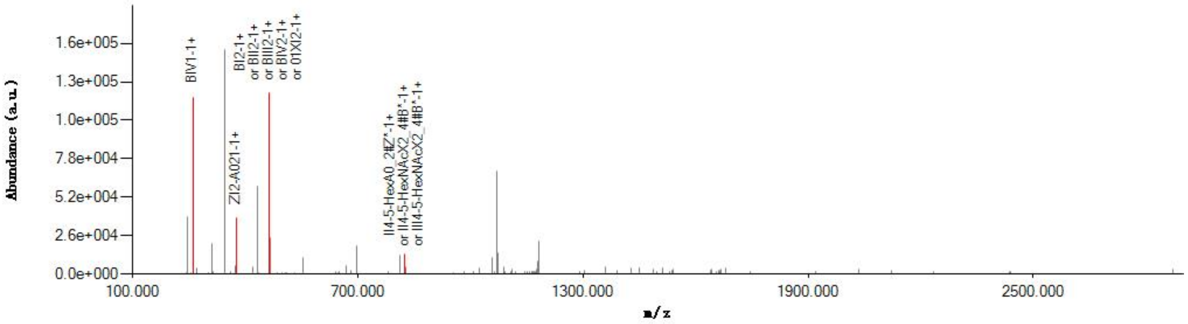


#### Graphical Fragmentation Map

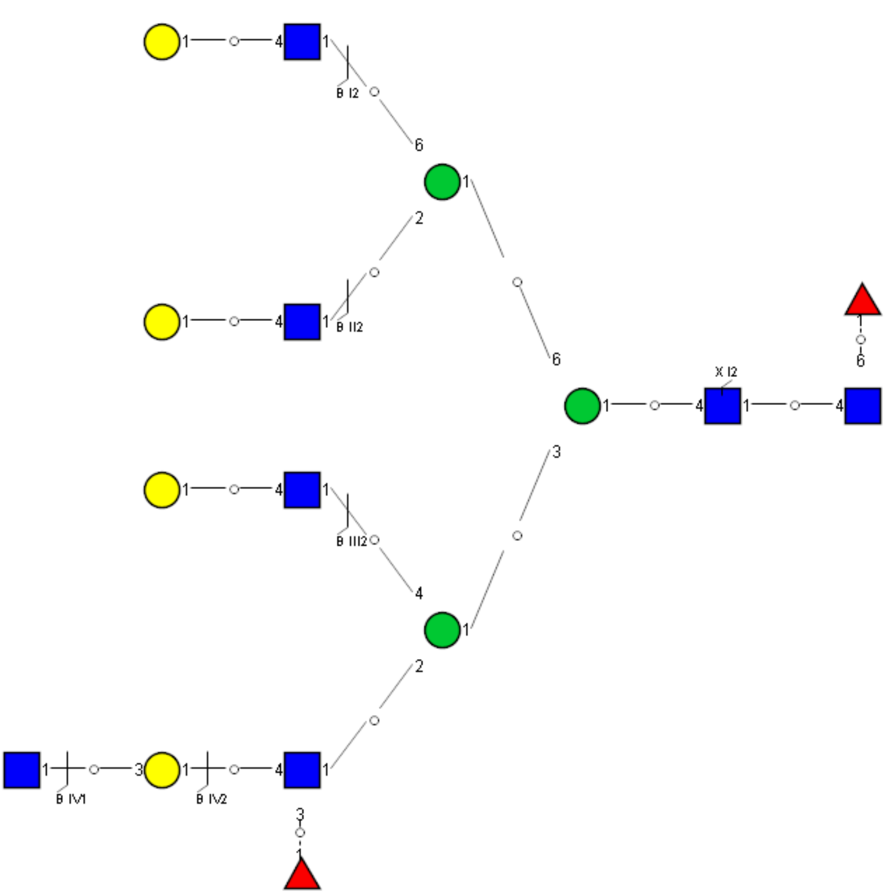

Spectrum Index: 39976(NoBHs:2)

#### Precursor iEF Map

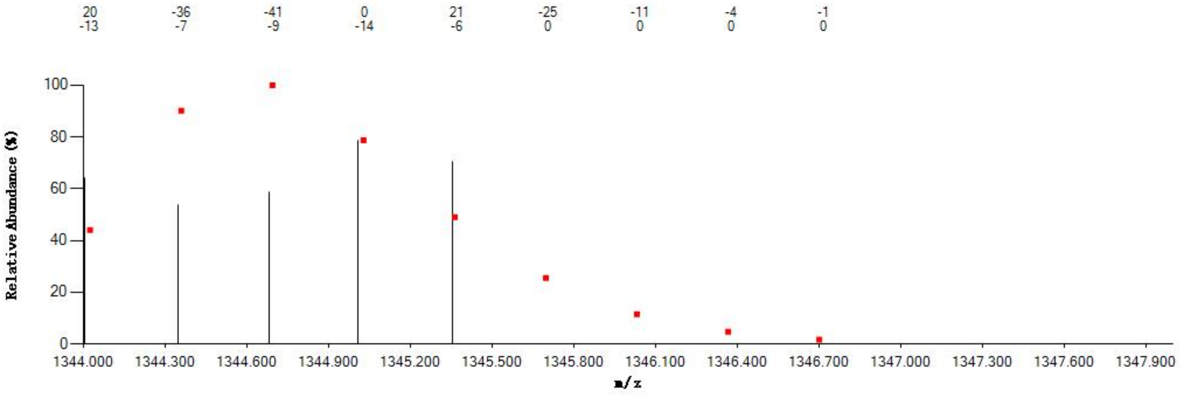


#### Annotated MS/MS Spectrum

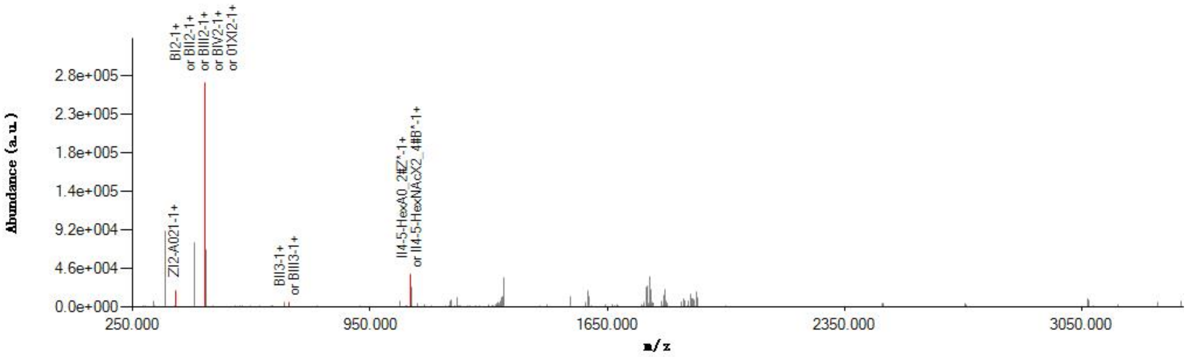


#### Graphical Fragmentation Map

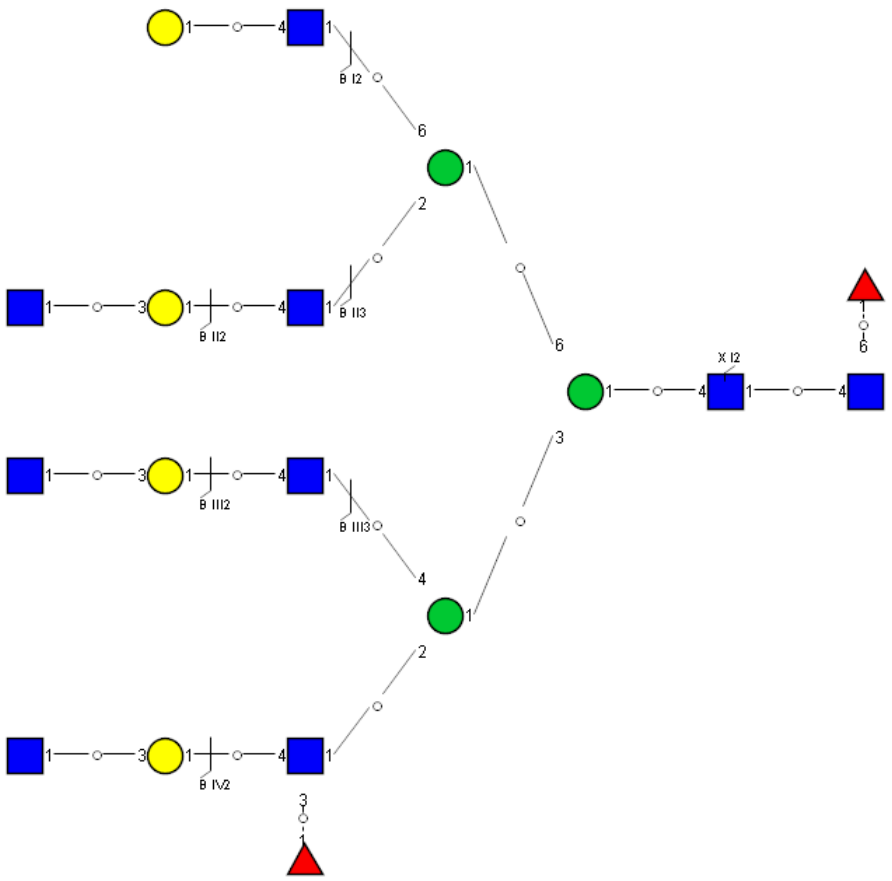

Spectrum Index: 39976(NoBHs:2)

#### Precursor iEF Map

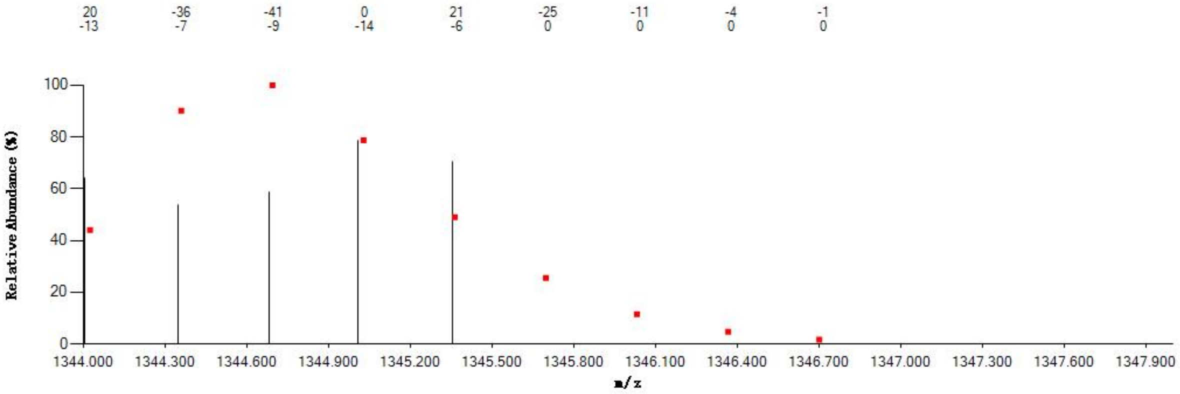


#### Annotated MS/MS Spectrum

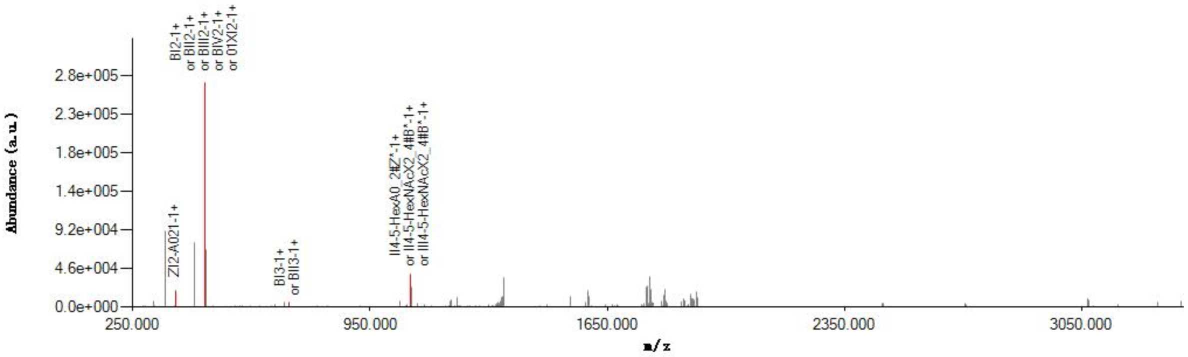


#### Graphical Fragmentation Map

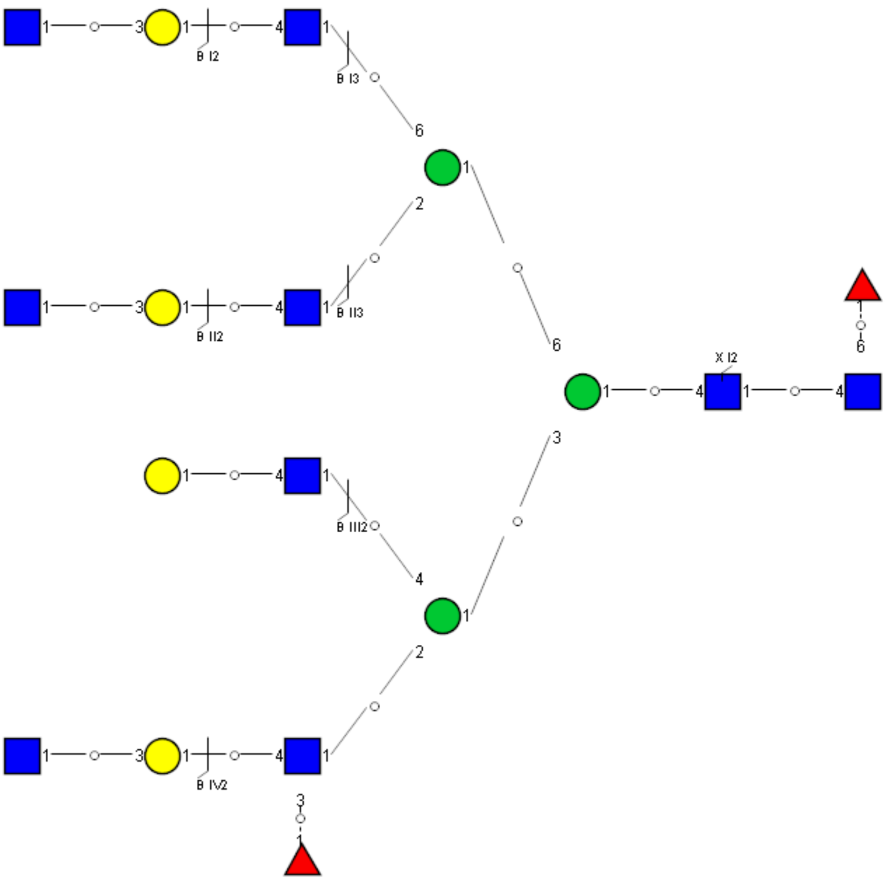

Spectrum Index: 40174(NoBHs:1)

Precursor iEF Map

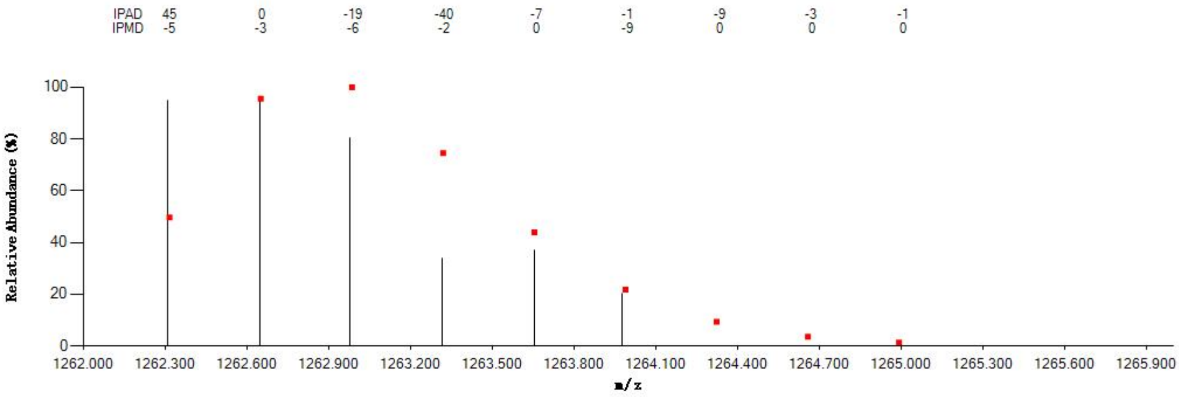

Annotated MS/MS Spectrum

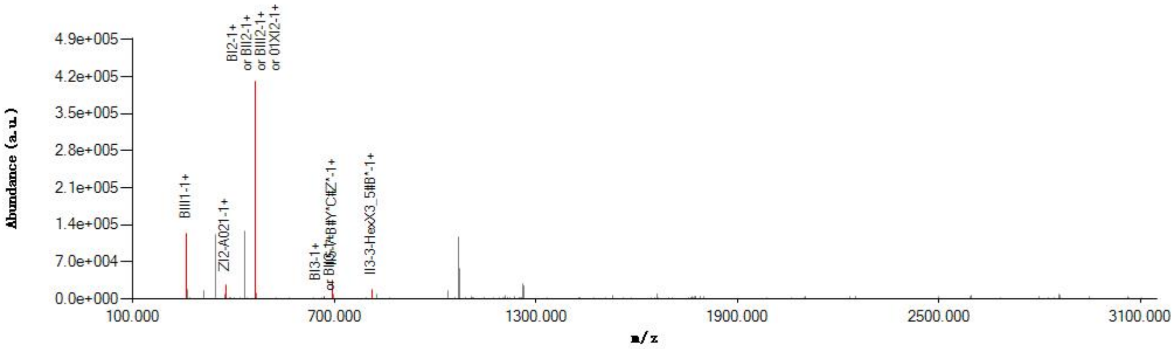

Graphical Fragmentation Map

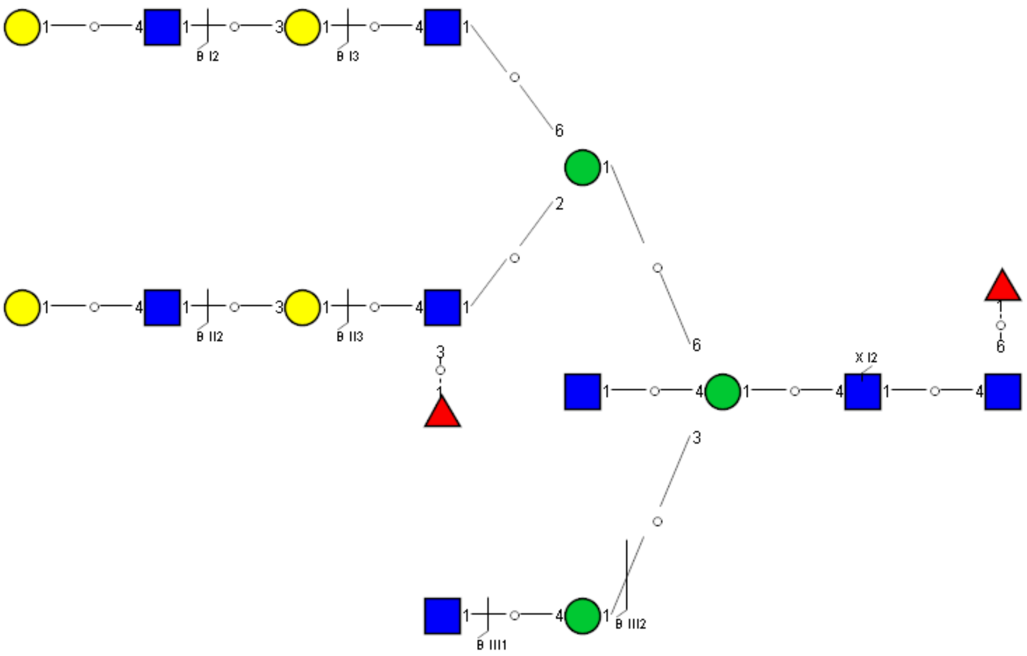

Spectrum Index: 40194(NoBHs:1)

#### Precursor iEF Map

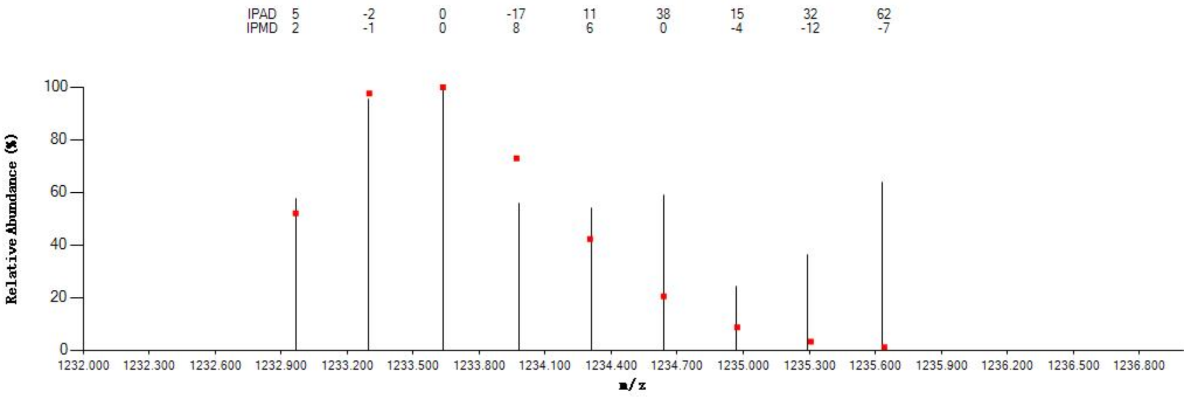


#### Annotated MS/MS Spectrum

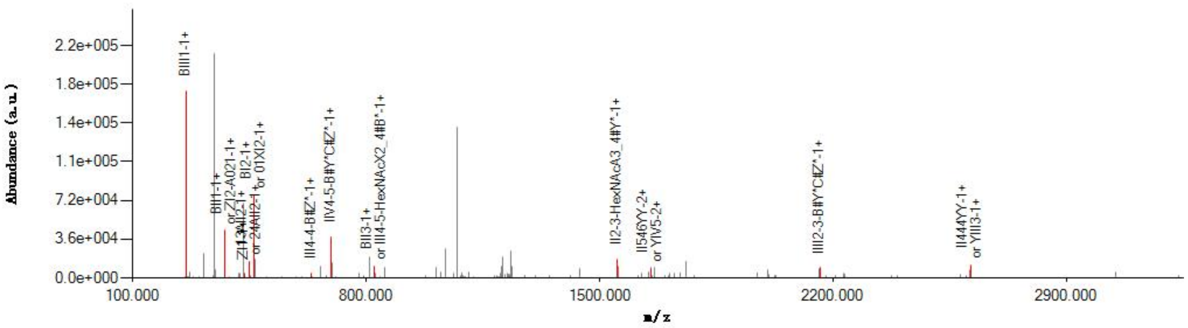


#### Graphical Fragmentation Map

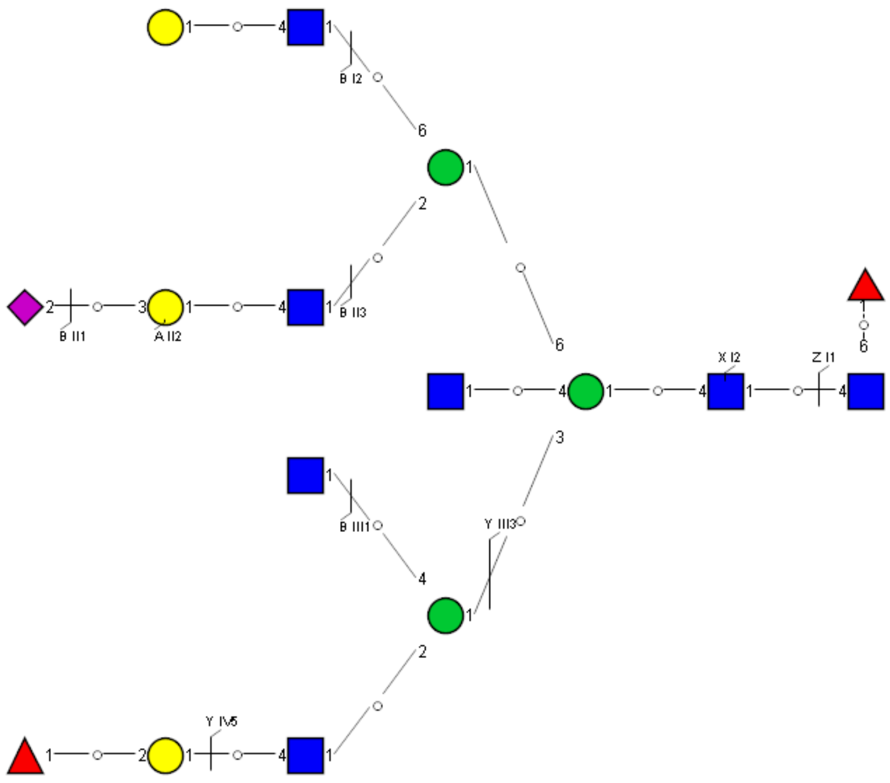

Spectrum Index: 40374(NoBHs:1)

#### Precursor iEF Map

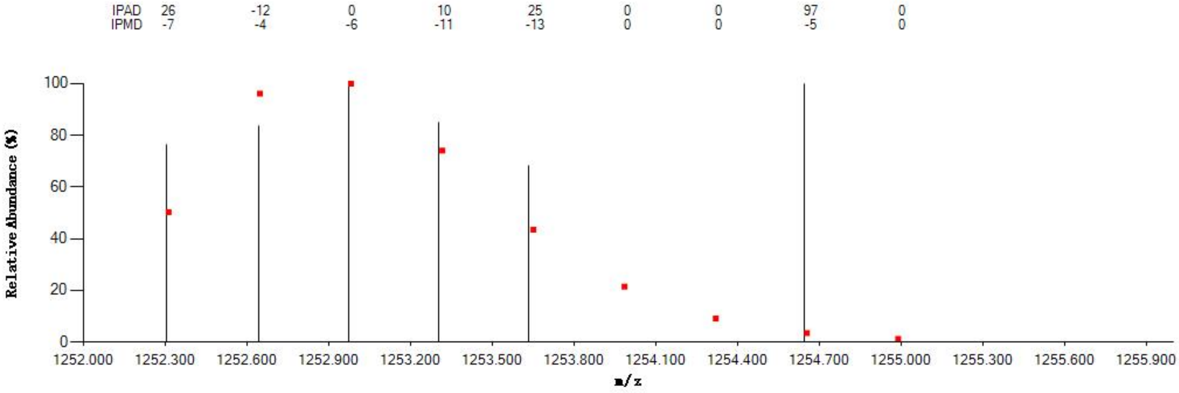


#### Annotated MS/MS Spectrum

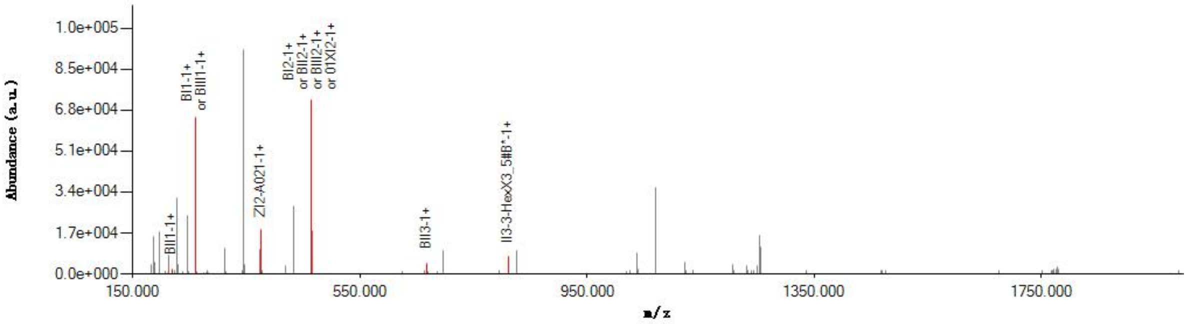


#### Graphical Fragmentation Map

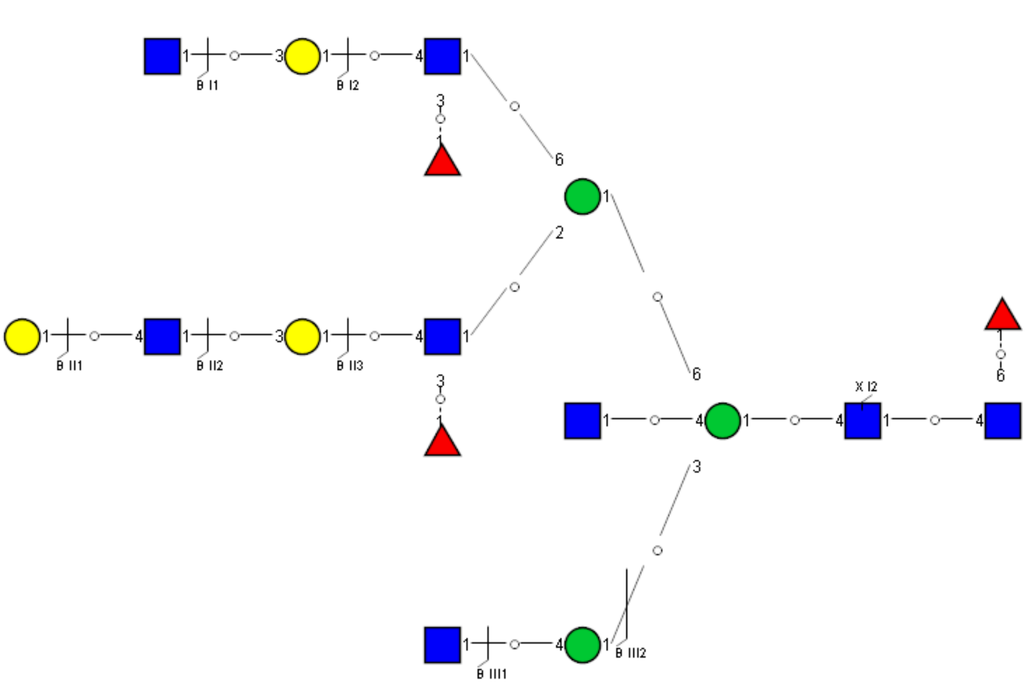

Spectrum Index: 40480(NoBHs:1)

#### Precursor iEF Map

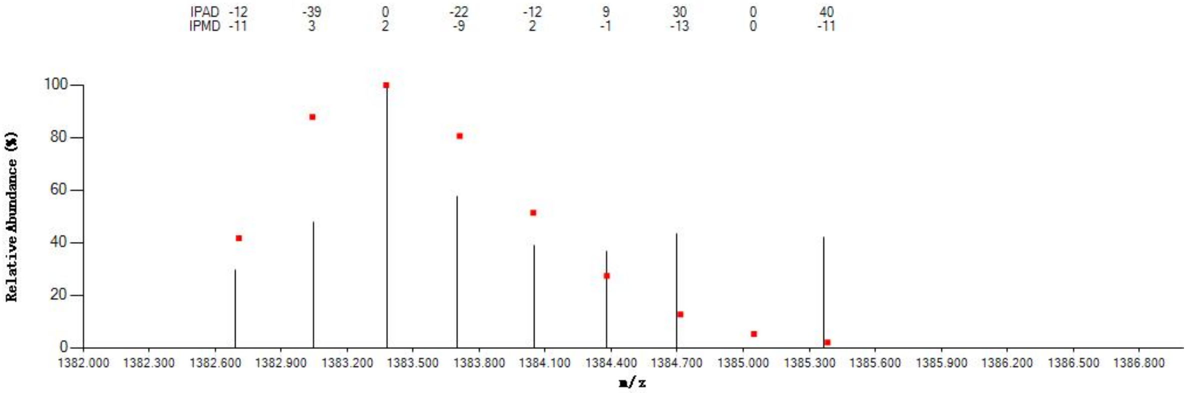


#### Annotated MS/MS Spectrum

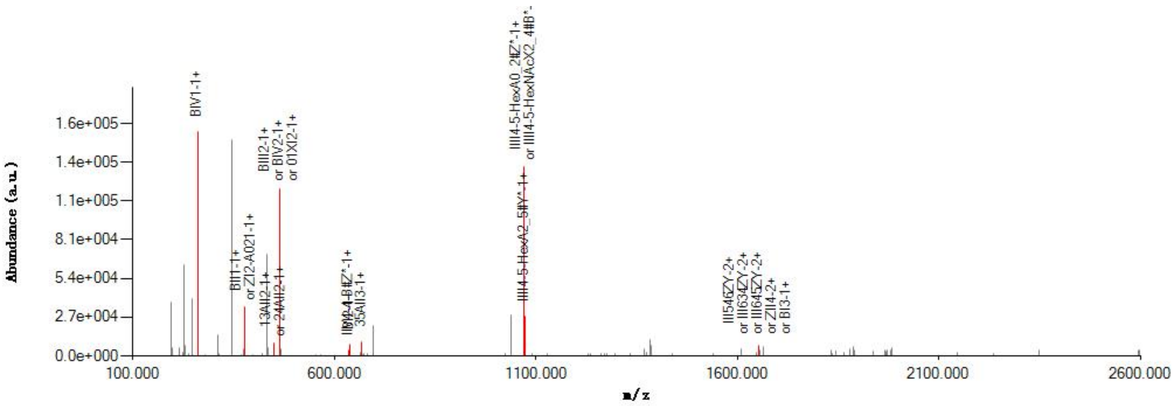


#### Graphical Fragmentation Map

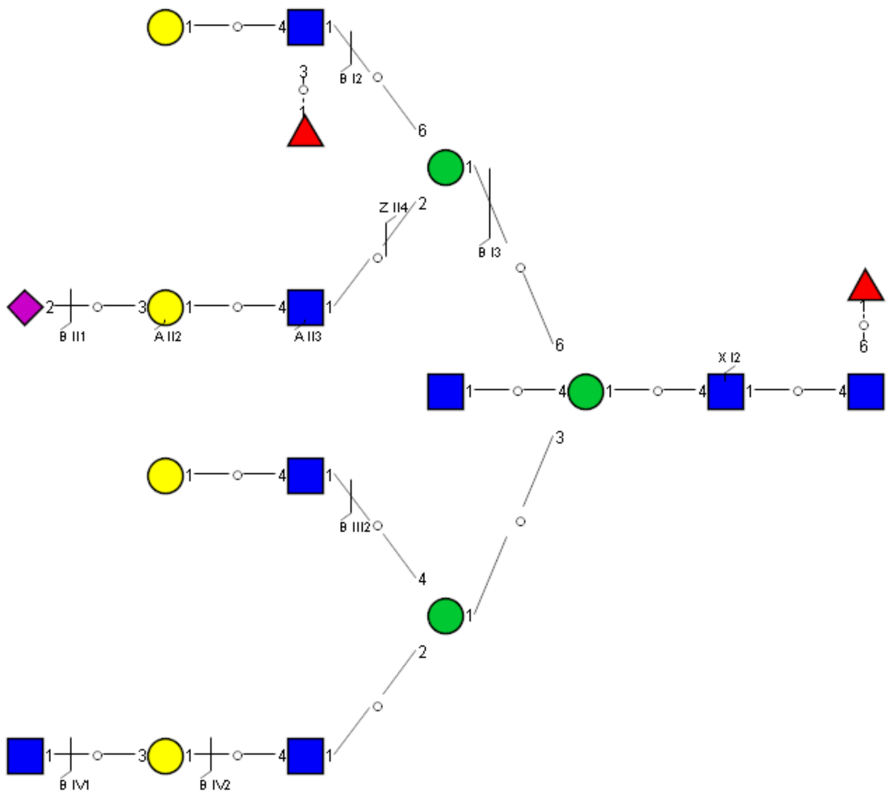

Spectrum Index: 40793(NoBHs:2)

#### Precursor iEF Map

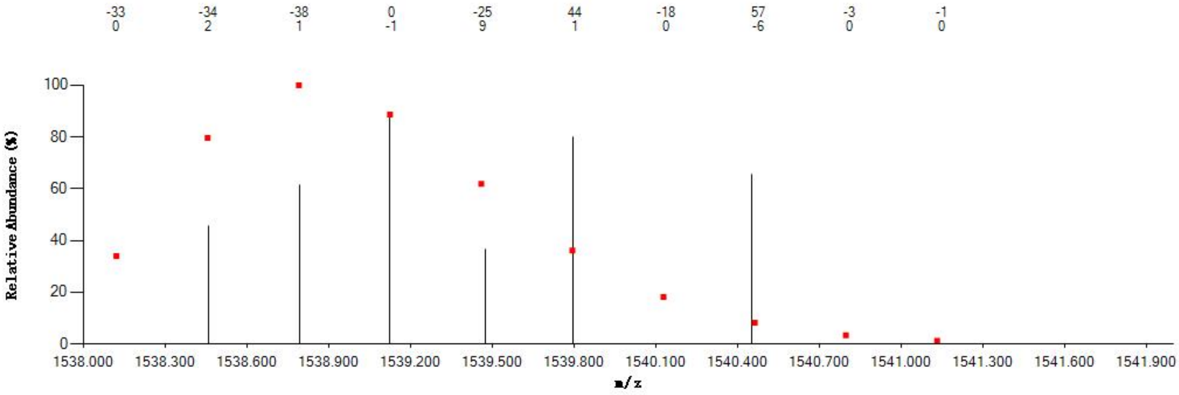


#### Annotated MS/MS Spectrum

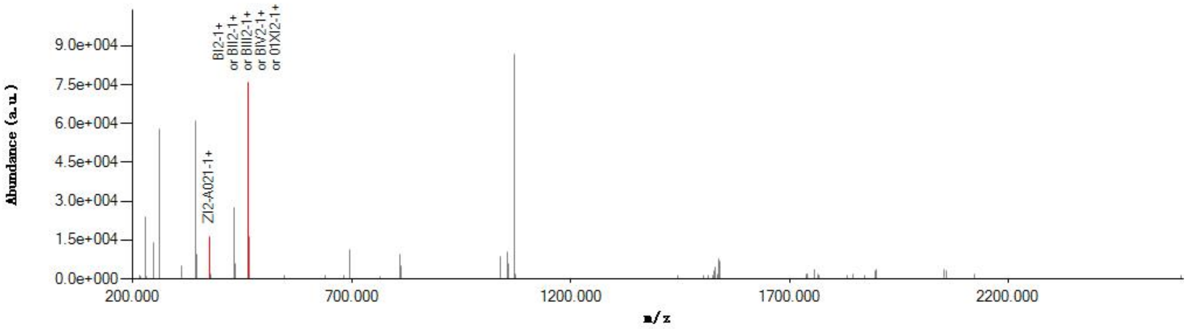


#### Graphical Fragmentation Map

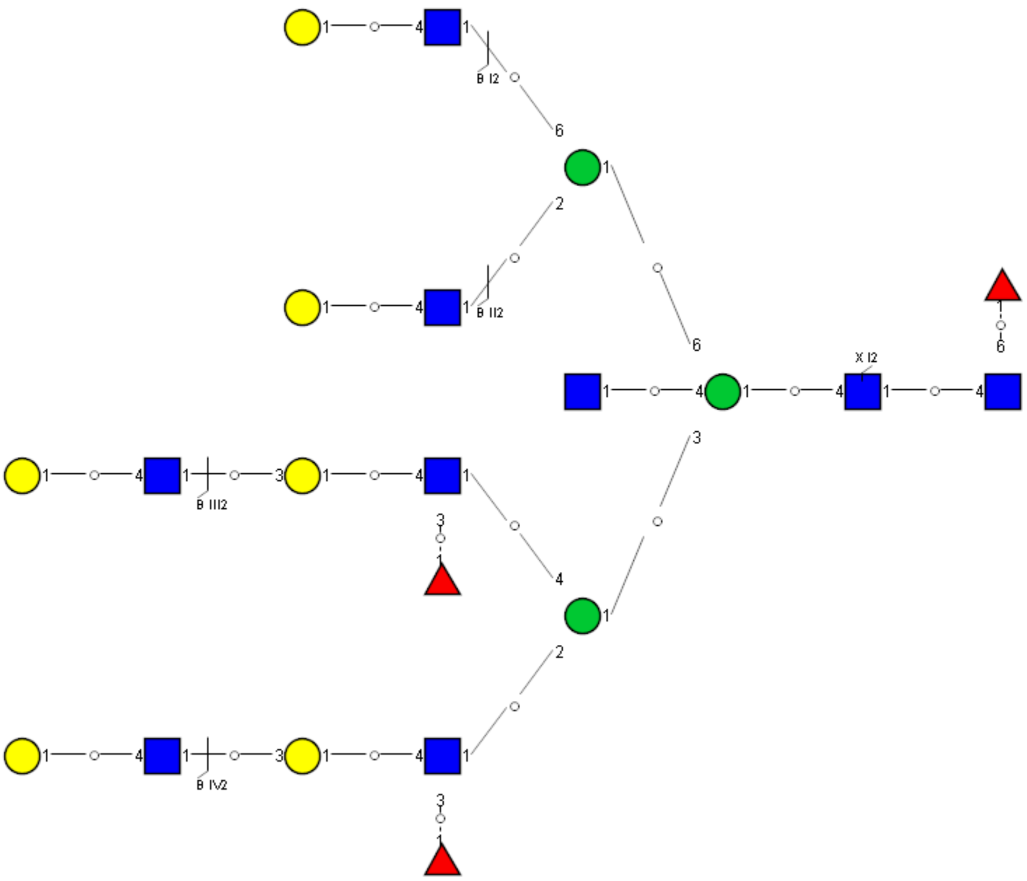

Spectrum Index: 40793(NoBHs:2)

#### Precursor iEF Map

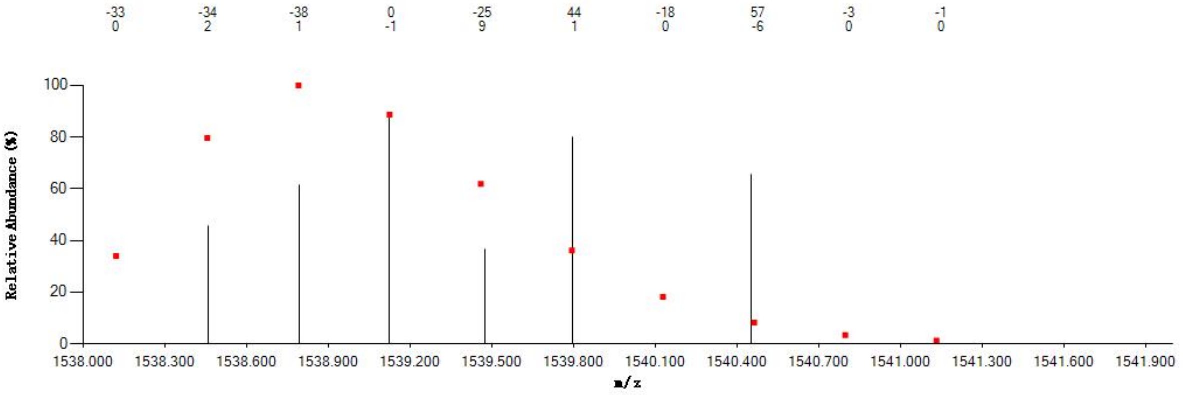


#### Annotated MS/MS Spectrum

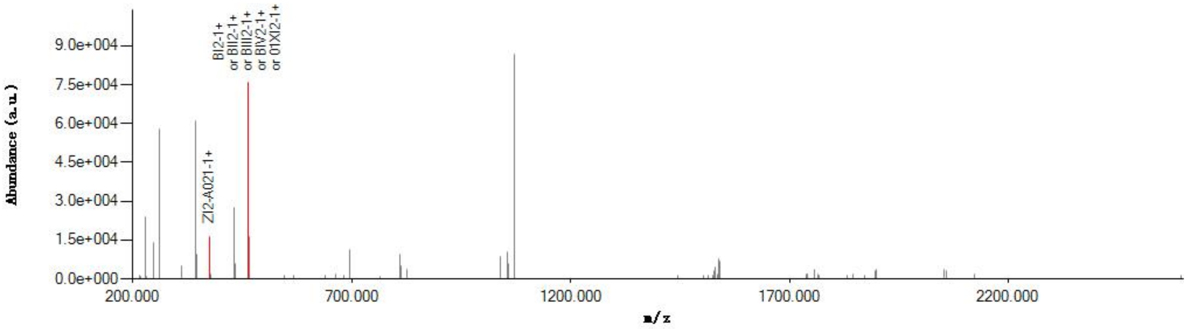


#### Graphical Fragmentation Map

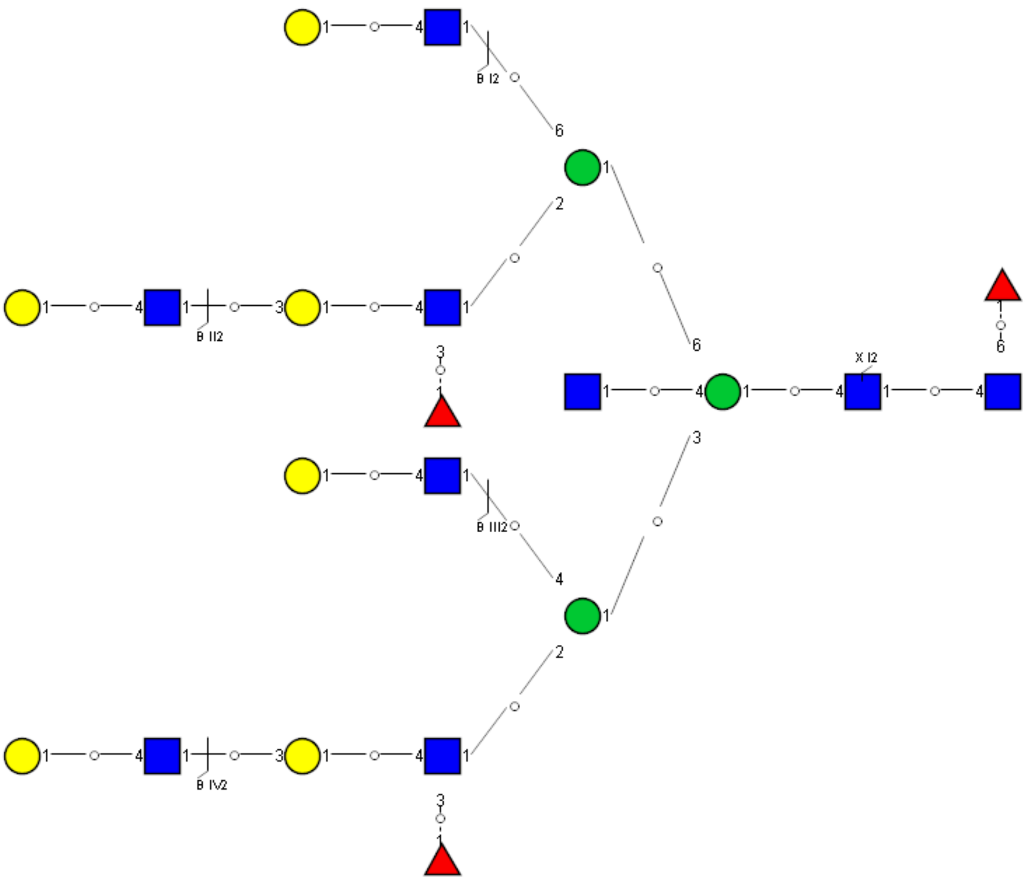

Spectrum Index: 41894(NoBHs:1)

#### Precursor iEF Map

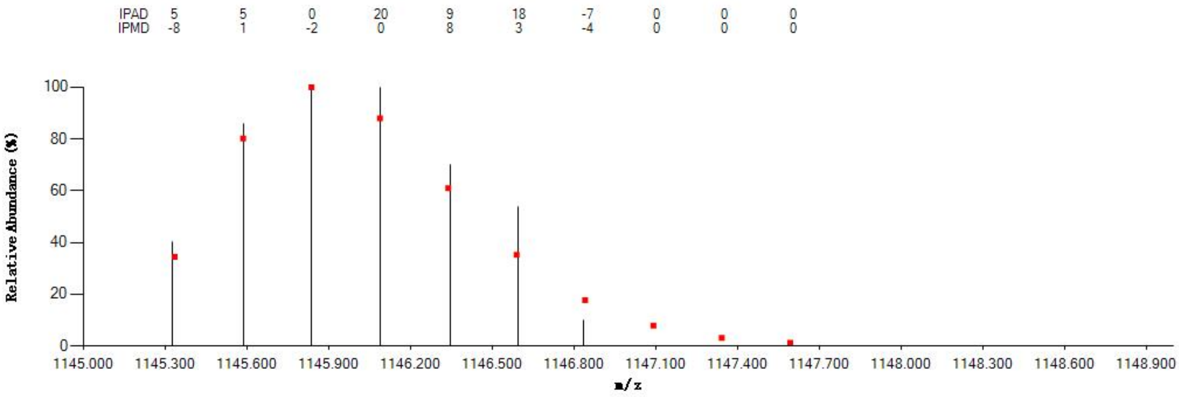


#### Annotated MS/MS Spectrum

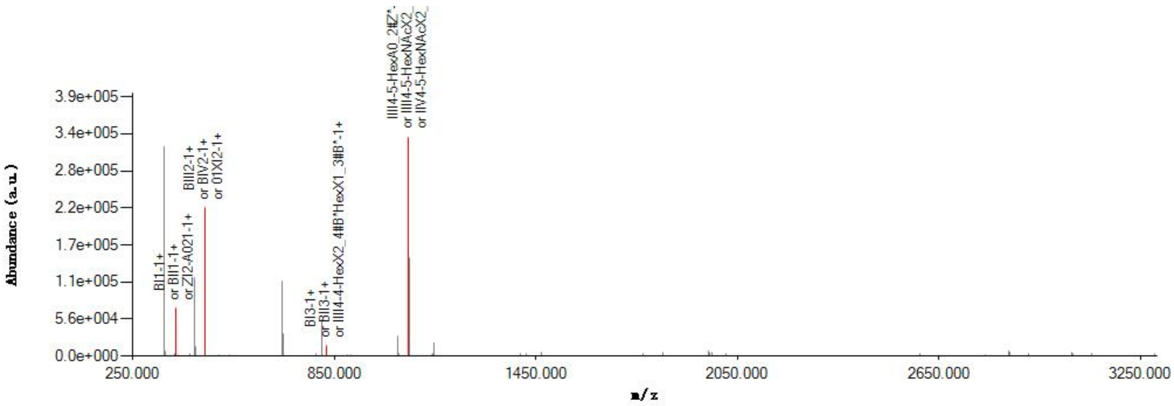


#### Graphical Fragmentation Map

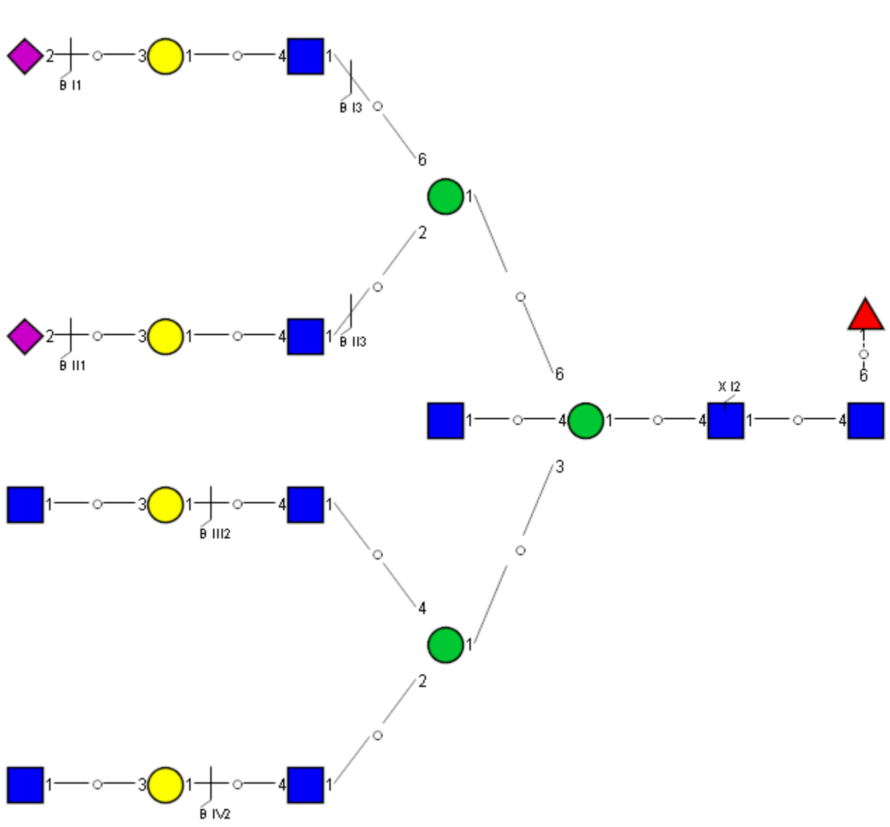

Spectrum Index: 41942(NoBHs:6)

#### Precursor iEF Map

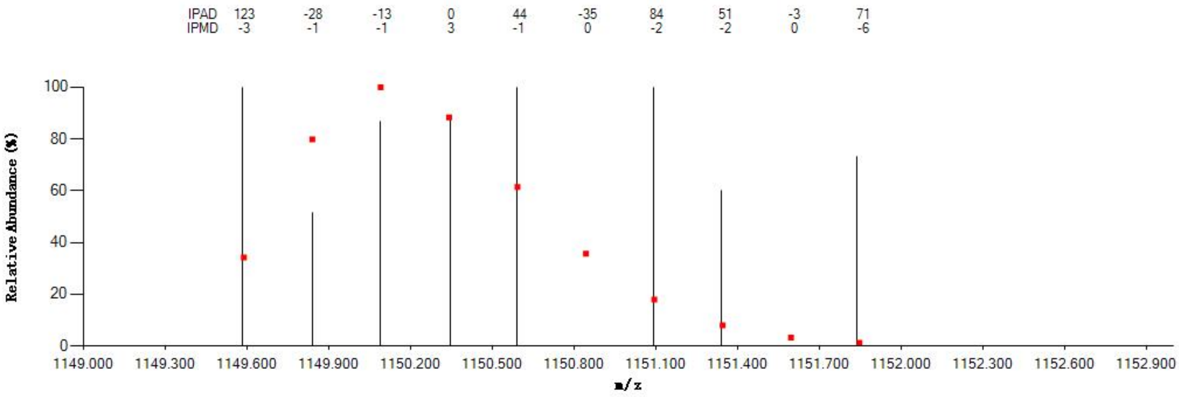


#### Annotated MS/MS Spectrum

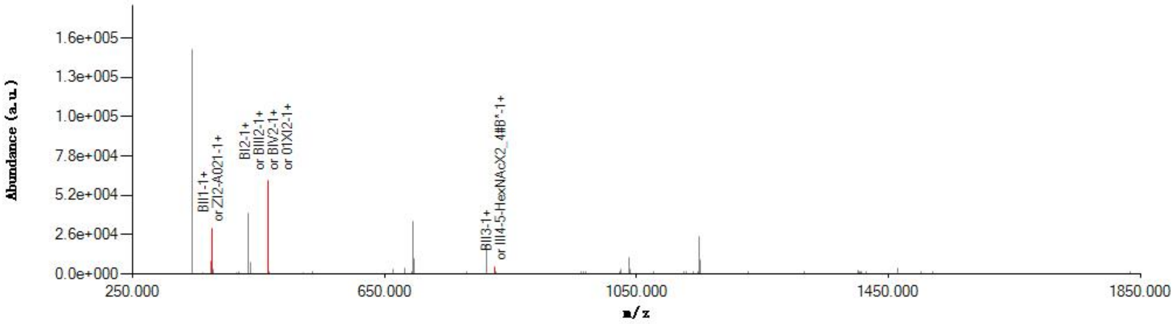


#### Graphical Fragmentation Map

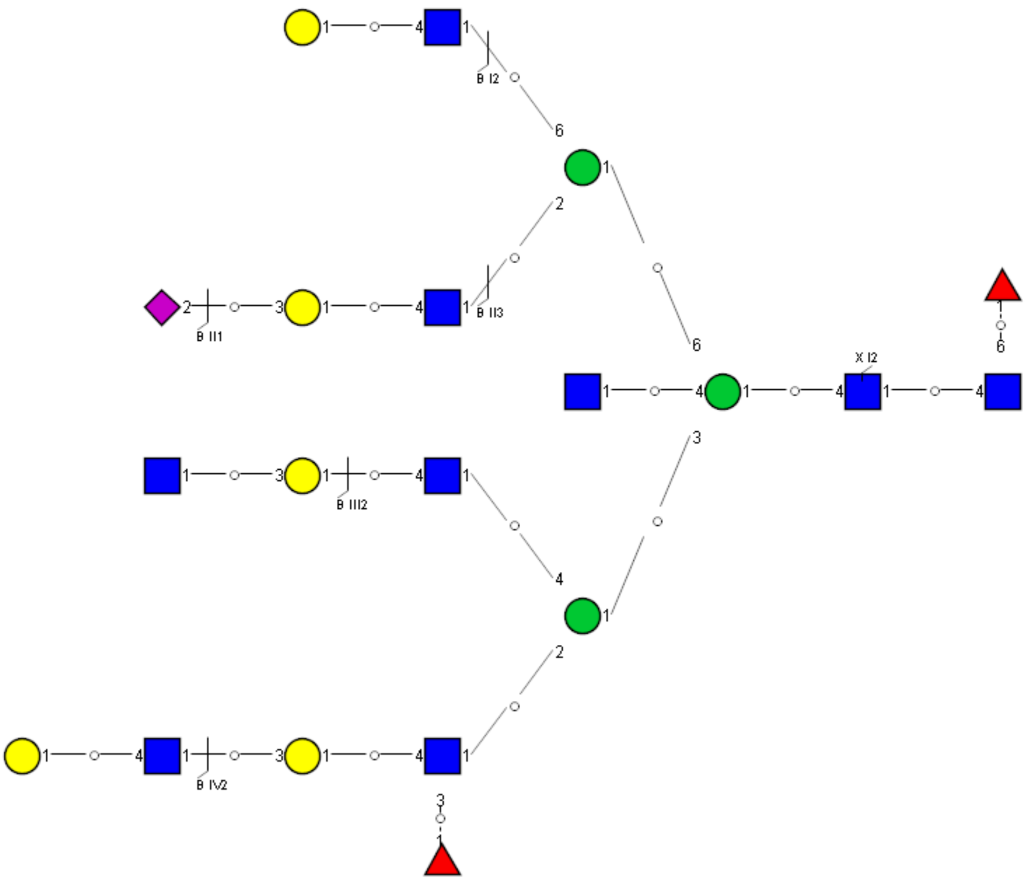

Spectrum Index: 41942(NoBHs:6)

#### Precursor iEF Map

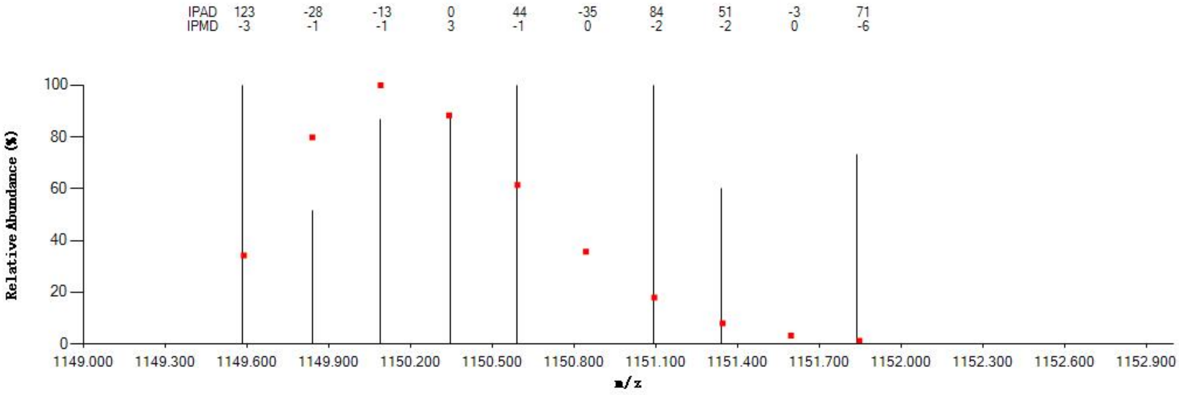

#### Annotated MS/MS Spectrum

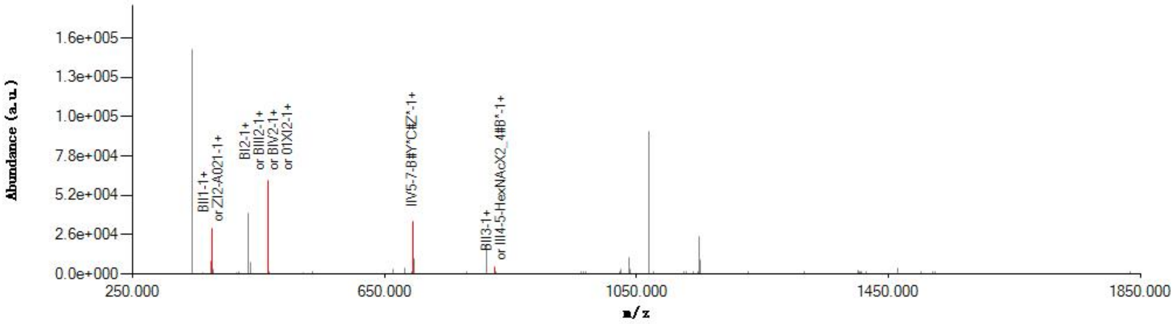

#### Graphical Fragmentation Map

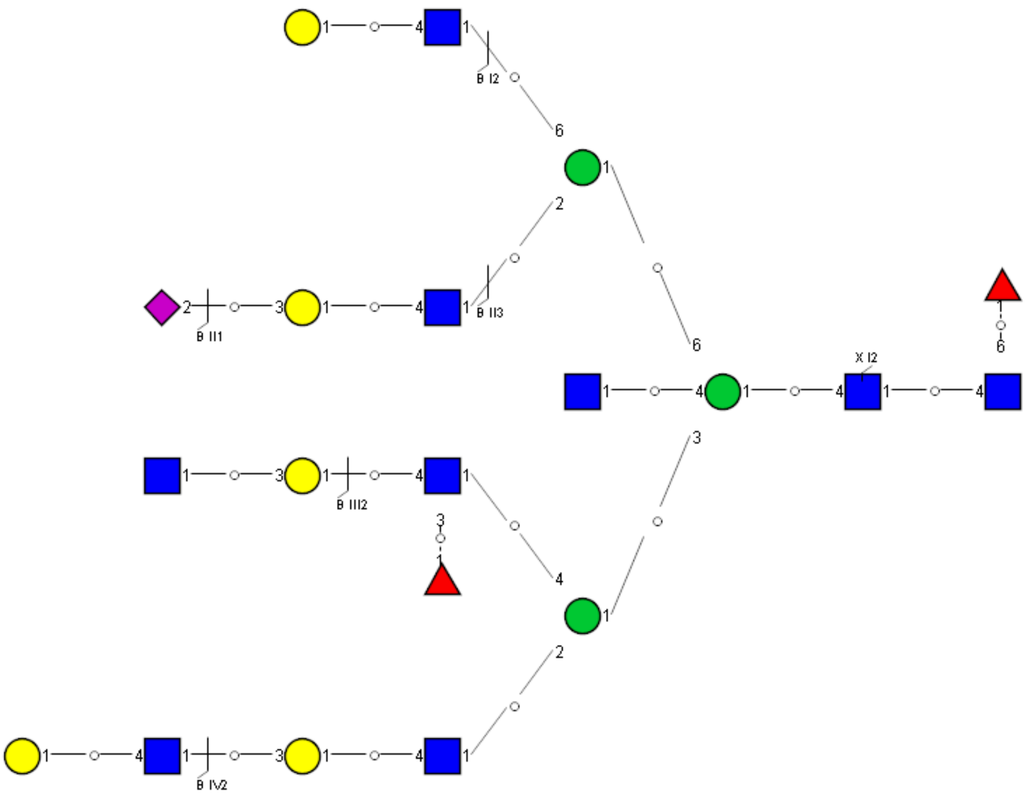

Spectrum Index: 41942(NoBHs:6)

#### Precursor iEF Map

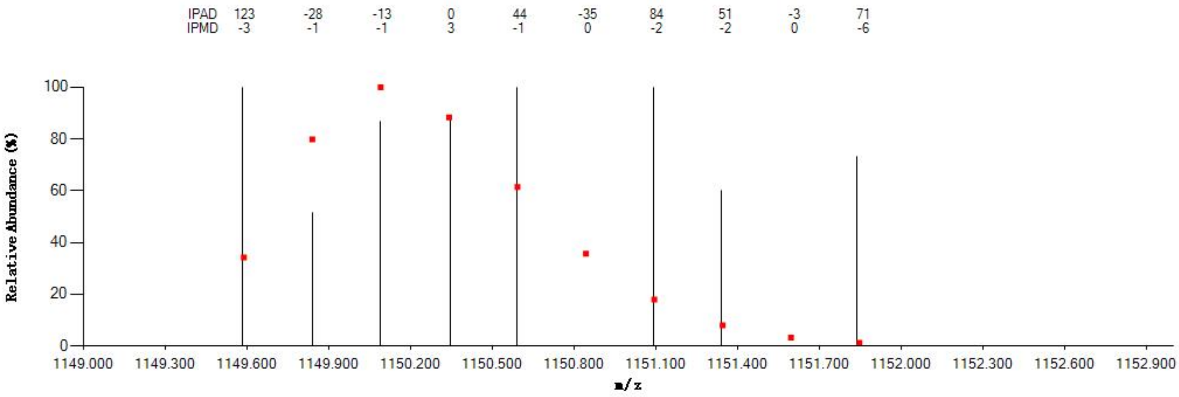


#### Annotated MS/MS Spectrum

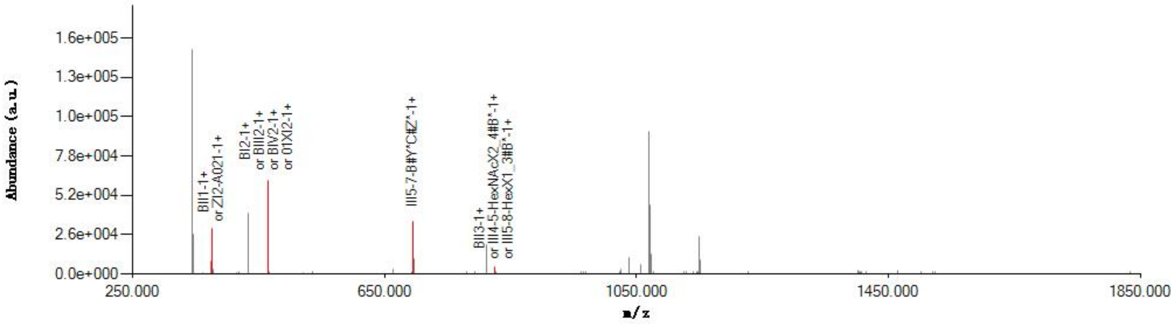


#### Graphical Fragmentation Map

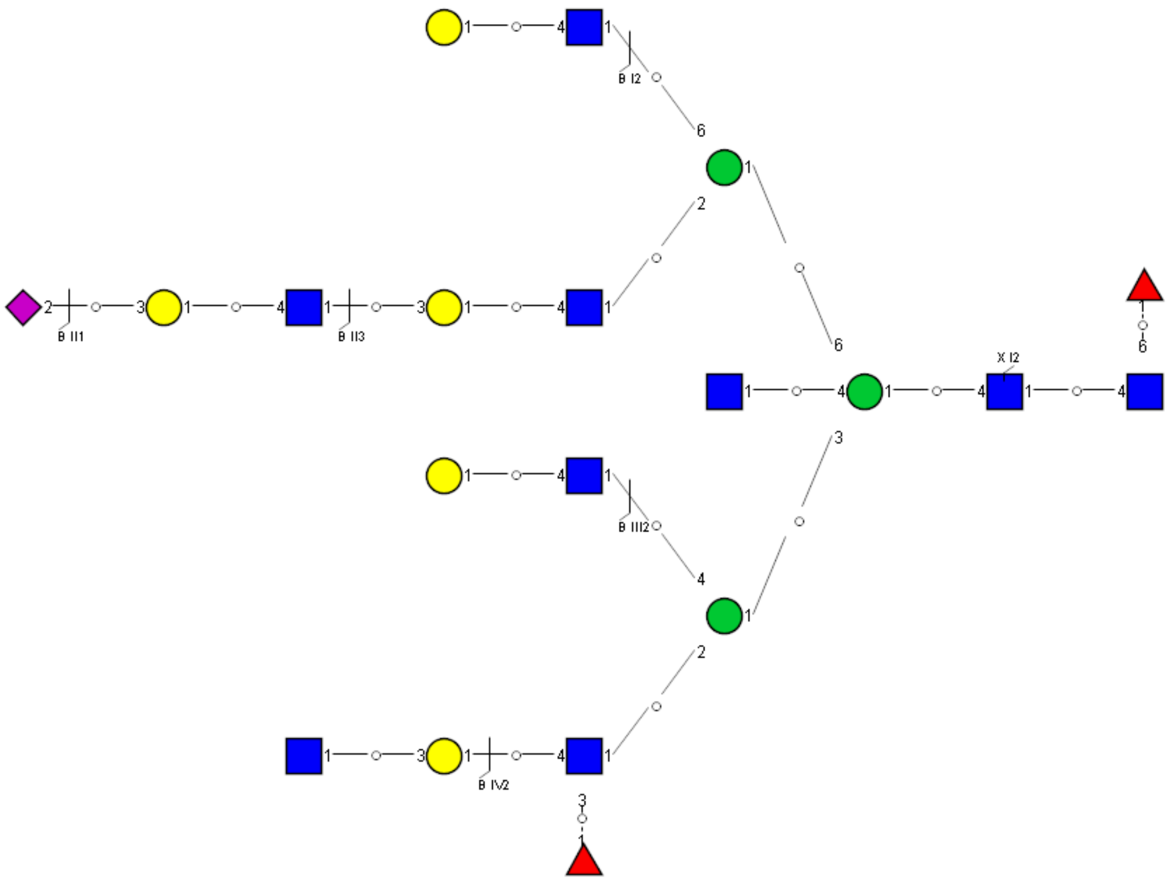

Spectrum Index: 41942(NoBHs:6)

#### Precursor iEF Map

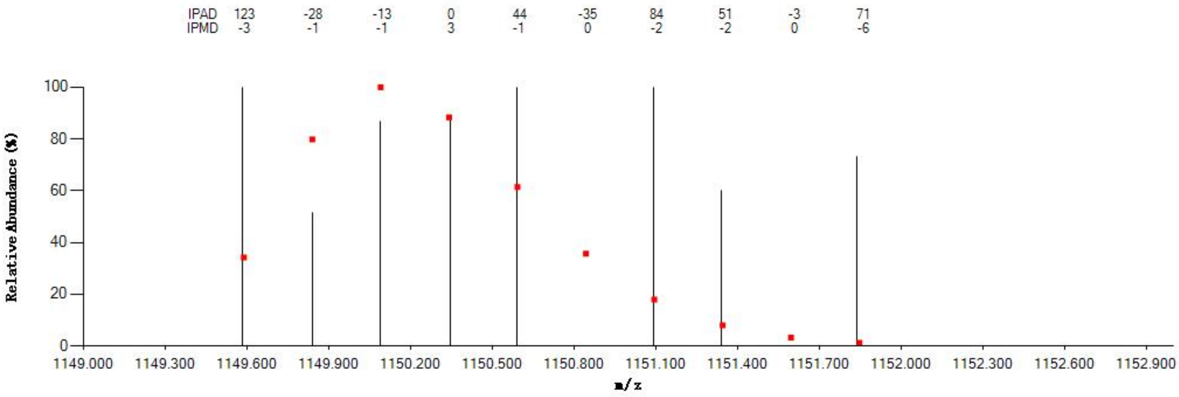

#### Annotated MS/MS Spectrum

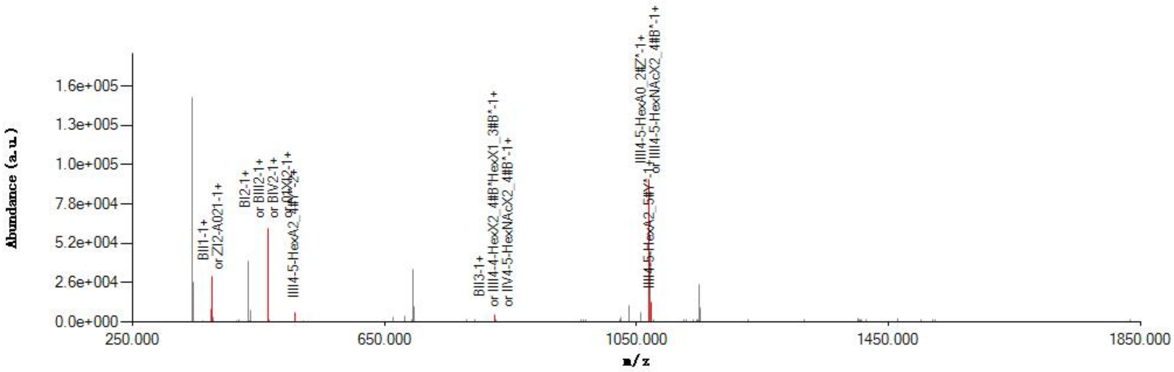

#### Graphical Fragmentation Map

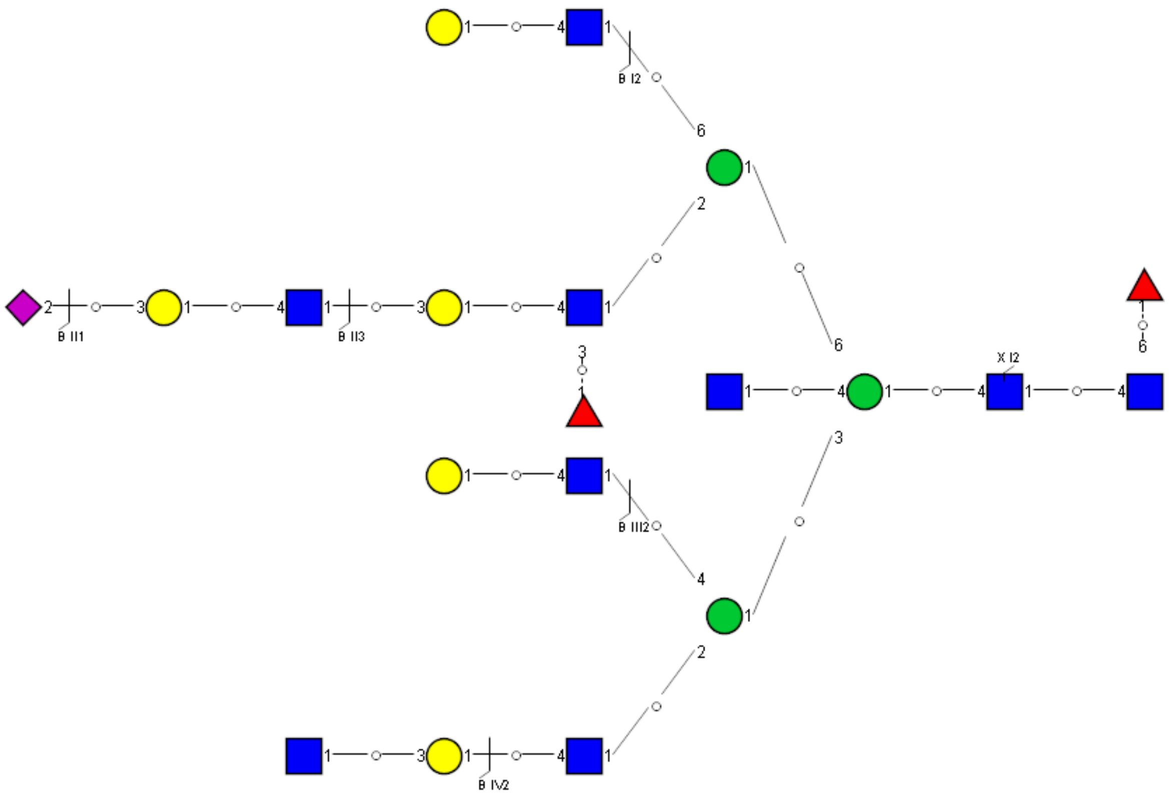

Spectrum Index: 41942(NoBHs:6)

#### Precursor iEF Map

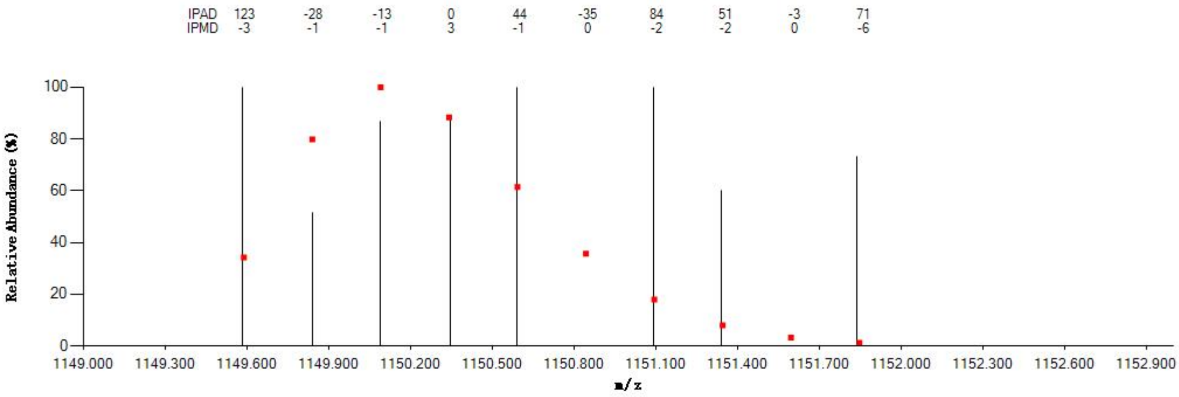


#### Annotated MS/MS Spectrum

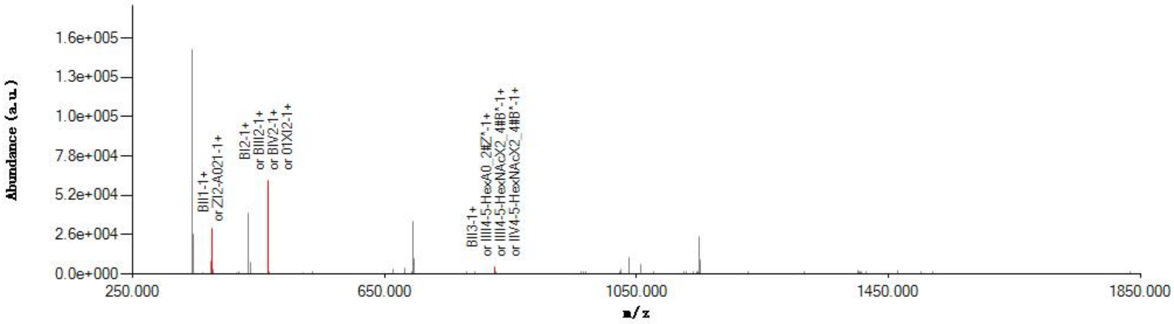


#### Graphical Fragmentation Map

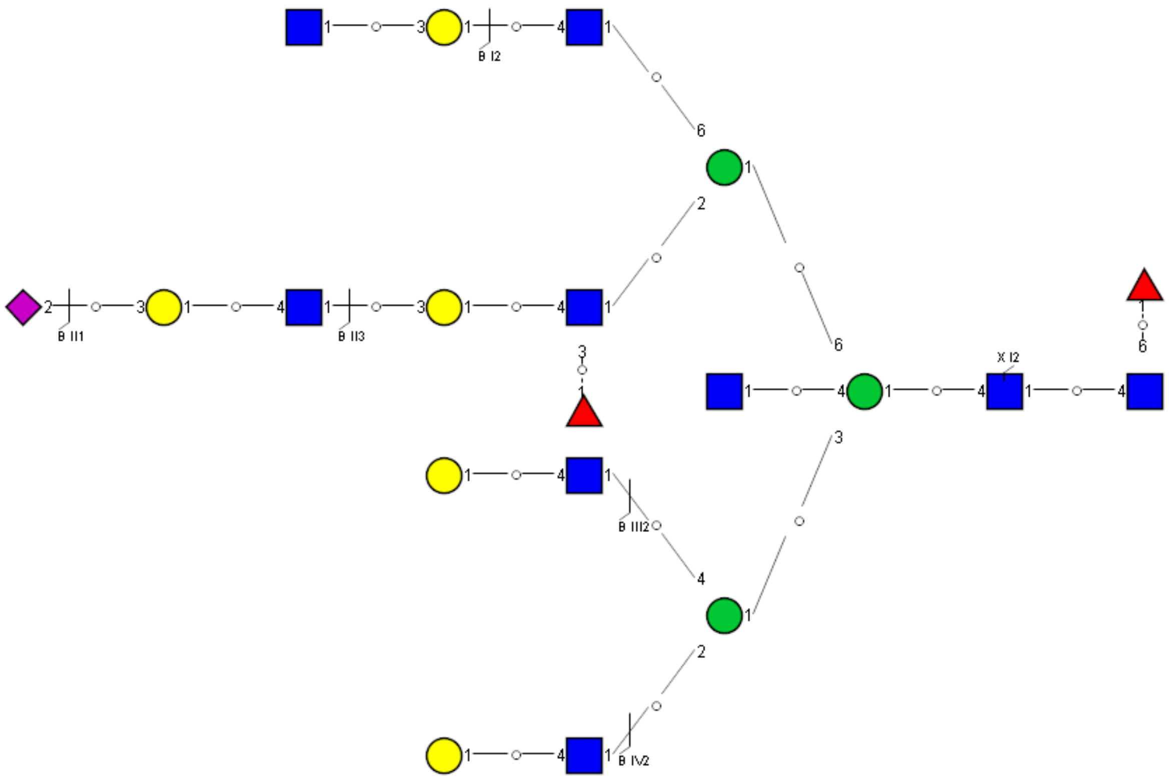

Spectrum Index: 41942(NoBHs:6)

#### Precursor iEF Map

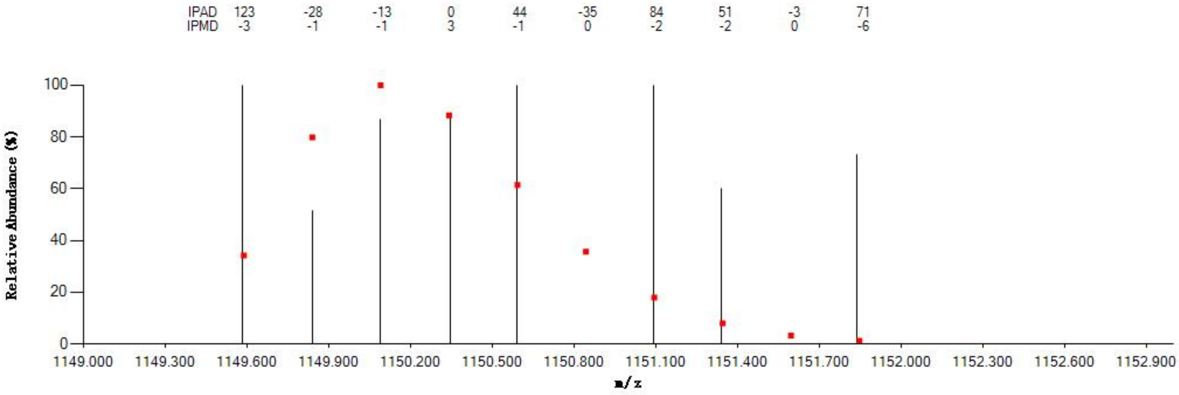

#### Annotated MS/MS Spectrum

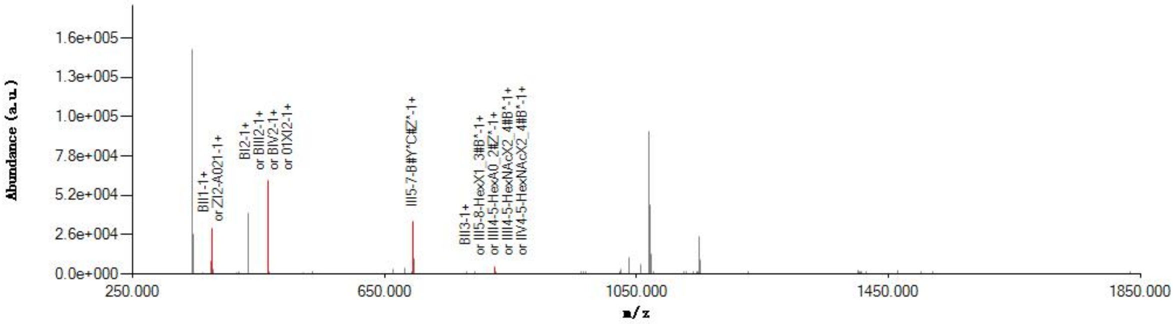

#### Graphical Fragmentation Map

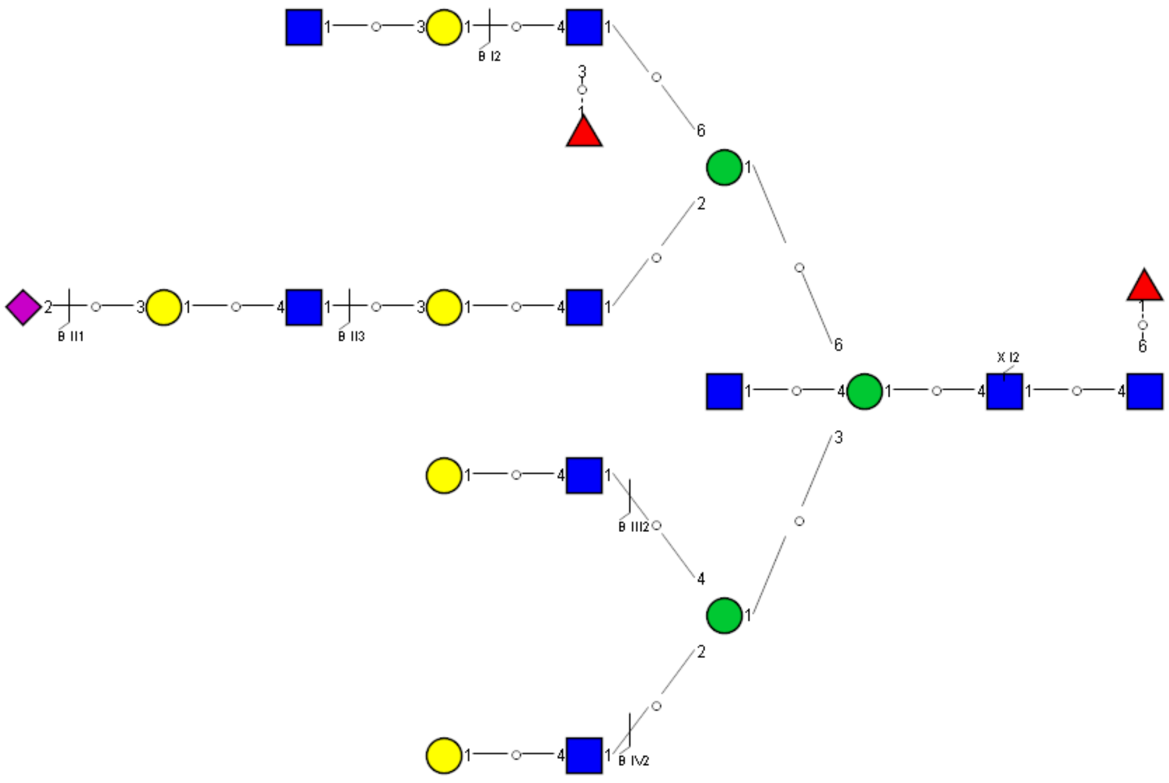

Spectrum Index: 42015(NoBHs:1)

#### Precursor iEF Map

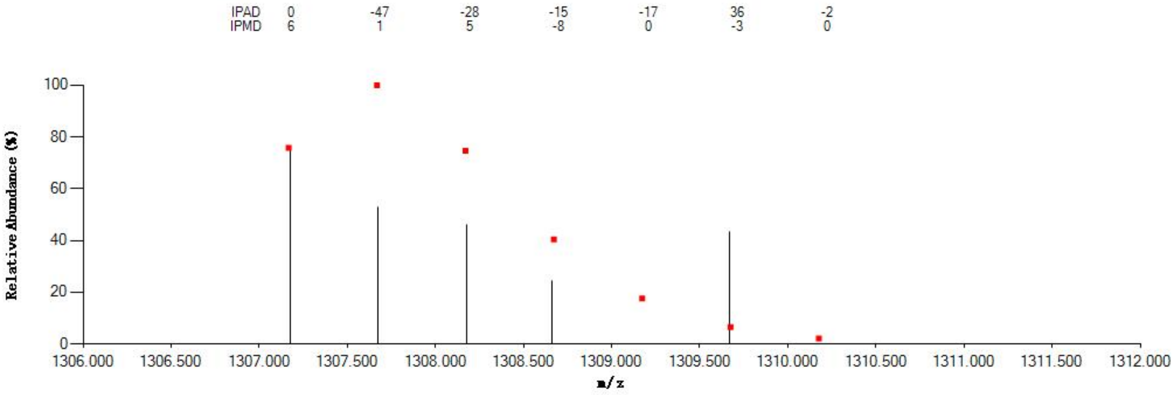


#### Annotated MS/MS Spectrum

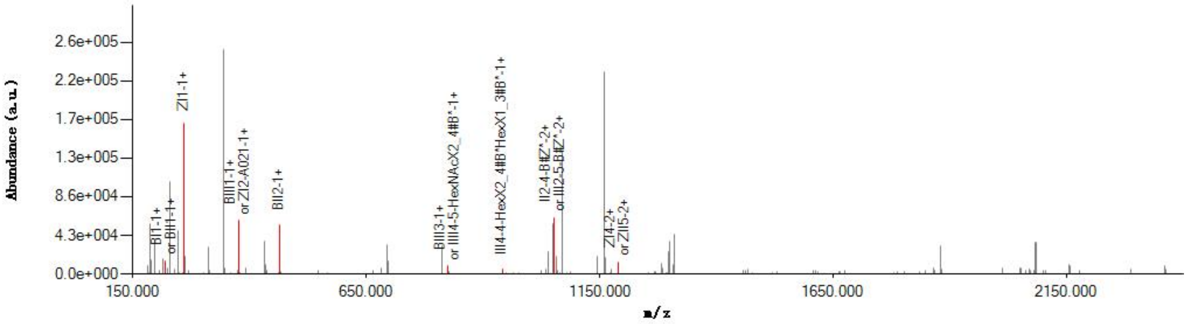


#### Graphical Fragmentation Map

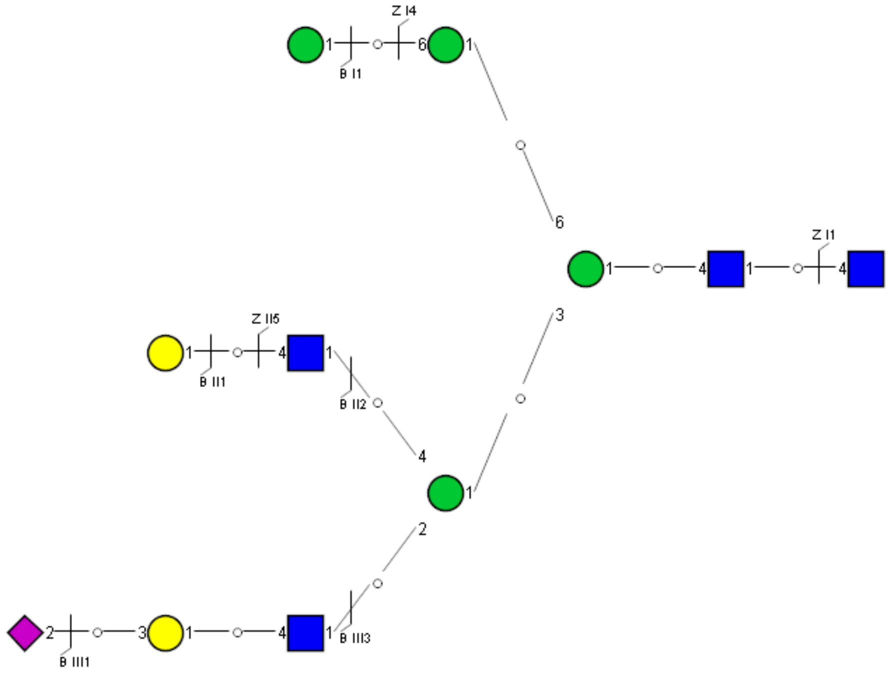

Spectrum Index: 27400(NoBHs:1)

#### Precursor iEF Map

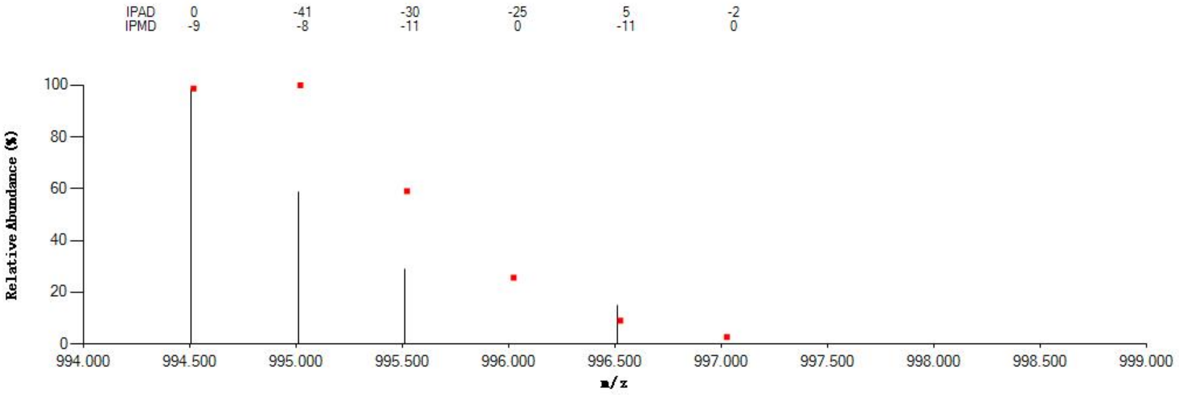


#### Annotated MS/MS Spectrum

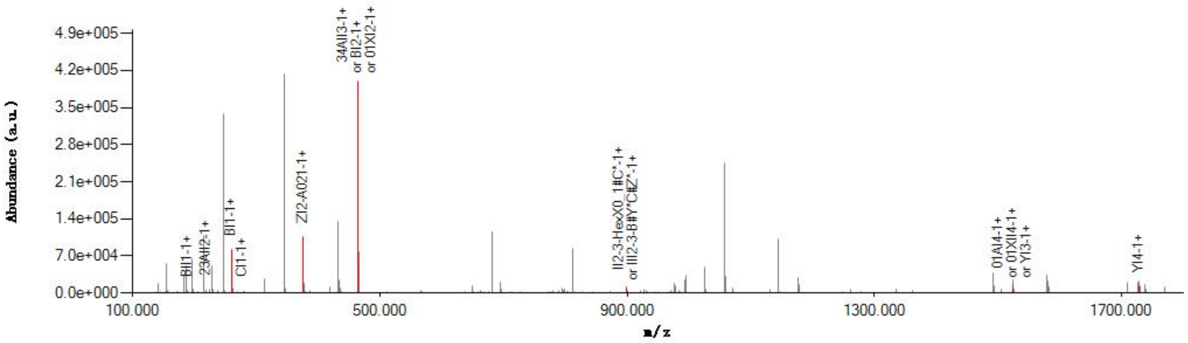


#### Graphical Fragmentation Map

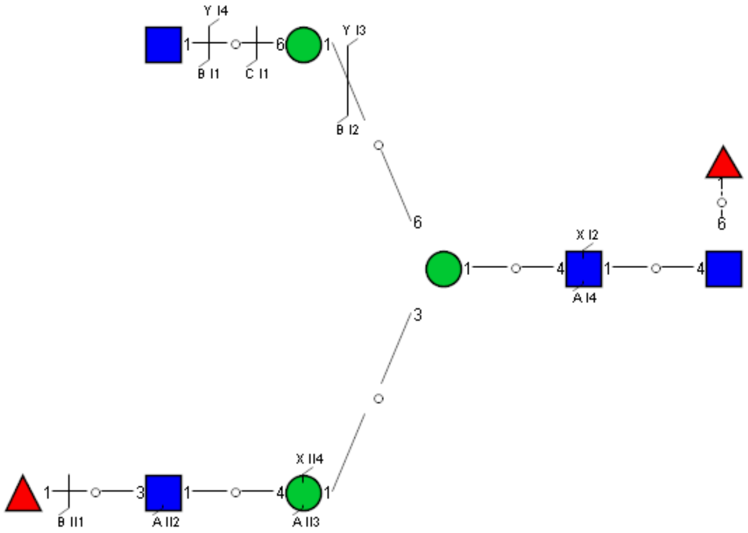

Spectrum Index: 27613(NoBHs:1)

#### Precursor iEF Map

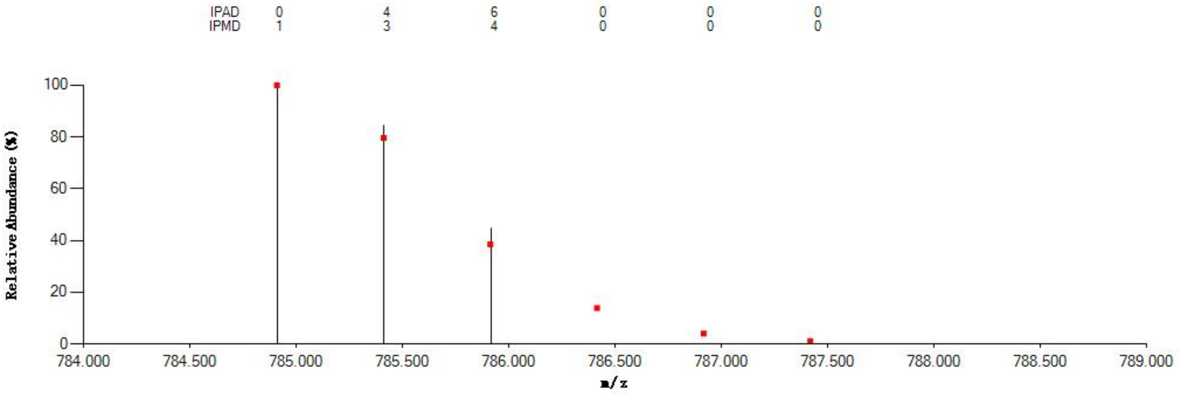


#### Annotated MS/MS Spectrum

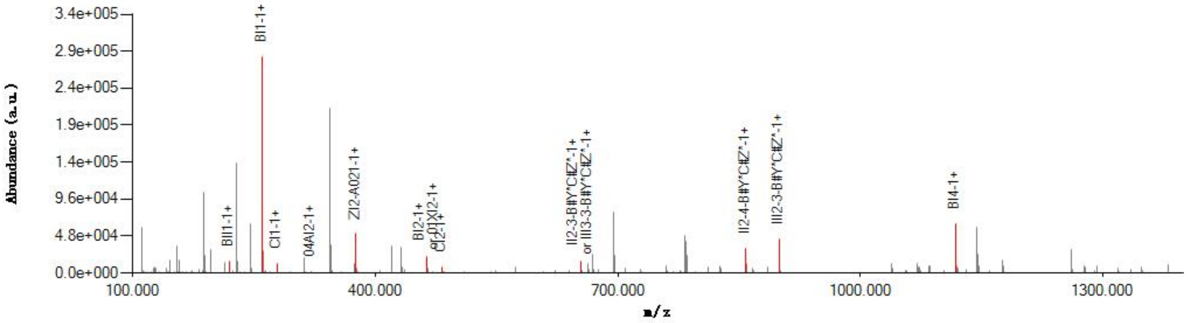


#### Graphical Fragmentation Map

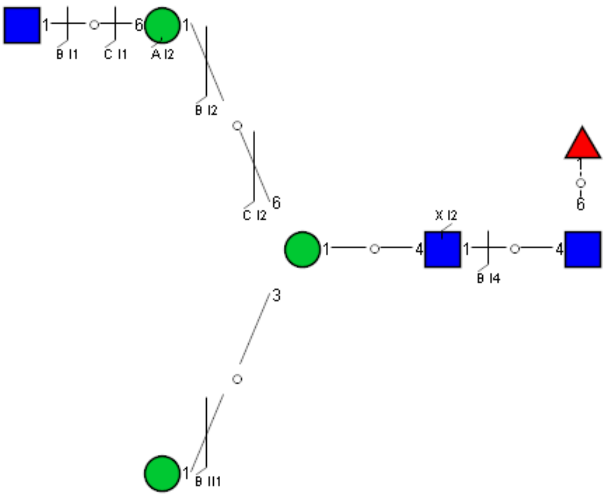

Spectrum Index: 28388(NoBHs:1)

#### Precursor iEF Map

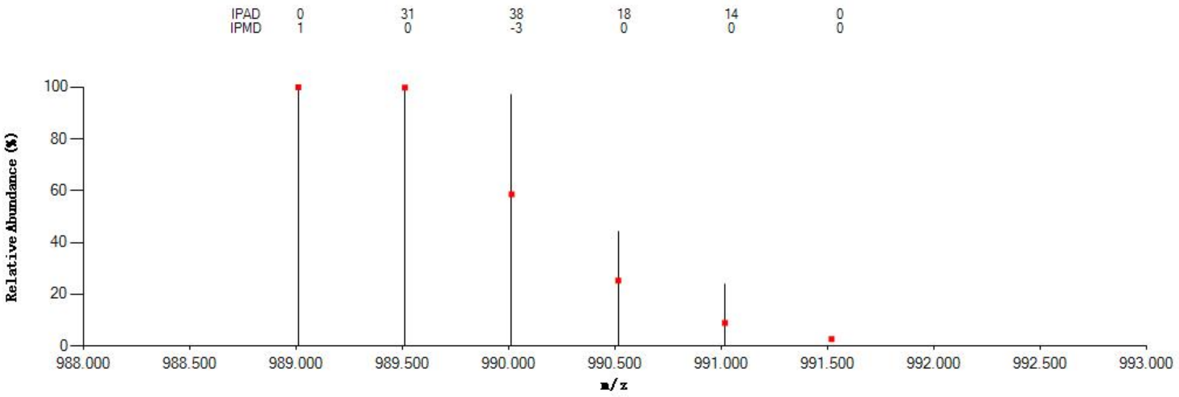


#### Annotated MS/MS Spectrum

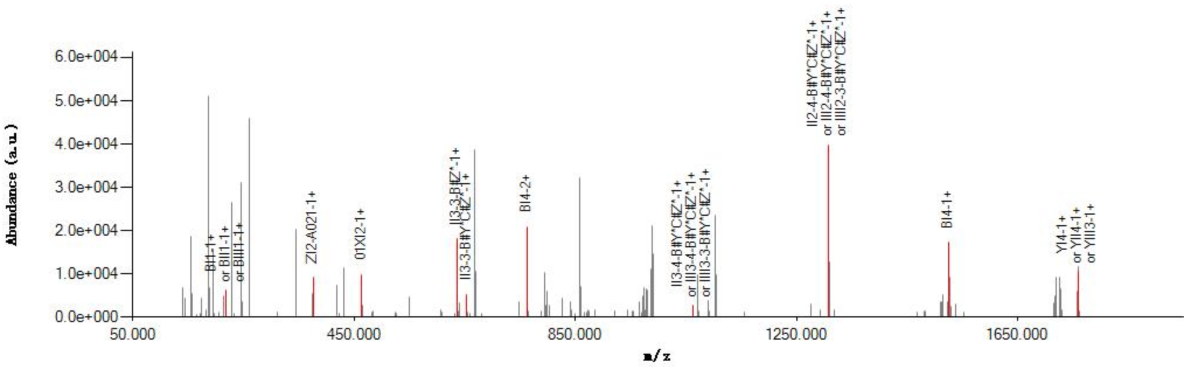


#### Graphical Fragmentation Map

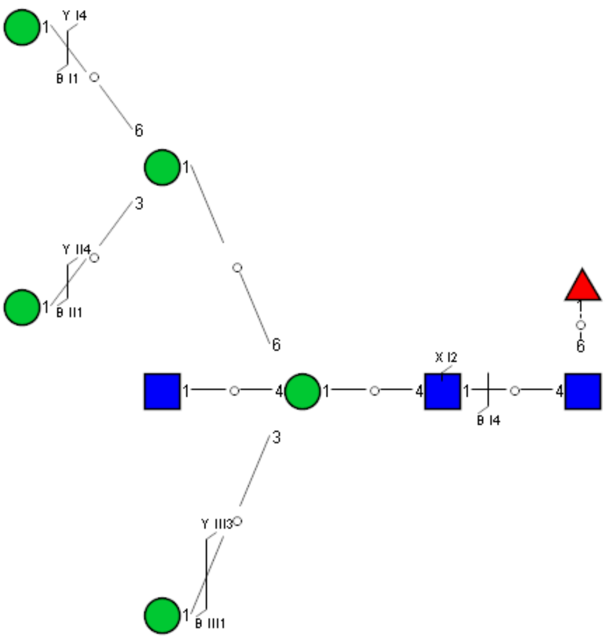

Spectrum Index: 28691(NoBHs:1)

#### Precursor iEF Map

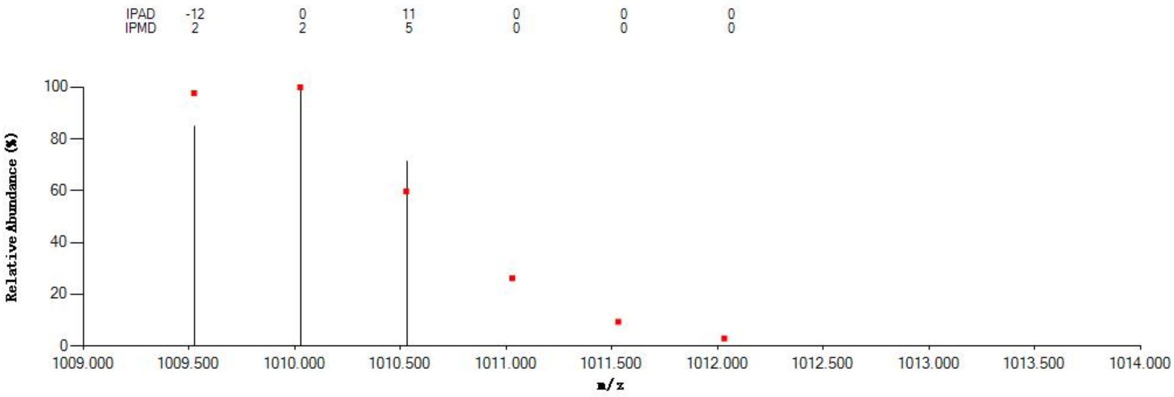


#### Annotated MS/MS Spectrum

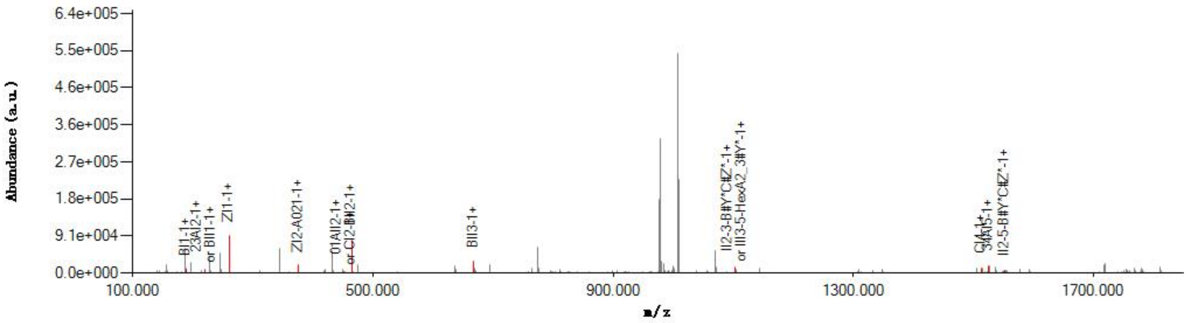


#### Graphical Fragmentation Map

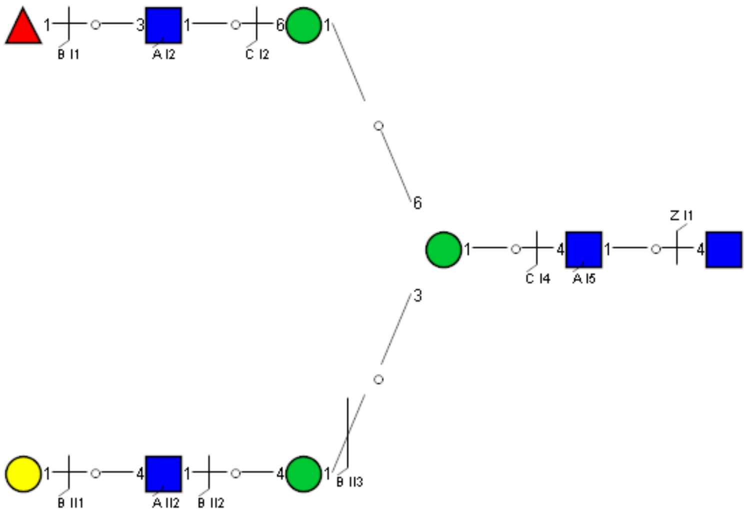

Spectrum Index: 29691(NoBHs:1)

#### Precursor iEF Map

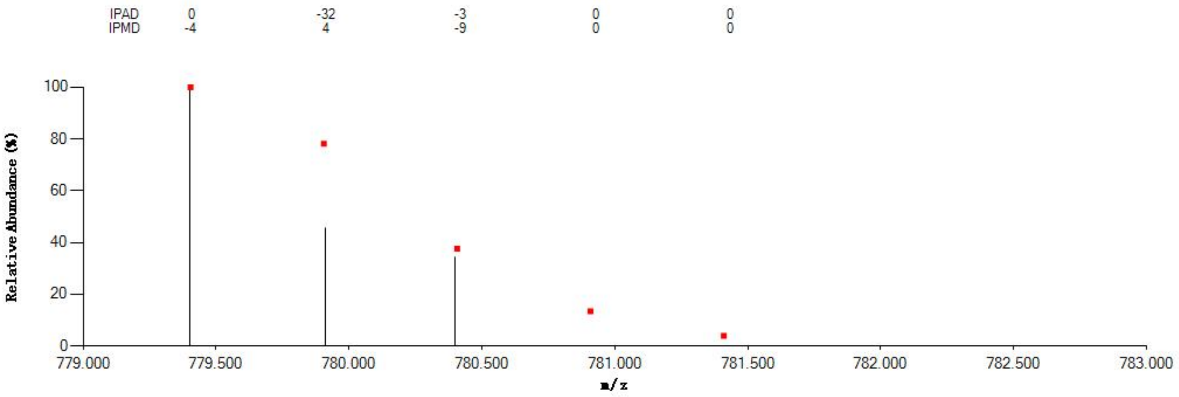

#### Annotated MS/MS Spectrum

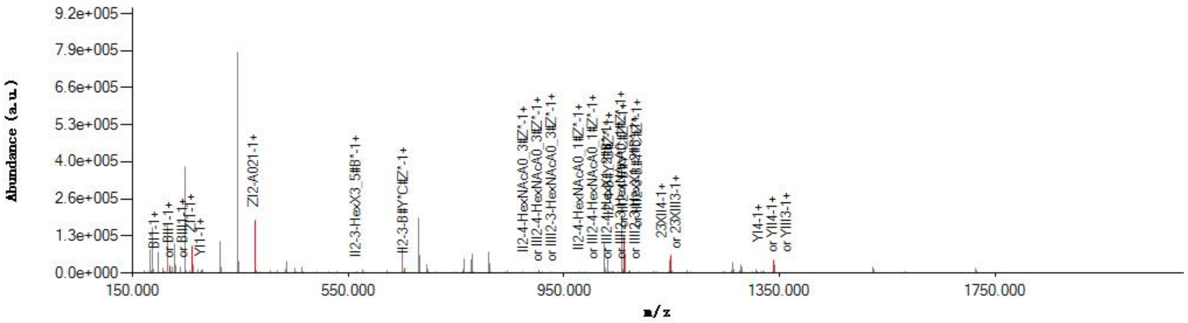

#### Graphical Fragmentation Map

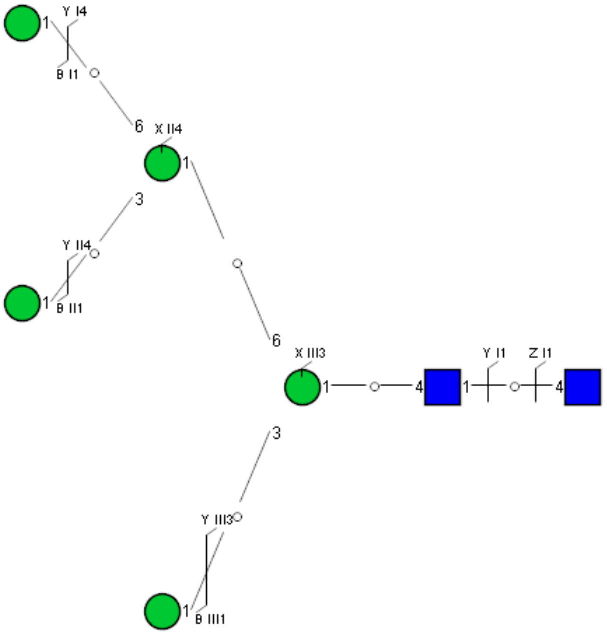

Spectrum Index: 30553(NoBHs:1)

Precursor iEF Map

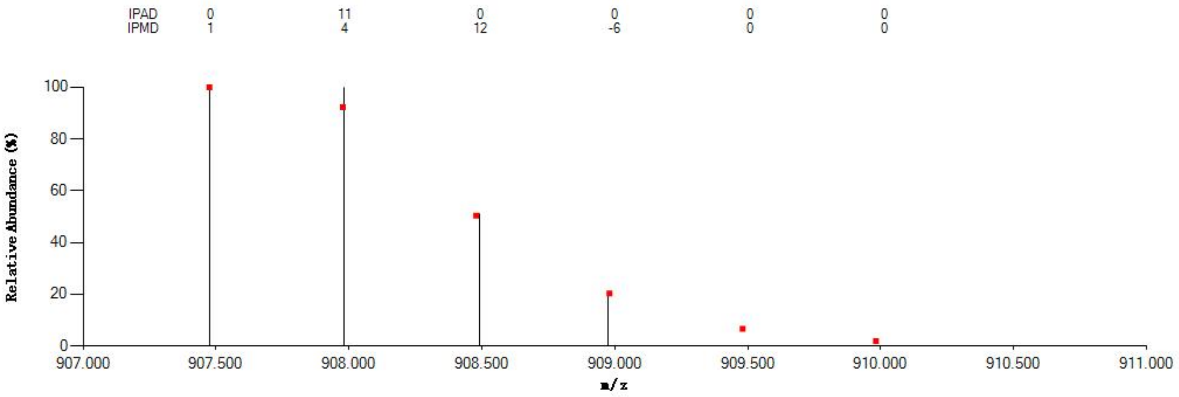


Annotated MS/MS Spectrum

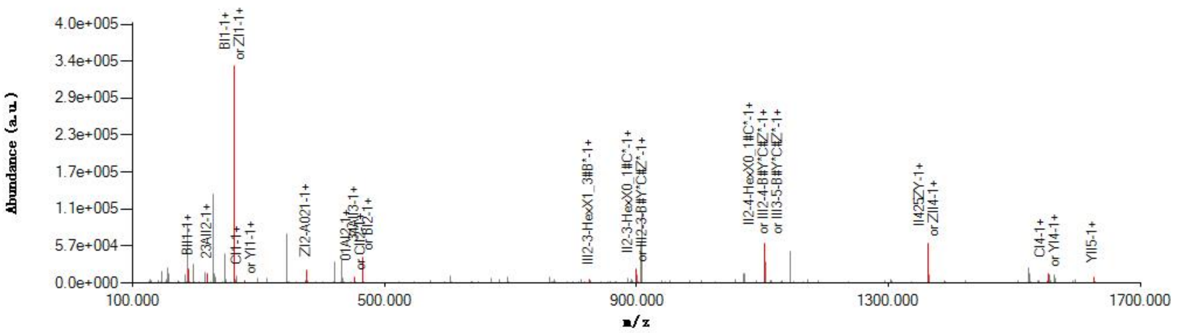


Graphical Fragmentation Map

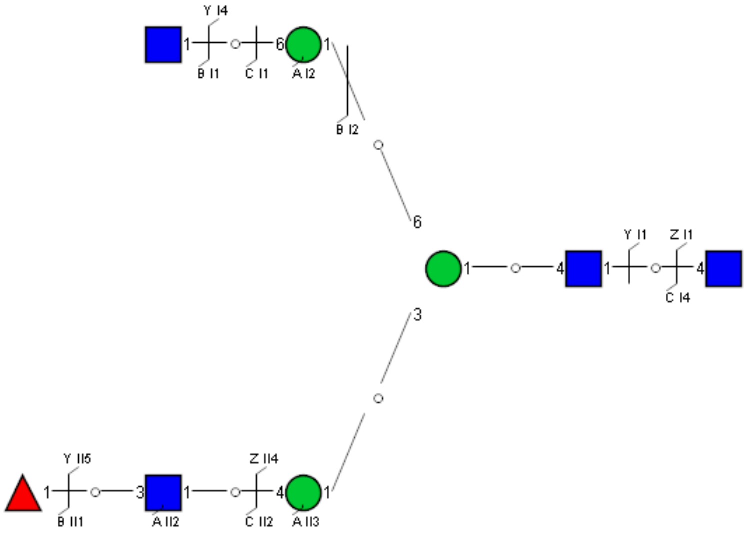

Spectrum Index: 31279(NoBHs:1)

#### Precursor iEF Map

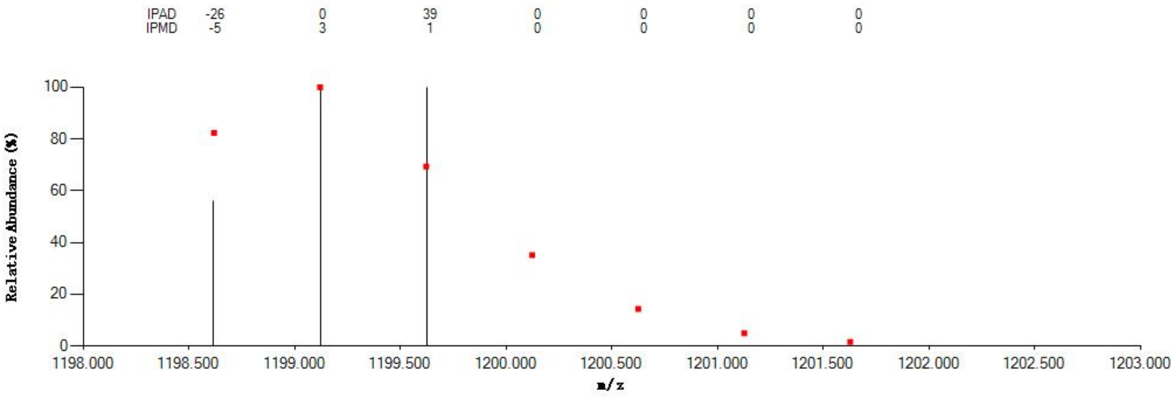

#### Annotated MS/MS Spectrum

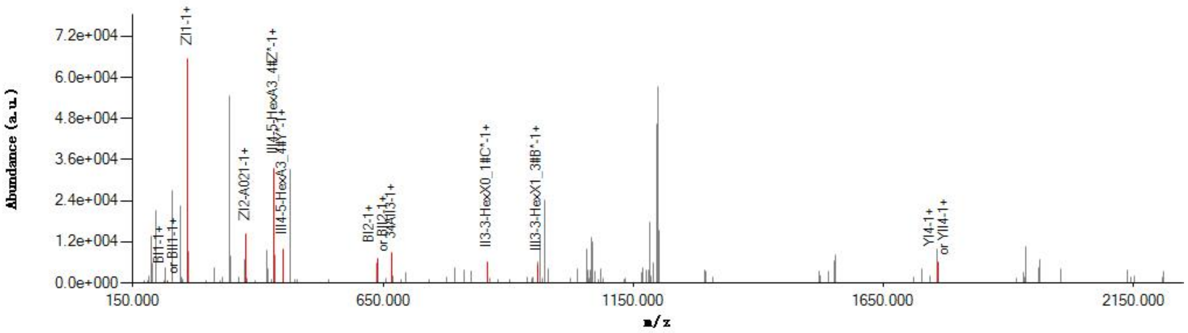

#### Graphical Fragmentation Map

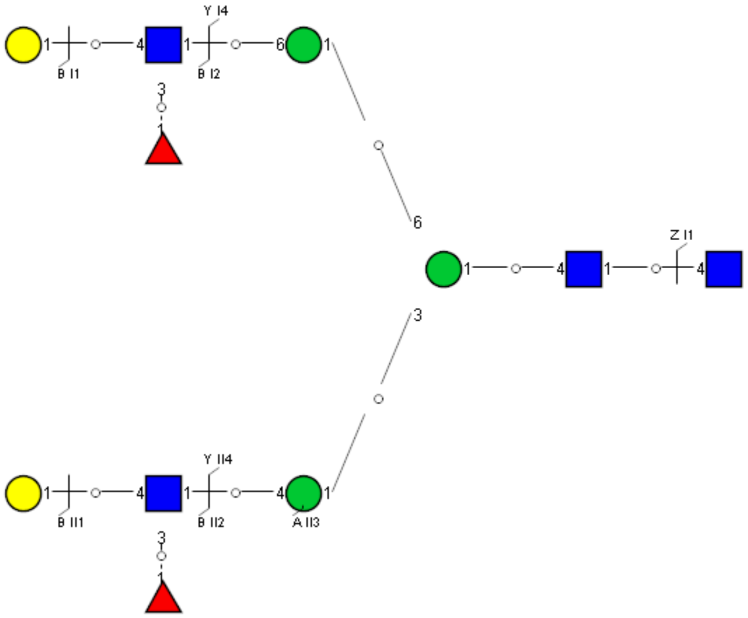

Spectrum Index: 31958(NoBHs:1)

#### Precursor iEF Map

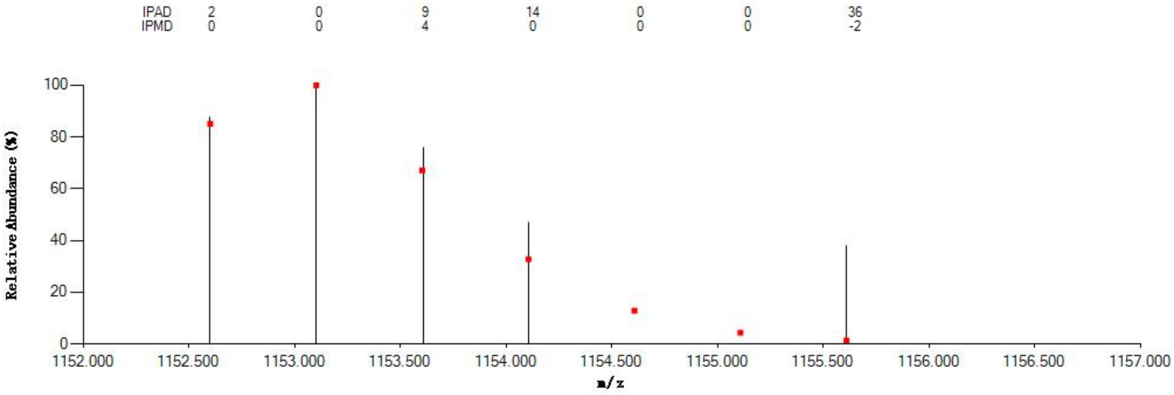


#### Annotated MS/MS Spectrum

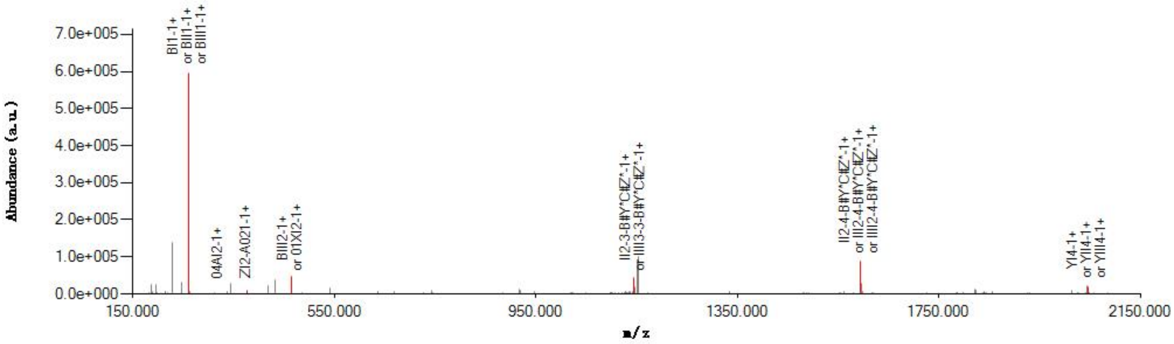


#### Graphical Fragmentation Map

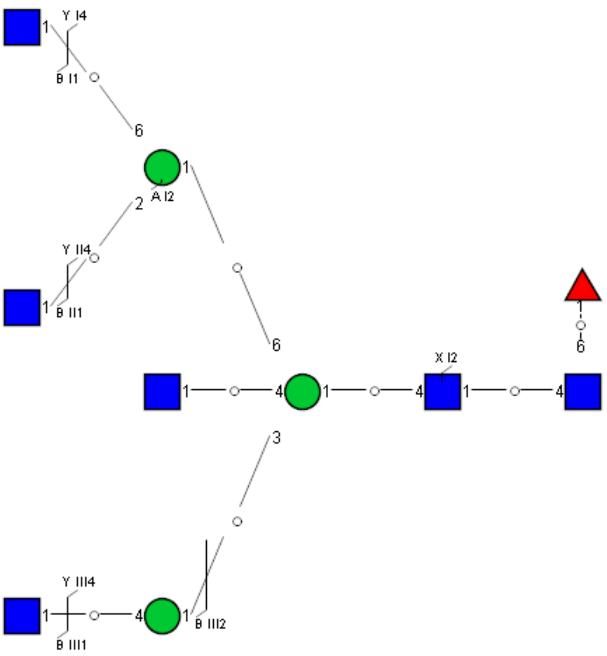

Spectrum Index: 32110(NoBHs:2)

#### Precursor iEF Map

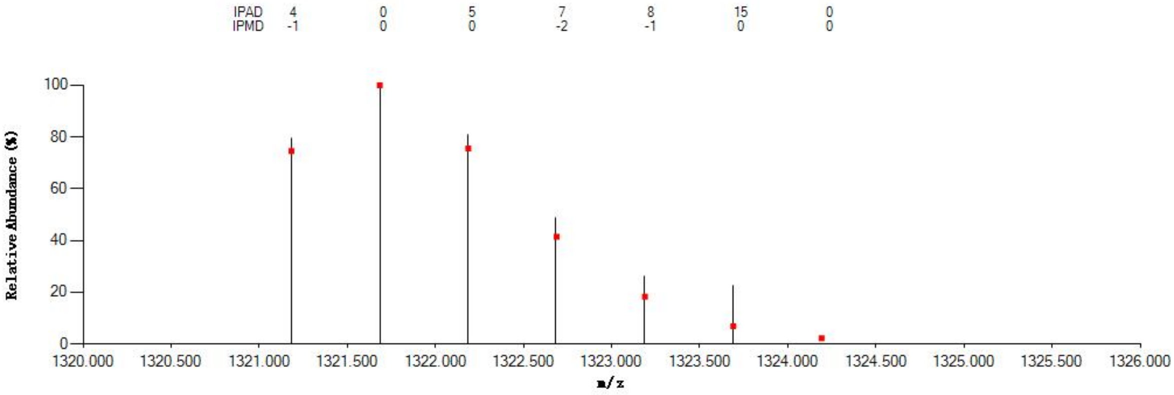

#### Annotated MS/MS Spectrum

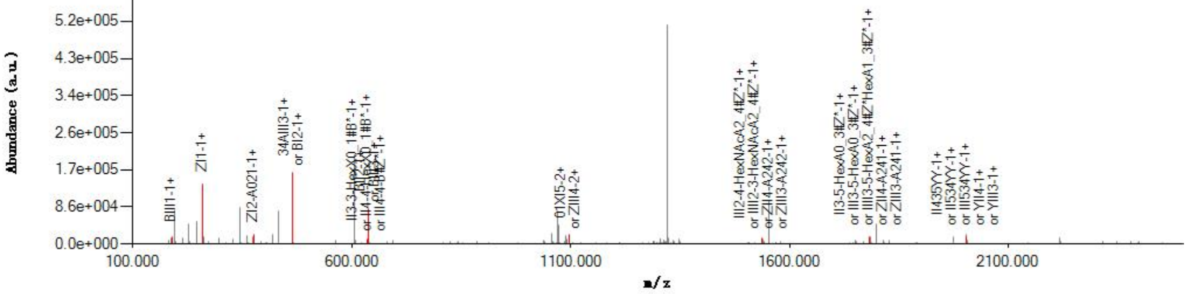

#### Graphical Fragmentation Map

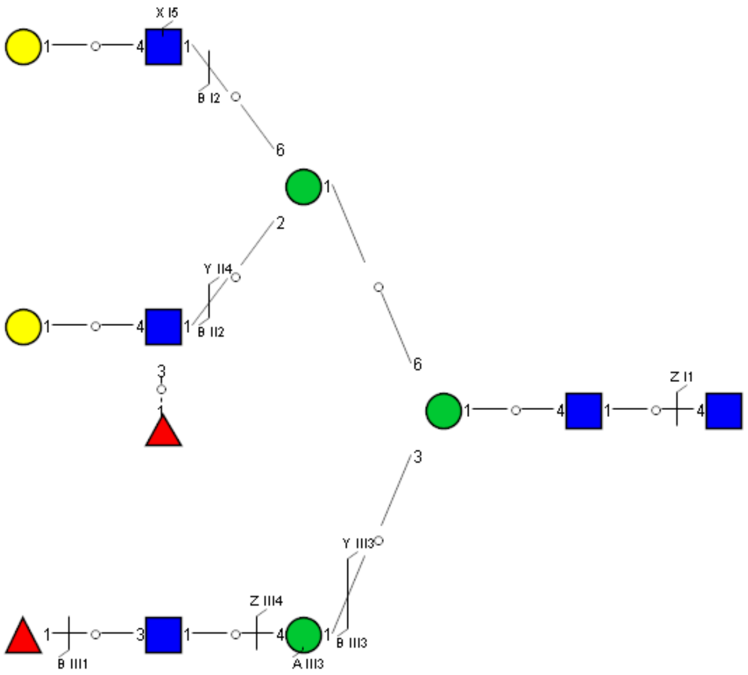

Spectrum Index: 32966(NoBHs:1)

#### Precursor iEF Map

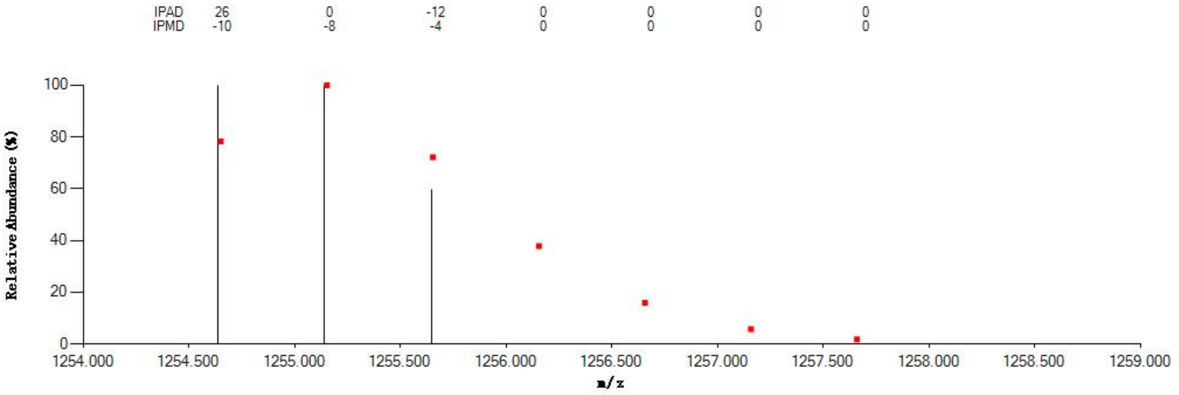


#### Annotated MS/MS Spectrum

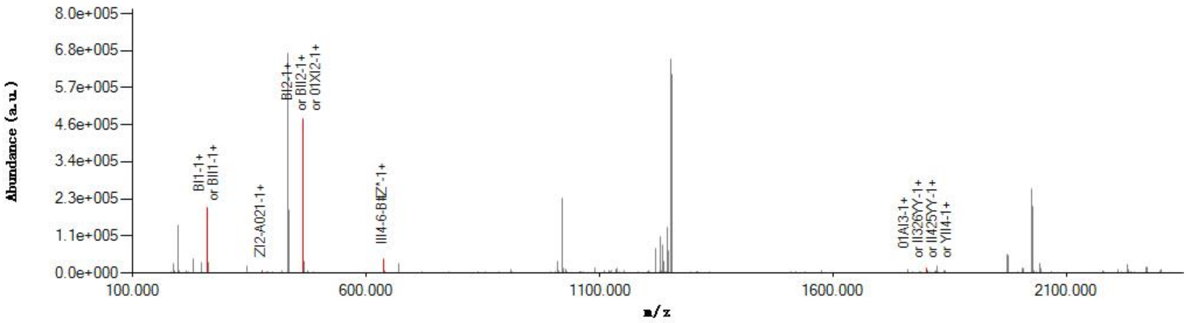


#### Graphical Fragmentation Map

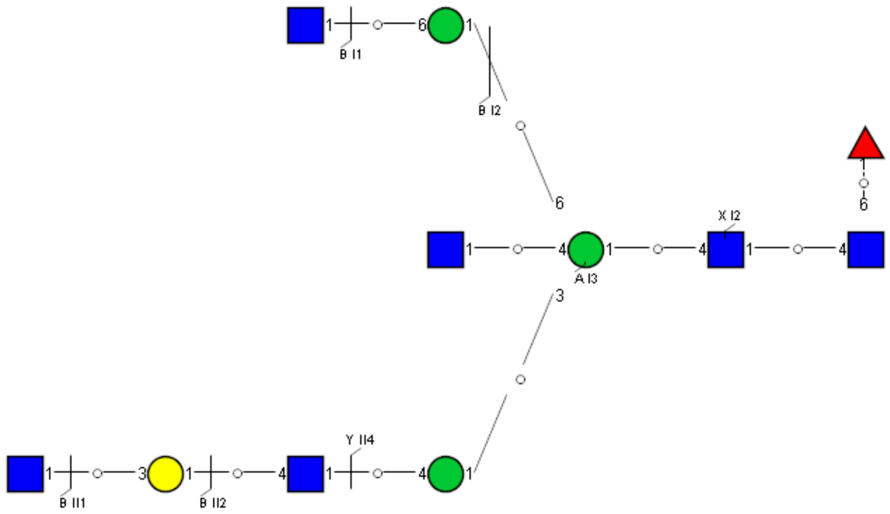

Spectrum Index: 32967(NoBHs:1)

#### Precursor iEF Map

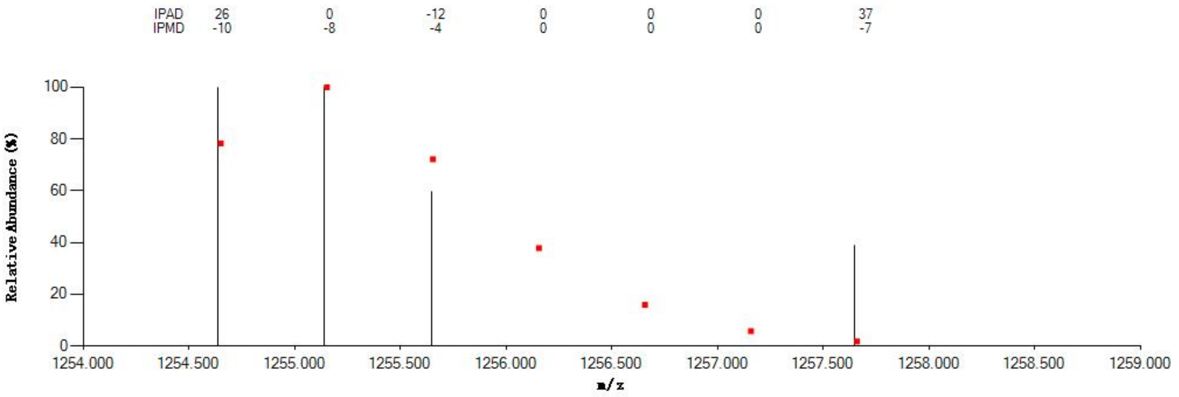


#### Annotated MS/MS Spectrum

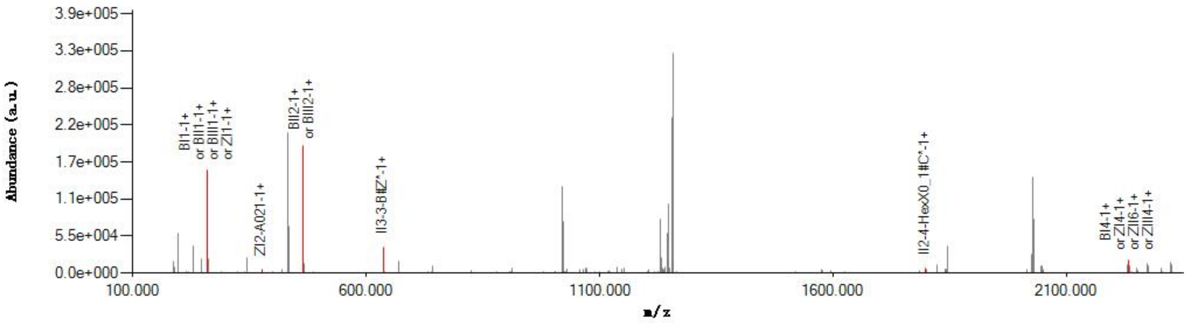


#### Graphical Fragmentation Map

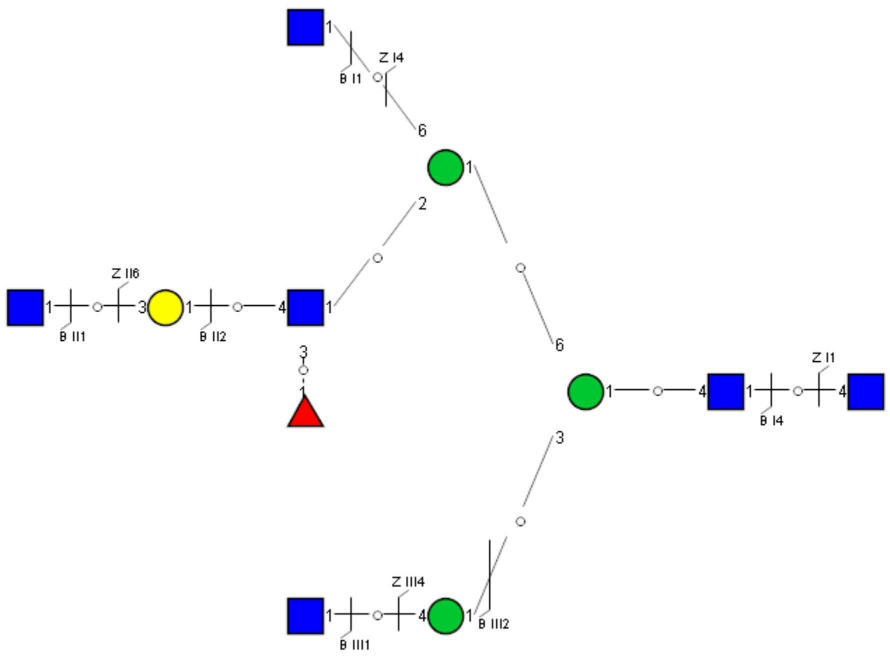

Spectrum Index: 34080(NoBHs:1)

#### Precursor iEF Map

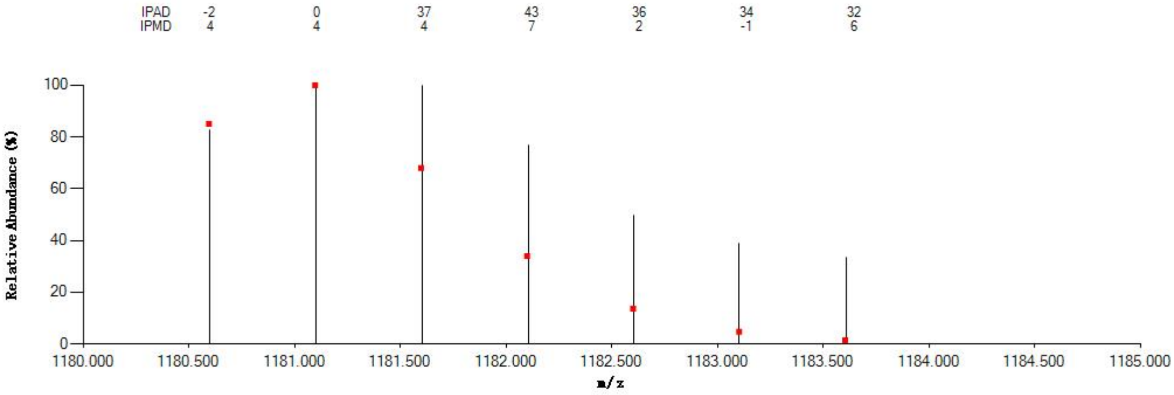

#### Annotated MS/MS Spectrum

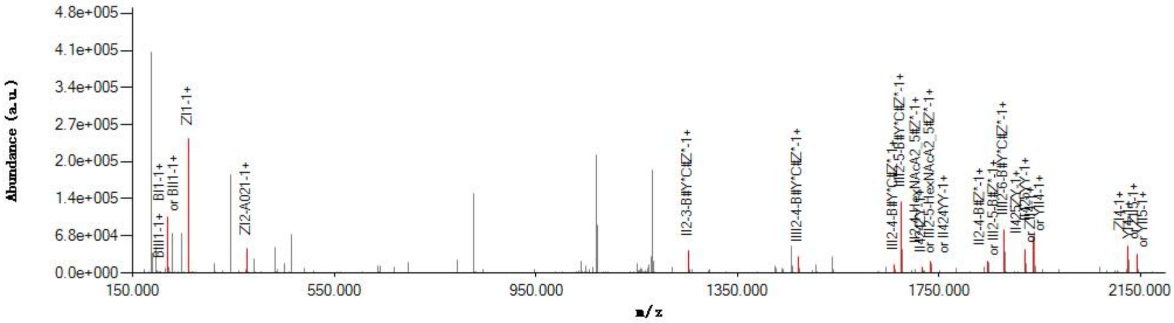

#### Graphical Fragmentation Map

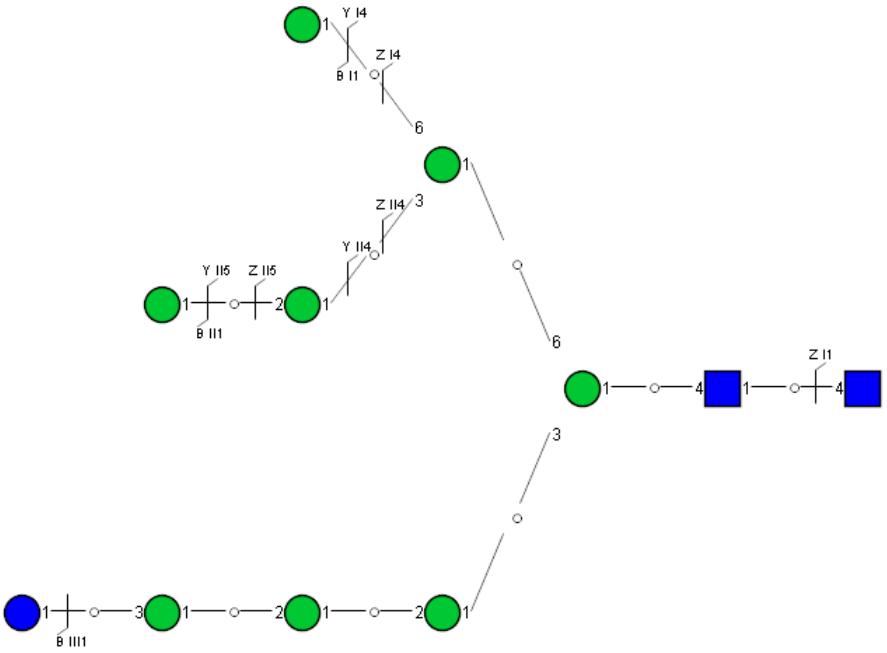

Spectrum Index: 34483(NoBHs:1)

#### Precursor iEF Map

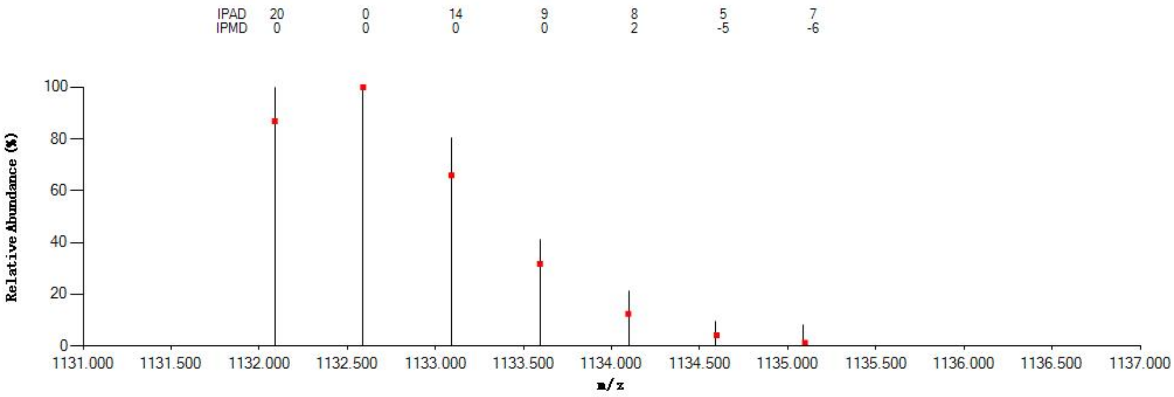


#### Annotated MS/MS Spectrum

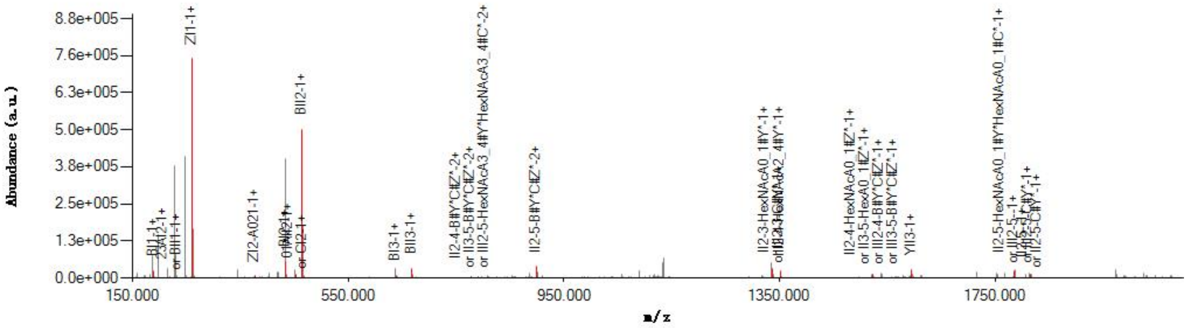


#### Graphical Fragmentation Map

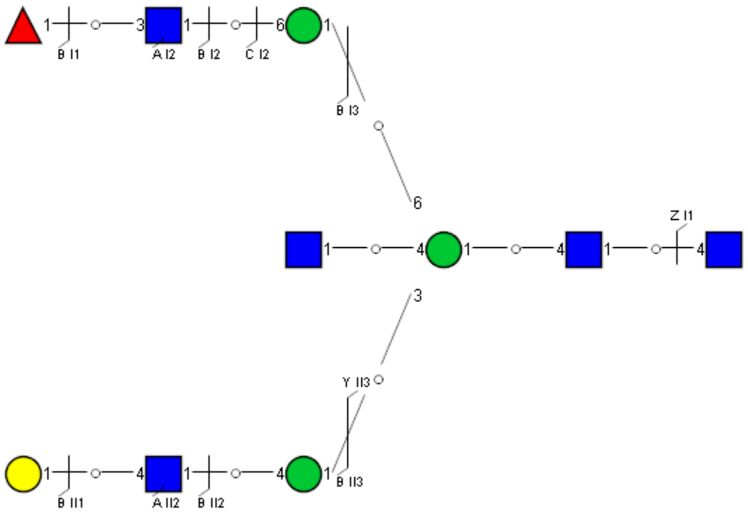

Spectrum Index: 35460(NoBHs:1)

#### Precursor iEF Map

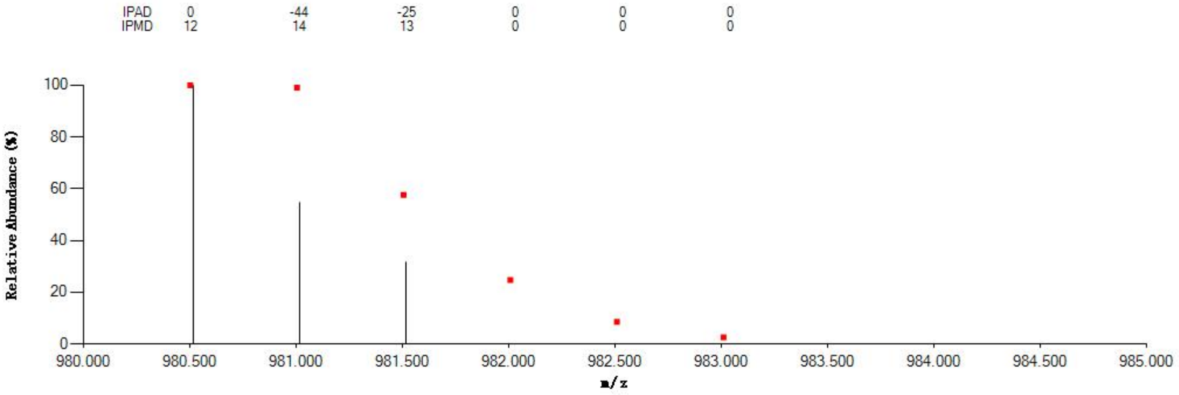

#### Annotated MS/MS Spectrum

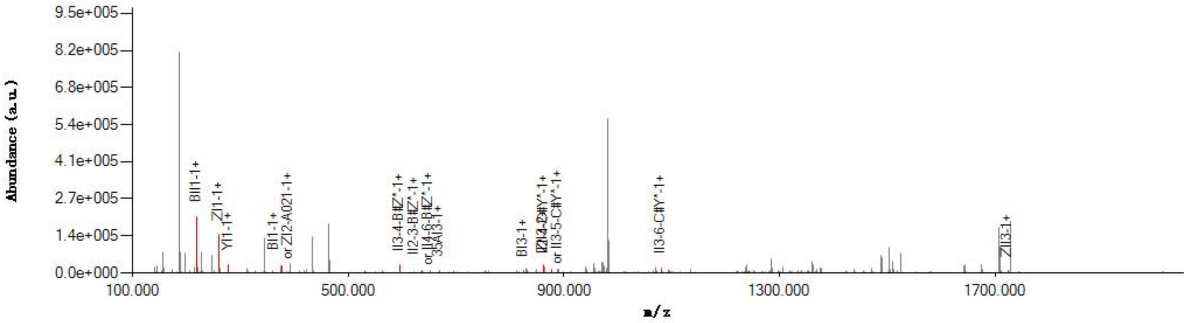

#### Graphical Fragmentation Map

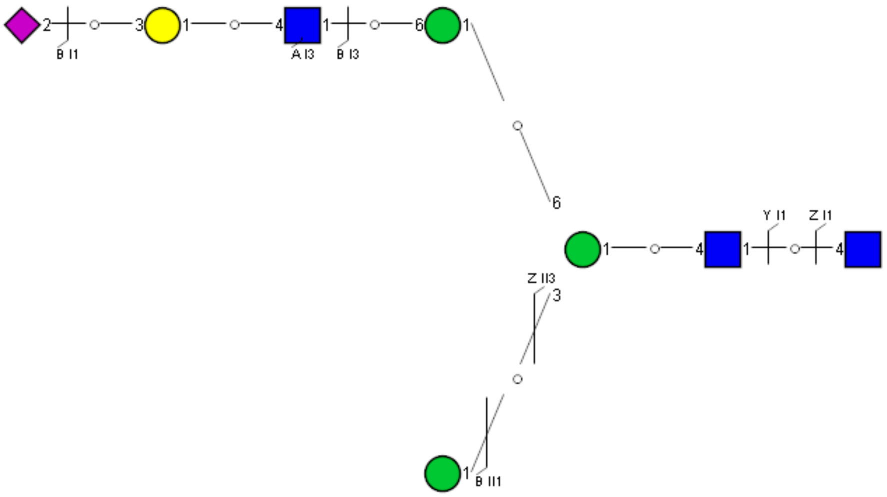

Spectrum Index: 36108(NoBHs:1)

#### Precursor iEF Map

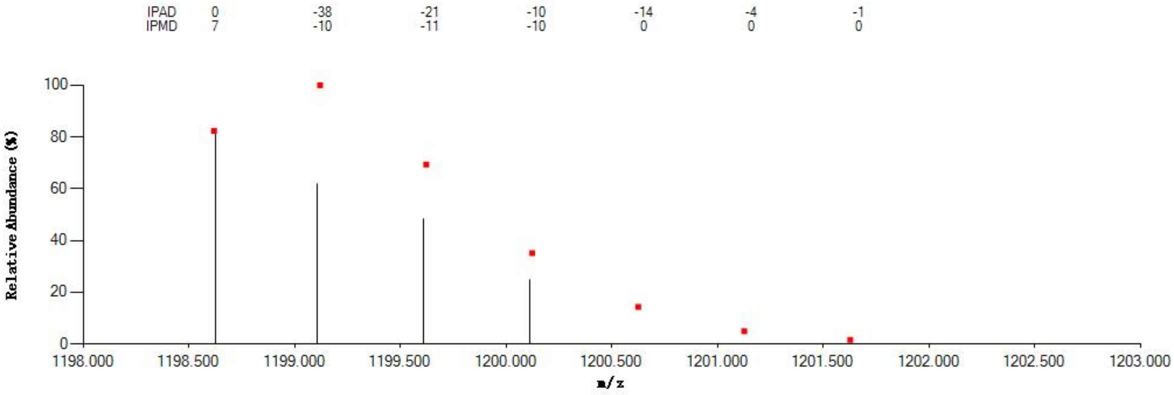

#### Annotated MS/MS Spectrum

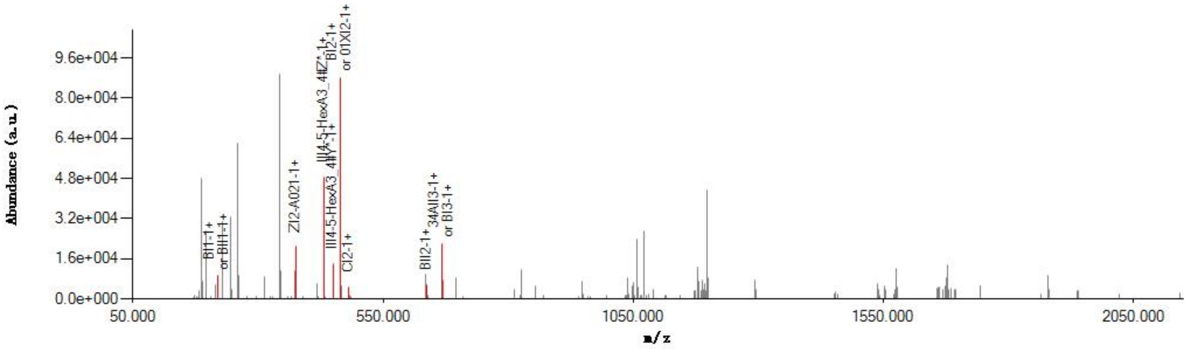

#### Graphical Fragmentation Map

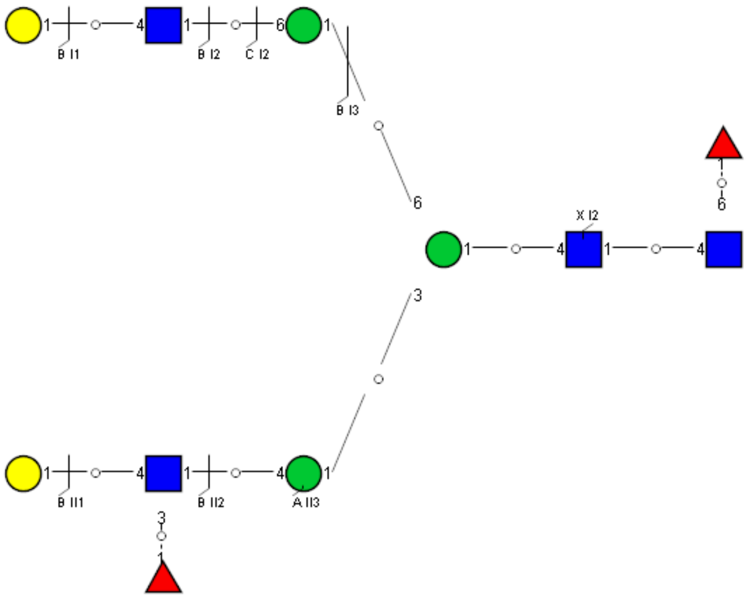

Spectrum Index: 36173(NoBHs:2)

#### Precursor iEF Map

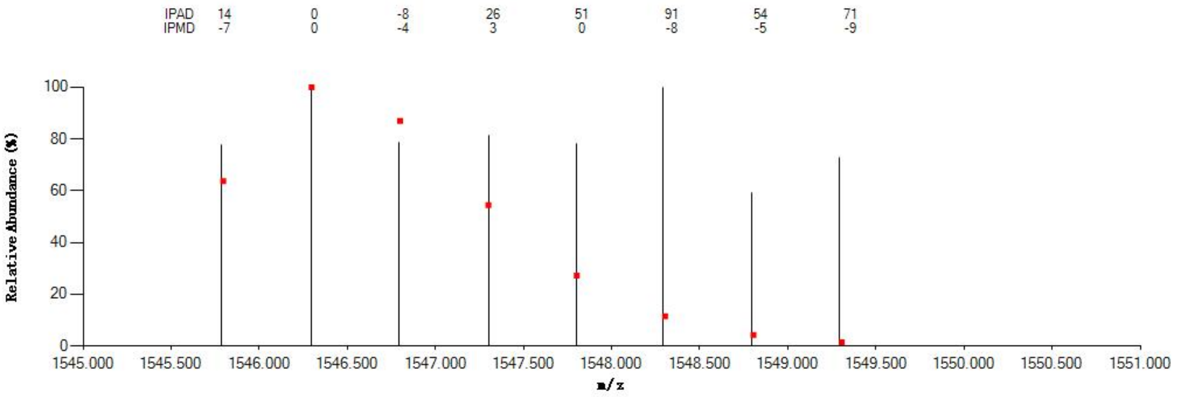


#### Annotated MS/MS Spectrum

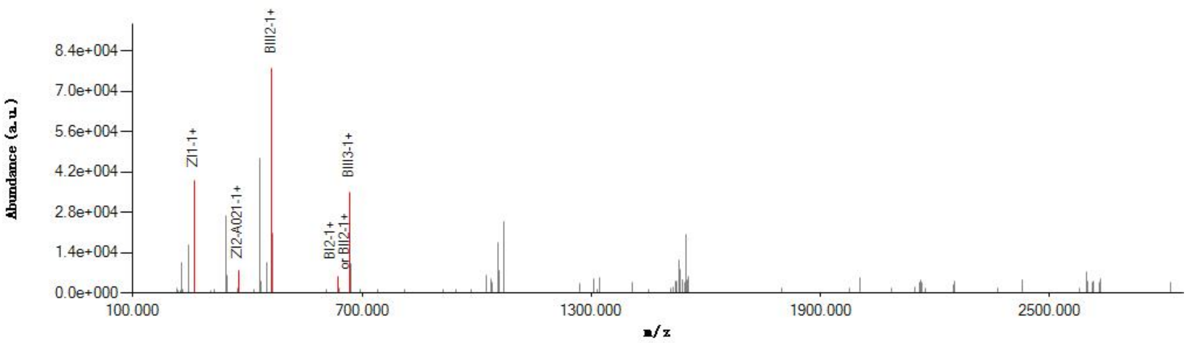


#### Graphical Fragmentation Map

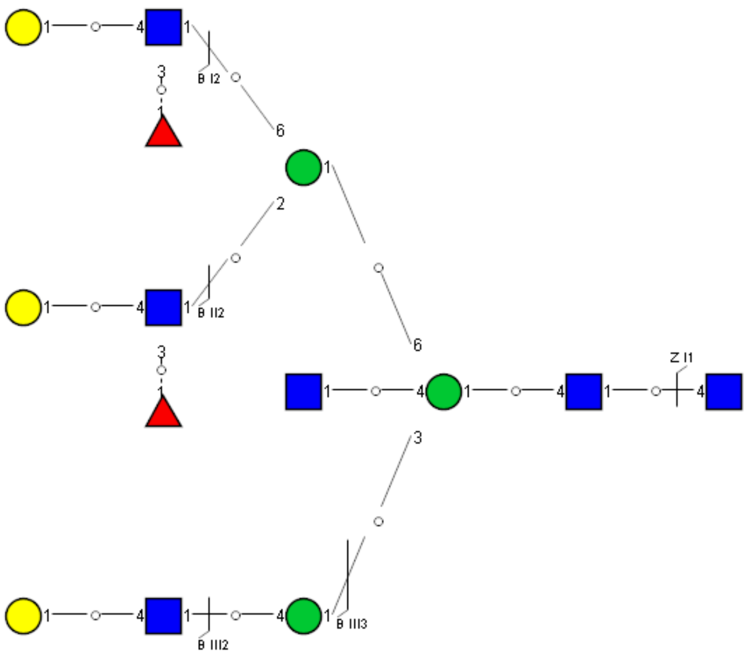

Spectrum Index: 36173(NoBHs:2)

#### Precursor iEF Map

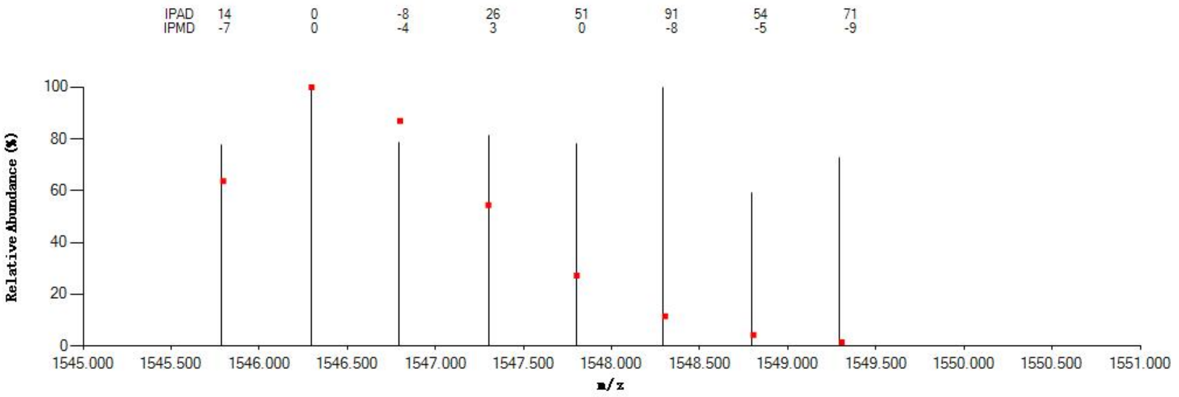


#### Annotated MS/MS Spectrum

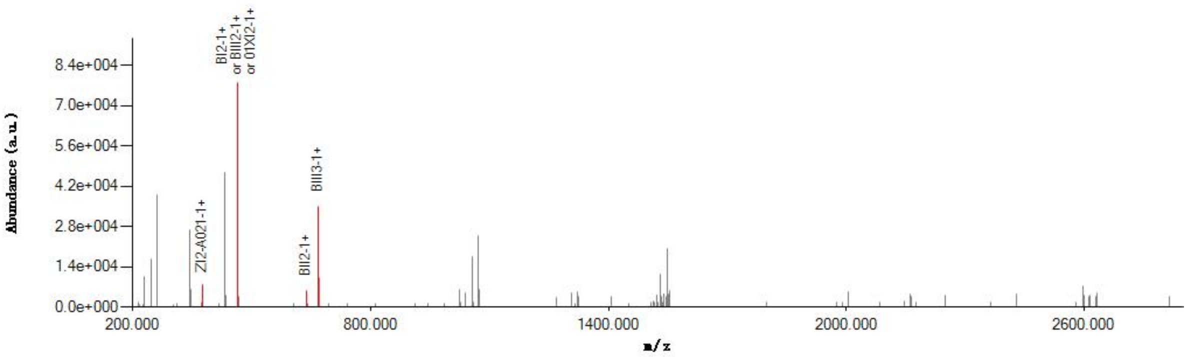


#### Graphical Fragmentation Map

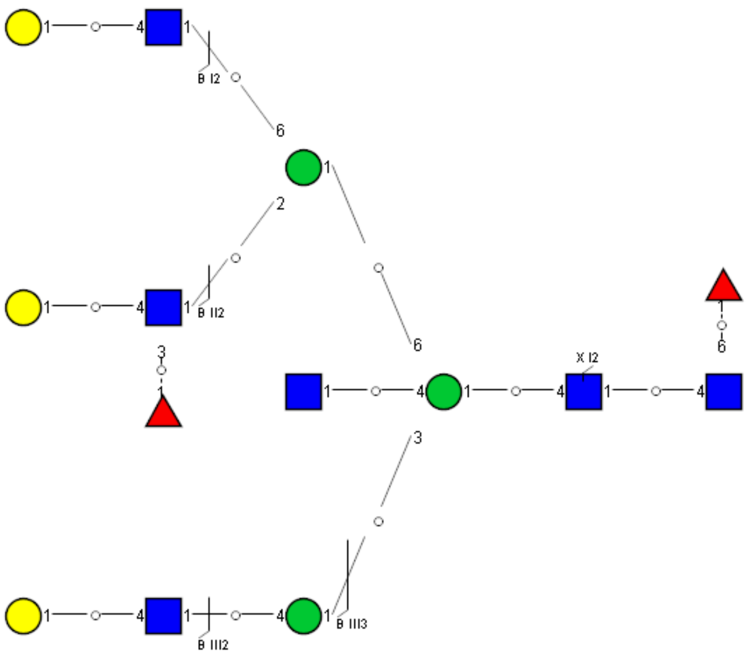

Spectrum Index: 36258(NoBHs:1)

#### Precursor iEF Map

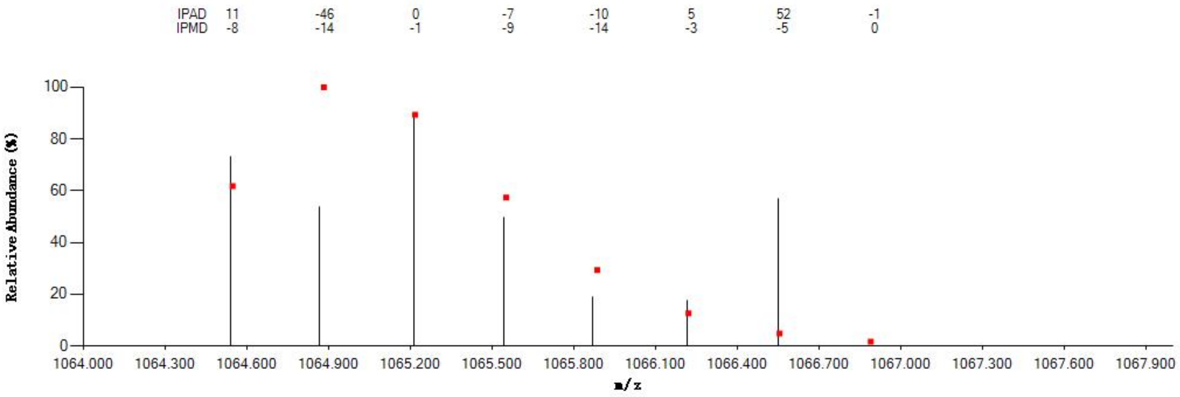


#### Annotated MS/MS Spectrum

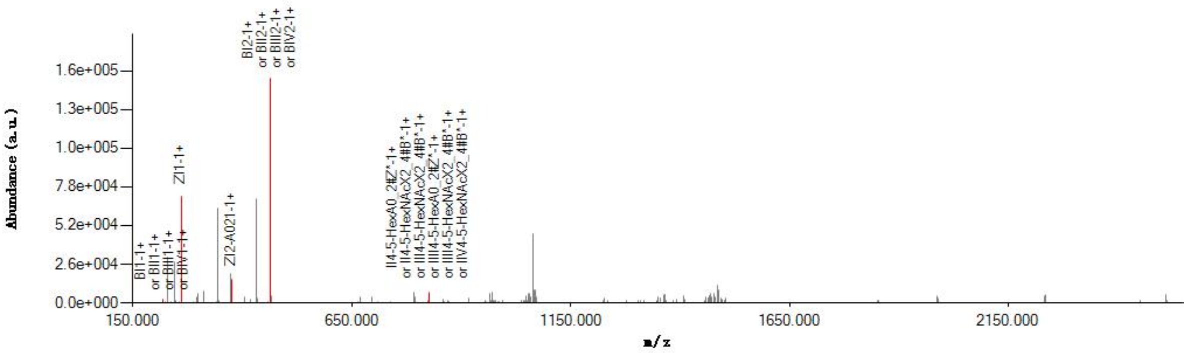


#### Graphical Fragmentation Map

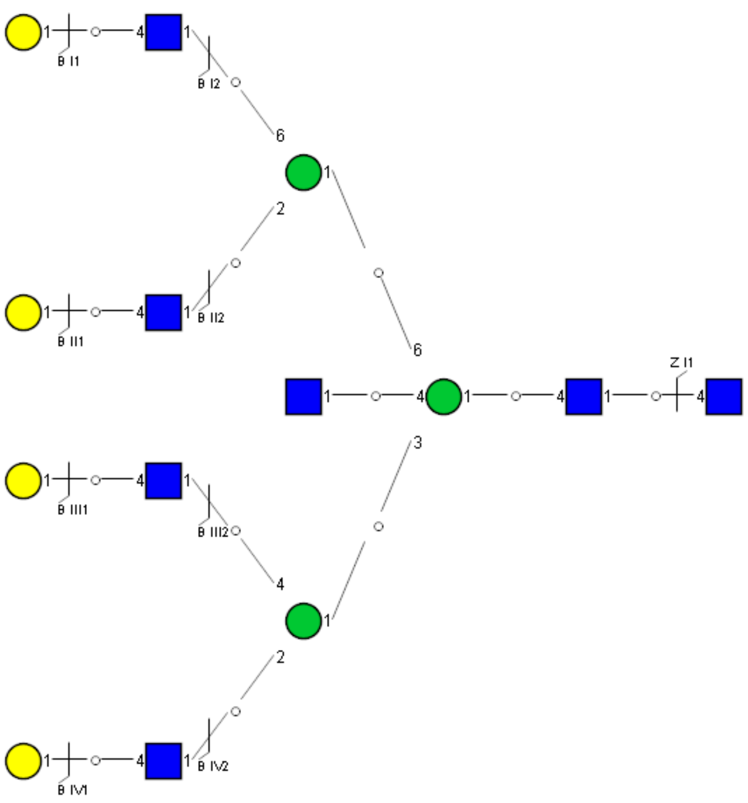

### Spectrum Index: 37426(NoBHs:1)

#### Precursor iEF Map

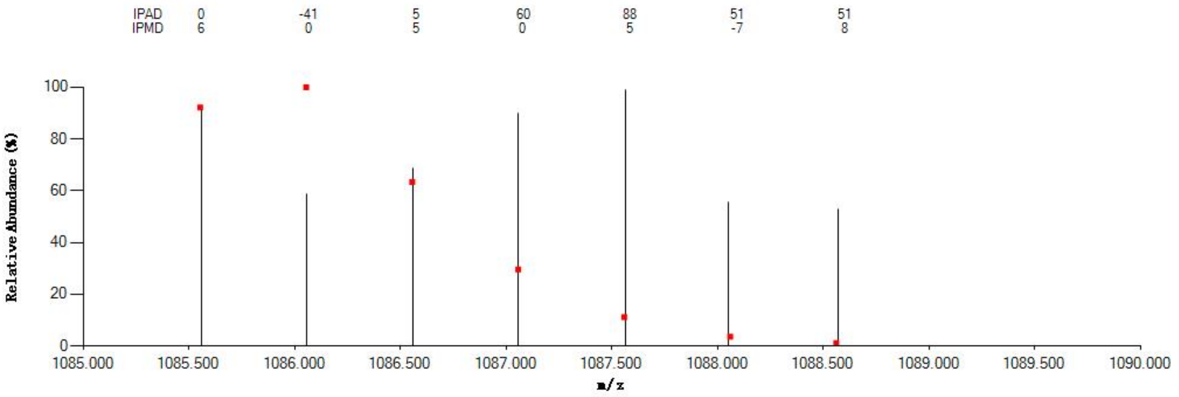


#### Annotated MS/MS Spectrum

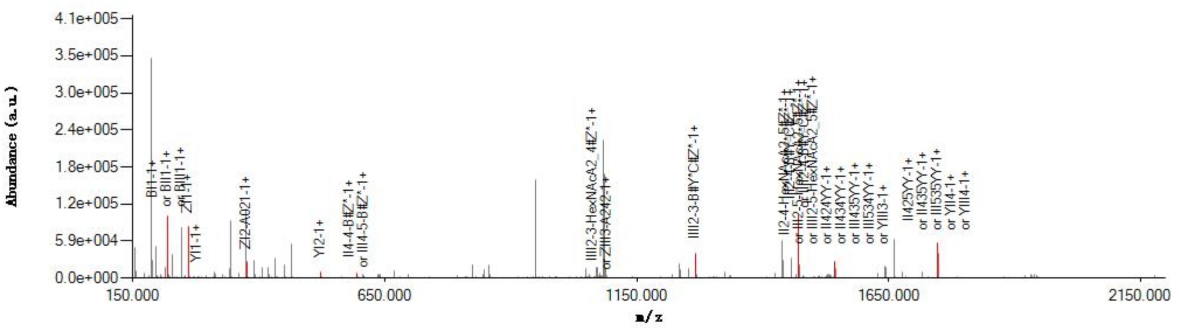


#### Graphical Fragmentation Map

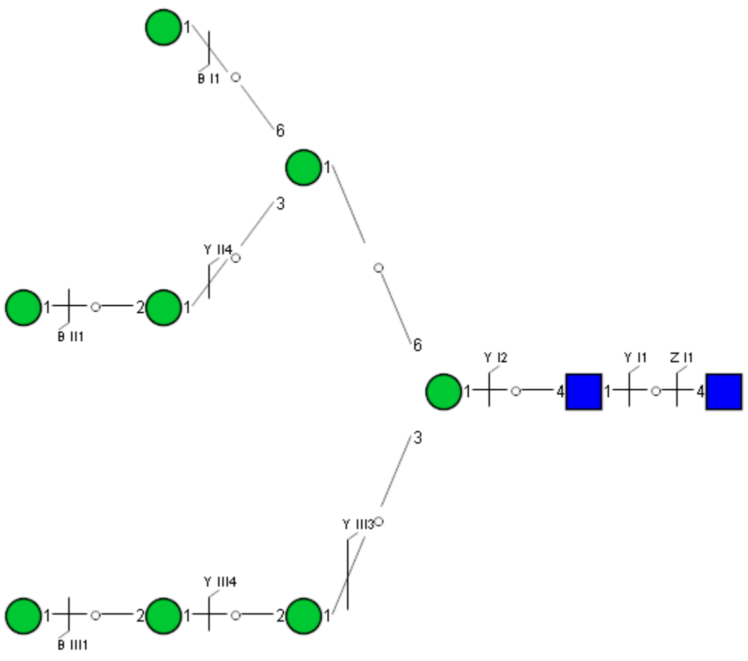

Spectrum Index: 37620(NoBHs:1)

#### Precursor iEF Map

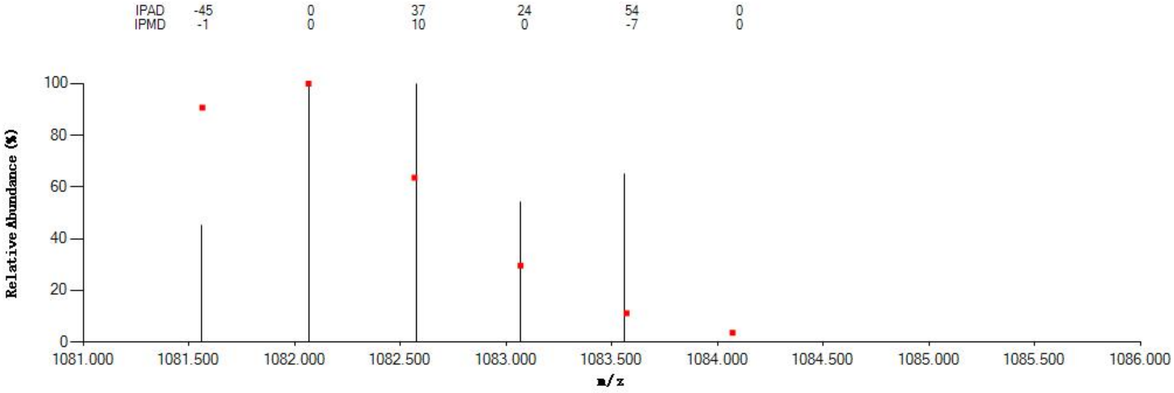


#### Annotated MS/MS Spectrum

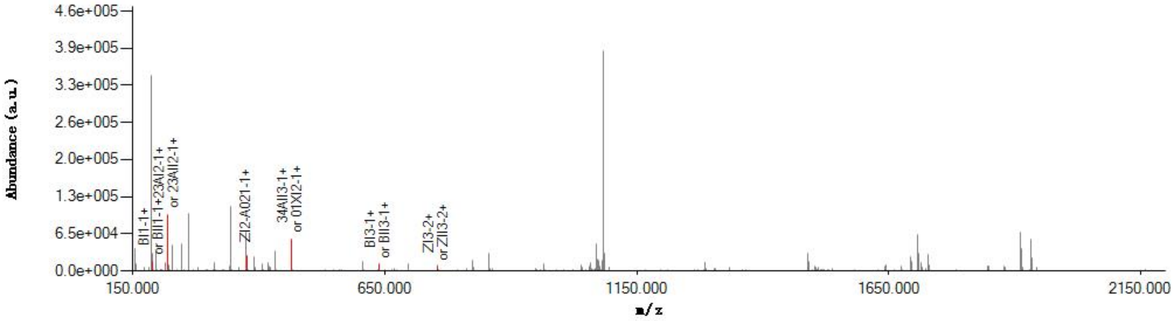


#### Graphical Fragmentation Map

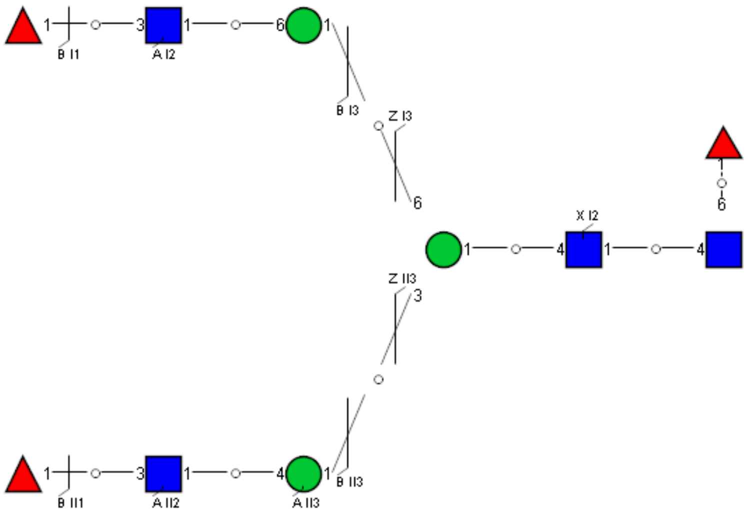

Spectrum Index: 37776(NoBHs:1)

#### Precursor iEF Map

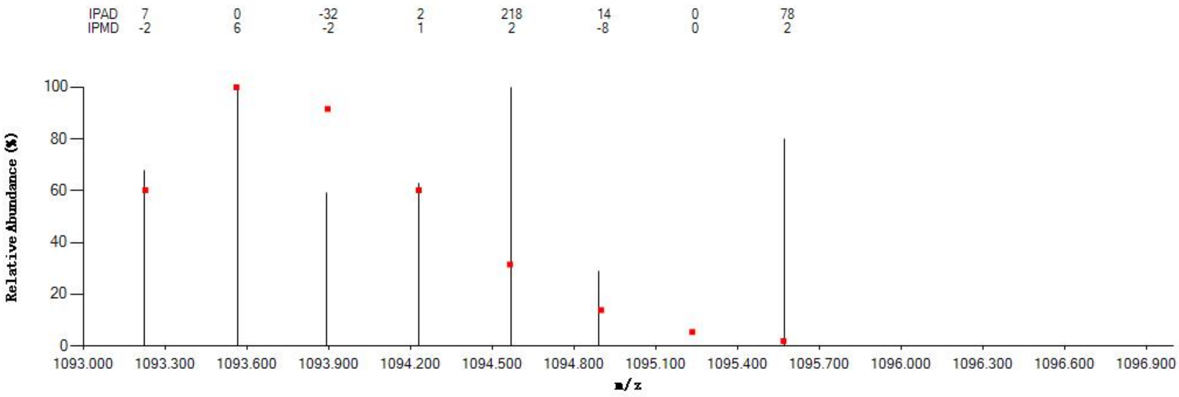


#### Annotated MS/MS Spectrum

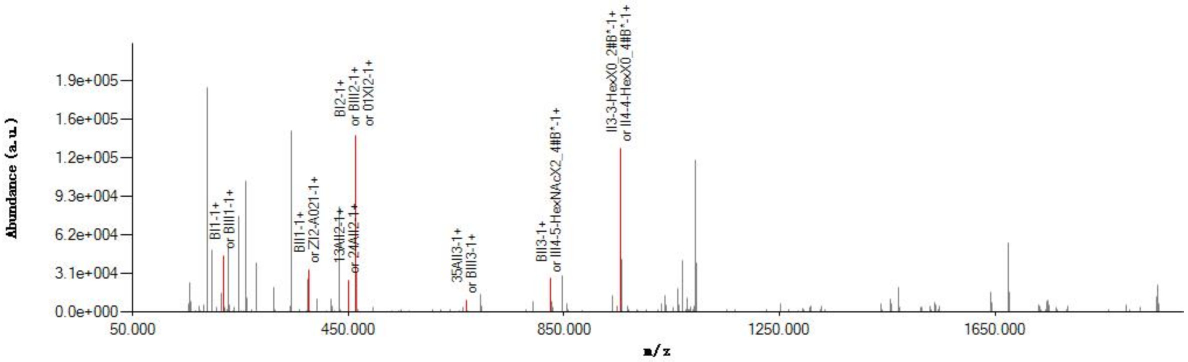


#### Graphical Fragmentation Map

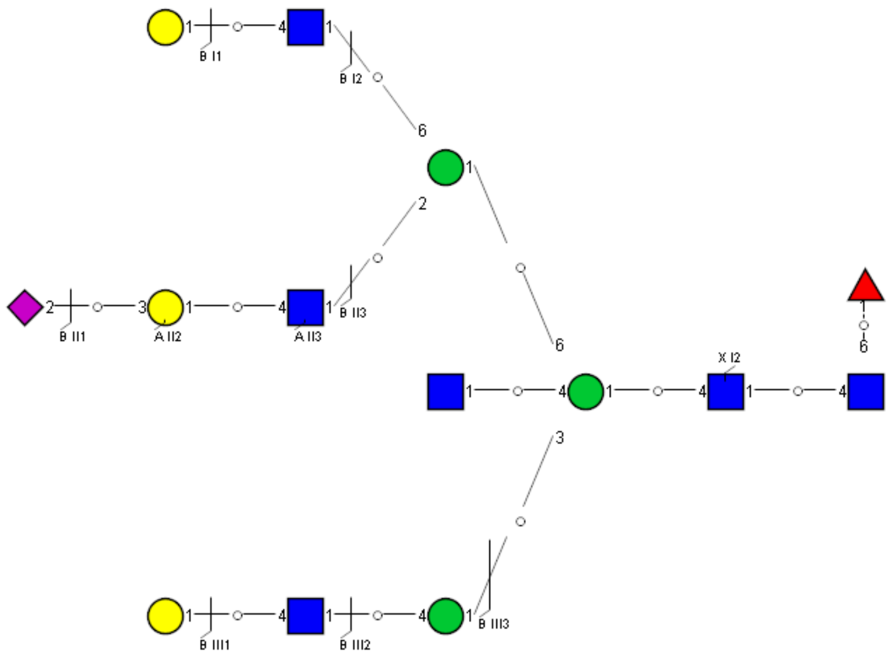

Spectrum Index: 38053(NoBHs:1)

#### Precursor iEF Map

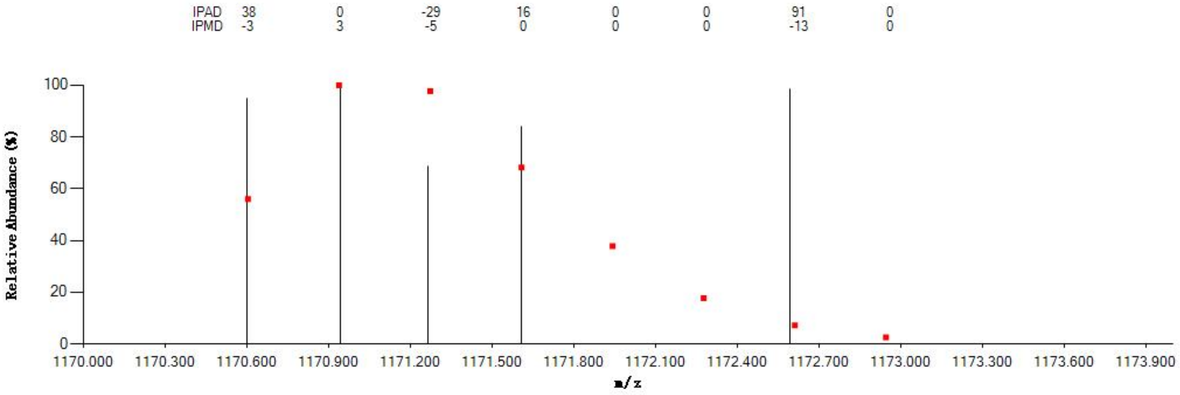


#### Annotated MS/MS Spectrum

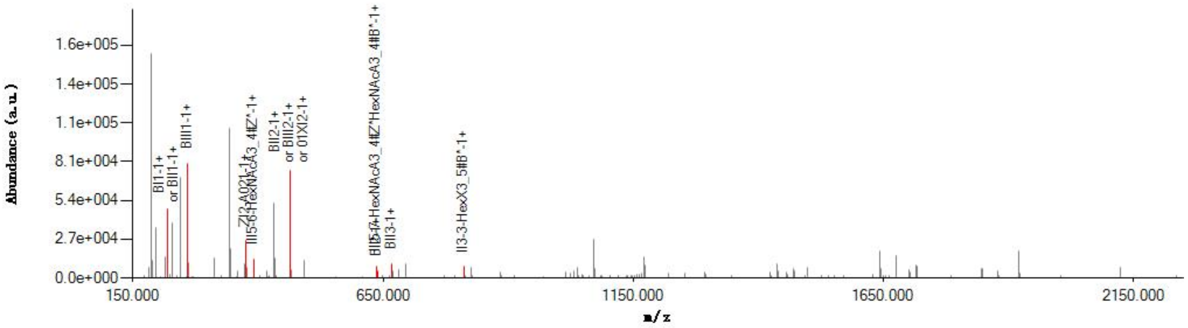


#### Graphical Fragmentation Map

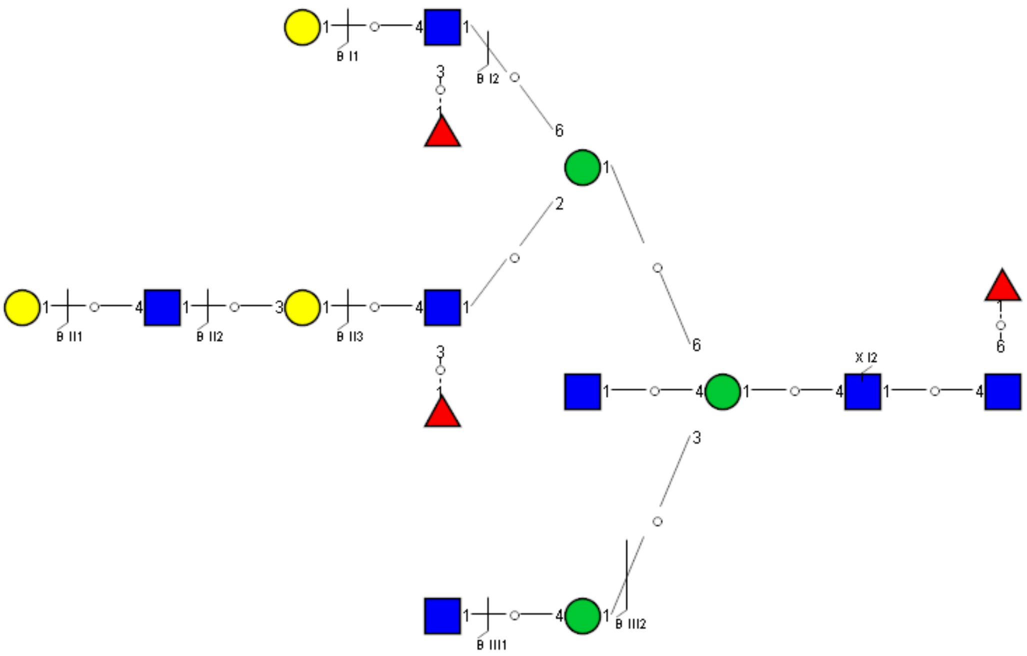

### Spectrum Index: 38236(NoBHs:2)

#### Precursor iEF Map

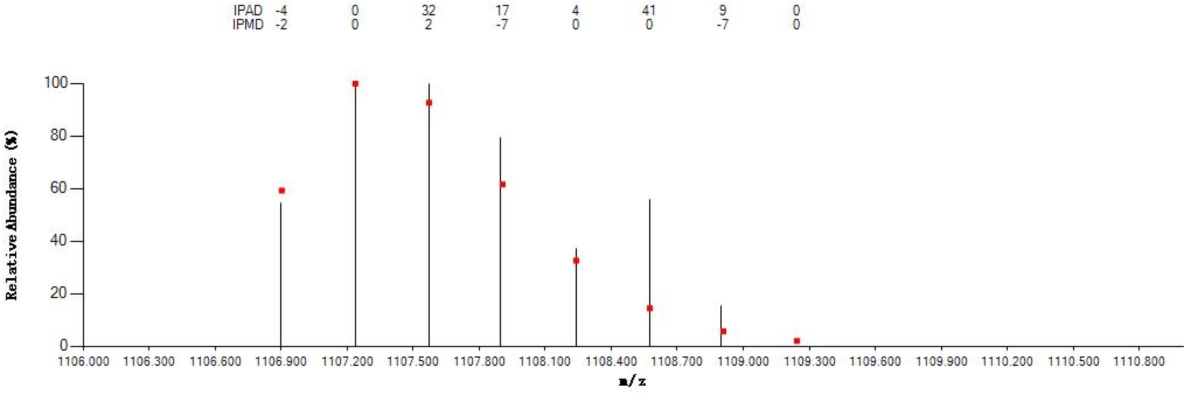


#### Annotated MS/MS Spectrum

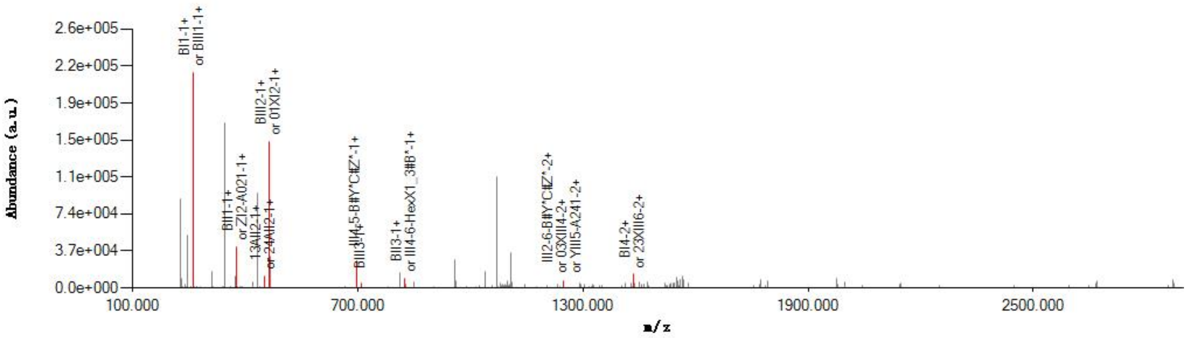


#### Graphical Fragmentation Map

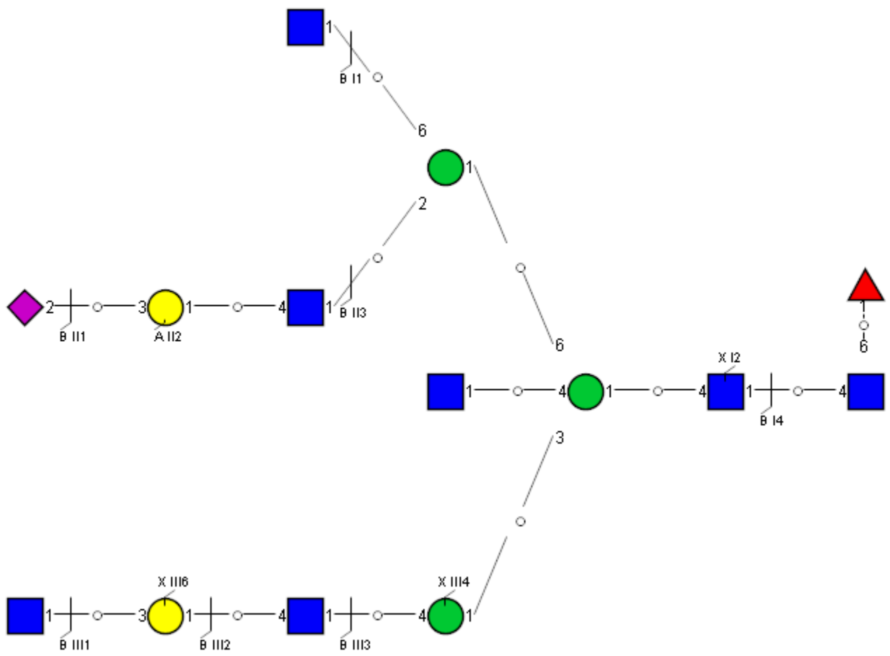

Spectrum Index: 38236(NoBHs:2)

#### Precursor iEF Map

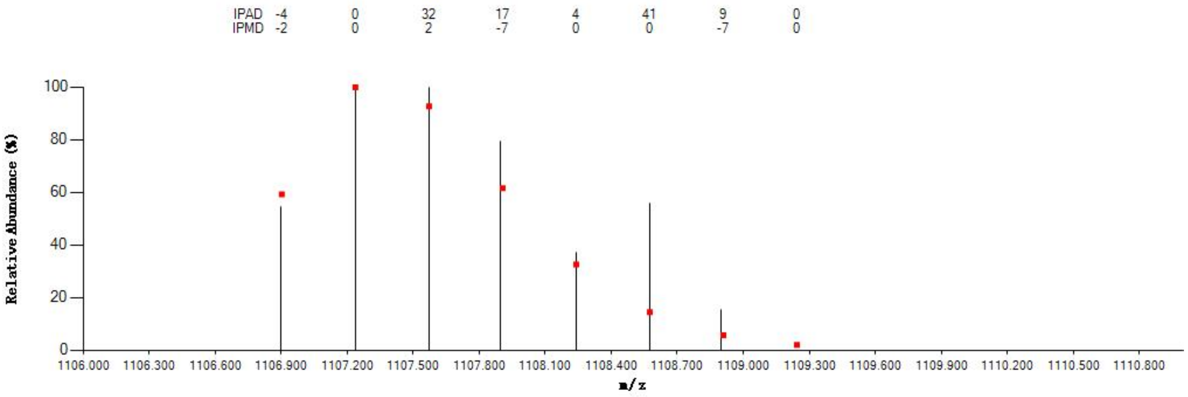

#### Annotated MS/MS Spectrum

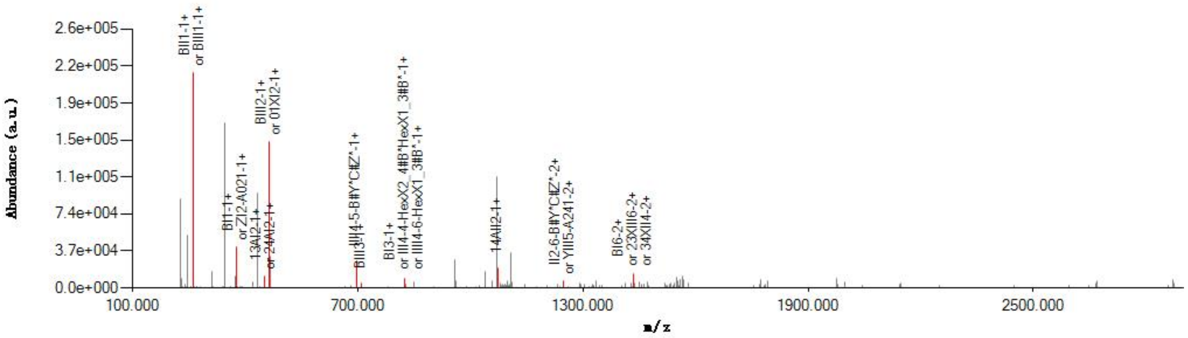

#### Graphical Fragmentation Map

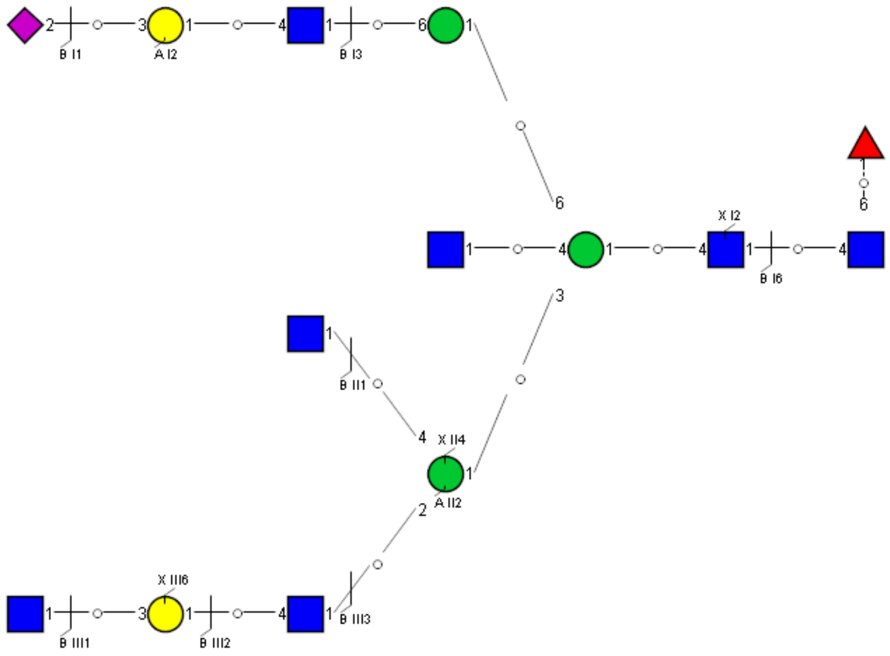

Spectrum Index: 38733(NoBHs:1)

#### Precursor iEF Map

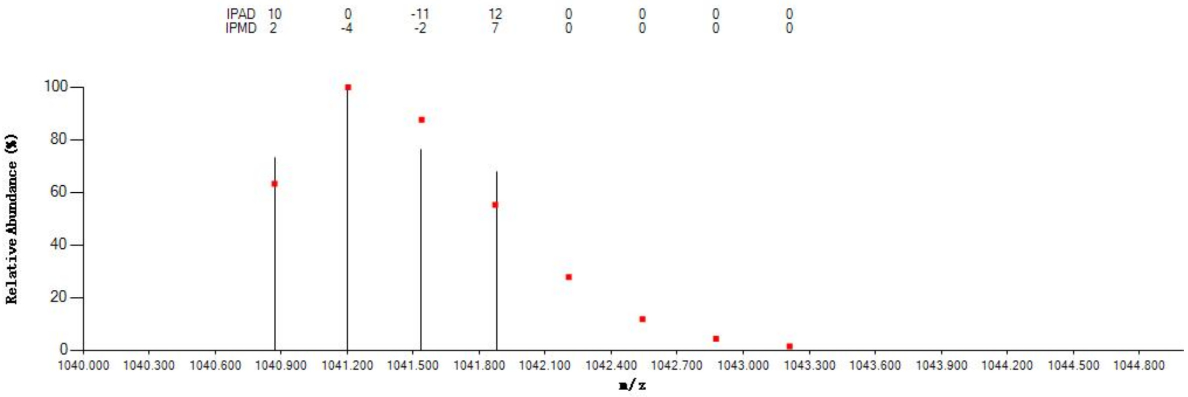


#### Annotated MS/MS Spectrum

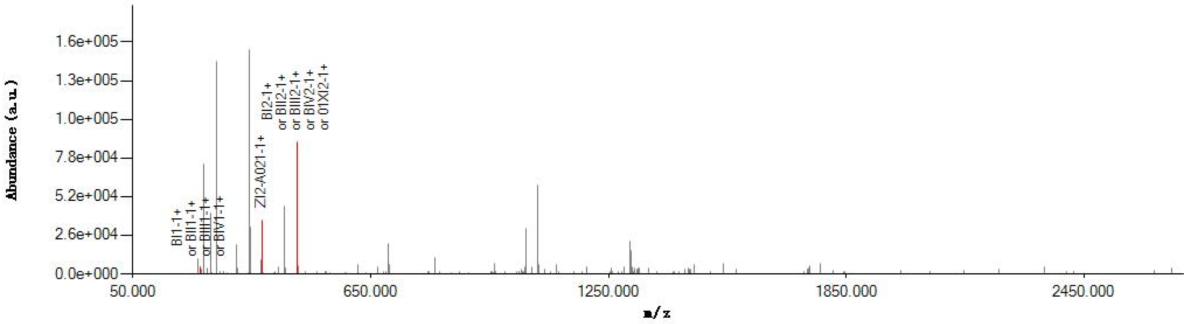


#### Graphical Fragmentation Map

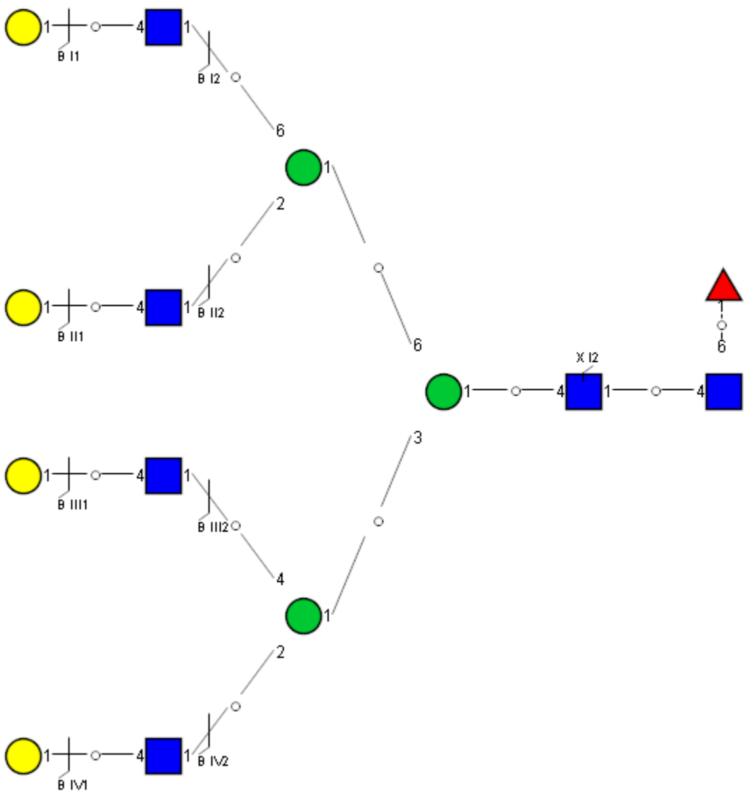

Spectrum Index: 38875(NoBHs:1)

#### Precursor iEF Map

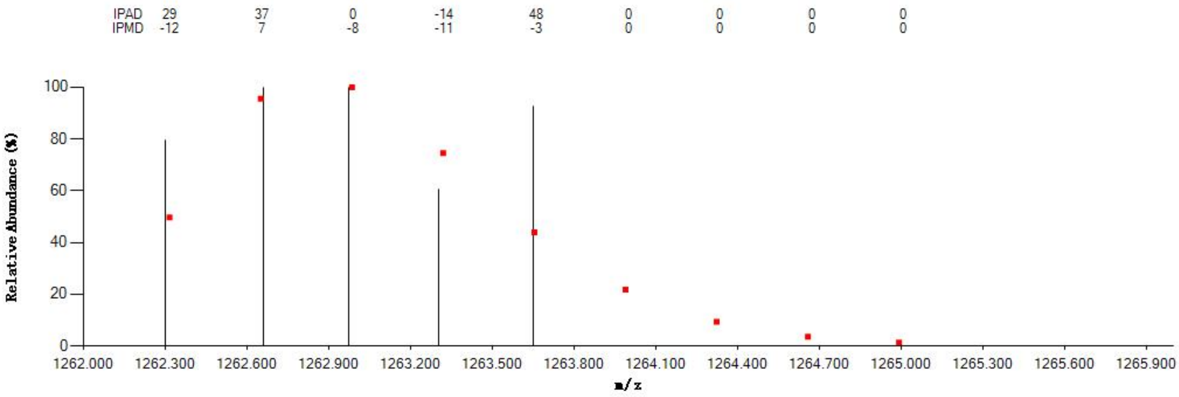


#### Annotated MS/MS Spectrum

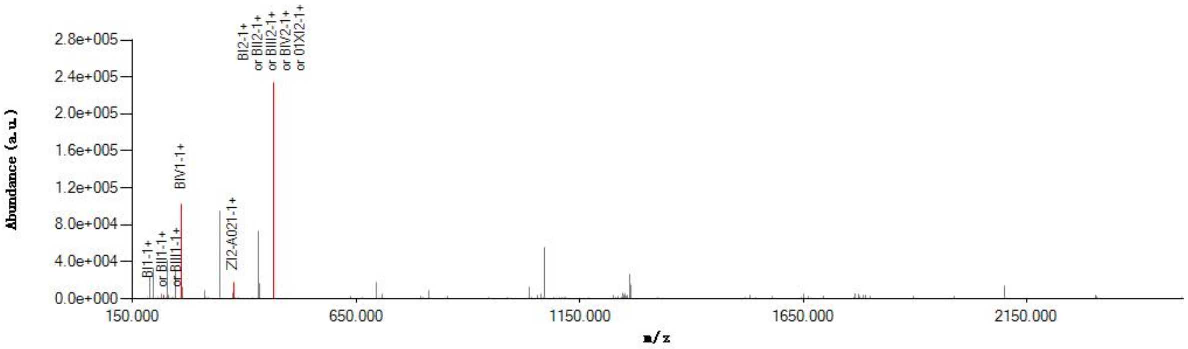


#### Graphical Fragmentation Map

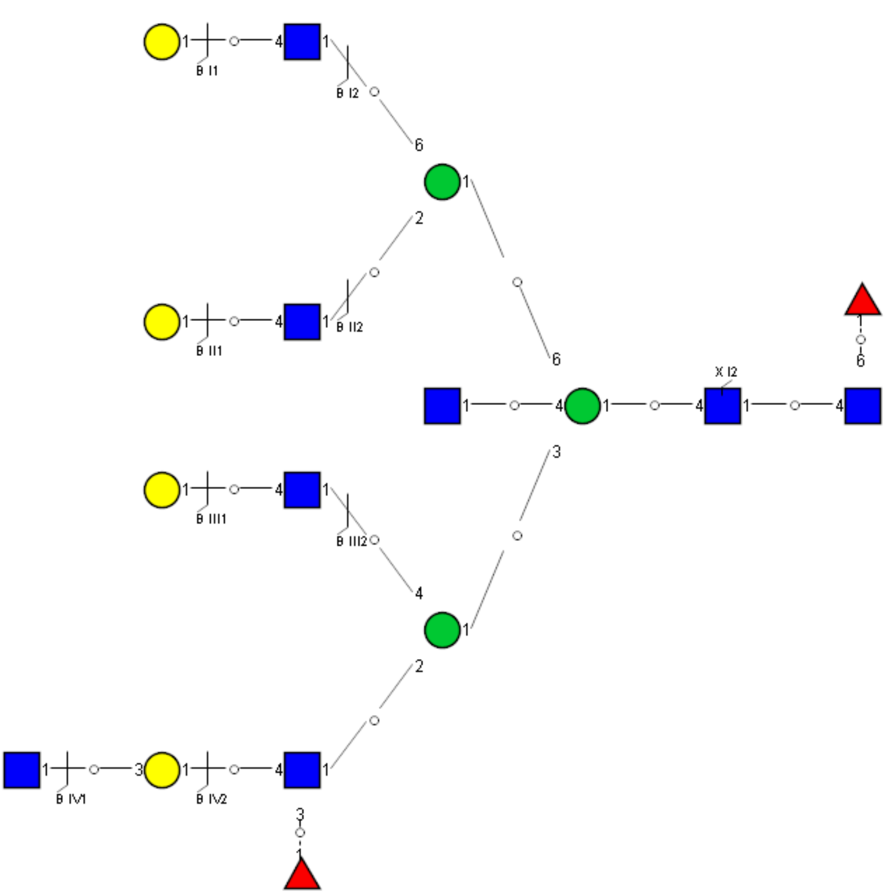

Spectrum Index: 39294(NoBHs:1)

#### Precursor iEF Map

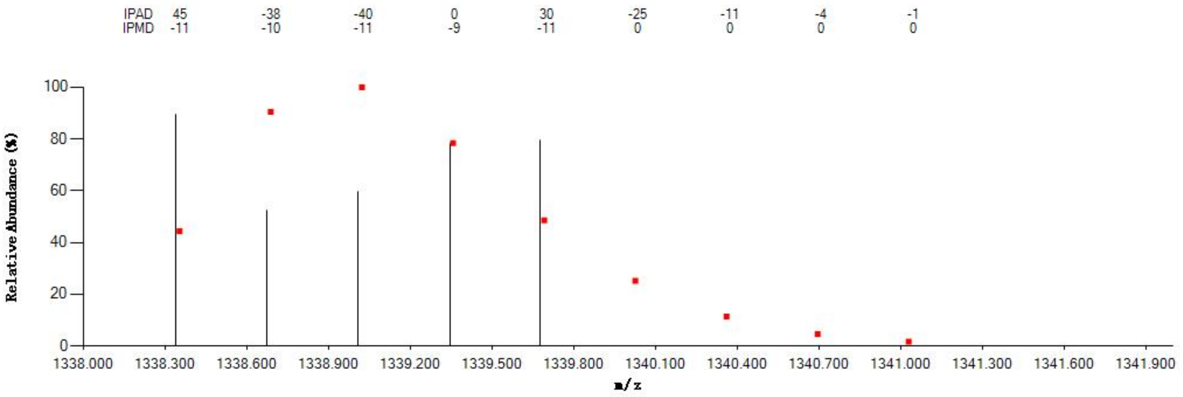


#### Annotated MS/MS Spectrum

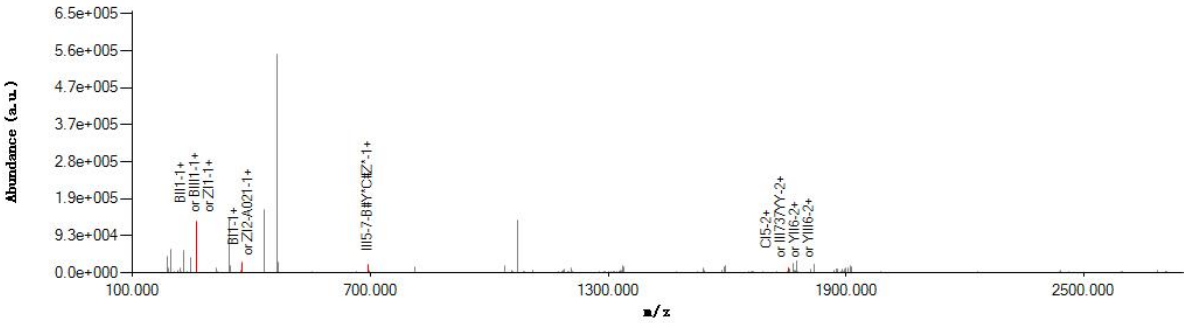


#### Graphical Fragmentation Map

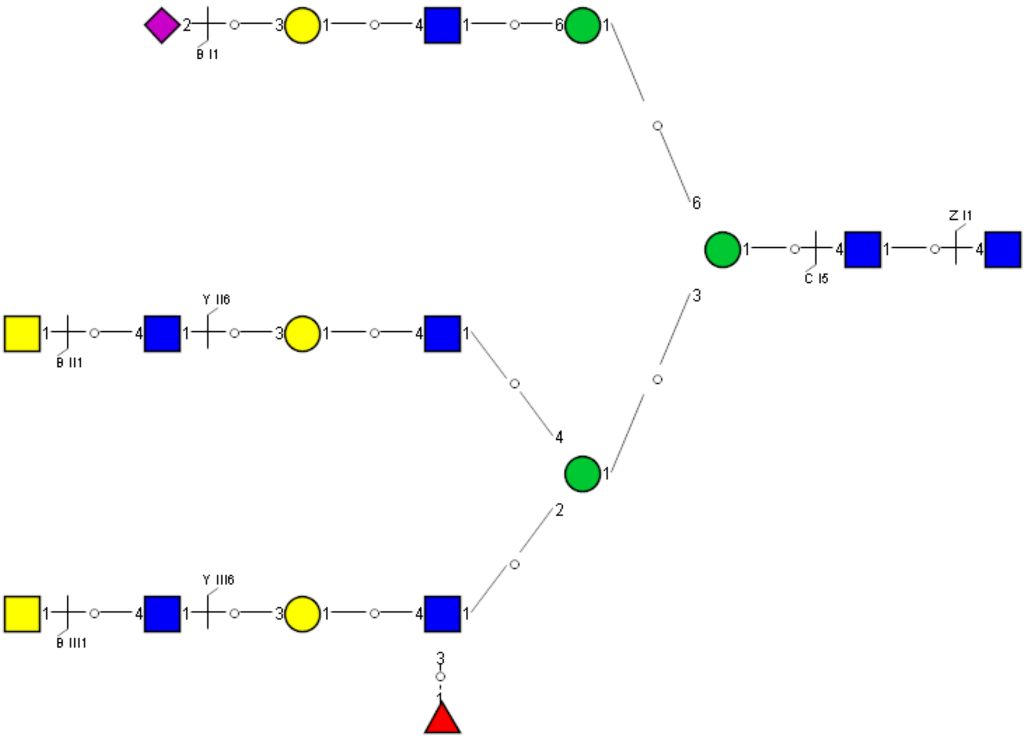

Spectrum Index: 39445(NoBHs:1)

#### Precursor iEF Map

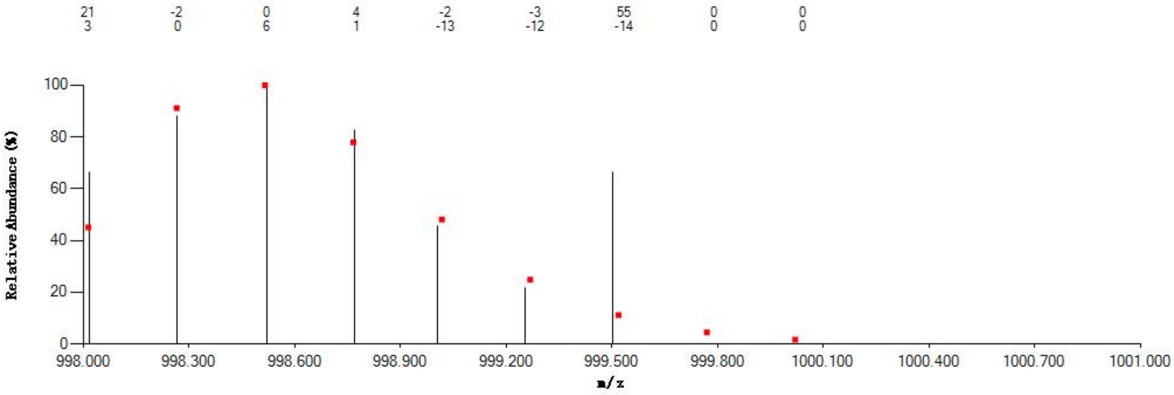


#### Annotated MS/MS Spectrum

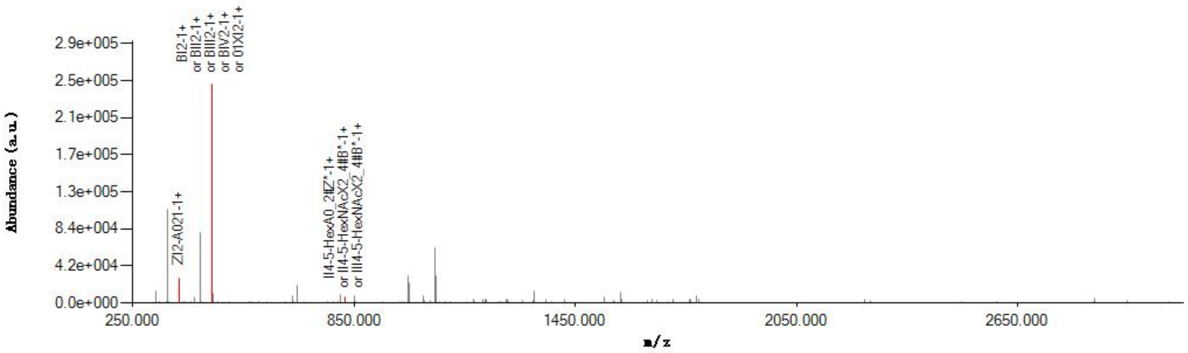


#### Graphical Fragmentation Map

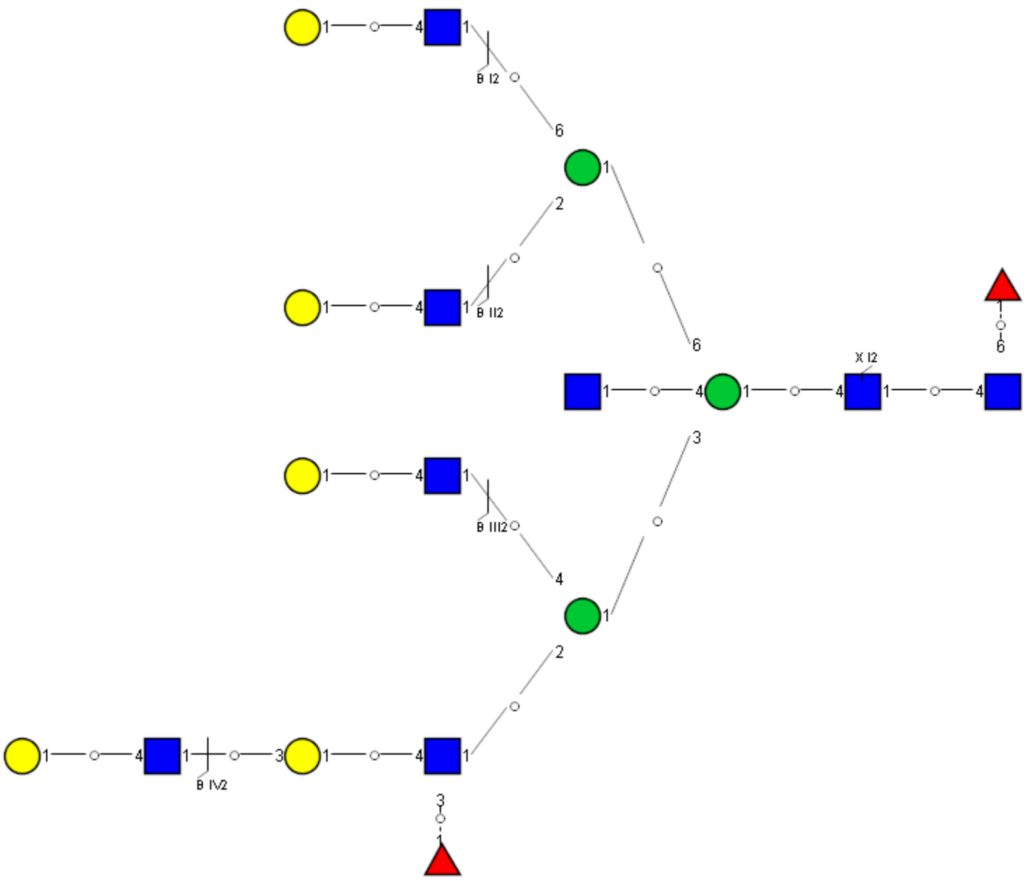

### Spectrum Index: 39548(NoBHs:1)

#### Precursor iEF Map

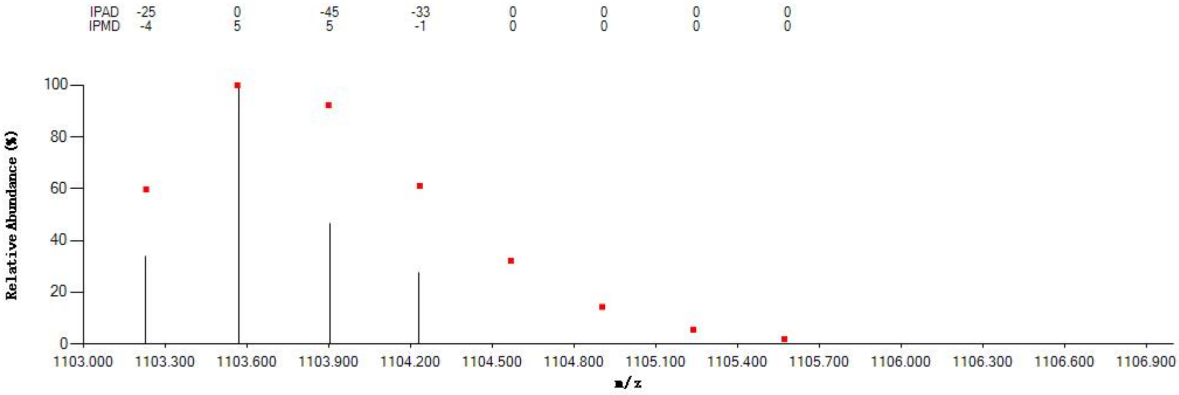


#### Annotated MS/MS Spectrum

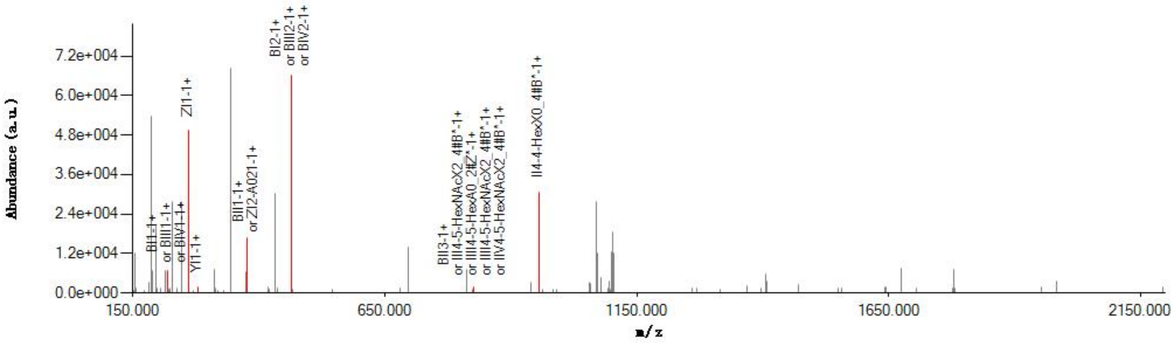


#### Graphical Fragmentation Map

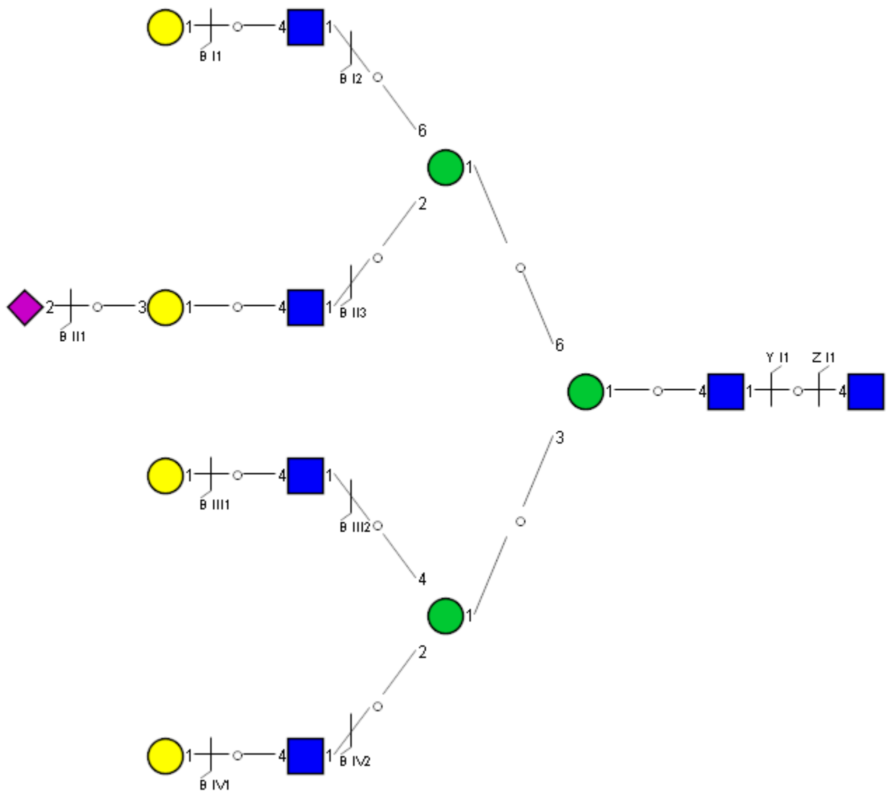

Spectrum Index: 39846(NoBHs:1)

#### Precursor iEF Map

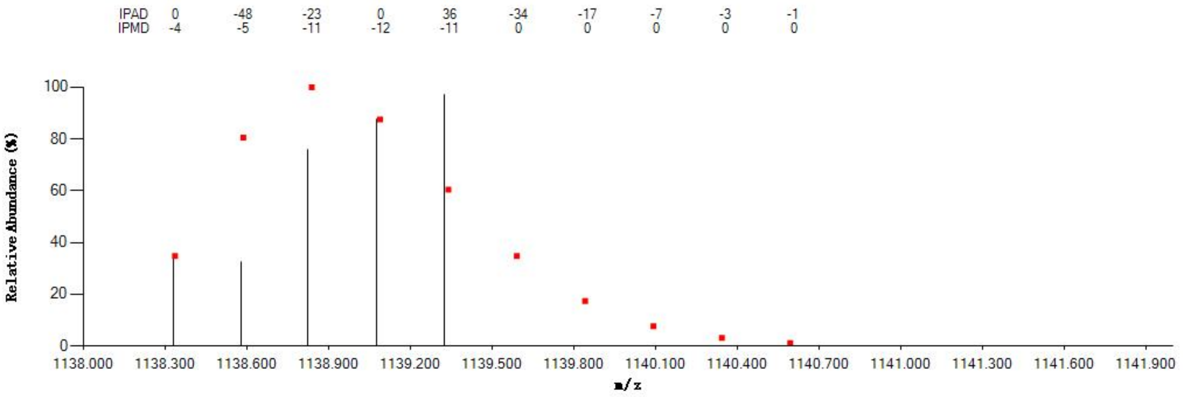


#### Annotated MS/MS Spectrum

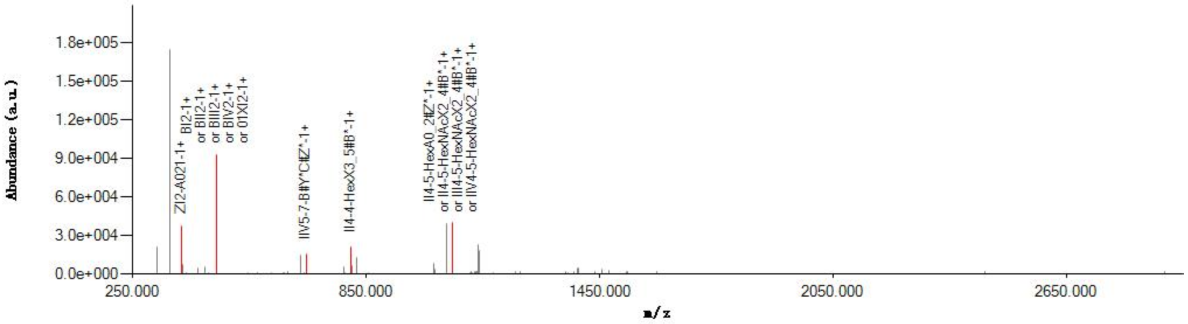


#### Graphical Fragmentation Map

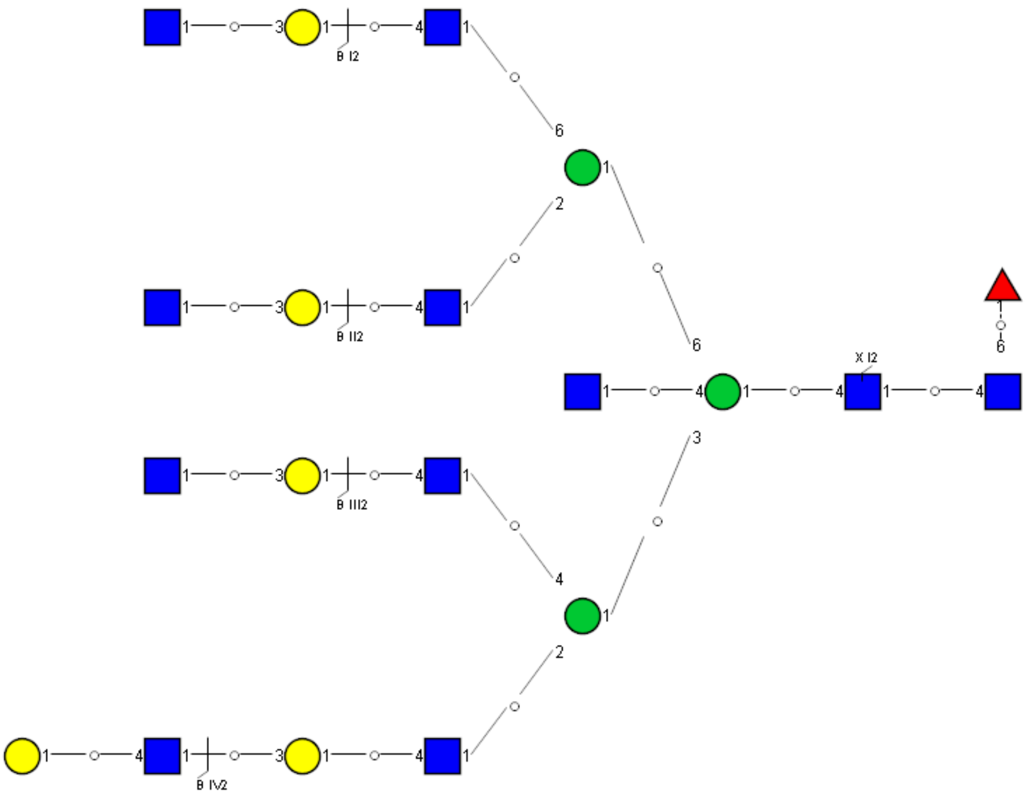

Spectrum Index: 40128(NoBHs:2)

#### Precursor iEF Map

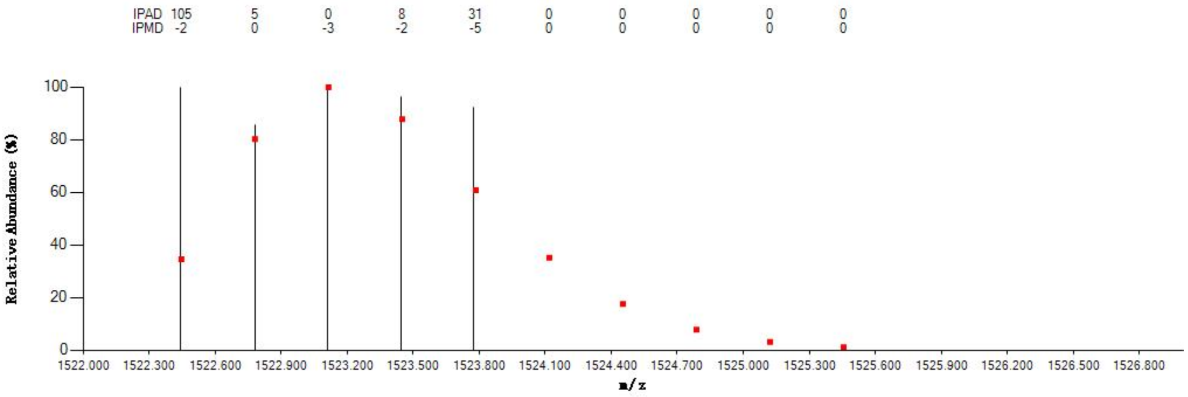


#### Annotated MS/MS Spectrum

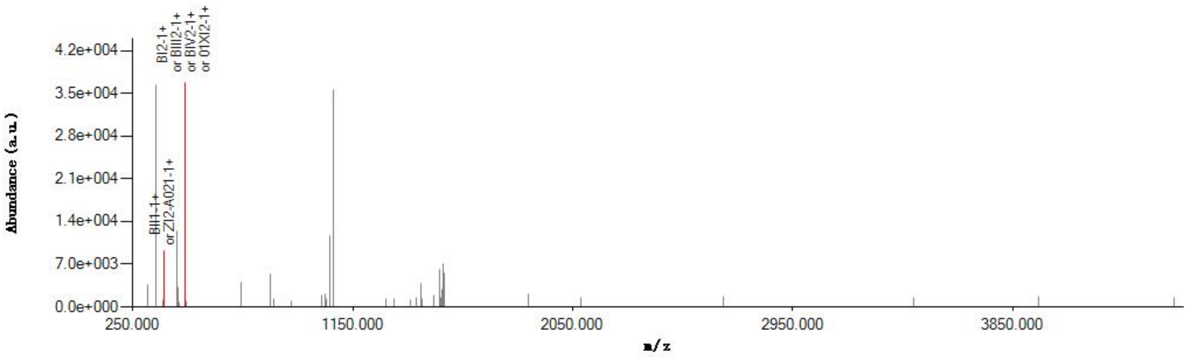


#### Graphical Fragmentation Map

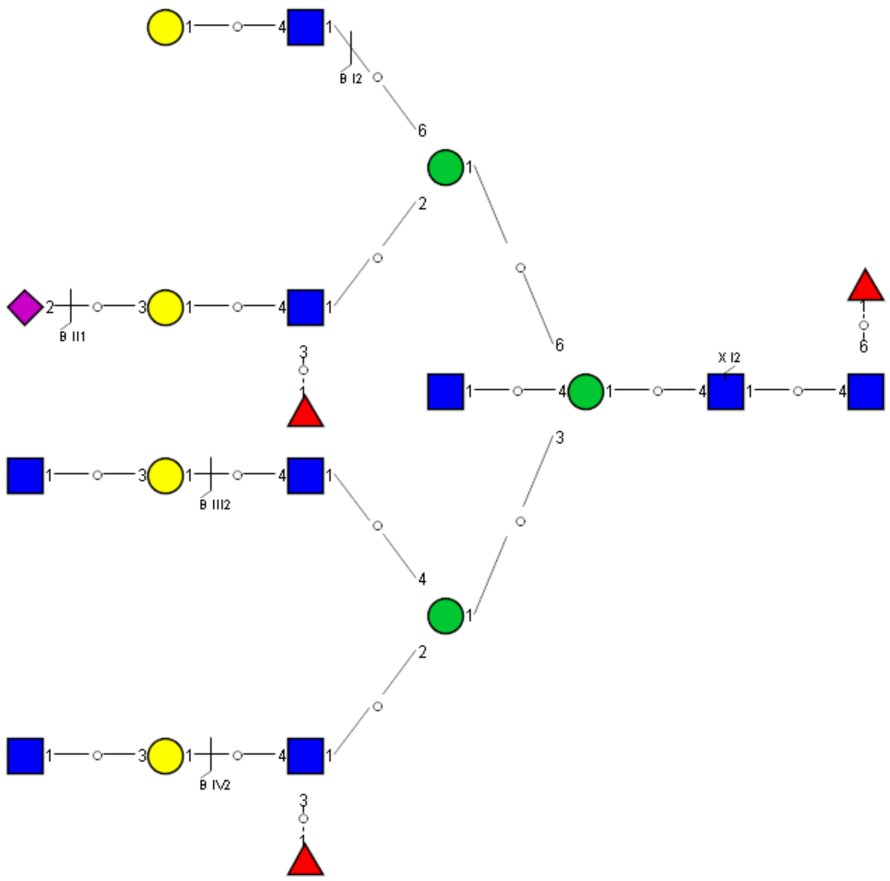

Spectrum Index: 40863(NoBHs:1)

#### Precursor iEF Map

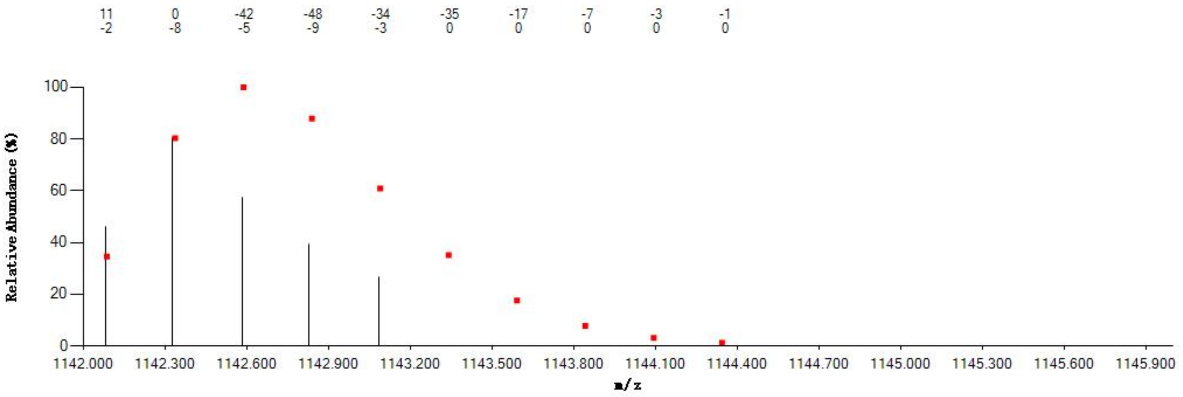


#### Annotated MS/MS Spectrum

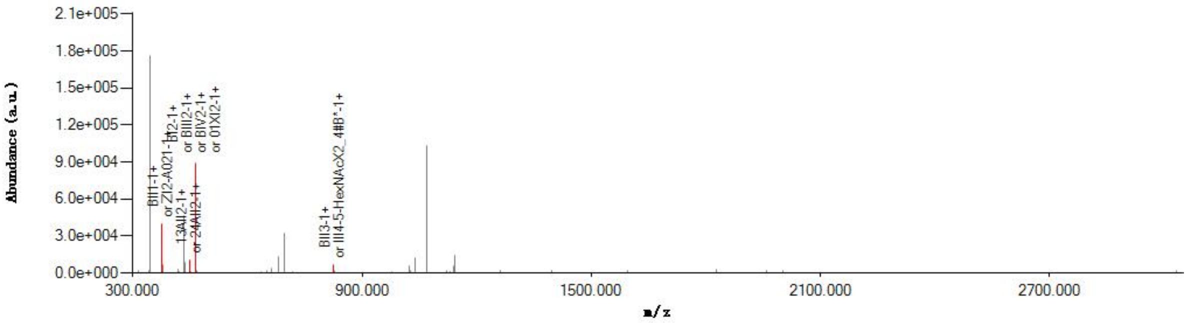


#### Graphical Fragmentation Map

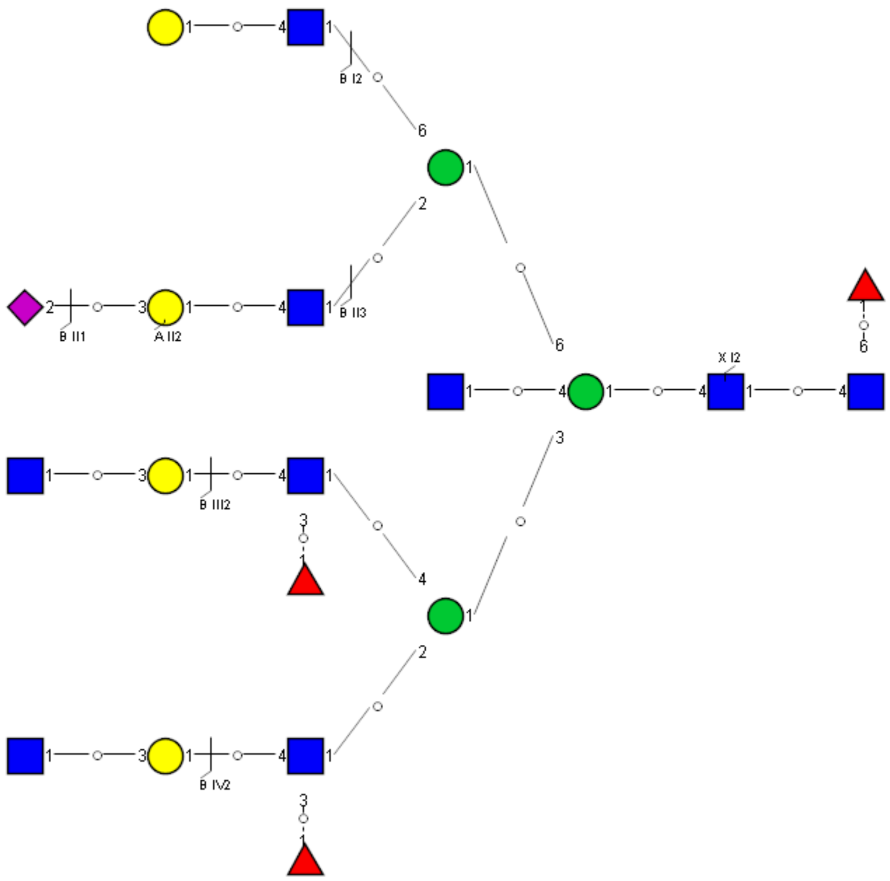

### Spectrum Index: 42120(NoBHs:1)

#### Precursor iEF Map

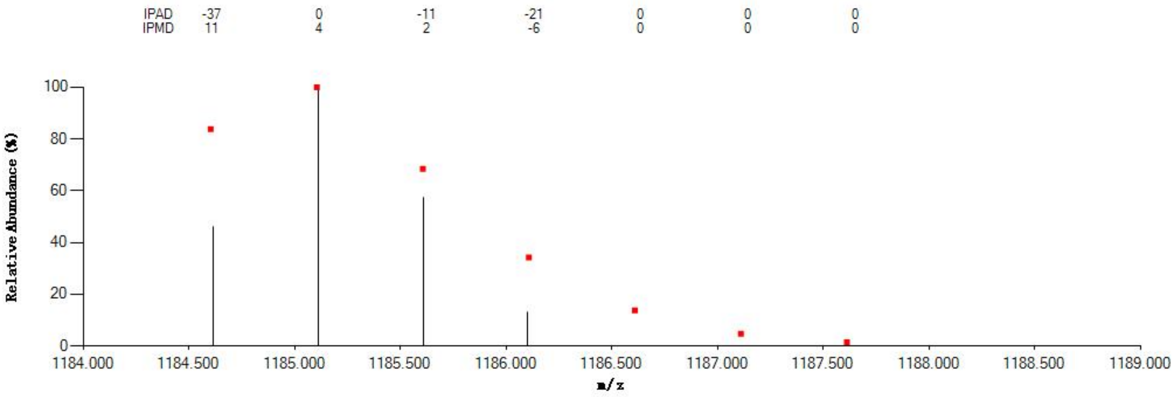


#### Annotated MS/MS Spectrum

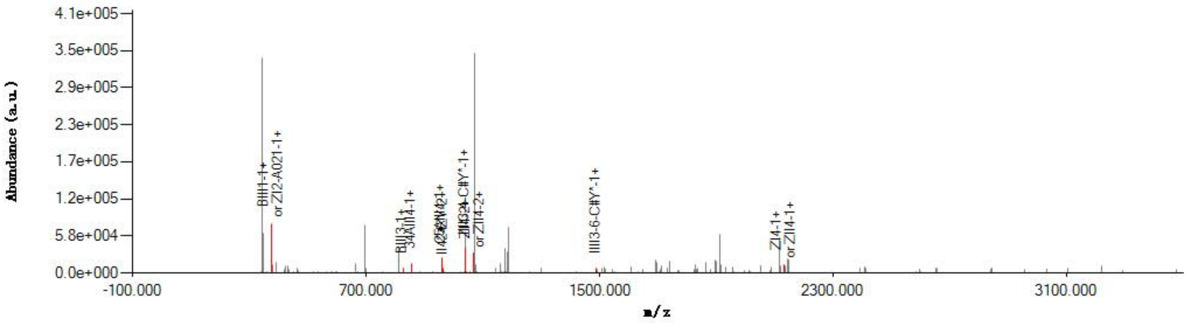


#### Graphical Fragmentation Map

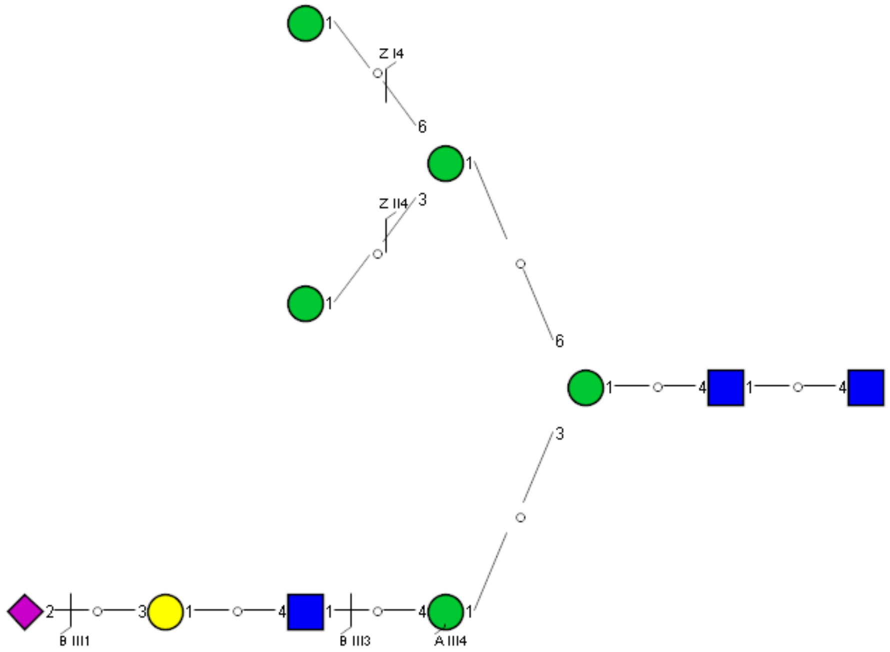
