## Supplemental Information 9 for "A draft of human N-glycans of glycoRNA"

Spectrum Index: 28981(NoBHs:1)

#### Precursor iEF Map

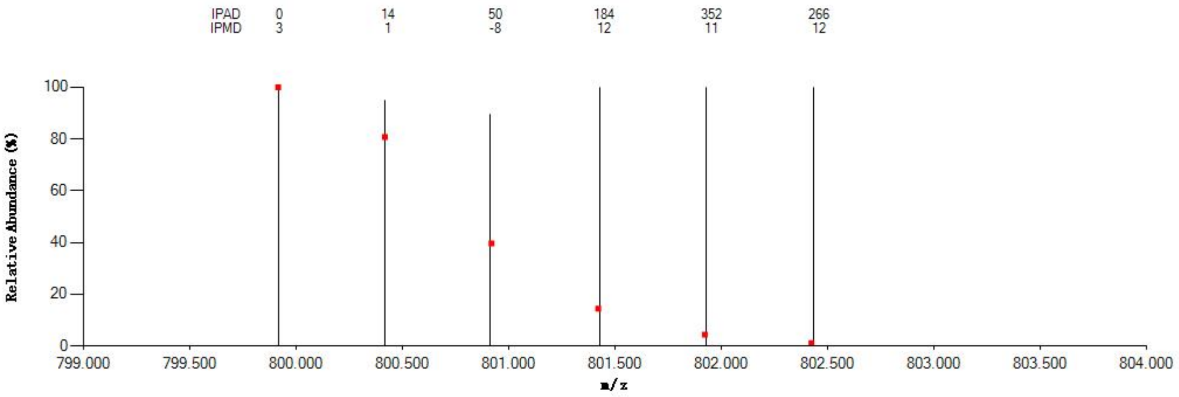


#### Annotated MS/MS Spectrum

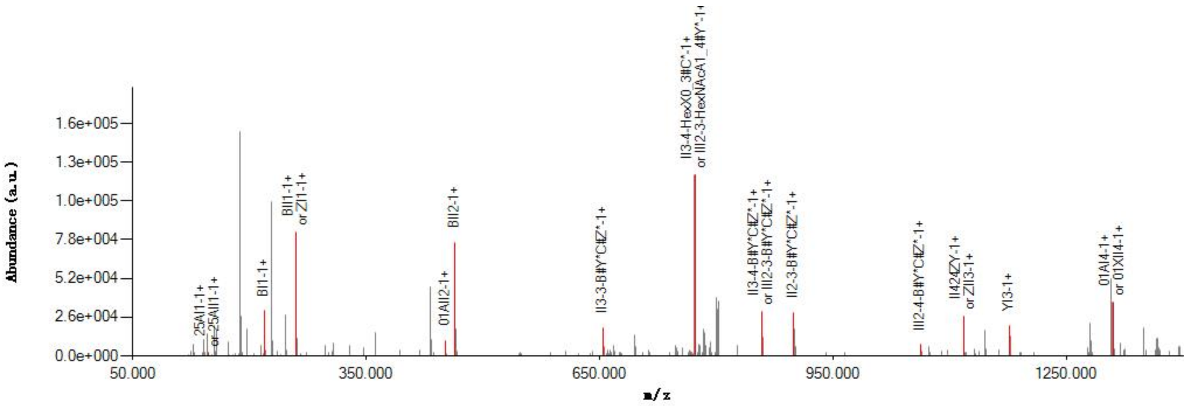


#### Graphical Fragmentation Map

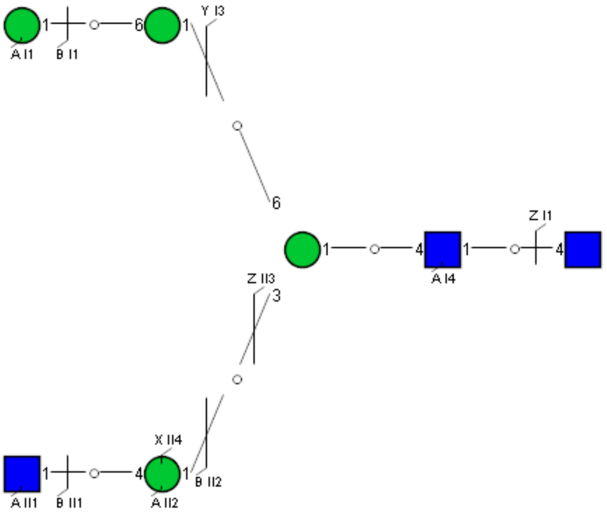

Spectrum Index: 29448(NoBHs:1)

#### Precursor iEF Map

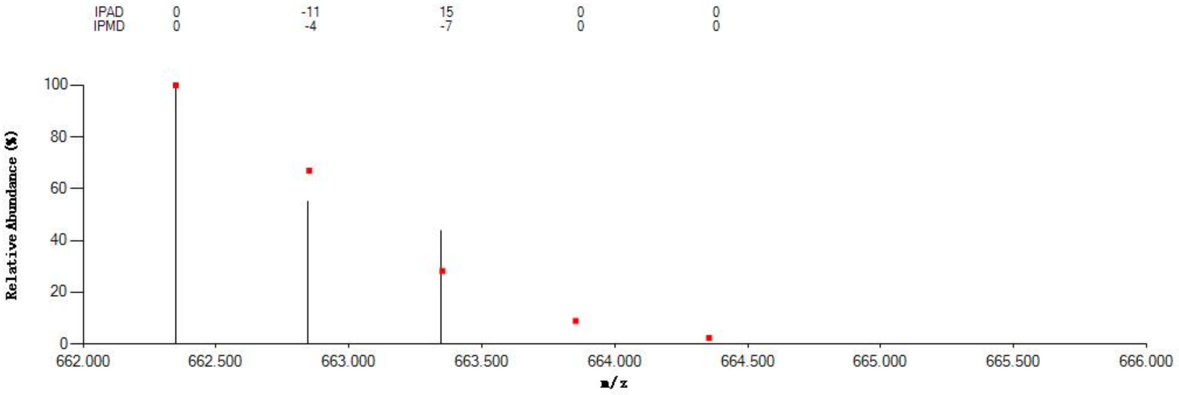

#### Annotated MS/MS Spectrum

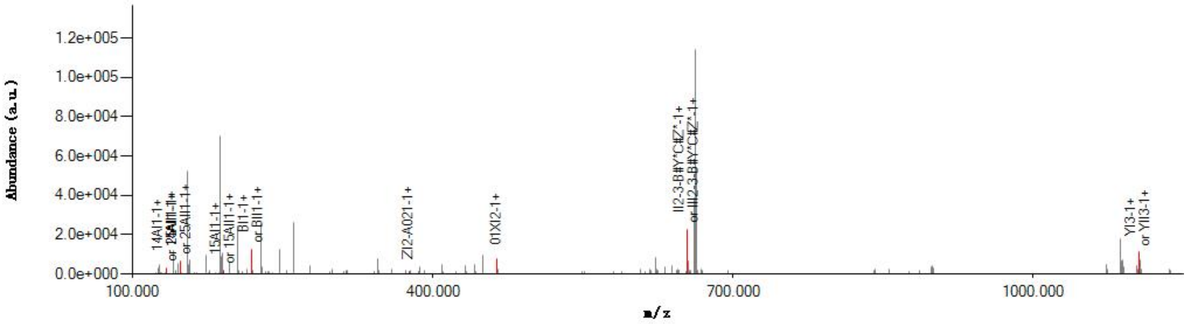

#### Graphical Fragmentation Map

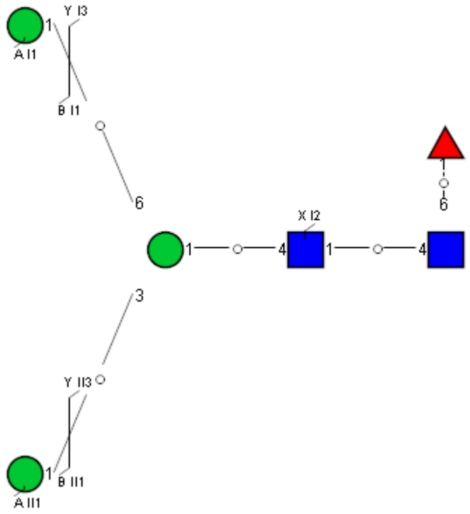

Spectrum Index: 29871(NoBHs:1)

#### Precursor iEF Map

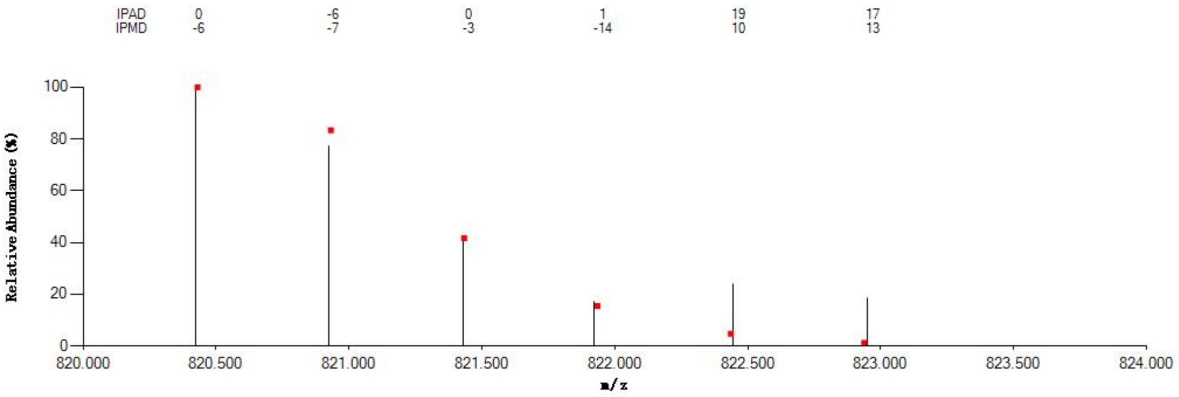


#### Annotated MS/MS Spectrum

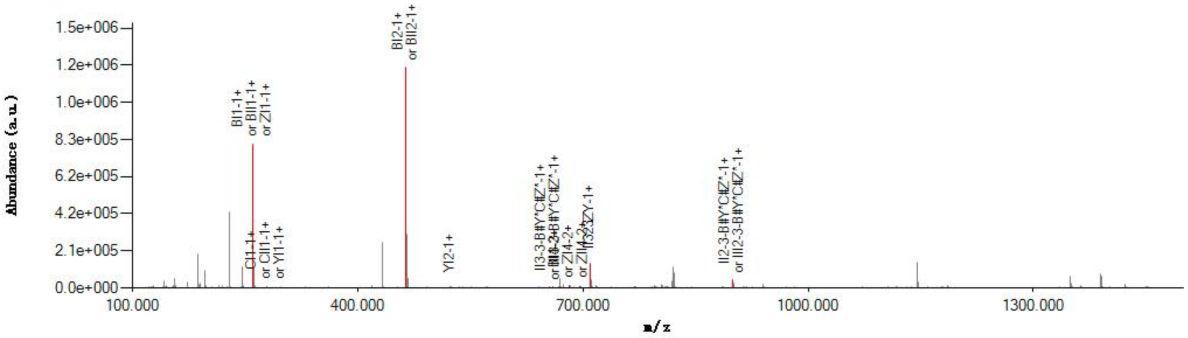


#### Graphical Fragmentation Map

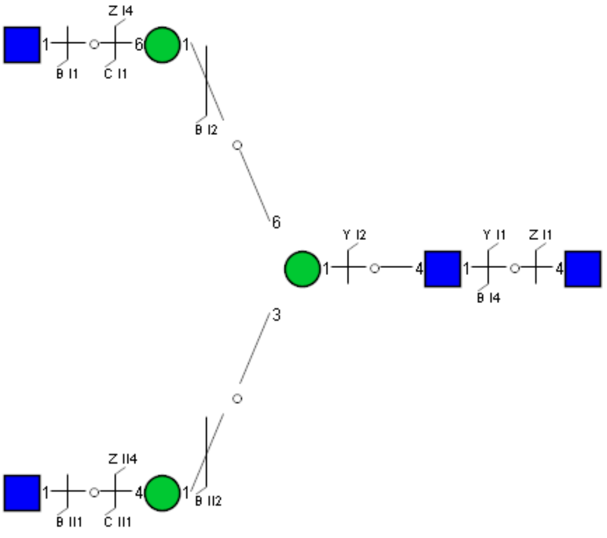

Spectrum Index: 30216(NoBHs:1)

#### Precursor iEF Map

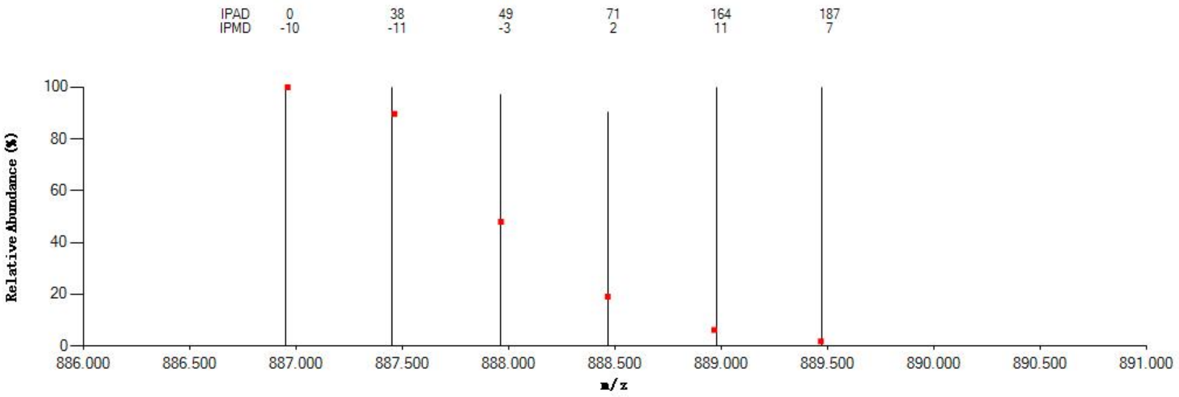

#### Annotated MS/MS Spectrum

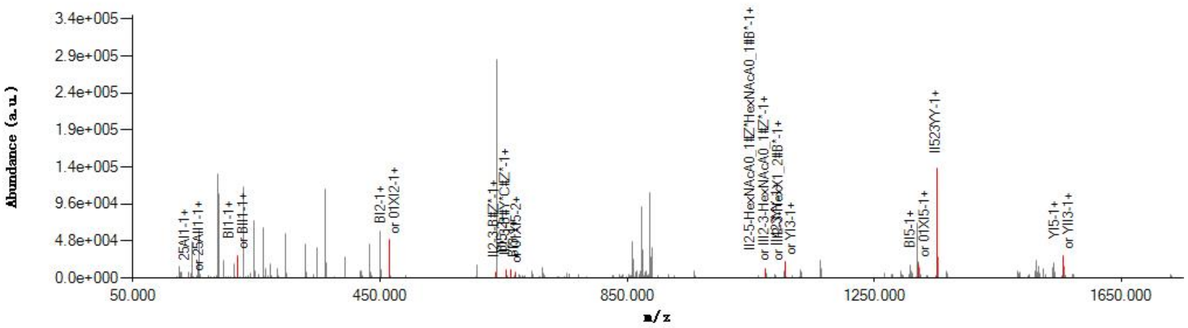

#### Graphical Fragmentation Map

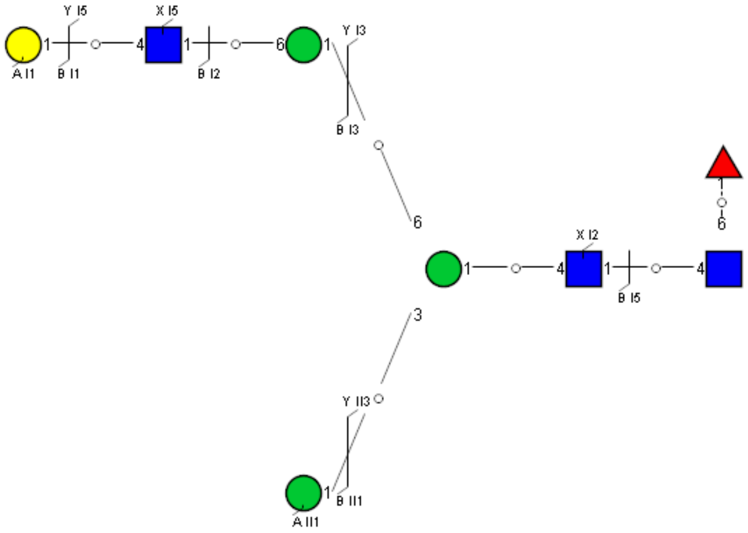

Spectrum Index: 30318(NoBHs:1)

#### Precursor iEF Map

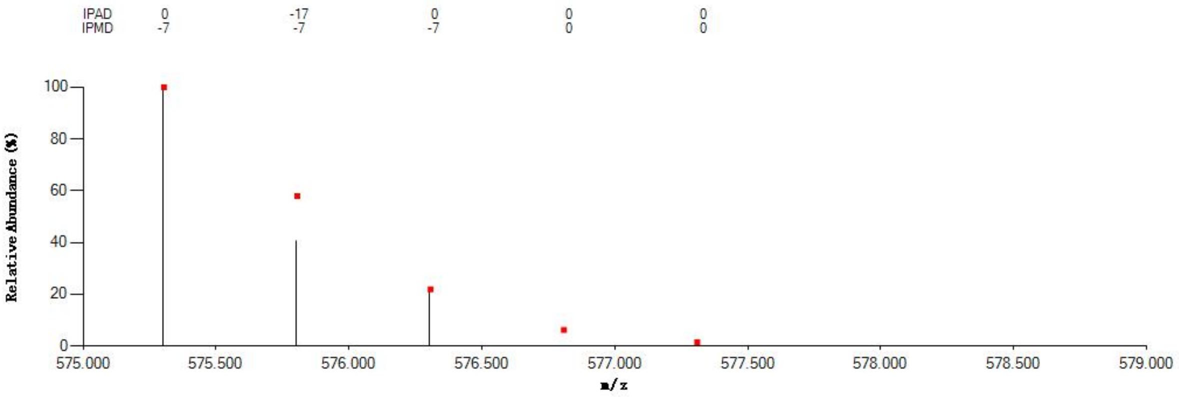


#### Annotated MS/MS Spectrum

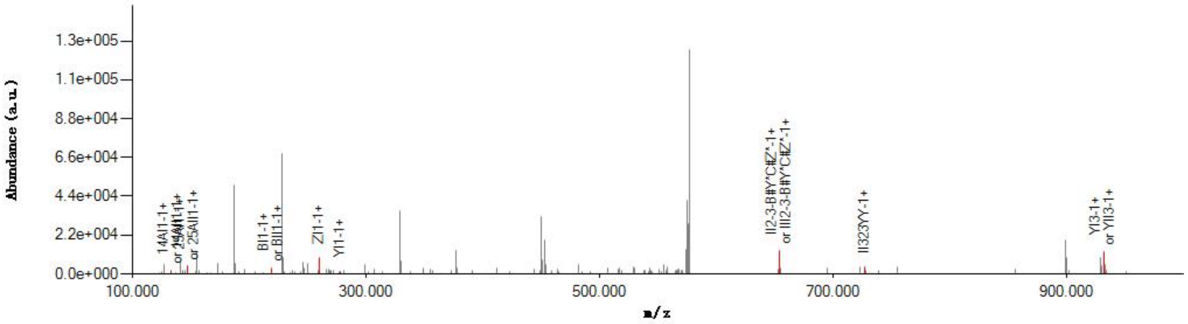


#### Graphical Fragmentation Map

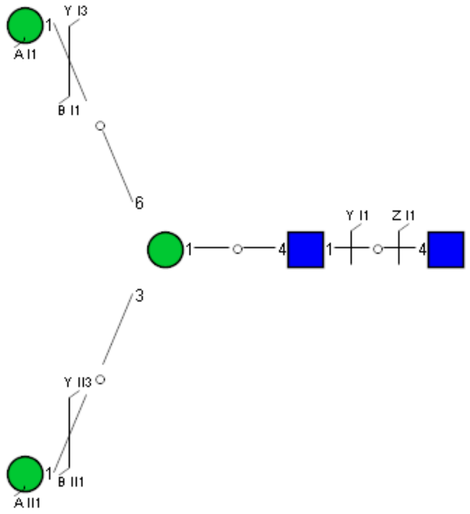

Spectrum Index: 30417(NoBHs:1)

#### Precursor iEF Map

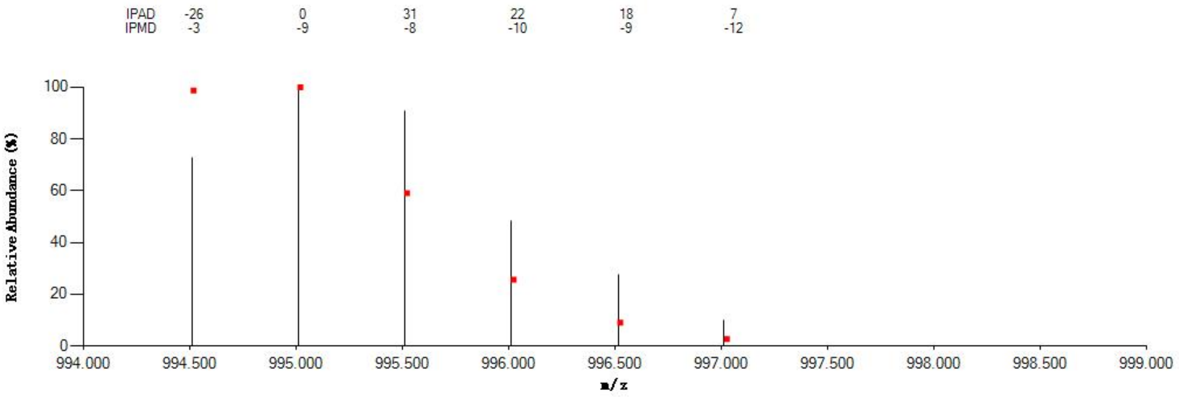


#### Annotated MS/MS Spectrum

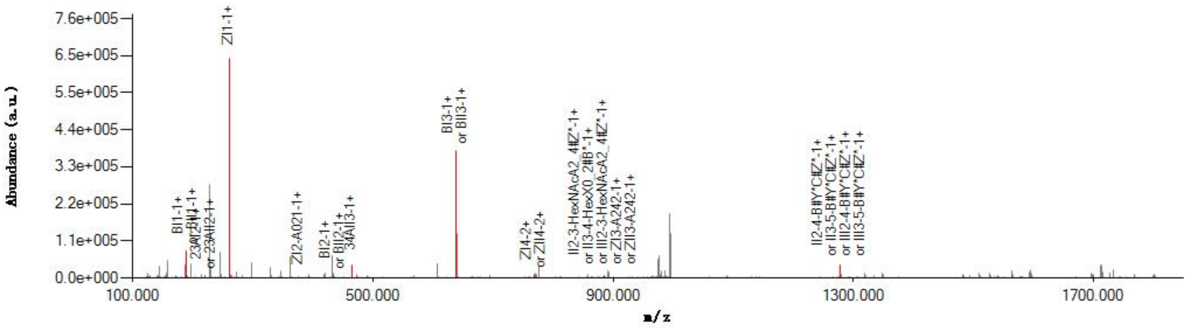


#### Graphical Fragmentation Map

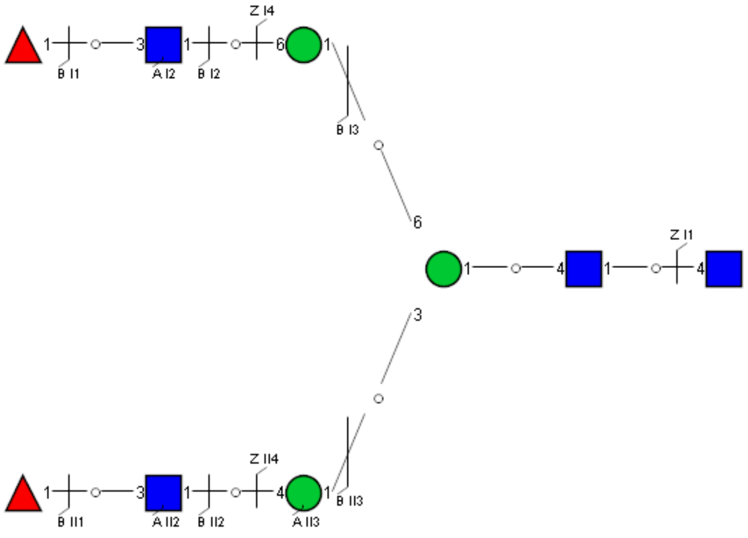

Spectrum Index: 31329(NoBHs:1)

#### Precursor iEF Map

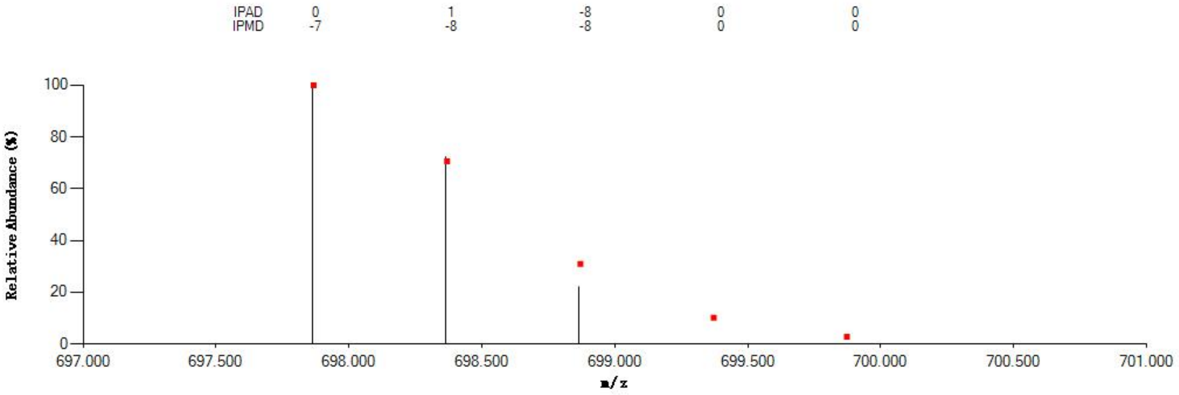


#### Annotated MS/MS Spectrum

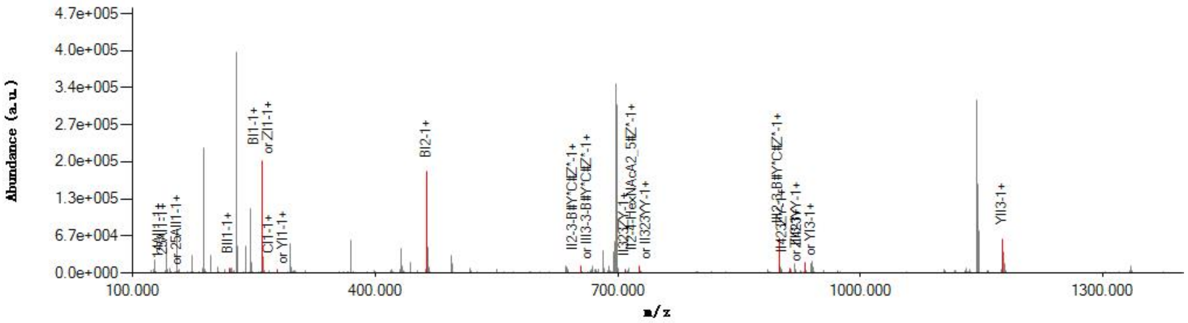


#### Graphical Fragmentation Map

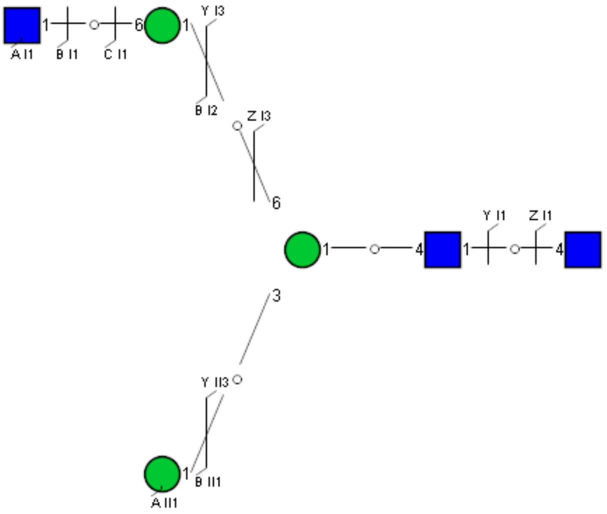

### Spectrum Index: 31501(NoBHs:1)

#### Precursor iEF Map

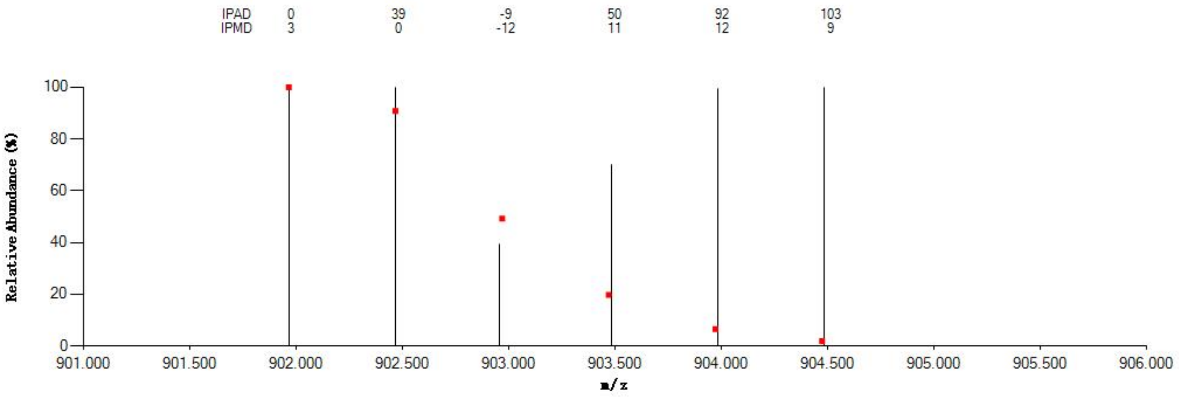


#### Annotated MS/MS Spectrum

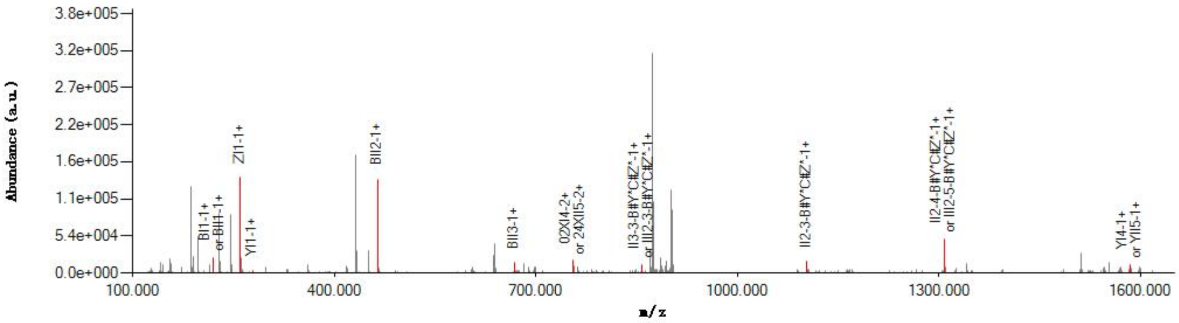


#### Graphical Fragmentation Map

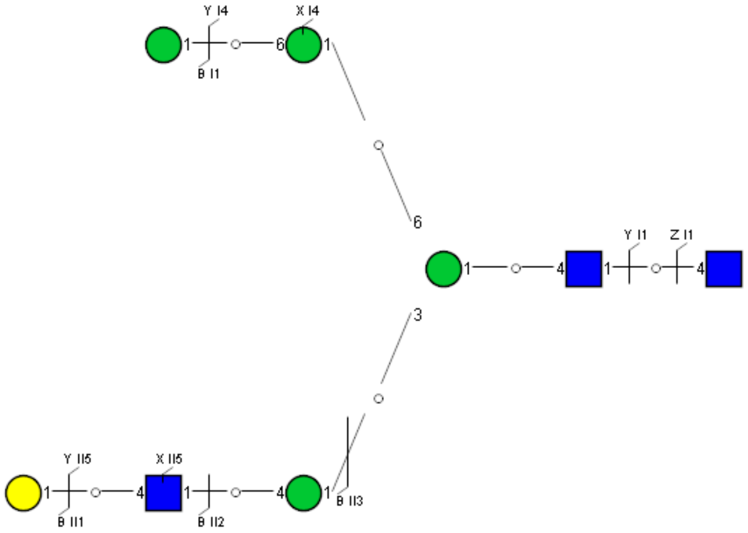

Spectrum Index: 31617(NoBHs:1)

#### Precursor iEF Map

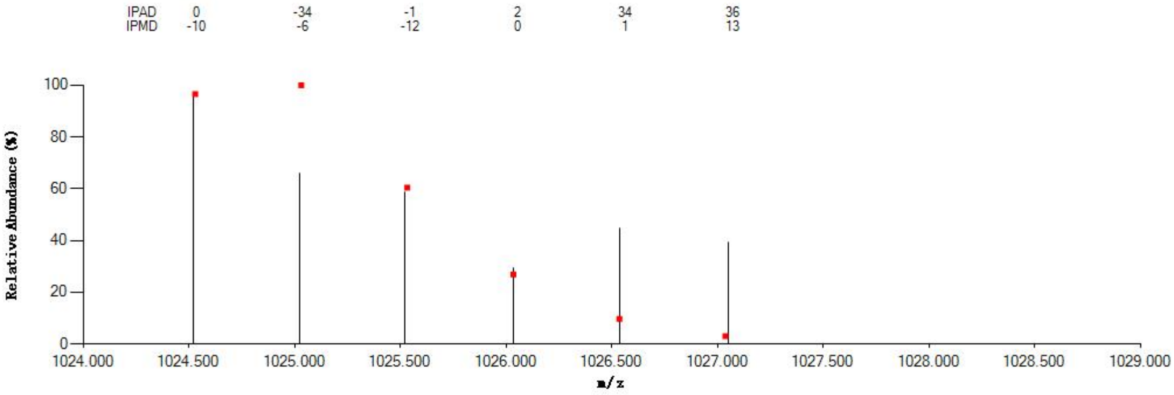


#### Annotated MS/MS Spectrum

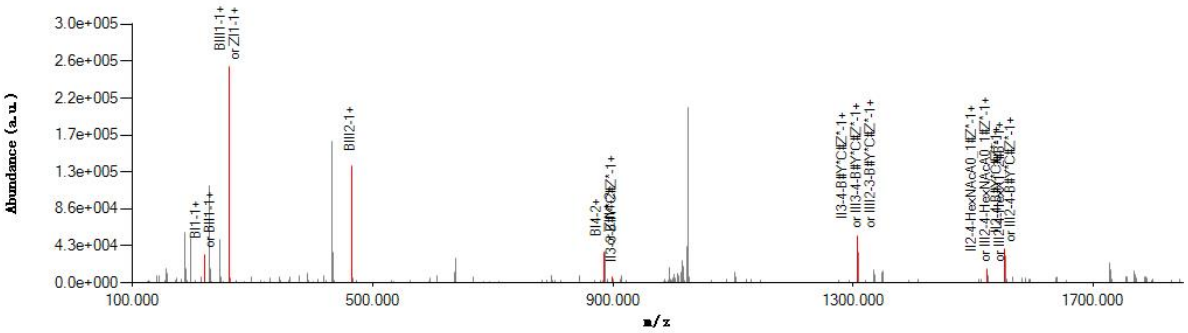


#### Graphical Fragmentation Map

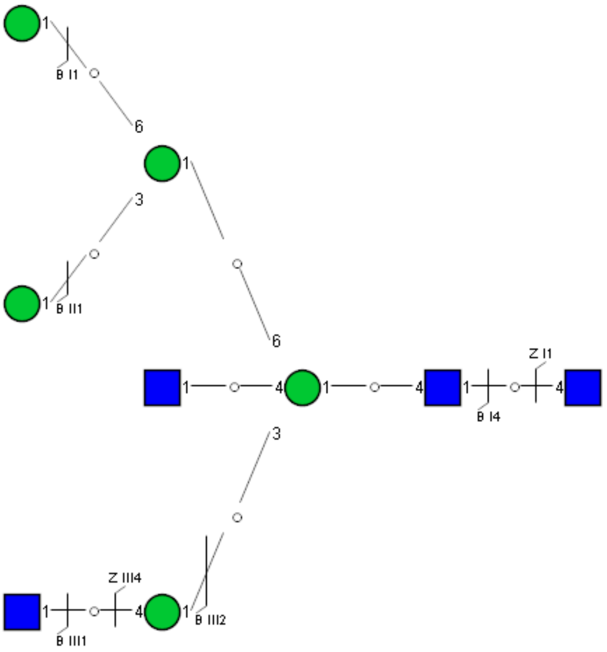

Spectrum Index: 31680(NoBHs:1)

#### Precursor iEF Map

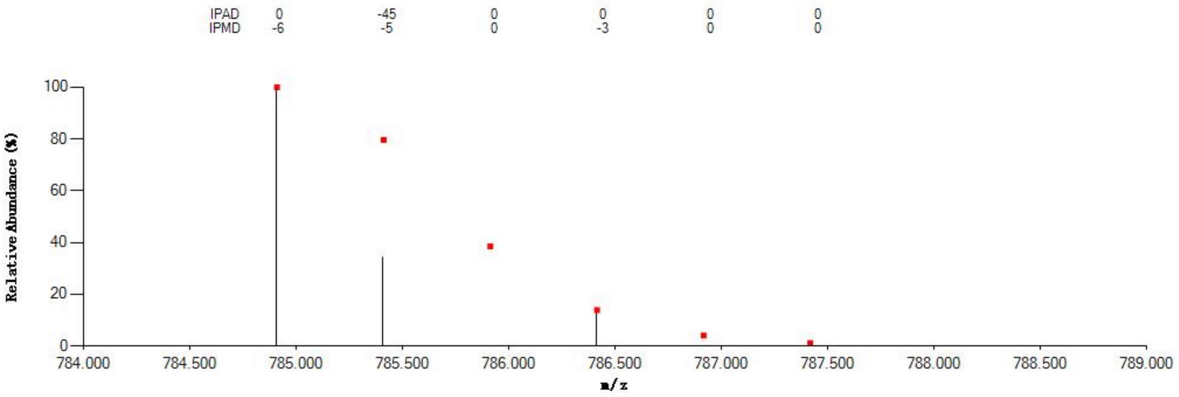


#### Annotated MS/MS Spectrum

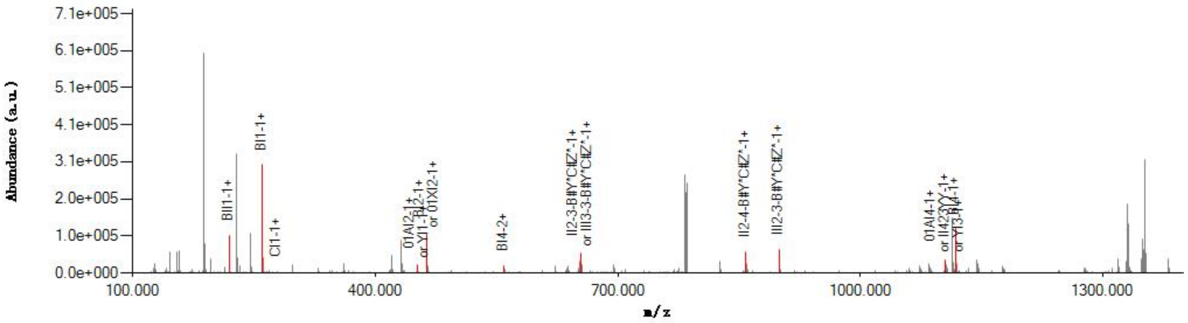


#### Graphical Fragmentation Map

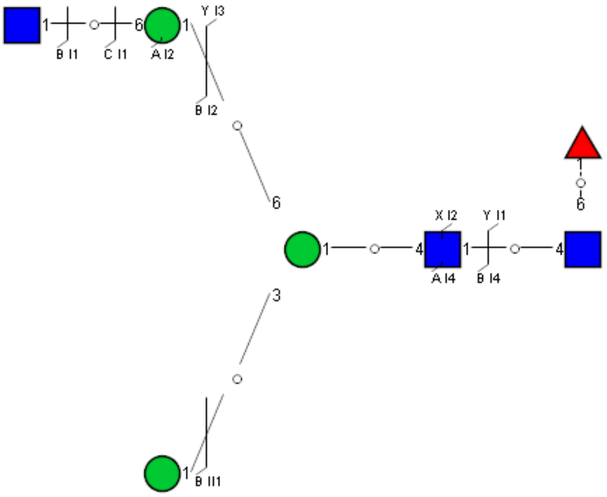

### Spectrum Index: 32118(NoBHs:1)

#### Precursor iEF Map

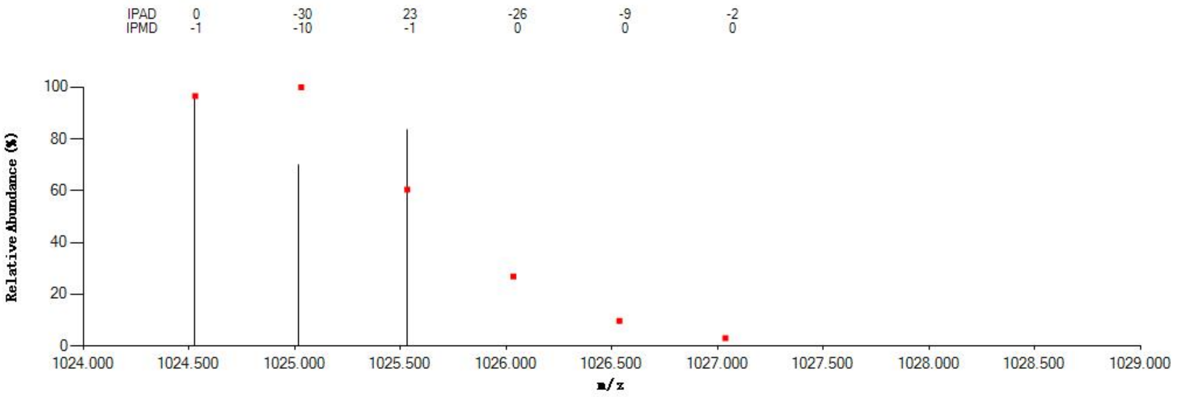


#### Annotated MS/MS Spectrum

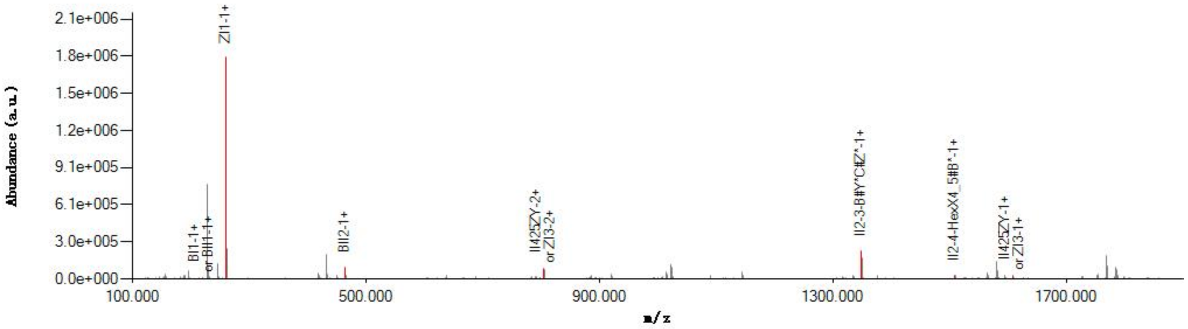


#### Graphical Fragmentation Map

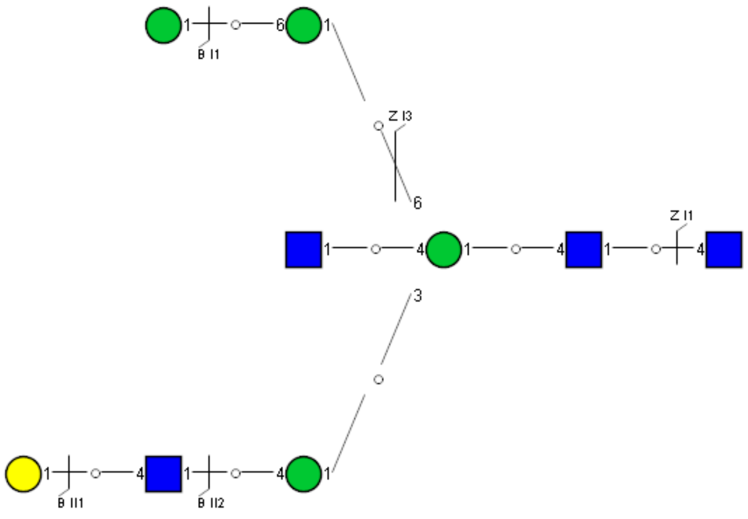

Spectrum Index: 32240(NoBHs:1)

#### Precursor iEF Map

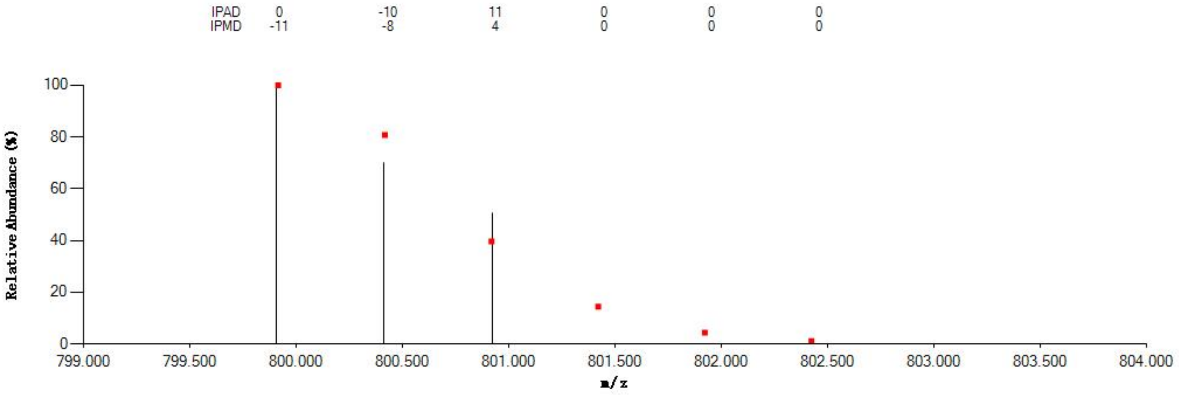


#### Annotated MS/MS Spectrum

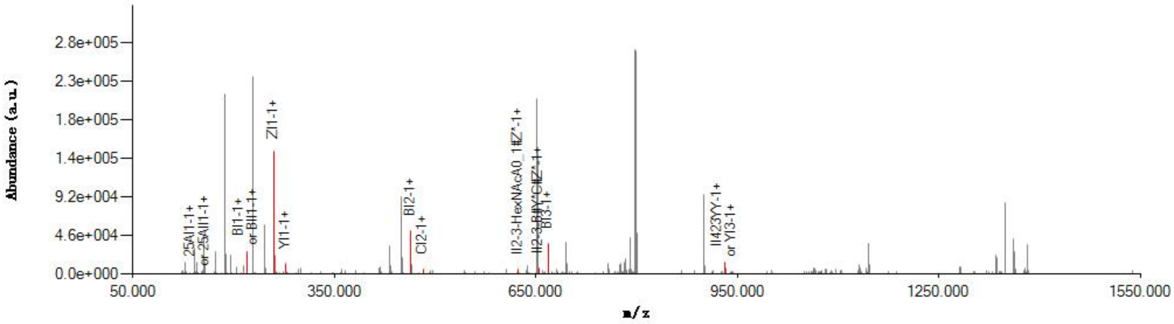


#### Graphical Fragmentation Map

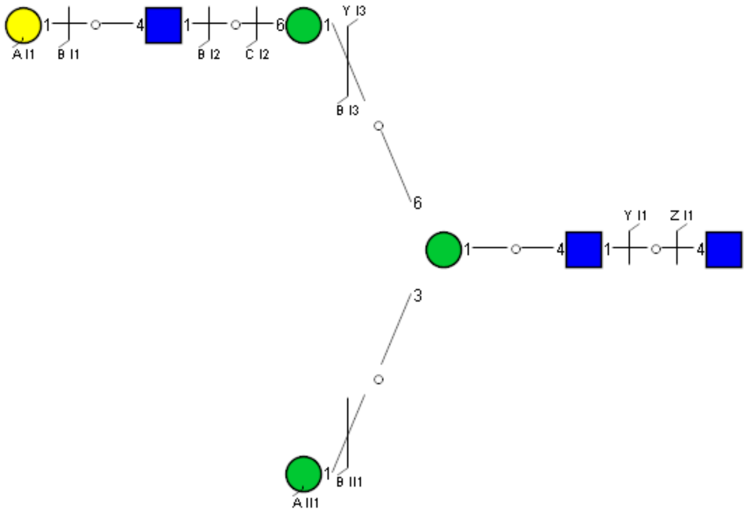

Spectrum Index: 32338(NoBHs:1)

#### Precursor iEF Map

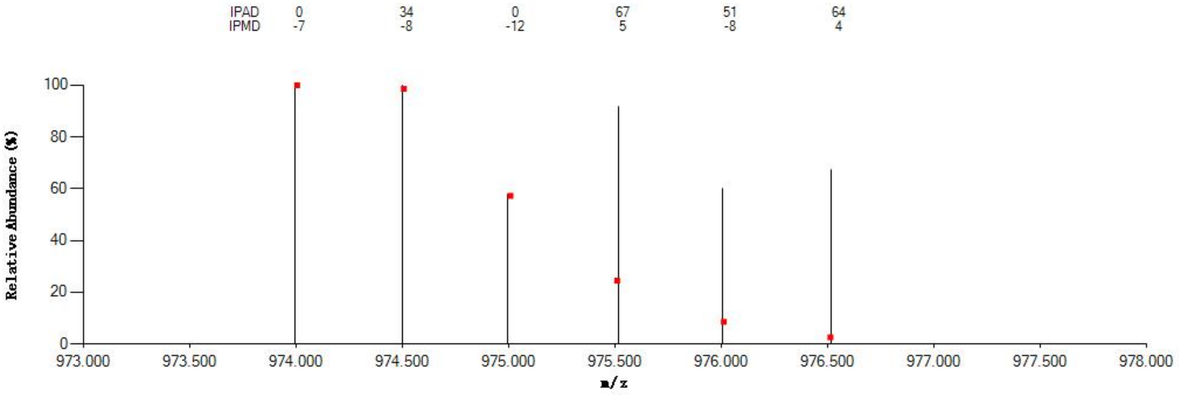


#### Annotated MS/MS Spectrum

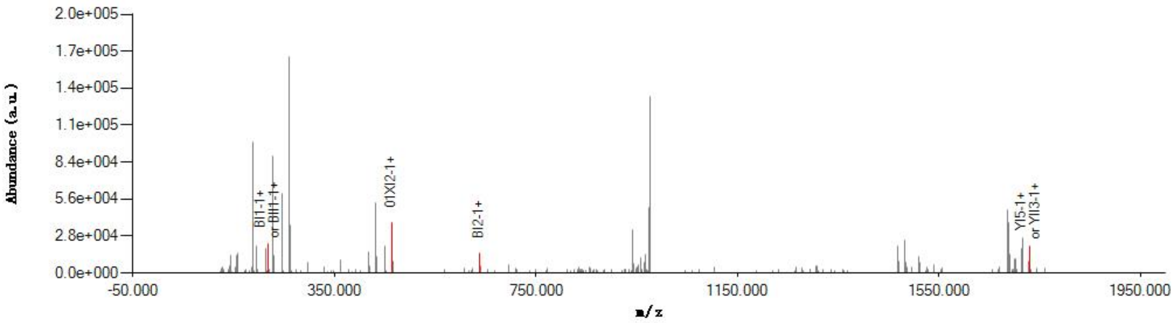


#### Graphical Fragmentation Map

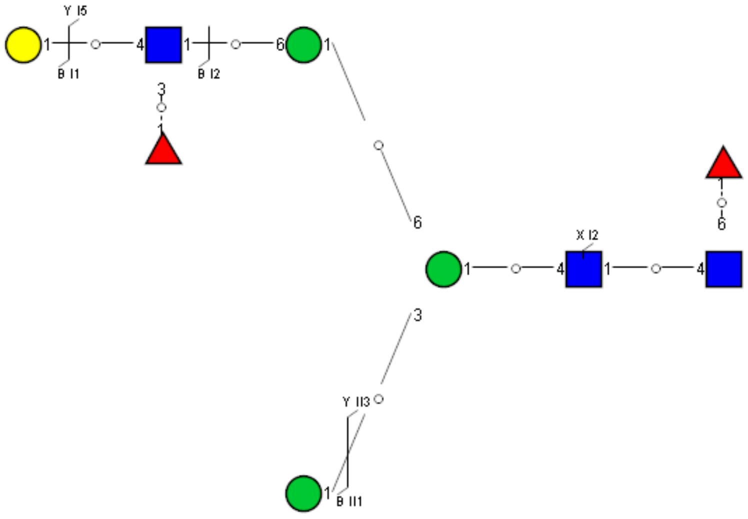

Spectrum Index: 32431(NoBHs:1)

#### Precursor iEF Map

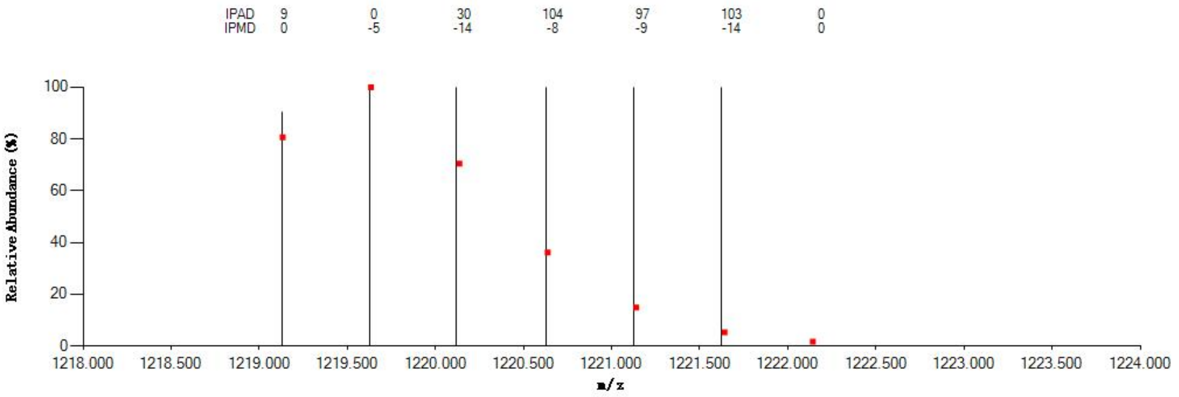


#### Annotated MS/MS Spectrum

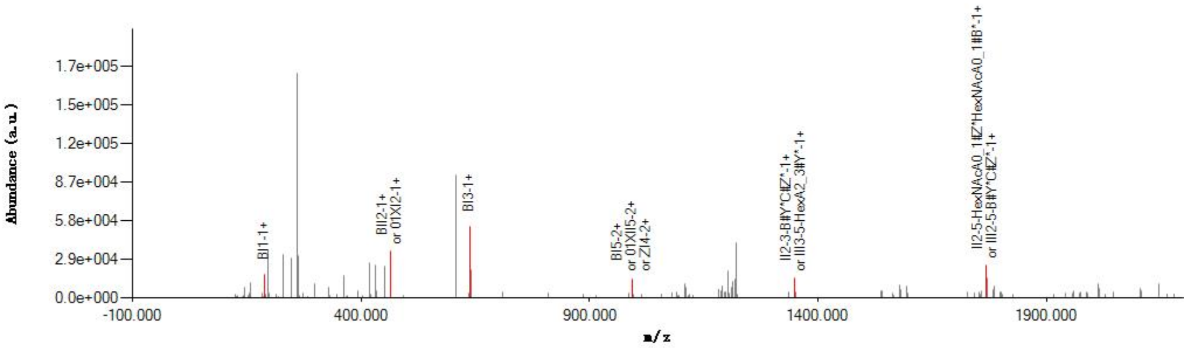


#### Graphical Fragmentation Map

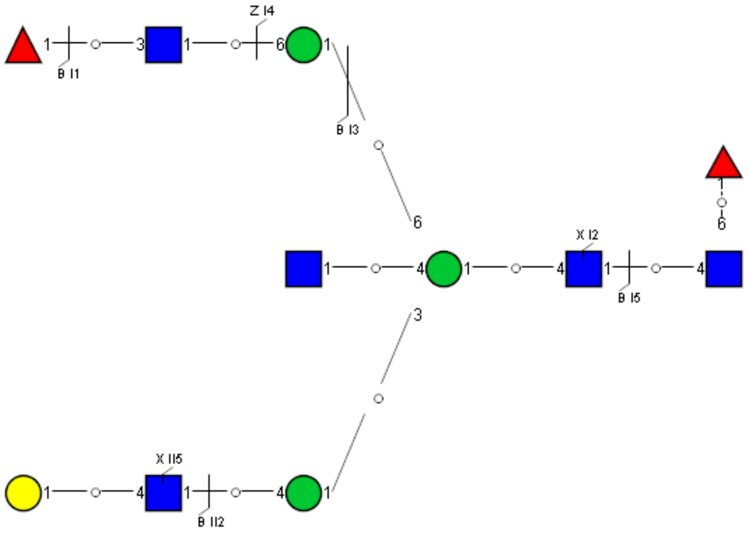

Spectrum Index: 32739(NoBHs:1)

#### Precursor iEF Map

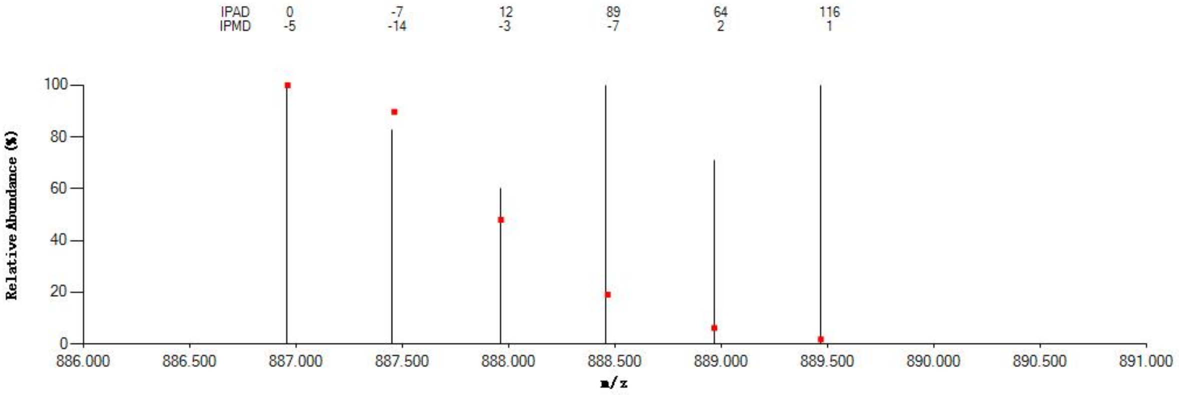

#### Annotated MS/MS Spectrum

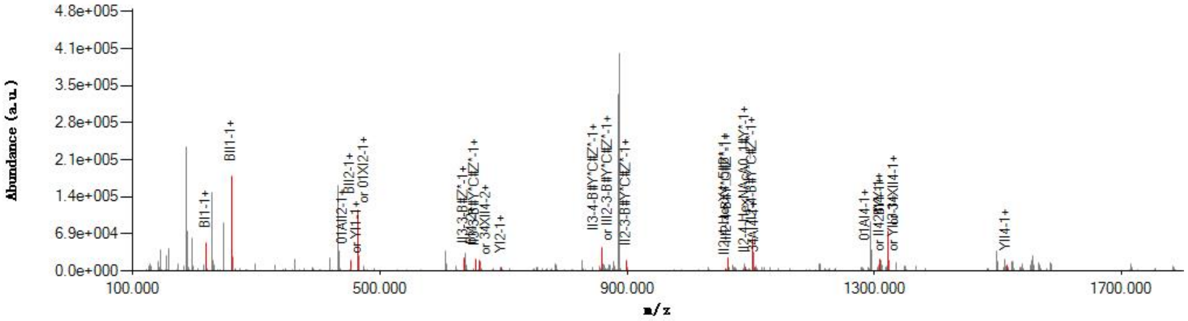

#### Graphical Fragmentation Map

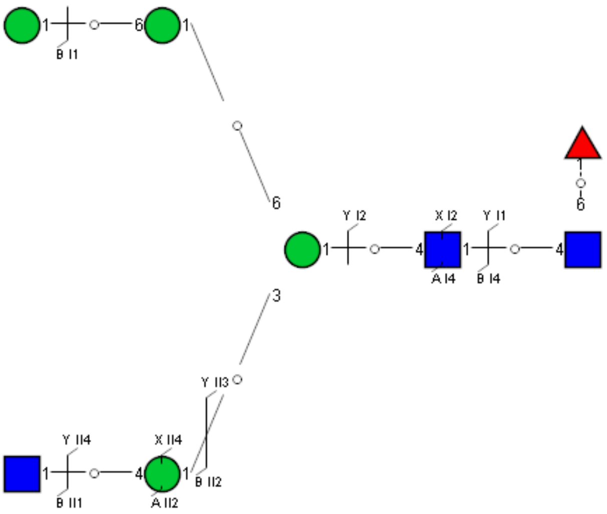

### Spectrum Index: 33013(NoBHs:1)

#### Precursor iEF Map

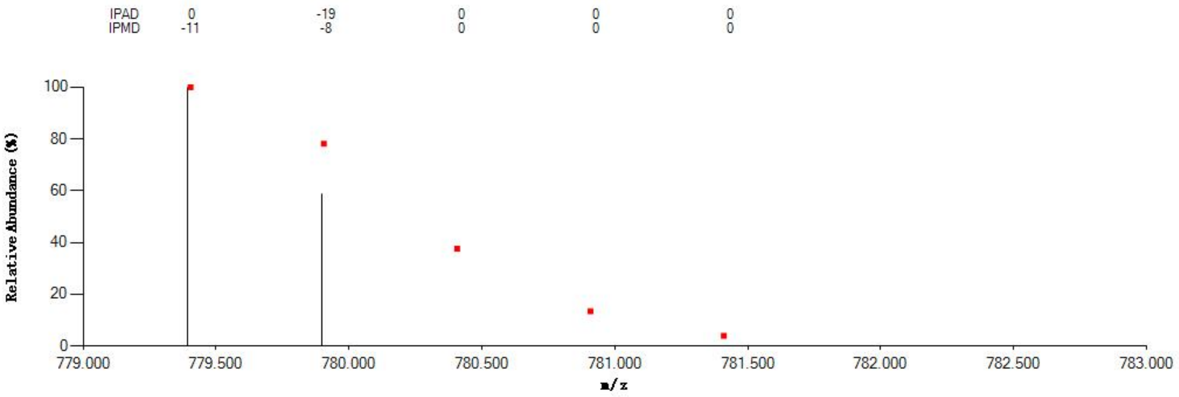


#### Annotated MS/MS Spectrum

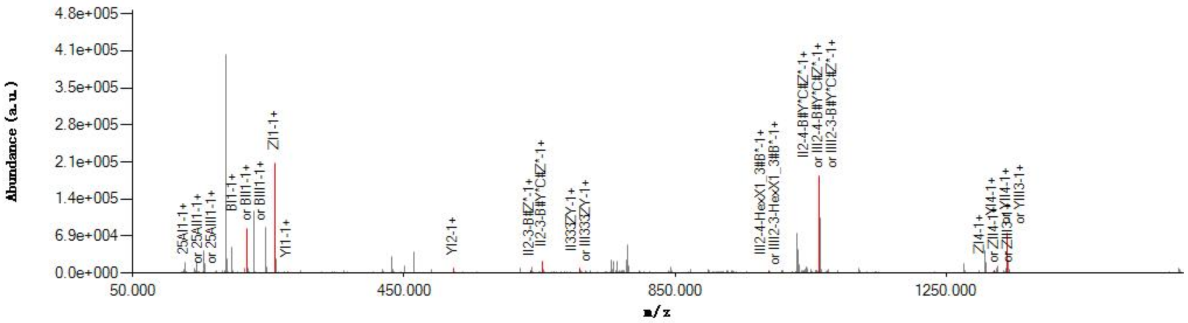


#### Graphical Fragmentation Map

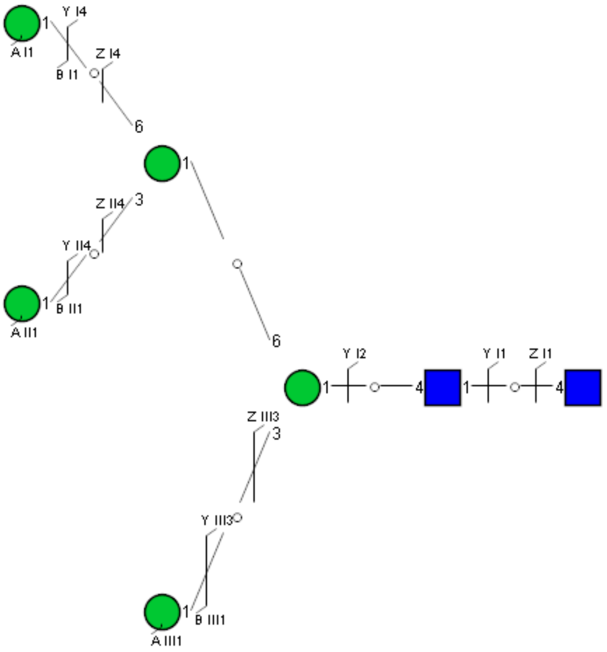

### Spectrum Index: 33061(NoBHs:1)

#### Precursor iEF Map

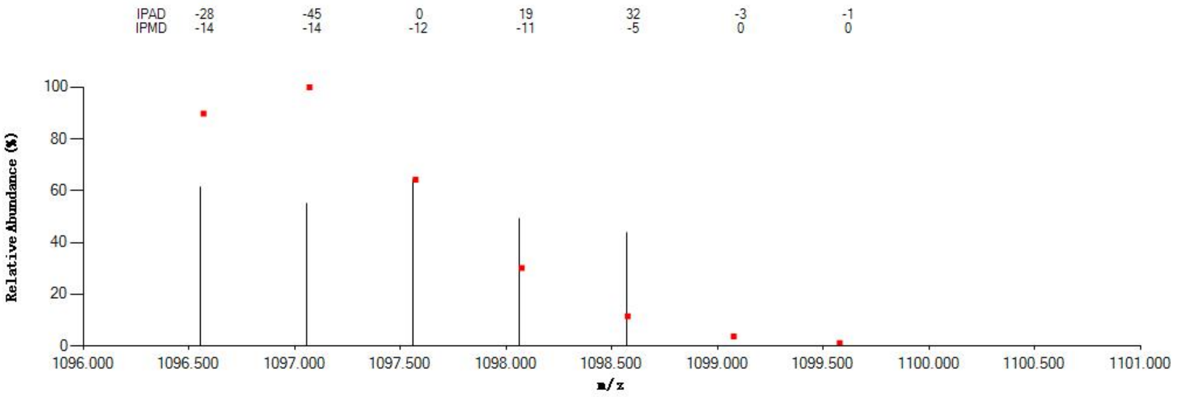


#### Annotated MS/MS Spectrum

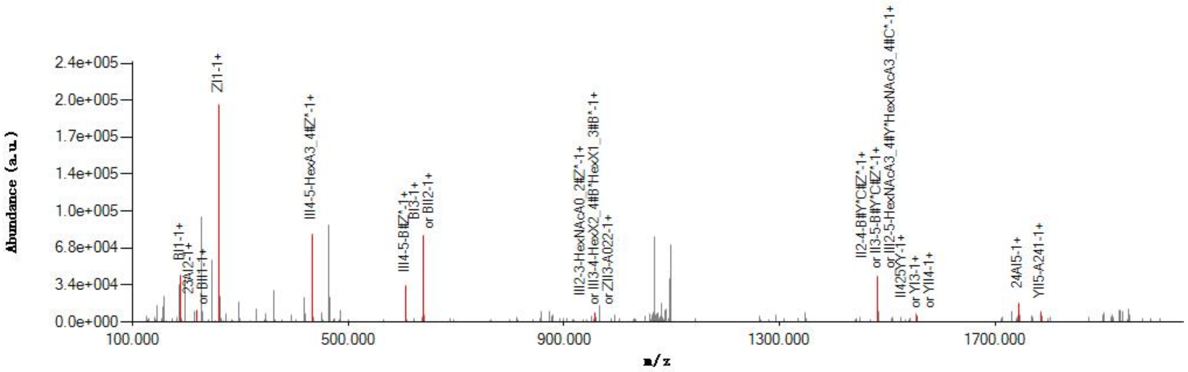


#### Graphical Fragmentation Map

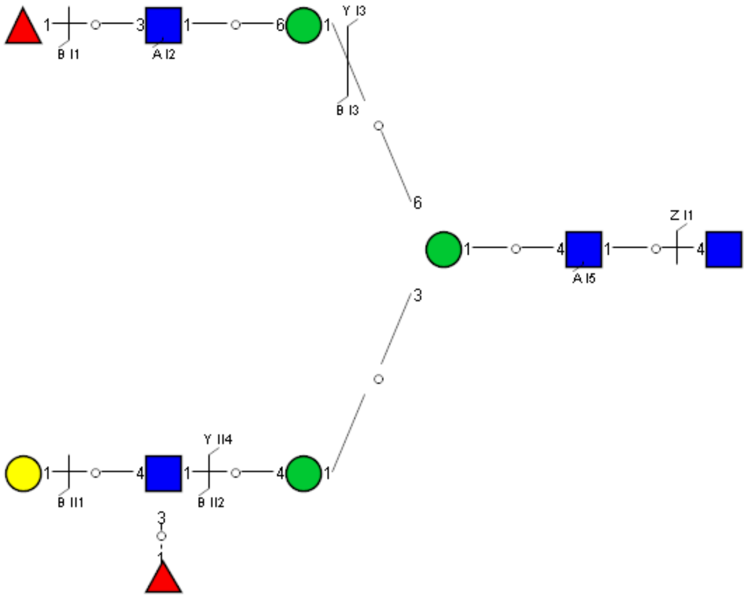

Spectrum Index: 33144(NoBHs:1)

Precursor iEF Map

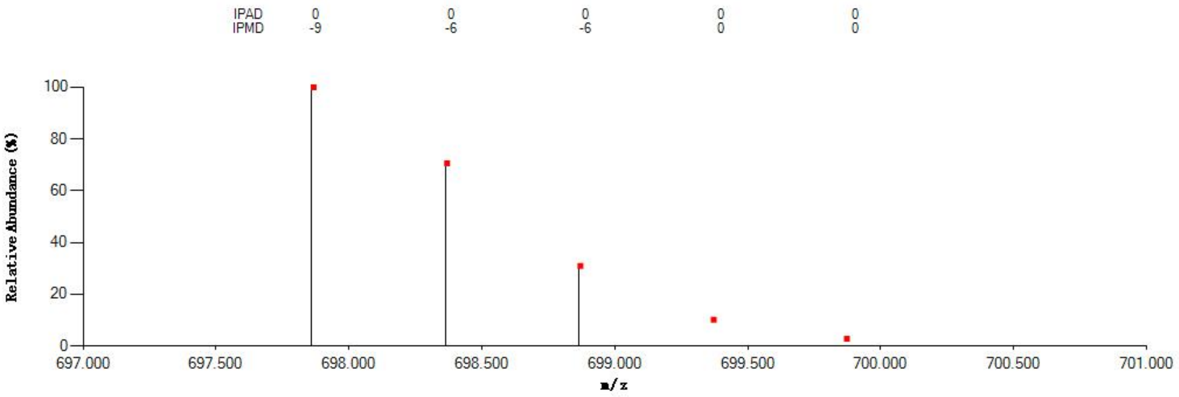


Annotated MS/MS Spectrum

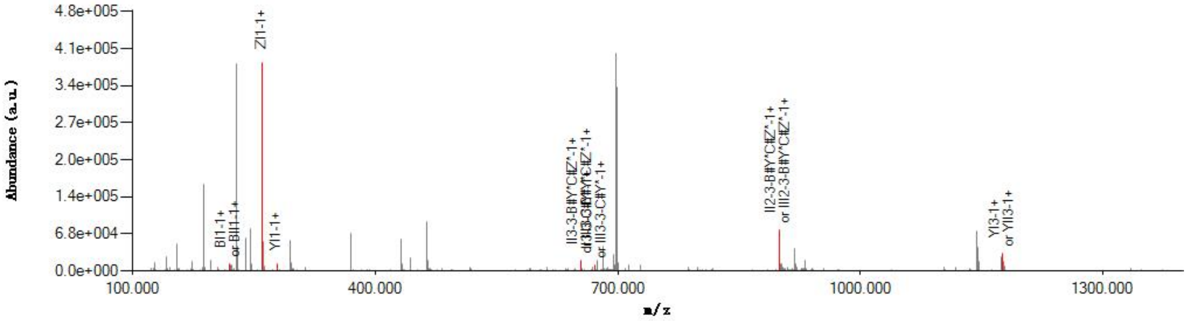


Graphical Fragmentation Map

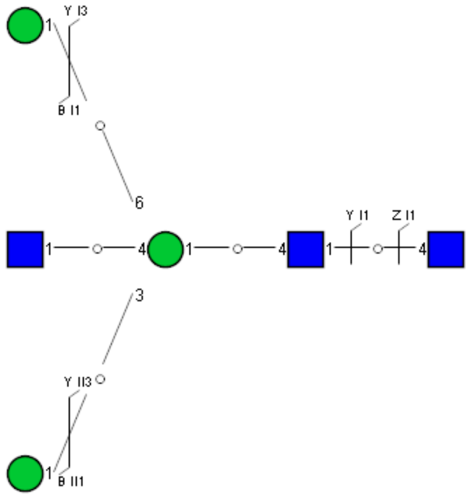

Spectrum Index: 33488(NoBHs:1)

#### Precursor iEF Map

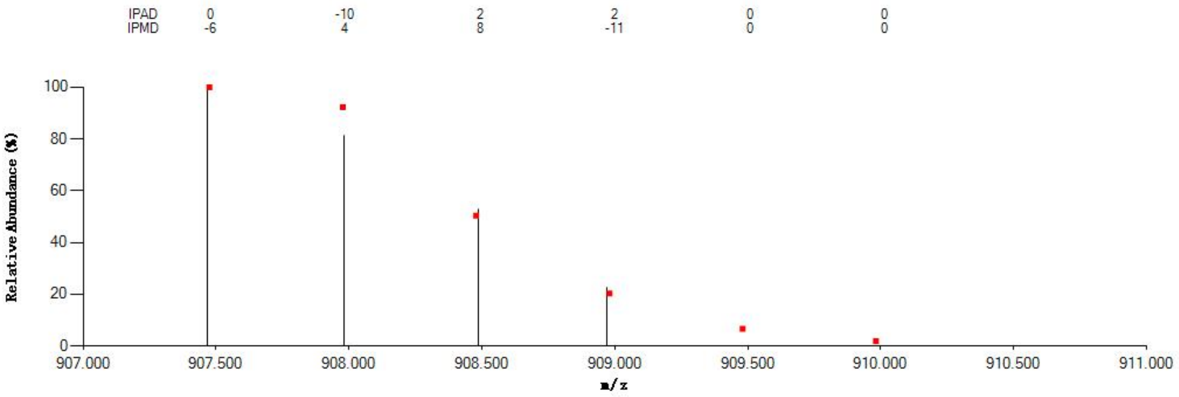

#### Annotated MS/MS Spectrum

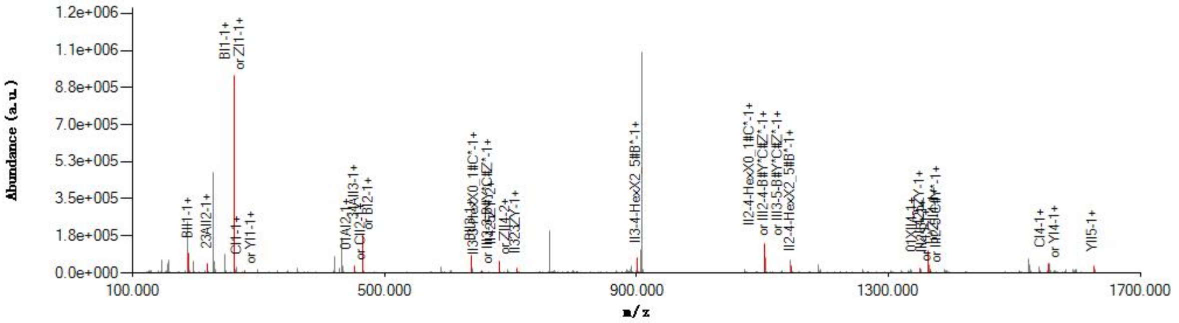

#### Graphical Fragmentation Map

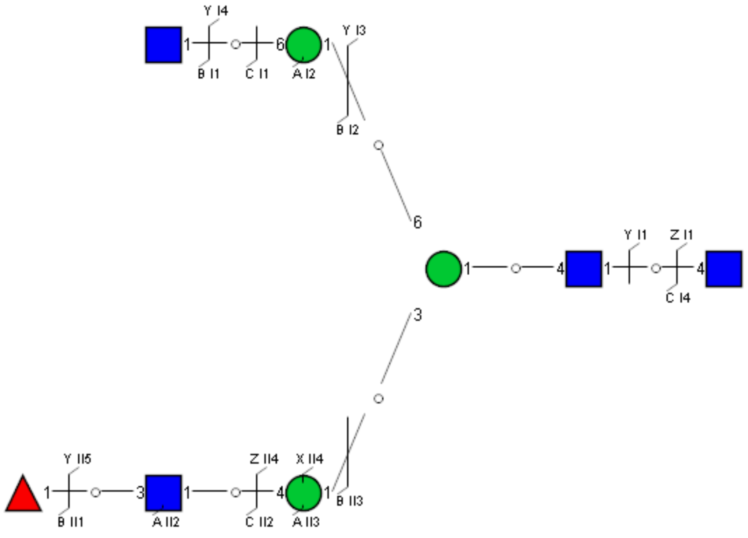

#### Spectrum Index: 33530(NoBHs:1)

#### Precursor iEF Map

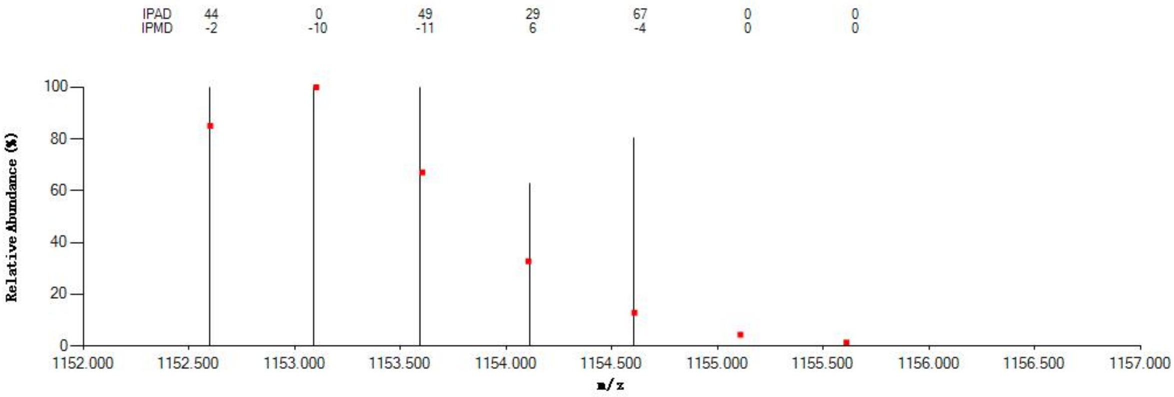


#### Annotated MS/MS Spectrum

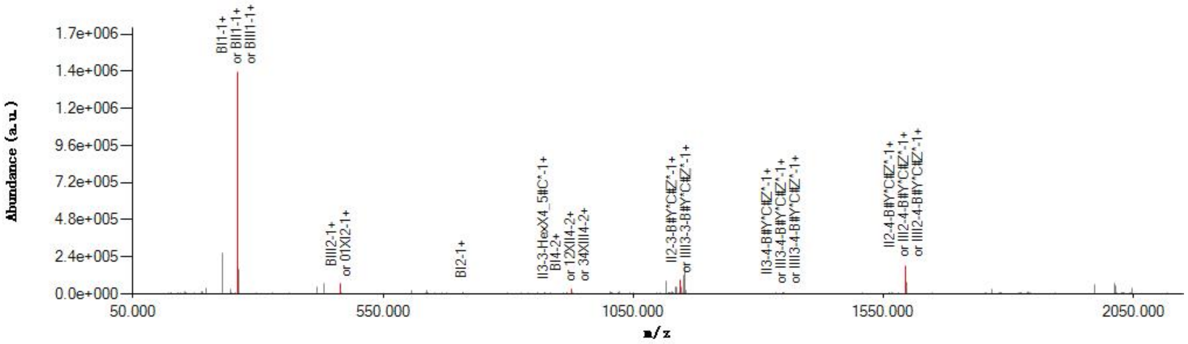


#### Graphical Fragmentation Map

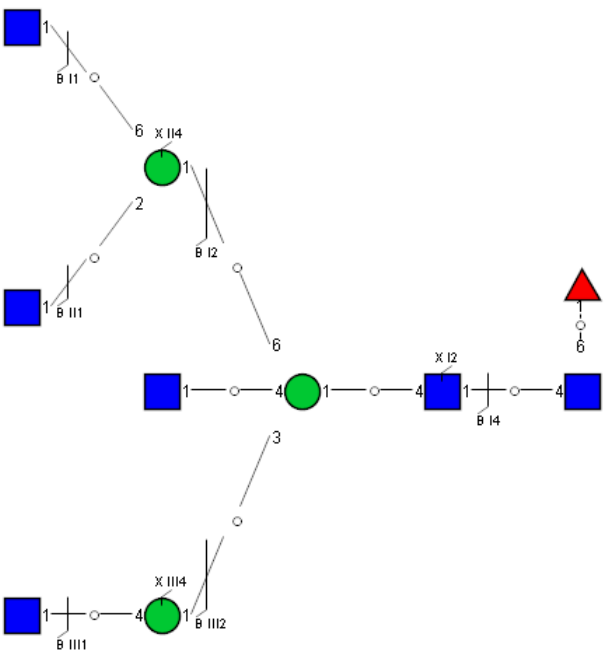

Spectrum Index: 33642(NoBHs:1)

#### Precursor iEF Map

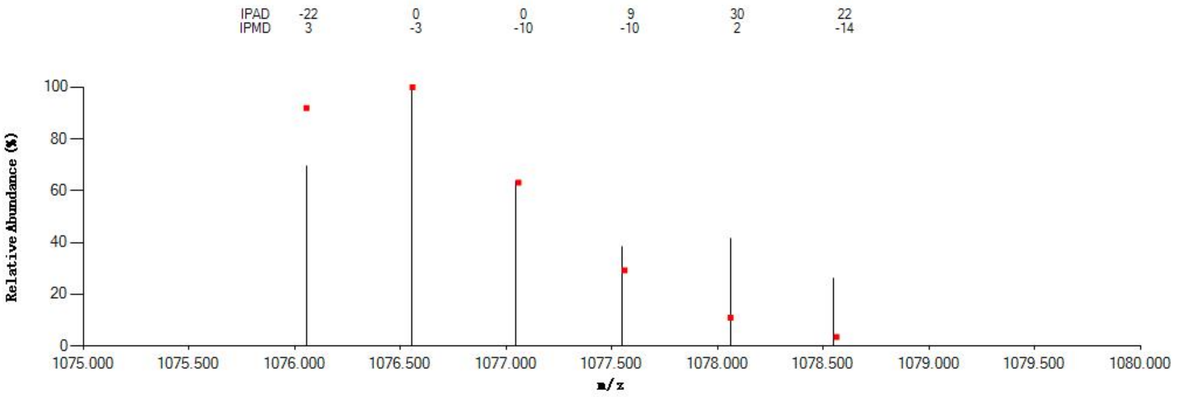


#### Annotated MS/MS Spectrum

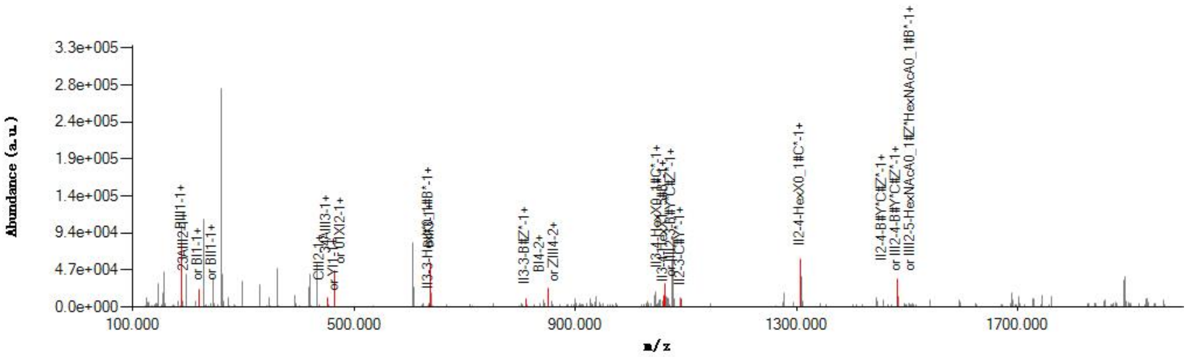


#### Graphical Fragmentation Map

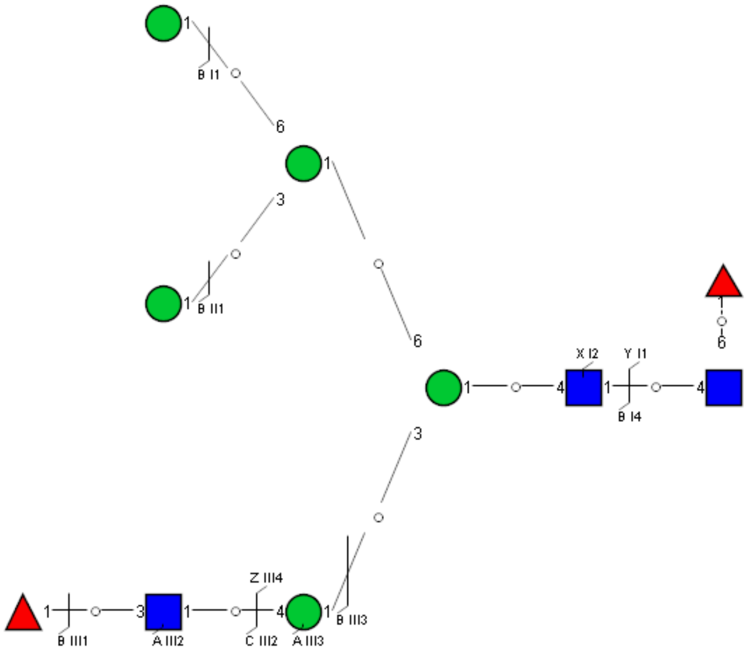

### Spectrum Index: 33659(NoBHs:1)

#### Precursor iEF Map

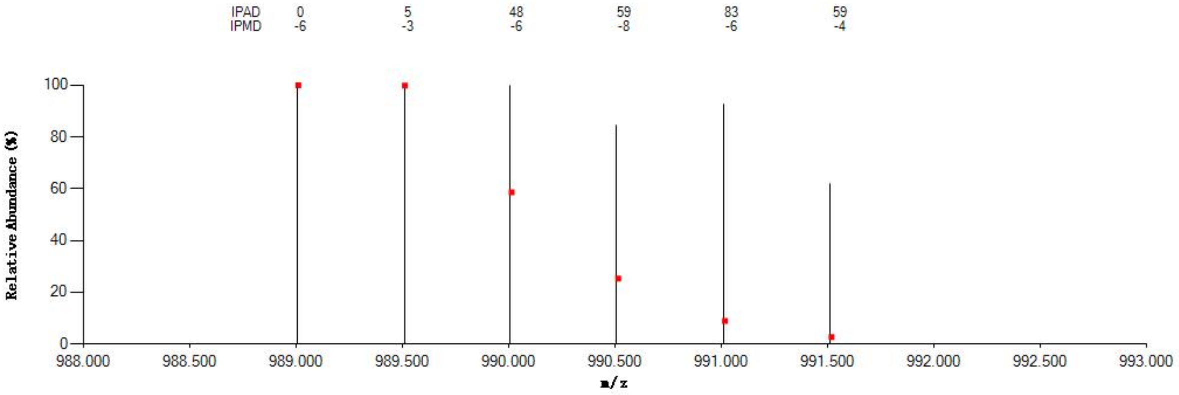


#### Annotated MS/MS Spectrum

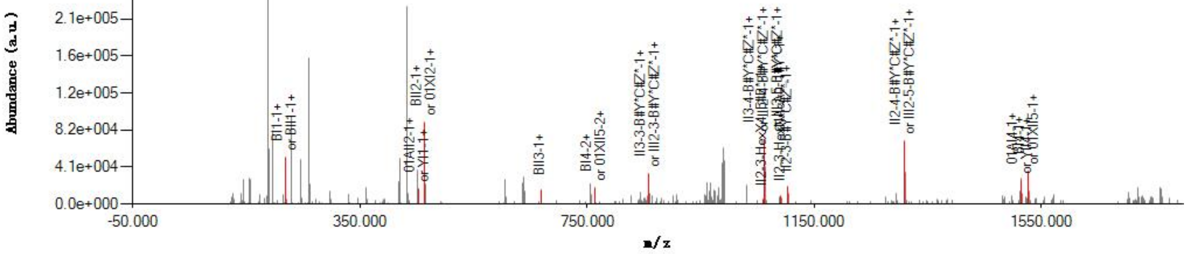


#### Graphical Fragmentation Map

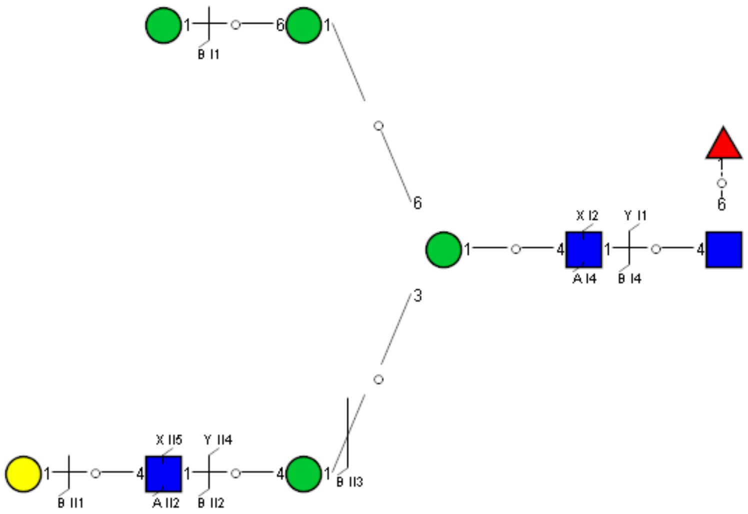

Spectrum Index: 33684(NoBHs:1)

#### Precursor iEF Map

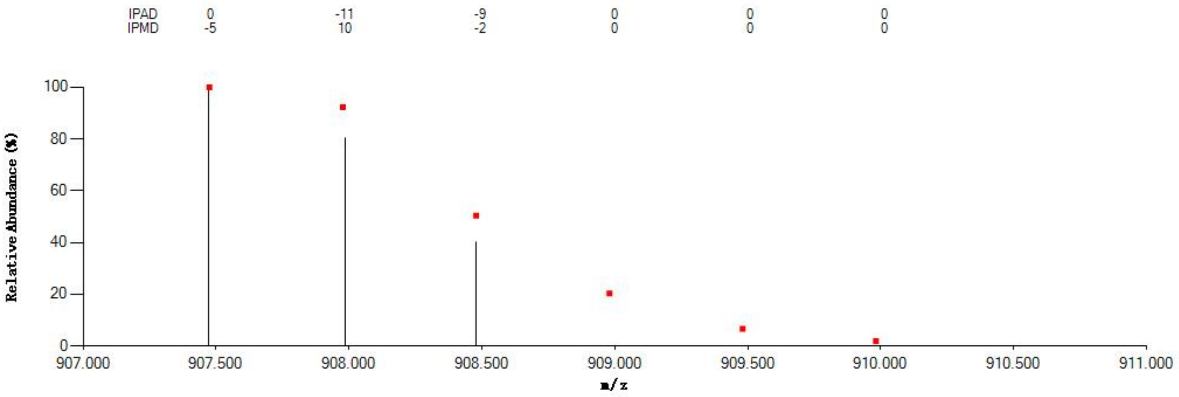


#### Annotated MS/MS Spectrum

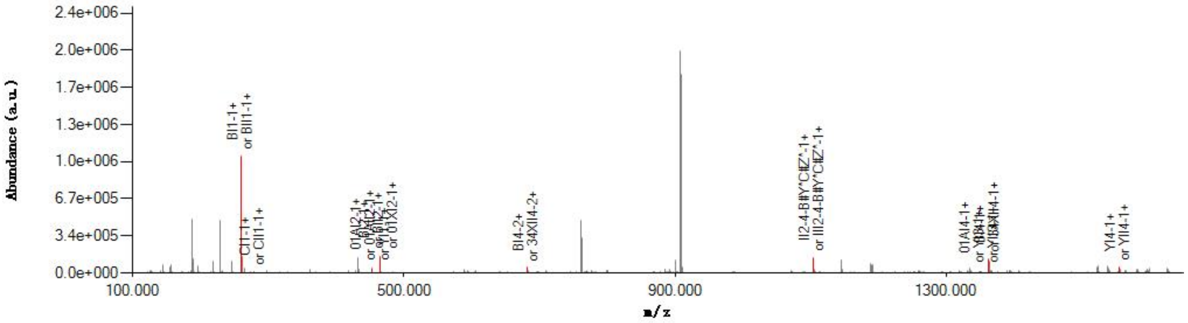


#### Graphical Fragmentation Map

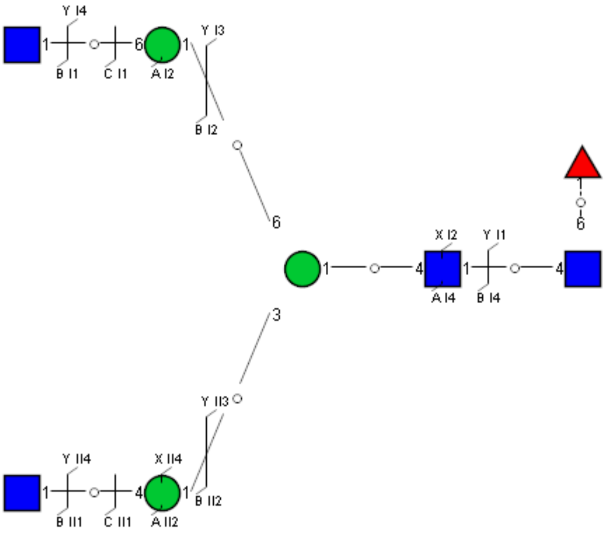

Spectrum Index: 33878(NoBHs:2)

#### Precursor iEF Map

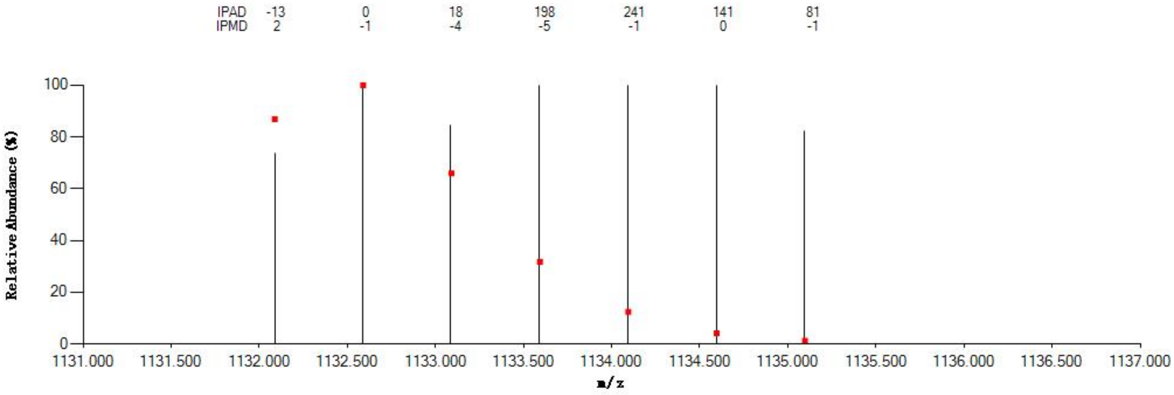


#### Annotated MS/MS Spectrum

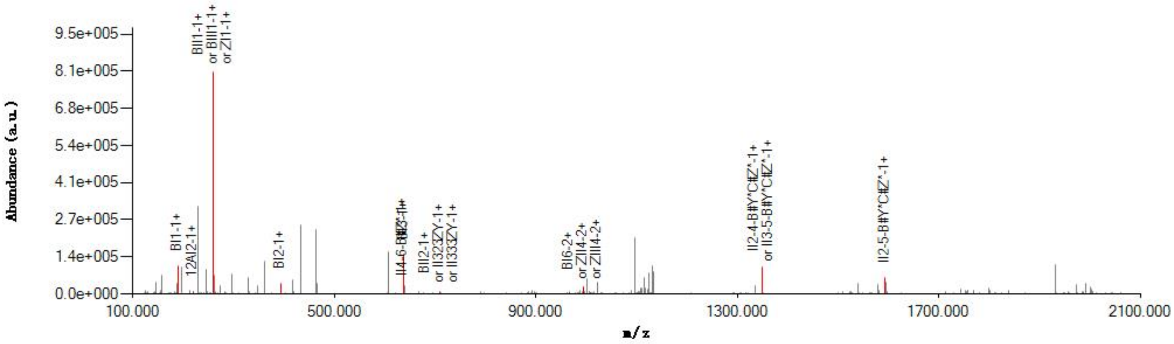


#### Graphical Fragmentation Map

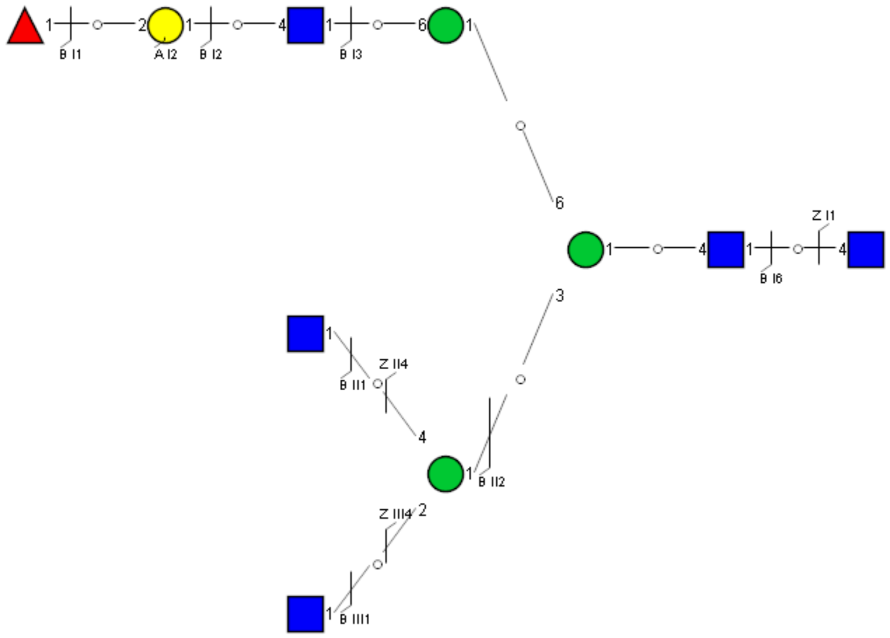

Spectrum Index: 34030(NoBHs:1)

#### Precursor iEF Map

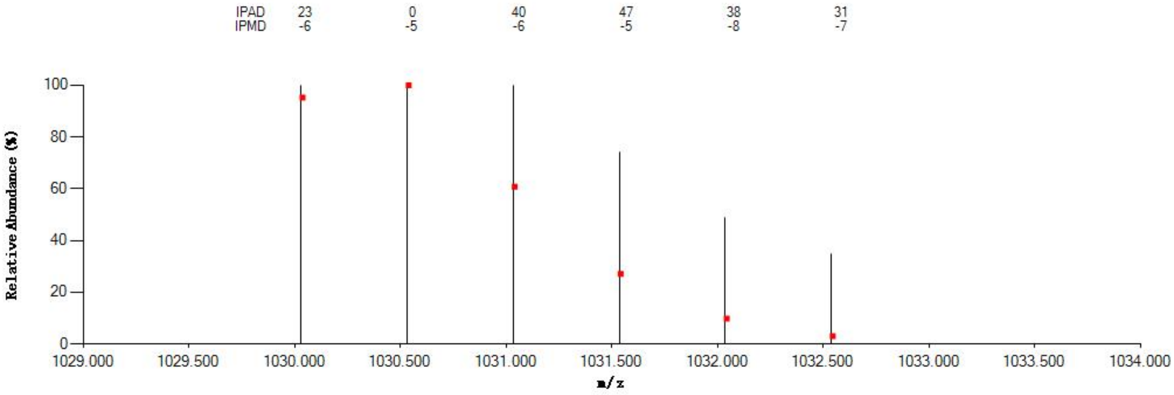

#### Annotated MS/MS Spectrum

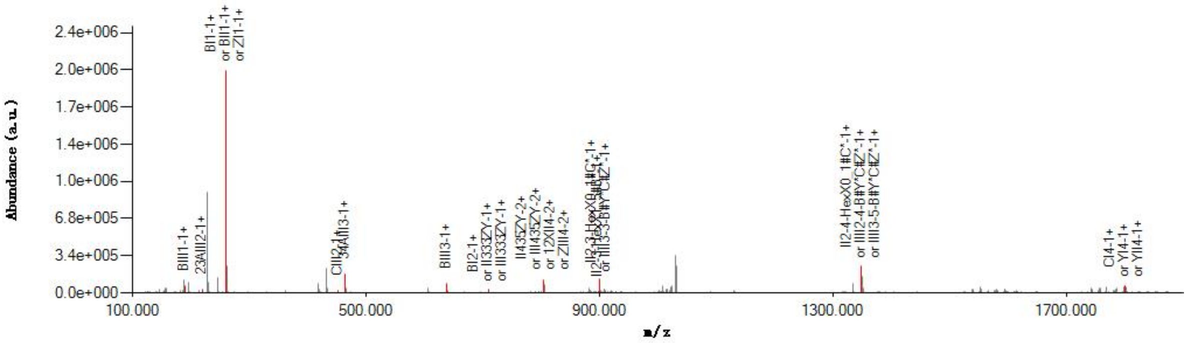

#### Graphical Fragmentation Map

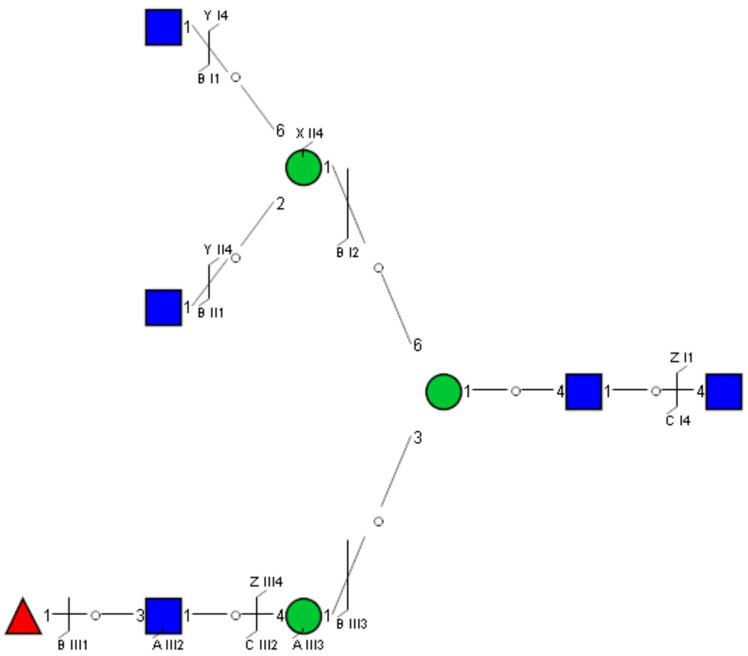

Spectrum Index: 34142(NoBHs:1)

#### Precursor iEF Map

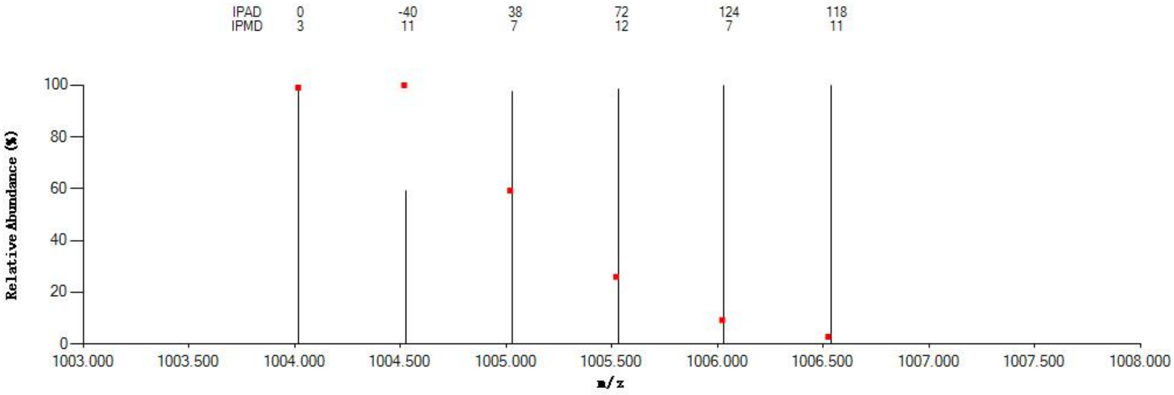

#### Annotated MS/MS Spectrum

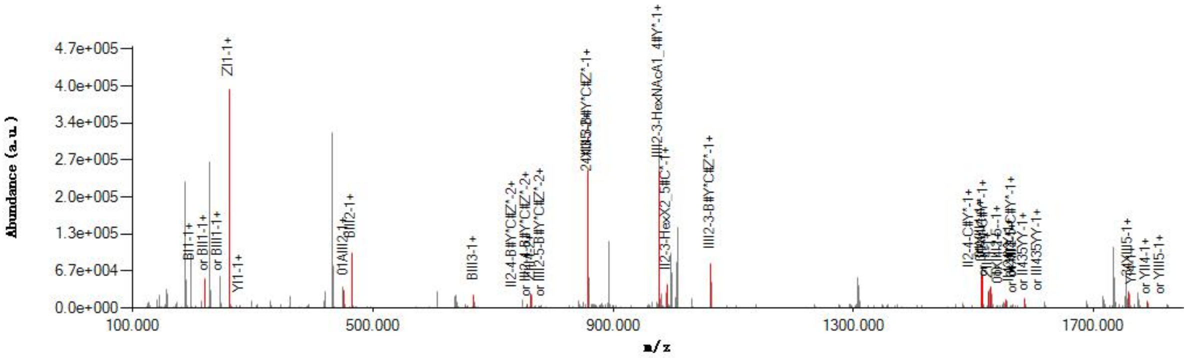

#### Graphical Fragmentation Map

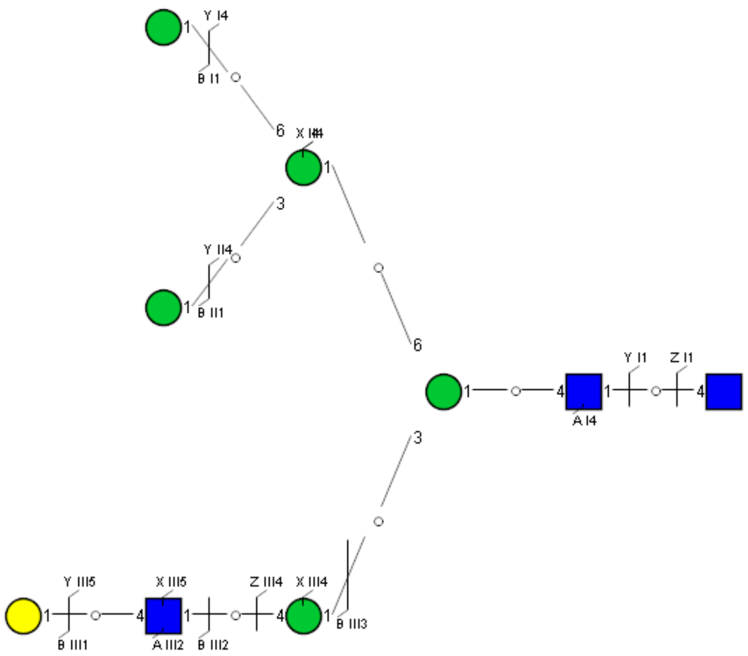

Spectrum Index: 34197(NoBHs:1)

#### Precursor iEF Map

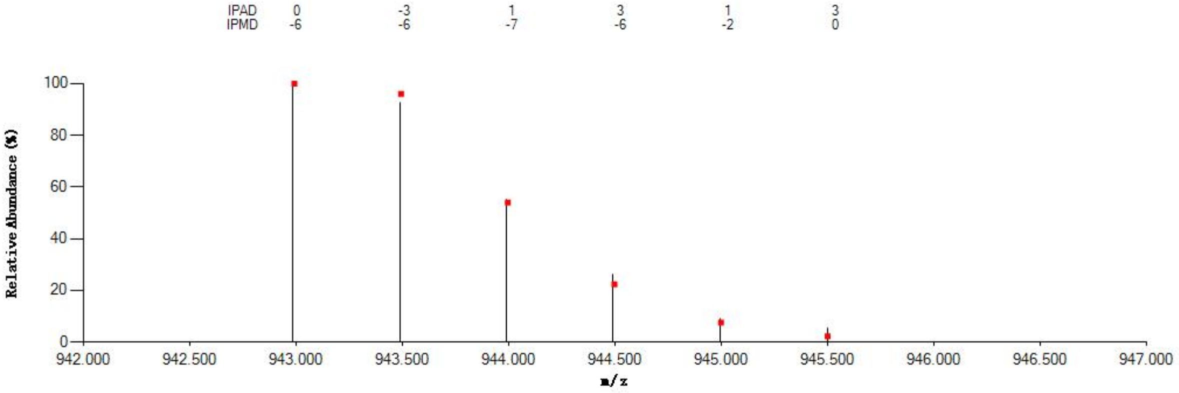


#### Annotated MS/MS Spectrum

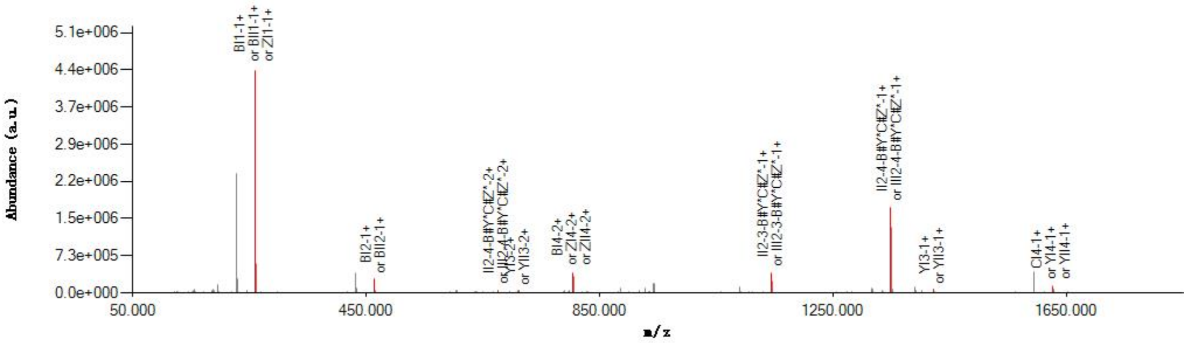


#### Graphical Fragmentation Map

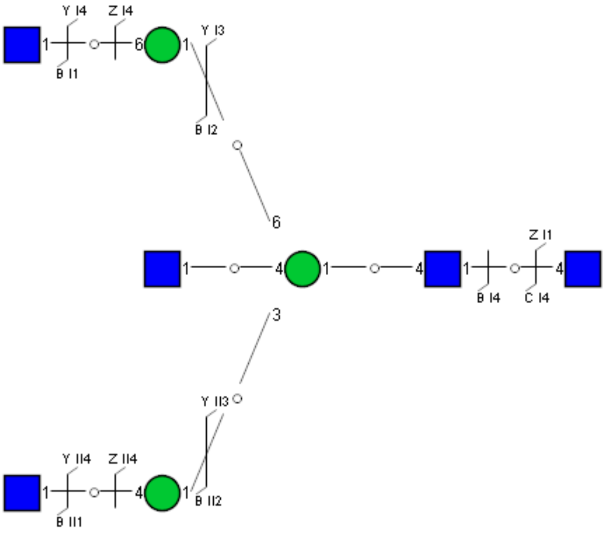

Spectrum Index: 34340(NoBHs:1)

#### Precursor iEF Map

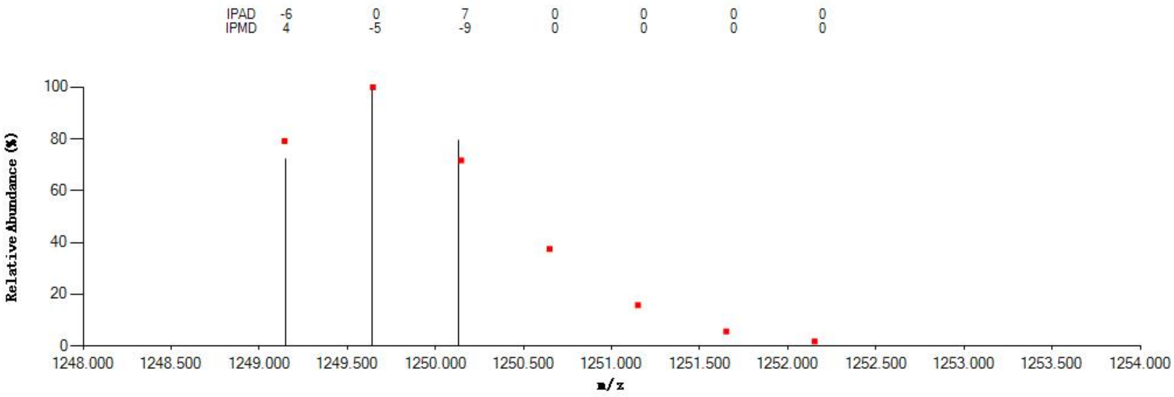


#### Annotated MS/MS Spectrum

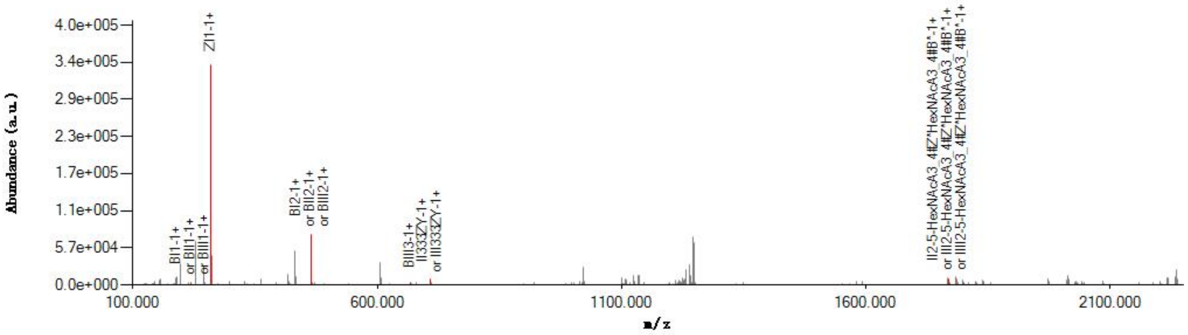


#### Graphical Fragmentation Map

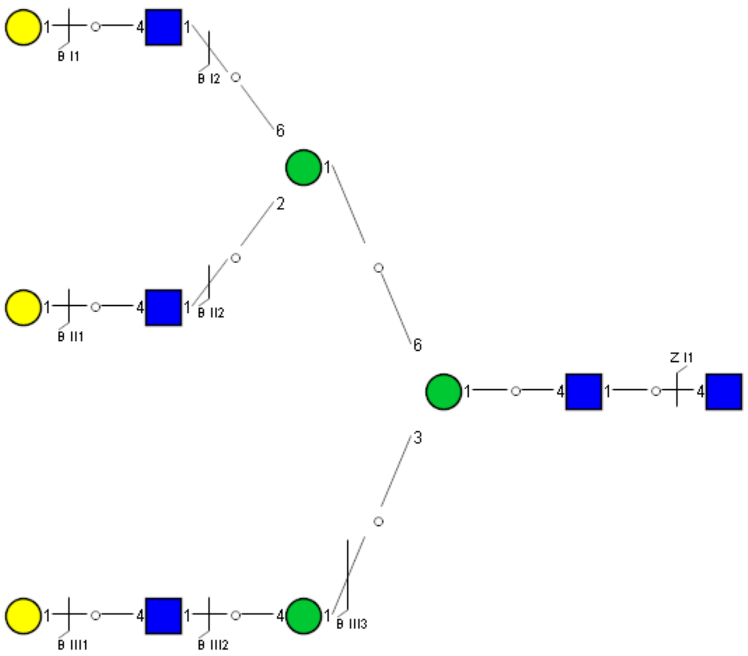

Spectrum Index: 34407(NoBHs:1)

#### Precursor iEF Map

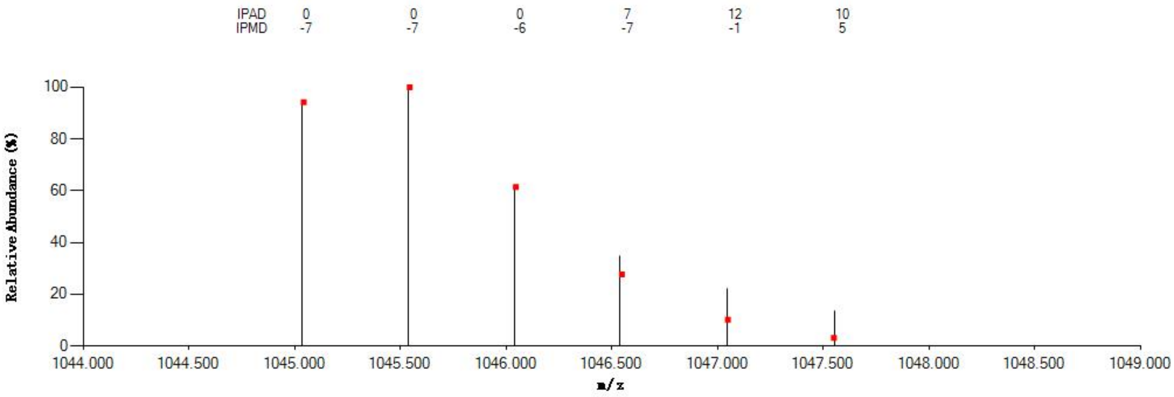

#### Annotated MS/MS Spectrum

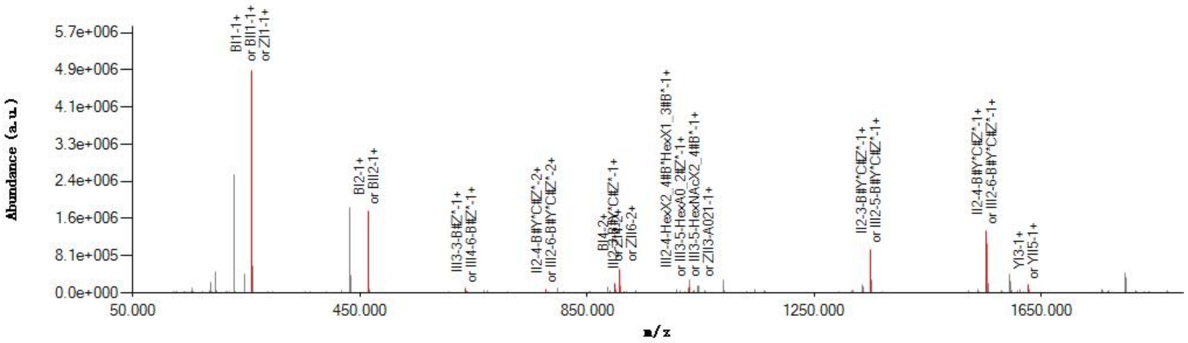

#### Graphical Fragmentation Map

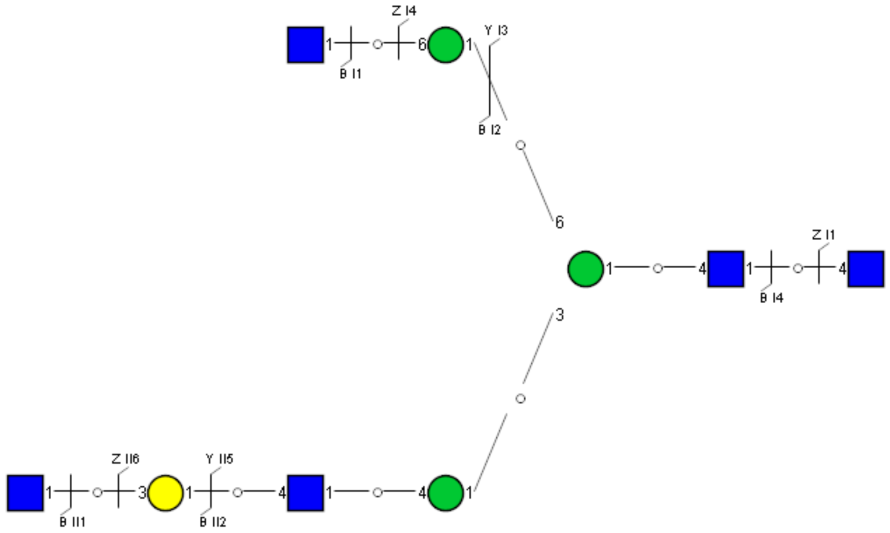

Spectrum Index: 34599(NoBHs:1)

#### Precursor iEF Map

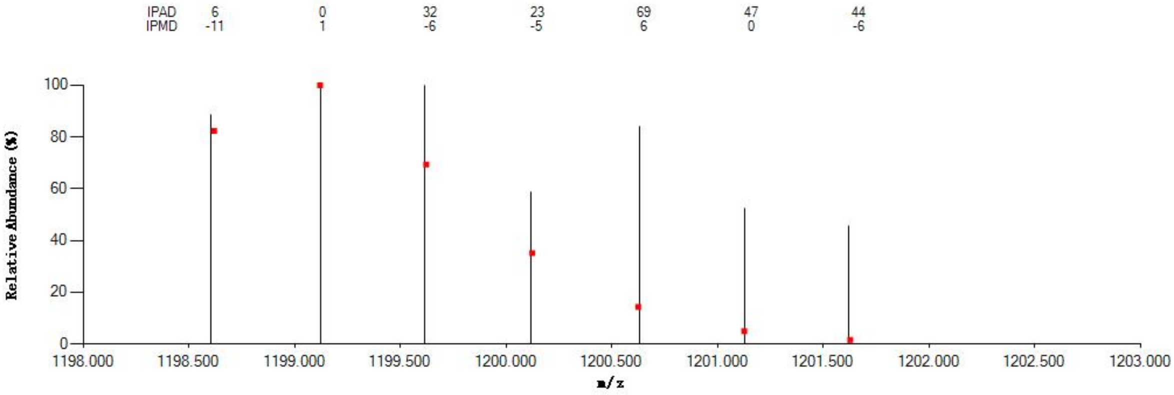


#### Annotated MS/MS Spectrum

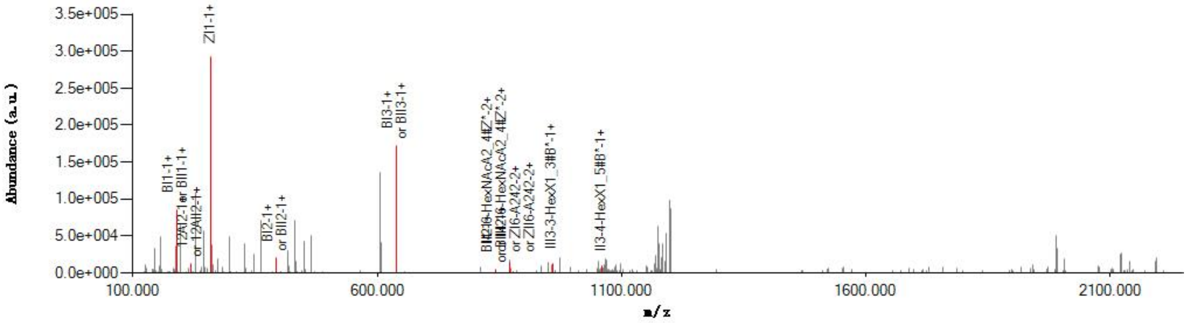


#### Graphical Fragmentation Map

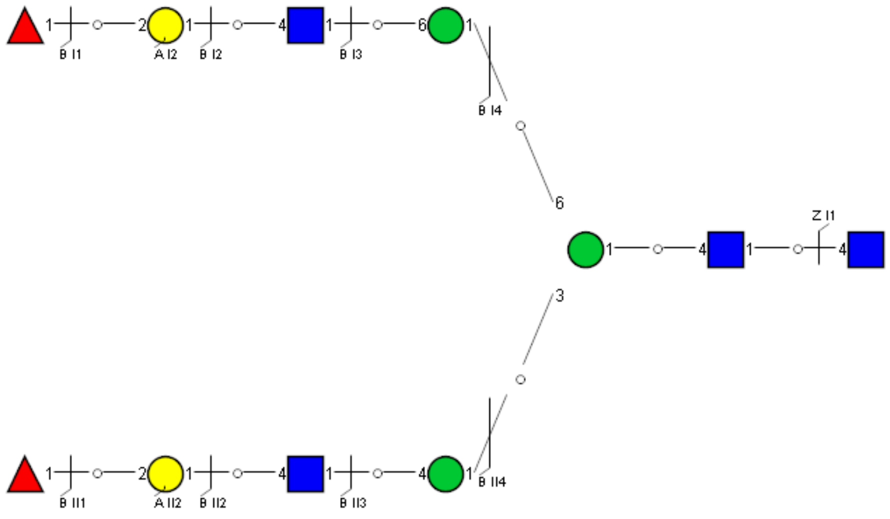

Spectrum Index: 34723(NoBHs:1)

Precursor iEF Map

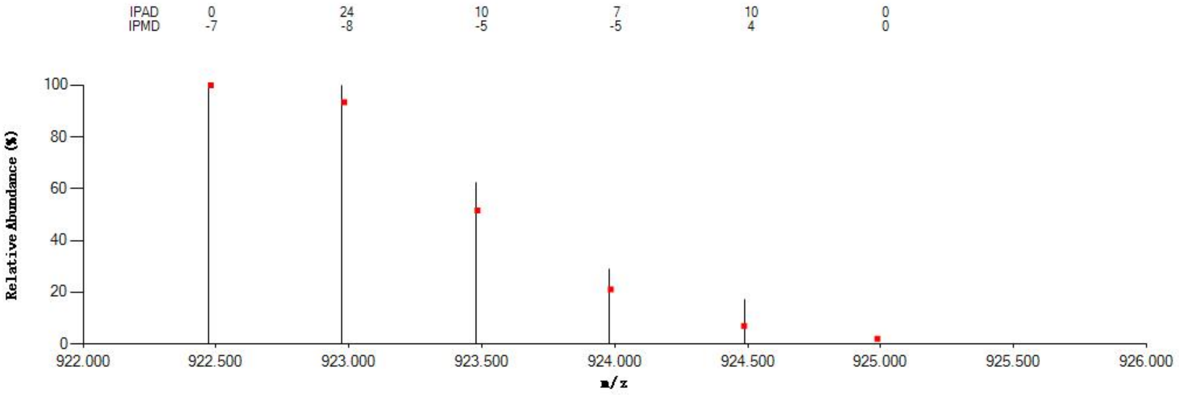


Annotated MS/MS Spectrum

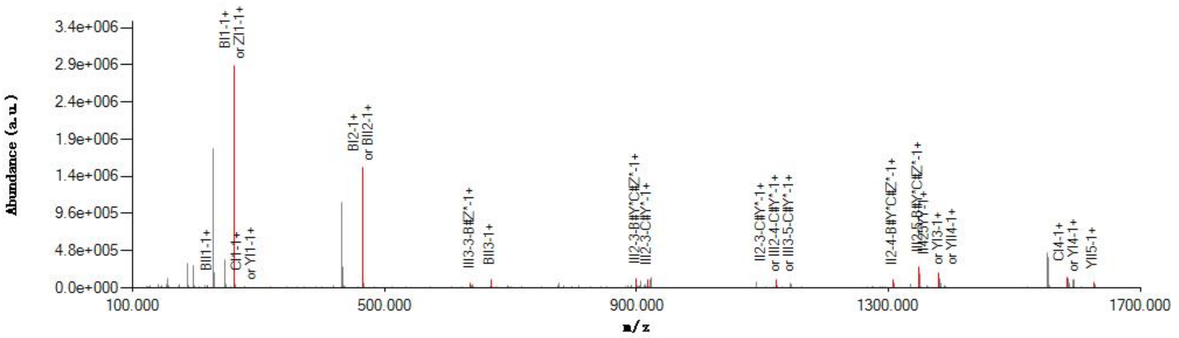


Graphical Fragmentation Map

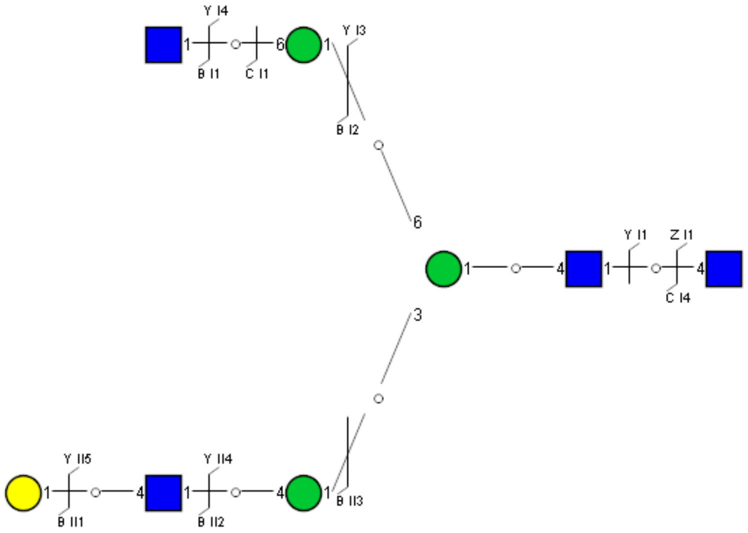

Spectrum Index: 34724(NoBHs:1)

#### Precursor iEF Map

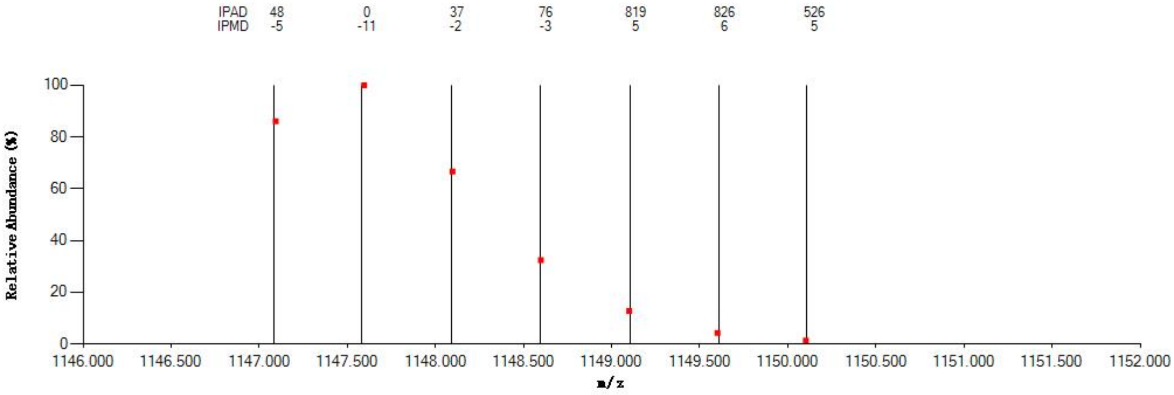


#### Annotated MS/MS Spectrum

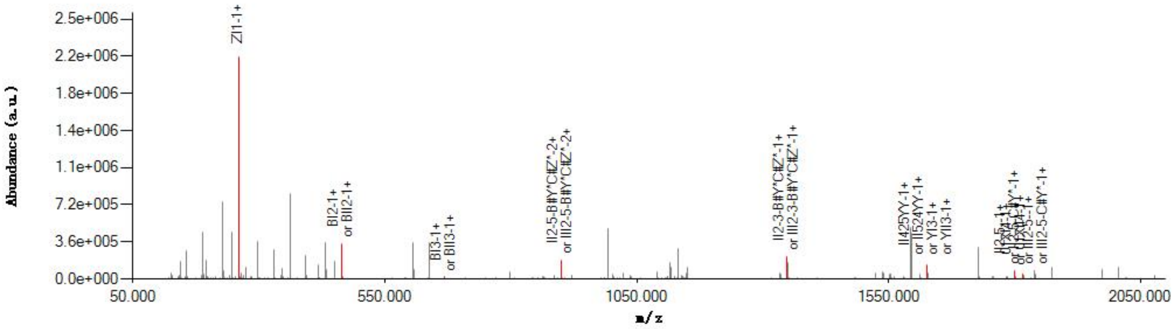


#### Graphical Fragmentation Map

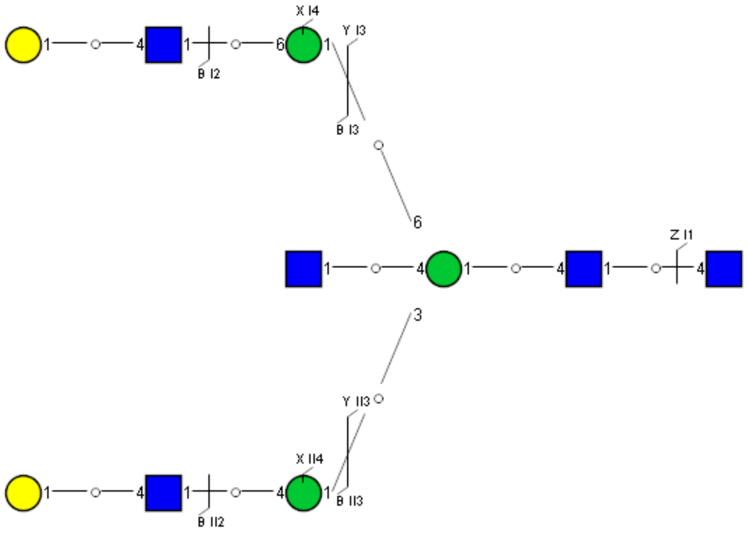

Spectrum Index: 34758(NoBHs:1)

#### Precursor iEF Map

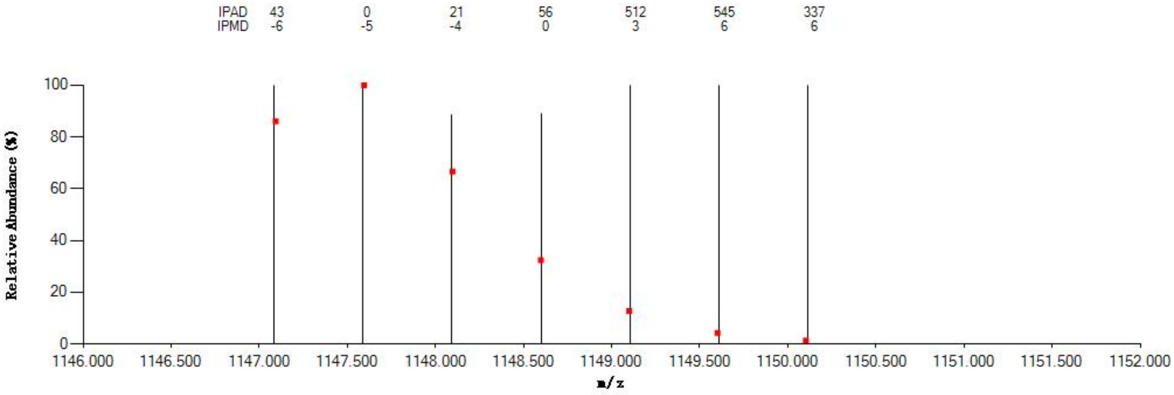

#### Annotated MS/MS Spectrum

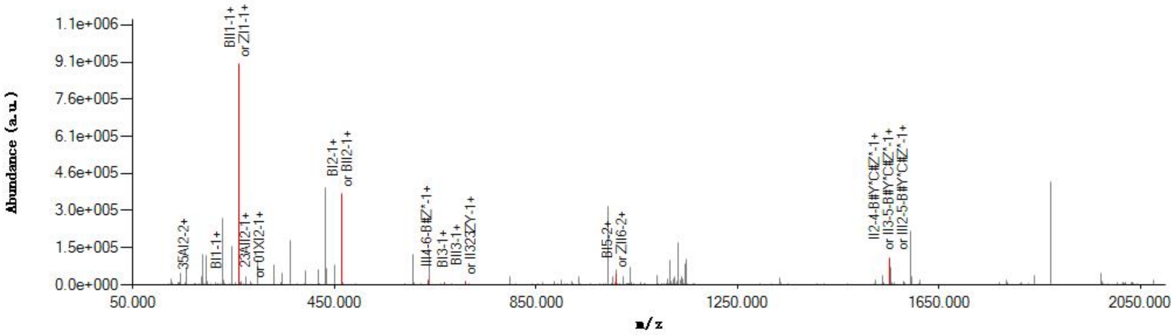

#### Graphical Fragmentation Map

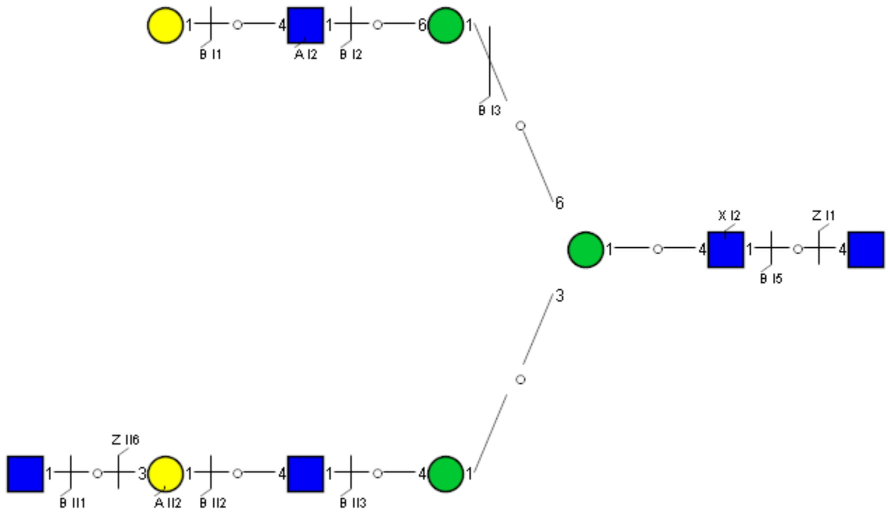

Spectrum Index: 34791(NoBHs:1)

#### Precursor iEF Map

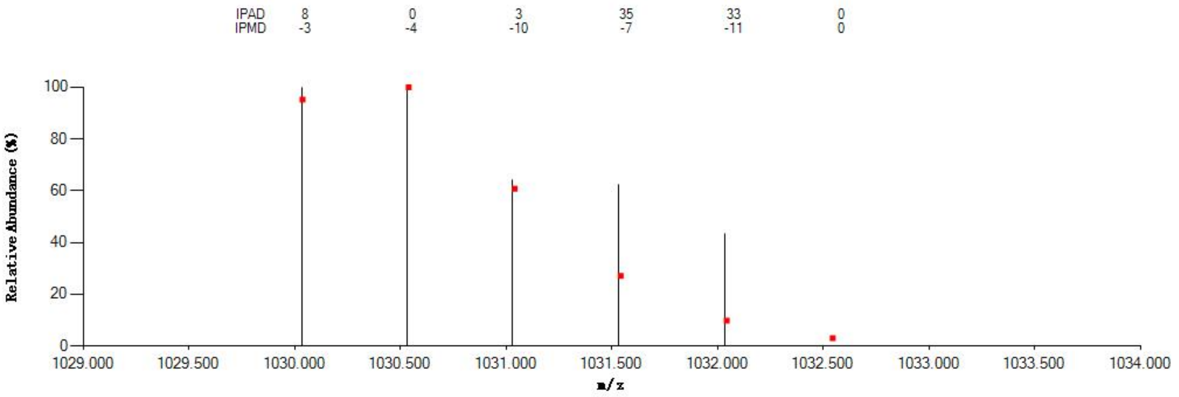


#### Annotated MS/MS Spectrum

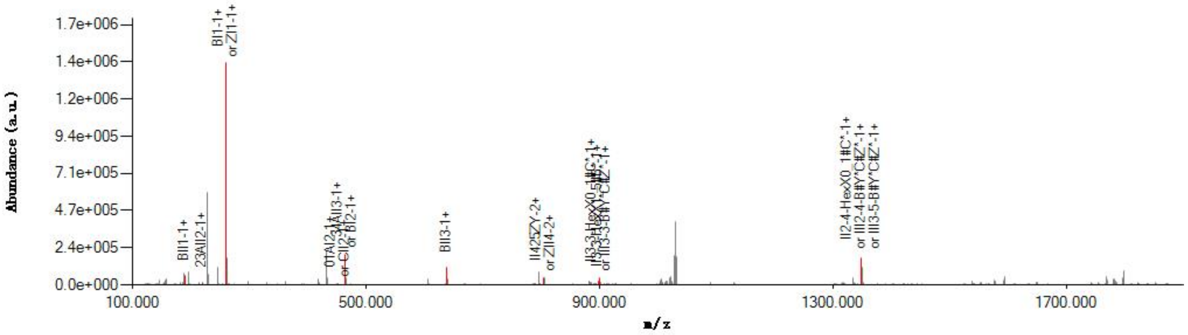


#### Graphical Fragmentation Map

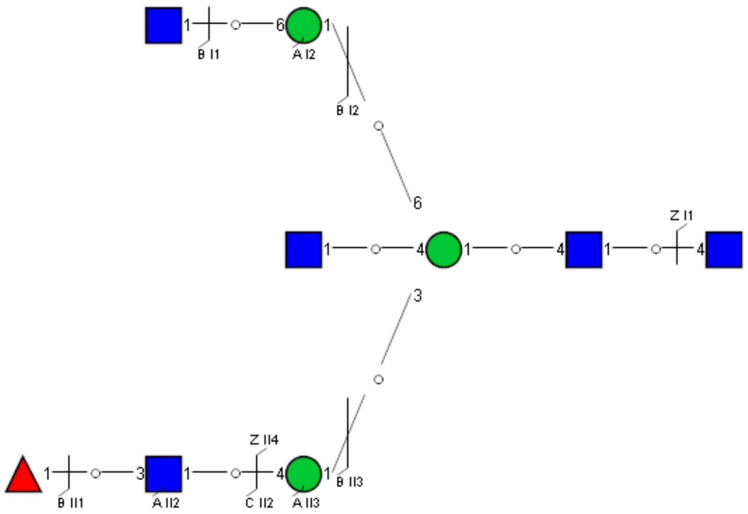

### Spectrum Index: 34837(NoBHs:1)

#### Precursor iEF Map

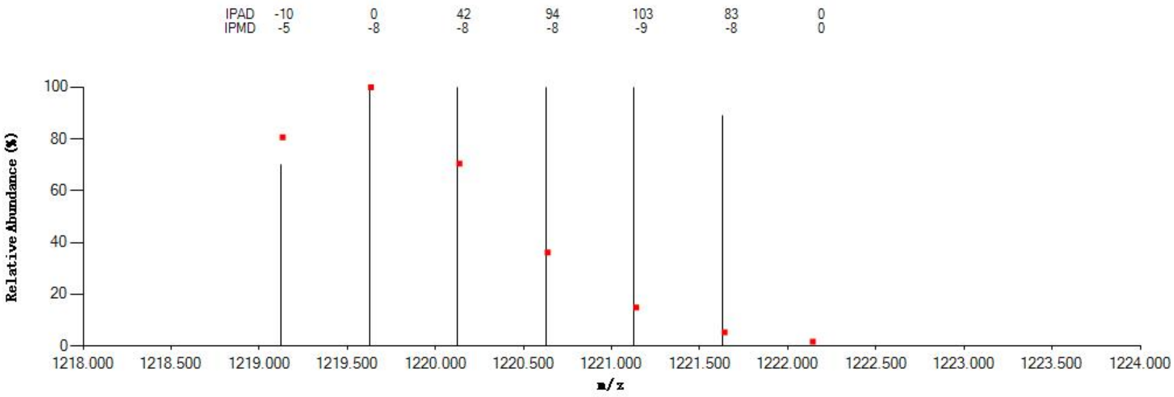


#### Annotated MS/MS Spectrum

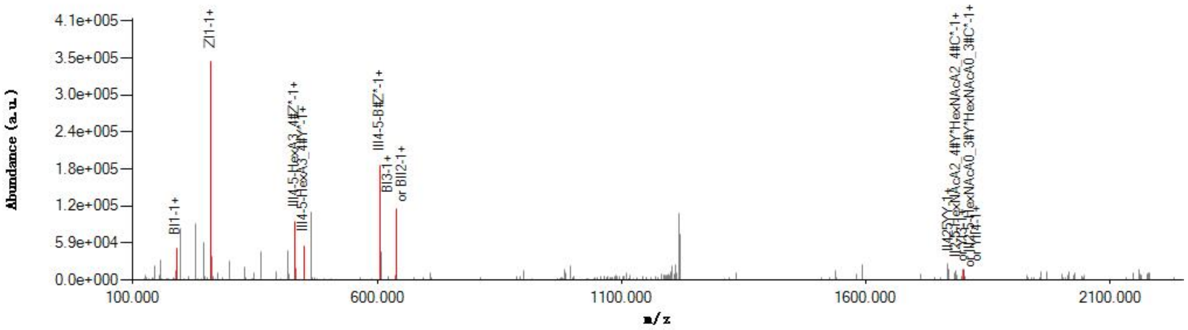


#### Graphical Fragmentation Map

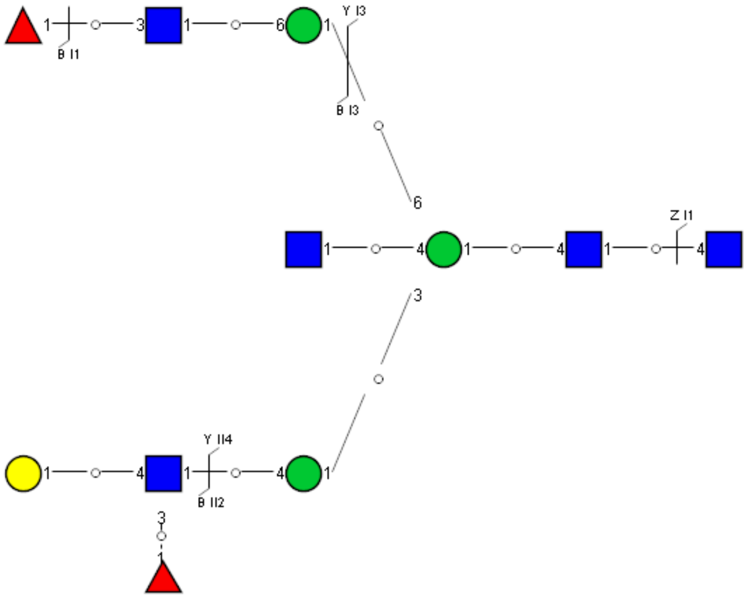

Spectrum Index: 34870(NoBHs:1)

#### Precursor iEF Map

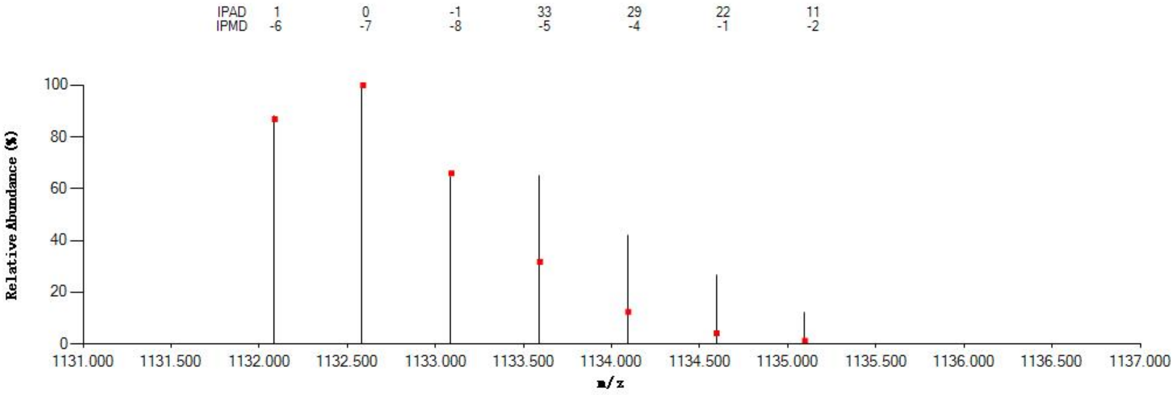


#### Annotated MS/MS Spectrum

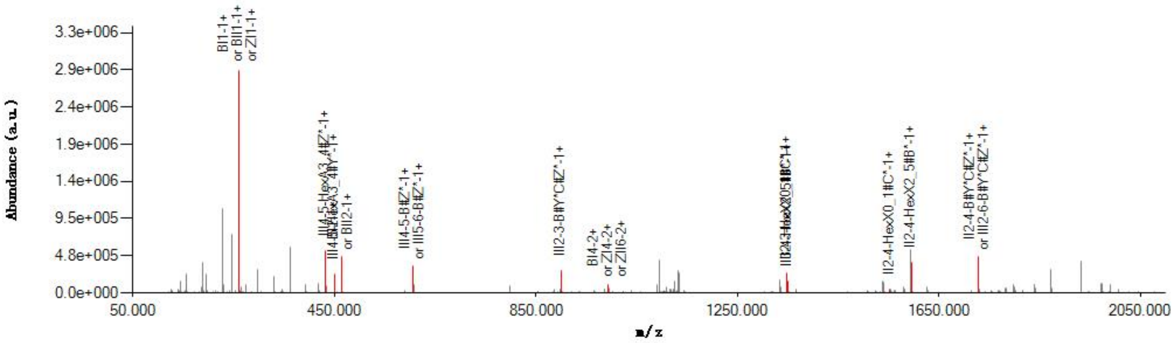


#### Graphical Fragmentation Map

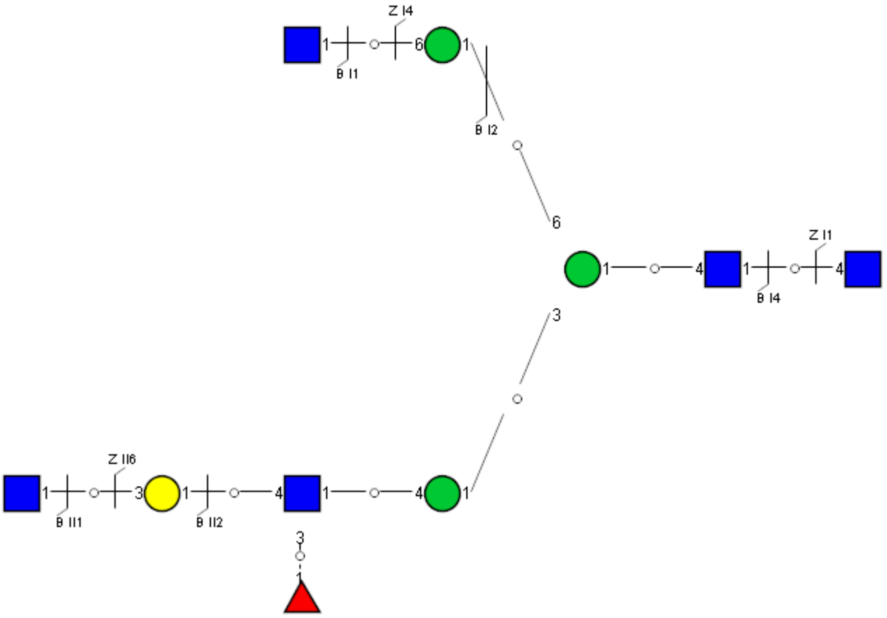

Spectrum Index: 34917(NoBHs:1)

#### Precursor iEF Map

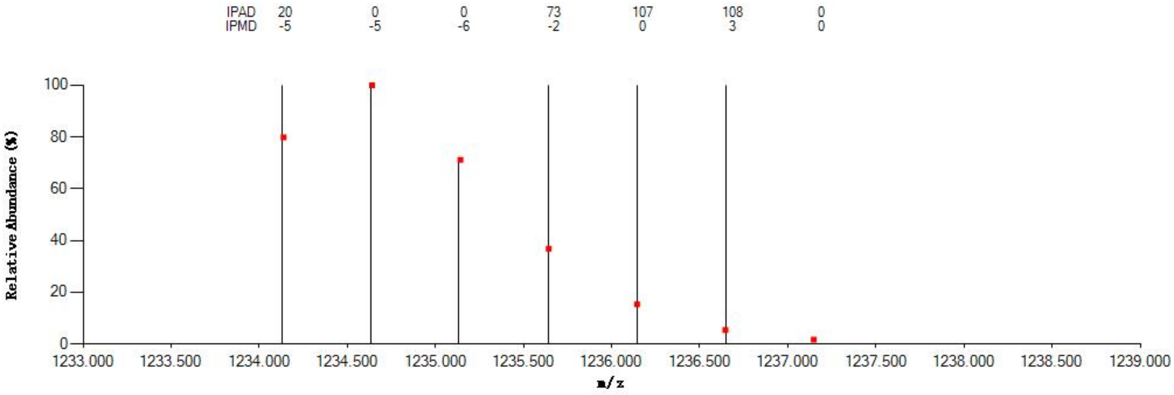


#### Annotated MS/MS Spectrum

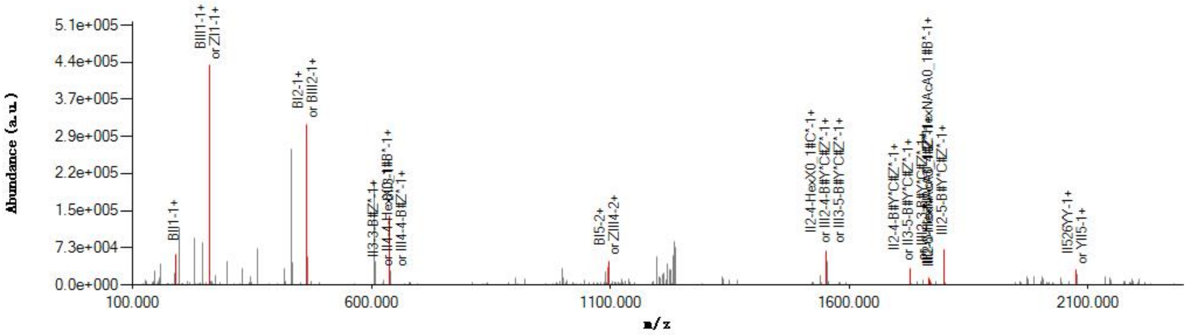


#### Graphical Fragmentation Map

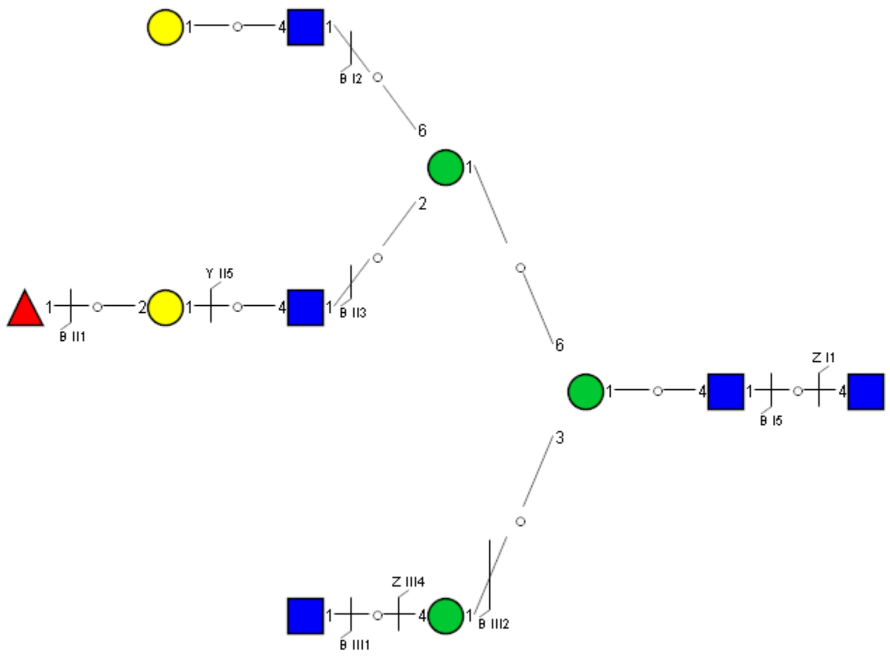

Spectrum Index: 34954(NoBHs:1)

#### Precursor iEF Map

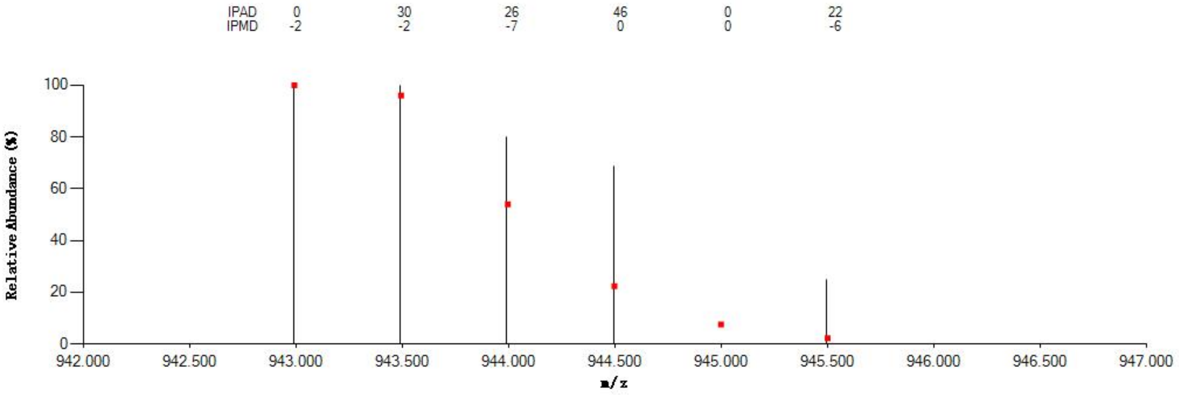

#### Annotated MS/MS Spectrum

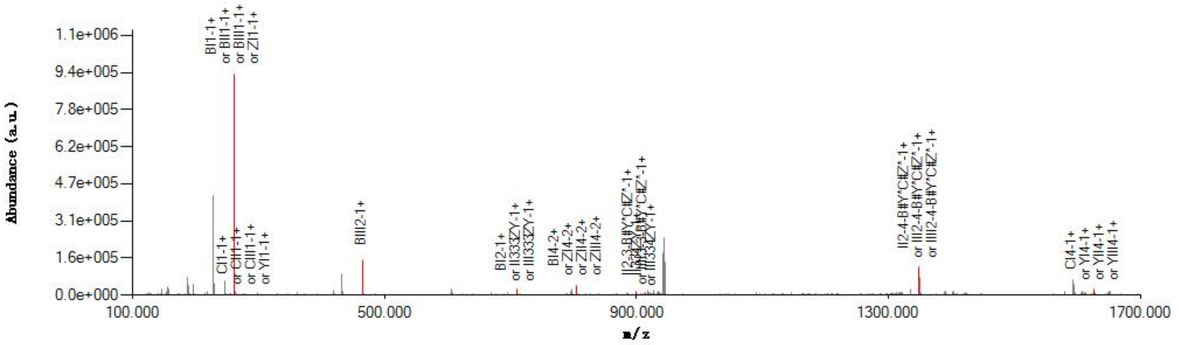

#### Graphical Fragmentation Map

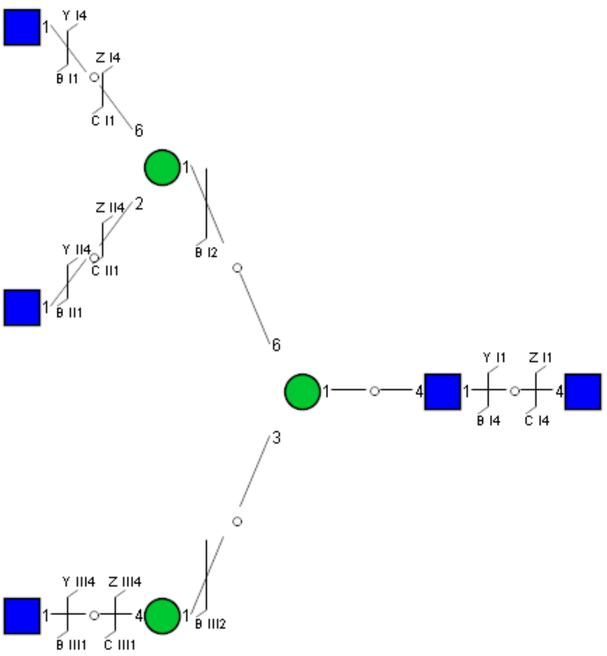

Spectrum Index: 34977(NoBHs:1)

#### Precursor iEF Map

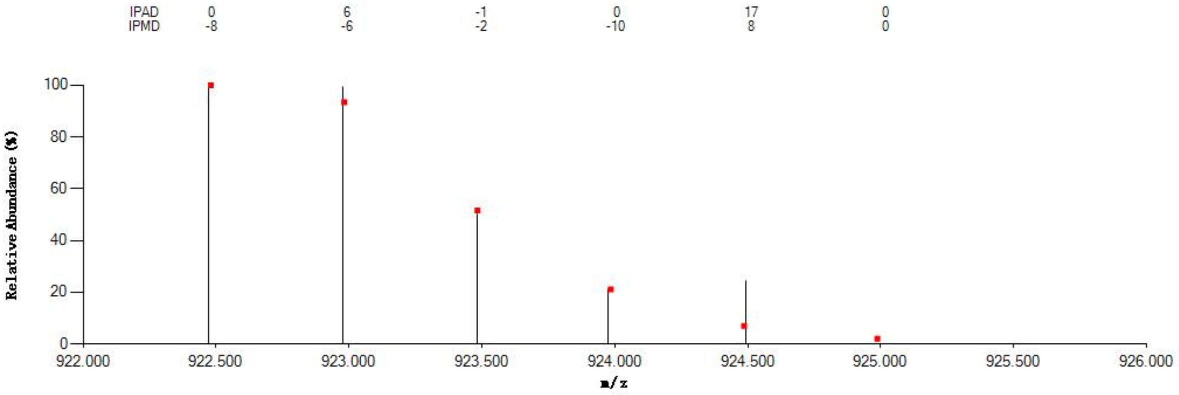


#### Annotated MS/MS Spectrum

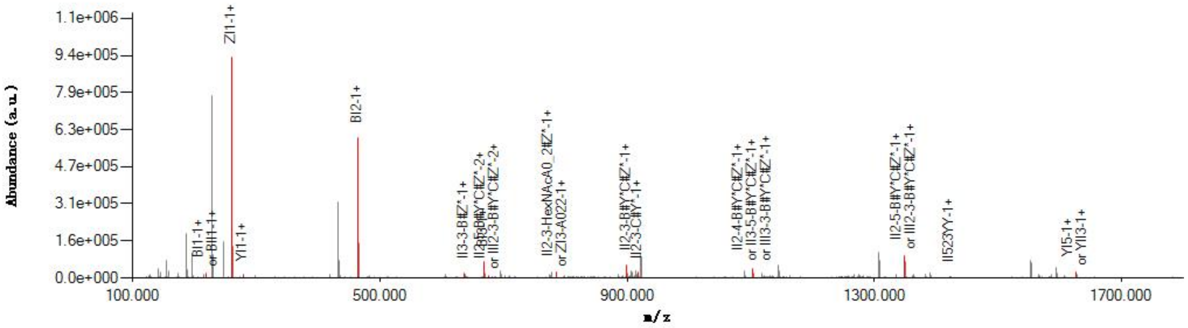


#### Graphical Fragmentation Map

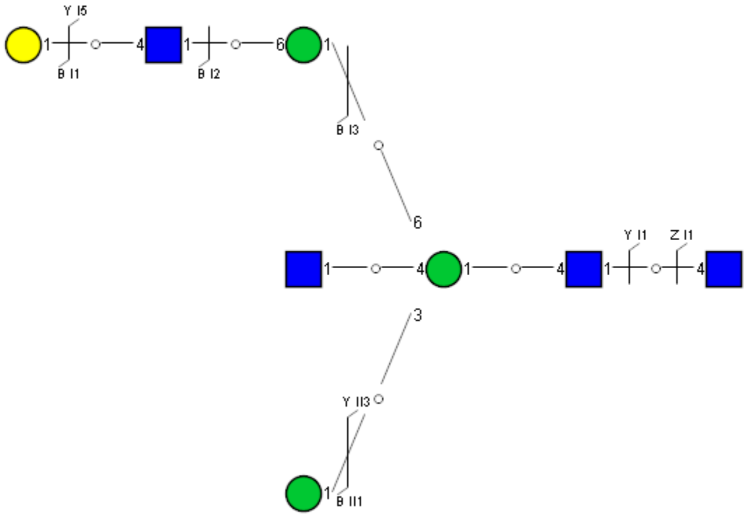

Spectrum Index: 35123(NoBHs:1)

#### Precursor iEF Map

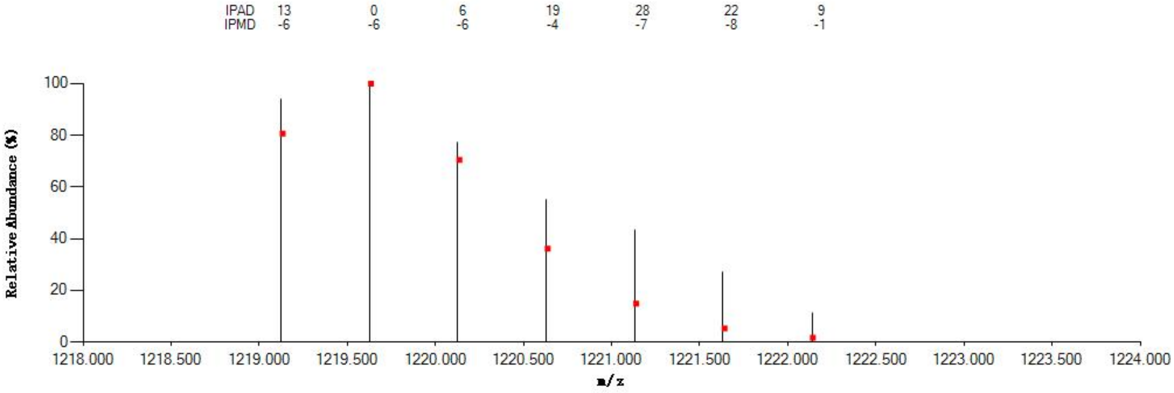


#### Annotated MS/MS Spectrum

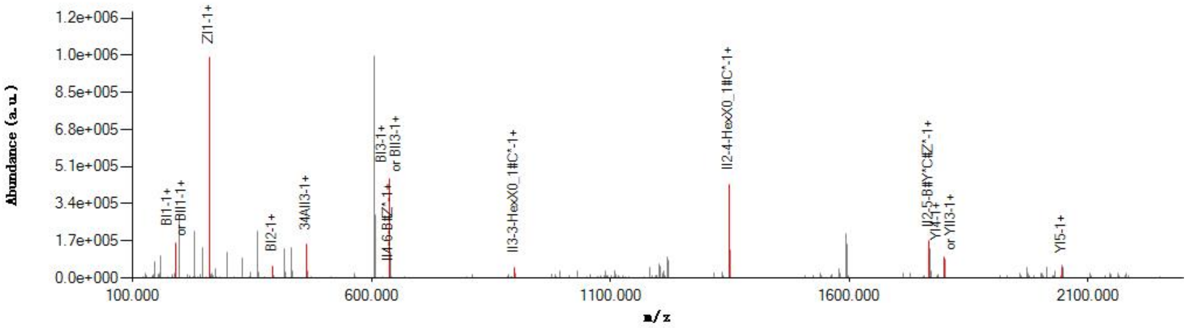


#### Graphical Fragmentation Map

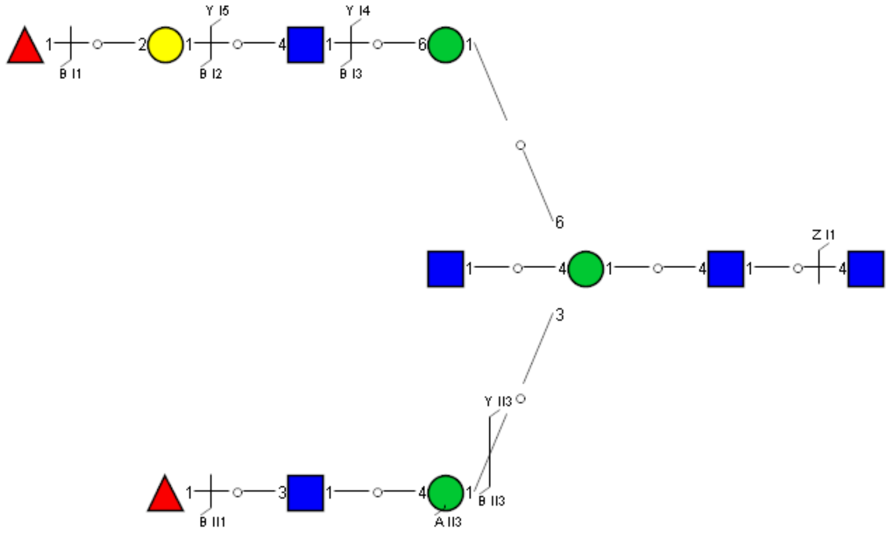

Spectrum Index: 35129(NoBHs:1)

#### Precursor iEF Map

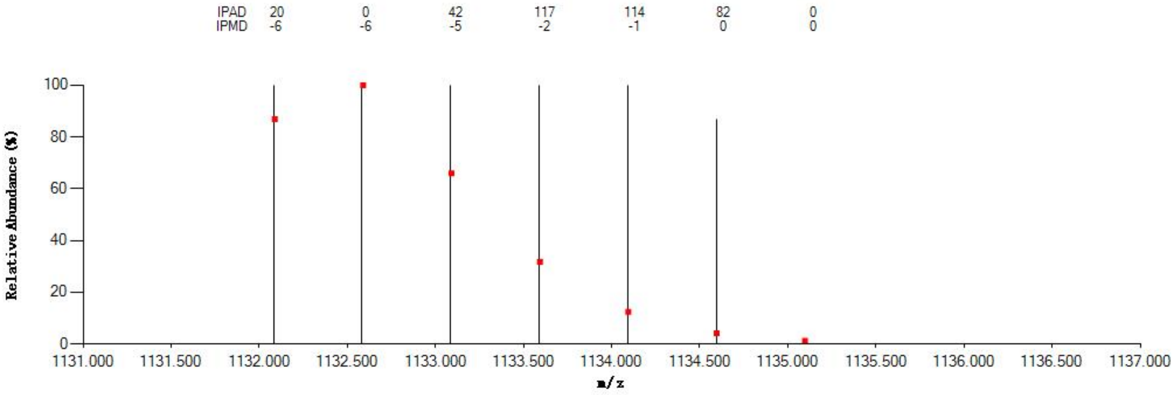


#### Annotated MS/MS Spectrum

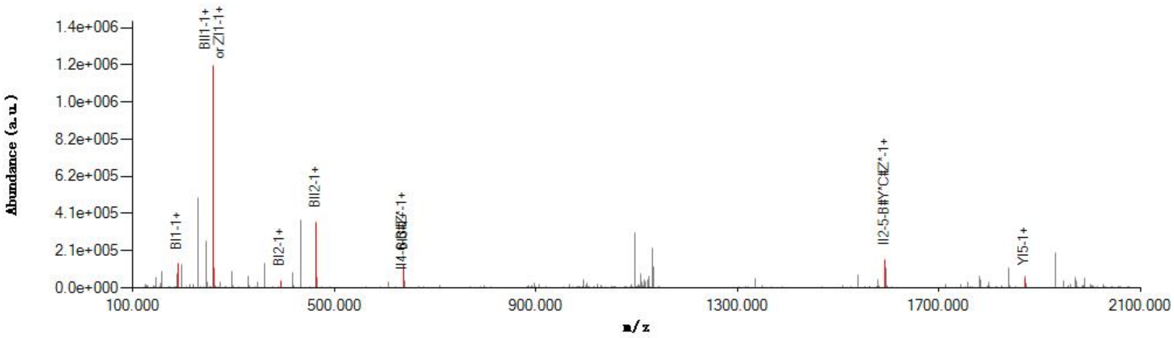


#### Graphical Fragmentation Map

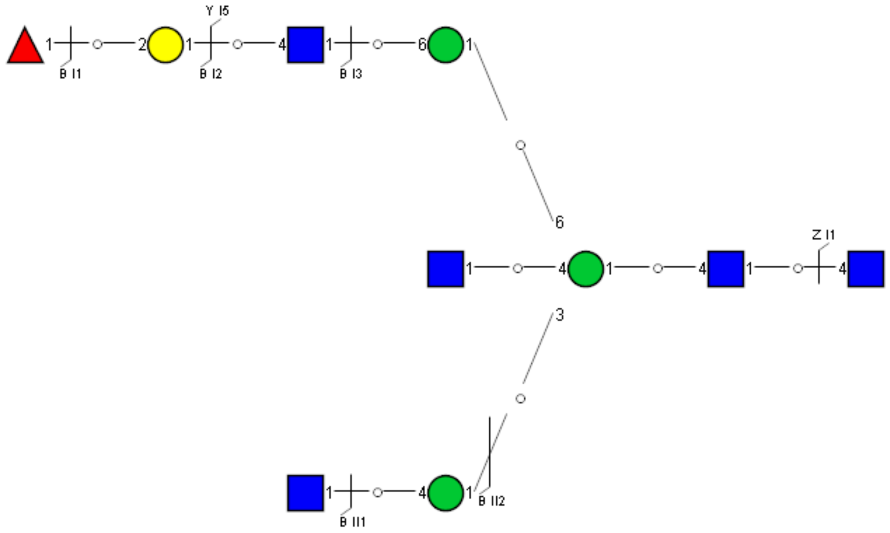

Spectrum Index: 35173(NoBHs:1)

#### Precursor iEF Map

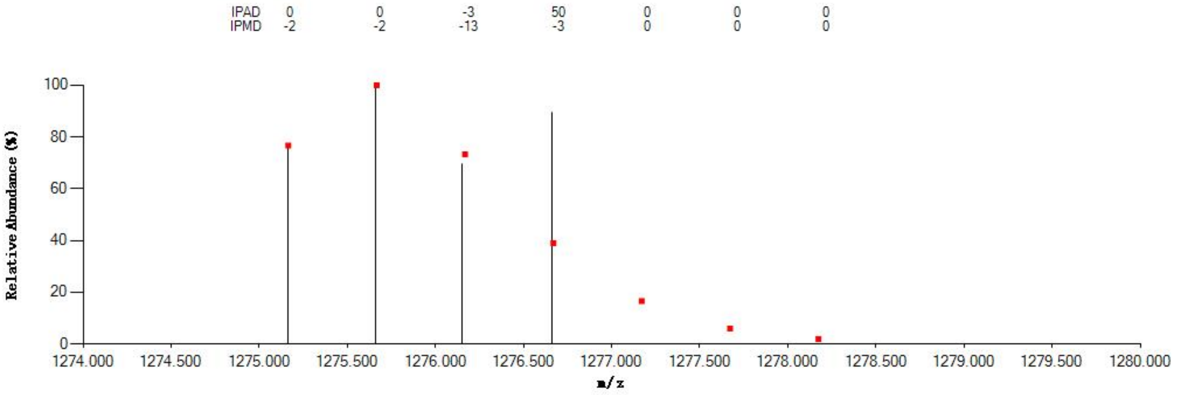


#### Annotated MS/MS Spectrum

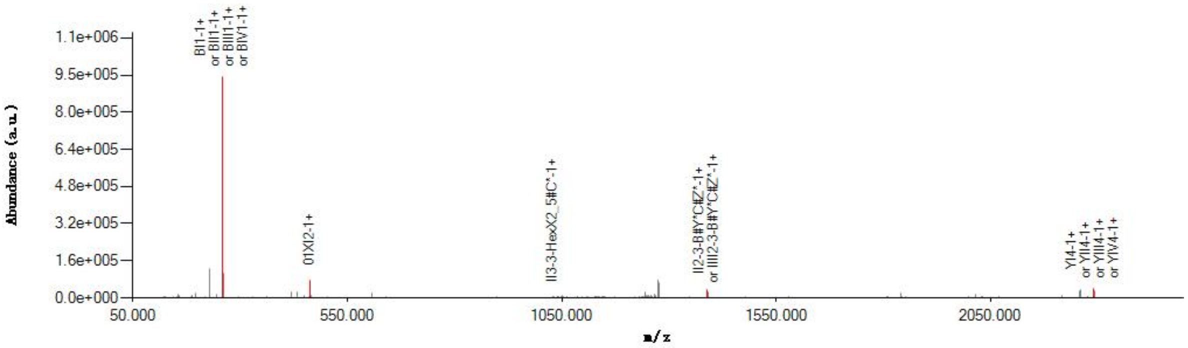


#### Graphical Fragmentation Map

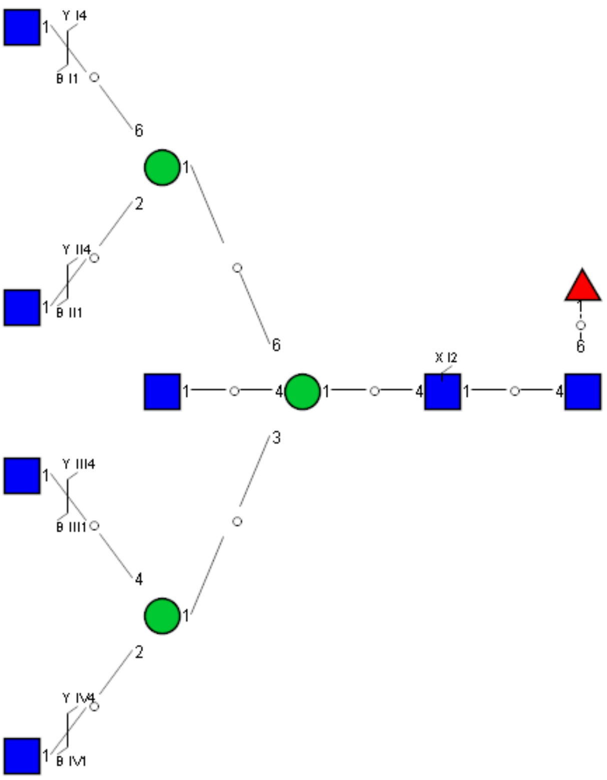

Spectrum Index: 35533(NoBHs:2)

#### Precursor iEF Map

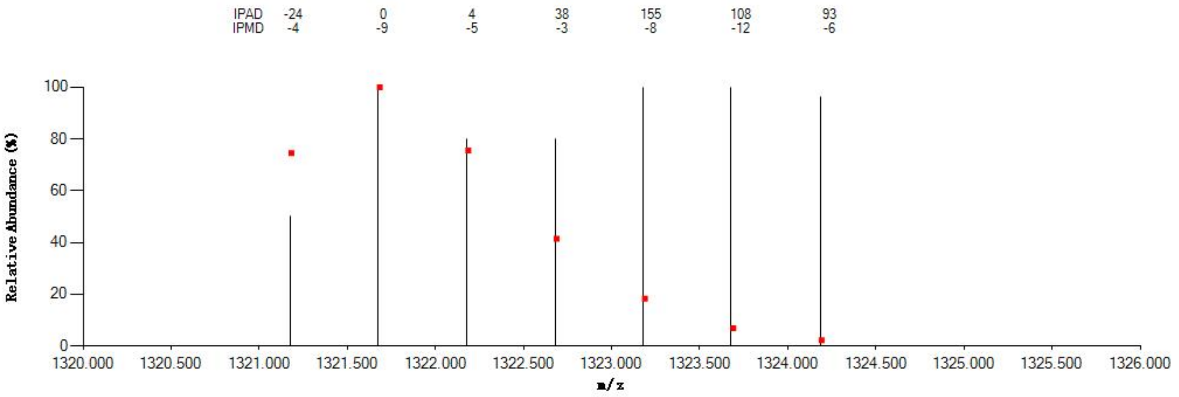


#### Annotated MS/MS Spectrum

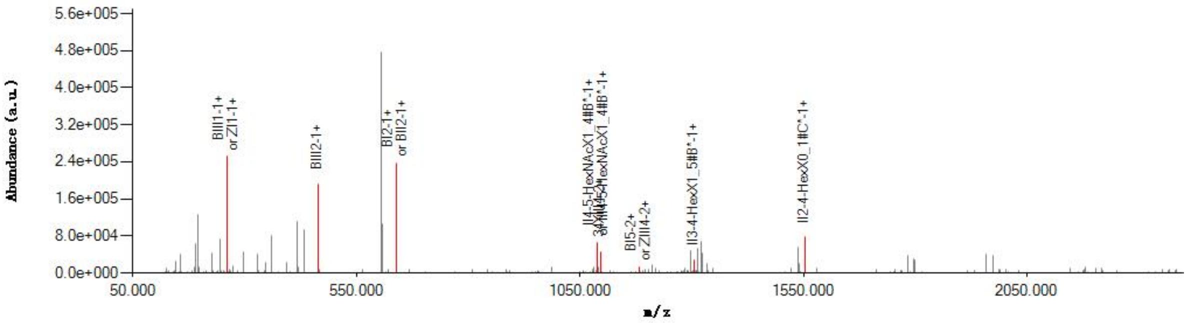


#### Graphical Fragmentation Map

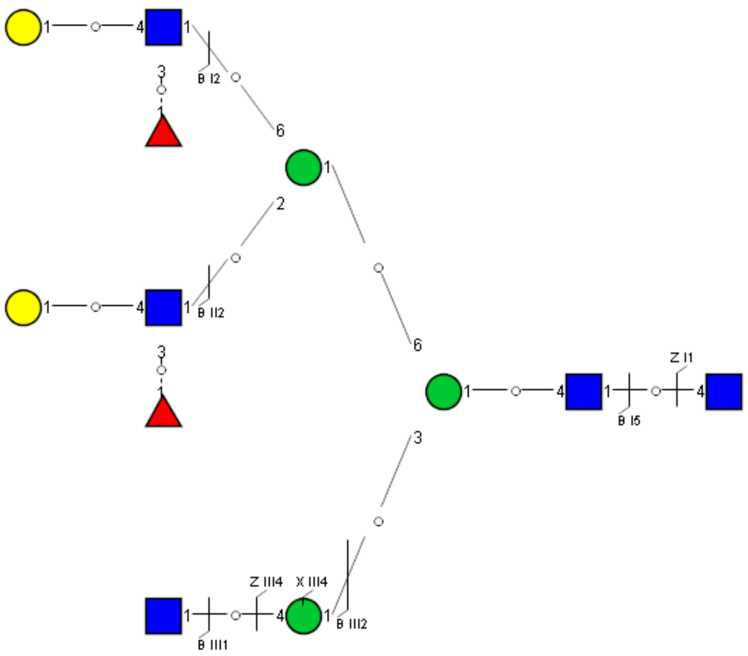

Spectrum Index: 35533(NoBHs:2)

#### Precursor iEF Map

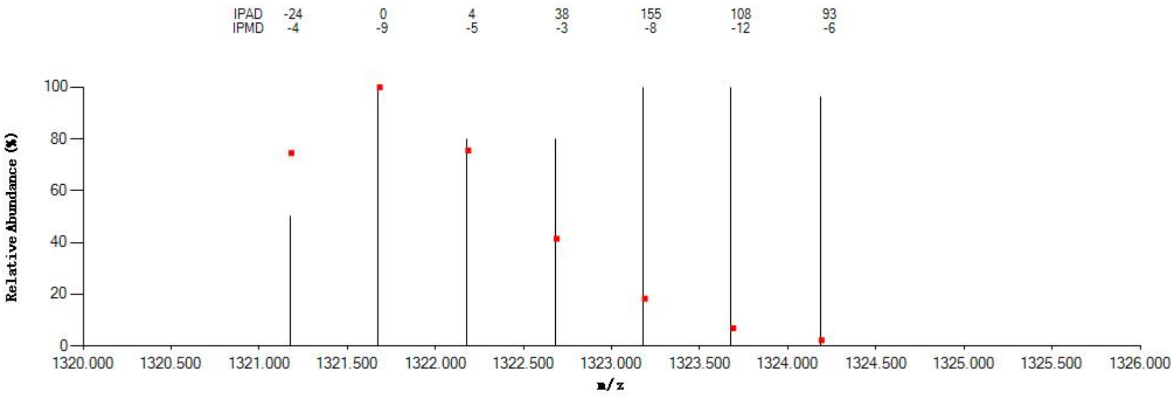


#### Annotated MS/MS Spectrum

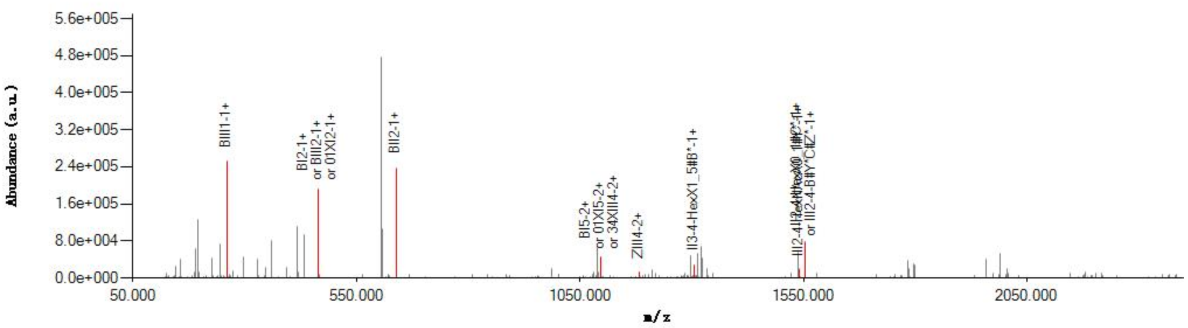


#### Graphical Fragmentation Map

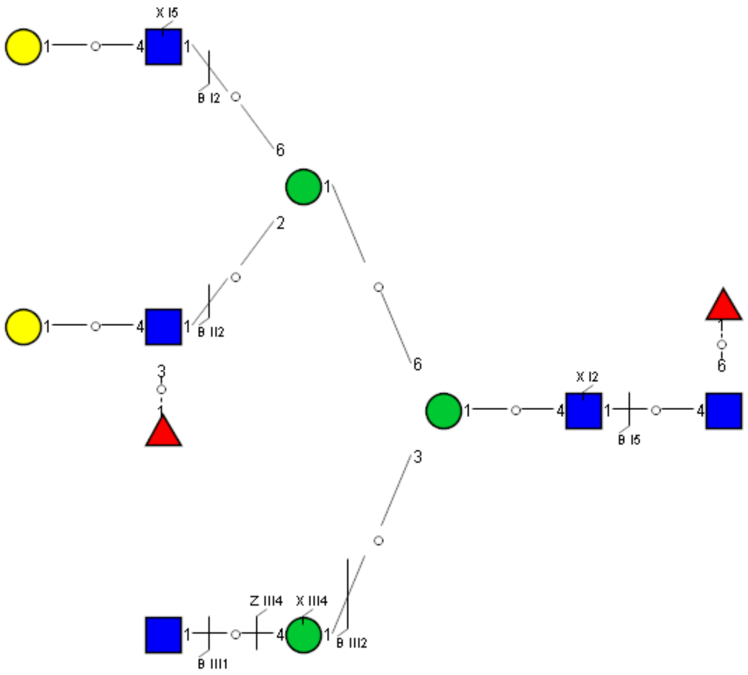

Spectrum Index: 35541(NoBHs:1)

#### Precursor iEF Map

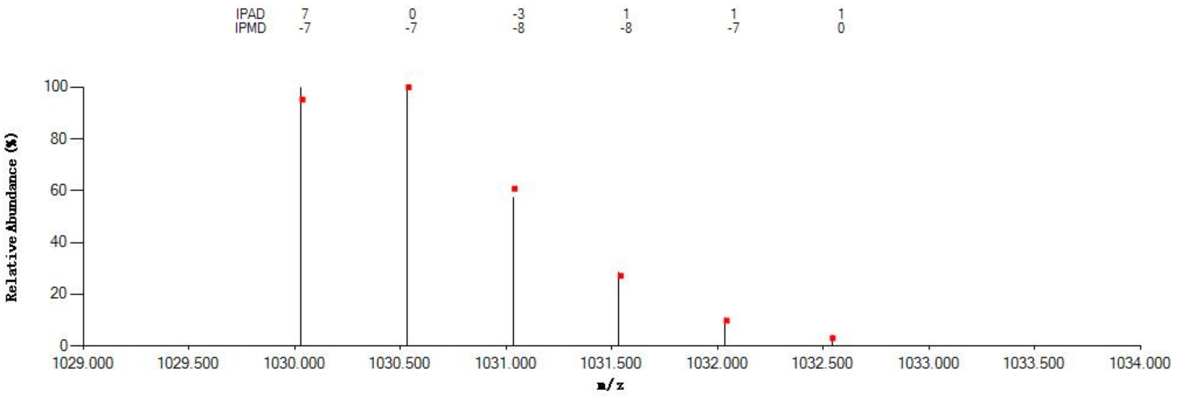


#### Annotated MS/MS Spectrum

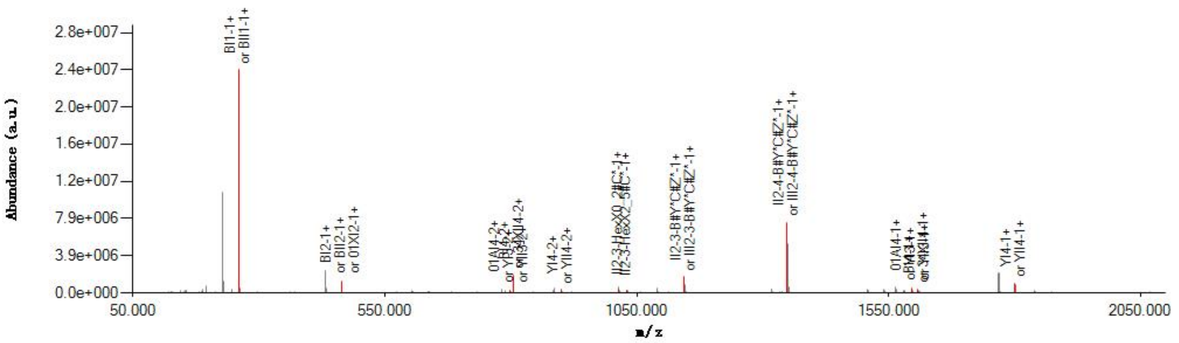


#### Graphical Fragmentation Map

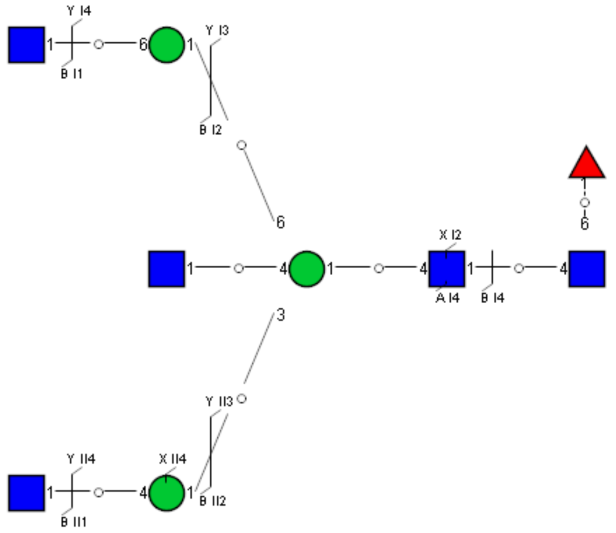

Spectrum Index: 35752(NoBHs:1)

#### Precursor iEF Map

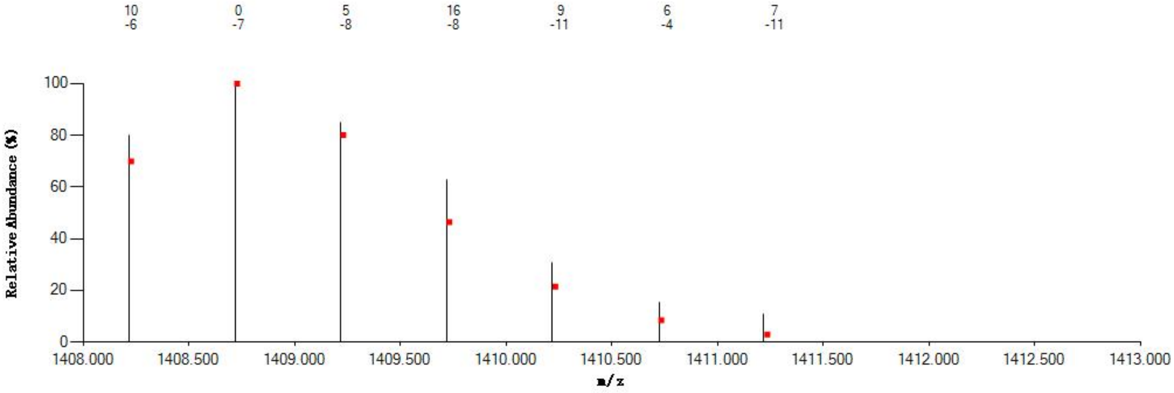

#### Annotated MS/MS Spectrum

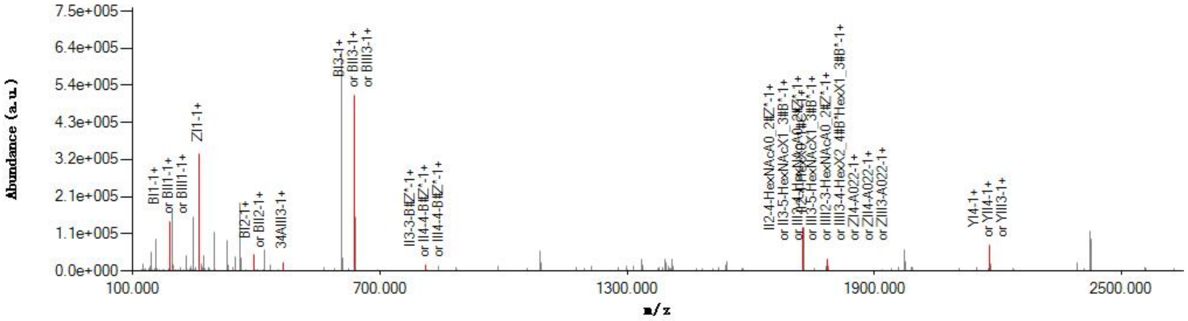

#### Graphical Fragmentation Map

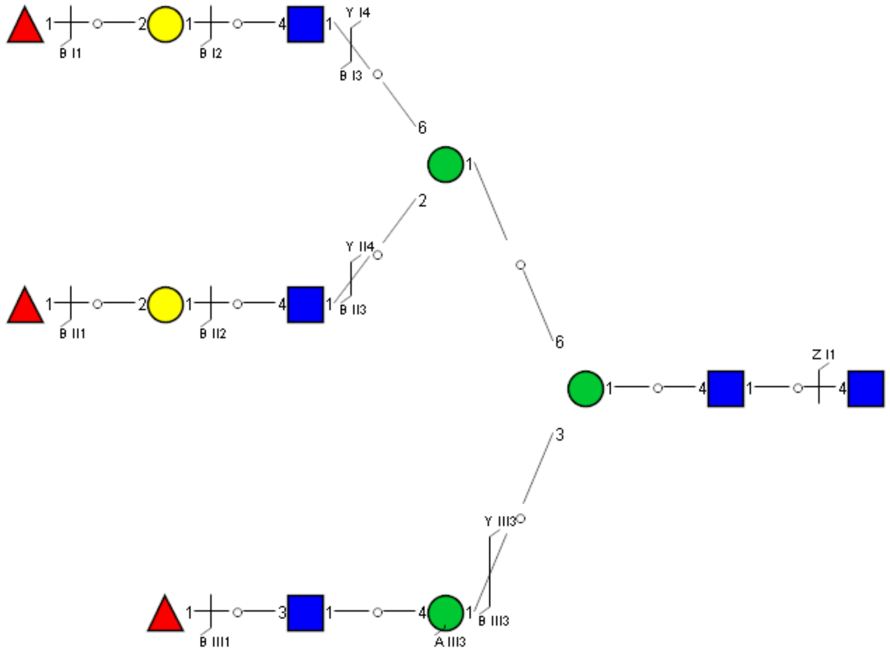

Spectrum Index: 35920(NoBHs:1)

#### Precursor iEF Map

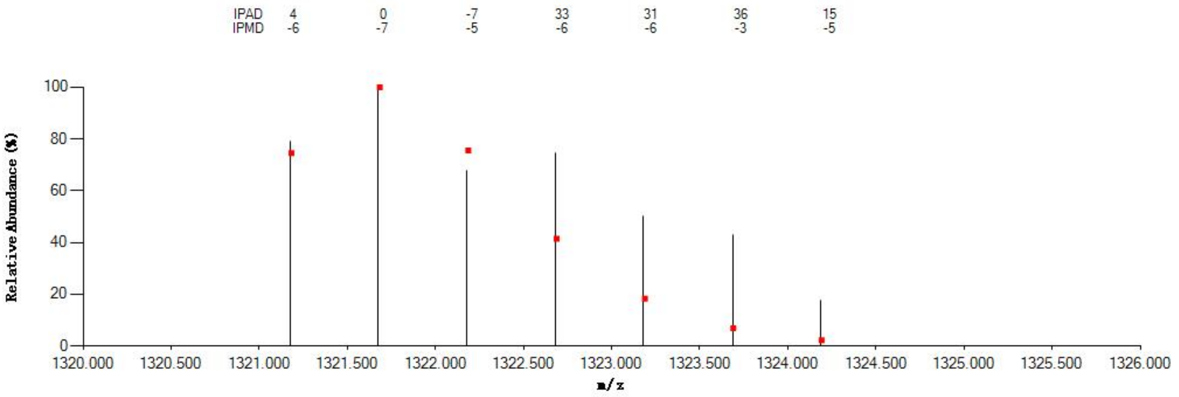


#### Annotated MS/MS Spectrum

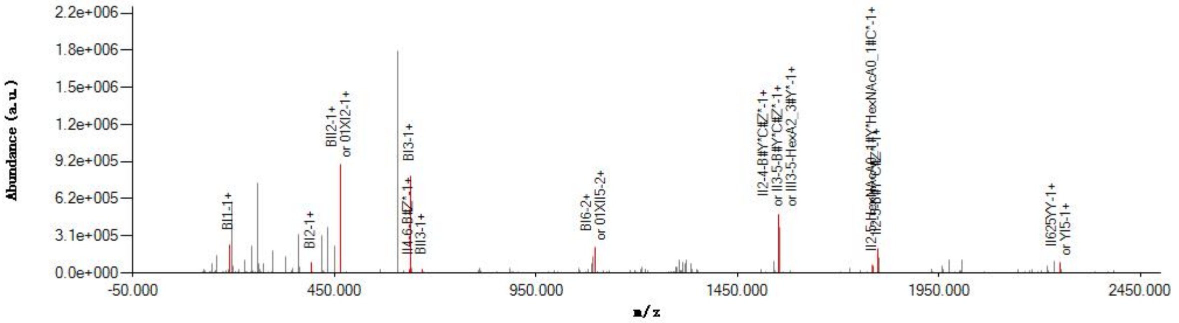


#### Graphical Fragmentation Map

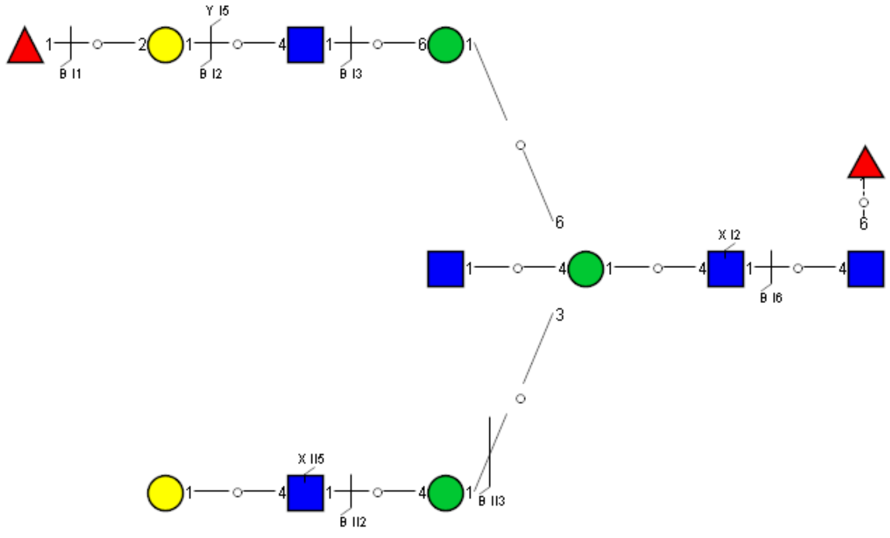

Spectrum Index: 36110(NoBHs:1)

#### Precursor iEF Map

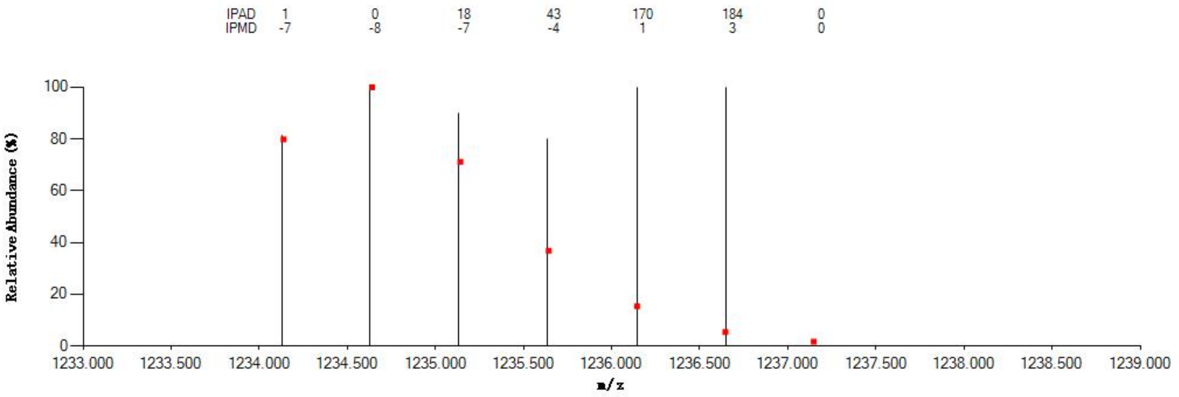


#### Annotated MS/MS Spectrum

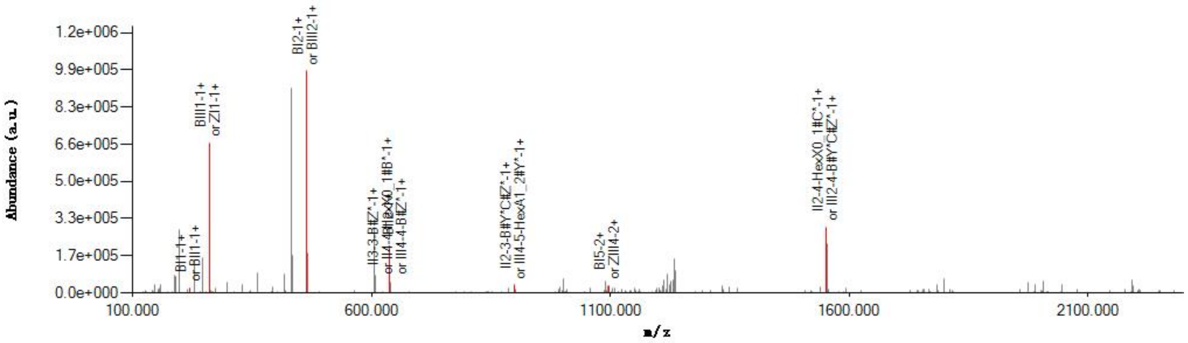


#### Graphical Fragmentation Map

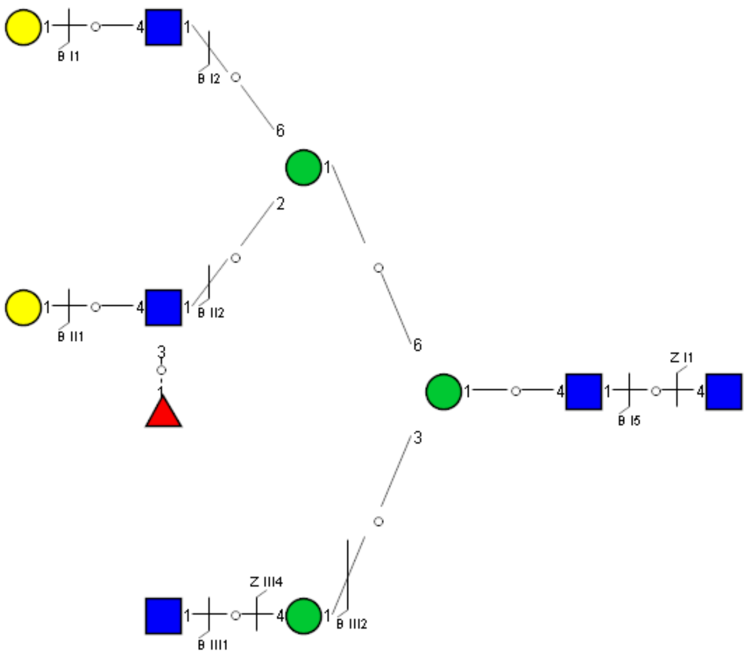

Spectrum Index: 36173(NoBHs:1)

#### Precursor iEF Map

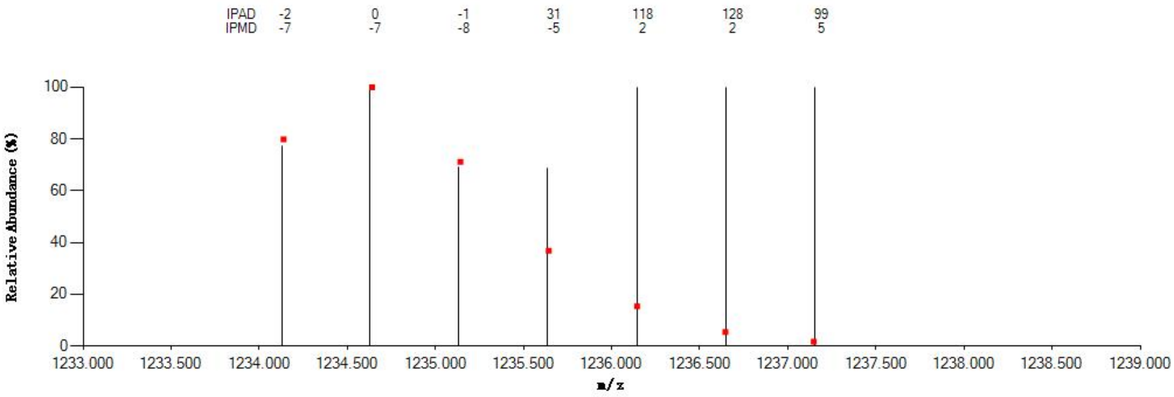


#### Annotated MS/MS Spectrum

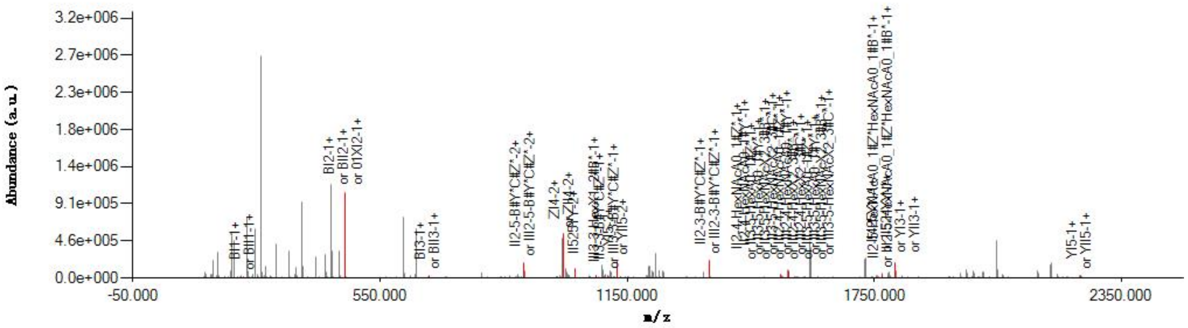


#### Graphical Fragmentation Map

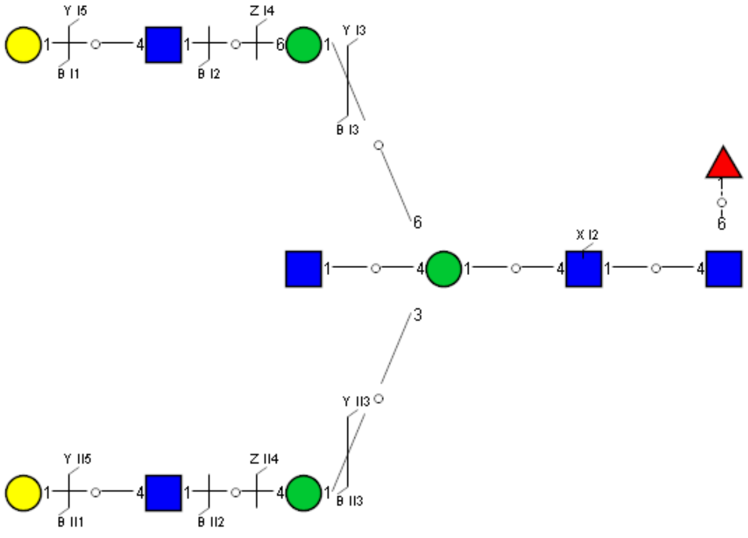

Spectrum Index: 36343(NoBHs:1)

#### Precursor iEF Map

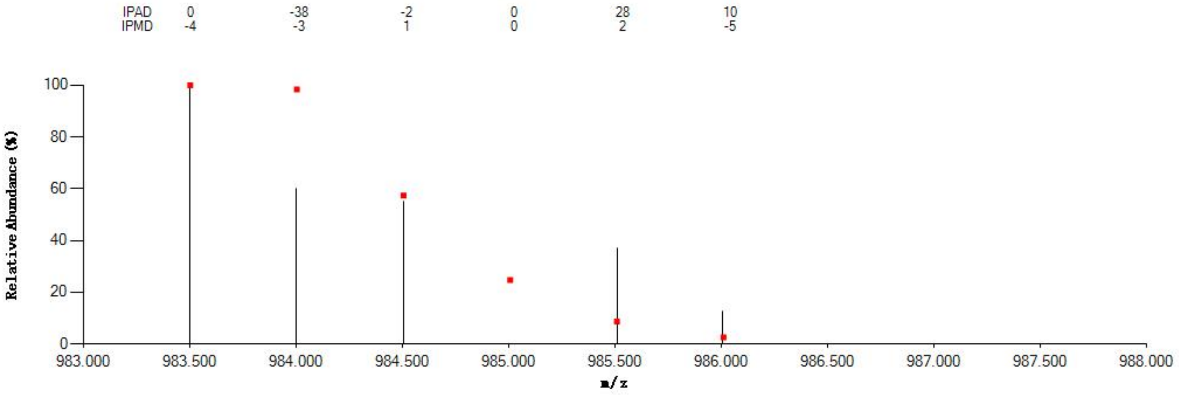

#### Annotated MS/MS Spectrum

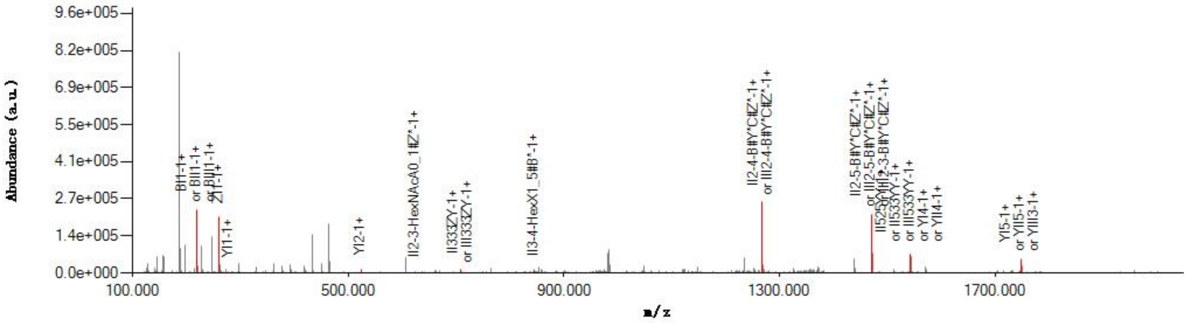

#### Graphical Fragmentation Map

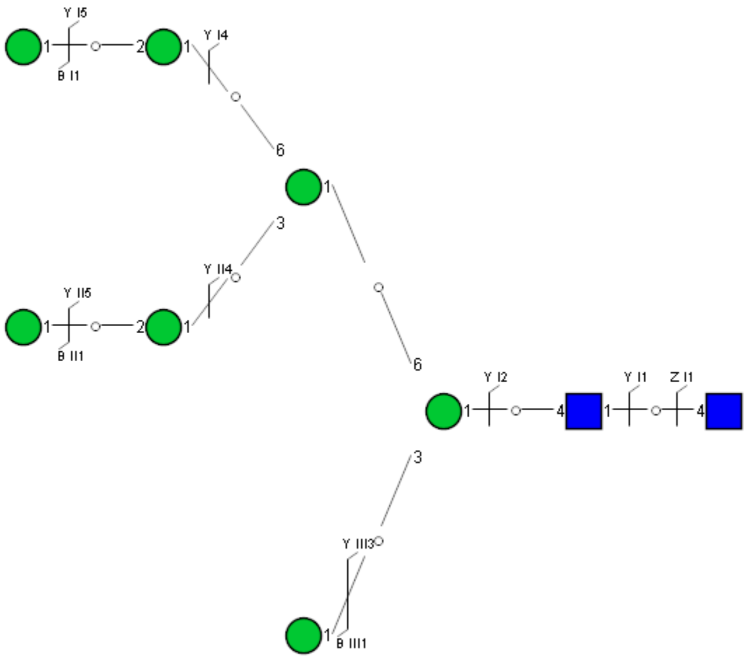

Spectrum Index: 36389(NoBHs:1)

#### Precursor iEF Map

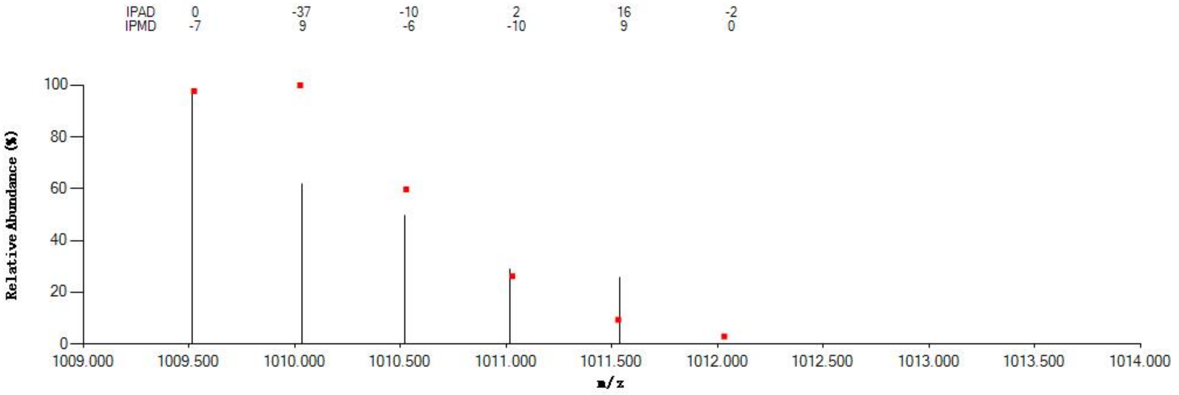


#### Annotated MS/MS Spectrum

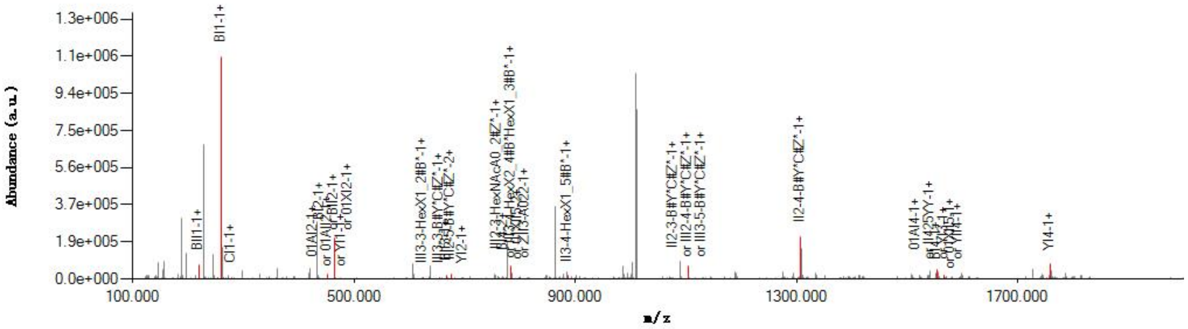


#### Graphical Fragmentation Map

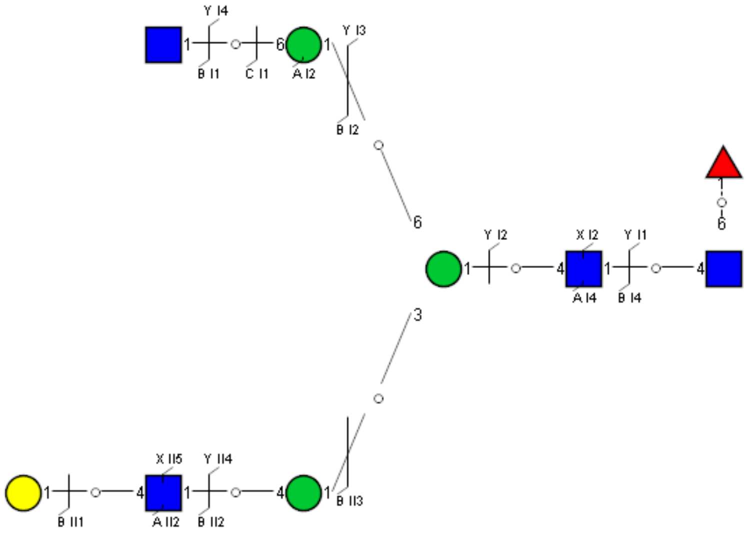

Spectrum Index: 36544(NoBHs:1)

#### Precursor iEF Map

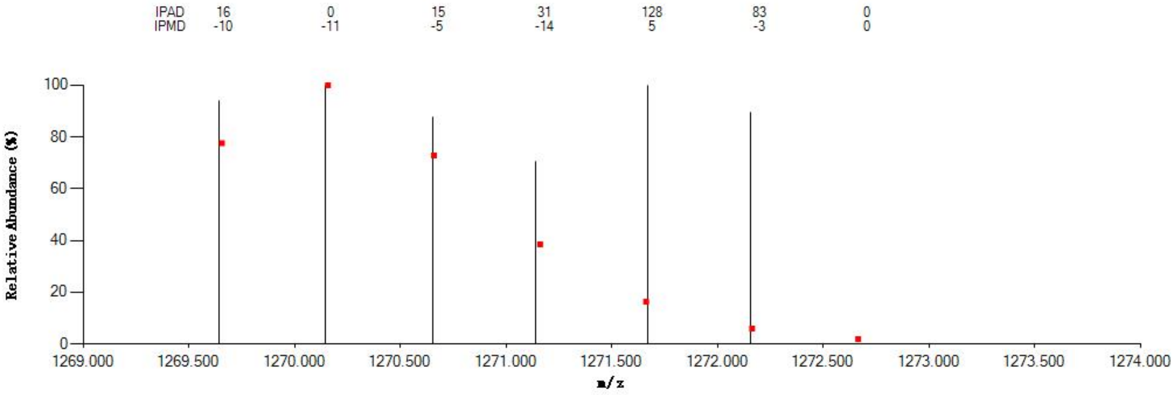

#### Annotated MS/MS Spectrum

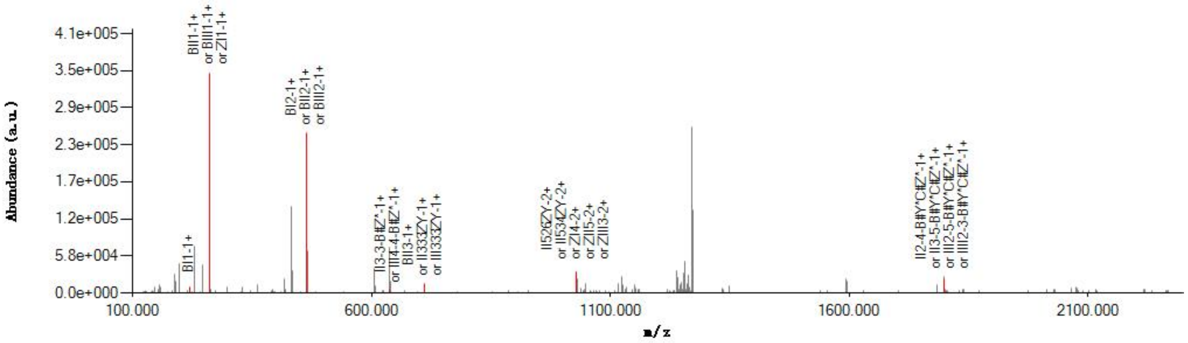

#### Graphical Fragmentation Map

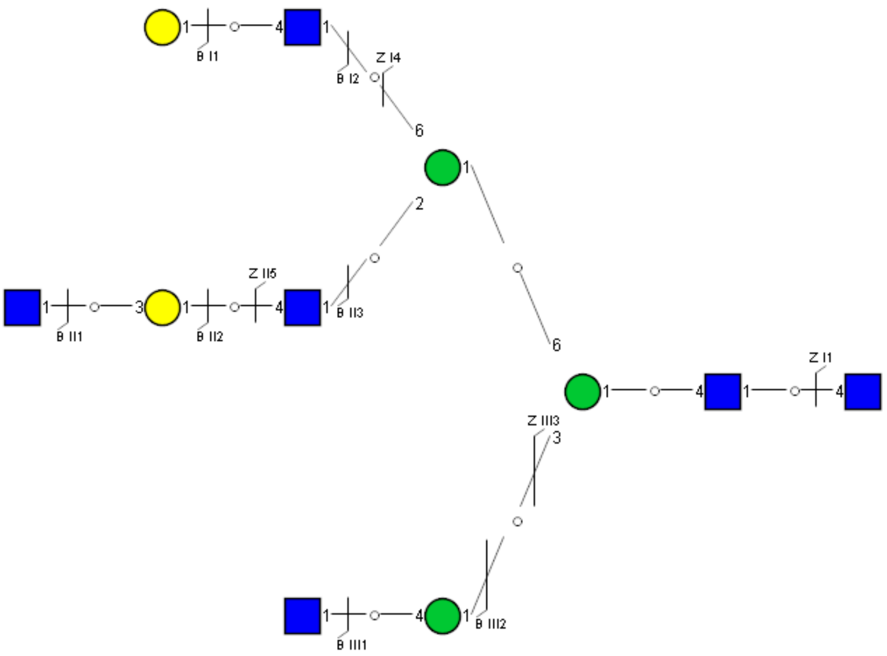

Spectrum Index: 36633(NoBHs:1)

#### Precursor iEF Map

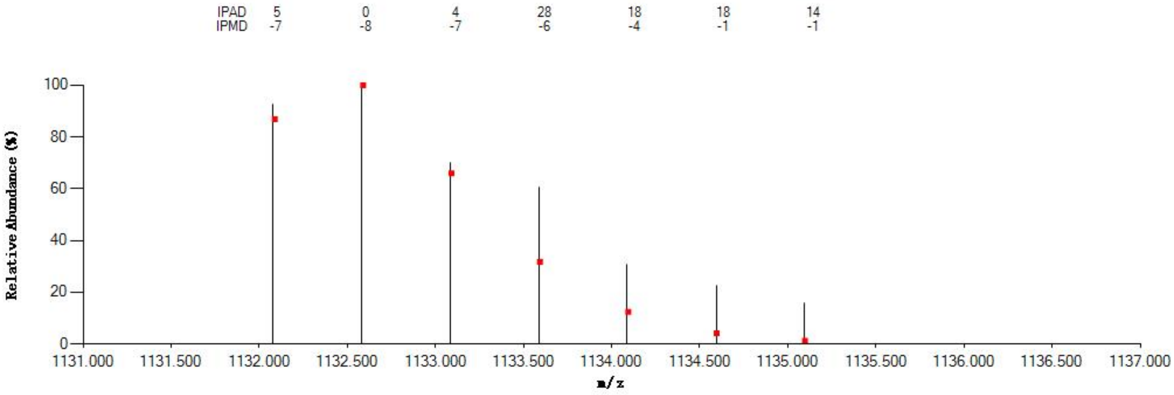

#### Annotated MS/MS Spectrum

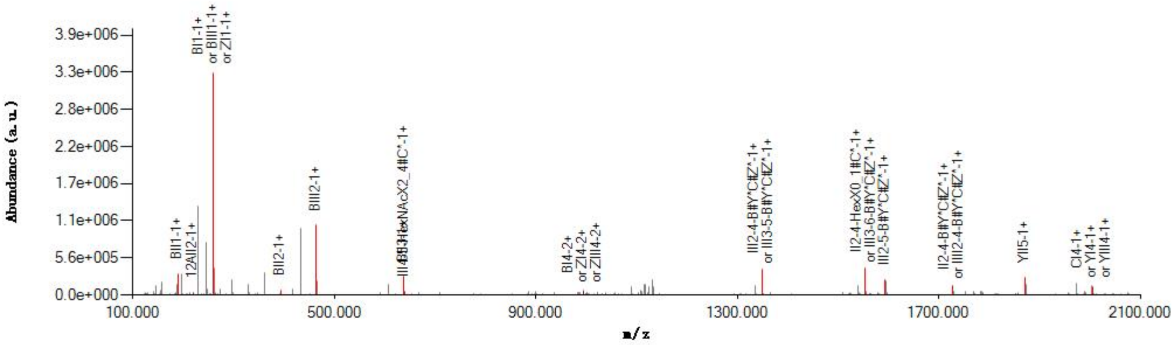

#### Graphical Fragmentation Map

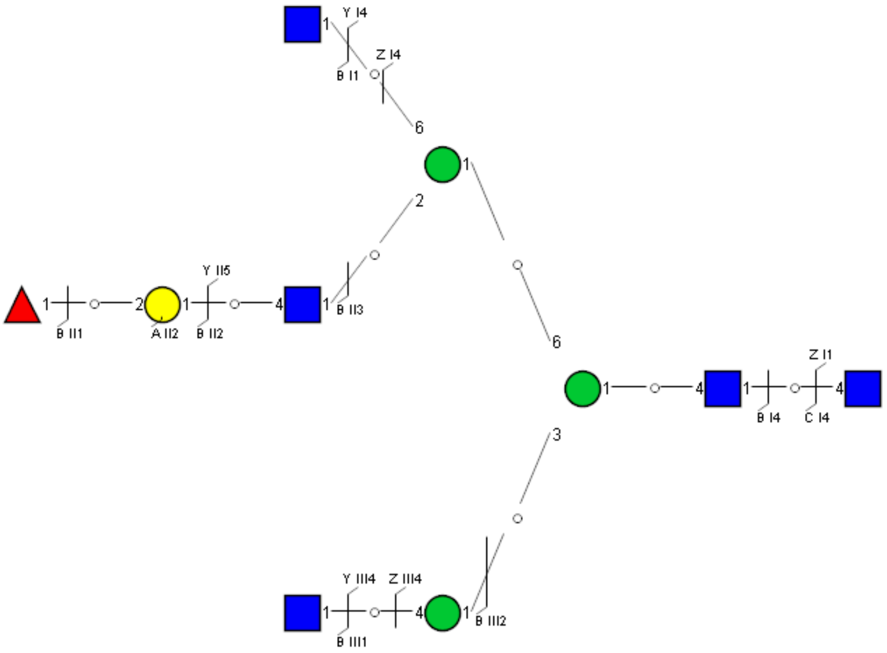

Spectrum Index: 36946(NoBHs:1)

#### Precursor iEF Map

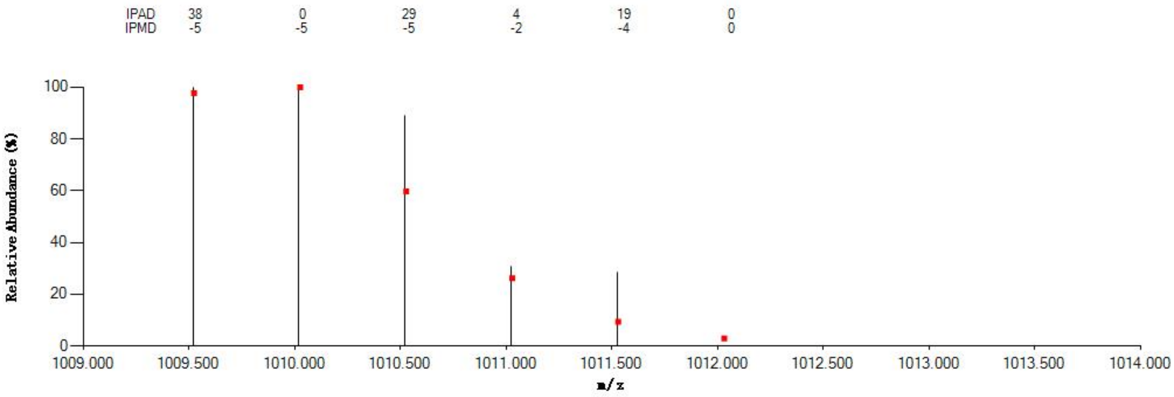

#### Annotated MS/MS Spectrum

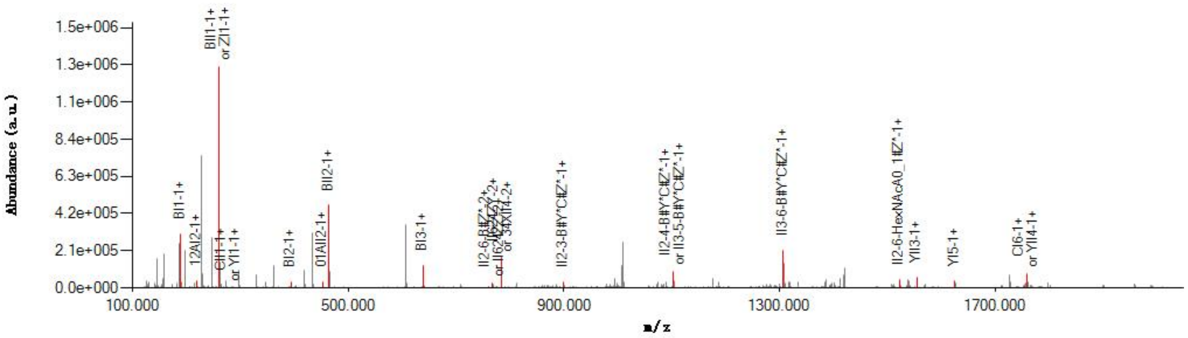

#### Graphical Fragmentation Map

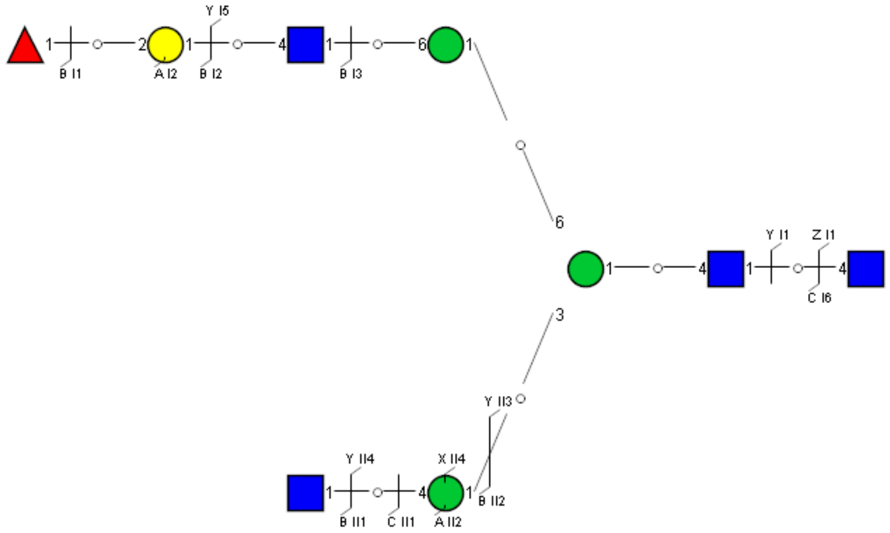

Spectrum Index: 37016(NoBHs:2)

#### Precursor iEF Map

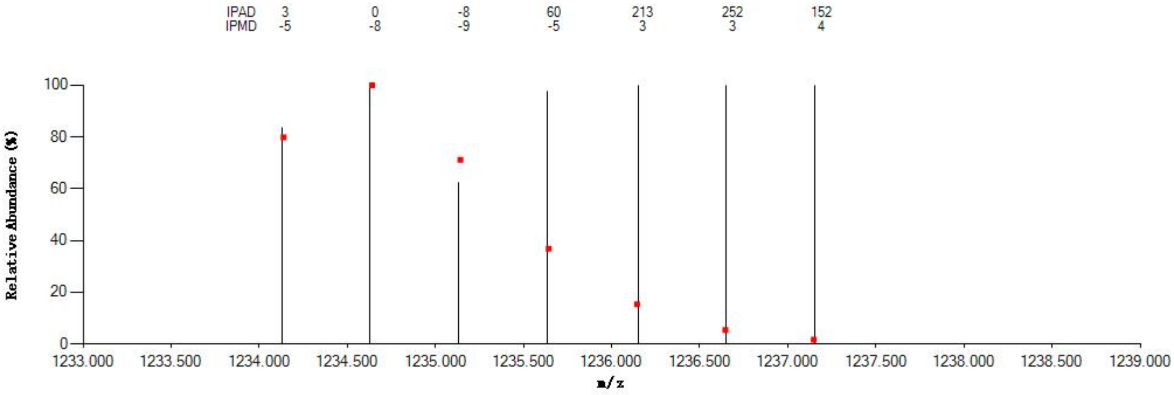


#### Annotated MS/MS Spectrum

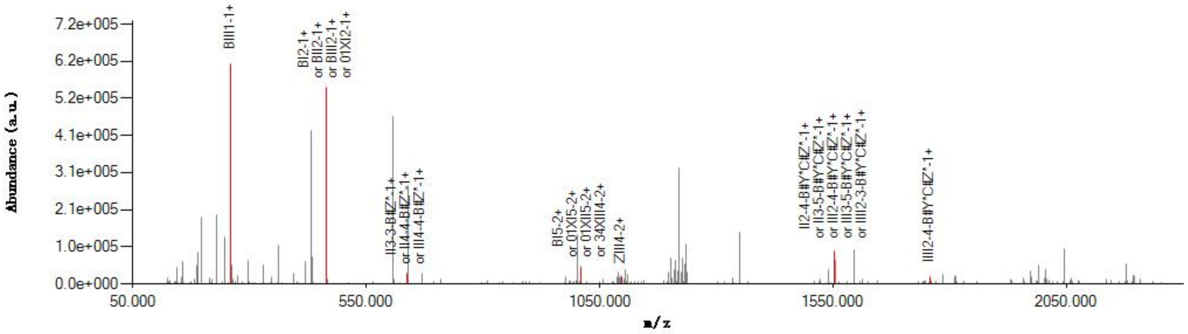


#### Graphical Fragmentation Map

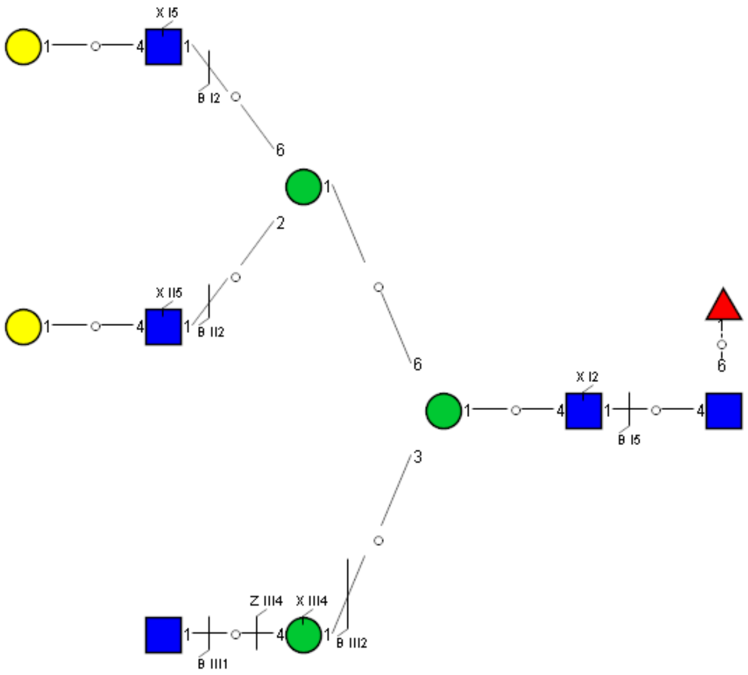

Spectrum Index: 37018(NoBHs:1)

#### Precursor iEF Map

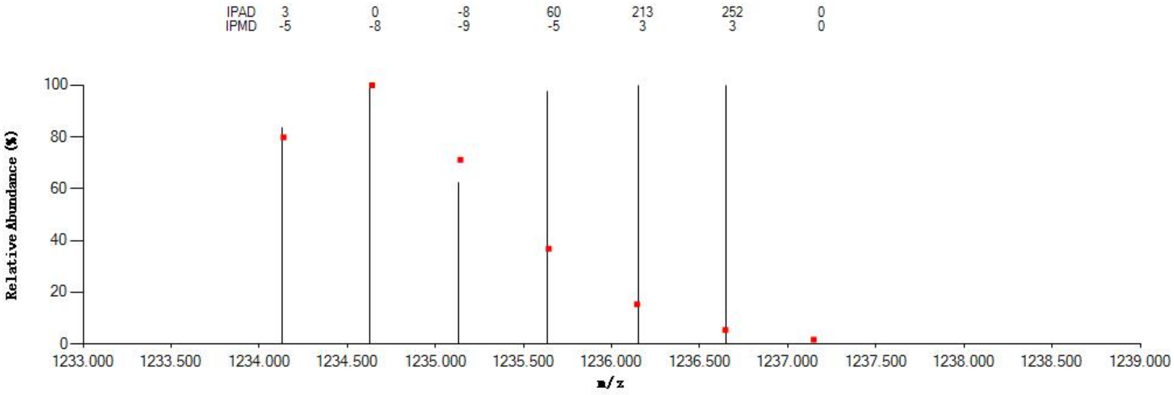


#### Annotated MS/MS Spectrum

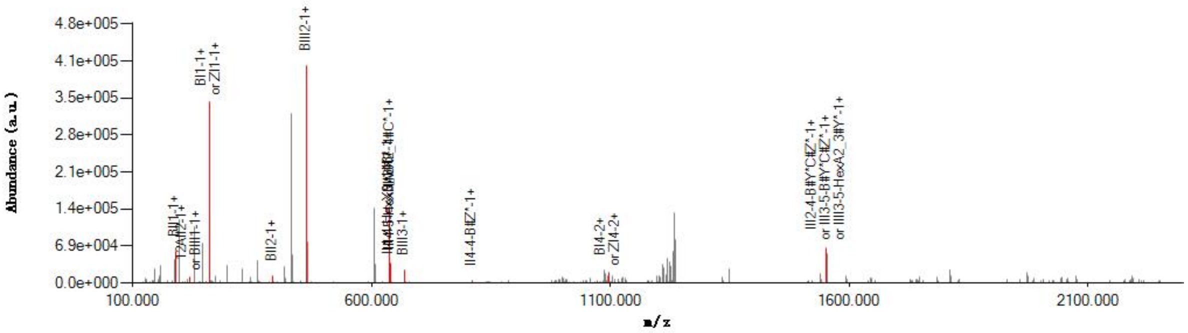


#### Graphical Fragmentation Map

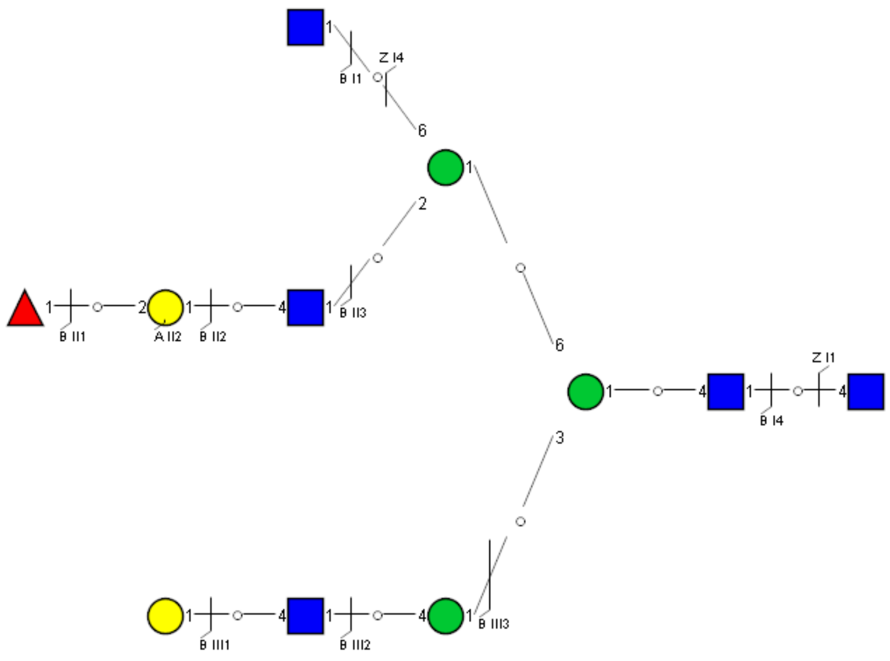

Spectrum Index: 37106(NoBHs:1)

#### Precursor iEF Map

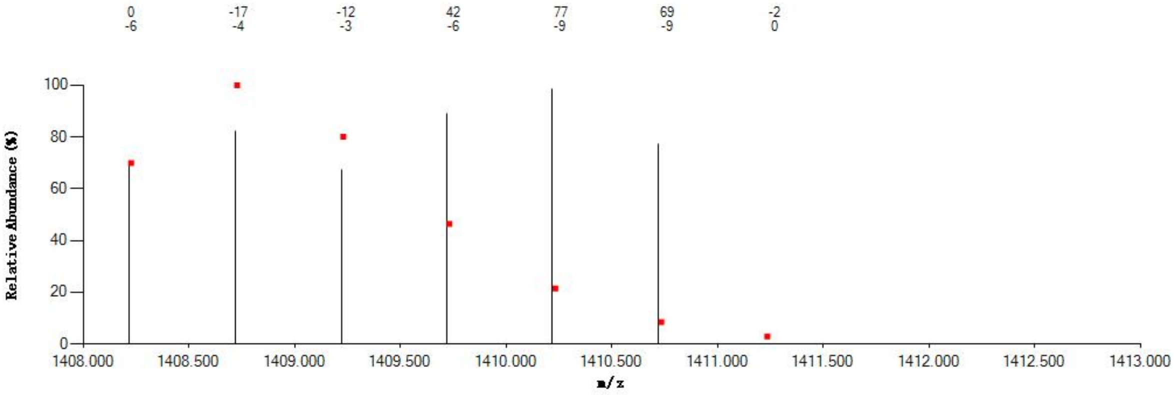

#### Annotated MS/MS Spectrum

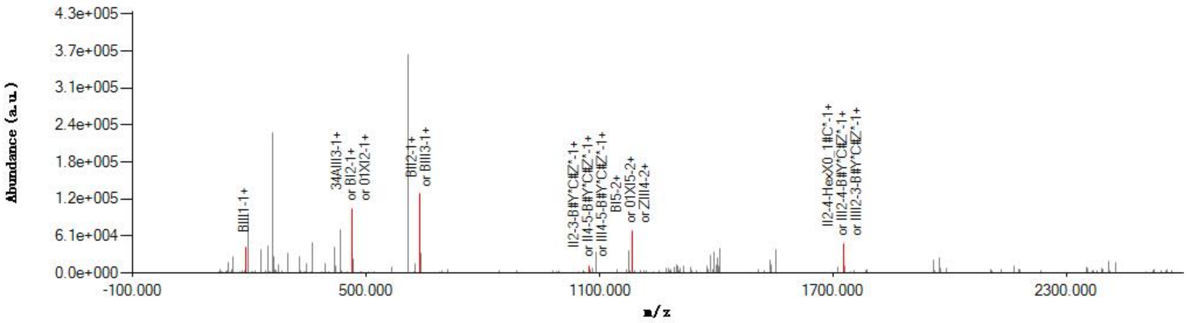

#### Graphical Fragmentation Map

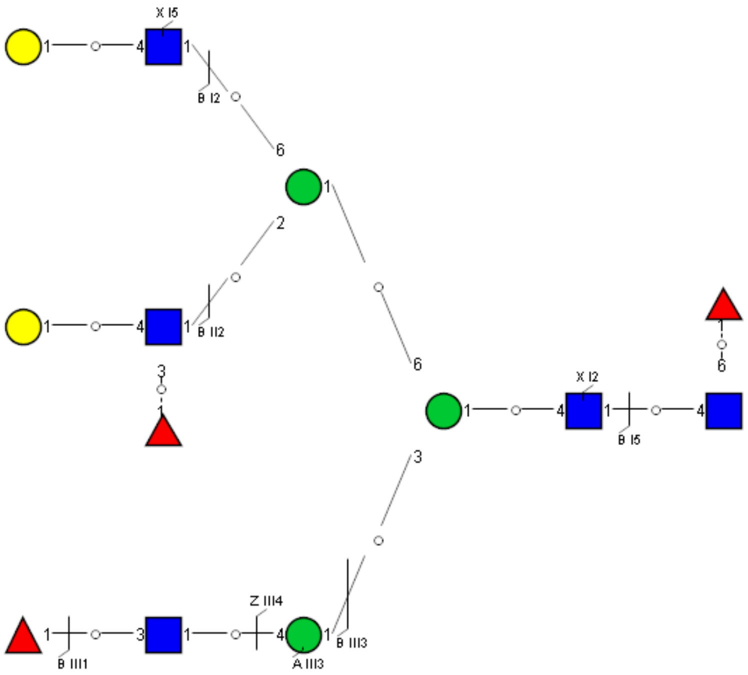

Spectrum Index: 37183(NoBHs:1)

#### Precursor iEF Map

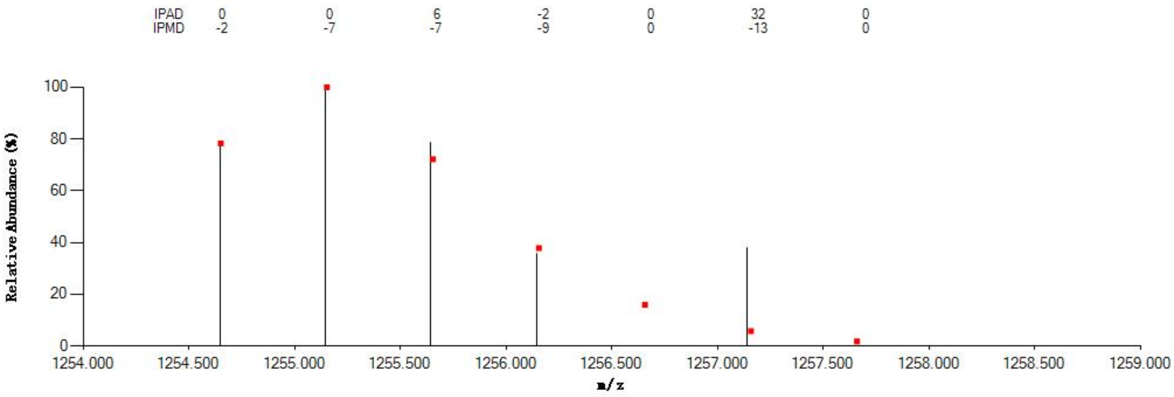

#### Annotated MS/MS Spectrum

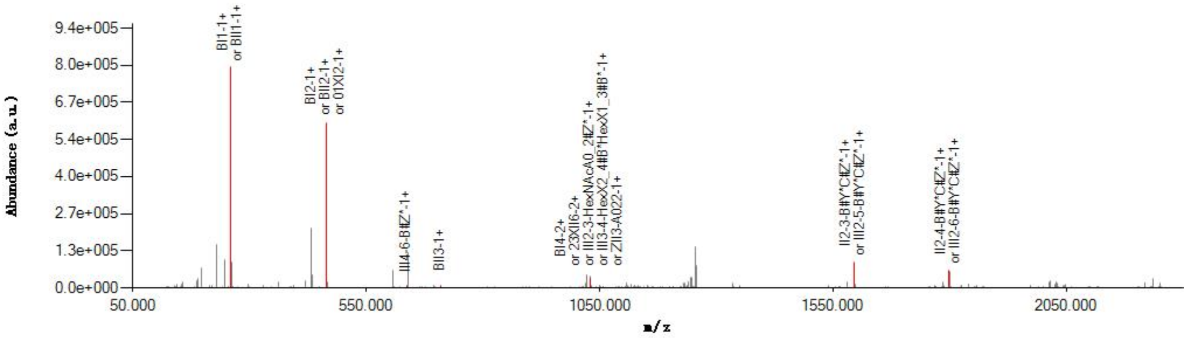

#### Graphical Fragmentation Map

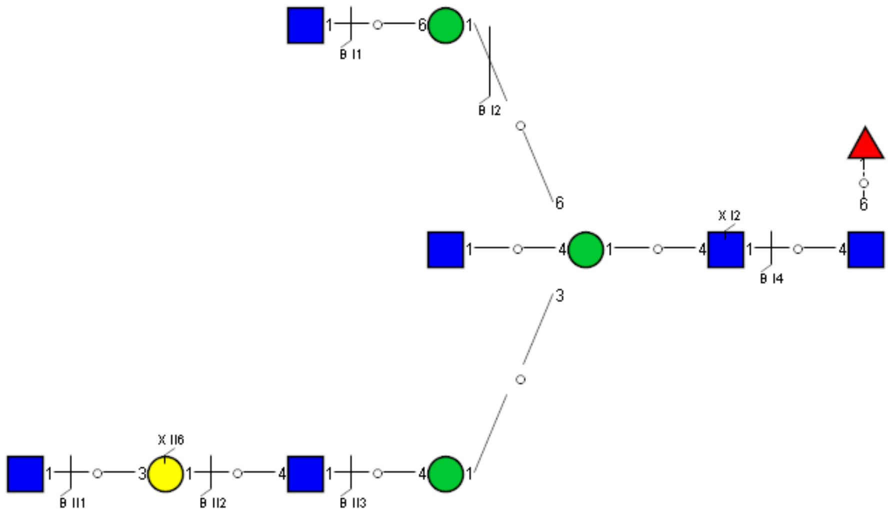

Spectrum Index: 37245(NoBHs:1)

#### Precursor iEF Map

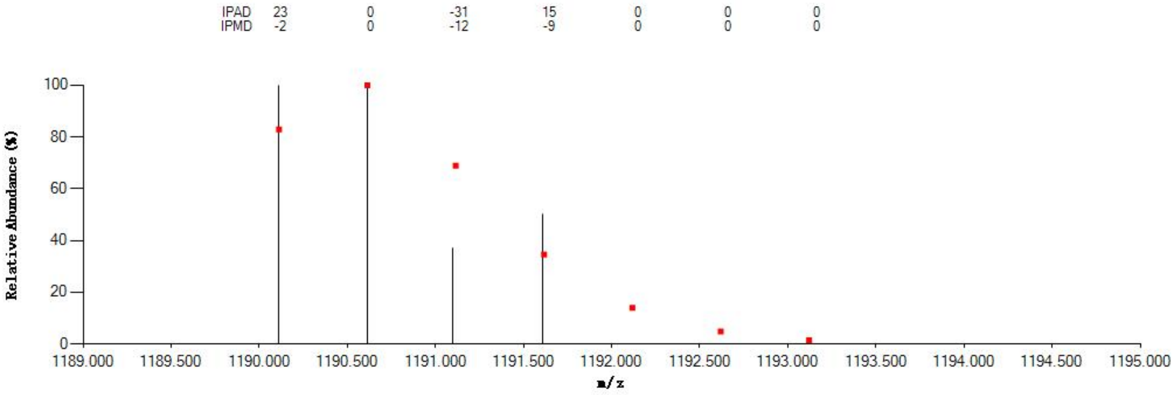


#### Annotated MS/MS Spectrum

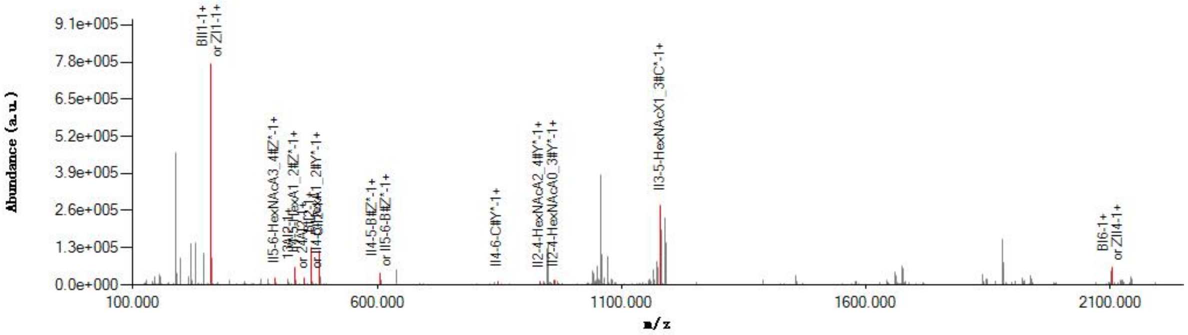


#### Graphical Fragmentation Map

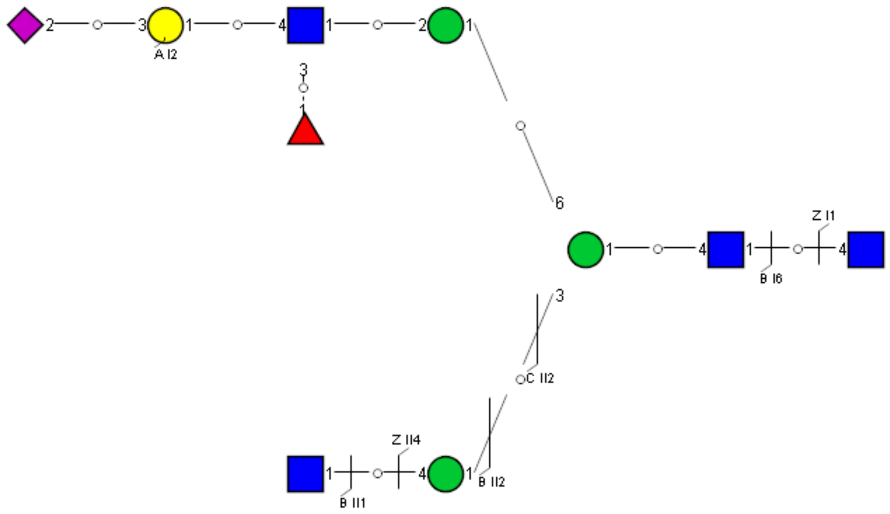

Spectrum Index: 37284(NoBHs:1)

#### Precursor iEF Map

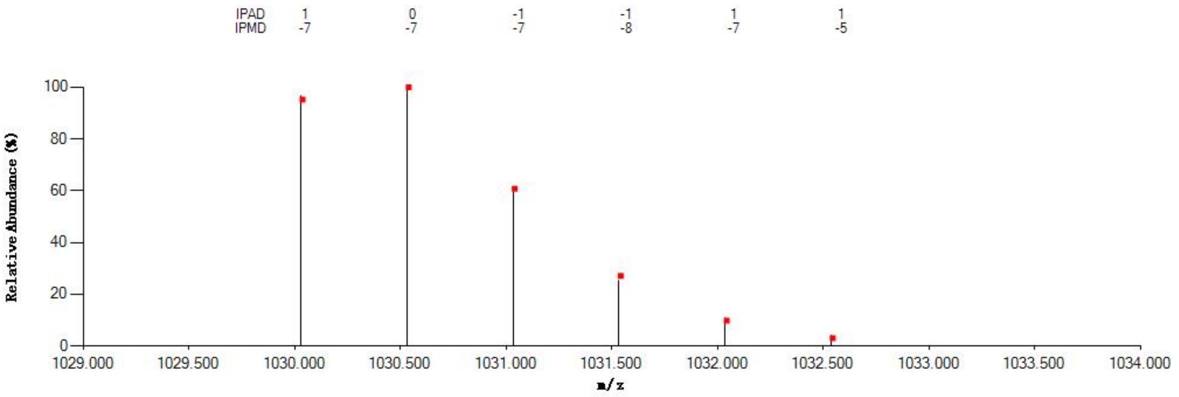


#### Annotated MS/MS Spectrum

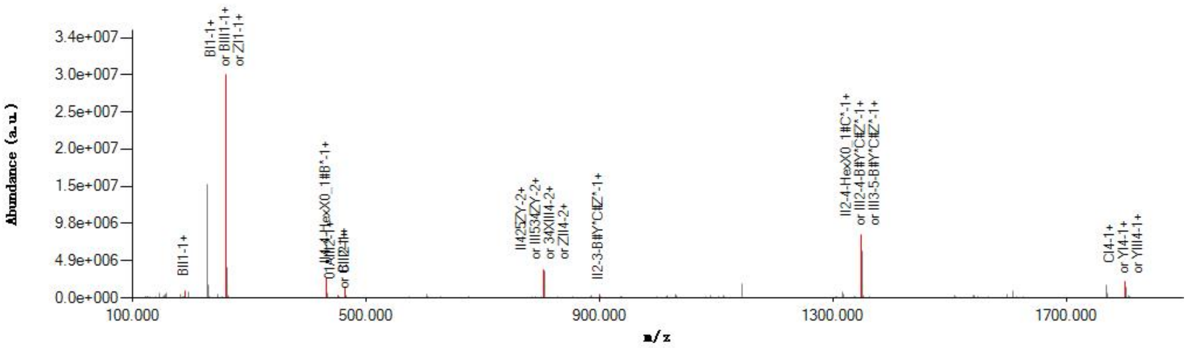


#### Graphical Fragmentation Map

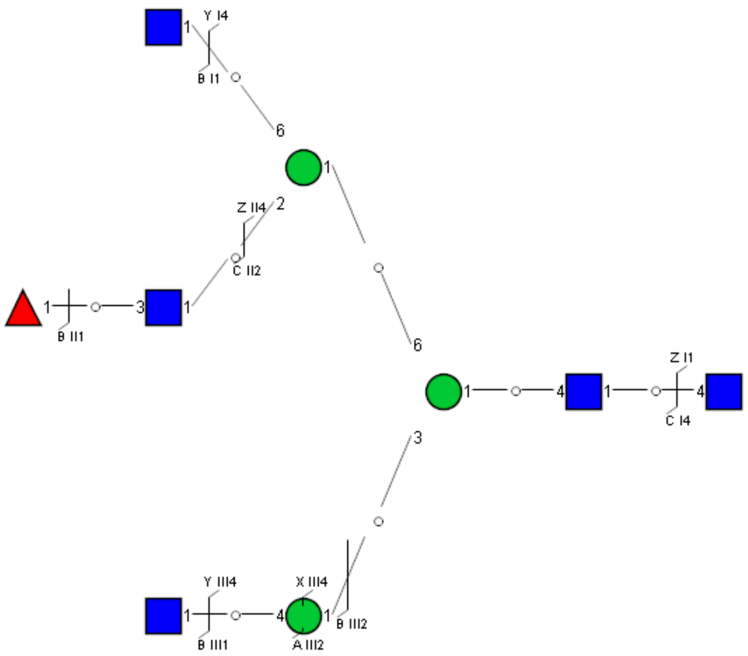

Spectrum Index: 37326(NoBHs:1)

Precursor iEF Map

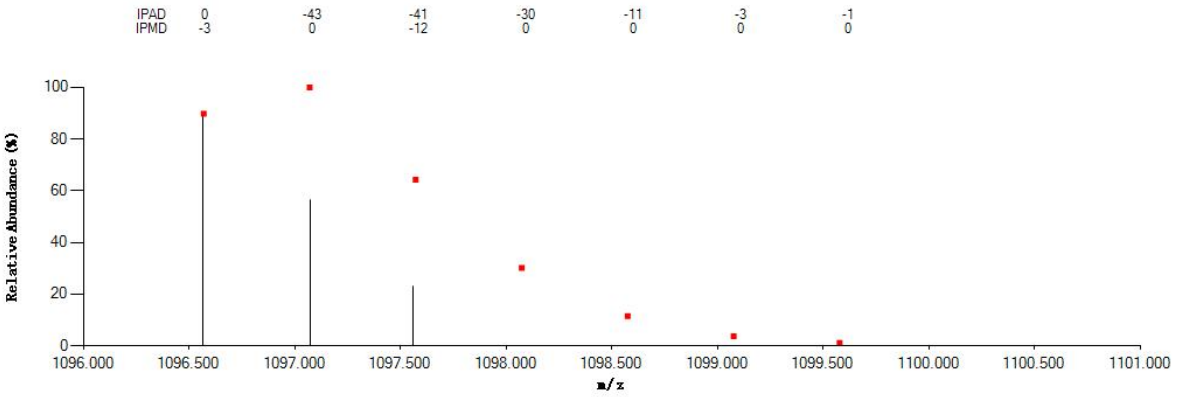


Annotated MS/MS Spectrum

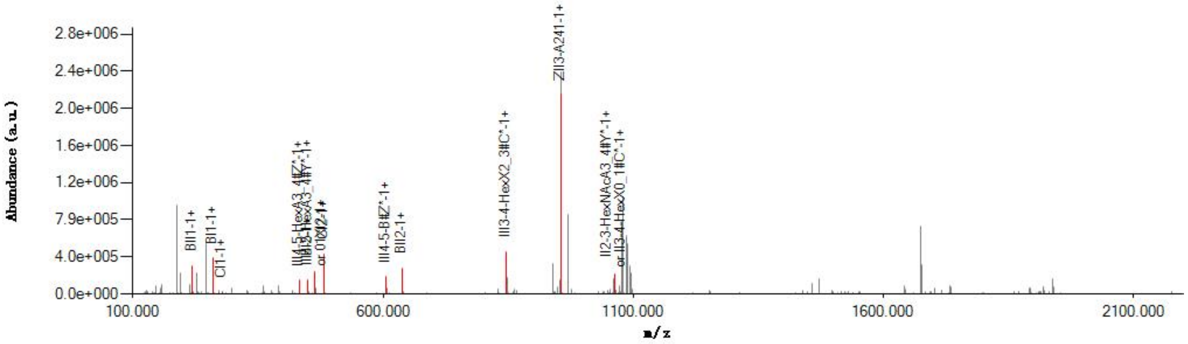


Graphical Fragmentation Map

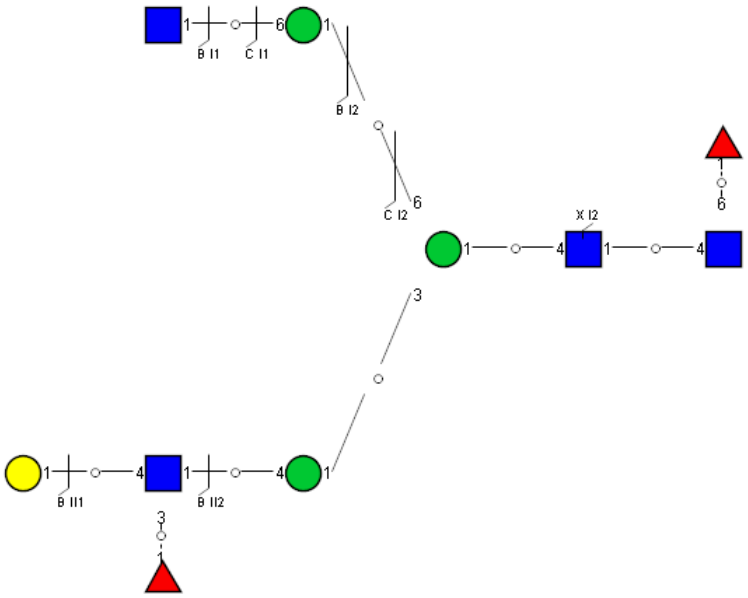

Spectrum Index: 37368(NoBHs:1)

#### Precursor iEF Map

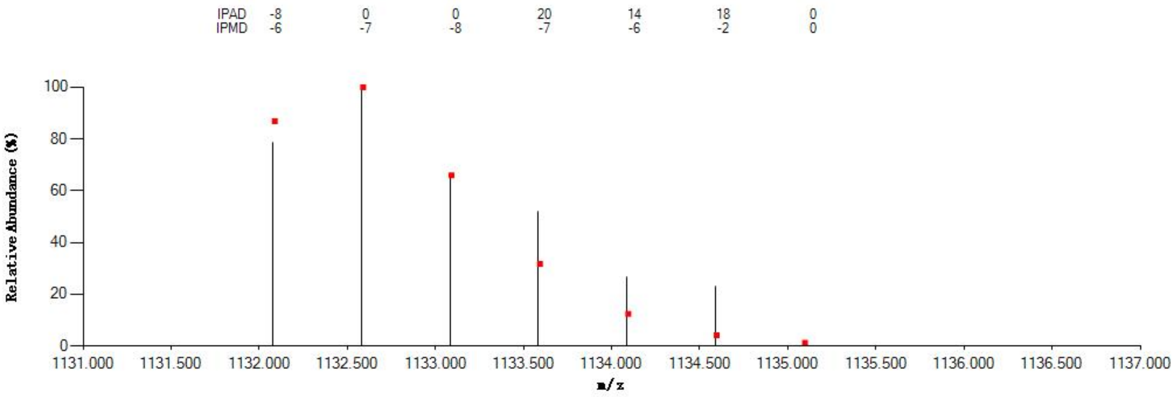


#### Annotated MS/MS Spectrum

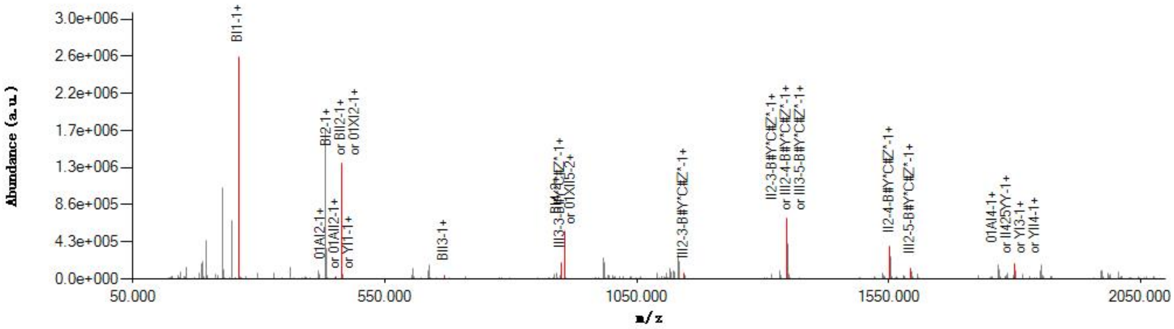


#### Graphical Fragmentation Map

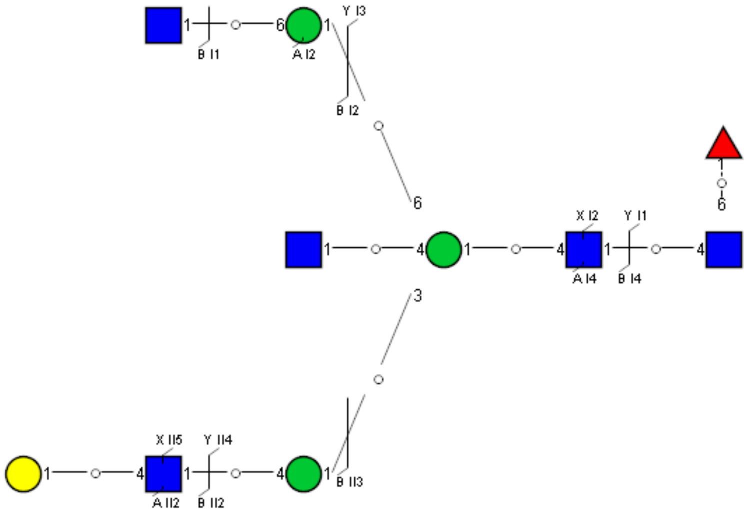

Spectrum Index: 37411(NoBHs:1)

#### Precursor iEF Map

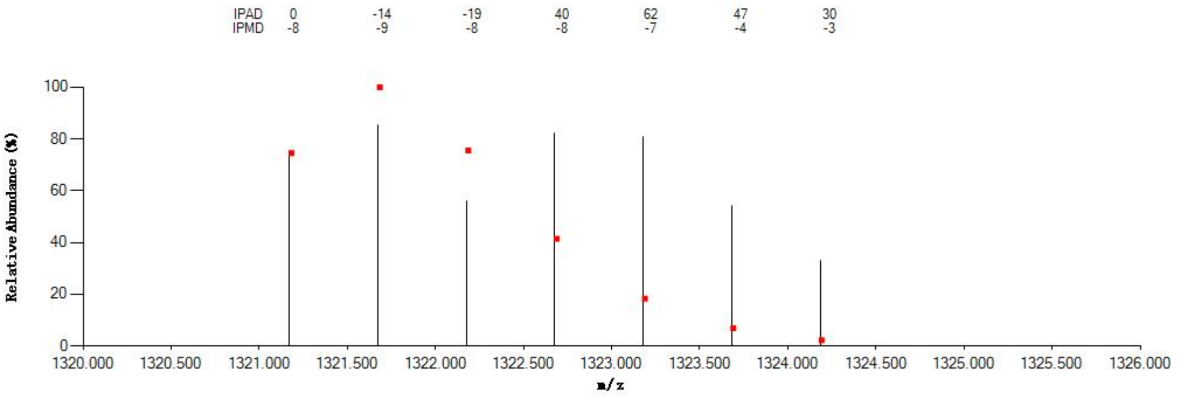


#### Annotated MS/MS Spectrum

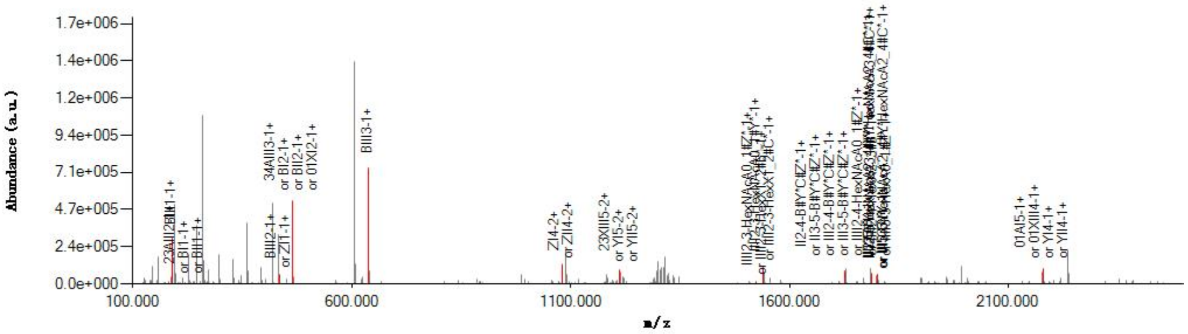


#### Graphical Fragmentation Map

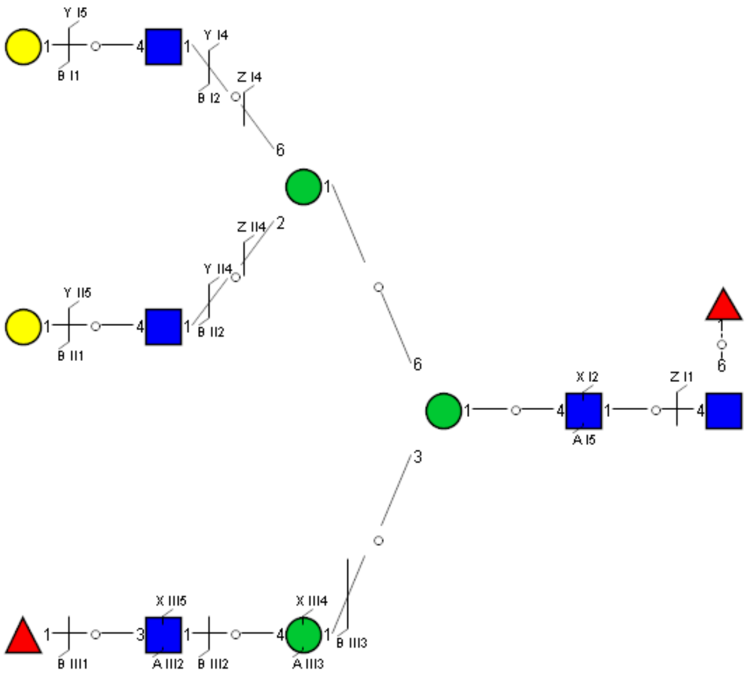

Spectrum Index: 37412(NoBHs:1)

#### Precursor iEF Map

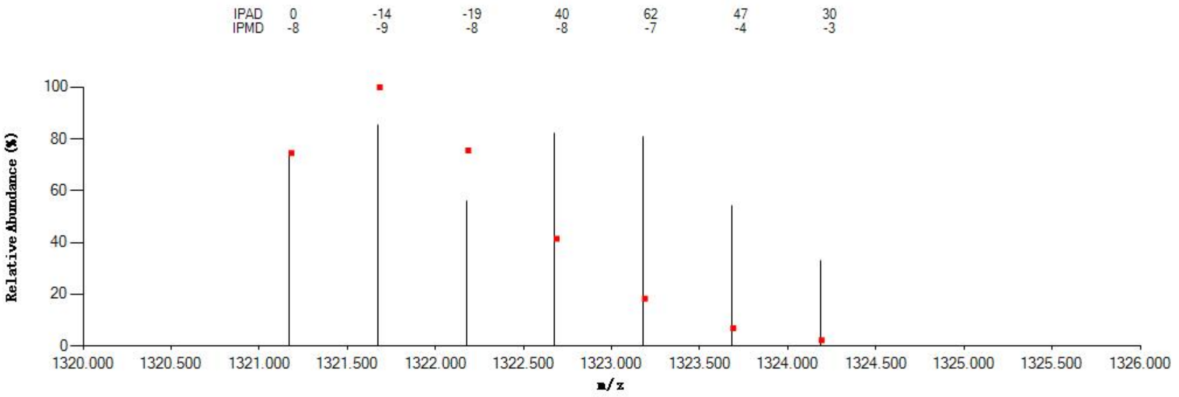


#### Annotated MS/MS Spectrum

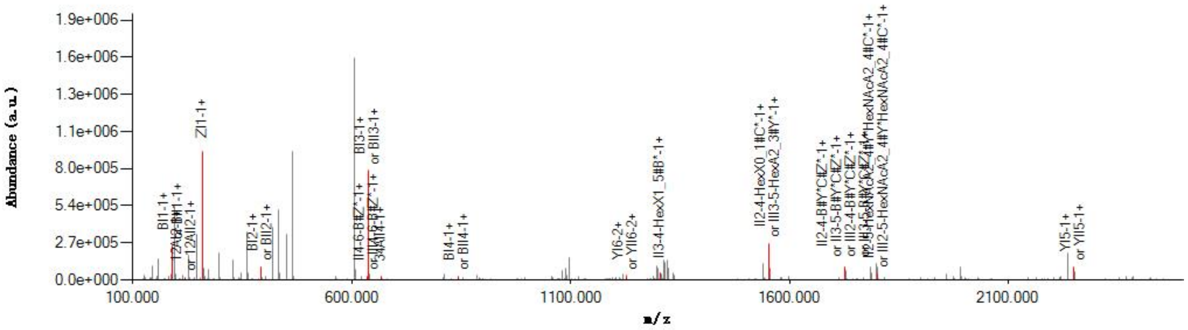


#### Graphical Fragmentation Map

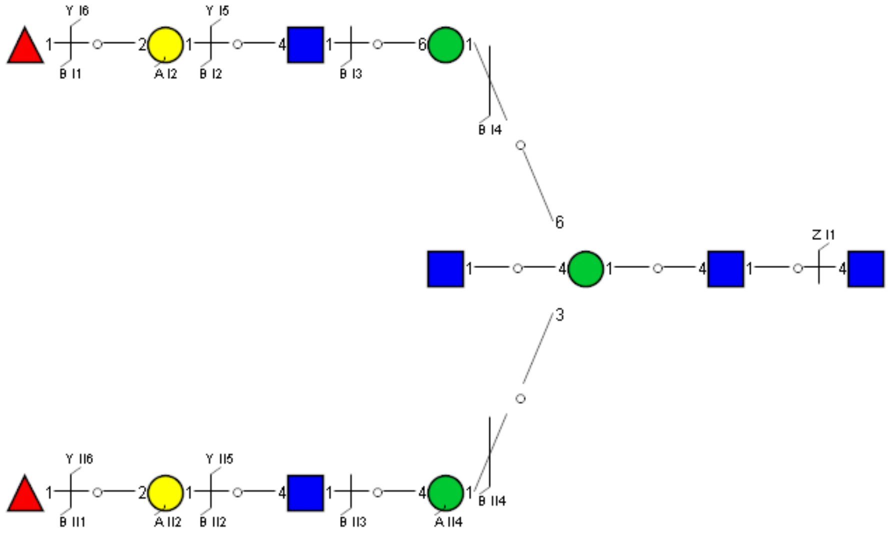

Spectrum Index: 37479(NoBHs:1)

#### Precursor iEF Map

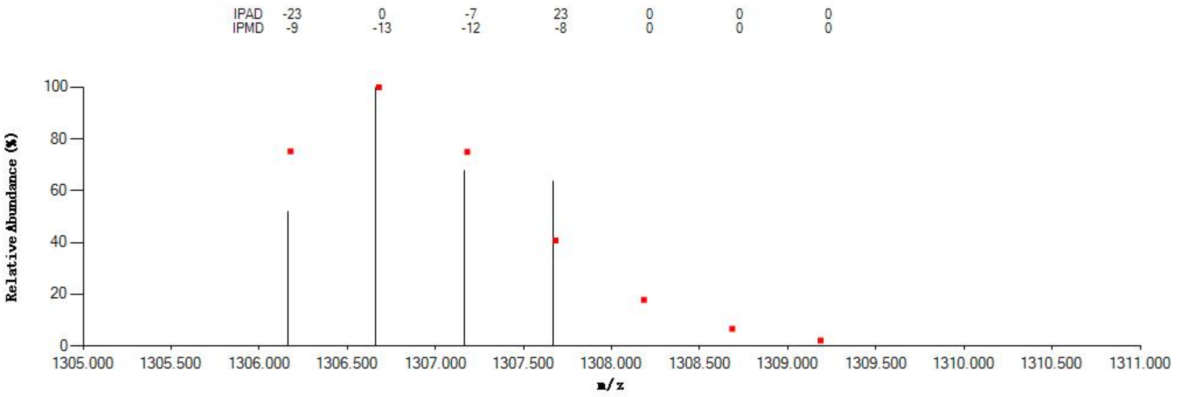


#### Annotated MS/MS Spectrum

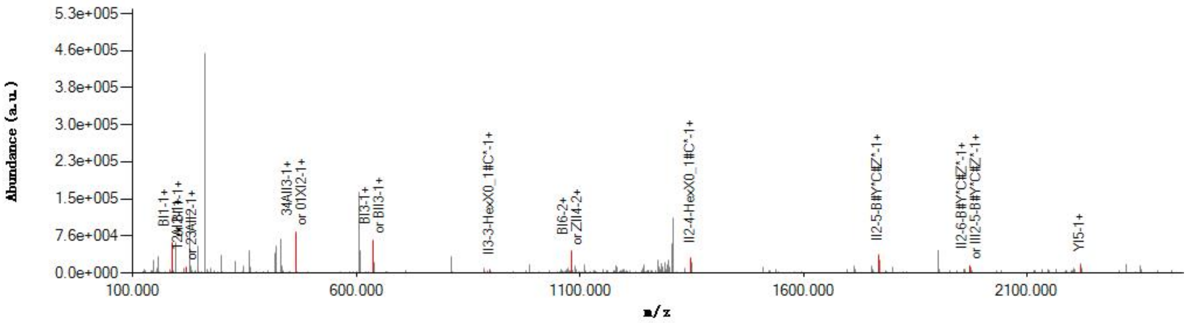


#### Graphical Fragmentation Map

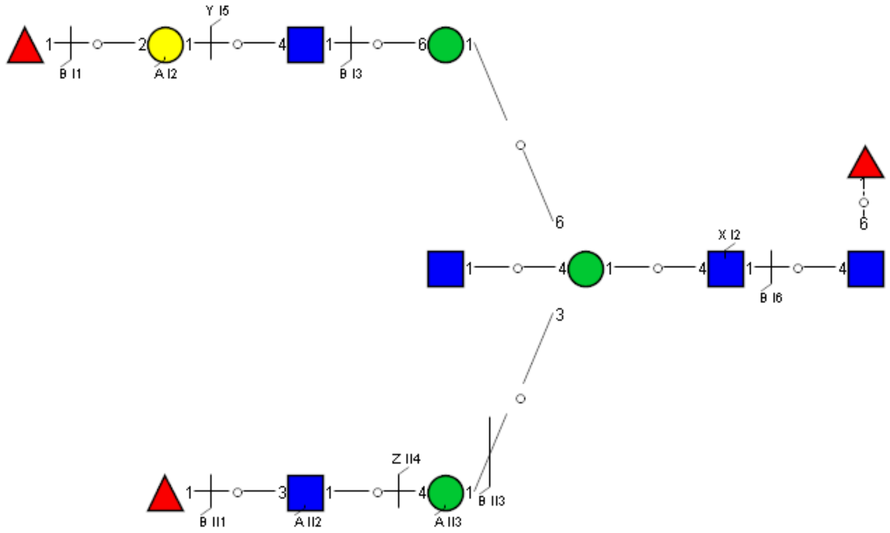

### Spectrum Index: 37516(NoBHs:1)

#### Precursor iEF Map

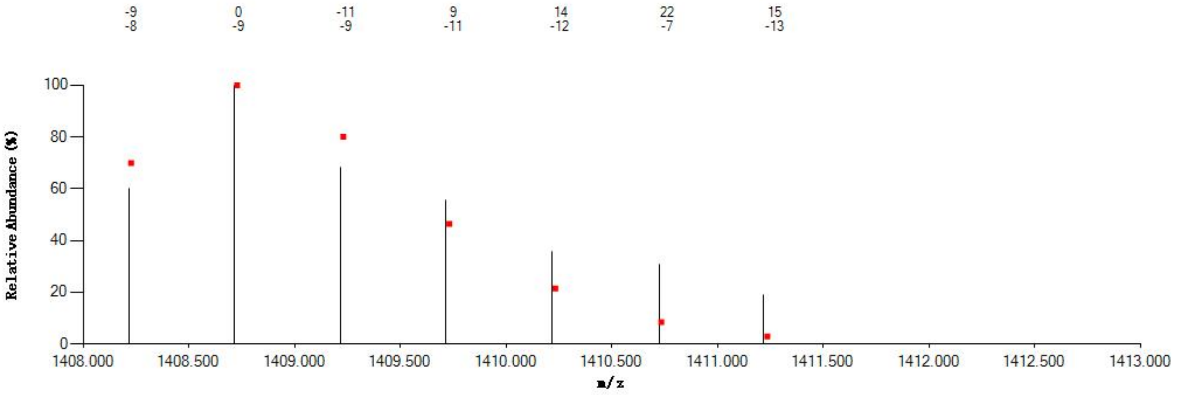


#### Annotated MS/MS Spectrum

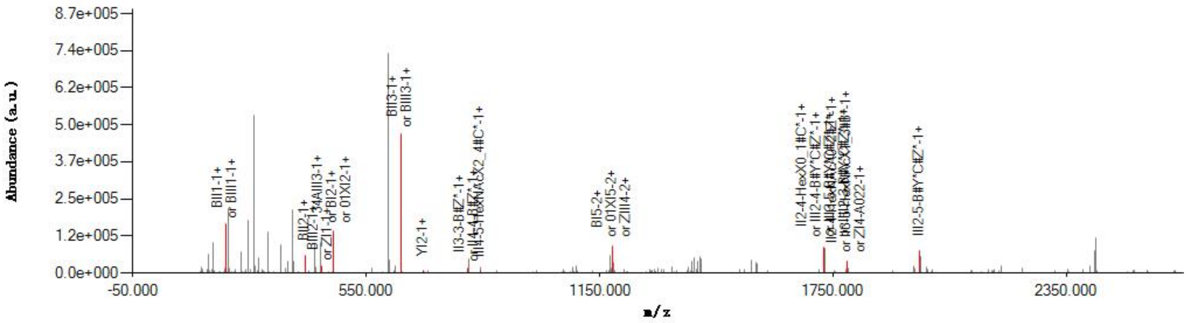


#### Graphical Fragmentation Map

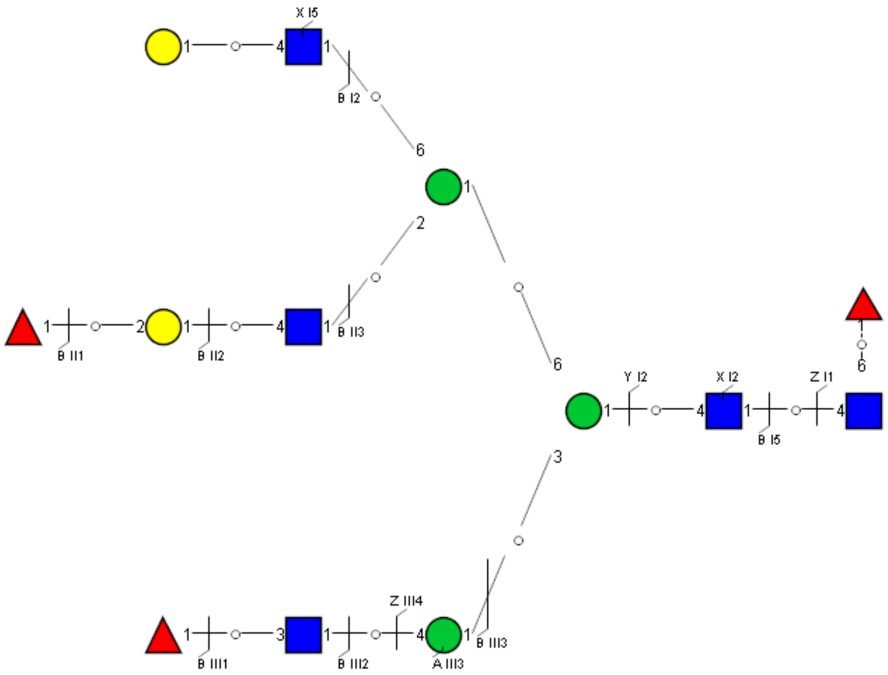

### Spectrum Index: 37540(NoBHs:1)

#### Precursor iEF Map

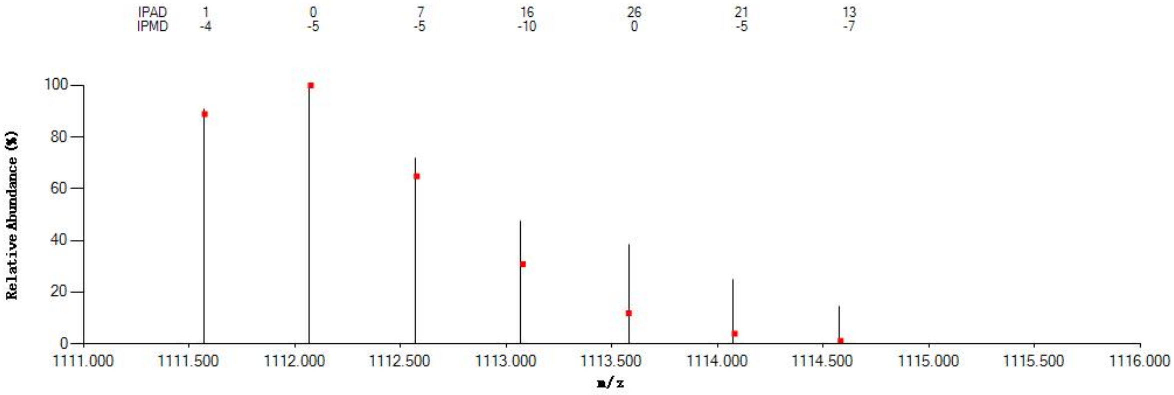

#### Annotated MS/MS Spectrum

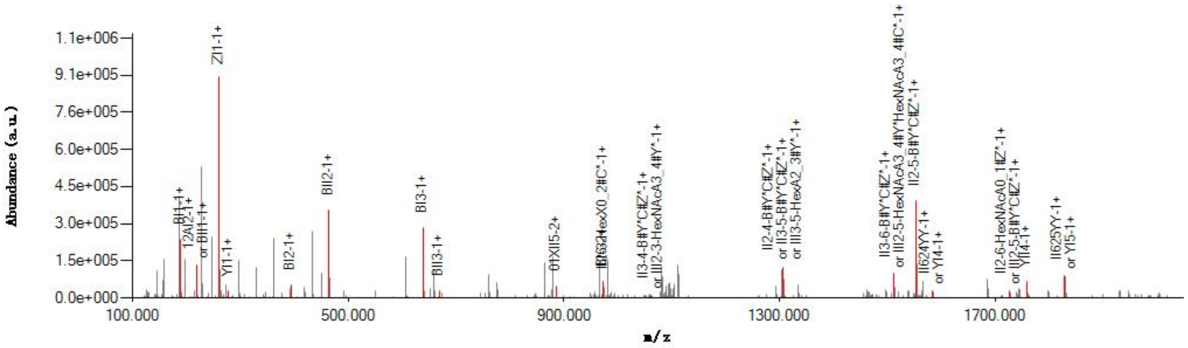

#### Graphical Fragmentation Map

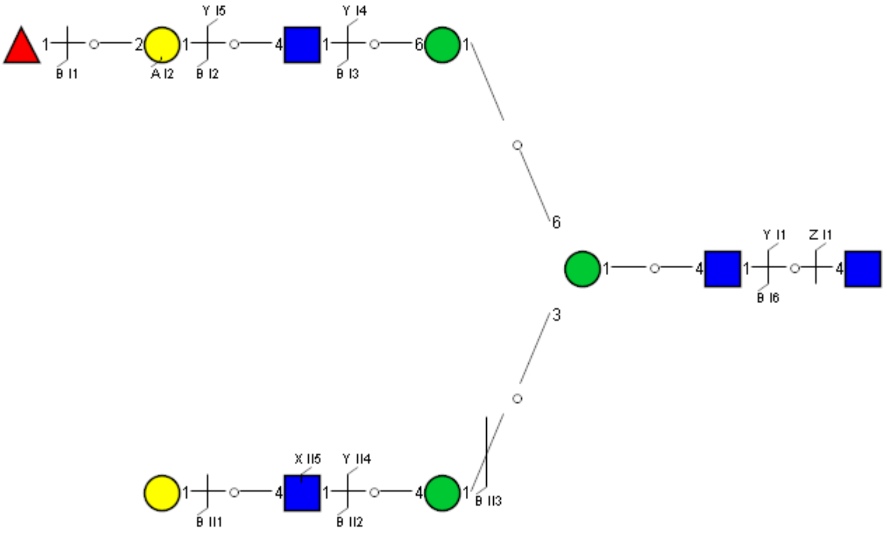

Spectrum Index: 37602(NoBHs:1)

#### Precursor iEF Map

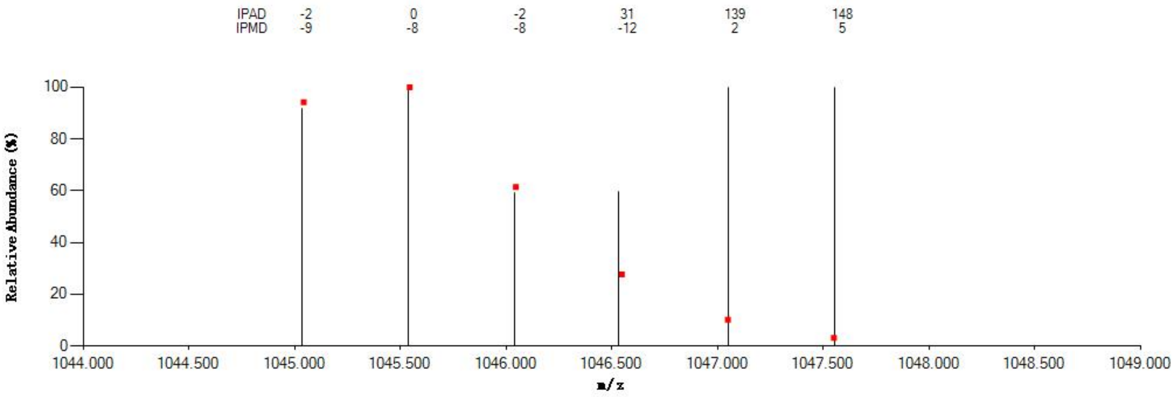


#### Annotated MS/MS Spectrum

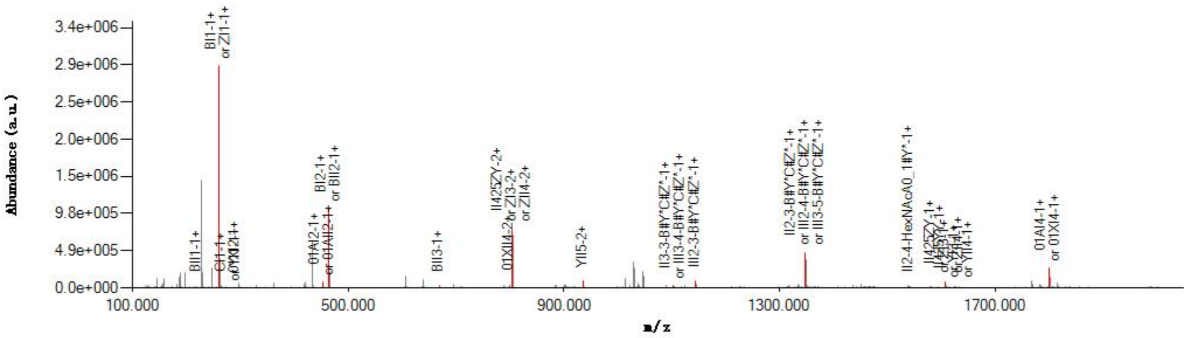


#### Graphical Fragmentation Map

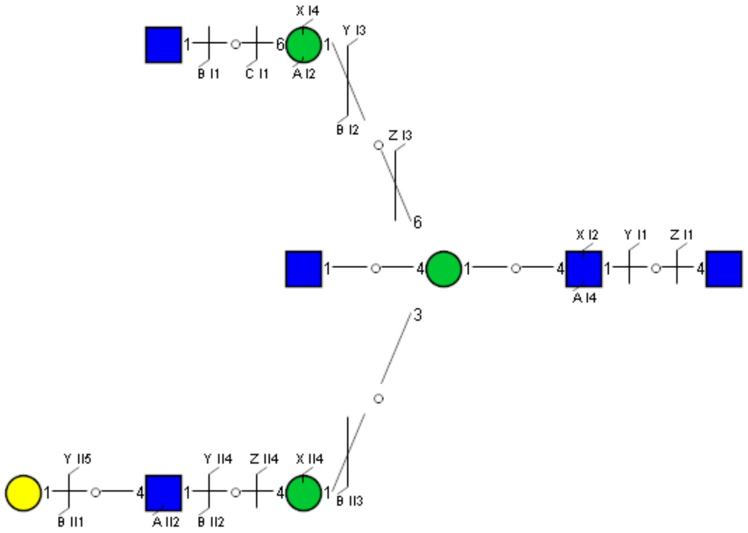

Spectrum Index: 37753(NoBHs:1)

#### Precursor iEF Map

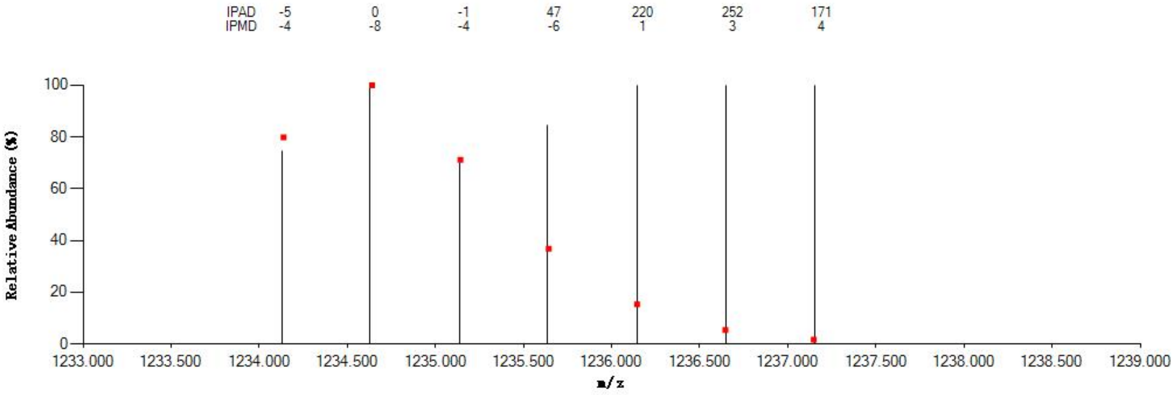

#### Annotated MS/MS Spectrum

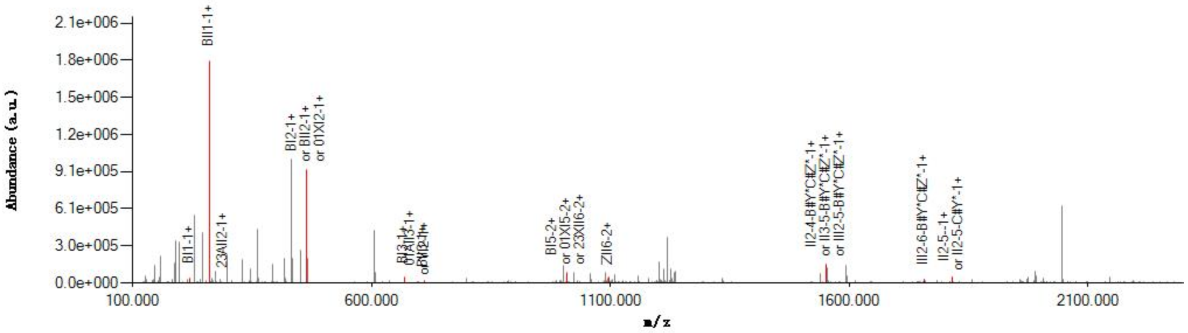

#### Graphical Fragmentation Map

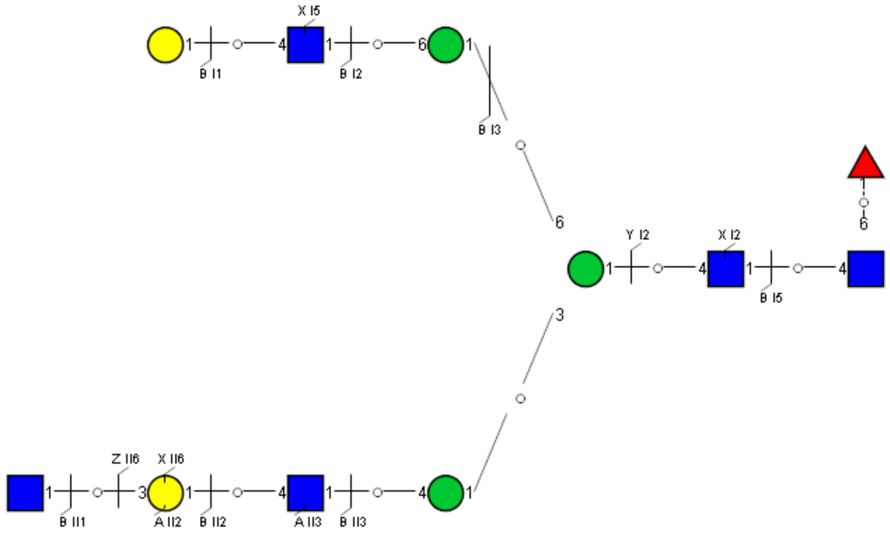

Spectrum Index: 37770(NoBHs:1)

#### Precursor iEF Map

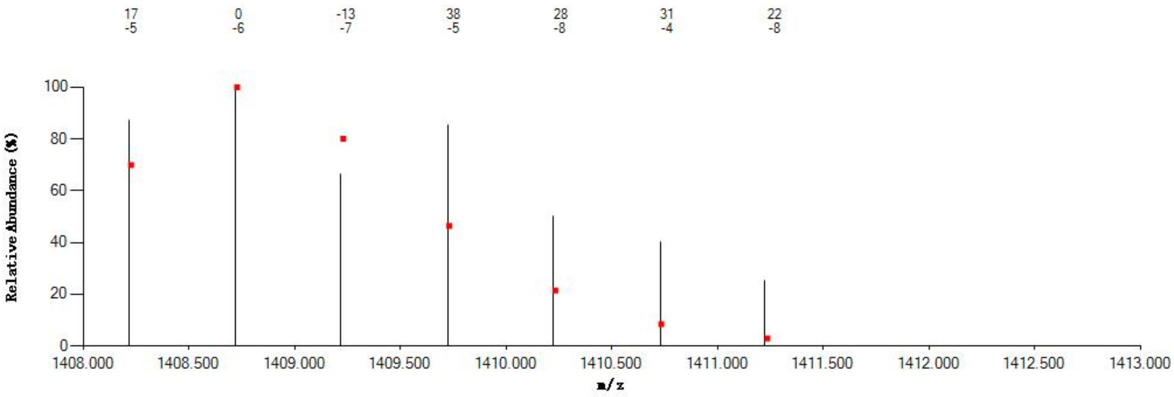


#### Annotated MS/MS Spectrum

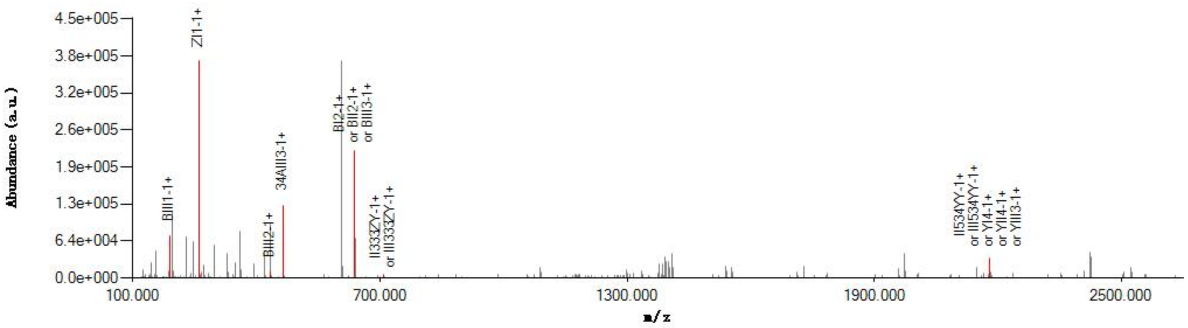


#### Graphical Fragmentation Map

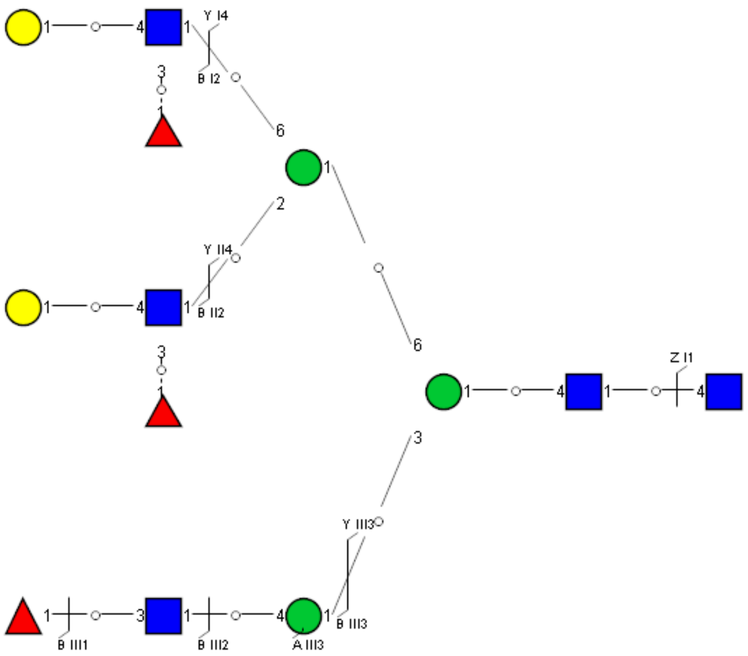

Spectrum Index: 37836(NoBHs:1)

#### Precursor iEF Map

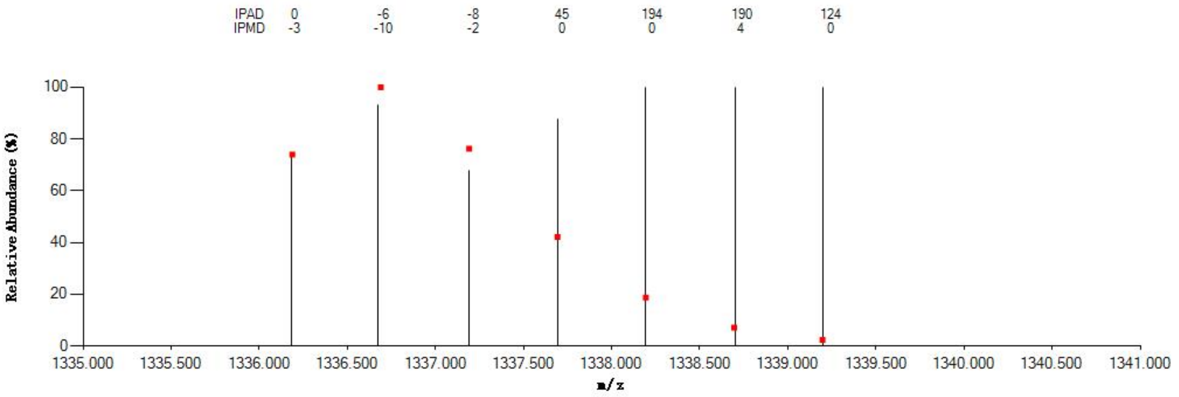


#### Annotated MS/MS Spectrum

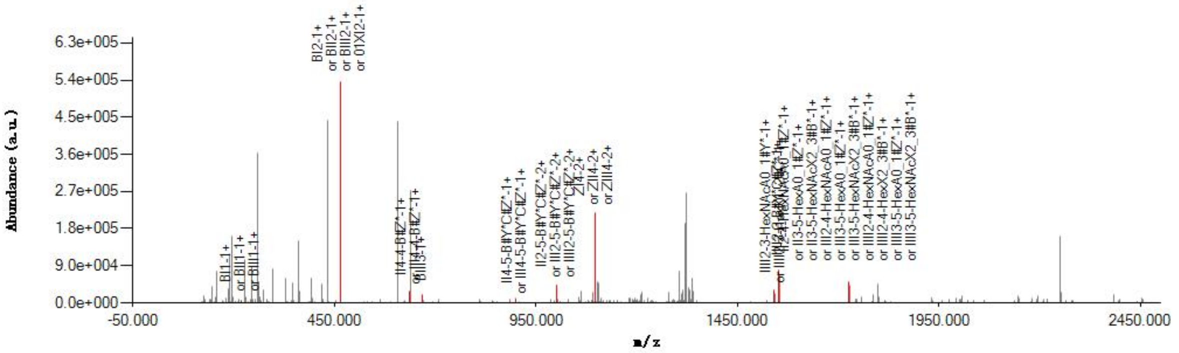


#### Graphical Fragmentation Map

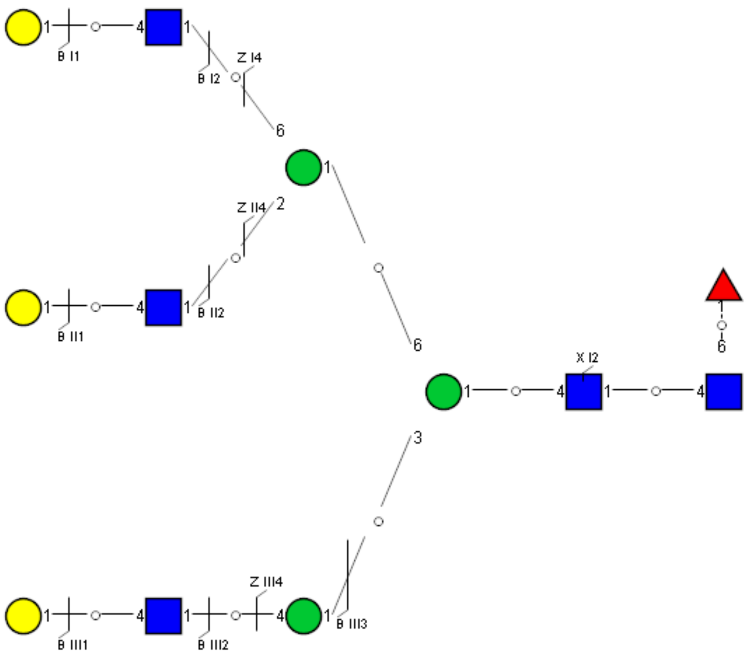

Spectrum Index: 37840(NoBHs:1)

#### Precursor iEF Map

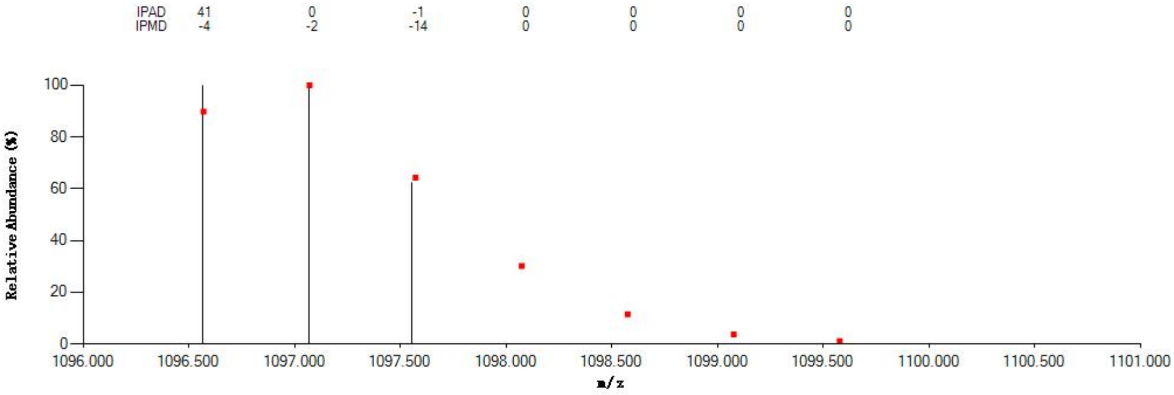


#### Annotated MS/MS Spectrum

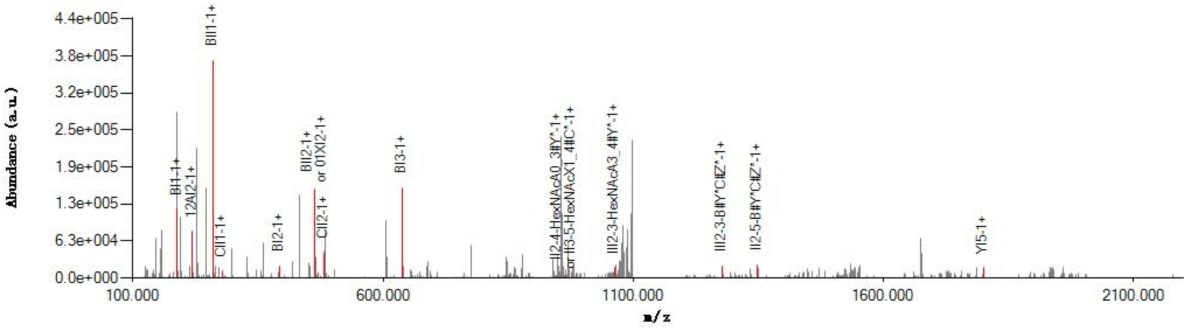


#### Graphical Fragmentation Map

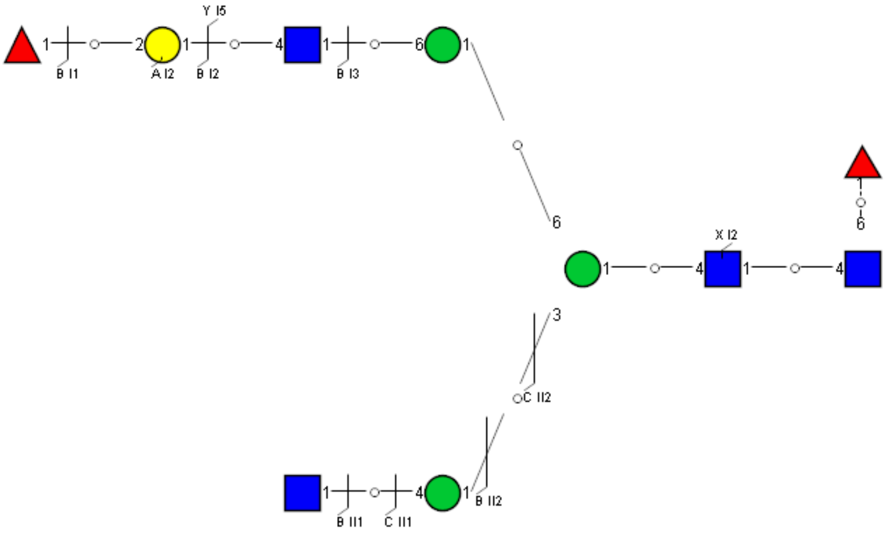

Spectrum Index: 37873(NoBHs:1)

#### Precursor iEF Map

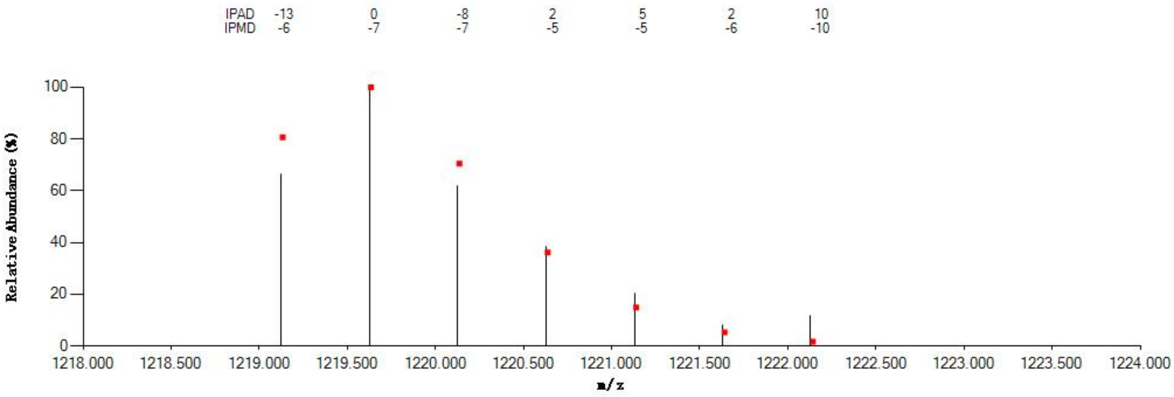


#### Annotated MS/MS Spectrum

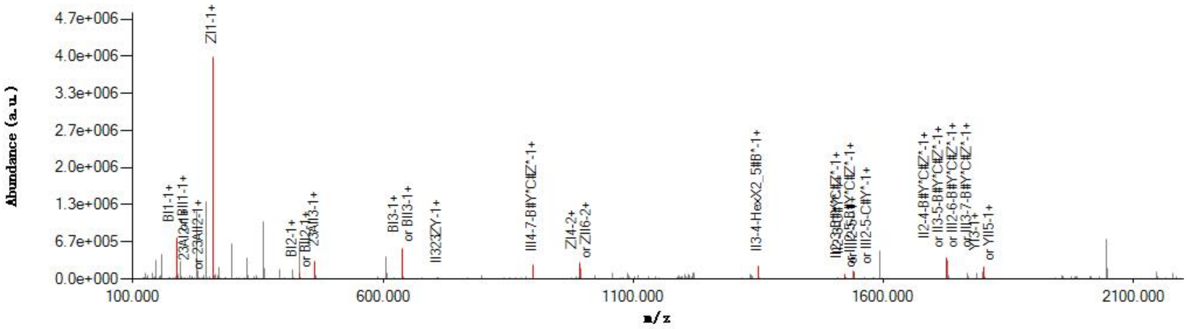


#### Graphical Fragmentation Map

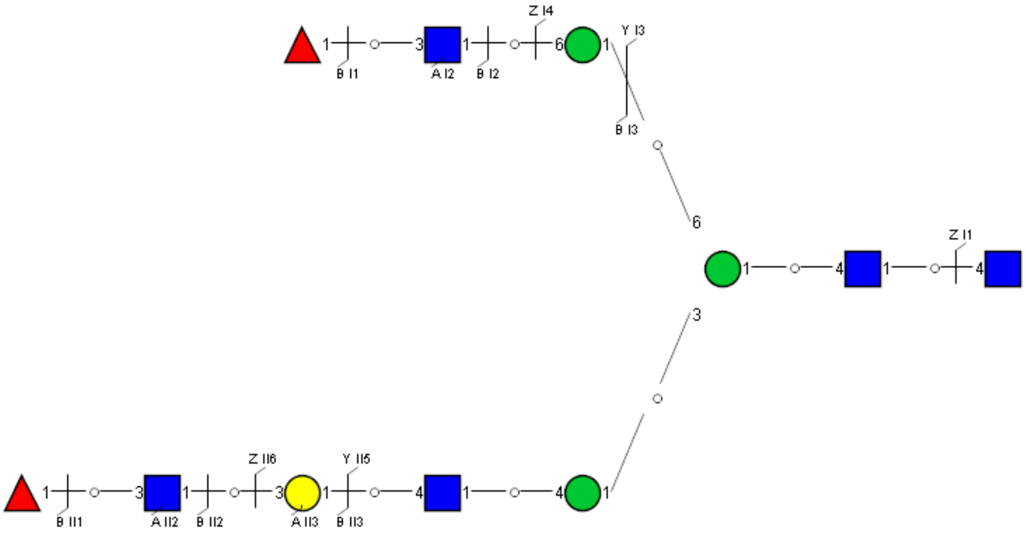

Spectrum Index: 37932(NoBHs:1)

#### Precursor iEF Map

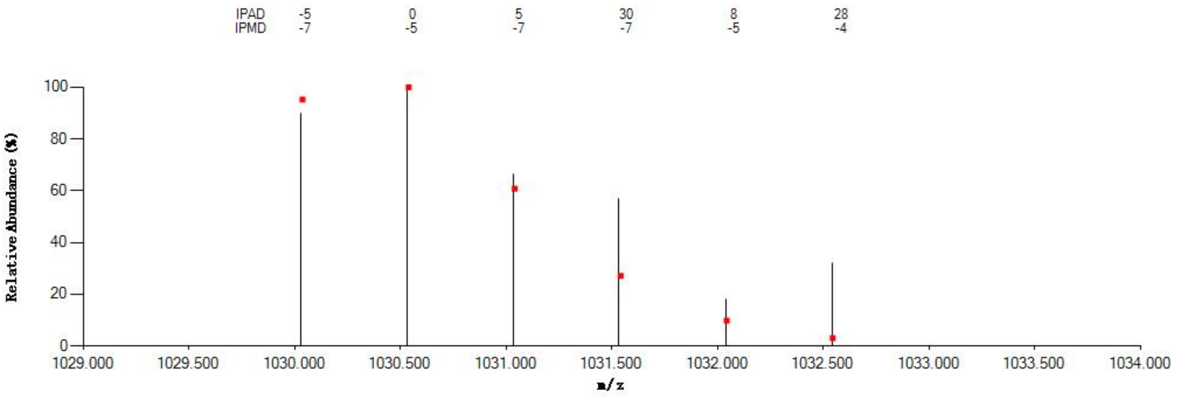


#### Annotated MS/MS Spectrum

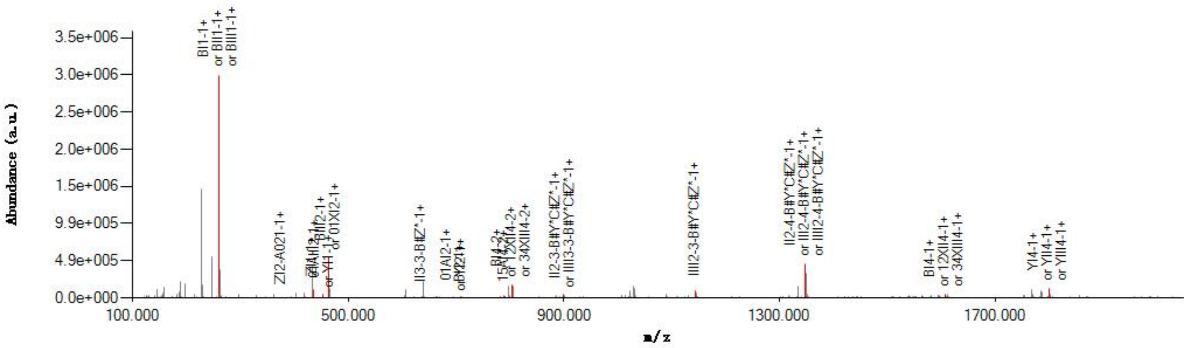


#### Graphical Fragmentation Map

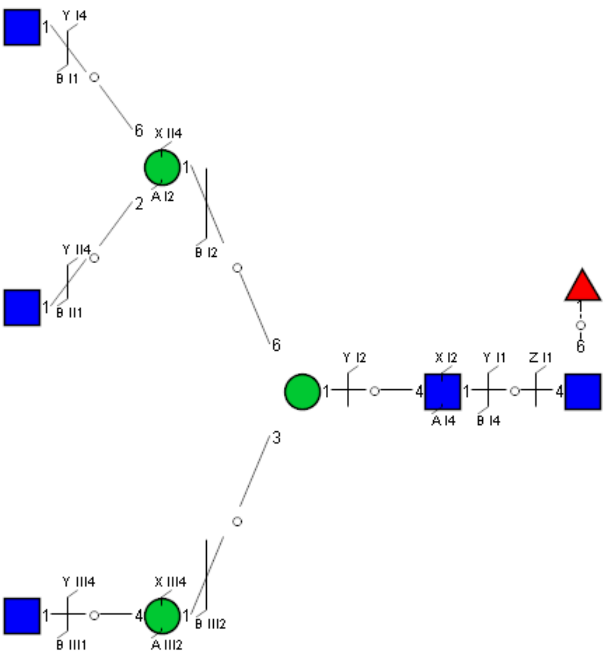

Spectrum Index: 37939(NoBHs:1)

#### Precursor iEF Map

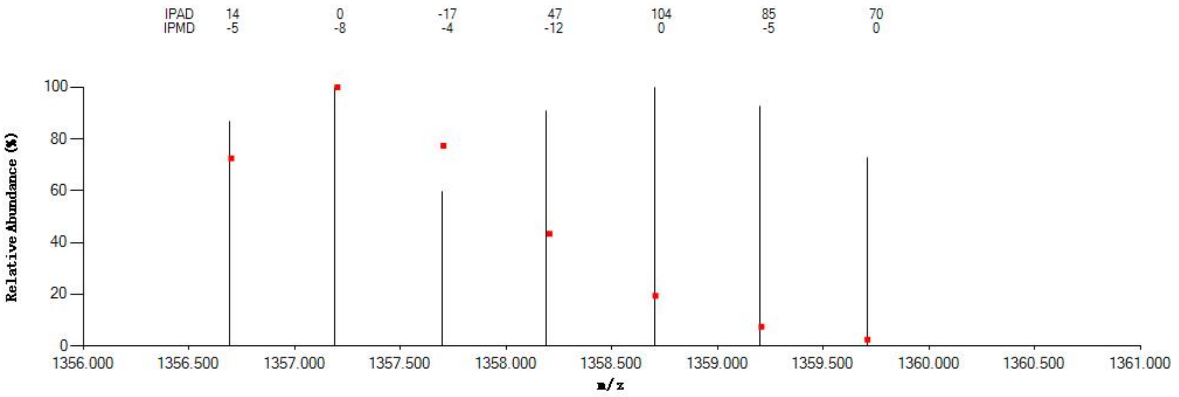


#### Annotated MS/MS Spectrum

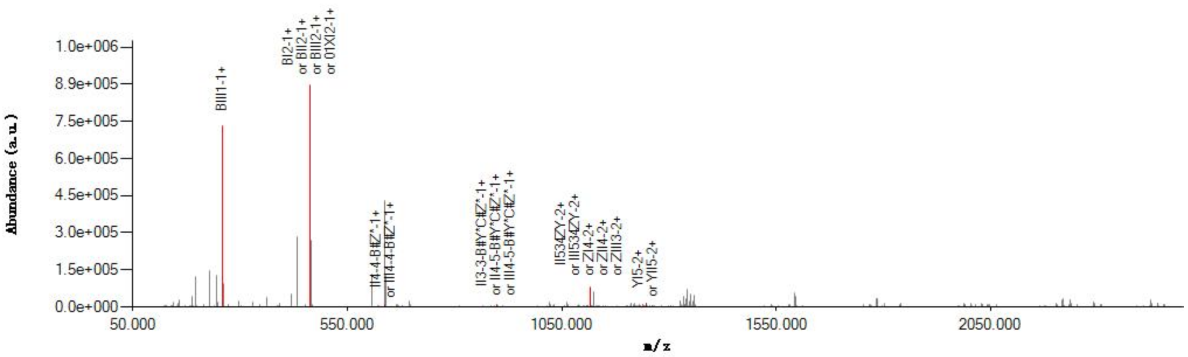


#### Graphical Fragmentation Map

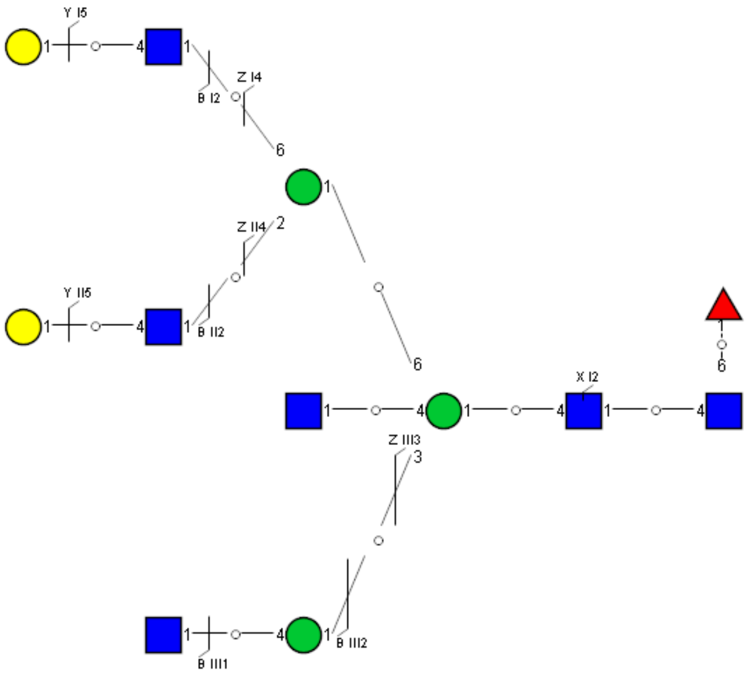

Spectrum Index: 38023(NoBHs:1)

#### Precursor iEF Map

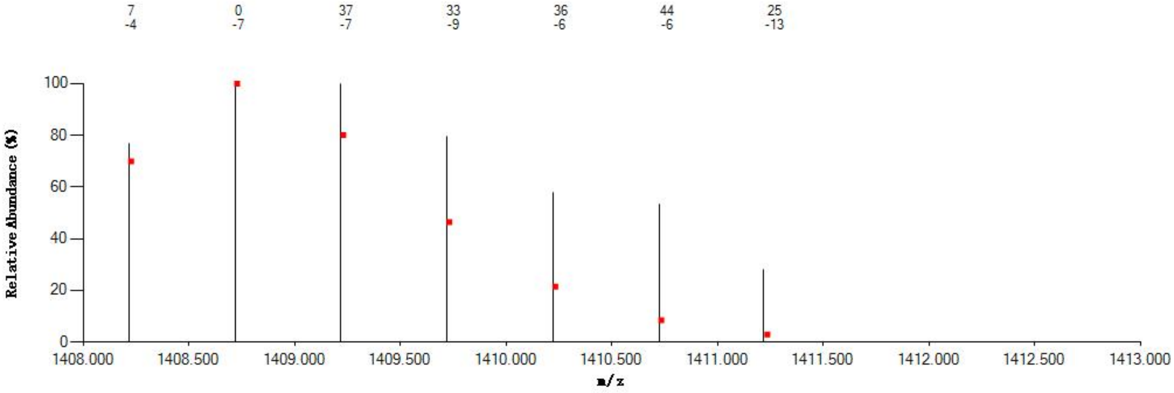


#### Annotated MS/MS Spectrum

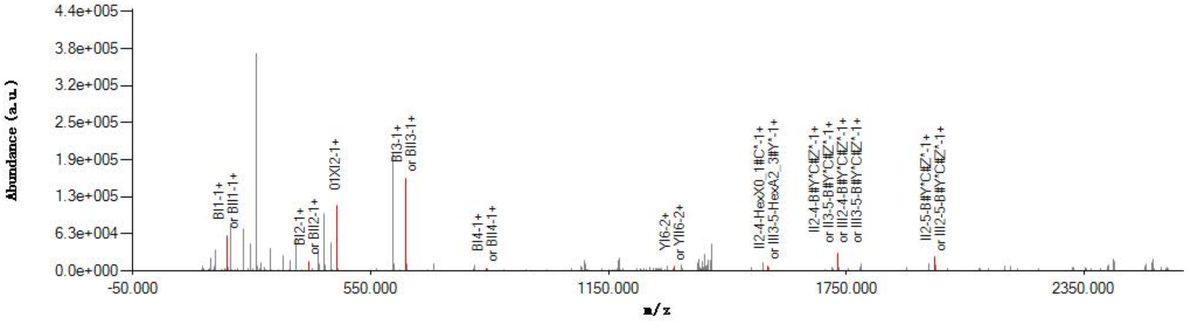


#### Graphical Fragmentation Map

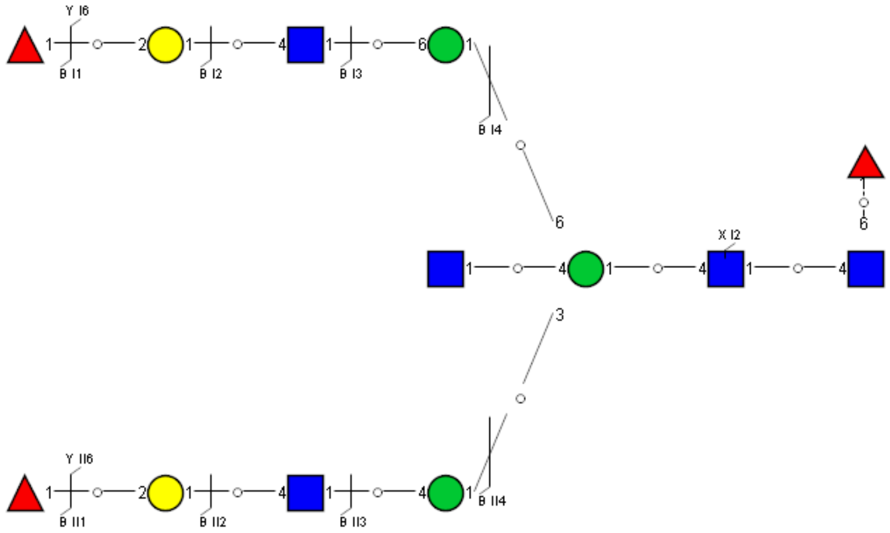

Spectrum Index: 38028(NoBHs:1)

Precursor iEF Map

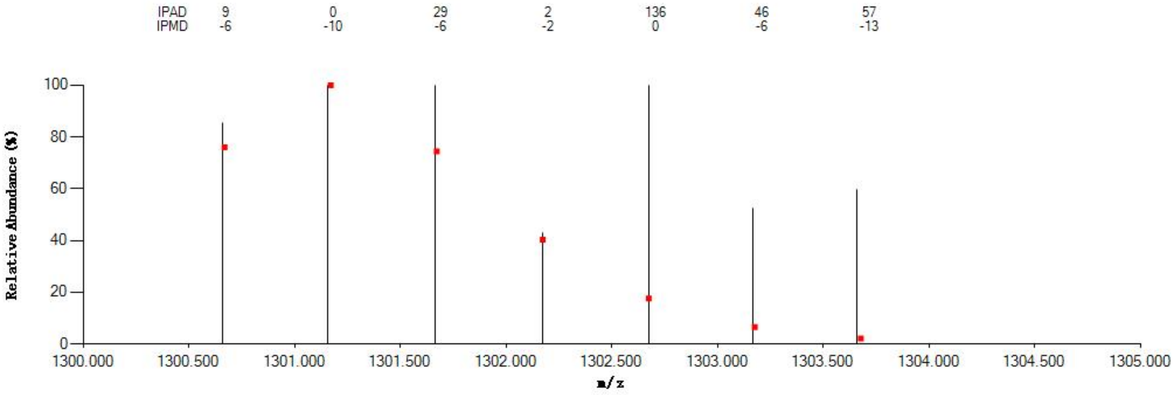


Annotated MS/MS Spectrum

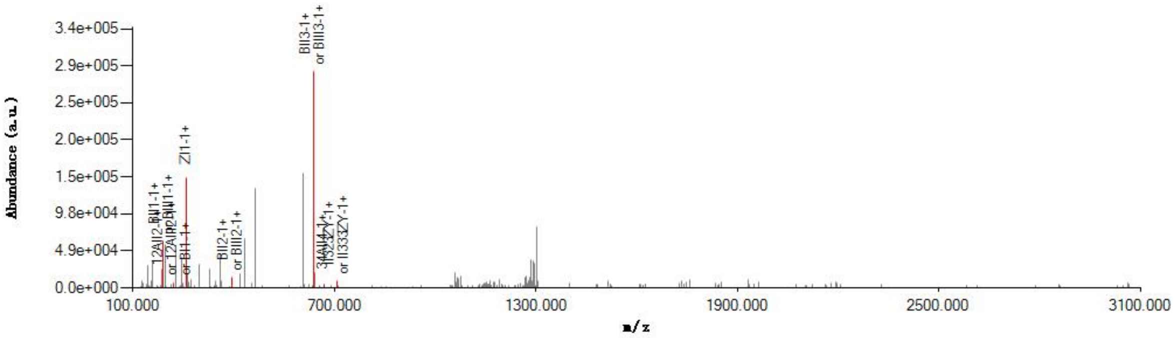


Graphical Fragmentation Map

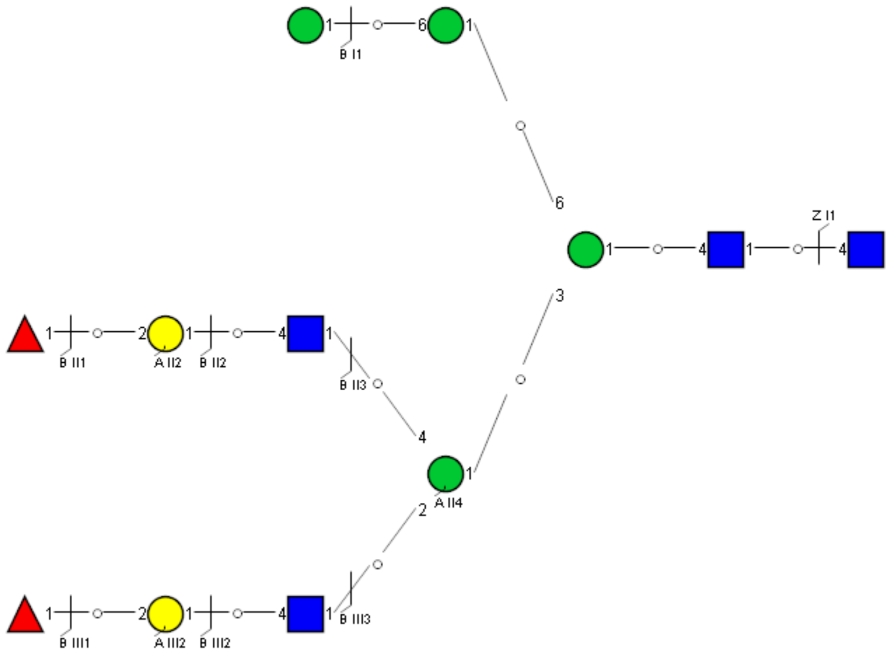

### Spectrum Index: 38125(NoBHs:1)

#### Precursor iEF Map

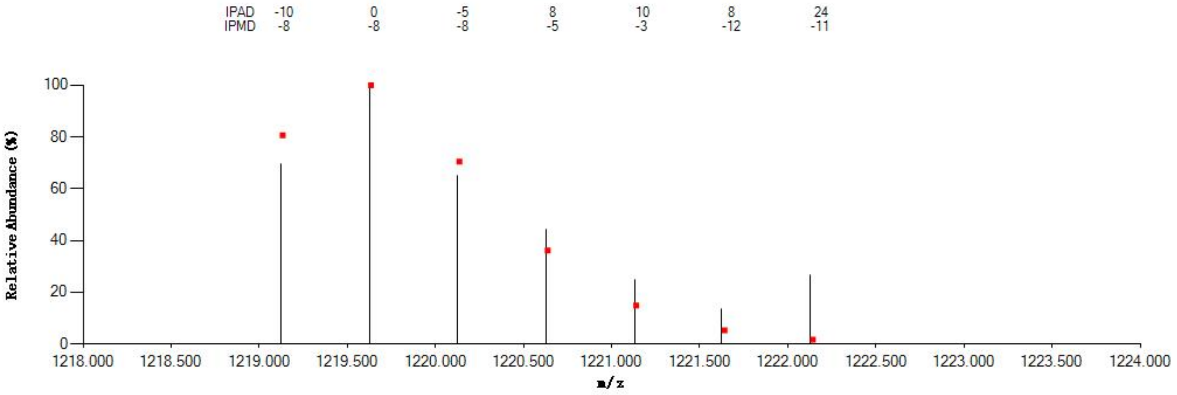


#### Annotated MS/MS Spectrum

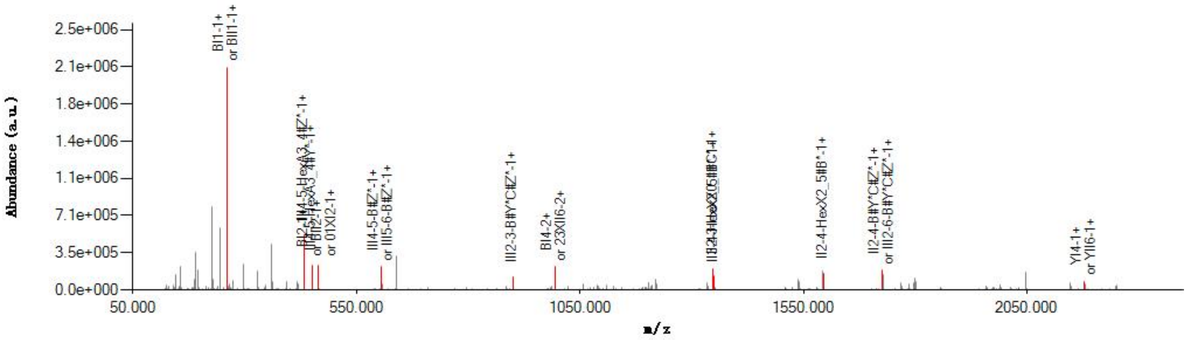


#### Graphical Fragmentation Map

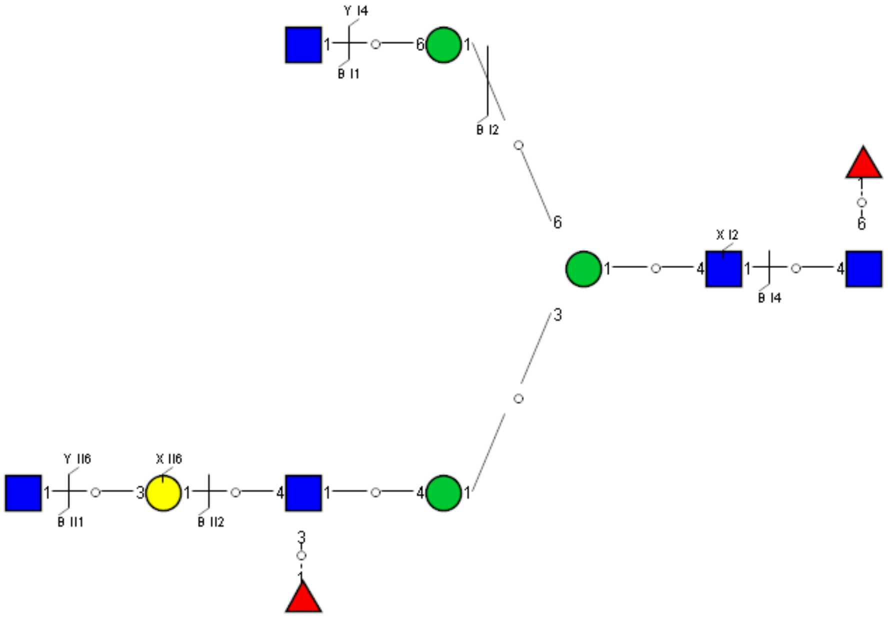

Spectrum Index: 38166(NoBHs:1)

#### Precursor iEF Map

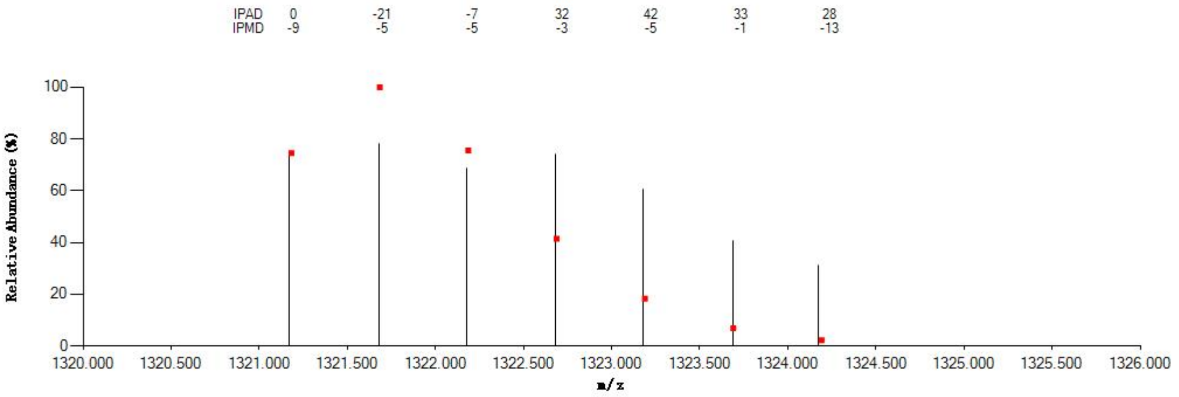


#### Annotated MS/MS Spectrum

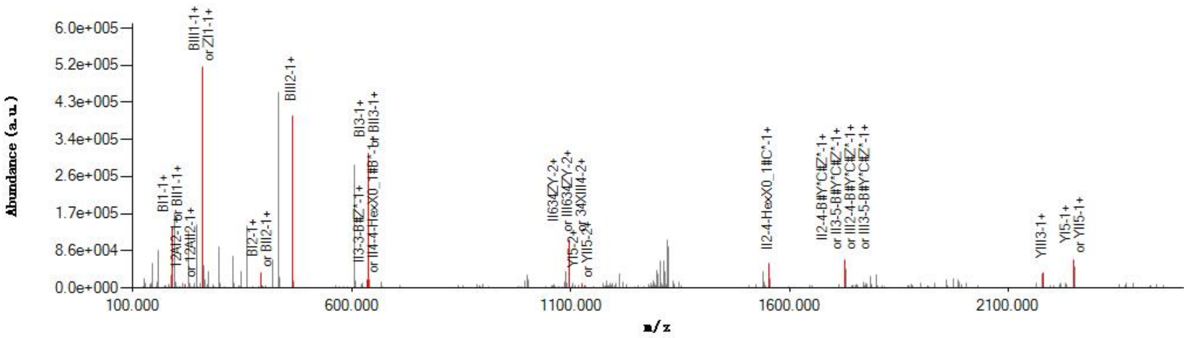


#### Graphical Fragmentation Map

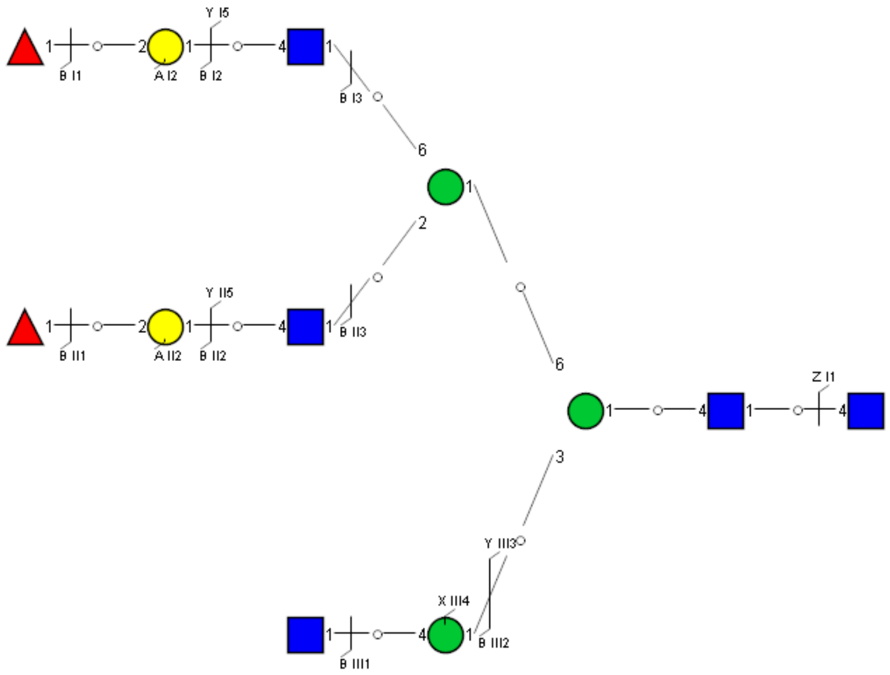

Spectrum Index: 38191(NoBHs:1)

#### Precursor iEF Map

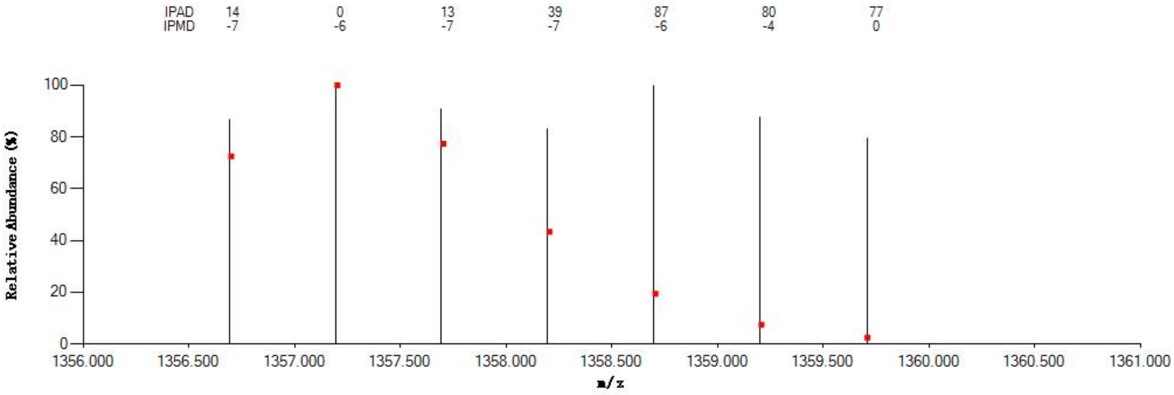


#### Annotated MS/MS Spectrum

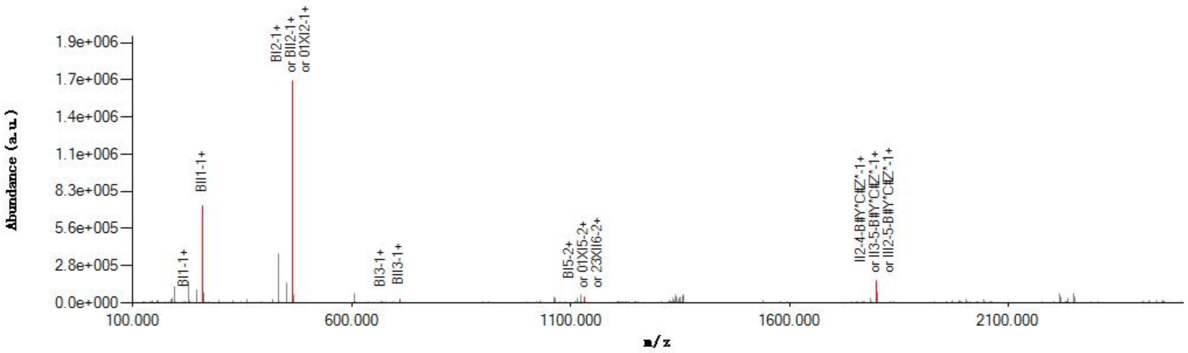


#### Graphical Fragmentation Map

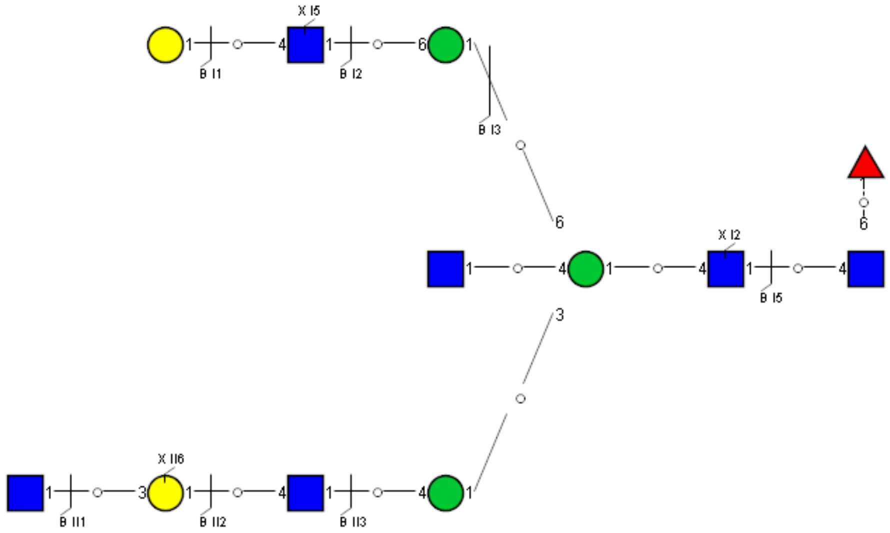

### Spectrum Index: 38243(NoBHs:1)

#### Precursor iEF Map

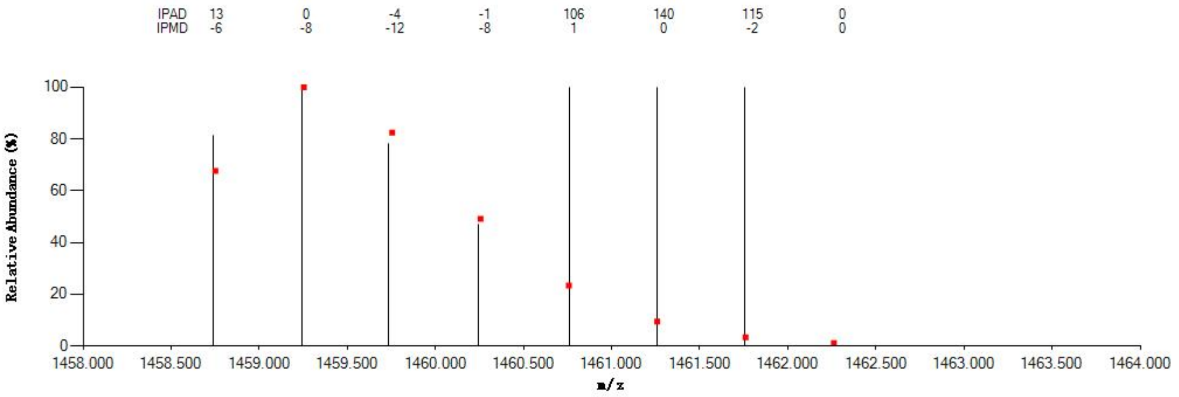


#### Annotated MS/MS Spectrum

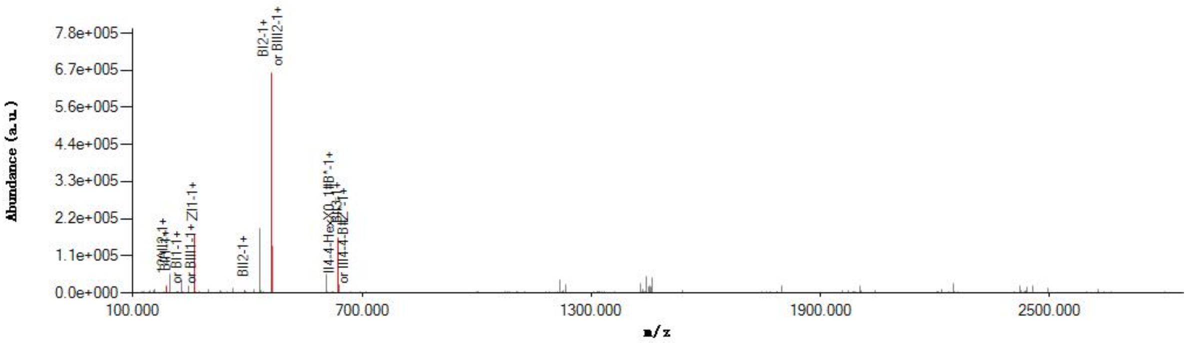


#### Graphical Fragmentation Map

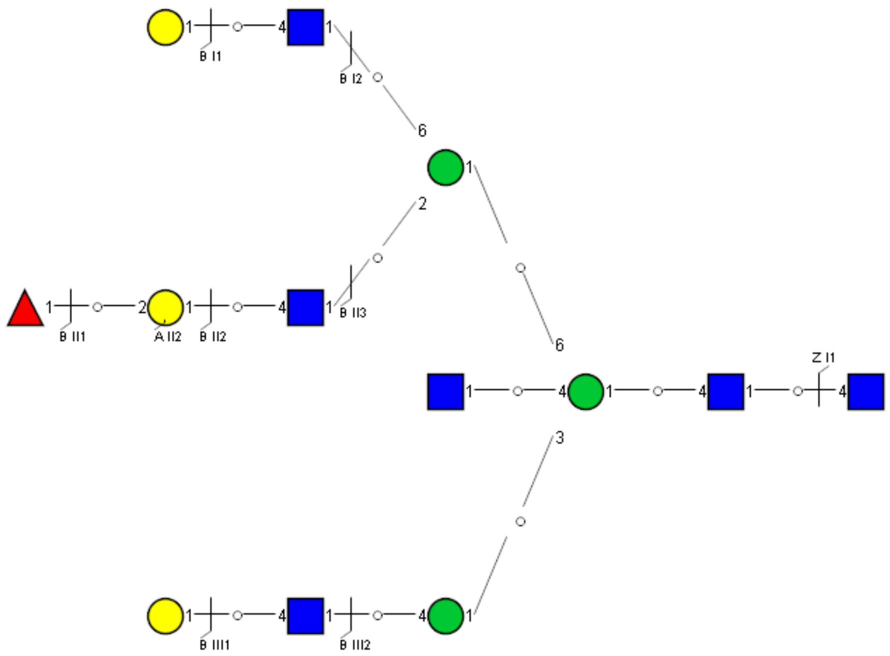

Spectrum Index: 38313(NoBHs:1)

#### Precursor iEF Map

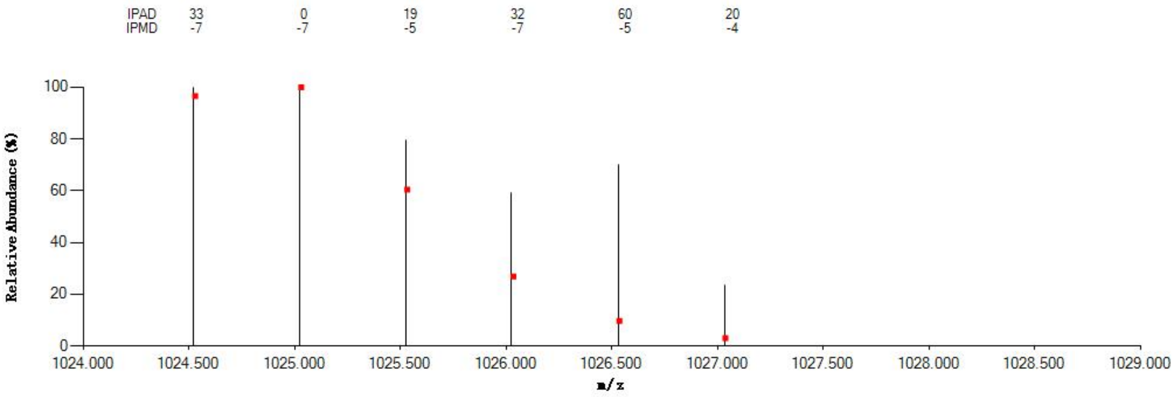


#### Annotated MS/MS Spectrum

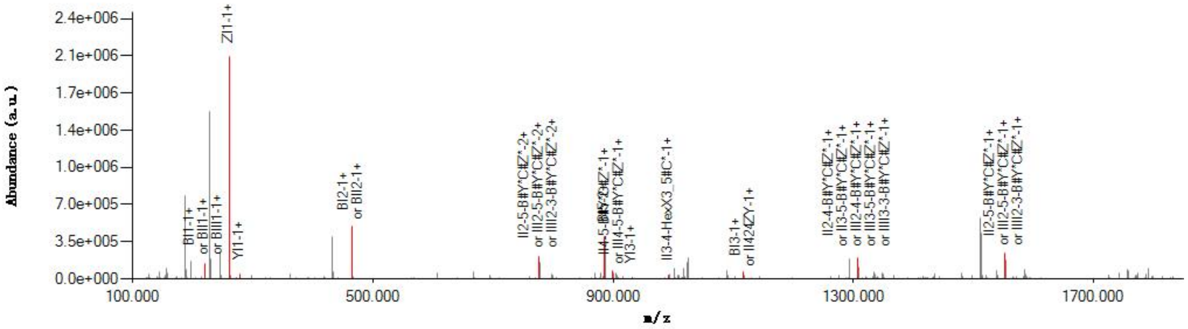


#### Graphical Fragmentation Map

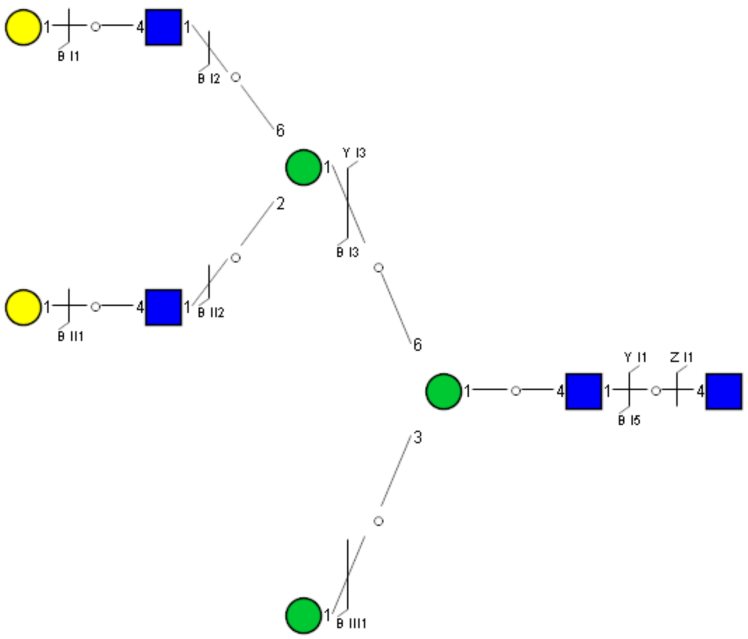

Spectrum Index: 38347(NoBHs:1)

#### Precursor iEF Map

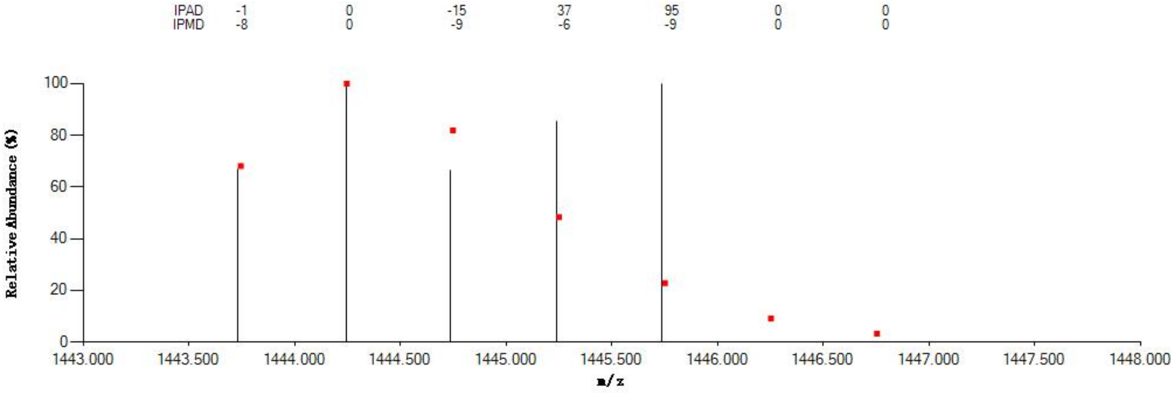


#### Annotated MS/MS Spectrum

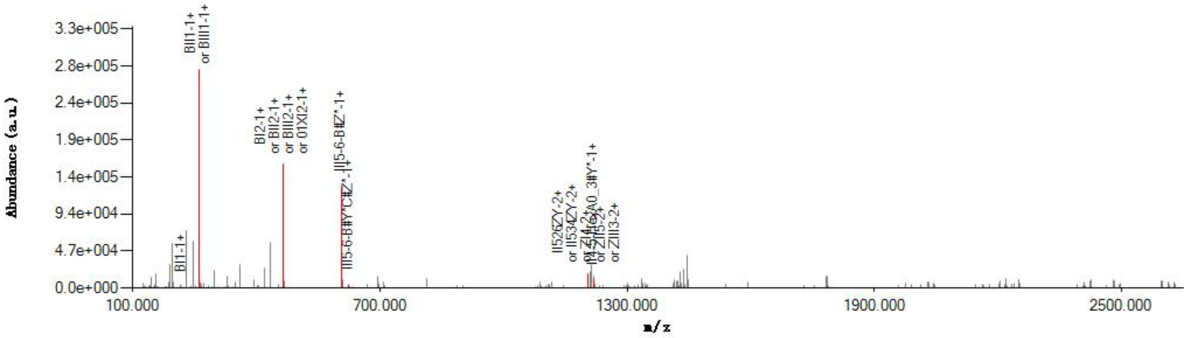


#### Graphical Fragmentation Map

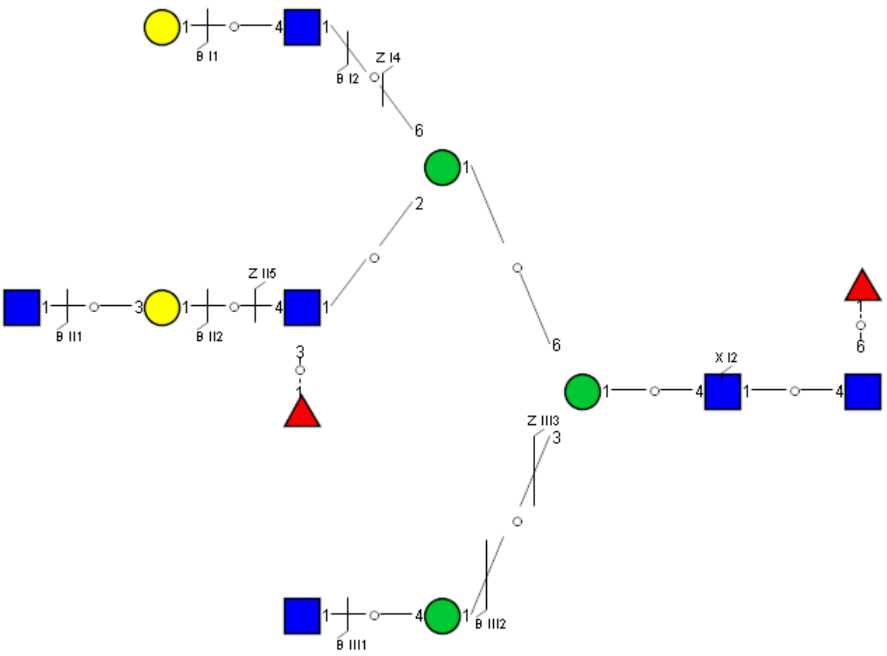

Spectrum Index: 38431(NoBHs:1)

#### Precursor iEF Map

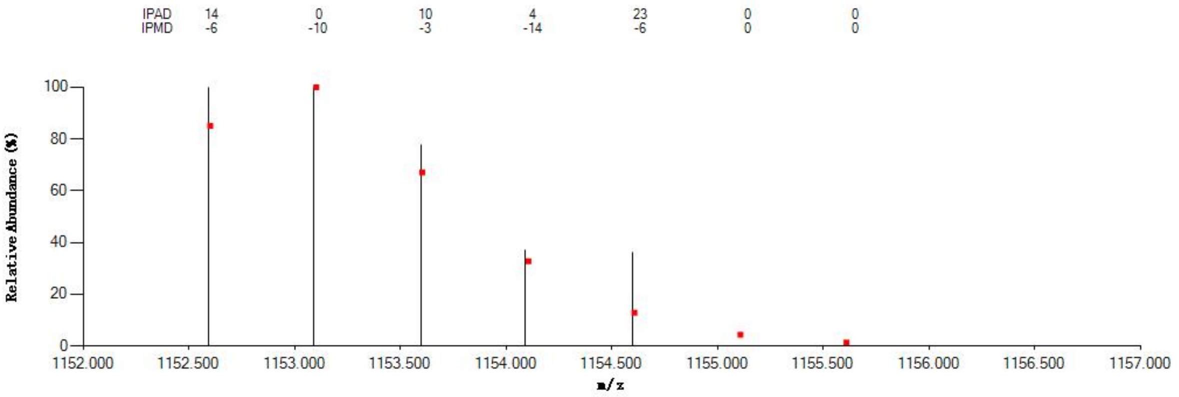


#### Annotated MS/MS Spectrum

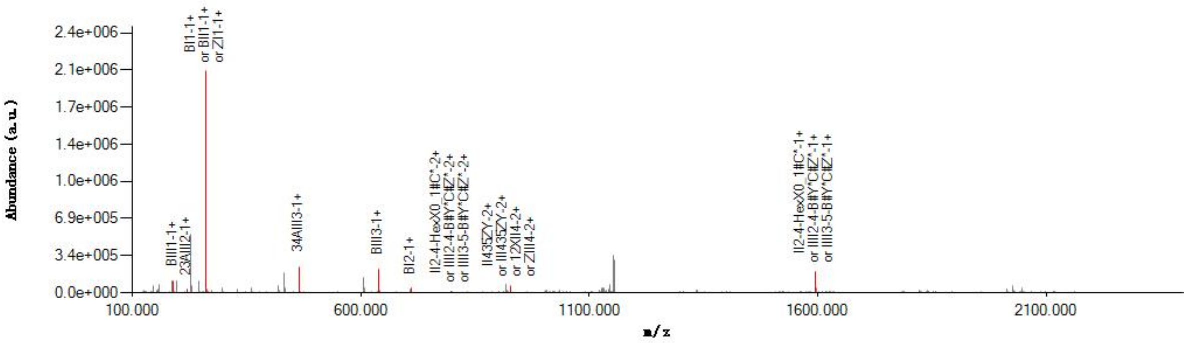


#### Graphical Fragmentation Map

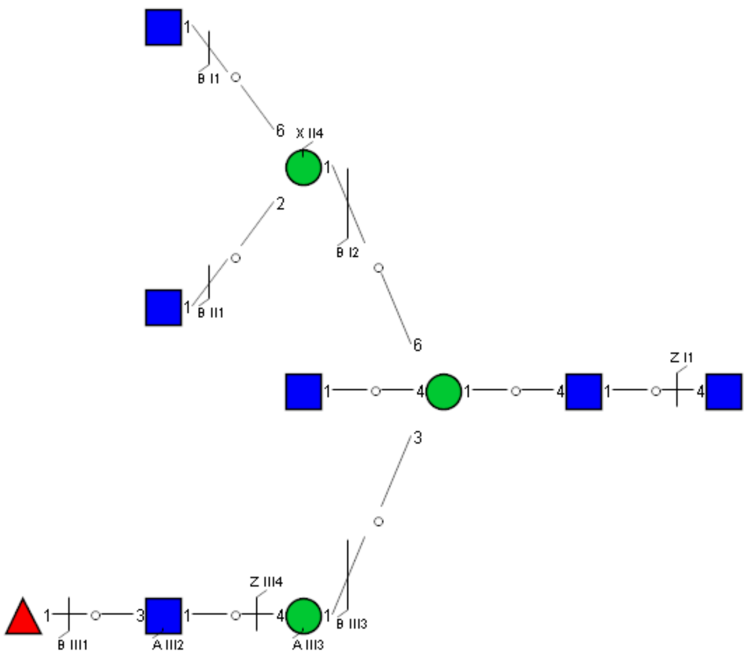

Spectrum Index: 38505(NoBHs:1)

#### Precursor iEF Map

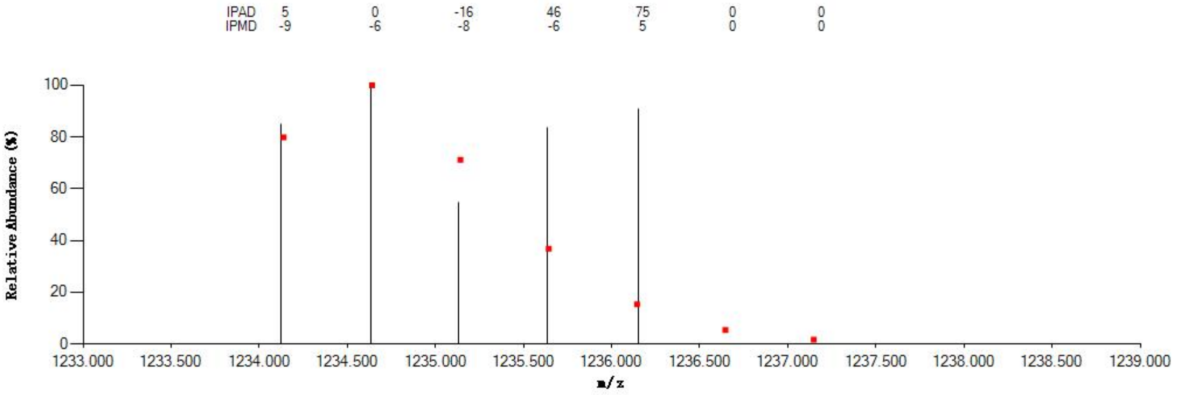


#### Annotated MS/MS Spectrum

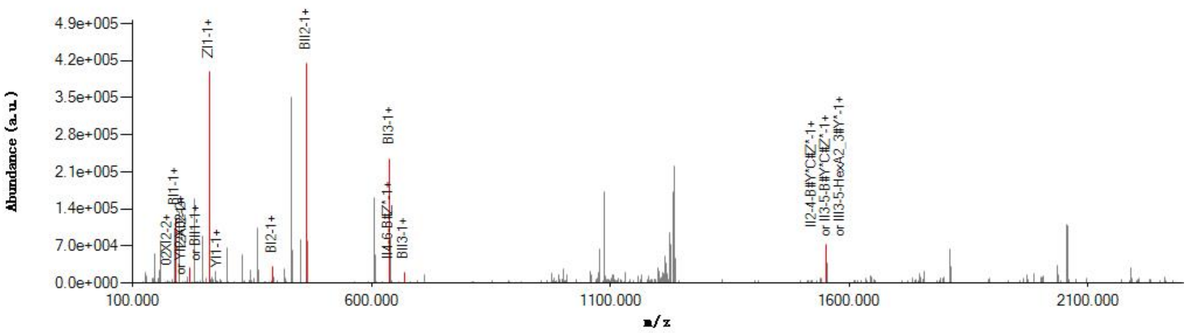


#### Graphical Fragmentation Map

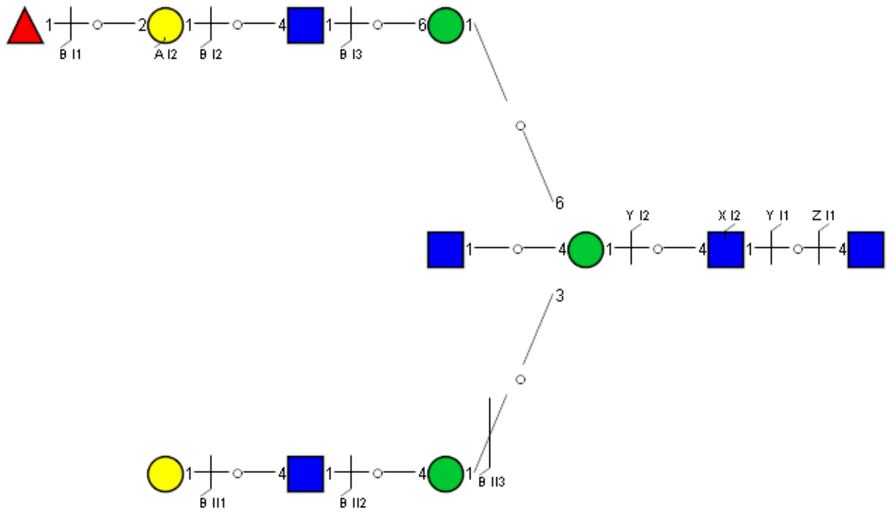

### Spectrum Index: 38660(NoBHs:1)

#### Precursor iEF Map

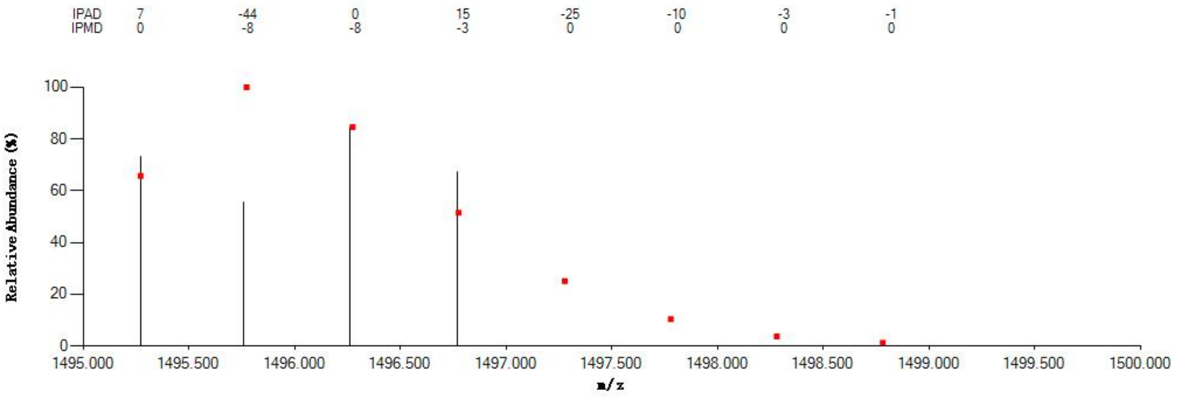


#### Annotated MS/MS Spectrum

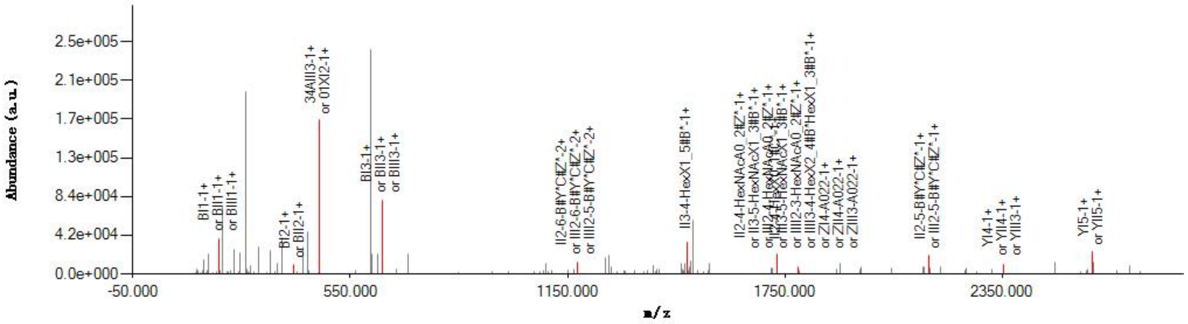


#### Graphical Fragmentation Map

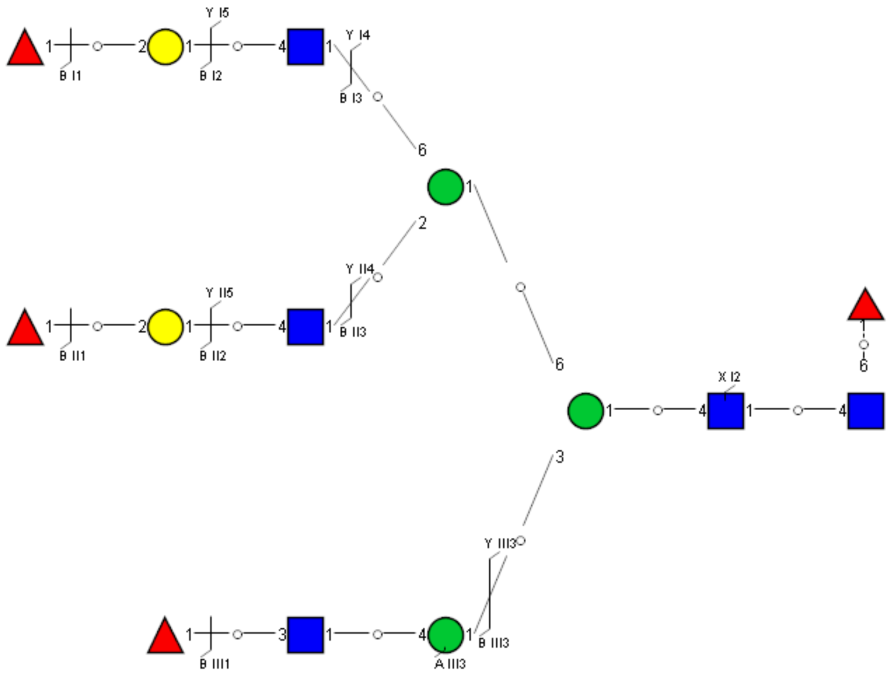

Spectrum Index: 38698(NoBHs:1)

#### Precursor iEF Map

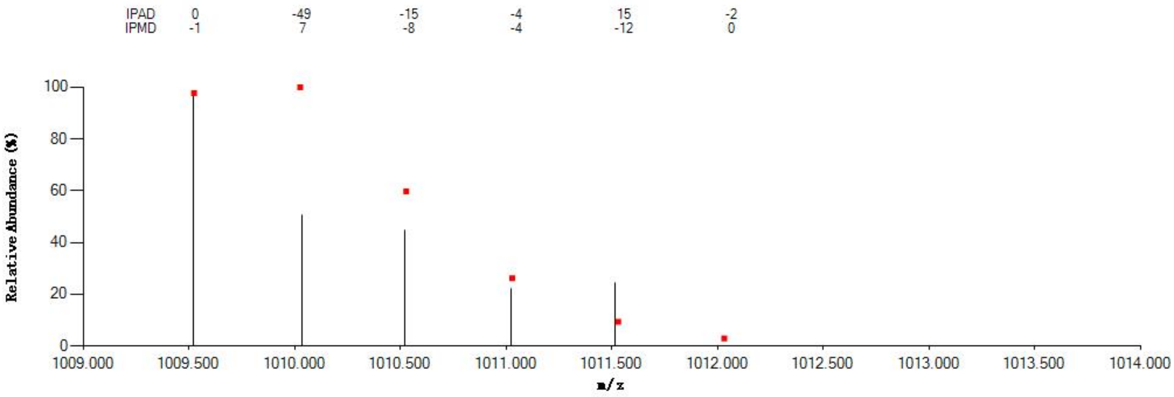


#### Annotated MS/MS Spectrum

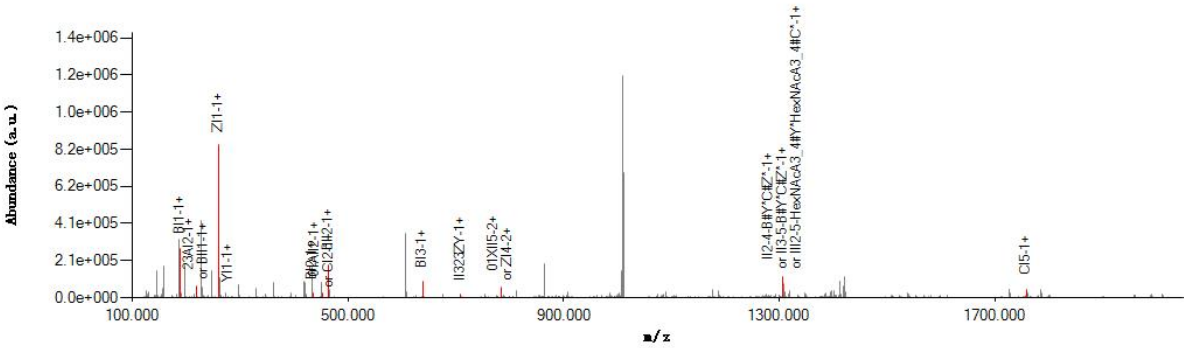


#### Graphical Fragmentation Map

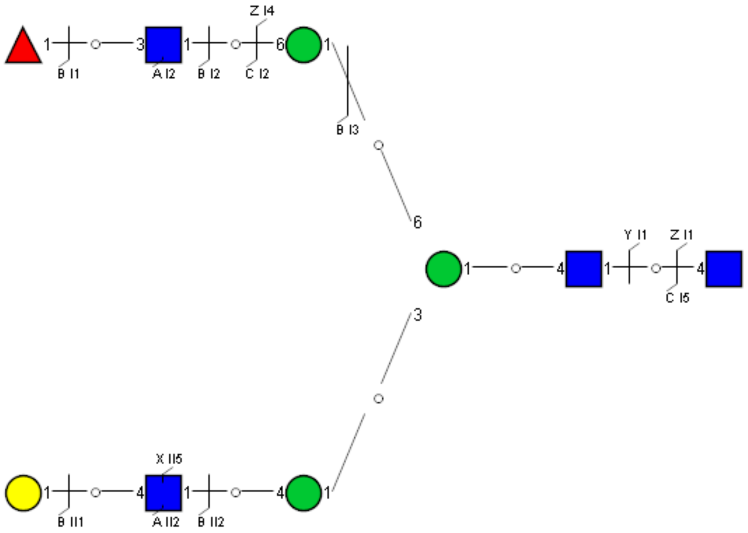

### Spectrum Index: 38758(NoBHs:1)

#### Precursor iEF Map

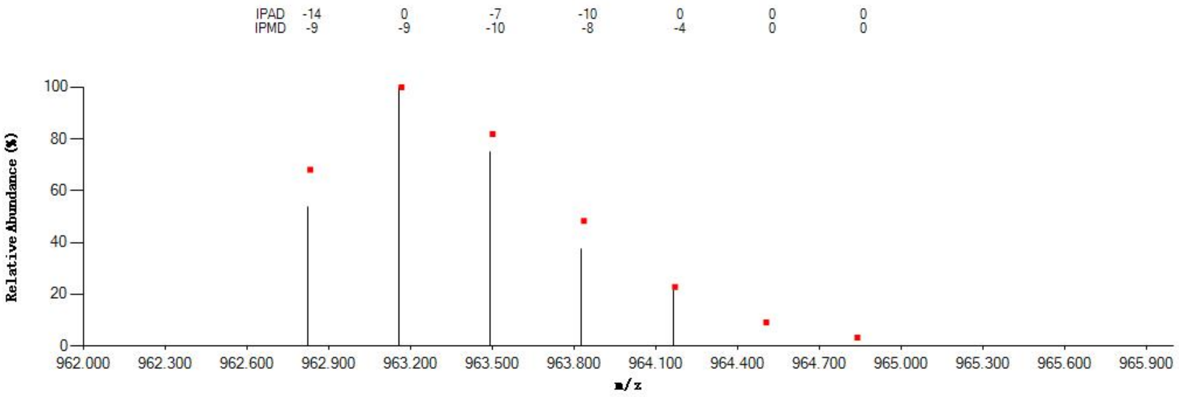


#### Annotated MS/MS Spectrum

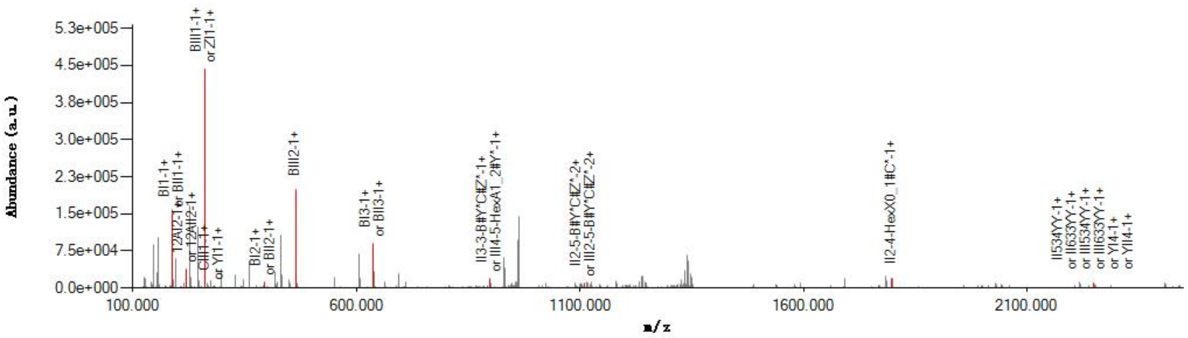


#### Graphical Fragmentation Map

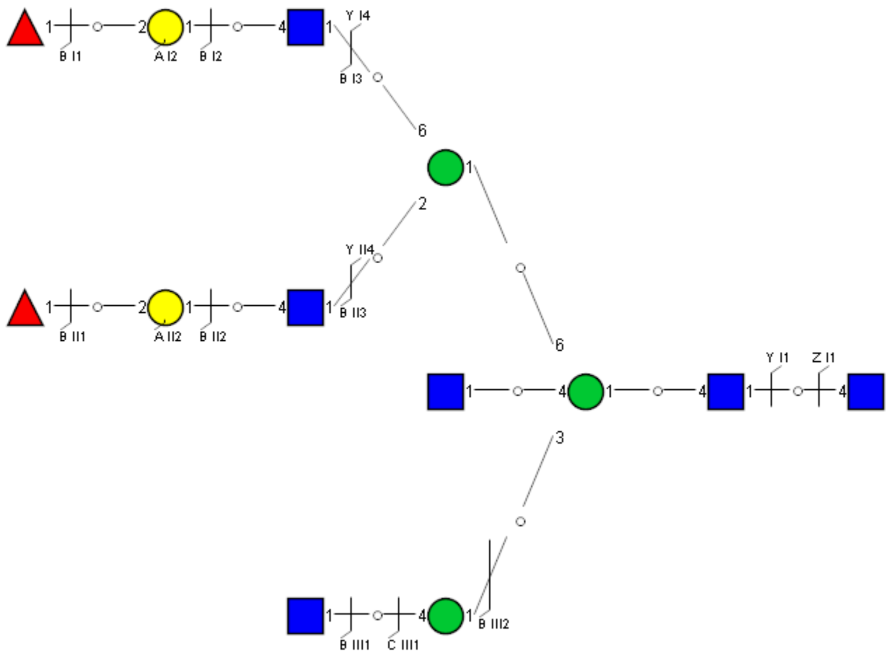

Spectrum Index: 38802(NoBHs:1)

#### Precursor iEF Map

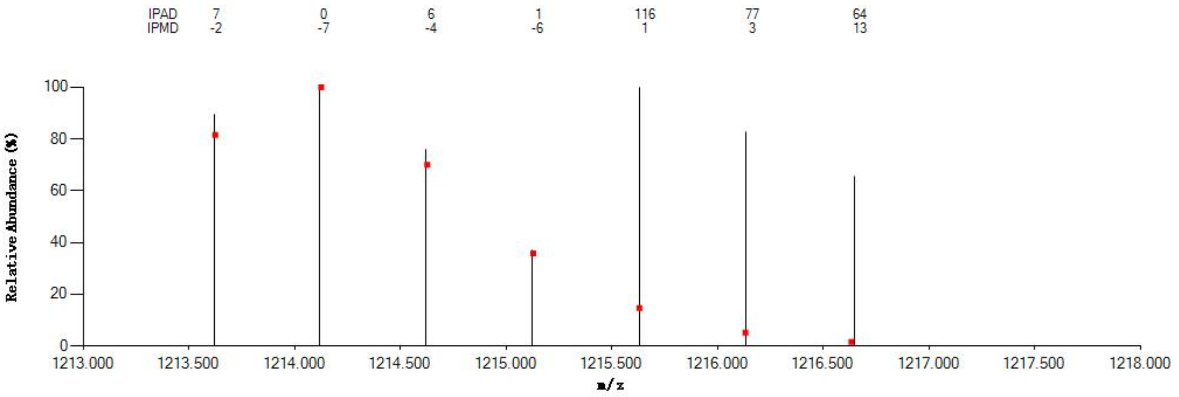


#### Annotated MS/MS Spectrum

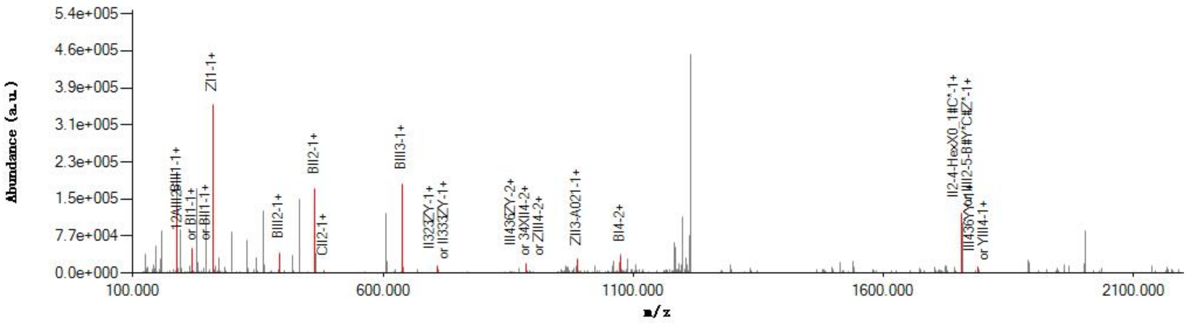


#### Graphical Fragmentation Map

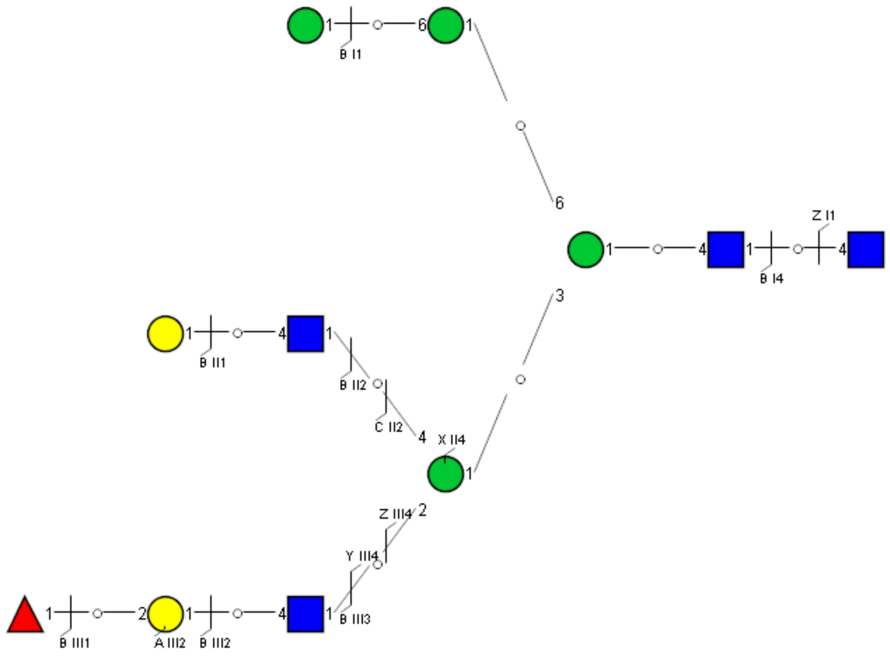

Spectrum Index: 39015(NoBHs:1)

#### Precursor iEF Map

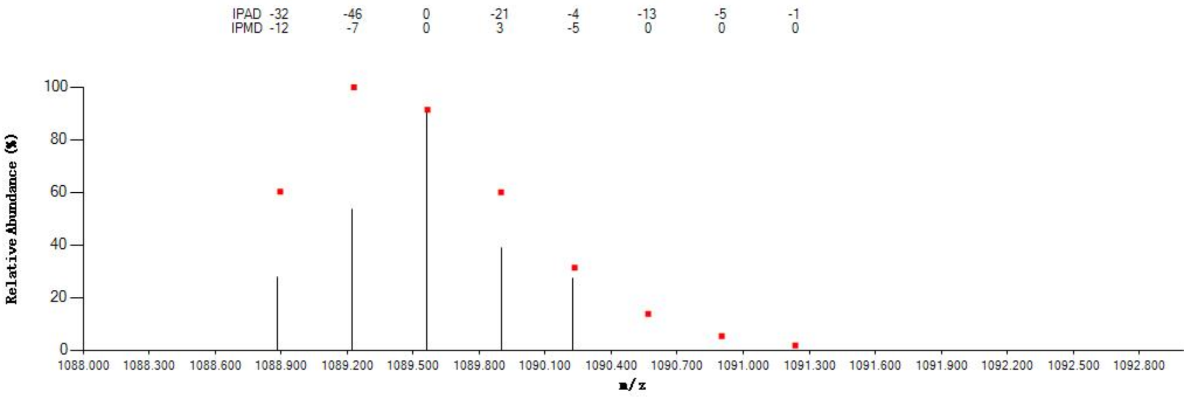


#### Annotated MS/MS Spectrum

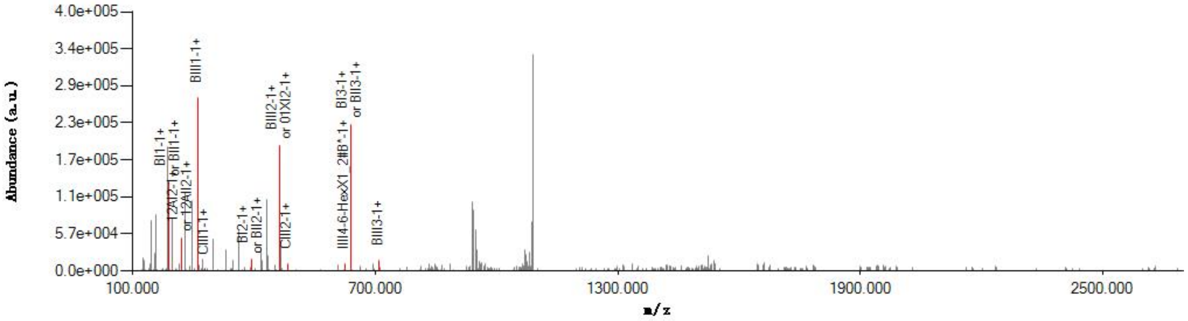


#### Graphical Fragmentation Map

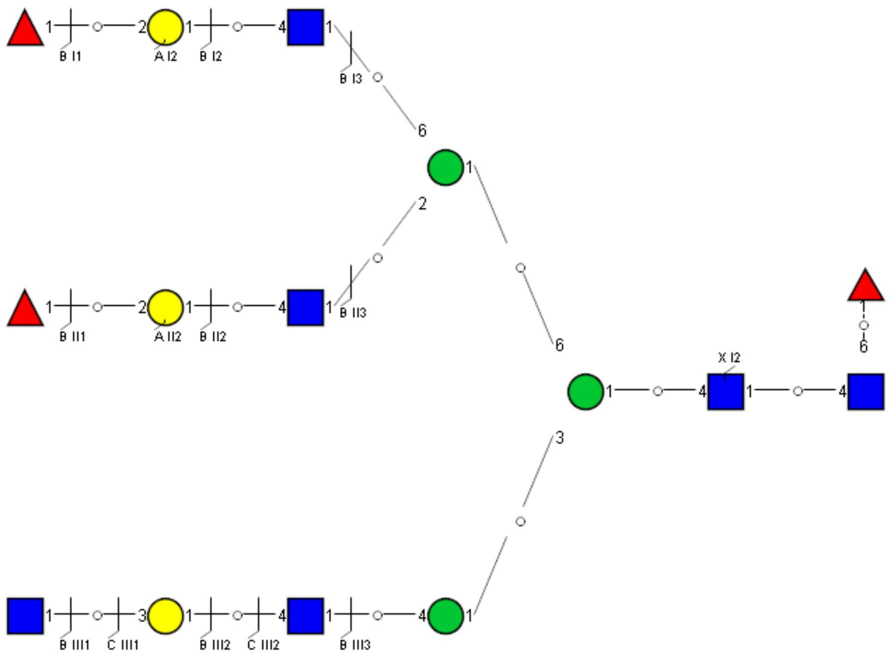

Spectrum Index: 39061(NoBHs:1)

#### Precursor iEF Map

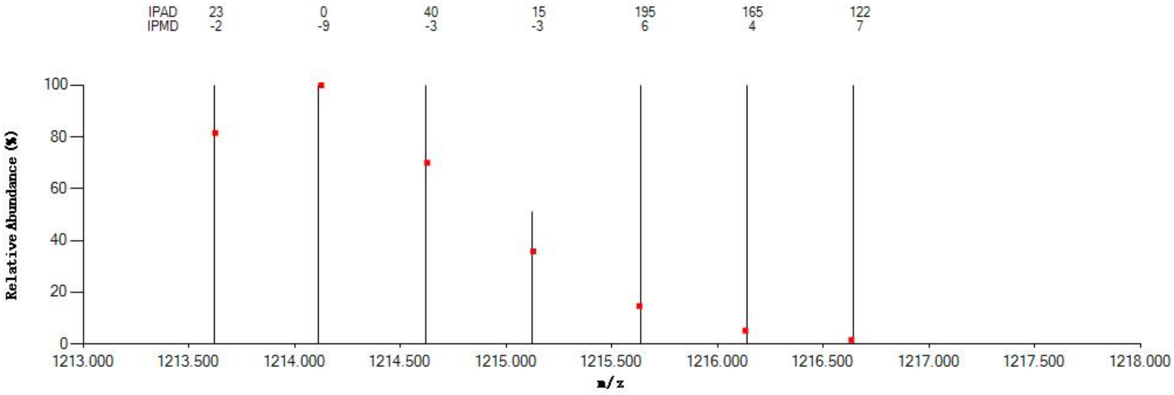


#### Annotated MS/MS Spectrum

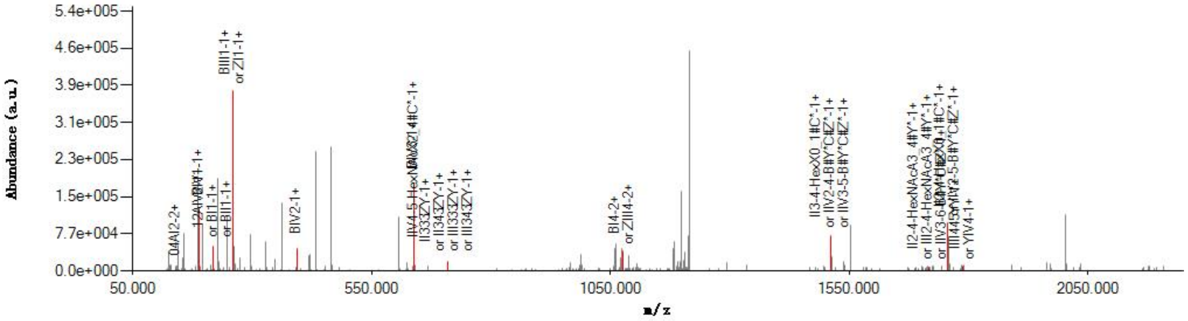


#### Graphical Fragmentation Map

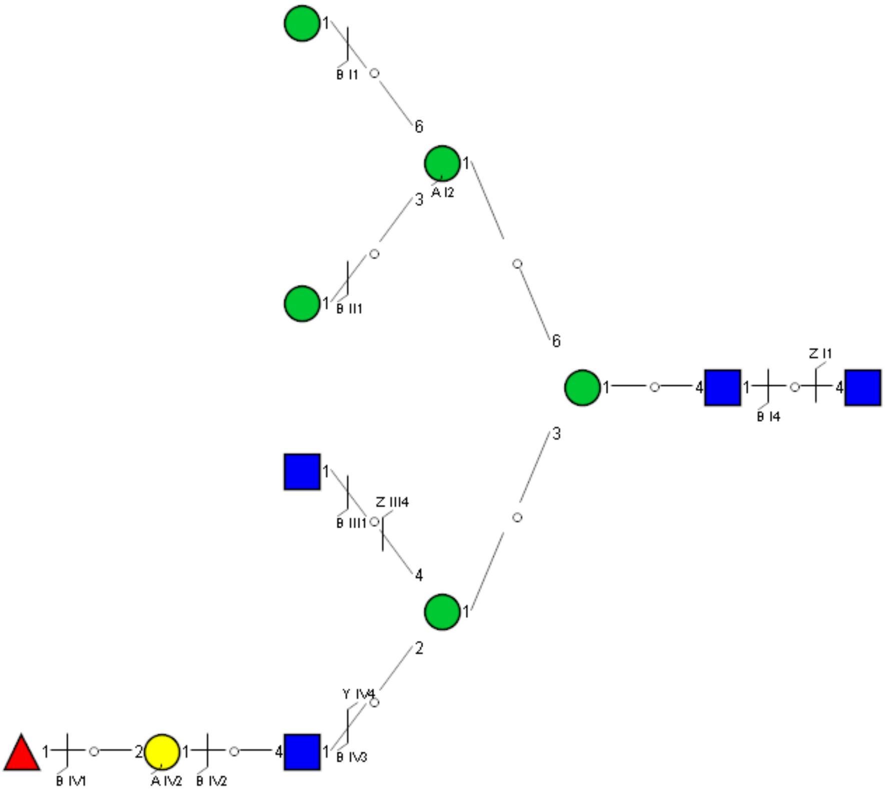

Spectrum Index: 39093(NoBHs:1)

#### Precursor iEF Map

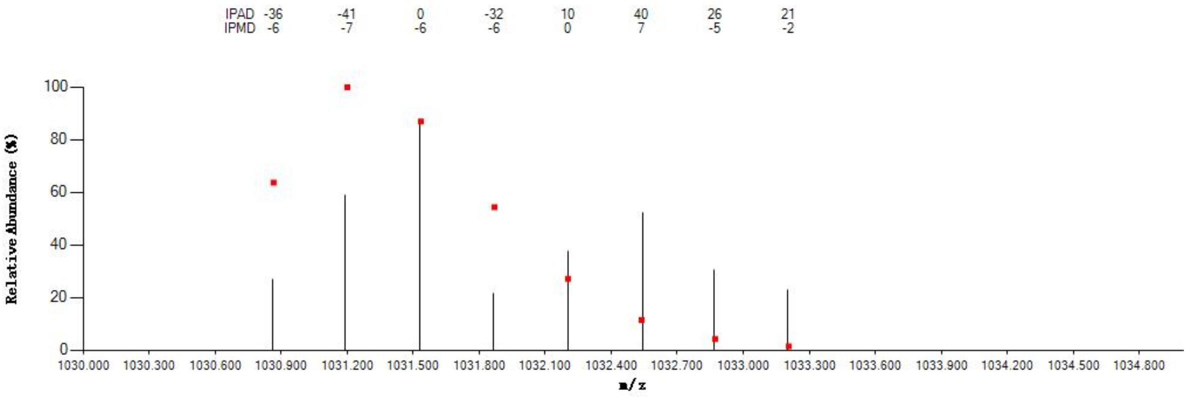


#### Annotated MS/MS Spectrum

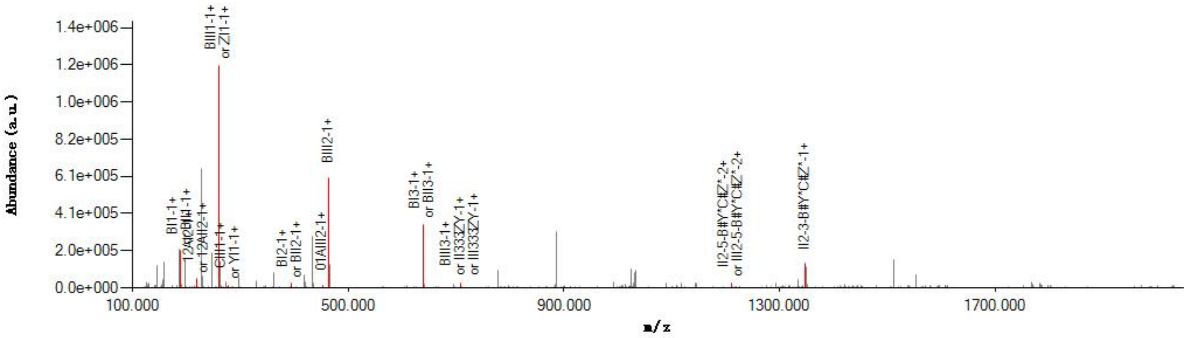


#### Graphical Fragmentation Map

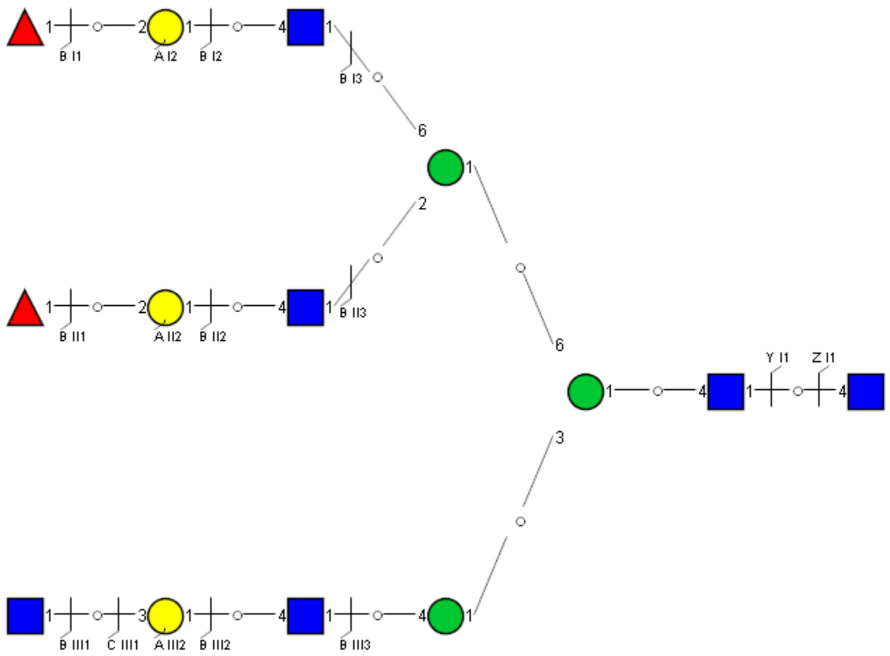

Spectrum Index: 39121(NoBHs:1)

#### Precursor iEF Map

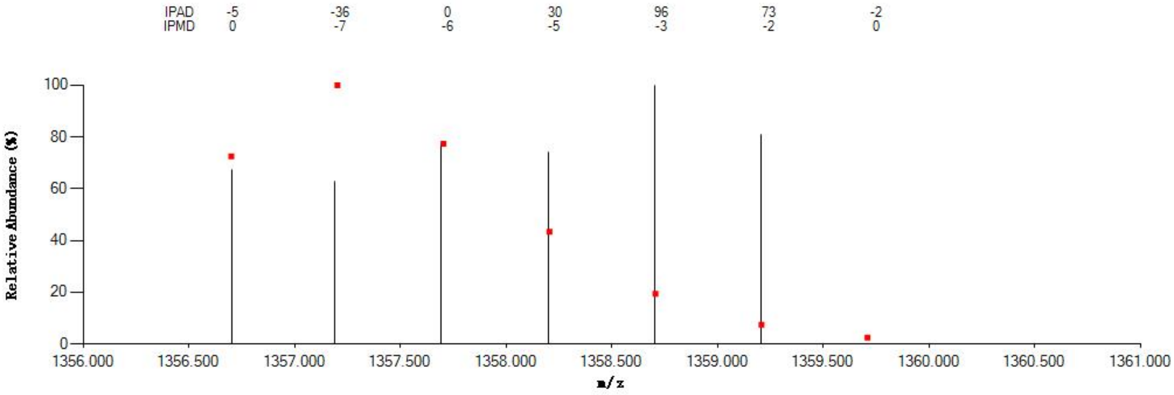

#### Annotated MS/MS Spectrum

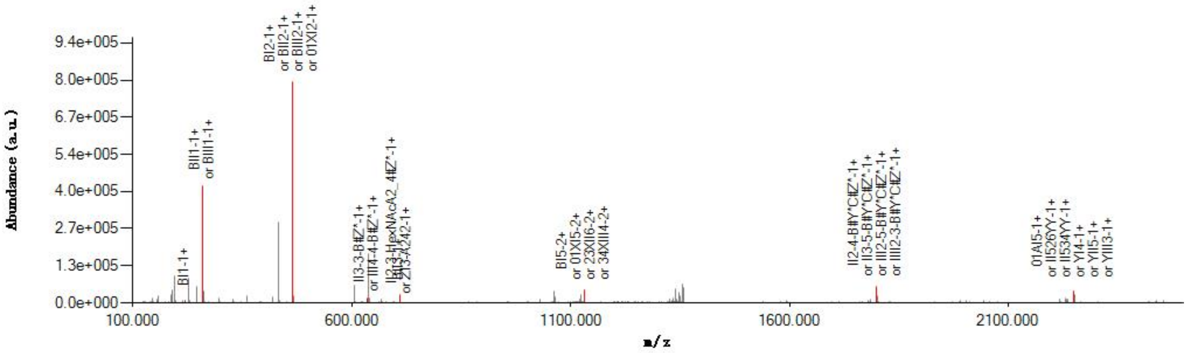

#### Graphical Fragmentation Map

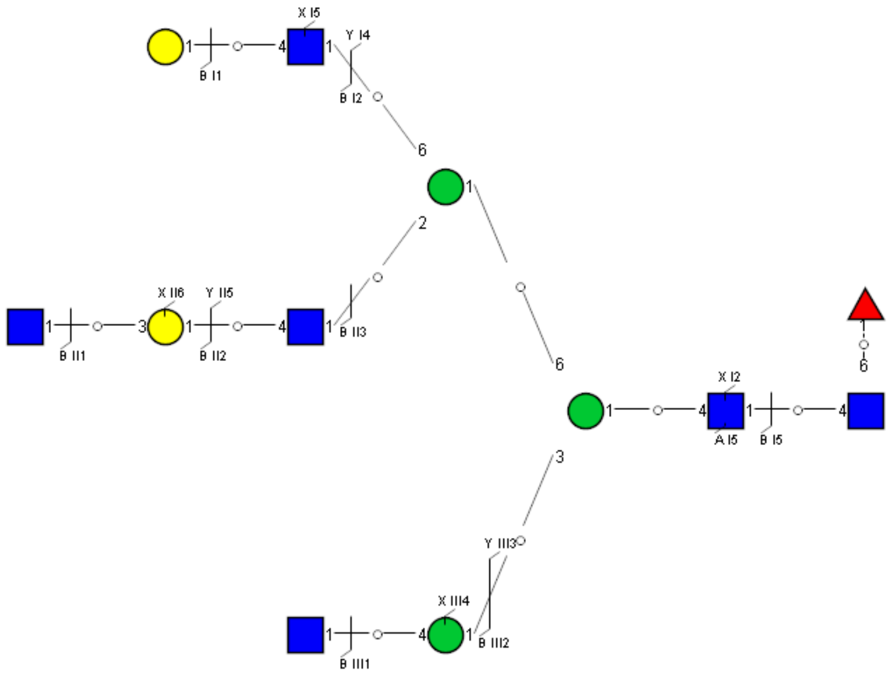

Spectrum Index: 39141(NoBHs:1)

Precursor iEF Map

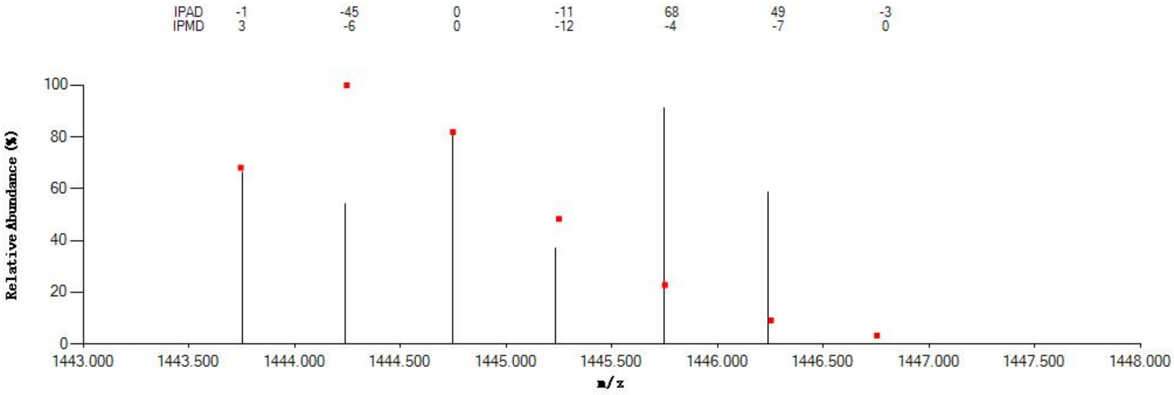


Annotated MS/MS Spectrum

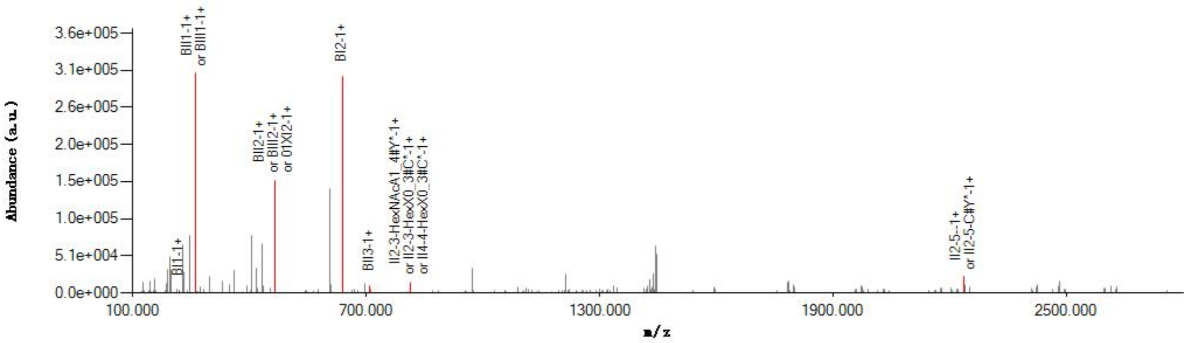


Graphical Fragmentation Map

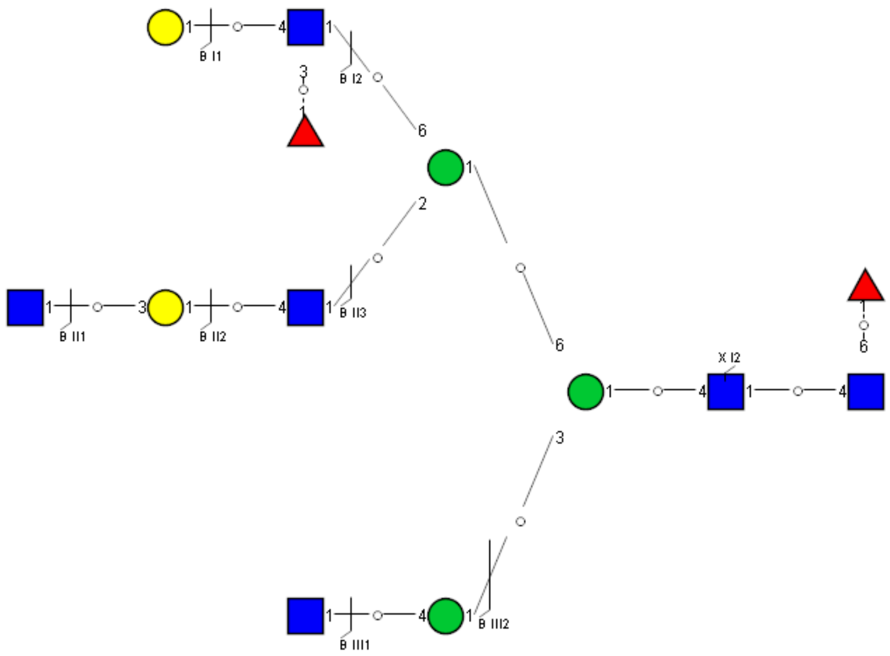

Spectrum Index: 39239(NoBHs:1)

#### Precursor iEF Map

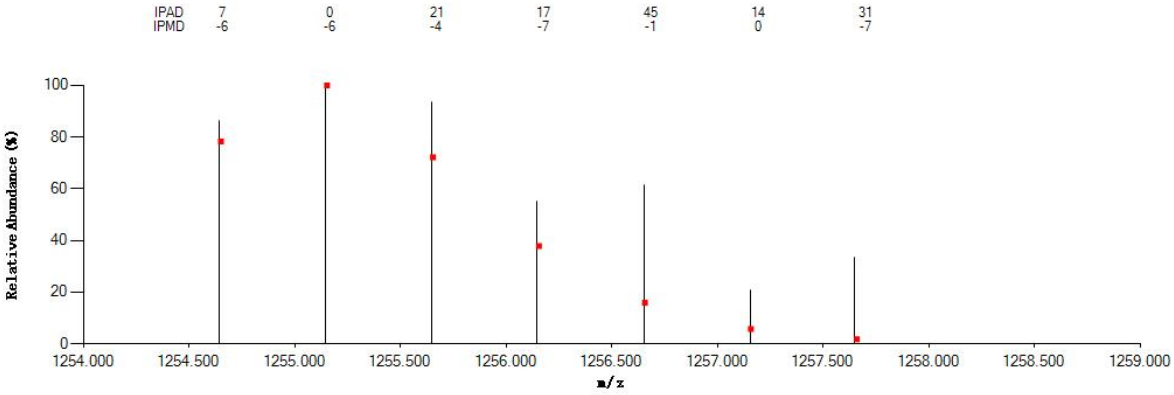

#### Annotated MS/MS Spectrum

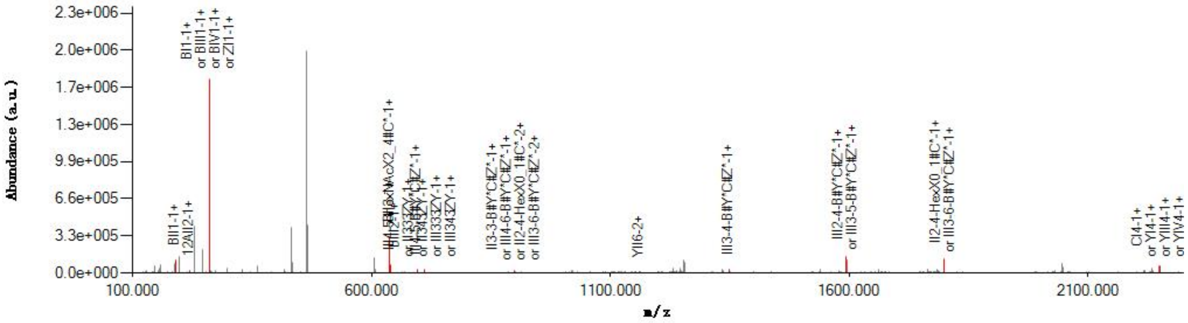

#### Graphical Fragmentation Map

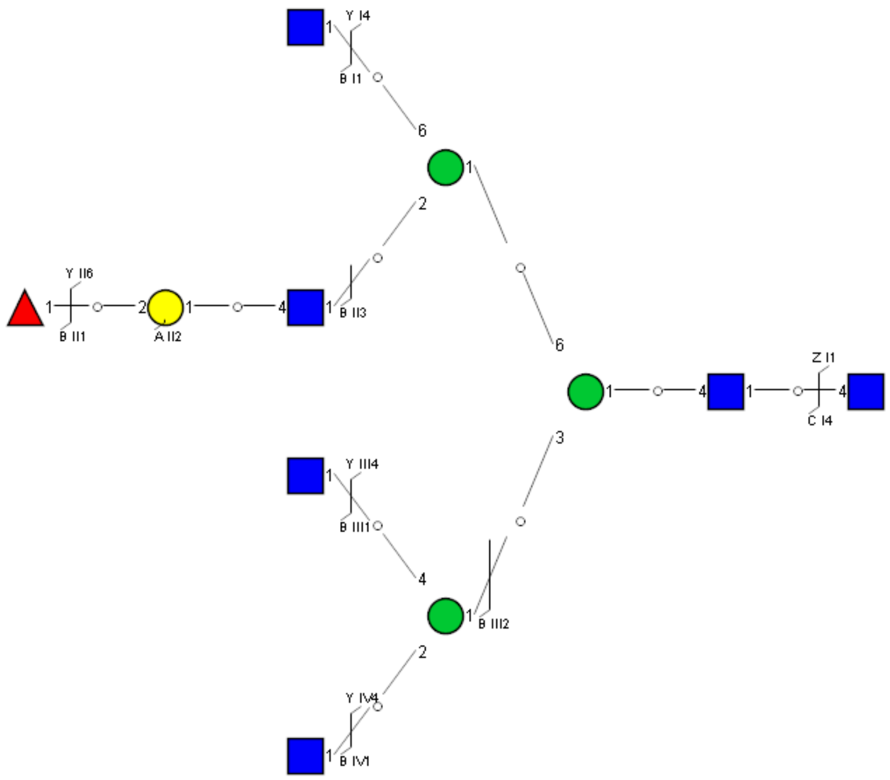

### Spectrum Index: 39298(NoBHs:1)

#### Precursor iEF Map

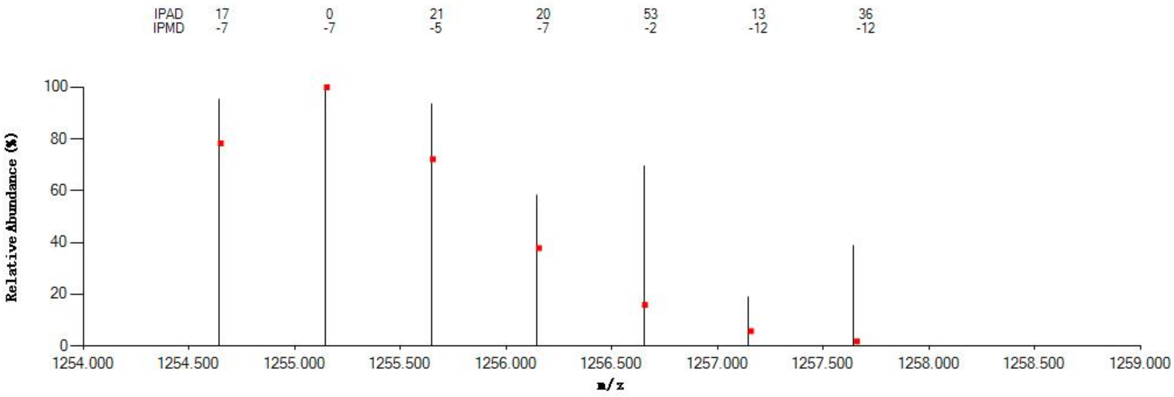


#### Annotated MS/MS Spectrum

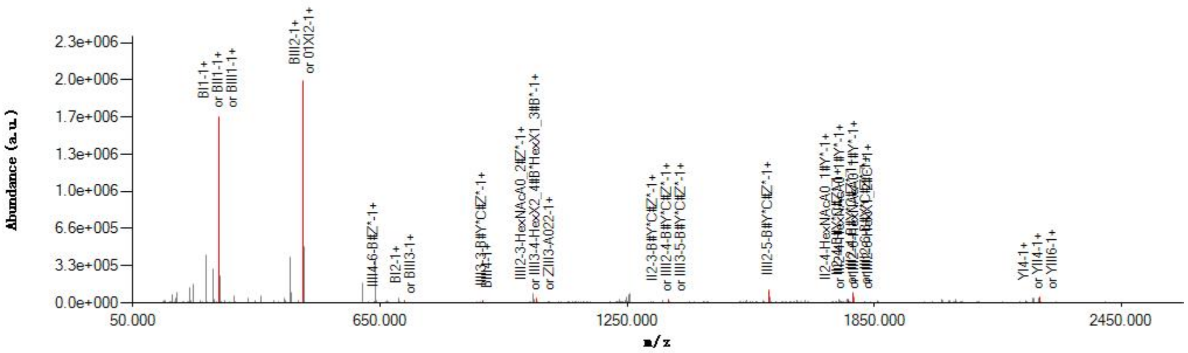


#### Graphical Fragmentation Map

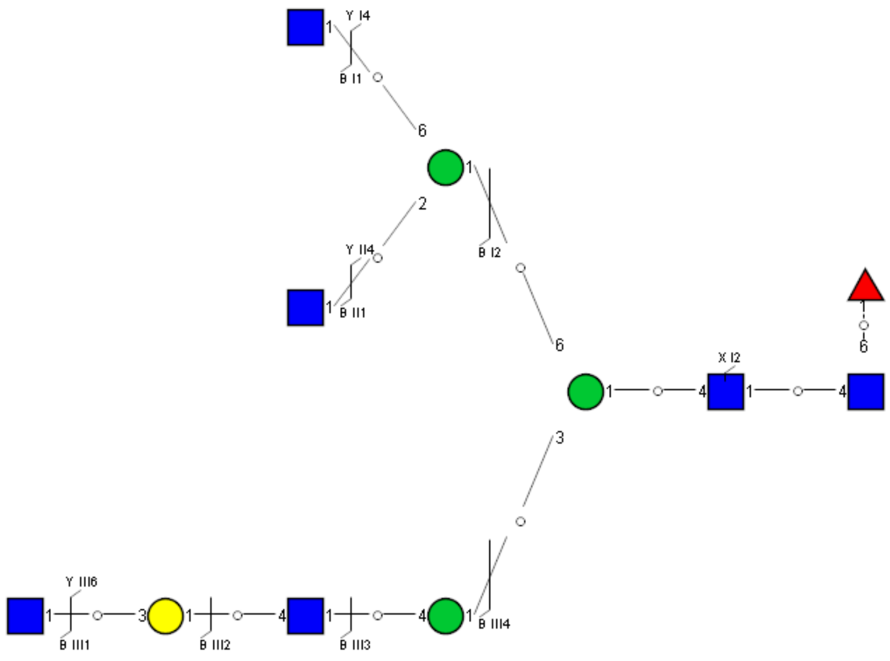

Spectrum Index: 39343(NoBHs:1)

#### Precursor iEF Map

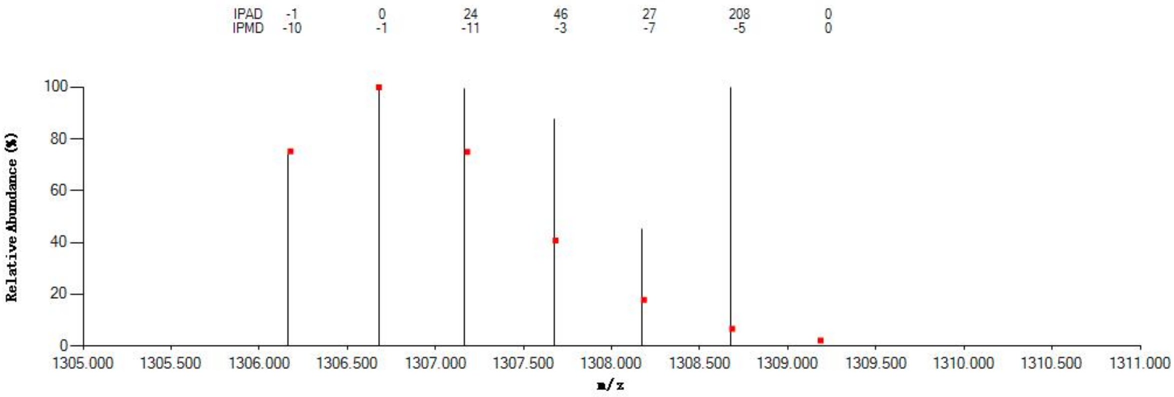


#### Annotated MS/MS Spectrum

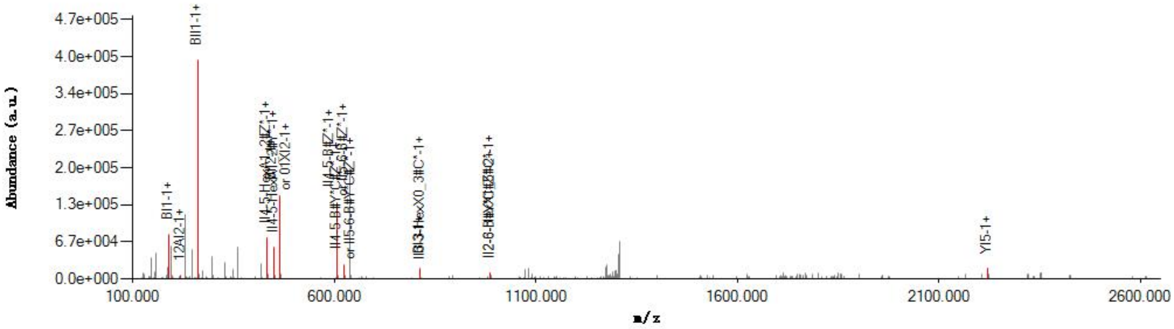


#### Graphical Fragmentation Map

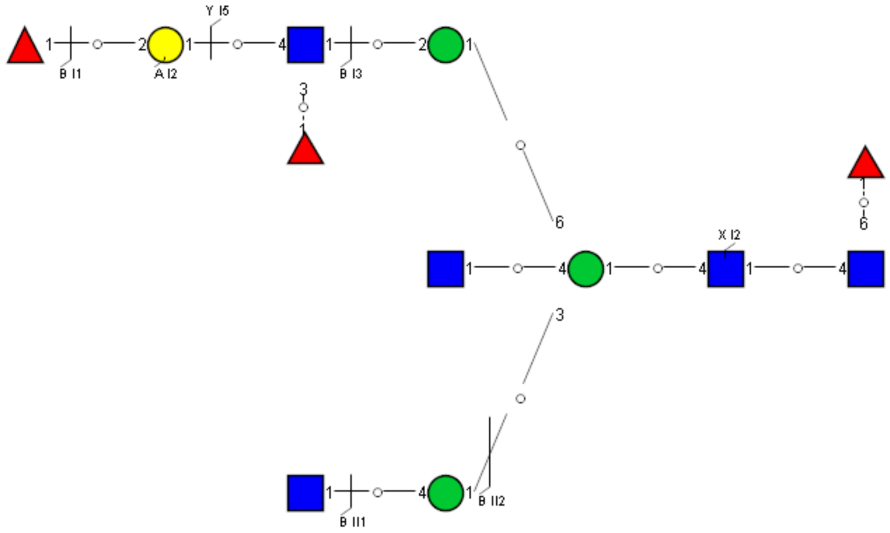

Spectrum Index: 39367(NoBHs:1)

#### Precursor iEF Map

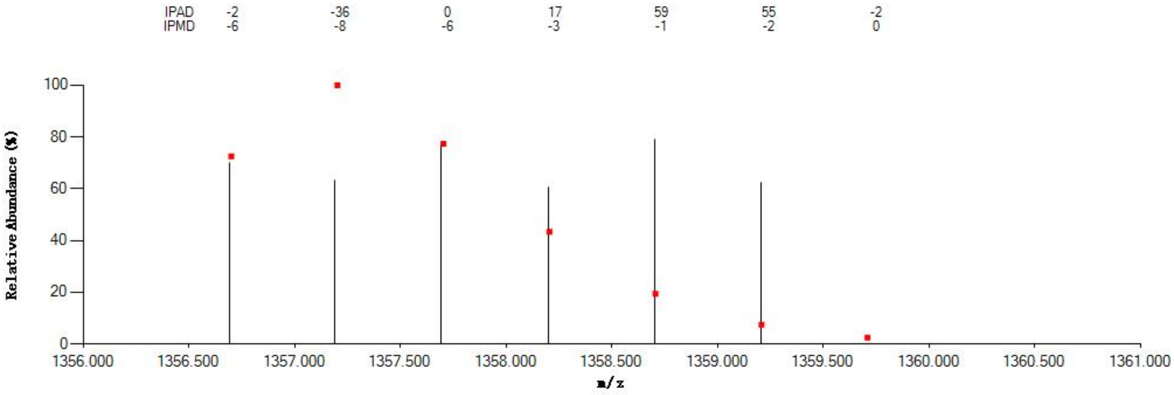


#### Annotated MS/MS Spectrum

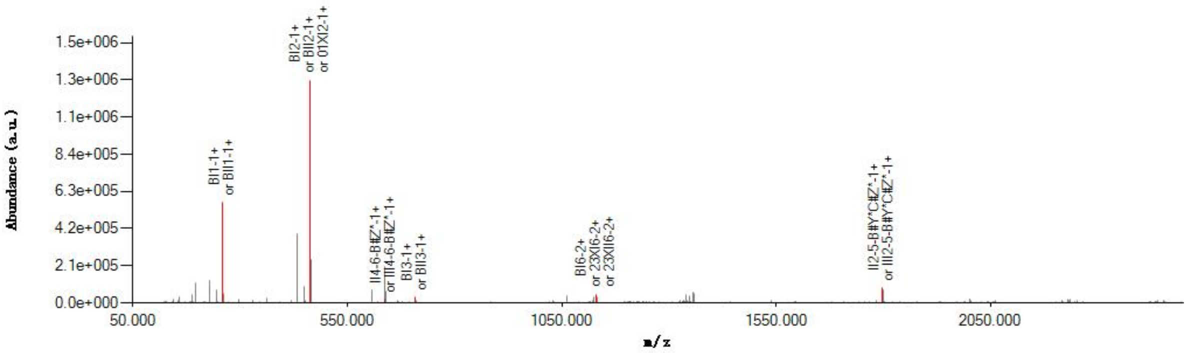


#### Graphical Fragmentation Map

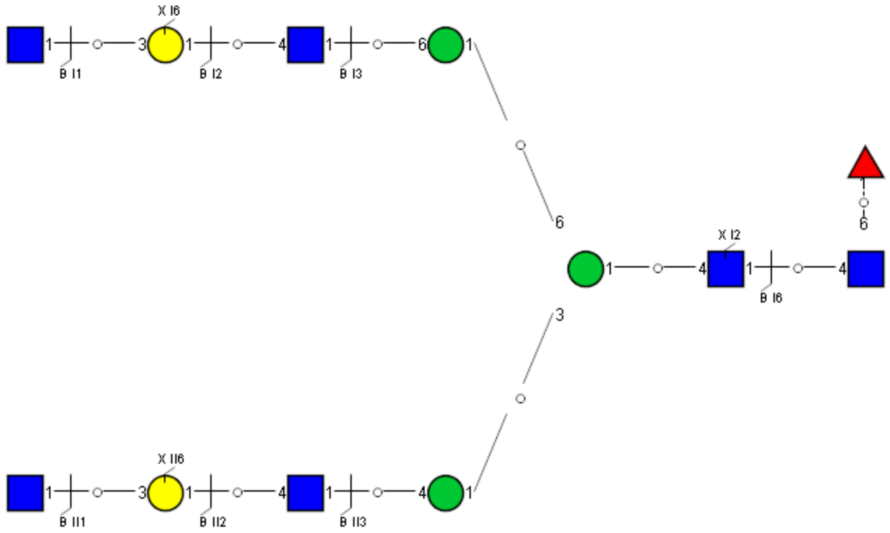

Spectrum Index: 39498(NoBHs:1)

#### Precursor iEF Map

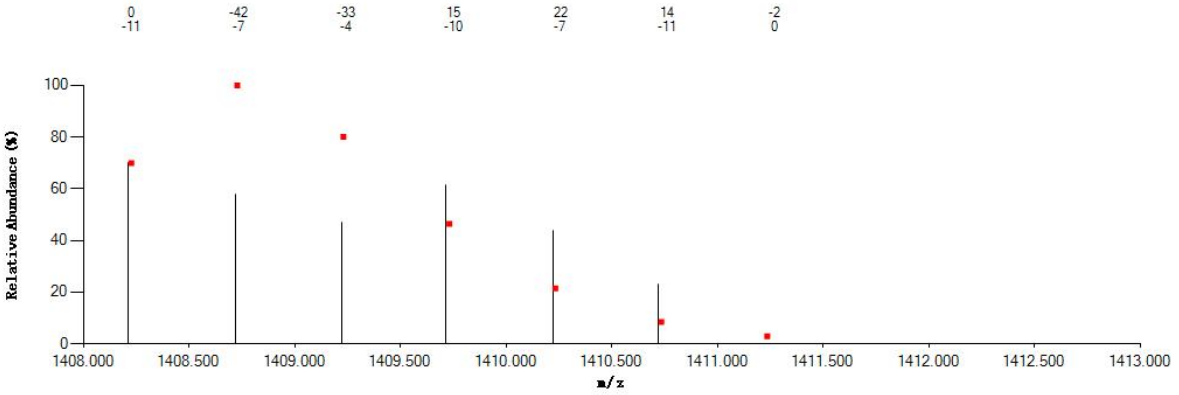


#### Annotated MS/MS Spectrum

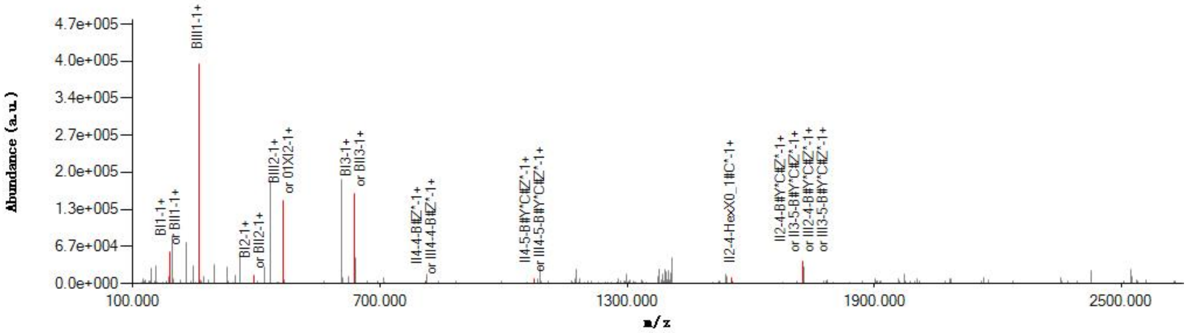


#### Graphical Fragmentation Map

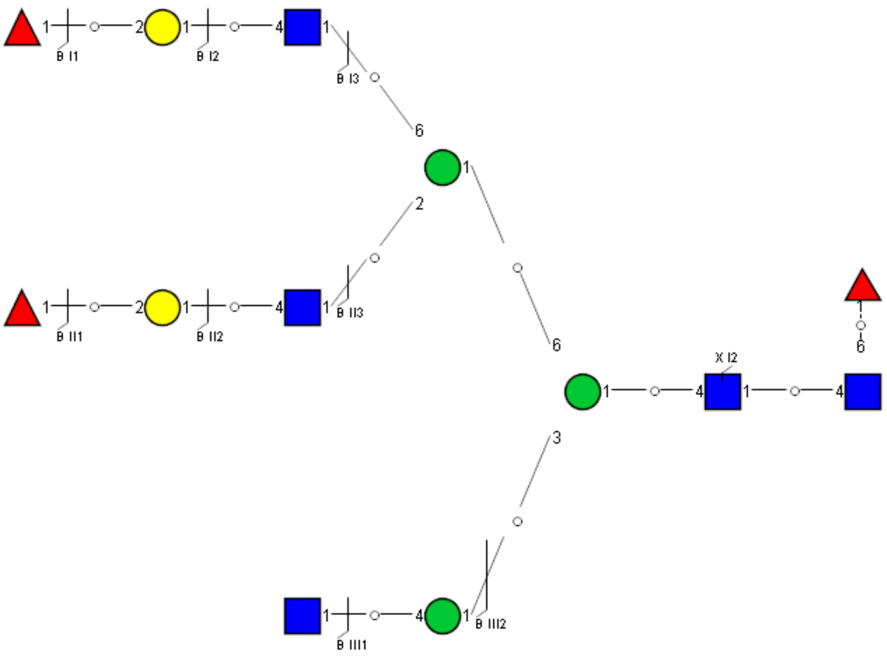

Spectrum Index: 39539(NoBHs:1)

#### Precursor iEF Map

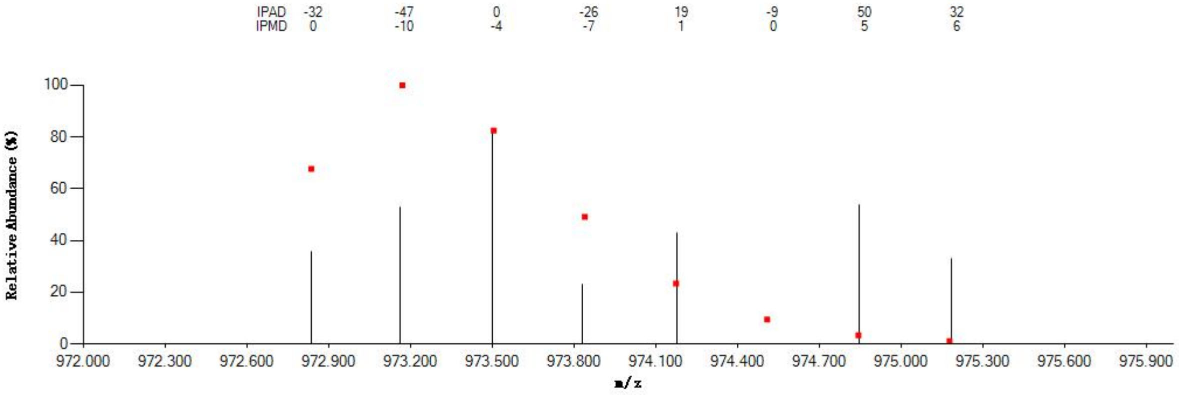


#### Annotated MS/MS Spectrum

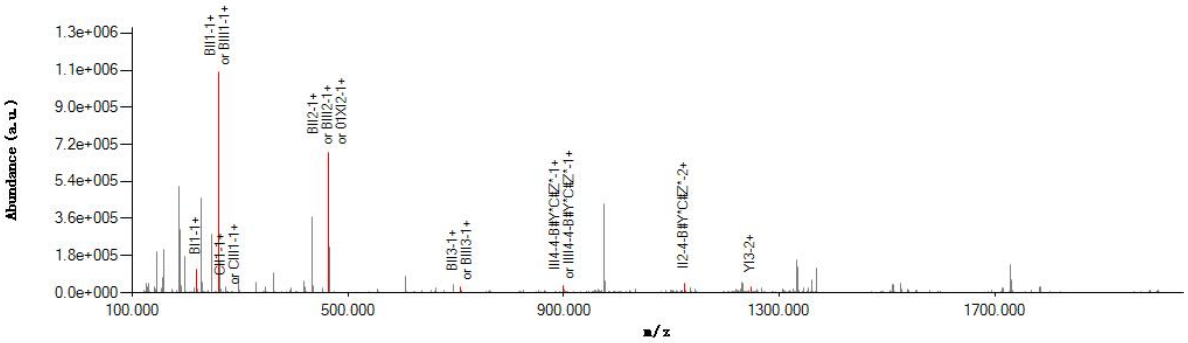


#### Graphical Fragmentation Map

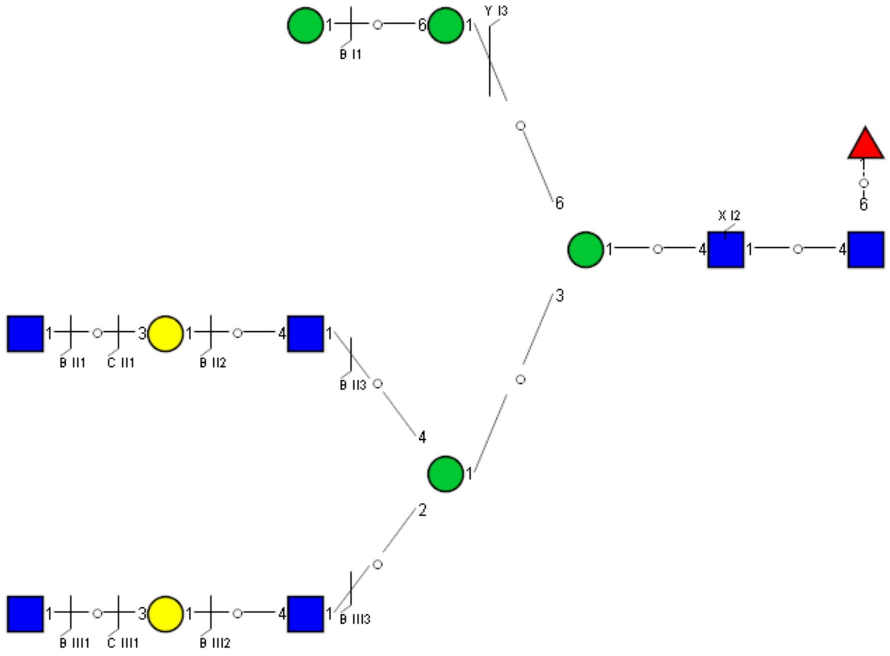

Spectrum Index: 39555(NoBHs:1)

#### Precursor iEF Map

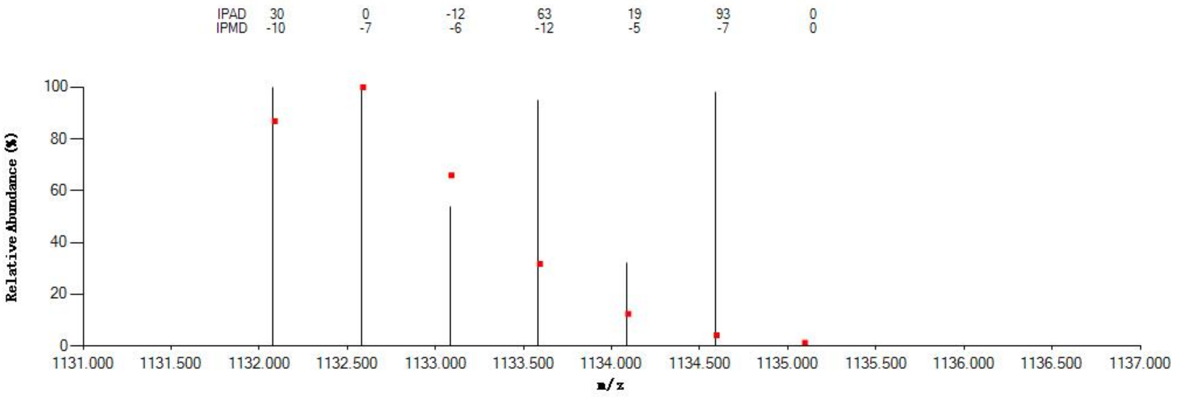

#### Annotated MS/MS Spectrum

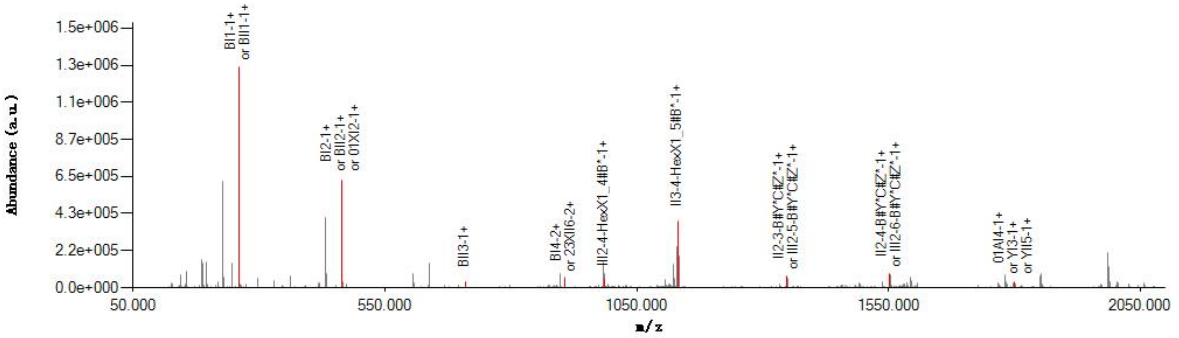

#### Graphical Fragmentation Map

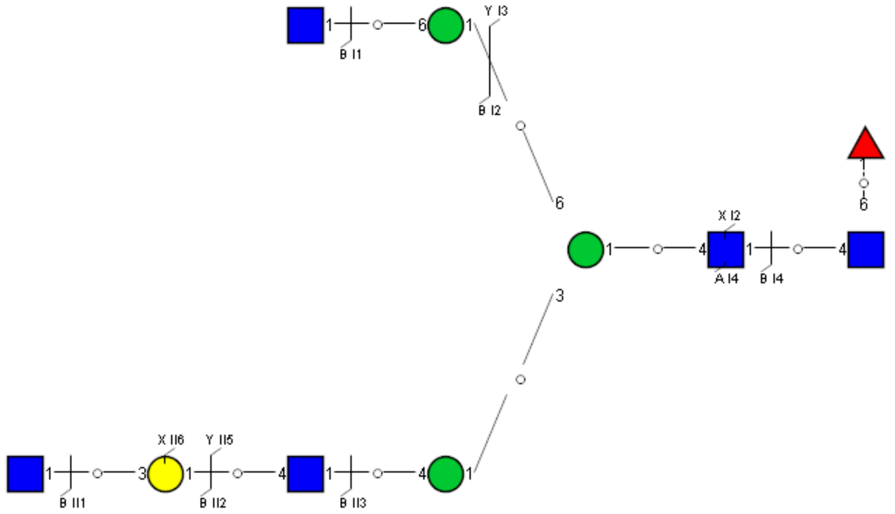

Spectrum Index: 39563(NoBHs:2)

#### Precursor iEF Map

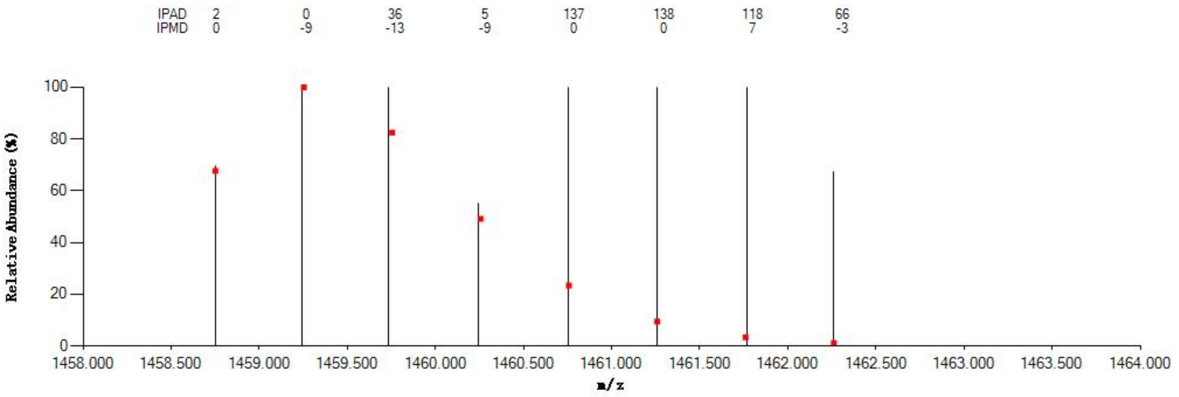


#### Annotated MS/MS Spectrum

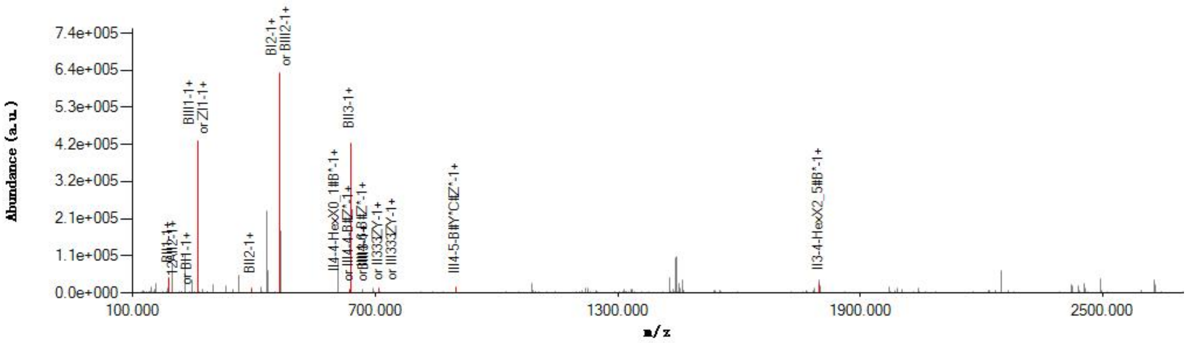


#### Graphical Fragmentation Map

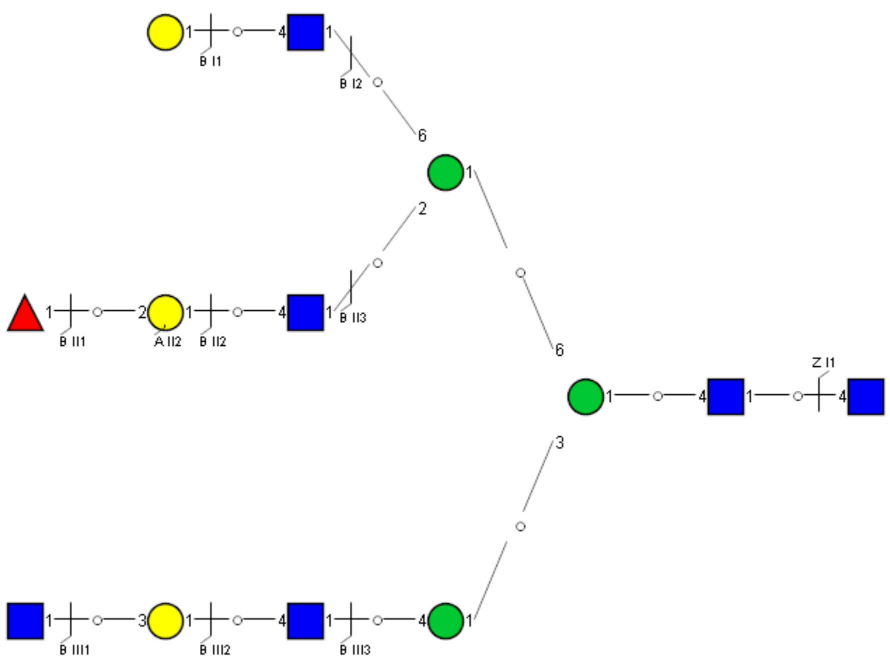

Spectrum Index: 39563(NoBHs:2)

#### Precursor iEF Map

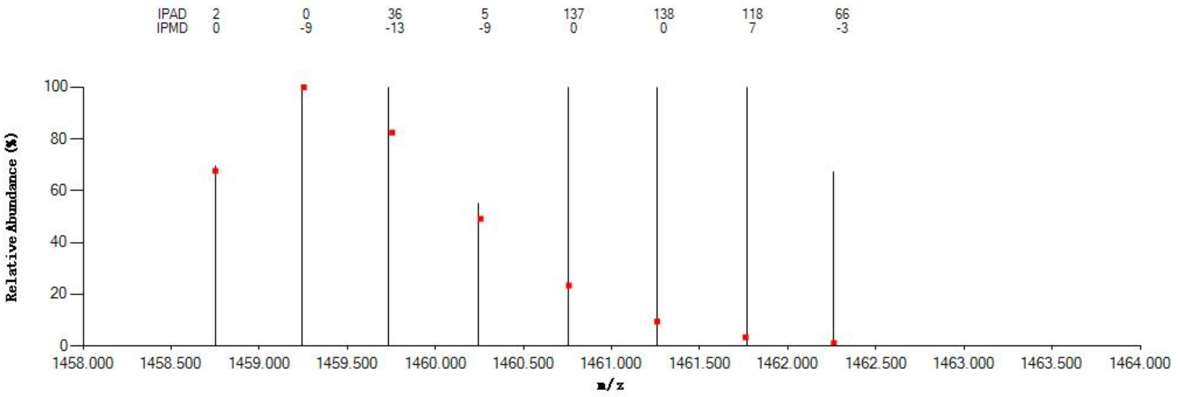


#### Annotated MS/MS Spectrum

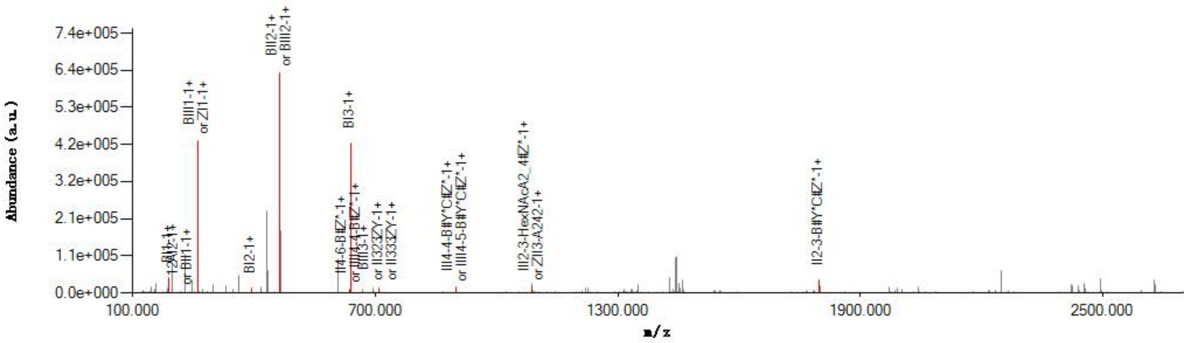


#### Graphical Fragmentation Map

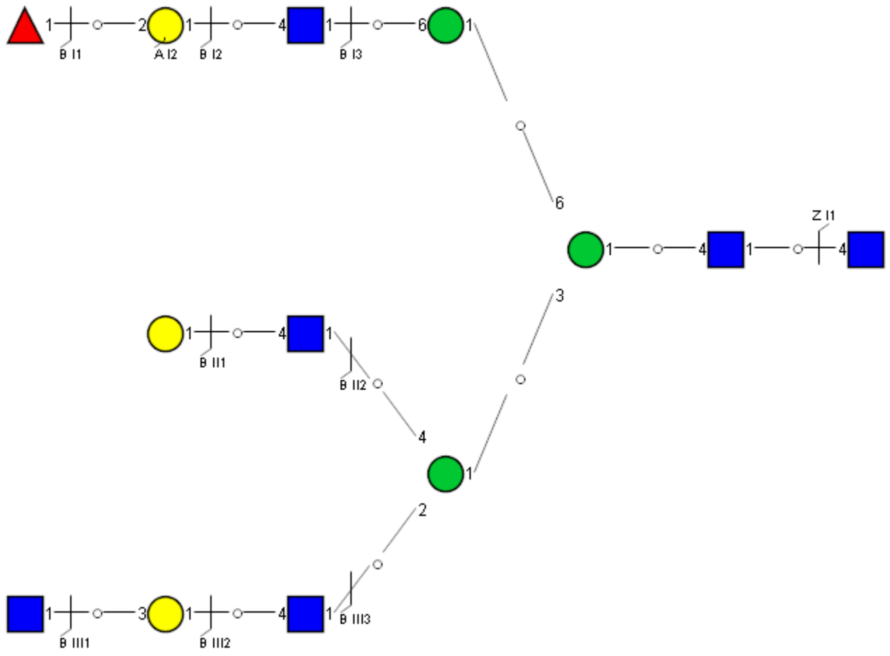

Spectrum Index: 39580(NoBHs:1)

#### Precursor iEF Map

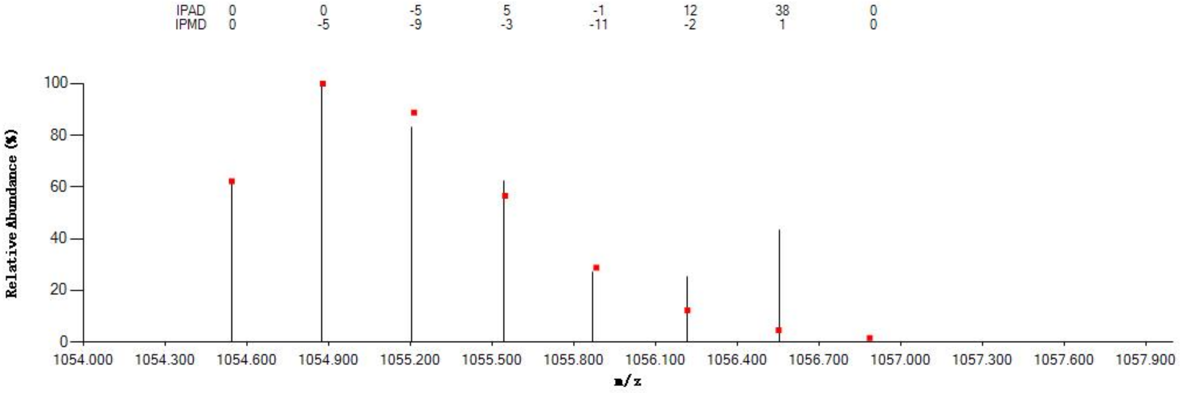


#### Annotated MS/MS Spectrum

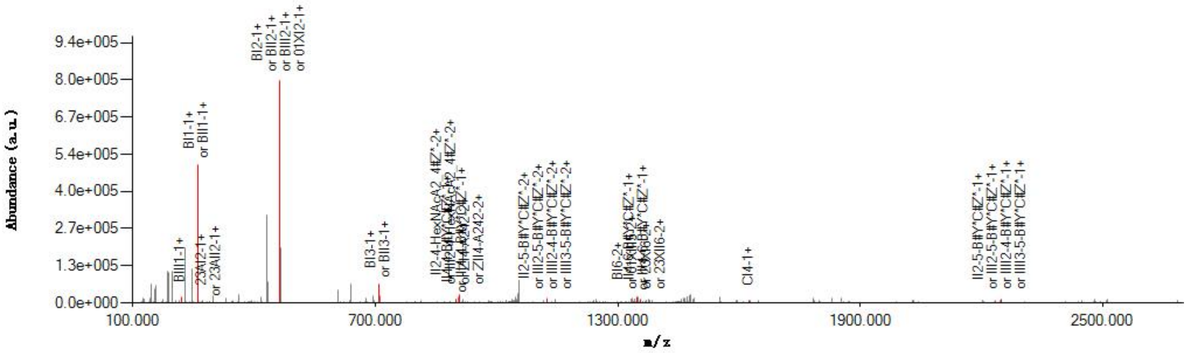


#### Graphical Fragmentation Map

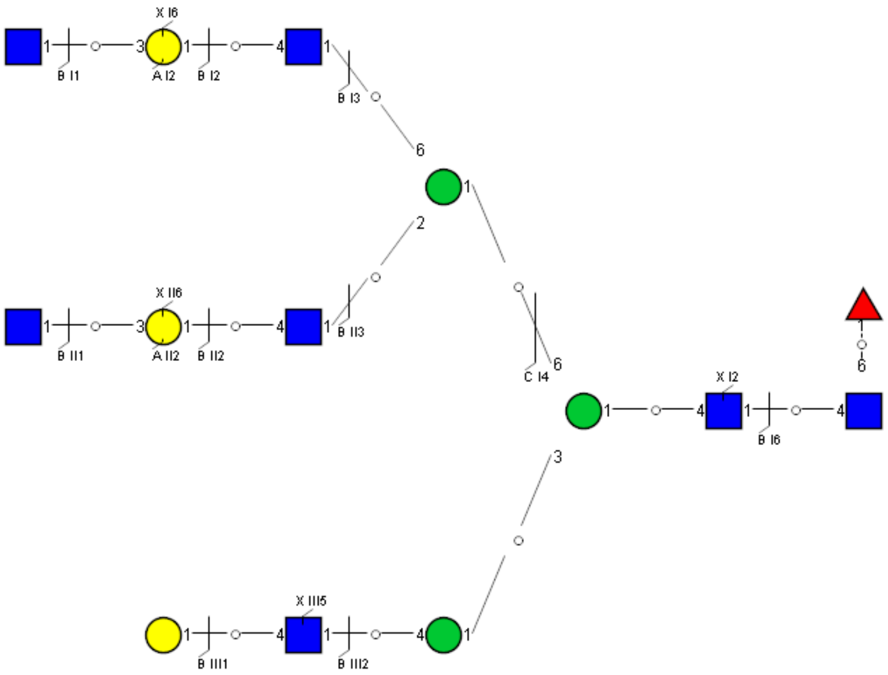

Spectrum Index: 39605(NoBHs:2)

#### Precursor iEF Map

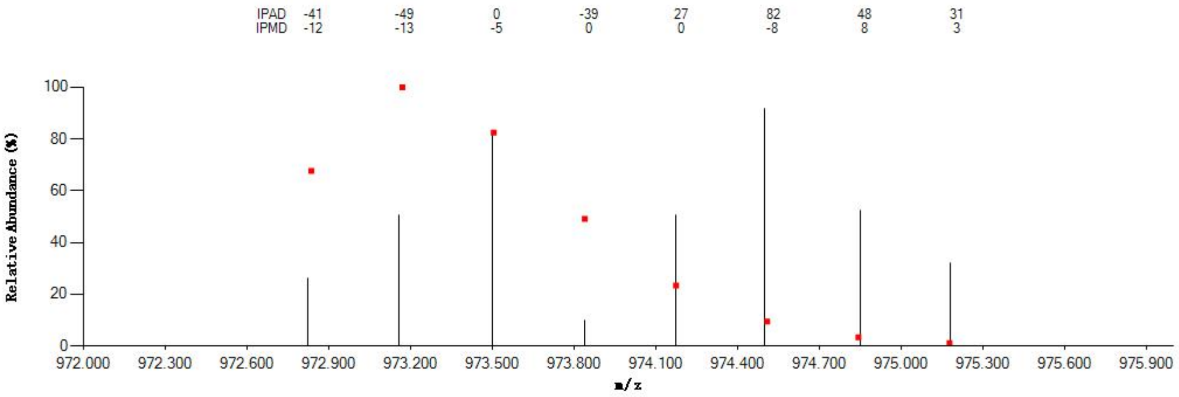


#### Annotated MS/MS Spectrum

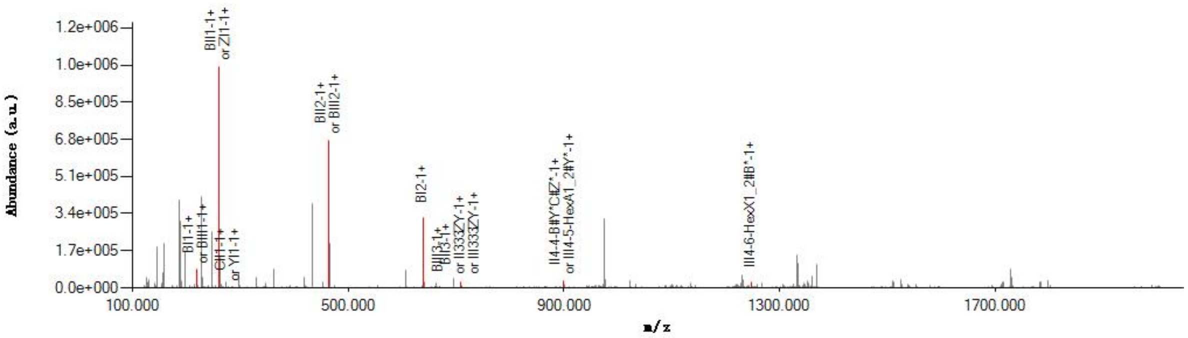


#### Graphical Fragmentation Map

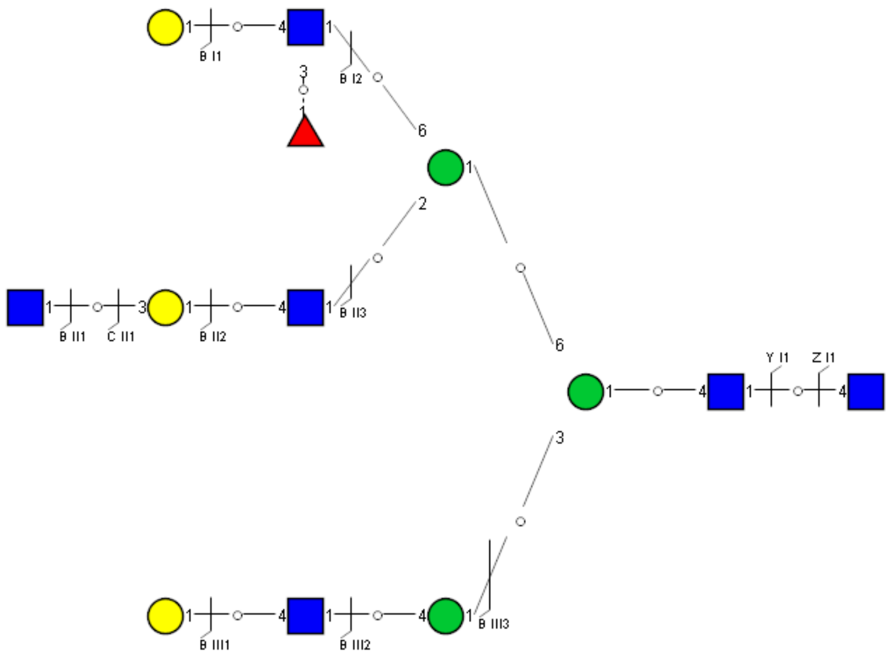

Spectrum Index: 39638(NoBHs:1)

#### Precursor iEF Map

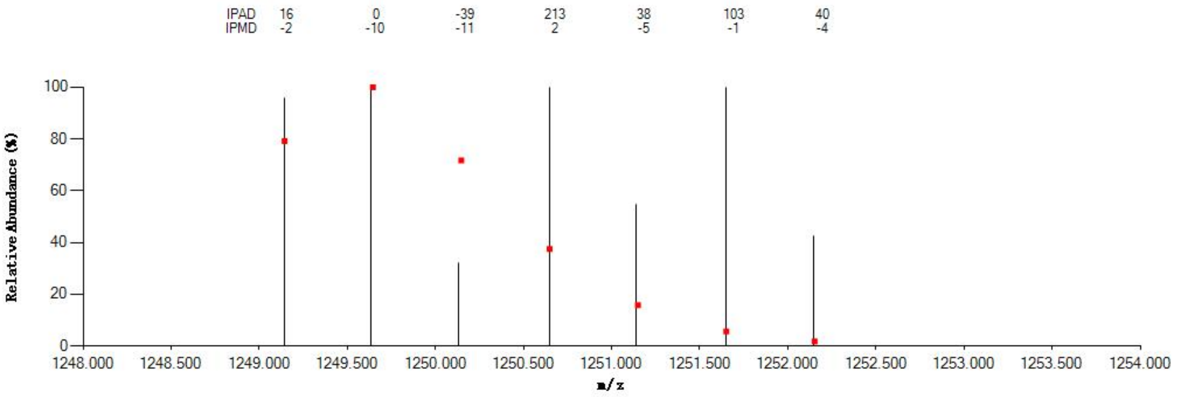

#### Annotated MS/MS Spectrum

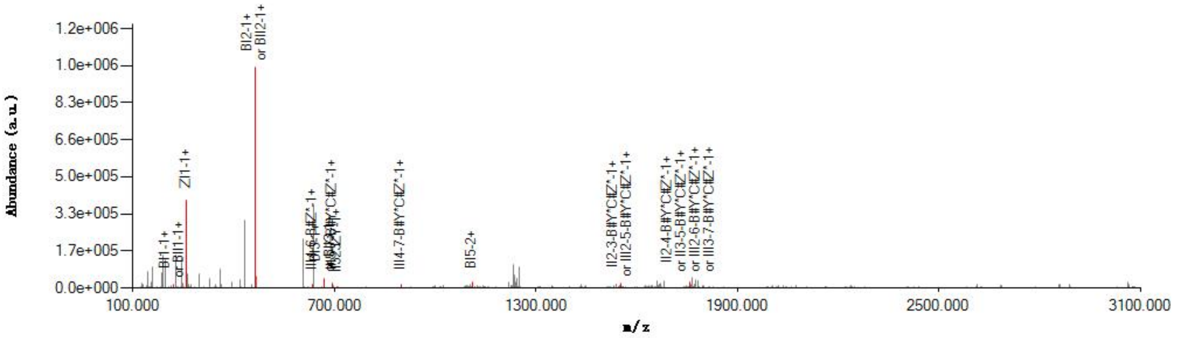

#### Graphical Fragmentation Map

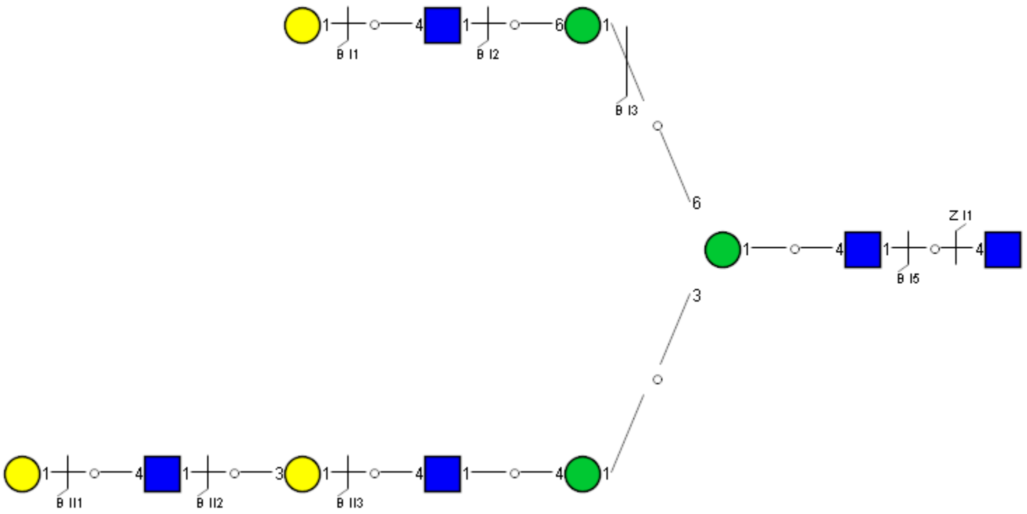

Spectrum Index: 39776(NoBHs:1)

Precursor iEF Map

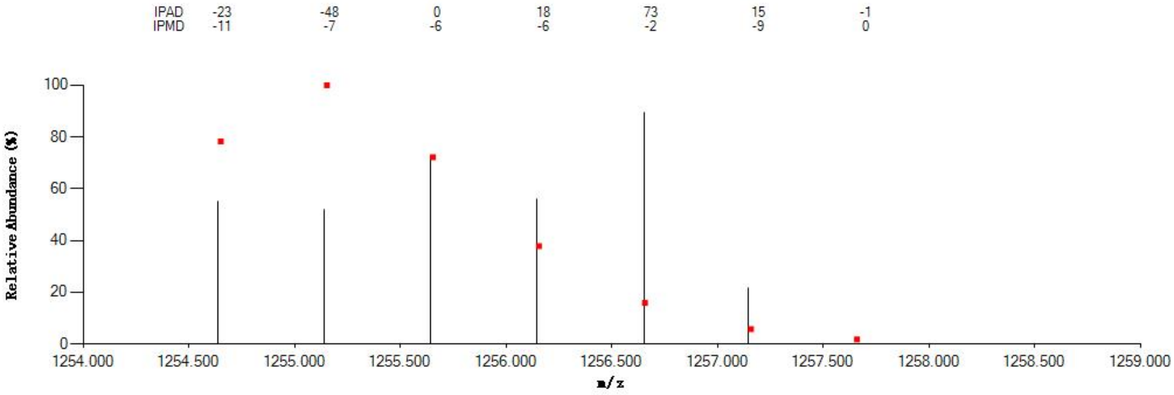

Annotated MS/MS Spectrum

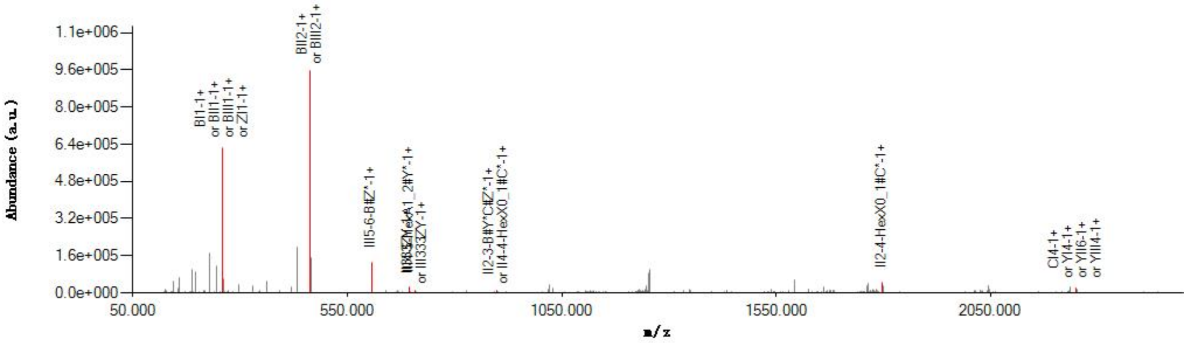

Graphical Fragmentation Map

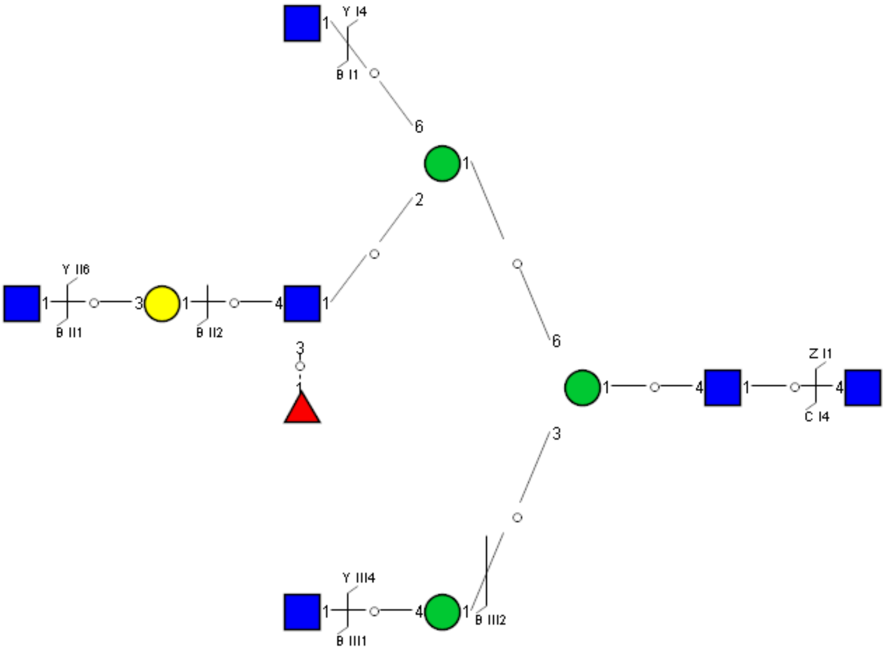

### Spectrum Index: 39796(NoBHs:1)

#### Precursor iEF Map

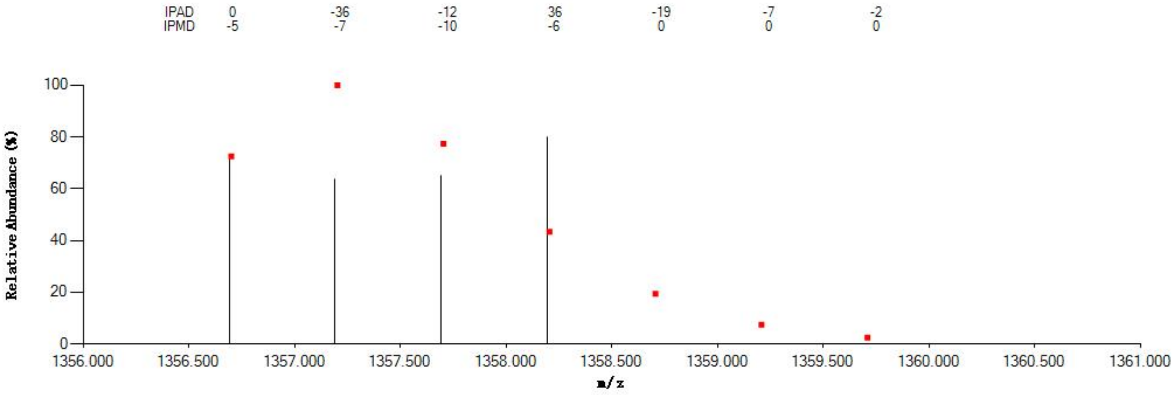


#### Annotated MS/MS Spectrum

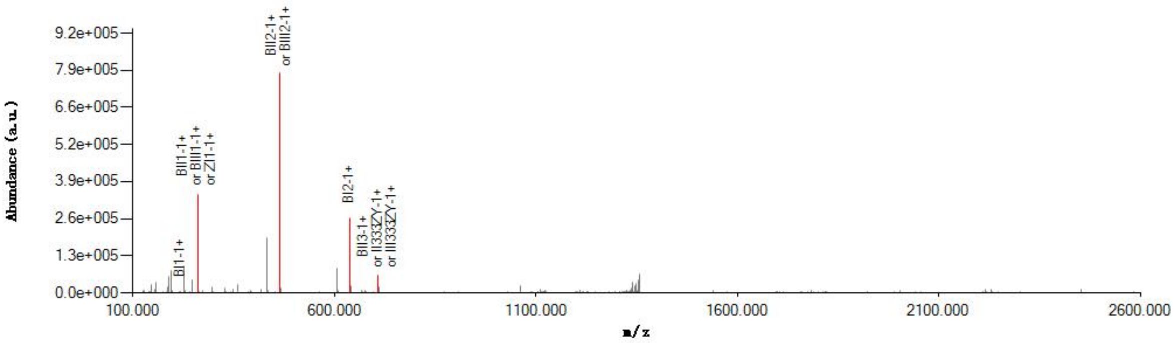


#### Graphical Fragmentation Map

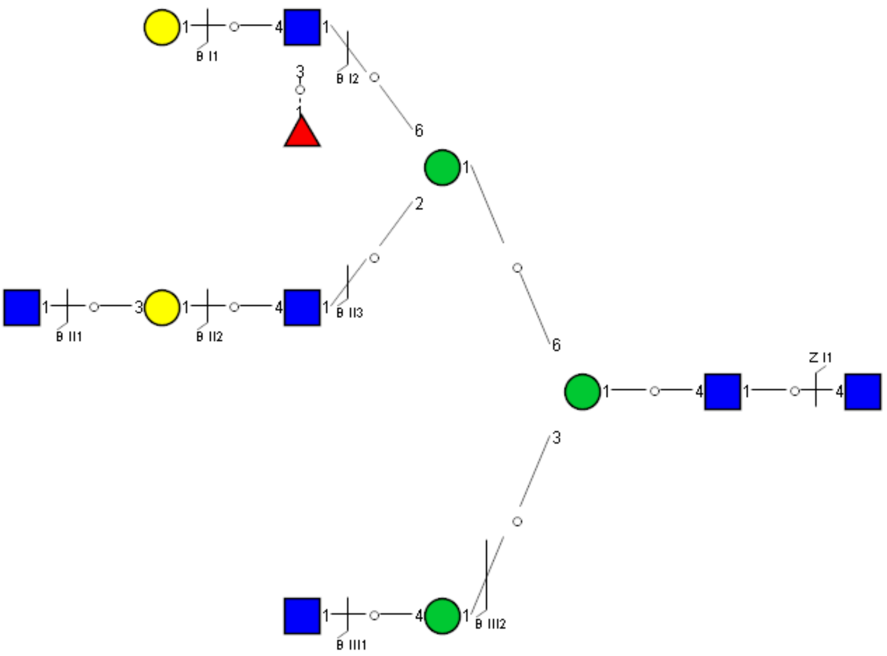

Spectrum Index: 39821(NoBHs:1)

#### Precursor iEF Map

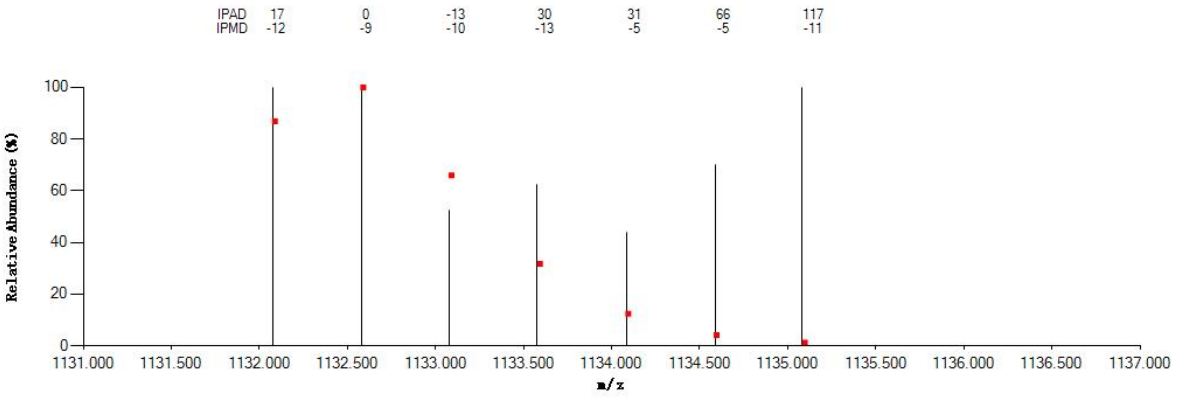


#### Annotated MS/MS Spectrum

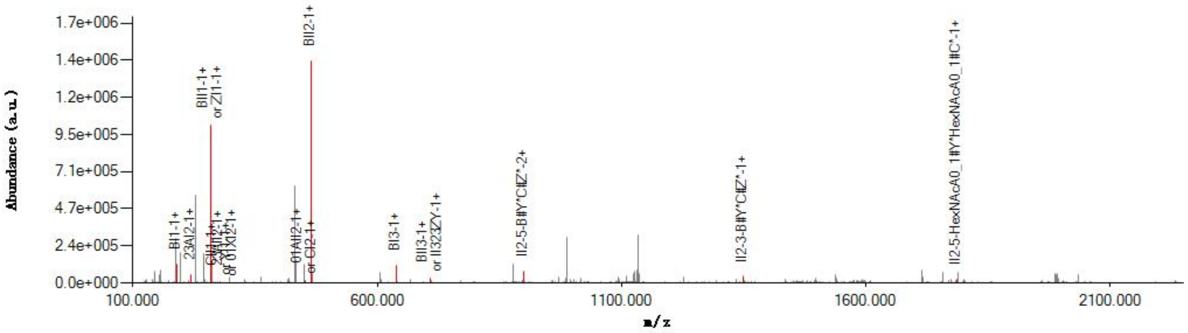


#### Graphical Fragmentation Map

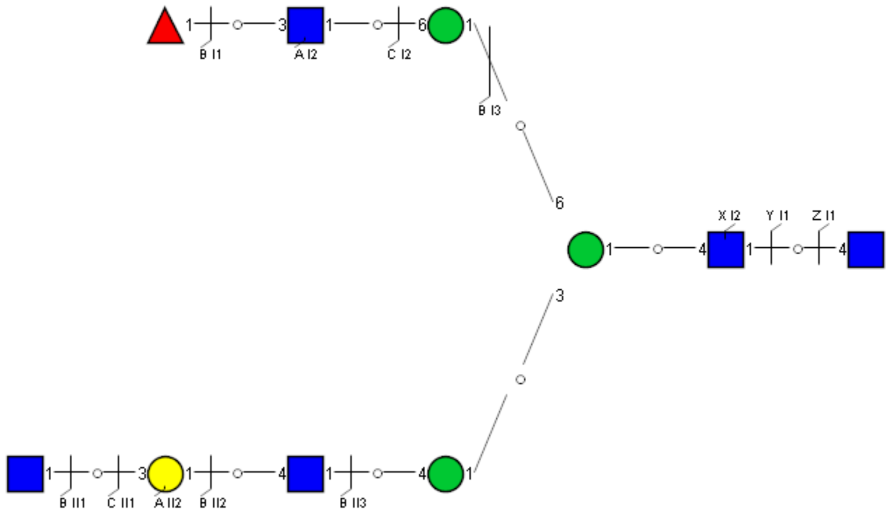

Spectrum Index: 39861(NoBHs:1)

#### Precursor iEF Map

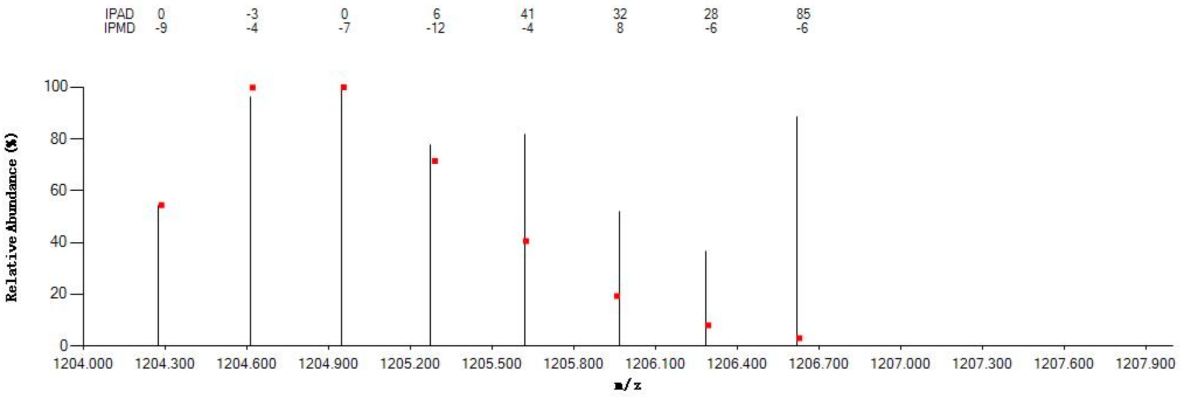


#### Annotated MS/MS Spectrum

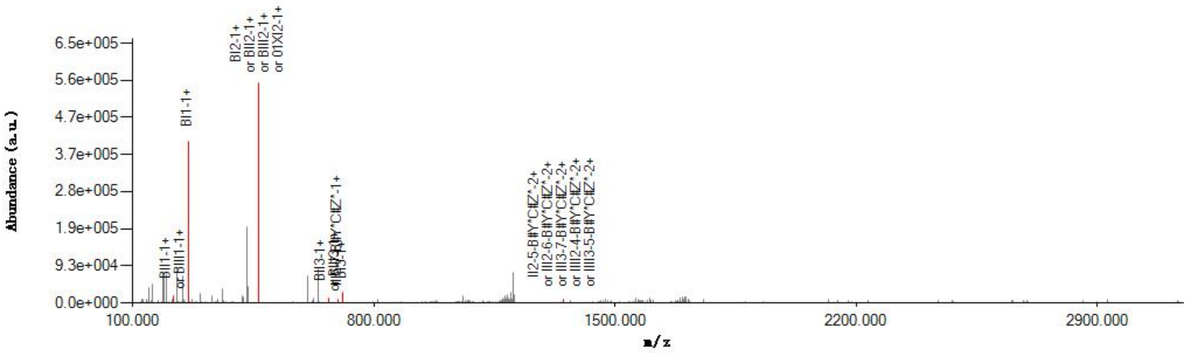


#### Graphical Fragmentation Map

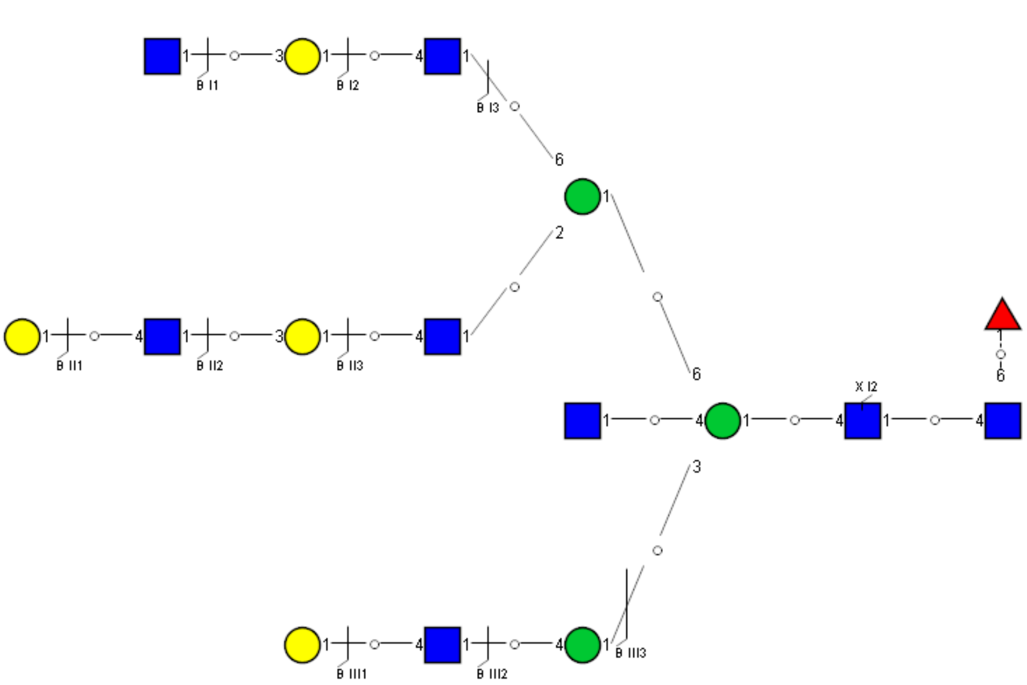

Spectrum Index: 39889(NoBHs:2)

#### Precursor iEF Map

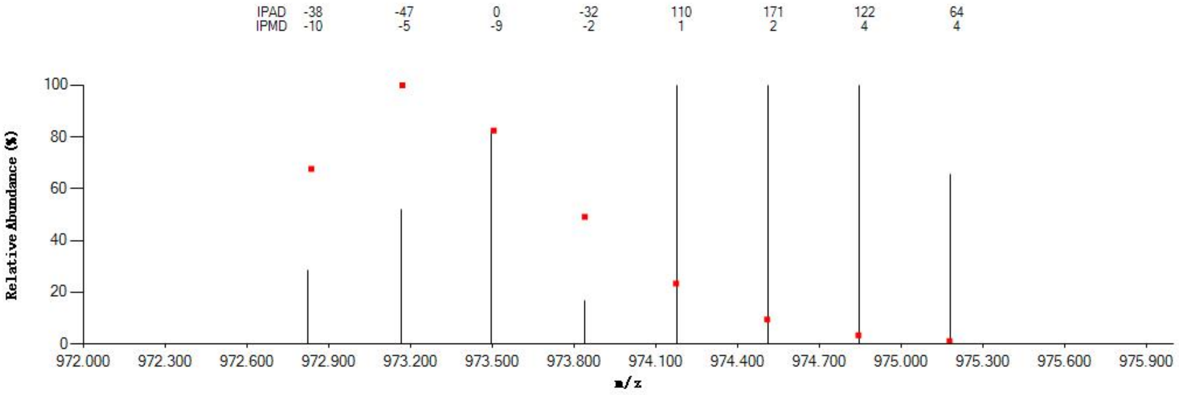


#### Annotated MS/MS Spectrum

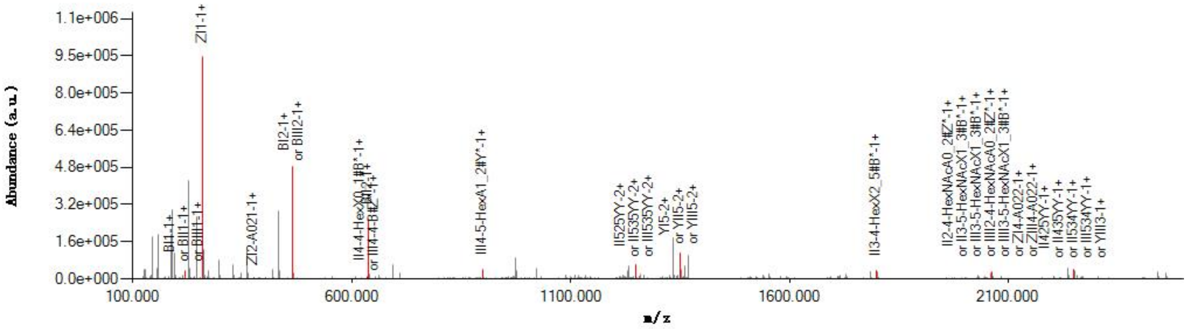


#### Graphical Fragmentation Map

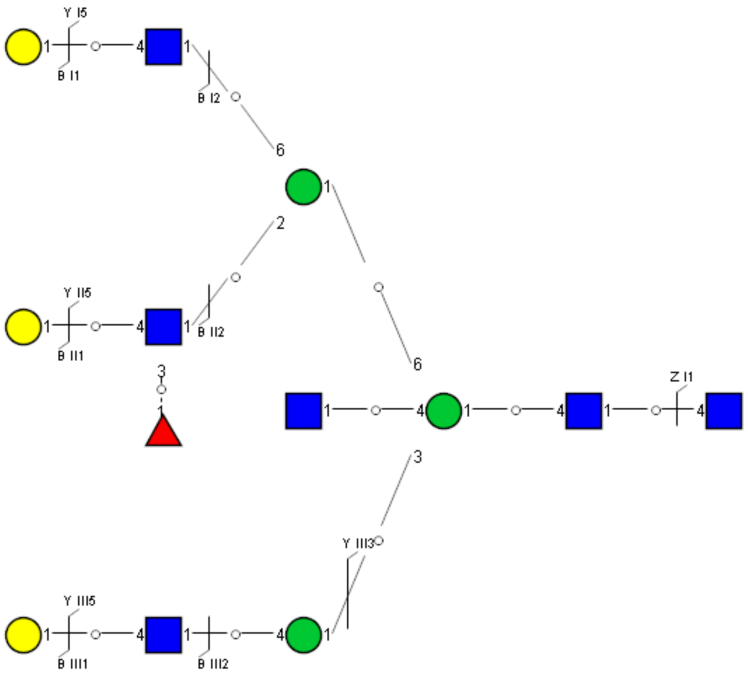

Spectrum Index: 39889(NoBHs:2)

#### Precursor iEF Map

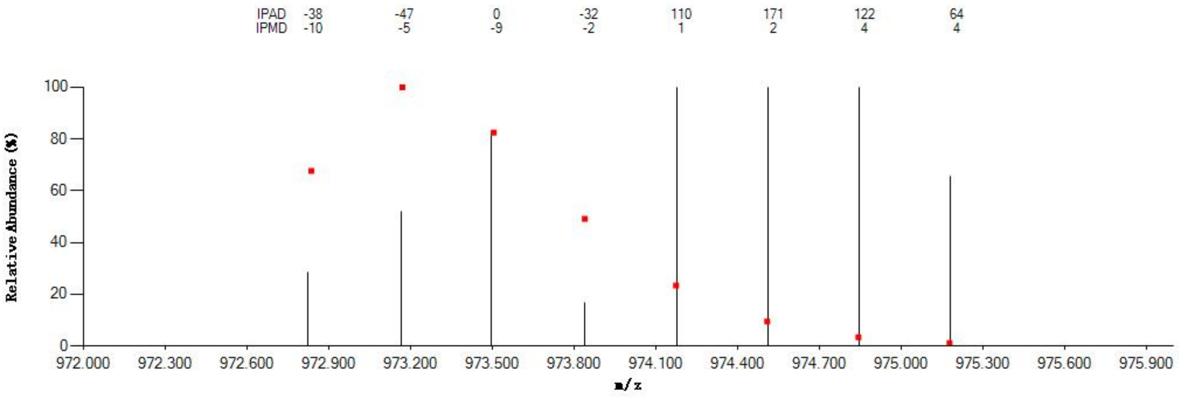


#### Annotated MS/MS Spectrum

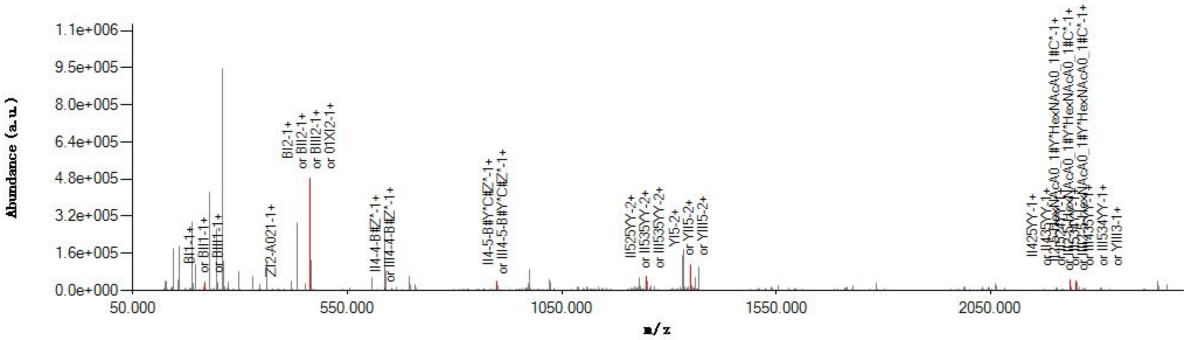


#### Graphical Fragmentation Map

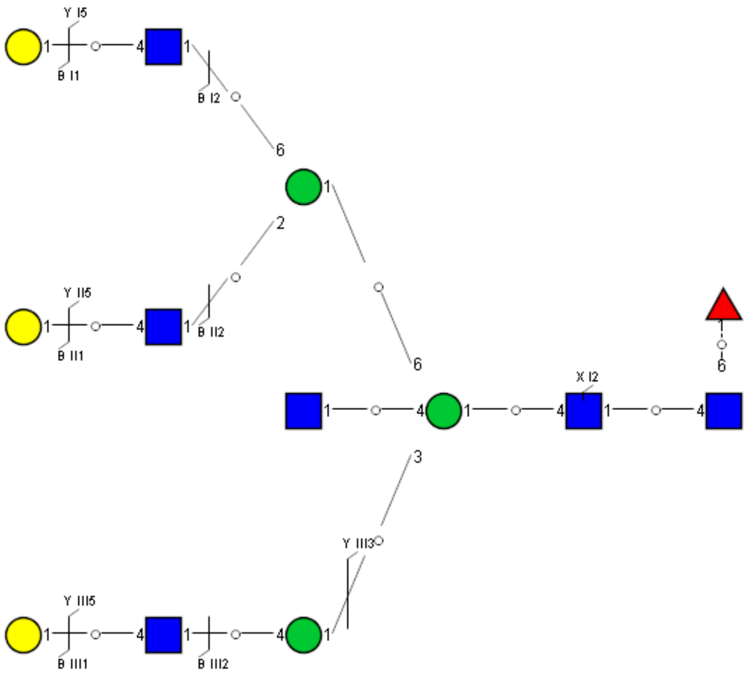

Spectrum Index: 39923(NoBHs:1)

#### Precursor iEF Map

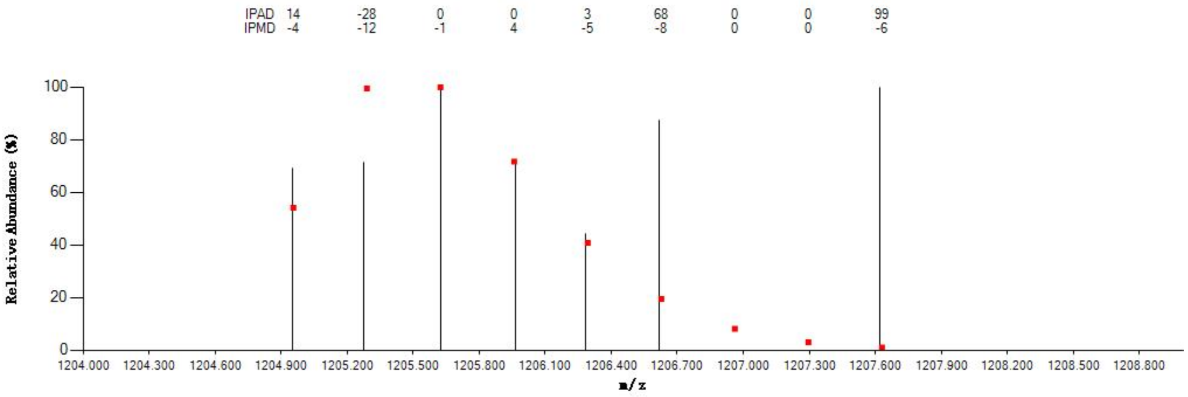


#### Annotated MS/MS Spectrum

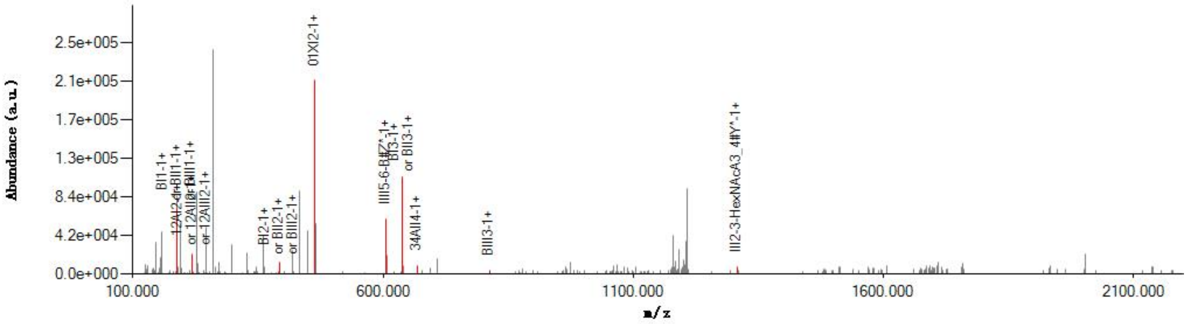


#### Graphical Fragmentation Map

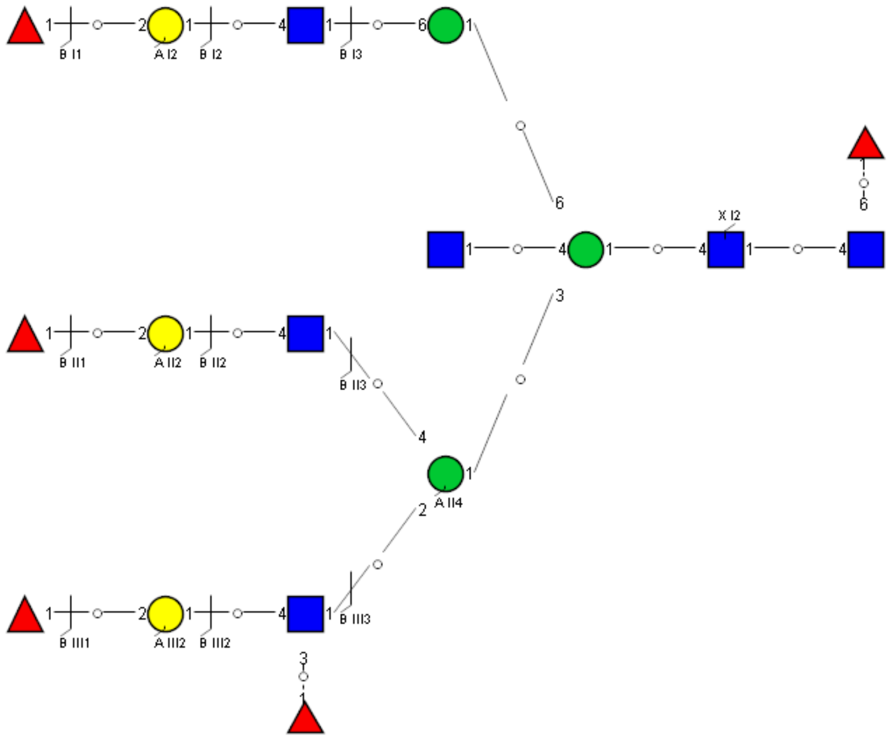

Spectrum Index: 39975(NoBHs:1)

#### Precursor iEF Map

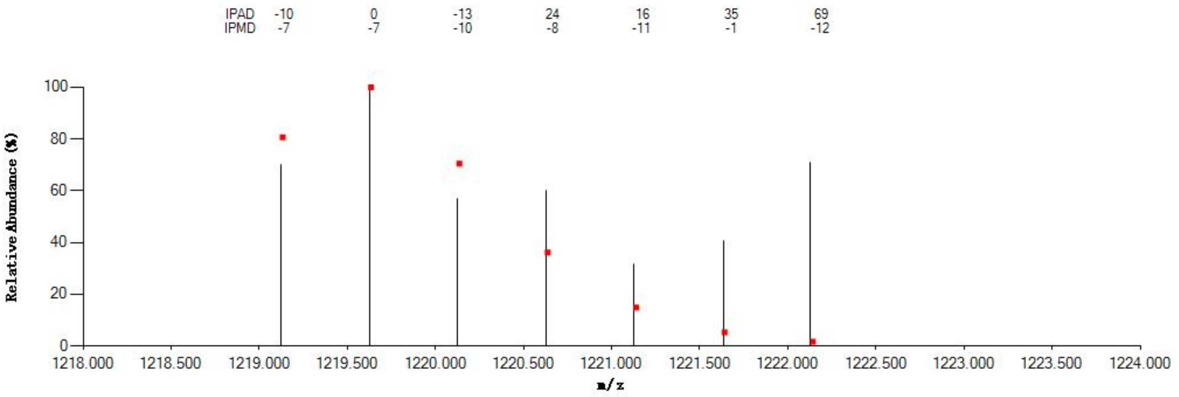

#### Annotated MS/MS Spectrum

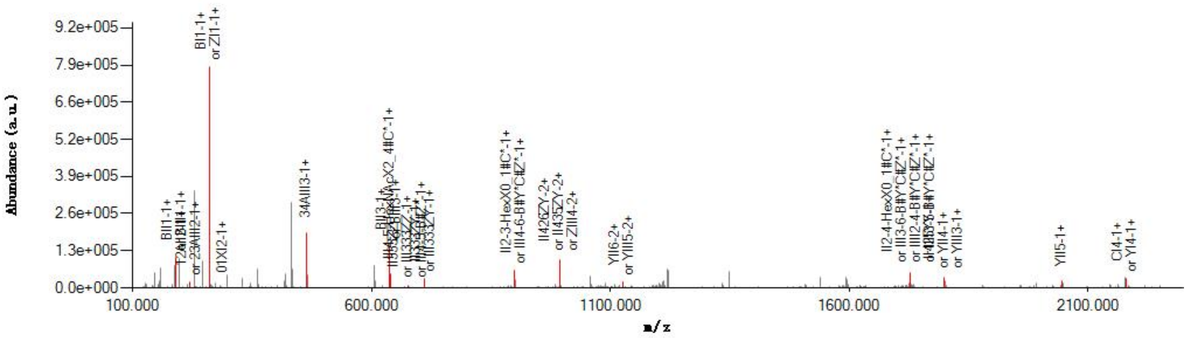

#### Graphical Fragmentation Map

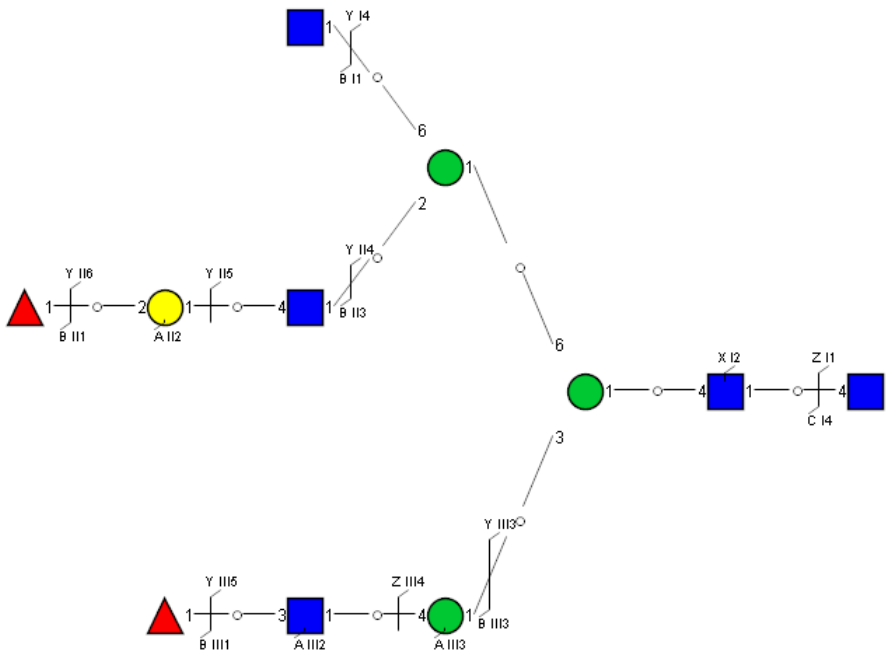

### Spectrum Index: 39981(NoBHs:1)

#### Precursor iEF Map

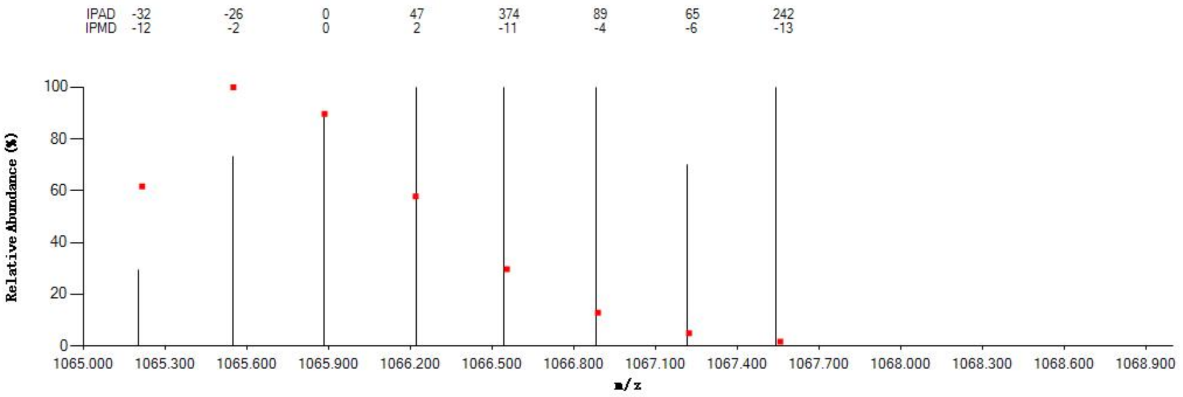


#### Annotated MS/MS Spectrum

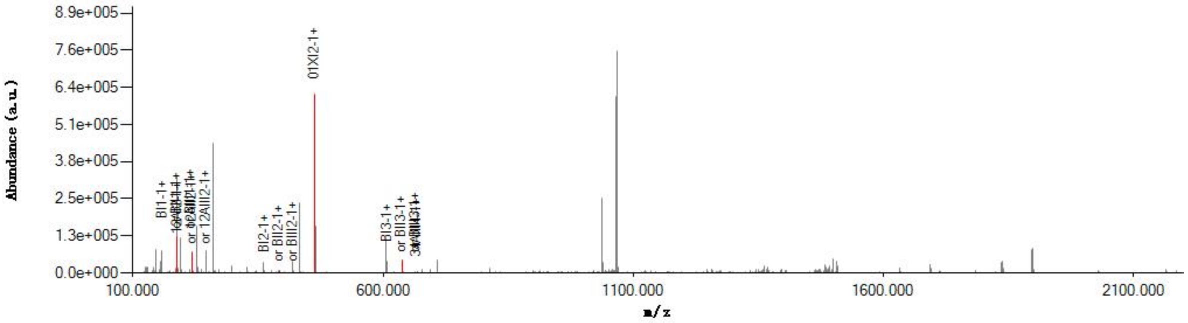


#### Graphical Fragmentation Map

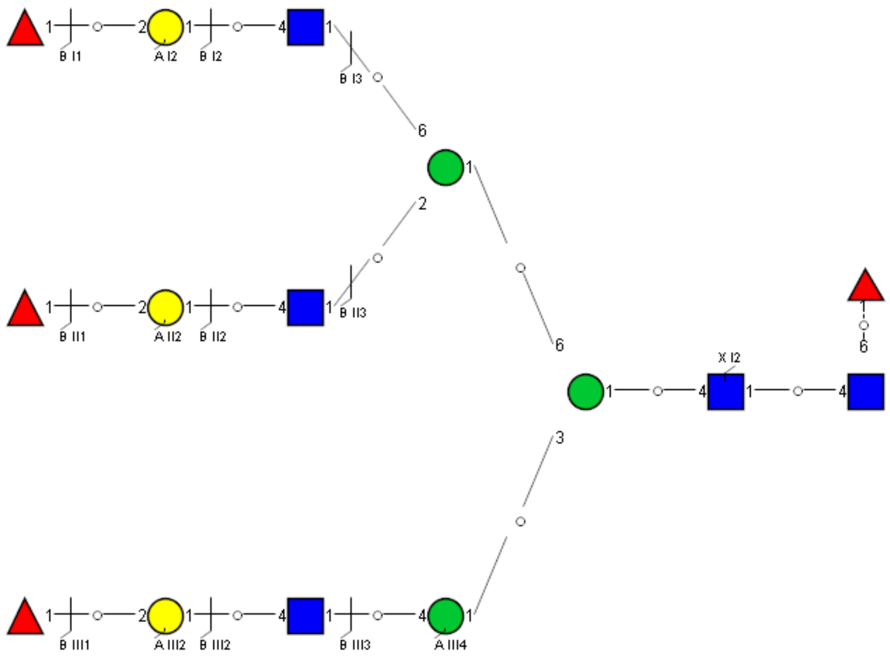

Spectrum Index: 40041(NoBHs:1)

#### Precursor iEF Map

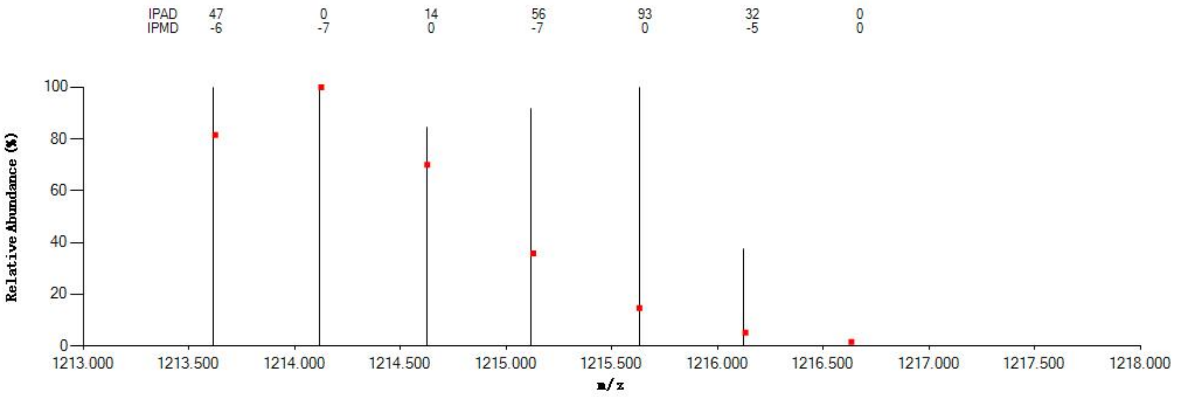


#### Annotated MS/MS Spectrum

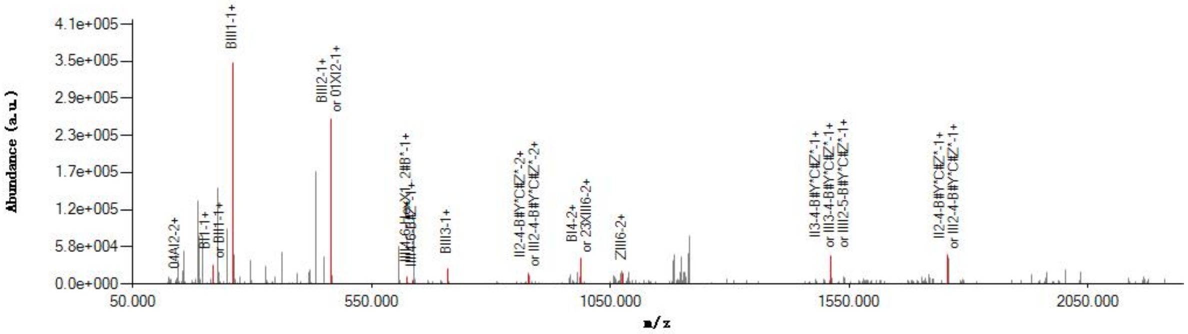


#### Graphical Fragmentation Map

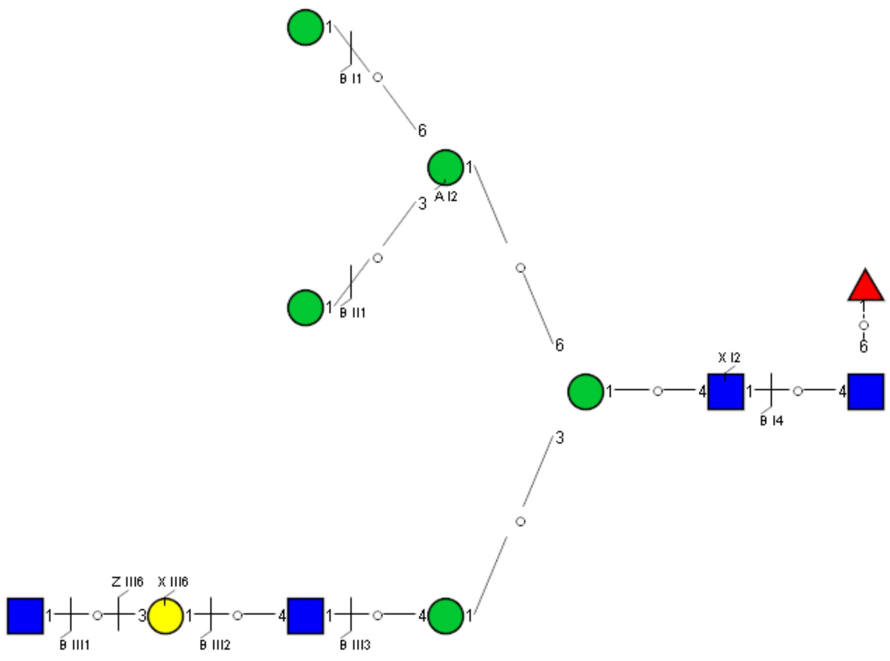

Spectrum Index: 40157(NoBHs:1)

#### Precursor iEF Map

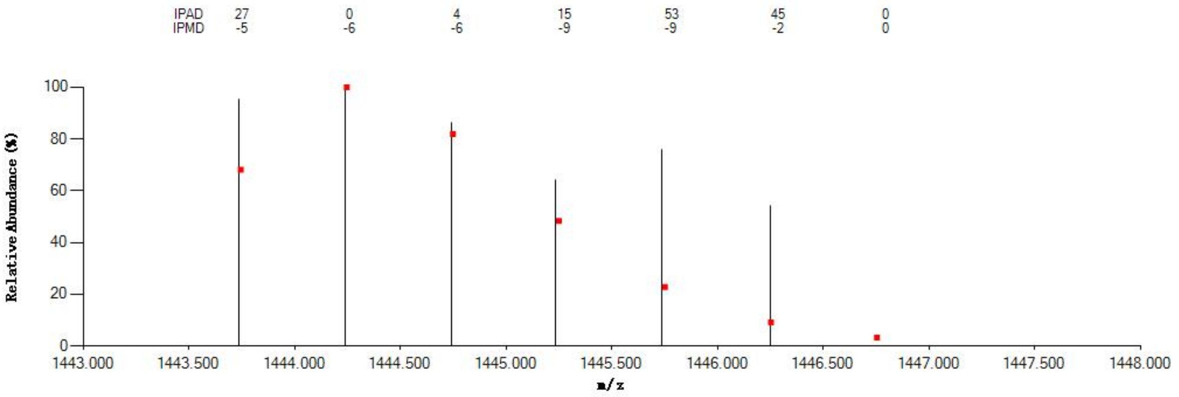


#### Annotated MS/MS Spectrum

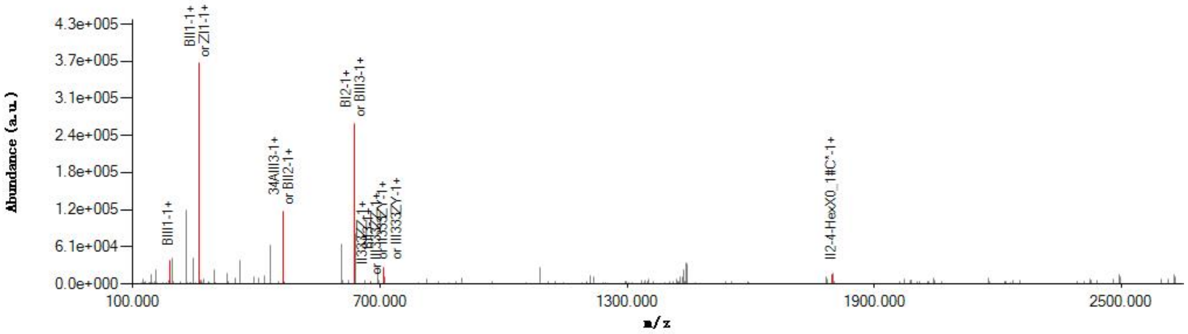


#### Graphical Fragmentation Map

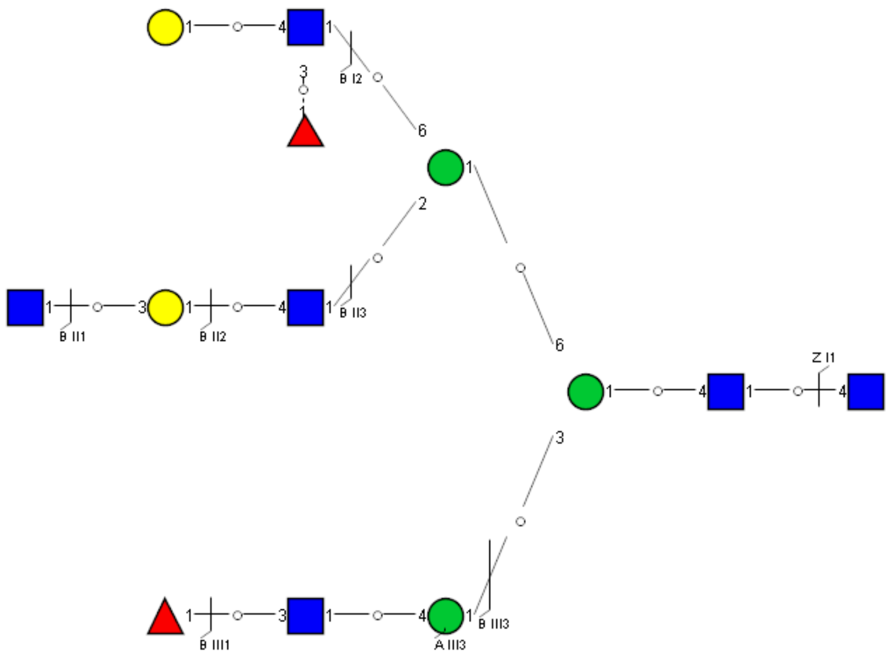

Spectrum Index: 40158(NoBHs:1)

#### Precursor iEF Map

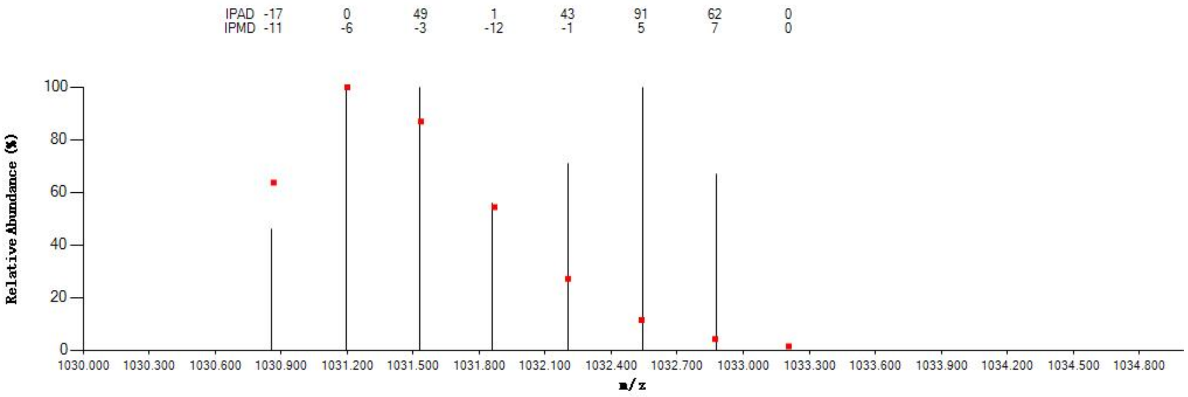


#### Annotated MS/MS Spectrum

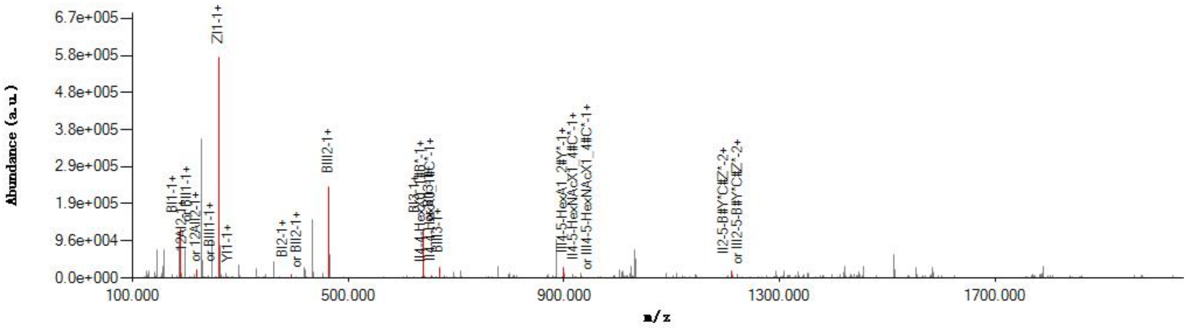


#### Graphical Fragmentation Map

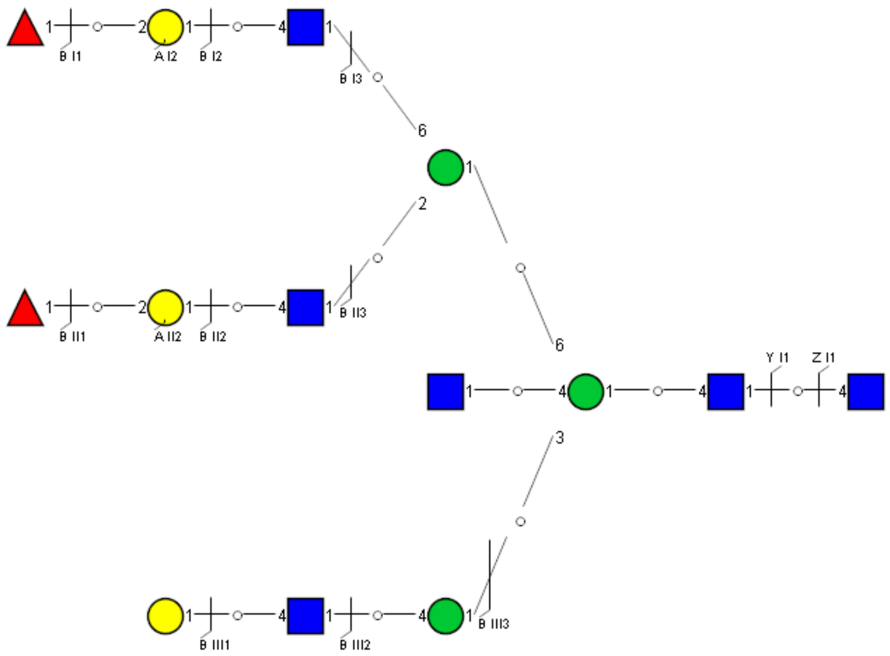

Spectrum Index: 40170(NoBHs:1)

Precursor iEF Map

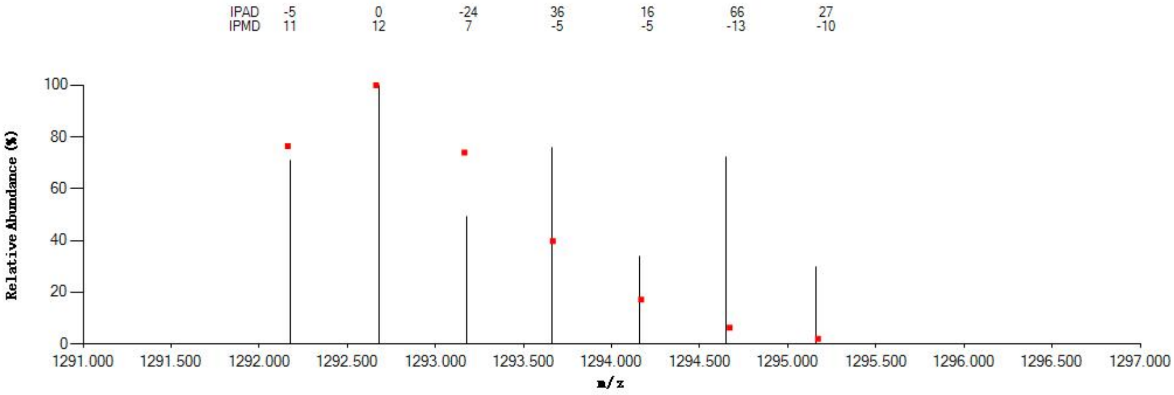


Annotated MS/MS Spectrum

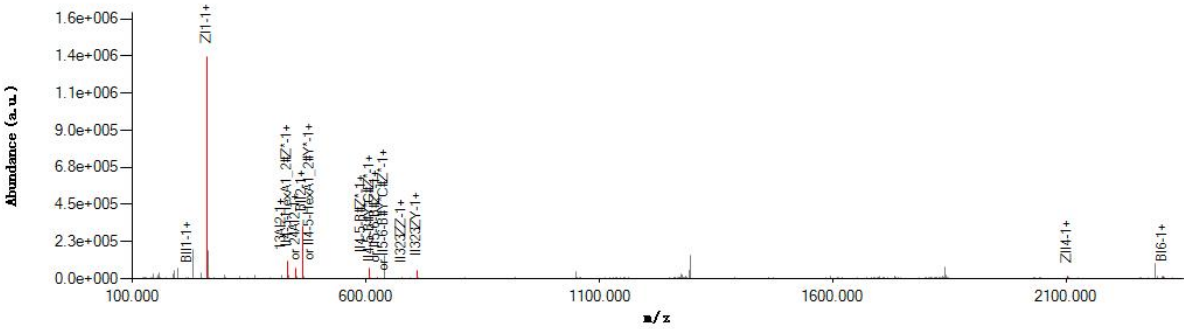


Graphical Fragmentation Map

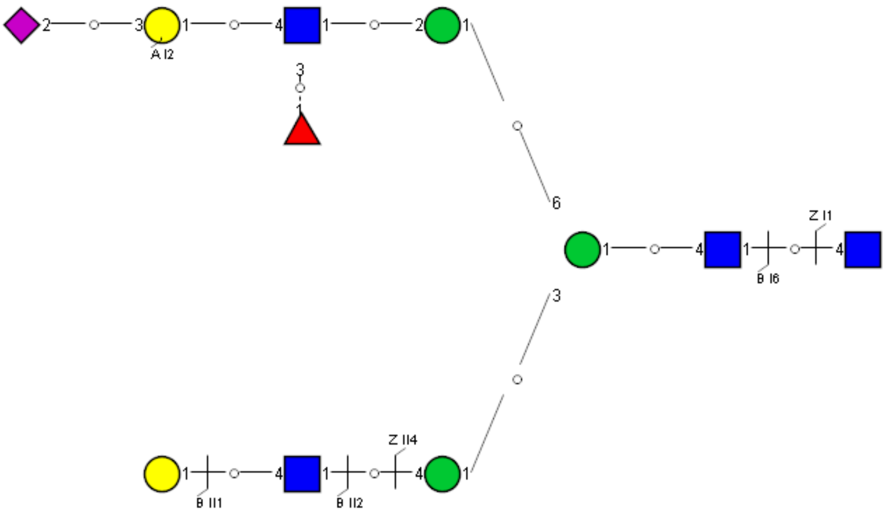

Spectrum Index: 40173(NoBHs:1)

Precursor iEF Map

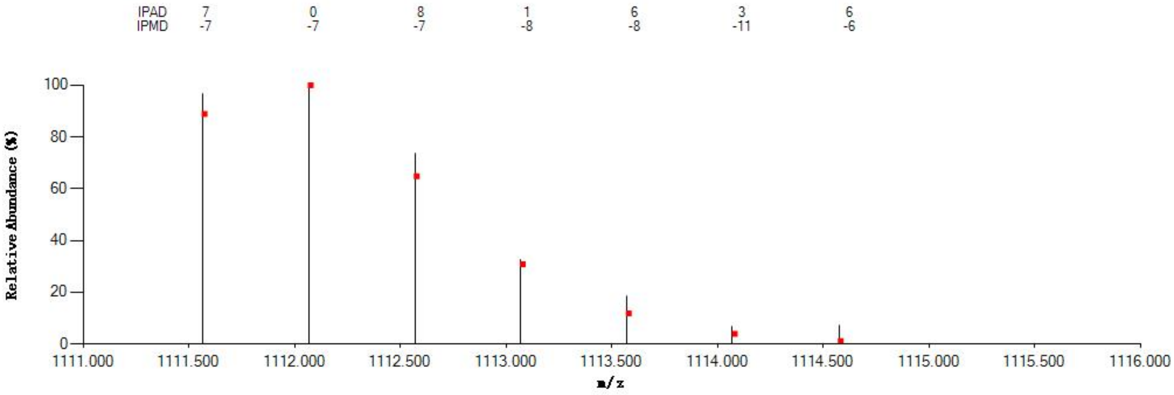

Annotated MS/MS Spectrum

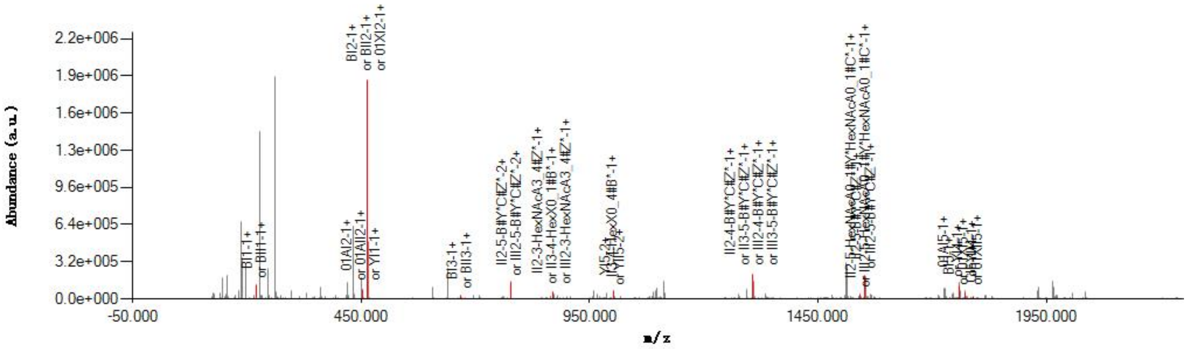

Graphical Fragmentation Map

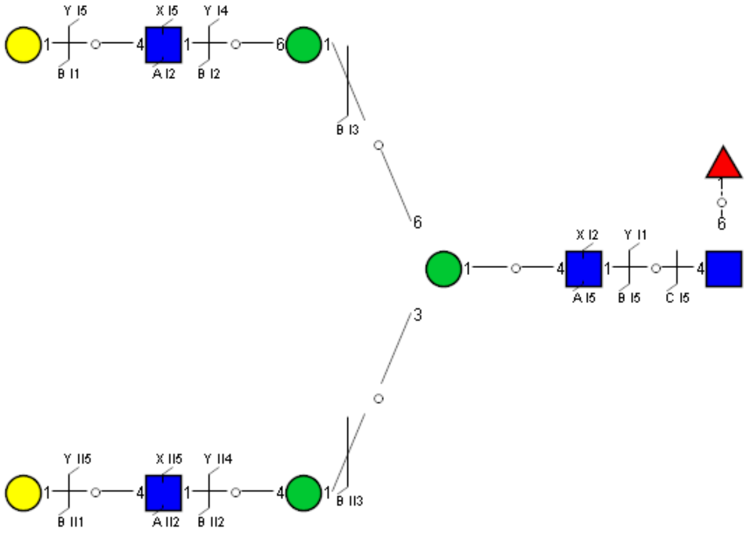

Spectrum Index: 40224(NoBHs:1)

#### Precursor iEF Map

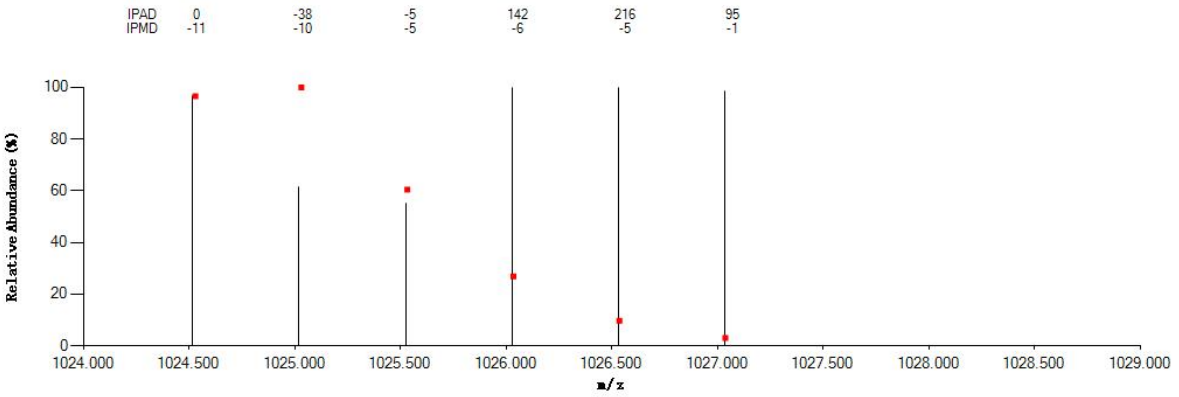


#### Annotated MS/MS Spectrum

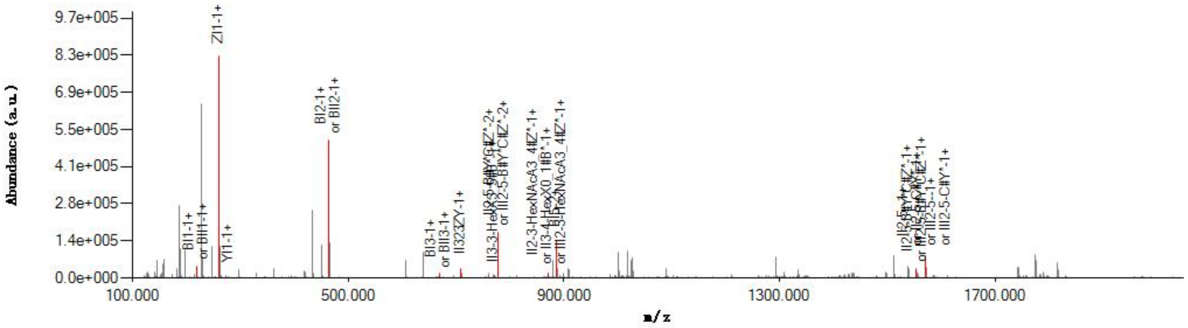


#### Graphical Fragmentation Map

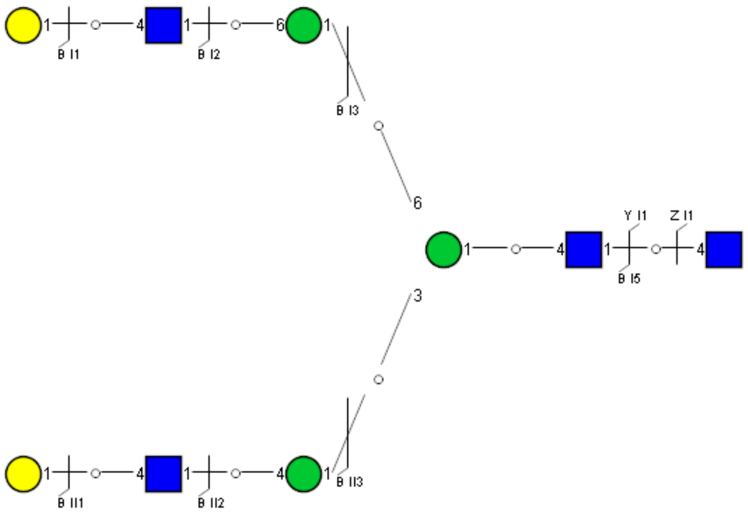

Spectrum Index: 40366(NoBHs:2)

#### Precursor iEF Map

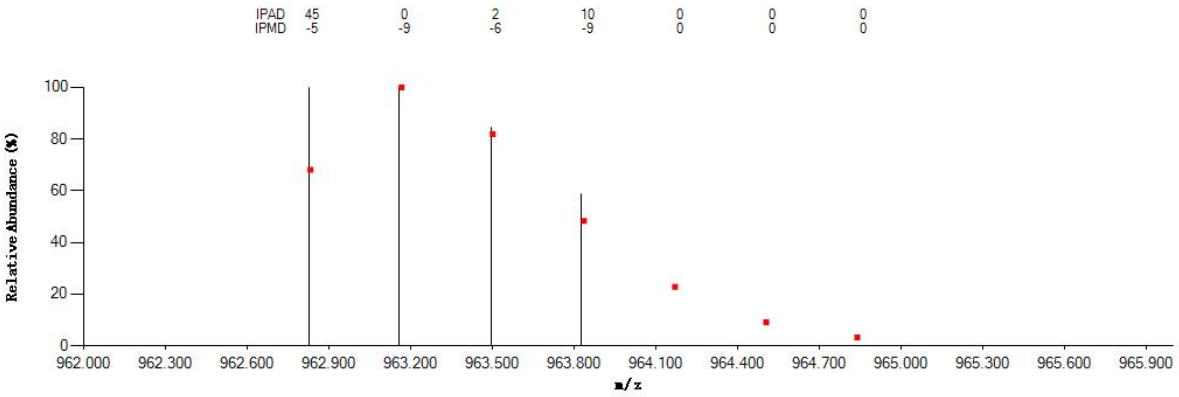


#### Annotated MS/MS Spectrum

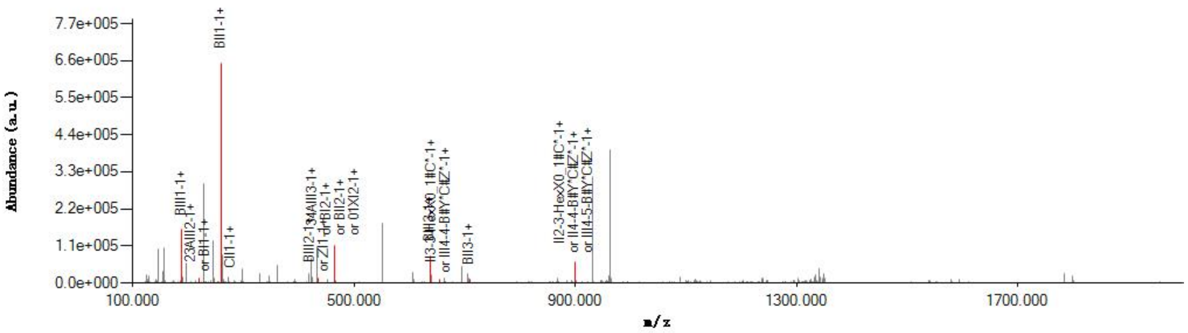


#### Graphical Fragmentation Map

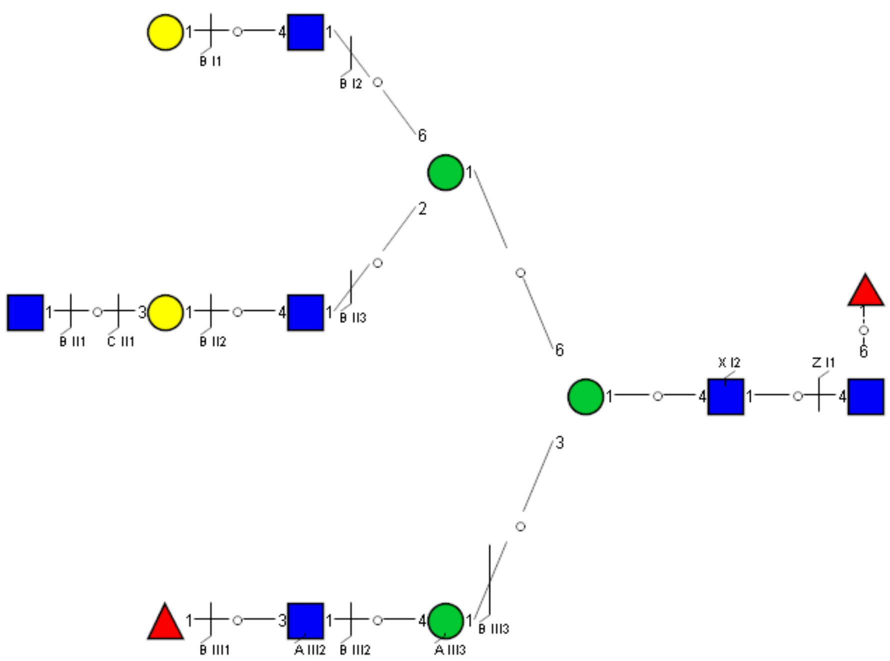

Spectrum Index: 40369(NoBHs:1)

#### Precursor iEF Map

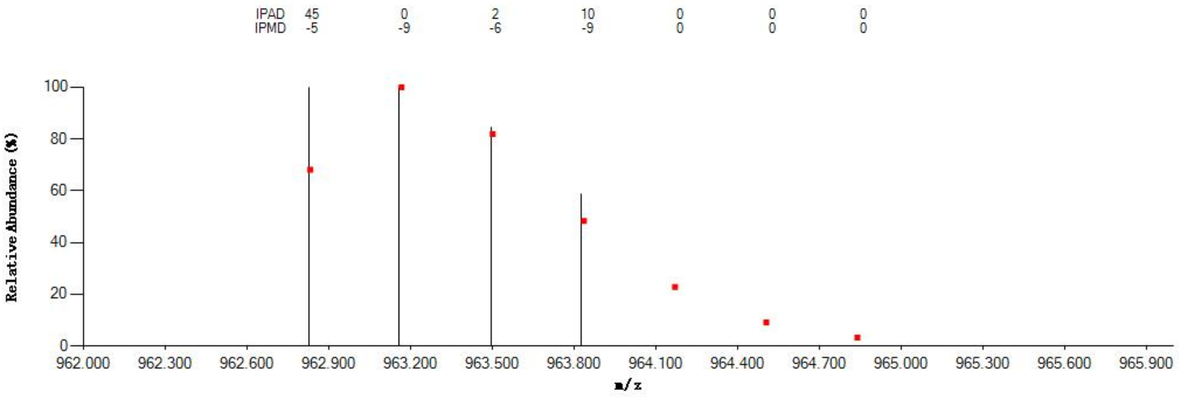


#### Annotated MS/MS Spectrum

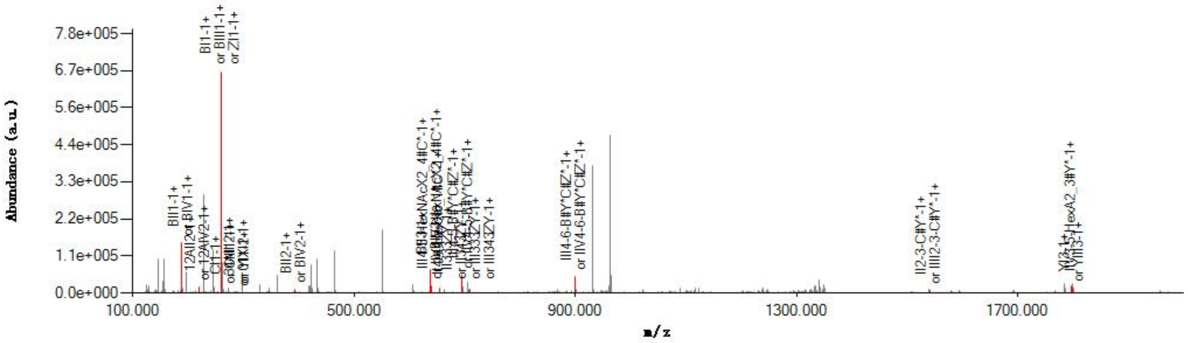


#### Graphical Fragmentation Map

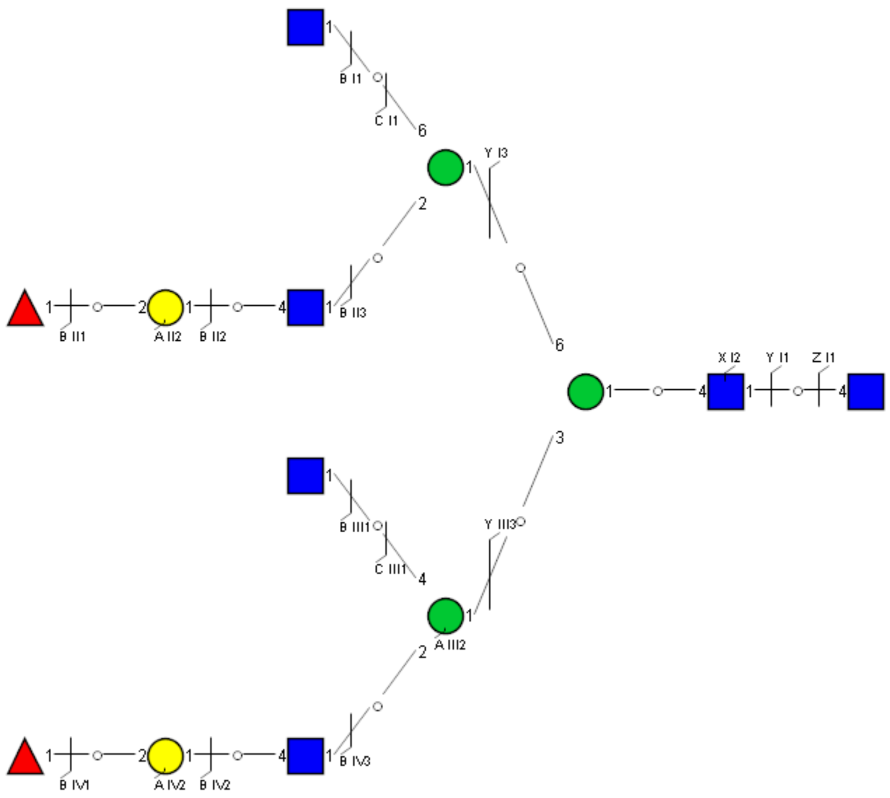

### Spectrum Index: 40489(NoBHs:1)

#### Precursor iEF Map

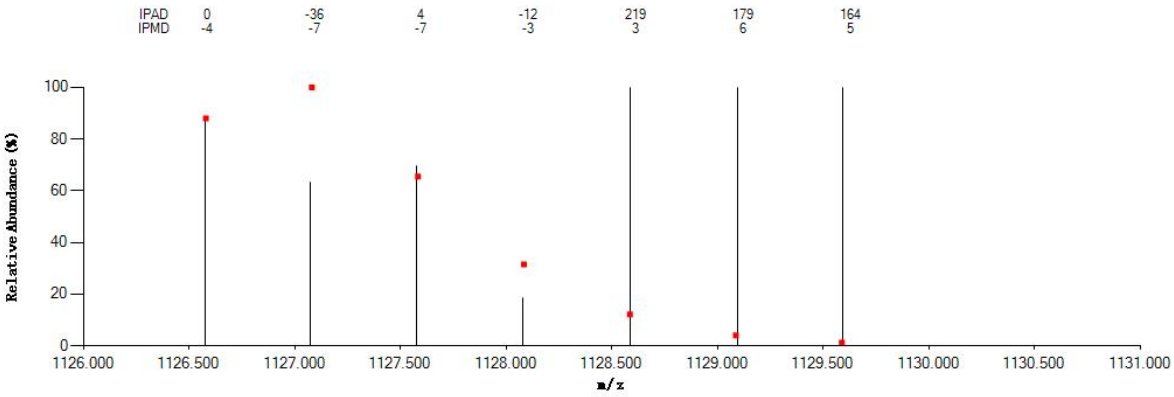


#### Annotated MS/MS Spectrum

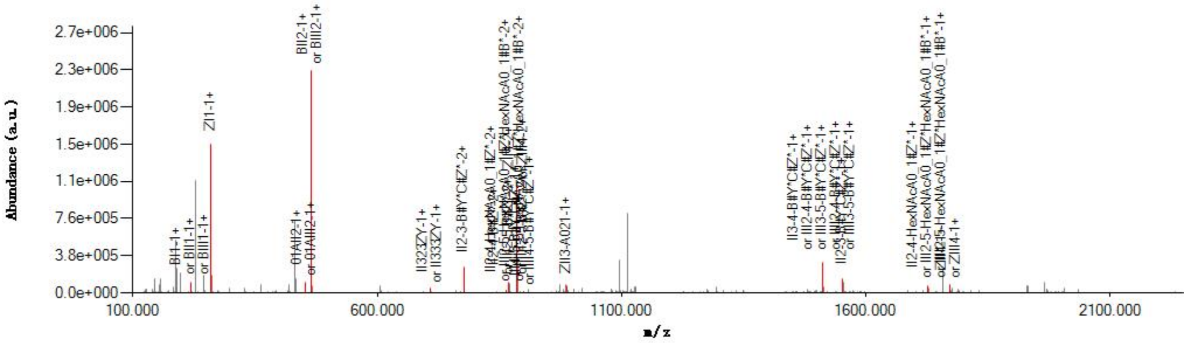


#### Graphical Fragmentation Map

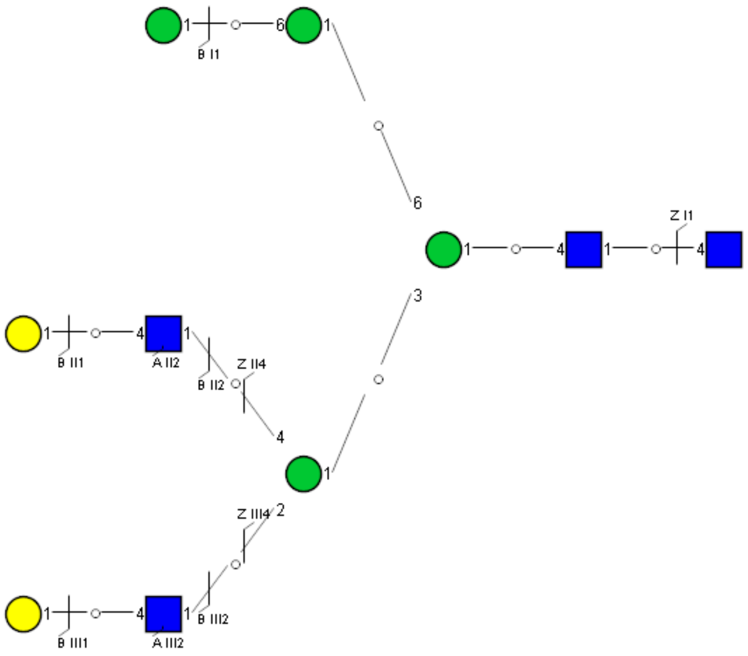

Spectrum Index: 40540(NoBHs:1)

#### Precursor iEF Map

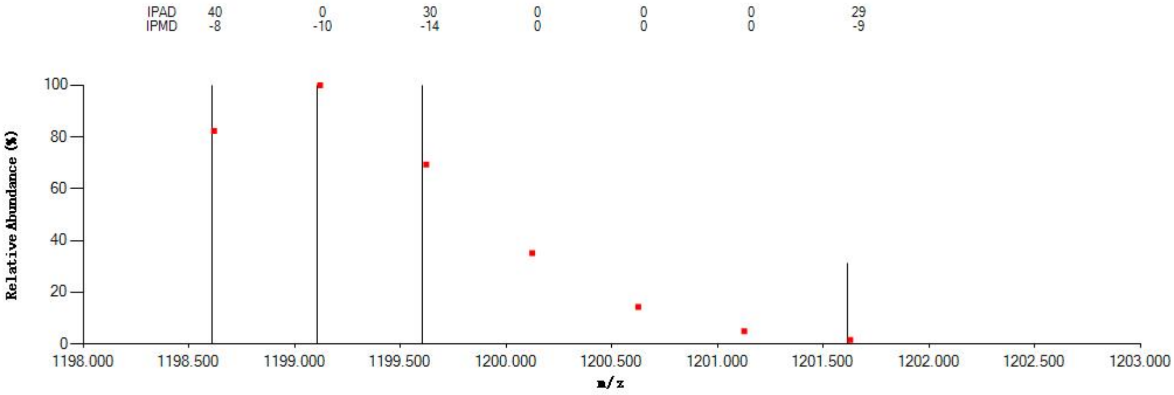

#### Annotated MS/MS Spectrum

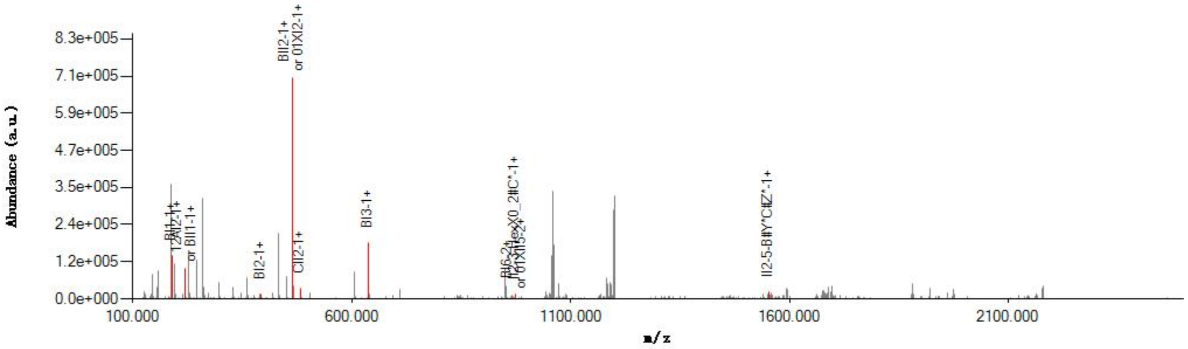

#### Graphical Fragmentation Map

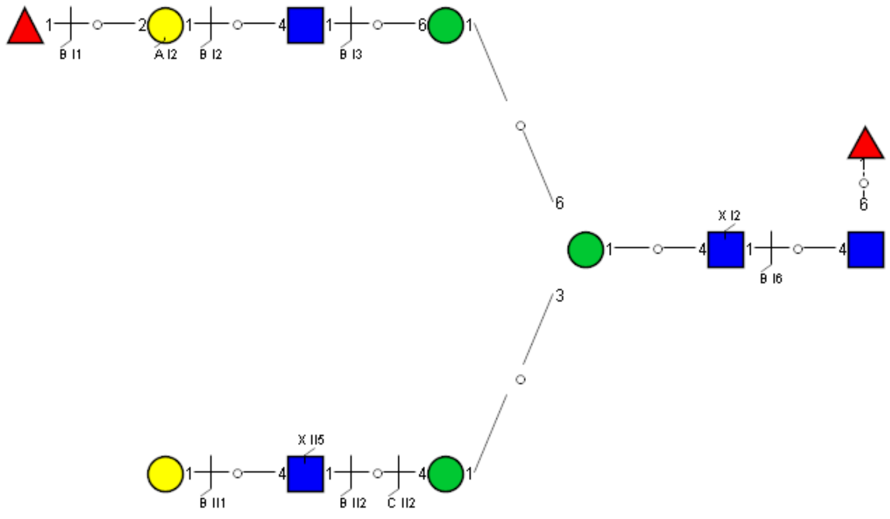

Spectrum Index: 40549(NoBHs:1)

#### Precursor iEF Map

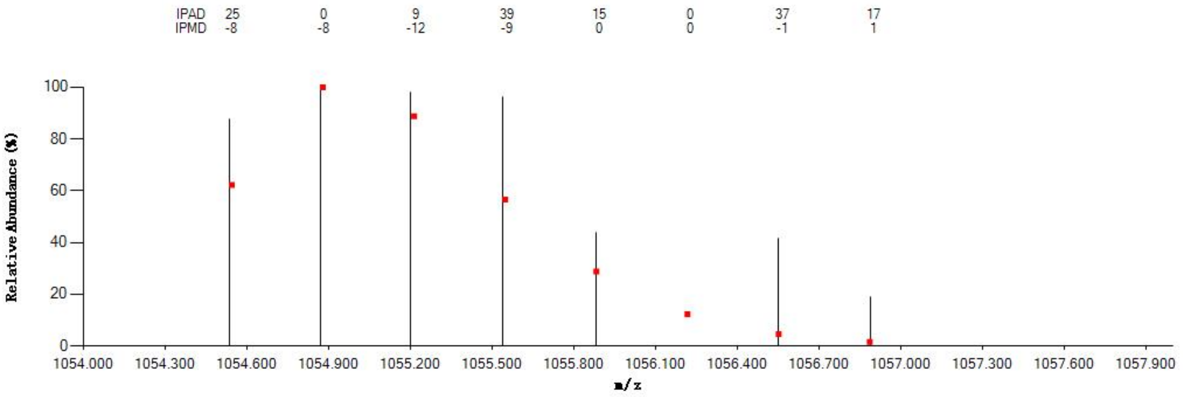


#### Annotated MS/MS Spectrum

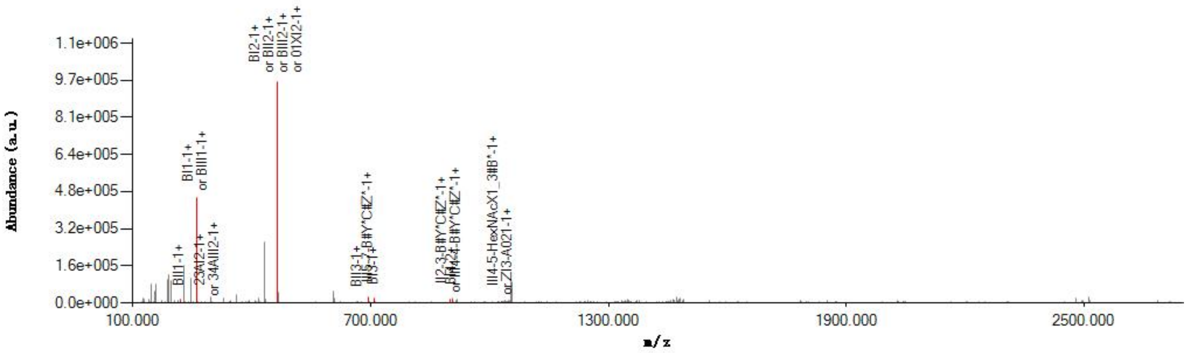


#### Graphical Fragmentation Map

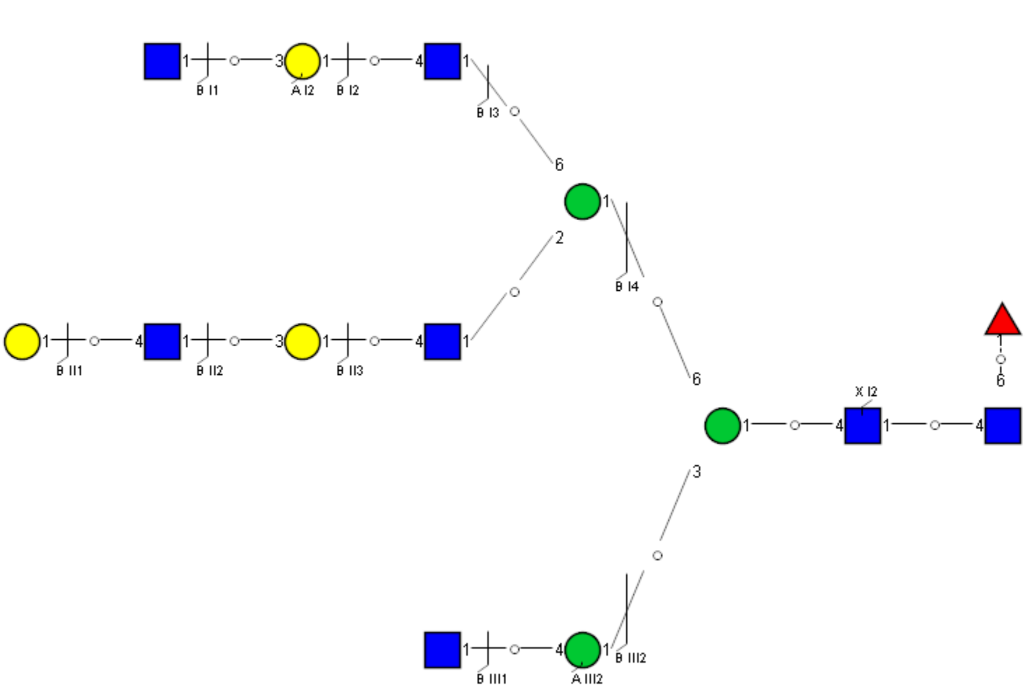

Spectrum Index: 41064(NoBHs:1)

#### Precursor iEF Map

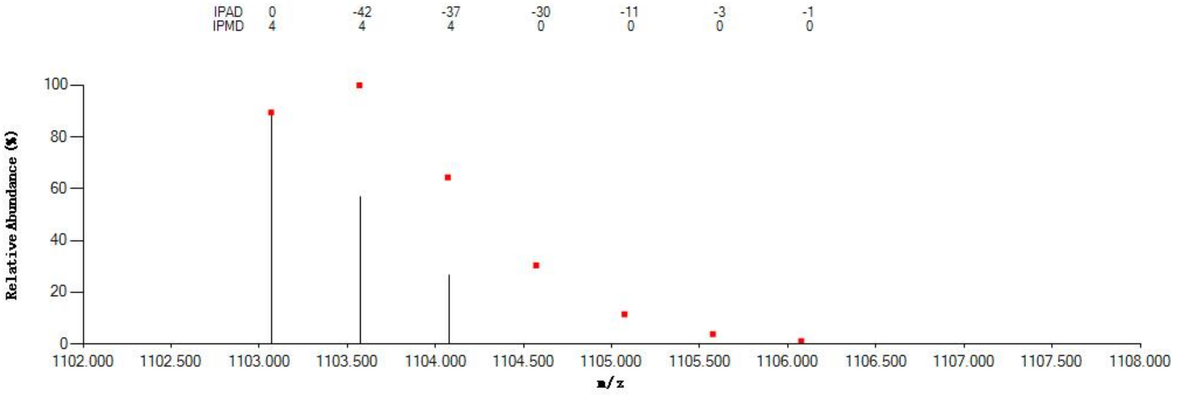


#### Annotated MS/MS Spectrum

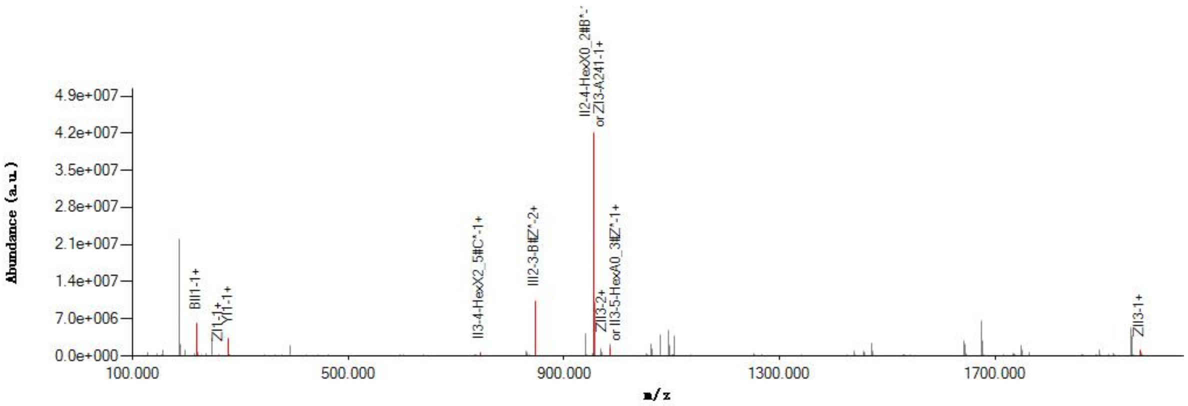


#### Graphical Fragmentation Map

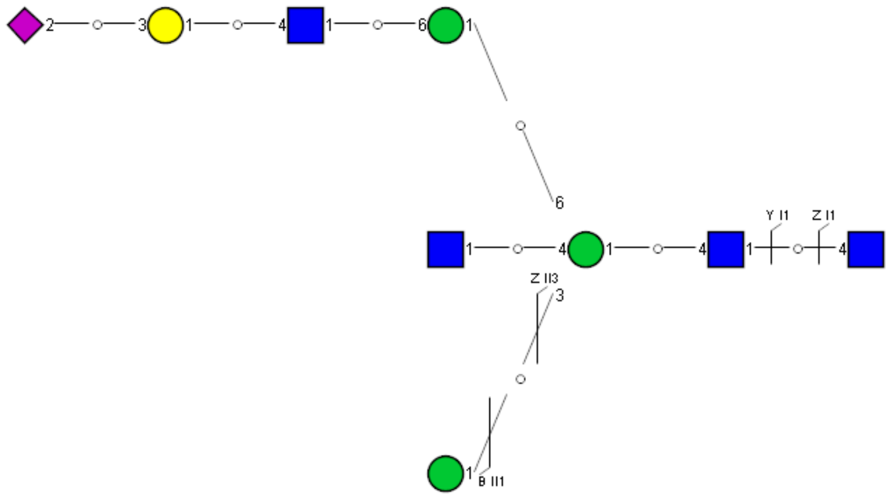

### Spectrum Index: 41505(NoBHs:1)

#### Precursor iEF Map

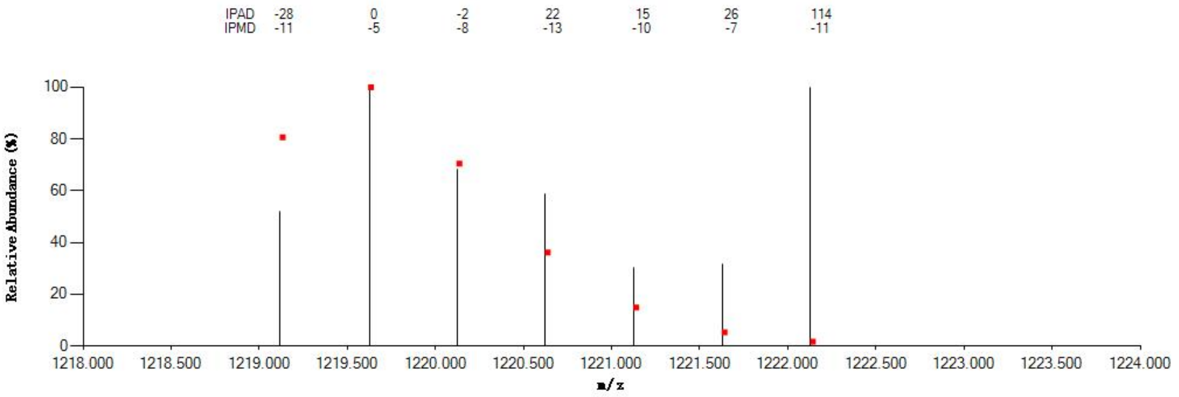


#### Annotated MS/MS Spectrum

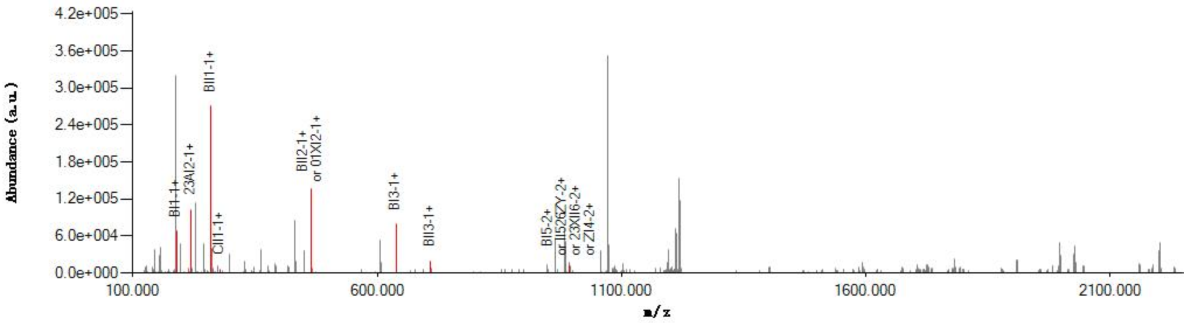


#### Graphical Fragmentation Map

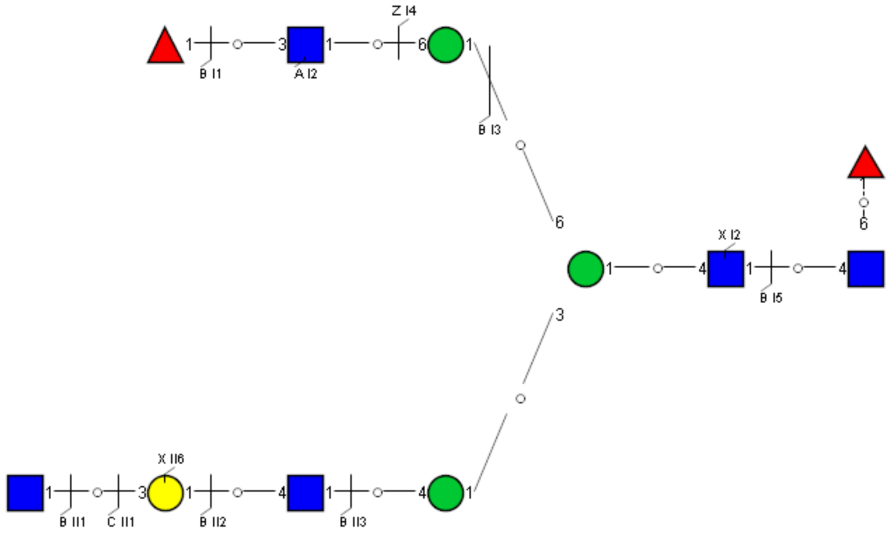

Spectrum Index: 41641(NoBHs:1)

#### Precursor iEF Map

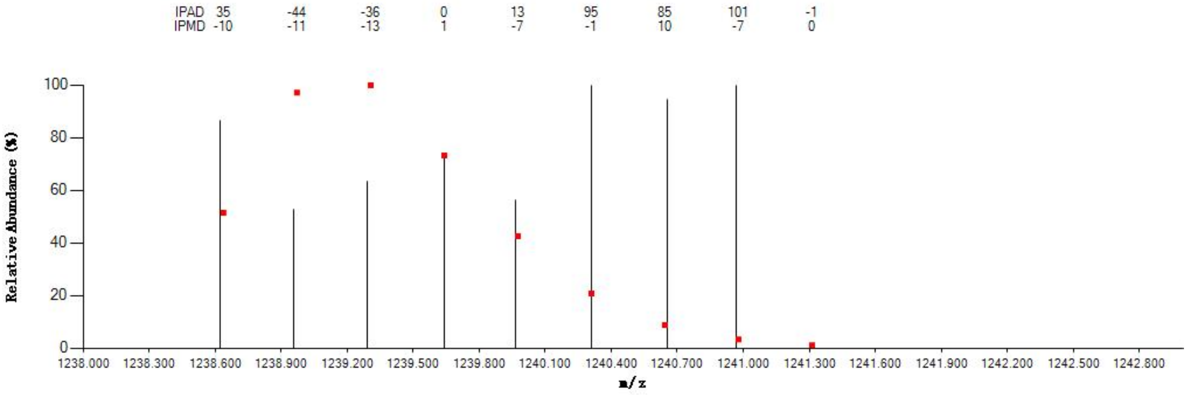


#### Annotated MS/MS Spectrum

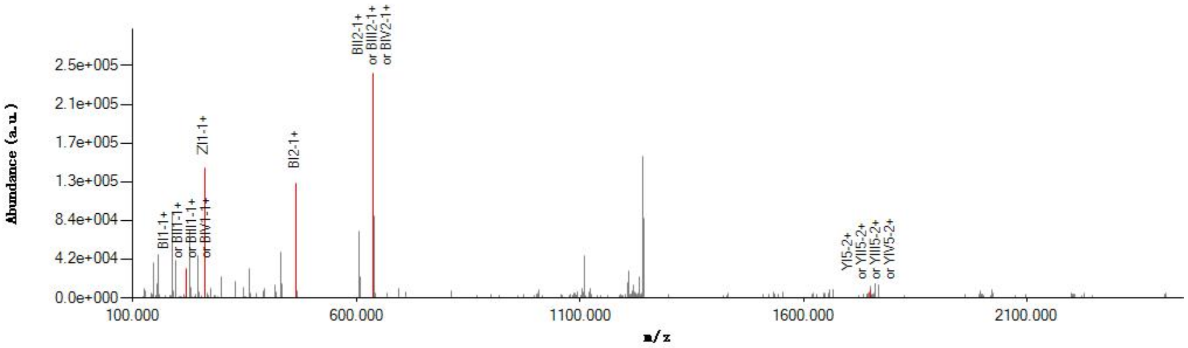


#### Graphical Fragmentation Map

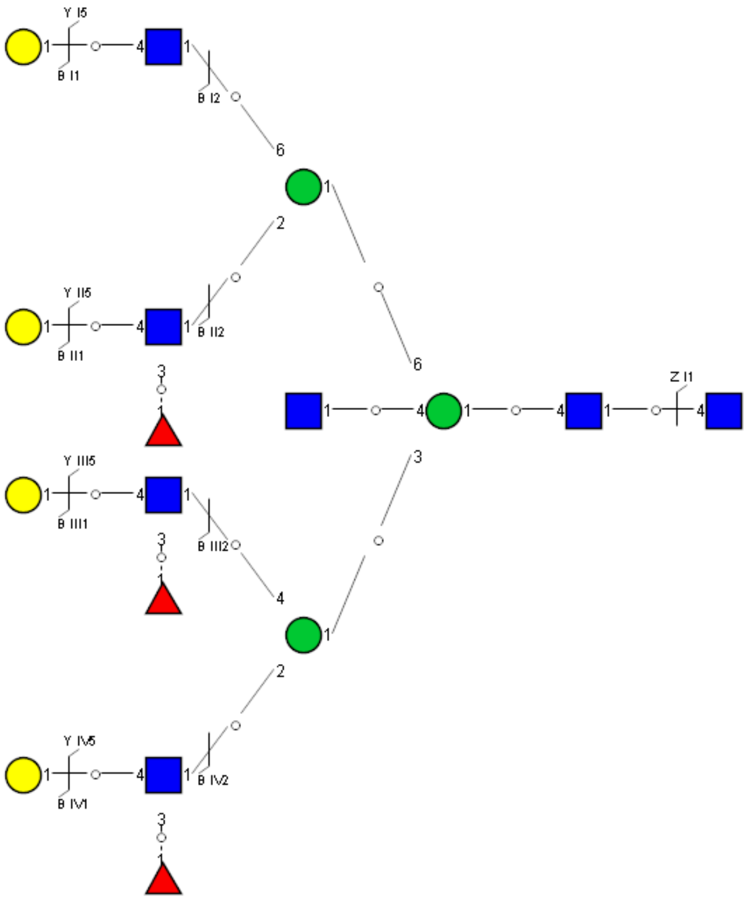

Spectrum Index: 41784(NoBHs:1)

#### Precursor iEF Map

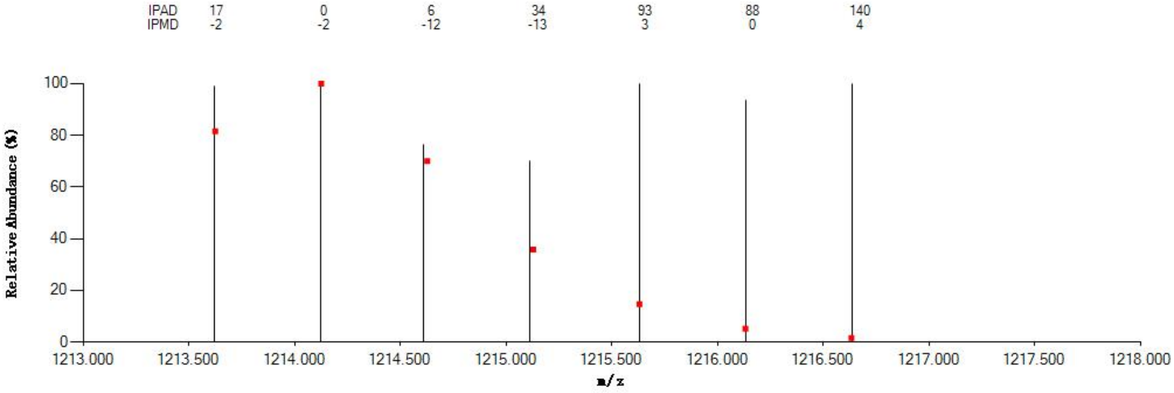


#### Annotated MS/MS Spectrum

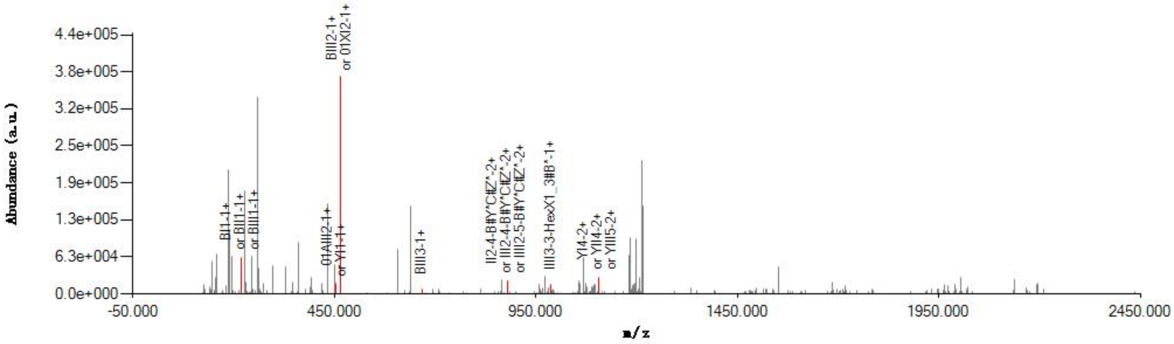


#### Graphical Fragmentation Map

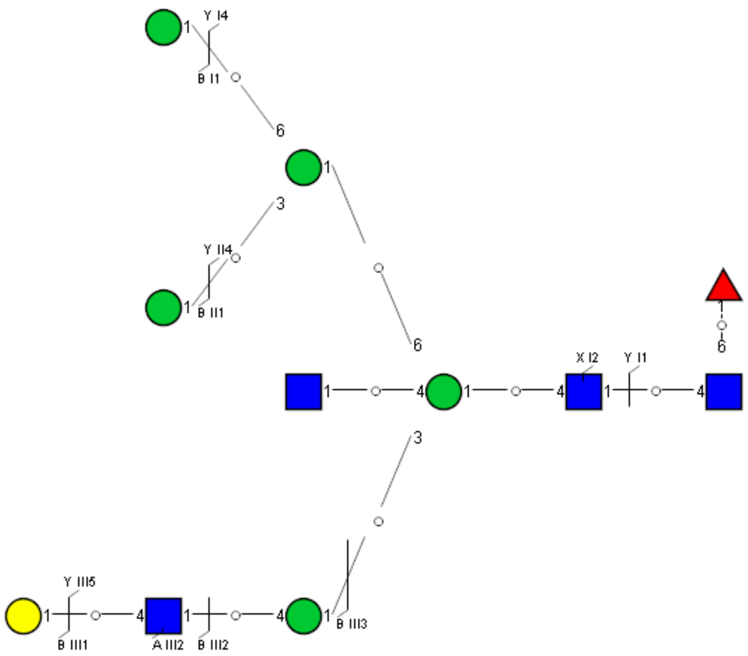

Spectrum Index: 41859(NoBHs:1)

#### Precursor iEF Map

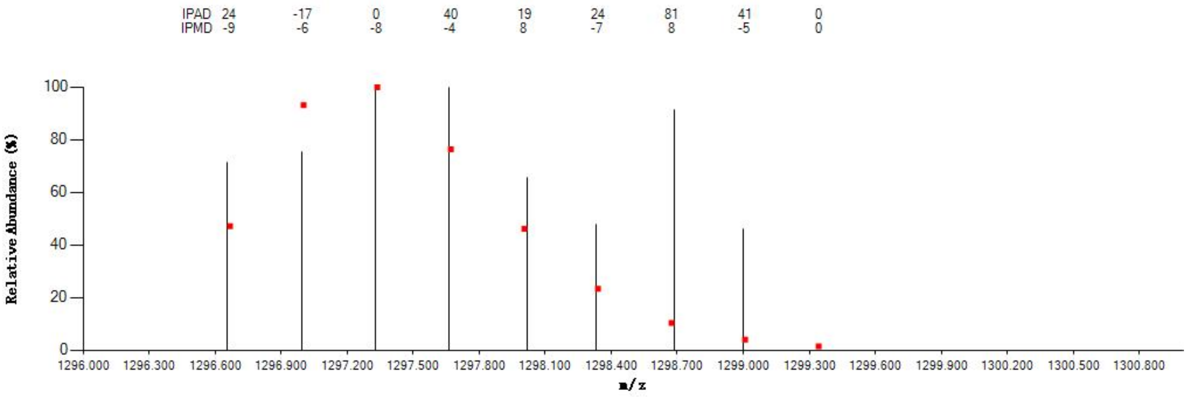


#### Annotated MS/MS Spectrum

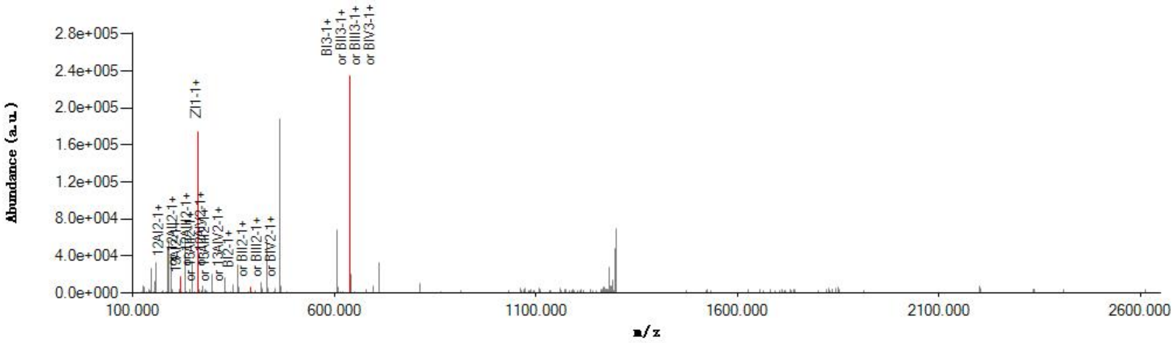


#### Graphical Fragmentation Map

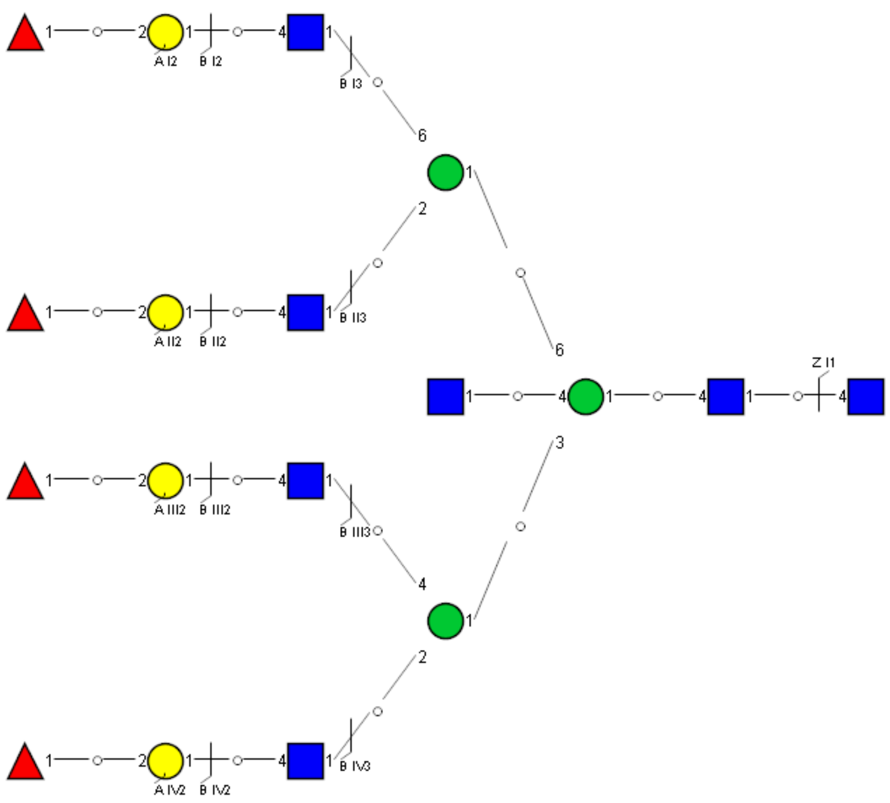

Spectrum Index: 42073(NoBHs:1)

#### Precursor iEF Map

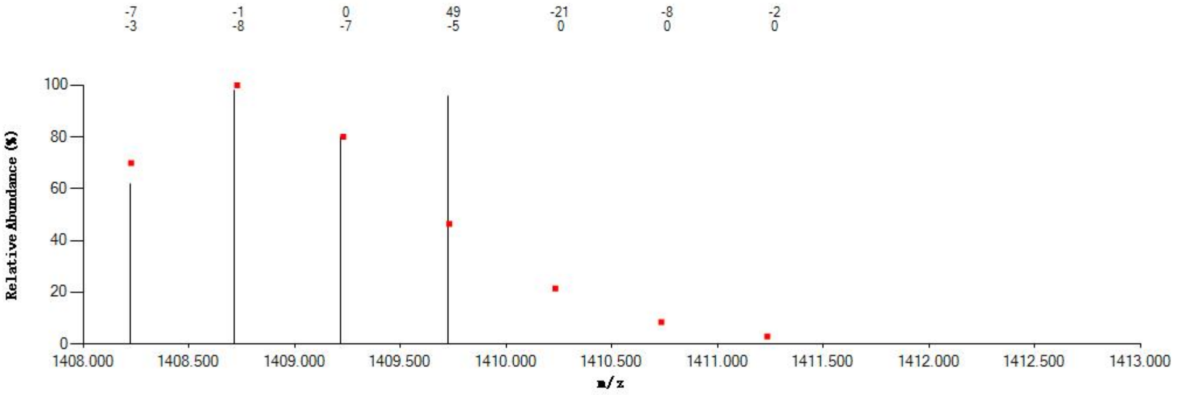


#### Annotated MS/MS Spectrum

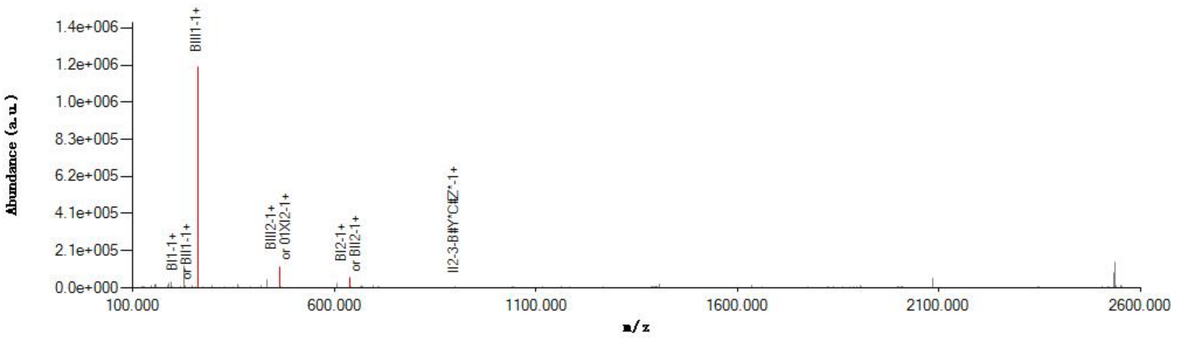


#### Graphical Fragmentation Map

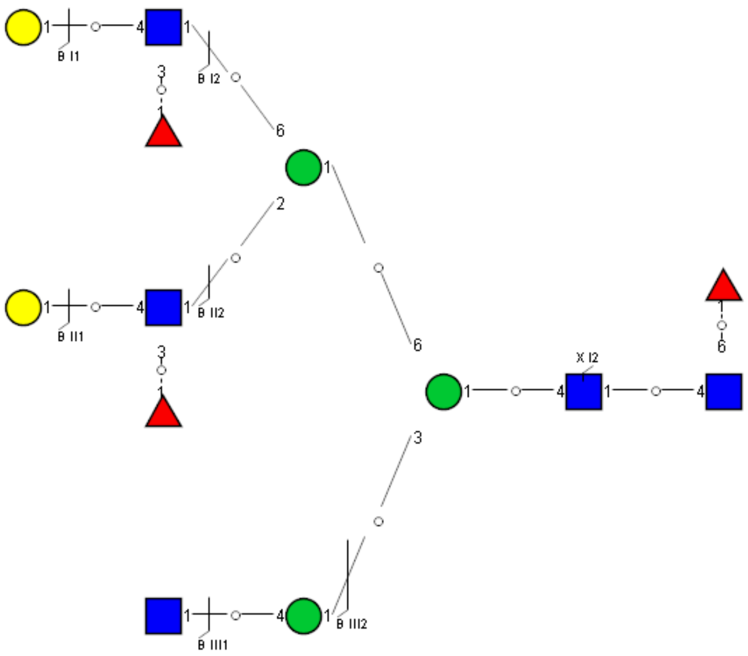

Spectrum Index: 42194(NoBHs:1)

#### Precursor iEF Map

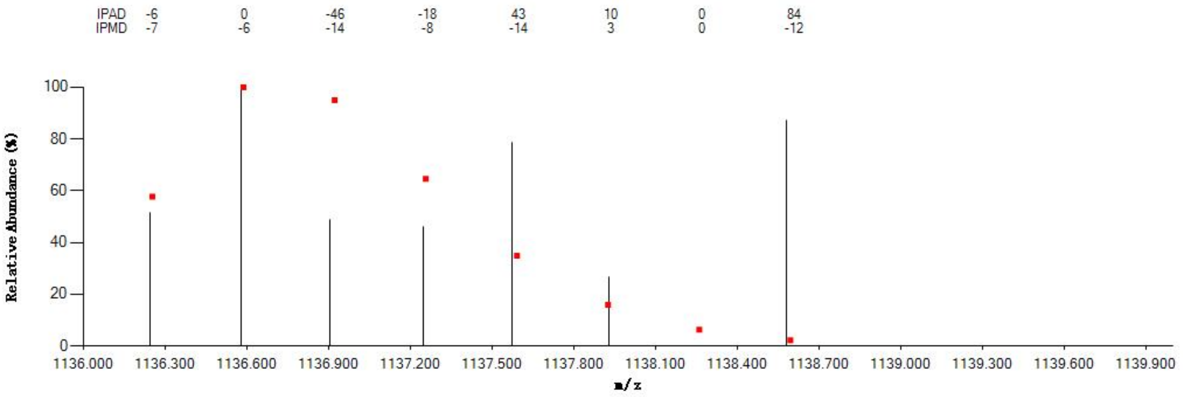


#### Annotated MS/MS Spectrum

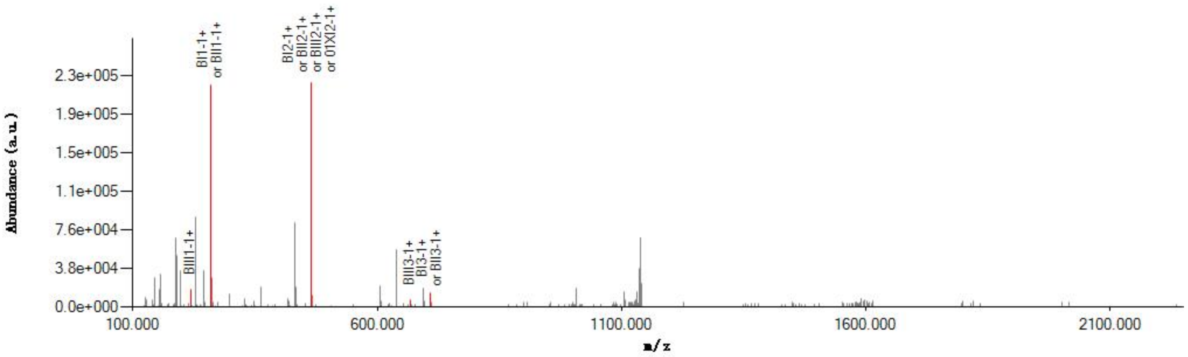


#### Graphical Fragmentation Map

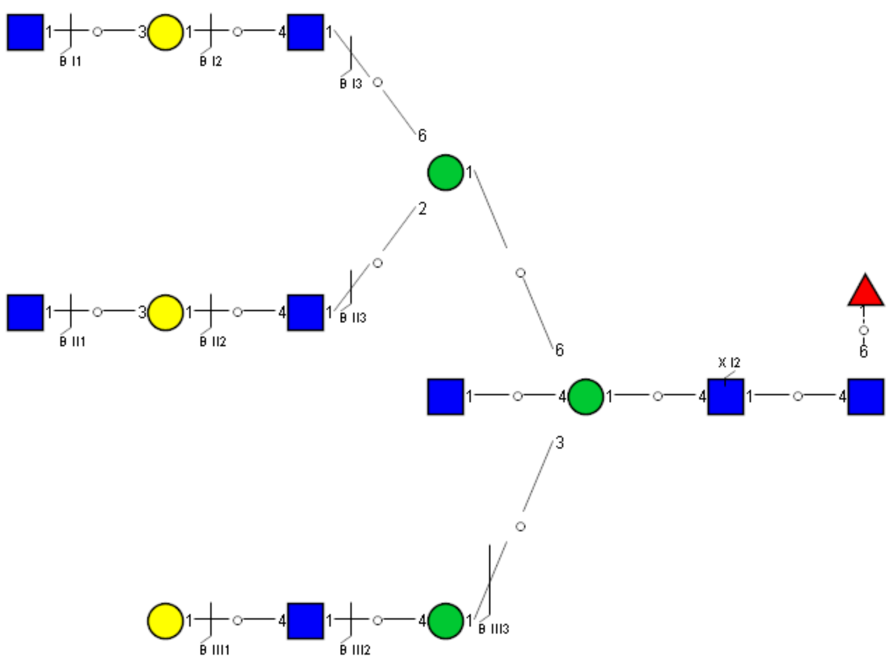

Spectrum Index: 42327(NoBHs:2)

#### Precursor iEF Map

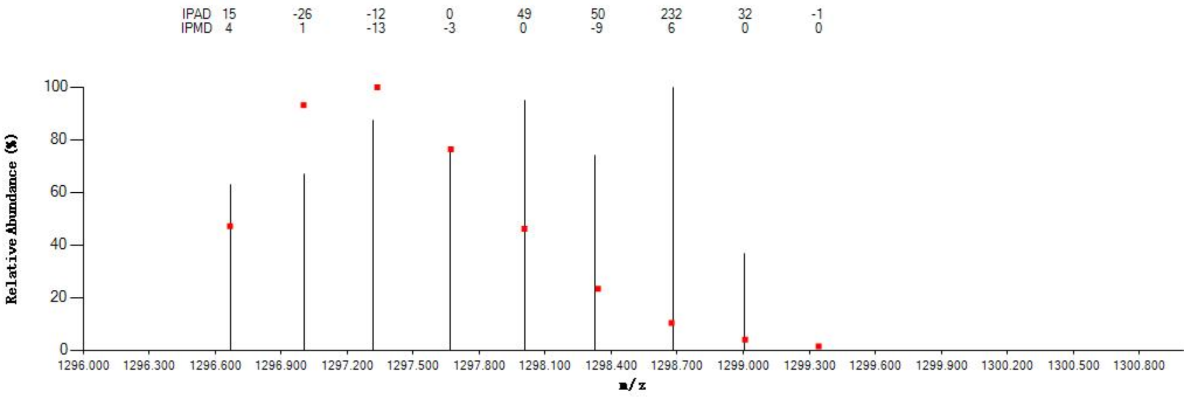


#### Annotated MS/MS Spectrum

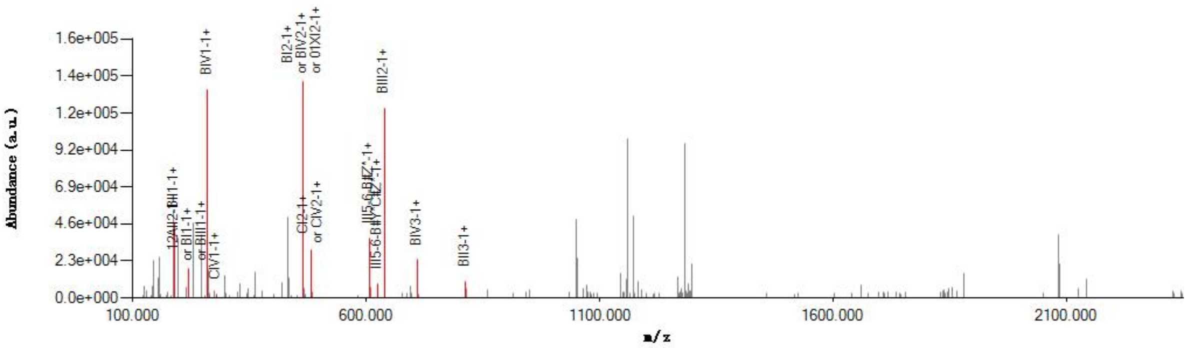


#### Graphical Fragmentation Map

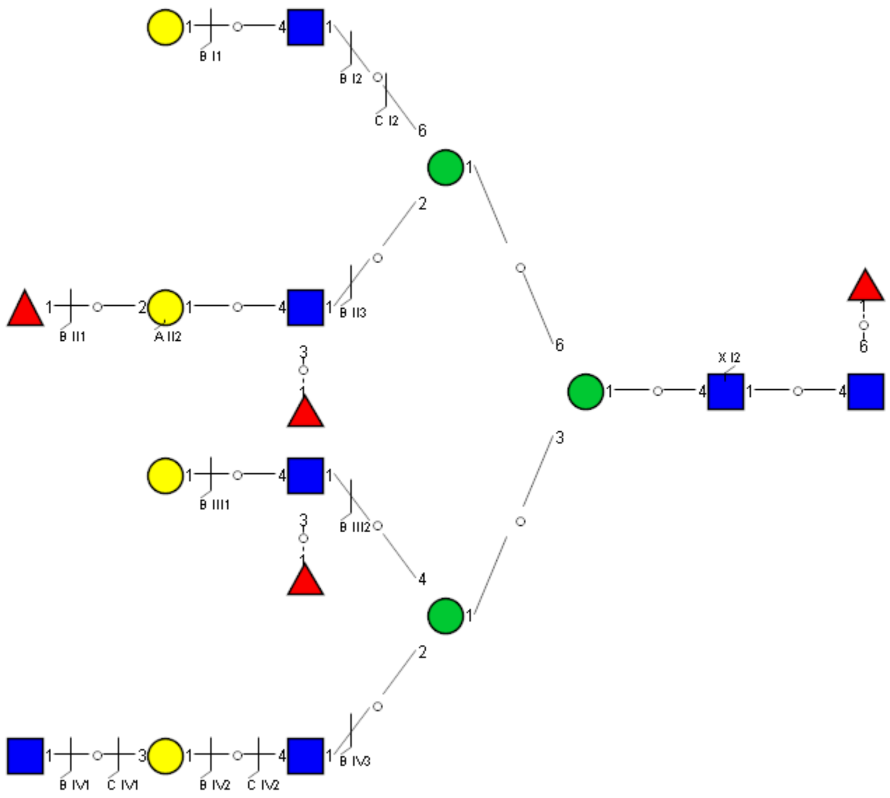

### Spectrum Index: 42327(NoBHs:2)

#### Precursor iEF Map

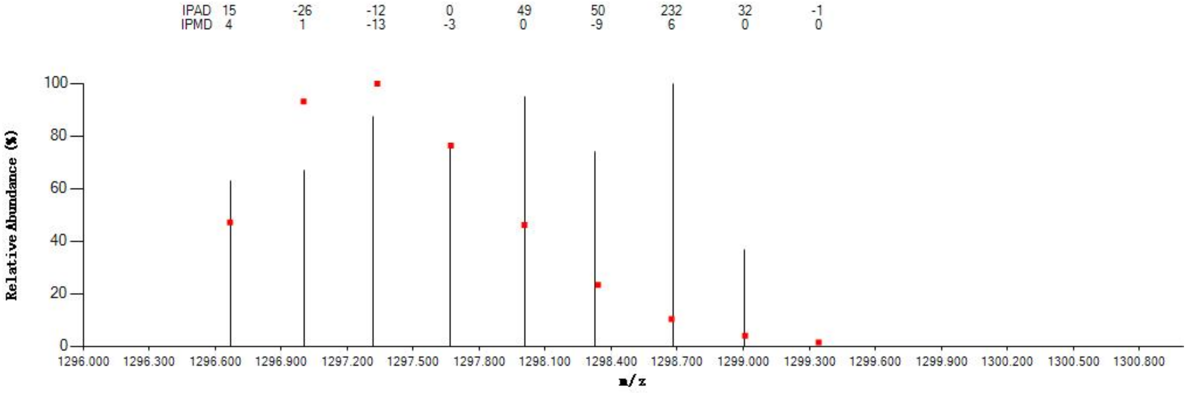


#### Annotated MS/MS Spectrum

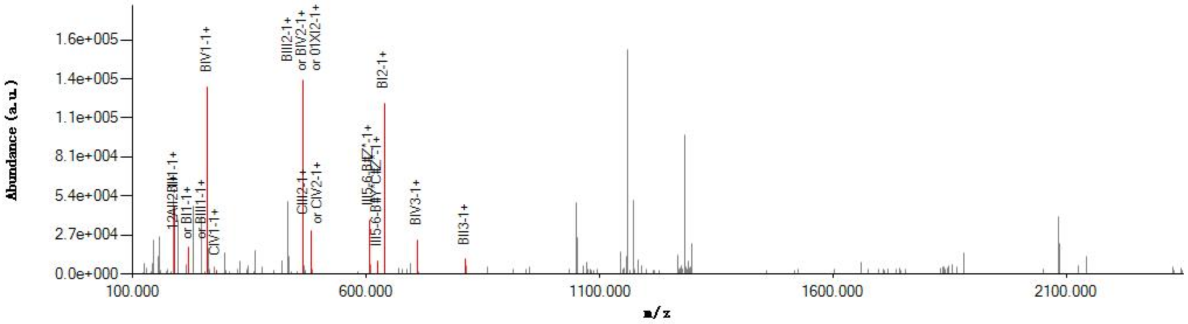


#### Graphical Fragmentation Map

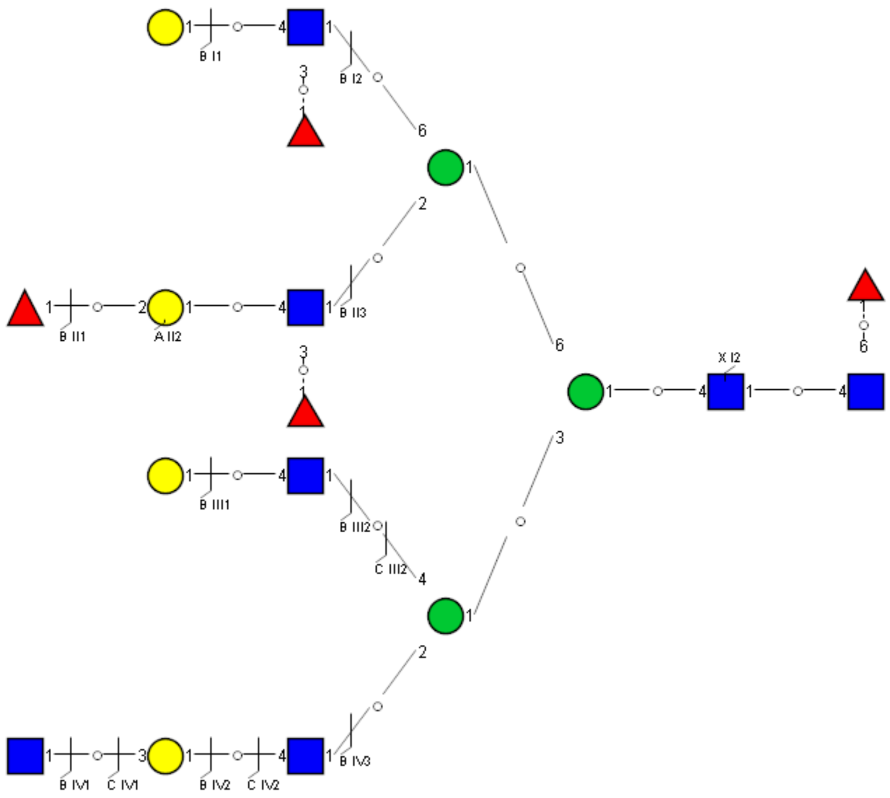

Spectrum Index: 42352(NoBHs:1)

#### Precursor iEF Map

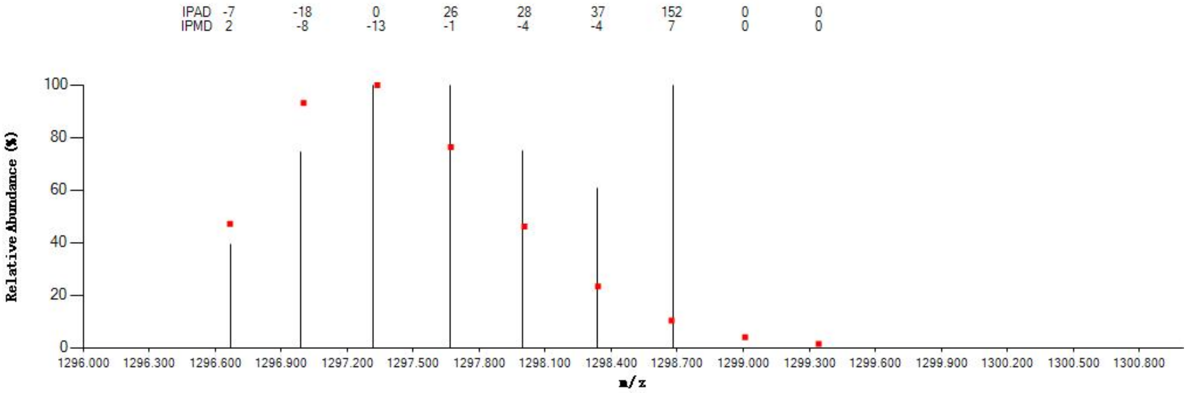


#### Annotated MS/MS Spectrum

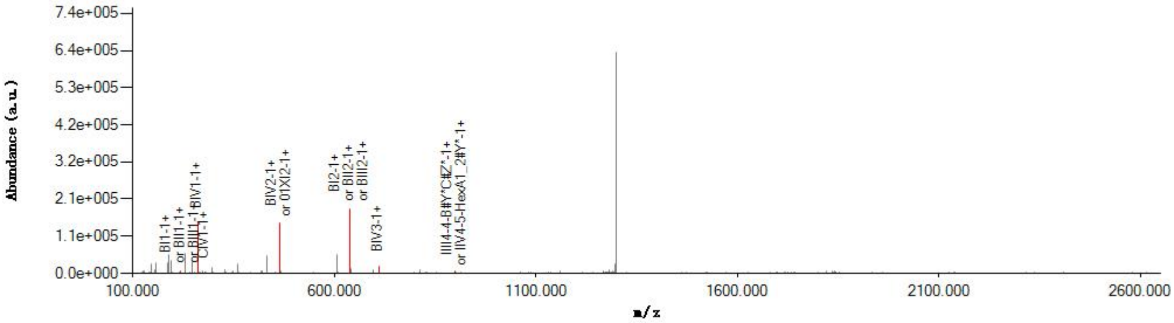


#### Graphical Fragmentation Map

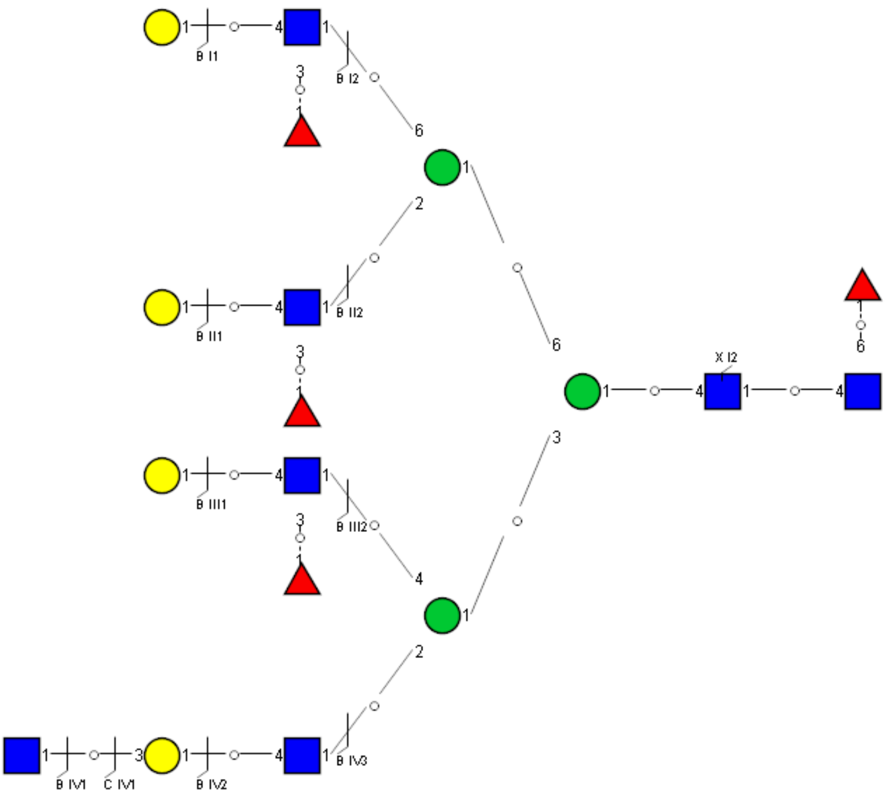

Spectrum Index: 42353(NoBHs:1)

#### Precursor iEF Map

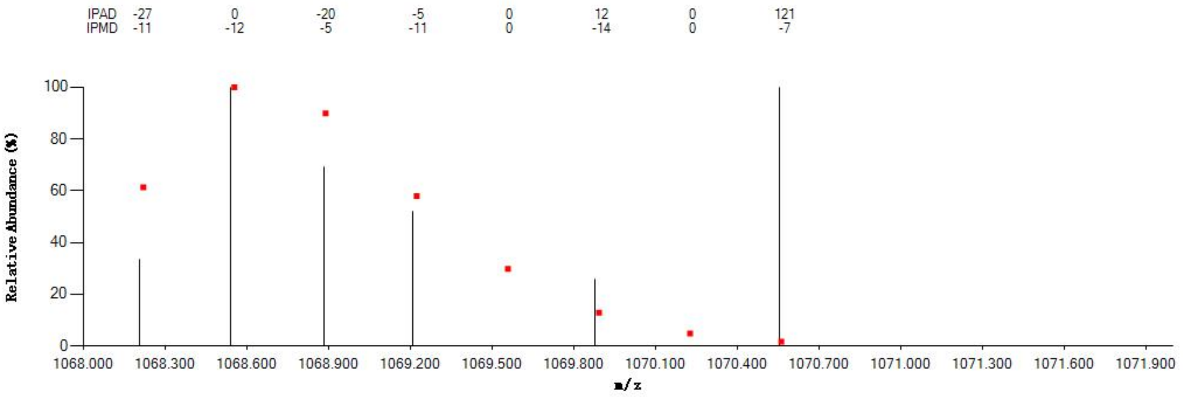


#### Annotated MS/MS Spectrum

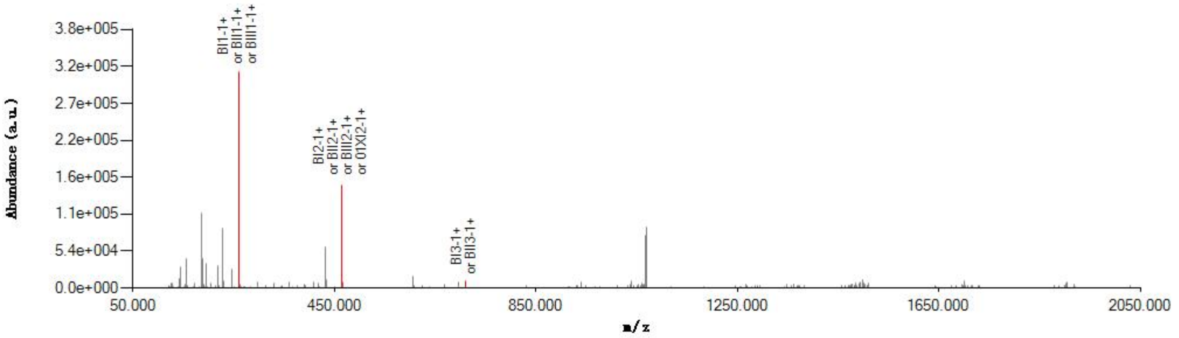


#### Graphical Fragmentation Map

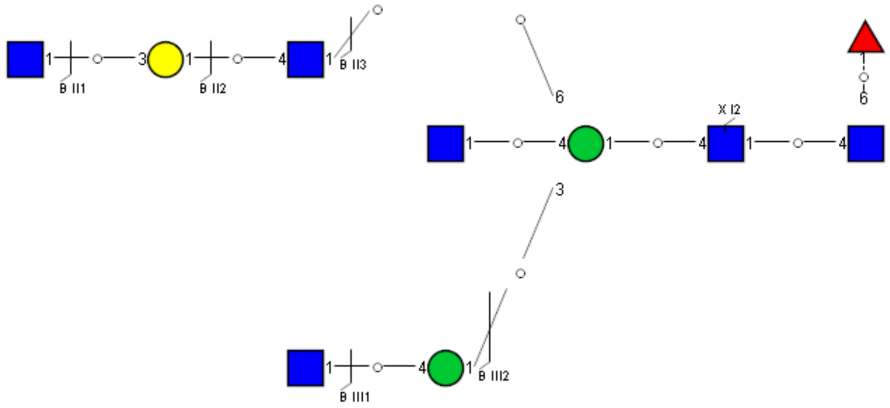

Spectrum Index: 42632(NoBHs:1)

#### Precursor iEF Map

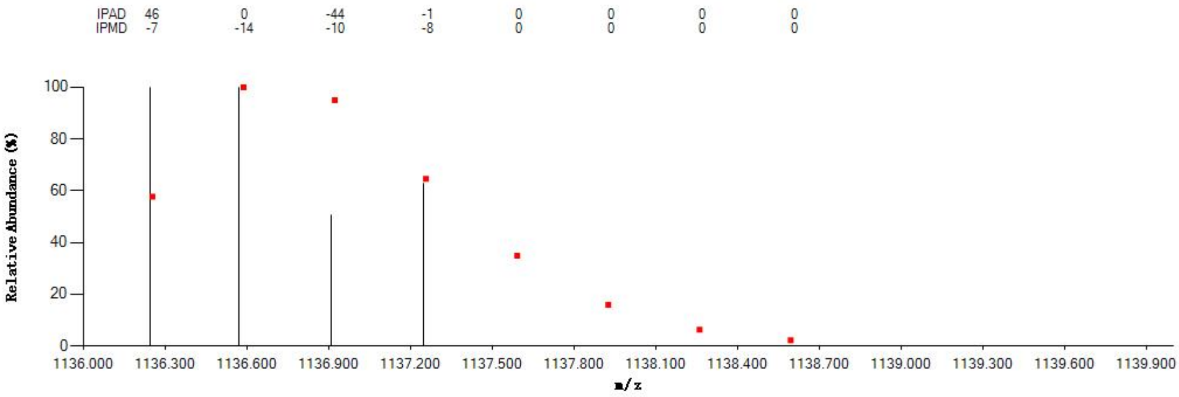


#### Annotated MS/MS Spectrum

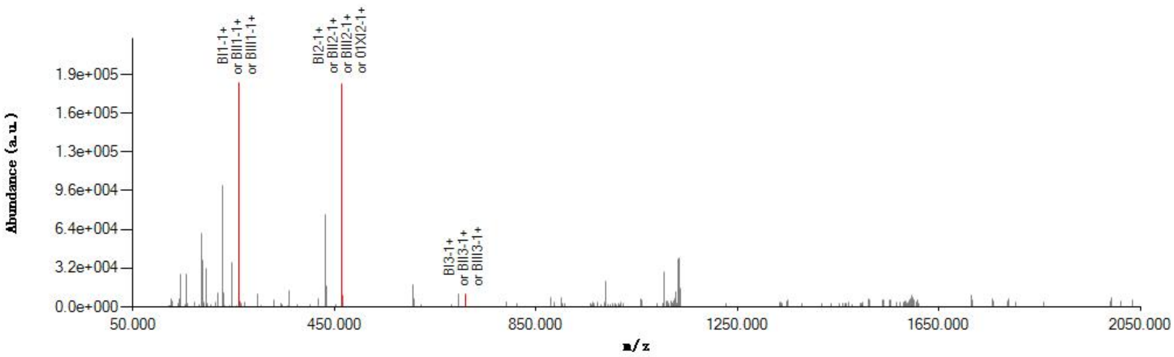


#### Graphical Fragmentation Map

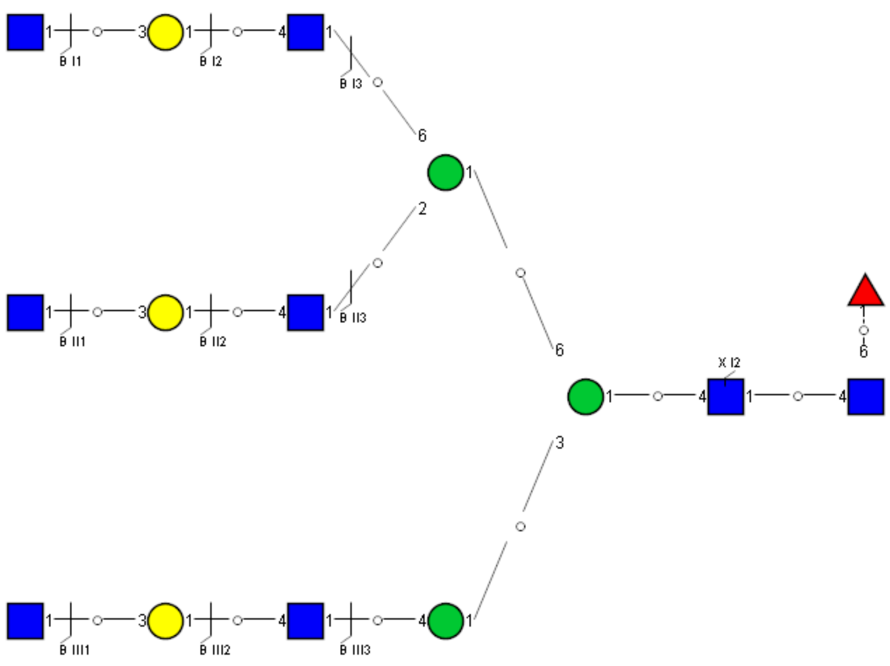

### Spectrum Index: 43082(NoBHs:1)

#### Precursor iEF Map

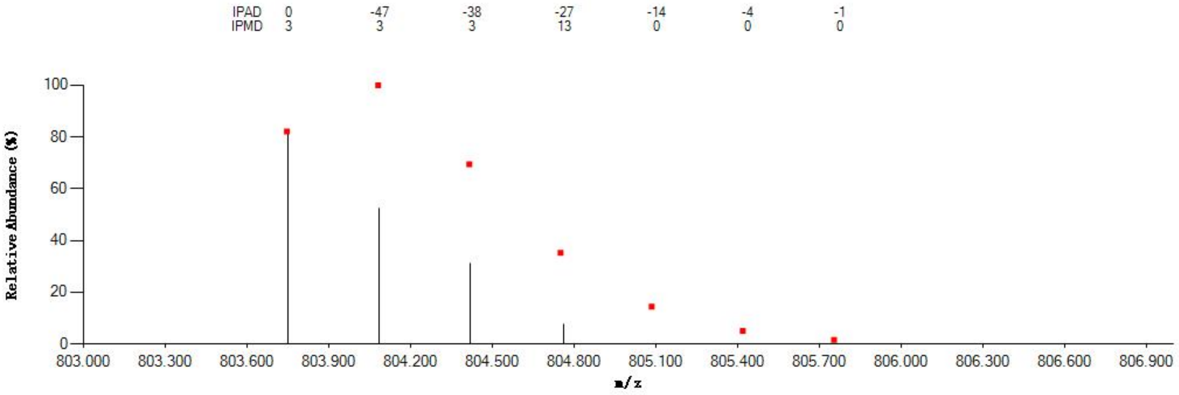


#### Annotated MS/MS Spectrum

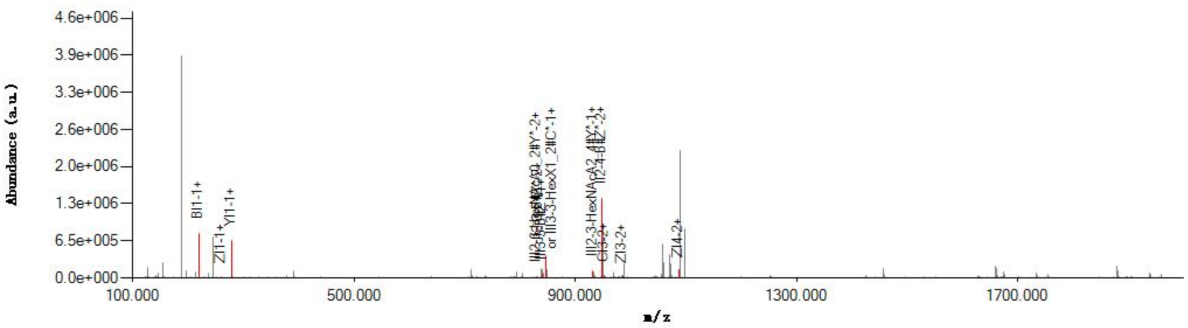


#### Graphical Fragmentation Map

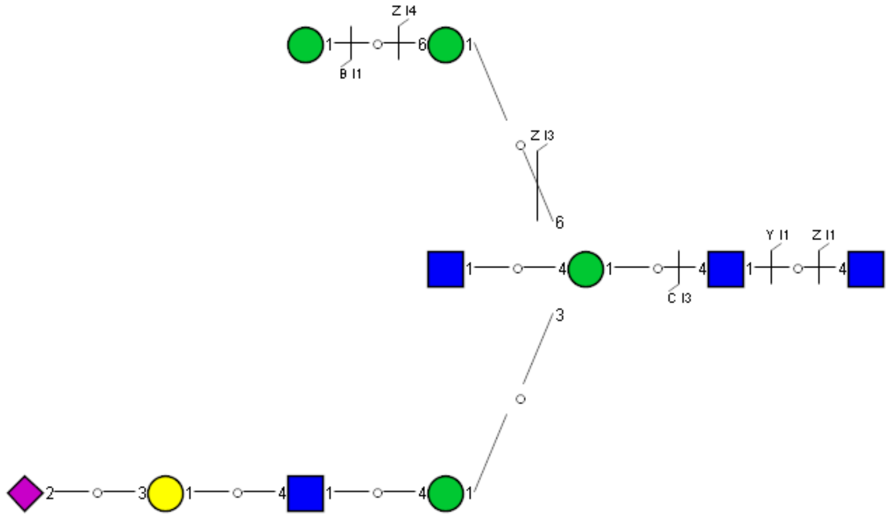

Spectrum Index: 43091(NoBHs:1)

#### Precursor iEF Map

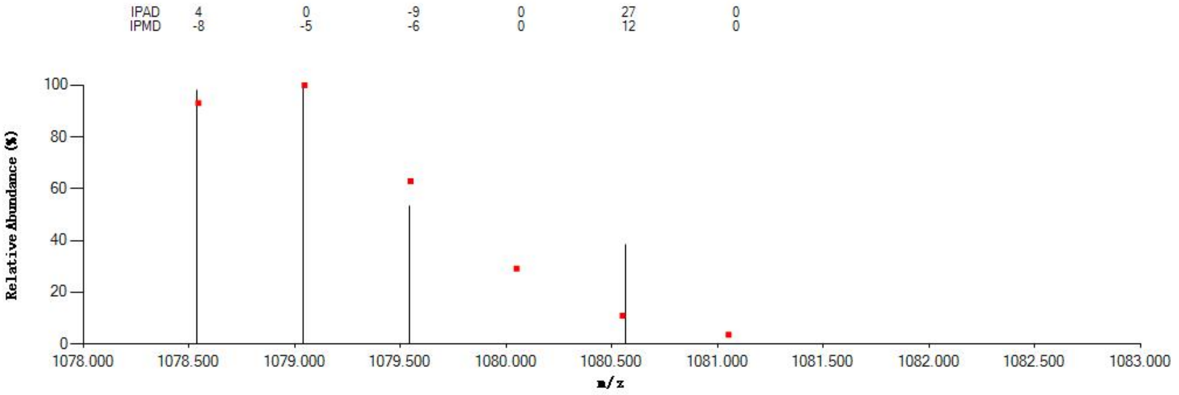


#### Annotated MS/MS Spectrum

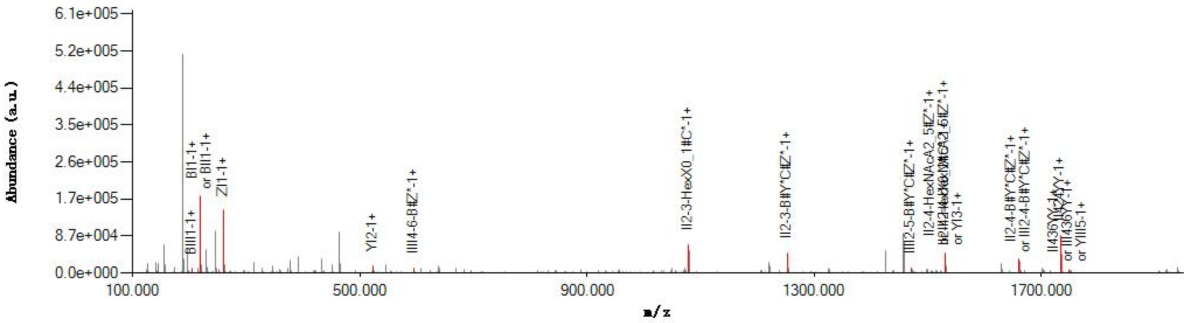


#### Graphical Fragmentation Map

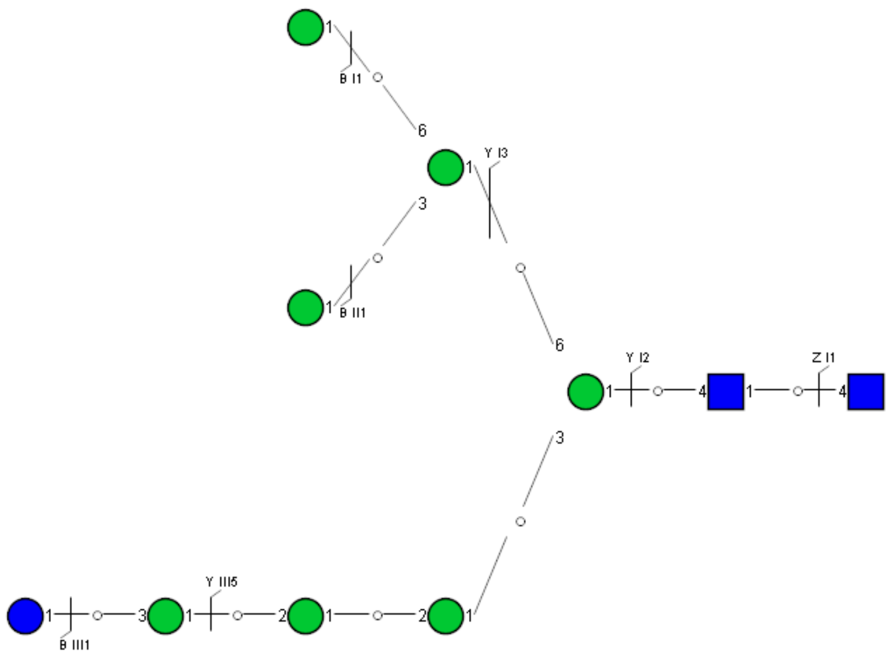

Spectrum Index: 43339(NoBHs:1)

#### Precursor iEF Map

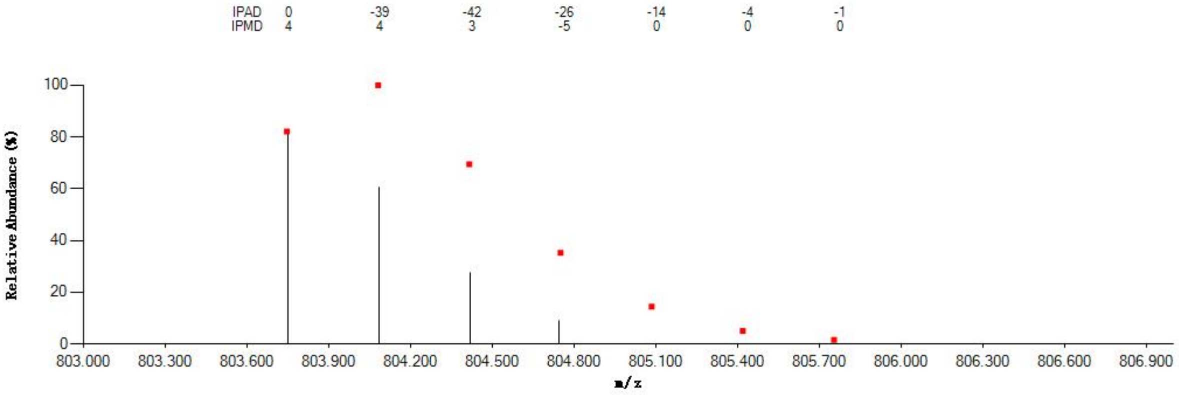


#### Annotated MS/MS Spectrum

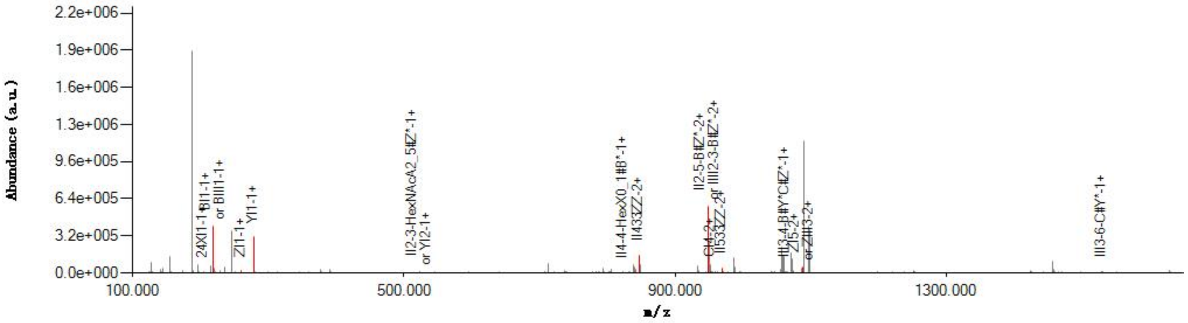


#### Graphical Fragmentation Map

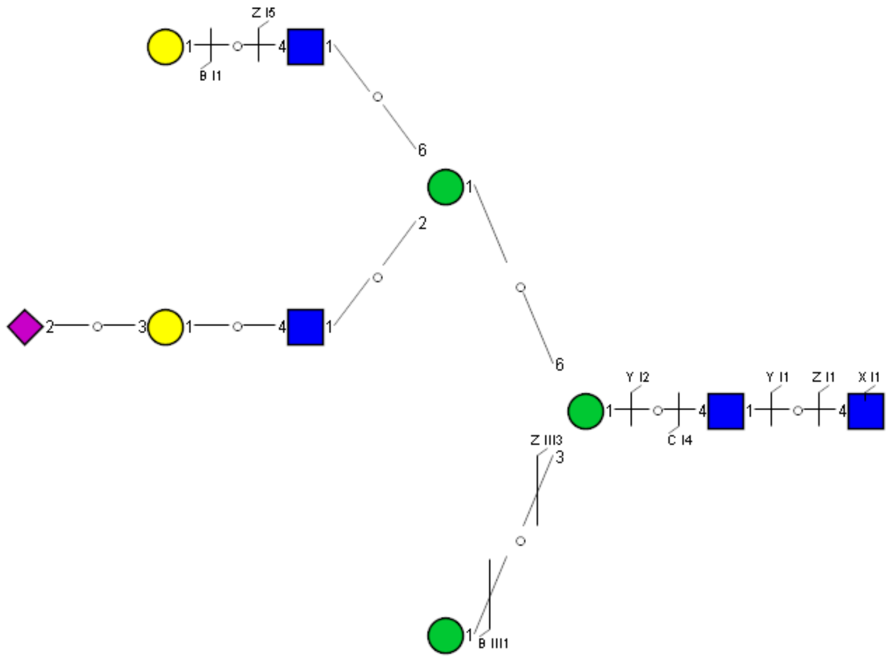

### Spectrum Index: 43477(NoBHs:1)

#### Precursor iEF Map

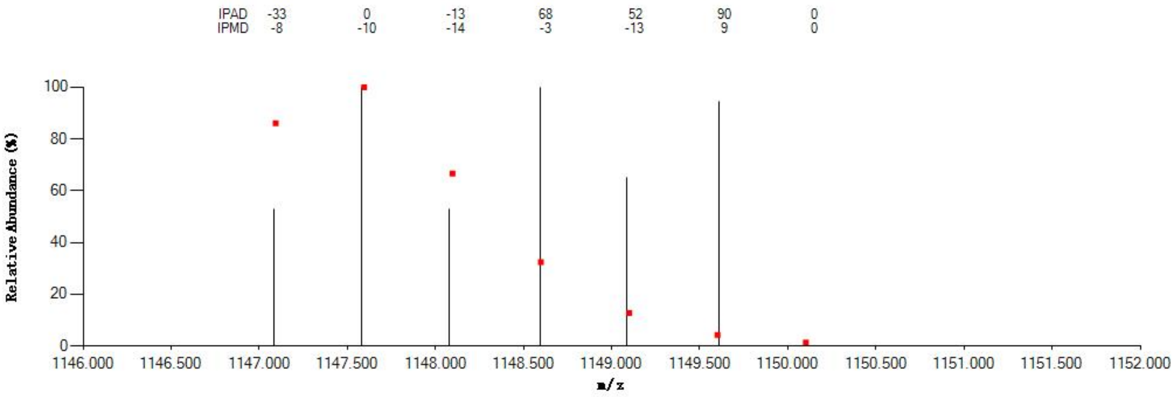


#### Annotated MS/MS Spectrum

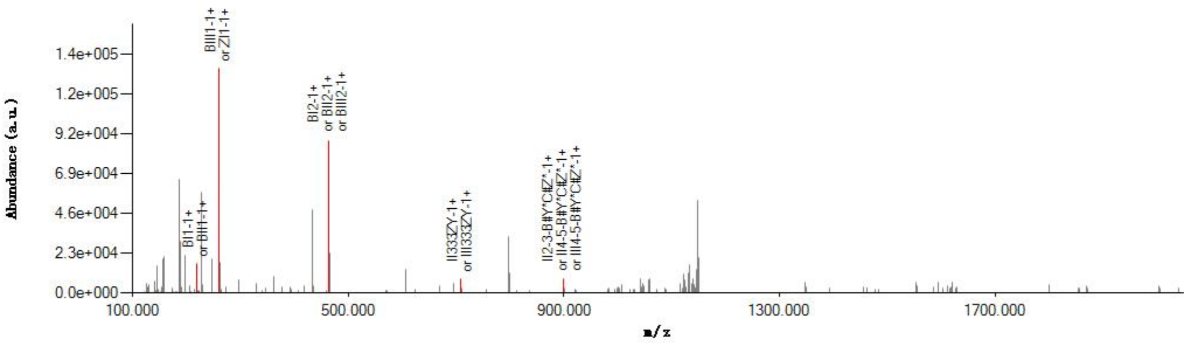


#### Graphical Fragmentation Map

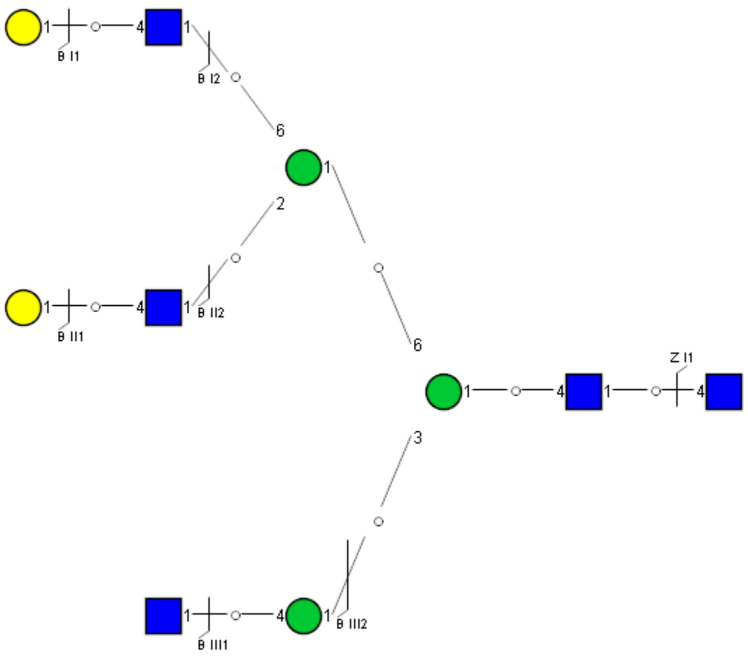

Spectrum Index: 43522(NoBHs:1)

#### Precursor iEF Map

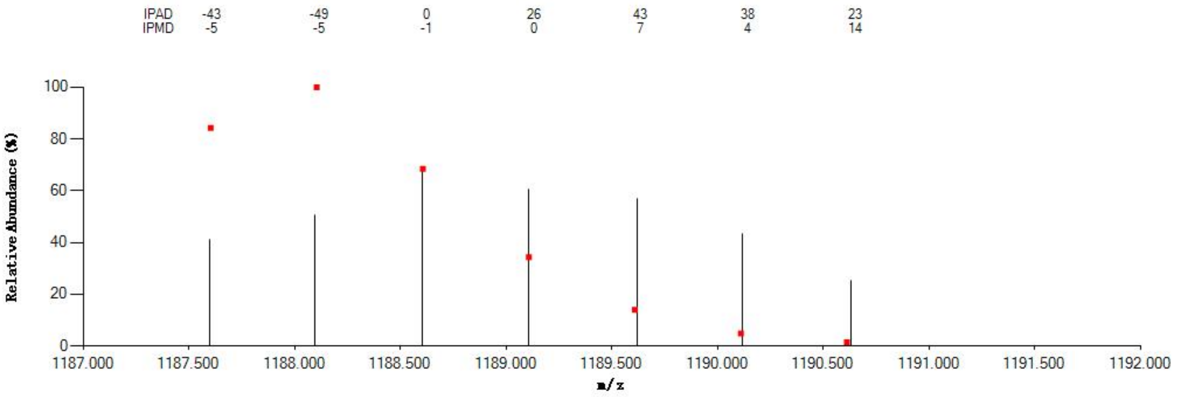

#### Annotated MS/MS Spectrum

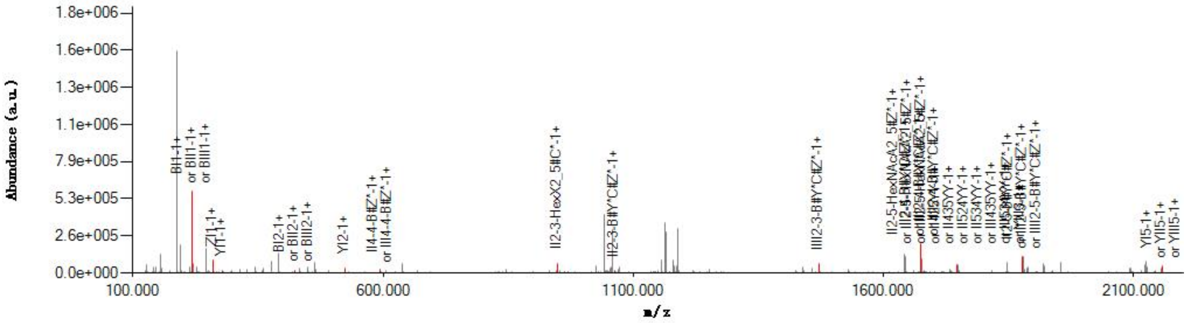

#### Graphical Fragmentation Map

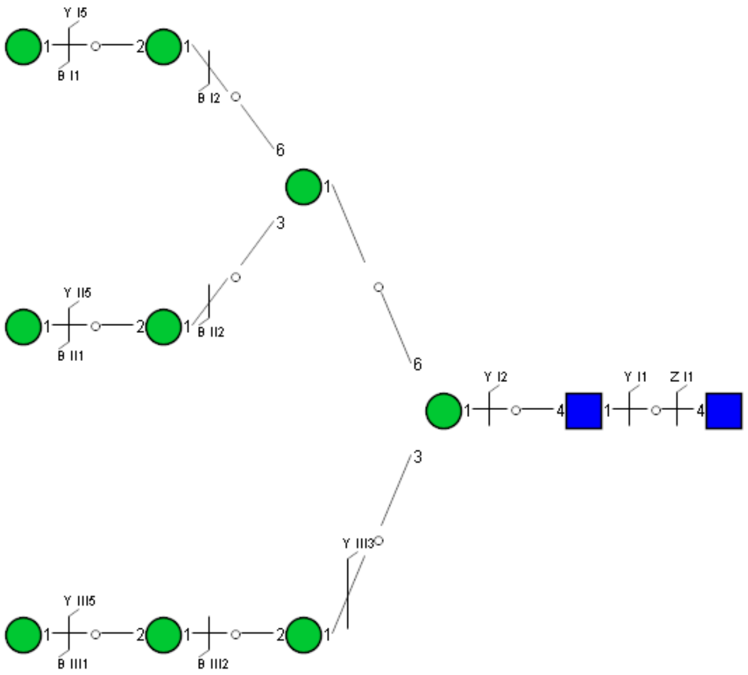

Spectrum Index: 44280(NoBHs:1)

#### Precursor iEF Map

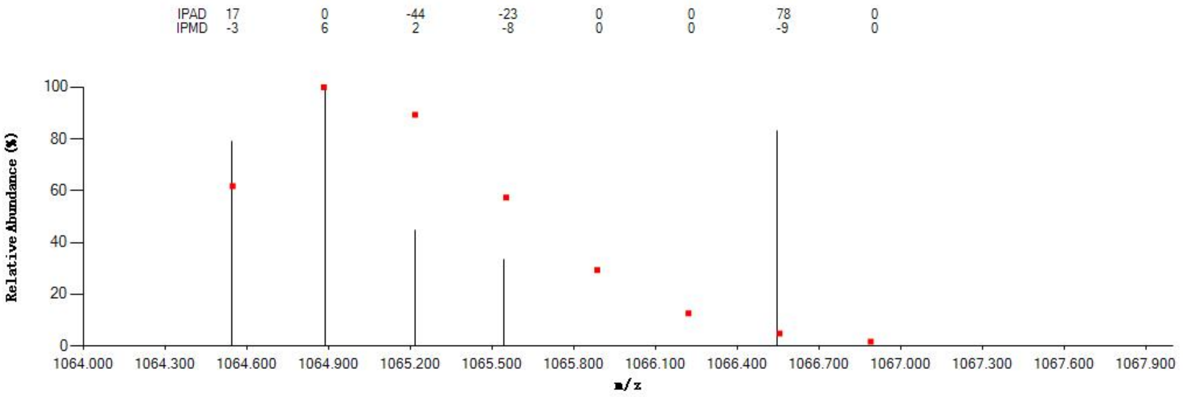

#### Annotated MS/MS Spectrum

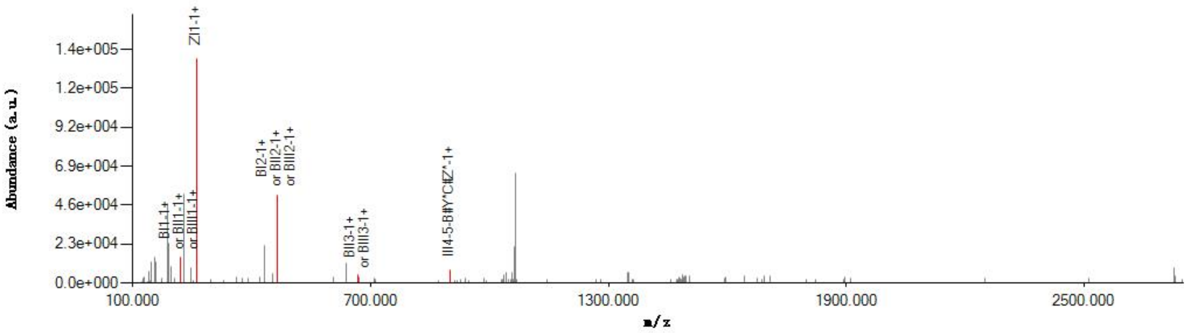

#### Graphical Fragmentation Map

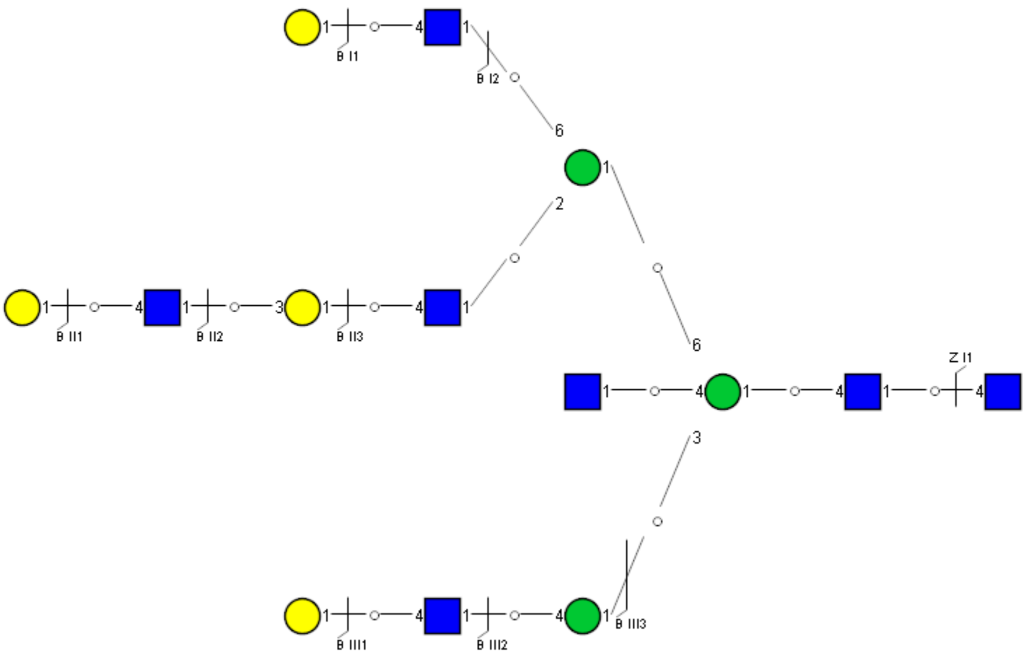

Spectrum Index: 45251(NoBHs:1)

#### Precursor iEF Map

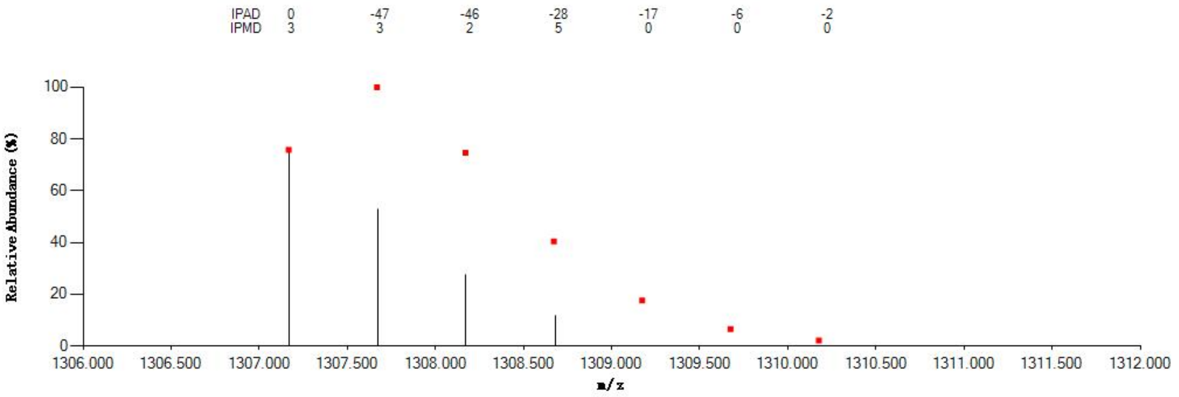


#### Annotated MS/MS Spectrum

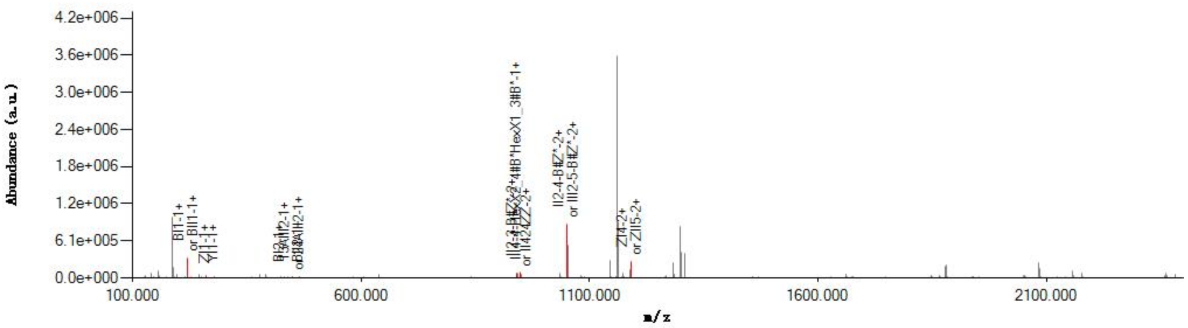


#### Graphical Fragmentation Map

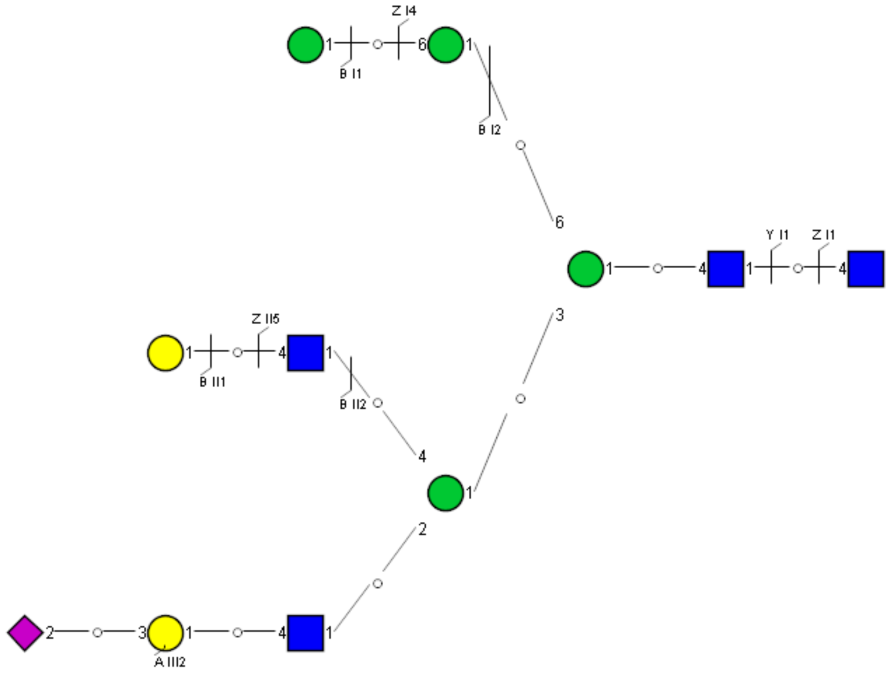

Spectrum Index: 45369(NoBHs:1)

#### Precursor iEF Map

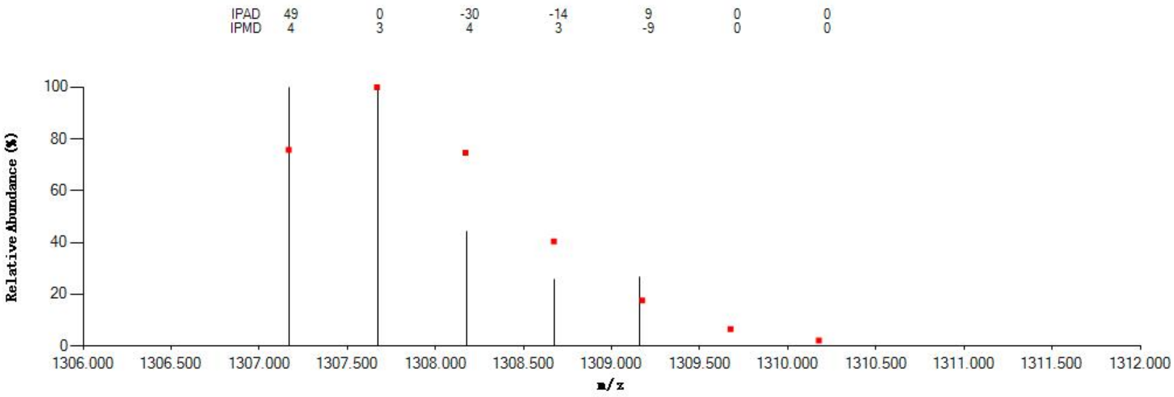


#### Annotated MS/MS Spectrum

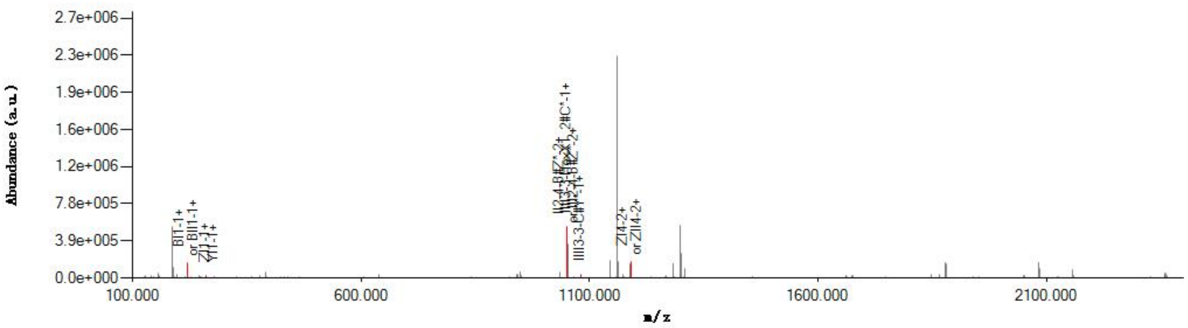


#### Graphical Fragmentation Map

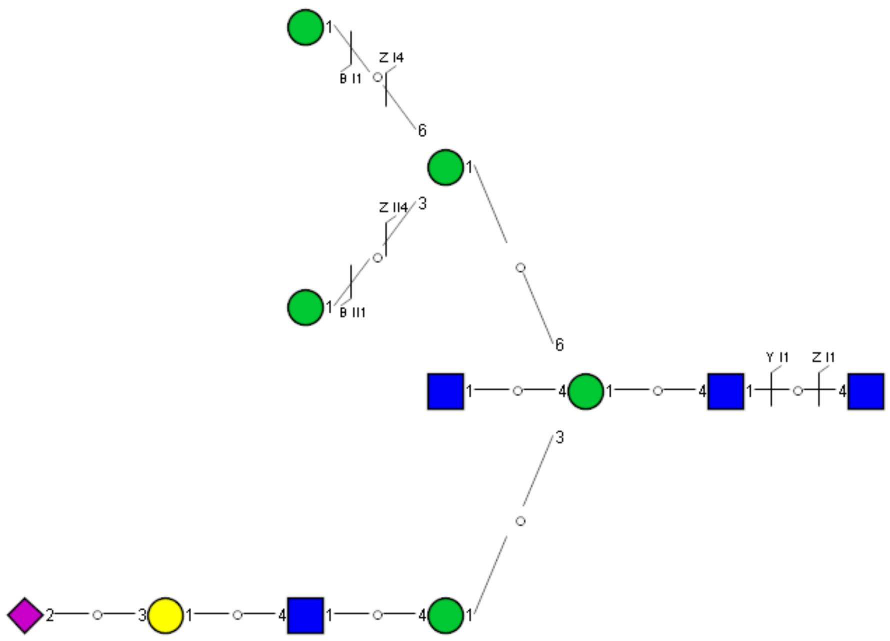

Spectrum Index: 46230(NoBHs:1)

#### Precursor iEF Map

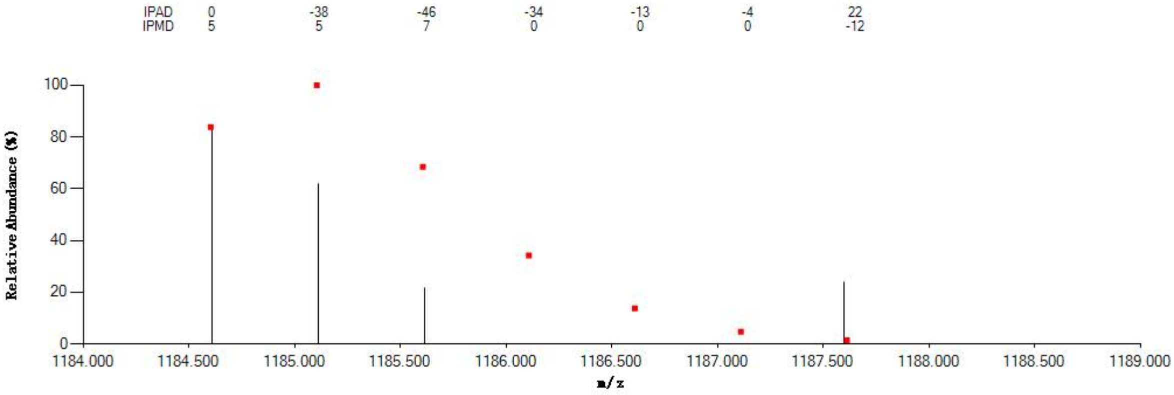


#### Annotated MS/MS Spectrum

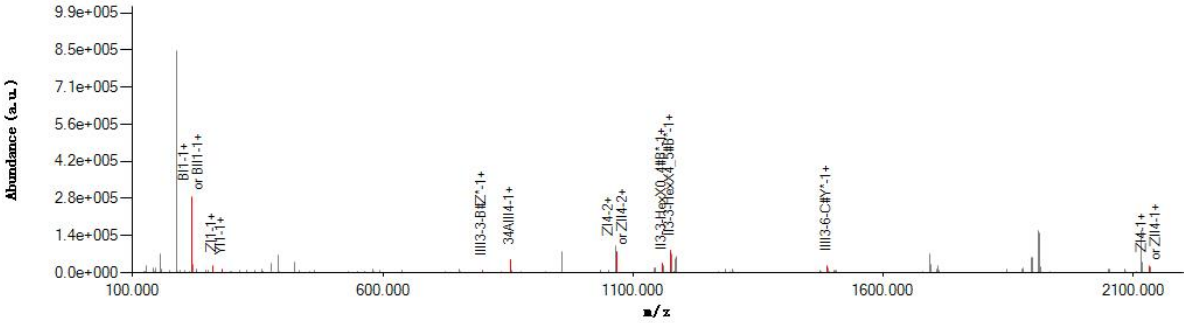


#### Graphical Fragmentation Map

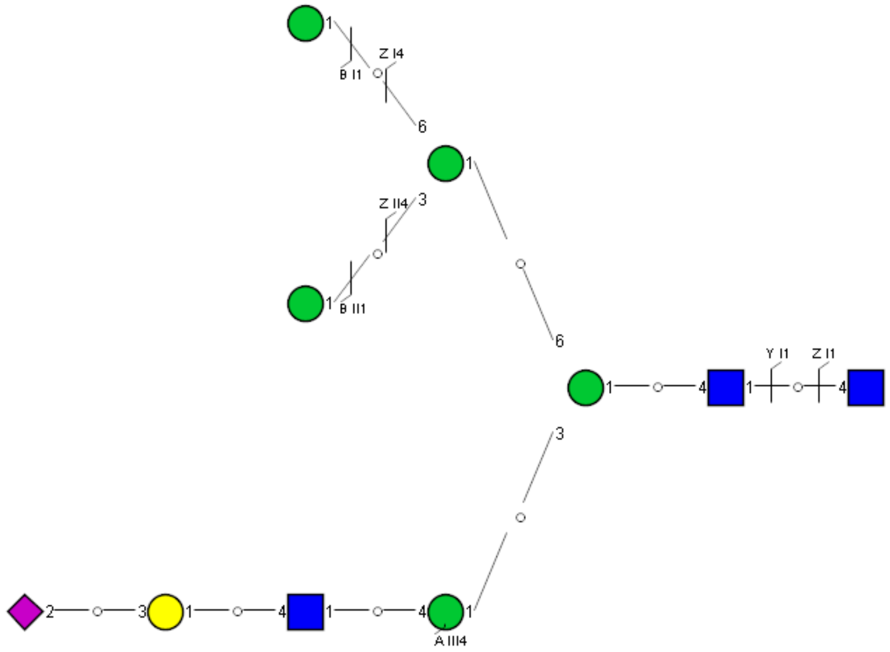

Spectrum Index: 49008(NoBHs:1)

#### Precursor iEF Map

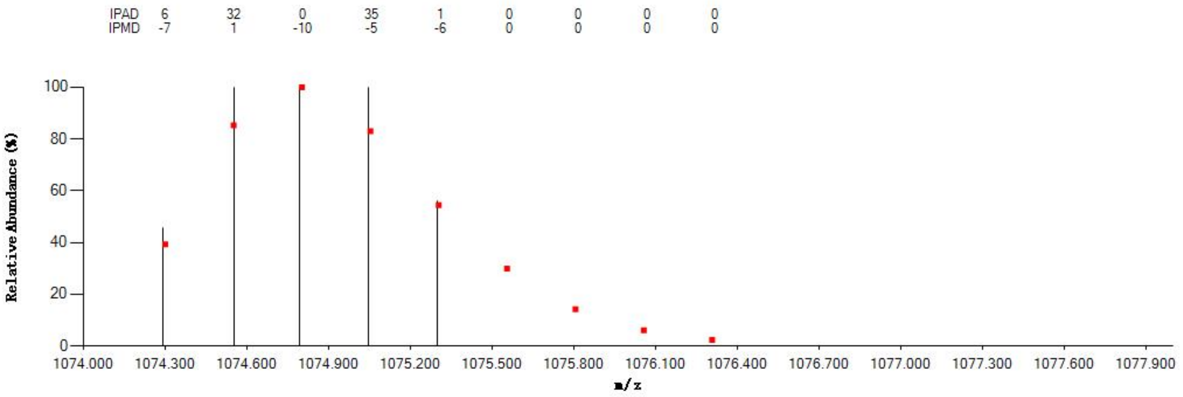

#### Annotated MS/MS Spectrum

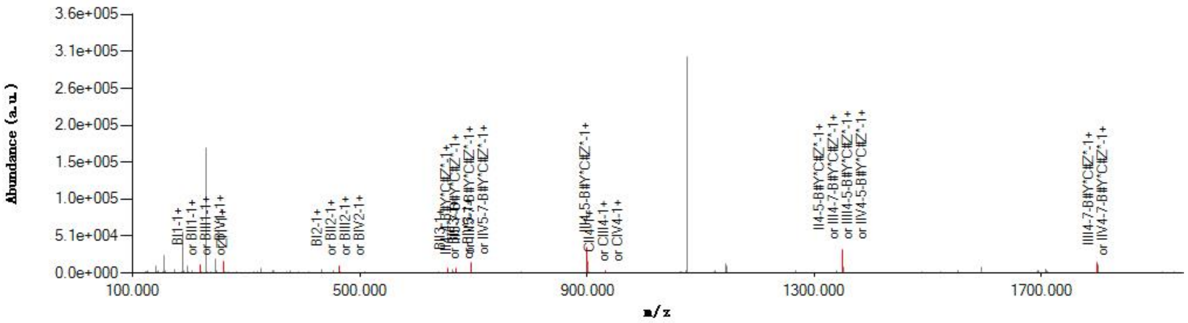

#### Graphical Fragmentation Map

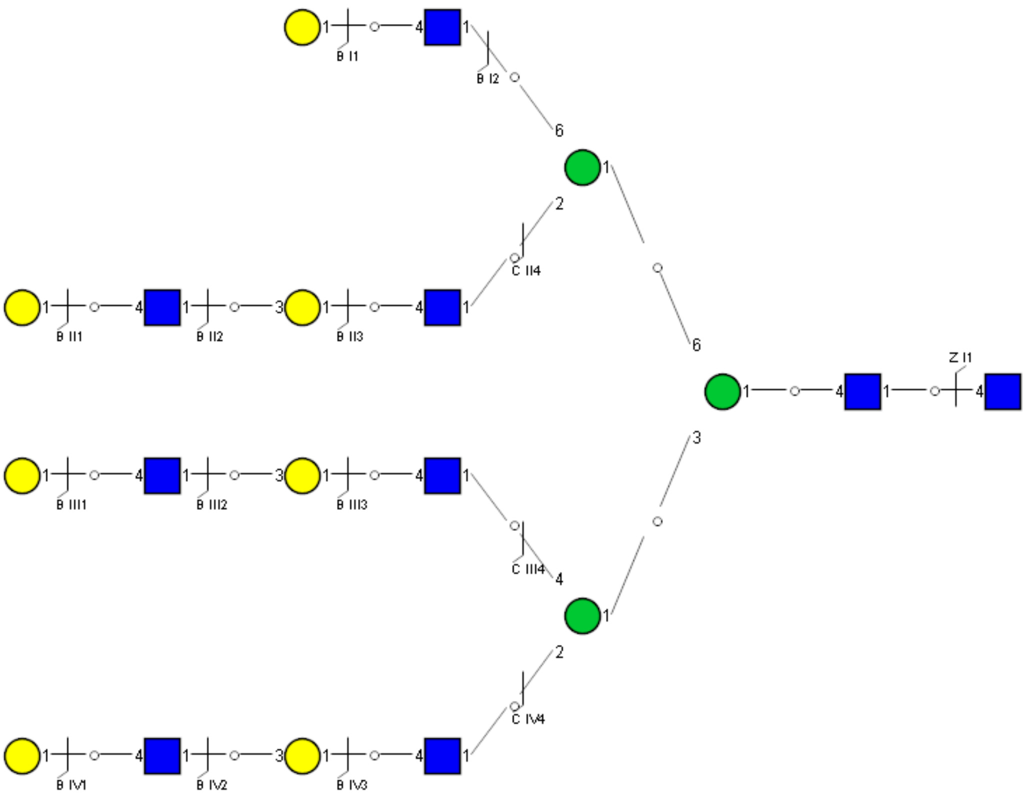

Spectrum Index: 60681(NoBHs:3)

#### Precursor iEF Map

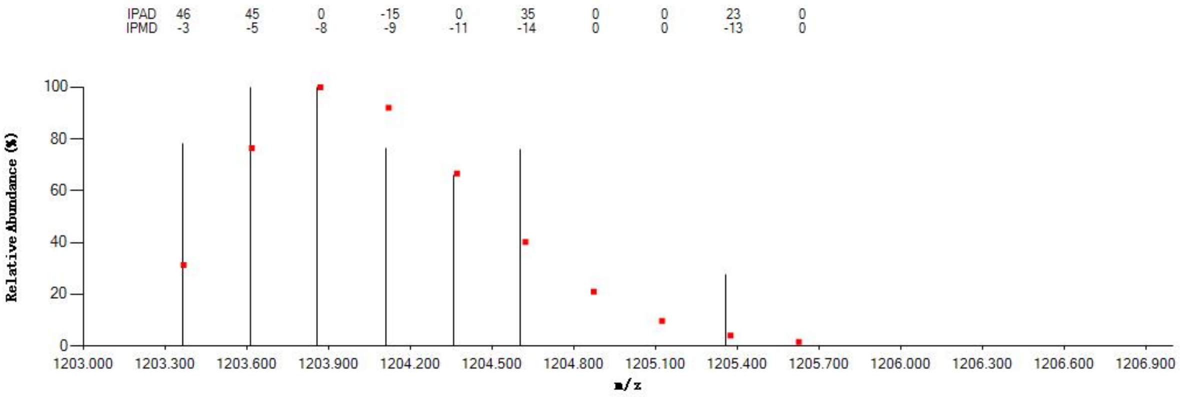


#### Annotated MS/MS Spectrum

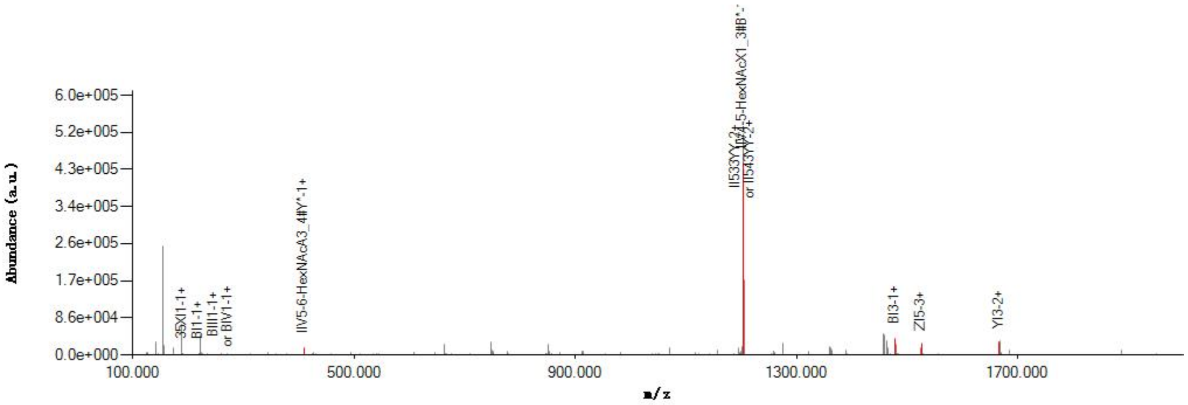


#### Graphical Fragmentation Map

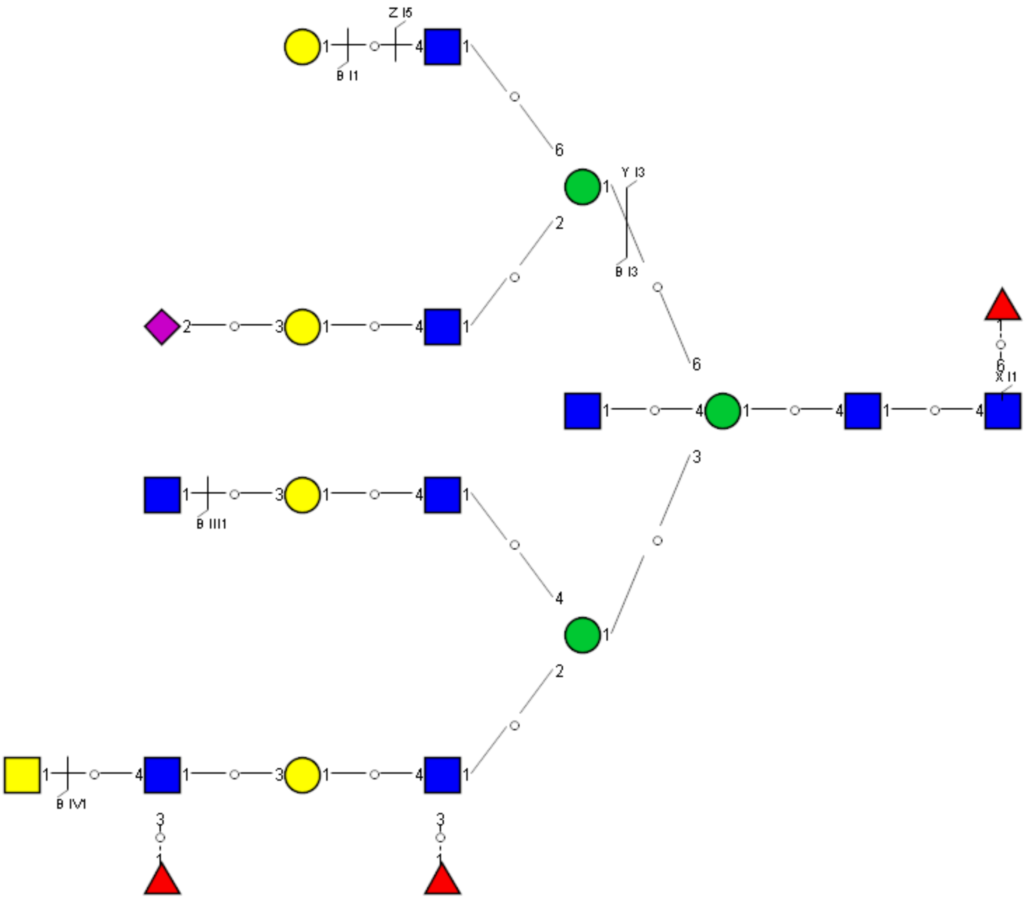

### Spectrum Index: 60681(NoBHs:3)

#### Precursor iEF Map

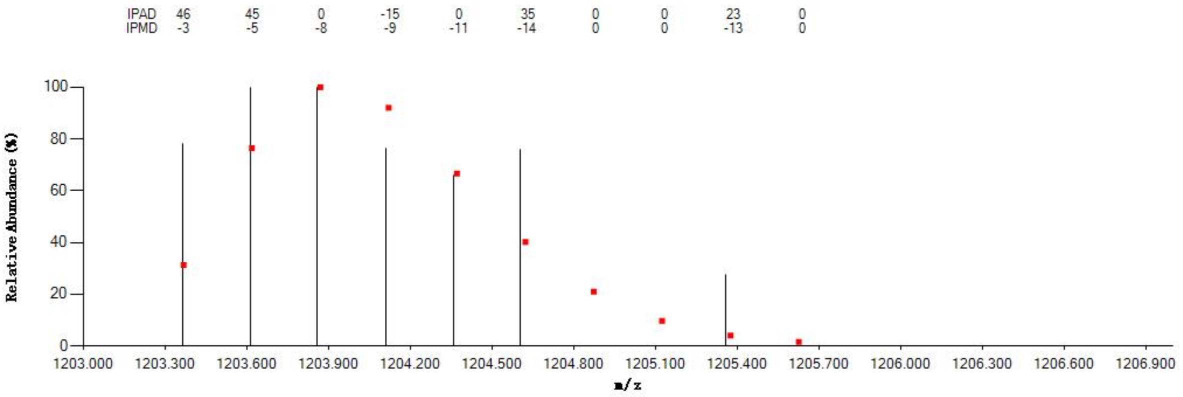


#### Annotated MS/MS Spectrum

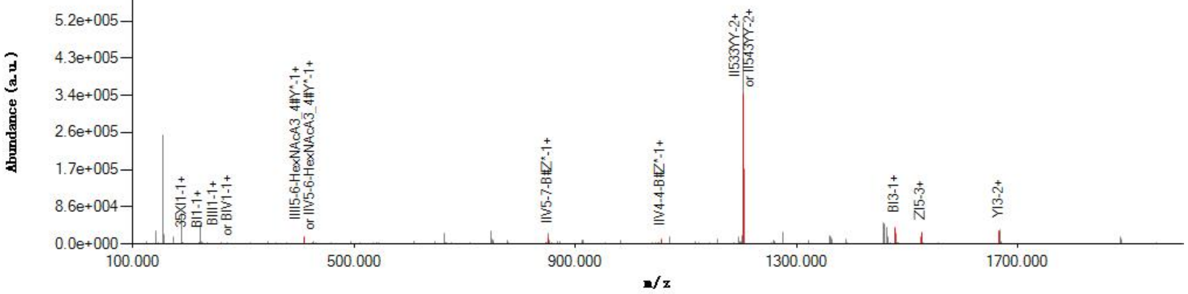


#### Graphical Fragmentation Map

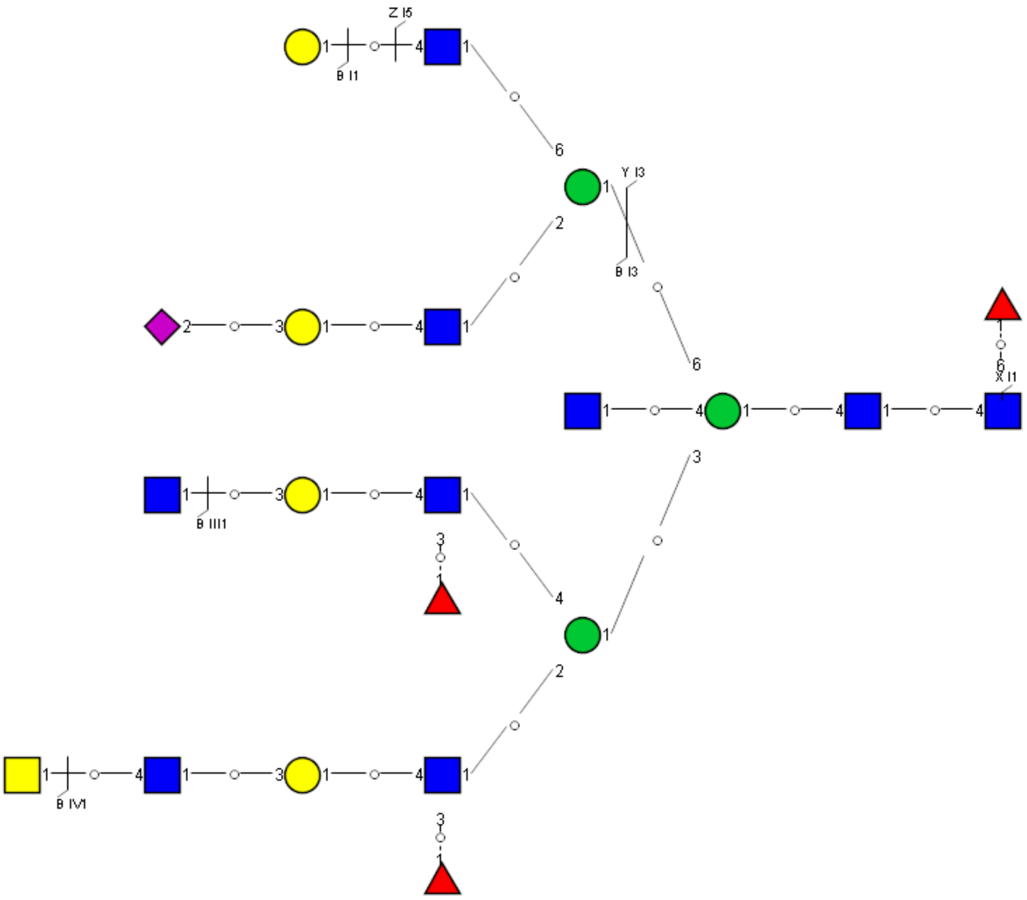

Spectrum Index: 60681(NoBHs:3)

#### Precursor iEF Map

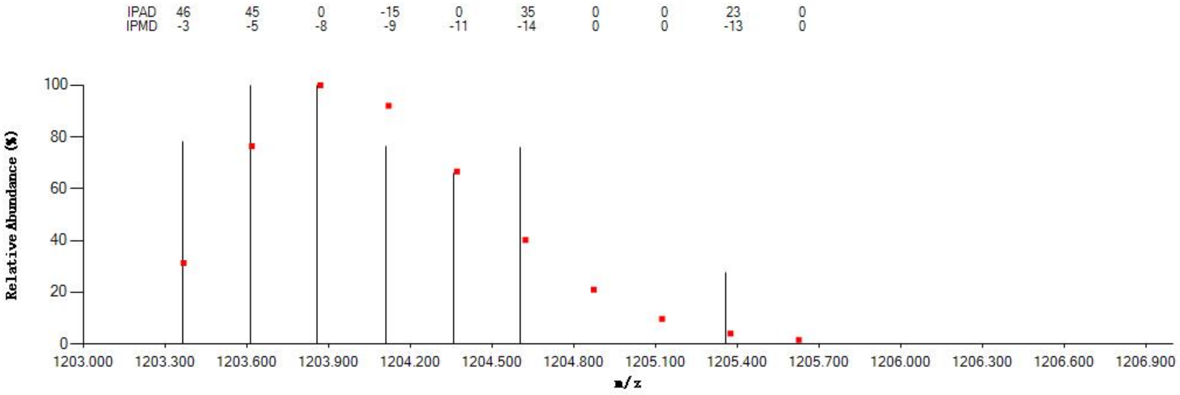


#### Annotated MS/MS Spectrum

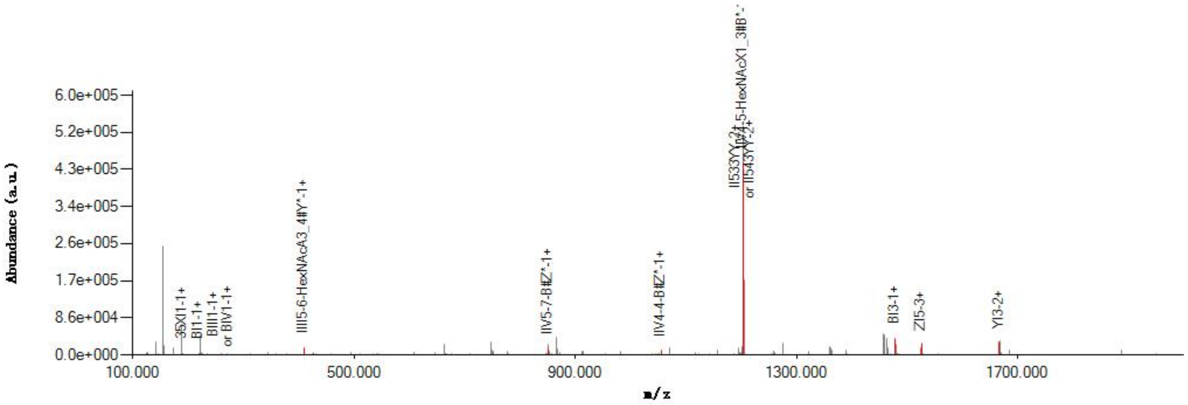


#### Graphical Fragmentation Map

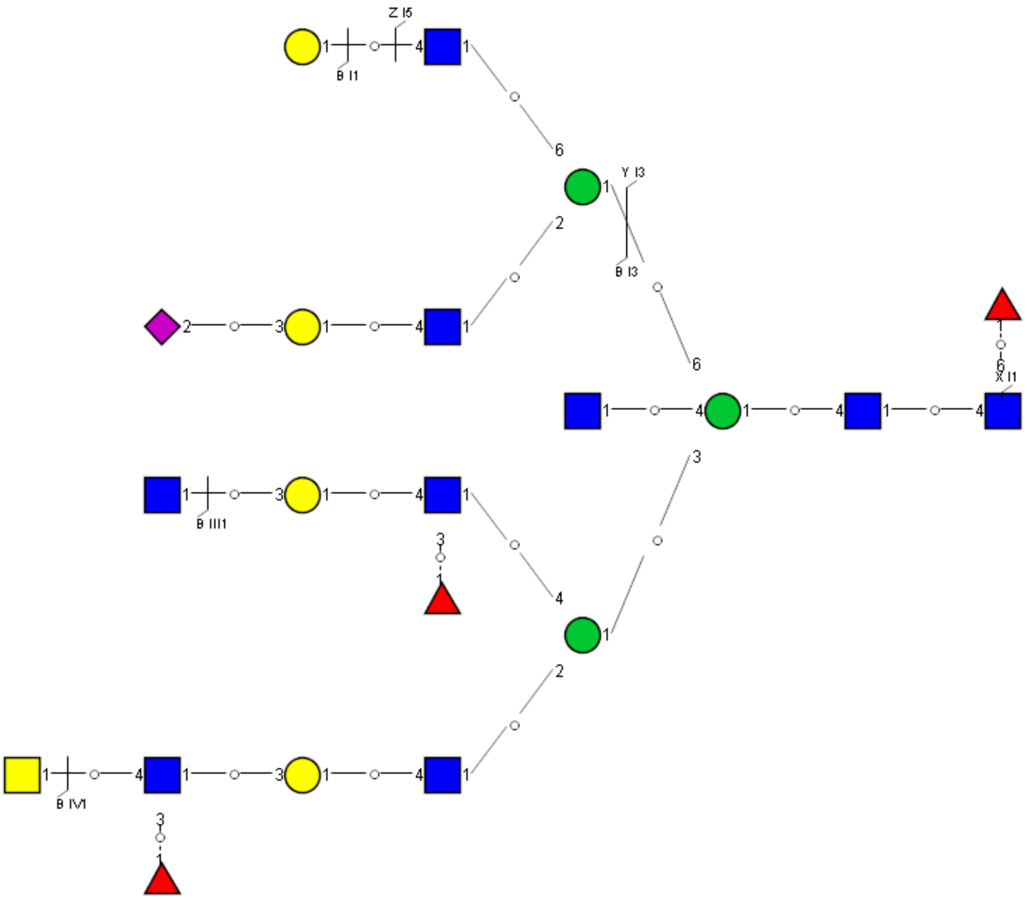

Spectrum Index: 65204(NoBHs:1)

#### Precursor iEF Map

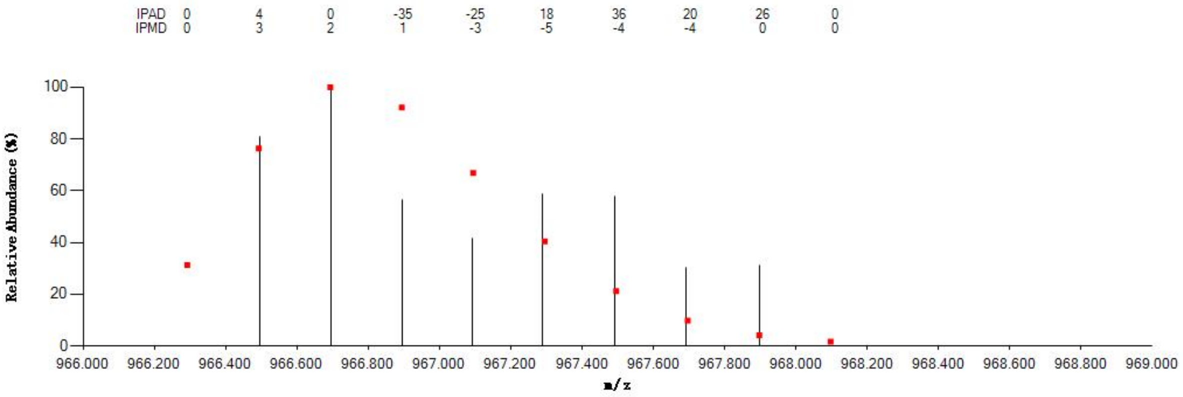


#### Annotated MS/MS Spectrum

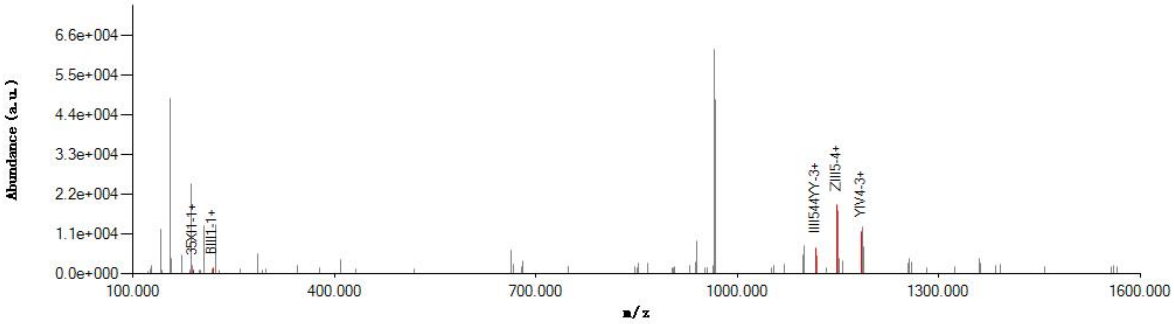


#### Graphical Fragmentation Map

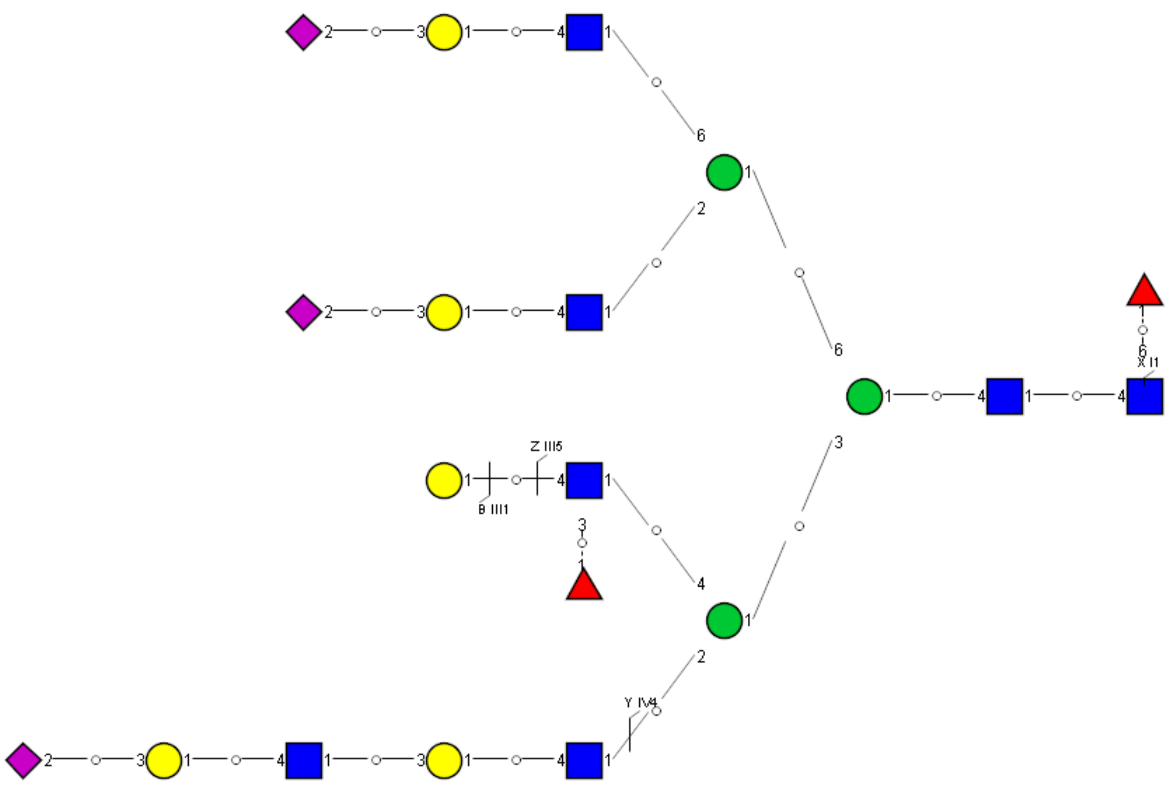

Spectrum Index: 65826(NoBHs:1)

#### Precursor iEF Map

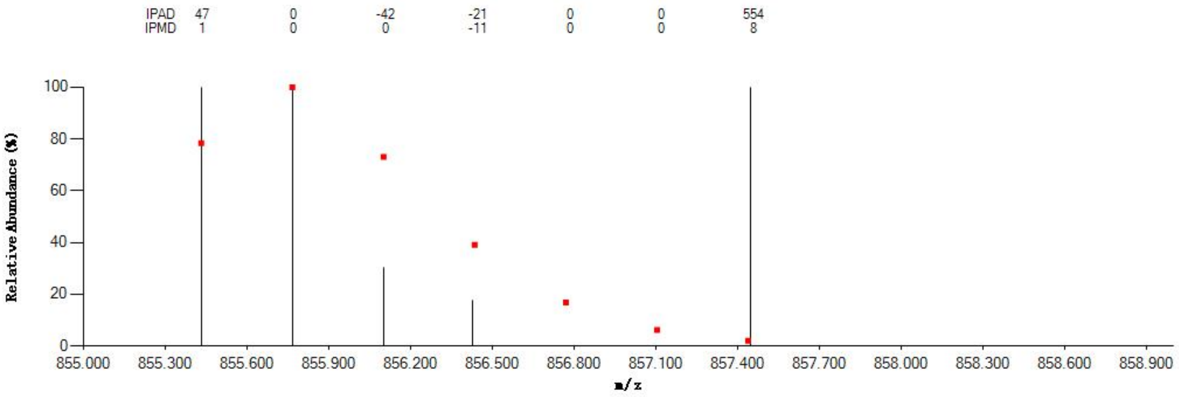


#### Annotated MS/MS Spectrum

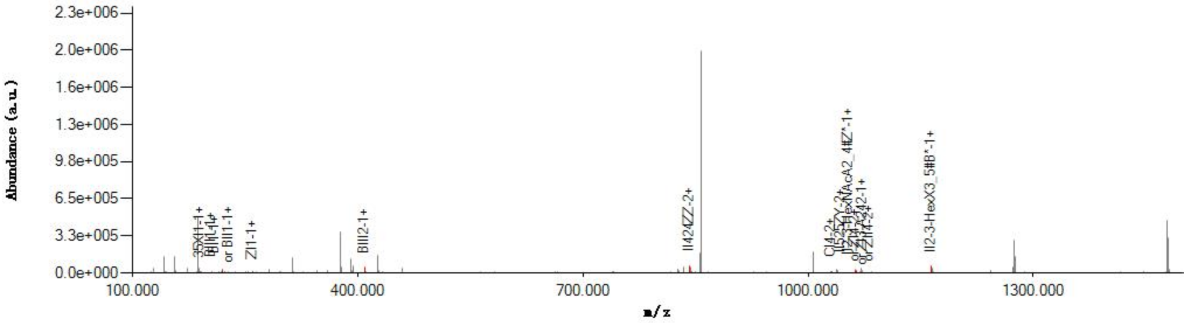


#### Graphical Fragmentation Map

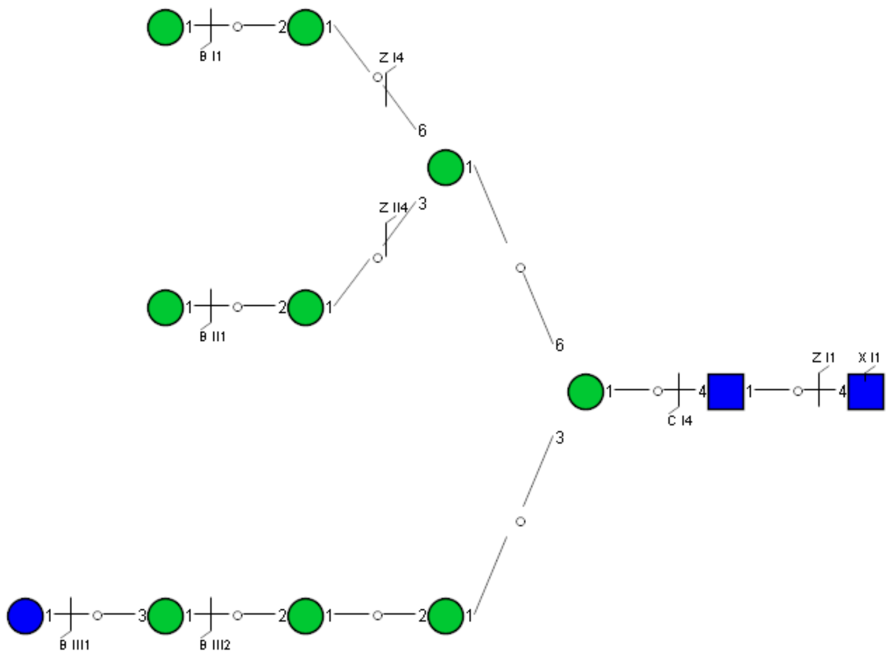

Spectrum Index: 66501(NoBHs:1)

#### Precursor iEF Map

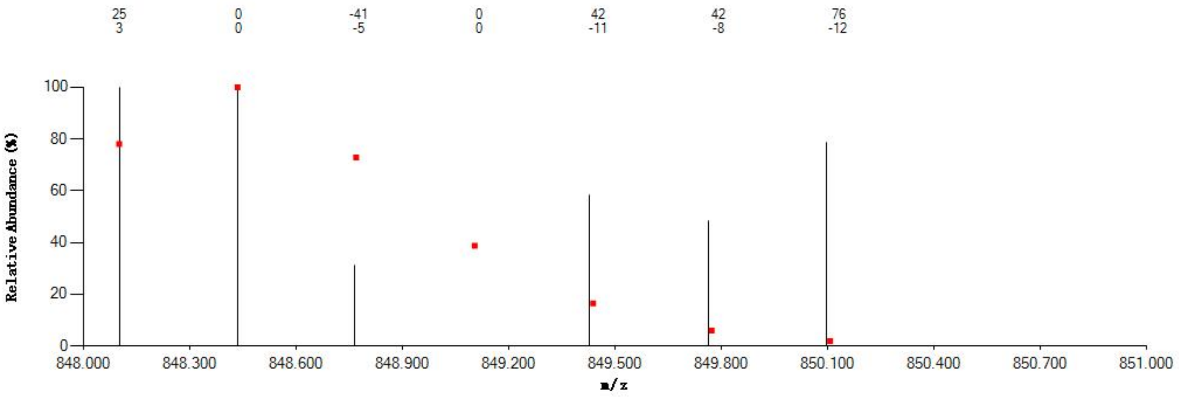


#### Annotated MS/MS Spectrum

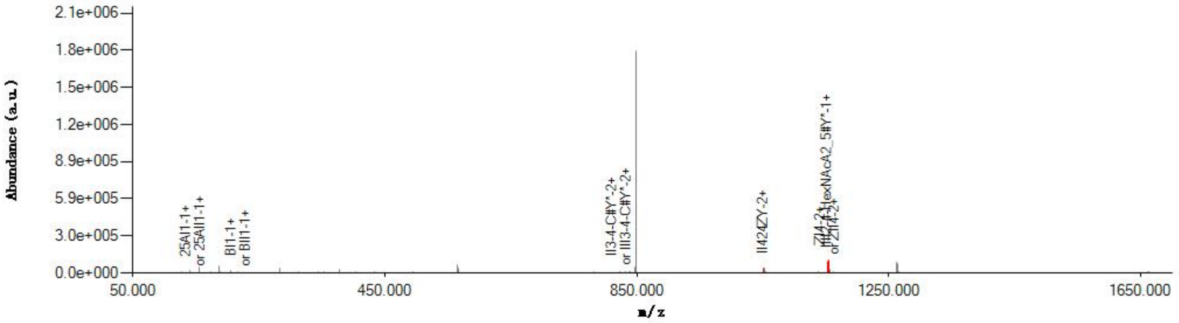


#### Graphical Fragmentation Map

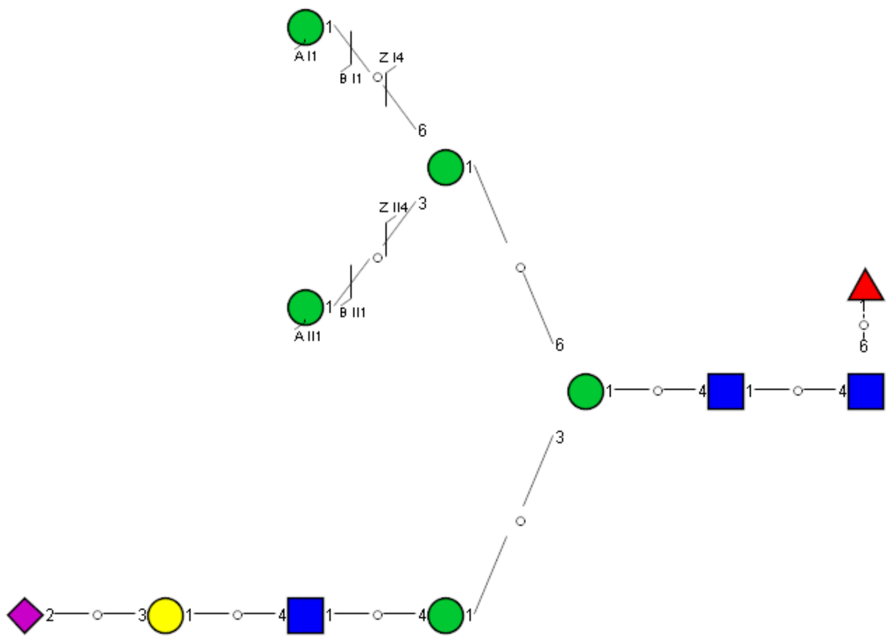

### Spectrum Index: 29476(NoBHs:1)

#### Precursor iEF Map

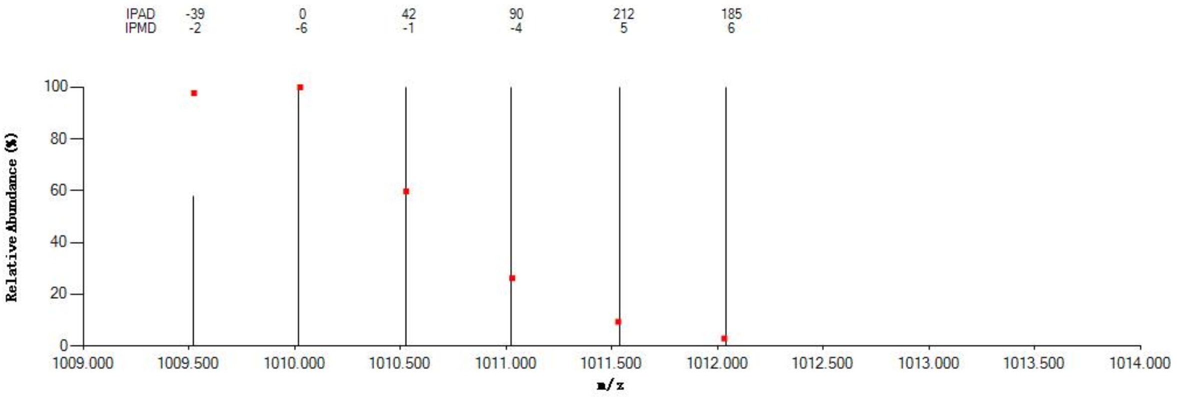


#### Annotated MS/MS Spectrum

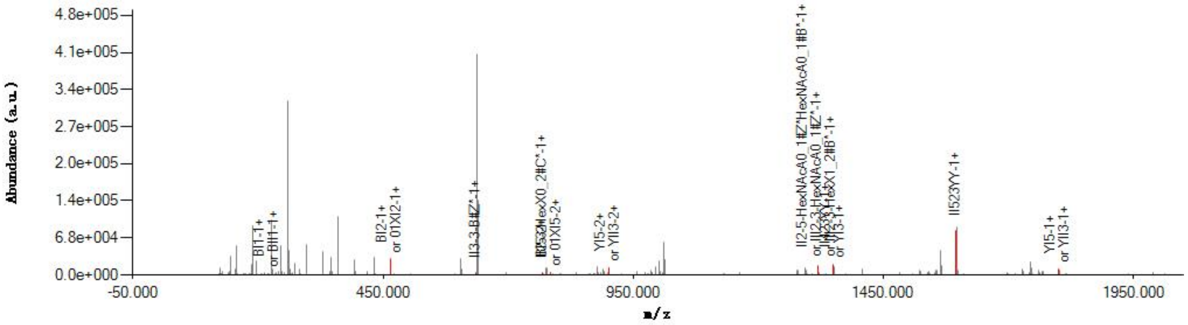


#### Graphical Fragmentation Map

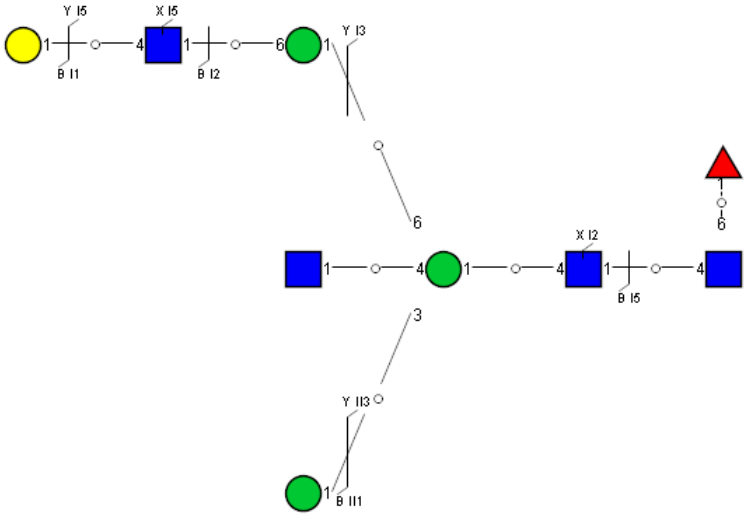

Spectrum Index: 30464(NoBHs:1)

#### Precursor iEF Map

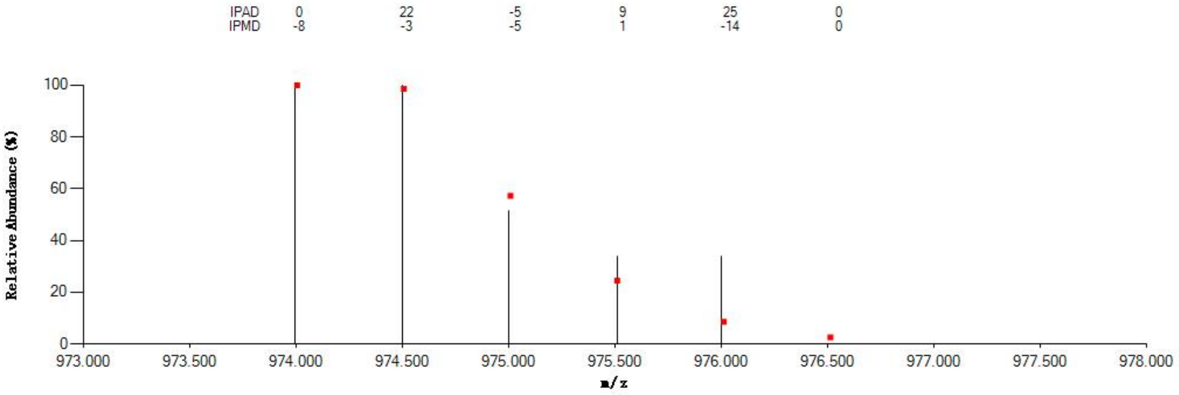


#### Annotated MS/MS Spectrum

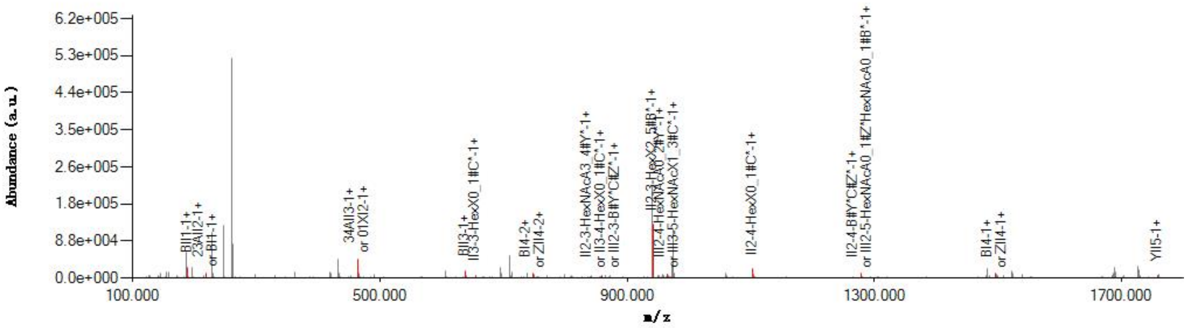


#### Graphical Fragmentation Map

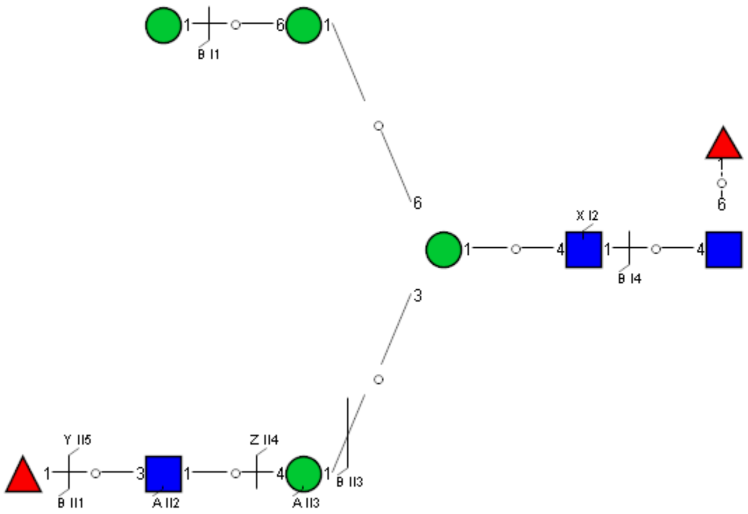

Spectrum Index: 31629(NoBHs:1)

#### Precursor iEF Map

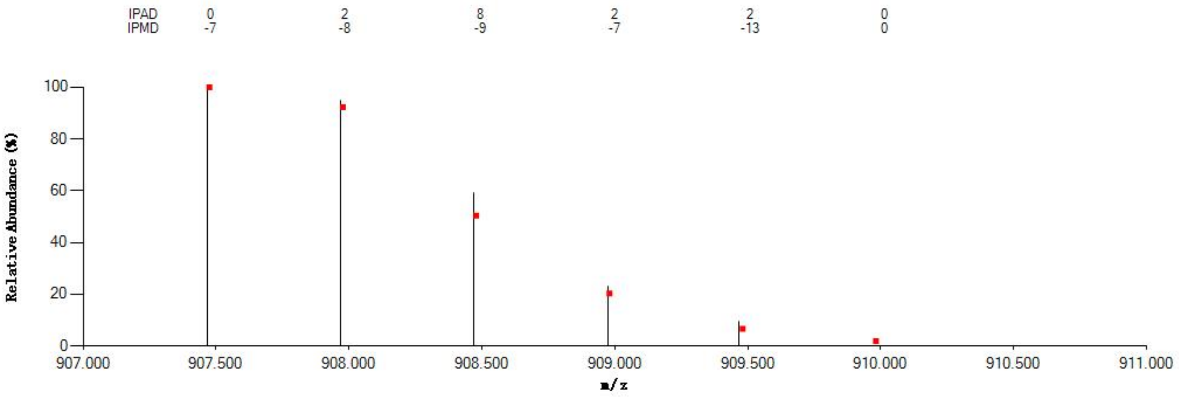


#### Annotated MS/MS Spectrum

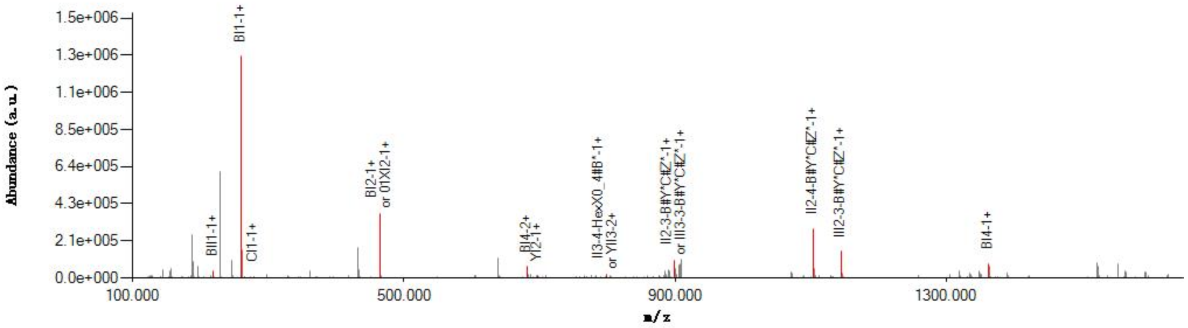


#### Graphical Fragmentation Map

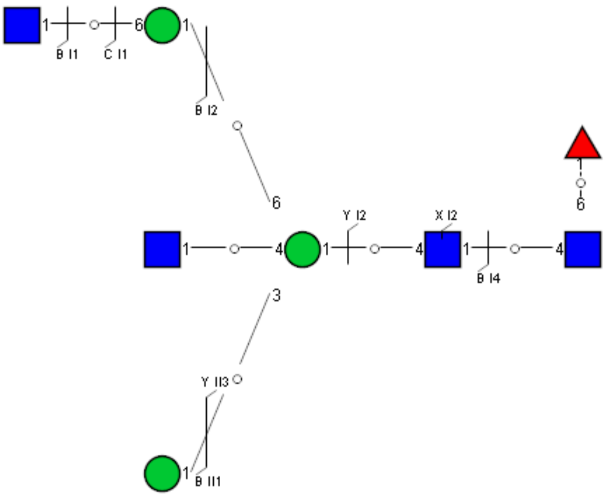

Spectrum Index: 31866(NoBHs:1)

#### Precursor iEF Map

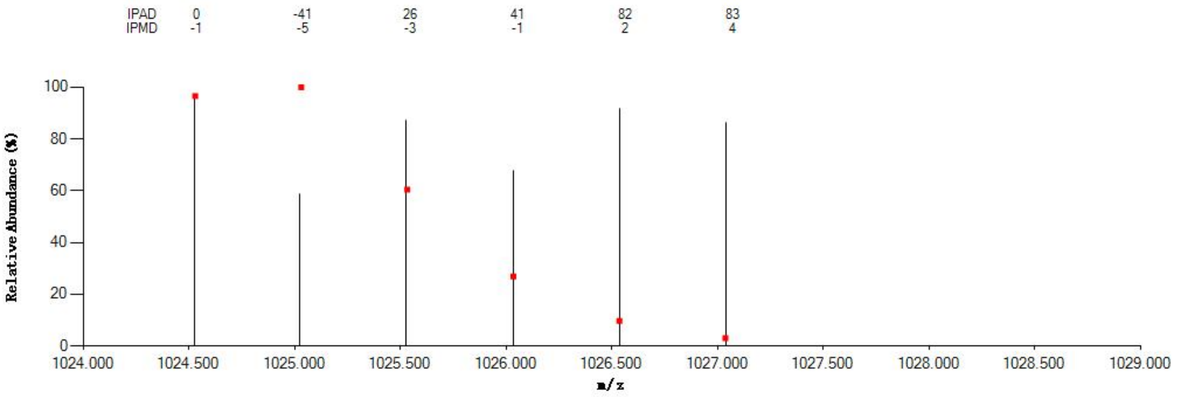


#### Annotated MS/MS Spectrum

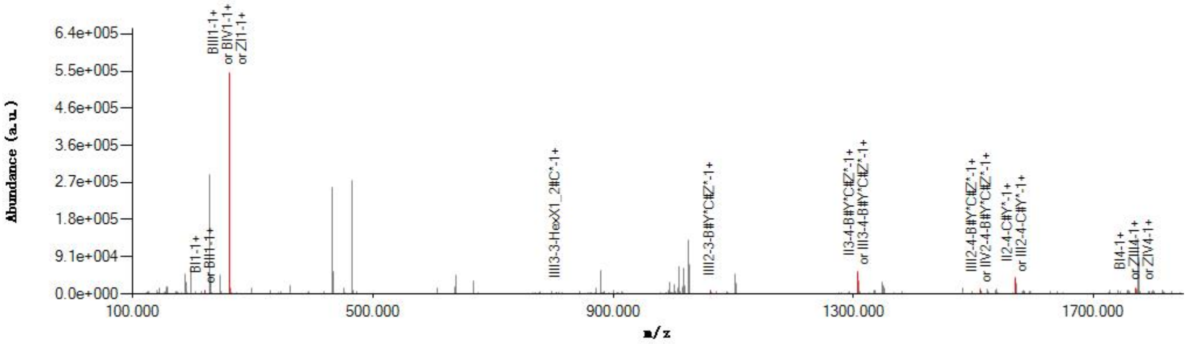


#### Graphical Fragmentation Map

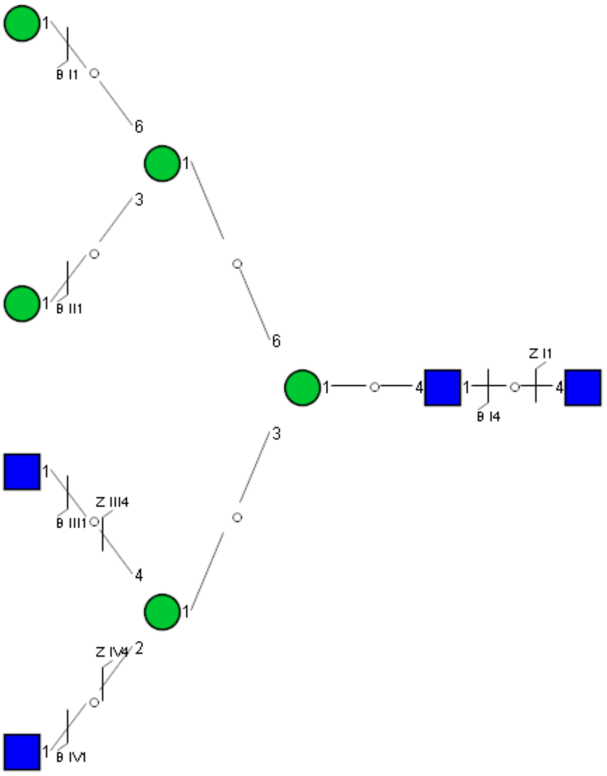

Spectrum Index: 33417(NoBHs:1)

#### Precursor iEF Map

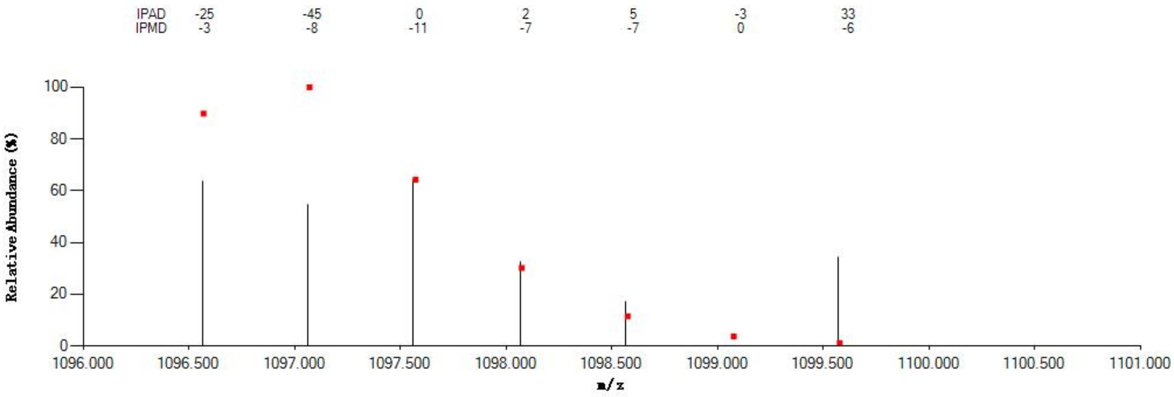


#### Annotated MS/MS Spectrum

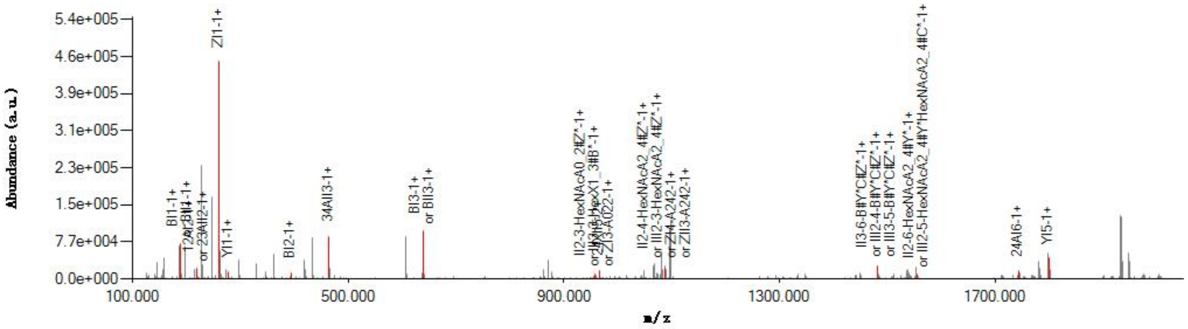


#### Graphical Fragmentation Map

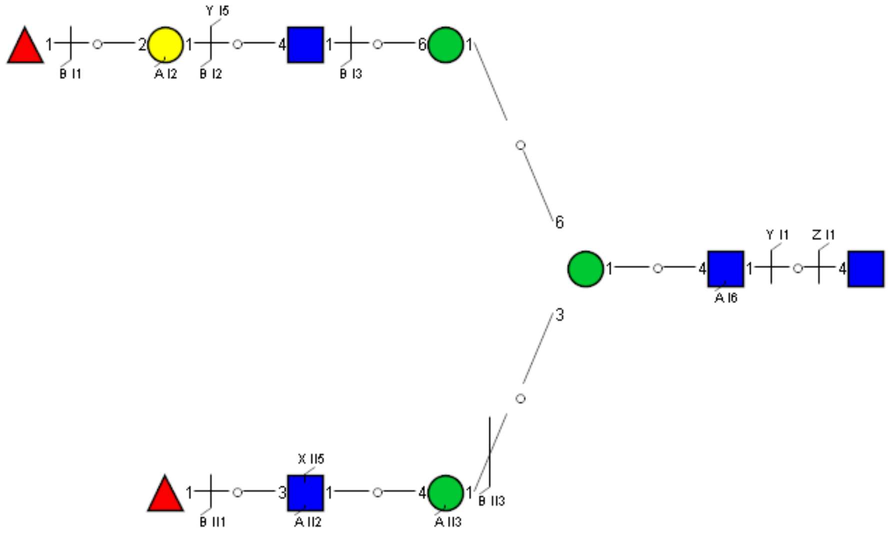

Spectrum Index: 34070(NoBHs:1)

#### Precursor iEF Map

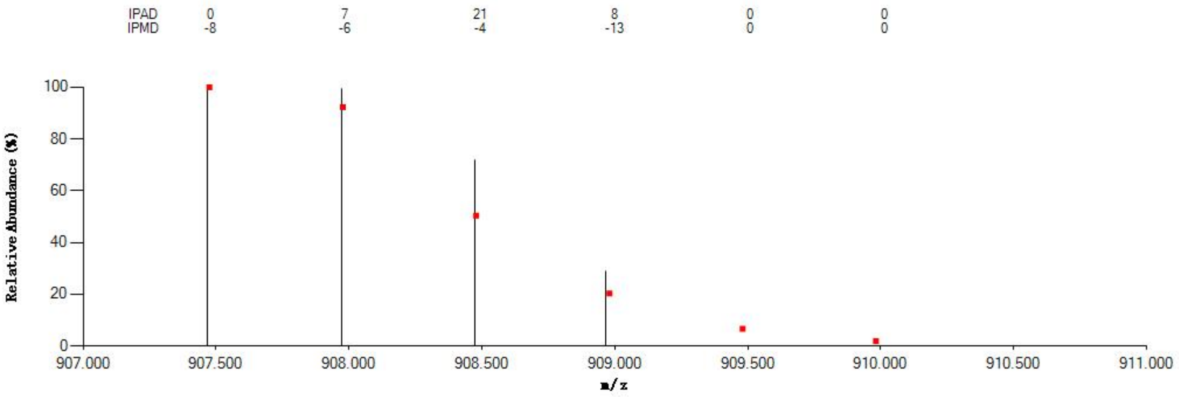


#### Annotated MS/MS Spectrum

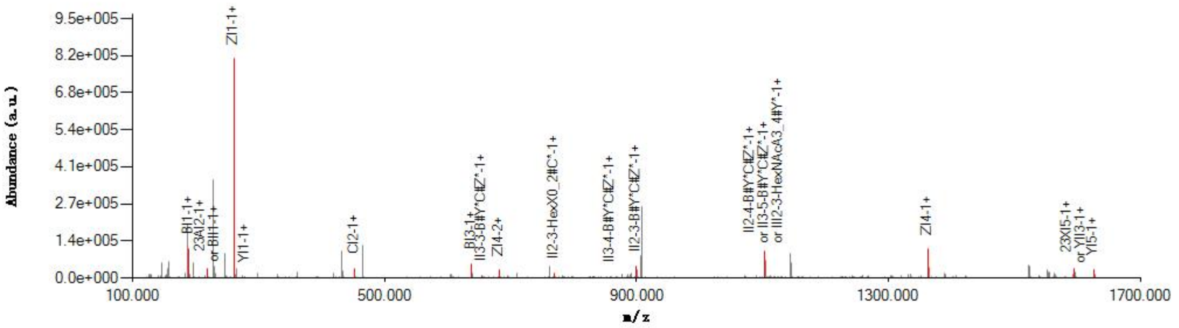


#### Graphical Fragmentation Map

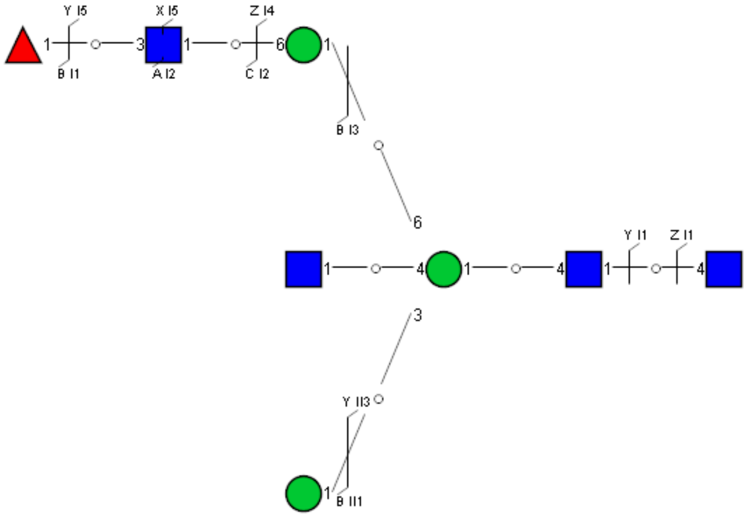

Spectrum Index: 34300(NoBHs:1)

#### Precursor iEF Map

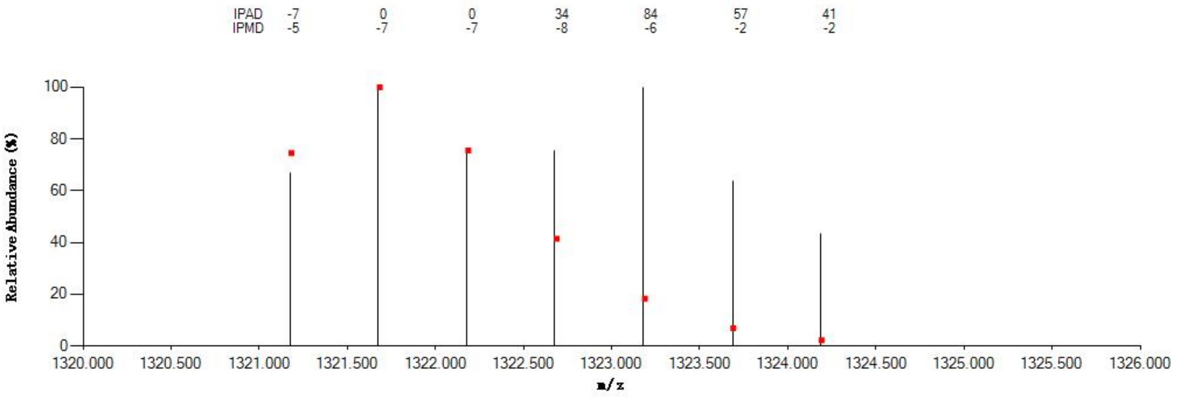


#### Annotated MS/MS Spectrum

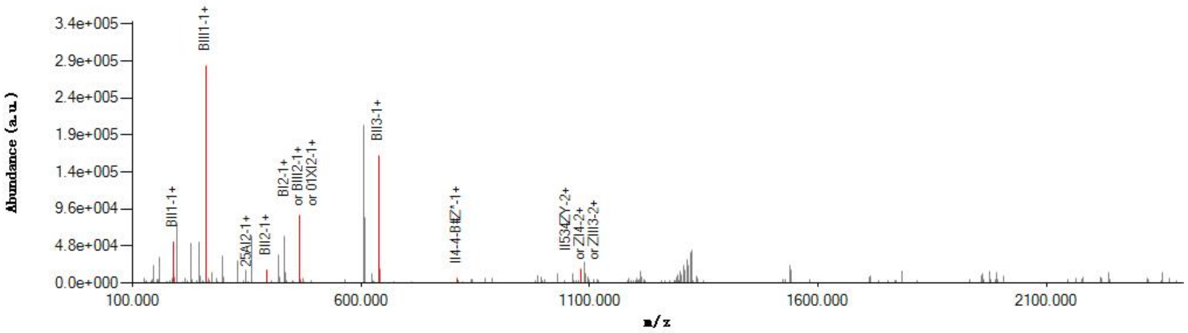


#### Graphical Fragmentation Map

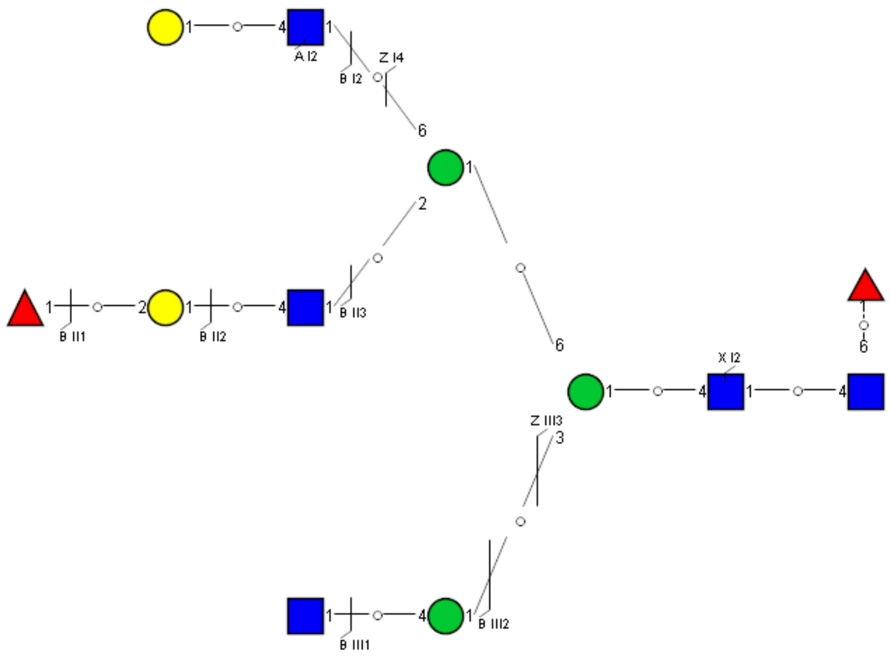

Spectrum Index: 35158(NoBHs:1)

#### Precursor iEF Map

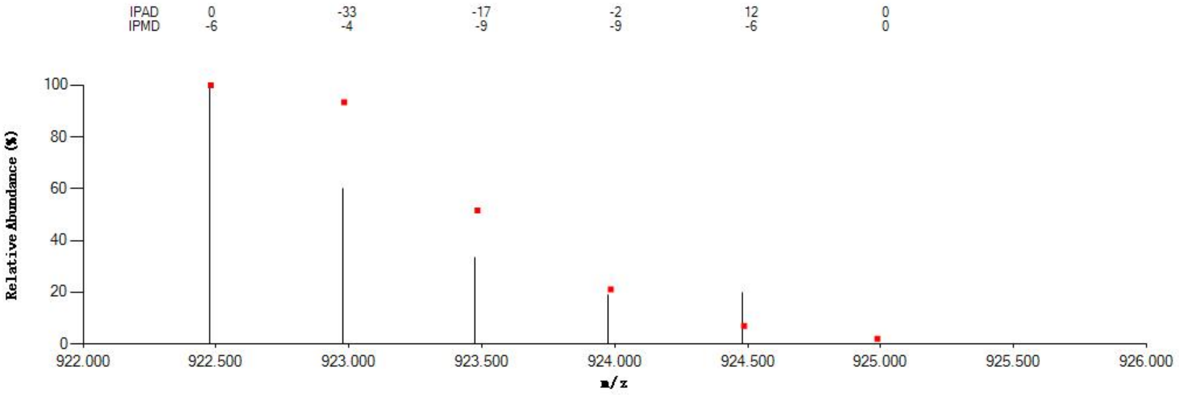


#### Annotated MS/MS Spectrum

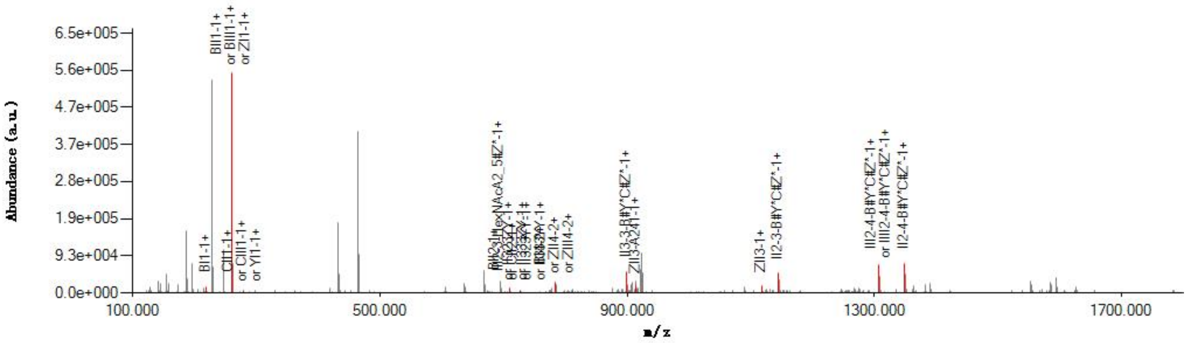


#### Graphical Fragmentation Map

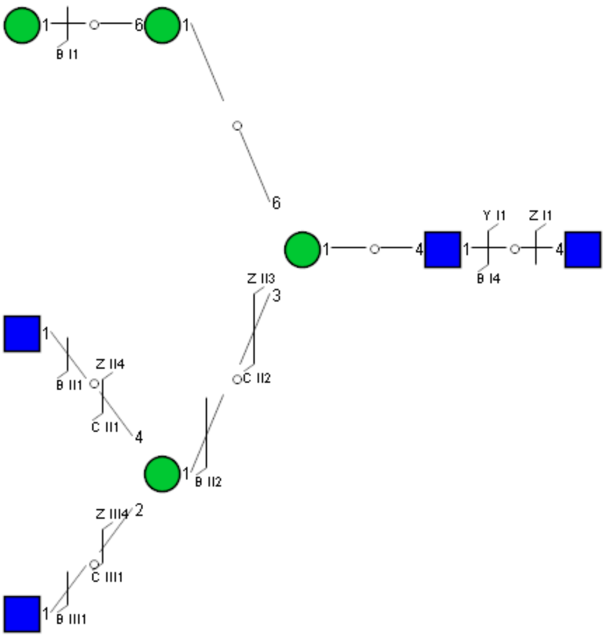

Spectrum Index: 35599(NoBHs:1)

#### Precursor iEF Map

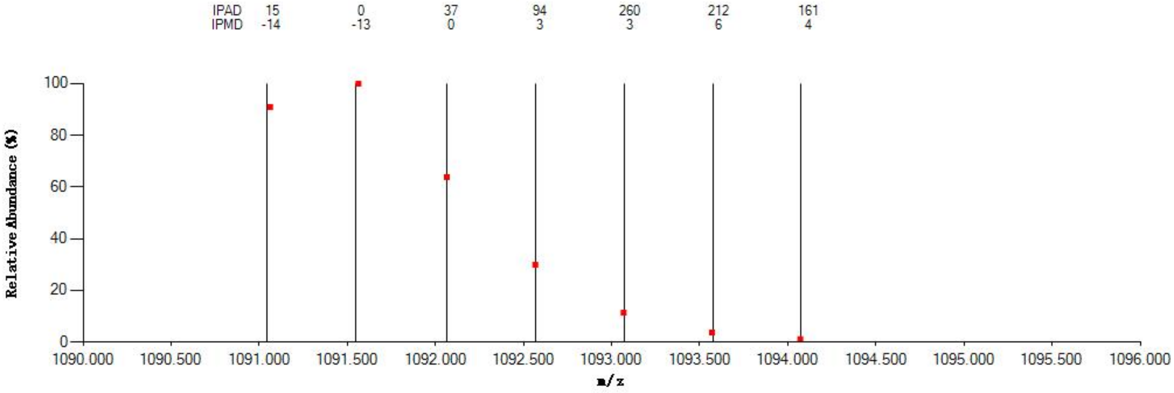


#### Annotated MS/MS Spectrum

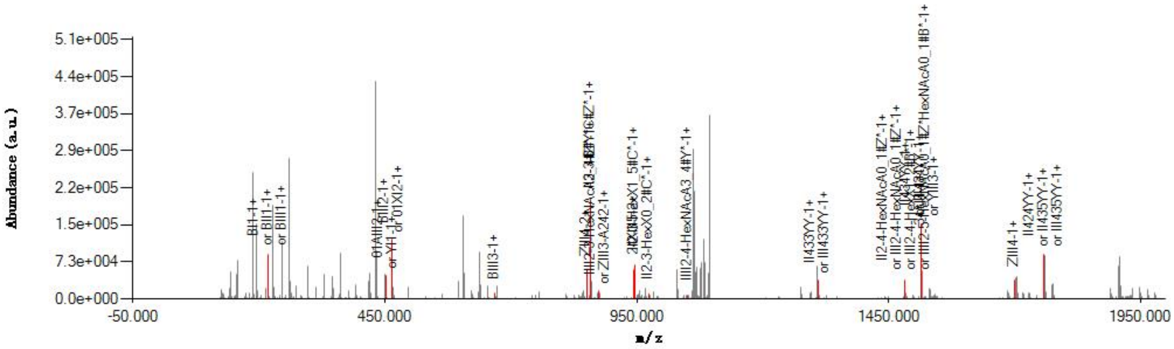


#### Graphical Fragmentation Map

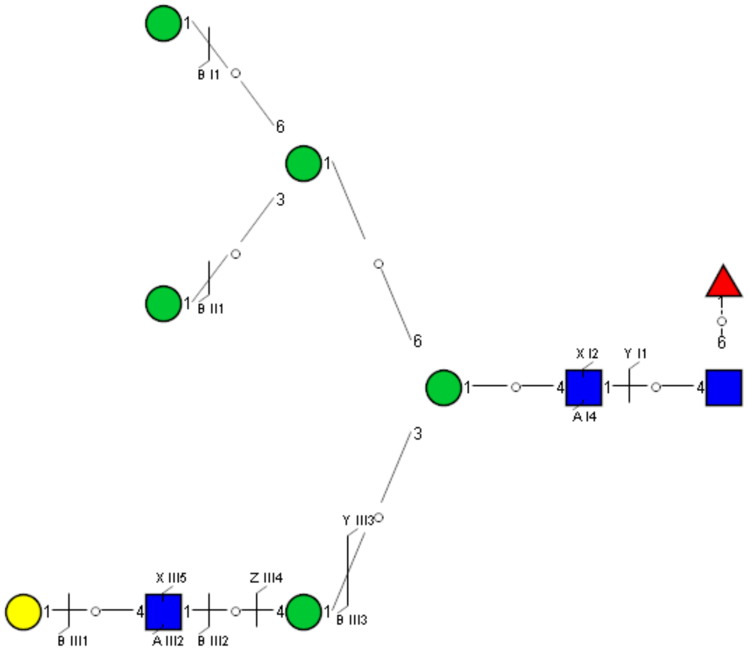

Spectrum Index: 36212(NoBHs:2)

#### Precursor iEF Map

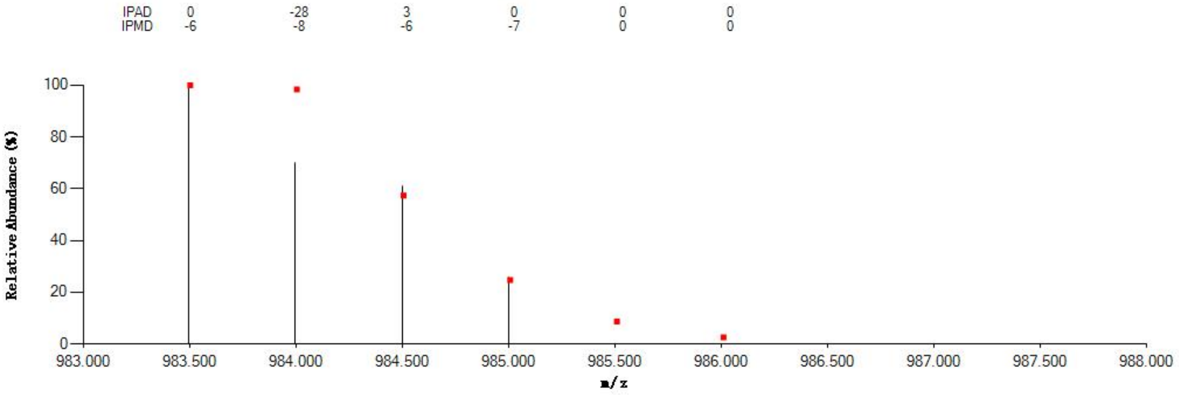


#### Annotated MS/MS Spectrum

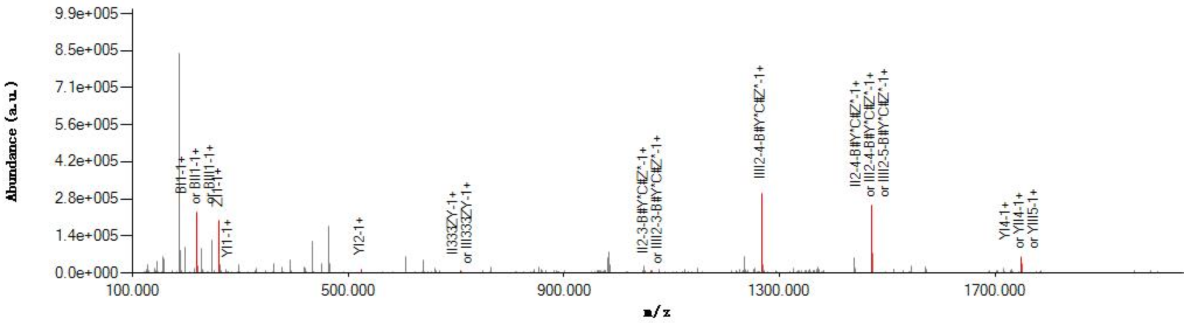


#### Graphical Fragmentation Map

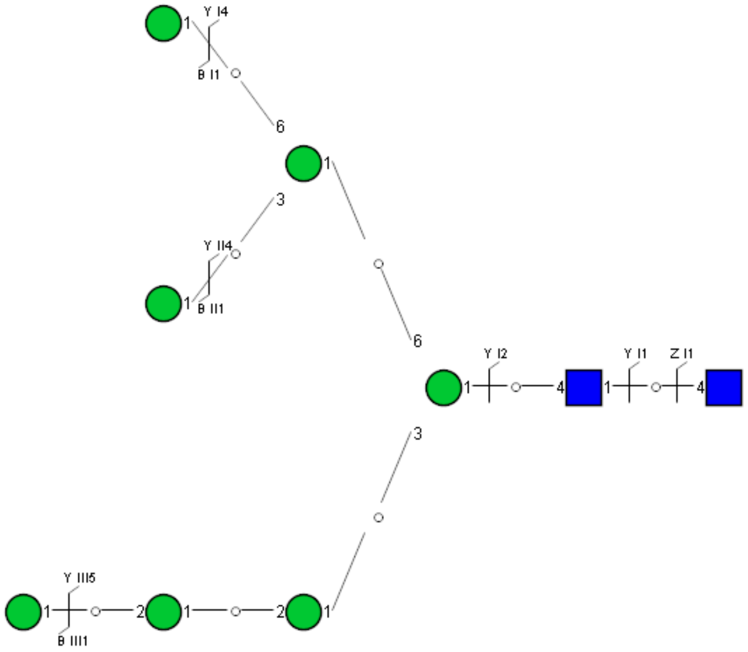

Spectrum Index: 36366(NoBHs:1)

#### Precursor iEF Map

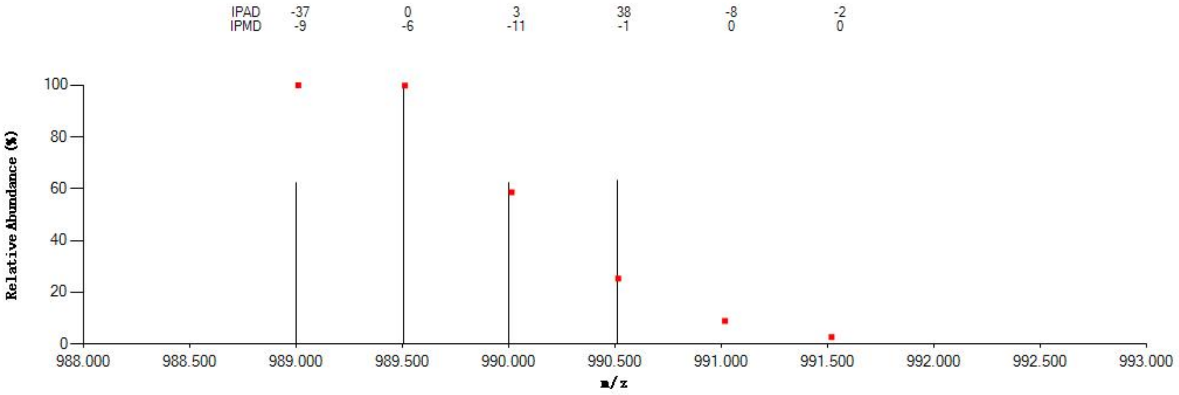


#### Annotated MS/MS Spectrum

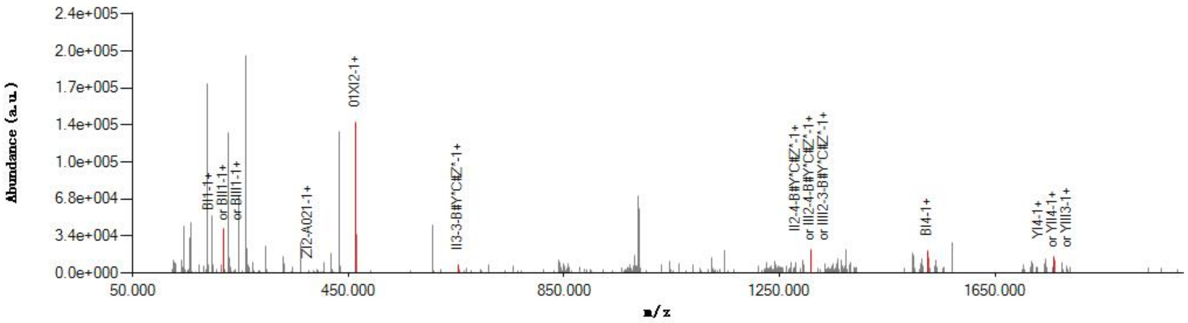


#### Graphical Fragmentation Map

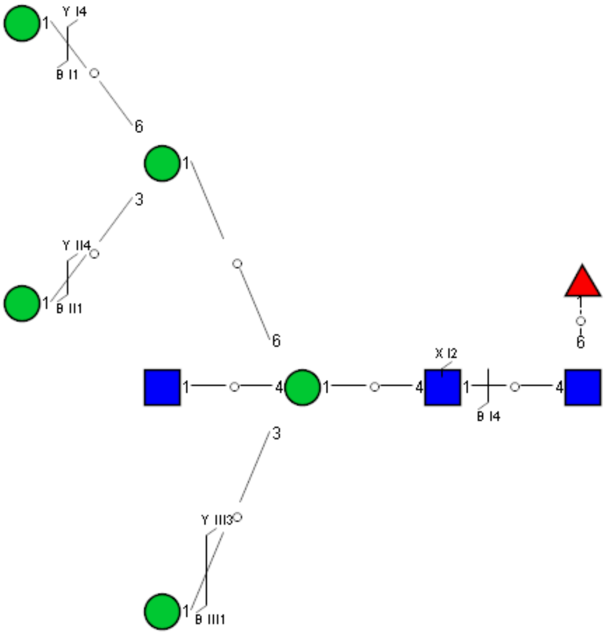

Spectrum Index: 36378(NoBHs:1)

#### Precursor iEF Map

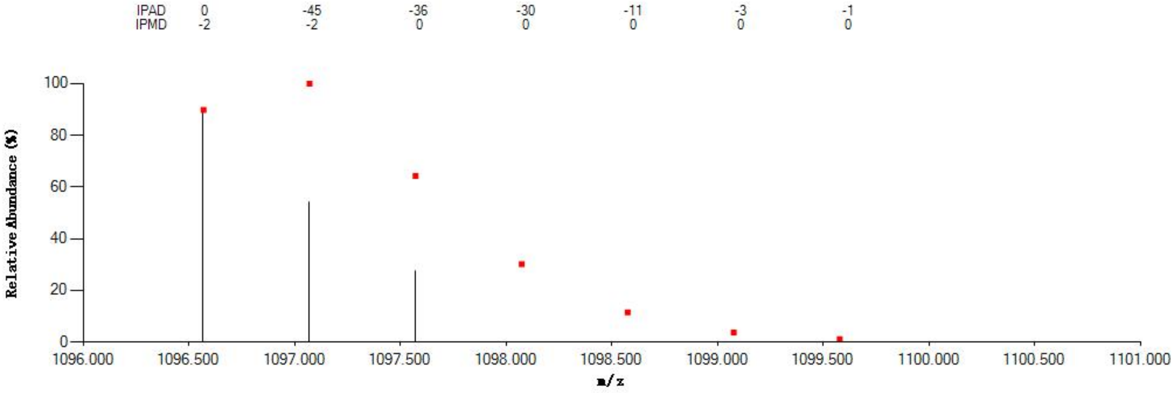


#### Annotated MS/MS Spectrum

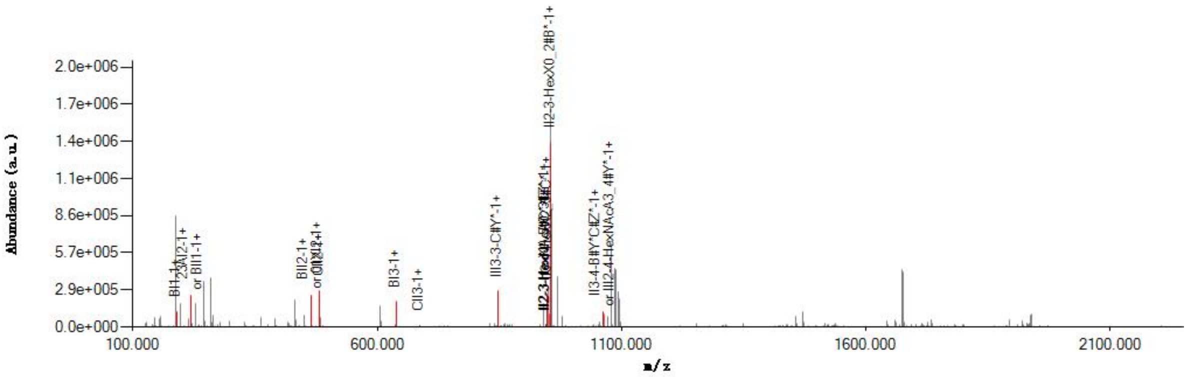


#### Graphical Fragmentation Map

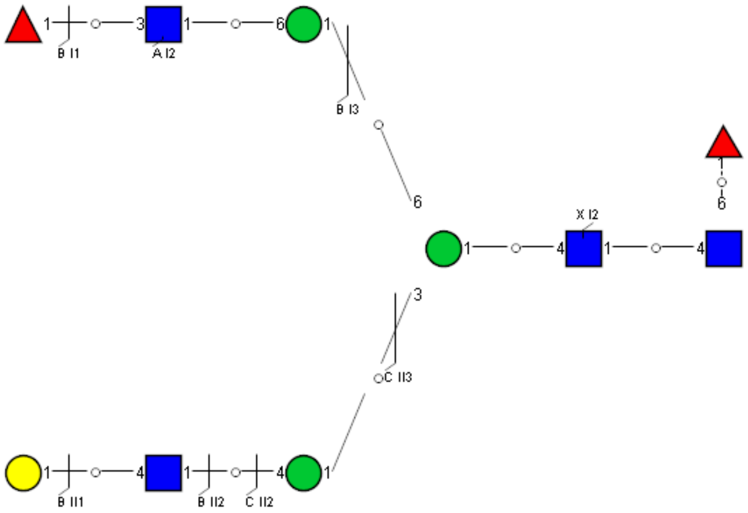

### Spectrum Index: 37406(NoBHs:1)

#### Precursor iEF Map

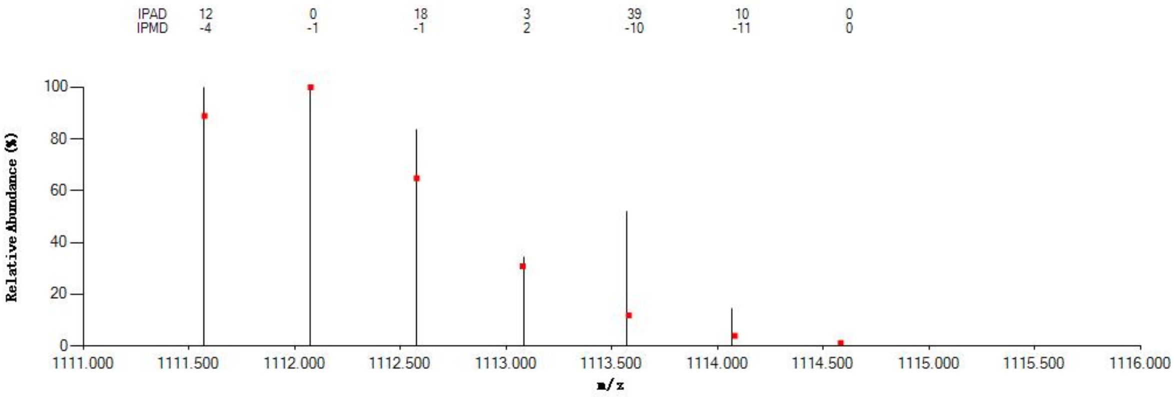


#### Annotated MS/MS Spectrum

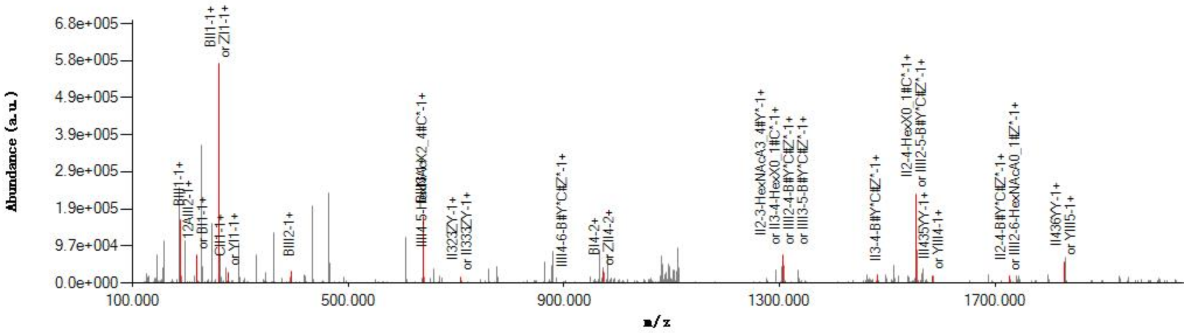


#### Graphical Fragmentation Map

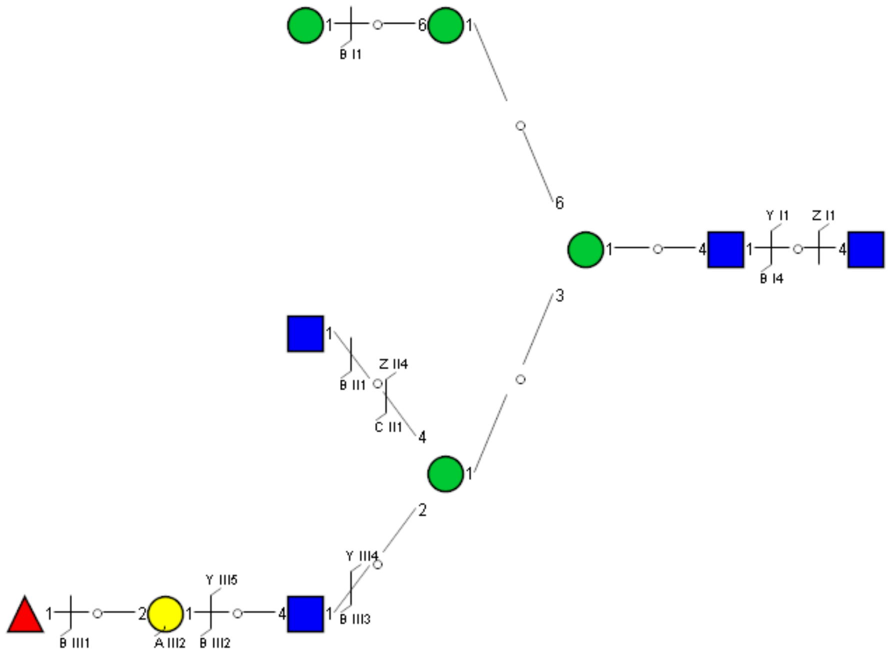

Spectrum Index: 37584(NoBHs:1)

#### Precursor iEF Map

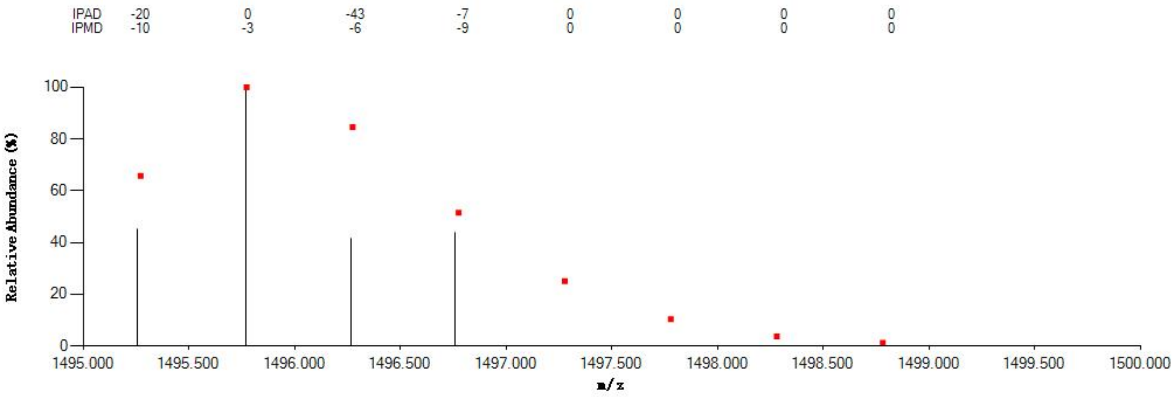


#### Annotated MS/MS Spectrum

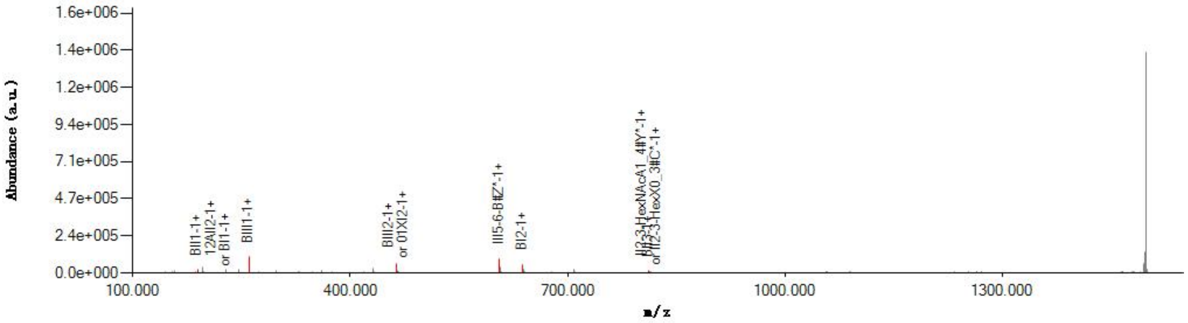


#### Graphical Fragmentation Map

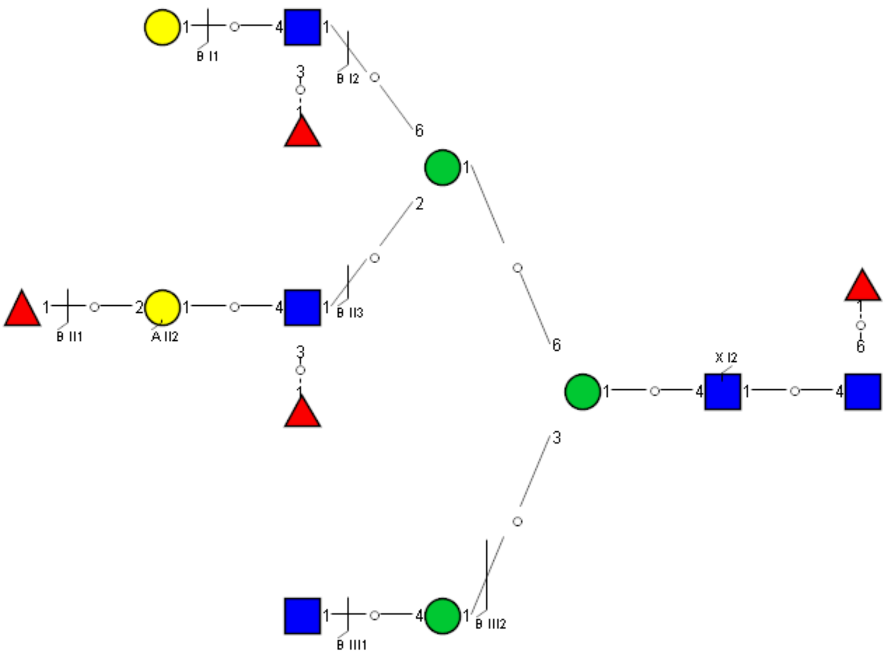

Spectrum Index: 37688(NoBHs:1)

#### Precursor iEF Map

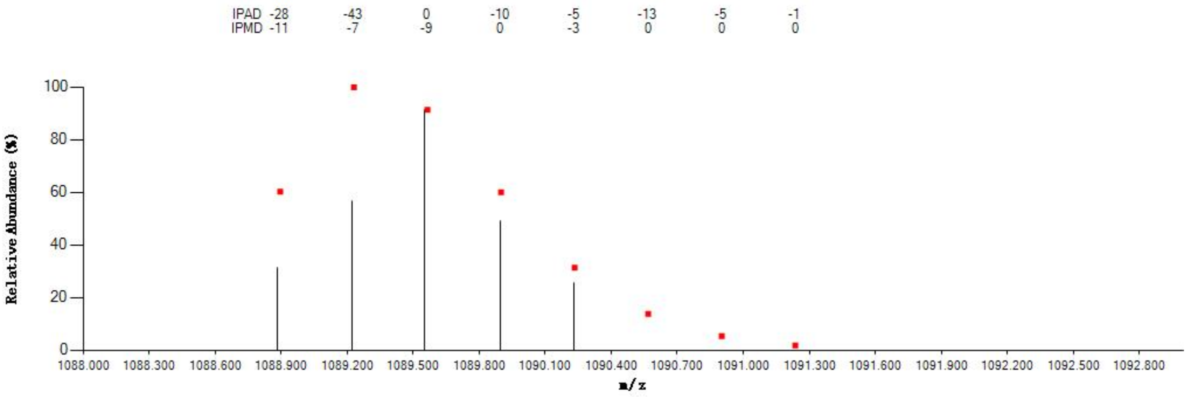


#### Annotated MS/MS Spectrum

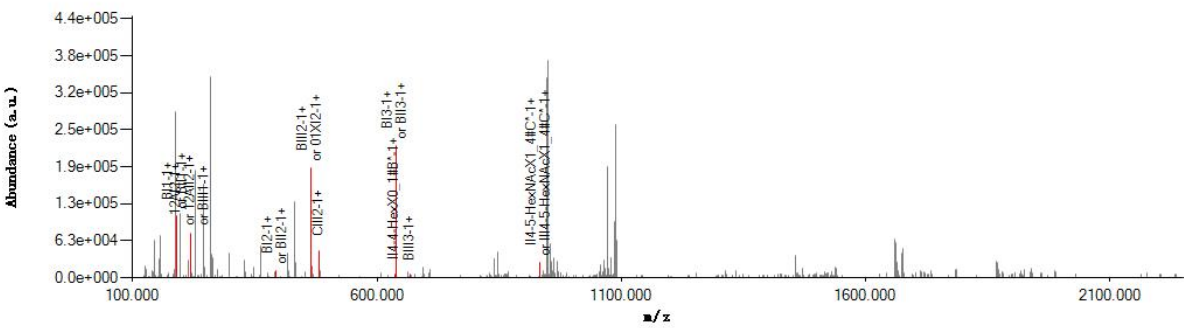


#### Graphical Fragmentation Map

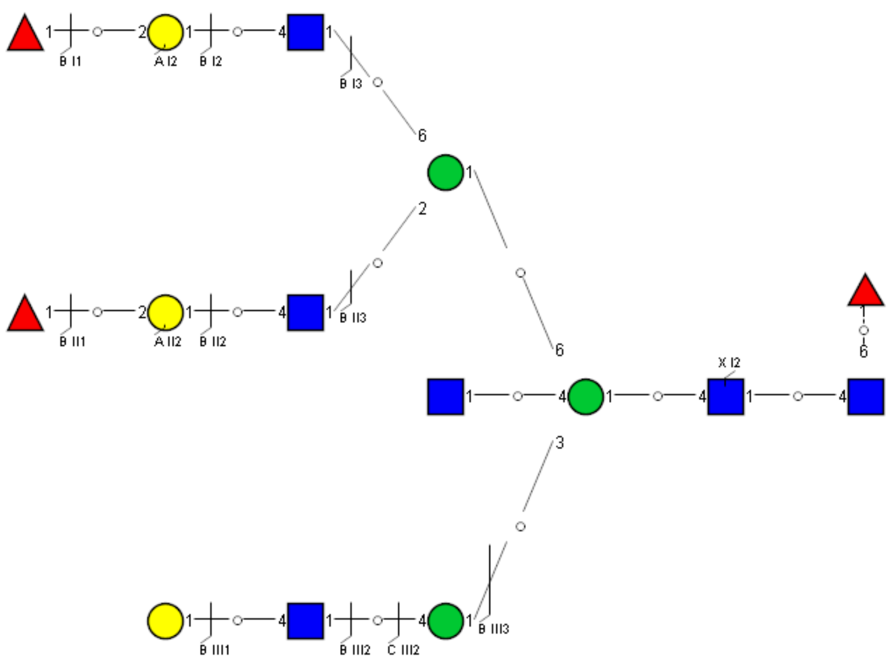

Spectrum Index: 37748(NoBHs:1)

#### Precursor iEF Map

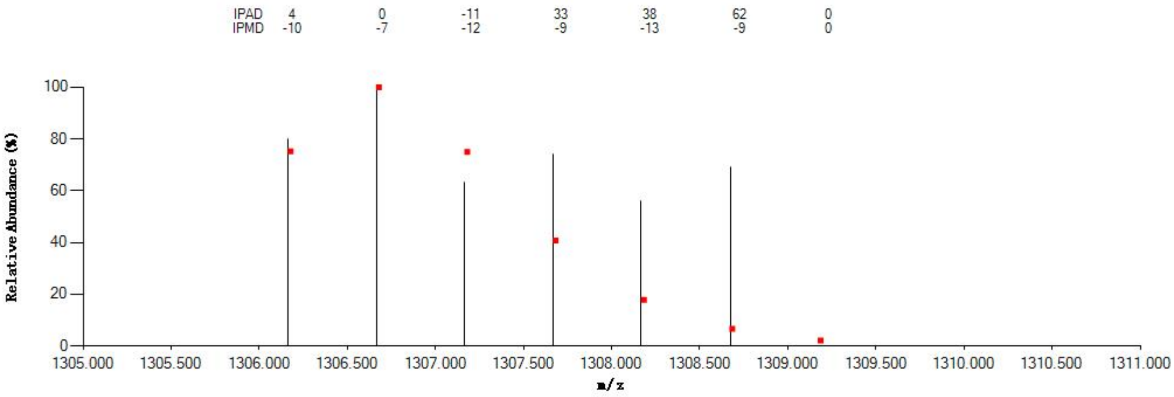


#### Annotated MS/MS Spectrum

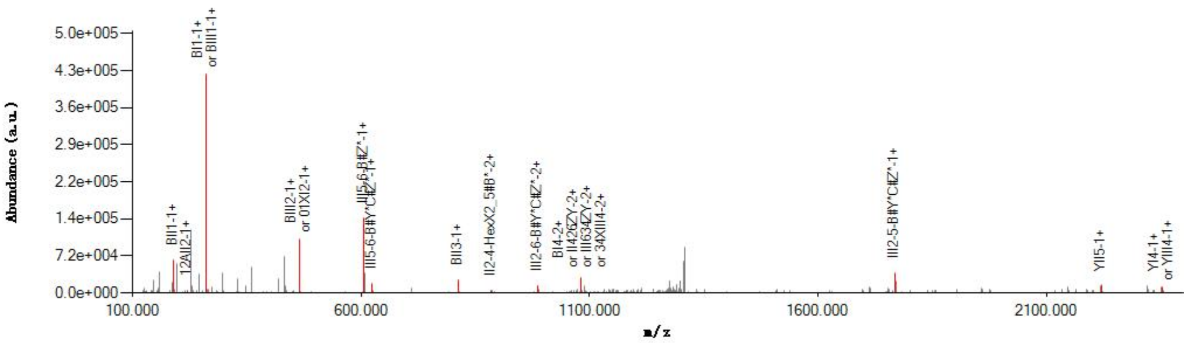


#### Graphical Fragmentation Map

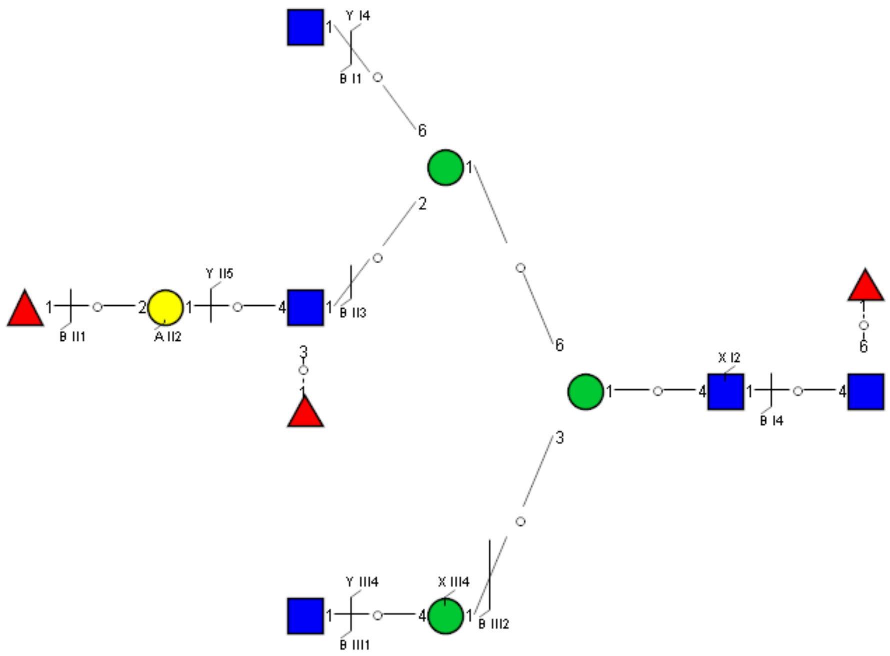

Spectrum Index: 37984(NoBHs:1)

#### Precursor iEF Map

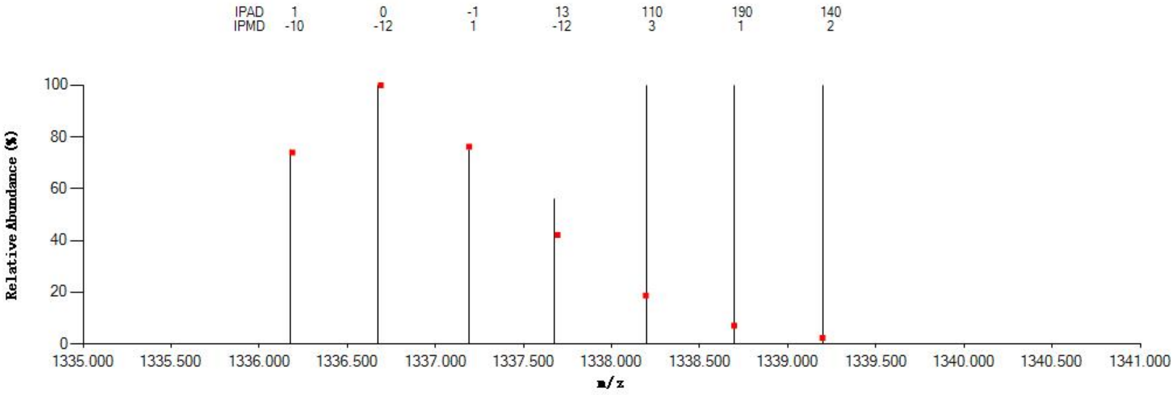


#### Annotated MS/MS Spectrum

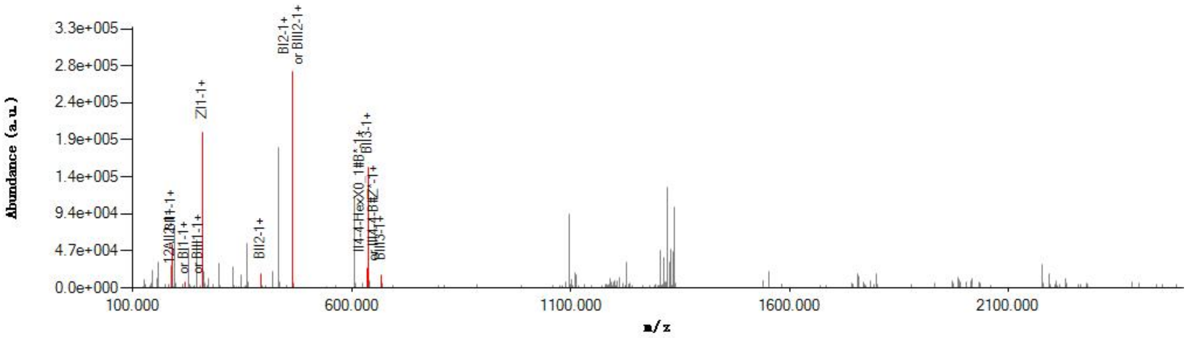


#### Graphical Fragmentation Map

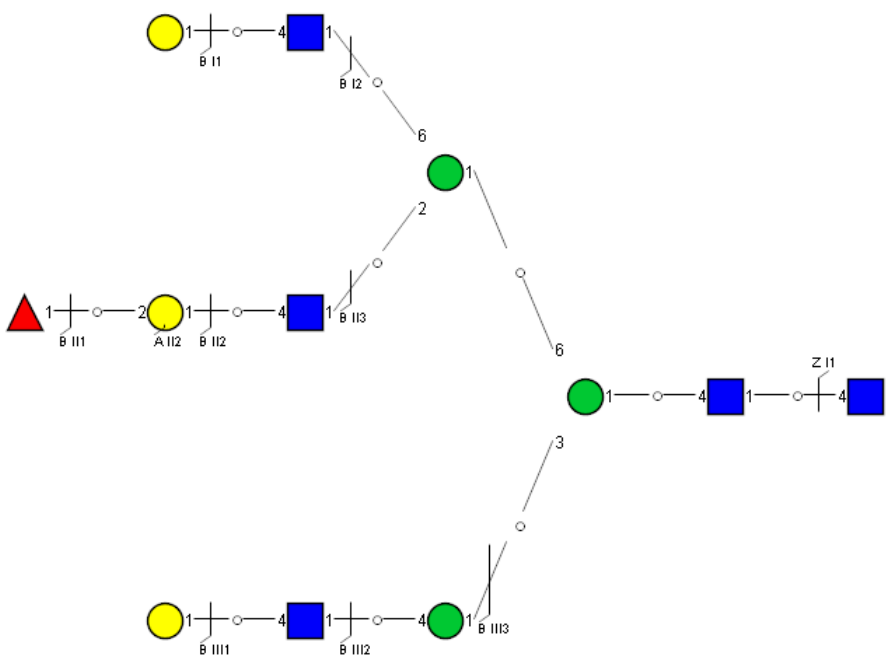

Spectrum Index: 37999(NoBHs:1)

#### Precursor iEF Map

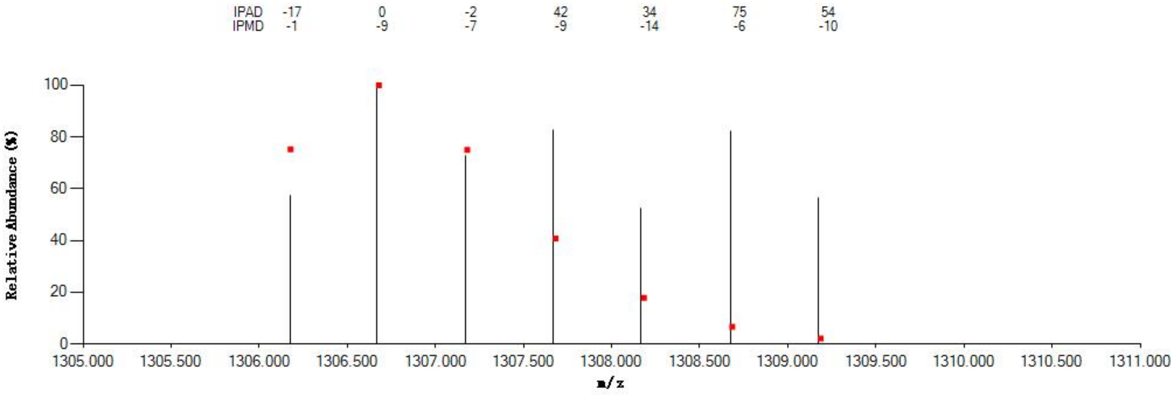


#### Annotated MS/MS Spectrum

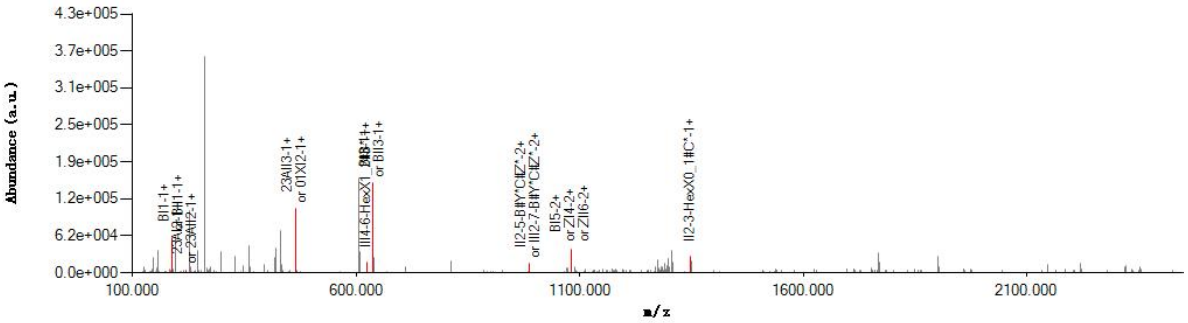


#### Graphical Fragmentation Map

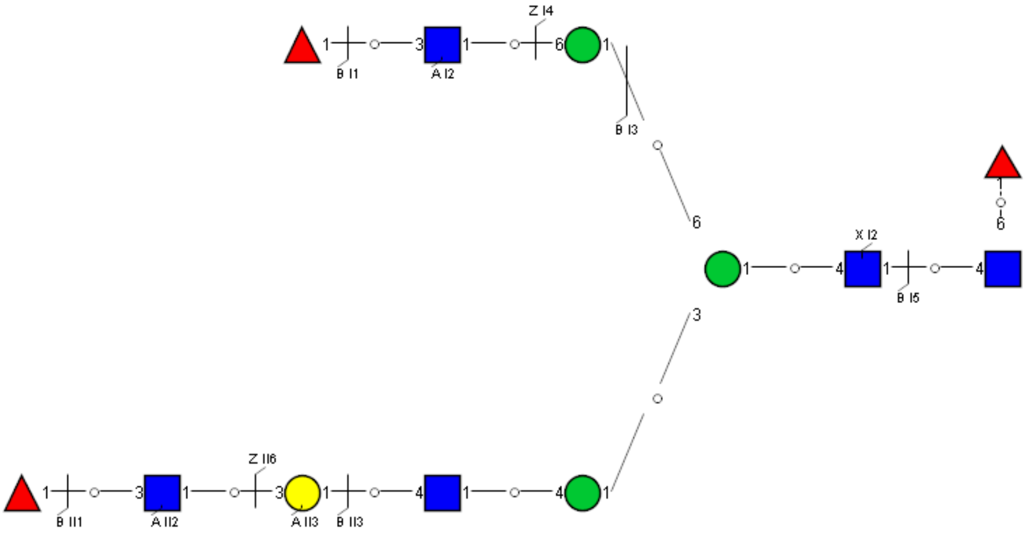

Spectrum Index: 38106(NoBHs:1)

#### Precursor iEF Map

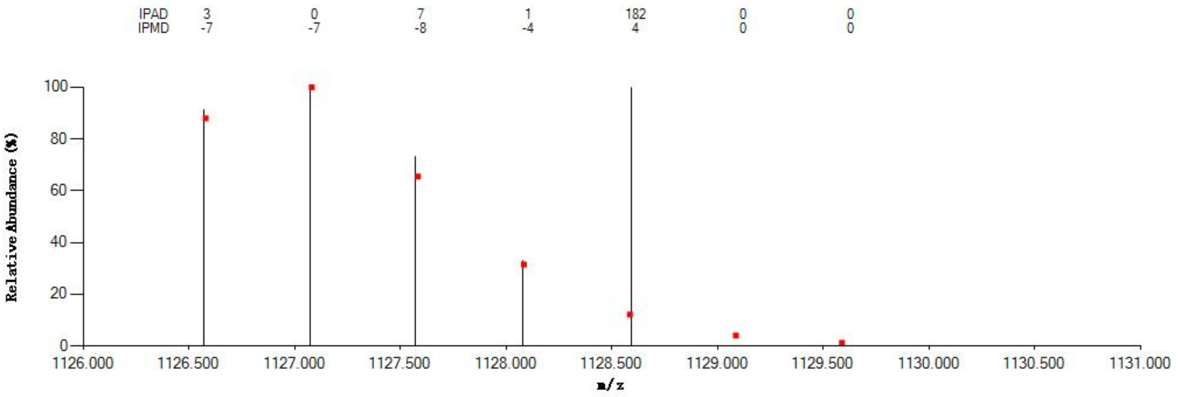

#### Annotated MS/MS Spectrum

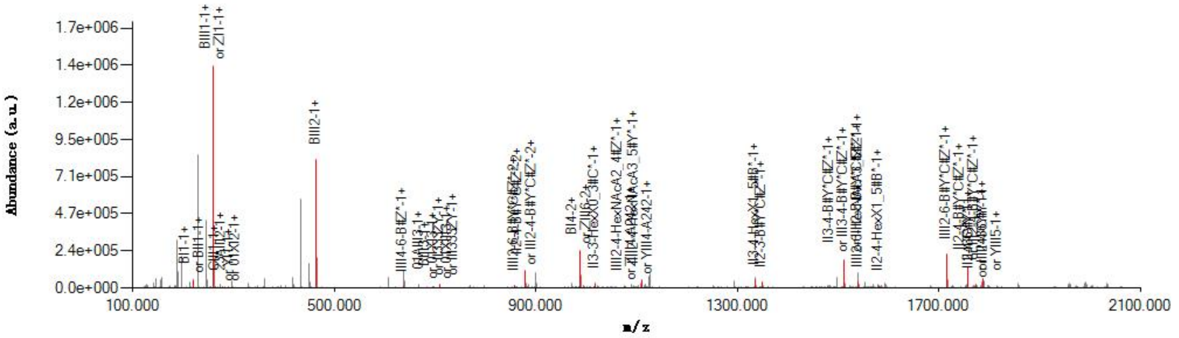

#### Graphical Fragmentation Map

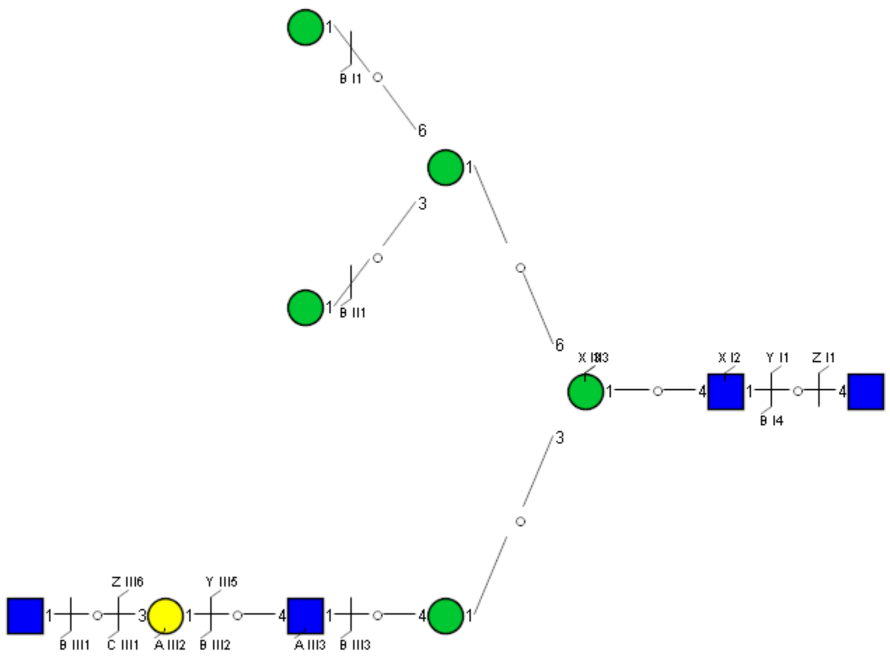

Spectrum Index: 38484(NoBHs:1)

#### Precursor iEF Map

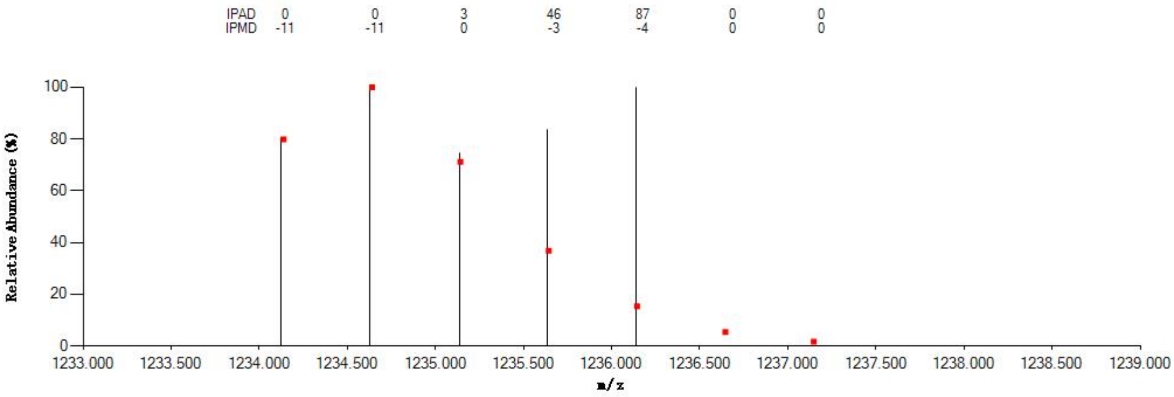


#### Annotated MS/MS Spectrum

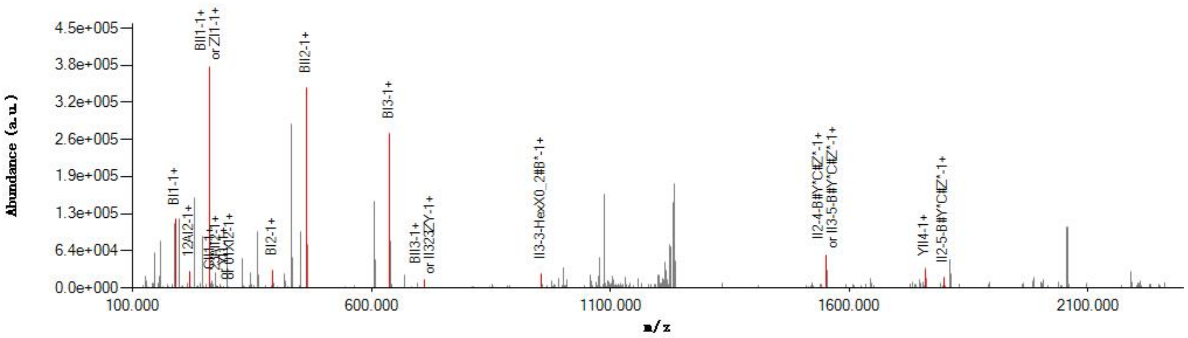


#### Graphical Fragmentation Map

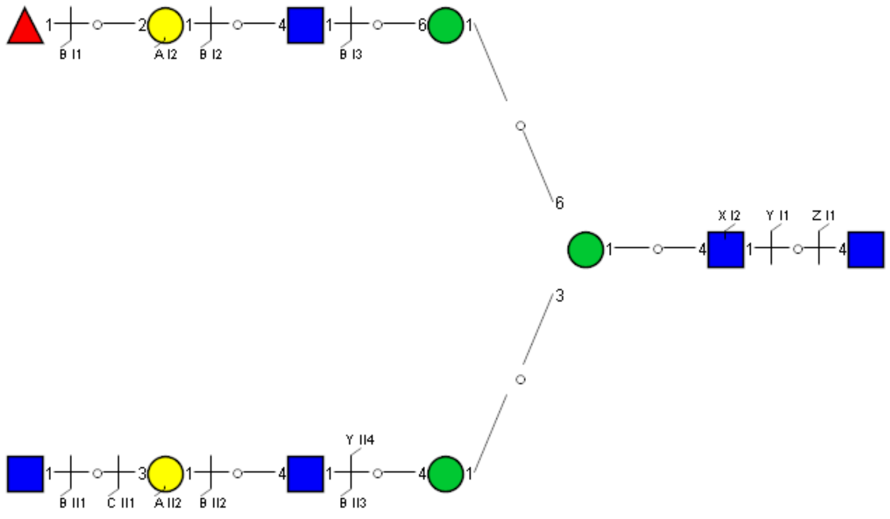

### Spectrum Index: 38498(NoBHs:1)

#### Precursor iEF Map

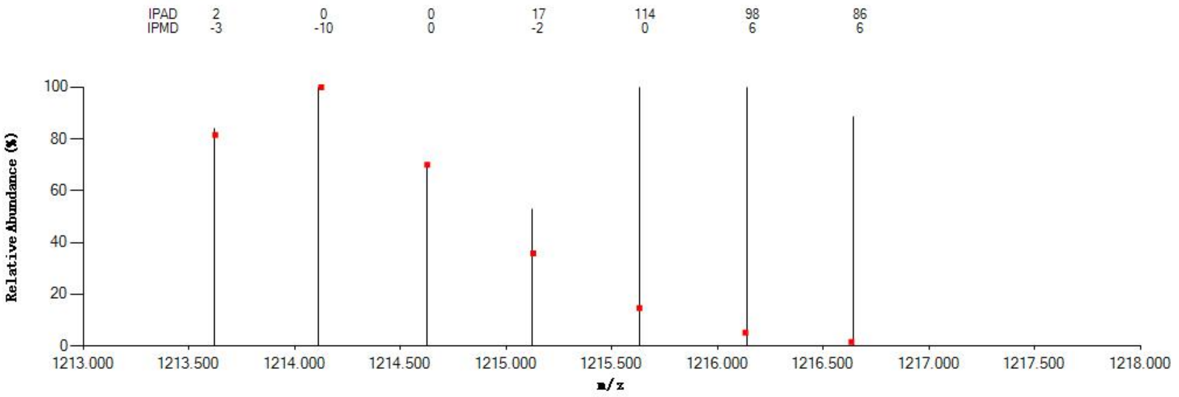


#### Annotated MS/MS Spectrum

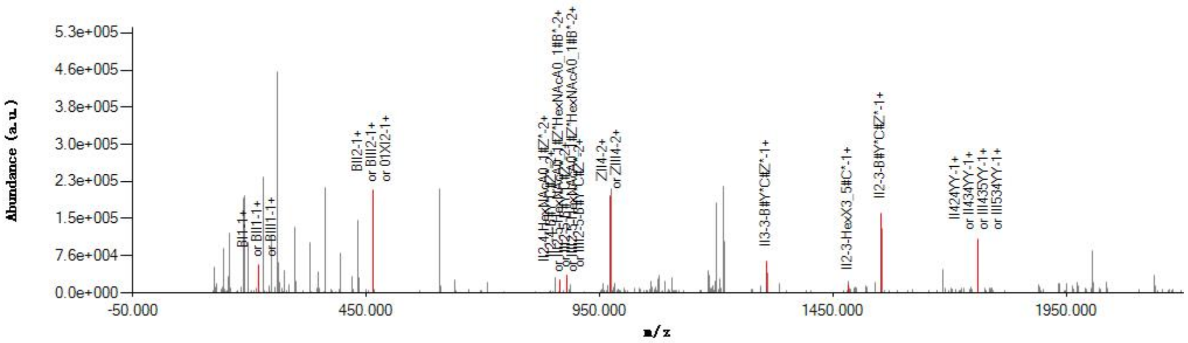


#### Graphical Fragmentation Map

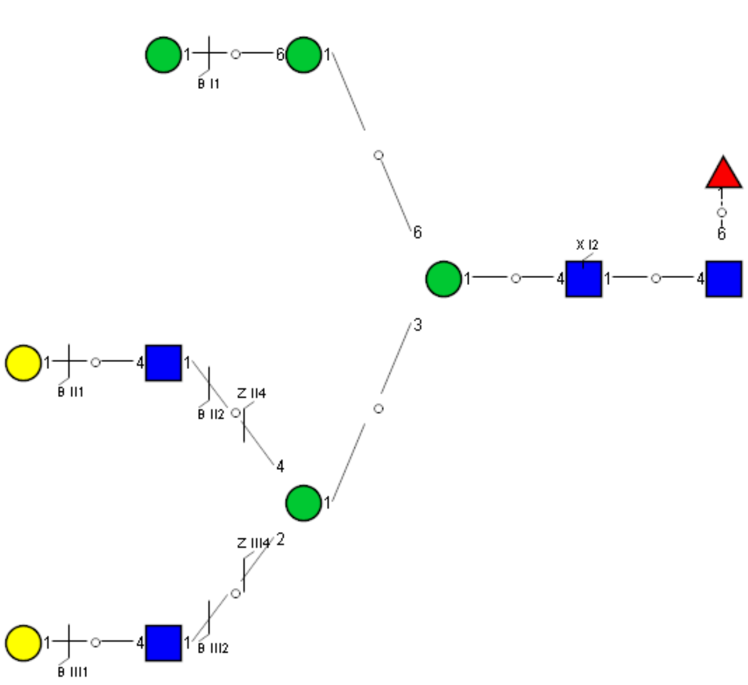

Spectrum Index: 38507(NoBHs:1)

#### Precursor iEF Map

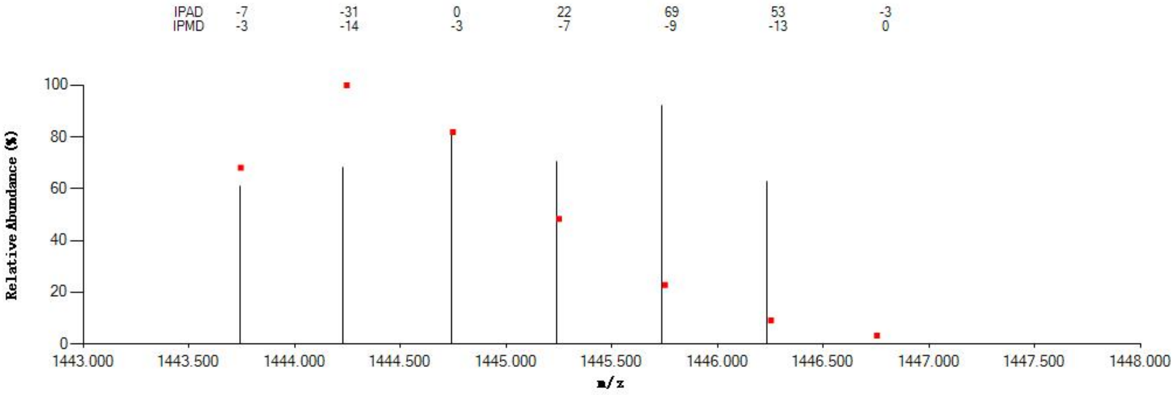


#### Annotated MS/MS Spectrum

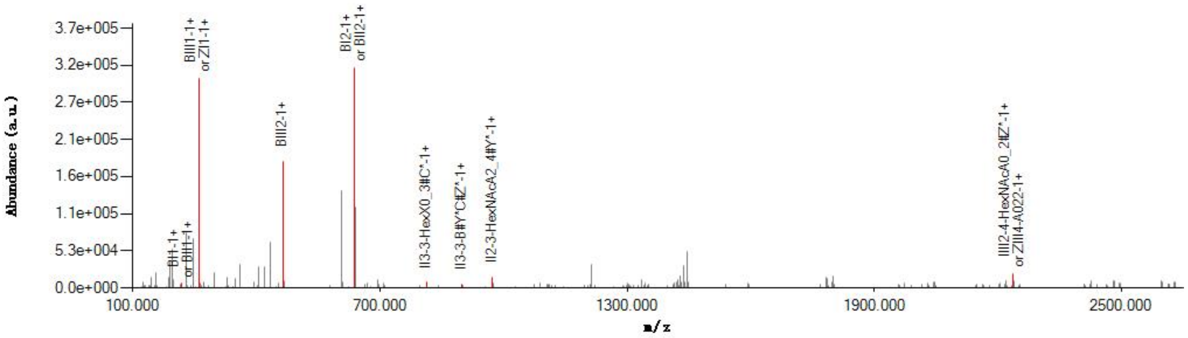


#### Graphical Fragmentation Map

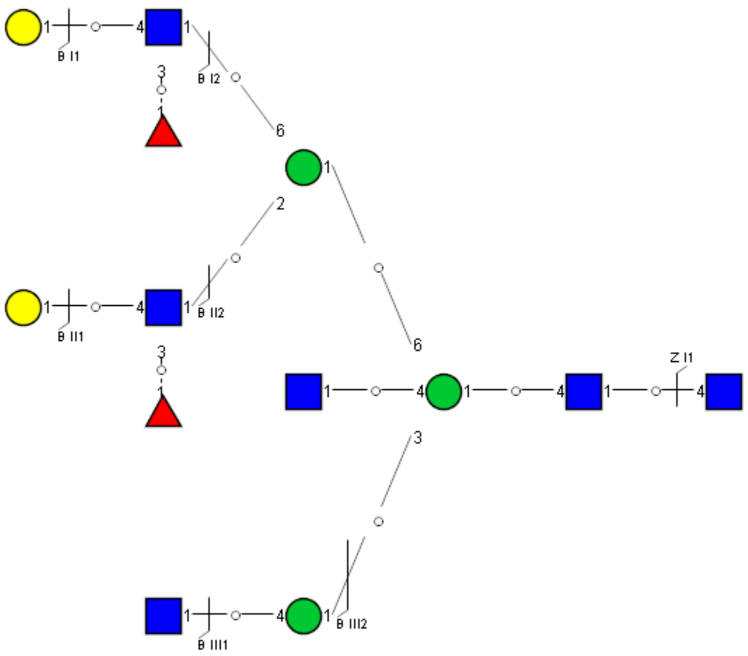

Spectrum Index: 38720(NoBHs:3)

#### Precursor iEF Map

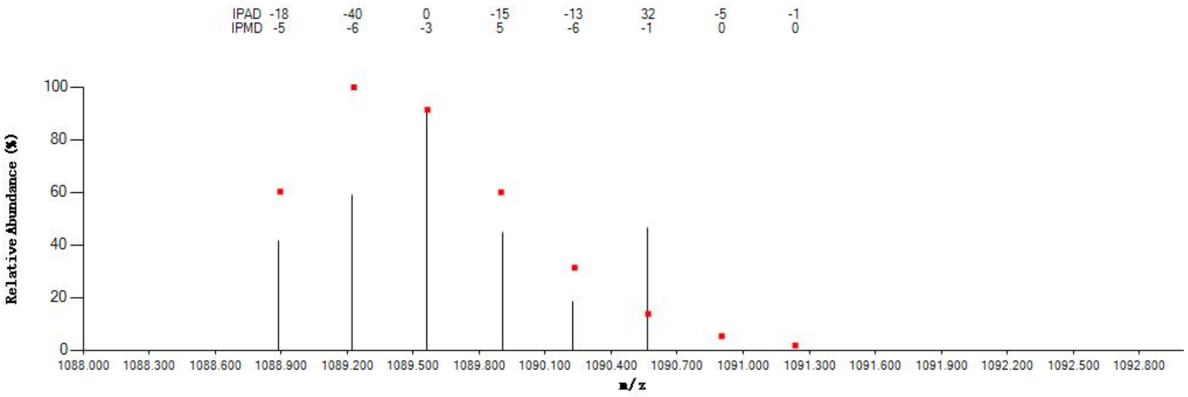


#### Annotated MS/MS Spectrum

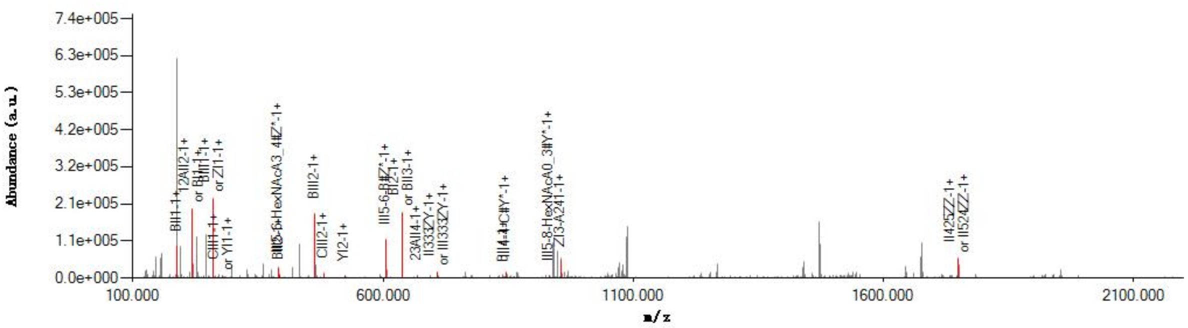


#### Graphical Fragmentation Map

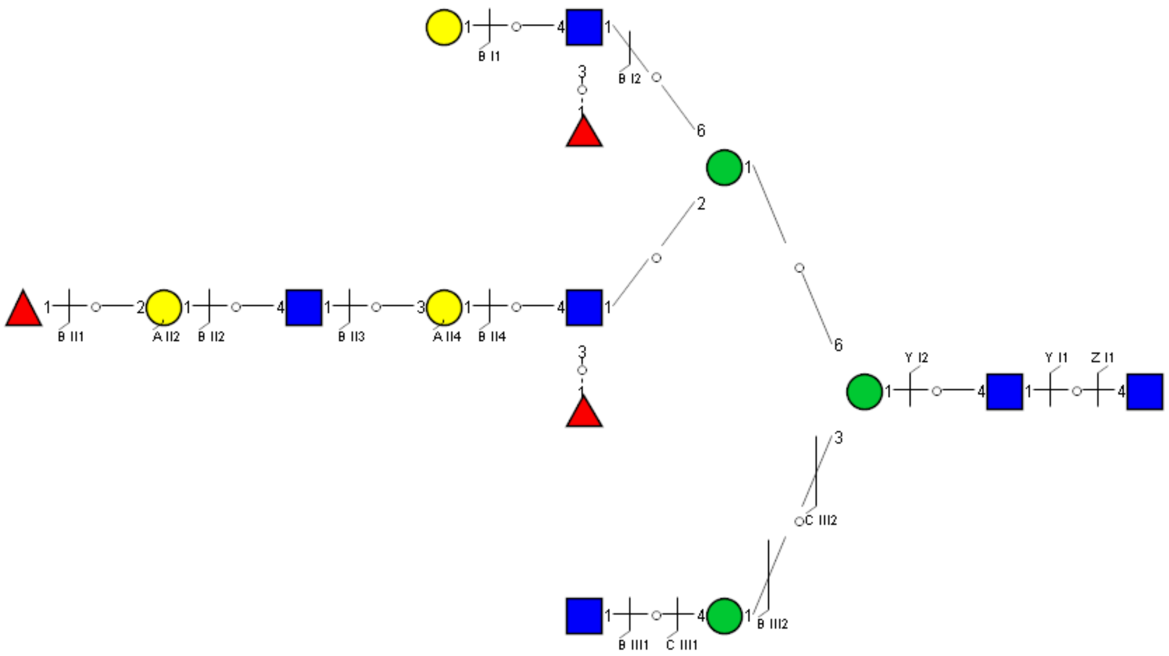

Spectrum Index: 38720(NoBHs:3)

#### Precursor iEF Map

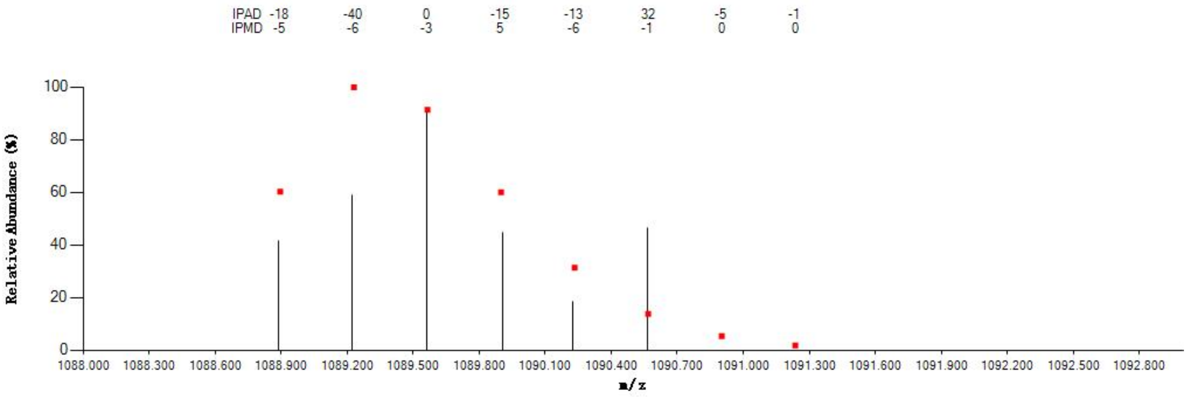


#### Annotated MS/MS Spectrum

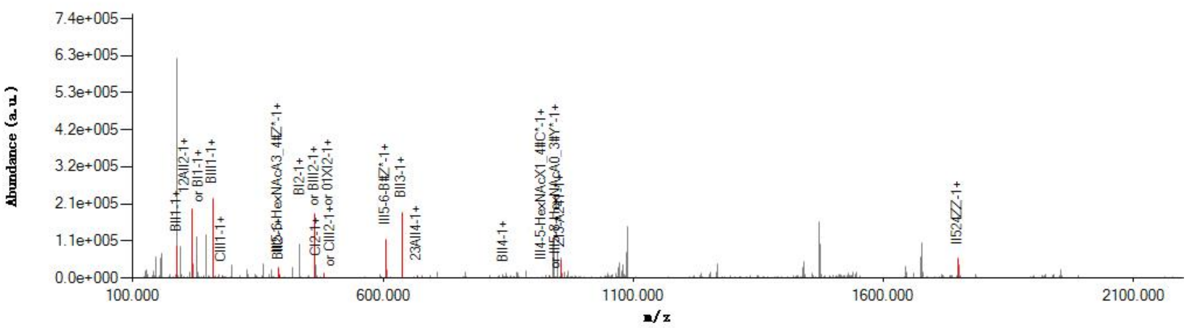


#### Graphical Fragmentation Map

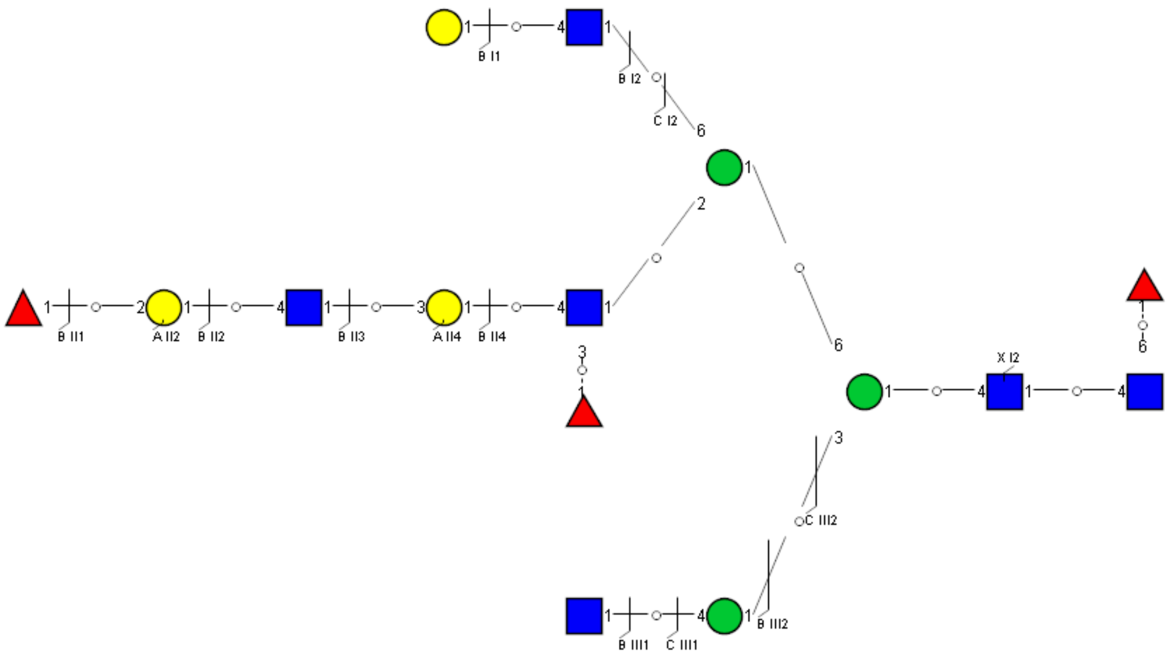

Spectrum Index: 38818(NoBHs:1)

#### Precursor iEF Map

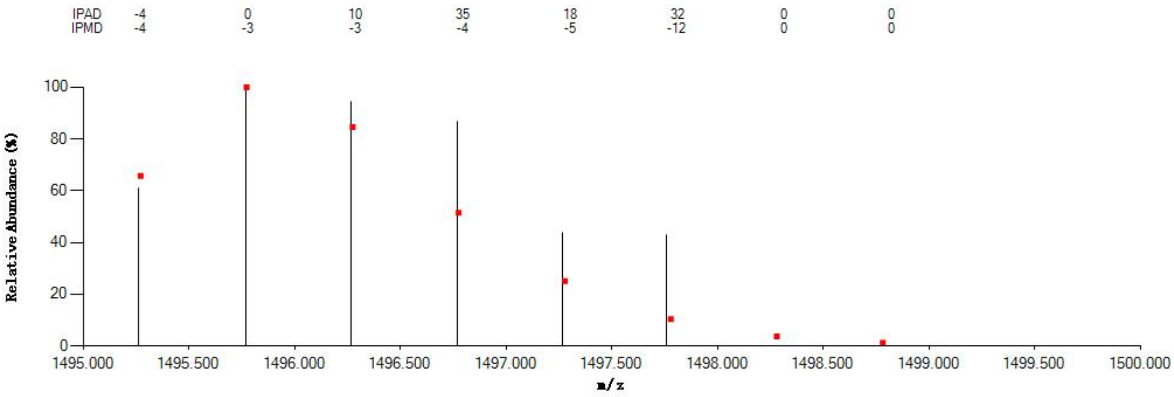


#### Annotated MS/MS Spectrum

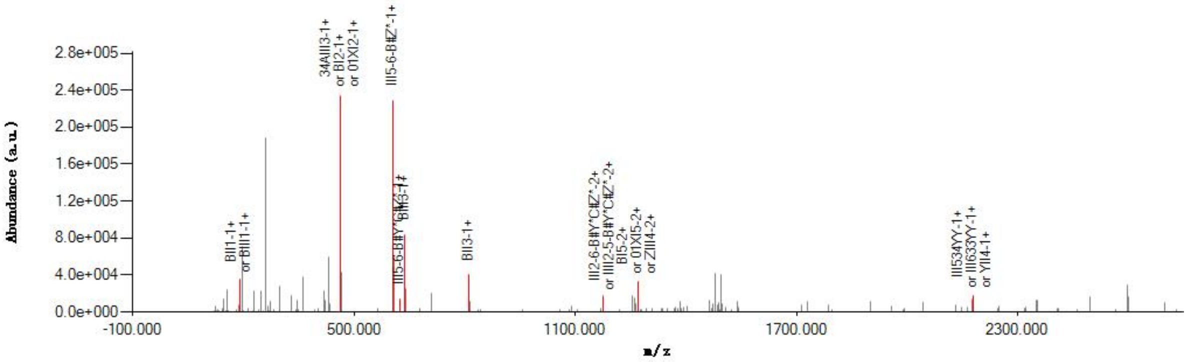


#### Graphical Fragmentation Map

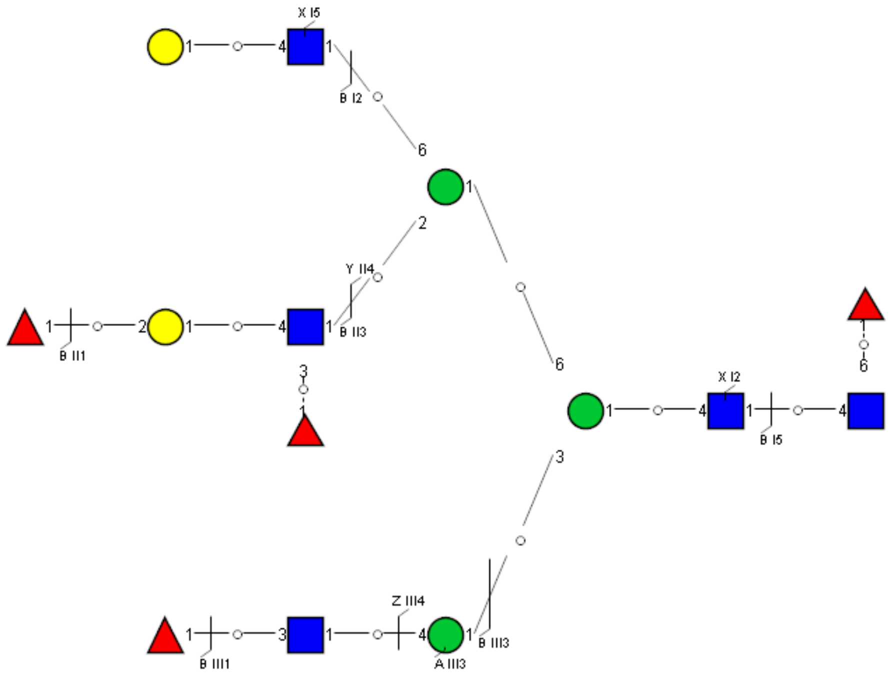

Spectrum Index: 39357(NoBHs:1)

#### Precursor iEF Map

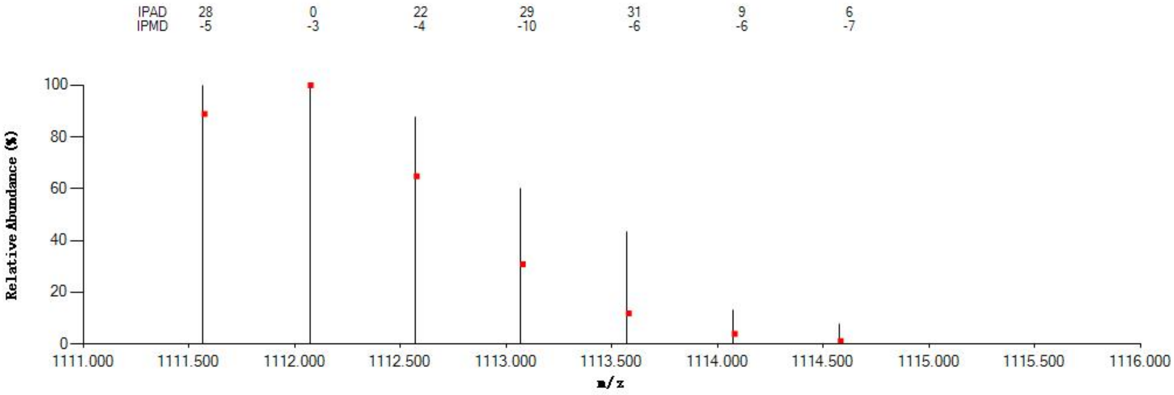


#### Annotated MS/MS Spectrum

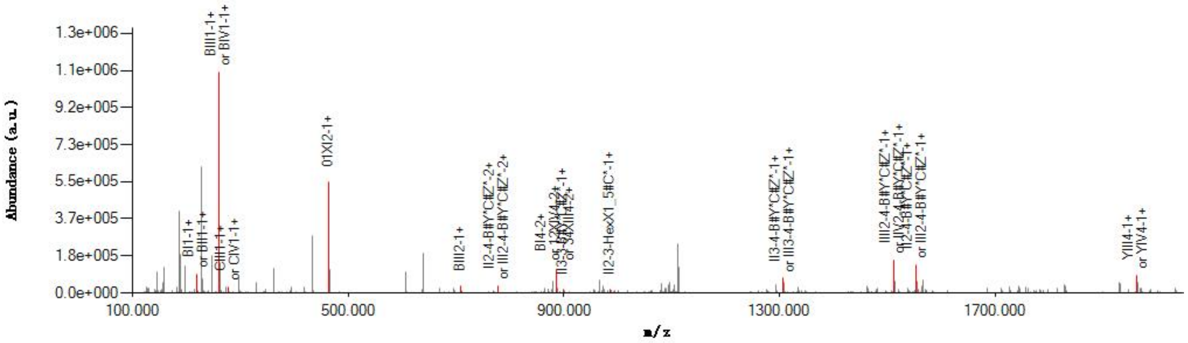


#### Graphical Fragmentation Map

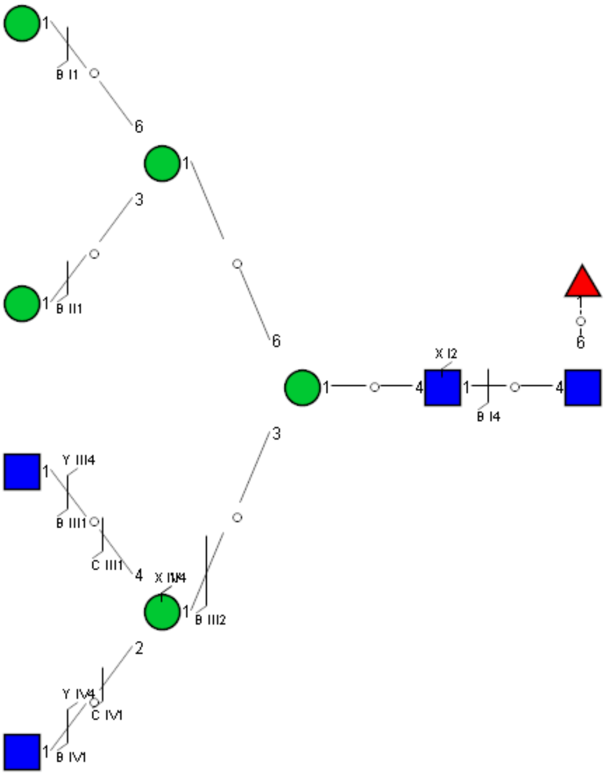

Spectrum Index: 39663(NoBHs:1)

#### Precursor iEF Map

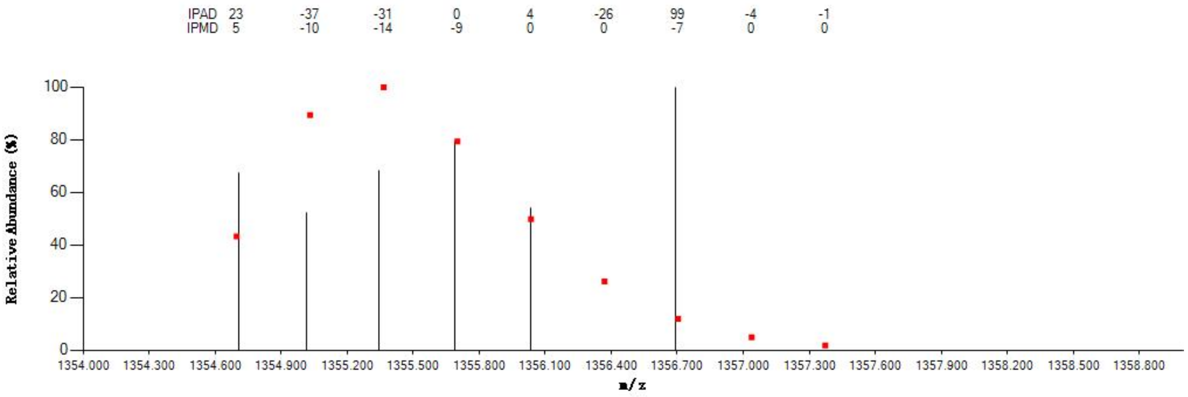


#### Annotated MS/MS Spectrum

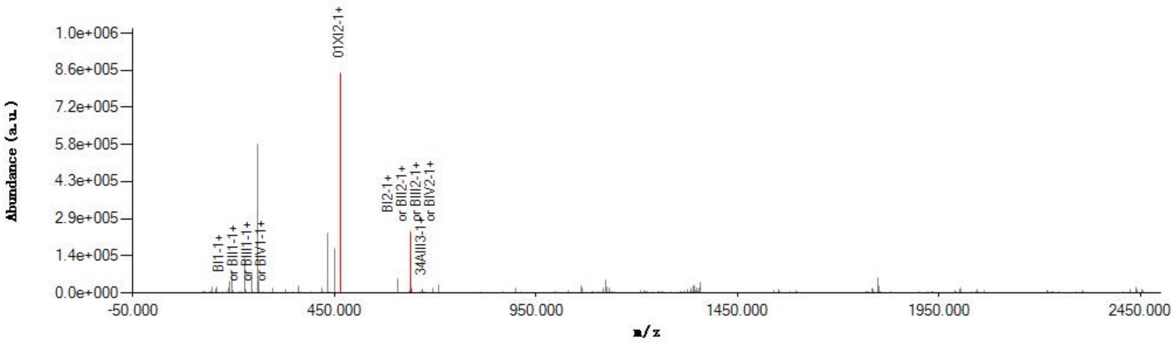


#### Graphical Fragmentation Map

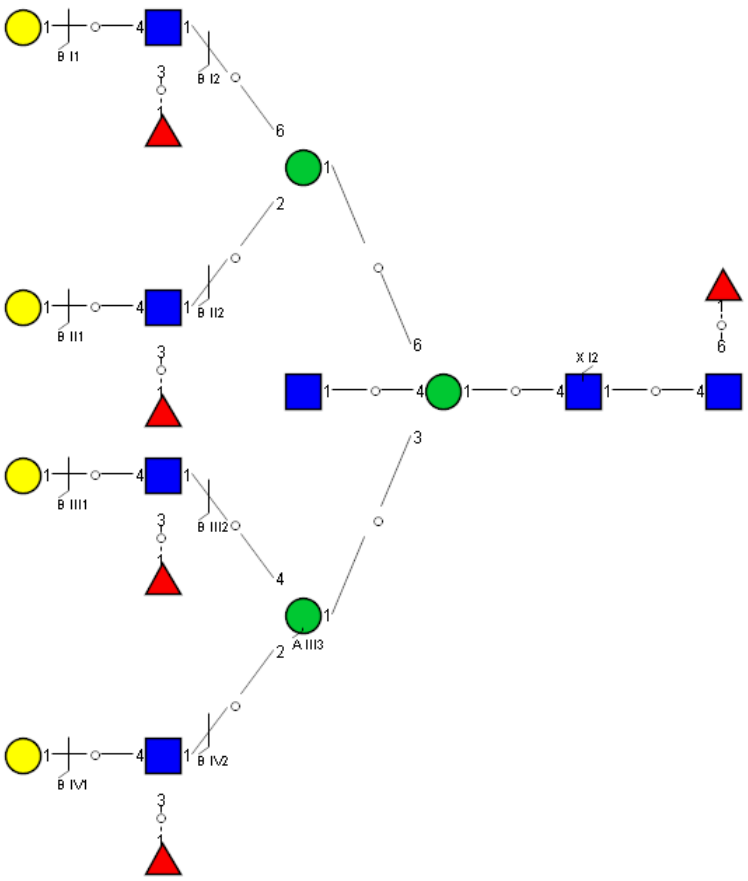

Spectrum Index: 39721(NoBHs:2)

#### Precursor iEF Map

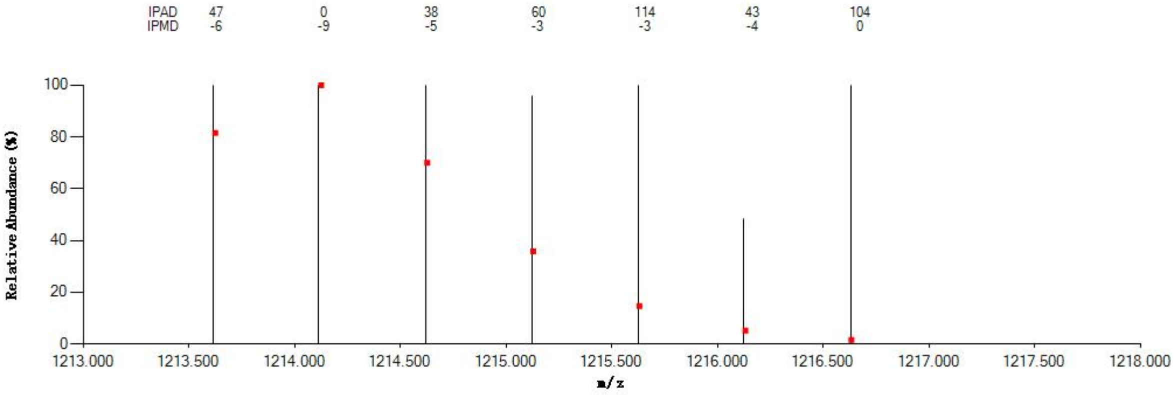


#### Annotated MS/MS Spectrum

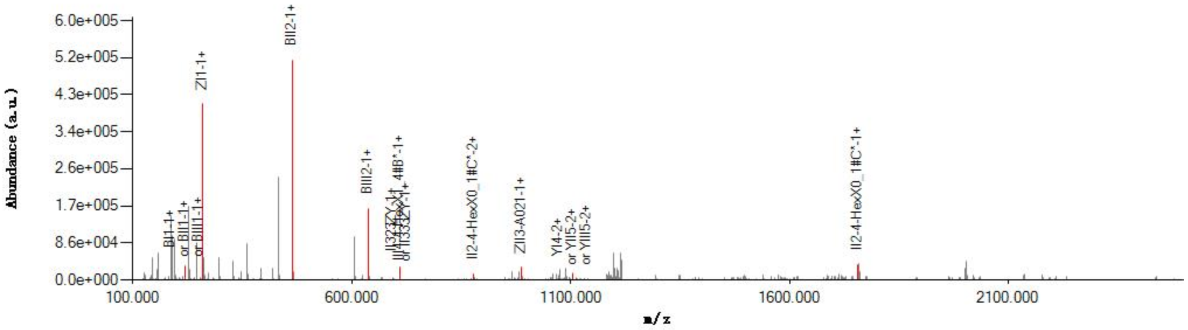


#### Graphical Fragmentation Map

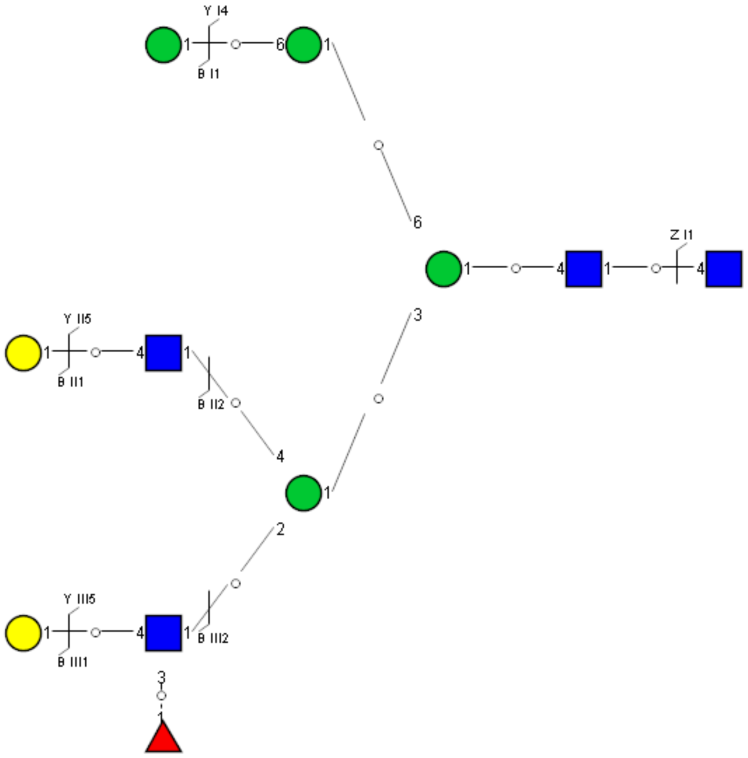

Spectrum Index: 39807(NoBHs:1)

#### Precursor iEF Map

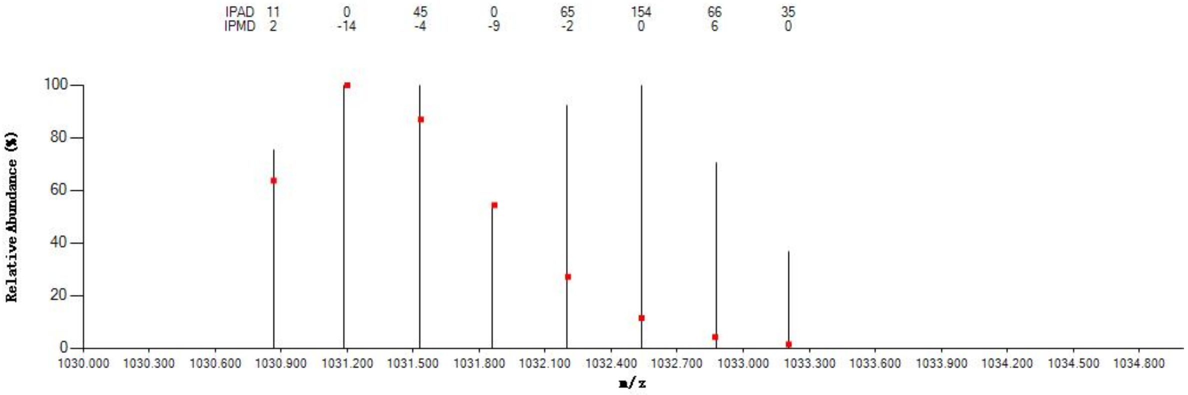


#### Annotated MS/MS Spectrum

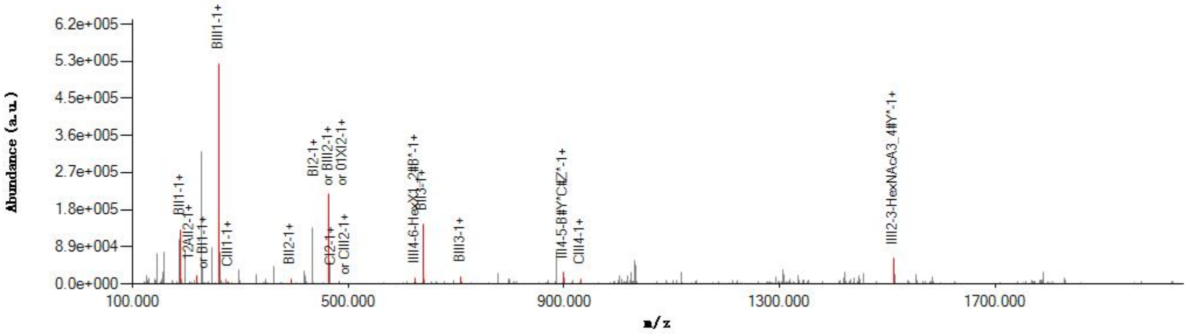


#### Graphical Fragmentation Map

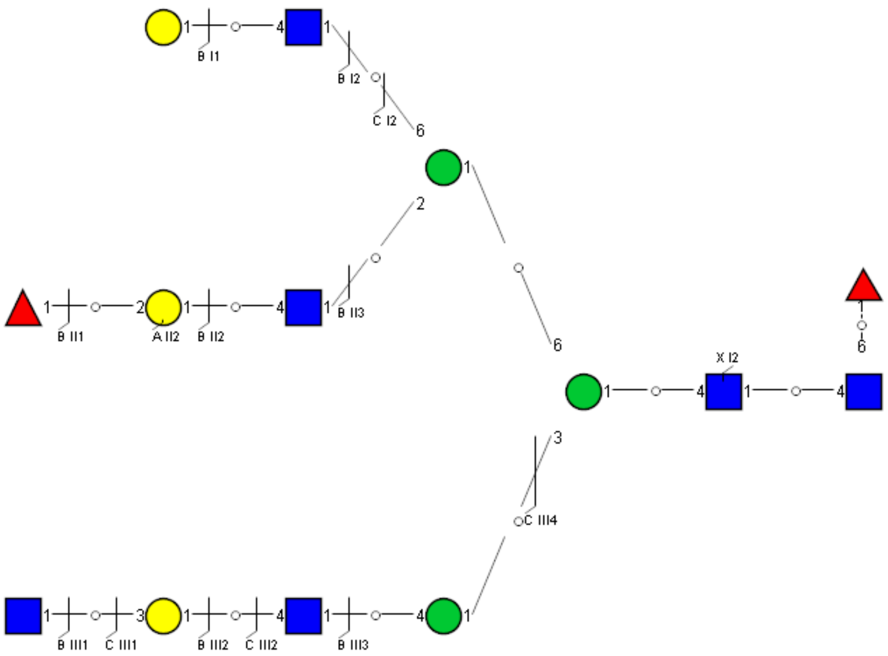

Spectrum Index: 39861(NoBHs:1)

#### Precursor iEF Map

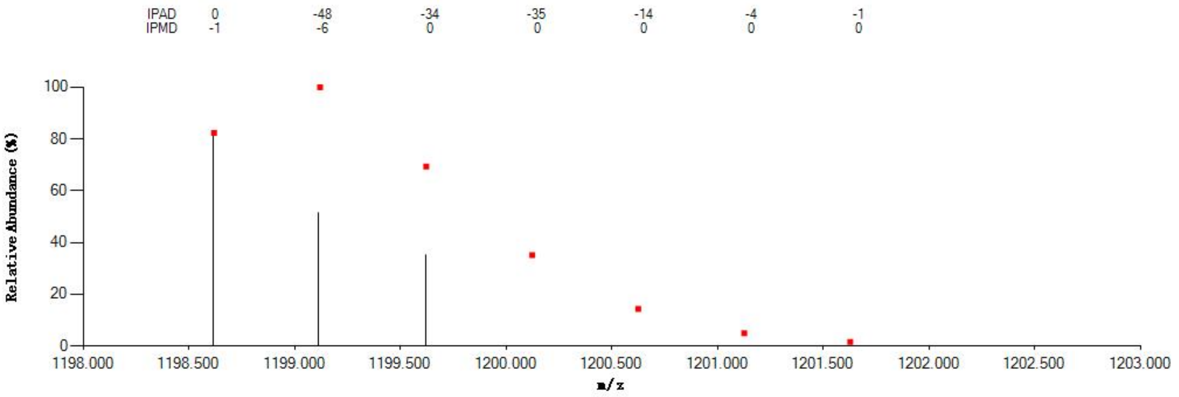

#### Annotated MS/MS Spectrum

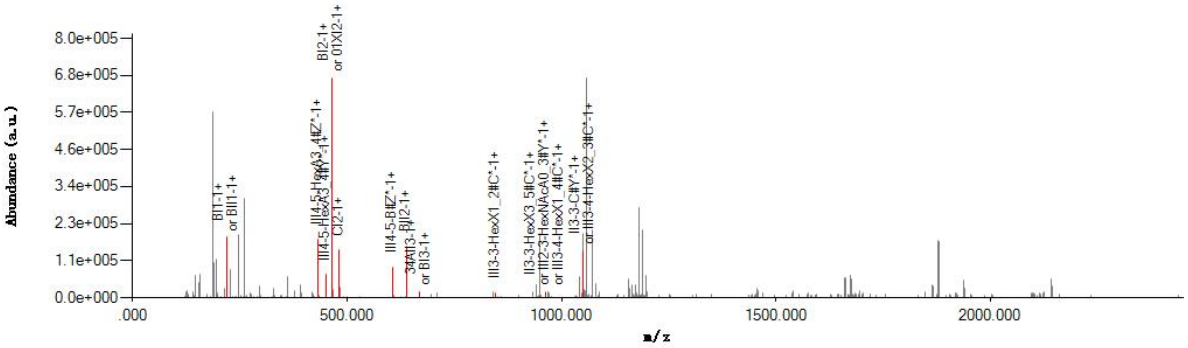

#### Graphical Fragmentation Map

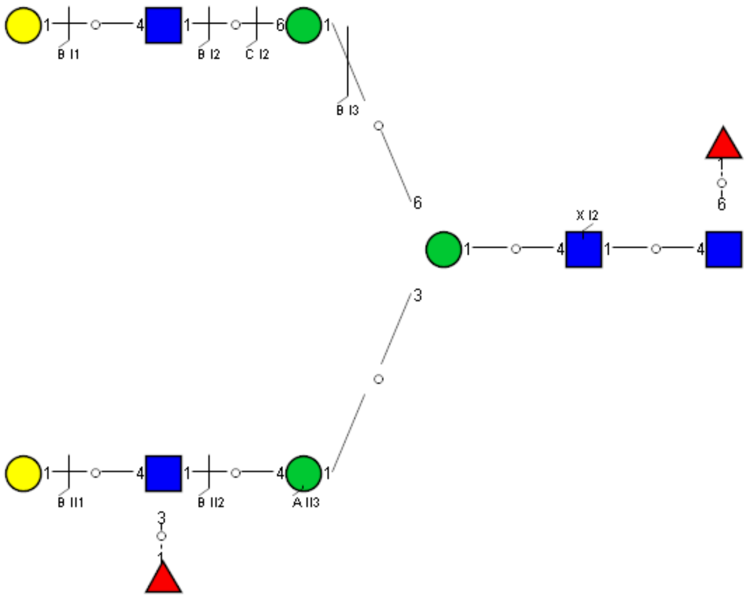

Spectrum Index: 39934(NoBHs:1)

#### Precursor iEF Map

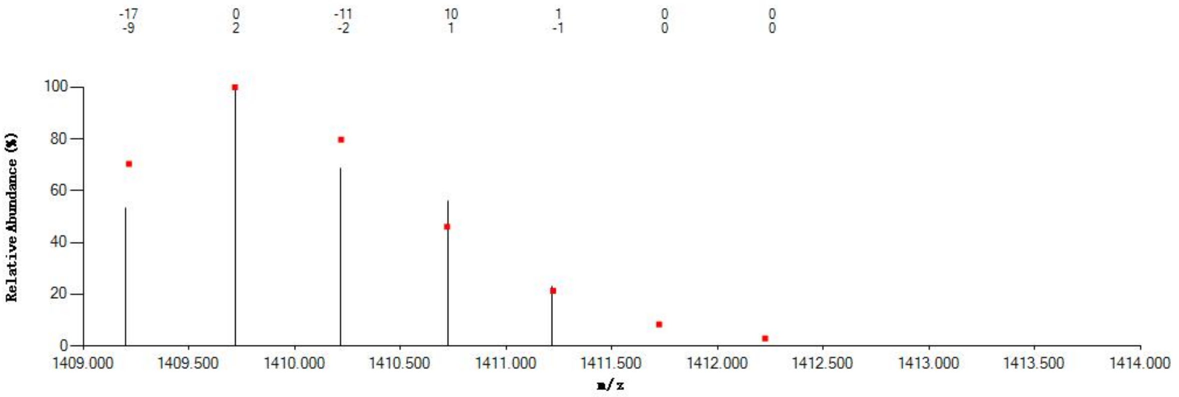


#### Annotated MS/MS Spectrum

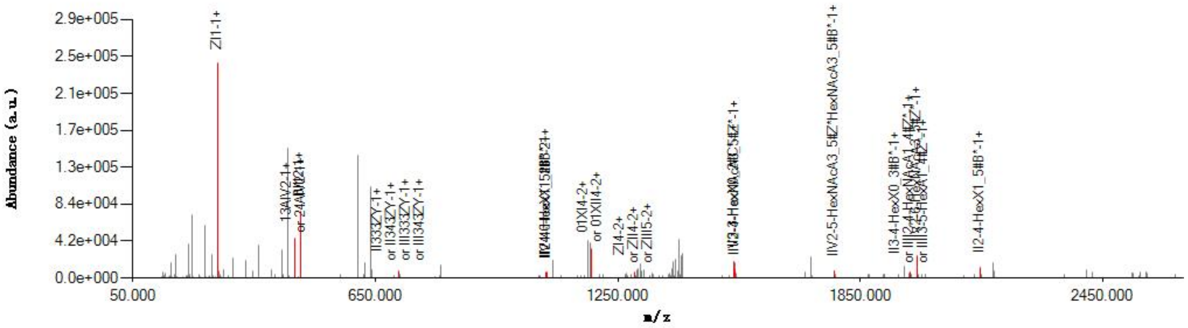


#### Graphical Fragmentation Map

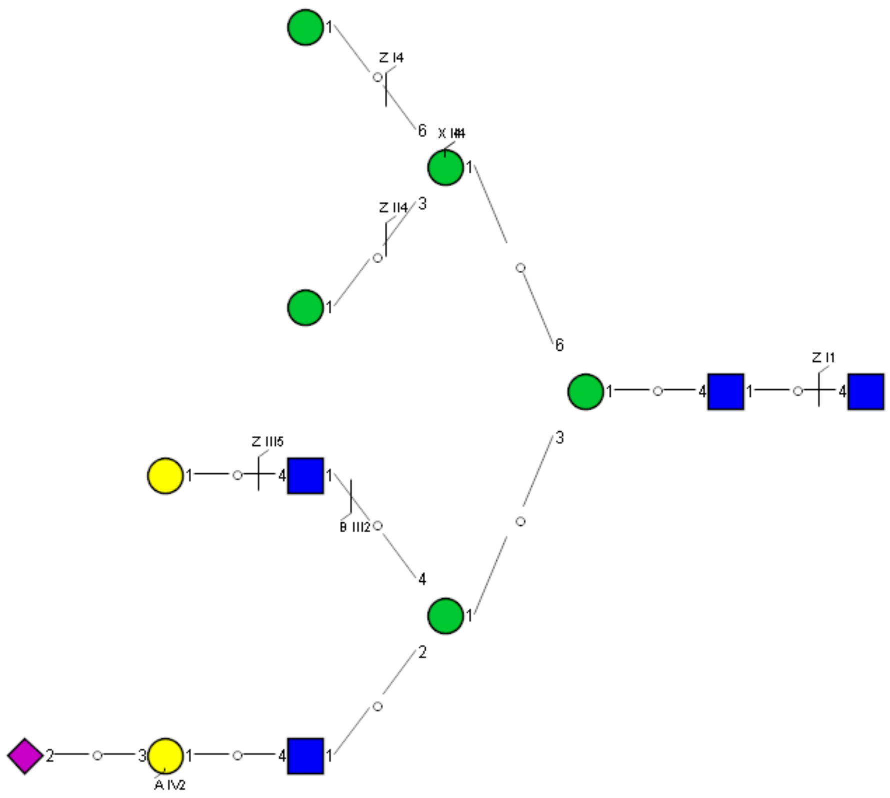

Spectrum Index: 40002(NoBHs:2)

#### Precursor iEF Map

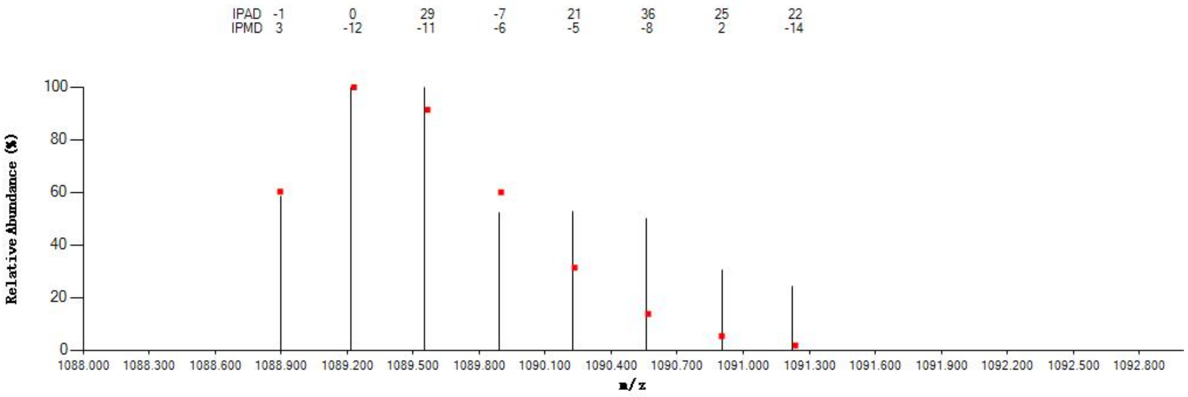


#### Annotated MS/MS Spectrum

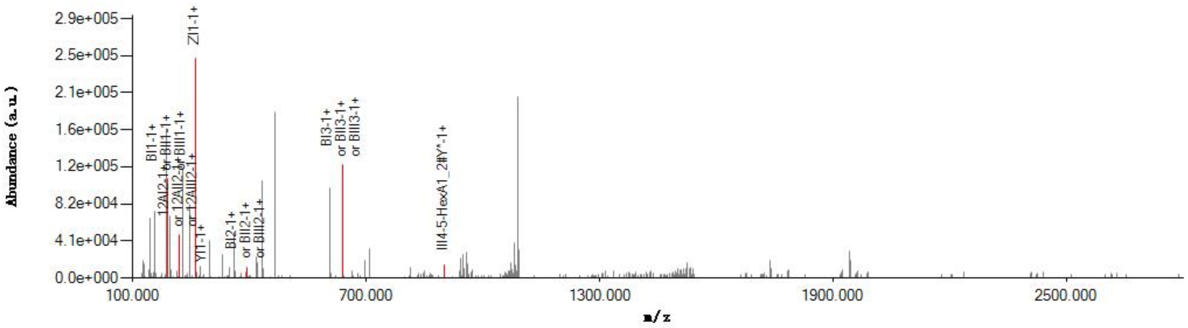


#### Graphical Fragmentation Map

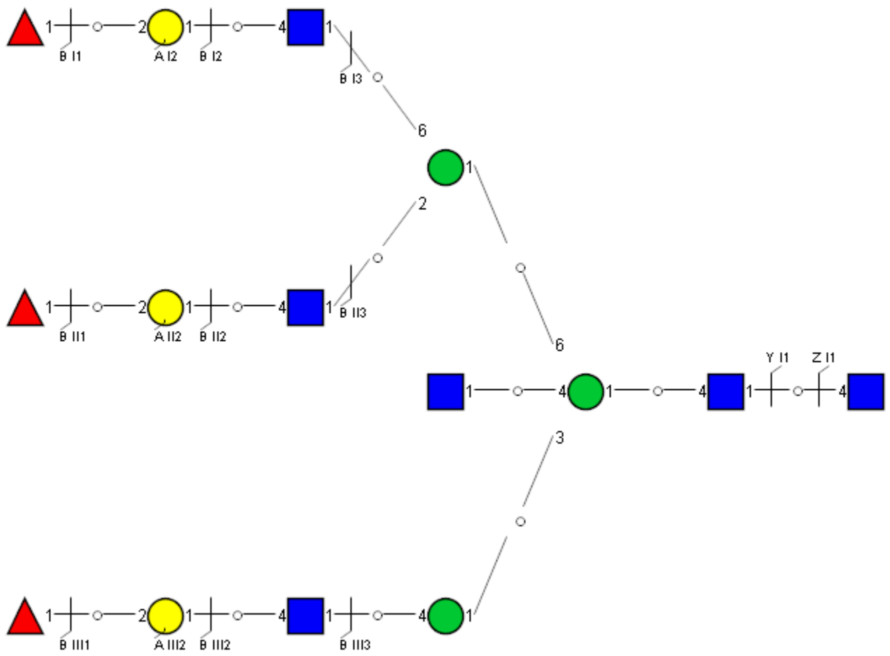

### Spectrum Index: 40313(NoBHs:3)

#### Precursor iEF Map

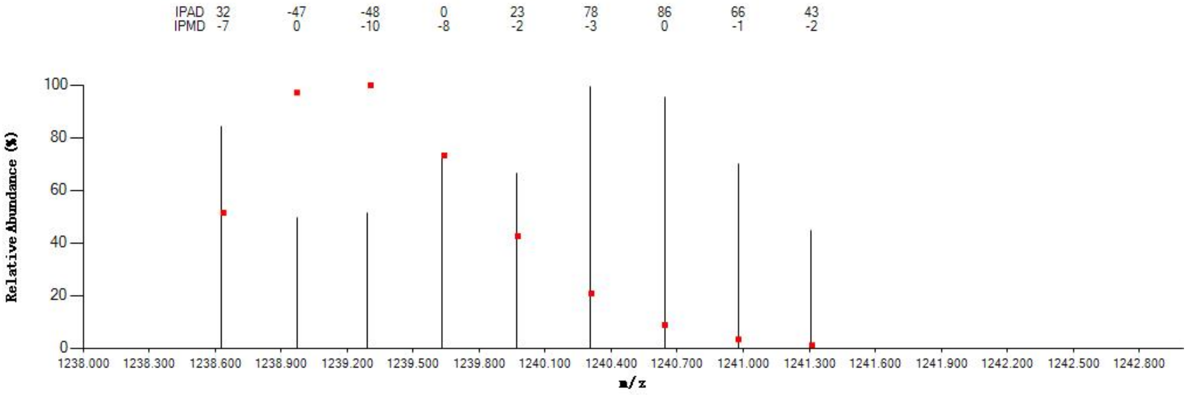


#### Annotated MS/MS Spectrum

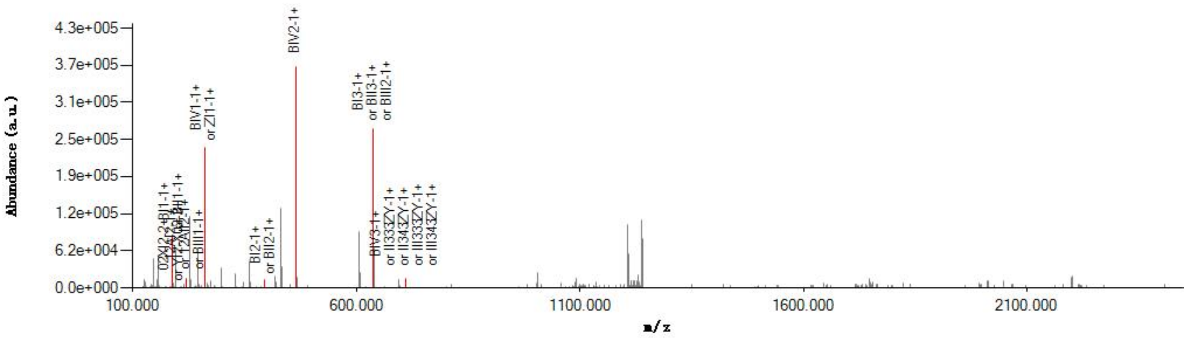


#### Graphical Fragmentation Map

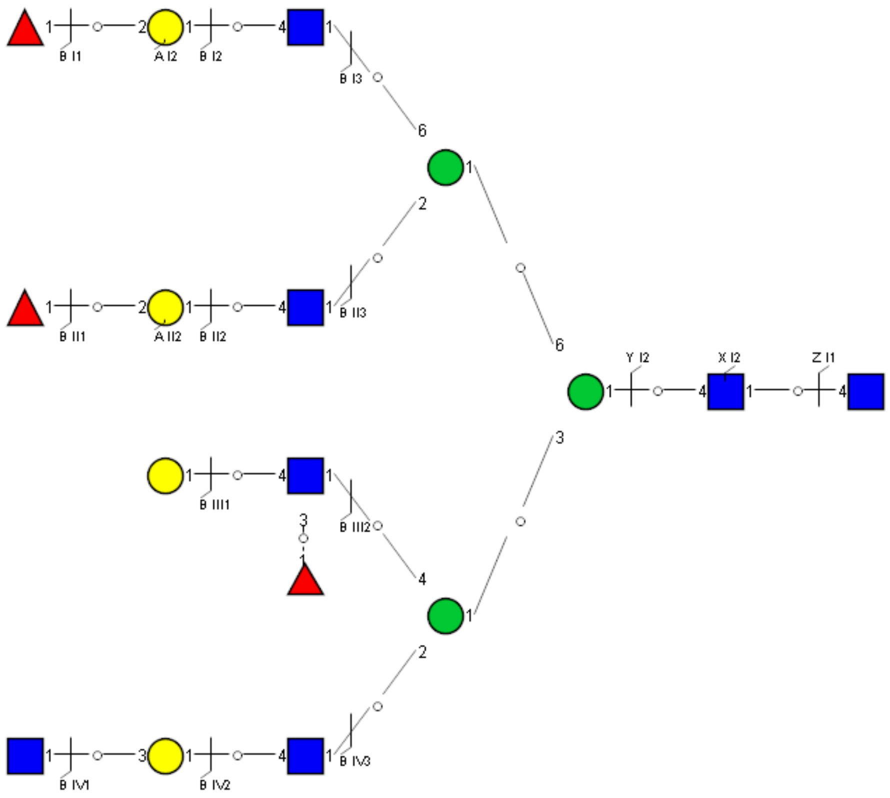

Spectrum Index: 40313(NoBHs:3)

#### Precursor iEF Map

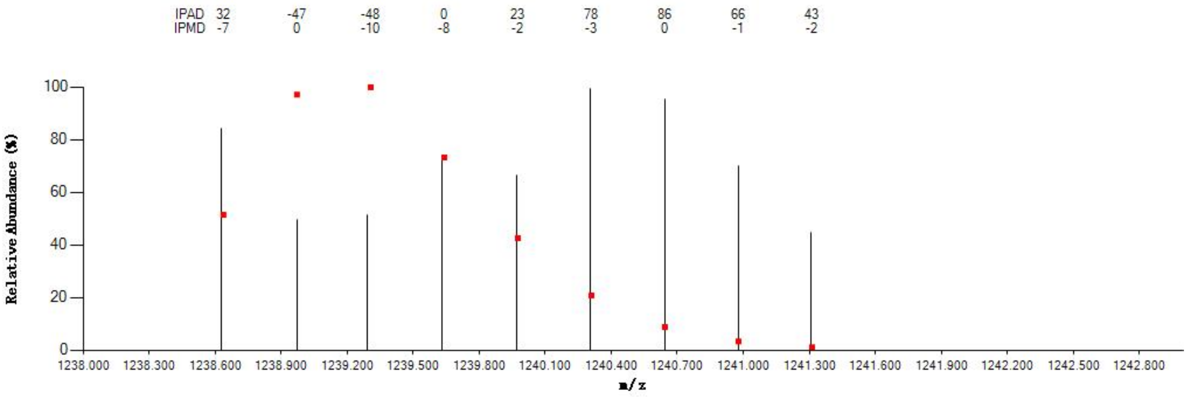


#### Annotated MS/MS Spectrum

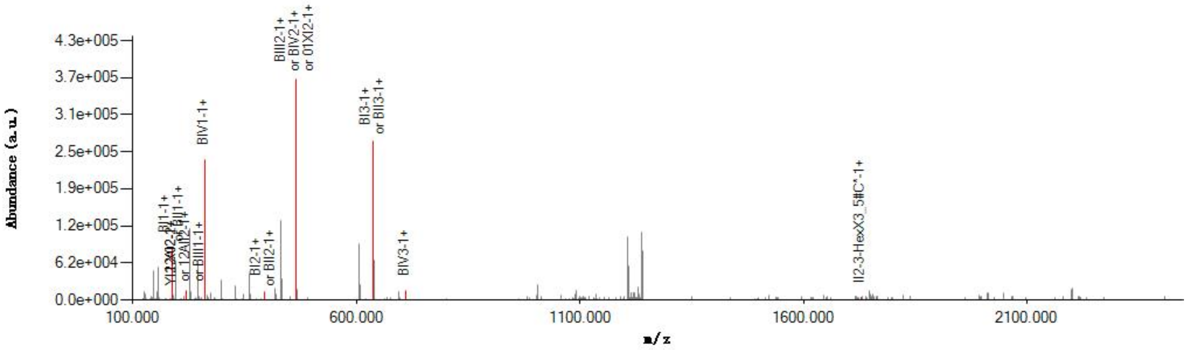


#### Graphical Fragmentation Map

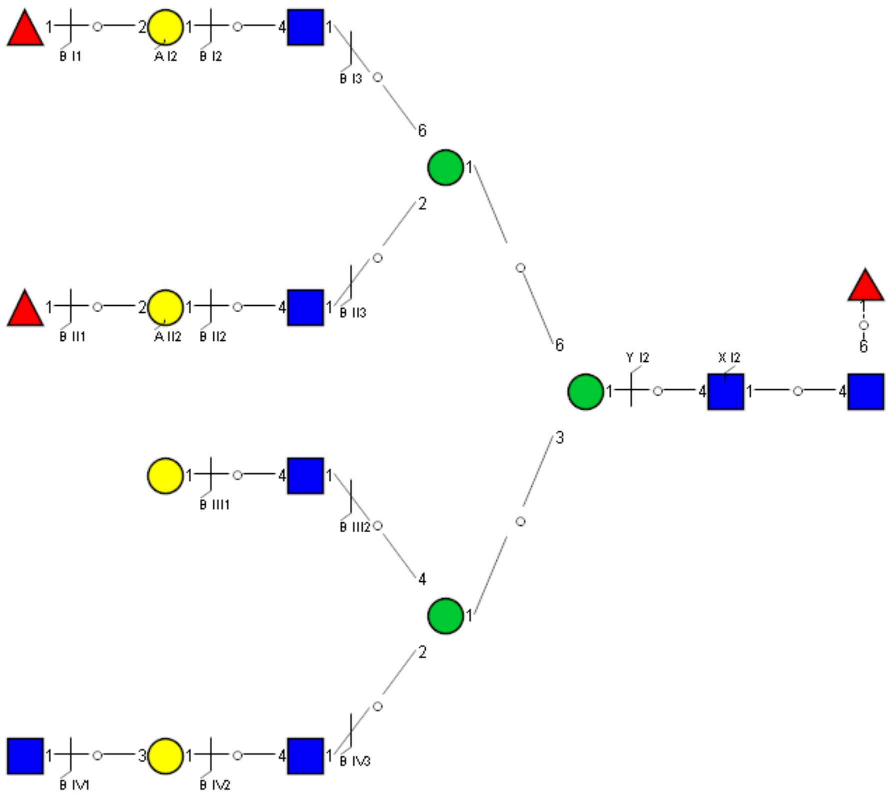

Spectrum Index: 40313(NoBHs:3)

#### Precursor iEF Map

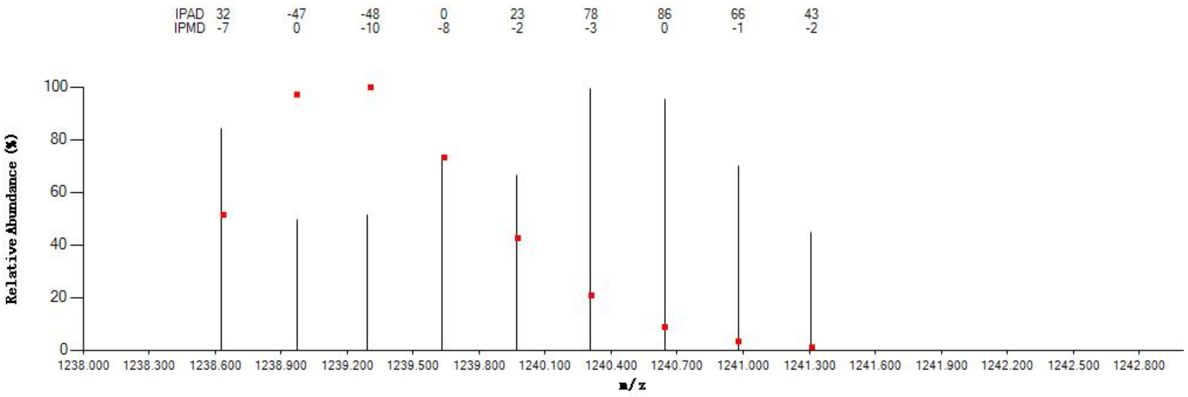


#### Annotated MS/MS Spectrum

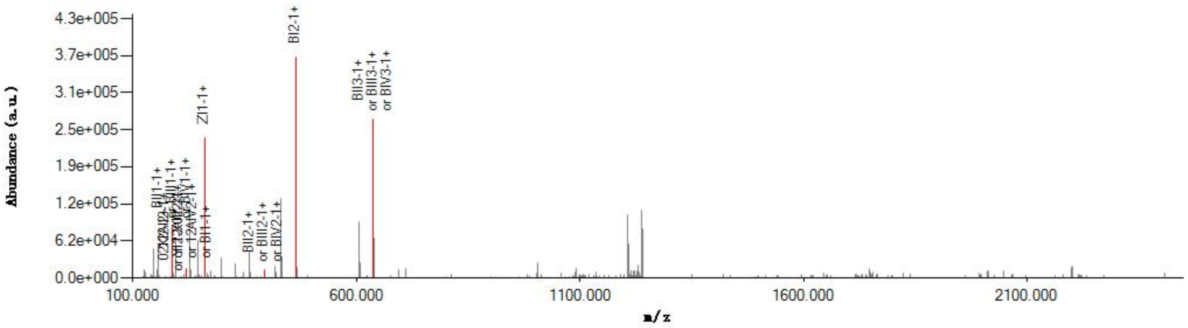


#### Graphical Fragmentation Map

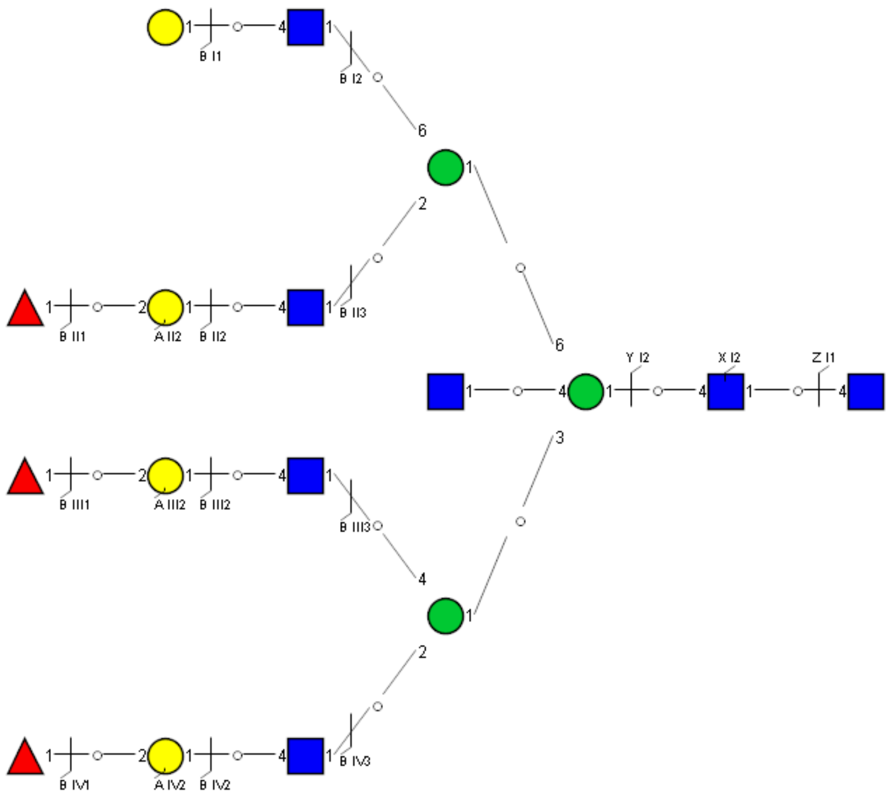

Spectrum Index: 40346(NoBHs:2)

#### Precursor iEF Map

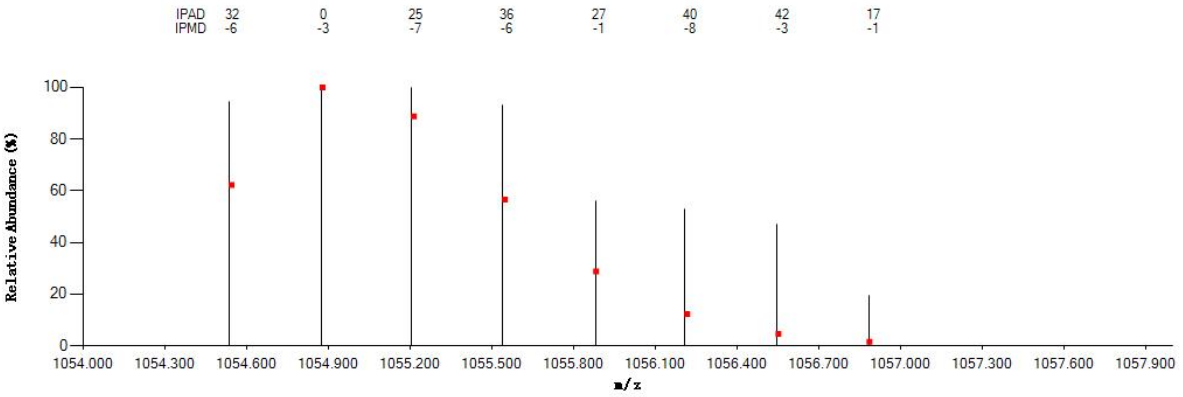

#### Annotated MS/MS Spectrum

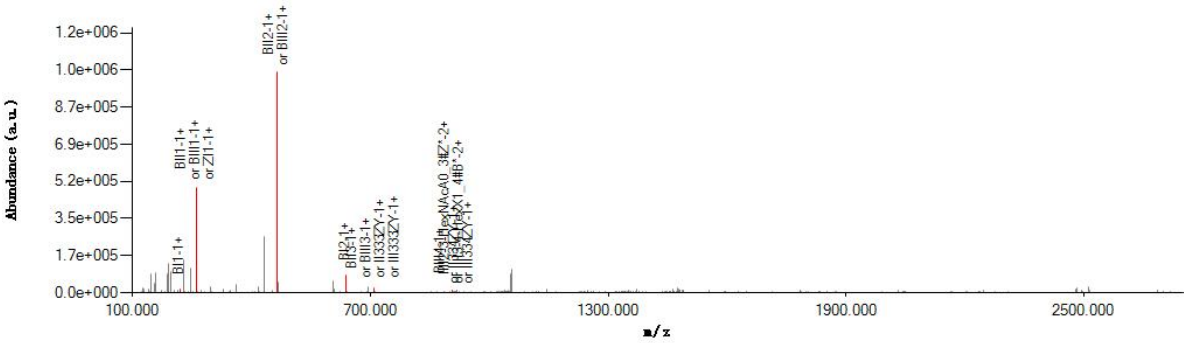

#### Graphical Fragmentation Map

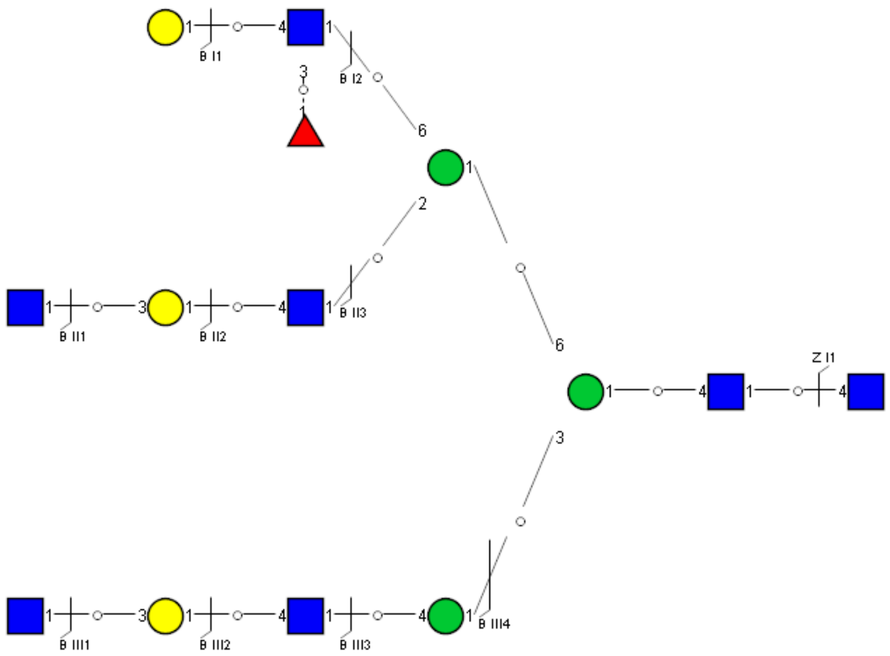

Spectrum Index: 40346(NoBHs:2)

#### Precursor iEF Map

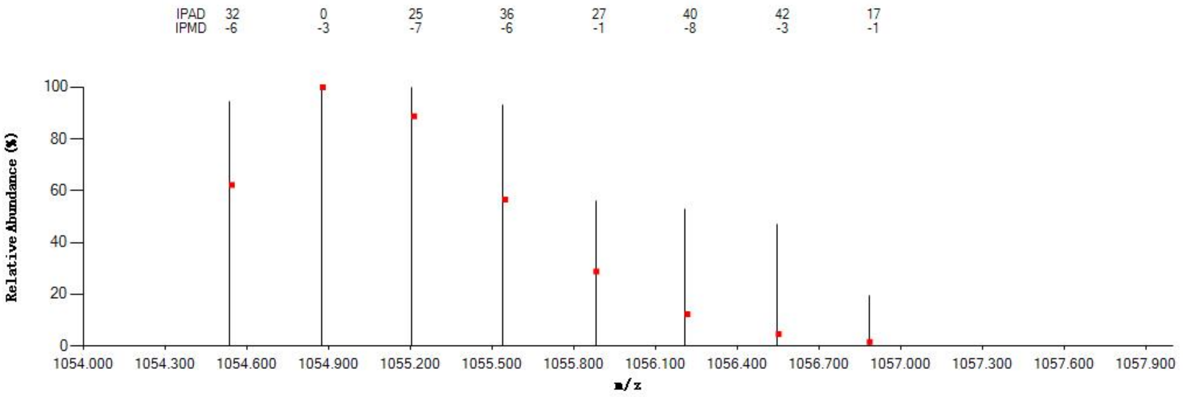


#### Annotated MS/MS Spectrum

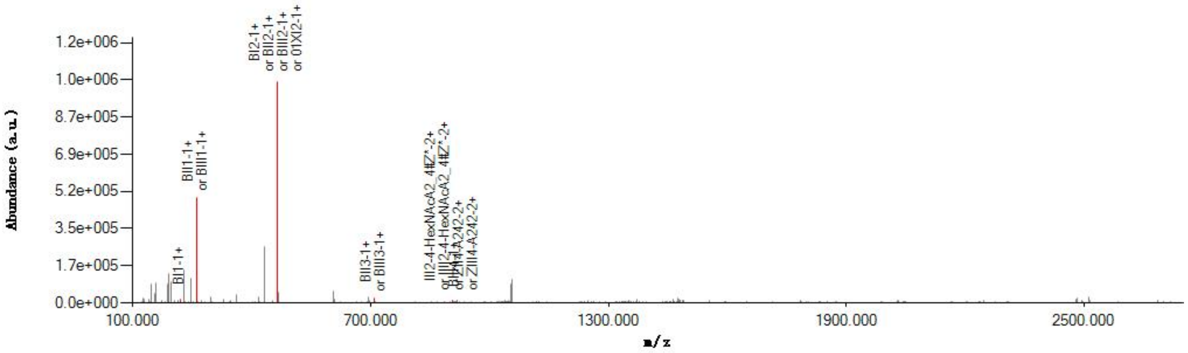


#### Graphical Fragmentation Map

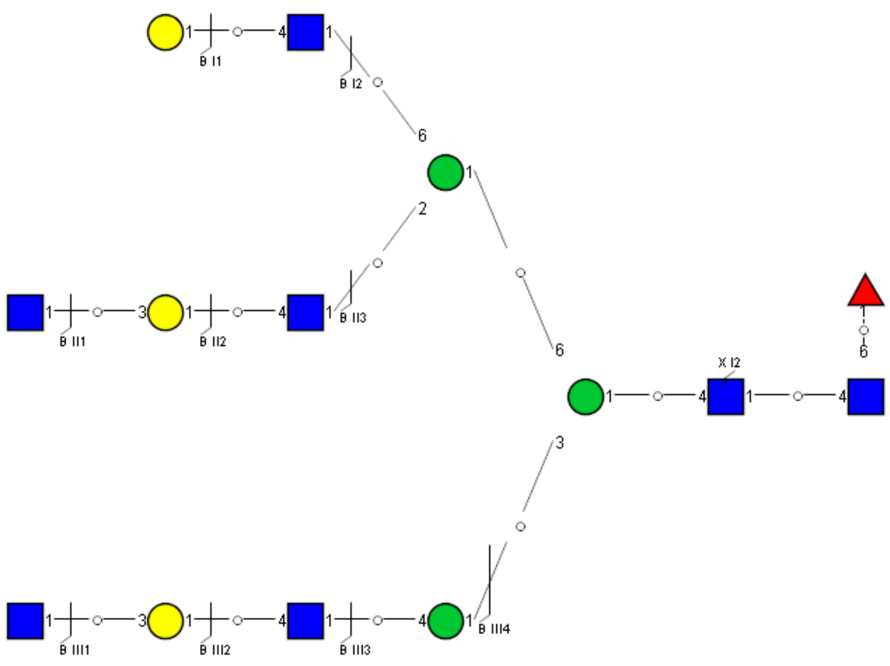

### Spectrum Index: 40892(NoBHs:1)

#### Precursor iEF Map

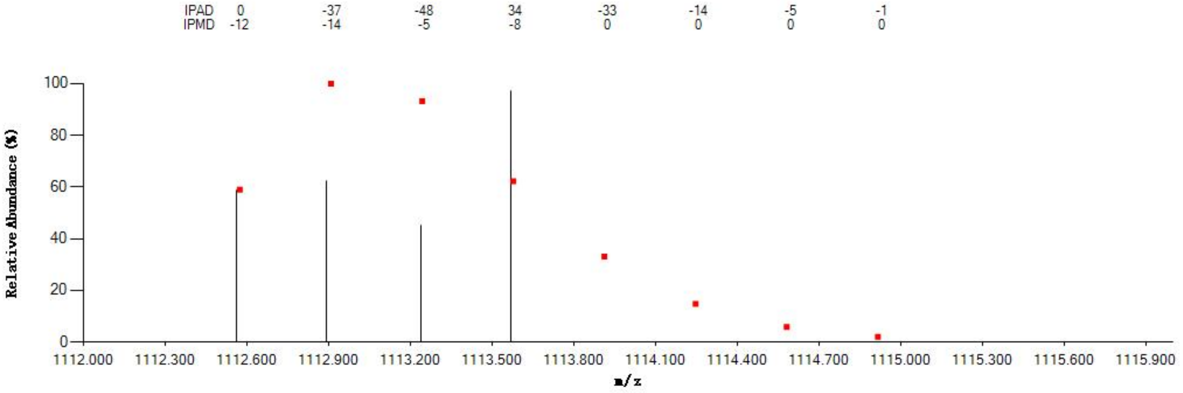


#### Annotated MS/MS Spectrum

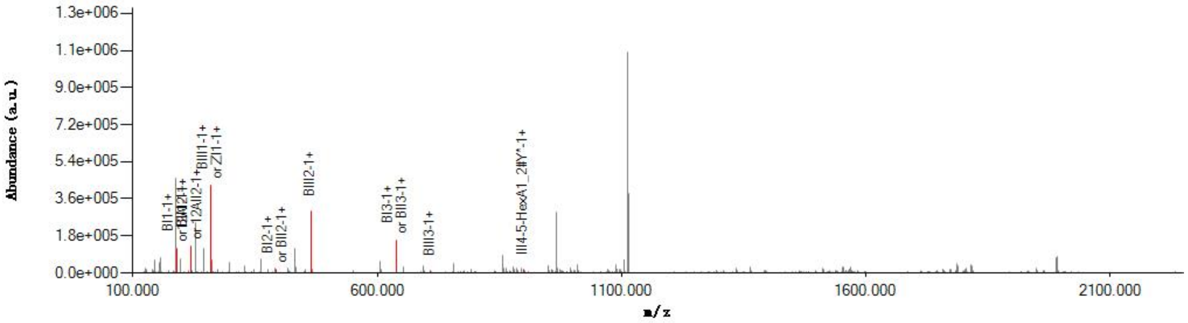


#### Graphical Fragmentation Map

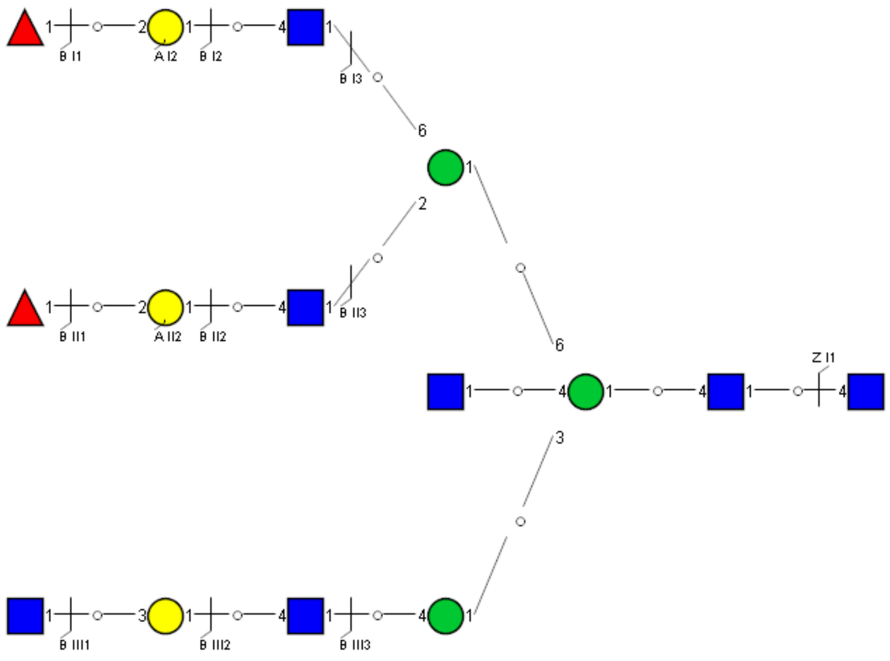

Spectrum Index: 41450(NoBHs:1)

Precursor iEF Map

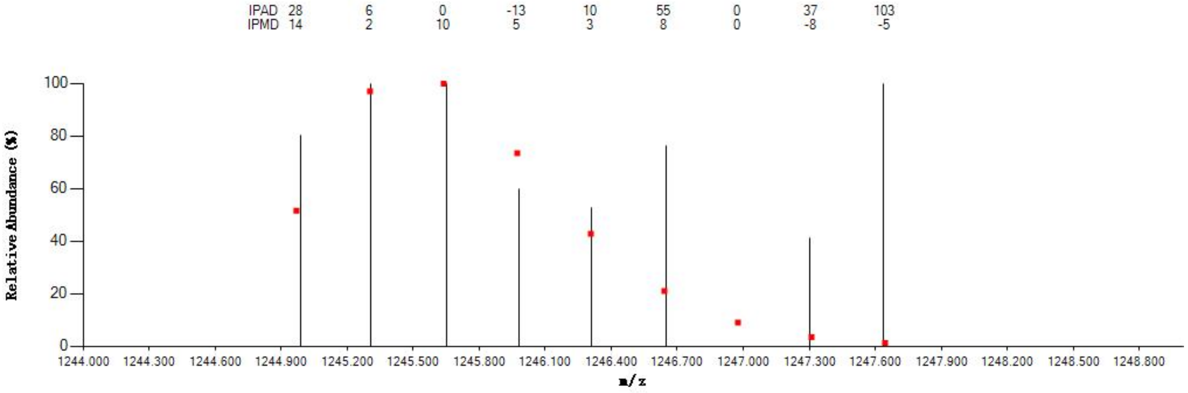


Annotated MS/MS Spectrum

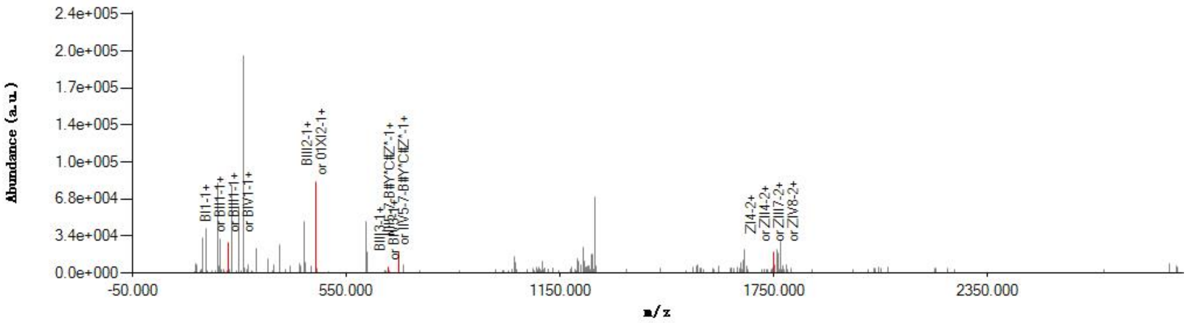


Graphical Fragmentation Map

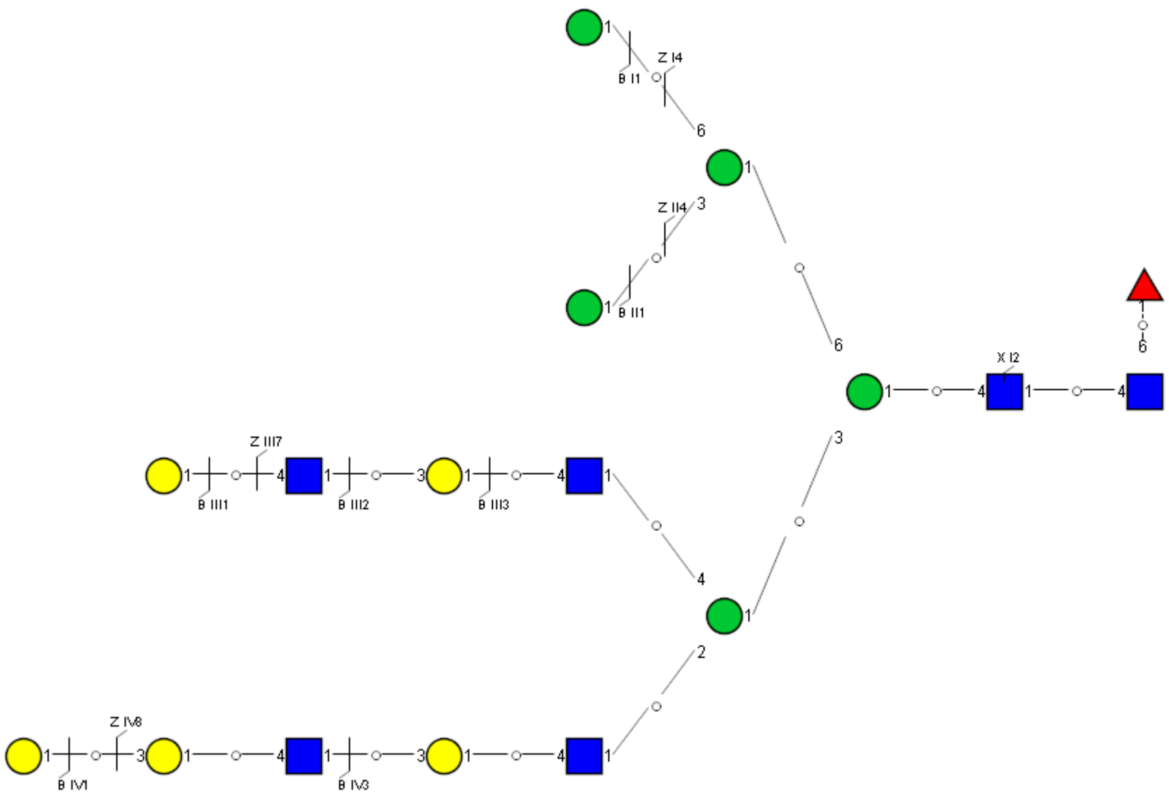

Spectrum Index: 41883(NoBHs:2)

#### Precursor iEF Map

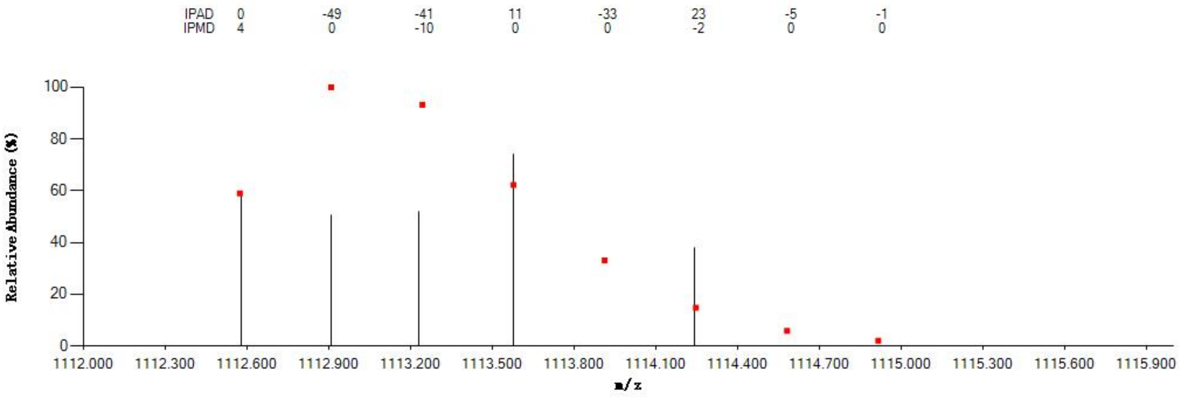


#### Annotated MS/MS Spectrum

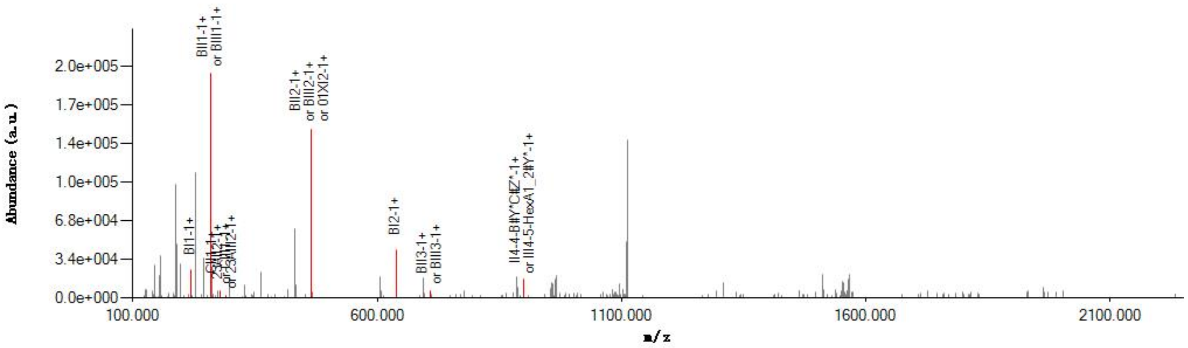


#### Graphical Fragmentation Map

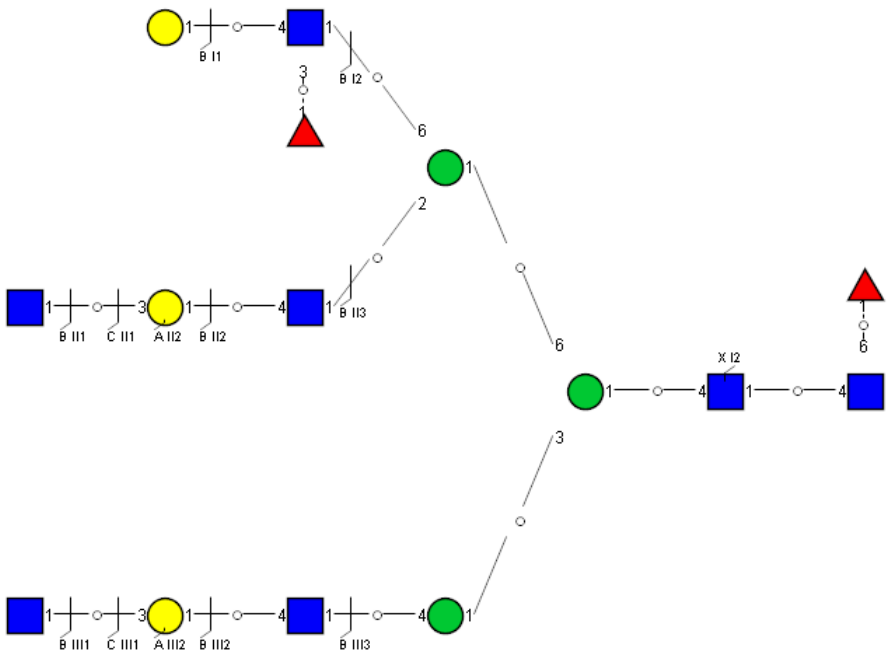

### Spectrum Index: 41883(NoBHs:2)

#### Precursor iEF Map

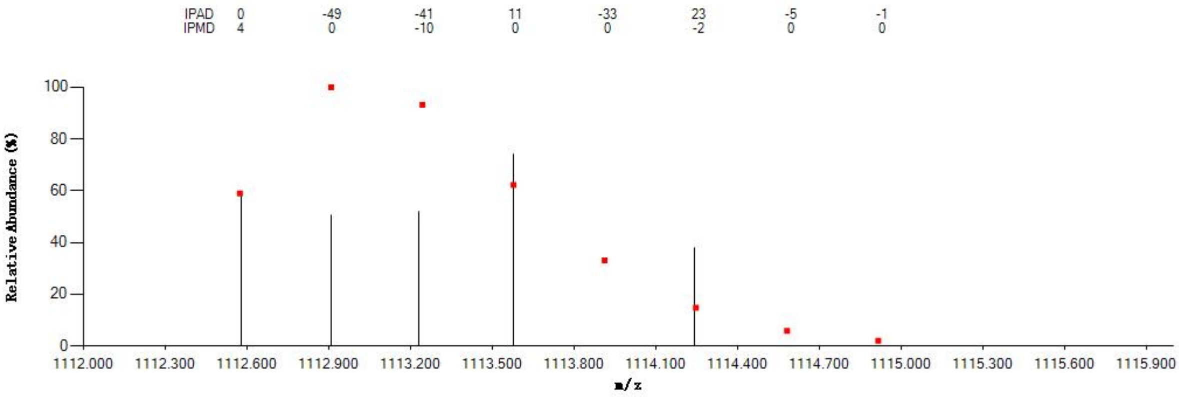


#### Annotated MS/MS Spectrum

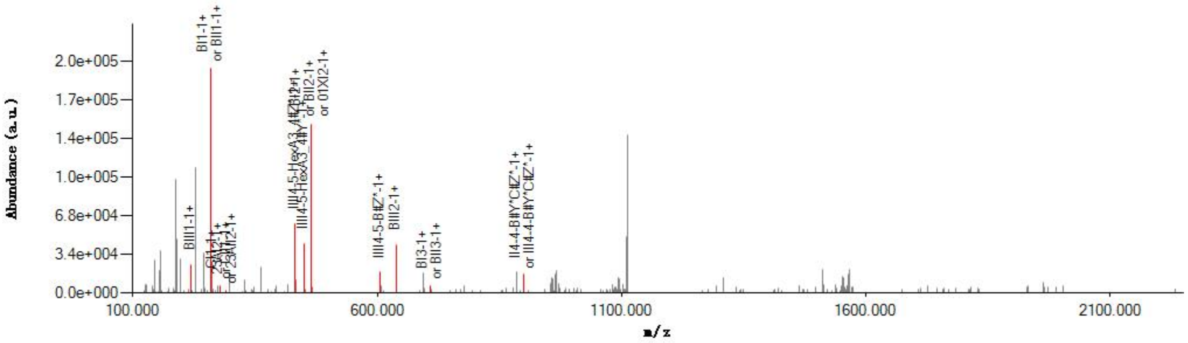


#### Graphical Fragmentation Map

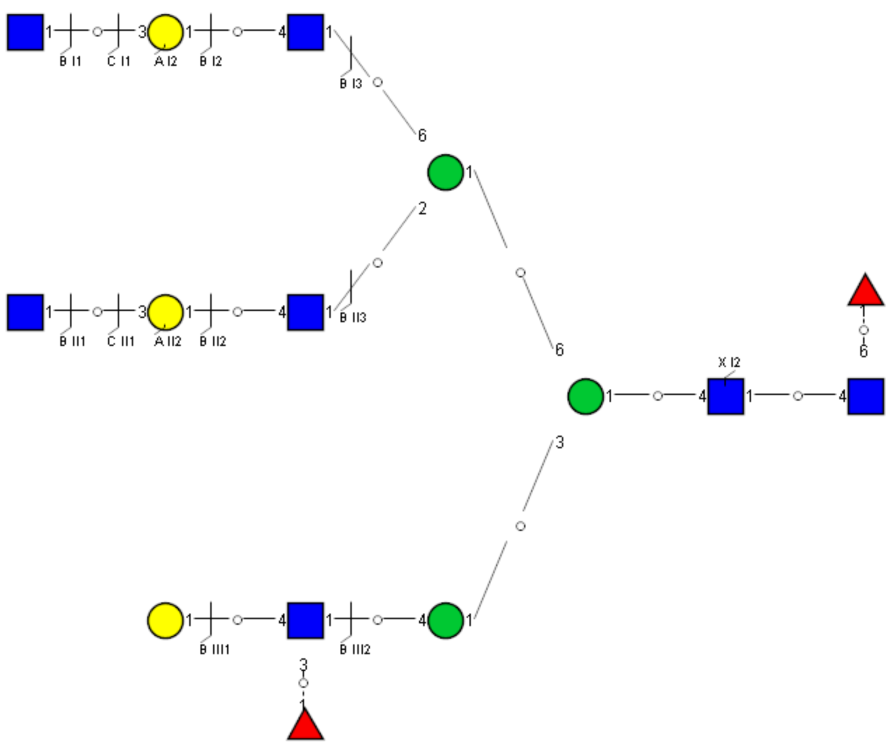

Spectrum Index: 42047(NoBHs:1)

#### Precursor iEF Map

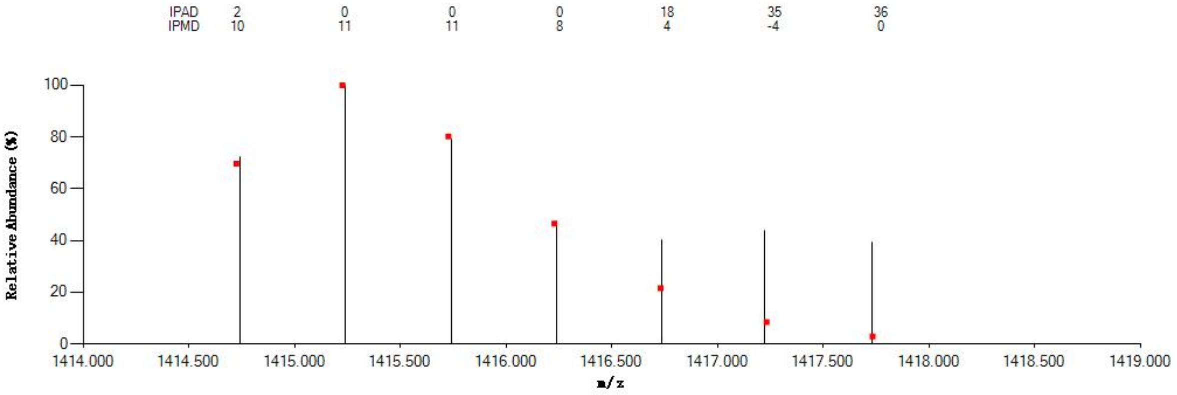


#### Annotated MS/MS Spectrum

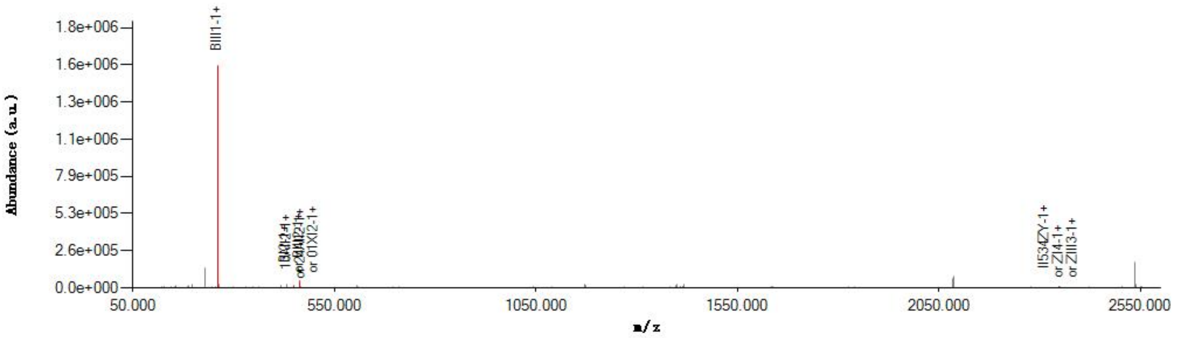


#### Graphical Fragmentation Map

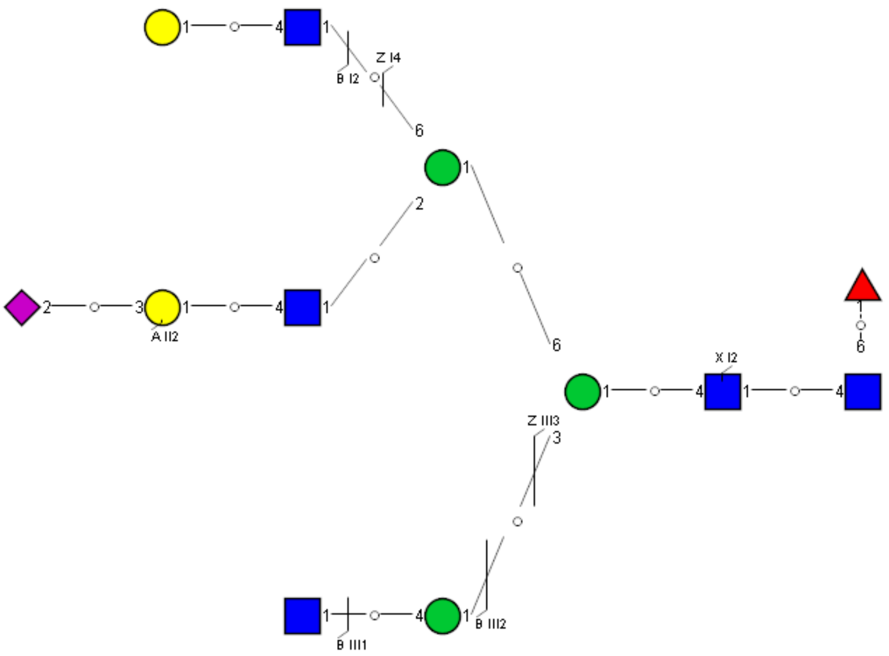

Spectrum Index: 42094(NoBHs:1)

#### Precursor iEF Map

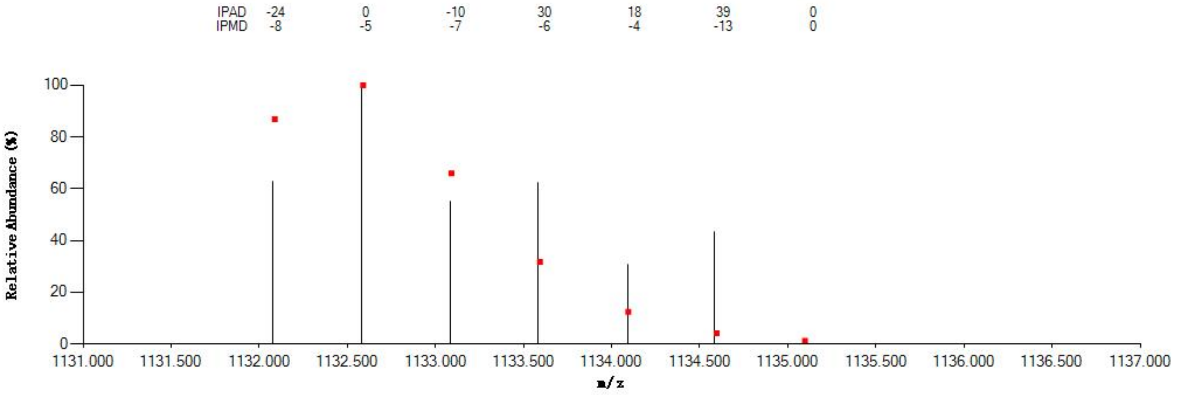


#### Annotated MS/MS Spectrum

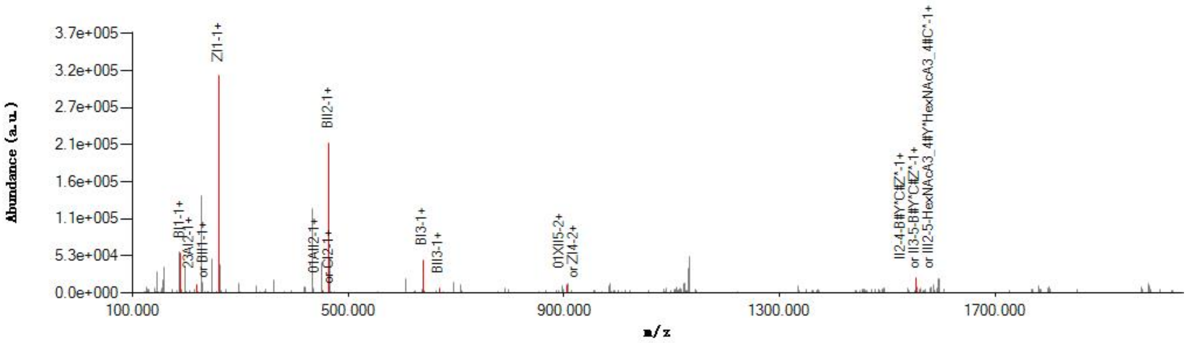


#### Graphical Fragmentation Map

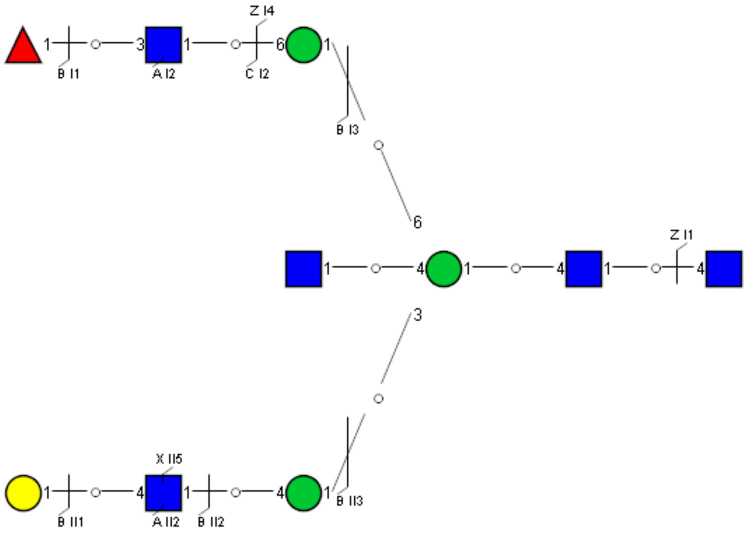

### Spectrum Index: 42163(NoBHs:1)

#### Precursor iEF Map

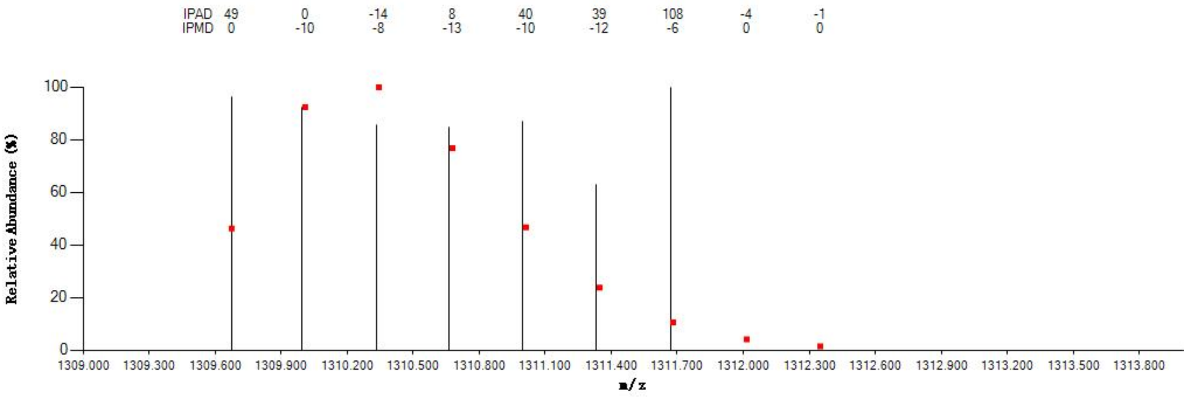


#### Annotated MS/MS Spectrum

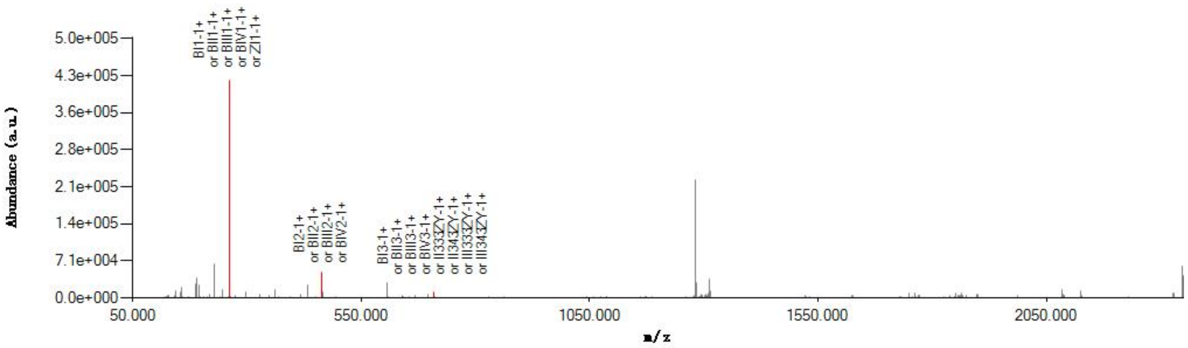


#### Graphical Fragmentation Map

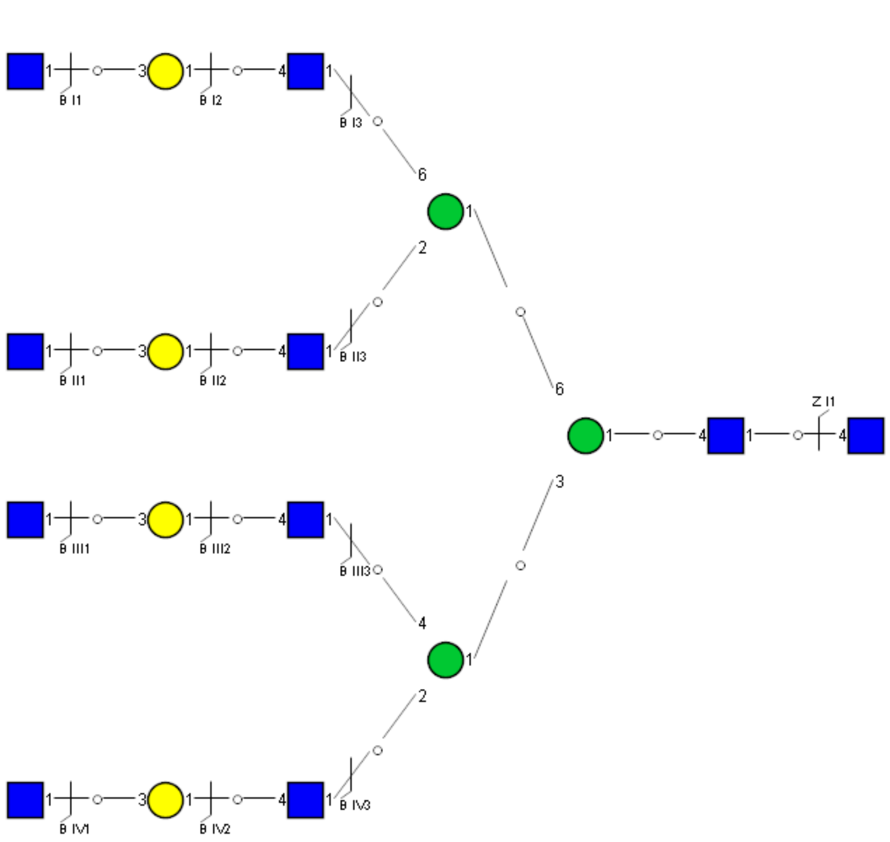

Spectrum Index: 49661(NoBHs:1)

#### Precursor iEF Map

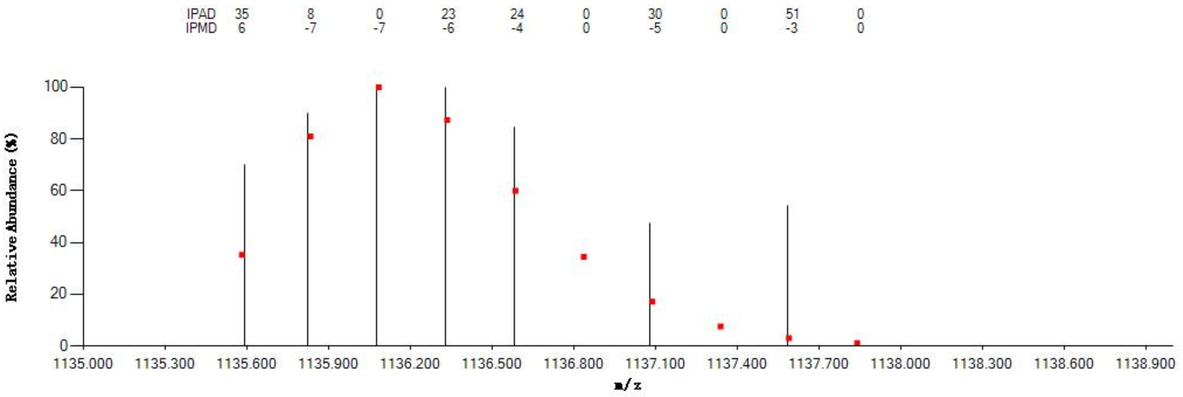

#### Annotated MS/MS Spectrum

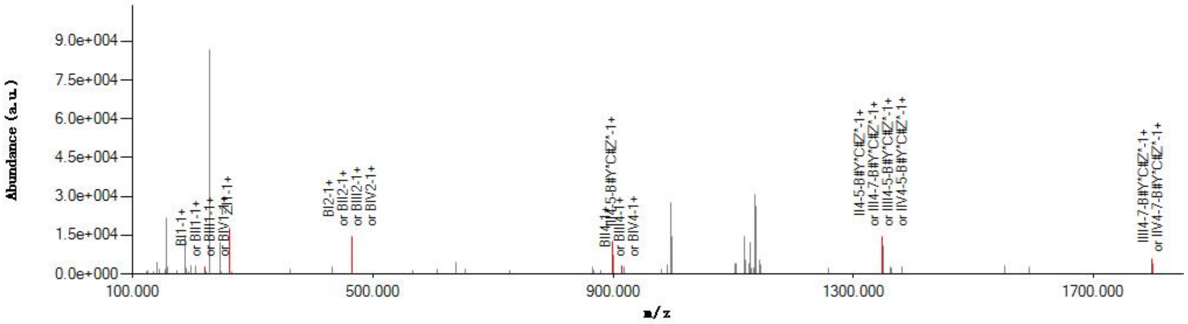

#### Graphical Fragmentation Map

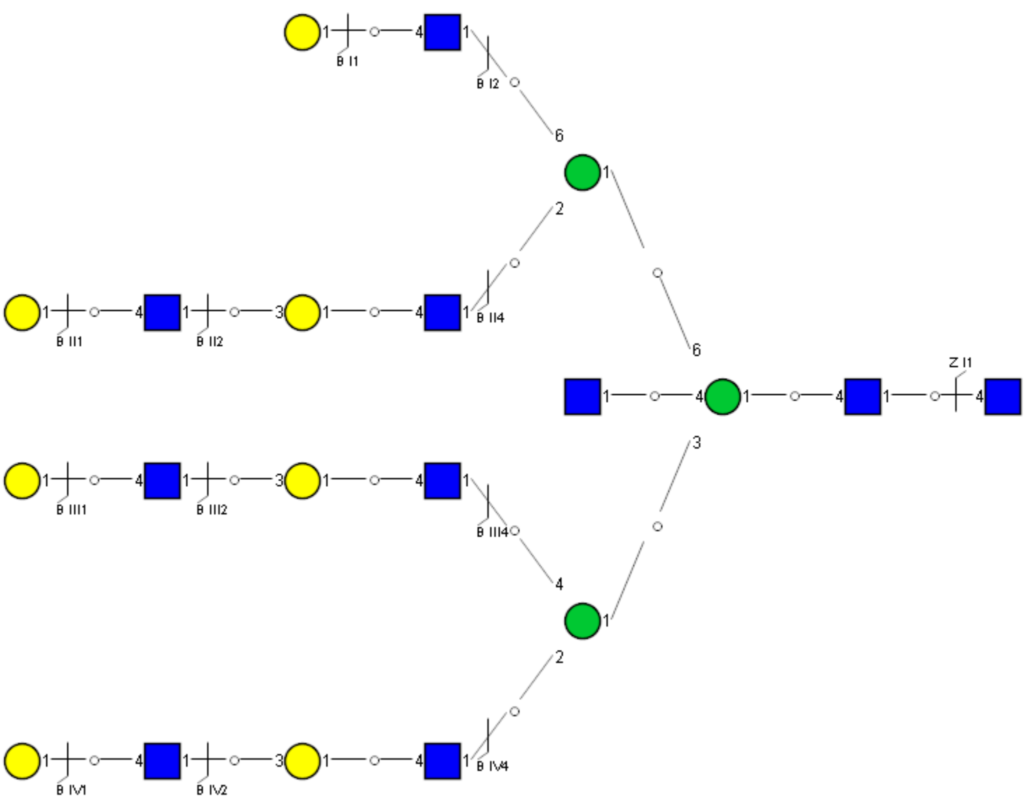

Spectrum Index: 60724(NoBHs:1)

Precursor iEF Map

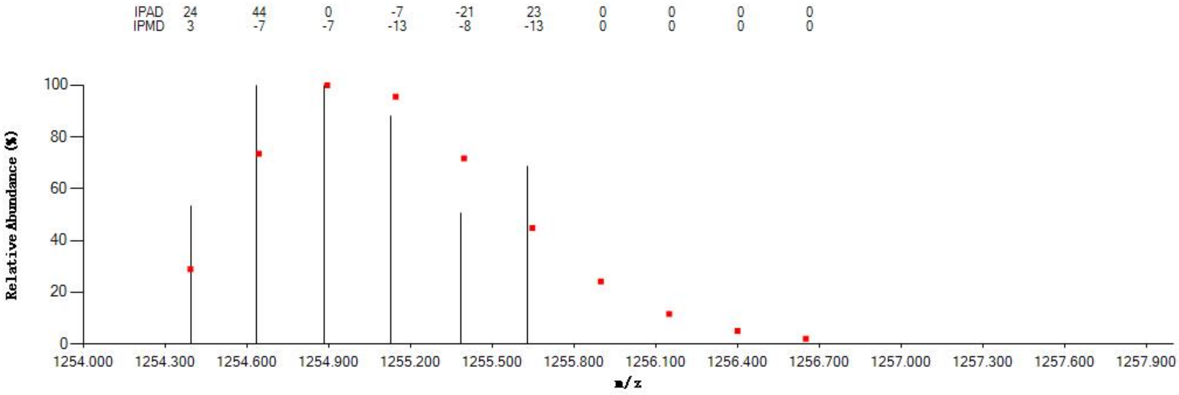

Annotated MS/MS Spectrum

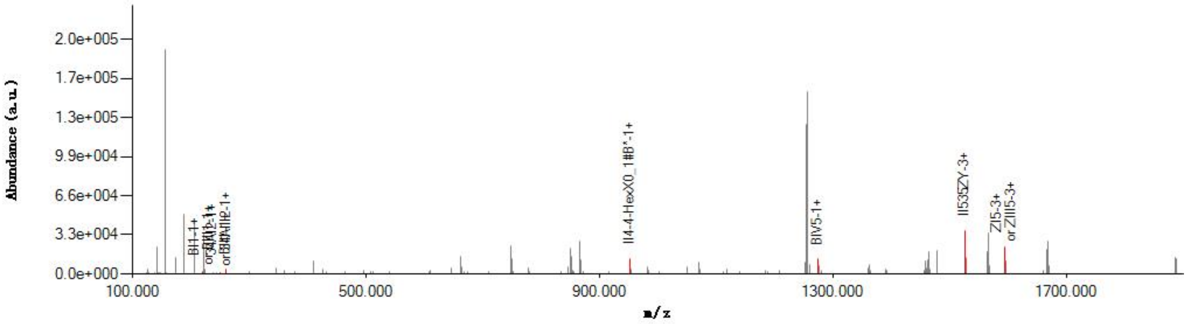

Graphical Fragmentation Map

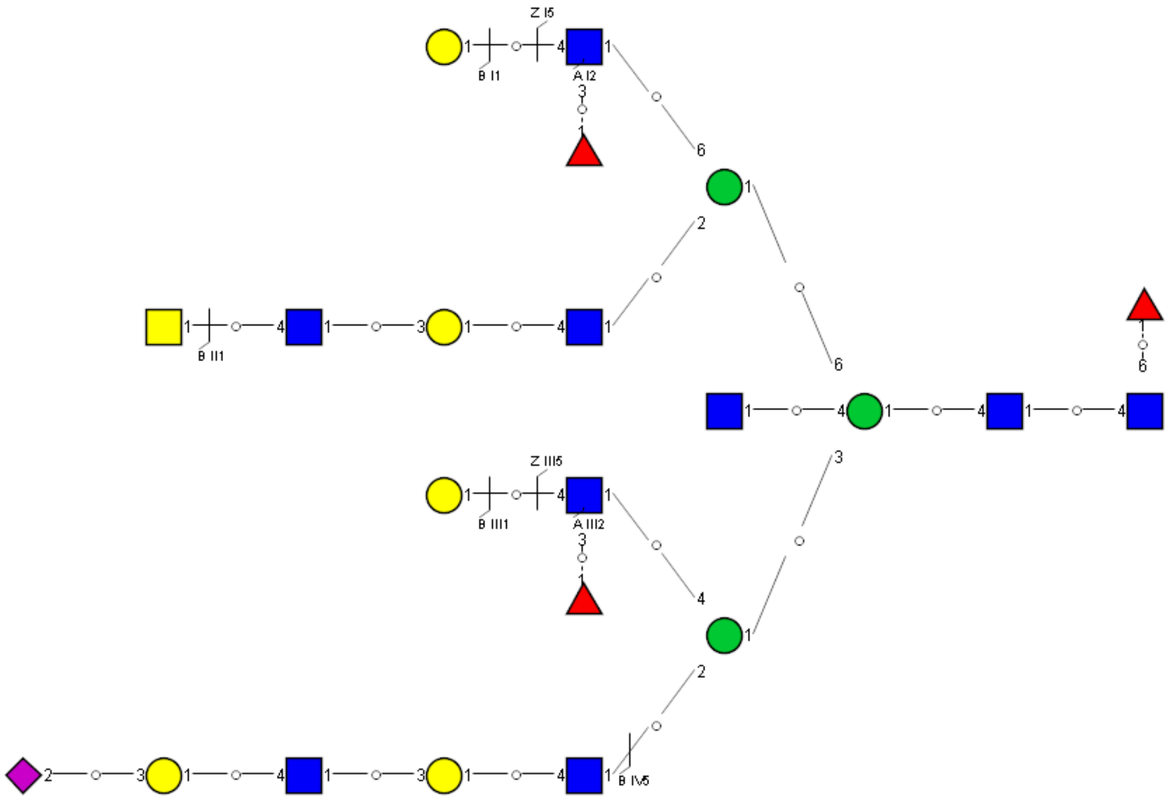

Spectrum Index: 30267(NoBHs:1)

#### Precursor iEF Map

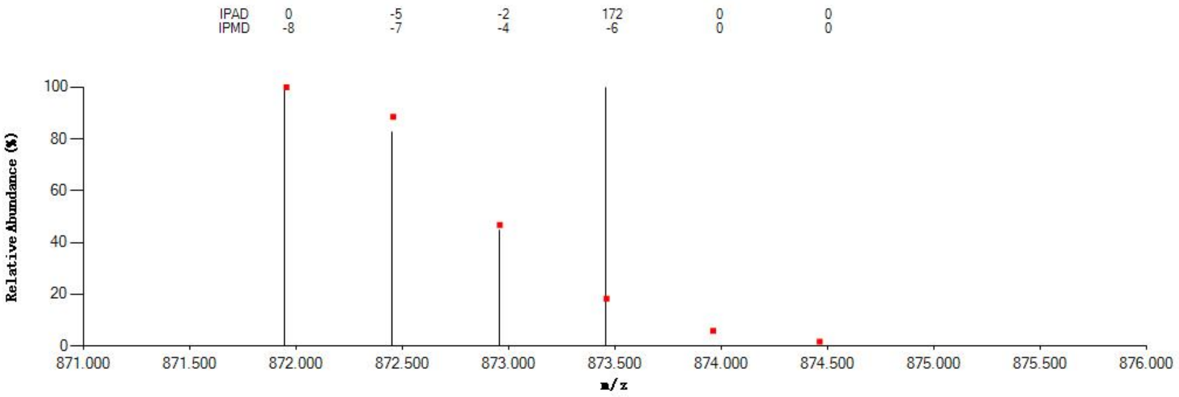


#### Annotated MS/MS Spectrum

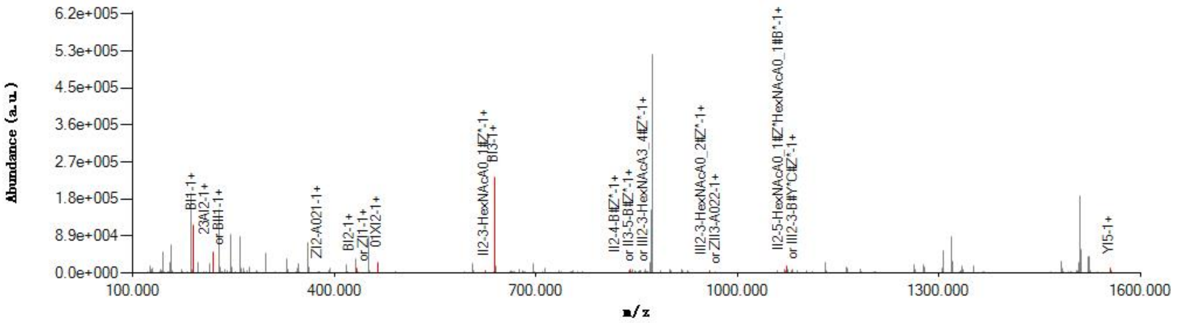


#### Graphical Fragmentation Map

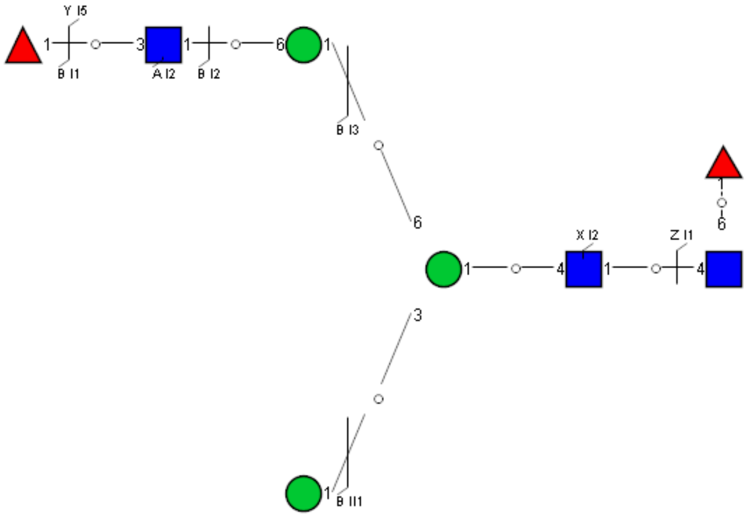

Spectrum Index: 31175(NoBHs:1)

#### Precursor iEF Map

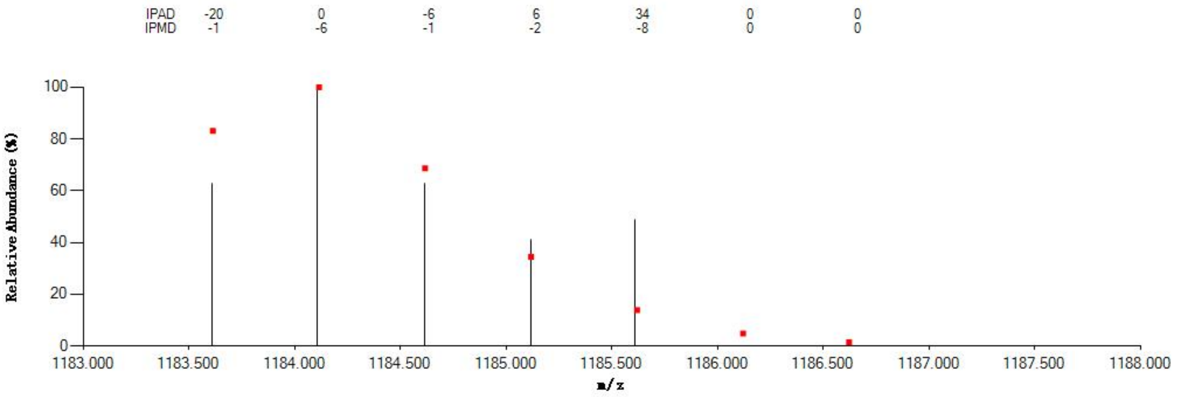


#### Annotated MS/MS Spectrum

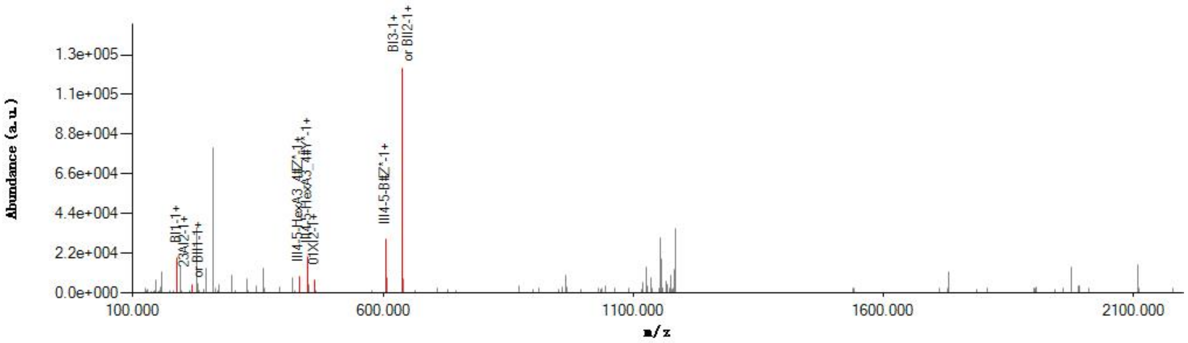


#### Graphical Fragmentation Map

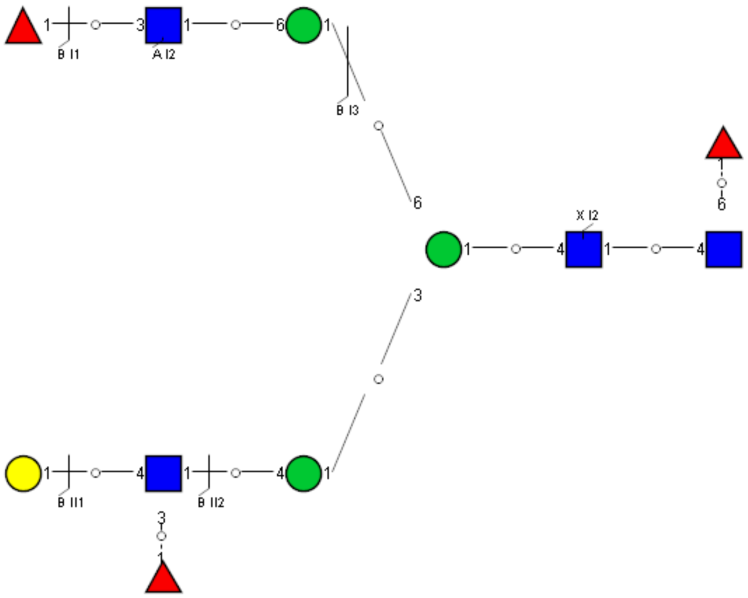

Spectrum Index: 31212(NoBHs:1)

Precursor iEF Map

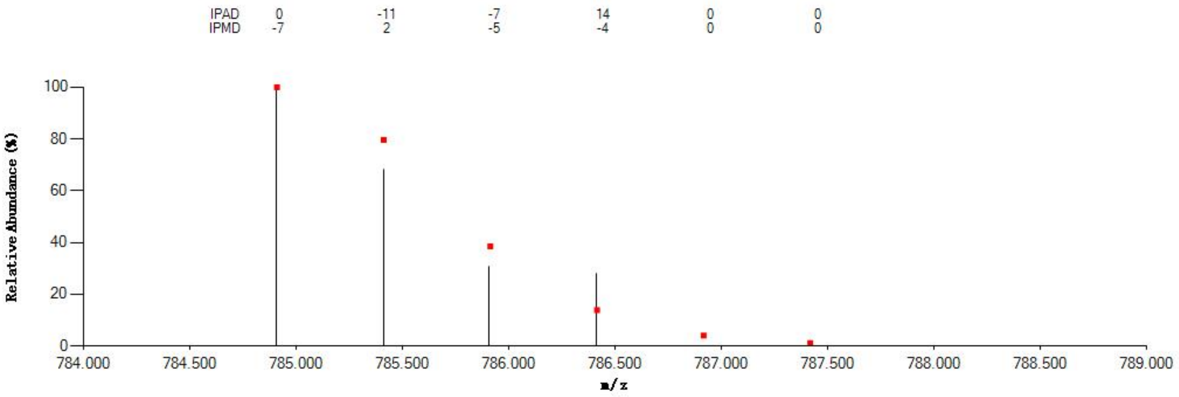


Annotated MS/MS Spectrum

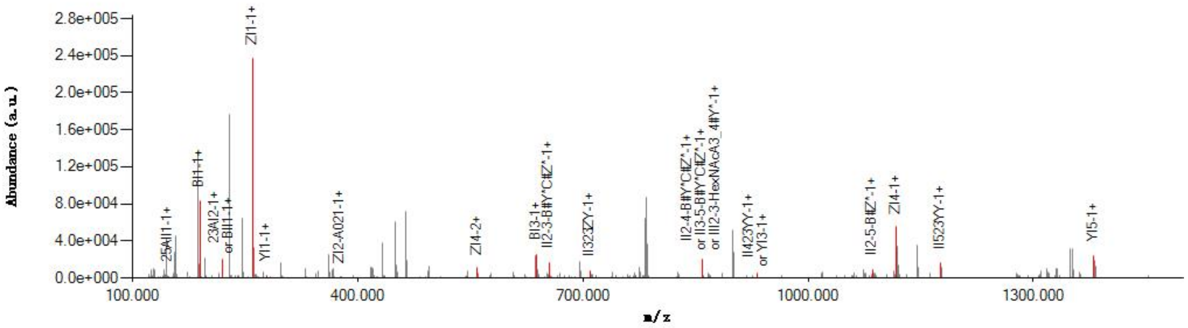


Graphical Fragmentation Map

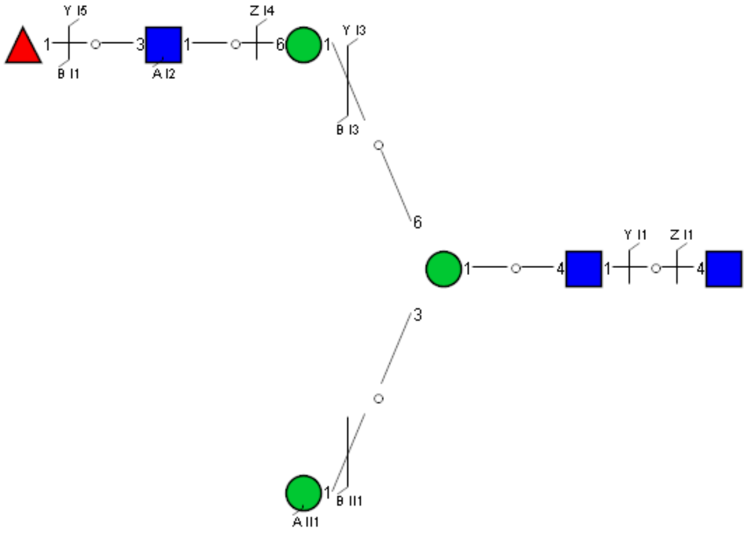

Spectrum Index: 32474(NoBHs:1)

#### Precursor iEF Map

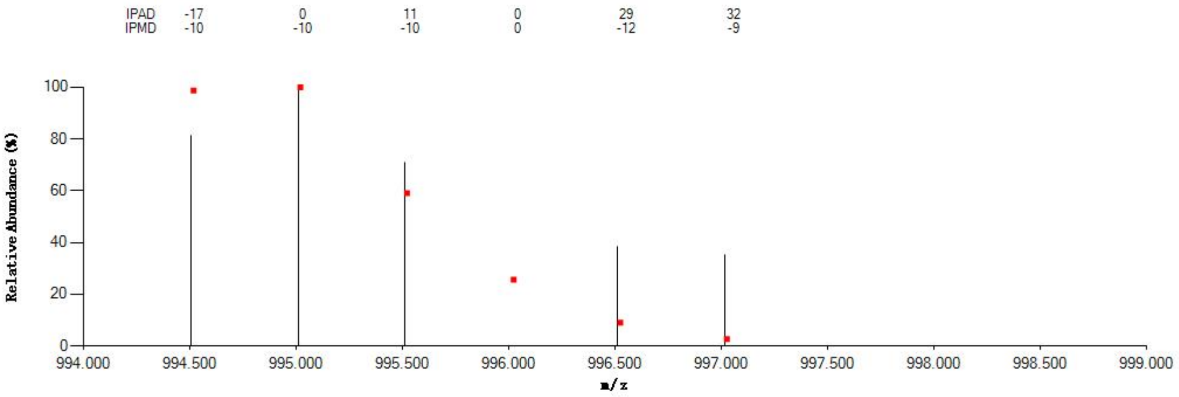


#### Annotated MS/MS Spectrum

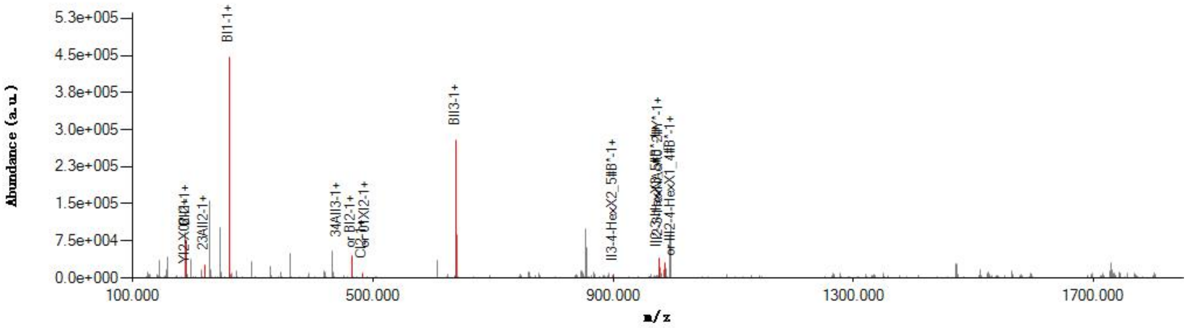


#### Graphical Fragmentation Map

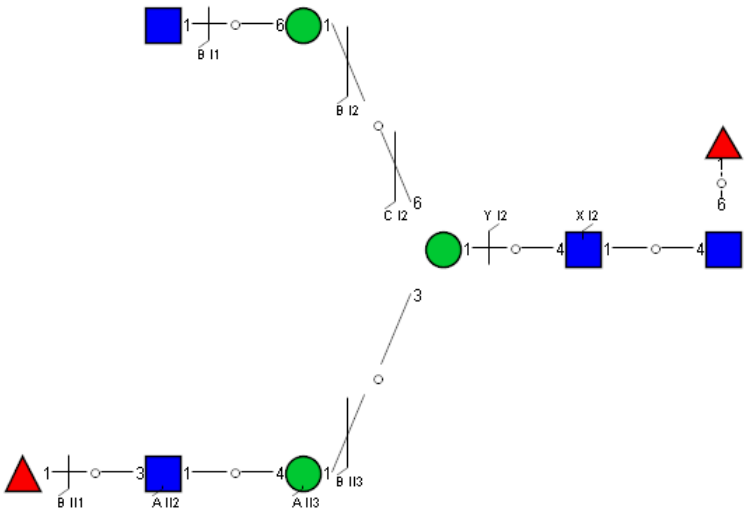

Spectrum Index: 32843(NoBHs:1)

#### Precursor iEF Map

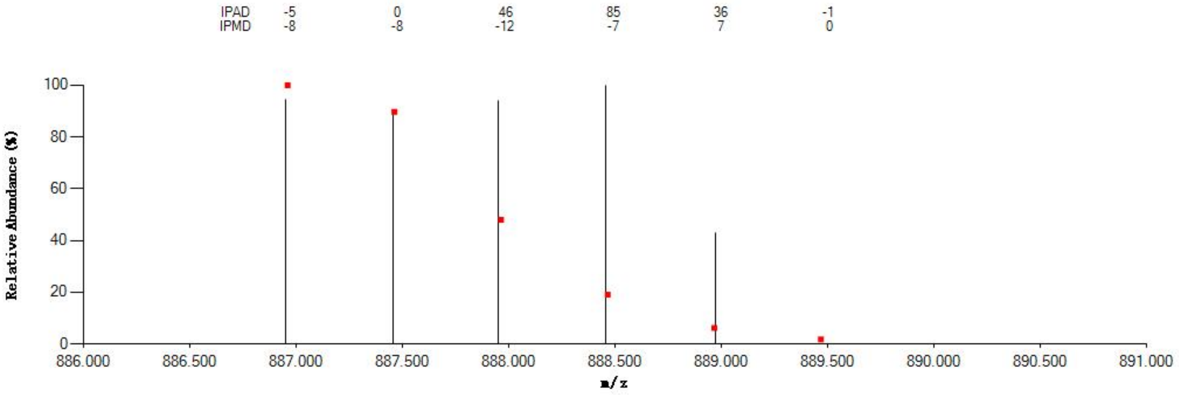


#### Annotated MS/MS Spectrum

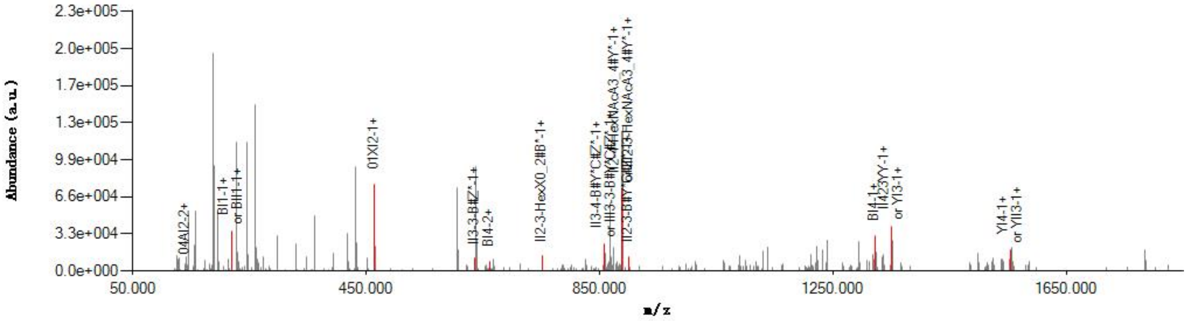


#### Graphical Fragmentation Map

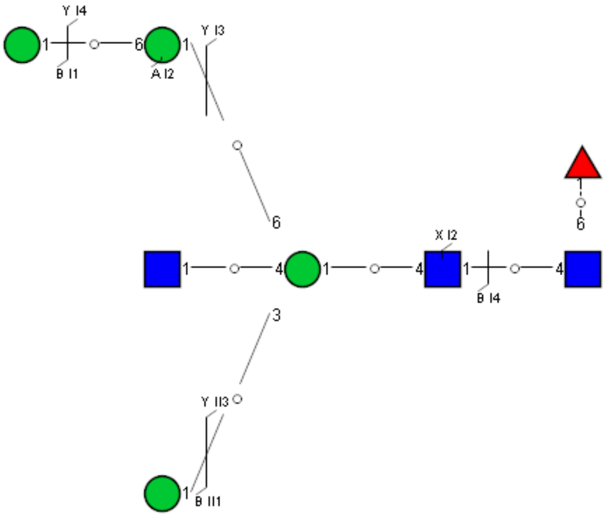

Spectrum Index: 33661(NoBHs:1)

#### Precursor iEF Map

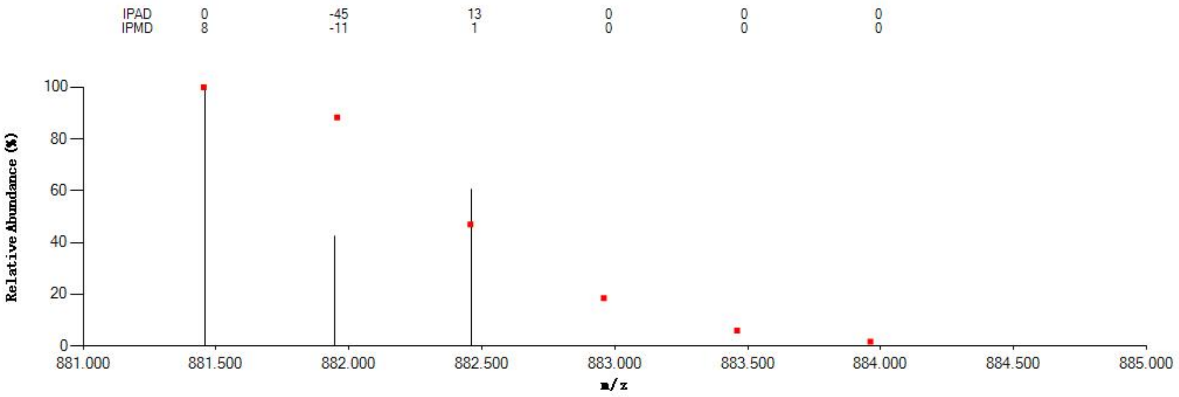


#### Annotated MS/MS Spectrum

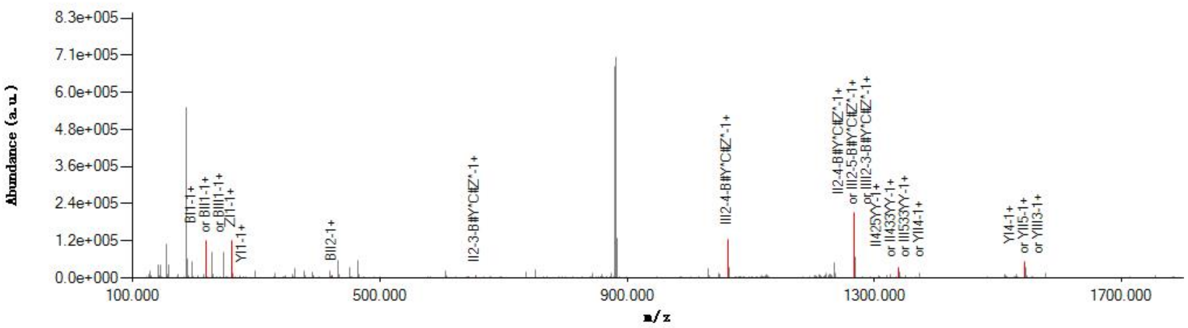


#### Graphical Fragmentation Map

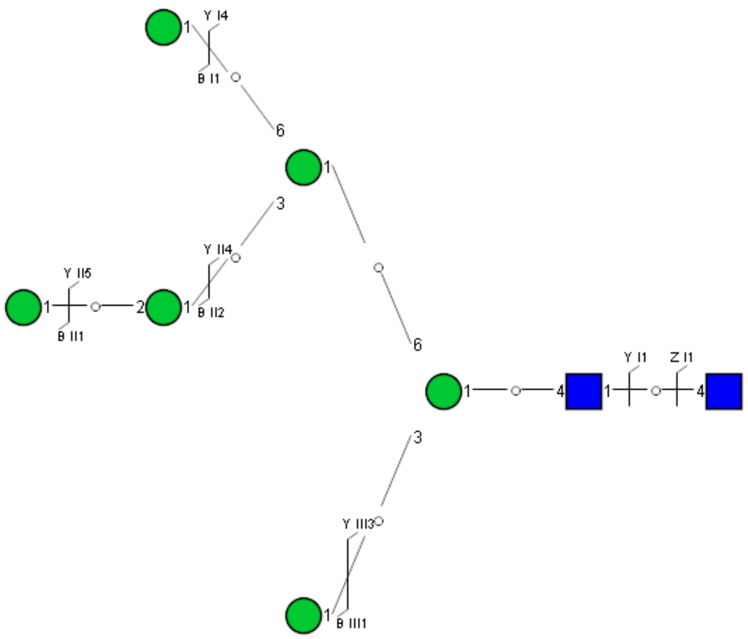

Spectrum Index: 34278(NoBHs:1)

#### Precursor iEF Map

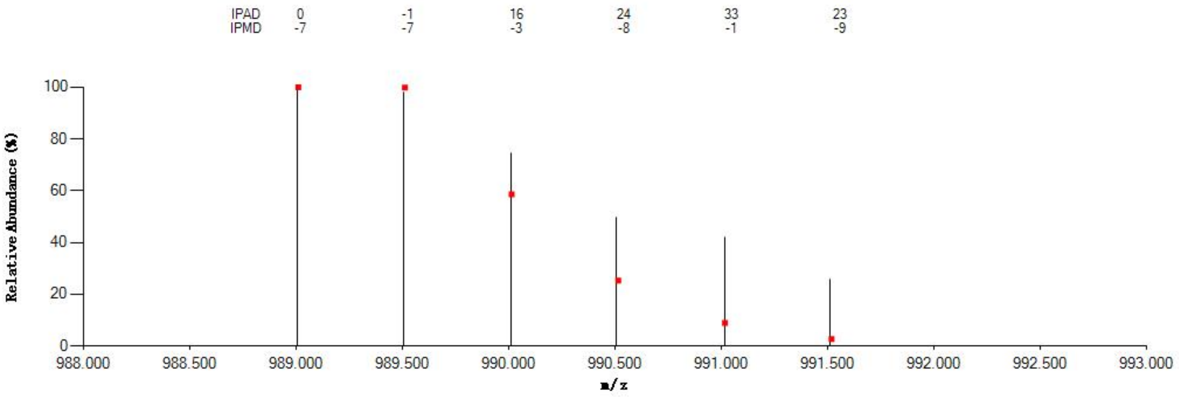


#### Annotated MS/MS Spectrum

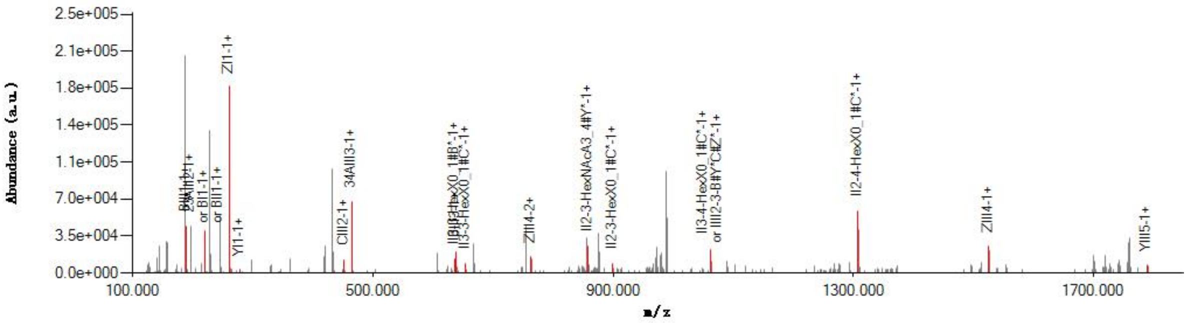


#### Graphical Fragmentation Map

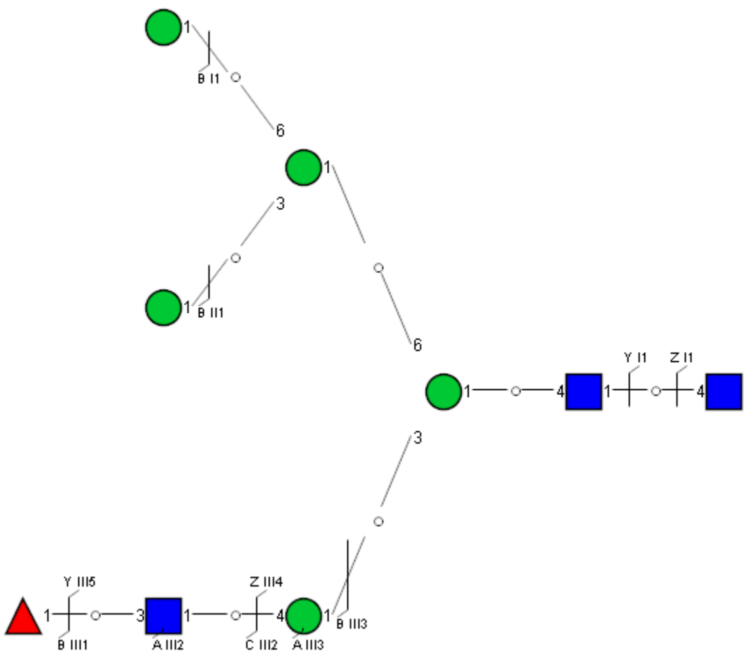

Spectrum Index: 34659(NoBHs:1)

#### Precursor iEF Map

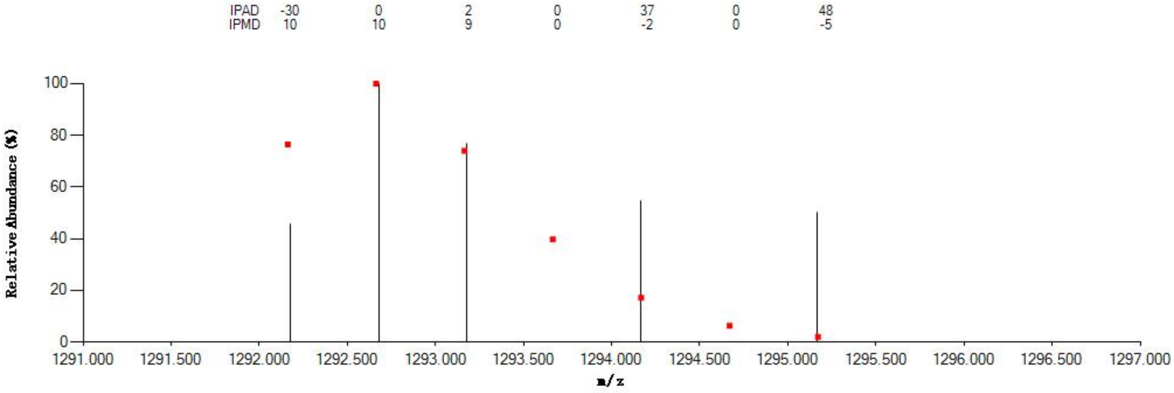


#### Annotated MS/MS Spectrum

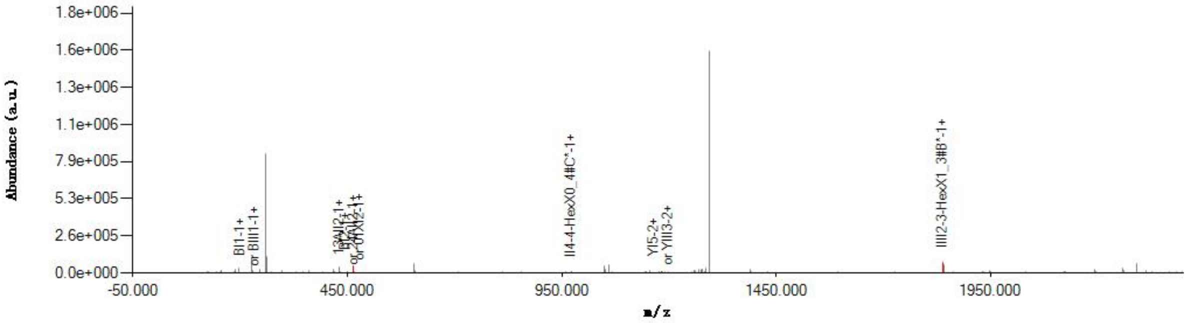


#### Graphical Fragmentation Map

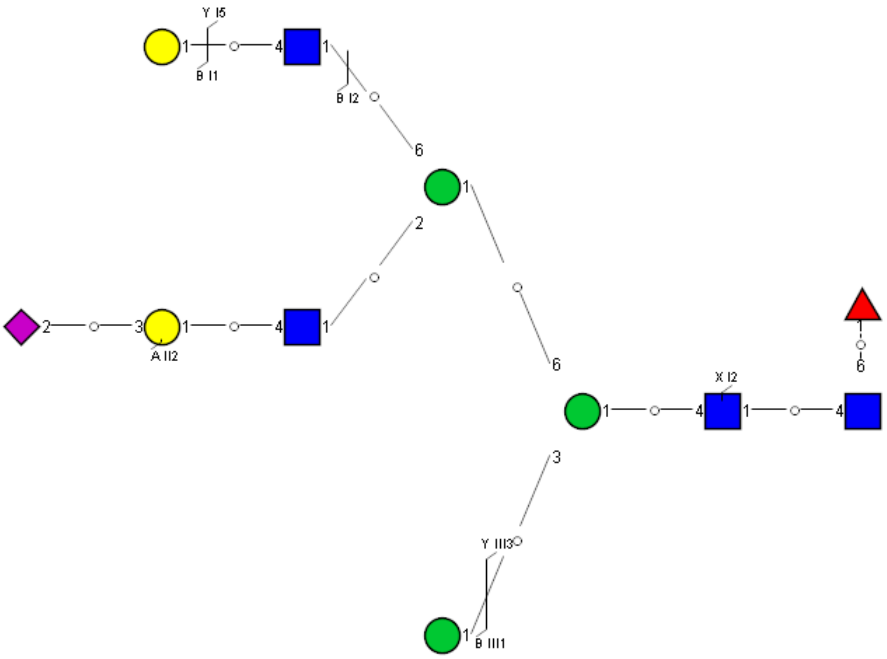

Spectrum Index: 35685(NoBHs:2)

#### Precursor iEF Map

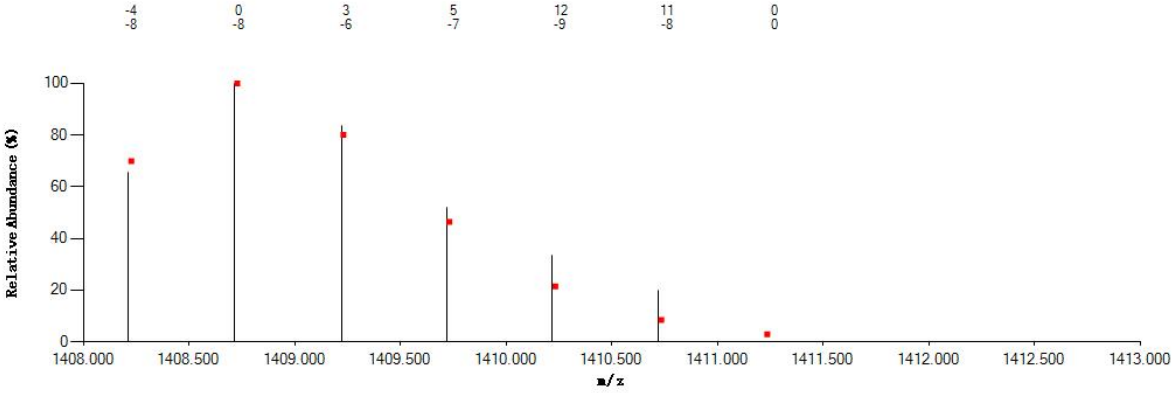

#### Annotated MS/MS Spectrum

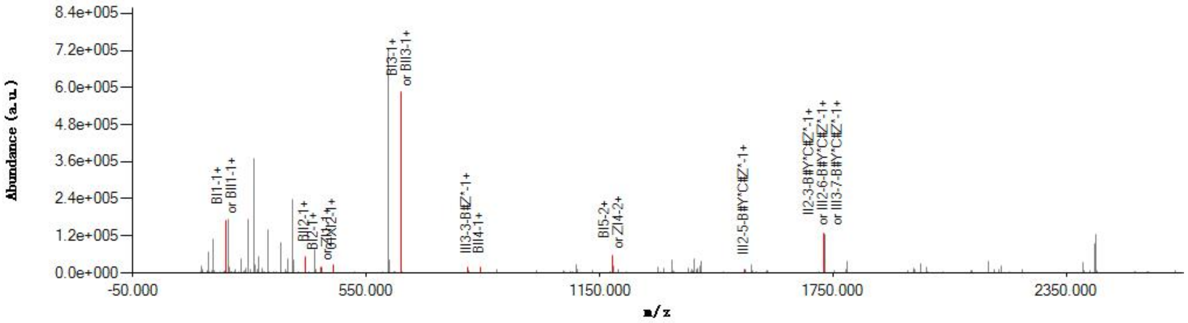

#### Graphical Fragmentation Map

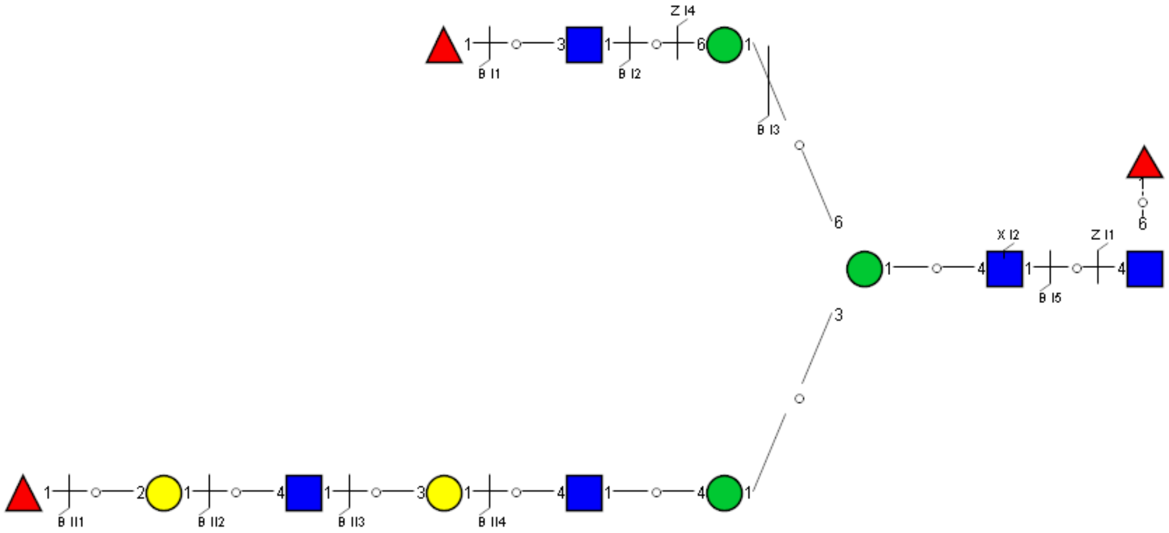

Spectrum Index: 35685(NoBHs:2)

#### Precursor iEF Map

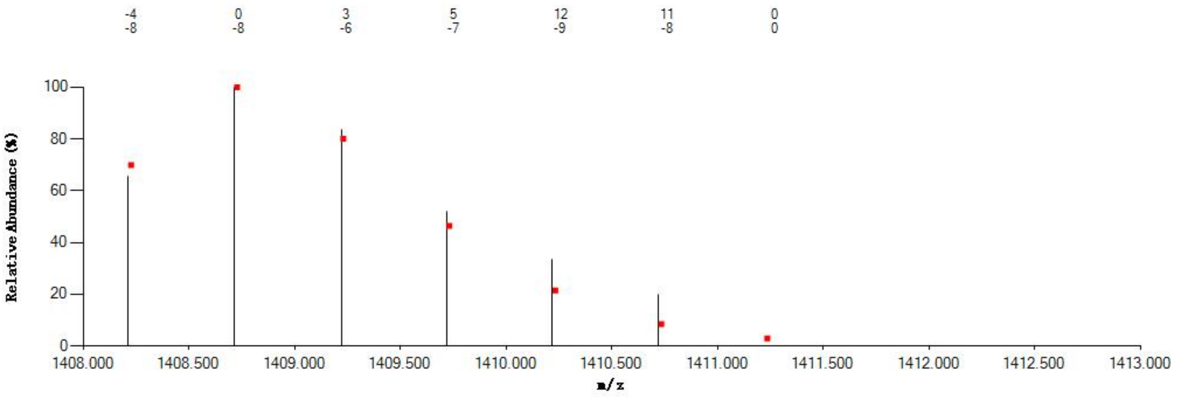


#### Annotated MS/MS Spectrum

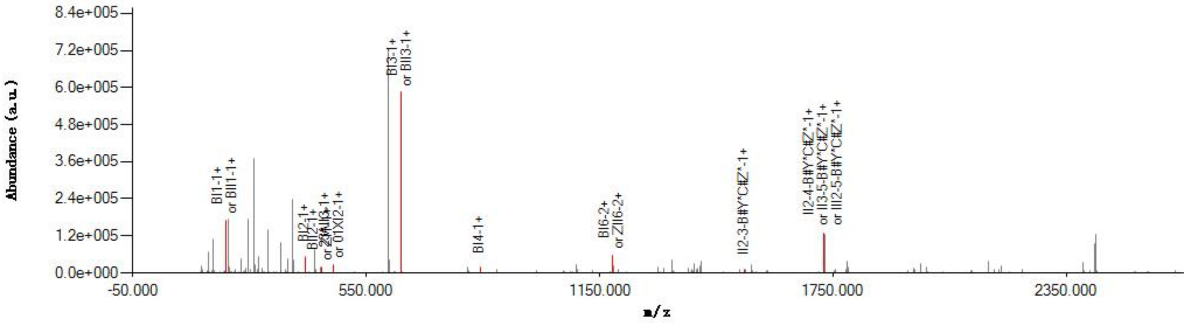


#### Graphical Fragmentation Map

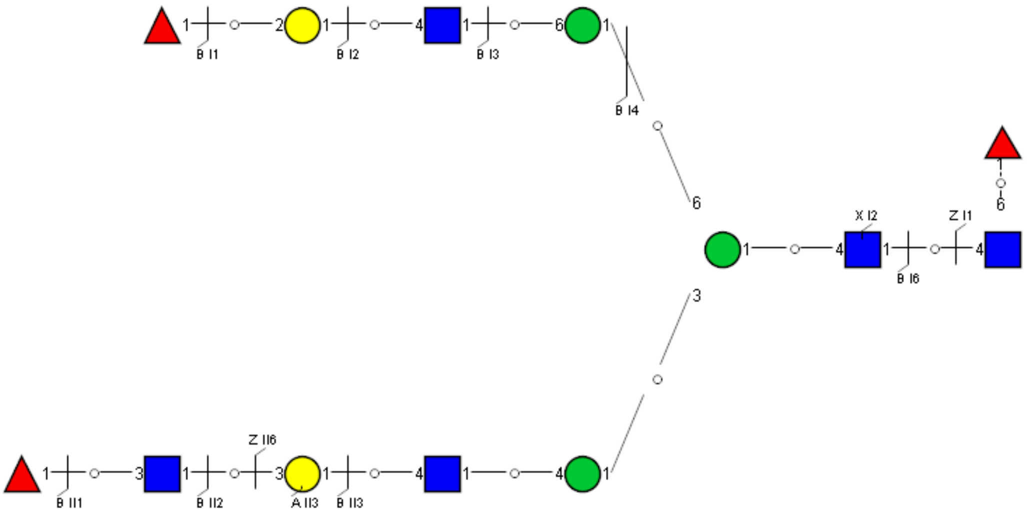

Spectrum Index: 35888(NoBHs:1)

#### Precursor iEF Map

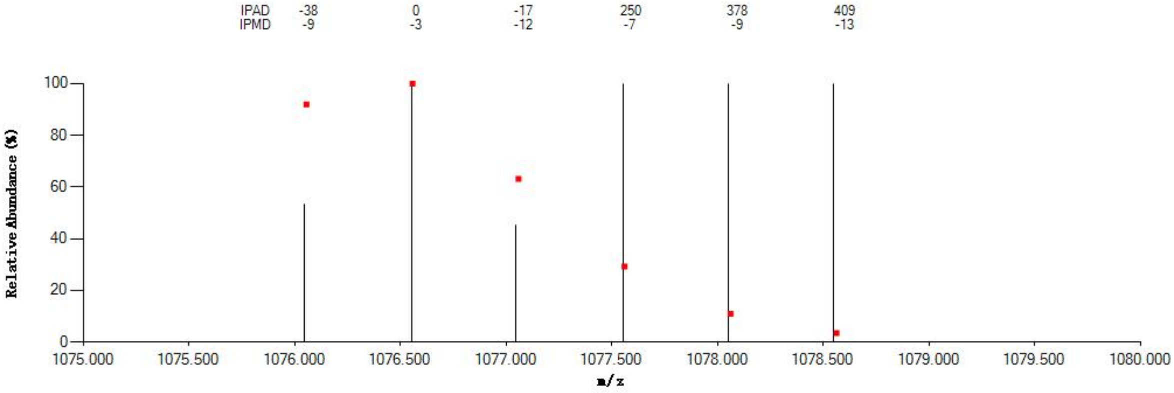


#### Annotated MS/MS Spectrum

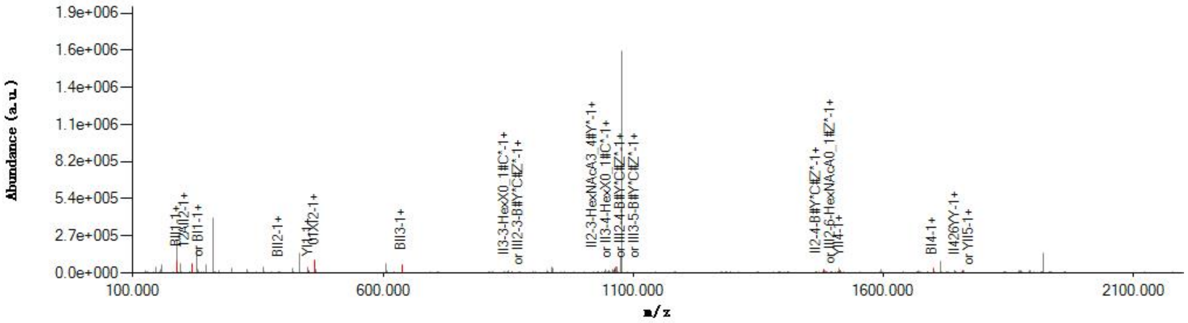


#### Graphical Fragmentation Map

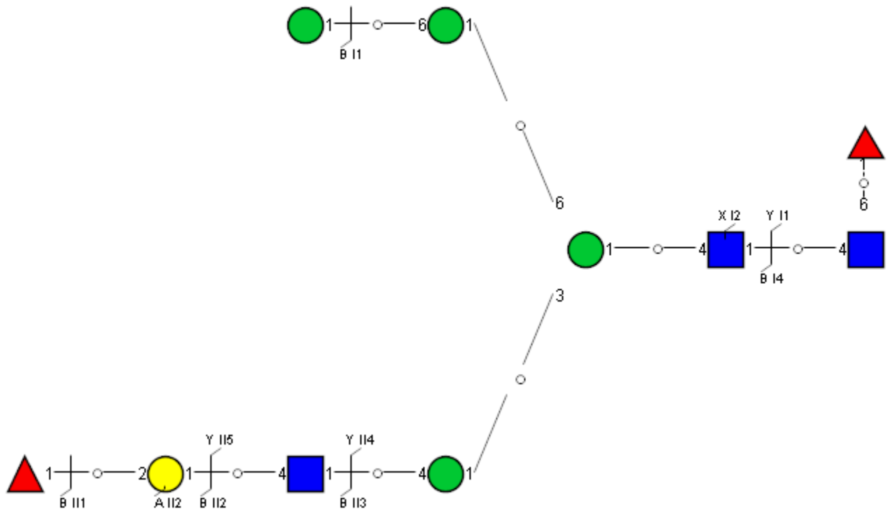

Spectrum Index: 36143(NoBHs:1)

#### Precursor iEF Map

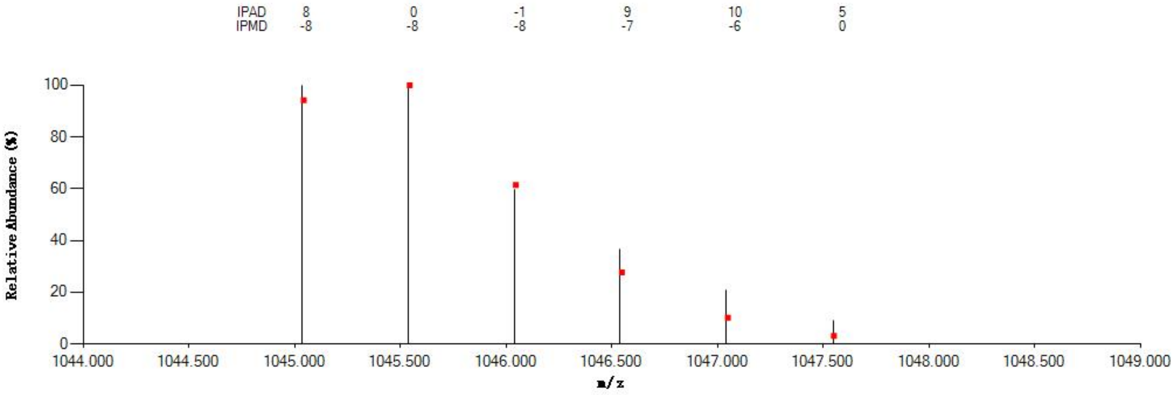


#### Annotated MS/MS Spectrum

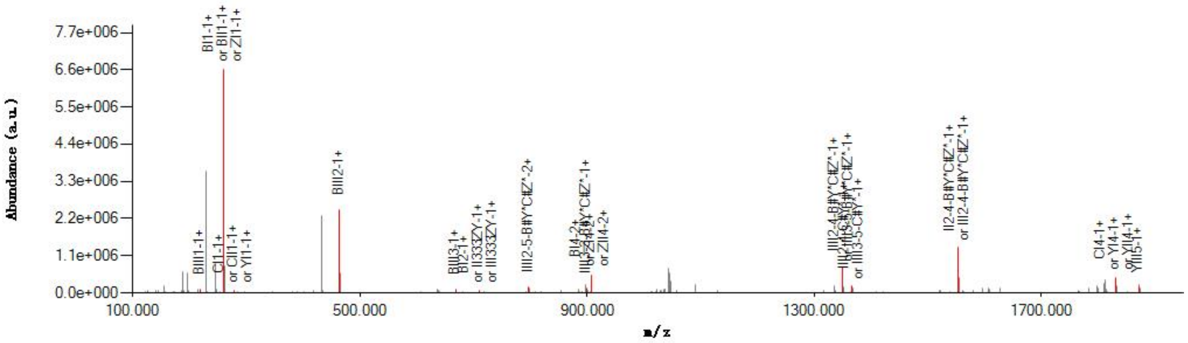


#### Graphical Fragmentation Map

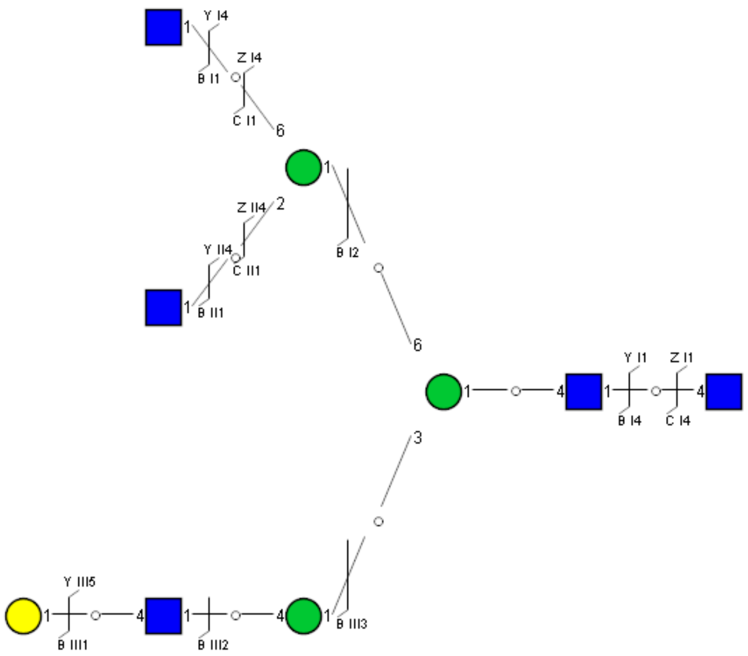

Spectrum Index: 36519(NoBHs:1)

#### Precursor iEF Map

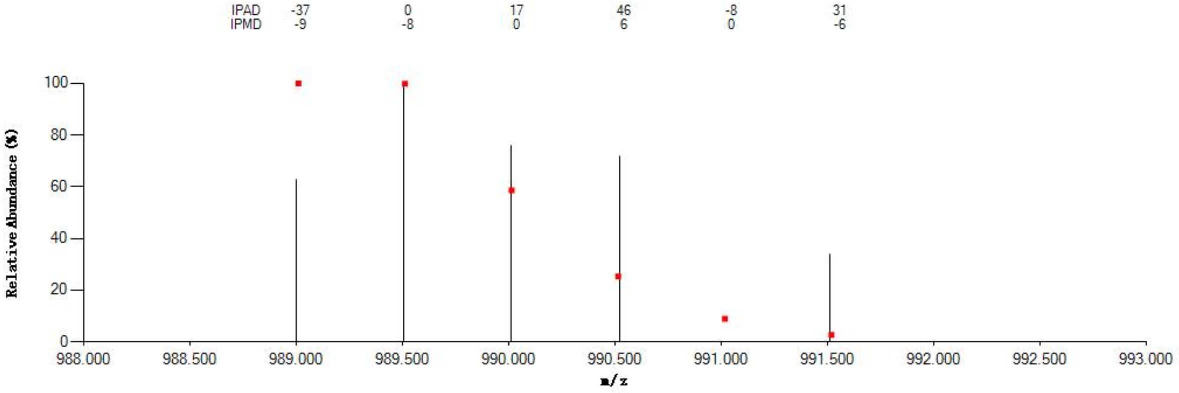


#### Annotated MS/MS Spectrum

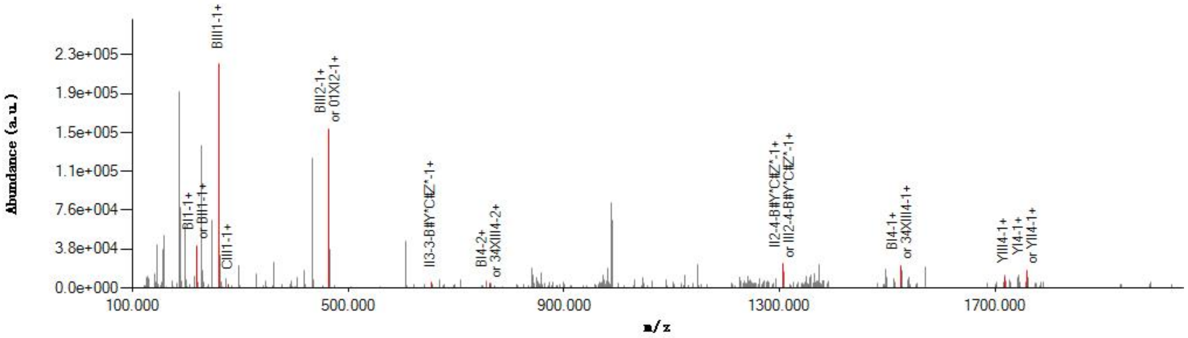


#### Graphical Fragmentation Map

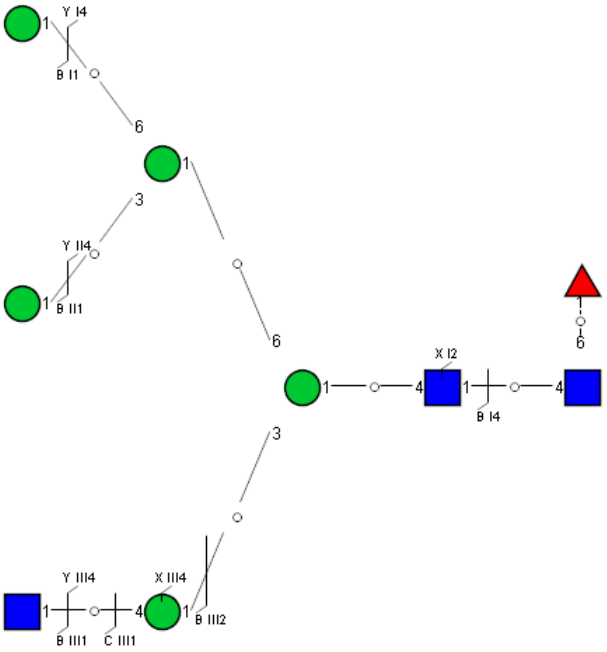

Spectrum Index: 36523(NoBHs:1)

#### Precursor iEF Map

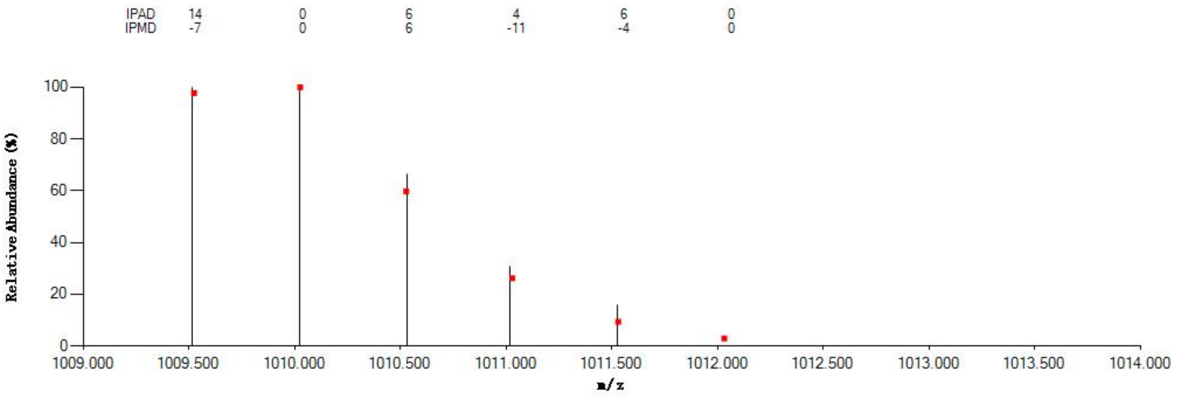


#### Annotated MS/MS Spectrum

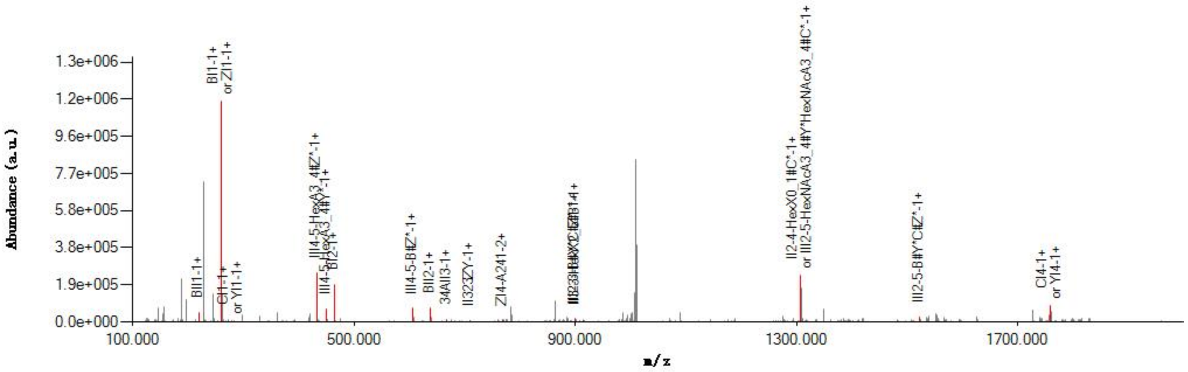


#### Graphical Fragmentation Map

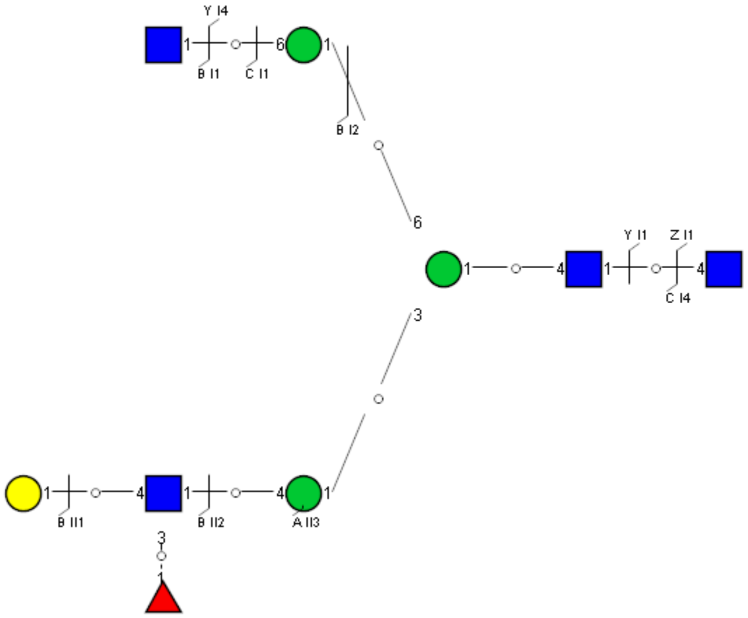

### Spectrum Index: 37361(NoBHs:1)

#### Precursor iEF Map

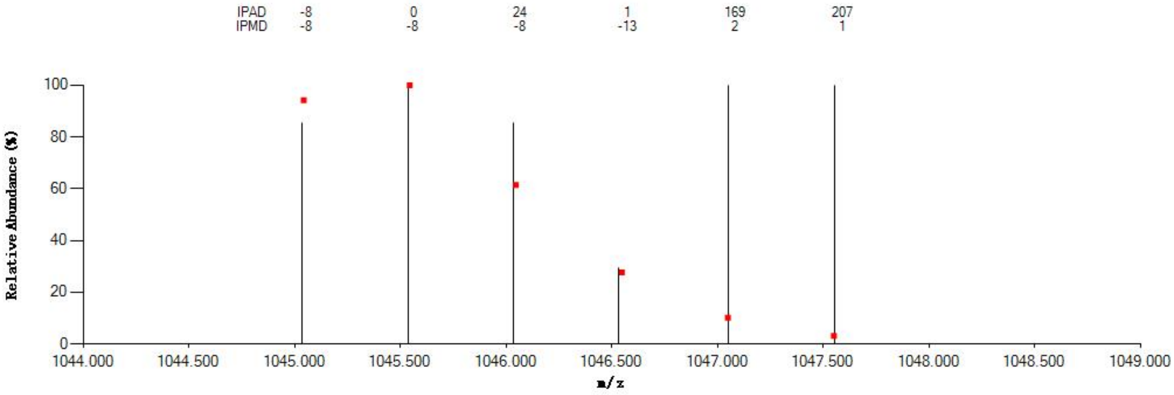


#### Annotated MS/MS Spectrum

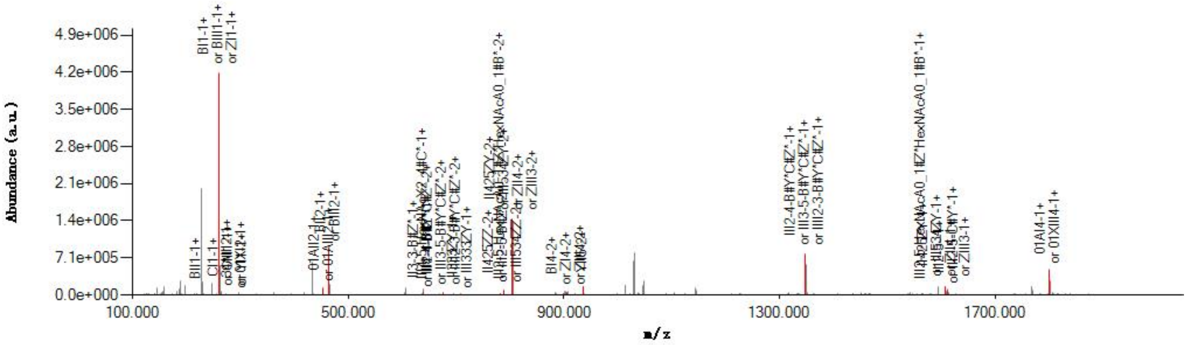


#### Graphical Fragmentation Map

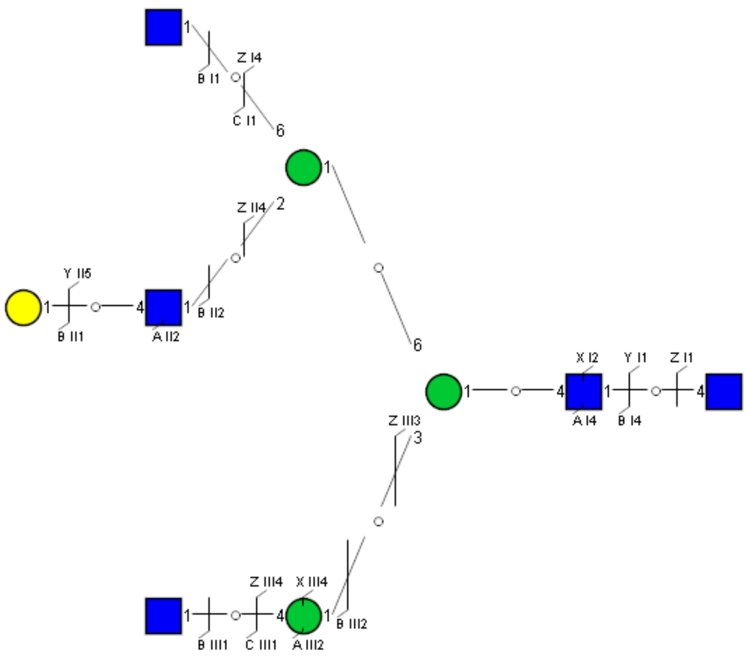

Spectrum Index: 37817(NoBHs:1)

#### Precursor iEF Map

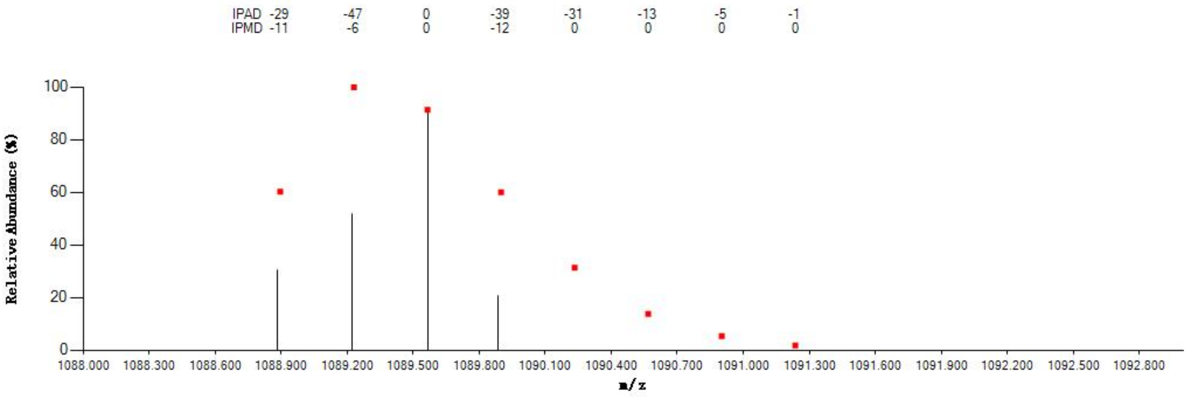


#### Annotated MS/MS Spectrum

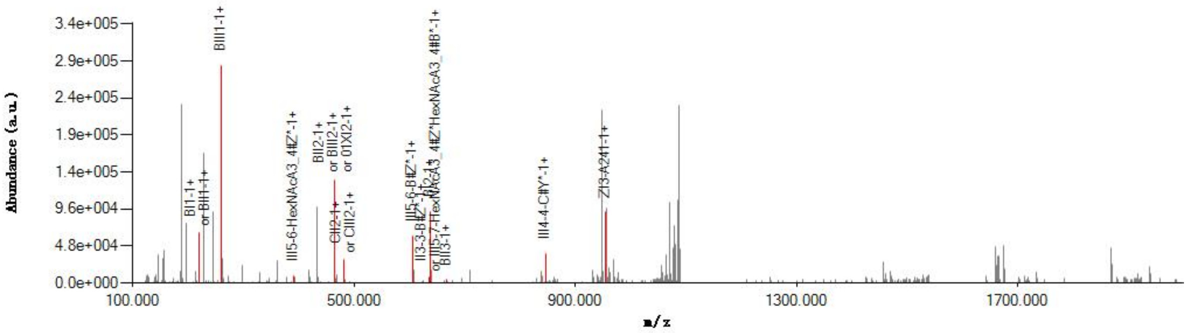


#### Graphical Fragmentation Map

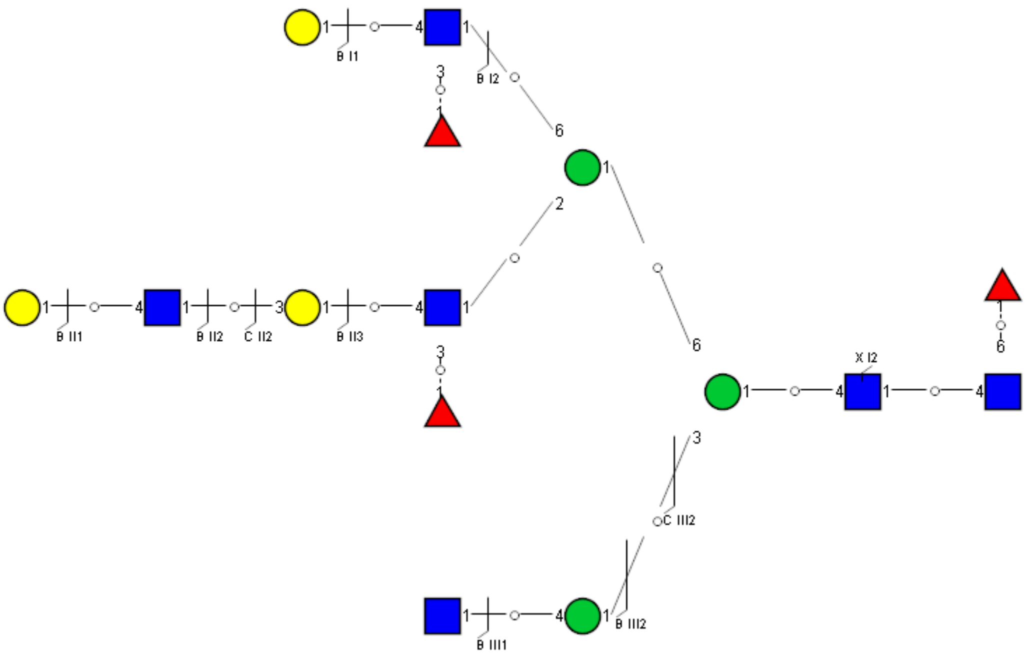

Spectrum Index: 37960(NoBHs:1)

#### Precursor iEF Map

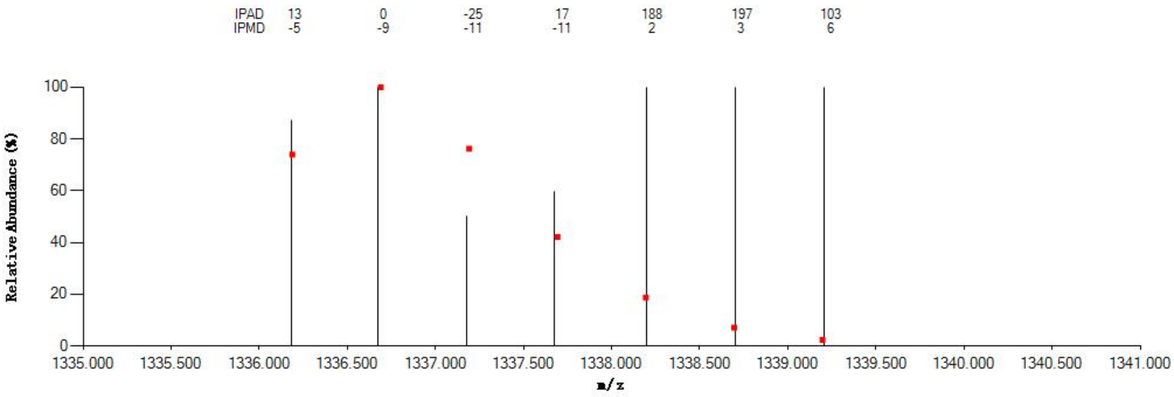


#### Annotated MS/MS Spectrum

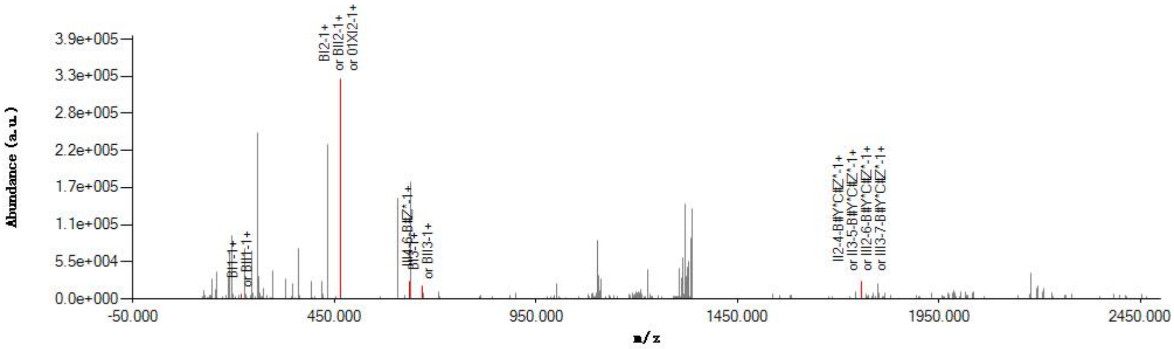


#### Graphical Fragmentation Map

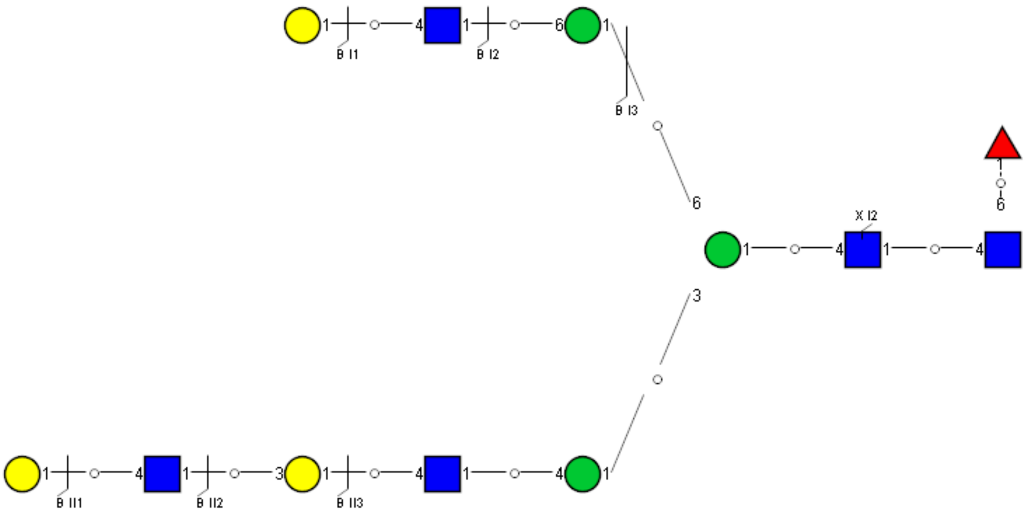

Spectrum Index: 38087(NoBHs:1)

#### Precursor iEF Map

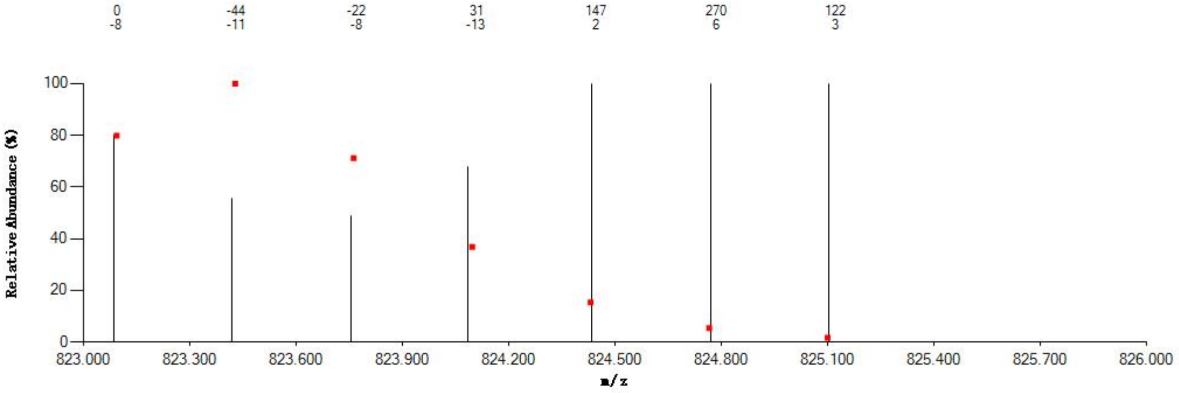

#### Annotated MS/MS Spectrum

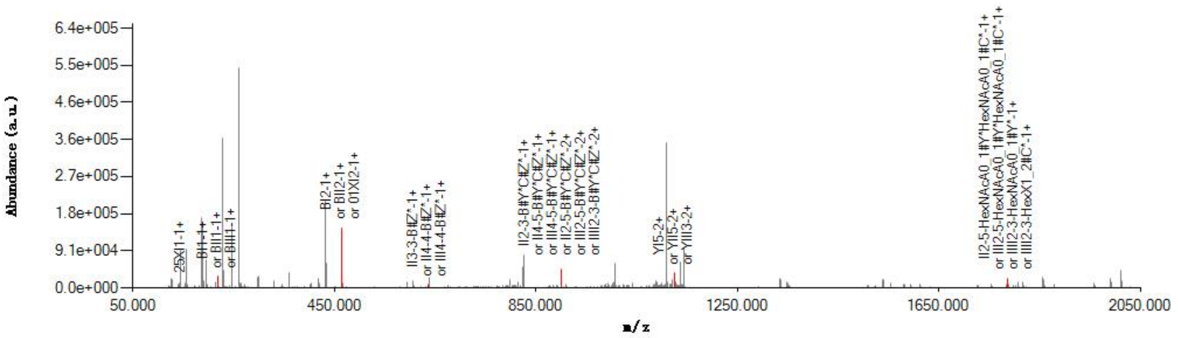

#### Graphical Fragmentation Map

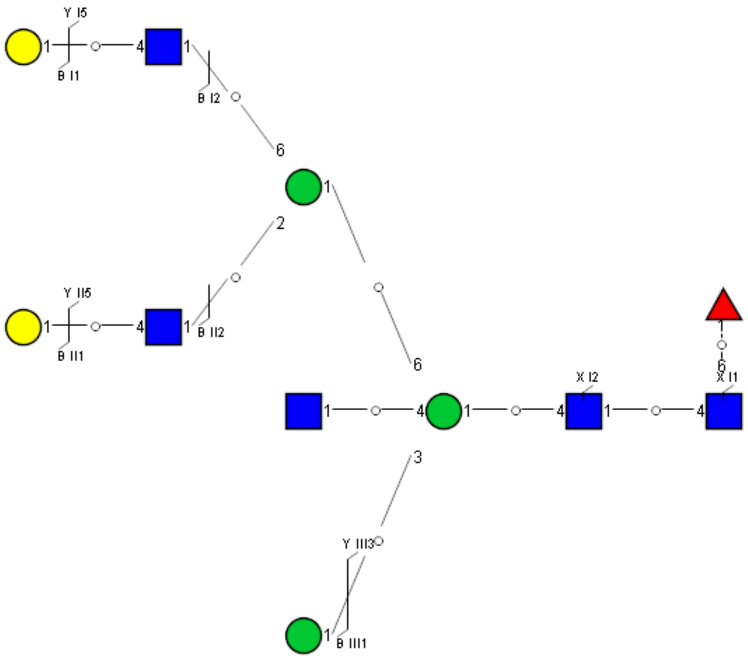

Spectrum Index: 38650(NoBHs:1)

#### Precursor iEF Map

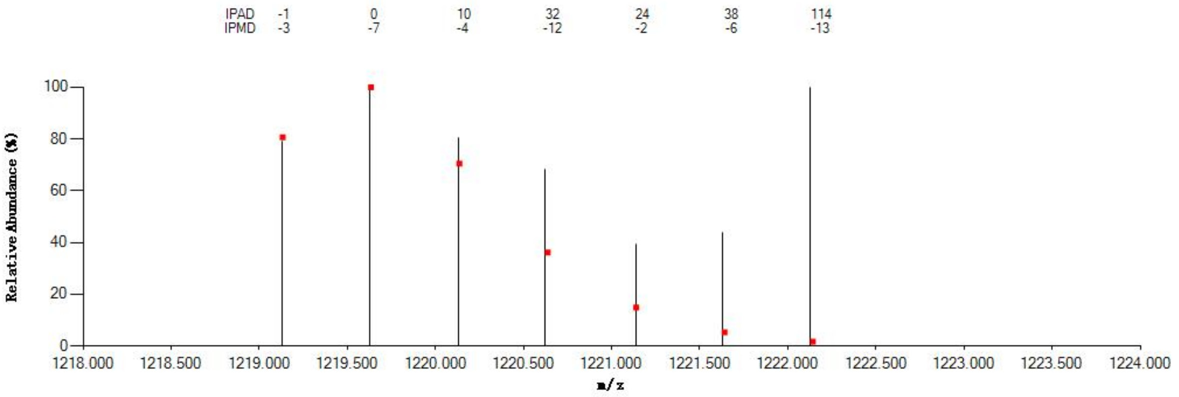


#### Annotated MS/MS Spectrum

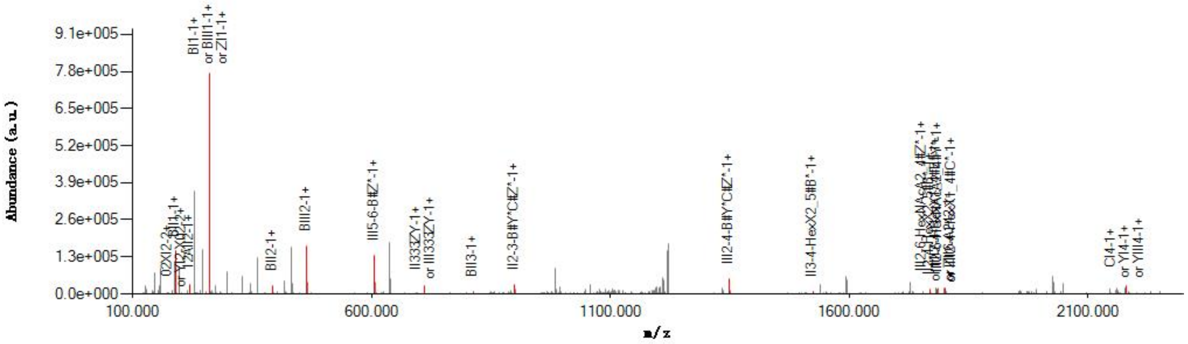


#### Graphical Fragmentation Map

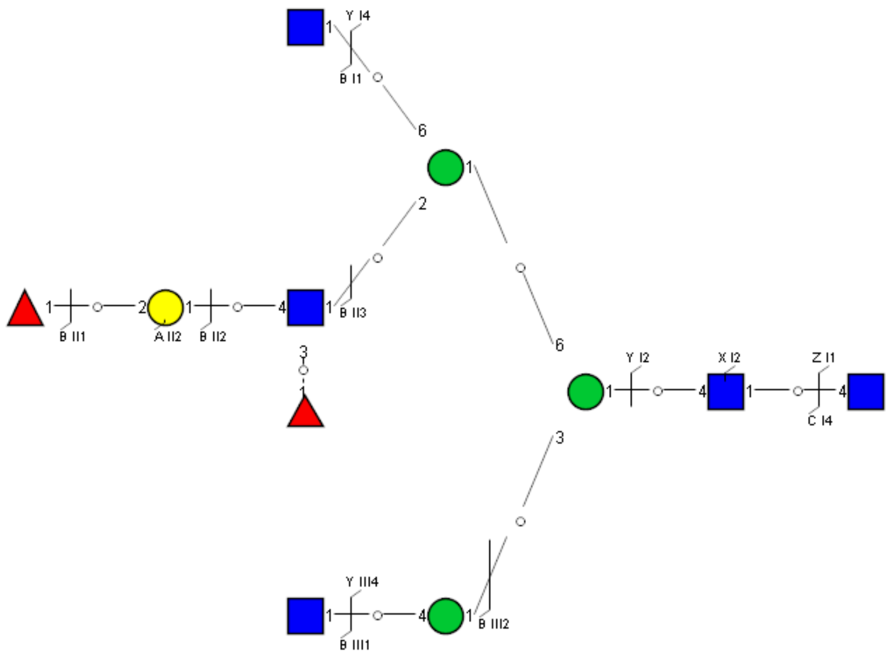

Spectrum Index: 38749(NoBHs:1)

#### Precursor iEF Map

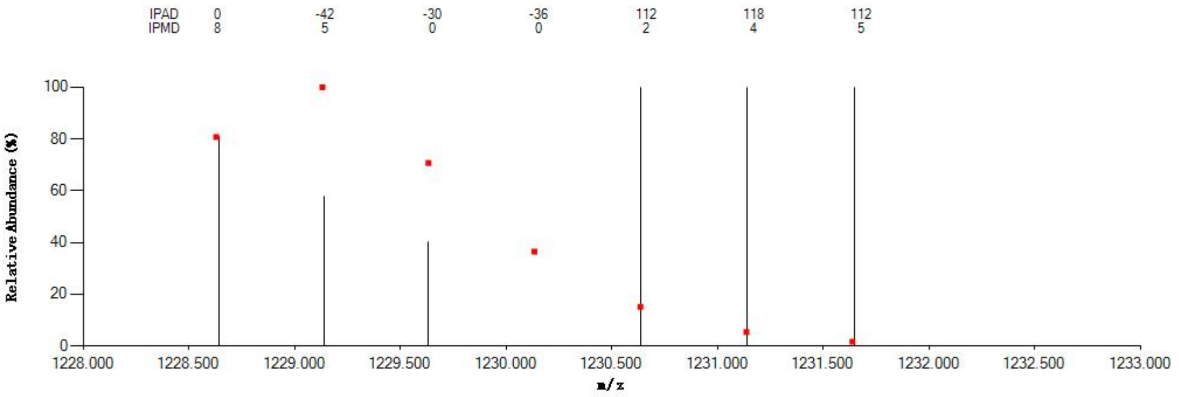


#### Annotated MS/MS Spectrum

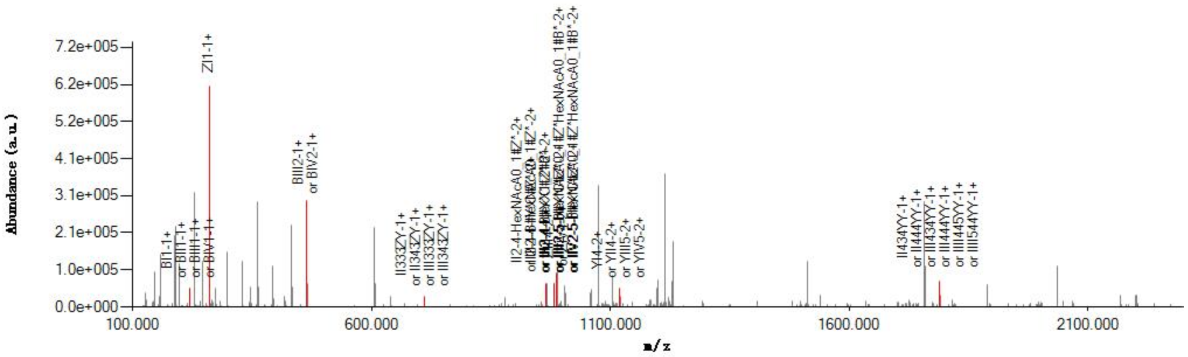


#### Graphical Fragmentation Map

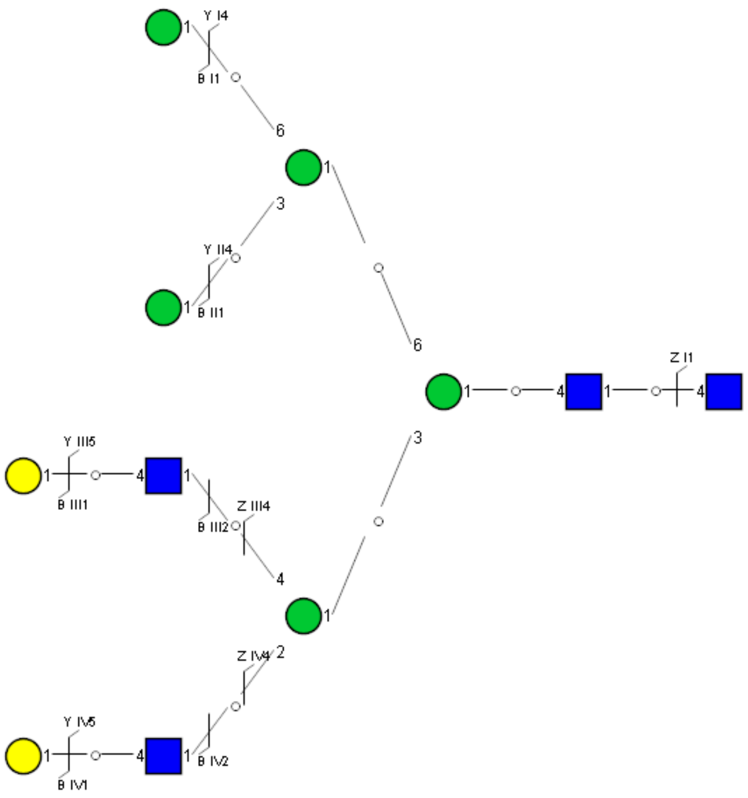

Spectrum Index: 38867(NoBHs:3)

#### Precursor iEF Map

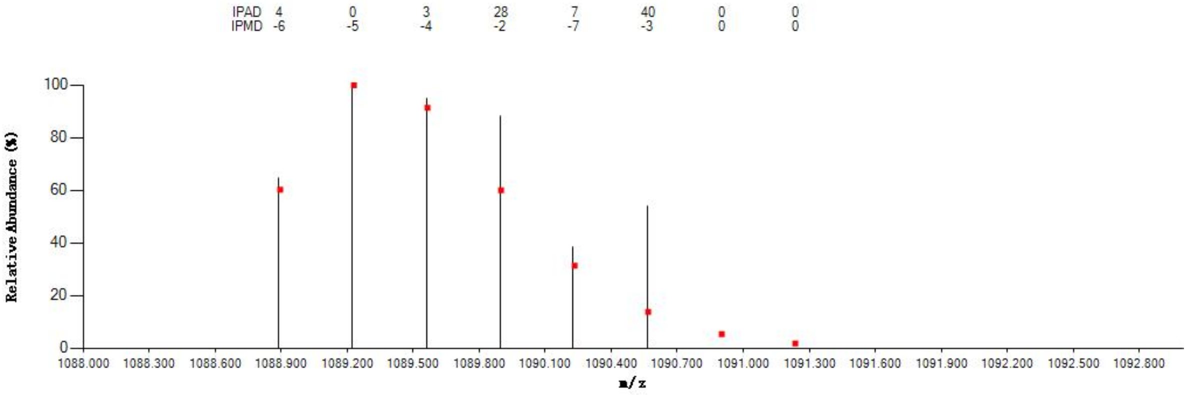


#### Annotated MS/MS Spectrum

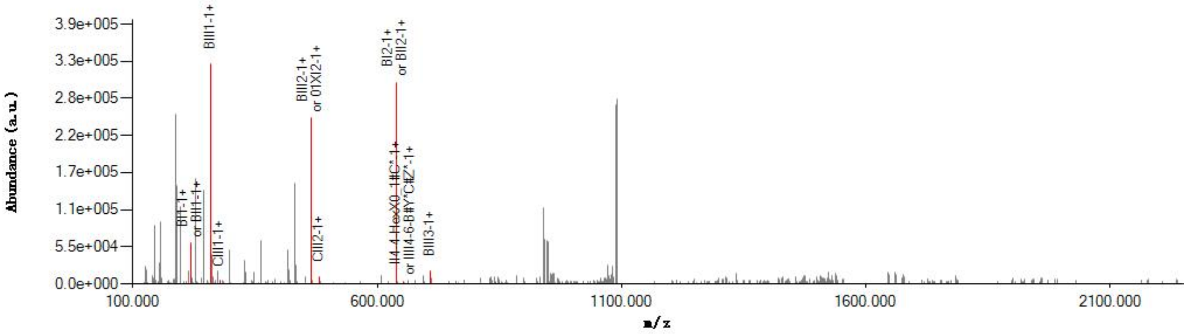


#### Graphical Fragmentation Map

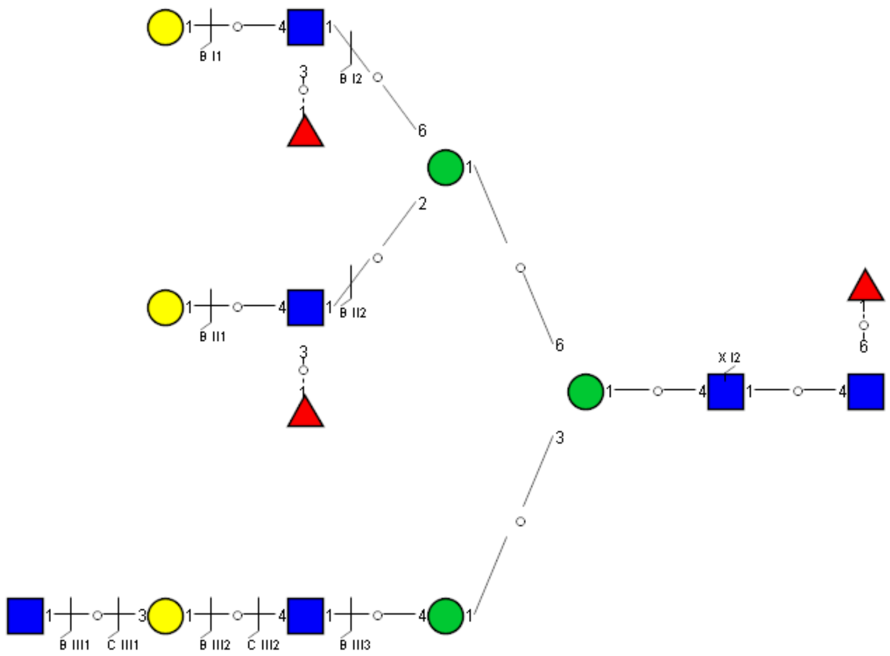

Spectrum Index: 38867(NoBHs:3)

#### Precursor iEF Map

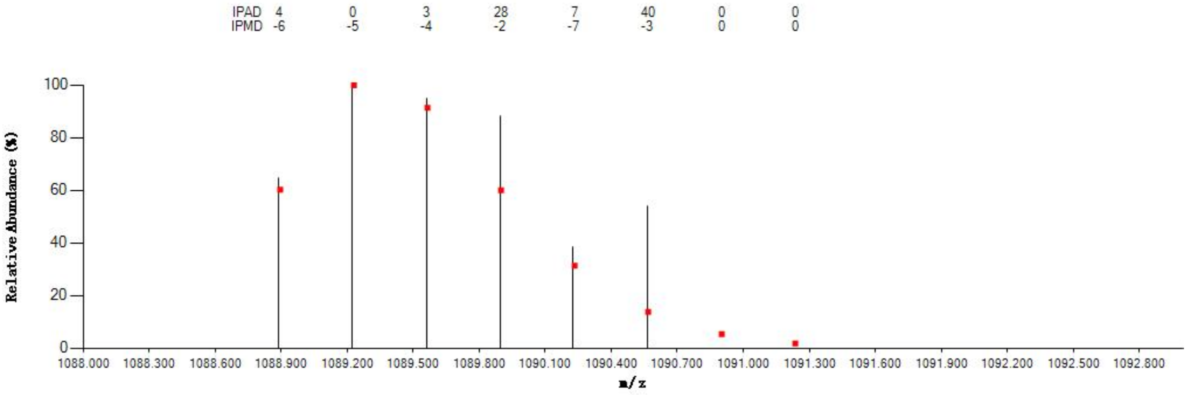


#### Annotated MS/MS Spectrum

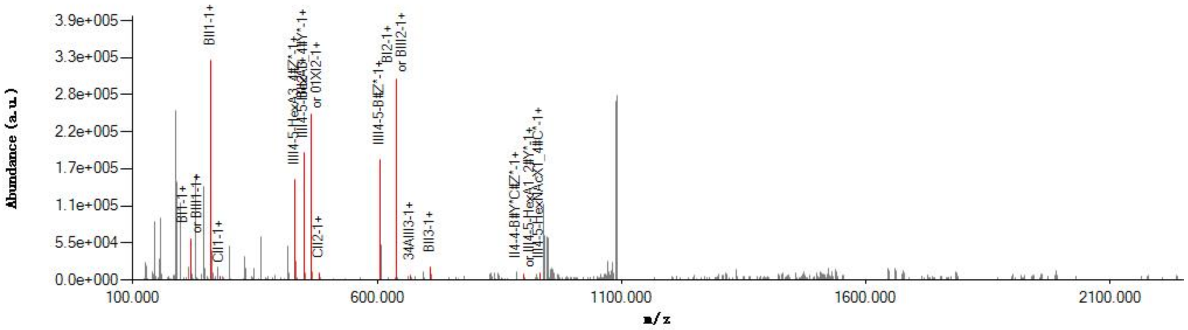


#### Graphical Fragmentation Map

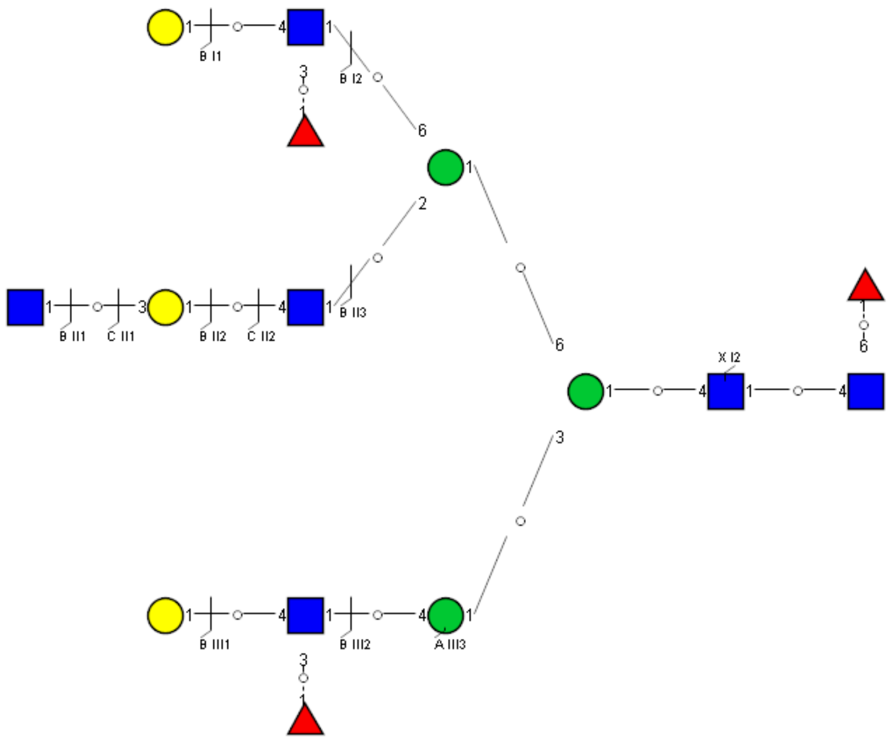

Spectrum Index: 38867(NoBHs:3)

#### Precursor iEF Map

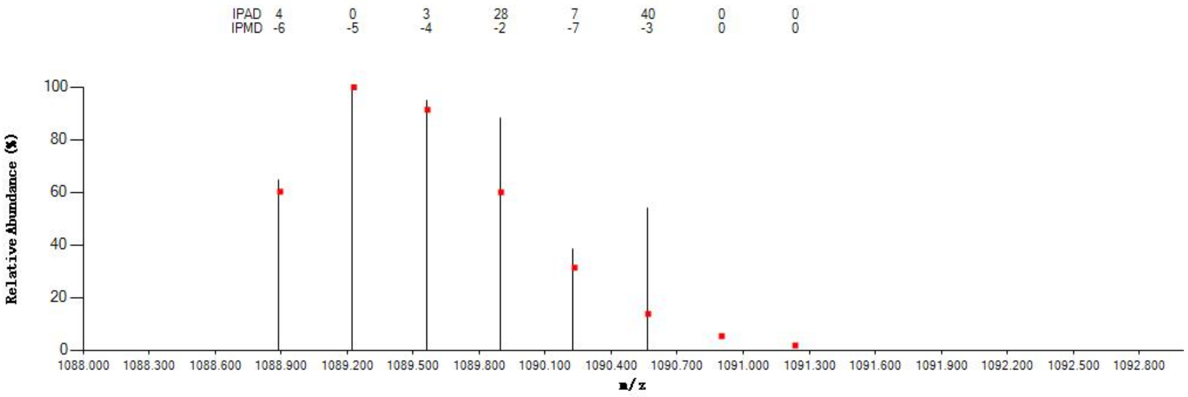

#### Annotated MS/MS Spectrum

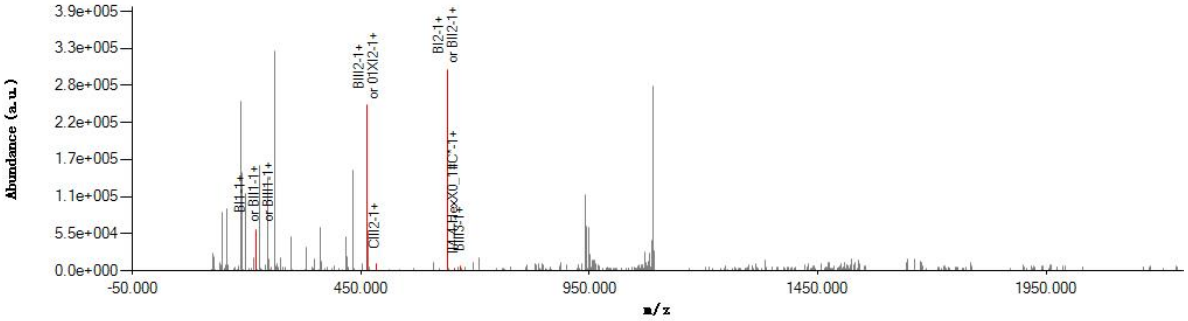

#### Graphical Fragmentation Map

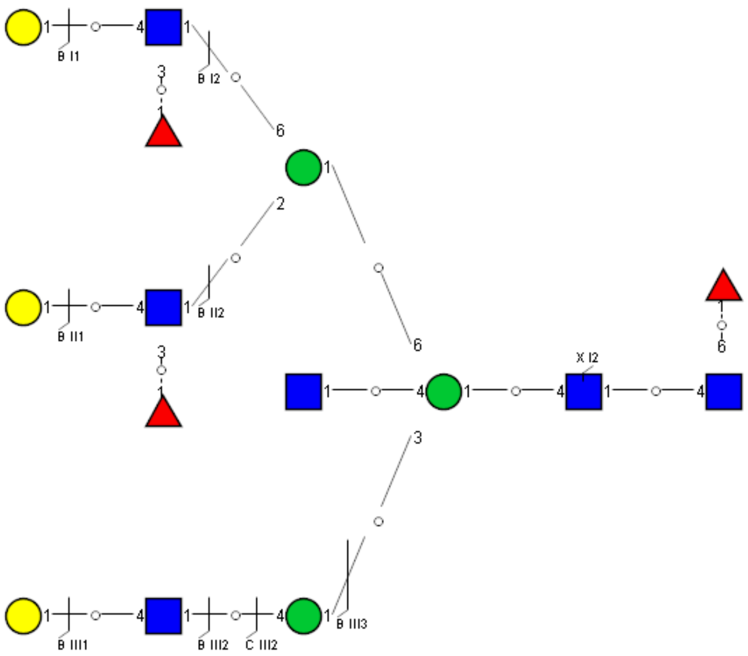

Spectrum Index: 39107(NoBHs:2)

#### Precursor iEF Map

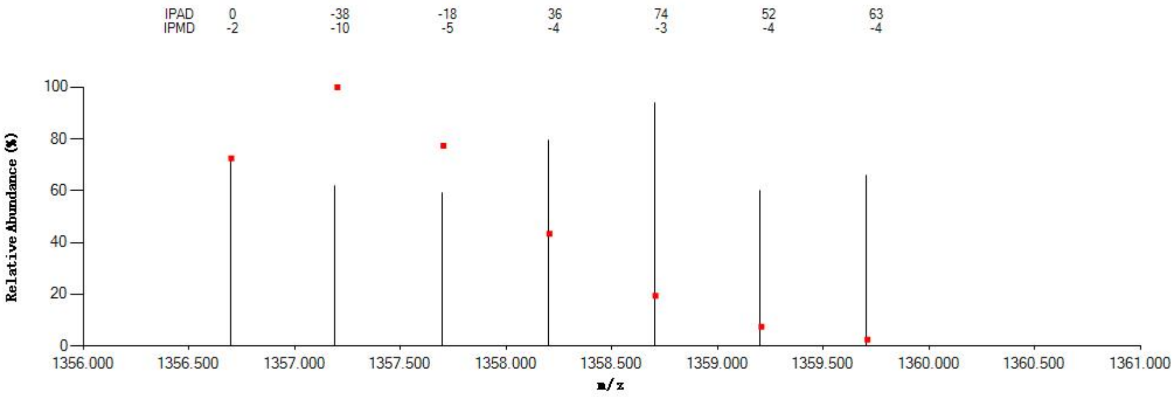

#### Annotated MS/MS Spectrum

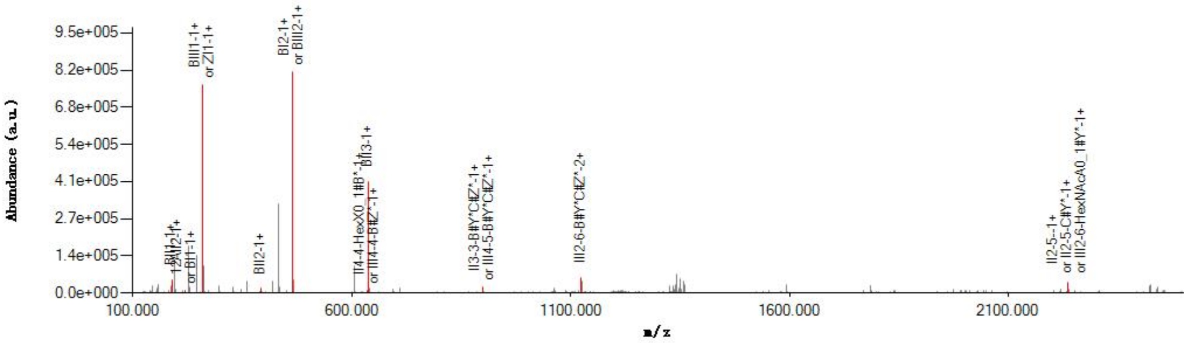

#### Graphical Fragmentation Map

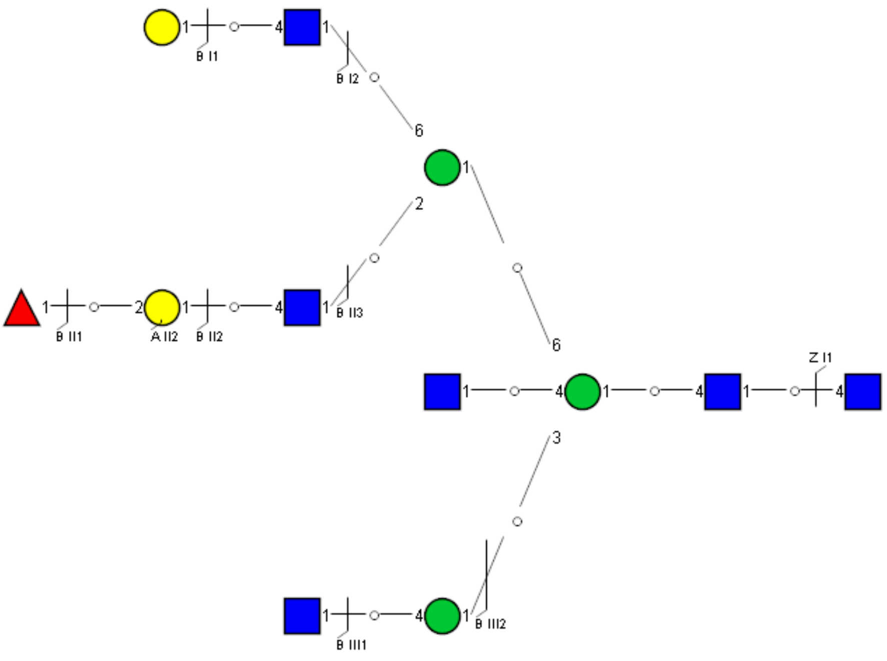

Spectrum Index: 39113(NoBHs:1)

#### Precursor iEF Map

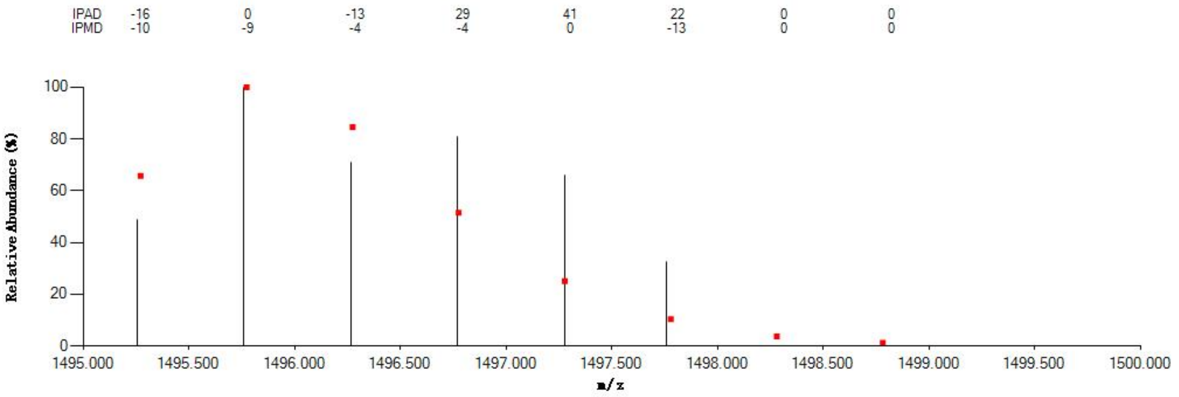


#### Annotated MS/MS Spectrum

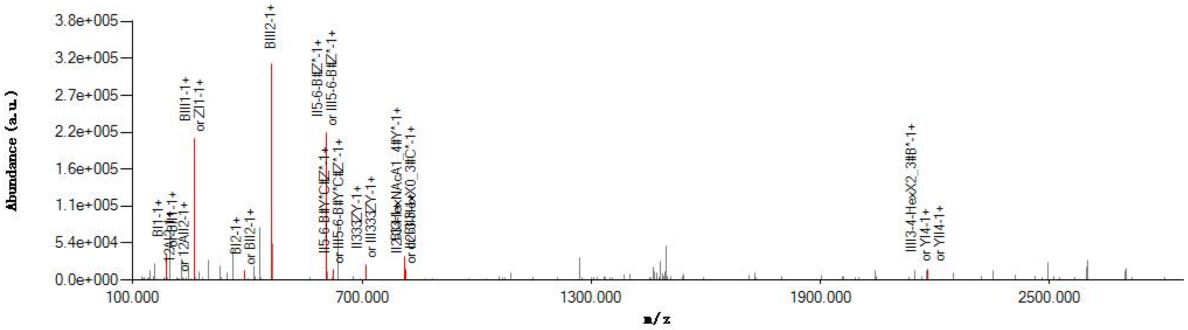


#### Graphical Fragmentation Map

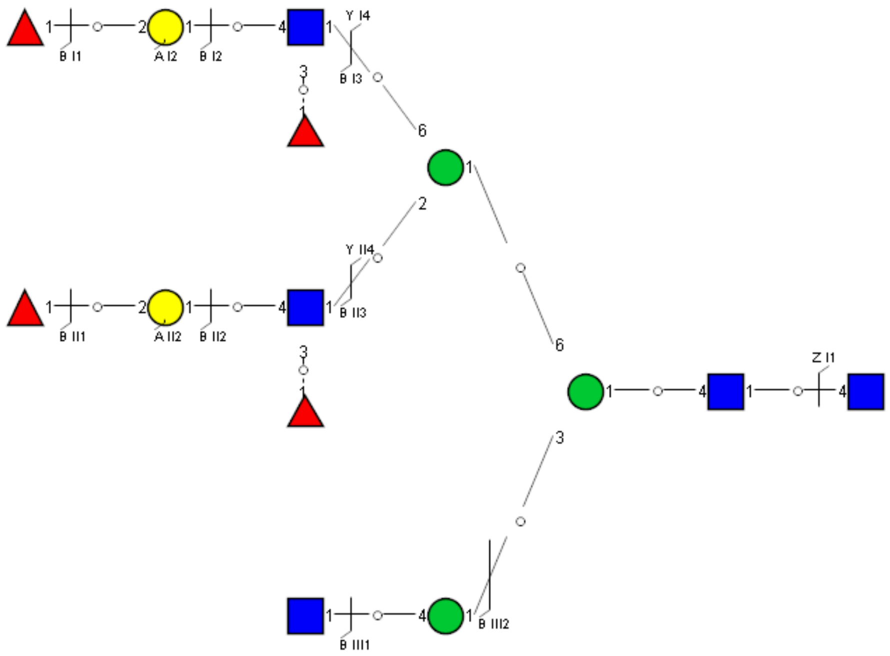

Spectrum Index: 39283(NoBHs:1)

#### Precursor iEF Map

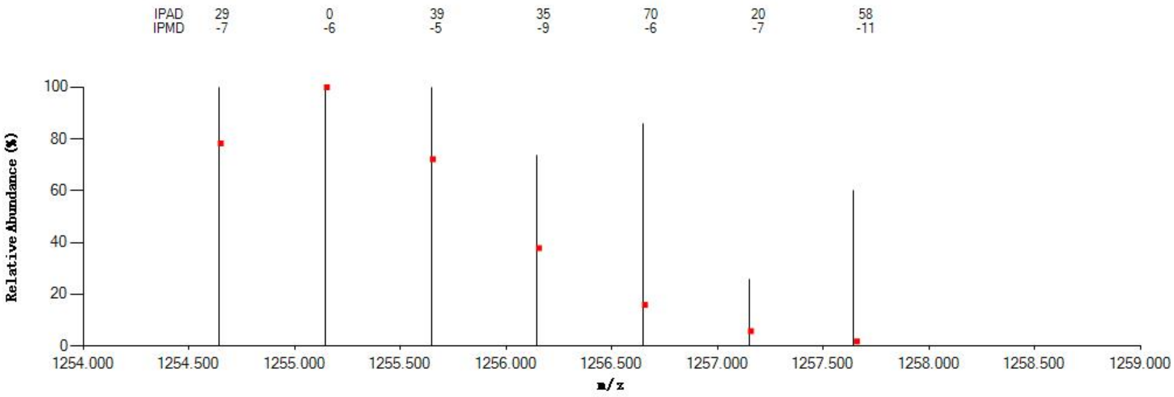


#### Annotated MS/MS Spectrum

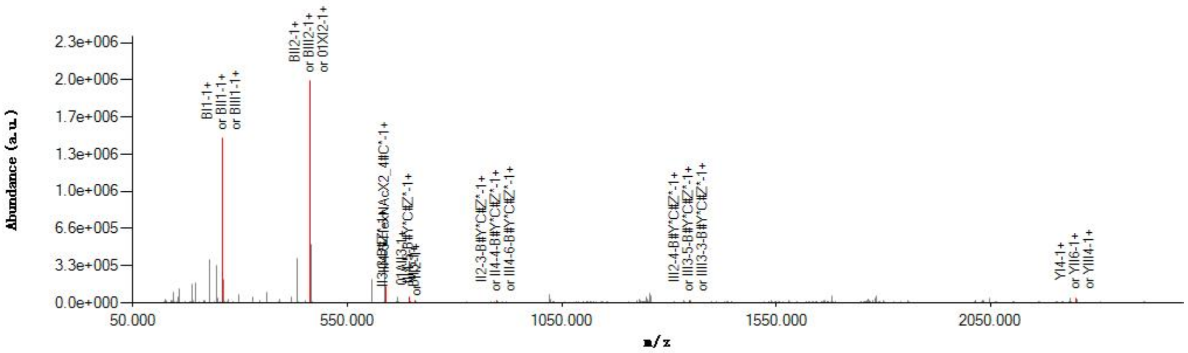


#### Graphical Fragmentation Map

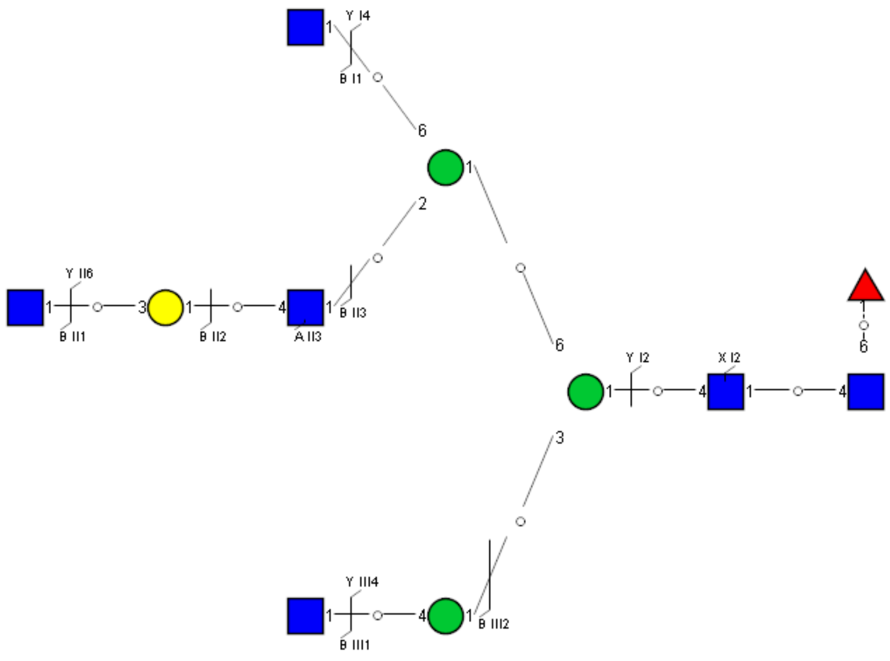

Spectrum Index: 39327(NoBHs:1)

#### Precursor iEF Map

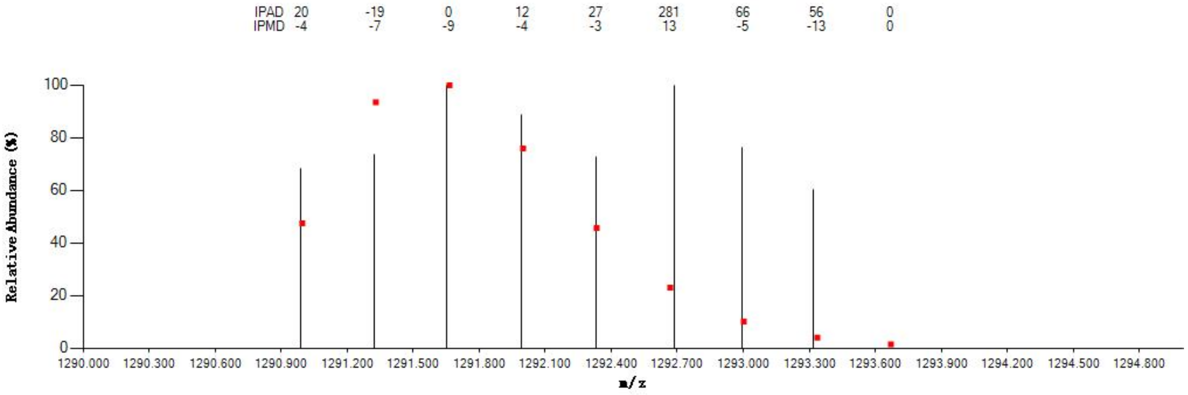


#### Annotated MS/MS Spectrum

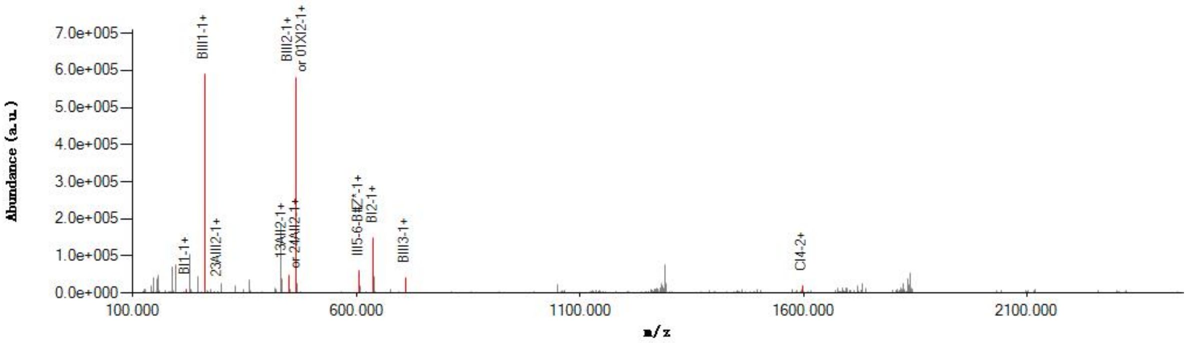


#### Graphical Fragmentation Map

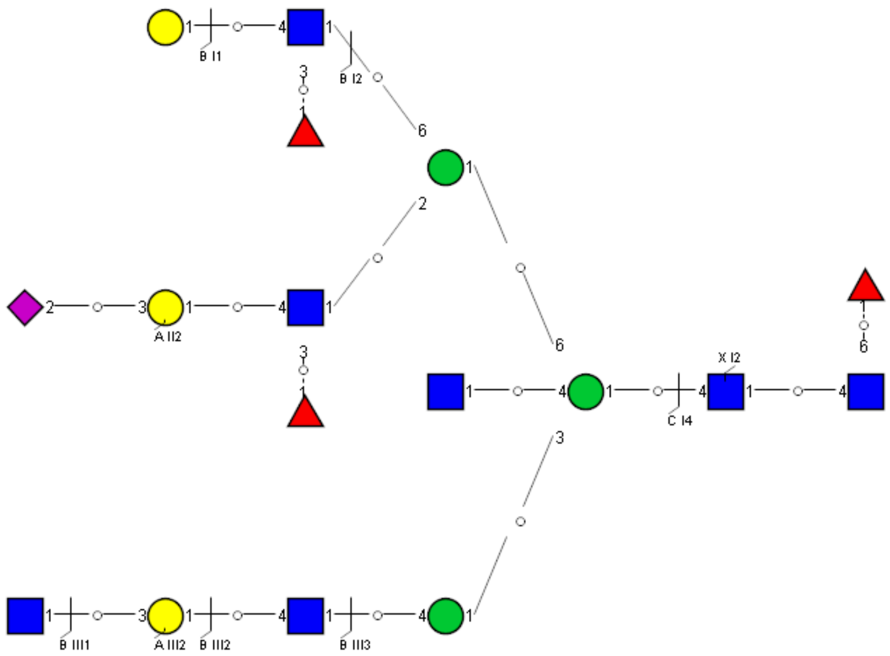

Spectrum Index: 39367(NoBHs:2)

Precursor iEF Map

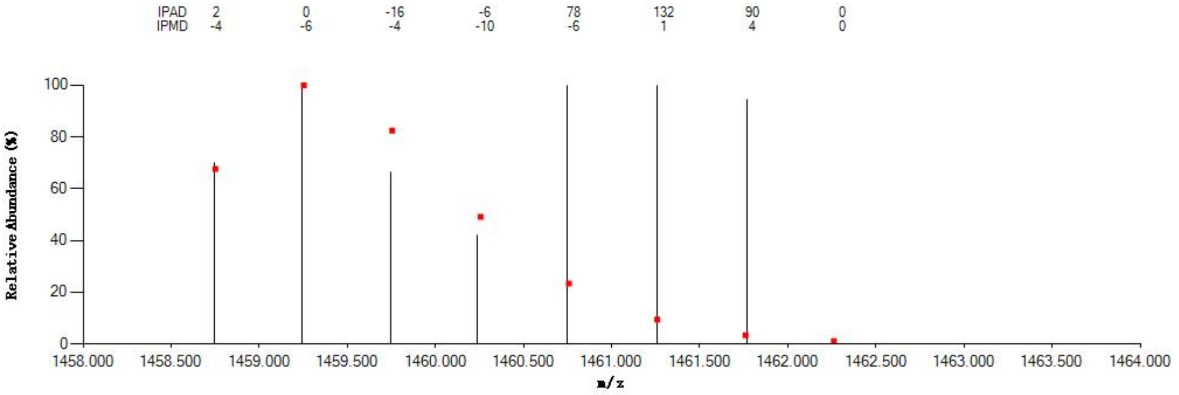


Annotated MS/MS Spectrum

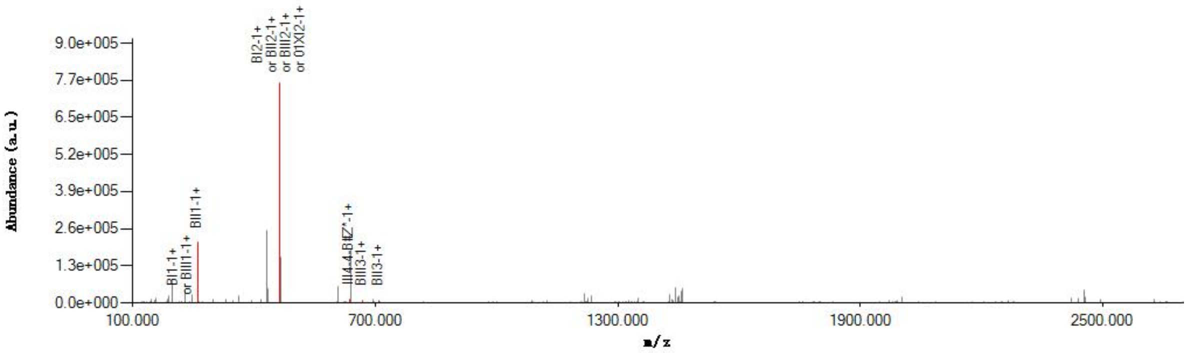


Graphical Fragmentation Map

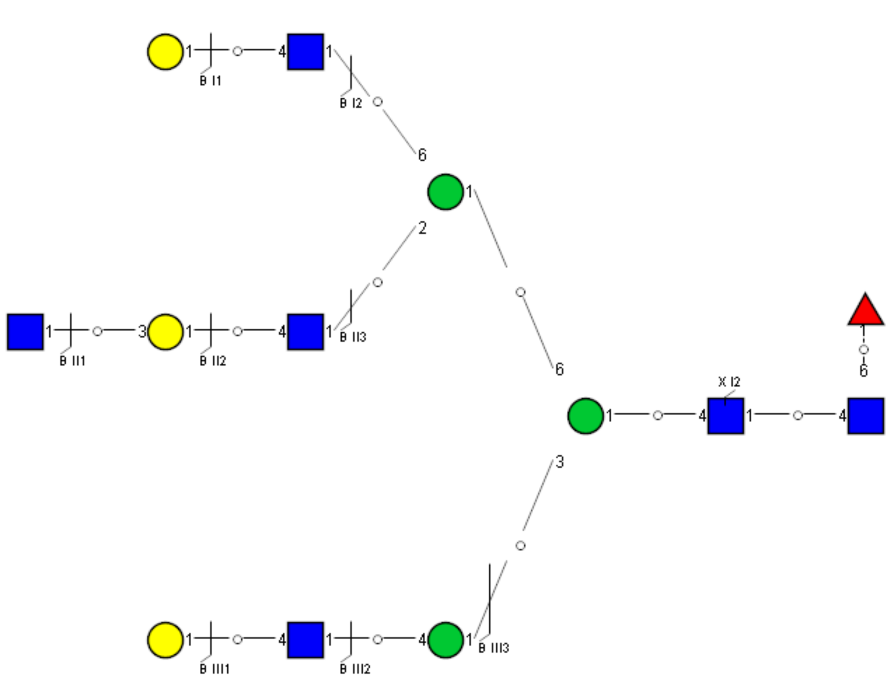

Spectrum Index: 39377(NoBHs:1)

#### Precursor iEF Map

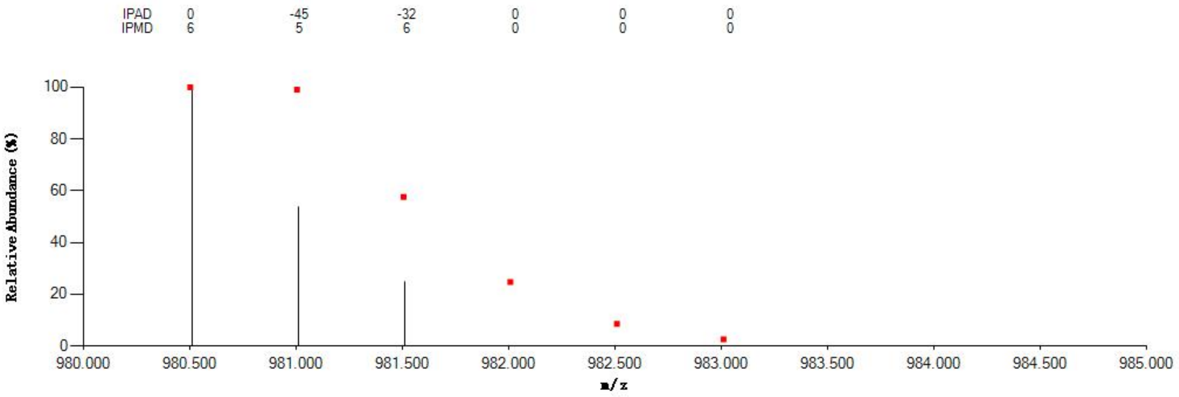


#### Annotated MS/MS Spectrum

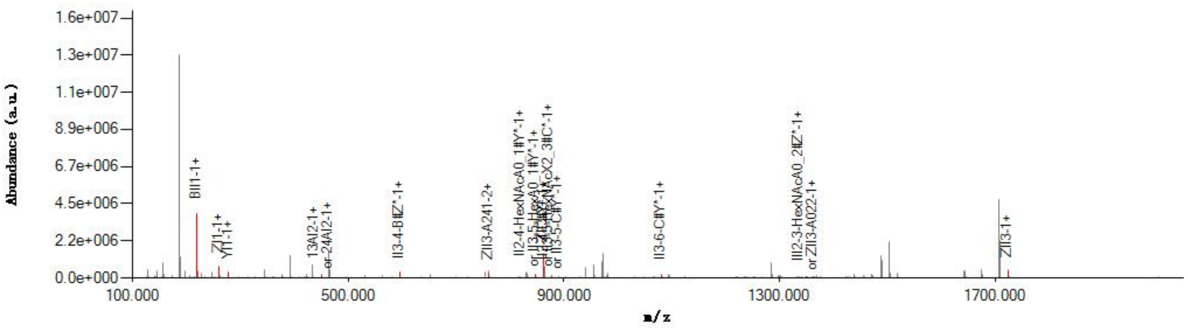


#### Graphical Fragmentation Map

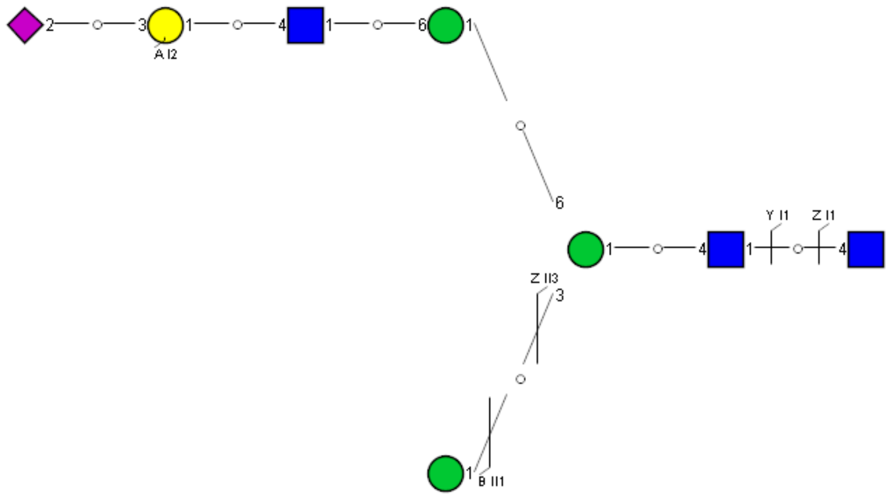

Spectrum Index: 39508(NoBHs:1)

#### Precursor iEF Map

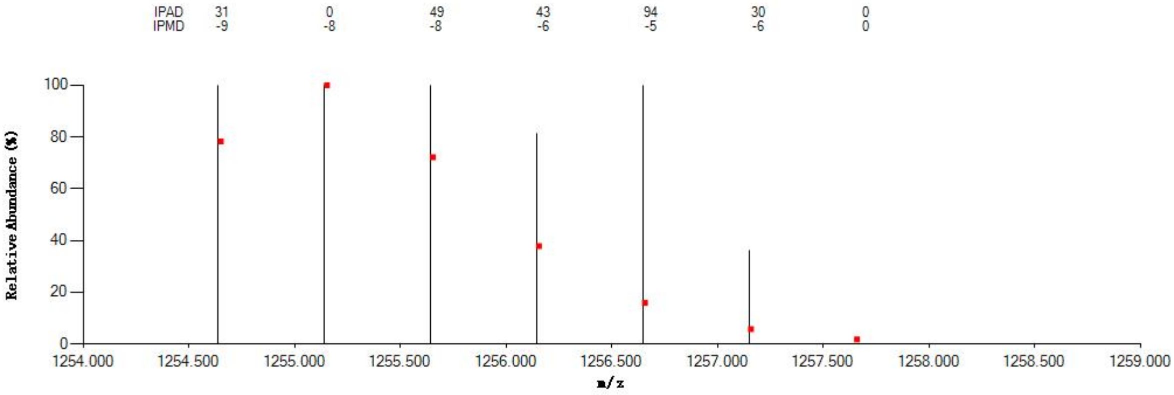


#### Annotated MS/MS Spectrum

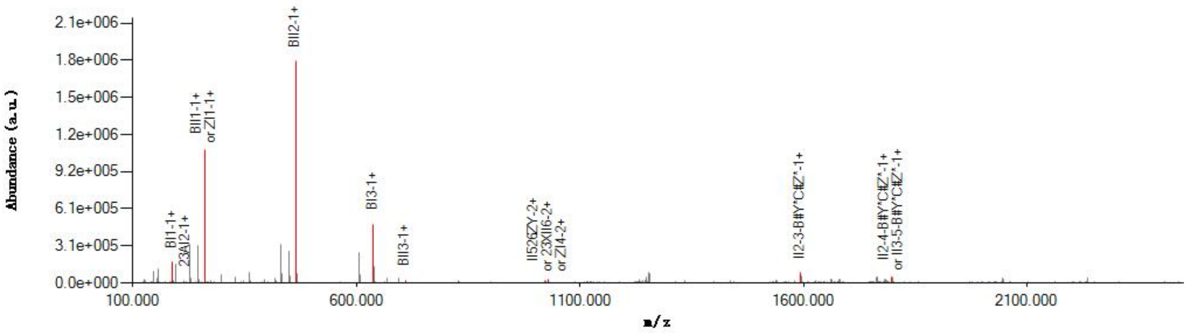


#### Graphical Fragmentation Map

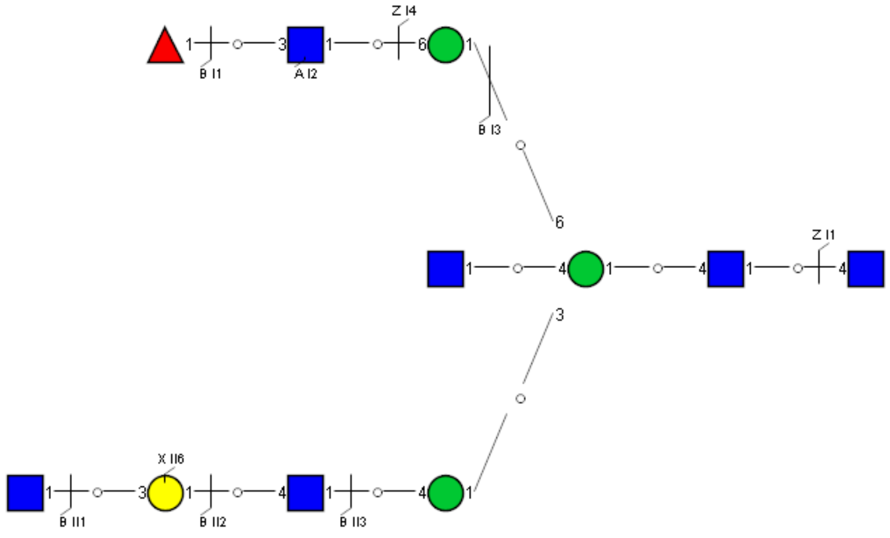

Spectrum Index: 39517(NoBHs:1)

#### Precursor iEF Map

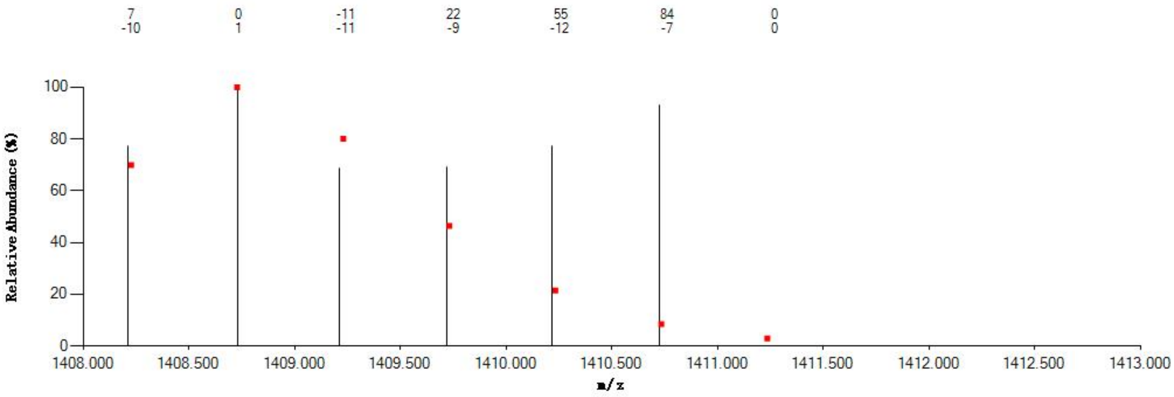


#### Annotated MS/MS Spectrum

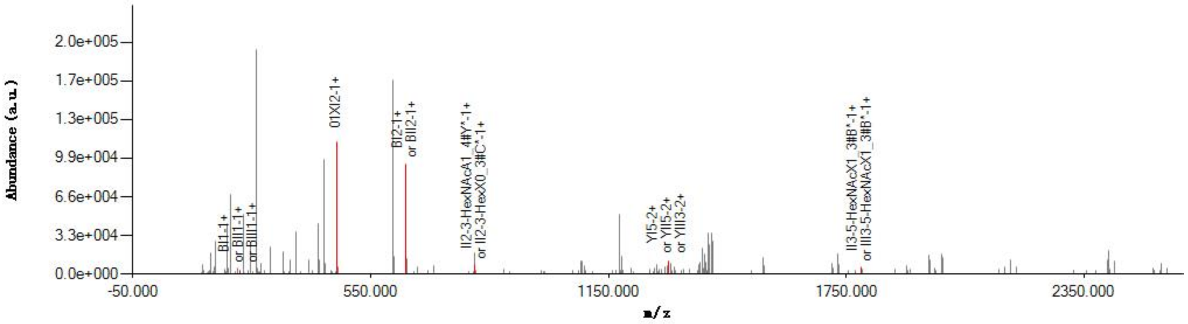


#### Graphical Fragmentation Map

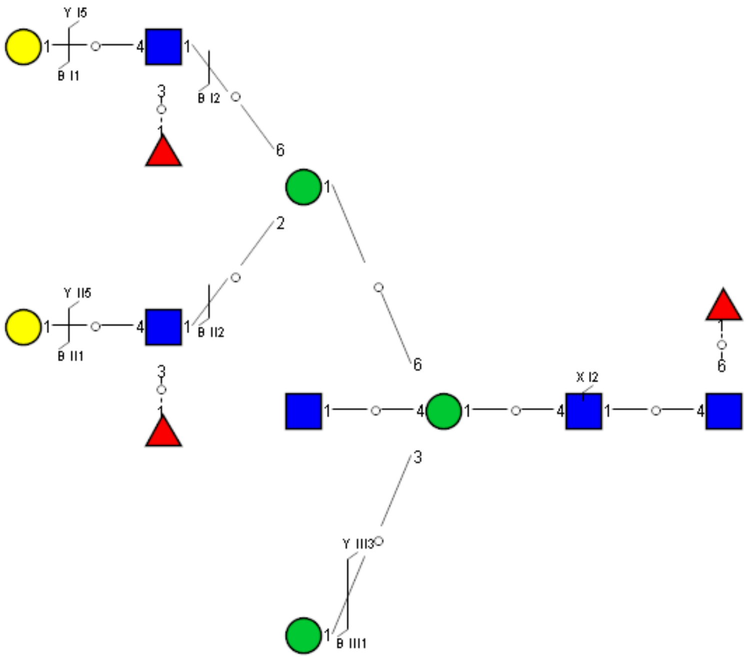

Spectrum Index: 39644(NoBHs:1)

#### Precursor iEF Map

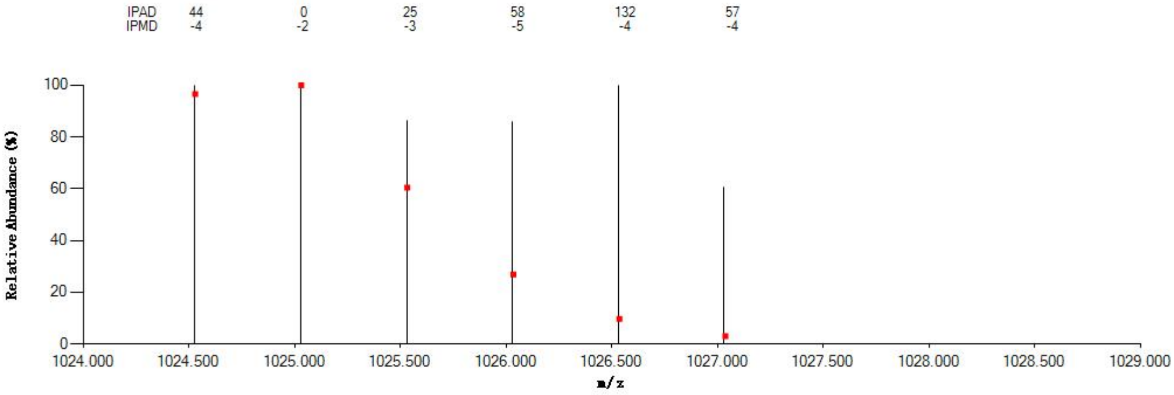


#### Annotated MS/MS Spectrum

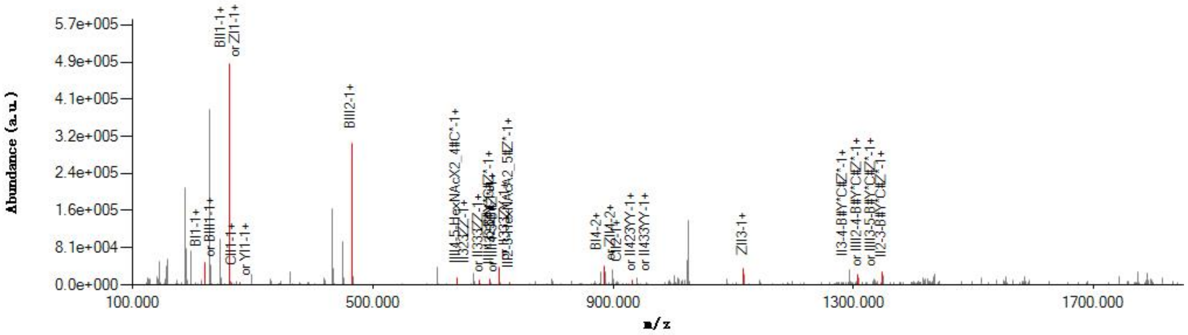


#### Graphical Fragmentation Map

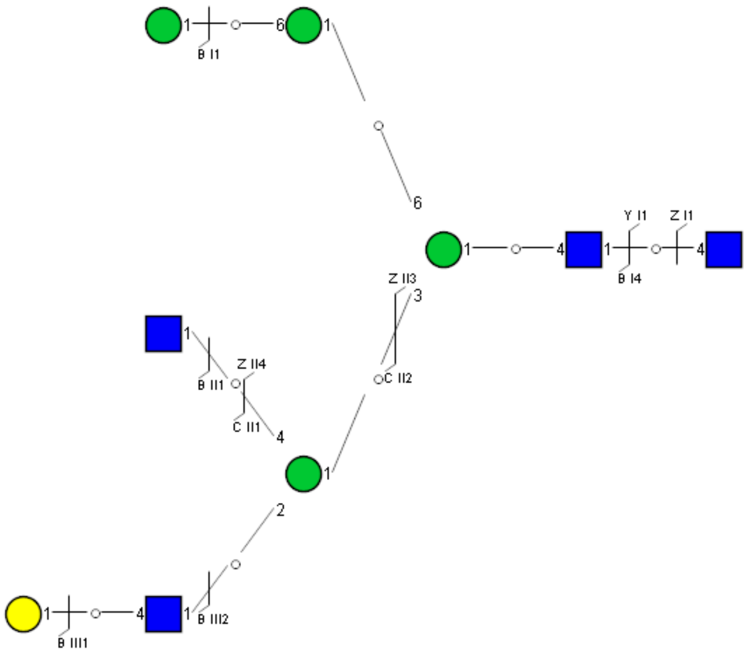

Spectrum Index: 39777(NoBHs:1)

Precursor iEF Map

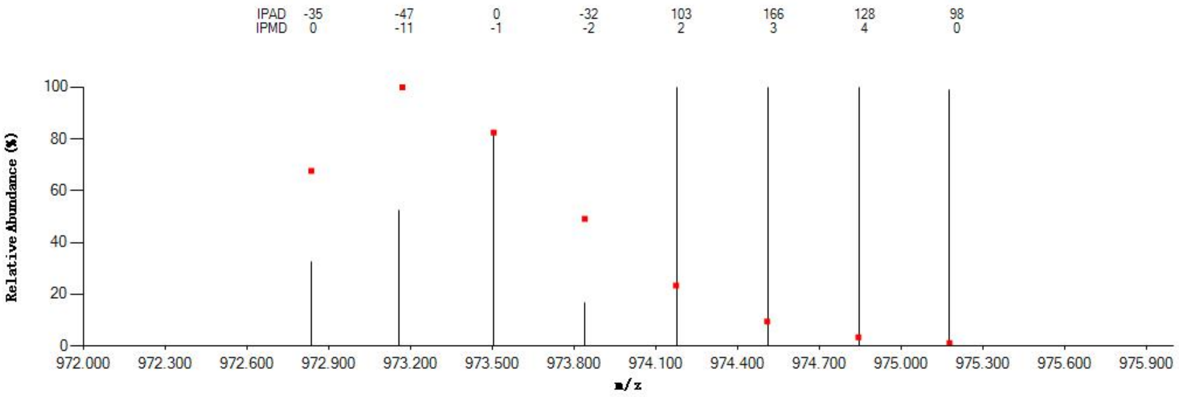

Annotated MS/MS Spectrum

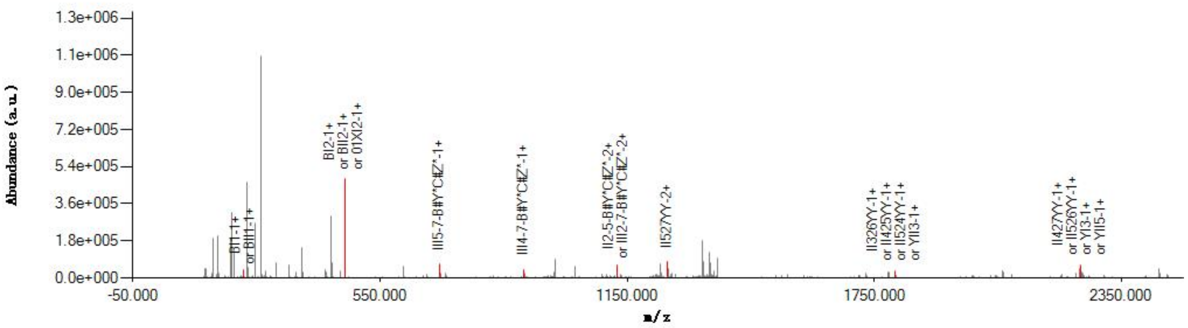

Graphical Fragmentation Map

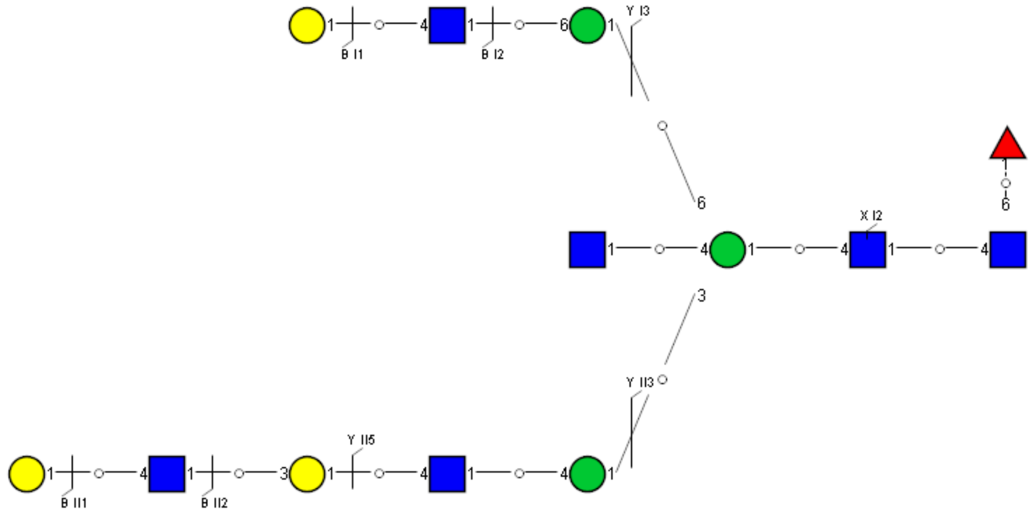

Spectrum Index: 39996(NoBHs:1)

#### Precursor iEF Map

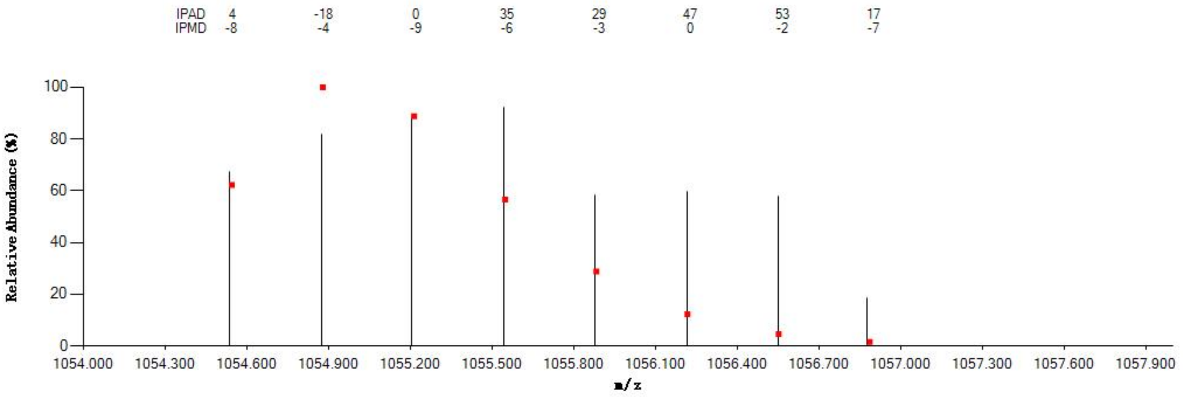


#### Annotated MS/MS Spectrum

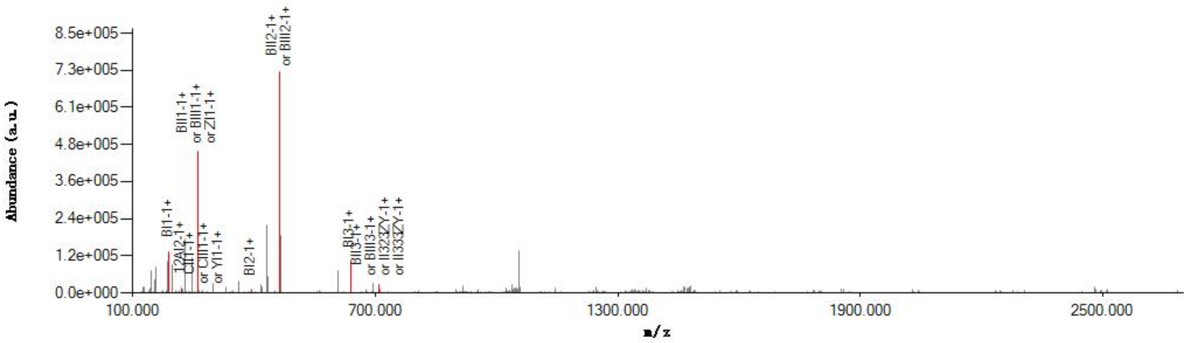


#### Graphical Fragmentation Map

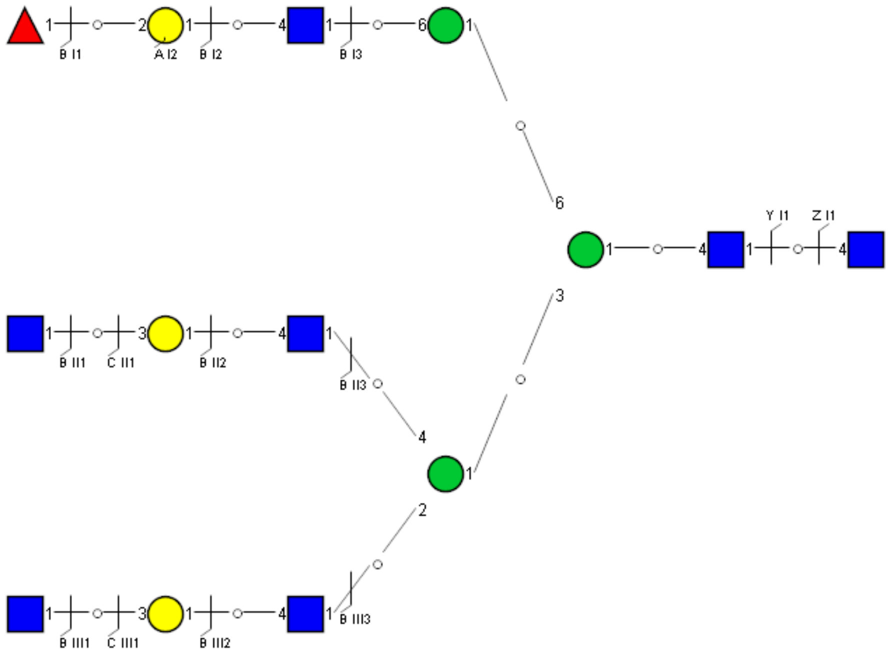

Spectrum Index: 40004(NoBHs:1)

#### Precursor iEF Map

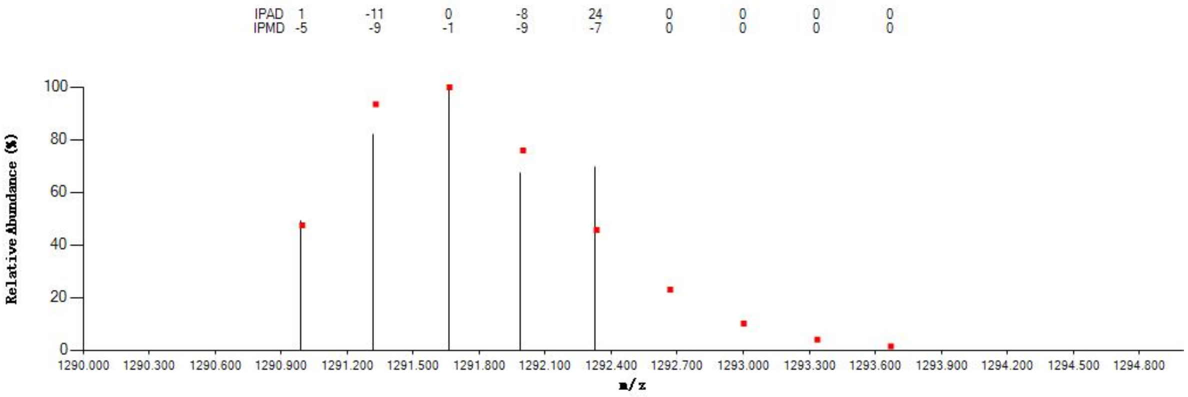


#### Annotated MS/MS Spectrum

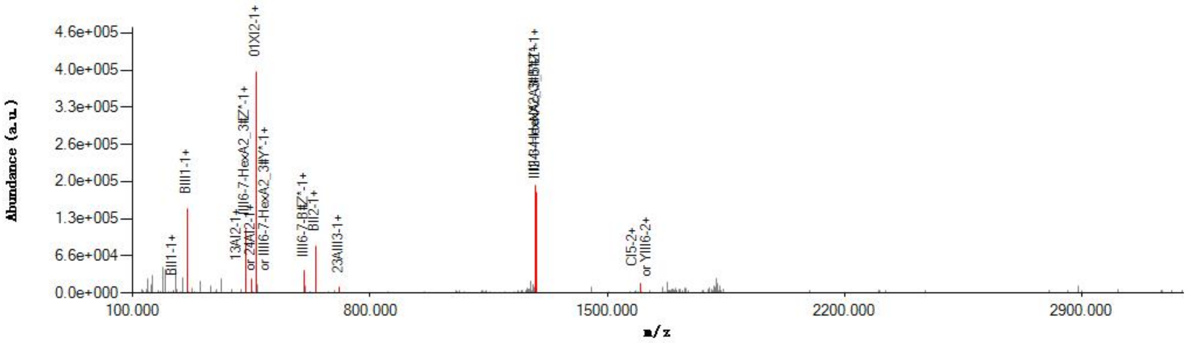


#### Graphical Fragmentation Map

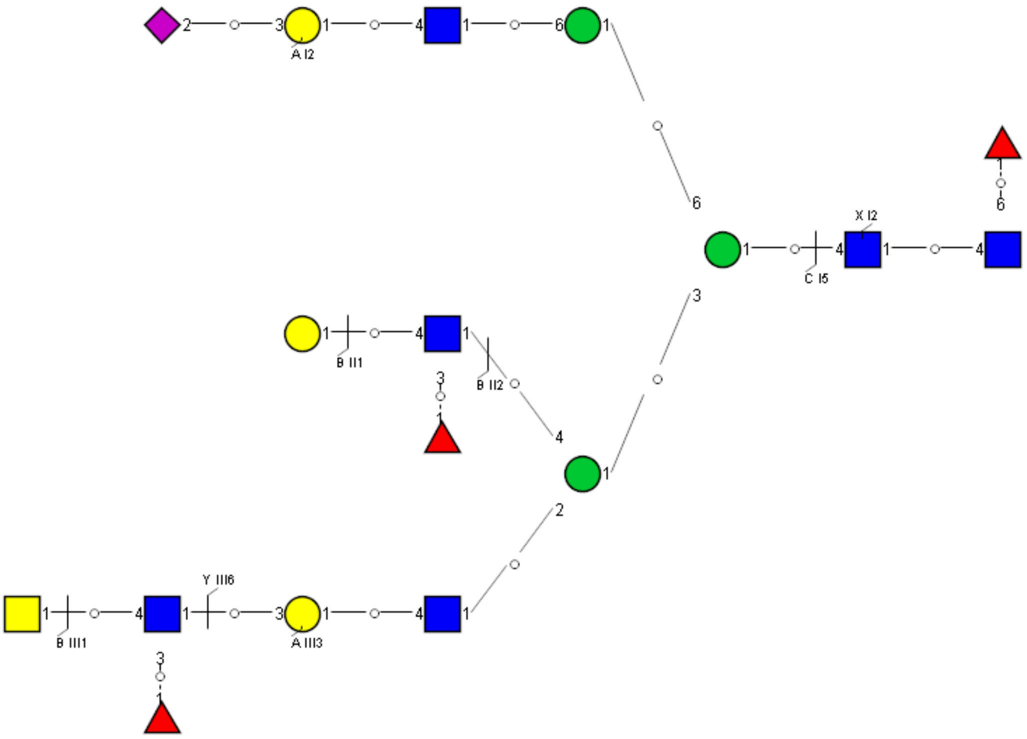

Spectrum Index: 40085(NoBHs:2)

#### Precursor iEF Map

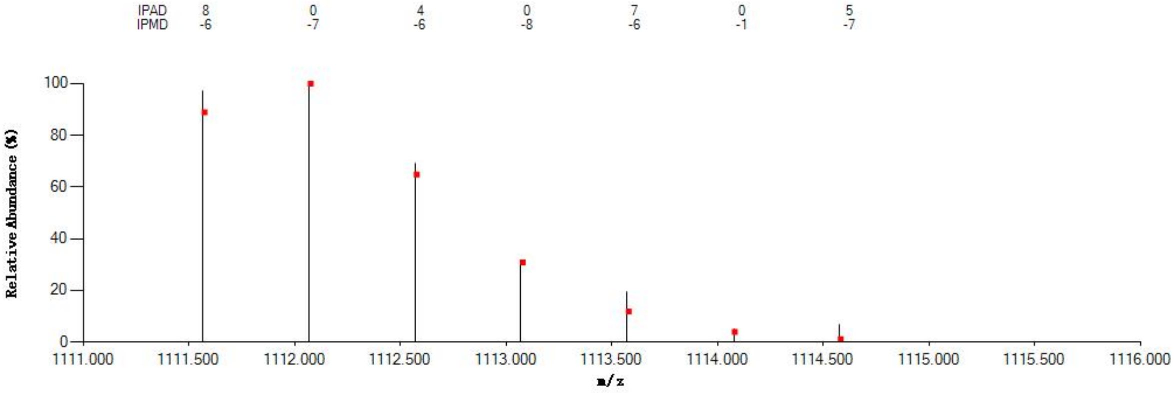


#### Annotated MS/MS Spectrum

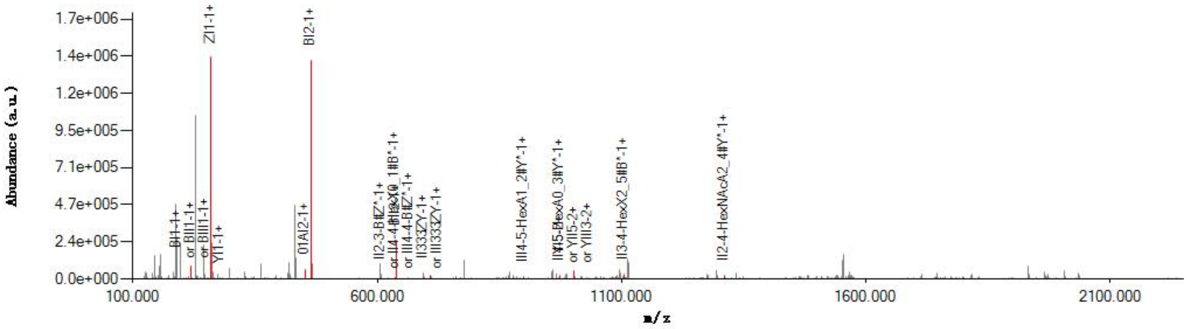


#### Graphical Fragmentation Map

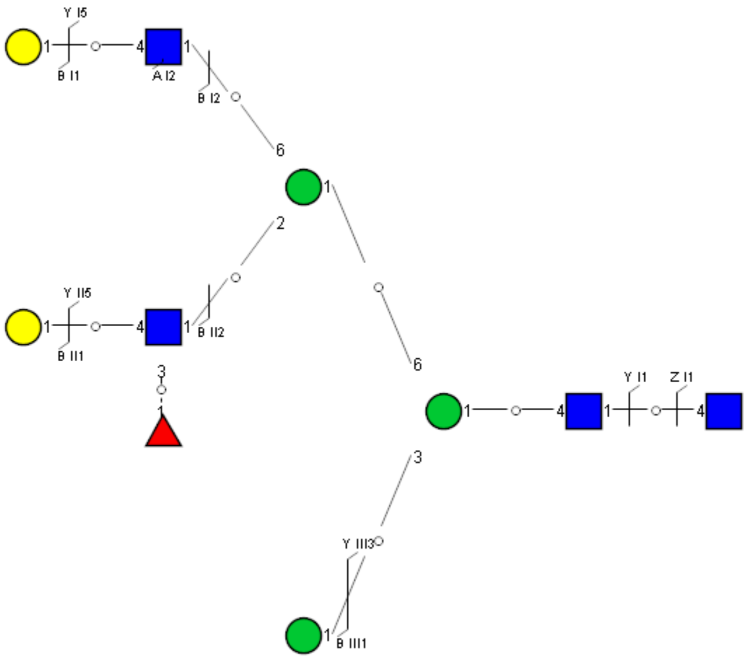

Spectrum Index: 40085(NoBHs:2)

#### Precursor iEF Map

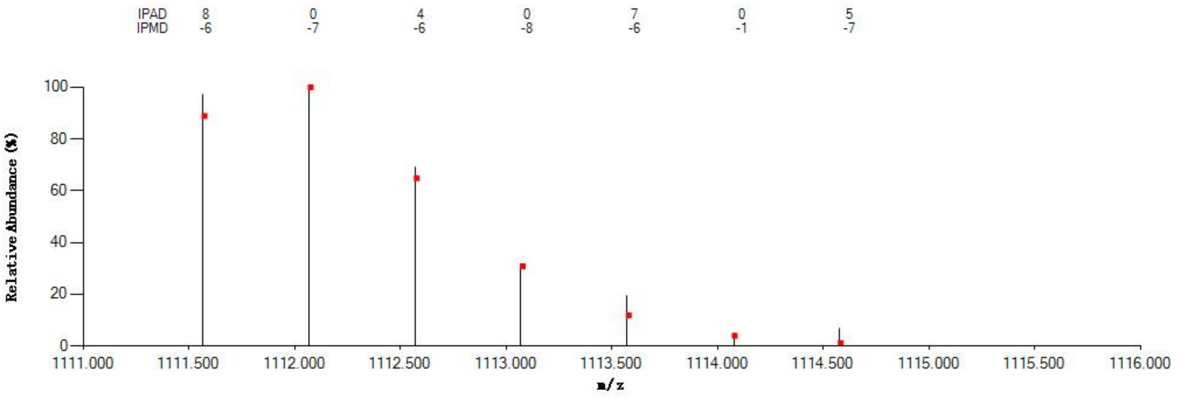


#### Annotated MS/MS Spectrum

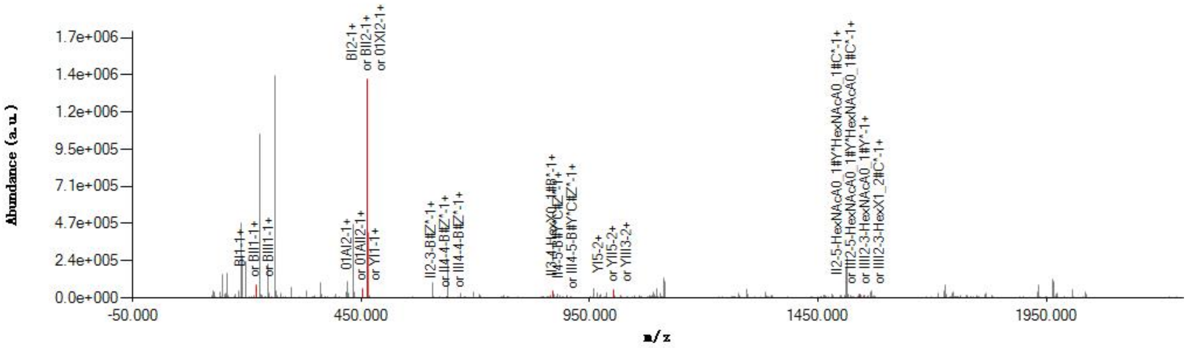


#### Graphical Fragmentation Map

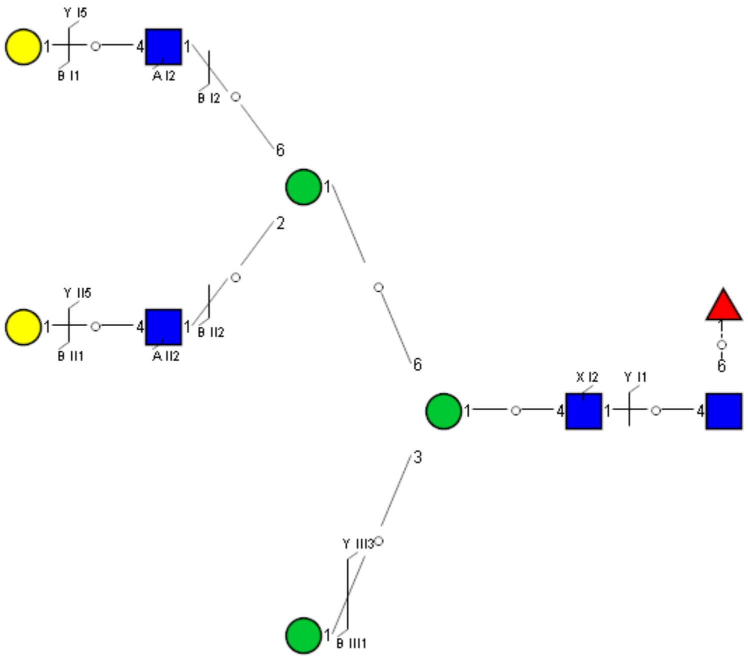

Spectrum Index: 40125(NoBHs:1)

#### Precursor iEF Map

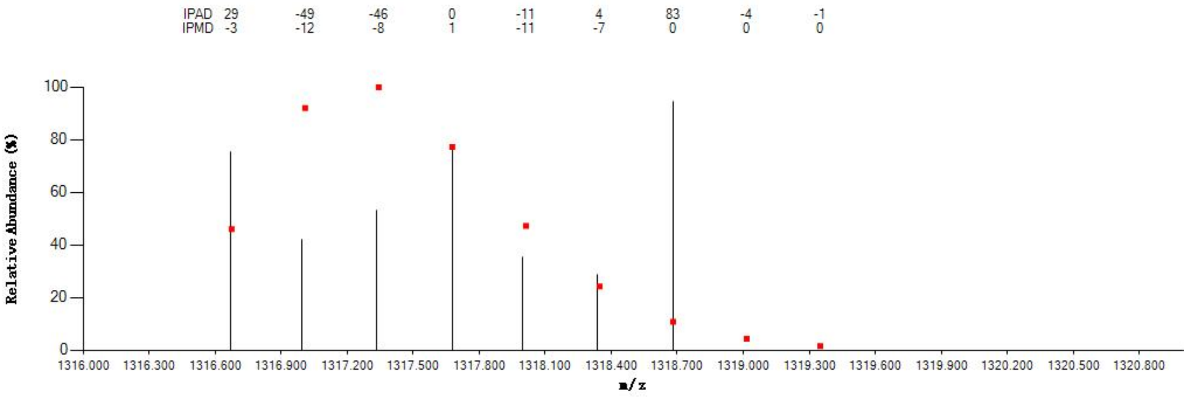


#### Annotated MS/MS Spectrum

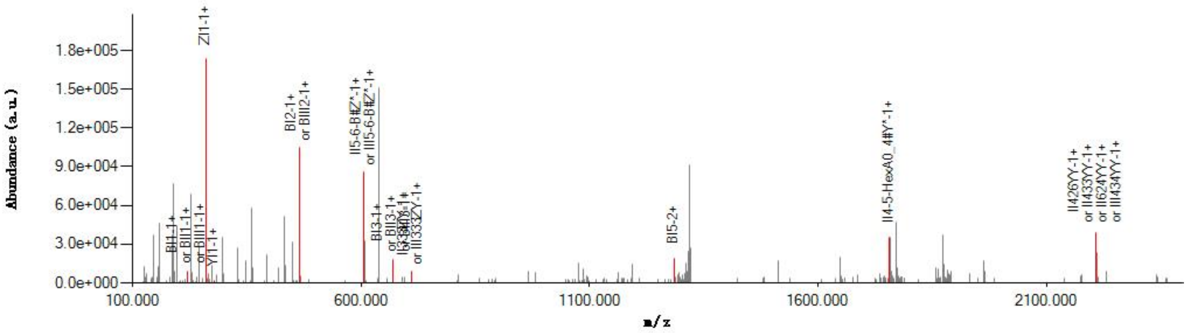


#### Graphical Fragmentation Map

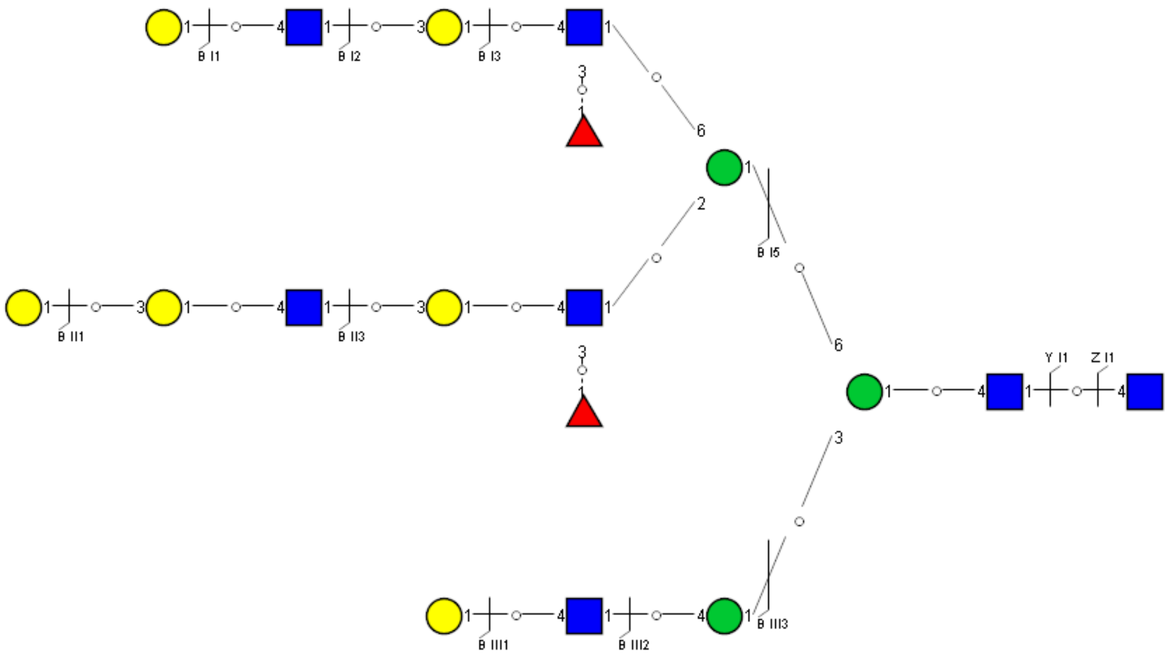

Spectrum Index: 40235(NoBHs:1)

#### Precursor iEF Map

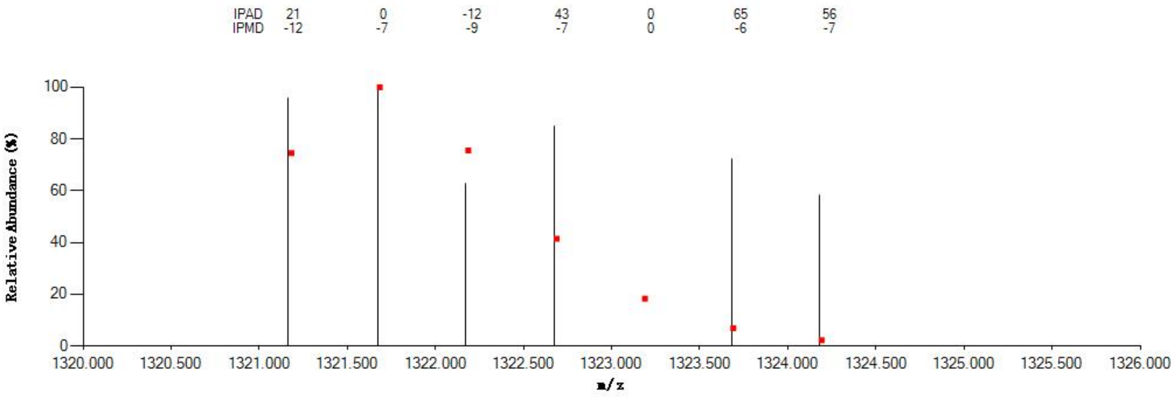


#### Annotated MS/MS Spectrum

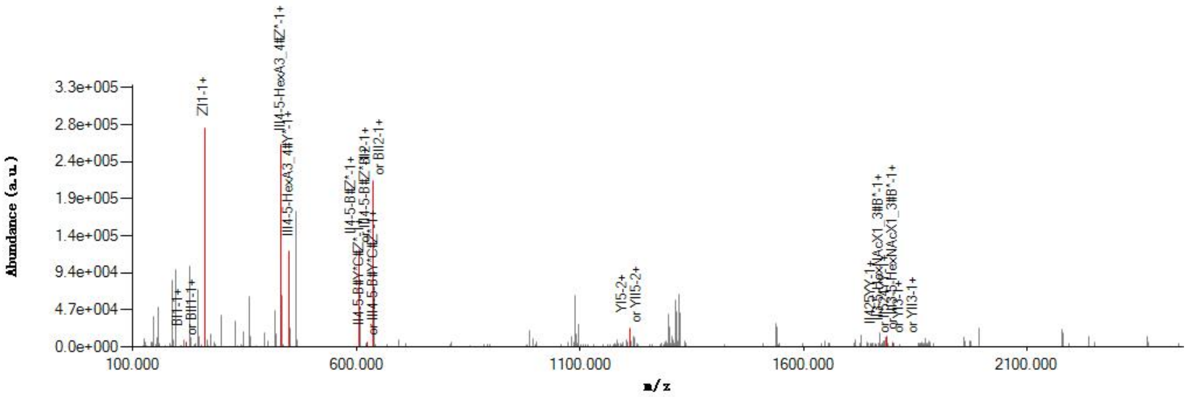


#### Graphical Fragmentation Map

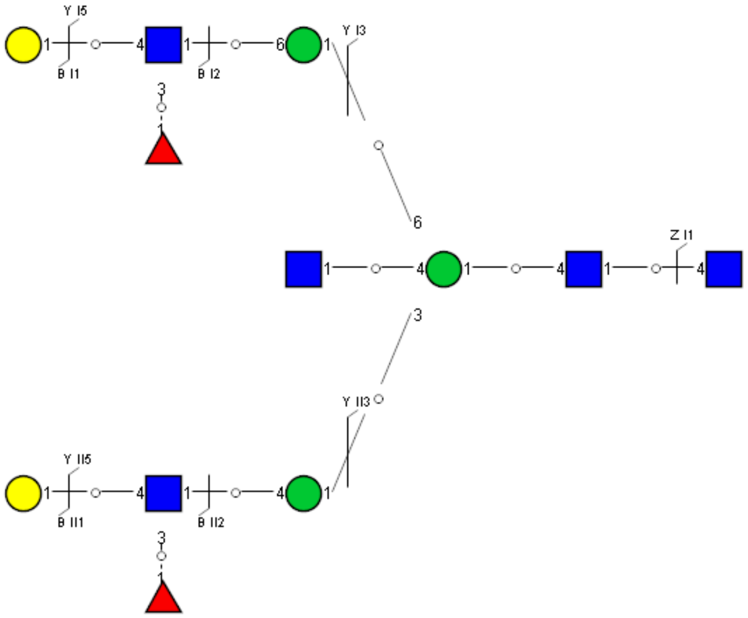

Spectrum Index: 40357(NoBHs:2)

#### Precursor iEF Map

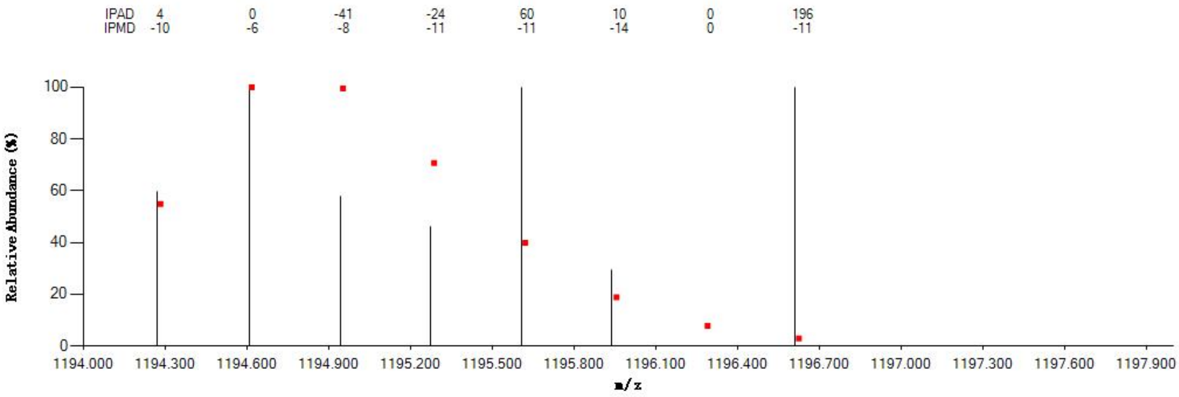


#### Annotated MS/MS Spectrum

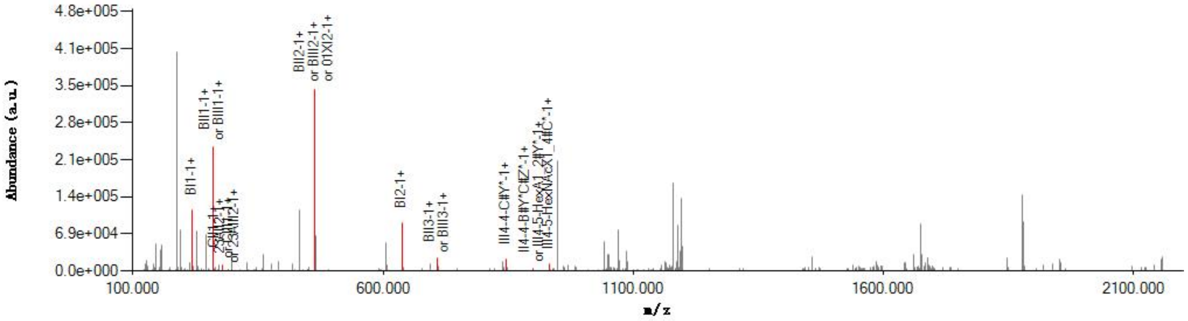


#### Graphical Fragmentation Map

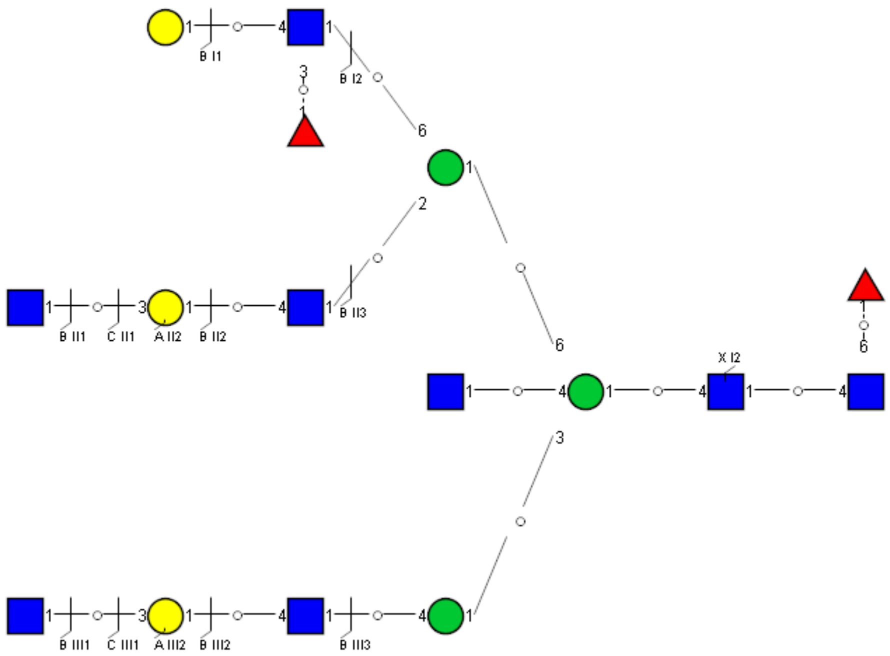

Spectrum Index: 40357(NoBHs:2)

#### Precursor iEF Map

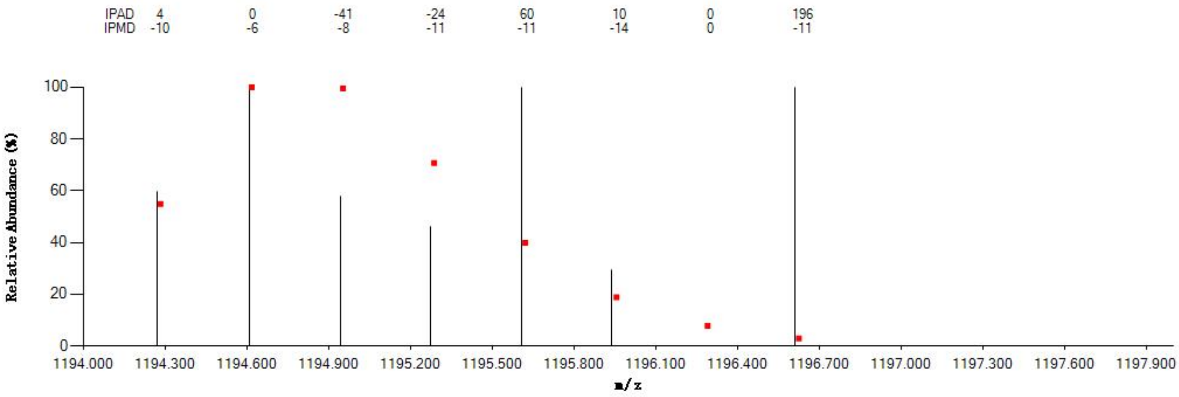

#### Annotated MS/MS Spectrum

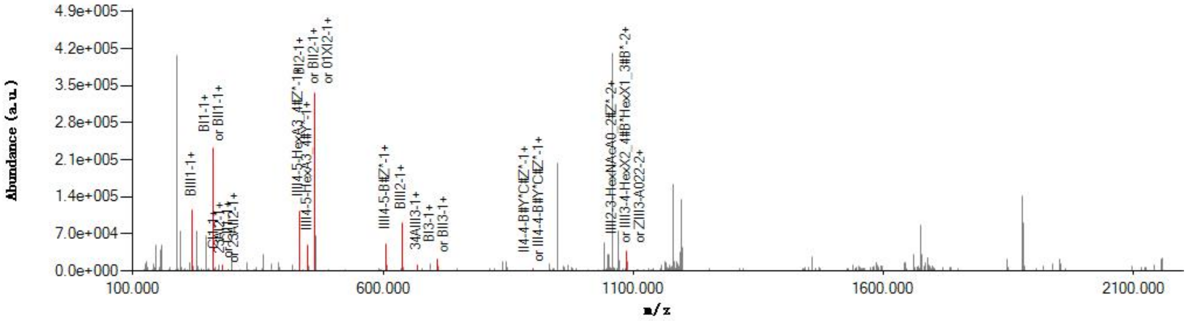

#### Graphical Fragmentation Map

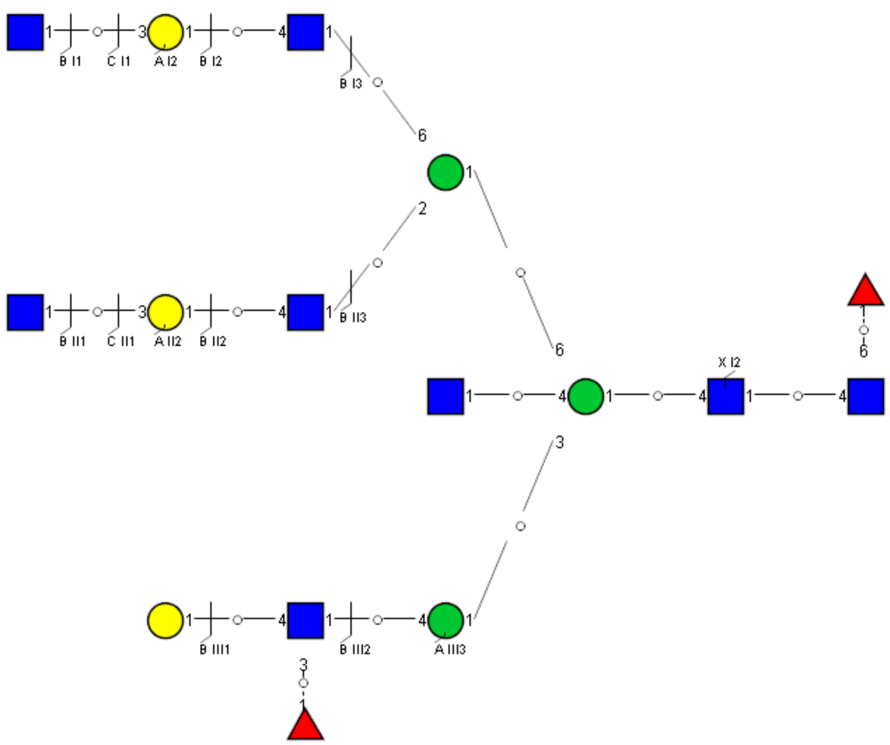

Spectrum Index: 40847(NoBHs:1)

#### Precursor iEF Map

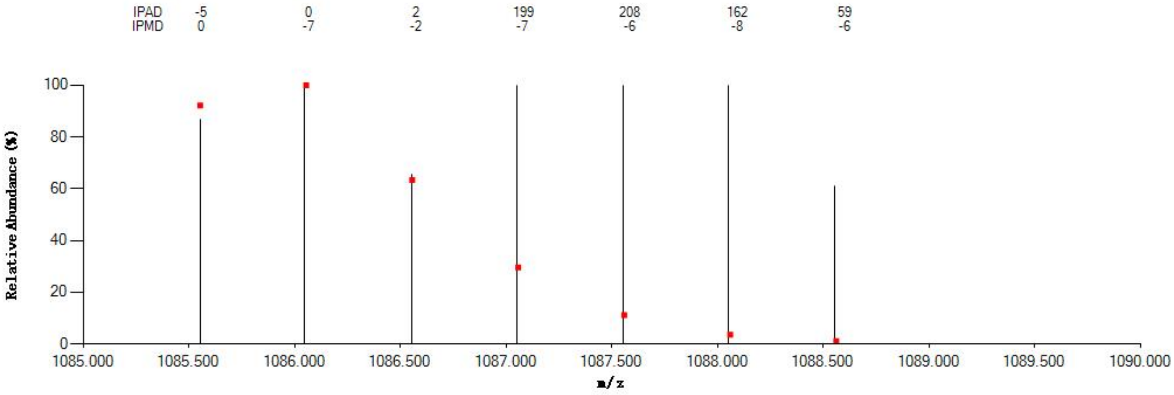

#### Annotated MS/MS Spectrum

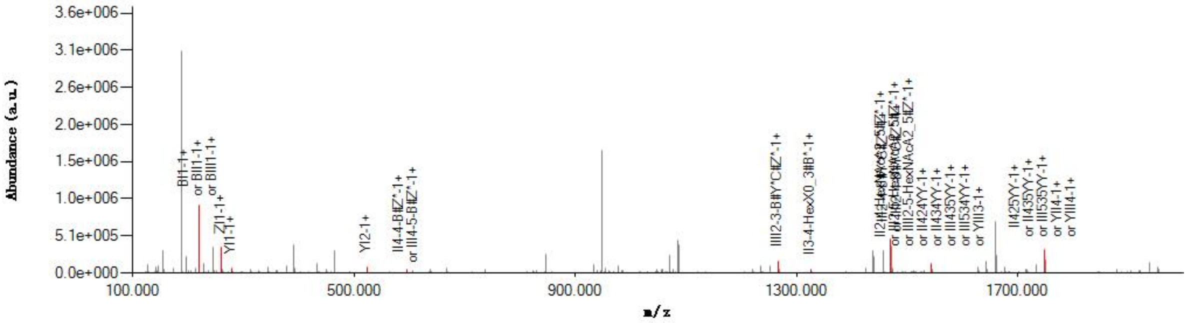

#### Graphical Fragmentation Map

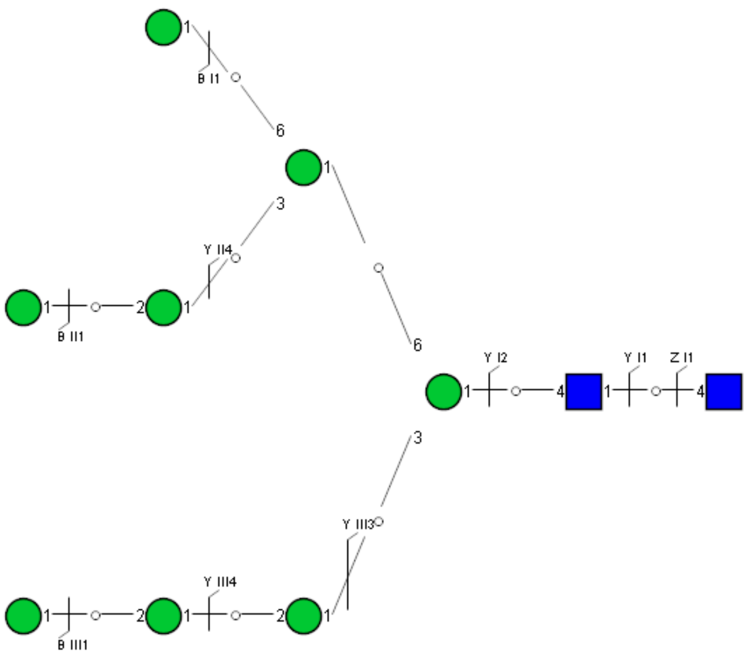

Spectrum Index: 41120(NoBHs:1)

#### Precursor iEF Map

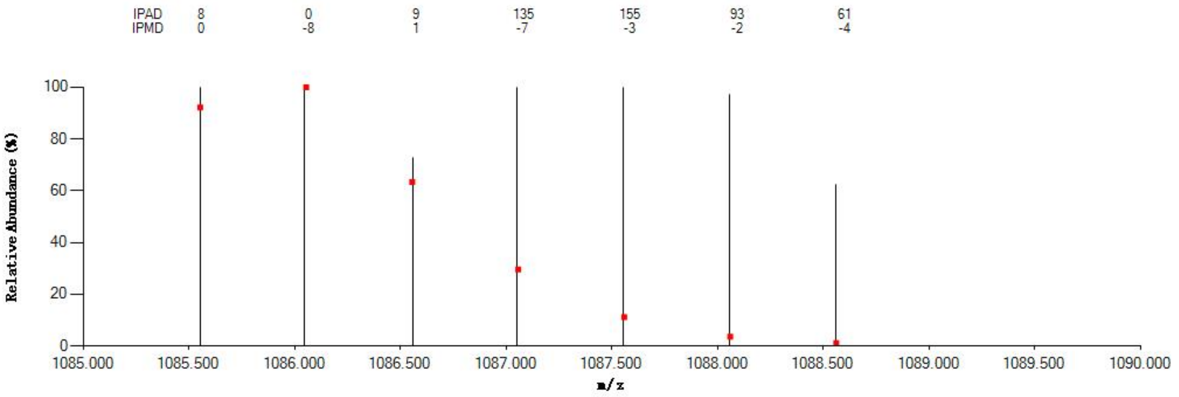


#### Annotated MS/MS Spectrum

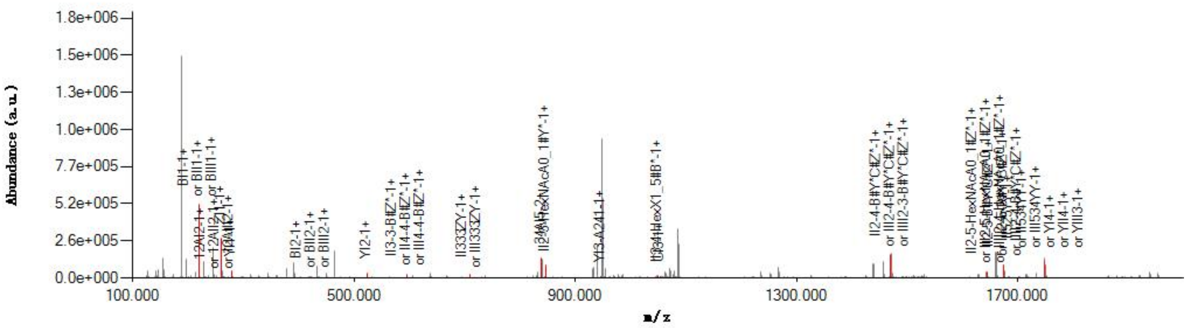


#### Graphical Fragmentation Map

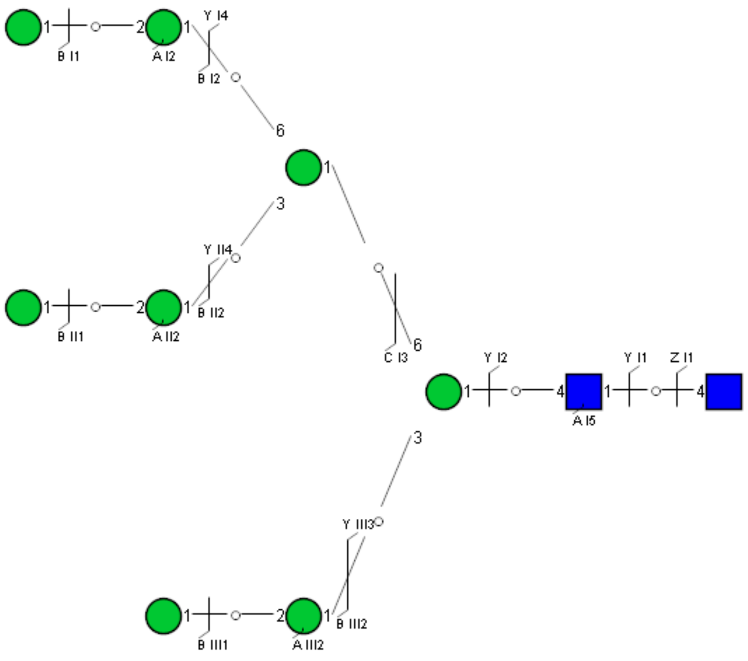

Spectrum Index: 41303(NoBHs:1)

#### Precursor iEF Map

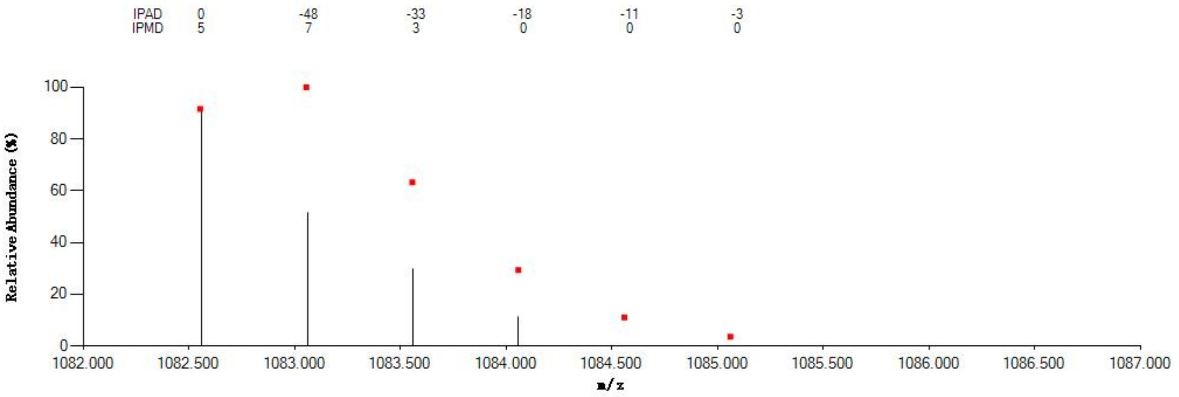


#### Annotated MS/MS Spectrum

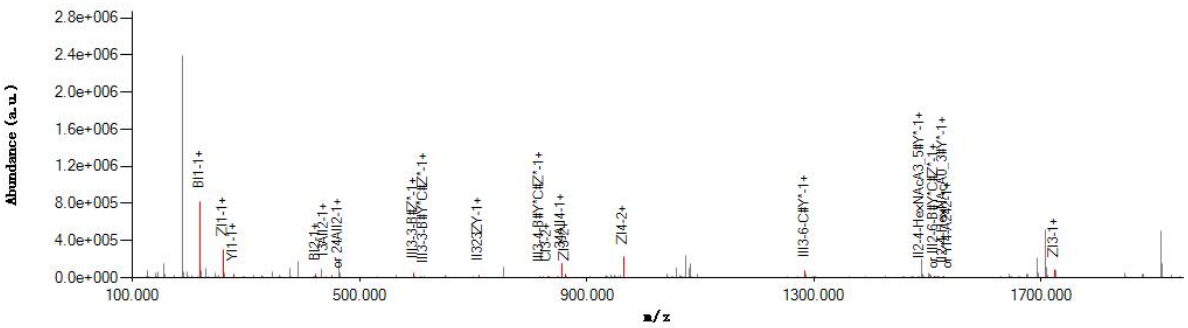


#### Graphical Fragmentation Map

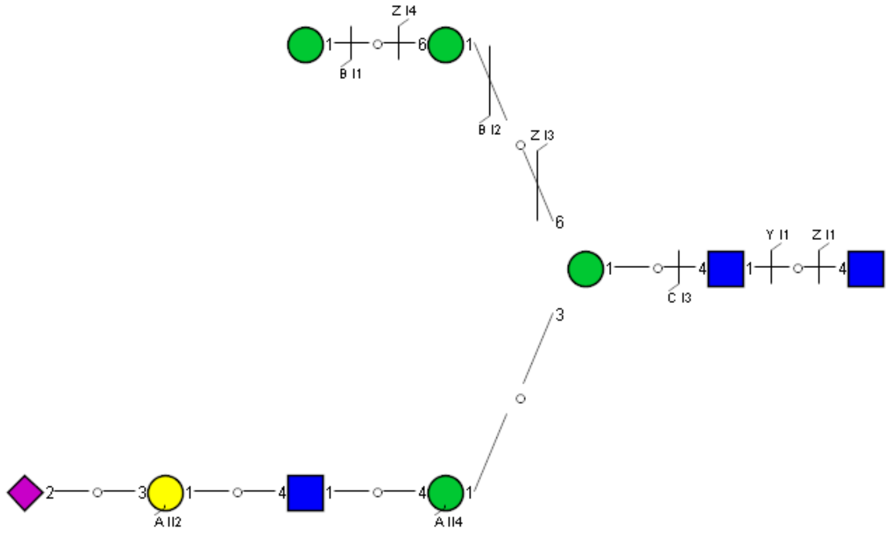

Spectrum Index: 41433(NoBHs:1)

#### Precursor iEF Map

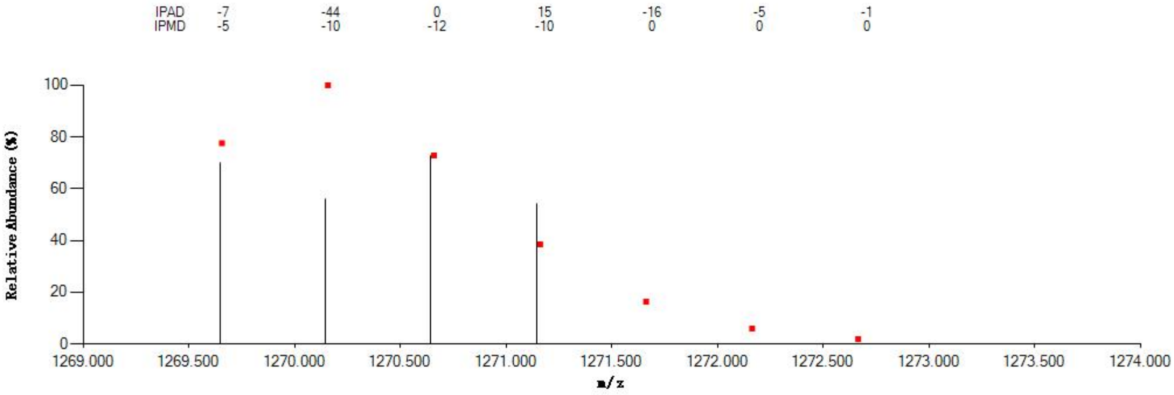


#### Annotated MS/MS Spectrum

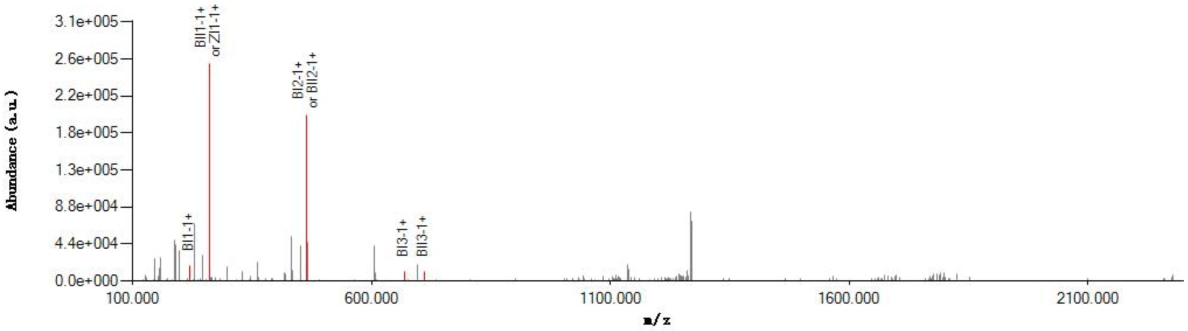


#### Graphical Fragmentation Map

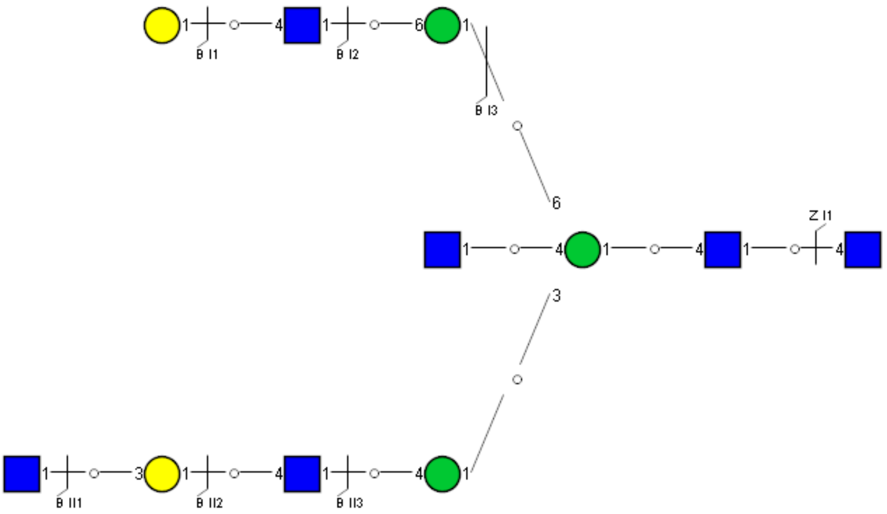

### Spectrum Index: 41616(NoBHs:1)

#### Precursor iEF Map

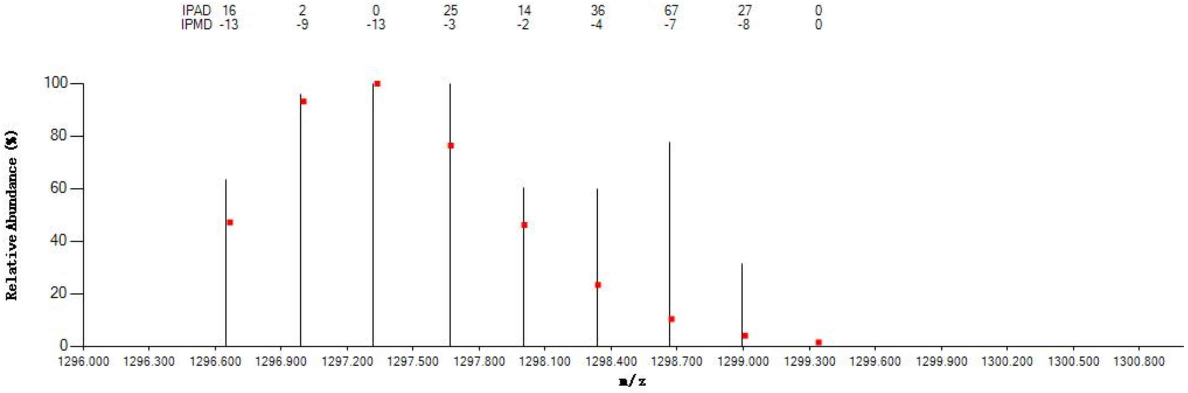


#### Annotated MS/MS Spectrum

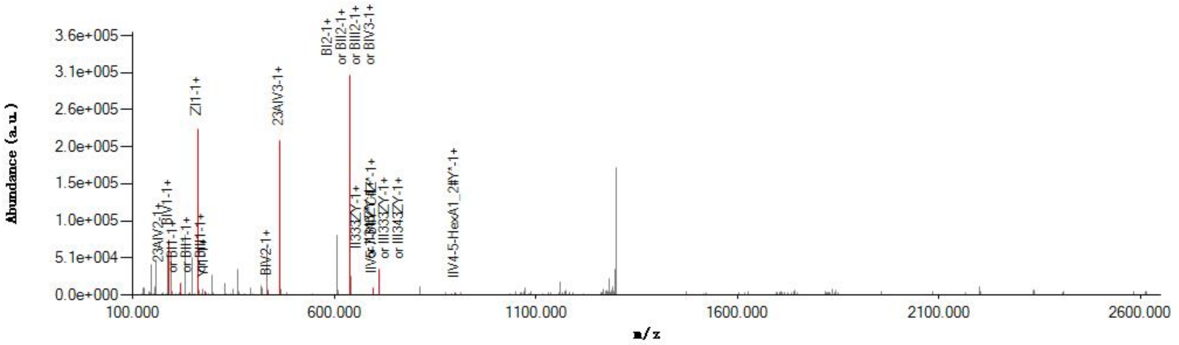


#### Graphical Fragmentation Map

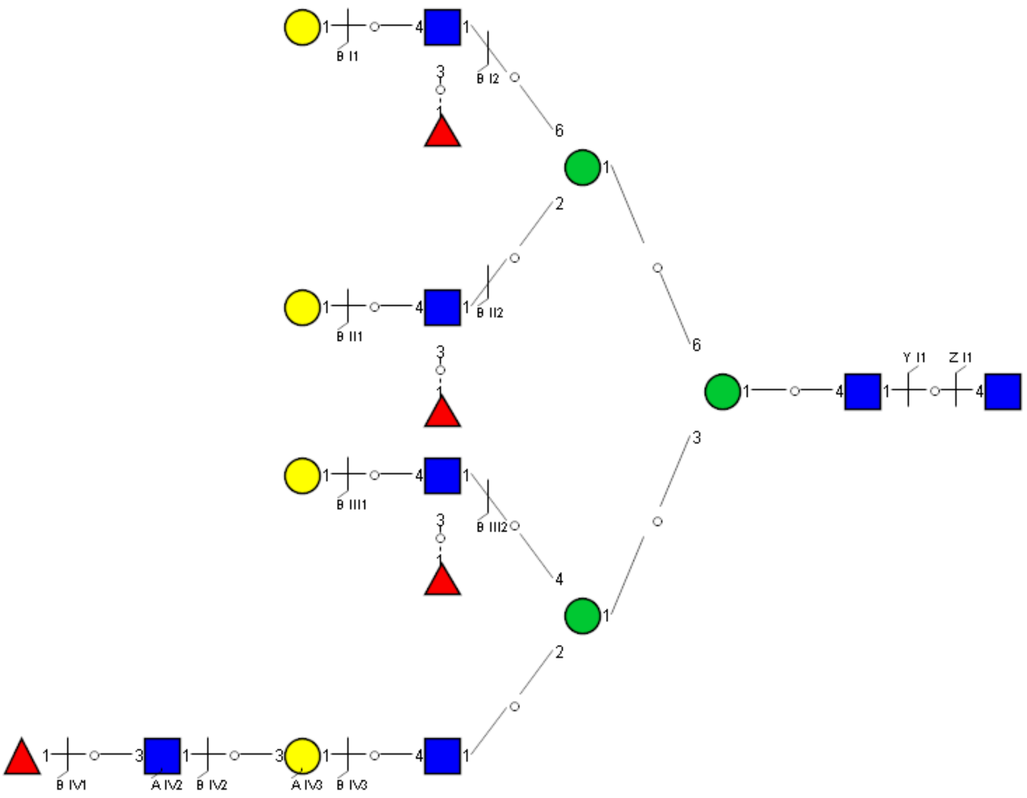

Spectrum Index: 41819(NoBHs:1)

Precursor iEF Map

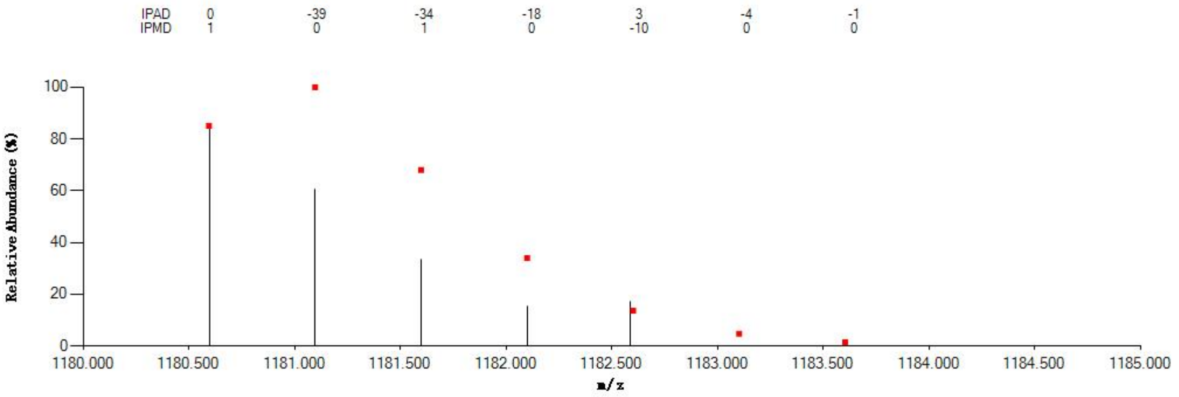

Annotated MS/MS Spectrum

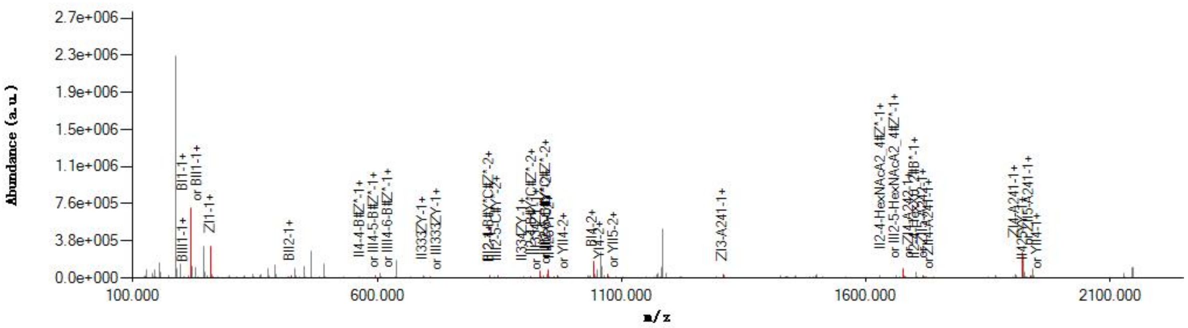

Graphical Fragmentation Map

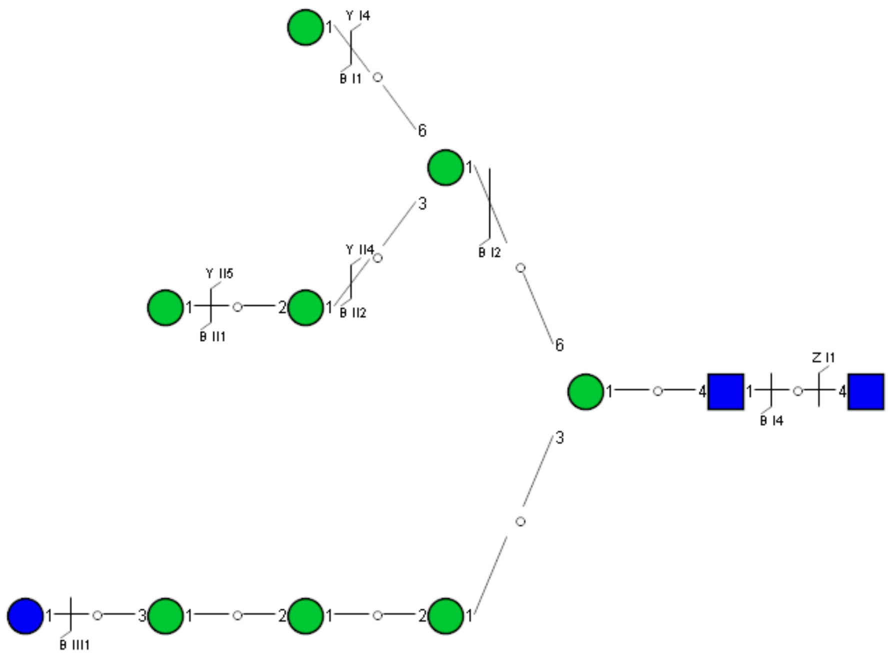

Spectrum Index: 41939(NoBHs:1)

#### Precursor iEF Map

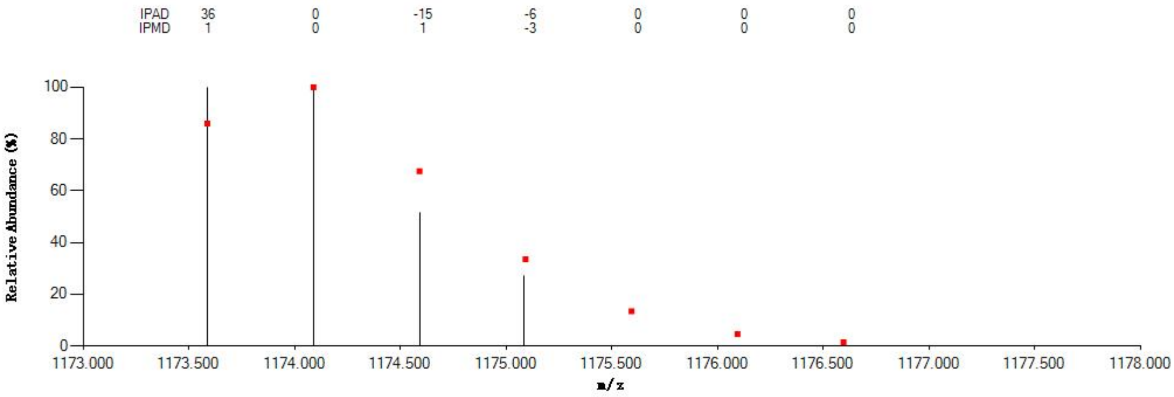


#### Annotated MS/MS Spectrum

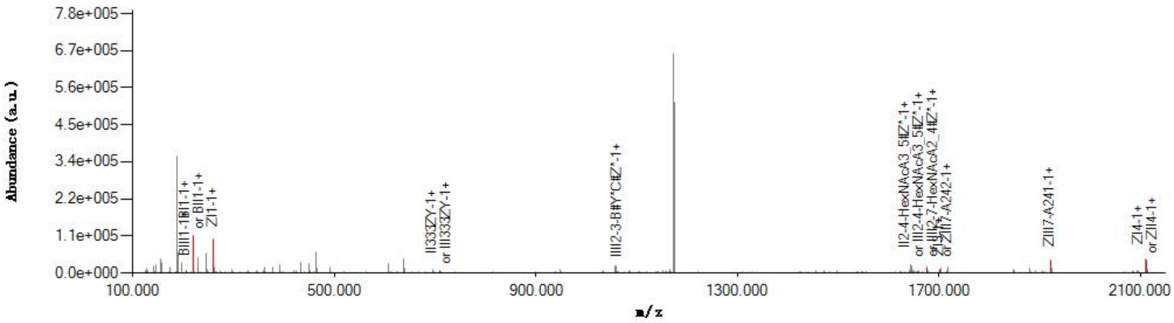


#### Graphical Fragmentation Map

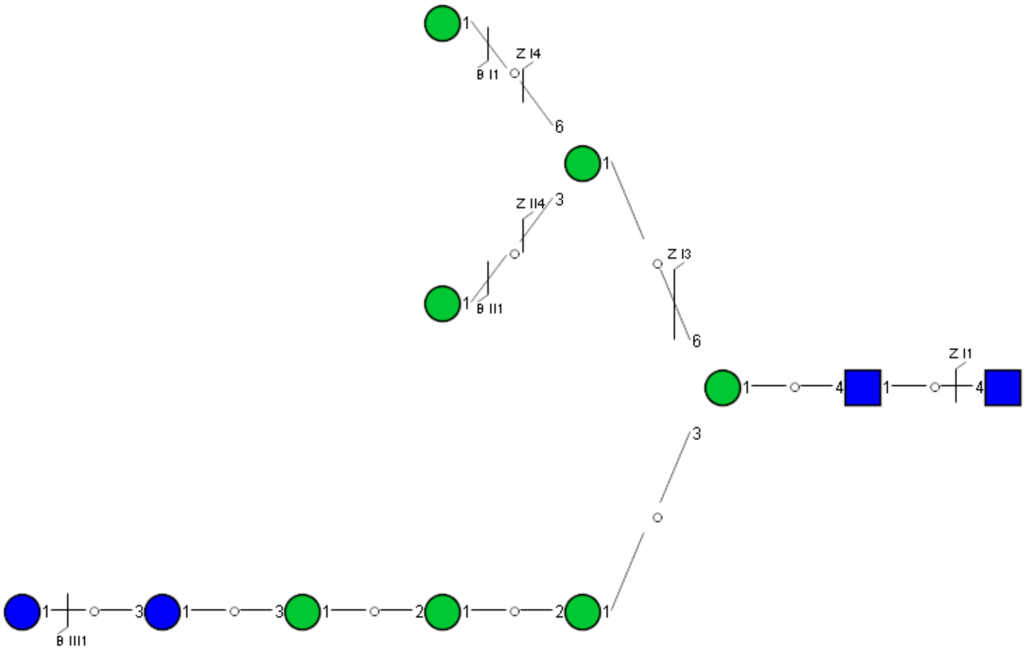

### Spectrum Index: 42097(NoBHs:1)

#### Precursor iEF Map

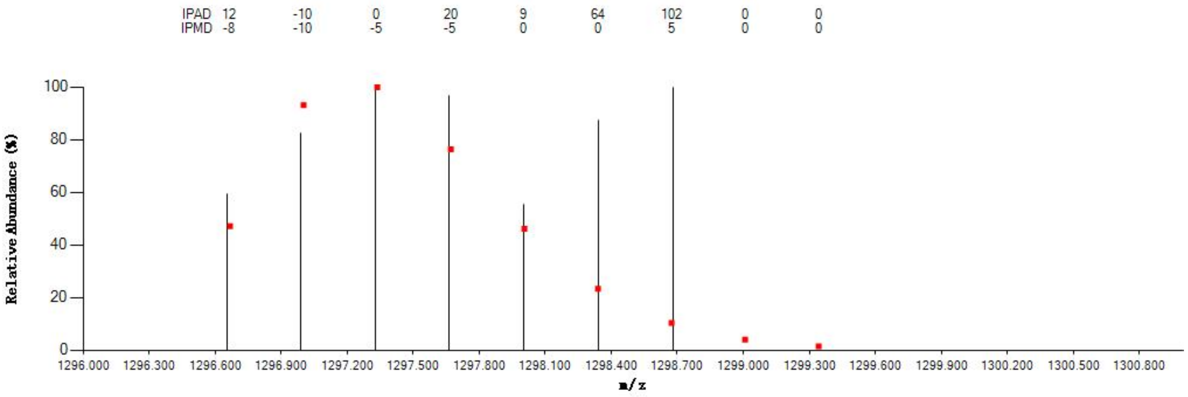


#### Annotated MS/MS Spectrum

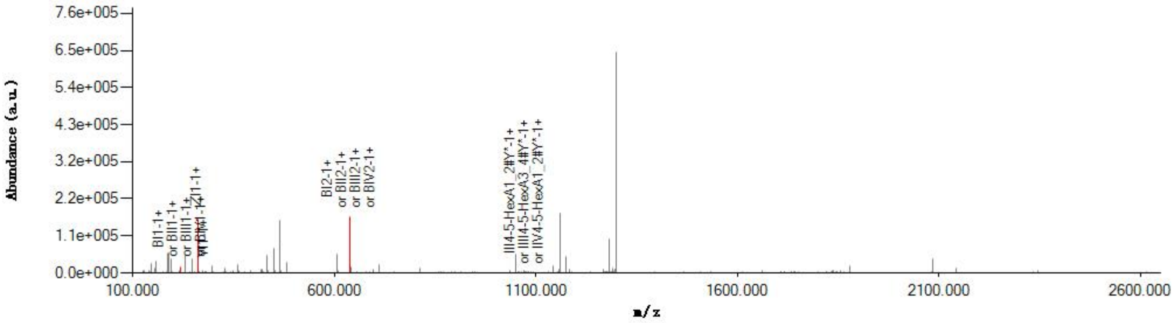


#### Graphical Fragmentation Map

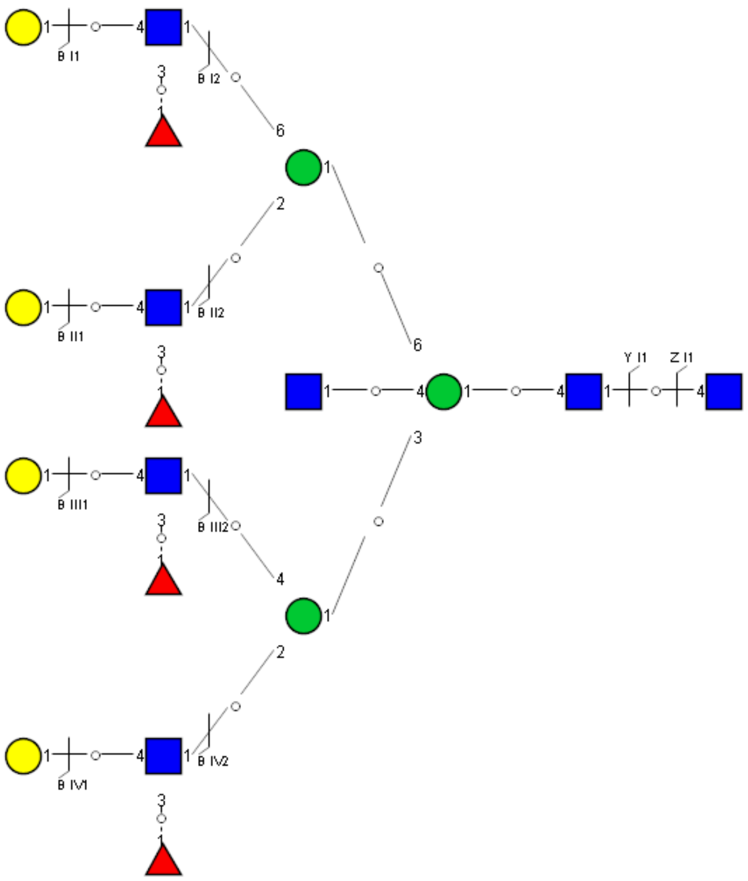

### Spectrum Index: 42209(NoBHs:2)

#### Precursor iEF Map

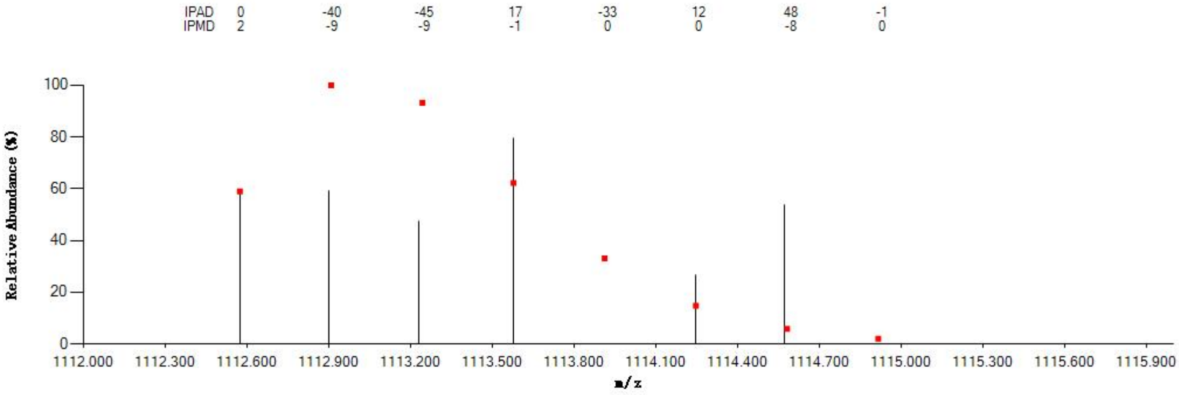


#### Annotated MS/MS Spectrum

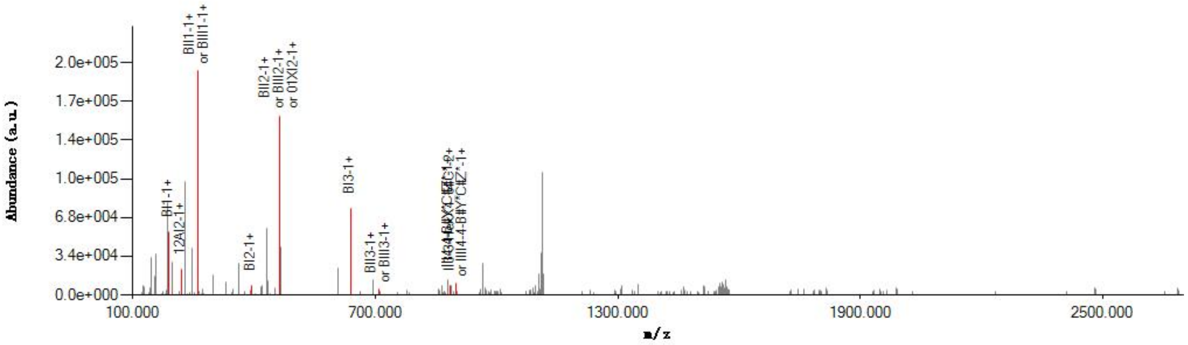


#### Graphical Fragmentation Map

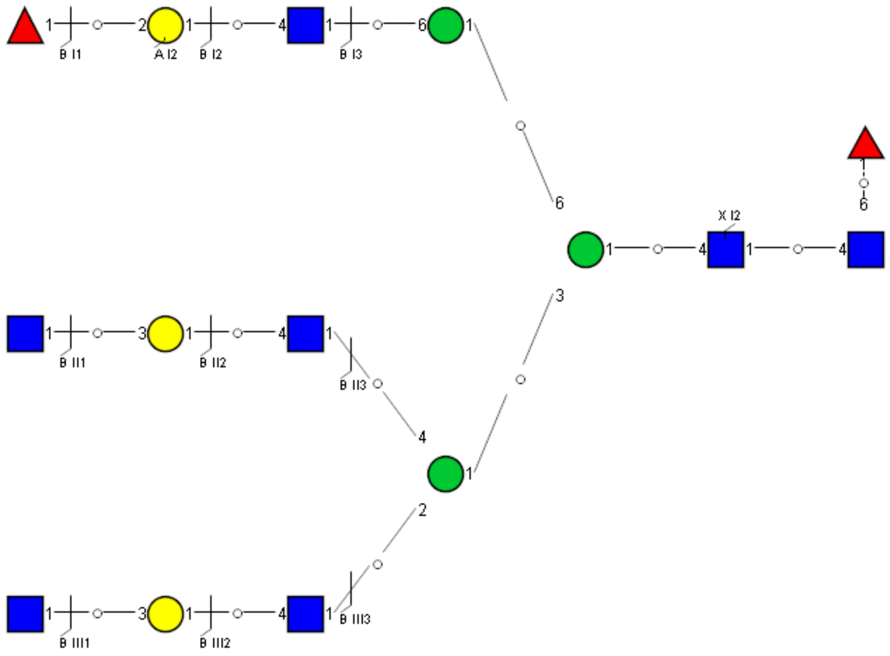

Spectrum Index: 42344(NoBHs:2)

#### Precursor iEF Map

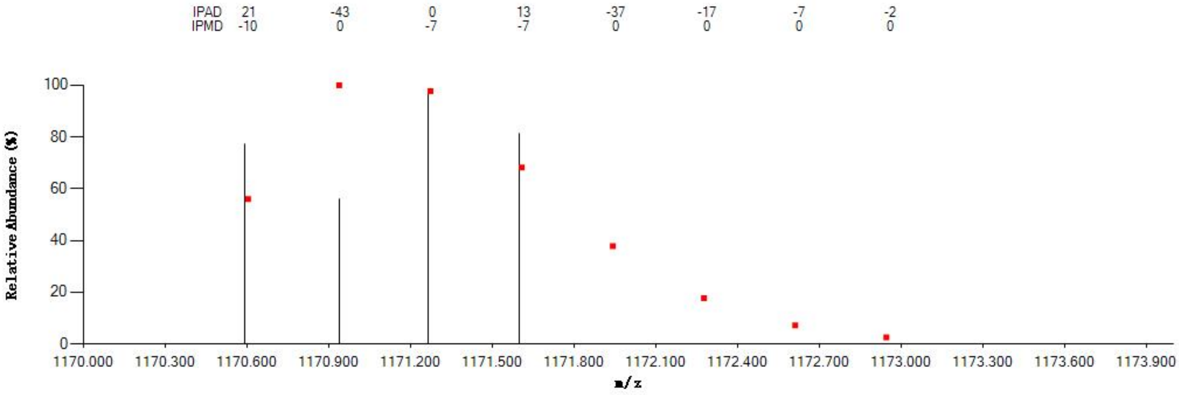


#### Annotated MS/MS Spectrum

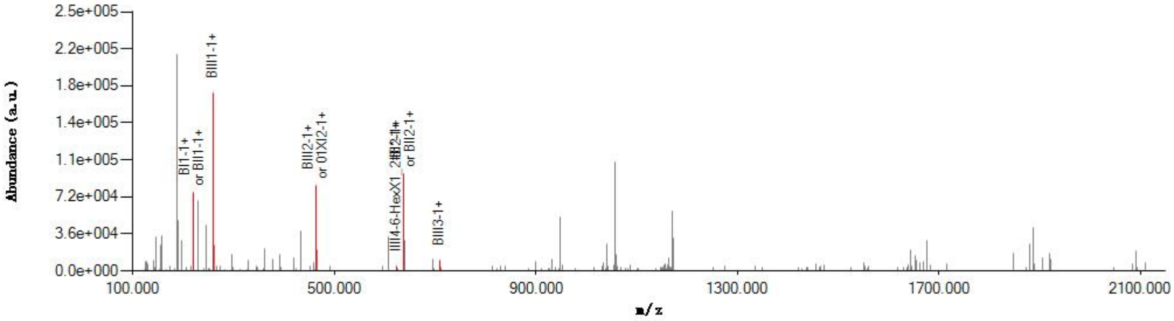


#### Graphical Fragmentation Map

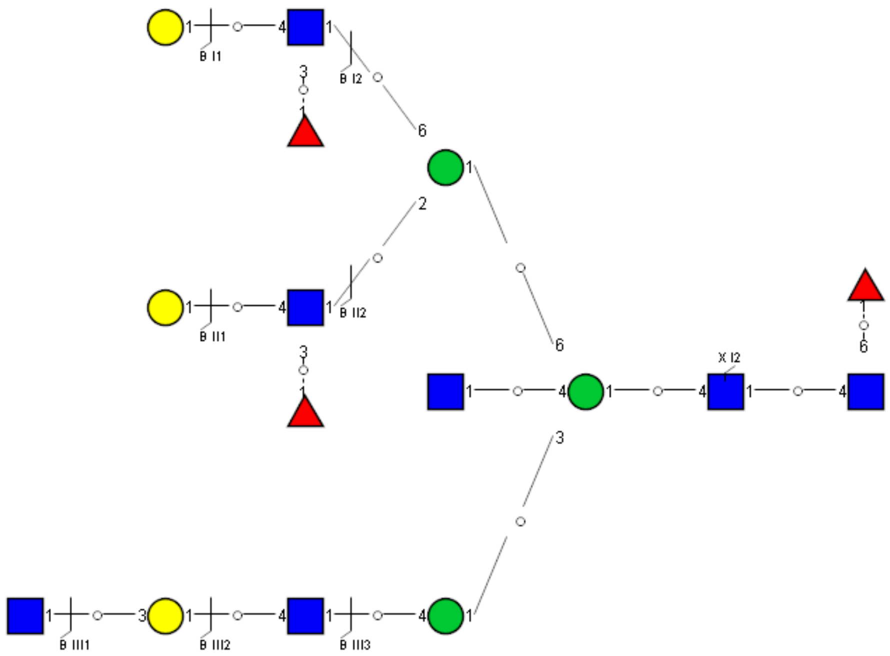

Spectrum Index: 42344(NoBHs:2)

Precursor iEF Map

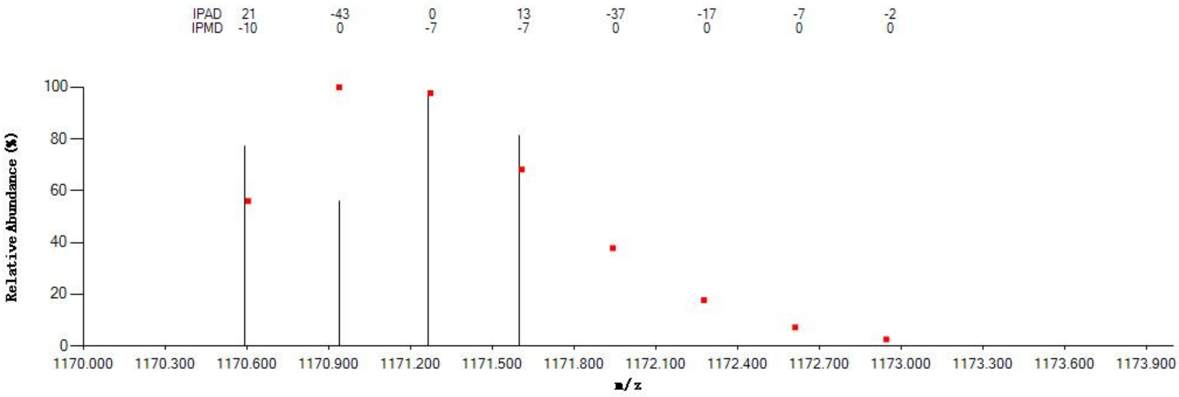


Annotated MS/MS Spectrum

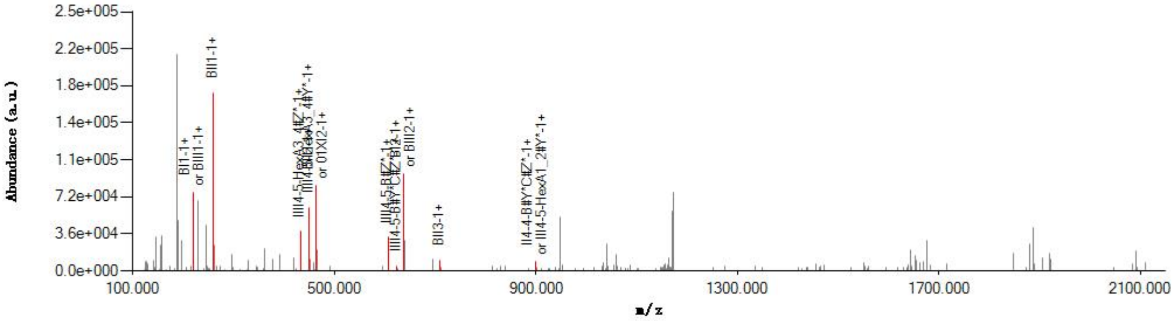


Graphical Fragmentation Map

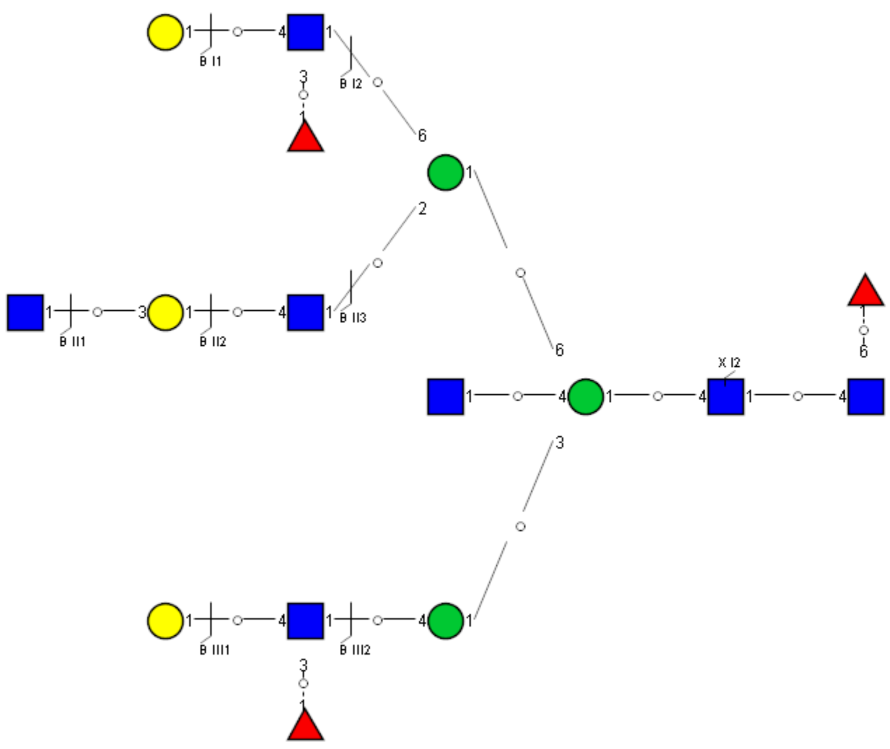

Spectrum Index: 65907(NoBHs:1)

Precursor iEF Map

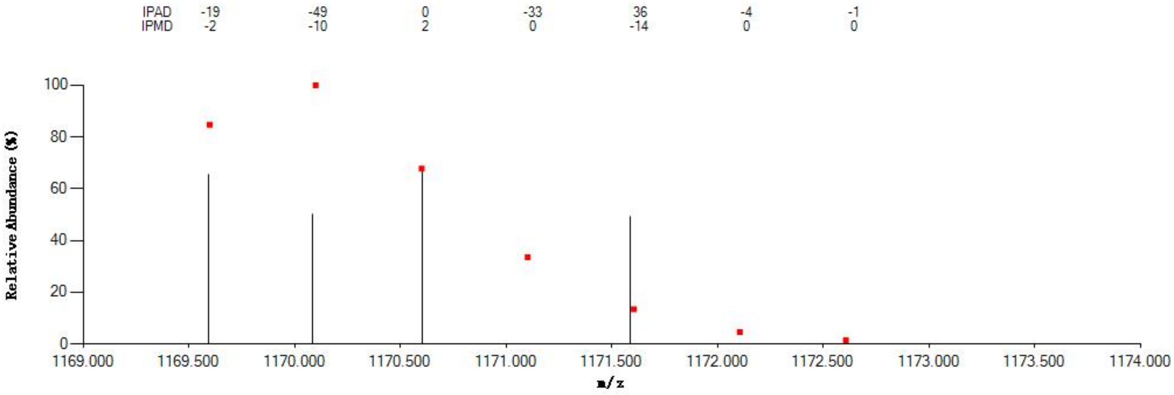


Annotated MS/MS Spectrum

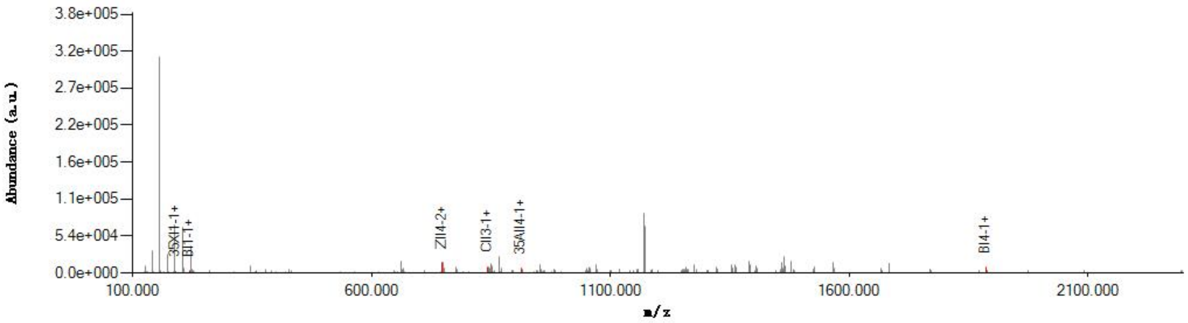


Graphical Fragmentation Map

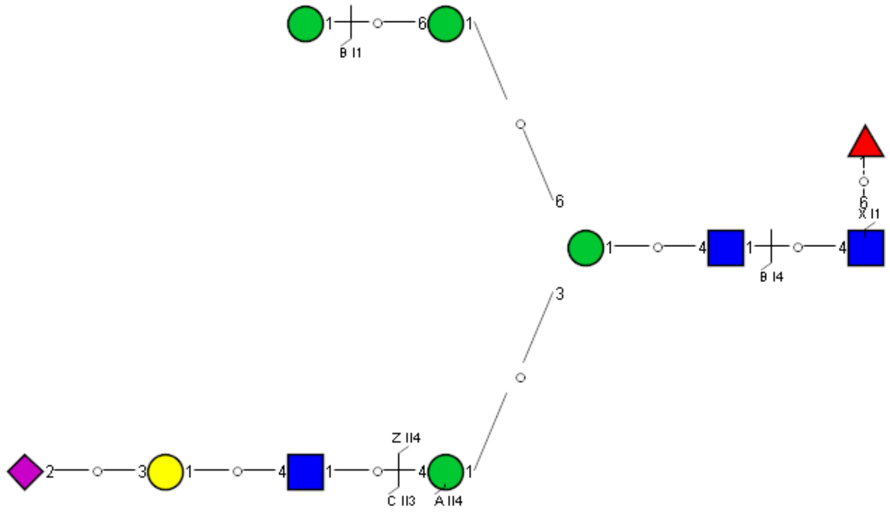
