## Supplemental Information 10 for "A draft of human N-glycans of glycoRNA"

### Spectrum Index: 28165(NoBHs:1)

#### Precursor iEF Map

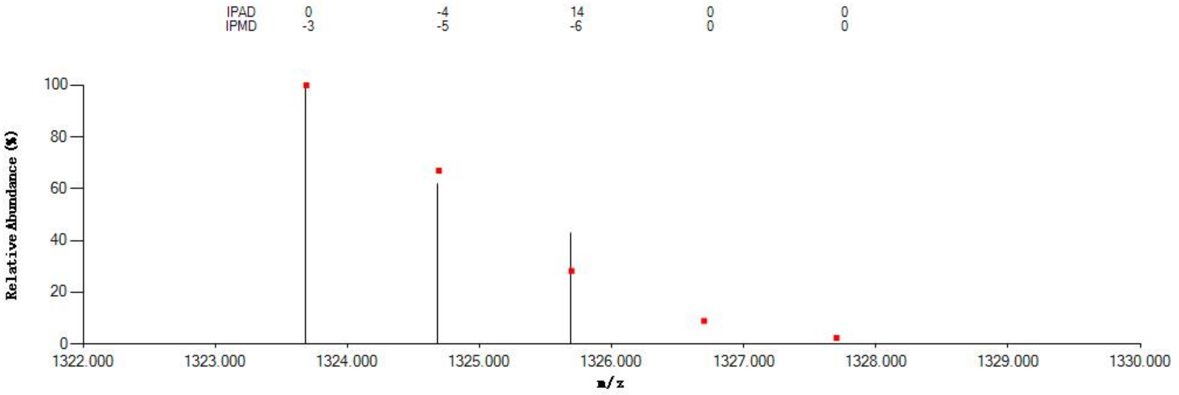


#### Annotated MS/MS Spectrum

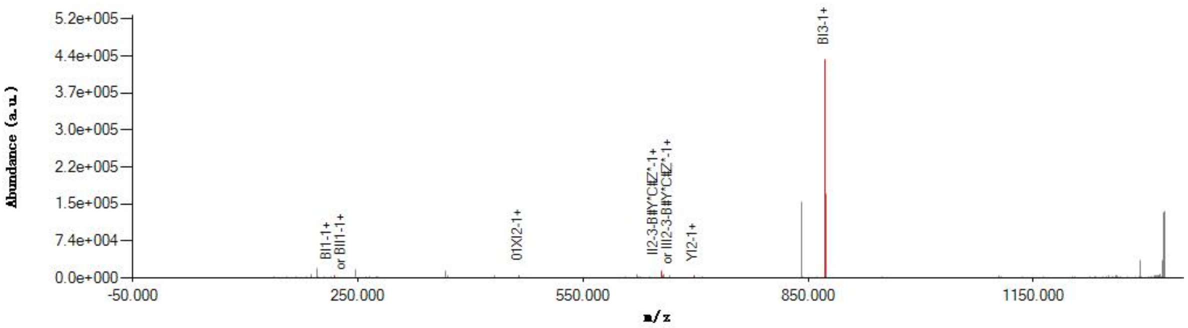


#### Graphical Fragmentation Map

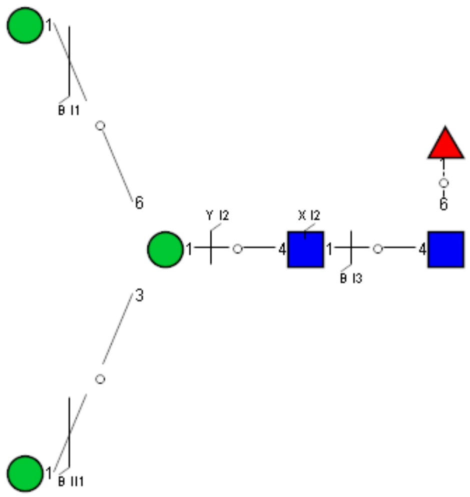

### Spectrum Index: 29360(NoBHs:1)

#### Precursor iEF Map

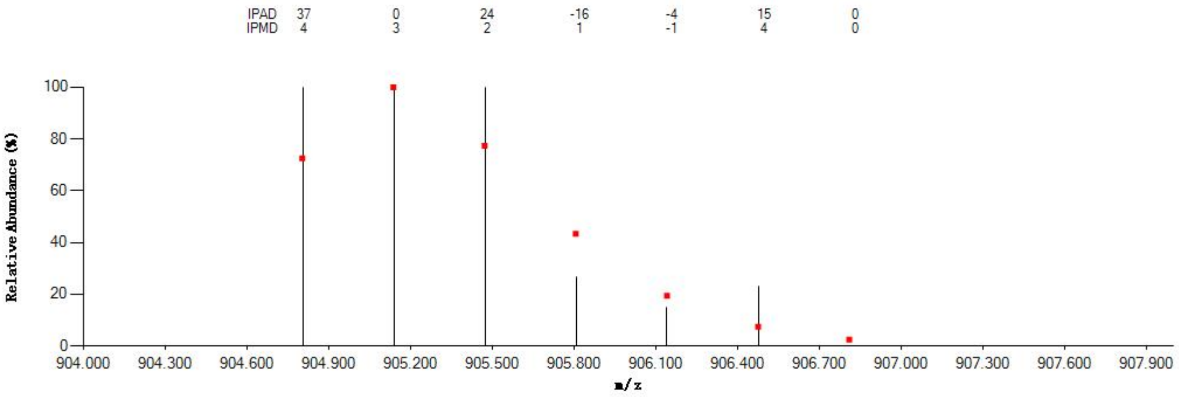


#### Annotated MS/MS Spectrum

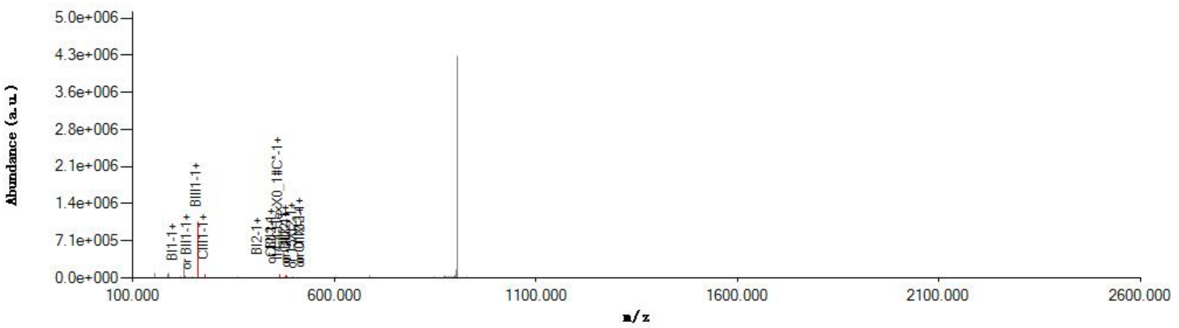


#### Graphical Fragmentation Map

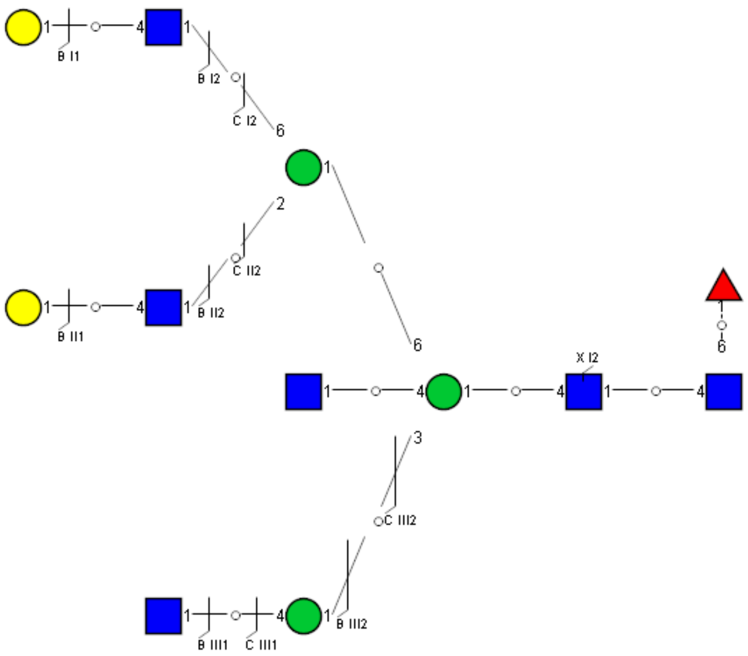

Spectrum Index: 29910(NoBHs:1)

#### Precursor iEF Map

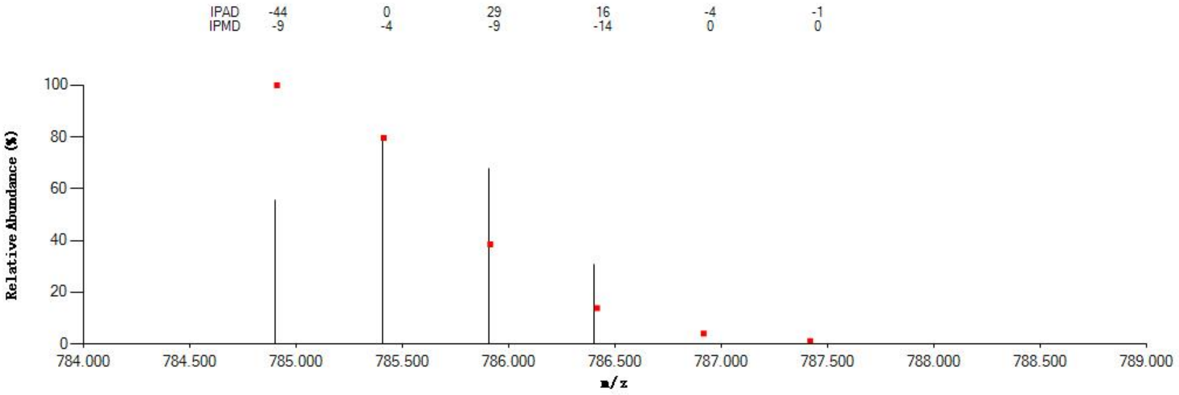


#### Annotated MS/MS Spectrum

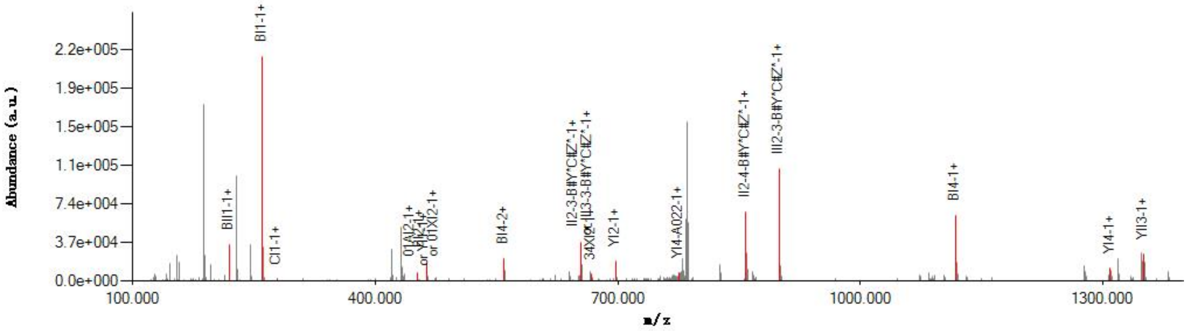


#### Graphical Fragmentation Map

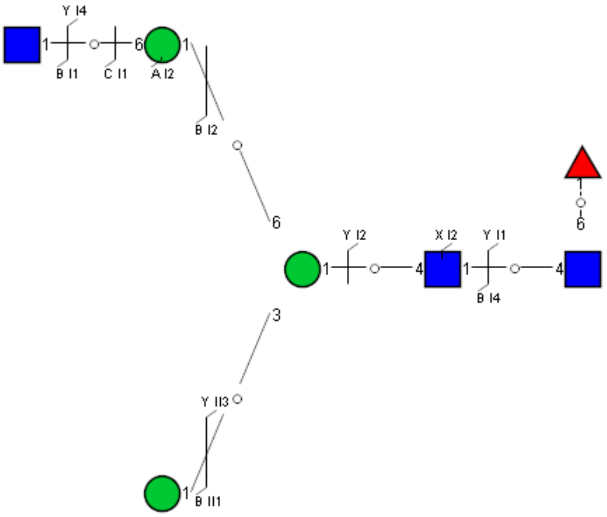

Spectrum Index: 31552(NoBHs:1)

#### Precursor iEF Map

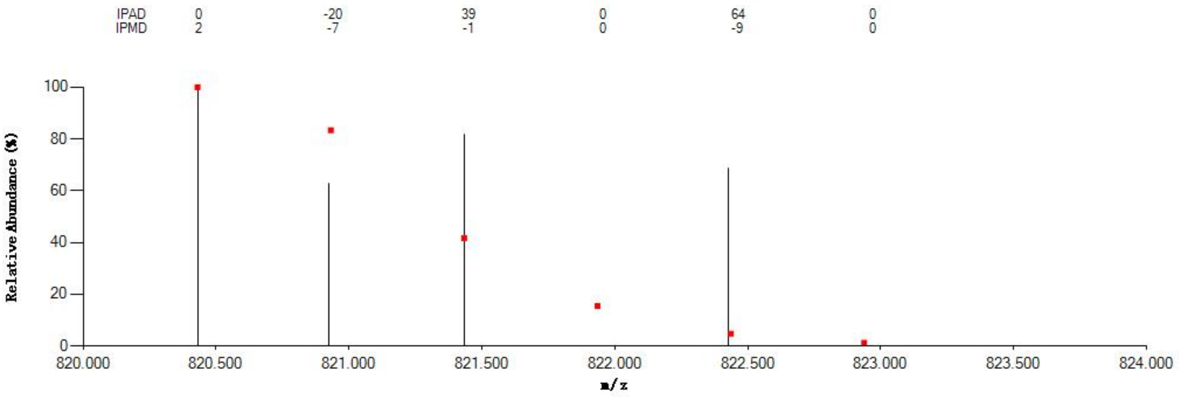


#### Annotated MS/MS Spectrum

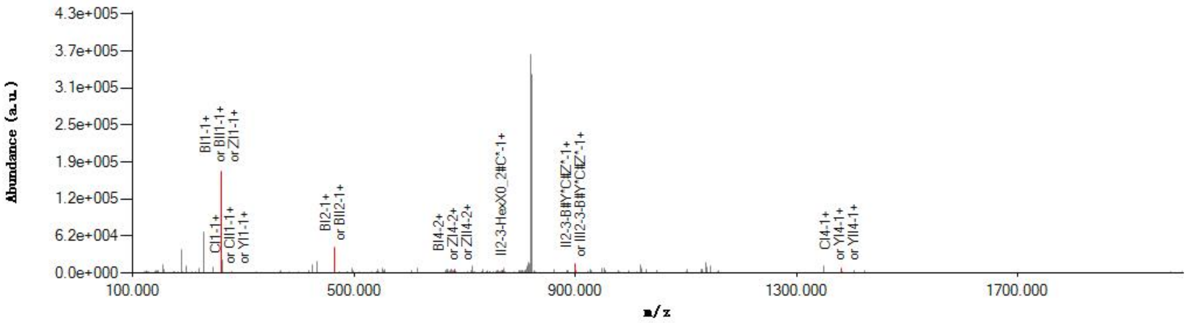


#### Graphical Fragmentation Map

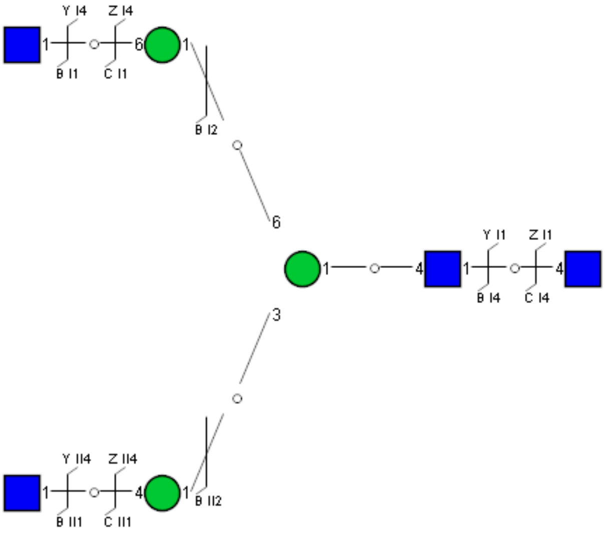

Spectrum Index: 32296(NoBHs:1)

#### Precursor iEF Map

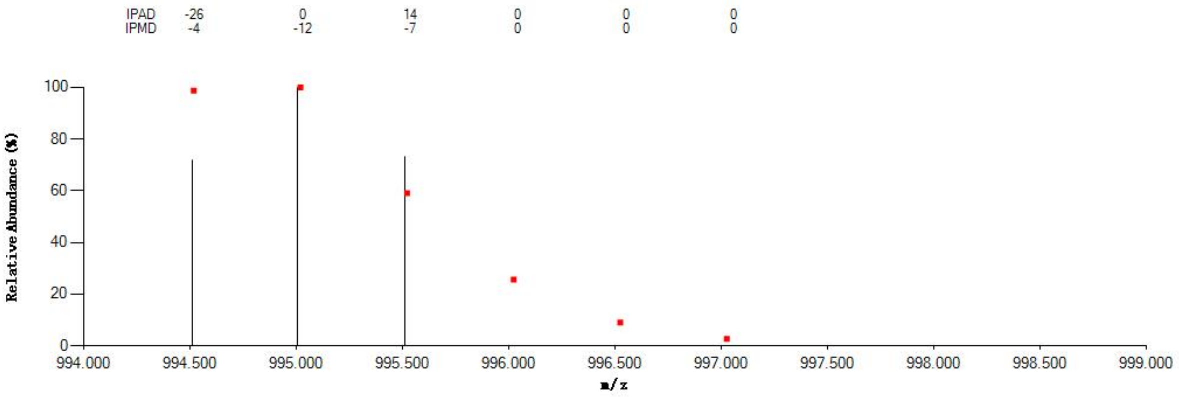


#### Annotated MS/MS Spectrum

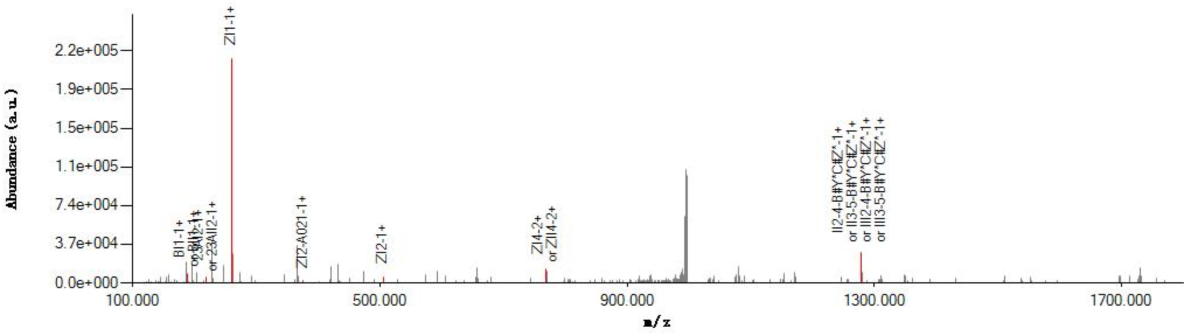


#### Graphical Fragmentation Map

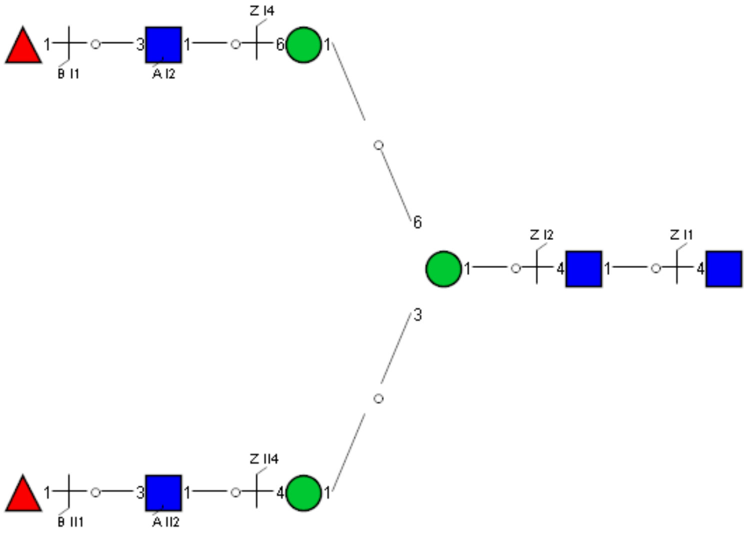

Spectrum Index: 33037(NoBHs:1)

#### Precursor iEF Map

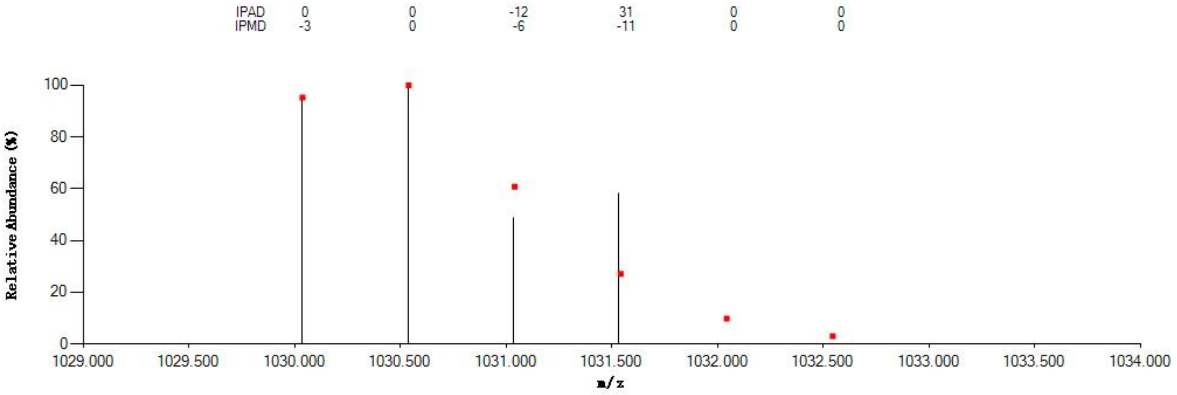


#### Annotated MS/MS Spectrum

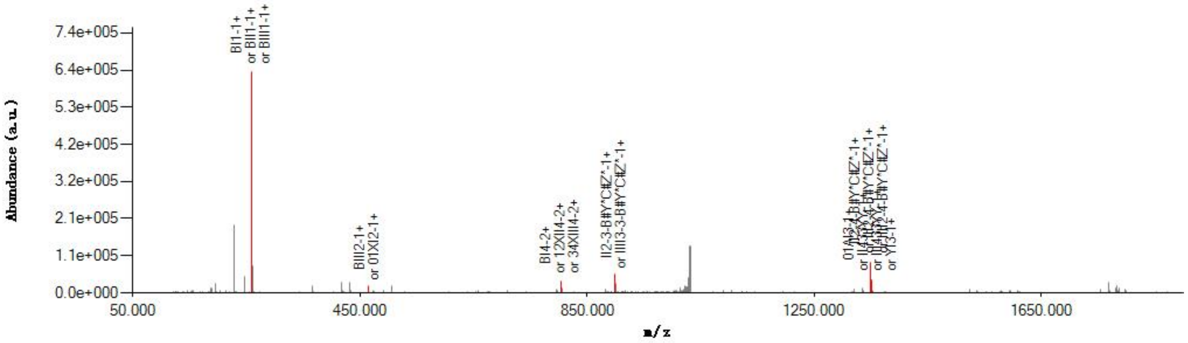


#### Graphical Fragmentation Map

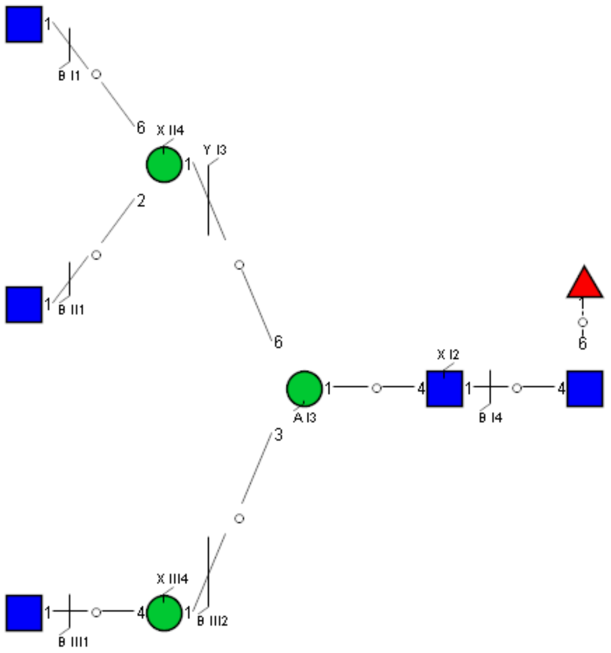

Spectrum Index: 33267(NoBHs:1)

#### Precursor iEF Map

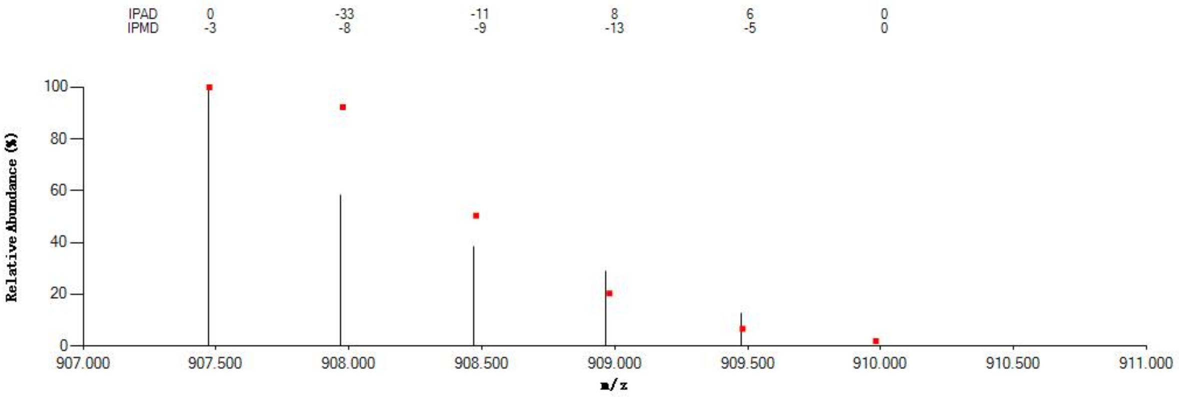

#### Annotated MS/MS Spectrum

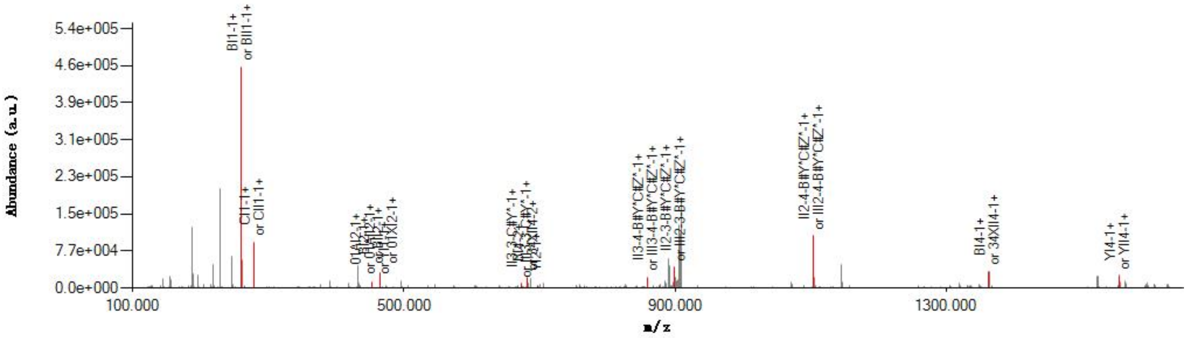

#### Graphical Fragmentation Map

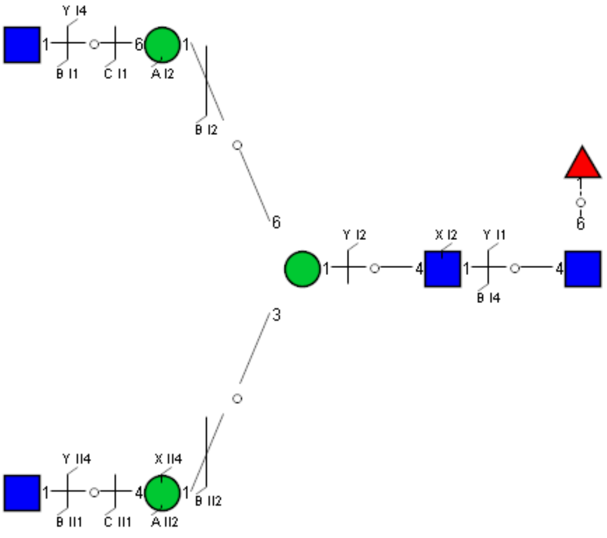

Spectrum Index: 33498(NoBHs:1)

#### Precursor iEF Map

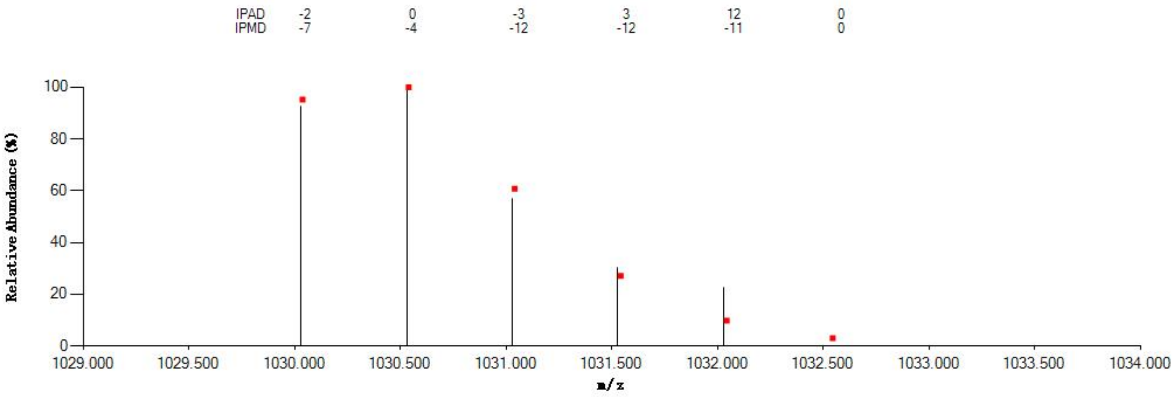


#### Annotated MS/MS Spectrum

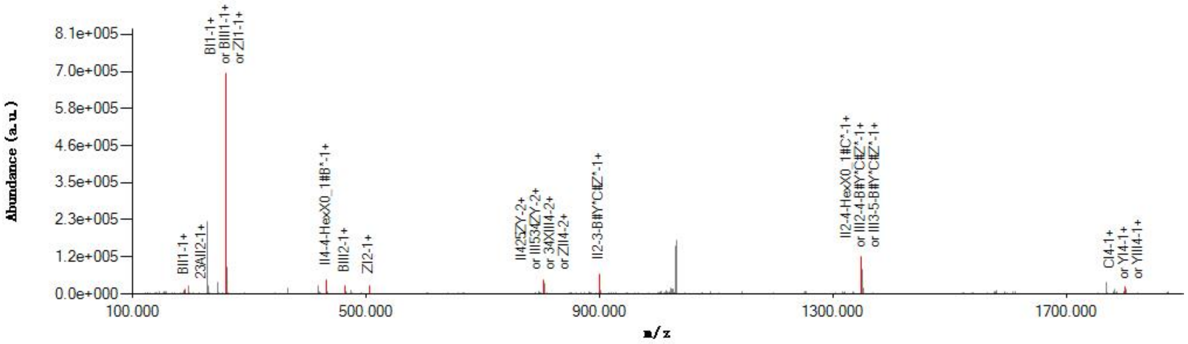


#### Graphical Fragmentation Map

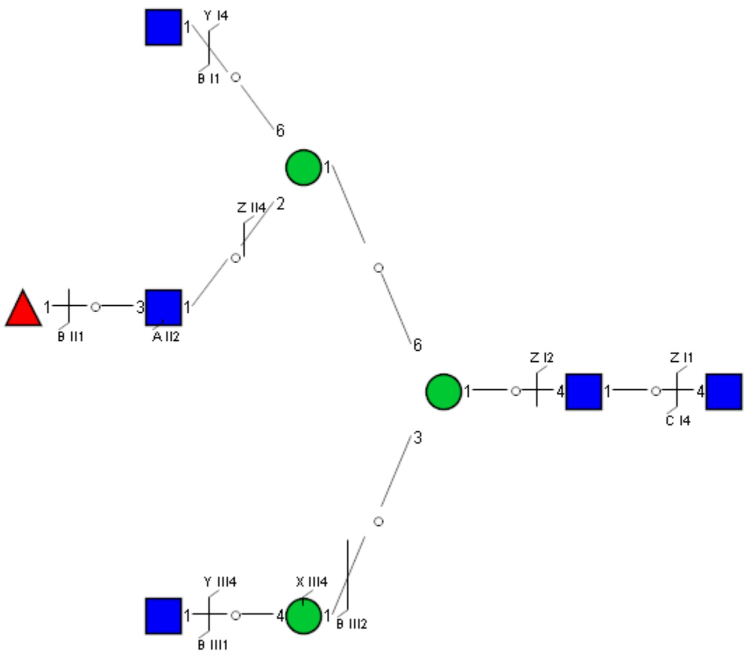

Spectrum Index: 33667(NoBHs:1)

#### Precursor iEF Map

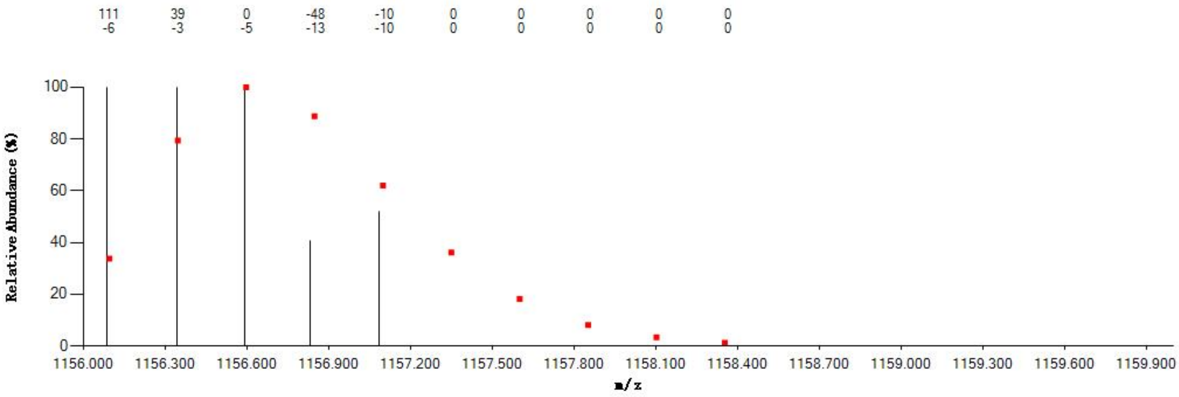


#### Annotated MS/MS Spectrum

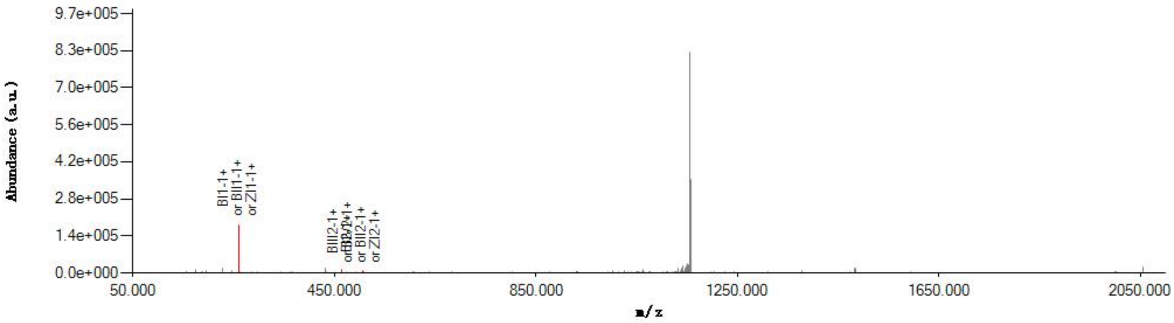


#### Graphical Fragmentation Map

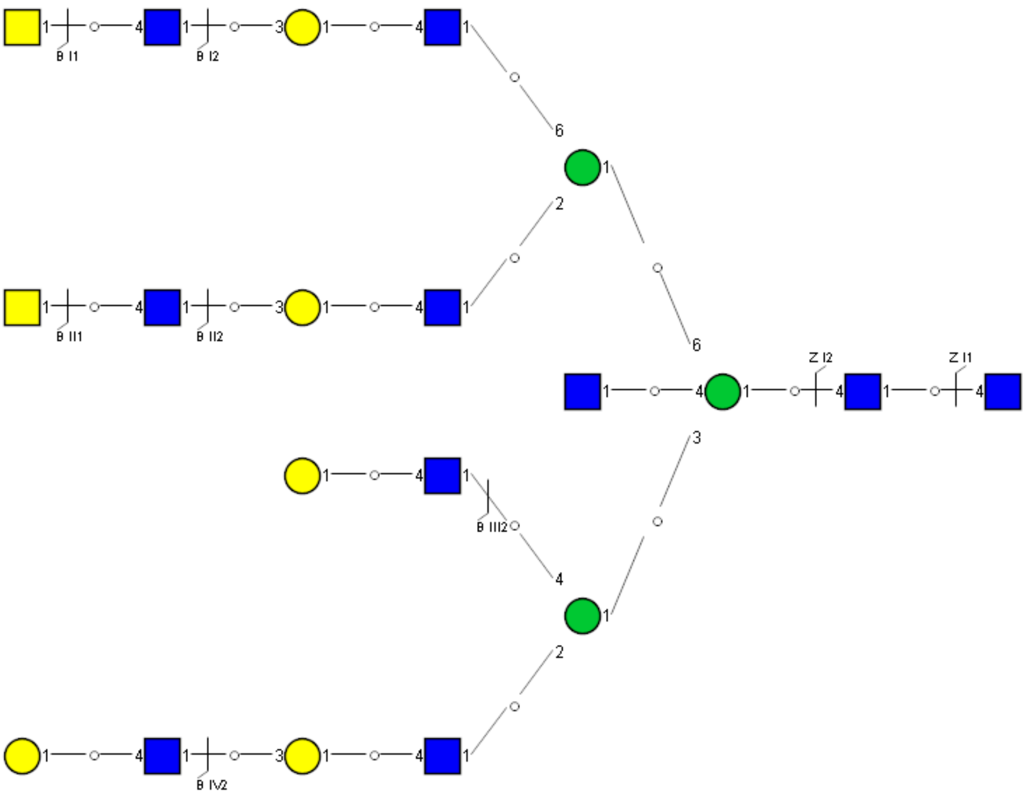

Spectrum Index: 33729(NoBHs:1)

#### Precursor iEF Map

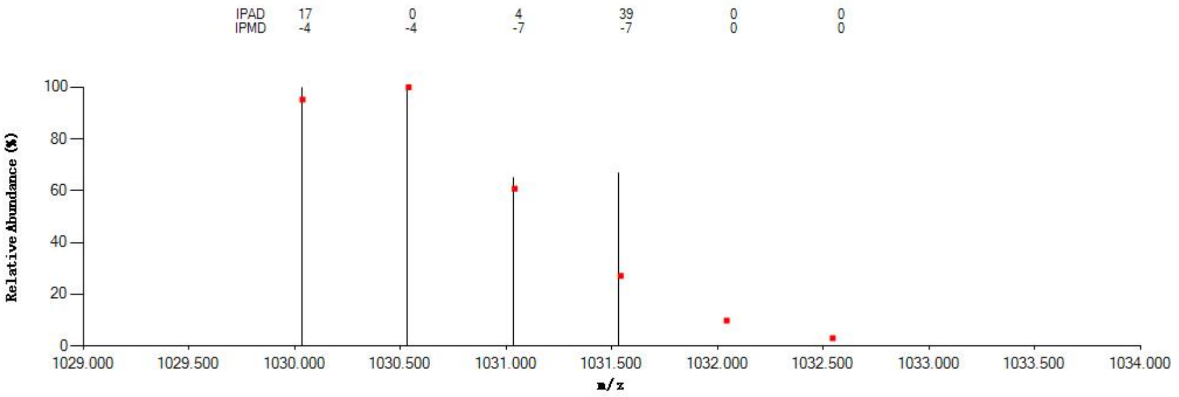


#### Annotated MS/MS Spectrum

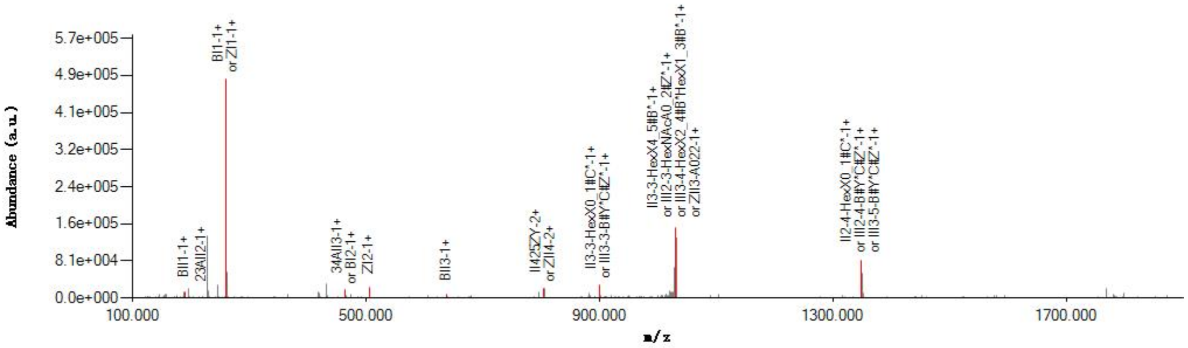


#### Graphical Fragmentation Map

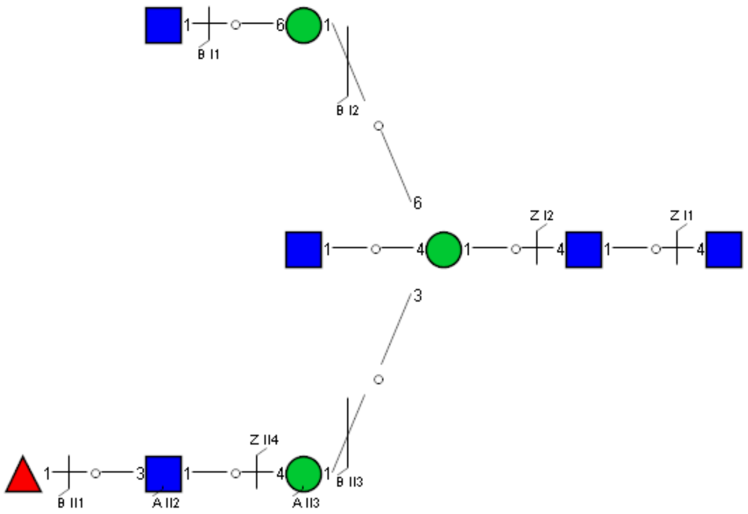

### Spectrum Index: 33857(NoBHs:1)

#### Precursor iEF Map

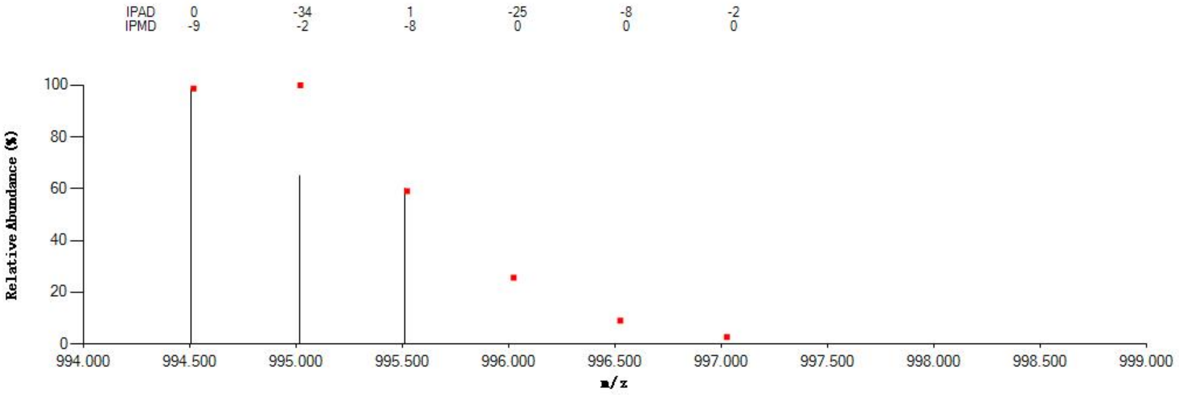


#### Annotated MS/MS Spectrum

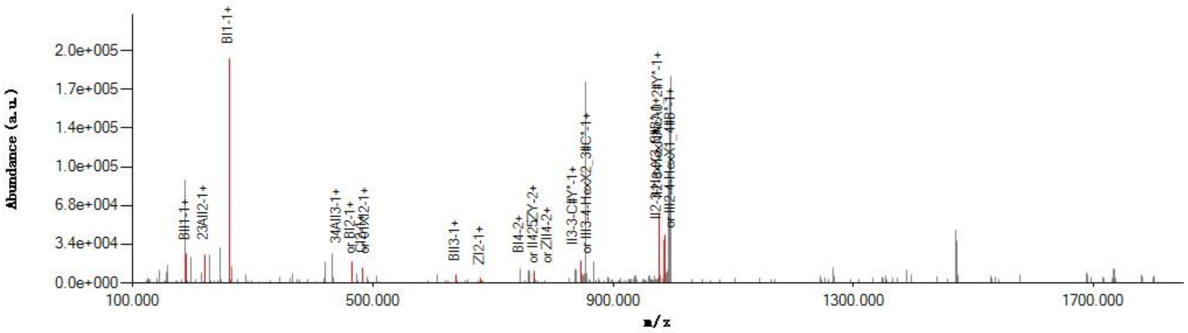


#### Graphical Fragmentation Map

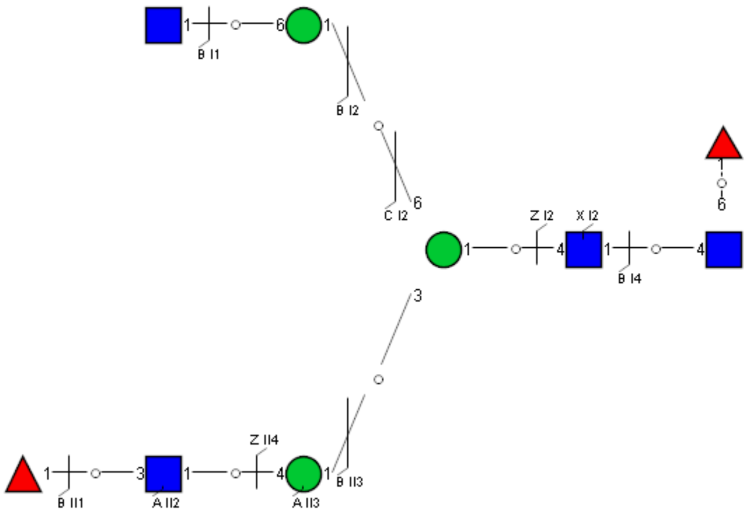

Spectrum Index: 33877(NoBHs:1)

#### Precursor iEF Map

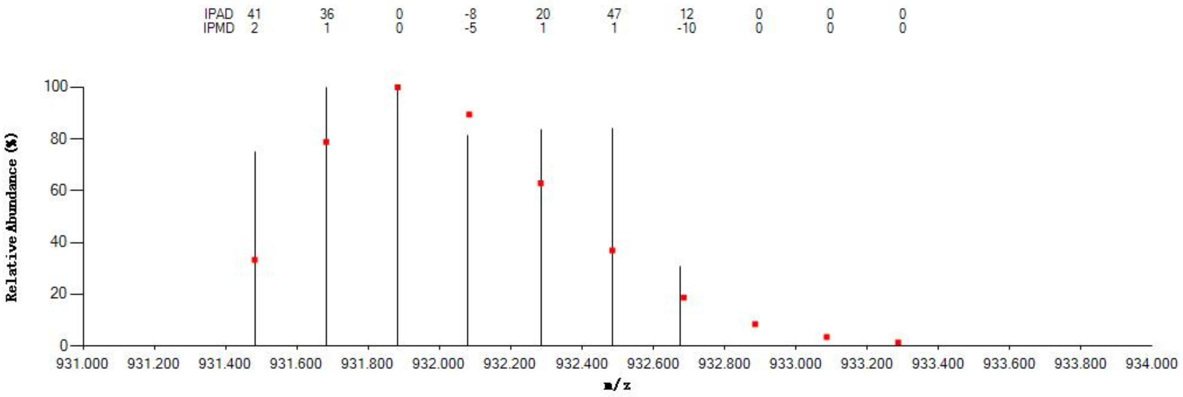

#### Annotated MS/MS Spectrum

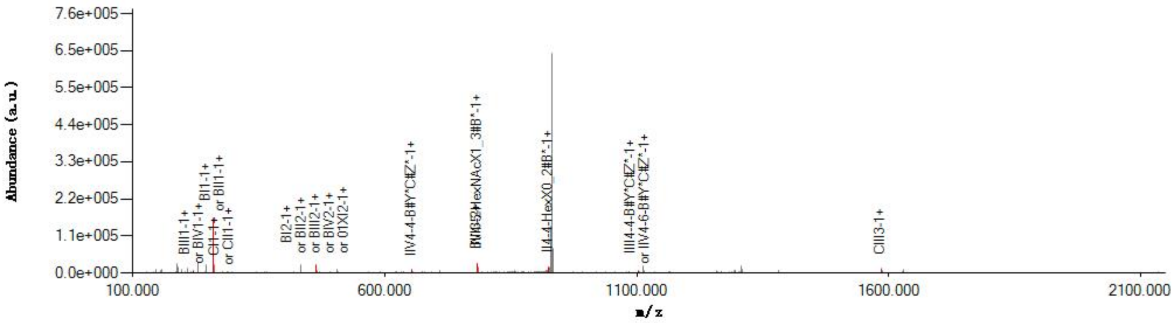

#### Graphical Fragmentation Map

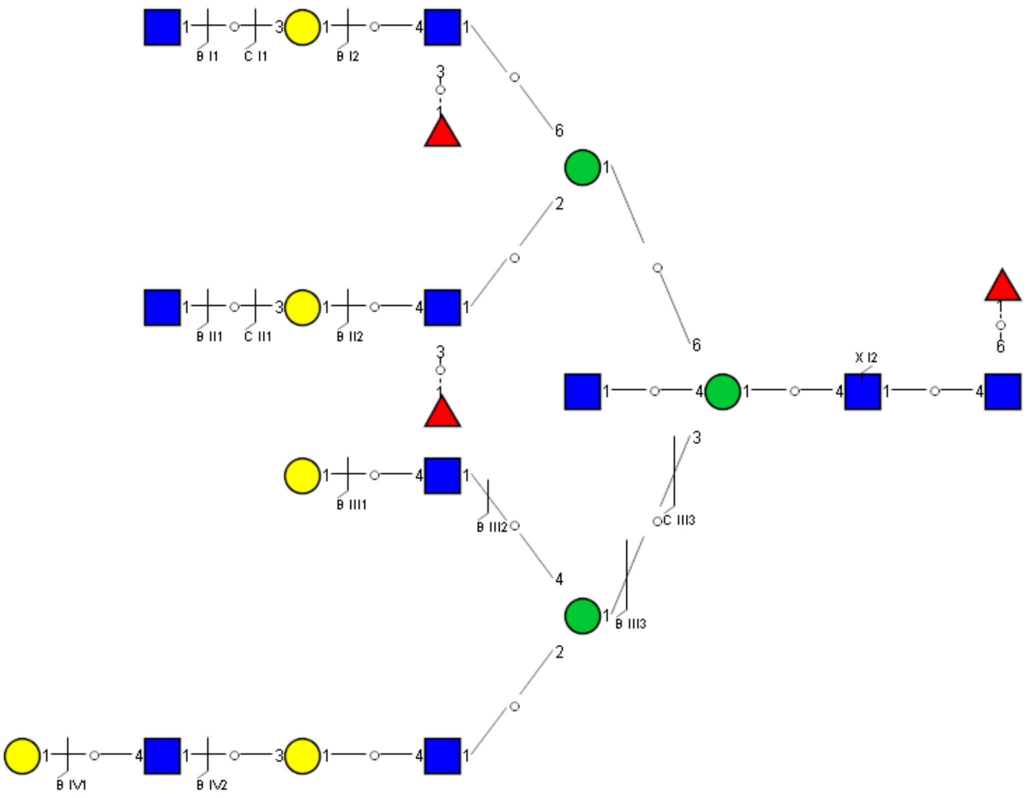

Spectrum Index: 33932(NoBHs:1)

#### Precursor iEF Map

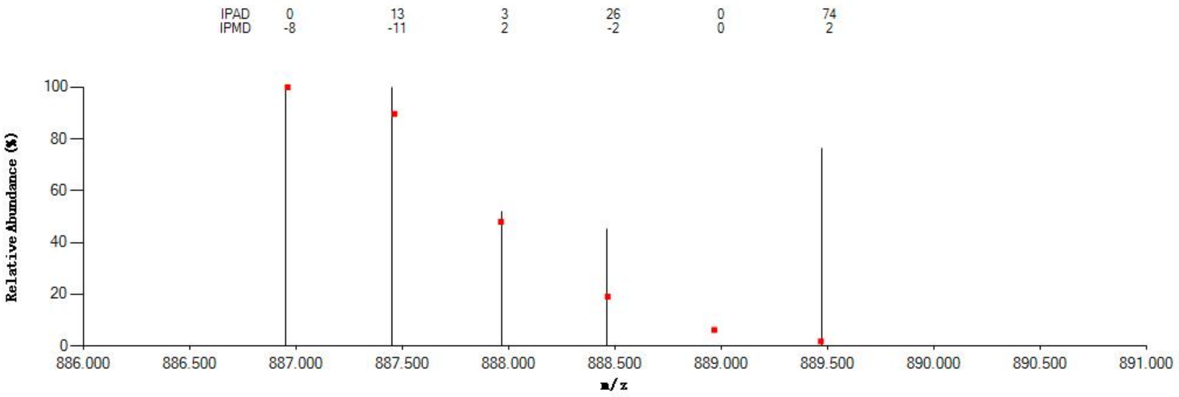


#### Annotated MS/MS Spectrum

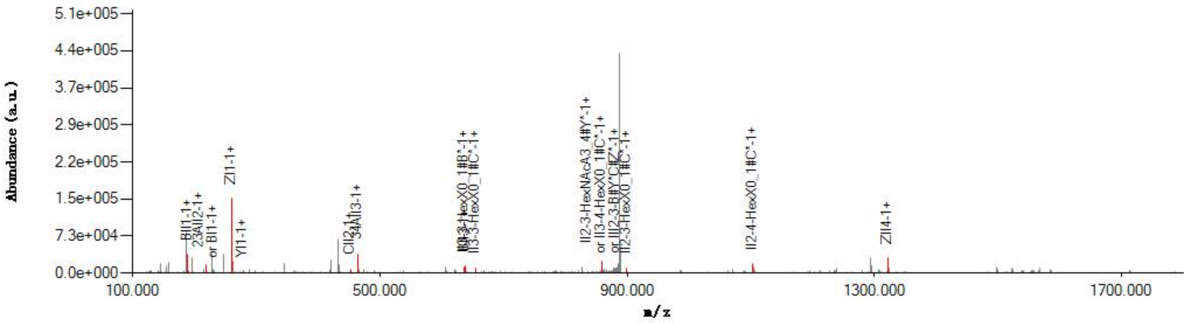


#### Graphical Fragmentation Map

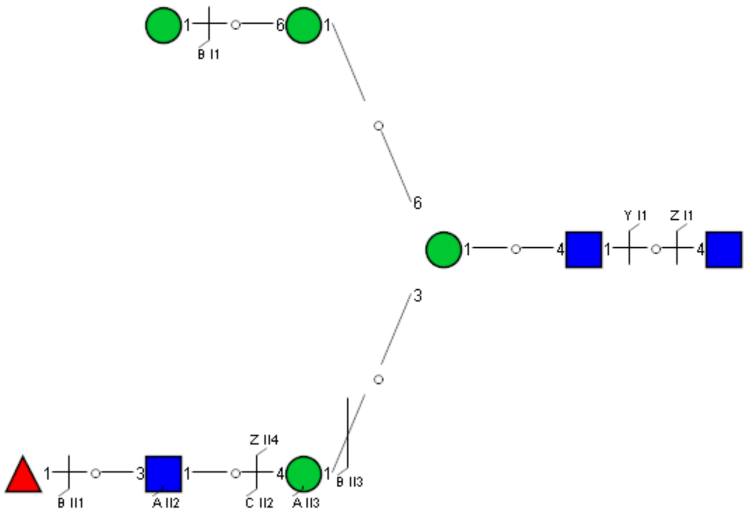

Spectrum Index: 33935(NoBHs:1)

Precursor iEF Map

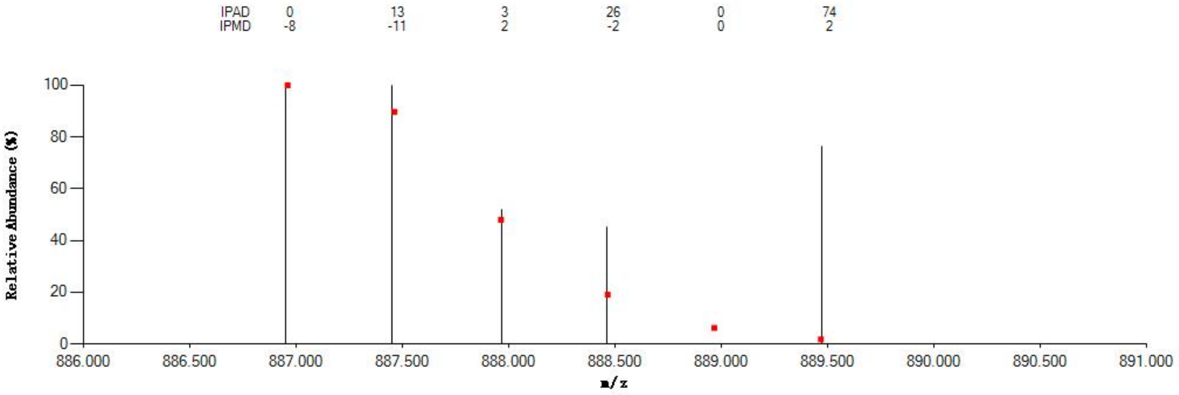


Annotated MS/MS Spectrum

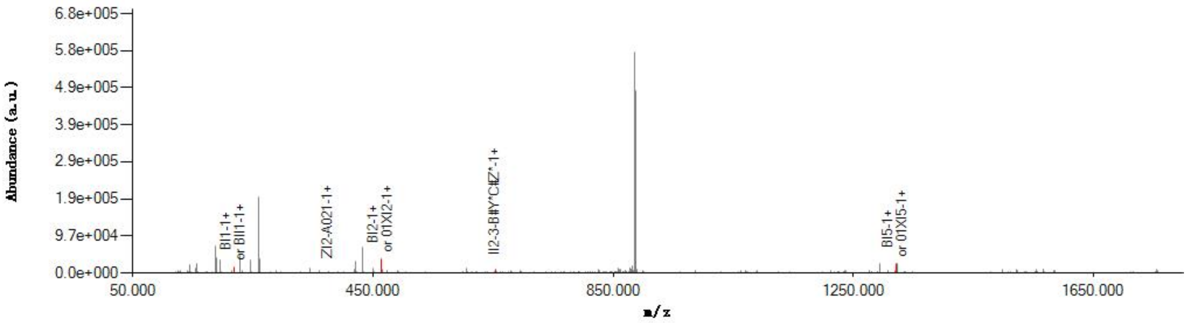


Graphical Fragmentation Map

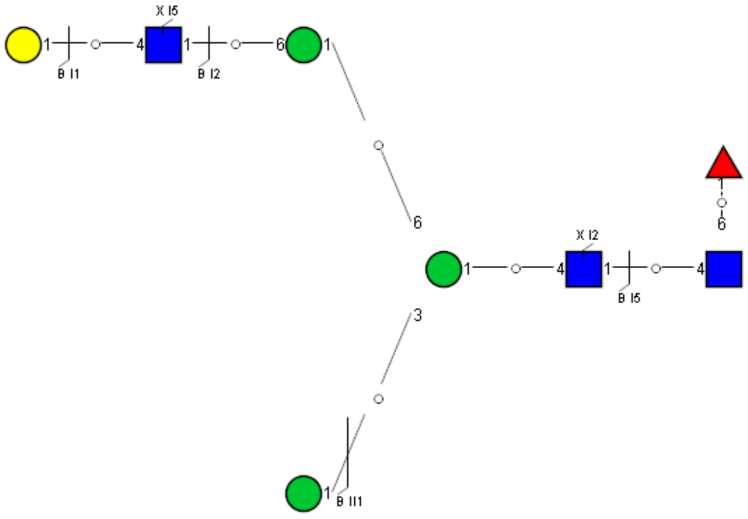

Spectrum Index: 33967(NoBHs:1)

#### Precursor iEF Map

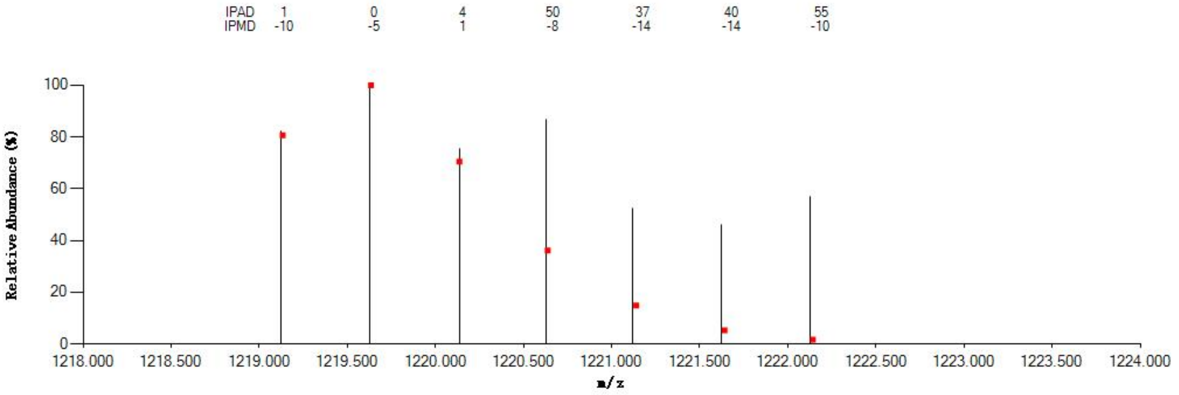

#### Annotated MS/MS Spectrum

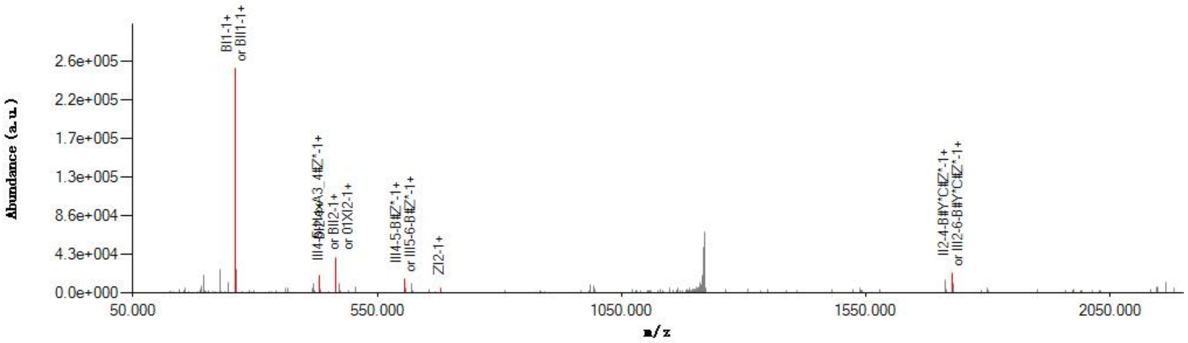

#### Graphical Fragmentation Map

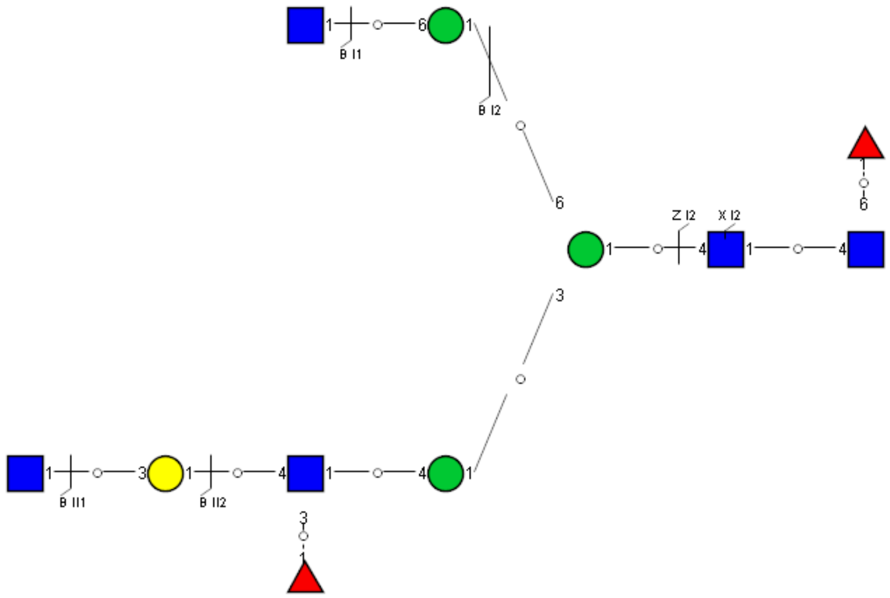

Spectrum Index: 34013(NoBHs:1)

#### Precursor iEF Map

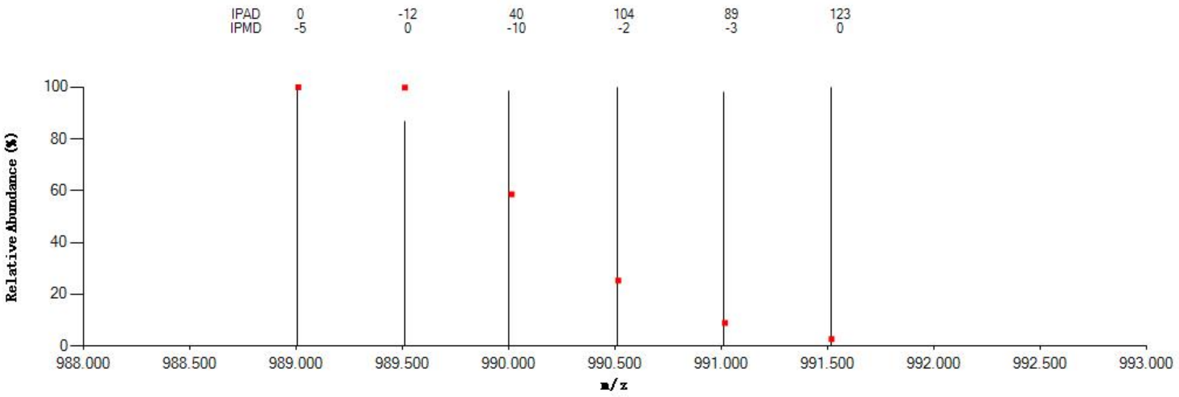


#### Annotated MS/MS Spectrum

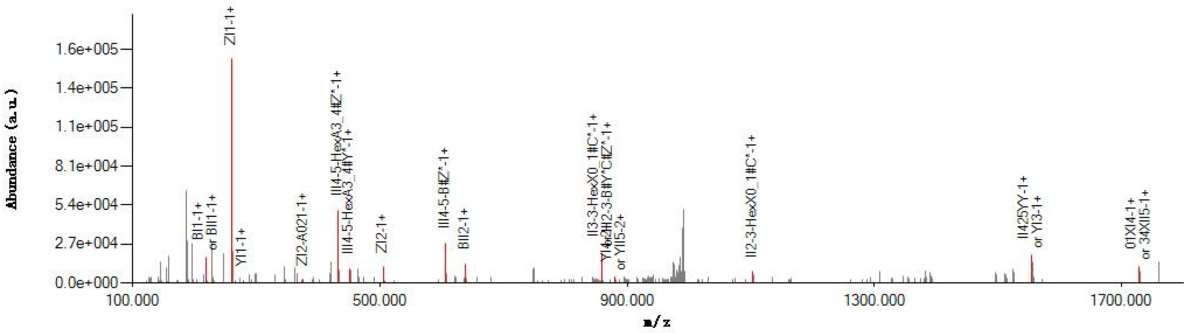


#### Graphical Fragmentation Map

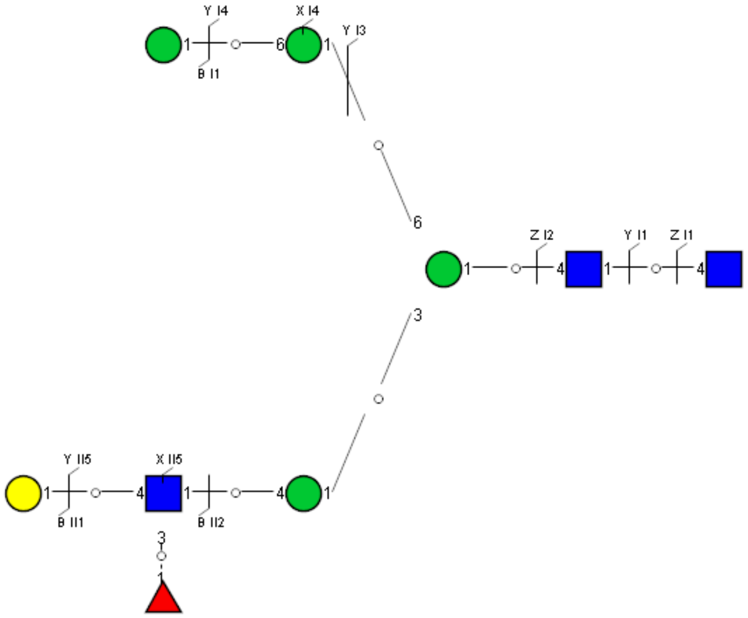

### Spectrum Index: 34026(NoBHs:1)

#### Precursor iEF Map

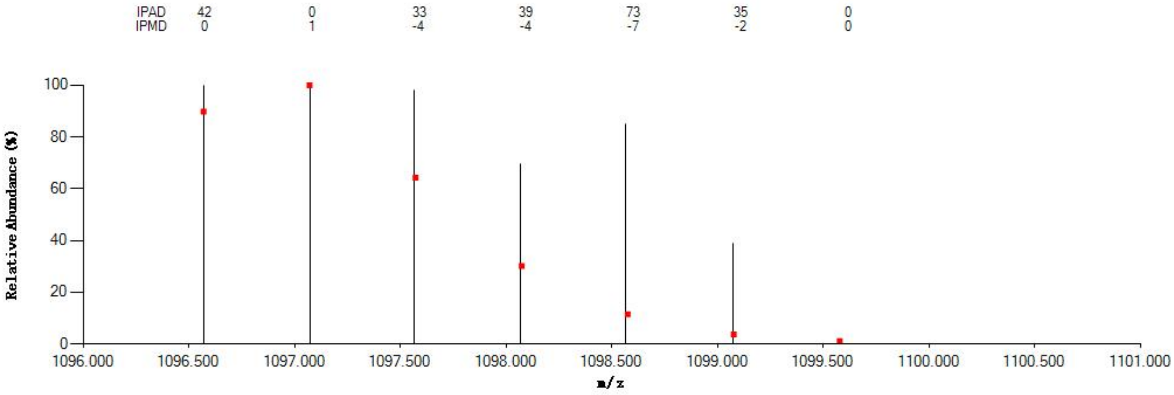


#### Annotated MS/MS Spectrum

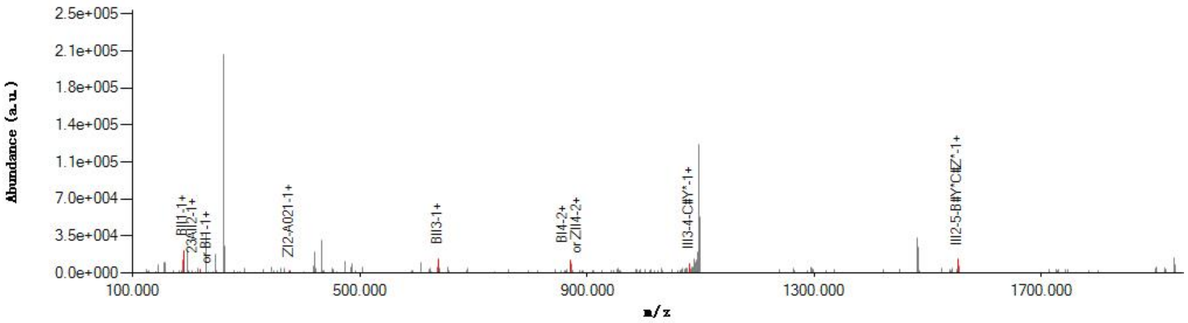


#### Graphical Fragmentation Map

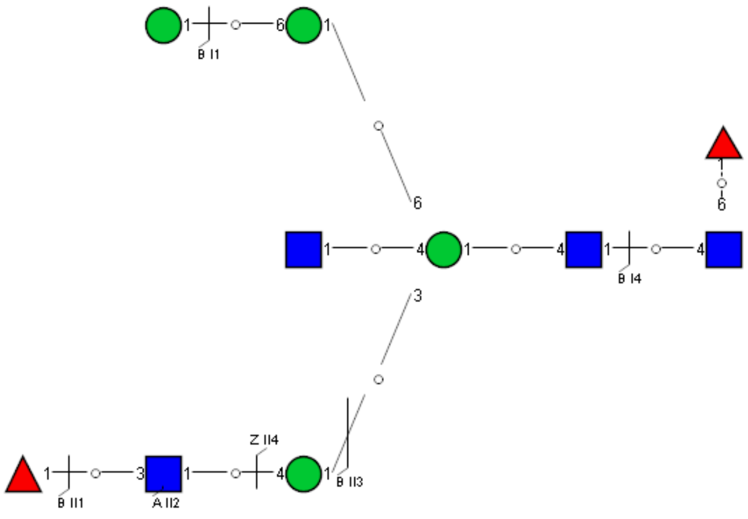

Spectrum Index: 34331(NoBHs:1)

#### Precursor iEF Map

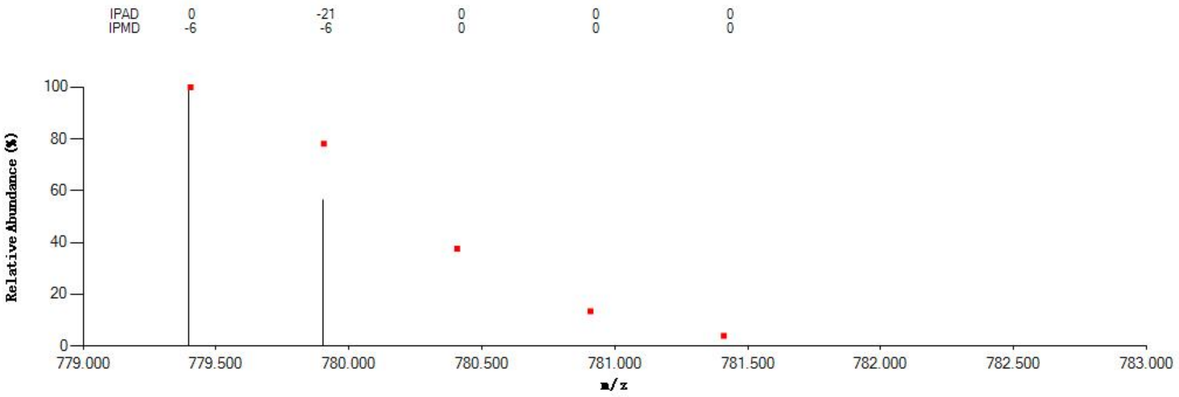

#### Annotated MS/MS Spectrum

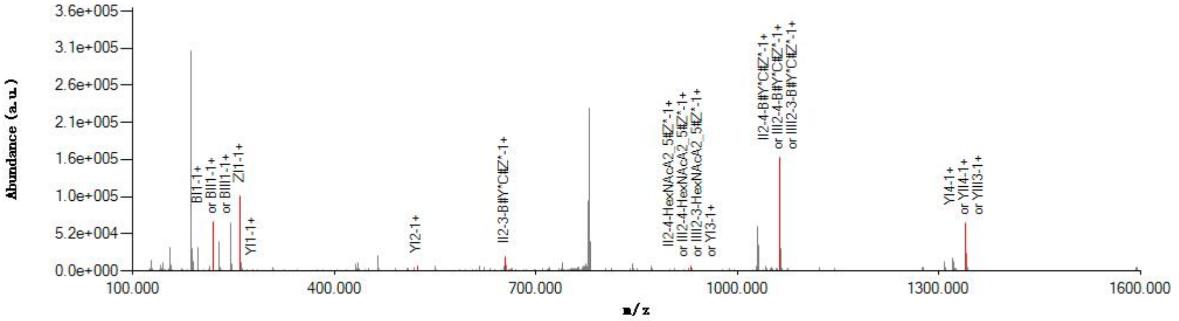

#### Graphical Fragmentation Map

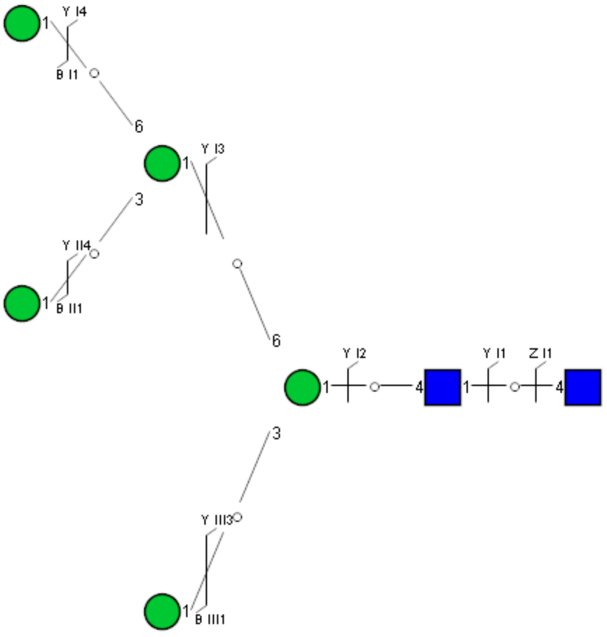

Spectrum Index: 34381(NoBHs:1)

#### Precursor iEF Map

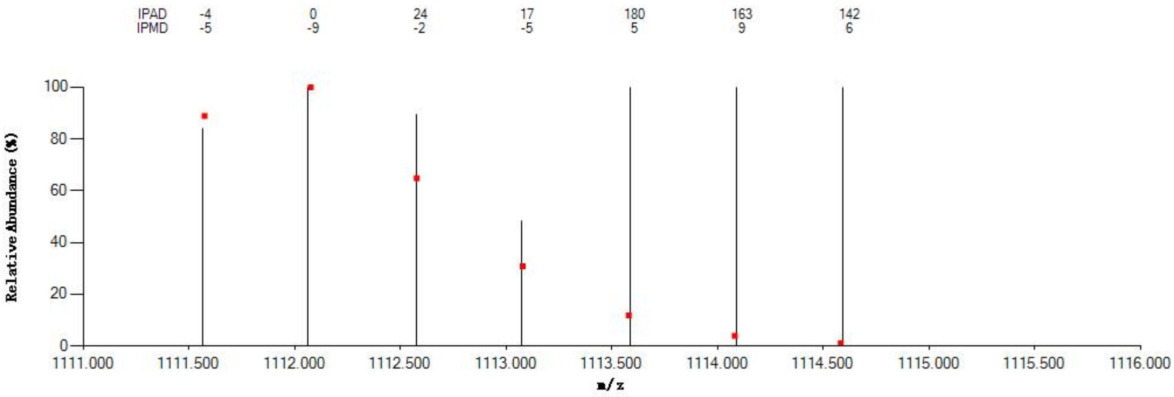


#### Annotated MS/MS Spectrum

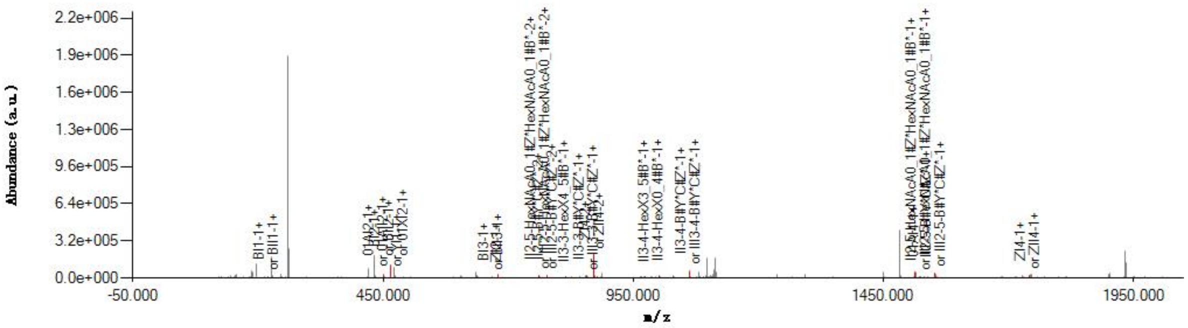


#### Graphical Fragmentation Map

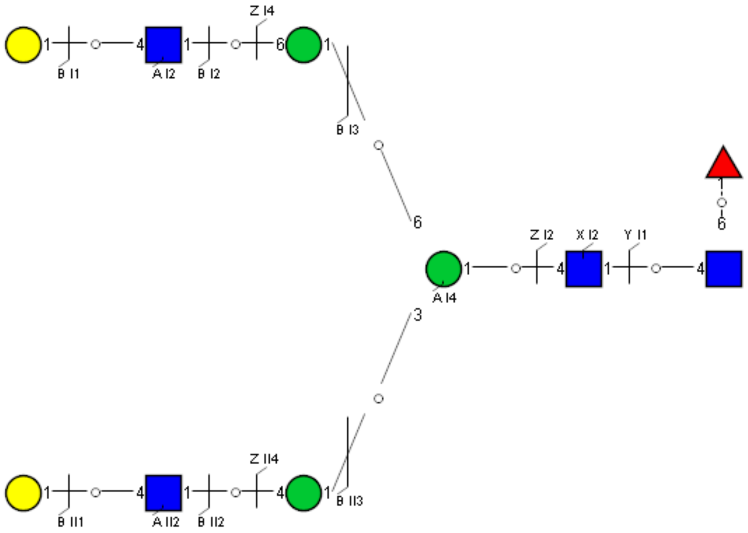

Spectrum Index: 34465(NoBHs:1)

#### Precursor iEF Map

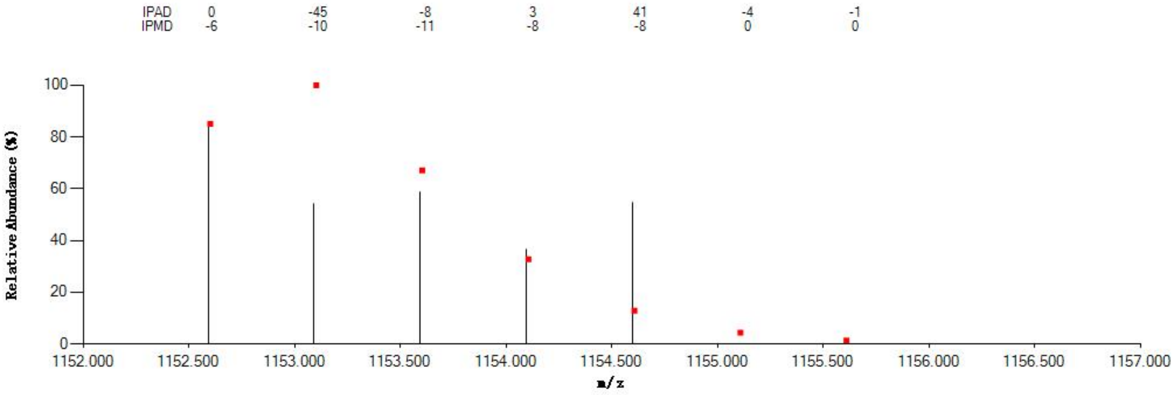


#### Annotated MS/MS Spectrum

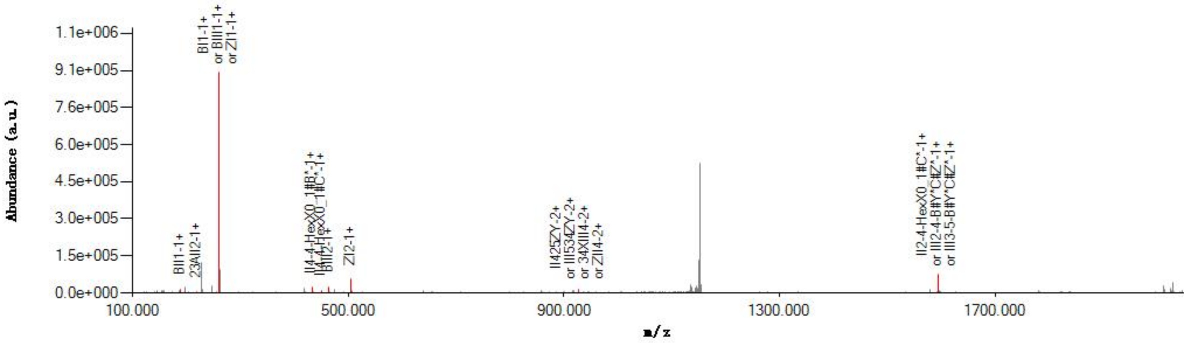


#### Graphical Fragmentation Map

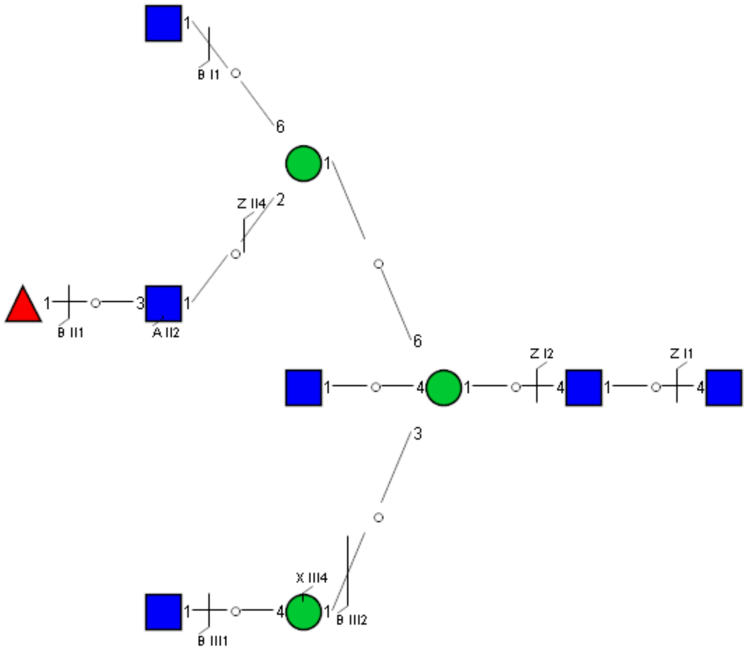

### Spectrum Index: 34487(NoBHs:1)

#### Precursor iEF Map

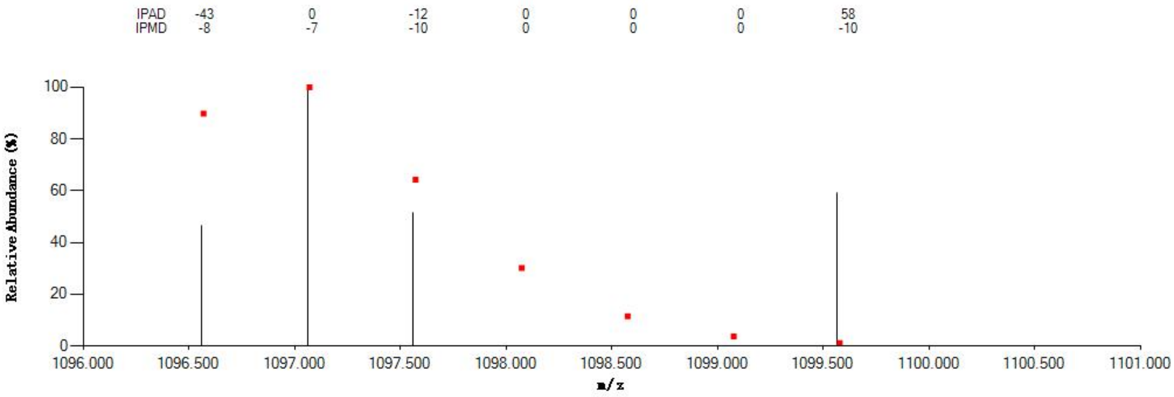

#### Annotated MS/MS Spectrum

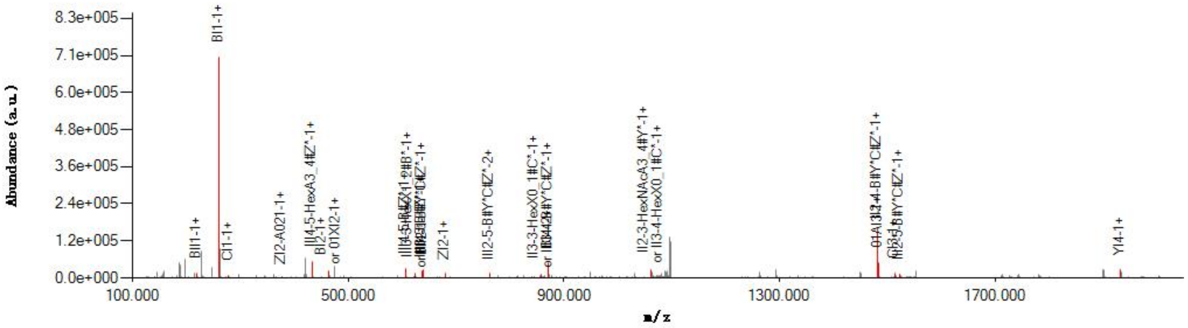

#### Graphical Fragmentation Map

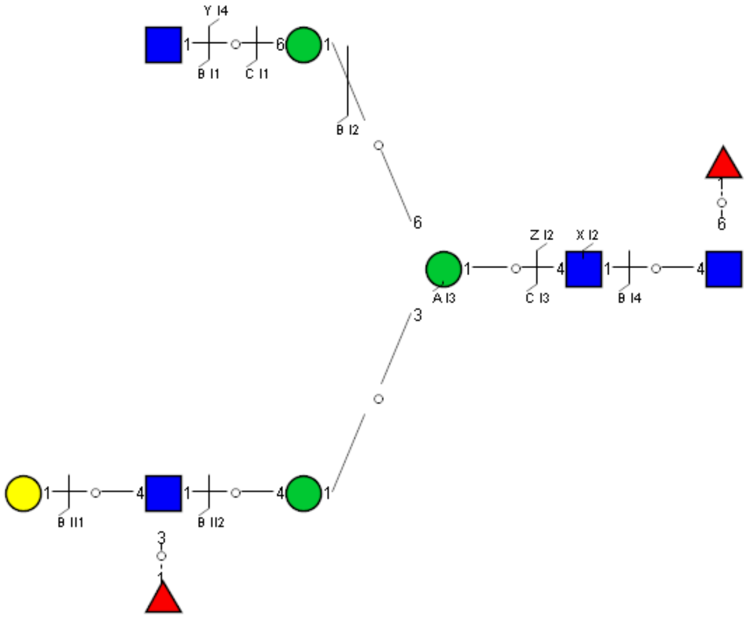

Spectrum Index: 34539(NoBHs:1)

#### Precursor iEF Map

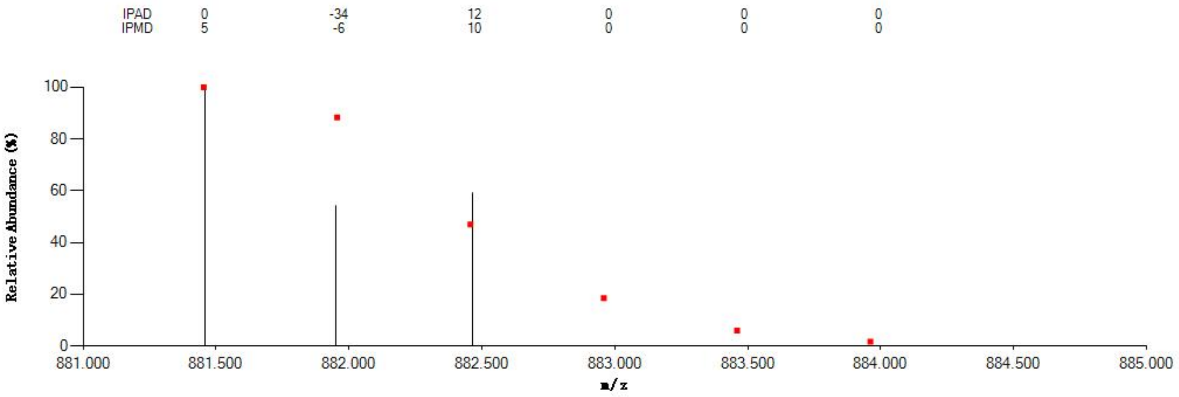


#### Annotated MS/MS Spectrum

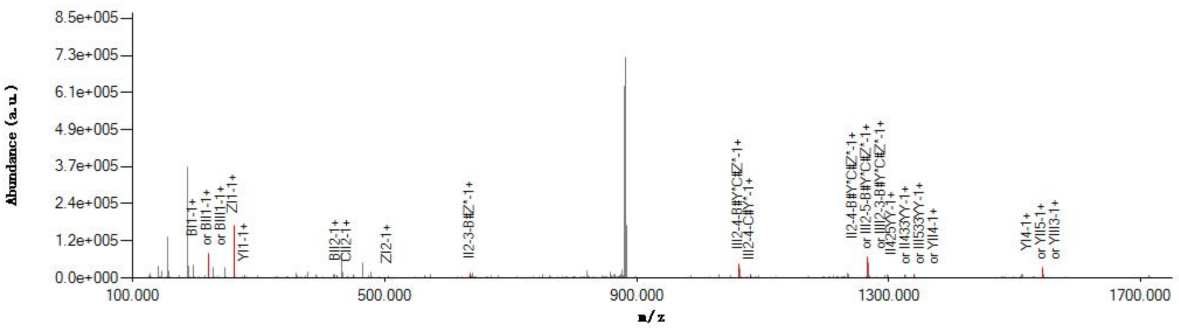


#### Graphical Fragmentation Map

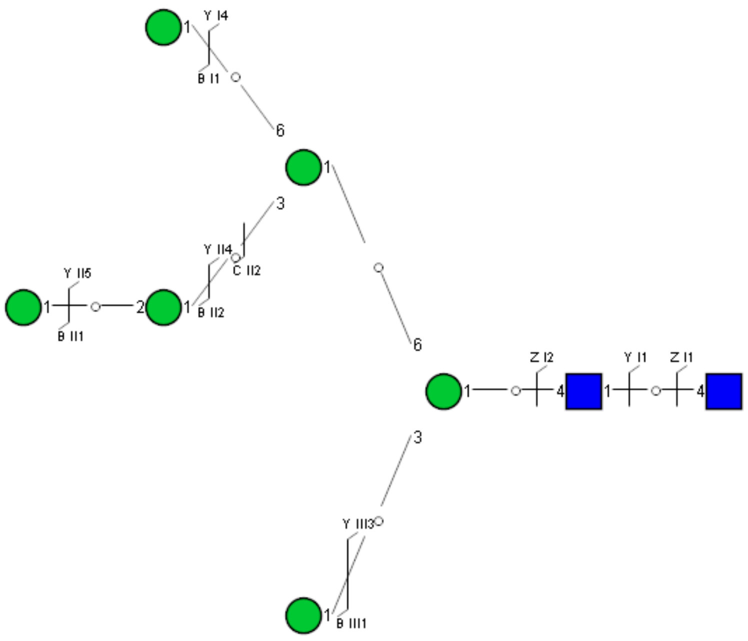

Spectrum Index: 34762(NoBHs:1)

#### Precursor iEF Map

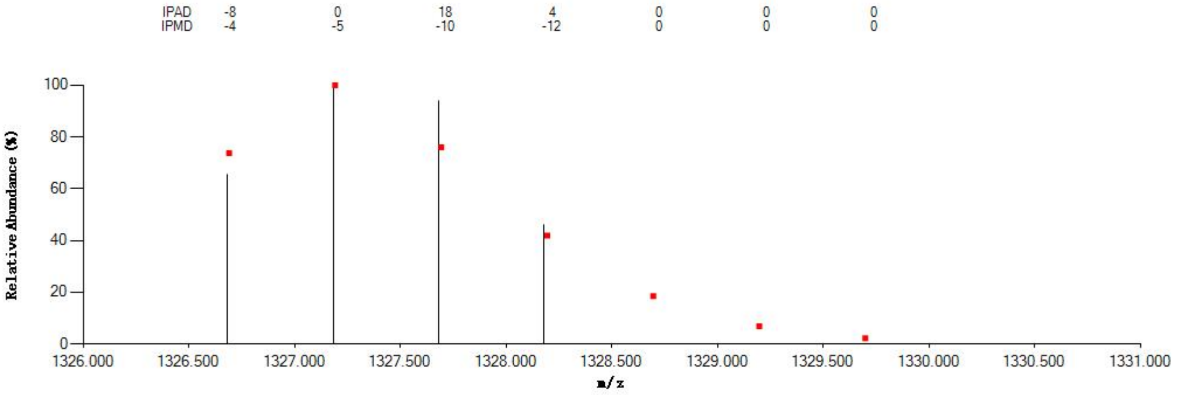


#### Annotated MS/MS Spectrum

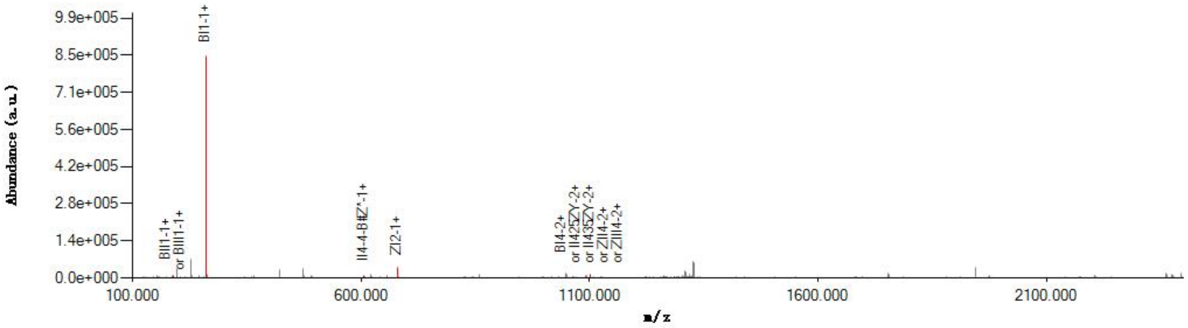


#### Graphical Fragmentation Map

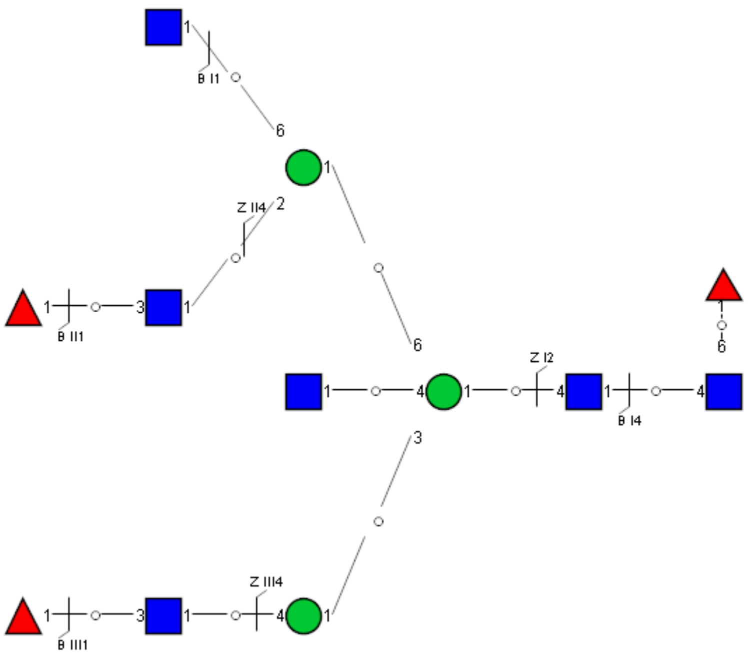

Spectrum Index: 35031(NoBHs:1)

#### Precursor iEF Map

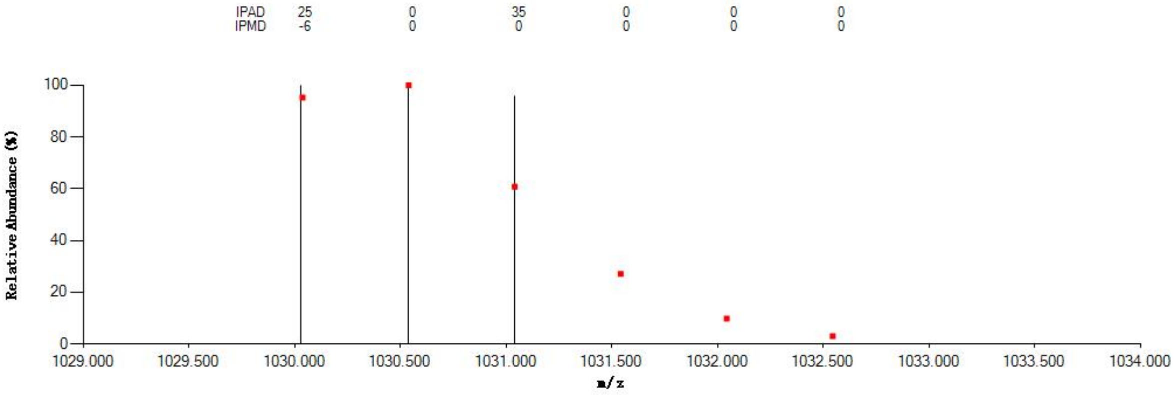

#### Annotated MS/MS Spectrum

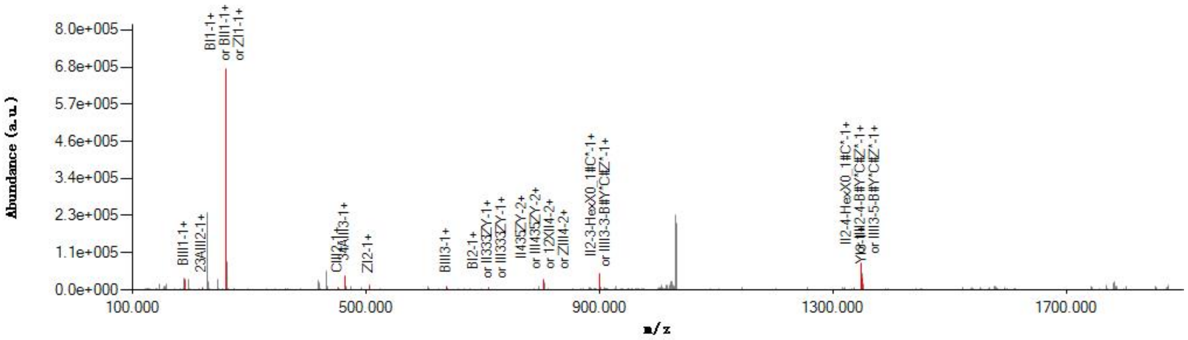

#### Graphical Fragmentation Map

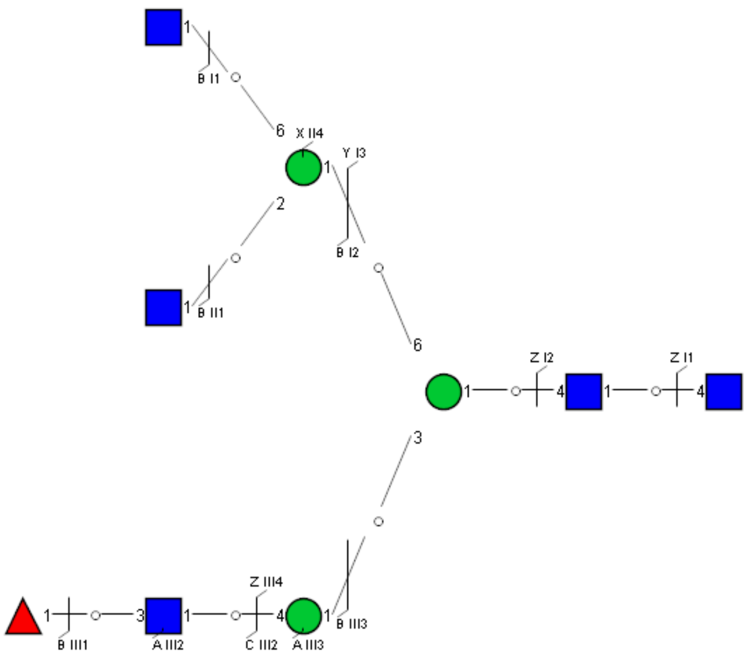

Spectrum Index: 35046(NoBHs:1)

#### Precursor iEF Map

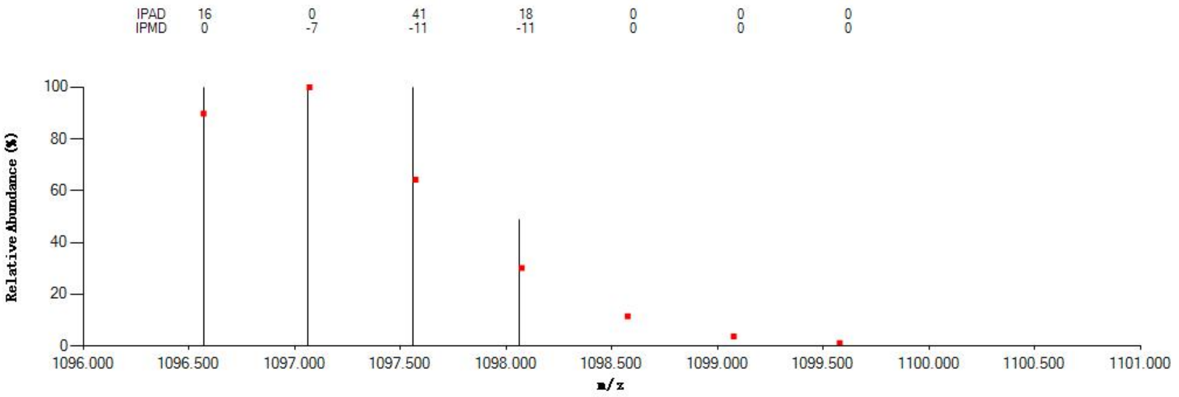


#### Annotated MS/MS Spectrum

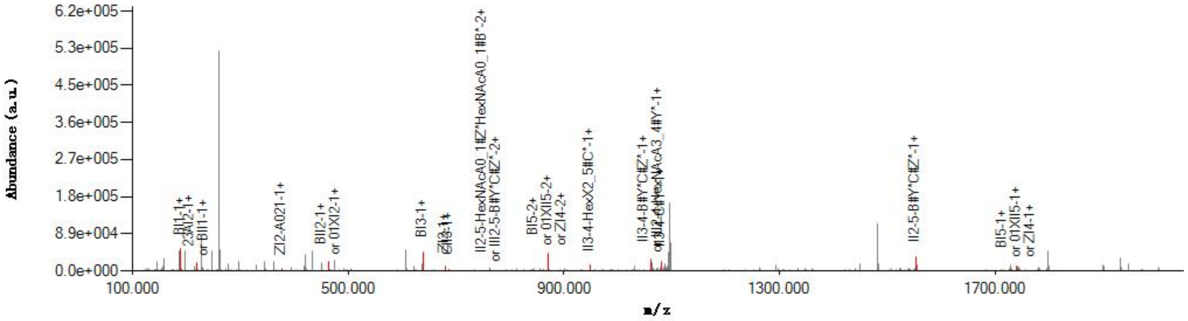


#### Graphical Fragmentation Map

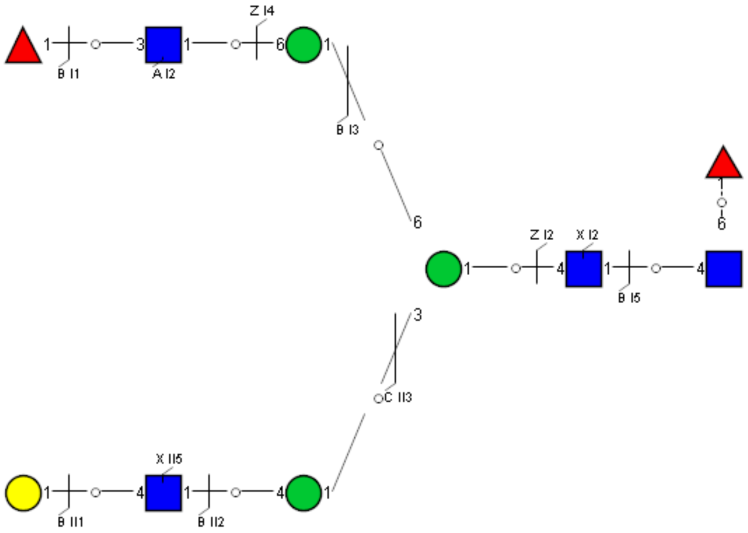

Spectrum Index: 35275(NoBHs:1)

#### Precursor iEF Map

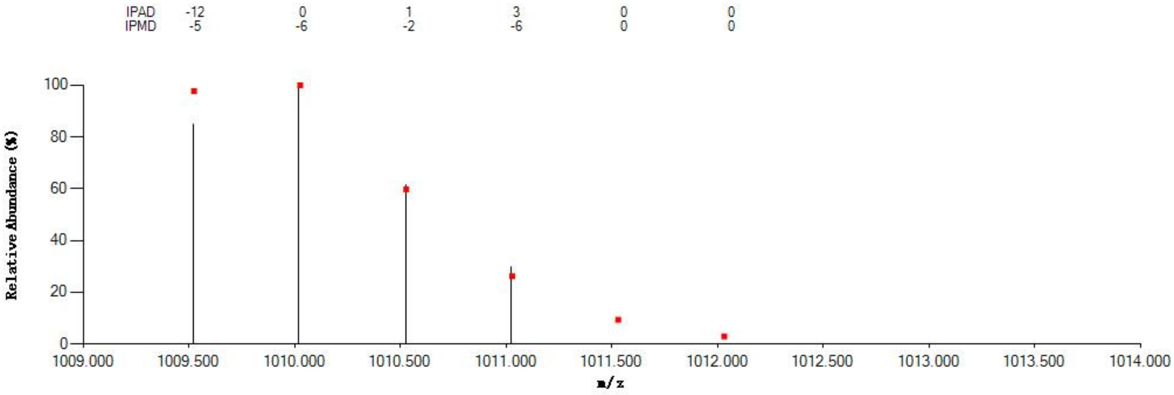

#### Annotated MS/MS Spectrum

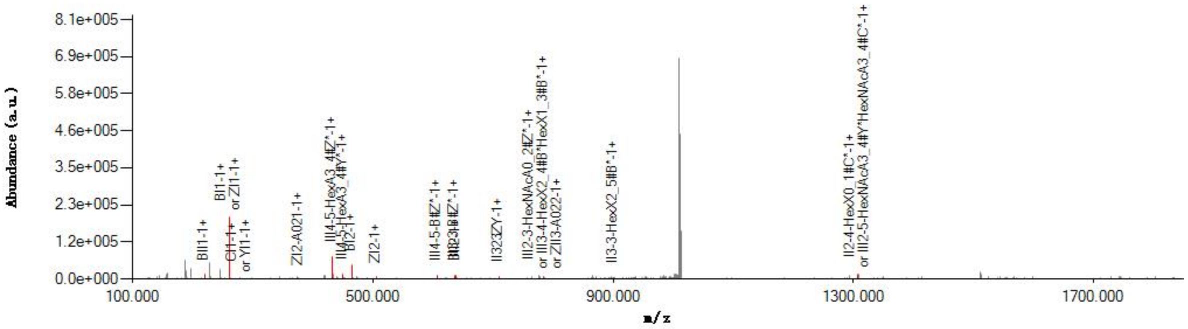

#### Graphical Fragmentation Map

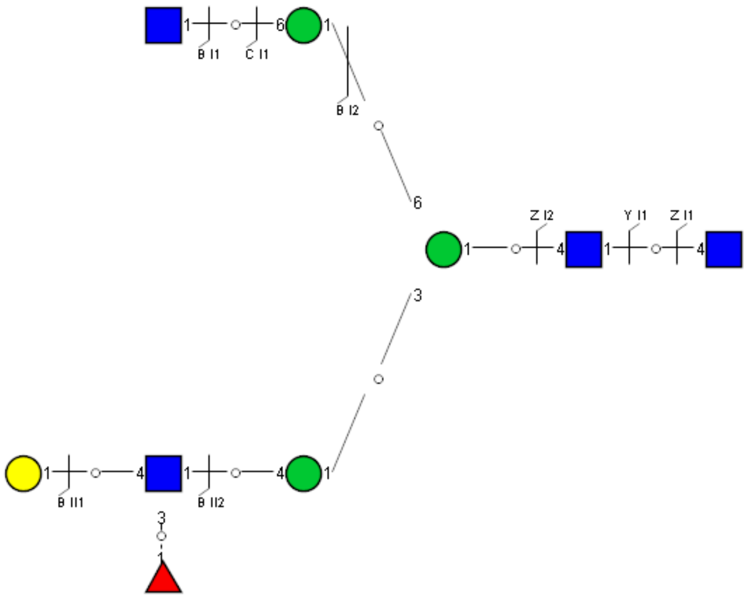

Spectrum Index: 35283(NoBHs:1)

#### Precursor iEF Map

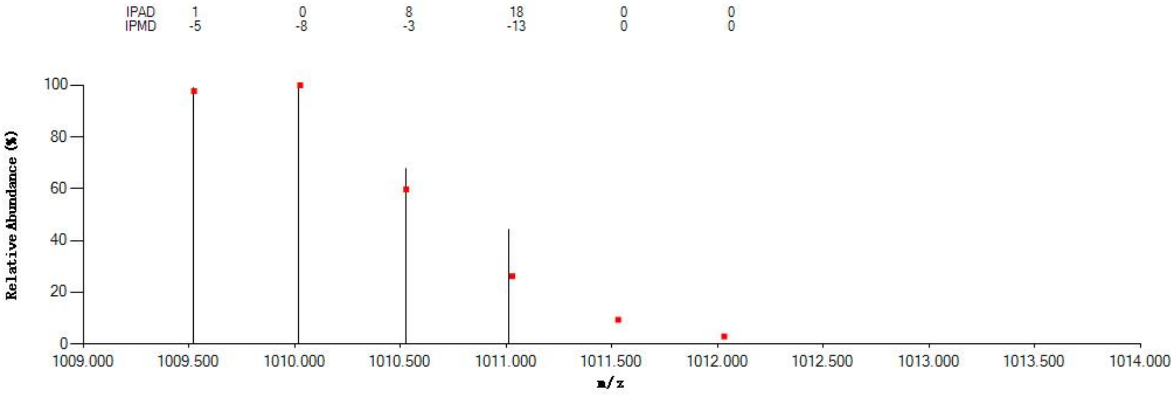


#### Annotated MS/MS Spectrum

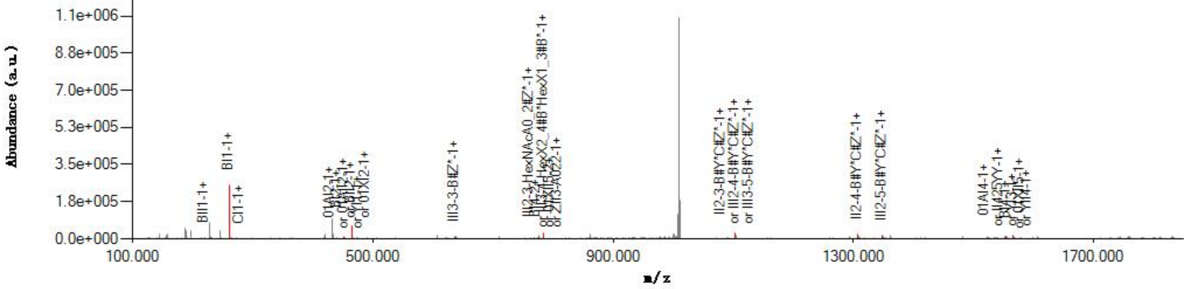


#### Graphical Fragmentation Map

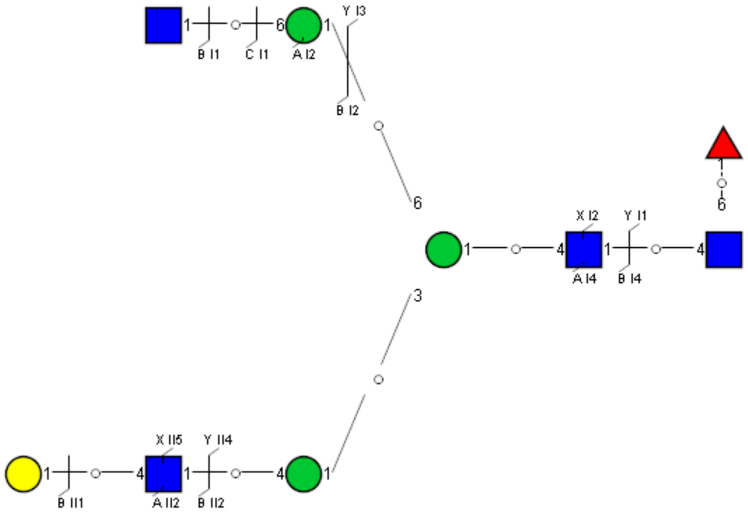

Spectrum Index: 35317(NoBHs:1)

#### Precursor iEF Map

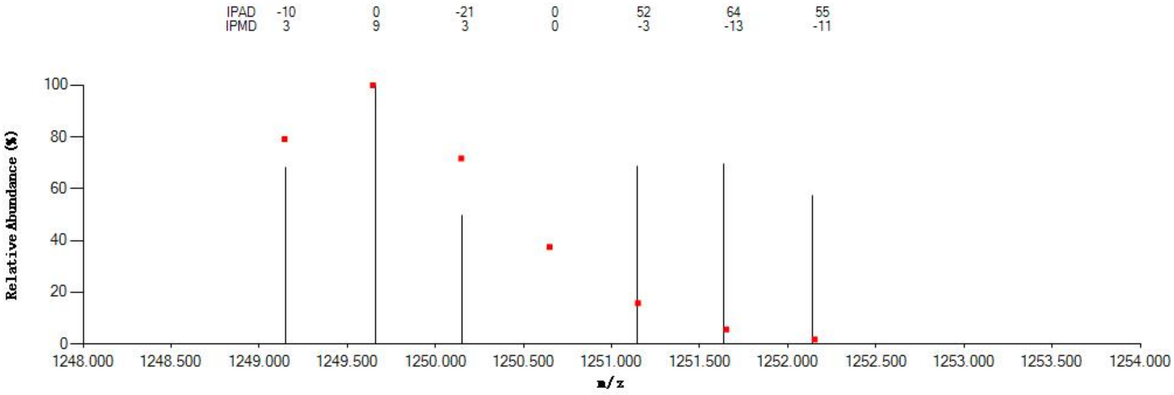


#### Annotated MS/MS Spectrum

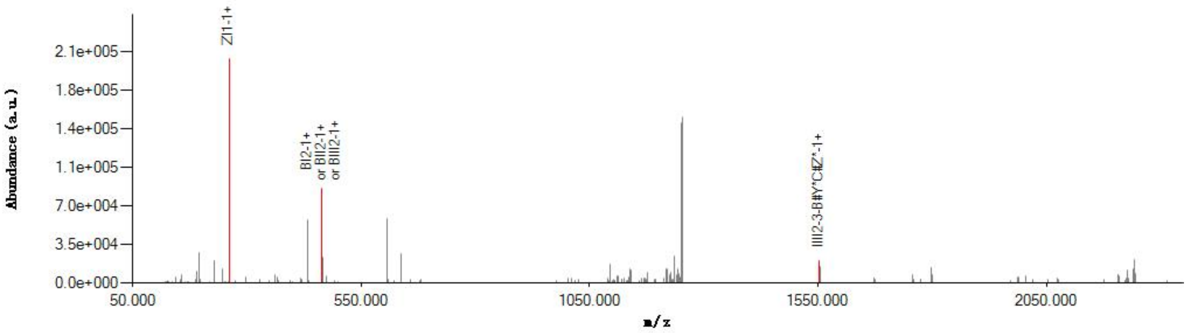


#### Graphical Fragmentation Map

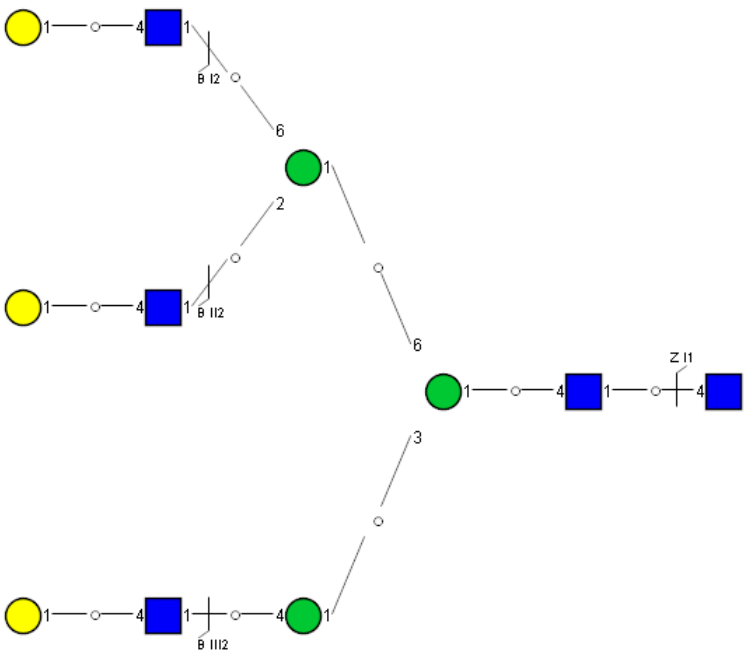

Spectrum Index: 35391(NoBHs:1)

#### Precursor iEF Map

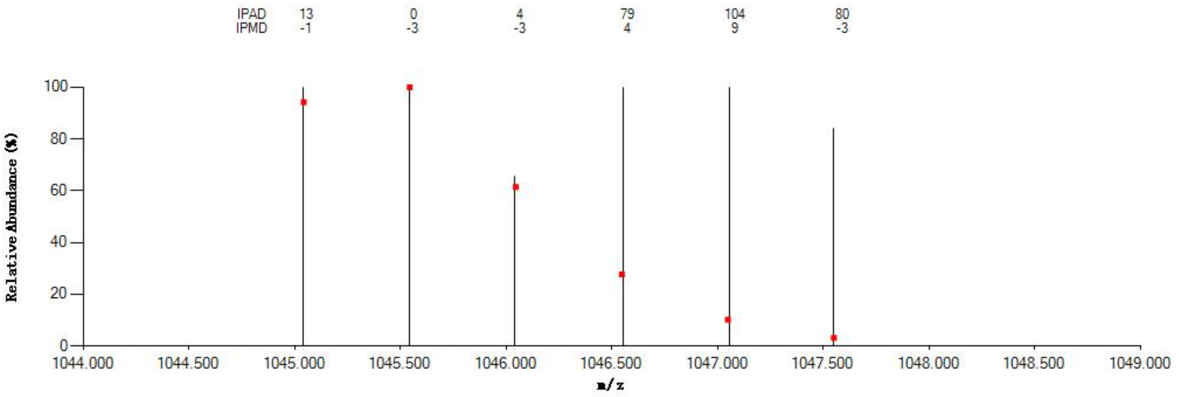

#### Annotated MS/MS Spectrum

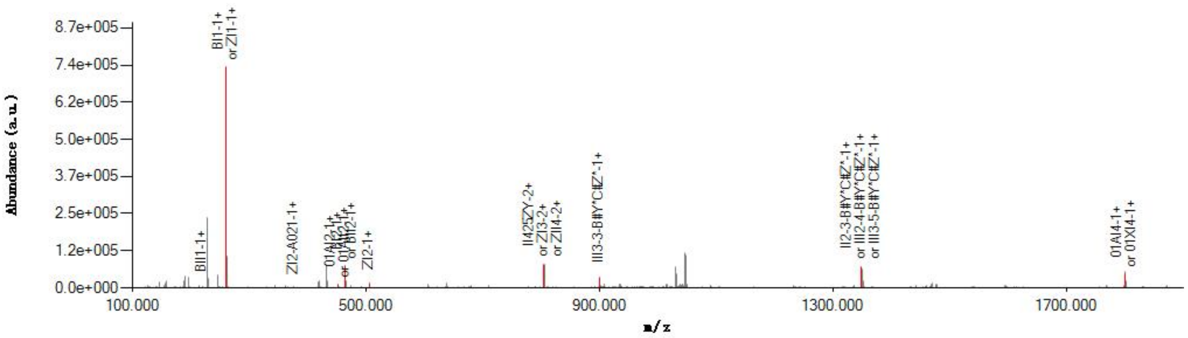

#### Graphical Fragmentation Map

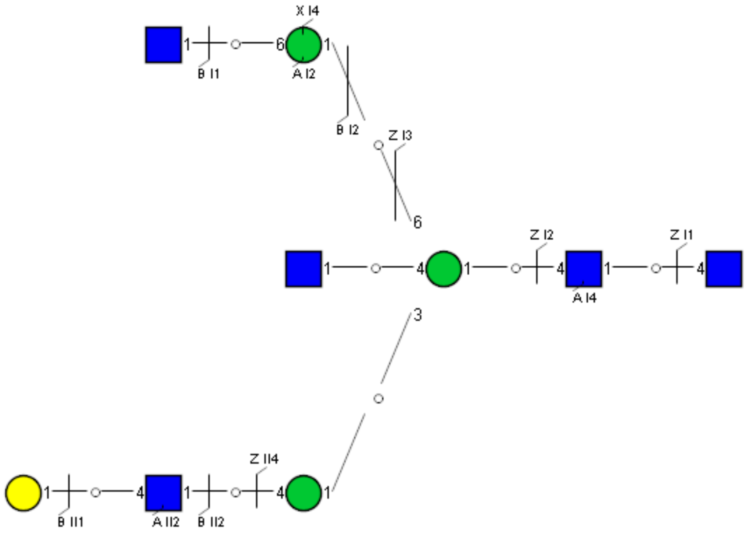

Spectrum Index: 35477(NoBHs:1)

#### Precursor iEF Map

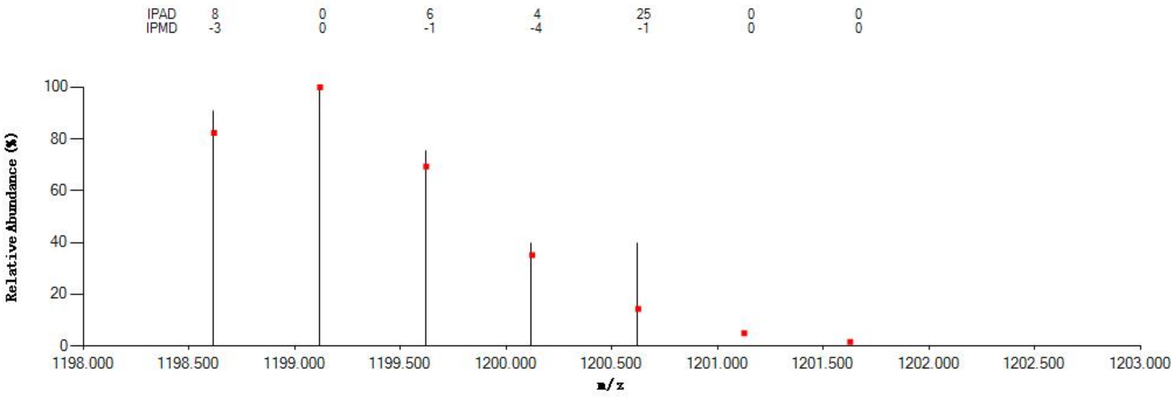


#### Annotated MS/MS Spectrum

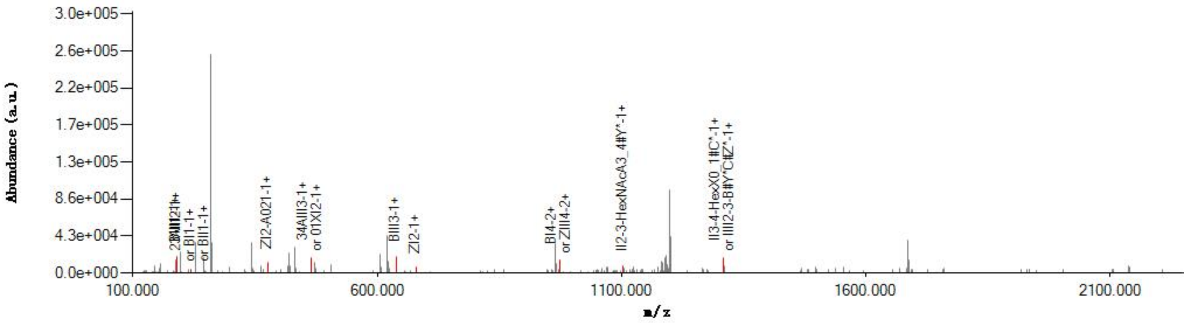


#### Graphical Fragmentation Map

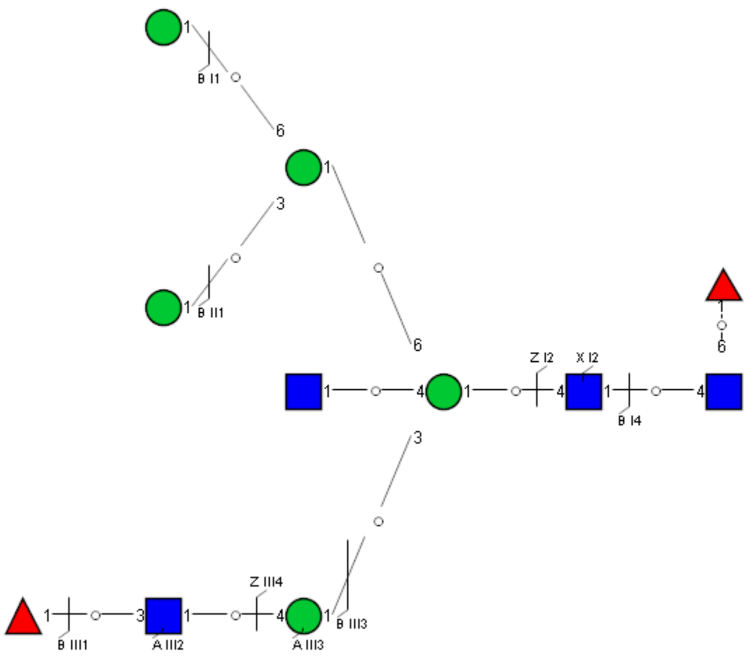

Spectrum Index: 35724(NoBHs:1)

#### Precursor iEF Map

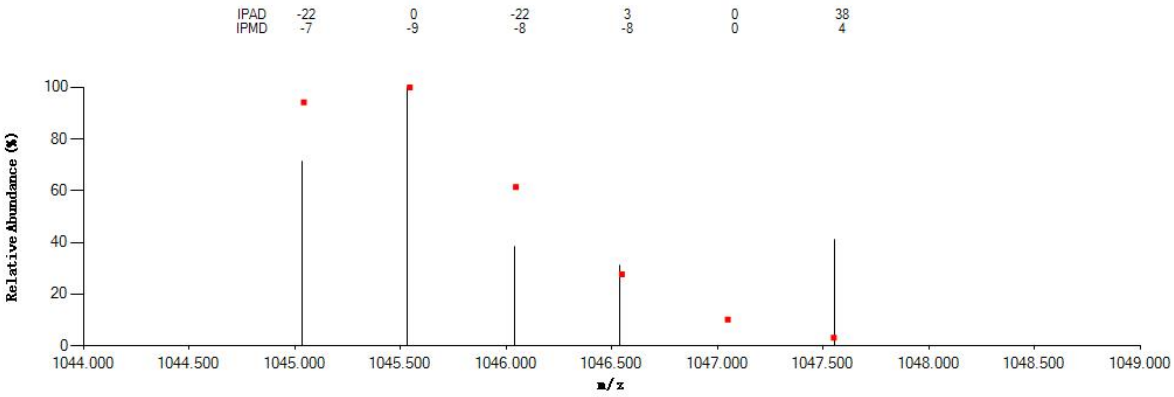


#### Annotated MS/MS Spectrum

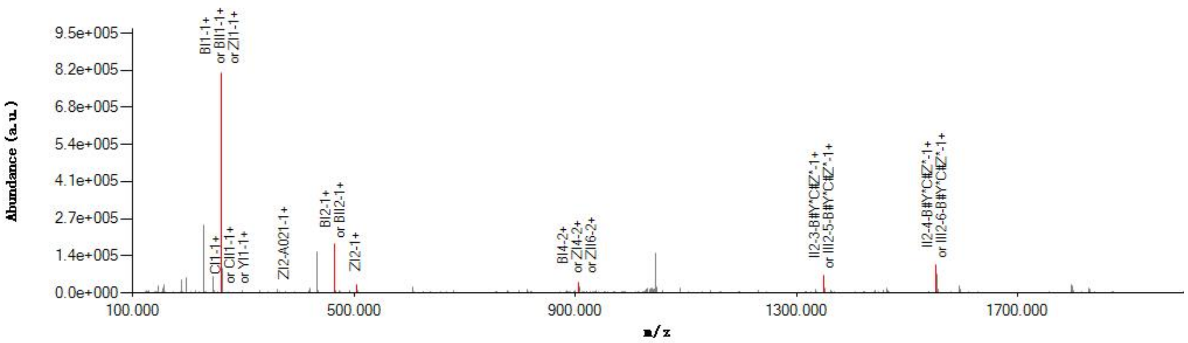


#### Graphical Fragmentation Map

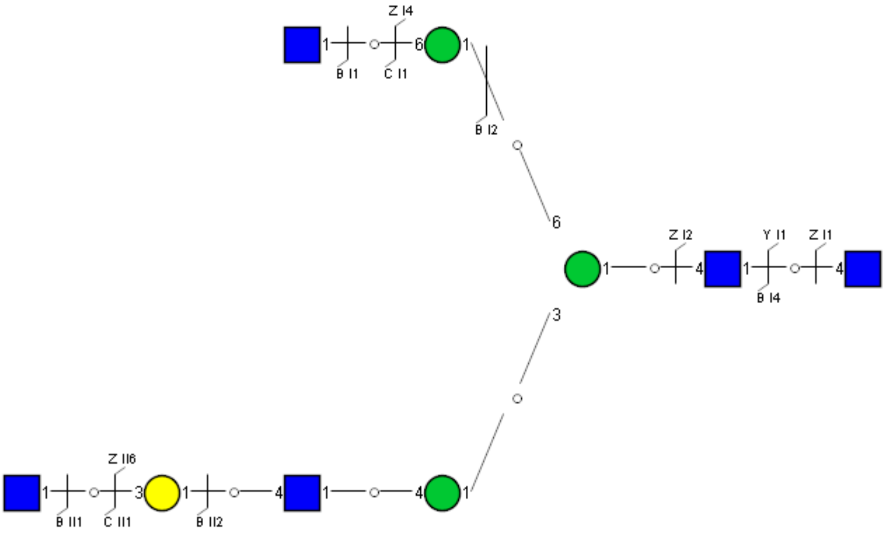

Spectrum Index: 35752(NoBHs:1)

#### Precursor iEF Map

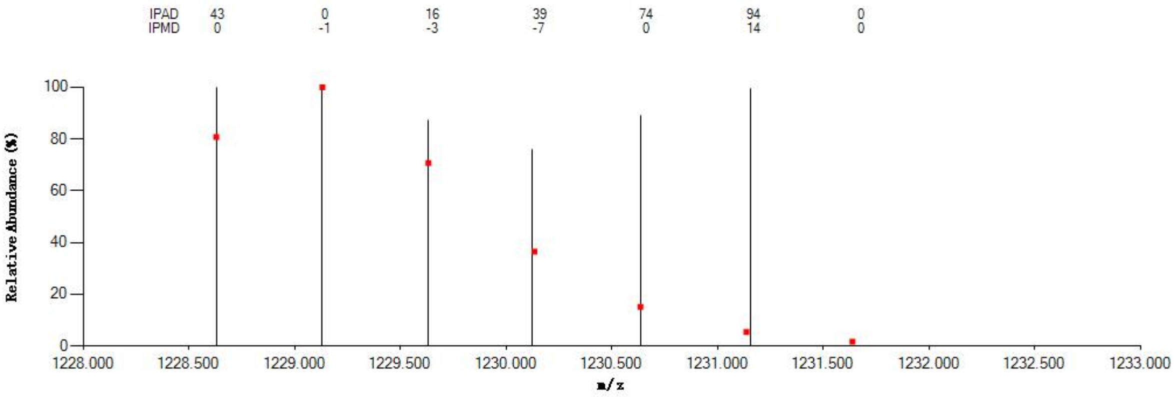


#### Annotated MS/MS Spectrum

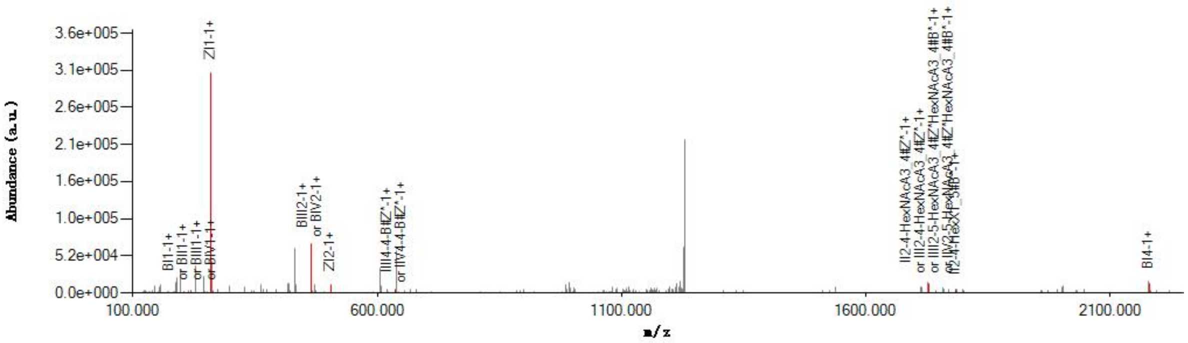


#### Graphical Fragmentation Map

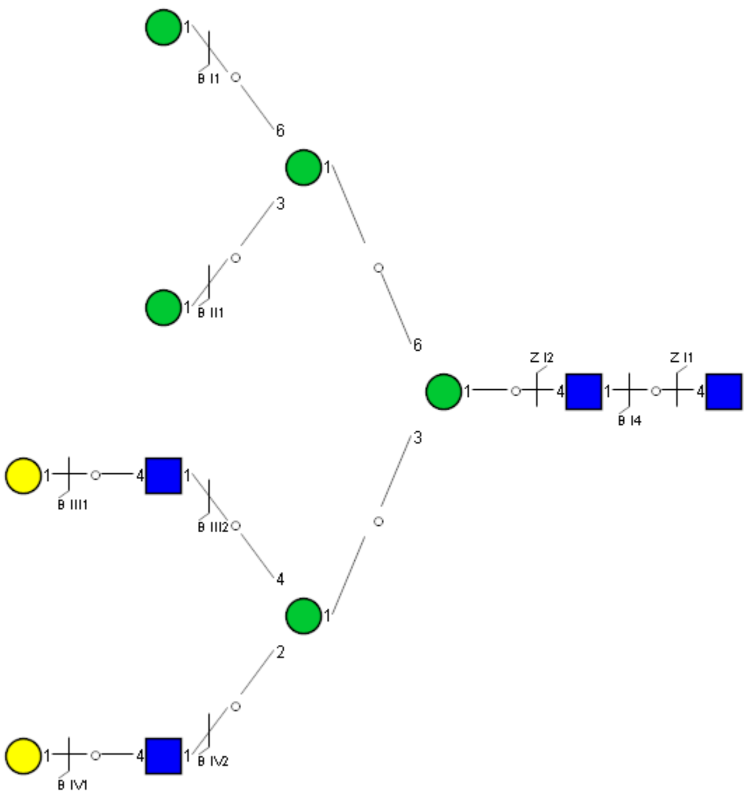

Spectrum Index: 35800(NoBHs:1)

#### Precursor iEF Map

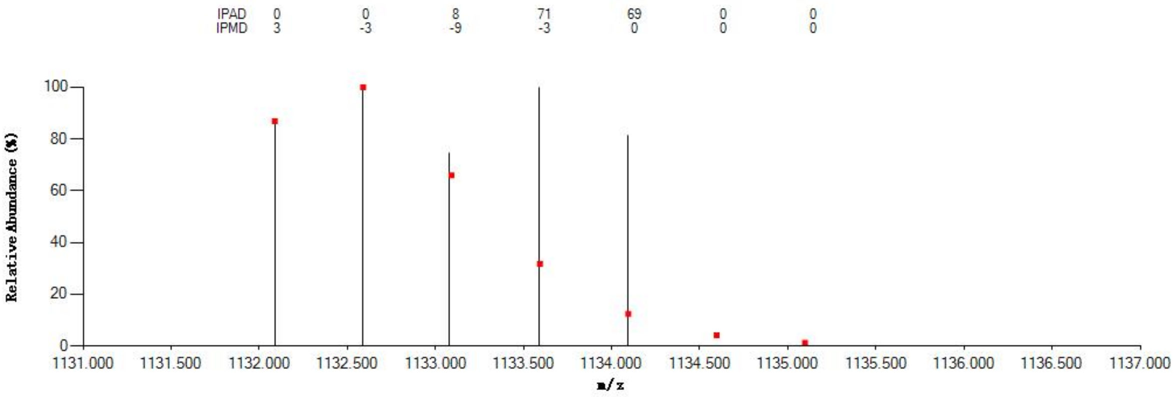


#### Annotated MS/MS Spectrum

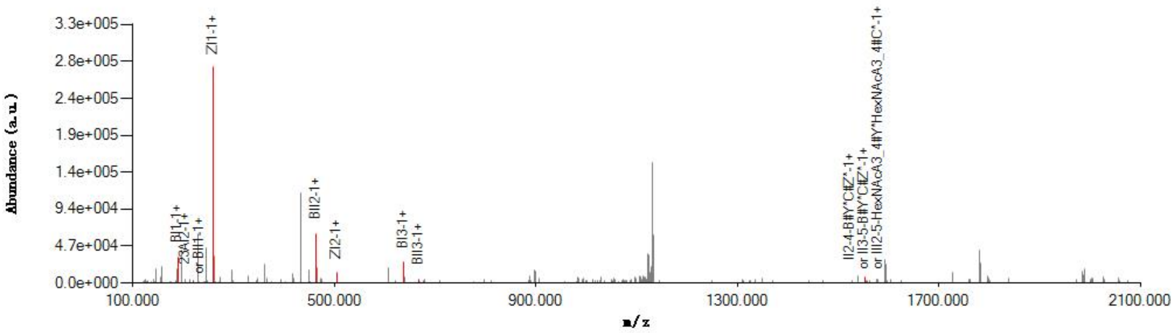


#### Graphical Fragmentation Map

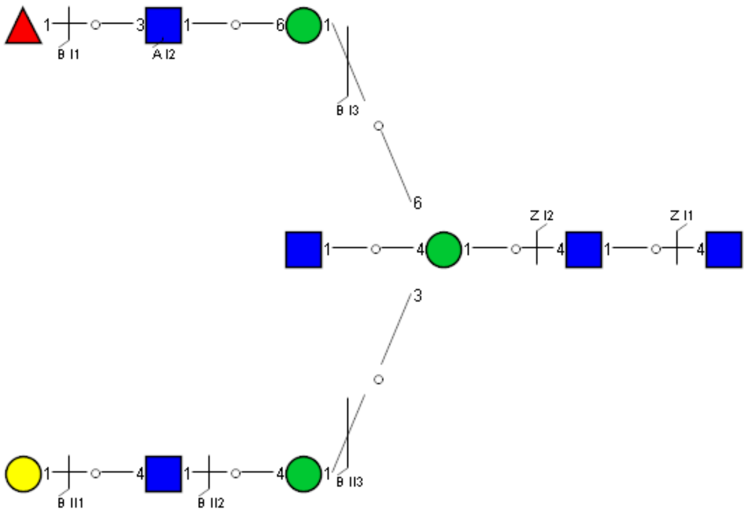

Spectrum Index: 35886(NoBHs:1)

Precursor iEF Map

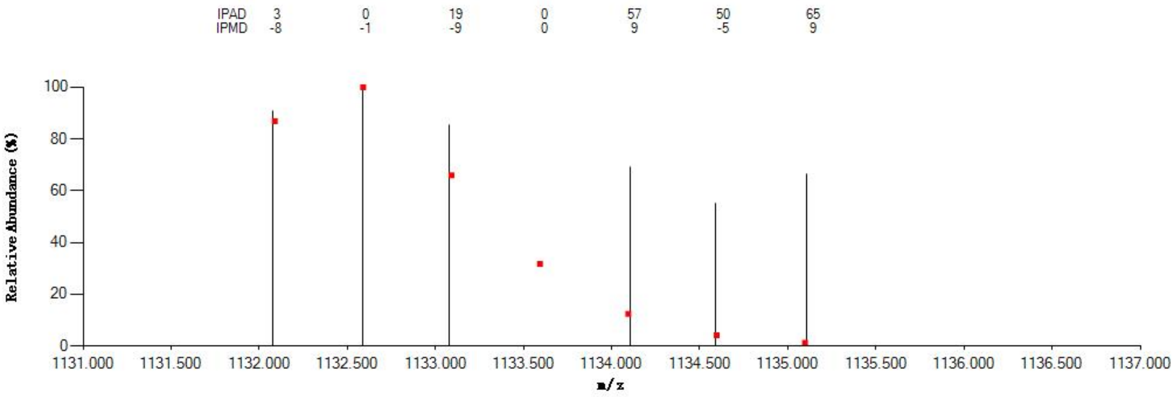


Annotated MS/MS Spectrum

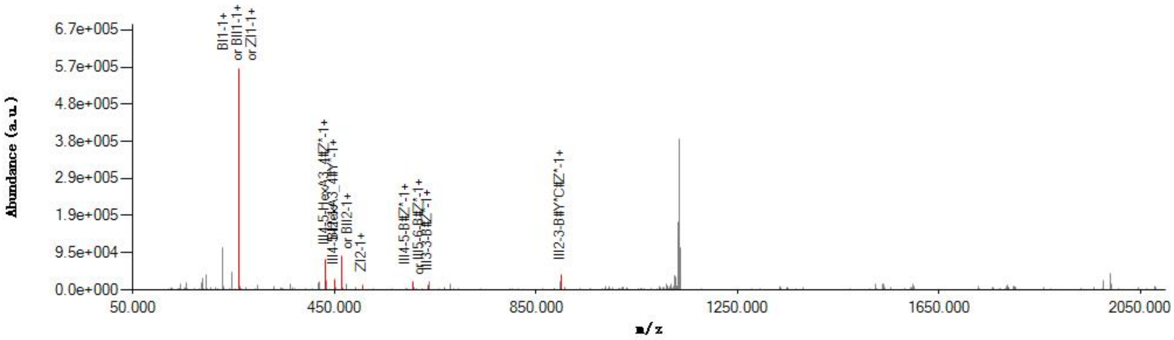


Graphical Fragmentation Map

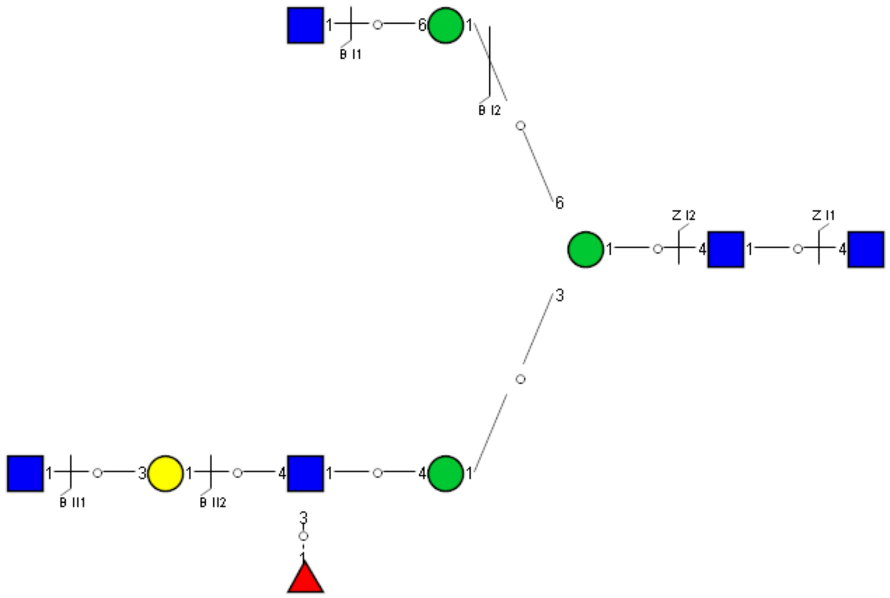

Spectrum Index: 35900(NoBHs:1)

#### Precursor iEF Map

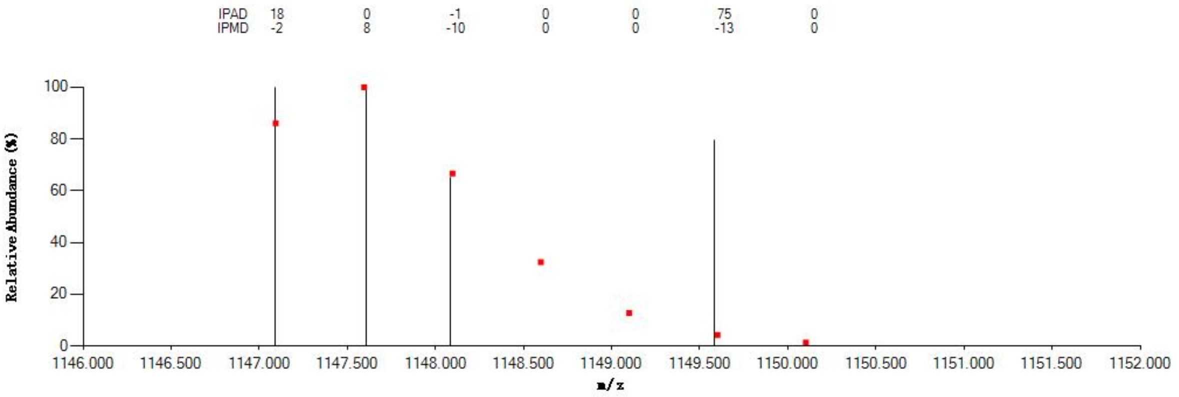

#### Annotated MS/MS Spectrum

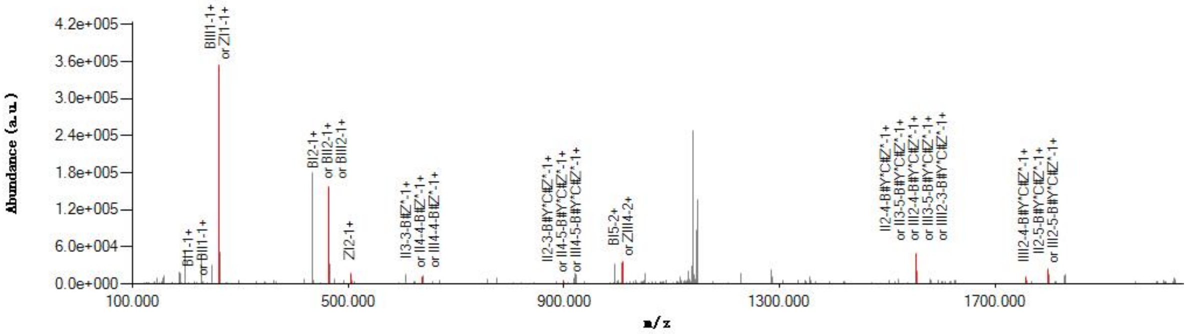

#### Graphical Fragmentation Map

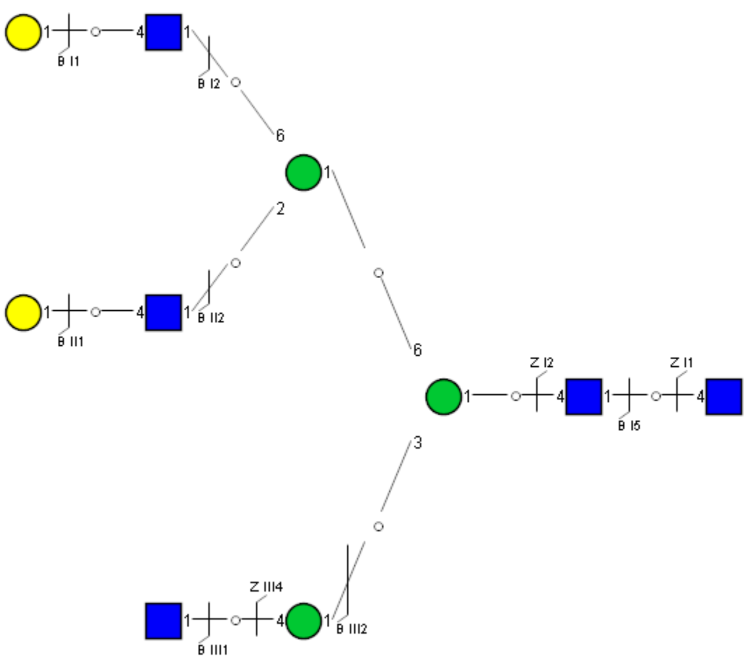

Spectrum Index: 35983(NoBHs:1)

#### Precursor iEF Map

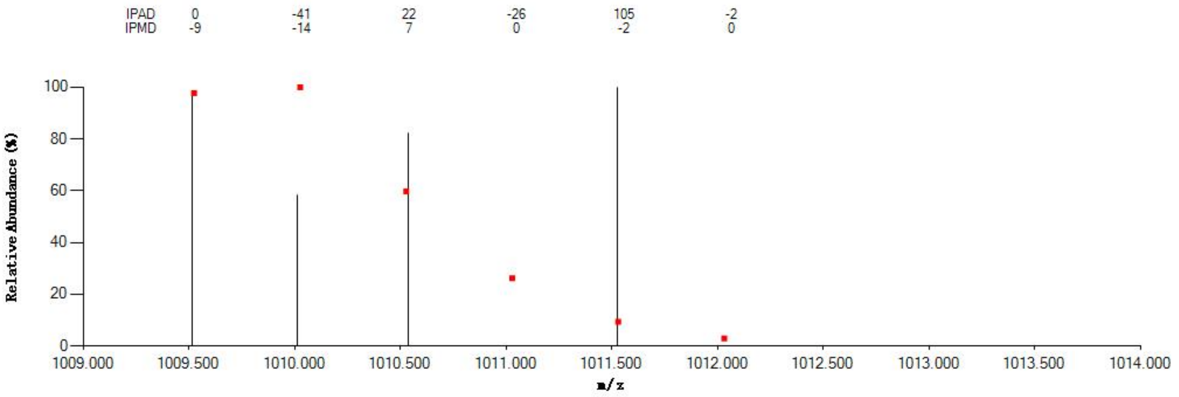


#### Annotated MS/MS Spectrum

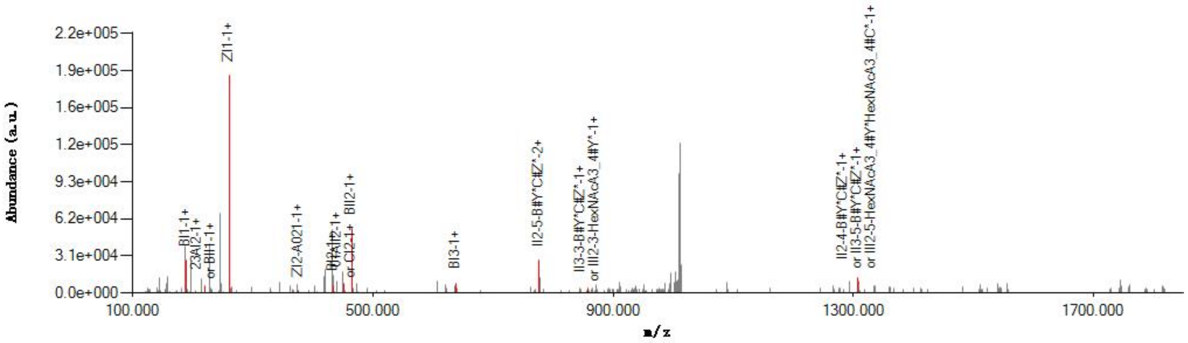


#### Graphical Fragmentation Map

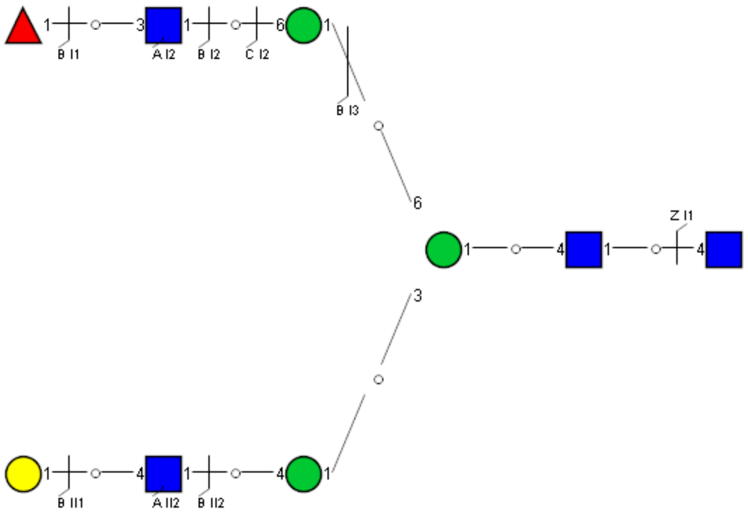

Spectrum Index: 36193(NoBHs:1)

#### Precursor iEF Map

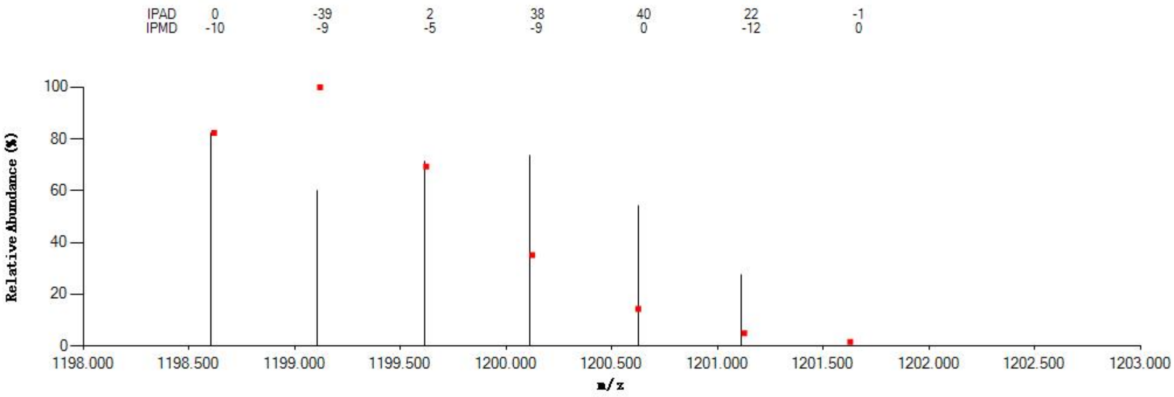


#### Annotated MS/MS Spectrum

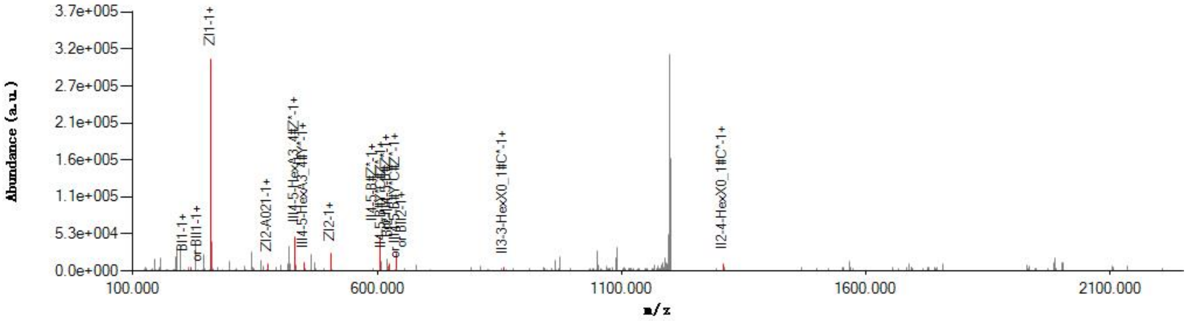


#### Graphical Fragmentation Map

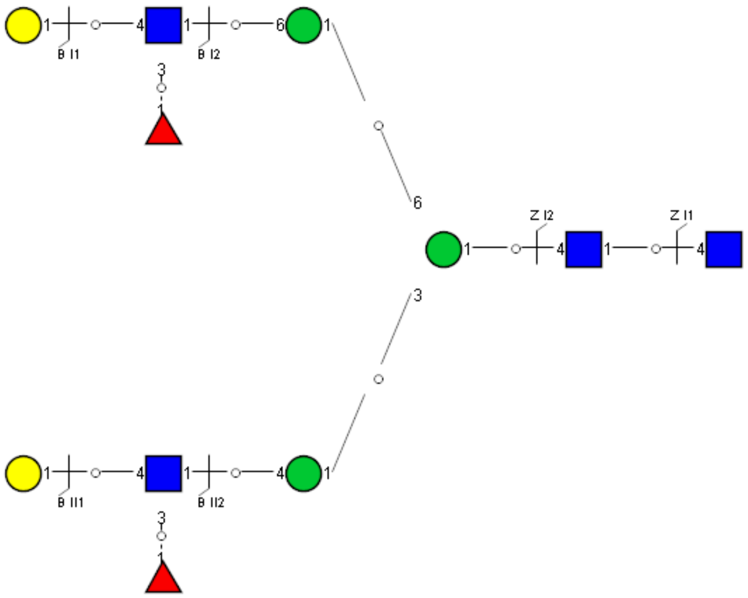

### Spectrum Index: 36200(NoBHs:1)

#### Precursor iEF Map

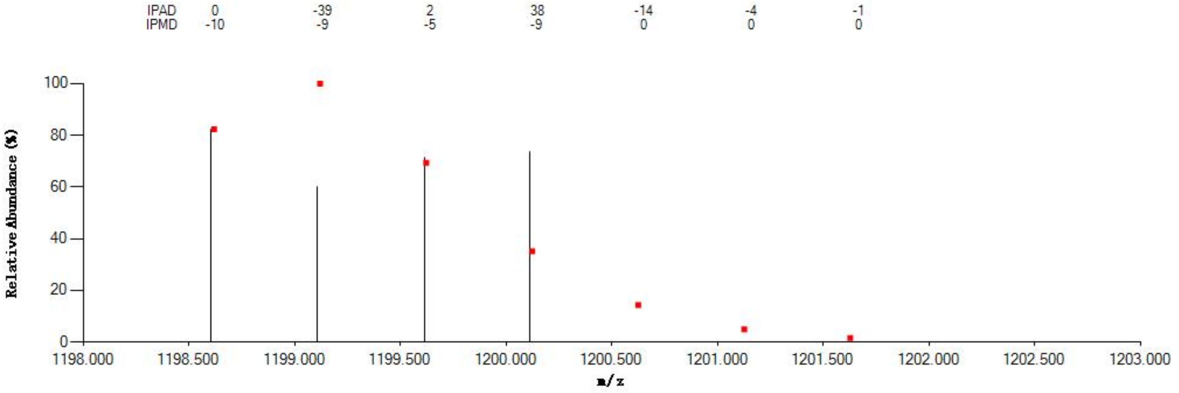


#### Annotated MS/MS Spectrum

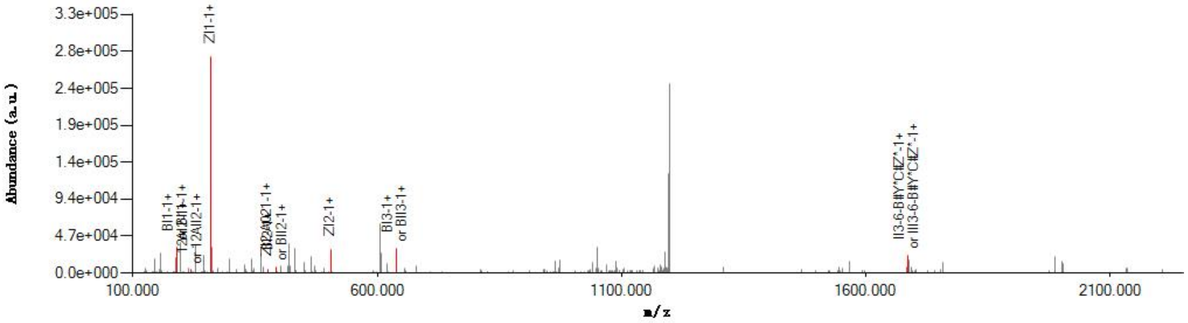


#### Graphical Fragmentation Map

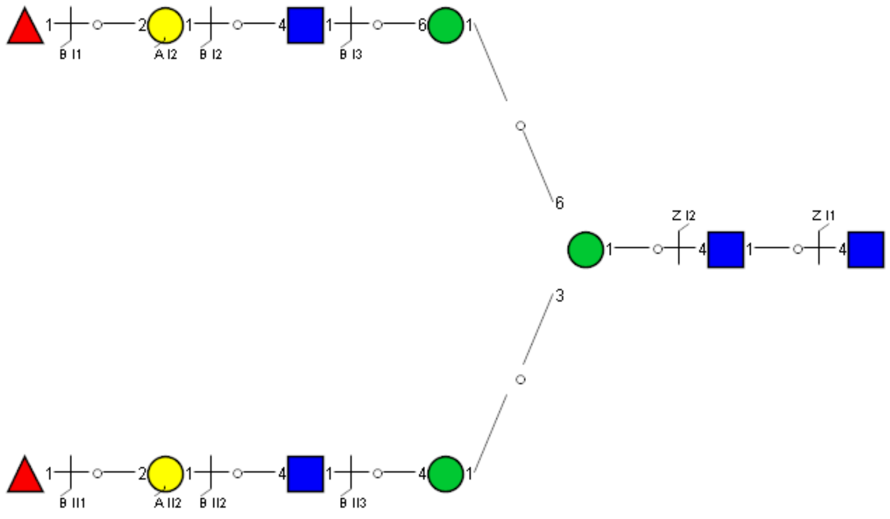

Spectrum Index: 36210(NoBHs:1)

#### Precursor iEF Map

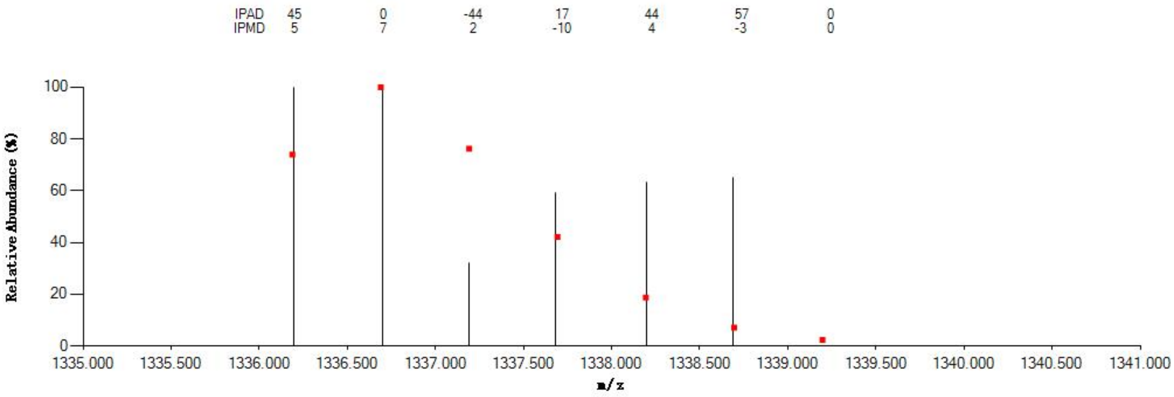


#### Annotated MS/MS Spectrum

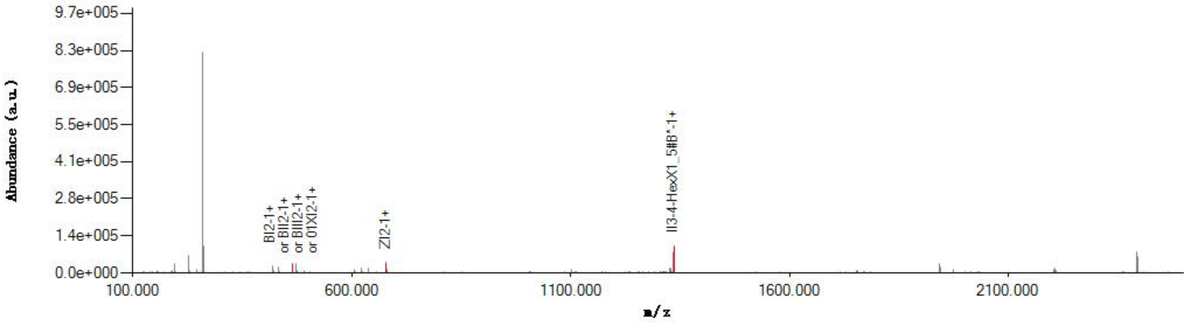


#### Graphical Fragmentation Map

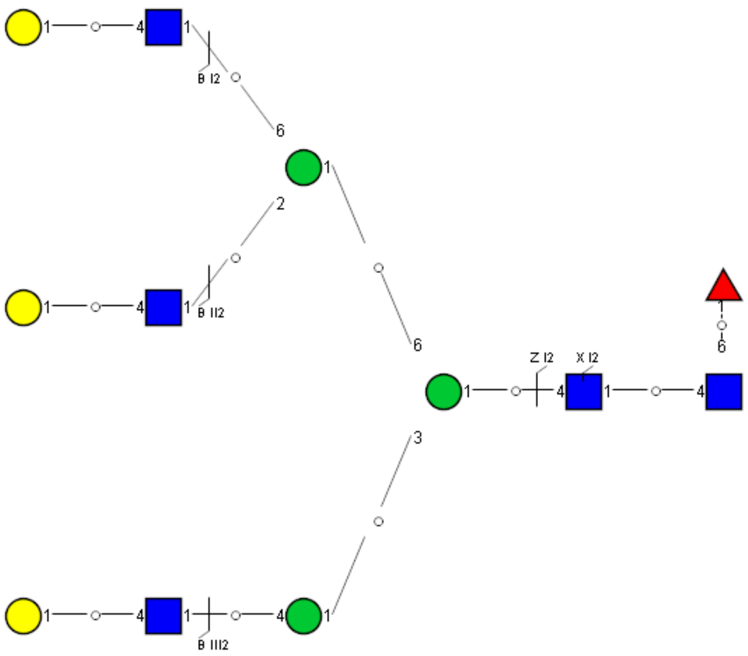

### Spectrum Index: 36237(NoBHs:1)

#### Precursor iEF Map

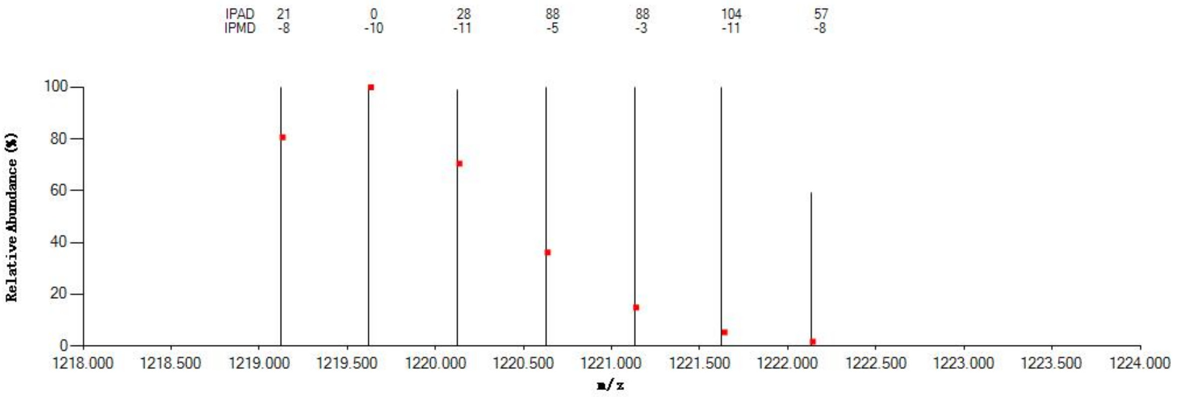


#### Annotated MS/MS Spectrum

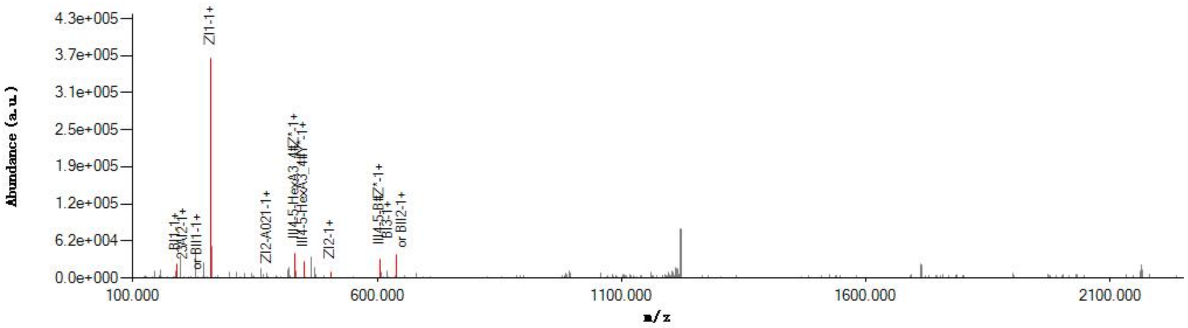


#### Graphical Fragmentation Map

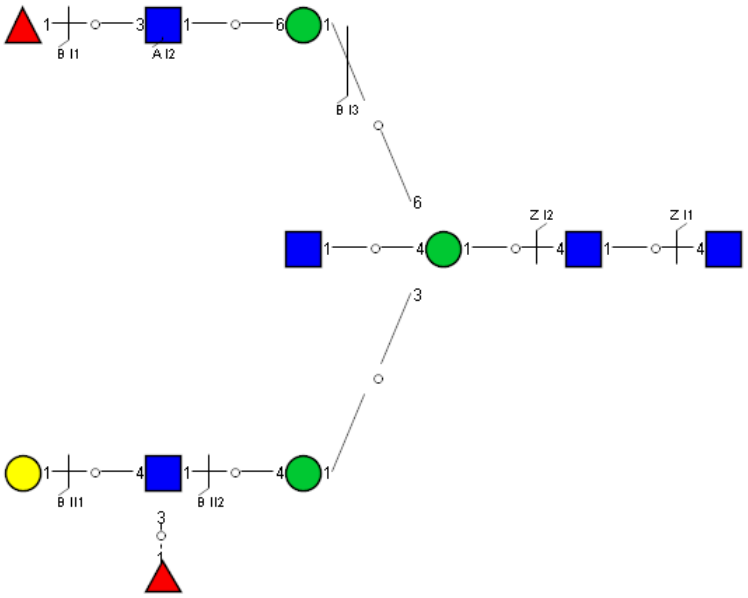

Spectrum Index: 36242(NoBHs:1)

#### Precursor iEF Map

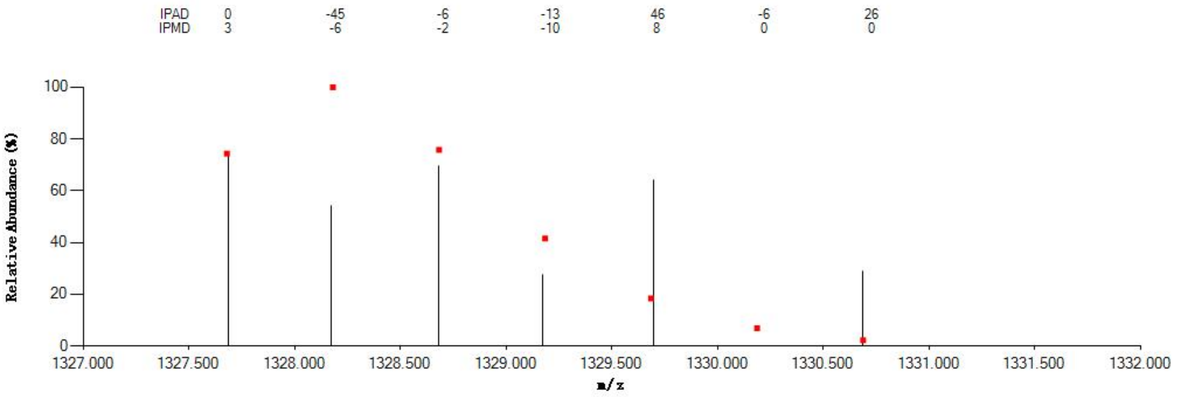


#### Annotated MS/MS Spectrum

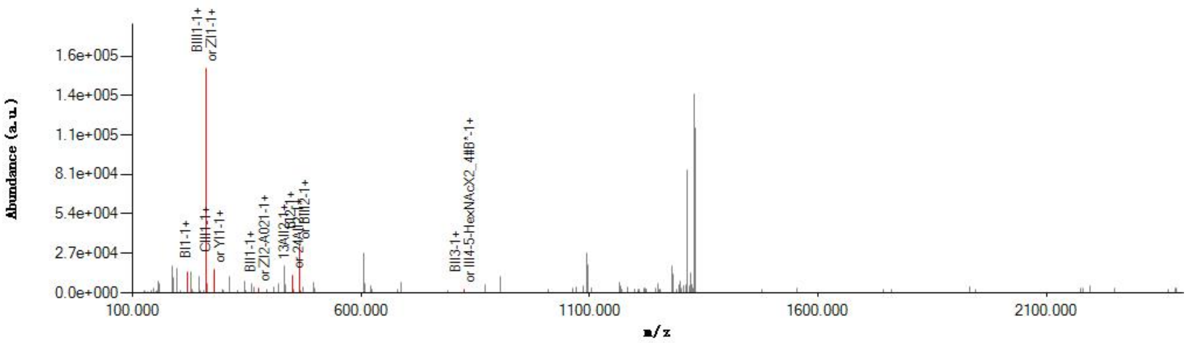


#### Graphical Fragmentation Map

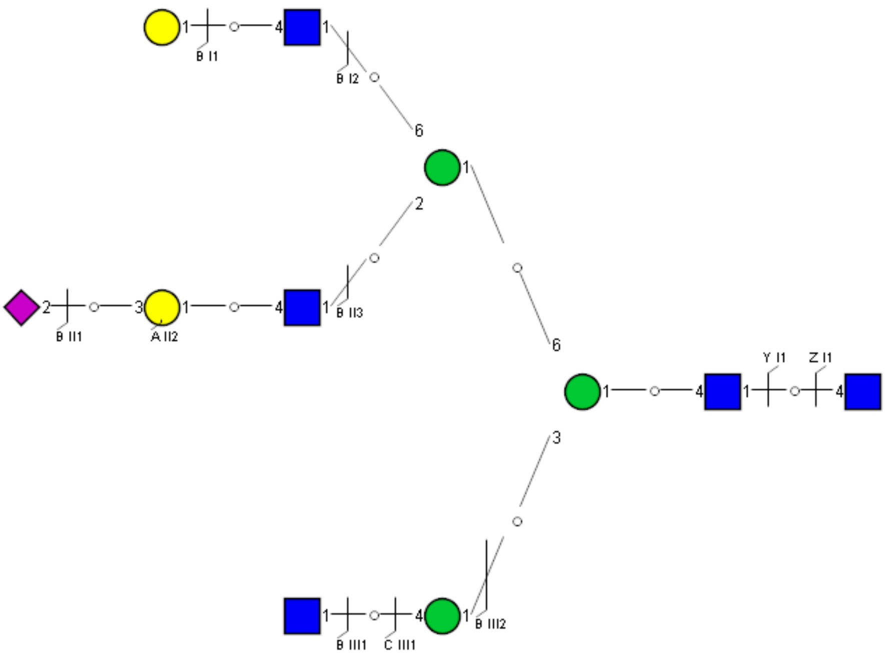

Spectrum Index: 36246(NoBHs:1)

#### Precursor iEF Map

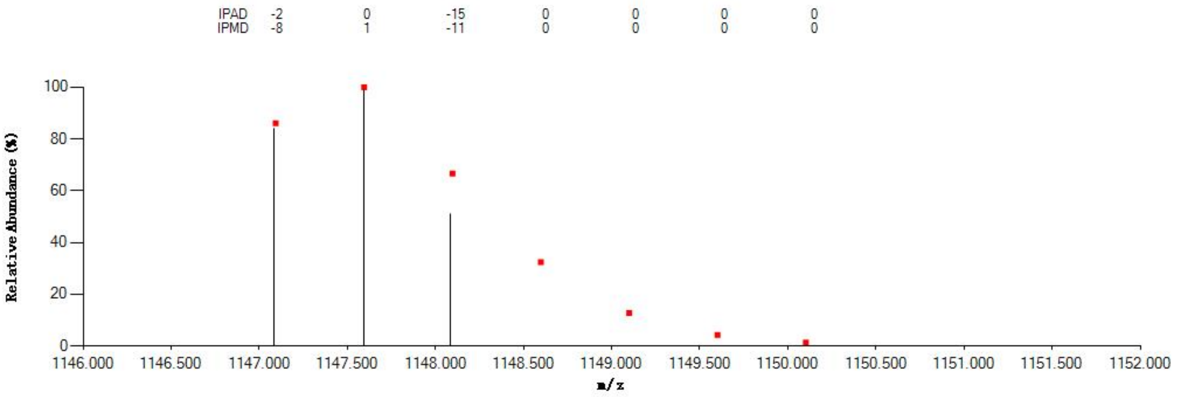


#### Annotated MS/MS Spectrum

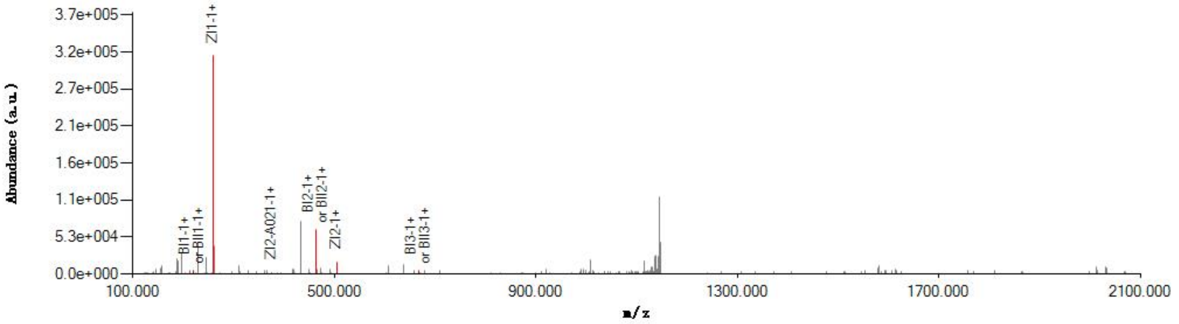


#### Graphical Fragmentation Map

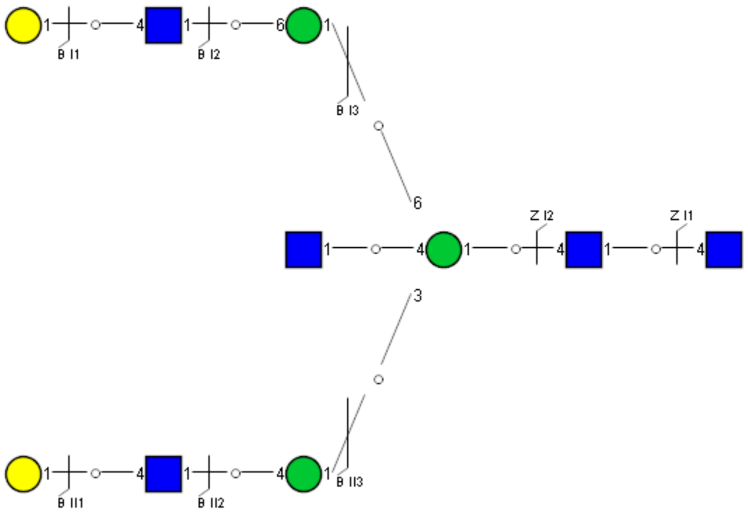

Spectrum Index: 36587(NoBHs:2)

#### Precursor iEF Map

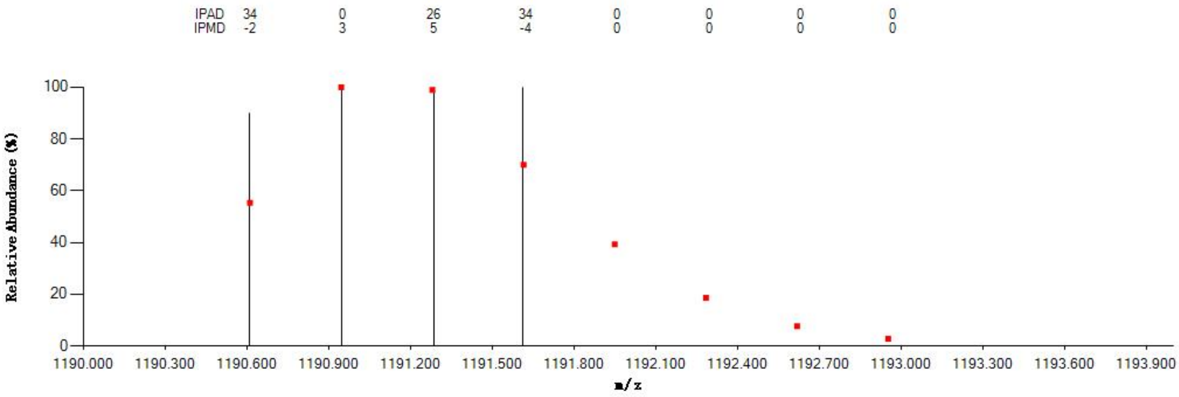


#### Annotated MS/MS Spectrum

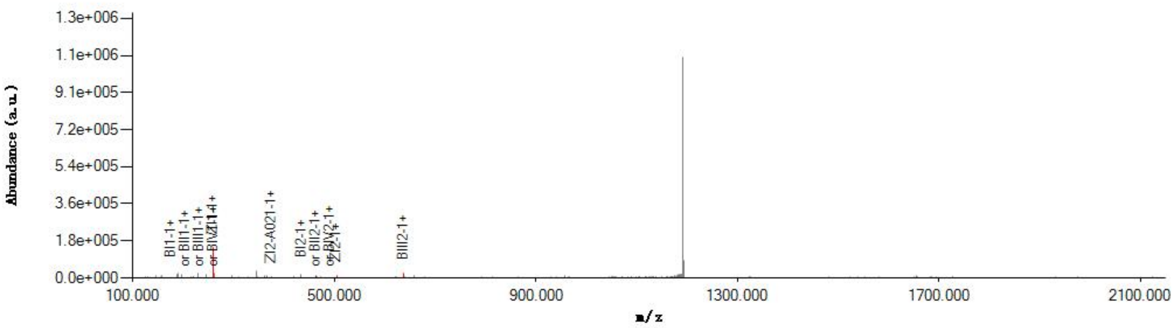


#### Graphical Fragmentation Map

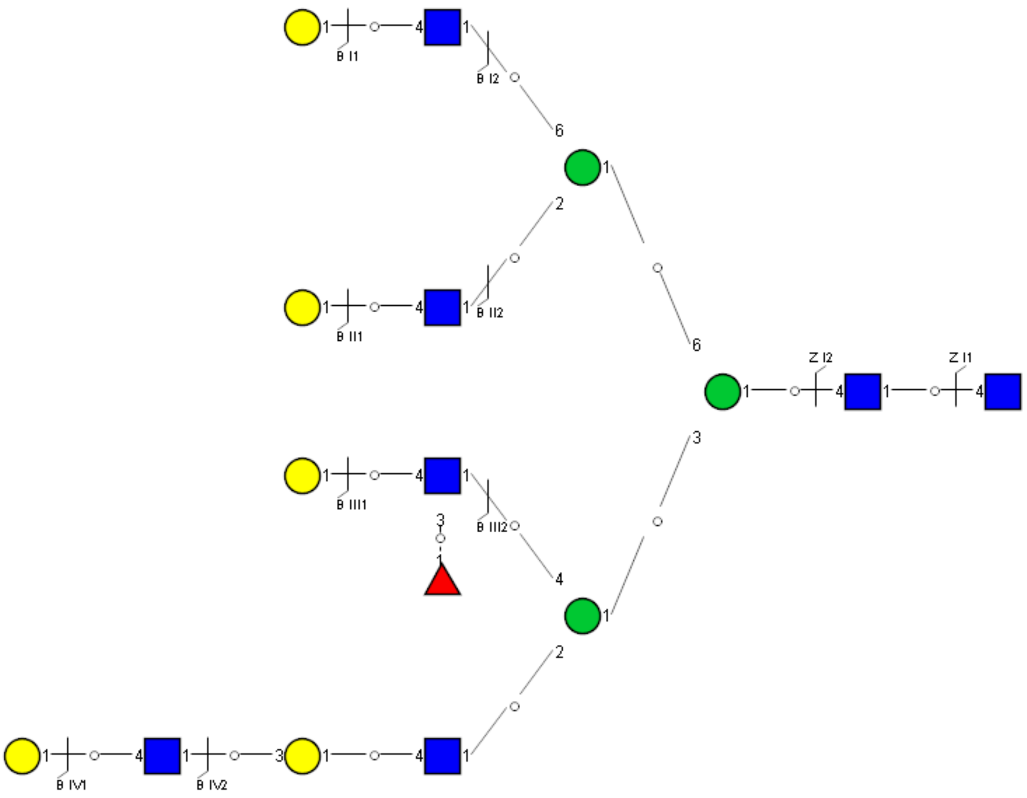

Spectrum Index: 36587(NoBHs:2)

#### Precursor iEF Map

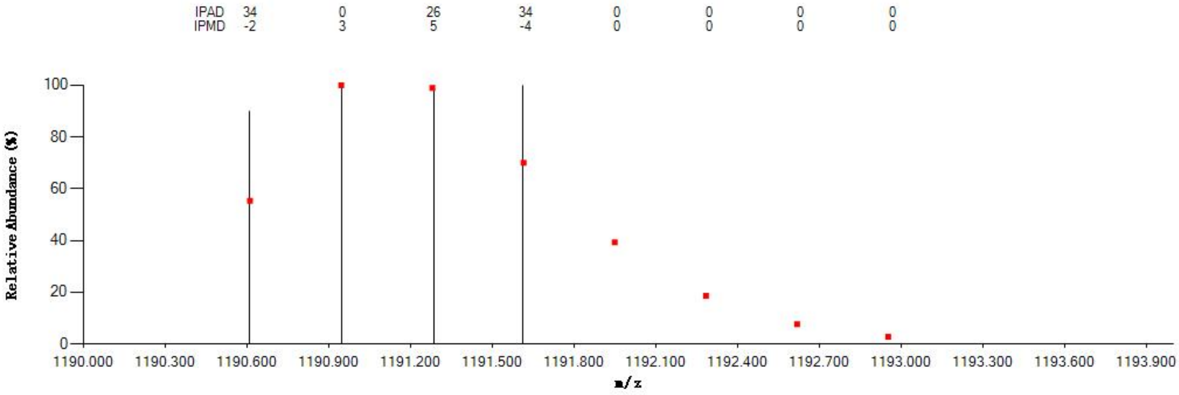


#### Annotated MS/MS Spectrum

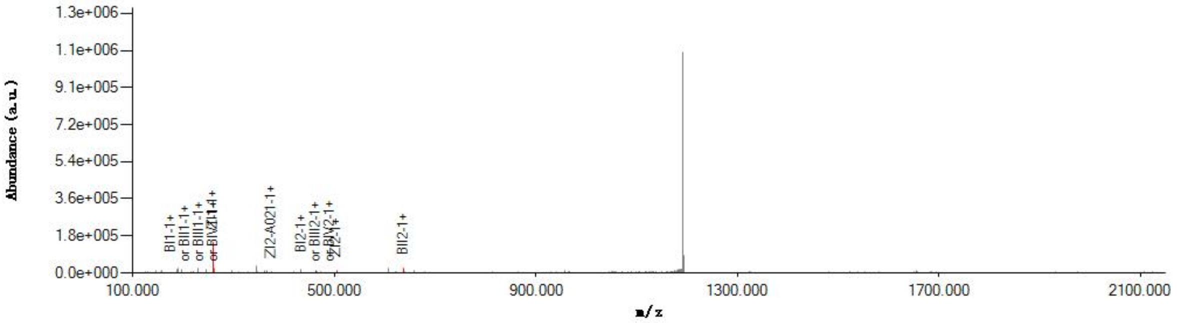


#### Graphical Fragmentation Map

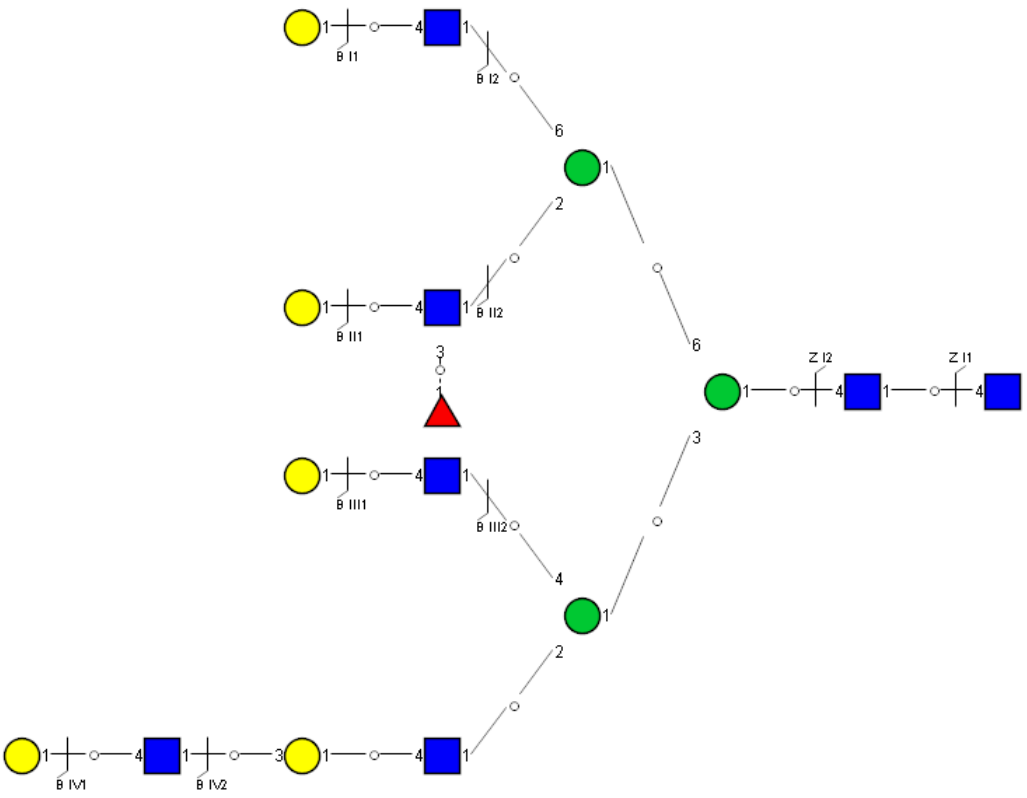

Spectrum Index: 36659(NoBHs:1)

#### Precursor iEF Map

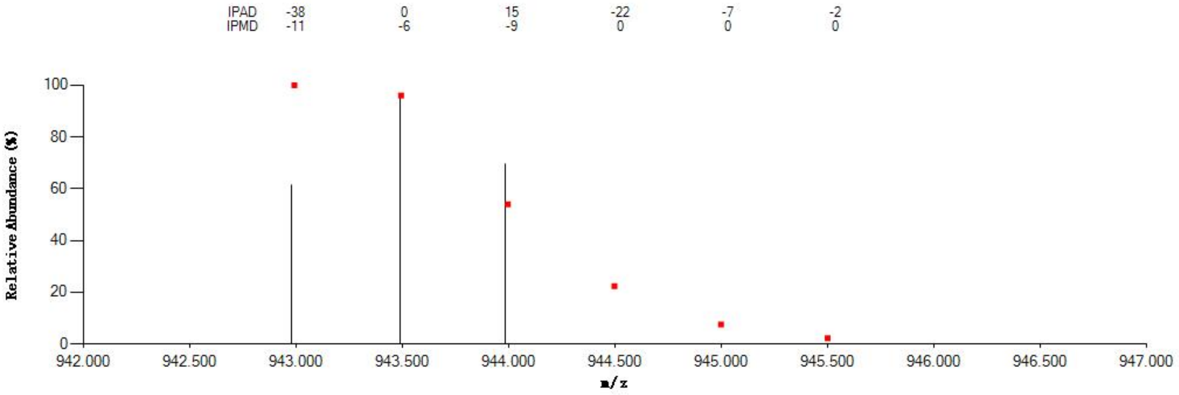


#### Annotated MS/MS Spectrum

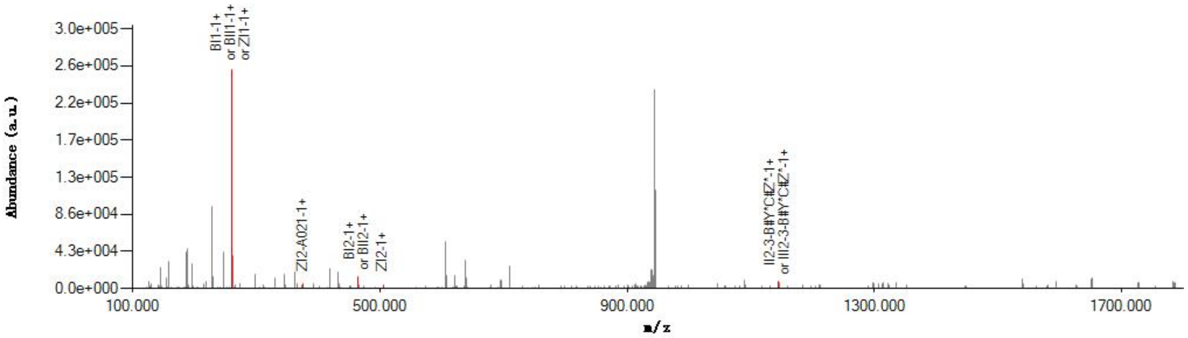


#### Graphical Fragmentation Map

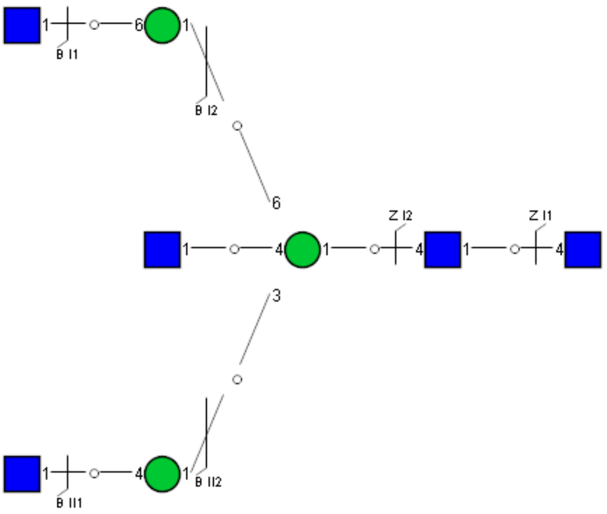

Spectrum Index: 36783(NoBHs:1)

#### Precursor iEF Map

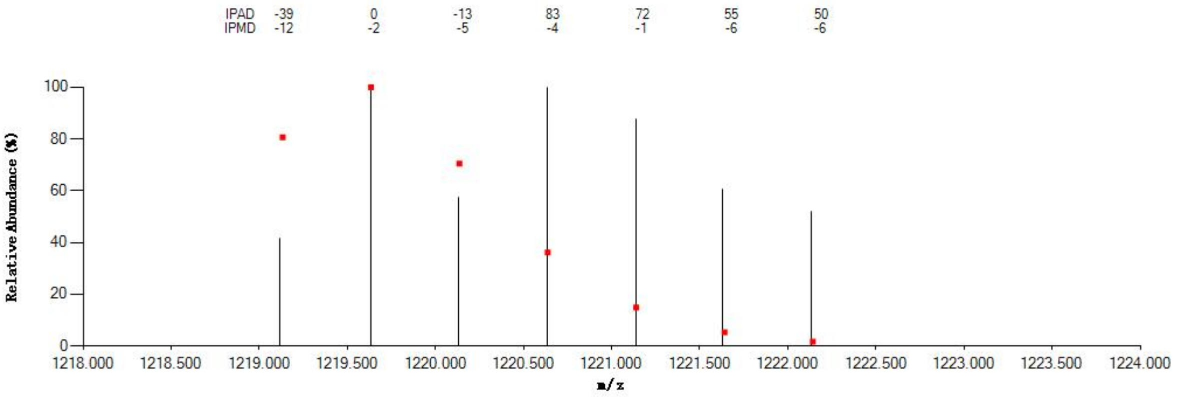


#### Annotated MS/MS Spectrum

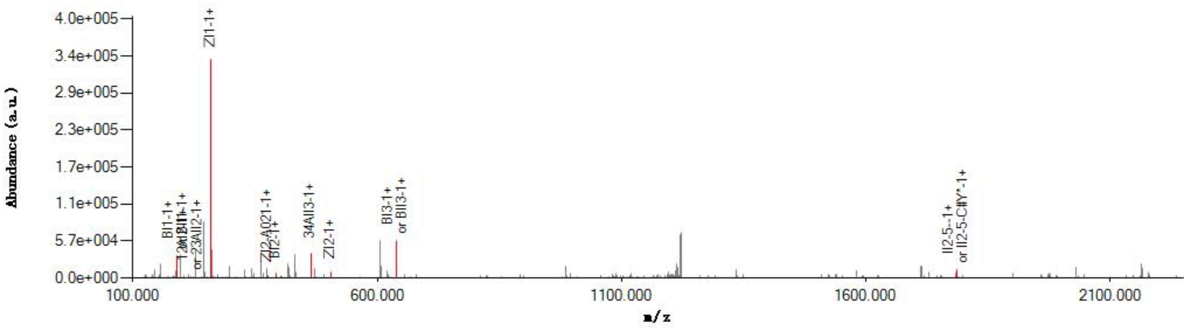


#### Graphical Fragmentation Map

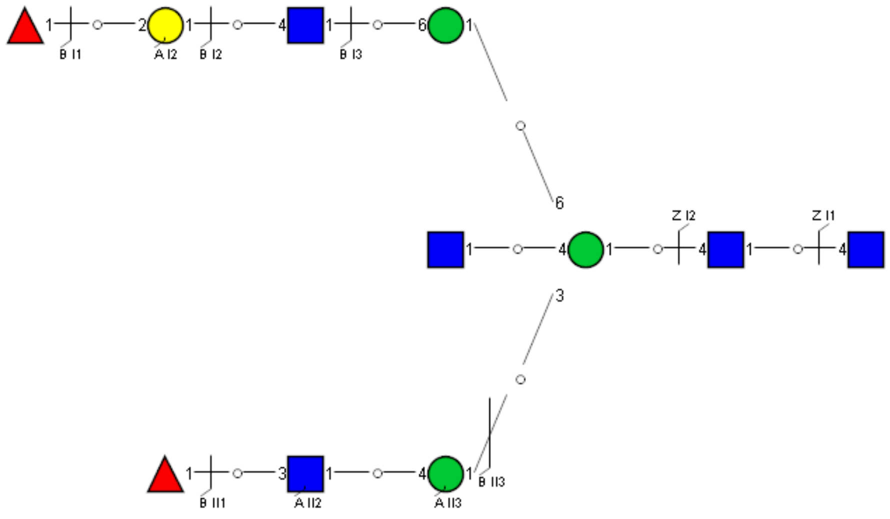

Spectrum Index: 36792(NoBHs:1)

#### Precursor iEF Map

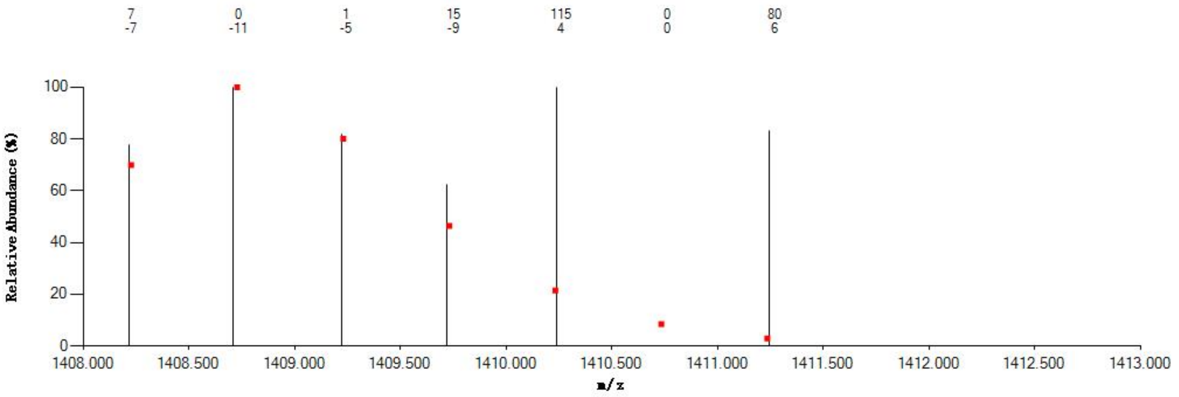


#### Annotated MS/MS Spectrum

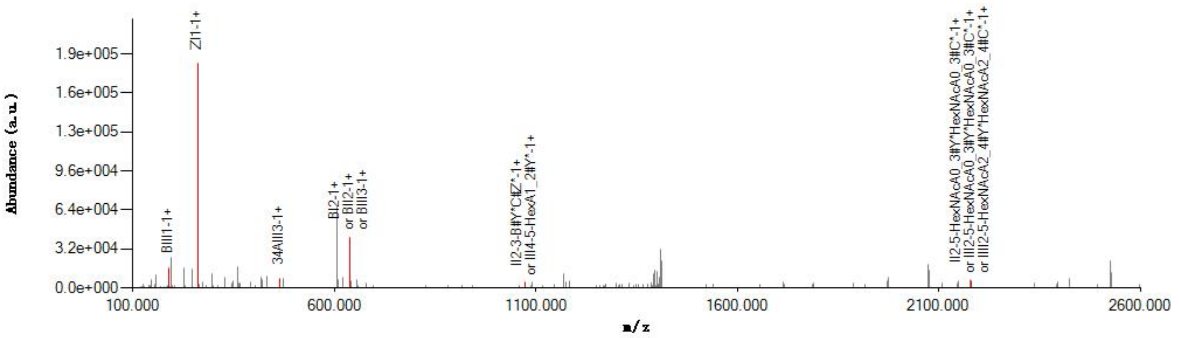


#### Graphical Fragmentation Map

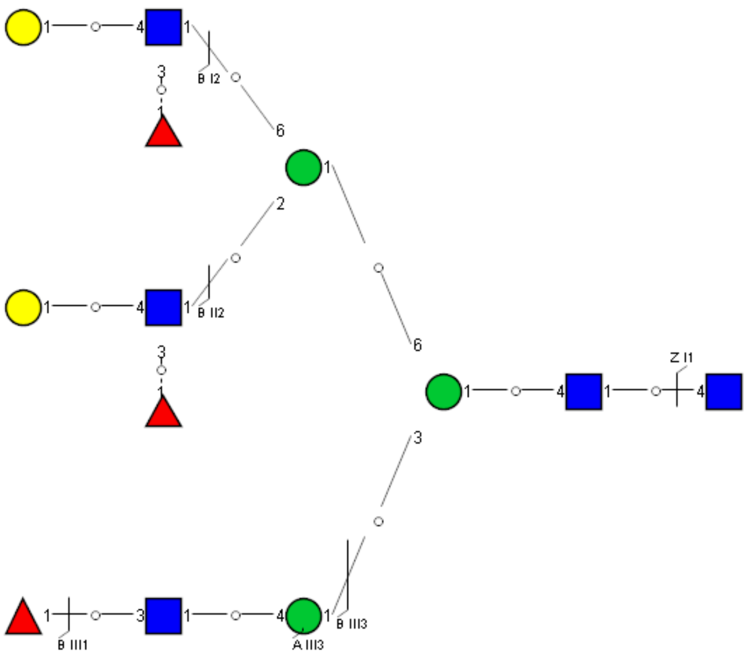

Spectrum Index: 36875(NoBHs:1)

#### Precursor iEF Map

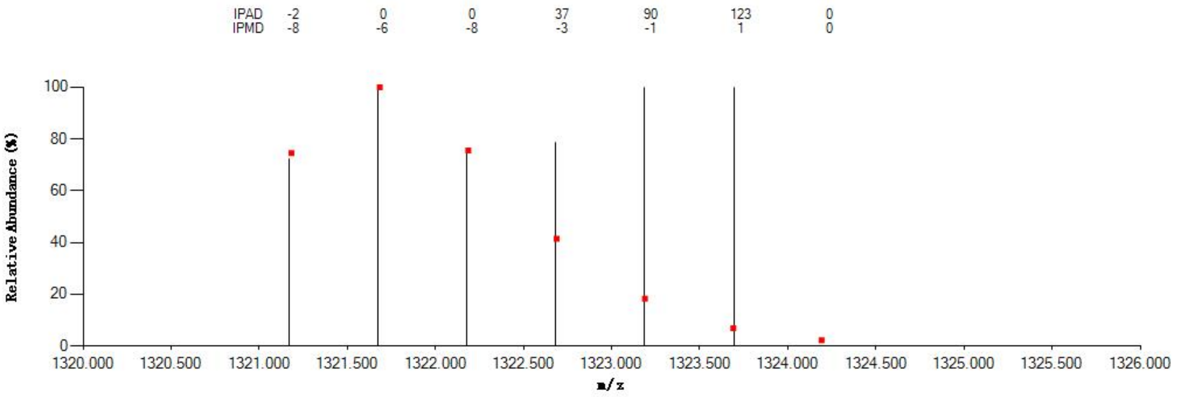


#### Annotated MS/MS Spectrum

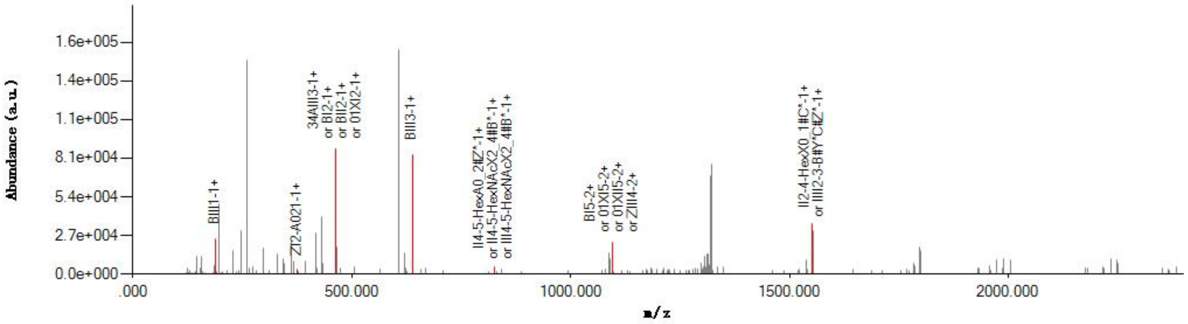


#### Graphical Fragmentation Map

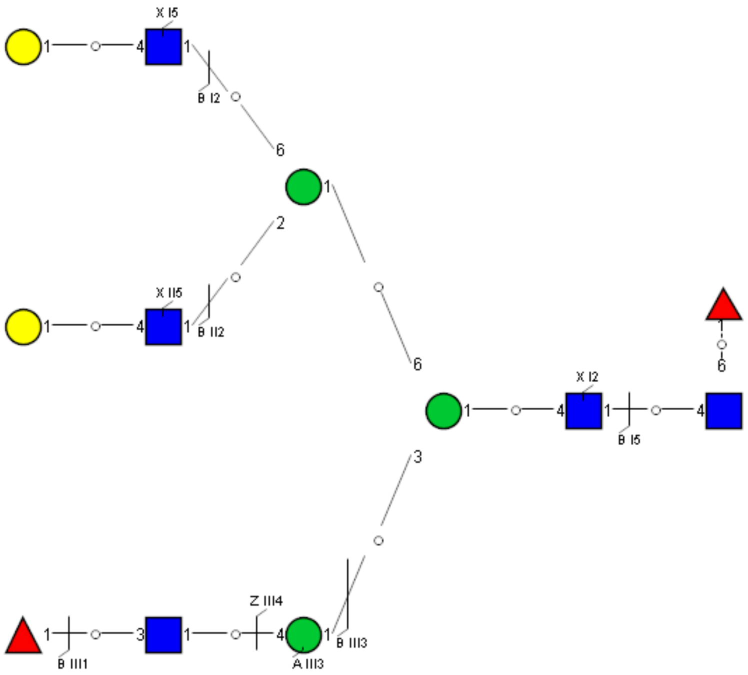

### Spectrum Index: 37185(NoBHs:1)

#### Precursor iEF Map

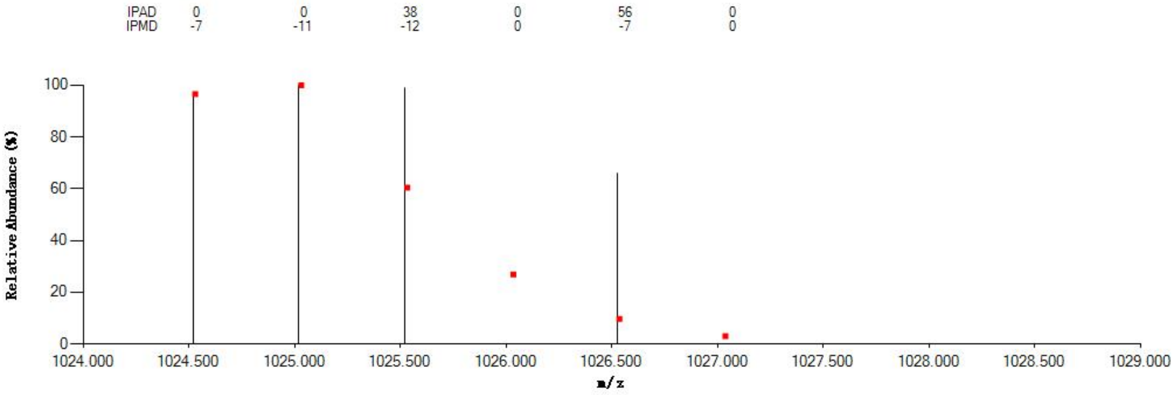


#### Annotated MS/MS Spectrum

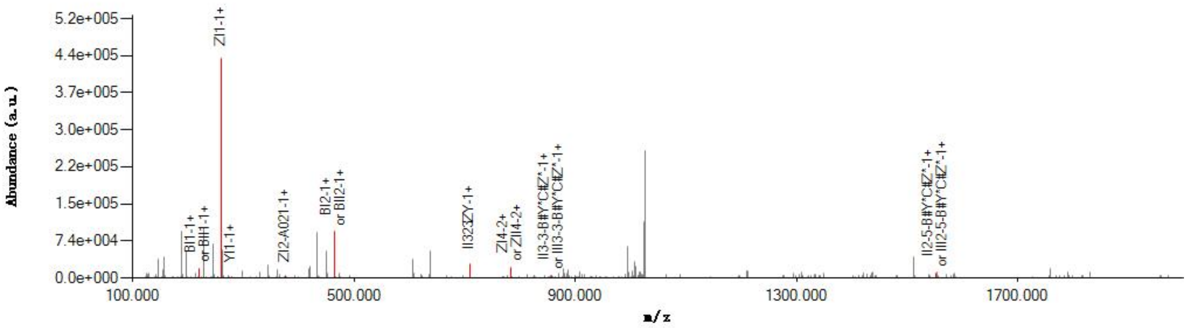


#### Graphical Fragmentation Map

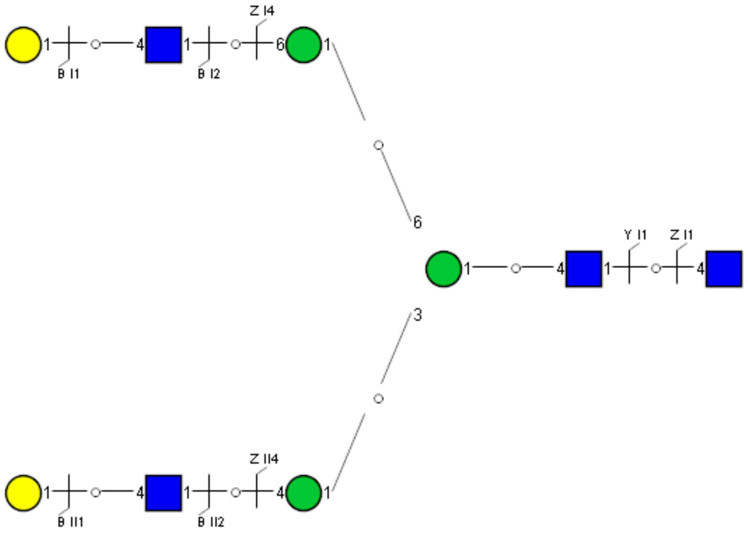

Spectrum Index: 37220(NoBHs:1)

#### Precursor iEF Map

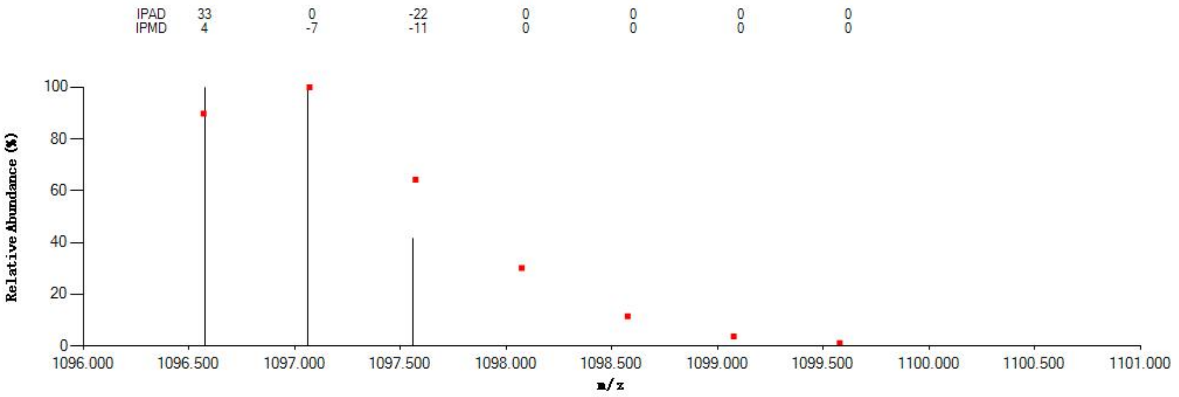


#### Annotated MS/MS Spectrum

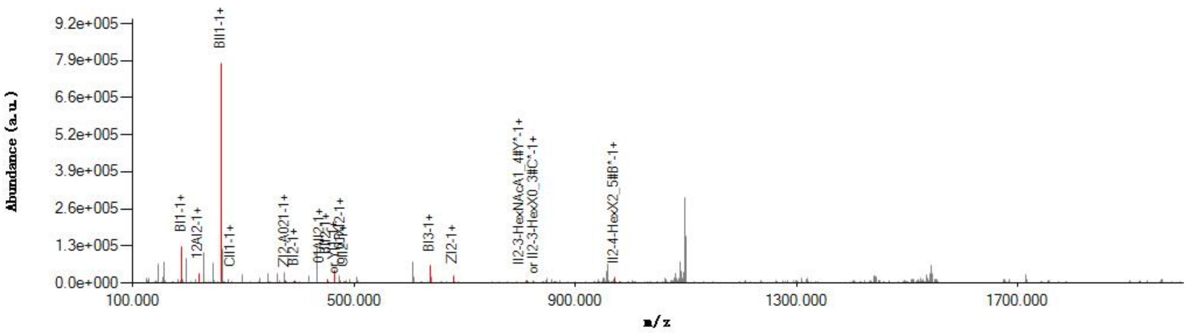


#### Graphical Fragmentation Map

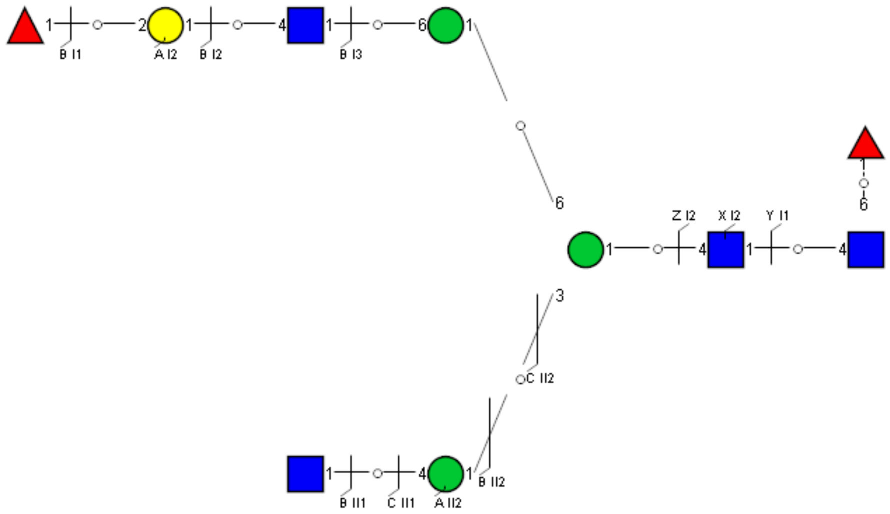

### Spectrum Index: 37319(NoBHs:1)

#### Precursor iEF Map

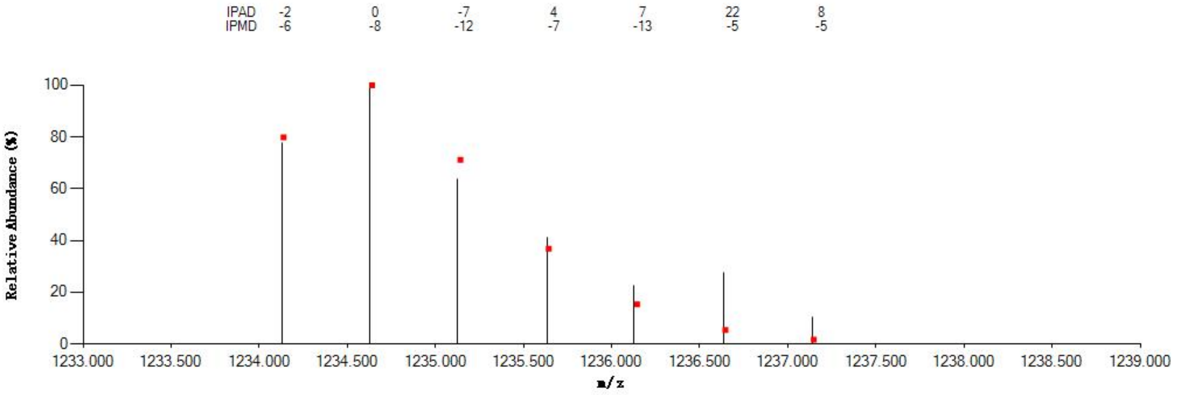


#### Annotated MS/MS Spectrum

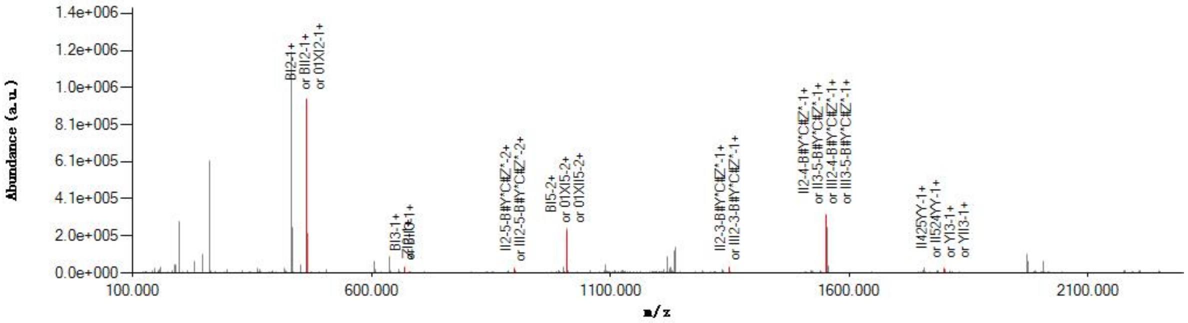


#### Graphical Fragmentation Map

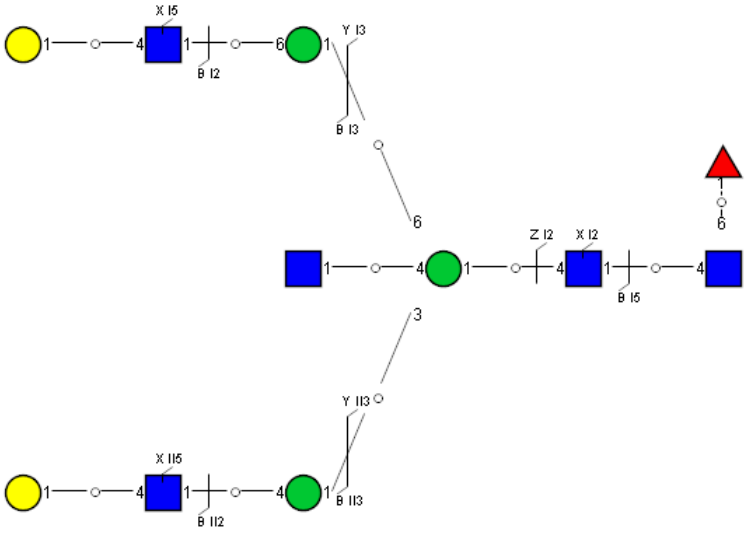

Spectrum Index: 37320(NoBHs:1)

#### Precursor iEF Map

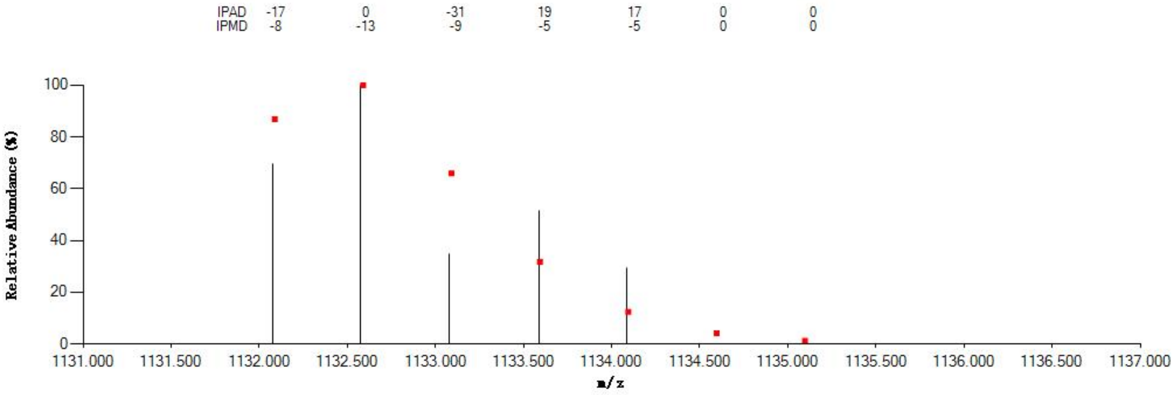


#### Annotated MS/MS Spectrum

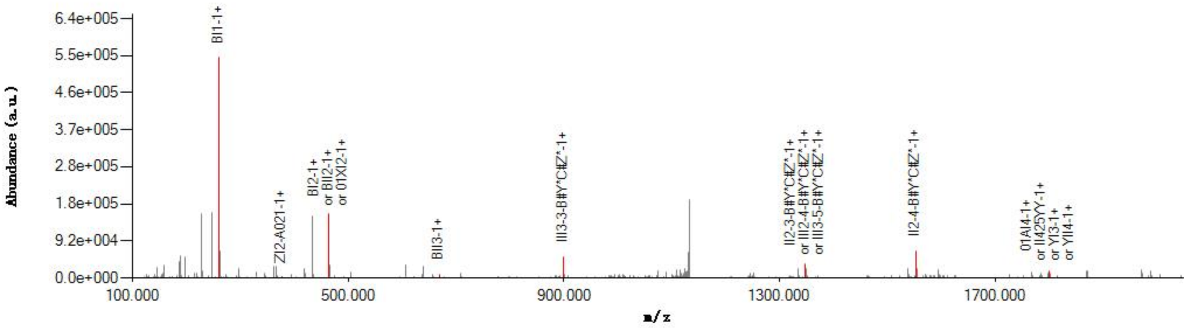


#### Graphical Fragmentation Map

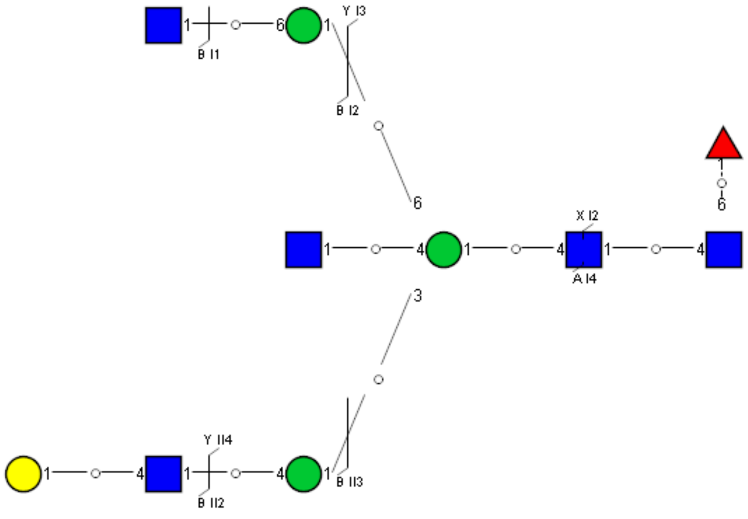

Spectrum Index: 37403(NoBHs:1)

#### Precursor iEF Map

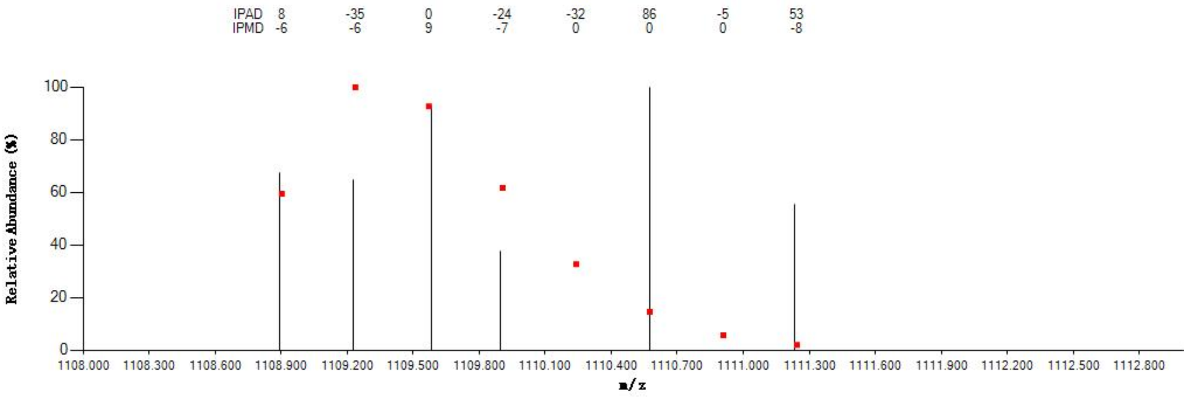


#### Annotated MS/MS Spectrum

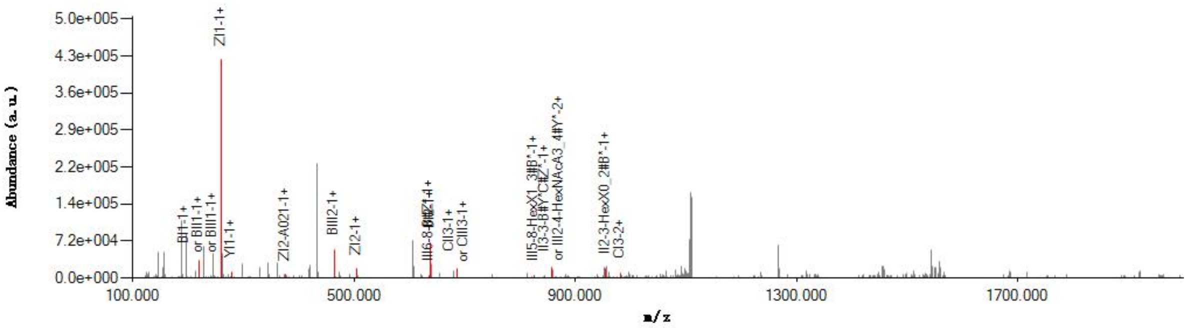


#### Graphical Fragmentation Map

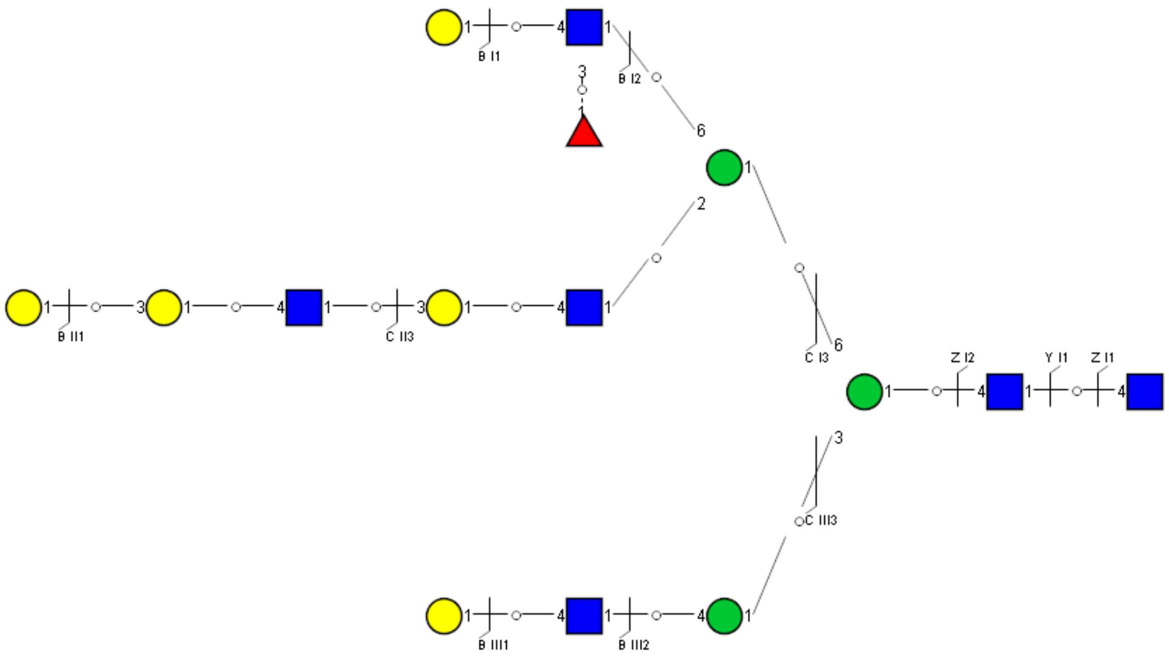

Spectrum Index: 37404(NoBHs:1)

#### Precursor iEF Map

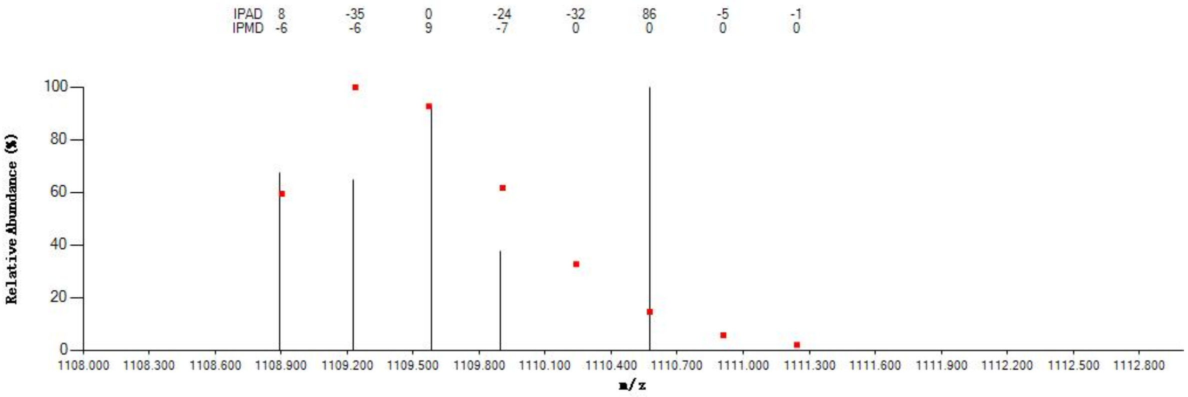


#### Annotated MS/MS Spectrum

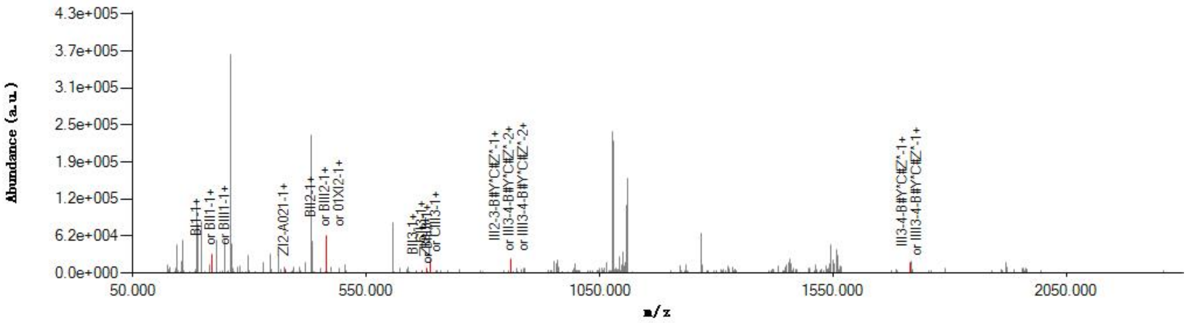


#### Graphical Fragmentation Map

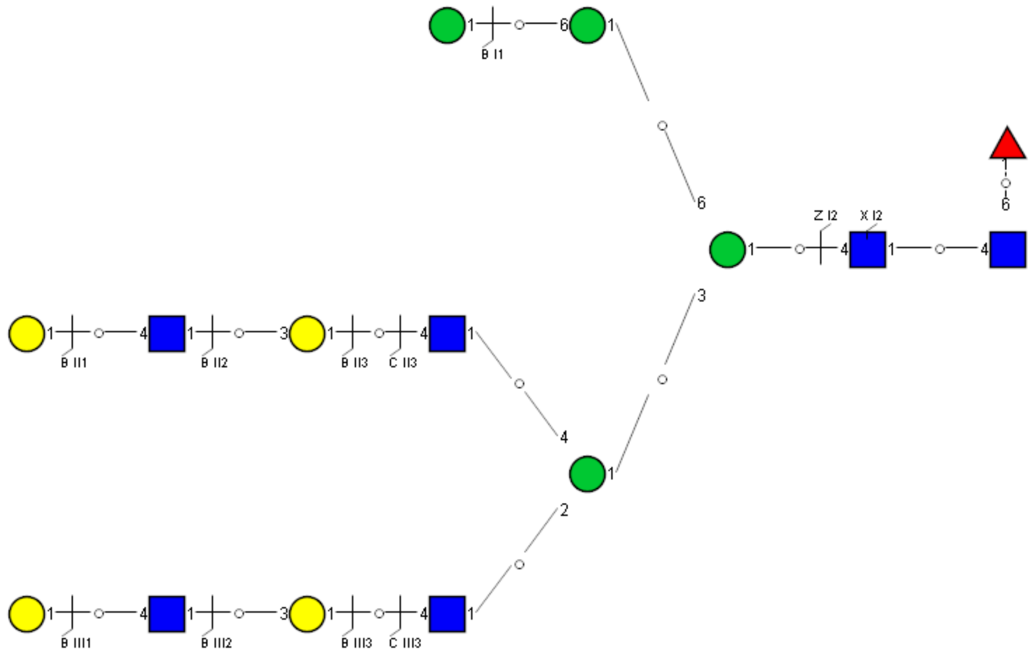

### Spectrum Index: 37456(NoBHs:1)

#### Precursor iEF Map

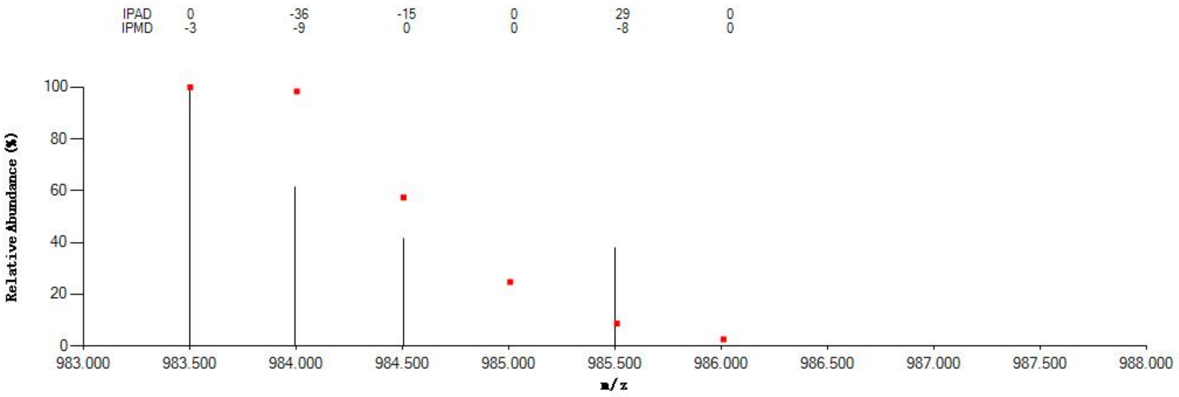

#### Annotated MS/MS Spectrum

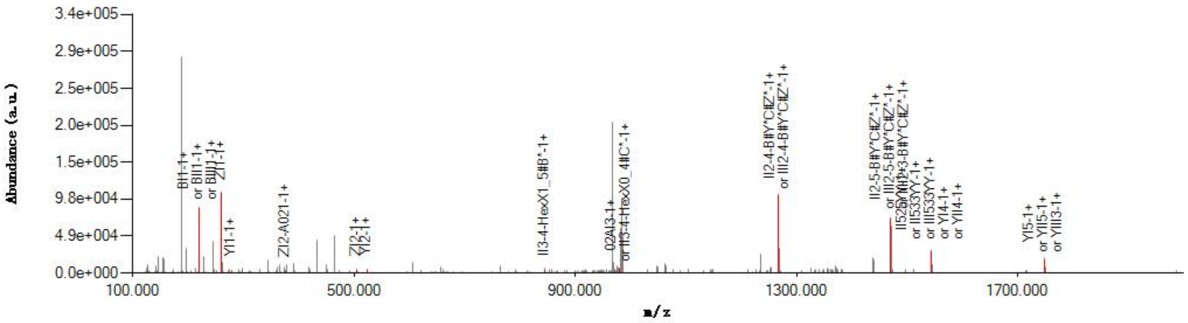

#### Graphical Fragmentation Map

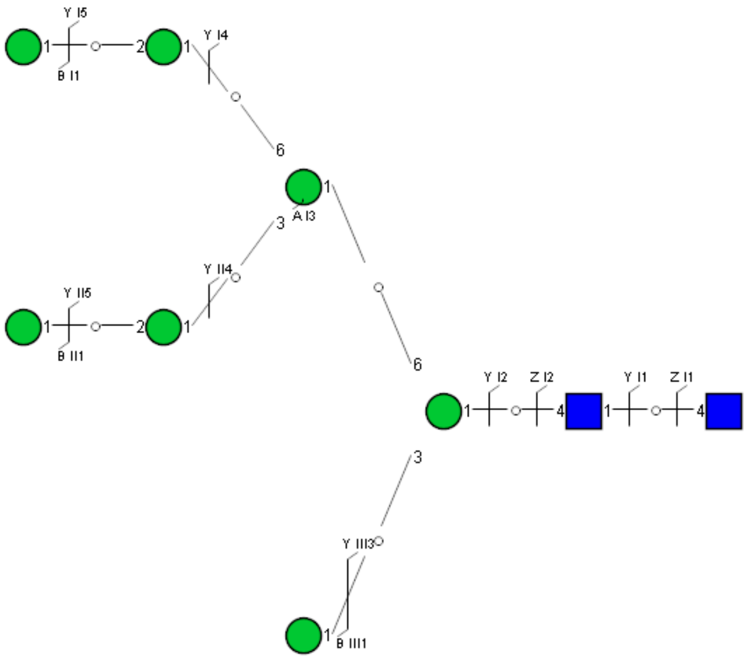

Spectrum Index: 37461(NoBHs:1)

Precursor iEF Map

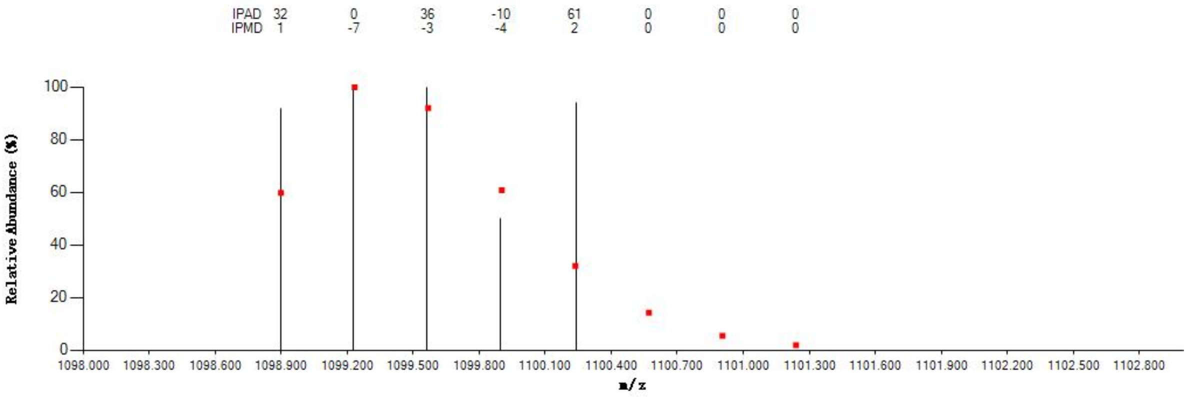

Annotated MS/MS Spectrum

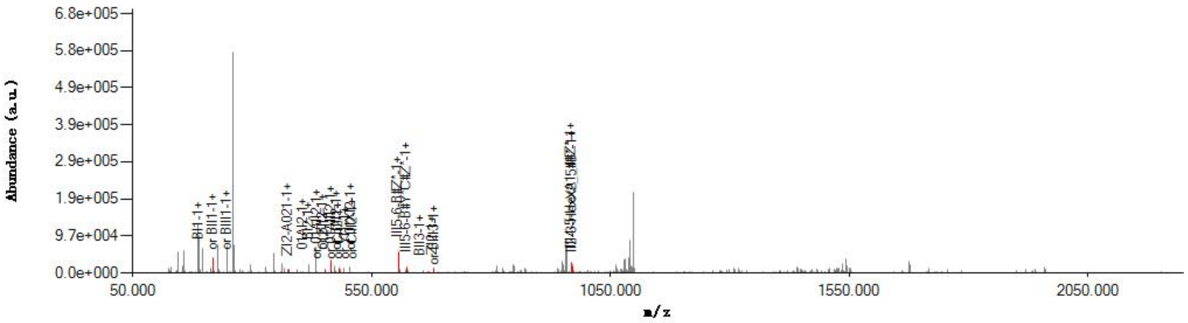

Graphical Fragmentation Map

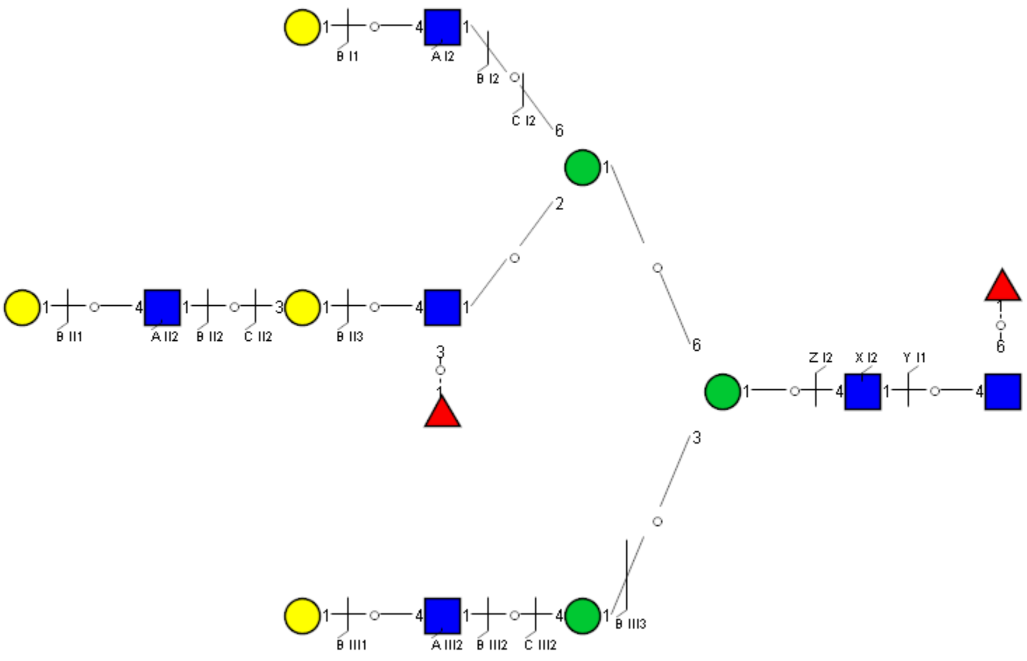

Spectrum Index: 37553(NoBHs:1)

#### Precursor iEF Map

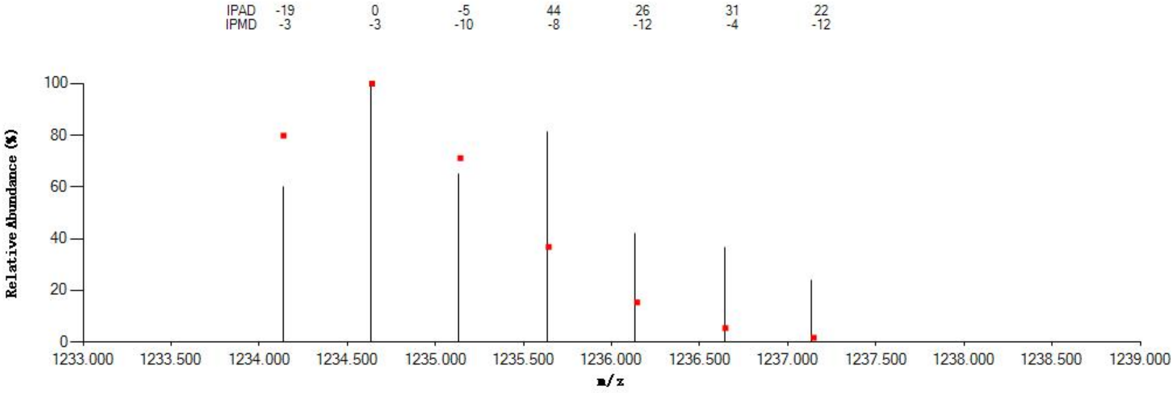


#### Annotated MS/MS Spectrum

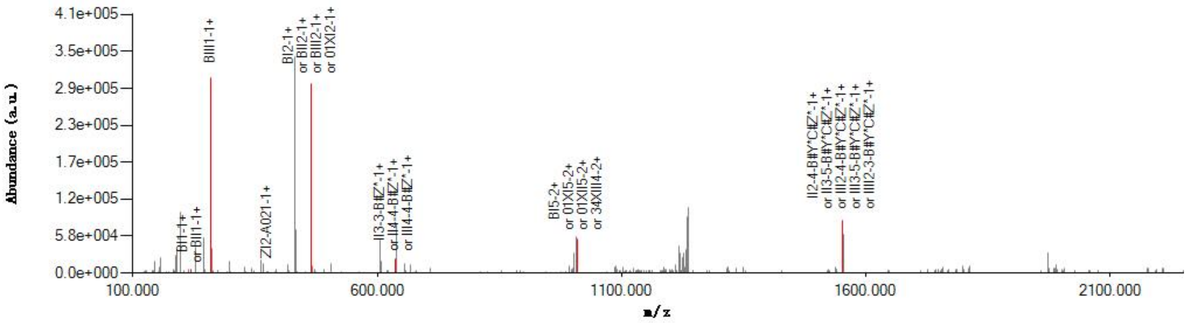


#### Graphical Fragmentation Map

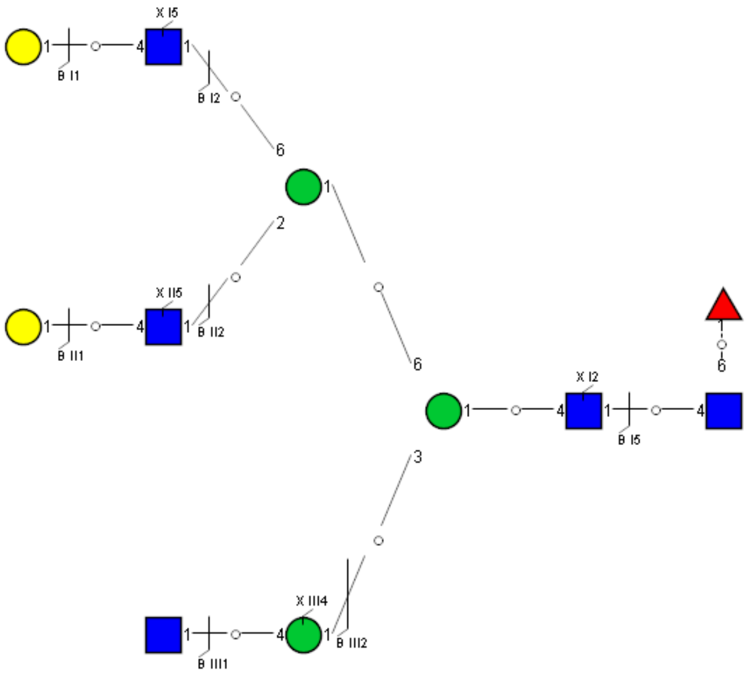

Spectrum Index: 37561(NoBHs:1)

#### Precursor iEF Map

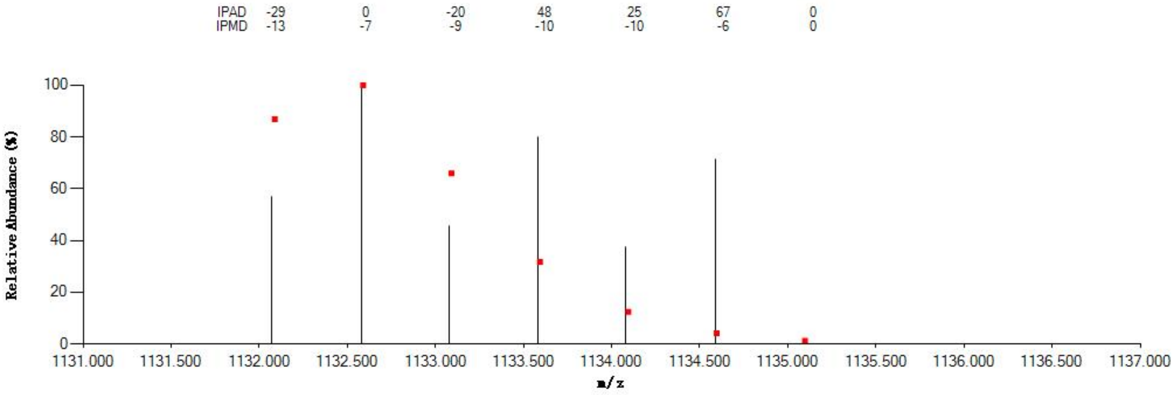


#### Annotated MS/MS Spectrum

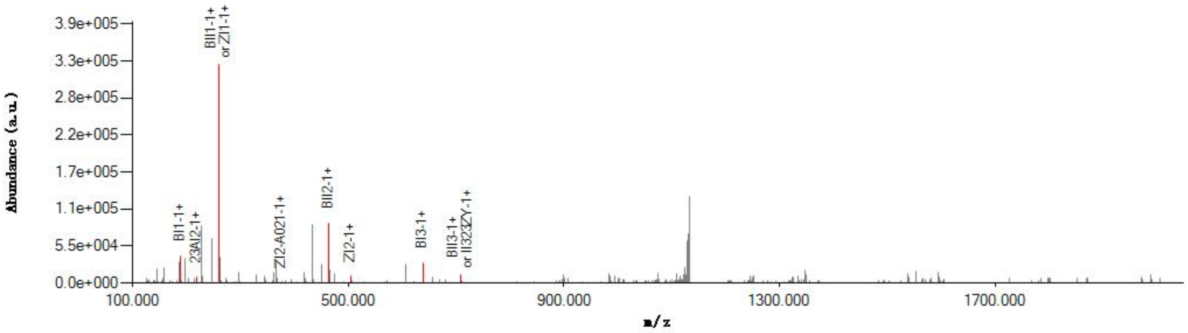


#### Graphical Fragmentation Map

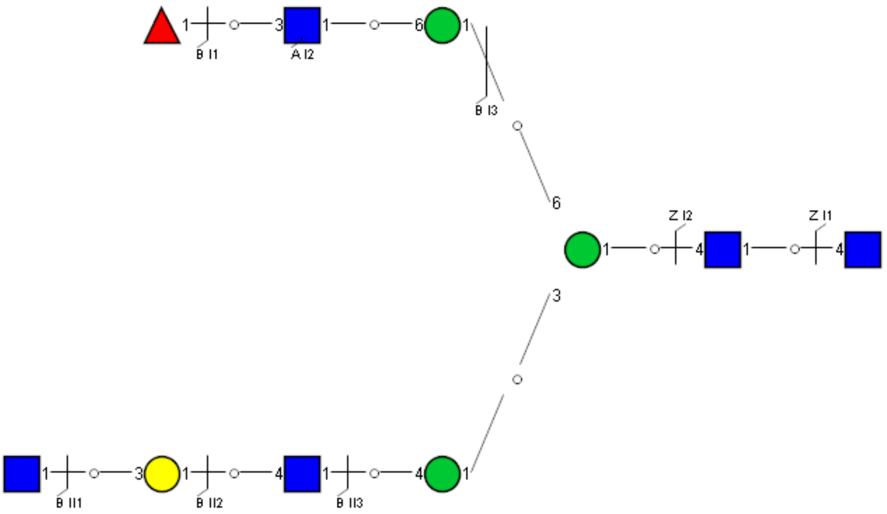

Spectrum Index: 37573(NoBHs:1)

#### Precursor iEF Map

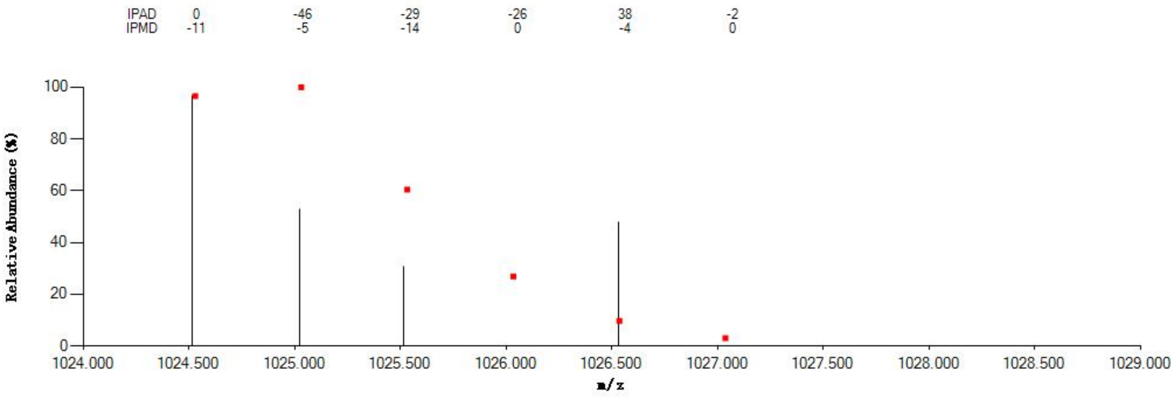


#### Annotated MS/MS Spectrum

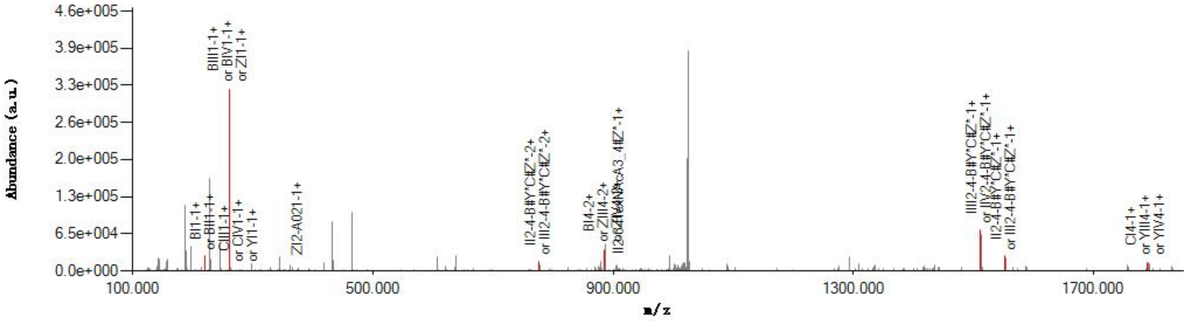


#### Graphical Fragmentation Map

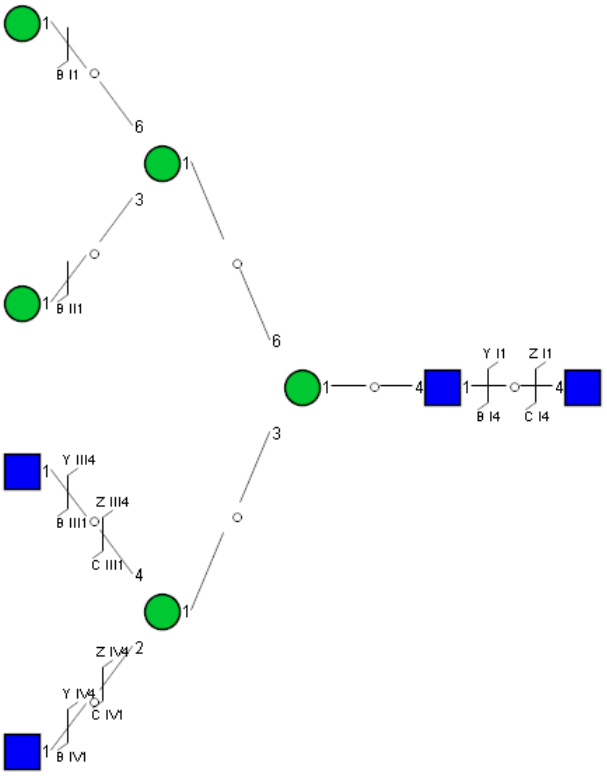

Spectrum Index: 37614(NoBHs:1)

#### Precursor iEF Map

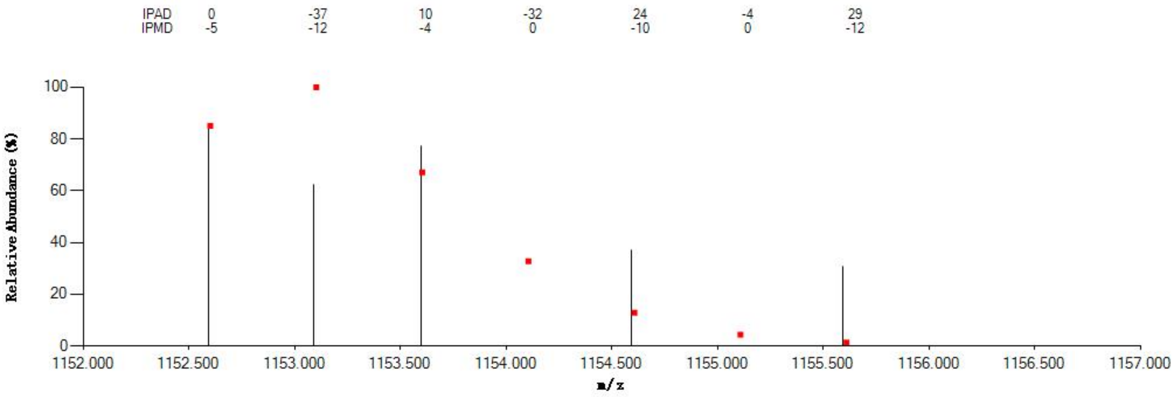


#### Annotated MS/MS Spectrum

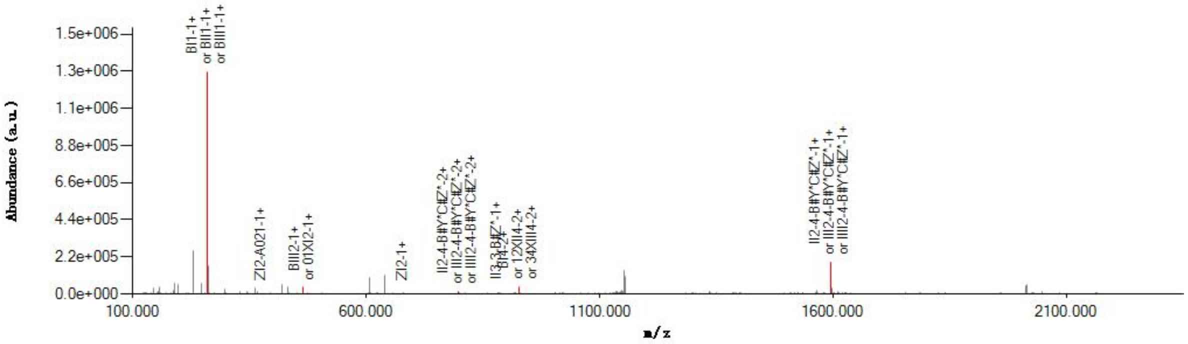


#### Graphical Fragmentation Map

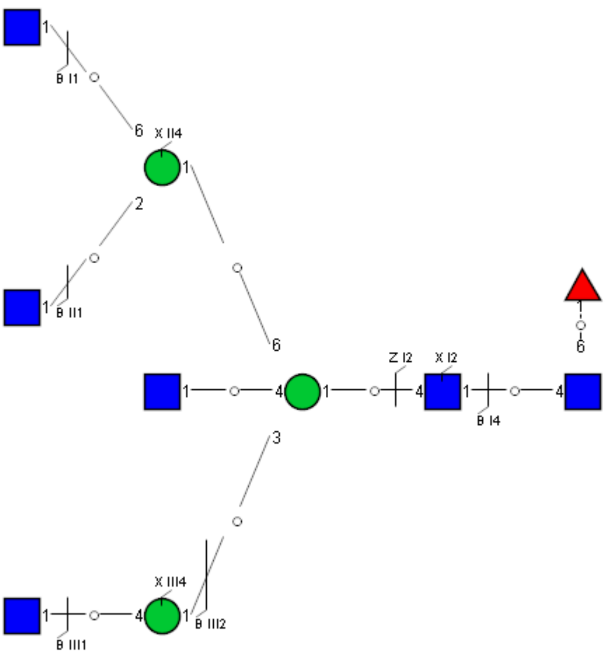

Spectrum Index: 38048(NoBHs:1)

#### Precursor iEF Map

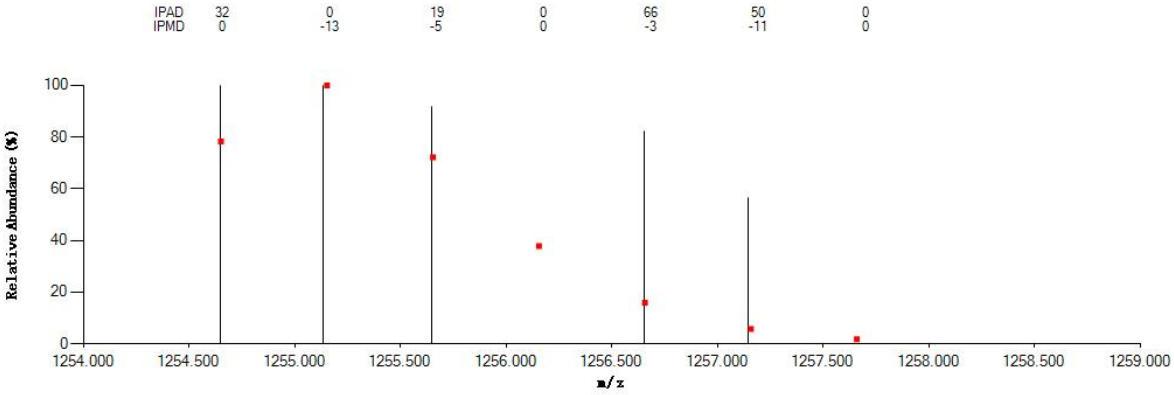


#### Annotated MS/MS Spectrum

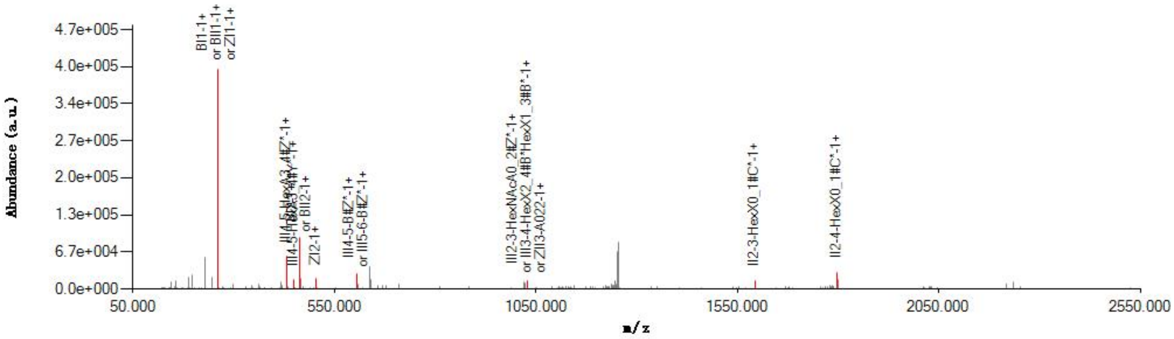


#### Graphical Fragmentation Map

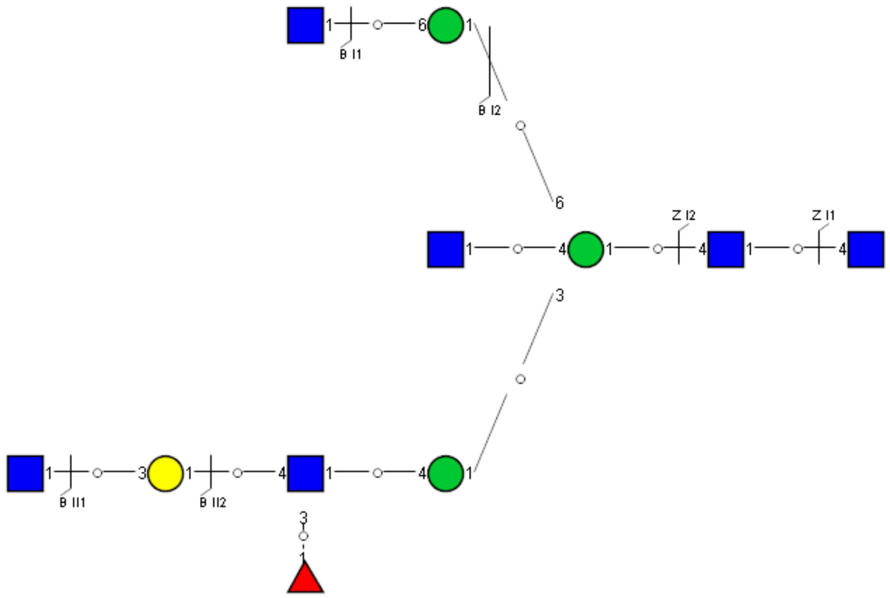

Spectrum Index: 38065(NoBHs:1)

#### Precursor iEF Map

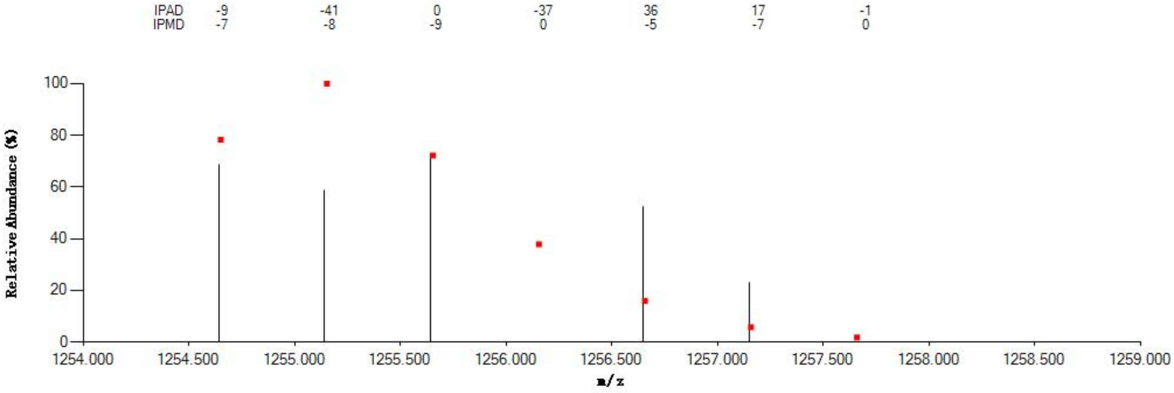


#### Annotated MS/MS Spectrum

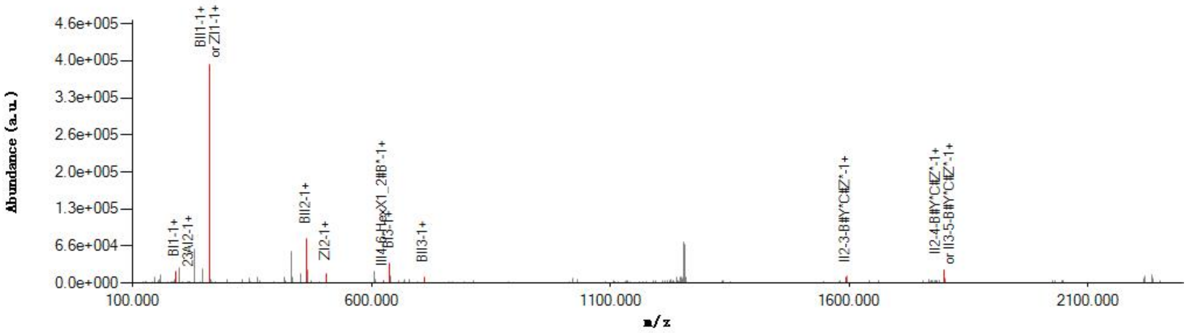


#### Graphical Fragmentation Map

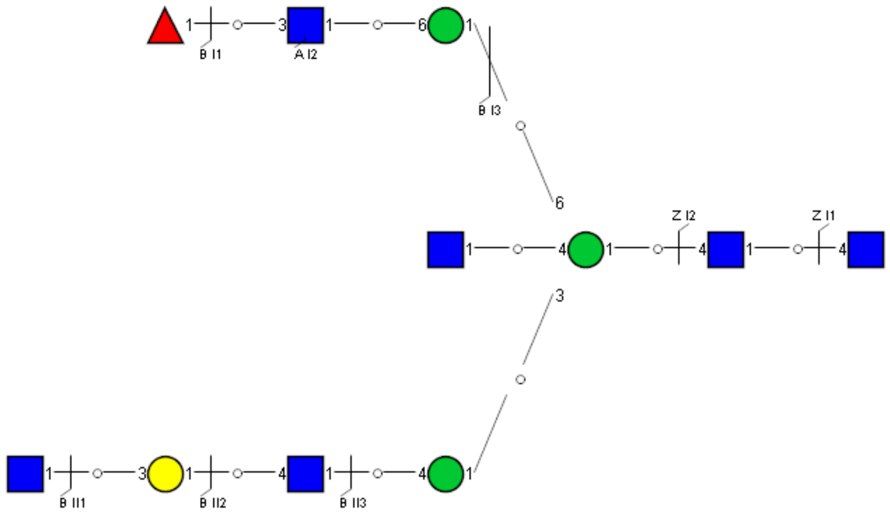

Spectrum Index: 38187(NoBHs:1)

#### Precursor iEF Map

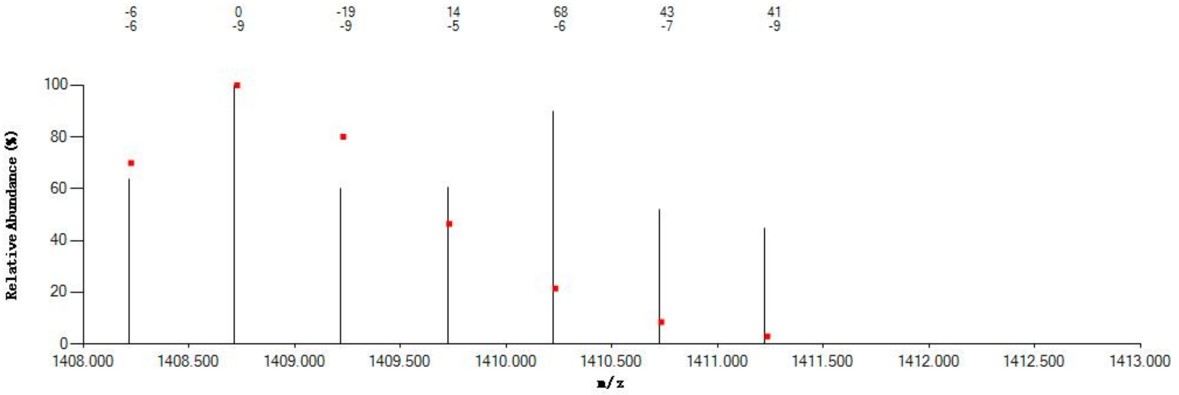

#### Annotated MS/MS Spectrum

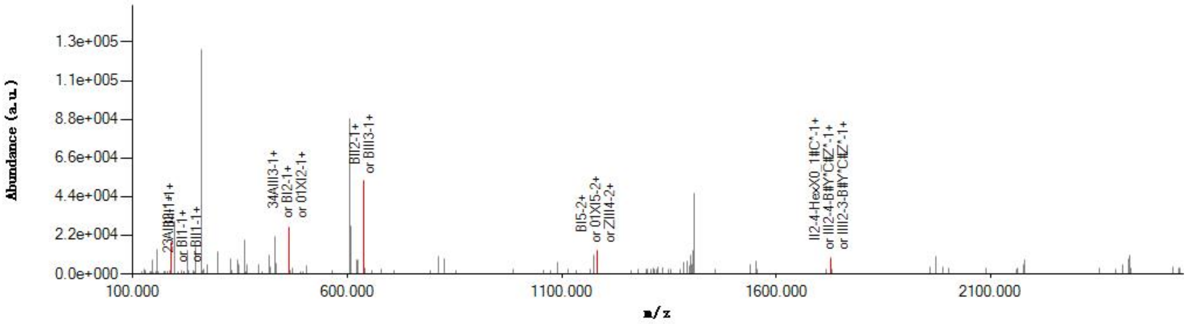

#### Graphical Fragmentation Map

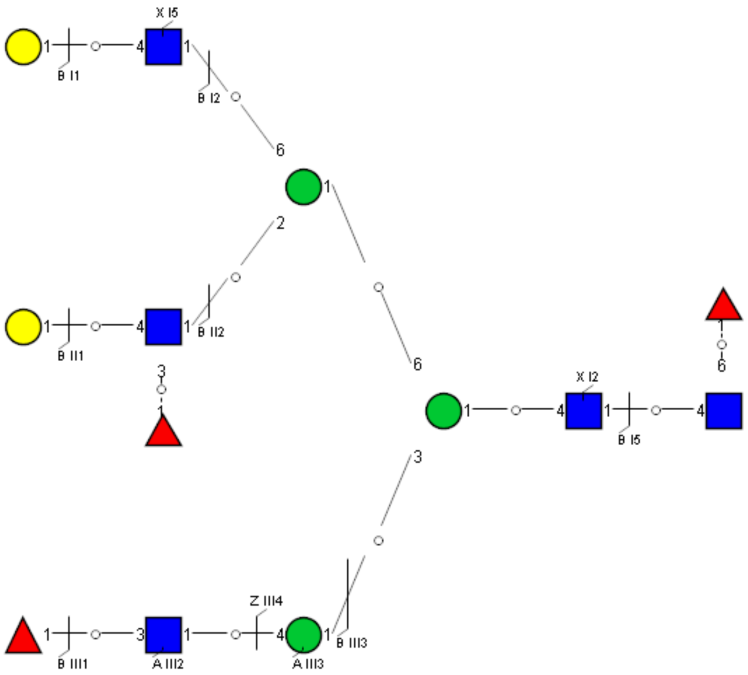

Spectrum Index: 38219(NoBHs:1)

#### Precursor iEF Map

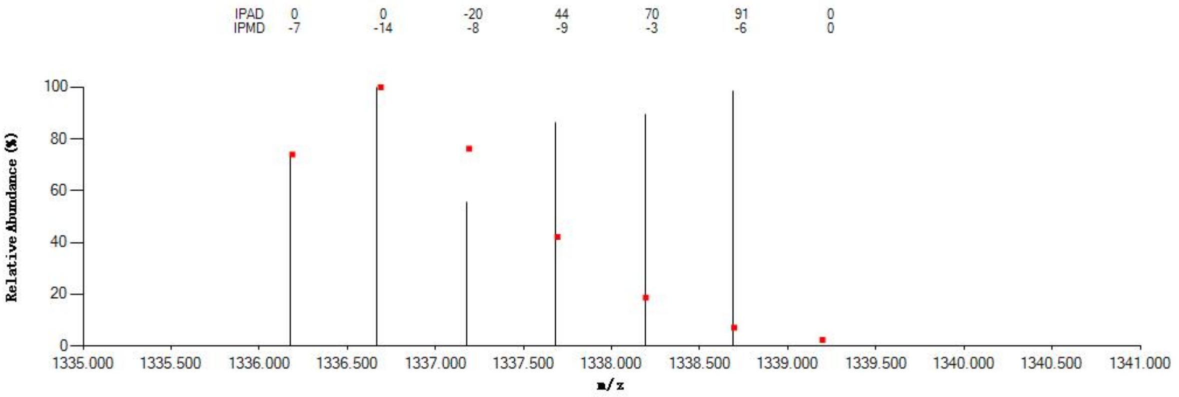


#### Annotated MS/MS Spectrum

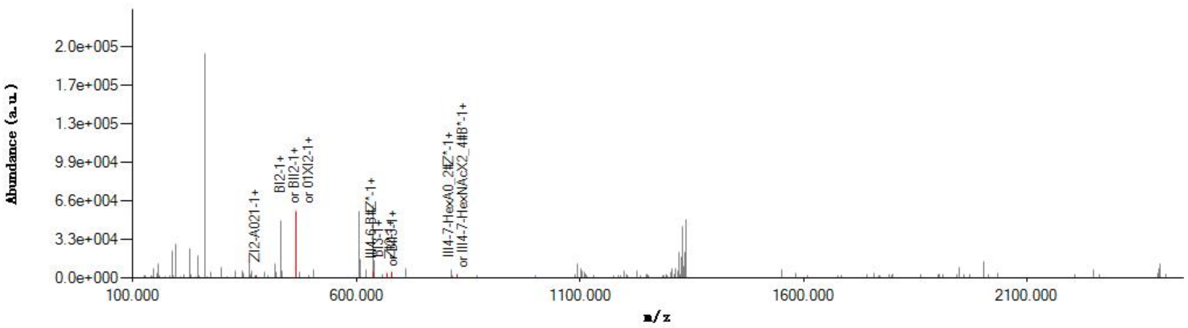


#### Graphical Fragmentation Map

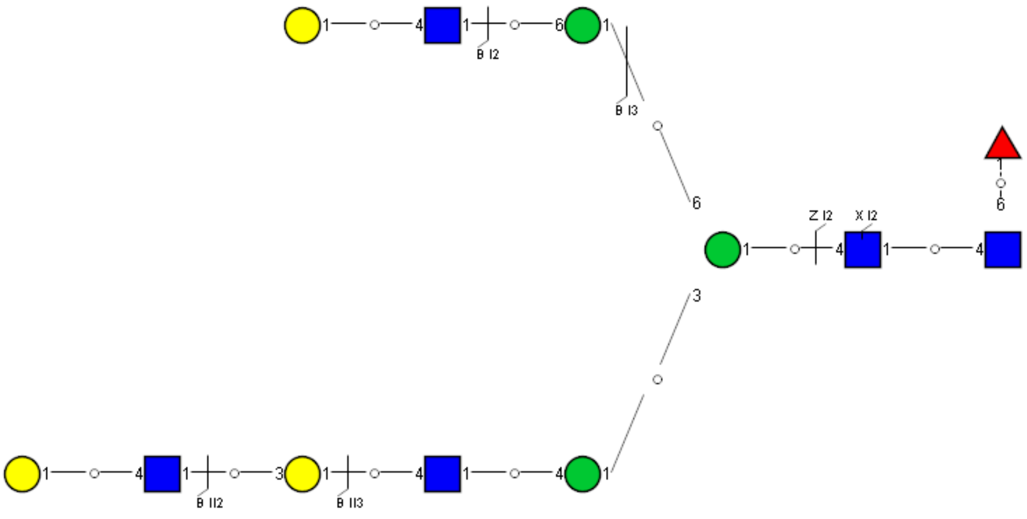

Spectrum Index: 38227(NoBHs:1)

#### Precursor iEF Map

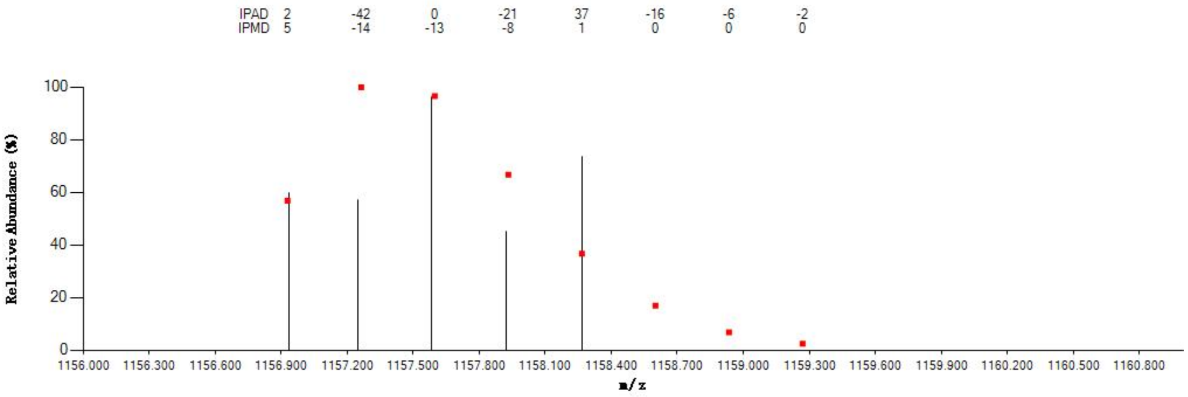


#### Annotated MS/MS Spectrum

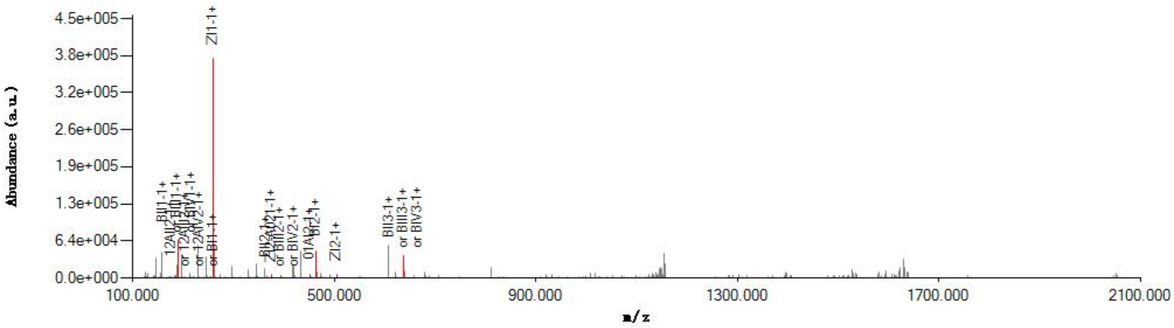


#### Graphical Fragmentation Map

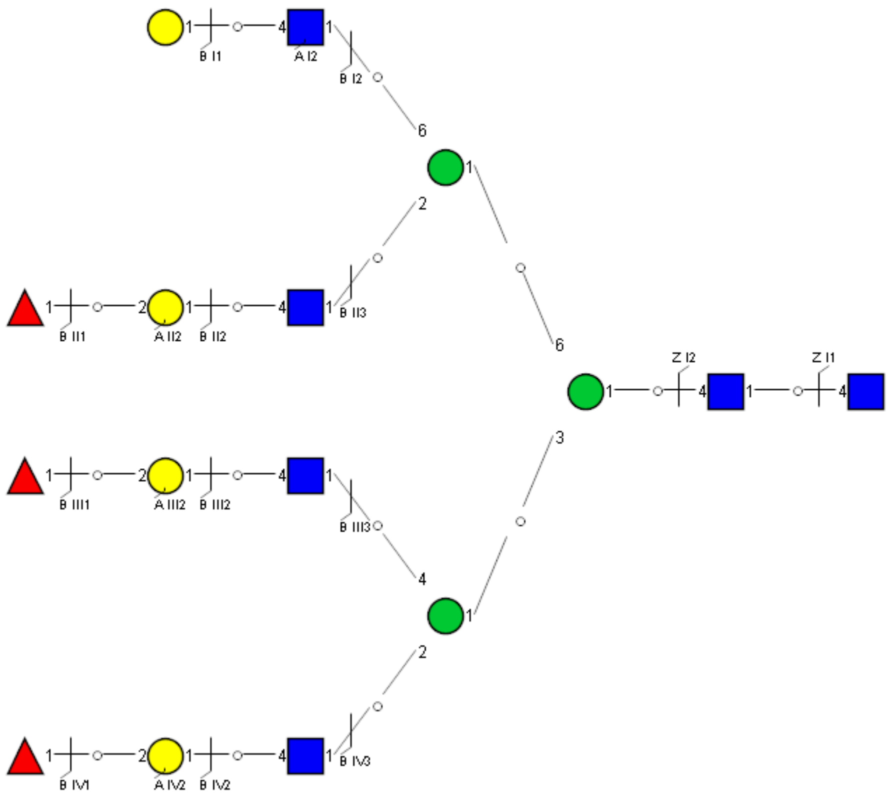

Spectrum Index: 38253(NoBHs:1)

#### Precursor iEF Map

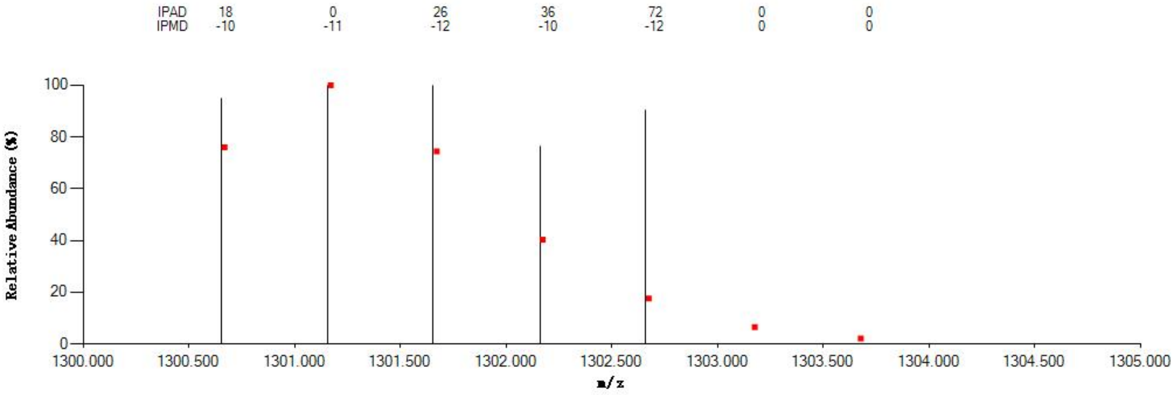


#### Annotated MS/MS Spectrum

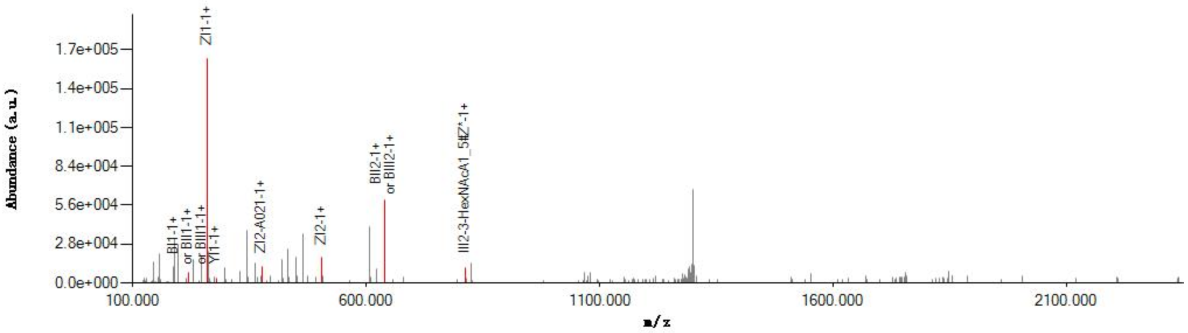


#### Graphical Fragmentation Map

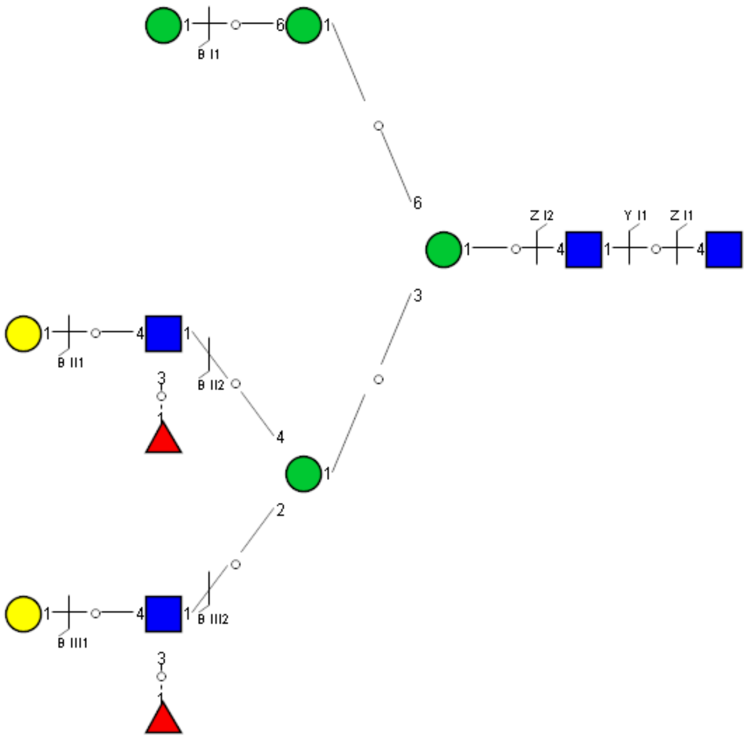

### Spectrum Index: 38524(NoBHs:1)

#### Precursor iEF Map

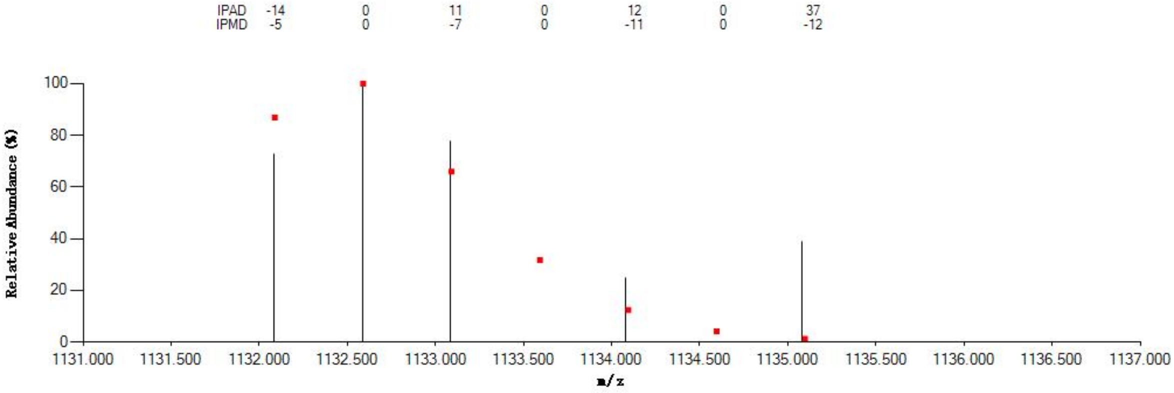


#### Annotated MS/MS Spectrum

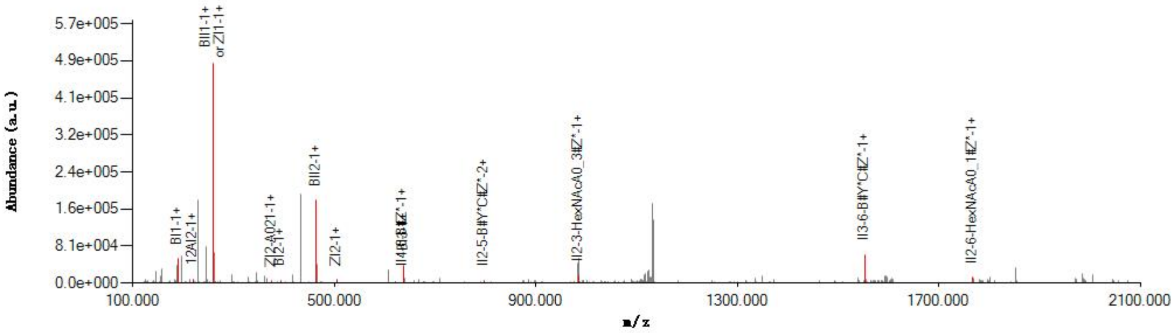


#### Graphical Fragmentation Map

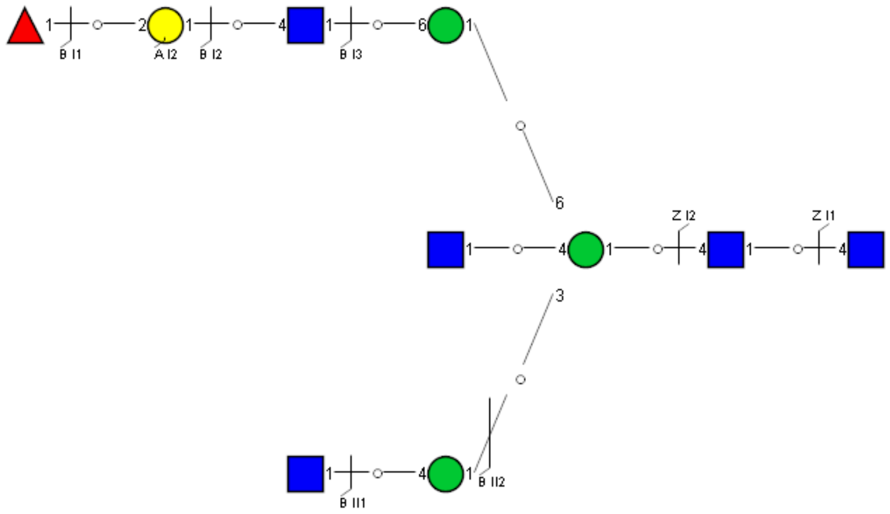

Spectrum Index: 38693(NoBHs:1)

#### Precursor iEF Map

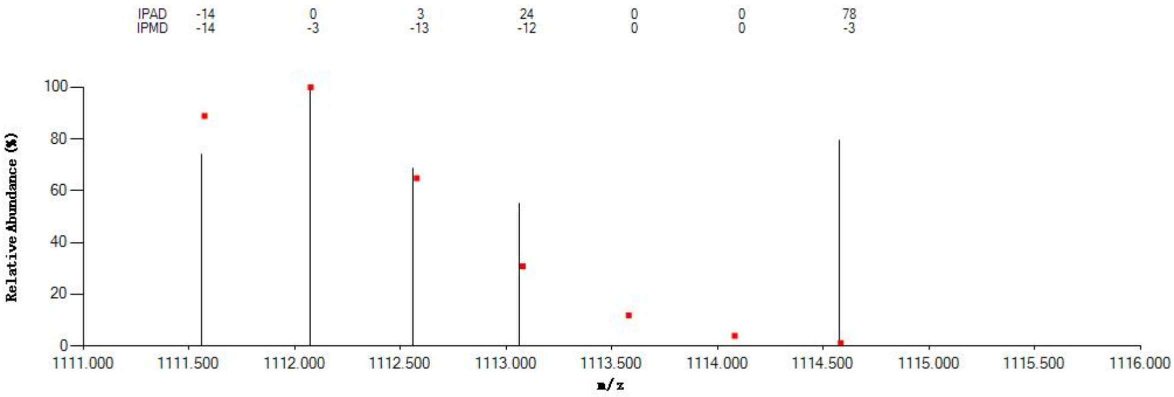


#### Annotated MS/MS Spectrum

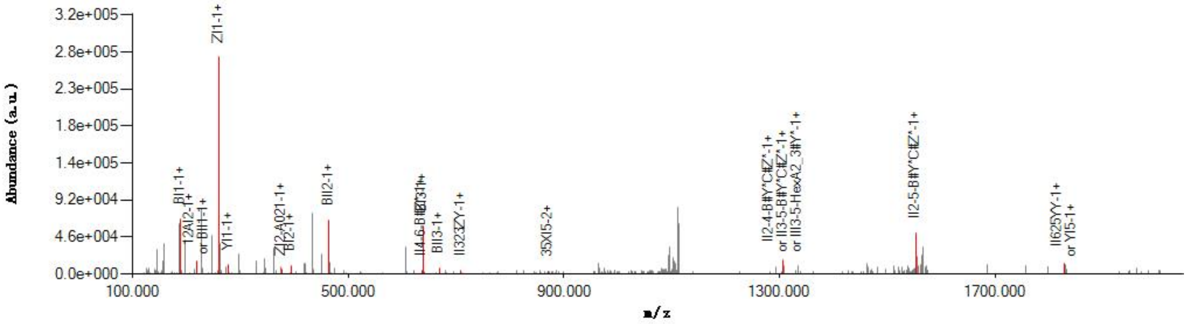


#### Graphical Fragmentation Map

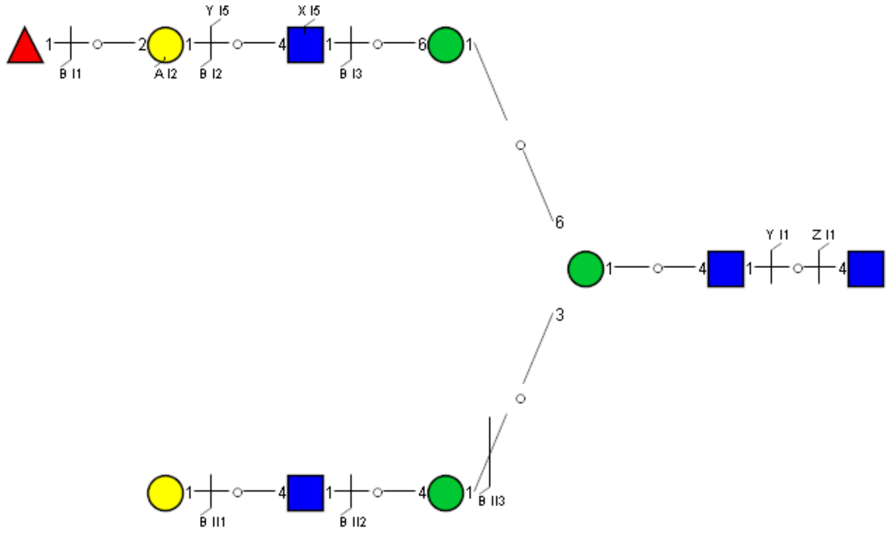

Spectrum Index: 38931(NoBHs:1)

#### Precursor iEF Map

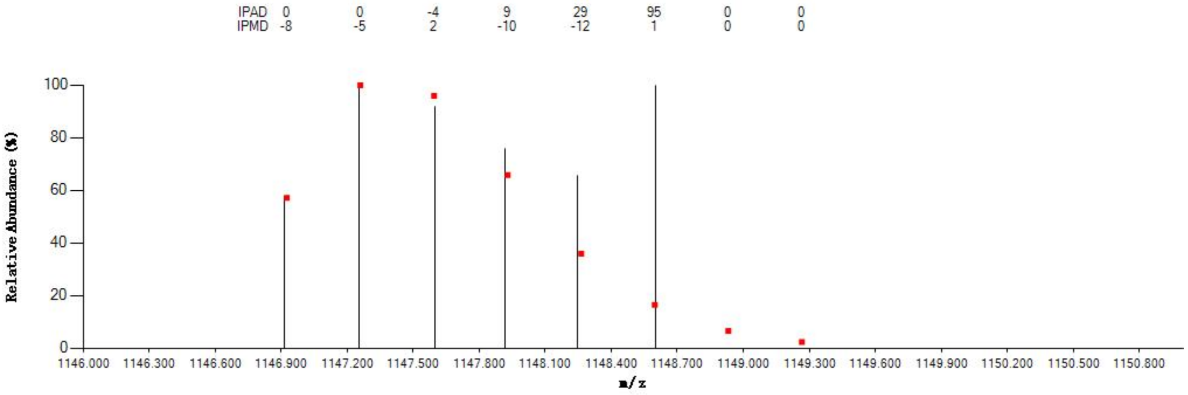

#### Annotated MS/MS Spectrum

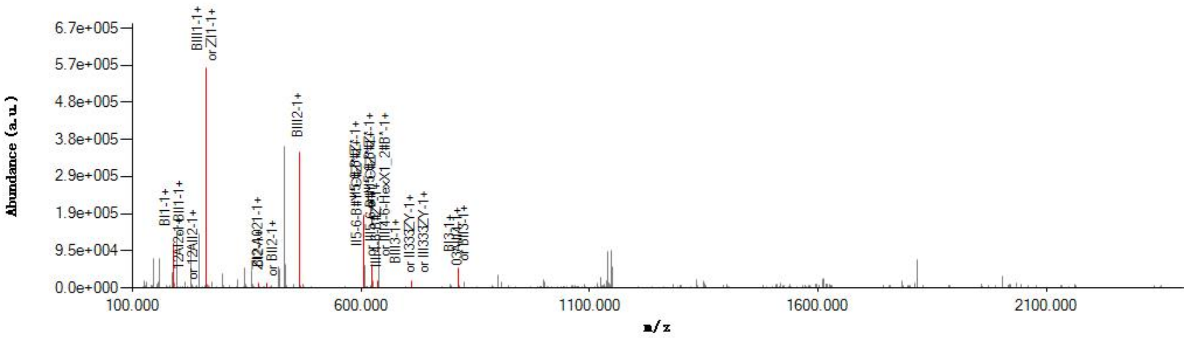

#### Graphical Fragmentation Map

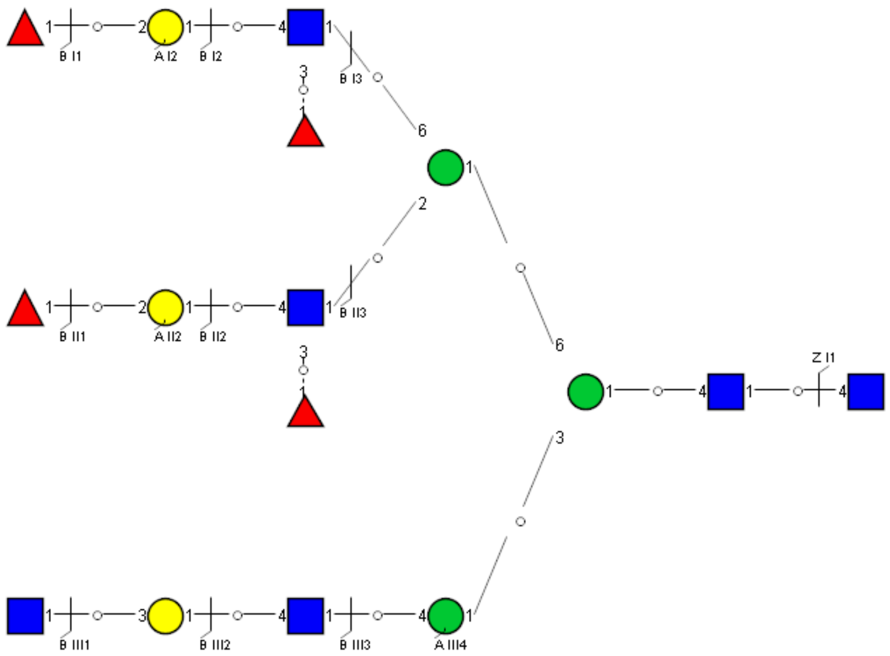

Spectrum Index: 38983(NoBHs:1)

#### Precursor iEF Map

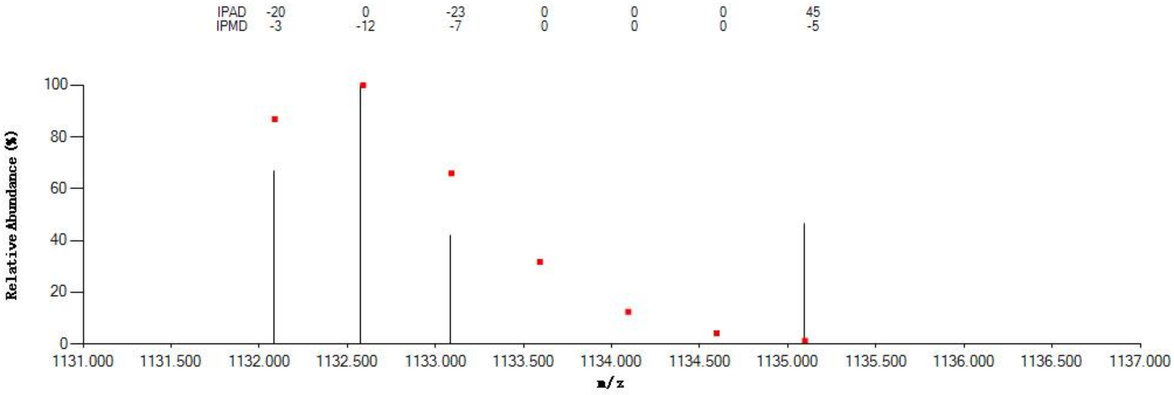


#### Annotated MS/MS Spectrum

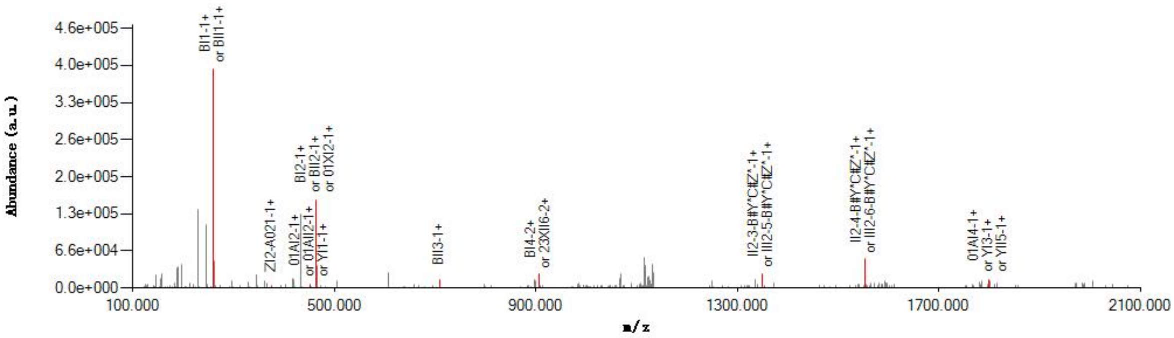


#### Graphical Fragmentation Map

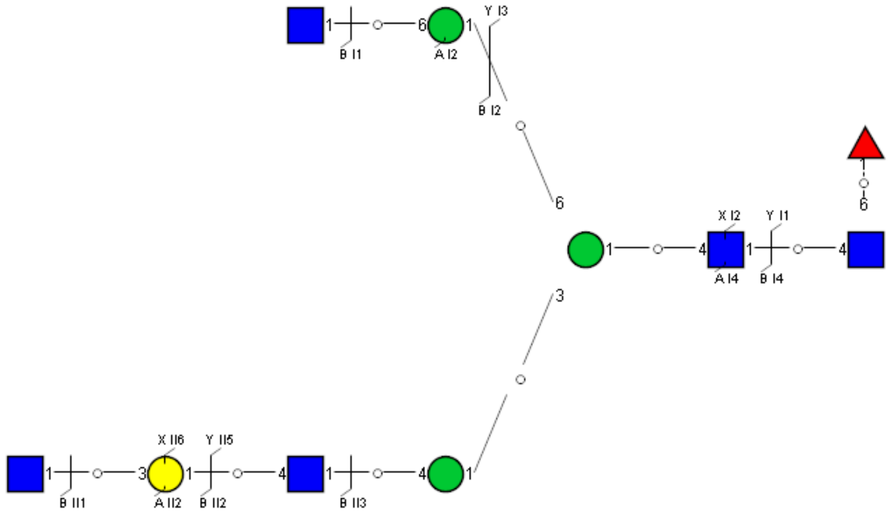

Spectrum Index: 38997(NoBHs:1)

#### Precursor iEF Map

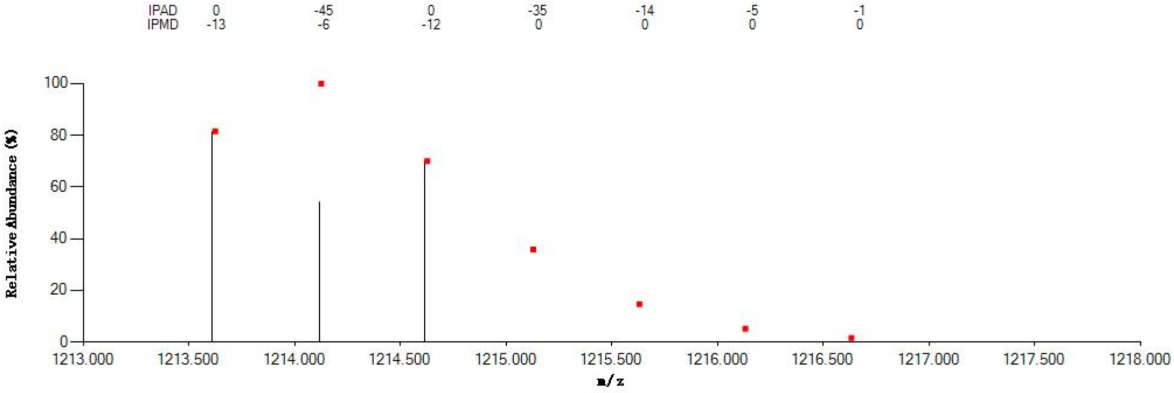


#### Annotated MS/MS Spectrum

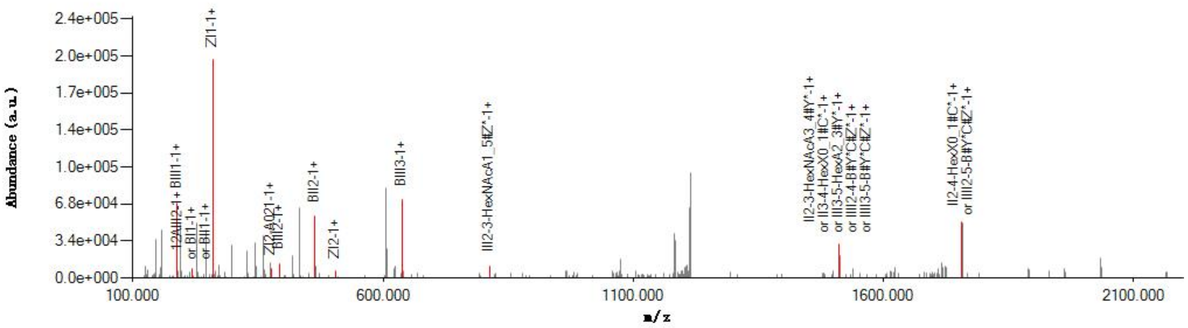


#### Graphical Fragmentation Map

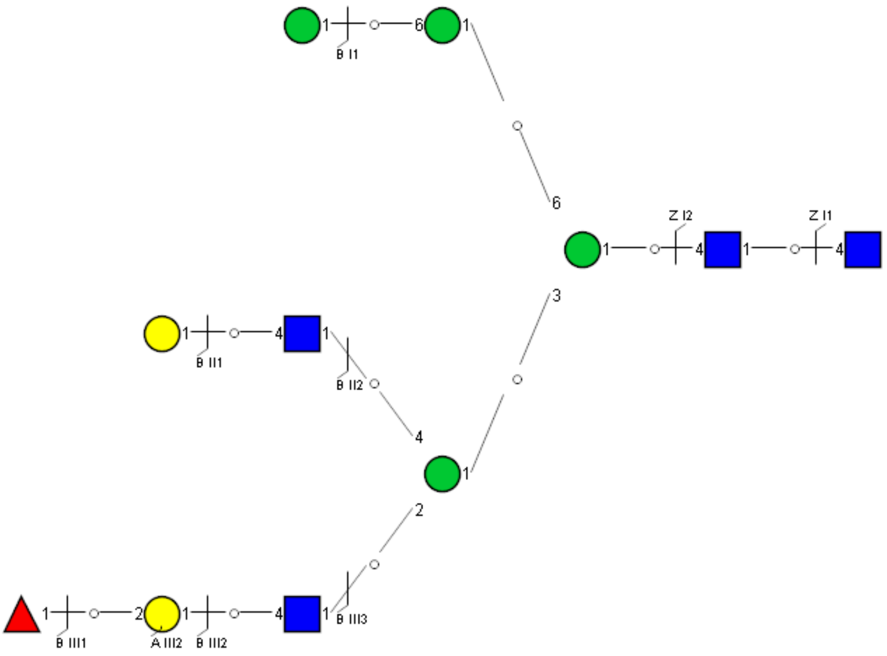

Spectrum Index: 39172(NoBHs:1)

#### Precursor iEF Map

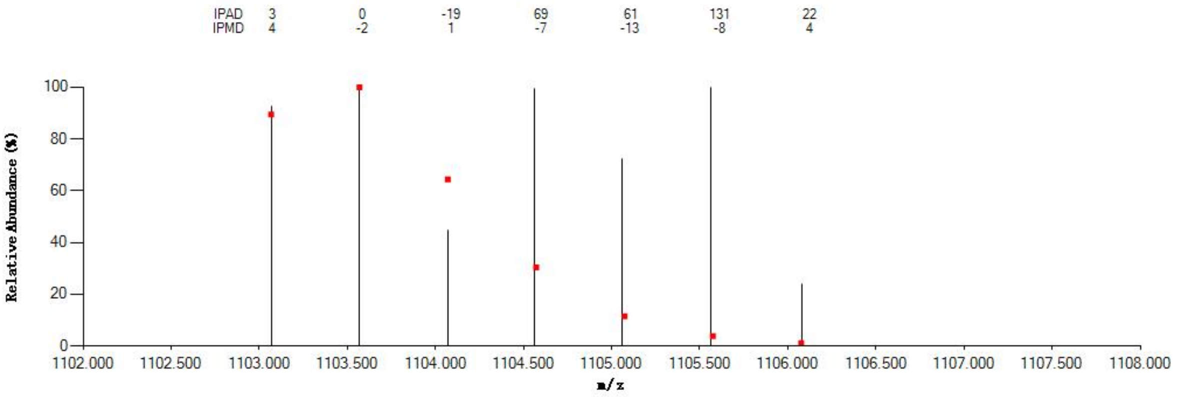


#### Annotated MS/MS Spectrum

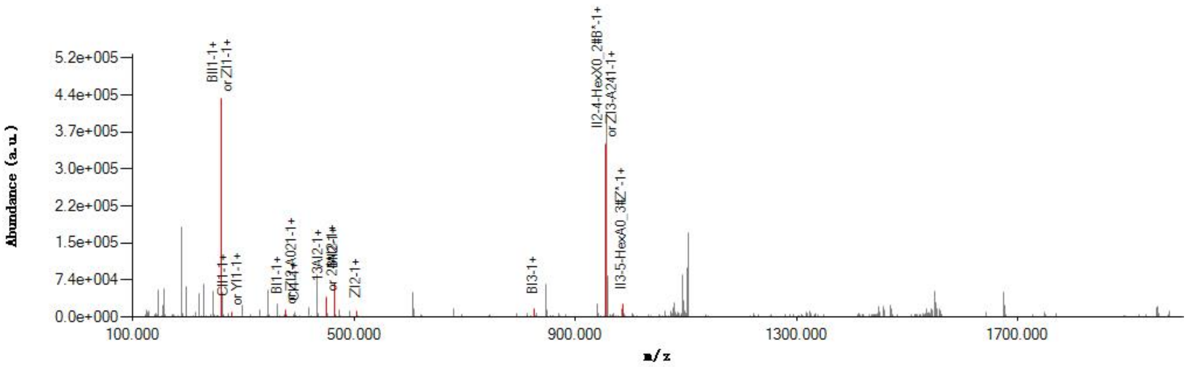


#### Graphical Fragmentation Map

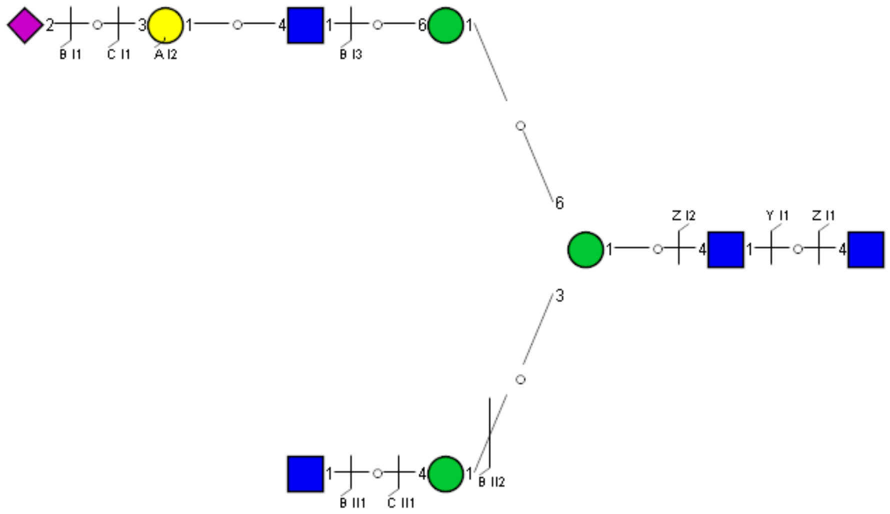

Spectrum Index: 39230(NoBHs:1)

#### Precursor iEF Map

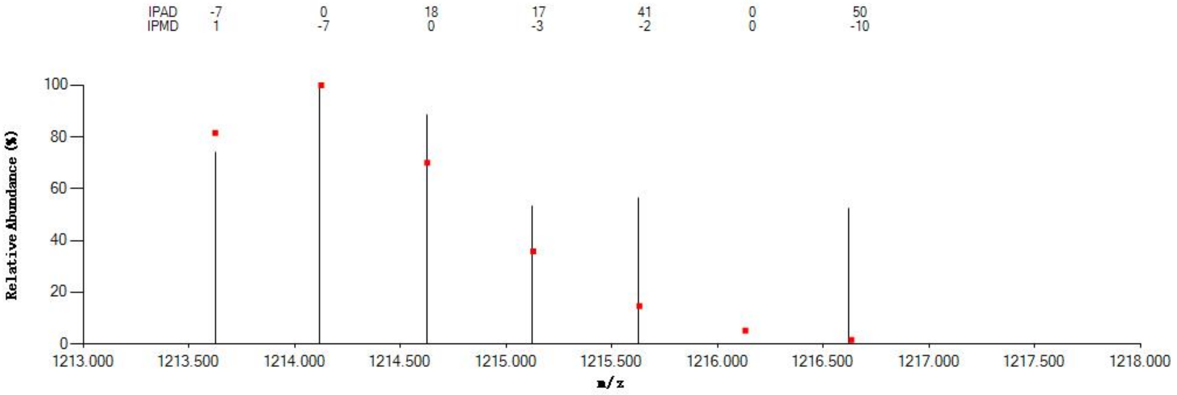


#### Annotated MS/MS Spectrum

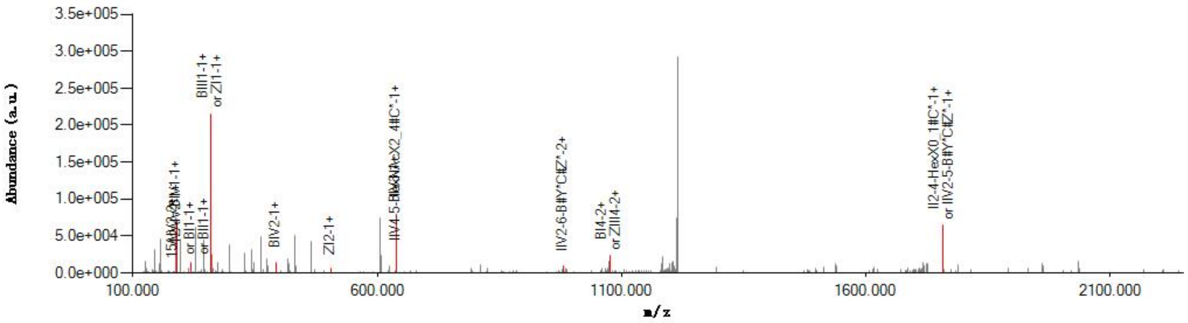


#### Graphical Fragmentation Map

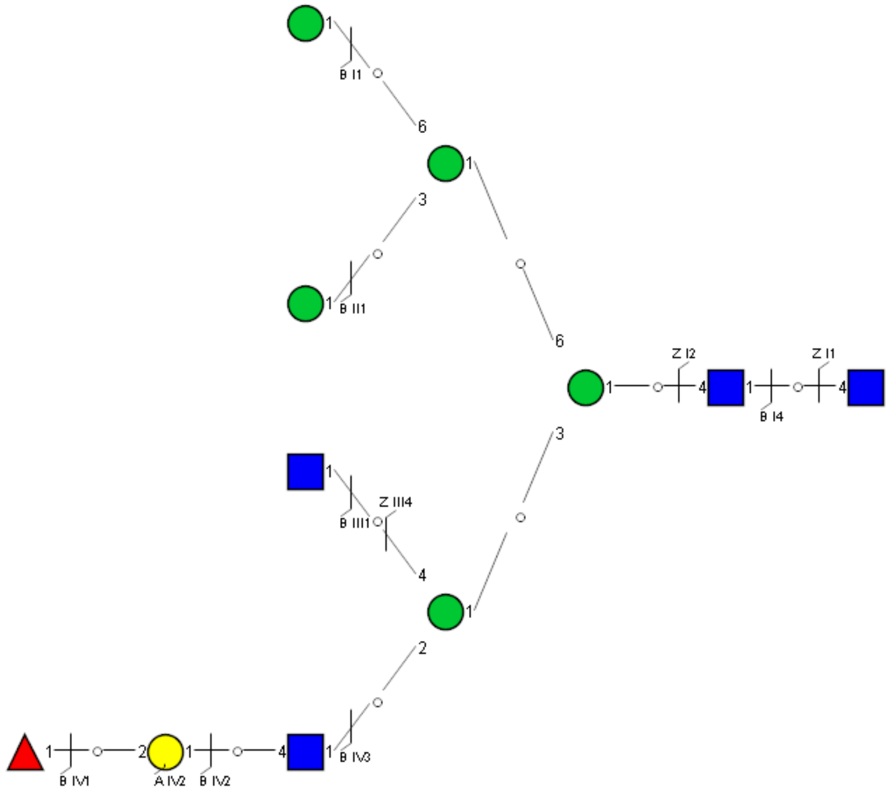

### Spectrum Index: 39298(NoBHs:1)

#### Precursor iEF Map

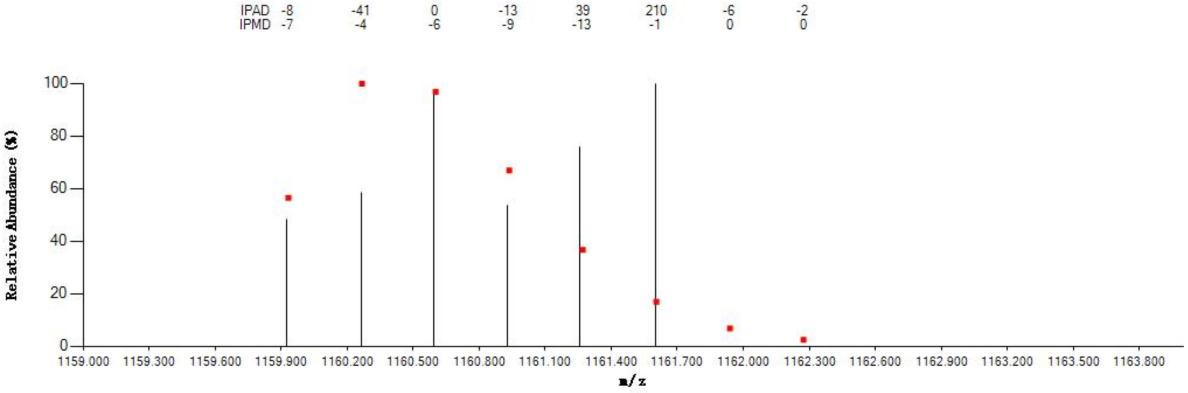


#### Annotated MS/MS Spectrum

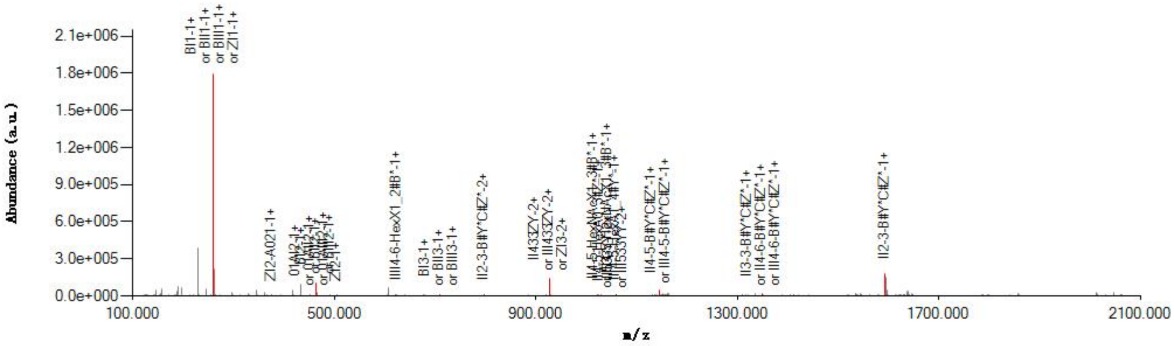


#### Graphical Fragmentation Map

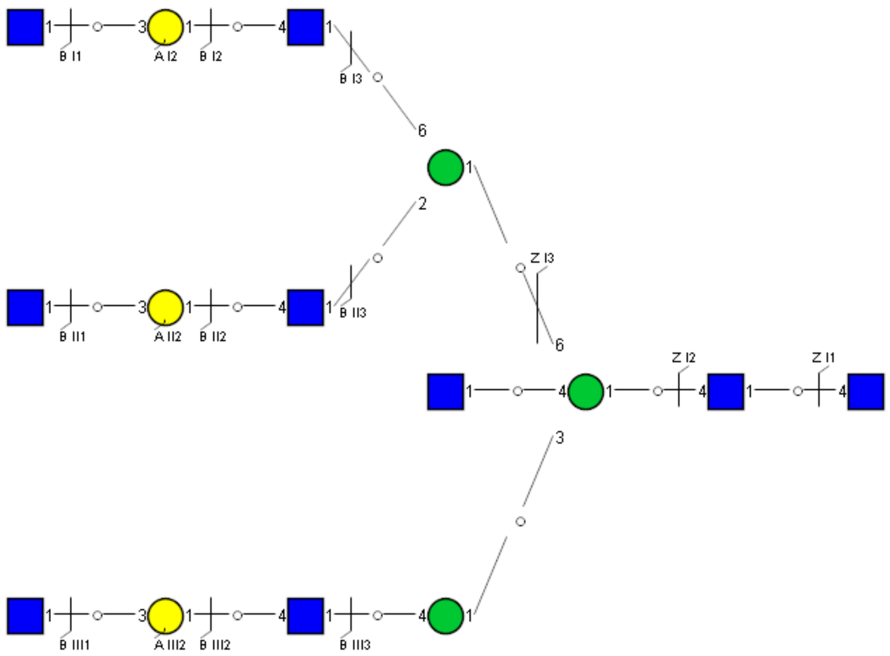

Spectrum Index: 39316(NoBHs:1)

#### Precursor iEF Map

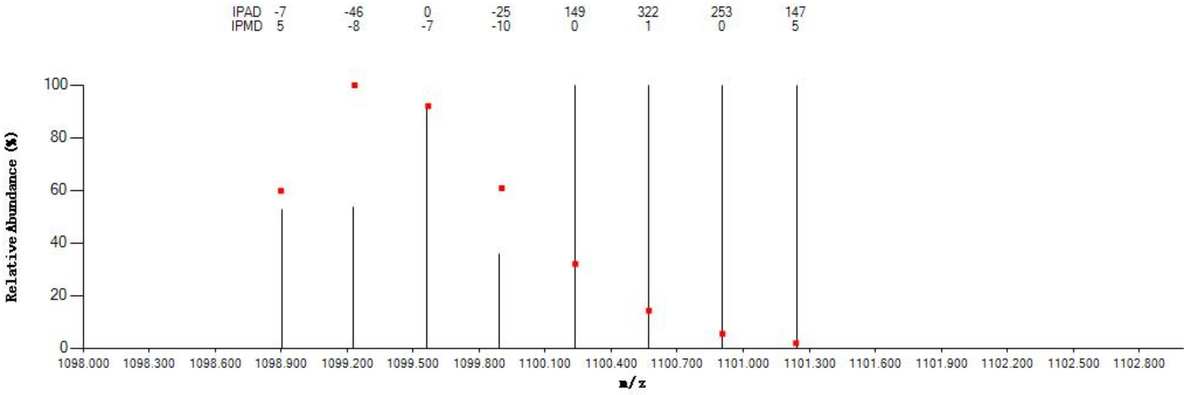


#### Annotated MS/MS Spectrum

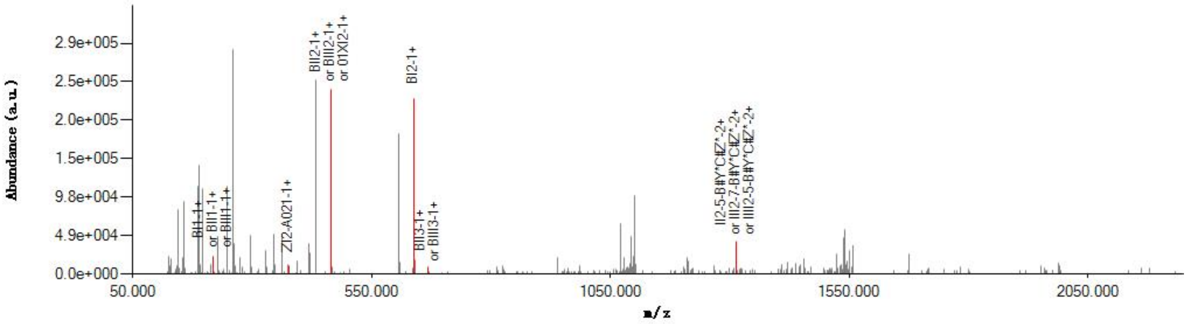


#### Graphical Fragmentation Map

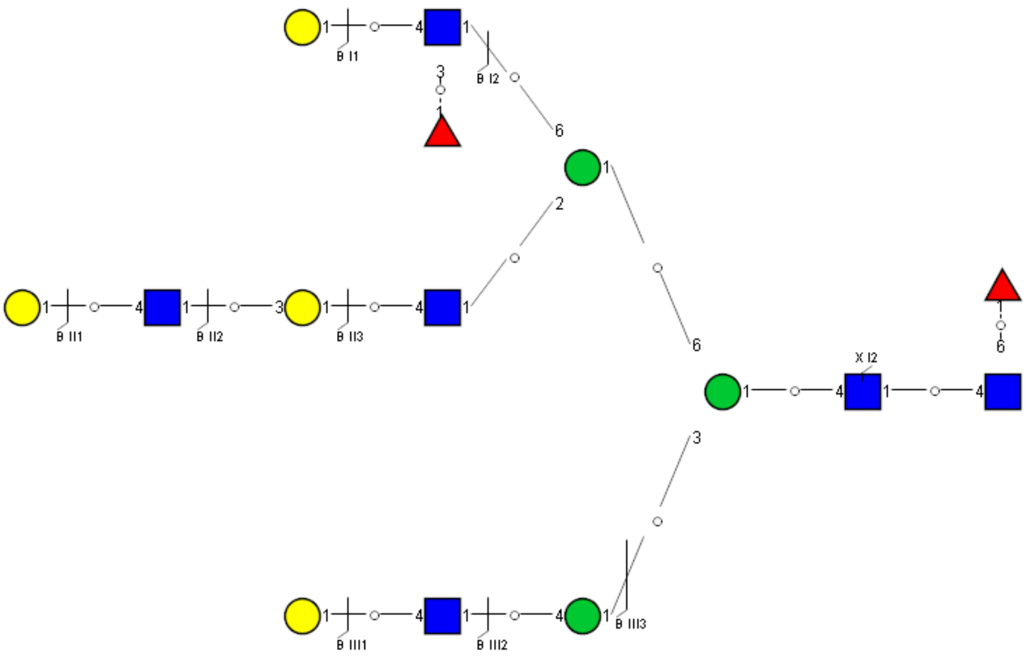

Spectrum Index: 39317(NoBHs:3)

#### Precursor iEF Map

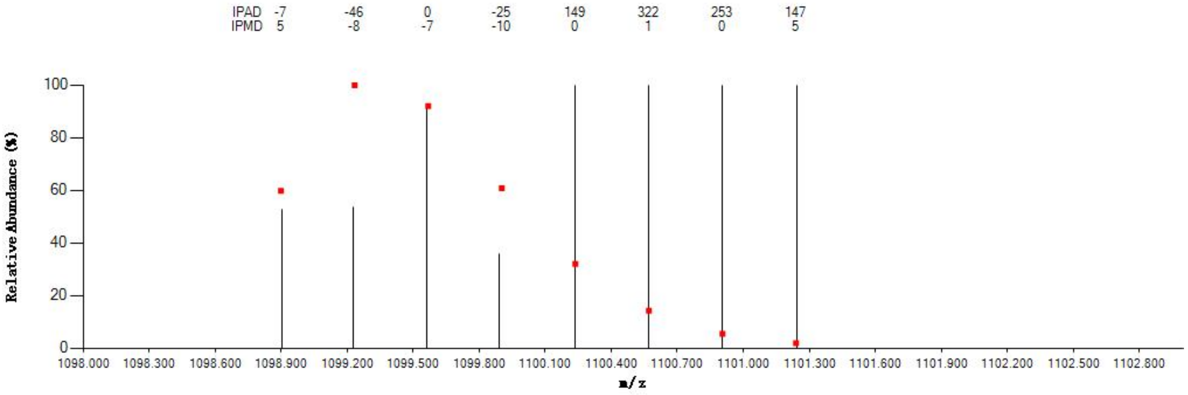


#### Annotated MS/MS Spectrum

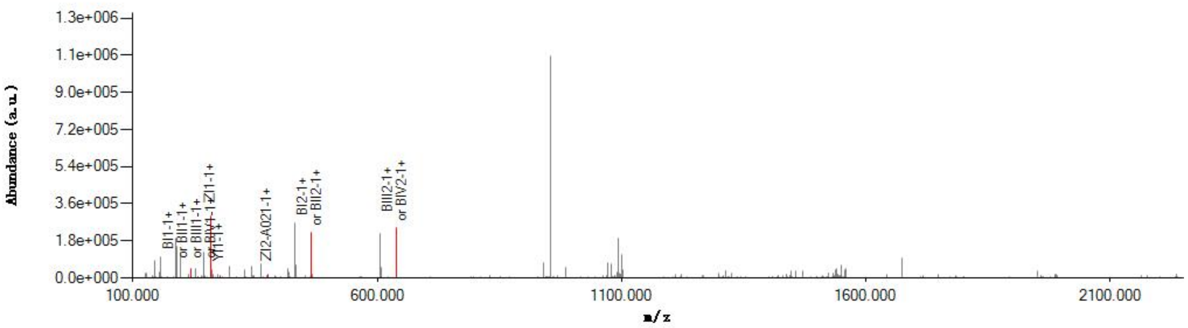


#### Graphical Fragmentation Map

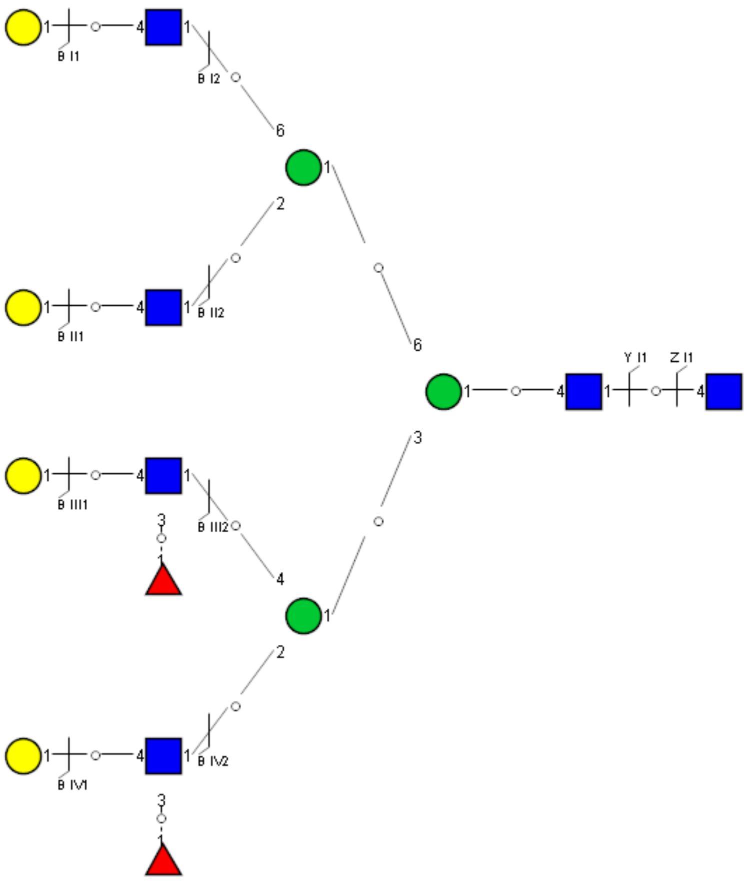

Spectrum Index: 39317(NoBHs:3)

#### Precursor iEF Map

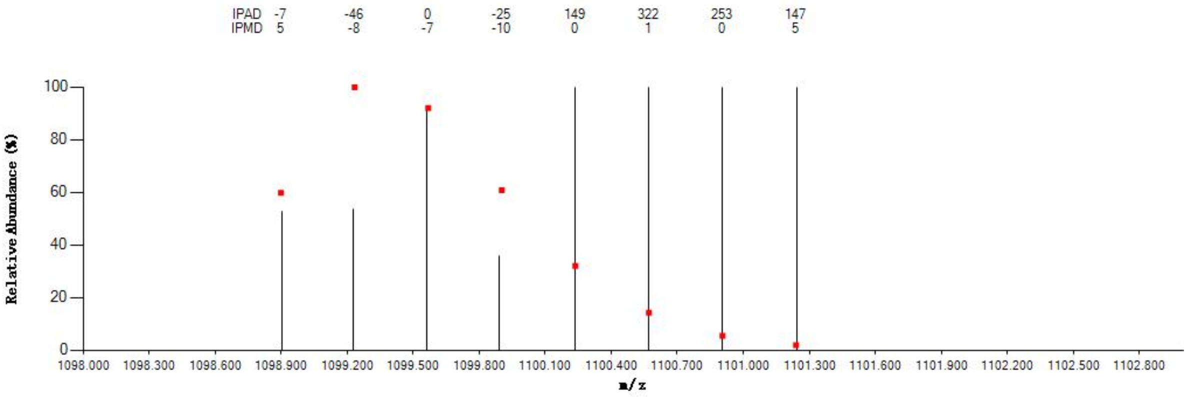


#### Annotated MS/MS Spectrum

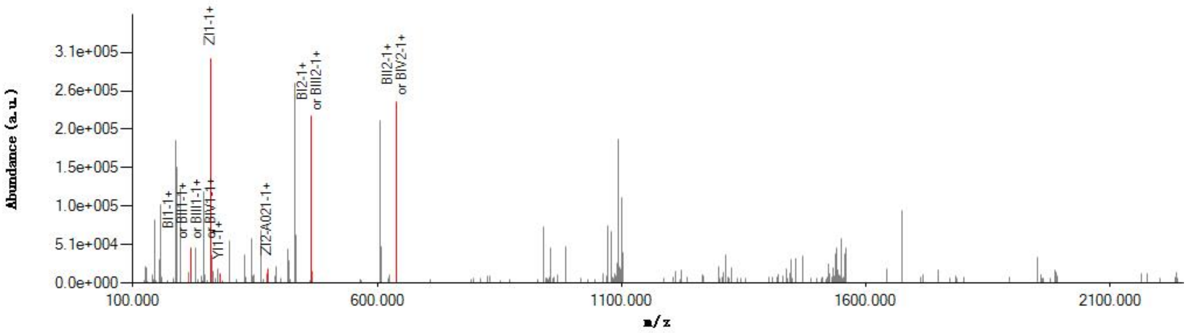


#### Graphical Fragmentation Map

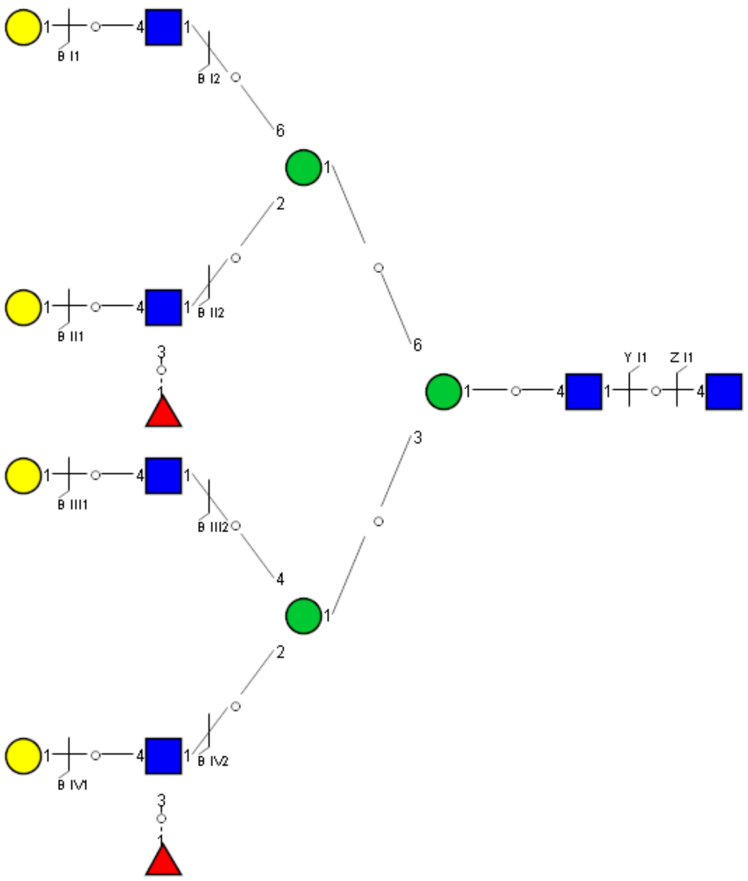

Spectrum Index: 39322(NoBHs:1)

Precursor iEF Map

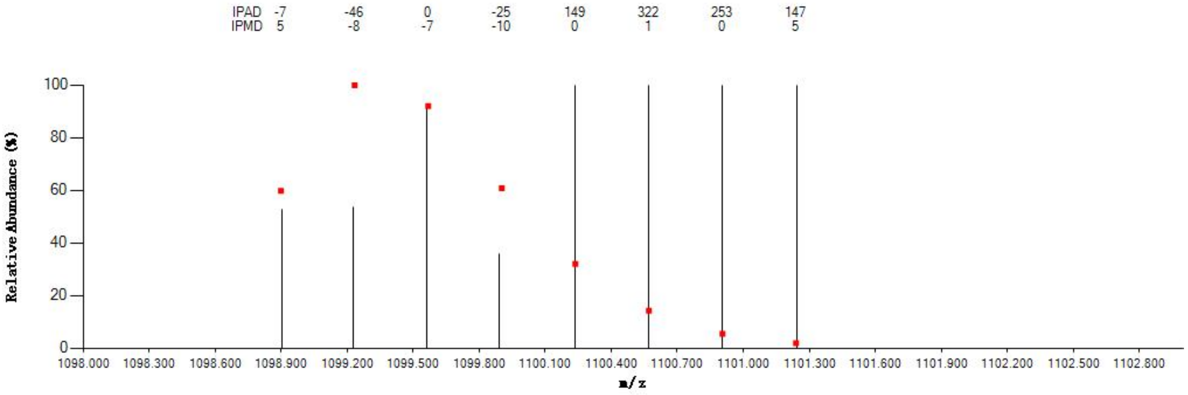

Annotated MS/MS Spectrum

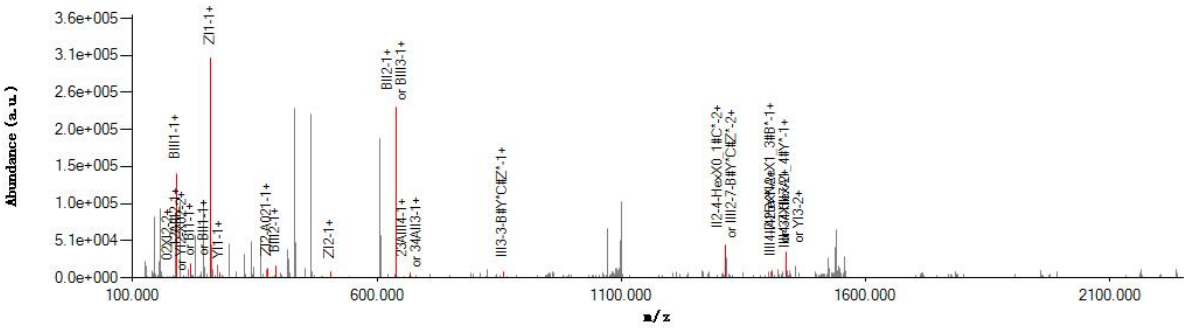

Graphical Fragmentation Map

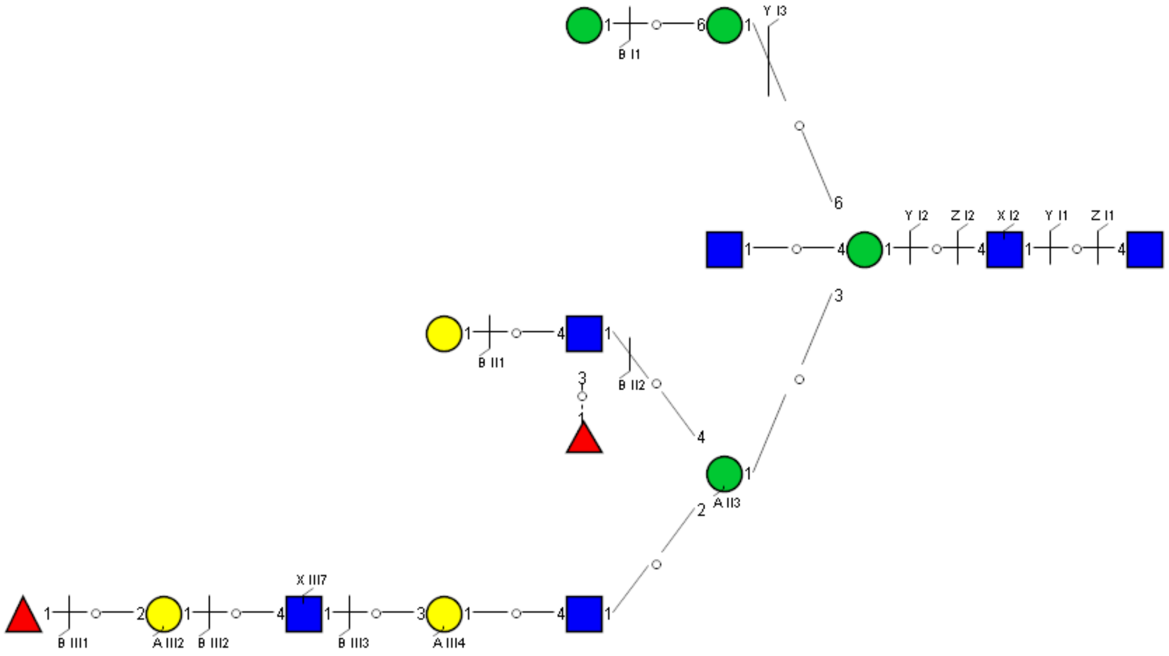

Spectrum Index: 39332(NoBHs:1)

#### Precursor iEF Map

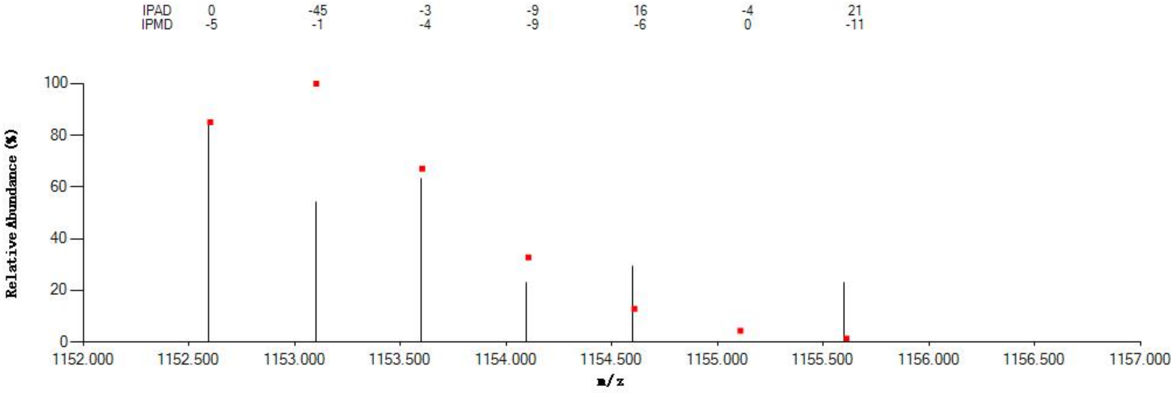


#### Annotated MS/MS Spectrum

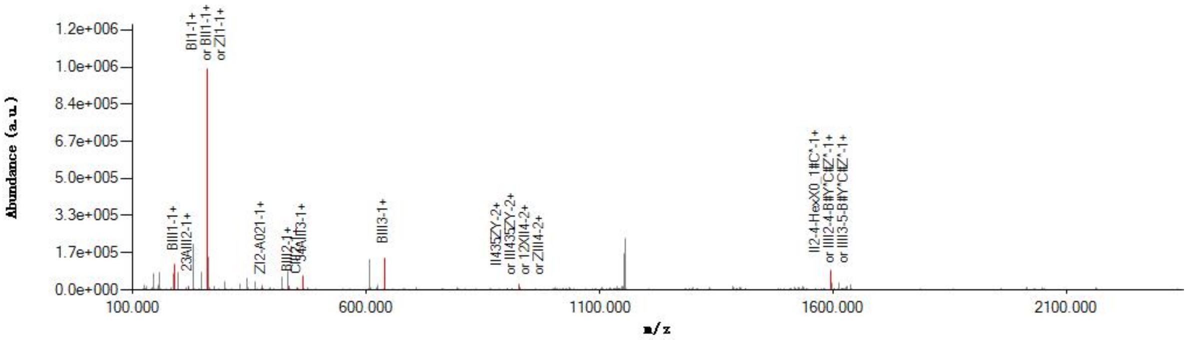


#### Graphical Fragmentation Map

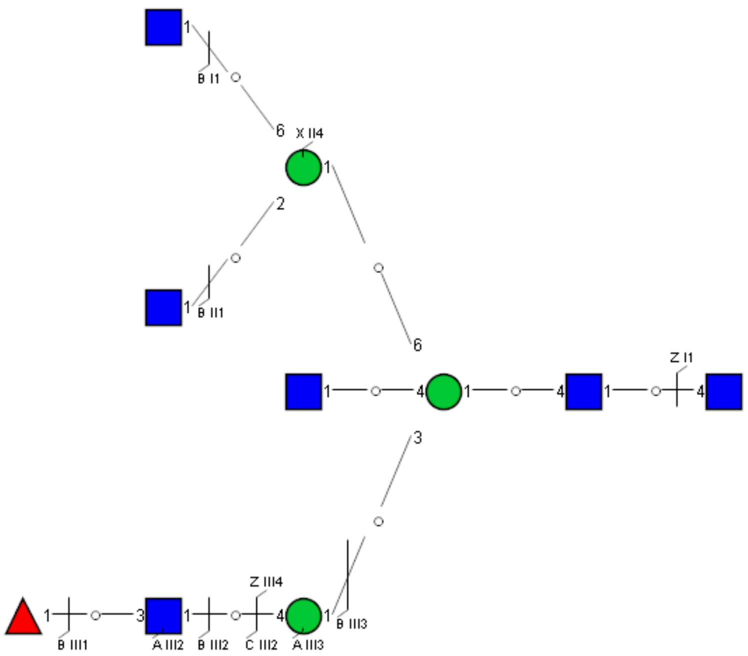

Spectrum Index: 39358(NoBHs:1)

#### Precursor iEF Map

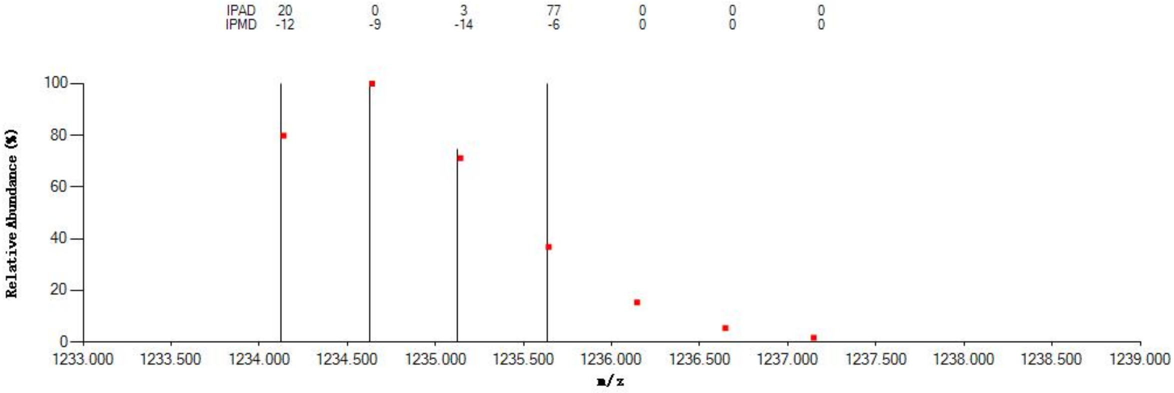


#### Annotated MS/MS Spectrum

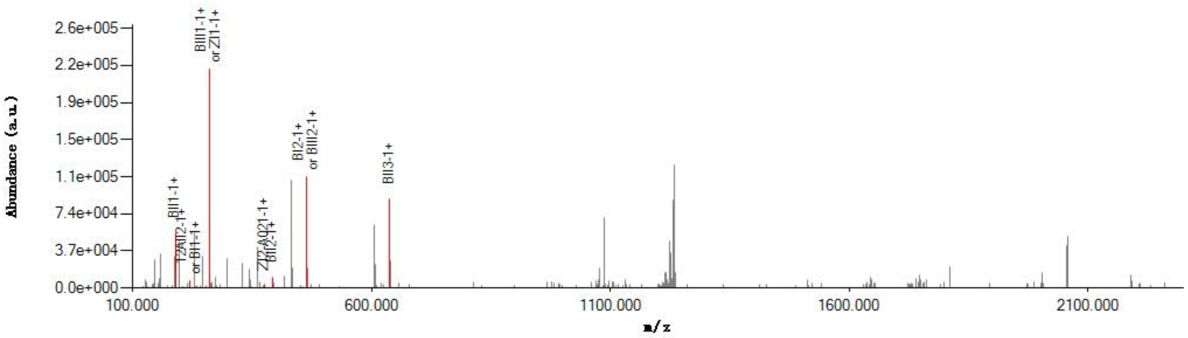


#### Graphical Fragmentation Map

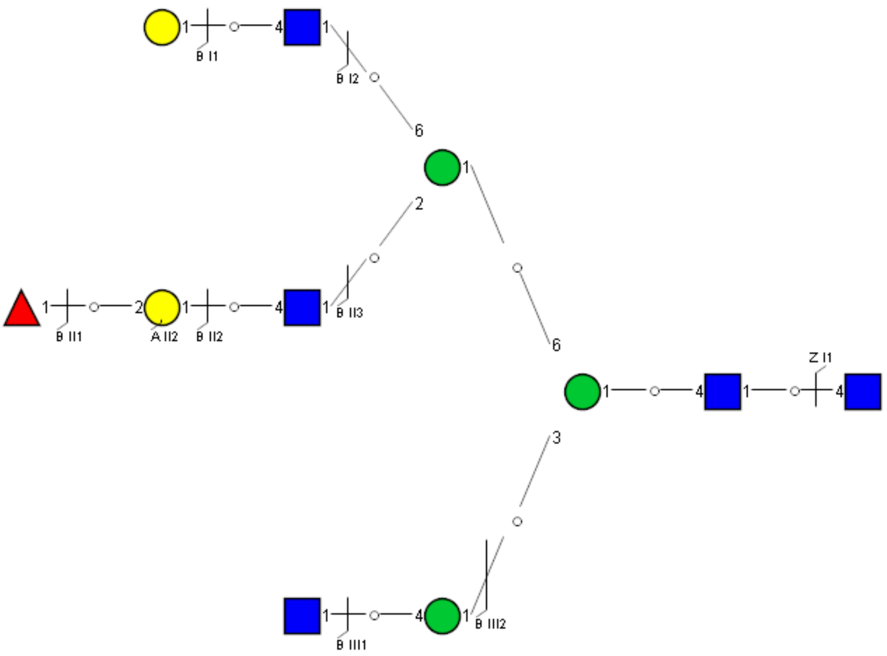

Spectrum Index: 39441(NoBHs:1)

#### Precursor iEF Map

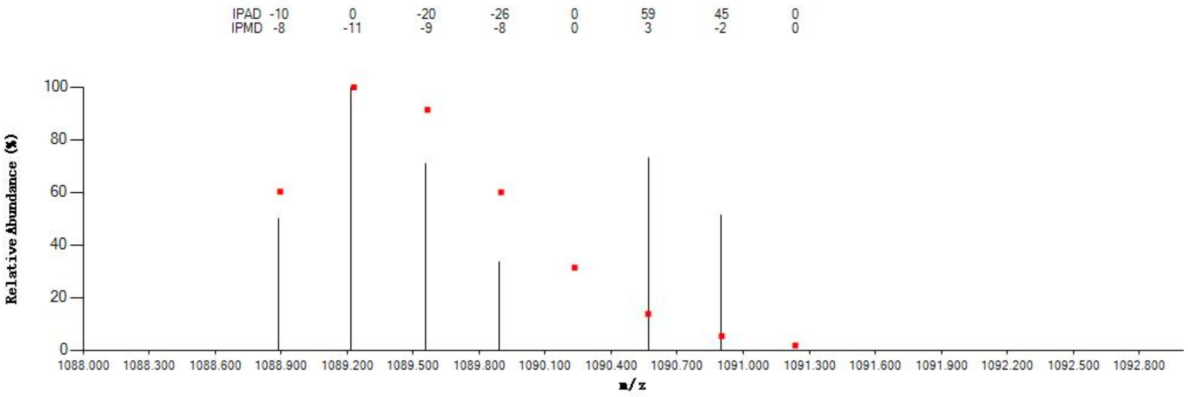


#### Annotated MS/MS Spectrum

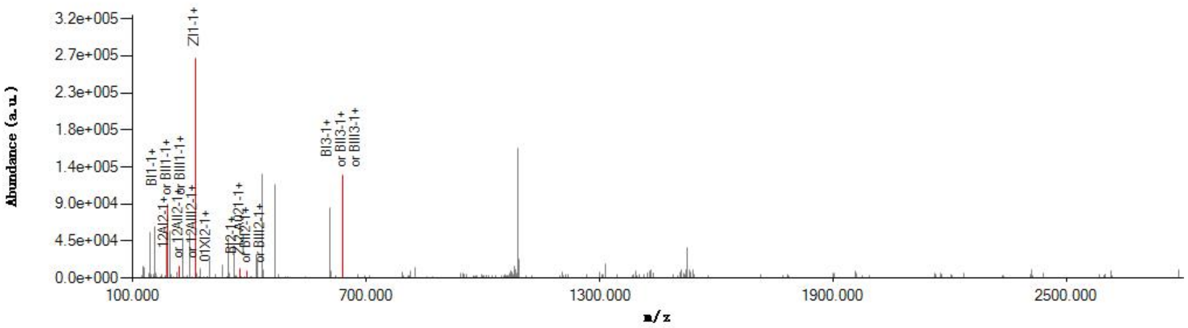


#### Graphical Fragmentation Map

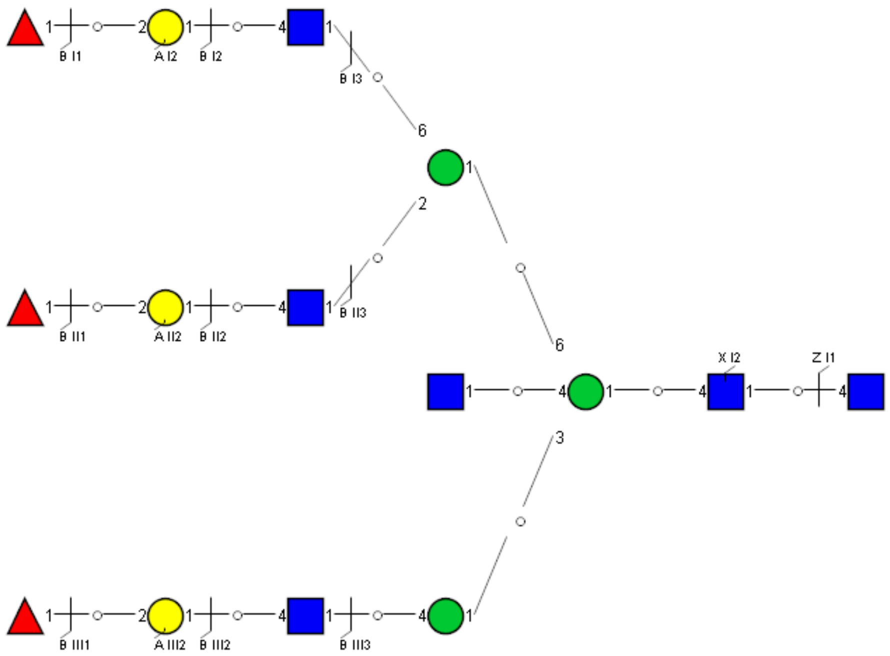

Spectrum Index: 39779(NoBHs:1)

#### Precursor iEF Map

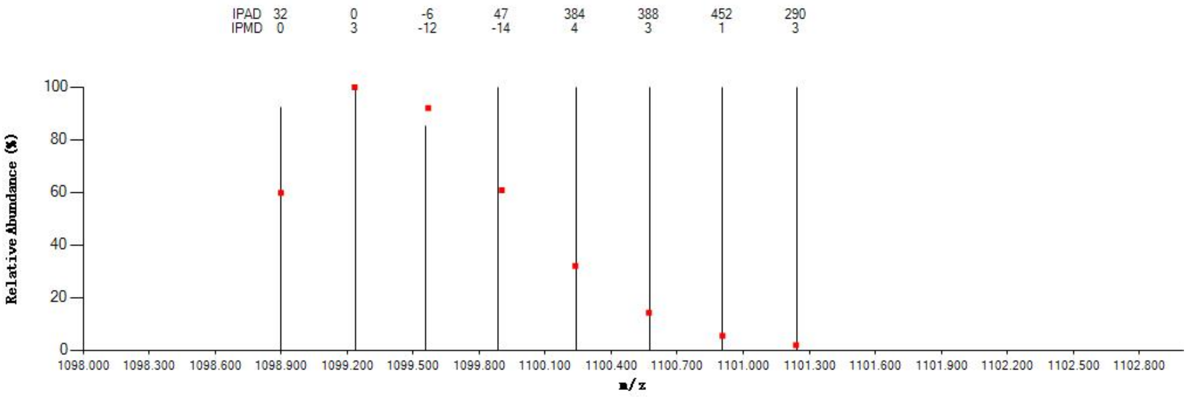


#### Annotated MS/MS Spectrum

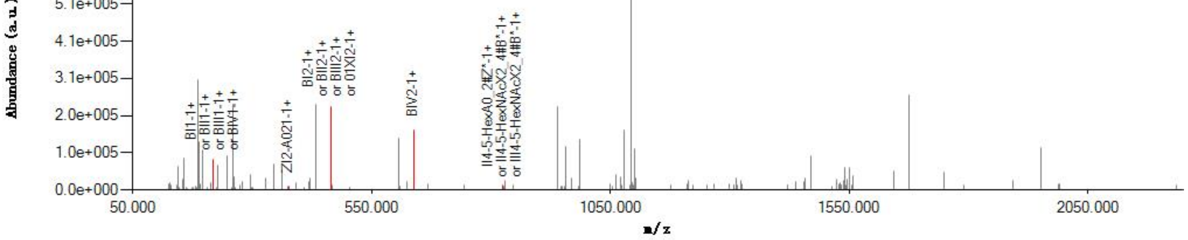


#### Graphical Fragmentation Map

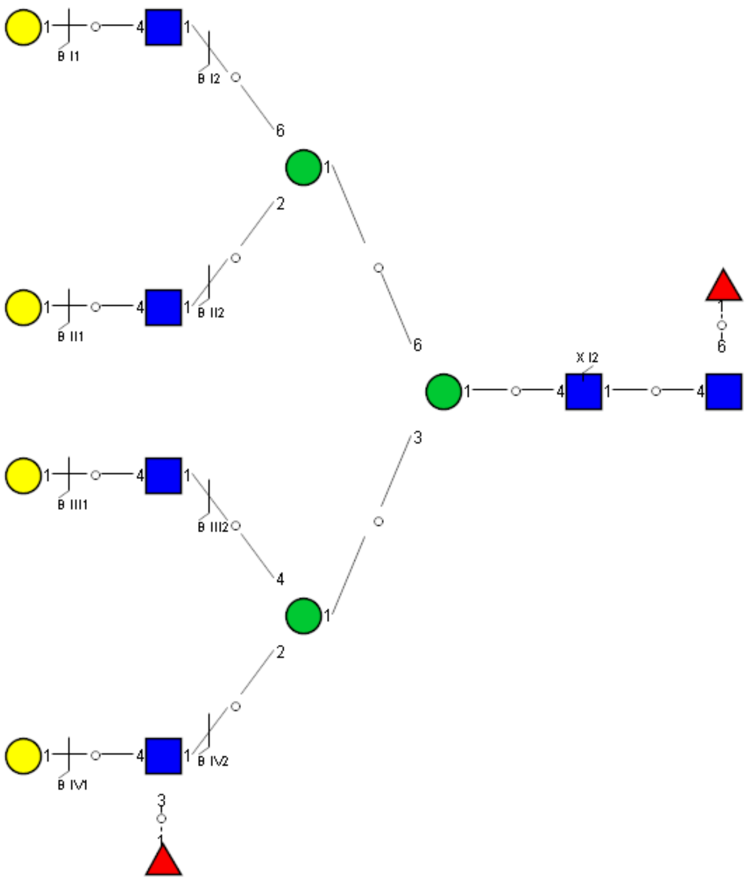

Spectrum Index: 39780(NoBHs:2)

#### Precursor iEF Map

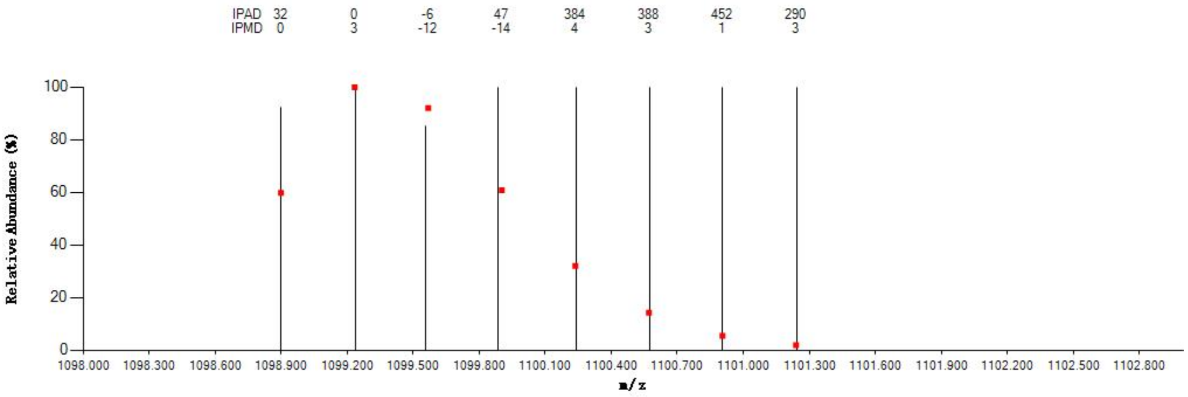

#### Annotated MS/MS Spectrum

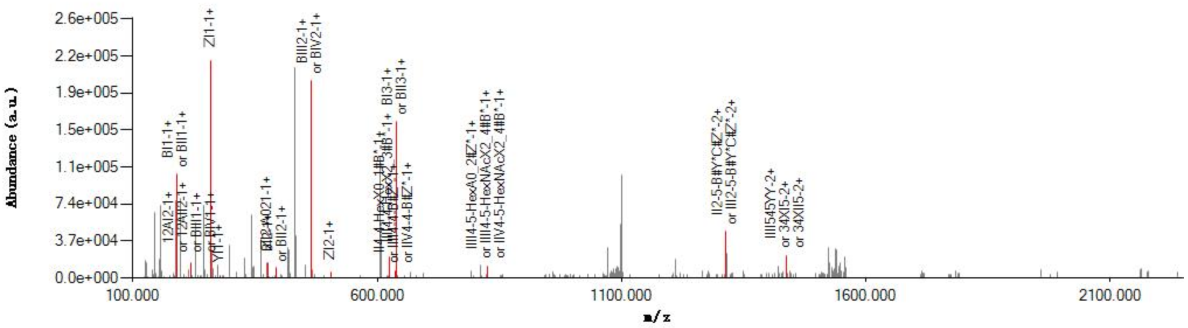

#### Graphical Fragmentation Map

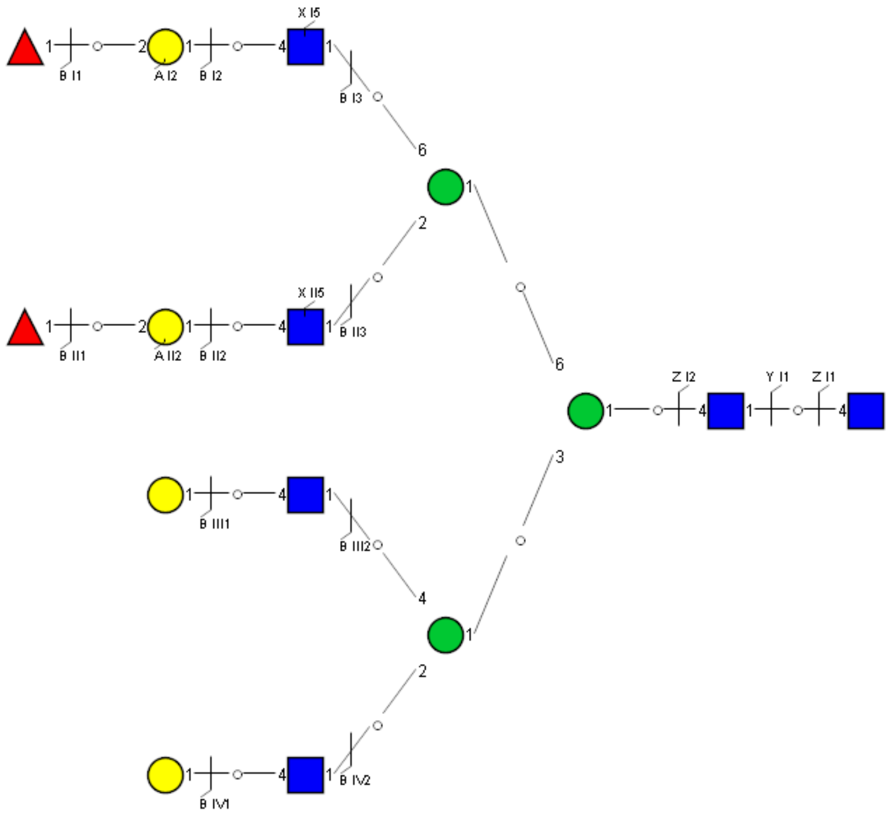

Spectrum Index: 39780(NoBHs:2)

#### Precursor iEF Map

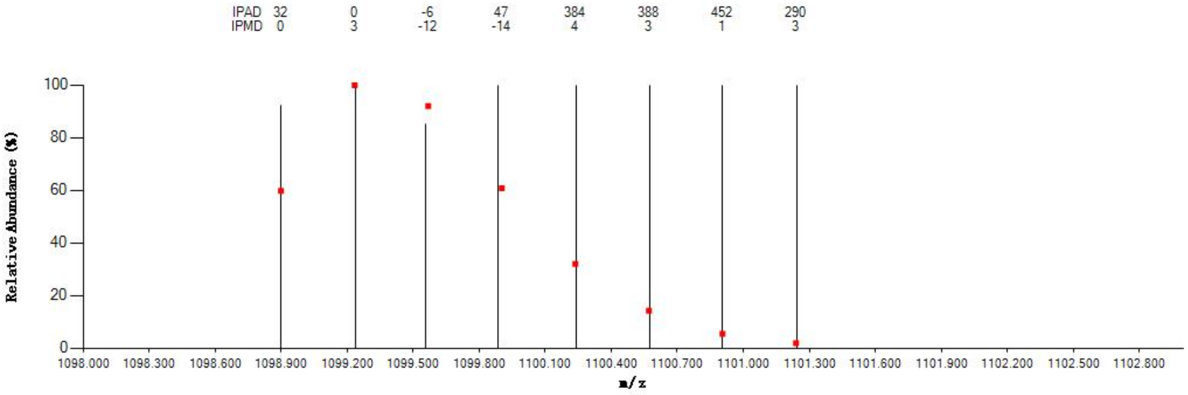

#### Annotated MS/MS Spectrum

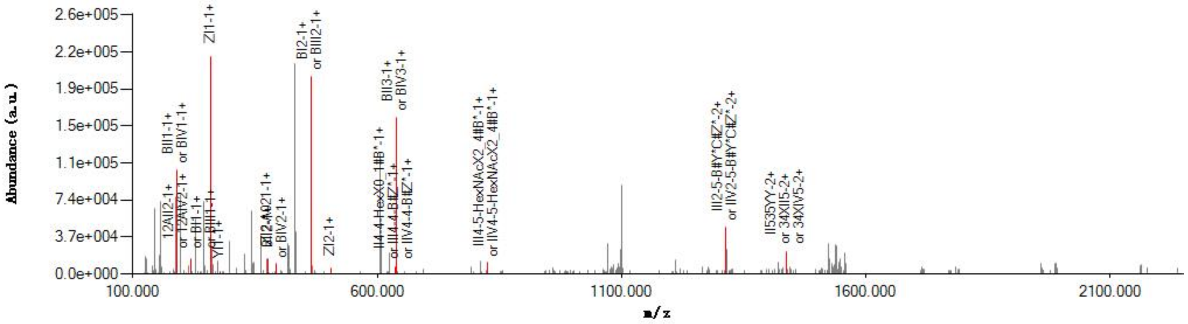

#### Graphical Fragmentation Map

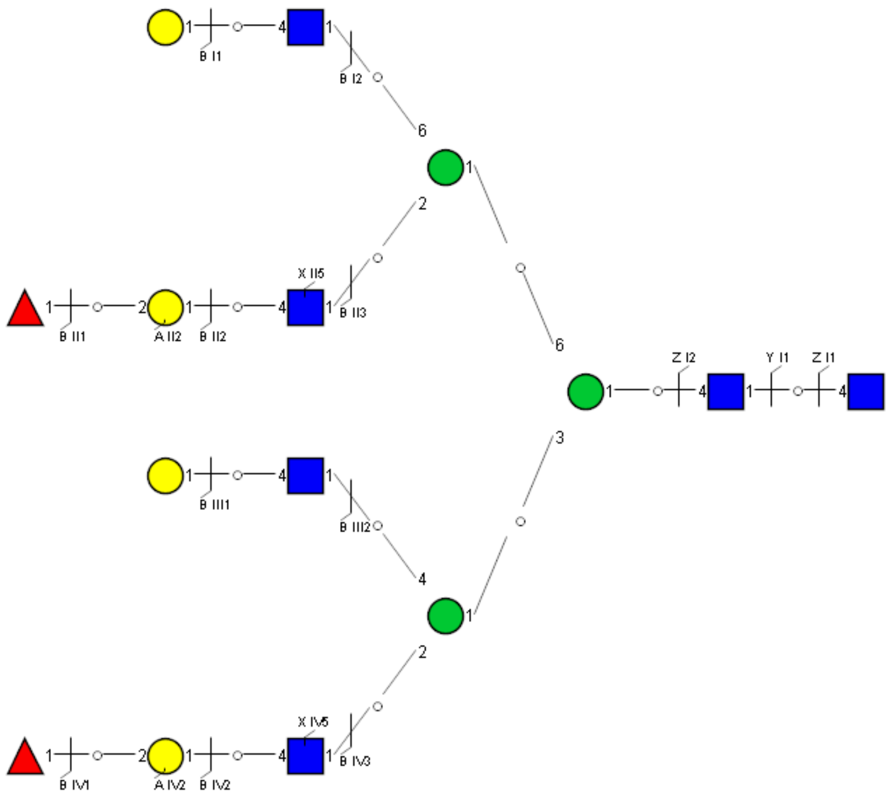

Spectrum Index: 39812(NoBHs:1)

#### Precursor iEF Map

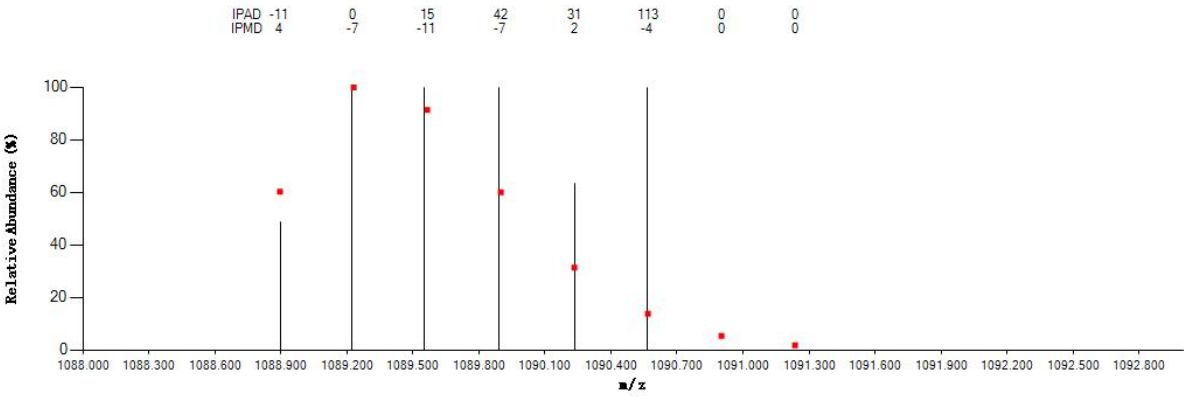


#### Annotated MS/MS Spectrum

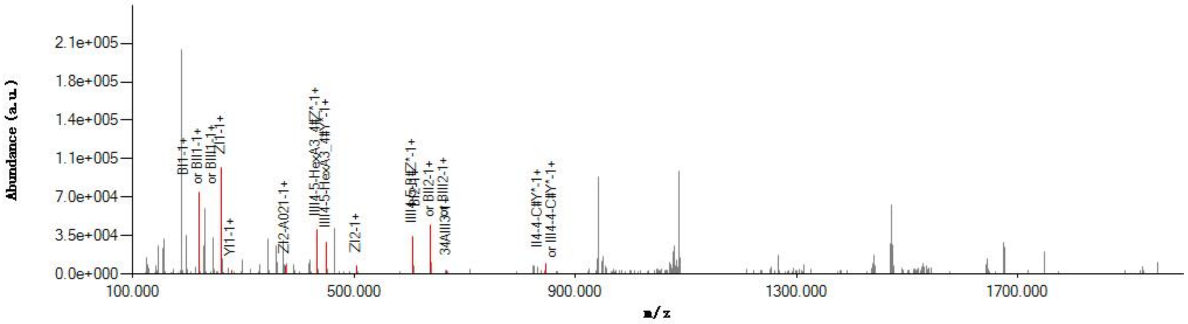


#### Graphical Fragmentation Map

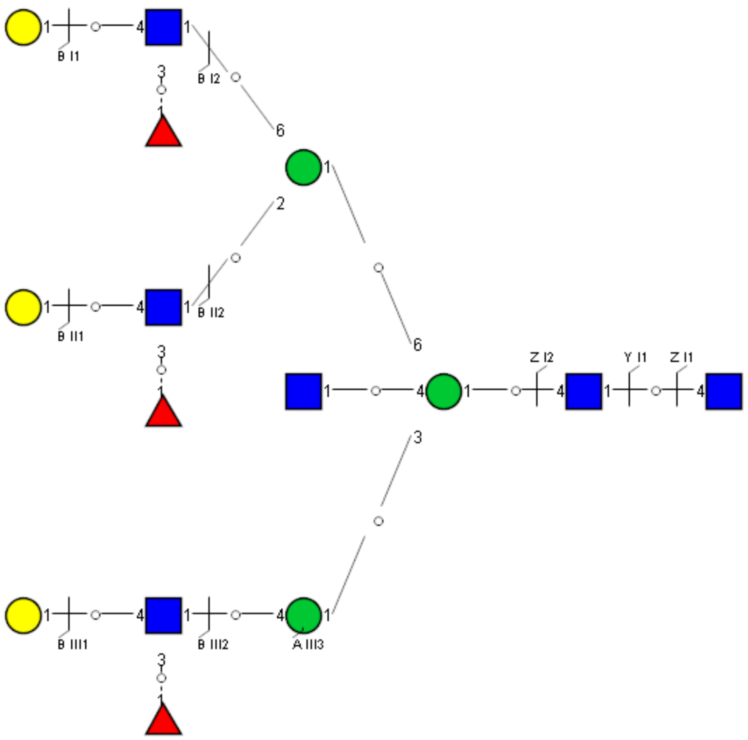

Spectrum Index: 39889(NoBHs:2)

#### Precursor iEF Map

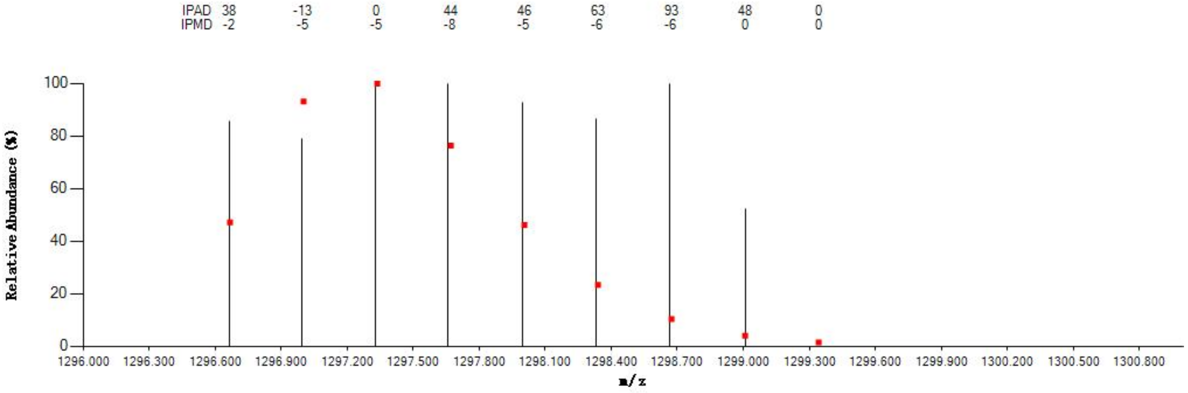

#### Annotated MS/MS Spectrum

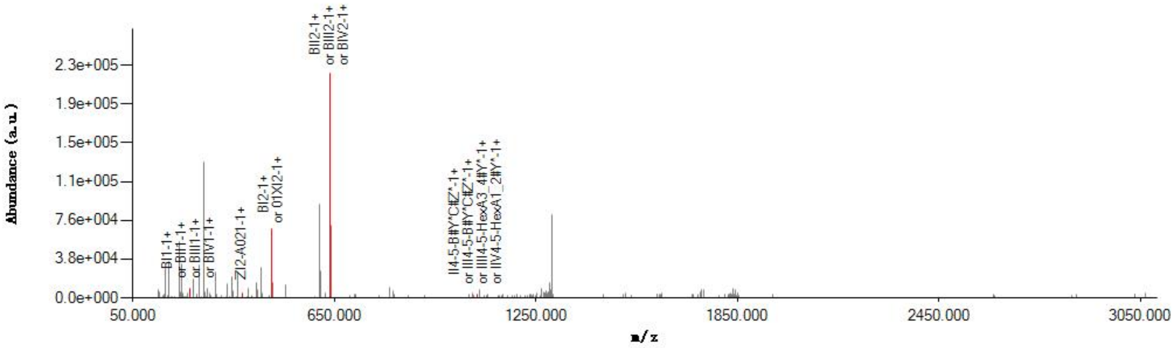

#### Graphical Fragmentation Map

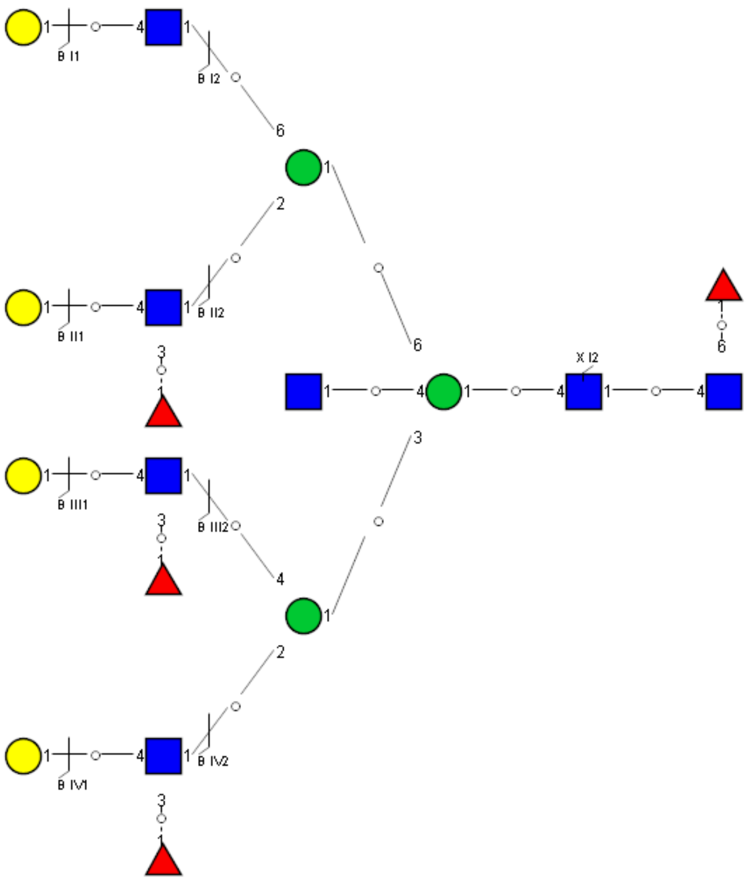

Spectrum Index: 39897(NoBHs:1)

#### Precursor iEF Map

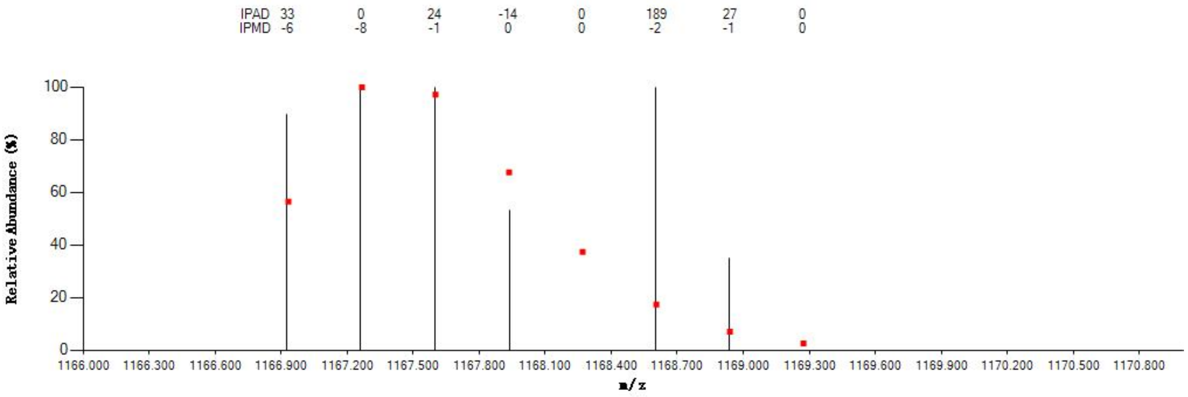


#### Annotated MS/MS Spectrum

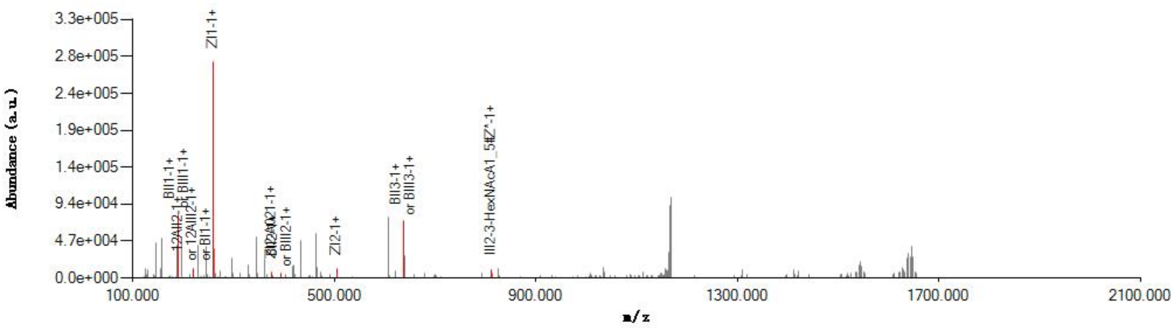


#### Graphical Fragmentation Map

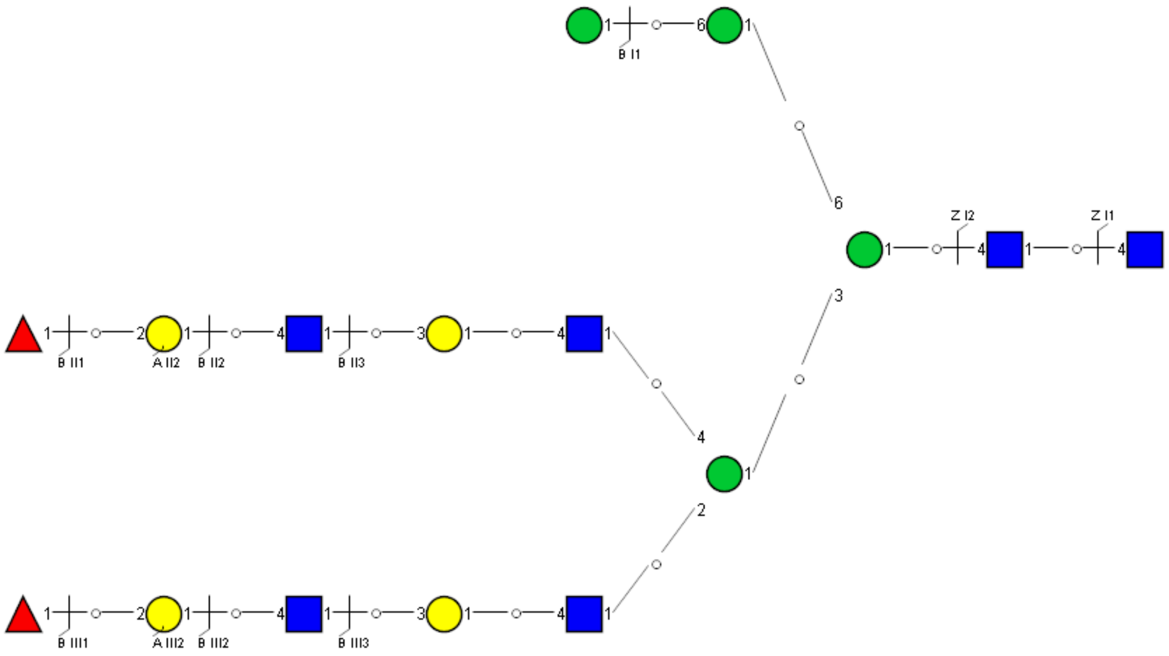

Spectrum Index: 39898(NoBHs:1)

#### Precursor iEF Map

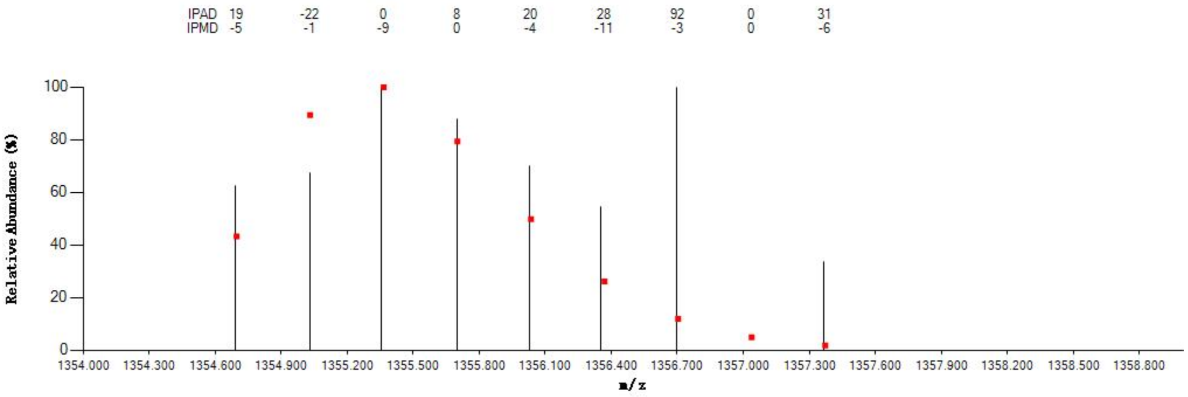


#### Annotated MS/MS Spectrum

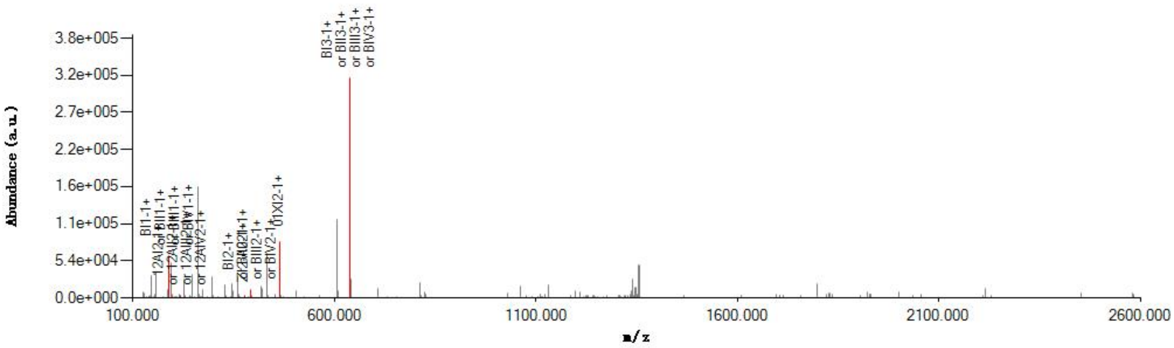


#### Graphical Fragmentation Map

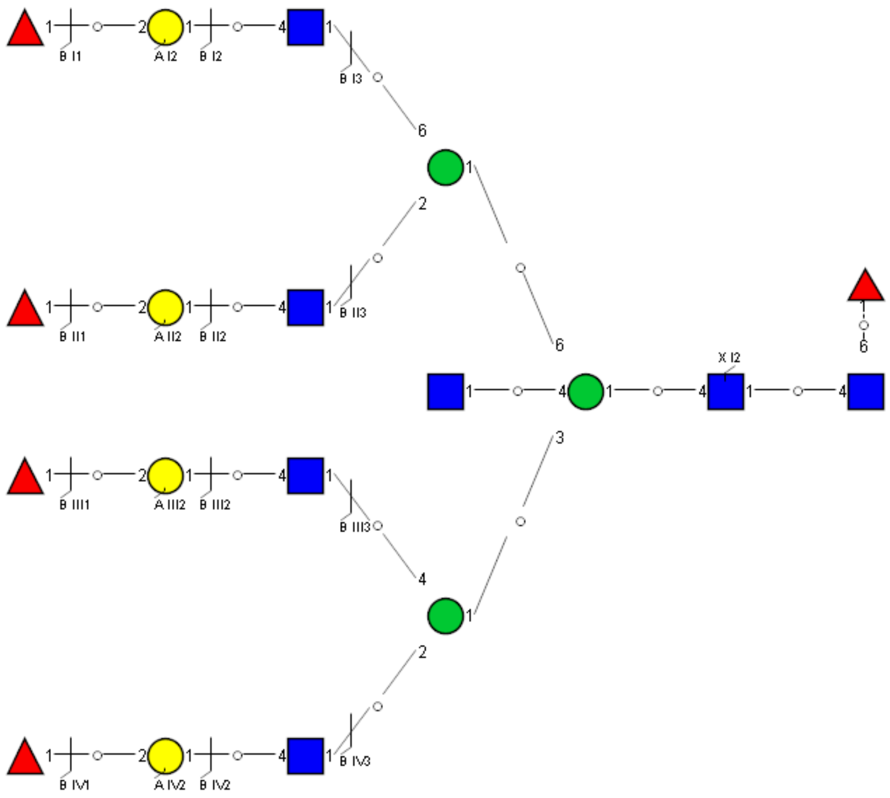

Spectrum Index: 39935(NoBHs:1)

#### Precursor iEF Map

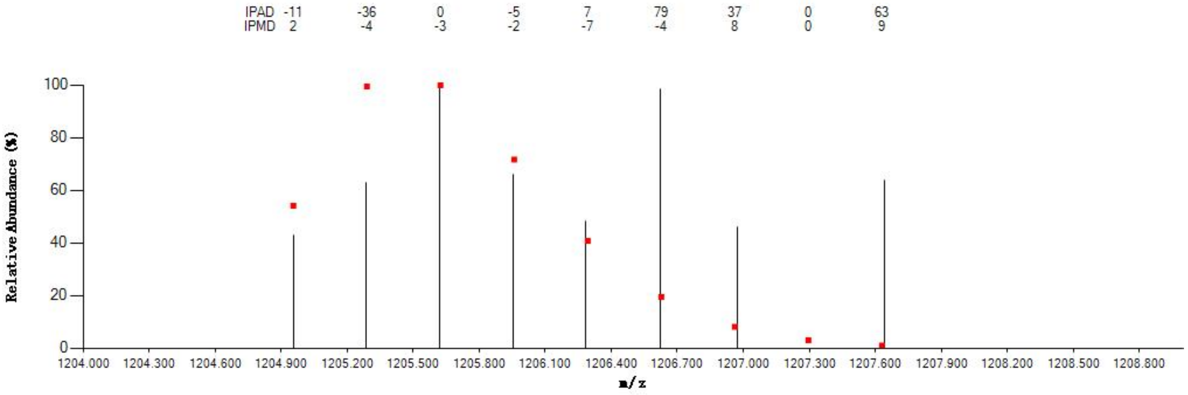

#### Annotated MS/MS Spectrum

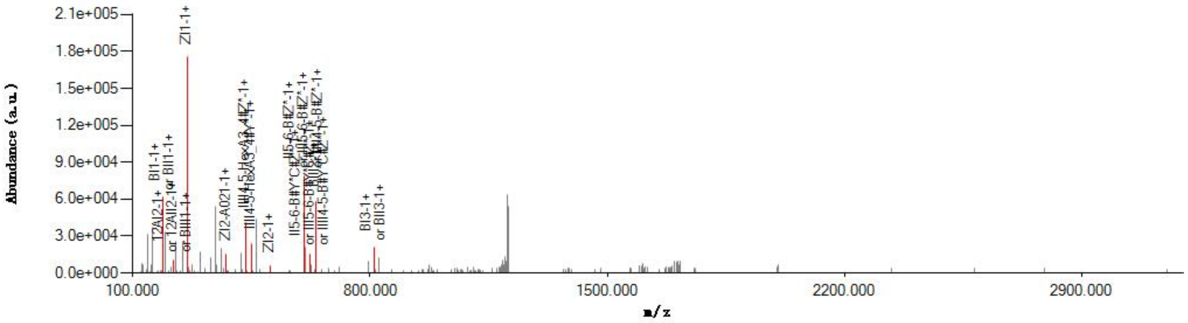

#### Graphical Fragmentation Map

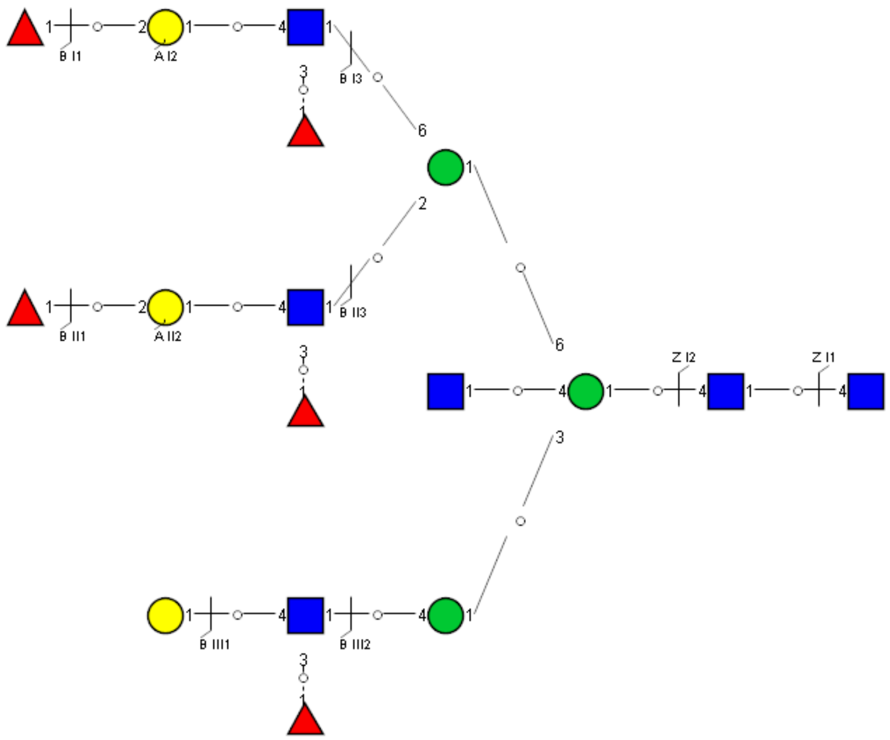

Spectrum Index: 39979(NoBHs:1)

Precursor iEF Map

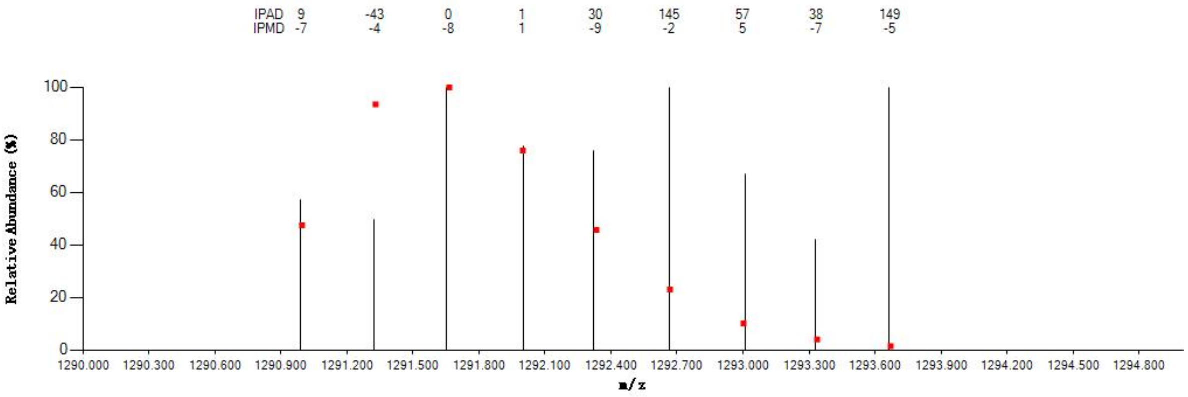


Annotated MS/MS Spectrum

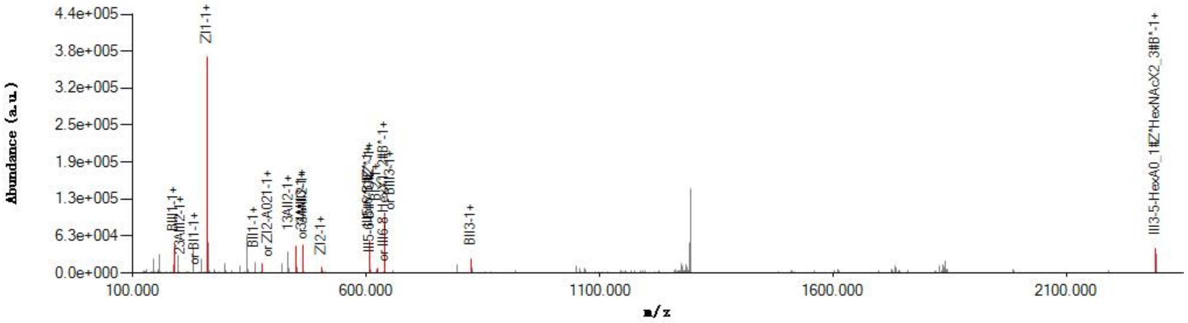


Graphical Fragmentation Map

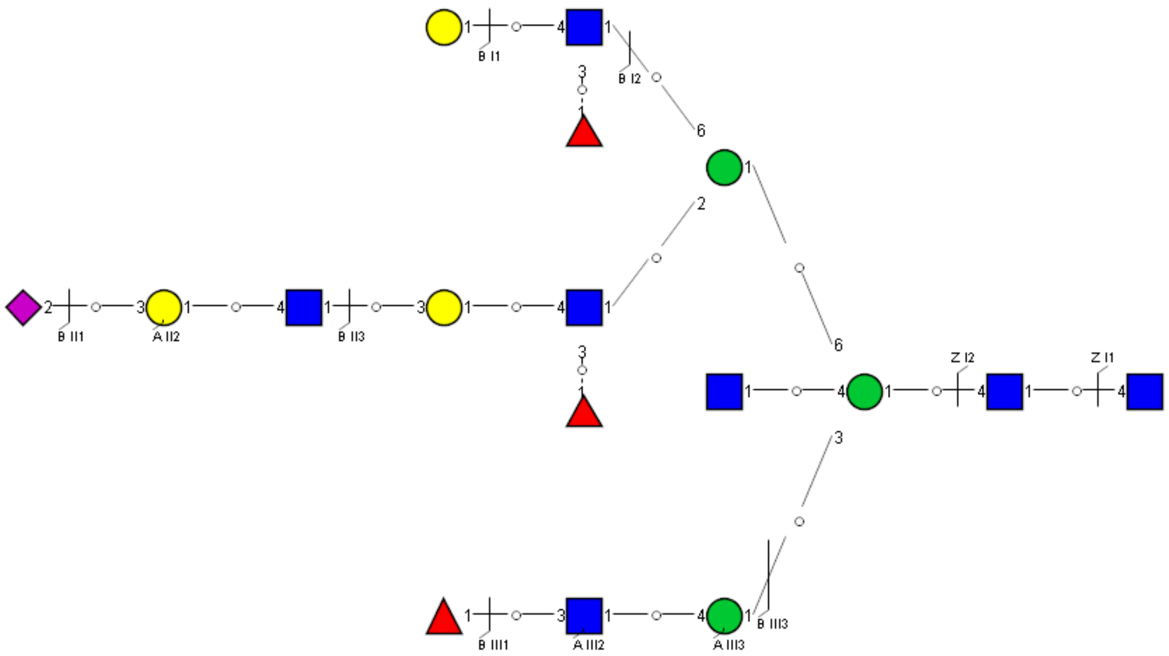

### Spectrum Index: 40013(NoBHs:4)

#### Precursor iEF Map

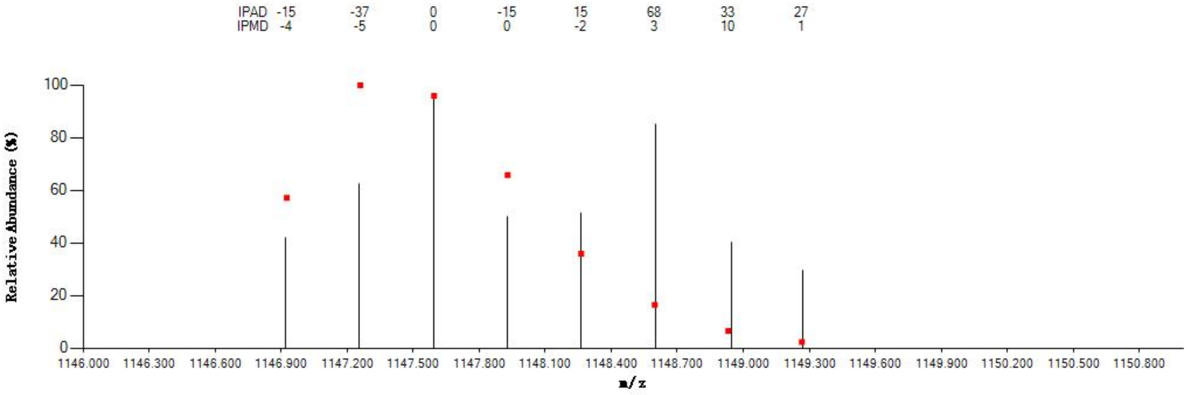


#### Annotated MS/MS Spectrum

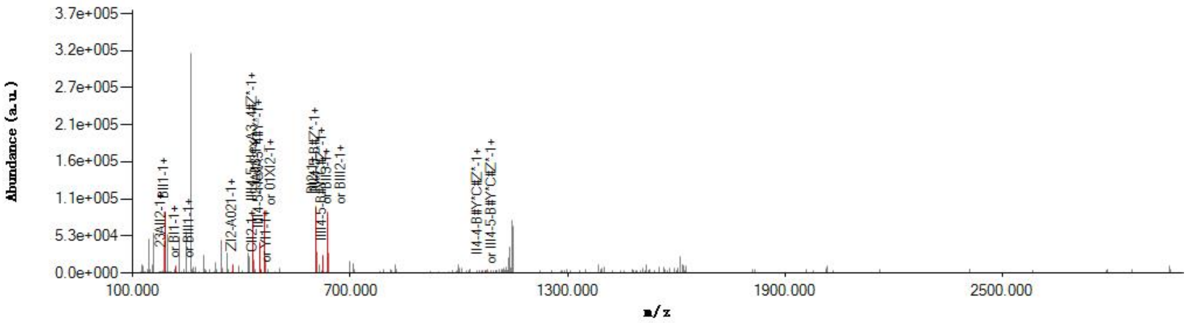


#### Graphical Fragmentation Map

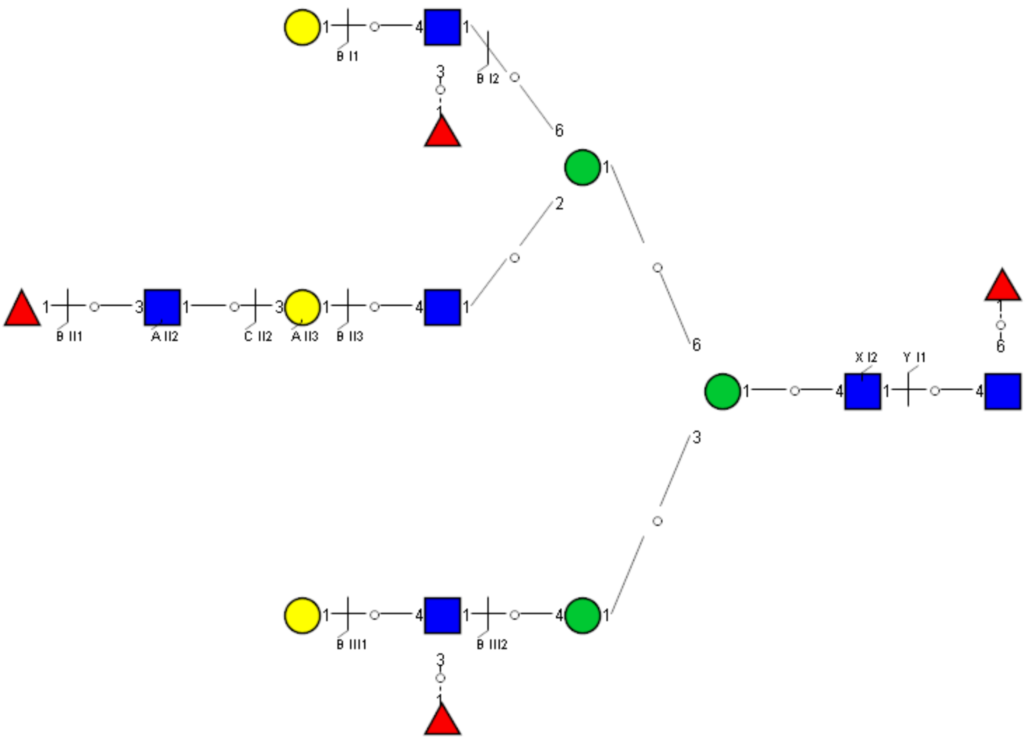

Spectrum Index: 40013(NoBHs:4)

#### Precursor iEF Map

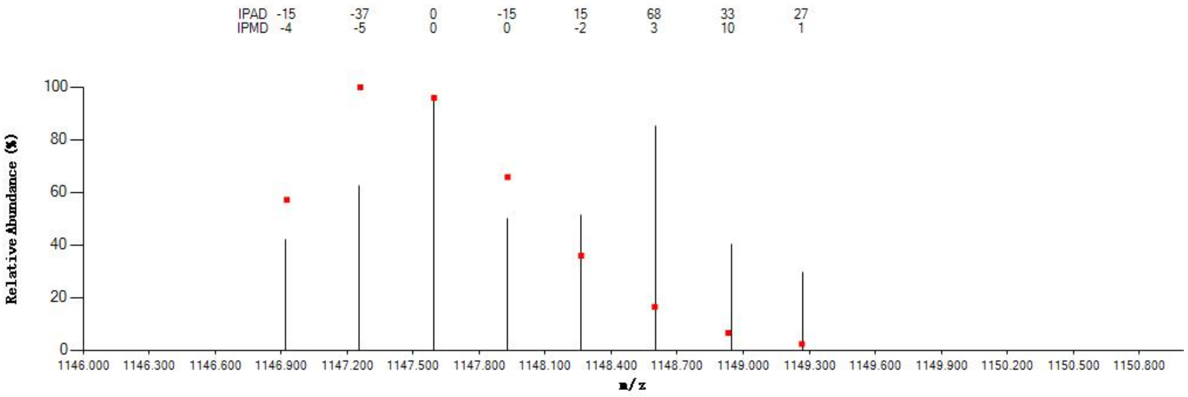

#### Annotated MS/MS Spectrum

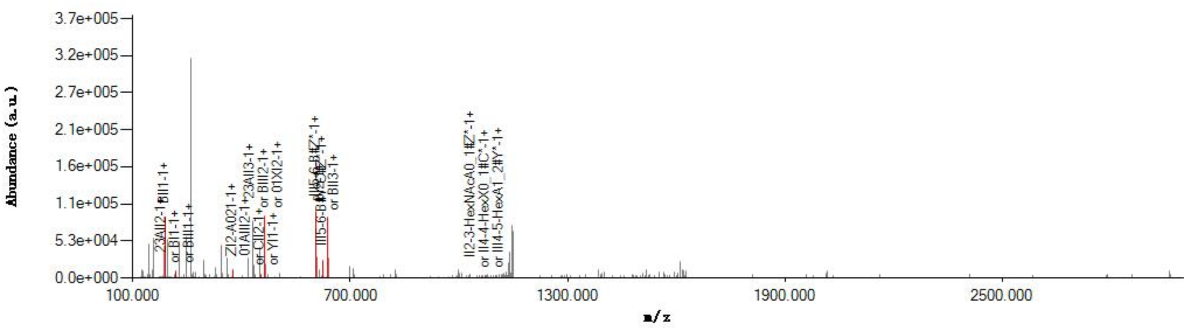

#### Graphical Fragmentation Map

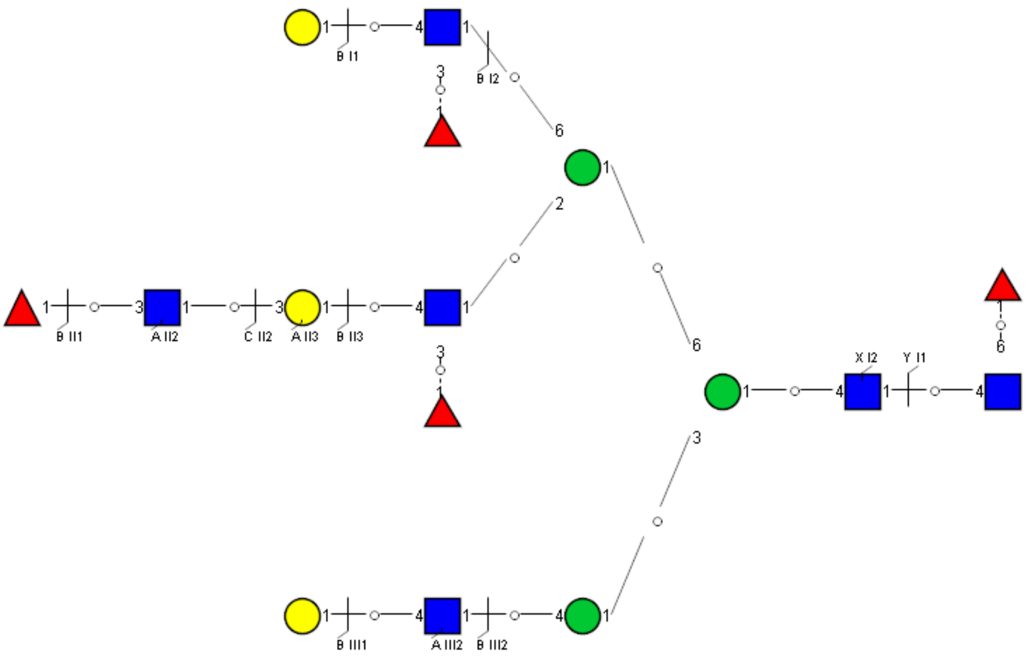

Spectrum Index: 40013(NoBHs:4)

#### Precursor iEF Map

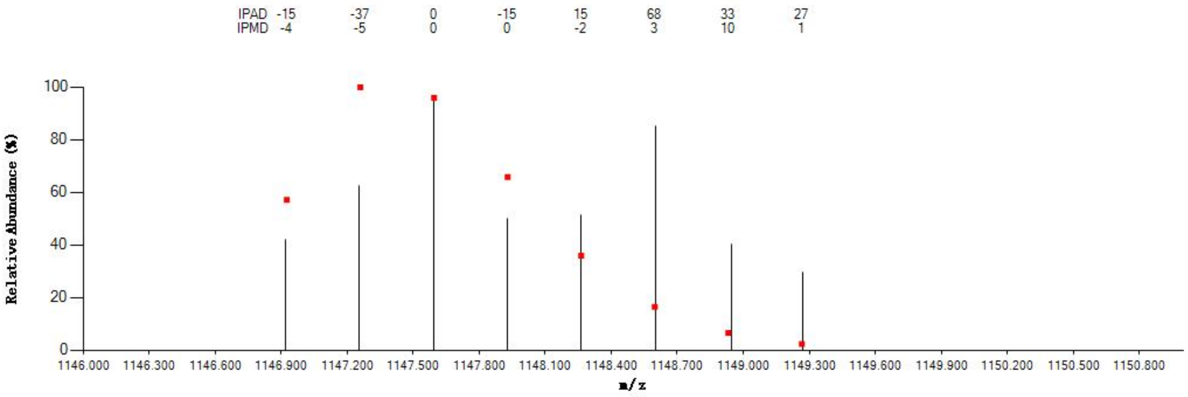

#### Annotated MS/MS Spectrum

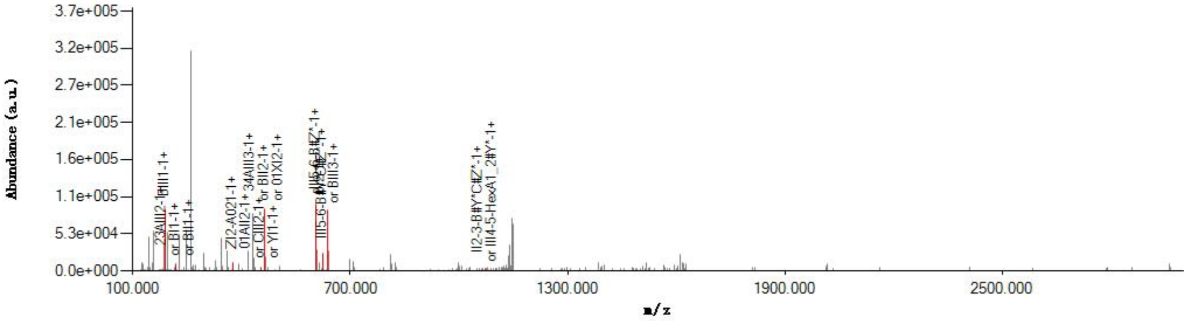

#### Graphical Fragmentation Map

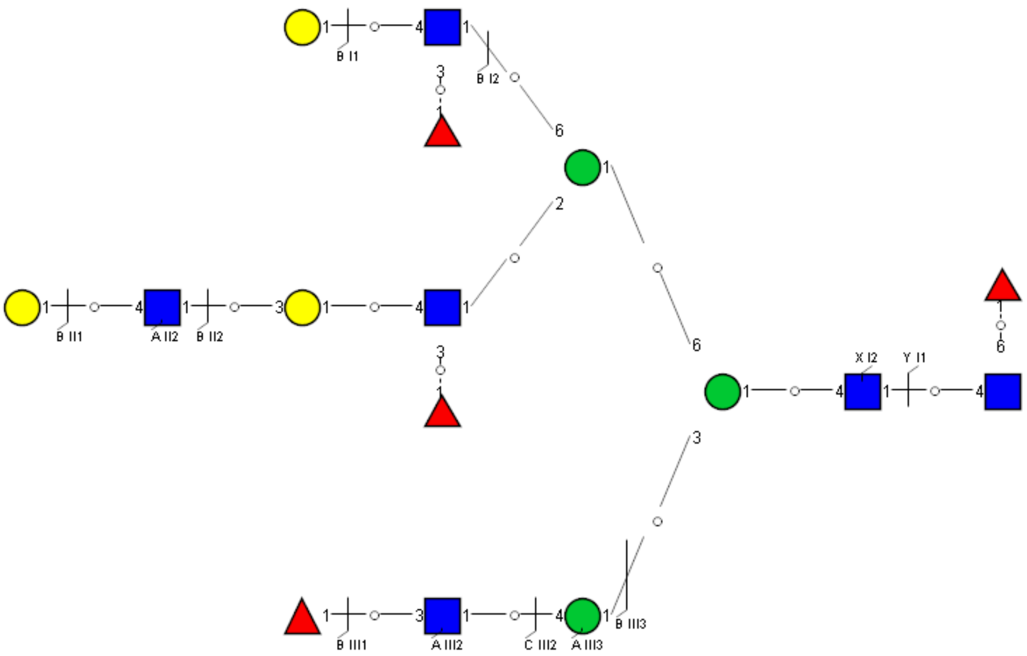

Spectrum Index: 40013(NoBHs:4)

#### Precursor iEF Map

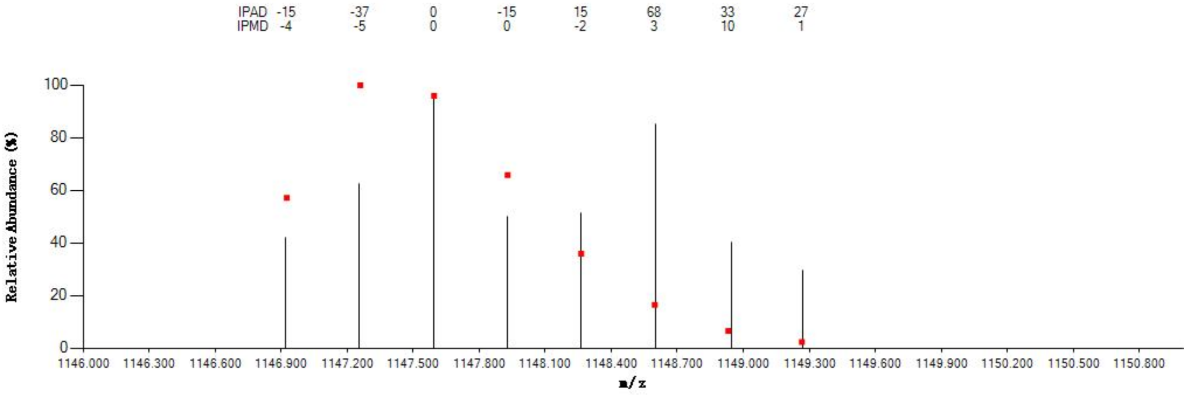


#### Annotated MS/MS Spectrum

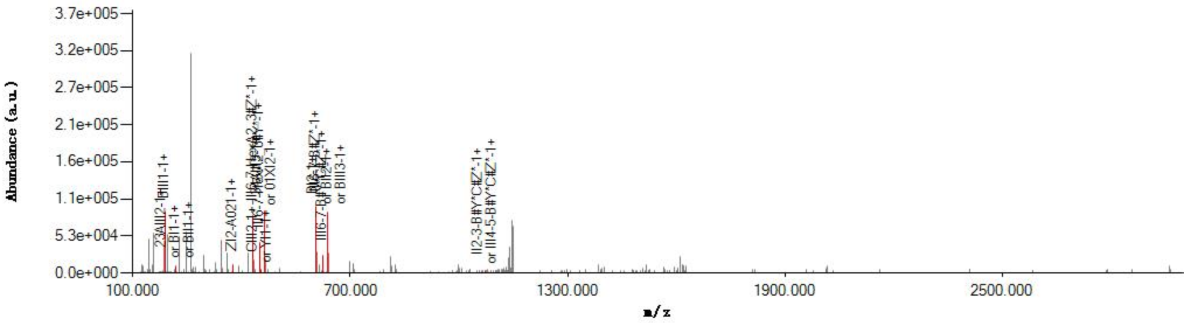


#### Graphical Fragmentation Map

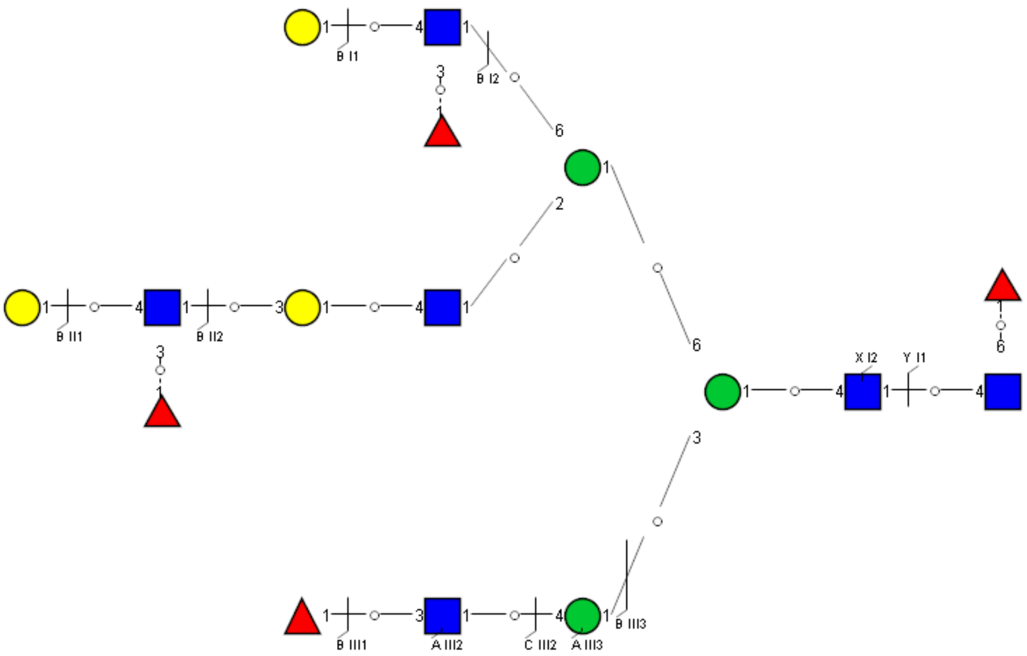

Spectrum Index: 40070(NoBHs:1)

#### Precursor iEF Map

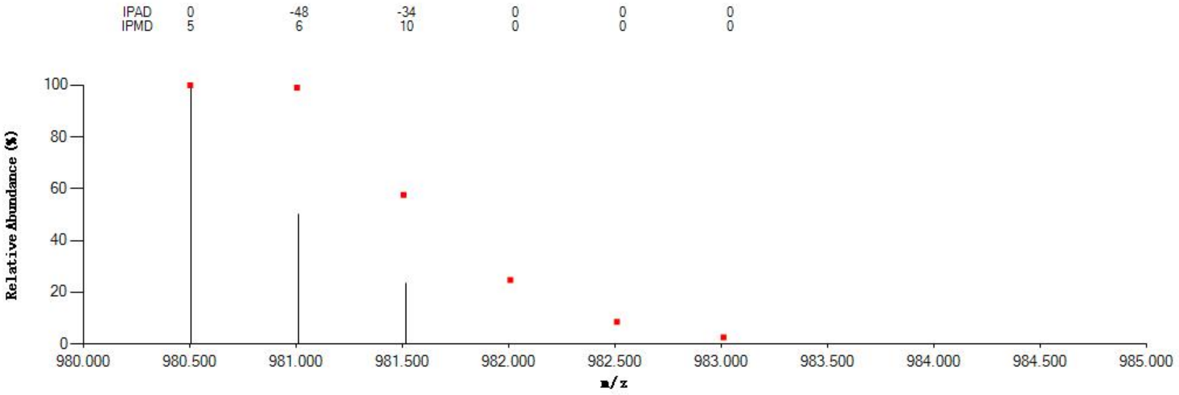


#### Annotated MS/MS Spectrum

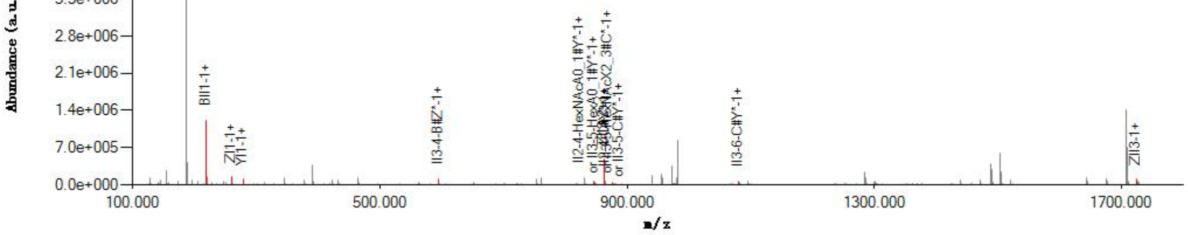


#### Graphical Fragmentation Map

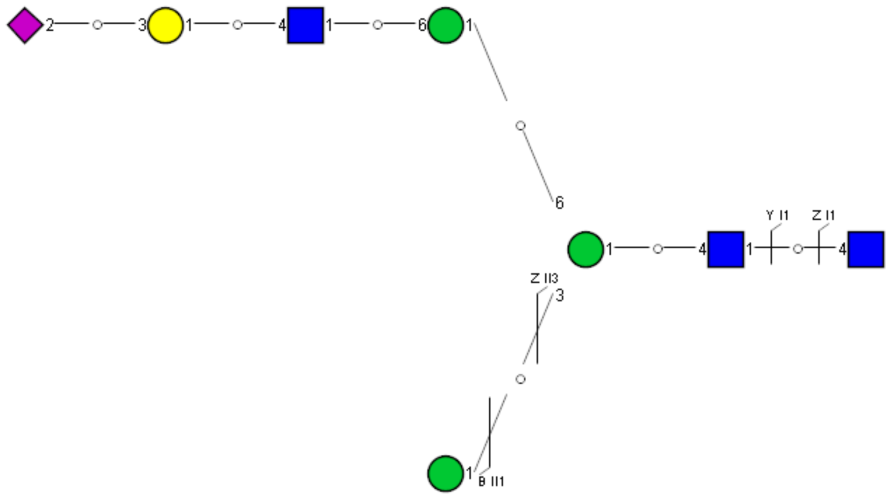

Spectrum Index: 40105(NoBHs:2)

#### Precursor iEF Map

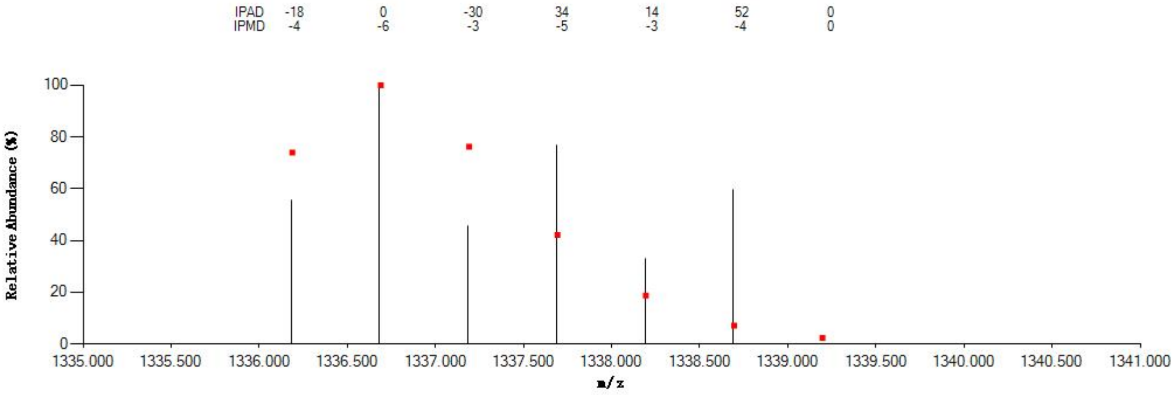


#### Annotated MS/MS Spectrum

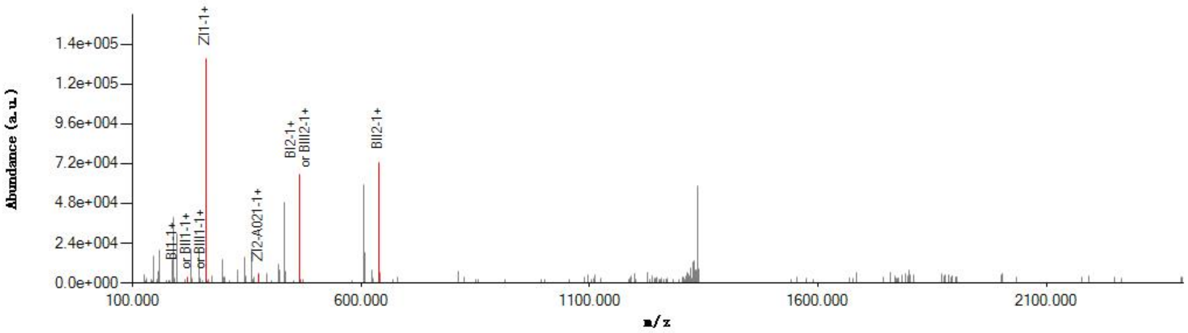


#### Graphical Fragmentation Map

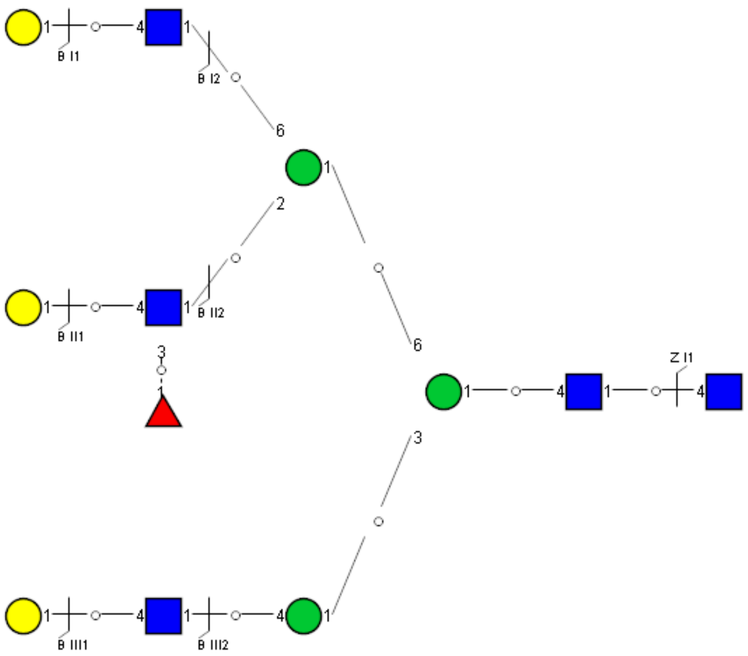

Spectrum Index: 40260(NoBHs:1)

#### Precursor iEF Map

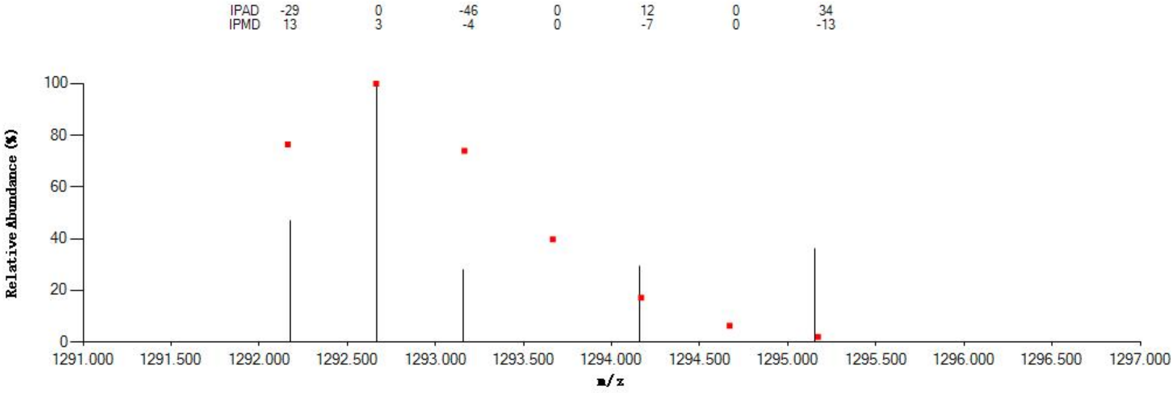


#### Annotated MS/MS Spectrum

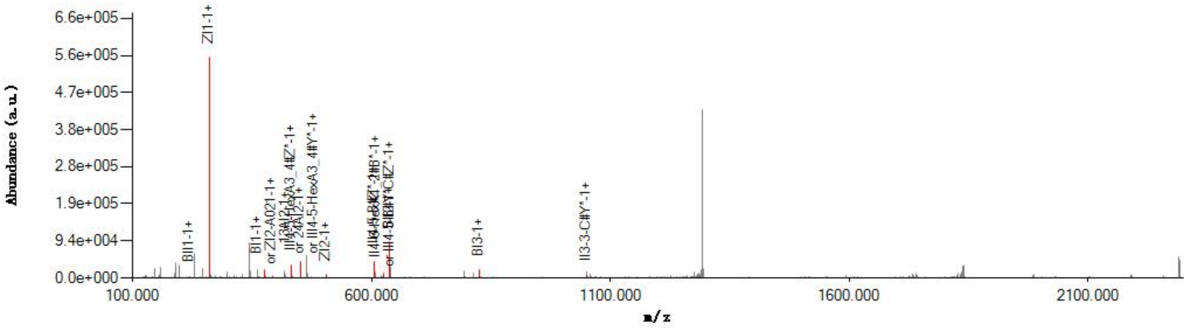


#### Graphical Fragmentation Map

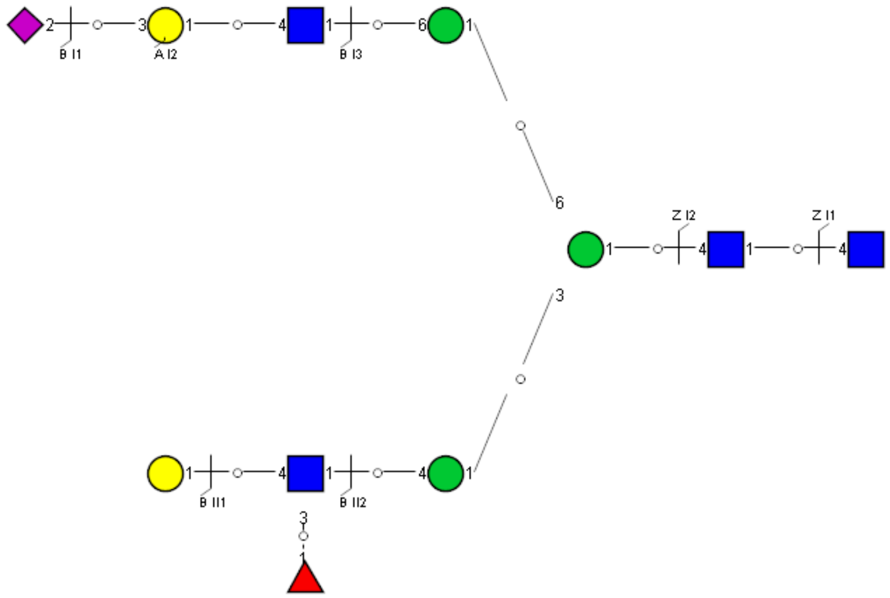

### Spectrum Index: 40269(NoBHs:1)

#### Precursor iEF Map

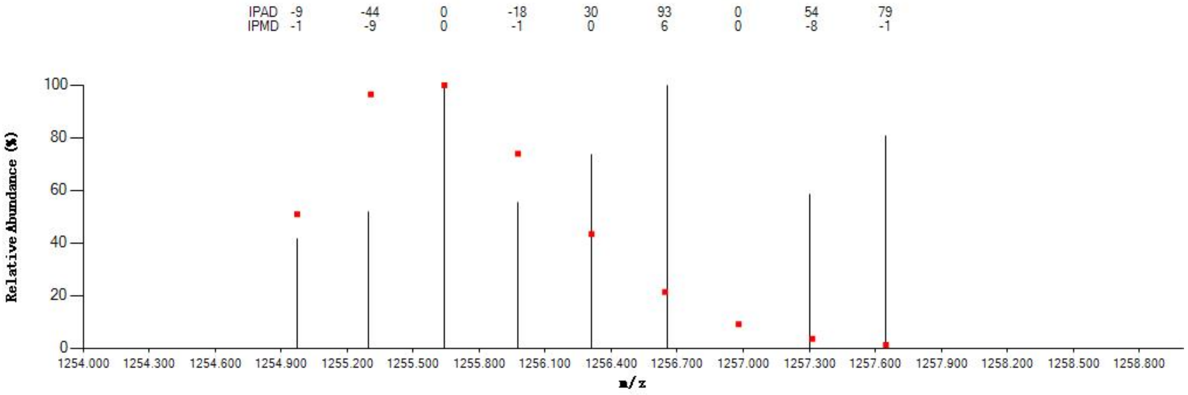

#### Annotated MS/MS Spectrum

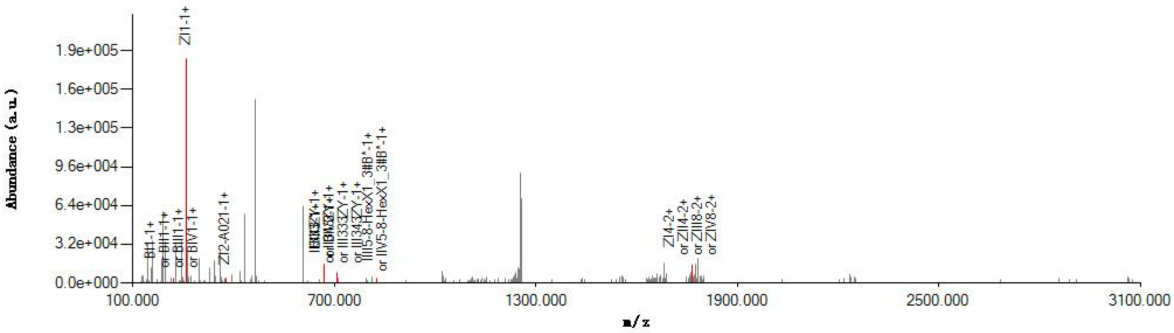

#### Graphical Fragmentation Map

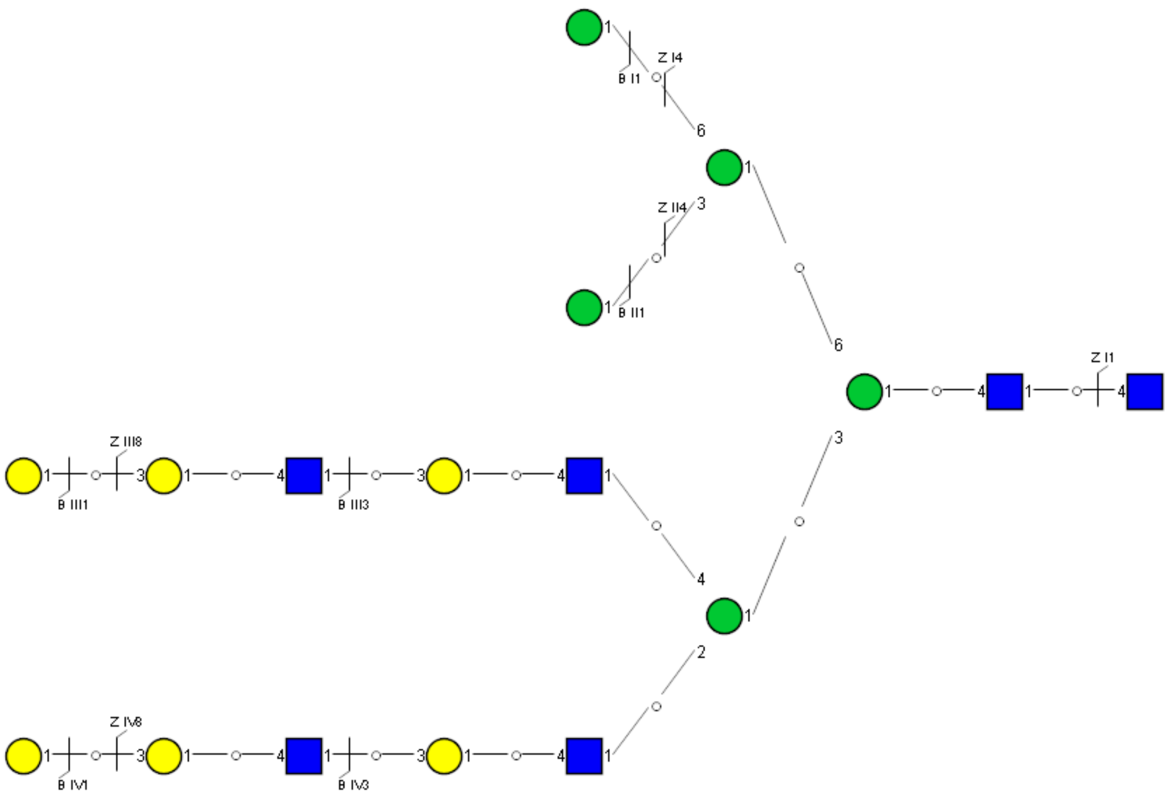

Spectrum Index: 40282(NoBHs:1)

#### Precursor iEF Map

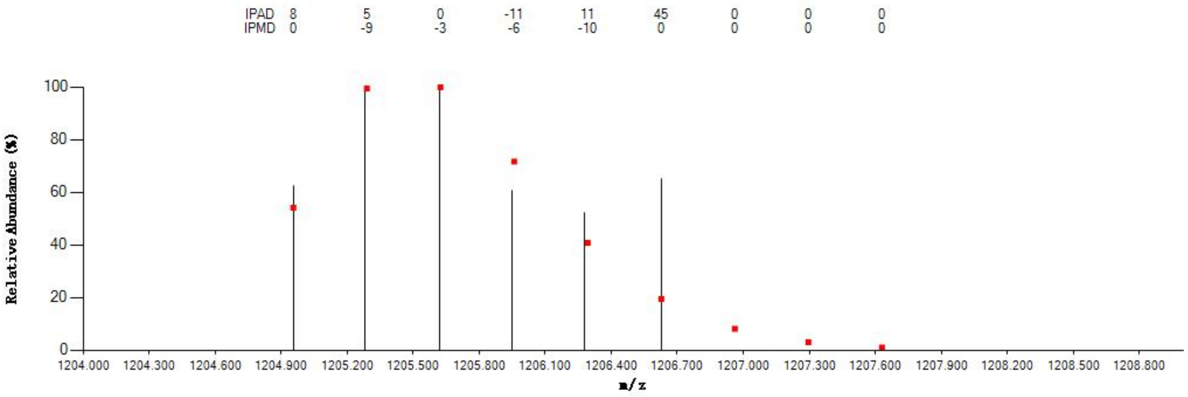

#### Annotated MS/MS Spectrum

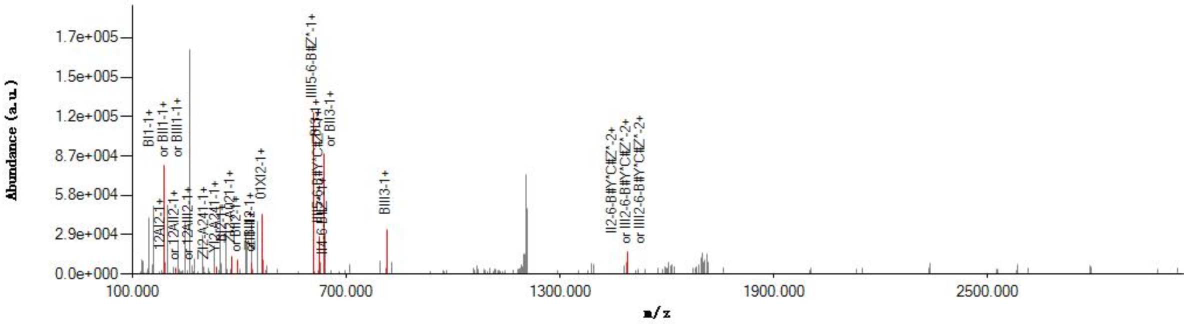

#### Graphical Fragmentation Map

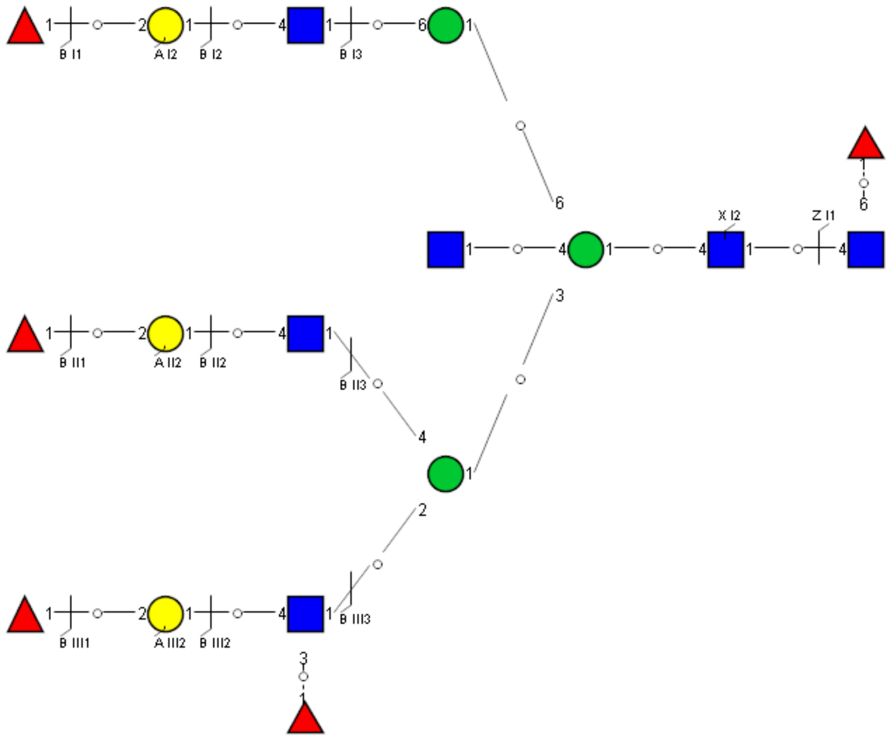

Spectrum Index: 40326(NoBHs:1)

#### Precursor iEF Map

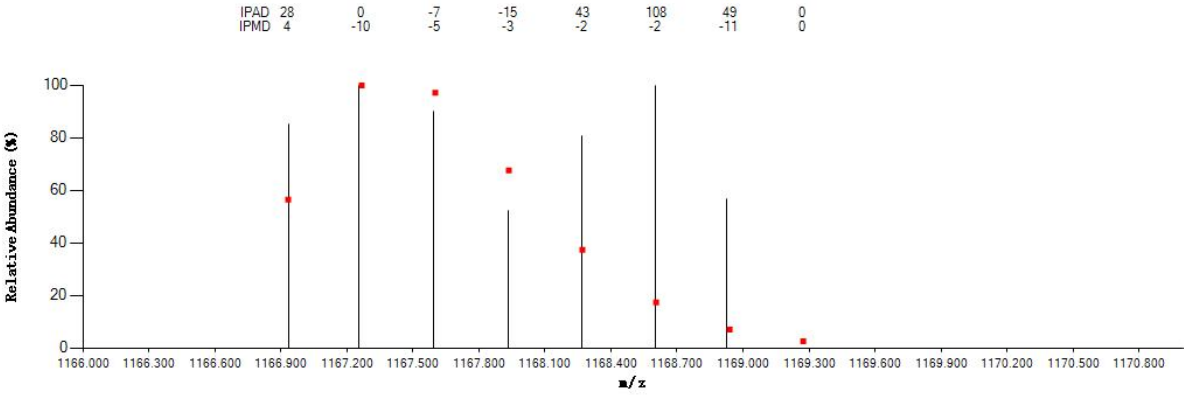


#### Annotated MS/MS Spectrum

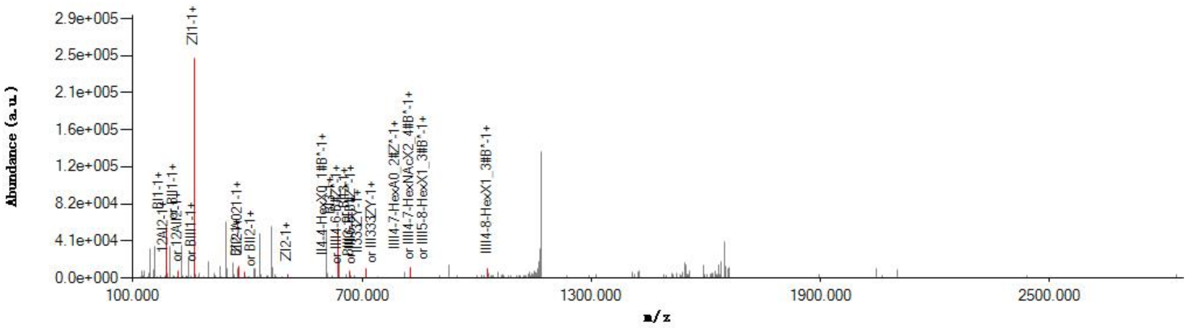


#### Graphical Fragmentation Map

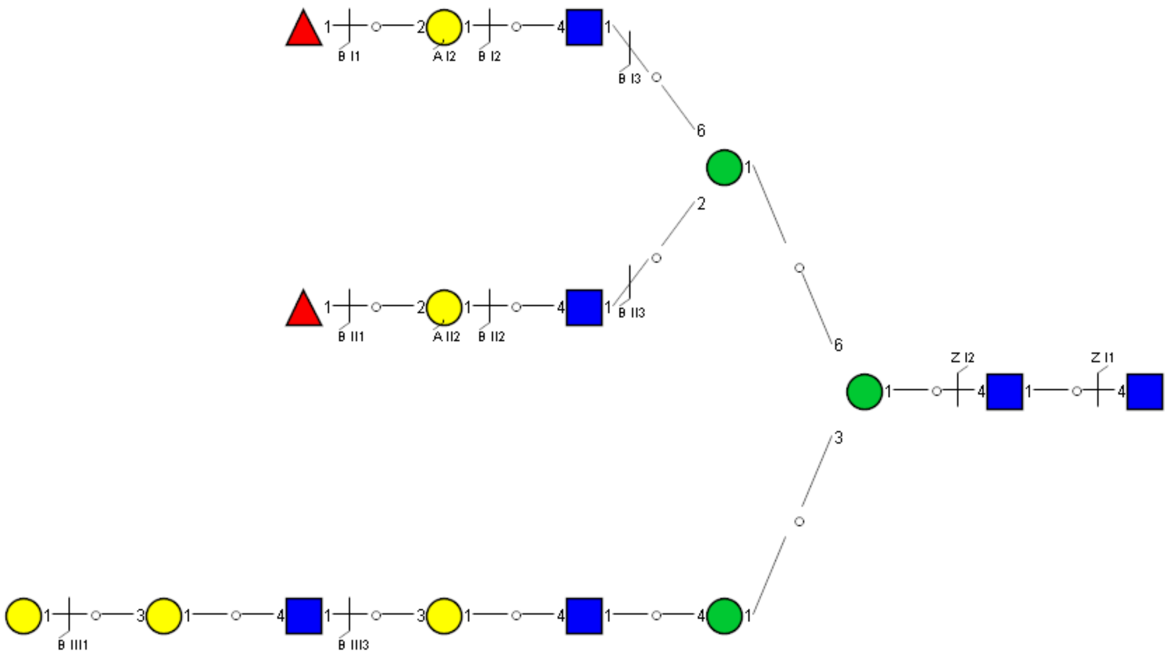

Spectrum Index: 40343(NoBHs:3)

#### Precursor iEF Map

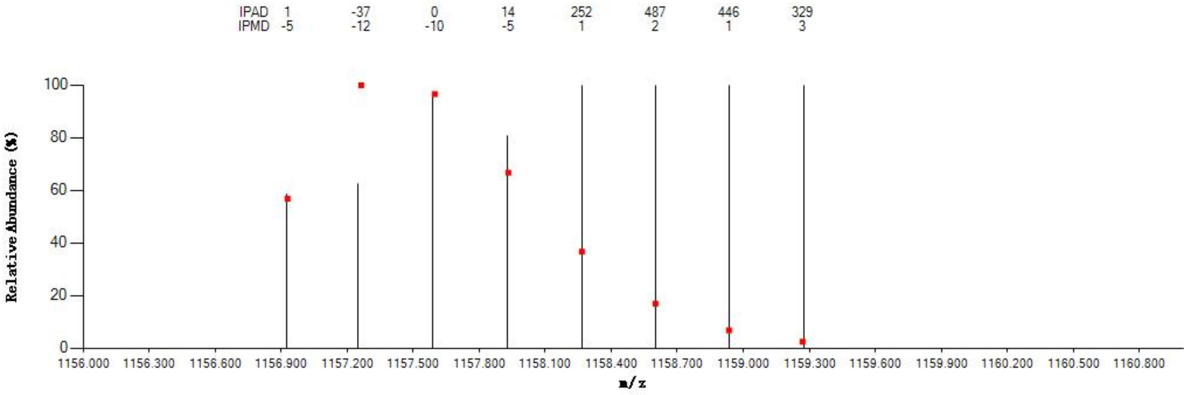


#### Annotated MS/MS Spectrum

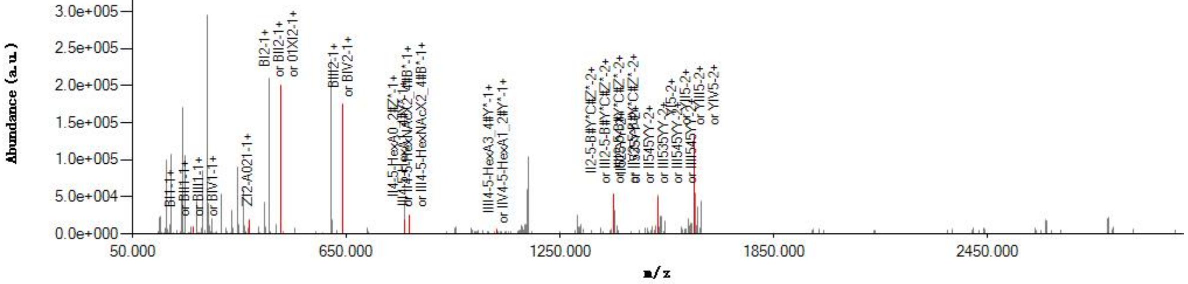


#### Graphical Fragmentation Map

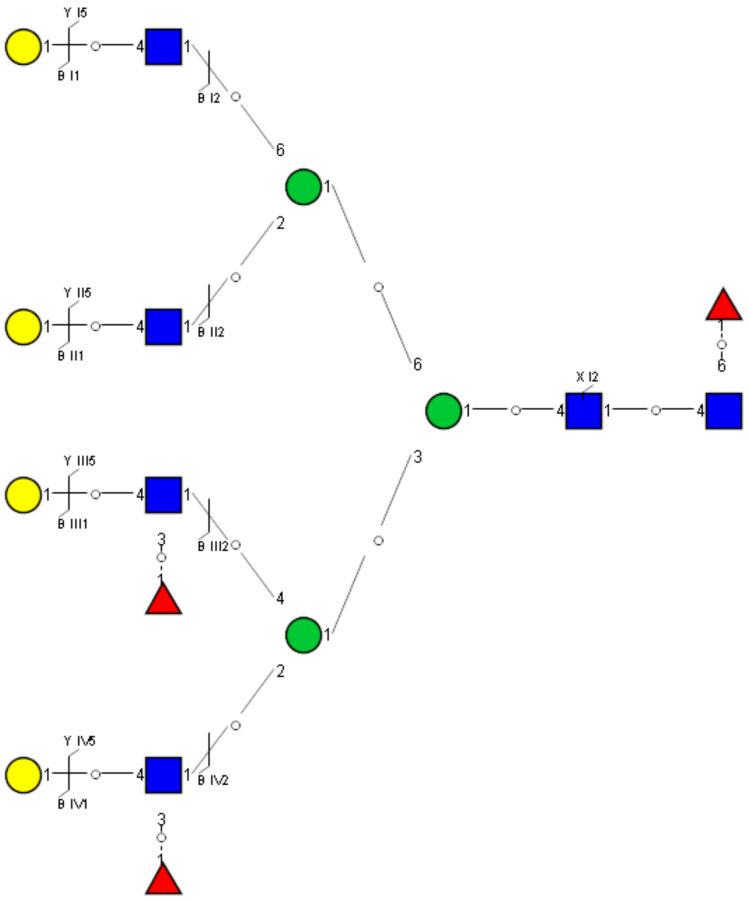

Spectrum Index: 40343(NoBHs:3)

#### Precursor iEF Map

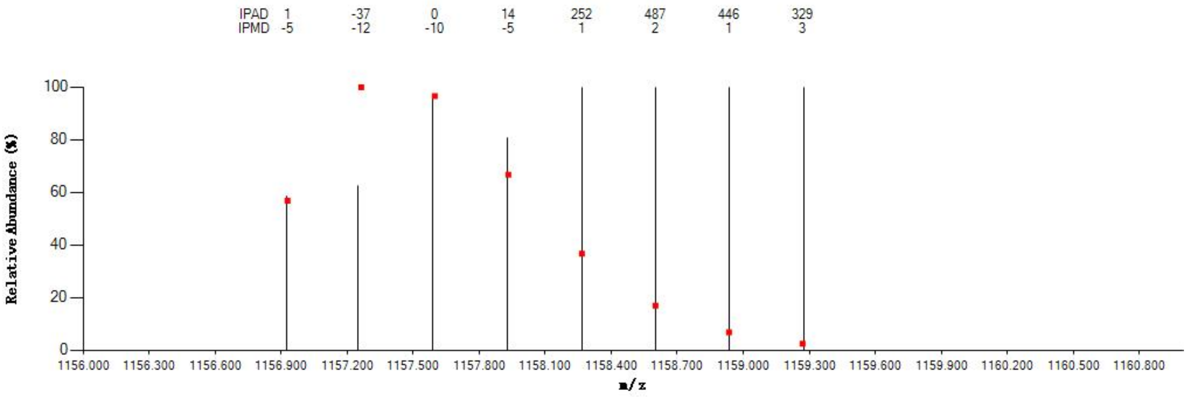

#### Annotated MS/MS Spectrum

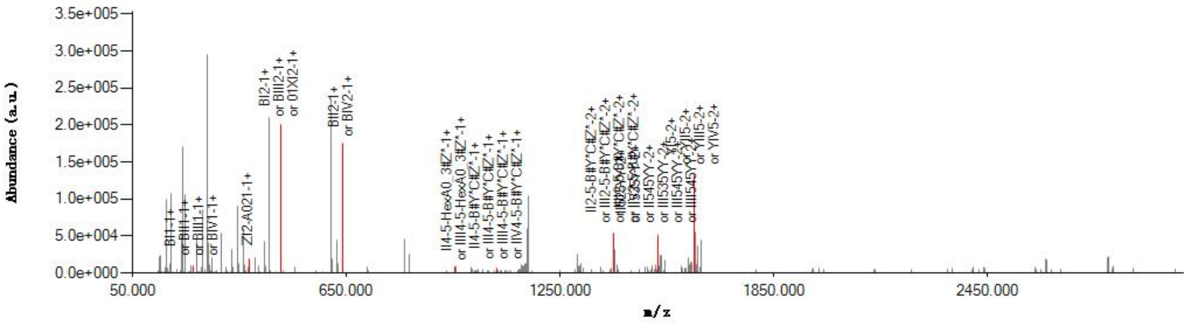

#### Graphical Fragmentation Map

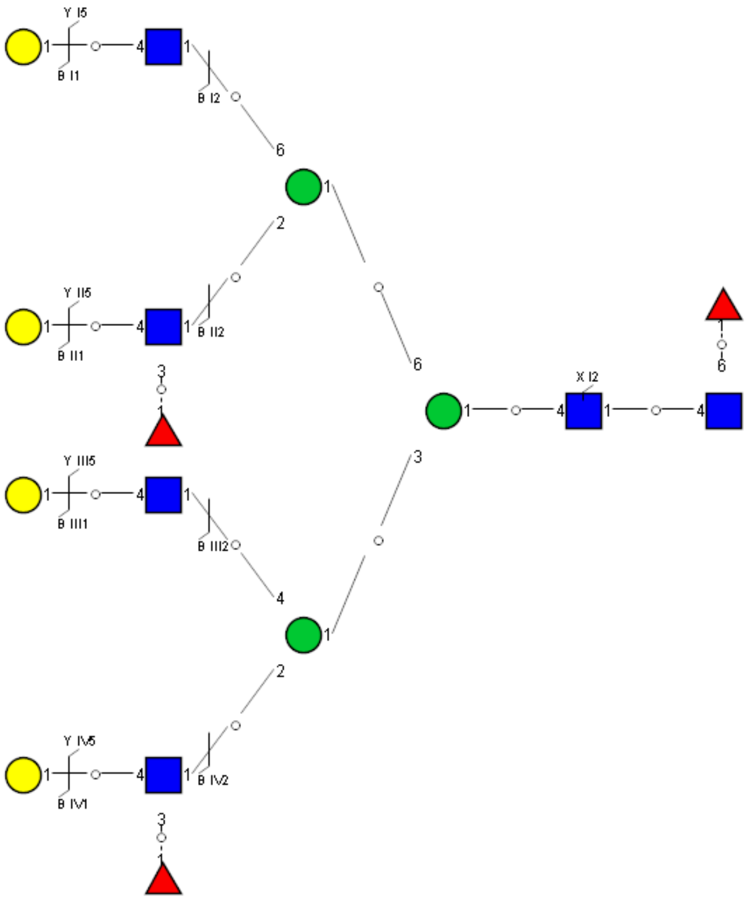

Spectrum Index: 40389(NoBHs:1)

#### Precursor iEF Map

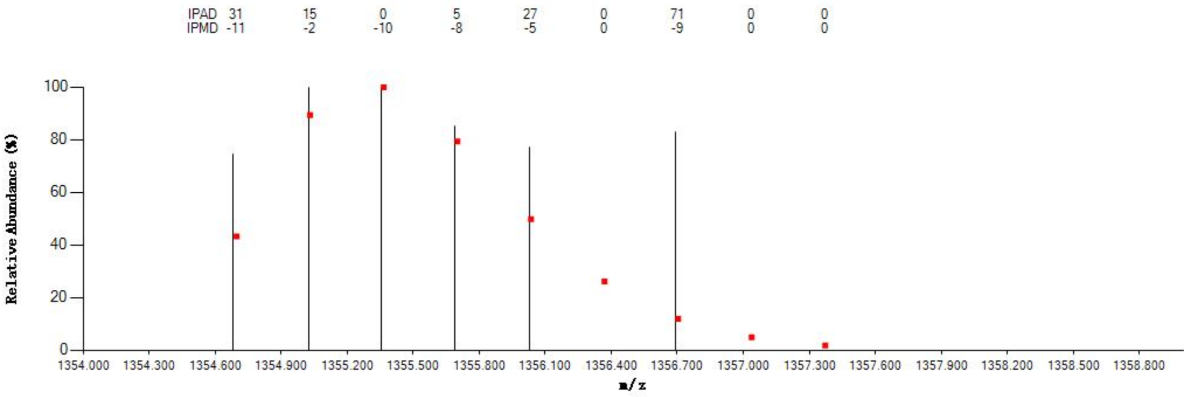


#### Annotated MS/MS Spectrum

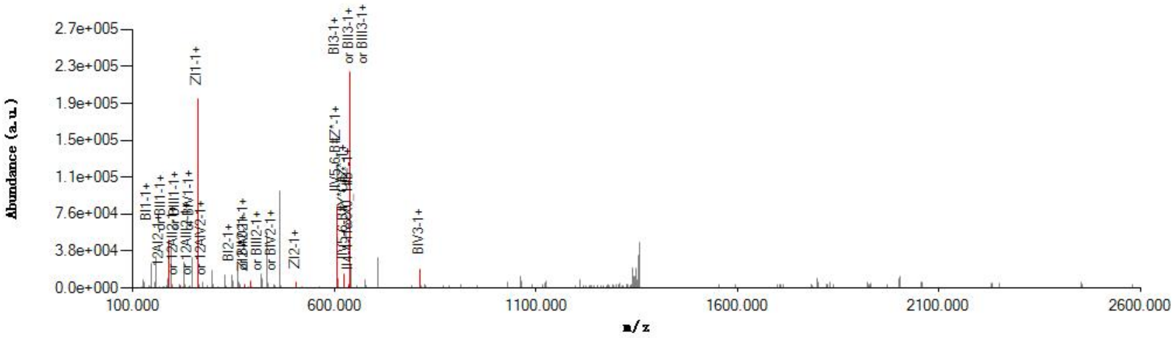


#### Graphical Fragmentation Map

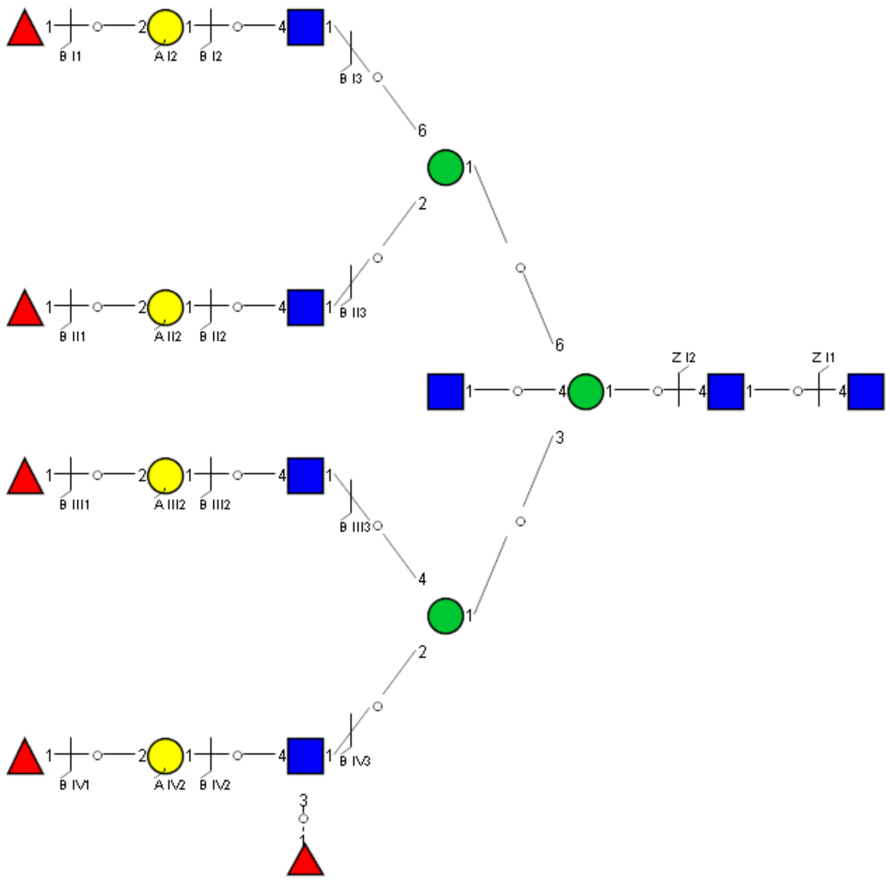

Spectrum Index: 40436(NoBHs:1)

#### Precursor iEF Map

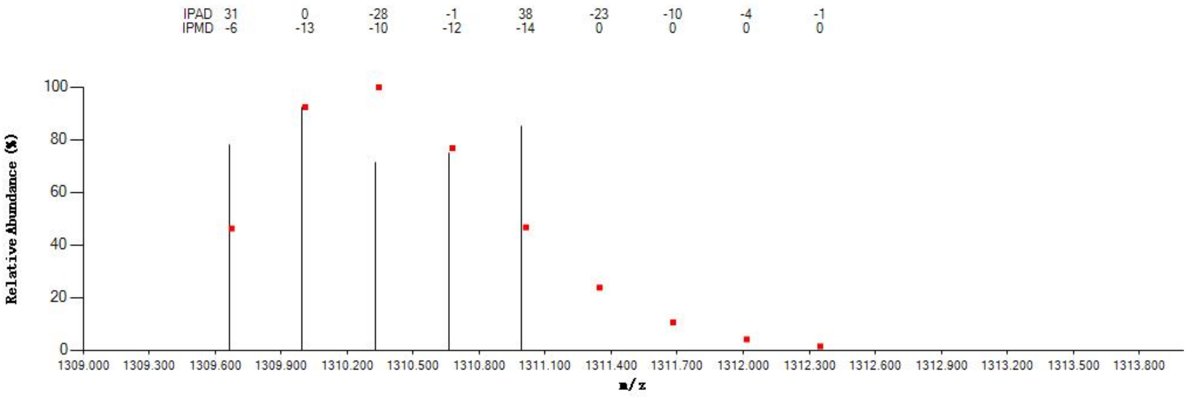


#### Annotated MS/MS Spectrum

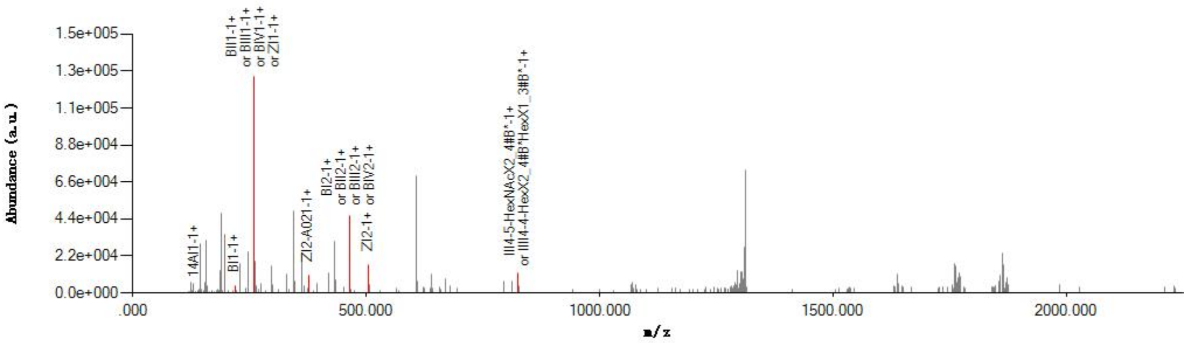


#### Graphical Fragmentation Map

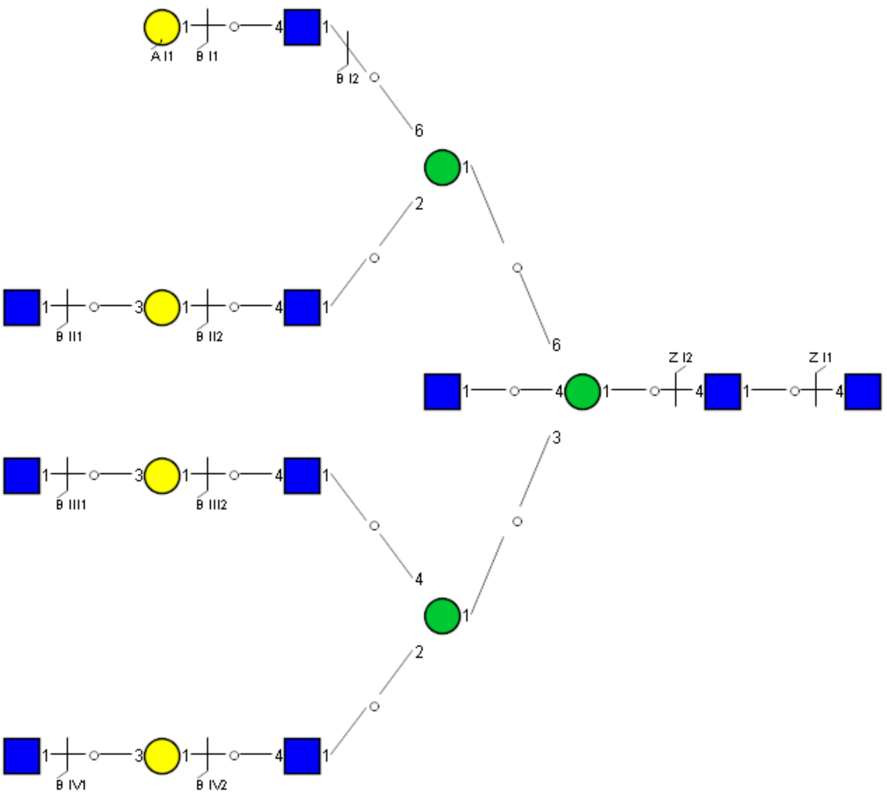

Spectrum Index: 40472(NoBHs:2)

#### Precursor iEF Map

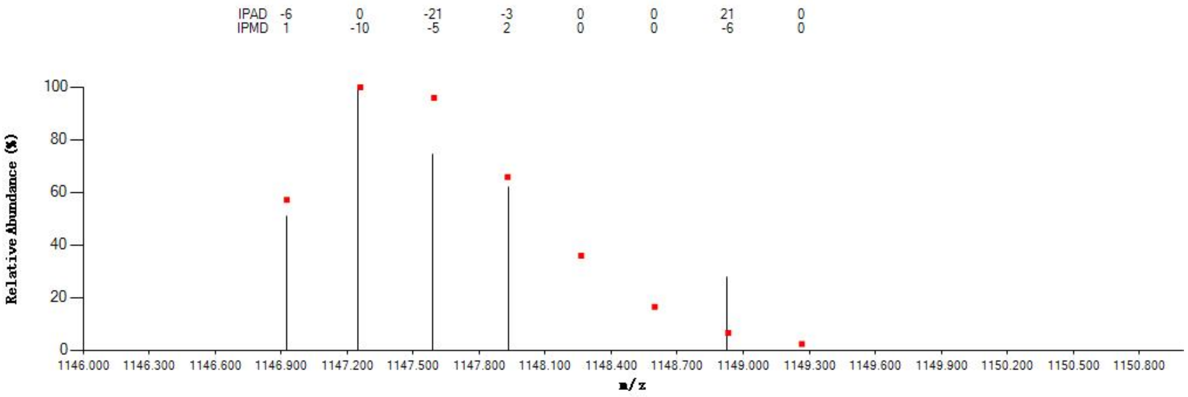


#### Annotated MS/MS Spectrum

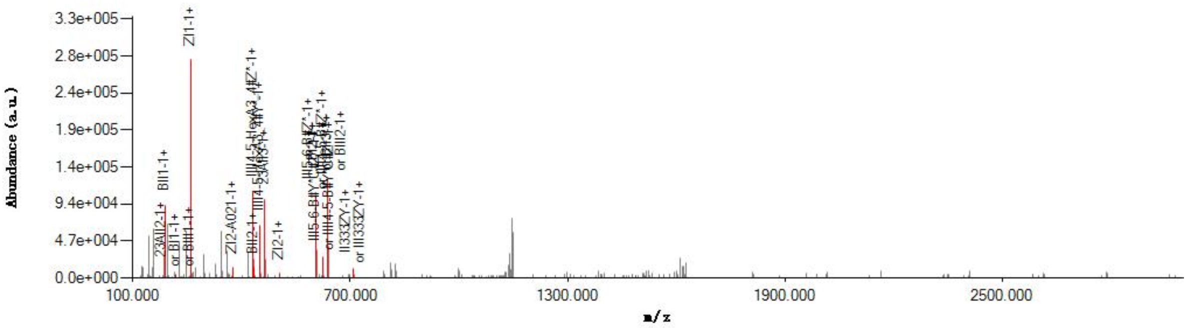


#### Graphical Fragmentation Map

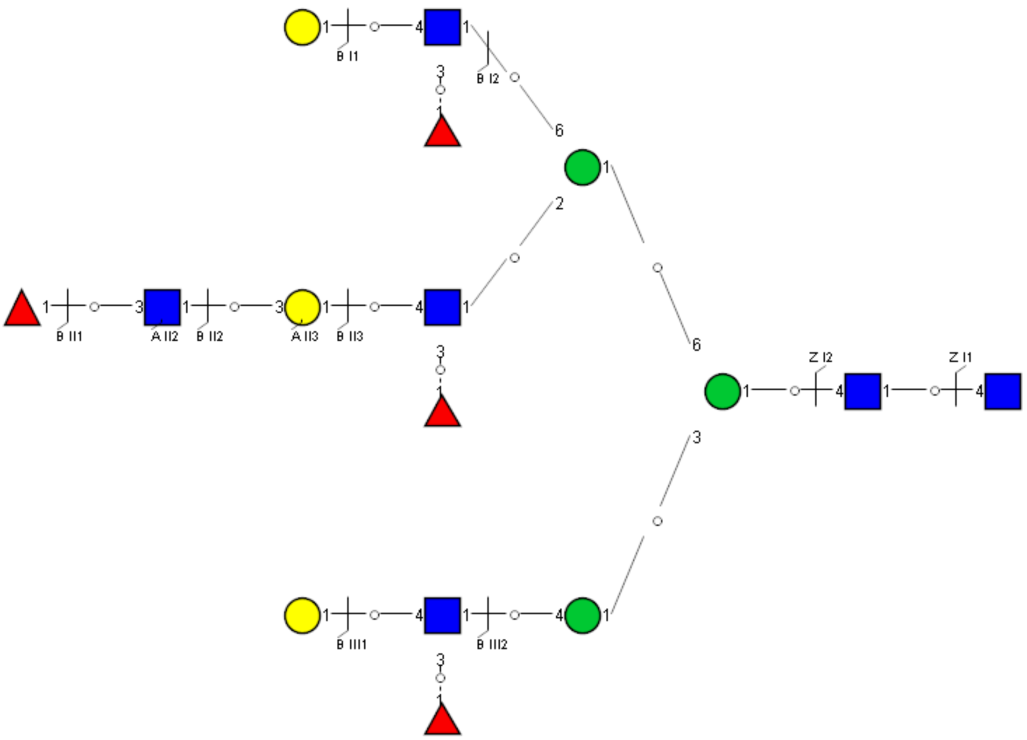

Spectrum Index: 40472(NoBHs:2)

#### Precursor iEF Map

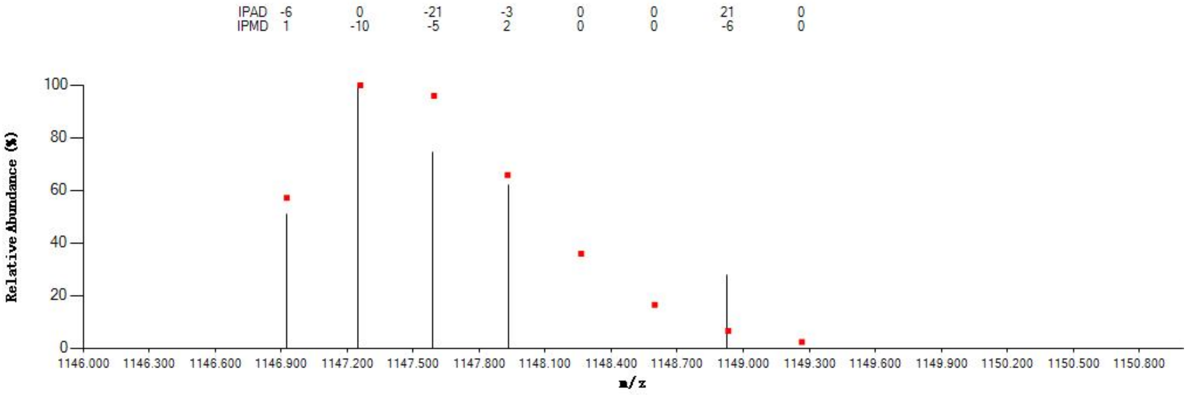


#### Annotated MS/MS Spectrum

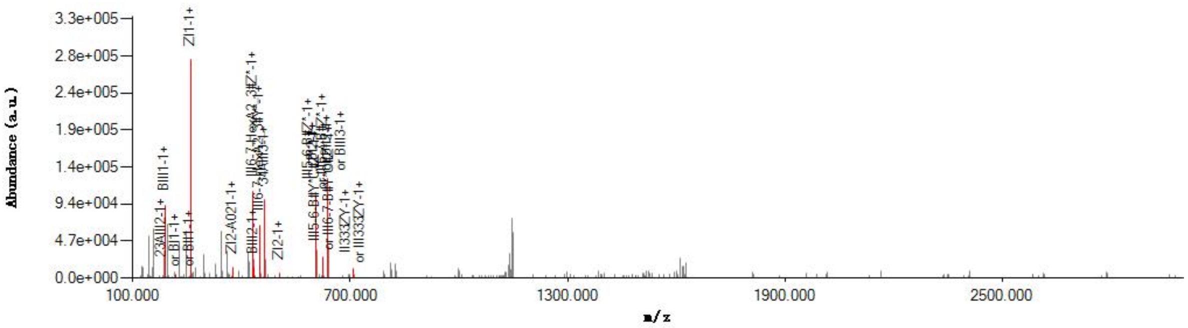


#### Graphical Fragmentation Map

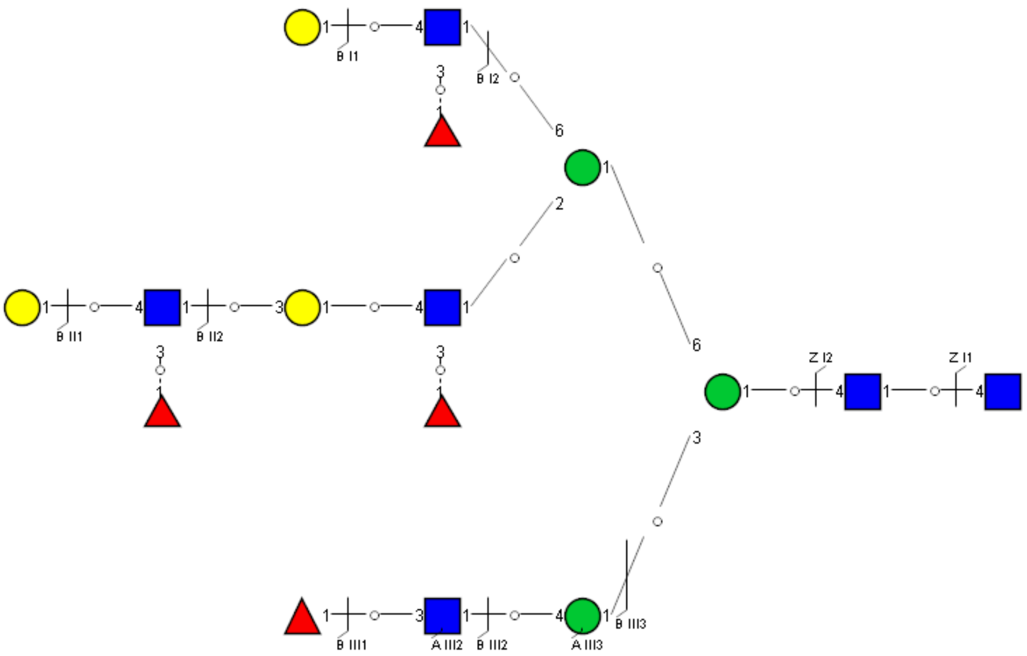

Spectrum Index: 40478(NoBHs:1)

#### Precursor iEF Map

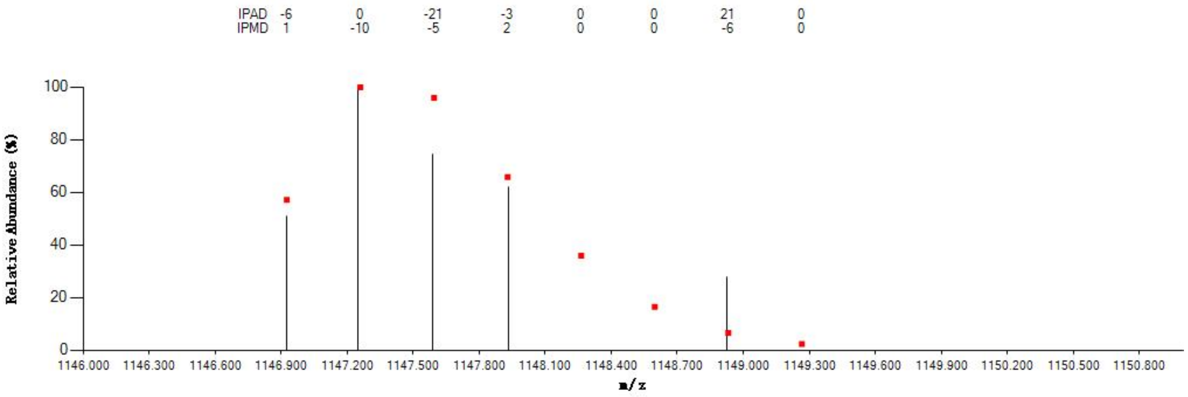


#### Annotated MS/MS Spectrum

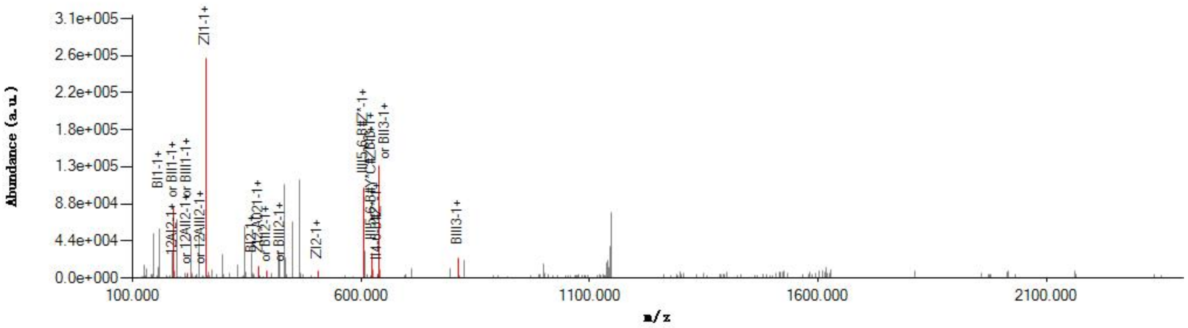


#### Graphical Fragmentation Map

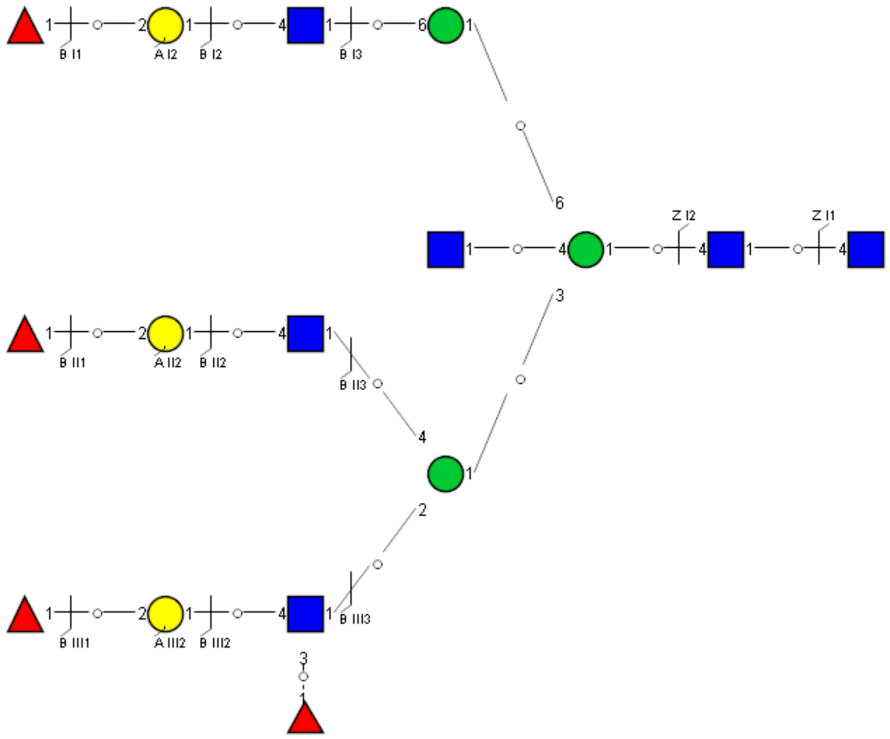

Spectrum Index: 40500(NoBHs:1)

#### Precursor iEF Map

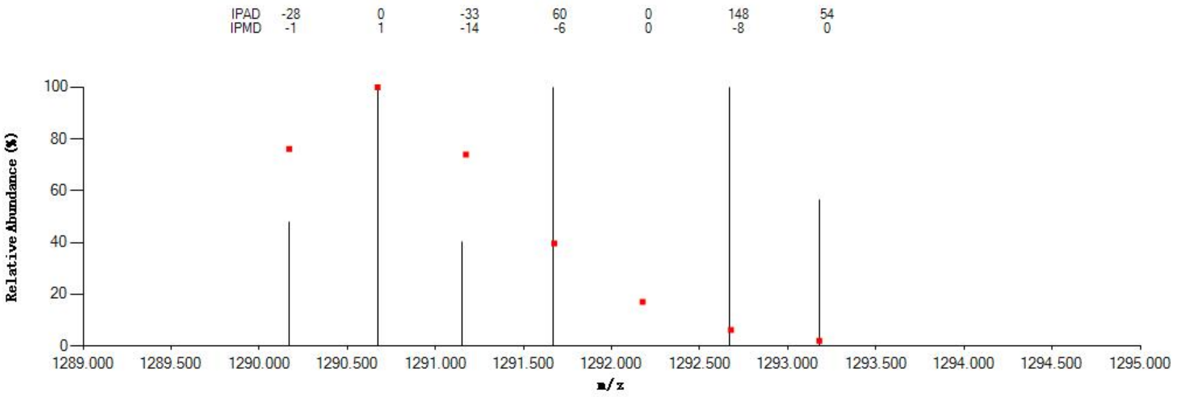

#### Annotated MS/MS Spectrum

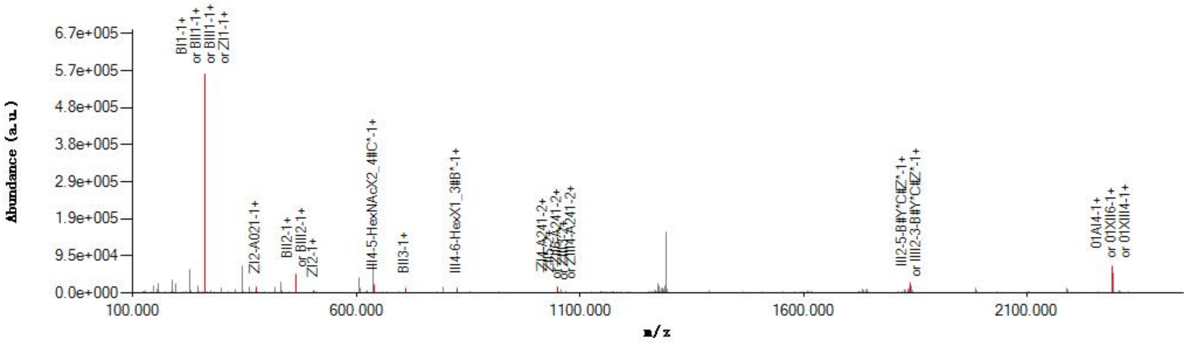

#### Graphical Fragmentation Map

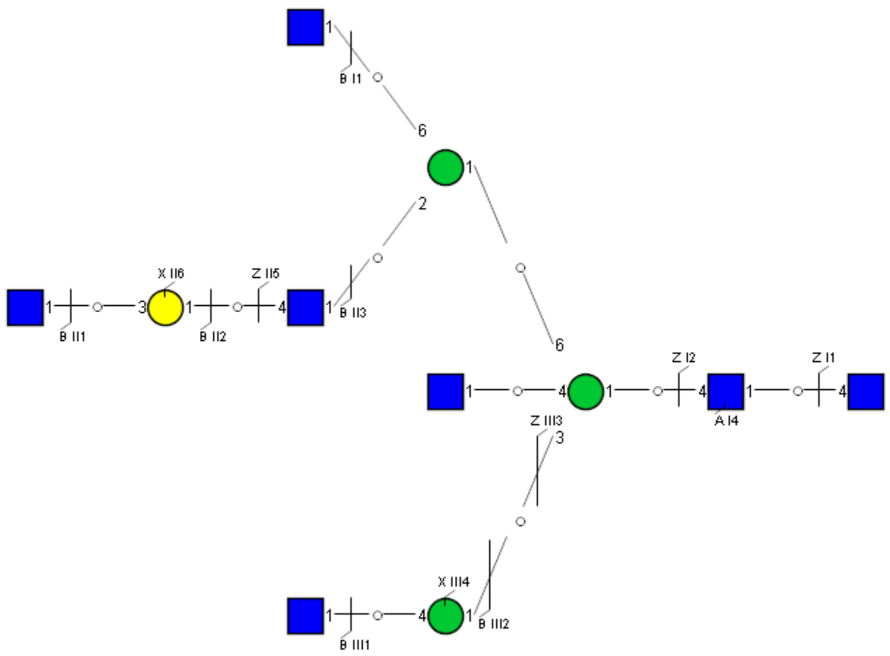

Spectrum Index: 40505(NoBHs:1)

#### Precursor iEF Map

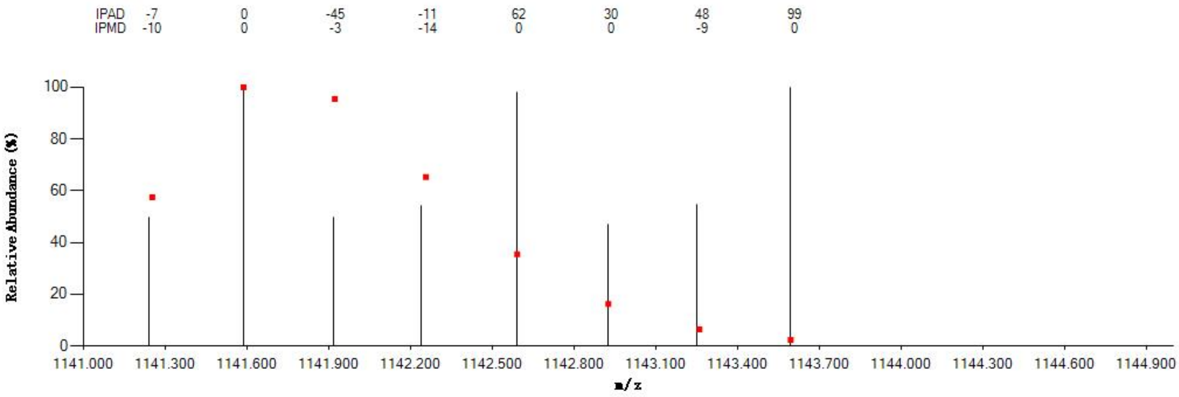


#### Annotated MS/MS Spectrum

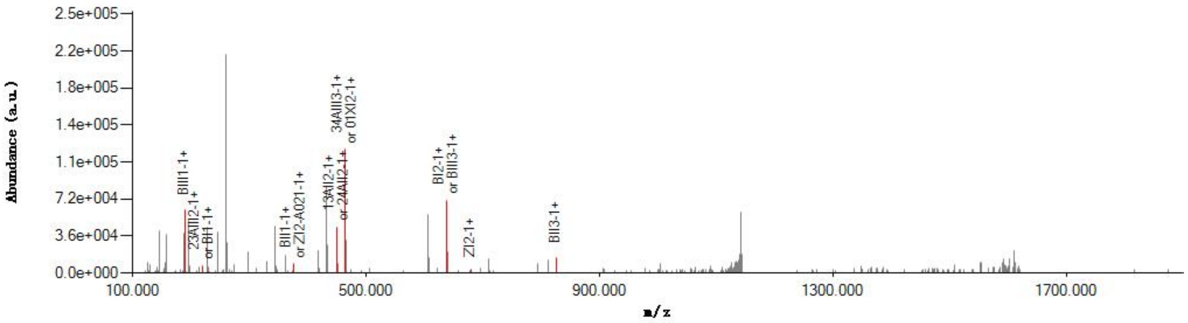


#### Graphical Fragmentation Map

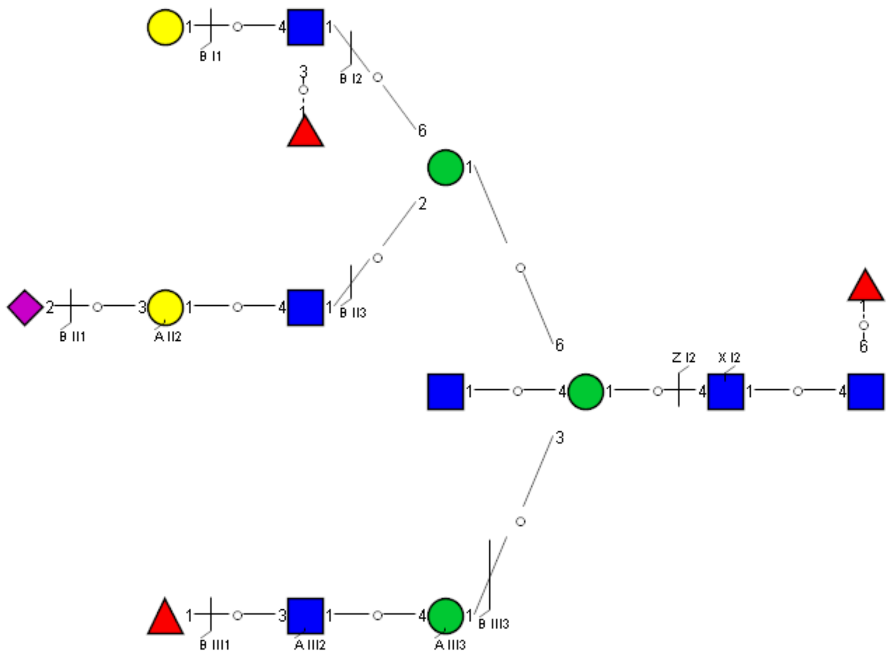

Spectrum Index: 40543(NoBHs:1)

#### Precursor iEF Map

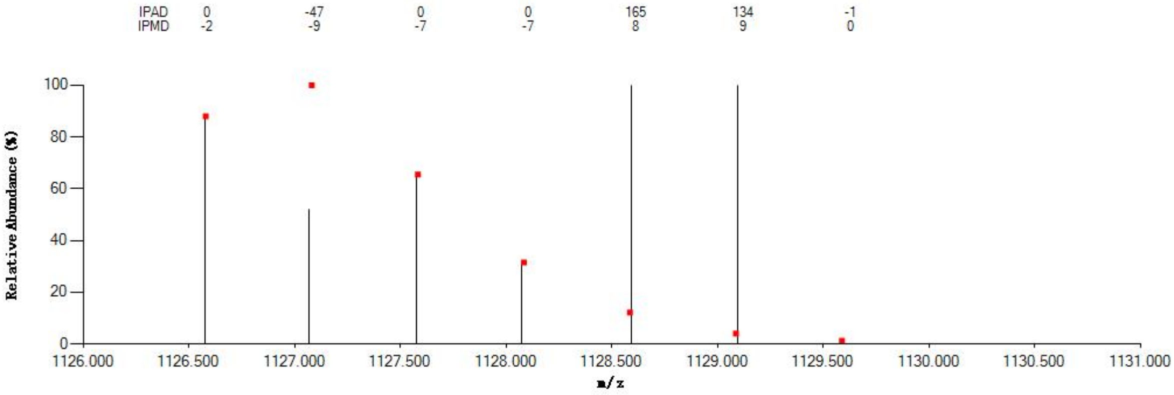


#### Annotated MS/MS Spectrum

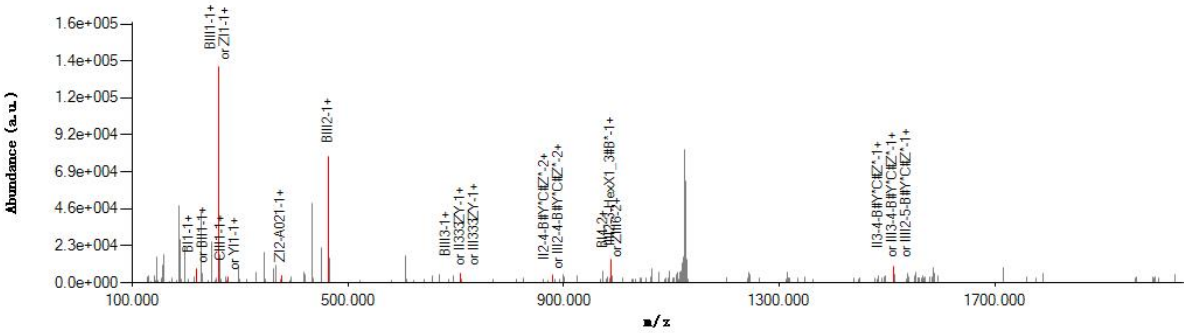


#### Graphical Fragmentation Map

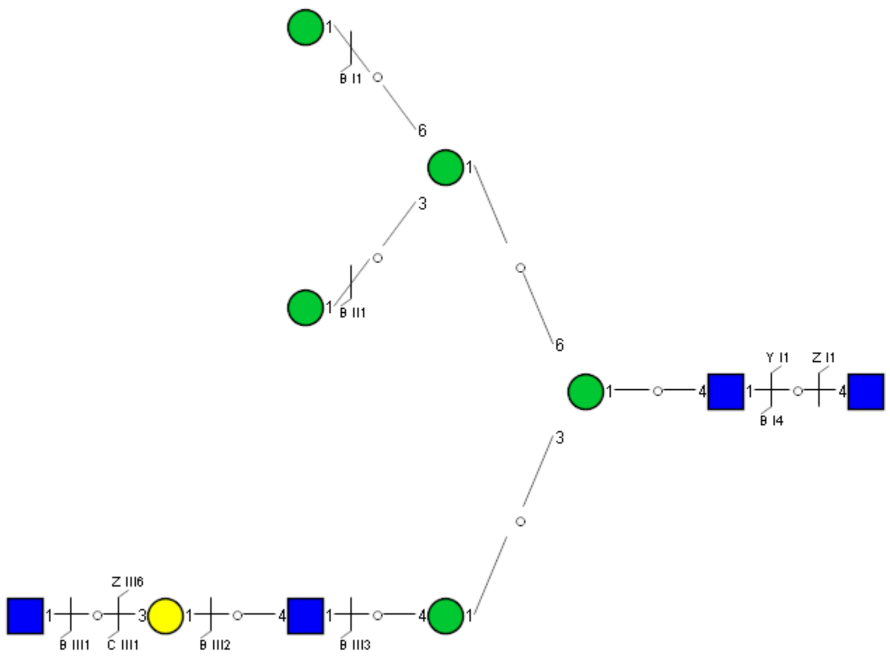

Spectrum Index: 40574(NoBHs:1)

#### Precursor iEF Map

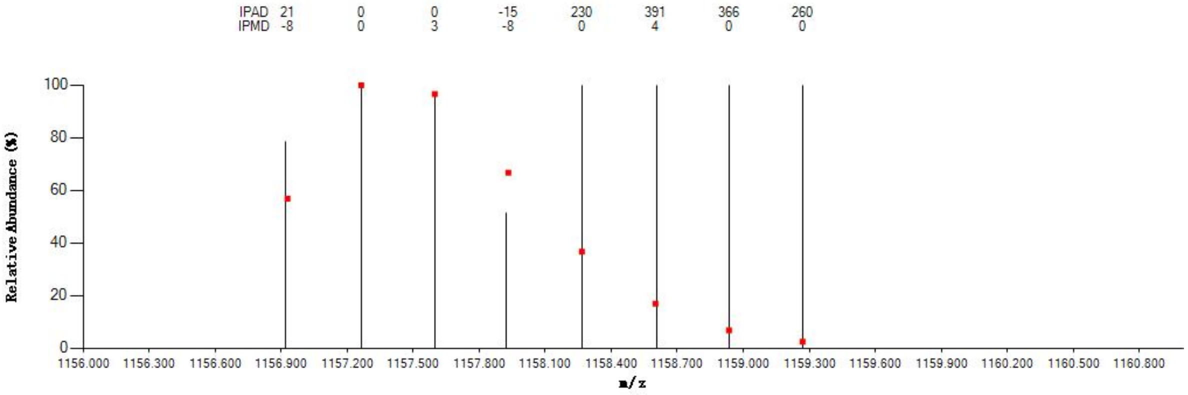


#### Annotated MS/MS Spectrum

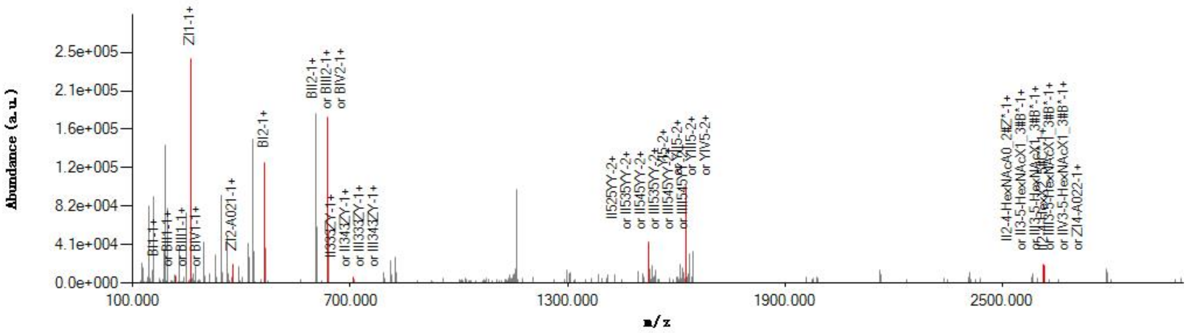


#### Graphical Fragmentation Map

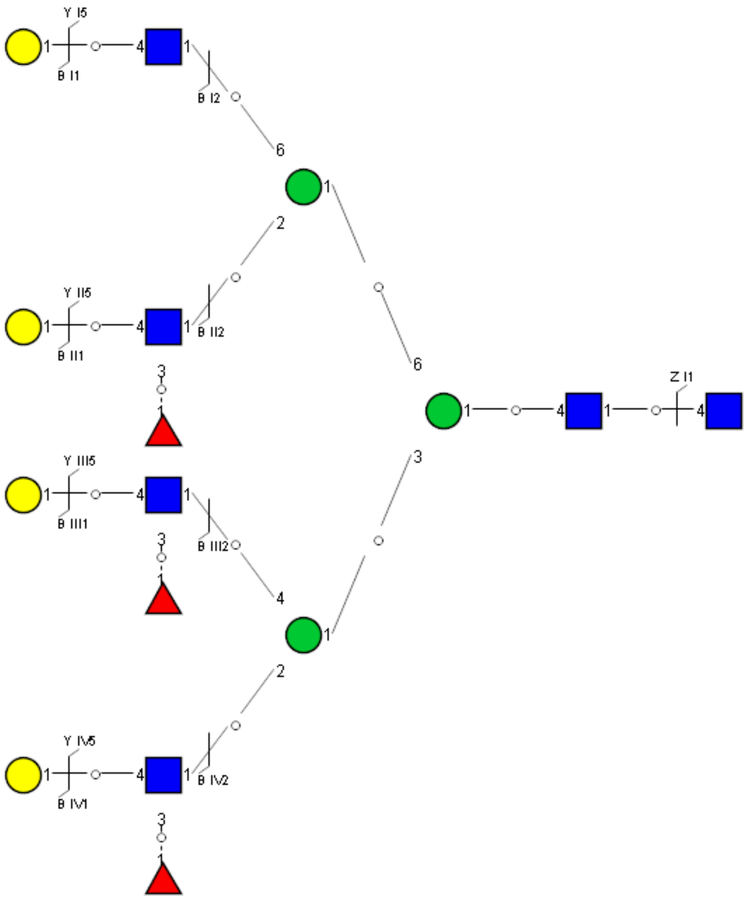

#### Spectrum Index: 40581(NoBHs:1)

##### Precursor iEF Map

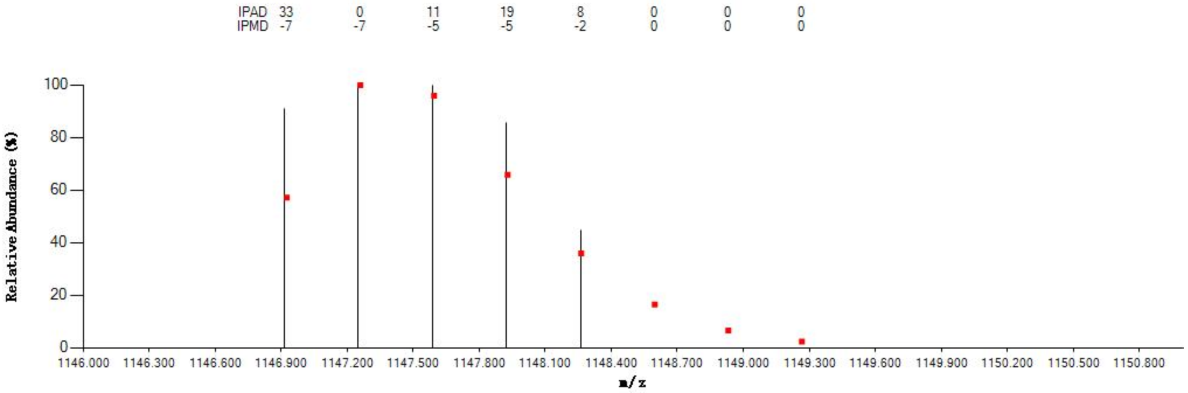


##### Annotated MS/MS Spectrum

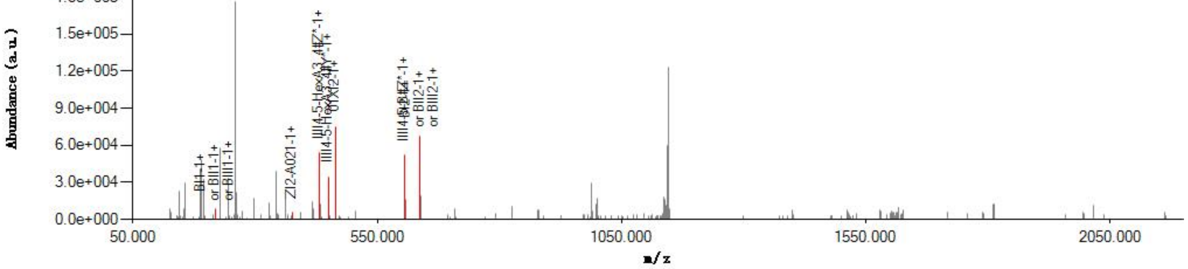


##### Graphical Fragmentation Map

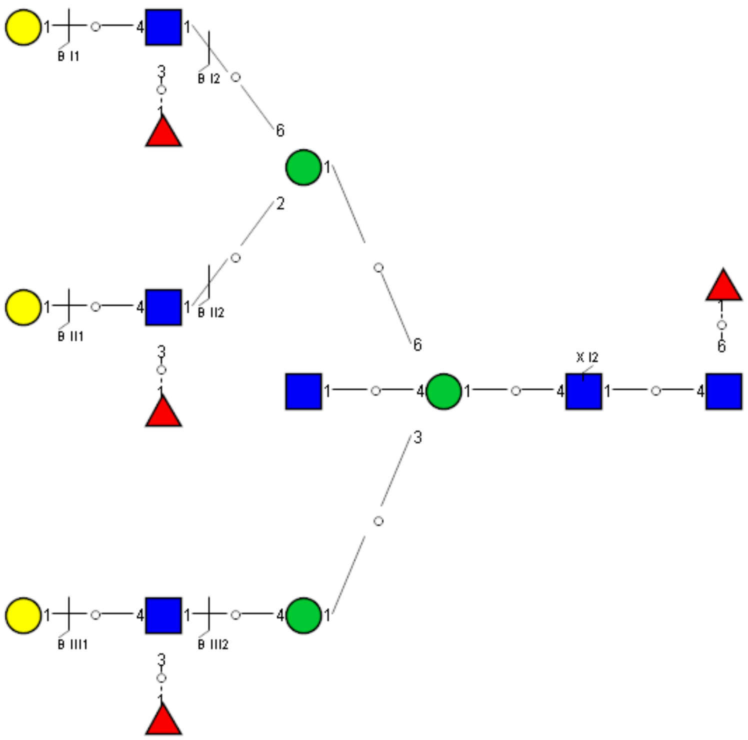

Spectrum Index: 40666(NoBHs:1)

#### Precursor iEF Map

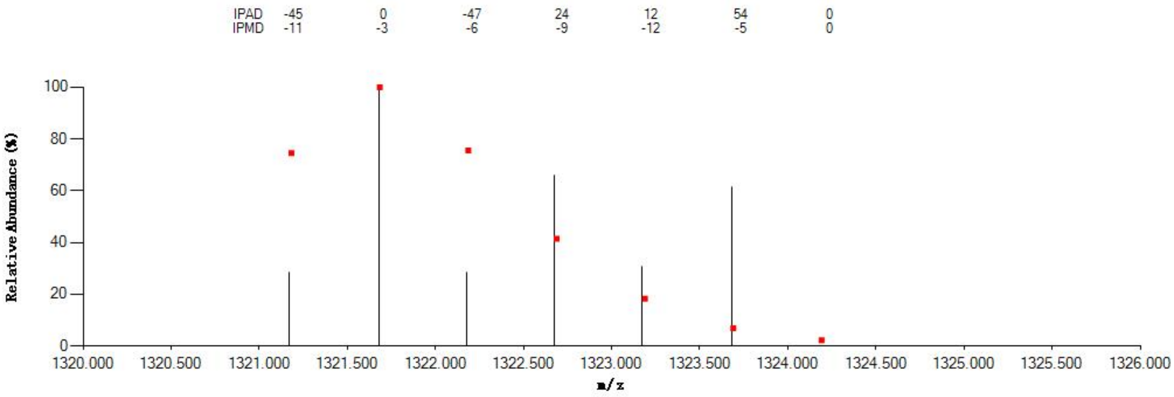


#### Annotated MS/MS Spectrum

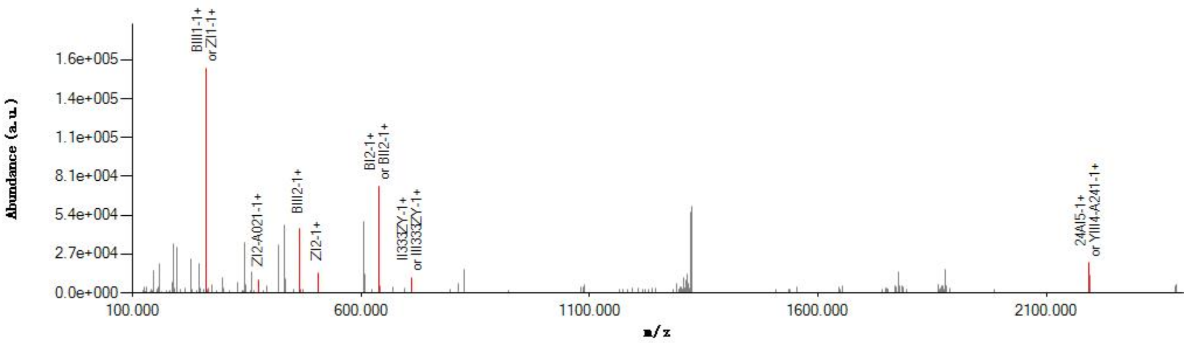


#### Graphical Fragmentation Map

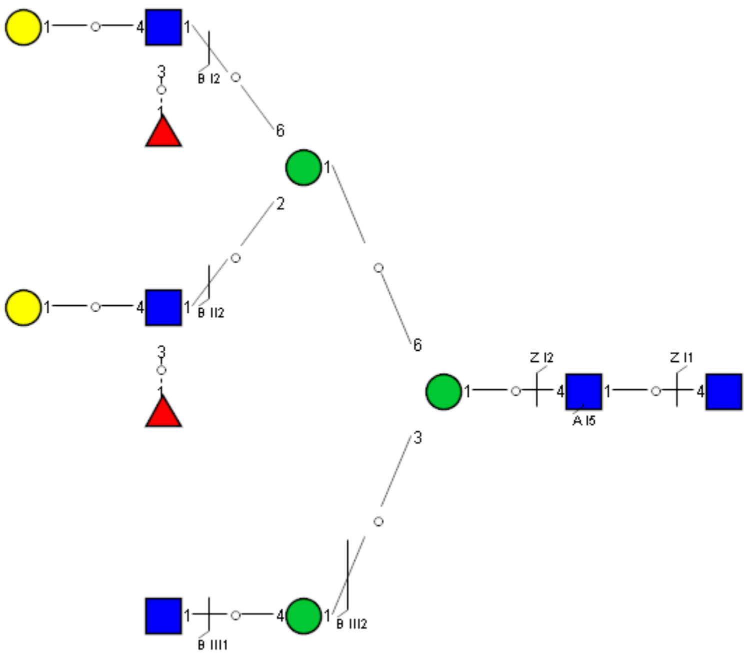

Spectrum Index: 40675(NoBHs:2)

#### Precursor iEF Map

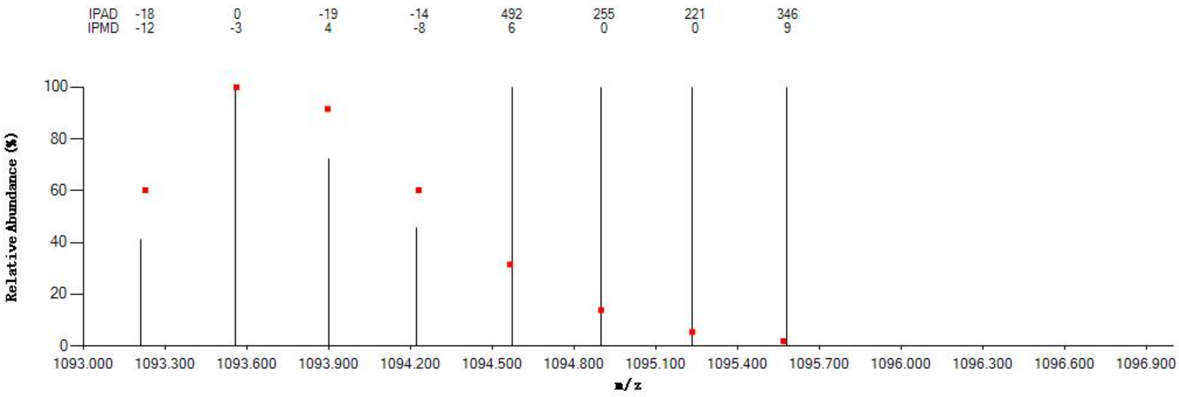

#### Annotated MS/MS Spectrum

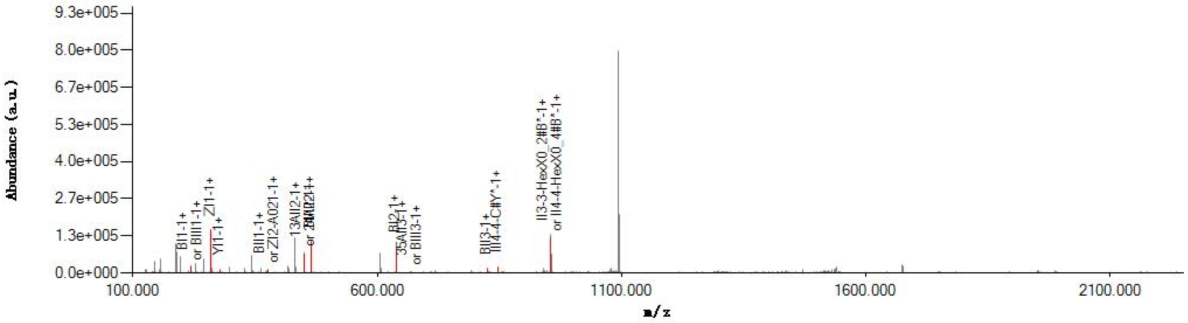

#### Graphical Fragmentation Map

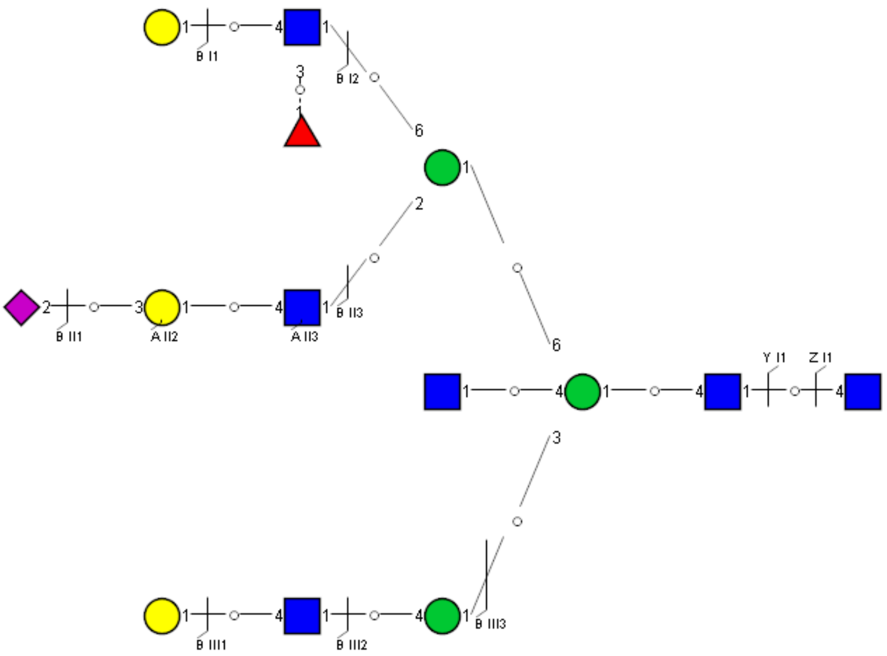

Spectrum Index: 40675(NoBHs:2)

#### Precursor iEF Map

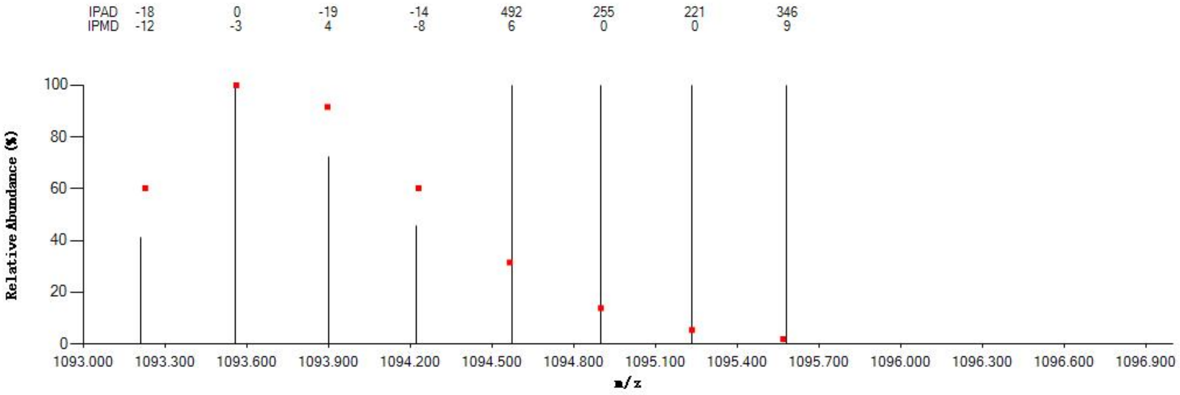


#### Annotated MS/MS Spectrum

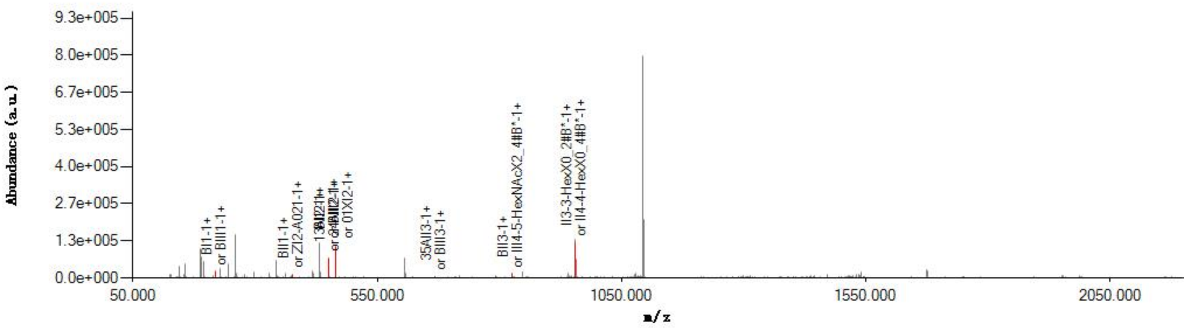


#### Graphical Fragmentation Map

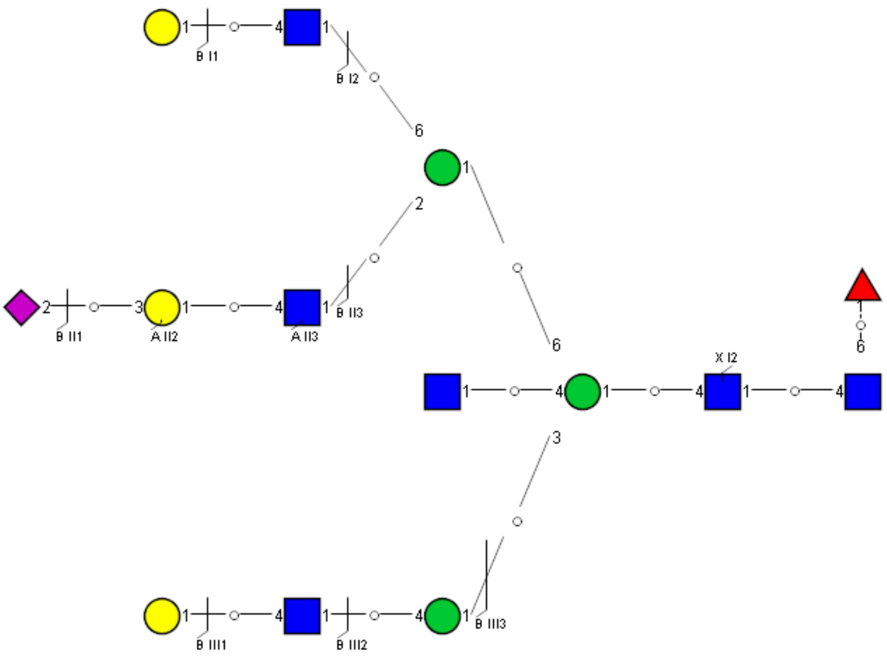

Spectrum Index: 40681(NoBHs:1)

#### Precursor iEF Map

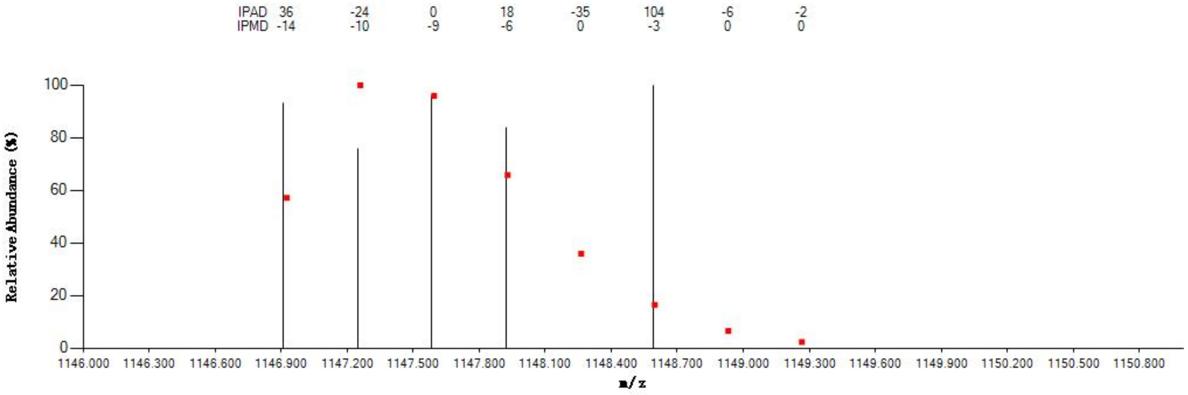

#### Annotated MS/MS Spectrum

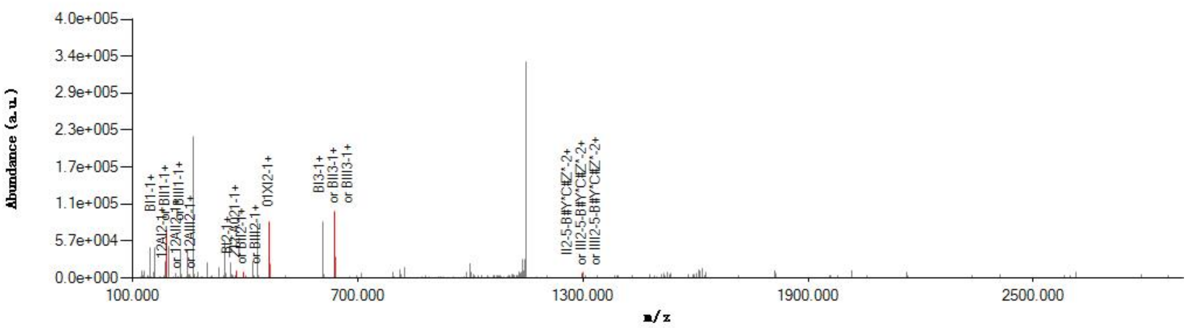

#### Graphical Fragmentation Map

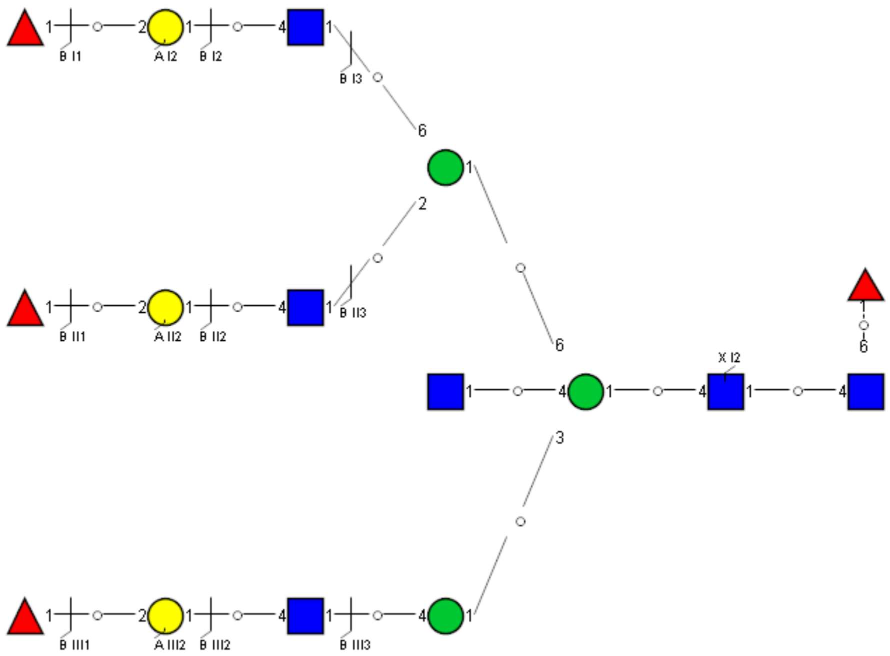

Spectrum Index: 40682(NoBHs:1)

#### Precursor iEF Map

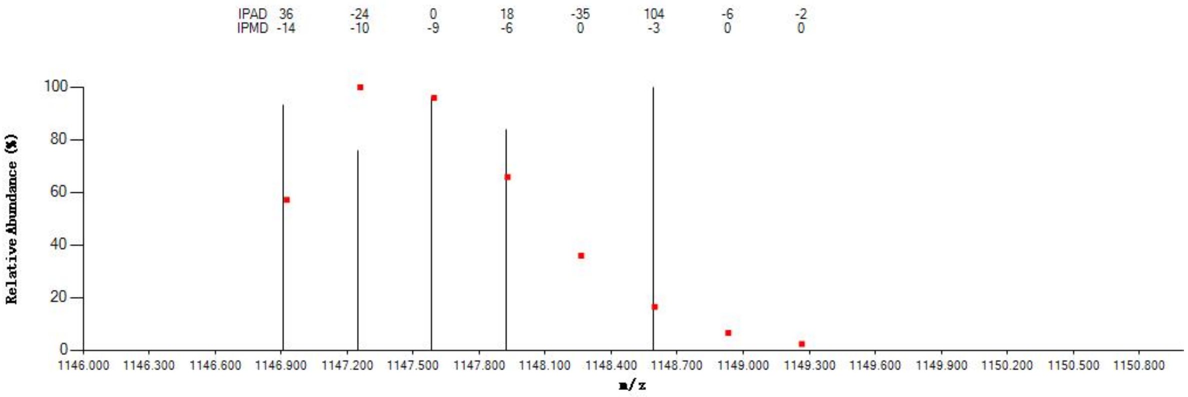


#### Annotated MS/MS Spectrum

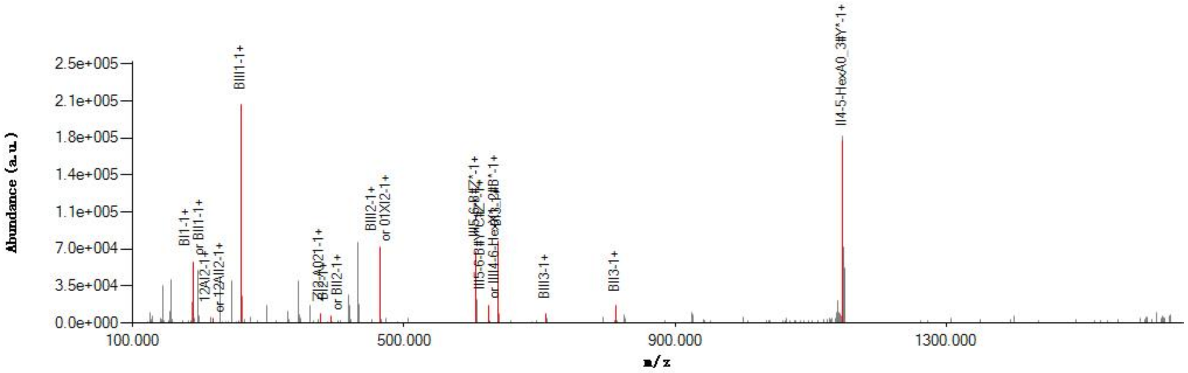


#### Graphical Fragmentation Map

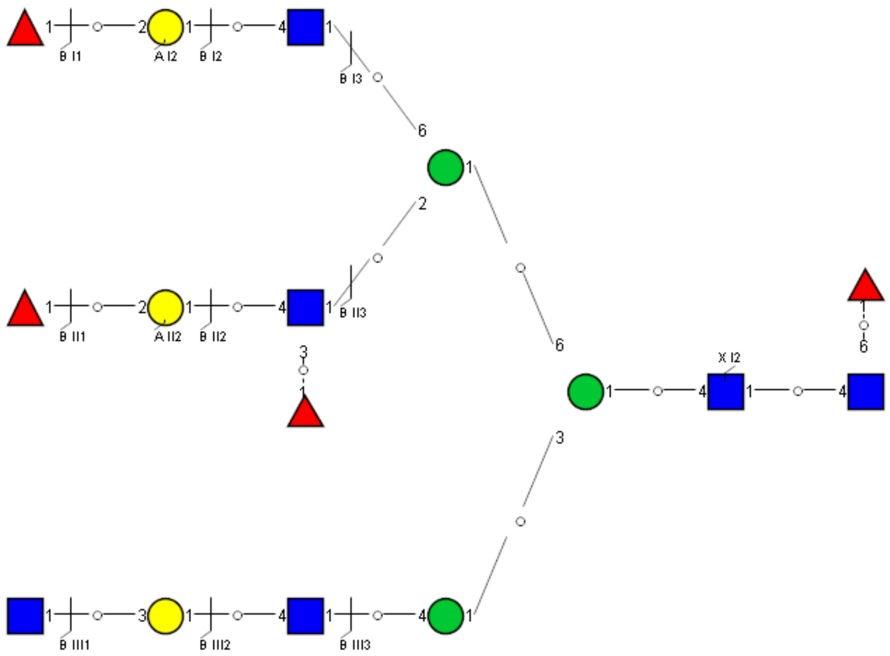

### Spectrum Index: 40757(NoBHs:3)

#### Precursor iEF Map

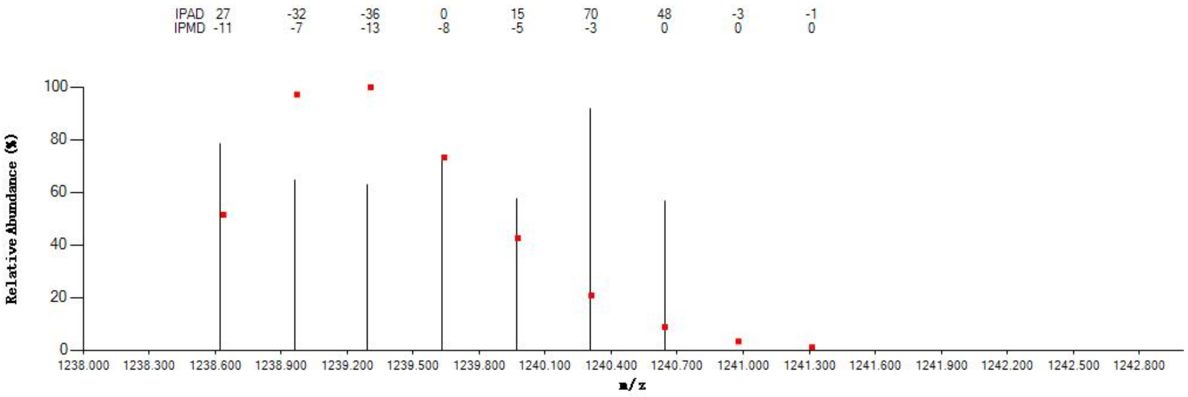


#### Annotated MS/MS Spectrum

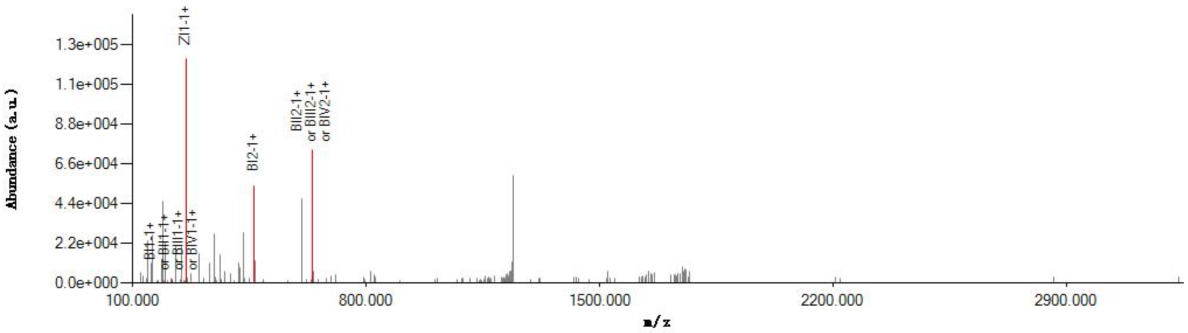


#### Graphical Fragmentation Map

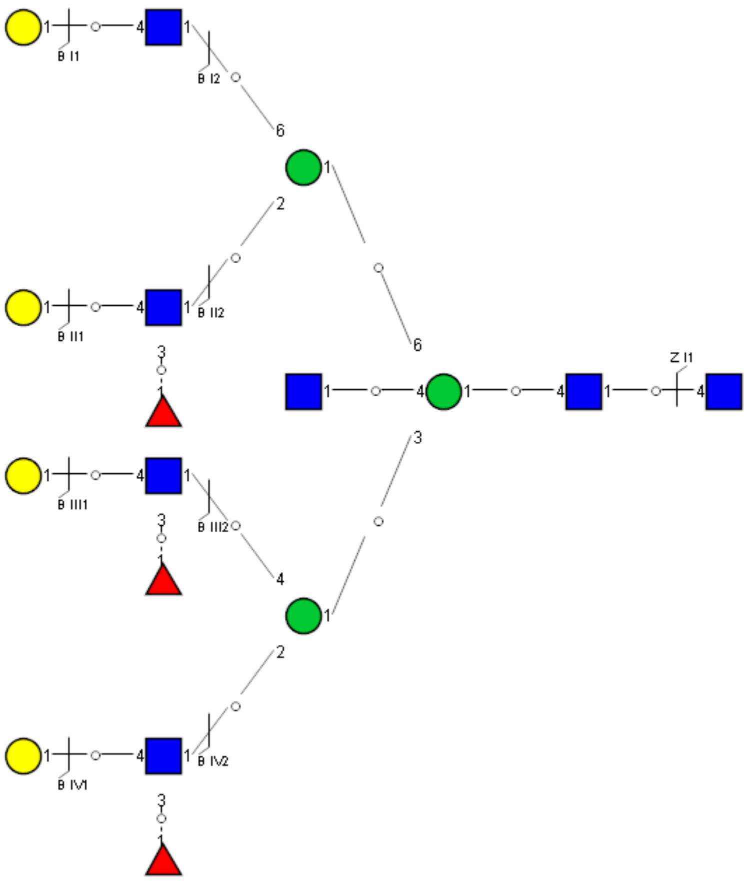

Spectrum Index: 40757(NoBHs:3)

#### Precursor iEF Map

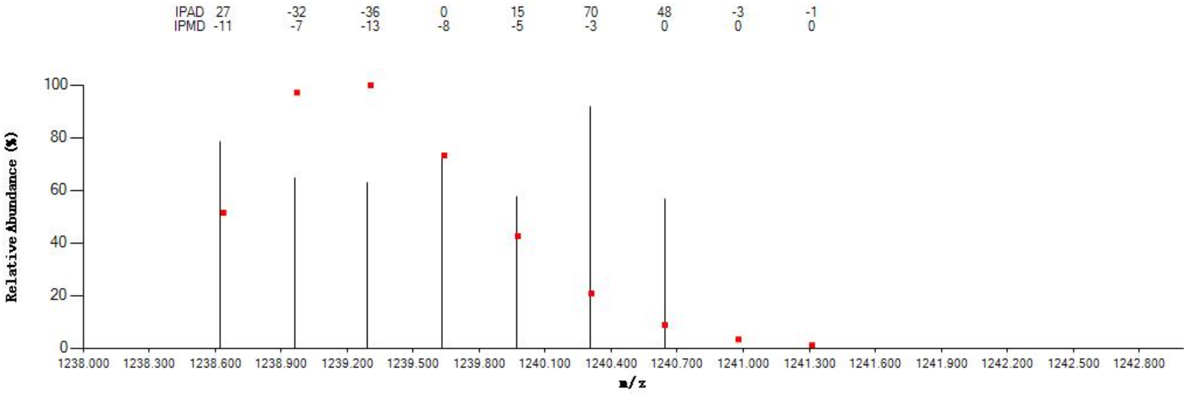


#### Annotated MS/MS Spectrum

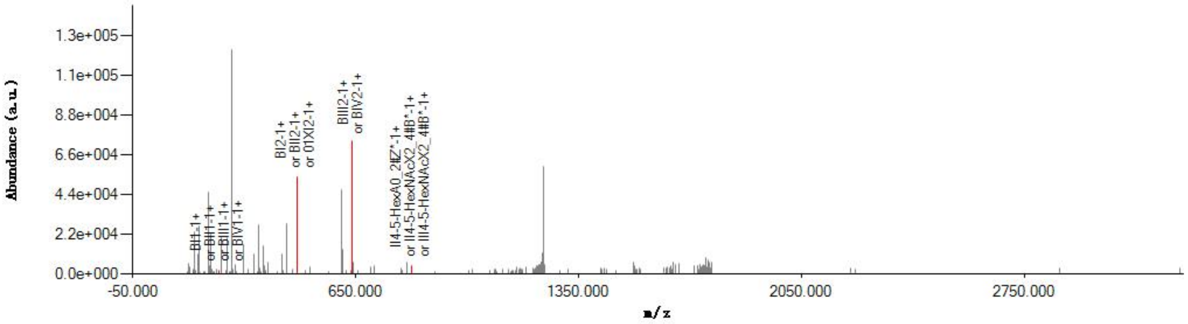


#### Graphical Fragmentation Map

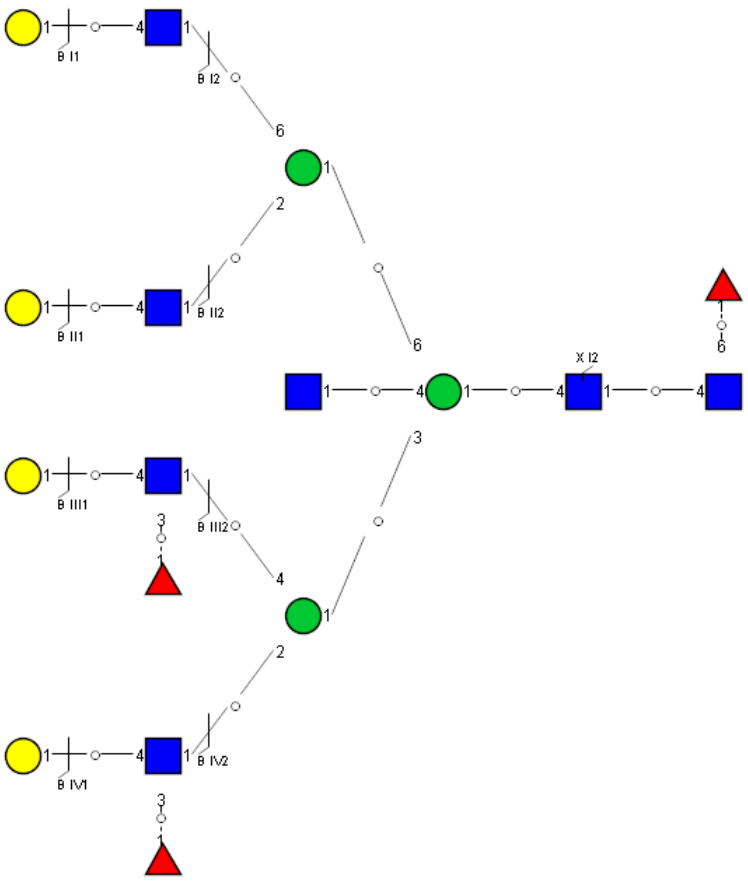

Spectrum Index: 40757(NoBHs:3)

Precursor iEF Map

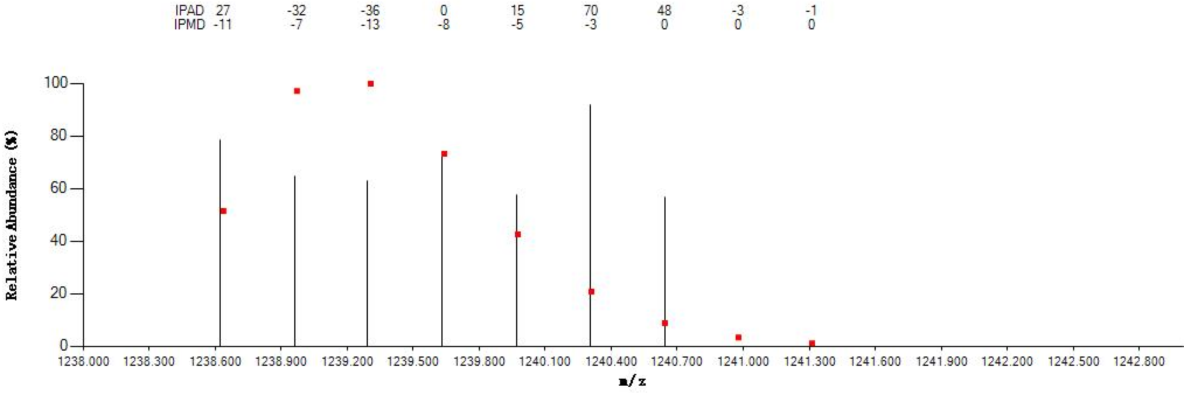


Annotated MS/MS Spectrum

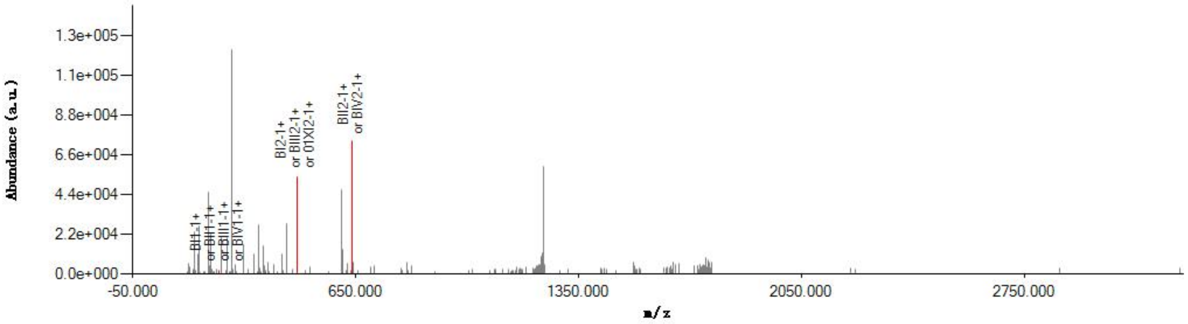


Graphical Fragmentation Map

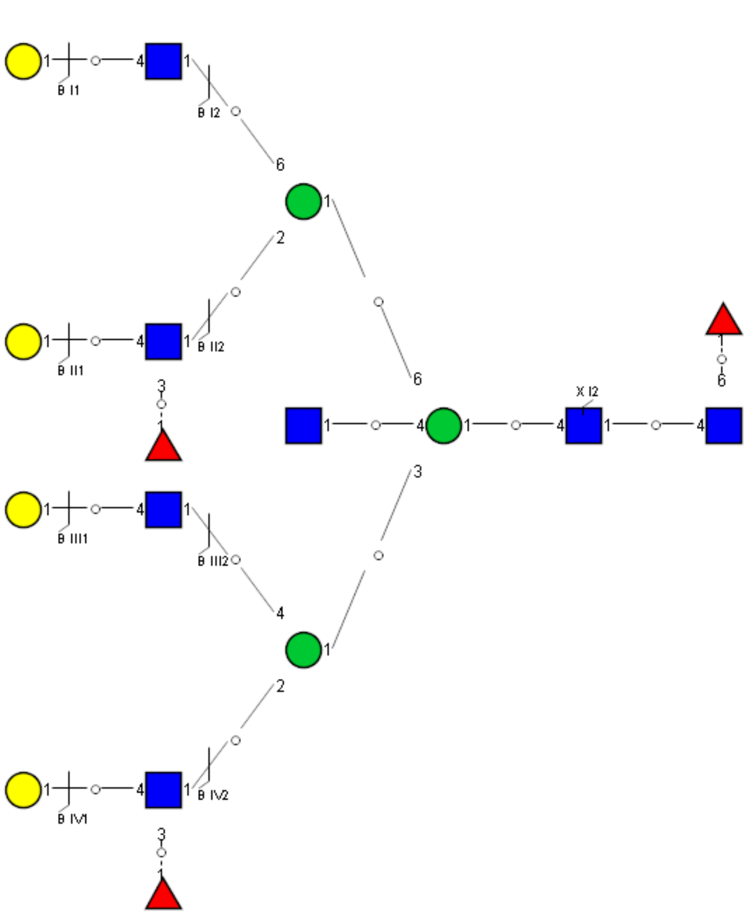

Spectrum Index: 40841(NoBHs:1)

#### Precursor iEF Map

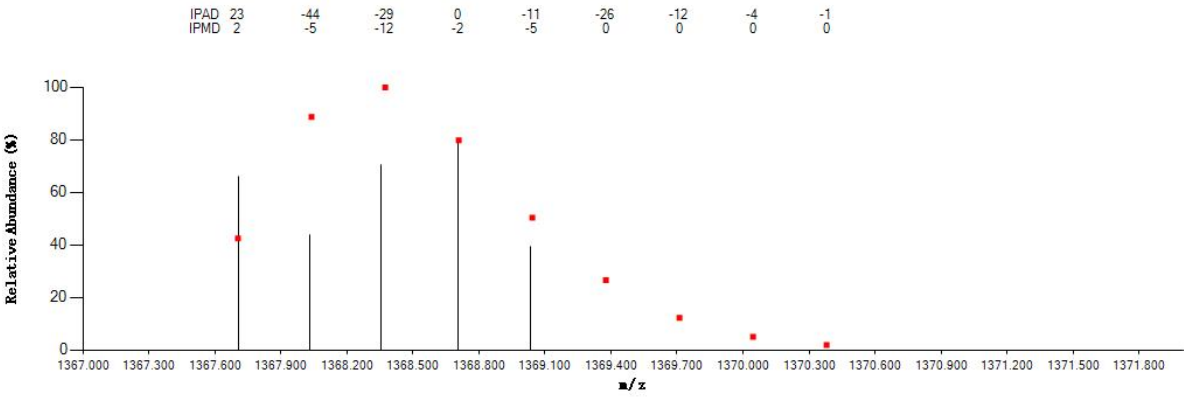


#### Annotated MS/MS Spectrum

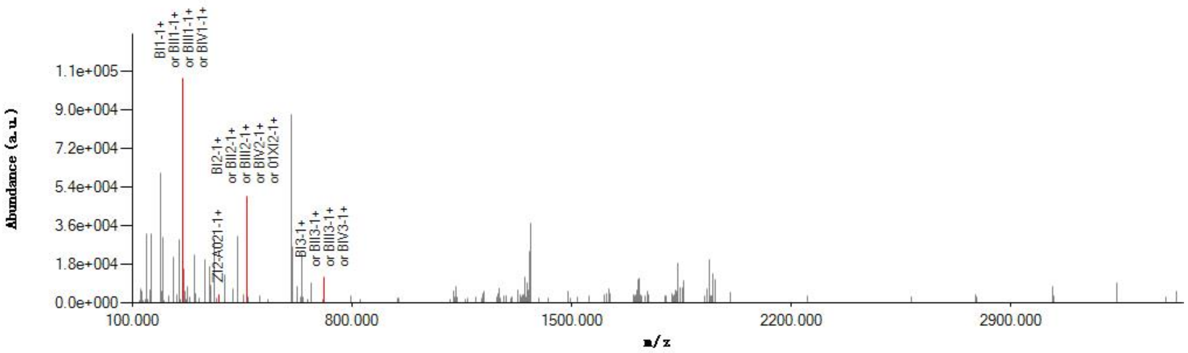


#### Graphical Fragmentation Map

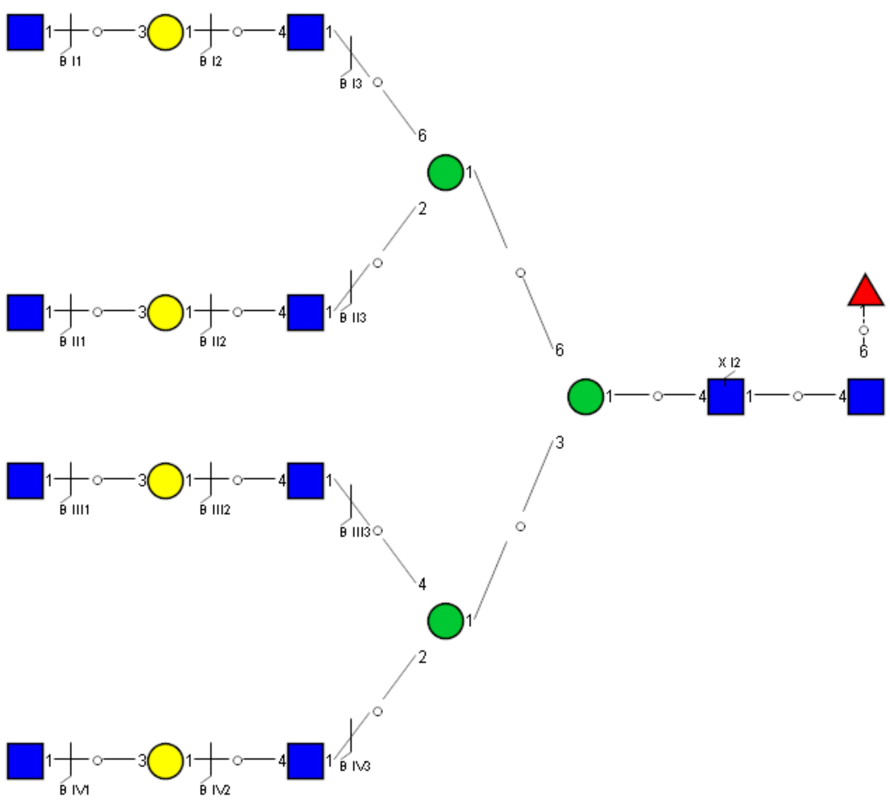

Spectrum Index: 40843(NoBHs:1)

#### Precursor iEF Map

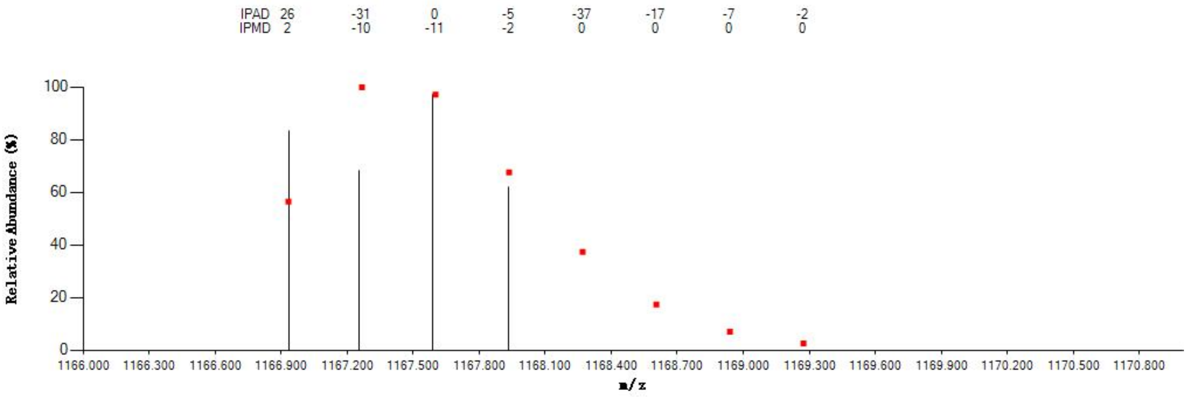


#### Annotated MS/MS Spectrum

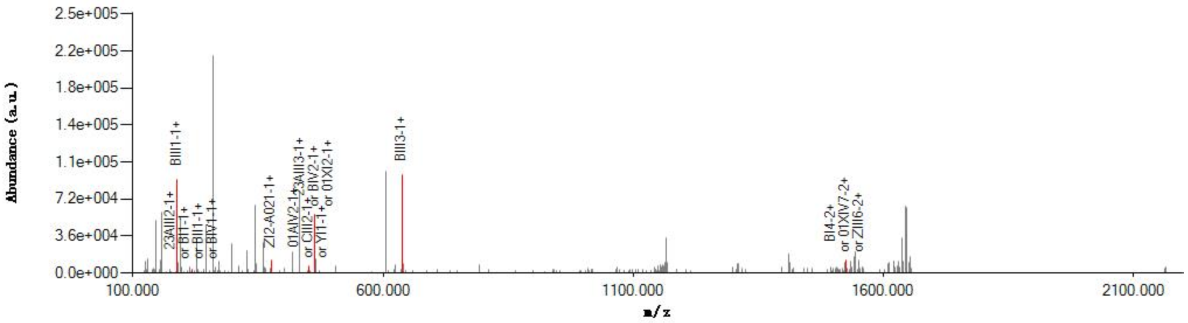


#### Graphical Fragmentation Map

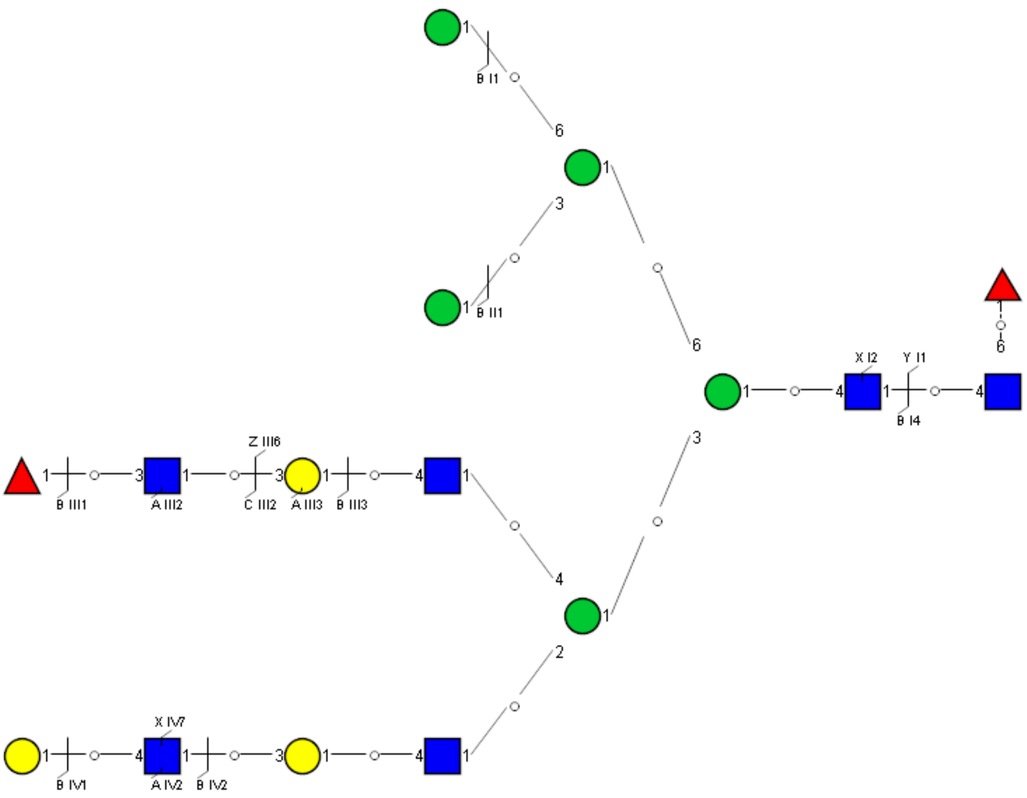

Spectrum Index: 40876(NoBHs:1)

#### Precursor iEF Map

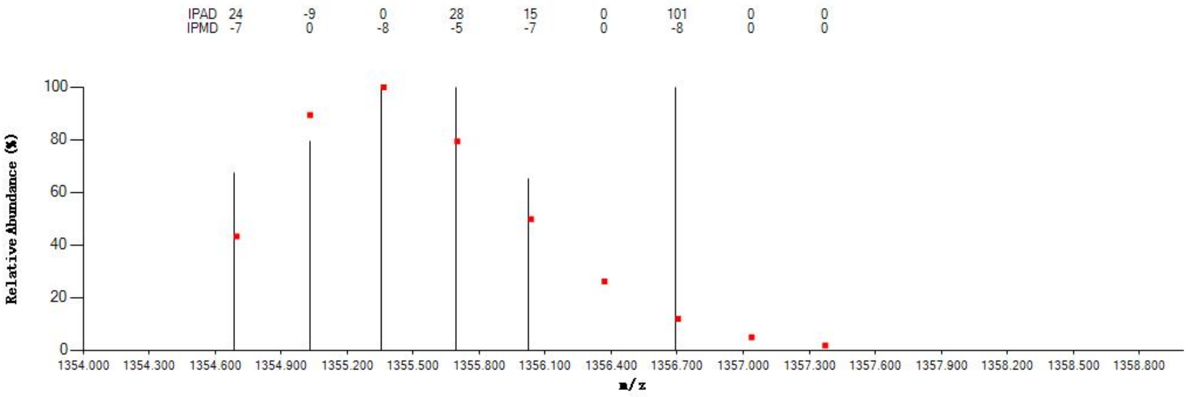


#### Annotated MS/MS Spectrum

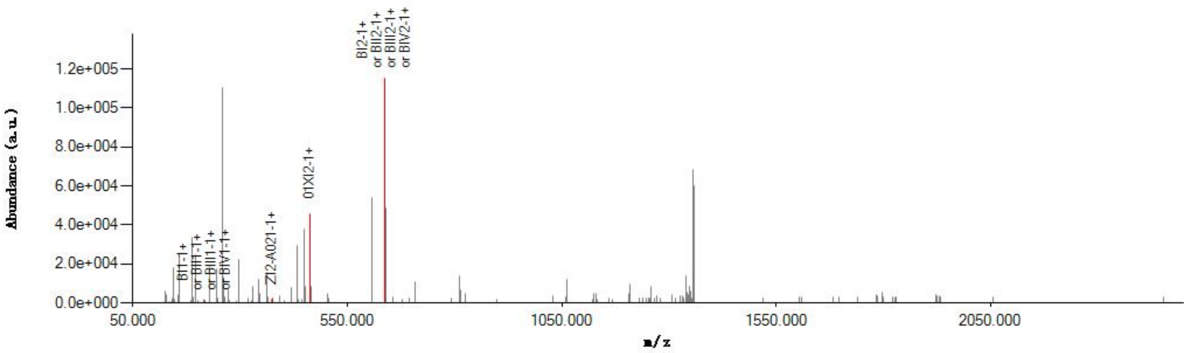


#### Graphical Fragmentation Map

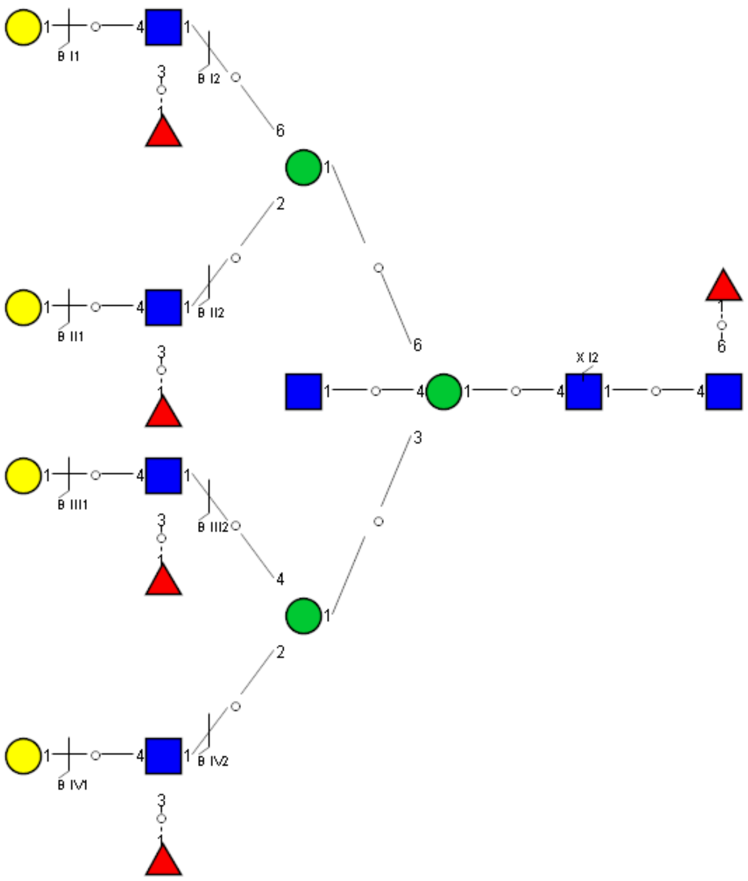

Spectrum Index: 40881(NoBHs:1)

#### Precursor iEF Map

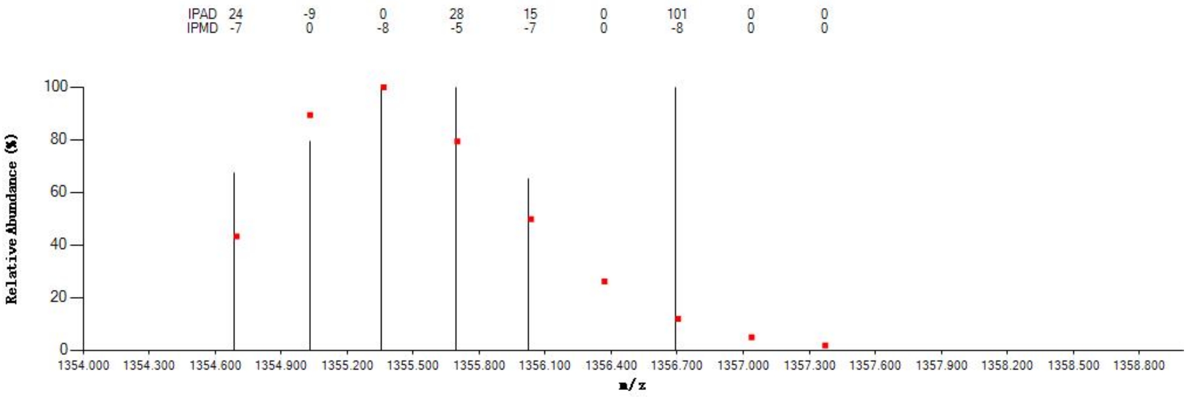


#### Annotated MS/MS Spectrum

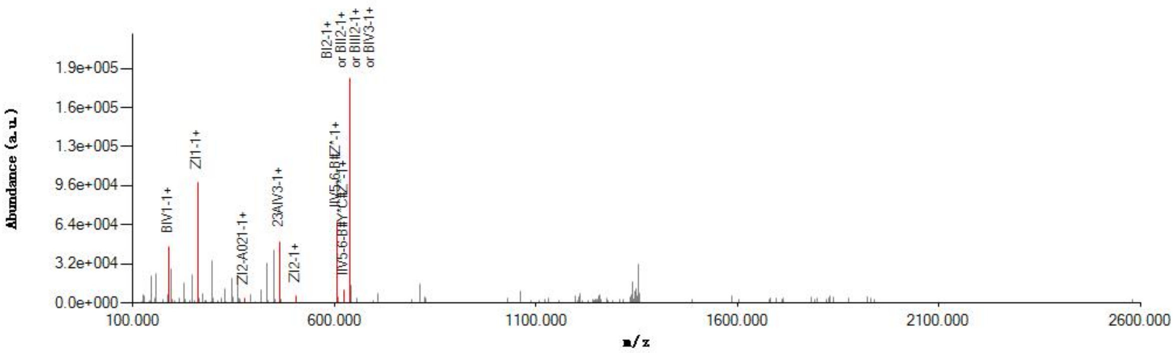


#### Graphical Fragmentation Map

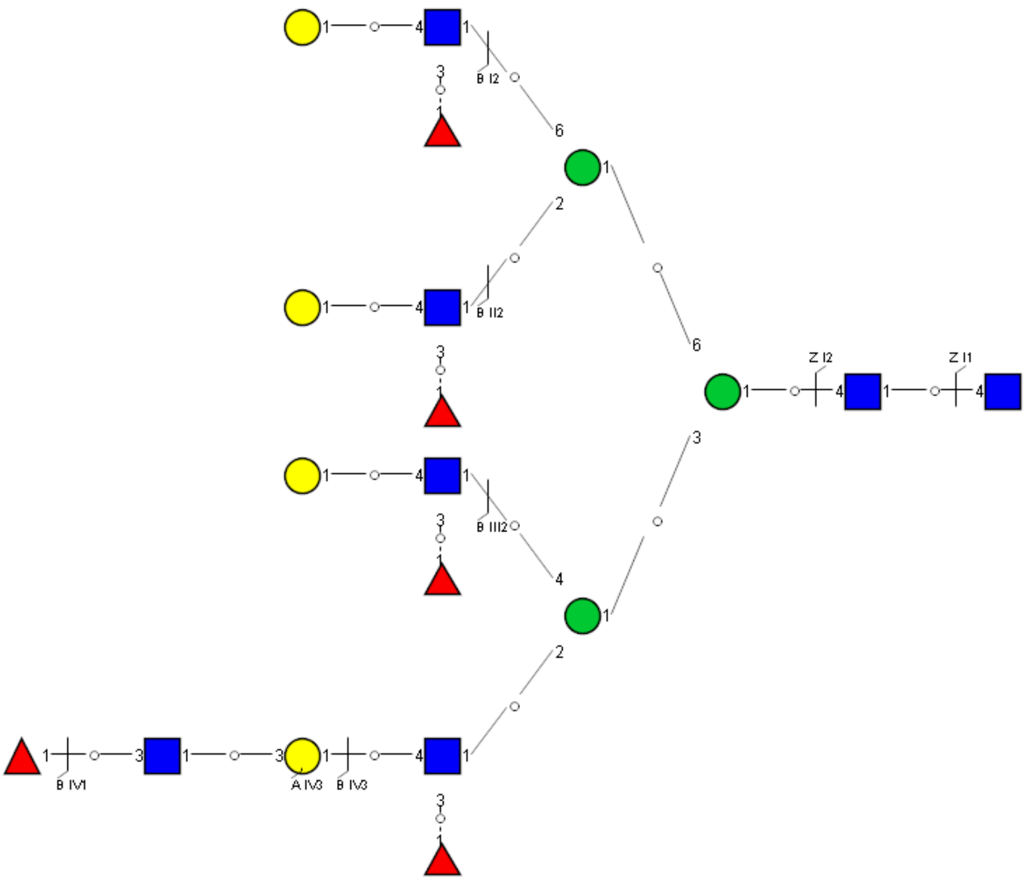

Spectrum Index: 41095(NoBHs:1)

#### Precursor iEF Map

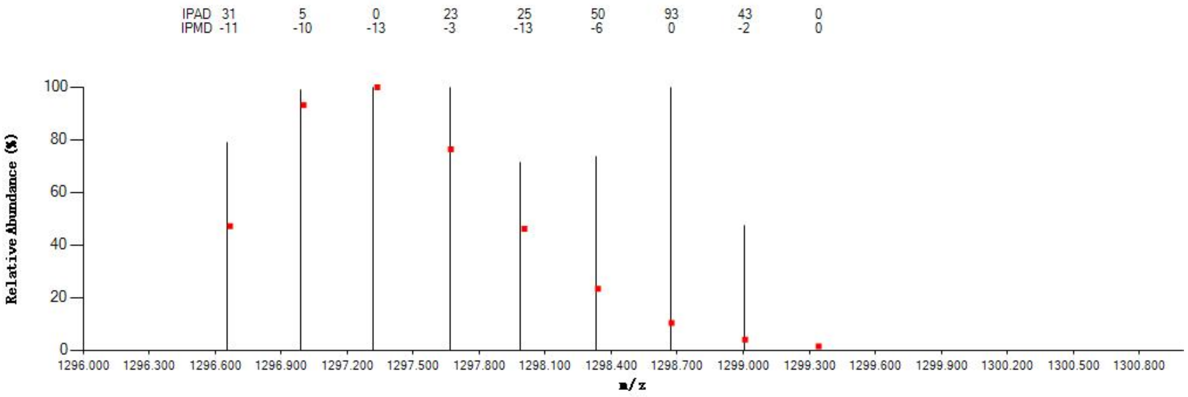


#### Annotated MS/MS Spectrum

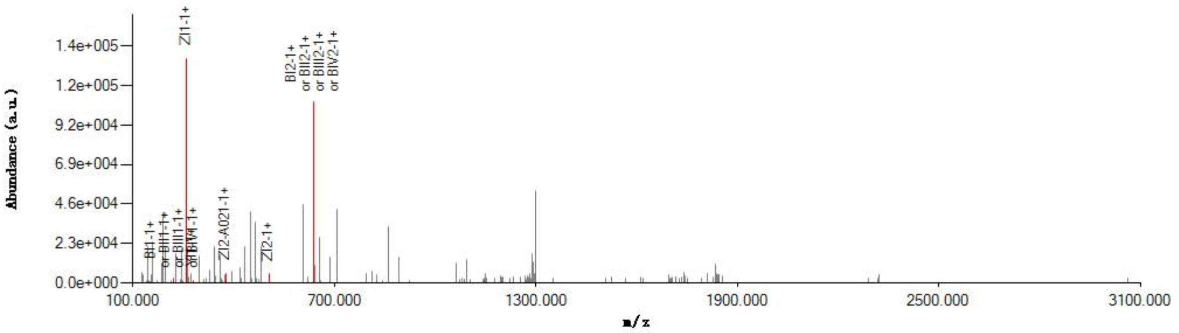


#### Graphical Fragmentation Map

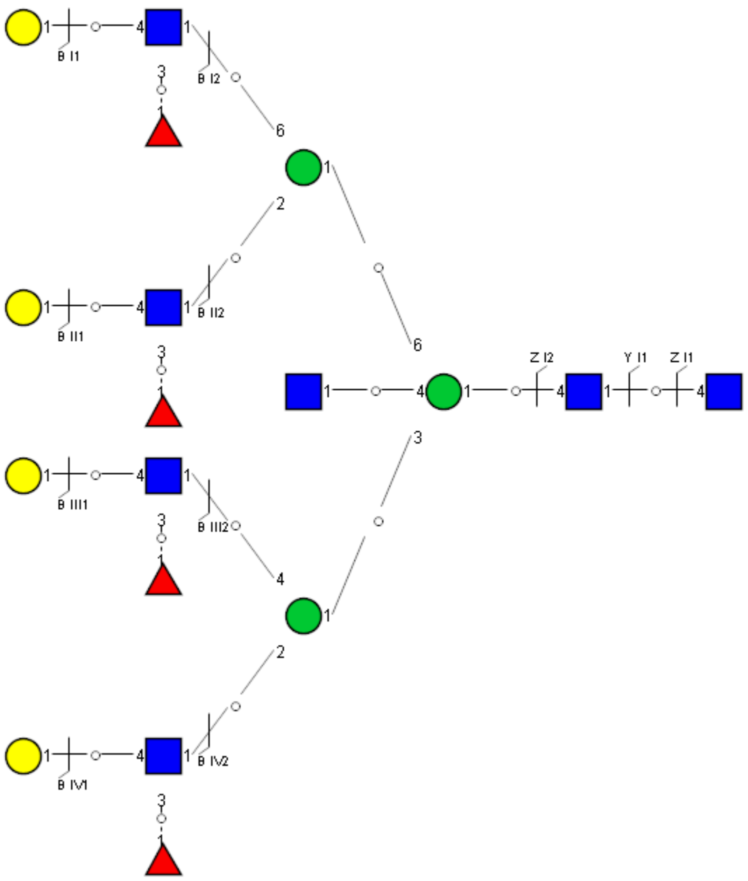

Spectrum Index: 41168(NoBHs:2)

#### Precursor iEF Map

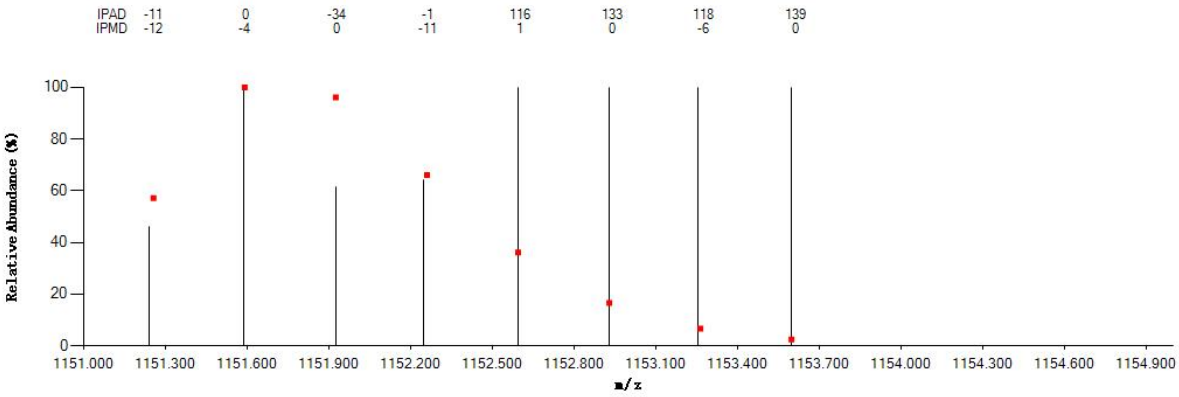


#### Annotated MS/MS Spectrum

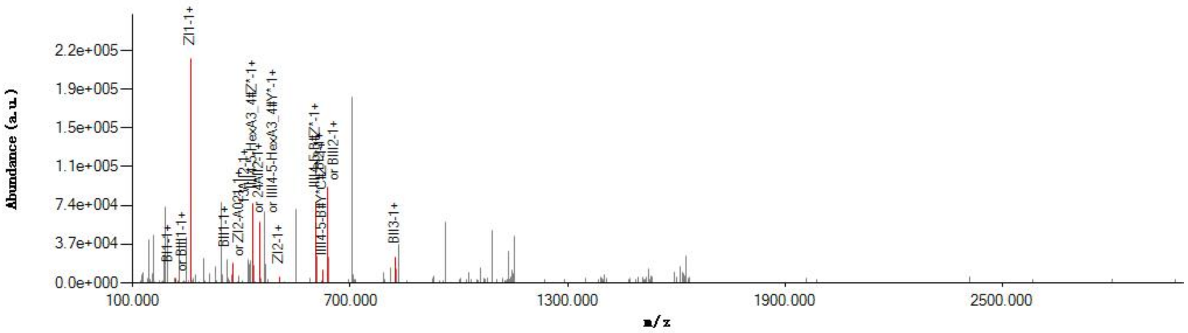


#### Graphical Fragmentation Map

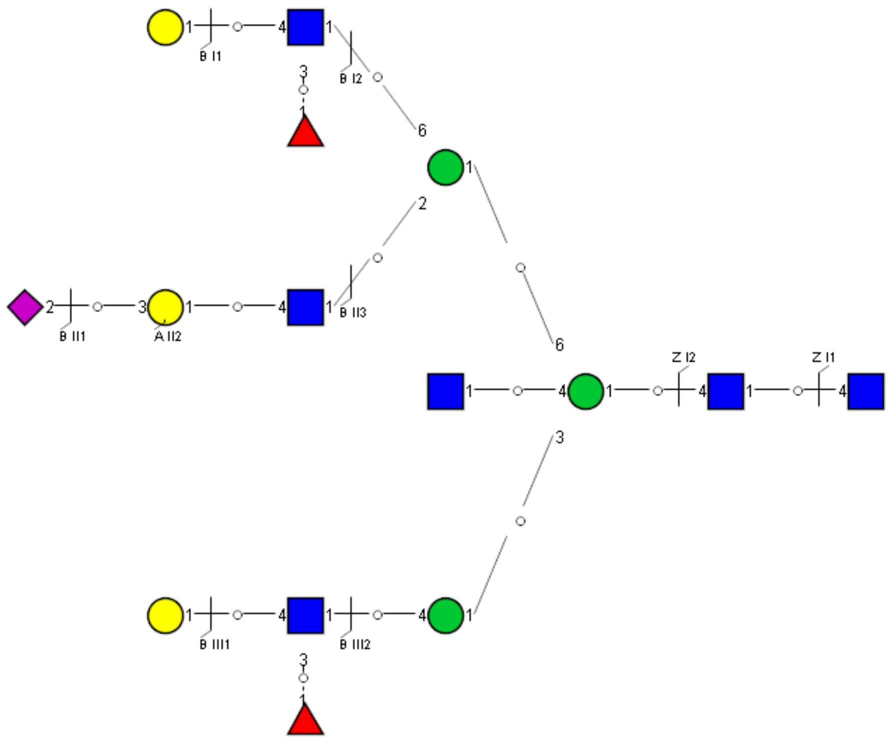

Spectrum Index: 41168(NoBHs:2)

#### Precursor iEF Map

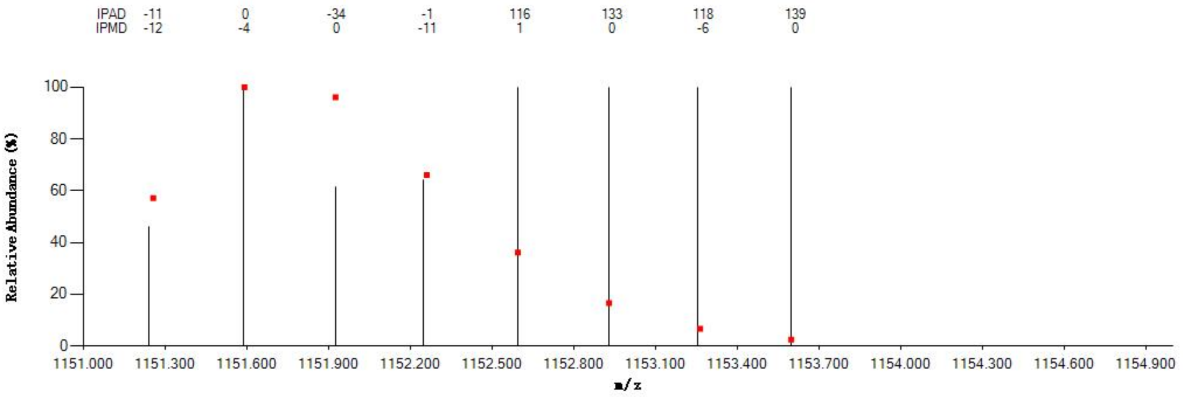


#### Annotated MS/MS Spectrum

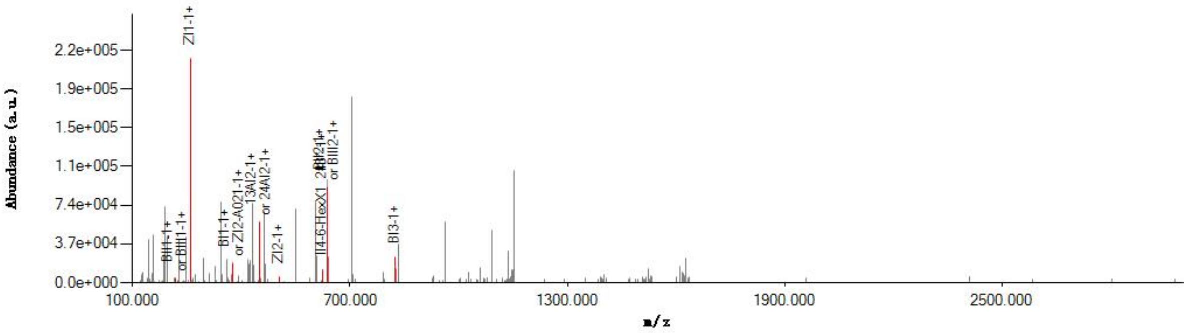


#### Graphical Fragmentation Map

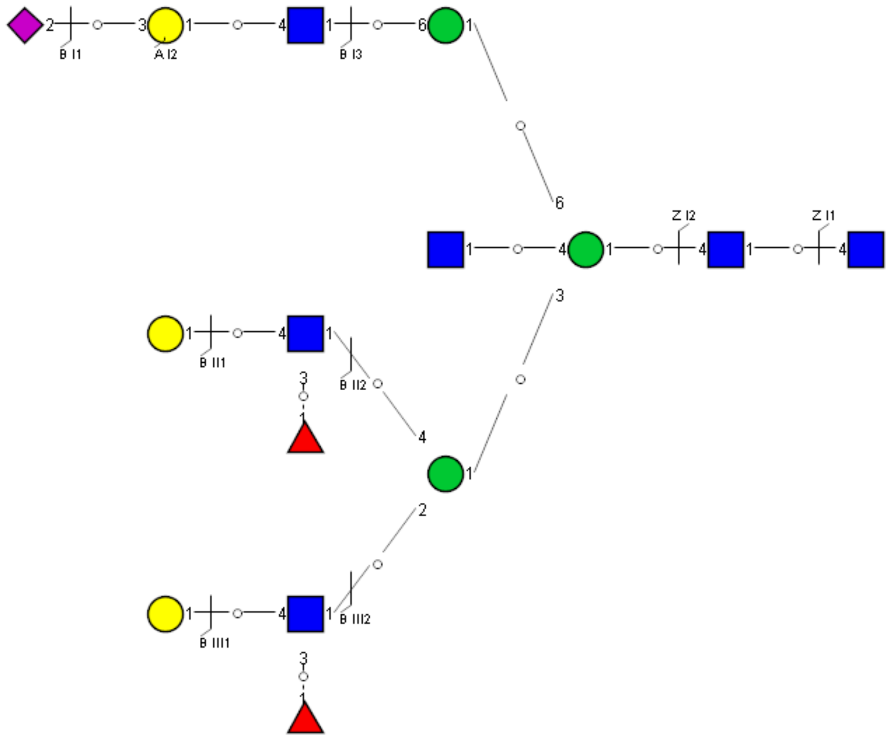

Spectrum Index: 41178(NoBHs:1)

#### Precursor iEF Map

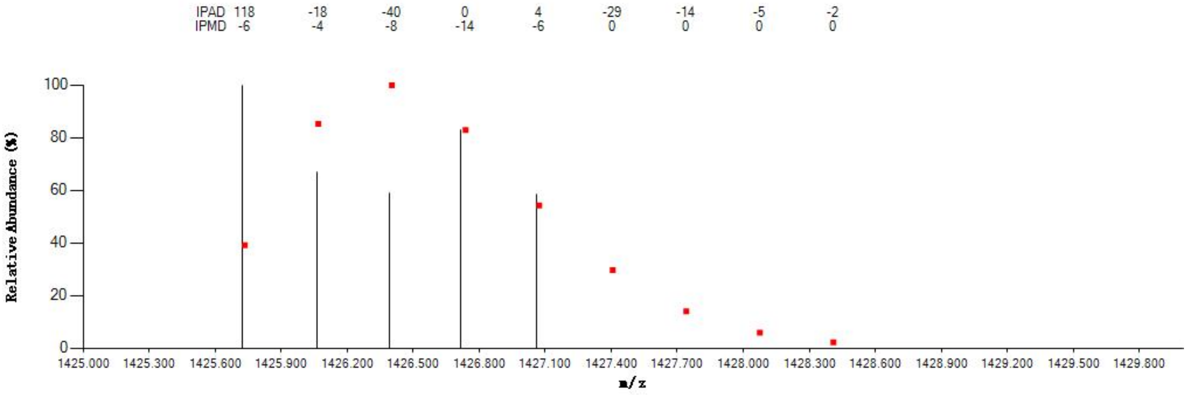

#### Annotated MS/MS Spectrum

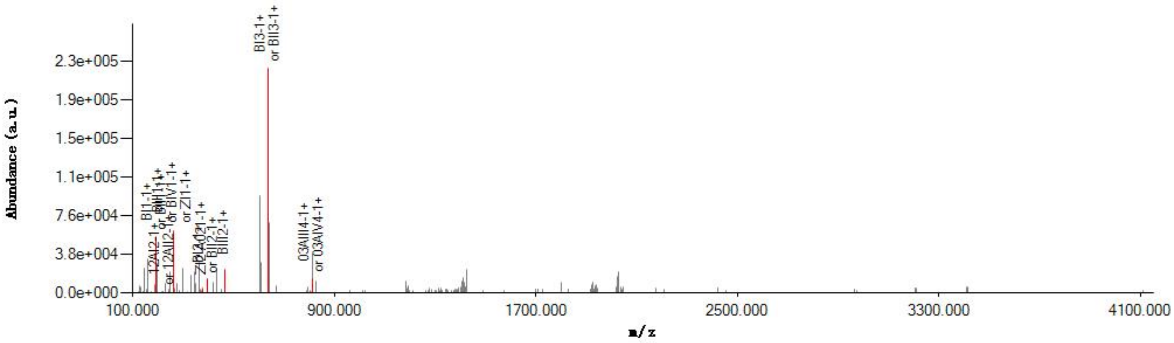

#### Graphical Fragmentation Map

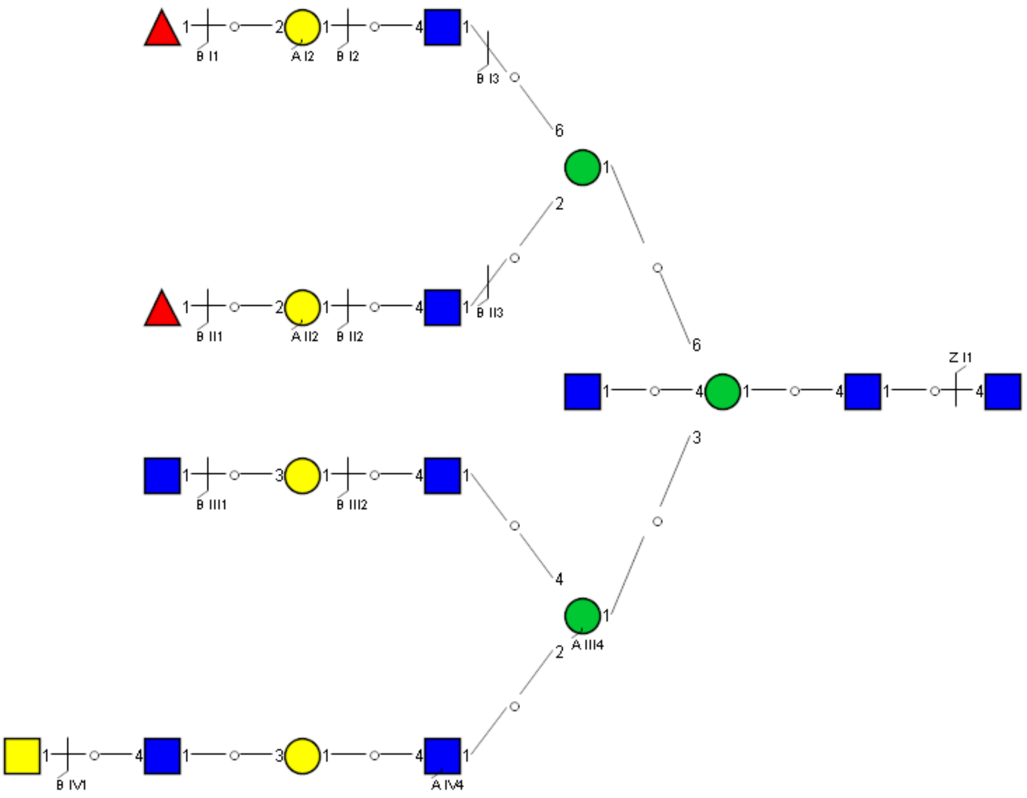

Spectrum Index: 41230(NoBHs:1)

#### Precursor iEF Map

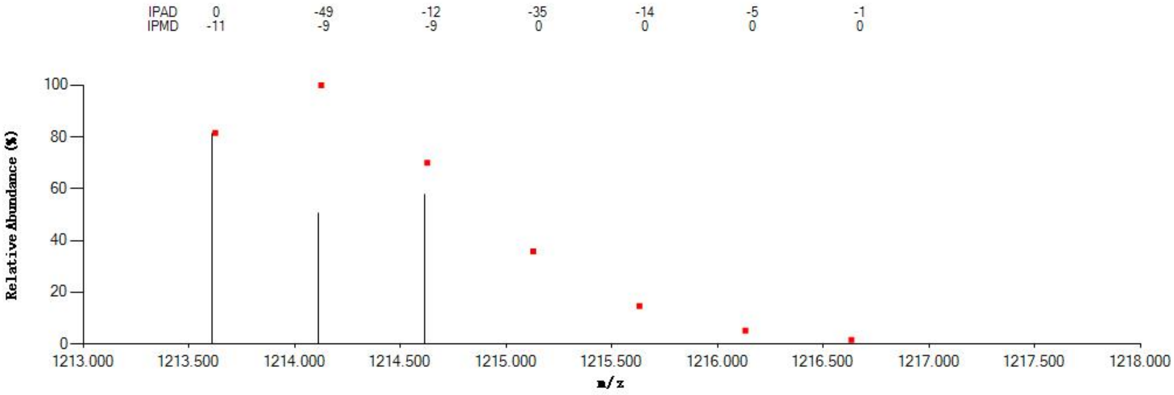


#### Annotated MS/MS Spectrum

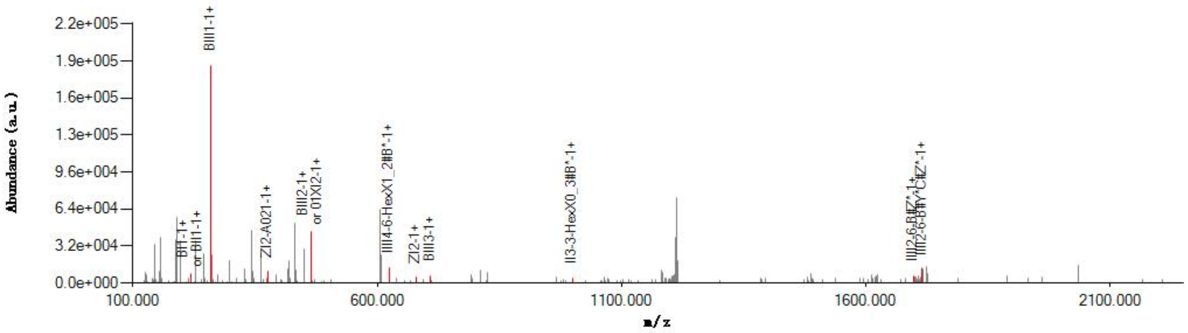


#### Graphical Fragmentation Map

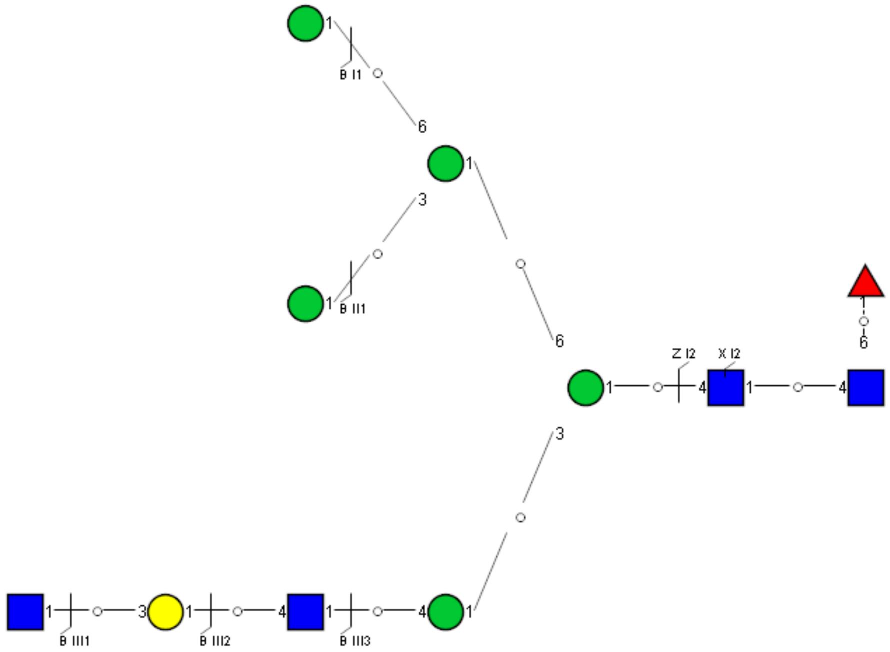

Spectrum Index: 41258(NoBHs:1)

#### Precursor iEF Map

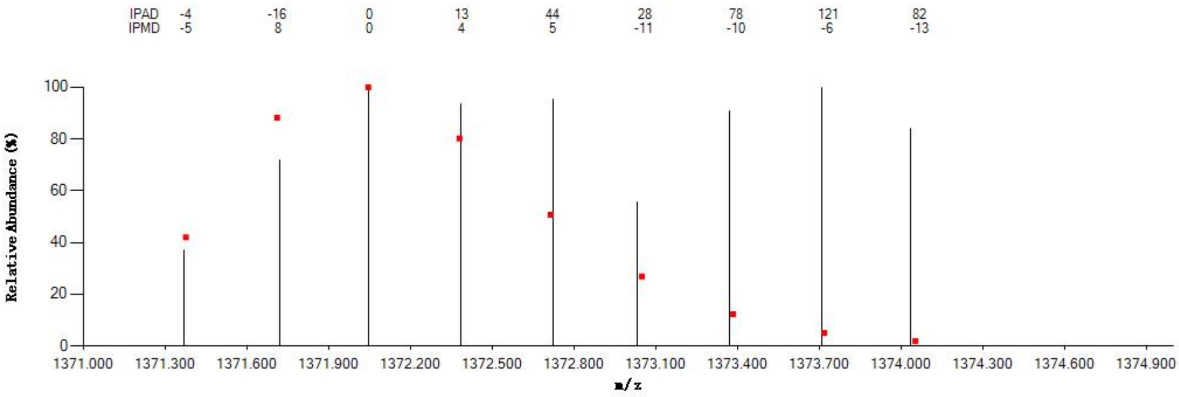


#### Annotated MS/MS Spectrum

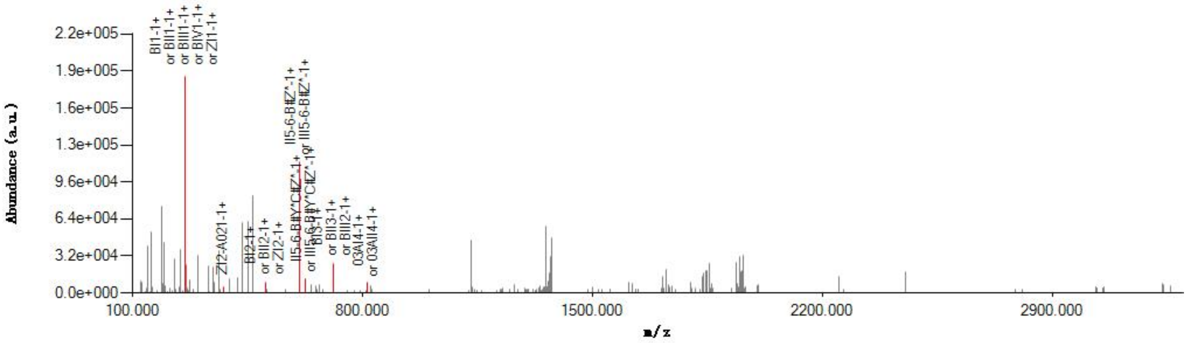


#### Graphical Fragmentation Map

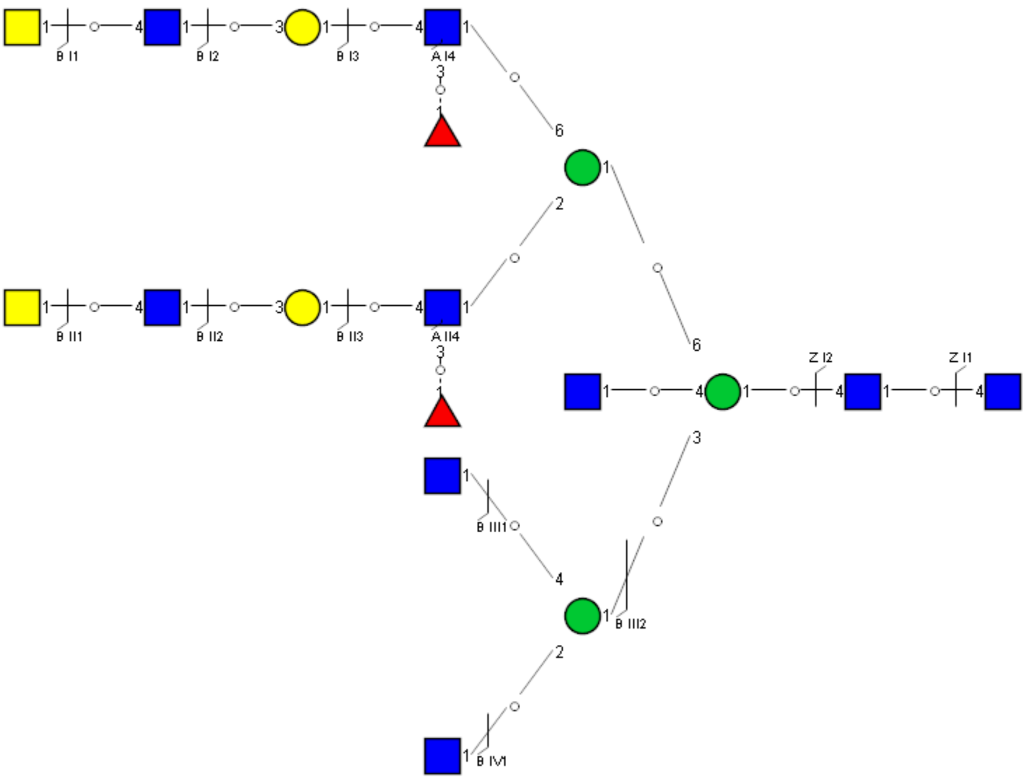

Spectrum Index: 41301(NoBHs:1)

#### Precursor iEF Map

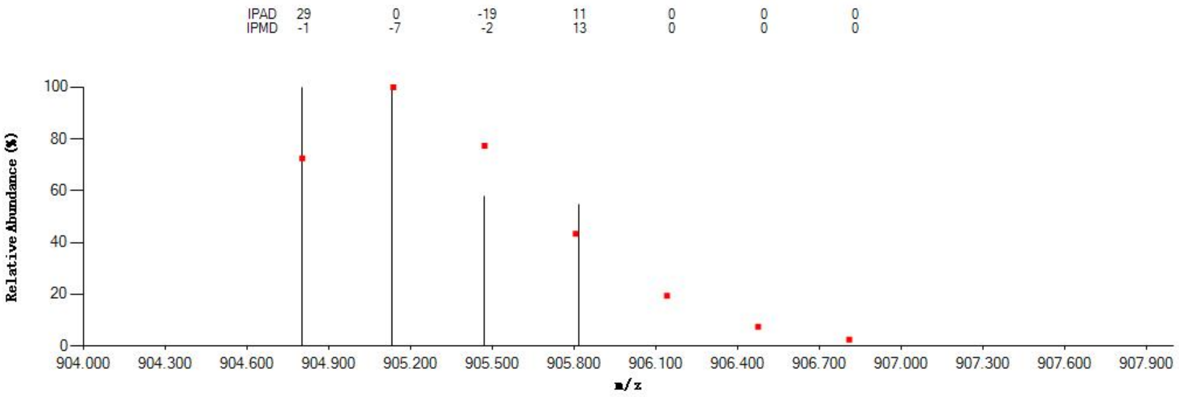

#### Annotated MS/MS Spectrum

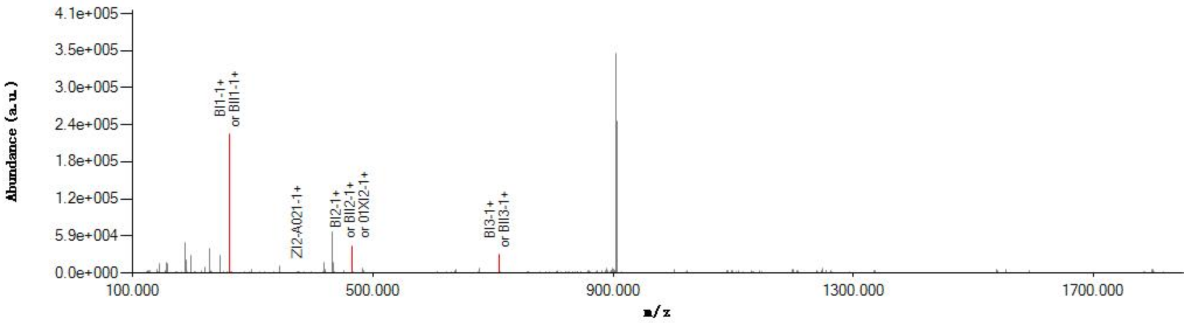

#### Graphical Fragmentation Map

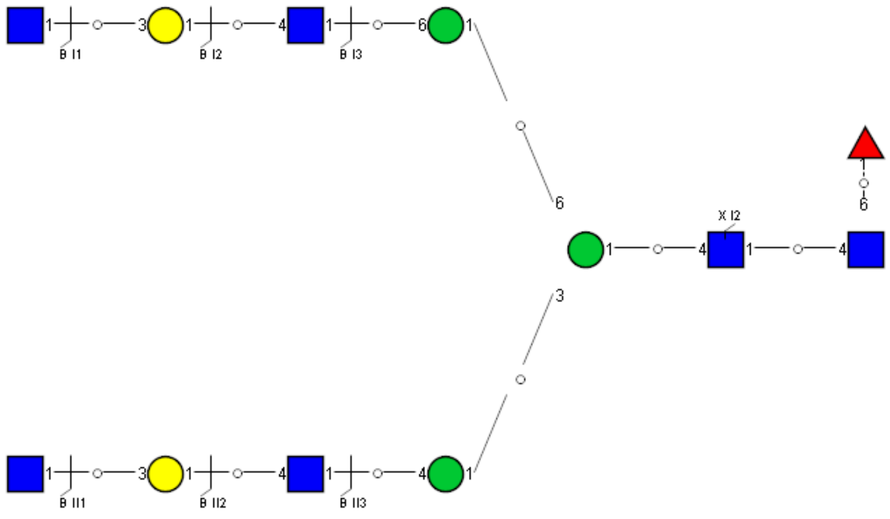

Spectrum Index: 41690(NoBHs:1)

#### Precursor iEF Map

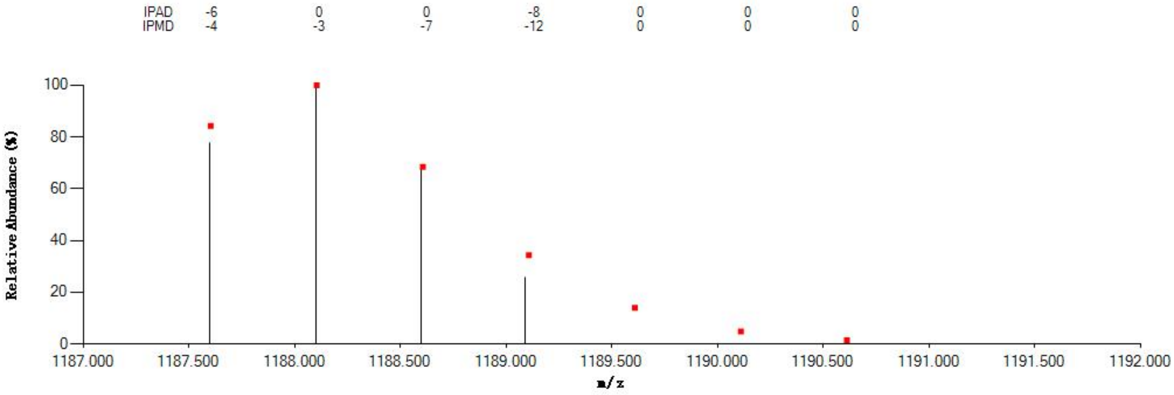


#### Annotated MS/MS Spectrum

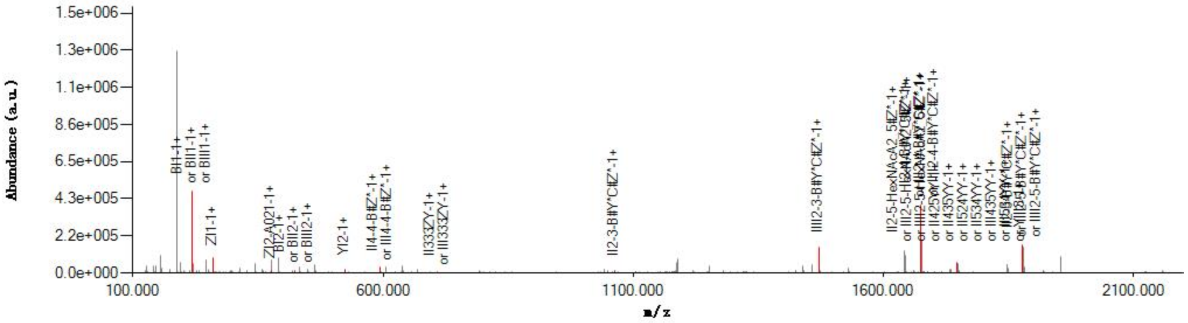


#### Graphical Fragmentation Map

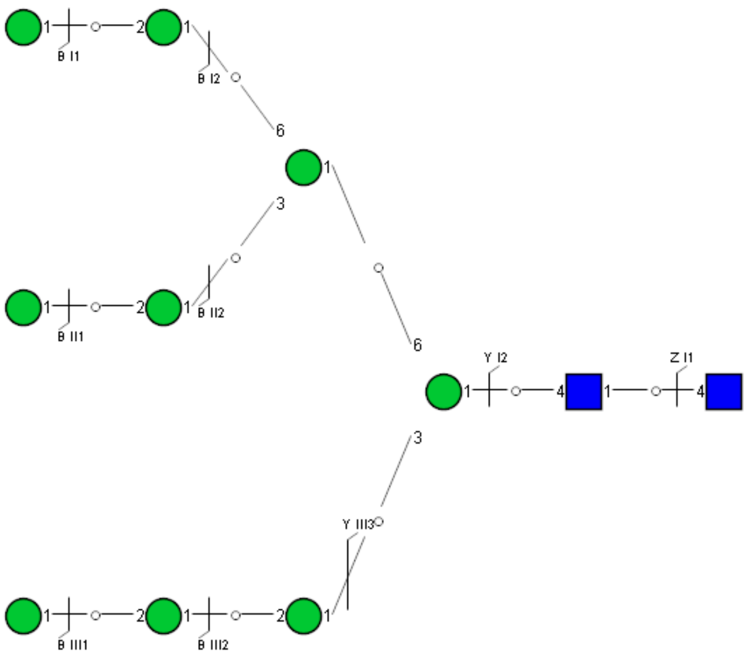

Spectrum Index: 41720(NoBHs:1)

#### Precursor iEF Map

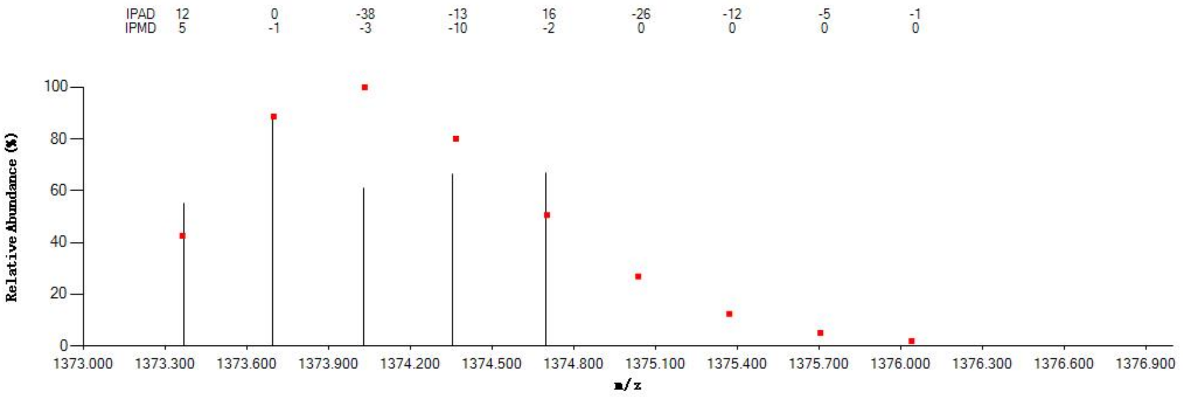

#### Annotated MS/MS Spectrum

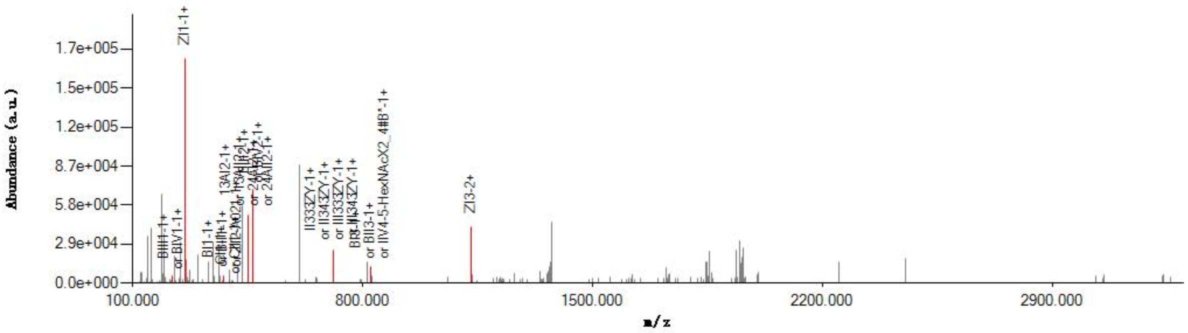

#### Graphical Fragmentation Map

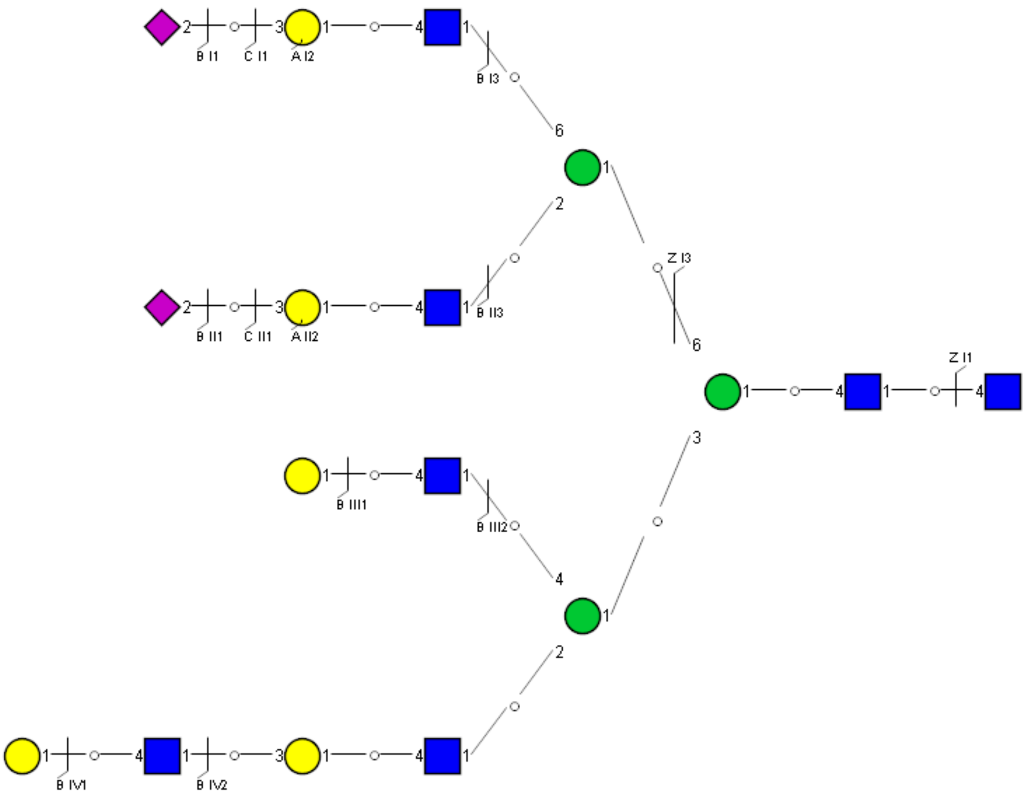

Spectrum Index: 42065(NoBHs:1)

#### Precursor iEF Map

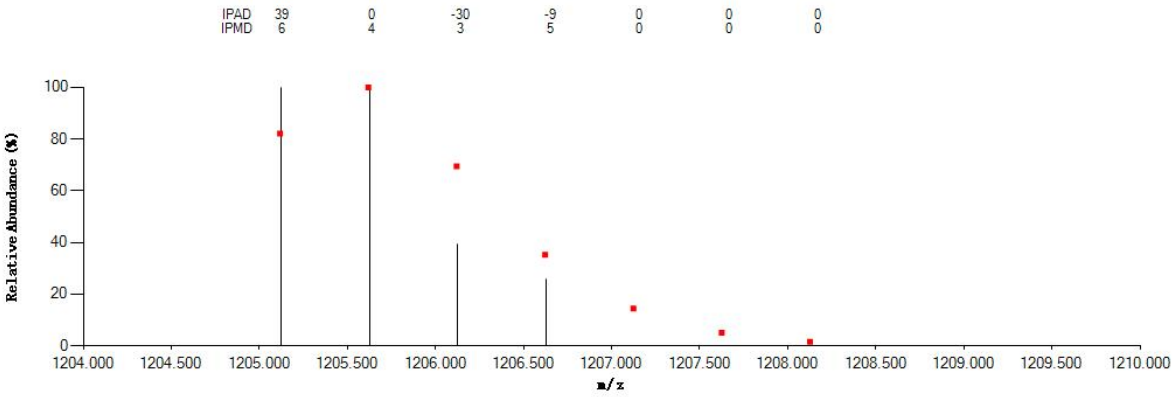


#### Annotated MS/MS Spectrum

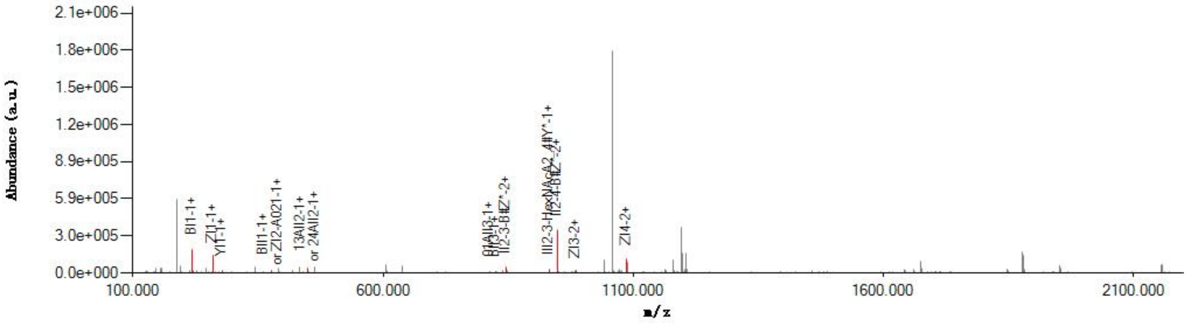


#### Graphical Fragmentation Map

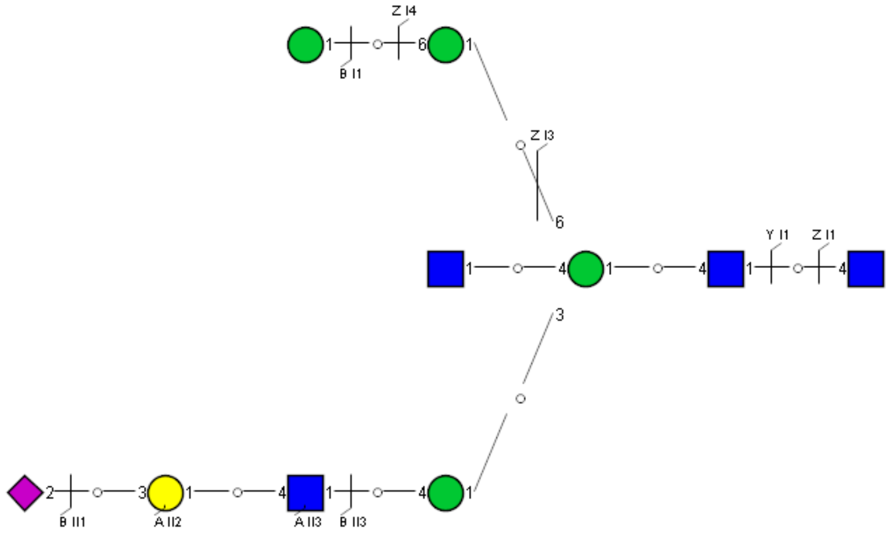

Spectrum Index: 42280(NoBHs:1)

#### Precursor iEF Map

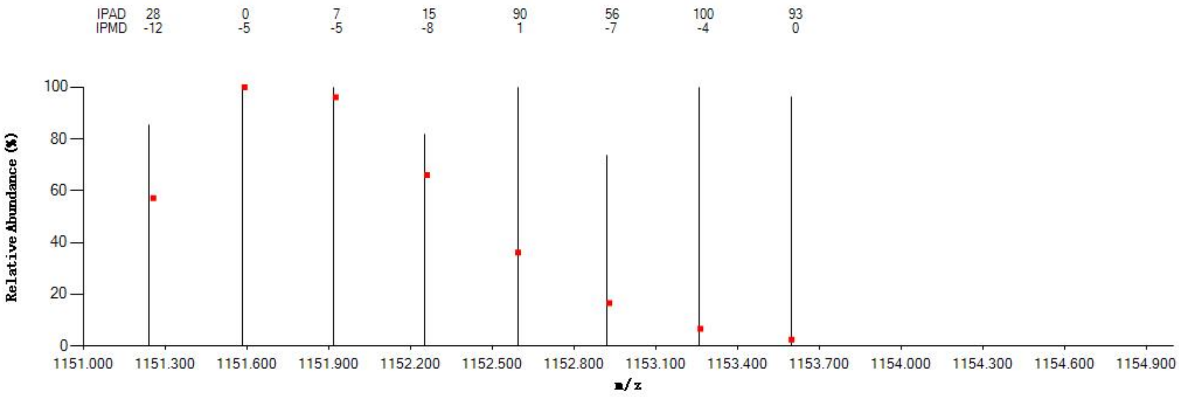


#### Annotated MS/MS Spectrum

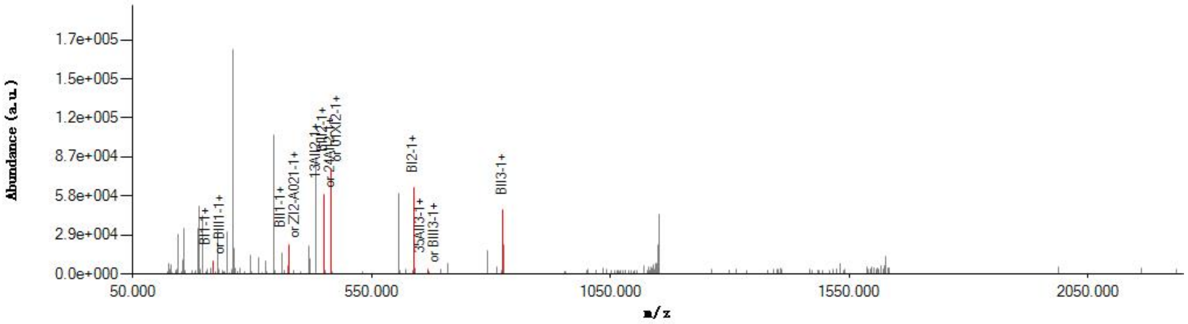


#### Graphical Fragmentation Map

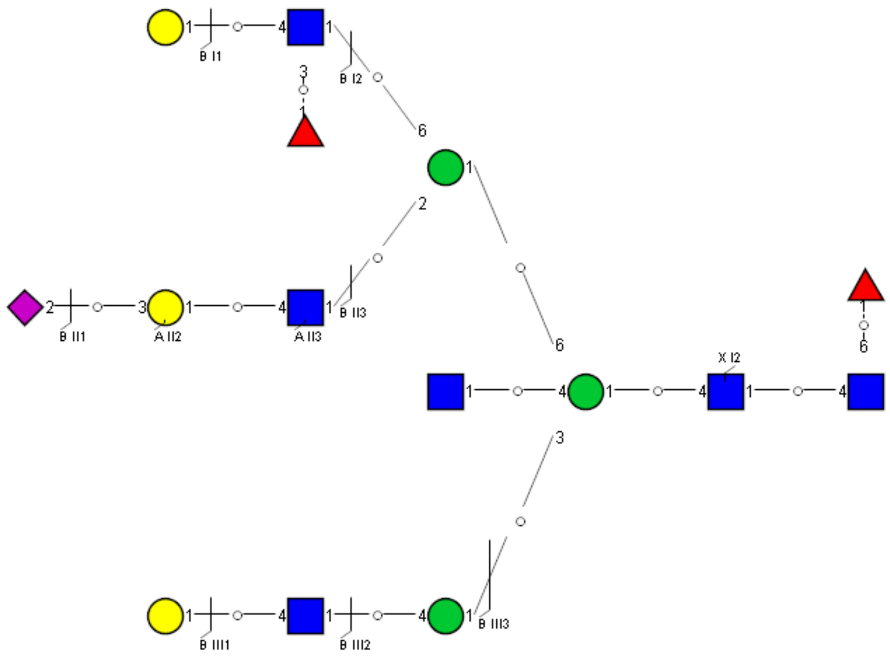

Spectrum Index: 42289(NoBHs:2)

#### Precursor iEF Map

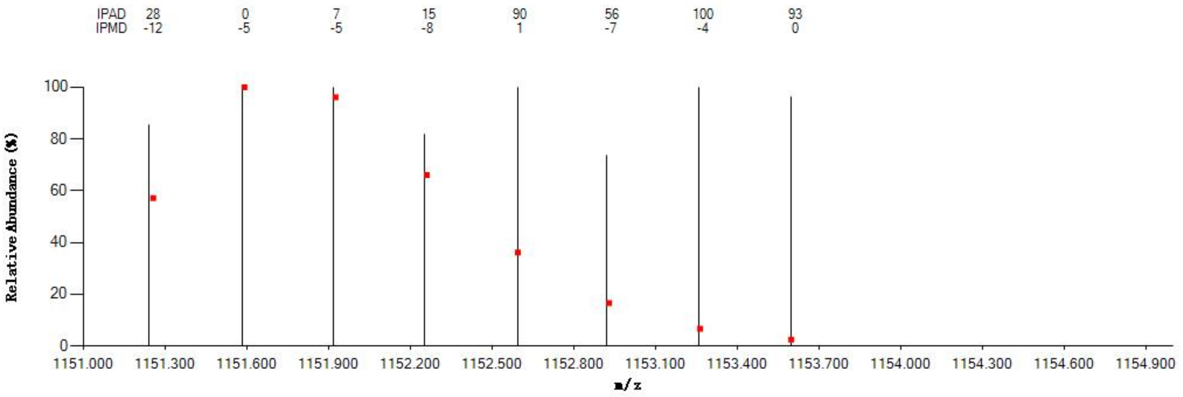

#### Annotated MS/MS Spectrum

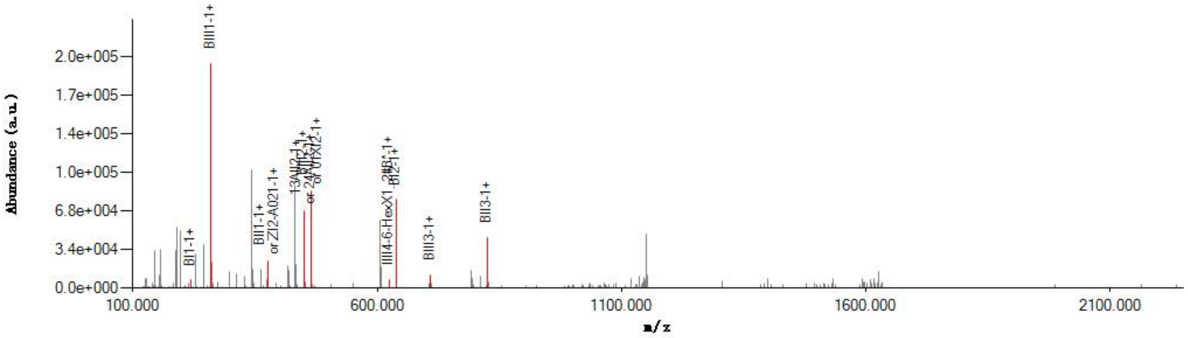

#### Graphical Fragmentation Map

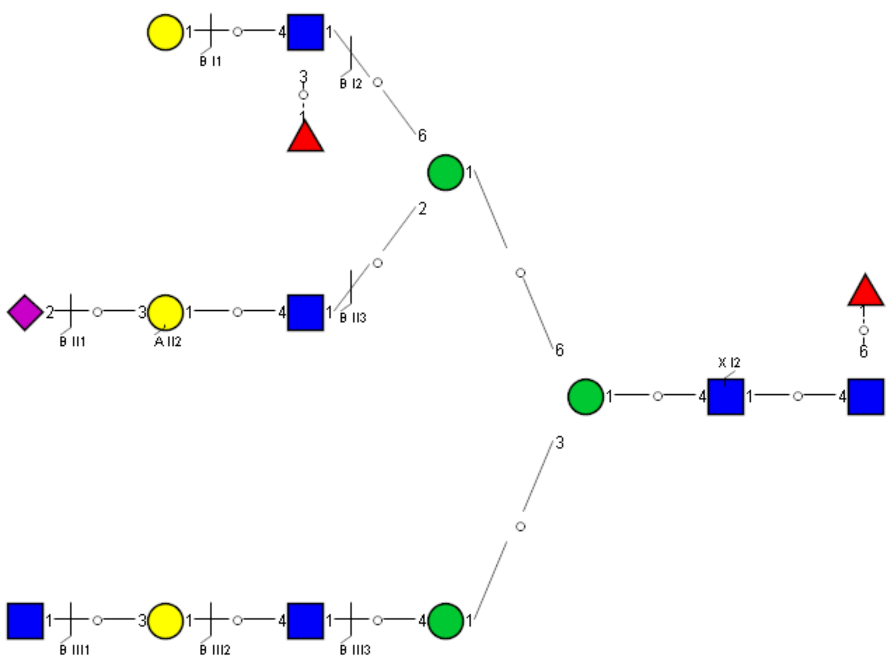

Spectrum Index: 42289(NoBHs:2)

Precursor iEF Map

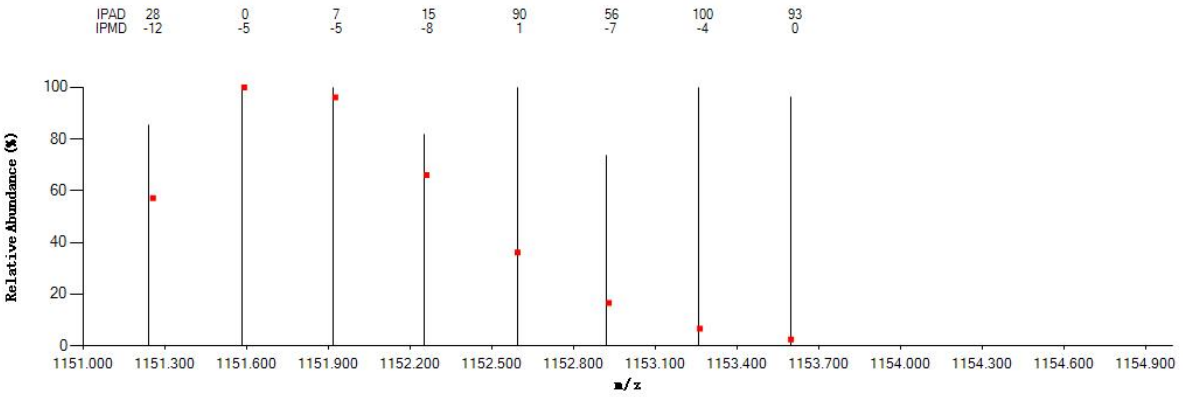


Annotated MS/MS Spectrum

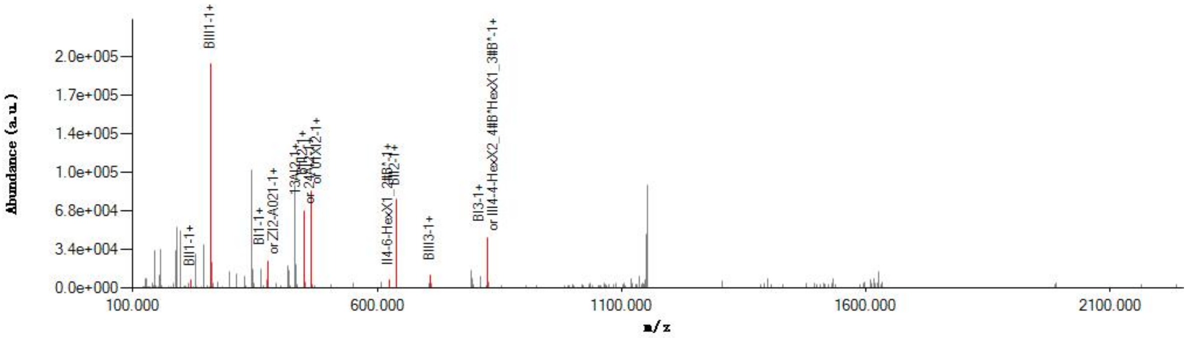


Graphical Fragmentation Map

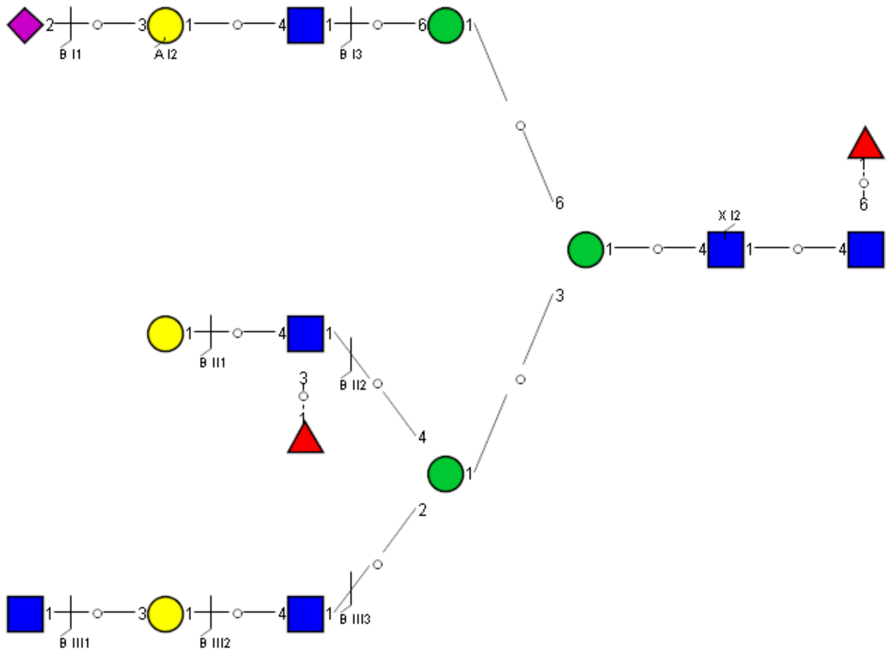

Spectrum Index: 42297(NoBHs:1)

#### Precursor iEF Map

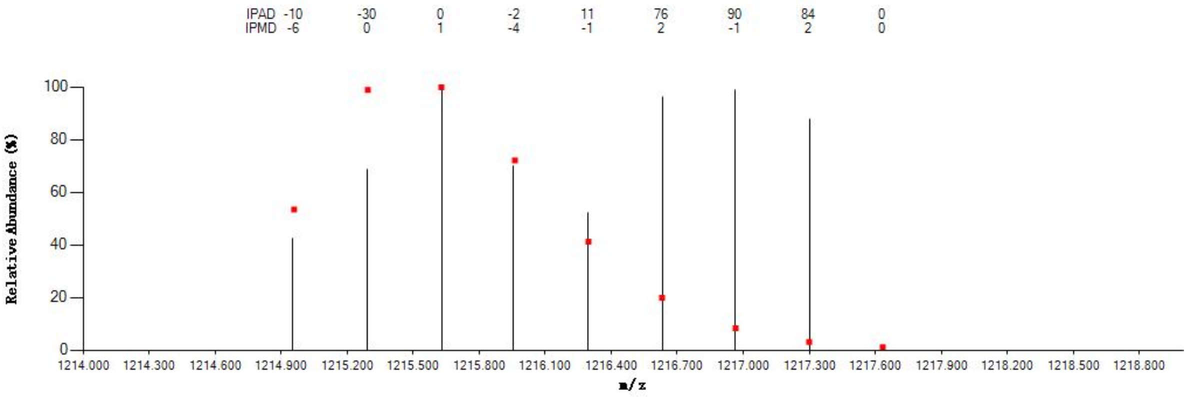

#### Annotated MS/MS Spectrum

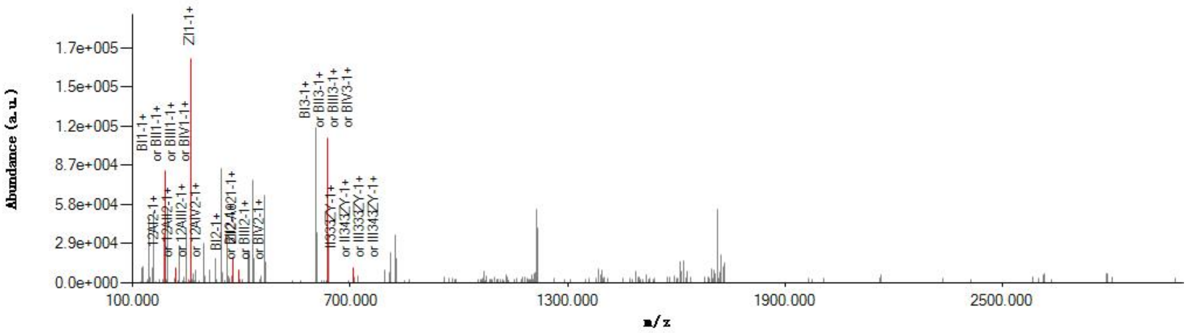

#### Graphical Fragmentation Map

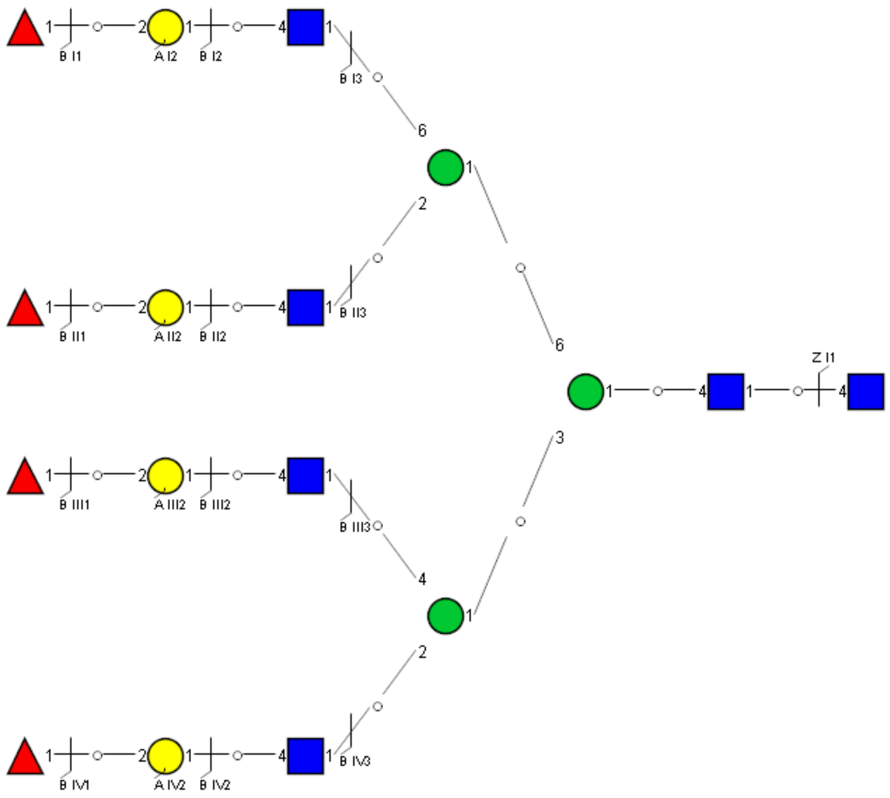

Spectrum Index: 42412(NoBHs:1)

#### Precursor iEF Map

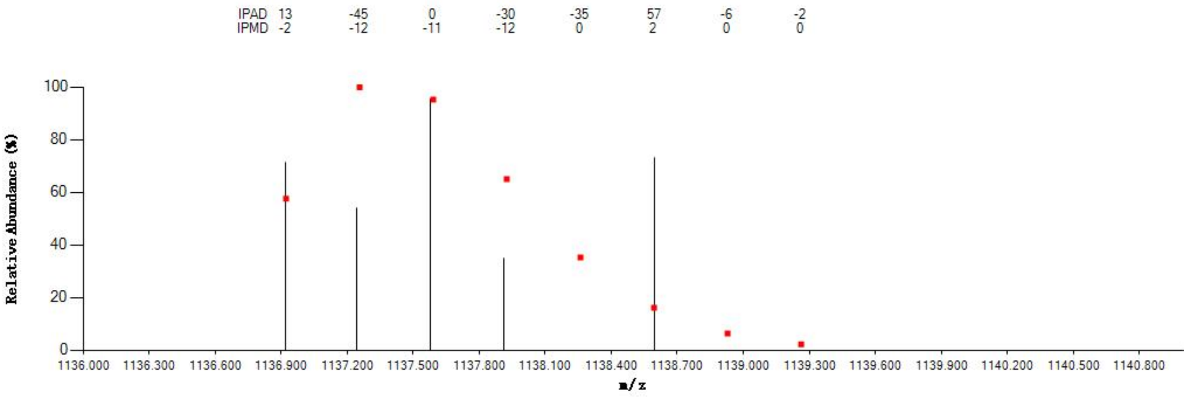


#### Annotated MS/MS Spectrum

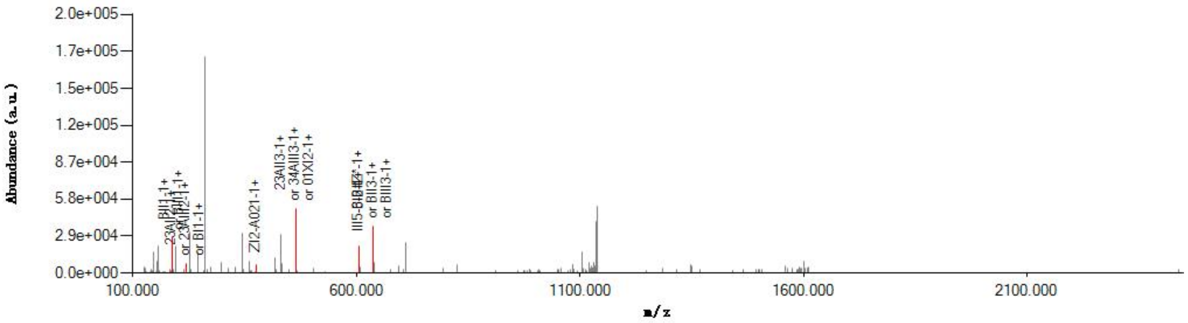


#### Graphical Fragmentation Map

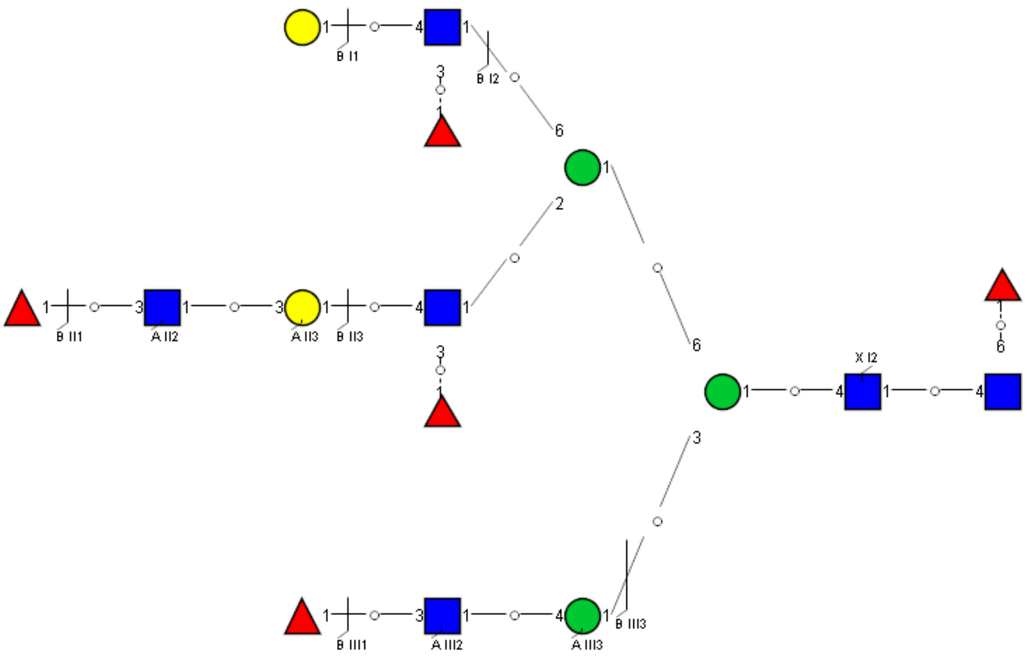

Spectrum Index: 42549(NoBHs:1)

#### Precursor iEF Map

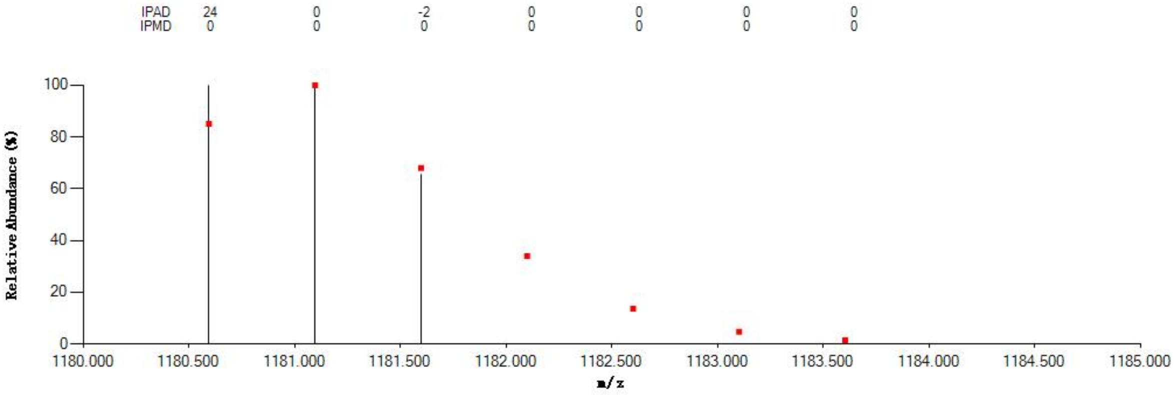


#### Annotated MS/MS Spectrum

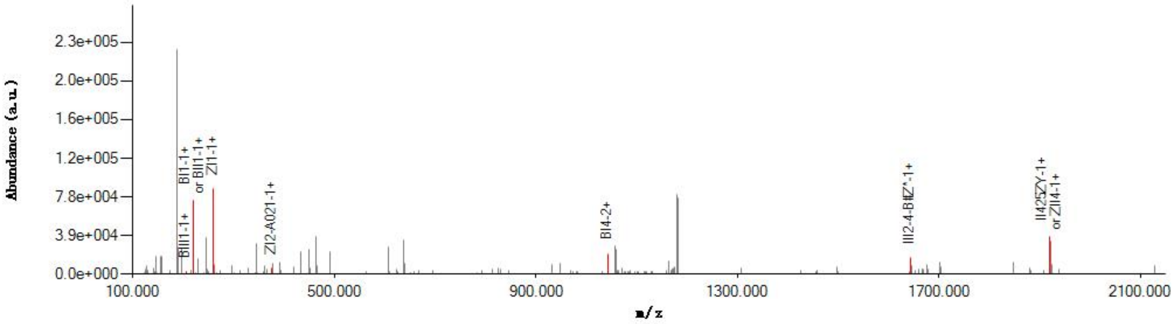


#### Graphical Fragmentation Map

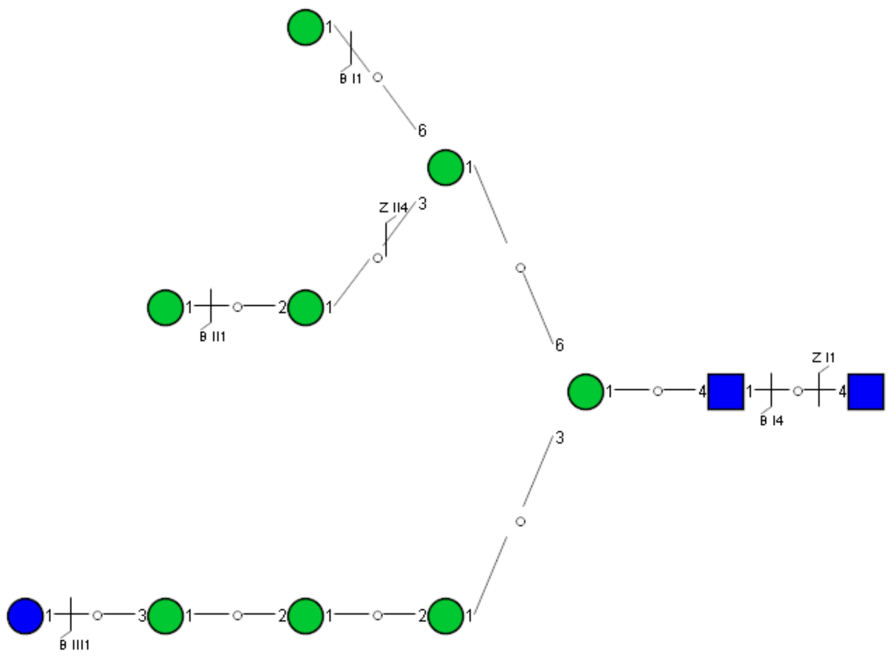

### Spectrum Index: 42614(NoBHs:1)

#### Precursor iEF Map

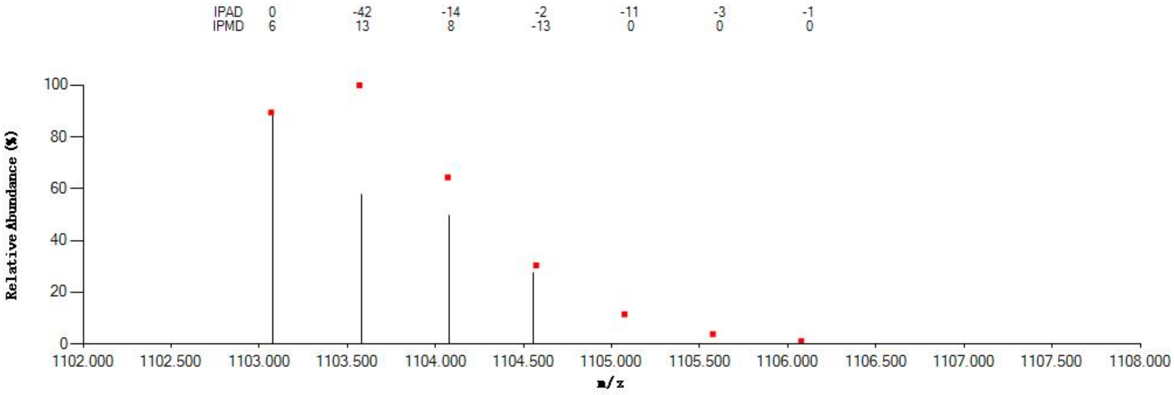

#### Annotated MS/MS Spectrum

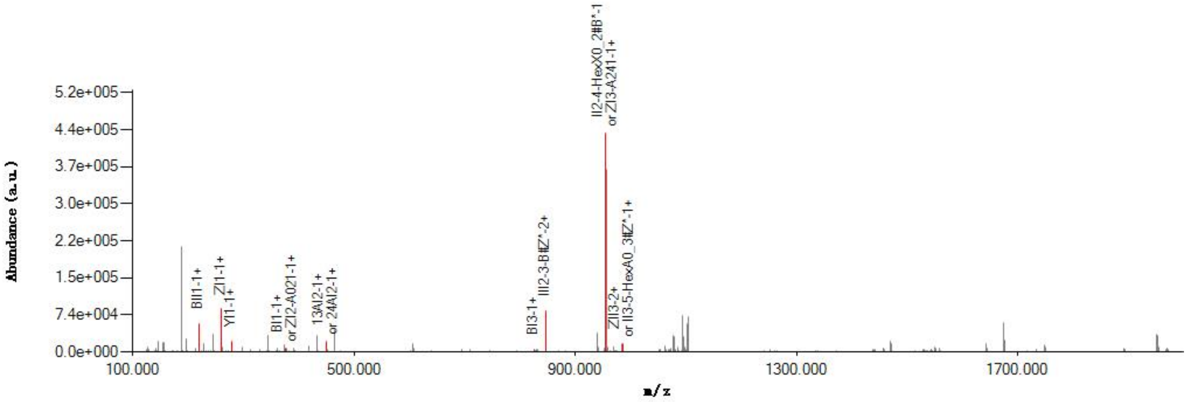

#### Graphical Fragmentation Map

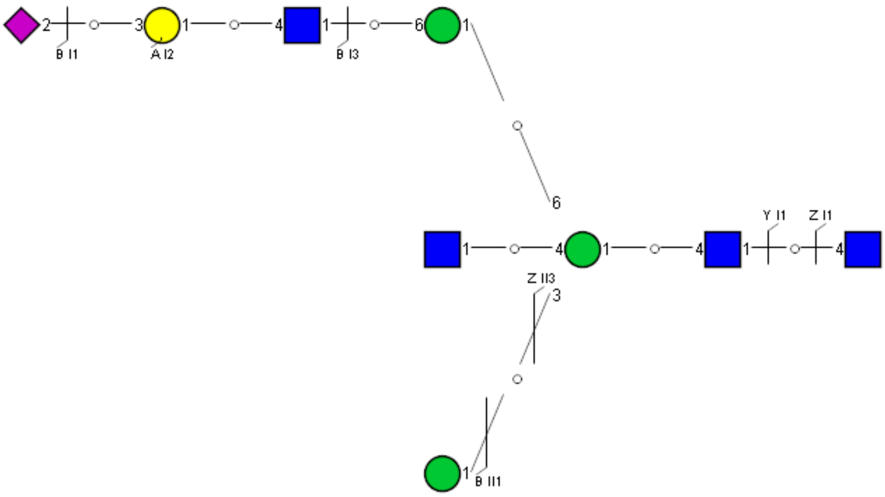

Spectrum Index: 42689(NoBHs:1)

Precursor iEF Map

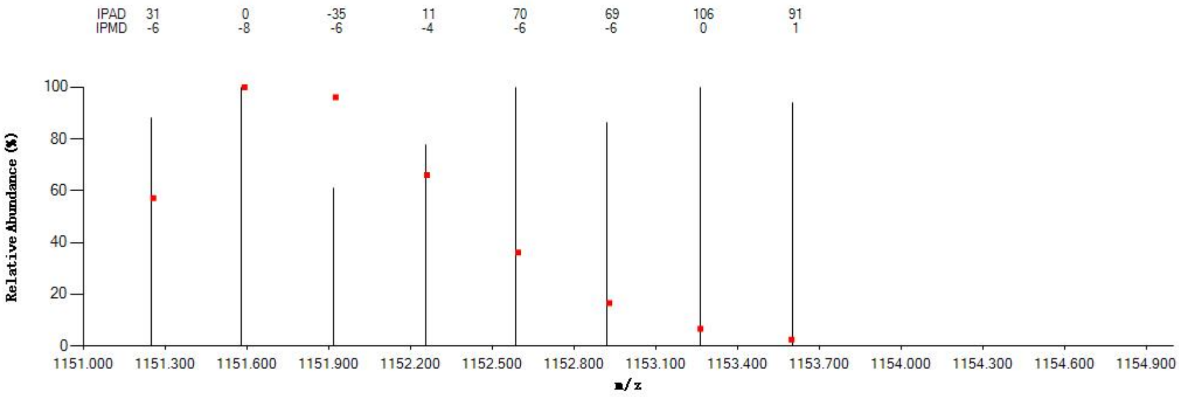

Annotated MS/MS Spectrum

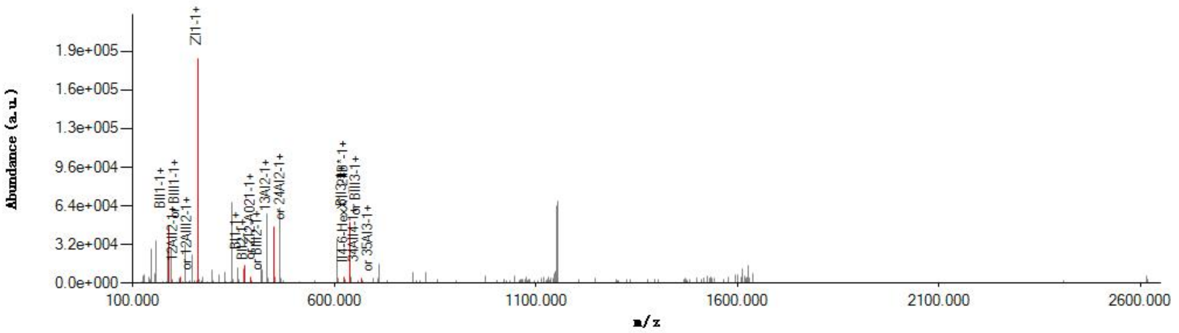

Graphical Fragmentation Map

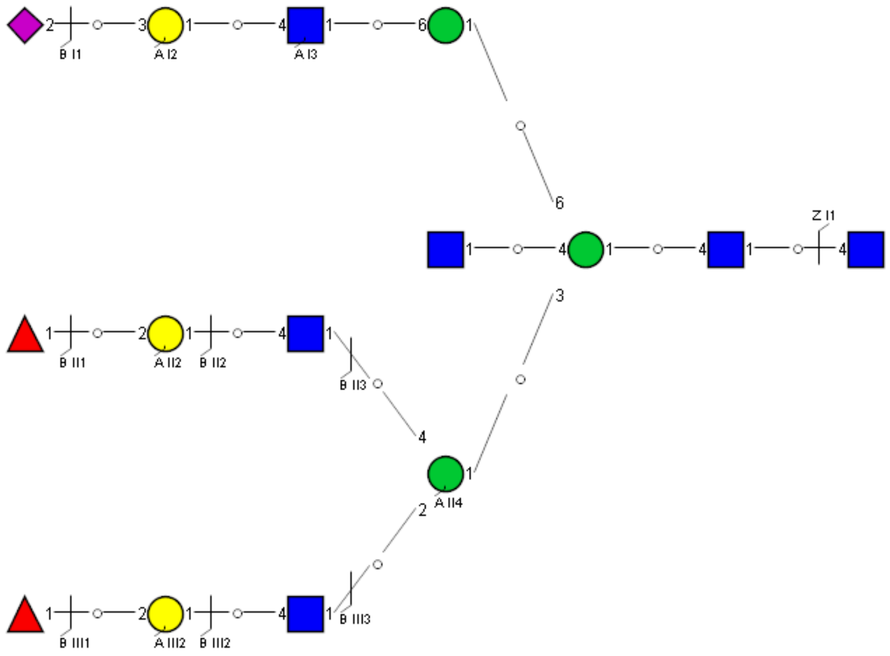

Spectrum Index: 42758(NoBHs:1)

#### Precursor iEF Map

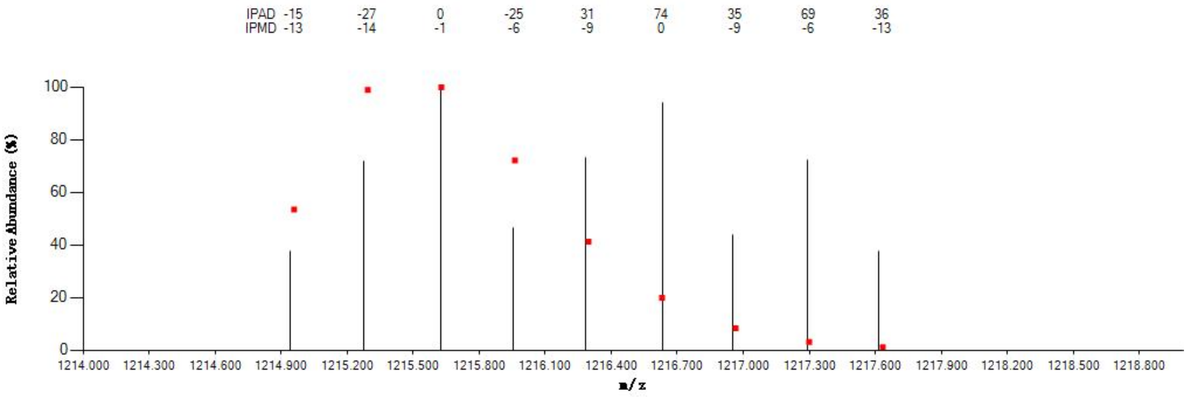


#### Annotated MS/MS Spectrum

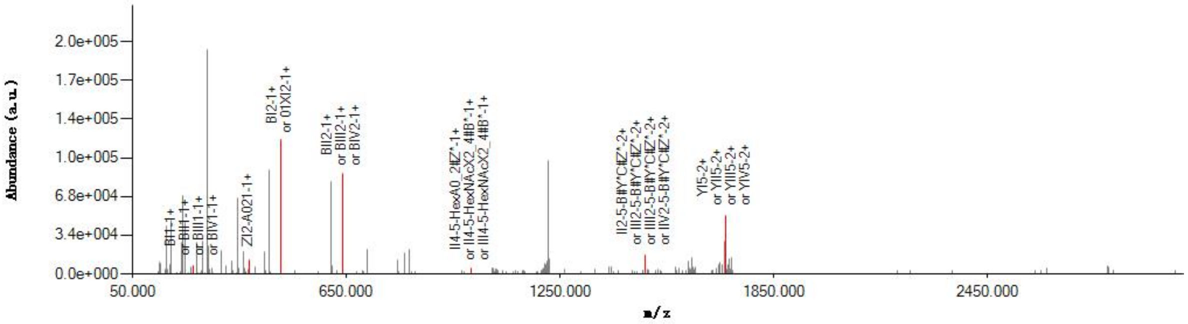


#### Graphical Fragmentation Map

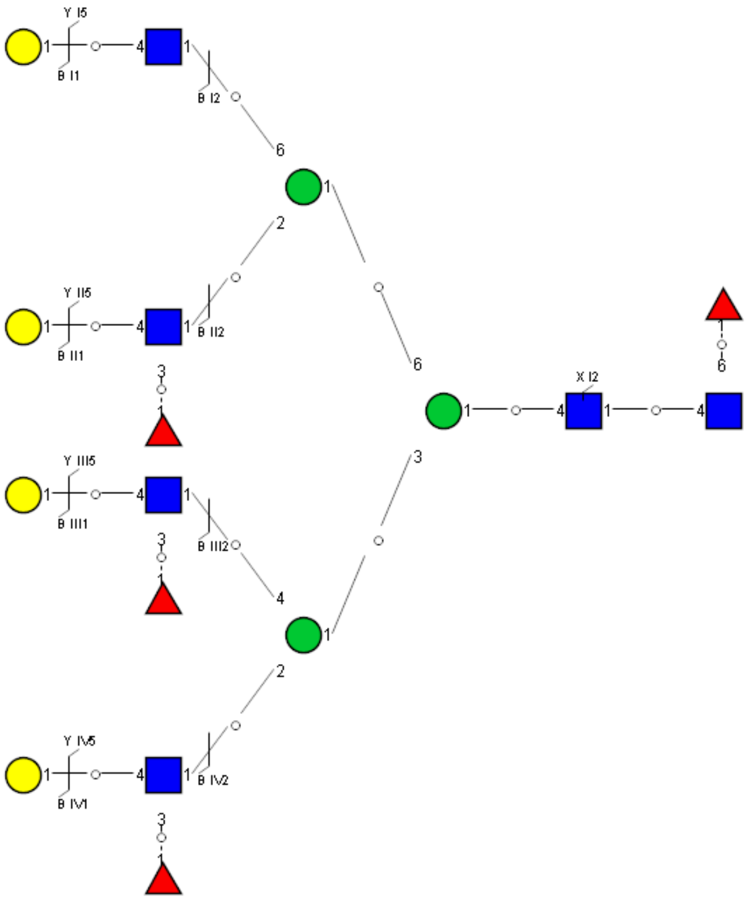

Spectrum Index: 42761(NoBHs:1)

#### Precursor iEF Map

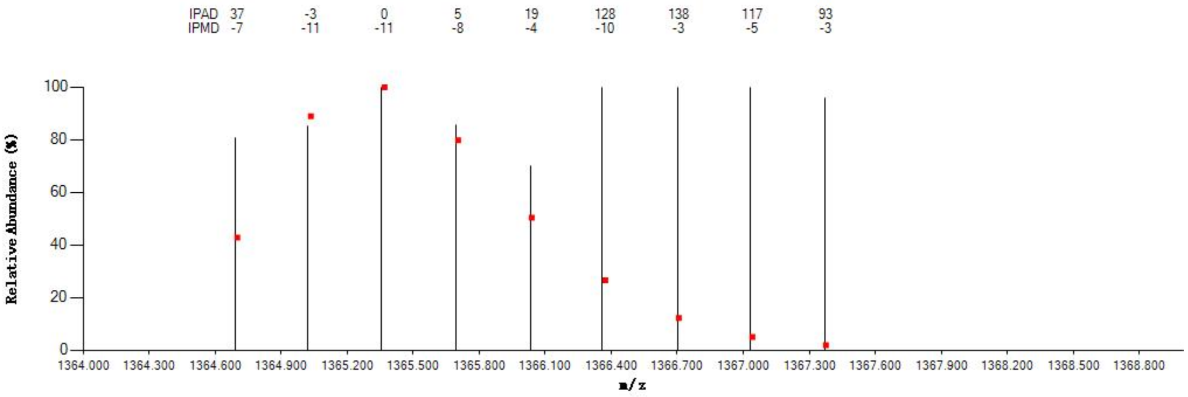


#### Annotated MS/MS Spectrum

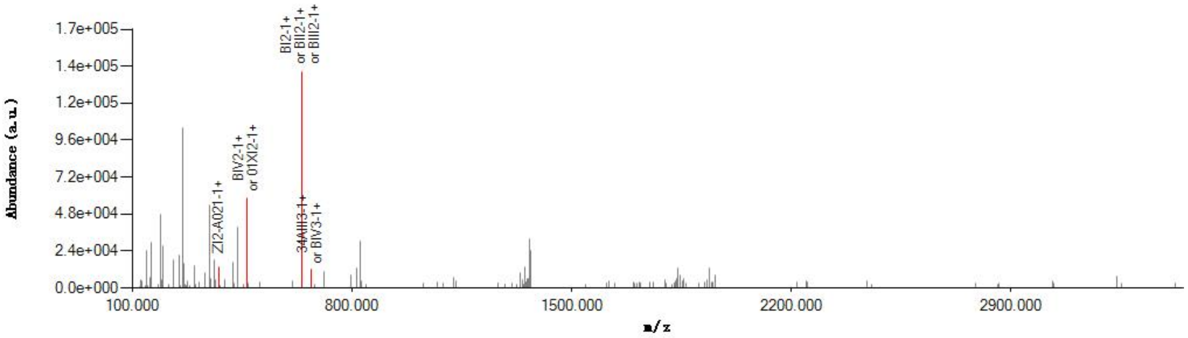


#### Graphical Fragmentation Map

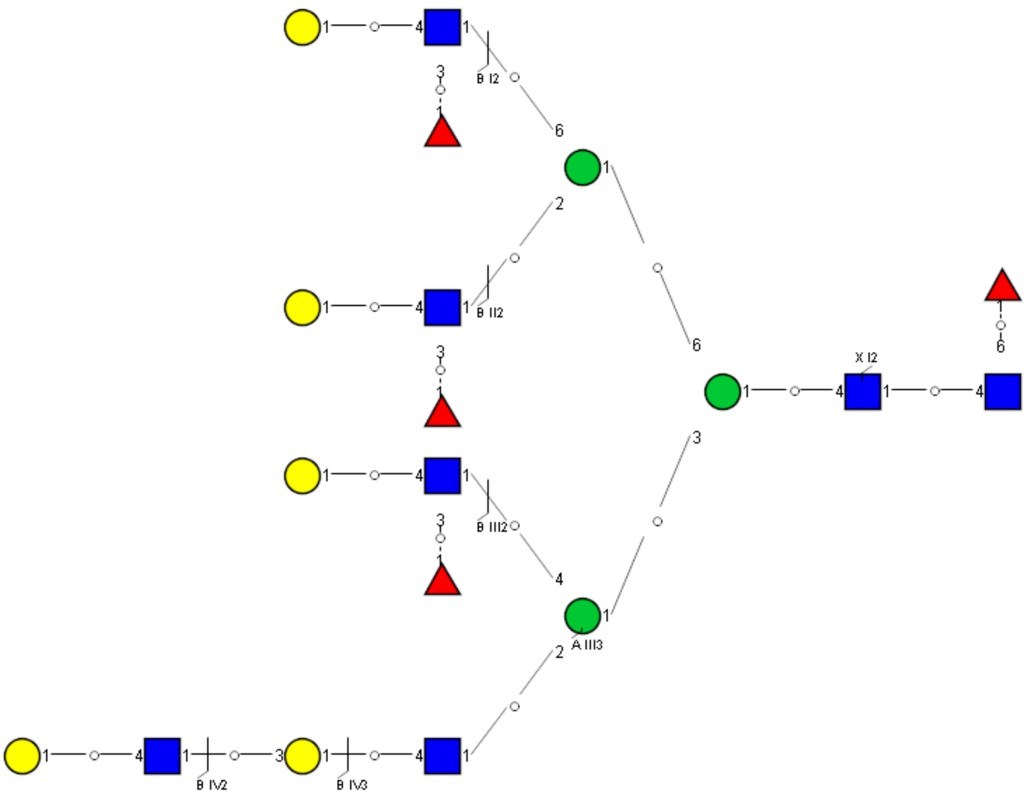

### Spectrum Index: 42764(NoBHs:2)

#### Precursor iEF Map

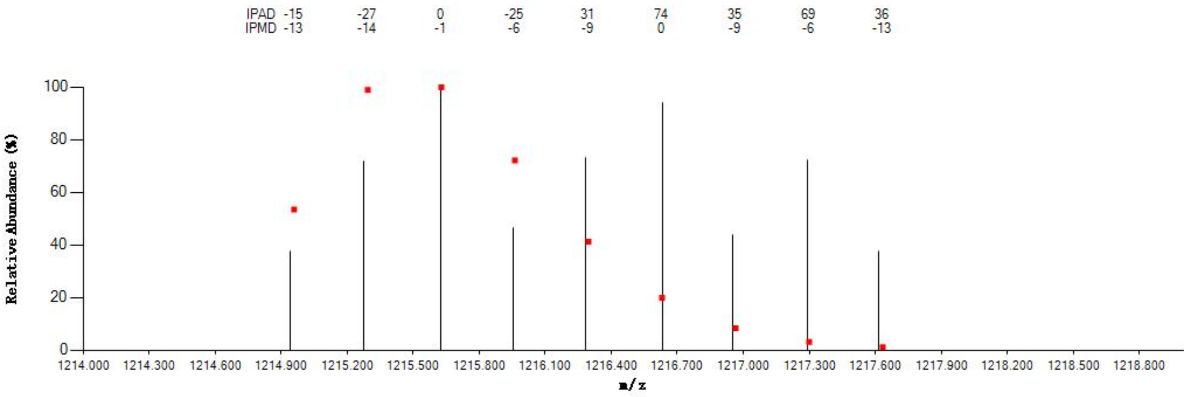


#### Annotated MS/MS Spectrum

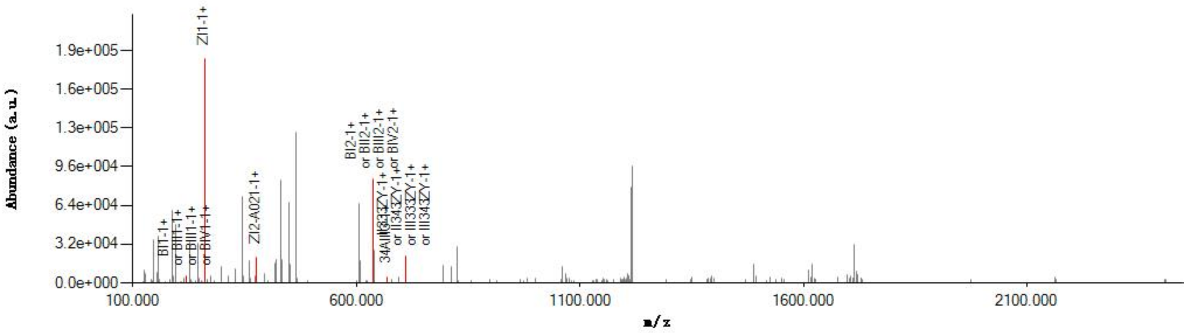


#### Graphical Fragmentation Map

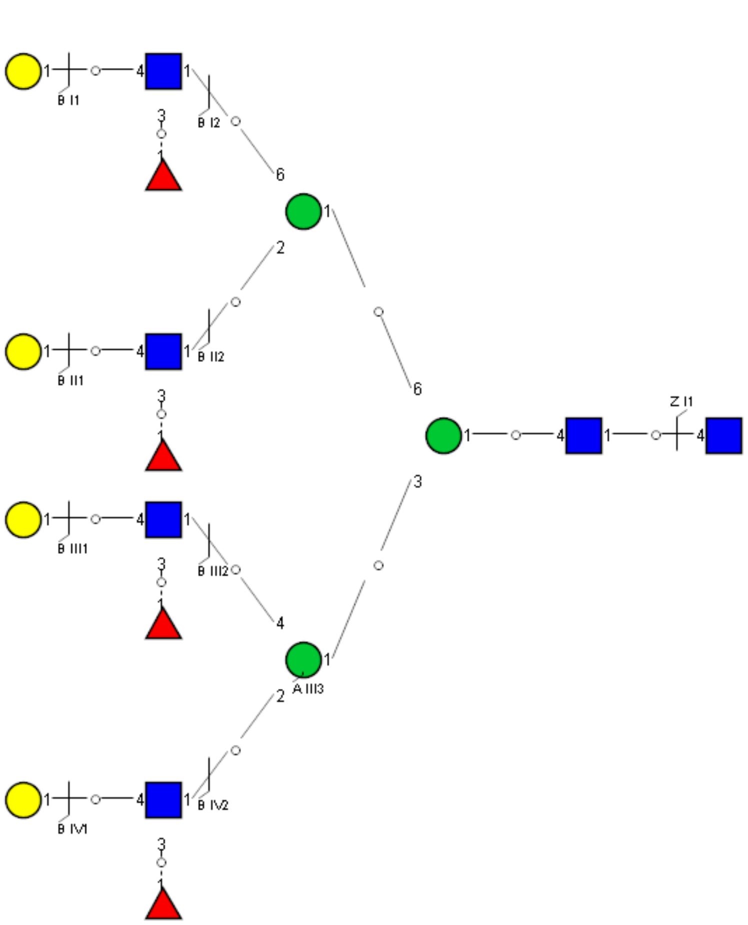

Spectrum Index: 42785(NoBHs:1)

#### Precursor iEF Map

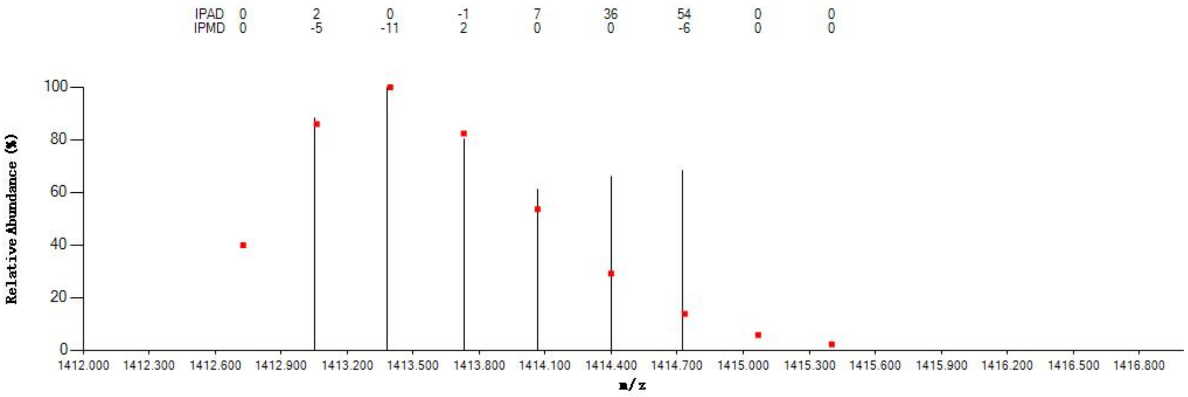


#### Annotated MS/MS Spectrum

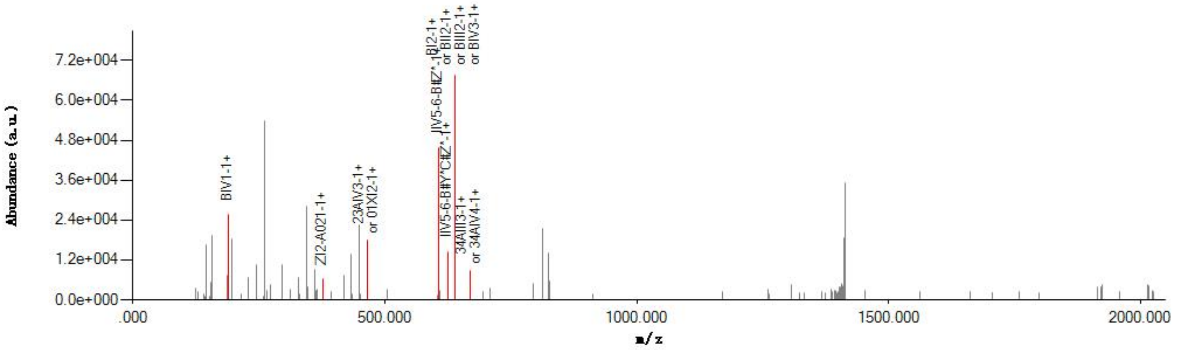


#### Graphical Fragmentation Map

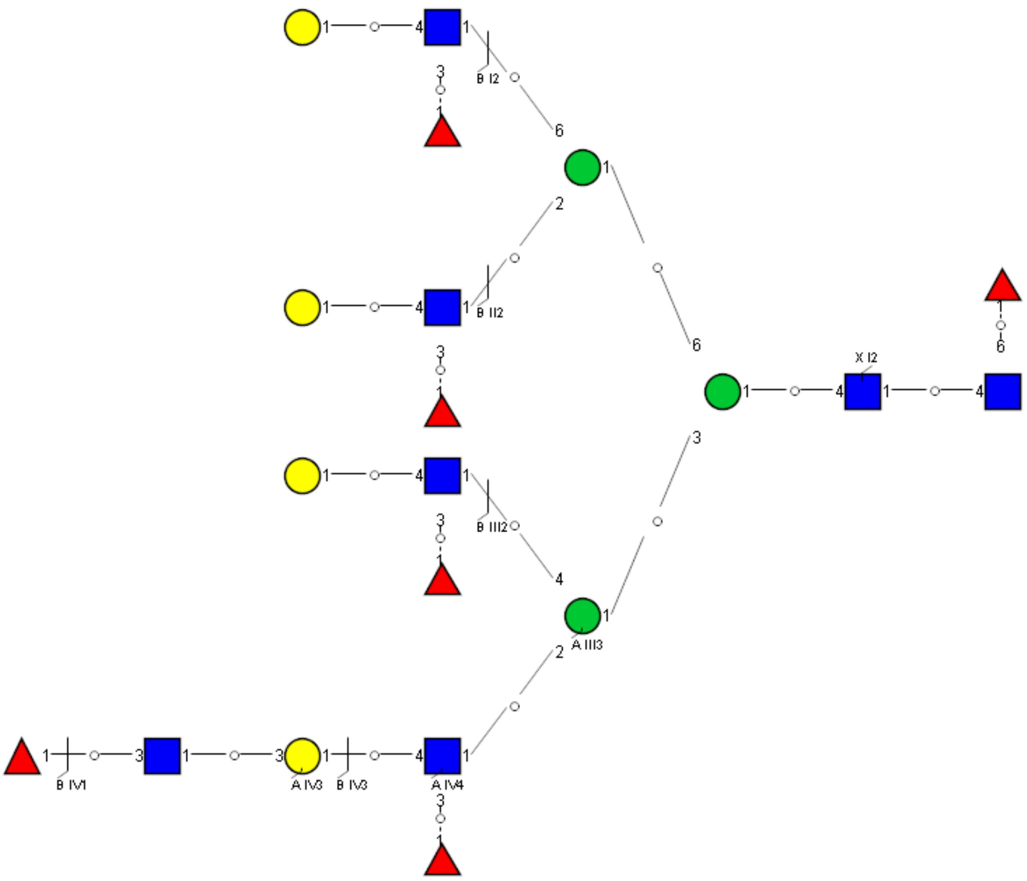

Spectrum Index: 42795(NoBHs:2)

#### Precursor iEF Map

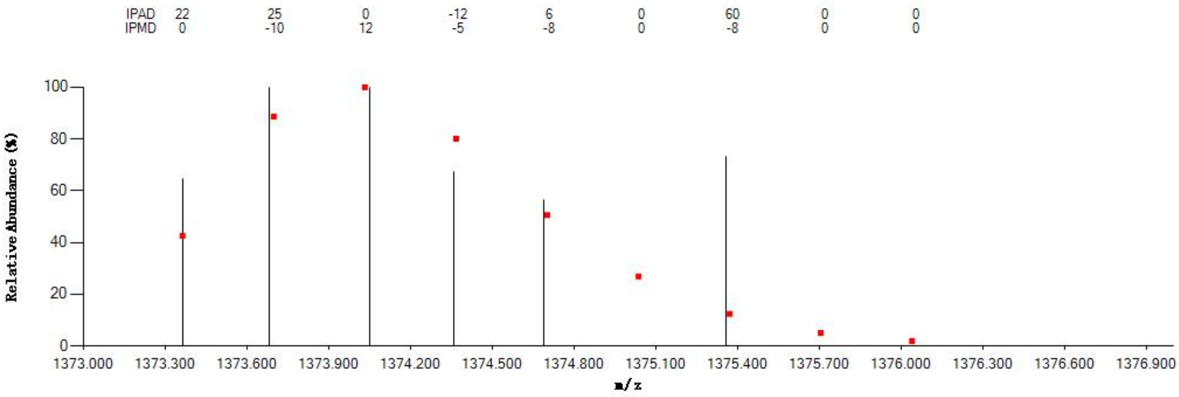

#### Annotated MS/MS Spectrum

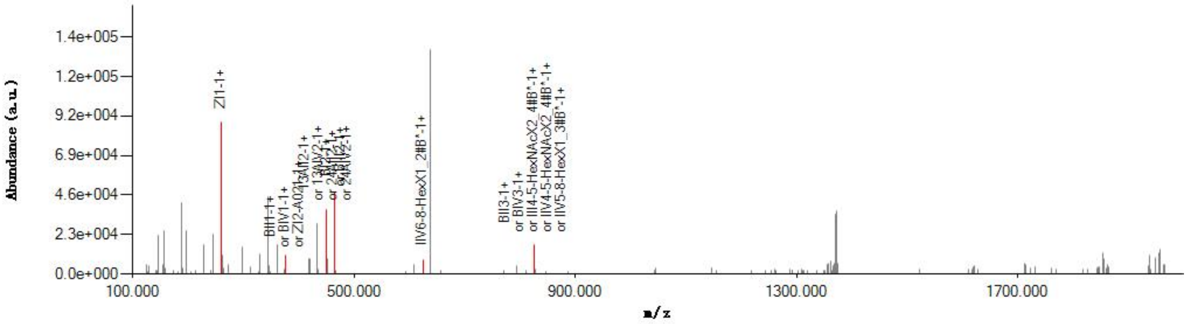

#### Graphical Fragmentation Map

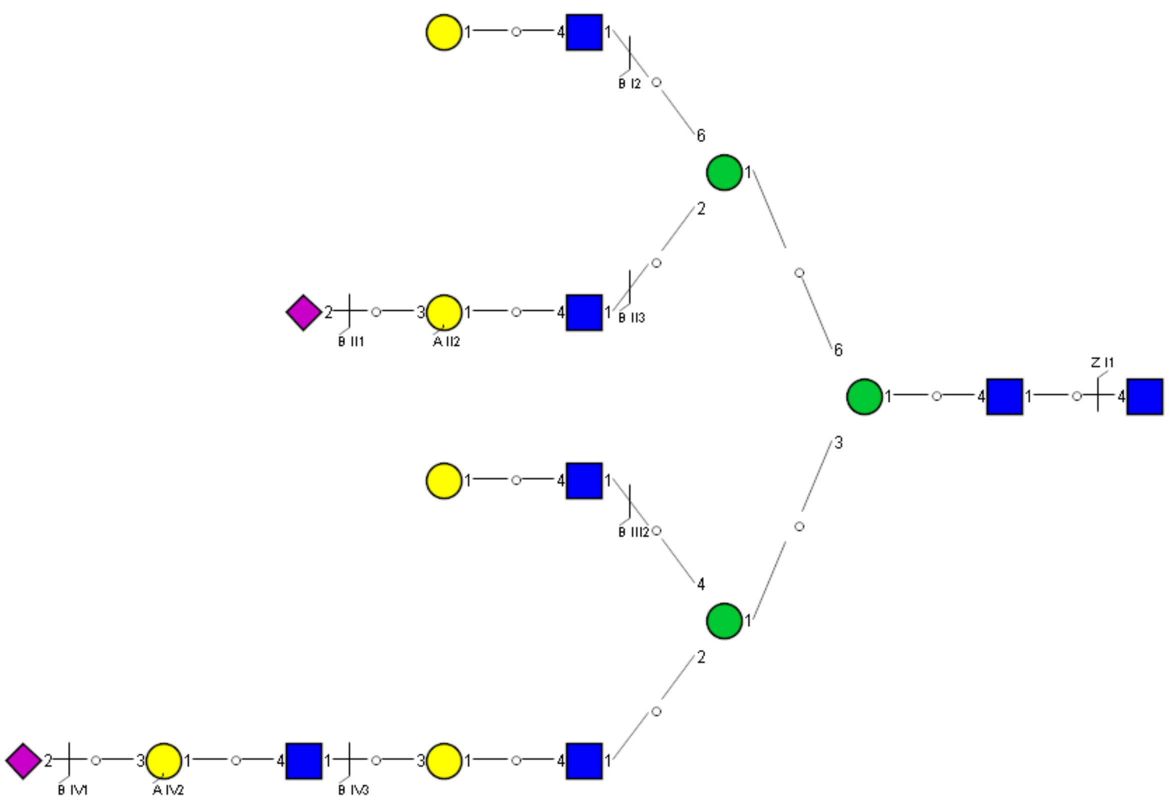

Spectrum Index: 42801(NoBHs:1)

#### Precursor iEF Map

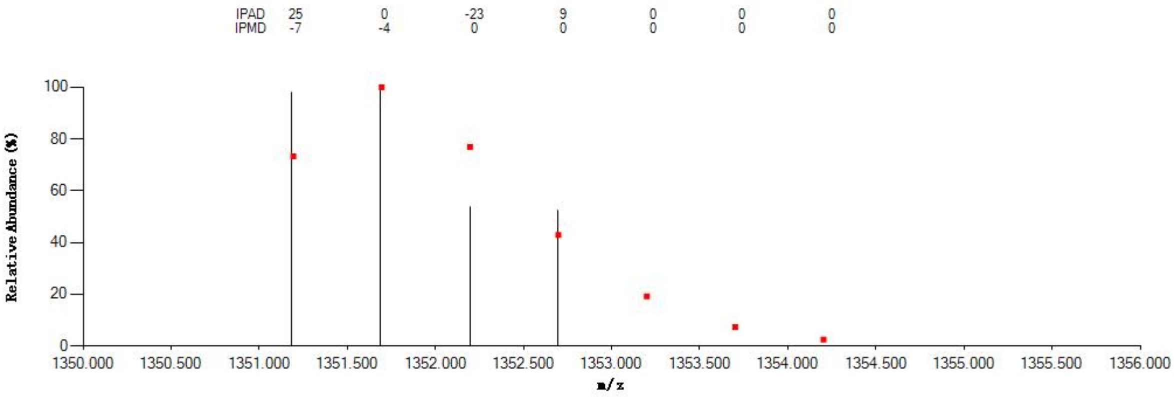


#### Annotated MS/MS Spectrum

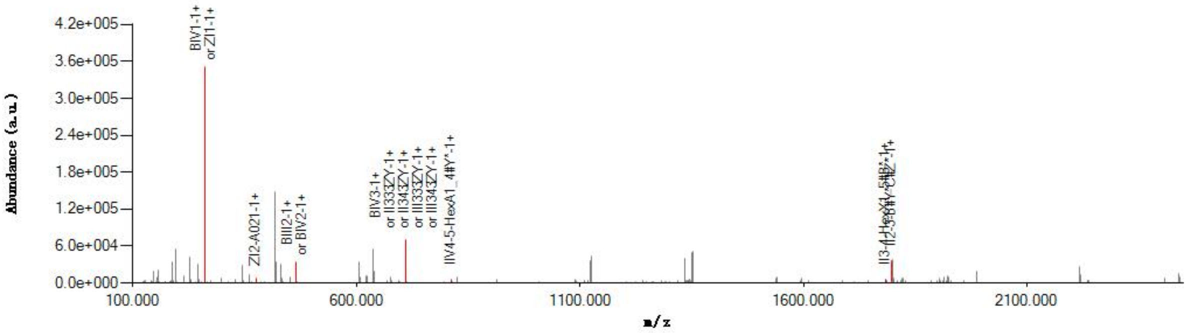


#### Graphical Fragmentation Map

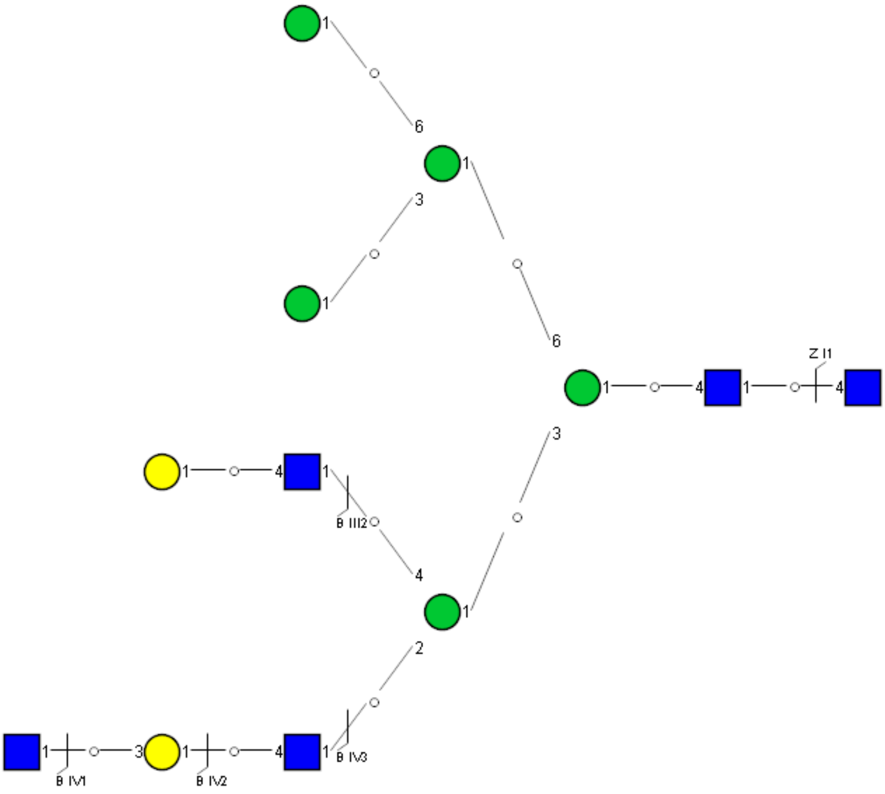

Spectrum Index: 42808(NoBHs:1)

#### Precursor iEF Map

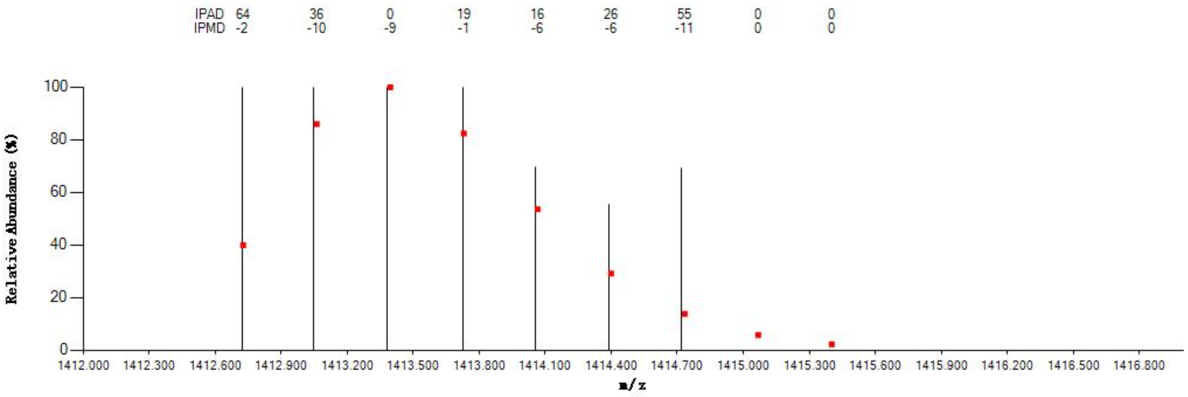


#### Annotated MS/MS Spectrum

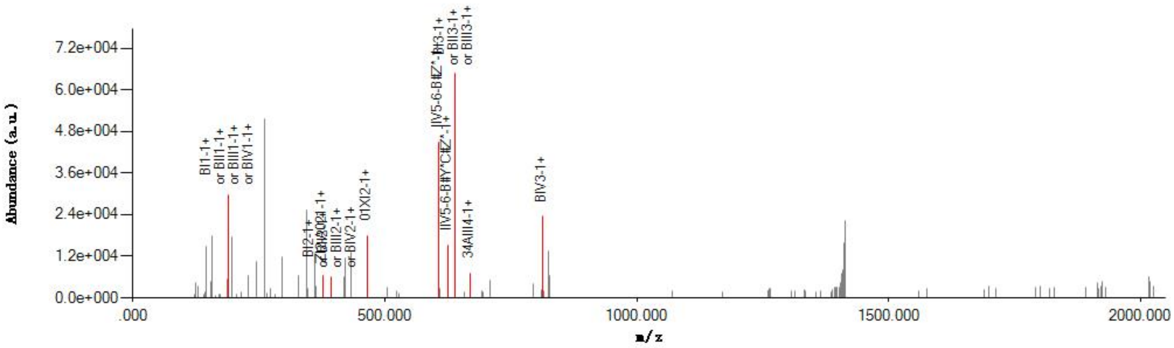


#### Graphical Fragmentation Map

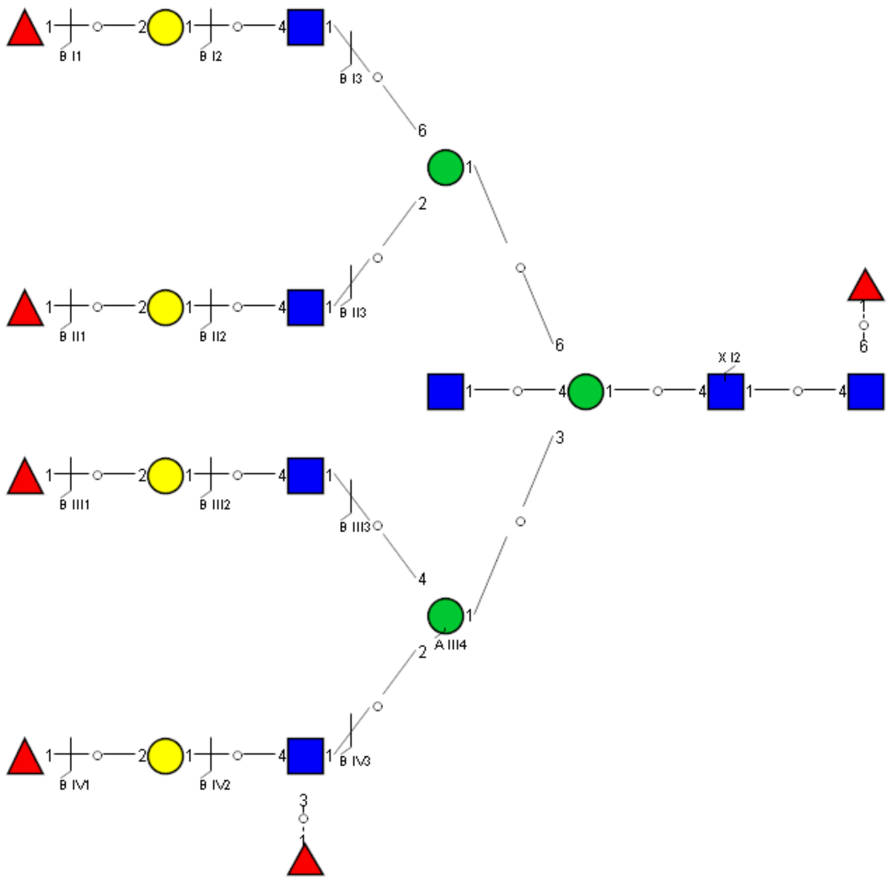

Spectrum Index: 42853(NoBHs:1)

#### Precursor iEF Map

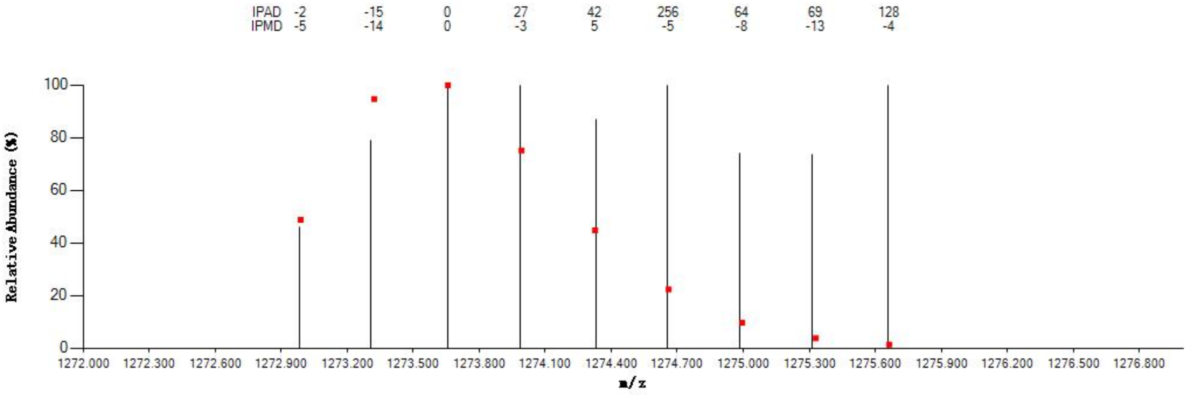


#### Annotated MS/MS Spectrum

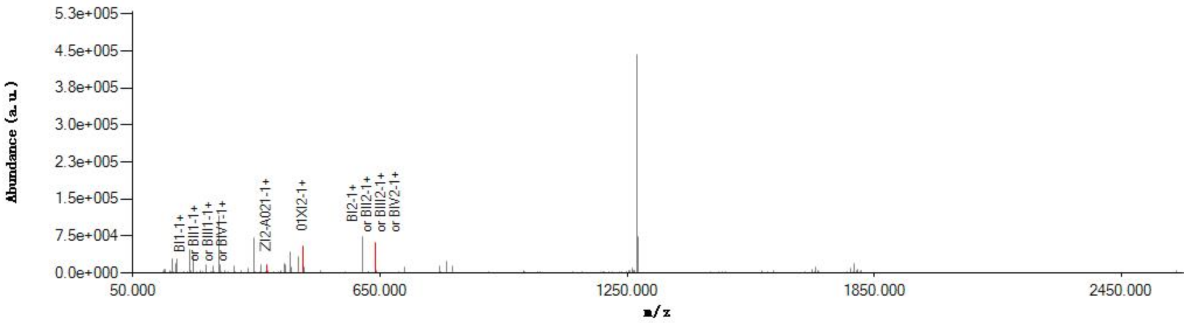


#### Graphical Fragmentation Map

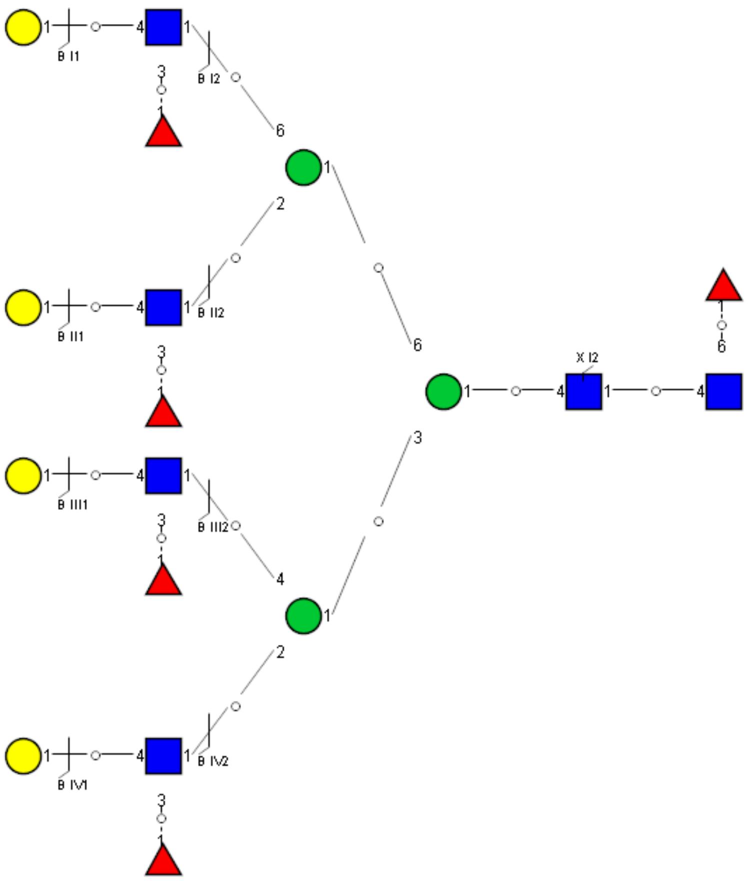

Spectrum Index: 42864(NoBHs:1)

#### Precursor iEF Map

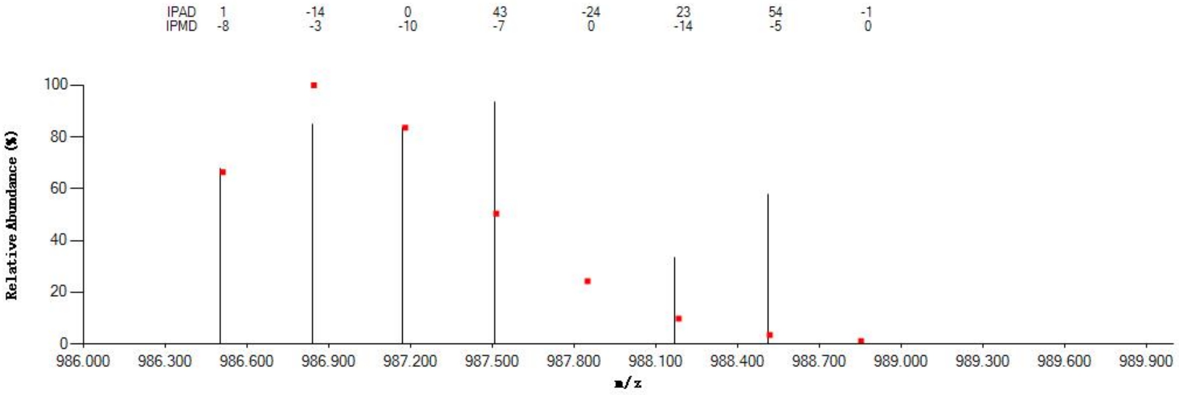


#### Annotated MS/MS Spectrum

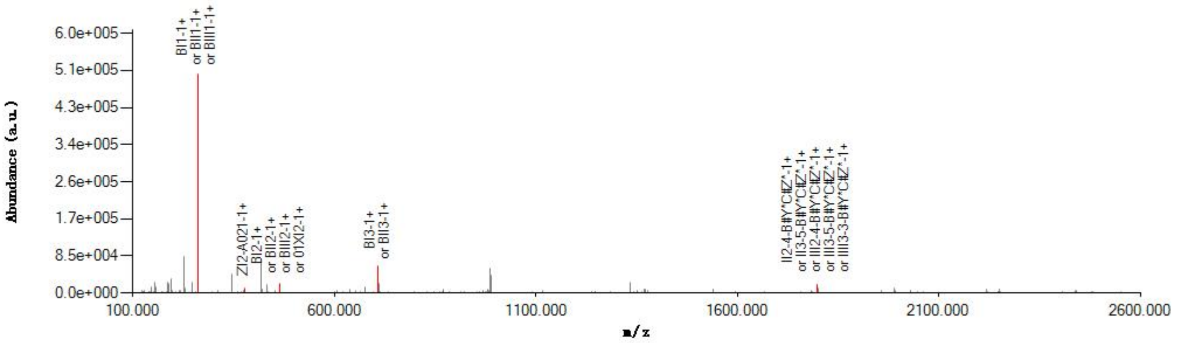


#### Graphical Fragmentation Map

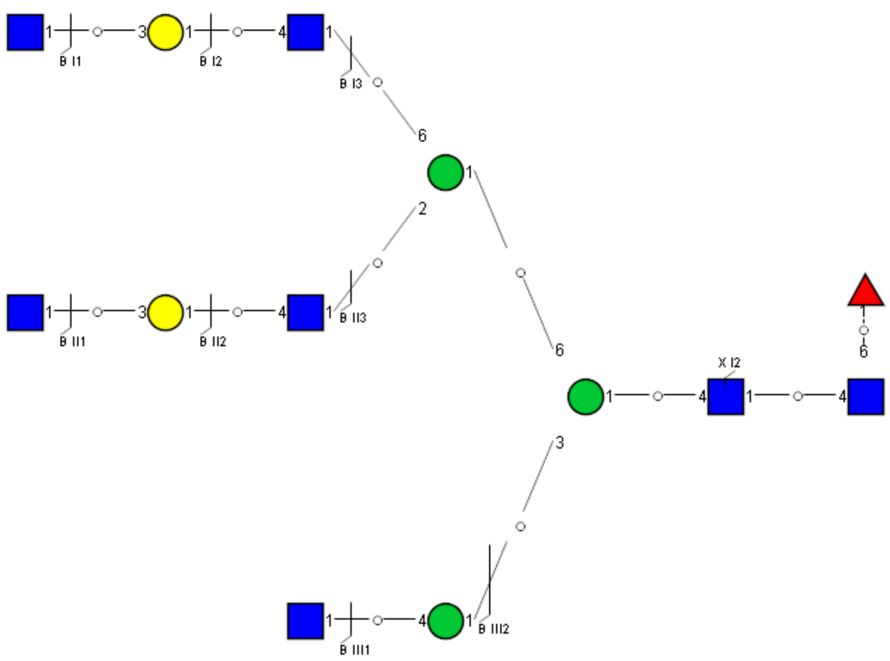

Spectrum Index: 43300(NoBHs:1)

Precursor iEF Map

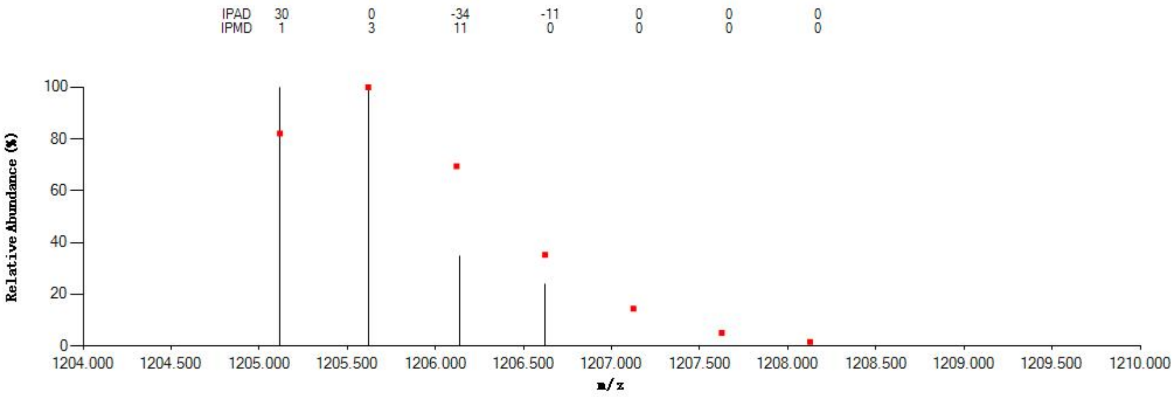


Annotated MS/MS Spectrum

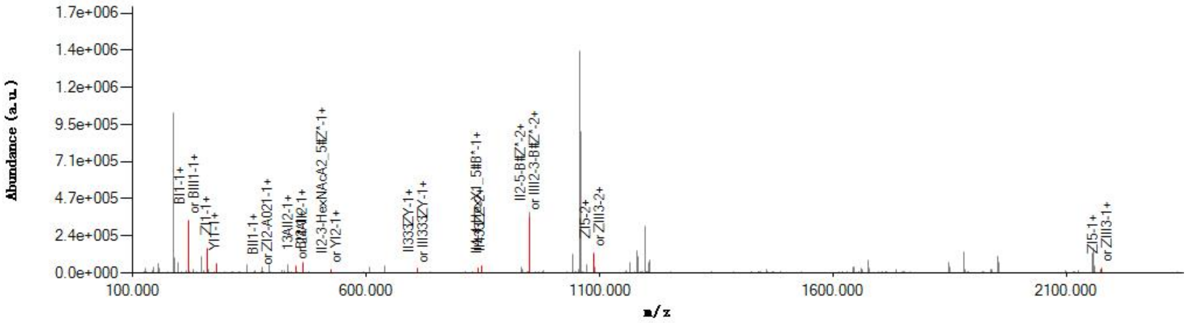


Graphical Fragmentation Map

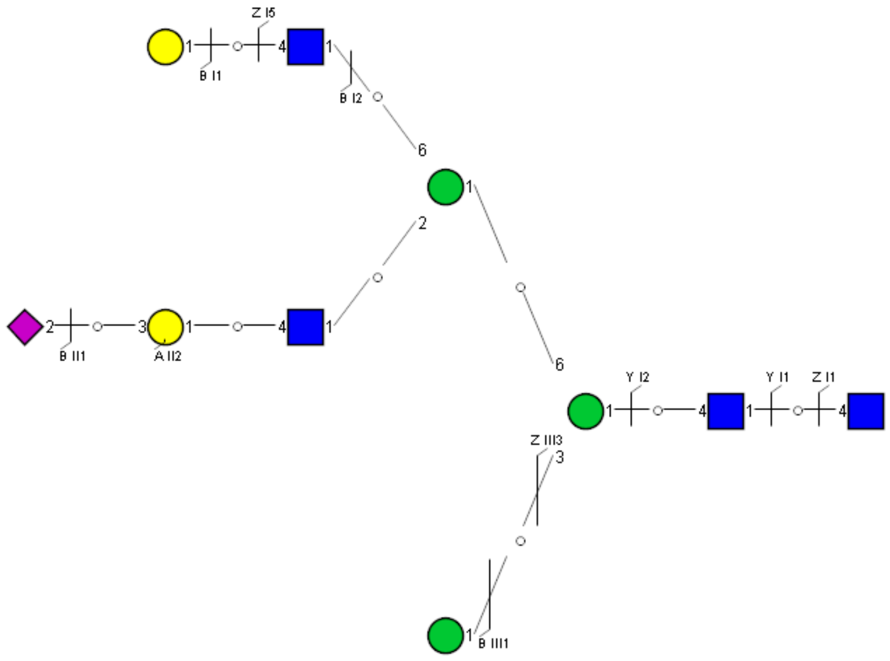

Spectrum Index: 43963(NoBHs:1)

#### Precursor iEF Map

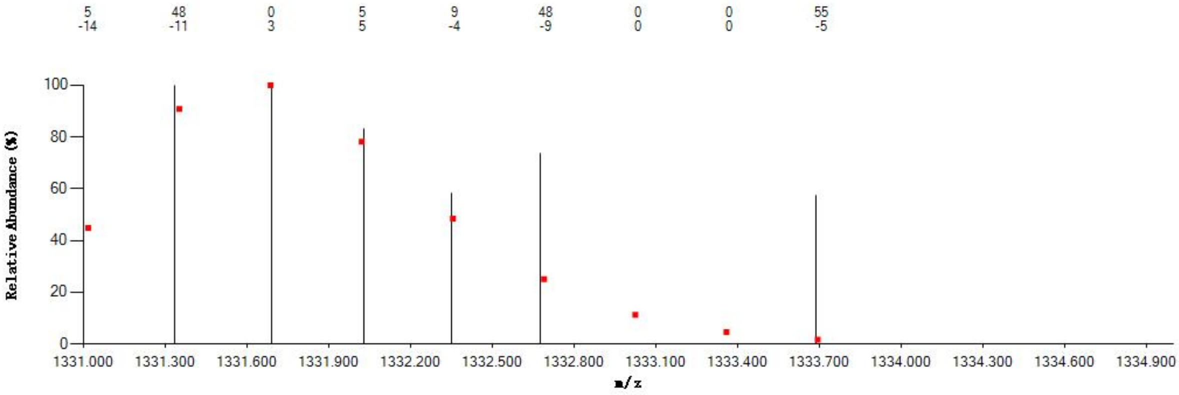


#### Annotated MS/MS Spectrum

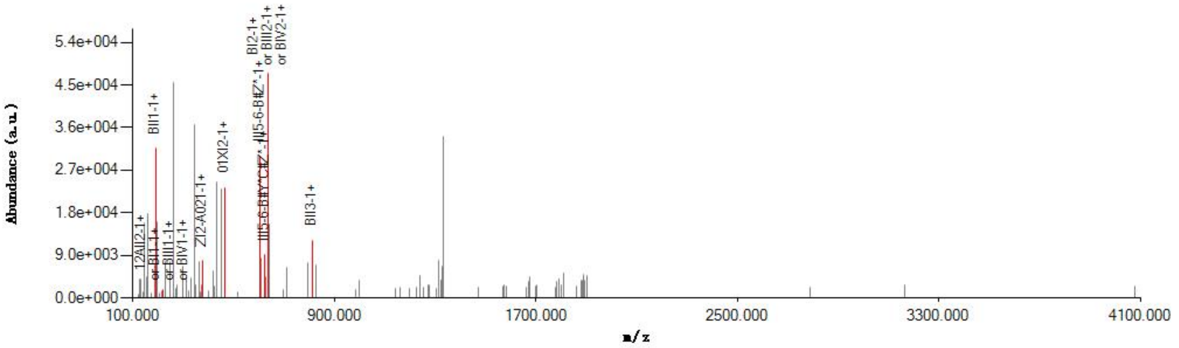


#### Graphical Fragmentation Map

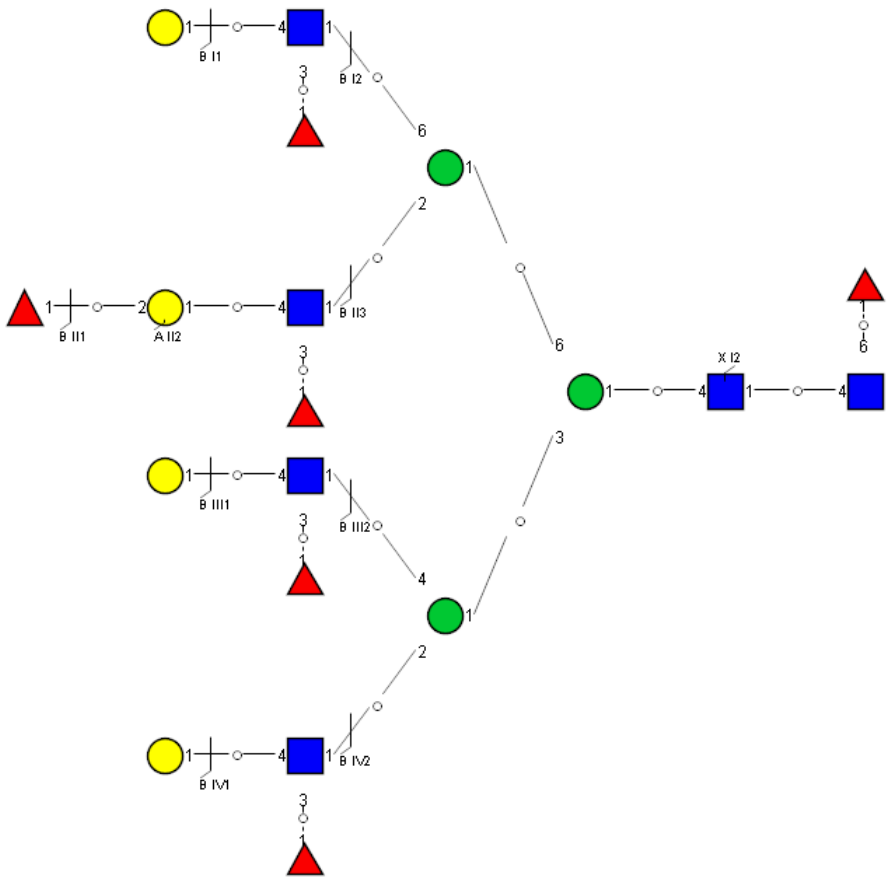

Spectrum Index: 45460(NoBHs:1)

#### Precursor iEF Map

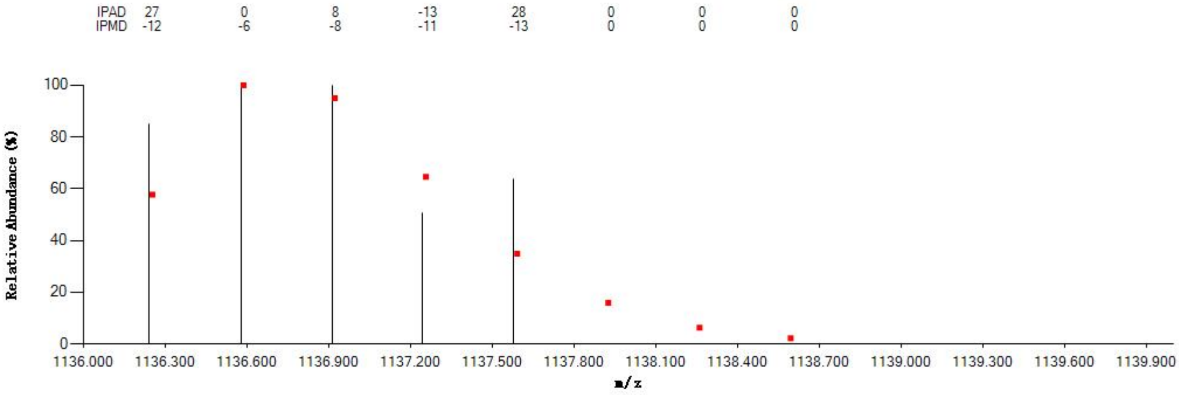


#### Annotated MS/MS Spectrum

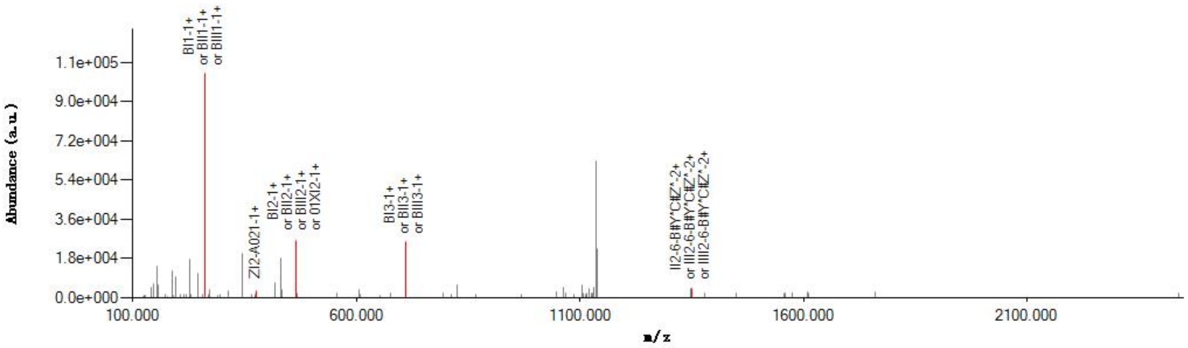


#### Graphical Fragmentation Map

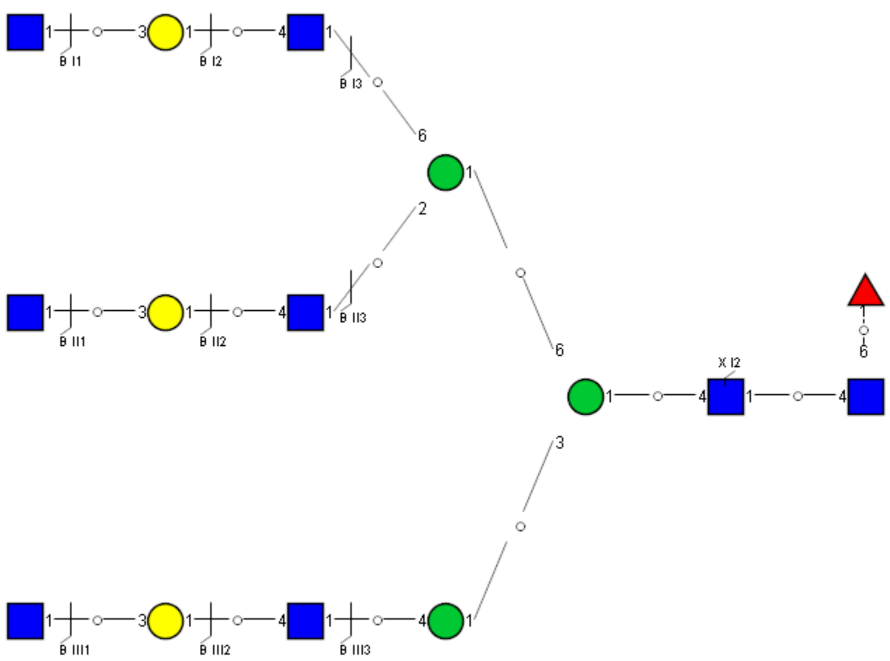

Spectrum Index: 47010(NoBHs:1)

Precursor iEF Map

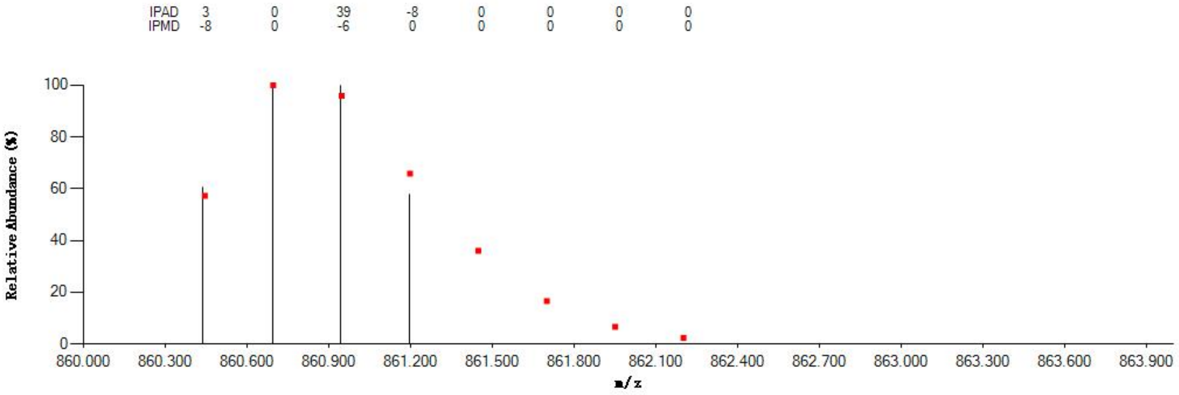


Annotated MS/MS Spectrum

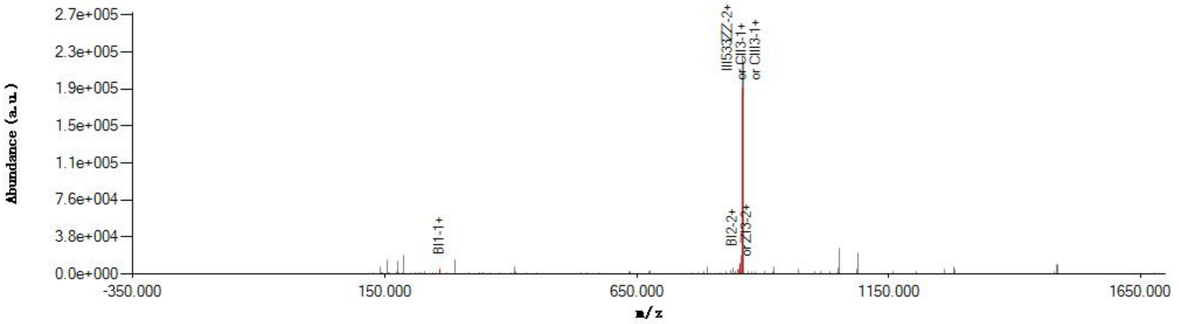


Graphical Fragmentation Map

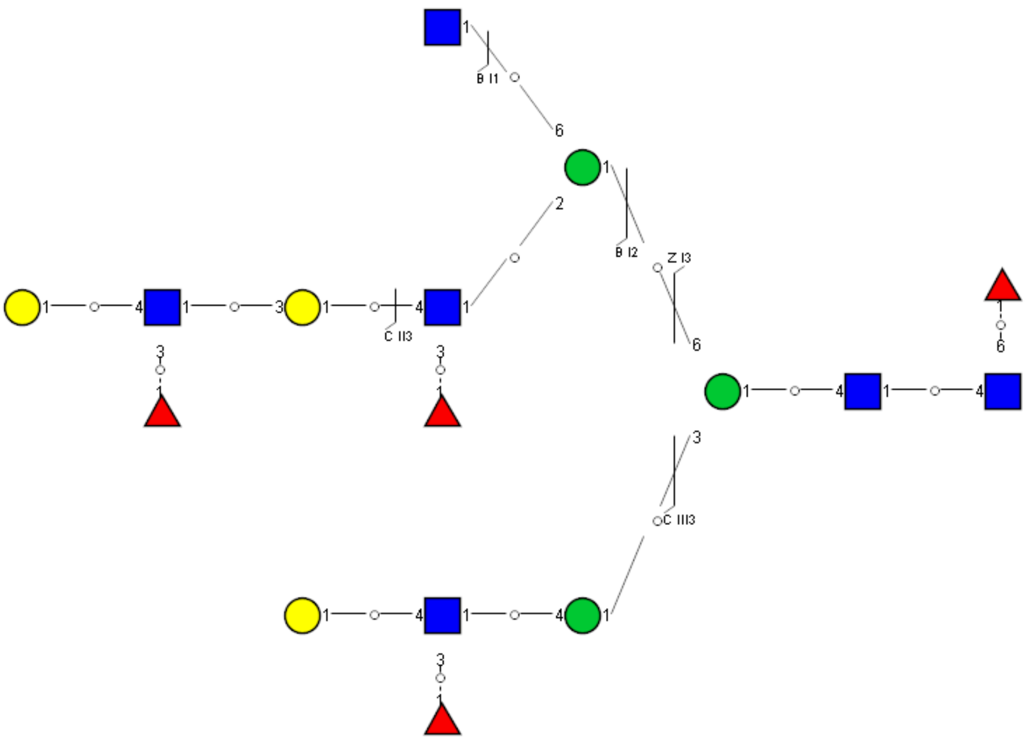

Spectrum Index: 25969(NoBHs:1)

#### Precursor iEF Map

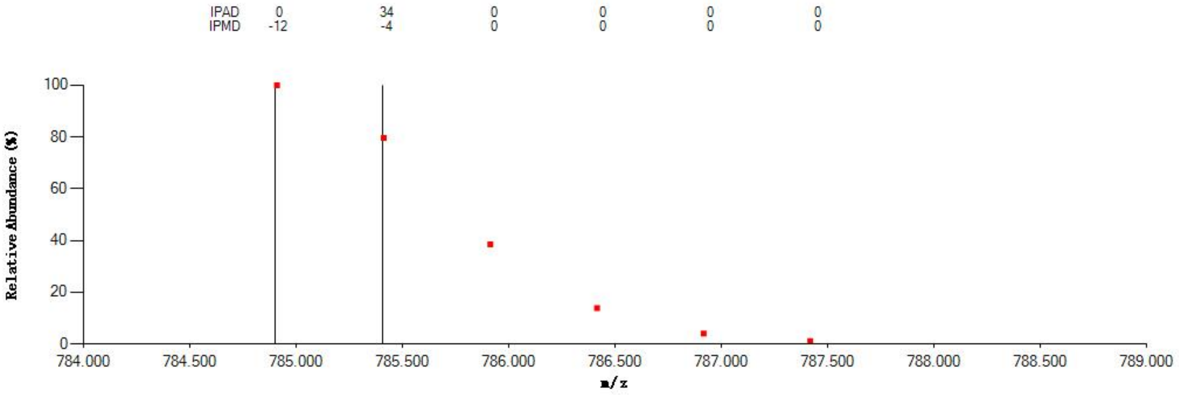


#### Annotated MS/MS Spectrum

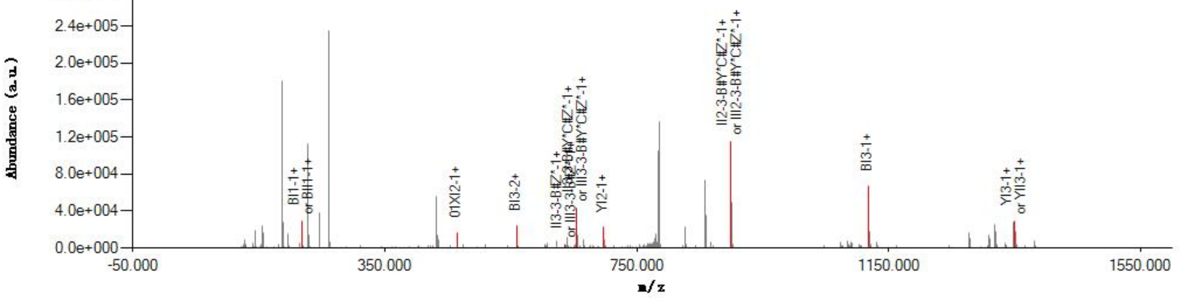


#### Graphical Fragmentation Map

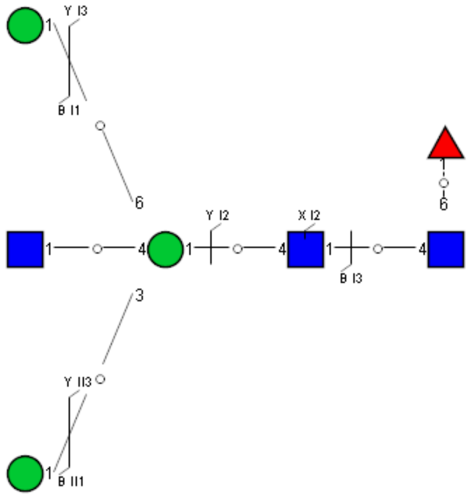

Spectrum Index: 27441(NoBHs:1)

#### Precursor iEF Map

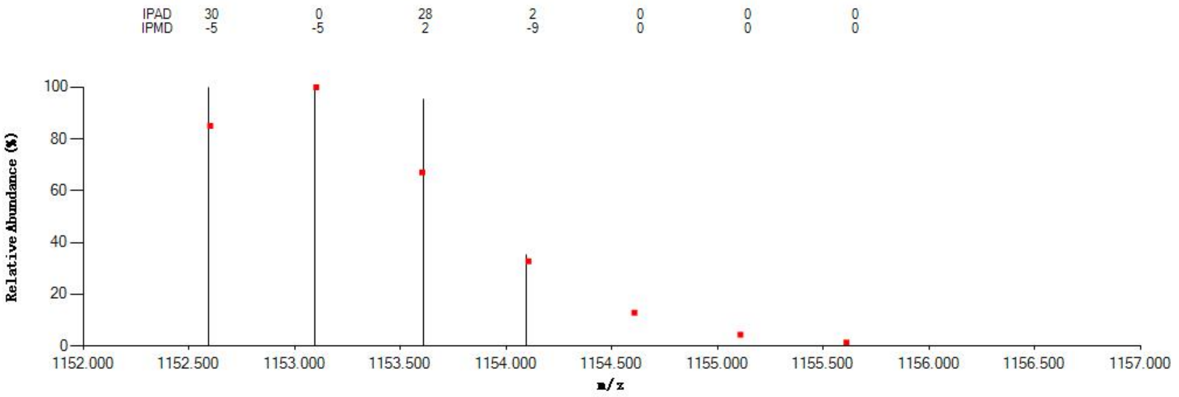


#### Annotated MS/MS Spectrum

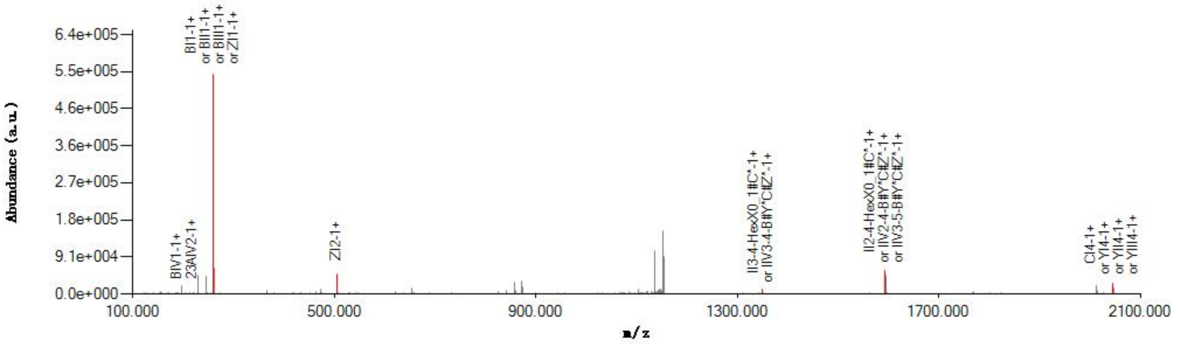


#### Graphical Fragmentation Map

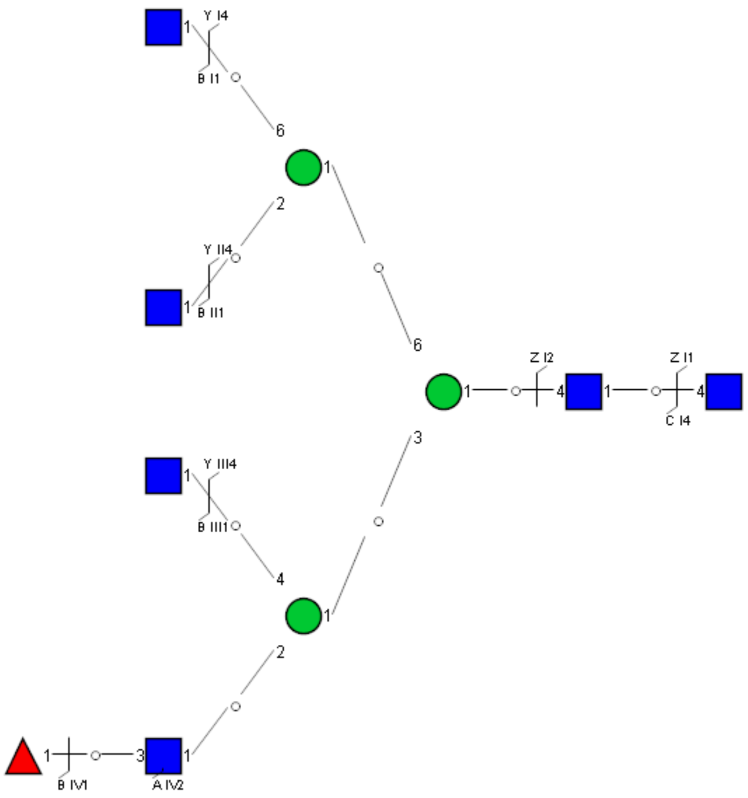

Spectrum Index: 27532(NoBHs:1)

#### Precursor iEF Map

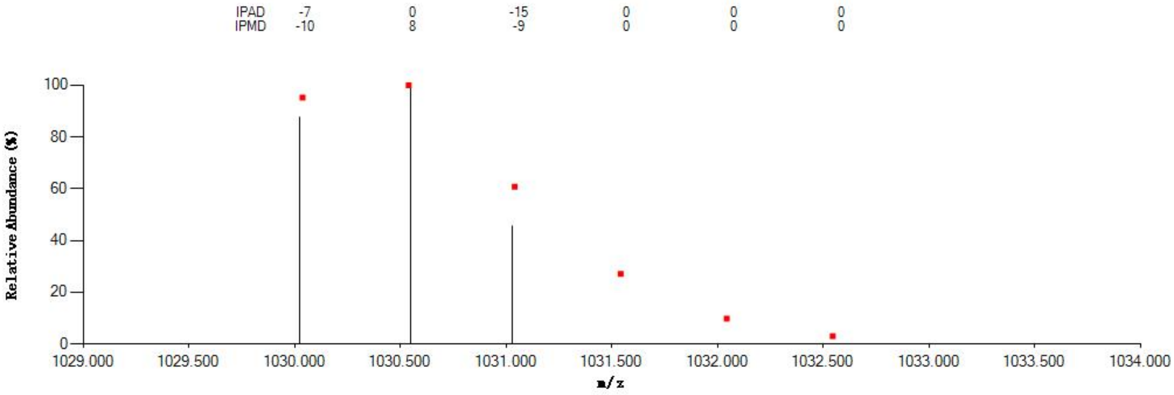


#### Annotated MS/MS Spectrum

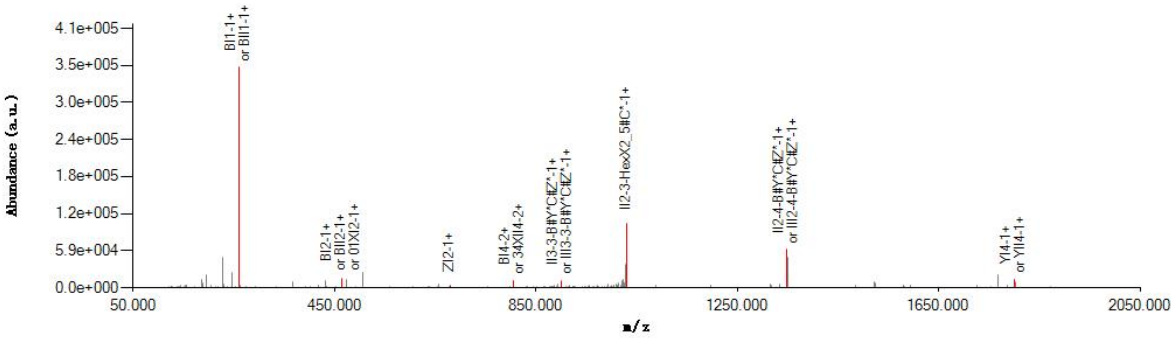


#### Graphical Fragmentation Map

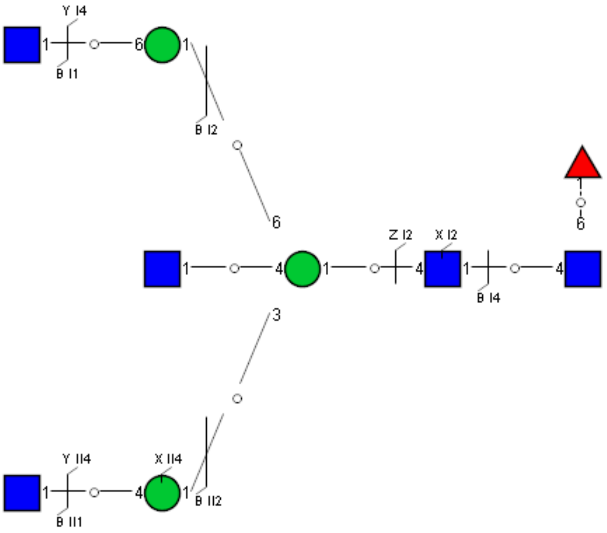

Spectrum Index: 27971(NoBHs:1)

#### Precursor iEF Map

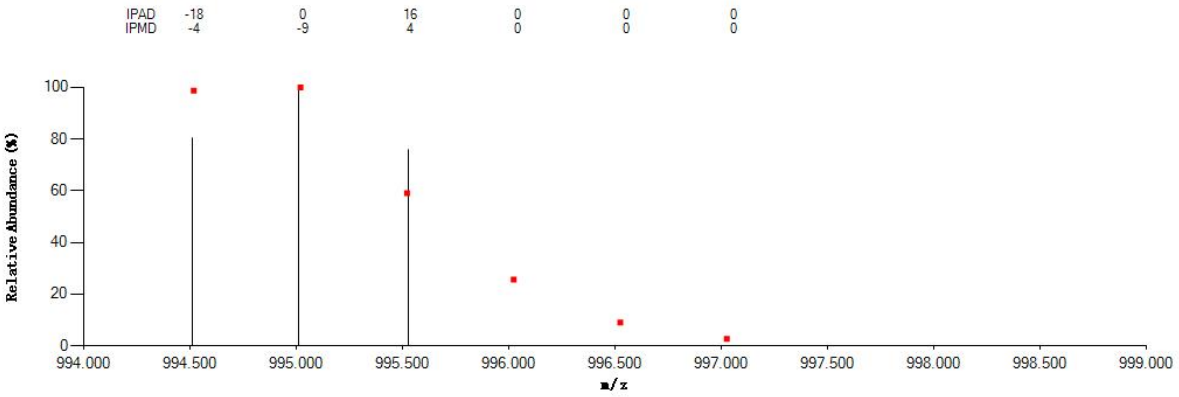


#### Annotated MS/MS Spectrum

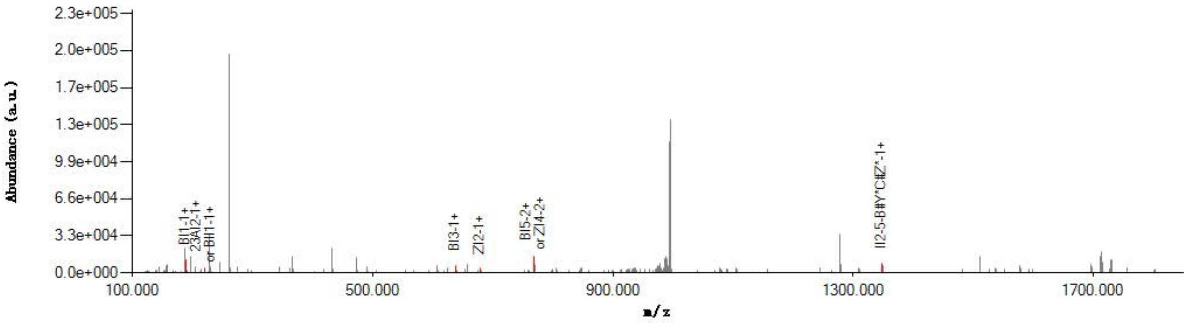


#### Graphical Fragmentation Map

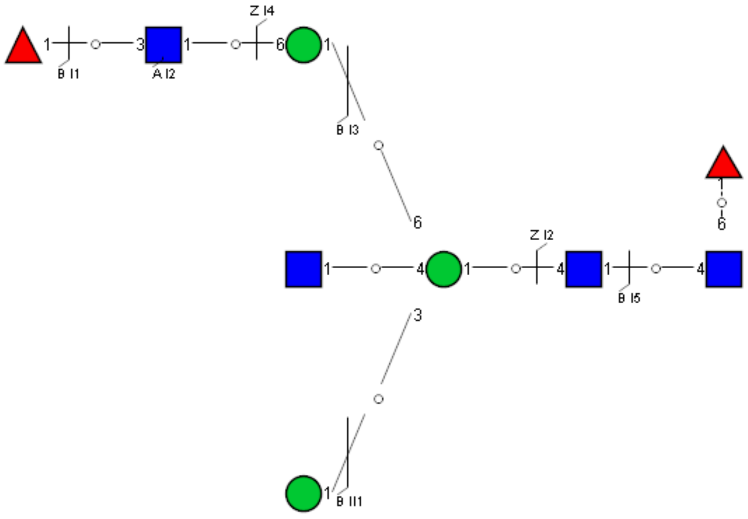

Spectrum Index: 28224(NoBHs:1)

#### Precursor iEF Map

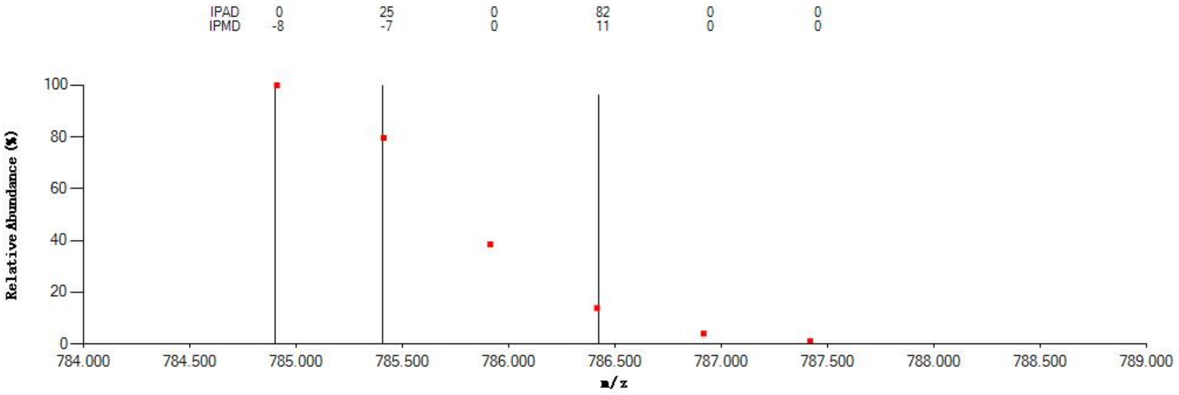

#### Annotated MS/MS Spectrum

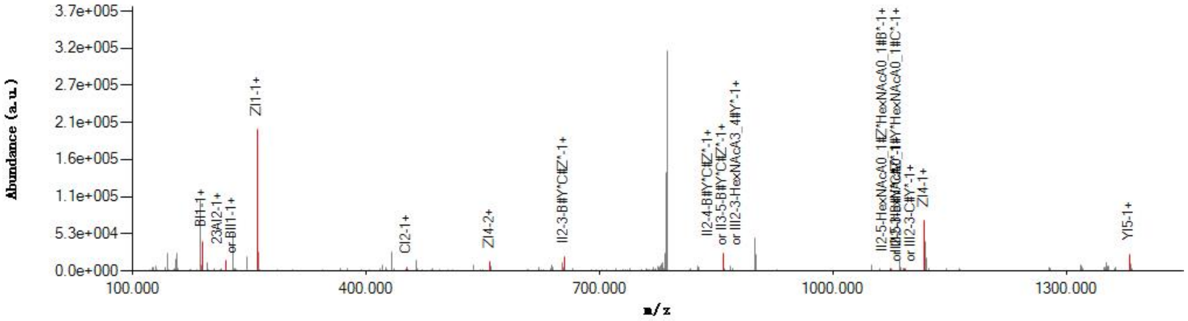

#### Graphical Fragmentation Map

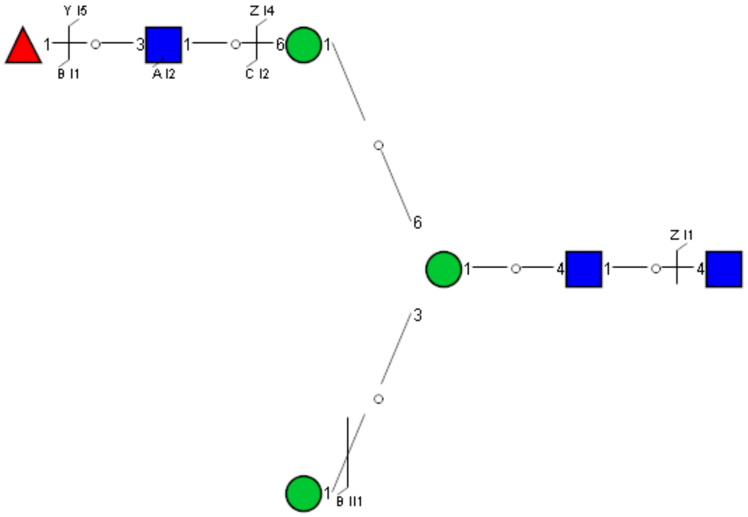

Spectrum Index: 28636(NoBHs:1)

Precursor iEF Map

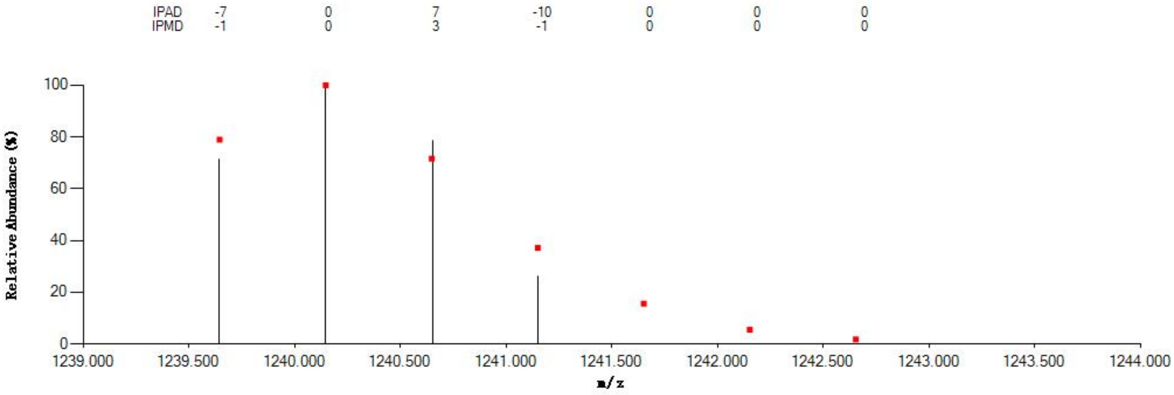


Annotated MS/MS Spectrum

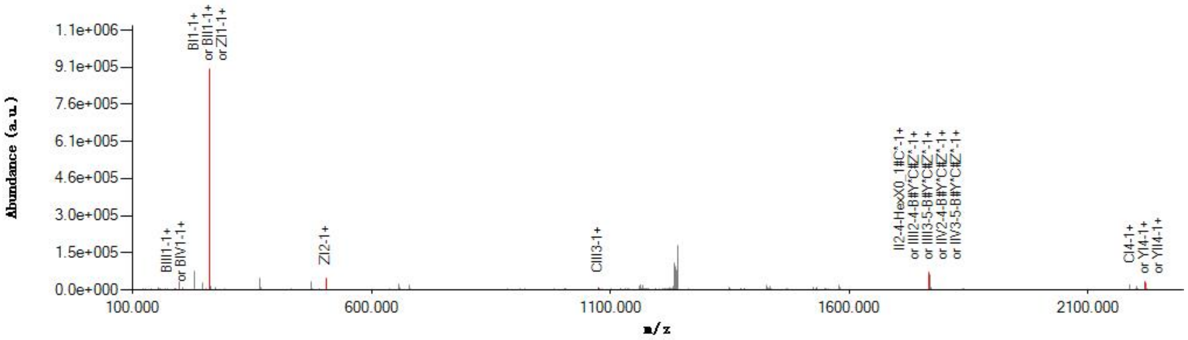


Graphical Fragmentation Map

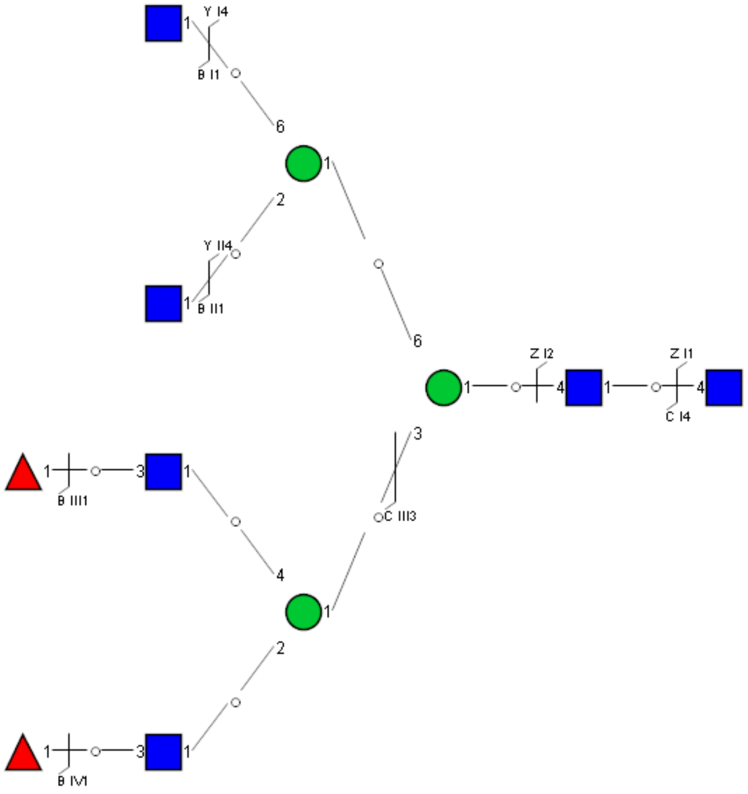

Spectrum Index: 29103(NoBHs:1)

#### Precursor iEF Map

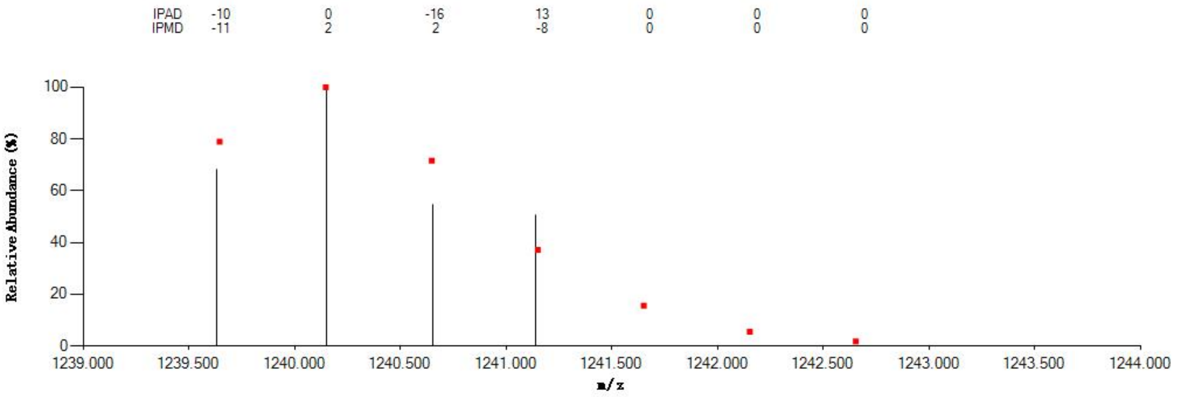


#### Annotated MS/MS Spectrum

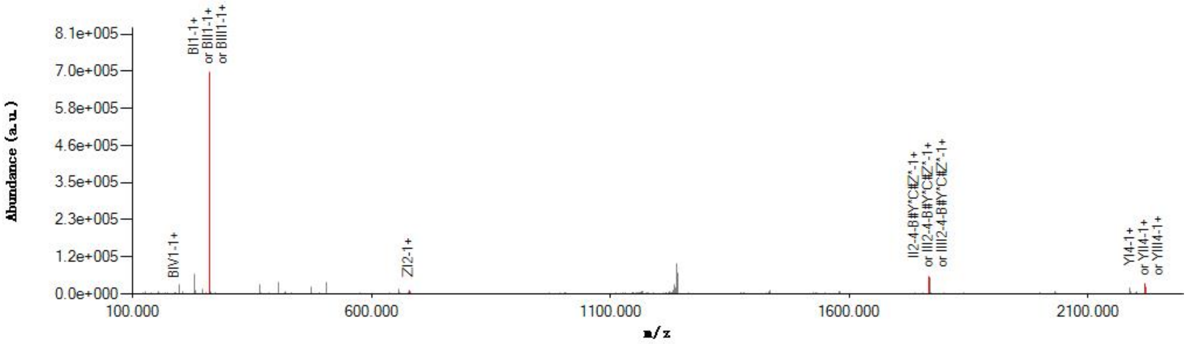


#### Graphical Fragmentation Map

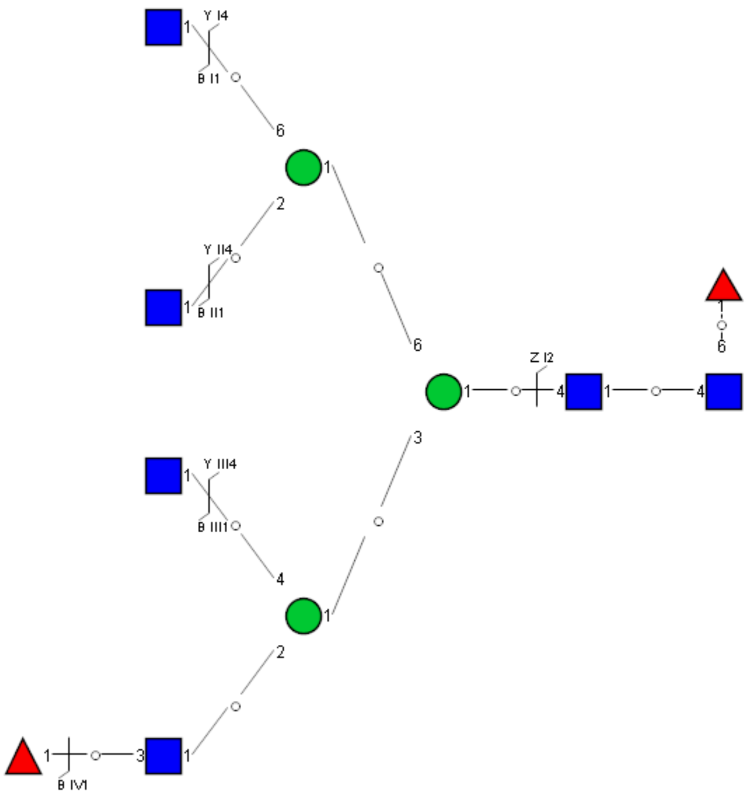

Spectrum Index: 29457(NoBHs:1)

#### Precursor iEF Map

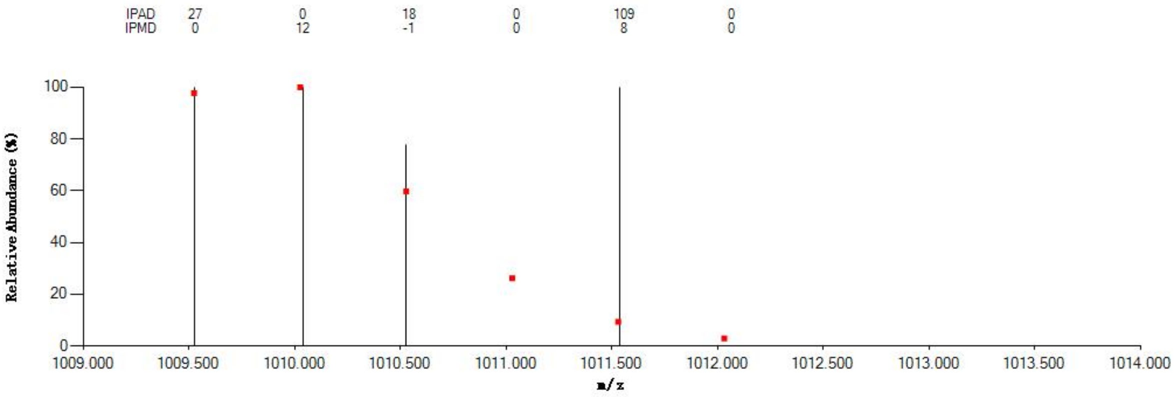

#### Annotated MS/MS Spectrum

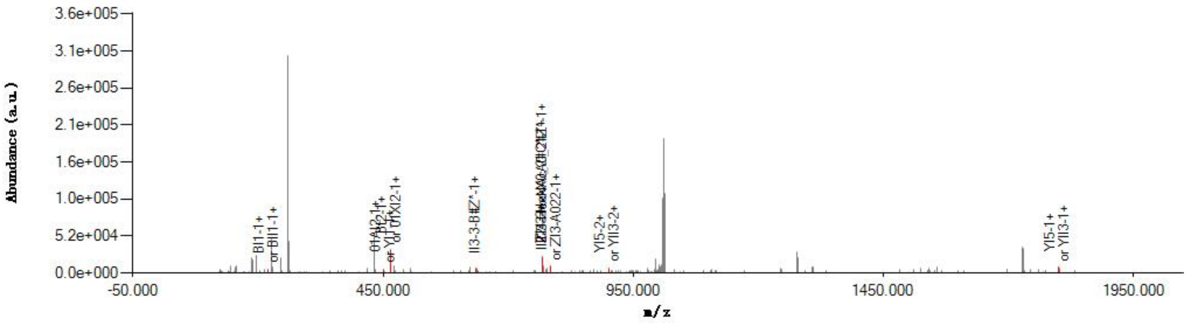

#### Graphical Fragmentation Map

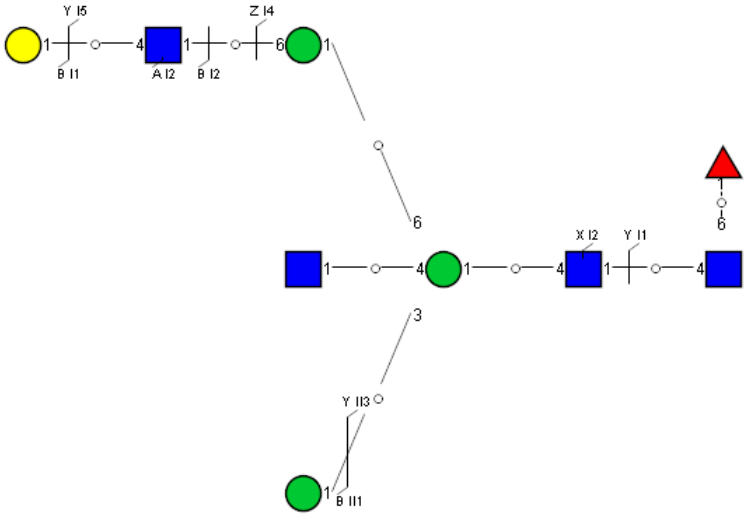

### Spectrum Index: 30709(NoBHs:1)

#### Precursor iEF Map

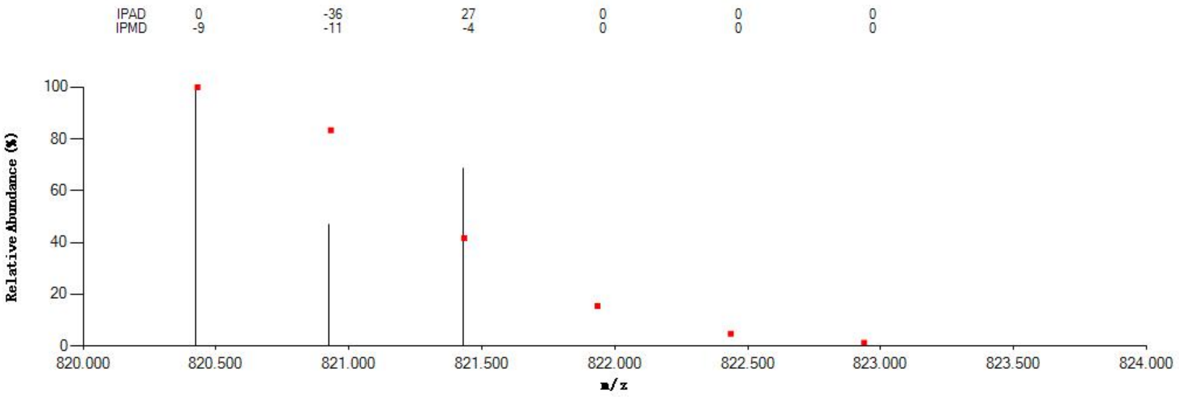


#### Annotated MS/MS Spectrum

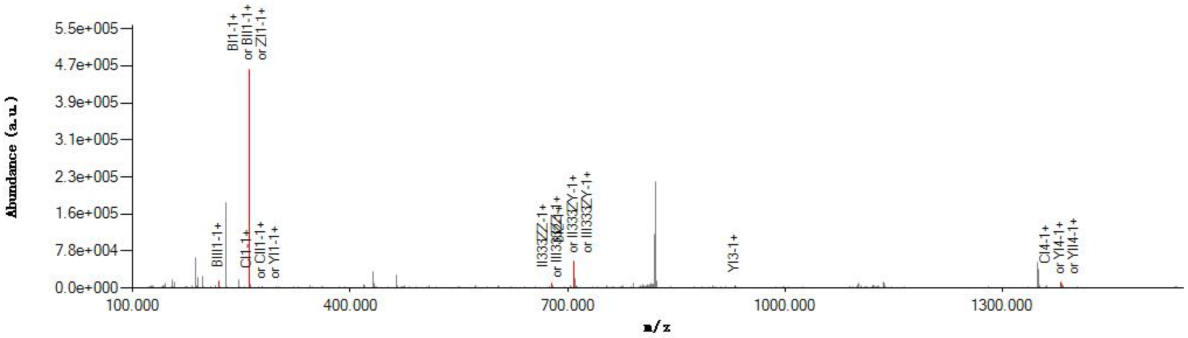


#### Graphical Fragmentation Map

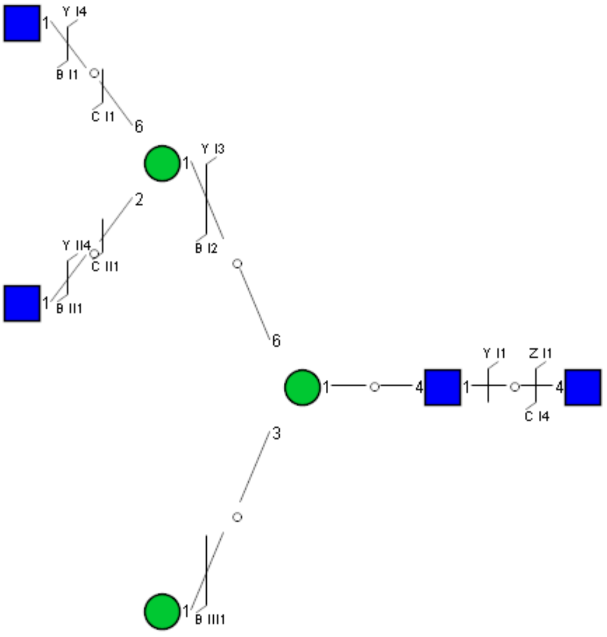

Spectrum Index: 30770(NoBHs:2)

#### Precursor iEF Map

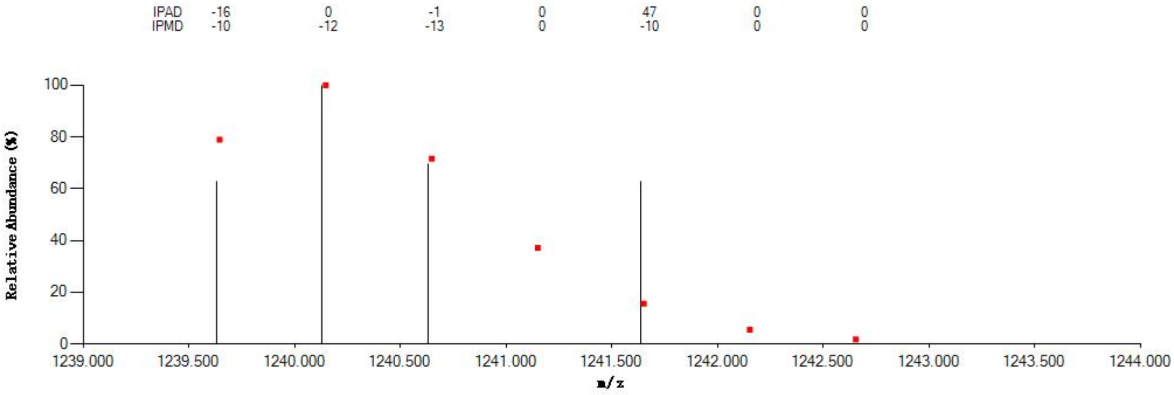


#### Annotated MS/MS Spectrum

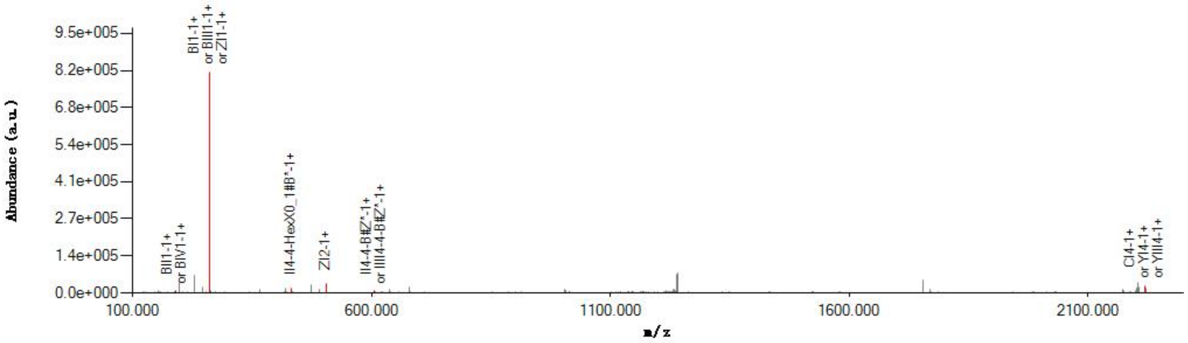


#### Graphical Fragmentation Map

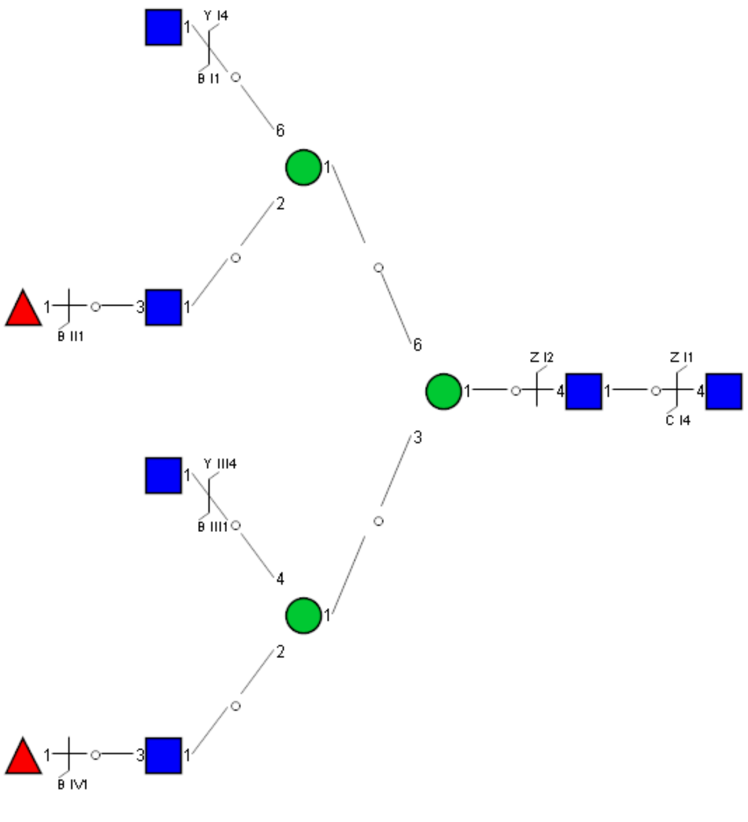

Spectrum Index: 31084(NoBHs:1)

#### Precursor iEF Map

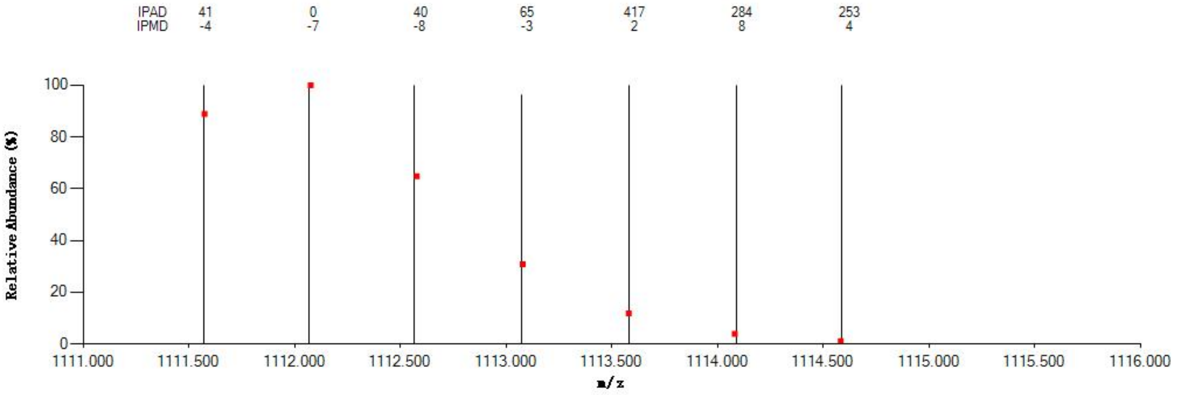


#### Annotated MS/MS Spectrum

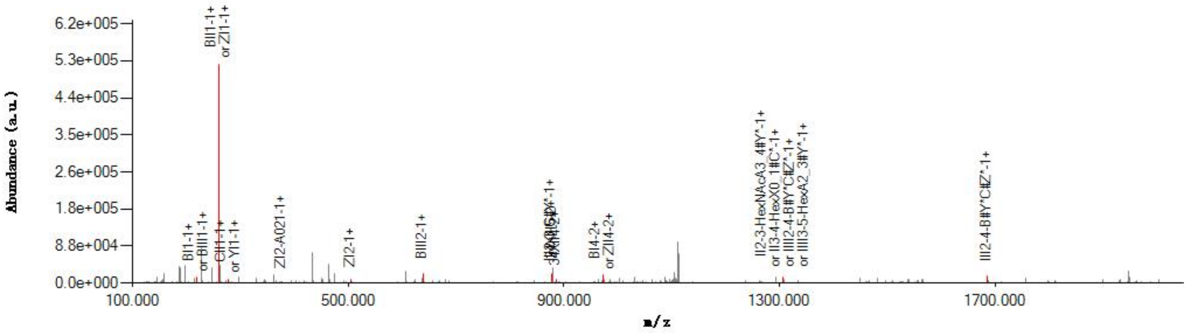


#### Graphical Fragmentation Map

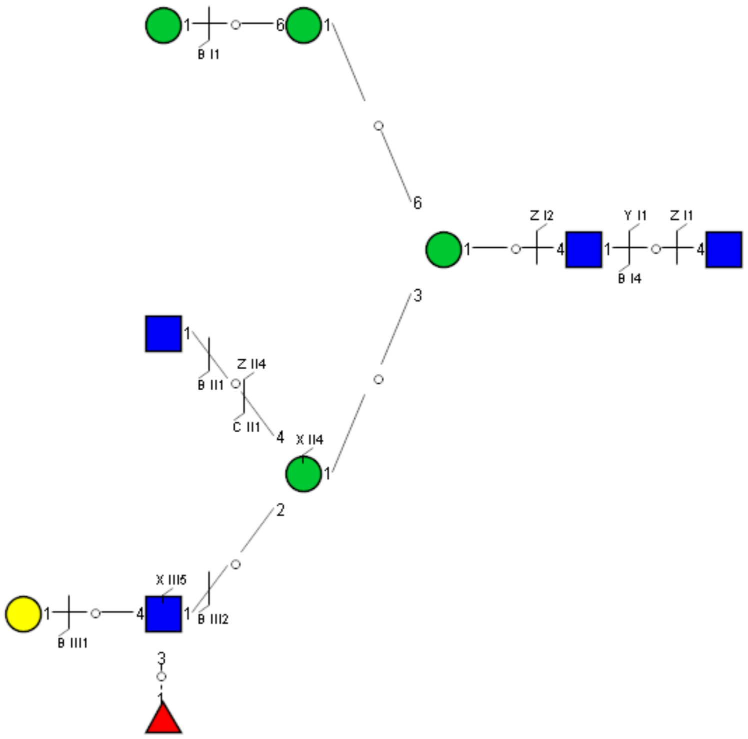

Spectrum Index: 31226(NoBHs:1)

#### Precursor iEF Map

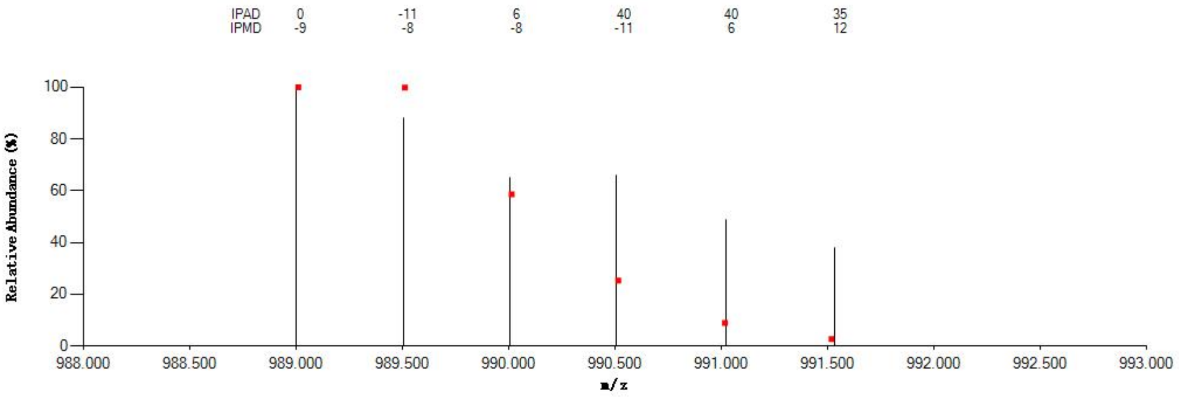

#### Annotated MS/MS Spectrum

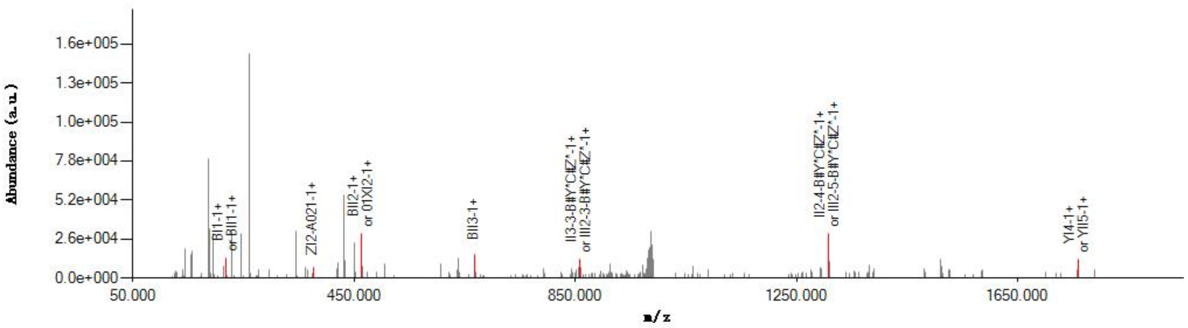

#### Graphical Fragmentation Map

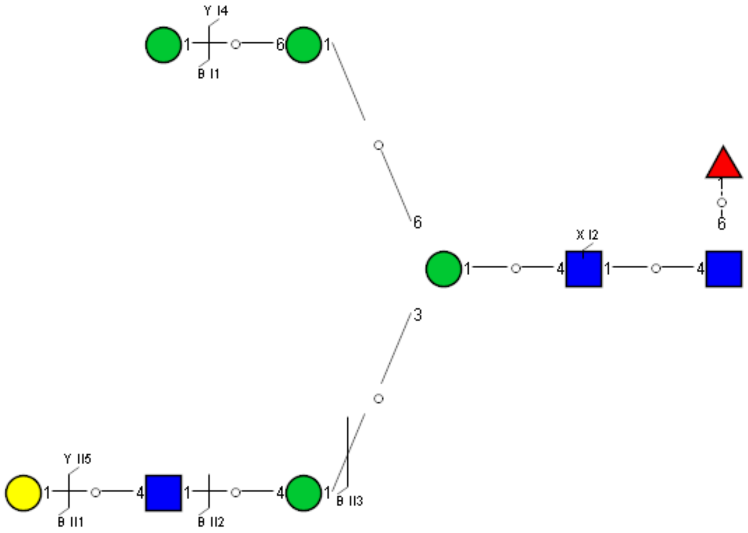

Spectrum Index: 32268(NoBHs:1)

#### Precursor iEF Map

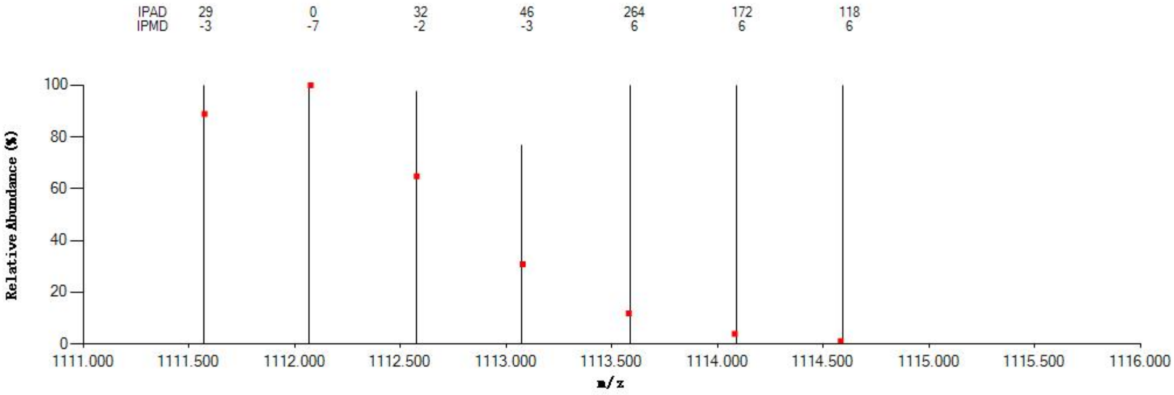


#### Annotated MS/MS Spectrum

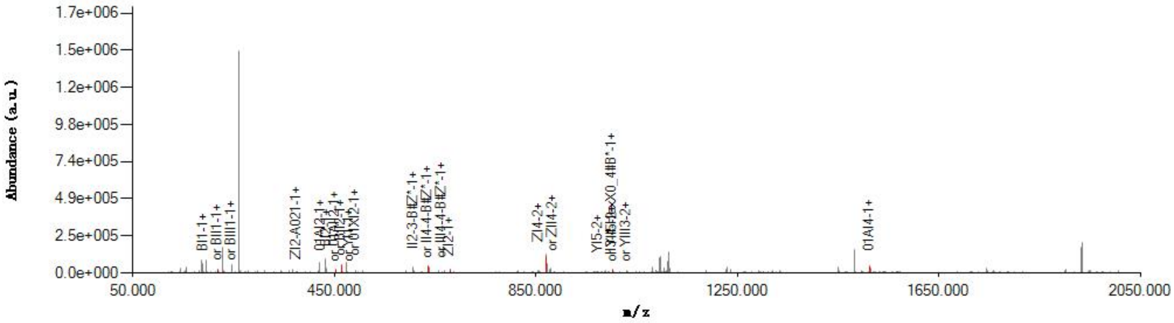


#### Graphical Fragmentation Map

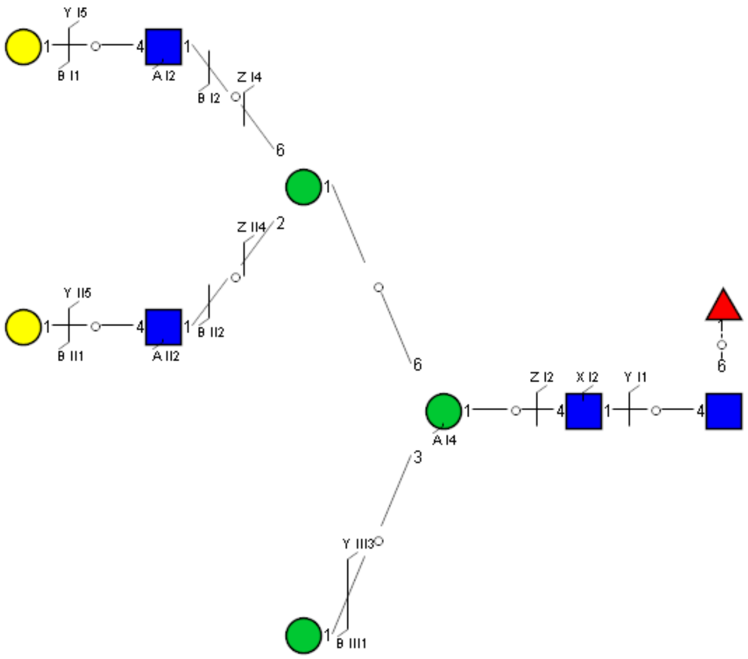

Spectrum Index: 32538(NoBHs:1)

#### Precursor iEF Map

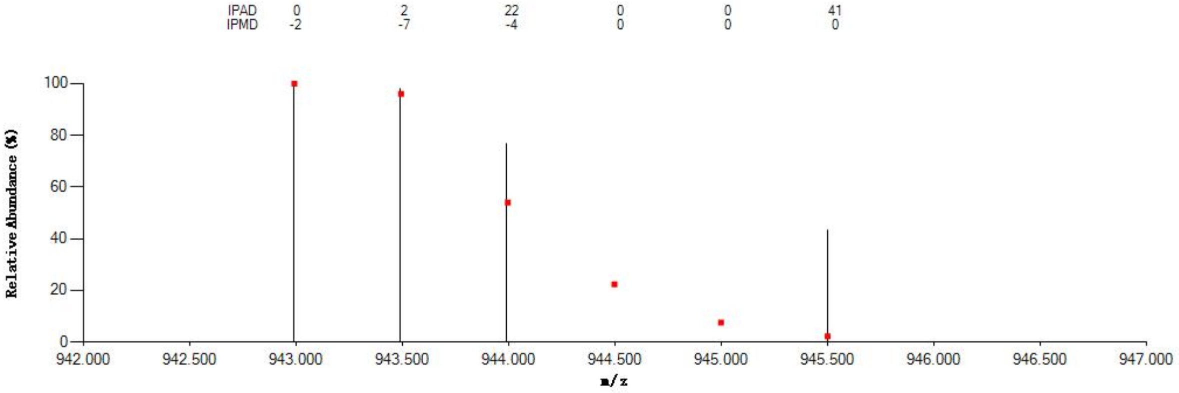


#### Annotated MS/MS Spectrum

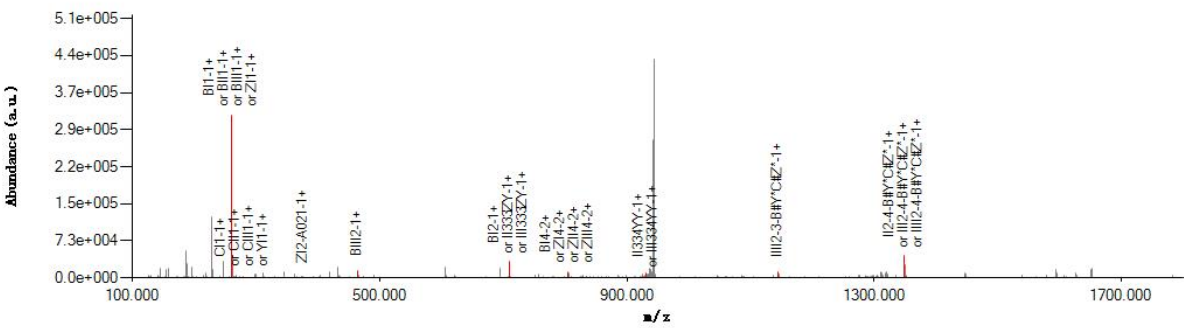


#### Graphical Fragmentation Map

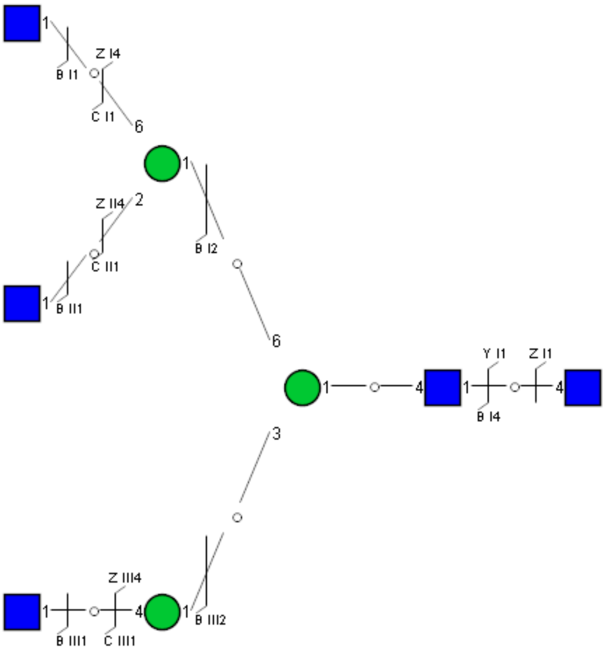

### Spectrum Index: 32546(NoBHs:1)

#### Precursor iEF Map

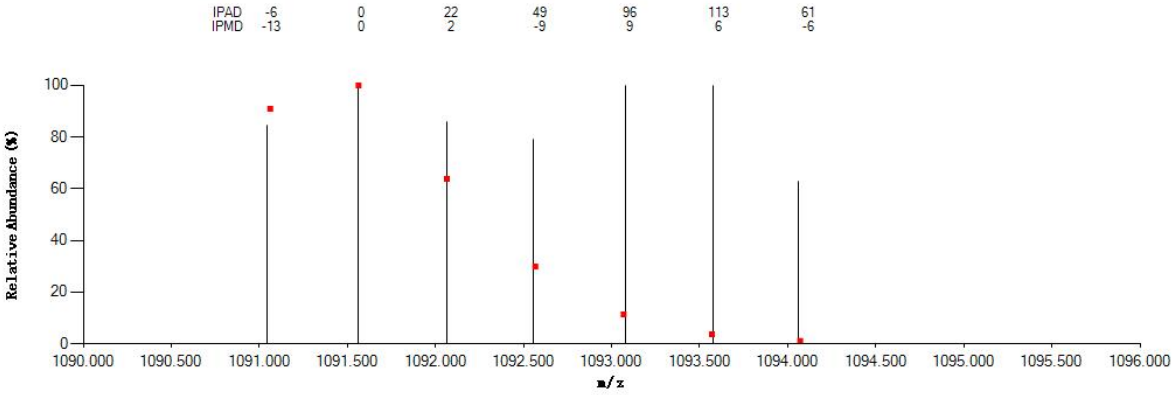


#### Annotated MS/MS Spectrum

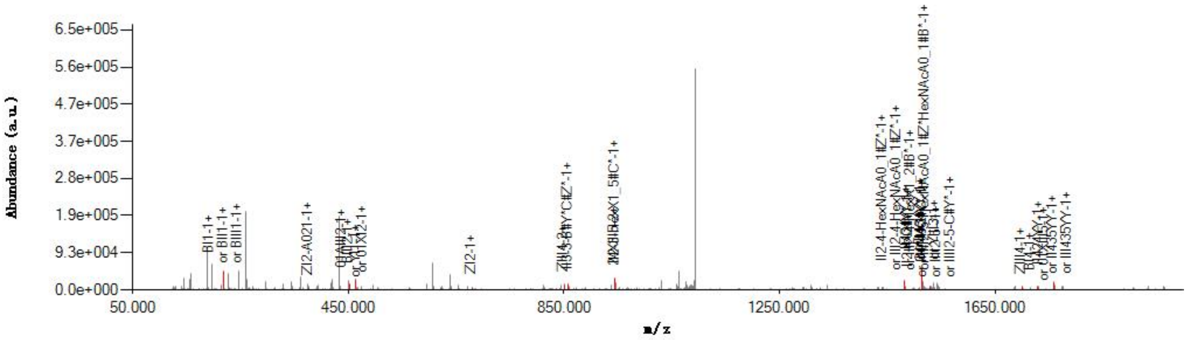


#### Graphical Fragmentation Map

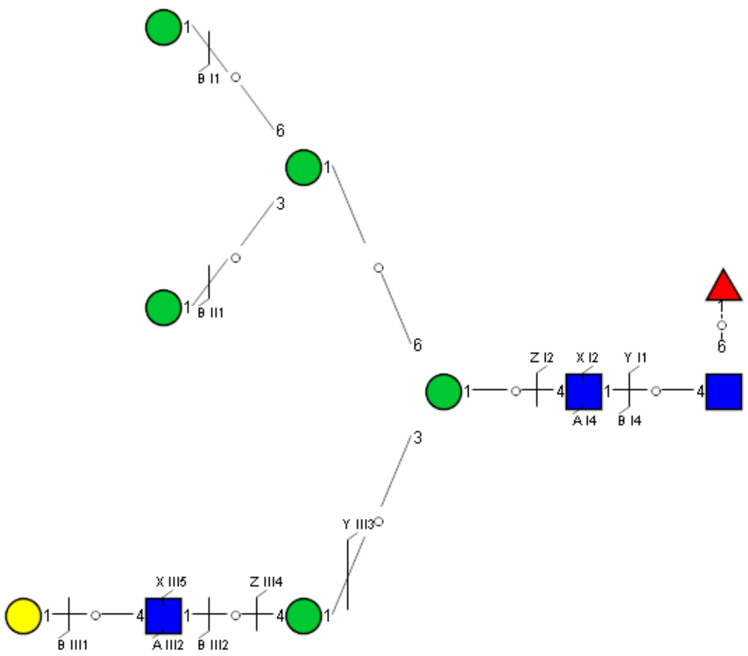

Spectrum Index: 33220(NoBHs:1)

#### Precursor iEF Map

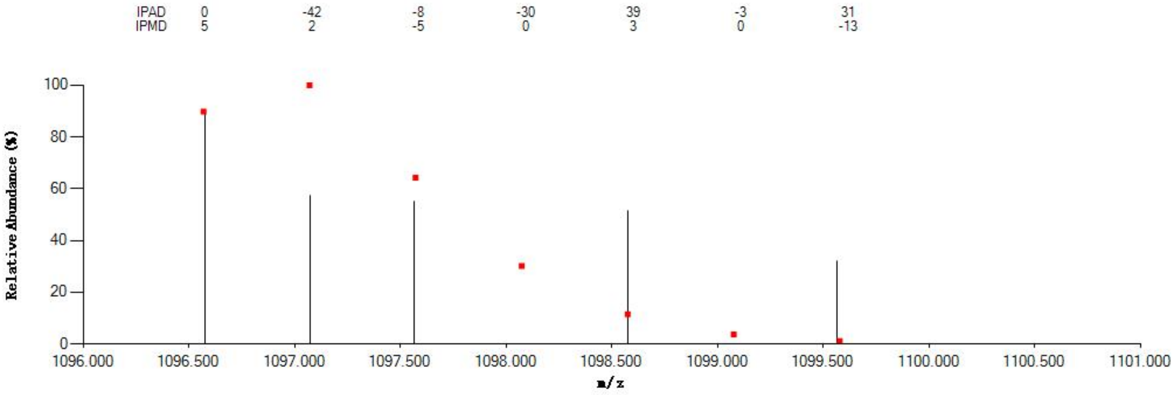


#### Annotated MS/MS Spectrum

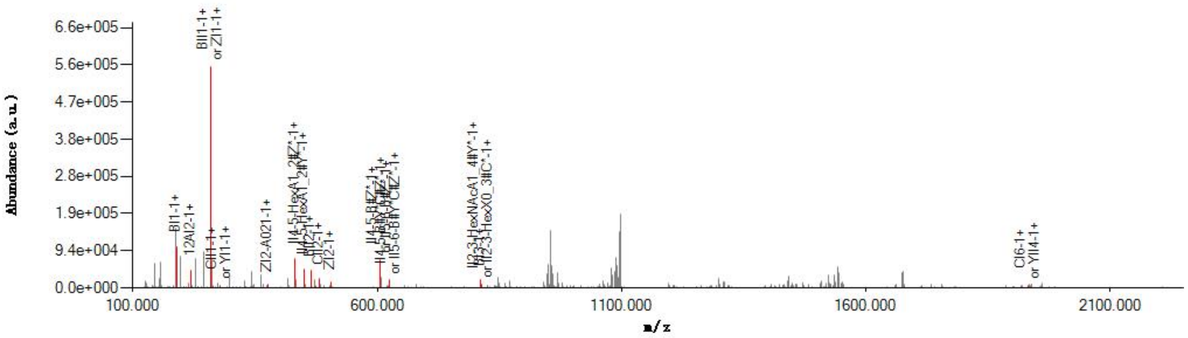


#### Graphical Fragmentation Map

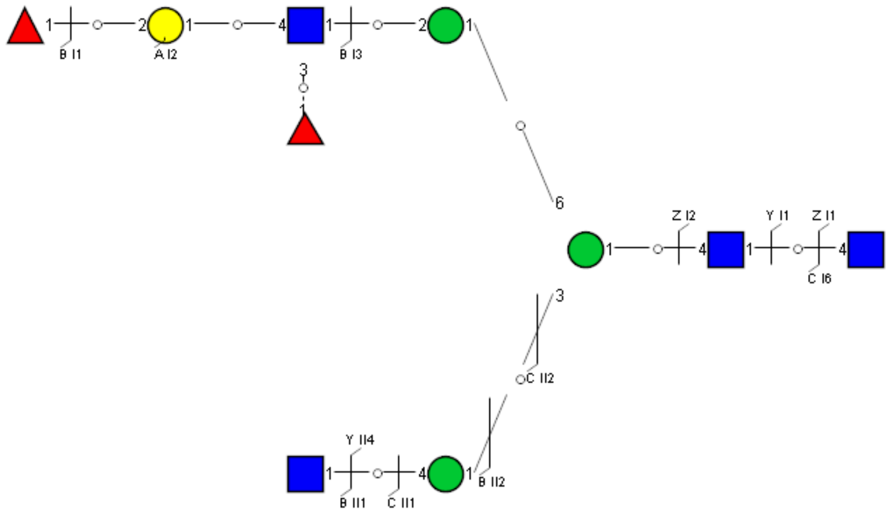

Spectrum Index: 33823(NoBHs:1)

#### Precursor iEF Map

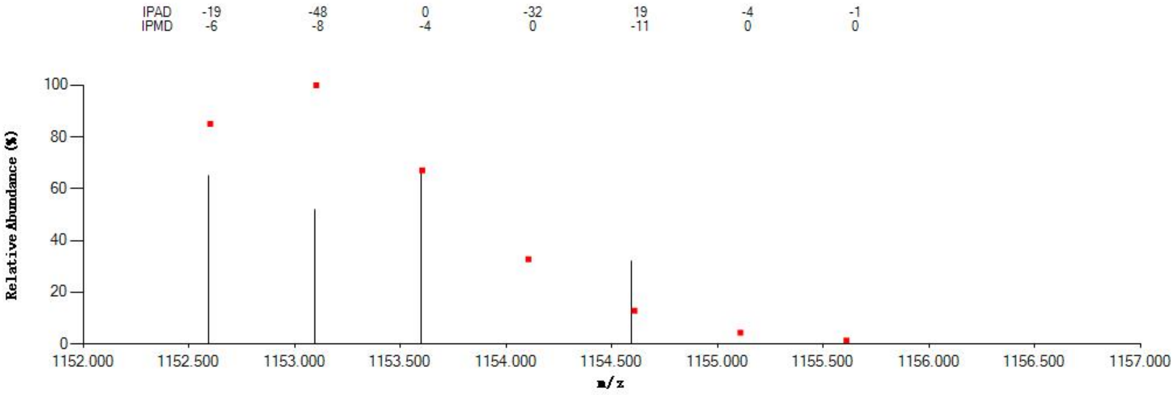

#### Annotated MS/MS Spectrum

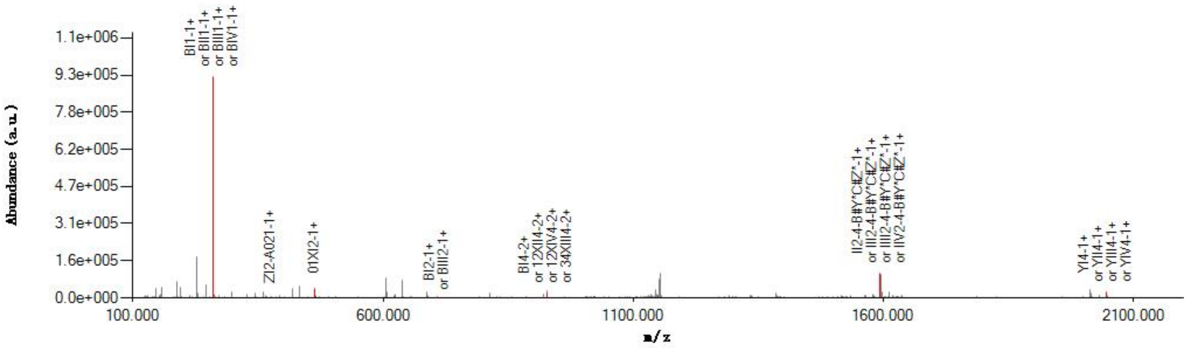

#### Graphical Fragmentation Map

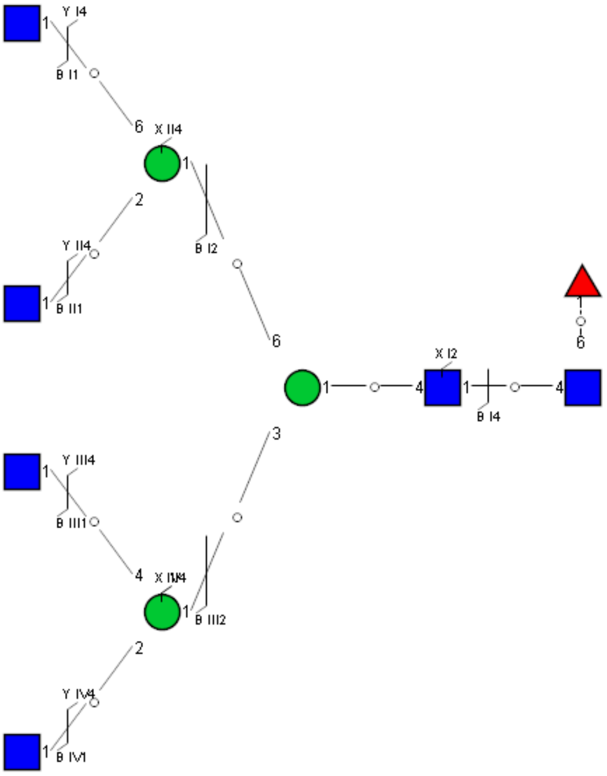

Spectrum Index: 34157(NoBHs:1)

#### Precursor iEF Map

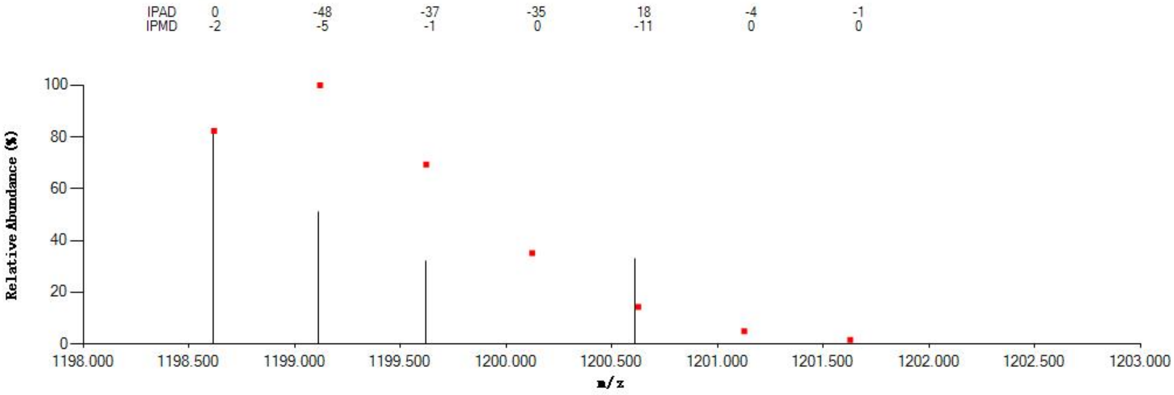


#### Annotated MS/MS Spectrum

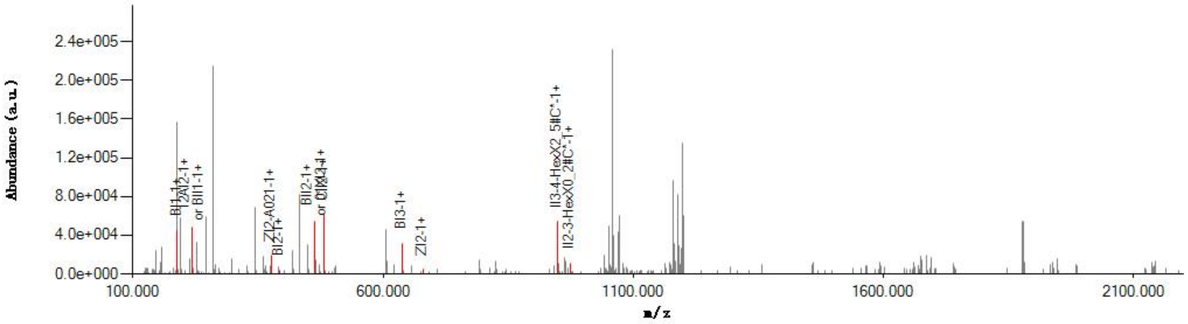


#### Graphical Fragmentation Map

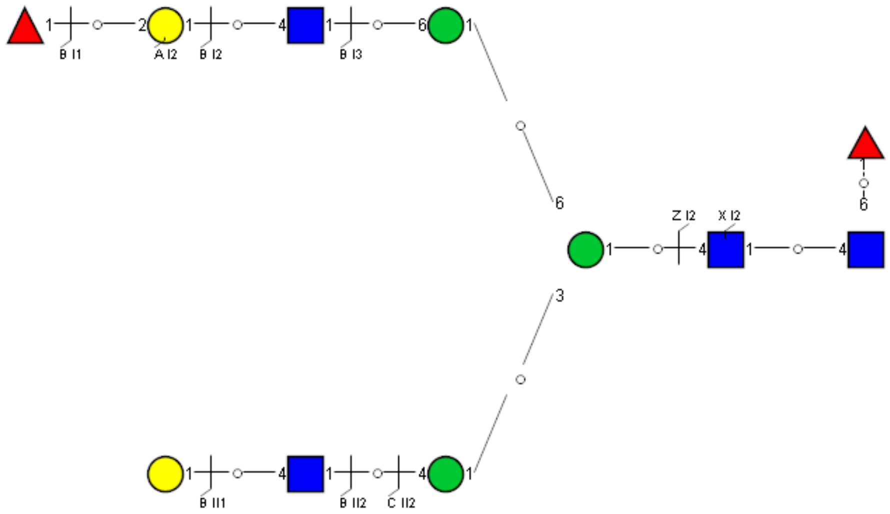

### Spectrum Index: 34549(NoBHs:1)

#### Precursor iEF Map

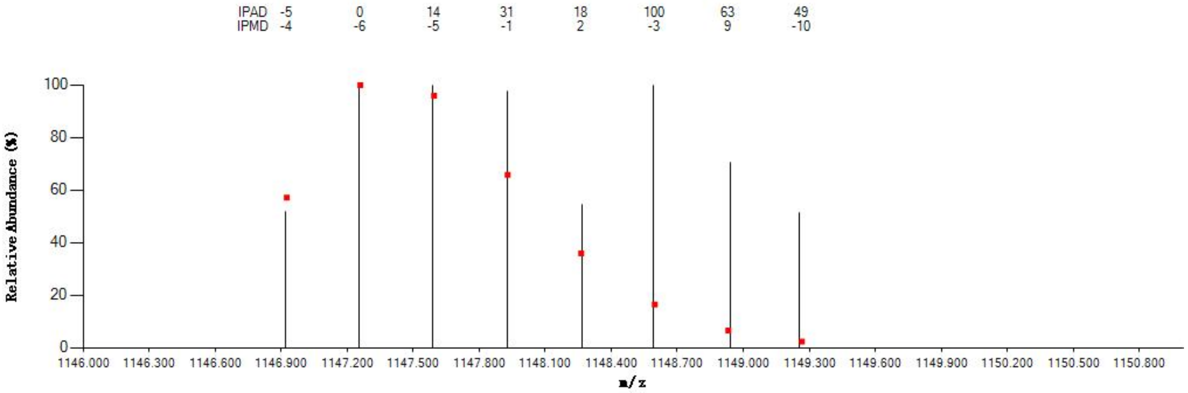


#### Annotated MS/MS Spectrum

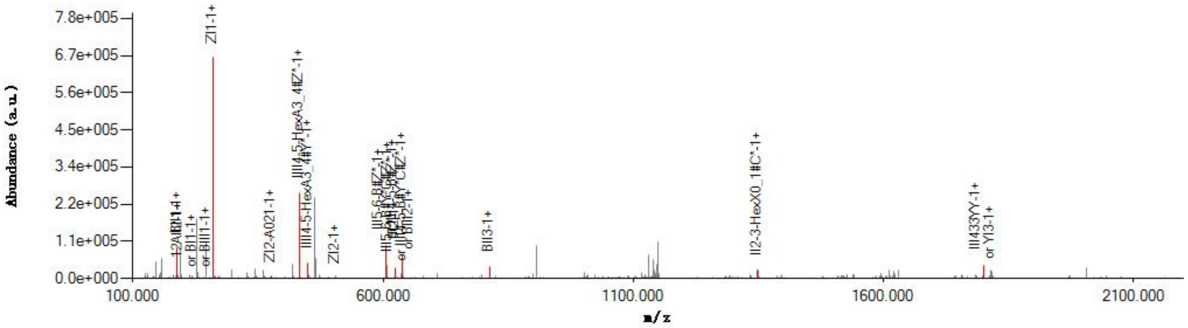


#### Graphical Fragmentation Map

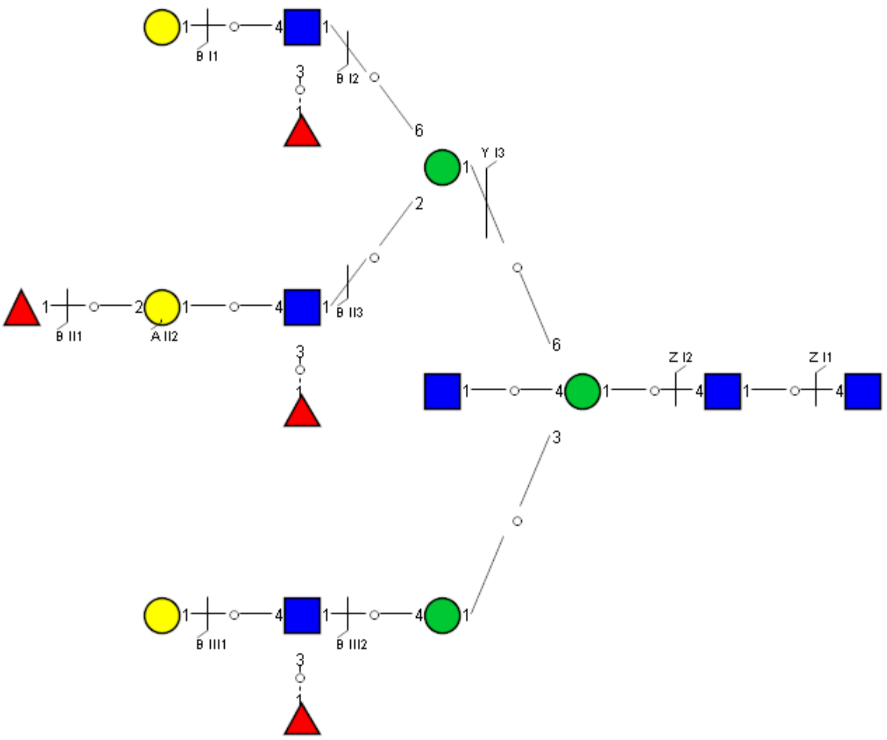

Spectrum Index: 34935(NoBHs:1)

#### Precursor iEF Map

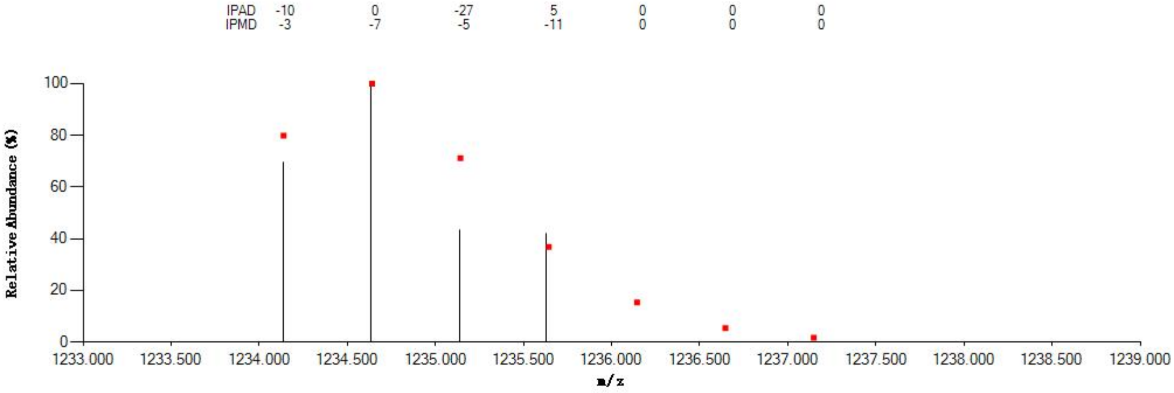


#### Annotated MS/MS Spectrum

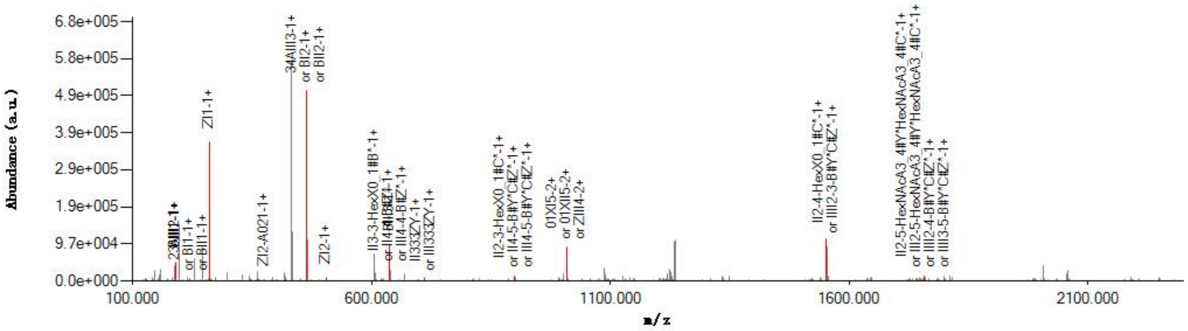


#### Graphical Fragmentation Map

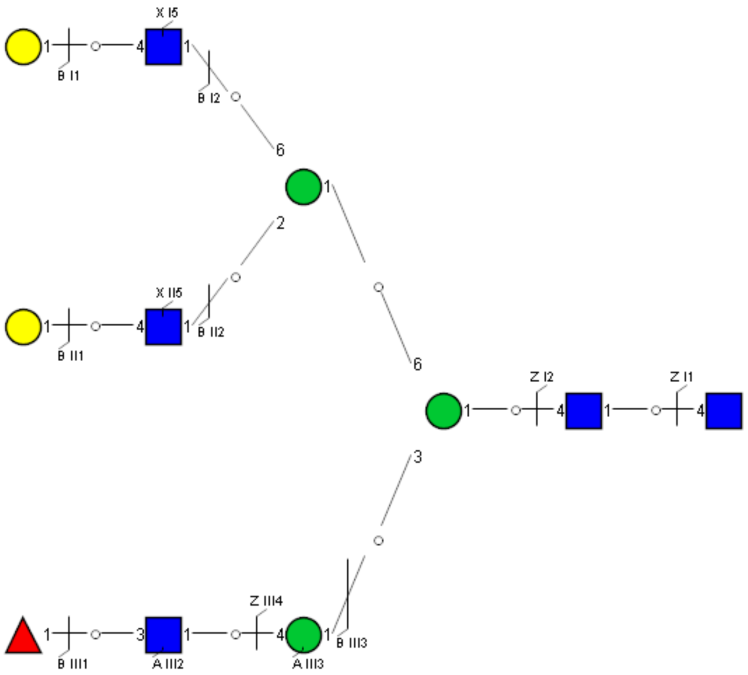

Spectrum Index: 35289(NoBHs:1)

#### Precursor iEF Map

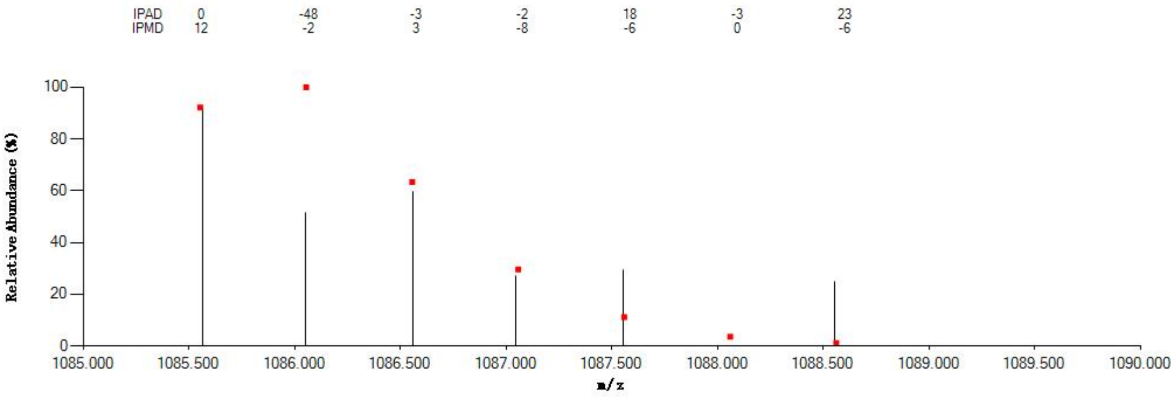


#### Annotated MS/MS Spectrum

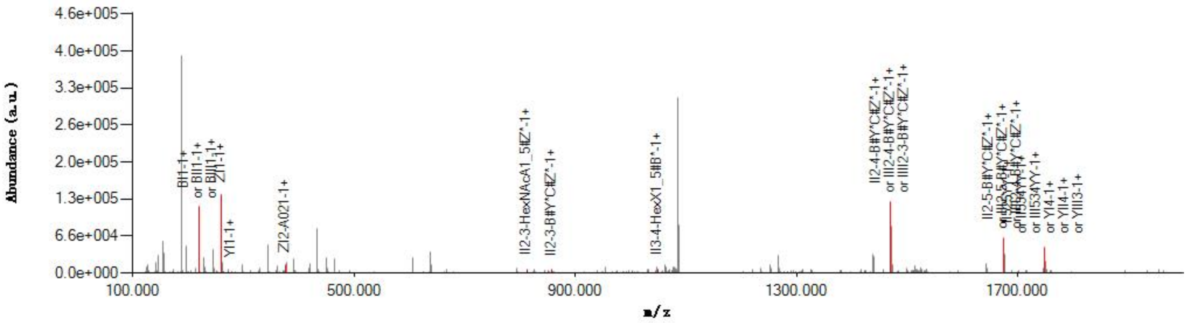


#### Graphical Fragmentation Map

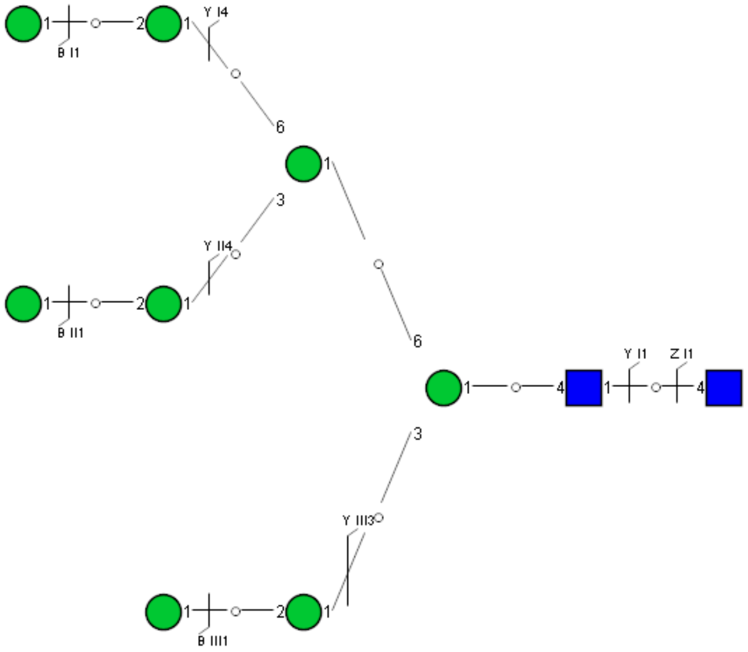

Spectrum Index: 36245(NoBHs:1)

#### Precursor iEF Map

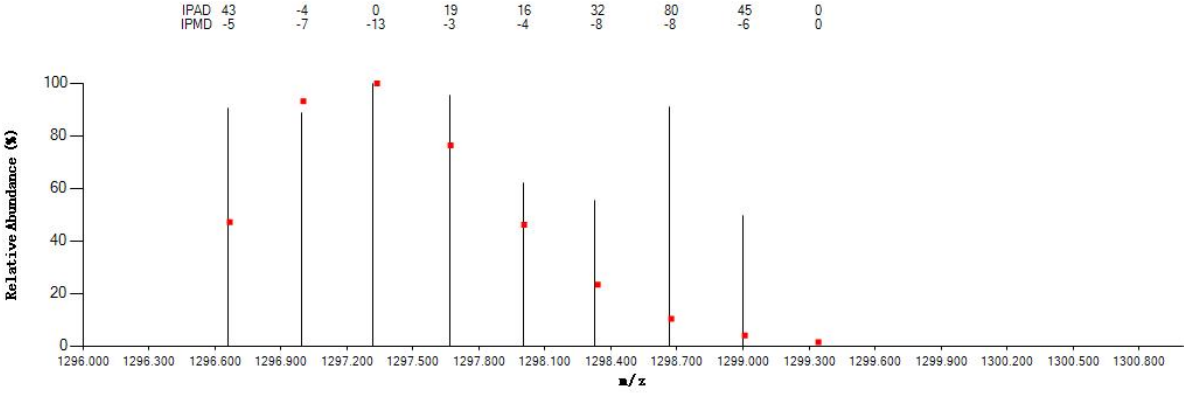


#### Annotated MS/MS Spectrum

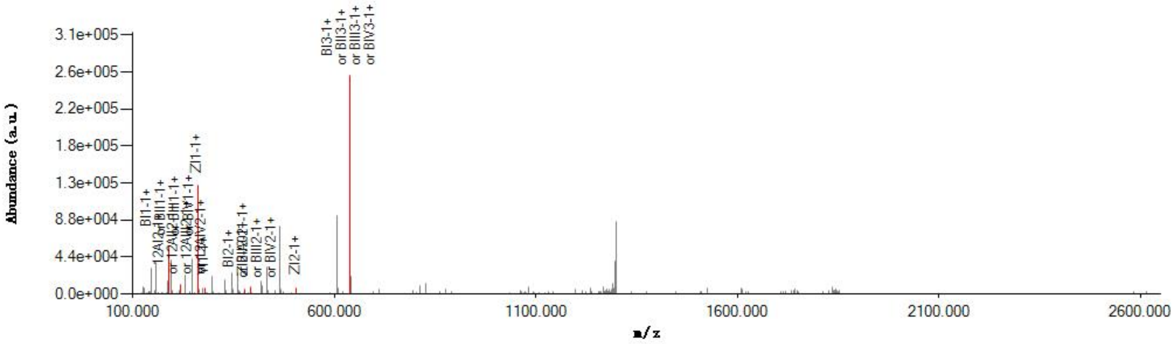


#### Graphical Fragmentation Map

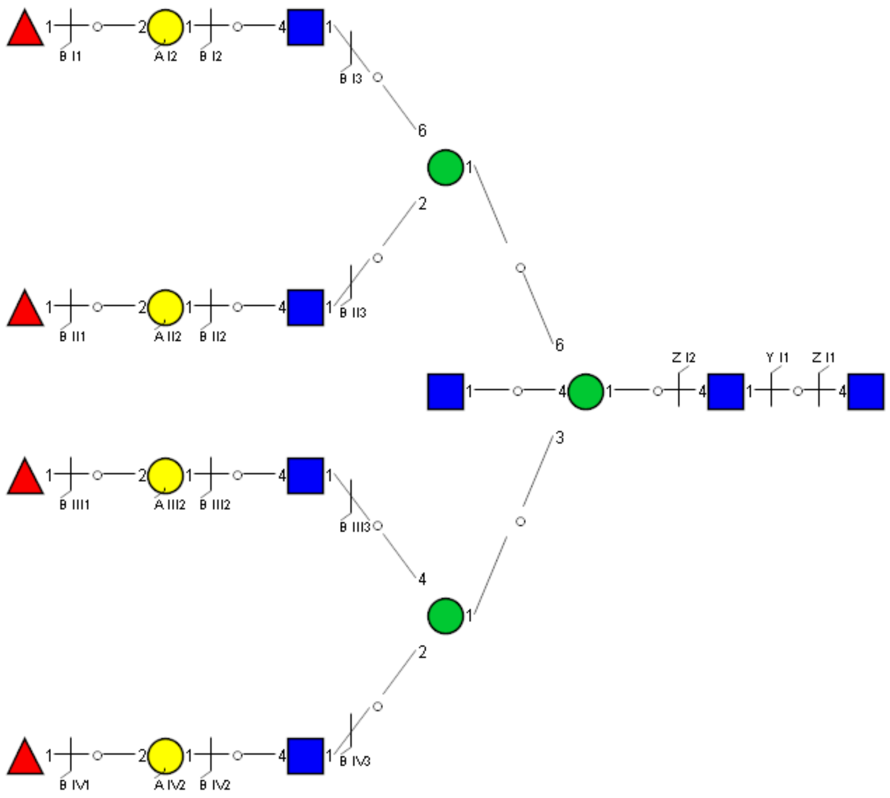

Spectrum Index: 38360(NoBHs:1)

#### Precursor iEF Map

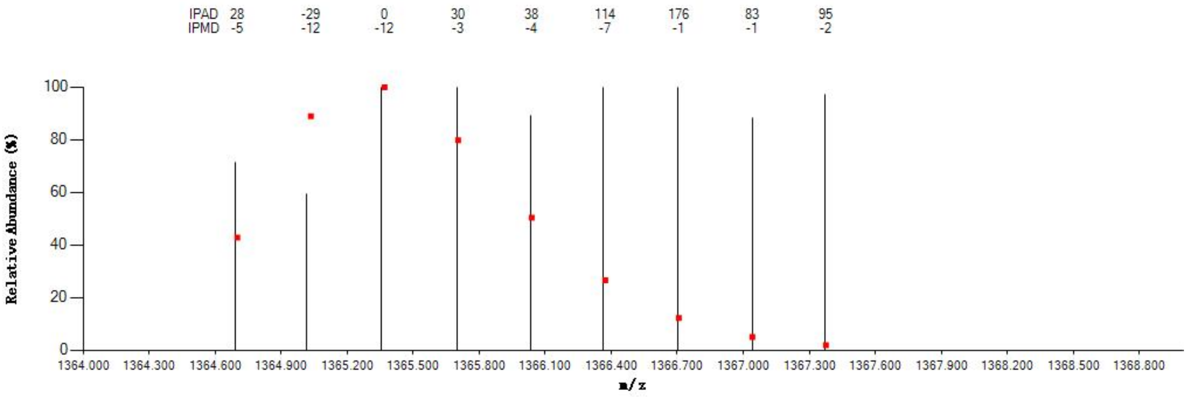


#### Annotated MS/MS Spectrum

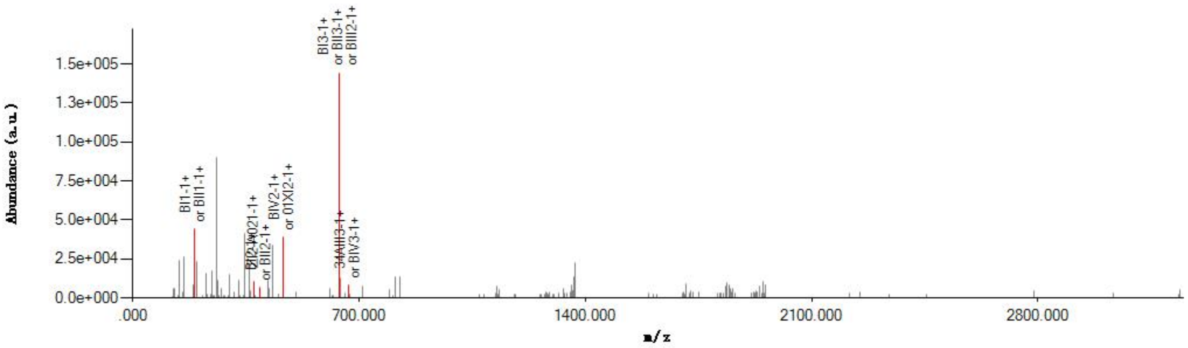


#### Graphical Fragmentation Map

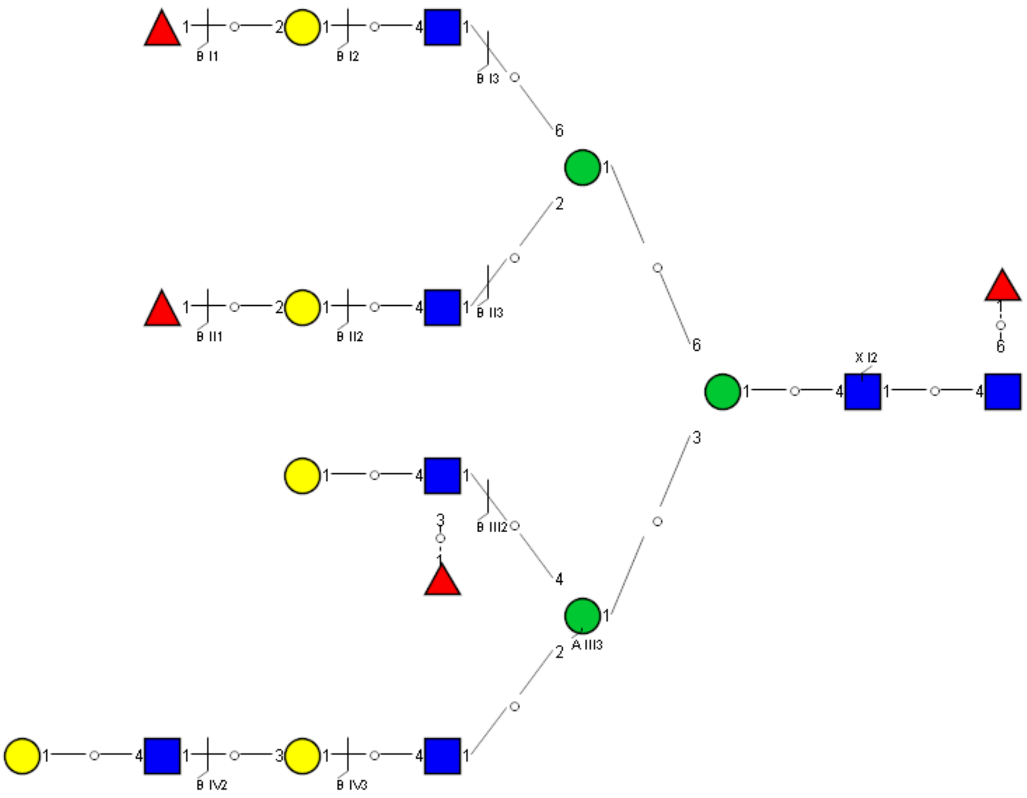

Spectrum Index: 38568(NoBHs:1)

#### Precursor iEF Map

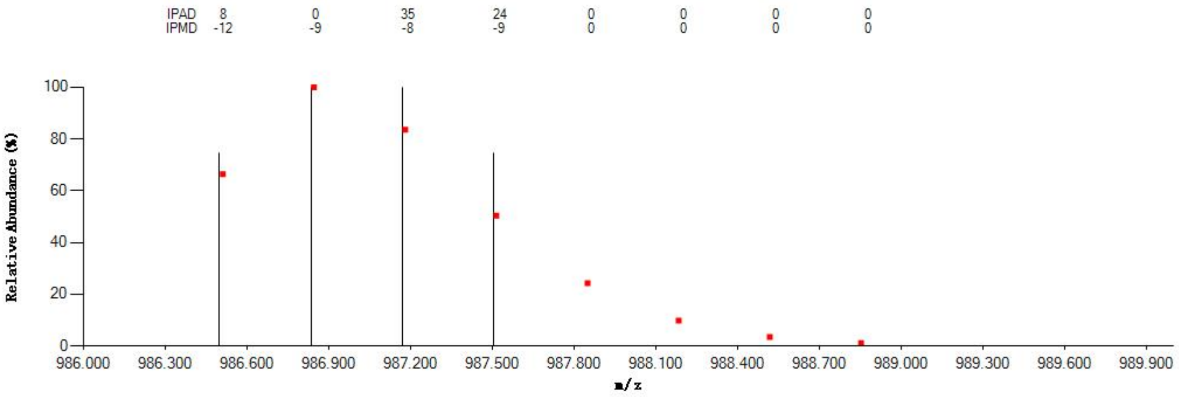


#### Annotated MS/MS Spectrum

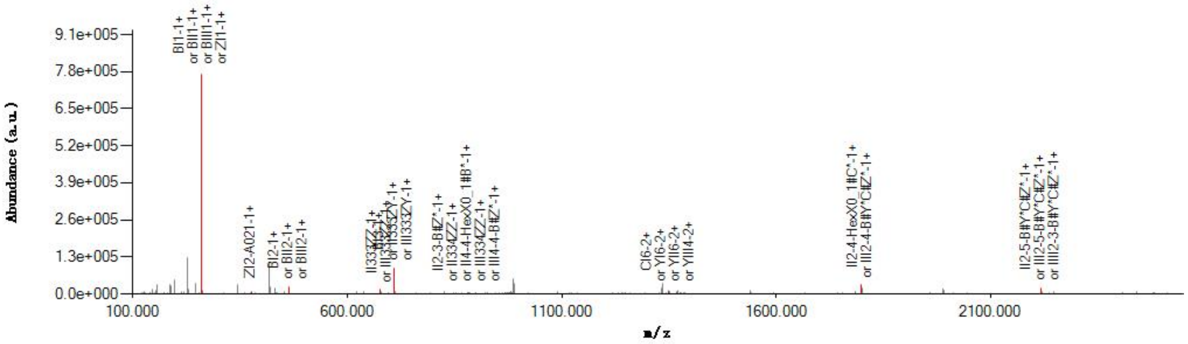


#### Graphical Fragmentation Map

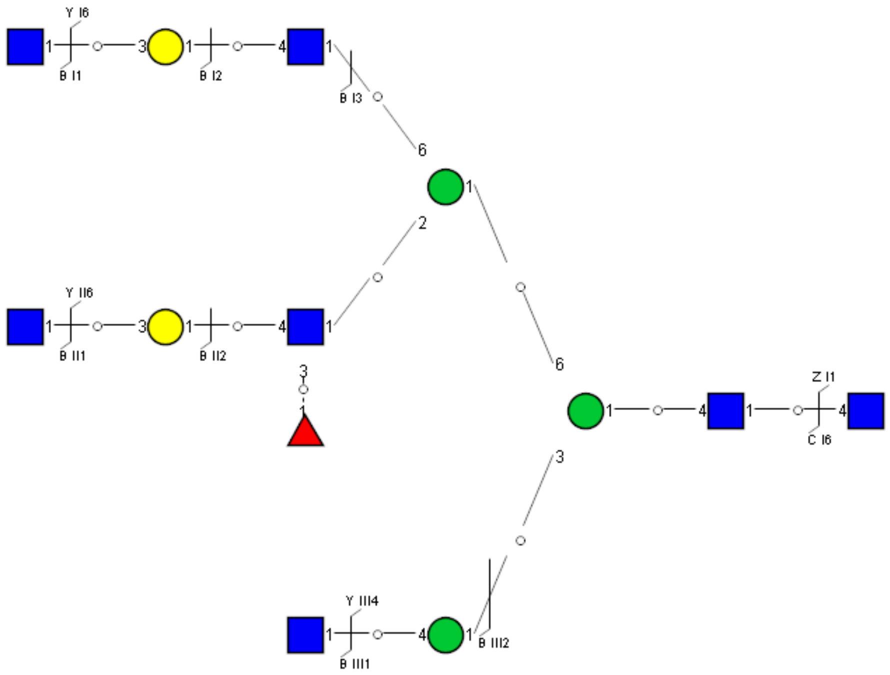

Spectrum Index: 41616(NoBHs:1)

#### Precursor iEF Map

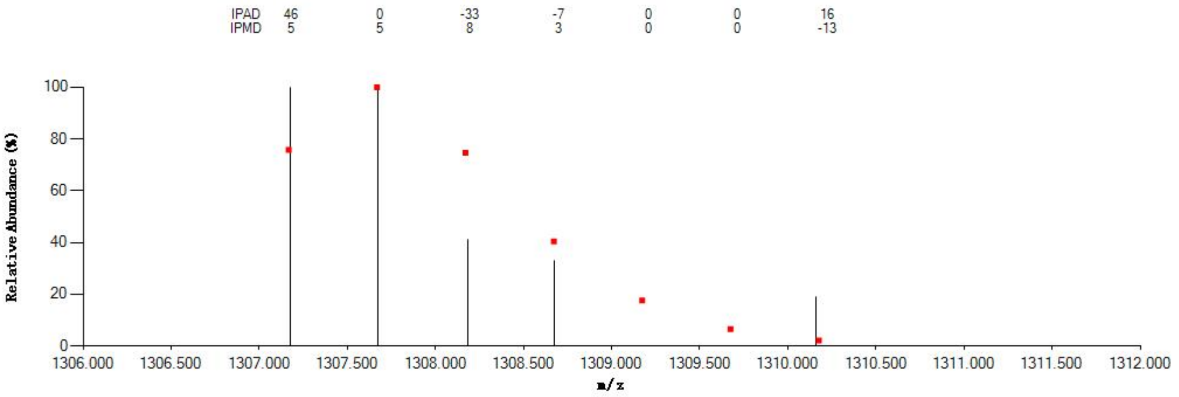


#### Annotated MS/MS Spectrum

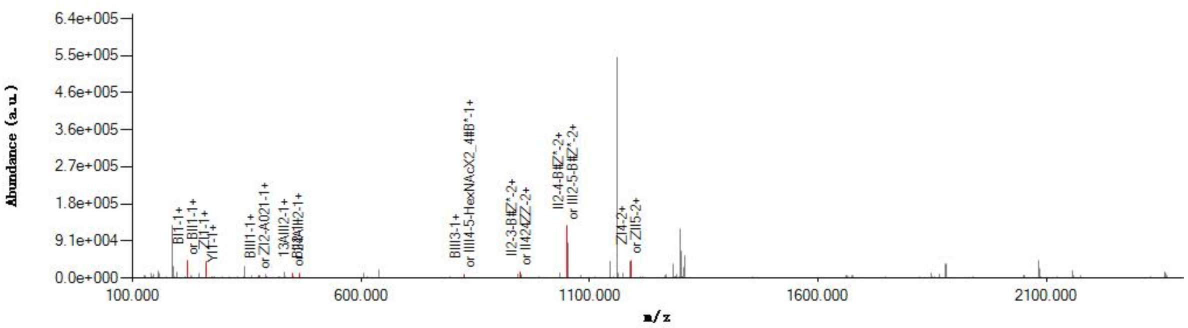


#### Graphical Fragmentation Map

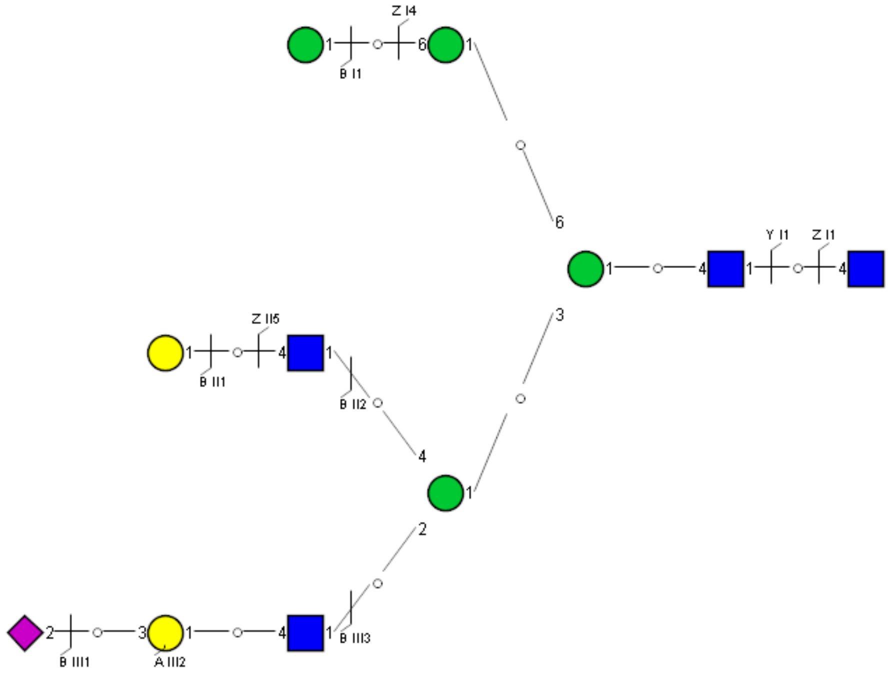

Spectrum Index: 41825(NoBHs:1)

#### Precursor iEF Map

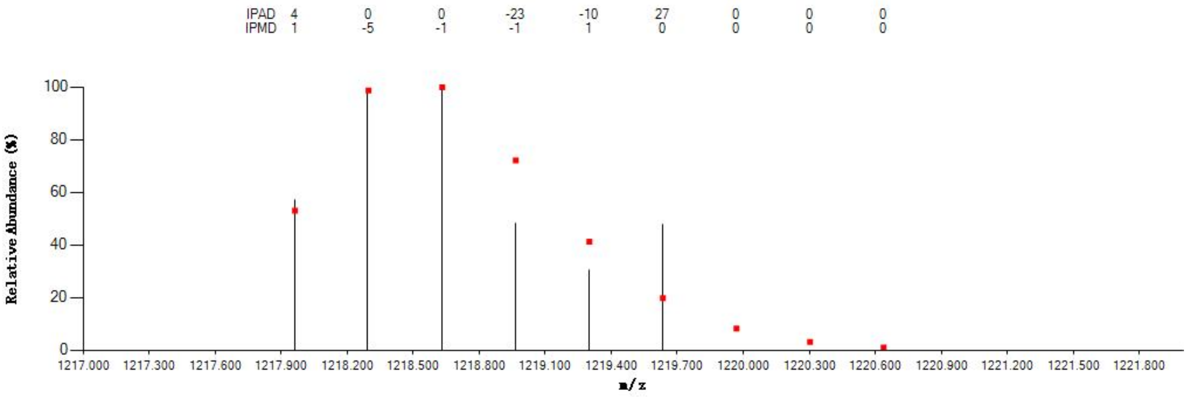


#### Annotated MS/MS Spectrum

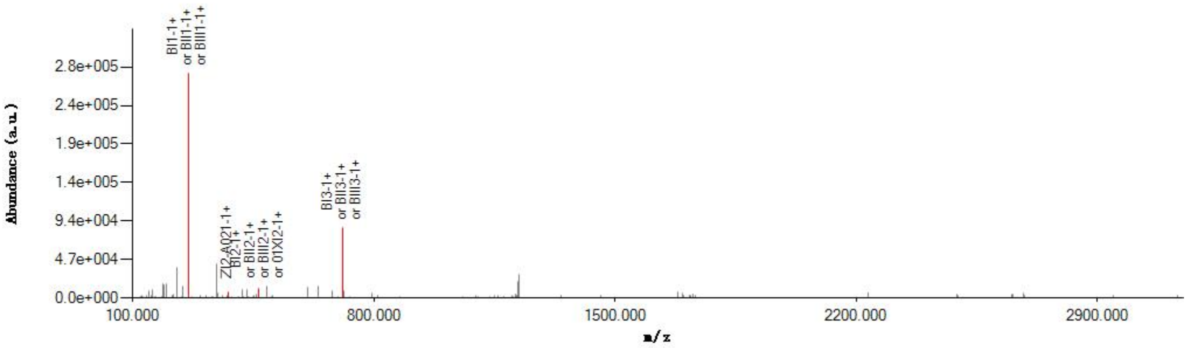


#### Graphical Fragmentation Map

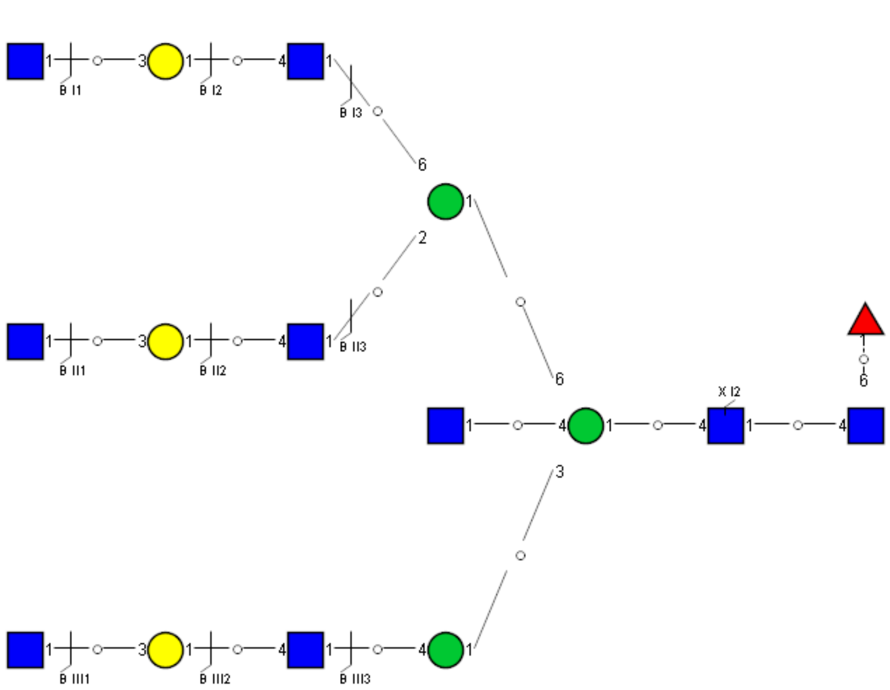

Spectrum Index: 42284(NoBHs:1)

#### Precursor iEF Map

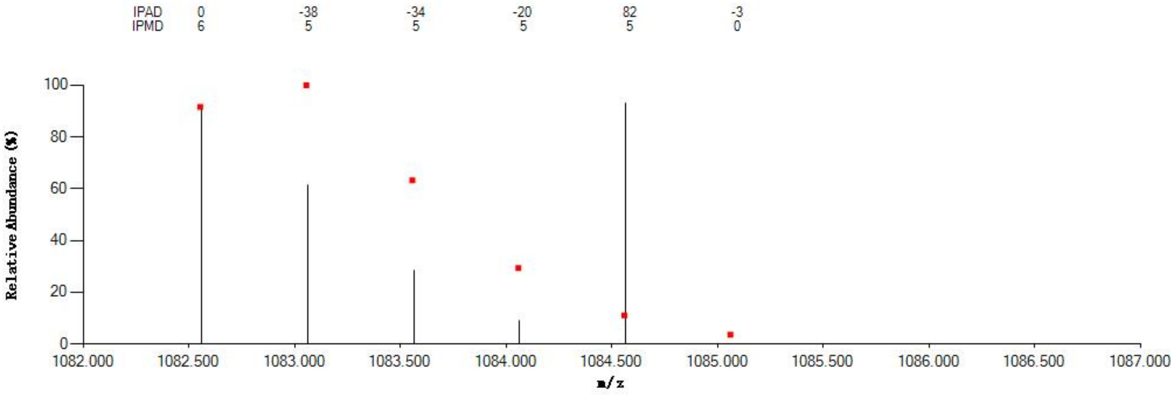


#### Annotated MS/MS Spectrum

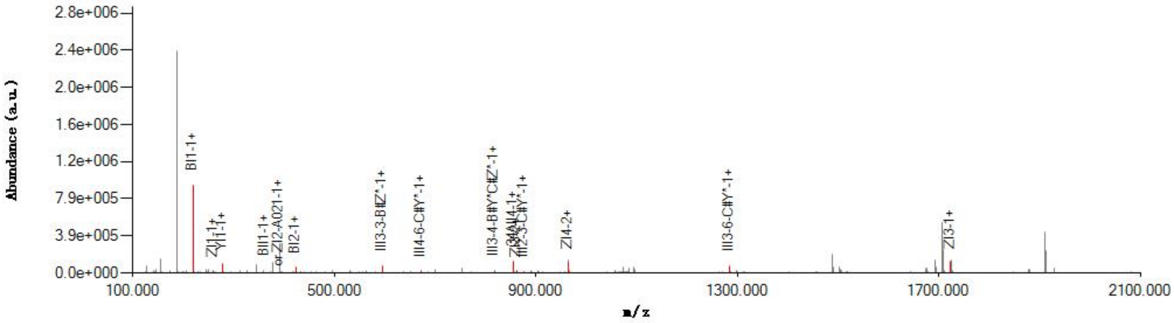


#### Graphical Fragmentation Map

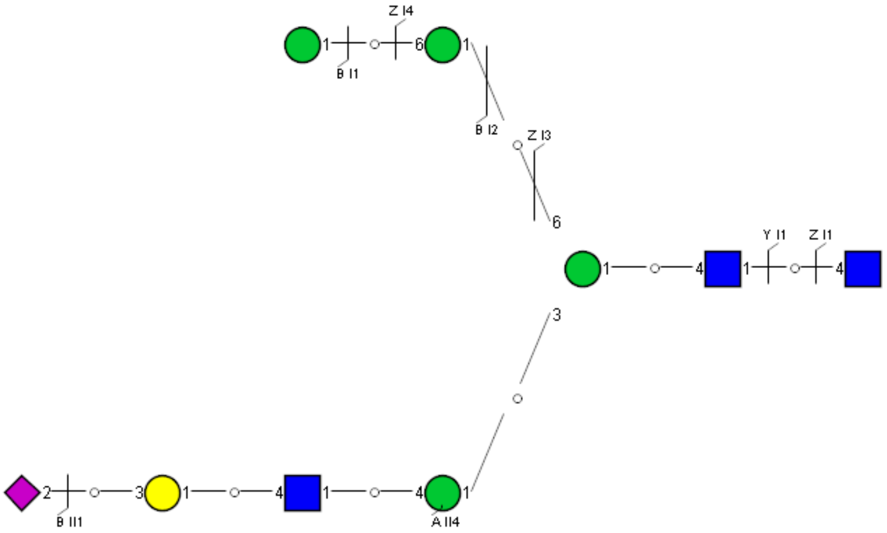

Spectrum Index: 28068(NoBHs:1)

#### Precursor iEF Map

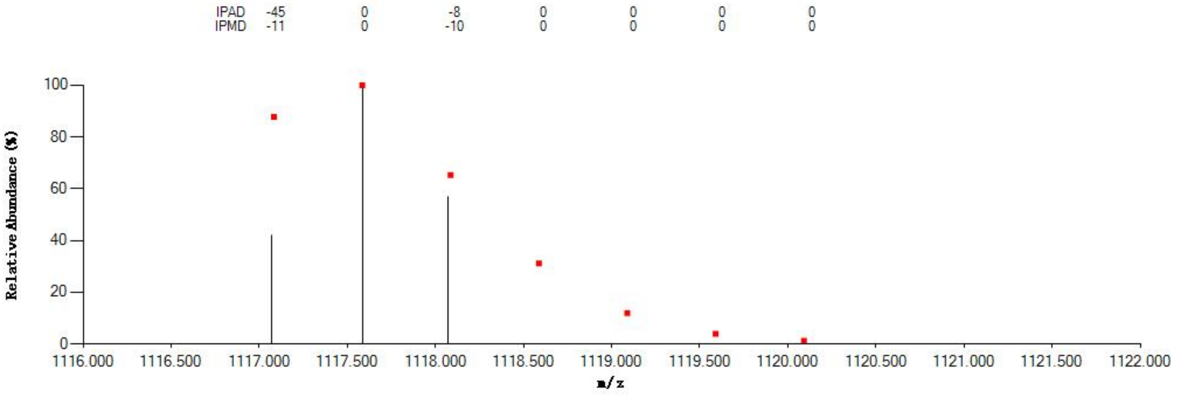

#### Annotated MS/MS Spectrum

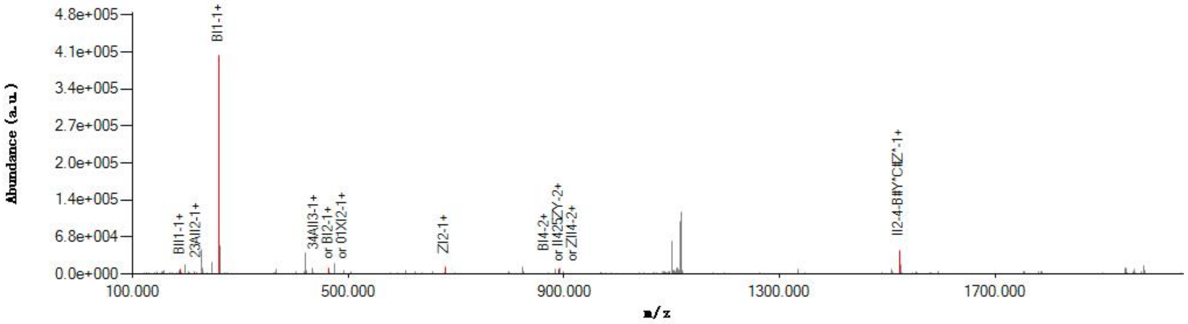

#### Graphical Fragmentation Map

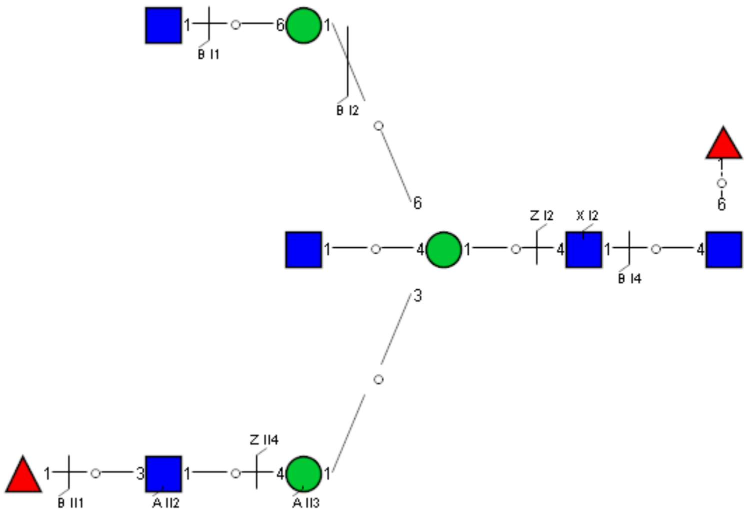

Spectrum Index: 30144(NoBHs:1)

Precursor iEF Map

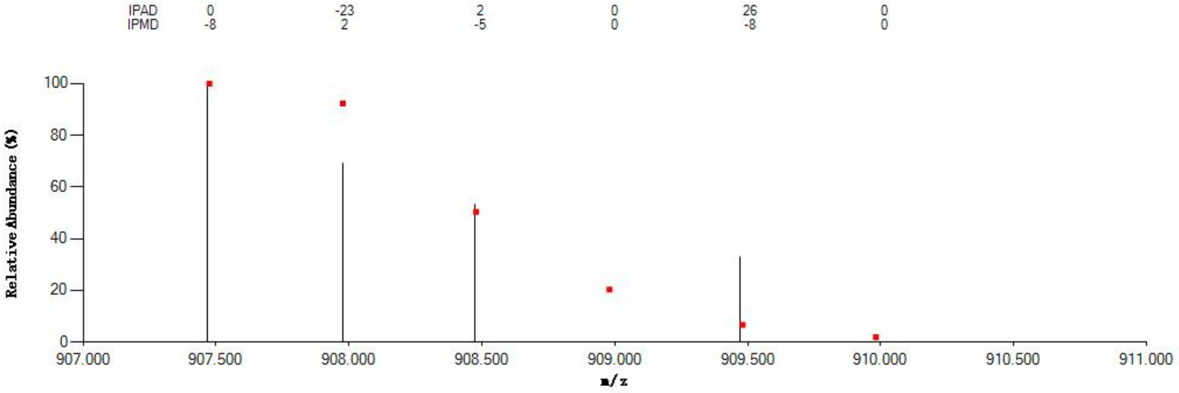


Annotated MS/MS Spectrum

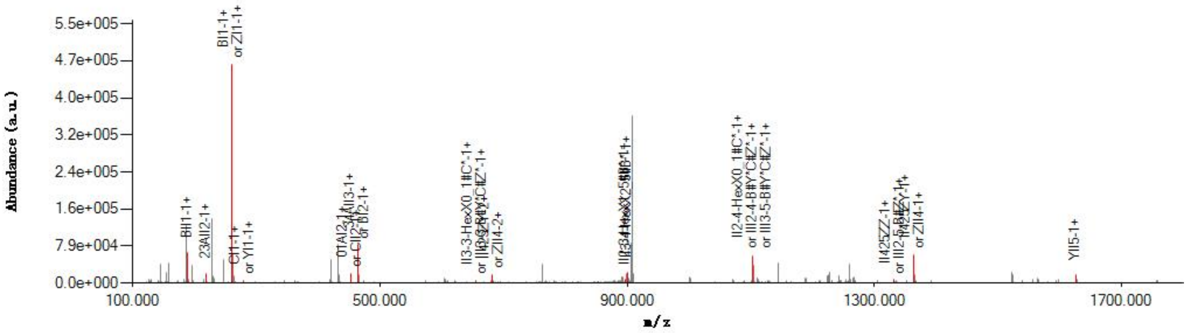


Graphical Fragmentation Map

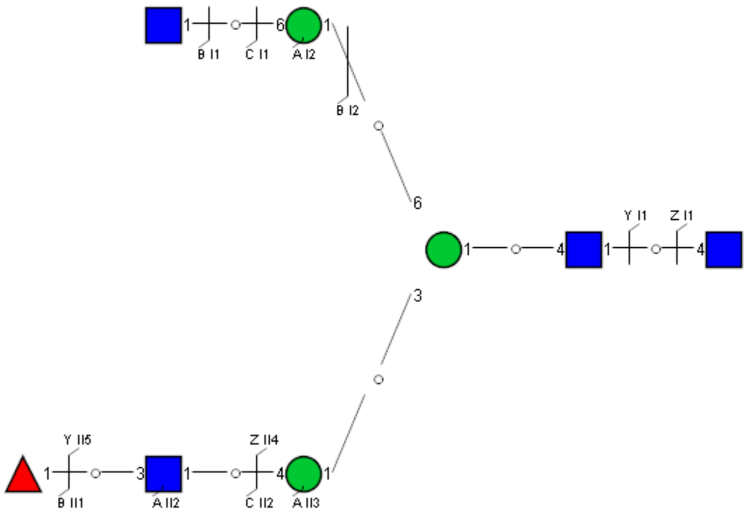

Spectrum Index: 30208(NoBHs:1)

#### Precursor iEF Map

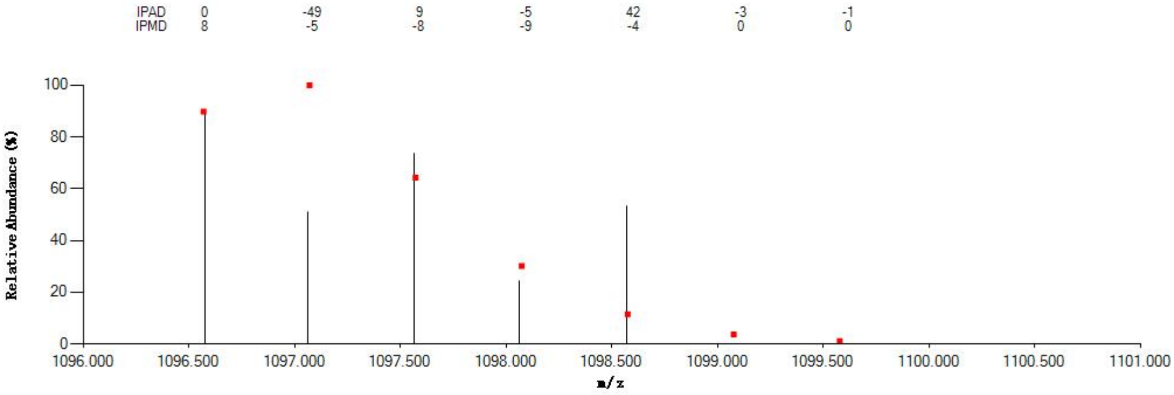


#### Annotated MS/MS Spectrum

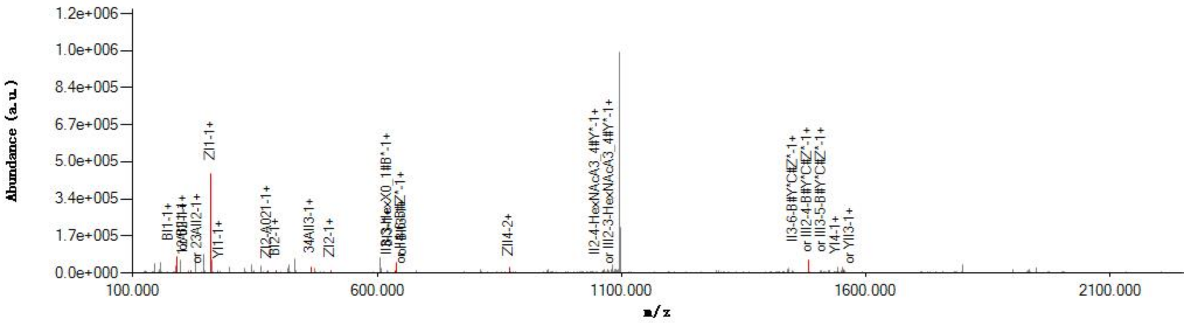


#### Graphical Fragmentation Map

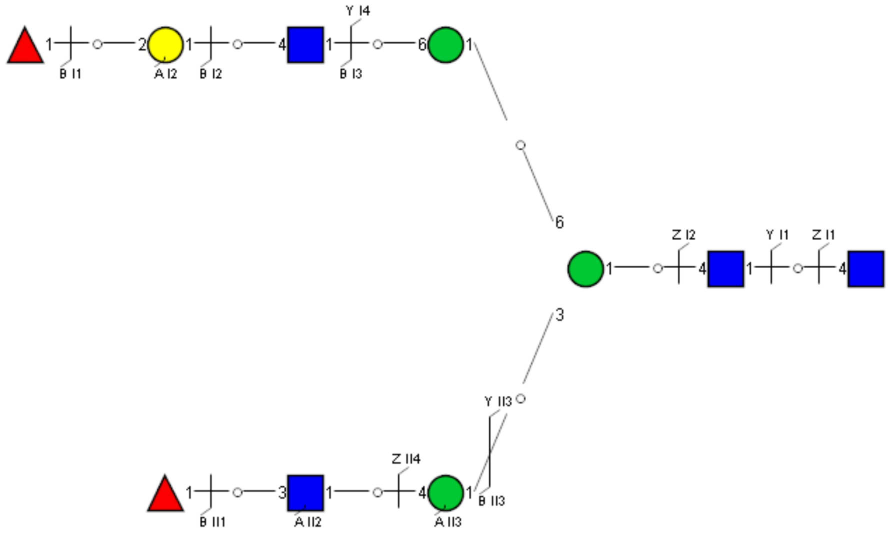

Spectrum Index: 30232(NoBHs:2)

#### Precursor iEF Map

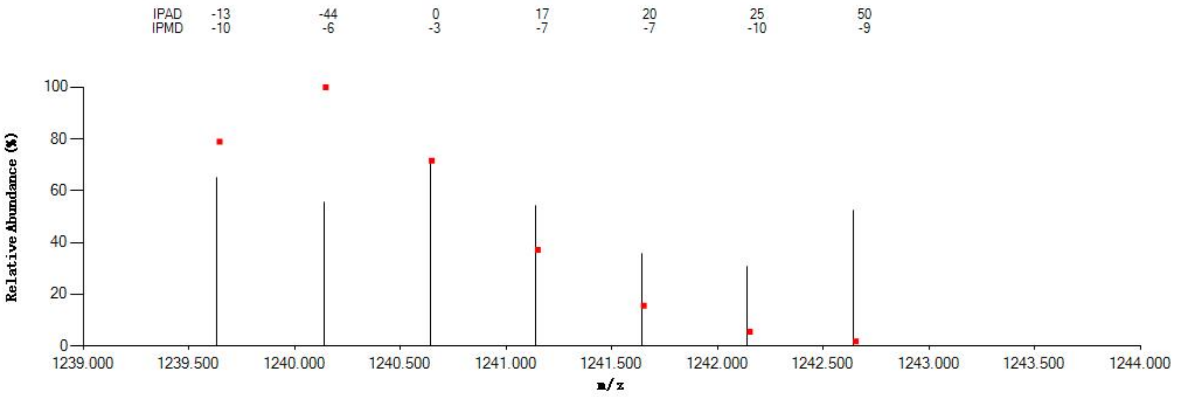


#### Annotated MS/MS Spectrum

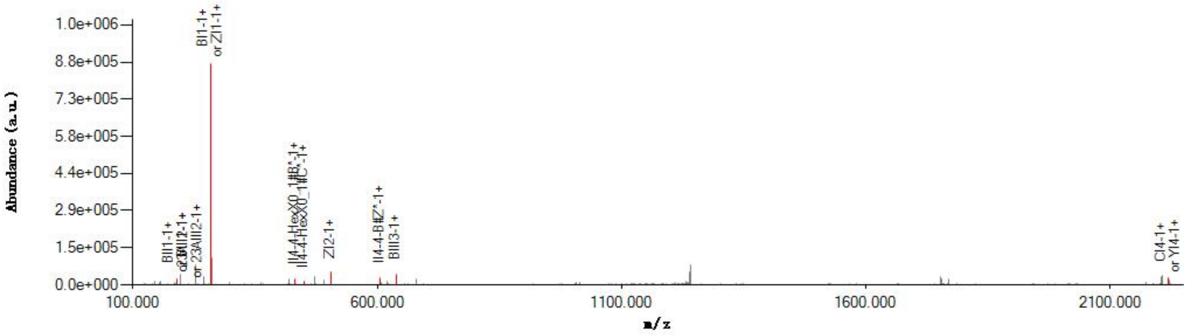


#### Graphical Fragmentation Map

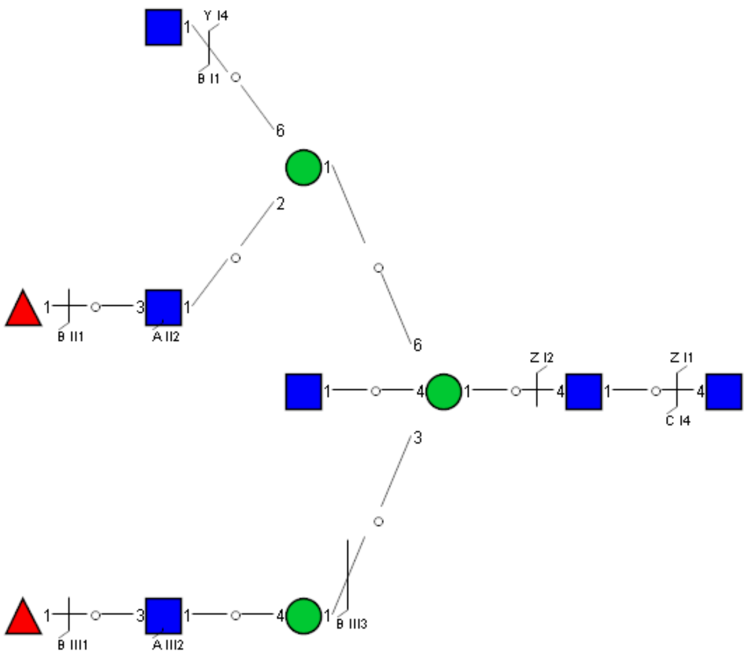

Spectrum Index: 30232(NoBHs:2)

#### Precursor iEF Map

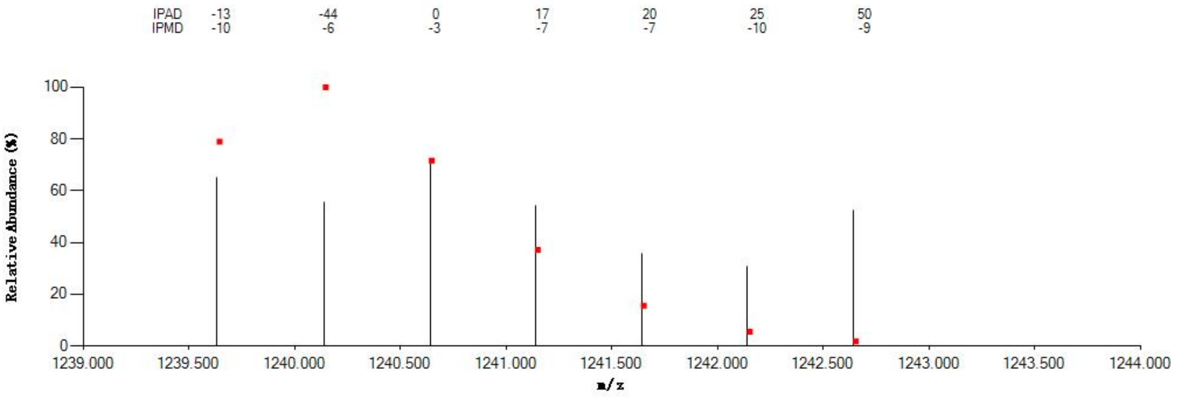


#### Annotated MS/MS Spectrum

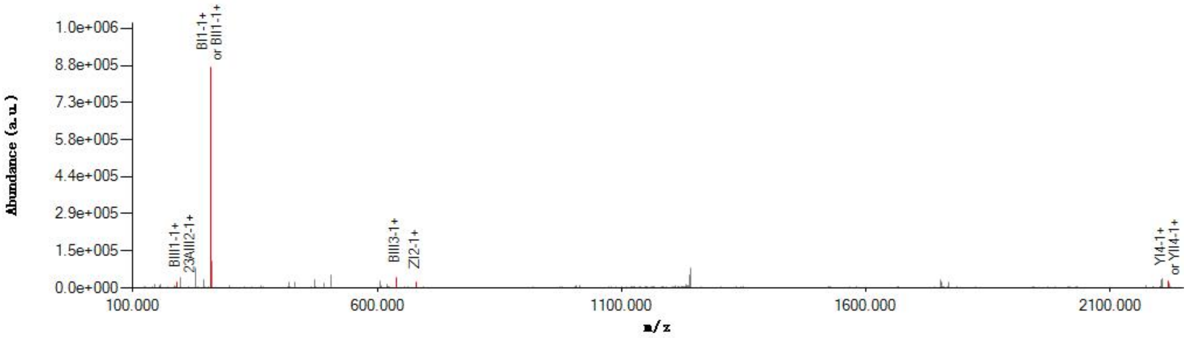


#### Graphical Fragmentation Map

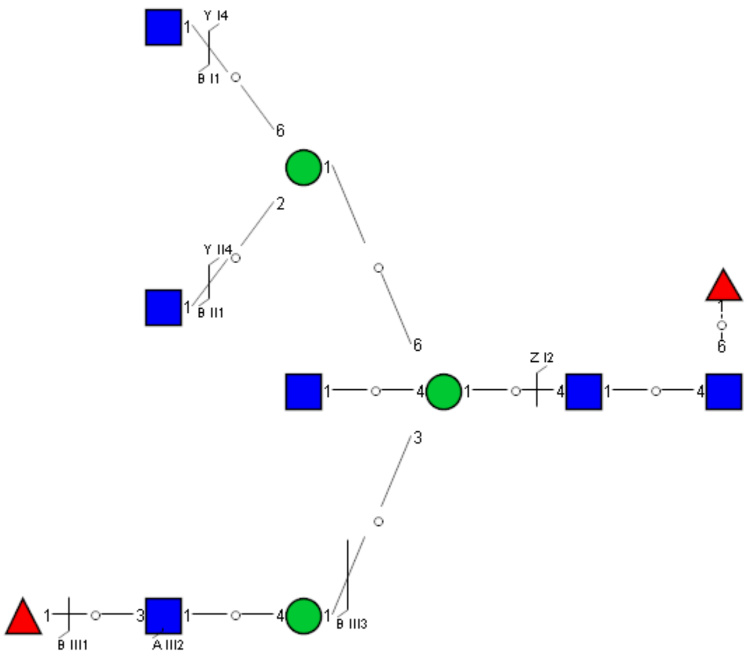

Spectrum Index: 30897(NoBHs:1)

#### Precursor iEF Map

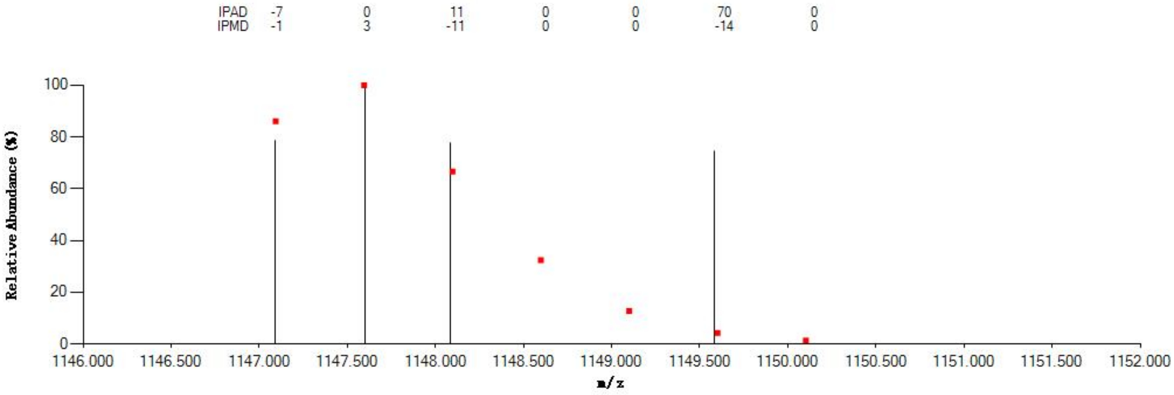


#### Annotated MS/MS Spectrum

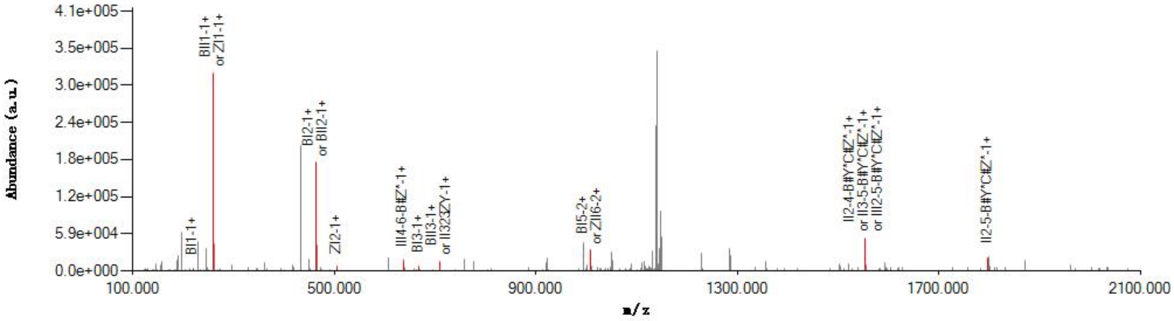


#### Graphical Fragmentation Map

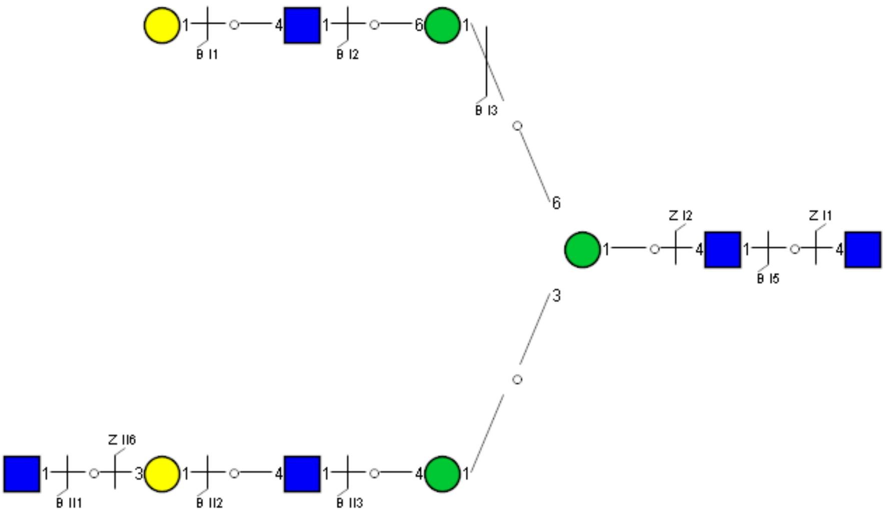

Spectrum Index: 31046(NoBHs:1)

#### Precursor iEF Map

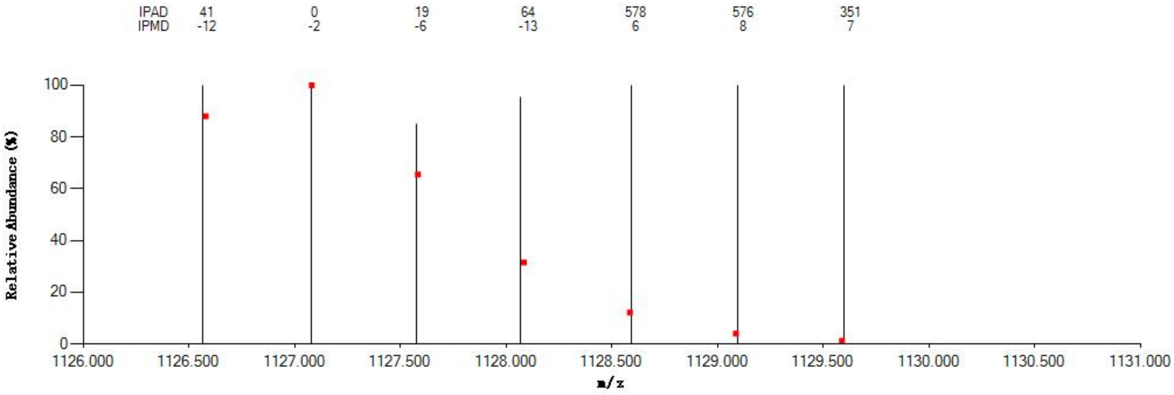


#### Annotated MS/MS Spectrum

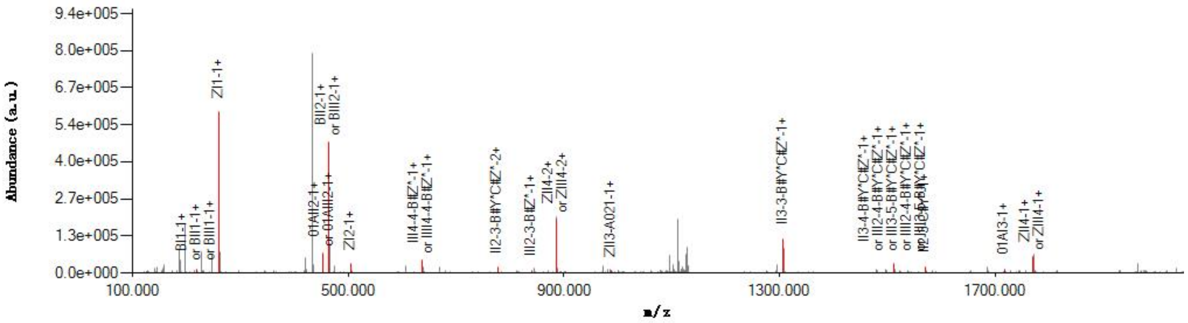


#### Graphical Fragmentation Map

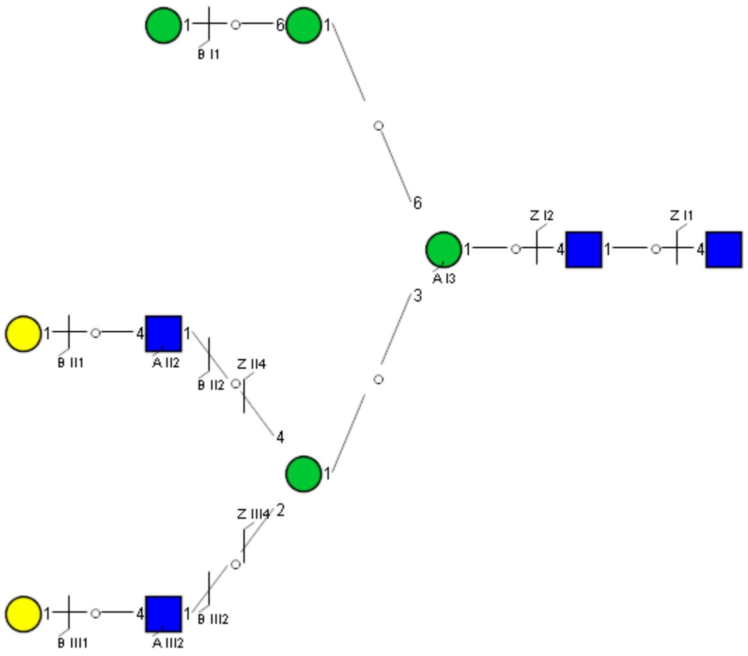

Spectrum Index: 31172(NoBHs:1)

#### Precursor iEF Map

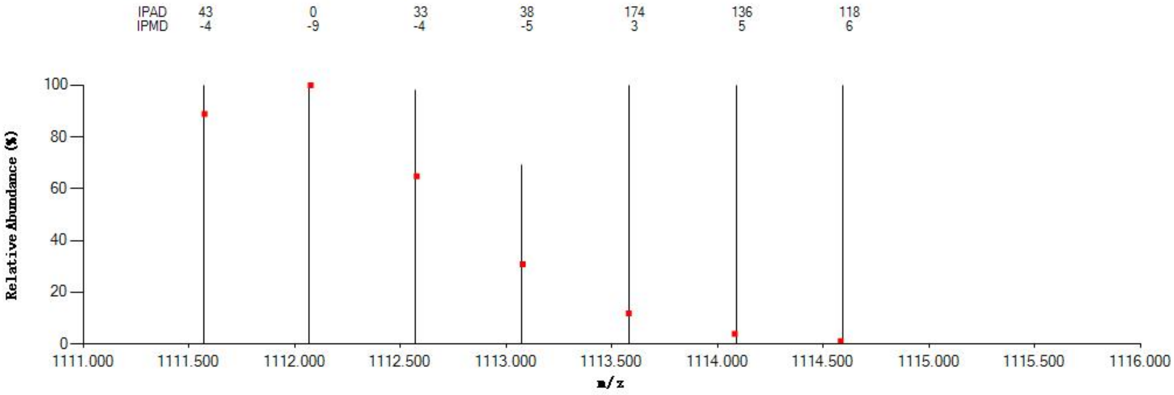


#### Annotated MS/MS Spectrum

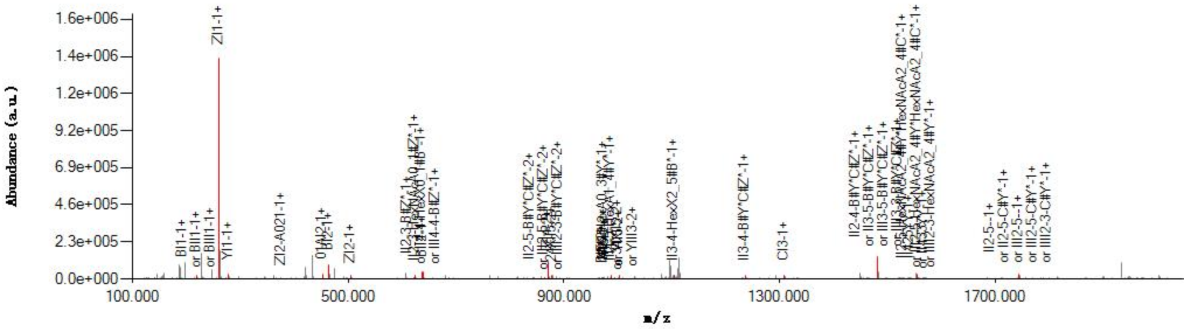


#### Graphical Fragmentation Map

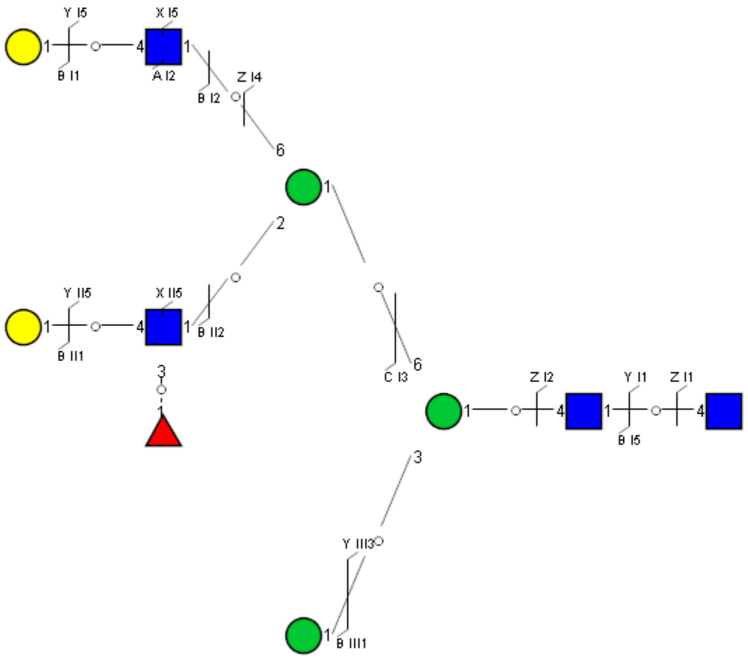

Spectrum Index: 33592(NoBHs:1)

#### Precursor iEF Map

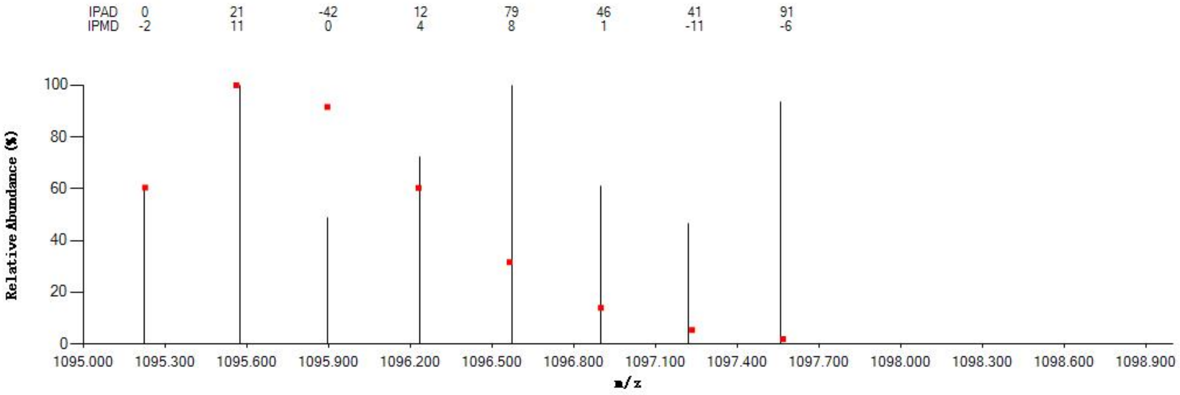


#### Annotated MS/MS Spectrum

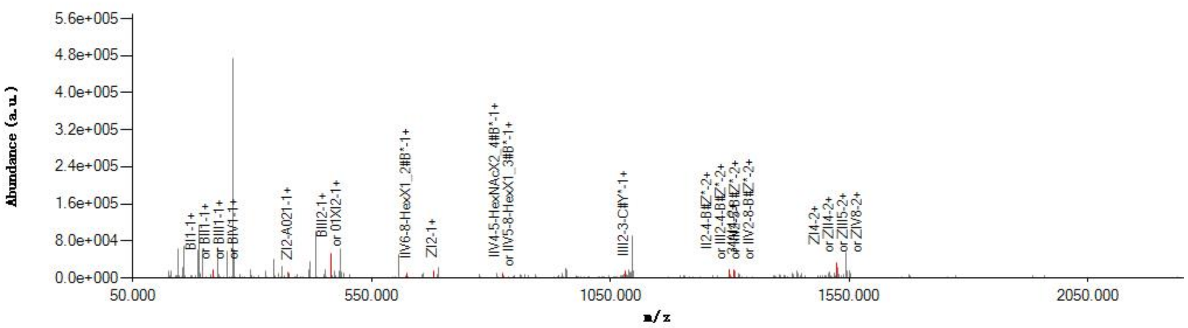


#### Graphical Fragmentation Map

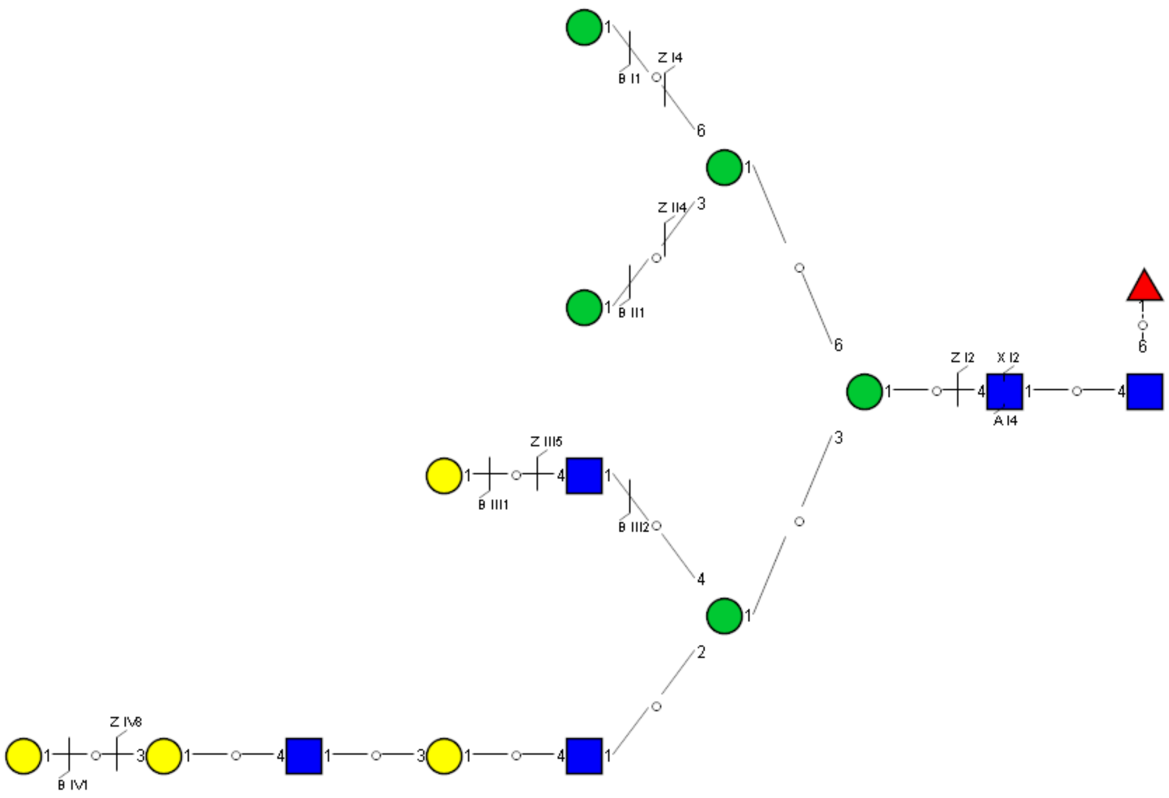

Spectrum Index: 34980(NoBHs:1)

#### Precursor iEF Map

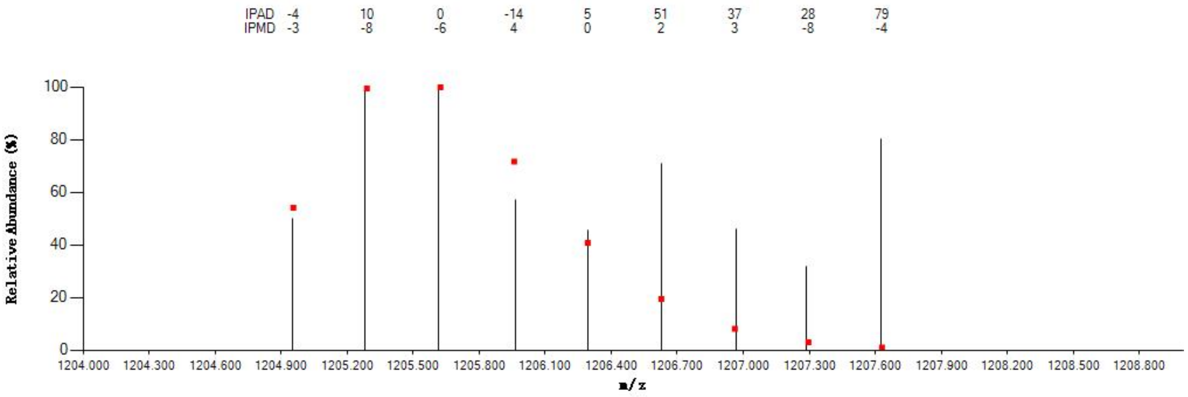


#### Annotated MS/MS Spectrum

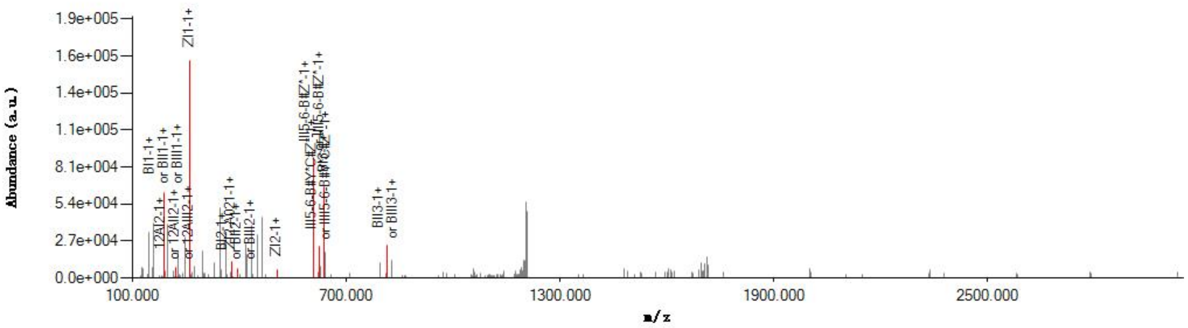


#### Graphical Fragmentation Map

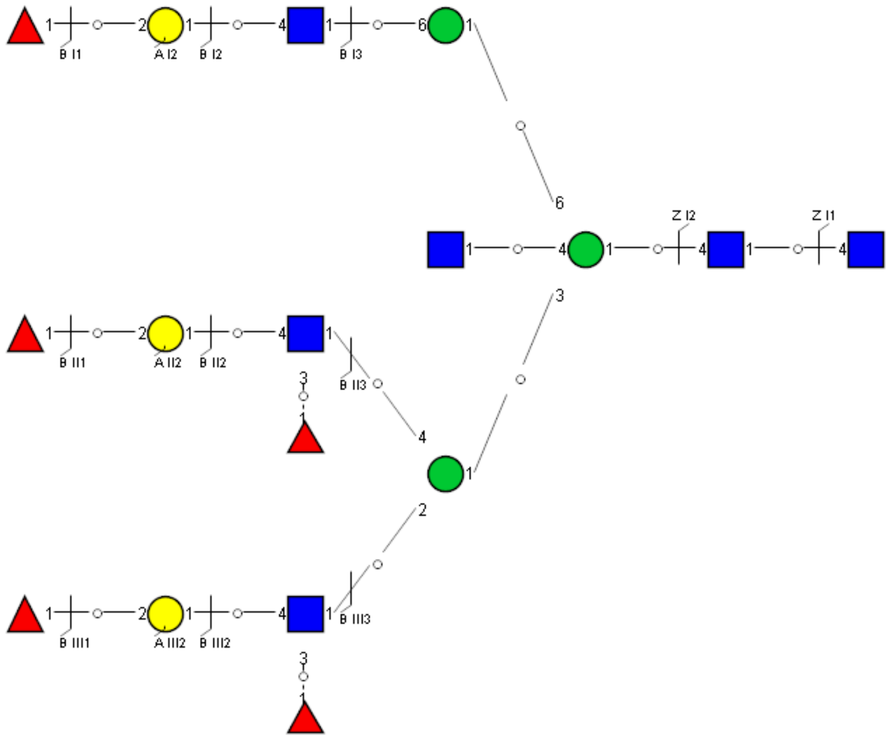

Spectrum Index: 35146(NoBHs:1)

#### Precursor iEF Map

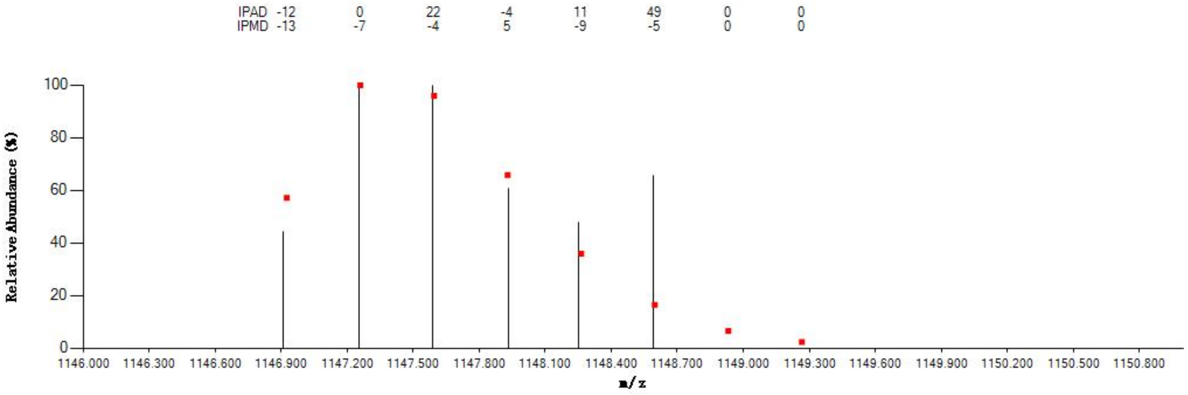


#### Annotated MS/MS Spectrum

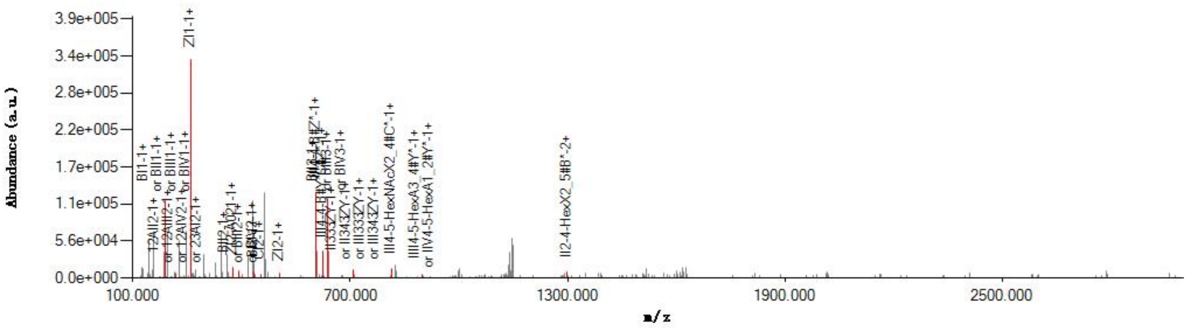


#### Graphical Fragmentation Map

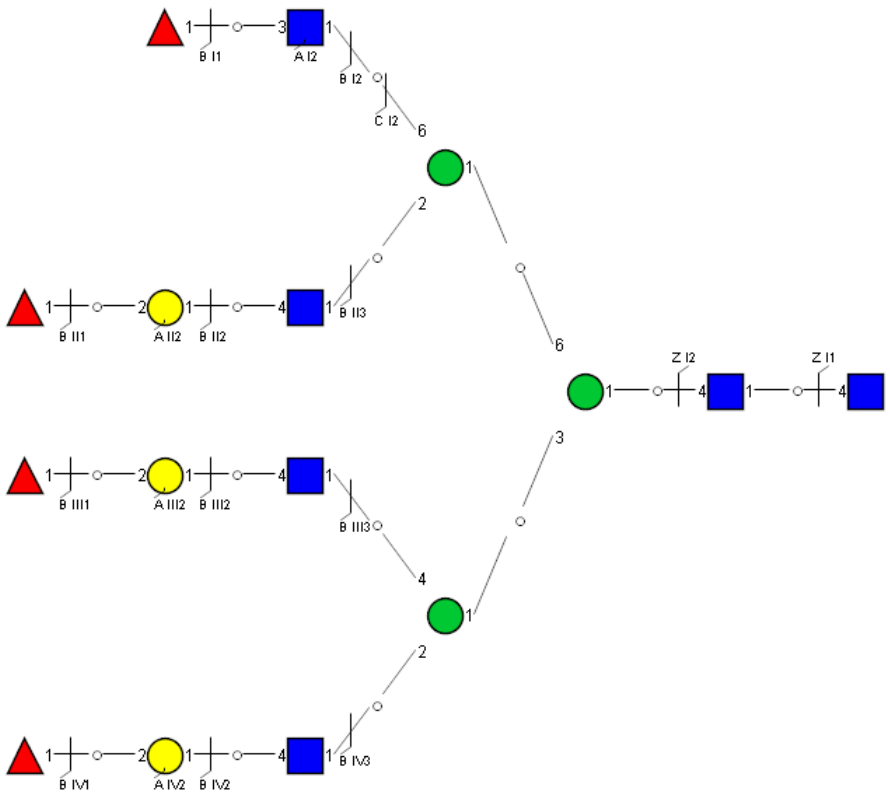

Spectrum Index: 35373(NoBHs:1)

#### Precursor iEF Map

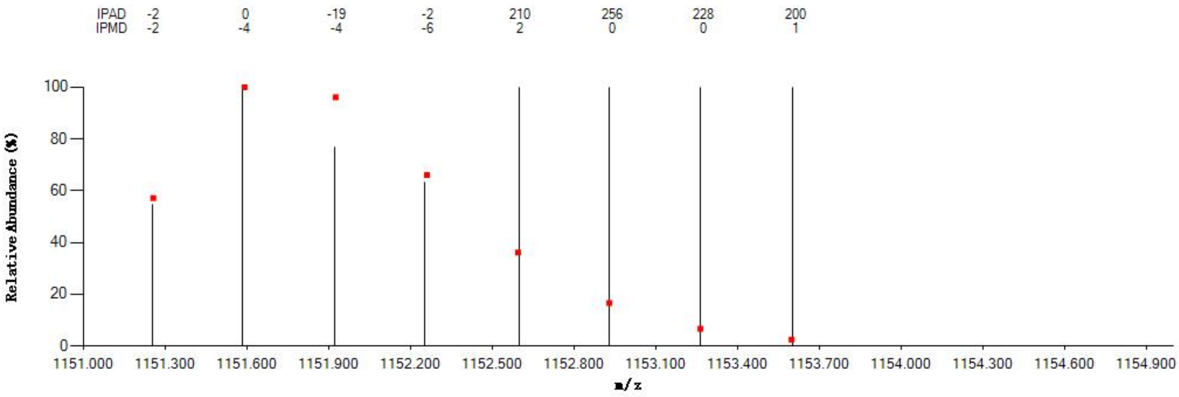

#### Annotated MS/MS Spectrum

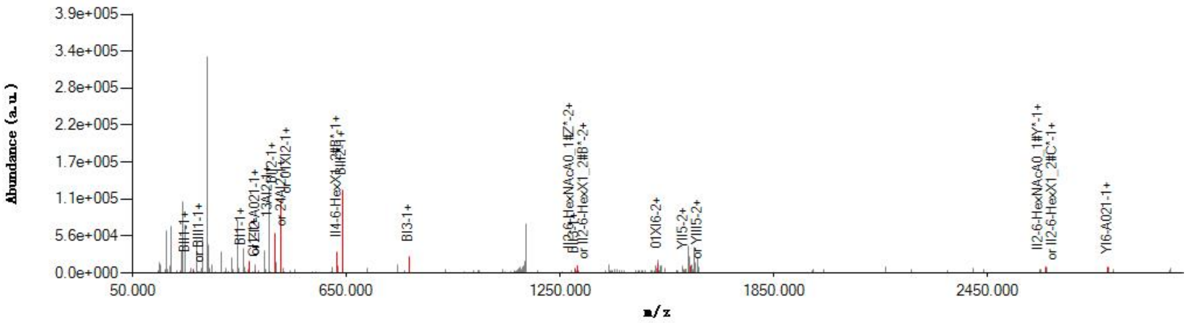

#### Graphical Fragmentation Map

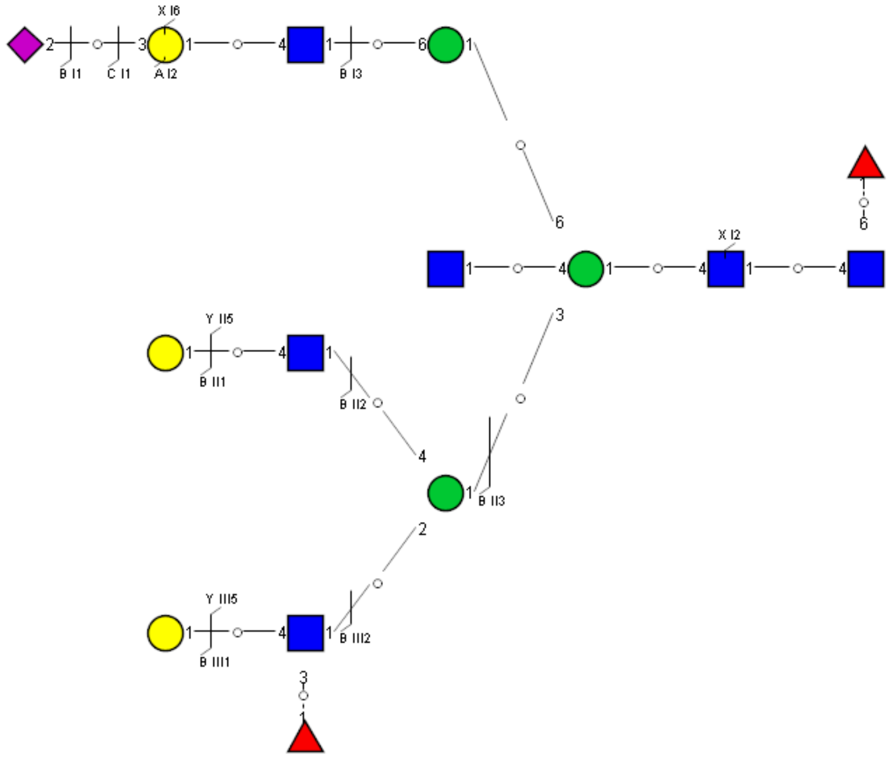

Spectrum Index: 35720(NoBHs:1)

#### Precursor iEF Map

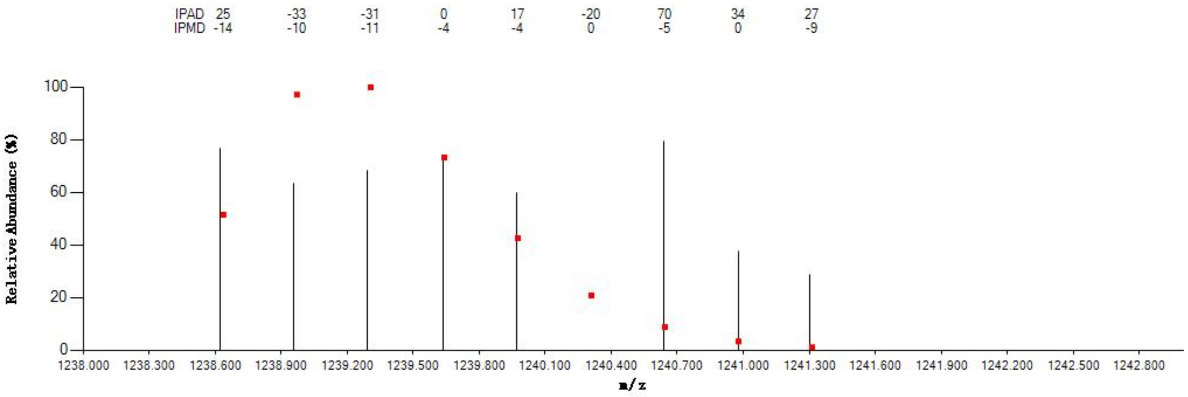


#### Annotated MS/MS Spectrum

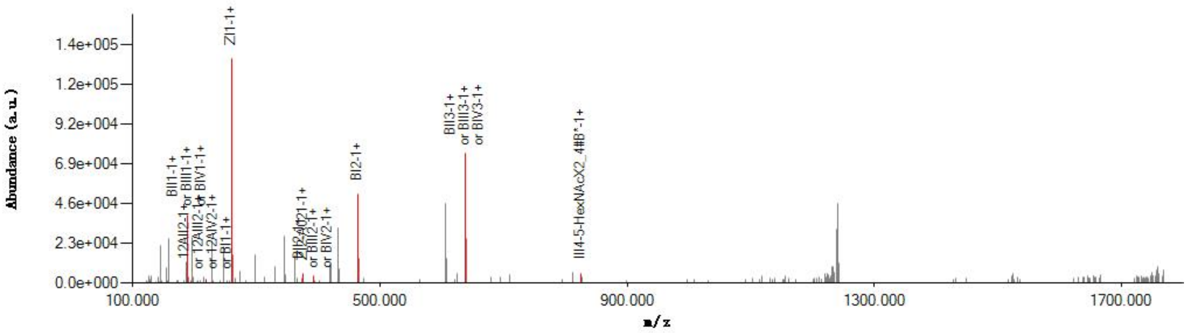


#### Graphical Fragmentation Map

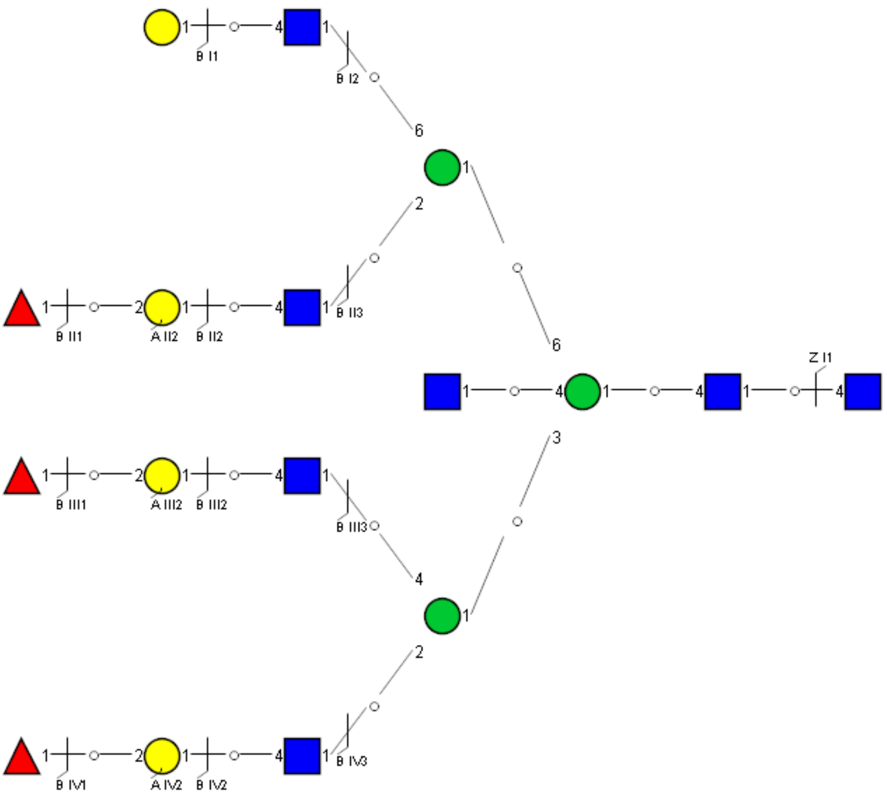

Spectrum Index: 37367(NoBHs:2)

#### Precursor iEF Map

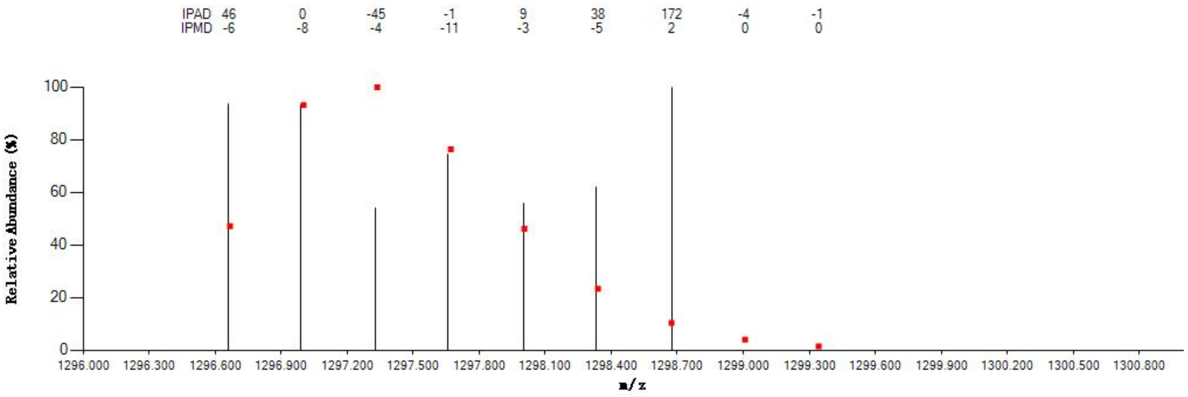


#### Annotated MS/MS Spectrum

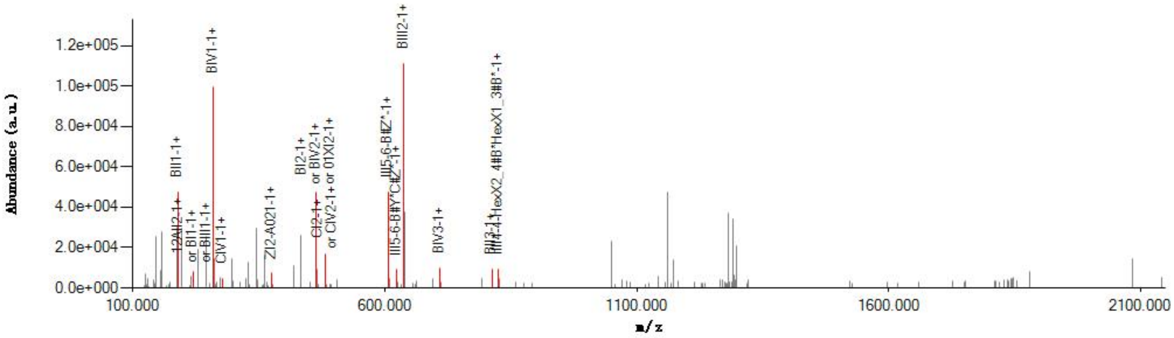


#### Graphical Fragmentation Map

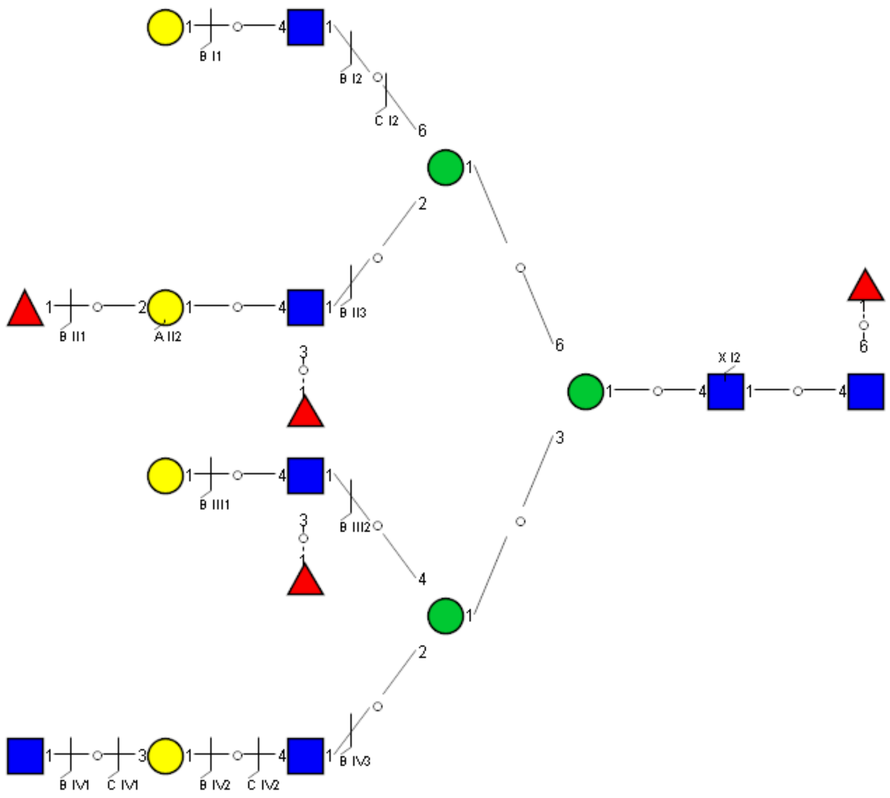

Spectrum Index: 37367(NoBHs:2)

#### Precursor iEF Map

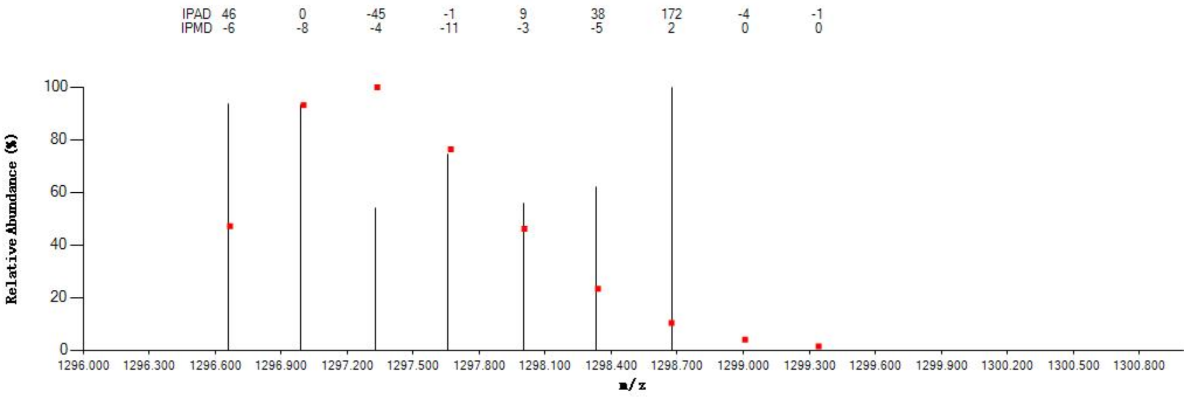


#### Annotated MS/MS Spectrum

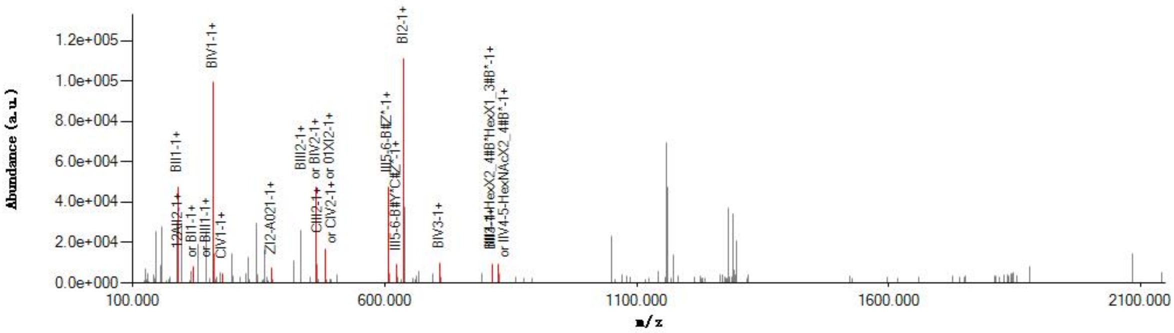


#### Graphical Fragmentation Map

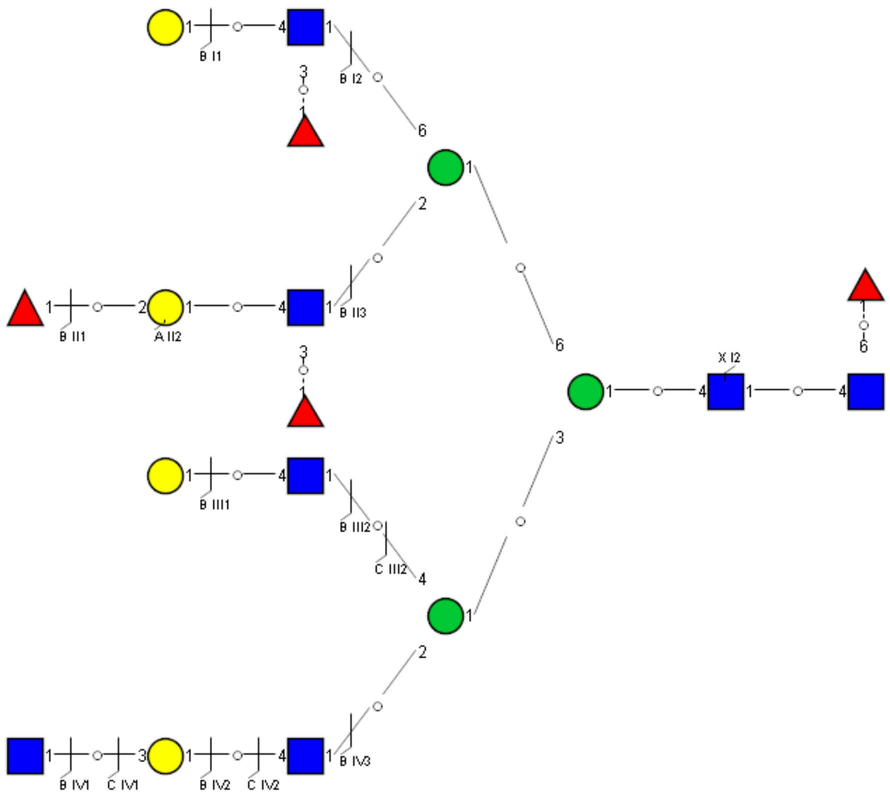

Spectrum Index: 37405(NoBHs:1)

#### Precursor iEF Map

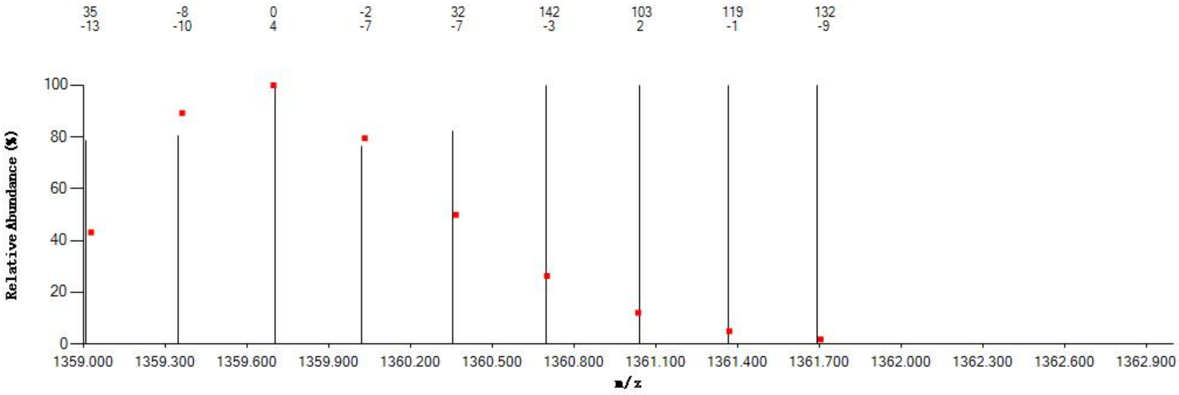


#### Annotated MS/MS Spectrum

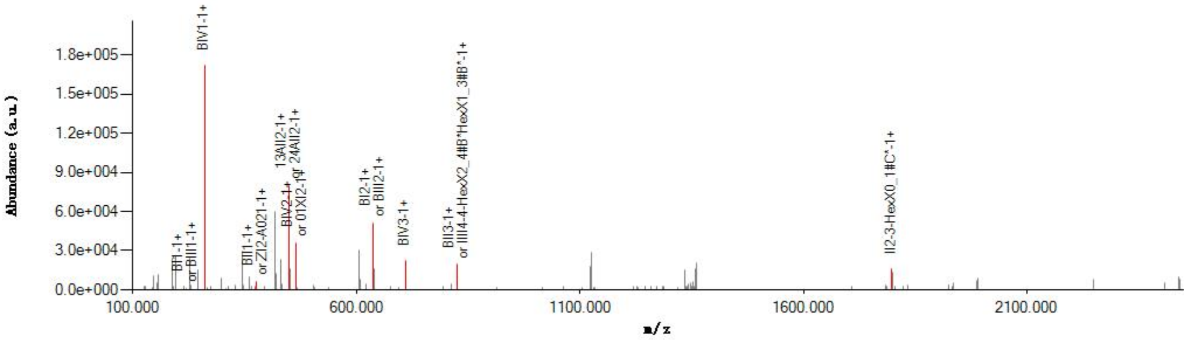


#### Graphical Fragmentation Map

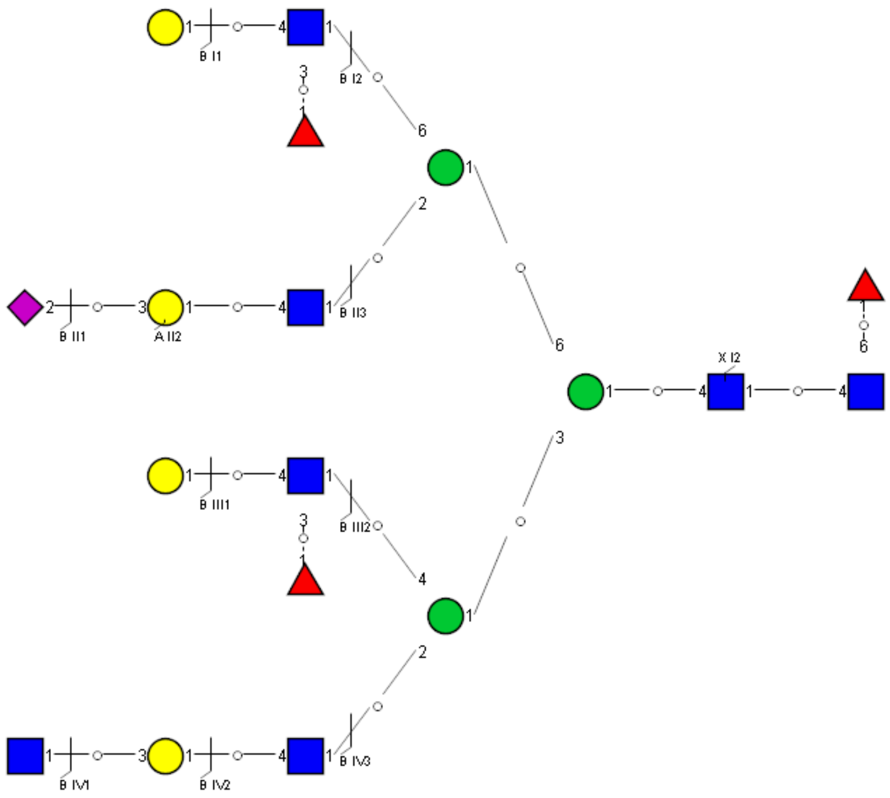

Spectrum Index: 37414(NoBHs:1)

#### Precursor iEF Map

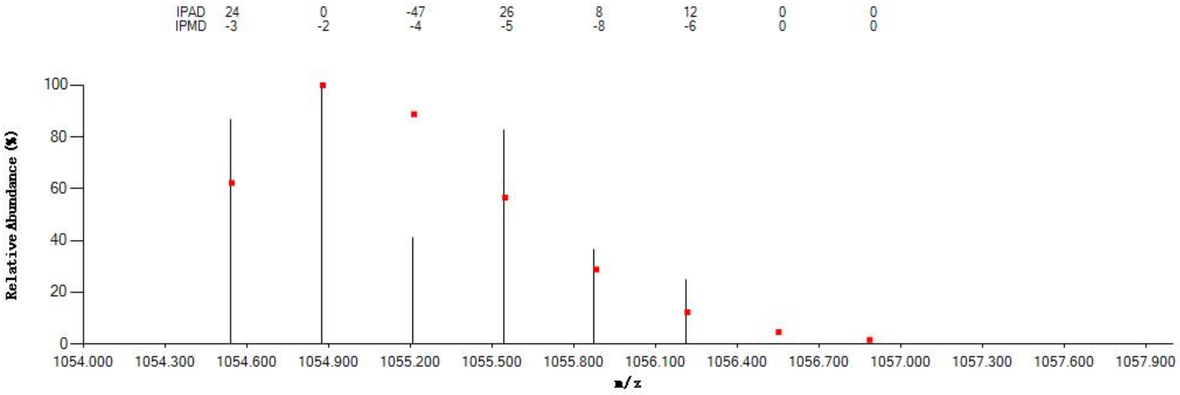


#### Annotated MS/MS Spectrum

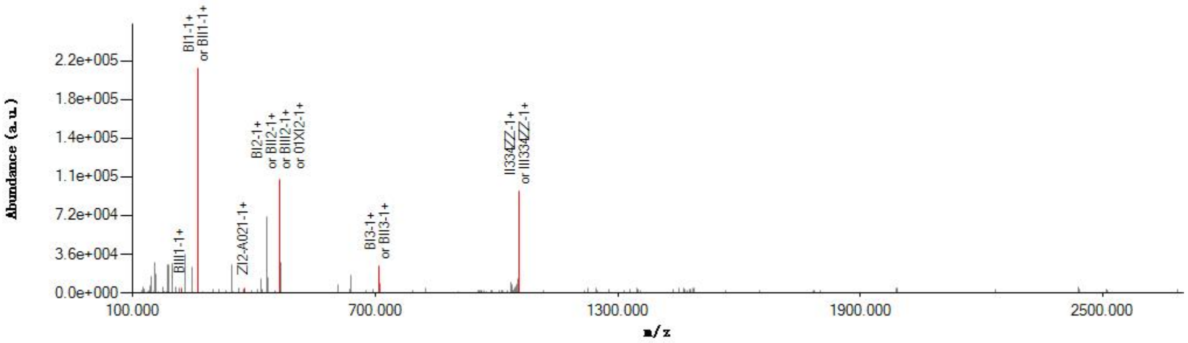


#### Graphical Fragmentation Map

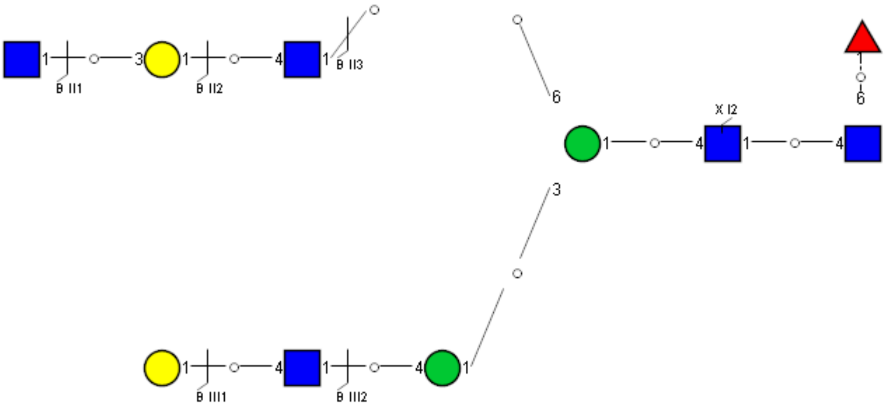
